# Nucleotide sequence analysis reveals the presence of PVY-Tam isolates affecting tamarillo in Colombia

**DOI:** 10.1101/2025.07.18.664922

**Authors:** Carolina Martínez-Moncayo, Tulio César Burbano-Lagos, Clara Ontañón, Inmaculada Ferriol, Juan José López-Moya, Mireia Uranga

## Abstract

Tamarillo or tree tomato (Solanum betaceum Cav.) is a fruit tree species of Andean origin with cultural and economic relevance in Colombia. However, the high incidence of complex viral diseases termed “virosis” in all tamarillo-growing regions of the country leads to huge production losses and seriously threatens its cultivation. The lack of effective treatments implies eradication as the only alternative in severe cases. In this work, we characterized the virome of eight tamarillo-growing locations across the Department of Nariño (Colombia). By in-depth sequence analysis of RNA libraries, we confirmed the presence of up to four different virus species belonging to the genera Torradovirus, Potyvirus and Polerovirus in symptomatic tamarillo plants. These results represent the first report of torradovirus infection in tamarillo. Additionally, we identified a novel isolate of potato virus Y-Tamarillo (PVY-Tam) in Nariño that could have originated in South America by a recent divergence of the PVY^N^ lineage. We propose that length variability in the P3N-PIPO protein, which in the case of PVY-Tam contains two premature stop codons not identified in other PVY isolates, might be involved with host-specific adaptations. Our findings broaden the knowledge of tamarillo virosis in the Andean region, and overall, worldwide, thus offering new possibilities for developing effective diagnostic and control strategies.

## INTRODUCTION

Tamarillo, or tree tomato (*Solanum betaceum Cav.*), is a fruit tree of Andean origin with economic relevance in Colombia and Ecuador, where it is cultivated for the fresh fruit market and food processing industry (Ramírez & Kallarackal, 2019). The fruit is egg-shaped with purple-red to golden-yellow skin, yellow-orange firm flesh and jelly with small seeds. Regarding its nutritional properties, it is relatively low in carbohydrates and a good source of dietary fibre, vitamins A, B_6_, C, and E, minerals (mainly potassium, phosphorus and magnesium) and a variety of functional bioactives with antioxidant activity like phenolics, anthocyanins, and carotenoids (Diep et al., 2020; Skinner & Hunter, 2013; Vasco et al., 2009). Although the exact origin of tamarillo is unknown, it is possibly native to South American countries since wild relatives are found in Colombia, Peru, Ecuador, Bolivia, Brazil and Chile (Morton, 2013; Duarte & Paull, 2015). In the late 19th century, tamarillo was globally introduced to Australia, New Zealand, Southeast Asia, and some countries in Europe and Africa (Bohs, 1995; Morton, 2013; Prohens & Nuez, 2001). Nowadays, it is commercially grown only in Colombia, Peru, Ecuador, New Zealand, and Australia (Diep et al., 2022; Ramírez & Kallarackal, 2019).

In Colombia, over 140,228 tons of tamarillo are produced annually in a total cultivated area of about 9,223 hectares distributed across 18 provinces, representing a relevant amount of the national fruit production (Buitrago, 2013). Tamarillo cultivation in Colombia is threatened by the high incidence of complex viral diseases known as “tamarillo virosis” (M. M. Jaramillo et al., 2012). Symptoms include mosaics, vein outgrowth, leaf deformation, yellowing, ringspots and severe reductions in fruit production and plant longevity. This disease was initially reported in 1991 in the department of Antioquia and has since rapidly spread to all tamarillo-growing regions of the country, including Nariño (Betancourt et al., 2003; Rodríguez, 2009), Cundinamarca (Álvarez, 2010; Cuspoca, 2007), and Antioquia (Ayala et al., 2010; Martínez et al., 2010), causing significant production losses ranging 50-80% annually. The lack of effective treatments implies eradication as the only alternative in severe virosis cases, which significantly reduces the crop’s lifespan and increases the economic costs by the need to replace orchards with healthy plants.

Several studies have reported that virosis in tamarillo can be caused by combinations of viruses from different genera, including members of *Alfamovirus* (alfalfa mosaic virus, AMV), *Cucumovirus* (cucumber mosaic virus, CMV), *Nepovirus* (tomato ringspot virus, ToRSV), *Polerovirus* (potato leafroll virus, PLRV), *Potexvirus* (potato aucuba mosaic virus, PAMV), *Potyvirus* (potato virus Y, PVY; tamarillo leaf malformation virus, TLMV; potato virus A, PVA), *Tobamovirus* (tobacco mosaic virus, TMV; tomato mosaic virus, ToMV), and *Tospovirus* (tomato spotted wilt virus, TSWV) (Eagles et al., 1994; Espinoza et al., 2017; Green et al., 2020; Gutiérrez et al., 2015a; Jaramillo et al., 2011; Vizuete et al., 2001). Some reports have identified up to eight viruses in a single infected plant, of which at least two were potyviruses able to establish sinergestic interactions with unrelated viruses (Gil et al., 2009; Gutiérrez et al., 2015b; Jaramillo et al., 2012).

In this work, we aimed to characterize the virome of symptomatic tamarillo plants cultivated across the Department of Nariño, Colombia. First, we generated RNA libraries from symptomatic plants collected in eight tamarillo-growing locations and submitted them to Next Generation Sequencing (NGS). Next, we studied the virus diversity of each sample using an innovative, web-based diagnostic tool named Genome Detective, confirming the presence of several viruses belonging to the genera *Torradovirus, Potyvirus*, and *Polerovirus*. Additionally, in-depth sequence analysis of the viral sequences enabled the identification of a novel isolate of potato virus Y-Tamarillo (PVY-Tam) that differs from those previously reported in the Andean region. We validated the presence of PVY-Tam in five of the eight sites using two virus-specific primer pairs. Phylogenetic analysis indicated that PVY-Tam originated in South America by a recent divergence of the PVY^N^ lineage. Moreover, length variations in the P3N-PIPO protein might be involved with host-specific adaptations of PVY-Tam. Overall, our findings contribute to a better understanding of tamarillo virosis in the Andean region, and overall, in other tamarillo-cultivating areas worldwide.

## MATERIALS AND METHODS

### Sample collection

The sampling for this study was conducted in eight tamarillo-growing orchards located in diverse altitudes (range 2000-2500 m) across the Department of Nariño (Colombia) (**Figure 1** and **Supplementary Table S1**). Each sample consisted of young apical shoots from tamarillo plants showing symptoms of viral infections (**Supplementary Table S2**), which were collected and transported in polypropylene coolers containing ice packs to the Molecular Biology Laboratory at the University of Nariño. Next, shoots were individually wrapped with paper towels, placed inside labeled paper bags, and immediately processed to minimize the risk of degradation. Part of the plant material was stored at –80°C for subsequent analyses.

**Figure 1.**
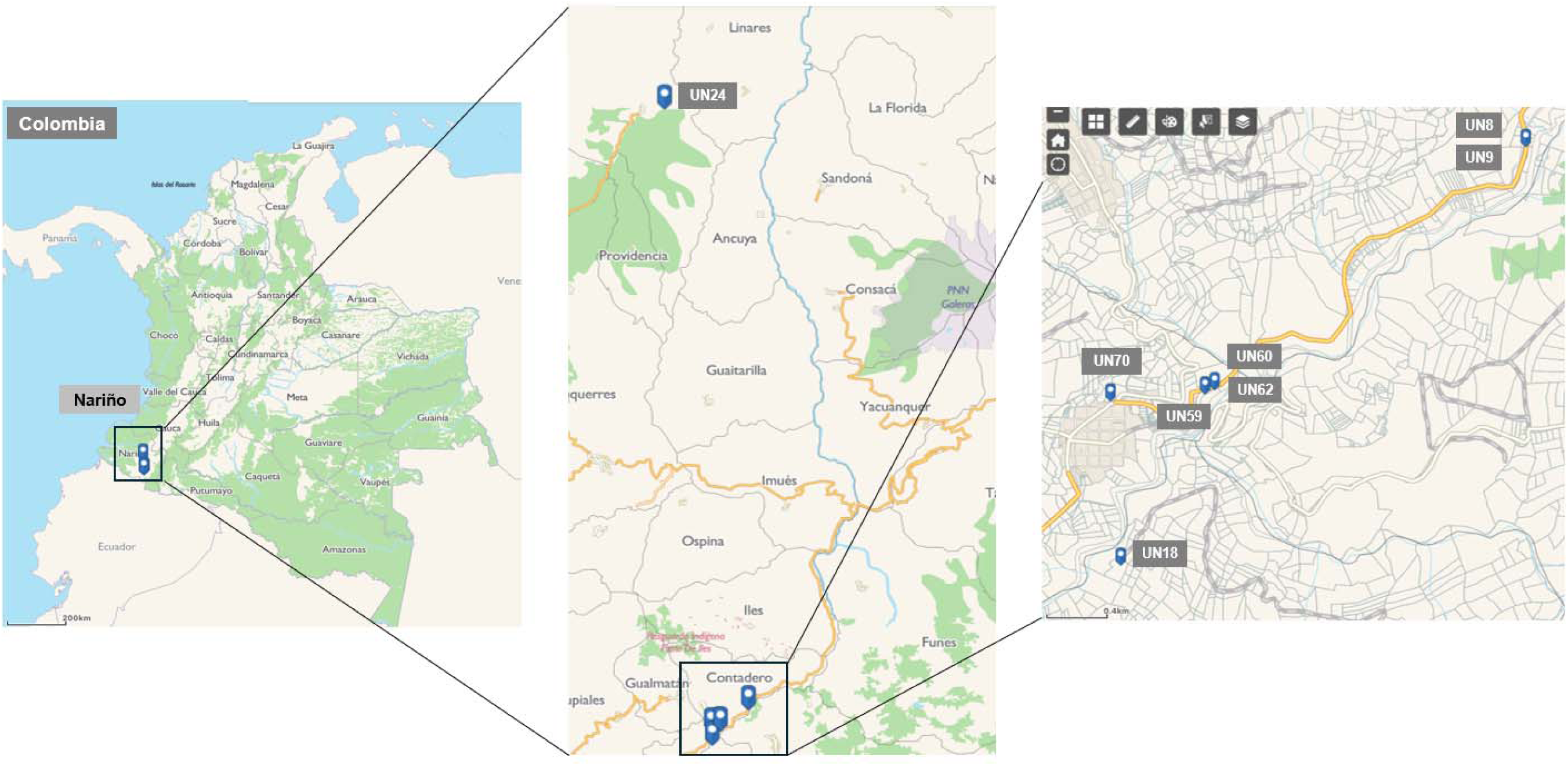
Geographic localization of tamarillo-growing orchards in Nariño (Colombia) analyzed in the study. Samples of tamarillo plants showing symptoms of virosis were collected in a total of eight locations distributed in an altitude range of 2000-2500 m across the department. All locations except UN24 were situated in Contadero region.

### Total RNA extraction and RNA-Seq library construction

Symptomatic shoots (50 mg) were frozen in liquid nitrogen and ground using a TissueLyser (Qiagen). Total RNA extraction was performed using TRIzol reagent (Life Technologies, USA) following the manufacturer’s instructions. RNA quality and concentration were estimated using a NanoDrop One (Thermo Fisher Scientific, USA) spectrophotometer and electrophoresis visualization on 1% agarose gels in 1X TAE buffer (40 mM Tris-acetate, 1 mM EDTA, pH 8.0) and staining with GelRed. Purified RNA samples were stored at –80°C until use. 200 ng of total RNA were used for RNA-Seq library construction. Messenger RNA (mRNA) was purified from total RNA using poly-T oligo-conjugated magnetic beads (New England Biolabs, USA), and ribosomal RNA (rRNA) depletion was performed using the TruSeq Stranded Total RNA kit with Ribo-Zero Plant (Illumina, USA), according to the manufacturer’s instructions. Fragmented RNA was employed for the synthesis of first-strand complementary DNA (cDNA) using random hexamer primers, followed by second-strand cDNA synthesis. Library construction was performed using the TruSeq RNA Sample Preparation kit (Illumina, USA) as described in (Zhao et al., 2024). Library quality and concentration were estimated using a Qubit (Thermo Fisher Scientific) fluorometer, and fragment size distribution was assessed by real-time PCR along with a Bioanalyzer. Finally, library sequencing was conducted at Novogene (USA) using the NovaSeq 6000sequencer platform (Illumina) to generate pooled, paired-end (read size: 50-150 bp) libraries.

### Identification of viral sequences using Genome Detective software

The obtained raw reads were submitted to the web-based software application Genome Detective (https://www.genomedetective.com/app/typingtool/virus/) to identify viral sequences. By using a novel reference-based linking alignment method that combines nucleotide and amino acid scores, Genome Detective allows a rapid and accurate assembly of RNA-seq reads into annotated viral contigs (Vilsker et al., 2019). For torradovirus samples, an additional analysis was performed calculating the nucleotide and amino acid p-distance of the RNA1 CG to GDD (Protease-Pol) conserved region from each contig to selected torradovirus reference sequences (**Table 1**) using the software MEGA 12 (Kumar et al., 2024).

**Table 1.**
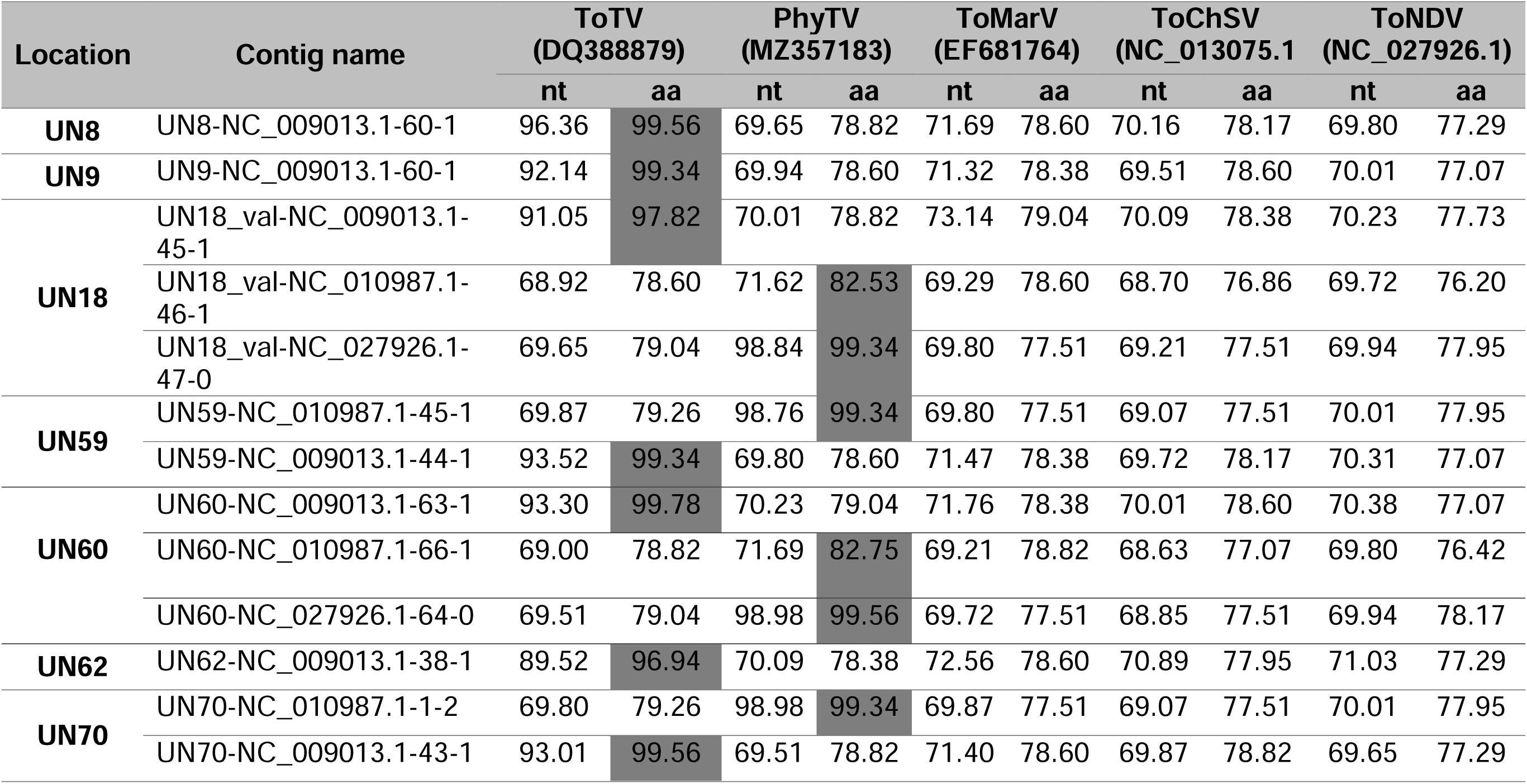
Nucleotide (nt) and amino acid (aa) sequence identities comparison of the Pro-Pol (CG to GDD) conserved region of each torradovirus-like RNA1 contig to the RNA1 reference genome of selected torradovirus species. Amino acid identities >80% in the Pro-Pol region are highlighted in grey. Accession numbers used for the Pro-Pol analysis: ToTV, tomato torrado virus (DQ388879); PhyTV, physalis torrado virus (MZ357183) ToMarV, tomato marchitez virus (EF681764); ToChSV, tomato chocolate spot virus (NC_013075.1); ToNDV, tomato necrotic dwarf virus (NCBI: NC_027926.1).

| Location | Contig name | ToTV<br>(DQ388879) |  | PhyTV<br>(MZ357183) |  | ToMarV<br>(EF681764) |  | ToChSV<br>(NC_013075.1) |  | ToNDV<br>(NC_027926.1) |  |
| --- | --- | --- | --- | --- | --- | --- | --- | --- | --- | --- | --- |
|  |  | nt | aa | nt | aa | nt | aa | nt | aa | nt | aa |
| **UN8** | UN8-NC_009013.1-60-1 | 96.36 | 99.56 | 69.65 | 78.82 | 71.69 | 78.60 | 70.16 | 78.17 | 69.80 | 77.29 |
| **UN9** | UN9-NC_009013.1-60-1 | 92.14 | 99.34 | 69.94 | 78.60 | 71.32 | 78.38 | 69.51 | 78.60 | 70.01 | 77.07 |
| **UN18** | UN18_val-NC_009013.1-45-1 | 91.05 | 97.82 | 70.01 | 78.82 | 73.14 | 79.04 | 70.09 | 78.38 | 70.23 | 77.73 |
|  | UN18_val-NC_010987.1-46-1 | 68.92 | 78.60 | 71.62 | 82.53 | 69.29 | 78.60 | 68.70 | 76.86 | 69.72 | 76.20 |
|  | UN18_val-NC_027926.1-47-0 | 69.65 | 79.04 | 98.84 | 99.34 | 69.80 | 77.51 | 69.21 | 77.51 | 69.94 | 77.95 |
| **UN59** | UN59-NC_010987.1-45-1 | 69.87 | 79.26 | 98.76 | 99.34 | 69.80 | 77.51 | 69.07 | 77.51 | 70.01 | 77.95 |
|  | UN59-NC_009013.1-44-1 | 93.52 | 99.34 | 69.80 | 78.60 | 71.47 | 78.38 | 69.72 | 78.17 | 70.31 | 77.07 |
| **UN60** | UN60-NC_009013.1-63-1 | 93.30 | 99.78 | 70.23 | 79.04 | 71.76 | 78.38 | 70.01 | 78.60 | 70.38 | 77.07 |
|  | UN60-NC_010987.1-66-1 | 69.00 | 78.82 | 71.69 | 82.75 | 69.21 | 78.82 | 68.63 | 77.07 | 69.80 | 76.42 |
|  | UN60-NC_027926.1-64-0 | 69.51 | 79.04 | 98.98 | 99.56 | 69.72 | 77.51 | 68.85 | 77.51 | 69.94 | 78.17 |
| **UN62** | UN62-NC_009013.1-38-1 | 89.52 | 96.94 | 70.09 | 78.38 | 72.56 | 78.60 | 70.89 | 77.95 | 71.03 | 77.29 |
| **UN70** | UN70-NC_010987.1-1-2 | 69.80 | 79.26 | 98.98 | 99.34 | 69.87 | 77.51 | 69.07 | 77.51 | 70.01 | 77.95 |
|  | UN70-NC_009013.1-43-1 | 93.01 | 99.56 | 69.51 | 78.82 | 71.40 | 78.60 | 69.87 | 78.82 | 69.65 | 77.29 |

### Validation of PVY-Tam by RT-PCR

Total RNA extraction was performed using TRIzol reagent (Life Technologies, USA) following the manufacturer’s instructions. A RT-PCR assay was performed to confirm the presence of PVY-Tam in the total RNA of the samples collected in the eight tamarillo-growing locations, as described by Martínez et al. (2014). The M-MLV Reverse Transcriptase kit and Taq DNA polymerase (both from Invitrogen, USA), were used for reverse transcription and PCR, respectively, following the manufacturer’s instructions, with a modification of the annealing temperature to 53 °C for primers PVY-T CP and PVY-T 3P. First, the consensus sequence of the polyprotein gene of PVY-Tam Nariño isolate was inferred by multiple alignment of the corresponding viral contigs from different locations. Then, specific primers were designed to amplify a 323-nucleotide (nt) fragment between the NIb protein and the coat protein (CP) regions, and a 404-nt fragment in the P3 protein region, respectively (**Supplementary Table S3**). RT-PCR products were visualized by electrophoresis on 1% agarose gels in 1X TAE buffer (40 mM Tris-acetate, 1 mM EDTA, pH 8.0) and stained with GelRed and then sent for Sanger sequencing to the National University of Colombia (Bogotá Campus).

### Phylogenetic analysis

The polyprotein gene of the PVY-Tam Nariño isolate was translated into its corresponding amino acid sequence and used to infer phylogenetic relationships with 19 selected potyvirus species belonging to the PVY clade (Gibbs & Ohshima, 2010a). Phylogenetic and geographic criteria were applied to select potyvirus species based on the most recently available phylogenetic tree generated by the International Committee on Taxonomy of Viruses (ICTV) *Potyviridae* Study Group (available at https://ictv.global/taxonomy/taxondetails?taxnode_id=202404562&taxon_name=Potyvirus), choosing one representative per main branch or node, with preference for species reported in South America. Additional phylogenetic analysis were performed with type members of the five major PVY clades (C, N, O, NTN and Chile) and a PVY-Tam isolate previously reported in Ecuador (Green et al., 2020; GenBank: MT380740). A list of the potyvirus species and PVY isolates used in the study is included in **Supplementary Table S4**. Tree construction was performed using the maximum likelihood method with 1000 bootstraps interactions available in MEGA12. CLUSTAL W with default parameters was used to generate multiple sequence alignments for the P3N-PIPO sequences.

## RESULTS

### Symptomatology of complex viral infections in tamarillo

The Department of Nariño has been previously reported to be affected by cases of tamarillo virosis (Betancourth et al., 2003; Rodríguez, 2009). To better understand the complex interactions between viral species in tamarillo, we selected a total of eight tamarillo-growing orchards located across the Nariño department (from now on, referred to as UN8 to UN70) (**Figure 1** and **Supplementary Table S1**). We observed viral infection symptoms in tamarillo plants collected in each location, including general mosaic with irregular mottling, chlorotic blisters, concentric ringspots, and deformation in leaves, as well as early breaking and pulp hardness in fruits (**Figure 2A** and **Supplementary Table S2**). Some plants also exhibited severe necrotic lesions in both leaves and fruits, possibly because of advanced viral infections or secondary co-infections with fungal pathogens. Conversely, tamarillo plants in UN24 showed a slightly different appearance consisting of partial chlorosis, mild leaf deformation and blistering.

**Figure 2.**
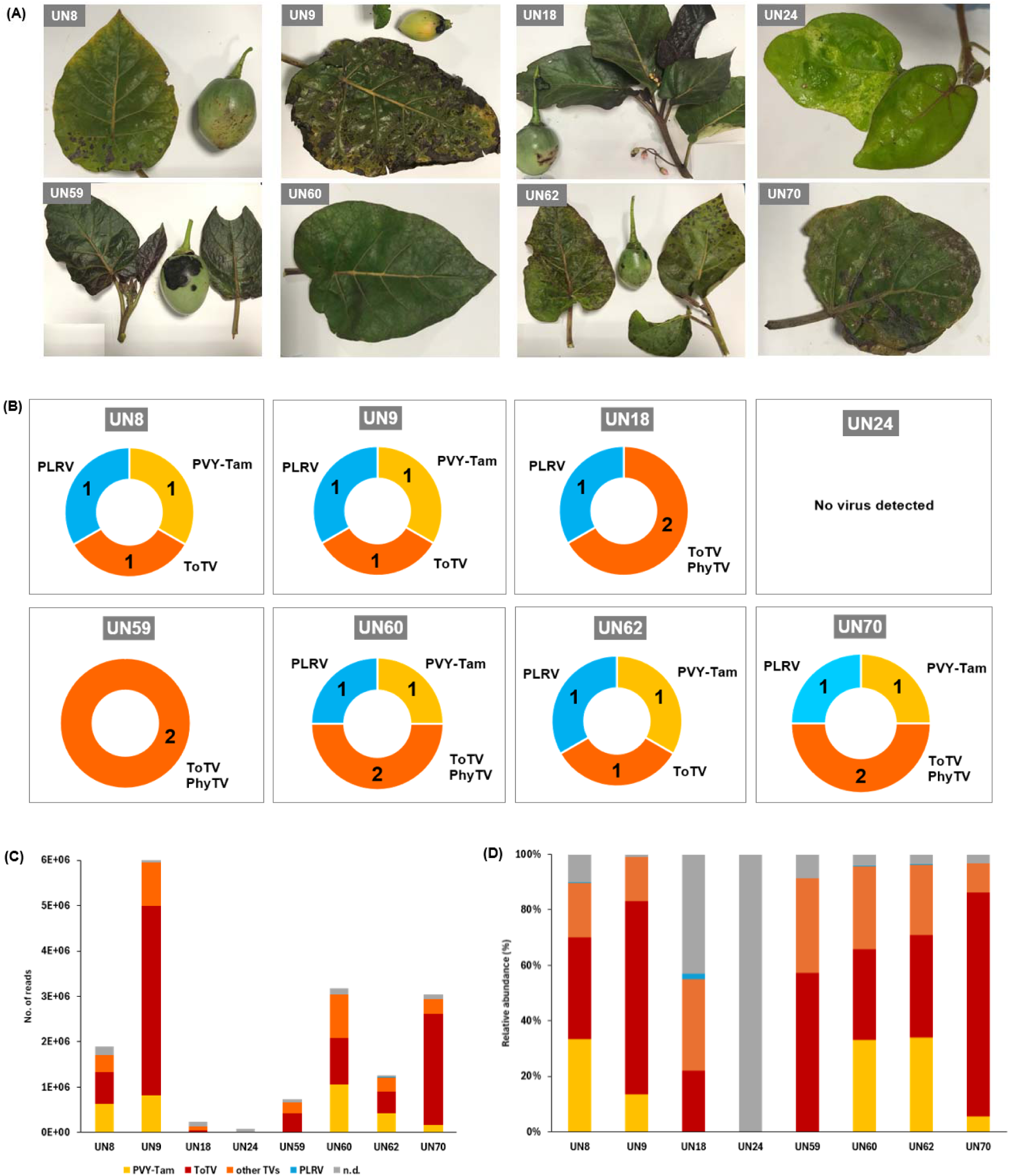
Characterization of virus diversity in tamarillo-growing orchards across Nariño. **(A)** Symptomatology of leaves and fruits from tamarillo plants collected in different geographic locations across Nariño. Symptoms include mosaicism, irregular mottling, chlorotic blisters, concentric ringspots and deformation in leaves, as well as early breaking and pulp hardness in fruits. **(B)** Ring chart representation of virus diversity in tamarillo plants showing virosis symptoms collected in the eight locations. The identified virus species are indicated with numbers and grouped to their corresponding genera (Potyvirus in yellow, Torradovirus in orange, and Polerovirus in blue). No viruses were detected on site UN24. **(C)** Number of RNA-seq reads and **(D)** relative abundance of the viral species identified in the eight geographic locations. The abundance percentage of each virus is normalized to the total numbers of RNA-seq reads counted in each sample. N.d., non-determined assignments.

### Characterization of virus diversity in tamarillo plants affected with virosis

To unravel the complex interactions between different viruses that can lead to tamarillo virosis, we generated RNA-seq libraries for symptomatic plants collected in the eight geographic locations. Analysis of RNA-seq reads using the Genome Detective software revealed that all tested samples, except for UN24, were positive for virus infections (**Supplementary e-Xtra Files**). Tomato torrado virus (ToTV) was found in all infected samples, where the assembled contigs showed > 90% amino acid sequence identities to its reference genome (**Supplementary Table S5**). Other assignments of members from the *Torradovirus* genera included tomato marchitez virus (ToMarV), tomato chocolate spot virus (ToChSV), and tomato necrotic dwarf virus (ToNDV), but amino acid identity levels were considerably lower (63.2-71%). To clarify the torradovirus assignments, we performed a Blastn analysis of the full-length RNA1 nucleotide contigs of each sample, which confirmed that ToTV and/or physalis torrado virus (PhyTV) were the only torradoviruses present in the different locations. The taxonomy criterion for members of the family *Secoviridae* is usually based on the conserved Pro-Pol region (from CG motif of the protease to the GDD motif of the Pol), and species demarcation is defined by amino acid sequence identities of >75% in the coat protein(s) (CP) or > 80% in the Pro-Pol region, respectively (Fuchs et al., 2022). However, CP cleavage sites are determined only for two torradoviruses (i.e. ToChSV and ToMarV) out of the twelve viral species recognized in this genus (Ferriol et al., 2016). Hence, for our study, we defined the species demarcation based on the Pro-Pol conserved region from RNA1. We identified seven RNA1 contigs for ToTV in samples from different locations. The Pro-Pol region for these sequences showed 68.92-96.36 % and 78.60-99.78% nucleotide and amino acid identity compared to that of ToTV isolate PRI-0301 (GenBank: DQ388879). Additionally, we found PhyTV in four out of eight locations. When using PhyTV isolate BPP2 (GenBank: MZ357183) as a reference, the Pro-Pol nucleotide and amino acid identities ranged between 69.51-98.98% and 78.38-99.56%, respectively (**Table 1**). Our findings constitute the very first evidence on the capacity of torradoviruses to infect tamarillo as a host species.

In addition to torradoviruses, we identified two viruses from unrelated genera in tamarillo plants: the potyvirus potato virus Y-Tamarillo (PVY-Tam) and the polerovirus potato leafroll virus (PLRV) appeared in five and six out of the eight locations, respectively (**Figure 2B**). Remarkably, five locations showed simultaneous infections with multiple viruses from these three different genera (*Potyvirus*, *Torradovirus* and *Polerovirus*). Regarding the total number of identified reads, UN9 contained by far the highest viral load of all samples (5.8×10^6^ reads), while the load of infection was significantly reduced in samples UN18 and UN59 where PVY-Tam was absent (1×10^5^ and 6×10^5^ reads, respectively) (**Figure 2C**). The predominant viral species in all infected samples was ToTV with a relative abundance of 34-86%, while PVY-Tam appeared in varying abundance (6-39.5%) and PLRV was marginally present (0.02-4%) (**Figure 2D**). Unfortunately, we could not determine the abundance of PhyTV in our samples since there were not assigned hits to the reference sequence of PhyTV (**Supplementary e-Xtra Files)**.

### Confirmation of a novel PVY-Tam isolate in Colombia

PVY is the most economically relevant virus worldwide, but there is limited knowledge of the predominant tamarillo-infecting PVY isolates in the Andean region and their effects on this crop. First, we determined the consensus nucleotide sequence of our PVY-Tam Nariño isolate by performing a multiple alignment of the viral contigs identified as this virus in all the PVY-positive locations. Then, we designed and validated two primer pairs to amplify specific regions between the viral proteins CP or within P3 (**Supplementary Table S3**). RT-PCR analysis of PVY-Tam CP and P3 using these primer pairs confirmed the presence of the virus in all the samples regarded as PVY-infected in the RNA-seq (**Supplementary Figure S1**), and we further validated the identity of PVY-Tam by Sanger sequencing of the RT-PCR products. Altogether, our results constitute the report of a novel PVY-Tam isolate present in Colombia.

### Unraveling PVY diversification in the Andean region

PVY belongs to a large clade of 19 virus species of the genus *Potyvirus* that are mostly present in South America, with 11 virus species being exclusively present on this continent (Gibbs & Ohshima, 2010). Phylogenetic analysis of the polyproteins from selected potyviruses of this clade confirmed that the PVY-Tam Nariño isolate is most closely related to another PVY-Tam previously reported in the neighboring region of Pichincha (Ecuador), about 350 km south of the Department of Nariño (Green et al., 2020) (**Figure 3A**). The two PVY-Tam isolates conform a well-defined clade with PVY and certain potyviruses originated in neighboring South American countries, such as alstroemeria mosaic virus (AlMV) from Ecuador and bidens mosaic virus (BiMV) from Brazil. In addition, our results pointed out that PVY-Tam is more distantly related to other potyviruses previously detected in cases of tamarillo virosis (e.g. colombian datura virus, CDV; tamarillo leaf malformation virus, TLMV; potato virus A, PVA).

**Figure 3.**
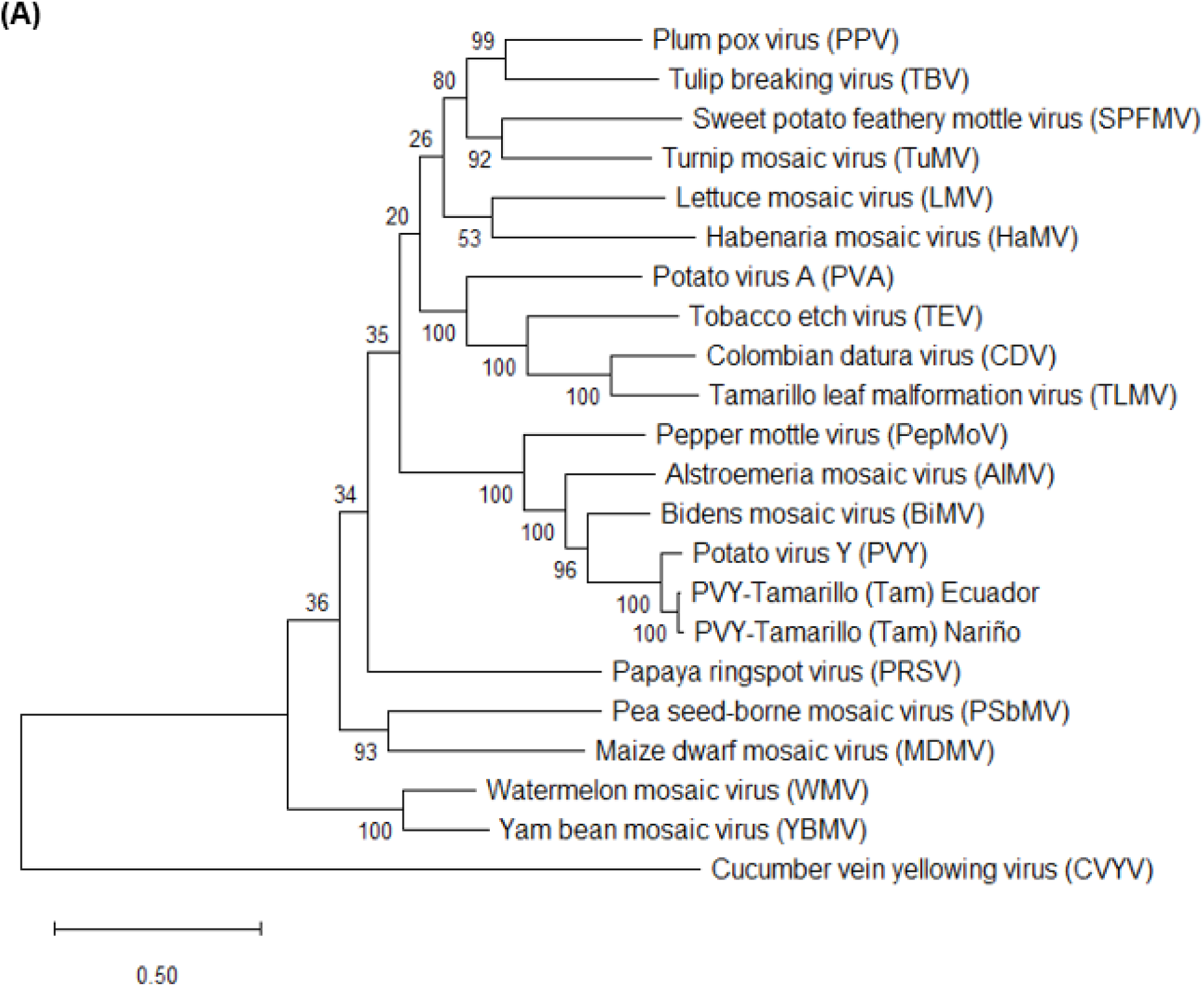
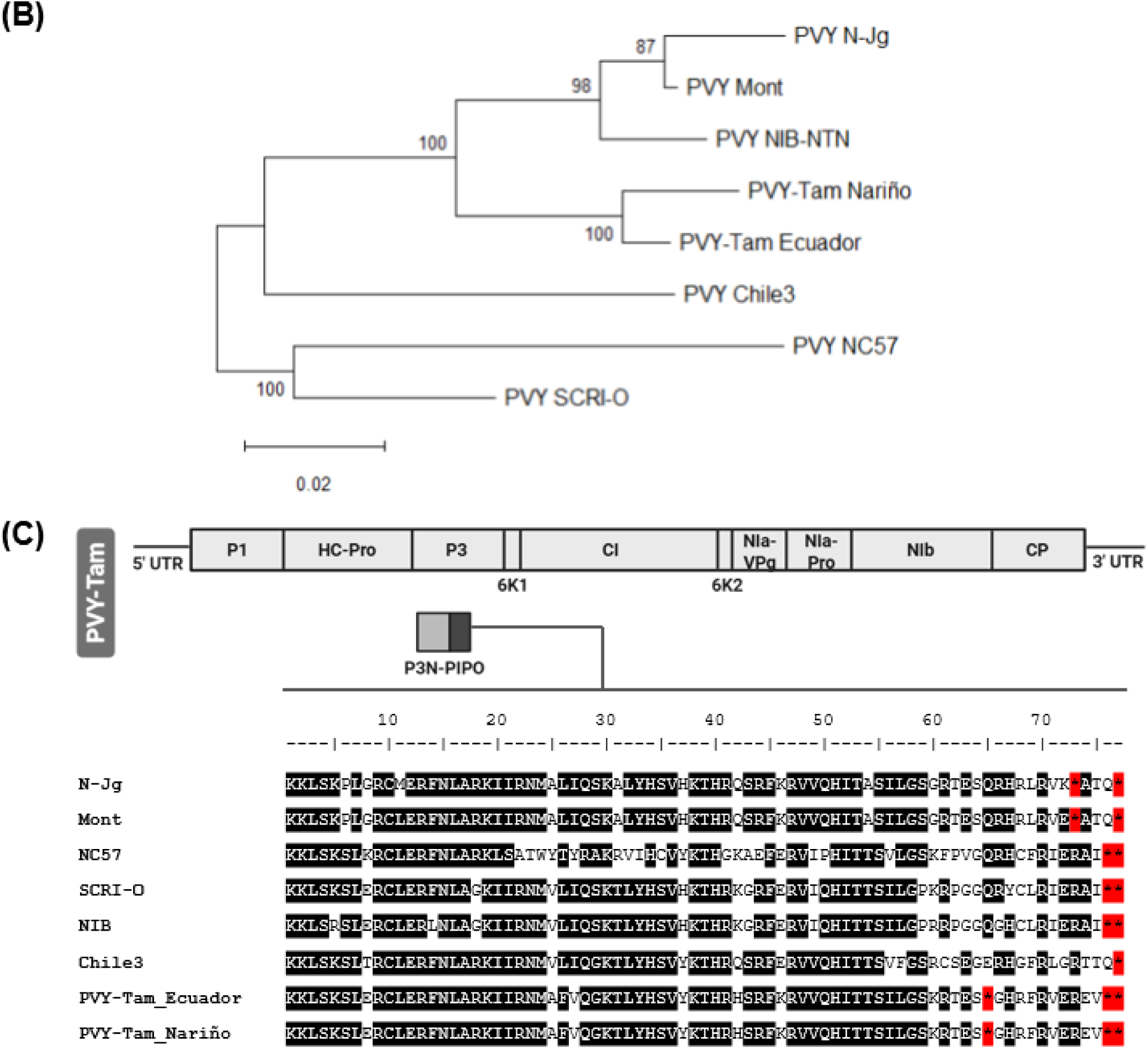
Phylogeny and molecular evolution of PVY-Tam. Maximum likelihood trees of the full-length polyprotein sequences for PVY-Tam and **(A)** 19 virus species of the genus *Potyvirus*; or **(B)** selected PVY isolates of the major clades. Trees are drawn to scale, with branch lengths measured in the number of substitutions per site and bootstrap values indicated for each node. The bar below indicates genetic distance. All positions containing gaps and missing data were eliminated. The tree from **Figure 3A** was rooted to the cucumber vein yellowing virus (CVYV, genus *Ipomovirus*). **(C)** Amino acid alignment of P3N-PIPO from PVY-Tam and selected PVY isolates of the major clades. Fully conserved residues are highlighted in black and stop codons are indicated as asterisks with red background.

PVY is classified into five major clades according to molecular, serological and biological properties (Singh et al., 2008). The C (common), O (ordinary) N (necrotic) and NTN (N x O recombinants) clades show a worldwide distribution and cause severe diseases in potato and other solanaceous crops, while the Chilean clade has been reported only in this country and is non-infectious in potato. To shed some light on the genetic diversity of PVY-Tam, we carried out additional phylogenetic analysis based on a selection of type members for each of the PVY clades: NC567 (C), SCRI-O (O), NIB (NTN), N-Jg (N-NA, North America), Mont (N-Eu, Europe) and Chile 3 (Chile). Comparison of the polyprotein sequences indicated that PVY-Tam isolates from Nariño and Ecuador conformed a separate clade within the PVY^N^ lineage that included the clades PVY^N-NA^, PVY^N-Eu^ and PVY^NTN^ (**Figure 3B**). To test whether the common occurrence of genetic recombination amongst PVY could be masking the geographical origin of PVY-Tam, we followed the approach described by Cuevas et al. (2012) to divide the PVY genomes into three recombination-free regions named R1 (310-2202 nt), R2 (2227-5628 nt) and R3 (5656-8991 nt). The phylogenetic analysis further confirmed that PVY-Tam isolates cluster within the PVY^N^ lineage in all three regions, thus discarding recombination events with PVY isolates from other clades (**Supplementary Figure S2**).

### Variability in P3N-PIPO length among PVY isolates

Next, we studied the variability of individual PVY proteins and their biological significance. Most proteins were highly conserved among all PVY representatives (>85% amino acid sequence similarities in most cases), except for P1 that is known for its variability among potyviruses, and P3N-PIPO (**Supplementary Table S5**). According to literature, P3N-PIPO encodes for a 76-amino acid (aa) protein in PVY, as it is determined by the UAA stop codon at position 77 (Dullemans et al., 2011). However, we observed that all our isolates carried additional stop codons leading to variations in the length of P3N-PIPO (**Figure 3C**): UGA at position 73 from PVY^N-NA^ and PVY^N-Eu^; UAA at position 76 for PVY^C^, PVY^O^ and PVY^NTN^; and CAA also at position 76 for PVY^Chile^. Intriguingly, the two PVY-Tam isolates presented the canonical UAA stop codon at position 77 plus two more UAAs at positions 65 and 76, leading to the shortest P3N-PIPO protein (64 aa) among the selected PVY isolates. These differences in the length of P3N-PIPO might have an influence on the host range and infectivity properties of the different PVY isolates.

## DISCUSSION

Viral diseases are the most devastating phytosanitary problem in tamarillo production in Colombia. The wide variety of symptoms is commonly associated with viral complexes from various genera that establish intricated yet unknown interactions, hindering the development of effective control or mitigation strategies. In this work, we used the innovative, web-based Genome Detective software that allows straightforward analysis of RNA-seq data to characterize the virome of tamarillo plants from diverse locations across the Department of Nariño, Colombia (**Figure 1**). Six out of the eight locations exhibited simultaneous infection with multiple viruses, where those belonging to the genus *Torradovirus* were the most prevalent (**Figure 2**). Torradoviruses are an emerging plant virus genus with a similar genome to that of picorna-like viruses in the *Secoviridae* family first described in the early 2000s asTomar the causal agents of a new disease in tomato (Van Der Vlugt et al., 2015). From the 12 species recognized by ICTV in the genus *Torradovirus*, many of them have been reported in South America including ToTV (Herrera-Vasquez et al., 2009; Verbeek & Dullemans, 2012a), ToChSV (Batuman et al., 2010; Verbeek et al., 2008), ToMarV (Verbeek et al., 2008), PhyTV (Corrales-Cabra et al., 2021a), tomato chocolate virus (ToChV) (Verbeek et al., 2010), cassava torrado-like virus (CTLV) (Carvajal-Yepes et al., 2014), and potato rugose stunting virus (PotRSV) (Alvarez-Quinto et al., 2023). We found that ToTV, the type member of the genus, was the prevailing virus species in all the virus-infected locations (**Figures 2C** and **2D**). This is the first evidence of ToTV in tamarillo plants in Colombia, since all previous reports corresponded to infections in tomato (Verbeek & Dullemans, 2012b). Additionally, based on sequence similarities of the torradovirus RNA1 Pro-Pol region we identified the presence of PhyTV in four out of the eight locations. PhyTV was first described in Colombia) in plants of cape gooseberry (*Physalis peruviana*) infected with several plant viruses, including PVY (Corrales-Cabra et al., 2021b). Altogether, our results provide strong evidence that tamarillo is a natural host of torradoviruses and open new possibilities for the identification of new viral species of this emerging genus in the upcoming years.

Potyviruses are frequent players in tamarillo virosis (Ayala et al., 2010; Gutiérrez et al., 2015; M. Jaramillo et al., 2011). PVY isolates infecting this crop (hereafter named PVY-Tam) were initially described in New Zealand (Chamberlain, 1954) and India (Bhargava, 1959), but later reports indicated that it is also widespread in tamarillo orchards in Ecuador (Vizuete, 1990). Based on serological and molecular analysis, the preliminary work by Jaramillo et al. (2011) provided the first evidence for PVY-Tam infection in the main tamarillo-growing regions of Colombia. In our study, we detected PVY-Tam in five out of the seven virus-infected locations, which interestingly showed more intense symptoms (e.g. leaf blistering and necrosis) and a burst in total viral load compared to that of the poty-free locations (**Figures 2A** and **2C**). It is well known that the potyviral sequence Pl/HC-Pro promotes the pathogenicity and accumulation of a broad range of unrelated viruses, leading to an enhanced severity of disease symptoms (Moreno & López-Moya, 2020; Pruss et al., 1997). We speculate that a similar PVY-Tam-associated synergism might lead to an increase in the accumulation of ToTV and other torradovirus partners, with negligible changes in the level of the potyvirus.

The agronomical relevance of PVY infection in many solanaceous crops has motivated an extensive study of its biological and genetic diversity. For many years, phylogenetic studies were mainly restricted to potato, pepper and tobacco as they are the three major crops affected by this virus (A. J. Gibbs et al., 2017). However, recent findings propose that other solanaceous species can allow the replication of host-specific isolates displaying extraordinary genetic diversity (Chikh-Ali et al., 2016; Green et al., 2017). Our phylogenetic analysis indicated that the PVY-Tam isolates found in the Andean region (i.e. Nariño and Ecuador) are closely related to other potyviruses infecting tamarillo, and they specifically cluster with some viruses first described in neighbouring countries (**Figure 3A**). These results support the hypothesis that South America is the centre of origin and diversification for the PVY clade (Quenouille et al., 2013). In accordance with the work by Green et al. (2020), we noticed that PVY-Tam forms a distinct clade within the PVY^N^ lineage that might have arisen by multiple mutations, recombination and gene rearrangement from a common geographical ancestor (**Figure 3B**). The short evolutionary distances depicted in our trees highlight that PVY diversification is very recent and is being significantly favored by human activities, mainly related to the cultivation and trade of potato tubers (Visser et al., 2012).

The P3N-PIPO protein is the product of a transcriptional slippage on a conserved GA6 motif on the P3 cistron and it shows variable lengths among potyviruses (Valli et al., 2024). It locates in the plasmodesmata, where it participates in the cell-to-cell movement of the virus in conjunction with other viral proteins like P3 and CI (Wei et al., 2010; Wen & Hajimorad, 2010). By comparing the P3N-PIPO sequences from all our PVY isolates, we found that both PVY-Tam isolates produce a 64-aa protein due to two premature stop codons at positions 65 and 76 (**Figure 3C**), which is the shortest protein we observed among all our PVY isolates (64 aa vs. 72-76 aa). These differences in the length of P3N-PIPO might be involved in host-specific adaptations and infectivity properties, as previously suggested by Cuevas et al. (2012). Nonetheless, additional experimental evidence and sequence analysis of P3N-PIPO from other potyviruses are required to shed light on the biological implications of this protein.

In summary, this study significantly contributes to the knowledge of complex viral infections in tamarillo in the Andean region and, overall, in other tamarillo-cultivating areas worldwide. We provide the first evidence on the capacity of torradoviruses to infect tamarillo, and we characterize a novel PVY-Tam isolate in Colombia that appears to carry host-driven adaptations. A deep understanding of the epidemiological variables influencing the establishment and expansion of tamarillo virosis in South America is crucial to developing effective and durable control methods for this disease. Further studies should address the interactions between different virus genera in the plant, with a special focus on the torradovirus-potyvirus synergism, and the biological implications of P3N-PIPO on host-driven adaptations of PVY.

## Supporting information

Genome Detective reports

## Acknowledgements & Funding

This project has received funding from the European Union’s Horizon 2020 research and innovation programme under grant agreement No 101000570 (VIRTIGATION). Work at Universidad de Nariño was funded by the Bécate-Nariño program for high-level human talent training under the postdoctoral fellowship modality (BPIN-2017000100028), coordinated by Fundación CeiBA with resources from the Sistema General de Regalías (SGR) allocated to the Gobernación de Nariño. Work at CRAG was funded by grant CEX2019-000902-S funded by MICIU/AEI/ 10.13039/501100011033 and by the CERCA Programme / Generalitat de Catalunya. The research lab also received funding by grant PID2022-139376OB-C33 funded by MICIU/AEI/ 10.13039/501100011033 by “ERDF/EU”. CO is recipient of fellowship PRE2020-094228 funded by MICIU/AEI /10.13039/501100011033 and by “ESF Investing in your future”. Work at ICA was funded by the CSIC (project No 202240I189).

## TABLES AND FIGURES

**Table 1**. Nucleotide and amino acid sequence identities comparison of the Pro-Pol (CG to GDD) conserved region of torradovirus-like RNA1 contigs to the RNA1 reference genome of selected torradovirus species.

**Figure 1**. Geographic localization of tamarillo-growing orchards in Nariño (Colombia) analyzed in the study.

**Figure 2**. Characterization of virus diversity in tamarillo-growing orchards across Nariño.

**Figure 3**. Phylogeny and molecular evolution of PVY-Tam.

## SUPPLEMENTARY DATA

**Supplementary Table S1.** Description of the geographic locations across Nariño (Colombia) in which tamarillo plants showing virosis symptoms were sampled.

**Supplementary Table S2.** Symptoms description of the tamarillo plants sampled in the different geographic locations.

**Supplementary Table S3.** Primers used for PVY-Tam diagnosis by RT-PCR.

**Supplementary Table S4.** List of the virus species belonging to the *Potyviridae* family and PVY isolates used in the phylogenetic analysis.

**Supplementary Table S5.** Identification of virus species in tamarillo-growing orchards across Nariño using Genome Detective.

**Supplementary Table S6.** Amino acid sequence identities of the individual proteins of PVY-Tam (Nariño) to those of type members from PVY clades and a PVY-Tam isolate previously reported in Ecuador.

**Supplementary Figure S1.** RT-PCR detection of PVY-Tam in infected plants from different geographic locations.

**Supplementary Figure S2.** Maximum likelihood trees of the R1, R2 and R3 regions of the nucleotide sequences of the polyprotein for PVY-Tam and selected PVY isolates of the major clades.

## SUPPLEMENTARY DATA

**Supplementary Table S1.**
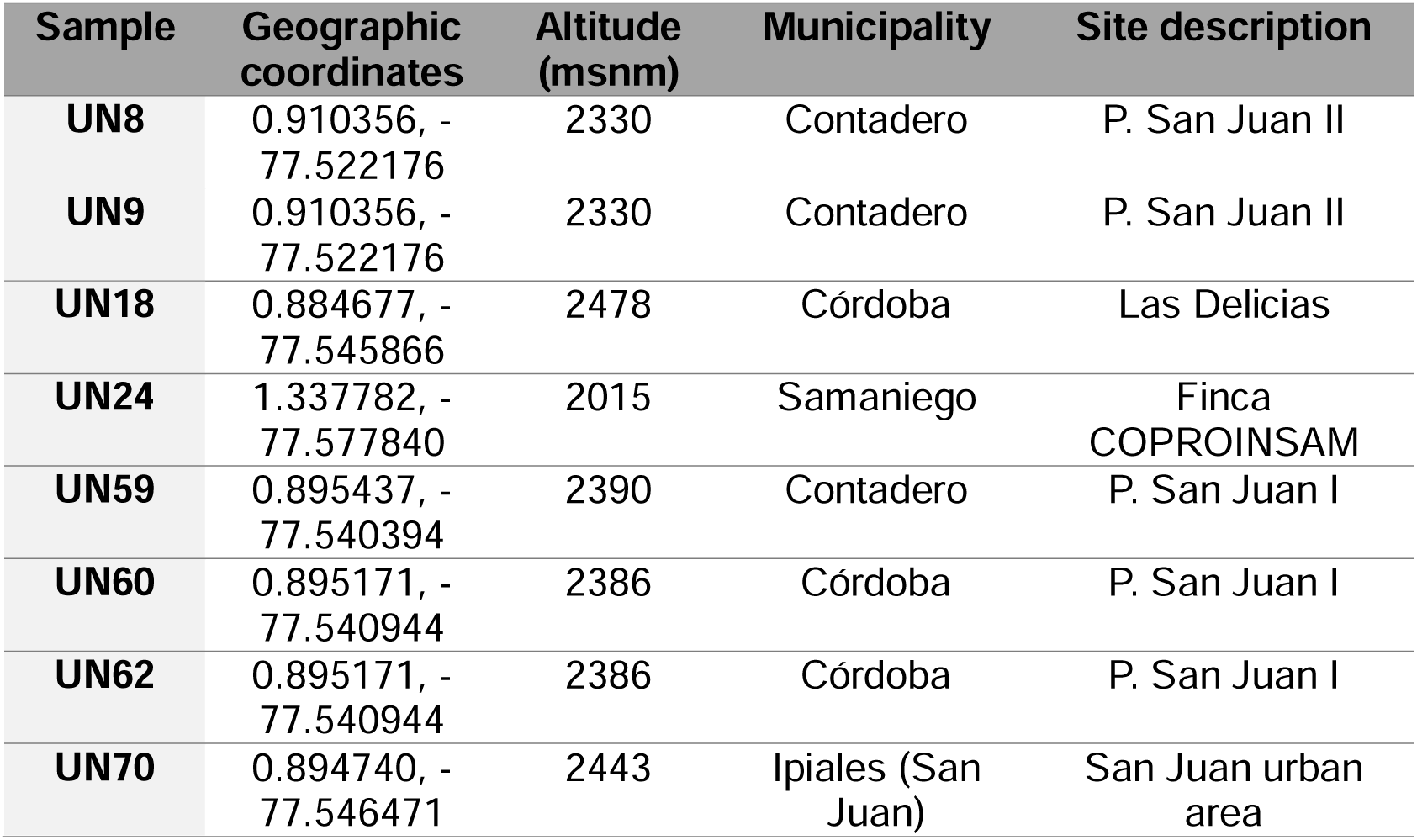
Description of the geographic locations across Nariño (Colombia) used in the study, in which samples of tamarillo plants showing symptoms of virosis were collected.

**Supplementary Table S2.**
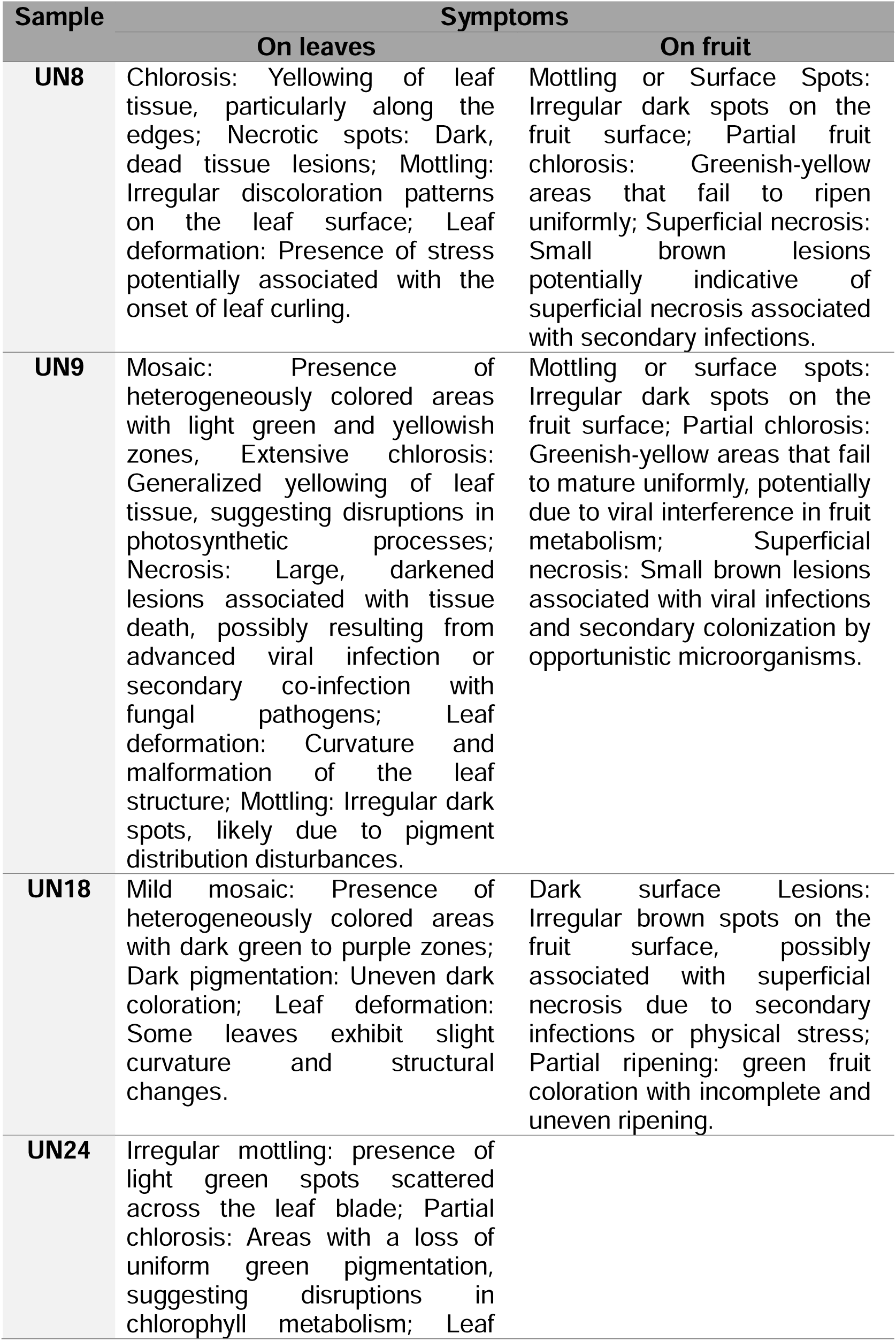
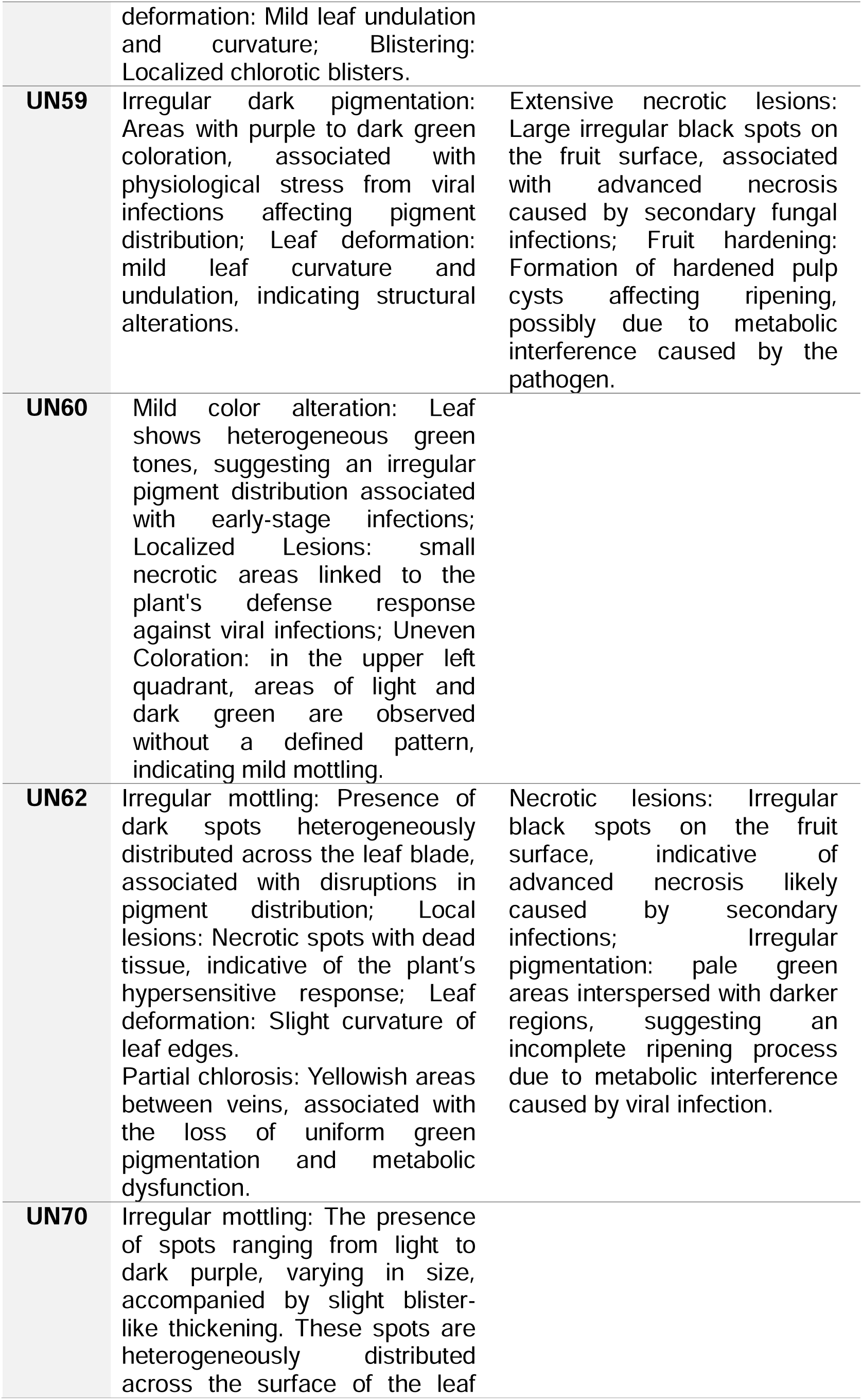
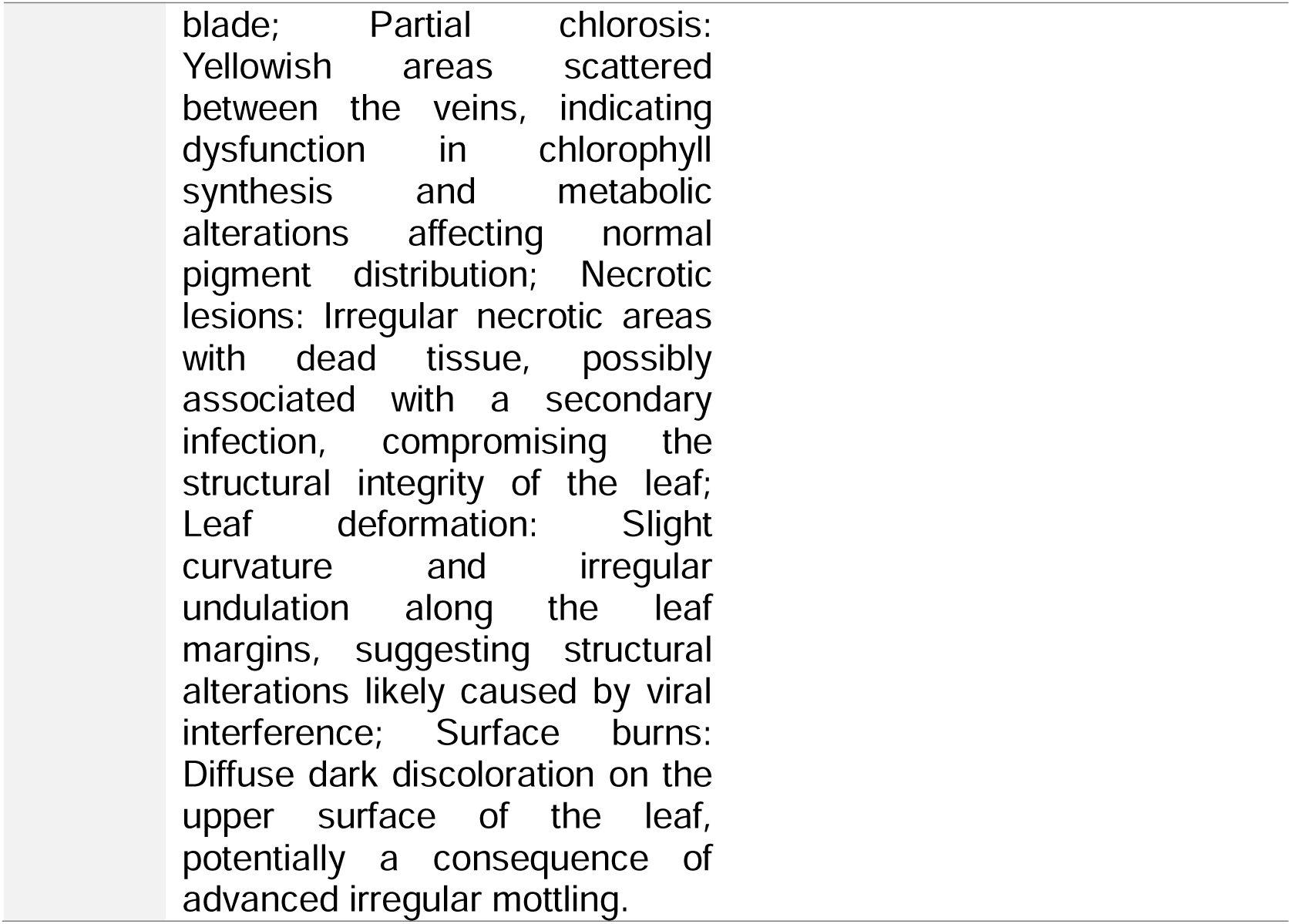
Symptoms description of the tamarillo plants sampled in the different geographic locations.

**Supplementary Table S3.**
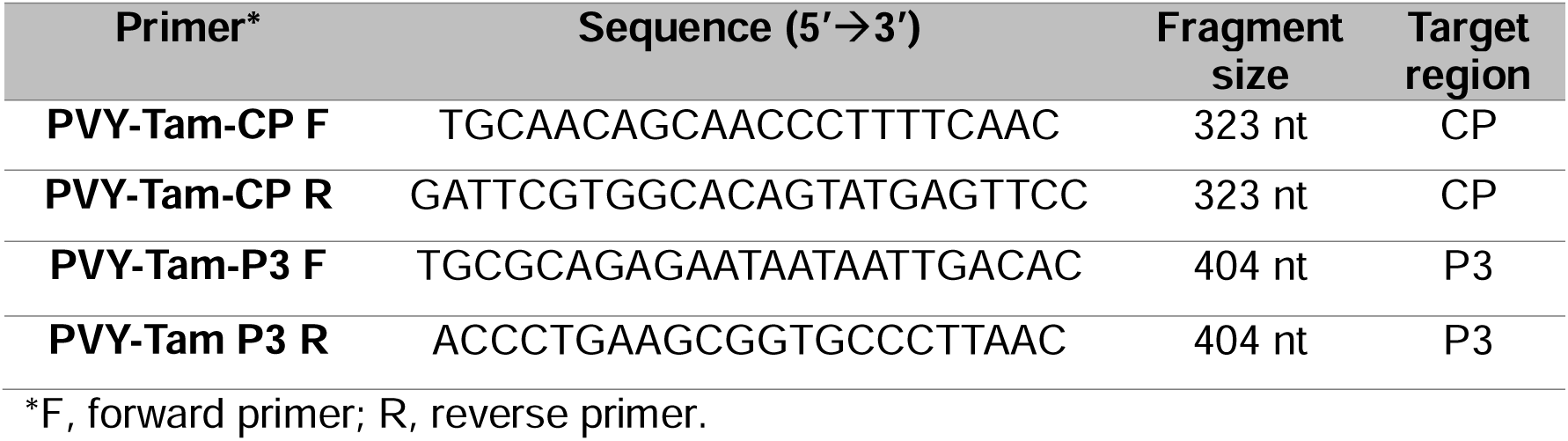
Primers used for PVY-Tam diagnosis by RT-PCR.

**Supplementary Table S4.**
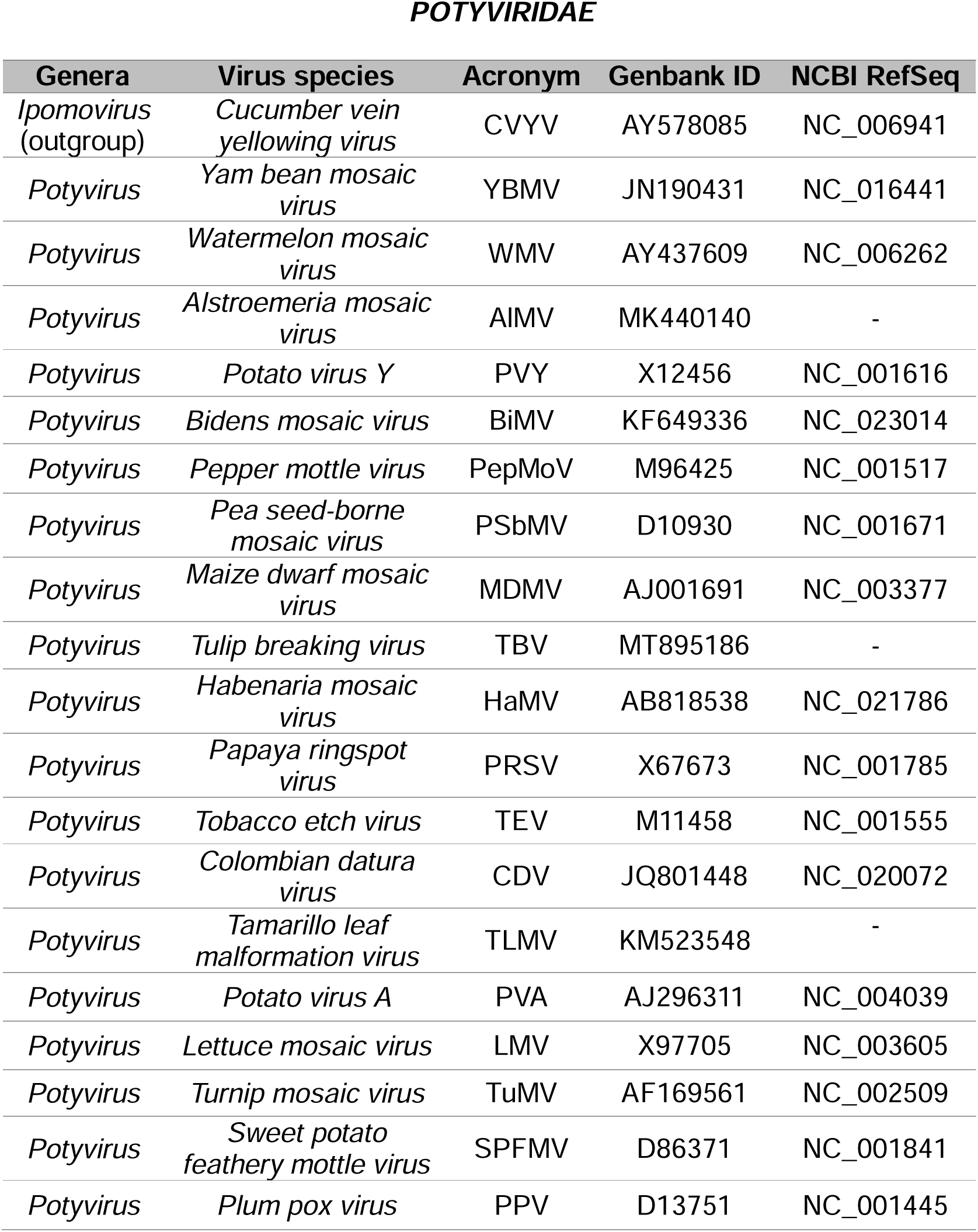
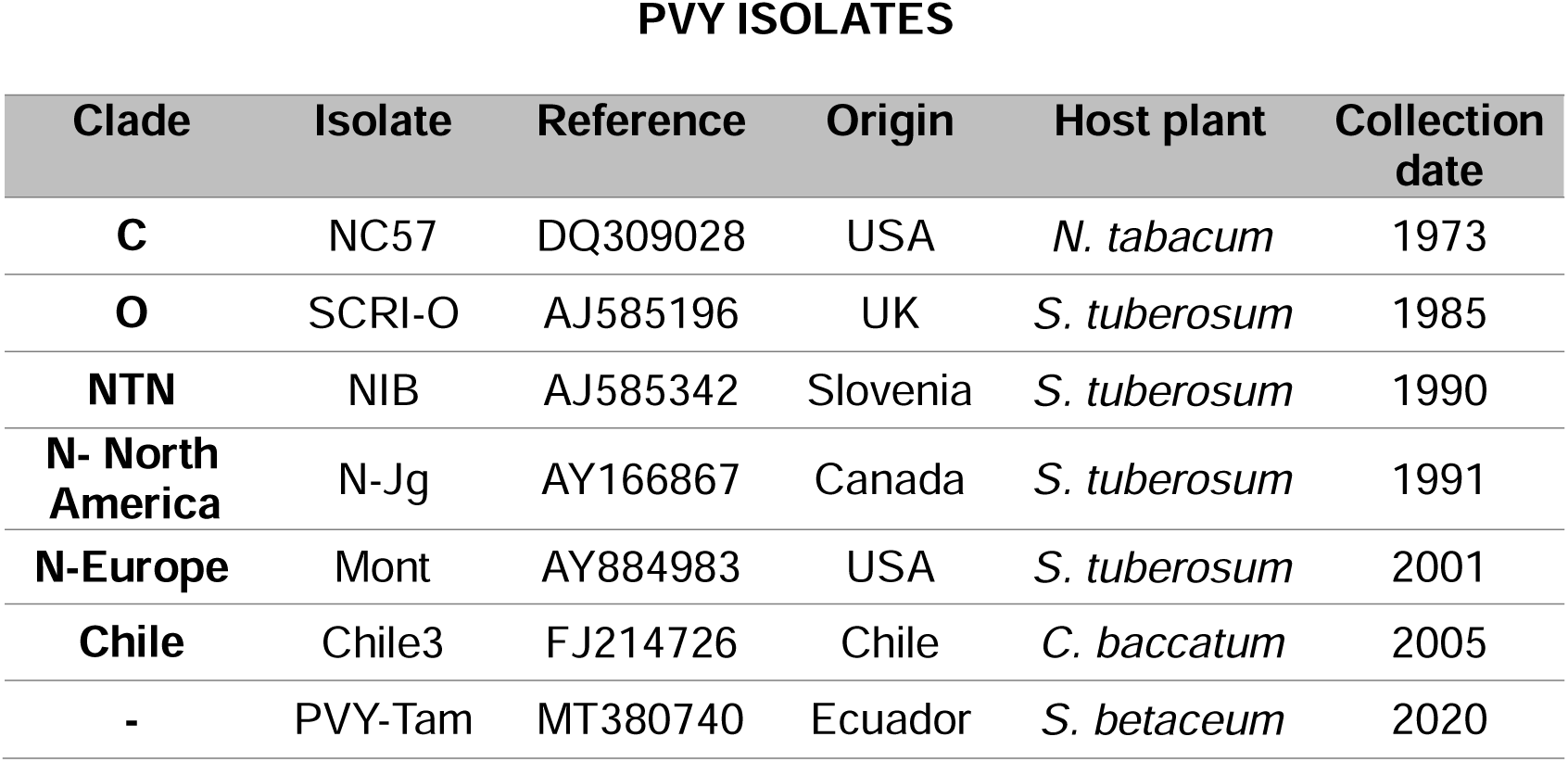
List of the virus species belonging to the *Potyviridae* family and PVY isolates used in the phylogenetic analysis.

**Supplementary Table S5.**
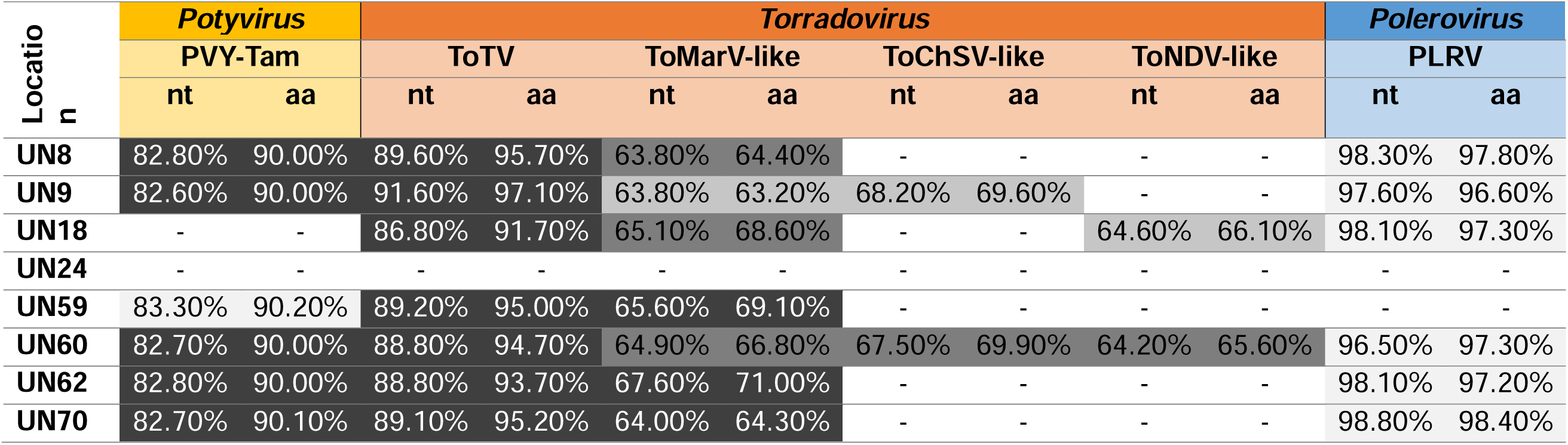
Identification of virus species in tamarillo-growing orchards across Nariño using Genome Detective software. Nucleotide (nt) and amino acid (aa) sequence identities correspond to an average of all the RNAs identified in different contigs for a given virus, as compared to their respective viral reference genomes. RNA-seq data that was filtered by number of reads, depth and coverage (indicated by colour gradation from highest to lowest intensity). UN24 is considered as negative control since no viruses are detected above the threshold. PVY-Tam, potato virus Y-Tamarillo (Genbank: MT380740); ToTV, tomato torrado virus (GB: DQ388879; DQ388880); ToMarV, tomato marchitez virus (GB: EF681764; EF681765); ToChSV, tomato chocolate spot virus (NCBI: NC_013075.1; NC_013076.1; ToNDV, tomato necrotic dwarf virus (NCBI: NC_027926.1; NC_027927.1); PLRV, potato leafroll virus (GB: D13954).

**Supplementary Table S6.**
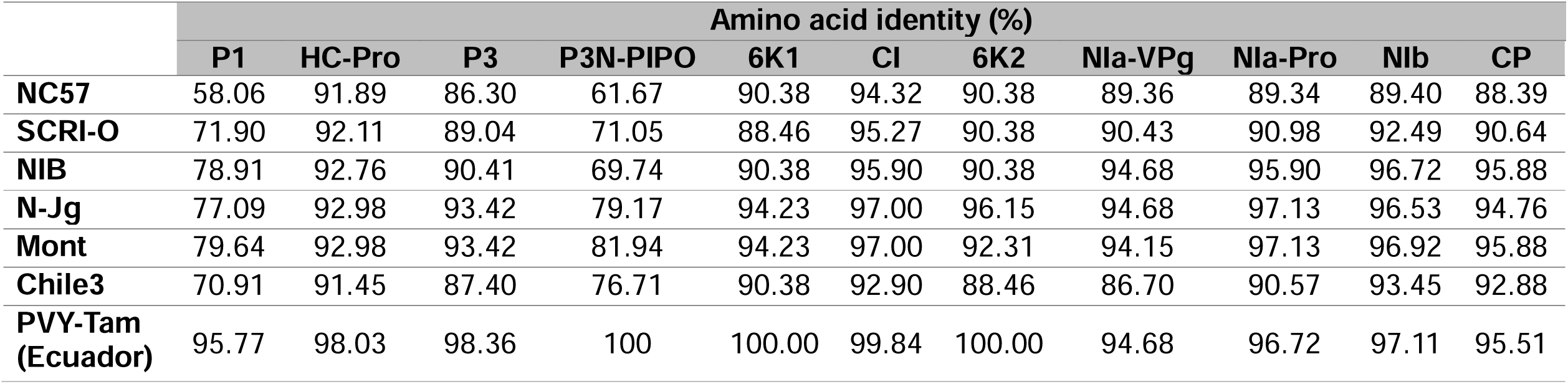
Amino acid sequence identities of the individual proteins of PVY-Tam (Nariño) to those of type members from the major PVY clades (C: NC57, O: SCRI-O, N: N-Jg and Mont, NTN: NIB, Chile: Chile3) and a PVY-Tam isolate previously reported in Ecuador (MT380740). Calculations were based on a two-sequence Blastp comparing the proteins from PVY-Tam and their respective from PVY isolates.

**Supplementary Figure S1.**
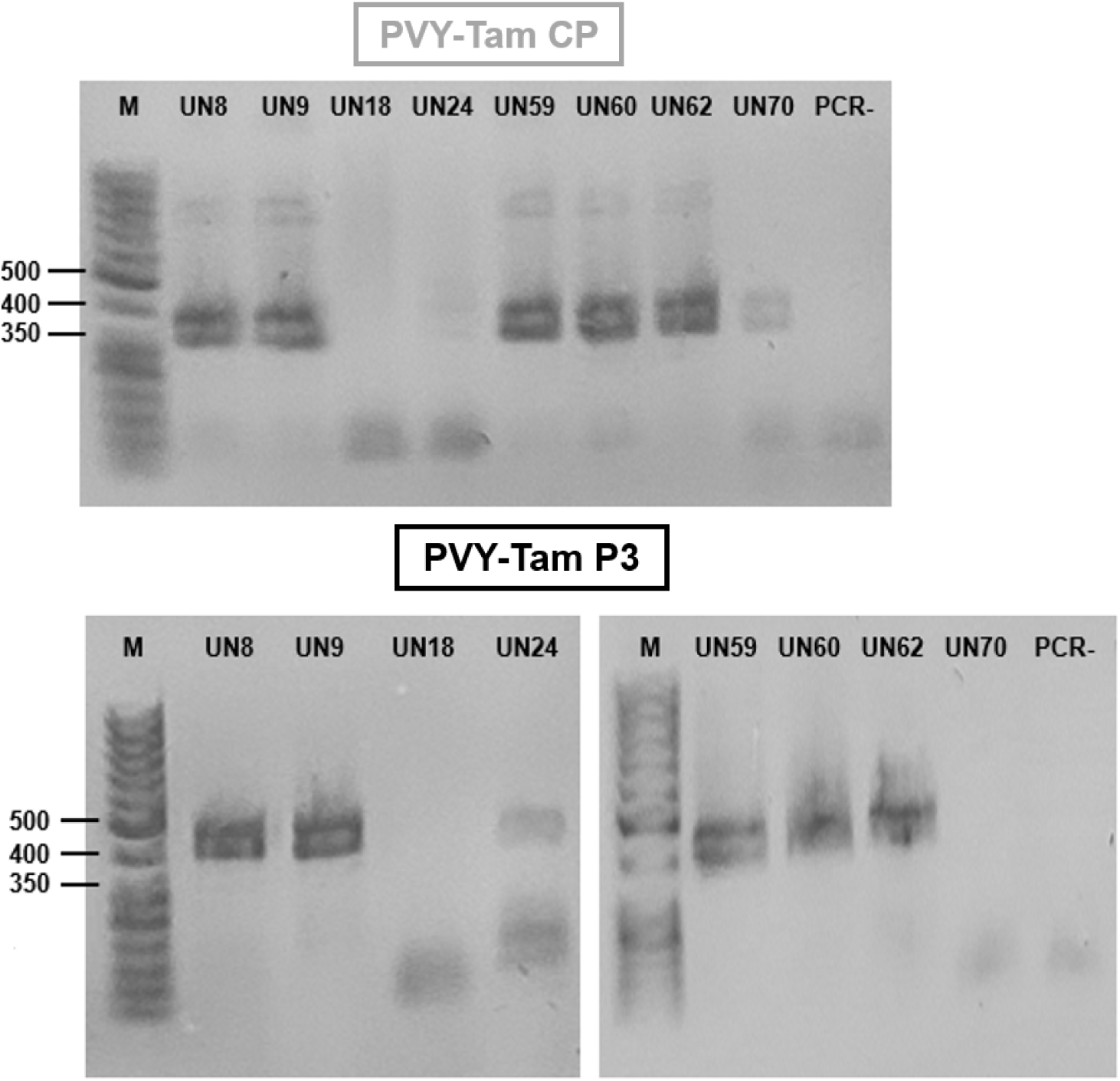
RT-PCR detection of PVY-Tam in infected plants from different geographical locations. The two sets of primer pairs corresponding to CP (left) and P3 (right) regions provide the same results. M, 50 bp DNA ladder with the length (bp) of some components indicated.

**Supplementary Figure S2.**
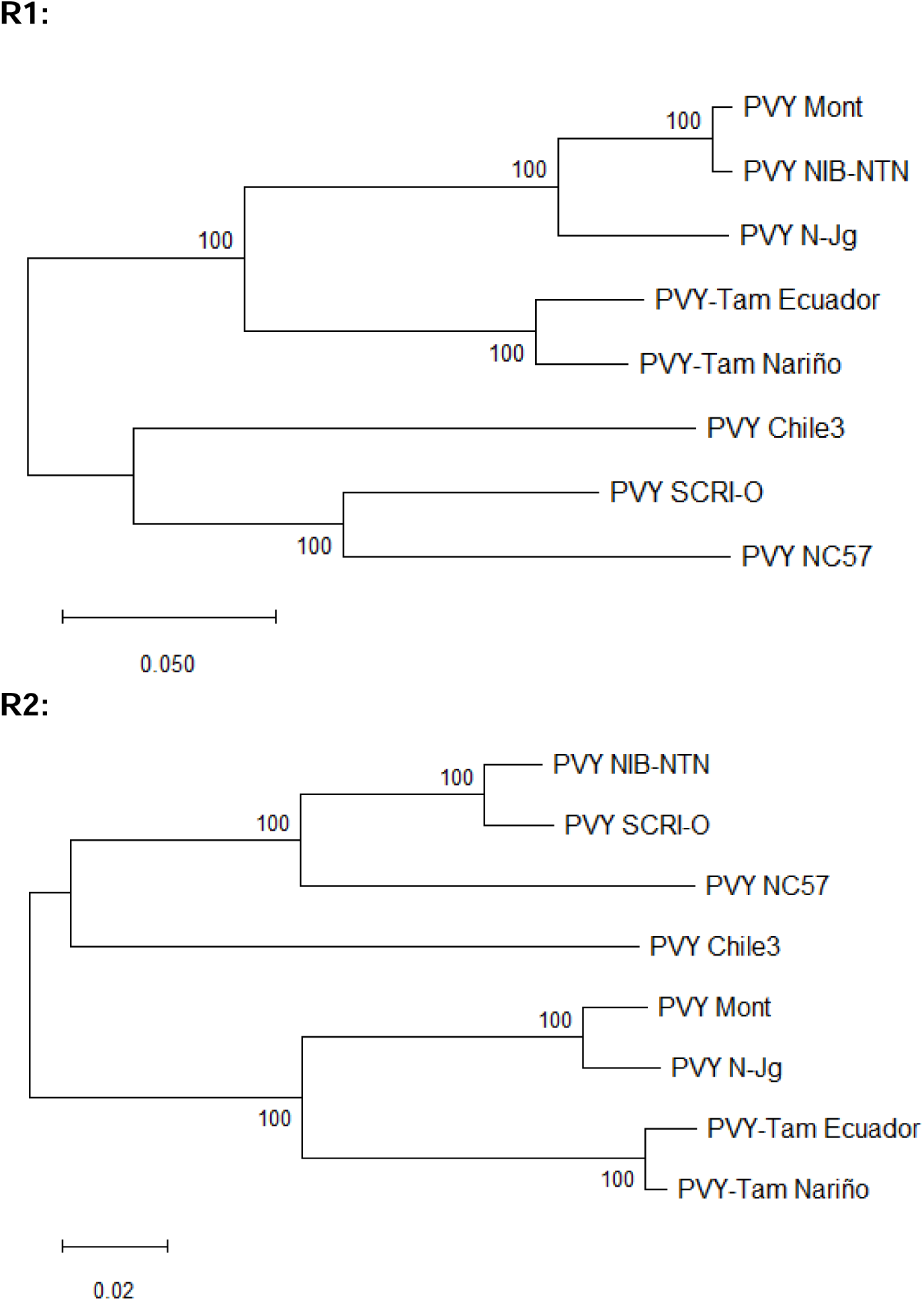
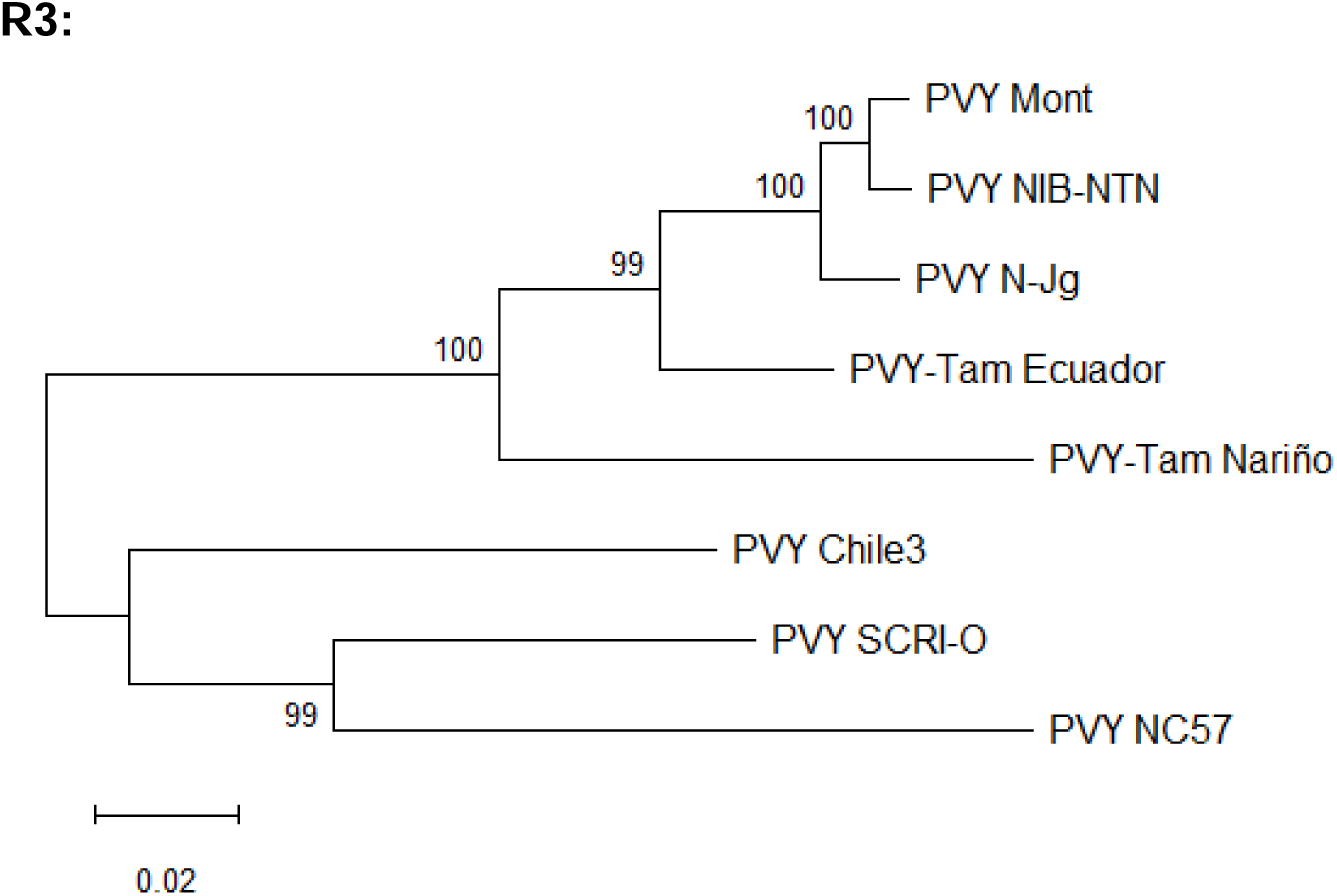
Maximum likelihood trees of the R1, R2 and R3 regions of the nucleotide sequences of the polyprotein for PVY-Tam and selected PVY isolates of the major clades. Trees are drawn to scale, with branch lengths measured in the number of substitutions per site and bootstrap values indicated above branches for each node. All positions containing gaps and missing data were eliminated.

## REFERENCES

1. Alvarez-Quinto, R., Amao, M., Muller, G., Fuentes, S., Grinstead, S., Fuentes-Bueno, I., Roenhorst, A., Westenberg, M., Botermans, M., Kreuze, J., & Mollov, D. (2023). Evidence that an Unnamed Isometric Virus Associated with Potato Rugose Disease in Peru Is a New Species of Genus Torradovirus. Phytopathology, 113(9), 1716–1728. 10.1094/PHYTO-11-22-0449-V

2. Ayala, M., González, E. P., Gutiérrez, P. A., Cotes, J. M., & Marín, M. A. (2010). Caracterización serológica y molecular de potyvirus asociados a la virosis del tomate de arbol en antioquia (colombia). Acta Biológica Colombiana, 15(3), 145–164. https://repositorio.unal.edu.co/handle/unal/28050

3. Batuman, O., Kuo, Y. W., Palmieri, M., Rojas, M. R., & Gilbertson, R. L. (2010). Tomato chocolate spot virus, a member of a new torradovirus species that causes a necrosis-associated disease of tomato in Guatemala. Archives of Virology, 155(6), 857–869. 10.1007/S00705-010-0653-9

4. Bhargava, K., & Joshi, R. (1959). A virus disease of tree tomato, Cyphomandra betacea Sendt. Due to Potato virus Y. American Journal of Potato Research, 36, 8.

5. Buitrago, D. (2013). Caracterización Socioeconómica de los productores de tomate de árbol en el departamento de Boyacá. In Vestigium Ire, 6, 55–64.

6. Carvajal-Yepes, M., Olaya, C., Lozano, I., Cuervo, M., Castaño, M., & Cuellar, W. J. (2014). Unraveling complex viral infections in cassava (Manihot esculenta Crantz) from colombia. Virus Research, 186, 76–86. 10.1016/j.virusres.2013.12.011

7. Chamberlain, E. (1954). Plant Viruses in New Zealand. In Department of Scientific and Industrial Research (Vol. 108).

8. Chikh-Ali, M., Vander Pol, D., Nikolaeva, O. V., Melzer, M. J., & Karasev, A. V. (2016). Biological and molecular characterization of a tomato isolate of potato virus Y (PVY) of the PVYC lineage. Archives of Virology, 161(12), 3561–3566. 10.1007/S00705-016-3071-9

9. Corrales-Cabra, E., Higuita, M., Hoyos, R., Gallo, Y., Marín, M., & Gutiérrez, P. (2021a). Prevalence of RNA viruses in seeds, plantlets, and adult plants of cape gooseberry (Physalis peruviana) in Antioquia (Colombia). Physiological and Molecular Plant Pathology, 116, 101715. 10.1016/J.PMPP.2021.101715

10. Corrales-Cabra, E., Higuita, M., Hoyos, R., Gallo, Y., Marín, M., & Gutiérrez, P. (2021b). Prevalence of RNA viruses in seeds, plantlets, and adult plants of cape gooseberry (Physalis peruviana) in Antioquia (Colombia). Physiological and Molecular Plant Pathology, 116, 101715. 10.1016/J.PMPP.2021.101715

11. Cuevas, J. M., Delaunay, A., Visser, J. C., Bellstedt, D. U., Jacquot, E., & Elena, S. F. (2012). Phylogeography and molecular evolution of potato virus Y. PLoS ONE, 7(5). 10.1371/JOURNAL.PONE.0037853

12. Dullemans, A. M., Cuperus, C., Verbeek, M., & van der Vlugt, R. A. A. (2011). Complete nucleotide sequence of a potato isolate of strain group C of potato virus Y from 1938. Archives of Virology, 156(3), 473–477. 10.1007/S00705-010-0853-3

13. Ferriol, I., Silva Junior, D. M., Nigg, J. C., Zamora-Macorra, E. J., & Falk, B. W. (2016). Identification of the cleavage sites of the RNA2-encoded polyproteins for two members of the genus Torradovirus by N-terminal sequencing of the virion capsid proteins. Virology, 498, 109–115. 10.1016/j.virol.2016.08.014

14. Fuchs, M., Hily, J. M., Petrzik, K., Sanfaçon, H., Thompson, J. R., van der Vlugt, R., & Wetzel, T. (2022). ICTV Virus Taxonomy Profile: Secoviridae 2022. Journal of General Virology, 103(12). 10.1099/JGV.0.001807

15. Gibbs, A. J., Ohshima, K., Yasaka, R., Mohammadi, M., Gibbs, M. J., & Jones, R. A. C. (2017). The phylogenetics of the global population of potato virus Y and its necrogenic recombinants. Virus Evolution, 3(1). 10.1093/VE/VEX002

16. Gibbs, A., & Ohshima, K. (2010a). Potyviruses and the digital revolution. Annual Review of Phytopathology, 48, 205–223. 10.1146/ANNUREV-PHYTO-073009-114404

17. Gibbs, A., & Ohshima, K. (2010b). Potyviruses and the digital revolution. Annual Review of Phytopathology, 48, 205–223. 10.1146/ANNUREV-PHYTO-073009-114404

18. Green, K. J., Chikh-Ali, M., Hamasaki, R. T., Melzer, M. J., & Karasev, A. V. (2017). Potato virus Y (PVY) isolates from Physalis peruviana are unable to systemically infect potato or pepper and form a distinct new lineage within the PVYC strain group. Phytopathology, 107(11), 1433–1439. 10.1094/PHYTO-04-17-0147-R

19. Green, K. J., Funke, C. N., Chojnacky, J., Alvarez-Quinto, R. A., Ochoa, J. B., Quito-Avila, D. F., & Karasev, A. V. (2020). Potato virus Y (PVY) isolates from solanum betaceum represent three novel recombinants within the PVYN strain group and are unable to systemically spread in potato. Phytopathology, 110(9), 1588–1596. 10.1094/PHYTO-04-20-0111-R

20. Gutiérrez, P. A., Alzate, J. F., & Marín Montoya, M. (2015). Genome sequence of a virus isolate from tamarillo (Solanum betaceum) in Colombia: evidence for a new potyvirus. Archives of Virology, 160(2), 557–560. 10.1007/S00705-014-2296-8

21. Herrera-V asquez, J. A., Alfaro-Fernández, A., Córdoba-Sellés, M. C., Cebrián, M. C., Font, M. I., & Jordá, C. (2009). First report of tomato torrado virus infecting tomato in single and mixed infections with Cucumber mosaic virus in panama. Plant Disease, 93(2), 198. 10.1094/PDIS-93-2-0198A

22. Jaramillo, M., Gutiérrez, P. A., Lagos, L. E., Cotes, J. M., & Marín, M. (2011). Detection of a complex of viruses in tamarillo (Solanum betaceum) orchards in the Andean region of Colombia. Tropical Plant Pathology, 36(3), 150–159. 10.1590/S1982-56762011000300003

23. Jaramillo, M. M., Álvarez, J. A., & Marín, M. (2012). Características de los virus asociados a la virosis del tomate de árbol (Solanum betaceum) en Colombia. Revista Lasallista de Investigación, 9(1), 115–127. https://dialnet.unirioja.es/servlet/articulo?codigo=4316856&info=resumen&idioma=E NG

24. Kumar, S., Stecher, G., Suleski, M., Sanderford, M., Sharma, S., & Tamura, K. (2024). MEGA12: Molecular Evolutionary Genetic Analysis version 12 for adaptive and green computing. Molecular Biology and Evolution, 41(12). 10.1093/MOLBEV/MSAE263

25. Martínez, J. E., Cotes, J. M., & Marín, M. (2010). Detección serológica y molecular de virus en áfidos asociados a cultivos de tomate de árbol con síntomas de virus en Antioquia y Nariño (Colombia). Revista de La Facultad de Ciencias Básicas, 6(2), 182–197. https://dialnet.unirioja.es/servlet/articulo?codigo=3662754&info=resumen&idioma=SPA

26. Moreno, A. B., & López-Moya, J. J. (2020). When viruses play team sports: Mixed infections in plants. Phytopathology, 110(1), 29–48. 10.1094/PHYTO-07-19-0250-FI/ASSET/IMAGES/LARGE/PHYTO-07-19-0250-FI_T2-1576622563041.JPEG

27. Pruss, G., Ge, X., Shi, X. M., Carrington, J. C., & Vance, V. B. (1997). Plant viral synergism: The potyviral genome encodes a broad-range pathogenicity enhancer that transactivates replication of heterologous viruses. Plant Cell, 9(6), 859–868. 10.1105/TPC.9.6.859

28. Quenouille, J., Vassilakos, N., & Moury, B. (2013). Potato virus Y: A major crop pathogen that has provided major insights into the evolution of viral pathogenicity. Molecular Plant Pathology, 14(5), 439–452. 10.1111/MPP.12024

29. Singh, R. P., Valkonen, J. P. T., Gray, S. M., Boonham, N., Jones, R. A. C., Kerlan, C., & Schubert, J. (2008). Discussion paper: The naming of Potato virus Y strains infecting potato. Archives of Virology, 153(1), 1–13. 10.1007/S00705-007-1059-1

30. Valli, A. A., Domingo-Calap, M. L., González de Prádena, A., García, J. A., Cui, H., Desbiez, C., & López-Moya, J. J. (2024). Reconceptualizing transcriptional slippage in plant RNA viruses. MBio, 15(10), e0212024. 10.1128/MBIO.02120-24

31. Van Der Vlugt, R. A. A., Verbeek, M., Dullemans, A. M., Wintermantel, W. M., Cuellar, W. J., Fox, A., & Thompson, J. R. (2015). Torradoviruses. Annual Review of Phytopathology, 53, 485–512. 10.1146/ANNUREV-PHYTO-080614-120021

32. Verbeek, M., & Dullemans, A. M. (2012a). First Report of Tomato torrado virus Infecting Tomato in Colombia. Plant Disease, 96(4), 592–592. 10.1094/PDIS-11-11-1000

33. Verbeek, M., & Dullemans, A. M. (2012b). First Report of Tomato torrado virus Infecting Tomato in Colombia. Plant Disease, 96(4), 592–592. 10.1094/PDIS-11-11-1000

34. Verbeek, M., Dullemans, A. M., Van Den Heuvel, J. F. J. M., Maris, P. C., & Van Der Vlugt, R. A. A. (2008). Tomato marchitez virus, a new plant picorna-like virus from tomato related to tomato torrado virus. Archives of Virology, 153(1), 127–134. 10.1007/S00705-007-1076-0

35. Verbeek, M., Dullemans, A., van den Heuvel, H., Maris, P., & van der Vlugt, R. (2010). Tomato chocolàte virus: A new plant virus infecting tomato and a proposed member of the genus Torradovirus. Archives of Virology, 155(5), 751–755. 10.1007/S00705-010-0640-1

36. Vilsker, M., Moosa, Y., Nooij, S., Fonseca, V., Ghysens, Y., Dumon, K., Pauwels, R., Alcantara, L. C., Vanden Eynden, E., Vandamme, A. M., Deforche, K., & De Oliveira, T. (2019). Genome Detective: An automated system for virus identification from high-throughput sequencing data. Bioinformatics, 35(5), 871–873. 10.1093/BIOINFORMATICS/BTY695

37. Visser, J. C., Bellstedt, D. U., & Pirie, M. D. (2012). The Recent Recombinant Evolution of a Major Crop Pathogen, Potato virus Y. PLoS ONE, 7(11). 10.1371/JOURNAL.PONE.0050631

38. Vizuete, B., Insuasti, M., Ochoa, J., & Ellis, M. (1990). Biological and serological characterization of tree tomato virus diseases in Ecuador. INIAP-Ohio State University.

39. Wei, T., Zhang, C., Hong, J., Xiong, R., Kasschau, K. D., Zhou, X., Carrington, J. C., & Wang, A. (2010). Formation of complexes at plasmodesmata for potyvirus intercellular movement is mediated by the viral protein P3N-PIPO. PLoS Pathogens, 6(6). 10.1371/JOURNAL.PPAT.1000962

40. Wen, R. H., & Hajimorad, M. R. (2010). Mutational analysis of the putative pipo of soybean mosaic virus suggests disruption of PIPO protein impedes movement. Virology, 400(1), 1–7. 10.1016/j.virol.2010.01.022

41. Zhao, Z., Cheng, J., Sun, W., Zhu, J., Lu, S., Feng, Y., Song, Z., Yang, Y., & Wu, X. (2024). The LINC01176-miR-218-5p-IL-36G Network is Responsible for the Pathogenesis of Psoriasis by Promoting Inflammation. *Clinical*, Cosmetic and Investigational Dermatology, 17, 1–12. 10.2147/CCID.S444265

