## Supplementary material for "Nucleotide sequence analysis reveals the presence of PVY-Tam isolates affecting tamarillo in Colombia": Suplementary Data

**SUPPLEMENTARY DATA**

**Supplementary Table S1.** Description of the geographic locations across Nariño (Colombia) used in the study, in which samples of tamarillo plants showing symptoms of virosis were collected.

| **Sample** | **Geographic coordinates** | **Altitude (msnm)** | **Municipality** | **Site description** |
| --- | --- | --- | --- | --- |
| **UN8** | 0.910356, -77.522176 | 2330 | Contadero | P. San Juan II |
| **UN9** | 0.910356, -77.522176 | 2330 | Contadero | P. San Juan II |
| **UN18** | 0.884677, -77.545866 | 2478 | Córdoba | Las Delicias |
| **UN24** | 1.337782, -77.577840 | 2015 | Samaniego | Finca COPROINSAM |
| **UN59** | 0.895437, -77.540394 | 2390 | Contadero | P. San Juan I |
| **UN60** | 0.895171, -77.540944 | 2386 | Córdoba | P. San Juan I |
| **UN62** | 0.895171, -77.540944 | 2386 | Córdoba | P. San Juan I |
| **UN70** | 0.894740, -77.546471 | 2443 | Ipiales (San Juan) | San Juan urban area |

**Supplementary Table S2.** Symptoms description of the tamarillo plants sampled in the different geographic locations.

| **Sample** | **Symptoms** | |
| --- | --- | --- |
|  | **On leaves** | **On fruit** |
| **UN8** | Chlorosis: Yellowing of leaf tissue, particularly along the edges; Necrotic spots: Dark, dead tissue lesions; Mottling: Irregular discoloration patterns on the leaf surface; Leaf deformation: Presence of stress potentially associated with the onset of leaf curling. | Mottling or Surface Spots: Irregular dark spots on the fruit surface; Partial fruit chlorosis: Greenish-yellow areas that fail to ripen uniformly; Superficial necrosis: Small brown lesions potentially indicative of superficial necrosis associated with secondary infections. |
| **UN9** | Mosaic: Presence of heterogeneously colored areas with light green and yellowish zones, Extensive chlorosis: Generalized yellowing of leaf tissue, suggesting disruptions in photosynthetic processes; Necrosis: Large, darkened lesions associated with tissue death, possibly resulting from advanced viral infection or secondary co-infection with fungal pathogens; Leaf deformation: Curvature and malformation of the leaf structure; Mottling: Irregular dark spots, likely due to pigment distribution disturbances. | Mottling or surface spots: Irregular dark spots on the fruit surface; Partial chlorosis: Greenish-yellow areas that fail to mature uniformly, potentially due to viral interference in fruit metabolism; Superficial necrosis: Small brown lesions associated with viral infections and secondary colonization by opportunistic microorganisms. |
| **UN18** | Mild mosaic: Presence of heterogeneously colored areas with dark green to purple zones; Dark pigmentation: Uneven dark coloration; Leaf deformation: Some leaves exhibit slight curvature and structural changes. | Dark surface Lesions: Irregular brown spots on the fruit surface, possibly associated with superficial necrosis due to secondary infections or physical stress; Partial ripening: green fruit coloration with incomplete and uneven ripening. |
| **UN24** | Irregular mottling: presence of light green spots scattered across the leaf blade; Partial chlorosis: Areas with a loss of uniform green pigmentation, suggesting disruptions in chlorophyll metabolism; Leaf deformation: Mild leaf undulation and curvature; Blistering: Localized chlorotic blisters. |  |
| **UN59** | Irregular dark pigmentation: Areas with purple to dark green coloration, associated with physiological stress from viral infections affecting pigment distribution; Leaf deformation: mild leaf curvature and undulation, indicating structural alterations. | Extensive necrotic lesions: Large irregular black spots on the fruit surface, associated with advanced necrosis caused by secondary fungal infections; Fruit hardening: Formation of hardened pulp cysts affecting ripening, possibly due to metabolic interference caused by the pathogen. |
| **UN60** | Mild color alteration: Leaf shows heterogeneous green tones, suggesting an irregular pigment distribution associated with early-stage infections; Localized Lesions: small necrotic areas linked to the plant's defense response against viral infections; Uneven Coloration: in the upper left quadrant, areas of light and dark green are observed without a defined pattern, indicating mild mottling. |  |
| **UN62** | Irregular mottling: Presence of dark spots heterogeneously distributed across the leaf blade, associated with disruptions in pigment distribution; Local lesions: Necrotic spots with dead tissue, indicative of the plant’s hypersensitive response; Leaf deformation: Slight curvature of leaf edges.  Partial chlorosis: Yellowish areas between veins, associated with the loss of uniform green pigmentation and metabolic dysfunction. | Necrotic lesions: Irregular black spots on the fruit surface, indicative of advanced necrosis likely caused by secondary infections; Irregular pigmentation: pale green areas interspersed with darker regions, suggesting an incomplete ripening process due to metabolic interference caused by viral infection. |
| **UN70** | Irregular mottling: The presence of spots ranging from light to dark purple, varying in size, accompanied by slight blister-like thickening. These spots are heterogeneously distributed across the surface of the leaf blade; Partial chlorosis: Yellowish areas scattered between the veins, indicating dysfunction in chlorophyll synthesis and metabolic alterations affecting normal pigment distribution; Necrotic lesions: Irregular necrotic areas with dead tissue, possibly associated with a secondary infection, compromising the structural integrity of the leaf; Leaf deformation: Slight curvature and irregular undulation along the leaf margins, suggesting structural alterations likely caused by viral interference; Surface burns: Diffuse dark discoloration on the upper surface of the leaf, potentially a consequence of advanced irregular mottling. |  |

**Supplementary Table S3.** Primers used for PVY-Tam diagnosis by RT-PCR.

| **Primer*** | **Sequence (5’**🡪**3’)** | **Fragment size** | **Target region** |
| --- | --- | --- | --- |
| **PVY-Tam-CP F** | TGCAACAGCAACCCTTTTCAAC | 323 nt | CP |
| **PVY-Tam-CP R** | GATTCGTGGCACAGTATGAGTTCC | 323 nt | CP |
| **PVY-Tam-P3 F** | TGCGCAGAGAATAATAATTGACAC | 404 nt | P3 |
| **PVY-Tam P3 R** | ACCCTGAAGCGGTGCCCTTAAC | 404 nt | P3 |

*F, forward primer; R, reverse primer.

**Supplementary Table S4.** List of the virus species belonging to the *Potyviridae* family and PVY isolates used in the phylogenetic analysis.

***POTYVIRIDAE***

| **Genera** | **Virus species** | **Acronym** | **Genbank ID** | **NCBI RefSeq** |
| --- | --- | --- | --- | --- |
| *Ipomovirus* (outgroup) | *Cucumber vein yellowing virus* | CVYV | AY578085 | NC_006941 |
| *Potyvirus* | *Yam bean mosaic virus* | YBMV | JN190431 | NC_016441 |
| *Potyvirus* | *Watermelon mosaic virus* | WMV | AY437609 | NC_006262 |
| *Potyvirus* | *Alstroemeria mosaic virus* | AlMV | MK440140 | - |
| *Potyvirus* | *Potato virus Y* | PVY | X12456 | NC_001616 |
| *Potyvirus* | *Bidens mosaic virus* | BiMV | KF649336 | NC_023014 |
| *Potyvirus* | *Pepper mottle virus* | PepMoV | M96425 | NC_001517 |
| *Potyvirus* | *Pea seed-borne mosaic virus* | PSbMV | D10930 | NC_001671 |
| *Potyvirus* | *Maize dwarf mosaic virus* | MDMV | AJ001691 | NC_003377 |
| *Potyvirus* | *Tulip breaking virus* | TBV | MT895186 | - |
| *Potyvirus* | *Habenaria mosaic virus* | HaMV | AB818538 | NC_021786 |
| *Potyvirus* | *Papaya ringspot virus* | PRSV | X67673 | NC_001785 |
| *Potyvirus* | *Tobacco etch virus* | TEV | M11458 | NC_001555 |
| *Potyvirus* | *Colombian datura virus* | CDV | JQ801448 | NC_020072 |
| *Potyvirus* | *Tamarillo leaf malformation virus* | TLMV | KM523548 | - |
| *Potyvirus* | *Potato virus A* | PVA | AJ296311 | NC_004039 |
| *Potyvirus* | *Lettuce mosaic virus* | LMV | X97705 | NC_003605 |
| *Potyvirus* | *Turnip mosaic virus* | TuMV | AF169561 | NC_002509 |
| *Potyvirus* | *Sweet potato feathery mottle virus* | SPFMV | D86371 | NC_001841 |
| *Potyvirus* | *Plum pox virus* | PPV | D13751 | NC_001445 |

**PVY ISOLATES**

| **Clade** | **Isolate** | **Reference** | **Origin** | **Host plant** | **Collection date** |
| --- | --- | --- | --- | --- | --- |
| **C** | NC57 | DQ309028 | USA | *N. tabacum* | 1973 |
| **O** | SCRI-O | AJ585196 | UK | *S. tuberosum* | 1985 |
| **NTN** | NIB | AJ585342 | Slovenia | *S. tuberosum* | 1990 |
| **N- North America** | N-Jg | AY166867 | Canada | *S. tuberosum* | 1991 |
| **N-Europe** | Mont | AY884983 | USA | *S. tuberosum* | 2001 |
| **Chile** | Chile3 | FJ214726 | Chile | *C. baccatum* | 2005 |
| **-** | PVY-Tam | MT380740 | Ecuador | *S. betaceum* | 2020 |

**Supplementary Table S5.** Identification of virus species in tamarillo-growing orchards across Nariño using Genome Detective software. Nucleotide (nt) and amino acid (aa) sequence identities correspond to an average of all the RNAs identified in different contigs for a given virus, as compared to their respective viral reference genomes. RNA-seq data that was filtered by number of reads, depth and coverage (indicated by colour gradation from highest to lowest intensity). UN24 is considered as negative control since no viruses are detected above the threshold. PVY-Tam, potato virus Y-Tamarillo (Genbank: MT380740); ToTV, tomato torrado virus (GB: DQ388879; DQ388880); ToMarV, tomato marchitez virus (GB: EF681764; EF681765); ToChSV, tomato chocolate spot virus (NCBI: NC_013075.1; NC_013076.1; ToNDV, tomato necrotic dwarf virus (NCBI: NC_027926.1; NC_027927.1); PLRV, potato leafroll virus (GB: D13954).

| **Location** | ***Potyvirus*** | | ***Torradovirus*** | | | | | | | | ***Polerovirus*** | |
| --- | --- | --- | --- | --- | --- | --- | --- | --- | --- | --- | --- | --- |
|  | **PVY-Tam** | | **ToTV** | | **ToMarV-like** | | **ToChSV-like** | | **ToNDV-like** | | **PLRV** | |
|  | **nt** | **aa** | **nt** | **aa** | **nt** | **aa** | **nt** | **aa** | **nt** | **aa** | **nt** | **aa** |
| **UN8** | 82.80% | 90.00% | 89.60% | 95.70% | 63.80% | 64.40% | - | - | - | - | 98.30% | 97.80% |
| **UN9** | 82.60% | 90.00% | 91.60% | 97.10% | 63.80% | 63.20% | 68.20% | 69.60% | - | - | 97.60% | 96.60% |
| **UN18** | - | - | 86.80% | 91.70% | 65.10% | 68.60% | - | - | 64.60% | 66.10% | 98.10% | 97.30% |
| **UN24** | - | - | - | - | - | - | - | - | - | - | - | - |
| **UN59** | 83.30% | 90.20% | 89.20% | 95.00% | 65.60% | 69.10% | - | - | - | - | - | - |
| **UN60** | 82.70% | 90.00% | 88.80% | 94.70% | 64.90% | 66.80% | 67.50% | 69.90% | 64.20% | 65.60% | 96.50% | 97.30% |
| **UN62** | 82.80% | 90.00% | 88.80% | 93.70% | 67.60% | 71.00% | - | - | - | - | 98.10% | 97.20% |
| **UN70** | 82.70% | 90.10% | 89.10% | 95.20% | 64.00% | 64.30% | - | - | - | - | 98.80% | 98.40% |

**Supplementary Table S6.** Amino acid sequence identities of the individual proteins of PVY-Tam (Nariño) to those of type members from the major PVY clades (C: NC57, O: SCRI-O, N: N-Jg and Mont, NTN: NIB, Chile: Chile3) and a PVY-Tam isolate previously reported in Ecuador (MT380740). Calculations were based on a two-sequence Blastp comparing the proteins from PVY-Tam and their respective from PVY isolates.

|  | **Amino acid identity (%)** | | | | | | | | | | |
| --- | --- | --- | --- | --- | --- | --- | --- | --- | --- | --- | --- |
|  | **P1** | **HC-Pro** | **P3** | **P3N-PIPO** | **6K1** | **CI** | **6K2** | **NIa-VPg** | **NIa-Pro** | **NIb** | **CP** |
| **NC57** | 58.06 | 91.89 | 86.30 | 61.67 | 90.38 | 94.32 | 90.38 | 89.36 | 89.34 | 89.40 | 88.39 |
| **SCRI-O** | 71.90 | 92.11 | 89.04 | 71.05 | 88.46 | 95.27 | 90.38 | 90.43 | 90.98 | 92.49 | 90.64 |
| **NIB** | 78.91 | 92.76 | 90.41 | 69.74 | 90.38 | 95.90 | 90.38 | 94.68 | 95.90 | 96.72 | 95.88 |
| **N-Jg** | 77.09 | 92.98 | 93.42 | 79.17 | 94.23 | 97.00 | 96.15 | 94.68 | 97.13 | 96.53 | 94.76 |
| **Mont** | 79.64 | 92.98 | 93.42 | 81.94 | 94.23 | 97.00 | 92.31 | 94.15 | 97.13 | 96.92 | 95.88 |
| **Chile3** | 70.91 | 91.45 | 87.40 | 76.71 | 90.38 | 92.90 | 88.46 | 86.70 | 90.57 | 93.45 | 92.88 |
| **PVY-Tam (Ecuador)** | 95.77 | 98.03 | 98.36 | 100 | 100.00 | 99.84 | 100.00 | 94.68 | 96.72 | 97.11 | 95.51 |

**Supplementary Figure S1.** RT-PCR detection of PVY-Tam in infected plants from different geographical locations. The two sets of primer pairs corresponding to CP (left) and P3 (right) regions provide the same results. M, 50 bp DNA ladder with the length (bp) of some components indicated.

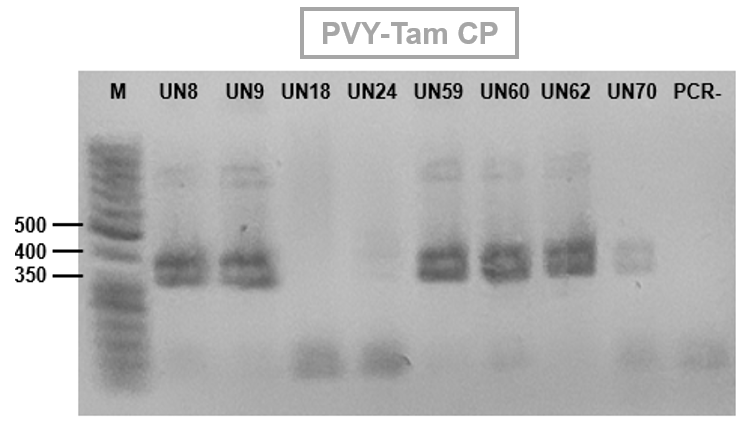

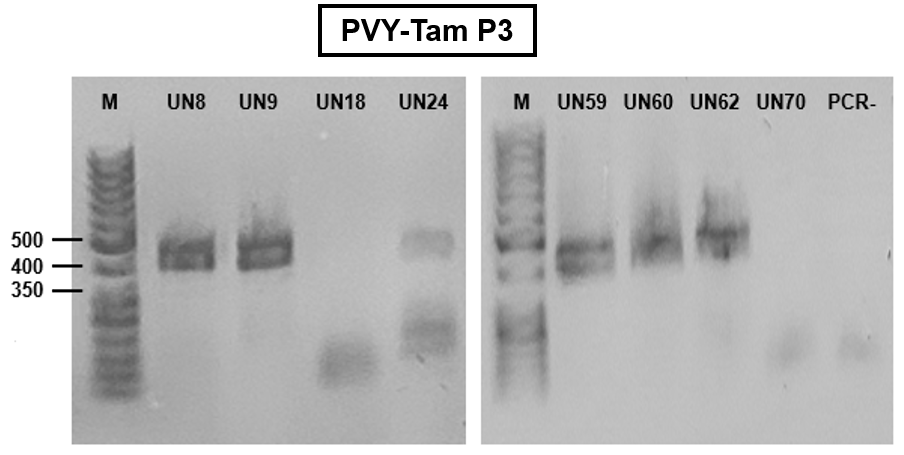

**Supplementary Figure S2.** Maximum likelihood trees of the R1, R2 and R3 regions of the nucleotide sequences of the polyprotein for PVY-Tam and selected PVY isolates of the major clades. Trees are drawn to scale, with branch lengths measured in the number of substitutions per site and bootstrap values indicated above branches for each node. All positions containing gaps and missing data were eliminated.

**R1:**

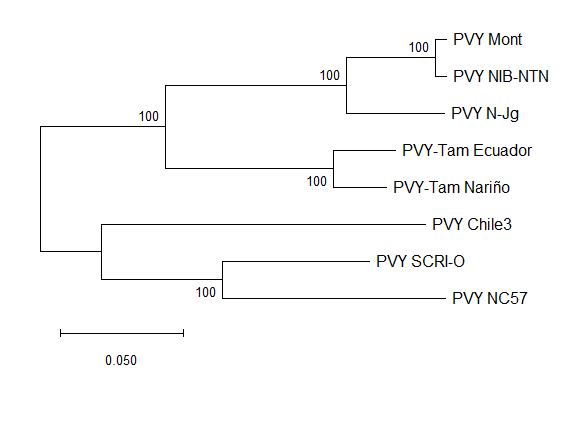

**R2:**

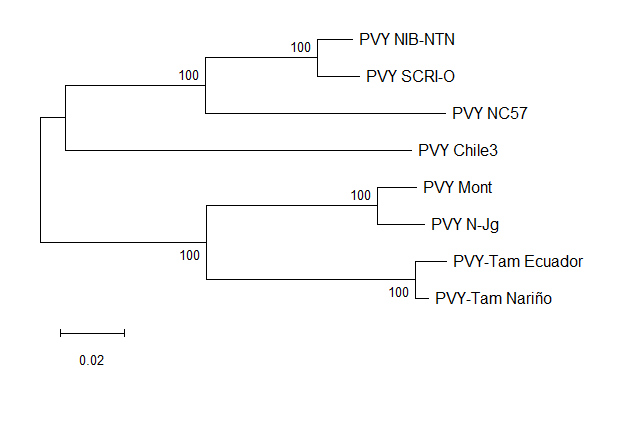

**R3:**

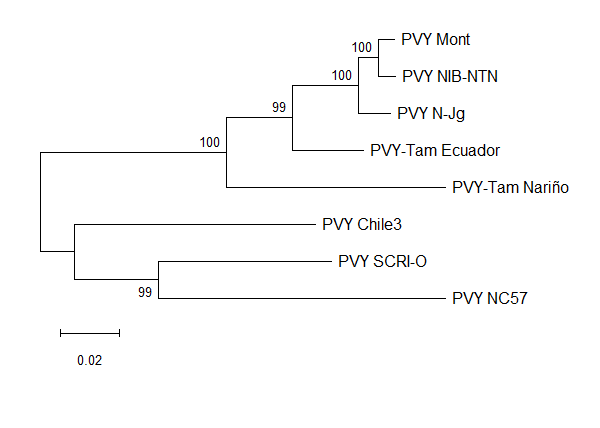
