## Supplementary material for "Nucleotide sequence analysis reveals the presence of PVY-Tam isolates affecting tamarillo in Colombia": Genome Detective reports

### Analysis of UN8

| | |
|---|---|
| **Technology** | Paired-end short reads |
| **Input Files** | UN8_R1.fq.gz (1.92 GB), UN8_R2.fq.gz (2 GB) |
| **Submitted On** | 2023-09-26 13:08:53 UTC |
| **Duration** | 3h 48m 49s |
| **Tool Version** | panviral2.64 |
| **Location** | https://www.genomedetective.com/db/ui/analysis/35ae2222-bf0b-41be-887f-70156779792e |

#### Statistics

| | |
|---|---|
| **Original Read Length** | 20 - 150 |
| **Trimmed Read Length** | 50 - 135 |

| | # Reads | % of Reads |
|---|---|---|
| **Input file** | 60259918 | 100.0% |
| **After QC** | 59835436 | 99.3% |
| **After filtering** | 4578960 | 7.6% |
| **Mapped back to contigs** | 1898442 | 3.2% |

#### Assignments

| Assignment | No. of Reads | Depth of Coverage | Identity | | Genome Coverage | |
|---|---|---|---|---|---|---|
| | | | NT | AA | | |
| Torradovirus lycopersici (4 segments out of 2) | 693606 | 3689.4 | 89.6% | 95.7% | 99.9% | |
| Torradovirus lycopersici (segment RNA 2) | 313618 | 7518.4 | 96.6% | 98.0% | 99.6% | 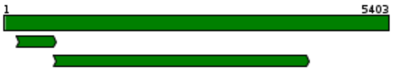  |
| Torradovirus lycopersici (segment RNA 1) | 186149 | 4265.8 | 94.3% | 98.8% | 99.4% | 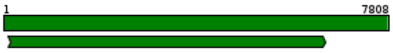  |
| Torradovirus lycopersici (segment RNA 1) | 114995 | 2254.7 | 90.5% | 97.8% | 87.0% | 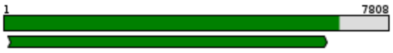  |
| Torradovirus lycopersici (segment RNA 1) | 78839 | 1527.7 | 78.6% | 89.9% | 86.9% | 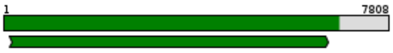  |
| Potato virus Y | 634281 | 8776.7 | 82.8% | 90.0% | 99.5% | 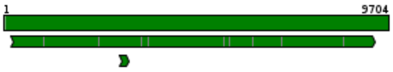  |
| Torradovirus marchitezum (2 segments out of 2) | 274810 | 4599.5 | 63.8% | 64.4% | 64.1% | |
| Torradovirus marchitezum (segment RNA 1) | 176957 | 3512.7 | 63.0% | 63.4% | 93.0% | 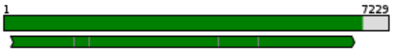  |

| Assignment | No. of Reads | Depth of Coverage | Identity | | Genome Coverage | |
|---|---|---|---|---|---|---|
| | | | NT | AA | | |
| Torradovirus marchitezum (segment RNA 2) | 97853 | 11498.6 | 69.3% | 72.8% | 21.6% | 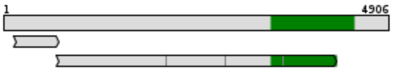 |
| Diachasmimorpha longicaudata entomopoxvirus (segment NC_043455.1) | 13895 | 1498.5 | 59.8% | 55.3% | 83.7% | 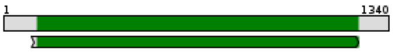 |
| Bracoviriform glomeratae (segment NC_043292.1) | 5742 | 2096.9 | 72.6% | 84.5% | 72.8% | 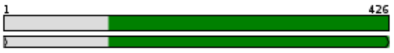 |
| Potato leafroll virus | 3146 | 72.1 | 98.3% | 97.8% | 97.3% | 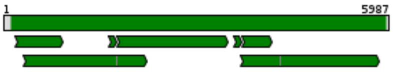 |
| Duamitovirus soch1 | 508 | 25.7 | 69.1% | 69.5% | 81.3% | 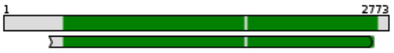 |
| Solendovirus venanicotianae | 430 | 21.1 | 79.8% | 76.8% | 30.0% | 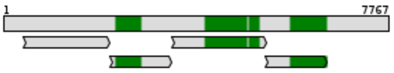 |
| Harvey murine sarcoma virus | 8 | 3.9 | 73.3% | 90.0% | 28.0% | 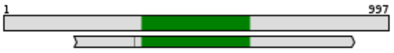 |

#### Discoveries

| Similar to | No. of Reads | Depth of Coverage | Identity | | Genome Coverage | |
|---|---|---|---|---|---|---|
| | | | NT | AA | | |
| Tomato chocolate spot virus (3 segments out of 2) | 101720 | 6454.9 | 65.8% | 66.0% | 15.3% | |
| Tomato chocolate spot virus (segment RNA2) | 50187 | 5116.5 | 65.2% | 70.5% | 24.5% | 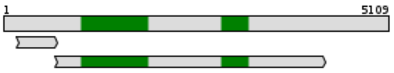 |
| Tomato chocolate spot virus (segment RNA 1) | 36615 | 15605.2 | 82.4% | 0.0% | 3.7% | 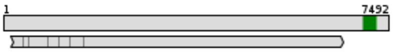 |
| Tomato chocolate spot virus (segment RNA2) | 14918 | 4272.8 | 56.2% | 51.9% | 7.7% | 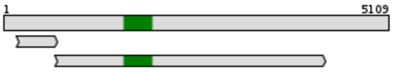 |
| Lausannevirus | 34961 | 23002.5 | 80.3% | 93.5% | 0.0% | 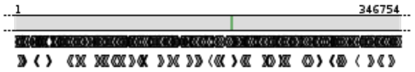 |
| Yellowstone lake phycodnavirus 1 | 29336 | 15316.6 | 82.1% | 93.1% | 0.1% | 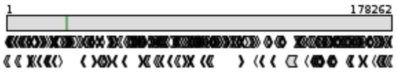 |
| Euproctis digramma nucleopolyhedrovirus | 26156 | 15809.8 | 79.5% | 92.0% | 0.1% | 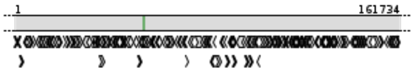 |
| Betabaculovirus clanastomosis | 22579 | 13042.5 | 83.6% | 86.7% | 0.1% | 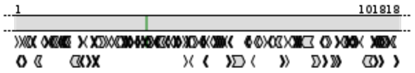 |
| Lausannevirus | 11975 | 7006.5 | 84.2% | 100.0% | 0.0% | 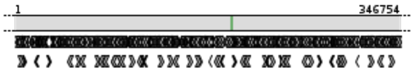 |

| Similar to | No. of Reads | Depth of Coverage | Identity | | Genome Coverage | |
|---|---|---|---|---|---|---|
| | | | NT | AA | | |
| Yellowstone lake phycodnavirus 1 | 11161 | 6804.1 | 83.0% | 90.2% | 0.1% | 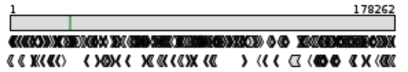 |
| Lausannevirus | 6209 | 3607.8 | 82.1% | 94.1% | 0.0% | 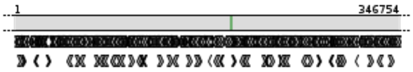 |
| Synechococcus phage S-CRM01 | 6209 | 3685.1 | 78.8% | 92.0% | 0.1% | 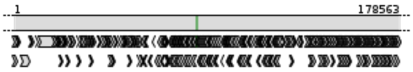 |
| Marseillevirus marseillevirus | 5547 | 3706.7 | 85.7% | 97.5% | 0.0% | 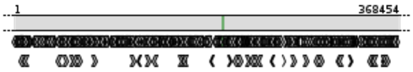 |
| Lausannevirus | 4798 | 3324.6 | 80.0% | 95.2% | 0.0% | 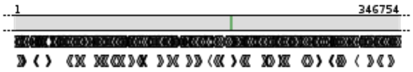 |
| Betabaculovirus disaccharalis | 2967 | 1794.9 | 82.4% | 92.9% | 0.1% | 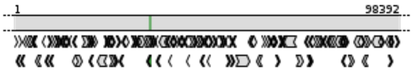 |
| Lausannevirus | 2902 | 1791.9 | 80.8% | 94.1% | 0.0% | 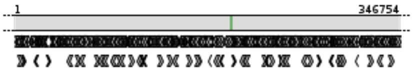 |
| Lausannevirus | 1319 | 631.3 | 83.8% | 95.3% | 0.0% | 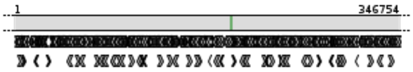 |
| Cladosporium fulvum T-1 virus | 1081 | 75.0 | 53.2% | 48.1% | 19.3% | 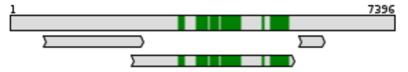 |
| Errantivirus | 891 | 139.9 | 53.1% | 40.8% | 9.3% | 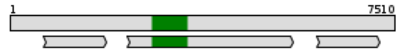 |
| Human gammaherpesvirus 8 (subtype: Could not assign) | 368 | 228.3 | 83.7% | 84.4% | 0.1% | 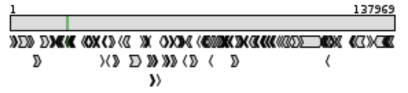 |
| Epiphyllum badnavirus 1 | 360 | 148.9 | 57.5% | 57.0% | 3.3% | 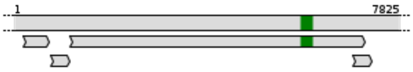 |
| Cassava brown streak virus | 308 | 70.8 | 60.9% | 58.0% | 5.9% | 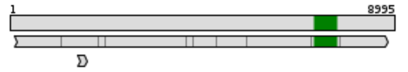 |
| Epiphyllum badnavirus 1 | 137 | 50.7 | 57.6% | 52.7% | 3.5% | 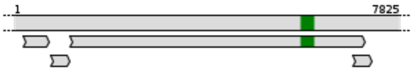 |
| Errantivirus | 133 | 22.2 | 52.4% | 46.2% | 9.5% | 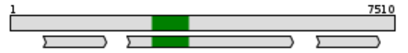 |
| Dioscovirus dioscoreae | 87 | 10.3 | 61.7% | 49.9% | 13.1% | 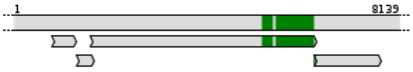 |
| Badnavirus occultipomeae | 84 | 29.0 | 51.1% | 41.3% | 4.0% | 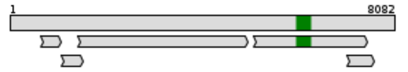 |

| Similar to | No. of Reads | Depth of Coverage | Identity | | Genome Coverage | |
|---|---|---|---|---|---|---|
| | | | NT | AA | | |
| Errantivirus | 73 | 22.1 | 54.6% | 45.3% | 5.1% | 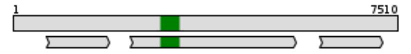  |
| Errantivirus | 64 | 20.6 | 56.9% | 47.5% | 4.7% | 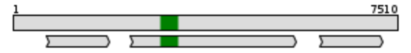  |
| Errantivirus | 59 | 18.4 | 53.1% | 42.4% | 5.0% | 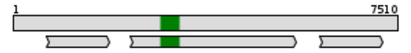  |
| Badnavirus maculacommelinae | 55 | 7.8 | 54.7% | 47.9% | 12.1% | 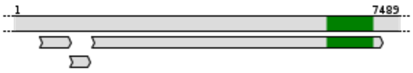  |
| Epiphyllum badnavirus 1 | 55 | 16.6 | 52.4% | 41.5% | 5.2% | 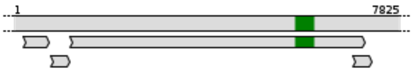  |
| Petuvirus venapetuniae | 50 | 7.5 | 51.7% | 43.1% | 11.4% | 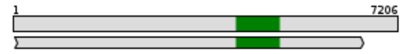  |
| Badnavirus occultipomeae | 40 | 11.3 | 55.0% | 40.7% | 4.5% | 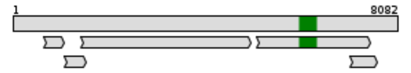  |
| Caulimovirus tessellobrassicae | 39 | 6.6 | 56.7% | 46.4% | 9.0% | 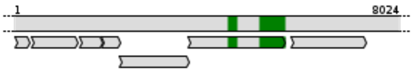  |
| Caulimovirus venafragariae | 39 | 11.4 | 64.1% | 67.6% | 5.4% | 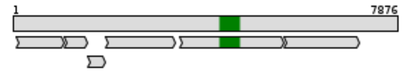  |
| Caulimovirus deformatiolamii | 27 | 5.8 | 54.5% | 45.0% | 7.5% | 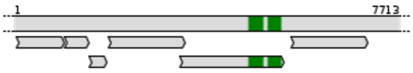  |
| Petuvirus venapetuniae | 25 | 5.8 | 55.2% | 49.5% | 7.6% | 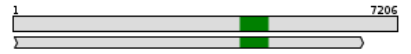  |
| Errantivirus | 23 | 7.6 | 55.4% | 47.5% | 4.8% | 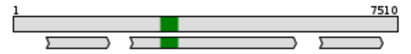  |
| Badnavirus maculaucubae | 20 | 8.4 | 57.2% | 57.5% | 3.7% | 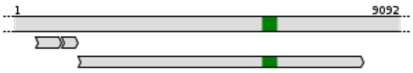  |
| Hubei noda-like virus 9 (2 segments out of 2) | 20 | 2.2 | 66.4% | 69.7% | 20.9% | |
| Hubei noda-like virus 9 (segment 1) | 11 | 1.9 | 66.4% | 69.9% | 21.6% | 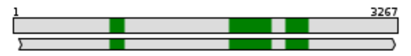  |
| Hubei noda-like virus 9 (segment 2) | 9 | 2.7 | 66.5% | 69.1% | 19.4% | 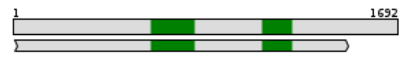  |
| Caulimovirus maculatractylodei | 18 | 4.7 | 56.7% | 47.0% | 6.3% | 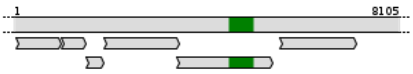  |
| Cavemovirus venamanihotis | 15 | 4.8 | 60.2% | 50.0% | 4.6% | 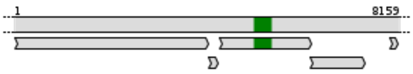  |

| Similar to | No. of Reads | Depth of Coverage | Identity | | Genome Coverage | |
|---|---|---|---|---|---|---|
| | | | NT | AA | | |
| Errantivirus | 14 | 5.4 | 56.4% | 48.6% | 4.2% | 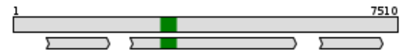 |
| Caulimovirus venafragariae | 14 | 6.4 | 60.9% | 61.9% | 3.2% | 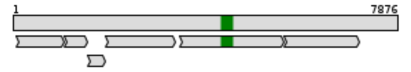 |
| Badnavirus maculasmallanthi | 13 | 4.7 | 51.1% | 41.2% | 4.0% | 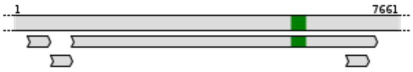 |
| Caulimovirus venafragariae | 9 | 4.2 | 63.6% | 63.5% | 3.2% | 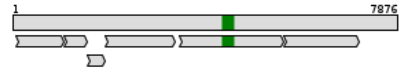 |
| Cavemovirus venamanihotis | 8 | 3.2 | 60.1% | 51.6% | 3.3% | 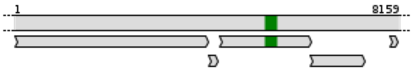 |
| Alphabaculovirus chofumiferanae | 7 | 4.5 | 78.7% | 80.0% | 0.1% | 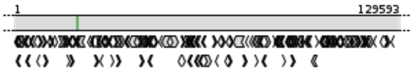 |
| Badnavirus volubetulae | 7 | 3.2 | 61.1% | 47.5% | 3.9% | 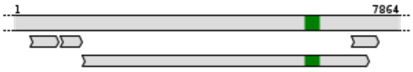 |
| Epiphyllum badnavirus 1 | 6 | 2.2 | 52.8% | 43.8% | 3.9% | 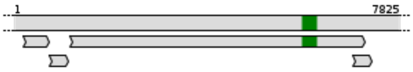 |
| Potato virus X | 4 | 2.3 | 96.6% | 95.6% | 3.7% | 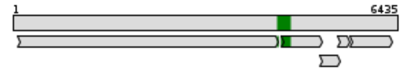 |
| Petuvirus venapetuniae | 4 | 2.4 | 68.4% | 69.3% | 3.1% | 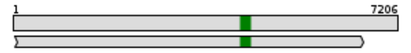 |
| Tomato associated geminivirus 1 | 4 | 1.8 | 63.7% | 59.0% | 9.0% | 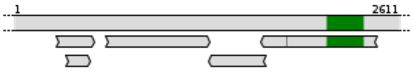 |
| Bracoviriform congregatae (segment Circle 7) | 3 | 2.5 | 75.6% | 86.7% | 0.5% | 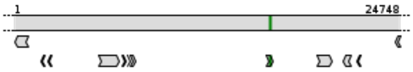 |
| Tobacco streak virus (segment RNA 1) | 2 | 1.0 | 74.3% | 80.0% | 7.7% | 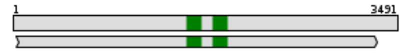 |
| Nepovirus aeonii (segment RNA2) | 2 | 1.1 | 69.0% | 66.7% | 6.0% | 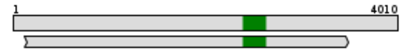 |
| Duamitovirus peex1 | 2 | 1.5 | 70.4% | 73.3% | 5.0% | 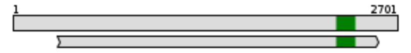 |
| Moloney murine leukemia virus | 2 | 1.3 | 81.2% | 78.3% | 2.7% | 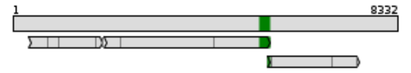 |
| Teseptimavirus EG1 | 2 | 1.2 | 75.1% | 98.6% | 0.5% | 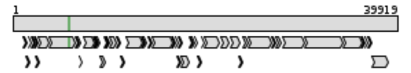 |
| Duamitovirus peex1 | 1 | 1.0 | 80.0% | 80.0% | 5.0% | 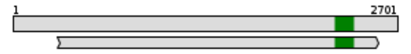 |

| Similar to | No. of Reads | Depth of Coverage | Identity | | Genome Coverage | |
|---|---|---|---|---|---|---|
| | | | NT | AA | | |
| Cyvirus cyprinidallo3 | 1 | 1.0 | 83.3% | 80.0% | 0.0% | 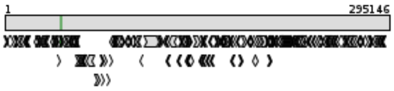  |
| Diplodia scrobiculata RNA virus 1 | 1 | 1.0 | 80.8% | 80.0% | 2.4% | 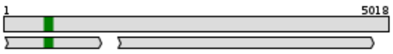  |

### NGS Details (UN8): Potato virus Y

#### Assembly

| | |
|---|---|
| **Coverage Length** | 9653 (1 contig(s)) |
| **Depth Of Coverage** | 8776.7 |
| **Number Of Reads** | 634281 |
| **Reads Per Million** | 10600.42 rpm (after QC) |
| **Ambiguities** | 0 |
| **Assembly Method** | de novo + reference guided assembly |
| **Consensus Caller** | Bcf Tools |

#### Coverage Map

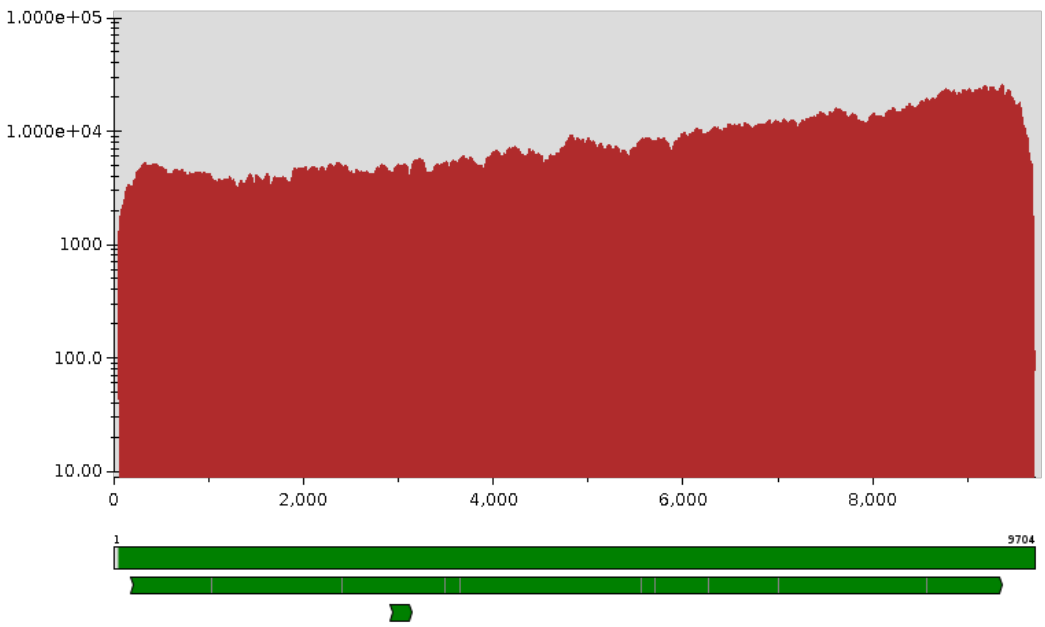

#### Assignment

| | |
|---|---|
| **Type** | Potato virus Y (Taxonomy ID: 12216) |
| **Reference Genome** | NC_001616.1 |
| **NT Identity (%)** | 82.7547 |
| **AA Identity (%)** | 90.0 |
| **Number Of Stop Codons** | 3 |
| **Number Of CDS** | 2 |

#### Alignment

| | |
|---|---|
| **Alignment Score** | 12530.0 (NT) + 19272.0 (AA) = 31802.0 |
| **Concordance (%)** | 78.3324 |

| **Alignment Method** | Global, seeded, nucleotide + amino acids (AGA) |
|---|---|

#### Genome Region

Sequence starts at position 52 and ends at position 9704 relative to NC_001616.1 reference sequence.

#### Alignment Detailed Statistics

| | Begin | End | Coverage | Score | Concordance | Matches | Identities | I/D/M/F* | Stop Codons |
|---|---|---|---|---|---|---|---|---|---|
| **NT** | **52** | **9704** | **99.5%** | **12530** | **65.2%** | **9642 (99.8%)** | **7985 (82.7%)** | **7/11** | |
| Mutations: | 75C>T, 79C>T, 83G>A, 85T>C, 87C>T, 89G>A, 90T>C, 93A>T, 94A>G, 95A>T, 100T>C, 101T>A, 102C>T, 104T>C, 107C>T, 110T>G, 112C>T, 114C>T, 115T>C, 116A>T, 117G>C, 119A>G, 120C>T, 121A>T, 125T>G, 126T>C, 127G>A, 128G>T, 131A>G, 132C>T, 139A>T, 139_140insG, 141C>T, 143C>T, 146C>T, 150C>T, 150_151insT, 154T>C, 156C>A, 158T>A, 160A>C, 164C>T, 168C>A, 169C>G, 171A>C, 172T>C, 173T>A, 174T>G, 177G>C, 178A>G, 179T>C, 180C>T, 181C>A, 182T>G, 183C>T, 193T>C, 194T>C, 196C>A, 202A>G, 208C>T, 209T>C, 210G>T, 211T>G, 220G>A, 221T>A, 223T>G, 226A>G, 235A>T, 241C>T, 247A>T, 251T>C, 255G>T, 256C>T, 258A>G, 262T>C, 263A>G, 265T>G, 267T>C, 273A>T, 277A>T, 280A>G, 283G>T, 286G>A, 287G>A, 289T>C, 293G>A, 294T>C, 301T>C, 304C>T, 305G>A, 306C>T, 309A>G, 311C>T, 316A>T, 317A>G, 318C>T, 319A>G, 322A>G, 331A>T, 339T>A, 340C>A, 342A>G, 347A>G, 348A>C, 349A>G, 350T>C, 355T>A, 356A>G, 357C>T, 364T>C, 365G>A, 366T>C, 367G>A, 368C>T, 369T>C, 373A>G, 376C>T, 384T>G, 388G>A, 391C>T, 394A>G, 400G>A, 406T>C, 408C>T, 409C>T, 412G>A, 415A>T, 416A>G, 417T>A, 424T>C, 427G>A, 433A>G, 436G>C, 437G>A, 440A>G, 441G>A, 443A>C, 444A>G, 445G>A, 448T>A, 454A>G, 456A>C, 460T>C, 461C>A, 469A>G, 475C>G, 481T>A, 502T>C, 505A>T, 511C>T, 517T>A, 520T>A, 523A>T, 531C>T, 535G>A, 536C>T, 539C>T, 540A>G, 541A>G, 549G>A, 550A>G, 556C>T, 559T>C, 562T>C, 568T>C, 571A>G, 576T>C, 580T>C, 585T>C, 586C>G, 593C>T, 594G>A, 595C>T, 597C>G, 599A>G, 601A>G, 602A>C, 603T>C, 604A>G, 613A>G, 616A>G, 619C>G, 622G>A, 625G>A, 628T>C, 631C>T, 635A>G, 637T>C, 644A>G, 646A>G, 655T>C, 661G>A, 663A>C, 664G>C, 667A>G, 679C>T, 682C>A, 683T>C, 685A>G, 691T>C, 699C>G, 703T>A, 706T>A, 719A>G, 721G>A, 722A>G, 724A>G, 725C>T, 727T>G, 730T>A, 731G>T, 734T>C, 735A>C, 736C>T, 737T>C, 738C>G, 739C>T, 744C>T, 745G>C, 749C>T, 751A>C, 754T>C, 765T>A, 770T>C, 773C>A, 774G>A, 775A>G, 776C>A, 790C>T, 801T>A, 813G>C, 814A>G, 816T>A, 818T>C, 823A>G, 825G>A, 826T>A, 833C>A, 835G>A, 841C>T, 851A>G, 853T>C, 859C>T, 860C>T, 862C>T, 863A>G, 865T>C, 867T>C, 869A>G, 871C>T, 872A>T, 874A>G, 876G>A, 879G>A, 886T>C, 898C>T, 903A>G, 906C>A, 907A>C, 909A>C, 910A>C, 912G>A, 913C>T, 922C>A, 931T>A, 938T>C, 939C>T, 940A>T, 943A>G, 952C>T, 967A>G, 973A>G, 976G>C, 980T>A, 982G>C, 991A>G, 994T>C, 999G>A, 1003T>C, 1007C>A, 1008A>G, 1009G>A, 1010A>G, 1012T>G, 1013A>G, 1018A>G, 1019A>C, 1021C>G, 1030C>T, 1039G>A, 1042T>C, 1045C>A, 1048C>A, 1051T>C, 1060G>A, 1063T>A, 1064C>T, 1069C>T, 1083G>A, 1093T>C, 1097T>G, 1099G>A, 1105C>T, 1114A>G, 1117T>G, 1123A>G, 1126T>A, 1129C>T, 1143G>A, 1147A>T, 1150T>A, 1154T>A, 1156G>A, 1165C>T, 1168T>C, 1172C>T, 1174T>A, 1201C>T, 1221G>A, 1229G>A, 1230T>C, 1231T>C, 1233G>A, 1234C>T, 1240G>A, 1246G>A, 1250T>C, 1252G>A, 1255T>C, 1257A>C, 1258A>G, 1267A>G, 1273T>C, 1276G>A, 1278A>G, 1283T>C, 1290C>T, 1291G>A, 1298G>A, 1299A>C, 1300C>T, 1306T>C, 1307A>G, 1309A>G, 1312T>C, 1316A>G, 1318T>A, 1327G>A, 1329T>C, 1333G>T, 1336A>G, 1343C>T, 1345A>G, 1354G>A, 1357G>T, 1360C>T, 1367C>T, 1369C>G, 1373C>A, 1378C>T, 1384G>A, 1385A>G, 1393A>G, 1396C>T, 1405G>T, 1408A>G, 1411G>A, 1415G>T, 1417A>T, 1420G>C, 1429T>C, 1432A>G, 1436G>A, 1438C>T, 1439T>C, 1441A>G, 1444T>C, 1453C>T, 1456G>A, 1468A>G, 1492A>G, 1499T>C, 1501G>A, 1504T>C, 1507G>A, 1510C>T, 1513A>G, 1516A>G, 1519A>T, 1522G>A, 1525C>T, 1528G>A, 1534C>T, 1540T>A, 1546C>T, 1555A>G, 1558T>C, 1561T>C, 1564A>C, 1567T>A, 1573C>T, 1576A>G, 1583T>C, 1588T>A, 1594G>A, 1600C>T, 1609G>A, 1618G>A, 1621C>T, 1624C>T, 1631C>T, 1639A>G, 1651C>T, 1654C>G, 1675T>C, 1693A>G, 1702T>C, 1707A>G, 1718G>A, 1726A>G, 1729C>T, 1732A>C, 1738T>C, 1742A>G, 1744C>T, 1747C>T, 1753T>C, 1758G>A, 1768G>A, 1786C>T, 1789A>G, 1790G>A, 1795C>T, 1805T>C, 1807A>G, 1813G>A, 1819G>A, 1825G>A, 1837C>T, 1840T>C, 1849A>G, 1858C>G, 1864A>G, 1870C>T, 1876T>C, 1882A>G, 1888T>A, 1891T>C, 1906T>C, 1915A>T, 1930T>C, 1936C>T, 1937A>G, 1939T>C, 1945A>G, 1951C>T, 1960A>T, 1978A>G, 1981T>A, 1984C>T, 1987T>C, 1996C>T, 2004T>A, 2020A>G, 2023G>A, 2026T>C, 2029G>T, 2032G>A, 2059T>C, 2068T>G, 2071C>T, 2072G>A, 2074G>A, 2087C>T, 2089G>A, 2101C>T, 2104G>A, 2107G>A, 2116G>A, 2122C>T, 2137C>T, 2158T>C, 2164C>T, 2170A>G, 2182T>C, 2188G>A, 2191C>A, 2208G>A, 2218T>C, 2224C>T, 2227A>T, 2242G>A, 2245C>T, 2254G>A, 2257T>C, 2260C>T, 2266C>G, 2272A>G, 2275G>A, 2293A>G, 2299C>T, 2302G>A, 2305G>A, 2314A>T, 2317T>C, 2326A>G, 2341G>A, 2347A>G, 2350T>C, 2356G>A, 2365T>C, 2371A>G, 2374A>G, 2383T>C, 2386A>T, 2389A>G, 2392T>C, 2395T>C, 2414A>G, 2415A>G, 2416T>G, 2419T>A, 2420A>T, 2440A>T, 2467T>C, 2476T>C, 2482G>A, 2491G>A, 2497G>T, 2503G>A, 2518A>G, 2521T>C, 2523A>G, 2533A>G, 2539G>A, 2545A>G, 2551G>A, 2554T>A, 2558C>T, 2561T>C, 2563G>C, 2572A>G, 2584T>C, 2587C>T, 2593G>C, 2599T>C, 2608T>C, 2620T>C, 2629G>A, 2632G>A, 2634G>A, 2638G>A, 2644T>A, 2653A>G, 2665T>C, 2678C>T, 2683A>G, 2689T>C, 2698A>C, 2707A>G, 2710G>A, 2716A>G, 2719A>G, 2722C>T, 2723G>A, 2725C>T, 2728A>G, 2734G>A, 2737T>A, 2746T>C, 2747G>A, 2749T>A, 2752A>T, 2755T>A, 2761C>T, 2764C>T, 2779T>C, 2785G>T, 2791C>T, 2795C>A, 2806C>T, 2816T>C, 2818G>C, 2822G>A, 2824G>A, 2827G>A, 2836T>C, 2839G>A, 2842T>C, 2863C>A, 2864C>T, 2882C>A, 2885A>C, 2886G>A, 2893C>T, 2895C>T, 2902C>T, 2905A>G, 2911T>C, 2923T>C, 2931A>G, 2940A>G, 2947T>C, 2962T>C, 2971_2972insA, 2989T>C, 2991C>T, 2994A>G, 2999G>A, 3000A>G, 3015C>T, 3021G>A, 3024A>T, 3025C>T, 3036C>T, 3039A>G, 3040A>C, 3042A>C, 3043G>A, 3052G>A, 3061A>G, 3082G>A, 3091C>T, 3099C>T, 3100C>A, 3104G>A, 3105T>G, 3106G>A, 3109C>T, 3112A>G, 3118T>C, 3121C>T, 3127A>G, 3132G>A, 3137C>A, 3138G>A, 3139A>G, 3151T>C, 3152T>A, 3154C>T, 3165G>A, 3169A>G, 3199C>T, 3203A>C, 3208T>C, 3214A>G, 3217C>T, 3218T>C, 3223C>A, 3225C>A, 3230G>A, 3235C>T, 3257T>C, 3262C>T, 3265C>T, 3266G>A, 3268A>G, 3271G>A, 3275G>A, 3284G>A, 3301G>A, 3315G>A, 3316A>G, 3318A>C, 3322C>T, 3325G>A, 3328G>A, 3337A>T, 3349G>A, 3355C>T, 3358C>T, 3359A>G, 3367A>G, 3370T>C, 3377T>C, 3379A>G, 3394A>G, 3397C>T, 3400T>G, 3406A>G, 3418C>T, 3421T>A, 3427T>C, 3433G>A, 3436A>T, 3439A>G, 3442C>T, 3445T>C, 3457G>A, 3460T>C, 3463T>A, 3469G>T, 3472G>A, 3496C>T, 3499G>A, 3505C>T, 3513T>G, 3520A>G, 3529A>G, 3532A>G, 3545A>G, 3548G>A, 3551T>C, 3554G>A, 3556C>T, 3583T>C, 3586T>C, 3589C>T, 3595C>T, 3607T>C, 3613T>C, 3616G>A, 3619T>C, 3623C>G, 3628C>T, 3632C>A, 3637T>C, 3638T>C, 3643A>G, 3661A>G, 3670G>A, 3682T>C, 3688G>A, 3697G>A, 3698A>G, 3706T>C, 3709T>C, 3713T>C, 3715G>A, 3717G>A, 3721G>A, 3731C>A, 3732G>A, 3736T>A, 3746C>T, 3748A>G, 3754A>G, 3763T>C, 3778A>G, 3781A>G, 3784C>T, 3796T>C, 3805A>T, 3811C>T, 3814A>G, 3829C>T, 3838T>C, 3844A>G, 3847A>C, 3856T>A, 3862G>T, 3871C>T, 3877C>A, 3883C>T, 3886A>G, 3892A>G, 3895C>T, 3898T>C, 3904A>G, 3905C>A, 3907G>A, 3913T>C, 3928G>A, 3931A>T, 3937G>A, 3955T>C, 3958G>A, 3961C>T, 3964A>G, 3967T>C, 3976A>G, 3979T>C, 3985A>T, 3997A>T, 4000G>A, 4006C>T, 4009T>G, 4012C>T, 4015A>G, 4024T>C, 4030A>G, 4033A>T, 4042G>A, 4052C>T, 4054G>A, 4057T>C, 4063T>A, 4066A>G, 4073A>G, 4078T>C, 4081C>T, 4084T>C, 4096C>T, 4108C>T, 4111A>T, 4117G>A, 4118C>T, 4120A>G, 4126C>T, 4128T>A, 4129C>T, 4132C>T, 4135T>C, 4147A>G, 4148T>C, 4151G>T, 4162C>T, 4166G>A, 4171A>C, 4180G>A, 4190C>T, 4198T>C, 4201C>T, 4210G>T, 4213C>T, 4219T>C, 4220C>T, 4223C>A, 4225G>A, 4231T>G, 4237T>C, 4252A>G, 4253T>C, 4255A>G, 4258A>G, 4270T>C, 4282A>G, 4285G>A, 4291A>G, 4303G>A, 4312C>T, 4319A>G, 4321A>G, 4324G>T, 4327G>A, 4336G>T, 4339T>C, 4348A>T, 4360A>G, 4366T>C, 4369T>A, 4375T>C, 4387T>C, 4395T>A, 4406G>A, 4411T>C, 4438T>C, 4444C>A, 4450C>T, 4456C>T, 4465C>T, 4475A>G, 4477G>T, 4480C>T, 4481A>T, 4483A>G, 4495C>T, 4507G>A, 4510C>T, 4513T>G, 4517C>T, 4528C>T, 4534A>G, 4540C>T, 4544G>T, 4545C>T, 4546G>A, 4548G>A, 4558T>C, 4564A>T, 4570C>T, 4576A>T, 4597G>A, 4600C>T, 4606C>T, 4609G>A, 4615G>A, 4618C>T, 4625T>C, 4627G>T, 4648C>T, 4651T>C, 4660G>A, 4663C>T, 4669T>A, 4679G>A, 4681G>C, 4684T>C, 4687T>G, 4699G>A, 4702A>G, 4708A>G, 4711G>A, 4712C>T, 4738A>G, 4744A>T, 4747A>G, 4748T>C, 4765T>C, 4796G>A, 4798C>T, 4803C>G, 4804T>C, 4807G>A, 4810G>T, 4813T>G, 4816T>C, 4831T>C, 4834C>T, 4840A>T, 4852A>G, 4855C>T, 4867T>C, 4870G>C, 4873T>A, 4882T>C, 4903A>T, 4909G>A, 4915T>C, 4918A>G, 4919T>C, 4921G>A, 4924T>C, 4927C>G, 4933T>C, 4936C>A, 4939G>A, 4942T>C, 4945C>T, 4963A>G, 4972T>C, 4990T>C, 5002A>G, 5023T>C, 5024T>C, 5029C>T, 5032T>C, 5038T>C, 5044A>T, 5053A>T, 5056G>A, 5057A>G, 5060A>G, 5071G>T, 5090C>A, 5095A>G, 5098G>T, 5100C>T, 5104A>G, 5107A>T, 5110T>C, 5113A>G, 5116G>A, 5119A>G, 5120G>T, 5122C>T, 5126A>G, 5137T>C, 5140C>T, 5143A>G, 5146G>A, 5161C>A, 5167A>G, 5173T>G, 5182G>A, 5183G>A, 5188C>T, 5191G>A, 5197A>G, 5200C>T, 5206C>T, 5209A>G, 5218C>T, 5230G>A, 5236C>T, 5239C>T, 5245C>T, 5263A>T, 5269C>T, 5272C>T, 5275C>A, 5278C>T, 5281A>C, 5284A>G, 5287T>C, 5288C>T, 5293A>T, 5294T>C, 5296G>A, 5302G>A, 5311T>A, 5314A>G, 5323A>T, 5347T>C, 5353C>T, 5365C>T, 5386T>C, 5395C>A, 5422A>G, 5425T>C, 5431C>T, 5437G>A, 5452T>C, 5458G>A, 5467T>C, 5470A>G, 5471C>T, 5473A>G, 5479A>G, 5482C>T, 5485A>C, 5488T>C, 5489T>C, 5491G>A, 5497T>C, 5506T>C, 5509G>A, 5512G>A, 5516T>C, 5530T>C, 5533G>A, 5536G>T, 5539G>A, 5548T>G, 5554C>T, 5560T>A, 5561G>A, 5563G>A, 5566G>C, 5569A>G, 5584C>G, 5593G>A, 5596G>A, 5597A>G, 5605C>A, 5617G>T, 5619C>T, 5620T>G, 5629G>A, 5630A>C, 5641C>T, 5650A>T, 5662A>T, 5665A>G, 5689A>G, 5693G>A, 5698G>A, 5713A>G, 5716G>C, 5719A>G, 5743C>A, 5755C>A, 5758T>C, 5764T>C, 5770A>G, 5779C>T, 5782T>C, 5791C>T, 5803C>T, 5836G>A, 5839A>G, 5845A>G, 5848A>G, 5853A>G, 5857T>C, 5860C>T, 5863A>T, 5869T>A, 5875T>C, 5884C>T, 5893C>T, 5894A>G, 5896C>G, 5899C>T, 5908G>A, 5911T>C, 5923G>A, 5938A>G, 5941C>T, 5944T>C, 5947T>C, 5959G>A, 5960C>G, 5961G>C, 5962G>C, 5974A>G, 5986T>A, 5989C>T, 5990A>G, 5998T>C, 6008A>C, 6010A>T, 6019A>T, 6022G>T, 6025A>T, 6030A>C, 6049C>A, 6058G>A, 6061A>G, 6064C>A, 6070T>C, 6072G>A, 6073T>C, 6074A>C, 6079G>A, 6081C>A, 6085A>T, 6091A>T, 6094C>T, 6111G>C, 6112T>C, 6118A>G, 6124G>A, 6127G>A, 6128A>G, 6130T>C, 6133T>C, 6136A>G, 6148C>T, 6157A>G, 6158G>A, 6163T>C, 6169A>G, 6170A>T, 6172A>G, 6178C>T, 6187A>G, 6190T>C, 6196G>A, 6203C>T, 6205C>T, 6208A>G, 6209C>T, 6211A>G, 6217G>A, 6229T>A, 6230G>A, 6238C>T, 6247G>A, 6253A>T, 6257G>A, 6258C>A, 6265G>A, 6274T>C, 6282A>G, 6286G>A, 6298C>T, 6301G>A, 6304A>G, 6307C>T, 6322C>T, 6328A>T, 6331T>G, 6334T>C, 6337G>A, 6338C>T, 6358T>C, 6361G>A, 6362G>A, 6367A>T, 6370G>A, 6376C>T, 6379T>G, 6385A>C, 6391A>T, 6394A>G, 6397C>T, 6401G>A, 6405C>T, 6412C>T, 6418A>G, 6421T>C, 6429A>T, 6436T>A, 6445G>A, 6450A>G, 6454C>G, 6460C>T, 6463T>A, 6469C>T, 6472G>A, 6475G>A, 6481T>C, 6485C>A, 6487C>T, 6490T>C, 6496C>T, 6502G>A, 6514T>C, 6517G>A, 6527C>A, 6532C>A, 6541G>A, 6544A>G, 6556C>T, 6559T>C, 6565G>A, 6569T>C, 6571G>A, 6586T>A, 6588C>T, 6598A>G, 6601A>G, 6604T>C, 6607T>C, 6616A>G, 6622C>T, 6631G>A, 6634G>A, 6635T>C, 6640T>A, 6646G>T, 6647A>G, 6652C>T, 6664C>T, 6670T>G, 6677A>G, 6679A>G, 6685C>T, 6691A>T, 6694C>T, 6715A>G, 6716G>A, 6719A>G, 6724A>G, 6734C>T, 6742G>A, 6745G>A, 6754C>T, 6760A>G, 6764A>T, 6766A>G, 6775T>A, 6782C>T, 6787A>T, 6790C>T, 6795C>T, 6796A>G, 6799C>A, 6802C>A, 6808C>T, 6811C>T, 6814C>T, 6823C>T, 6829A>G, 6847C>T, 6853A>C, 6857A>G, 6862G>A, 6871A>G, 6875G>A, 6876T>C, 6877C>A, 6880G>A, 6883T>A, 6889T>G, 6901C>T, 6916C>T, 6919G>A, 6925A>G, 6928T>C, 6934C>A, 6949G>C, 6955C>T, 6958A>G, 6976A>G, 6980C>T, 6985C>T, 6997A>T, 7000G>T, 7001G>A, 7003G>A, 7012T>C, 7015G>A, 7018T>C, 7032T>A, 7039C>T, 7040T>C, 7042G>A, 7048A>G, 7060T>A, 7063C>A, 7078C>T, 7082T>C, 7084A>G, 7087A>G, 7090C>A, 7096T>C, 7099A>G, 7102T>C, 7111G>A, 7117A>G, 7125C>A, 7129A>G, 7132T>C, 7133C>T, 7135G>A, 7141G>A, 7144T>C, 7145G>T, 7147A>G, 7150G>A, 7159A>T, 7160_7165delGAGGCA, 7171C>T, 7189G>T, 7198A>G, 7201C>T, 7205C>T, 7210T>C, 7216T>A, 7219G>A, 7225C>A, 7228G>A, 7240G>A, 7243T>C, 7249A>G, 7252T>G, 7258T>A, 7261T>C, 7264T>A, 7267C>G, 7270G>A, 7273_7274insT, 7274C>G, 7275G>T, 7281A>T, 7284T>C, 7296C>T, 7311C>T, 7314C>T, 7317G>A, 7321T>A, 7322delG, 7327C>T, 7330C>T, 7333C>T, 7334A>C, 7342T>C, 7343G>A, 7344C>A, 7345A>C, 7354T>C, 7363A>G, 7369T>C, 7372C>T, 7378G>T, 7381C>T, 7396A>T, 7399C>T, 7405C>T, 7407G>T, 7408T>G, 7414G>T, 7415T>G, 7420A>G, 7429C>T, 7432T>C, 7441T>C, 7459G>A, 7462A>G, 7468A>T, 7471C>A, 7477A>G, 7480C>T, 7486G>T, 7495T>C, 7498A>G, 7501T>C, 7503T>C, 7504G>A, 7513T>A, 7519C>T, 7522A>G, 7531G>A, 7537G>A, 7540C>T, 7546T>C, 7570G>A, 7573G>A, 7579G>A, 7588T>C, 7595C>T, 7604T>C, 7606G>A, 7615T>A, 7624T>C, 7633C>T, 7636C>T, 7639T>C, 7648T>C, 7666A>G, 7672T>C, 7678A>T, 7684G>A, 7690T>C, 7696T>C, 7702T>G, 7709G>A, 7713A>G, 7714A>G, 7715C>T, 7726T>C, 7735G>A, 7744A>G, 7747C>T, 7762C>T, 7768G>A, 7771T>C, 7780A>G, 7783A>C, 7786T>C, 7793C>T, 7795A>G, 7798C>T, 7804T>C, 7812C>T, 7834A>G, 7837C>T, 7844G>T, 7846G>A, 7855G>A, 7861G>A, 7870A>G, 7873C>T, 7876T>A, 7879G>A, 7882T>A, 7883G>A, 7894A>T, 7897T>C, 7921C>T, 7927G>A, 7936T>C, 7939T>C, 7942T>C, 7966G>A, 7969T>C, 7975T>C, 7987C>T, 7990T>C, 8005T>C, 8032A>G, 8038C>T, 8041C>T, 8043G>A, 8080G>A, 8089G>A, 8095G>A, 8098G>A, 8107C>T, 8110T>C, 8116T>C, 8128A>G, 8146T>C, 8151A>G, 8152C>T, 8164G>A, 8178G>A, 8182G>A, 8191G>A, 8197C>T, 8203C>T, 8208G>A, 8213C>T, 8215G>A, 8218A>G, 8221C>T, 8224G>A, 8239A>G, 8254G>A, 8260T>C, 8263A>G, 8281T>C, 8284A>G, 8294C>G, 8296A>G, 8299G>A, 8302C>T, 8317T>C, 8320C>T, 8326T>A, 8335G>A, 8338C>T, 8349C>T, 8353A>G, 8362C>T, 8365A>G, 8368C>T, 8380C>T, 8383A>G, 8389A>G, 8404T>C, 8408G>T, 8413A>G, 8419G>T, 8434T>C, 8440T>C, 8452G>A, 8458A>G, 8460G>A, 8464A>G, 8467G>A, 8479G>A, 8480G>A, 8482T>A, 8497G>A, 8498C>T, 8500A>G, 8502G>A, 8503A>G, 8506C>T, 8515A>T, 8524C>T, 8527A>C, 8528T>C, 8530A>G, 8539G>C, 8546C>T, 8547T>G, 8552T>A, 8563A>G, 8574C>G, 8585A>G, 8598G>A, 8602C>T, 8603A>T, 8604A>C, 8605C>T, 8620A>G, 8622C>A, 8625A>G, 8626G>A, 8629G>A, 8632C>T, 8638C>T, 8641G>A, 8647C>T, 8648C>T, 8649C>T, 8650G>C, 8653C>T, 8656A>G, 8658G>A, 8664A>C, 8665T>G, 8674T>G, 8679C>T, 8680A>T, 8683C>A, 8692G>A, 8695A>T, 8707G>A, 8708A>C, 8716G>A, 8722C>T, 8725G>A, 8731A>G, 8737A>G, 8743C>T, 8745C>A, 8746A>G, 8752G>A, 8755A>T, 8770C>T, 8771T>C, 8776A>G, 8779T>C, 8782G>A, 8791T>C, 8797A>G, 8800A>G, 8803A>G, 8809T>C, 8812T>C, 8818T>C, 8824G>A, 8827A>G, 8848G>A, 8854T>C, 8856A>G, 8857G>A, 8863G>A, 8865G>A, 8867A>C, 8869G>T, 8872A>G, 8883G>A, 8888A>T, 8929C>T, 8954G>A, 8959C>T, 8962A>T, 8983G>A, 8984A>G, 8986T>C, 8988A>T, 8998G>A, 9004G>A, 9007G>A, 9016C>T, 9037C>A, 9085A>T, 9097C>T, 9131A>G, 9136A>T, 9145G>T, 9149A>G, 9150T>G, 9151G>A, 9152G>A, 9157A>G, 9160G>T, 9163T>C, 9184G>A, 9196A>G, 9214G>A, 9217A>G, 9220G>A, 9244A>T, 9247G>A, 9256C>T, 9260C>T, 9262T>C, 9274G>A, 9280C>T, 9289C>T, 9298A>C, 9304G>A, 9334C>T, 9364C>G, 9377_9378insT, 9387C>T, 9404G>A, 9411C>T, 9412A>G, 9414A>T, 9437C>T, 9454A>G, 9458T>C, 9467G>A, 9470_9471insT, 9482G>A, 9483C>T, 9498T>C, 9516T>G, 9517delA, 9533G>A, 9534G>T, 9548A>T, 9551C>T, 9553_9554delTA, 9556A>T, 9557T>A, 9558A>T, 9574T>A, 9581G>T, 9582T>G, 9583G>C, 9584_9585insG, 9590C>T, 9591T>C, 9607C>T, 9610T>C, 9615delC, 9620A>G, 9648A>G, 9674T>C, 9690C>T | | | | | | | | |

| | Begin | End | Coverage | Score | Concordance | Matches | Identities | I/D/M/F* | Stop Codons |
|---|---|---|---|---|---|---|---|---|---|
| **NT** | **52** | **9704** | **99.5%** | **12530** | **65.2%** | **9642 (99.8%)** | **7985 (82.7%)** | **7/11** | |
| **CDS** | | | | | | | | | |
| PVYgp1 | 1 | 3064 | 100% | 19042 | 90.2% | 3062 (99.9%) | 2771 (90.4%) | 2/2/3/3 | 1 |
| Protein mutations: | Y4Q (194T>C 196C>A), C9L (209T>C 210G>T 211T>G), F13M (221T>A 223T>G), S23P (251T>C), C24F (255G>T 256C>T), E25G (258A>G), I27V (263A>G 265T>G), V28A (267T>C), E30V (273A>T), A35T (287G>A 289T>C), V37T (293G>A 294T>C), A41I (305G>A 306C>T), D42G (309A>G), E44D (316A>T), T45V (317A>G 318C>T 319A>G), L52Q (339T>A 340C>A), K53R (342A>G), K55A (347A>G 348A>C 349A>G), Y56H (350T>C), T58V (356A>G 357C>T), V61T (365G>A 366T>C 367G>A), L62S (368C>T 369T>C), F67C (384T>G), A75V (408C>T 409C>T), M78E (416A>G 417T>A), E85K (437G>A), R86E (440A>G 441G>A), K87R (443A>C 444A>G 445G>A), D88E (448T>A), E91A (456A>C), H93N (461C>A), D111E (517T>A), S116F (531C>T), P118S (536C>T), Q119W (539C>T 540A>G 541A>G), R122K (549G>A 550A>G), M131T (576T>C), V134A (585T>C 586C>G), R137Y (593C>T 594G>A 595C>T), P138R (597C>G), I139V (599A>G 601A>G), I140P (602A>C 603T>C 604A>G), M147I (625G>A), I151V (635A>G 637T>C), I154V (644A>G 646A>G), E160A (663A>C 664G>C), H166Q (682C>A), T172S (699C>G), H174Q (706T>A), K179E (719A>G 721G>A), I180V (722A>G 724A>G), A183S (731G>T), Y184P (734T>C 735A>C 736C>T), S185R (737T>C 738C>G 739C>T), A187V (744C>T 745G>C), R189C (749C>T 751A>C), M194K (765T>A), R197K (773C>A 774G>A 775A>G), M206K (801T>A), G210A (813G>C 814A>G), L211H (816T>A), R214Q (825G>A 826T>A), N223D (851A>G 853T>C), R226C (860C>T 862C>T), T227A (863A>G 865T>C), I228T (867T>C), N229D (869A>G 871C>T), I230L (872A>T 874A>G), R231Q (876G>A), R232K (879G>A), N240S (903A>G), T241N (906C>A 907A>C), K242T (909A>C 910A>C), S243N (912G>A 913C>T), S252L (938T>C 939C>T 940A>T), L266I (980T>A 982G>C), R272K (999G>A), Q275R (1007C>A 1008A>G 1009G>A), S276G (1010A>G 1012T>G), I277V (1013A>G), N279Q (1019A>C 1021C>G), D288E (1048C>A), R300Q (1083G>A), S305A (1097T>G 1099G>A), R320K (1143G>A), L324I (1154T>A 1156G>A), S346N (1221G>A), V349T (1229G>A 1230T>C 1231T>C), S350N (1233G>A 1234C>T), K358T (1257A>C 1258A>G), N365S (1278A>G), A369V (1290C>T 1291G>A), D372T (1298G>A 1299A>C 1300C>T), I375V (1307A>G 1309A>G), N378E (1316A>G 1318T>A), I382T (1329T>C), L397I (1373C>A), I401V (1385A>G), E407D (1405G>T), A411S (1415G>T 1417A>T), V418I (1436G>A 1438C>T), E508G (1707A>G), A512T (1718G>A), I520V (1742A>G 1744C>T), S525N (1758G>A), V536I (1790G>A), I585V (1937A>G 1939T>C), F607Y (2004T>A), I628M (2068T>G), V630I (2072G>A 2074G>A), R675K (2208G>A), D694E (2266C>G), N744G (2414A>G 2415A>G 2416T>G), S746C (2420A>T), K780R (2523A>G), R817K (2634G>A), V847I (2723G>A 2725C>T), A855T (2747G>A 2749T>A), H871N (2795C>A), V880I (2822G>A 2824G>A), P900T (2882C>A), S901H (2885A>C 2886G>A), T904M (2895C>T), N916S (2931A>G), N919S (2940A>G), E929_N930insX (2971_2972insA), T936I (2991C>T), H937R (2994A>G), E939R (2999G>A 3000A>G), S944F (3015C>T), R946Q (3021G>A), Y947F (3024A>T 3025C>T), T951I (3036C>T), E952G (3039A>G 3040A>C), K953T (3042A>C 3043G>A), A972V (3099C>T 3100C>A), V974R (3104G>A 3105T>G 3106G>A), S983N (3132G>A), R985K (3137C>A 3138G>A 3139A>G), F990I (3152T>A 3154C>T), C994Y (3165G>A), F1012L (3218T>C), T1014N (3225C>A), V1016I (3230G>A), V1028M (3266G>A 3268A>G), V1031M (3275G>A), A1034T (3284G>A), R1044K (3315G>A 3316A>G), E1045A (3318A>C), M1059V (3359A>G), D1072E (3400T>G), V1110G (3513T>G), M1121V (3545A>G), A1122T (3548G>A), V1124I (3554G>A 3556C>T), L1147V (3623C>G), L1150M (3632C>A), Y1152H (3638T>C), I1172V (3698A>G), S1178N (3717G>A), R1183K (3731C>A 3732G>A), I1297V (4073A>G), F1315Y (4128T>A 4129C>T), A1323S (4151G>T), V1328I (4166G>A), L1347I (4223C>A 4225G>A), I1379V (4319A>G 4321A>G), F1404Y (4395T>A), V1408I (4406G>A), M1431V (4475A>G 4477G>T), T1433S (4481A>T 4483A>G), A1454L (4544G>T 4545C>T 4546G>A), R1455K (4548G>A), V1499I (4679G>A 4681G>C), V1538I (4796G>A 4798C>T), T1540S (4803C>G 4804T>C), S1625G (5057A>G), T1626A (5060A>G), L1636I (5090C>A), A1639V (5100C>T), E1641D (5107A>T), V1646F (5120G>T 5122C>T), I1648V (5126A>G), V1667I (5183G>A), A1793T (5561G>A 5563G>A), I1805V (5597A>G), N1807K (5605C>A), A1812V (5619C>T 5620T>G), I1816L (5630A>C), V1837I (5693G>A), K1890R (5853A>G), I1904V (5894A>G 5896C>G), R1926A (5960C>G 5961G>C 5962G>C), I1936V (5990A>G), E1945D (6019A>T), K1949T (6030A>C), D1955E (6049C>A), M1958I (6058G>A), S1963N (6072G>A 6073T>C), N1964H (6074A>C), T1966N (6081C>A), C1976S (6111G>C 6112T>C), I1982V (6128A>G 6130T>C), V1992I (6158G>A), T1996S (6170A>T 6172A>G), L2007F (6203C>T 6205C>T), V2016I (6230G>A), A2025K (6257G>A 6258C>A), K2033R (6282A>G), A2060T (6362G>A), V2073I (6401G>A), A2074V (6405C>T), Y2082F (6429A>T), Q2089R (6450A>G), H2101N (6485C>A 6487C>T), L2115I (6527C>A), T2135I (6588C>T), Y2151H (6635T>C), I2155V (6647A>G), I2165V (6677A>G 6679A>G), D2178N (6716G>A), N2179D (6719A>G), I2194L (6764A>T 6766A>G), A2204V (6795C>T 6796A>G), H2205Q (6799C>A), N2225D (6857A>G), V2231T (6875G>A 6876T>C 6877C>A), D2250E (6934C>A), V2273I (7001G>A 7003G>A), F2283Y (7032T>A), T2314K (7125C>A), A2321S (7145G>T 7147A>G), E2326_A2327del (7160_7165delGAGGCA), D2344E (7216T>A), D2358E (7258T>A), D2363_R2364insX (7273_7274insT), R2364V (7274C>G 7275G>T), H2366L (7281A>T), L2367S (7284T>C), S2371L (7296C>T), S2376F (7311C>T), T2377I (7314C>T), C2378Y (7317G>A), N2379K (7321T>A), K2384Q (7334A>C), A2387N (7343G>A 7344C>A 7345A>C), S2408M (7407G>T 7408T>G), C2411G (7415T>G), L2440S (7503T>C 7504G>A), D2509N (7709G>A), K2510R (7713A>G 7714A>G), T2543I (7812C>T), V2554L (7844G>T 7846G>A), V2567I (7883G>A), S2620N (8043G>A), N2656S (8151A>G 8152C>T), R2665K (8178G>A), R2675K (8208G>A), P2704A (8294C>G 8296A>G), S2722F (8349C>T), A2742S (8408G>T), M2756I (8452G>A), R2759K (8460G>A), A2766T (8480G>A 8482T>A), R2773K (8502G>A 8503A>G), E2777D (8515A>T), E2785D (8539G>C), L2788C (8546C>T 8547T>G), S2790T (8552T>A), A2797G (8574C>G), I2801V (8585A>G), G2805E (8598G>A), N2807S (8603A>T 8604A>C 8605C>T), P2813Q (8622C>A), E2814G (8625A>G 8626G>A), P2822F (8648C>T 8649C>T 8650G>C), G2825E (8658G>A), D2827A (8664A>C 8665T>G), A2832V (8679C>T 8680A>T), T2854K (8745C>A 8746A>G), E2891G (8856A>G 8857G>A), R2894Q (8865G>A), M2895L (8867A>C 8869G>T), G2900E (8883G>A), T2902S (8888A>T), V2924I (8954G>A), N2934D (8984A>G 8986T>C), E2935V (8988A>T), I2983V (9131A>G), M2989G (9149A>G 9150T>G 9151G>A), G2990S (9152G>A), P3026S (9260C>T 9262T>C), Q3038H (9298A>C) | | | | | | | | |

| | Begin | End | Coverage | Score | Concordance | Matches | Identities | I/D/M/F* | Stop Codons |
|---|---|---|---|---|---|---|---|---|---|
| **NT** | **52** | **9704** | **99.5%** | **12530** | **65.2%** | **9642 (99.8%)** | **7985 (82.7%)** | **7/11** | |

Codon mutations:

ACT3ACC (193T>C), TAC4CAA (194T>C 196C>A), TCA6TCG (202A>G), ATC8ATT (208C>T), TGT9CTG (209T>C 210G>T 211T>G), TCG12TCA (220G>A), TTT13ATG (221T>A 223T>G), GAA14GAG (226A>G), CTA17CTT (235A>T), TAC19TAT (241C>T), CCA21CCT (247A>T), TCT23CCT (251T>C), TGC24TTT (255G>T 256C>T), GAG25GGG (258A>G), CAT26CAC (262T>C), ATT27GTG (263A>G 265T>G), GTG28GCG (267T>C), GAA30GTA (273A>T), CGA31CGT (277A>T), GAA32GAG (280A>G), GTG33GTT (283G>T), CCG34CCA (286G>A), GCT35ACC (287G>A 289T>C), GTT37ACT (293G>A 294T>C), CCT39CCC (301T>C), TTC40TTT (304C>T), GCA41ATA (305G>A 306C>T), GAT42GGT (309A>G), CTG43TTG (311C>T), GAA44GAT (316A>T), ACA45GTG (317A>G 318C>T 319A>G), CAA46CAG (322A>G), GCA49GCT (331A>T), CTC52CAA (339T>A 340C>A), AAG53AGG (342A>G), AAA55GCG (347A>G 348A>C 349A>G), TAT56CAT (350T>C), GCT57GCA (355T>A), ACT58GTT (356A>G 357C>T), CGT60CGC (364T>C), GTG61ACA (365G>A 366T>C 367G>A), CTC62TCC (368C>T 369T>C), AAA63AAG (373A>G), AAC64AAT (376C>T), TTT67TGT (384T>G), ACG68ACA (388G>A), TAC69TAT (391C>T), CGA70CGG (394A>G), AAG72AAA (400G>A), GAT74GAC (406T>C), GCC75GTT (408C>T 409C>T), CAG76CAA (412G>A), ATA77ATT (415A>T), ATG78GAG (416A>G 417T>A), ATT80ATC (424T>C), CAG81CAA (427G>A), AAA83AAG (433A>G), CTG84CTC (436G>C), GAG85AAG (437G>A), AGG86GAG (440A>G 441G>A), AAG87CGA (443A>C 444A>G 445G>A), GAT88GAA (448T>A), GAA90GAG (454A>G), GAA91GCA (456A>C), TAT92TAC (460T>C), CAC93AAC (461C>A), CAA95CAG (469A>G), GCC97GCG (475C>G), CCT99CCA (481T>A), ACT106ACC (502T>C), ATA107ATT (505A>T), GGC109GGT (511C>T), GAT111GAA (517T>A), CCT112CCA (520T>A), CCA113CCT (523A>T), TCT116TTT (531C>T), GAG117GAA (535G>A), CCA118TCA (536C>T), CAA119TGG (539C>T 540A>G 541A>G), AGA122AAG (549G>A 550A>G), ATC124ATT (556C>T), ATT125ATC (559T>C), CAT126CAC (562T>C), ACT128ACC (568T>C), CCA129CCG (571A>G), ATG131ACG (576T>C), CGT132CGC (580T>C), GTC134GCG (585T>C 586C>G), CGC137TAT (593C>T 594G>A 595C>T), CCC138CGC (597C>G), ATA139GTG (599A>G 601A>G), ATA140CCG (602A>C 603T>C 604A>G), ACA143ACG (613A>G), GAA144GAG (616A>G), GGC145GGG (619C>G), CAG146CAA (622G>A), ATG147ATA (625G>A), AAT148AAC (628T>C), CAC149CAT (631C>T), ATT151GTC (635A>G 637T>C), ATA154GTG (644A>G 646A>G), ATT157ATC (655T>C), TCG159TCA (661G>A), GAG160GCC (663A>C 664G>C), AAA161AAG (667A>G), GTC165GTT (679C>T), CAC166CAA (682C>A), TTA167CTG (683T>C 685A>G), AGT169AGC (691T>C), ACC172AGC (699C>G), ACT173ACA (703T>A), CAT174CAA (706T>A), AAG179GAA (719A>G 721G>A), ATA180GTG (722A>G 724A>G), CTT181TTG (725C>T 727T>G), GGT182GGA (730T>A), GCA183TCA (731G>T), TAC184CCT (734T>C 735A>C 736C>T), TCC185CGT (737T>C 738C>G 739C>T), GCG187GTC (744C>T 745G>C), CGA189TGC (749C>T 751A>C), ACT190ACC (754T>C), ATG194AAG (765T>A), TTG196CTG (770T>C), CGA197AAG (773C>A 774G>A 775A>G), CGG198AGG (776C>A), TTC202TTT (790C>T), ATG206AAG (801T>A), GGA210GCG (813G>C 814A>G), CTT211CAT (816T>A), TTG212CTG (818T>C), CAA213CAG (823A>G), CGT214CAA (825G>A 826T>A), CGG217AGA (833C>A 835G>A), GAC219GAT (841C>T), AAT223GAC (851A>G 853T>C), GTC225GTT (859C>T), CGC226TGT (860C>T 862C>T), ACT227GCC (863A>G 865T>C), ATC228ACC (867T>C), AAC229GAT (869A>G 871C>T), ATA230TTG (872A>T 874A>G), CGA231CAA (876G>A), AGG232AAG (879G>A), GAT234GAC (886T>C), ATC238ATT (898C>T), AAC240AGC (903A>G), ACA241AAC (906C>A 907A>C), AAA242ACC (909A>C 910A>C), AGC243AAT (912G>A 913C>T), GGC246GGA (922C>A), GGT249GGA (931T>A), TCA252CTT (938T>C 939C>T 940A>T), GGA253GGG (943A>G), TTC256TTT (952C>T), TCA261TCG (967A>G), GAA263GAG (973A>G), GGG264GGC (976G>C), TTG266ATC (980T>A 982G>C), GCA269GCG (991A>G), CGT270CGC (994T>C), AGA272AAA (999G>A), GTT273GTC (1003T>C), CAG275AGA (1007C>A 1008A>G 1009G>A), AGT276GGG (1010A>G 1012T>G), ATT277GTT (1013A>G), TTA278TTG (1018A>G), AAC279CAG (1019A>C 1021C>G), ATC282ATT (1030C>T), TCG285TCA (1039G>A), AAT286AAC (1042T>C), GCC287GCA (1045C>A), GAC288GAA (1048C>A), AAT289AAC (1051T>C), AAG292AAA (1060G>A), GGT293GGA (1063T>A), CTG294TTG (1064C>T), GAC295GAT (1069C>T), CGA300CAA (1083G>A), TAT303TAC (1093T>C), TCG305GCA (1097T>G 1099G>A), CAC307CAT (1105C>T), GTA310GTG (1114A>G), GCT311GCG (1117T>G), TTA313TTG (1123A>G), CCT314CCA (1126T>A), GTC315GTT (1129C>T), AGG320AAG (1143G>A), GTA321GTT (1147A>T), GCT322GCA (1150T>A), TTG324ATA (1154T>A 1156G>A), CAC327CAT (1165C>T), AGT328AGC (1168T>C), CTT330TTA (1172C>T 1174T>A), ACC339ACT (1201C>T), AGC346AAC (1221G>A), GTT349ACC (1229G>A 1230T>C 1231T>C), AGC350AAT (1233G>A 1234C>T), CTG352CTA (1240G>A), AAG354AAA (1246G>A), TTG356CTA (1250T>C 1252G>A), CAT357CAC (1255T>C), AAA358ACG (1257A>C 1258A>G), AGA361AGG (1267A>G), GGT363GGC (1273T>C), TTG364TTA (1276G>A), AAT365AGT (1278A>G), TTG367CTG (1283T>C), GCG369GTA (1290C>T 1291G>A), GAC372ACT (1298G>A 1299A>C 1300C>T), TTT374TTC (1306T>C), ATA375GTG (1307A>G 1309A>G), CAT376CAC (1312T>C), AAT378GAA (1316A>G 1318T>A), TTG381TTA (1327G>A), ATA382ACA (1329T>C), GCG383GCT (1333G>T), TTA384TTG (1336A>G), CTA387TTG (1343C>T 1345A>G), CCG390CCA (1354G>A), GTG391GTT (1357G>T), GAC392GAT (1360C>T), CTC395TTG (1367C>T 1369C>G), CTT397ATT (1373C>A), TTC398TTT (1378C>T), GAG400GAA (1384G>A), ATA401GTA (1385A>G), AAA403AAG (1393A>G), TCC404TCT (1396C>T), GAG407GAT (1405G>T), AAA408AAG (1408A>G), CAG409CAA (1411G>A), GCA411TCT (1415G>T 1417A>T), CCG412CCC (1420G>C), AAT415AAC (1429T>C), TTA416TTG (1432A>G), GTC418ATT (1436G>A 1438C>T), TTA419CTG (1439T>C 1441A>G), AAT420AAC (1444T>C), TTC423TTT (1453C>T), CTG424CTA (1456G>A), GAA428GAG (1468A>G), GTA436GTG (1492A>G), TTG439CTA (1499T>C 1501G>A), AGT440AGC (1504T>C), TTG441TTA (1507G>A), CTC442CTT (1510C>T), GAA443GAG (1513A>G), TTA444TTG (1516A>G), GCA445GCT (1519A>T), AGG446AGA (1522G>A), TTC447TTT (1525C>T), CAG448CAA (1528G>A), AAC450AAT (1534C>T), ACT452ACA (1540T>A), AAC454AAT (1546C>T), AAA457AAG (1555A>G), GGT458GGC (1558T>C), GAT459GAC (1561T>C), ATA460ATC (1564A>C), TCT461TCA (1567T>A), TTC463TTT (1573C>T), AGA464AGG (1576A>G), TTA467CTA (1583T>C), TCT468TCA (1588T>A), AAG470AAA (1594G>A), AAC472AAT (1600C>T), CTG475CTA (1609G>A), TCG478TCA (1618G>A), TGC479TGT (1621C>T), GAC480GAT (1624C>T), CTG483TTG (1631C>T), AAA485AAG (1639A>G), TTC489TTT (1651C>T), CTC490CTG (1654C>G), CAT497CAC (1675T>C), TCA503TCG (1693A>G), TTT506TTC (1702T>C), GAA508GGA (1707A>G), GCA512ACA (1718G>A), GGA514GGG (1726A>G), TAC515TAT (1729C>T), TCA516TCC (1732A>C), TAT518TAC (1738T>C), ATC520GTT (1742A>G 1744C>T), CGC521CGT (1747C>T), CAT523CAC (1753T>C), AGT525AAT (1758G>A), AGG528AGA (1768G>A), AAC534AAT (1786C>T), TTA535TTG (1789A>G), GTT536ATT (1790G>A), GTC537GTT (1795C>T), TTA541CTG (1805T>C 1807A>G), GAG543GAA (1813G>A), AGG545AGA (1819G>A), AAG547AAA (1825G>A), GAC551GAT (1837C>T), TAT552TAC (1840T>C), CAA555CAG (1849A>G), GTC558GTG (1858C>G), AAA560AAG (1864A>G), TGC562TGT (1870C>T), AGT564AGC (1876T>C), AAA566AAG (1882A>G), GGT568GGA (1888T>A), AAT569AAC (1891T>C), TGT574TGC (1906T>C), ACA577ACT (1915A>T), GGT582GGC (1930T>C), GCC584GCT (1936C>T), ATT585GTC (1937A>G 1939T>C), TCA587TCG (1945A>G), TTC589TTT (1951C>T), CCA592CCT (1960A>T), GTA598GTG (1978A>G), ATT599ATA (1981T>A), GGC600GGT (1984C>T), AAT601AAC (1987T>C), GAC604GAT (1996C>T), TTT607TAT (2004T>A), AAA612AAG (2020A>G), GGG613GGA (2023G>A), GAT614GAC (2026T>C), TCG615TCT (2029G>T), GAG616GAA (2032G>A), TAT625TAC (2059T>C), ATT628ATG (2068T>G), AAC629AAT (2071C>T), GTG630ATA (2072G>A 2074G>A), CTG635TTA (2087C>T 2089G>A), AGC639AGT (2101C>T), GAG640GAA (2104G>A), GAG641GAA (2107G>A), AAG644AAA (2116G>A), TTC646TTT (2122C>T), CGC651CGT (2137C>T), CTT658CTC (2158T>C), ACC660ACT (2164C>T), CCA662CCG (2170A>G), GAT666GAC (2182T>C), GCG668GCA (2188G>A), ACC669ACA (2191C>A), AGA675AAA (2208G>A), TAT678TAC (2218T>C), GAC680GAT (2224C>T), GTA681GTT (2227A>T), TTG686TTA (2242G>A), CCC687CCT (2245C>T), TTG690TTA (2254G>A), GTT691GTC (2257T>C), GAC692GAT (2260C>T), GAC694GAG (2266C>G), CAA696CAG (2272A>G), ACG697ACA (2275G>A), TCA703TCG (2293A>G), GGC705GGT (2299C>T), TCG706TCA (2302G>A), CAG707CAA (2305G>A), GGA710GGT (2314A>T), TAT711TAC (2317T>C), CTA714CTG (2326A>G), GTG719GTA (2341G>A), CAA721CAG (2347A>G), CTT722CTC (2350T>C), TTG724TTA (2356G>A), AAT727AAC (2365T>C), GAA729GAG (2371A>G), TTA730TTG (2374A>G), GAT733GAC (2383T>C), ATA734ATT (2386A>T), AAA735AAG (2389A>G), CAT736CAC (2392T>C), TAT737TAC (2395T>C), AAT744GGG (2414A>G 2415A>G 2416T>G), GCT745GCA (2419T>A), AGC746TGC (2420A>T), ACA752ACT (2440A>T), GTT761GTC (2467T>C), TCT764TCC (2476T>C), TCG766TCA (2482G>A), CTG769CTA (2491G>A), CTG771CTT (2497G>T), TTG773TTA (2503G>A), AGA778AGG (2518A>G), CCT779CCC (2521T>C), AAG780AGG (2523A>G), AGA783AGG (2533A>G), TTG785TTA (2539G>A), TTA787TTG (2545A>G), GAG789GAA (2551G>A), CCT790CCA (2554T>A), CTG792TTG (2558C>T), TTG793CTC (2561T>C 2563G>C), TCA796TCG (2572A>G), CCT800CCC (2584T>C), GGC801GGT (2587C>T), CTG803CTC (2593G>C), GCT805GCC (2599T>C), AAT808AAC (2608T>C), TTT812TTC (2620T>C), GCG815GCA (2629G>A), GTG816GTA (2632G>A), AGG817AAG (2634G>A), TTG818TTA (2638G>A), ATT820ATA (2644T>A), AAA823AAG (2653A>G), GCT827GCC (2665T>C), CTA832TTA (2678C>T), CTA833CTG (2683A>G), GCT835GCC (2689T>C), CTA838CTC (2698A>C), TCA841TCG (2707A>G), GCG842GCA (2710G>A), GAA844GAG (2716A>G), ACA845ACG (2719A>G), CTC846CTT (2722C>T), GTC847ATT (2723G>A 2725C>T), GCA848GCG (2728A>G), AGG850AGA (2734G>A), ATT851ATA (2737T>A), GAT854GAC (2746T>C), GCT855ACA (2747G>A 2749T>A), GCA856GCT (2752A>T), GCT857GCA (2755T>A), GAC859GAT (2761C>T), CTC860CTT (2764C>T), TGT865TGC (2779T>C), GGG867GGT (2785G>T), AAC869AAT (2791C>T), CAT871AAT (2795C>A), TAC874TAT (2806C>T), TTG878CTC (2816T>C 2818G>C), GTG880ATA (2822G>A 2824G>A), TTG881TTA (2827G>A), GTT884GTC (2836T>C), AAG885AAA (2839G>A), AAT886AAC (2842T>C), ACC893ACA (2863C>A), CTA894TTA (2864C>T), CCA900ACA (2882C>A), AGT901CAT (2885A>C 2886G>A), AAC903AAT (2893C>T), ACG904ATG (2895C>T), GTC906GTT (2902C>T), GTA907GTG (2905A>G), ATT909ATC (2911T>C), AAT913AAC (2923T>C), AAT916AGT (2931A>G), AAC919AGC (2940A>G), GCT921GCC (2947T>C), ACT926ACC (2962T>C), GAA929_AAT930insA-- (2971_2972insA), GGT935GGC (2989T>C), ACT936ATT (2991C>T), CAT937CGT (2994A>G), GAG939AGG (2999G>A 3000A>G), TCC944TTC (3015C>T), CGG946CAG (3021G>A), TAC947TTT (3024A>T 3025C>T), ACA951ATA (3036C>T), GAA952GGC (3039A>G 3040A>C), AAG953ACA (3042A>C 3043G>A), TTG956TTA (3052G>A), TTA959TTG (3061A>G), GCG966GCA (3082G>A), GGC969GGT (3091C>T), GCC972GTA (3099C>T 3100C>A), GTG974AGA (3104G>A 3105T>G 3106G>A), GTC975GTT (3109C>T), AAA976AAG (3112A>G), ACT978ACC (3118T>C), GCC979GCT (3121C>T), GGA981GGG (3127A>G), AGC983AAC (3132G>A), CGA985AAG (3137C>A 3138G>A 3139A>G), TAT989TAC (3151T>C), TTC990ATT (3152T>A 3154C>T), TGT994TAT (3165G>A), GTA995GTG (3169A>G), ATC1005ATT (3199C>T), AGG1007CGG (3203A>C), CGT1008CGC (3208T>C), CCA1010CCG (3214A>G), ACC1011ACT (3217C>T), TTT1012CTT (3218T>C), GTC1013GTA (3223C>A), ACT1014AAT (3225C>A), GTT1016ATT (3230G>A), AAC1017AAT (3235C>T), TTA1025CTA (3257T>C), ACC1026ACT (3262C>T), AGC1027AGT (3265C>T), GTA1028ATG (3266G>A 3268A>G), GTG1029GTA (3271G>A), GTG1031ATG (3275G>A), GCA1034ACA (3284G>A), CAG1039CAA (3301G>A), AGA1044AAG (3315G>A 3316A>G), GAA1045GCA (3318A>C), ATC1046ATT (3322C>T), GAG1047GAA (3325G>A), TTG1048TTA (3328G>A), ATA1051ATT (3337A>T), GAG1055GAA (3349G>A), GTC1057GTT (3355C>T), TGC1058TGT (3358C>T), ATG1059GTG (3359A>G), CTA1061CTG (3367A>G), TAT1062TAC (3370T>C), TTA1065CTG (3377T>C 3379A>G), GAA1070GAG (3394A>G), CGC1071CGT (3397C>T), GAT1072GAG (3400T>G), ACA1074ACG (3406A>G), TAC1078TAT (3418C>T), ATT1079ATA (3421T>A), TAT1081TAC (3427T>C), AAG1083AAA (3433G>A), TCA1084TCT (3436A>T), GTA1085GTG (3439A>G), AAC1086AAT (3442C>T), CCT1087CCC (3445T>C), CAG1091CAA (3457G>A), TTT1092TTC (3460T>C), GCT1093GCA (3463T>A), GCG1095GCT (3469G>T), CAG1096CAA (3472G>A), CAC1104CAT (3496C>T), CAG1105CAA (3499G>A), TCC1107TCT (3505C>T), GTT1110GGT (3513T>G), AAA1112AAG (3520A>G), GAA1115GAG (3529A>G), CAA1116CAG (3532A>G), ATG1121GTG (3545A>G), GCT1122ACT (3548G>A), TTA1123CTA (3551T>C), GTC1124ATT (3554G>A 3556C>T), AGT1133AGC (3583T>C), GAT1134GAC (3586T>C), TGC1135TGT (3589C>T), TTC1137TTT (3595C>T), AAT1141AAC (3607T>C), TTT1143TTC (3613T>C), AAG1144AAA (3616G>A), GGT1145GGC (3619T>C), CTT1147GTT (3623C>G), TCC1148TCT (3628C>T), CTG1150ATG (3632C>A), GAT1151GAC (3637T>C), TAT1152CAT (3638T>C), GAA1153GAG (3643A>G), TTA1159TTG (3661A>G), GTG1162GTA (3670G>A), TTT1166TTC (3682T>C), GAG1168GAA (3688G>A), GAG1171GAA (3697G>A), ATT1172GTT (3698A>G), GAT1174GAC (3706T>C), TTT1175TTC (3709T>C), TTG1177CTA (3713T>C 3715G>A), AGT1178AAT (3717G>A), GAG1179GAA (3721G>A), CGA1183AAA (3731C>A 3732G>A), ACT1184ACA (3736T>A), CTA1188TTG (3746C>T 3748A>G), ACA1190ACG (3754A>G), AGT1193AGC (3763T>C), CGA1198CGG (3778A>G), CAA1199CAG (3781A>G), ATC1200ATT (3784C>T), CAT1204CAC (3796T>C), CCA1207CCT (3805A>T), TAC1209TAT (3811C>T), AGA1210AGG (3814A>G), TTC1215TTT (3829C>T), TTT1218TTC (3838T>C), AGA1220AGG (3844A>G), GCA1221GCC (3847A>C), GTT1224GTA (3856T>A), GTG1226GTT (3862G>T), GAC1229GAT (3871C>T), GCC1231GCA (3877C>A), AGC1233AGT (3883C>T), GAA1234GAG (3886A>G), CTA1236CTG (3892A>G), GAC1237GAT (3895C>T), TTT1238TTC (3898T>C), GTA1240GTG (3904A>G), CGG1241AGA (3905C>A 3907G>A), GCT1243GCC (3913T>C), AAG1248AAA (3928G>A), TCA1249TCT (3931A>T), GGG1251GGA (3937G>A), AGT1257AGC (3955T>C), GTG1258GTA (3958G>A), GCC1259GCT (3961C>T), GGA1260GGG (3964A>G), TCT1261TCC (3967T>C), TTA1264TTG (3976A>G), ATT1265ATC (3979T>C), CCA1267CCT (3985A>T), CTA1271CTT (3997A>T), GCG1272GCA (4000G>A), AAC1274AAT (4006C>T), GTT1275GTG (4009T>G), TTC1276TTT (4012C>T), AAA1277AAG (4015A>G), TCT1280TCC (4024T>C), GAA1282GAG (4030A>G), CCA1283CCT (4033A>T), AAG1286AAA (4042G>A), CTG1290TTA (4052C>T 4054G>A), CGT1291CGC (4057T>C), CGT1293CGA (4063T>A), GGA1294GGG (4066A>G), ATA1297GTA (4073A>G), TTT1298TTC (4078T>C), GGC1299GGT (4081C>T), TCT1300TCC (4084T>C), TCC1304TCT (4096C>T), AGC1308AGT (4108C>T), GGA1309GGT (4111A>T), GCG1311GCA (4117G>A), CTA1312TTG (4118C>T 4120A>G), TAC1314TAT (4126C>T), TTC1315TAT (4128T>A 4129C>T), GCC1316GCT (4132C>T), AAT1317AAC (4135T>C), CAA1321CAG (4147A>G), TTA1322CTA (4148T>C), GCT1323TCT (4151G>T), AAC1326AAT (4162C>T), GTA1328ATA (4166G>A), ATA1329ATC (4171A>C), GAG1332GAA (4180G>A), CTG1336TTG (4190C>T), CCT1338CCC (4198T>C), TCC1339TCT (4201C>T), GCG1342GCT (4210G>T), TTC1343TTT (4213C>T), AGT1345AGC (4219T>C), CTG1346TTG (4220C>T), CTG1347ATA (4223C>A 4225G>A), GTT1349GTG (4231T>G), CAT1351CAC (4237T>C), GTA1356GTG (4252A>G), TTA1357CTG (4253T>C 4255A>G), AAA1358AAG (4258A>G), ACT1362ACC (4270T>C), AGA1366AGG (4282A>G), GAG1367GAA (4285G>A), GAA1369GAG (4291A>G), CAG1373CAA (4303G>A), GTC1376GTT (4312C>T), ATA1379GTG (4319A>G 4321A>G), GTG1380GTT (4324G>T), GAG1381GAA (4327G>A), CTG1384CTT (4336G>T), TCT1385TCC (4339T>C), TCA1388TCT (4348A>T), GCA1392GCG (4360A>G), GGT1394GGC (4366T>C), TCT1395TCA (4369T>A), ACT1397ACC (4375T>C), GTT1401GTC (4387T>C), TTT1404TAT (4395T>A), GTA1408ATA (4406G>A), CTT1409CTC (4411T>C), GTT1418GTC (4438T>C), ACC1420ACA (4444C>A), GCC1422GCT (4450C>T), CTC1424CTT (4456C>T), GAC1427GAT (4465C>T), ATG1431GTT (4475A>G 4477G>T), GTC1432GTT (4480C>T), ACA1433TCG (4481A>T 4483A>G), GGC1437GGT (4495C>T), AAG1441AAA (4507G>A), CAC1442CAT (4510C>T), GGT1443GGG (4513T>G), CTA1445TTA (4517C>T), GTC1448GTT (4528C>T), AAA1450AAG (4534A>G), ACC1452ACT (4540C>T), GCG1454TTA (4544G>T 4545C>T 4546G>A), AGA1455AAA (4548G>A), TTT1458TTC (4558T>C), GTA1460GTT (4564A>T), ACC1462ACT (4570C>T), ATA1464ATT (4576A>T), TTG1471TTA (4597G>A), GAC1472GAT (4600C>T), GAC1474GAT (4606C>T), GTG1475GTA (4609G>A), GTG1477GTA (4615G>A), GAC1478GAT (4618C>T), TTG1481CTT (4625T>C 4627G>T), GAC1488GAT (4648C>T), ATT1489ATC (4651T>C), AGG1492AGA (4660G>A), AGC1493AGT (4663C>T), GCT1495GCA (4669T>A), GTG1499ATC (4679G>A 4681G>C), AGT1500AGC (4684T>C), GTT1501GTG (4687T>G), GAG1505GAA (4699G>A), AGA1506AGG (4702A>G), CAA1508CAG (4708A>G), AGG1509AGA (4711G>A), CTG1510TTG (4712C>T), AAA1518AAG (4738A>G), GTA1520GTT (4744A>T), GCA1521GCG (4747A>G), TTG1522CTG (4748T>C), ACT1527ACC (4765T>C), GTC1538ATT (4796G>A 4798C>T), ACT1540AGC (4803C>G 4804T>C), GAG1541GAA (4807G>A), GCG1542GCT (4810G>T), GCT1543GCG (4813T>G), CTT1544CTC (4816T>C), TAT1549TAC

| | | | | | | | | | |
|---|---|---|---|---|---|---|---|---|---|
| | (4831T>C), AAC1550AAT (4834C>T), CCA1552CCT (4840A>T), GGA1556GGG (4852A>G), GGC1557GGT (4855C>T), AGT1561AGC (4867T>C), CTG1562CTC (4870G>C), ATT1563ATA (4873T>A), TGT1566TGC (4882T>C), ACA1573ACT (4903A>T), CAG1575CAA (4909G>A), TTT1577TTC (4915T>C), GAA1578GAG (4918A>G), TTG1579CTA (4919T>C 4921G>A), AGT1580AGC (4924T>C), CCC1581CCG (4927C>G), TTT1583TTC (4933T>C), ATC1584ATA (4936C>A), CAG1585CAA (4939G>A), AAT1586AAC (4942T>C), TTC1587TTT (4945C>T), TCA1593TCG (4963A>G), CCT1596CCC (4972T>C), CTT1602CTC (4990T>C), AAA1606AAG (5002A>G), CCT1613CCC (5023T>C), TTG1614CTG (5024T>C), TGC1615TGT (5029C>T), GAT1616GAC (5032T>C), TCT1618TCC (5038T>C), CCA1620CCT (5044A>T), GCA1623GCT (5053A>T), TCG1624TCA (5056G>A), AGT1625GGT (5057A>G), ACT1626GCT (5060A>G), TCG1629TCT (5071G>T), CTT1636ATT (5090C>A), GGA1637GGG (5095A>G), GTG1638GTT (5098G>T), GCC1639GTC (5100C>T), TTA1640TTG (5104A>G), GAA1641GAT (5107A>T), ATT1642ATC (5110T>C), CCA1643CCG (5113A>G), AAG1644AAA (5116G>A), CAA1645CAG (5119A>G), GTC1646TTT (5120G>T 5122C>T), ATT1648GTT (5126A>G), CAT1651CAC (5137T>C), ATC1652ATT (5140C>T), AAA1653AAG (5143A>G), GAG1654GAA (5146G>A), CTC1659CTA (5161C>A), GAA1661GAG (5167A>G), CTT1663CTG (5173T>G), ACG1666ACA (5182G>A), GTT1667ATT (5183G>A), GTC1668GTT (5188C>T), AAG1669AAA (5191G>A), AAA1671AAG (5197A>G), GAC1672GAT (5200C>T), TGC1674TGT (5206C>T), TTA1675TTG (5209A>G), AGC1678AGT (5218C>T), TCG1682TCA (5230G>A), ATC1684ATT (5236C>T), AGC1685AGT (5239C>T), ATC1687ATT (5245C>T), ACA1693ACT (5263A>T), CTC1695CTT (5269C>T), TTC1696TTT (5272C>T), GCC1697GCA (5275C>A), ATC1698ATT (5278C>T), CCA1699CCC (5281A>C), AGA1700AGG (5284A>G), ACT1701ACC (5287T>C), CTA1702TTA (5288C>T), ATA1703ATT (5293A>T), TTG1704CTA (5294T>C 5296G>A), GAG1706GAA (5302G>A), CTT1709CTA (5311T>A), GAA1710GAG (5314A>G), CGA1713CGT (5323A>T), AGT1721AGC (5347T>C), ATC1723ATT (5353C>T), TGC1727TGT (5365C>T), GTT1734GTC (5386T>C), ACC1737ACA (5395C>A), AAA1746AAG (5422A>G), GAT1747GAC (5425T>C), ACC1749ACT (5431C>T), GAG1751GAA (5437G>A), CTT1756CTC (5452T>C), AAG1758AAA (5458G>A), AGT1761AGC (5467T>C), CAA1762CAG (5470A>G), CTA1763TTG (5471C>T 5473A>G), GAA1765GAG (5479A>G), TTC1766TTT (5482C>T), TCA1767TCC (5485A>C), AAT1768AAC (5488T>C), TTG1769CTA (5489T>C 5491G>A), GGT1771GGC (5497T>C), TGT1774TGC (5506T>C), GAG1775GAA (5509G>A), GAG1776GAA (5512G>A), TTA1778CTA (5516T>C), TAT1782TAC (5530T>C), GAG1783GAA (5533G>A), TCG1784TCT (5536G>T), TTG1785TTA (5539G>A), GTT1788GTG (5548T>G), CAC1790CAT (5554C>T), GCT1792GCA (5560T>A), GCG1793ACA (5561G>A 5563G>A), ACG1794ACC (5566G>C), TCA1795TCG (5569A>G), CTC1800CTG (5584C>G), AAG1803AAA (5593G>A), GGG1804GGA (5596G>A), ATT1805GTT (5597A>G), AAC1807AAA (5605C>A), GTG1811GTT (5617G>T), GCT1812GTG (5619C>T 5620T>G), TTG1815TTA (5629G>A), ATC1816CTC (5630A>C), GGC1819GGT (5641C>T), GCA1822GCT (5650A>T), ATA1826ATT (5662A>T), GGA1827GGG (5665A>G), CAA1835CAG (5689A>G), GTT1837ATT (5693G>A), GAG1838GAA (5698G>A), CAA1843CAG (5713A>G), GGG1844GGC (5716G>C), AAA1845AAG (5719A>G), GCC1853GCA (5743C>A), CGC1857CGA (5755C>A), CAT1858CAC (5758T>C), CGT1860CGC (5764T>C), AAA1862AAG (5770A>G), GGC1865GGT (5779C>T), TTT1866TTC (5782T>C), GAC1869GAT (5791C>T), GAC1873GAT (5803C>T), AGG1884AGA (5836G>A), AAA1885AAG (5839A>G), GGA1887GGG (5845A>G), AAA1888AAG (5848A>G), AAA1890AGA (5853A>G), GGT1891GGC (5857T>C), ACC1892ACT (5860C>T), ACA1893ACT (5863A>T), GGT1895GGA (5869T>A), GGT1897GGC (5875T>C), AGC1900AGT (5884C>T), TTC1903TTT (5893C>T), ATC1904GTG (5894A>G 5896C>G), AAC1905AAT (5899C>T), GGG1908GGA (5908G>A), TTT1909TTC (5911T>C), GAG1913GAA (5923G>A), CAA1918CAG (5938A>G), TTC1919TTT (5941C>T), GTT1920GTC (5944T>C), GAT1921GAC (5947T>C), GGG1925GGA (5959G>A), CGG1926GCC (5960C>G 5961G>C 5962G>C), GAA1930GAG (5974A>G), GCT1934GCA (5986T>A), GAC1935GAT (5989C>T), ATT1936GTT (5990A>G), GAT1938GAC (5998T>C), AGA1942CGT (6008A>C 6010A>T), GAA1945GAT (6019A>T), GTG1946GTT (6022G>T), CGA1947CGT (6025A>T), AAA1949ACA (6030A>C), GAC1955GAA (6049C>A), ATG1958ATA (6058G>A), CAA1959CAG (6061A>G), GCC1960GCA (6064C>A), GGT1962GGC (6070T>C), AGT1963AAC (6072G>A 6073T>C), AAC1964CAC (6074A>C), ACG1965ACA (6079G>A), ACC1966AAC (6081C>A), ATA1967ATT (6085A>T), GCA1969GCT (6091A>T), TAC1970TAT (6094C>T), TGT1976TCC (6111G>C 6112T>C), AAA1978AAG (6118A>G), TTG1980TTA (6124G>A), AAG1981AAA (6127G>A), ATT1982GTC (6128A>G 6130T>C), GAT1983GAC (6133T>C), TTA1984TTG (6136A>G), AAC1988AAT (6148C>T), AAA1991AAG (6157A>G), GTT1992ATT (6158G>A), TGT1993TGC (6163T>C), AAA1995AAG (6169A>G), ACA1996TCG (6170A>T 6172A>G), GGC1998GGT (6178C>T), AAA2001AAG (6187A>G), TTT2002TTC (6190T>C), GAG2004GAA (6196G>A), CTC2007TTT (6203C>T 6205C>T), GAA2008GAG (6208A>G), CTA2009TTG (6209C>T 6211A>G), CAG2011CAA (6217G>A), GCT2015GCA (6229T>A), GTA2016ATA (6230G>A), GTC2018GTT (6238C>T), AAG2021AAA (6247G>A), ATA2023ATT (6253A>T), GCA2025AAA (6257G>A 6258C>A), GAG2027GAA (6265G>A), CAT2030CAC (6274T>C), AAA2033AGA (6282A>G), TCG2034TCA (6286G>A), GGC2038GGT (6298C>T), TTG2039TTA (6301G>A), AGA2040AGG (6304A>G), GAC2041GAT (6307C>T), GCC2046GCT (6322C>T), ACA2048ACT (6328A>T), GTT2049GTG (6331T>G), TGT2050TGC (6334T>C), AGG2051AGA (6337G>A), CTG2052TTG (6338C>T), TAT2058TAC (6358T>C), GGG2059GGA (6361G>A), GCA2060ACA (6362G>A), TCA2061TCT (6367A>T), GAG2062GAA (6370G>A), TAC2064TAT (6376C>T), GGT2065GGG (6379T>G), GGA2067GGC (6385A>C), GGA2069GGT (6391A>T), GCA2070GCG (6394A>G), TAC2071TAT (6397C>T), GTA2073ATA (6401G>A), GCG2074GTG (6405C>T), CAC2076CAT (6412C>T), TTA2078TTG (6418A>G), TTT2079TTC (6421T>C), TAC2082TTC (6429A>T), GGT2084GGA (6436T>A), GAG2087GAA (6445G>A), CAA2089CGA (6450A>G), TCC2090TCG (6454C>G), CAC2092CAT (6460C>T), GGT2093GGA (6463T>A), TTC2095TTT (6469C>T), AGG2096AGA (6472G>A), GTG2097GTA (6475G>A), AAT2099AAC (6481T>C), CAC2101AAT (6485C>A 6487C>T), AGT2102AGC (6490T>C), AGC2104AGT (6496C>T), CTG2106CTA (6502G>A), GGT2110GGC (6514T>C), AGG2111AGA (6517G>A), CTC2115ATC (6527C>A), ATC2116ATA (6532C>A), CCG2119CCA (6541G>A), AAA2120AAG (6544A>G), GTC2124GTT (6556C>T), TTT2125TTC (6559T>C), CAG2127CAA (6565G>A), TTG2129CTA (6569T>C 6571G>A), CCT2134CCA (6586T>A), ACA2135ATA (6588C>T), GAA2138GAG (6598A>G), AGA2139AGG (6601A>G), ATT2140ATC (6604T>C), TGT2141TGC (6607T>C), GGA2144GGG (6616A>G), AAC2146AAT (6622C>T), GAG2149GAA (6631G>A), AAG2150AAA (6634G>A), TAT2151CAT (6635T>C), GCT2152GCA (6640T>A), TCG2154TCT (6646G>T), ATC2155GTC (6647A>G), ATC2156ATT (6652C>T), AGC2160AGT (6664C>T), ACT2162ACG (6670T>G), ATA2165GTG (6677A>G 6679A>G), GGC2167GGT (6685C>T), ACA2169ACT (6691A>T), TTC2170TTT (6694C>T), ACA2177ACG (6715A>G), GAT2178AAT (6716G>A), AAT2179GAT (6719A>G), GGA2180GGG (6724A>G), CTA2184TTA (6734C>T), GTG2186GTA (6742G>A), GTG2187GTA (6745G>A), GCC2190GCT (6754C>T), GGA2192GGG (6760A>G), ATA2194TTG (6764A>T 6766A>G), ATT2197ATA (6775T>A), CTG2200TTG (6782C>T), GCA2201GCT (6787A>T), AAC2202AAT (6790C>T), GCA2204GTG (6795C>T 6796A>G), CAC2205CAA (6799C>A), ACC2206ACA (6802C>A), AAC2208AAT (6808C>T), TAC2209TAT (6811C>T), TAC2210TAT (6814C>T), TTC2213TTT (6823C>T), GAA2215GAG (6829A>G), TAC2221TAT (6847C>T), CGA2223CGC (6853A>C), AAT2225GAT (6857A>G), GAG2226GAA (6862G>A), GAA2229GAG (6871A>G), GTC2231ACA (6875G>A 6876T>C 6877C>A), AAG2232AAA (6880G>A), TCT2233TCA (6883T>A), GTT2235GTG (6889T>G), GAC2239GAT (6901C>T), GGC2244GGT (6916C>T), CCG2245CCA (6919G>A), AAA2247AAG (6925A>G), CTT2248CTC (6928T>C), GAC2250GAA (6934C>A), GGG2255GGC (6949G>C), TTC2257TTT (6955C>T), AAA2258AAG (6958A>G), CAA2264CAG (6976A>G), CTA2266TTA (6980C>T), ATC2267ATT (6985C>T), GTA2271GTT (6997A>T), GTG2272GTT (7000G>T), GTG2273ATA (7001G>A 7003G>A), GCT2276GCC (7012T>C), AAG2277AAA (7015G>A), CAT2278CAC (7018T>C), TTT2283TAT (7032T>A), GCC2285GCT (7039C>T), TTG2286CTA (7040T>C 7042G>A), GGA2288GGG (7048A>G), GCT2292GCA (7060T>A), GTC2293GTA (7063C>A), AGC2298AGT (7078C>T), TTA2300CTG (7082T>C 7084A>G), GTA2301GTG (7087A>G), ACC2302ACA (7090C>A), CAT2304CAC (7096T>C), GTA2305GTG (7099A>G), GTT2306GTC (7102T>C), GAG2309GAA (7111G>A), CGA2311CGG (7117A>G), ACA2314AAA (7125C>A), GAA2315GAG (7129A>G), TTT2316TTC (7132T>C), CTG2317TTA (7133C>T 7135G>A), GTG2319GTA (7141G>A), GAT2320GAC (7144T>C), GCA2321TCG (7145G>T 7147A>G), GAG2322GAA (7150G>A), GCA2325GCT (7159A>T), GAG2326_GCA2327del (7160_7165delGAGGCA), TTC2329TTT (7171C>T), GCG2335GCT (7189G>T), AAA2338AAG (7198A>G), AGC2339AGT (7201C>T), CTA2341TTA (7205C>T), AAT2342AAC (7210T>C), GAT2344GAA (7216T>A), GCG2345GCA (7219G>A), ATC2347ATA (7225C>A), AAG2348AAA (7228G>A), AAG2352AAA (7240G>A), TAT2353TAC (7243T>C), AAA2355AAG (7249A>G), CCT2356CCG (7252T>G), GAT2358GAA (7258T>A), GTT2359GTC (7261T>C), GGT2360GGA (7264T>A), GTC2361GTG (7267C>G), GTG2362GTA (7270G>A), GAT2363_CGG2364insT-- (7273_7274insT), CGG2364GTG (7274C>G 7275G>T), CAT2366CTT (7281A>T), TTG2367TCG (7284T>C), TCA2371TTA (7296C>T), TCT2376TTT (7311C>T), ACC2377ATC (7314C>T), TGC2378TAC (7317G>A), AAT2379AAA (7321T>A), GTG2380-TG (7322delG), CAC2381CAT (7327C>T), GGC2382GGT (7330C>T), TTC2383TTT (7333C>T), AAG2384CAG (7334A>C), TGT2386TGC (7342T>C), GCA2387AAC (7343G>A 7344C>A 7345A>C), ACT2390ACC (7354T>C), CAA2393CAG (7363A>G), ATT2395ATC (7369T>C), TTC2396TTT (7372C>T), GCG2398GCT (7378G>T), CTC2399CTT (7381C>T), GCA2404GCT (7396A>T), GTC2405GTT (7399C>T), GCC2407GCT (7405C>T), AGT2408ATG (7407G>T 7408T>G), GGG2410GGT (7414G>T), TGC2411GGC (7415T>G), AAA2412AAG (7420A>G), GAC2415GAT (7429C>T), TAT2416TAC (7432T>C), CAT2419CAC (7441T>C), AAG2425AAA (7459G>A), GAA2426GAG (7462A>G), ATA2428ATT (7468A>T), GTC2429GTA (7471C>A), CAA2431CAG (7477A>G), AGC2432AGT (7480C>T), CTG2434CTT (7486G>T), TAT2437TAC (7495T>C), AAA2438AAG (7498A>G), GGT2439GGC (7501T>C), TTG2440TCA (7503T>C 7504G>A), ATT2443ATA (7513T>A), AAC2445AAT (7519C>T), GGA2446GGG (7522A>G), AAG2449AAA (7531G>A), GAG2451GAA (7537G>A), CTC2452CTT (7540C>T), TGT2454TGC (7546T>C), AAG2462AAA (7570G>A), ACG2463ACA (7573G>A), ACG2465ACA (7579G>A), GCT2468GCC (7588T>C), CTA2471TTA (7595C>T), TTG2474CTA (7604T>C 7606G>A), GGT2477GGA (7615T>A), TGT2480TGC (7624T>C), GAC2483GAT (7633C>T), TTC2484TTT (7636C>T), AAT2485AAC (7639T>C), TTT2488TTC (7648T>C), GAA2494GAG (7666A>G), TGT2496TGC (7672T>C), ACA2498ACT (7678A>T), GGG2500GGA (7684G>A), ACT2502ACC (7690T>C), TTT2504TTC (7696T>C), GGT2506GGG (7702T>G), GAT2509AAT (7709G>A), AAA2510AGG (7713A>G 7714A>G), CTG2511TTG (7715C>T), CGT2514CGC (7726T>C), GAG2517GAA (7735G>A), GTA2520GTG (7744A>G), TAC2521TAT (7747C>T), GGC2526GGT (7762C>T), CAG2528CAA (7768G>A), TTT2529TTC (7771T>C), TCA2532TCG (7780A>G), CTA2533CTC (7783A>C), ACT2534ACC (7786T>C), CTA2537TTG (7793C>T 7795A>G), ATC2538ATT (7798C>T), GCT2540GCC (7804T>C), ACC2543ATC (7812C>T), GAA2550GAG (7834A>G), GAC2551GAT (7837C>T), GTG2554TTA (7844G>T 7846G>A), CAG2557CAA (7855G>A), CTG2559CTA (7861G>A), TTA2562TTG (7870A>G), TAC2563TAT (7873C>T), ACT2564ACA (7876T>A), GAG2565GAA (7879G>A), ATT2566ATA (7882T>A), GTT2567ATT (7883G>A), CCA2570CCT (7894A>T), ATT2571ATC (7897T>C), GTC2579GTT (7921C>T), AAG2581AAA (7927G>A), GGT2584GGC (7936T>C), AAT2585AAC (7939T>C), AAT2586AAC (7942T>C), GTG2594GTA (7966G>A), GAT2595GAC (7969T>C), TCT2597TCC (7975T>C), GTC2601GTT (7987C>T), CTT2602CTC (7990T>C), GCT2607GCC (8005T>C), GAA2616GAG (8032A>G), ATC2618ATT (8038C>T), GAC2619GAT (8041C>T), AGC2620AAC (8043G>A), TTG2632TTA (8080G>A), GTG2635GTA (8089G>A), CCG2637CCA (8095G>A), GAG2638GAA (8098G>A), AGC2641AGT (8107C>T), ATT2642ATC (8110T>C), GAT2644GAC (8116T>C), CAA2648CAG (8128A>G), GGT2654GGC (8146T>C), AAC2656AGT (8151A>G 8152C>T), TCG2660TCA (8164G>A), AGG2665AAG (8178G>A), AAG2666AAA (8182G>A), TTG2669TTA (8191G>A), TTC2671TTT (8197C>T), TCC2673TCT (8203C>T), AGA2675AAA (8208G>A), CTG2677TTA (8213C>T 8215G>A), CTA2678CTG (8218A>G), ATC2679ATT (8221C>T), GAG2680GAA (8224G>A), CCA2685CCG (8239A>G), GAG2690GAA (8254G>A), ATT2692ATC (8260T>C), GTA2693GTG (8263A>G), GAT2699GAC (8281T>C), AGA2700AGG (8284A>G), CCA2704GCG (8294C>G 8296A>G), GAG2705GAA (8299G>A), CAC2706CAT (8302C>T), ATT2711ATC (8317T>C), TGC2712TGT (8320C>T), GCT2714GCA (8326T>A), GAG2717GAA (8335G>A), TCC2718TCT (8338C>T), TCT2722TTT (8349C>T), GAA2723GAG (8353A>G), CAC2726CAT (8362C>T), CAA2727CAG (8365A>G), ATC2728ATT (8368C>T), TAC2732TAT (8380C>T), TCA2733TCG (8383A>G), TTA2735TTG (8389A>G), CCT2740CCC (8404T>C), GCA2742TCA (8408G>T), ACA2743ACG (8413A>G), GCG2745GCT (8419G>T), GCT2750GCC (8434T>C), TAT2752TAC (8440T>C), ATG2756ATA (8452G>A), CTA2758CTG (8458A>G), AGG2759AAG (8460G>A), AAA2760AAG (8464A>G), CTG2761CTA (8467G>A), AGG2765AGA (8479G>A), GCT2766ACA (8480G>A 8482T>A), GAG2771GAA (8497G>A), CTA2772TTG (8498C>T 8500A>G), AGA2773AAG (8502G>A 8503A>G), GCC2774GCT (8506C>T), GAA2777GAT (8515A>T), GTC2780GTT (8524C>T), GCA2781GCC (8527A>C), TTA2782CTG (8528T>C 8530A>G), GAG2785GAC (8539G>C), CTT2788TGT (8546C>T 8547T>G), TCT2790ACT (8552T>A), GTA2793GTG (8563A>G), GCA2797GGA (8574C>G), ATT2801GTT (8585A>G), GGA2805GAA (8598G>A), AGC2806AGT (8602C>T), AAC2807TCT (8603A>T 8604A>C 8605C>T), AAA2812AAG (8620A>G), CCA2813CAA (8622C>A), GAG2814GGA (8625A>G 8626G>A), CAG2815CAA (8629G>A), GGC2816GGT (8632C>T), ATC2818ATT (8638C>T), CAG2819CAA (8641G>A), AAC2821AAT (8647C>T), CCG2822TTC (8648C>T 8649C>T 8650G>C), AAC2823AAT (8653C>T), AAA2824AAG (8656A>G), GGA2825GAA (8658G>A), GAT2827GCG (8664A>C 8665T>G), GTT2830GTG (8674T>G), GCA2832GTT (8679C>T 8680A>T), GGC2833GGA (8683C>A), GGG2836GGA (8692G>A), ACA2837ACT (8695A>T), CCG2841CCA (8707G>A), AGA2842CGA (8708A>C), AAG2844AAA (8716G>A), ATC2846ATT (8722C>T), ACG2847ACA (8725G>A), AAA2849AAG (8731A>G), AGA2851AGG (8737A>G), CCC2853CCT (8743C>T), ACA2854AAG (8745C>A 8746A>G), AAG2856AAA (8752G>A), GGA2857GGT (8755A>T), AAC2862AAT (8770C>T), TTA2863CTA (8771T>C), GAA2864GAG (8776A>G), CAT2865CAC (8779T>C), TTG2866TTA (8782G>A), TAT2869TAC (8791T>C), CCA2871CCG (8797A>G), CAA2872CAG (8800A>G), CAA2873CAG (8803A>G), GAT2875GAC (8809T>C), ATT2876ATC (8812T>C), AAT2878AAC (8818T>C), CGG2880CGA (8824G>A), GCA2881GCG (8827A>G), ACG2888ACA (8848G>A), TAT2890TAC (8854T>C), GAG2891GGA (8856A>G 8857G>A), GTG2893GTA (8863G>A), CGG2894CAG (8865G>A), ATG2895CTT (8867A>C 8869G>T), GCA2896GCG (8872A>G), GGA2900GAA (8883G>A), ACT2902TCT (8888A>T), TGC2915TGT (8929C>T), GTC2924ATC (8954G>A), AAC2925AAT (8959C>T), GGA2926GGT (8962A>T), GGG2933GGA (8983G>A), AAT2934GAC (8984A>G 8986T>C), GAA2935GTA (8988A>T), GAG2938GAA (8998G>A), CCG2940CCA (9004G>A), TTG2941TTA (9007G>A), ATC2944ATT (9016C>T), ACC2951ACA (9037C>A), ATA2967ATT (9085A>T), AAC2971AAT (9097C>T), ATT2983GTT (9131A>G), CGA2984CGT (9136A>T), CGG2987CGT (9145G>T), ATG2989GGA (9149A>G 9150T>G 9151G>A), GGT2990AGT (9152G>A), TTA2991TTG (9157A>G), GCG2992GCT (9160G>T), CGT2993CGC (9163T>C), GAG3000GAA (9184G>A), CGA3004CGG (9196A>G), AGG3010AGA (9214G>A), GAA3011GAG (9217A>G), GCG3012GCA (9220G>A), GCA3020GCT (9244A>T), TTG3021TTA (9247G>A), GCC3024GCT (9256C>T), CCT3026TCC (9260C>T 9262T>C), GGG3030GGA (9274G>A), GAC3032GAT (9280C>T), ATC3035ATT (9289C>T), CAA3038CAC (9298A>C), GAG3040GAA (9304G>A), GTC3050GTT (9334C>T), GTC3060GTG (9364C>G) | | | | | | | | |
| PVYgp2 | 1 | 75 | 100% | 230 | 43.7% | 75 (98.7%) | 55 (72.4%) | 1/0/2/1 | 2 |
| Protein mutations: | R17_K17insX (2971_2972insA), Y23H (2989T>C 2991C>T), R26K (2999G>A 3000A>G), H35Y (3025C>T), K40H (3040A>C 3042A>C), G41S (3043G>A), E44K (3052G>A), I47V (3061A>G), V54I (3082G>A), P57S (3091C>T), P60T (3100C>A), G61E (3104G>A 3105T>G), G62S (3106G>A), Q63* (3109C>T), R64G (3112A>G), C66R (3118T>C), L67F (3121C>T), I69V (3127A>G), A72E (3137C>A 3138G>A), I73V (3139A>G) | | | | | | | | |
| Codon mutations: | TTA2CTA (2923T>C), AAA4AAG (2931A>G), GAA7GAG (2940A>G), TTG10CTG (2947T>C), TTG15CTG (2962T>C), AGA17_AAA17insAA- (2971_2972insA), TAC23CAT (2989T>C 2991C>T), TCA24TCG (2994A>G), AGA26AAG (2999G>A 3000A>G), ATC31ATT (3015C>T), TCG33TCA (3021G>A), GTA34GTT (3024A>T), CAT35TAT (3025C>T), CAC38CAT (3036C>T), AGA39AGG (3039A>G), AAA40CAC (3040A>C 3042A>C), GGC41AGC (3043G>A), GAA44AAA (3052G>A), ATA47GTA (3061A>G), GTT54ATT (3082G>A), CCG57TCG (3091C>T), CGC59CGT (3099C>T), CCA60ACA (3100C>A), GGT61GAG (3104G>A 3105T>G), GGT62AGT (3106G>A), CAA63TAA (3109C>T), AGG64GGG (3112A>G), TGC66CGC (3118T>C), CTC67TTC (3121C>T), ATT69GTT (3127A>G), GAG70GAA (3132G>A), GCG72GAA (3137C>A 3138G>A), ATT73GTT (3139A>G) | | | | | | | | |

| | Begin | End | Coverage | Score | Concordance | Matches | Identities | I/D/M/F* | Stop Codons |
|---|---|---|---|---|---|---|---|---|---|
| **NT** | **52** | **9704** | **99.5%** | **12530** | **65.2%** | **9642 (99.8%)** | **7985 (82.7%)** | **7/11** | |

**Proteins**

| | Begin | End | Coverage | Score | Concordance | Matches | Identities | I/D/M/F* | Stop Codons |
|---|---|---|---|---|---|---|---|---|---|
| polyprotein (NP_056759.1) | 1 | 3064 | 100% | 19042 | 90.2% | 3062 (99.9%) | 2771 (90.4%) | 2/2/3/3 | 1 |
| Protein mutations: | Y4Q (194T>C 196C>A), C9L (209T>C 210G>T 211T>G), F13M (221T>A 223T>G), S23P (251T>C), C24F (255G>T 256C>T), E25G (258A>G), I27V (263A>G 265T>G), V28A (267T>C), E30V (273A>T), A35T (287G>A 289T>C), V37T (293G>A 294T>C), A41I (305G>A 306C>T), D42G (309A>G), E44D (316A>T), T45V (317A>G 318C>T 319A>G), L52Q (339T>A 340C>A), K53R (342A>G), K55A (347A>G 348A>C 349A>G), Y56H (350T>C), T58V (356A>G 357C>T), V61T (365G>A 366T>C 367G>A), L62S (368C>T 369T>C), F67C (384T>G), A75V (408C>T 409C>T), M78E (416A>G 417T>A), E85K (437G>A), R86E (440A>G 441G>A), K87R (443A>C 444A>G 445G>A), D88E (448T>A), E91A (456A>C), H93N (461C>A), D111E (517T>A), S116F (531C>T), P118S (536C>T), Q119W (539C>T 540A>G 541A>G), R122K (549G>A 550A>G), M131T (576T>C), V134A (585T>C 586C>G), R137Y (593C>T 594G>A 595C>T), P138R (597C>G), I139V (599A>G 601A>G), I140P (602A>C 603T>C 604A>G), M147I (625G>A), I151V (635A>G 637T>C), I154V (644A>G 646A>G), E160A (663A>C 664G>C), H166Q (682C>A), T172S (699C>G), H174Q (706T>A), K179E (719A>G 721G>A), I180V (722A>G 724A>G), A183S (731G>T), Y184P (734T>C 735A>C 736C>T), S185R (737T>C 738C>G 739C>T), A187V (744C>T 745G>C), R189C (749C>T 751A>C), M194K (765T>A), R197K (773C>A 774G>A 775A>G), M206K (801T>A), G210A (813G>C 814A>G), L211H (816T>A), R214Q (825G>A 826T>A), N223D (851A>G 853T>C), R226C (860C>T 862C>T), T227A (863A>G 865T>C), I228T (867T>C), N229D (869A>G 871C>T), I230L (872A>T 874A>G), R231Q (876G>A), R232K (879G>A), N240S (903A>G), T241N (906C>A 907A>C), K242T (909A>C 910A>C), S243N (912G>A 913C>T), S252L (938T>C 939C>T 940A>T), L266I (980T>A 982G>C), R272K (999G>A), Q275R (1007C>A 1008A>G 1009G>A), S276G (1010A>G 1012T>G), I277V (1013A>G), N279Q (1019A>C 1021C>G), D288E (1048C>A), R300Q (1083G>A), S305A (1097T>G 1099G>A), R320K (1143G>A), L324I (1154T>A 1156G>A), S346N (1221G>A), V349T (1229G>A 1230T>C 1231T>C), S350N (1233G>A 1234C>T), K358T (1257A>C 1258A>G), N365S (1278A>G), A369V (1290C>T 1291G>A), D372T (1298G>A 1299A>C 1300C>T), I375V (1307A>G 1309A>G), N378E (1316A>G 1318T>A), I382T (1329T>C), L397I (1373C>A), I401V (1385A>G), E407D (1405G>T), A411S (1415G>T 1417A>T), V418I (1436G>A 1438C>T), E508G (1707A>G), A512T (1718G>A), I520V (1742A>G 1744C>T), S525N (1758G>A), V536I (1790G>A), I585V (1937A>G 1939T>C), F607Y (2004T>A), I628M (2068T>G), V630I (2072G>A 2074G>A), R675K (2208G>A), D694E (2266C>G), N744G (2414A>G 2415A>G 2416T>G), S746C (2420A>T), K780R (2523A>G), R817K (2634G>A), V847I (2723G>A 2725C>T), A855T (2747G>A 2749T>A), H871N (2795C>A), V880I (2822G>A 2824G>A), P900T (2882C>A), S901H (2885A>C 2886G>A), T904M (2895C>T), N916S (2931A>G), N919S (2940A>G), E929_N930insX (2971_2972insA), T936I (2991C>T), H937R (2994A>G), E939R (2999G>A 3000A>G), S944F (3015C>T), R946Q (3021G>A), Y947F (3024A>T 3025C>T), T951I (3036C>T), E952G (3039A>G 3040A>C), K953T (3042A>C 3043G>A), A972V (3099C>T 3100C>A), V974R (3104G>A 3105T>G 3106G>A), S983N (3132G>A), R985K (3137C>A 3138G>A 3139A>G), F990I (3152T>A 3154C>T), C994Y (3165G>A), F1012L (3218T>C), T1014N (3225C>A), V1016I (3230G>A), V1028M (3266G>A 3268A>G), V1031M (3275G>A), A1034T (3284G>A), R1044K (3315G>A 3316A>G), E1045A (3318A>C), M1059V (3359A>G), D1072E (3400T>G), V1110G (3513T>G), M1121V (3545A>G), A1122T (3548G>A), V1124I (3554G>A 3556C>T), L1147V (3623C>G), L1150M (3632C>A), Y1152H (3638T>C), I1172V (3698A>G), S1178N (3717G>A), R1183K (3731C>A 3732G>A), I1297V (4073A>G), F1315Y (4128T>A 4129C>T), A1323S (4151G>T), V1328I (4166G>A), L1347I (4223C>A 4225G>A), I1379V (4319A>G 4321A>G), F1404Y (4395T>A), V1408I (4406G>A), M1431V (4475A>G 4477G>T), T1433S (4481A>T 4483A>G), A1454L (4544G>T 4545C>T 4546G>A), R1455K (4548G>A), V1499I (4679G>A 4681G>C), V1538I (4796G>A 4798C>T), T1540S (4803C>G 4804T>C), S1625G (5057A>G), T1626A (5060A>G), L1636I (5090C>A), A1639V (5100C>T), E1641D (5107A>T), V1646F (5120G>T 5122C>T), I1648V (5126A>G), V1667I (5183G>A), A1793T (5561G>A 5563G>A), I1805V (5597A>G), N1807K (5605C>A), A1812V (5619C>T 5620T>G), I1816L (5630A>C), V1837I (5693G>A), K1890R (5853A>G), I1904V (5894A>G 5896C>G), R1926A (5960C>G 5961G>C 5962G>C), I1936V (5990A>G), E1945D (6019A>T), K1949T (6030A>C), D1955E (6049C>A), M1958I (6058G>A), S1963N (6072G>A 6073T>C), N1964H (6074A>C), T1966N (6081C>A), C1976S (6111G>C 6112T>C), I1982V (6128A>G 6130T>C), V1992I (6158G>A), T1996S (6170A>T 6172A>G), L2007F (6203C>T 6205C>T), V2016I (6230G>A), A2025K (6257G>A 6258C>A), K2033R (6282A>G), A2060T (6362G>A), V2073I (6401G>A), A2074V (6405C>T), Y2082F (6429A>T), Q2089R (6450A>G), H2101N (6485C>A 6487C>T), L2115I (6527C>A), T2135I (6588C>T), Y2151H (6635T>C), I2155V (6647A>G), I2165V (6677A>G 6679A>G), D2178N (6716G>A), N2179D (6719A>G), I2194L (6764A>T 6766A>G), A2204V (6795C>T 6796A>G), H2205Q (6799C>A), N2225D (6857A>G), V2231T (6875G>A 6876T>C 6877C>A), D2250E (6934C>A), V2273I (7001G>A 7003G>A), F2283Y (7032T>A), T2314K (7125C>A), A2321S (7145G>T 7147A>G), E2326_A2327del (7160_7165delGAGGCA), D2344E (7216T>A), D2358E (7258T>A), D2363_R2364insX (7273_7274insT), R2364V (7274C>G 7275G>T), H2366L (7281A>T), L2367S (7284T>C), S2371L (7296C>T), S2376F (7311C>T), T2377I (7314C>T), C2378Y (7317G>A), N2379K (7321T>A), K2384Q (7334A>C), A2387N (7343G>A 7344C>A 7345A>C), S2408M (7407G>T 7408T>G), C2411G (7415T>G), L2440S (7503T>C 7504G>A), D2509N (7709G>A), K2510R (7713A>G 7714A>G), T2543I (7812C>T), V2554L (7844G>T 7846G>A), V2567I (7883G>A), S2620N (8043G>A), N2656S (8151A>G 8152C>T), R2665K (8178G>A), R2675K (8208G>A), P2704A (8294C>G 8296A>G), S2722F (8349C>T), A2742S (8408G>T), M2756I (8452G>A), R2759K (8460G>A), A2766T (8480G>A 8482T>A), R2773K (8502G>A 8503A>G), E2777D (8515A>T), E2785D (8539G>C), L2788C (8546C>T 8547T>G), S2790T (8552T>A), A2797G (8574C>G), I2801V (8585A>G), G2805E (8598G>A), N2807S (8603A>T 8604A>C 8605C>T), P2813Q (8622C>A), E2814G (8625A>G 8626G>A), P2822F (8648C>T 8649C>T 8650G>C), G2825E (8658G>A), D2827A (8664A>C 8665T>G), A2832V (8679C>T 8680A>T), T2854K (8745C>A 8746A>G), E2891G (8856A>G 8857G>A), R2894Q (8865G>A), M2895L (8867A>C 8869G>T), G2900E (8883G>A), T2902S (8888A>T), V2924I (8954G>A), N2934D (8984A>G 8986T>C), E2935V (8988A>T), I2983V (9131A>G), M2989G (9149A>G 9150T>G 9151G>A), G2990S (9152G>A), P3026S (9260C>T 9262T>C), Q3038H (9298A>C) | | | | | | | | |

| | Begin | End | Coverage | Score | Concordance | Matches | Identities | I/D/M/F* | Stop Codons |
|---|---|---|---|---|---|---|---|---|---|
| **NT** | **52** | **9704** | **99.5%** | **12530** | **65.2%** | **9642 (99.8%)** | **7985 (82.7%)** | **7/11** | |

Codon mutations:

ACT3ACC (193T>C), TAC4CAA (194T>C 196C>A), TCA6TCG (202A>G), ATC8ATT (208C>T), TGT9CTG (209T>C 210G>T 211T>G), TCG12TCA (220G>A), TTT13ATG (221T>A 223T>G), GAA14GAG (226A>G), CTA17CTT (235A>T), TAC19TAT (241C>T), CCA21CCT (247A>T), TCT23CCT (251T>C), TGC24TTT (255G>T 256C>T), GAG25GGG (258A>G), CAT26CAC (262T>C), ATT27GTG (263A>G 265T>G), GTG28GCG (267T>C), GAA30GTA (273A>T), CGA31CGT (277A>T), GAA32GAG (280A>G), GTG33GTT (283G>T), CCG34CCA (286G>A), GCT35ACC (287G>A 289T>C), GTT37ACT (293G>A 294T>C), CCT39CCC (301T>C), TTC40TTT (304C>T), GCA41ATA (305G>A 306C>T), GAT42GGT (309A>G), CTG43TTG (311C>T), GAA44GAT (316A>T), ACA45GTG (317A>G 318C>T 319A>G), CAA46CAG (322A>G), GCA49GCT (331A>T), CTC52CAA (339T>A 340C>A), AAG53AGG (342A>G), AAA55GCG (347A>G 348A>C 349A>G), TAT56CAT (350T>C), GCT57GCA (355T>A), ACT58GTT (356A>G 357C>T), CGT60CGC (364T>C), GTG61ACA (365G>A 366T>C 367G>A), CTC62TCC (368C>T 369T>C), AAA63AAG (373A>G), AAC64AAT (376C>T), TTT67TGT (384T>G), ACG68ACA (388G>A), TAC69TAT (391C>T), CGA70CGG (394A>G), AAG72AAA (400G>A), GAT74GAC (406T>C), GCC75GTT (408C>T 409C>T), CAG76CAA (412G>A), ATA77ATT (415A>T), ATG78GAG (416A>G 417T>A), ATT80ATC (424T>C), CAG81CAA (427G>A), AAA83AAG (433A>G), CTG84CTC (436G>C), GAG85AAG (437G>A), AGG86GAG (440A>G 441G>A), AAG87CGA (443A>C 444A>G 445G>A), GAT88GAA (448T>A), GAA90GAG (454A>G), GAA91GCA (456A>C), TAT92TAC (460T>C), CAC93AAC (461C>A), CAA95CAG (469A>G), GCC97GCG (475C>G), CCT99CCA (481T>A), ACT106ACC (502T>C), ATA107ATT (505A>T), GGC109GGT (511C>T), GAT111GAA (517T>A), CCT112CCA (520T>A), CCA113CCT (523A>T), TCT116TTT (531C>T), GAG117GAA (535G>A), CCA118TCA (536C>T), CAA119TGG (539C>T 540A>G 541A>G), AGA122AAG (549G>A 550A>G), ATC124ATT (556C>T), ATT125ATC (559T>C), CAT126CAC (562T>C), ACT128ACC (568T>C), CCA129CCG (571A>G), ATG131ACG (576T>C), CGT132CGC (580T>C), GTC134GCG (585T>C 586C>G), CGC137TAT (593C>T 594G>A 595C>T), CCC138CGC (597C>G), ATA139GTG (599A>G 601A>G), ATA140CCG (602A>C 603T>C 604A>G), ACA143ACG (613A>G), GAA144GAG (616A>G), GGC145GGG (619C>G), CAG146CAA (622G>A), ATG147ATA (625G>A), AAT148AAC (628T>C), CAC149CAT (631C>T), ATT151GTC (635A>G 637T>C), ATA154GTG (644A>G 646A>G), ATT157ATC (655T>C), TCG159TCA (661G>A), GAG160GCC (663A>C 664G>C), AAA161AAG (667A>G), GTC165GTT (679C>T), CAC166CAA (682C>A), TTA167CTG (683T>C 685A>G), AGT169AGC (691T>C), ACC172AGC (699C>G), ACT173ACA (703T>A), CAT174CAA (706T>A), AAG179GAA (719A>G 721G>A), ATA180GTG (722A>G 724A>G), CTT181TTG (725C>T 727T>G), GGT182GGA (730T>A), GCA183TCA (731G>T), TAC184CCT (734T>C 735A>C 736C>T), TCC185CGT (737T>C 738C>G 739C>T), GCG187GTC (744C>T 745G>C), CGA189TGC (749C>T 751A>C), ACT190ACC (754T>C), ATG194AAG (765T>A), TTG196CTG (770T>C), CGA197AAG (773C>A 774G>A 775A>G), CGG198AGG (776C>A), TTC202TTT (790C>T), ATG206AAG (801T>A), GGA210GCG (813G>C 814A>G), CTT211CAT (816T>A), TTG212CTG (818T>C), CAA213CAG (823A>G), CGT214CAA (825G>A 826T>A), CGG217AGA (833C>A 835G>A), GAC219GAT (841C>T), AAT223GAC (851A>G 853T>C), GTC225GTT (859C>T), CGC226TGT (860C>T 862C>T), ACT227GCC (863A>G 865T>C), ATC228ACC (867T>C), AAC229GAT (869A>G 871C>T), ATA230TTG (872A>T 874A>G), CGA231CAA (876G>A), AGG232AAG (879G>A), GAT234GAC (886T>C), ATC238ATT (898C>T), AAC240AGC (903A>G), ACA241AAC (906C>A 907A>C), AAA242ACC (909A>C 910A>C), AGC243AAT (912G>A 913C>T), GGC246GGA (922C>A), GGT249GGA (931T>A), TCA252CTT (938T>C 939C>T 940A>T), GGA253GGG (943A>G), TTC256TTT (952C>T), TCA261TCG (967A>G), GAA263GAG (973A>G), GGG264GGC (976G>C), TTG266ATC (980T>A 982G>C), GCA269GCG (991A>G), CGT270CGC (994T>C), AGA272AAA (999G>A), GTT273GTC (1003T>C), CAG275AGA (1007C>A 1008A>G 1009G>A), AGT276GGG (1010A>G 1012T>G), ATT277GTT (1013A>G), TTA278TTG (1018A>G), AAC279CAG (1019A>C 1021C>G), ATC282ATT (1030C>T), TCG285TCA (1039G>A), AAT286AAC (1042T>C), GCC287GCA (1045C>A), GAC288GAA (1048C>A), AAT289AAC (1051T>C), AAG292AAA (1060G>A), GGT293GGA (1063T>A), CTG294TTG (1064C>T), GAC295GAT (1069C>T), CGA300CAA (1083G>A), TAT303TAC (1093T>C), TCG305GCA (1097T>G 1099G>A), CAC307CAT (1105C>T), GTA310GTG (1114A>G), GCT311GCG (1117T>G), TTA313TTG (1123A>G), CCT314CCA (1126T>A), GTC315GTT (1129C>T), AGG320AAG (1143G>A), GTA321GTT (1147A>T), GCT322GCA (1150T>A), TTG324ATA (1154T>A 1156G>A), CAC327CAT (1165C>T), AGT328AGC (1168T>C), CTT330TTA (1172C>T 1174T>A), ACC339ACT (1201C>T), AGC346AAC (1221G>A), GTT349ACC (1229G>A 1230T>C 1231T>C), AGC350AAT (1233G>A 1234C>T), CTG352CTA (1240G>A), AAG354AAA (1246G>A), TTG356CTA (1250T>C 1252G>A), CAT357CAC (1255T>C), AAA358ACG (1257A>C 1258A>G), AGA361AGG (1267A>G), GGT363GGC (1273T>C), TTG364TTA (1276G>A), AAT365AGT (1278A>G), TTG367CTG (1283T>C), GCG369GTA (1290C>T 1291G>A), GAC372ACT (1298G>A 1299A>C 1300C>T), TTT374TTC (1306T>C), ATA375GTG (1307A>G 1309A>G), CAT376CAC (1312T>C), AAT378GAA (1316A>G 1318T>A), TTG381TTA (1327G>A), ATA382ACA (1329T>C), GCG383GCT (1333G>T), TTA384TTG (1336A>G), CTA387TTG (1343C>T 1345A>G), CCG390CCA (1354G>A), GTG391GTT (1357G>T), GAC392GAT (1360C>T), CTC395TTG (1367C>T 1369C>G), CTT397ATT (1373C>A), TTC398TTT (1378C>T), GAG400GAA (1384G>A), ATA401GTA (1385A>G), AAA403AAG (1393A>G), TCC404TCT (1396C>T), GAG407GAT (1405G>T), AAA408AAG (1408A>G), CAG409CAA (1411G>A), GCA411TCT (1415G>T 1417A>T), CCG412CCC (1420G>C), AAT415AAC (1429T>C), TTA416TTG (1432A>G), GTC418ATT (1436G>A 1438C>T), TTA419CTG (1439T>C 1441A>G), AAT420AAC (1444T>C), TTC423TTT (1453C>T), CTG424CTA (1456G>A), GAA428GAG (1468A>G), GTA436GTG (1492A>G), TTG439CTA (1499T>C 1501G>A), AGT440AGC (1504T>C), TTG441TTA (1507G>A), CTC442CTT (1510C>T), GAA443GAG (1513A>G), TTA444TTG (1516A>G), GCA445GCT (1519A>T), AGG446AGA (1522G>A), TTC447TTT (1525C>T), CAG448CAA (1528G>A), AAC450AAT (1534C>T), ACT452ACA (1540T>A), AAC454AAT (1546C>T), AAA457AAG (1555A>G), GGT458GGC (1558T>C), GAT459GAC (1561T>C), ATA460ATC (1564A>C), TCT461TCA (1567T>A), TTC463TTT (1573C>T), AGA464AGG (1576A>G), TTA467CTA (1583T>C), TCT468TCA (1588T>A), AAG470AAA (1594G>A), AAC472AAT (1600C>T), CTG475CTA (1609G>A), TCG478TCA (1618G>A), TGC479TGT (1621C>T), GAC480GAT (1624C>T), CTG483TTG (1631C>T), AAA485AAG (1639A>G), TTC489TTT (1651C>T), CTC490CTG (1654C>G), CAT497CAC (1675T>C), TCA503TCG (1693A>G), TTT506TTC (1702T>C), GAA508GGA (1707A>G), GCA512ACA (1718G>A), GGA514GGG (1726A>G), TAC515TAT (1729C>T), TCA516TCC (1732A>C), TAT518TAC (1738T>C), ATC520GTT (1742A>G 1744C>T), CGC521CGT (1747C>T), CAT523CAC (1753T>C), AGT525AAT (1758G>A), AGG528AGA (1768G>A), AAC534AAT (1786C>T), TTA535TTG (1789A>G), GTT536ATT (1790G>A), GTC537GTT (1795C>T), TTA541CTG (1805T>C 1807A>G), GAG543GAA (1813G>A), AGG545AGA (1819G>A), AAG547AAA (1825G>A), GAC551GAT (1837C>T), TAT552TAC (1840T>C), CAA555CAG (1849A>G), GTC558GTG (1858C>G), AAA560AAG (1864A>G), TGC562TGT (1870C>T), AGT564AGC (1876T>C), AAA566AAG (1882A>G), GGT568GGA (1888T>A), AAT569AAC (1891T>C), TGT574TGC (1906T>C), ACA577ACT (1915A>T), GGT582GGC (1930T>C), GCC584GCT (1936C>T), ATT585GTC (1937A>G 1939T>C), TCA587TCG (1945A>G), TTC589TTT (1951C>T), CCA592CCT (1960A>T), GTA598GTG (1978A>G), ATT599ATA (1981T>A), GGC600GGT (1984C>T), AAT601AAC (1987T>C), GAC604GAT (1996C>T), TTT607TAT (2004T>A), AAA612AAG (2020A>G), GGG613GGA (2023G>A), GAT614GAC (2026T>C), TCG615TCT (2029G>T), GAG616GAA (2032G>A), TAT625TAC (2059T>C), ATT628ATG (2068T>G), AAC629AAT (2071C>T), GTG630ATA (2072G>A 2074G>A), CTG635TTA (2087C>T 2089G>A), AGC639AGT (2101C>T), GAG640GAA (2104G>A), GAG641GAA (2107G>A), AAG644AAA (2116G>A), TTC646TTT (2122C>T), CGC651CGT (2137C>T), CTT658CTC (2158T>C), ACC660ACT (2164C>T), CCA662CCG (2170A>G), GAT666GAC (2182T>C), GCG668GCA (2188G>A), ACC669ACA (2191C>A), AGA675AAA (2208G>A), TAT678TAC (2218T>C), GAC680GAT (2224C>T), GTA681GTT (2227A>T), TTG686TTA (2242G>A), CCC687CCT (2245C>T), TTG690TTA (2254G>A), GTT691GTC (2257T>C), GAC692GAT (2260C>T), GAC694GAG (2266C>G), CAA696CAG (2272A>G), ACG697ACA (2275G>A), TCA703TCG (2293A>G), GGC705GGT (2299C>T), TCG706TCA (2302G>A), CAG707CAA (2305G>A), GGA710GGT (2314A>T), TAT711TAC (2317T>C), CTA714CTG (2326A>G), GTG719GTA (2341G>A), CAA721CAG (2347A>G), CTT722CTC (2350T>C), TTG724TTA (2356G>A), AAT727AAC (2365T>C), GAA729GAG (2371A>G), TTA730TTG (2374A>G), GAT733GAC (2383T>C), ATA734ATT (2386A>T), AAA735AAG (2389A>G), CAT736CAC (2392T>C), TAT737TAC (2395T>C), AAT744GGG (2414A>G 2415A>G 2416T>G), GCT745GCA (2419T>A), AGC746TGC (2420A>T), ACA752ACT (2440A>T), GTT761GTC (2467T>C), TCT764TCC (2476T>C), TCG766TCA (2482G>A), CTG769CTA (2491G>A), CTG771CTT (2497G>T), TTG773TTA (2503G>A), AGA778AGG (2518A>G), CCT779CCC (2521T>C), AAG780AGG (2523A>G), AGA783AGG (2533A>G), TTG785TTA (2539G>A), TTA787TTG (2545A>G), GAG789GAA (2551G>A), CCT790CCA (2554T>A), CTG792TTG (2558C>T), TTG793CTC (2561T>C 2563G>C), TCA796TCG (2572A>G), CCT800CCC (2584T>C), GGC801GGT (2587C>T), CTG803CTC (2593G>C), GCT805GCC (2599T>C), AAT808AAC (2608T>C), TTT812TTC (2620T>C), GCG815GCA (2629G>A), GTG816GTA (2632G>A), AGG817AAG (2634G>A), TTG818TTA (2638G>A), ATT820ATA (2644T>A), AAA823AAG (2653A>G), GCT827GCC (2665T>C), CTA832TTA (2678C>T), CTA833CTG (2683A>G), GCT835GCC (2689T>C), CTA838CTC (2698A>C), TCA841TCG (2707A>G), GCG842GCA (2710G>A), GAA844GAG (2716A>G), ACA845ACG (2719A>G), CTC846CTT (2722C>T), GTC847ATT (2723G>A 2725C>T), GCA848GCG (2728A>G), AGG850AGA (2734G>A), ATT851ATA (2737T>A), GAT854GAC (2746T>C), GCT855ACA (2747G>A 2749T>A), GCA856GCT (2752A>T), GCT857GCA (2755T>A), GAC859GAT (2761C>T), CTC860CTT (2764C>T), TGT865TGC (2779T>C), GGG867GGT (2785G>T), AAC869AAT (2791C>T), CAT871AAT (2795C>A), TAC874TAT (2806C>T), TTG878CTC (2816T>C 2818G>C), GTG880ATA (2822G>A 2824G>A), TTG881TTA (2827G>A), GTT884GTC (2836T>C), AAG885AAA (2839G>A), AAT886AAC (2842T>C), ACC893ACA (2863C>A), CTA894TTA (2864C>T), CCA900ACA (2882C>A), AGT901CAT (2885A>C 2886G>A), AAC903AAT (2893C>T), ACG904ATG (2895C>T), GTC906GTT (2902C>T), GTA907GTG (2905A>G), ATT909ATC (2911T>C), AAT913AAC (2923T>C), AAT916AGT (2931A>G), AAC919AGC (2940A>G), GCT921GCC (2947T>C), ACT926ACC (2962T>C), GAA929_AAT930insA-- (2971_2972insA), GGT935GGC (2989T>C), ACT936ATT (2991C>T), CAT937CGT (2994A>G), GAG939AGG (2999G>A 3000A>G), TCC944TTC (3015C>T), CGG946CAG (3021G>A), TAC947TTT (3024A>T 3025C>T), ACA951ATA (3036C>T), GAA952GGC (3039A>G 3040A>C), AAG953ACA (3042A>C 3043G>A), TTG956TTA (3052G>A), TTA959TTG (3061A>G), GCG966GCA (3082G>A), GGC969GGT (3091C>T), GCC972GTA (3099C>T 3100C>A), GTG974AGA (3104G>A 3105T>G 3106G>A), GTC975GTT (3109C>T), AAA976AAG (3112A>G), ACT978ACC (3118T>C), GCC979GCT (3121C>T), GGA981GGG (3127A>G), AGC983AAC (3132G>A), CGA985AAG (3137C>A 3138G>A 3139A>G), TAT989TAC (3151T>C), TTC990ATT (3152T>A 3154C>T), TGT994TAT (3165G>A), GTA995GTG (3169A>G), ATC1005ATT (3199C>T), AGG1007CGG (3203A>C), CGT1008CGC (3208T>C), CCA1010CCG (3214A>G), ACC1011ACT (3217C>T), TTT1012CTT (3218T>C), GTC1013GTA (3223C>A), ACT1014AAT (3225C>A), GTT1016ATT (3230G>A), AAC1017AAT (3235C>T), TTA1025CTA (3257T>C), ACC1026ACT (3262C>T), AGC1027AGT (3265C>T), GTA1028ATG (3266G>A 3268A>G), GTG1029GTA (3271G>A), GTG1031ATG (3275G>A), GCA1034ACA (3284G>A), CAG1039CAA (3301G>A), AGA1044AAG (3315G>A 3316A>G), GAA1045GCA (3318A>C), ATC1046ATT (3322C>T), GAG1047GAA (3325G>A), TTG1048TTA (3328G>A), ATA1051ATT (3337A>T), GAG1055GAA (3349G>A), GTC1057GTT (3355C>T), TGC1058TGT (3358C>T), ATG1059GTG (3359A>G), CTA1061CTG (3367A>G), TAT1062TAC (3370T>C), TTA1065CTG (3377T>C 3379A>G), GAA1070GAG (3394A>G), CGC1071CGT (3397C>T), GAT1072GAG (3400T>G), ACA1074ACG (3406A>G), TAC1078TAT (3418C>T), ATT1079ATA (3421T>A), TAT1081TAC (3427T>C), AAG1083AAA (3433G>A), TCA1084TCT (3436A>T), GTA1085GTG (3439A>G), AAC1086AAT (3442C>T), CCT1087CCC (3445T>C), CAG1091CAA (3457G>A), TTT1092TTC (3460T>C), GCT1093GCA (3463T>A), GCG1095GCT (3469G>T), CAG1096CAA (3472G>A), CAC1104CAT (3496C>T), CAG1105CAA (3499G>A), TCC1107TCT (3505C>T), GTT1110GGT (3513T>G), AAA1112AAG (3520A>G), GAA1115GAG (3529A>G), CAA1116CAG (3532A>G), ATG1121GTG (3545A>G), GCT1122ACT (3548G>A), TTA1123CTA (3551T>C), GTC1124ATT (3554G>A 3556C>T), AGT1133AGC (3583T>C), GAT1134GAC (3586T>C), TGC1135TGT (3589C>T), TTC1137TTT (3595C>T), AAT1141AAC (3607T>C), TTT1143TTC (3613T>C), AAG1144AAA (3616G>A), GGT1145GGC (3619T>C), CTT1147GTT (3623C>G), TCC1148TCT (3628C>T), CTG1150ATG (3632C>A), GAT1151GAC (3637T>C), TAT1152CAT (3638T>C), GAA1153GAG (3643A>G), TTA1159TTG (3661A>G), GTG1162GTA (3670G>A), TTT1166TTC (3682T>C), GAG1168GAA (3688G>A), GAG1171GAA (3697G>A), ATT1172GTT (3698A>G), GAT1174GAC (3706T>C), TTT1175TTC (3709T>C), TTG1177CTA (3713T>C 3715G>A), AGT1178AAT (3717G>A), GAG1179GAA (3721G>A), CGA1183AAA (3731C>A 3732G>A), ACT1184ACA (3736T>A), CTA1188TTG (3746C>T 3748A>G), ACA1190ACG (3754A>G), AGT1193AGC (3763T>C), CGA1198CGG (3778A>G), CAA1199CAG (3781A>G), ATC1200ATT (3784C>T), CAT1204CAC (3796T>C), CCA1207CCT (3805A>T), TAC1209TAT (3811C>T), AGA1210AGG (3814A>G), TTC1215TTT (3829C>T), TTT1218TTC (3838T>C), AGA1220AGG (3844A>G), GCA1221GCC (3847A>C), GTT1224GTA (3856T>A), GTG1226GTT (3862G>T), GAC1229GAT (3871C>T), GCC1231GCA (3877C>A), AGC1233AGT (3883C>T), GAA1234GAG (3886A>G), CTA1236CTG (3892A>G), GAC1237GAT (3895C>T), TTT1238TTC (3898T>C), GTA1240GTG (3904A>G), CGG1241AGA (3905C>A 3907G>A), GCT1243GCC (3913T>C), AAG1248AAA (3928G>A), TCA1249TCT (3931A>T), GGG1251GGA (3937G>A), AGT1257AGC (3955T>C), GTG1258GTA (3958G>A), GCC1259GCT (3961C>T), GGA1260GGG (3964A>G), TCT1261TCC (3967T>C), TTA1264TTG (3976A>G), ATT1265ATC (3979T>C), CCA1267CCT (3985A>T), CTA1271CTT (3997A>T), GCG1272GCA (4000G>A), AAC1274AAT (4006C>T), GTT1275GTG (4009T>G), TTC1276TTT (4012C>T), AAA1277AAG (4015A>G), TCT1280TCC (4024T>C), GAA1282GAG (4030A>G), CCA1283CCT (4033A>T), AAG1286AAA (4042G>A), CTG1290TTA (4052C>T 4054G>A), CGT1291CGC (4057T>C), CGT1293CGA (4063T>A), GGA1294GGG (4066A>G), ATA1297GTA (4073A>G), TTT1298TTC (4078T>C), GGC1299GGT (4081C>T), TCT1300TCC (4084T>C), TCC1304TCT (4096C>T), AGC1308AGT (4108C>T), GGA1309GGT (4111A>T), GCG1311GCA (4117G>A), CTA1312TTG (4118C>T 4120A>G), TAC1314TAT (4126C>T), TTC1315TAT (4128T>A 4129C>T), GCC1316GCT (4132C>T), AAT1317AAC (4135T>C), CAA1321CAG (4147A>G), TTA1322CTA (4148T>C), GCT1323TCT (4151G>T), AAC1326AAT (4162C>T), GTA1328ATA (4166G>A), ATA1329ATC (4171A>C), GAG1332GAA (4180G>A), CTG1336TTG (4190C>T), CCT1338CCC (4198T>C), TCC1339TCT (4201C>T), GCG1342GCT (4210G>T), TTC1343TTT (4213C>T), AGT1345AGC (4219T>C), CTG1346TTG (4220C>T), CTG1347ATA (4223C>A 4225G>A), GTT1349GTG (4231T>G), CAT1351CAC (4237T>C), GTA1356GTG (4252A>G), TTA1357CTG (4253T>C 4255A>G), AAA1358AAG (4258A>G), ACT1362ACC (4270T>C), AGA1366AGG (4282A>G), GAG1367GAA (4285G>A), GAA1369GAG (4291A>G), CAG1373CAA (4303G>A), GTC1376GTT (4312C>T), ATA1379GTG (4319A>G 4321A>G), GTG1380GTT (4324G>T), GAG1381GAA (4327G>A), CTG1384CTT (4336G>T), TCT1385TCC (4339T>C), TCA1388TCT (4348A>T), GCA1392GCG (4360A>G), GGT1394GGC (4366T>C), TCT1395TCA (4369T>A), ACT1397ACC (4375T>C), GTT1401GTC (4387T>C), TTT1404TAT (4395T>A), GTA1408ATA (4406G>A), CTT1409CTC (4411T>C), GTT1418GTC (4438T>C), ACC1420ACA (4444C>A), GCC1422GCT (4450C>T), CTC1424CTT (4456C>T), GAC1427GAT (4465C>T), ATG1431GTT (4475A>G 4477G>T), GTC1432GTT (4480C>T), ACA1433TCG (4481A>T 4483A>G), GGC1437GGT (4495C>T), AAG1441AAA (4507G>A), CAC1442CAT (4510C>T), GGT1443GGG (4513T>G), CTA1445TTA (4517C>T), GTC1448GTT (4528C>T), AAA1450AAG (4534A>G), ACC1452ACT (4540C>T), GCG1454TTA (4544G>T 4545C>T 4546G>A), AGA1455AAA (4548G>A), TTT1458TTC (4558T>C), GTA1460GTT (4564A>T), ACC1462ACT (4570C>T), ATA1464ATT (4576A>T), TTG1471TTA (4597G>A), GAC1472GAT (4600C>T), GAC1474GAT (4606C>T), GTG1475GTA (4609G>A), GTG1477GTA (4615G>A), GAC1478GAT (4618C>T), TTG1481CTT (4625T>C 4627G>T), GAC1488GAT (4648C>T), ATT1489ATC (4651T>C), AGG1492AGA (4660G>A), AGC1493AGT (4663C>T), GCT1495GCA (4669T>A), GTG1499ATC (4679G>A 4681G>C), AGT1500AGC (4684T>C), GTT1501GTG (4687T>G), GAG1505GAA (4699G>A), AGA1506AGG (4702A>G), CAA1508CAG (4708A>G), AGG1509AGA (4711G>A), CTG1510TTG (4712C>T), AAA1518AAG (4738A>G), GTA1520GTT (4744A>T), GCA1521GCG (4747A>G), TTG1522CTG (4748T>C), ACT1527ACC (4765T>C), GTC1538ATT (4796G>A 4798C>T), ACT1540AGC (4803C>G 4804T>C), GAG1541GAA (4807G>A), GCG1542GCT (4810G>T), GCT1543GCG (4813T>G), CTT1544CTC (4816T>C), TAT1549TAC

| | | | | | | | | | |
|---|---|---|---|---|---|---|---|---|---|
| | (4831T>C), AAC1550AAT (4834C>T), CCA1552CCT (4840A>T), GGA1556GGG (4852A>G), GGC1557GGT (4855C>T), AGT1561AGC (4867T>C), CTG1562CTC (4870G>C), ATT1563ATA (4873T>A), TGT1566TGC (4882T>C), ACA1573ACT (4903A>T), CAG1575CAA (4909G>A), TTT1577TTC (4915T>C), GAA1578GAG (4918A>G), TTG1579CTA (4919T>C 4921G>A), AGT1580AGC (4924T>C), CCC1581CCG (4927C>G), TTT1583TTC (4933T>C), ATC1584ATA (4936C>A), CAG1585CAA (4939G>A), AAT1586AAC (4942T>C), TTC1587TTT (4945C>T), TCA1593TCG (4963A>G), CCT1596CCC (4972T>C), CTT1602CTC (4990T>C), AAA1606AAG (5002A>G), CCT1613CCC (5023T>C), TTG1614CTG (5024T>C), TGC1615TGT (5029C>T), GAT1616GAC (5032T>C), TCT1618TCC (5038T>C), CCA1620CCT (5044A>T), GCA1623GCT (5053A>T), TCG1624TCA (5056G>A), AGT1625GGT (5057A>G), ACT1626GCT (5060A>G), TCG1629TCT (5071G>T), CTT1636ATT (5090C>A), GGA1637GGG (5095A>G), GTG1638GTT (5098G>T), GCC1639GTC (5100C>T), TTA1640TTG (5104A>G), GAA1641GAT (5107A>T), ATT1642ATC (5110T>C), CCA1643CCG (5113A>G), AAG1644AAA (5116G>A), CAA1645CAG (5119A>G), GTC1646TTT (5120G>T 5122C>T), ATT1648GTT (5126A>G), CAT1651CAC (5137T>C), ATC1652ATT (5140C>T), AAA1653AAG (5143A>G), GAG1654GAA (5146G>A), CTC1659CTA (5161C>A), GAA1661GAG (5167A>G), CTT1663CTG (5173T>G), ACG1666ACA (5182G>A), GTT1667ATT (5183G>A), GTC1668GTT (5188C>T), AAG1669AAA (5191G>A), AAA1671AAG (5197A>G), GAC1672GAT (5200C>T), TGC1674TGT (5206C>T), TTA1675TTG (5209A>G), AGC1678AGT (5218C>T), TCG1682TCA (5230G>A), ATC1684ATT (5236C>T), AGC1685AGT (5239C>T), ATC1687ATT (5245C>T), ACA1693ACT (5263A>T), CTC1695CTT (5269C>T), TTC1696TTT (5272C>T), GCC1697GCA (5275C>A), ATC1698ATT (5278C>T), CCA1699CCC (5281A>C), AGA1700AGG (5284A>G), ACT1701ACC (5287T>C), CTA1702TTA (5288C>T), ATA1703ATT (5293A>T), TTG1704CTA (5294T>C 5296G>A), GAG1706GAA (5302G>A), CTT1709CTA (5311T>A), GAA1710GAG (5314A>G), CGA1713CGT (5323A>T), AGT1721AGC (5347T>C), ATC1723ATT (5353C>T), TGC1727TGT (5365C>T), GTT1734GTC (5386T>C), ACC1737ACA (5395C>A), AAA1746AAG (5422A>G), GAT1747GAC (5425T>C), ACC1749ACT (5431C>T), GAG1751GAA (5437G>A), CTT1756CTC (5452T>C), AAG1758AAA (5458G>A), AGT1761AGC (5467T>C), CAA1762CAG (5470A>G), CTA1763TTG (5471C>T 5473A>G), GAA1765GAG (5479A>G), TTC1766TTT (5482C>T), TCA1767TCC (5485A>C), AAT1768AAC (5488T>C), TTG1769CTA (5489T>C 5491G>A), GGT1771GGC (5497T>C), TGT1774TGC (5506T>C), GAG1775GAA (5509G>A), GAG1776GAA (5512G>A), TTA1778CTA (5516T>C), TAT1782TAC (5530T>C), GAG1783GAA (5533G>A), TCG1784TCT (5536G>T), TTG1785TTA (5539G>A), GTT1788GTG (5548T>G), CAC1790CAT (5554C>T), GCT1792GCA (5560T>A), GCG1793ACA (5561G>A 5563G>A), ACG1794ACC (5566G>C), TCA1795TCG (5569A>G), CTC1800CTG (5584C>G), AAG1803AAA (5593G>A), GGG1804GGA (5596G>A), ATT1805GTT (5597A>G), AAC1807AAA (5605C>A), GTG1811GTT (5617G>T), GCT1812GTG (5619C>T 5620T>G), TTG1815TTA (5629G>A), ATC1816CTC (5630A>C), GGC1819GGT (5641C>T), GCA1822GCT (5650A>T), ATA1826ATT (5662A>T), GGA1827GGG (5665A>G), CAA1835CAG (5689A>G), GTT1837ATT (5693G>A), GAG1838GAA (5698G>A), CAA1843CAG (5713A>G), GGG1844GGC (5716G>C), AAA1845AAG (5719A>G), GCC1853GCA (5743C>A), CGC1857CGA (5755C>A), CAT1858CAC (5758T>C), CGT1860CGC (5764T>C), AAA1862AAG (5770A>G), GGC1865GGT (5779C>T), TTT1866TTC (5782T>C), GAC1869GAT (5791C>T), GAC1873GAT (5803C>T), AGG1884AGA (5836G>A), AAA1885AAG (5839A>G), GGA1887GGG (5845A>G), AAA1888AAG (5848A>G), AAA1890AGA (5853A>G), GGT1891GGC (5857T>C), ACC1892ACT (5860C>T), ACA1893ACT (5863A>T), GGT1895GGA (5869T>A), GGT1897GGC (5875T>C), AGC1900AGT (5884C>T), TTC1903TTT (5893C>T), ATC1904GTG (5894A>G 5896C>G), AAC1905AAT (5899C>T), GGG1908GGA (5908G>A), TTT1909TTC (5911T>C), GAG1913GAA (5923G>A), CAA1918CAG (5938A>G), TTC1919TTT (5941C>T), GTT1920GTC (5944T>C), GAT1921GAC (5947T>C), GGG1925GGA (5959G>A), CGG1926GCC (5960C>G 5961G>C 5962G>C), GAA1930GAG (5974A>G), GCT1934GCA (5986T>A), GAC1935GAT (5989C>T), ATT1936GTT (5990A>G), GAT1938GAC (5998T>C), AGA1942CGT (6008A>C 6010A>T), GAA1945GAT (6019A>T), GTG1946GTT (6022G>T), CGA1947CGT (6025A>T), AAA1949ACA (6030A>C), GAC1955GAA (6049C>A), ATG1958ATA (6058G>A), CAA1959CAG (6061A>G), GCC1960GCA (6064C>A), GGT1962GGC (6070T>C), AGT1963AAC (6072G>A 6073T>C), AAC1964CAC (6074A>C), ACG1965ACA (6079G>A), ACC1966AAC (6081C>A), ATA1967ATT (6085A>T), GCA1969GCT (6091A>T), TAC1970TAT (6094C>T), TGT1976TCC (6111G>C 6112T>C), AAA1978AAG (6118A>G), TTG1980TTA (6124G>A), AAG1981AAA (6127G>A), ATT1982GTC (6128A>G 6130T>C), GAT1983GAC (6133T>C), TTA1984TTG (6136A>G), AAC1988AAT (6148C>T), AAA1991AAG (6157A>G), GTT1992ATT (6158G>A), TGT1993TGC (6163T>C), AAA1995AAG (6169A>G), ACA1996TCG (6170A>T 6172A>G), GGC1998GGT (6178C>T), AAA2001AAG (6187A>G), TTT2002TTC (6190T>C), GAG2004GAA (6196G>A), CTC2007TTT (6203C>T 6205C>T), GAA2008GAG (6208A>G), CTA2009TTG (6209C>T 6211A>G), CAG2011CAA (6217G>A), GCT2015GCA (6229T>A), GTA2016ATA (6230G>A), GTC2018GTT (6238C>T), AAG2021AAA (6247G>A), ATA2023ATT (6253A>T), GCA2025AAA (6257G>A 6258C>A), GAG2027GAA (6265G>A), CAT2030CAC (6274T>C), AAA2033AGA (6282A>G), TCG2034TCA (6286G>A), GGC2038GGT (6298C>T), TTG2039TTA (6301G>A), AGA2040AGG (6304A>G), GAC2041GAT (6307C>T), GCC2046GCT (6322C>T), ACA2048ACT (6328A>T), GTT2049GTG (6331T>G), TGT2050TGC (6334T>C), AGG2051AGA (6337G>A), CTG2052TTG (6338C>T), TAT2058TAC (6358T>C), GGG2059GGA (6361G>A), GCA2060ACA (6362G>A), TCA2061TCT (6367A>T), GAG2062GAA (6370G>A), TAC2064TAT (6376C>T), GGT2065GGG (6379T>G), GGA2067GGC (6385A>C), GGA2069GGT (6391A>T), GCA2070GCG (6394A>G), TAC2071TAT (6397C>T), GTA2073ATA (6401G>A), GCG2074GTG (6405C>T), CAC2076CAT (6412C>T), TTA2078TTG (6418A>G), TTT2079TTC (6421T>C), TAC2082TTC (6429A>T), GGT2084GGA (6436T>A), GAG2087GAA (6445G>A), CAA2089CGA (6450A>G), TCC2090TCG (6454C>G), CAC2092CAT (6460C>T), GGT2093GGA (6463T>A), TTC2095TTT (6469C>T), AGG2096AGA (6472G>A), GTG2097GTA (6475G>A), AAT2099AAC (6481T>C), CAC2101AAT (6485C>A 6487C>T), AGT2102AGC (6490T>C), AGC2104AGT (6496C>T), CTG2106CTA (6502G>A), GGT2110GGC (6514T>C), AGG2111AGA (6517G>A), CTC2115ATC (6527C>A), ATC2116ATA (6532C>A), CCG2119CCA (6541G>A), AAA2120AAG (6544A>G), GTC2124GTT (6556C>T), TTT2125TTC (6559T>C), CAG2127CAA (6565G>A), TTG2129CTA (6569T>C 6571G>A), CCT2134CCA (6586T>A), ACA2135ATA (6588C>T), GAA2138GAG (6598A>G), AGA2139AGG (6601A>G), ATT2140ATC (6604T>C), TGT2141TGC (6607T>C), GGA2144GGG (6616A>G), AAC2146AAT (6622C>T), GAG2149GAA (6631G>A), AAG2150AAA (6634G>A), TAT2151CAT (6635T>C), GCT2152GCA (6640T>A), TCG2154TCT (6646G>T), ATC2155GTC (6647A>G), ATC2156ATT (6652C>T), AGC2160AGT (6664C>T), ACT2162ACG (6670T>G), ATA2165GTG (6677A>G 6679A>G), GGC2167GGT (6685C>T), ACA2169ACT (6691A>T), TTC2170TTT (6694C>T), ACA2177ACG (6715A>G), GAT2178AAT (6716G>A), AAT2179GAT (6719A>G), GGA2180GGG (6724A>G), CTA2184TTA (6734C>T), GTG2186GTA (6742G>A), GTG2187GTA (6745G>A), GCC2190GCT (6754C>T), GGA2192GGG (6760A>G), ATA2194TTG (6764A>T 6766A>G), ATT2197ATA (6775T>A), CTG2200TTG (6782C>T), GCA2201GCT (6787A>T), AAC2202AAT (6790C>T), GCA2204GTG (6795C>T 6796A>G), CAC2205CAA (6799C>A), ACC2206ACA (6802C>A), AAC2208AAT (6808C>T), TAC2209TAT (6811C>T), TAC2210TAT (6814C>T), TTC2213TTT (6823C>T), GAA2215GAG (6829A>G), TAC2221TAT (6847C>T), CGA2223CGC (6853A>C), AAT2225GAT (6857A>G), GAG2226GAA (6862G>A), GAA2229GAG (6871A>G), GTC2231ACA (6875G>A 6876T>C 6877C>A), AAG2232AAA (6880G>A), TCT2233TCA (6883T>A), GTT2235GTG (6889T>G), GAC2239GAT (6901C>T), GGC2244GGT (6916C>T), CCG2245CCA (6919G>A), AAA2247AAG (6925A>G), CTT2248CTC (6928T>C), GAC2250GAA (6934C>A), GGG2255GGC (6949G>C), TTC2257TTT (6955C>T), AAA2258AAG (6958A>G), CAA2264CAG (6976A>G), CTA2266TTA (6980C>T), ATC2267ATT (6985C>T), GTA2271GTT (6997A>T), GTG2272GTT (7000G>T), GTG2273ATA (7001G>A 7003G>A), GCT2276GCC (7012T>C), AAG2277AAA (7015G>A), CAT2278CAC (7018T>C), TTT2283TAT (7032T>A), GCC2285GCT (7039C>T), TTG2286CTA (7040T>C 7042G>A), GGA2288GGG (7048A>G), GCT2292GCA (7060T>A), GTC2293GTA (7063C>A), AGC2298AGT (7078C>T), TTA2300CTG (7082T>C 7084A>G), GTA2301GTG (7087A>G), ACC2302ACA (7090C>A), CAT2304CAC (7096T>C), GTA2305GTG (7099A>G), GTT2306GTC (7102T>C), GAG2309GAA (7111G>A), CGA2311CGG (7117A>G), ACA2314AAA (7125C>A), GAA2315GAG (7129A>G), TTT2316TTC (7132T>C), CTG2317TTA (7133C>T 7135G>A), GTG2319GTA (7141G>A), GAT2320GAC (7144T>C), GCA2321TCG (7145G>T 7147A>G), GAG2322GAA (7150G>A), GCA2325GCT (7159A>T), GAG2326_GCA2327del (7160_7165delGAGGCA), TTC2329TTT (7171C>T), GCG2335GCT (7189G>T), AAA2338AAG (7198A>G), AGC2339AGT (7201C>T), CTA2341TTA (7205C>T), AAT2342AAC (7210T>C), GAT2344GAA (7216T>A), GCG2345GCA (7219G>A), ATC2347ATA (7225C>A), AAG2348AAA (7228G>A), AAG2352AAA (7240G>A), TAT2353TAC (7243T>C), AAA2355AAG (7249A>G), CCT2356CCG (7252T>G), GAT2358GAA (7258T>A), GTT2359GTC (7261T>C), GGT2360GGA (7264T>A), GTC2361GTG (7267C>G), GTG2362GTA (7270G>A), GAT2363_CGG2364insT-- (7273_7274insT), CGG2364GTG (7274C>G 7275G>T), CAT2366CTT (7281A>T), TTG2367TCG (7284T>C), TCA2371TTA (7296C>T), TCT2376TTT (7311C>T), ACC2377ATC (7314C>T), TGC2378TAC (7317G>A), AAT2379AAA (7321T>A), GTG2380-TG (7322delG), CAC2381CAT (7327C>T), GGC2382GGT (7330C>T), TTC2383TTT (7333C>T), AAG2384CAG (7334A>C), TGT2386TGC (7342T>C), GCA2387AAC (7343G>A 7344C>A 7345A>C), ACT2390ACC (7354T>C), CAA2393CAG (7363A>G), ATT2395ATC (7369T>C), TTC2396TTT (7372C>T), GCG2398GCT (7378G>T), CTC2399CTT (7381C>T), GCA2404GCT (7396A>T), GTC2405GTT (7399C>T), GCC2407GCT (7405C>T), AGT2408ATG (7407G>T 7408T>G), GGG2410GGT (7414G>T), TGC2411GGC (7415T>G), AAA2412AAG (7420A>G), GAC2415GAT (7429C>T), TAT2416TAC (7432T>C), CAT2419CAC (7441T>C), AAG2425AAA (7459G>A), GAA2426GAG (7462A>G), ATA2428ATT (7468A>T), GTC2429GTA (7471C>A), CAA2431CAG (7477A>G), AGC2432AGT (7480C>T), CTG2434CTT (7486G>T), TAT2437TAC (7495T>C), AAA2438AAG (7498A>G), GGT2439GGC (7501T>C), TTG2440TCA (7503T>C 7504G>A), ATT2443ATA (7513T>A), AAC2445AAT (7519C>T), GGA2446GGG (7522A>G), AAG2449AAA (7531G>A), GAG2451GAA (7537G>A), CTC2452CTT (7540C>T), TGT2454TGC (7546T>C), AAG2462AAA (7570G>A), ACG2463ACA (7573G>A), ACG2465ACA (7579G>A), GCT2468GCC (7588T>C), CTA2471TTA (7595C>T), TTG2474CTA (7604T>C 7606G>A), GGT2477GGA (7615T>A), TGT2480TGC (7624T>C), GAC2483GAT (7633C>T), TTC2484TTT (7636C>T), AAT2485AAC (7639T>C), TTT2488TTC (7648T>C), GAA2494GAG (7666A>G), TGT2496TGC (7672T>C), ACA2498ACT (7678A>T), GGG2500GGA (7684G>A), ACT2502ACC (7690T>C), TTT2504TTC (7696T>C), GGT2506GGG (7702T>G), GAT2509AAT (7709G>A), AAA2510AGG (7713A>G 7714A>G), CTG2511TTG (7715C>T), CGT2514CGC (7726T>C), GAG2517GAA (7735G>A), GTA2520GTG (7744A>G), TAC2521TAT (7747C>T), GGC2526GGT (7762C>T), CAG2528CAA (7768G>A), TTT2529TTC (7771T>C), TCA2532TCG (7780A>G), CTA2533CTC (7783A>C), ACT2534ACC (7786T>C), CTA2537TTG (7793C>T 7795A>G), ATC2538ATT (7798C>T), GCT2540GCC (7804T>C), ACC2543ATC (7812C>T), GAA2550GAG (7834A>G), GAC2551GAT (7837C>T), GTG2554TTA (7844G>T 7846G>A), CAG2557CAA (7855G>A), CTG2559CTA (7861G>A), TTA2562TTG (7870A>G), TAC2563TAT (7873C>T), ACT2564ACA (7876T>A), GAG2565GAA (7879G>A), ATT2566ATA (7882T>A), GTT2567ATT (7883G>A), CCA2570CCT (7894A>T), ATT2571ATC (7897T>C), GTC2579GTT (7921C>T), AAG2581AAA (7927G>A), GGT2584GGC (7936T>C), AAT2585AAC (7939T>C), AAT2586AAC (7942T>C), GTG2594GTA (7966G>A), GAT2595GAC (7969T>C), TCT2597TCC (7975T>C), GTC2601GTT (7987C>T), CTT2602CTC (7990T>C), GCT2607GCC (8005T>C), GAA2616GAG (8032A>G), ATC2618ATT (8038C>T), GAC2619GAT (8041C>T), AGC2620AAC (8043G>A), TTG2632TTA (8080G>A), GTG2635GTA (8089G>A), CCG2637CCA (8095G>A), GAG2638GAA (8098G>A), AGC2641AGT (8107C>T), ATT2642ATC (8110T>C), GAT2644GAC (8116T>C), CAA2648CAG (8128A>G), GGT2654GGC (8146T>C), AAC2656AGT (8151A>G 8152C>T), TCG2660TCA (8164G>A), AGG2665AAG (8178G>A), AAG2666AAA (8182G>A), TTG2669TTA (8191G>A), TTC2671TTT (8197C>T), TCC2673TCT (8203C>T), AGA2675AAA (8208G>A), CTG2677TTA (8213C>T 8215G>A), CTA2678CTG (8218A>G), ATC2679ATT (8221C>T), GAG2680GAA (8224G>A), CCA2685CCG (8239A>G), GAG2690GAA (8254G>A), ATT2692ATC (8260T>C), GTA2693GTG (8263A>G), GAT2699GAC (8281T>C), AGA2700AGG (8284A>G), CCA2704GCG (8294C>G 8296A>G), GAG2705GAA (8299G>A), CAC2706CAT (8302C>T), ATT2711ATC (8317T>C), TGC2712TGT (8320C>T), GCT2714GCA (8326T>A), GAG2717GAA (8335G>A), TCC2718TCT (8338C>T), TCT2722TTT (8349C>T), GAA2723GAG (8353A>G), CAC2726CAT (8362C>T), CAA2727CAG (8365A>G), ATC2728ATT (8368C>T), TAC2732TAT (8380C>T), TCA2733TCG (8383A>G), TTA2735TTG (8389A>G), CCT2740CCC (8404T>C), GCA2742TCA (8408G>T), ACA2743ACG (8413A>G), GCG2745GCT (8419G>T), GCT2750GCC (8434T>C), TAT2752TAC (8440T>C), ATG2756ATA (8452G>A), CTA2758CTG (8458A>G), AGG2759AAG (8460G>A), AAA2760AAG (8464A>G), CTG2761CTA (8467G>A), AGG2765AGA (8479G>A), GCT2766ACA (8480G>A 8482T>A), GAG2771GAA (8497G>A), CTA2772TTG (8498C>T 8500A>G), AGA2773AAG (8502G>A 8503A>G), GCC2774GCT (8506C>T), GAA2777GAT (8515A>T), GTC2780GTT (8524C>T), GCA2781GCC (8527A>C), TTA2782CTG (8528T>C 8530A>G), GAG2785GAC (8539G>C), CTT2788TGT (8546C>T 8547T>G), TCT2790ACT (8552T>A), GTA2793GTG (8563A>G), GCA2797GGA (8574C>G), ATT2801GTT (8585A>G), GGA2805GAA (8598G>A), AGC2806AGT (8602C>T), AAC2807TCT (8603A>T 8604A>C 8605C>T), AAA2812AAG (8620A>G), CCA2813CAA (8622C>A), GAG2814GGA (8625A>G 8626G>A), CAG2815CAA (8629G>A), GGC2816GGT (8632C>T), ATC2818ATT (8638C>T), CAG2819CAA (8641G>A), AAC2821AAT (8647C>T), CCG2822TTC (8648C>T 8649C>T 8650G>C), AAC2823AAT (8653C>T), AAA2824AAG (8656A>G), GGA2825GAA (8658G>A), GAT2827GCG (8664A>C 8665T>G), GTT2830GTG (8674T>G), GCA2832GTT (8679C>T 8680A>T), GGC2833GGA (8683C>A), GGG2836GGA (8692G>A), ACA2837ACT (8695A>T), CCG2841CCA (8707G>A), AGA2842CGA (8708A>C), AAG2844AAA (8716G>A), ATC2846ATT (8722C>T), ACG2847ACA (8725G>A), AAA2849AAG (8731A>G), AGA2851AGG (8737A>G), CCC2853CCT (8743C>T), ACA2854AAG (8745C>A 8746A>G), AAG2856AAA (8752G>A), GGA2857GGT (8755A>T), AAC2862AAT (8770C>T), TTA2863CTA (8771T>C), GAA2864GAG (8776A>G), CAT2865CAC (8779T>C), TTG2866TTA (8782G>A), TAT2869TAC (8791T>C), CCA2871CCG (8797A>G), CAA2872CAG (8800A>G), CAA2873CAG (8803A>G), GAT2875GAC (8809T>C), ATT2876ATC (8812T>C), AAT2878AAC (8818T>C), CGG2880CGA (8824G>A), GCA2881GCG (8827A>G), ACG2888ACA (8848G>A), TAT2890TAC (8854T>C), GAG2891GGA (8856A>G 8857G>A), GTG2893GTA (8863G>A), CGG2894CAG (8865G>A), ATG2895CTT (8867A>C 8869G>T), GCA2896GCG (8872A>G), GGA2900GAA (8883G>A), ACT2902TCT (8888A>T), TGC2915TGT (8929C>T), GTC2924ATC (8954G>A), AAC2925AAT (8959C>T), GGA2926GGT (8962A>T), GGG2933GGA (8983G>A), AAT2934GAC (8984A>G 8986T>C), GAA2935GTA (8988A>T), GAG2938GAA (8998G>A), CCG2940CCA (9004G>A), TTG2941TTA (9007G>A), ATC2944ATT (9016C>T), ACC2951ACA (9037C>A), ATA2967ATT (9085A>T), AAC2971AAT (9097C>T), ATT2983GTT (9131A>G), CGA2984CGT (9136A>T), CGG2987CGT (9145G>T), ATG2989GGA (9149A>G 9150T>G 9151G>A), GGT2990AGT (9152G>A), TTA2991TTG (9157A>G), GCG2992GCT (9160G>T), CGT2993CGC (9163T>C), GAG3000GAA (9184G>A), CGA3004CGG (9196A>G), AGG3010AGA (9214G>A), GAA3011GAG (9217A>G), GCG3012GCA (9220G>A), GCA3020GCT (9244A>T), TTG3021TTA (9247G>A), GCC3024GCT (9256C>T), CCT3026TCC (9260C>T 9262T>C), GGG3030GGA (9274G>A), GAC3032GAT (9280C>T), ATC3035ATT (9289C>T), CAA3038CAC (9298A>C), GAG3040GAA (9304G>A), GTC3050GTT (9334C>T), GTC3060GTG (9364C>G) | | | | | | | | |
| P1 protein (NP_734243.1) | 1 | 284 | 100% | 1394 | 72.7% | 284 (100%) | 203 (71.5%) | 0/0/0/0 | 0 |

| | Begin | End | Coverage | Score | Concordance | Matches | Identities | I/D/M/F* | Stop Codons |
|---|---|---|---|---|---|---|---|---|---|
| **NT** | **52** | **9704** | **99.5%** | **12530** | **65.2%** | **9642 (99.8%)** | **7985 (82.7%)** | **7/11** | |
| Protein mutations: | Y4Q (194T>C 196C>A), C9L (209T>C 210G>T 211T>G), F13M (221T>A 223T>G), S23P (251T>C), C24F (255G>T 256C>T), E25G (258A>G), I27V (263A>G 265T>G), V28A (267T>C), E30V (273A>T), A35T (287G>A 289T>C), V37T (293G>A 294T>C), A41I (305G>A 306C>T), D42G (309A>G), E44D (316A>T), T45V (317A>G 318C>T 319A>G), L52Q (339T>A 340C>A), K53R (342A>G), K55A (347A>G 348A>C 349A>G), Y56H (350T>C), T58V (356A>G 357C>T), V61T (365G>A 366T>C 367G>A), L62S (368C>T 369T>C), F67C (384T>G), A75V (408C>T 409C>T), M78E (416A>G 417T>A), E85K (437G>A), R86E (440A>G 441G>A), K87R (443A>C 444A>G 445G>A), D88E (448T>A), E91A (456A>C), H93N (461C>A), D111E (517T>A), S116F (531C>T), P118S (536C>T), Q119W (539C>T 540A>G 541A>G), R122K (549G>A 550A>G), M131T (576T>C), V134A (585T>C 586C>G), R137Y (593C>T 594G>A 595C>T), P138R (597C>G), I139V (599A>G 601A>G), I140P (602A>C 603T>C 604A>G), M147I (625G>A), I151V (635A>G 637T>C), I154V (644A>G 646A>G), E160A (663A>C 664G>C), H166Q (682C>A), T172S (699C>G), H174Q (706T>A), K179E (719A>G 721G>A), I180V (722A>G 724A>G), A183S (731G>T), Y184P (734T>C 735A>C 736C>T), S185R (737T>C 738C>G 739C>T), A187V (744C>T 745G>C), R189C (749C>T 751A>C), M194K (765T>A), R197K (773C>A 774G>A 775A>G), M206K (801T>A), G210A (813G>C 814A>G), L211H (816T>A), R214Q (825G>A 826T>A), N223D (851A>G 853T>C), R226C (860C>T 862C>T), T227A (863A>G 865T>C), I228T (867T>C), N229D (869A>G 871C>T), I230L (872A>T 874A>G), R231Q (876G>A), R232K (879G>A), N240S (903A>G), T241N (906C>A 907A>C), K242T (909A>C 910A>C), S243N (912G>A 913C>T), S252L (938T>C 939C>T 940A>T), L266I (980T>A 982G>C), R272K (999G>A), Q275R (1007C>A 1008A>G 1009G>A), S276G (1010A>G 1012T>G), I277V (1013A>G), N279Q (1019A>C 1021C>G) | | | | | | | | |
| Codon mutations: | ACT3ACC (193T>C), TAC4CAA (194T>C 196C>A), TCA6TCG (202A>G), ATC8ATT (208C>T), TGT9CTG (209T>C 210G>T 211T>G), TCG12TCA (220G>A), TTT13ATG (221T>A 223T>G), GAA14GAG (226A>G), CTA17CTT (235A>T), TAC19TAT (241C>T), CCA21CCT (247A>T), TCT23CCT (251T>C), TGC24TTT (255G>T 256C>T), GAG25GGG (258A>G), CAT26CAC (262T>C), ATT27GTG (263A>G 265T>G), GTG28GCG (267T>C), GAA30GTA (273A>T), CGA31CGT (277A>T), GAA32GAG (280A>G), GTG33GTT (283G>T), CCG34CCA (286G>A), GCT35ACC (287G>A 289T>C), GTT37ACT (293G>A 294T>C), CCT39CCC (301T>C), TTC40TTT (304C>T), GCA41ATA (305G>A 306C>T), GAT42GGT (309A>G), CTG43TTG (311C>T), GAA44GAT (316A>T), ACA45GTG (317A>G 318C>T 319A>G), CAA46CAG (322A>G), GCA49GCT (331A>T), CTC52CAA (339T>A 340C>A), AAG53AGG (342A>G), AAA55GCG (347A>G 348A>C 349A>G), TAT56CAT (350T>C), GCT57GCA (355T>A), ACT58GTT (356A>G 357C>T), CGT60CGC (364T>C), GTG61ACA (365G>A 366T>C 367G>A), CTC62TCC (368C>T 369T>C), AAA63AAG (373A>G), AAC64AAT (376C>T), TTT67TGT (384T>G), ACG68ACA (388G>A), TAC69TAT (391C>T), CGA70CGG (394A>G), AAG72AAA (400G>A), GAT74GAC (406T>C), GCC75GTT (408C>T 409C>T), CAG76CAA (412G>A), ATA77ATT (415A>T), ATG78GAG (416A>G 417T>A), ATT80ATC (424T>C), CAG81CAA (427G>A), AAA83AAG (433A>G), CTG84CTC (436G>C), GAG85AAG (437G>A), AGG86GAG (440A>G 441G>A), AAG87CGA (443A>C 444A>G 445G>A), GAT88GAA (448T>A), GAA90GAG (454A>G), GAA91GCA (456A>C), TAT92TAC (460T>C), CAC93AAC (461C>A), CAA95CAG (469A>G), GCC97GCG (475C>G), CCT99CCA (481T>A), ACT106ACC (502T>C), ATA107ATT (505A>T), GGC109GGT (511C>T), GAT111GAA (517T>A), CCT112CCA (520T>A), CCA113CCT (523A>T), TCT116TTT (531C>T), GAG117GAA (535G>A), CCA118TCA (536C>T), CAA119TGG (539C>T 540A>G 541A>G), AGA122AAG (549G>A 550A>G), ATC124ATT (556C>T), ATT125ATC (559T>C), CAT126CAC (562T>C), ACT128ACC (568T>C), CCA129CCG (571A>G), ATG131ACG (576T>C), CGT132CGC (580T>C), GTC134GCG (585T>C 586C>G), CGC137TAT (593C>T 594G>A 595C>T), CCC138CGC (597C>G), ATA139GTG (599A>G 601A>G), ATA140CCG (602A>C 603T>C 604A>G), ACA143ACG (613A>G), GAA144GAG (616A>G), GGC145GGG (619C>G), CAG146CAA (622G>A), ATG147ATA (625G>A), AAT148AAC (628T>C), CAC149CAT (631C>T), ATT151GTC (635A>G 637T>C), ATA154GTG (644A>G 646A>G), ATT157ATC (655T>C), TCG159TCA (661G>A), GAG160GCC (663A>C 664G>C), AAA161AAG (667A>G), GTC165GTT (679C>T), CAC166CAA (682C>A), TTA167CTG (683T>C 685A>G), AGT169AGC (691T>C), ACC172AGC (699C>G), ACT173ACA (703T>A), CAT174CAA (706T>A), AAG179GAA (719A>G 721G>A), ATA180GTG (722A>G 724A>G), CTT181TTG (725C>T 727T>G), GGT182GGA (730T>A), GCA183TCA (731G>T), TAC184CCT (734T>C 735A>C 736C>T), TCC185CGT (737T>C 738C>G 739C>T), GCG187GTC (744C>T 745G>C), CGA189TGC (749C>T 751A>C), ACT190ACC (754T>C), ATG194AAG (765T>A), TTG196CTG (770T>C), CGA197AAG (773C>A 774G>A 775A>G), CGG198AGG (776C>A), TTC202TTT (790C>T), ATG206AAG (801T>A), GGA210GCG (813G>C 814A>G), CTT211CAT (816T>A), TTG212CTG (818T>C), CAA213CAG (823A>G), CGT214CAA (825G>A 826T>A), CGG217AGA (833C>A 835G>A), GAC219GAT (841C>T), AAT223GAC (851A>G 853T>C), GTC225GTT (859C>T), CGC226TGT (860C>T 862C>T), ACT227GCC (863A>G 865T>C), ATC228ACC (867T>C), AAC229GAT (869A>G 871C>T), ATA230TTG (872A>T 874A>G), CGA231CAA (876G>A), AGG232AAG (879G>A), GAT234GAC (886T>C), ATC238ATT (898C>T), AAC240AGC (903A>G), ACA241AAC (906C>A 907A>C), AAA242ACC (909A>C 910A>C), AGC243AAT (912G>A 913C>T), GGC246GGA (922C>A), GGT249GGA (931T>A), TCA252CTT (938T>C 939C>T 940A>T), GGA253GGG (943A>G), TTC256TTT (952C>T), TCA261TCG (967A>G), GAA263GAG (973A>G), GGG264GGC (976G>C), TTG266ATC (980T>A 982G>C), GCA269GCG (991A>G), CGT270CGC (994T>C), AGA272AAA (999G>A), GTT273GTC (1003T>C), CAG275AGA (1007C>A 1008A>G 1009G>A), AGT276GGG (1010A>G 1012T>G), ATT277GTT (1013A>G), TTA278TTG (1018A>G), AAC279CAG (1019A>C 1021C>G), ATC282ATT (1030C>T) | | | | | | | | |
| HC-Pro protein (NP_734244.1) | 1 | 456 | 100% | 3144 | 95.7% | 456 (100%) | 425 (93.2%) | 0/0/0/0 | 0 |
| Protein mutations: | D4E (1048C>A), R16Q (1083G>A), S21A (1097T>G 1099G>A), R36K (1143G>A), L40I (1154T>A 1156G>A), S62N (1221G>A), V65T (1229G>A 1230T>C 1231T>C), S66N (1233G>A 1234C>T), K74T (1257A>C 1258A>G), N81S (1278A>G), A85V (1290C>T 1291G>A), D88T (1298G>A 1299A>C 1300C>T), I91V (1307A>G 1309A>G), N94E (1316A>G 1318T>A), I98T (1329T>C), L113I (1373C>A), I117V (1385A>G), E123D (1405G>T), A127S (1415G>T 1417A>T), V134I (1436G>A 1438C>T), E224G (1707A>G), A228T (1718G>A), I236V (1742A>G 1744C>T), S241N (1758G>A), V252I (1790G>A), I301V (1937A>G 1939T>C), F323Y (2004T>A), I344M (2068T>G), V346I (2072G>A 2074G>A), R391K (2208G>A), D410E (2266C>G) | | | | | | | | |
| Codon mutations: | TCG1TCA (1039G>A), AAT2AAC (1042T>C), GCC3GCA (1045C>A), GAC4GAA (1048C>A), AAT5AAC (1051T>C), AAG8AAA (1060G>A), GGT9GGA (1063T>A), CTG10TTG (1064C>T), GAC11GAT (1069C>T), CGA16CAA (1083G>A), TAT19TAC (1093T>C), TCG21GCA (1097T>G 1099G>A), CAC23CAT (1105C>T), GTA26GTG (1114A>G), GCT27GCG (1117T>G), TTA29TTG (1123A>G), CCT30CCA (1126T>A), GTC31GTT (1129C>T), AGG36AAG (1143G>A), GTA37GTT (1147A>T), GCT38GCA (1150T>A), TTG40ATA (1154T>A 1156G>A), CAC43CAT (1165C>T), AGT44AGC (1168T>C), CTT46TTA (1172C>T 1174T>A), ACC55ACT (1201C>T), AGC62AAC (1221G>A), GTT65ACC (1229G>A 1230T>C 1231T>C), AGC66AAT (1233G>A 1234C>T), CTG68CTA (1240G>A), AAG70AAA (1246G>A), TTG72CTA (1250T>C 1252G>A), CAT73CAC (1255T>C), AAA74ACG (1257A>C 1258A>G), AGA77AGG (1267A>G), GGT79GGC (1273T>C), TTG80TTA (1276G>A), AAT81AGT (1278A>G), TTG83CTG (1283T>C), GCG85GTA (1290C>T 1291G>A), GAC88ACT (1298G>A 1299A>C 1300C>T), TTT90TTC (1306T>C), ATA91GTG (1307A>G 1309A>G), CAT92CAC (1312T>C), AAT94GAA (1316A>G 1318T>A), TTG97TTA (1327G>A), ATA98ACA (1329T>C), GCG99GCT (1333G>T), TTA100TTG (1336A>G), CTA103TTG (1343C>T 1345A>G), CCG106CCA (1354G>A), GTG107GTT (1357G>T), GAC108GAT (1360C>T), CTC111TTG (1367C>T 1369C>G), CTT113ATT (1373C>A), TTC114TTT (1378C>T), GAG116GAA (1384G>A), ATA117GTA (1385A>G), AAA119AAG (1393A>G), TCC120TCT (1396C>T), GAG123GAT (1405G>T), AAA124AAG (1408A>G), CAG125CAA (1411G>A), GCA127TCT (1415G>T 1417A>T), CCG128CCC (1420G>C), AAT131AAC (1429T>C), TTA132TTG (1432A>G), GTC134ATT (1436G>A 1438C>T), TTA135CTG (1439T>C 1441A>G), AAT136AAC (1444T>C), TTC139TTT (1453C>T), CTG140CTA (1456G>A), GAA144GAG (1468A>G), GTA152GTG (1492A>G), TTG155CTA (1499T>C 1501G>A), AGT156AGC (1504T>C), TTG157TTA (1507G>A), CTC158CTT (1510C>T), GAA159GAG (1513A>G), TTA160TTG (1516A>G), GCA161GCT (1519A>T), AGG162AGA (1522G>A), TTC163TTT (1525C>T), CAG164CAA (1528G>A), AAC166AAT (1534C>T), ACT168ACA (1540T>A), AAC170AAT (1546C>T), AAA173AAG (1555A>G), GGT174GGC (1558T>C), GAT175GAC (1561T>C), ATA176ATC (1564A>C), TCT177TCA (1567T>A), TTC179TTT (1573C>T), AGA180AGG (1576A>G), TTA183CTA (1583T>C), TCT184TCA (1588T>A), AAG186AAA (1594G>A), AAC188AAT (1600C>T), CTG191CTA (1609G>A), TCG194TCA (1618G>A), TGC195TGT (1621C>T), GAC196GAT (1624C>T), CTG199TTG (1631C>T), AAA201AAG (1639A>G), TTC205TTT (1651C>T), CTC206CTG (1654C>G), CAT213CAC (1675T>C), TCA219TCG (1693A>G), TTT222TTC (1702T>C), GAA224GGA (1707A>G), GCA228ACA (1718G>A), GGA230GGG (1726A>G), TAC231TAT (1729C>T), TCA232TCC (1732A>C), TAT234TAC (1738T>C), ATC236GTT (1742A>G 1744C>T), CGC237CGT (1747C>T), CAT239CAC (1753T>C), AGT241AAT (1758G>A), AGG244AGA (1768G>A), AAC250AAT (1786C>T), TTA251TTG (1789A>G), GTT252ATT (1790G>A), GTC253GTT (1795C>T), TTA257CTG (1805T>C 1807A>G), GAG259GAA (1813G>A), AGG261AGA (1819G>A), AAG263AAA (1825G>A), GAC267GAT (1837C>T), TAT268TAC (1840T>C), CAA271CAG (1849A>G), GTC274GTG (1858C>G), AAA276AAG (1864A>G), TGC278TGT (1870C>T), AGT280AGC (1876T>C), AAA282AAG (1882A>G), GGT284GGA (1888T>A), AAT285AAC (1891T>C), TGT290TGC (1906T>C), ACA293ACT (1915A>T), GGT298GGC (1930T>C), GCC300GCT (1936C>T), ATT301GTC (1937A>G 1939T>C), TCA303TCG (1945A>G), TTC305TTT (1951C>T), CCA308CCT (1960A>T), GTA314GTG (1978A>G), ATT315ATA (1981T>A), GGC316GGT (1984C>T), AAT317AAC (1987T>C), GAC320GAT (1996C>T), TTT323TAT (2004T>A), AAA328AAG (2020A>G), GGG329GGA (2023G>A), GAT330GAC (2026T>C), TCG331TCT (2029G>T), GAG332GAA (2032G>A), TAT341TAC (2059T>C), ATT344ATG (2068T>G), AAC345AAT (2071C>T), GTG346ATA (2072G>A 2074G>A), CTG351TTA (2087C>T 2089G>A), AGC355AGT (2101C>T), GAG356GAA (2104G>A), GAG357GAA (2107G>A), AAG360AAA (2116G>A), TTC362TTT (2122C>T), CGC367CGT (2137C>T), CTT374CTC (2158T>C), ACC376ACT (2164C>T), CCA378CCG (2170A>G), GAT382GAC (2182T>C), GCG384GCA (2188G>A), ACC385ACA (2191C>A), AGA391AAA (2208G>A), TAT394TAC (2218T>C), CAG396GAT (2224C>T), GTA397GTT (2227A>T), TTG402TTA (2242G>A), CCC403CCT (2245C>T), TTG406TTA (2254G>A), GTT407GTC (2257T>C), GAC408GAT (2260C>T), GAC410GAG (2266C>G), CAA412CAG (2272A>G), ACG413ACA (2275G>A), TCA419TCG (2293A>G), GGC421GGT (2299C>T), TCG422TCA (2302G>A), CAG423CAA (2305G>A), GGA426GGT (2314A>T), TAT427TAC (2317T>C), CTA430CTG (2326A>G), GTG435GTA (2341G>A), CAA437CAG (2347A>G), CTT438CTC (2350T>C), TTG440TTA (2356G>A), AAT443AAC (2365T>C), GAA445GAG (2371A>G), TTA446TTG (2374A>G), GAT449GAC (2383T>C), ATA450ATT (2386A>T), AAA451AAG (2389A>G), CAT452CAC (2392T>C), TAT453TAC (2395T>C) | | | | | | | | |
| P3 protein (NP_734251.1) | 1 | 365 | 100% | 2043 | 84.5% | 365 (99.7%) | 327 (89.3%) | 1/0/1/1 | 0 |
| Protein mutations: | N4G (2414A>G 2415A>G 2416T>G), S6C (2420A>T), K40R (2523A>G), R77K (2634G>A), V107I (2723G>A 2725C>T), A115T (2747G>A 2749T>A), H131N (2795C>A), V140I (2822G>A 2824G>A), P160T (2882C>A), S161H (2885A>C 2886G>A), T164M (2895C>T), N176S (2931A>G), N179S (2940A>G), E189_N190insX (2971_2972insA), T196I (2991C>T), H197R (2994A>G), E199R (2999G>A 3000A>G), S204F (3015C>T), R206Q (3021G>A), Y207F (3024A>T 3025C>T), T211I (3036C>T), E212G (3039A>G 3040A>C), K213T (3042A>C 3043G>A), A232V (3099C>T 3100C>A), V234R (3104G>A 3105T>G 3106G>A), S243N (3132G>A), R245K (3137C>A 3138G>A 3139A>G), F250I (3152T>A 3154C>T), C254Y (3165G>A), F272L (3218T>C), T274N (3225C>A), V276I (3230G>A), V288M (3266G>A 3268A>G), V291M (3275G>A), A294T (3284G>A), R304K (3315G>A 3316A>G), E305A (3318A>C), M319V (3359A>G), D332E (3400T>G) | | | | | | | | |

| | Begin | End | Coverage | Score | Concordance | Matches | Identities | I/D/M/F* | Stop Codons |
|---|---|---|---|---|---|---|---|---|---|
| **NT** | **52** | **9704** | **99.5%** | **12530** | **65.2%** | **9642 (99.8%)** | **7985 (82.7%)** | **7/11** | |
| Codon mutations: | AAT4GGG (2414A>G 2415A>G 2416T>G), GCT5GCA (2419T>A), AGC6TGC (2420A>T), ACA12ACT (2440A>T), GTT21GTC (2467T>C), TCT24TCC (2476T>C), TCG26TCA (2482G>A), CTG29CTA (2491G>A), CTG31CTT (2497G>T), TTG33TTA (2503G>A), AGA38AGG (2518A>G), CCT39CCC (2521T>C), AAG40AGG (2523A>G), AGA43AGG (2533A>G), TTG45TTA (2539G>A), TTA47TTG (2545A>G), GAG49GAA (2551G>A), CCT50CCA (2554T>A), CTG52TTG (2558C>T), TTG53CTC (2561T>C 2563G>C), TCA56TCG (2572A>G), CCT60CCC (2584T>C), GGC61GGT (2587C>T), CTG63CTC (2593G>C), GCT65GCC (2599T>C), AAT68AAC (2608T>C), TTT72TTC (2620T>C), GCG75GCA (2629G>A), GTG76GTA (2632G>A), AGG77AAG (2634G>A), TTG78TTA (2638G>A), ATT80ATA (2644T>A), AAA83AAG (2653A>G), GCT87GCC (2665T>C), CTA92TTA (2678C>T), CTA93CTG (2683A>G), GCT95GCC (2689T>C), CTA98CTC (2698A>C), TCA101TCG (2707A>G), GCG102GCA (2710G>A), GAA104GAG (2716A>G), ACA105ACG (2719A>G), CTC106CTT (2722C>T), GTC107ATT (2723G>A 2725C>T), GCA108GCG (2728A>G), AGG110AGA (2734G>A), ATT111ATA (2737T>A), GAT114GAC (2746T>C), GCT115ACA (2747G>A 2749T>A), GCA116GCT (2752A>T), GCT117GCA (2755T>A), GAC119GAT (2761C>T), CTC120CTT (2764C>T), TGT125TGC (2779T>C), GGG127GGT (2785G>T), AAC129AAT (2791C>T), CAT131AAT (2795C>A), TAC134TAT (2806C>T), TTG138CTC (2816T>C 2818G>C), GTG140ATA (2822G>A 2824G>A), TTG141TTA (2827G>A), GTT144GTC (2836T>C), AAG145AAA (2839G>A), AAT146AAC (2842T>C), ACC153ACA (2863C>A), CTA154TTA (2864C>T), CCA160ACA (2882C>A), AGT161CAT (2885A>C 2886G>A), AAC163AAT (2893C>T), ACG164ATG (2895C>T), GTC166GTT (2902C>T), GTA167GTG (2905A>G), ATT169ATC (2911T>C), AAT173AAC (2923T>C), AAT176AGT (2931A>G), AAC179AGC (2940A>G), GCT181GCC (2947T>C), ACT186ACC (2962T>C), GAA189_AAT190insA-- (2971_2972insA), GGT195GGC (2989T>C), ACT196ATT (2991C>T), CAT197CGT (2994A>G), GAG199AGG (2999G>A 3000A>G), TCC204TTC (3015C>T), CGG206CAG (3021G>A), TAC207TTT (3024A>T 3025C>T), ACA211ATA (3036C>T), GAA212GGC (3039A>G 3040A>C), AAG213ACA (3042A>C 3043G>A), TTG216TTA (3052G>A), TTA219TTG (3061A>G), GCG226GCA (3082G>A), GGC229GGT (3091C>T), GCC232GTA (3099C>T 3100C>A), GTG234AGA (3104G>A 3105T>G 3106G>A), GTC235GTT (3109C>T), AAA236AAG (3112A>G), ACT238ACC (3118T>C), GCC239GCT (3121C>T), GGA241GGG (3127A>G), AGC243AAC (3132G>A), CGA245AAG (3137C>A 3138G>A 3139A>G), TAT249TAC (3151T>C), TTC250ATT (3152T>A 3154C>T), TGT254TAT (3165G>A), GTA255GTG (3169A>G), ATC265ATT (3199C>T), AGG267CGG (3203A>C), CGT268CGC (3208T>C), CCA270CCG (3214A>G), ACC271ACT (3217C>T), TTT272CTT (3218T>C), GTC273GTA (3223C>A), ACT274AAT (3225C>A), GTT276ATT (3230G>A), AAC277AAT (3235C>T), TTA285CTA (3257T>C), ACC286ACT (3262C>T), AGC287AGT (3265C>T), GTA288ATG (3266G>A 3268A>G), GTG289GTA (3271G>A), GTG291ATG (3275G>A), GCA294ACA (3284G>A), CAG299CAA (3301G>A), AGA304AAG (3315G>A 3316A>G), GAA305GCA (3318A>C), ATC306ATT (3322C>T), GAG307GAA (3325G>A), TTG308TTA (3328G>A), ATA311ATT (3337A>T), GAG315GAA (3349G>A), GTC317GTT (3355C>T), TGC318TGT (3358C>T), ATG319GTG (3359A>G), CTA321CTG (3367A>G), TAT322TAC (3370T>C), TTA325CTG (3377T>C 3379A>G), GAA330GAG (3394A>G), CGC331CGT (3397C>T), GAT332GAG (3400T>G), ACA334ACG (3406A>G), TAC338TAT (3418C>T), ATT339ATA (3421T>A), TAT341TAC (3427T>C), AAG343AAA (3433G>A), TCA344TCT (3436A>T), GTA345GTG (3439A>G), AAC346AAT (3442C>T), CCT347CCC (3445T>C), CAG351CAA (3457G>A), TTT352TTC (3460T>C), GCT353GCA (3463T>A), GCG355GCT (3469G>T), CAG356CAA (3472G>A), CAC364CAT (3496C>T), CAG365CAA (3499G>A) | | | | | | | | |
| 6K1 protein (NP_734245.1) | 1 | 52 | 100% | 305 | 90.5% | 52 (100%) | 45 (86.5%) | 0/0/0/0 | 0 |
| Protein mutations: | V5G (3513T>G), M16V (3545A>G), A17T (3548G>A), V19I (3554G>A 3556C>T), L42V (3623C>G), L45M (3632C>A), Y47H (3638T>C) | | | | | | | | |
| Codon mutations: | TCC2TCT (3505C>T), GTT5GGT (3513T>G), AAA7AAG (3520A>G), GAA10GAG (3529A>G), CAA11CAG (3532A>G), ATG16GTG (3545A>G), GCT17ACT (3548G>A), TTA18CTA (3551T>C), GTC19ATT (3554G>A 3556C>T), AGT28AGC (3583T>C), GAT29GAC (3586T>C), TGC30TGT (3589C>T), TTC32TTT (3595C>T), AAT36AAC (3607T>C), TTT38TTC (3613T>C), AAG39AAA (3616G>A), GGT40GGC (3619T>C), CTT42GTT (3623C>G), TCC43TCT (3628C>T), CTG45ATG (3632C>A), GAT46GAC (3637T>C), TAT47CAT (3638T>C), GAA48GAG (3643A>G) | | | | | | | | |
| CI protein (NP_734246.1) | 1 | 634 | 100% | 4190 | 97.9% | 634 (100%) | 608 (95.9%) | 0/0/0/0 | 0 |
| Protein mutations: | I15V (3698A>G), S21N (3717G>A), R26K (3731C>A 3732G>A), I140V (4073A>G), F158Y (4128T>A 4129C>T), A166S (4151G>T), V171I (4166G>A), L190I (4223C>A 4225G>A), I222V (4319A>G 4321A>G), F247Y (4395T>A), V251I (4406G>A), M274V (4475A>G 4477G>T), T276S (4481A>T 4483A>G), A297L (4544G>T 4545C>T 4546G>A), R298K (4548G>A), V342I (4679G>A 4681G>C), V381I (4796G>A 4798C>T), T383S (4803C>G 4804T>C), S468G (5057A>G), T469A (5060A>G), L479I (5090C>A), A482V (5100C>T), E484D (5107A>T), V489F (5120G>T 5122C>T), I491V (5126A>G), V510I (5183G>A) | | | | | | | | |
| Codon mutations: | TTA2TTG (3661A>G), GTG5GTA (3670G>A), TTT9TTC (3682T>C), GAG11GAA (3688G>A), GAG14GAA (3697G>A), ATT15GTT (3698A>G), GAT17GAC (3706T>C), TTT18TTC (3709T>C), TTG20CTA (3713T>C 3715G>A), AGT21AAT (3717G>A), GAG22GAA (3721G>A), CGA26AAA (3731C>A 3732G>A), ACT27ACA (3736T>A), CTA31TTG (3746C>T 3748A>G), ACA33ACG (3754A>G), AGT36AGC (3763T>C), CGA41CGG (3778A>G), CAA42CAG (3781A>G), ATC43ATT (3784C>T), CAT47CAC (3796T>C), CCA50CCT (3805A>T), TAC52TAT (3811C>T), AGA53AGG (3814A>G), TTC58TTT (3829C>T), TTT61TTC (3838T>C), AGA63AGG (3844A>G), GCA64GCC (3847A>C), GTT67GTA (3856T>A), GTG69GTT (3862G>T), GAC72GAT (3871C>T), GCC74GCA (3877C>A), AGC76AGT (3883C>T), GAA77GAG (3886A>G), CTA79CTG (3892A>G), GAC80GAT (3895C>T), TTT81TTC (3898T>C), GTA83GTG (3904A>G), CGG84AGA (3905C>A 3907G>A), GCT86GCC (3913T>C), AAG91AAA (3928G>A), TCA92TCT (3931A>T), GGG94GGA (3937G>A), AGT100AGC (3955T>C), GTG101GTA (3958G>A), GCC102GCT (3961C>T), GGA103GGG (3964A>G), TCT104TCC (3967T>C), TTA107TTG (3976A>G), ATT108ATC (3979T>C), CCA110CCT (3985A>T), CTA114CTT (3997A>T), GCG115GCA (4000G>A), AAC117AAT (4006C>T), GTT118GTG (4009T>G), TTC119TTT (4012C>T), AAA120AAG (4015A>G), TCT123TCC (4024T>C), GAA125GAG (4030A>G), CCA126CCT (4033A>T), AAG129AAA (4042G>A), CTG133TTA (4052C>T 4054G>A), CGT134CGC (4057T>C), CGT136CGA (4063T>A), GGA137GGG (4066A>G), ATA140GTA (4073A>G), TTT141TTC (4078T>C), GGC142GGT (4081C>T), TCT143TCC (4084T>C), TCC147TCT (4096C>T), AGC151AGT (4108C>T), GGA152GGT (4111A>T), GCG154GCA (4117G>A), CTA155TTG (4118C>T 4120A>G), TAC157TAT (4126C>T), TTC158TAT (4128T>A 4129C>T), GCC159GCT (4132C>T), AAT160AAC (4135T>C), CAA164CAG (4147A>G), TTA165CTA (4148T>C), GCT166TCT (4151G>T), AAC169AAT (4162C>T), GTA171ATA (4166G>A), ATA172ATC (4171A>C), GAG175GAA (4180G>A), CTG179TTG (4190C>T), CCT181CCC (4198T>C), TCC182TCT (4201C>T), GCG185GCT (4210G>T), TTC186TTT (4213C>T), AGT188AGC (4219T>C), CTG189TTG (4220C>T), CTG190ATA (4223C>A 4225G>A), GTT192GTG (4231T>G), CAT194CAC (4237T>C), GTA199GTG (4252A>G), TTA200CTG (4253T>C 4255A>G), AAA201AAG (4258A>G), ACT205ACC (4270T>C), AGA209AGG (4282A>G), GAG210GAA (4285G>A), GAA212GAG (4291A>G), CAG216CAA (4303G>A), GTC219GTT (4312C>T), ATA222GTG (4319A>G 4321A>G), GTG223GTT (4324G>T), GAG224GAA (4327G>A), CTG227CTT (4336G>T), TCT228TCC (4339T>C), TCA231TCT (4348A>T), GCA235GCG (4360A>G), GGT237GGC (4366T>C), TCT238TCA (4369T>A), ACT240ACC (4375T>C), GTT244GTC (4387T>C), TTT247TAT (4395T>A), GTA251ATA (4406G>A), CTT252CTC (4411T>C), GTT261GTC (4438T>C), ACC263ACA (4444C>A), GCC265GCT (4450C>T), CTC267CTT (4456C>T), GAC270GAT (4465C>T), ATG274GTT (4475A>G 4477G>T), GTC275GTT (4480C>T), ACA276TCG (4481A>T 4483A>G), GGC280GGT (4495C>T), AAG284AAA (4507G>A), CAC285CAT (4510C>T), GGT286GGG (4513T>G), CTA288TTA (4517C>T), GTC291GTT (4528C>T), AAA293AAG (4534A>G), ACC295ACT (4540C>T), GCG297TTA (4544G>T 4545C>T 4546G>A), AGA298AAA (4548G>A), TTT301TTC (4558T>C), GTA303GTT (4564A>T), ACC305ACT (4570C>T), ATA307ATT (4576A>T), TTG314TTA (4597G>A), GAC315GAT (4600C>T), GAC317GAT (4606C>T), GTG318GTA (4609G>A), GTG320GTA (4615G>A), GAC321GAT (4618C>T), TTG324CTT (4625T>C 4627G>T), GAC331GAT (4648C>T), ATT332ATC (4651T>C), AGG335AGA (4660G>A), AGC336AGT (4663C>T), GCT338GCA (4669T>A), GTG342ATC (4679G>A 4681G>C), AGT343AGC (4684T>C), GTT344GTG (4687T>G), GAG348GAA (4699G>A), AGA349AGG (4702A>G), CAA351CAG (4708A>G), AGG352AGA (4711G>A), CTG353TTG (4712C>T), AAA361AAG (4738A>G), GTA363GTT (4744A>T), GCA364GCG (4747A>G), TTG365CTG (4748T>C), ACT370ACC (4765T>C), GTC381ATT (4796G>A 4798C>T), ACT383AGC (4803C>G 4804T>C), GAG384GAA (4807G>A), GCG385GCT (4810G>T), GCT386GCG (4813T>G), CTT387CTC (4816T>C), TAT392TAC (4831T>C), AAC393AAT (4834C>T), CCA395CCT (4840A>T), GGA399GGG (4852A>G), GGC400GGT (4855C>T), AGT404AGC (4867T>C), CTG405CTC (4870G>C), ATT406ATA (4873T>A), TGT409TGC (4882T>C), ACA416ACT (4903A>T), CAG418CAA (4909G>A), TTT420TTC (4915T>C), GAA421GAG (4918A>G), TTG422CTA (4919T>C 4921G>A), AGT423AGC (4924T>C), CCC424CCG (4927C>G), TTT426TTC (4933T>C), ATC427ATA (4936C>A), CAG428CAA (4939G>A), AAT429AAC (4942T>C), TTC430TTT (4945C>T), TCA436TCG (4963A>G), CCT439CCC (4972T>C), CTT445CTC (4990T>C), AAA449AAG (5002A>G), CCT456CCC (5023T>C), TTG457CTG (5024T>C), TGC458TGT (5029C>T), GAT459GAC (5032T>C), TCT461TCC (5038T>C), CCA463CCT (5044A>T), GCA466GCT (5053A>T), TCG467TCA (5056G>A), AGT468GGT (5057A>G), ACT469GCT (5060A>G), TCG472TCT (5071G>T), CTT479ATT (5090C>A), GGA480GGG (5095A>G), GTG481GTT (5098G>T), GCC482GTC (5100C>T), TTA483TTG (5104A>G), GAA484GAT (5107A>T), ATT485ATC (5110T>C), CCA486CCG (5113A>G), AAG487AAA (5116G>A), CAA488CAG (5119A>G), GTC489TTT (5120G>T 5122C>T), ATT491GTT (5126A>G), CAT494CAC (5137T>C), ATC495ATT (5140C>T), AAA496AAG (5143A>G), GAG497GAA (5146G>A), CTC502CTA (5161C>A), GAA504GAG (5167A>G), CTT506CTG (5173T>G), ACG509ACA (5182G>A), GTT510ATT (5183G>A), GTC511GTT (5188C>T), AAG512AAA (5191G>A), AAA514AAG (5197A>G), GAC515GAT (5200C>T), TGC517TGT (5206C>T), TTA518TTG (5209A>G), AGC521AGT (5218C>T), TCG525TCA (5230G>A), ATC527ATT (5236C>T), AGC528AGT (5239C>T), ATC530ATT (5245C>T), ACA536ACT (5263A>T), CTC538CTT (5269C>T), TTC539TTT (5272C>T), GCC540GCA (5275C>A), ATC541ATT (5278C>T), CCA542CCC (5281A>C), AGA543AGG (5284A>G), ACT544ACC (5287T>C), CTA545TTA (5288C>T), ATA546ATT (5293A>T), TTG547CTA (5294T>C 5296G>A), GAG549GAA (5302G>A), CTT552CTA (5311T>A), GAA553GAG (5314A>G), CGA556CGT (5323A>T), AGT564AGC (5347T>C), ATC566ATT (5353C>T), TGC570TGT (5365C>T), GTT577GTC (5386T>C), ACC580ACA (5395C>A), AAA589AAG (5422A>G), GAT590GAC (5425T>C), ACC592ACT (5431C>T), GAG594GAA (5437G>A), CTT599CTC (5452T>C), AAG601AAA (5458G>A), AGT604AGC (5467T>C), CAA605CAG (5470A>G), CTA606TTG (5471C>T 5473A>G), GAA608GAG (5479A>G), TTC609TTT (5482C>T), TCA610TCC (5485A>C), AAT611AAC (5488T>C), TTG612CTA (5489T>C 5491G>A), GGT614GGC (5497T>C), TGT617TGC (5506T>C), GAG618GAA (5509G>A), GAG619GAA (5512G>A), TTA621CTA (5516T>C), TAT625TAC (5530T>C), GAG626GAA (5533G>A), TCG627TCT (5536G>T), TTG628TTA (5539G>A), GTT631GTG (5548T>G), CAC633CAT (5554C>T) | | | | | | | | |
| 6K2 protein (NP_734247.1) | 1 | 52 | 100% | 303 | 93.5% | 52 (100%) | 46 (88.5%) | 0/0/0/0 | 0 |
| Protein mutations: | A2T (5561G>A 5563G>A), I14V (5597A>G), N16K (5605C>A), A21V (5619C>T 5620T>G), I25L (5630A>C), V46I (5693G>A) | | | | | | | | |
| Codon mutations: | GCT1GCA (5560T>A), GCG2ACA (5561G>A 5563G>A), ACG3ACC (5566G>C), TCA4TCG (5569A>G), CTC9CTG (5584C>G), AAG12AAA (5593G>A), GGG13GGA (5596G>A), ATT14GTT (5597A>G), AAC16AAA (5605C>A), GTG20GTT (5617G>T), GCT21GTG (5619C>T 5620T>G), TTG24TTA (5629G>A), ATC25CTC (5630A>C), GGC28GGT (5641C>T), GCA31GCT (5650A>T), ATA35ATT (5662A>T), GGA36GGG (5665A>G), CAA44CAG (5689A>G), GTT46ATT (5693G>A), GAG47GAA (5698G>A), CAA52CAG (5713A>G) | | | | | | | | |
| NIa-VPg protein (NP_734252.1) | 1 | 188 | 100% | 1227 | 93.3% | 188 (100%) | 170 (90.4%) | 0/0/0/0 | 0 |
| Protein mutations: | K47R (5853A>G), I61V (5894A>G 5896C>G), R83A (5960C>G 5961G>C 5962G>C), I93V (5990A>G), E102D (6019A>T), K106T (6030A>C), D112E (6049C>A), M115I (6058G>A), S120N (6072G>A 6073T>C), N121H (6074A>C), T123N (6081C>A), C133S (6111G>C 6112T>C), I139V (6128A>G 6130T>C), V149I (6158G>A), T153S (6170A>T 6172A>G), L164F (6203C>T 6205C>T), V173I (6230G>A), A182K (6257G>A 6258C>A) | | | | | | | | |

| | Begin | End | Coverage | Score | Concordance | Matches | Identities | I/D/M/F* | Stop Codons |
|---|---|---|---|---|---|---|---|---|---|
| **NT** | **52** | **9704** | **99.5%** | **12530** | **65.2%** | **9642 (99.8%)** | **7985 (82.7%)** | **7/11** | |
| Codon mutations: | GGG1GGC (5716G>C), AAA2AAG (5719A>G), GCC10GCA (5743C>A), CGC14CGA (5755C>A), CAT15CAC (5758T>C), CGT17CGC (5764T>C), AAA19AAG (5770A>G), GGC22GGT (5779C>T), TTT23TTC (5782T>C), GAC26GAT (5791C>T), GAC30GAT (5803C>T), AGG41AGA (5836G>A), AAA42AAG (5839A>G), GGA44GGG (5845A>G), AAA45AAG (5848A>G), AAA47AGA (5853A>G), GGT48GGC (5857T>C), ACC49ACT (5860C>T), ACA50ACT (5863A>T), GGT52GGA (5869T>A), GGT54GGC (5875T>C), AGC57AGT (5884C>T), TTC60TTT (5893C>T), ATC61GTG (5894A>G 5896C>G), AAC62AAT (5899C>T), GGG65GGA (5908G>A), TTT66TTC (5911T>C), GAG70GAA (5923G>A), CAA75CAG (5938A>G), TTC76TTT (5941C>T), GTT77GTC (5944T>C), GAT78GAC (5947T>C), GGG82GGA (5959G>A), CGG83GCC (5960C>G 5961G>C 5962G>C), GAA87GAG (5974A>G), GCT91GCA (5986T>A), GAC92GAT (5989C>T), ATT93GTT (5990A>G), GAT95GAC (5998T>C), AGA99CGT (6008A>C 6010A>T), GAA102GAT (6019A>T), GTG103GTT (6022G>T), CGA104CGT (6025A>T), AAA106ACA (6030A>C), GAC112GAA (6049C>A), ATG115ATA (6058G>A), CAA116CAG (6061A>G), GCC117GCA (6064C>A), GGT119GGC (6070T>C), AGT120AAC (6072G>A 6073T>C), AAC121CAC (6074A>C), ACG122ACA (6079G>A), ACC123AAC (6081C>A), ATA124ATT (6085A>T), GCA126GCT (6091A>T), TAC127TAT (6094C>T), TGT133TCC (6111G>C 6112T>C), AAA135AAG (6118A>G), TTG137TTA (6124G>A), AAG138AAA (6127G>A), ATT139GTC (6128A>G 6130T>C), GAT140GAC (6133T>C), TTA141TTG (6136A>G), AAC145AAT (6148C>T), AAA148AAG (6157A>G), GTT149ATT (6158G>A), TGT150TGC (6163T>C), AAA152AAG (6169A>G), ACA153TCG (6170A>T 6172A>G), GGC155GGT (6178C>T), AAA158AAG (6187A>G), TTT159TTC (6190T>C), GAG161GAA (6196G>A), CTC164TTT (6203C>T 6205C>T), GAA165GAG (6208A>G), CTA166TTG (6209C>T 6211A>G), CAG168CAA (6217G>A), GCT172GCA (6229T>A), GTA173ATA (6230G>A), GTC175GTT (6238C>T), AAG178AAA (6247G>A), ATA180ATT (6253A>T), GCA182AAA (6257G>A 6258C>A), GAG184GAA (6265G>A), CAT187CAC (6274T>C) | | | | | | | | |
| NIa-Pro protein (NP_734248.1) | 1 | 244 | 100% | 1654 | 93.7% | 244 (100%) | 223 (91.4%) | 0/0/0/0 | 0 |
| Protein mutations: | K2R (6282A>G), A29T (6362G>A), V42I (6401G>A), A43V (6405C>T), Y51F (6429A>T), Q58R (6450A>G), H70N (6485C>A 6487C>T), L84I (6527C>A), T104I (6588C>T), Y120H (6635T>C), I124V (6647A>G), I134V (6677A>G 6679A>G), D147N (6716G>A), N148D (6719A>G), I163L (6764A>T 6766A>G), A173V (6795C>T 6796A>G), H174Q (6799C>A), N194D (6857A>G), V200T (6875G>A 6876T>C 6877C>A), D219E (6934C>A), V242I (7001G>A 7003G>A) | | | | | | | | |
| Codon mutations: | AAA2AGA (6282A>G), TCG3TCA (6286G>A), GGC7GGT (6298C>T), TTG8TTA (6301G>A), AGA9AGG (6304A>G), GAC10GAT (6307C>T), GCC15GCT (6322C>T), ACA17ACT (6328A>T), GTT18GTG (6331T>G), TGT19TGC (6334T>C), AGG20AGA (6337G>A), CTG21TTG (6338C>T), TAT27TAC (6358T>C), GGG28GGA (6361G>A), GCA29ACA (6362G>A), TCA30TCT (6367A>T), GAG31GAA (6370G>A), TAC33TAT (6376C>T), GGT34GGG (6379T>G), GGA36GGC (6385A>C), GGA38GGT (6391A>T), GCA39GCG (6394A>G), TAC40TAT (6397C>T), GTA42ATA (6401G>A), GCG43GTG (6405C>T), CAC45CAT (6412C>T), TTA47TTG (6418A>G), TTT48TTC (6421T>C), TAC51TTC (6429A>T), GGT53GGA (6436T>A), GAG56GAA (6445G>A), CAA58CGA (6450A>G), TCC59TCG (6454C>G), CAC61CAT (6460C>T), GGT62GGA (6463T>A), TTC64TTT (6469C>T), AGG65AGA (6472G>A), GTG66GTA (6475G>A), AAT68AAC (6481T>C), CAC70AAT (6485C>A 6487C>T), AGT71AGC (6490T>C), AGC73AGT (6496C>T), CTG75CTA (6502G>A), GGT79GGC (6514T>C), AGG80AGA (6517G>A), CTC84ATC (6527C>A), ATC85ATA (6532C>A), CCG88CCA (6541G>A), AAA89AAG (6544A>G), GTC93GTT (6556C>T), TTT94TTC (6559T>C), CAG96CAA (6565G>A), TTG98CTA (6569T>C 6571G>A), CCT103CCA (6586T>A), ACA104ATA (6588C>T), GAA107GAG (6598A>G), AGA108AGG (6601A>G), ATT109ATC (6604T>C), TGT110TGC (6607T>C), GGA113GGG (6616A>G), AAC115AAT (6622C>T), GAG118GAA (6631G>A), AAG119AAA (6634G>A), TAT120CAT (6635T>C), GCT121GCA (6640T>A), TCG123TCT (6646G>T), ATC124GTC (6647A>G), ATC125ATT (6652C>T), AGC129AGT (6664C>T), ACT131ACG (6670T>G), ATA134GTG (6677A>G 6679A>G), GGC136GGT (6685C>T), ACA138ACT (6691A>T), TTC139TTT (6694C>T), ACA146ACG (6715A>G), GAT147AAT (6716G>A), AAT148GAT (6719A>G), GGA149GGG (6724A>G), CTA153TTA (6734C>T), GTG155GTA (6742G>A), GTG156GTA (6745G>A), GCC159GCT (6754C>T), GGA161GGG (6760A>G), ATA163TTG (6764A>T 6766A>G), ATT166ATA (6775T>A), CTG169TTG (6782C>T), GCA170GCT (6787A>T), AAC171AAT (6790C>T), GCA173GTG (6795C>T 6796A>G), CAC174CAA (6799C>A), ACC175ACA (6802C>A), AAC177AAT (6808C>T), TAC178TAT (6811C>T), TAC179TAT (6814C>T), TTC182TTT (6823C>T), GAA184GAG (6829A>G), TAC190TAT (6847C>T), CGA192CGC (6853A>C), AAT194GAT (6857A>G), GAG195GAA (6862G>A), GAA198GAG (6871A>G), GTC200ACA (6875G>A 6876T>C 6877C>A), AAG201AAA (6880G>A), TCT202TCA (6883T>A), GTT204GTG (6889T>G), GAC208GAT (6901C>T), GGC213GGT (6916C>T), CCG214CCA (6919G>A), AAA216AAG (6925A>G), CTT217CTC (6928T>C), GAC219GAA (6934C>A), GGG224GGC (6949G>C), TTC226TTT (6955C>T), AAA227AAG (6958A>G), CAA233CAG (6976A>G), CTA235TTA (6980C>T), ATC236ATT (6985C>T), GTA240GTT (6997A>T), GTG241GTT (7000G>T), GTG242ATA (7001G>A 7003G>A) | | | | | | | | |
| NIb protein (NP_734249.1) | 1 | 521 | 100% | 3129 | 86.1% | 519 (99.4%) | 480 (92.0%) | 1/2/2/2 | 0 |
| Protein mutations: | F8Y (7032T>A), T39K (7125C>A), A46S (7145G>T 7147A>G), E51_A52del (7160_7165delGAGGCA), D69E (7216T>A), D83E (7258T>A), D88_R89insX (7273_7274insT), R89V (7274C>G 7275G>T), H91L (7281A>T), L92S (7284T>C), S96L (7296C>T), S101F (7311C>T), T102I (7314C>T), C103Y (7317G>A), N104K (7321T>A), K109Q (7334A>C), A112N (7343G>A 7344C>A 7345A>C), S133M (7407G>T 7408T>G), C136G (7415T>G), L165S (7503T>C 7504G>A), D234N (7709G>A), K235R (7713A>G 7714A>G), T268I (7812C>T), V279L (7844G>T 7846G>A), V292I (7883G>A), S345N (8043G>A), N381S (8151A>G 8152C>T), R390K (8178G>A), R400K (8208G>A), P429A (8294C>G 8296A>G), S447F (8349C>T), A467S (8408G>T), M481I (8452G>A), R484K (8460G>A), A491T (8480G>A 8482T>A), R498K (8502G>A 8503A>G), E502D (8515A>T), E510D (8539G>C), L513C (8546C>T 8547T>G), S515T (8552T>A) | | | | | | | | |
| Codon mutations: | GCT1GCC (7012T>C), AAG2AAA (7015G>A), CAT3CAC (7018T>C), TTT8TAT (7032T>A), GCC10GCT (7039C>T), TTG11CTA (7040T>C 7042G>A), GGA13GGG (7048A>G), GCT17GCA (7060T>A), GTC18GTA (7063C>A), AGC23AGT (7078C>T), TTA25CTG (7082T>C 7084A>G), GTA26GTG (7087A>G), ACC27ACA (7090C>A), CAT29CAC (7096T>C), GTA30GTG (7099A>G), GTT31GTC (7102T>C), GAG34GAA (7111G>A), CGA36CGG (7117A>G), ACA39AAA (7125C>A), GAA40GAG (7129A>G), TTT41TTC (7132T>C), CTG42TTA (7133C>T 7135G>A), GTG44GTA (7141G>A), GAT45GAC (7144T>C), GCA46TCG (7145G>T 7147A>G), GAG47GAA (7150G>A), GCA50GCT (7159A>T), GAG51_GCA52del (7160_7165delGAGGCA), TTC54TTT (7171C>T), GCG60GCT (7189G>T), AAA63AAG (7198A>G), AGC64AGT (7201C>T), CTA66TTA (7205C>T), AAT67AAC (7210T>C), GAT69GAA (7216T>A), GCG70GCA (7219G>A), ATC72ATA (7225C>A), AAG73AAA (7228G>A), AAG77AAA (7240G>A), TAT78TAC (7243T>C), AAA80AAG (7249A>G), CCT81CCG (7252T>G), GAT83GAA (7258T>A), GTT84GTC (7261T>C), GGT85GGA (7264T>A), GTC86GTG (7267C>G), GTG87GTA (7270G>A), GAT88_CGG89insT-- (7273_7274insT), CGG89GTG (7274C>G 7275G>T), CAT91CTT (7281A>T), TTG92TCG (7284T>C), TCA96TTA (7296C>T), TCT101TTT (7311C>T), ACC102ATC (7314C>T), TGC103TAC (7317G>A), AAT104AAA (7321T>A), GTG105-TG (7322delG), CAC106CAT (7327C>T), GGC107GGT (7330C>T), TTC108TTT (7333C>T), AAG109CAG (7334A>C), TGT111TGC (7342T>C), GCA112AAC (7343G>A 7344C>A 7345A>C), ACT115ACC (7354T>C), CAA118CAG (7363A>G), ATT120ATC (7369T>C), TTC121TTT (7372C>T), GCG123GCT (7378G>T), CTC124CTT (7381C>T), GCA129GCT (7396A>T), GTC130GTT (7399C>T), GCC132GCT (7405C>T), AGT133ATG (7407G>T 7408T>G), GGG135GGT (7414G>T), TGC136GGC (7415T>G), AAA137AAG (7420A>G), GAC140GAT (7429C>T), TAT141TAC (7432T>C), CAT144CAC (7441T>C), AAG150AAA (7459G>A), GAA151GAG (7462A>G), ATA153ATT (7468A>T), GTC154GTA (7471C>A), CAA156CAG (7477A>G), AGC157AGT (7480C>T), CTG159CTT (7486G>T), TAT162TAC (7495T>C), AAA163AAG (7498A>G), GGT164GGC (7501T>C), TTG165TCA (7503T>C 7504G>A), ATT168ATA (7513T>A), AAC170AAT (7519C>T), GGA171GGG (7522A>G), AAG174AAA (7531G>A), GAG176GAA (7537G>A), CTC177CTT (7540C>T), TGT179TGC (7546T>C), AAG187AAA (7570G>A), ACG188ACA (7573G>A), ACG190ACA (7579G>A), GCT193GCC (7588T>C), CTA196TTA (7595C>T), TTG199CTA (7604T>C 7606G>A), GGT202GGA (7615T>A), TGT205TGC (7624T>C), GAC208GAT (7633C>T), TTC209TTT (7636C>T), AAT210AAC (7639T>C), TTT213TTC (7648T>C), GAA219GAG (7666A>G), TGT221TGC (7672T>C), ACA223ACT (7678A>T), GGG225GGA (7684G>A), ACT227ACC (7690T>C), TTT229TTC (7696T>C), GGT231GGG (7702T>G), GAT234AAT (7709G>A), AAA235AGG (7713A>G 7714A>G), CTG236TTG (7715C>T), CGT239CGC (7726T>C), GAG242GAA (7735G>A), GTA245GTG (7744A>G), TAC246TAT (7747C>T), GGC251GGT (7762C>T), CAG253CAA (7768G>A), TTT254TTC (7771T>C), TCA257TCG (7780A>G), CTA258CTC (7783A>C), ACT259ACC (7786T>C), CTA262TTG (7793C>T 7795A>G), ATC263ATT (7798C>T), GCT265GCC (7804T>C), ACC268ATC (7812C>T), GAA275GAG (7834A>G), GAC276GAT (7837C>T), GTG279TTA (7844G>T 7846G>A), CAG282CAA (7855G>A), CTG284CTA (7861G>A), TTA287TTG (7870A>G), TAC288TAT (7873C>T), ACT289ACA (7876T>A), GAG290GAA (7879G>A), ATT291ATA (7882T>A), GTT292ATT (7883G>A), CCA295CCT (7894A>T), ATT296ATC (7897T>C), GTC304GTT (7921C>T), AAG306AAA (7927G>A), GGT309GGC (7936T>C), AAT310AAC (7939T>C), AAT311AAC (7942T>C), GTG319GTA (7966G>A), GAT320GAC (7969T>C), TCT322TCC (7975T>C), GTC326GTT (7987C>T), CTT327CTC (7990T>C), GCT332GCC (8005T>C), GAA341GAG (8032A>G), ATC343ATT (8038C>T), GAC344GAT (8041C>T), AGC345AAC (8043G>A), TTG357TTA (8080G>A), GTG360GTA (8089G>A), CCG362CCA (8095G>A), GAG363GAA (8098G>A), AGC366AGT (8107C>T), ATT367ATC (8110T>C), GAT369GAC (8116T>C), CAA373CAG (8128A>G), GGT379GGC (8146T>C), AAC381AGT (8151A>G 8152C>T), TCG385TCA (8164G>A), AGG390AAG (8178G>A), AAG391AAA (8182G>A), TTG394TTA (8191G>A), TTC396TTT (8197C>T), TCC398TCT (8203C>T), AGA400AAA (8208G>A), CTG402TTA (8213C>T 8215G>A), CTA403CTG (8218A>G), ATC404ATT (8221C>T), GAG405GAA (8224G>A), CCA410CCG (8239A>G), GAG415GAA (8254G>A), ATT417ATC (8260T>C), GTA418GTG (8263A>G), GAT424GAC (8281T>C), AGA425AGG (8284A>G), CCA429GCG (8294C>G 8296A>G), GAG430GAA (8299G>A), CAC431CAT (8302C>T), ATT436ATC (8317T>C), TGC437TGT (8320C>T), GCT439GCA (8326T>A), GAG442GAA (8335G>A), TCC443TCT (8338C>T), TCT447TTT (8349C>T), GAA448GAG (8353A>G), CAC451CAT (8362C>T), CAA452CAG (8365A>G), ATC453ATT (8368C>T), TAC457TAT (8380C>T), TCA458TCG (8383A>G), TTA460TTG (8389A>G), CCT465CCC (8404T>C), GCA467TCA (8408G>T), ACA468ACG (8413A>G), GCG470GCT (8419G>T), GCT475GCC (8434T>C), TAT477TAC (8440T>C), ATG481ATA (8452G>A), CTA483CTG (8458A>G), AGG484AAG (8460G>A), AAA485AAG (8464A>G), CTG486CTA (8467G>A), AGG490AGA (8479G>A), GCT491ACA (8480G>A 8482T>A), GAG496GAA (8497G>A), CTA497TTG (8498C>T 8500A>G), AGA498AAG (8502G>A 8503A>G), GCC499GCT (8506C>T), GAA502GAT (8515A>T), GTC505GTT (8524C>T), GCA506GCC (8527A>C), TTA507CTG (8528T>C 8530A>G), GAG510GAC (8539G>C), CTT513TGT (8546C>T 8547T>G), TCT515ACT (8552T>A), GTA518GTG (8563A>G) | | | | | | | | |
| coat protein (NP_734250.1) | 1 | 267 | 100% | 1652 | 90.6% | 267 (100%) | 243 (91.0%) | 0/0/0/0 | 0 |
| Protein mutations: | A1G (8574C>G), I5V (8585A>G), G9E (8598G>A), N11S (8603A>T 8604A>C 8605C>T), P17Q (8622C>A), E18G (8625A>G 8626G>A), P26F (8648C>T 8649C>T 8650G>C), G29E (8658G>A), D31A (8664A>C 8665T>G), A36V (8679C>T 8680A>T), T58K (8745C>A 8746A>G), E95G (8856A>G 8857G>A), R98Q (8865G>A), M99L (8867A>C 8869G>T), G104E (8883G>A), T106S (8888A>T), V128I (8954G>A), N138D (8984A>G 8986T>C), E139V (8988A>T), I187V (9131A>G), M193G (9149A>G 9150T>G 9151G>A), G194S (9152G>A), P230S (9260C>T 9262T>C), Q242H (9298A>C) | | | | | | | | |
| Codon mutations: | GCA1GGA (8574C>G), ATT5GTT (8585A>G), GGA9GAA (8598G>A), AGC10AGT (8602C>T), AAC11TCT (8603A>T 8604A>C 8605C>T), AAA16AAG (8620A>G), CCA17CAA (8622C>A), GAG18GGA (8625A>G 8626G>A), CAG19CAA (8629G>A), GGC20GGT (8632C>T), ATC22ATT (8638C>T), CAG23CAA (8641G>A), AAC25AAT (8647C>T), CCG26TTC (8648C>T 8649C>T 8650G>C), AAC27AAT (8653C>T), AAA28AAG (8656A>G), GGA29GAA (8658G>A), GAT31GCG (8664A>C 8665T>G), GTT34GTG (8674T>G), GCA36GTT (8679C>T 8680A>T), GGC37GGA (8683C>A), GGG40GGA (8692G>A), ACA41ACT (8695A>T), CCG45CCA (8707G>A), AGA46CGA (8708A>C), AAG48AAA (8716G>A), ATC50ATT (8722C>T), ACG51ACA (8725G>A), AAA53AAG (8731A>G), AGA55AGG (8737A>G), CCC57CCT (8743C>T), ACA58AAG (8745C>A 8746A>G), AAG60AAA (8752G>A), GGA61GGT (8755A>T), AAC66AAT (8770C>T), TTA67CTA (8771T>C), GAA68GAG (8776A>G), CAT69CAC (8779T>C), TTG70TTA (8782G>A), TAT73TAC (8791T>C), CCA75CCG (8797A>G), CAA76CAG (8800A>G), CAA77CAG (8803A>G), GAT79GAC (8809T>C), ATT80ATC (8812T>C), AAT82AAC (8818T>C), CGG84CGA (8824G>A), GCA85GCG (8827A>G), ACG92ACA (8848G>A), TAT94TAC (8854T>C), GAG95GGA (8856A>G 8857G>A), GTG97GTA (8863G>A), CGG98CAG (8865G>A), ATG99CTT (8867A>C 8869G>T), GCA100GCG (8872A>G), GGA104GAA (8883G>A), ACT106TCT (8888A>T), TGC119TGT (8929C>T), GTC128ATC (8954G>A), AAC129AAT (8959C>T), GGA130GGT (8962A>T), GGG137GGA (8983G>A), AAT138GAC (8984A>G 8986T>C), GAA139GTA (8988A>T), GAG142GAA (8998G>A), CCG144CCA (9004G>A), TTG145TTA (9007G>A), ATC148ATT (9016C>T), ACC155ACA (9037C>A), ATA171ATT (9085A>T), AAC175AAT (9097C>T), ATT187GTT (9131A>G), CGA188CGT (9136A>T), CGG191CGT (9145G>T), ATG193GGA (9149A>G 9150T>G 9151G>A), GGT194AGT (9152G>A), TTA195TTG (9157A>G), GCG196GCT (9160G>T), CGT197CGC (9163T>C), GAG204GAA (9184G>A), CGA208CGG (9196A>G), AGG214AGA (9214G>A), GAA215GAG (9217A>G), GCG216GCA (9220G>A), GCA224GCT (9244A>T), TTG225TTA (9247G>A), GCC228GCT (9256C>T), CCT230TCC (9260C>T 9262T>C), GGG234GGA (9274G>A), GAC236GAT (9280C>T), ATC239ATT (9289C>T), CAA242CAC (9298A>C), GAG244GAA (9304G>A), GTC254GTT (9334C>T), GTC264GTG (9364C>G) | | | | | | | | |

| | Begin | End | Coverage | Score | Concordance | Matches | Identities | I/D/M/F* | Stop Codons |
|---|---|---|---|---|---|---|---|---|---|
| **NT** | **52** | **9704** | **99.5%** | **12530** | **65.2%** | **9642 (99.8%)** | **7985 (82.7%)** | **7/11** | |
| PIPO (YP_006393460.1) | 1 | 75 | 100% | 230 | 43.7% | 75 (98.7%) | 55 (72.4%) | 1/0/2/1 | 2 |
| Protein mutations: | R17_K17insX (2971_2972insA), Y23H (2989T>C 2991C>T), R26K (2999G>A 3000A>G), H35Y (3025C>T), K40H (3040A>C 3042A>C), G41S (3043G>A), E44K (3052G>A), I47V (3061A>G), V54I (3082G>A), P57S (3091C>T), P60T (3100C>A), G61E (3104G>A 3105T>G), G62S (3106G>A), Q63* (3109C>T), R64G (3112A>G), C66R (3118T>C), L67F (3121C>T), I69V (3127A>G), A72E (3137C>A 3138G>A), I73V (3139A>G) | | | | | | | | |
| Codon mutations: | TTA2CTA (2923T>C), AAA4AAG (2931A>G), GAA7GAG (2940A>G), TTG10CTG (2947T>C), TTG15CTG (2962T>C), AGA17_AAA17insAA- (2971_2972insA), TAC23CAT (2989T>C 2991C>T), TCA24TCG (2994A>G), AGA26AAG (2999G>A 3000A>G), ATC31ATT (3015C>T), TCG33TCA (3021G>A), GTA34GTT (3024A>T), CAT35TAT (3025C>T), CAC38CAT (3036C>T), AGA39AGG (3039A>G), AAA40CAC (3040A>C 3042A>C), GGC41AGC (3043G>A), GAA44AAA (3052G>A), ATA47GTA (3061A>G), GTT54ATT (3082G>A), CCG57TCG (3091C>T), CGC59CGT (3099C>T), CCA60ACA (3100C>A), GGT61GAG (3104G>A 3105T>G), GGT62AGT (3106G>A), CAA63TAA (3109C>T), AGG64GGG (3112A>G), TGC66CGC (3118T>C), CTC67TTC (3121C>T), ATT69GTT (3127A>G), GAG70GAA (3132G>A), GCG72GAA (3137C>A 3138G>A), ATT73GTT (3139A>G) | | | | | | | | |

*: Inserts / Deletes / Misaligned / Frameshifts

#### Analysis details

This analysis was performed with panviral2.64

### NGS Details (UN8): Torradovirus lycopersici (segment RNA 1)

#### Assembly

| | |
|---|---|
| **Coverage Length** | 6786 (1 contig(s)) |
| **Depth Of Coverage** | 1527.7 |
| **Number Of Reads** | 78839 |
| **Reads Per Million** | 1317.60 rpm (after QC) |
| **Ambiguities** | 0 |
| **Assembly Method** | de novo + reference guided assembly |
| **Consensus Caller** | Bcf Tools |

#### Coverage Map

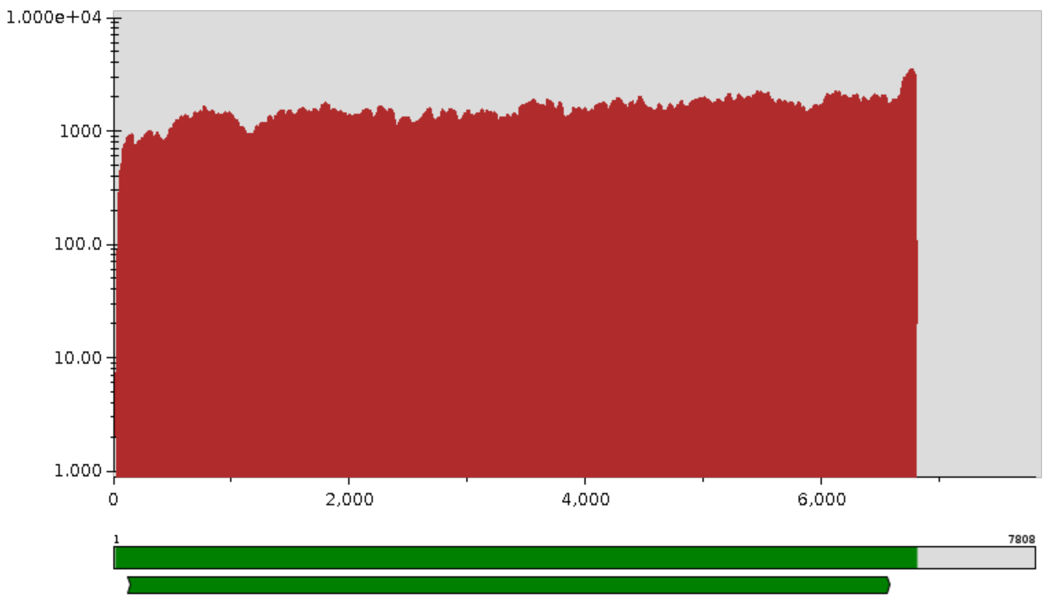

#### Assignment

| | |
|---|---|
| **Type** | Torradovirus lycopersici (Taxonomy ID: 3048378) |
| **Reference Genome** | NC_009013.1 |
| **NT Identity (%)** | 78.5609 |
| **AA Identity (%)** | 89.9027 |
| **Number Of Stop Codons** | 1 |
| **Number Of CDS** | 1 |

#### Alignment

| | |
|---|---|
| **Alignment Score** | 7777.0 (NT) + 13837.0 (AA) = 21614.0 |
| **Concordance (%)** | 75.4916 |

| Alignment Method | Global, seeded, nucleotide + amino acids (AGA) |
|---|---|

#### Genome Region

Sequence starts at position 21 and ends at position 6806 relative to NC_009013.1 reference sequence.

#### Alignment Detailed Statistics

| | Begin | End | Coverage | Score | Concordance | Matches | Identities | I/D/M/F* | Stop Codons |
|---|---|---|---|---|---|---|---|---|---|
| **NT** | **21** | **6806** | **86.9%** | **7777** | **57.4%** | **6785 (99.6%)** | **5350 (78.5%)** | **25/1** | |
| Mutations: | 65A>G, 72C>T, 77A>T, 78A>G, 83C>T, 83_84insTACTTTATATTGTTTTGTGCT, 88T>C, 90C>G, 92T>C, 93T>A, 101G>A, 102G>C, 103C>A, 104A>G, 115T>C, 118C>T, 121G>A, 122A>T, 124G>A, 128C>T, 130C>T, 134T>C, 135T>C, 136C>T, 138A>G, 140T>C, 142A>C, 144T>C, 145T>A, 147C>T, 148T>G, 151A>G, 155T>A, 157C>T, 159C>T, 161A>G, 178T>G, 187A>T, 196T>A, 197G>C, 198C>A, 199T>A, 208T>C, 211T>C, 214C>T, 220C>T, 223T>A, 226G>T, 247A>C, 253A>G, 256T>C, 262T>A, 265C>T, 271A>G, 275T>C, 280G>A, 283T>C, 286A>G, 289T>G, 292A>T, 301T>A, 302T>A, 307C>T, 310T>C, 311A>T, 313C>T, 315G>A, 316C>T, 322T>C, 324A>G, 325C>T, 328T>A, 334T>C, 337A>G, 340G>A, 343A>G, 349G>A, 358T>A, 359T>C, 364T>A, 367A>G, 370C>T, 373C>T, 376C>A, 382T>A, 394C>T, 397A>C, 400C>T, 409A>G, 412A>C, 415T>C, 427T>G, 433G>A, 439T>C, 440G>A, 443A>G, 451A>G, 457G>A, 473A>G, 485G>A, 487C>T, 493T>C, 496G>A, 502G>T, 526T>C, 553T>A, 559C>T, 568A>T, 573G>A, 578C>T, 586A>T, 595G>A, 599A>T, 601T>A, 604C>T, 606T>C, 607A>T, 615A>G, 618C>T, 619A>T, 628C>T, 640T>C, 650T>C, 652A>G, 655C>G, 659T>C, 661A>G, 664G>C, 673C>T, 676A>G, 694A>G, 707C>T, 715G>A, 718A>G, 719T>G, 721C>T, 722A>G, 727A>T, 730C>T, 739T>G, 748G>T, 751A>T, 754A>G, 758C>T, 760C>A, 763A>T, 766C>T, 772G>C, 775T>A, 778T>A, 781C>A, 790C>T, 793G>A, 796T>A, 799A>T, 802T>G, 805T>C, 806A>C, 808A>T, 817T>A, 820A>C, 821T>A, 826C>T, 829G>A, 832A>C, 835A>C, 838A>G, 842C>T, 844G>A, 847C>T, 853G>A, 862T>A, 865T>C, 877T>C, 881T>C, 883G>T, 886T>C, 889C>A, 895C>T, 898G>A, 901T>C, 913T>C, 949A>T, 958T>A, 964T>A, 967T>C, 976A>T, 977G>A, 979T>A, 988T>A, 991T>C, 1001T>C, 1003G>T, 1004T>G, 1006T>A, 1009G>A, 1018C>T, 1019A>G, 1020C>T, 1033G>A, 1054A>C, 1060T>C, 1063C>A, 1064T>C, 1066G>T, 1072T>C, 1078A>G, 1081C>T, 1093T>C, 1096C>A, 1109C>T, 1111A>G, 1129A>G, 1135T>A, 1141C>T, 1147T>C, 1159A>G, 1165C>A, 1180G>A, 1183A>T, 1186G>A, 1189A>C, 1192A>C, 1195A>T, 1196A>T, 1198C>T, 1204T>G, 1207T>G, 1219G>C, 1225C>T, 1237T>A, 1240C>T, 1243G>T, 1246C>T, 1249A>G, 1255G>A, 1258A>T, 1264T>C, 1285T>A, 1291T>C, 1300G>A, 1309C>A, 1318G>A, 1319C>T, 1321G>A, 1327C>T, 1336T>G, 1337T>C, 1339G>T, 1343C>A, 1345C>G, 1351A>T, 1354G>A, 1357A>C, 1361T>C, 1363G>T, 1367A>C, 1372C>G, 1378T>C, 1381G>A, 1390A>T, 1405A>C, 1408T>C, 1411T>C, 1426T>A, 1432G>T, 1450T>A, 1451C>T, 1462G>T, 1465A>T, 1471A>G, 1480C>T, 1481A>G, 1486G>A, 1495A>T, 1501C>T, 1507C>A, 1510T>A, 1516A>T, 1522C>A, 1525G>A, 1528C>A, 1534A>T, 1553C>A, 1555C>G, 1568C>A, 1570A>G, 1573T>C, 1576A>T, 1582C>T, 1585A>C, 1588C>A, 1591C>T, 1600A>C, 1607G>T, 1609T>A, 1612G>A, 1618T>G, 1624C>A, 1643A>C, 1645A>G, 1649C>A, 1657T>G, 1660T>G, 1663G>A, 1669C>T, 1681T>A, 1684G>A, 1687T>C, 1691A>G, 1692C>T, 1693T>A, 1699T>A, 1702C>T, 1703A>G, 1705T>A, 1706T>G, 1708T>C, 1714T>G, 1715G>A, 1716A>G, 1726G>C, 1729T>G, 1732G>A, 1735C>T, 1737A>G, 1738A>G, 1739A>C, 1741A>T, 1744C>A, 1755C>T, 1756A>C, 1762G>A, 1769A>G, 1770G>A, 1771C>T, 1777T>C, 1780T>C, 1781T>C, 1783G>A, 1786G>T, 1810G>A, 1813T>G, 1819A>C, 1825C>A, 1828A>T, 1831A>C, 1837G>C, 1839A>T, 1844G>A, 1849A>C, 1850T>A, 1852A>C, 1858G>A, 1876G>C, 1877T>A, 1879A>T, 1885G>A, 1889A>G, 1891A>G, 1894A>T, 1906T>A, 1907A>C, 1908T>C, 1909G>A, 1911T>G, 1915T>C, 1918C>T, 1921A>G, 1924T>C, 1927C>A, 1936T>C, 1937G>A, 1939T>A, 1942G>A, 1945T>G, 1948C>G, 1951G>A, 1957T>G, 1960A>C, 1978G>A, 1984C>T, 1994T>C, 1996G>A, 2002G>T, 2005A>G, 2008T>C, 2014A>G, 2016A>T, 2020T>C, 2023T>C, 2044A>G, 2050T>C, 2053C>T, 2056T>A, 2062C>T, 2071A>G, 2082T>A, 2092A>T, 2099T>C, 2101C>T, 2110A>G, 2111A>T, 2113A>C, 2119C>T, 2120A>C, 2128T>C, 2130A>C, 2131T>A, 2134C>T, 2140G>C, 2141T>C, 2143T>A, 2146A>G, 2149T>A, 2155T>C, 2158T>C, 2160A>C, 2164C>T, 2176G>C, 2177A>G, 2179A>T, 2182G>A, 2188C>T, 2194T>A, 2200T>C, 2224C>A, 2227C>T, 2230T>G, 2240A>T, 2241A>G, 2245G>T, 2248T>C, 2250G>A, 2251C>A, 2257C>T, 2260T>G, 2263G>A, 2270C>T, 2272G>A, 2281A>C, 2282A>T, 2287G>A, 2290C>T, 2296C>T, 2299A>T, 2305T>A, 2311G>T, 2314C>T, 2318C>T, 2320G>A, 2323A>T, 2326C>T, 2327G>A, 2329G>C, 2332G>A, 2335G>A, 2341T>A, 2344C>A, 2347C>T, 2350A>G, 2356C>T, 2359G>A, 2365A>T, 2368A>T, 2371T>C, 2375A>C, 2377G>T, 2380T>C, 2387C>A, 2389C>A, 2392A>G, 2404G>A, 2407T>C, 2409G>A, 2410G>A, 2416C>T, 2476A>G, 2479G>A, 2485C>T, 2488G>A, 2491A>G, 2494T>G, 2497G>T, 2500T>C, 2503G>A, 2506T>A, 2509C>G, 2510C>T, 2512T>G, 2515T>C, 2516T>C, 2524A>C, 2536A>T, 2542G>T, 2554A>T, 2557C>T, 2563T>G, 2571A>G, 2572A>G, 2575C>A, 2587T>C, 2600A>C, 2602A>G, 2611T>C, 2614C>A, 2617C>T, 2626T>C, 2629C>T, 2639C>A, 2650G>A, 2662C>T, 2677G>A, 2678C>T, 2683A>G, 2692G>A, 2695T>A, 2704G>T, 2710A>G, 2716A>C, 2726G>A, 2731C>T, 2743A>C, 2744T>C, 2761C>G, 2765T>C, 2767G>C, 2768A>C, 2776T>G, 2788C>T, 2800C>A, 2806C>T, 2809C>T, 2812G>T, 2815C>A, 2819G>A, 2821G>A, 2827A>T, 2830G>A, 2842T>C, 2845A>C, 2848C>T, 2852C>A, 2854T>A, 2860C>G, 2863A>T, 2864A>G, 2865C>T, 2866T>G, 2869C>T, 2875A>T, 2878A>C, 2882G>T, 2884C>G, 2890A>C, 2908C>T, 2911T>C, 2926G>T, 2929C>T, 2930T>C, 2932G>A, 2935G>A, 2941A>G, 2947T>A, 2950C>G, 2953G>T, 2956C>G, 2959T>C, 2965A>T, 2971C>T, 2974T>C, 2980C>G, 2986T>G, 2992T>C, 2998A>G, 3011C>A, 3025T>C, 3031T>A, 3034G>A, 3037C>A, 3038A>T, 3046G>T, 3049A>C, 3052T>C, 3064A>T, 3073G>A, 3079T>C, 3083C>A, 3085A>G, 3101T>C, 3103G>T, 3115G>T, 3124C>A, 3136T>G, 3146A>T, 3148G>C, 3160C>T, 3166T>C, 3169C>A, 3178G>A, 3184T>A, 3196T>C, 3202T>C, 3226T>C, 3238T>G, 3239C>T, 3241T>A, 3242C>T, 3244T>G, 3247G>A, 3253C>T, 3256G>A, 3265A>G, 3274C>T, 3277T>A, 3286T>A, 3298T>A, 3310A>G, 3319C>T, 3325G>C, 3328T>C, 3331A>T, 3334A>C, 3346A>G, 3349T>G, 3352T>C, 3356C>A, 3365T>C, 3371G>A, 3373T>C, 3374C>T, 3376T>G, 3379G>A, 3382T>A, 3385T>G, 3388G>T, 3394A>G, 3400A>C, 3406A>G, 3412C>T, 3413C>G, 3416C>T, 3418C>G, 3421T>G, 3428C>T, 3430A>G, 3431C>A, 3439G>A, 3445G>T, 3457A>G, 3466G>A, 3474T>A, 3475T>C, 3476G>C, 3481C>A, 3484A>T, 3492C>A, 3505C>A, 3508G>T, 3514C>A, 3520A>G, 3523A>C, 3526G>A, 3529G>C, 3530C>A, 3532A>G, 3534G>A, 3541C>T, 3542T>C, 3544G>C, 3553T>A, 3559G>A, 3565A>T, 3580T>C, 3583C>A, 3586T>C, 3589A>G, 3591C>T, 3592T>C, 3593T>C, 3594C>A, 3602C>A, 3604T>A, 3607G>A, 3610A>G, 3613T>C, 3622G>A, 3625C>T, 3628G>C, 3629G>T, 3637A>C, 3640A>G, 3643A>G, 3646C>T, 3652A>G, 3658A>C, 3661A>G, 3664T>C, 3673G>A, 3676C>T, 3679T>C, 3682G>C, 3684G>A, 3685G>A, 3686C>T, 3688A>G, 3691C>T, 3693A>T, 3700G>A, 3701C>T, 3703A>G, 3706T>G, 3709A>G, 3710G>T, 3712A>T, 3713A>G, 3715T>G, 3716T>C, 3722G>A, 3724G>A, 3729A>G, 3733C>G, 3742T>G, 3752A>G, 3758G>A, 3763A>T, 3766G>A, 3776G>A, 3778G>T, 3781G>T, 3782C>T, 3784C>G, 3790T>A, 3793T>C, 3802T>C, 3808T>C, 3814C>T, 3817G>A, 3824T>C, 3841G>T, 3847G>A, 3850C>T, 3856G>A, 3868C>T, 3871T>C, 3883G>A, 3889G>A, 3895A>C, 3901A>C, 3907A>T, 3910T>C, 3911C>A, 3916C>G, 3920C>T, 3922C>G, 3925A>G, 3928G>A, 3934T>C, 3949C>T, 3955G>A, 3958T>A, 3959T>C, 3961G>A, 3973T>C, 3982A>T, 3995C>T, 4000C>T, 4003C>A, 4006C>A, 4009A>G, 4015C>A, 4018A>G, 4028T>C, 4030G>A, 4033G>T, 4034T>C, 4039A>G, 4045T>G, 4051G>C, 4055C>A, 4063C>A, 4067T>C, 4069G>A, 4075A>G, 4078T>A, 4081T>A, 4084C>T, 4093C>T, 4096T>C, 4118C>T, 4123T>C, 4129C>T, 4147C>G, 4157C>T, 4160C>G, 4162A>T, 4189T>A, 4192T>G, 4195A>G, 4202C>T, 4204G>A, 4205T>C, 4207G>A, 4210T>A, 4213T>C, 4214G>A, 4219C>T, 4220A>C, 4222A>G, 4225A>T, 4228C>T, 4237A>G, 4246T>C, 4249T>A, 4252C>T, 4255T>C, 4282C>T, 4288C>A, 4294A>G, 4303T>C, 4309T>G, 4318C>T, 4327A>C, 4335C>A, 4336T>A, 4339T>C, 4342T>C, 4348C>T, 4351C>A, 4354T>C, 4358A>C, 4360A>C, 4363T>C, 4366G>A, 4367C>T, 4369T>G, 4372T>G, 4381T>C, 4384A>T, 4387C>T, 4396T>C, 4399T>C, 4405G>A, 4414T>G, 4417T>C, 4418A>C, 4420G>C, 4421T>C, 4426G>T, 4429A>G, 4432T>C, 4438T>A, 4441A>T, 4444A>G, 4450A>T, 4453C>T, 4456C>T, 4459T>C, 4462C>T, 4465C>T, 4468C>T, 4495T>G, 4501A>T, 4507T>C, 4510T>A, 4513A>G, 4514T>C, 4522A>G, 4528T>C, 4534A>T, 4537T>A, 4540A>G, 4546A>T, 4549G>C, 4552C>T, 4555A>G, 4558C>A, 4570C>T, 4576A>T, 4582G>T, 4585A>T, 4588C>A, 4600C>T, 4606A>T, 4615T>A, 4618C>A, 4623A>G, 4624A>T, 4625T>C, 4627G>C, 4636T>C, 4637A>T, 4639T>G, 4648T>A, 4651T>A, 4660A>G, 4661T>C, 4663G>A, 4678T>C, 4684A>C, 4696C>A, 4702G>A, 4705C>T, 4714T>C, 4716G>A, 4717C>T, 4718A>C, 4719G>A, 4732A>G, 4735C>T, 4736T>C, 4738G>T, 4741T>A, 4745C>T, 4747A>G, 4753T>C, 4756A>T, 4759C>T, 4762A>G, 4765T>C, 4768T>C, 4771G>A, 4774G>A, 4786A>G, 4789C>T, 4801T>C, 4802C>T, 4813A>G, 4816T>C, 4825A>G, 4828A>C, 4834C>T, 4840A>T, 4843G>A, 4846T>C, 4850G>A, 4851T>C, 4867T>C, 4868C>T, 4870C>G, 4879G>A, 4885C>T, 4888A>T, 4891C>T, 4894G>T, 4897A>C, 4898G>A, 4903A>G, 4909A>C, 4915C>T, 4921G>T, 4927G>A, 4930T>G, 4936A>C, 4945G>T, 4947C>A, 4948A>T, 4954T>G, 4958A>C, 4963T>C, 4969G>A, 4975G>A, 4981C>T, 4984A>T, 4987C>T, 4988A>G, 4990C>A, 4999T>G, 5002A>T, 5005T>C, 5017G>A, 5020G>A, 5023C>T, 5029G>T, 5030G>A, 5035G>A, 5038T>A, 5041A>T, 5044A>C, 5050A>T, 5059G>T, 5062C>T, 5072C>T, 5074T>A, 5079T>C, 5080T>A, 5083A>T, 5089A>C, 5090G>T, 5092T>A, 5098G>T, 5101T>A, 5102T>C, 5105C>G, 5107C>T, 5110A>T, 5113C>T, 5114A>G, 5116T>A, 5119T>C, 5120G>A, 5122T>A, 5123C>T, 5131T>C, 5132T>A, 5141G>A, 5143G>T, 5146T>C, 5149T>A, 5152T>C, 5155T>A, 5159C>A, 5173C>T, 5176C>T, 5177T>G, 5179G>C, 5180C>A, 5182A>G, 5185A>T, 5194A>C, 5197G>T, 5200G>A, 5206C>T, 5207A>T, 5209C>A, 5212C>A, 5218T>C, 5227C>T, 5233T>C, 5239A>C, 5242G>A, 5257A>T, 5260A>G, 5263G>A, 5266C>A, 5269C>A, 5276A>C, 5278G>A, 5290G>T, 5293T>C, 5296T>C, 5302T>A, 5314T>C, 5326C>T, 5329C>T, 5336A>G, 5341T>C, 5342G>A, 5344A>C, 5347C>A, 5353G>A, 5357A>T, 5359C>T, 5360A>C, 5362A>C, 5366C>A, 5368T>C, 5369G>A, 5371G>T, 5374T>A, 5377T>C, 5386C>G, 5389T>C, 5392G>A, 5399A>G, 5401T>G, 5404T>A, 5407A>T, 5410G>T, 5413C>T, 5416T>A, 5419A>T, 5420T>C, 5422G>A, 5429C>T, 5431G>A, 5434C>T, 5437T>G, 5440C>A, 5443A>G, 5446T>C, 5449T>A, 5452T>C, 5459G>A, 5461T>C, 5462G>A, 5467G>C, 5479C>T, 5482A>T, 5485A>T, 5486G>A, 5491T>G, 5492A>C, 5494A>T, 5498C>T, 5500C>G, 5503T>C, 5506A>T, 5509T>C, 5512A>T, 5515A>T, 5524C>T, 5527G>A, 5530T>C, 5531C>T, 5533T>G, 5534C>G, 5537A>C, 5543G>T, 5548T>C, 5551A>C, 5558G>C, 5566T>A, 5569A>T, 5575A>G, 5578T>C, 5582A>G, 5584T>G, 5591G>T, 5593C>A, 5602G>A, 5603A>C, 5604C>T, 5608A>G, 5611A>T, 5613A>T, 5617A>T, 5620T>A, 5626A>T, 5632T>C, 5638T>G, 5641T>C, 5656G>A, 5659A>G, 5662G>A, 5664G>A, 5665A>T, 5672C>T, 5673T>G, 5674A>T, 5675C>A, 5676A>G, 5677G>T, 5686G>A, 5687C>A, 5689C>A, 5690T>C, 5692G>T, 5707A>T, 5710T>A, 5711T>C, 5716G>T, 5719A>G, 5734T>C, 5737T>C, 5739G>A, 5743G>A, 5746T>A, 5749G>A, 5751T>G, 5753C>T, 5767G>T, 5773G>C, 5774A>G, 5776A>G, 5779T>C, 5784T>C, 5785T>A, 5786T>C, 5788A>T, 5789G>T, 5791C>G, 5794A>G, 5797A>G, 5799A>G, 5800G>T, 5804T>A, 5805T>C, 5806C>T, 5812T>C, 5815C>G, 5818C>T, 5826A>T, 5830C>T, 5836A>C, 5843G>A, 5845G>A, 5846A>T, 5848G>C, 5849T>C, 5850T>A, 5851G>A, 5857T>G, 5860C>A, 5864C>T, 5866T>G, 5872C>A, 5878G>T, 5881T>C, 5884T>C, 5887C>T, 5890C>T, 5894A>C, 5902G>A, 5907A>T, 5914T>C, 5923T>C, 5929G>A, 5942C>T, 5943A>T, 5944C>T, 5950G>A, 5953A>T, 5955A>G, 5956C>T, 5965G>T, 5968T>G, 5971C>T, 5974T>C, 5975A>C, 5977A>T, 5980G>A, 5983T>C, 5986T>C, 5989C>T, 5990T>G, 5992A>C, 5998A>G, 6002A>G, 6003C>T, 6004C>T, 6009G>A, 6010G>A, 6013C>A, 6016A>G, 6022T>A, 6028C>T, 6031A>T, 6033C>T, 6034A>C, 6035G>T, 6040T>A, 6043G>A, 6044T>A, 6046A>G, 6049G>A, 6050T>C, 6055T>G, 6059G>A, 6062G>A, 6064T>C, 6067C>T, 6070T>C, 6079C>G, 6085T>C, 6088G>A, 6089C>T, 6092C>T, 6094T>A, 6097T>A, 6100T>G, 6103G>A, 6106G>A, 6112A>G, 6121A>T, 6124C>A, 6125A>C, 6129A>G, 6130G>T, 6139T>C, 6142A>G, 6145T>C, 6148T>C, 6150G>C, 6152G>T, 6154C>G, 6160G>A, 6163C>G, 6175G>A, 6177A>G, 6184A>G, 6185A>G, 6186C>A, 6187T>G, 6189A>G, 6193C>T, 6196A>G, 6199C>T, 6202G>C, 6208A>G, 6214T>G, 6215C>G, 6217G>T, 6218A>C, 6221G>T, 6222A>C, 6223A>T, 6225G>A, 6226G>A, 6227A>G, 6229C>G, 6230A>G, 6232A>G, 6235G>A, 6237A>G, 6238A>T, 6239G>A, 6240T>C, 6241G>A, 6242A>G, 6244A>G, 6251T>C, 6253A>C, 6256C>A, 6262A>T, 6263C>T, 6266T>A, 6268G>T, 6273G>T, 6274C>G, 6283G>A, 6287C>T, 6289T>G, 6291T>A, 6292T>A, 6294A>G, 6296G>C, 6298A>C, 6301T>A, 6302T>A, 6303T>C, 6304A>G, 6307C>A, 6308T>C, 6310G>T, 6311G>T, 6312A>C, 6313T>C, 6316C>T, 6317A>G, 6319A>G, 6322G>A, 6323A>G, 6325C>G, 6328G>A, 6334A>G, 6337T>C, 6344C>T, 6364T>C, 6371G>A, 6373A>T, 6382C>T, 6383A>G, 6385C>G, 6397C>A, 6404T>C, 6405G>A, 6412C>T, 6413C>G, 6414G>T, 6415A>G, 6416T>A, 6417C>A, 6418A>C, 6421A>T, 6433G>T, 6436C>T, 6437T>A, 6445C>T, 6448G>A, 6449A>T, 6455T>G, 6457C>A, 6458T>A, 6461C>T, 6463T>G, 6472C>T, 6478G>T, 6479C>T, 6481T>A, 6485C>A, 6490A>C, 6491C>A, 6494C>G, 6495A>T, 6496G>A, 6497A>C, 6499A>T, 6500A>G, 6501T>A, 6502T>G, 6503G>A, 6505T>C, 6514G>T, 6515T>C, 6517C>A, 6518T>C, 6520C>T, 6529A>T, 6541G>A, 6544G>A, 6545C>T, 6547T>G, 6548T>C, 6552C>T, 6553A>G, 6556A>C, 6562A>T, 6566G>A, 6567G>A, 6568C>T, 6571C>T, 6574A>G, 6587G>A, 6591T>A, 6592A>T, 6593A>T, 6595T>C, 6602G>A, 6603C>T, 6647_6648insGTTA, 6669G>A, 6670C>T, 6675C>T, 6685A>G, 6692A>C, 6715C>T, 6716G>A, 6721A>T, 6727T>C, 6742A>T, 6752T>C, 6759G>A, 6766delC, 6776C>A, 6784C>T, 6786T>C, 6787G>A | | | | | | | | |

**CDS**

| | | | | | | | | | |
|---|---|---|---|---|---|---|---|---|---|
| ToTV_sRNA1gp1 | 1 | 2159 | 100% | 13837 | 91.8% | 2159 (100%) | 1941 (89.9%) | 0/0/0/0 | 1 |

| | Begin | End | Coverage | Score | Concordance | Matches | Identities | I/D/M/F* | Stop Codons |
|---|---|---|---|---|---|---|---|---|---|
| **NT** | **21** | **6806** | **86.9%** | **7777** | **57.4%** | **6785 (99.6%)** | **5350 (78.5%)** | **25/1** | |
| Protein mutations: | M6L (122A>T 124G>A), P8S (128C>T 130C>T), F10P (134T>C 135T>C 136C>T), N11S (138A>G), S12P (140T>C 142A>C), V13A (144T>C 145T>A), T14M (147C>T 148T>G), C17S (155T>A 157C>T), A18V (159C>T), T19A (161A>G), A31Q (197G>C 198C>A 199T>A), S66T (302T>A), T69S (311A>T 313C>T), S70N (315G>A 316C>T), N73S (324A>G 325C>T), A112T (440G>A), S113G (443A>G), N123D (473A>G), D127N (485G>A 487C>T), S156N (573G>A), T165S (599A>T 601T>A), I167T (606T>C 607A>T), N170S (615A>G), T171I (618C>T 619A>T), D183E (655C>G), S205A (719T>G 721C>T), M206V (722A>G), S239T (821T>A), V291I (977G>A 979T>A), S300A (1004T>G 1006T>A), T305V (1019A>G 1020C>T), T364S (1196A>T 1198C>T), D367E (1207T>G), E415D (1351A>T), I459V (1481A>G), A501S (1607G>T 1609T>A), T529V (1691A>G 1692C>T 1693T>A), T533A (1703A>G 1705T>A), S534A (1706T>G 1708T>C), E537R (1715G>A 1716A>G), K544R (1737A>G 1738A>G), T550I (1755C>T 1756A>C), S555D (1769A>G 1770G>A 1771C>T), Y578F (1839A>T), A580T (1844G>A), S582T (1850T>A 1852A>C), S591T (1877T>A 1879A>T), I595V (1889A>G 1891A>G), D600E (1906T>A), M601P (1907A>C 1908T>C 1909G>A), L602W (1911T>G), A611T (1937G>A 1939T>A), Y637F (2016A>T), F659Y (2082T>A), S665P (2099T>C 2101C>T), T669S (2111A>T 2113A>C), N675T (2130A>C 2131T>A), Y679Q (2141T>C 2143T>A), E685A (2160A>C), I691V (2177A>G 2179A>T), N712C (2240A>T 2241A>G), E713D (2245G>T), S715K (2250G>A 2251C>A), M726L (2282A>T), V741I (2327G>A 2329G>C), R768K (2409G>A 2410G>A), Q822R (2571A>G 2572A>G), G874S (2726G>A), V905I (2819G>A 2821G>A), T920V (2864A>G 2865C>T 2866T>G), A926S (2882G>T 2884C>G), T978S (3038A>T), Q993K (3083C>A 3085A>G), T1014S (3146A>T 3148G>C), D1089N (3371G>A 3373T>C), Q1103E (3413C>G), F1123Y (3474T>A 3475T>C), V1124L (3476G>C), A1129D (3492C>A), R1143K (3534G>A), A1162V (3591C>T 3592T>C), S1163Q (3593T>C 3594C>A), A1175S (3629G>T), R1193K (3684G>A 3685G>A), Y1196F (3693A>T), A1202S (3710G>T 3712A>T), I1203V (3713A>G 3715T>G), V1206I (3722G>A 3724G>A), H1208R (3729A>G), K1216E (3752A>G), V1218I (3758G>A), V1224I (3776G>A 3778G>T), P1352A (4160C>G 4162A>T), I1362M (4192T>G), G1370S (4214G>A), P1410Q (4335C>A 4336T>A), E1506G (4623A>G 4624A>T), T1511S (4637A>T 4639T>G), S1537N (4716G>A 4717C>T), S1538H (4718A>C 4719G>A), V1582T (4850G>A 4851T>C), V1598I (4898G>A), E1610D (4936A>C), T1614N (4947C>A 4948A>T), N1618H (4958A>C), T1628A (4988A>G 4990C>A), V1642M (5030G>A), I1658T (5079T>C 5080T>A), A1662S (5090G>T 5092T>A), P1667A (5105C>G 5107C>T), N1670E (5114A>G 5116T>A), V1672I (5120G>A 5122T>A), S1676T (5132T>A), V1679I (5141G>A 5143G>T), S1691A (5177T>G 5179G>C), Q1697H (5197G>T), T1701S (5207A>T 5209C>A), S1744G (5336A>G), V1746I (5342G>A 5344A>C), T1751S (5357A>T 5359C>T), K1752H (5360A>C 5362A>C), H1754N (5366C>A 5368T>C), V1755I (5369G>A 5371G>T), I1765V (5399A>G 5401T>G), V1785I (5459G>A 5461T>C), V1786I (5462G>A), D1794N (5486G>A), T1796P (5492A>C 5494A>T), Q1800H (5506A>T), L1810V (5534C>G), K1811Q (5537A>C), A1813S (5543G>T), E1818Q (5558G>C), I1826V (5582A>G 5584T>G), A1829S (5591G>T 5593C>A), T1833L (5603A>C 5604C>T), Y1836F (5613A>T), R1853N (5664G>A 5665A>T), L1856C (5672C>T 5673T>G 5674A>T), Q1857S (5675C>A 5676A>G 5677G>T), Y1869H (5711T>C), R1878K (5739G>A), L1882W (5751T>G), M1887I (5767G>T), K1889N (5773G>C), I1890V (5774A>G 5776A>G), I1893T (5784T>C 5785T>A), V1895L (5789G>T 5791C>G), E1898G (5799A>G 5800G>T), F1900T (5804T>A 5805T>C 5806C>T), Y1907F (5826A>T), V1913I (5843G>A 5845G>A), M1914F (5846A>T 5848G>C), L1915Q (5849T>C 5850T>A 5851G>A), D1918E (5860C>A), N1930H (5894A>C), Y1934F (5907A>T), H1946F (5942C>T 5943A>T 5944C>T), E1949D (5953A>T), N1950S (5955A>G 5956C>T), S1962A (5990T>G 5992A>C), T1966V (6002A>G 6003C>T 6004C>T), G1968E (6009G>A 6010G>A), T1976I (6033C>T 6034A>C), A1977S (6035G>T), S1980T (6044T>A 6046A>G), F1982L (6050T>C), D1985N (6059G>A), V1986I (6062G>A 6064T>C), H1997Q (6097T>A), I2002M (6112A>G), M2007L (6125A>C), E2008G (6129A>G 6130G>T), C2015S (6150G>C), V2016L (6152G>T 6154C>G), E2024G (6177A>G), T2027E (6185A>G 6186C>A 6187T>G), H2028R (6189A>G), M2032I (6202G>C), D2036E (6214T>G), L2037V (6215C>G 6217G>T), T2038P (6218A>C), E2039S (6221G>T 6222A>C 6223A>T), G2040E (6225G>A 6226G>A), I2041V (6227A>G 6229C>G), I2042V (6230A>G 6232A>G), K2044S (6237A>G 6238A>T), V2045T (6239G>A 6240T>C 6241G>A), I2046V (6242A>G 6244A>G), S2054T (6266T>A 6268G>T), S2056M (6273G>T 6274C>G), V2062E (6291T>A 6292T>A), D2063G (6294A>G), V2064L (6296G>C 6298A>C), L2066T (6302T>A 6303T>C 6304A>G), D2069S (6311G>T 6312A>C 6313T>C), I2071V (6317A>G 6319A>G), I2073V (6323A>G 6325C>G), A2089T (6371G>A 6373A>T), I2093V (6383A>G 6385C>G), C2100H (6404T>C 6405G>A), R2103V (6413C>G 6414G>T 6415A>G), S2104N (6416T>A 6417C>A 6418A>C), C2111S (6437T>A), T2115S (6449A>T), S2117A (6455T>G 6457C>A), S2118T (6458T>A), H2127N (6485C>A), R2128S (6490A>C), H2129N (6491C>A), Q2130V (6494C>G 6495A>T 6496G>A), T2131P (6497A>C 6499A>T), I2132E (6500A>G 6501T>A 6502T>G), D2133N (6503G>A 6505T>C), S2137P (6515T>C 6517C>A), F2138L (6518T>C 6520C>T), T2149M (6552C>T 6553A>G), G2154N (6566G>A 6567G>A 6568C>T) | | | | | | | | |

| | Begin | End | Coverage | Score | Concordance | Matches | Identities | I/D/M/F* | Stop Codons |
|---|---|---|---|---|---|---|---|---|---|
| NT | 21 | 6806 | 86.9% | 7777 | 57.4% | 6785 (99.6%) | 5350 (78.5%) | 25/1 | |

Codon mutations:

TTT3TTC (115T>C), TCC4TCT (118C>T), AAG5AAA (121G>A), ATG6TTA (122A>T 124G>A), CCC8TCT (128C>T 130C>T), TTC10CCT (134T>C 135T>C 136C>T), AAC11AGC (138A>G), TCA12CCC (140T>C 142A>C), GTT13GCA (144T>C 145T>A), ACT14ATG (147C>T 148T>G), GAA15GAG (151A>G), TGC17AGT (155T>A 157C>T), GCT18GTT (159C>T), ACC19GCC (161A>G), TCT24TCG (178T>G), TCA27TCT (187A>T), ACT30ACA (196T>A), GCT31CAA (197G>C 198C>A 199T>A), AGT34AGC (208T>C), AAT35AAC (211T>C), TTC36TTT (214C>T), CGC38CGT (220C>T), ACT39ACA (223T>A), CTG40CTT (226G>T), TCA47TCC (247A>C), CAA49CAG (253A>G), ATT50ATC (256T>C), TCT52TCA (262T>A), CAC53CAT (265C>T), GAA55GAG (271A>G), TTG57CTG (275T>C), AAG58AAA (280G>A), CCT59CCC (283T>C), TCA60TCG (286A>G), GTT61GTG (289T>G), ACA62ACT (292A>T), GCT65GCA (301T>A), TCT66ACT (302T>A), TCC67TCT (307C>T), TTT68TTC (310T>C), ACC69TCT (311A>T 313C>T), AGC70AAT (315G>A 316C>T), TGT72TGC (322T>C), AAC73AGT (324A>G 325C>T), TCT74TCA (328T>A), ACT76ACC (334T>C), AAA77AAG (337A>G), TTG78TTA (340G>A), TTA79TTG (343A>G), AAG81AAA (349G>A), ACT84ACA (358T>A), TTA85CTA (359T>C), ATT86ATA (364T>A), GAA87GAG (367A>G), CCC88CCT (370C>T), TTC89TTT (373C>T), ATC90ATA (376C>A), GCT92GCA (382T>A), GTC96GTT (394C>T), GCA97GCC (397A>C), TCC98TCT (400C>T), AAA101AAG (409A>G), TCA102TCC (412A>C), ATT103ATC (415T>C), GTT107GTG (427T>G), AAG109AAA (433G>A), GTT111GTC (439T>C), GCA112ACA (440G>A), AGC113GGC (443A>G), AAA115AAG (451A>G), AAG117AAA (457G>A), AAC123GAC (473A>G), GAC127AAT (485G>A 487C>T), TCT129TCC (493T>C), GAG130GAA (496G>A), GTG132GTT (502G>T), CTT140CTC (526T>C), ATT149ATA (553T>A), TTC151TTT (559C>T), CCA154CCT (568A>T), AGT156AAT (573G>A), CTA158TTA (578C>T), GGA160GGT (586A>T), CAG163CAA (595G>A), ACT165TCA (599A>T 601T>A), CAC166CAT (604C>T), ATA167ACT (606T>C 607A>T), AAT170AGT (615A>G), ACA171ATT (618C>T 619A>T), AAC174AAT (628C>T), GCT178GCC (640T>C), TTA182CTG (650T>C 652A>G), GAC183GAG (655C>G), TTA185CTG (659T>C 661A>G), CCG186CCC (664G>C), TCC189TCT (673C>T), CAA190CAG (676A>G), GCA196GCG (694A>G), CTA201TTA (707C>T), GGG203GGA (715G>A), GAA204GAG (718A>G), TCC205GCT (719T>G 721C>T), ATG206GTG (722A>G), TCA207TCT (727A>T), CAC208CAT (730C>T), CTT211CTG (739T>G), GTG214GTT (748G>T), GTA215GTT (751A>T), GCA216GCG (754A>G), CTC218TTA (758C>T 760C>A), ATA219ATT (763A>T), GCC220GCT (766C>T), GGG222GGC (772G>C), ATT223ATA (775T>A), TCT224TCA (778T>A), ACC225ACA (781C>A), TTC228TTT (790C>T), GTG229GTA (793G>A), GCT230GCA (796T>A), GTA231GTT (799A>T), CCT232CCG (802T>G), GGT233GGC (805T>C), AGA234CGT (806A>C 808A>T), GGT237GGA (817T>A), CTA238CTC (820A>C), TCC239ACC (821T>A), AAC240AAT (826C>T), CCG241CCA (829G>A), CTA242CTC (832A>C), TCA243TCC (835A>C), AAA244AAG (838A>G), CTG246TTA (842C>T 844G>A), TAC247TAT (847C>T), GCG249GCA (853G>A), GGT252GGA (862T>A), GCT253GCC (865T>C), AAT257AAC (877T>C), TTG259CTT (881T>C 883G>T), TTT260TTC (886T>C), ACC261ACA (889C>A), TTC263TTT (895C>T), AGG264AGA (898G>A), AAT265AAC (901T>C), TGT269TGC (913T>C), ATA281ATT (949A>T), ATT284ATA (958T>A), GGT286GGA (964T>A), TTT287TTC (967T>C), CCA290CCT (976A>T), GTT291ATA (977G>A 979T>A), GCT294GCA (988T>A), ATT295ATC (991T>C), TTG299CTT (1001T>C 1003G>T), TCT300GCA (1004T>G 1006T>A), ACG301ACA (1009G>A), TTC304TTT (1018C>T), ACG305GTG (1019A>G 1020C>T), GAG309GAA (1033G>A), CCA316CCC (1054A>C), CAT318CAC (1060T>C), CGC319CGA (1063C>A), TTG320CTT (1064T>C 1066G>T), AAT322AAC (1072T>C), GCA324GCG (1078A>G), AAC325AAT (1081C>T), ACT329ACC (1093T>C), ATC330ATA (1096C>A), CTA335TTG (1109C>T 1111A>G), AAA341AAG (1129A>G), TCT343TCA (1135T>A), TAC345TAT (1141C>T), GCT347GCC (1147T>C), GTA351GTG (1159A>G), GCC353GCA (1165C>A), AGG358AGA (1180G>A), GTA359GTT (1183A>T), ACG360ACA (1186G>A), GCA361GCC (1189A>C), GCA362GCC (1192A>C), ATA363ATT (1195A>T), ACC364TCT (1196A>T 1198C>T), CTT366CTG (1204T>G), GAT367GAG (1207T>G), GGG371GGC (1219G>C), AAC373AAT (1225C>T), ACT377ACA (1237T>A), GGC378GGT (1240C>T), GTG379GTT (1243G>T), GGC380GGT (1246C>T), CAA381CAG (1249A>G), AGG383AGA (1255G>A), GCA384GCT (1258A>T), CCT386CCC (1264T>C), GGT393GGA (1285T>A), AGT395AGC (1291T>C), GGG398GGA (1300G>A), ACC401ACA (1309C>A), CGG404CGA (1318G>A), CTG405TTA (1319C>T 1321G>A), ATC407ATT (1327C>T), GTT410GTG (1336T>G), TTG411CTT (1337T>C 1339G>T), CGC413AGG (1343C>A 1345C>G), GAA415GAT (1351A>T), GAG416GAA (1354G>A), CCA417CCC (1357A>C), TTG419CTT (1361T>C 1363G>T), AGG421CGG (1367A>C), CTC422CTG (1372C>G), GCT424GCC (1378T>C), GTG425GTA (1381G>A), CGA428CGT (1390A>T), TCA433TCC (1405A>C), AAT434AAC (1408T>C), TAT435TAC (1411T>C), GCT440GCA (1426T>A), CTG442CTT (1432G>T), GCT448GCA (1450T>A), CTG449TTG (1451C>T), GGG452GGT (1462G>T), GCA453GCT (1465A>T), CAA455CAG (1471A>G), GAC458GAT (1480C>T), ATT459GTT (1481A>G), AAG460AAA (1486G>A), ATA463ATT (1495A>T), ATC465ATT (1501C>T), TCC467TCA (1507C>A), ACT468ACA (1510T>A), CCA470CCT (1516A>T), CCC472CCA (1522C>A), TTG473TTA (1525G>A), CCC474CCA (1528C>A), GCA476GCT (1534A>T), CGC483AGG (1553C>A 1555C>G), CGA488AGG (1568C>A 1570A>G), TAT489TAC (1573T>C), ATA490ATT (1576A>T), GCC492GCT (1582C>T), ACA493ACC (1585A>C), TCC494TCA (1588C>A), AAC495AAT (1591C>T), TCA498TCC (1600A>C), GCT501TCA (1607G>T 1609T>A), CAG502CAA (1612G>A), GGT504GGG (1618T>G), ACC506ACA (1624C>A), AGA513CGG (1643A>C 1645A>G), CGG515AGG (1649C>A), GTT517GTG (1657T>G), CTT518CTG (1660T>G), GTG519GTA (1663G>A), TGC521TGT (1669C>T), GGT525GGA (1681T>A), GAG526GAA (1684G>A), TTT527TTC (1687T>C), ACT529GTA (1691A>G 1692C>T 1693T>A), GCT531GCA (1699T>A), CCC532CCT (1702C>T), ACT533GCA (1703A>G 1705T>A), TCT534GCC (1706T>G 1708T>C), CTT536CTG (1714T>G), GAA537AGA (1715G>A 1716A>G), GTG540GTC (1726G>C), GTT541GTG (1729T>G), GAG542GAA (1732G>A), AGC543AGT (1735C>T), AAA544AGG (1737A>G 1738A>G), AGA545CGT (1739A>C 1741A>T), CCC546CCA (1744C>A), ACA550ATC (1755C>T 1756A>C), AGG552AGA (1762G>A), AGC555GAT (1769A>G 1770G>A 1771C>T), GAT557GAC (1777T>C), GAT558GAC (1780T>C), TTG559CTA (1781T>C 1783G>A), CTG560CTT (1786G>T), AAG568AAA (1810G>A), GTT569GTG (1813T>G), GCA571GCC (1819A>C), ATC573ATA (1825C>A), TCA574TCT (1828A>T), GGA575GGC (1831A>C), CTG577CTC (1837G>C), TAT578TTT (1839A>T), GCC580ACC (1844G>A), ACA581ACC (1849A>C), TCA582ACC (1850T>A 1852A>C), AAG584AAA (1858G>A), GTG590GTC (1876G>C), TCA591ACT (1877T>A 1879A>T), GAG593GAA (1885G>A), ATA595GTG (1889A>G 1891A>G), ATA596ATT (1894A>T), GAT600GAA (1906T>A), ATG601CCA (1907A>C 1908T>C 1909G>A), TTG602TGG (1911T>G), AAT603AAC (1915T>C), ATC604ATT (1918C>T), CAA605CAG (1921A>G), GAT606GAC (1924T>C), ACC607ACA (1927C>A), GAT610GAC (1936T>C), GCT611ACA (1937G>A 1939T>A), AAG612AAA (1942G>A), CTT613CTG (1945T>G), CCC614CCG (1948C>G), GTG615GTA (1951G>A), GTT617GTG (1957T>G), GTA618GTC (1960A>C), GTG624GTA (1978G>A), TAC626TAT (1984C>T), TTG630CTA (1994T>C 1996G>A), GTG632GTT (2002G>T), GAA633GAG (2005A>G), GCT634GCC (2008T>C), AAA636AAG (2014A>G), TAT637TTT (2016A>T), GCT638GCC (2020T>C), TAT639TAC (2023T>C), GCA646GCG (2044A>G), TAT648TAC (2050T>C), TTC649TTT (2053C>T), CCT650CCA (2056T>A), GAC652GAT (2062C>T), AAA655AAG (2071A>G), TTT659TAT (2082T>A), CTA662CTT (2092A>T), TCC665CCT (2099T>C 2101C>T), GAA668GAG (2110A>G), ACA669TCC (2111A>T 2113A>C), ACC671ACT (2119C>T), AGG672CGG (2120A>C), GTT674GTC (2128T>C), AAT675ACA (2130A>C 2131T>A), GAC676GAT (2134C>T), CTG678CTC (2140G>C), TAT679CAA (2141T>C 2143T>A), CAA680CAG (2146A>G), GGT681GGA (2149T>A), TGT683TGC (2155T>C), AAT684AAC (2158T>C), GAG685GCG (2160A>C), AAC686AAT (2164C>T), CTG690CTC (2176G>C), ATA691GTT (2177A>G 2179A>T), AAG692AAA (2182G>A), ATC694ATT (2188C>T), GCT696GCA (2194T>A), TAT698TAC (2200T>C), ATC706ATA (2224C>A), TAC707TAT (2227C>T), GCT708GCG (2230T>G), AAT712TGT (2240A>T 2241A>G), GAG713GAT (2245G>T), CGT714CGC (2248T>C), AGC715AAA (2250G>A 2251C>A), GAC717GAT (2257C>T), GTT718GTG (2260T>G), CAG719CAA (2263G>A), CTG722TTA (2270C>T 2272G>A), GGA725GGC (2281A>C), ATG726TTG (2282A>T), GCG727GCA (2287G>A), CAC728CAT (2290C>T), GCC730GCT (2296C>T), ATA731ATT (2299A>T), GCT733GCA (2305T>A), GTG735GTT (2311G>T), GAC736GAT (2314C>T), CTG738TTA (2318C>T 2320G>A), ATA739ATT (2323A>T), TGC740TGT (2326C>T), GTG741ATC (2327G>A 2329G>C), GAG742GAA (2332G>A), CAG743CAA (2335G>A), CCT745CCA (2341T>A), GCC746GCA (2344C>A), CAC747CAT (2347C>T), GTA748GTG (2350A>G), TTC750TTT (2356C>T), TTG751TTA (2359G>A), ACA753ACT (2365A>T), GCA754GCT (2368A>T), TTT755TTC (2371T>C), AGG757CGT (2375A>C 2377G>T), TAT758TAC (2380T>C), CGC761AGA (2387C>A 2389C>A), AGA762AGG (2392A>G), CAG766CAA (2404G>A), CCT767CCC (2407T>C), AGG768AAA (2409G>A 2410G>A), TCC770TCT (2416C>T), TTA790TTG (2476A>G), CCG791CCA (2479G>A), ATC793ATT (2485C>T), TTG794TTA (2488G>A), AGA795AGG (2491A>G), GTT796GTG (2494T>G), CTG797CTT (2497G>T), ATT798ATC (2500T>C), AAG799AAA (2503G>A), GCT800GCA (2506T>A), GGC801GGG (2509C>G), CTT802TTG (2510C>T 2512T>G), ATT803ATC (2515T>C), TTA804CTA (2516T>C), GCA806GCC (2524A>C), GCA810GCT (2536A>T), GGG812GGT (2542G>T), GCA816GCT (2554A>T), TTC817TTT (2557C>T), GCT819GCG (2563T>G), CAA822CGG (2571A>G 2572A>G), TCC823TCA (2575C>A), CCT827CCC (2587T>C), AGA832CGG (2600A>C 2602A>G), ATT835ATC (2611T>C), ACC836ACA (2614C>A), AAC837AAT (2617C>T), AAT840AAC (2626T>C), AGC841AGT (2629C>T), CGG845AGG (2639C>A), AAG848AAA (2650G>A), ATC852ATT (2662C>T), TTG857TTA (2677G>A), CTG858TTG (2678C>T), GCA859GCG (2683A>G), AAG862AAA (2692G>A), GGT863GGA (2695T>A), CTG866CTT (2704G>T), AAA868AAG (2710A>G), GCA870GCC (2716A>C), GGC874AGC (2726G>A), TTC875TTT (2731C>T), GGA879GGC (2743A>C), TTG880CTG (2744T>C), GTC885GTG (2761C>G), TTG887CTC (2765T>C 2767G>C), AGA888CGA (2768A>C), GGT890GGG (2776T>G), TAC894TAT (2788C>T), ACC898ACA (2800C>A), GTC900GTT (2806C>T), TGC901TGT (2809C>T), TCG902TCT (2812G>T), GGC903GGA (2815C>A), GTG905ATA (2819G>A 2821G>A), ACA907ACT (2827A>T), GTG908GTA (2830G>A), TTT912TTC (2842T>C), TCA913TCC (2845A>C), AGC914AGT (2848C>T), CGT916AGA (2852C>A 2854T>A), GGC918GGG (2860C>G), ACA919ACT (2863A>T), ACT920GTG (2864A>G 2865C>T 2866T>G), TTC921TTT (2869C>T), ATA923ATT (2875A>T), ATA924ATC (2878A>C), GCC926TCG (2882G>T 2884C>G), TCA928TCC (2890A>C), TAC934TAT (2908C>T), AAT935AAC (2911T>C), CGG940CGT (2926G>T), TTC941TTT (2929C>T), TTG942CTA (2930T>C 2932G>A), AAG943AAA (2935G>A), CAA945CAG (2941A>G), ATT947ATA (2947T>A), GTC948GTG (2950C>G), CTG949CTT (2953G>T), CTC950CTG (2956C>G), AAT951AAC (2959T>C), GGA953GGT (2965A>T), CCC955CCT (2971C>T), GAT956GAC (2974T>C), CCC958CCG (2980C>G), CCT960CCG (2986T>G), ATT962ATC (2992T>C), AAA964AAG (2998A>G), CGG969AGG (3011C>A), TGT973TGC (3025T>C), TCT975TCA (3031T>A), AAG976AAA (3034G>A), GGC977GGA (3037C>A), ACT978TCT (3038A>T), GGG980GGT (3046G>T), GTA981GTC (3049A>C), TGT982TGC (3052T>C), GTA986GTT (3064A>T), AAG989AAA (3073G>A), GCT991GCC (3079T>C), CAA993AAG (3083C>A 3085A>G), TTG999CTT (3101T>C 3103G>T), GTG1003GTT (3115G>T), TCC1006TCA (3124C>A), CTT1010CTG (3136T>G), ACG1014TCC (3146A>T 3148G>C), GAC1018GAT (3160C>T), AGT1020AGC (3166T>C), TCC1021TCA (3169C>A), CTG1024CTA (3178G>A), TCT1026TCA (3184T>A), TAT1030TAC (3196T>C), TAT1032TAC (3202T>C), GAT1040GAC (3226T>C), CTT1044CTG (3238T>G), CTT1045TTA (3239C>T 3241T>A), CTT1046TTG (3242C>T 3244T>G), TTG1047TTA (3247G>A), AAC1049AAT (3253C>T), GTG1050GTA (3256G>A), AAA1053AAG (3265A>G), TGC1056TGT (3274C>T), ATT1057ATA (3277T>A), ATT1060ATA (3286T>A), TCT1064TCA (3298T>A), GAA1068GAG (3310A>G), CAC1071CAT (3319C>T), GGG1073GGC (3325G>C), TTT1074TTC (3328T>C), GTA1075GTT (3331A>T), GCA1076GCC (3334A>C), ACA1080ACG (3346A>G), GCT1081GCG (3349T>G), ATT1082ATC (3352T>C), CGA1084AGA (3356C>A), TTG1087CTG (3365T>C), GAT1089AAC (3371G>A 3373T>C), CTT1090TTG (3374C>T 3376T>G), CCG1091CCA (3379G>A), ACT1092ACA (3382T>A), GGT1093GGG (3385T>G), CCG1094CCT (3388G>T), AAA1096AAG (3394A>G), GCA1098GCC (3400A>C), GTA1100GTG (3406A>G), TGC1102TGT (3412C>T), CAG1103GAG (3413C>G), CTC1104TTG (3416C>T 3418C>G), CTT1105CTG (3421T>G), CTA1108TTG (3428C>T 3430A>G), CGA1109AGA (3431C>A), AGG1111AGA (3439G>A), GCG1113GCT (3445G>T), GAA1117GAG (3457A>G), CAG1120CAA (3466G>A), TTT1123TAC (3474T>A 3475T>C), GTT1124CTT (3476G>C), GGC1125GGA (3481C>A), ACA1126ACT (3484A>T), GCT1129GAT (3492C>A), GCC1133GCA (3505C>A), ACG1134ACT (3508G>T), CCC1136CCA (3514C>A), AAA1138AAG (3520A>G), ACA1139ACC (3523A>C), ACG1140ACA (3526G>A), CTG1141CTC (3529G>C), CGA1142AGG (3530C>A 3532A>G), AGG1143AAG (3534G>A), GGC1145GGT (3541C>T), TTG1146CTC (3542T>C 3544G>C), GCT1149GCA (3553T>A), GGG1151GGA (3559G>A), GCA1153GCT (3565A>T), TCT1158TCC (3580T>C), ATC1159ATA (3583C>A), ATT1160ATC (3586T>C), TCA1161TCG (3589A>G), GCT1162GTC (3591C>T 3592T>C), TCA1163CAA (3593T>C 3594C>A), CGT1166AGA (3602C>A 3604T>A), GGG1167GGA (3607G>A), GAA1168GAG (3610A>G), GGT1169GGC (3613T>C), CCG1172CCA (3622G>A), TAC1173TAT (3625C>T), GTG1174GTC (3628G>C), GCT1175TCT (3629G>T), ATA1177ATC (3637A>C), CAA1178CAG (3640A>G), AAA1179AAG (3643A>G), TAC1180TAT (3646C>T), GAA1182GAG (3652A>G), GCA1184GCC (3658A>C), CAA1185CAG (3661A>G), AAT1186AAC (3664T>C), GAG1189GAA (3673G>A), GAC1190GAT (3676C>T), ATT1191ATC (3679T>C), GCG1192GCC (3682G>C), AGG1193AAA (3684G>A 3685G>A), CTA1194TTG (3686C>T 3688A>G), GCC1195GCT (3691C>T), TAT1196TTT (3693A>T), GGG1198GGA (3700G>A), CTA1199TTG (3701C>T 3703A>G), CGT1200CGG (3706T>G), CAA1201CAG (3709A>G), GCA1202TCT (3710G>T 3712A>T), ATT1203GTG (3713A>G 3715T>G), TTG1204CTG (3716T>C), GTG1206ATA (3722G>A 3724G>A), CAC1208CGC (3729A>G), TCC1209TCG (3733C>G), GTT1212GTG (3742T>G), AAG1216GAG (3752A>G), GTC1218ATC (3758G>A), ACA1219ACT (3763A>T), CAG1220CAA (3766G>A), GTG1224ATT (3776G>A 3778G>T), GTG1225GTT (3781G>T), CTC1226TTG (3782C>T 3784C>G), GGT1228GGA (3790T>A), GTT1229GTC (3793T>C), TTT1232TTC (3802T>C), TAT1234TAC (3808T>C), GAC1236GAT (3814C>T), GGG1237GGA (3817G>A), TTG1240CTG (3824T>C), GGG1245GGT (3841G>T), CCG1247CCA (3847G>A), TAC1248TAT (3850C>T), AAG1250AAA (3856G>A), GGC1254GGT (3868C>T), ACT1255ACC (3871T>C), GAG1259GAA (3883G>A), GTG1261GTA (3889G>A), CCA1263CCC (3895A>C), GGA1265GGC (3901A>C), GGA1267GGT (3907A>T), GAT1268GAC (3910T>C), CGA1269AGA (3911C>A), GTC1270GTG (3916C>G), CTC1272TTG (3920C>T 3922C>G), AAA1273AAG (3925A>G), AGG1274AGA (3928G>A), ACT1276ACC (3934T>C), GAC1281GAT (3949C>T), GAG1283GAA (3955G>A), GCT1284GCA (3958T>A), TTG1285CTA (3959T>C 3961G>A), ATT1289ATC (3973T>C), GGA1292GGT (3982A>T), CTG1297TTG (3995C>T), GTC1298GTT (4000C>T), ACC1299ACA (4003C>A), ACC1300ACA (4006C>A), CAA1301CAG (4009A>G), GCC1303GCA (4015C>A), AAA1304AAG (4018A>G), TTG1308CTA (4028T>C 4030G>A), CCG1309CCT (4033G>T), TTG1310CTG (4034T>C), GAA1311GAG (4039A>G), GTT1313GTG (4045T>G), GGG1315GGC (4051G>C), CGG1317AGG (4055C>A), ACC1319ACA (4063C>A), TTG1321CTA (4067T>C 4069G>A), GAA1323GAG (4075A>G), ATT1324ATA (4078T>A), CTT1325CTA (4081T>A), CCC1326CCT (4084C>T), TAC1329TAT (4093C>T), AAT1330AAC (4096T>C), CTG1338TTG (4118C>T), GAT1339GAC (4123T>C), TCC1341TCT (4129C>T), TCC1347TCG (4147C>G), CTG1351TTG (4157C>T), CCA1352GCT (4160C>G 4162A>T), GGT1361GGA (4189T>A), ATT1362ATG (4192T>G), GAA1363GAG (4195A>G), CTG1366TTA (4202C>T 4204G>A), TTG1367CTA (4205T>C 4207G>A), GCT1368GCA (4210T>A), AAT1369AAC (4213T>C), GGC1370AGC (4214G>A), TTC1371TTT (4219C>T), AGA1372CGG (4220A>C 4222A>G), GCA1373GCT (4225A>T), GTC1374GTT (4228C>T), ACA1377ACG (4237A>G), TCT1380TCC (4246T>C), GCT1381GCA (4249T>A), GAC1382GAT (4252C>T), TAT1383TAC (4255T>C), ATC1392ATT (4282C>T), GCC1394GCA (4288C>A), CAA1396CAG (4294A>G), TGT1399TGC (4303T>C), CTT1401CTG (4309T>G), GAC1404GAT (4318C>T), GGA1407GGC (4327A>C), CCT1410CAA (4335C>A 4336T>A), GGT1411GGC (4339T>C), TCT1412TCC (4342T>C), GAC1414GAT (4348C>T), TCC1415TCA (4351C>A), AAT1416AAC (4354T>C), AGA1418CGC (4358A>C 4360A>C), CAT1419CAC (4363T>C), GTG1420GTA (4366G>A), CTT1421TTG (4367C>T 4369T>G), CTT1422CTG (4372T>G), GCT1425GCC (4381T>C), TCA1426TCT (4384A>T), TGC1427TGT (4387C>T), ACT1430ACC (4396T>C), ATT1431ATC (4399T>C), GAG1433GAA (4405G>A), GTT1436GTG (4414T>G), TTT1437TTC (4417T>C), AGG1438CGC (4418A>C 4420G>C), TTG1439CTG (4421T>C), GTG1440GTT (4426G>T), GGA1441GGG (4429A>G), GGT1442GGC (4432T>C), CCT1444CCA (4438T>A), TCA1445TCT (4441A>T), GGA1446GGG (4444A>G), GCA1448GCT (4450A>T), CTC1449CTT (4453C>T), ACC1450ACT (4456C>T), GTT1451GTC (4459T>C),

ATC1452ATT (4462C>T), TTC1453TTT (4465C>T), AAC1454AAT (4468C>T), CGT1463CGG (4495T>G), GCA1465GCT (4501A>T), ATT1467ATC (4507T>C), TCT1468TCA (4510T>A), CTA1469CTG (4513A>G), TTG1470CTG (4514T>C), AGA1472AGG (4522A>G), CAT1474CAC (4528T>C), GCA1476GCT (4534A>T), GCT1477GCA (4537T>A), CAA1478CAG (4540A>G), ATA1480ATT (4546A>T), GGG1481GGC (4549G>C), TGC1482TGT (4552C>T), AAA1483AAG (4555A>G), CCC1484CCA (4558C>A), AAC1488AAT (4570C>T), CTA1490CTT (4576A>T), GTG1492GTT (4582G>T), GCA1493GCT (4585A>T), GTC1494GTA (4588C>A), GAC1498GAT (4600C>T), CTA1500CTT (4606A>T), GTT1503GTA (4615T>A), CCC1504CCA (4618C>A), GAA1506GGT (4623A>G 4624A>T), TTG1507CTC (4625T>C 4627G>C), TAT1510TAC (4636T>C), ACT1511TCG (4637A>T 4639T>G), GCT1514GCA (4648T>A), ATT1515ATA (4651T>A), GAA1518GAG (4660A>G), TTG1519CTA (4661T>C 4663G>A), GTT1524GTC (4678T>C), ATA1526ATC (4684A>C), ATC1530ATA (4696C>A), AAG1532AAA (4702G>A), AAC1533AAT (4705C>T), GTT1536GTC (4714T>C), AGC1537AAT (4716G>A 4717C>T), AGT1538CAT (4718A>C 4719G>A), CTA1542CTG (4732A>G), GAC1543GAT (4735C>T), TTG1544CTT (4736T>C 4738G>T), TCT1545TCA (4741T>A), CTA1547TTG (4745C>T 4747A>G), TTT1549TTC (4753T>C), CTA1550CTT (4756A>T), AGC1551AGT (4759C>T), AGA1552AGG (4762A>G), GGT1553GGC (4765T>C), TTT1554TTC (4768T>C), AAG1555AAA (4771G>A), AGG1556AGA (4774G>A), GGA1560GGG (4786A>G), TAC1561TAT (4789C>T), CCT1565CCC (4801T>C), CTG1566TTG (4802C>T), GTA1569GTG (4813A>G), TCT1570TCC (4816T>C), GAA1573GAG (4825A>G), CCA1574CCC (4828A>C), TAC1576TAT (4834C>T), ATA1578ATT (4840A>T), AGG1579AGA (4843G>A), CCT1580CCC (4846T>C), GTT1582ACT (4850G>A 4851T>C), GCT1587GCC (4867T>C), CTC1588TTG (4868C>T 4870C>G), TTG1591TTA (4879G>A), AAC1593AAT (4885C>T), ATA1594ATT (4888A>T), GAC1595GAT (4891C>T), ACG1596ACT (4894G>T), GGA1597GGC (4897A>C), GTT1598ATT (4898G>A), AGA1599AGG (4903A>G), GCA1601GCC (4909A>C), CAC1603CAT (4915C>T), GGG1605GGT (4921G>T), CAG1607CAA (4927G>A), GTT1608GTG (4930T>G), GAA1610GAC (4936A>C), GTG1613GTT (4945G>T), ACA1614AAT (4947C>A 4948A>T), GTT1616GTG (4954T>G), AAC1618CAC (4958A>C), GCT1619GCC (4963T>C), AAG1621AAA (4969G>A), CGG1623CGA (4975G>A), TTC1625TTT (4981C>T), CCA1626CCT (4984A>T), GCC1627GCT (4987C>T), ACC1628GCA (4988A>G 4990C>A), CCT1631CCG (4999T>G), ACA1632ACT (5002A>T), TAT1633TAC (5005T>C), GAG1637GAA (5017G>A), CAG1638CAA (5020G>A), GAC1639GAT (5023C>T), CTG1641CTT (5029G>T), GTG1642ATG (5030G>A), GAG1643GAA (5035G>A), GTT1644GTA (5038T>A), ACA1645ACT (5041A>T), GGA1646GGC (5044A>C), CCA1648CCT (5050A>T), GGG1651GGT (5059G>T), CTC1652CTT (5062C>T), CTT1656TTA (5072C>T 5074T>A), ATT1658ACA (5079T>C 5080T>A), GCA1659GCT (5083A>T), TCA1661TCC (5089A>C), GCT1662TCA (5090G>T 5092T>A), GTG1664GTT (5098G>T), CCT1665CCA (5101T>A), TTG1666CTG (5102T>C), CCC1667GCT (5105C>G 5107C>T), CCA1668CCT (5110A>T), GGC1669GGT (5113C>T), AAT1670GAA (5114A>G 5116T>A), ACT1671ACC (5119T>C), GTT1672ATA (5120G>A 5122T>A), CTG1673TTG (5123C>T), TTT1675TTC (5131T>C), TCT1676ACT (5132T>A), GTG1679ATT (5141G>A 5143G>T), CAT1680CAC (5146T>C), ACT1681ACA (5149T>A), TTT1682TTC (5152T>C), GCT1683GCA (5155T>A), CGA1685AGA (5159C>A), TGC1689TGT (5173C>T), TCC1690TCT (5176C>T), TCG1691GCC (5177T>G 5179G>C), CGA1692AGG (5180C>A 5182A>G), ACA1693ACT (5185A>T), GCA1696GCC (5194A>C), CAG1697CAT (5197G>T), CAG1698CAA (5200G>A), GAC1700GAT (5206C>T), ACC1701TCA (5207A>T 5209C>A), ACC1702ACA (5212C>A), GTT1704GTC (5218T>C), AAC1707AAT (5227C>T), ACT1709ACC (5233T>C), GCA1711GCC (5239A>C), AAG1712AAA (5242G>A), GTA1717GTT (5257A>T), AGA1718AGG (5260A>G), GGG1719GGA (5263G>A), CCC1720CCA (5266C>A), ACC1721ACA (5269C>A), AGG1724CGA (5276A>C 5278G>A), GGG1728GGT (5290G>T), CTT1729CTC (5293T>C), ATT1730ATC (5296T>C), CCT1732CCA (5302T>A), GCT1736GCC (5314T>C), GAC1740GAT (5326C>T), CAC1741CAT (5329C>T), AGC1744GGC (5336A>G), ATT1745ATC (5341T>C), GTA1746ATC (5342G>A 5344A>C), ACC1747ACA (5347C>A), GGG1749GGA (5353G>A), ACC1751TCT (5357A>T 5359C>T), AAA1752CAC (5360A>C 5362A>C), CAT1754AAC (5366C>A 5368T>C), GTG1755ATT (5369G>A 5371G>T), GTT1756GTA (5374T>A), TTT1757TTC (5377T>C), GGC1760GGG (5386C>G), AAT1761AAC (5389T>C), GGG1762GGA (5392G>A), ATT1765GTG (5399A>G 5401T>G), GGT1766GGA (5404T>A), CCA1767CCT (5407A>T), GTG1768GTT (5410G>T), TGC1769TGT (5413C>T), GCT1770GCA (5416T>A), GCA1771GCT (5419A>T), TTG1772CTA (5420T>C 5422G>A), CTG1775TTA (5429C>T 5431G>A), TCC1776TCT (5434C>T), ACT1777ACG (5437T>G), GGC1778GGA (5440C>A), CAA1779CAG (5443A>G), TAT1780TAC (5446T>C), TCT1781TCA (5449T>A), TCT1782TCC (5452T>C), GTT1785ATC (5459G>A 5461T>C), GTT1786ATT (5462G>A), GTG1787GTC (5467G>C), ACC1791ACT (5479C>T), ATA1792ATT (5482A>T), GCA1793GCT (5485A>T), GAT1794AAT (5486G>A), GTT1795GTG (5491T>G), ACA1796CCT (5492A>C 5494A>T), CTC1798TTG (5498C>T 5500C>G), AGT1799AGC (5503T>C), CAA1800CAT (5506A>T), TAT1801TAC (5509T>C), CCA1802CCT (5512A>T), GGA1803GGT (5515A>T), GCC1806GCT (5524C>T), AAG1807AAA (5527G>A), TAT1808TAC (5530T>C), CTT1809TTG (5531C>T 5533T>G), CTG1810GTG (5534C>G), AAA1811CAA (5537A>C), GCT1813TCT (5543G>T), GAT1814GAC (5548T>C), ACA1815ACC (5551A>C), GAA1818CAA (5558G>C), CTT1820CTA (5566T>A), GCA1821GCT (5569A>T), ACA1823ACG (5575A>G), TGT1824TGC (5578T>C), ATT1826GTG (5582A>G 5584T>G), GCC1829TCA (5591G>T 5593C>A), GAG1832GAA (5602G>A), ACA1833CTA (5603A>C 5604C>T), CCA1834CCG (5608A>G), GCA1835GCT (5611A>T), TAC1836TTC (5613A>T), ATA1837ATT (5617A>T), CCT1838CCA (5620T>A), GGA1840GGT (5626A>T), TTT1842TTC (5632T>C), CTT1844CTG (5638T>G), GGT1845GGC (5641T>C), GTG1850GTA (5656G>A), CAA1851CAG (5659A>G), GGG1852GGA (5662G>A), AGA1853AAT (5664G>A 5665A>T), CTA1856TGT (5672C>T 5673T>G 5674A>T), CAG1857AGT (5675C>A 5676A>G 5677G>T), CAG1860CAA (5686G>A), CGC1861AGA (5687C>A 5689C>A), TTG1862CTT (5690T>C 5692G>T), GGA1867GGT (5707A>T), CCT1868CCA (5710T>A), TAT1869CAT (5711T>C), GGG1870GGT (5716G>T), GGA1871GGG (5719A>G), ACT1876ACC (5734T>C), ACT1877ACC (5737T>C), AGA1878AAA (5739G>A), GAG1879GAA (5743G>A), CTT1880CTA (5746T>A), AAG1881AAA (5749G>A), TTG1882TGG (5751T>G), CTG1883TTG (5753C>T), ATG1887ATT (5767G>T), AAG1889AAC (5773G>C), ATA1890GTG (5774A>G 5776A>G), TAT1891TAC (5779T>C), ATT1893ACA (5784T>C 5785T>A), TTA1894CTT (5786T>C 5788A>T), GTC1895TTG (5789G>T 5791C>G), CAA1896CAG (5794A>G), AGA1897AGG (5797A>G), GAG1898GGT (5799A>G 5800G>T), TTC1900ACT (5804T>A 5805T>C 5806C>T), ACT1902ACC (5812T>C), CTC1903CTG (5815C>G), TAC1904TAT (5818C>T), TAT1907TTT (5826A>T), CTC1908CTT (5830C>T), TCA1910TCC (5836A>C), GTG1913ATA (5843G>A 5845G>A), ATG1914TTC (5846A>T 5848G>C), TTG1915CAA (5849T>C 5850T>A 5851G>A), GTT1917GTG (5857T>G), GAC1918GAA (5860C>A), CTT1920TTG (5864C>T 5866T>G), CTC1922CTA (5872C>A), GGG1924GGT (5878G>T), TTT1925TTC (5881T>C), TTT1926TTC (5884T>C), CCC1927CCT (5887C>T), CGC1928CGT (5890C>T), AAT1930CAT (5894A>C), GTG1932GTA (5902G>A), TAC1934TTC (5907A>T), CTT1936CTC (5914T>C), TTT1939TTC (5923T>C), CTG1941CTA (5929G>A), CAC1946TTT (5942C>T 5943A>T 5944C>T), AAG1948AAA (5950G>A), GAA1949GAT (5953A>T), AAC1950AGT (5955A>G 5956C>T), ACG1953ACT (5965G>T), GTT1954GTG (5968T>G), TAC1955TAT (5971C>T), TTT1956TTC (5974T>C), AGA1957CGT (5975A>C 5977A>T), AAG1958AAA (5980G>A), CCT1959CCC (5983T>C), TTT1960TTC (5986T>C), CTC1961CTT (5989C>T), TCA1962GCC (5990T>G 5992A>C), AAA1964AAG (5998A>G), ACC1966GTT (6002A>G 6003C>T 6004C>T), GGG1968GAA (6009G>A 6010G>A), TCC1969TCA (6013C>A), AAA1970AAG (6016A>G), ATT1972ATA (6022T>A), TCC1974TCT (6028C>T), GCA1975GCT (6031A>T), ACA1976ATC (6033C>T 6034A>C), GCT1977TCT (6035G>T), GCT1978GCA (6040T>A), GAG1979GAA (6043G>A), TCA1980ACG (6044T>A 6046A>G), TTG1981TTA (6049G>A), TTT1982CTT (6050T>C), GGT1983GGG (6055T>G), GAT1985AAT (6059G>A), GTT1986ATC (6062G>A 6064T>C), TCC1987TCT (6067C>T), GCT1988GCC (6070T>C), CTC1991CTG (6079C>G), AGT1993AGC (6085T>C), AGG1994AGA (6088G>A), CTA1995TTA (6089C>T), CTT1996TTA (6092C>T 6094T>A), CAT1997CAA (6097T>A), CTT1998CTG (6100T>G), CAG1999CAA (6103G>A), AAG2000AAA (6106G>A), ATA2002ATG (6112A>G), TCA2005TCT (6121A>T), TCC2006TCA (6124C>A), ATG2007CTG (6125A>C), GAG2008GGT (6129A>G 6130G>T), TTT2011TTC (6139T>C), AAA2012AAG (6142A>G), ATT2013ATC (6145T>C), TAT2014TAC (6148T>C), TGT2015TCT (6150G>C), GTC2016TTG (6152G>T 6154C>G), CAG2018CAA (6160G>A), GGC2019GGG (6163C>G), AAG2023AAA (6175G>A), GAA2024GGA (6177A>G), GTA2026GTG (6184A>G), ACT2027GAG (6185A>G 6186C>A 6187T>G), CAC2028CGC (6189A>G), TTC2029TTT (6193C>T), CAA2030CAG (6196A>G), CGC2031CGT (6199C>T), ATG2032ATC (6202G>C), CAA2034CAG (6208A>G), GAT2036GAG (6214T>G), CTG2037GTT (6215C>G 6217G>T), ACA2038CCA (6218A>C), GAA2039TCT (6221G>T 6222A>C 6223A>T), GGG2040GAA (6225G>A 6226G>A), ATC2041GTG (6227A>G 6229C>G), ATA2042GTG (6230A>G 6232A>G), GAG2043GAA (6235G>A), AAA2044AGT (6237A>G 6238A>T), GTG2045ACA (6239G>A 6240T>C 6241G>A), ATA2046GTG (6242A>G 6244A>G), TTA2049CTC (6251T>C 6253A>C), ACC2050ACA (6256C>A), ACA2052ACT (6262A>T), CTG2053TTG (6263C>T), TCG2054ACT (6266T>A 6268G>T), AGC2056ATG (6273G>T 6274C>G), GTG2059GTA (6283G>A), CTT2061TTG (6287C>T 6289T>G), GTT2062GAA (6291T>A 6292T>A), GAT2063GGT (6294A>G), GTA2064CTC (6296G>C 6298A>C), CCT2065CCA (6301T>A), TTA2066ACG (6302T>A 6303T>C 6304A>G), GGC2067GGA (6307C>A), TTG2068CTT (6308T>C 6310G>T), GAT2069TCC (6311G>T 6312A>C 6313T>C), AAC2070AAT (6316C>T), ATA2071GTG (6317A>G 6319A>G), GAG2072GAA (6322G>A), ATC2073GTG (6323A>G 6325C>G), CAG2074CAA (6328G>A), AAA2076AAG (6334A>G), GGT2077GGC (6337T>C), CTA2080TTA (6344C>T), AAT2086AAC (6364T>C), GCA2089ACT (6371G>A 6373A>T), GCC2092GCT (6382C>T), ATC2093GTG (6383A>G 6385C>G), CTC2097CTA (6397C>A), TGT2100CAT (6404T>C 6405G>A), AAC2102AAT (6412C>T), CGA2103GTG (6413C>G 6414G>T 6415A>G), TCA2104AAC (6416T>A 6417C>A 6418A>C), ACA2105ACT (6421A>T), GTG2109GTT (6433G>T), TTC2110TTT (6436C>T), TGT2111AGT (6437T>A), TAC2113TAT (6445C>T), ACG2114ACA (6448G>A), ACT2115TCT (6449A>T), TCC2117GCA (6455T>G 6457C>A), TCT2118ACT (6458T>A), CTT2119TTG (6461C>T 6463T>G), GCC2122GCT (6472C>T), GTG2124GTT (6478G>T), CTT2125TTA (6479C>T 6481T>A), CAT2127AAT (6485C>A), AGA2128AGC (6490A>C), CAT2129AAT (6491C>A), CAG2130GTA (6494C>G 6495A>T 6496G>A), ACA2131CCT (6497A>C 6499A>T), ATT2132GAG (6500A>G 6501T>A 6502T>G), GAT2133AAC (6503G>A 6505T>C), CCG2136CCT (6514G>T), TCC2137CCA (6515T>C 6517C>A), TTC2138CTT (6518T>C 6520C>T), ACA2141ACT (6529A>T), GTG2145GTA (6541G>A), TTG2146TTA (6544G>A), CTT2147TTG (6545C>T 6547T>G), TTG2148CTG (6548T>C), ACA2149ATG (6552C>T 6553A>G), CCA2150CCC (6556A>C), CTA2152CTT (6562A>T), GGC2154AAT (6566G>A 6567G>A 6568C>T), TAC2155TAT (6571C>T), AAA2156AAG (6574A>G)

#### Proteins

| | | | | | | | | | |
|---|---|---|---|---|---|---|---|---|---|
| polyprotein (YP_001039627.1) | 1 | 2159 | 100% | 13837 | 91.8% | 2159 (100%) | 1941 (89.9%) | 0/0/0/0 | 1 |
| Protein mutations: | M6L (122A>T 124G>A), P8S (128C>T 130C>T), F10P (134T>C 135T>C 136C>T), N11S (138A>G), S12P (140T>C 142A>C), V13A (144T>C 145T>A), T14M (147C>T 148T>G), C17S (155T>A 157C>T), A18V (159C>T), T19A (161A>G), A31Q (197G>C 198C>A 199T>A), S66T (302T>A), T69S (311A>T 313C>T), S70N (315G>A 316C>T), N73S (324A>G 325C>T), A112T (440G>A), S113G (443A>G), N123D (473A>G), D127N (485G>A 487C>T), S156N (573G>A), T165S (599A>T 601T>A), I167T (606T>C 607A>T), N170S (615A>G), T171I (618C>T 619A>T), D183E (655C>G), S205A (719T>G 721C>T), M206V (722A>G), S239T (821T>A), V291I (977G>A 979T>A), S300A (1004T>G 1006T>A), T305V (1019A>G 1020C>T), T364S (1196A>T 1198C>T), D367E (1207T>G), E415D (1351A>T), I459V (1481A>G), A501S (1607G>T 1609T>A), T529V (1691A>G 1692C>T 1693T>A), T533A (1703A>G 1705T>A), S534A (1706T>G 1708T>C), E537R (1715G>A 1716A>G), K544R (1737A>G 1738A>G), T550I (1755C>T 1756A>C), S555D (1769A>G 1770G>A 1771C>T), Y578F (1839A>T), A580T (1844G>A), S582T (1850T>A 1852A>C), S591T (1877T>A 1879A>T), I595V (1889A>G 1891A>G), D600E (1906T>A), M601P (1907A>C 1908T>C 1909G>A), L602W (1911T>G), A611T (1937G>A 1939T>A), Y637F (2016A>T), F659Y (2082T>A), S665P (2099T>C 2101C>T), T669S (2111A>T 2113A>C), N675T (2130A>C 2131T>A), Y679Q (2141T>C 2143T>A), E685A (2160A>C), I691V (2177A>G 2179A>T), N712C (2240A>T 2241A>G), E713D (2245G>T), S715K (2250G>A 2251C>A), M726L (2282A>T), V741I (2327G>A 2329G>C), R768K (2409G>A 2410G>A), Q822R (2571A>G 2572A>G), G874S (2726G>A), V905I (2819G>A 2821G>A), T920V (2864A>G 2865C>T 2866T>G), A926S (2882G>T 2884C>G), T978S (3038A>T), Q993K (3083C>A 3085A>G), T1014S (3146A>T 3148G>C), D1089N (3371G>A 3373T>C), Q1103E (3413C>G), F1123Y (3474T>A 3475T>C), V1124L (3476G>C), A1129D (3492C>A), R1143K (3534G>A), A1162V (3591C>T 3592T>C), S1163Q (3593T>C 3594C>A), A1175S (3629G>T), R1193K (3684G>A 3685G>A), Y1196F (3693A>T), A1202S (3710G>T 3712A>T), I1203V (3713A>G 3715T>G), V1206I (3722G>A 3724G>A), H1208R (3729A>G), K1216E (3752A>G), V1218I (3758G>A), V1224I (3776G>A 3778G>T), P1352A (4160C>G 4162A>T), I1362M (4192T>G), G1370S (4214G>A), P1410Q (4335C>A 4336T>A), E1506G (4623A>G 4624A>T), T1511S (4637A>T 4639T>G), S1537N (4716G>A 4717C>T), S1538H (4718A>C 4719G>A), V1582T (4850G>A 4851T>C), V1598I (4898G>A), E1610D (4936A>C), T1614N (4947C>A 4948A>T), N1618H (4958A>C), T1628A (4988A>G 4990C>A), V1642M (5030G>A), I1658T (5079T>C 5080T>A), A1662S (5090G>T 5092T>A), P1667A (5105C>G 5107C>T), N1670E (5114A>G 5116T>A), V1672I (5120G>A 5122T>A), S1676T (5132T>A), V1679I (5141G>A 5143G>T), S1691A (5177T>G 5179G>C), Q1697H (5197G>T), T1701S (5207A>T 5209C>A), S1744G (5336A>G), V1746I (5342G>A 5344A>C), T1751S (5357A>T 5359C>T), K1752H (5360A>C 5362A>C), H1754N (5366C>A 5368T>C), V1755I (5369G>A 5371G>T), I1765V (5399A>G 5401T>G), V1785I (5459G>A 5461T>C), V1786I (5462G>A), D1794N (5486G>A), T1796P (5492A>C 5494A>T), Q1800H (5506A>T), L1810V (5534C>G), K1811Q (5537A>C), A1813S (5543G>T), E1818Q (5558G>C), I1826V (5582A>G 5584T>G), A1829S (5591G>T 5593C>A), T1833L (5603A>C 5604C>T), Y1836F (5613A>T), R1853N (5664G>A 5665A>T), L1856C (5672C>T 5673T>G 5674A>T), Q1857S (5675C>A 5676A>G 5677G>T), Y1869H (5711T>C), R1878K (5739G>A), L1882W (5751T>G), M1887I (5767G>T), K1889N (5773G>C), I1890V (5774A>G 5776A>G), I1893T (5784T>C 5785T>A), V1895L (5789G>T 5791C>G), E1898G (5799A>G 5800G>T), F1900T (5804T>A 5805T>C 5806C>T), Y1907F (5826A>T), V1913I (5843G>A 5845G>A), M1914F (5846A>T 5848G>C), L1915Q (5849T>C 5850T>A 5851G>A), D1918E (5860C>A), N1930H (5894A>C), Y1934F (5907A>T), H1946F (5942C>T 5943A>T 5944C>T), E1949D (5953A>T), N1950S (5955A>G 5956C>T), S1962A (5990T>G 5992A>C), T1966V (6002A>G 6003C>T 6004C>T), G1968E (6009G>A 6010G>A), T1976I (6033C>T 6034A>C), A1977S (6035G>T), S1980T (6044T>A 6046A>G), F1982L (6050T>C), D1985N (6059G>A), V1986I (6062G>A 6064T>C), H1997Q (6097T>A), I2002M (6112A>G), M2007L (6125A>C), E2008G (6129A>G 6130G>T), C2015S (6150G>C), V2016L (6152G>T 6154C>G), E2024G (6177A>G), T2027E (6185A>G 6186C>A 6187T>G), H2028R (6189A>G), M2032I (6202G>C), D2036E (6214T>G), L2037V (6215C>G 6217G>T), T2038P (6218A>C), E2039S (6221G>T 6222A>C 6223A>T), G2040E (6225G>A 6226G>A), I2041V (6227A>G 6229C>G), I2042V (6230A>G 6232A>G), K2044S (6237A>G 6238A>T), V2045T (6239G>A 6240T>C 6241G>A), I2046V (6242A>G 6244A>G), S2054T (6266T>A 6268G>T), S2056M (6273G>T 6274C>G), V2062E (6291T>A 6292T>A), D2063G (6294A>G), V2064L (6296G>C 6298A>C), L2066T (6302T>A 6303T>C 6304A>G), D2069S (6311G>T 6312A>C 6313T>C), I2071V (6317A>G 6319A>G), I2073V (6323A>G 6325C>G), A2089T (6371G>A 6373A>T), I2093V (6383A>G 6385C>G), C2100H (6404T>C 6405G>A), R2103V (6413C>G 6414G>T 6415A>G), S2104N (6416T>A 6417C>A 6418A>C), C2111S (6437T>A), T2115S (6449A>T), S2117A (6455T>G 6457C>A), S2118T (6458T>A), H2127N (6485C>A), R2128S (6490A>C), H2129N (6491C>A), Q2130V (6494C>G 6495A>T 6496G>A), T2131P (6497A>C 6499A>T), I2132E (6500A>G 6501T>A 6502T>G), D2133N (6503G>A 6505T>C), S2137P (6515T>C 6517C>A), F2138L (6518T>C 6520C>T), T2149M (6552C>T 6553A>G), G2154N (6566G>A 6567G>A 6568C>T) | | | | | | | | |

| | Begin | End | Coverage | Score | Concordance | Matches | Identities | I/D/M/F* | Stop Codons |
|---|---|---|---|---|---|---|---|---|---|
| **NT** | **21** | **6806** | **86.9%** | **7777** | **57.4%** | **6785 (99.6%)** | **5350 (78.5%)** | **25/1** | |

Codon mutations:

TTT3TTC (115T>C), TCC4TCT (118C>T), AAG5AAA (121G>A), ATG6TTA (122A>T 124G>A), CCC8TCT (128C>T 130C>T), TTC10CCT (134T>C 135T>C 136C>T), AAC11AGC (138A>G), TCA12CCC (140T>C 142A>C), GTT13GCA (144T>C 145T>A), ACT14ATG (147C>T 148T>G), GAA15GAG (151A>G), TGC17AGT (155T>A 157C>T), GCT18GTT (159C>T), ACC19GCC (161A>G), TCT24TCG (178T>G), TCA27TCT (187A>T), ACT30ACA (196T>A), GCT31CAA (197G>C 198C>A 199T>A), AGT34AGC (208T>C), AAT35AAC (211T>C), TTC36TTT (214C>T), CGC38CGT (220C>T), ACT39ACA (223T>A), CTG40CTT (226G>T), TCA47TCC (247A>C), CAA49CAG (253A>G), ATT50ATC (256T>C), TCT52TCA (262T>A), CAC53CAT (265C>T), GAA55GAG (271A>G), TTG57CTG (275T>C), AAG58AAA (280G>A), CCT59CCC (283T>C), TCA60TCG (286A>G), GTT61GTG (289T>G), ACA62ACT (292A>T), GCT65GCA (301T>A), TCT66ACT (302T>A), TCC67TCT (307C>T), TTT68TTC (310T>C), ACC69TCT (311A>T 313C>T), AGC70AAT (315G>A 316C>T), TGT72TGC (322T>C), AAC73AGT (324A>G 325C>T), TCT74TCA (328T>A), ACT76ACC (334T>C), AAA77AAG (337A>G), TTG78TTA (340G>A), TTA79TTG (343A>G), AAG81AAA (349G>A), ACT84ACA (358T>A), TTA85CTA (359T>C), ATT86ATA (364T>A), GAA87GAG (367A>G), CCC88CCT (370C>T), TTC89TTT (373C>T), ATC90ATA (376C>A), GCT92GCA (382T>A), GTC96GTT (394C>T), GCA97GCC (397A>C), TCC98TCT (400C>T), AAA101AAG (409A>G), TCA102TCC (412A>C), ATT103ATC (415T>C), GTT107GTG (427T>G), AAG109AAA (433G>A), GTT111GTC (439T>C), GCA112ACA (440G>A), AGC113GGC (443A>G), AAA115AAG (451A>G), AAG117AAA (457G>A), AAC123GAC (473A>G), GAC127AAT (485G>A 487C>T), TCT129TCC (493T>C), GAG130GAA (496G>A), GTG132GTT (502G>T), CTT140CTC (526T>C), ATT149ATA (553T>A), TTC151TTT (559C>T), CCA154CCT (568A>T), AGT156AAT (573G>A), CTA158TTA (578C>T), GGA160GGT (586A>T), CAG163CAA (595G>A), ACT165TCA (599A>T 601T>A), CAC166CAT (604C>T), ATA167ACT (606T>C 607A>T), AAT170AGT (615A>G), ACA171ATT (618C>T 619A>T), AAC174AAT (628C>T), GCT178GCC (640T>C), TTA182CTG (650T>C 652A>G), GAC183GAG (655C>G), TTA185CTG (659T>C 661A>G), CCG186CCC (664G>C), TCC189TCT (673C>T), CAA190CAG (676A>G), GCA196GCG (694A>G), CTA201TTA (707C>T), GGG203GGA (715G>A), GAA204GAG (718A>G), TCC205GCT (719T>G 721C>T), ATG206GTG (722A>G), TCA207TCT (727A>T), CAC208CAT (730C>T), CTT211CTG (739T>G), GTG214GTT (748G>T), GTA215GTT (751A>T), GCA216GCG (754A>G), CTC218TTA (758C>T 760C>A), ATA219ATT (763A>T), GCC220GCT (766C>T), GGG222GGC (772G>C), ATT223ATA (775T>A), TCT224TCA (778T>A), ACC225ACA (781C>A), TTC228TTT (790C>T), GTG229GTA (793G>A), GCT230GCA (796T>A), GTA231GTT (799A>T), CCT232CCG (802T>G), GGT233GGC (805T>C), AGA234CGT (806A>C 808A>T), GGT237GGA (817T>A), CTA238CTC (820A>C), TCC239ACC (821T>A), AAC240AAT (826C>T), CCG241CCA (829G>A), CTA242CTC (832A>C), TCA243TCC (835A>C), AAA244AAG (838A>G), CTG246TTA (842C>T 844G>A), TAC247TAT (847C>T), GCG249GCA (853G>A), GGT252GGA (862T>A), GCT253GCC (865T>C), AAT257AAC (877T>C), TTG259CTT (881T>C 883G>T), TTT260TTC (886T>C), ACC261ACA (889C>A), TTC263TTT (895C>T), AGG264AGA (898G>A), AAT265AAC (901T>C), TGT269TGC (913T>C), ATA281ATT (949A>T), ATT284ATA (958T>A), GGT286GGA (964T>A), TTT287TTC (967T>C), CCA290CCT (976A>T), GTT291ATA (977G>A 979T>A), GCT294GCA (988T>A), ATT295ATC (991T>C), TTG299CTT (1001T>C 1003G>T), TCT300GCA (1004T>G 1006T>A), ACG301ACA (1009G>A), TTC304TTT (1018C>T), ACG305GTG (1019A>G 1020C>T), GAG309GAA (1033G>A), CCA316CCC (1054A>C), CAT318CAC (1060T>C), CGC319CGA (1063C>A), TTG320CTT (1064T>C 1066G>T), AAT322AAC (1072T>C), GCA324GCG (1078A>G), AAC325AAT (1081C>T), ACT329ACC (1093T>C), ATC330ATA (1096C>A), CTA335TTG (1109C>T 1111A>G), AAA341AAG (1129A>G), TCT343TCA (1135T>A), TAC345TAT (1141C>T), GCT347GCC (1147T>C), GTA351GTG (1159A>G), GCC353GCA (1165C>A), AGG358AGA (1180G>A), GTA359GTT (1183A>T), ACG360ACA (1186G>A), GCA361GCC (1189A>C), GCA362GCC (1192A>C), ATA363ATT (1195A>T), ACC364TCT (1196A>T 1198C>T), CTT366CTG (1204T>G), GAT367GAG (1207T>G), GGG371GGC (1219G>C), AAC373AAT (1225C>T), ACT377ACA (1237T>A), GGC378GGT (1240C>T), GTG379GTT (1243G>T), GGC380GGT (1246C>T), CAA381CAG (1249A>G), AGG383AGA (1255G>A), GCA384GCT (1258A>T), CCT386CCC (1264T>C), GGT393GGA (1285T>A), AGT395AGC (1291T>C), GGG398GGA (1300G>A), ACC401ACA (1309C>A), CGG404CGA (1318G>A), CTG405TTA (1319C>T 1321G>A), ATC407ATT (1327C>T), GTT410GTG (1336T>G), TTG411CTT (1337T>C 1339G>T), CGC413AGG (1343C>A 1345C>G), GAA415GAT (1351A>T), GAG416GAA (1354G>A), CCA417CCC (1357A>C), TTG419CTT (1361T>C 1363G>T), AGG421CGG (1367A>C), CTC422CTG (1372C>G), GCT424GCC (1378T>C), GTG425GTA (1381G>A), CGA428CGT (1390A>T), TCA433TCC (1405A>C), AAT434AAC (1408T>C), TAT435TAC (1411T>C), GCT440GCA (1426T>A), CTG442CTT (1432G>T), GCT448GCA (1450T>A), CTG449TTG (1451C>T), GGG452GGT (1462G>T), GCA453GCT (1465A>T), CAA455CAG (1471A>G), GAC458GAT (1480C>T), ATT459GTT (1481A>G), AAG460AAA (1486G>A), ATA463ATT (1495A>T), ATC465ATT (1501C>T), TCC467TCA (1507C>A), ACT468ACA (1510T>A), CCA470CCT (1516A>T), CCC472CCA (1522C>A), TTG473TTA (1525G>A), CCC474CCA (1528C>A), GCA476GCT (1534A>T), CGC483AGG (1553C>A 1555C>G), CGA488AGG (1568C>A 1570A>G), TAT489TAC (1573T>C), ATA490ATT (1576A>T), GCC492GCT (1582C>T), ACA493ACC (1585A>C), TCC494TCA (1588C>A), AAC495AAT (1591C>T), TCA498TCC (1600A>C), GCT501TCA (1607G>T 1609T>A), CAG502CAA (1612G>A), GGT504GGG (1618T>G), ACC506ACA (1624C>A), AGA513CGG (1643A>C 1645A>G), CGG515AGG (1649C>A), GTT517GTG (1657T>G), CTT518CTG (1660T>G), GTG519GTA (1663G>A), TGC521TGT (1669C>T), GGT525GGA (1681T>A), GAG526GAA (1684G>A), TTT527TTC (1687T>C), ACT529GTA (1691A>G 1692C>T 1693T>A), GCT531GCA (1699T>A), CCC532CCT (1702C>T), ACT533GCA (1703A>G 1705T>A), TCT534GCC (1706T>G 1708T>C), CTT536CTG (1714T>G), GAA537AGA (1715G>A 1716A>G), GTG540GTC (1726G>C), GTT541GTG (1729T>G), GAG542GAA (1732G>A), AGC543AGT (1735C>T), AAA544AGG (1737A>G 1738A>G), AGA545CGT (1739A>C 1741A>T), CCC546CCA (1744C>A), ACA550ATC (1755C>T 1756A>C), AGG552AGA (1762G>A), AGC555GAT (1769A>G 1770G>A 1771C>T), GAT557GAC (1777T>C), GAT558GAC (1780T>C), TTG559CTA (1781T>C 1783G>A), CTG560CTT (1786G>T), AAG568AAA (1810G>A), GTT569GTG (1813T>G), GCA571GCC (1819A>C), ATC573ATA (1825C>A), TCA574TCT (1828A>T), GGA575GGC (1831A>C), CTG577CTC (1837G>C), TAT578TTT (1839A>T), GCC580ACC (1844G>A), ACA581ACC (1849A>C), TCA582ACC (1850T>A 1852A>C), AAG584AAA (1858G>A), GTG590GTC (1876G>C), TCA591ACT (1877T>A 1879A>T), GAG593GAA (1885G>A), ATA595GTG (1889A>G 1891A>G), ATA596ATT (1894A>T), GAT600GAA (1906T>A), ATG601CCA (1907A>C 1908T>C 1909G>A), TTG602TGG (1911T>G), AAT603AAC (1915T>C), ATC604ATT (1918C>T), CAA605CAG (1921A>G), GAT606GAC (1924T>C), ACC607ACA (1927C>A), GAT610GAC (1936T>C), GCT611ACA (1937G>A 1939T>A), AAG612AAA (1942G>A), CTT613CTG (1945T>G), CCC614CCG (1948C>G), GTG615GTA (1951G>A), GTT617GTG (1957T>G), GTA618GTC (1960A>C), GTG624GTA (1978G>A), TAC626TAT (1984C>T), TTG630CTA (1994T>C 1996G>A), GTG632GTT (2002G>T), GAA633GAG (2005A>G), GCT634GCC (2008T>C), AAA636AAG (2014A>G), TAT637TTT (2016A>T), GCT638GCC (2020T>C), TAT639TAC (2023T>C), GCA646GCG (2044A>G), TAT648TAC (2050T>C), TTC649TTT (2053C>T), CCT650CCA (2056T>A), GAC652GAT (2062C>T), AAA655AAG (2071A>G), TTT659TAT (2082T>A), CTA662CTT (2092A>T), TCC665CCT (2099T>C 2101C>T), GAA668GAG (2110A>G), ACA669TCC (2111A>T 2113A>C), ACC671ACT (2119C>T), AGG672CGG (2120A>C), GTT674GTC (2128T>C), AAT675ACA (2130A>C 2131T>A), GAC676GAT (2134C>T), CTG678CTC (2140G>C), TAT679CAA (2141T>C 2143T>A), CAA680CAG (2146A>G), GGT681GGA (2149T>A), TGT683TGC (2155T>C), AAT684AAC (2158T>C), GAG685GCG (2160A>C), AAC686AAT (2164C>T), CTG690CTC (2176G>C), ATA691GTT (2177A>G 2179A>T), AAG692AAA (2182G>A), ATC694ATT (2188C>T), GCT696GCA (2194T>A), TAT698TAC (2200T>C), ATC706ATA (2224C>A), TAC707TAT (2227C>T), GCT708GCG (2230T>G), AAT712TGT (2240A>T 2241A>G), GAG713GAT (2245G>T), CGT714CGC (2248T>C), AGC715AAA (2250G>A 2251C>A), GAC717GAT (2257C>T), GTT718GTG (2260T>G), CAG719CAA (2263G>A), CTG722TTA (2270C>T 2272G>A), GGA725GGC (2281A>C), ATG726TTG (2282A>T), GCG727GCA (2287G>A), CAC728CAT (2290C>T), GCC730GCT (2296C>T), ATA731ATT (2299A>T), GCT733GCA (2305T>A), GTG735GTT (2311G>T), GAC736GAT (2314C>T), CTG738TTA (2318C>T 2320G>A), ATA739ATT (2323A>T), TGC740TGT (2326C>T), GTG741ATC (2327G>A 2329G>C), GAG742GAA (2332G>A), CAG743CAA (2335G>A), CCT745CCA (2341T>A), GCC746GCA (2344C>A), CAC747CAT (2347C>T), GTA748GTG (2350A>G), TTC750TTT (2356C>T), TTG751TTA (2359G>A), ACA753ACT (2365A>T), GCA754GCT (2368A>T), TTT755TTC (2371T>C), AGG757CGT (2375A>C 2377G>T), TAT758TAC (2380T>C), CGC761AGA (2387C>A 2389C>A), AGA762AGG (2392A>G), CAG766CAA (2404G>A), CCT767CCC (2407T>C), AGG768AAA (2409G>A 2410G>A), TCC770TCT (2416C>T), TTA790TTG (2476A>G), CCG791CCA (2479G>A), ATC793ATT (2485C>T), TTG794TTA (2488G>A), AGA795AGG (2491A>G), GTT796GTG (2494T>G), CTG797CTT (2497G>T), ATT798ATC (2500T>C), AAG799AAA (2503G>A), GCT800GCA (2506T>A), GGC801GGG (2509C>G), CTT802TTG (2510C>T 2512T>G), ATT803ATC (2515T>C), TTA804CTA (2516T>C), GCA806GCC (2524A>C), GCA810GCT (2536A>T), GGG812GGT (2542G>T), GCA816GCT (2554A>T), TTC817TTT (2557C>T), GCT819GCG (2563T>G), CAA822CGG (2571A>G 2572A>G), TCC823TCA (2575C>A), CCT827CCC (2587T>C), AGA832CGG (2600A>C 2602A>G), ATT835ATC (2611T>C), ACC836ACA (2614C>A), AAC837AAT (2617C>T), AAT840AAC (2626T>C), AGC841AGT (2629C>T), CGG845AGG (2639C>A), AAG848AAA (2650G>A), ATC852ATT (2662C>T), TTG857TTA (2677G>A), CTG858TTG (2678C>T), GCA859GCG (2683A>G), AAG862AAA (2692G>A), GGT863GGA (2695T>A), CTG866CTT (2704G>T), AAA868AAG (2710A>G), GCA870GCC (2716A>C), GGC874AGC (2726G>A), TTC875TTT (2731C>T), GGA879GGC (2743A>C), TTG880CTG (2744T>C), GTC885GTG (2761C>G), TTG887CTC (2765T>C 2767G>C), AGA888CGA (2768A>C), GGT890GGG (2776T>G), TAC894TAT (2788C>T), ACC898ACA (2800C>A), GTC900GTT (2806C>T), TGC901TGT (2809C>T), TCG902TCT (2812G>T), GGC903GGA (2815C>A), GTG905ATA (2819G>A 2821G>A), ACA907ACT (2827A>T), GTG908GTA (2830G>A), TTT912TTC (2842T>C), TCA913TCC (2845A>C), AGC914AGT (2848C>T), CGT916AGA (2852C>A 2854T>A), GGC918GGG (2860C>G), ACA919ACT (2863A>T), ACT920GTG (2864A>G 2865C>T 2866T>G), TTC921TTT (2869C>T), ATA923ATT (2875A>T), ATA924ATC (2878A>C), GCC926TCG (2882G>T 2884C>G), TCA928TCC (2890A>C), TAC934TAT (2908C>T), AAT935AAC (2911T>C), CGG940CGT (2926G>T), TTC941TTT (2929C>T), TTG942CTA (2930T>C 2932G>A), AAG943AAA (2935G>A), CAA945CAG (2941A>G), ATT947ATA (2947T>A), GTC948GTG (2950C>G), CTG949CTT (2953G>T), CTC950CTG (2956C>G), AAT951AAC (2959T>C), GGA953GGT (2965A>T), CCC955CCT (2971C>T), GAT956GAC (2974T>C), CCC958CCG (2980C>G), CCT960CCG (2986T>G), ATT962ATC (2992T>C), AAA964AAG (2998A>G), CGG969AGG (3011C>A), TGT973TGC (3025T>C), TCT975TCA (3031T>A), AAG976AAA (3034G>A), GGC977GGA (3037C>A), ACT978TCT (3038A>T), GGG980GGT (3046G>T), GTA981GTC (3049A>C), TGT982TGC (3052T>C), GTA986GTT (3064A>T), AAG989AAA (3073G>A), GCT991GCC (3079T>C), CAA993AAG (3083C>A 3085A>G), TTG999CTT (3101T>C 3103G>T), GTG1003GTT (3115G>T), TCC1006TCA (3124C>A), CTT1010CTG (3136T>G), ACG1014TCC (3146A>T 3148G>C), GAC1018GAT (3160C>T), AGT1020AGC (3166T>C), TCC1021TCA (3169C>A), CTG1024CTA (3178G>A), TCT1026TCA (3184T>A), TAT1030TAC (3196T>C), TAT1032TAC (3202T>C), GAT1040GAC (3226T>C), CTT1044CTG (3238T>G), CTT1045TTA (3239C>T 3241T>A), CTT1046TTG (3242C>T 3244T>G), TTG1047TTA (3247G>A), AAC1049AAT (3253C>T), GTG1050GTA (3256G>A), AAA1053AAG (3265A>G), TGC1056TGT (3274C>T), ATT1057ATA (3277T>A), ATT1060ATA (3286T>A), TCT1064TCA (3298T>A), GAA1068GAG (3310A>G), CAC1071CAT (3319C>T), GGG1073GGC (3325G>C), TTT1074TTC (3328T>C), GTA1075GTT (3331A>T), GCA1076GCC (3334A>C), ACA1080ACG (3346A>G), GCT1081GCG (3349T>G), ATT1082ATC (3352T>C), CGA1084AGA (3356C>A), TTG1087CTG (3365T>C), GAT1089AAC (3371G>A 3373T>C), CTT1090TTG (3374C>T 3376T>G), CCG1091CCA (3379G>A), ACT1092ACA (3382T>A), GGT1093GGG (3385T>G), CCG1094CCT (3388G>T), AAA1096AAG (3394A>G), GCA1098GCC (3400A>C), GTA1100GTG (3406A>G), TGC1102TGT (3412C>T), CAG1103GAG (3413C>G), CTC1104TTG (3416C>T 3418C>G), CTT1105CTG (3421T>G), CTA1108TTG (3428C>T 3430A>G), CGA1109AGA (3431C>A), AGG1111AGA (3439G>A), GCG1113GCT (3445G>T), GAA1117GAG (3457A>G), CAG1120CAA (3466G>A), TTT1123TAC (3474T>A 3475T>C), GTT1124CTT (3476G>C), GGC1125GGA (3481C>A), ACA1126ACT (3484A>T), GCT1129GAT (3492C>A), GCC1133GCA (3505C>A), ACG1134ACT (3508G>T), CCC1136CCA (3514C>A), AAA1138AAG (3520A>G), ACA1139ACC (3523A>C), ACG1140ACA (3526G>A), CTG1141CTC (3529G>C), CGA1142AGG (3530C>A 3532A>G), AGG1143AAG (3534G>A), GGC1145GGT (3541C>T), TTG1146CTC (3542T>C 3544G>C), GCT1149GCA (3553T>A), GGG1151GGA (3559G>A), GCA1153GCT (3565A>T), TCT1158TCC (3580T>C), ATC1159ATA (3583C>A), ATT1160ATC (3586T>C), TCA1161TCG (3589A>G), GCT1162GTC (3591C>T 3592T>C), TCA1163CAA (3593T>C 3594C>A), CGT1166AGA (3602C>A 3604T>A), GGG1167GGA (3607G>A), GAA1168GAG (3610A>G), GGT1169GGC (3613T>C), CCG1172CCA (3622G>A), TAC1173TAT (3625C>T), GTG1174GTC (3628G>C), GCT1175TCT (3629G>T), ATA1177ATC (3637A>C), CAA1178CAG (3640A>G), AAA1179AAG (3643A>G), TAC1180TAT (3646C>T), GAA1182GAG (3652A>G), GCA1184GCC (3658A>C), CAA1185CAG (3661A>G), AAT1186AAC (3664T>C), GAG1189GAA (3673G>A), GAC1190GAT (3676C>T), ATT1191ATC (3679T>C), GCG1192GCC (3682G>C), AGG1193AAA (3684G>A 3685G>A), CTA1194TTG (3686C>T 3688A>G), GCC1195GCT (3691C>T), TAT1196TTT (3693A>T), GGG1198GGA (3700G>A), CTA1199TTG (3701C>T 3703A>G), CGT1200CGG (3706T>G), CAA1201CAG (3709A>G), GCA1202TCT (3710G>T 3712A>T), ATT1203GTG (3713A>G 3715T>G), TTG1204CTG (3716T>C), GTG1206ATA (3722G>A 3724G>A), CAC1208CGC (3729A>G), TCC1209TCG (3733C>G), GTT1212GTG (3742T>G), AAG1216GAG (3752A>G), GTC1218ATC (3758G>A), ACA1219ACT (3763A>T), CAG1220CAA (3766G>A), GTG1224ATT (3776G>A 3778G>T), GTG1225GTT (3781G>T), CTC1226TTG (3782C>T 3784C>G), GGT1228GGA (3790T>A), GTT1229GTC (3793T>C), TTT1232TTC (3802T>C), TAT1234TAC (3808T>C), GAC1236GAT (3814C>T), GGG1237GGA (3817G>A), TTG1240CTG (3824T>C), GGG1245GGT (3841G>T), CCG1247CCA (3847G>A), TAC1248TAT (3850C>T), AAG1250AAA (3856G>A), GGC1254GGT (3868C>T), ACT1255ACC (3871T>C), GAG1259GAA (3883G>A), GTG1261GTA (3889G>A), CCA1263CCC (3895A>C), GGA1265GGC (3901A>C), GGA1267GGT (3907A>T), GAT1268GAC (3910T>C), CGA1269AGA (3911C>A), GTC1270GTG (3916C>G), CTC1272TTG (3920C>T 3922C>G), AAA1273AAG (3925A>G), AGG1274AGA (3928G>A), ACT1276ACC (3934T>C), GAC1281GAT (3949C>T), GAG1283GAA (3955G>A), GCT1284GCA (3958T>A), TTG1285CTA (3959T>C 3961G>A), ATT1289ATC (3973T>C), GGA1292GGT (3982A>T), CTG1297TTG (3995C>T), GTC1298GTT (4000C>T), ACC1299ACA (4003C>A), ACC1300ACA (4006C>A), CAA1301CAG (4009A>G), GCC1303GCA (4015C>A), AAA1304AAG (4018A>G), TTG1308CTA (4028T>C 4030G>A), CCG1309CCT (4033G>T), TTG1310CTG (4034T>C), GAA1311GAG (4039A>G), GTT1313GTG (4045T>G), GGG1315GGC (4051G>C), CGG1317AGG (4055C>A), ACC1319ACA (4063C>A), TTG1321CTA (4067T>C 4069G>A), GAA1323GAG (4075A>G), ATT1324ATA (4078T>A), CTT1325CTA (4081T>A), CCC1326CCT (4084C>T), TAC1329TAT (4093C>T), AAT1330AAC (4096T>C), CTG1338TTG (4118C>T), GAT1339GAC (4123T>C), TCC1341TCT (4129C>T), TCC1347TCG (4147C>G), CTG1351TTG (4157C>T), CCA1352GCT (4160C>G 4162A>T), GGT1361GGA (4189T>A), ATT1362ATG (4192T>G), GAA1363GAG (4195A>G), CTG1366TTA (4202C>T 4204G>A), TTG1367CTA (4205T>C 4207G>A), GCT1368GCA (4210T>A), AAT1369AAC (4213T>C), GGC1370AGC (4214G>A), TTC1371TTT (4219C>T), AGA1372CGG (4220A>C 4222A>G), GCA1373GCT (4225A>T), GTC1374GTT (4228C>T), ACA1377ACG (4237A>G), TCT1380TCC (4246T>C), GCT1381GCA (4249T>A), GAC1382GAT (4252C>T), TAT1383TAC (4255T>C), ATC1392ATT (4282C>T), GCC1394GCA (4288C>A), CAA1396CAG (4294A>G), TGT1399TGC (4303T>C), CTT1401CTG (4309T>G), GAC1404GAT (4318C>T), GGA1407GGC (4327A>C), CCT1410CAA (4335C>A 4336T>A), GGT1411GGC (4339T>C), TCT1412TCC (4342T>C), GAC1414GAT (4348C>T), TCC1415TCA (4351C>A), AAT1416AAC (4354T>C), AGA1418CGC (4358A>C 4360A>C), CAT1419CAC (4363T>C), GTG1420GTA (4366G>A), CTT1421TTG (4367C>T 4369T>G), CTT1422CTG (4372T>G), GCT1425GCC (4381T>C), TCA1426TCT (4384A>T), TGC1427TGT (4387C>T), ACT1430ACC (4396T>C), ATT1431ATC (4399T>C), GAG1433GAA (4405G>A), GTT1436GTG (4414T>G), TTT1437TTC (4417T>C), AGG1438CGC (4418A>C 4420G>C), TTG1439CTG (4421T>C), GTG1440GTT (4426G>T), GGA1441GGG (4429A>G), GGT1442GGC (4432T>C), CCT1444CCA (4438T>A), TCA1445TCT (4441A>T), GGA1446GGG (4444A>G), GCA1448GCT (4450A>T), CTC1449CTT (4453C>T), ACC1450ACT (4456C>T), GTT1451GTC (4459T>C),

ATC1452ATT (4462C>T), TTC1453TTT (4465C>T), AAC1454AAT (4468C>T), CGT1463CGG (4495T>G), GCA1465GCT (4501A>T), ATT1467ATC (4507T>C), TCT1468TCA (4510T>A), CTA1469CTG (4513A>G), TTG1470CTG (4514T>C), AGA1472AGG (4522A>G), CAT1474CAC (4528T>C), GCA1476GCT (4534A>T), GCT1477GCA (4537T>A), CAA1478CAG (4540A>G), ATA1480ATT (4546A>T), GGG1481GGC (4549G>C), TGC1482TGT (4552C>T), AAA1483AAG (4555A>G), CCC1484CCA (4558C>A), AAC1488AAT (4570C>T), CTA1490CTT (4576A>T), GTG1492GTT (4582G>T), GCA1493GCT (4585A>T), GTC1494GTA (4588C>A), GAC1498GAT (4600C>T), CTA1500CTT (4606A>T), GTT1503GTA (4615T>A), CCC1504CCA (4618C>A), GAA1506GGT (4623A>G 4624A>T), TTG1507CTC (4625T>C 4627G>C), TAT1510TAC (4636T>C), ACT1511TCG (4637A>T 4639T>G), GCT1514GCA (4648T>A), ATT1515ATA (4651T>A), GAA1518GAG (4660A>G), TTG1519CTA (4661T>C 4663G>A), GTT1524GTC (4678T>C), ATA1526ATC (4684A>C), ATC1530ATA (4696C>A), AAG1532AAA (4702G>A), AAC1533AAT (4705C>T), GTT1536GTC (4714T>C), AGC1537AAT (4716G>A 4717C>T), AGT1538CAT (4718A>C 4719G>A), CTA1542CTG (4732A>G), GAC1543GAT (4735C>T), TTG1544CTT (4736T>C 4738G>T), TCT1545TCA (4741T>A), CTA1547TTG (4745C>T 4747A>G), TTT1549TTC (4753T>C), CTA1550CTT (4756A>T), AGC1551AGT (4759C>T), AGA1552AGG (4762A>G), GGT1553GGC (4765T>C), TTT1554TTC (4768T>C), AAG1555AAA (4771G>A), AGG1556AGA (4774G>A), GGA1560GGG (4786A>G), TAC1561TAT (4789C>T), CCT1565CCC (4801T>C), CTG1566TTG (4802C>T), GTA1569GTG (4813A>G), TCT1570TCC (4816T>C), GAA1573GAG (4825A>G), CCA1574CCC (4828A>C), TAC1576TAT (4834C>T), ATA1578ATT (4840A>T), AGG1579AGA (4843G>A), CCT1580CCC (4846T>C), GTT1582ACT (4850G>A 4851T>C), GCT1587GCC (4867T>C), CTC1588TTG (4868C>T 4870C>G), TTG1591TTA (4879G>A), AAC1593AAT (4885C>T), ATA1594ATT (4888A>T), GAC1595GAT (4891C>T), ACG1596ACT (4894G>T), GGA1597GGC (4897A>C), GTT1598ATT (4898G>A), AGA1599AGG (4903A>G), GCA1601GCC (4909A>C), CAC1603CAT (4915C>T), GGG1605GGT (4921G>T), CAG1607CAA (4927G>A), GTT1608GTG (4930T>G), GAA1610GAC (4936A>C), GTG1613GTT (4945G>T), ACA1614AAT (4947C>A 4948A>T), GTT1616GTG (4954T>G), AAC1618CAC (4958A>C), GCT1619GCC (4963T>C), AAG1621AAA (4969G>A), CGG1623CGA (4975G>A), TTC1625TTT (4981C>T), CCA1626CCT (4984A>T), GCC1627GCT (4987C>T), ACC1628GCA (4988A>G 4990C>A), CCT1631CCG (4999T>G), ACA1632ACT (5002A>T), TAT1633TAC (5005T>C), GAG1637GAA (5017G>A), CAG1638CAA (5020G>A), GAC1639GAT (5023C>T), CTG1641CTT (5029G>T), GTG1642ATG (5030G>A), GAG1643GAA (5035G>A), GTT1644GTA (5038T>A), ACA1645ACT (5041A>T), GGA1646GGC (5044A>C), CCA1648CCT (5050A>T), GGG1651GGT (5059G>T), CTC1652CTT (5062C>T), CTT1656TTA (5072C>T 5074T>A), ATT1658ACA (5079T>C 5080T>A), GCA1659GCT (5083A>T), TCA1661TCC (5089A>C), GCT1662TCA (5090G>T 5092T>A), GTG1664GTT (5098G>T), CCT1665CCA (5101T>A), TTG1666CTG (5102T>C), CCC1667GCT (5105C>G 5107C>T), CCA1668CCT (5110A>T), GGC1669GGT (5113C>T), AAT1670GAA (5114A>G 5116T>A), ACT1671ACC (5119T>C), GTT1672ATA (5120G>A 5122T>A), CTG1673TTG (5123C>T), TTT1675TTC (5131T>C), TCT1676ACT (5132T>A), GTG1679ATT (5141G>A 5143G>T), CAT1680CAC (5146T>C), ACT1681ACA (5149T>A), TTT1682TTC (5152T>C), GCT1683GCA (5155T>A), CGA1685AGA (5159C>A), TGC1689TGT (5173C>T), TCC1690TCT (5176C>T), TCG1691GCC (5177T>G 5179G>C), CGA1692AGG (5180C>A 5182A>G), ACA1693ACT (5185A>T), GCA1696GCC (5194A>C), CAG1697CAT (5197G>T), CAG1698CAA (5200G>A), GAC1700GAT (5206C>T), ACC1701TCA (5207A>T 5209C>A), ACC1702ACA (5212C>A), GTT1704GTC (5218T>C), AAC1707AAT (5227C>T), ACT1709ACC (5233T>C), GCA1711GCC (5239A>C), AAG1712AAA (5242G>A), GTA1717GTT (5257A>T), AGA1718AGG (5260A>G), GGG1719GGA (5263G>A), CCC1720CCA (5266C>A), ACC1721ACA (5269C>A), AGG1724CGA (5276A>C 5278G>A), GGG1728GGT (5290G>T), CTT1729CTC (5293T>C), ATT1730ATC (5296T>C), CCT1732CCA (5302T>A), GCT1736GCC (5314T>C), GAC1740GAT (5326C>T), CAC1741CAT (5329C>T), AGC1744GGC (5336A>G), ATT1745ATC (5341T>C), GTA1746ATC (5342G>A 5344A>C), ACC1747ACA (5347C>A), GGG1749GGA (5353G>A), ACC1751TCT (5357A>T 5359C>T), AAA1752CAC (5360A>C 5362A>C), CAT1754AAC (5366C>A 5368T>C), GTG1755ATT (5369G>A 5371G>T), GTT1756GTA (5374T>A), TTT1757TTC (5377T>C), GGC1760GGG (5386C>G), AAT1761AAC (5389T>C), GGG1762GGA (5392G>A), ATT1765GTG (5399A>G 5401T>G), GGT1766GGA (5404T>A), CCA1767CCT (5407A>T), GTG1768GTT (5410G>T), TGC1769TGT (5413C>T), GCT1770GCA (5416T>A), GCA1771GCT (5419A>T), TTG1772CTA (5420T>C 5422G>A), CTG1775TTA (5429C>T 5431G>A), TCC1776TCT (5434C>T), ACT1777ACG (5437T>G), GGC1778GGA (5440C>A), CAA1779CAG (5443A>G), TAT1780TAC (5446T>C), TCT1781TCA (5449T>A), TCT1782TCC (5452T>C), GTT1785ATC (5459G>A 5461T>C), GTT1786ATT (5462G>A), GTG1787GTC (5467G>C), ACC1791ACT (5479C>T), ATA1792ATT (5482A>T), GCA1793GCT (5485A>T), GAT1794AAT (5486G>A), GTT1795GTG (5491T>G), ACA1796CCT (5492A>C 5494A>T), CTC1798TTG (5498C>T 5500C>G), AGT1799AGC (5503T>C), CAA1800CAT (5506A>T), TAT1801TAC (5509T>C), CCA1802CCT (5512A>T), GGA1803GGT (5515A>T), GCC1806GCT (5524C>T), AAG1807AAA (5527G>A), TAT1808TAC (5530T>C), CTT1809TTG (5531C>T 5533T>G), CTG1810GTG (5534C>G), AAA1811CAA (5537A>C), GCT1813TCT (5543G>T), GAT1814GAC (5548T>C), ACA1815ACC (5551A>C), GAA1818CAA (5558G>C), CTT1820CTA (5566T>A), GCA1821GCT (5569A>T), ACA1823ACG (5575A>G), TGT1824TGC (5578T>C), ATT1826GTG (5582A>G 5584T>G), GCC1829TCA (5591G>T 5593C>A), GAG1832GAA (5602G>A), ACA1833CTA (5603A>C 5604C>T), CCA1834CCG (5608A>G), GCA1835GCT (5611A>T), TAC1836TTC (5613A>T), ATA1837ATT (5617A>T), CCT1838CCA (5620T>A), GGA1840GGT (5626A>T), TTT1842TTC (5632T>C), CTT1844CTG (5638T>G), GGT1845GGC (5641T>C), GTG1850GTA (5656G>A), CAA1851CAG (5659A>G), GGG1852GGA (5662G>A), AGA1853AAT (5664G>A 5665A>T), CTA1856TGT (5672C>T 5673T>G 5674A>T), CAG1857AGT (5675C>A 5676A>G 5677G>T), CAG1860CAA (5686G>A), CGC1861AGA (5687C>A 5689C>A), TTG1862CTT (5690T>C 5692G>T), GGA1867GGT (5707A>T), CCT1868CCA (5710T>A), TAT1869CAT (5711T>C), GGG1870GGT (5716G>T), GGA1871GGG (5719A>G), ACT1876ACC (5734T>C), ACT1877ACC (5737T>C), AGA1878AAA (5739G>A), GAG1879GAA (5743G>A), CTT1880CTA (5746T>A), AAG1881AAA (5749G>A), TTG1882TGG (5751T>G), CTG1883TTG (5753C>T), ATG1887ATT (5767G>T), AAG1889AAC (5773G>C), ATA1890GTG (5774A>G 5776A>G), TAT1891TAC (5779T>C), ATT1893ACA (5784T>C 5785T>A), TTA1894CTT (5786T>C 5788A>T), GTC1895TTG (5789G>T 5791C>G), CAA1896CAG (5794A>G), AGA1897AGG (5797A>G), GAG1898GGT (5799A>G 5800G>T), TTC1900ACT (5804T>A 5805T>C 5806C>T), ACT1902ACC (5812T>C), CTC1903CTG (5815C>G), TAC1904TAT (5818C>T), TAT1907TTT (5826A>T), CTC1908CTT (5830C>T), TCA1910TCC (5836A>C), GTG1913ATA (5843G>A 5845G>A), ATG1914TTC (5846A>T 5848G>C), TTG1915CAA (5849T>C 5850T>A 5851G>A), GTT1917GTG (5857T>G), GAC1918GAA (5860C>A), CTT1920TTG (5864C>T 5866T>G), CTC1922CTA (5872C>A), GGG1924GGT (5878G>T), TTT1925TTC (5881T>C), TTT1926TTC (5884T>C), CCC1927CCT (5887C>T), CGC1928CGT (5890C>T), AAT1930CAT (5894A>C), GTG1932GTA (5902G>A), TAC1934TTC (5907A>T), CTT1936CTC (5914T>C), TTT1939TTC (5923T>C), CTG1941CTA (5929G>A), CAC1946TTT (5942C>T 5943A>T 5944C>T), AAG1948AAA (5950G>A), GAA1949GAT (5953A>T), AAC1950AGT (5955A>G 5956C>T), ACG1953ACT (5965G>T), GTT1954GTG (5968T>G), TAC1955TAT (5971C>T), TTT1956TTC (5974T>C), AGA1957CGT (5975A>C 5977A>T), AAG1958AAA (5980G>A), CCT1959CCC (5983T>C), TTT1960TTC (5986T>C), CTC1961CTT (5989C>T), TCA1962GCC (5990T>G 5992A>C), AAA1964AAG (5998A>G), ACC1966GTT (6002A>G 6003C>T 6004C>T), GGG1968GAA (6009G>A 6010G>A), TCC1969TCA (6013C>A), AAA1970AAG (6016A>G), ATT1972ATA (6022T>A), TCC1974TCT (6028C>T), GCA1975GCT (6031A>T), ACA1976ATC (6033C>T 6034A>C), GCT1977TCT (6035G>T), GCT1978GCA (6040T>A), GAG1979GAA (6043G>A), TCA1980ACG (6044T>A 6046A>G), TTG1981TTA (6049G>A), TTT1982CTT (6050T>C), GGT1983GGG (6055T>G), GAT1985AAT (6059G>A), GTT1986ATC (6062G>A 6064T>C), TCC1987TCT (6067C>T), GCT1988GCC (6070T>C), CTC1991CTG (6079C>G), AGT1993AGC (6085T>C), AGG1994AGA (6088G>A), CTA1995TTA (6089C>T), CTT1996TTA (6092C>T 6094T>A), CAT1997CAA (6097T>A), CTT1998CTG (6100T>G), CAG1999CAA (6103G>A), AAG2000AAA (6106G>A), ATA2002ATG (6112A>G), TCA2005TCT (6121A>T), TCC2006TCA (6124C>A), ATG2007CTG (6125A>C), GAG2008GGT (6129A>G 6130G>T), TTT2011TTC (6139T>C), AAA2012AAG (6142A>G), ATT2013ATC (6145T>C), TAT2014TAC (6148T>C), TGT2015TCT (6150G>C), GTC2016TTG (6152G>T 6154C>G), CAG2018CAA (6160G>A), GGC2019GGG (6163C>G), AAG2023AAA (6175G>A), GAA2024GGA (6177A>G), GTA2026GTG (6184A>G), ACT2027GAG (6185A>G 6186C>A 6187T>G), CAC2028CGC (6189A>G), TTC2029TTT (6193C>T), CAA2030CAG (6196A>G), CGC2031CGT (6199C>T), ATG2032ATC (6202G>C), CAA2034CAG (6208A>G), GAT2036GAG (6214T>G), CTG2037GTT (6215C>G 6217G>T), ACA2038CCA (6218A>C), GAA2039TCT (6221G>T 6222A>C 6223A>T), GGG2040GAA (6225G>A 6226G>A), ATC2041GTG (6227A>G 6229C>G), ATA2042GTG (6230A>G 6232A>G), GAG2043GAA (6235G>A), AAA2044AGT (6237A>G 6238A>T), GTG2045ACA (6239G>A 6240T>C 6241G>A), ATA2046GTG (6242A>G 6244A>G), TTA2049CTC (6251T>C 6253A>C), ACC2050ACA (6256C>A), ACA2052ACT (6262A>T), CTG2053TTG (6263C>T), TCG2054ACT (6266T>A 6268G>T), AGC2056ATG (6273G>T 6274C>G), GTG2059GTA (6283G>A), CTT2061TTG (6287C>T 6289T>G), GTT2062GAA (6291T>A 6292T>A), GAT2063GGT (6294A>G), GTA2064CTC (6296G>C 6298A>C), CCT2065CCA (6301T>A), TTA2066ACG (6302T>A 6303T>C 6304A>G), GGC2067GGA (6307C>A), TTG2068CTT (6308T>C 6310G>T), GAT2069TCC (6311G>T 6312A>C 6313T>C), AAC2070AAT (6316C>T), ATA2071GTG (6317A>G 6319A>G), GAG2072GAA (6322G>A), ATC2073GTG (6323A>G 6325C>G), CAG2074CAA (6328G>A), AAA2076AAG (6334A>G), GGT2077GGC (6337T>C), CTA2080TTA (6344C>T), AAT2086AAC (6364T>C), GCA2089ACT (6371G>A 6373A>T), GCC2092GCT (6382C>T), ATC2093GTG (6383A>G 6385C>G), CTC2097CTA (6397C>A), TGT2100CAT (6404T>C 6405G>A), AAC2102AAT (6412C>T), CGA2103GTG (6413C>G 6414G>T 6415A>G), TCA2104AAC (6416T>A 6417C>A 6418A>C), ACA2105ACT (6421A>T), GTG2109GTT (6433G>T), TTC2110TTT (6436C>T), TGT2111AGT (6437T>A), TAC2113TAT (6445C>T), ACG2114ACA (6448G>A), ACT2115TCT (6449A>T), TCC2117GCA (6455T>G 6457C>A), TCT2118ACT (6458T>A), CTT2119TTG (6461C>T 6463T>G), GCC2122GCT (6472C>T), GTG2124GTT (6478G>T), CTT2125TTA (6479C>T 6481T>A), CAT2127AAT (6485C>A), AGA2128AGC (6490A>C), CAT2129AAT (6491C>A), CAG2130GTA (6494C>G 6495A>T 6496G>A), ACA2131CCT (6497A>C 6499A>T), ATT2132GAG (6500A>G 6501T>A 6502T>G), GAT2133AAC (6503G>A 6505T>C), CCG2136CCT (6514G>T), TCC2137CCA (6515T>C 6517C>A), TTC2138CTT (6518T>C 6520C>T), ACA2141ACT (6529A>T), GTG2145GTA (6541G>A), TTG2146TTA (6544G>A), CTT2147TTG (6545C>T 6547T>G), TTG2148CTG (6548T>C), ACA2149ATG (6552C>T 6553A>G), CCA2150CCC (6556A>C), CTA2152CTT (6562A>T), GGC2154AAT (6566G>A 6567G>A 6568C>T), TAC2155TAT (6571C>T), AAA2156AAG (6574A>G)

*: Inserts / Deletes / Misaligned / Frameshifts

#### Analysis details

This analysis was performed with panviral2.64

### NGS Details (UN8): Torradovirus lycopersici (segment RNA 1)

#### Assembly

| | |
|---|---|
| **Coverage Length** | 6792 (1 contig(s)) |
| **Depth Of Coverage** | 2254.7 |
| **Number Of Reads** | 114995 |
| **Reads Per Million** | 1921.85 rpm (after QC) |
| **Ambiguities** | 0 |
| **Assembly Method** | de novo + reference guided assembly |
| **Consensus Caller** | Bcf Tools |

#### Coverage Map

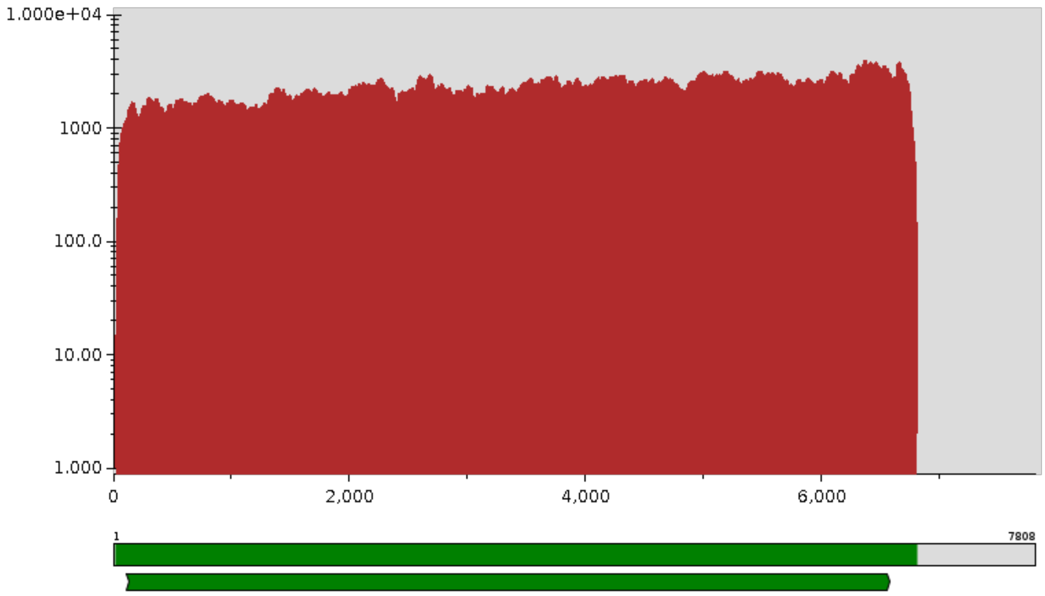

#### Assignment

| | |
|---|---|
| **Type** | Torradovirus lycopersici (Taxonomy ID: 3048378) |
| **Reference Genome** | NC_009013.1 |
| **NT Identity (%)** | 90.4769 |
| **AA Identity (%)** | 97.8231 |
| **Number Of Stop Codons** | 1 |
| **Number Of CDS** | 1 |

#### Alignment

| | |
|---|---|
| **Alignment Score** | 10966.0 (NT) + 14853.0 (AA) = 25819.0 |
| **Concordance (%)** | 90.1407 |

| Alignment Method | Global, seeded, nucleotide + amino acids (AGA) |
|---|---|

#### Genome Region

Sequence starts at position 22 and ends at position 6813 relative to NC_009013.1 reference sequence.

#### Alignment Detailed Statistics

| | Begin | End | Coverage | Score | Concordance | Matches | Identities | I/D/M/F* | Stop Codons |
|---|---|---|---|---|---|---|---|---|---|
| **NT** | **22** | **6813** | **87.0%** | **10966** | **80.8%** | **6791 (99.9%)** | **6147 (90.5%)** | **3/1** | |
| Mutations: | 22T>A, 69C>T, 76_77insT, 83C>T, 83_84insT, 85T>C, 87C>T, 95A>G, 96C>T, 98T>C, 99T>C, 100T>C, 101G>A, 102G>A, 115T>C, 127C>T, 128C>T, 130C>T, 135T>C, 148T>C, 151A>G, 158G>T, 161A>G, 175T>C, 178T>C, 184T>C, 190G>A, 199T>A, 202T>C, 205T>A, 220C>T, 224C>T, 253A>G, 259T>C, 262T>A, 265C>T, 338T>C, 340G>A, 349G>A, 361A>G, 367A>G, 370C>T, 376C>T, 385C>T, 388T>C, 409A>G, 421T>C, 433G>A, 436T>A, 451A>G, 454T>C, 472T>C, 487C>T, 490A>G, 502G>A, 559C>T, 577A>G, 578C>T, 583T>A, 637A>C, 655C>T, 676A>G, 706T>C, 707C>T, 709A>G, 727A>G, 760C>T, 781C>T, 790C>T, 799A>G, 805T>C, 818C>T, 820A>G, 832A>T, 835A>C, 841T>C, 842C>T, 847C>T, 883G>A, 886T>C, 892G>A, 901T>C, 913T>C, 925C>T, 938C>T, 952A>G, 955C>T, 964T>G, 970G>A, 973T>C, 977G>A, 980T>C, 997C>T, 1006T>C, 1012C>T, 1013T>C, 1018C>T, 1033G>A, 1063C>T, 1064T>C, 1078A>T, 1081C>T, 1093T>C, 1105A>G, 1108T>C, 1109C>T, 1165C>T, 1180G>A, 1196A>G, 1231G>A, 1240C>T, 1255G>A, 1267T>C, 1276A>G, 1300G>A, 1303A>G, 1319C>T, 1324C>T, 1360G>A, 1361T>C, 1366C>T, 1408T>C, 1411T>C, 1417T>C, 1423T>C, 1426T>C, 1429C>T, 1430C>T, 1441C>T, 1450T>C, 1451C>T, 1480C>T, 1481A>G, 1489T>C, 1522C>T, 1552C>T, 1555C>T, 1570A>G, 1573T>C, 1582C>T, 1585A>T, 1591C>T, 1594T>C, 1607G>T, 1612G>A, 1618T>G, 1619C>T, 1624C>T, 1630C>T, 1639T>C, 1642G>A, 1645A>G, 1669C>T, 1672G>A, 1681T>A, 1684G>T, 1699T>C, 1705T>C, 1737A>G, 1738A>G, 1750A>T, 1756A>G, 1765T>G, 1781T>C, 1783G>A, 1784C>T, 1789A>G, 1810G>A, 1825C>T, 1834A>G, 1867G>A, 1879A>G, 1888A>G, 1910T>C, 1912G>A, 1960A>G, 1966A>G, 1984C>T, 1990G>A, 1993G>A, 2009T>C, 2023T>C, 2056T>C, 2077C>A, 2082T>A, 2092A>G, 2107A>G, 2128T>C, 2138C>T, 2143T>C, 2149T>C, 2155T>C, 2174C>T, 2179A>C, 2182G>A, 2194T>A, 2200T>C, 2215G>A, 2218C>T, 2224C>T, 2236A>G, 2241A>G, 2250G>A, 2251C>T, 2257C>T, 2263G>A, 2275T>C, 2287G>A, 2290C>T, 2314C>T, 2320G>A, 2323A>T, 2347C>T, 2350A>G, 2356C>T, 2359G>A, 2362C>T, 2377G>A, 2380T>C, 2392A>G, 2407T>C, 2409G>A, 2443T>C, 2446G>A, 2479G>A, 2485C>T, 2497G>A, 2500T>C, 2503G>A, 2509C>T, 2512T>C, 2542G>A, 2548C>T, 2557C>T, 2620G>A, 2626T>C, 2650G>A, 2665T>C, 2675T>C, 2683A>G, 2692G>A, 2702C>T, 2710A>G, 2719T>C, 2725T>C, 2731C>T, 2734C>T, 2743A>G, 2744T>C, 2752A>G, 2773A>G, 2785C>T, 2788C>T, 2809C>T, 2821G>T, 2827A>G, 2842T>C, 2860C>T, 2866T>A, 2869C>T, 2872A>T, 2893T>A, 2896C>A, 2926G>A, 2929C>T, 2930T>C, 2941A>G, 2944T>C, 2950C>T, 2951C>T, 2980C>T, 2983G>A, 2998A>G, 3007T>C, 3016T>C, 3028T>C, 3031T>C, 3034G>A, 3040T>C, 3058A>G, 3064A>G, 3073G>A, 3079T>C, 3109T>C, 3115G>A, 3145A>G, 3151C>T, 3163T>C, 3169C>T, 3172T>C, 3202T>C, 3241T>C, 3253C>T, 3256G>A, 3262C>T, 3283T>C, 3289T>C, 3298T>C, 3307A>G, 3319C>T, 3358A>T, 3406A>C, 3418C>A, 3421T>C, 3463A>G, 3466G>A, 3481C>T, 3484A>T, 3508G>A, 3559G>A, 3568A>G, 3574T>C, 3577A>G, 3586T>C, 3591C>T, 3625C>T, 3628G>A, 3646C>T, 3661A>G, 3664T>C, 3673G>A, 3676C>T, 3686C>T, 3706T>C, 3716T>C, 3721T>A, 3739A>T, 3751A>G, 3754G>A, 3758G>A, 3766G>A, 3781G>A, 3805C>T, 3808T>C, 3814C>T, 3817G>C, 3824T>C, 3835T>C, 3844T>C, 3857T>C, 3859G>A, 3868C>T, 3871T>C, 3883G>A, 3898T>C, 3907A>T, 3916C>T, 3925A>G, 3928G>A, 3949C>T, 3955G>A, 3961G>A, 3970A>G, 3973T>C, 4034T>C, 4066T>C, 4075A>G, 4084C>T, 4100T>C, 4118C>T, 4129C>T, 4147C>A, 4159G>A, 4162A>T, 4168A>G, 4186A>T, 4195A>G, 4207G>A, 4210T>C, 4228C>T, 4234T>C, 4243T>C, 4273A>G, 4282C>A, 4288C>T, 4291T>C, 4300T>C, 4318C>T, 4324T>C, 4339T>C, 4343C>T, 4348C>T, 4354T>C, 4396T>C, 4408C>T, 4414T>G, 4417T>C, 4420G>A, 4459T>C, 4462C>T, 4465C>T, 4468C>T, 4471T>C, 4474T>C, 4486T>C, 4489T>C, 4504T>C, 4510T>A, 4511C>T, 4514T>C, 4519A>G, 4543T>C, 4555A>G, 4570C>T, 4574C>T, 4576A>G, 4588C>T, 4591T>C, 4612A>G, 4625T>C, 4630T>C, 4648T>C, 4660A>G, 4666G>A, 4672G>C, 4679A>G, 4684A>T, 4687G>A, 4696C>T, 4702G>A, 4705C>T, 4714T>C, 4720T>C, 4723C>T, 4730C>T, 4735C>T, 4747A>G, 4756A>C, 4765T>G, 4772A>C, 4774G>A, 4780T>C, 4781C>T, 4786A>G, 4795A>G, 4802C>T, 4807A>G, 4813A>G, 4816T>C, 4834C>T, 4858T>C, 4861C>T, 4864T>C, 4903A>G, 4924T>C, 4946A>G, 4960C>T, 4975G>A, 4993A>C, 4999T>C, 5012T>C, 5017G>A, 5023C>T, 5044A>C, 5059G>A, 5071A>G, 5074T>C, 5077T>C, 5079T>C, 5086T>C, 5102T>C, 5104G>A, 5119T>C, 5120G>A, 5122T>C, 5123C>T, 5128T>C, 5132T>A, 5140A>G, 5146T>C, 5161A>G, 5167T>C, 5170C>T, 5176C>A, 5209C>T, 5244G>A, 5245A>G, 5251A>G, 5287A>G, 5290G>A, 5302T>C, 5332C>T, 5354G>A, 5356G>A, 5365A>T, 5392G>A, 5413C>T, 5419A>C, 5429C>T, 5437T>C, 5464T>C, 5470A>G, 5491T>G, 5492A>T, 5497T>C, 5518T>C, 5533T>C, 5537A>C, 5560A>G, 5563G>A, 5575A>T, 5578T>C, 5581A>G, 5590A>G, 5591G>T, 5599T>A, 5603A>G, 5608A>G, 5611A>G, 5617A>C, 5623A>G, 5635C>T, 5641T>A, 5644T>C, 5647T>C, 5656G>T, 5668T>C, 5674A>G, 5677G>A, 5678T>C, 5686G>A, 5690T>C, 5702G>A, 5704G>A, 5710T>C, 5716G>A, 5719A>C, 5731C>T, 5734T>C, 5737T>C, 5739G>A, 5746T>C, 5750T>C, 5752G>A, 5758C>T, 5773G>A, 5782A>G, 5784T>C, 5786T>C, 5797A>G, 5806C>T, 5836A>T, 5849T>C, 5850T>A, 5851G>A, 5869G>A, 5875A>G, 5878G>C, 5884T>C, 5887C>A, 5890C>T, 5902G>T, 5908C>T, 5909T>C, 5914T>C, 5923T>C, 5927C>T, 5932C>T, 5955A>G, 5965G>A, 5998A>G, 6004C>T, 6016A>G, 6028C>T, 6031A>T, 6033C>T, 6035G>T, 6055T>A, 6067C>A, 6070T>C, 6094T>C, 6100T>C, 6103G>A, 6106G>A, 6109C>T, 6112A>T, 6131T>C, 6139T>C, 6143A>C, 6145T>C, 6154C>T, 6157C>T, 6163C>T, 6166C>T, 6167C>T, 6187T>C, 6193C>T, 6196A>G, 6202G>A, 6208A>G, 6211A>G, 6217G>A, 6223A>G, 6226G>A, 6230A>G, 6232A>G, 6235G>A, 6244A>T, 6253A>G, 6256C>A, 6265G>A, 6268G>A, 6273G>A, 6274C>T, 6295T>C, 6296G>T, 6298A>G, 6304A>G, 6308T>C, 6315A>T, 6319A>T, 6322G>A, 6323A>G, 6327A>G, 6331T>C, 6344C>T, 6358T>C, 6382C>T, 6385C>T, 6406T>C, 6412C>T, 6414G>A, 6427A>G, 6442C>G, 6445C>T, 6448G>A, 6451T>C, 6457C>A, 6466T>C, 6484A>C, 6490A>G, 6493T>C, 6500A>T, 6517C>T, 6532C>T, 6545C>T, 6568C>T, 6571C>T, 6587G>A, 6643A>G, 6670C>T, 6685A>G, 6715C>T, 6721A>T, 6742A>T, 6766delC, 6786T>C, 6798_6799insA | | | | | | | | |

##### CDS

| | Begin | End | Coverage | Score | Concordance | Matches | Identities | I/D/M/F* | Stop Codons |
|---|---|---|---|---|---|---|---|---|---|
| ToTV_sRNA1gp1 | 1 | 2159 | 100% | 14853 | 98.6% | 2159 (100%) | 2112 (97.8%) | 0/0/0/0 | 1 |
| Protein mutations: | P8S (128C>T 130C>T), F10S (135T>C), A18S (158G>T), T19A (161A>G), D159E (583T>A), V291I (977G>A), T364A (1196A>G), I459V (1481A>G), A501S (1607G>T), E526D (1684G>T), K544R (1737A>G 1738A>G), F659Y (2082T>A), N712S (2241A>G), S715N (2250G>A 2251C>T), R768K (2409G>A), A1162V (3591C>T), V1218I (3758G>A), P1413S (4343C>T), I1525V (4679A>G), T1614A (4946A>G), I1658T (5079T>C), V1672I (5120G>A 5122T>C), S1676T (5132T>A), R1713K (5244G>A 5245A>G), V1750I (5354G>A 5356G>A), T1796S (5492A>T), K1811Q (5537A>C), A1829S (5591G>T), T1833A (5603A>G), V1866I (5702G>A 5704G>A), R1878K (5739G>A), I1893T (5784T>C), L1915Q (5849T>C 5850T>A 5851G>A), N1950S (5955A>G), T1976I (6033C>T), A1977S (6035G>T), I2013L (6143A>C 6145T>C), M2032I (6202G>A), I2042V (6230A>G 6232A>G), S2056N (6273G>A 6274C>T), V2064L (6296G>T 6298A>G), N2070I (6315A>T), I2073V (6323A>G), Q2074R (6327A>G), R2103Q (6414G>A), I2132F (6500A>T), L2147F (6545C>T) | | | | | | | | |

| | Begin | End | Coverage | Score | Concordance | Matches | Identities | I/D/M/F* | Stop Codons |
|---|---|---|---|---|---|---|---|---|---|
| **NT** | **22** | **6813** | **87.0%** | **10966** | **80.8%** | **6791 (99.9%)** | **6147 (90.5%)** | **3/1** | |
| Codon mutations: | TTT3TTC (115T>C), TTC7TTT (127C>T), CCC8TCT (128C>T 130C>T), TTC10TCC (135T>C), ACT14ACC (148T>C), GAA15GAG (151A>G), GCT18TCT (158G>T), ACC19GCC (161A>G), GGT23GGC (175T>C), TCT24TCC (178T>C), TTT26TTC (184T>C), GAG28GAA (190G>A), GCT31GCA (199T>A), AGT32AGC (202T>C), ATT33ATA (205T>A), CGC38CGT (220C>T), CTG40TTG (224C>T), CAA49CAG (253A>G), TCT51TCC (259T>C), TCT52TCA (262T>A), CAC53CAT (265C>T), TTG78CTA (338T>C 340G>A), AAG81AAA (349G>A), TTA85TTG (361A>G), GAA87GAG (367A>G), CCC88CCT (370C>T), ATC90ATT (376C>T), TAC93TAT (385C>T), TCT94TCC (388T>C), AAA101AAG (409A>G), GAT105GAC (421T>C), AAG109AAA (433G>A), ATT110ATA (436T>A), AAA115AAG (451A>G), GAT116GAC (454T>C), TTT122TTC (472T>C), GAC127GAT (487C>T), AAA128AAG (490A>G), GTG132GTA (502G>A), TTC151TTT (559C>T), GTA157GTG (577A>G), CTA158TTA (578C>T), GAT159GAA (583T>A), TCA177TCC (637A>C), GAC183GAT (655C>T), CAA190CAG (676A>G), GTT200GTC (706T>C), CTA201TTG (707C>T 709A>G), TCA207TCG (727A>G), CTC218CTT (760C>T), ACC225ACT (781C>T), TTC228TTT (790C>T), GTA231GTG (799A>G), GGT233GGC (805T>C), CTA238TTG (818C>T 820A>G), CTA242CTT (832A>T), TCA243TCC (835A>C), ATT245ATC (841T>C), CTG246TTG (842C>T), TAC247TAT (847C>T), TTG259TTA (883G>A), TTT260TTC (886T>C), CTG262CTA (892G>A), AAT265AAC (901T>C), TGT269TGC (913T>C), GCC273GCT (925C>T), CTG278TTG (938C>T), GTA282GTG (952A>G), GAC283GAT (955C>T), GGT286GGG (964T>G), AAG288AAA (970G>A), AAT289AAC (973T>C), GTT291ATT (977G>A), TTG292CTG (980T>C), GCC297GCT (997C>T), TCT300TCC (1006T>C), GAC302GAT (1012C>T), TTA303CTA (1013T>C), TTC304TTT (1018C>T), GAG309GAA (1033G>A), CGC319CGT (1063C>T), TTG320CTG (1064T>C), GCA324GCT (1078A>T), AAC325AAT (1081C>T), ACT329ACC (1093T>C), CAA333CAG (1105A>G), CAT334CAC (1108T>C), CTA335TTA (1109C>T), GCC353GCT (1165C>T), AGG358AGA (1180G>A), ACC364GCC (1196A>G), CAG375CAA (1231G>A), GGC378GGT (1240C>T), AGG383AGA (1255G>A), TTT387TTC (1267T>C), CAA390CAG (1276A>G), GGG398GGA (1300G>A), AAA399AAG (1303A>G), CTG405TTG (1319C>T), TTC406TTT (1324C>T), AAG418AAA (1360G>A), TTG419CTG (1361T>C), AAC420AAT (1366C>T), AAT434AAC (1408T>C), TAT435TAC (1411T>C), CAT437CAC (1417T>C), ACT439ACC (1423T>C), GCT440GCC (1426T>C), ATC441ATT (1429C>T), CTG442TTG (1430C>T), GAC445GAT (1441C>T), GCT448GCC (1450T>C), CTG449TTG (1451C>T), GAC458GAT (1480C>T), ATT459GTT (1481A>G), GAT461GAC (1489T>C), CCC472CCT (1522C>T), GGC482GGT (1552C>T), CGC483CGT (1555C>T), CGA488CGG (1570A>G), TAT489TAC (1573T>C), GCC492GCT (1582C>T), ACA493ACT (1585A>T), AAC495AAT (1591C>T), CTT496CTC (1594T>C), GCT501TCT (1607G>T), CAG502CAA (1612G>A), GGT504GGG (1618T>G), CTA505TTA (1619C>T), ACC506ACT (1624C>T), CCC508CCT (1630C>T), TTT511TTC (1639T>C), GAG512GAA (1642G>A), AGA513AGG (1645A>G), TGC521TGT (1669C>T), AGG522AGA (1672G>A), GGT525GGA (1681T>A), GAG526GAT (1684G>T), GCT531GCC (1699T>C), ACT533ACC (1705T>C), AAA544AGG (1737A>G 1738A>G), GCA548GCT (1750A>T), ACA550ACG (1756A>G), GGT553GGG (1765T>G), TTG559CTA (1781T>C 1783G>A), CTG560TTG (1784C>T), GAA561GAG (1789A>G), AAG568AAA (1810G>A), ATC573ATT (1825C>T), AAA576AAG (1834A>G), CAG587CAA (1867G>A), TCA591TCG (1879A>G), GAA594GAG (1888A>G), TTG602CTA (1910T>C 1912G>A), GTA618GTG (1960A>G), GAA620GAG (1966A>G), TAC626TAT (1984C>T), CAG628CAA (1990G>A), GAG629GAA (1993G>A), TTG635CTG (2009T>C), TAT639TAC (2023T>C), CCT650CCC (2056T>C), GCC657GCA (2077C>A), TTT659TAT (2082T>A), CTA662CTG (2092A>G), AAA667AAG (2107A>G), GTT674GTC (2128T>C), CTG678TTG (2138C>T), TAT679TAC (2143T>C), GGT681GGC (2149T>C), TGT683TGC (2155T>C), CTG690TTG (2174C>T), ATA691ATC (2179A>C), AAG692AAA (2182G>A), GCT696GCA (2194T>A), TAT698TAC (2200T>C), AAG703AAA (2215G>A), AGC704AGT (2218C>T), ATC706ATT (2224C>T), ACA710ACG (2236A>G), AAT712AGT (2241A>G), AGC715AAT (2250G>A 2251C>T), GAC717GAT (2257C>T), CAG719CAA (2263G>A), CCT723CCC (2275T>C), GCG727GCA (2287G>A), CAC728CAT (2290C>T), GAC736GAT (2314C>T), CTG738CTA (2320G>A), ATA739ATT (2323A>T), CAC747CAT (2347C>T), GTA748GTG (2350A>G), TTC750TTT (2356C>T), TTG751TTA (2359G>A), TAC752TAT (2362C>T), AGG757AGA (2377G>A), TAT758TAC (2380T>C), AGA762AGG (2392A>G), CCT767CCC (2407T>C), AGG768AAG (2409G>A), ATT779ATC (2443T>C), GTG780GTA (2446G>A), CCG791CCA (2479G>A), ATC793ATT (2485C>T), CTG797CTA (2497G>A), ATT798ATC (2500T>C), AAG799AAA (2503G>A), GGC801GGT (2509C>T), CTT802CTC (2512T>C), GGG812GGA (2542G>A), GTC814GTT (2548C>T), TTC817TTT (2557C>T), GAG838GAA (2620G>A), AAT840AAC (2626T>C), AAG848AAA (2650G>A), TTT853TTC (2665T>C), TTG857CTG (2675T>C), GCA859GCG (2683A>G), AAG862AAA (2692G>A), CTG866TTG (2702C>T), AAA868AAG (2710A>G), AGT871AGC (2719T>C), GAT873GAC (2725T>C), TTC875TTT (2731C>T), ATC876ATT (2734C>T), GGA879GGG (2743A>G), TTG880CTG (2744T>C), AAA882AAG (2752A>G), CTA889CTG (2773A>G), GTC893GTT (2785C>T), TAC894TAT (2788C>T), TGC901TGT (2809C>T), GTG905GTT (2821G>T), ACA907ACG (2827A>G), TTT912TTC (2842T>C), GGC918GGT (2860C>T), ACT920ACA (2866T>A), TTC921TTT (2869C>T), TCA922TCT (2872A>T), ATT929ATA (2893T>A), TCC930TCA (2896C>A), CGG940CGA (2926G>A), TTC941TTT (2929C>T), TTG942CTG (2930T>C), CAA945CAG (2941A>G), GAT946GAC (2944T>C), GTC948GTT (2950C>T), CTG949TTG (2951C>T), CCC958CCT (2980C>T), AAG959AAA (2983G>A), AAA964AAG (2998A>G), CCT967CCC (3007T>C), GAT970GAC (3016T>C), TTT974TTC (3028T>C), TCT975TCC (3031T>C), AAG976AAA (3034G>A), ACT978ACC (3040T>C), AGA984AGG (3058A>G), GTA986GTG (3064A>G), AAG989AAA (3073G>A), GCT991GCC (3079T>C), TTT1001TTC (3109T>C), GTG1003GTA (3115G>A), GTA1013GTG (3145A>G), TAC1015TAT (3151C>T), TCT1019TCC (3163T>C), TCC1021TCT (3169C>T), TTT1022TTC (3172T>C), TAT1032TAC (3202T>C), CTT1045CTC (3241T>C), AAC1049AAT (3253C>T), GTG1050GTA (3256G>A), GAC1052GAT (3262C>T), GGT1059GGC (3283T>C), CAT1061CAC (3289T>C), TCT1064TCC (3298T>C), GAA1067GAG (3307A>G), CAC1071CAT (3319C>T), CGA1084CGT (3358A>T), GTA1100GTC (3406A>C), CTC1104CTA (3418C>A), CTT1105CTC (3421T>C), AAA1119AAG (3463A>G), CAG1120CAA (3466G>A), GGC1125GGT (3481C>T), ACA1126ACT (3484A>T), ACG1134ACA (3508G>A), GGG1151GGA (3559G>A), GAA1154GAG (3568A>G), GCT1156GCC (3574T>C), CCA1157CCG (3577A>G), ATT1160ATC (3586T>C), GCT1162GTT (3591C>T), TAC1173TAT (3625C>T), GTG1174GTA (3628G>A), TAC1180TAT (3646C>T), CAA1185CAG (3661A>G), AAT1186AAC (3664T>C), GAG1189GAA (3673G>A), GAC1190GAT (3676C>T), CTA1194TTA (3686C>T), CGT1200CGC (3706T>C), TTG1204CTG (3716T>C), CCT1205CCA (3721T>A), CGA1211CGT (3739A>T), GGA1215GGG (3751A>G), AAG1216AAA (3754G>A), GTC1218ATC (3758G>A), CAG1220CAA (3766G>A), GTG1225GTA (3781G>A), GAC1233GAT (3805C>T), TAT1234TAC (3808T>C), GAC1236GAT (3814C>T), GGG1237GGC (3817G>C), TTG1240CTG (3824T>C), TCT1243TCC (3835T>C), TAT1246TAC (3844T>C), TTG1251CTA (3857T>C 3859G>A), GGC1254GGT (3868C>T), ACT1255ACC (3871T>C), GAG1259GAA (3883G>A), AGT1264AGC (3898T>C), GGA1267GGT (3907A>T), GTC1270GTT (3916C>T), AAA1273AAG (3925A>G), AGG1274AGA (3928G>A), GAC1281GAT (3949C>T), GAG1283GAA (3955G>A), TTG1285TTA (3961G>A), GAA1288GAG (3970A>G), ATT1289ATC (3973T>C), TTG1310CTG (4034T>C), CGT1320CGC (4066T>C), GAA1323GAG (4075A>G), CCC1326CCT (4084C>T), TTG1332CTG (4100T>C), CTG1338TTG (4118C>T), TCC1341TCT (4129C>T), TCC1347TCA (4147C>A), CTG1351CTA (4159G>A), CCA1352CCT (4162A>T), AAA1354AAG (4168A>G), GGA1360GGT (4186A>T), GAA1363GAG (4195A>G), TTG1367TTA (4207G>A), GCT1368GCC (4210T>C), GTC1374GTT (4228C>T), GAT1376GAC (4234T>C), TTT1379TTC (4243T>C), AGA1389AGG (4273A>G), ATC1392ATA (4282C>A), GCC1394GCT (4288C>T), TTT1395TTC (4291T>C), TTT1398TTC (4300T>C), GAC1404GAT (4318C>T), TAT1406TAC (4324T>C), GGT1411GGC (4339T>C), CCA1413TCA (4343C>T), GAC1414GAT (4348C>T), AAT1416AAC (4354T>C), ACT1430ACC (4396T>C), AAC1434AAT (4408C>T), GTT1436GTG (4414T>G), TTT1437TTC (4417T>C), AGG1438AGA (4420G>A), GTT1451GTC (4459T>C), ATC1452ATT (4462C>T), TTC1453TTT (4465C>T), AAC1454AAT (4468C>T), TCT1455TCC (4471T>C), CTT1456CTC (4474T>C), TTT1460TTC (4486T>C), TAT1461TAC (4489T>C), TTT1466TTC (4504T>C), TCT1468TCA (4510T>A), CTA1469TTA (4511C>T), TTG1470CTG (4514T>C), AGA1471AGG (4519A>G), GCT1479GCC (4543T>C), AAA1483AAG (4555A>G), AAC1488AAT (4570C>T), CTA1490TTG (4574C>T 4576A>G), GTC1494GTT (4588C>T), TAT1495TAC (4591T>C), GCA1502GCG (4612A>G), TTG1507CTG (4625T>C), CAT1508CAC (4630T>C), GCT1514GCC (4648T>C), GAA1518GAG (4660A>G), GAG1520GAA (4666G>A), GTG1522GTC (4672G>C), ATT1525GTT (4679A>G), ATA1526ATT (4684A>T), AAG1527AAA (4687G>A), ATC1530ATT (4696C>T), AAG1532AAA (4702G>A), AAC1533AAT (4705C>T), GTT1536GTC (4714T>C), AGT1538AGC (4720T>C), TCC1539TCT (4723C>T), CTA1542TTA (4730C>T), GAC1543GAT (4735C>T), CTA1547CTG (4747A>G), CTA1550CTC (4756A>C), GGT1553GGG (4765T>G), AGG1556CGA (4772A>C 4774G>A), CGT1558CGC (4780T>C), CTA1559TTA (4781C>T), GGA1560GGG (4786A>G), CAA1563CAG (4795A>G), CTG1566TTG (4802C>T), AAA1567AAG (4807A>G), GTA1569GTG (4813A>G), TCT1570TCC (4816T>C), TAC1576TAT (4834C>T), TGT1584TGC (4858T>C), CCC1585CCT (4861C>T), GAT1586GAC (4864T>C), AGA1599AGG (4903A>G), CCT1606CCC (4924T>C), ACA1614GCA (4946A>G), AAC1618AAT (4960C>T), CGG1623CGA (4975G>A), ACA1629ACC (4993A>C), CCT1631CCC (4999T>C), TTG1636CTG (5012T>C), GAG1637GAA (5017G>A), GAC1639GAT (5023C>T), GGA1646GGC (5044A>C), GGG1651GGA (5059G>A), GAA1655GAG (5071A>G), CTT1656CTC (5074T>C), CAT1657CAC (5077T>C), ATT1658ACT (5079T>C), GCT1660GCC (5086T>C), TTG1666CTA (5102T>C 5104G>A), ACT1671ACC (5119T>C), GTT1672ATC (5120G>A 5122T>C), CTG1673TTG (5123C>T), AAT1674AAC (5128T>C), TCT1676ACT (5132T>A), GGA1678GGG (5140A>G), CAT1680CAC (5146T>C), CGA1685CGG (5161A>G), AGT1687AGC (5167T>C), TTC1688TTT (5170C>T), TCC1690TCA (5176C>A), ACC1701ACT (5209C>T), AGA1713AAG (5244G>A 5245A>G), ACA1715ACG (5251A>G), GAA1727GAG (5287A>G), GGG1728GGA (5290G>A), CCT1732CCC (5302T>C), ATC1742ATT (5332C>T), GTG1750ATA (5354G>A 5356G>A), CCA1753CCT (5365A>T), GGG1762GGA (5392G>A), TGC1769TGT (5413C>T), GCA1771GCC (5419A>C), CTG1775TTG (5429C>T), ACT1777ACC (5437T>C), GTT1786GTC (5464T>C), AGA1788AGG (5470A>G), GTT1795GTG (5491T>G), ACA1796TCA (5492A>T), GAT1797GAC (5497T>C), GGT1804GGC (5518T>C), CTT1809CTC (5533T>C), AAA1811CAA (5537A>C), GAA1818GAG (5560A>G), GAG1819GAA (5563G>A), ACA1823ACT (5575A>T), TGT1824TGC (5578T>C), AAA1825AAG (5581A>G), CAA1828CAG (5590A>G), GCC1829TCC (5591G>T), GGT1831GGA (5599T>A), ACA1833GCA (5603A>G), CCA1834CCG (5608A>G), GCA1835GCG (5611A>G), ATA1837ATC (5617A>C), CAA1839CAG (5623A>G), TCC1843TCT (5635C>T), GGT1845GGA (5641T>A), AAT1846AAC (5644T>C), TTT1847TTC (5647T>C), GTG1850GTT (5656G>T), ATT1854ATC (5668T>C), CTA1856CTG (5674A>G), CAG1857CAA (5677G>A), TTG1858CTG (5678T>C), CAG1860CAA (5686G>A), TTG1862CTG (5690T>C), GTG1866ATA (5702G>A 5704G>A), CCT1868CCC (5710T>C), GGG1870GGA (5716G>A), GGA1871GGC (5719A>C), CAC1875CAT (5731C>T), ACT1876ACC (5734T>C), ACT1877ACC (5737T>C), AGA1878AAA (5739G>A), CTT1880CTC (5746T>C), TTG1882CTA (5750T>C 5752G>A), CTC1884CTT (5758C>T), AAG1889AAA (5773G>A), CAA1892CAG (5782A>G), ATT1893ACT (5784T>C), TTA1894CTA (5786T>C), AGA1897AGG (5797A>G), TTC1900TTT (5806C>T), TCA1910TCT (5836A>T), TTG1915CAA (5849T>C 5850T>A 5851G>A), AGG1921AGA (5869G>A), CAA1923CAG (5875A>G), GGG1924GGC (5878G>C), TTT1926TTC (5884T>C), CCC1927CCA (5887C>A), CGC1928CGT (5890C>T), GTG1932GTT (5902G>T), TAC1934TAT (5908C>T), TTG1935CTG (5909T>C), CTT1936CTC (5914T>C), TTT1939TTC (5923T>C), CTG1941TTG (5927C>T), AGC1942AGT (5932C>T), AAC1950AGC (5955A>G), ACG1953ACA (5965G>A), AAA1964AAG (5998A>G), ACC1966ACT (6004C>T), AAA1970AAG (6016A>G), TCC1974TCT (6028C>T), GCA1975GCT (6031A>T), ACA1976ATA (6033C>T), GCT1977TCT (6035G>T), GGT1983GGA (6055T>A), TCC1987TCA (6067C>A), GCT1988GCC (6070T>C), CTT1996CTC (6094T>C), CTT1998CTC (6100T>C), CAG1999CAA (6103G>A), AAG2000AAA (6106G>A), CCC2001CCT (6109C>T), ATA2002ATT (6112A>T), TTG2009CTG (6131T>C), TTT2011TTC (6139T>C), ATT2013CTC (6143A>C 6145T>C), GTC2016GTT (6154C>T), ATC2017ATT (6157C>T), GGC2019GGT (6163C>T), CAC2020CAT (6166C>T), CTG2021TTG (6167C>T), ACT2027ACC (6187T>C), TTC2029TTT (6193C>T), CAA2030CAG (6196A>G), ATG2032ATA (6202G>A), CAA2034CAG (6208A>G), CAA2035CAG (6211A>G), CTG2037CTA (6217G>A), GAA2039GAG (6223A>G), GGG2040GGA (6226G>A), ATA2042GTG (6230A>G 6232A>G), GAG2043GAA (6235G>A), ATA2046ATT (6244A>T), TTA2049TTG (6253A>G), ACC2050ACA (6256C>A), CTG2053CTA (6265G>A), TCG2054TCA (6268G>A), AGC2056AAT (6273G>A 6274C>T), GAT2063GAC (6295T>C), GTA2064TTG (6296G>T 6298A>G), TTA2066TTG (6304A>G), TTG2068CTG (6308T>C), AAC2070ATC (6315A>T), ATA2071ATT (6319A>T), GAG2072GAA (6322G>A), ATC2073GTC (6323A>G), CAG2074CGG (6327A>G), GAT2075GAC (6331T>C), CTA2080TTA (6344C>T), AAT2084AAC (6358T>C), GCC2092GCT (6382C>T), ATC2093ATT (6385C>T), TGT2100TGC (6406T>C), AAC2102AAT (6412C>T), CGA2103CAA (6414G>A), AAA2107AAG (6427A>G), CGC2112CGG (6442C>G), TAC2113TAT (6445C>T), ACG2114ACA (6448G>A), ACT2115ACC (6451T>C), TCC2117TCA (6457C>A), GTT2120GTC (6466T>C), GCA2126GCC (6484A>C), AGA2128AGG (6490A>G), CAT2129CAC (6493T>C), ATT2132TTT (6500A>T), TCC2137TCT (6517C>T), CAC2142CAT (6532C>T), CTT2147TTT (6545C>T), GGC2154GGT (6568C>T), TAC2155TAT (6571C>T) | | | | | | | | |

#### Proteins

| | Begin | End | Coverage | Score | Concordance | Matches | Identities | I/D/M/F* | Stop Codons |
|---|---|---|---|---|---|---|---|---|---|
| polyprotein (YP_001039627.1) | 1 | 2159 | 100% | 14853 | 98.6% | 2159 (100%) | 2112 (97.8%) | 0/0/0/0 | 1 |
| Protein mutations: | P8S (128C>T 130C>T), F10S (135T>C), A18S (158G>T), T19A (161A>G), D159E (583T>A), V291I (977G>A), T364A (1196A>G), I459V (1481A>G), A501S (1607G>T), E526D (1684G>T), K544R (1737A>G 1738A>G), F659Y (2082T>A), N712S (2241A>G), S715N (2250G>A 2251C>T), R768K (2409G>A), A1162V (3591C>T), V1218I (3758G>A), P1413S (4343C>T), I1525V (4679A>G), T1614A (4946A>G), I1658T (5079T>C), V1672I (5120G>A 5122T>C), S1676T (5132T>A), R1713K (5244G>A 5245A>G), V1750I (5354G>A 5356G>A), T1796S (5492A>T), K1811Q (5537A>C), A1829S (5591G>T), T1833A (5603A>G), V1866I (5702G>A 5704G>A), R1878K (5739G>A), I1893T (5784T>C), L1915Q (5849T>C 5850T>A 5851G>A), N1950S (5955A>G), T1976I (6033C>T), A1977S (6035G>T), I2013L (6143A>C 6145T>C), M2032I (6202G>A), I2042V (6230A>G 6232A>G), S2056N (6273G>A 6274C>T), V2064L (6296G>T 6298A>G), N2070I (6315A>T), I2073V (6323A>G), Q2074R (6327A>G), R2103Q (6414G>A), I2132F (6500A>T), L2147F (6545C>T) | | | | | | | | |

| | Begin | End | Coverage | Score | Concordance | Matches | Identities | I/D/M/F* | Stop Codons |
|---|---|---|---|---|---|---|---|---|---|
| **NT** | **22** | **6813** | **87.0%** | **10966** | **80.8%** | **6791 (99.9%)** | **6147 (90.5%)** | **3/1** | |

Codon mutations:

TTT3TTC (115T>C), TTC7TTT (127C>T), CCC8TCT (128C>T 130C>T), TTC10TCC (135T>C), ACT14ACC (148T>C), GAA15GAG (151A>G), GCT18TCT (158G>T), ACC19GCC (161A>G), GGT23GGC (175T>C), TCT24TCC (178T>C), TTT26TTC (184T>C), GAG28GAA (190G>A), GCT31GCA (199T>A), AGT32AGC (202T>C), ATT33ATA (205T>A), CGC38CGT (220C>T), CTG40TTG (224C>T), CAA49CAG (253A>G), TCT51TCC (259T>C), TCT52TCA (262T>A), CAC53CAT (265C>T), TTG78CTA (338T>C 340G>A), AAG81AAA (349G>A), TTA85TTG (361A>G), GAA87GAG (367A>G), CCC88CCT (370C>T), ATC90ATT (376C>T), TAC93TAT (385C>T), TCT94TCC (388T>C), AAA101AAG (409A>G), GAT105GAC (421T>C), AAG109AAA (433G>A), ATT110ATA (436T>A), AAA115AAG (451A>G), GAT116GAC (454T>C), TTT122TTC (472T>C), GAC127GAT (487C>T), AAA128AAG (490A>G), GTG132GTA (502G>A), TTC151TTT (559C>T), GTA157GTG (577A>G), CTA158TTA (578C>T), GAT159GAA (583T>A), TCA177TCC (637A>C), GAC183GAT (655C>T), CAA190CAG (676A>G), GTT200GTC (706T>C), CTA201TTG (707C>T 709A>G), TCA207TCG (727A>G), CTC218CTT (760C>T), ACC225ACT (781C>T), TTC228TTT (790C>T), GTA231GTG (799A>G), GGT233GGC (805T>C), CTA238TTG (818C>T 820A>G), CTA242CTT (832A>T), TCA243TCC (835A>C), ATT245ATC (841T>C), CTG246TTG (842C>T), TAC247TAT (847C>T), TTG259TTA (883G>A), TTT260TTC (886T>C), CTG262CTA (892G>A), AAT265AAC (901T>C), TGT269TGC (913T>C), GCC273GCT (925C>T), CTG278TTG (938C>T), GTA282GTG (952A>G), GAC283GAT (955C>T), GGT286GGG (964T>G), AAG288AAA (970G>A), AAT289AAC (973T>C), GTT291ATT (977G>A), TTG292CTG (980T>C), GCC297GCT (997C>T), TCT300TCC (1006T>C), GAC302GAT (1012C>T), TTA303CTA (1013T>C), TTC304TTT (1018C>T), GAG309GAA (1033G>A), CGC319CGT (1063C>T), TTG320CTG (1064T>C), GCA324GCT (1078A>T), AAC325AAT (1081C>T), ACT329ACC (1093T>C), CAA333CAG (1105A>G), CAT334CAC (1108T>C), CTA335TTA (1109C>T), GCC353GCT (1165C>T), AGG358AGA (1180G>A), ACC364GCC (1196A>G), CAG375CAA (1231G>A), GGC378GGT (1240C>T), AGG383AGA (1255G>A), TTT387TTC (1267T>C), CAA390CAG (1276A>G), GGG398GGA (1300G>A), AAA399AAG (1303A>G), CTG405TTG (1319C>T), TTC406TTT (1324C>T), AAG418AAA (1360G>A), TTG419CTG (1361T>C), AAC420AAT (1366C>T), AAT434AAC (1408T>C), TAT435TAC (1411T>C), CAT437CAC (1417T>C), ACT439ACC (1423T>C), GCT440GCC (1426T>C), ATC441ATT (1429C>T), CTG442TTG (1430C>T), GAC445GAT (1441C>T), GCT448GCC (1450T>C), CTG449TTG (1451C>T), GAC458GAT (1480C>T), ATT459GTT (1481A>G), GAT461GAC (1489T>C), CCC472CCT (1522C>T), GGC482GGT (1552C>T), CGC483CGT (1555C>T), CGA488CGG (1570A>G), TAT489TAC (1573T>C), GCC492GCT (1582C>T), ACA493ACT (1585A>T), AAC495AAT (1591C>T), CTT496CTC (1594T>C), GCT501TCT (1607G>T), CAG502CAA (1612G>A), GGT504GGG (1618T>G), CTA505TTA (1619C>T), ACC506ACT (1624C>T), CCC508CCT (1630C>T), TTT511TTC (1639T>C), GAG512GAA (1642G>A), AGA513AGG (1645A>G), TGC521TGT (1669C>T), AGG522AGA (1672G>A), GGT525GGA (1681T>A), GAG526GAT (1684G>T), GCT531GCC (1699T>C), ACT533ACC (1705T>C), AAA544AGG (1737A>G 1738A>G), GCA548GCT (1750A>T), ACA550ACG (1756A>G), GGT553GGG (1765T>G), TTG559CTA (1781T>C 1783G>A), CTG560TTG (1784C>T), GAA561GAG (1789A>G), AAG568AAA (1810G>A), ATC573ATT (1825C>T), AAA576AAG (1834A>G), CAG587CAA (1867G>A), TCA591TCG (1879A>G), GAA594GAG (1888A>G), TTG602CTA (1910T>C 1912G>A), GTA618GTG (1960A>G), GAA620GAG (1966A>G), TAC626TAT (1984C>T), CAG628CAA (1990G>A), GAG629GAA (1993G>A), TTG635CTG (2009T>C), TAT639TAC (2023T>C), CCT650CCC (2056T>C), GCC657GCA (2077C>A), TTT659TAT (2082T>A), CTA662CTG (2092A>G), AAA667AAG (2107A>G), GTT674GTC (2128T>C), CTG678TTG (2138C>T), TAT679TAC (2143T>C), GGT681GGC (2149T>C), TGT683TGC (2155T>C), CTG690TTG (2174C>T), ATA691ATC (2179A>C), AAG692AAA (2182G>A), GCT696GCA (2194T>A), TAT698TAC (2200T>C), AAG703AAA (2215G>A), AGC704AGT (2218C>T), ATC706ATT (2224C>T), ACA710ACG (2236A>G), AAT712AGT (2241A>G), AGC715AAT (2250G>A 2251C>T), GAC717GAT (2257C>T), CAG719CAA (2263G>A), CCT723CCC (2275T>C), GCG727GCA (2287G>A), CAC728CAT (2290C>T), GAC736GAT (2314C>T), CTG738CTA (2320G>A), ATA739ATT (2323A>T), CAC747CAT (2347C>T), GTA748GTG (2350A>G), TTC750TTT (2356C>T), TTG751TTA (2359G>A), TAC752TAT (2362C>T), AGG757AGA (2377G>A), TAT758TAC (2380T>C), AGA762AGG (2392A>G), CCT767CCC (2407T>C), AGG768AAG (2409G>A), ATT779ATC (2443T>C), GTG780GTA (2446G>A), CCG791CCA (2479G>A), ATC793ATT (2485C>T), CTG797CTA (2497G>A), ATT798ATC (2500T>C), AAG799AAA (2503G>A), GGC801GGT (2509C>T), CTT802CTC (2512T>C), GGG812GGA (2542G>A), GTC814GTT (2548C>T), TTC817TTT (2557C>T), GAG838GAA (2620G>A), AAT840AAC (2626T>C), AAG848AAA (2650G>A), TTT853TTC (2665T>C), TTG857CTG (2675T>C), GCA859GCG (2683A>G), AAG862AAA (2692G>A), CTG866TTG (2702C>T), AAA868AAG (2710A>G), AGT871AGC (2719T>C), GAT873GAC (2725T>C), TTC875TTT (2731C>T), ATC876ATT (2734C>T), GGA879GGG (2743A>G), TTG880CTG (2744T>C), AAA882AAG (2752A>G), CTA889CTG (2773A>G), GTC893GTT (2785C>T), TAC894TAT (2788C>T), TGC901TGT (2809C>T), GTG905GTT (2821G>T), ACA907ACG (2827A>G), TTT912TTC (2842T>C), GGC918GGT (2860C>T), ACT920ACA (2866T>A), TTC921TTT (2869C>T), TCA922TCT (2872A>T), ATT929ATA (2893T>A), TCC930TCA (2896C>A), CGG940CGA (2926G>A), TTC941TTT (2929C>T), TTG942CTG (2930T>C), CAA945CAG (2941A>G), GAT946GAC (2944T>C), GTC948GTT (2950C>T), CTG949TTG (2951C>T), CCC958CCT (2980C>T), AAG959AAA (2983G>A), AAA964AAG (2998A>G), CCT967CCC (3007T>C), GAT970GAC (3016T>C), TTT974TTC (3028T>C), TCT975TCC (3031T>C), AAG976AAA (3034G>A), ACT978ACC (3040T>C), AGA984AGG (3058A>G), GTA986GTG (3064A>G), AAG989AAA (3073G>A), GCT991GCC (3079T>C), TTT1001TTC (3109T>C), GTG1003GTA (3115G>A), GTA1013GTG (3145A>G), TAC1015TAT (3151C>T), TCT1019TCC (3163T>C), TCC1021TCT (3169C>T), TTT1022TTC (3172T>C), TAT1032TAC (3202T>C), CTT1045CTC (3241T>C), AAC1049AAT (3253C>T), GTG1050GTA (3256G>A), GAC1052GAT (3262C>T), GGT1059GGC (3283T>C), CAT1061CAC (3289T>C), TCT1064TCC (3298T>C), GAA1067GAG (3307A>G), CAC1071CAT (3319C>T), CGA1084CGT (3358A>T), GTA1100GTC (3406A>C), CTC1104CTA (3418C>A), CTT1105CTC (3421T>C), AAA1119AAG (3463A>G), CAG1120CAA (3466G>A), GGC1125GGT (3481C>T), ACA1126ACT (3484A>T), ACG1134ACA (3508G>A), GGG1151GGA (3559G>A), GAA1154GAG (3568A>G), GCT1156GCC (3574T>C), CCA1157CCG (3577A>G), ATT1160ATC (3586T>C), GCT1162GTT (3591C>T), TAC1173TAT (3625C>T), GTG1174GTA (3628G>A), TAC1180TAT (3646C>T), CAA1185CAG (3661A>G), AAT1186AAC (3664T>C), GAG1189GAA (3673G>A), GAC1190GAT (3676C>T), CTA1194TTA (3686C>T), CGT1200CGC (3706T>C), TTG1204CTG (3716T>C), CCT1205CCA (3721T>A), CGA1211CGT (3739A>T), GGA1215GGG (3751A>G), AAG1216AAA (3754G>A), GTC1218ATC (3758G>A), CAG1220CAA (3766G>A), GTG1225GTA (3781G>A), GAC1233GAT (3805C>T), TAT1234TAC (3808T>C), GAC1236GAT (3814C>T), GGG1237GGC (3817G>C), TTG1240CTG (3824T>C), TCT1243TCC (3835T>C), TAT1246TAC (3844T>C), TTG1251CTA (3857T>C 3859G>A), GGC1254GGT (3868C>T), ACT1255ACC (3871T>C), GAG1259GAA (3883G>A), AGT1264AGC (3898T>C), GGA1267GGT (3907A>T), GTC1270GTT (3916C>T), AAA1273AAG (3925A>G), AGG1274AGA (3928G>A), GAC1281GAT (3949C>T), GAG1283GAA (3955G>A), TTG1285TTA (3961G>A), GAA1288GAG (3970A>G), ATT1289ATC (3973T>C), TTG1310CTG (4034T>C), CGT1320CGC (4066T>C), GAA1323GAG (4075A>G), CCC1326CCT (4084C>T), TTG1332CTG (4100T>C), CTG1338TTG (4118C>T), TCC1341TCT (4129C>T), TCC1347TCA (4147C>A), CTG1351CTA (4159G>A), CCA1352CCT (4162A>T), AAA1354AAG (4168A>G), GGA1360GGT (4186A>T), GAA1363GAG (4195A>G), TTG1367TTA (4207G>A), GCT1368GCC (4210T>C), GTC1374GTT (4228C>T), GAT1376GAC (4234T>C), TTT1379TTC (4243T>C), AGA1389AGG (4273A>G), ATC1392ATA (4282C>A), GCC1394GCT (4288C>T), TTT1395TTC (4291T>C), TTT1398TTC (4300T>C), GAC1404GAT (4318C>T), TAT1406TAC (4324T>C), GGT1411GGC (4339T>C), CCA1413TCA (4343C>T), GAC1414GAT (4348C>T), AAT1416AAC (4354T>C), ACT1430ACC (4396T>C), AAC1434AAT (4408C>T), GTT1436GTG (4414T>G), TTT1437TTC (4417T>C), AGG1438AGA (4420G>A), GTT1451GTC (4459T>C), ATC1452ATT (4462C>T), TTC1453TTT (4465C>T), AAC1454AAT (4468C>T), TCT1455TCC (4471T>C), CTT1456CTC (4474T>C), TTT1460TTC (4486T>C), TAT1461TAC (4489T>C), TTT1466TTC (4504T>C), TCT1468TCA (4510T>A), CTA1469TTA (4511C>T), TTG1470CTG (4514T>C), AGA1471AGG (4519A>G), GCT1479GCC (4543T>C), AAA1483AAG (4555A>G), AAC1488AAT (4570C>T), CTA1490TTG (4574C>T 4576A>G), GTC1494GTT (4588C>T), TAT1495TAC (4591T>C), GCA1502GCG (4612A>G), TTG1507CTG (4625T>C), CAT1508CAC (4630T>C), GCT1514GCC (4648T>C), GAA1518GAG (4660A>G), GAG1520GAA (4666G>A), GTG1522GTC (4672G>C), ATT1525GTT (4679A>G), ATA1526ATT (4684A>T), AAG1527AAA (4687G>A), ATC1530ATT (4696C>T), AAG1532AAA (4702G>A), AAC1533AAT (4705C>T), GTT1536GTC (4714T>C), AGT1538AGC (4720T>C), TCC1539TCT (4723C>T), CTA1542TTA (4730C>T), GAC1543GAT (4735C>T), CTA1547CTG (4747A>G), CTA1550CTC (4756A>C), GGT1553GGG (4765T>G), AGG1556CGA (4772A>C 4774G>A), CGT1558CGC (4780T>C), CTA1559TTA (4781C>T), GGA1560GGG (4786A>G), CAA1563CAG (4795A>G), CTG1566TTG (4802C>T), AAA1567AAG (4807A>G), GTA1569GTG (4813A>G), TCT1570TCC (4816T>C), TAC1576TAT (4834C>T), TGT1584TGC (4858T>C), CCC1585CCT (4861C>T), GAT1586GAC (4864T>C), AGA1599AGG (4903A>G), CCT1606CCC (4924T>C), ACA1614GCA (4946A>G), AAC1618AAT (4960C>T), CGG1623CGA (4975G>A), ACA1629ACC (4993A>C), CCT1631CCC (4999T>C), TTG1636CTG (5012T>C), GAG1637GAA (5017G>A), GAC1639GAT (5023C>T), GGA1646GGC (5044A>C), GGG1651GGA (5059G>A), GAA1655GAG (5071A>G), CTT1656CTC (5074T>C), CAT1657CAC (5077T>C), ATT1658ACT (5079T>C), GCT1660GCC (5086T>C), TTG1666CTA (5102T>C 5104G>A), ACT1671ACC (5119T>C), GTT1672ATC (5120G>A 5122T>C), CTG1673TTG (5123C>T), AAT1674AAC (5128T>C), TCT1676ACT (5132T>A), GGA1678GGG (5140A>G), CAT1680CAC (5146T>C), CGA1685CGG (5161A>G), AGT1687AGC (5167T>C), TTC1688TTT (5170C>T), TCC1690TCA (5176C>A), ACC1701ACT (5209C>T), AGA1713AAG (5244G>A 5245A>G), ACA1715ACG (5251A>G), GAA1727GAG (5287A>G), GGG1728GGA (5290G>A), CCT1732CCC (5302T>C), ATC1742ATT (5332C>T), GTG1750ATA (5354G>A 5356G>A), CCA1753CCT (5365A>T), GGG1762GGA (5392G>A), TGC1769TGT (5413C>T), GCA1771GCC (5419A>C), CTG1775TTG (5429C>T), ACT1777ACC (5437T>C), GTT1786GTC (5464T>C), AGA1788AGG (5470A>G), GTT1795GTG (5491T>G), ACA1796TCA (5492A>T), GAT1797GAC (5497T>C), GGT1804GGC (5518T>C), CTT1809CTC (5533T>C), AAA1811CAA (5537A>C), GAA1818GAG (5560A>G), GAG1819GAA (5563G>A), ACA1823ACT (5575A>T), TGT1824TGC (5578T>C), AAA1825AAG (5581A>G), CAA1828CAG (5590A>G), GCC1829TCC (5591G>T), GGT1831GGA (5599T>A), ACA1833GCA (5603A>G), CCA1834CCG (5608A>G), GCA1835GCG (5611A>G), ATA1837ATC (5617A>C), CAA1839CAG (5623A>G), TCC1843TCT (5635C>T), GGT1845GGA (5641T>A), AAT1846AAC (5644T>C), TTT1847TTC (5647T>C), GTG1850GTT (5656G>T), ATT1854ATC (5668T>C), CTA1856CTG (5674A>G), CAG1857CAA (5677G>A), TTG1858CTG (5678T>C), CAG1860CAA (5686G>A), TTG1862CTG (5690T>C), GTG1866ATA (5702G>A 5704G>A), CCT1868CCC (5710T>C), GGG1870GGA (5716G>A), GGA1871GGC (5719A>C), CAC1875CAT (5731C>T), ACT1876ACC (5734T>C), ACT1877ACC (5737T>C), AGA1878AAA (5739G>A), CTT1880CTC (5746T>C), TTG1882CTA (5750T>C 5752G>A), CTC1884CTT (5758C>T), AAG1889AAA (5773G>A), CAA1892CAG (5782A>G), ATT1893ACT (5784T>C), TTA1894CTA (5786T>C), AGA1897AGG (5797A>G), TTC1900TTT (5806C>T), TCA1910TCT (5836A>T), TTG1915CAA (5849T>C 5850T>A 5851G>A), AGG1921AGA (5869G>A), CAA1923CAG (5875A>G), GGG1924GGC (5878G>C), TTT1926TTC (5884T>C), CCC1927CCA (5887C>A), CGC1928CGT (5890C>T), GTG1932GTT (5902G>T), TAC1934TAT (5908C>T), TTG1935CTG (5909T>C), CTT1936CTC (5914T>C), TTT1939TTC (5923T>C), CTG1941TTG (5927C>T), AGC1942AGT (5932C>T), AAC1950AGC (5955A>G), ACG1953ACA (5965G>A), AAA1964AAG (5998A>G), ACC1966ACT (6004C>T), AAA1970AAG (6016A>G), TCC1974TCT (6028C>T), GCA1975GCT (6031A>T), ACA1976ATA (6033C>T), GCT1977TCT (6035G>T), GGT1983GGA (6055T>A), TCC1987TCA (6067C>A), GCT1988GCC (6070T>C), CTT1996CTC (6094T>C), CTT1998CTC (6100T>C), CAG1999CAA (6103G>A), AAG2000AAA (6106G>A), CCC2001CCT (6109C>T), ATA2002ATT (6112A>T), TTG2009CTG (6131T>C), TTT2011TTC (6139T>C), ATT2013CTC (6143A>C 6145T>C), GTC2016GTT (6154C>T), ATC2017ATT (6157C>T), GGC2019GGT (6163C>T), CAC2020CAT (6166C>T), CTG2021TTG (6167C>T), ACT2027ACC (6187T>C), TTC2029TTT (6193C>T), CAA2030CAG (6196A>G), ATG2032ATA (6202G>A), CAA2034CAG (6208A>G), CAA2035CAG (6211A>G), CTG2037CTA (6217G>A), GAA2039GAG (6223A>G), GGG2040GGA (6226G>A), ATA2042GTG (6230A>G 6232A>G), GAG2043GAA (6235G>A), ATA2046ATT (6244A>T), TTA2049TTG (6253A>G), ACC2050ACA (6256C>A), CTG2053CTA (6265G>A), TCG2054TCA (6268G>A), AGC2056AAT (6273G>A 6274C>T), GAT2063GAC (6295T>C), GTA2064TTG (6296G>T 6298A>G), TTA2066TTG (6304A>G), TTG2068CTG (6308T>C), AAC2070ATC (6315A>T), ATA2071ATT (6319A>T), GAG2072GAA (6322G>A), ATC2073GTC (6323A>G), CAG2074CGG (6327A>G), GAT2075GAC (6331T>C), CTA2080TTA (6344C>T), AAT2084AAC (6358T>C), GCC2092GCT (6382C>T), ATC2093ATT (6385C>T), TGT2100TGC (6406T>C), AAC2102AAT (6412C>T), CGA2103CAA (6414G>A), AAA2107AAG (6427A>G), CGC2112CGG (6442C>G), TAC2113TAT (6445C>T), ACG2114ACA (6448G>A), ACT2115ACC (6451T>C), TCC2117TCA (6457C>A), GTT2120GTC (6466T>C), GCA2126GCC (6484A>C), AGA2128AGG (6490A>G), CAT2129CAC (6493T>C), ATT2132TTT (6500A>T), TCC2137TCT (6517C>T), CAC2142CAT (6532C>T), CTT2147TTT (6545C>T), GGC2154GGT (6568C>T), TAC2155TAT (6571C>T)

*: Inserts / Deletes / Misaligned / Frameshifts

#### Analysis details

This analysis was performed with panviral2.64

### NGS Details (UN8): Torradovirus lycopersici (segment RNA 1)

#### Assembly

| | |
|---|---|
| **Coverage Length** | 7760 (1 contig(s)) |
| **Depth Of Coverage** | 4265.8 |
| **Number Of Reads** | 186149 |
| **Reads Per Million** | 3111.02 rpm (after QC) |
| **Ambiguities** | 0 |
| **Assembly Method** | de novo + reference guided assembly |
| **Consensus Caller** | Bcf Tools |

#### Coverage Map

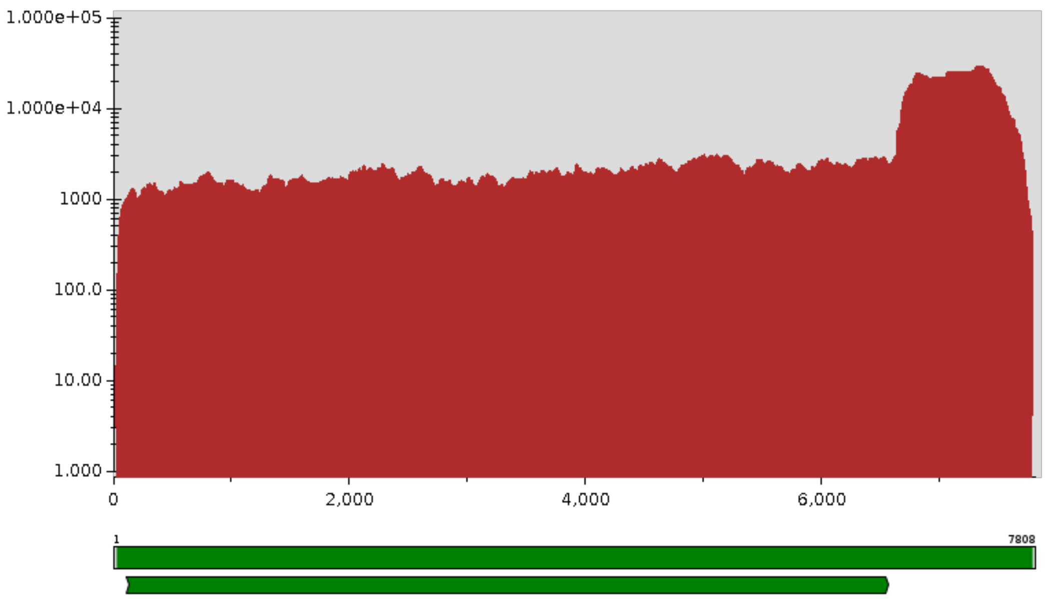

#### Assignment

| | |
|---|---|
| **Type** | Torradovirus lycopersici (Taxonomy ID: 3048378) |
| **Reference Genome** | NC_009013.1 |
| **NT Identity (%)** | 94.2857 |
| **AA Identity (%)** | 98.8421 |
| **Number Of Stop Codons** | 1 |
| **Number Of CDS** | 1 |

#### Alignment

| | |
|---|---|
| **Alignment Score** | 12464.0 (NT) + 14956.0 (AA) = 27420.0 |
| **Concordance (%)** | 93.7051 |

| Alignment Method | Global, seeded, nucleotide + amino acids (AGA) |
|---|---|

#### Genome Region

Sequence starts at position 27 and ends at position 7786 relative to NC_009013.1 reference sequence.

#### Alignment Detailed Statistics

| | Begin | End | Coverage | Score | Concordance | Matc |
|---|---|---|---|---|---|---|
| **NT** | **27** | **7786** | **99.4%** | **12464** | **87.8%** | **7359 (9** |
| Mutations: | 28C>T, 76_77insT, 87C>T, 115T>C, 127C>T, 128C>T, 130C>T, 135T>C, 158G>A, 160T>C, 161A>G, 187A>G, 193T>C, 208T>C, 220C>T, 224C>T, 328T>C, 334T>C, 340G>A, 349G>A, 373C>T, 376C><br>925C>T, 952A>G, 964T>G, 970G>A, 997C>T, 1006T>C, 1033G>A, 1060T>C, 1196A>G, 1207T>C, 1222G>A, 1234C>T, 1240C>T, 1255G>A, 1276A>G, 1300G>A, 1351A>G, 1372C>T, 1411T>C, 1417T<br>1789A>G, 1897A>G, 1910T>C, 1954T>C, 1960A>G, 1984C>T, 2014A>G, 2042G>A, 2092A>G, 2138C>T, 2149T>C, 2155T>C, 2174C>T, 2200T>C, 2227C>T, 2251C>T, 2257C>T, 2263G>A, 2287G>A,<br>2512T>C, 2516T>C, 2620G>A, 2662C>T, 2665T>C, 2728C>T, 2734C>T, 2743A>G, 2765T>C, 2773A>G, 2809C>T, 2812G>A, 2827A>G, 2869C>T, 2881T>C, 2914C>T, 2935G>A, 2950C>T, 2951C>T,<br>3451T>C, 3499A>G, 3534G>A, 3538T>C, 3541C>T, 3580T>C, 3583C>T, 3610A>G, 3613T>C, 3625C>T, 3628G>A, 3643A>G, 3676C>T, 3685G>A, 3701C>T, 3710G>A, 3739A>T, 3754G>A, 3784C>T,<br>4129C>T, 4150C>T, 4157C>T, 4159G>A, 4168A>G, 4210T>C, 4219C>T, 4294A>G, 4351C>T, 4357C>T, 4360A>G, 4393T>C, 4417T>C, 4421T>C, 4477C>T, 4543T>C, 4591T>C, 4636T>C, 4640C>T, 4<br>4813A>G, 4816T>C, 4843G>A, 4898G>A, 4906G>A, 4969G>A, 4988A>G, 4993A>T, 5005T>C, 5012T>C, 5029G>A, 5065C>T, 5074T>C, 5077T>C, 5079T>C, 5102T>C, 5118C>T, 5119T>C, 5120G>A,<br>5369G>A, 5371G>A, 5386C>T, 5392G>A, 5407A>G, 5437T>C, 5440C>T, 5461T>C, 5471T>C, 5482A>G, 5533T>A, 5572T>C, 5578T>C, 5584T>C, 5617A>T, 5641T>C, 5647T>C, 5695T>C, 5746T>C, 5<br>6049G>A, 6055T>A, 6073T>C, 6079C>T, 6091A>G, 6092C>A, 6106G>A, 6118T>C, 6121A>G, 6148T>C, 6178A>G, 6184A>G, 6193C>T, 6196A>G, 6200A>T, 6205C>T, 6247G>A, 6256C>A, 6262A>G,<br>6721A>T, 6742A>T, 6766delC, 6786T>C, 6798_6799insA, 6806T>G, 6807_6832delAACGGAGAGTACTGACTTTAACTAGT, 6835G>T, 6838_6850delGTCCGGCTCCATT, 6854A>T, 6858C>T, 6859C>A,<br>6912_6916delCGTGT, 6918C>T, 6921T>G, 6923C>G, 6924T>A, 6927_6941delGTTAGAAAGTTCCTT, 6947A>C, 6948G>T, 6949_6972delGTTTACAAAGTACGGGGAGCACTC, 6976G>A, 6984T>C, 69<br>7043T>C, 7045T>G, 7047_7060delGCCGTGATGAAAGA, 7063A>T, 7069C>A,<br>7071_7272delTCCCCCTTATTGGGAGGCTGAAACTACACATATGTAGTGGGTTTGACTGAGTCCTATAATCAGTCCGTTTGAAATTCGATAATTTTCCGTAGCTTGCGTCAAGCTGCTCACGTTAGGGGTGT<br>7292G>T, 7293A>T, 7297_7298insC, 7299G>A, 7311T>A, 7318G>T, 7320G>T, 7331A>T, 7333G>A, 7343C>A, 7344C>G, 7346A>T, 7349_7350insC, 7354_7355insGAGCACTGCCATTA, 7358A>C, 736<br>7398C>A, 7399G>A, 7400A>T, 7404C>T, 7407A>T, 7408G>T, 7423A>T, 7425G>A, 7435T>C, 7436G>A, 7444C>T, 7445A>G, 7457A>G, 7458T>C, 7460G>A, 7498C>T, 7503C>T, 7505A>G, 7511_7512<br>7585T>C, 7589T>A, 7600T>G, 7601T>A, 7602T>G, 7603_7604insC, 7610C>T, 7612G>A, 7615A>G, 7616A>C, 7617A>T, 7618C>T, 7625_7631delATTGCTT, 7637G>T, 7639T>A, 7641T>A, 7644_7650<br>7698G>A, 7717A>C, 7719T>A, 7723T>C, 7739T>A, 7742_7743insC, 7748A>G, 7749T>C, 7751T>C, 7752G>A, 7754_7755insG, 7756_7757insT, 7762C>T, 7763A>C, 7764A>T, 7767C>T, 7773G>T, 777 | | | | | |
| **CDS** | | | | | | |
| ToTV_sRNA1gp1 | 1 | 2159 | 100% | 14956 | 99.2% | 2159 (1 |
| Protein mutations: | P8S (128C>T 130C>T), F10S (135T>C), A18T (158G>A 160T>C), T19A (161A>G), T364A (1196A>G), A646T (2042G>A), R768K (2409G>A), V773I (2423G>A), R1143K (3534G>A), A1202T (3710G>A),<br>V1672I (5120G>A), T1681N (5148C>A), R1713K (5244G>A), V1755I (5369G>A 5371G>A), I1792M (5482A>G), N1947S (5946A>G), T1976I (6033C>T), L1996I (6092C>A), M2032L (6200A>T), A2089V ( | | | | | |
| Codon mutations: | TTT3TTC (115T>C), TTC7TTT (127C>T), CCC8TCT (128C>T 130C>T), TTC10TCC (135T>C), GCT18ACC (158G>A 160T>C), ACC19GCC (161A>G), TCA27TCG (187A>G), CTT29CTC (193T>C), AGT<br>(340G>A), AAG81AAA (349G>A), TTC89TTT (373C>T), ATC90ATT (376C>T), AGC113AGT (445C>T), GAC127GAT (487C>T), GTG132GTA (502G>A), TCT145TCC (541T>C), TTC151TTT (559C>T), C<br>CCA210CCG (736A>G), CTG246TTG (842C>T), CAA258CAG (880A>G), GCC273GCT (925C>T), GTA282GTG (952A>G), GGT286GGG (964T>G), AAG288AAA (970G>A), GCC297GCT (997C>T), TC<br>GAG372GAA (1222G>A), CAC376CAT (1234C>T), GGC378GGT (1240C>T), AGG383AGA (1255G>A), CAA390CAG (1276A>G), GGG398GGA (1300G>A), GAA415GAG (1351A>G), CTC422CTT (137<br>(1543G>A), AAA481AAG (1549A>G), GGC482GGT (1552C>T), AAC495AAT (1591C>T), CAG502CAA (1612G>A), GAT516GAC (1654T>C), TGC521TGT (1669C>T), AGC555AGT (1771C>T), TTG559<br>GTA618GTG (1960A>G), TAC626TAT (1984C>T), AAA636AAG (2014A>G), GCA646ACA (2042G>A), CTA662CTG (2092A>G), CTG678TTG (2138C>T), GGT681GGC (2149T>C), TGT683TGC (2155T<br>(2257C>T), CAG719CAA (2263G>A), GCG727GCA (2287G>A), CAC728CAT (2290C>T), CTG738TTG (2318C>T), TTC750TTT (2356C>T), TTG751TTA (2359G>A), TAT758TAC (2380T>C), AGG768A<br>GTT796GTC (2494T>C), GGC801GGT (2509C>T), CTT802CTC (2512T>C), TTA804CTA (2516T>C), GAG838GAA (2620G>A), ATC852ATT (2662C>T), TTT853TTC (2665T>C), GGC874GGT (2728C><br>(2809C>T), TCG902TCA (2812G>A), ACA907ACG (2827A>G), TTC921TTT (2869C>T), CAT925CAC (2881T>C), GCC936GCT (2914C>T), AAG943AAA (2935G>A), GTC948GTT (2950C>T), CTG949T<br>TTT1001TTC (3109T>C), TCT1019TCC (3163T>C), TCC1021TCT (3169C>T), AAA1023AAG (3175A>G), CTG1024TTG (3176C>T), TCT1029TCA (3193T>A), AAA1053AAG (3265A>G), CTC1104CTT (<br>GGC1145GGT (3541C>T), TCT1158TCC (3580T>C), ATC1159ATT (3583C>T), GAA1168GAG (3610A>G), GGT1169GGC (3613T>C), TAC1173TAT (3625C>T), GTG1174GTA (3628G>A), AAA1179AA<br>CGA1211CGT (3739A>T), AAG1216AAA (3754G>A), CTC1226CTT (3784C>T), GAC1236GAT (3814C>T), GAG1262GAA (3892G>A), GAC1281GAT (3949C>T), TTG1285CTG (3959T>C), AAA1304AA<br>(4084C>T), CAT1328CAC (4090T>C), TAC1329TAT (4093C>T), TCC1341TCT (4129C>T), CAC1348CAT (4150C>T), CTG1351TTA (4157C>T 4159G>A), AAA1354AAG (4168A>G), GCT1368GCC (421<br>AGA1418AGG (4360A>G), TAT1429TAC (4393T>C), TTT1437TTC (4417T>C), TTG1439CTG (4421T>C), CTC1457CTT (4477C>T), GCT1479GCC (4543T>C), TAT1495TAC (4591T>C), TAT1510TAC (<br>(4696C>T), GAC1531GAT (4699C>T), AGC1551AGT (4759C>T), TTT1554TTC (4768T>C), AGG1556AGA (4774G>A), CAC1557CAT (4777C>T), CTA1559CTG (4783A>G), GGA1560GGG (4786A>G),<br>(4898G>A), GAG1600GAA (4906G>A), AAG1621AAA (4969G>A), ACC1628GCC (4988A>G), ACA1629ACT (4993A>T), TAT1633TAC (5005T>C), TTG1636CTG (5012T>C), CTG1641CTA (5029G>A),<br>(5102T>C), ACT1671ATC (5118C>T 5119T>C), GTT1672ATT (5120G>A), CTG1673CTC (5125G>C), CAT1680CAC (5146T>C), ACT1681AAT (5148C>A), CAG1698CAA (5200G>A), GTT1704GTG (52<br>ATT1745ATA (5341T>A), GTA1746GTG (5344A>G), ACC1747ACT (5347C>T), GTG1755ATA (5369G>A 5371G>A), GGC1760GGT (5386C>T), GGG1762GGA (5392G>A), CCA1767CCG (5407A>G),<br>(5482A>G), CTT1809CTA (5533T>A), GAT1822GAC (5572T>C), TGT1824TGC (5578T>C), ATT1826ATC (5584T>C), ATA1837ATT (5617A>T), GGT1845GGC (5641T>C), TTT1847TTC (5647T>C), CC<br>(5797A>G), TAT1907TAC (5827T>C), GGG1924GGA (5878G>A), CCC1927CCT (5887C>T), CAG1944CAA (5938G>A), CAC1946CAT (5944C>T), AAT1947AGT (5946A>G), TTT1960TTC (5986T>C),<br>(6073T>C), CTC1991CTT (6079C>T), CTA1995CTG (6091A>G), CTT1996ATT (6092C>A), AAG2000AAA (6106G>A), TGT2004TGC (6118T>C), TCA2005TCG (6121A>G), TAT2014TAC (6148T>C), G<br>(6200A>T), TAC2033TAT (6205C>T), TTG2047TTA (6247G>A), ACC2050ACA (6256C>A), ACA2052ACG (6262A>G), CTG2053CTA (6265G>A), GTA2064GTG (6298A>G), GAT2069GAC (6313T>C),<br>(6493T>C) | | | | | |
| **Proteins** | | | | | | |
| polyprotein (YP_001039627.1) | 1 | 2159 | 100% | 14956 | 99.2% | 2159 (1 |
| Protein mutations: | P8S (128C>T 130C>T), F10S (135T>C), A18T (158G>A 160T>C), T19A (161A>G), T364A (1196A>G), A646T (2042G>A), R768K (2409G>A), V773I (2423G>A), R1143K (3534G>A), A1202T (3710G>A),<br>V1672I (5120G>A), T1681N (5148C>A), R1713K (5244G>A), V1755I (5369G>A 5371G>A), I1792M (5482A>G), N1947S (5946A>G), T1976I (6033C>T), L1996I (6092C>A), M2032L (6200A>T), A2089V ( | | | | | |
| Codon mutations: | TTT3TTC (115T>C), TTC7TTT (127C>T), CCC8TCT (128C>T 130C>T), TTC10TCC (135T>C), GCT18ACC (158G>A 160T>C), ACC19GCC (161A>G), TCA27TCG (187A>G), CTT29CTC (193T>C), AGT<br>(340G>A), AAG81AAA (349G>A), TTC89TTT (373C>T), ATC90ATT (376C>T), AGC113AGT (445C>T), GAC127GAT (487C>T), GTG132GTA (502G>A), TCT145TCC (541T>C), TTC151TTT (559C>T), C<br>CCA210CCG (736A>G), CTG246TTG (842C>T), CAA258CAG (880A>G), GCC273GCT (925C>T), GTA282GTG (952A>G), GGT286GGG (964T>G), AAG288AAA (970G>A), GCC297GCT (997C>T), TC<br>GAG372GAA (1222G>A), CAC376CAT (1234C>T), GGC378GGT (1240C>T), AGG383AGA (1255G>A), CAA390CAG (1276A>G), GGG398GGA (1300G>A), GAA415GAG (1351A>G), CTC422CTT (137<br>(1543G>A), AAA481AAG (1549A>G), GGC482GGT (1552C>T), AAC495AAT (1591C>T), CAG502CAA (1612G>A), GAT516GAC (1654T>C), TGC521TGT (1669C>T), AGC555AGT (1771C>T), TTG559<br>GTA618GTG (1960A>G), TAC626TAT (1984C>T), AAA636AAG (2014A>G), GCA646ACA (2042G>A), CTA662CTG (2092A>G), CTG678TTG (2138C>T), GGT681GGC (2149T>C), TGT683TGC (2155T<br>(2257C>T), CAG719CAA (2263G>A), GCG727GCA (2287G>A), CAC728CAT (2290C>T), CTG738TTG (2318C>T), TTC750TTT (2356C>T), TTG751TTA (2359G>A), TAT758TAC (2380T>C), AGG768A<br>GTT796GTC (2494T>C), GGC801GGT (2509C>T), CTT802CTC (2512T>C), TTA804CTA (2516T>C), GAG838GAA (2620G>A), ATC852ATT (2662C>T), TTT853TTC (2665T>C), GGC874GGT (2728C><br>(2809C>T), TCG902TCA (2812G>A), ACA907ACG (2827A>G), TTC921TTT (2869C>T), CAT925CAC (2881T>C), GCC936GCT (2914C>T), AAG943AAA (2935G>A), GTC948GTT (2950C>T), CTG949T<br>TTT1001TTC (3109T>C), TCT1019TCC (3163T>C), TCC1021TCT (3169C>T), AAA1023AAG (3175A>G), CTG1024TTG (3176C>T), TCT1029TCA (3193T>A), AAA1053AAG (3265A>G), CTC1104CTT (<br>GGC1145GGT (3541C>T), TCT1158TCC (3580T>C), ATC1159ATT (3583C>T), GAA1168GAG (3610A>G), GGT1169GGC (3613T>C), TAC1173TAT (3625C>T), GTG1174GTA (3628G>A), AAA1179AA<br>CGA1211CGT (3739A>T), AAG1216AAA (3754G>A), CTC1226CTT (3784C>T), GAC1236GAT (3814C>T), GAG1262GAA (3892G>A), GAC1281GAT (3949C>T), TTG1285CTG (3959T>C), AAA1304AA<br>(4084C>T), CAT1328CAC (4090T>C), TAC1329TAT (4093C>T), TCC1341TCT (4129C>T), CAC1348CAT (4150C>T), CTG1351TTA (4157C>T 4159G>A), AAA1354AAG (4168A>G), GCT1368GCC (421<br>AGA1418AGG (4360A>G), TAT1429TAC (4393T>C), TTT1437TTC (4417T>C), TTG1439CTG (4421T>C), CTC1457CTT (4477C>T), GCT1479GCC (4543T>C), TAT1495TAC (4591T>C), TAT1510TAC (<br>(4696C>T), GAC1531GAT (4699C>T), AGC1551AGT (4759C>T), TTT1554TTC (4768T>C), AGG1556AGA (4774G>A), CAC1557CAT (4777C>T), CTA1559CTG (4783A>G), GGA1560GGG (4786A>G),<br>(4898G>A), GAG1600GAA (4906G>A), AAG1621AAA (4969G>A), ACC1628GCC (4988A>G), ACA1629ACT (4993A>T), TAT1633TAC (5005T>C), TTG1636CTG (5012T>C), CTG1641CTA (5029G>A),<br>(5102T>C), ACT1671ATC (5118C>T 5119T>C), GTT1672ATT (5120G>A), CTG1673CTC (5125G>C), CAT1680CAC (5146T>C), ACT1681AAT (5148C>A), CAG1698CAA (5200G>A), GTT1704GTG (52<br>ATT1745ATA (5341T>A), GTA1746GTG (5344A>G), ACC1747ACT (5347C>T), GTG1755ATA (5369G>A 5371G>A), GGC1760GGT (5386C>T), GGG1762GGA (5392G>A), CCA1767CCG (5407A>G),<br>(5482A>G), CTT1809CTA (5533T>A), GAT1822GAC (5572T>C), TGT1824TGC (5578T>C), ATT1826ATC (5584T>C), ATA1837ATT (5617A>T), GGT1845GGC (5641T>C), TTT1847TTC (5647T>C), CC<br>(5797A>G), TAT1907TAC (5827T>C), GGG1924GGA (5878G>A), CCC1927CCT (5887C>T), CAG1944CAA (5938G>A), CAC1946CAT (5944C>T), AAT1947AGT (5946A>G), TTT1960TTC (5986T>C),<br>(6073T>C), CTC1991CTT (6079C>T), CTA1995CTG (6091A>G), CTT1996ATT (6092C>A), AAG2000AAA (6106G>A), TGT2004TGC (6118T>C), TCA2005TCG (6121A>G), TAT2014TAC (6148T>C), G<br>(6200A>T), TAC2033TAT (6205C>T), TTG2047TTA (6247G>A), ACC2050ACA (6256C>A), ACA2052ACG (6262A>G), CTG2053CTA (6265G>A), GTA2064GTG (6298A>G), GAT2069GAC (6313T>C),<br>(6493T>C) | | | | | |

*: Inserts / Deletes / Misaligned / Frameshifts

#### Analysis details

This analysis was performed with panviral2.64

### NGS Details (UN8): Torradovirus lycopersici (segment RNA 2)

#### Assembly

| | |
|---|---|
| **Coverage Length** | 5382 (1 contig(s)) |
| **Depth Of Coverage** | 7518.4 |
| **Number Of Reads** | 313618 |
| **Reads Per Million** | 5241.34 rpm (after QC) |
| **Ambiguities** | 0 |
| **Assembly Method** | read mapping against reference + variant calling |
| **Consensus Caller** | Bcf Tools |

#### Coverage Map

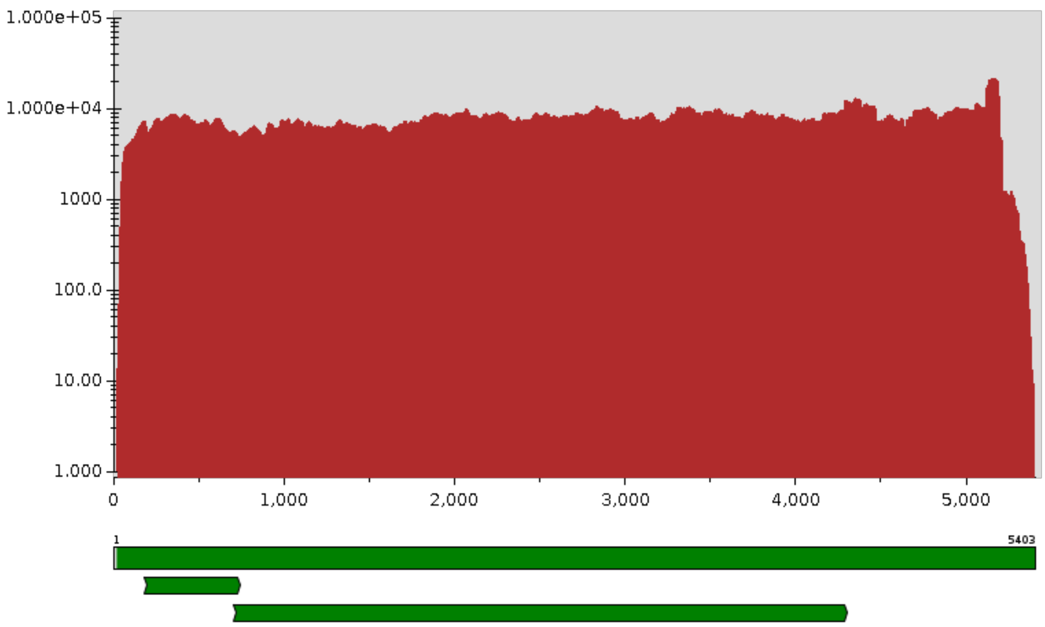

#### Assignment

| | |
|---|---|
| **Type** | Torradovirus lycopersici (Taxonomy ID: 3048378) |
| **Reference Genome** | NC_009032.1 |
| **NT Identity (%)** | 96.5633 |
| **AA Identity (%)** | 97.9813 |
| **Number Of Stop Codons** | 2 |
| **Number Of CDS** | 2 |

#### Alignment

| | |
|---|---|
| **Alignment Score** | 10018.0 (NT) + 9318.0 (AA) = 19336.0 |
| **Concordance (%)** | 95.4817 |

| Alignment Method | Global, seeded, nucleotide + amino acids (AGA) |
|---|---|

#### Genome Region

Sequence starts at position 22 and ends at position 5403 relative to NC_009032.1 reference sequence.

#### Alignment Detailed Statistics

| | Begin | End | Coverage | Score | Concordance | Matches | Identities | I/D/M/F* | Stop Codons |
|---|---|---|---|---|---|---|---|---|---|
| **NT** | **22** | **5403** | **99.6%** | **10018** | **93.1%** | **5382 (99.9%)** | **5198 (96.6%)** | **1/0** | |
| Mutations: | 89G>A, 90C>T, 97C>T, 107G>A, 111C>A, 137T>C, 144A>G, 161A>G, 224G>A, 277C>T, 280T>C, 283C>T, 295T>C, 301A>C, 313C>T, 343T>C, 401T>C, 433C>T, 440C>T, 448C>A, 460T>C, 481C>A, 487A>G, 507G>A, 583A>T, 586A>G, 589A>G, 654T>A, 679G>A, 696T>C, 721C>T, 733A>G, 748A>G, 750A>G, 797T>C, 806C>T, 827A>G, 844A>T, 866A>G, 965T>C, 968A>G, 1010T>G, 1061T>C, 1085G>A, 1106T>C, 1109C>T, 1139G>A, 1226A>G, 1271T>C, 1277A>G, 1300C>T, 1316C>T, 1337T>C, 1418G>A, 1421A>G, 1457C>T, 1490G>A, 1502C>T, 1508G>A, 1547C>T, 1557C>T, 1608T>C, 1619G>A, 1637G>A, 1646T>C, 1651A>G, 1662C>T, 1685T>C, 1694T>C, 1724C>T, 1757A>G, 1796T>A, 1859C>T, 1863C>T, 1878A>G, 1892C>T, 1967G>A, 1970A>G, 1994T>C, 2072G>A, 2084G>A, 2108A>G, 2128G>A, 2140A>C, 2159T>A, 2189C>T, 2205A>G, 2249T>C, 2263A>C, 2270A>G, 2285T>C, 2303A>G, 2312T>C, 2321T>C, 2324G>A, 2375C>T, 2429G>A, 2439C>T, 2465T>C, 2471C>T, 2501A>C, 2503G>A, 2519A>G, 2570G>A, 2573T>C, 2612C>T, 2630C>T, 2660T>C, 2708A>G, 2720C>T, 2732T>C, 2792C>T, 2819G>A, 2840T>C, 2841C>T, 2885T>C, 2939A>G, 2969C>T, 3003T>C, 3050G>A, 3122T>C, 3128T>C, 3131G>A, 3168C>T, 3215T>C, 3236C>T, 3265G>A, 3269T>C, 3320T>C, 3323T>C, 3335C>T, 3368A>G, 3386A>G, 3413C>T, 3422A>G, 3431T>C, 3437C>T, 3446T>C, 3470T>C, 3512T>C, 3514T>C, 3543T>A, 3551C>T, 3560T>A, 3569A>G, 3595A>G, 3626A>G, 3638A>G, 3662A>T, 3674G>A, 3716C>T, 3722C>T, 3743G>A, 3767C>T, 3773C>T, 3800C>T, 3827C>T, 3893T>C, 3933A>G, 3995C>T, 4013G>C, 4040C>T, 4046C>T, 4059A>G, 4061A>G, 4069A>G, 4100A>T, 4112C>T, 4130C>T, 4190A>G, 4224C>T, 4250C>T, 4457A>G, 4518G>A, 4556A>G, 4590G>A, 4592T>C, 4812C>T, 5180T>G, 5192C>T, 5206C>T, 5218C>T, 5235_5236insT, 5249A>G, 5262A>T | | | | | | | | |

##### CDS

| | Begin | End | Coverage | Score | Concordance | Matches | Identities | I/D/M/F* | Stop Codons |
|---|---|---|---|---|---|---|---|---|---|
| ToTV_sRNA2gp1 | 1 | 188 | 100% | 1186 | 98.1% | 188 (100%) | 184 (97.9%) | 0/0/0/0 | 1 |
| Protein mutations: | A15T (224G>A), R109K (507G>A), L158Q (654T>A), L172P (696T>C) | | | | | | | | |
| Codon mutations: | GCA15ACA (224G>A), GGC32GGT (277C>T), TCT33TCC (280T>C), GGC34GGT (283C>T), AGT38AGC (295T>C), CCA40CCC (301A>C), TTC44TTT (313C>T), ATT54ATC (343T>C), TTA74CTA (401T>C), GGC84GGT (433C>T), CTA87TTA (440C>T), GGC89GGA (448C>A), CAT93CAC (460T>C), ACC100ACA (481C>A), AGA102AGG (487A>G), AGG109AAG (507G>A), ACA134ACT (583A>T), CAA135CAG (586A>G), TTA136TTG (589A>G), CTA158CAA (654T>A), TTG166TTA (679G>A), CTA172CCA (696T>C), GGC180GGT (721C>T), GAA184GAG (733A>G) | | | | | | | | |
| ToTV_sRNA2gp2 | 1 | 1199 | 100% | 8132 | 98.2% | 1199 (100%) | 1175 (98.0%) | 0/0/0/0 | 1 |
| Protein mutations: | A7V (721C>T), K11R (733A>G), K16R (748A>G), S17G (750A>G), Y48F (844A>T), A200V (1300C>T), H286Y (1557C>T), Y303H (1608T>C), K317R (1651A>G), H388Y (1863C>T), I393V (1878A>G), R476K (2128G>A), K480T (2140A>C), M502V (2205A>G), E521A (2263A>C), R601K (2503G>A), R855K (3265G>A), V938A (3514T>C), S948T (3543T>A), D965G (3595A>G), T1078A (3933A>G), I1120V (4059A>G 4061A>G), N1123S (4069A>G), H1175Y (4224C>T) | | | | | | | | |
| Codon mutations: | GCT7GTT (721C>T), AAG11AGG (733A>G), AAA16AGA (748A>G), AGT17GGT (750A>G), GTT32GTC (797T>C), GGC35GGT (806C>T), AAA42AAG (827A>G), TAT48TTT (844A>T), CAA55CAG (866A>G), CAT88CAC (965T>C), GGA89GGG (968A>G), GTT103GTG (1010T>G), AGT120AGC (1061T>C), CTG128CTA (1085G>A), TTT135TTC (1106T>C), TCC136TCT (1109C>T), CCG146CCA (1139G>A), TCA175TCG (1226A>G), ATT190ATC (1271T>C), TCA192TCG (1277A>G), GCT200GTT (1300C>T), CAC205CAT (1316C>T), GAT212GAC (1337T>C), GTG239GTA (1418G>A), GGA240GGG (1421A>G), AGC252AGT (1457C>T), CGG263CGA (1490G>A), CTC267CTT (1502C>T), GTG269GTA (1508G>A), ACC282ACT (1547C>T), CAC286TAC (1557C>T), TAC303CAC (1608T>C), GAG306GAA (1619G>A), TTG312TTA (1637G>A), GTT315GTC (1646T>C), AAG317AGG (1651A>G), CTG321TTG (1662C>T), GAT328GAC (1685T>C), GGT331GGC (1694T>C), CCC341CCT (1724C>T), CAA352CAG (1757A>G), GTT365GTA (1796T>A), ACC386ACT (1859C>T), CAC388TAC (1863C>T), ATA393GTA (1878A>G), TAC397TAT (1892C>T), GAG422GAA (1967G>A), CTA423CTG (1970A>G), TAT431TAC (1994T>C), GAG457GAA (2072G>A), GTG461GTA (2084G>A), AGA469AGG (2108A>G), AGG476AAG (2128G>A), AAA480ACA (2140A>C), ACT486ACA (2159T>A), TTC496TTT (2189C>T), ATG502GTG (2205A>G), GCT516GCC (2249T>C), GAA521GCA (2263A>C), GTA523GTG (2270A>G), GTT528GTC (2285T>C), CAA534CAG (2303A>G), AAT537AAC (2312T>C), GGT540GGC (2321T>C), GCG541GCA (2324G>A), TTC558TTT (2375C>T), GAG576GAA (2429G>A), CTG580TTG (2439C>T), GGT588GGC (2465T>C), TCC590TCT (2471C>T), CTA600CTC (2501A>C), AGG601AAG (2503G>A), GCA606GCG (2519A>G), TCG623TCA (2570G>A), GGT624GGC (2573T>C), AAC637AAT (2612C>T), ATC643ATT (2630C>T), CAT653CAC (2660T>C), ACA669ACG (2708A>G), ATC673ATT (2720C>T), GTT677GTC (2732T>C), CAC697CAT (2792C>T), GAG706GAA (2819G>A), CAT713CAC (2840T>C), CTG714TTG (2841C>T), CGT728CGC (2885T>C), CGA746CGG (2939A>G), TAC756TAT (2969C>T), TTA768CTA (3003T>C), ACG783ACA (3050G>A), ATT807ATC (3122T>C), ATT809ATC (3128T>C), GAG810GAA (3131G>A), CTG823TTG (3168C>T), AAT838AAC (3215T>C), AAC845AAT (3236C>T), AGG855AAG (3265G>A), TGT856TGC (3269T>C), AGT873AGC (3320T>C), TCT874TCC (3323T>C), CCC878CCT (3335C>T), ACA889ACG (3368A>G), GTA895GTG (3386A>G), GTC904GTT (3413C>T), ACA907ACG (3422A>G), CTT910CTC (3431T>C), TGC912TGT (3437C>T), GAT915GAC (3446T>C), TCT923TCC (3470T>C), GCT937GCC (3512T>C), GTG938GCG (3514T>C), TCT948ACT (3543T>A), CGC950CGT (3551C>T), ATT953ATA (3560T>A), TTA956TTG (3569A>G), GAT965GGT (3595A>G), CAA975CAG (3626A>G), AGA979AGG (3638A>G), GTA987GTT (3662A>T), GTG991GTA (3674G>A), TGC1005TGT (3716C>T), CCC1007CCT (3722C>T), CAG1014CAA (3743G>A), ATC1022ATT (3767C>T), TGC1024TGT (3773C>T), TTC1033TTT (3800C>T), TAC1042TAT (3827C>T), GAT1064GAC (3893T>C), ACC1078GCC (3933A>G), AAC1098AAT (3995C>T), GTG1104GTC (4013G>C), CGC1113CGT (4040C>T), ACC1115ACT (4046C>T), ATA1120GTG (4059A>G 4061A>G), AAT1123AGT (4069A>G), CGA1133CGT (4100A>T), CTC1137CTT (4112C>T), GCC1143GCT (4130C>T), CAA1163CAG (4190A>G), CAT1175TAT (4224C>T), ACC1183ACT (4250C>T) | | | | | | | | |

##### Proteins

| | Begin | End | Coverage | Score | Concordance | Matches | Identities | I/D/M/F* | Stop Codons |
|---|---|---|---|---|---|---|---|---|---|
| Orf1 (YP_001040017.1) | 1 | 188 | 100% | 1186 | 98.1% | 188 (100%) | 184 (97.9%) | 0/0/0/0 | 1 |
| Protein mutations: | A15T (224G>A), R109K (507G>A), L158Q (654T>A), L172P (696T>C) | | | | | | | | |
| Codon mutations: | GCA15ACA (224G>A), GGC32GGT (277C>T), TCT33TCC (280T>C), GGC34GGT (283C>T), AGT38AGC (295T>C), CCA40CCC (301A>C), TTC44TTT (313C>T), ATT54ATC (343T>C), TTA74CTA (401T>C), GGC84GGT (433C>T), CTA87TTA (440C>T), GGC89GGA (448C>A), CAT93CAC (460T>C), ACC100ACA (481C>A), AGA102AGG (487A>G), AGG109AAG (507G>A), ACA134ACT (583A>T), CAA135CAG (586A>G), TTA136TTG (589A>G), CTA158CAA (654T>A), TTG166TTA (679G>A), CTA172CCA (696T>C), GGC180GGT (721C>T), GAA184GAG (733A>G) | | | | | | | | |
| polyprotein (YP_001040018.1) | 1 | 1199 | 100% | 8132 | 98.2% | 1199 (100%) | 1175 (98.0%) | 0/0/0/0 | 1 |
| Protein mutations: | A7V (721C>T), K11R (733A>G), K16R (748A>G), S17G (750A>G), Y48F (844A>T), A200V (1300C>T), H286Y (1557C>T), Y303H (1608T>C), K317R (1651A>G), H388Y (1863C>T), I393V (1878A>G), R476K (2128G>A), K480T (2140A>C), M502V (2205A>G), E521A (2263A>C), R601K (2503G>A), R855K (3265G>A), V938A (3514T>C), S948T (3543T>A), D965G (3595A>G), T1078A (3933A>G), I1120V (4059A>G 4061A>G), N1123S (4069A>G), H1175Y (4224C>T) | | | | | | | | |
| Codon mutations: | GCT7GTT (721C>T), AAG11AGG (733A>G), AAA16AGA (748A>G), AGT17GGT (750A>G), GTT32GTC (797T>C), GGC35GGT (806C>T), AAA42AAG (827A>G), TAT48TTT (844A>T), CAA55CAG (866A>G), CAT88CAC (965T>C), GGA89GGG (968A>G), GTT103GTG (1010T>G), AGT120AGC (1061T>C), CTG128CTA (1085G>A), TTT135TTC (1106T>C), TCC136TCT (1109C>T), CCG146CCA (1139G>A), TCA175TCG (1226A>G), ATT190ATC (1271T>C), TCA192TCG (1277A>G), GCT200GTT (1300C>T), CAC205CAT (1316C>T), GAT212GAC (1337T>C), GTG239GTA (1418G>A), GGA240GGG (1421A>G), AGC252AGT (1457C>T), CGG263CGA (1490G>A), CTC267CTT (1502C>T), GTG269GTA (1508G>A), ACC282ACT (1547C>T), CAC286TAC (1557C>T), TAC303CAC (1608T>C), GAG306GAA (1619G>A), TTG312TTA (1637G>A), GTT315GTC (1646T>C), AAG317AGG (1651A>G), CTG321TTG (1662C>T), GAT328GAC (1685T>C), GGT331GGC (1694T>C), CCC341CCT (1724C>T), CAA352CAG (1757A>G), GTT365GTA (1796T>A), ACC386ACT (1859C>T), CAC388TAC (1863C>T), ATA393GTA (1878A>G), TAC397TAT (1892C>T), GAG422GAA (1967G>A), CTA423CTG (1970A>G), TAT431TAC (1994T>C), GAG457GAA (2072G>A), GTG461GTA (2084G>A), AGA469AGG (2108A>G), AGG476AAG (2128G>A), AAA480ACA (2140A>C), ACT486ACA (2159T>A), TTC496TTT (2189C>T), ATG502GTG (2205A>G), GCT516GCC (2249T>C), GAA521GCA (2263A>C), GTA523GTG (2270A>G), GTT528GTC (2285T>C), CAA534CAG (2303A>G), AAT537AAC (2312T>C), GGT540GGC (2321T>C), GCG541GCA (2324G>A), TTC558TTT (2375C>T), GAG576GAA (2429G>A), CTG580TTG (2439C>T), GGT588GGC (2465T>C), TCC590TCT (2471C>T), CTA600CTC (2501A>C), AGG601AAG (2503G>A), GCA606GCG (2519A>G), TCG623TCA (2570G>A), GGT624GGC (2573T>C), AAC637AAT (2612C>T), ATC643ATT (2630C>T), CAT653CAC (2660T>C), ACA669ACG (2708A>G), ATC673ATT (2720C>T), GTT677GTC (2732T>C), CAC697CAT (2792C>T), GAG706GAA (2819G>A), CAT713CAC (2840T>C), CTG714TTG (2841C>T), CGT728CGC (2885T>C), CGA746CGG (2939A>G), TAC756TAT (2969C>T), TTA768CTA (3003T>C), ACG783ACA (3050G>A), ATT807ATC (3122T>C), ATT809ATC (3128T>C), GAG810GAA (3131G>A), CTG823TTG (3168C>T), AAT838AAC (3215T>C), AAC845AAT (3236C>T), AGG855AAG (3265G>A), TGT856TGC (3269T>C), AGT873AGC (3320T>C), TCT874TCC (3323T>C), CCC878CCT (3335C>T), ACA889ACG (3368A>G), GTA895GTG (3386A>G), GTC904GTT (3413C>T), ACA907ACG (3422A>G), CTT910CTC (3431T>C), TGC912TGT (3437C>T), GAT915GAC (3446T>C), TCT923TCC (3470T>C), GCT937GCC (3512T>C), GTG938GCG (3514T>C), TCT948ACT (3543T>A), CGC950CGT (3551C>T), ATT953ATA (3560T>A), TTA956TTG (3569A>G), GAT965GGT (3595A>G), CAA975CAG (3626A>G), AGA979AGG (3638A>G), GTA987GTT (3662A>T), GTG991GTA (3674G>A), TGC1005TGT (3716C>T), CCC1007CCT (3722C>T), CAG1014CAA (3743G>A), ATC1022ATT (3767C>T), TGC1024TGT (3773C>T), TTC1033TTT (3800C>T), TAC1042TAT (3827C>T), GAT1064GAC (3893T>C), ACC1078GCC (3933A>G), AAC1098AAT (3995C>T), GTG1104GTC (4013G>C), CGC1113CGT (4040C>T), ACC1115ACT (4046C>T), ATA1120GTG (4059A>G 4061A>G), AAT1123AGT (4069A>G), CGA1133CGT (4100A>T), CTC1137CTT (4112C>T), GCC1143GCT (4130C>T), CAA1163CAG (4190A>G), CAT1175TAT (4224C>T), ACC1183ACT (4250C>T) | | | | | | | | |

*: Inserts / Deletes / Misaligned / Frameshifts

#### Analysis details

This analysis was performed with panviral2.64

### NGS Details (UN8): Torradovirus marchitezum (segment RNA 2)

#### Assembly

| | |
|---|---|
| **Coverage Length** | 1059 (1 contig(s)) |
| **Depth Of Coverage** | 11498.6 |
| **Number Of Reads** | 97853 |
| **Reads Per Million** | 1635.37 rpm (after QC) |
| **Ambiguities** | 0 |
| **Assembly Method** | de novo + reference guided assembly |
| **Consensus Caller** | Bcf Tools |

#### Coverage Map

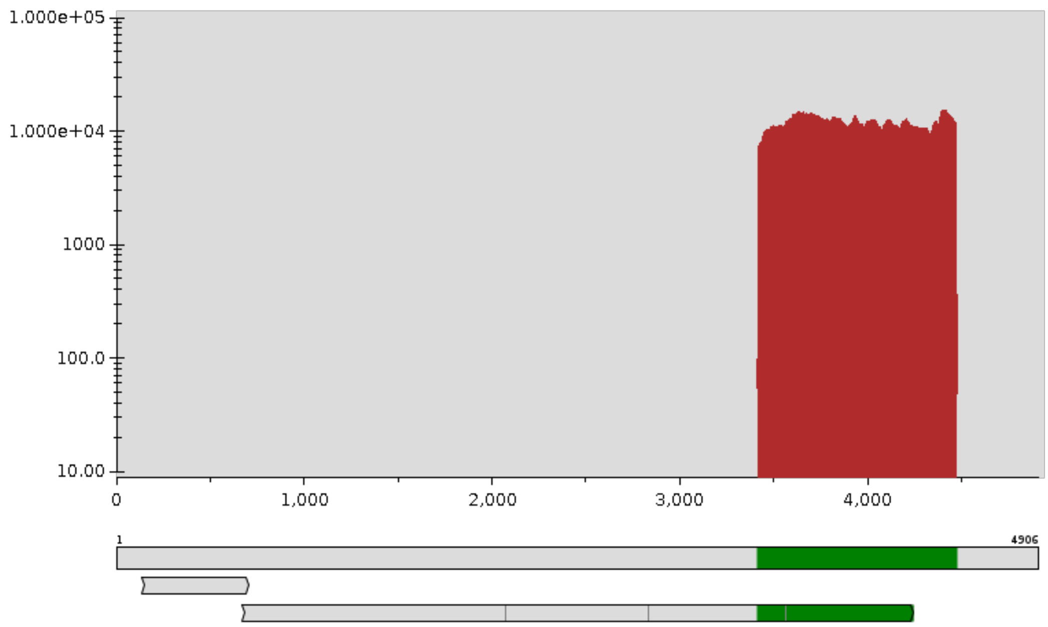

#### Assignment

| | |
|---|---|
| **Type** | Torradovirus marchitezum (Taxonomy ID: 3048376) |
| **Reference Genome** | NC_010988.1 |
| **NT Identity (%)** | 69.2669 |
| **AA Identity (%)** | 72.7599 |
| **Number Of Stop Codons** | 1 |
| **Number Of CDS** | 2 |

#### Alignment

| | |
|---|---|
| **Alignment Score** | 771.0 (NT) + 1569.0 (AA) = 2340.0 |
| **Concordance (%)** | 57.1289 |

| Alignment Method | Global, seeded, nucleotide + amino acids (AGA) |
|---|---|

#### Genome Region

Sequence starts at position 3416 and ends at position 4474 relative to NC_010988.1 reference sequence.

#### Alignment Detailed Statistics

| | Begin | End | Coverage | Score | Concordance | Matches | Identities | I/D/M/F* | Stop Codons |
|---|---|---|---|---|---|---|---|---|---|
| **NT** | **3416** | **4474** | **21.6%** | **771** | **36.8%** | **1054 (98.6%)** | **737 (68.9%)** | **10/5** | |

Mutations: 3427G>T, 3436A>T, 3442C>G, 3445A>G, 3448C>T, 3449T>G, 3451A>C, 3457T>C, 3460C>T, 3461C>T, 3463T>G, 3466C>T, 3476G>A, 3478T>C, 3481A>G, 3482T>A, 3484A>T, 3493G>A, 3497A>C, 3506T>C, 3514G>A, 3517C>A, 3520C>T, 3521A>G, 3528A>T, 3529G>C, 3532C>T, 3533C>G, 3535G>T, 3538G>T, 3544C>G, 3544_3545insAAGAAG, 3550T>C, 3551C>A, 3552C>A, 3554C>G, 3555C>A, 3556C>G, 3559G>A, 3560A>G, 3561A>T, 3564C>G, 3568A>G, 3569A>G, 3570G>T, 3572A>G, 3573T>C, 3574T>A, 3577G>T, 3578T>G, 3579C>G, 3580C>A, 3581A>C, 3583A>T, 3584A>C, 3586A>G, 3589G>A, 3595T>C, 3598C>T, 3604T>A, 3610A>T, 3619C>T, 3622C>T, 3623A>T, 3624T>C, 3625G>T, 3627G>C, 3628T>C, 3638G>A, 3639A>C, 3641G>A, 3642A>G, 3643A>C, 3644A>C, 3646A>C, 3647T>C, 3650T>G, 3655A>T, 3656A>C, 3661T>A, 3667A>T, 3670A>G, 3676C>T, 3677A>G, 3680A>C, 3682G>A, 3685T>C, 3686G>T, 3687A>C, 3689G>C, 3690G>C, 3691G>C, 3692C>A, 3693A>C, 3700T>C, 3712A>C, 3715C>A, 3718T>G, 3724A>C, 3730A>C, 3739T>C, 3740A>T, 3742A>T, 3751T>C, 3760A>T, 3763T>C, 3767T>C, 3769G>T, 3770A>G, 3772T>G, 3775T>C, 3776T>A, 3778A>C, 3779A>C, 3781T>C, 3782G>A, 3784G>C, 3787A>C, 3790C>T, 3793A>G, 3796A>C, 3800T>G, 3801C>G, 3803A>T, 3804G>C, 3806C>A, 3807C>A, 3811T>C, 3812G>A, 3817A>T, 3826A>T, 3832T>G, 3838T>G, 3841T>C, 3842G>A, 3860C>T, 3862A>T, 3863G>T, 3864C>A, 3868G>T, 3869C>G, 3871T>G, 3872A>G, 3873A>C, 3874T>A, 3875A>G, 3877A>G, 3880A>G, 3881A>G, 3882A>C, 3883T>A, 3884T>C, 3890G>C, 3892A>T, 3901T>C, 3904A>T, 3907C>T, 3908A>T, 3910A>C, 3917G>A, 3922T>C, 3925A>C, 3931T>A, 3934G>T, 3935G>A, 3937A>C, 3940A>T, 3943T>C, 3945C>T, 3946G>T, 3949C>T, 3952A>C, 3955C>T, 3956A>G, 3957C>T, 3958T>G, 3967T>C, 3971G>A, 3973G>A, 3982A>T, 3983A>C, 3986C>A, 3989C>T, 3994A>T, 3996G>A, 3998A>G, 3999T>C, 4000G>A, 4001A>C, 4003G>A, 4009C>T, 4010T>G, 4011C>A, 4012A>C, 4014G>A, 4015C>T, 4019C>A, 4020A>G, 4021A>T, 4024C>T, 4027C>T, 4030C>T, 4031A>G, 4033G>T, 4036A>T, 4037C>T, 4039A>G, 4045T>C, 4047G>A, 4048G>A, 4049C>T, 4051T>G, 4054A>T, 4055A>C, 4056A>T, 4060G>T, 4061A>T, 4063C>G, 4066A>C, 4070T>C, 4072G>A, 4075T>A, 4078T>C, 4079T>G, 4080C>T, 4081C>A, 4083C>A, 4085A>T, 4088G>A, 4090G>T, 4093T>C, 4102G>A, 4105A>C, 4106C>A, 4108T>G, 4109A>G, 4111A>T, 4112T>A, 4113C>A, 4114T>G, 4117T>A, 4120A>G, 4122T>C, 4123T>A, 4129C>T, 4131A>G, 4135G>A, 4138G>A, 4139T>C, 4141G>T, 4154T>C, 4155C>T, 4156T>G, 4159C>T, 4162C>T, 4180C>T, 4181A>G, 4192T>G, 4195C>T, 4198C>T, 4201T>C, 4204G>A, 4213A>T, 4219T>C, 4222T>A, 4225G>A, 4228A>G, 4229C>A, 4231T>A, 4234T>C, 4237T>C, 4238G>T, 4242T>A, 4243T>C, 4248A>C, 4252T>A, 4279A>T, 4279_4280insT, 4280A>T, 4280_4281insC, 4283A>G, 4286C>T, 4287T>A, 4288T>C, 4291C>A, 4294C>T, 4306G>A, 4309G>A, 4312G>T, 4313C>A, 4313_4314insG, 4315A>G, 4316A>T, 4318T>C, 4319G>T, 4320G>A, 4323T>G, 4324T>A, 4328T>C, 4330G>A, 4333A>G, 4335G>T, 4336A>C, 4338A>G, 4339_4343delAGGGA, 4345A>T, 4346G>C, 4349A>C, 4364C>T, 4365T>C, 4369A>G, 4372A>G, 4373G>A, 4392T>C, 4393C>T, 4419_4420insG, 4422A>G, 4426G>A, 4430G>A, 4437C>T, 4441T>C, 4449A>T, 4451T>G

##### CDS

| | Begin | End | Coverage | Score | Concordance | Matches | Identities | I/D/M/F* | Stop Codons |
|---|---|---|---|---|---|---|---|---|---|
| ToMarV_RNA2gp2 | 916 | 1192 | 23.2% | 1569 | 78.4% | 277 (99.3%) | 203 (72.8%) | 2/0/0/0 | 1 |

Protein mutations: S927A (3449T>G 3451A>C), D936N (3476G>A 3478T>C), S938T (3482T>A 3484A>T), I943L (3497A>C), D949E (3517C>A), I951V (3521A>G), E953V (3528A>T 3529G>C), Q955D (3533C>G 3535G>T), E956D (3538G>T), G958_K959insKK (3544_3545insAAGAAG), P961K (3551C>A 3552C>A), P962E (3554C>G 3555C>A 3556C>G), K964V (3560A>G 3561A>T), A965G (3564C>G), S967V (3569A>G 3570G>T), I968A (3572A>G 3573T>C 3574T>A), S970G (3578T>G 3579C>G 3580C>A), I971L (3581A>C 3583A>T), M985S (3623A>T 3624T>C 3625G>T), G986A (3627G>C 3628T>C), D990T (3638G>A 3639A>C), E991S (3641G>A 3642A>G 3643A>C), L994V (3650T>G), I996L (3656A>C), I1003V (3677A>G), E1006S (3686G>T 3687A>C), G1007P (3689G>C 3690G>C 3691G>C), H1008T (3692C>A 3693A>C), T1024S (3740A>T 3742A>T), N1034E (3770A>G 3772T>G), S1036T (3776T>A 3778A>C), I1037L (3779A>C 3781T>C), V1038I (3782G>A 3784G>C), S1044G (3800T>G 3801C>G), P1046N (3806C>A 3807C>A), V1048I (3812G>A), F1056L (3838T>G), A1058T (3842G>A), P1064S (3860C>T 3862A>T), A1065Y (3863G>T 3864C>A), L1067V (3869C>G 3871T>G), N1068A (3872A>G 3873A>C 3874T>A), K1069E (3875A>G 3877A>G), N1071A (3881A>G 3882A>C 3883T>A), Y1072H (3884T>C), V1074L (3890G>C 3892A>T), T1080S (3908A>T 3910A>C), D1083N (3917G>A), A1089T (3935G>A 3937A>C), T1092I (3945C>T 3946G>T), T1096V (3956A>G 3957C>T 3958T>G), E1101K (3971G>A 3973G>A), H1107Y (3989C>T), R1109K (3996G>A), M1110A (3998A>G 3999T>C 4000G>A), S1114D (4010T>G 4011C>A 4012A>C), S1115N (4014G>A 4015C>T), Q1117S (4019C>A 4020A>G 4021A>T), M1121V (4031A>G 4033G>T), R1126K (4047G>A 4048G>A), N1129L (4055A>C 4056A>T), I1131L (4061A>T 4063C>G), S1137V (4079T>G 4080C>T 4081C>A), A1138D (4083C>A), I1139L (4085A>T), V1140I (4088G>A 4090G>T), L1146M (4106C>A 4108T>G), I1147V (4109A>G 4111A>T), S1148K (4112T>A 4113C>A 4114T>G), L1151P (4122T>C 4123T>A), K1154R (4131A>G), S1162L (4154T>C 4155C>T 4156T>G), N1171D (4181A>G), A1190S (4238G>T), I1191N (4242T>A 4243T>C)

Codon mutations: GGG919GGT (3427G>T), ACA922ACT (3436A>T), GGC924GGG (3442C>G), AAA925AAG (3445A>G), AGC926AGT (3448C>T), TCA927GCC (3449T>G 3451A>C), GAT929GAC (3457T>C), TTC930TTT (3460C>T), CTT931TTG (3461C>T 3463T>G), TTC932TTT (3466C>T), GAT936AAC (3476G>A 3478T>C), TTA937TTG (3481A>G), TCA938ACT (3482T>A 3484A>T), AAG941AAA (3493G>A), ATT943CTT (3497A>C), TTG946CTG (3506T>C), AAG948AAA (3514G>A), GAC949GAA (3517C>A), AAC950AAT (3520C>T), ATT951GTT (3521A>G), GAG953GTC (3528A>T 3529G>C), TAC954TAT (3532C>T), CAG955GAT (3533C>G 3535G>T), GAG956GAT (3538G>T), GGC958GGG (3544C>G), GGC958_AAA959insAAGAAG (3544_3545insAAGAAG), GAT960GAC (3550T>C), CCG961AAG (3551C>A 3552C>A), CCC962GAG (3554C>G 3555C>A 3556C>G), AAG963AAA (3559G>A), AAG964GTG (3560A>G 3561A>T), GCT965GGT (3564C>G), CAA966CAG (3568A>G), AGT967GTT (3569A>G 3570G>T), ATT968GCA (3572A>G 3573T>C 3574T>A), CTG969CTT (3577G>T), TCC970GGA (3578T>G 3579C>G 3580C>A), ATA971CTT (3581A>C 3583A>T), AGA972CGG (3584A>C 3586A>G), GAG973GAA (3589G>A), TTT975TTC (3595T>C), TCC976TCT (3598C>T), GGT978GGA (3604T>A), GTA980GTT (3610A>T), TTC983TTT (3619C>T), TGC984TGT (3622C>T), ATG985TCT (3623A>T 3624T>C 3625G>T), GGT986GCC (3627G>C 3628T>C), GAC990ACC (3638G>A 3639A>C), GAA991AGC (3641G>A 3642A>G 3643A>C), AGA992CGC (3644A>C 3646A>C), TTG993CTG (3647T>C), TTG994GTG (3650T>G), GTA995GTT (3655A>T), ATT996CTT (3656A>C), CCT997CCA (3661T>A), GCA999GCT (3667A>T), CCA1000CCG (3670A>G), TCC1002TCT (3676C>T), ATA1003GTA (3677A>G), AGG1004CGA (3680A>C 3682G>A), TTT1005TTC (3685T>C), GAA1006TCA (3686G>T 3687A>C), GGG1007CCC (3689G>C 3690G>C 3691G>C), CAC1008ACC (3692C>A 3693A>C), CCT1010CCC (3700T>C), GCA1014GCC (3712A>C), ATC1015ATA (3715C>A), ACT1016ACG (3718T>G), CCA1018CCC (3724A>C), ATA1020ATC (3730A>C), TGT1023TGC (3739T>C), ACA1024TCT (3740A>T 3742A>T), TGT1027TGC (3751T>C), TCA1030TCT (3760A>T), GGT1031GGC (3763T>C), TTG1033CTT (3767T>C 3769G>T), AAT1034GAG (3770A>G 3772T>G), TAT1035TAC (3775T>C), TCA1036ACC (3776T>A 3778A>C), ATT1037CTC (3779A>C 3781T>C), GTG1038ATC (3782G>A 3784G>C), ATA1039ATC (3787A>C), CAC1040CAT (3790C>T), AGA1041AGG (3793A>G), GTA1042GTC (3796A>C), TCC1044GGC (3800T>G 3801C>G), AGT1045TCT (3803A>T 3804G>C), CCT1046AAT (3806C>A 3807C>A), AAT1047AAC (3811T>C), GTT1048ATT (3812G>A), GGA1049GGT (3817A>T), CTA1052CTT (3826A>T), GTT1054GTG (3832T>G), TTT1056TTG (3838T>G), GAT1057GAC (3841T>C), GCC1058ACC (3842G>A), CCA1064TCT (3860C>T 3862A>T), GCT1065TAT (3863G>T 3864C>A), GGG1066GGT (3868G>T), CTT1067GTG (3869C>G 3871T>G), AAT1068GCA (3872A>G 3873A>C 3874T>A), AAA1069GAG (3875A>G 3877A>G), GGA1070GGG (3880A>G), AAT1071GCA (3881A>G 3882A>C 3883T>A), TAT1072CAT (3884T>C), GTA1074CTT (3890G>C 3892A>T), GGT1077GGC (3901T>C), GGA1078GGT (3904A>T), GGC1079GGT (3907C>T), ACA1080TCC (3908A>T 3910A>C), GAT1083AAT (3917G>A), TTT1084TTC (3922T>C), TCA1085TCC (3925A>C), GGT1087GGA (3931T>A), GTG1088GTT (3934G>T), GCA1089ACC (3935G>A 3937A>C), ACA1090ACT (3940A>T), AAT1091AAC (3943T>C), ACG1092ATT (3945C>T 3946G>T), TTC1093TTT (3949C>T), TCA1094TCC (3952A>C), TTC1095TTT (3955C>T), ACT1096GTG (3956A>G 3957C>T 3958T>G), GAT1099GAC (3967T>C), GAG1101AAA (3971G>A 3973G>A), CCA1104CCT (3982A>T), AGG1105CGG (3983A>C), CGG1106AGG (3986C>A), CAT1107TAT (3989C>T), ACA1108ACT (3994A>T), AGG1109AAG (3996G>A), ATG1110GCA (3998A>G 3999T>C 4000G>A), AGG1111CGA (4001A>C 4003G>A), TTC1113TTT (4009C>T), TCA1114GAC (4010T>G 4011C>A 4012A>C), AGC1115AAT (4014G>A 4015C>T), CAA1117AGT (4019C>A 4020A>G 4021A>T), TCC1118TCT (4024C>T), CGC1119CGT (4027C>T), ATC1120ATT (4030C>T), ATG1121GTT (4031A>G 4033G>T), TCA1122TCT (4036A>T), CTA1123TTG (4037C>T 4039A>G), GAT1125GAC (4045T>C), AGG1126AAA (4047G>A 4048G>A), CTT1127TTG (4049C>T 4051T>G), GGA1128GGT (4054A>T), AAT1129CTT (4055A>C 4056A>T), CTG1130CTT (4060G>T), ATC1131TTG (4061A>T 4063C>G), ATA1132ATC (4066A>C), TTG1134CTA (4070T>C 4072G>A), CCT1135CCA (4075T>A), CCT1136CCC (4078T>C), TCC1137GTA (4079T>G 4080C>T 4081C>A), GCC1138GAC (4083C>A), ATA1139TTA (4085A>T), GTG1140ATT (4088G>A 4090G>T), AGT1141AGC (4093T>C), GAG1144GAA (4102G>A), ATA1145ATC (4105A>C), CTT1146ATG (4106C>A 4108T>G), ATA1147GTT (4109A>G 4111A>T), TCT1148AAG (4112T>A 4113C>A 4114T>G), CCT1149CCA (4117T>A), GGA1150GGG (4120A>G), CTT1151CCA (4122T>C 4123T>A), TTC1153TTT (4129C>T), AAG1154AGG (4131A>G), TTG1155TTA (4135G>A), GAG1156GAA (4138G>A), TTG1157CTT (4139T>C 4141G>T), TCT1162CTG (4154T>C 4155C>T 4156T>G), GCC1163GCT (4159C>T), AAC1164AAT (4162C>T), GGC1170GGT (4180C>T), AAT1171GAT (4181A>G), ACT1174ACG (4192T>G), CAC1175CAT (4195C>T), ACC1176ACT (4198C>T), TAT1177TAC (4201T>C), CAG1178CAA (4204G>A), TCA1181TCT (4213A>T), TTT1183TTC (4219T>C), TCT1184TCA (4222T>A), GAG1185GAA (4225G>A), CTA1186CTG (4228A>G), CGT1187AGA (4229C>A 4231T>A), GAT1188GAC (4234T>C), TTT1189TTC (4237T>C), GCG1190TCG (4238G>T), ATT1191AAC (4242T>A 4243T>C)

##### Proteins

| | Begin | End | Coverage | Score | Concordance | Matches | Identities | I/D/M/F* | Stop Codons |
|---|---|---|---|---|---|---|---|---|---|
| polyprotein (YP_001976149.1) | 916 | 1192 | 23.2% | 1569 | 78.4% | 277 (99.3%) | 203 (72.8%) | 2/0/0/0 | 1 |

Protein mutations: S927A (3449T>G 3451A>C), D936N (3476G>A 3478T>C), S938T (3482T>A 3484A>T), I943L (3497A>C), D949E (3517C>A), I951V (3521A>G), E953V (3528A>T 3529G>C), Q955D (3533C>G 3535G>T), E956D (3538G>T), G958_K959insKK (3544_3545insAAGAAG), P961K (3551C>A 3552C>A), P962E (3554C>G 3555C>A 3556C>G), K964V (3560A>G 3561A>T), A965G (3564C>G), S967V (3569A>G 3570G>T), I968A (3572A>G 3573T>C 3574T>A), S970G (3578T>G 3579C>G 3580C>A), I971L (3581A>C 3583A>T), M985S (3623A>T 3624T>C 3625G>T), G986A (3627G>C 3628T>C), D990T (3638G>A 3639A>C), E991S (3641G>A 3642A>G 3643A>C), L994V (3650T>G), I996L (3656A>C), I1003V (3677A>G), E1006S (3686G>T 3687A>C), G1007P (3689G>C 3690G>C 3691G>C), H1008T (3692C>A 3693A>C), T1024S (3740A>T 3742A>T), N1034E (3770A>G 3772T>G), S1036T (3776T>A 3778A>C), I1037L (3779A>C 3781T>C), V1038I (3782G>A 3784G>C), S1044G (3800T>G 3801C>G), P1046N (3806C>A 3807C>A), V1048I (3812G>A), F1056L (3838T>G), A1058T (3842G>A), P1064S (3860C>T 3862A>T), A1065Y (3863G>T 3864C>A), L1067V (3869C>G 3871T>G), N1068A (3872A>G 3873A>C 3874T>A), K1069E (3875A>G 3877A>G), N1071A (3881A>G 3882A>C 3883T>A), Y1072H (3884T>C), V1074L (3890G>C 3892A>T), T1080S (3908A>T 3910A>C), D1083N (3917G>A), A1089T (3935G>A 3937A>C), T1092I (3945C>T 3946G>T), T1096V (3956A>G 3957C>T 3958T>G), E1101K (3971G>A 3973G>A), H1107Y (3989C>T), R1109K (3996G>A), M1110A (3998A>G 3999T>C 4000G>A), S1114D (4010T>G 4011C>A 4012A>C), S1115N (4014G>A 4015C>T), Q1117S (4019C>A 4020A>G 4021A>T), M1121V (4031A>G 4033G>T), R1126K (4047G>A 4048G>A), N1129L (4055A>C 4056A>T), I1131L (4061A>T 4063C>G), S1137V (4079T>G 4080C>T 4081C>A), A1138D (4083C>A), I1139L (4085A>T), V1140I (4088G>A 4090G>T), L1146M (4106C>A 4108T>G), I1147V (4109A>G 4111A>T), S1148K (4112T>A 4113C>A 4114T>G), L1151P (4122T>C 4123T>A), K1154R (4131A>G), S1162L (4154T>C 4155C>T 4156T>G), N1171D (4181A>G), A1190S (4238G>T), I1191N (4242T>A 4243T>C)

| | Begin | End | Coverage | Score | Concordance | Matches | Identities | I/D/M/F* | Stop Codons |
|---|---|---|---|---|---|---|---|---|---|
| **NT** | **3416** | **4474** | **21.6%** | **771** | **36.8%** | **1054 (98.6%)** | **737 (68.9%)** | **10/5** | |
| Codon mutations: | GGG919GGT (3427G>T), ACA922ACT (3436A>T), GGC924GGG (3442C>G), AAA925AAG (3445A>G), AGC926AGT (3448C>T), TCA927GCC (3449T>G 3451A>C), GAT929GAC (3457T>C), TTC930TTT (3460C>T), CTT931TTG (3461C>T 3463T>G), TTC932TTT (3466C>T), GAT936AAC (3476G>A 3478T>C), TTA937TTG (3481A>G), TCA938ACT (3482T>A 3484A>T), AAG941AAA (3493G>A), ATT943CTT (3497A>C), TTG946CTG (3506T>C), AAG948AAA (3514G>A), GAC949GAA (3517C>A), AAC950AAT (3520C>T), ATT951GTT (3521A>G), GAG953GTC (3528A>T 3529G>C), TAC954TAT (3532C>T), CAG955GAT (3533C>G 3535G>T), GAG956GAT (3538G>T), GGC958GGG (3544C>G), GGC958_AAA959insAAGAAG (3544_3545insAAGAAG), GAT960GAC (3550T>C), CCG961AAG (3551C>A 3552C>A), CCC962GAG (3554C>G 3555C>A 3556C>G), AAG963AAA (3559G>A), AAG964GTG (3560A>G 3561A>T), GCT965GGT (3564C>G), CAA966CAG (3568A>G), AGT967GTT (3569A>G 3570G>T), ATT968GCA (3572A>G 3573T>C 3574T>A), CTG969CTT (3577G>T), TCC970GGA (3578T>G 3579C>G 3580C>A), ATA971CTT (3581A>C 3583A>T), AGA972CGG (3584A>C 3586A>G), GAG973GAA (3589G>A), TTT975TTC (3595T>C), TCC976TCT (3598C>T), GGT978GGA (3604T>A), GTA980GTT (3610A>T), TTC983TTT (3619C>T), TGC984TGT (3622C>T), ATG985TCT (3623A>T 3624T>C 3625G>T), GGT986GCC (3627G>C 3628T>C), GAC990ACC (3638G>A 3639A>C), GAA991AGC (3641G>A 3642A>G 3643A>C), AGA992CGC (3644A>C 3646A>C), TTG993CTG (3647T>C), TTG994GTG (3650T>G), GTA995GTT (3655A>T), ATT996CTT (3656A>C), CCT997CCA (3661T>A), GCA999GCT (3667A>T), CCA1000CCG (3670A>G), TCC1002TCT (3676C>T), ATA1003GTA (3677A>G), AGG1004CGA (3680A>C 3682G>A), TTT1005TTC (3685T>C), GAA1006TCA (3686G>T 3687A>C), GGG1007CCC (3689G>C 3690G>C 3691G>C), CAC1008ACC (3692C>A 3693A>C), CCT1010CCC (3700T>C), GCA1014GCC (3712A>C), ATC1015ATA (3715C>A), ACT1016ACG (3718T>G), CCA1018CCC (3724A>C), ATA1020ATC (3730A>C), TGT1023TGC (3739T>C), ACA1024TCT (3740A>T 3742A>T), TGT1027TGC (3751T>C), TCA1030TCT (3760A>T), GGT1031GGC (3763T>C), TTG1033CTT (3767T>C 3769G>T), AAT1034GAG (3770A>G 3772T>G), TAT1035TAC (3775T>C), TCA1036ACC (3776T>A 3778A>C), ATT1037CTC (3779A>C 3781T>C), GTG1038ATC (3782G>A 3784G>C), ATA1039ATC (3787A>C), CAC1040CAT (3790C>T), AGA1041AGG (3793A>G), GTA1042GTC (3796A>C), TCC1044GGC (3800T>G 3801C>G), AGT1045TCT (3803A>T 3804G>C), CCT1046AAT (3806C>A 3807C>A), AAT1047AAC (3811T>C), GTT1048ATT (3812G>A), GGA1049GGT (3817A>T), CTA1052CTT (3826A>T), GTT1054GTG (3832T>G), TTT1056TTG (3838T>G), GAT1057GAC (3841T>C), GCC1058ACC (3842G>A), CCA1064TCT (3860C>T 3862A>T), GCT1065TAT (3863G>T 3864C>A), GGG1066GGT (3868G>T), CTT1067GTG (3869C>G 3871T>G), AAT1068GCA (3872A>G 3873A>C 3874T>A), AAA1069GAG (3875A>G 3877A>G), GGA1070GGG (3880A>G), AAT1071GCA (3881A>G 3882A>C 3883T>A), TAT1072CAT (3884T>C), GTA1074CTT (3890G>C 3892A>T), GGT1077GGC (3901T>C), GGA1078GGT (3904A>T), GGC1079GGT (3907C>T), ACA1080TCC (3908A>T 3910A>C), GAT1083AAT (3917G>A), TTT1084TTC (3922T>C), TCA1085TCC (3925A>C), GGT1087GGA (3931T>A), GTG1088GTT (3934G>T), GCA1089ACC (3935G>A 3937A>C), ACA1090ACT (3940A>T), AAT1091AAC (3943T>C), ACG1092ATT (3945C>T 3946G>T), TTC1093TTT (3949C>T), TCA1094TCC (3952A>C), TTC1095TTT (3955C>T), ACT1096GTG (3956A>G 3957C>T 3958T>G), GAT1099GAC (3967T>C), GAG1101AAA (3971G>A 3973G>A), CCA1104CCT (3982A>T), AGG1105CGG (3983A>C), CGG1106AGG (3986C>A), CAT1107TAT (3989C>T), ACA1108ACT (3994A>T), AGG1109AAG (3996G>A), ATG1110GCA (3998A>G 3999T>C 4000G>A), AGG1111CGA (4001A>C 4003G>A), TTC1113TTT (4009C>T), TCA1114GAC (4010T>G 4011C>A 4012A>C), AGC1115AAT (4014G>A 4015C>T), CAA1117AGT (4019C>A 4020A>G 4021A>T), TCC1118TCT (4024C>T), CGC1119CGT (4027C>T), ATC1120ATT (4030C>T), ATG1121GTT (4031A>G 4033G>T), TCA1122TCT (4036A>T), CTA1123TTG (4037C>T 4039A>G), GAT1125GAC (4045T>C), AGG1126AAA (4047G>A 4048G>A), CTT1127TTG (4049C>T 4051T>G), GGA1128GGT (4054A>T), AAT1129CTT (4055A>C 4056A>T), CTG1130CTT (4060G>T), ATC1131TTG (4061A>T 4063C>G), ATA1132ATC (4066A>C), TTG1134CTA (4070T>C 4072G>A), CCT1135CCA (4075T>A), CCT1136CCC (4078T>C), TCC1137GTA (4079T>G 4080C>T 4081C>A), GCC1138GAC (4083C>A), ATA1139TTA (4085A>T), GTG1140ATT (4088G>A 4090G>T), AGT1141AGC (4093T>C), GAG1144GAA (4102G>A), ATA1145ATC (4105A>C), CTT1146ATG (4106C>A 4108T>G), ATA1147GTT (4109A>G 4111A>T), TCT1148AAG (4112T>A 4113C>A 4114T>G), CCT1149CCA (4117T>A), GGA1150GGG (4120A>G), CTT1151CCA (4122T>C 4123T>A), TTC1153TTT (4129C>T), AAG1154AGG (4131A>G), TTG1155TTA (4135G>A), GAG1156GAA (4138G>A), TTG1157CTT (4139T>C 4141G>T), TCT1162CTG (4154T>C 4155C>T 4156T>G), GCC1163GCT (4159C>T), AAC1164AAT (4162C>T), GGC1170GGT (4180C>T), AAT1171GAT (4181A>G), ACT1174ACG (4192T>G), CAC1175CAT (4195C>T), ACC1176ACT (4198C>T), TAT1177TAC (4201T>C), CAG1178CAA (4204G>A), TCA1181TCT (4213A>T), TTT1183TTC (4219T>C), TCT1184TCA (4222T>A), GAG1185GAA (4225G>A), CTA1186CTG (4228A>G), CGT1187AGA (4229C>A 4231T>A), GAT1188GAC (4234T>C), TTT1189TTC (4237T>C), GCG1190TCG (4238G>T), ATT1191AAC (4242T>A 4243T>C) | | | | | | | | |
| Coat protein B (YP_001976156.1) | 194 | 244 | 20.9% | 263 | 74.9% | 51 (96.2%) | 38 (71.7%) | 2/0/0/0 | 0 |
| Protein mutations: | S205A (3449T>G 3451A>C), D214N (3476G>A 3478T>C), S216T (3482T>A 3484A>T), I221L (3497A>C), D227E (3517C>A), I229V (3521A>G), E231V (3528A>T 3529G>C), Q233D (3533C>G 3535G>T), E234D (3538G>T), G236_K237insKK (3544_3545insAAGAAG), P239K (3551C>A 3552C>A), P240E (3554C>G 3555C>A 3556C>G), K242V (3560A>G 3561A>T), A243G (3564C>G) | | | | | | | | |
| Codon mutations: | GGG197GGT (3427G>T), ACA200ACT (3436A>T), GGC202GGG (3442C>G), AAA203AAG (3445A>G), AGC204AGT (3448C>T), TCA205GCC (3449T>G 3451A>C), GAT207GAC (3457T>C), TTC208TTT (3460C>T), CTT209TTG (3461C>T 3463T>G), TTC210TTT (3466C>T), GAT214AAC (3476G>A 3478T>C), TTA215TTG (3481A>G), TCA216ACT (3482T>A 3484A>T), AAG219AAA (3493G>A), ATT221CTT (3497A>C), TTG224CTG (3506T>C), AAG226AAA (3514G>A), GAC227GAA (3517C>A), AAC228AAT (3520C>T), ATT229GTT (3521A>G), GAG231GTC (3528A>T 3529G>C), TAC232TAT (3532C>T), CAG233GAT (3533C>G 3535G>T), GAG234GAT (3538G>T), GGC236GGG (3544C>G), GGC236_AAA237insAAGAAG (3544_3545insAAGAAG), GAT238GAC (3550T>C), CCG239AAG (3551C>A 3552C>A), CCC240GAG (3554C>G 3555C>A 3556C>G), AAG241AAA (3559G>A), AAG242GTG (3560A>G 3561A>T), GCT243GGT (3564C>G), CAA244CAG (3568A>G) | | | | | | | | |
| Coat protein C (YP_001976157.1) | 1 | 225 | 100% | 1305 | 79.1% | 225 (100%) | 164 (72.9%) | 0/0/0/0 | 0 |
| Protein mutations: | S1V (3569A>G 3570G>T), I2A (3572A>G 3573T>C 3574T>A), S4G (3578T>G 3579C>G 3580C>A), I5L (3581A>C 3583A>T), M19S (3623A>T 3624T>C 3625G>T), G20A (3627G>C 3628T>C), D24T (3638G>A 3639A>C), E25S (3641G>A 3642A>G 3643A>C), L28V (3650T>G), I30L (3656A>C), I37V (3677A>G), E40S (3686G>T 3687A>C), G41P (3689G>C 3690G>C 3691G>C), H42T (3692C>A 3693A>C), T58S (3740A>T 3742A>T), N68E (3770A>G 3772T>G), S70T (3776T>A 3778A>C), I71L (3779A>C 3781T>C), V72I (3782G>A 3784G>C), S78G (3800T>G 3801C>G), P80N (3806C>A 3807C>A), V82I (3812G>A), F90L (3838T>G), A92T (3842G>A), P98S (3860C>T 3862A>T), A99Y (3863G>T 3864C>A), L101V (3869C>G 3871T>G), N102A (3872A>G 3873A>C 3874T>A), K103E (3875A>G 3877A>G), N105A (3881A>G 3882A>C 3883T>A), Y106H (3884T>C), V108L (3890G>C 3892A>T), T114S (3908A>T 3910A>C), D117N (3917G>A), A123T (3935G>A 3937A>C), T126I (3945C>T 3946G>T), T130V (3956A>G 3957C>T 3958T>G), E135K (3971G>A 3973G>A), H141Y (3989C>T), R143K (3996G>A), M144A (3998A>G 3999T>C 4000G>A), S148D (4010T>G 4011C>A 4012A>C), S149N (4014G>A 4015C>T), Q151S (4019C>A 4020A>G 4021A>T), M155V (4031A>G 4033G>T), R160K (4047G>A 4048G>A), N163L (4055A>C 4056A>T), I165L (4061A>T 4063C>G), S171V (4079T>G 4080C>T 4081C>A), A172D (4083C>A), I173L (4085A>T), V174I (4088G>A 4090G>T), L180M (4106C>A 4108T>G), I181V (4109A>G 4111A>T), S182K (4112T>A 4113C>A 4114T>G), L185P (4122T>C 4123T>A), K188R (4131A>G), S196L (4154T>C 4155C>T 4156T>G), N205D (4181A>G), A224S (4238G>T), I225N (4242T>A 4243T>C) | | | | | | | | |
| Codon mutations: | AGT1GTT (3569A>G 3570G>T), ATT2GCA (3572A>G 3573T>C 3574T>A), CTG3CTT (3577G>T), TCC4GGA (3578T>G 3579C>G 3580C>A), ATA5CTT (3581A>C 3583A>T), AGA6CGG (3584A>C 3586A>G), GAG7GAA (3589G>A), TTT9TTC (3595T>C), TCC10TCT (3598C>T), GGT12GGA (3604T>A), GTA14GTT (3610A>T), TTC17TTT (3619C>T), TGC18TGT (3622C>T), ATG19TCT (3623A>T 3624T>C 3625G>T), GGT20GCC (3627G>C 3628T>C), GAC24ACC (3638G>A 3639A>C), GAA25AGC (3641G>A 3642A>G 3643A>C), AGA26CGC (3644A>C 3646A>C), TTG27CTG (3647T>C), TTG28GTG (3650T>G), GTA29GTT (3655A>T), ATT30CTT (3656A>C), CCT31CCA (3661T>A), GCA33GCT (3667A>T), CCA34CCG (3670A>G), TCC36TCT (3676C>T), ATA37GTA (3677A>G), AGG38CGA (3680A>C 3682G>A), TTT39TTC (3685T>C), GAA40TCA (3686G>T 3687A>C), GGG41CCC (3689G>C 3690G>C 3691G>C), CAC42ACC (3692C>A 3693A>C), CCT44CCC (3700T>C), GCA48GCC (3712A>C), ATC49ATA (3715C>A), ACT50ACG (3718T>G), CCA52CCC (3724A>C), ATA54ATC (3730A>C), TGT57TGC (3739T>C), ACA58TCT (3740A>T 3742A>T), TGT61TGC (3751T>C), TCA64TCT (3760A>T), GGT65GGC (3763T>C), TTG67CTT (3767T>C 3769G>T), AAT68GAG (3770A>G 3772T>G), TAT69TAC (3775T>C), TCA70ACC (3776T>A 3778A>C), ATT71CTC (3779A>C 3781T>C), GTG72ATC (3782G>A 3784G>C), ATA73ATC (3787A>C), CAC74CAT (3790C>T), AGA75AGG (3793A>G), GTA76GTC (3796A>C), TCC78GGC (3800T>G 3801C>G), AGT79TCT (3803A>T 3804G>C), CCT80AAT (3806C>A 3807C>A), AAT81AAC (3811T>C), GTT82ATT (3812G>A), GGA83GGT (3817A>T), CTA86CTT (3826A>T), GTT88GTG (3832T>G), TTT90TTG (3838T>G), GAT91GAC (3841T>C), GCC92ACC (3842G>A), CCA98TCT (3860C>T 3862A>T), GCT99TAT (3863G>T 3864C>A), GGG100GGT (3868G>T), CTT101GTG (3869C>G 3871T>G), AAT102GCA (3872A>G 3873A>C 3874T>A), AAA103GAG (3875A>G 3877A>G), GGA104GGG (3880A>G), AAT105GCA (3881A>G 3882A>C 3883T>A), TAT106CAT (3884T>C), GTA108CTT (3890G>C 3892A>T), GGT111GGC (3901T>C), GGA112GGT (3904A>T), GGC113GGT (3907C>T), ACA114TCC (3908A>T 3910A>C), GAT117AAT (3917G>A), TTT118TTC (3922T>C), TCA119TCC (3925A>C), GGT121GGA (3931T>A), GTG122GTT (3934G>T), GCA123ACC (3935G>A 3937A>C), ACA124ACT (3940A>T), AAT125AAC (3943T>C), ACG126ATT (3945C>T 3946G>T), TTC127TTT (3949C>T), TCA128TCC (3952A>C), TTC129TTT (3955C>T), ACT130GTG (3956A>G 3957C>T 3958T>G), GAT133GAC (3967T>C), GAG135AAA (3971G>A 3973G>A), CCA138CCT (3982A>T), AGG139CGG (3983A>C), CGG140AGG (3986C>A), CAT141TAT (3989C>T), ACA142ACT (3994A>T), AGG143AAG (3996G>A), ATG144GCA (3998A>G 3999T>C 4000G>A), AGG145CGA (4001A>C 4003G>A), TTC147TTT (4009C>T), TCA148GAC (4010T>G 4011C>A 4012A>C), AGC149AAT (4014G>A 4015C>T), CAA151AGT (4019C>A 4020A>G 4021A>T), TCC152TCT (4024C>T), CGC153CGT (4027C>T), ATC154ATT (4030C>T), ATG155GTT (4031A>G 4033G>T), TCA156TCT (4036A>T), CTA157TTG (4037C>T 4039A>G), GAT159GAC (4045T>C), AGG160AAA (4047G>A 4048G>A), CTT161TTG (4049C>T 4051T>G), GGA162GGT (4054A>T), AAT163CTT (4055A>C 4056A>T), CTG164CTT (4060G>T), ATC165TTG (4061A>T 4063C>G), ATA166ATC (4066A>C), TTG168CTA (4070T>C 4072G>A), CCT169CCA (4075T>A), CCT170CCC (4078T>C), TCC171GTA (4079T>G 4080C>T 4081C>A), GCC172GAC (4083C>A), ATA173TTA (4085A>T), GTG174ATT (4088G>A 4090G>T), AGT175AGC (4093T>C), GAG178GAA (4102G>A), ATA179ATC (4105A>C), CTT180ATG (4106C>A 4108T>G), ATA181GTT (4109A>G 4111A>T), TCT182AAG (4112T>A 4113C>A 4114T>G), CCT183CCA (4117T>A), GGA184GGG (4120A>G), CTT185CCA (4122T>C 4123T>A), TTC187TTT (4129C>T), AAG188AGG (4131A>G), TTG189TTA (4135G>A), GAG190GAA (4138G>A), TTG191CTT (4139T>C 4141G>T), TCT196CTG (4154T>C 4155C>T 4156T>G), GCC197GCT (4159C>T), AAC198AAT (4162C>T), GGC204GGT (4180C>T), AAT205GAT (4181A>G), ACT208ACG (4192T>G), CAC209CAT (4195C>T), ACC210ACT (4198C>T), TAT211TAC (4201T>C), CAG212CAA (4204G>A), TCA215TCT (4213A>T), TTT217TTC (4219T>C), TCT218TCA (4222T>A), GAG219GAA (4225G>A), CTA220CTG (4228A>G), CGT221AGA (4229C>A 4231T>A), GAT222GAC (4234T>C), TTT223TTC (4237T>C), GCG224TCG (4238G>T), ATT225AAC (4242T>A 4243T>C) | | | | | | | | |

*: Inserts / Deletes / Misaligned / Frameshifts

#### Analysis details

This analysis was performed with panviral2.64

### NGS Details (UN8): Torradovirus marchitezum (segment RNA 1)

#### Assembly

| | |
|---|---|
| **Coverage Length** | 6723 (1 contig(s)) |
| **Depth Of Coverage** | 3512.7 |
| **Number Of Reads** | 176957 |
| **Reads Per Million** | 2957.39 rpm (after QC) |
| **Ambiguities** | 0 |
| **Assembly Method** | de novo + reference guided assembly |
| **Consensus Caller** | Bcf Tools |

#### Coverage Map

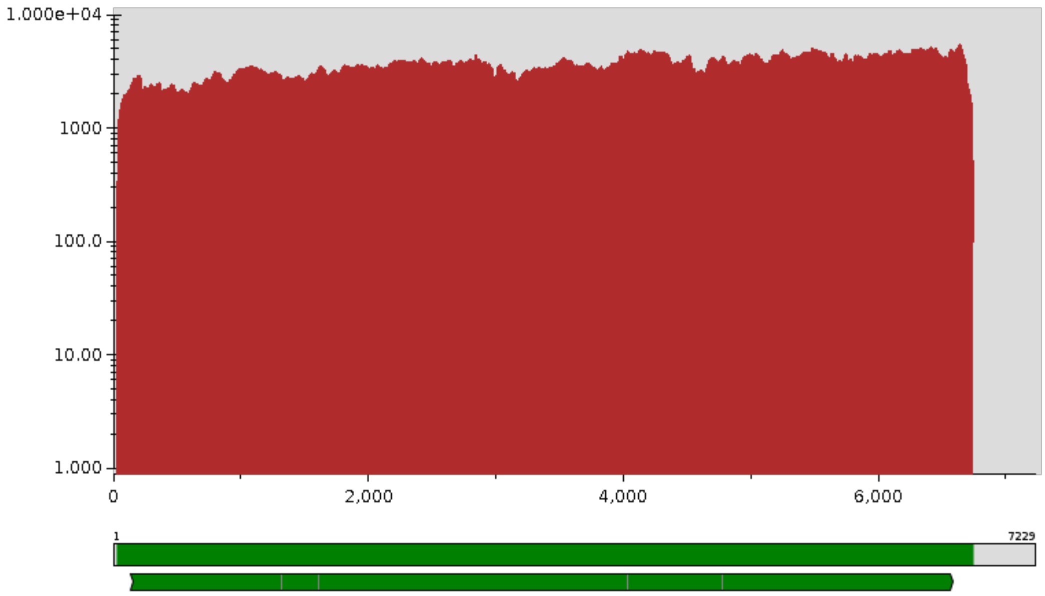

#### Assignment

| | |
|---|---|
| **Type** | Torradovirus marchitezum (Taxonomy ID: 3048376) |
| **Reference Genome** | NC_010987.1 |
| **NT Identity (%)** | 62.9646 |
| **AA Identity (%)** | 63.3737 |
| **Number Of Stop Codons** | 0 |
| **Number Of CDS** | 1 |

#### Alignment

| | |
|---|---|
| **Alignment Score** | 3297.0 (NT) + 9863.0 (AA) = 13160.0 |
| **Concordance (%)** | 47.0084 |

| | |
|---|---|
| **Alignment Method** | Global, seeded, nucleotide + amino acids (AGA) |

#### Genome Region

Sequence starts at position 25 and ends at position 6747 relative to NC_010987.1 reference sequence.

#### Alignment Detailed Statistics

| | Begin | End | Coverage | Score | Concordance | Matches | Identities | I/D/M/F* | Stop Codons |
|---|---|---|---|---|---|---|---|---|---|
| **NT** | **25** | **6747** | **93.0%** | **3297** | **25.1%** | **6658 (98.4%)** | **4218 (62.4%)** | **41/65** | |

| | Begin | End | Coverage | Score | Concordance | Matches | Identities | I/D/M/F* | Stop Codons |
|---|---|---|---|---|---|---|---|---|---|
| **NT** | **25** | **6747** | **93.0%** | **3297** | **25.1%** | **6658 (98.4%)** | **4218 (62.4%)** | **41/65** | |

Mutations:

32T>C, 37G>A, 40C>T, 59A>C, 61T>C, 62T>A, 63G>C, 66A>C, 67T>A, 68T>A, 70G>A, 71A>C, 74C>T, 75T>G, 76T>C, 77G>T, 79A>T, 81A>C, 82C>T, 85T>C, 87C>A,
89_99delAACATCTTCTT, 108A>T, 112G>C, 114C>T, 115C>T, 116A>C, 118T>C, 119A>T, 122T>G, 125A>T, 128C>A, 129C>A, 130T>A, 133_134delAT, 136T>A, 146T>A, 160T>G,
161C>G, 162T>A, 163C>G, 164C>T, 165_188delAAGGTTGCTGGTGTTGATGACGCC, 189A>C, 190G>C, 192T>C, 195T>G, 200C>T, 201A>T, 203C>A, 204A>C, 217G>C, 218T>A,
222C>G, 227A>T, 228A>T, 229A>C, 234A>T, 239C>T, 240A>T, 241A>C, 242A>T, 243T>C, 251A>G, 252G>A, 254G>T, 255G>C, 257T>C, 258A>T, 259G>C, 260T>C, 264A>G,
268C>A, 269C>T, 271C>A, 272T>A, 273A>G, 275G>T, 283A>C, 284G>C, 288T>G, 289C>T, 296T>C, 297C>A, 299T>A, 306C>G, 308G>C, 314A>G, 315T>C, 317T>A, 320G>T,
322C>T, 323T>G, 325A>C, 326T>A, 328T>C, 329G>A, 330T>G, 332C>A, 338T>A, 343T>C, 344G>C, 345G>T, 347C>T, 350T>A, 353T>C, 355G>A, 357T>G, 359A>T, 360C>G,
362T>A, 363A>G, 365T>A, 367A>G, 374G>A, 377C>T, 380A>G, 385G>C, 386T>A, 387G>A, 388C>A, 390C>A, 392T>A, 395A>G, 398T>A, 401A>C, 404T>C, 405C>A, 407T>C,
410A>G, 412C>G, 413G>T, 416C>T, 417T>G, 418C>A, 422C>T, 423T>G, 425G>T, 426T>A, 427C>A, 428C>A, 431A>T, 441T>A, 446C>A, 447A>G, 448A>T, 449G>A, 453A>C,
455G>T, 458T>A, 459T>C, 461A>T, 465T>A, 468T>G, 471G>T, 472A>T, 473G>A, 477T>G, 479G>C, 480T>A, 481C>A, 482C>G, 483A>G, 485T>C, 489G>T, 491T>C, 493A>T,
494A>C, 495T>C, 502T>A, 503T>C, 505C>A, 506T>A, 510G>T, 512G>A, 516A>G, 518T>A, 522A>G, 523G>T, 527A>G, 528G>T, 529A>C, 530T>A, 531G>T, 542T>G, 543G>T,
545A>T, 546T>A, 547A>T, 548T>A, 550G>C, 555C>T, 557T>A, 560A>T, 567A>T, 569A>G, 570A>G, 572A>C, 573T>A, 575G>T, 578A>G, 579G>C, 581G>A, 582C>T, 584T>A,
587A>G, 589T>A, 593T>A, 596T>C, 600A>G, 601G>T, 602T>A, 611T>A, 612G>A, 613A>C, 615G>C, 616G>C, 618G>T, 621T>A, 622C>A, 626C>T, 627A>G, 628C>T, 629T>A,
632C>T, 633A>G, 634A>T, 635A>T, 637A>C, 639A>T, 641C>G, 646T>C, 647G>A, 648A>G, 650A>T, 653C>T, 659C>T, 660A>T, 662G>T, 665T>C, 668A>T, 669A>T, 670G>C,
671T>A, 672C>T, 674A>G, 677C>T, 697A>C, 702G>T, 704C>T, 706A>G, 707A>G, 708C>A, 709G>A, 710A>G, 713C>T, 714A>T, 717A>C, 718T>A, 720G>C, 725C>G, 731T>A,
732A>G, 733G>C, 734T>A, 735A>G, 737G>A, 746G>T, 750T>A, 751C>G, 752A>T, 753G>T, 755T>A, 756T>A, 758A>G, 770G>T, 771T>A, 772C>G, 773T>C, 776G>A, 777T>C,
779G>T, 780A>T, 782G>A, 786A>T, 789A>T, 791A>G, 803G>T, 806T>C, 809G>A, 812G>T, 815G>A, 818G>A, 821C>T, 822A>G, 827A>T, 830G>T, 836T>G, 848C>T, 849T>C,
851G>T, 852A>T, 853G>C, 854C>A, 861T>C, 866C>A, 872T>C, 873C>T, 875C>G, 878T>C, 881G>T, 882A>G, 884A>T, 896T>G, 899G>A, 911A>G, 914A>G, 918A>C, 919C>A,
920T>G, 921C>T, 923T>G, 927C>A, 929T>G, 932T>C, 938A>G, 944C>T, 947C>A, 950C>T, 953G>A, 956T>C, 974A>G, 978A>G, 980A>G, 983C>T, 984G>A, 988C>T, 989C>T,
995A>C, 998C>T, 1010G>A, 1019T>G, 1024G>A, 1028A>T, 1031G>C, 1032C>T, 1034G>A, 1037C>T, 1040T>C, 1046G>T, 1050G>C, 1059C>A, 1060A>C, 1064G>A, 1068G>T,
1069A>C, 1070T>A, 1072C>A, 1073A>G, 1076G>T, 1077T>G, 1078G>C, 1082T>A, 1085T>A, 1086G>T, 1088A>T, 1090C>T, 1100G>A, 1103C>T, 1109C>A, 1113A>C, 1114A>C,
1115G>T, 1118T>A, 1121C>T, 1124T>A, 1126C>T, 1127C>T, 1130A>G, 1134A>C, 1136C>A, 1140C>T, 1142G>A, 1144G>A, 1145G>A, 1148A>G, 1149G>C, 1151A>G, 1154T>C,
1155C>G, 1160G>A, 1163C>A, 1178A>C, 1193G>A, 1199C>T, 1200A>T, 1215A>T, 1216G>C, 1226T>A, 1227G>A, 1232G>A, 1233T>C, 1238A>T, 1242G>T, 1244T>G, 1245T>G,
1246A>T, 1253A>G, 1254A>G, 1255G>C, 1256T>C, 1257T>A, 1259T>C, 1260A>C, 1261G>A, 1262G>A, 1265C>T, 1266A>T, 1267T>C, 1268G>T, 1271C>T, 1280G>A, 1284C>A,
1286T>G, 1292A>G, 1298T>C, 1304A>G, 1313C>T, 1316G>A, 1317T>G, 1319C>T, 1325G>A, 1337T>A, 1340A>G, 1342C>G, 1350T>C, 1352A>T, 1355T>C, 1367G>T, 1368T>C,
1370G>C, 1374A>G, 1375G>C, 1385G>A, 1391A>G, 1394C>G, 1397T>C, 1400G>C, 1401C>T, 1403A>G, 1409A>G, 1418G>A, 1421G>A, 1445T>A, 1448C>T, 1457T>A, 1458A>G,
1460C>G, 1463G>T, 1472C>T, 1478A>T, 1481A>T, 1482T>C, 1484G>T, 1487A>G, 1496T>A, 1505C>T, 1508A>G, 1511T>C, 1514T>A, 1517A>G, 1529T>C, 1538A>T, 1541A>C,
1544A>G, 1547A>T, 1553C>T, 1554T>C, 1556A>T, 1565C>T, 1571T>C, 1577C>T, 1583C>A, 1589T>C, 1592C>T, 1595C>A, 1598C>A, 1604T>C, 1607A>C, 1616A>C, 1623C>T,
1628C>A, 1631A>T, 1637T>A, 1640C>T, 1643A>G, 1649T>G, 1652T>A, 1655A>G, 1661A>T, 1667T>A, 1673G>A, 1674C>A, 1679A>G, 1682G>A, 1685C>T, 1688C>T, 1689C>T,
1691G>A, 1694G>T, 1700T>C, 1702T>G, 1703G>A, 1708A>T, 1715G>T, 1719T>G, 1720C>A, 1721C>T, 1722A>C, 1724T>A, 1725G>A, 1727A>T, 1729A>C, 1733T>A, 1736G>A,
1738G>C, 1739A>C, 1742T>C, 1745C>A, 1746A>G, 1747G>T, 1748A>T, 1752A>G, 1754C>T, 1755A>G, 1757A>G, 1763G>A, 1781T>C, 1787T>A, 1790C>T, 1793G>A, 1794A>G,
1795A>C, 1799G>A, 1800A>T, 1806A>G, 1807G>A, 1811T>C, 1814T>C, 1818G>C, 1820G>A, 1829A>G, 1832C>A, 1835A>G, 1838T>C, 1840A>G, 1844A>G, 1847T>C, 1848G>T,
1849C>A, 1850A>T, 1853A>G, 1856A>G, 1861A>G, 1862G>C, 1866T>C, 1868G>T, 1870T>A, 1872G>A, 1874A>T, 1875G>A, 1878G>A, 1879A>C, 1883T>A, 1886C>A, 1887A>T,
1889A>G, 1890A>G, 1891G>C, 1893C>G, 1894C>A, 1895T>C, 1899A>G, 1901A>T, 1910T>C, 1911C>A, 1913A>T, 1917T>G, 1921C>T, 1922A>T, 1928T>A, 1929G>A, 1932A>C,
1934A>G, 1936C>A, 1937G>T, 1941G>T, 1943G>C, 1944G>C, 1946T>A, 1948G>A, 1949G>T, 1952T>C, 1953_1955delAGA, 1958G>A, 1960G>A, 1961C>A, 1962A>G, 1963C>T,
1964C>T, 1965A>T, 1966A>T, 1969A>C, 1970A>T, 1971G>C, 1972A>C, 1980A>G, 1981T>C, 1983T>A, 1984T>C, 1985G>T, 1986G>T, 1987T>A, 1988G>T, 1989C>G, 1990C>A,
1991C>G, 1992A>C, 1993C>A, 1994T>G, 1995G>A, 1996T>C, 1997T>A, 1998G>A, 2000G>T, 2002G>C, 2003T>A, 2007A>G, 2008A>C, 2012T>A, 2014G>A, 2016G>A, 2018G>T,
2019A>G, 2021C>T, 2024T>C, 2025G>T, 2027A>C, 2028A>C, 2029G>A, 2030A>G, 2033G>A, 2039G>A, 2042A>T, 2048A>C, 2049T>C, 2051G>C, 2060A>T, 2066G>A, 2069C>G,
2073T>A, 2076G>A, 2079C>A, 2080C>A, 2081T>A, 2082G>T, 2083A>T, 2084G>T, 2087A>G, 2088C>T, 2093T>C, 2096T>A, 2102C>T, 2104A>T, 2105T>C, 2106C>T, 2107A>C,
2108T>C, 2117C>T, 2120G>C, 2127T>G, 2128C>T, 2129T>G, 2132T>A, 2133G>T, 2134A>C, 2136G>T, 2137A>C, 2141A>T, 2142C>G, 2143A>C, 2144T>C, 2148G>C, 2152T>A,
2156C>T, 2157A>G, 2159C>A, 2161A>T, 2162A>G, 2166A>G, 2167G>T, 2169G>A, 2170T>A, 2171C>T, 2172A>G, 2174C>T, 2175A>T, 2181T>G, 2182A>T, 2183T>C, 2185G>A,
2193G>A, 2194A>G, 2196G>C, 2199G>C, 2201A>G, 2208C>G, 2209G>C, 2210A>T, 2219A>G, 2221A>G, 2237C>T, 2241A>G, 2243A>C, 2249C>T, 2252T>C, 2268G>A, 2269C>G,
2270A>C, 2271T>A, 2279C>T, 2285G>A, 2286A>G, 2289A>T, 2290A>T, 2291A>G, 2294A>G, 2296C>A, 2297A>T, 2300G>C, 2303G>A, 2304A>C, 2305C>G, 2306A>C, 2307G>A,
2309G>T, 2310A>C, 2312G>T, 2313A>G, 2314G>A, 2316A>C, 2317C>A, 2318T>A, 2321A>G, 2322T>A, 2323T>A, 2324T>A, 2327A>G, 2328C>T, 2331T>A, 2332C>A, 2333T>G,
2334T>G, 2335G>C, 2336C>G, 2337A>C, 2340G>C, 2342T>G, 2345A>T, 2350T>C, 2351T>A, 2354C>T, 2357C>A, 2360C>T, 2362T>C, 2363A>G, 2369C>T, 2372G>A, 2381A>G,
2385T>A, 2386T>G, 2387T>A, 2390T>A, 2396T>C, 2397T>C, 2399A>G, 2405A>T, 2408A>T, 2415A>C, 2417A>C, 2426C>T, 2428A>G, 2429C>G, 2435A>T, 2436T>G,
2438_2439insGGT, 2440A>C, 2441A>T, 2444G>C, 2445C>A, 2446C>A, 2447A>G, 2451G>A, 2452A>G, 2454T>A, 2455C>A, 2456A>T, 2462C>T, 2464A>T, 2465A>C, 2466A>G,
2467T>A, 2468T>A, 2472T>A, 2473A>T, 2474C>G, 2475C>A, 2476A>T, 2478A>G, 2479A>G, 2480A>T, 2484G>A, 2485T>A, 2486T>G, 2489A>C, 2490T>A, 2492T>A, 2495C>G,
2497A>T, 2498G>C, 2499G>C, 2501T>G, 2503C>T, 2504A>G, 2508T>A, 2509G>A, 2510G>T, 2511A>G, 2512G>A, 2514C>T, 2516T>G, 2521A>T, 2522T>C, 2523G>A, 2524C>T,
2525A>C, 2526C>T, 2530G>A, 2531A>G, 2537C>T, 2546A>C, 2549T>A, 2556A>T, 2558A>G, 2561G>A, 2564T>A, 2573A>C, 2574G>A, 2575T>C, 2576T>C, 2577T>A, 2579T>G,
2580G>A, 2581G>A, 2582A>T, 2584G>C, 2586A>G, 2591A>T, 2594A>T, 2597T>C, 2598T>A, 2600G>T, 2601G>A, 2603G>T, 2607T>C, 2608G>T, 2609G>C, 2610C>A, 2613A>G,
2614G>T, 2615T>A, 2616A>G, 2617A>C, 2618C>A, 2619T>A, 2623T>C, 2627C>T, 2628A>T, 2629A>T, 2630T>A, 2633A>T, 2634T>G, 2635C>A, 2636G>T, 2645A>G, 2648T>C,
2651G>A, 2660A>G, 2672A>C, 2678T>C, 2684G>A, 2687C>T, 2690G>A, 2693T>G, 2696T>G, 2698A>C, 2699G>T, 2699_2700insCAT, 2700C>T, 2703A>G, 2704G>C, 2708T>A,
2709C>T, 2710T>C, 2711T>C, 2712C>T, 2714C>G, 2715G>C, 2717T>A, 2720T>G, 2724A>G, 2725G>C, 2726T>G, 2727T>G, 2729T>A, 2730C>G, 2732A>C, 2733T>A, 2734C>A,
2735A>C, 2736T>A, 2737T>A, 2738G>C, 2739G>C, 2740C>A, 2741A>T, 2742C>G, 2743A>T, 2744A>G, 2747T>C, 2747_2748insCGA, 2753T>A, 2755C>A, 2756T>A, 2757G>C,
2759A>G, 2771A>T, 2775C>G, 2777G>A, 2780C>T, 2783C>G, 2784A>C, 2786G>C, 2792T>C, 2798G>T, 2801G>T, 2804A>G, 2808C>A, 2812G>C, 2822G>C, 2825C>T, 2828T>C,
2834C>T, 2837C>G, 2840C>T, 2843G>T, 2858A>T, 2859A>C, 2861G>C, 2864C>T, 2867A>G, 2870T>A, 2876A>C, 2879C>T, 2880C>A, 2881A>G, 2882C>A, 2885T>C, 2886G>A,
2887C>T, 2888T>G, 2889C>A, 2891A>G, 2897T>C, 2903A>T, 2907A>T, 2909A>C, 2915C>T, 2920C>A, 2921C>G, 2922A>C, 2927T>C, 2928C>A, 2929G>T, 2930A>T, 2932C>G,
2936A>T, 2939T>G, 2942A>G, 2944A>T, 2945C>T, 2948C>T, 2953G>C, 2954A>C, 2955G>C, 2964A>T, 2966T>G, 2969T>A, 2975C>T, 2981C>T, 2982A>G, 2984C>G, 2988C>T,
2994C>A, 2996C>G, 2999T>A, 3002T>G, 3017A>C, 3020G>A, 3021C>T, 3023T>A, 3026C>T, 3029C>A, 3030C>A, 3032T>A, 3038C>T, 3044A>T, 3045A>C, 3047A>C, 3050G>A,
3056G>T, 3062C>T, 3064T>A, 3066A>T, 3068A>C, 3078C>A, 3079A>T, 3080A>G, 3083A>C, 3090T>G, 3091G>T, 3092T>G, 3095A>G, 3101T>A, 3104G>T, 3107T>A, 3109C>A,
3110A>G, 3116A>C, 3117C>A, 3118G>A, 3119T>G, 3120C>A, 3121T>A, 3122T>A, 3123G>T, 3124G>C, 3131A>G, 3135T>G, 3137T>G, 3140C>T, 3143G>A, 3144A>T, 3145T>A,
3146G>T, 3167A>C, 3173A>C, 3175T>A, 3176T>A, 3182G>C, 3183G>A, 3184A>C, 3190A>G, 3192C>T, 3194G>A, 3195A>G, 3197C>T, 3200C>G, 3201T>A, 3202C>A, 3203C>T,
3204A>G, 3205G>C, 3206T>A, 3210A>G, 3211A>C, 3212G>T, 3213A>C, 3216T>A, 3221T>A, 3225C>G, 3226A>T, 3227A>T, 3228T>G, 3230T>A, 3236A>G, 3245A>T, 3249G>A,
3251G>T, 3254A>G, 3260T>A, 3263C>T, 3269C>T, 3272G>T, 3273T>C, 3275G>T, 3278A>C, 3279C>T, 3281G>A, 3284A>G, 3289G>C, 3293T>G, 3299T>C, 3305A>G, 3308T>G,
3309G>T, 3310T>G, 3311G>C, 3320C>A, 3323C>A, 3326C>T, 3335A>T, 3343G>A, 3344C>T, 3345G>A, 3346T>C, 3347A>C, 3350A>T, 3353T>A, 3356A>G, 3357C>A, 3358G>A,
3359T>G, 3362A>T, 3364T>A, 3365T>C, 3366A>G, 3368C>G, 3369T>G, 3377T>C, 3380T>A, 3386T>A, 3387A>G, 3389C>G, 3391A>T, 3392C>T, 3395G>A, 3398A>T, 3402C>T,
3404A>G, 3407G>A, 3416A>C, 3419T>G, 3422G>A, 3425G>C, 3426G>A, 3428C>T, 3433C>T, 3434A>G, 3437A>T, 3443A>G, 3452T>A, 3453A>C, 3455T>G, 3456C>T, 3462T>C,
3463C>A, 3464A>C, 3465A>C, 3467T>G, 3470A>G, 3471A>T, 3472G>C, 3475G>A, 3476A>G, 3479A>G, 3480A>G, 3481C>A, 3483C>A, 3484A>G, 3485G>A, 3486C>T, 3488T>C,
3497A>G, 3500C>A, 3503A>G, 3507T>G, 3508A>C, 3509C>T, 3511A>T, 3513C>T, 3515T>G, 3518A>C, 3521A>T, 3524T>A, 3527A>G, 3528C>T, 3529A>T, 3533G>A, 3534T>A,
3539C>T, 3542C>T, 3548T>G, 3551C>A, 3557G>A, 3560C>A, 3563T>C, 3566G>T, 3567C>A, 3568G>A, 3569G>A, 3570A>C, 3571A>G, 3572A>C, 3575C>T, 3577A>C, 3578A>C,
3579T>C, 3587A>G, 3590A>T, 3593C>T, 3596A>G, 3597C>G, 3598C>A, 3602A>T, 3603G>C, 3608A>T, 3611A>C, 3620T>C, 3624A>T, 3626A>T, 3628T>C, 3629T>C, 3630C>A,
3632T>A, 3641A>G, 3644T>A, 3647T>A, 3650T>G, 3656C>T, 3671G>T, 3674A>C, 3675A>G, 3676T>A, 3680A>G, 3683C>T, 3697G>C, 3698T>C, 3699G>A, 3700G>A, 3701A>C,
3707T>G, 3710T>G, 3711G>C, 3712A>C, 3713C>A, 3719C>A, 3722A>G, 3725A>G, 3728A>C, 3731C>T, 3734A>G, 3735A>G, 3736A>G, 3740C>T, 3743G>A, 3744T>G, 3745G>C,
3746C>A, 3749G>C, 3750C>A, 3752G>T, 3755A>T, 3759A>G, 3761C>G, 3762A>C, 3764G>T, 3765C>A, 3766G>A, 3767T>G, 3769G>T, 3770C>G, 3774A>C, 3775A>G, 3776G>A,
3777A>G, 3779C>G, 3782T>C, 3788A>G, 3789C>A, 3790G>A, 3791T>A, 3794A>T, 3795T>G, 3796G>C, 3797T>A, 3798G>A, 3799A>T, 3800A>T, 3803G>A, 3804G>A, 3827A>G,
3838G>T, 3839T>C, 3845C>T, 3847A>T, 3853G>C, 3854C>A, 3860G>A, 3861C>T, 3869A>G, 3872T>A, 3875C>T, 3878A>T, 3884C>T, 3887C>T, 3894A>T, 3898G>A, 3899G>A,
3900A>C, 3902G>T, 3905G>T, 3906A>T, 3907T>C, 3908G>T, 3914G>A, 3917A>G, 3927C>G, 3930T>G, 3933A>C, 3934C>A, 3935T>A, 3936G>C, 3937G>C, 3938C>A, 3941A>C,
3942G>T, 3943G>C, 3944A>G, 3945G>C, 3950A>G, 3953G>T, 3954G>C, 3955A>T, 3956A>T, 3965A>G, 3966G>A, 3971T>A, 3977A>C, 3983A>T, 3996C>T, 3998A>G, 4001T>C,
4002G>T, 4003T>C, 4008A>G, 4010T>A, 4014A>C, 4019C>A, 4021T>C, 4022C>A, 4023C>T, 4024A>T, 4028G>T, 4031T>C, 4034G>T, 4040C>A, 4046A>G, 4049C>T, 4052C>G,
4064C>G, 4065C>T, 4067C>G, 4079A>G, 4088C>G, 4094A>G, 4101A>C, 4103G>C, 4106C>G, 4109T>C, 4112G>A, 4115A>C, 4116C>T, 4118T>G, 4130C>T, 4137T>C, 4142C>G,
4143A>C, 4148G>A, 4151T>C, 4154T>C, 4160T>C, 4163C>T, 4166A>T, 4169C>T, 4170A>T, 4171C>G, 4172A>T, 4182T>G, 4183T>C, 4184G>A, 4190T>C, 4192C>T, 4193T>A,
4194A>C, 4196A>C, 4199A>T, 4205A>G, 4208T>A, 4214T>C, 4215G>A, 4220T>A, 4223A>G, 4226T>C, 4227T>C, 4228C>A, 4239T>G, 4242T>C, 4247A>T, 4249A>G, 4251G>A,
4253G>C, 4259A>G, 4260G>T, 4262T>A, 4265G>A, 4268A>T, 4271C>T, 4273T>A, 4280T>C, 4286T>C, 4292T>C, 4295C>G, 4316T>C, 4317G>A, 4319T>C, 4325T>A, 4331G>A,
4344T>C, 4346G>A, 4354A>C, 4355A>T, 4357A>T, 4358C>T, 4361C>T, 4364A>T, 4365T>G, 4366C>A, 4367A>T, 4368A>G, 4376C>G, 4385T>G, 4388C>A, 4395C>A, 4397A>G,
4400T>C, 4401G>A, 4404C>T, 4406T>A, 4418A>C, 4424T>C, 4430C>T, 4433A>C, 4436G>C, 4439C>T, 4443G>A, 4445T>C, 4448A>G, 4449G>T, 4455A>C, 4457G>C, 4458T>C,
4460G>A, 4463T>G, 4466G>T, 4469C>A, 4475A>C, 4478A>T, 4481A>T, 4487A>T, 4488C>T, 4490A>G, 4493G>T, 4499C>A, 4502C>T, 4508T>A, 4509C>T, 4511C>G, 4514C>A,
4530C>A, 4535T>C, 4538C>G, 4548T>C, 4553A>G, 4556A>G, 4562C>A, 4566A>G, 4568T>C, 4569G>A, 4570C>A, 4571T>A, 4576G>A, 4577G>A, 4578G>T, 4581A>G, 4583A>G,
4586A>C, 4587G>A, 4592A>G, 4597G>A, 4601T>C, 4611C>T, 4613A>G, 4622T>A, 4625T>A, 4631A>T, 4637C>T, 4640T>C, 4641C>T, 4643T>G, 4649T>G, 4652A>T, 4656T>A,
4658A>G, 4662C>T, 4664C>G, 4665C>A, 4666A>G, 4673T>C, 4674T>G, 4675C>A, 4679G>A, 4683A>T, 4684A>C, 4685T>C, 4688A>T, 4692C>A, 4693A>G, 4694T>A, 4697G>A,
4698T>C, 4700A>G, 4704C>G, 4705T>C, 4706G>T, 4709C>T, 4712T>C, 4715A>C, 4716A>G, 4731C>T, 4733T>G, 4739A>G, 4742G>A, 4751T>A, 4752A>G, 4754T>G, 4755G>A,
4758G>A, 4759T>C, 4761C>A, 4768A>T, 4769A>G, 4784A>C, 4787T>C, 4793G>C, 4799A>G, 4805T>C, 4808A>G, 4817T>A, 4818C>T, 4820T>G, 4826C>T, 4829C>A, 4835T>A,
4838T>A, 4844G>A, 4854A>G, 4868G>T, 4871T>C, 4877A>T, 4880A>G, 4883T>A, 4886C>A, 4887C>G, 4888C>T, 4889A>G, 4892T>A, 4901C>T, 4902A>G, 4903C>T, 4904T>A,
4907G>A, 4910C>T, 4914A>T, 4915T>A, 4916G>T, 4923G>A, 4925G>T, 4934A>C, 4937A>C, 4938C>A, 4940T>A, 4946T>C, 4955C>T, 4958T>G, 4960G>A, 4961T>G, 4963T>C,
4964G>C, 4965G>A, 4973C>A, 4975A>G, 4976G>T, 4977C>T, 4982C>A, 4983T>A, 4984T>A, 4985A>T, 4993A>T, 4994G>T, 4995A>G, 4996C>G, 4997G>A, 5000G>T, 5001T>C,
5003G>C, 5004G>A, 5006T>G, 5007G>C, 5009A>G, 5012G>A, 5013G>A, 5014G>A, 5015T>C, 5016T>C, 5019A>C, 5020G>C, 5021A>C, 5022G>A, 5023C>T, 5024T>C, 5025G>A,
5031T>A, 5033T>G, 5036T>C, 5037T>A, 5039C>T, 5042T>C, 5044T>A, 5045G>T, 5049G>C, 5051G>T, 5055C>A, 5056A>G, 5057G>T, 5058G>C, 5064A>T, 5066T>G, 5067G>A,
5070A>G, 5075A>C, 5081G>A, 5085T>G, 5086C>A, 5087A>G, 5091G>A, 5092C>T, 5093C>T, 5094C>G, 5095T>G, 5097A>G, 5098C>T, 5099A>T, 5100A>T, 5102T>G, 5103T>A,
5104G>T, 5105T>G, 5108A>G, 5109A>C, 5111G>A, 5112G>T, 5115A>G, 5117A>T, 5120A>T, 5121G>A, 5122C>A, 5123T>A, 5125T>A, 5127T>A, 5129C>A, 5132T>C, 5135G>C,
5138A>T, 5141G>A, 5142G>C, 5143A>C, 5144C>T, 5147A>T, 5150T>G, 5153G>A, 5159C>T, 5160_5162delACA, 5172C>G, 5173G>A, 5174T>G, 5176A>G, 5178C>G, 5180G>C,
5181A>C, 5184T>G, 5185G>A, 5186G>A, 5192A>G, 5195A>T, 5196A>G, 5198C>T, 5199A>G, 5201T>A, 5202G>T, 5203G>T, 5207T>C, 5211T>A, 5212C>G, 5213A>T, 5216A>T,
5219T>G, 5226G>T, 5228C>A, 5229C>A, 5232T>C, 5234C>A, 5235A>C, 5236C>A, 5237G>A, 5241G>T, 5242A>C, 5243T>A, 5244G>T, 5245A>C, 5249G>T, 5250T>A, 5255T>C,
5256A>C, 5258T>G, 5264C>T, 5267T>C, 5270A>T, 5273T>A, 5276A>C, 5277A>C, 5278A>G, 5279G>C, 5281A>G, 5282G>A, 5283T>A, 5285C>A, 5287A>C, 5288T>A, 5294T>G,
5300T>G, 5303A>T, 5309C>T, 5318C>T, 5333G>A, 5339G>C, 5346A>T, 5349T>G, 5354T>A, 5357A>C, 5358A>C, 5359T>A, 5364G>C, 5365C>A, 5366C>T, 5370A>C, 5371A>G,
5372G>C, 5375T>G, 5376G>A, 5377T>A, 5378G>T, 5379T>C, 5383C>A, 5386G>C, 5387A>T, 5390A>T, 5391C>G, 5395T>G, 5399A>T, 5403A>T, 5404A>C, 5409G>A, 5410T>C,
5411T>C, 5417G>T, 5423A>C, 5429A>T, 5435T>A, 5436G>A, 5438C>A, 5440G>C, 5441C>T, 5447T>C, 5456G>A, 5457C>T, 5460A>T, 5462G>T, 5465C>A, 5468A>G, 5471A>C,
5474A>T, 5475A>C, 5476G>A, 5477G>T, 5483T>C, 5484T>A, 5486T>G, 5488T>C, 5489G>C, 5490G>A, 5491A>C, 5492G>A, 5496A>C, 5498A>T, 5501A>T, 5504T>G, 5505A>C,
5507A>C, 5508T>C, 5510G>A, 5513C>A, 5516A>C, 5519T>G, 5521G>C, 5522A>T, 5523A>G, 5524A>G, 5525T>G, 5531T>G, 5534C>T, 5537C>G, 5540T>C, 5546T>C, 5549T>C,
5555C>T, 5560T>C, 5564G>A, 5567T>C, 5570T>G, 5573T>C, 5574C>A, 5576A>G, 5582G>T, 5583C>G, 5584C>A, 5585T>A, 5586C>G, 5588T>C, 5588_5589insACC, 5589G>T,
5590G>C, 5592G>C, 5597A>G, 5598G>T, 5602C>G, 5603T>A, 5606A>G, 5612T>C, 5613G>A, 5617C>A, 5618G>T, 5619C>G, 5620T>G, 5622_5624delCAC, 5626G>C, 5627T>C,
5629G>C, 5635T>A, 5636G>C, 5642A>T, 5642_5643insCACAGT, 5644G>C, 5645G>A, 5649G>T, 5650A>C, 5652T>G, 5654G>T, 5657C>A, 5658A>C, 5659A>T, 5660C>T,
5663C>A, 5664C>A, 5666C>G, 5669T>A, 5670A>G, 5672A>T, 5674A>C, 5676G>T, 5677G>C, 5678T>C, 5681T>C, 5682T>C, 5684A>C, 5685G>T, 5687A>T, 5688A>G, 5691C>A,

| | |
|---|---|
| | 5695C>T, 5696A>G, 5700A>G, 5701T>C, 5702G>T, 5705G>T, 5706A>T, 5714T>C, 5715G>A, 5716T>C, 5717G>A, 5718G>A, 5720A>T, 5723A>T, 5726A>T, 5727C>G, 5729T>A, 5735A>T, 5740G>C, 5741A>C, 5742A>C, 5744T>G, 5746T>G, 5747T>C, 5750C>A, 5753A>G, 5754C>A, 5757G>A, 5759T>C, 5762T>G, 5763G>A, 5766A>C, 5767G>A, 5769T>G, 5770T>C, 5771G>T, 5772C>T, 5774A>G, 5775C>G, 5777C>G, 5778T>A, 5779A>C, 5780T>A, 5783G>A, 5784G>T, 5785A>C, 5786G>T, 5787C>G, 5788A>T, 5789G>C, 5790G>A, 5792A>T, 5798A>T, 5799G>A, 5800A>C, 5801G>T, 5802T>C, 5804A>T, 5805A>G, 5806T>C, 5807T>C, 5808C>G, 5809A>C, 5812A>G, 5813G>A, 5814A>G, 5815A>G, 5816T>A, 5817A>G, 5818C>T, 5819A>C, 5823C>G, 5825A>G, 5828T>A, 5829C>T, 5837C>T, 5843G>A, 5844C>T, 5846G>C, 5847A>T, 5848G>C, 5849T>C, 5850A>C, 5851T>A, 5852G>A, 5853G>C, 5855A>C, 5856A>G, 5859G>T, 5860T>G, 5861G>T, 5862C>G, 5865C>A, 5866A>T, 5867A>G, 5868C>A, 5870G>A, 5871A>C, 5873A>T, 5876T>G, 5880G>C, 5882T>G, 5883A>G, 5885G>A, 5886C>A, 5887T>G, 5891G>A, 5895T>G, 5897T>A, 5898T>G, 5900C>A, 5901C>T, 5902C>T, 5903A>C, 5905A>T, 5906A>G, 5908A>G, 5909A>G, 5911C>G, 5912T>C, 5913A>G, 5914C>T, 5915C>T, 5917T>A, 5918T>A, 5919C>A, 5921G>A, 5922A>C, 5924G>A, 5925C>A, 5927C>G, 5928A>T, 5929A>T, 5930A>G, 5931A>T, 5932A>C, 5933T>C, 5936T>C, 5937G>A, 5939T>G, 5940G>A, 5943G>C, 5944C>A, 5945T>A, 5946G>C, 5948A>G, 5949T>A, 5950G>T, 5951T>G, 5952C>T, 5953T>C, 5956C>A, 5958T>A, 5959T>C, 5961A>C, 5965C>T, 5966A>T, 5967A>G, 5968G>C, 5969G>C, 5972_5973insCCATGTTGTGTCACA, 5978G>T, 5979C>A, 5981C>A, 5983T>C, 5985G>T, 5986T>G, 5987T>G, 5988T>C, 5990C>A, 5992A>C, 5993A>T, 5997A>G, 5999A>C, 6000C>A, 6003T>G, 6005G>C, 6007G>T, 6008T>G, 6009T>A, 6011C>A, 6013C>A, 6015T>G, 6017G>T, 6020A>G, 6022T>A, 6024T>C, 6025G>C, 6026T>A, 6029T>A, 6030_6032delGAA, 6033A>T, 6034G>C, 6039_6050delGTTGCTTCAAGG, 6051T>C, 6056A>T, 6059G>A, 6061G>C, 6062T>A, 6063C>G, 6065A>G, 6068C>T, 6070C>T, 6074C>G, 6078T>C, 6079T>C, 6080G>T, 6083T>A, 6086T>A, 6087T>C, 6088C>T, 6089C>A, 6092A>T, 6093T>G, 6097A>G, 6102T>A, 6104A>G, 6108G>C, 6109A>G, 6110A>C, 6112G>A, 6113A>T, 6114C>A, 6115A>G, 6116C>T, 6117A>G, 6118C>G, 6121G>A, 6122A>G, 6126A>T, 6127T>A, 6128G>C, 6129A>T, 6130G>C, 6131T>A, 6135C>A, 6136A>C, 6137G>A, 6139G>C, 6140T>A, 6141A>C, 6146G>A, 6153C>T, 6155G>T, 6157A>G, 6161A>G, 6164T>C, 6165C>A, 6167A>G, 6169T>G, 6170A>T, 6171A>C, 6173C>G, 6174T>G, 6175T>A, 6176C>A, 6177C>G, 6178A>G, 6179A>G, 6180G>T, 6181T>A, 6182A>T, 6183A>T, 6186A>G, 6187C>A, 6188A>T, 6190G>A, 6191T>A, 6193A>C, 6194G>C, 6198T>G, 6200A>G, 6201A>G, 6203G>C, 6204A>T, 6205A>G, 6206G>C, 6209T>C, 6210G>A, 6211A>G, 6212A>T, 6215T>G, 6219T>A, 6221T>G, 6222C>G, 6223A>G, 6224G>C, 6226A>T, 6227G>T, 6228A>T, 6229A>G, 6230G>T, 6231A>G, 6233A>G, 6234T>C, 6236C>G, 6237A>G, 6238C>A, 6239C>A, 6240A>C, 6241C>G, 6243C>A, 6245A>C, 6246C>T, 6248A>G, 6249A>G, 6251C>G, 6252A>C, 6253A>G, 6254A>C, 6255G>A, 6256T>C, 6258T>A, 6260C>T, 6261T>G, 6262T>C, 6263G>T, 6270G>T, 6271A>C, 6272G>C, 6273G>A, 6274A>C, 6276A>T, 6277G>C, 6278T>A, 6280A>T, 6281T>G, 6282C>A, 6284G>C, 6285A>G, 6286C>A, 6287T>G, 6288G>A, 6290G>T, 6294T>A, 6295G>C, 6297A>G, 6299A>G, 6300G>T, 6302T>C, 6303T>A, 6305T>A, 6306C>G, 6308G>T, 6310T>G, 6314C>T, 6315C>T, 6316A>T, 6317G>T, 6318A>G, 6319G>A, 6320G>T, 6321_6323delATA, 6324A>C, 6326C>T, 6328C>T, 6330C>A, 6332G>A, 6333A>G, 6334A>C, 6336T>G, 6338C>G, 6339A>G, 6340G>A, 6341A>C, 6342G>T, 6344G>C, 6345C>A, 6348G>C, 6349A>C, 6350A>T, 6353T>G, 6359T>C, 6361A>C, 6362T>C, 6364T>G, 6366C>G, 6368T>A, 6369C>A, 6370C>T, 6372G>A, 6374A>T, 6376G>A, 6377A>T, 6378A>G, 6379T>C, 6383C>T, 6389T>C, 6392C>T, 6394T>C, 6402A>G, 6404A>G, 6405A>T, 6407A>G, 6411T>G, 6412C>A, 6413T>A, 6416G>C, 6420G>A, 6421A>G, 6423T>C, 6424C>A, 6427A>G, 6428T>G, 6429A>T, 6430G>C, 6431C>A, 6432C>A, 6433G>A, 6434A>G, 6435G>T, 6436A>C, 6437T>A, 6438A>C, 6439G>A, 6440T>A, 6442A>G, 6444T>A, 6445C>A, 6446C>G, 6448A>T, 6449T>C, 6450T>A, 6453G>T, 6455G>T, 6456T>G, 6458A>C, 6460C>A, 6462C>G, 6463C>A, 6464C>A, 6465A>G, 6467C>T, 6468A>G, 6471C>A, 6473G>A, 6474G>T, 6475G>C, 6476G>A, 6477T>A, 6478G>A, 6479T>C, 6480T>C, 6482C>G, 6483A>C, 6488C>A, 6489C>G, 6490T>C, 6494C>T, 6495G>T, 6496T>A, 6497G>T, 6498C>T, 6503T>A, 6505G>A, 6507C>A, 6509A>G, 6510G>C, 6511G>T, 6514A>C, 6515G>A, 6517A>G, 6518A>T, 6519G>C, 6520A>T, 6521T>A, 6522C>G, 6524T>G, 6525C>T, 6526C>A, 6528G>T, 6529T>A, 6531C>G, 6532G>A, 6533G>T, 6534A>T, 6535G>C, 6536G>C, 6537T>A, 6539T>A, 6540T>G, 6541C>A, 6543A>T, 6545G>T, 6546G>T, 6548T>G, 6549T>G, 6550T>A, 6552C>T, 6554T>G, 6555A>T, 6556A>T, 6558A>C, 6560C>A, 6561T>A, 6562G>T, 6563T>A, 6566A>T, 6572T>A, 6576A>G, 6582T>G, 6583A>T, 6588G>T, 6590G>C, 6594T>G, 6596A>G, 6597G>T, 6598T>C, 6599C>G, 6600A>G, 6602G>A, 6603C>T, 6605T>A, 6608delT, 6612T>C, 6615A>T, 6618T>G, 6619G>C, 6620T>A, 6621G>C, 6622C>A, 6623G>T, 6624A>G, 6626A>T, 6629C>T, 6643G>A, 6647_6648insACAAACAT, 6649T>A, 6652A>G, 6654C>T, 6655G>T, 6657T>G, 6658A>T, 6659G>A, 6660G>T, 6661A>C, 6663G>A, 6664G>C, 6667T>G, 6669G>A, 6672A>C, 6683C>T, 6687C>T, 6688T>C, 6692A>G, 6695A>G, 6696G>A, 6715T>C |

**CDS**

| | | | | | | | | | |
|---|---|---|---|---|---|---|---|---|---|
| ToMarV_RNA1gp1 | 1 | 2152 | 100% | 9863 | 66.5% | 2135 (98.7%) | 1360 (62.9%) | 11/17/0/0 | 0 |

| | Begin | End | Coverage | Score | Concordance | Matches | Identities | I/D/M/F* | Stop Codons |
|---|---|---|---|---|---|---|---|---|---|
| **NT** | **25** | **6747** | **93.0%** | **3297** | **25.1%** | **6658 (98.4%)** | **4218 (62.4%)** | **41/65** | |

Protein mutations:

F7W (160T>G 161C>G), K9_A16del (165_188delAAGGTTGCTGGTGTTGATGACGCC), S17P (189A>C 190G>C), S18P (192T>C), S19A (195T>G), T21S (201A>T 203C>A), T22P (204A>C), G26A (217G>C 218T>A), L28V (222C>G), N30S (228A>T 229A>C), I32F (234A>T), K34S (240A>T 241A>C 242A>T), F35L (243T>C), A38T (252G>A 254G>T), V39L (255G>C 257T>C), I42V (264A>G), T43N (268C>A 269C>T), T44K (271C>A 272T>A), M45V (273A>G 275G>T), K48T (283A>C 284G>C), S50V (288T>G 289C>T), L53I (297C>A 299T>A), L56V (306C>G 308G>C), S59P (315T>C 317T>A), T61M (322C>T 323T>G), D62A (325A>C 326T>A), V63A (328T>C 329G>A), S64A (330T>G 332C>A), V68A (343T>C 344G>C), A69S (345G>T 347C>T), S72N (355G>A), S73A (357T>G 359A>T), L74V (360C>G 362T>A), N75E (363A>G 365T>A), K76R (367A>G), G82A (385G>C 386T>A), A83K (387G>A 388C>A), L84I (390C>A 392T>A), I85M (395A>G), D86E (398T>A), L89I (405C>A 407T>C), A91G (412C>G 413G>T), S93D (417T>G 418C>A), L95V (423T>G 425G>T), S96K (426T>A 427C>A 428C>A), S101T (441T>A), K103V (447A>G 448A>T 449G>A), M105L (453A>C 455G>T), L109M (465T>A), F110V (468T>G), E111L (471G>T 472A>T 473G>A), L113V (477T>G 479G>C), S114K (480T>A 481C>A 482C>G), N115D (483A>G 485T>C), V117F (489G>T 491T>C), K118I (493A>T 494A>C), F121Y (502T>A 503T>C), A122E (505C>A 506T>A), V124L (510G>T 512G>A), N126E (516A>G 518T>A), S128V (522A>G 523G>T), D130S (528G>T 529A>C 530T>A), V131L (531G>T), V135F (543G>T 545A>T), Y136I (546T>A 547A>T 548T>A), G137A (550G>C), I143L (567A>T 569A>G), T144A (570A>G 572A>C), L145I (573T>A 575G>T), V147L (579G>C 581G>A), F150Y (589T>A), S154V (600A>G 601G>T 602T>A), E158T (612G>A 613A>C), G159P (615G>C 616G>C), V160F (618G>T), S161N (621T>A 622C>A), T163V (627A>G 628C>T 629T>A), K165V (633A>G 634A>T 635A>T), N166T (637A>C), I167L (639A>T 641C>G), L169S (646T>C 647G>A), I170V (648A>G 650A>T), M174F (660A>T 662G>T), K186T (697A>C), A188S (702G>T 704C>T), E189G (706A>G 707A>G), R190K (708C>A 709G>A 710A>G), T192S (714A>T), M193Q (717A>C 718T>A), V194L (720G>C), S198A (732A>G 733G>C 734T>A), M199V (735A>G 737G>A), V205L (753G>T 755T>A), S206T (756T>A 758A>G), M214L (780A>T 782G>A), T216S (786A>T), I217L (789A>T 791A>G), I228V (822A>G), T248A (882A>G 884A>T), T260Q (918A>C 919C>A 920T>G), I280V (978A>G 980A>G), G282S (984G>A), T283I (988C>T 989C>T), S295N (1024G>A), E304Q (1050G>C), Q307T (1059C>A 1060A>C), D310S (1068G>T 1069A>C 1070T>A), A311E (1072C>A 1073A>G), C313A (1077T>G 1078G>C), D314E (1082T>A), A316S (1086G>T 1088A>T), T317M (1090C>T), K325P (1113A>C 1114A>C 1115G>T), T329I (1126C>T 1127C>T), N332Q (1134A>C 1136C>A), R335K (1144G>A 1145G>A), E337Q (1149G>C 1151A>G), L339V (1155C>G), M354L (1200A>T), A363T (1227G>A), E366D (1238A>T), V368L (1242G>T 1244T>G), Y369V (1245T>G 1246A>T), S372A (1254A>G 1255G>C 1256T>C), C373S (1257T>A 1259T>C), R374Q (1260A>C 1261G>A 1262G>A), M376S (1266A>T 1267T>C 1268G>T), S393A (1317T>G 1319C>T), S401C (1342C>G), R412A (1374A>G 1375G>C), I440V (1458A>G 1460C>G), M521R (1702T>G 1703G>A), E523V (1708A>T), E525D (1715G>T), S527D (1719T>G 1720C>A 1721C>T), T528P (1722A>C 1724T>A), E529N (1725G>A 1727A>T), D530A (1729A>C), G533A (1738G>C 1739A>C), R536V (1746A>G 1747G>T 1748A>T), N538D (1752A>G 1754C>T), I539V (1755A>G 1757A>G), N552A (1794A>G 1795A>C), T554S (1800A>T), S556D (1806A>G 1807G>A), E560Q (1818G>C 1820G>A), Q567R (1840A>G), A570Y (1848G>T 1849C>A 1850A>T), K574S (1861A>G 1862G>C), F577Y (1870T>A), E578N (1872G>A 1874A>T), A579T (1875G>A), E580T (1878G>A 1879A>C), I583L (1887A>T 1889A>G), S584A (1890A>G 1891G>C), P585D (1893C>G 1894C>A 1895T>C), I587V (1899A>G 1901A>T), Q591N (1911C>A 1913A>T), S593A (1917T>G), A594V (1921C>T 1922A>T), D596E (1928T>A), V597M (1929G>A), I598L (1932A>C 1934A>G), A599D (1936C>A 1937G>T), V601F (1941G>T 1943G>C), D602Q (1944G>C 1946T>A), G603D (1948G>A 1949G>T), R605del (1953_1955delAGA), R607Q (1960G>A 1961C>A), T608V (1962A>G 1963C>T 1964C>T), K609L (1965A>T 1966A>T), Q610P (1969A>C 1970A>T), D611P (1971G>C 1972A>C), I614A (1980A>G 1981T>C), L615T (1983T>A 1984T>C 1985G>T), V616Y (1986G>T 1987T>A 1988G>T), P617E (1989C>G 1990C>A 1991C>G), T618Q (1992A>C 1993C>A 1994T>G), V619T (1995G>A 1996T>C 1997T>A), V620I (1998G>A 2000G>T), S621T (2002G>C 2003T>A), N623A (2007A>G 2008A>C), D624E (2012T>A), R625K (2014G>A), V626I (2016G>A 2018G>T), I627V (2019A>G 2021C>T), A629S (2025G>T 2027A>C), R630Q (2028A>C 2029G>A 2030A>G), L645M (2073T>A), D646N (2076G>A), P647K (2079C>A 2080C>A 2081T>A), E648F (2082G>T 2083A>T 2084G>T), L650F (2088C>T), H655L (2104A>T 2105T>C), H656S (2106C>T 2107A>C 2108T>C), M660I (2120G>C), S663V (2127T>G 2128C>T 2129T>G), D665S (2133G>T 2134A>C), D666S (2136G>T 2137A>C), E667D (2141A>T), H668A (2142C>G 2143A>C 2144T>C), E670Q (2148G>C), I671N (2152T>A), N673E (2157A>G 2159C>A), K674M (2161A>T 2162A>G), R676V (2166A>G 2167G>T), V677N (2169G>A 2170T>A 2171C>T), N678D (2172A>G 2174C>T), M679L (2175A>T), Y681V (2181T>G 2182A>T 2183T>C), R682K (2185G>A), D685S (2193G>A 2194A>G), A686P (2196G>C), E687Q (2199G>C 2201A>G), R690A (2208C>G 2209G>C 2210A>T), Q694R (2221A>G), I701V (2241A>G 2243A>C), A710S (2268G>A 2269C>G 2270A>C), S711T (2271T>A), K716E (2286A>G), K717L (2289A>T 2290A>T 2291A>G), A719D (2296C>A 2297A>T), T722R (2304A>C 2305C>G 2306A>C), E723N (2307G>A 2309G>T), M724L (2310A>C 2312G>T), R725E (2313A>G 2314G>A), T726Q (2316A>C 2317C>A 2318T>A), F728K (2322T>A 2323T>A 2324T>A), H730Y (2328C>T), S731K (2331T>A 2332C>A 2333T>G), C732A (2334T>G 2335G>C 2336C>G), I733L (2337A>C), D734Q (2340G>C 2342T>G), V737A (2350T>C 2351T>A), I741T (2362T>C 2363A>G), F749R (2385T>A 2386T>G 2387T>A), N763R (2428A>G 2429C>G), S766A (2436T>G), S766_K767insG (2438_2439insGGT), K767T (2440A>C 2441A>T), E768D (2444G>C), P769K (2445C>A 2446C>A 2447A>G), E771R (2451G>A 2452A>G), S772N (2454T>A 2455C>A 2456A>T), K775I (2464A>T 2465A>C), I776E (2466A>G 2467T>A 2468T>A), Y778M (2472T>A 2473A>T 2474C>G), H779I (2475C>A 2476A>T), K780G (2478A>G 2479A>G 2480A>T), V782K (2484G>A 2485T>A 2486T>G), E783D (2489A>C), Y784K (2490T>A 2492T>A), I785M (2495C>G), K786I (2497A>T 2498G>C), D787Q (2499G>C 2501T>G), T788M (2503C>T 2504A>G), W790N (2508T>A 2509G>A 2510G>T), S791D (2511A>G 2512G>A), Y794F (2521A>T 2522T>C), A795I (2523G>A 2524C>T 2525A>C), R797K (2530G>A 2531A>G), I806L (2556A>T 2558A>G), M807I (2561G>A), V812T (2574G>A 2575T>C 2576T>C), F813M (2577T>A 2579T>G), G814N (2580G>A 2581G>A 2582A>T), G815A (2584G>C), I816V (2586A>G), L820I (2598T>A 2600G>T), A821T (2601G>A 2603G>T), W823L (2607T>C 2608G>T 2609G>C), Q824K (2610C>A), S825V (2613A>G 2614G>T 2615T>A), N826A (2616A>G 2617A>C 2618C>A), S827T (2619T>A), F828S (2623T>C), N830L (2628A>T 2629A>T 2630T>A), S832D (2634T>G 2635C>A 2636G>T), N852K (2696T>G), K853T (2698A>C 2699G>T), K853_L854insH (2699_2700insCAT), L854F (2700C>T), R855A (2703A>G 2704G>C), N856K (2708T>A), L857S (2709C>T 2710T>C 2711T>C), V859L (2715G>C 2717T>A), S862A (2724A>G 2725G>C 2726T>G), S863A (2727T>G 2729T>A), Q864D (2730C>G 2732A>C), S865N (2733T>A 2734C>A 2735A>C), L866N (2736T>A 2737T>A 2738G>C), A867H (2739G>C 2740C>A 2741A>T), Q868V (2742C>G 2743A>T 2744A>G), D869_W870insR (2747_2748insCGA), A872E (2755C>A 2756T>A), E873Q (2757G>C 2759A>G), Q879E (2775C>G 2777G>A), K882H (2784A>C 2786G>C), L890I (2808C>A), G891A (2812G>C), M907L (2859A>C 2861G>C), H914R (2880C>A 2881A>G 2882C>A), A916M (2886G>A 2887C>T 2888T>G), T923S (2907A>T 2909A>C), A927E (2920C>A 2921C>G), N928H (2922A>C), R930I (2928C>A 2929G>T 2930A>T), S931C (2932C>G), K932N (2936A>T), Y935F (2944A>T 2945C>T), R938T (2953G>C 2954A>C), E939Q (2955G>C), I942L (2964A>T 2966T>G), I948V (2982A>G 2984C>G), P961S (3021C>T 3023T>A), E972D (3056G>T), F975Y (3064T>A), T976S (3066A>T 3068A>C), Q980M (3078C>A 3079A>T 3080A>G), C984V (3090T>G 3091G>T 3092T>G), T990K (3109C>A 3110A>G), R993K (3117C>A 3118G>A 3119T>G), L994K (3120C>A 3121T>A 3122T>A), G995S (3123G>T 3124G>C), F999V (3135T>G 3137T>G), M1002Y (3144A>T 3145T>A 3146G>T), V1012E (3175T>A 3176T>A), E1015T (3183G>A 3184A>C), E1017G (3190A>G), N1019D (3195A>G 3197C>T), S1021N (3201T>A 3202C>A 3203C>T), S1022A (3204A>G 3205G>C 3206T>A), K1024A (3210A>G 3211A>C 3212G>T), I1025L (3213A>C), C1026S (3216T>A), Q1029V (3225C>G 3226A>T 3227A>T), S1030A (3228T>G 3230T>A), G1037S (3249G>A 3251G>T), S1050T (3289G>C), V1057C (3309G>T 3310T>G 3311G>C), G1068D (3343G>A 3344C>T), V1069T (3345G>A 3346T>C 3347A>C), R1073K (3357C>A 3358G>A 3359T>G), F1075Y (3364T>A 3365T>C), I1076V (3366A>G 3368C>G), S1077A (3369T>G), I1083V (3387A>G 3389C>G), Y1084F (3391A>T 3392C>T), E1086D (3398A>T), V1096I (3426G>A 3428C>T), A1098V (3433C>T 3434A>G), D1104E (3452T>A), I1105L (3453A>C 3455T>G), S1108H (3462T>C 3463C>A 3464A>C), I1109L (3465A>C 3467T>G), R1112K (3475G>A 3476A>G), T1114E (3480A>G 3481C>A), Q1115R (3483C>A 3484A>G 3485G>A), L1116F (3486C>T 3488T>C), N1120K (3500C>A), Y1123A (3507T>G 3508A>C 3509C>T), Y1124F (3511A>T), Q1130L (3528C>T 3529A>T), L1132M (3534T>A), R1143K (3567C>A 3568G>A 3569G>A), K1144R (3570A>C 3571A>G 3572A>C), Q1146P (3577A>C 3578A>C), P1153D (3597C>G 3598C>A), E1155Q (3603G>C), T1162S (3624A>T 3626A>T), V1163A (3628T>C 3629T>C), H1164K (3630C>A 3632T>A), D1169E (3647T>A), M1179E (3675A>G 3676T>A), C1186S (3697G>C 3698T>C), G1187N (3699G>A 3700G>A 3701A>C), D1189E (3707T>G), D1190E (3710T>G), D1191P (3711G>C 3712A>C 3713C>A), N1199G (3735A>G 3736A>G), C1202A (3744T>G 3745G>C 3746C>A), L1204I (3750C>A 3752G>T), I1207V (3759A>G 3761C>G), M1208L (3762A>C 3764G>T), R1209K (3765C>A 3766G>A 3767T>G), S1210M (3769G>T 3770C>G), K1212R (3774A>C 3775A>G 3776G>A), I1213V (3777A>G 3779C>G), R1217K (3789C>A 3790G>A 3791T>A), C1219A (3795T>G 3796G>C 3797T>A), E1220I (3798G>A 3799A>T 3800A>T), D1222N (3804G>A), C1233F (3838G>T 3839T>C), Y1236F (3847A>T), G1238A (3853G>C 3854C>A), M1252L (3894A>T), G1253E (3898G>A 3899G>A), M1254L (3900A>C 3902G>T), M1256S (3906A>T 3907T>C 3908G>T), Q1263E (3927C>G), S1264A (3930T>G), T1265Q (3933A>C 3934C>A 3935T>A), G1266P (3936G>C 3937G>C 3938C>A), E1267D (3941A>C), G1268S (3942G>T 3943G>C 3944A>G), E1269Q (3945G>C), E1272L (3954G>C 3955A>T 3956A>T), D1276N (3966G>A), E1281D (3983A>T), V1288S (4002G>T 4003T>C), I1290V (4008A>G 4010T>A), K1292Q (4014A>C), I1294T (4021T>C 4022C>A), H1295F (4023C>T 4024A>T), T1344C (4170A>T 4171C>G 4172A>T), L1348A (4182T>G 4183T>C 4184G>A), A1351V (4192C>T 4193T>A), I1352L (4194A>C 4196A>C), D1359N (4215G>A), S1363H (4227T>C 4228C>A), L1367V (4239T>G), N1370S (4249A>G), G1371S (4251G>A 4253G>C), A1374S (4260G>T 4262T>A), V1378E (4273T>A), V1393I (4317G>A 4319T>C), E1405A (4354A>C 4355A>T), Y1406F (4357A>T 4358C>T), S1409D (4365T>G 4366C>A 4367A>T), K1410E (4368A>G), D1415E (4385T>G), A1421T (4401G>A), D1435N (4443G>A 4445T>C), V1437L (4449G>T), I1476V (4566A>G 4568T>C), A1477K (4569G>A 4570C>A 4571T>A), R1479K (4576G>A 4577G>A), A1480S (4578G>T), I1481V (4581A>G 4583A>G), V1483I (4587G>A), S1486N (4597G>A), L1506M (4656T>A 4658A>G), Q1509R (4665C>A 4666A>G), S1512D (4674T>G 4675C>A), N1515S (4683A>T 4684A>C 4685T>C), H1518R (4692C>A 4693A>G 4694T>A), L1522A (4704C>G 4705T>C 4706G>T), I1526V (4716A>G), N1538E (4752A>G 4754T>G), E1539K (4755G>A), V1540T (4758G>A 4759T>C), Q1541K (4761C>A), Q1543L (4768A>T 4769A>G), H1563Q (4829C>A), I1572V (4854A>G), M1576I (4868G>T), P1583V (4887C>G 4888C>T 4889A>G), T1588V (4902A>G 4903C>T 4904T>A), M1592Y (4914A>T 4915T>A 4916G>T), V1595I (4923G>A 4925G>T), R1607Q (4960G>A 4961T>G), V1608A (4963T>C 4964G>C), A1609T (4965G>A), D1611E (4973C>A), K1612S (4975A>G 4976G>T), L1613F (4977C>T), L1615N (4983T>A 4984T>A 4985A>T), Q1618L (4993A>T 4994G>T), T1619G (4995A>G 4996C>G 4997G>A), D1622K (5004G>A 5006T>G), E1623Q (5007G>C 5009A>G), G1625N (5013G>A 5014G>A 5015T>C), F1626L (5016T>C), R1627P (5019A>C 5020G>C 5021A>C), A1628I (5022G>A 5023C>T 5024T>C), V1629M (5025G>A), F1631M (5031T>A 5033T>G), S1633T (5037T>A 5039C>T), L1635Y (5044T>A 5045G>T), V1637L (5049G>C 5051G>T), Q1639S (5055C>A 5056A>G 5057G>T), E1640Q (5058G>C), I1642L (5064A>T 5066T>G), A1643T (5067G>A), K1644E (5070A>G), S1649E (5085T>G 5086C>A 5087A>G), A1651I (5091G>A 5092C>T 5093C>T), L1652G (5094C>G 5095T>G), T1653V (5097A>G 5098C>T 5099A>T), I1654L (5100A>T 5102T>G), C1655M (5103T>A 5104G>T 5105T>G), M1657L (5109A>C 5111G>A), A1658S (5112G>T), K1659D (5115A>G 5117A>T), A1661K (5121G>A 5122C>A 5123T>A), I1662N (5125T>A), S1663T (5127T>A 5129C>A), D1668P (5142G>C 5143A>C 5144C>T), T1674del (5160_5162delACA), R1678E (5172C>G 5173G>A 5174T>G), D1679G (5176A>G), L1680V (5178C>G 5180G>C), N1681H (5181A>C), W1682E (5184T>G 5185G>A 5186G>A), N1686D (5196A>G 5198C>T), I1687V (5199A>G 5201T>A), G1688F (5202G>T 5203G>T), A1696S (5226G>T 5228C>A), H1697N (5229C>A), Y1698Q (5232T>C 5234C>A), T1699Q (5235A>C 5236C>A 5237G>A), D1701S (5241G>T 5242A>C 5243T>A), E1702S (5244G>T 5245A>C), Y1704N (5250T>A), I1706L (5256A>C 5258T>G), K1713R (5277A>C 5278A>G 5279G>C), K1714R (5281A>G 5282G>A), S1715T (5283T>A 5285C>A), N1716T (5287A>C 5288T>A), M1731I (5333G>A), M1736L (5346A>T), S1737A (5349T>G), M1740Q (5358A>C 5359T>A), A1742H (5364G>C 5365C>A 5366C>T), K1744R (5370A>C 5371A>G 5372G>C), N1745K (5375T>G), V1746N (5376G>A 5377T>A 5378G>T), A1748E (5383C>A), G1749A (5386G>C 5387A>T), H1751D (5391C>G), V1752G (5395T>G), N1755S (5403A>T 5404A>C), V1757T (5409G>A 5410T>C 5411T>C), V1766I (5436G>A 5438C>A), G1767A (5440G>C 5441C>T), M1774F (5460A>T 5462G>T), R1779H (5475A>C 5476G>A 5477G>T), C1782R (5484T>A 5486T>G), V1783A (5488T>C 5489G>C), E1784T (5490G>A 5491A>C 5492G>A), I1786L (5496A>C 5498A>T), I1793M (5519T>G), G1794A (5521G>C 5522A>T), N1795G (5523A>G 5524A>G 5525T>G), V1807A (5560T>C), F1810L (5570T>G), Q1812K (5574C>A 5576A>G), P1815E (5583C>G 5584C>A 5585T>A), H1816D (5586C>G 5588T>C), H1816_G1817insT (5588_5589insACC), G1817S (5589G>T 5590G>C), D1818H (5592G>C), V1820L (5598G>T), A1821G (5602C>G 5603T>A), A1825T (5613G>A), S1826Y (5617C>A 5618G>T), L1827G (5619C>G 5620T>G), H1828del (5622_5624delCAC), S1829T (5626G>C 5627T>C), S1830T (5629G>C), M1832N (5635T>A 5636G>C), Q1834H (5642A>T), Q1834_G1835insHS (5642_5643insCACAGT), G1835A (5644G>C 5645G>A), E1837S (5649G>T 5650A>C), L1838V (5652T>G 5654G>T), N1840L (5658A>C 5659A>T 5660C>T), I1844V (5670A>G 5672A>T), H1845P (5674A>C), G1846S (5676G>T 5677G>C 5678T>C), A1849S (5685G>T 5687A>T), K1850E (5688A>G), Q1851K (5691C>A), T1852M (5695C>T 5696A>G), M1854A (5700A>G 5701T>C 5702G>T), M1856L (5706A>T), V1859T (5715G>A 5716T>C 5717G>A), V1860I (5718G>A 5720A>T), H1863E (5727C>G 5729T>A), G1867A (5740G>C 5741A>C), N1868Q (5742A>C 5744T>G), F1869C (5746T>G 5747T>C), Q1872K (5754C>A), D1873N (5757G>A 5759T>C), E1875K (5763G>A), S1876H (5766A>C 5767G>A), L1877A (5769T>G 5770T>C 5771G>T), H1879E (5775C>G 5777C>G), Y1880T (5778T>A 5779A>C 5780T>A), E1882S (5784G>T 5785A>C 5786G>T), Q1883V (5787C>G 5788A>T 5789G>C), G1884S (5790G>A 5792A>T), E1887T (5799G>A 5800A>C 5801G>T), I1889A (5805A>G 5806T>C 5807T>C), Q1890A (5808C>G 5809A>C), K1891R (5812A>G 5813G>A), N1892G (5814A>G 5815A>G 5816T>A), T1893V (5817A>G 5818C>T 5819A>C), L1895V (5823C>G 5825A>G), L1902F (5844C>T 5846G>C), M1904Q (5850A>C 5851T>A 5852G>A), E1905H (5853G>C 5855A>C), N1906D (5856A>G), V1907C (5859G>T 5860T>G 5861G>T), Q1908E (5862C>G), Q1909M (5865C>A 5866A>T 5867A>G), L1910I (5868C>A 5870G>A), I1911L (5871A>C 5873A>T), D1912E (5876T>G), V1914L (5880G>C 5882T>G), K1915E (5883A>G 5885G>A), L1916S (5886C>A 5887T>G), F1919V (5895T>G 5897T>A), F1920V (5898T>G 5900C>A), P1921F (5901C>T 5902C>T 5903A>C), K1922M (5905A>T 5906A>G), E1923G (5908A>G 5909A>G), T1924S (5911C>G 5912T>C), T1925V (5913A>G 5914C>T 5915C>T), I1926K (5917T>A 5918T>A), Q1927K (5919C>A 5921G>A), K1928Q (5922A>C 5924G>A), L1929M (5925C>A 5927C>G), K1930L (5928A>T 5929A>T 5930A>G), N1931S (5931A>T 5932A>C 5933T>C), V1933M (5937G>A 5939T>G), D1934N (5940G>A), A1935Q (5943G>C 5944C>A 5945T>A), E1936Q (5946G>C 5948A>G), C1937M (5949T>A 5950G>T 5951T>G), L1938S (5952C>T 5953T>C), T1939K (5956C>A), F1940T (5958T>A 5959T>C), K1941Q (5961A>C), A1942V (5965C>T 5966A>T), R1943A (5967A>G 5968G>C 5969G>C), S1944_F1945insPCCVT (5972_5973insCCATGTTGTGTCACA), R1946S (5978G>T), H1947K (5979C>A 5981C>A), V1948A (5983T>C), V1949W (5985G>T 5986T>G 5987T>G), F1950L (5988T>C 5990C>A), K1951T (5992A>C 5993A>T), K1953D (5997A>G 5999A>C), L1954I (6000C>A), L1955V (6003T>G 6005G>C), S1956M (6007G>T 6008T>G), S1957T (6009T>A 6011C>A), T1958K (6013C>A), W1959G (6015T>G 6017G>T), M1961K (6022T>A), C1962P (6024T>C 6025G>C 6026T>A), E1964del (6030_6032delGAA), V1967_R1970del (6039_6050delGTTGCTTCAAGG), S1971P (6051T>C), S1974T (6061G>C 6062T>A), L1975V (6063C>G 6065A>G), P1977L (6070C>T), L1980P (6078T>C 6079T>C 6080G>T), S1983L (6087T>C 6088C>T 6089C>A), L1985V (6093T>G), K1986R (6097A>G), L1988M (6102T>A 6104A>G), E1990R (6108G>C 6109A>G 6110A>C), R1991N (6112G>A 6113A>T), H1992S (6114C>A 6115A>G 6116C>T), T1993G (6117A>G 6118C>G), R1994K (6121G>A 6122A>G), M1996Y (6126A>T 6127T>A

6128G>C), Q1999T (6135C>A 6136A>C 6137G>A), S2000T (6139G>C 6140T>A), M2001L (6141A>C), L2005F (6153C>T 6155G>T), K2006R (6157A>G), I2007M (6161A>G), L2009M (6165C>A 6167A>G), L2010C (6169T>G 6170A>T), N2011Q (6171A>C 6173C>G), F2012E (6174T>G 6175T>A 6176C>A), Q2013G (6177C>G 6178A>G 6179A>G), V2014Y (6180G>T 6181T>A 6182A>T), I2015L (6183A>T), T2016D (6186A>G 6187C>A 6188A>T), S2017K (6190G>A 6191T>A), E2018A (6193A>C 6194G>C), L2020V (6198T>G 6200A>G), K2021D (6201A>G 6203G>C), K2022C (6204A>T 6205A>G 6206G>C), E2024S (6210G>A 6211A>G 6212A>T), D2025E (6215T>G), F2027M (6219T>A 6221T>G), Q2028G (6222C>G 6223A>G 6224G>C), E2029V (6226A>T 6227G>T), K2030C (6228A>T 6229A>G 6230G>T), I2031V (6231A>G 6233A>G), S2032P (6234T>C 6236C>G), T2033E (6237A>G 6238C>A 6239C>A), T2034R (6240A>C 6241C>G), L2035I (6243C>A 6245A>C), I2037V (6249A>G 6251C>G), K2038R (6252A>C 6253A>G 6254A>C), V2039T (6255G>A 6256T>C), F2040I (6258T>A 6260C>T), L2041A (6261T>G 6262T>C 6263G>T), E2044S (6270G>T 6271A>C 6272G>C), E2045T (6273G>A 6274A>C), Y2047L (6280A>T 6281T>G), Q2048N (6282C>A 6284G>C), T2049E (6285A>G 6286C>A 6287T>G), E2050N (6288G>A 6290G>T), C2052T (6294T>A 6295G>C), I2053V (6297A>G 6299A>G), D2054Y (6300G>T 6302T>C), S2055T (6303T>A 6305T>A), Q2056D (6306C>G 6308G>T), V2057G (6310T>G), Q2059F (6315C>T 6316A>T 6317G>T), R2060D (6318A>G 6319G>A 6320G>T), I2061del (6321_6323delATA), N2062H (6324A>C 6326C>T), S2063F (6328C>T), Q2064K (6330C>A 6332G>A), K2065A (6333A>G 6334A>C), F2066V (6336T>G 6338C>G), R2067D (6339A>G 6340G>A 6341A>C), V2068F (6342G>T 6344G>C), Q2069K (6345C>A), E2070P (6348G>C 6349A>C 6350A>T), Y2074S (6361A>C 6362T>C), L2075W (6364T>G), H2076E (6366C>G 6368T>A), P2077I (6369C>A 6370C>T), E2078N (6372G>A 6374A>T), G2079D (6376G>A 6377A>T), I2080A (6378A>G 6379T>C), V2085A (6394T>C), I2088V (6402A>G 6404A>G), I2089L (6405A>T 6407A>G), S2091E (6411T>G 6412C>A 6413T>A), E2094R (6420G>A 6421A>G), S2095H (6423T>C 6424C>A), Y2096W (6427A>G 6428T>G), R2098K (6432C>A 6433G>A 6434A>G), D2099S (6435G>T 6436A>C 6437T>A), S2100Q (6438A>C 6439G>A 6440T>A), N2101S (6442A>G), S2102K (6444T>A 6445C>A 6446C>G), Y2103F (6448A>T 6449T>C), S2104T (6450T>A), V2105F (6453G>T 6455G>T), S2106A (6456T>G 6458A>C), T2107N (6460C>A), P2108E (6462C>G 6463C>A 6464C>A), I2109V (6465A>G 6467C>T), K2110E (6468A>G), L2111I (6471C>A 6473G>A), G2112S (6474G>T 6475G>C 6476G>A), C2113N (6477T>A 6478G>A 6479T>C), F2114L (6480T>C 6482C>G), I2115L (6483A>C), F2116L (6488C>A), L2117A (6489C>G 6490T>C), V2119Y (6495G>T 6496T>A 6497G>T), D2121E (6503T>A), S2122N (6505G>A), Q2123K (6507C>A 6509A>G), G2124L (6510G>C 6511G>T), K2125T (6514A>C 6515G>A), E2126G (6517A>G 6518A>T), D2127L (6519G>C 6520A>T 6521T>A), H2128E (6522C>G 6524T>G), P2129Y (6525C>T 6526C>A), V2130Y (6528G>T 6529T>A), R2131D (6531C>G 6532G>A 6533G>T), R2132S (6534A>T 6535G>C 6536G>C), F2133I (6537T>A 6539T>A), S2134E (6540T>G 6541C>A), T2135S (6543A>T 6545G>T), V2136L (6546G>T 6548T>G), F2137D (6549T>G 6550T>A), K2139L (6555A>T 6556A>T), N2140Q (6558A>C 6560C>A), C2141I (6561T>A 6562G>T 6563T>A), E2142D (6566A>T), T2146A (6576A>G), Y2148V (6582T>G 6583A>T), E2150Y (6588G>T 6590G>C), *2152G (6594T>G 6596A>G)

| | Begin | End | Coverage | Score | Concordance | Matches | Identities | I/D/M/F* | Stop Codons |
|---|---|---|---|---|---|---|---|---|---|
| **NT** | **25** | **6747** | **93.0%** | **3297** | **25.1%** | **6658 (98.4%)** | **4218 (62.4%)** | **41/65** | |

GCT2GCA (146T>A), TTC7TGG (160T>G 161C>G), TCC8AGT (162T>A 163C>G 164C>T), AAG9_GCC16del (165_188delAAGGTTGCTGGTGTTGATGACGCC), AGT17CCT (189A>C 190G>C), TCT18CCT (192T>C), TCT19GCT (195T>G), TCC20TCT (200C>T), ACC21TCA (201A>T 203C>A), ACT22CCT (204A>C), GGT26GCA (217G>C 218T>A), CTT28GTT (222C>G), TCA29TCT (227A>T), AAT30TCT (228A>T 229A>C), ATT32TTT (234A>T), TCC33TCT (239C>T), AAA34TCT (240A>T 241A>C 242A>T), TTT35CTT (243T>C), AAA37AAG (251A>G), GCG38ACT (252G>A 254G>T), GTT39CTC (255G>C 257T>C), AGT40TCC (258A>T 259G>C 260T>C), ATT42GTT (264A>G), ACC43AAT (268C>A 269C>T), ACT44AAA (271C>A 272T>A), ATG45GTT (273A>G 275G>T), AAG48ACC (283A>C 284G>C), TCA50GTA (288T>G 289C>T), CAT52CAC (296T>C), CTT53ATA (297C>A 299T>A), CTG56GTC (306C>G 308G>C), CCA58CCG (314A>G), TCT59CCA (315T>C 317T>A), GTG60GTT (320G>T), ACT61ATG (322C>T 323T>G), GAT62GCA (325A>C 326T>A), GTG63GCA (328T>C 329G>A), TCC64GCA (330T>G 332C>A), TCT66TCA (338T>A), GTG68GCC (343T>C 344G>C), GCC69TCT (345G>T 347C>T), ACT70ACA (350T>A), TGT71TGC (353T>C), AGT72AAT (355G>A), TCA73GCT (357T>G 359A>T), CTT74GTA (360C>G 362T>A), AAT75GAA (363A>G 365T>A), AAA76AGA (367A>G), TTG78TTA (374G>A), GAC79GAT (377C>T), AAA80AAG (380A>G), GGT82GCA (385G>C 386T>A), GCA83AAA (387G>A 388C>A), CTT84ATA (390C>A 392T>A), ATA85ATG (395A>G), GAT86GAA (398T>A), CCA87CCC (401A>C), TTT88TTC (404T>C), CTT89ATC (405C>A 407T>C), AAA90AAG (410A>G), GCG91GGT (412C>G 413G>T), TAC92TAT (416C>T), TCT93GAT (417T>G 418C>A), TTC94TTT (422C>T), TTG95GTT (423T>G 425G>T), TCC96AAA (426T>A 427C>A 428C>A), ACA97ACT (431A>T), TCT101ACT (441T>A), ATC102ATA (446C>A), AAG103GTA (447A>G 448A>T 449G>A), ATG105CTT (453A>C 455G>T), GTT106GTA (458T>A), TTA107CTT (459T>C 461A>T), TTG109ATG (465T>A), TTT110GTT (468T>G), GAG111TTA (471G>T 472A>T 473G>A), TTG113GTC (477T>G 479G>C), TCC114AAG (480T>A 481C>A 482C>G), AAT115GAC (483A>G 485T>C), GTT117TTC (489G>T 491T>C), AAA118ATC (493A>T 494A>C), TTA119CTA (495T>C), TTT121TAC (502T>A 503T>C), GCT122GAA (505C>A 506T>A), GTG124TTA (510G>T 512G>A), AAT126GAA (516A>G 518T>A), AGT128GTT (522A>G 523G>T), GAA129GAG (527A>G), GAT130TCA (528G>T 529A>C 530T>A), GTA131TTA (531G>T), GTT134GTG (542T>G), GTA135TTT (543G>T 545A>T), TAT136ATA (546T>A 547A>T 548T>A), GGT137GCT (550G>C), CTT139TTA (555C>T 557T>A), GTA140GTT (560A>T), ATA143TTG (567A>T 569A>G), ACA144GCC (570A>G 572A>C), TTG145ATT (573T>A 575G>T), TTA146TTG (578A>G), GTG147CTA (579G>C 581G>A), CTT148TTA (582C>T 584T>A), CTA149CTG (587A>G), TTT150TAT (589T>A), GTT151GTA (593T>A), TGT152TGC (596T>C), AGT154GTA (600A>G 601G>T 602T>A), GTT157GTA (611T>A), GAG158ACG (612G>A 613A>C), GGT159CCT (615G>C 616G>C), GTT160TTT (618G>T), TCT161AAT (621T>A 622C>A), AAC162AAT (626C>T), ACT163GTA (627A>G 628C>T 629T>A), GTC164GTT (632C>T), AAA165GTT (633A>G 634A>T 635A>T), AAT166ACT (637A>C), ATC167TTG (639A>T 641C>G), TTG169TCA (646T>C 647G>A), ATA170GTT (648A>G 650A>T), GTC171GTT (653C>T), AAC173AAT (659C>T), ATG174TTT (660A>T 662G>T), TTT175TTC (665T>C), TCA176TCT (668A>T), AGT177TCA (669A>T 670G>C 671T>A), CTA178TTG (672C>T 674A>G), TAC179TAT (677C>T), AAA186ACA (697A>C), GCC188TCT (702G>T 704C>T), GAA189GGG (706A>G 707A>G), CGA190AAG (708C>A 709G>A 710A>G), TTC191TTT (713C>T), ACC192TCC (714A>T), ATG193CAG (717A>C 718T>A), GTG194CTG (720G>C), GCC195GCG (725C>G), GCT197GCA (731T>A), AGT198GCA (732A>G 733G>C 734T>A), ATG199GTA (735A>G 737G>A), GGG202GGT (746G>T), TCA204AGT (750T>A 751C>G 752A>T), GTT205TTA (753G>T 755T>A), TCA206ACG (756T>A 758A>G), ACG210ACT (770G>T), TCT211AGC (771T>A 772C>G 773T>C), CAG212CAA (776G>A), TTG213CTT (777T>C 779G>T), ATG214TTA (780A>T 782G>A), ACA216TCA (786A>T), ATA217TTG (789A>T 791A>G), GGG221GGT (803G>T), ATT222ATC (806T>C), TCG223TCA (809G>A), ACG224ACT (812G>T), TTG225TTA (815G>A), GTG226GTA (818G>A), TTC227TTT (821C>T), ATT228GTT (822A>G), GCA229GCT (827A>T), GTG230GTT (830G>T), GGT232GGG (836T>G), GGC236GGT (848C>T), TTG237CTT (849T>C 851G>T), AGC238TCA (852A>T 853G>C 854C>A), TTG241CTG (861T>C), TCC242TCA (866C>A), ATT244ATC (872T>C), CTC245TTG (873C>T 875C>G), TAT246TAC (878T>C), TCG247TCT (881G>T), ACA248GCT (882A>G 884A>T), GCT252GCG (896T>G), CAG253CAA (899G>A), CAA257CAG (911A>G), CTA258CTG (914A>G), ACT260CAG (918A>C 919C>A 920T>G), CTT261TTG (921C>T 923T>G), CGT263AGG (927C>A 929T>G), AAT264AAC (932T>C), AAA266AAG (938A>G), TGC268TGT (944C>T), ACC269ACA (947C>A), TCC270TCT (950C>T), CAG271CAA (953G>A), GCT272GCC (956T>C), GAA278GAG (974A>G), ATA280GTG (978A>G 980A>G), GTC281GTT (983C>T), GGC282AGC (984G>A), ACC283ATT (988C>T 989C>T), GGA285GGC (995A>C), TTC286TTT (998C>T), GTG290GTA (1010G>A), GCT293GCG (1019T>G), AGT295AAT (1024G>A), GCA296GCT (1028A>T), ACG297ACC (1031G>C), CTG298TTA (1032C>T 1034G>A), TCC299TCT (1037C>T), ACT300ACC (1040T>C), CTG302CTT (1046G>T), GAG304CAG (1050G>C), CAG307ACG (1059C>A 1060A>C), GAG308GAA (1064G>A), GAT310TCA (1068G>T 1069A>C 1070T>A), GCA311GAG (1072C>A 1073A>G), GTG312GTT (1076G>T), TGT313GCT (1077T>G 1078G>C), GAT314GAA (1082T>A), CCT315CCA (1085T>A), GCA316TCT (1086G>T 1088A>T), ACG317ATG (1090C>T), GAG320GAA (1100G>A), AAC321AAT (1103C>T), GCC323GCA (1109C>A), AAG325CCT (1113A>C 1114A>C 1115G>T), GCT326GCA (1118T>A), TTC327TTT (1121C>T), CCT328CCA (1124T>A), ACC329ATT (1126C>T 1127C>T), AAA330AAG (1130A>G), AAC332CAA (1134A>C 1136C>A), CTG334TTA (1140C>T 1142G>A), AGG335AAA (1144G>A 1145G>A), GAA336GAG (1148A>G), GAA337CAG (1149G>C 1151A>G), GCT338GCC (1154T>C), CTC339GTC (1155C>G), AAG340AAA (1160G>A), ATC341ATA (1163C>A), GCA346GCC (1178A>C), GCG351GCA (1193G>A), TTC353TTT (1199C>T), ATG354TTG (1200A>T), AGT359TCT (1215A>T 1216G>C), ATT362ATA (1226T>A), GCG363ACG (1227G>A), CAG364CAA (1232G>A), TTA365CTA (1233T>C), GAA366GAT (1238A>T), GTT368TTG (1242G>T 1244T>G), TAT369GTT (1245T>G 1246A>T), GAA371GAG (1253A>G), AGT372GCC (1254A>G 1255G>C 1256T>C), TGT373AGC (1257T>A 1259T>C), AGG374CAA (1260A>C 1261G>A 1262G>A), CAC375CAT (1265C>T), ATG376TCT (1266A>T 1267T>C 1268G>T), GGC377GGT (1271C>T), CAG380CAA (1280G>A), CGT382AGG (1284C>A 1286T>G), GAA384GAG (1292A>G), TTT386TTC (1298T>C), GTA388GTG (1304A>G), TTC391TTT (1313C>T), GGG392GGA (1316G>A), TCC393GCT (1317T>G 1319C>T), GGG395GGA (1325G>A), TCT399TCA (1337T>A), ACA400ACG (1340A>G), TCC401TGC (1342C>G), TTA404CTT (1350T>C 1352A>T), TTT405TTC (1355T>C), GTG409GTT (1367G>T), TTG410CTC (1368T>C 1370G>C), AGA412GCA (1374A>G 1375G>C), GAG415GAA (1385G>A), AAA417AAG (1391A>G), CTC418CTG (1394C>G), AAT419AAC (1397T>C), CGG420CGC (1400G>C), CTA421TTG (1401C>T 1403A>G), GCA423GCG (1409A>G), AAG426AAA (1418G>A), AGG427AGA (1421G>A), GCT435GCA (1445T>A), CAC436CAT (1448C>T), GCT439GCA (1457T>A), ATC440GTG (1458A>G 1460C>G), CTG441CTT (1463G>T), GAC444GAT (1472C>T), GGA446GGT (1478A>T), GCA447GCT (1481A>T), TTG448CTT (1482T>C 1484G>T), CGA449CGG (1487A>G), GCT452GCA (1496T>A), TGC455TGT (1505C>T), CAA456CAG (1508A>G), GAT457GAC (1511T>C), ATT458ATA (1514T>A), AAA459AAG (1517A>G), GAT463GAC (1529T>C), TCA466TCT (1538A>T), ACA467ACC (1541A>C), CAA468CAG (1544A>G), CCA469CCT (1547A>T), CCC471CCT (1553C>T), TTA472CTT (1554T>C 1556A>T), GCC475GCT (1565C>T), GTT477GTC (1571T>C), GAC479GAT (1577C>T), GGC481GGA (1583C>A), CAT483CAC (1589T>C), TTC484TTT (1592C>T), ACC485ACA (1595C>A), TCC486TCA (1598C>A), TAT488TAC (1604T>C), ATA489ATC (1607A>C), ACA492ACC (1616A>C), CTG495TTG (1623C>T), ATC496ATA (1628C>A), TCA497TCT (1631A>T), CCT499CCA (1637T>A), GCC500GCT (1640C>T), CAA501CAG (1643A>G), GGT503GGG (1649T>G), CTT504CTA (1652T>A), ACA505ACG (1655A>G), CCA507CCT (1661A>T), GCT509GCA (1667T>A), GAG511GAA (1673G>A), CGG512AGG (1674C>A), AGA513AGG (1679A>G), AGG514AGA (1682G>A), GAC515GAT (1685C>T), ATC516ATT (1688C>T), CTG517TTA (1689C>T 1691G>A), GTG518GTT (1694G>T), TGT520TGC (1700T>C), ATG521AGA (1702T>G 1703G>A), GAG523GTG (1708A>T), GAG525GAT (1715G>T), TCC527GAT (1719T>G 1720C>A 1721C>T), ACT528CCA (1722A>C 1724T>A), GAA529AAT (1725G>A 1727A>T), GAT530GCT (1729A>C), CCT531CCA (1733T>A), ACG532ACA (1736G>A), GGA533GCC (1738G>C 1739A>C), CAT534CAC (1742T>C), CTC535CTA (1745C>A), AGA536GTT (1746A>G 1747G>T 1748A>T), AAC538GAT (1752A>G 1754C>T), ATA539GTG (1755A>G 1757A>G), GAG541GAA (1763G>A), GCT547GCC (1781T>C), ACT549ACA (1787T>A), CAC550CAT (1790C>T), AGG551AGA (1793G>A), AAT552GCT (1794A>G 1795A>C), TTG553TTA (1799G>A), ACC554TCC (1800A>T), AGT556GAT (1806A>G 1807G>A), GAT557GAC (1811T>C), CTT558CTC (1814T>C), GAG560CAA (1818G>C 1820G>A), GTA563GTG (1829A>G), GCC564GCA (1832C>A), AAA565AAG (1835A>G), TGT566TGC (1838T>C), CAG567CGG (1840A>G), GTA568GTG (1844A>G), CAT569CAC (1847T>C), GCA570TAT (1848G>T 1849C>A 1850A>T), GAA571GAG (1853A>G), GTA572GTG (1856A>G), AAG574AGC (1861A>G 1862G>C), TTG576CTT (1866T>C 1868G>T), TTT577TAT (1870T>A), GAA578AAT (1872G>A 1874A>T), GCT579ACT (1875G>A), GAA580ACA (1878G>A 1879A>C), TCT581TCA (1883T>A), GGC582GGA (1886C>A), ATA583TTG (1887A>T 1889A>G), AGT584GCT (1890A>G 1891G>C), CCT585GAC (1893C>G 1894C>A 1895T>C), ATA587GTT (1899A>G 1901A>T), GTT590GTC (1910T>C), CAA591AAT (1911C>A 1913A>T), TCT593GCT (1917T>G), GCA594GTT (1921C>T 1922A>T), GAT596GAA (1928T>A), GTG597ATG (1929G>A), ATA598CTG (1932A>C 1934A>G), GCG599GAT (1936C>A 1937G>T), GTG601TTC (1941G>T 1943G>C), GAT602CAA (1944G>C 1946T>A), GGG603GAT (1948G>A 1949G>T), GCT604GCC (1952T>C), AGA605del (1953_1955delAGA), TTG606TTA (1958G>A), CGC607CAA (1960G>A 1961C>A), ACC608GTT (1962A>G 1963C>T 1964C>T), AAG609TTG (1965A>T 1966A>T), CAA610CCT (1969A>C 1970A>T), GAT611CCT (1971G>C 1972A>C), ATA614GCA (1980A>G 1981T>C), TTG615ACT (1983T>A 1984T>C 1985G>T), GTG616TAT (1986G>T 1987T>A 1988G>T), CCC617GAG (1989C>G 1990C>A 1991C>G), ACT618CAG (1992A>C 1993C>A 1994T>G), GTT619ACA (1995G>A 1996T>C 1997T>A), GTG620ATT (1998G>A 2000G>T), AGT621ACA (2002G>C 2003T>A), AAT623GCT (2007A>G 2008A>C), GAT624GAA (2012T>A), AGA625AAA (2014G>A), GTG626ATT (2016G>A 2018G>T), ATC627GTT (2019A>G 2021C>T), TAT628TAC (2024T>C), GCA629TCC (2025G>T 2027A>C), AGA630CAG (2028A>C 2029G>A 2030A>G), GAG631GAA (2033G>A), ACG633ACA (2039G>A), GTA634GTT (2042A>T), GCA636GCC (2048A>C), TTG637CTC (2049T>C 2051G>C), GCA640GCT (2060A>T), CAG642CAA (2066G>A), GGC643GGG (2069C>G), TTG645ATG (2073T>A), GAC646AAC (2076G>A), CCT647AAA (2079C>A 2080C>A 2081T>A), GAG648TTT (2082G>T 2083A>T 2084G>T), GAA649GAG (2087A>G), CTT650TTT (2088C>T), TTT651TTC (2093T>C), CCT652CCA (2096T>A), GAC654GAT (2102C>T), CAT655CTC (2104A>T 2105T>C), CAT656TCC (2106C>T 2107A>C 2108T>C), GCC659GCT (2117C>T), ATG660ATC (2120G>C), TCT663GTG (2127T>G 2128C>T 2129T>G), CTT664CTA (2132T>A), GAT665TCT (2133G>T 2134A>C), GAT666TCT (2136G>T 2137A>C), GAA667GAT (2141A>T), CAT668GCC (2142C>G 2143A>C 2144T>C), GAG670CAG (2148G>C), ATT671AAT (2152T>A), TTC672TTT (2156C>T), AAC673GAA (2157A>G 2159C>A), AAA674ATG (2161A>T 2162A>G), AGA676GTA (2166A>G 2167G>T), GTC677AAT (2169G>A 2170T>A 2171C>T), AAC678GAT (2172A>G 2174C>T), ATG679TTG (2175A>T), TAT681GTC (2181T>G 2182A>T 2183T>C), AGA682AAA (2185G>A), GAT685AGT (2193G>A 2194A>G), GCT686CCT (2196G>C), GAA687CAG (2199G>C 2201A>G), CGA690GCT (2208C>G 2209G>C 2210A>T), GTA693GTG (2219A>G), CAA694CGA (2221A>G), GAC699GAT (2237C>T), ATA701GTC (2241A>G 2243A>C), CAC703CAT (2249C>T), TTT704TTC (2252T>C), GCA710AGC (2268G>A 2269C>G 2270A>C), TCA711ACA (2271T>A), ATC713ATT (2279C>T), GAG715GAA (2285G>A), AAG716GAG (2286A>G), AAA717TTG (2289A>T 2290A>T 2291A>G), CTA718CTG (2294A>G), GCA719GAT (2296C>A 2297A>T), GTG720GTC (2300G>C), CAG721CAA (2303G>A), ACA722CGC (2304A>C 2305C>G 2306A>C), GAG723AAT (2307G>A 2309G>T), ATG724CTT (2310A>C 2312G>T), AGG725GAG (2313A>G 2314G>A), ACT726CAA (2316A>C 2317C>A 2318T>A), GGA727GGG (2321A>G), TTT728AAA (2322T>A 2323T>A 2324T>A), GCA729GCG (2327A>G), CAT730TAT (2328C>T), TCT731AAG (2331T>A 2332C>A 2333T>G), TGC732GCG (2334T>G 2335G>C 2336C>G), ATT733CTT (2337A>C), GAT734CAG (2340G>C 2342T>G), GCA735GCT (2345A>T), GTT737GCA (2350T>C 2351T>A), GAC738GAT (2354C>T), ACC739ACA (2357C>A), CTC740CTT (2360C>T), ATA741ACG (2362T>C 2363A>G), ATC743ATT (2369C>T), GAG744GAA (2372G>A), CCA747CCG (2381A>G), TTT749AGA (2385T>A 2386T>G 2387T>A), GTT750GTA (2390T>A), TTT752TTC (2396T>C), TTA753CTG (2397T>C 2399A>G), ACA755ACT (2405A>T), GCA756GCT (2408A>T), AGA759CGC (2415A>C 2417A>C), TGC762TGT (2426C>T), AAC763AGG (2428A>G 2429C>G), GTA765GTT (2435A>T), TCA766GCA (2436T>G), TCA766_AAA767insGGT (2438_2439insGGT), AAA767ACT (2440A>C 2441A>T), GAG768GAC (2444G>C), CCA769AAG (2445C>A 2446C>A 2447A>G), GAA771AGA (2451G>A 2452A>G), TCA772AAT (2454T>A 2455C>A 2456A>T), ATC774ATT (2462C>T), AAA775ATC (2464A>T 2465A>C), ATT776GAA (2466A>G 2467T>A 2468T>A), TAC778ATG (2472T>A 2473A>T 2474C>G), CAT779ATT (2475C>A 2476A>T), AAA780GGT (2478A>G 2479A>G 2480A>T), GTT782AAG (2484G>A 2485T>A 2486T>G), GAA783GAC (2489A>C), TAT784AAA (2490T>A 2492T>A), ATC785ATG (2495C>G), AAG786ATC (2497A>T 2498G>C), GAT787CAG (2499G>C 2501T>G), ACA788ATG (2503C>T 2504A>G), TGG790AAT (2508T>A 2509G>A 2510G>T), AGC791GAC (2511A>G 2512G>A), CTT792TTG (2514C>T 2516T>G), TAT794TTC (2521A>T 2522T>C), GCA795ATC (2523G>A 2524C>T 2525A>C), CTG796TTG (2526C>T), AGA797AAG (2530G>A 2531A>G), CTC799CTT (2537C>T), GCA802GCC (2546A>C), GGT803GGA (2549T>A), ATA806TTG (2556A>T 2558A>G), ATG807ATA (2561G>A), GCT808GCA (2564T>A), GGA811GGC (2573A>C), GTT812ACC (2574G>A 2575T>C 2576T>C), TTT813ATG (2577T>A 2579T>G), GGA814AAT (2580G>A 2581G>A 2582A>T), GGC815GCC (2584G>C), ATT816GTT (2586A>G), ACA817ACT (2591A>T), GCA818GCT (2594A>T), TTT819TTC (2597T>C), TTG820ATT (2598T>A 2600G>T), GCG821ACT (2601G>A 2603G>T), TGG823CTC (2607T>C 2608G>T 2609G>C), CAG824AAG (2610C>A), AGT825GTA (2613A>G 2614G>T 2615T>A), AAC826GCA (2616A>G 2617A>C 2618C>A), TCT827ACT (2619T>A), TTC828TCC (2623T>C), CCC829CCT (2627C>T), AAT830TTA (2628A>T 2629A>T 2630T>A), GCA831GCT (2633A>T), TCG832GAT (2634T>G 2635C>A 2636G>T), GGA835GGG (2645A>G), GGT836GGC (2648T>C), GTG837GTA (2651G>A), GAA840GAG (2660A>G), ATA844ATC (2672A>C), AGT846AGC (2678T>C), AAG848AAA (2684G>A), AAC849AAT (2687C>T), AAG850AAA (2690G>A), GGT851GGG (2693T>G), AAT852AAG (2696T>G), AAG853ACT (2698A>C 2699G>T), AAG853_CTC854insCAT (2699_2700insCAT), CTC854TTC (2700C>T), AGA855GCA (2703A>G 2704G>C), AAT856AAA (2708T>A), CTT857TCC (2709C>T 2710T>C 2711T>C), CTC858TTG (2712C>T 2714C>G), GTT859CTA (2715G>C 2717T>A), GGT860GGG (2720T>G), AGT862GCG (2724A>G 2725G>C 2726T>G), TCT863GCA (2727T>G 2729T>A), CAA864GAC (2730C>G 2732A>C), TCA865AAC (2733T>A 2734C>A 2735A>C), TTG866AAC (2736T>A 2737T>A 2738G>C), GCA867CAT (2739G>C 2740C>A 2741A>T), CAA868GTG (2742C>G 2743A>T 2744A>G), GAT869GAC (2747T>C), GAT869_TGG870insCGA (2747_2748insCGA), GCT871GCA (2753T>A), GCT872GAA (2755C>A 2756T>A), GAA873CAG (2757G>C 2759A>G), GTA877GTT (2771A>T), CAG879GAA (2775C>G 2777G>A), AGC880AGT (2780C>T), CTC881CTG (2783C>G), AAG882CAC (2784A>C 2786G>C), AAT884AAC (2792T>C), GTG886GTT (2798G>T), GTG887GTT (2801G>T), TTA888TTG (2804A>G), CTT890ATT (2808C>A), GGT891GCT (2812G>C), GTG894GTC (2822G>C), TAC895TAT (2825C>T), TTT896TTC (2828T>C), GGC898GGT (2834C>T), ACC899ACG (2837C>G), TAC900TAT (2840C>T), GTG901GTT (2843G>T), ATA906ATT (2858A>T), ATG907CTC (2859A>C 2861G>C), ACC908ACT (2864C>T), GTA909GTG (2867A>G), GCT910GCA (2870T>A), GCA912GCC (2876A>C), TTC913TTT (2879C>T), CAC914AGA (2880C>A 2881A>G 2882C>A), AAT915AAC (2885T>C), GCT916ATG (2886G>A 2887C>T 2888T>G), CGA917AGG (2889C>A 2891A>G), GGT919GGC (2897T>C), CCA921CCT (2903A>T), ACA923TCC (2907A>T 2909A>C), ATC925ATT (2915C>T), GCC927GAG (2920C>A 2921C>G), AAT928CAT (2922A>C), TCT929TCC (2927T>C), CGA930ATT (2928C>A 2929G>T 2930A>T), TCT931TGT (2932C>G), AAA932AAT (2936A>T), GTT933GTG (2939T>G), CAA934CAG (2942A>G), TAC935TTT (2944A>T 2945C>T), AAC936AAT (2948C>T), AGA938ACC (2953G>C 2954A>C), GAA939CAA (2955G>C), ATT942TTG (2964A>T 2966T>G), ATT943ATA (2969T>A), GGC945GGT (2975C>T), GAC947GAT (2981C>T), ATC948GTG (2982A>G 2984C>G), CTG950TTG (2988C>T), CGC952AGG (2994C>A 2996C>G), GTT953GTA (2999T>A), GGT954GGG (3002T>G), CCA959CCC (3017A>C), AAG960AAA (3020G>A), CCT961TCA (3021C>T 3023T>A), GAC962GAT (3026C>T), ATC963ATA (3029C>A), CGT964AGA (3030C>A 3032T>A), CAC966CAT (3038C>T), CCA968CCT (3044A>T), AGA969CGC (3045A>C 3047A>C), AGG970AGA (3050G>A), GAG972GAT (3056G>T),

Codon mutations:

TGC974TGT (3062C>T), TTC975TAC (3064T>A), ACA976TCC (3066A>T 3068A>C), CAA980ATG (3078C>A 3079A>T 3080A>G), GGA981GGC (3083A>C), TGT984GTG (3090T>G 3091G>T 3092T>G), AGA985AGG (3095A>G), GTT987GTA (3101T>A), GCG988GCT (3104G>T), TCT989TCA (3107T>A), ACA990AAG (3109C>A 3110A>G), CCA992CCC (3116A>C), CGT993AAG (3117C>A 3118G>A 3119T>G), CTT994AAA (3120C>A 3121T>A 3122T>A), GGC995TCC (3123G>T 3124G>C), TTA997TTG (3131A>G), TTT999GTG (3135T>G 3137T>G), CTC1000CTT (3140C>T), AAG1001AAA (3143G>A), ATG1002TAT (3144A>T 3145T>A 3146G>T), GGA1009GGC (3167A>C), ACA1011ACC (3173A>C), GTT1012GAA (3175T>A 3176T>A), GTG1014GTC (3182G>C), GAA1015ACA (3183G>A 3184A>C), GAA1017GGA (3190A>G), CTG1018TTA (3192C>T 3194G>A), AAC1019GAT (3195A>G 3197C>T), TCC1020TCG (3200C>G), TCC1021AAT (3201T>A 3202C>A 3203C>T), AGT1022GCA (3204A>G 3205G>C 3206T>A), AAG1024GCT (3210A>G 3211A>C 3212G>T), ATT1025CTT (3213A>C), TGC1026AGC (3216T>A), TCT1027TCA (3221T>A), CAA1029GTT (3225C>G 3226A>T 3227A>T), TCT1030GCA (3228T>G 3230T>A), GAA1032GAG (3236A>G), ATA1035ATT (3245A>T), GGG1037AGT (3249G>A 3251G>T), GAA1038GAG (3254A>G), GGT1040GGA (3260T>A), GAC1041GAT (3263C>T), GGC1043GGT (3269C>T), ACG1044ACT (3272G>T), TTG1045CTT (3273T>C 3275G>T), CTA1046CTC (3278A>C), CTG1047TTA (3279C>T 3281G>A), TTA1048TTG (3284A>G), AGT1050ACT (3289G>C), GTT1051GTG (3293T>G), AAT1053AAC (3299T>C), CAA1055CAG (3305A>G), CCT1056CCG (3308T>G), GTG1057TGC (3309G>T 3310T>G 3311G>C), GGC1060GGA (3320C>A), ATC1061ATA (3323C>A), CAC1062CAT (3326C>T), TCA1065TCT (3335A>T), GGC1068GAT (3343G>A 3344C>T), GTA1069ACC (3345G>A 3346T>C 3347A>C), GCA1070GCT (3350A>T), GCT1071GCA (3353T>A), GAA1072GAG (3356A>G), CGT1073AAG (3357C>A 3358G>A 3359T>G), GGA1074GGT (3362A>T), TTT1075TAC (3364T>A 3365T>C), ATC1076GTG (3366A>G 3368C>G), TCT1077GCT (3369T>G), AAT1079AAC (3377T>C), GCT1080GCA (3380T>A), GCT1082GCA (3386T>A), ATC1083GTG (3387A>G 3389C>G), TAC1084TTT (3391A>T 3392C>T), AGG1085AGA (3395G>A), GAA1086GAT (3398A>T), CTA1088TTG (3402C>T 3404A>G), GAG1089GAA (3407G>A), CCA1092CCC (3416A>C), ACT1093ACG (3419T>G), GGG1094GGA (3422G>A), CCG1095CCC (3425G>C), GTC1096ATT (3426G>A 3428C>T), GCA1098GTG (3433C>T 3434A>G), GCA1099GCT (3437A>T), GTA1101GTG (3443A>G), GAT1104GAA (3452T>A), ATT1105CTG (3453A>C 3455T>G), CTG1106TTG (3456C>T), TCA1108CAC (3462T>C 3463C>A 3464A>C), ATT1109CTG (3465A>C 3467T>G), AGA1110AGG (3470A>G), AGC1111TCC (3471A>T 3472G>C), AGA1112AAG (3475G>A 3476A>G), GAA1113GAG (3479A>G), ACA1114GAA (3480A>G 3481C>A), CAG1115AGA (3483C>A 3484A>G 3485G>A), CTT1116TTC (3486C>T 3488T>C), GAA1119GAG (3497A>G), AAC1120AAA (3500C>A), CAA1121CAG (3503A>G), TAC1123GCT (3507T>G 3508A>C 3509C>T), TAC1124TTC (3511A>T), CTT1125TTG (3513C>T 3515T>G), GGA1126GGC (3518A>C), ACA1127ACT (3521A>T), GTT1128GTA (3524T>A), CCA1129CCG (3527A>G), CAG1130TTG (3528C>T 3529A>T), GAG1131GAA (3533G>A), TTG1132ATG (3534T>A), GCC1133GCT (3539C>T), GCC1134GCT (3542C>T), GTT1136GTG (3548T>G), CCC1137CCA (3551C>A), AAG1139AAA (3557G>A), ACC1140ACA (3560C>A), ACT1141ACC (3563T>C), CTG1142CTT (3566G>T), CGG1143AAA (3567C>A 3568G>A 3569G>A), AAA1144CGC (3570A>C 3571A>G 3572A>C), AGC1145AGT (3575C>T), CAA1146CCC (3577A>C 3578A>C), TTG1147CTG (3579T>C), GAA1149GAG (3587A>G), GCA1150GCT (3590A>T), TTC1151TTT (3593C>T), GGA1152GGG (3596A>G), CCT1153GAT (3597C>G 3598C>A), GCA1154GCT (3602A>T), GAG1155CAG (3603G>C), ACA1156ACT (3608A>T), GCA1157GCC (3611A>C), ATT1160ATC (3620T>C), ACA1162TCT (3624A>T 3626A>T), GTT1163GCC (3628T>C 3629T>C), CAT1164AAA (3630C>A 3632T>A), AGA1167AGG (3641A>G), GGT1168GGA (3644T>A), GAT1169GAA (3647T>A), GGT1170GGG (3650T>G), GAC1172GAT (3656C>T), GGG1177GGT (3671G>T), GTA1178GTC (3674A>C), ATG1179GAG (3675A>G 3676T>A), AAA1180AAG (3680A>G), TAC1181TAT (3683C>T), TGT1186TCC (3697G>C 3698T>C), GGA1187AAC (3699G>A 3700G>A 3701A>C), GAT1189GAG (3707T>G), GAT1190GAG (3710T>G), GAC1191CCA (3711G>C 3712A>C 3713C>A), GCC1193GCA (3719C>A), AAA1194AAG (3722A>G), CTA1195CTG (3725A>G), GCA1196GCC (3728A>C), TTC1197TTT (3731C>T), GAA1198GAG (3734A>G), AAT1199GGT (3735A>G 3736A>G), CTC1200CTT (3740C>T), AAG1201AAA (3743G>A), TGC1202GCA (3744T>G 3745G>C 3746C>A), TCG1203TCC (3749G>C), CTG1204ATT (3750C>A 3752G>T), CTA1205CTT (3755A>T), ATC1207GTG (3759A>G 3761C>G), ATG1208CTT (3762A>C 3764G>T), CGT1209AAG (3765C>A 3766G>A 3767T>G), AGC1210ATG (3769G>T 3770C>G), AAG1212CGA (3774A>C 3775A>G 3776G>A), ATC1213GTG (3777A>G 3779C>G), CCT1214CCC (3782T>C), GGA1216GGG (3788A>G), CGT1217AAA (3789C>A 3790G>A 3791T>A), CCA1218CCT (3794A>T), TGT1219GCA (3795T>G 3796G>C 3797T>A), GAA1220ATT (3798G>A 3799A>T 3800A>T), AGG1221AGA (3803G>A), GAT1222AAT (3804G>A), GGA1229GGG (3827A>G), TGT1233TTC (3838G>T 3839T>C), TAC1235TAT (3845C>T), TAT1236TTT (3847A>T), GGC1238GCA (3853G>C 3854C>A), GAG1240GAA (3860G>A), CTG1241TTG (3861C>T), ACA1243ACG (3869A>G), TCT1244TCA (3872T>A), TGC1245TGT (3875C>T), GGA1246GGT (3878A>T), CCC1248CCT (3884C>T), TTC1249TTT (3887C>T), ATG1252TTG (3894A>T), GGG1253GAA (3898G>A 3899G>A), ATG1254CTT (3900A>C 3902G>T), GGG1255GGT (3905G>T), ATG1256TCT (3906A>T 3907T>C 3908G>T), AAG1258AAA (3914G>A), AGA1259AGG (3917A>G), CAA1263GAA (3927C>G), TCC1264GCC (3930T>G), ACT1265CAA (3933A>C 3934C>A 3935T>A), GGC1266CCA (3936G>C 3937G>C 3938C>A), GAA1267GAC (3941A>C), GGA1268TCG (3942G>T 3943G>C 3944A>G), GAG1269CAG (3945G>C), AGA1270AGG (3950A>G), GTG1271GTT (3953G>T), GAA1272CTT (3954G>C 3955A>T 3956A>T), AGA1275AGG (3965A>G), GAC1276AAC (3966G>A), ACT1277ACA (3971T>A), GTA1279GTC (3977A>C), GAA1281GAT (3983A>T), CTA1286TTG (3996C>T 3998A>G), GAT1287GAC (4001T>C), GTG1288TCG (4002G>T 4003T>C), ATT1290GTA (4008A>G 4010T>A), AAA1292CAA (4014A>C), GGC1293GGA (4019C>A), ATC1294ACA (4021T>C 4022C>A), CAT1295TTT (4023C>T 4024A>T), GTG1296GTT (4028G>T), GAT1297GAC (4031T>C), CTG1298CTT (4034G>T), ACC1300ACA (4040C>A), CAA1302CAG (4046A>G), TGC1303TGT (4049C>T), GCC1304GCG (4052C>G), CGC1308CGG (4064C>G), CTC1309TTG (4065C>T 4067C>G), AAA1313AAG (4079A>G), GGC1316GGG (4088C>G), AGA1318AGG (4094A>G), AGG1321CGC (4101A>C 4103G>C), CTC1322CTG (4106C>G), TTT1323TTC (4109T>C), GAG1324GAA (4112G>A), ATA1325ATC (4115A>C), CTT1326TTG (4116C>T 4118T>G), TAC1330TAT (4130C>T), TTG1333CTG (4137T>C), GTC1334GTG (4142C>G), AGG1335CGG (4143A>C), AAG1336AAA (4148G>A), TAT1337TAC (4151T>C), TTT1338TTC (4154T>C), GAT1340GAC (4160T>C), TTC1341TTT (4163C>T), TCA1342TCT (4166A>T), GCC1343GCT (4169C>T), ACA1344TGT (4170A>T 4171C>G 4172A>T), TTG1348GCA (4182T>G 4183T>C 4184G>A), AAT1350AAC (4190T>C), GCT1351GTA (4192C>T 4193T>A), ATA1352CTC (4194A>C 4196A>C), CCA1353CCT (4199A>T), AAA1355AAG (4205A>G), GTT1356GTA (4208T>A), ATT1358ATC (4214T>C), GAT1359AAT (4215G>A), CCT1360CCA (4220T>A), ACA1361ACG (4223A>G), AGT1362AGC (4226T>C), TCT1363CAT (4227T>C 4228C>A), TTG1367GTG (4239T>G), TTG1368CTG (4242T>C), GCA1369GCT (4247A>T), AAT1370AGT (4249A>G), GGG1371AGC (4251G>A 4253G>C), AGA1373AGG (4259A>G), GCT1374TCA (4260G>T 4262T>A), GTG1375GTA (4265G>A), TCA1376TCT (4268A>T), GAC1377GAT (4271C>T), GTG1378GAG (4273T>A), TTT1380TTC (4280T>C), GCT1382GCC (4286T>C), TAT1384TAC (4292T>C), TCC1385TCG (4295C>G), CCT1392CCC (4316T>C), GTT1393ATC (4317G>A 4319T>C), GCT1395GCA (4325T>A), CAG1397CAA (4331G>A), TTG1402CTA (4344T>C 4346G>A), GAA1405GCT (4354A>C 4355A>T), TAC1406TTT (4357A>T 4358C>T), TAC1407TAT (4361C>T), GGA1408GGT (4364A>T), TCA1409GAT (4365T>G 4366C>A 4367A>T), AAG1410GAG (4368A>G), GGC1412GGG (4376C>G), GAT1415GAG (4385T>G), TCC1416TCA (4388C>A), CGA1419AGG (4395C>A 4397A>G), CAT1420CAC (4400T>C), GCA1421ACA (4401G>A), CTT1422TTA (4404C>T 4406T>A), GCA1426GCC (4418A>C), TGT1428TGC (4424T>C), TAC1430TAT (4430C>T), ACA1431ACC (4433A>C), CTG1432CTC (4436G>C), TGC1433TGT (4439C>T), GAT1435AAC (4443G>A 4445T>C), AAA1436AAG (4448A>G), GTG1437TTG (4449G>T), AGG1439CGC (4455A>C 4457G>C), TTG1440CTA (4458T>C 4460G>A), GTT1441GTG (4463T>G), GGG1442GGT (4466G>T), GGC1443GGA (4469C>A), CCA1445CCC (4475A>C), TCA1446TCT (4478A>T), GGA1447GGT (4481A>T), GCA1449GCT (4487A>T), CTA1450TTG (4488C>T 4490A>G), ACG1451ACT (4493G>T), ATC1453ATA (4499C>A), TTC1454TTT (4502C>T), TCT1456TCA (4508T>A), CTC1457TTG (4509C>T 4511C>G), CTC1458CTA (4514C>A), CGA1464AGA (4530C>A), TAT1465TAC (4535T>C), GCC1466GCG (4538C>G), TTG1470CTG (4548T>C), TTA1471TTG (4553A>G), AGA1472AGG (4556A>G), CCC1474CCA (4562C>A), ATT1476GTC (4566A>G 4568T>C), GCT1477AAA (4569G>A 4570C>A 4571T>A), AGG1479AAA (4576G>A 4577G>A), GCT1480TCT (4578G>T), ATA1481GTG (4581A>G 4583A>G), GGA1482GGC (4586A>C), GTT1483ATT (4587G>A), AAA1484AAG (4592A>G), AGT1486AAT (4597G>A), GAT1487GAC (4601T>C), CTA1491TTG (4611C>T 4613A>G), GCT1494GCA (4622T>A), GTT1495GTA (4625T>A), GGA1497GGT (4631A>T), GAC1499GAT (4637C>T), AAT1500AAC (4640T>C), CTT1501TTG (4641C>T 4643T>G), GCT1503GCG (4649T>G), GTA1504GTT (4652A>T), TTA1506ATG (4656T>A 4658A>G), CTC1508TTG (4662C>T 4664C>G), CAG1509AGG (4665C>A 4666A>G), TAT1511TAC (4673T>C), TCT1512GAT (4674T>G 4675C>A), CTG1513CTA (4679G>A), AAT1515TCC (4683A>T 4684A>C 4685T>C), ATA1516ATT (4688A>T), CAT1518AGA (4692C>A 4693A>G 4694T>A), GAG1519GAA (4697G>A), TTA1520CTG (4698T>C 4700A>G), CTG1522GCT (4704C>G 4705T>C 4706G>T), GTC1523GTT (4709C>T), AAT1524AAC (4712T>C), GTA1525GTC (4715A>C), ATC1526GTC (4716A>G), CTT1531TTG (4731C>T 4733T>G), AAA1533AAG (4739A>G), TCG1534TCA (4742G>A), GTT1537GTA (4751T>A), AAT1538GAG (4752A>G 4754T>G), GAG1539AAG (4755G>A), GTA1540ACA (4758G>A 4759T>C), CAA1541AAA (4761C>A), CAA1543CTG (4768A>T 4769A>G), CTA1548CTC (4784A>C), ACT1549ACC (4787T>C), CTG1551CTC (4793G>C), AGA1553AGG (4799A>G), TTT1555TTC (4805T>C), AAA1556AAG (4808A>G), GCT1559GCA (4817T>A), CTT1560TTG (4818C>T 4820T>G), TAC1562TAT (4826C>T), CAC1563CAA (4829C>A), GCT1565GCA (4835T>A), CCT1566CCA (4838T>A), AAG1568AAA (4844G>A), ATC1572GTC (4854A>G), ATG1576ATT (4868G>T), TAT1577TAC (4871T>C), ATA1579ATT (4877A>T), AGA1580AGG (4880A>G), CCT1581CCA (4883T>A), GCC1582GCA (4886C>A), CCA1583GTG (4887C>G 4888C>T 4889A>G), GGT1584GGA (4892T>A), GAC1587GAT (4901C>T), ACT1588GTA (4902A>G 4903C>T 4904T>A), CAG1589CAA (4907G>A), GCC1590GCT (4910C>T), ATG1592TAT (4914A>T 4915T>A 4916G>T), GTG1595ATT (4923G>A 4925G>T), GGA1598GGC (4934A>C), ATA1599ATC (4937A>C), CGT1600AGA (4938C>A 4940T>A), GCT1602GCC (4946T>C), CAC1605CAT (4955C>T), GGT1606GGG (4958T>G), CGT1607CAG (4960G>A 4961T>G), GTG1608GCC (4963T>C 4964G>C), GCT1609ACT (4965G>A), GAC1611GAA (4973C>A), AAG1612AGT (4975A>G 4976G>T), CTT1613TTT (4977C>T), GTC1614GTA (4982C>A), TTA1615AAT (4983T>A 4984T>A 4985A>T), CAG1618CTT (4993A>T 4994G>T), ACG1619GGA (4995A>G 4996C>G 4997G>A), GCG1620GCT (5000G>T), TTG1621CTC (5001T>C 5003G>C), GAT1622AAG (5004G>A 5006T>G), GAA1623CAG (5007G>C 5009A>G), AGG1624AGA (5012G>A), GGT1625AAC (5013G>A 5014G>A 5015T>C), TTC1626CTC (5016T>C), AGA1627CCC (5019A>C 5020G>C 5021A>C), GCT1628ATC (5022G>A 5023C>T 5024T>C), GTG1629ATG (5025G>A), TTT1631ATG (5031T>A 5033T>G), CCT1632CCC (5036T>C), TCC1633ACT (5037T>A 5039C>T), TAT1634TAC (5042T>C), TTG1635TAT (5044T>A 5045G>T), GTG1637CTT (5049G>C 5051G>T), CAG1639AGT (5055C>A 5056A>G 5057G>T), GAA1640CAA (5058G>C), ATT1642TTG (5064A>T 5066T>G), GCA1643ACA (5067G>A), AAG1644GAG (5070A>G), GTA1645GTC (5075A>C), GGG1647GGA (5081G>A), TCA1649GAG (5085T>G 5086C>A 5087A>G), GCC1651ATT (5091G>A 5092C>T 5093C>T), CTG1652GGG (5094C>G 5095T>G), ACA1653GTT (5097A>G 5098C>T 5099A>T), ATT1654TTG (5100A>T 5102T>G), TGT1655ATG (5103T>A 5104G>T 5105T>G), GAA1656GAG (5108A>G), ATG1657CTA (5109A>C 5111G>A), GCA1658TCA (5112G>T), AAA1659GAT (5115A>G 5117A>T), GCA1660GCT (5120A>T), GCT1661AAA (5121G>A 5122C>A 5123T>A), ATT1662AAT (5125T>A), TCC1663ACA (5127T>A 5129C>A), TAT1664TAC (5132T>C), ACG1665ACC (5135G>C), CCA1666CCT (5138A>T), TTG1667TTA (5141G>A), GAC1668CCT (5142G>C 5143A>C 5144C>T), GCA1669GCT (5147A>T), GGT1670GGG (5150T>G), GAG1671GAA (5153G>A), ATC1673ATT (5159C>T), ACA1674del (5160_5162delACA), CGT1678GAG (5172C>G 5173G>A 5174T>G), GAT1679GGT (5176A>G), CTG1680GTC (5178C>G 5180G>C), AAT1681CAT (5181A>C), TGG1682GAA (5184T>G 5185G>A 5186G>A), GCA1684GCG (5192A>G), CCA1685CCT (5195A>T), AAC1686GAT (5196A>G 5198C>T), ATT1687GTA (5199A>G 5201T>A), GGT1688TTT (5202G>T 5203G>T), TTT1689TTC (5207T>C), TCA1691AGT (5211T>A 5212C>G 5213A>T), GCA1692GCT (5216A>T), CGT1693CGG (5219T>G), GCC1696TCA (5226G>T 5228C>A), CAC1697AAC (5229C>A), TAC1698CAA (5232T>C 5234C>A), ACG1699CAA (5235A>C 5236C>A 5237G>A), GAT1701TCA (5241G>T 5242A>C 5243T>A), GAA1702TCA (5244G>T 5245A>C), GGG1703GGT (5249G>T), TAC1704AAC (5250T>A), ATT1705ATC (5255T>C), ATT1706CTG (5256A>C 5258T>G), AAC1708AAT (5264C>T), TGT1709TGC (5267T>C), ACA1710ACT (5270A>T), GGT1711GGA (5273T>A), GCA1712GCC (5276A>C), AAG1713CGC (5277A>C 5278A>G 5279G>C), AAG1714AGA (5281A>G 5282G>A), TCC1715ACA (5283T>A 5285C>A), AAT1716ACA (5287A>C 5288T>A), GTT1718GTG (5294T>G), GGT1720GGG (5300T>G), CCA1721CCT (5303A>T), AAC1723AAT (5309C>T), GAC1726GAT (5318C>T), ATG1731ATA (5333G>A), CCG1733CCC (5339G>C), ATG1736TTG (5346A>T), TCA1737GCA (5349T>G), GCT1738GCA (5354T>A), ATA1739ATC (5357A>C), ATG1740CAG (5358A>C 5359T>A), GCC1742CAT (5364G>C 5365C>A 5366C>T), AAG1744CGC (5370A>C 5371A>G 5372G>C), AAT1745AAG (5375T>G), GTG1746AAT (5376G>A 5377T>A 5378G>T), TTG1747CTG (5379T>C), GCA1748GAA (5383C>A), GGA1749GCT (5386G>C 5387A>T), GGA1750GGT (5390A>T), CAT1751GAT (5391C>G), GTA1752GGA (5395T>G), GCA1753GCT (5399A>T), AAT1755TCT (5403A>T 5404A>C), GTT1757ACC (5409G>A 5410T>C 5411T>C), GTG1759GTT (5417G>T), GGA1761GGC (5423A>C), GGA1763GGT (5429A>T), GCT1765GCA (5435T>A), GTC1766ATA (5436G>A 5438C>A), GGC1767GCT (5440G>C 5441C>T), ATT1769ATC (5447T>C), GCG1772GCA (5456G>A), CTG1773TTG (5457C>T), ATG1774TTT (5460A>T 5462G>T), GCC1775GCA (5465C>A), TTA1776TTG (5468A>G), GCA1777GCC (5471A>C), ACA1778ACT (5474A>T), AGG1779CAT (5475A>C 5476G>A 5477G>T), TAT1781TAC (5483T>C), TGT1782AGG (5484T>A 5486T>G), GTG1783GCC (5488T>C 5489G>C), GAG1784ACA (5490G>A 5491A>C 5492G>A), ATA1786CTT (5496A>C 5498A>T), ATA1787ATT (5501A>T), GTT1788GTG (5504T>G), AGA1789CGC (5505A>C 5507A>C), TTG1790CTA (5508T>C 5510G>A), CGC1791CGA (5513C>A), ACA1792ACC (5516A>C), ATT1793ATG (5519T>G), GGA1794GCT (5521G>C 5522A>T), AAT1795GGG (5523A>G 5524A>G 5525T>G), CTT1797CTG (5531T>G), GAC1798GAT (5534C>T), CTC1799CTG (5537C>G), AAT1800AAC (5540T>C), TAT1802TAC (5546T>C), CCT1803CCC (5549T>C), GGC1805GGT (5555C>T), GTG1807GCG (5560T>C), CAG1808CAA (5564G>A), TAT1809TAC (5567T>C), TTT1810TTG (5570T>G), CTT1811CTC (5573T>C), CAA1812AAG (5574C>A 5576A>G), GTG1814GTT (5582G>T), CCT1815GAA (5583C>G 5584C>A 5585T>A), CAT1816GAC (5586C>G 5588T>C), CAT1816_GGA1817insACC (5588_5589insACC), GGA1817TCA (5589G>T 5590G>C), GAC1818CAC (5592G>C), AAA1819AAG (5597A>G), GTG1820TTG (5598G>T), GCT1821GGA (5602C>G 5603T>A), CAA1822CAG (5606A>G), GGT1824GGC (5612T>C), GCA1825ACA (5613G>A), TCG1826TAT (5617C>A 5618G>T), CTC1827GGC (5619C>G 5620T>G), CAC1828del (5622_5624delCAC), AGT1829ACC (5626G>C 5627T>C), AGT1830ACT (5629G>C), ATG1832AAC (5635T>A 5636G>C), CAA1834CAT (5642A>T), CAA1834_GGG1835insCACAGT (5642_5643insCACAGT), GGG1835GCA (5644G>C 5645G>A), GAA1837TCA (5649G>T 5650A>C), TTG1838GTT (5652T>G 5654G>T), GGC1839GGA (5657C>A), AAC1840CTT (5658A>C 5659A>T 5660C>T), CTC1841CTA (5663C>A), CGC1842AGG (5664C>A 5666C>G), ATT1843ATA (5669T>A), ATA1844GTT (5670A>G 5672A>T), CAT1845CCT (5674A>C), GGT1846TCC (5676G>T 5677G>C 5678T>C), GAT1847GAC (5681T>C), TTA1848CTC (5682T>C 5684A>C), GCA1849TCT (5685G>T 5687A>T), AAA1850GAA (5688A>G), CAG1851AAG (5691C>A), ACA1852ATG (5695C>T 5696A>G), ATG1854GCT (5700A>G 5701T>C 5702G>T), CGG1855CGT (5705G>T), ATG1856TTG (5706A>T), TAT1858TAC (5714T>C), GTG1859ACA (5715G>A 5716T>C 5717G>A), GTA1860ATT (5718G>A 5720A>T), GGA1861GGT (5723A>T), CCA1862CCT (5726A>T), CAT1863GAA (5727C>G 5729T>A), GGA1865GGT (5735A>T), GGA1867GCC (5740G>C 5741A>C), AAT1868CAG (5742A>C 5744T>G), TTT1869TGC (5746T>G 5747T>C), TCC1870TCA (5750C>A), ACA1871ACG (5753A>G), CAG1872AAG (5754C>A), GAT1873AAC (5757G>A 5759T>C), CTT1874CTG (5762T>G), GAG1875AAG (5763G>A), AGT1876CAT (5766A>C 5767G>A), TTG1877GCT (5769T>G 5770T>C 5771G>T), CTA1878TTG (5772C>T 5774A>G), CAC1879GAG (5775C>G 5777C>G), TAT1880ACA (5778T>A 5779A>C 5780T>A), TTG1881TTA (5783G>A), GAG1882TCT (5784G>T 5785A>C 5786G>T), CAG1883GTC (5787C>G 5788A>T 5789G>C), GGA1884AGT (5790G>A 5792A>T), GCA1886GCT (5798A>T), GAG1887ACT (5799G>A 5800A>C 5801G>T), TTA1888CTT (5802T>C 5804A>T), ATT1889GCC (5805A>G 5806T>C 5807T>C), CAA1890GCA (5808C>G 5809A>C), AAG1891AGA (5812A>G 5813G>A), AAT1892GGA (5814A>G 5815A>G 5816T>A), ACA1893GTC (5817A>G 5818C>T 5819A>C), CTA1895GTG (5823C>G 5825A>G), ACT1896ACA (5828T>A), CTG1897TTG (5829C>T), TTC1899TTT (5837C>T), GAG1901GAA (5843G>A), CTG1902TTC (5844C>T 5846G>C), AGT1903TCC (5847A>T 5848G>C 5849T>C), ATG1904CAA (5850A>C 5851T>A 5852G>A), GAA1905CAC (5853G>C 5855A>C), AAT1906GAT (5856A>G), GTG1907TGT (5859G>T 5860T>G 5861G>T), CAA1908GAA (5862C>G), CAA1909ATG (5865C>A 5866A>T 5867A>G), CTG1910ATA (5868C>A 5870G>A), ATA1911CTT (5871A>C 5873A>T), GAT1912GAG (5876T>G), GTT1914CTG (5880G>C 5882T>G), AAG1915GAA (5883A>G 5885G>A), CTT1916AGT (5886C>A 5887T>G), CAG1917CAA (5891G>A), TTT1919GTA (5895T>G 5897T>A), TTC1920GTA (5898T>G 5900C>A), CCA1921TTC (5901C>T 5902C>T 5903A>C), AAA1922ATG (5905A>T 5906A>G), GAA1923GGG (5908A>G 5909A>G), ACT1924AGC

(5911C>G 5912T>C), ACC1925GTT (5913A>G 5914C>T 5915C>T), ATT1926AAA (5917T>A 5918T>A), CAG1927AAA (5919C>A 5921G>A), AAG1928CAA (5922A>C 5924G>A), CTC1929ATG (5925C>A 5927C>G), AAA1930TTG (5928A>T 5929A>T 5930A>G), AAT1931TCC (5931A>T 5932A>C 5933T>C), TTT1932TTC (5936T>C), GTT1933ATG (5937G>A 5939T>G), GAT1934AAT (5940G>A), GCT1935CAA (5943G>C 5944C>A 5945T>A), GAA1936CAG (5946G>C 5948A>G), TGT1937ATG (5949T>A 5950G>T 5951T>G), CTA1938TCA (5952C>T 5953T>C), ACA1939AAA (5956C>A), TTC1940ACC (5958T>A 5959T>C), AAA1941CAA (5961A>C), GCA1942GTT (5965C>T 5966A>T), AGG1943GCC (5967A>G 5968G>C 5969G>C), AGC1944_TTT1945insCCATGTTGTGTCACA (5972_5973insCCATGTTGTGTCACA), AGG1946AGT (5978G>T), CAC1947AAA (5979C>A 5981C>A), GTA1948GCA (5983T>C), GTT1949TGG (5985G>T 5986T>G 5987T>G), TTC1950CTA (5988T>C 5990C>A), AAA1951ACT (5992A>C 5993A>T), AAA1953GAC (5997A>G 5999A>C), CTT1954ATT (6000C>A), TTG1955GTC (6003T>G 6005G>C), AGT1956ATG (6007G>T 6008T>G), TCC1957ACA (6009T>A 6011C>A), ACA1958AAA (6013C>A), TGG1959GGT (6015T>G 6017G>T), AAA1960AAG (6020A>G), ATG1961AAG (6022T>A), TGT1962CCA (6024T>C 6025G>C 6026T>A), GGT1963GGA (6029T>A), GAA1964del (6030_6032delGAA), AGT1965TCT (6033A>T 6034G>C), GTT1967_AGG1970del (6039_6050delGTTGCTTCAAGG), TCT1971CCT (6051T>C), GCA1972GCT (6056A>T), GAG1973GAA (6059G>A), AGT1974ACA (6061G>C 6062T>A), CTA1975GTG (6063C>G 6065A>G), TTC1976TTT (6068C>T), CCT1977CTT (6070C>T), GGC1978GGG (6074C>G), TTG1980CCT (6078T>C 6079T>C 6080G>T), TCT1981TCA (6083T>A), GCT1982GCA (6086T>A), TCC1983CTA (6087T>C 6088C>T 6089C>A), GTA1984GTT (6092A>T), TTG1985GTG (6093T>G), AAA1986AGA (6097A>G), TTA1988ATG (6102T>A 6104A>G), GAA1990CGC (6108G>C 6109A>G 6110A>C), AGA1991AAT (6112G>A 6113A>T), CAC1992AGT (6114C>A 6115A>G 6116C>T), ACA1993GGA (6117A>G 6118C>G), AGA1994AAG (6121G>A 6122A>G), ATG1996TAC (6126A>T 6127T>A 6128G>C), AGT1997TCA (6129A>T 6130G>C 6131T>A), CAG1999ACA (6135C>A 6136A>C 6137G>A), AGT2000ACA (6139G>C 6140T>A), ATG2001CTG (6141A>C), GAG2002GAA (6146G>A), CTG2005TTT (6153C>T 6155G>T), AAA2006AGA (6157A>G), ATA2007ATG (6161A>G), TAT2008TAC (6164T>C), CTA2009ATG (6165C>A 6167A>G), TTA2010TGT (6169T>G 6170A>T), AAC2011CAG (6171A>C 6173C>G), TTC2012GAA (6174T>G 6175T>A 6176C>A), CAA2013GGG (6177C>G 6178A>G 6179A>G), GTA2014TAT (6180G>T 6181T>A 6182A>T), ATA2015TTA (6183A>T), ACA2016GAT (6186A>G 6187C>A 6188A>T), AGT2017AAA (6190G>A 6191T>A), GAG2018GCC (6193A>C 6194G>C), TTA2020GTG (6198T>G 6200A>G), AAG2021GAC (6201A>G 6203G>C), AAG2022TGC (6204A>T 6205A>G 6206G>C), TTT2023TTC (6209T>C), GAA2024AGT (6210G>A 6211A>G 6212A>T), GAT2025GAG (6215T>G), TTT2027ATG (6219T>A 6221T>G), CAG2028GGC (6222C>G 6223A>G 6224G>C), GAG2029GTT (6226A>T 6227G>T), AAG2030TGT (6228A>T 6229A>G 6230G>T), ATA2031GTG (6231A>G 6233A>G), TCC2032CCG (6234T>C 6236C>G), ACC2033GAA (6237A>G 6238C>A 6239C>A), ACC2034CGC (6240A>C 6241C>G), CTA2035ATC (6243C>A 6245A>C), CTA2036TTG (6246C>T 6248A>G), ATC2037GTG (6249A>G 6251C>G), AAA2038CGC (6252A>C 6253A>G 6254A>C), GTA2039ACA (6255G>A 6256T>C), TTC2040ATT (6258T>A 6260C>T), TTG2041GCT (6261T>G 6262T>C 6263G>T), GAG2044TCC (6270G>T 6271A>C 6272G>C), GAG2045ACG (6273G>A 6274A>C), AGT2046TCA (6276A>T 6277G>C 6278T>A), TAT2047TTG (6280A>T 6281T>G), CAG2048AAC (6282C>A 6284G>C), ACT2049GAG (6285A>G 6286C>A 6287T>G), GAG2050AAT (6288G>A 6290G>T), TGT2052ACT (6294T>A 6295G>C), ATA2053GTG (6297A>G 6299A>G), GAT2054TAC (6300G>T 6302T>C), TCT2055ACA (6303T>A 6305T>A), CAG2056GAT (6306C>G 6308G>T), GTG2057GGG (6310T>G), CTC2058CTT (6314C>T), CAG2059TTT (6315C>T 6316A>T 6317G>T), AGG2060GAT (6318A>G 6319G>A 6320G>T), ATA2061del (6321_6323delATA), AAC2062CAT (6324A>C 6326C>T), TCC2063TTC (6328C>T), CAG2064AAA (6330C>A 6332G>A), AAA2065GCA (6333A>G 6334A>C), TTC2066GTG (6336T>G 6338C>G), AGA2067GAC (6339A>G 6340G>A 6341A>C), GTG2068TTC (6342G>T 6344G>C), CAG2069AAG (6345C>A), GAA2070CCT (6348G>C 6349A>C 6350A>T), GGT2071GGG (6353T>G), TTT2073TTC (6359T>C), TAT2074TCC (6361A>C 6362T>C), TTG2075TGG (6364T>G), CAT2076GAA (6366C>G 6368T>A), CCT2077ATT (6369C>A 6370C>T), GAA2078AAT (6372G>A 6374A>T), GGA2079GAT (6376G>A 6377A>T), ATC2080GCC (6378A>G 6379T>C), AAC2081AAT (6383C>T), AAT2083AAC (6389T>C), GCC2084GCT (6392C>T), GTA2085GCA (6394T>C), ATA2088GTG (6402A>G 6404A>G), ATA2089TTG (6405A>T 6407A>G), TCT2091GAA (6411T>G 6412C>A 6413T>A), CTG2092CTC (6416G>C), GAG2094AGG (6420G>A 6421A>G), TCC2095CAC (6423T>C 6424C>A), TAT2096TGG (6427A>G 6428T>G), AGC2097TCA (6429A>T 6430G>C 6431C>A), CGA2098AAG (6432C>A 6433G>A 6434A>G), GAT2099TCA (6435G>T 6436A>C 6437T>A), AGT2100CAA (6438A>C 6439G>A 6440T>A), AAC2101AGC (6442A>G), TCC2102AAG (6444T>A 6445C>A 6446C>G), TAT2103TTC (6448A>T 6449T>C), TCT2104ACT (6450T>A), GTG2105TTT (6453G>T 6455G>T), TCA2106GCC (6456T>G 6458A>C), ACT2107AAT (6460C>A), CCC2108GAA (6462C>G 6463C>A 6464C>A), ATC2109GTT (6465A>G 6467C>T), AAA2110GAA (6468A>G), CTG2111ATA (6471C>A 6473G>A), GGG2112TCA (6474G>T 6475G>C 6476G>A), TGT2113AAC (6477T>A 6478G>A 6479T>C), TTC2114CTG (6480T>C 6482C>G), ATT2115CTT (6483A>C), TTC2116TTA (6488C>A), CTT2117GCT (6489C>G 6490T>C), GTC2118GTT (6494C>T), GTG2119TAT (6495G>T 6496T>A 6497G>T), CTA2120TTA (6498C>T), GAT2121GAA (6503T>A), AGT2122AAT (6505G>A), CAA2123AAG (6507C>A 6509A>G), GGC2124CTC (6510G>C 6511G>T), AAG2125ACA (6514A>C 6515G>A), GAA2126GGT (6517A>G 6518A>T), GAT2127CTA (6519G>C 6520A>T 6521T>A), CAT2128GAG (6522C>G 6524T>G), CCT2129TAT (6525C>T 6526C>A), GTT2130TAT (6528G>T 6529T>A), CGG2131GAT (6531C>G 6532G>A 6533G>T), AGG2132TCC (6534A>T 6535G>C 6536G>C), TTT2133ATA (6537T>A 6539T>A), TCA2134GAA (6540T>G 6541C>A), ACG2135TCT (6543A>T 6545G>T), GTT2136TTG (6546G>T 6548T>G), TTC2137GAC (6549T>G 6550T>A), CTT2138TTG (6552C>T 6554T>G), AAA2139TTA (6555A>T 6556A>T), AAC2140CAA (6558A>C 6560C>A), TGT2141ATA (6561T>A 6562G>T 6563T>A), GAA2142GAT (6566A>T), ATT2144ATA (6572T>A), ACA2146GCA (6576A>G), TAC2148GTC (6582T>G 6583A>T), GAG2150TAC (6588G>T 6590G>C), TGA2152GGG (6594T>G 6596A>G)

#### Proteins

| | | | | | | | | | |
|---|---|---|---|---|---|---|---|---|---|
| polyprotein (YP_001976147.1) | 1 | 2152 | 100% | 9863 | 66.5% | 2135 (98.7%) | 1360 (62.9%) | 11/17/0/0 | 0 |

| | Begin | End | Coverage | Score | Concordance | Matches | Identities | I/D/M/F* | Stop Codons |
|---|---|---|---|---|---|---|---|---|---|
| NT | 25 | 6747 | 93.0% | 3297 | 25.1% | 6658 (98.4%) | 4218 (62.4%) | 41/65 | |

Protein mutations:

F7W (160T>G 161C>G), K9_A16del (165_188delAAGGTTGCTGGTGTTGATGACGCC), S17P (189A>C 190G>C), S18P (192T>C), S19A (195T>G), T21S (201A>T 203C>A), T22P (204A>C), G26A (217G>C 218T>A), L28V (222C>G), N30S (228A>T 229A>C), I32F (234A>T), K34S (240A>T 241A>C 242A>T), F35L (243T>C), A38T (252G>A 254G>T), V39L (255G>C 257T>C), I42V (264A>G), T43N (268C>A 269C>T), T44K (271C>A 272T>A), M45V (273A>G 275G>T), K48T (283A>C 284G>C), S50V (288T>G 289C>T), L53I (297C>A 299T>A), L56V (306C>G 308G>C), S59P (315T>C 317T>A), T61M (322C>T 323T>G), D62A (325A>C 326T>A), V63A (328T>C 329G>A), S64A (330T>G 332C>A), V68A (343T>C 344G>C), A69S (345G>T 347C>T), S72N (355G>A), S73A (357T>G 359A>T), L74V (360C>G 362T>A), N75E (363A>G 365T>A), K76R (367A>G), G82A (385G>C 386T>A), A83K (387G>A 388C>A), L84I (390C>A 392T>A), I85M (395A>G), D86E (398T>A), L89I (405C>A 407T>C), A91G (412C>G 413G>T), S93D (417T>G 418C>A), L95V (423T>G 425G>T), S96K (426T>A 427C>A 428C>A), S101T (441T>A), K103V (447A>G 448A>T 449G>A), M105L (453A>C 455G>T), L109M (465T>A), F110V (468T>G), E111L (471G>T 472A>T 473G>A), L113V (477T>G 479G>C), S114K (480T>A 481C>A 482C>G), N115D (483A>G 485T>C), V117F (489G>T 491T>C), K118I (493A>T 494A>C), F121Y (502T>A 503T>C), A122E (505C>A 506T>A), V124L (510G>T 512G>A), N126E (516A>G 518T>A), S128V (522A>G 523G>T), D130S (528G>T 529A>C 530T>A), V131L (531G>T), V135F (543G>T 545A>T), Y136I (546T>A 547A>T 548T>A), G137A (550G>C), I143L (567A>T 569A>G), T144A (570A>G 572A>C), L145I (573T>A 575G>T), V147L (579G>C 581G>A), F150Y (589T>A), S154V (600A>G 601G>T 602T>A), E158T (612G>A 613A>C), G159P (615G>C 616G>C), V160F (618G>T), S161N (621T>A 622C>A), T163V (627A>G 628C>T 629T>A), K165V (633A>G 634A>T 635A>T), N166T (637A>C), I167L (639A>T 641C>G), L169S (646T>C 647G>A), I170V (648A>G 650A>T), M174F (660A>T 662G>T), K186T (697A>C), A188S (702G>T 704C>T), E189G (706A>G 707A>G), R190K (708C>A 709G>A 710A>G), T192S (714A>T), M193Q (717A>C 718T>A), V194L (720G>C), S198A (732A>G 733G>C 734T>A), M199V (735A>G 737G>A), V205L (753G>T 755T>A), S206T (756T>A 758A>G), M214L (780A>T 782G>A), T216S (786A>T), I217L (789A>T 791A>G), I228V (822A>G), T248A (882A>G 884A>T), T260Q (918A>C 919C>A 920T>G), I280V (978A>G 980A>G), G282S (984G>A), T283I (988C>T 989C>T), S295N (1024G>A), E304Q (1050G>C), Q307T (1059C>A 1060A>C), D310S (1068G>T 1069A>C 1070T>A), A311E (1072C>A 1073A>G), C313A (1077T>G 1078G>C), D314E (1082T>A), A316S (1086G>T 1088A>T), T317M (1090C>T), K325P (1113A>C 1114A>C 1115G>T), T329I (1126C>T 1127C>T), N332Q (1134A>C 1136C>A), R335K (1144G>A 1145G>A), E337Q (1149G>C 1151A>G), L339V (1155C>G), M354L (1200A>T), A363T (1227G>A), E366D (1238A>T), V368L (1242G>T 1244T>G), Y369V (1245T>G 1246A>T), S372A (1254A>G 1255G>C 1256T>C), C373S (1257T>A 1259T>C), R374Q (1260A>C 1261G>A 1262G>A), M376S (1266A>T 1267T>C 1268G>T), S393A (1317T>G 1319C>T), S401C (1342C>G), R412A (1374A>G 1375G>C), I440V (1458A>G 1460C>G), M521R (1702T>G 1703G>A), E523V (1708A>T), E525D (1715G>T), S527D (1719T>G 1720C>A 1721C>T), T528P (1722A>C 1724T>A), E529N (1725G>A 1727A>T), D530A (1729A>C), G533A (1738G>C 1739A>C), R536V (1746A>G 1747G>T 1748A>T), N538D (1752A>G 1754C>T), I539V (1755A>G 1757A>G), N552A (1794A>G 1795A>C), T554S (1800A>T), S556D (1806A>G 1807G>A), E560Q (1818G>C 1820G>A), Q567R (1840A>G), A570Y (1848G>T 1849C>A 1850A>T), K574S (1861A>G 1862G>C), F577Y (1870T>A), E578N (1872G>A 1874A>T), A579T (1875G>A), E580T (1878G>A 1879A>C), I583L (1887A>T 1889A>G), S584A (1890A>G 1891G>C), P585D (1893C>G 1894C>A 1895T>C), I587V (1899A>G 1901A>T), Q591N (1911C>A 1913A>T), S593A (1917T>G), A594V (1921C>T 1922A>T), D596E (1928T>A), V597M (1929G>A), I598L (1932A>C 1934A>G), A599D (1936C>A 1937G>T), V601F (1941G>T 1943G>C), D602Q (1944G>C 1946T>A), G603D (1948G>A 1949G>T), R605del (1953_1955delAGA), R607Q (1960G>A 1961C>A), T608V (1962A>G 1963C>T 1964C>T), K609L (1965A>T 1966A>T), Q610P (1969A>C 1970A>T), D611P (1971G>C 1972A>C), I614A (1980A>G 1981T>C), L615T (1983T>A 1984T>C 1985G>T), V616Y (1986G>T 1987T>A 1988G>T), P617E (1989C>G 1990C>A 1991C>G), T618Q (1992A>C 1993C>A 1994T>G), V619T (1995G>A 1996T>C 1997T>A), V620I (1998G>A 2000G>T), S621T (2002G>C 2003T>A), N623A (2007A>G 2008A>C), D624E (2012T>A), R625K (2014G>A), V626I (2016G>A 2018G>T), I627V (2019A>G 2021C>T), A629S (2025G>T 2027A>C), R630Q (2028A>C 2029G>A 2030A>G), L645M (2073T>A), D646N (2076G>A), P647K (2079C>A 2080C>A 2081T>A), E648F (2082G>T 2083A>T 2084G>T), L650F (2088C>T), H655L (2104A>T 2105T>C), H656S (2106C>T 2107A>C 2108T>C), M660I (2120G>C), S663V (2127T>G 2128C>T 2129T>G), D665S (2133G>T 2134A>C), D666S (2136G>T 2137A>C), E667D (2141A>T), H668A (2142C>G 2143A>C 2144T>C), E670Q (2148G>C), I671N (2152T>A), N673E (2157A>G 2159C>A), K674M (2161A>T 2162A>G), R676V (2166A>G 2167G>T), V677N (2169G>A 2170T>A 2171C>T), N678D (2172A>G 2174C>T), M679L (2175A>T), Y681V (2181T>G 2182A>T 2183T>C), R682K (2185G>A), D685S (2193G>A 2194A>G), A686P (2196G>C), E687Q (2199G>C 2201A>G), R690A (2208C>G 2209G>C 2210A>T), Q694R (2221A>G), I701V (2241A>G 2243A>C), A710S (2268G>A 2269C>G 2270A>C), S711T (2271T>A), K716E (2286A>G), K717L (2289A>T 2290A>T 2291A>G), A719D (2296C>A 2297A>T), T722R (2304A>C 2305C>G 2306A>C), E723N (2307G>A 2309G>T), M724L (2310A>C 2312G>T), R725E (2313A>G 2314G>A), T726Q (2316A>C 2317C>A 2318T>A), F728K (2322T>A 2323T>A 2324T>A), H730Y (2328C>T), S731K (2331T>A 2332C>A 2333T>G), C732A (2334T>G 2335G>C 2336C>G), I733L (2337A>C), D734Q (2340G>C 2342T>G), V737A (2350T>C 2351T>A), I741T (2362T>C 2363A>G), F749R (2385T>A 2386T>G 2387T>A), N763R (2428A>G 2429C>G), S766A (2436T>G), S766_K767insG (2438_2439insGGT), K767T (2440A>C 2441A>T), E768D (2444G>C), P769K (2445C>A 2446C>A 2447A>G), E771R (2451G>A 2452A>G), S772N (2454T>A 2455C>A 2456A>T), K775I (2464A>T 2465A>C), I776E (2466A>G 2467T>A 2468T>A), Y778M (2472T>A 2473A>T 2474C>G), H779I (2475C>A 2476A>T), K780G (2478A>G 2479A>G 2480A>T), V782K (2484G>A 2485T>A 2486T>G), E783D (2489A>C), Y784K (2490T>A 2492T>A), I785M (2495C>G), K786I (2497A>T 2498G>C), D787Q (2499G>C 2501T>G), T788M (2503C>T 2504A>G), W790N (2508T>A 2509G>A 2510G>T), S791D (2511A>G 2512G>A), Y794F (2521A>T 2522T>C), A795I (2523G>A 2524C>T 2525A>C), R797K (2530G>A 2531A>G), I806L (2556A>T 2558A>G), M807I (2561G>A), V812T (2574G>A 2575T>C 2576T>C), F813M (2577T>A 2579T>G), G814N (2580G>A 2581G>A 2582A>T), G815A (2584G>C), I816V (2586A>G), L820I (2598T>A 2600G>T), A821T (2601G>A 2603G>T), W823L (2607T>C 2608G>T 2609G>C), Q824K (2610C>A), S825V (2613A>G 2614G>T 2615T>A), N826A (2616A>G 2617A>C 2618C>A), S827T (2619T>A), F828S (2623T>C), N830L (2628A>T 2629A>T 2630T>A), S832D (2634T>G 2635C>A 2636G>T), N852K (2696T>G), K853T (2698A>C 2699G>T), K853_L854insH (2699_2700insCAT), L854F (2700C>T), R855A (2703A>G 2704G>C), N856K (2708T>A), L857S (2709C>T 2710T>C 2711T>C), V859L (2715G>C 2717T>A), S862A (2724A>G 2725G>C 2726T>G), S863A (2727T>G 2729T>A), Q864D (2730C>G 2732A>C), S865N (2733T>A 2734C>A 2735A>C), L866N (2736T>A 2737T>A 2738G>C), A867H (2739G>C 2740C>A 2741A>T), Q868V (2742C>G 2743A>T 2744A>G), D869_W870insR (2747_2748insCGA), A872E (2755C>A 2756T>A), E873Q (2757G>C 2759A>G), Q879E (2775C>G 2777G>A), K882H (2784A>C 2786G>C), L890I (2808C>A), G891A (2812G>C), M907L (2859A>C 2861G>C), H914R (2880C>A 2881A>G 2882C>A), A916M (2886G>A 2887C>T 2888T>G), T923S (2907A>T 2909A>C), A927E (2920C>A 2921C>G), N928H (2922A>C), R930I (2928C>A 2929G>T 2930A>T), S931C (2932C>G), K932N (2936A>T), Y935F (2944A>T 2945C>T), R938T (2953G>C 2954A>C), E939Q (2955G>C), I942L (2964A>T 2966T>G), I948V (2982A>G 2984C>G), P961S (3021C>T 3023T>A), E972D (3056G>T), F975Y (3064T>A), T976S (3066A>T 3068A>C), Q980M (3078C>A 3079A>T 3080A>G), C984V (3090T>G 3091G>T 3092T>G), T990K (3109C>A 3110A>G), R993K (3117C>A 3118G>A 3119T>G), L994K (3120C>A 3121T>A 3122T>A), G995S (3123G>T 3124G>C), F999V (3135T>G 3137T>G), M1002Y (3144A>T 3145T>A 3146G>T), V1012E (3175T>A 3176T>A), E1015T (3183G>A 3184A>C), E1017G (3190A>G), N1019D (3195A>G 3197C>T), S1021N (3201T>A 3202C>A 3203C>T), S1022A (3204A>G 3205G>C 3206T>A), K1024A (3210A>G 3211A>C 3212G>T), I1025L (3213A>C), C1026S (3216T>A), Q1029V (3225C>G 3226A>T 3227A>T), S1030A (3228T>G 3230T>A), G1037S (3249G>A 3251G>T), S1050T (3289G>C), V1057C (3309G>T 3310T>G 3311G>C), G1068D (3343G>A 3344C>T), V1069T (3345G>A 3346T>C 3347A>C), R1073K (3357C>A 3358G>A 3359T>G), F1075Y (3364T>A 3365T>C), I1076V (3366A>G 3368C>G), S1077A (3369T>G), I1083V (3387A>G 3389C>G), Y1084F (3391A>T 3392C>T), E1086D (3398A>T), V1096I (3426G>A 3428C>T), A1098V (3433C>T 3434A>G), D1104E (3452T>A), I1105L (3453A>C 3455T>G), S1108H (3462T>C 3463C>A 3464A>C), I1109L (3465A>C 3467T>G), R1112K (3475G>A 3476A>G), T1114E (3480A>G 3481C>A), Q1115R (3483C>A 3484A>G 3485G>A), L1116F (3486C>T 3488T>C), N1120K (3500C>A), Y1123A (3507T>G 3508A>C 3509C>T), Y1124F (3511A>T), Q1130L (3528C>T 3529A>T), L1132M (3534T>A), R1143K (3567C>A 3568G>A 3569G>A), K1144R (3570A>C 3571A>G 3572A>C), Q1146P (3577A>C 3578A>C), P1153D (3597C>G 3598C>A), E1155Q (3603G>C), T1162S (3624A>T 3626A>T), V1163A (3628T>C 3629T>C), H1164K (3630C>A 3632T>A), D1169E (3647T>A), M1179E (3675A>G 3676T>A), C1186S (3697G>C 3698T>C), G1187N (3699G>A 3700G>A 3701A>C), D1189E (3707T>G), D1190E (3710T>G), D1191P (3711G>C 3712A>C 3713C>A), N1199G (3735A>G 3736A>G), C1202A (3744T>G 3745G>C 3746C>A), L1204I (3750C>A 3752G>T), I1207V (3759A>G 3761C>G), M1208L (3762A>C 3764G>T), R1209K (3765C>A 3766G>A 3767T>G), S1210M (3769G>T 3770C>G), K1212R (3774A>C 3775A>G 3776G>A), I1213V (3777A>G 3779C>G), R1217K (3789C>A 3790G>A 3791T>A), C1219A (3795T>G 3796G>C 3797T>A), E1220I (3798G>A 3799A>T 3800A>T), D1222N (3804G>A), C1233F (3838G>T 3839T>C), Y1236F (3847A>T), G1238A (3853G>C 3854C>A), M1252L (3894A>T), G1253E (3898G>A 3899G>A), M1254L (3900A>C 3902G>T), M1256S (3906A>T 3907T>C 3908G>T), Q1263E (3927C>G), S1264A (3930T>G), T1265Q (3933A>C 3934C>A 3935T>A), G1266P (3936G>C 3937G>C 3938C>A), E1267D (3941A>C), G1268S (3942G>T 3943G>C 3944A>G), E1269Q (3945G>C), E1272L (3954G>C 3955A>T 3956A>T), D1276N (3966G>A), E1281D (3983A>T), V1288S (4002G>T 4003T>C), I1290V (4008A>G 4010T>A), K1292Q (4014A>C), I1294T (4021T>C 4022C>A), H1295F (4023C>T 4024A>T), T1344C (4170A>T 4171C>G 4172A>T), L1348A (4182T>G 4183T>C 4184G>A), A1351V (4192C>T 4193T>A), I1352L (4194A>C 4196A>C), D1359N (4215G>A), S1363H (4227T>C 4228C>A), L1367V (4239T>G), N1370S (4249A>G), G1371S (4251G>A 4253G>C), A1374S (4260G>T 4262T>A), V1378E (4273T>A), V1393I (4317G>A 4319T>C), E1405A (4354A>C 4355A>T), Y1406F (4357A>T 4358C>T), S1409D (4365T>G 4366C>A 4367A>T), K1410E (4368A>G), D1415E (4385T>G), A1421T (4401G>A), D1435N (4443G>A 4445T>C), V1437L (4449G>T), I1476V (4566A>G 4568T>C), A1477K (4569G>A 4570C>A 4571T>A), R1479K (4576G>A 4577G>A), A1480S (4578G>T), I1481V (4581A>G 4583A>G), V1483I (4587G>A), S1486N (4597G>A), L1506M (4656T>A 4658A>G), Q1509R (4665C>A 4666A>G), S1512D (4674T>G 4675C>A), N1515S (4683A>T 4684A>C 4685T>C), H1518R (4692C>A 4693A>G 4694T>A), L1522A (4704C>G 4705T>C 4706G>T), I1526V (4716A>G), N1538E (4752A>G 4754T>G), E1539K (4755G>A), V1540T (4758G>A 4759T>C), Q1541K (4761C>A), Q1543L (4768A>T 4769A>G), H1563Q (4829C>A), I1572V (4854A>G), M1576I (4868G>T), P1583V (4887C>G 4888C>T 4889A>G), T1588V (4902A>G 4903C>T 4904T>A), M1592Y (4914A>T 4915T>A 4916G>T), V1595I (4923G>A 4925G>T), R1607Q (4960G>A 4961T>G), V1608A (4963T>C 4964G>C), A1609T (4965G>A), D1611E (4973C>A), K1612S (4975A>G 4976G>T), L1613F (4977C>T), L1615N (4983T>A 4984T>A 4985A>T), Q1618L (4993A>T 4994G>T), T1619G (4995A>G 4996C>G 4997G>A), D1622K (5004G>A 5006T>G), E1623Q (5007G>C 5009A>G), G1625N (5013G>A 5014G>A 5015T>C), F1626L (5016T>C), R1627P (5019A>C 5020G>C 5021A>C), A1628I (5022G>A 5023C>T 5024T>C), V1629M (5025G>A), F1631M (5031T>A 5033T>G), S1633T (5037T>A 5039C>T), L1635Y (5044T>A 5045G>T), V1637L (5049G>C 5051G>T), Q1639S (5055C>A 5056A>G 5057G>T), E1640Q (5058G>C), I1642L (5064A>T 5066T>G), A1643T (5067G>A), K1644E (5070A>G), S1649E (5085T>G 5086C>A 5087A>G), A1651I (5091G>A 5092C>T 5093C>T), L1652G (5094C>G 5095T>G), T1653V (5097A>G 5098C>T 5099A>T), I1654L (5100A>T 5102T>G), C1655M (5103T>A 5104G>T 5105T>G), M1657L (5109A>C 5111G>A), A1658S (5112G>T), K1659D (5115A>G 5117A>T), A1661K (5121G>A 5122C>A 5123T>A), I1662N (5125T>A), S1663T (5127T>A 5129C>A), D1668P (5142G>C 5143A>C 5144C>T), T1674del (5160_5162delACA), R1678E (5172C>G 5173G>A 5174T>G), D1679G (5176A>G), L1680V (5178C>G 5180G>C), N1681H (5181A>C), W1682E (5184T>G 5185G>A 5186G>A), N1686D (5196A>G 5198C>T), I1687V (5199A>G 5201T>A), G1688F (5202G>T 5203G>T), A1696S (5226G>T 5228C>A), H1697N (5229C>A), Y1698Q (5232T>C 5234C>A), T1699Q (5235A>C 5236C>A 5237G>A), D1701S (5241G>T 5242A>C 5243T>A), E1702S (5244G>T 5245A>C), Y1704N (5250T>A), I1706L (5256A>C 5258T>G), K1713R (5277A>C 5278A>G 5279G>C), K1714R (5281A>G 5282G>A), S1715T (5283T>A 5285C>A), N1716T (5287A>C 5288T>A), M1731I (5333G>A), M1736L (5346A>T), S1737A (5349T>G), M1740Q (5358A>C 5359T>A), A1742H (5364G>C 5365C>A 5366C>T), K1744R (5370A>C 5371A>G 5372G>C), N1745K (5375T>G), V1746N (5376G>A 5377T>A 5378G>T), A1748E (5383C>A), G1749A (5386G>C 5387A>T), H1751D (5391C>G), V1752G (5395T>G), N1755S (5403A>T 5404A>C), V1757T (5409G>A 5410T>C 5411T>C), V1766I (5436G>A 5438C>A), G1767A (5440G>C 5441C>T), M1774F (5460A>T 5462G>T), R1779H (5475A>C 5476G>A 5477G>T), C1782R (5484T>A 5486T>G), V1783A (5488T>C 5489G>C), E1784T (5490G>A 5491A>C 5492G>A), I1786L (5496A>C 5498A>T), I1793M (5519T>G), G1794A (5521G>C 5522A>T), N1795G (5523A>G 5524A>G 5525T>G), V1807A (5560T>C), F1810L (5570T>G), Q1812K (5574C>A 5576A>G), P1815E (5583C>G 5584C>A 5585T>A), H1816D (5586C>G 5588T>C), H1816_G1817insT (5588_5589insACC), G1817S (5589G>T 5590G>C), D1818H (5592G>C), V1820L (5598G>T), A1821G (5602C>G 5603T>A), A1825T (5613G>A), S1826Y (5617C>A 5618G>T), L1827G (5619C>G 5620T>G), H1828del (5622_5624delCAC), S1829T (5626G>C 5627T>C), S1830T (5629G>C), M1832N (5635T>A 5636G>C), Q1834H (5642A>T), Q1834_G1835insHS (5642_5643insCACAGT), G1835A (5644G>C 5645G>A), E1837S (5649G>T 5650A>C), L1838V (5652T>G 5654G>T), N1840L (5658A>C 5659A>T 5660C>T), I1844V (5670A>G 5672A>T), H1845P (5674A>C), G1846S (5676G>T 5677G>C 5678T>C), A1849S (5685G>T 5687A>T), K1850E (5688A>G), Q1851K (5691C>A), T1852M (5695C>T 5696A>G), M1854A (5700A>G 5701T>C 5702G>T), M1856L (5706A>T), V1859T (5715G>A 5716T>C 5717G>A), V1860I (5718G>A 5720A>T), H1863E (5727C>G 5729T>A), G1867A (5740G>C 5741A>C), N1868Q (5742A>C 5744T>G), F1869C (5746T>G 5747T>C), Q1872K (5754C>A), D1873N (5757G>A 5759T>C), E1875K (5763G>A), S1876H (5766A>C 5767G>A), L1877A (5769T>G 5770T>C 5771G>T), H1879E (5775C>G 5777C>G), Y1880T (5778T>A 5779A>C 5780T>A), E1882S (5784G>T 5785A>C 5786G>T), Q1883V (5787C>G 5788A>T 5789G>C), G1884S (5790G>A 5792A>T), E1887T (5799G>A 5800A>C 5801G>T), I1889A (5805A>G 5806T>C 5807T>C), Q1890A (5808C>G 5809A>C), K1891R (5812A>G 5813G>A), N1892G (5814A>G 5815A>G 5816T>A), T1893V (5817A>G 5818C>T 5819A>C), L1895V (5823C>G 5825A>G), L1902F (5844C>T 5846G>C), M1904Q (5850A>C 5851T>A 5852G>A), E1905H (5853G>C 5855A>C), N1906D (5856A>G), V1907C (5859G>T 5860T>G 5861G>T), Q1908E (5862C>G), Q1909M (5865C>A 5866A>T 5867A>G), L1910I (5868C>A 5870G>A), I1911L (5871A>C 5873A>T), D1912E (5876T>G), V1914L (5880G>C 5882T>G), K1915E (5883A>G 5885G>A), L1916S (5886C>A 5887T>G), F1919V (5895T>G 5897T>A), F1920V (5898T>G 5900C>A), P1921F (5901C>T 5902C>T 5903A>C), K1922M (5905A>T 5906A>G), E1923G (5908A>G 5909A>G), T1924S (5911C>G 5912T>C), T1925V (5913A>G 5914C>T 5915C>T), I1926K (5917T>A 5918T>A), Q1927K (5919C>A 5921G>A), K1928Q (5922A>C 5924G>A), L1929M (5925C>A 5927C>G), K1930L (5928A>T 5929A>T 5930A>G), N1931S (5931A>T 5932A>C 5933T>C), V1933M (5937G>A 5939T>G), D1934N (5940G>A), A1935Q (5943G>C 5944C>A 5945T>A), E1936Q (5946G>C 5948A>G), C1937M (5949T>A 5950G>T 5951T>G), L1938S (5952C>T 5953T>C), T1939K (5956C>A), F1940T (5958T>A 5959T>C), K1941Q (5961A>C), A1942V (5965C>T 5966A>T), R1943A (5967A>G 5968G>C 5969G>C), S1944_F1945insPCCVT (5972_5973insCCATGTTGTGTCACA), R1946S (5978G>T), H1947K (5979C>A 5981C>A), V1948A (5983T>C), V1949W (5985G>T 5986T>G 5987T>G), F1950L (5988T>C 5990C>A), K1951T (5992A>C 5993A>T), K1953D (5997A>G 5999A>C), L1954I (6000C>A), L1955V (6003T>G 6005G>C), S1956M (6007G>T 6008T>G), S1957T (6009T>A 6011C>A), T1958K (6013C>A), W1959G (6015T>G 6017G>T), M1961K (6022T>A), C1962P (6024T>C 6025G>C 6026T>A), E1964del (6030_6032delGAA), V1967_R1970del (6039_6050delGTTGCTTCAAGG), S1971P (6051T>C), S1974T (6061G>C 6062T>A), L1975V (6063C>G 6065A>G), P1977L (6070C>T), L1980P (6078T>C 6079T>C 6080G>T), S1983L (6087T>C 6088C>T 6089C>A), L1985V (6093T>G), K1986R (6097A>G), L1988M (6102T>A 6104A>G), E1990R (6108G>C 6109A>G 6110A>C), R1991N (6112G>A 6113A>T), H1992S (6114C>A 6115A>G 6116C>T), T1993G (6117A>G 6118C>G), R1994K (6121G>A 6122A>G), M1996Y (6126A>T 6127T>A

6128G>C), Q1999T (6135C>A 6136A>C 6137G>A), S2000T (6139G>C 6140T>A), M2001L (6141A>C), L2005F (6153C>T 6155G>T), K2006R (6157A>G), I2007M (6161A>G), L2009M (6165C>A 6167A>G), L2010C (6169T>G 6170A>T), N2011Q (6171A>C 6173C>G), F2012E (6174T>G 6175T>A 6176C>A), Q2013G (6177C>G 6178A>G 6179A>G), V2014Y (6180G>T 6181T>A 6182A>T), I2015L (6183A>T), T2016D (6186A>G 6187C>A 6188A>T), S2017K (6190G>A 6191T>A), E2018A (6193A>C 6194G>C), L2020V (6198T>G 6200A>G), K2021D (6201A>G 6203G>C), K2022C (6204A>T 6205A>G 6206G>C), E2024S (6210G>A 6211A>G 6212A>T), D2025E (6215T>G), F2027M (6219T>A 6221T>G), Q2028G (6222C>G 6223A>G 6224G>C), E2029V (6226A>T 6227G>T), K2030C (6228A>T 6229A>G 6230G>T), I2031V (6231A>G 6233A>G), S2032P (6234T>C 6236C>G), T2033E (6237A>G 6238C>A 6239C>A), T2034R (6240A>C 6241C>G), L2035I (6243C>A 6245A>C), I2037V (6249A>G 6251C>G), K2038R (6252A>C 6253A>G 6254A>C), V2039T (6255G>A 6256T>C), F2040I (6258T>A 6260C>T), L2041A (6261T>G 6262T>C 6263G>T), E2044S (6270G>T 6271A>C 6272G>C), E2045T (6273G>A 6274A>C), Y2047L (6280A>T 6281T>G), Q2048N (6282C>A 6284G>C), T2049E (6285A>G 6286C>A 6287T>G), E2050N (6288G>A 6290G>T), C2052T (6294T>A 6295G>C), I2053V (6297A>G 6299A>G), D2054Y (6300G>T 6302T>C), S2055T (6303T>A 6305T>A), Q2056D (6306C>G 6308G>T), V2057G (6310T>G), Q2059F (6315C>T 6316A>T 6317G>T), R2060D (6318A>G 6319G>A 6320G>T), I2061del (6321_6323delATA), N2062H (6324A>C 6326C>T), S2063F (6328C>T), Q2064K (6330C>A 6332G>A), K2065A (6333A>G 6334A>C), F2066V (6336T>G 6338C>G), R2067D (6339A>G 6340G>A 6341A>C), V2068F (6342G>T 6344G>C), Q2069K (6345C>A), E2070P (6348G>C 6349A>C 6350A>T), Y2074S (6361A>C 6362T>C), L2075W (6364T>G), H2076E (6366C>G 6368T>A), P2077I (6369C>A 6370C>T), E2078N (6372G>A 6374A>T), G2079D (6376G>A 6377A>T), I2080A (6378A>G 6379T>C), V2085A (6394T>C), I2088V (6402A>G 6404A>G), I2089L (6405A>T 6407A>G), S2091E (6411T>G 6412C>A 6413T>A), E2094R (6420G>A 6421A>G), S2095H (6423T>C 6424C>A), Y2096W (6427A>G 6428T>G), R2098K (6432C>A 6433G>A 6434A>G), D2099S (6435G>T 6436A>C 6437T>A), S2100Q (6438A>C 6439G>A 6440T>A), N2101S (6442A>G), S2102K (6444T>A 6445C>A 6446C>G), Y2103F (6448A>T 6449T>C), S2104T (6450T>A), V2105F (6453G>T 6455G>T), S2106A (6456T>G 6458A>C), T2107N (6460C>A), P2108E (6462C>G 6463C>A 6464C>A), I2109V (6465A>G 6467C>T), K2110E (6468A>G), L2111I (6471C>A 6473G>A), G2112S (6474G>T 6475G>C 6476G>A), C2113N (6477T>A 6478G>A 6479T>C), F2114L (6480T>C 6482C>G), I2115L (6483A>C), F2116L (6488C>A), L2117A (6489C>G 6490T>C), V2119Y (6495G>T 6496T>A 6497G>T), D2121E (6503T>A), S2122N (6505G>A), Q2123K (6507C>A 6509A>G), G2124L (6510G>C 6511G>T), K2125T (6514A>C 6515G>A), E2126G (6517A>G 6518A>T), D2127L (6519G>C 6520A>T 6521T>A), H2128E (6522C>G 6524T>G), P2129Y (6525C>T 6526C>A), V2130Y (6528G>T 6529T>A), R2131D (6531C>G 6532G>A 6533G>T), R2132S (6534A>T 6535G>C 6536G>C), F2133I (6537T>A 6539T>A), S2134E (6540T>G 6541C>A), T2135S (6543A>T 6545G>T), V2136L (6546G>T 6548T>G), F2137D (6549T>G 6550T>A), K2139L (6555A>T 6556A>T), N2140Q (6558A>C 6560C>A), C2141I (6561T>A 6562G>T 6563T>A), E2142D (6566A>T), T2146A (6576A>G), Y2148V (6582T>G 6583A>T), E2150Y (6588G>T 6590G>C), *2152G (6594T>G 6596A>G)

| | Begin | End | Coverage | Score | Concordance | Matches | Identities | I/D/M/F* | Stop Codons |
|---|---|---|---|---|---|---|---|---|---|
| **NT** | **25** | **6747** | **93.0%** | **3297** | **25.1%** | **6658 (98.4%)** | **4218 (62.4%)** | **41/65** | |

GCT2GCA (146T>A), TTC7TGG (160T>G 161C>G), TCC8AGT (162T>A 163C>G 164C>T), AAG9_GCC16del (165_188delAAGGTTGCTGGTGTTGATGACGCC), AGT17CCT (189A>C 190G>C), TCT18CCT (192T>C), TCT19GCT (195T>G), TCC20TCT (200C>T), ACC21TCA (201A>T 203C>A), ACT22CCT (204A>C), GGT26GCA (217G>C 218T>A), CTT28GTT (222C>G), TCA29TCT (227A>T), AAT30TCT (228A>T 229A>C), ATT32TTT (234A>T), TCC33TCT (239C>T), AAA34TCT (240A>T 241A>C 242A>T), TTT35CTT (243T>C), AAA37AAG (251A>G), GCG38ACT (252G>A 254G>T), GTT39CTC (255G>C 257T>C), AGT40TCC (258A>T 259G>C 260T>C), ATT42GTT (264A>G), ACC43AAT (268C>A 269C>T), ACT44AAA (271C>A 272T>A), ATG45GTT (273A>G 275G>T), AAG48ACC (283A>C 284G>C), TCA50GTA (288T>G 289C>T), CAT52CAC (296T>C), CTT53ATA (297C>A 299T>A), CTG56GTC (306C>G 308G>C), CCA58CCG (314A>G), TCT59CCA (315T>C 317T>A), GTG60GTT (320G>T), ACT61ATG (322C>T 323T>G), GAT62GCA (325A>C 326T>A), GTG63GCA (328T>C 329G>A), TCC64GCA (330T>G 332C>A), TCT66TCA (338T>A), GTG68GCC (343T>C 344G>C), GCC69TCT (345G>T 347C>T), ACT70ACA (350T>A), TGT71TGC (353T>C), AGT72AAT (355G>A), TCA73GCT (357T>G 359A>T), CTT74GTA (360C>G 362T>A), AAT75GAA (363A>G 365T>A), AAA76AGA (367A>G), TTG78TTA (374G>A), GAC79GAT (377C>T), AAA80AAG (380A>G), GGT82GCA (385G>C 386T>A), GCA83AAA (387G>A 388C>A), CTT84ATA (390C>A 392T>A), ATA85ATG (395A>G), GAT86GAA (398T>A), CCA87CCC (401A>C), TTT88TTC (404T>C), CTT89ATC (405C>A 407T>C), AAA90AAG (410A>G), GCG91GGT (412C>G 413G>T), TAC92TAT (416C>T), TCT93GAT (417T>G 418C>A), TTC94TTT (422C>T), TTG95GTT (423T>G 425G>T), TCC96AAA (426T>A 427C>A 428C>A), ACA97ACT (431A>T), TCT101ACT (441T>A), ATC102ATA (446C>A), AAG103GTA (447A>G 448A>T 449G>A), ATG105CTT (453A>C 455G>T), GTT106GTA (458T>A), TTA107CTT (459T>C 461A>T), TTG109ATG (465T>A), TTT110GTT (468T>G), GAG111TTA (471G>T 472A>T 473G>A), TTG113GTC (477T>G 479G>C), TCC114AAG (480T>A 481C>A 482C>G), AAT115GAC (483A>G 485T>C), GTT117TTC (489G>T 491T>C), AAA118ATC (493A>T 494A>C), TTA119CTA (495T>C), TTT121TAC (502T>A 503T>C), GCT122GAA (505C>A 506T>A), GTG124TTA (510G>T 512G>A), AAT126GAA (516A>G 518T>A), AGT128GTT (522A>G 523G>T), GAA129GAG (527A>G), GAT130TCA (528G>T 529A>C 530T>A), GTA131TTA (531G>T), GTT134GTG (542T>G), GTA135TTT (543G>T 545A>T), TAT136ATA (546T>A 547A>T 548T>A), GGT137GCT (550G>C), CTT139TTA (555C>T 557T>A), GTA140GTT (560A>T), ATA143TTG (567A>T 569A>G), ACA144GCC (570A>G 572A>C), TTG145ATT (573T>A 575G>T), TTA146TTG (578A>G), GTG147CTA (579G>C 581G>A), CTT148TTA (582C>T 584T>A), CTA149CTG (587A>G), TTT150TAT (589T>A), GTT151GTA (593T>A), TGT152TGC (596T>C), AGT154GTA (600A>G 601G>T 602T>A), GTT157GTA (611T>A), GAG158ACG (612G>A 613A>C), GGT159CCT (615G>C 616G>C), GTT160TTT (618G>T), TCT161AAT (621T>A 622C>A), AAC162AAT (626C>T), ACT163GTA (627A>G 628C>T 629T>A), GTC164GTT (632C>T), AAA165GTT (633A>G 634A>T 635A>T), AAT166ACT (637A>C), ATC167TTG (639A>T 641C>G), TTG169TCA (646T>C 647G>A), ATA170GTT (648A>G 650A>T), GTC171GTT (653C>T), AAC173AAT (659C>T), ATG174TTT (660A>T 662G>T), TTT175TTC (665T>C), TCA176TCT (668A>T), AGT177TCA (669A>T 670G>C 671T>A), CTA178TTG (672C>T 674A>G), TAC179TAT (677C>T), AAA186ACA (697A>C), GCC188TCT (702G>T 704C>T), GAA189GGG (706A>G 707A>G), CGA190AAG (708C>A 709G>A 710A>G), TTC191TTT (713C>T), ACC192TCC (714A>T), ATG193CAG (717A>C 718T>A), GTG194CTG (720G>C), GCC195GCG (725C>G), GCT197GCA (731T>A), AGT198GCA (732A>G 733G>C 734T>A), ATG199GTA (735A>G 737G>A), GGG202GGT (746G>T), TCA204AGT (750T>A 751C>G 752A>T), GTT205TTA (753G>T 755T>A), TCA206ACG (756T>A 758A>G), ACG210ACT (770G>T), TCT211AGC (771T>A 772C>G 773T>C), CAG212CAA (776G>A), TTG213CTT (777T>C 779G>T), ATG214TTA (780A>T 782G>A), ACA216TCA (786A>T), ATA217TTG (789A>T 791A>G), GGG221GGT (803G>T), ATT222ATC (806T>C), TCG223TCA (809G>A), ACG224ACT (812G>T), TTG225TTA (815G>A), GTG226GTA (818G>A), TTC227TTT (821C>T), ATT228GTT (822A>G), GCA229GCT (827A>T), GTG230GTT (830G>T), GGT232GGG (836T>G), GGC236GGT (848C>T), TTG237CTT (849T>C 851G>T), AGC238TCA (852A>T 853G>C 854C>A), TTG241CTG (861T>C), TCC242TCA (866C>A), ATT244ATC (872T>C), CTC245TTG (873C>T 875C>G), TAT246TAC (878T>C), TCG247TCT (881G>T), ACA248GCT (882A>G 884A>T), GCT252GCG (896T>G), CAG253CAA (899G>A), CAA257CAG (911A>G), CTA258CTG (914A>G), ACT260CAG (918A>C 919C>A 920T>G), CTT261TTG (921C>T 923T>G), CGT263AGG (927C>A 929T>G), AAT264AAC (932T>C), AAA266AAG (938A>G), TGC268TGT (944C>T), ACC269ACA (947C>A), TCC270TCT (950C>T), CAG271CAA (953G>A), GCT272GCC (956T>C), GAA278GAG (974A>G), ATA280GTG (978A>G 980A>G), GTC281GTT (983C>T), GGC282AGC (984G>A), ACC283ATT (988C>T 989C>T), GGA285GGC (995A>C), TTC286TTT (998C>T), GTG290GTA (1010G>A), GCT293GCG (1019T>G), AGT295AAT (1024G>A), GCA296GCT (1028A>T), ACG297ACC (1031G>C), CTG298TTA (1032C>T 1034G>A), TCC299TCT (1037C>T), ACT300ACC (1040T>C), CTG302CTT (1046G>T), GAG304CAG (1050G>C), CAG307ACG (1059C>A 1060A>C), GAG308GAA (1064G>A), GAT310TCA (1068G>T 1069A>C 1070T>A), GCA311GAG (1072C>A 1073A>G), GTG312GTT (1076G>T), TGT313GCT (1077T>G 1078G>C), GAT314GAA (1082T>A), CCT315CCA (1085T>A), GCA316TCT (1086G>T 1088A>T), ACG317ATG (1090C>T), GAG320GAA (1100G>A), AAC321AAT (1103C>T), GCC323GCA (1109C>A), AAG325CCT (1113A>C 1114A>C 1115G>T), GCT326GCA (1118T>A), TTC327TTT (1121C>T), CCT328CCA (1124T>A), ACC329ATT (1126C>T 1127C>T), AAA330AAG (1130A>G), AAC332CAA (1134A>C 1136C>A), CTG334TTA (1140C>T 1142G>A), AGG335AAA (1144G>A 1145G>A), GAA336GAG (1148A>G), GAA337CAG (1149G>C 1151A>G), GCT338GCC (1154T>C), CTC339GTC (1155C>G), AAG340AAA (1160G>A), ATC341ATA (1163C>A), GCA346GCC (1178A>C), GCG351GCA (1193G>A), TTC353TTT (1199C>T), ATG354TTG (1200A>T), AGT359TCT (1215A>T 1216G>C), ATT362ATA (1226T>A), GCG363ACG (1227G>A), CAG364CAA (1232G>A), TTA365CTA (1233T>C), GAA366GAT (1238A>T), GTT368TTG (1242G>T 1244T>G), TAT369GTT (1245T>G 1246A>T), GAA371GAG (1253A>G), AGT372GCC (1254A>G 1255G>C 1256T>C), TGT373AGC (1257T>A 1259T>C), AGG374CAA (1260A>C 1261G>A 1262G>A), CAC375CAT (1265C>T), ATG376TCT (1266A>T 1267T>C 1268G>T), GGC377GGT (1271C>T), CAG380CAA (1280G>A), CGT382AGG (1284C>A 1286T>G), GAA384GAG (1292A>G), TTT386TTC (1298T>C), GTA388GTG (1304A>G), TTC391TTT (1313C>T), GGG392GGA (1316G>A), TCC393GCT (1317T>G 1319C>T), GGG395GGA (1325G>A), TCT399TCA (1337T>A), ACA400ACG (1340A>G), TCC401TGC (1342C>G), TTA404CTT (1350T>C 1352A>T), TTT405TTC (1355T>C), GTG409GTT (1367G>T), TTG410CTC (1368T>C 1370G>C), AGA412GCA (1374A>G 1375G>C), GAG415GAA (1385G>A), AAA417AAG (1391A>G), CTC418CTG (1394C>G), AAT419AAC (1397T>C), CGG420CGC (1400G>C), CTA421TTG (1401C>T 1403A>G), GCA423GCG (1409A>G), AAG426AAA (1418G>A), AGG427AGA (1421G>A), GCT435GCA (1445T>A), CAC436CAT (1448C>T), GCT439GCA (1457T>A), ATC440GTG (1458A>G 1460C>G), CTG441CTT (1463G>T), GAC444GAT (1472C>T), GGA446GGT (1478A>T), GCA447GCT (1481A>T), TTG448CTT (1482T>C 1484G>T), CGA449CGG (1487A>G), GCT452GCA (1496T>A), TGC455TGT (1505C>T), CAA456CAG (1508A>G), GAT457GAC (1511T>C), ATT458ATA (1514T>A), AAA459AAG (1517A>G), GAT463GAC (1529T>C), TCA466TCT (1538A>T), ACA467ACC (1541A>C), CAA468CAG (1544A>G), CCA469CCT (1547A>T), CCC471CCT (1553C>T), TTA472CTT (1554T>C 1556A>T), GCC475GCT (1565C>T), GTT477GTC (1571T>C), GAC479GAT (1577C>T), GGC481GGA (1583C>A), CAT483CAC (1589T>C), TTC484TTT (1592C>T), ACC485ACA (1595C>A), TCC486TCA (1598C>A), TAT488TAC (1604T>C), ATA489ATC (1607A>C), ACA492ACC (1616A>C), CTG495TTG (1623C>T), ATC496ATA (1628C>A), TCA497TCT (1631A>T), CCT499CCA (1637T>A), GCC500GCT (1640C>T), CAA501CAG (1643A>G), GGT503GGG (1649T>G), CTT504CTA (1652T>A), ACA505ACG (1655A>G), CCA507CCT (1661A>T), GCT509GCA (1667T>A), GAG511GAA (1673G>A), CGG512AGG (1674C>A), AGA513AGG (1679A>G), AGG514AGA (1682G>A), GAC515GAT (1685C>T), ATC516ATT (1688C>T), CTG517TTA (1689C>T 1691G>A), GTG518GTT (1694G>T), TGT520TGC (1700T>C), ATG521AGA (1702T>G 1703G>A), GAG523GTG (1708A>T), GAG525GAT (1715G>T), TCC527GAT (1719T>G 1720C>A 1721C>T), ACT528CCA (1722A>C 1724T>A), GAA529AAT (1725G>A 1727A>T), GAT530GCT (1729A>C), CCT531CCA (1733T>A), ACG532ACA (1736G>A), GGA533GCC (1738G>C 1739A>C), CAT534CAC (1742T>C), CTC535CTA (1745C>A), AGA536GTT (1746A>G 1747G>T 1748A>T), AAC538GAT (1752A>G 1754C>T), ATA539GTG (1755A>G 1757A>G), GAG541GAA (1763G>A), GCT547GCC (1781T>C), ACT549ACA (1787T>A), CAC550CAT (1790C>T), AGG551AGA (1793G>A), AAT552GCT (1794A>G 1795A>C), TTG553TTA (1799G>A), ACC554TCC (1800A>T), AGT556GAT (1806A>G 1807G>A), GAT557GAC (1811T>C), CTT558CTC (1814T>C), GAG560CAA (1818G>C 1820G>A), GTA563GTG (1829A>G), GCC564GCA (1832C>A), AAA565AAG (1835A>G), TGT566TGC (1838T>C), CAG567CGG (1840A>G), GTA568GTG (1844A>G), CAT569CAC (1847T>C), GCA570TAT (1848G>T 1849C>A 1850A>T), GAA571GAG (1853A>G), GTA572GTG (1856A>G), AAG574AGC (1861A>G 1862G>C), TTG576CTT (1866T>C 1868G>T), TTT577TAT (1870T>A), GAA578AAT (1872G>A 1874A>T), GCT579ACT (1875G>A), GAA580ACA (1878G>A 1879A>C), TCT581TCA (1883T>A), GGC582GGA (1886C>A), ATA583TTG (1887A>T 1889A>G), AGT584GCT (1890A>G 1891G>C), CCT585GAC (1893C>G 1894C>A 1895T>C), ATA587GTT (1899A>G 1901A>T), GTT590GTC (1910T>C), CAA591AAT (1911C>A 1913A>T), TCT593GCT (1917T>G), GCA594GTT (1921C>T 1922A>T), GAT596GAA (1928T>A), GTG597ATG (1929G>A), ATA598CTG (1932A>C 1934A>G), GCG599GAT (1936C>A 1937G>T), GTG601TTC (1941G>T 1943G>C), GAT602CAA (1944G>C 1946T>A), GGG603GAT (1948G>A 1949G>T), GCT604GCC (1952T>C), AGA605del (1953_1955delAGA), TTG606TTA (1958G>A), CGC607CAA (1960G>A 1961C>A), ACC608GTT (1962A>G 1963C>T 1964C>T), AAG609TTG (1965A>T 1966A>T), CAA610CCT (1969A>C 1970A>T), GAT611CCT (1971G>C 1972A>C), ATA614GCA (1980A>G 1981T>C), TTG615ACT (1983T>A 1984T>C 1985G>T), GTG616TAT (1986G>T 1987T>A 1988G>T), CCC617GAG (1989C>G 1990C>A 1991C>G), ACT618CAG (1992A>C 1993C>A 1994T>G), GTT619ACA (1995G>A 1996T>C 1997T>A), GTG620ATT (1998G>A 2000G>T), AGT621ACA (2002G>C 2003T>A), AAT623GCT (2007A>G 2008A>C), GAT624GAA (2012T>A), AGA625AAA (2014G>A), GTG626ATT (2016G>A 2018G>T), ATC627GTT (2019A>G 2021C>T), TAT628TAC (2024T>C), GCA629TCC (2025G>T 2027A>C), AGA630CAG (2028A>C 2029G>A 2030A>G), GAG631GAA (2033G>A), ACG633ACA (2039G>A), GTA634GTT (2042A>T), GCA636GCC (2048A>C), TTG637CTC (2049T>C 2051G>C), GCA640GCT (2060A>T), CAG642CAA (2066G>A), GGC643GGG (2069C>G), TTG645ATG (2073T>A), GAC646AAC (2076G>A), CCT647AAA (2079C>A 2080C>A 2081T>A), GAG648TTT (2082G>T 2083A>T 2084G>T), GAA649GAG (2087A>G), CTT650TTT (2088C>T), TTT651TTC (2093T>C), CCT652CCA (2096T>A), GAC654GAT (2102C>T), CAT655CTC (2104A>T 2105T>C), CAT656TCC (2106C>T 2107A>C 2108T>C), GCC659GCT (2117C>T), ATG660ATC (2120G>C), TCT663GTG (2127T>G 2128C>T 2129T>G), CTT664CTA (2132T>A), GAT665TCT (2133G>T 2134A>C), GAT666TCT (2136G>T 2137A>C), GAA667GAT (2141A>T), CAT668GCC (2142C>G 2143A>C 2144T>C), GAG670CAG (2148G>C), ATT671AAT (2152T>A), TTC672TTT (2156C>T), AAC673GAA (2157A>G 2159C>A), AAA674ATG (2161A>T 2162A>G), AGA676GTA (2166A>G 2167G>T), GTC677AAT (2169G>A 2170T>A 2171C>T), AAC678GAT (2172A>G 2174C>T), ATG679TTG (2175A>T), TAT681GTC (2181T>G 2182A>T 2183T>C), AGA682AAA (2185G>A), GAT685AGT (2193G>A 2194A>G), GCT686CCT (2196G>C), GAA687CAG (2199G>C 2201A>G), CGA690GCT (2208C>G 2209G>C 2210A>T), GTA693GTG (2219A>G), CAA694CGA (2221A>G), GAC699GAT (2237C>T), ATA701GTC (2241A>G 2243A>C), CAC703CAT (2249C>T), TTT704TTC (2252T>C), GCA710AGC (2268G>A 2269C>G 2270A>C), TCA711ACA (2271T>A), ATC713ATT (2279C>T), GAG715GAA (2285G>A), AAG716GAG (2286A>G), AAA717TTG (2289A>T 2290A>T 2291A>G), CTA718CTG (2294A>G), GCA719GAT (2296C>A 2297A>T), GTG720GTC (2300G>C), CAG721CAA (2303G>A), ACA722CGC (2304A>C 2305C>G 2306A>C), GAG723AAT (2307G>A 2309G>T), ATG724CTT (2310A>C 2312G>T), AGG725GAG (2313A>G 2314G>A), ACT726CAA (2316A>C 2317C>A 2318T>A), GGA727GGG (2321A>G), TTT728AAA (2322T>A 2323T>A 2324T>A), GCA729GCG (2327A>G), CAT730TAT (2328C>T), TCT731AAG (2331T>A 2332C>A 2333T>G), TGC732GCG (2334T>G 2335G>C 2336C>G), ATT733CTT (2337A>C), GAT734CAG (2340G>C 2342T>G), GCA735GCT (2345A>T), GTT737GCA (2350T>C 2351T>A), GAC738GAT (2354C>T), ACC739ACA (2357C>A), CTC740CTT (2360C>T), ATA741ACG (2362T>C 2363A>G), ATC743ATT (2369C>T), GAG744GAA (2372G>A), CCA747CCG (2381A>G), TTT749AGA (2385T>A 2386T>G 2387T>A), GTT750GTA (2390T>A), TTT752TTC (2396T>C), TTA753CTG (2397T>C 2399A>G), ACA755ACT (2405A>T), GCA756GCT (2408A>T), AGA759CGC (2415A>C 2417A>C), TGC762TGT (2426C>T), AAC763AGG (2428A>G 2429C>G), GTA765GTT (2435A>T), TCA766GCA (2436T>G), TCA766_AAA767insGGT (2438_2439insGGT), AAA767ACT (2440A>C 2441A>T), GAG768GAC (2444G>C), CCA769AAG (2445C>A 2446C>A 2447A>G), GAA771AGA (2451G>A 2452A>G), TCA772AAT (2454T>A 2455C>A 2456A>T), ATC774ATT (2462C>T), AAA775ATC (2464A>T 2465A>C), ATT776GAA (2466A>G 2467T>A 2468T>A), TAC778ATG (2472T>A 2473A>T 2474C>G), CAT779ATT (2475C>A 2476A>T), AAA780GGT (2478A>G 2479A>G 2480A>T), GTT782AAG (2484G>A 2485T>A 2486T>G), GAA783GAC (2489A>C), TAT784AAA (2490T>A 2492T>A), ATC785ATG (2495C>G), AAG786ATC (2497A>T 2498G>C), GAT787CAG (2499G>C 2501T>G), ACA788ATG (2503C>T 2504A>G), TGG790AAT (2508T>A 2509G>A 2510G>T), AGC791GAC (2511A>G 2512G>A), CTT792TTG (2514C>T 2516T>G), TAT794TTC (2521A>T 2522T>C), GCA795ATC (2523G>A 2524C>T 2525A>C), CTG796TTG (2526C>T), AGA797AAG (2530G>A 2531A>G), CTC799CTT (2537C>T), GCA802GCC (2546A>C), GGT803GGA (2549T>A), ATA806TTG (2556A>T 2558A>G), ATG807ATA (2561G>A), GCT808GCA (2564T>A), GGA811GGC (2573A>C), GTT812ACC (2574G>A 2575T>C 2576T>C), TTT813ATG (2577T>A 2579T>G), GGA814AAT (2580G>A 2581G>A 2582A>T), GGC815GCC (2584G>C), ATT816GTT (2586A>G), ACA817ACT (2591A>T), GCA818GCT (2594A>T), TTT819TTC (2597T>C), TTG820ATT (2598T>A 2600G>T), GCG821ACT (2601G>A 2603G>T), TGG823CTC (2607T>C 2608G>T 2609G>C), CAG824AAG (2610C>A), AGT825GTA (2613A>G 2614G>T 2615T>A), AAC826GCA (2616A>G 2617A>C 2618C>A), TCT827ACT (2619T>A), TTC828TCC (2623T>C), CCC829CCT (2627C>T), AAT830TTA (2628A>T 2629A>T 2630T>A), GCA831GCT (2633A>T), TCG832GAT (2634T>G 2635C>A 2636G>T), GGA835GGG (2645A>G), GGT836GGC (2648T>C), GTG837GTA (2651G>A), GAA840GAG (2660A>G), ATA844ATC (2672A>C), AGT846AGC (2678T>C), AAG848AAA (2684G>A), AAC849AAT (2687C>T), AAG850AAA (2690G>A), GGT851GGG (2693T>G), AAT852AAG (2696T>G), AAG853ACT (2698A>C 2699G>T), AAG853_CTC854insCAT (2699_2700insCAT), CTC854TTC (2700C>T), AGA855GCA (2703A>G 2704G>C), AAT856AAA (2708T>A), CTT857TCC (2709C>T 2710T>C 2711T>C), CTC858TTG (2712C>T 2714C>G), GTT859CTA (2715G>C 2717T>A), GGT860GGG (2720T>G), AGT862GCG (2724A>G 2725G>C 2726T>G), TCT863GCA (2727T>G 2729T>A), CAA864GAC (2730C>G 2732A>C), TCA865AAC (2733T>A 2734C>A 2735A>C), TTG866AAC (2736T>A 2737T>A 2738G>C), GCA867CAT (2739G>C 2740C>A 2741A>T), CAA868GTG (2742C>G 2743A>T 2744A>G), GAT869GAC (2747T>C), GAT869_TGG870insCGA (2747_2748insCGA), GCT871GCA (2753T>A), GCT872GAA (2755C>A 2756T>A), GAA873CAG (2757G>C 2759A>G), GTA877GTT (2771A>T), CAG879GAA (2775C>G 2777G>A), AGC880AGT (2780C>T), CTC881CTG (2783C>G), AAG882CAC (2784A>C 2786G>C), AAT884AAC (2792T>C), GTG886GTT (2798G>T), GTG887GTT (2801G>T), TTA888TTG (2804A>G), CTT890ATT (2808C>A), GGT891GCT (2812G>C), GTG894GTC (2822G>C), TAC895TAT (2825C>T), TTT896TTC (2828T>C), GGC898GGT (2834C>T), ACC899ACG (2837C>G), TAC900TAT (2840C>T), GTG901GTT (2843G>T), ATA906ATT (2858A>T), ATG907CTC (2859A>C 2861G>C), ACC908ACT (2864C>T), GTA909GTG (2867A>G), GCT910GCA (2870T>A), GCA912GCC (2876A>C), TTC913TTT (2879C>T), CAC914AGA (2880C>A 2881A>G 2882C>A), AAT915AAC (2885T>C), GCT916ATG (2886G>A 2887C>T 2888T>G), CGA917AGG (2889C>A 2891A>G), GGT919GGC (2897T>C), CCA921CCT (2903A>T), ACA923TCC (2907A>T 2909A>C), ATC925ATT (2915C>T), GCC927GAG (2920C>A 2921C>G), AAT928CAT (2922A>C), TCT929TCC (2927T>C), CGA930ATT (2928C>A 2929G>T 2930A>T), TCT931TGT (2932C>G), AAA932AAT (2936A>T), GTT933GTG (2939T>G), CAA934CAG (2942A>G), TAC935TTT (2944A>T 2945C>T), AAC936AAT (2948C>T), AGA938ACC (2953G>C 2954A>C), GAA939CAA (2955G>C), ATT942TTG (2964A>T 2966T>G), ATT943ATA (2969T>A), GGC945GGT (2975C>T), GAC947GAT (2981C>T), ATC948GTG (2982A>G 2984C>G), CTG950TTG (2988C>T), CGC952AGG (2994C>A 2996C>G), GTT953GTA (2999T>A), GGT954GGG (3002T>G), CCA959CCC (3017A>C), AAG960AAA (3020G>A), CCT961TCA (3021C>T 3023T>A), GAC962GAT (3026C>T), ATC963ATA (3029C>A), CGT964AGA (3030C>A 3032T>A), CAC966CAT (3038C>T), CCA968CCT (3044A>T), AGA969CGC (3045A>C 3047A>C), AGG970AGA (3050G>A), GAG972GAT (3056G>T),

Codon mutations:

TGC974TGT (3062C>T), TTC975TAC (3064T>A), ACA976TCC (3066A>T 3068A>C), CAA980ATG (3078C>A 3079A>T 3080A>G), GGA981GGC (3083A>C), TGT984GTG (3090T>G 3091G>T 3092T>G), AGA985AGG (3095A>G), GTT987GTA (3101T>A), GCG988GCT (3104G>T), TCT989TCA (3107T>A), ACA990AAG (3109C>A 3110A>G), CCA992CCC (3116A>C), CGT993AAG (3117C>A 3118G>A 3119T>G), CTT994AAA (3120C>A 3121T>A 3122T>A), GGC995TCC (3123G>T 3124G>C), TTA997TTG (3131A>G), TTT999GTG (3135T>G 3137T>G), CTC1000CTT (3140C>T), AAG1001AAA (3143G>A), ATG1002TAT (3144A>T 3145T>A 3146G>T), GGA1009GGC (3167A>C), ACA1011ACC (3173A>C), GTT1012GAA (3175T>A 3176T>A), GTG1014GTC (3182G>C), GAA1015ACA (3183G>A 3184A>C), GAA1017GGA (3190A>G), CTG1018TTA (3192C>T 3194G>A), AAC1019GAT (3195A>G 3197C>T), TCC1020TCG (3200C>G), TCC1021AAT (3201T>A 3202C>A 3203C>T), AGT1022GCA (3204A>G 3205G>C 3206T>A), AAG1024GCT (3210A>G 3211A>C 3212G>T), ATT1025CTT (3213A>C), TGC1026AGC (3216T>A), TCT1027TCA (3221T>A), CAA1029GTT (3225C>G 3226A>T 3227A>T), TCT1030GCA (3228T>G 3230T>A), GAA1032GAG (3236A>G), ATA1035ATT (3245A>T), GGG1037AGT (3249G>A 3251G>T), GAA1038GAG (3254A>G), GGT1040GGA (3260T>A), GAC1041GAT (3263C>T), GGC1043GGT (3269C>T), ACG1044ACT (3272G>T), TTG1045CTT (3273T>C 3275G>T), CTA1046CTC (3278A>C), CTG1047TTA (3279C>T 3281G>A), TTA1048TTG (3284A>G), AGT1050ACT (3289G>C), GTT1051GTG (3293T>G), AAT1053AAC (3299T>C), CAA1055CAG (3305A>G), CCT1056CCG (3308T>G), GTG1057TGC (3309G>T 3310T>G 3311G>C), GGC1060GGA (3320C>A), ATC1061ATA (3323C>A), CAC1062CAT (3326C>T), TCA1065TCT (3335A>T), GGC1068GAT (3343G>A 3344C>T), GTA1069ACC (3345G>A 3346T>C 3347A>C), GCA1070GCT (3350A>T), GCT1071GCA (3353T>A), GAA1072GAG (3356A>G), CGT1073AAG (3357C>A 3358G>A 3359T>G), GGA1074GGT (3362A>T), TTT1075TAC (3364T>A 3365T>C), ATC1076GTG (3366A>G 3368C>G), TCT1077GCT (3369T>G), AAT1079AAC (3377T>C), GCT1080GCA (3380T>A), GCT1082GCA (3386T>A), ATC1083GTG (3387A>G 3389C>G), TAC1084TTT (3391A>T 3392C>T), AGG1085AGA (3395G>A), GAA1086GAT (3398A>T), CTA1088TTG (3402C>T 3404A>G), GAG1089GAA (3407G>A), CCA1092CCC (3416A>C), ACT1093ACG (3419T>G), GGG1094GGA (3422G>A), CCG1095CCC (3425G>C), GTC1096ATT (3426G>A 3428C>T), GCA1098GTG (3433C>T 3434A>G), GCA1099GCT (3437A>T), GTA1101GTG (3443A>G), GAT1104GAA (3452T>A), ATT1105CTG (3453A>C 3455T>G), CTG1106TTG (3456C>T), TCA1108CAC (3462T>C 3463C>A 3464A>C), ATT1109CTG (3465A>C 3467T>G), AGA1110AGG (3470A>G), AGC1111TCC (3471A>T 3472G>C), AGA1112AAG (3475G>A 3476A>G), GAA1113GAG (3479A>G), ACA1114GAA (3480A>G 3481C>A), CAG1115AGA (3483C>A 3484A>G 3485G>A), CTT1116TTC (3486C>T 3488T>C), GAA1119GAG (3497A>G), AAC1120AAA (3500C>A), CAA1121CAG (3503A>G), TAC1123GCT (3507T>G 3508A>C 3509C>T), TAC1124TTC (3511A>T), CTT1125TTG (3513C>T 3515T>G), GGA1126GGC (3518A>C), ACA1127ACT (3521A>T), GTT1128GTA (3524T>A), CCA1129CCG (3527A>G), CAG1130TTG (3528C>T 3529A>T), GAG1131GAA (3533G>A), TTG1132ATG (3534T>A), GCC1133GCT (3539C>T), GCC1134GCT (3542C>T), GTT1136GTG (3548T>G), CCC1137CCA (3551C>A), AAG1139AAA (3557G>A), ACC1140ACA (3560C>A), ACT1141ACC (3563T>C), CTG1142CTT (3566G>T), CGG1143AAA (3567C>A 3568G>A 3569G>A), AAA1144CGC (3570A>C 3571A>G 3572A>C), AGC1145AGT (3575C>T), CAA1146CCC (3577A>C 3578A>C), TTG1147CTG (3579T>C), GAA1149GAG (3587A>G), GCA1150GCT (3590A>T), TTC1151TTT (3593C>T), GGA1152GGG (3596A>G), CCT1153GAT (3597C>G 3598C>A), GCA1154GCT (3602A>T), GAG1155CAG (3603G>C), ACA1156ACT (3608A>T), GCA1157GCC (3611A>C), ATT1160ATC (3620T>C), ACA1162TCT (3624A>T 3626A>T), GTT1163GCC (3628T>C 3629T>C), CAT1164AAA (3630C>A 3632T>A), AGA1167AGG (3641A>G), GGT1168GGA (3644T>A), GAT1169GAA (3647T>A), GGT1170GGG (3650T>G), GAC1172GAT (3656C>T), GGG1177GGT (3671G>T), GTA1178GTC (3674A>C), ATG1179GAG (3675A>G 3676T>A), AAA1180AAG (3680A>G), TAC1181TAT (3683C>T), TGT1186TCC (3697G>C 3698T>C), GGA1187AAC (3699G>A 3700G>A 3701A>C), GAT1189GAG (3707T>G), GAT1190GAG (3710T>G), GAC1191CCA (3711G>C 3712A>C 3713C>A), GCC1193GCA (3719C>A), AAA1194AAG (3722A>G), CTA1195CTG (3725A>G), GCA1196GCC (3728A>C), TTC1197TTT (3731C>T), GAA1198GAG (3734A>G), AAT1199GGT (3735A>G 3736A>G), CTC1200CTT (3740C>T), AAG1201AAA (3743G>A), TGC1202GCA (3744T>G 3745G>C 3746C>A), TCG1203TCC (3749G>C), CTG1204ATT (3750C>A 3752G>T), CTA1205CTT (3755A>T), ATC1207GTG (3759A>G 3761C>G), ATG1208CTT (3762A>C 3764G>T), CGT1209AAG (3765C>A 3766G>A 3767T>G), AGC1210ATG (3769G>T 3770C>G), AAG1212CGA (3774A>C 3775A>G 3776G>A), ATC1213GTG (3777A>G 3779C>G), CCT1214CCC (3782T>C), GGA1216GGG (3788A>G), CGT1217AAA (3789C>A 3790G>A 3791T>A), CCA1218CCT (3794A>T), TGT1219GCA (3795T>G 3796G>C 3797T>A), GAA1220ATT (3798G>A 3799A>T 3800A>T), AGG1221AGA (3803G>A), GAT1222AAT (3804G>A), GGA1229GGG (3827A>G), TGT1233TTC (3838G>T 3839T>C), TAC1235TAT (3845C>T), TAT1236TTT (3847A>T), GGC1238GCA (3853G>C 3854C>A), GAG1240GAA (3860G>A), CTG1241TTG (3861C>T), ACA1243ACG (3869A>G), TCT1244TCA (3872T>A), TGC1245TGT (3875C>T), GGA1246GGT (3878A>T), CCC1248CCT (3884C>T), TTC1249TTT (3887C>T), ATG1252TTG (3894A>T), GGG1253GAA (3898G>A 3899G>A), ATG1254CTT (3900A>C 3902G>T), GGG1255GGT (3905G>T), ATG1256TCT (3906A>T 3907T>C 3908G>T), AAG1258AAA (3914G>A), AGA1259AGG (3917A>G), CAA1263GAA (3927C>G), TCC1264GCC (3930T>G), ACT1265CAA (3933A>C 3934C>A 3935T>A), GGC1266CCA (3936G>C 3937G>C 3938C>A), GAA1267GAC (3941A>C), GGA1268TCG (3942G>T 3943G>C 3944A>G), GAG1269CAG (3945G>C), AGA1270AGG (3950A>G), GTG1271GTT (3953G>T), GAA1272CTT (3954G>C 3955A>T 3956A>T), AGA1275AGG (3965A>G), GAC1276AAC (3966G>A), ACT1277ACA (3971T>A), GTA1279GTC (3977A>C), GAA1281GAT (3983A>T), CTA1286TTG (3996C>T 3998A>G), GAT1287GAC (4001T>C), GTG1288TCG (4002G>T 4003T>C), ATT1290GTA (4008A>G 4010T>A), AAA1292CAA (4014A>C), GGC1293GGA (4019C>A), ATC1294ACA (4021T>C 4022C>A), CAT1295TTT (4023C>T 4024A>T), GTG1296GTT (4028G>T), GAT1297GAC (4031T>C), CTG1298CTT (4034G>T), ACC1300ACA (4040C>A), CAA1302CAG (4046A>G), TGC1303TGT (4049C>T), GCC1304GCG (4052C>G), CGC1308CGG (4064C>G), CTC1309TTG (4065C>T 4067C>G), AAA1313AAG (4079A>G), GGC1316GGG (4088C>G), AGA1318AGG (4094A>G), AGG1321CGC (4101A>C 4103G>C), CTC1322CTG (4106C>G), TTT1323TTC (4109T>C), GAG1324GAA (4112G>A), ATA1325ATC (4115A>C), CTT1326TTG (4116C>T 4118T>G), TAC1330TAT (4130C>T), TTG1333CTG (4137T>C), GTC1334GTG (4142C>G), AGG1335CGG (4143A>C), AAG1336AAA (4148G>A), TAT1337TAC (4151T>C), TTT1338TTC (4154T>C), GAT1340GAC (4160T>C), TTC1341TTT (4163C>T), TCA1342TCT (4166A>T), GCC1343GCT (4169C>T), ACA1344TGT (4170A>T 4171C>G 4172A>T), TTG1348GCA (4182T>G 4183T>C 4184G>A), AAT1350AAC (4190T>C), GCT1351GTA (4192C>T 4193T>A), ATA1352CTC (4194A>C 4196A>C), CCA1353CCT (4199A>T), AAA1355AAG (4205A>G), GTT1356GTA (4208T>A), ATT1358ATC (4214T>C), GAT1359AAT (4215G>A), CCT1360CCA (4220T>A), ACA1361ACG (4223A>G), AGT1362AGC (4226T>C), TCT1363CAT (4227T>C 4228C>A), TTG1367GTG (4239T>G), TTG1368CTG (4242T>C), GCA1369GCT (4247A>T), AAT1370AGT (4249A>G), GGG1371AGC (4251G>A 4253G>C), AGA1373AGG (4259A>G), GCT1374TCA (4260G>T 4262T>A), GTG1375GTA (4265G>A), TCA1376TCT (4268A>T), GAC1377GAT (4271C>T), GTG1378GAG (4273T>A), TTT1380TTC (4280T>C), GCT1382GCC (4286T>C), TAT1384TAC (4292T>C), TCC1385TCG (4295C>G), CCT1392CCC (4316T>C), GTT1393ATC (4317G>A 4319T>C), GCT1395GCA (4325T>A), CAG1397CAA (4331G>A), TTG1402CTA (4344T>C 4346G>A), GAA1405GCT (4354A>C 4355A>T), TAC1406TTT (4357A>T 4358C>T), TAC1407TAT (4361C>T), GGA1408GGT (4364A>T), TCA1409GAT (4365T>G 4366C>A 4367A>T), AAG1410GAG (4368A>G), GGC1412GGG (4376C>G), GAT1415GAG (4385T>G), TCC1416TCA (4388C>A), CGA1419AGG (4395C>A 4397A>G), CAT1420CAC (4400T>C), GCA1421ACA (4401G>A), CTT1422TTA (4404C>T 4406T>A), GCA1426GCC (4418A>C), TGT1428TGC (4424T>C), TAC1430TAT (4430C>T), ACA1431ACC (4433A>C), CTG1432CTC (4436G>C), TGC1433TGT (4439C>T), GAT1435AAC (4443G>A 4445T>C), AAA1436AAG (4448A>G), GTG1437TTG (4449G>T), AGG1439CGC (4455A>C 4457G>C), TTG1440CTA (4458T>C 4460G>A), GTT1441GTG (4463T>G), GGG1442GGT (4466G>T), GGC1443GGA (4469C>A), CCA1445CCC (4475A>C), TCA1446TCT (4478A>T), GGA1447GGT (4481A>T), GCA1449GCT (4487A>T), CTA1450TTG (4488C>T 4490A>G), ACG1451ACT (4493G>T), ATC1453ATA (4499C>A), TTC1454TTT (4502C>T), TCT1456TCA (4508T>A), CTC1457TTG (4509C>T 4511C>G), CTC1458CTA (4514C>A), CGA1464AGA (4530C>A), TAT1465TAC (4535T>C), GCC1466GCG (4538C>G), TTG1470CTG (4548T>C), TTA1471TTG (4553A>G), AGA1472AGG (4556A>G), CCC1474CCA (4562C>A), ATT1476GTC (4566A>G 4568T>C), GCT1477AAA (4569G>A 4570C>A 4571T>A), AGG1479AAA (4576G>A 4577G>A), GCT1480TCT (4578G>T), ATA1481GTG (4581A>G 4583A>G), GGA1482GGC (4586A>C), GTT1483ATT (4587G>A), AAA1484AAG (4592A>G), AGT1486AAT (4597G>A), GAT1487GAC (4601T>C), CTA1491TTG (4611C>T 4613A>G), GCT1494GCA (4622T>A), GTT1495GTA (4625T>A), GGA1497GGT (4631A>T), GAC1499GAT (4637C>T), AAT1500AAC (4640T>C), CTT1501TTG (4641C>T 4643T>G), GCT1503GCG (4649T>G), GTA1504GTT (4652A>T), TTA1506ATG (4656T>A 4658A>G), CTC1508TTG (4662C>T 4664C>G), CAG1509AGG (4665C>A 4666A>G), TAT1511TAC (4673T>C), TCT1512GAT (4674T>G 4675C>A), CTG1513CTA (4679G>A), AAT1515TCC (4683A>T 4684A>C 4685T>C), ATA1516ATT (4688A>T), CAT1518AGA (4692C>A 4693A>G 4694T>A), GAG1519GAA (4697G>A), TTA1520CTG (4698T>C 4700A>G), CTG1522GCT (4704C>G 4705T>C 4706G>T), GTC1523GTT (4709C>T), AAT1524AAC (4712T>C), GTA1525GTC (4715A>C), ATC1526GTC (4716A>G), CTT1531TTG (4731C>T 4733T>G), AAA1533AAG (4739A>G), TCG1534TCA (4742G>A), GTT1537GTA (4751T>A), AAT1538GAG (4752A>G 4754T>G), GAG1539AAG (4755G>A), GTA1540ACA (4758G>A 4759T>C), CAA1541AAA (4761C>A), CAA1543CTG (4768A>T 4769A>G), CTA1548CTC (4784A>C), ACT1549ACC (4787T>C), CTG1551CTC (4793G>C), AGA1553AGG (4799A>G), TTT1555TTC (4805T>C), AAA1556AAG (4808A>G), GCT1559GCA (4817T>A), CTT1560TTG (4818C>T 4820T>G), TAC1562TAT (4826C>T), CAC1563CAA (4829C>A), GCT1565GCA (4835T>A), CCT1566CCA (4838T>A), AAG1568AAA (4844G>A), ATC1572GTC (4854A>G), ATG1576ATT (4868G>T), TAT1577TAC (4871T>C), ATA1579ATT (4877A>T), AGA1580AGG (4880A>G), CCT1581CCA (4883T>A), GCC1582GCA (4886C>A), CCA1583GTG (4887C>G 4888C>T 4889A>G), GGT1584GGA (4892T>A), GAC1587GAT (4901C>T), ACT1588GTA (4902A>G 4903C>T 4904T>A), CAG1589CAA (4907G>A), GCC1590GCT (4910C>T), ATG1592TAT (4914A>T 4915T>A 4916G>T), GTG1595ATT (4923G>A 4925G>T), GGA1598GGC (4934A>C), ATA1599ATC (4937A>C), CGT1600AGA (4938C>A 4940T>A), GCT1602GCC (4946T>C), CAC1605CAT (4955C>T), GGT1606GGG (4958T>G), CGT1607CAG (4960G>A 4961T>G), GTG1608GCC (4963T>C 4964G>C), GCT1609ACT (4965G>A), GAC1611GAA (4973C>A), AAG1612AGT (4975A>G 4976G>T), CTT1613TTT (4977C>T), GTC1614GTA (4982C>A), TTA1615AAT (4983T>A 4984T>A 4985A>T), CAG1618CTT (4993A>T 4994G>T), ACG1619GGA (4995A>G 4996C>G 4997G>A), GCG1620GCT (5000G>T), TTG1621CTC (5001T>C 5003G>C), GAT1622AAG (5004G>A 5006T>G), GAA1623CAG (5007G>C 5009A>G), AGG1624AGA (5012G>A), GGT1625AAC (5013G>A 5014G>A 5015T>C), TTC1626CTC (5016T>C), AGA1627CCC (5019A>C 5020G>C 5021A>C), GCT1628ATC (5022G>A 5023C>T 5024T>C), GTG1629ATG (5025G>A), TTT1631ATG (5031T>A 5033T>G), CCT1632CCC (5036T>C), TCC1633ACT (5037T>A 5039C>T), TAT1634TAC (5042T>C), TTG1635TAT (5044T>A 5045G>T), GTG1637CTT (5049G>C 5051G>T), CAG1639AGT (5055C>A 5056A>G 5057G>T), GAA1640CAA (5058G>C), ATT1642TTG (5064A>T 5066T>G), GCA1643ACA (5067G>A), AAG1644GAG (5070A>G), GTA1645GTC (5075A>C), GGG1647GGA (5081G>A), TCA1649GAG (5085T>G 5086C>A 5087A>G), GCC1651ATT (5091G>A 5092C>T 5093C>T), CTG1652GGG (5094C>G 5095T>G), ACA1653GTT (5097A>G 5098C>T 5099A>T), ATT1654TTG (5100A>T 5102T>G), TGT1655ATG (5103T>A 5104G>T 5105T>G), GAA1656GAG (5108A>G), ATG1657CTA (5109A>C 5111G>A), GCA1658TCA (5112G>T), AAA1659GAT (5115A>G 5117A>T), GCA1660GCT (5120A>T), GCT1661AAA (5121G>A 5122C>A 5123T>A), ATT1662AAT (5125T>A), TCC1663ACA (5127T>A 5129C>A), TAT1664TAC (5132T>C), ACG1665ACC (5135G>C), CCA1666CCT (5138A>T), TTG1667TTA (5141G>A), GAC1668CCT (5142G>C 5143A>C 5144C>T), GCA1669GCT (5147A>T), GGT1670GGG (5150T>G), GAG1671GAA (5153G>A), ATC1673ATT (5159C>T), ACA1674del (5160_5162delACA), CGT1678GAG (5172C>G 5173G>A 5174T>G), GAT1679GGT (5176A>G), CTG1680GTC (5178C>G 5180G>C), AAT1681CAT (5181A>C), TGG1682GAA (5184T>G 5185G>A 5186G>A), GCA1684GCG (5192A>G), CCA1685CCT (5195A>T), AAC1686GAT (5196A>G 5198C>T), ATT1687GTA (5199A>G 5201T>A), GGT1688TTT (5202G>T 5203G>T), TTT1689TTC (5207T>C), TCA1691AGT (5211T>A 5212C>G 5213A>T), GCA1692GCT (5216A>T), CGT1693CGG (5219T>G), GCC1696TCA (5226G>T 5228C>A), CAC1697AAC (5229C>A), TAC1698CAA (5232T>C 5234C>A), ACG1699CAA (5235A>C 5236C>A 5237G>A), GAT1701TCA (5241G>T 5242A>C 5243T>A), GAA1702TCA (5244G>T 5245A>C), GGG1703GGT (5249G>T), TAC1704AAC (5250T>A), ATT1705ATC (5255T>C), ATT1706CTG (5256A>C 5258T>G), AAC1708AAT (5264C>T), TGT1709TGC (5267T>C), ACA1710ACT (5270A>T), GGT1711GGA (5273T>A), GCA1712GCC (5276A>C), AAG1713CGC (5277A>C 5278A>G 5279G>C), AAG1714AGA (5281A>G 5282G>A), TCC1715ACA (5283T>A 5285C>A), AAT1716ACA (5287A>C 5288T>A), GTT1718GTG (5294T>G), GGT1720GGG (5300T>G), CCA1721CCT (5303A>T), AAC1723AAT (5309C>T), GAC1726GAT (5318C>T), ATG1731ATA (5333G>A), CCG1733CCC (5339G>C), ATG1736TTG (5346A>T), TCA1737GCA (5349T>G), GCT1738GCA (5354T>A), ATA1739ATC (5357A>C), ATG1740CAG (5358A>C 5359T>A), GCC1742CAT (5364G>C 5365C>A 5366C>T), AAG1744CGC (5370A>C 5371A>G 5372G>C), AAT1745AAG (5375T>G), GTG1746AAT (5376G>A 5377T>A 5378G>T), TTG1747CTG (5379T>C), GCA1748GAA (5383C>A), GGA1749GCT (5386G>C 5387A>T), GGA1750GGT (5390A>T), CAT1751GAT (5391C>G), GTA1752GGA (5395T>G), GCA1753GCT (5399A>T), AAT1755TCT (5403A>T 5404A>C), GTT1757ACC (5409G>A 5410T>C 5411T>C), GTG1759GTT (5417G>T), GGA1761GGC (5423A>C), GGA1763GGT (5429A>T), GCT1765GCA (5435T>A), GTC1766ATA (5436G>A 5438C>A), GGC1767GCT (5440G>C 5441C>T), ATT1769ATC (5447T>C), GCG1772GCA (5456G>A), CTG1773TTG (5457C>T), ATG1774TTT (5460A>T 5462G>T), GCC1775GCA (5465C>A), TTA1776TTG (5468A>G), GCA1777GCC (5471A>C), ACA1778ACT (5474A>T), AGG1779CAT (5475A>C 5476G>A 5477G>T), TAT1781TAC (5483T>C), TGT1782AGG (5484T>A 5486T>G), GTG1783GCC (5488T>C 5489G>C), GAG1784ACA (5490G>A 5491A>C 5492G>A), ATA1786CTT (5496A>C 5498A>T), ATA1787ATT (5501A>T), GTT1788GTG (5504T>G), AGA1789CGC (5505A>C 5507A>C), TTG1790CTA (5508T>C 5510G>A), CGC1791CGA (5513C>A), ACA1792ACC (5516A>C), ATT1793ATG (5519T>G), GGA1794GCT (5521G>C 5522A>T), AAT1795GGG (5523A>G 5524A>G 5525T>G), CTT1797CTG (5531T>G), GAC1798GAT (5534C>T), CTC1799CTG (5537C>G), AAT1800AAC (5540T>C), TAT1802TAC (5546T>C), CCT1803CCC (5549T>C), GGC1805GGT (5555C>T), GTG1807GCG (5560T>C), CAG1808CAA (5564G>A), TAT1809TAC (5567T>C), TTT1810TTG (5570T>G), CTT1811CTC (5573T>C), CAA1812AAG (5574C>A 5576A>G), GTG1814GTT (5582G>T), CCT1815GAA (5583C>G 5584C>A 5585T>A), CAT1816GAC (5586C>G 5588T>C), CAT1816_GGA1817insACC (5588_5589insACC), GGA1817TCA (5589G>T 5590G>C), GAC1818CAC (5592G>C), AAA1819AAG (5597A>G), GTG1820TTG (5598G>T), GCT1821GGA (5602C>G 5603T>A), CAA1822CAG (5606A>G), GGT1824GGC (5612T>C), GCA1825ACA (5613G>A), TCG1826TAT (5617C>A 5618G>T), CTC1827GGC (5619C>G 5620T>G), CAC1828del (5622_5624delCAC), AGT1829ACC (5626G>C 5627T>C), AGT1830ACT (5629G>C), ATG1832AAC (5635T>A 5636G>C), CAA1834CAT (5642A>T), CAA1834_GGG1835insCACAGT (5642_5643insCACAGT), GGG1835GCA (5644G>C 5645G>A), GAA1837TCA (5649G>T 5650A>C), TTG1838GTT (5652T>G 5654G>T), GGC1839GGA (5657C>A), AAC1840CTT (5658A>C 5659A>T 5660C>T), CTC1841CTA (5663C>A), CGC1842AGG (5664C>A 5666C>G), ATT1843ATA (5669T>A), ATA1844GTT (5670A>G 5672A>T), CAT1845CCT (5674A>C), GGT1846TCC (5676G>T 5677G>C 5678T>C), GAT1847GAC (5681T>C), TTA1848CTC (5682T>C 5684A>C), GCA1849TCT (5685G>T 5687A>T), AAA1850GAA (5688A>G), CAG1851AAG (5691C>A), ACA1852ATG (5695C>T 5696A>G), ATG1854GCT (5700A>G 5701T>C 5702G>T), CGG1855CGT (5705G>T), ATG1856TTG (5706A>T), TAT1858TAC (5714T>C), GTG1859ACA (5715G>A 5716T>C 5717G>A), GTA1860ATT (5718G>A 5720A>T), GGA1861GGT (5723A>T), CCA1862CCT (5726A>T), CAT1863GAA (5727C>G 5729T>A), GGA1865GGT (5735A>T), GGA1867GCC (5740G>C 5741A>C), AAT1868CAG (5742A>C 5744T>G), TTT1869TGC (5746T>G 5747T>C), TCC1870TCA (5750C>A), ACA1871ACG (5753A>G), CAG1872AAG (5754C>A), GAT1873AAC (5757G>A 5759T>C), CTT1874CTG (5762T>G), GAG1875AAG (5763G>A), AGT1876CAT (5766A>C 5767G>A), TTG1877GCT (5769T>G 5770T>C 5771G>T), CTA1878TTG (5772C>T 5774A>G), CAC1879GAG (5775C>G 5777C>G), TAT1880ACA (5778T>A 5779A>C 5780T>A), TTG1881TTA (5783G>A), GAG1882TCT (5784G>T 5785A>C 5786G>T), CAG1883GTC (5787C>G 5788A>T 5789G>C), GGA1884AGT (5790G>A 5792A>T), GCA1886GCT (5798A>T), GAG1887ACT (5799G>A 5800A>C 5801G>T), TTA1888CTT (5802T>C 5804A>T), ATT1889GCC (5805A>G 5806T>C 5807T>C), CAA1890GCA (5808C>G 5809A>C), AAG1891AGA (5812A>G 5813G>A), AAT1892GGA (5814A>G 5815A>G 5816T>A), ACA1893GTC (5817A>G 5818C>T 5819A>C), CTA1895GTG (5823C>G 5825A>G), ACT1896ACA (5828T>A), CTG1897TTG (5829C>T), TTC1899TTT (5837C>T), GAG1901GAA (5843G>A), CTG1902TTC (5844C>T 5846G>C), AGT1903TCC (5847A>T 5848G>C 5849T>C), ATG1904CAA (5850A>C 5851T>A 5852G>A), GAA1905CAC (5853G>C 5855A>C), AAT1906GAT (5856A>G), GTG1907TGT (5859G>T 5860T>G 5861G>T), CAA1908GAA (5862C>G), CAA1909ATG (5865C>A 5866A>T 5867A>G), CTG1910ATA (5868C>A 5870G>A), ATA1911CTT (5871A>C 5873A>T), GAT1912GAG (5876T>G), GTT1914CTG (5880G>C 5882T>G), AAG1915GAA (5883A>G 5885G>A), CTT1916AGT (5886C>A 5887T>G), CAG1917CAA (5891G>A), TTT1919GTA (5895T>G 5897T>A), TTC1920GTA (5898T>G 5900C>A), CCA1921TTC (5901C>T 5902C>T 5903A>C), AAA1922ATG (5905A>T 5906A>G), GAA1923GGG (5908A>G 5909A>G), ACT1924AGC

| | | | | | | | | | |
|---|---|---|---|---|---|---|---|---|---|
| | (5911C>G 5912T>C), ACC1925GTT (5913A>G 5914C>T 5915C>T), ATT1926AAA (5917T>A 5918T>A), CAG1927AAA (5919C>A 5921G>A), AAG1928CAA (5922A>C 5924G>A), CTC1929ATG (5925C>A 5927C>G), AAA1930TTG (5928A>T 5929A>T 5930A>G), AAT1931TCC (5931A>T 5932A>C 5933T>C), TTT1932TTC (5936T>C), GTT1933ATG (5937G>A 5939T>G), GAT1934AAT (5940G>A), GCT1935CAA (5943G>C 5944C>A 5945T>A), GAA1936CAG (5946G>C 5948A>G), TGT1937ATG (5949T>A 5950G>T 5951T>G), CTA1938TCA (5952C>T 5953T>C), ACA1939AAA (5956C>A), TTC1940ACC (5958T>A 5959T>C), AAA1941CAA (5961A>C), GCA1942GTT (5965C>T 5966A>T), AGG1943GCC (5967A>G 5968G>C 5969G>C), AGC1944_TTT1945insCCATGTTGTGTCACA (5972_5973insCCATGTTGTGTCACA), AGG1946AGT (5978G>T), CAC1947AAA (5979C>A 5981C>A), GTA1948GCA (5983T>C), GTT1949TGG (5985G>T 5986T>G 5987T>G), TTC1950CTA (5988T>C 5990C>A), AAA1951ACT (5992A>C 5993A>T), AAA1953GAC (5997A>G 5999A>C), CTT1954ATT (6000C>A), TTG1955GTC (6003T>G 6005G>C), AGT1956ATG (6007G>T 6008T>G), TCC1957ACA (6009T>A 6011C>A), ACA1958AAA (6013C>A), TGG1959GGT (6015T>G 6017G>T), AAA1960AAG (6020A>G), ATG1961AAG (6022T>A), TGT1962CCA (6024T>C 6025G>C 6026T>A), GGT1963GGA (6029T>A), GAA1964del (6030_6032delGAA), AGT1965TCT (6033A>T 6034G>C), GTT1967_AGG1970del (6039_6050delGTTGCTTCAAGG), TCT1971CCT (6051T>C), GCA1972GCT (6056A>T), GAG1973GAA (6059G>A), AGT1974ACA (6061G>C 6062T>A), CTA1975GTG (6063C>G 6065A>G), TTC1976TTT (6068C>T), CCT1977CTT (6070C>T), GGC1978GGG (6074C>G), TTG1980CCT (6078T>C 6079T>C 6080G>T), TCT1981TCA (6083T>A), GCT1982GCA (6086T>A), TCC1983CTA (6087T>C 6088C>T 6089C>A), GTA1984GTT (6092A>T), TTG1985GTG (6093T>G), AAA1986AGA (6097A>G), TTA1988ATG (6102T>A 6104A>G), GAA1990CGC (6108G>C 6109A>G 6110A>C), AGA1991AAT (6112G>A 6113A>T), CAC1992AGT (6114C>A 6115A>G 6116C>T), ACA1993GGA (6117A>G 6118C>G), AGA1994AAG (6121G>A 6122A>G), ATG1996TAC (6126A>T 6127T>A 6128G>C), AGT1997TCA (6129A>T 6130G>C 6131T>A), CAG1999ACA (6135C>A 6136A>C 6137G>A), AGT2000ACA (6139G>C 6140T>A), ATG2001CTG (6141A>C), GAG2002GAA (6146G>A), CTG2005TTT (6153C>T 6155G>T), AAA2006AGA (6157A>G), ATA2007ATG (6161A>G), TAT2008TAC (6164T>C), CTA2009ATG (6165C>A 6167A>G), TTA2010TGT (6169T>G 6170A>T), AAC2011CAG (6171A>C 6173C>G), TTC2012GAA (6174T>G 6175T>A 6176C>A), CAA2013GGG (6177C>G 6178A>G 6179A>G), GTA2014TAT (6180G>T 6181T>A 6182A>T), ATA2015TTA (6183A>T), ACA2016GAT (6186A>G 6187C>A 6188A>T), AGT2017AAA (6190G>A 6191T>A), GAG2018GCC (6193A>C 6194G>C), TTA2020GTG (6198T>G 6200A>G), AAG2021GAC (6201A>G 6203G>C), AAG2022TGC (6204A>T 6205A>G 6206G>C), TTT2023TTC (6209T>C), GAA2024AGT (6210G>A 6211A>G 6212A>T), GAT2025GAG (6215T>G), TTT2027ATG (6219T>A 6221T>G), CAG2028GGC (6222C>G 6223A>G 6224G>C), GAG2029GTT (6226A>T 6227G>T), AAG2030TGT (6228A>T 6229A>G 6230G>T), ATA2031GTG (6231A>G 6233A>G), TCC2032CCG (6234T>C 6236C>G), ACC2033GAA (6237A>G 6238C>A 6239C>A), ACC2034CGC (6240A>C 6241C>G), CTA2035ATC (6243C>A 6245A>C), CTA2036TTG (6246C>T 6248A>G), ATC2037GTG (6249A>G 6251C>G), AAA2038CGC (6252A>C 6253A>G 6254A>C), GTA2039ACA (6255G>A 6256T>C), TTC2040ATT (6258T>A 6260C>T), TTG2041GCT (6261T>G 6262T>C 6263G>T), GAG2044TCC (6270G>T 6271A>C 6272G>C), GAG2045ACG (6273G>A 6274A>C), AGT2046TCA (6276A>T 6277G>C 6278T>A), TAT2047TTG (6280A>T 6281T>G), CAG2048AAC (6282C>A 6284G>C), ACT2049GAG (6285A>G 6286C>A 6287T>G), GAG2050AAT (6288G>A 6290G>T), TGT2052ACT (6294T>A 6295G>C), ATA2053GTG (6297A>G 6299A>G), GAT2054TAC (6300G>T 6302T>C), TCT2055ACA (6303T>A 6305T>A), CAG2056GAT (6306C>G 6308G>T), GTG2057GGG (6310T>G), CTC2058CTT (6314C>T), CAG2059TTT (6315C>T 6316A>T 6317G>T), AGG2060GAT (6318A>G 6319G>A 6320G>T), ATA2061del (6321_6323delATA), AAC2062CAT (6324A>C 6326C>T), TCC2063TTC (6328C>T), CAG2064AAA (6330C>A 6332G>A), AAA2065GCA (6333A>G 6334A>C), TTC2066GTG (6336T>G 6338C>G), AGA2067GAC (6339A>G 6340G>A 6341A>C), GTG2068TTC (6342G>T 6344G>C), CAG2069AAG (6345C>A), GAA2070CCT (6348G>C 6349A>C 6350A>T), GGT2071GGG (6353T>G), TTT2073TTC (6359T>C), TAT2074TCC (6361A>C 6362T>C), TTG2075TGG (6364T>G), CAT2076GAA (6366C>G 6368T>A), CCT2077ATT (6369C>A 6370C>T), GAA2078AAT (6372G>A 6374A>T), GGA2079GAT (6376G>A 6377A>T), ATC2080GCC (6378A>G 6379T>C), AAC2081AAT (6383C>T), AAT2083AAC (6389T>C), GCC2084GCT (6392C>T), GTA2085GCA (6394T>C), ATA2088GTG (6402A>G 6404A>G), ATA2089TTG (6405A>T 6407A>G), TCT2091GAA (6411T>G 6412C>A 6413T>A), CTG2092CTC (6416G>C), GAG2094AGG (6420G>A 6421A>G), TCC2095CAC (6423T>C 6424C>A), TAT2096TGG (6427A>G 6428T>G), AGC2097TCA (6429A>T 6430G>C 6431C>A), CGA2098AAG (6432C>A 6433G>A 6434A>G), GAT2099TCA (6435G>T 6436A>C 6437T>A), AGT2100CAA (6438A>C 6439G>A 6440T>A), AAC2101AGC (6442A>G), TCC2102AAG (6444T>A 6445C>A 6446C>G), TAT2103TTC (6448A>T 6449T>C), TCT2104ACT (6450T>A), GTG2105TTT (6453G>T 6455G>T), TCA2106GCC (6456T>G 6458A>C), ACT2107AAT (6460C>A), CCC2108GAA (6462C>G 6463C>A 6464C>A), ATC2109GTT (6465A>G 6467C>T), AAA2110GAA (6468A>G), CTG2111ATA (6471C>A 6473G>A), GGG2112TCA (6474G>T 6475G>C 6476G>A), TGT2113AAC (6477T>A 6478G>A 6479T>C), TTC2114CTG (6480T>C 6482C>G), ATT2115CTT (6483A>C), TTC2116TTA (6488C>A), CTT2117GCT (6489C>G 6490T>C), GTC2118GTT (6494C>T), GTG2119TAT (6495G>T 6496T>A 6497G>T), CTA2120TTA (6498C>T), GAT2121GAA (6503T>A), AGT2122AAT (6505G>A), CAA2123AAG (6507C>A 6509A>G), GGC2124CTC (6510G>C 6511G>T), AAG2125ACA (6514A>C 6515G>A), GAA2126GGT (6517A>G 6518A>T), GAT2127CTA (6519G>C 6520A>T 6521T>A), CAT2128GAG (6522C>G 6524T>G), CCT2129TAT (6525C>T 6526C>A), GTT2130TAT (6528G>T 6529T>A), CGG2131GAT (6531C>G 6532G>A 6533G>T), AGG2132TCC (6534A>T 6535G>C 6536G>C), TTT2133ATA (6537T>A 6539T>A), TCA2134GAA (6540T>G 6541C>A), ACG2135TCT (6543A>T 6545G>T), GTT2136TTG (6546G>T 6548T>G), TTC2137GAC (6549T>G 6550T>A), CTT2138TTG (6552C>T 6554T>G), AAA2139TTA (6555A>T 6556A>T), AAC2140CAA (6558A>C 6560C>A), TGT2141ATA (6561T>A 6562G>T 6563T>A), GAA2142GAT (6566A>T), ATT2144ATA (6572T>A), ACA2146GCA (6576A>G), TAC2148GTC (6582T>G 6583A>T), GAG2150TAC (6588G>T 6590G>C), TGA2152GGG (6594T>G 6596A>G) | | | | | | | | |
| helicase (YP_001976154.1) | 1 | 98 | 100% | 654 | 97.5% | 98 (100%) | 95 (96.9%) | 0/0/0/0 | 0 |
| Protein mutations: | S5C (1342C>G), R16A (1374A>G 1375G>C), I44V (1458A>G 1460C>G) | | | | | | | | |
| Codon mutations: | TCT3TCA (1337T>A), ACA4ACG (1340A>G), TCC5TGC (1342C>G), TTA8CTT (1350T>C 1352A>T), TTT9TTC (1355T>C), GTG13GTT (1367G>T), TTG14CTC (1368T>C 1370G>C), AGA16GCA (1374A>G 1375G>C), GAG19GAA (1385G>A), AAA21AAG (1391A>G), CTC22CTG (1394C>G), AAT23AAC (1397T>C), CGG24CGC (1400G>C), CTA25TTG (1401C>T 1403A>G), GCA27GCG (1409A>G), AAG30AAA (1418G>A), AGG31AGA (1421G>A), GCT39GCA (1445T>A), CAC40CAT (1448C>T), GCT43GCA (1457T>A), ATC44GTG (1458A>G 1460C>G), CTG45CTT (1463G>T), GAC48GAT (1472C>T), GGA50GGT (1478A>T), GCA51GCT (1481A>T), TTG52CTT (1482T>C 1484G>T), CGA53CGG (1487A>G), GCT56GCA (1496T>A), TGC59TGT (1505C>T), CAA60CAG (1508A>G), GAT61GAC (1511T>C), ATT62ATA (1514T>A), AAA63AAG (1517A>G), GAT67GAC (1529T>C), TCA70TCT (1538A>T), ACA71ACC (1541A>C), CAA72CAG (1544A>G), CCA73CCT (1547A>T), CCC75CCT (1553C>T), TTA76CTT (1554T>C 1556A>T), GCC79GCT (1565C>T), GTT81GTC (1571T>C), GAC83GAT (1577C>T), GGC85GGA (1583C>A), CAT87CAC (1589T>C), TTC88TTT (1592C>T), ACC89ACA (1595C>A), TCC90TCA (1598C>A), TAT92TAC (1604T>C), ATA93ATC (1607A>C), ACA96ACC (1616A>C) | | | | | | | | |
| RNA-dependent RNA polymerase (YP_001976153.1) | 1 | 249 | 100% | 1552 | 88.4% | 249 (100%) | 210 (84.3%) | 0/0/0/0 | 0 |
| Protein mutations: | T40C (4170A>T 4171C>G 4172A>T), L44A (4182T>G 4183T>C 4184G>A), A47V (4192C>T 4193T>A), I48L (4194A>C 4196A>C), D55N (4215G>A), S59H (4227T>C 4228C>A), L63V (4239T>G), N66S (4249A>G), G67S (4251G>A 4253G>C), A70S (4260G>T 4262T>A), V74E (4273T>A), V89I (4317G>A 4319T>C), E101A (4354A>C 4355A>T), Y102F (4357A>T 4358C>T), S105D (4365T>G 4366C>A 4367A>T), K106E (4368A>G), D111E (4385T>G), A117T (4401G>A), D131N (4443G>A 4445T>C), V133L (4449G>T), I172V (4566A>G 4568T>C), A173K (4569G>A 4570C>A 4571T>A), R175K (4576G>A 4577G>A), A176S (4578G>T), I177V (4581A>G 4583A>G), V179I (4587G>A), S182N (4597G>A), L202M (4656T>A 4658A>G), Q205R (4665C>A 4666A>G), S208D (4674T>G 4675C>A), N211S (4683A>T 4684A>C 4685T>C), H214R (4692C>A 4693A>G 4694T>A), L218A (4704C>G 4705T>C 4706G>T), I222V (4716A>G), N234E (4752A>G 4754T>G), E235K (4755G>A), V236T (4758G>A 4759T>C), Q237K (4761C>A), Q239L (4768A>T 4769A>G) | | | | | | | | |
| Codon mutations: | CGC4CGG (4064C>G), CTC5TTG (4065C>T 4067C>G), AAA9AAG (4079A>G), GGC12GGG (4088C>G), AGA14AGG (4094A>G), AGG17CGC (4101A>C 4103G>C), CTC18CTG (4106C>G), TTT19TTC (4109T>C), GAG20GAA (4112G>A), ATA21ATC (4115A>C), CTT22TTG (4116C>T 4118T>G), TAC26TAT (4130C>T), TTG29CTG (4137T>C), GTC30GTG (4142C>G), AGG31CGG (4143A>C), AAG32AAA (4148G>A), TAT33TAC (4151T>C), TTT34TTC (4154T>C), GAT36GAC (4160T>C), TTC37TTT (4163C>T), TCA38TCT (4166A>T), GCC39GCT (4169C>T), ACA40TGT (4170A>T 4171C>G 4172A>T), TTG44GCA (4182T>G 4183T>C 4184G>A), AAT46AAC (4190T>C), GCT47GTA (4192C>T 4193T>A), ATA48CTC (4194A>C 4196A>C), CCA49CCT (4199A>T), AAA51AAG (4205A>G), GTT52GTA (4208T>A), ATT54ATC (4214T>C), GAT55AAT (4215G>A), CCT56CCA (4220T>A), ACA57ACG (4223A>G), AGT58AGC (4226T>C), TCT59CAT (4227T>C 4228C>A), TTG63GTG (4239T>G), TTG64CTG (4242T>C), GCA65GCT (4247A>T), AAT66AGT (4249A>G), GGG67AGC (4251G>A 4253G>C), AGA69AGG (4259A>G), GCT70TCA (4260G>T 4262T>A), GTG71GTA (4265G>A), TCA72TCT (4268A>T), GAC73GAT (4271C>T), GTG74GAG (4273T>A), TTT76TTC (4280T>C), GCT78GCC (4286T>C), TAT80TAC (4292T>C), TCC81TCG (4295C>G), CCT88CCC (4316T>C), GTT89ATC (4317G>A 4319T>C), GCT91GCA (4325T>A), CAG93CAA (4331G>A), TTG98CTA (4344T>C 4346G>A), GAA101GCT (4354A>C 4355A>T), TAC102TTT (4357A>T 4358C>T), TAC103TAT (4361C>T), GGA104GGT (4364A>T), TCA105GAT (4365T>G 4366C>A 4367A>T), AAG106GAG (4368A>G), GGC108GGG (4376C>G), GAT111GAG (4385T>G), TCC112TCA (4388C>A), CGA115AGG (4395C>A 4397A>G), CAT116CAC (4400T>C), GCA117ACA (4401G>A), CTT118TTA (4404C>T 4406T>A), GCA122GCC (4418A>C), TGT124TGC (4424T>C), TAC126TAT (4430C>T), ACA127ACC (4433A>C), CTG128CTC (4436G>C), TGC129TGT (4439C>T), GAT131AAC (4443G>A 4445T>C), AAA132AAG (4448A>G), GTG133TTG (4449G>T), AGG135CGC (4455A>C 4457G>C), TTG136CTA (4458T>C 4460G>A), GTT137GTG (4463T>G), GGG138GGT (4466G>T), GGC139GGA (4469C>A), CCA141CCC (4475A>C), TCA142TCT (4478A>T), GGA143GGT (4481A>T), GCA145GCT (4487A>T), CTA146TTG (4488C>T 4490A>G), ACG147ACT (4493G>T), ATC149ATA (4499C>A), TTC150TTT (4502C>T), TCT152TCA (4508T>A), CTC153TTG (4509C>T 4511C>G), CTC154CTA (4514C>A), CGA160AGA (4530C>A), TAT161TAC (4535T>C), GCC162GCG (4538C>G), TTG166CTG (4548T>C), TTA167TTG (4553A>G), AGA168AGG (4556A>G), CCC170CCA (4562C>A), ATT172GTC (4566A>G 4568T>C), GCT173AAA (4569G>A 4570C>A 4571T>A), AGG175AAA (4576G>A 4577G>A), GCT176TCT (4578G>T), ATA177GTG (4581A>G 4583A>G), GGA178GGC (4586A>C), GTT179ATT (4587G>A), AAA180AAG (4592A>G), AGT182AAT (4597G>A), GAT183GAC (4601T>C), CTA187TTG (4611C>T 4613A>G), GCT190GCA (4622T>A), GTT191GTA (4625T>A), GGA193GGT (4631A>T), GAC195GAT (4637C>T), AAT196AAC (4640T>C), CTT197TTG (4641C>T 4643T>G), GCT199GCG (4649T>G), GTA200GTT (4652A>T), TTA202ATG (4656T>A 4658A>G), CTC204TTG (4662C>T 4664C>G), CAG205AGG (4665C>A 4666A>G), TAT207TAC (4673T>C), TCT208GAT (4674T>G 4675C>A), CTG209CTA (4679G>A), AAT211TCC (4683A>T 4684A>C 4685T>C), ATA212ATT (4688A>T), CAT214AGA (4692C>A 4693A>G 4694T>A), GAG215GAA (4697G>A), TTA216CTG (4698T>C 4700A>G), CTG218GCT (4704C>G 4705T>C 4706G>T), GTC219GTT (4709C>T), AAT220AAC (4712T>C), GTA221GTC (4715A>C), ATC222GTC (4716A>G), CTT227TTG (4731C>T 4733T>G), AAA229AAG (4739A>G), TCG230TCA (4742G>A), GTT233GTA (4751T>A), AAT234GAG (4752A>G 4754T>G), GAG235AAG (4755G>A), GTA236ACA (4758G>A 4759T>C), CAA237AAA (4761C>A), CAA239CTG (4768A>T 4769A>G), CTA244CTC (4784A>C), ACT245ACC (4787T>C), CTG247CTC (4793G>C), AGA249AGG (4799A>G) | | | | | | | | |

*: Inserts / Deletes / Misaligned / Frameshifts

#### Analysis details

This analysis was performed with panviral2.64

### NGS Details (UN8): Diachasmimorpha longicaudata entomopoxvirus (segment NC_043455.1)

#### Assembly

| | |
|---|---|
| **Coverage Length** | 1121 (1 contig(s)) |
| **Depth Of Coverage** | 1498.5 |
| **Number Of Reads** | 13895 |
| **Reads Per Million** | 232.22 rpm (after QC) |
| **Ambiguities** | 0 |
| **Assembly Method** | de novo + reference guided assembly |
| **Consensus Caller** | Bcf Tools |

#### Coverage Map

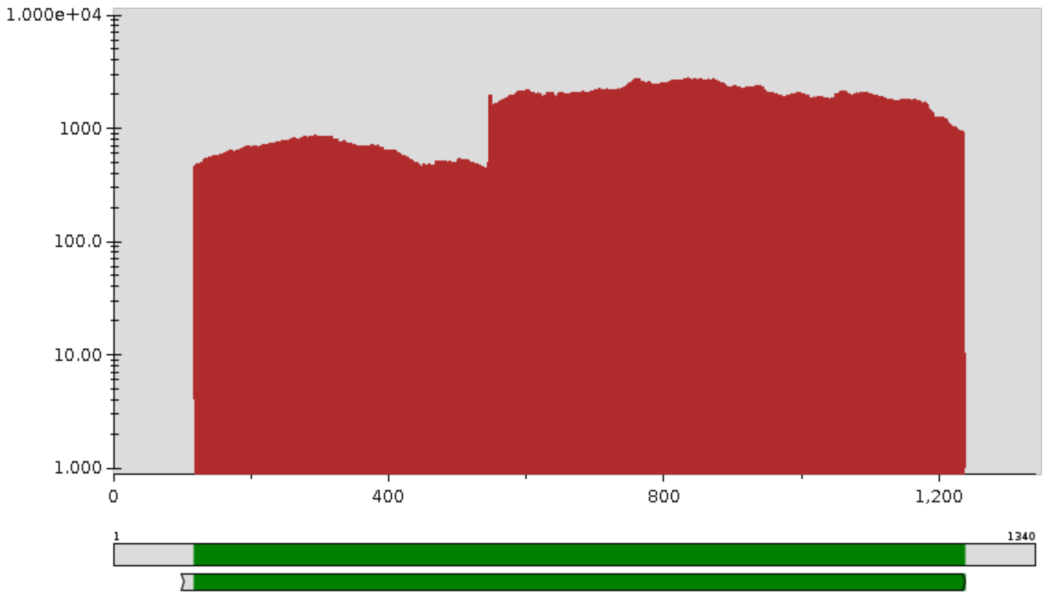

#### Assignment

| | |
|---|---|
| **Type** | Diachasmimorpha longicaudata entomopoxvirus (Taxonomy ID: 109981) |
| **Reference Genome** | NC_043455.1 |
| **NT Identity (%)** | 59.7681 |
| **AA Identity (%)** | 55.3476 |
| **Number Of Stop Codons** | 0 |
| **Number Of CDS** | 1 |

#### Alignment

| | |
|---|---|
| **Alignment Score** | 420.0 (NT) + 1460.0 (AA) = 1880.0 |
| **Concordance (%)** | 40.1366 |

| Alignment Method | Global, seeded, nucleotide + amino acids (AGA) |
|---|---|

#### Genome Region

Sequence starts at position 118 and ends at position 1238 relative to NC_043455.1 reference sequence.

#### Alignment Detailed Statistics

| | Begin | End | Coverage | Score | Concordance | Matches | Identities | I/D/M/F* | Stop Codons |
|---|---|---|---|---|---|---|---|---|---|
| **NT** | **118** | **1238** | **83.7%** | **420** | **18.9%** | **1118 (99.5%)** | **670 (59.6%)** | **3/3** | |
| Mutations: | 118T>G, 119G>A, 120A>G, 121A>T, 126A>C, 127C>T, 128C>G, 130T>G, 136C>G, 137C>A, 139C>T, 142G>A, 143A>G, 145A>T, 146A>G, 148T>C, 149A>C, 151C>T, 154A>T, 156A>G, 157G>A, 160T>G, 163A>T, 166T>C, 167T>C, 168C>A, 169T>G, 171G>A, 172T>C, 175A>G, 178T>C, 187T>A, 190A>T, 191A>G, 193T>C, 203A>C, 204A>G, 208C>T, 209A>G, 211T>A, 214T>G, 217A>T, 218T>A, 219G>T, 223T>A, 226T>G, 230A>C, 231A>G, 232A>T, 238G>C, 241A>T, 243T>C, 244C>T, 247A>G, 256G>C, 259A>T, 262A>C, 271A>C, 272G>T, 274A>C, 276C>T, 277A>G, 278T>A, 279A>T, 283A>T, 284A>C, 286C>T, 288G>C, 289T>C, 294T>G, 295A>C, 298G>A, 299C>G, 300A>T, 301A>T, 302A>G, 304A>G, 310T>C, 311T>A, 312C>A, 313T>A, 320A>G, 322T>G, 323A>G, 325C>A, 331T>A, 337C>A, 341A>T, 343C>T, 346A>T, 352T>G, 355A>G, 356T>C, 358G>T, 361T>A, 362C>G, 363T>C, 364C>T, 368G>A, 370T>G, 371C>G, 374C>A, 375G>A, 376T>G, 379A>G, 380T>A, 382G>T, 383C>T, 384A>T, 385A>G, 386A>G, 394G>A, 395A>G, 400T>C, 401C>A, 403A>T, 404_406delTAT, 409T>C, 410T>A, 412T>A, 413A>C, 415A>G, 416T>G, 417G>C, 418T>A, 421A>T, 423T>C, 424T>G, 438C>A, 442T>C, 443A>G, 445C>G, 446A>G, 447A>G, 448G>T, 452A>G, 453G>A, 454C>T, 455C>A, 456A>T, 457G>C, 458G>A, 459A>G, 462C>A, 463T>G, 467A>G, 469_470insCAT, 470A>G, 471A>G, 474C>T, 475C>A, 478G>A, 482T>G, 484G>C, 485A>T, 486T>C, 500C>A, 503A>G, 505G>C, 506A>T, 507T>G, 511T>C, 512C>A, 514A>G, 515C>A, 517C>T, 519C>A, 520C>A, 521C>A, 523A>G, 525A>G, 528G>C, 530A>T, 532C>A, 533G>C, 534A>G, 535T>A, 540A>G, 541A>G, 544A>C, 547T>C, 550A>G, 553A>G, 554G>T, 559A>T, 560A>T, 562A>G, 574T>C, 577G>A, 581T>C, 583G>C, 584A>T, 585T>C, 586A>T, 587G>A, 588A>G, 589T>A, 590A>G, 591A>G, 595T>C, 596T>A, 597T>A, 602A>C, 604A>G, 607A>T, 608C>T, 610A>T, 616T>A, 620G>C, 622A>G, 623T>C, 628T>G, 629G>C, 630A>C, 632A>C, 633G>C, 634T>A, 635C>A, 637T>A, 638G>A, 646T>G, 647A>G, 648T>G, 650T>G, 652A>T, 655A>C, 658A>T, 661A>C, 664C>T, 665G>A, 667T>G, 671T>C, 674A>G, 675G>A, 678T>C, 680A>C, 682C>T, 683A>G, 685T>G, 687C>T, 688A>T, 689T>A, 692C>A, 693A>G, 694G>A, 695G>A, 696T>A, 697C>G, 700T>C, 705G>A, 706A>C, 707G>A, 709T>G, 715A>G, 717A>G, 718A>G, 724G>T, 727A>G, 730A>G, 731A>C, 732A>G, 735C>A, 739A>G, 740T>C, 742G>C, 745A>T, 748C>T, 754C>T, 759G>A, 765A>T, 770A>G, 772A>C, 778A>C, 779A>G, 781G>T, 784A>G, 785A>G, 787T>A, 790T>G, 792T>G, 793T>G, 796A>G, 797G>C, 798C>T, 799A>T, 810T>G, 820T>C, 823C>G, 824C>A, 825A>C, 826C>T, 829A>G, 830A>G, 831G>C, 833C>A, 838T>C, 841A>G, 843C>G, 844A>T, 845C>G, 847A>C, 850A>C, 854T>G, 855G>T, 859T>C, 862A>T, 863C>A, 865T>G, 866A>C, 869C>A, 871A>G, 878G>T, 879T>G, 880A>G, 885T>C, 886G>T, 889A>T, 890T>A, 891G>A, 892T>A, 893T>A, 895A>G, 896A>C, 897C>G, 898A>T, 900A>G, 901T>C, 902A>C, 904A>T, 905A>G, 908T>C, 909T>A, 915C>T, 919A>T, 920A>G, 921G>C, 922C>A, 924T>C, 925C>T, 934T>C, 938T>G, 939C>A, 940C>T, 943A>G, 944C>A, 946A>T, 947G>A, 948A>C, 950C>A, 952T>A, 956T>A, 958T>C, 966A>G, 967A>G, 970A>G, 973T>C, 974A>C, 977G>T, 978A>C, 982T>C, 983A>T, 984A>C, 986A>T, 989A>C, 991A>T, 992A>G, 994A>G, 998C>A, 1000T>C, 1001T>A, 1007A>G, 1012A>C, 1013T>C, 1015A>T, 1018A>T, 1019A>C, 1021A>T, 1024C>T, 1033T>A, 1036A>G, 1040A>G, 1042A>C, 1045T>C, 1046T>C, 1048A>T, 1051T>C, 1054C>T, 1057T>C, 1063T>C, 1064T>C, 1070C>A, 1071A>C, 1073A>C, 1075T>A, 1076A>C, 1077G>C, 1087T>C, 1088A>C, 1090A>G, 1103A>C, 1105A>T, 1111T>A, 1112A>C, 1120C>A, 1121C>A, 1123A>G, 1126A>G, 1129G>T, 1130A>G, 1141T>C, 1145A>G, 1147A>C, 1150T>C, 1151G>A, 1153A>G, 1156C>T, 1159A>T, 1160T>G, 1161T>A, 1163C>A, 1164C>G, 1165A>G, 1167A>T, 1172G>T, 1173C>T, 1177A>C, 1178T>A, 1181G>C, 1183A>G, 1185C>A, 1186G>A, 1192T>C, 1195T>C, 1196A>G, 1197C>T, 1199A>G, 1200A>T, 1204A>C, 1207T>G, 1210A>G, 1211A>C, 1213G>C, 1218A>C, 1219G>C, 1223A>G, 1227T>C, 1232T>C, 1234G>C, 1235A>C, 1237A>G | | | | | | | | |
| **CDS** | | | | | | | | | |
| FLA14_p101 | 7 | 380 | 98.4% | 1460 | 59.1% | 373 (99.5%) | 207 (55.2%) | 1/1/0/0 | 0 |
| Protein mutations: | E7S (119G>A 120A>G 121A>T), D9A (126A>C 127C>T), H10E (128C>G 130T>G), L13I (137C>A 139C>T), K15D (143A>G 145A>T), N16D (146A>G 148T>C), I17L (149A>C 151C>T), K19R (156A>G 157G>A), S23Q (167T>C 168C>A 169T>G), C24Y (171G>A 172T>C), T31A (191A>G 193T>C), K35R (203A>C 204A>G), I37V (209A>G 211T>A), F38L (214T>G), C40I (218T>A 219G>T), K44R (230A>C 231A>G 232A>T), V48A (243T>C 244C>T), A58S (272G>T 274A>C), T59M (276C>T 277A>G), Y60I (278T>A 279A>T), I62L (284A>C 286C>T), S63T (288G>C 289T>C), L65C (294T>G 295A>C), Q67V (299C>G 300A>T 301A>T), I68V (302A>G 304A>G), S71K (311T>A 312C>A 313T>A), N74E (320A>G 322T>G), I75V (323A>G 325C>A), T81S (341A>T 343C>T), L88A (362C>G 363T>C 364C>T), A90T (368G>A 370T>G), Q91E (371C>G), R92K (374C>A 375G>A 376T>G), L94I (380T>A 382G>T), Q95L (383C>T 384A>T 385A>G), T96A (386A>G), N99D (395A>G), L101I (401C>A 403A>T), Y102del (404_406delTAT), F104I (410T>A 412T>A), K105Q (413A>C 415A>G), C106A (416T>G 417G>C 418T>A), Q107H (421A>T), V108A (423T>C 424T>G), T113K (438C>A), I115V (443A>G 445C>G), K116G (446A>G 447A>G 448G>T), S118D (452A>G 453G>A 454C>T), Q119I (455C>A 456A>T 457G>C), E120R (458G>A 459A>G), T121K (462C>A 463T>G), K123E (467A>G), K123_K124insH (469_470insCAT), K124G (470A>G 471A>G), A125V (474C>T 475C>A), L128V (482T>G 484G>C), I129S (485A>T 486T>C), M135V (503A>G 505G>C), I136C (506A>T 507T>G), L138M (512C>A 514A>G), L139I (515C>A 517C>T), T140K (519C>A 520C>A), K142R (525A>G), S143T (528G>C), I144L (530A>T 532C>A), D145R (533G>C 534A>G 535T>A), K147R (540A>G 541A>G), I151M (553A>G), V152F (554G>T), I154L (560A>T 562A>G), I162S (584A>T 585T>C 586A>T), D163R (587G>A 588A>G 589T>A), N164G (590A>G 591A>G), L166K (596T>A 597T>A), K168Q (602A>C 604A>G), Q170Y (608C>T 610A>T), E174Q (620G>C 622A>G), F175L (623T>C), F176L (628T>G), E177P (629G>C 630A>C), S178P (632A>C 633G>C 634T>A), H179K (635C>A 637T>A), V180I (638G>A), I183G (647A>G 648T>G), L184V (650T>G 652A>T), L185F (655A>C), V189M (665G>A 667T>G), S191P (671T>C), R192E (674A>G 675G>A), V193A (678T>C), I194L (680A>C 682C>T), N195E (683A>G 685T>G), T196I (687C>T 688A>T), S197T (689T>A), Q198R (692C>A 693A>G 694G>A), V199K (695G>A 696T>A 697C>G), R202N (705G>A 706A>C), D203K (707G>A 709T>G), K206R (717A>G 718A>G), N211R (731A>C 732A>G), A212D (735C>A), R220K (759G>A), Y222F (765A>T), I224V (770A>G 772A>C), K227D (779A>G 781G>T), N229E (785A>G 787T>A), D230E (790T>G), F231W (792T>G 793T>G), A233L (797G>C 798C>T 799A>T), F237C (810T>G), D241E (823C>G), H242T (824C>A 825A>C 826C>T), S244A (830A>G 831G>C), L245I (833C>A), T248S (843C>G 844A>T), L249V (845C>G 847A>C), C252V (854T>G 855G>T), Q257K (869C>A 871A>G), V260W (878G>T 879T>G 880A>G), M262T (885T>C 886G>T), E263D (889A>T), C264K (890T>A 891G>A 892T>A), L265M (893T>A 895A>G), T266R (896A>C 897C>G 898A>T), N267S (900A>G 901T>C), N269D (905A>G), F270H (908T>C 909T>A), A272V (915C>T), S274A (920A>G 921G>C 922C>A), I275T (924T>C 925C>T), S280D (938T>G 939C>A 940C>T), Q282N (944C>A 946A>T), E283T (947G>A 948A>C), F286I (956T>A 958T>C), K289R (966A>G 967A>G), D293S (977G>T 978A>C), K295S (983A>T 984A>C), T296S (986A>T), I298V (992A>G 994A>G), L300I (998C>A 1000T>C), S301T (1001T>A), N303D (1007A>G), I314V (1040A>G 1042A>C), H324T (1070C>A 1071A>C), N325Q (1073A>C 1075T>A), R326P (1076A>C 1077G>C), I330L (1088A>C 1090A>G), I344V (1130A>G), I349V (1145A>G 1147A>C), E351K (1151G>A 1153A>G), E353D (1159A>T), L354E (1160T>G 1161T>A), P355R (1163C>A 1164C>G 1165A>G), K356M (1167A>T), A358F (1172G>T 1173C>T), E359D (1177A>C), L360I (1178T>A), E361Q (1181G>C 1183A>G), T362K (1185C>A 1186G>A), T366V (1196A>G 1197C>T), K367V (1199A>G 1200A>T), D369E (1207T>G), M371L (1211A>C 1213G>C), E373A (1218A>C 1219G>C), I375V (1223A>G), V376A (1227T>C), I379L (1235A>C 1237A>G) | | | | | | | | |

| | Begin | End | Coverage | Score | Concordance | Matches | Identities | I/D/M/F* | Stop Codons |
|---|---|---|---|---|---|---|---|---|---|
| **NT** | **118** | **1238** | **83.7%** | **420** | **18.9%** | **1118 (99.5%)** | **670 (59.6%)** | **3/3** | |

Codon mutations: GCT6..G (118T>G), GAA7AGT (119G>A 120A>G 121A>T), GAC9GCT (126A>C 127C>T), CAT10GAG (128C>G 130T>G), GGC12GGG (136C>G), CTC13ATT (137C>A 139C>T), AAG14AAA (142G>A), AAA15GAT (143A>G 145A>T), AAT16GAC (146A>G 148T>C), ATC17CTT (149A>C 151C>T), CTA18CTT (154A>T), AAG19AGA (156A>G 157G>A), GGT20GGG (160T>G), ATA21ATT (163A>T), TAT22TAC (166T>C), TCT23CAG (167T>C 168C>A 169T>G), TGT24TAC (171G>A 172T>C), GGA25GGG (175A>G), TTT26TTC (178T>C), CCT29CCA (187T>A), TCA30TCT (190A>T), ACT31GCC (191A>G 193T>C), AAA35CGA (203A>C 204A>G), GCC36GCT (208C>T), ATT37GTA (209A>G 211T>A), TTT38TTG (214T>G), CCA39CCT (217A>T), TGT40ATT (218T>A 219G>T), ATT41ATA (223T>A), TCT42TCG (226T>G), AAA44CGT (230A>C 231A>G 232A>T), GTG46GTC (238G>C), ATA47ATT (241A>T), GTC48GCT (243T>C 244C>T), CAA49CAG (247A>G), TCG52TCC (256G>C), GGA53GGT (259A>T), ACA54ACC (262A>C), ACA57ACC (271A>C), GCA58TCC (272G>T 274A>C), ACA59ATG (276C>T 277A>G), TAT60ATT (278T>A 279A>T), GCA61GCT (283A>T), ATC62CTT (284A>C 286C>T), AGT63ACC (288G>C 289T>C), TTA65TGC (294T>G 295A>C), CAG66CAA (298G>A), CAA67GTT (299C>G 300A>T 301A>T), ATA68GTG (302A>G 304A>G), ACT70ACC (310T>C), TCT71AAA (311T>A 312C>A 313T>A), AAT74GAG (320A>G 322T>G), ATC75GTA (323A>G 325C>A), GCT77GCA (331T>A), ATC79ATA (337C>A), ACC81TCT (341A>T 343C>T), CCA82CCT (346A>T), CGT84CGG (352T>G), GAA85GAG (355A>G), TTG86CTT (356T>C 358G>T), GCT87GCA (361T>A), CTC88GCT (362C>G 363T>C 364C>T), GCT90ACG (368G>A 370T>G), CAA91GAA (371C>G), CGT92AAG (374C>A 375G>A 376T>G), GTA93GTG (379A>G), TTG94ATT (380T>A 382G>T), CAA95TTG (383C>T 384A>T 385A>G), ACA96GCA (386A>G), GGG98GGA (394G>A), AAT99GAT (395A>G), TAT100TAC (400T>C), CTA101ATT (401C>A 403A>T), TAT102del (404_406delTAT), AAT103AAC (409T>C), TTT104ATA (410T>A 412T>A), AAA105CAG (413A>C 415A>G), TGT106GCA (416T>G 417G>C 418T>A), CAA107CAT (421A>T), GTT108GCG (423T>C 424T>G), ACA113AAA (438C>A), AGT114AGC (442T>C), ATC115GTG (443A>G 445C>G), AAG116GGT (446A>G 447A>G 448G>T), AGC118GAT (452A>G 453G>A 454C>T), CAG119ATC (455C>A 456A>T 457G>C), GAA120AGA (458G>A 459A>G), ACT121AAG (462C>A 463T>G), AAG123GAG (467A>G), AAG123_AAA124insCAT (469_470insCAT), AAA124GGA (470A>G 471A>G), GCC125GTA (474C>T 475C>A), CAG126CAA (478G>A), TTG128GTC (482T>G 484G>C), ATT129TCT (485A>T 486T>C), CGA134AGA (500C>A), ATG135GTC (503A>G 505G>C), ATT136TGT (506A>T 507T>G), GAT137GAC (511T>C), CTA138ATG (512C>A 514A>G), CTC139ATT (515C>A 517C>T), ACC140AAA (519C>A 520C>A), CGA141AGG (521C>A 523A>G), AAA142AGA (525A>G), AGT143ACT (528G>C), ATC144TTA (530A>T 532C>A), GAT145CGA (533G>C 534A>G 535T>A), AAA147AGG (540A>G 541A>G), GCA148GCC (544A>C), ATT149ATC (547T>C), AAA150AAG (550A>G), ATA151ATG (553A>G), GTT152TTT (554G>T), GTA153GTT (559A>T), ATA154TTG (560A>T 562A>G), GAT158GAC (574T>C), GAG159GAA (577G>A), TTG161CTC (581T>C 583G>C), ATA162TCT (584A>T 585T>C 586A>T), GAT163AGA (587G>A 588A>G 589T>A), AAT164GGT (590A>G 591A>G), TTT165TTC (595T>C), TTG166AAG (596T>A 597T>A), AAA168CAG (602A>C 604A>G), ATA169ATT (607A>T), CAA170TAT (608C>T 610A>T), ATT172ATA (616T>A), GAA174CAG (620G>C 622A>G), TTT175CTT (623T>C), TTT176TTG (628T>G), GAA177CCA (629G>C 630A>C), AGT178CCA (632A>C 633G>C 634T>A), CAT179AAA (635C>A 637T>A), GTC180ATC (638G>A), GTT182GTG (646T>G), ATT183GGT (647A>G 648T>G), TTA184GTT (650T>G 652A>T), TTA185TTC (655A>C), TCA186TCT (658A>T), GCA187GCC (661A>C), ACC188ACT (664C>T), GTT189ATG (665G>A 667T>G), TCA191CCA (671T>C), AGG192GAG (674A>G 675G>A), GTT193GCT (678T>C), ATC194CTT (680A>C 682C>T), AAT195GAG (683A>G 685T>G), ACA196ATT (687C>T 688A>T), TCT197ACT (689T>A), CAG198AGA (692C>A 693A>G 694G>A), GTC199AAG (695G>A 696T>A 697C>G), TTT200TTC (700T>C), AGA202AAC (705G>A 706A>C), GAT203AAG (707G>A 709T>G), GTA205GTG (715A>G), AAA206AGG (717A>G 718A>G), CTG208CTT (724G>T), GTA209GTG (727A>G), AAA210AAG (730A>G), AAT211CGT (731A>C 732A>G), GCT212GAT (735C>A), GAA213GAG (739A>G), TTG214CTC (740T>C 742G>C), ACA215ACT (745A>T), CTC216CTT (748C>T), GGC218GGT (754C>T), AGG220AAG (759G>A), TAT222TTT (765A>T), ATA224GTC (770A>G 772A>C), GTA226GTC (778A>C), AAG227GAT (779A>G 781G>T), AAA228AAG (784A>G), AAT229GAA (785A>G 787T>A), GAT230GAG (790T>G), TTT231TGG (792T>G 793T>G), AAA232AAG (796A>G), GCA233CTT (797G>C 798C>T 799A>T), TTT237TGT (810T>G), TAT240TAC (820T>C), GAC241GAG (823C>G), CAC242ACT (824C>A 825A>C 826C>T), TTA243TTG (829A>G), AGC244GCC (830A>G 831G>C), CTC245ATC (833C>A), ACT246ACC (838T>C), CAA247CAG (841A>G), ACA248AGT (843C>G 844A>T), CTA249GTC (845C>G 847A>C), ATA250ATC (850A>C), TGT252GTT (854T>G 855G>T), AAT253AAC (859T>C), ACA254ACT (862A>T), CGT255AGG (863C>A 865T>G), AGA256CGA (866A>C), CAA257AAG (869C>A 871A>G), GTA260TGG (878G>T 879T>G 880A>G), ATG262ACT (885T>C 886G>T), GAA263GAT (889A>T), TGT264AAA (890T>A 891G>A 892T>A), TTA265ATG (893T>A 895A>G), ACA266CGT (896A>C 897C>G 898A>T), AAT267AGC (900A>G 901T>C), AGA268CGT (902A>C 904A>T), AAT269GAT (905A>G), TTC270CAC (908T>C 909T>A), GCA272GTA (915C>T), TCA273TCT (919A>T), AGC274GCA (920A>G 921G>C 922C>A), ATC275ACT (924T>C 925C>T), GAT278GAC (934T>C), TCC280GAT (938T>G 939C>A 940C>T), CAA281CAG (943A>G), CAA282AAT (944C>A 946A>T), GAA283ACA (947G>A 948A>C), CGT284AGA (950C>A 952T>A), TTT286ATC (956T>A 958T>C), AAA289AGG (966A>G 967A>G), GAA290GAG (970A>G), TTT291TTC (973T>C), AGA292CGA (974A>C), GAT293TCT (977G>T 978A>C), GGT294GGC (982T>C), AAA295TCA (983A>T 984A>C), ACT296TCT (986A>T), AGA297CGT (989A>C 991A>T), ATA298GTG (992A>G 994A>G), CTT300ATC (998C>A 1000T>C), TCA301ACA (1001T>A), AAT303GAT (1007A>G), CTA304CTC (1012A>C), TTA305CTT (1013T>C 1015A>T), GCA306GCT (1018A>T), AGA307CGT (1019A>C 1021A>T), GGC308GGT (1024C>T), GTT311GTA (1033T>A), CAA312CAG (1036A>G), ATA314GTC (1040A>G 1042A>C), TCT315TCC (1045T>C), TTA316CTT (1046T>C 1048A>T), GTT317GTC (1051T>C), ATC318ATT (1054C>T), AAT319AAC (1057T>C), GAT321GAC (1063T>C), TTG322CTG (1064T>C), CAT324ACT (1070C>A 1071A>C), AAT325CAA (1073A>C 1075T>A), AGA326CCA (1076A>C 1077G>C), TAT329TAC (1087T>C), ATA330CTG (1088A>C 1090A>G), AGA335CGT (1103A>C 1105A>T), GGT337GGA (1111T>A), AGA338CGA (1112A>C), GGC340GGA (1120C>A), CGA341AGG (1121C>A 1123A>G), AAA342AAG (1126A>G), GGG343GGT (1129G>T), ATT344GTT (1130A>G), AAT347AAC (1141T>C), ATA349GTC (1145A>G 1147A>C), ACT350ACC (1150T>C), GAA351AAG (1151G>A 1153A>G), GAC352GAT (1156C>T), GAA353GAT (1159A>T), TTA354GAA (1160T>G 1161T>A), CCA355AGG (1163C>A 1164C>G 1165A>G), AAG356ATG (1167A>T), GCT358TTT (1172G>T 1173C>T), GAA359GAC (1177A>C), TTA360ATA (1178T>A), GAA361CAG (1181G>C 1183A>G), ACG362AAA (1185C>A 1186G>A), TAT364TAC (1192T>C), AAT365AAC (1195T>C), ACC366GTC (1196A>G 1197C>T), AAA367GTA (1199A>G 1200A>T), ATA368ATC (1204A>C), GAT369GAG (1207T>G), GAA370GAG (1210A>G), ATG371CTC (1211A>C 1213G>C), GAG373GCC (1218A>C 1219G>C), ATT375GTT (1223A>G), GTT376GCT (1227T>C), TTG378CTC (1232T>C 1234G>C), ATA379CTG (1235A>C 1237A>G)

#### Proteins

| | Begin | End | Coverage | Score | Concordance | Matches | Identities | I/D/M/F* | Stop Codons |
|---|---|---|---|---|---|---|---|---|---|
| hypothetical protein (YP_009666148.1) | 7 | 380 | 98.4% | 1460 | 59.1% | 373 (99.5%) | 207 (55.2%) | 1/1/0/0 | 0 |

Protein mutations: E7S (119G>A 120A>G 121A>T), D9A (126A>C 127C>T), H10E (128C>G 130T>G), L13I (137C>A 139C>T), K15D (143A>G 145A>T), N16D (146A>G 148T>C), I17L (149A>C 151C>T), K19R (156A>G 157G>A), S23Q (167T>C 168C>A 169T>G), C24Y (171G>A 172T>C), T31A (191A>G 193T>C), K35R (203A>C 204A>G), I37V (209A>G 211T>A), F38L (214T>G), C40I (218T>A 219G>T), K44R (230A>C 231A>G 232A>T), V48A (243T>C 244C>T), A58S (272G>T 274A>C), T59M (276C>T 277A>G), Y60I (278T>A 279A>T), I62L (284A>C 286C>T), S63T (288G>C 289T>C), L65C (294T>G 295A>C), Q67V (299C>G 300A>T 301A>T), I68V (302A>G 304A>G), S71K (311T>A 312C>A 313T>A), N74E (320A>G 322T>G), I75V (323A>G 325C>A), T81S (341A>T 343C>T), L88A (362C>G 363T>C 364C>T), A90T (368G>A 370T>G), Q91E (371C>G), R92K (374C>A 375G>A 376T>G), L94I (380T>A 382G>T), Q95L (383C>T 384A>T 385A>G), T96A (386A>G), N99D (395A>G), L101I (401C>A 403A>T), Y102del (404_406delTAT), F104I (410T>A 412T>A), K105Q (413A>C 415A>G), C106A (416T>G 417G>C 418T>A), Q107H (421A>T), V108A (423T>C 424T>G), T113K (438C>A), I115V (443A>G 445C>G), K116G (446A>G 447A>G 448G>T), S118D (452A>G 453G>A 454C>T), Q119I (455C>A 456A>T 457G>C), E120R (458G>A 459A>G), T121K (462C>A 463T>G), K123E (467A>G), K123_K124insH (469_470insCAT), K124G (470A>G 471A>G), A125V (474C>T 475C>A), L128V (482T>G 484G>C), I129S (485A>T 486T>C), M135V (503A>G 505G>C), I136C (506A>T 507T>G), L138M (512C>A 514A>G), L139I (515C>A 517C>T), T140K (519C>A 520C>A), K142R (525A>G), S143T (528G>C), I144L (530A>T 532C>A), D145R (533G>C 534A>G 535T>A), K147R (540A>G 541A>G), I151M (553A>G), V152F (554G>T), I154L (560A>T 562A>G), I162S (584A>T 585T>C 586A>T), D163R (587G>A 588A>G 589T>A), N164G (590A>G 591A>G), L166K (596T>A 597T>A), K168Q (602A>C 604A>G), Q170Y (608C>T 610A>T), E174Q (620G>C 622A>G), F175L (623T>C), F176L (628T>G), E177P (629G>C 630A>C), S178P (632A>C 633G>C 634T>A), H179K (635C>A 637T>A), V180I (638G>A), I183G (647A>G 648T>G), L184V (650T>G 652A>T), L185F (655A>C), V189M (665G>A 667T>G), S191P (671T>C), R192E (674A>G 675G>A), V193A (678T>C), I194L (680A>C 682C>T), N195E (683A>G 685T>G), T196I (687C>T 688A>T), S197T (689T>A), Q198R (692C>A 693A>G 694G>A), V199K (695G>A 696T>A 697C>G), R202N (705G>A 706A>C), D203K (707G>A 709T>G), K206R (717A>G 718A>G), N211R (731A>C 732A>G), A212D (735C>A), R220K (759G>A), Y222F (765A>T), I224V (770A>G 772A>C), K227D (779A>G 781G>T), N229E (785A>G 787T>A), D230E (790T>G), F231W (792T>G 793T>G), A233L (797G>C 798C>T 799A>T), F237C (810T>G), D241E (823C>G), H242T (824C>A 825A>C 826C>T), S244A (830A>G 831G>C), L245I (833C>A), T248S (843C>G 844A>T), L249V (845C>G 847A>C), C252V (854T>G 855G>T), Q257K (869C>A 871A>G), V260W (878G>T 879T>G 880A>G), M262T (885T>C 886G>T), E263D (889A>T), C264K (890T>A 891G>A 892T>A), L265M (893T>A 895A>G), T266R (896A>C 897C>G 898A>T), N267S (900A>G 901T>C), N269D (905A>G), F270H (908T>C 909T>A), A272V (915C>T), S274A (920A>G 921G>C 922C>A), I275T (924T>C 925C>T), S280D (938T>G 939C>A 940C>T), Q282N (944C>A 946A>T), E283T (947G>A 948A>C), F286I (956T>A 958T>C), K289R (966A>G 967A>G), D293S (977G>T 978A>C), K295S (983A>T 984A>C), T296S (986A>T), I298V (992A>G 994A>G), L300I (998C>A 1000T>C), S301T (1001T>A), N303D (1007A>G), I314V (1040A>G 1042A>C), H324T (1070C>A 1071A>C), N325Q (1073A>C 1075T>A), R326P (1076A>C 1077G>C), I330L (1088A>C 1090A>G), I344V (1130A>G), I349V (1145A>G 1147A>C), E351K (1151G>A 1153A>G), E353D (1159A>T), L354E (1160T>G 1161T>A), P355R (1163C>A 1164C>G 1165A>G), K356M (1167A>T), A358F (1172G>T 1173C>T), E359D (1177A>C), L360I (1178T>A), E361Q (1181G>C 1183A>G), T362K (1185C>A 1186G>A), T366V (1196A>G 1197C>T), K367V (1199A>G 1200A>T), D369E (1207T>G), M371L (1211A>C 1213G>C), E373A (1218A>C 1219G>C), I375V (1223A>G), V376A (1227T>C), I379L (1235A>C 1237A>G)

| | Begin | End | Coverage | Score | Concordance | Matches | Identities | I/D/M/F* | Stop Codons |
|---|---|---|---|---|---|---|---|---|---|
| **NT** | **118** | **1238** | **83.7%** | **420** | **18.9%** | **1118 (99.5%)** | **670 (59.6%)** | **3/3** | |
| Codon mutations: | GCT6..G (118T>G), GAA7AGT (119G>A 120A>G 121A>T), GAC9GCT (126A>C 127C>T), CAT10GAG (128C>G 130T>G), GGC12GGG (136C>G), CTC13ATT (137C>A 139C>T), AAG14AAA (142G>A), AAA15GAT (143A>G 145A>T), AAT16GAC (146A>G 148T>C), ATC17CTT (149A>C 151C>T), CTA18CTT (154A>T), AAG19AGA (156A>G 157G>A), GGT20GGG (160T>G), ATA21ATT (163A>T), TAT22TAC (166T>C), TCT23CAG (167T>C 168C>A 169T>G), TGT24TAC (171G>A 172T>C), GGA25GGG (175A>G), TTT26TTC (178T>C), CCT29CCA (187T>A), TCA30TCT (190A>T), ACT31GCC (191A>G 193T>C), AAA35CGA (203A>C 204A>G), GCC36GCT (208C>T), ATT37GTA (209A>G 211T>A), TTT38TTG (214T>G), CCA39CCT (217A>T), TGT40ATT (218T>A 219G>T), ATT41ATA (223T>A), TCT42TCG (226T>G), AAA44CGT (230A>C 231A>G 232A>T), GTG46GTC (238G>C), ATA47ATT (241A>T), GTC48GCT (243T>C 244C>T), CAA49CAG (247A>G), TCG52TCC (256G>C), GGA53GGT (259A>T), ACA54ACC (262A>C), ACA57ACC (271A>C), GCA58TCC (272G>T 274A>C), ACA59ATG (276C>T 277A>G), TAT60ATT (278T>A 279A>T), GCA61GCT (283A>T), ATC62CTT (284A>C 286C>T), AGT63ACC (288G>C 289T>C), TTA65TGC (294T>G 295A>C), CAG66CAA (298G>A), CAA67GTT (299C>G 300A>T 301A>T), ATA68GTG (302A>G 304A>G), ACT70ACC (310T>C), TCT71AAA (311T>A 312C>A 313T>A), AAT74GAG (320A>G 322T>G), ATC75GTA (323A>G 325C>A), GCT77GCA (331T>A), ATC79ATA (337C>A), ACC81TCT (341A>T 343C>T), CCA82CCT (346A>T), CGT84CGG (352T>G), GAA85GAG (355A>G), TTG86CTT (356T>C 358G>T), GCT87GCA (361T>A), CTC88GCT (362C>G 363T>C 364C>T), GCT90ACG (368G>A 370T>G), CAA91GAA (371C>G), CGT92AAG (374C>A 375G>A 376T>G), GTA93GTG (379A>G), TTG94ATT (380T>A 382G>T), CAA95TTG (383C>T 384A>T 385A>G), ACA96GCA (386A>G), GGG98GGA (394G>A), AAT99GAT (395A>G), TAT100TAC (400T>C), CTA101ATT (401C>A 403A>T), TAT102del (404_406delTAT), AAT103AAC (409T>C), TTT104ATA (410T>A 412T>A), AAA105CAG (413A>C 415A>G), TGT106GCA (416T>G 417G>C 418T>A), CAA107CAT (421A>T), GTT108GCG (423T>C 424T>G), ACA113AAA (438C>A), AGT114AGC (442T>C), ATC115GTG (443A>G 445C>G), AAG116GGT (446A>G 447A>G 448G>T), AGC118GAT (452A>G 453G>A 454C>T), CAG119ATC (455C>A 456A>T 457G>C), GAA120AGA (458G>A 459A>G), ACT121AAG (462C>A 463T>G), AAG123GAG (467A>G), AAG123_AAA124insCAT (469_470insCAT), AAA124GGA (470A>G 471A>G), GCC125GTA (474C>T 475C>A), CAG126CAA (478G>A), TTG128GTC (482T>G 484G>C), ATT129TCT (485A>T 486T>C), CGA134AGA (500C>A), ATG135GTC (503A>G 505G>C), ATT136TGT (506A>T 507T>G), GAT137GAC (511T>C), CTA138ATG (512C>A 514A>G), CTC139ATT (515C>A 517C>T), ACC140AAA (519C>A 520C>A), CGA141AGG (521C>A 523A>G), AAA142AGA (525A>G), AGT143ACT (528G>C), ATC144TTA (530A>T 532C>A), GAT145CGA (533G>C 534A>G 535T>A), AAA147AGG (540A>G 541A>G), GCA148GCC (544A>C), ATT149ATC (547T>C), AAA150AAG (550A>G), ATA151ATG (553A>G), GTT152TTT (554G>T), GTA153GTT (559A>T), ATA154TTG (560A>T 562A>G), GAT158GAC (574T>C), GAG159GAA (577G>A), TTG161CTC (581T>C 583G>C), ATA162TCT (584A>T 585T>C 586A>T), GAT163AGA (587G>A 588A>G 589T>A), AAT164GGT (590A>G 591A>G), TTT165TTC (595T>C), TTG166AAG (596T>A 597T>A), AAA168CAG (602A>C 604A>G), ATA169ATT (607A>T), CAA170TAT (608C>T 610A>T), ATT172ATA (616T>A), GAA174CAG (620G>C 622A>G), TTT175CTT (623T>C), TTT176TTG (628T>G), GAA177CCA (629G>C 630A>C), AGT178CCA (632A>C 633G>C 634T>A), CAT179AAA (635C>A 637T>A), GTC180ATC (638G>A), GTT182GTG (646T>G), ATT183GGT (647A>G 648T>G), TTA184GTT (650T>G 652A>T), TTA185TTC (655A>C), TCA186TCT (658A>T), GCA187GCC (661A>C), ACC188ACT (664C>T), GTT189ATG (665G>A 667T>G), TCA191CCA (671T>C), AGG192GAG (674A>G 675G>A), GTT193GCT (678T>C), ATC194CTT (680A>C 682C>T), AAT195GAG (683A>G 685T>G), ACA196ATT (687C>T 688A>T), TCT197ACT (689T>A), CAG198AGA (692C>A 693A>G 694G>A), GTC199AAG (695G>A 696T>A 697C>G), TTT200TTC (700T>C), AGA202AAC (705G>A 706A>C), GAT203AAG (707G>A 709T>G), GTA205GTG (715A>G), AAA206AGG (717A>G 718A>G), CTG208CTT (724G>T), GTA209GTG (727A>G), AAA210AAG (730A>G), AAT211CGT (731A>C 732A>G), GCT212GAT (735C>A), GAA213GAG (739A>G), TTG214CTC (740T>C 742G>C), ACA215ACT (745A>T), CTC216CTT (748C>T), GGC218GGT (754C>T), AGG220AAG (759G>A), TAT222TTT (765A>T), ATA224GTC (770A>G 772A>C), GTA226GTC (778A>C), AAG227GAT (779A>G 781G>T), AAA228AAG (784A>G), AAT229GAA (785A>G 787T>A), GAT230GAG (790T>G), TTT231TGG (792T>G 793T>G), AAA232AAG (796A>G), GCA233CTT (797G>C 798C>T 799A>T), TTT237TGT (810T>G), TAT240TAC (820T>C), GAC241GAG (823C>G), CAC242ACT (824C>A 825A>C 826C>T), TTA243TTG (829A>G), AGC244GCC (830A>G 831G>C), CTC245ATC (833C>A), ACT246ACC (838T>C), CAA247CAG (841A>G), ACA248AGT (843C>G 844A>T), CTA249GTC (845C>G 847A>C), ATA250ATC (850A>C), TGT252GTT (854T>G 855G>T), AAT253AAC (859T>C), ACA254ACT (862A>T), CGT255AGG (863C>A 865T>G), AGA256CGA (866A>C), CAA257AAG (869C>A 871A>G), GTA260TGG (878G>T 879T>G 880A>G), ATG262ACT (885T>C 886G>T), GAA263GAT (889A>T), TGT264AAA (890T>A 891G>A 892T>A), TTA265ATG (893T>A 895A>G), ACA266CGT (896A>C 897C>G 898A>T), AAT267AGC (900A>G 901T>C), AGA268CGT (902A>C 904A>T), AAT269GAT (905A>G), TTC270CAC (908T>C 909T>A), GCA272GTA (915C>T), TCA273TCT (919A>T), AGC274GCA (920A>G 921G>C 922C>A), ATC275ACT (924T>C 925C>T), GAT278GAC (934T>C), TCC280GAT (938T>G 939C>A 940C>T), CAA281CAG (943A>G), CAA282AAT (944C>A 946A>T), GAA283ACA (947G>A 948A>C), CGT284AGA (950C>A 952T>A), TTT286ATC (956T>A 958T>C), AAA289AGG (966A>G 967A>G), GAA290GAG (970A>G), TTT291TTC (973T>C), AGA292CGA (974A>C), GAT293TCT (977G>T 978A>C), GGT294GGC (982T>C), AAA295TCA (983A>T 984A>C), ACT296TCT (986A>T), AGA297CGT (989A>C 991A>T), ATA298GTG (992A>G 994A>G), CTT300ATC (998C>A 1000T>C), TCA301ACA (1001T>A), AAT303GAT (1007A>G), CTA304CTC (1012A>C), TTA305CTT (1013T>C 1015A>T), GCA306GCT (1018A>T), AGA307CGT (1019A>C 1021A>T), GGC308GGT (1024C>T), GTT311GTA (1033T>A), CAA312CAG (1036A>G), ATA314GTC (1040A>G 1042A>C), TCT315TCC (1045T>C), TTA316CTT (1046T>C 1048A>T), GTT317GTC (1051T>C), ATC318ATT (1054C>T), AAT319AAC (1057T>C), GAT321GAC (1063T>C), TTG322CTG (1064T>C), CAT324ACT (1070C>A 1071A>C), AAT325CAA (1073A>C 1075T>A), AGA326CCA (1076A>C 1077G>C), TAT329TAC (1087T>C), ATA330CTG (1088A>C 1090A>G), AGA335CGT (1103A>C 1105A>T), GGT337GGA (1111T>A), AGA338CGA (1112A>C), GGC340GGA (1120C>A), CGA341AGG (1121C>A 1123A>G), AAA342AAG (1126A>G), GGG343GGT (1129G>T), ATT344GTT (1130A>G), AAT347AAC (1141T>C), ATA349GTC (1145A>G 1147A>C), ACT350ACC (1150T>C), GAA351AAG (1151G>A 1153A>G), GAC352GAT (1156C>T), GAA353GAT (1159A>T), TTA354GAA (1160T>G 1161T>A), CCA355AGG (1163C>A 1164C>G 1165A>G), AAG356ATG (1167A>T), GCT358TTT (1172G>T 1173C>T), GAA359GAC (1177A>C), TTA360ATA (1178T>A), GAA361CAG (1181G>C 1183A>G), ACG362AAA (1185C>A 1186G>A), TAT364TAC (1192T>C), AAT365AAC (1195T>C), ACC366GTC (1196A>G 1197C>T), AAA367GTA (1199A>G 1200A>T), ATA368ATC (1204A>C), GAT369GAG (1207T>G), GAA370GAG (1210A>G), ATG371CTC (1211A>C 1213G>C), GAG373GCC (1218A>C 1219G>C), ATT375GTT (1223A>G), GTT376GCT (1227T>C), TTG378CTC (1232T>C 1234G>C), ATA379CTG (1235A>C 1237A>G) | | | | | | | | |

*: Inserts / Deletes / Misaligned / Frameshifts

#### Analysis details

This analysis was performed with panviral2.64

### NGS Details (UN8): Bracoviriform glomeratae (segment NC_043292.1)

#### Assembly

| | |
|---|---|
| **Coverage Length** | 310 (1 contig(s)) |
| **Depth Of Coverage** | 2096.9 |
| **Number Of Reads** | 5742 |
| **Reads Per Million** | 95.96 rpm (after QC) |
| **Ambiguities** | 0 |
| **Assembly Method** | de novo + reference guided assembly |
| **Consensus Caller** | Bcf Tools |

#### Coverage Map

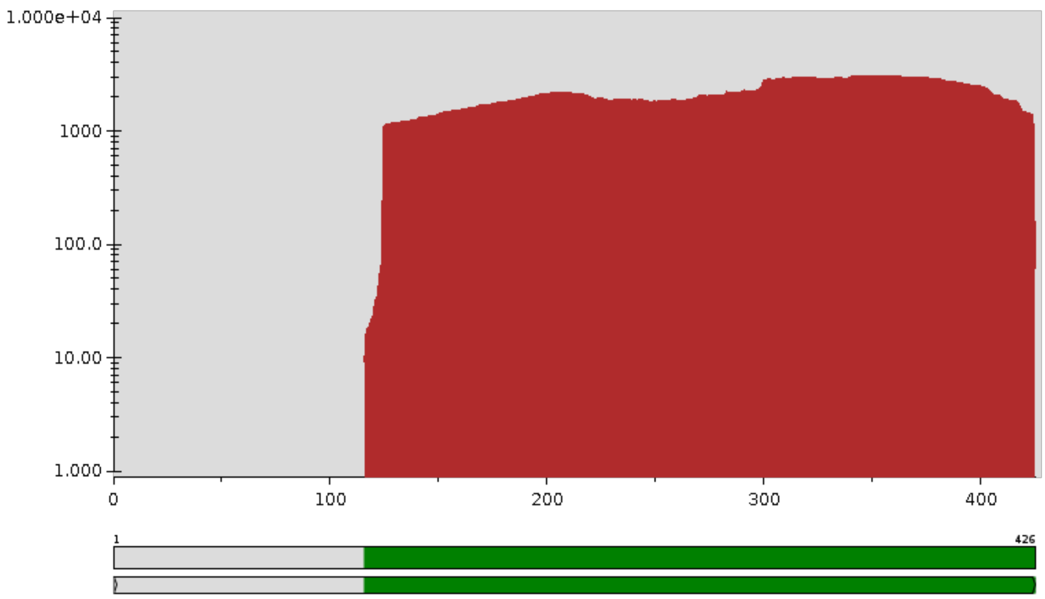

#### Assignment

| | |
|---|---|
| **Type** | Bracoviriform glomeratae (Taxonomy ID: 257816) |
| **Reference Genome** | NC_043292.1 |
| **NT Identity (%)** | 72.5806 |
| **AA Identity (%)** | 84.466 |
| **Number Of Stop Codons** | 1 |
| **Number Of CDS** | 1 |

#### Alignment

| | |
|---|---|
| **Alignment Score** | 280.0 (NT) + 567.0 (AA) = 847.0 |
| **Concordance (%)** | 66.3273 |

| Alignment Method | Global, seeded, nucleotide + amino acids (AGA) |
|---|---|

#### Genome Region

Sequence starts at position 117 and ends at position 426 relative to NC_043292.1 reference sequence.

#### Alignment Detailed Statistics

| | Begin | End | Coverage | Score | Concordance | Matches | Identities | I/D/M/F* | Stop Codons |
|---|---|---|---|---|---|---|---|---|---|
| **NT** | **117** | **426** | **72.8%** | **280** | **45.2%** | **310 (100%)** | **225 (72.6%)** | **0/0** | |
| Mutations: | 126G>T, 129C>A, 132A>G, 136A>G, 137C>G, 138G>C, 150A>C, 153A>G, 155C>G, 156G>A, 159C>A, 166C>A, 168T>G, 169T>C, 171T>C, 172C>A, 174C>G, 177A>G, 183T>A, 184C>A, 189T>C, 198A>G, 201A>C, 207C>T, 214A>G, 219C>T, 222T>C, 223C>A, 225T>G, 228G>T, 235C>A, 237T>G, 243A>T, 246C>G, 255C>T, 261A>G, 262T>C, 264A>G, 265G>A, 273A>G, 278T>C, 279T>C, 284A>G, 285T>A, 288T>G, 289T>C, 300C>T, 301C>T, 303C>G, 309C>T, 312T>G, 315C>T, 322G>T, 329T>C, 333T>C, 336C>T, 339A>G, 345A>C, 347A>G, 348A>G, 349C>A, 351T>A, 360C>T, 363C>T, 366C>T, 375C>T, 378C>T, 384A>C, 385T>C, 387G>C, 390A>G, 391C>A, 392A>G, 393T>A, 394A>C, 396A>G, 399C>A, 400C>A, 402T>G, 404T>C, 405G>T, 406A>C, 414T>G, 416A>T, 420A>T | | | | | | | | |
| **CDS** | | | | | | | | | |
| FK954_p501 | 40 | 142 | 72.5% | 567 | 85.3% | 103 (100%) | 87 (84.5%) | 0/0/0/0 | 1 |
| Protein mutations: | L42F (126G>T), T46G (136A>G 137C>G 138G>C), A52G (155C>G 156G>A), Y57H (169T>C 171T>C), T72A (214A>G), V89I (265G>A), I93T (278T>C 279T>C), D95G (284A>G 285T>A), A108S (322G>T), I110T (329T>C), K116R (347A>G 348A>G), H131R (391C>A 392A>G 393T>A), K132Q (394A>C 396A>G), M135T (404T>C 405G>T), I136L (406A>C), Y139F (416A>T) | | | | | | | | |
| Codon mutations: | TTG42TTT (126G>T), GGC43GGA (129C>A), AAA44AAG (132A>G), ACG46GGC (136A>G 137C>G 138G>C), GGA50GGC (150A>C), AAA51AAG (153A>G), GCG52GGA (155C>G 156G>A), GGC53GGA (159C>A), CGT56AGG (166C>A 168T>G), TAT57CAC (169T>C 171T>C), CGC58AGG (172C>A 174C>G), AAA59AAG (177A>G), CTT61CTA (183T>A), CGA62AGA (184C>A), GAT63GAC (189T>C), CAA66CAG (198A>G), GGA67GGC (201A>C), ACC69ACT (207C>T), ACT72GCT (214A>G), ATC73ATT (219C>T), CGT74CGC (222T>C), CGT75AGG (223C>A 225T>G), CTG76CTT (228G>T), CGT79AGG (235C>A 237T>G), GGA81GGT (243A>T), GTC82GTG (246C>G), ATC85ATT (255C>T), GGA87GGG (261A>G), TTA88CTG (262T>C 264A>G), GTC89ATC (265G>A), GAA91GAG (273A>G), ATT93ACC (278T>C 279T>C), GAT95GGA (284A>G 285T>A), GTT96GTG (288T>G), TTG97CTG (289T>C), TTC100TTT (300C>T), CTC101TTG (301C>T 303C>G), AAC103AAT (309C>T), GTT104GTG (312T>G), ATC105ATT (315C>T), GCT108TCT (322G>T), ATC110ACC (329T>C), TAT111TAC (333T>C), ACC112ACT (336C>T), GAA113GAG (339A>G), GCA115GCC (345A>C), AAA116AGG (347A>G 348A>G), CGT117AGA (349C>A 351T>A), GTC120GTT (360C>T), ACC121ACT (363C>T), GCC122GCT (366C>T), GTC125GTT (375C>T), GTC126GTT (378C>T), GCA128GCC (384A>C), TTG129CTC (385T>C 387G>C), AAA130AAG (390A>G), CAT131AGA (391C>A 392A>G 393T>A), AAA132CAG (394A>C 396A>G), GGC133GGA (399C>A), CGT134AGG (400C>A 402T>G), ATG135ACT (404T>C 405G>T), ATC136CTC (406A>C), GGT138GGG (414T>G), TAT139TTT (416A>T), GGA140GGT (420A>T) | | | | | | | | |
| **Proteins** | | | | | | | | | |
| putative histone 4 (YP_009665791.1) | 40 | 142 | 72.5% | 567 | 85.3% | 103 (100%) | 87 (84.5%) | 0/0/0/0 | 1 |
| Protein mutations: | L42F (126G>T), T46G (136A>G 137C>G 138G>C), A52G (155C>G 156G>A), Y57H (169T>C 171T>C), T72A (214A>G), V89I (265G>A), I93T (278T>C 279T>C), D95G (284A>G 285T>A), A108S (322G>T), I110T (329T>C), K116R (347A>G 348A>G), H131R (391C>A 392A>G 393T>A), K132Q (394A>C 396A>G), M135T (404T>C 405G>T), I136L (406A>C), Y139F (416A>T) | | | | | | | | |
| Codon mutations: | TTG42TTT (126G>T), GGC43GGA (129C>A), AAA44AAG (132A>G), ACG46GGC (136A>G 137C>G 138G>C), GGA50GGC (150A>C), AAA51AAG (153A>G), GCG52GGA (155C>G 156G>A), GGC53GGA (159C>A), CGT56AGG (166C>A 168T>G), TAT57CAC (169T>C 171T>C), CGC58AGG (172C>A 174C>G), AAA59AAG (177A>G), CTT61CTA (183T>A), CGA62AGA (184C>A), GAT63GAC (189T>C), CAA66CAG (198A>G), GGA67GGC (201A>C), ACC69ACT (207C>T), ACT72GCT (214A>G), ATC73ATT (219C>T), CGT74CGC (222T>C), CGT75AGG (223C>A 225T>G), CTG76CTT (228G>T), CGT79AGG (235C>A 237T>G), GGA81GGT (243A>T), GTC82GTG (246C>G), ATC85ATT (255C>T), GGA87GGG (261A>G), TTA88CTG (262T>C 264A>G), GTC89ATC (265G>A), GAA91GAG (273A>G), ATT93ACC (278T>C 279T>C), GAT95GGA (284A>G 285T>A), GTT96GTG (288T>G), TTG97CTG (289T>C), TTC100TTT (300C>T), CTC101TTG (301C>T 303C>G), AAC103AAT (309C>T), GTT104GTG (312T>G), ATC105ATT (315C>T), GCT108TCT (322G>T), ATC110ACC (329T>C), TAT111TAC (333T>C), ACC112ACT (336C>T), GAA113GAG (339A>G), GCA115GCC (345A>C), AAA116AGG (347A>G 348A>G), CGT117AGA (349C>A 351T>A), GTC120GTT (360C>T), ACC121ACT (363C>T), GCC122GCT (366C>T), GTC125GTT (375C>T), GTC126GTT (378C>T), GCA128GCC (384A>C), TTG129CTC (385T>C 387G>C), AAA130AAG (390A>G), CAT131AGA (391C>A 392A>G 393T>A), AAA132CAG (394A>C 396A>G), GGC133GGA (399C>A), CGT134AGG (400C>A 402T>G), ATG135ACT (404T>C 405G>T), ATC136CTC (406A>C), GGT138GGG (414T>G), TAT139TTT (416A>T), GGA140GGT (420A>T) | | | | | | | | |

*: Inserts / Deletes / Misaligned / Frameshifts

#### Analysis details

This analysis was performed with panviral2.64

#### NGS Details (UN8): Potato leafroll virus

##### Assembly

| | |
|---|---|
| **Coverage Length** | 5827 (1 contig(s)) |
| **Depth Of Coverage** | 72.1 |
| **Number Of Reads** | 3146 |
| **Reads Per Million** | 52.58 rpm (after QC) |
| **Ambiguities** | 0 |
| **Assembly Method** | read mapping against reference + variant calling |
| **Consensus Caller** | Bcf Tools |

##### Coverage Map

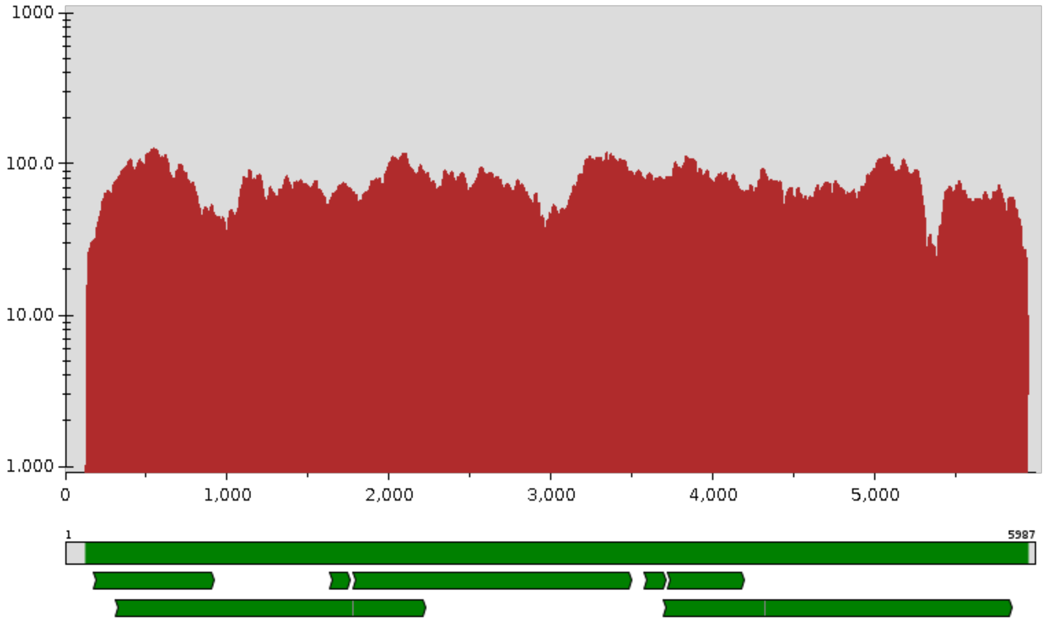

##### Assignment

| | |
|---|---|
| **Type** | Potato leafroll virus (Taxonomy ID: 12045) |
| **Reference Genome** | NC_001747.1 |
| **NT Identity (%)** | 98.3173 |
| **AA Identity (%)** | 97.8212 |
| **Number Of Stop Codons** | 9 |
| **Number Of CDS** | 8 |

##### Alignment

| | |
|---|---|
| **Alignment Score** | 11226.0 (NT) + 20943.0 (AA) = 32169.0 |
| **Concordance (%)** | 97.2882 |

| Alignment Method | Global, seeded, nucleotide + amino acids (AGA) |
|---|---|

#### Genome Region

Sequence starts at position 122 and ends at position 5948 relative to NC_001747.1 reference sequence.

#### Alignment Detailed Statistics

| | Begin | End | Coverage | Score | Concordance | Matches | Identities | I/D/M/F* | Stop Codons |
|---|---|---|---|---|---|---|---|---|---|
| **NT** | **122** | **5948** | **97.3%** | **11226** | **96.6%** | **5822 (99.9%)** | **5726 (98.2%)** | **2/5** | |
| Mutations: | 656G>A, 665A>G, 699A>C, 822C>T, 825T>G, 903G>A, 931A>G, 958G>A, 1036G>A, 1126G>A, 1132T>G, 1147C>T, 1171C>T, 1204A>G, 1234A>T, 1300T>C, 1319G>A, 1327C>T, 1356G>A, 1577G>T, 1593G>A, 1611A>G, 1716A>G, 1754C>T, 1779A>G, 1894C>A, 1909A>C, 1915G>A, 1957C>A, 2007T>C, 2051G>A, 2154C>T, 2265A>G, 2283G>A, 2313A>G, 2316T>G, 2470G>A, 2555C>T, 2586A>G, 2640C>T, 2649C>T, 2729A>G, 2849C>T, 2856G>A, 2877C>T, 2898T>C, 2931T>C, 3000T>C, 3033C>T, 3036C>T, 3069C>T, 3117C>T, 3171A>G, 3186T>C, 3321C>T, 3378G>T, 3379A>G, 3387C>T, 3614T>A, 3619C>T, 3652T>A, 3721T>G, 3748_3750delTGC, 3788T>G, 3830C>T, 3939A>T, 4031A>G, 4094G>A, 4113G>A, 4166C>T, 4172T>C, 4208C>T, 4214T>G, 4658G>A, 4661T>C, 4733C>G, 4817T>C, 4883C>T, 4886T>C, 4913A>T, 4949G>A, 4964A>G, 4979C>T, 5038C>T, 5062C>T, 5071A>G, 5072C>T, 5129G>A, 5162C>T, 5189C>T, 5199C>T, 5216A>C, 5268A>G, 5689_5690insGA, 5692_5693delGT, 5741T>G, 5748A>G, 5781G>A, 5925T>C | | | | | | | | |
| **CDS** | | | | | | | | | |
| PLRVgp1 | 1 | 248 | 100% | 1715 | 99.0% | 248 (100%) | 245 (98.8%) | 0/0/0/0 | 1 |
| Protein mutations: | S161N (656G>A), Q164R (665A>G), Q175H (699A>C) | | | | | | | | |
| Codon mutations: | AGT161AAT (656G>A), CAA164CGA (665A>G), CAA175CAC (699A>C), CGC216CGT (822C>T), GCT217GCG (825T>G), AAG243AAA (903G>A) | | | | | | | | |
| PLRVgp2 | 1 | 1063 | 100% | 7327 | 98.2% | 1063 (100%) | 1041 (97.9%) | 0/0/0/0 | 1 |
| Protein mutations: | V117I (656G>A), K120E (665A>G), N131T (699A>C), A172V (822C>T), L173R (825T>G), R199K (903G>A), M273I (1126G>A), A338T (1319G>A), G350E (1356G>A), V424F (1577G>T), R429K (1593G>A), D435G (1611A>G), E470G (1716A>G), Q530K (1894C>A), E537K (1915G>A), Q551K (1957C>A), G582E (2051G>A), A722T (2470G>A), A750V (2555C>T), N808S (2729A>G), P848L (2849C>T), I1025V (3379A>G) | | | | | | | | |
| Codon mutations: | GTA117ATA (656G>A), AAA120GAA (665A>G), AAC131ACC (699A>C), GCG172GTG (822C>T), CTA173CGA (825T>G), AGG199AAG (903G>A), GAA208GAG (931A>G), CAG217CAA (958G>A), TTG243TTA (1036G>A), ATG273ATA (1126G>A), ACT275ACG (1132T>G), TTC280TTT (1147C>T), TCC288TCT (1171C>T), CTA299CTG (1204A>G), ACA309ACT (1234A>T), CAT331CAC (1300T>C), GCC338ACC (1319G>A), CAC340CAT (1327C>T), GGA350GAA (1356G>A), GTC424TTC (1577G>T), AGA429AAA (1593G>A), GAC435GGC (1611A>G), GAG470GGG (1716A>G), CTA483TTA (1754C>T), GCA491GCG (1779A>G), CAA530AAA (1894C>A), AGA535CGA (1909A>C), GAA537AAA (1915G>A), CAA551AAA (1957C>A), TTT567TTC (2007T>C), GGA582GAA (2051G>A), CAC616CAT (2154C>T), TCA653TCG (2265A>G), GCG659GCA (2283G>A), ACA669ACG (2313A>G), GTT670GTG (2316T>G), GCA722ACA (2470G>A), GCC750GTC (2555C>T), CTA760CTG (2586A>G), CAC778CAT (2640C>T), AGC781AGT (2649C>T), AAT808AGT (2729A>G), CCA848CTA (2849C>T), GTG850GTA (2856G>A), CAC857CAT (2877C>T), ACT864ACC (2898T>C), TAT875TAC (2931T>C), CGT898CGC (3000T>C), AAC909AAT (3033C>T), TCC910TCT (3036C>T), GCC921GCT (3069C>T), TCC937TCT (3117C>T), GCA955GCG (3171A>G), GAT960GAC (3186T>C), AAC1005AAT (3321C>T), GTG1024GTT (3378G>T), ATT1025GTT (3379A>G), AAC1027AAT (3387C>T) | | | | | | | | |
| PLRVgp3 | 1 | 640 | 100% | 4199 | 97.8% | 640 (100%) | 623 (97.3%) | 0/0/0/0 | 1 |
| Protein mutations: | V117I (656G>A), K120E (665A>G), N131T (699A>C), A172V (822C>T), L173R (825T>G), R199K (903G>A), M273I (1126G>A), A338T (1319G>A), G350E (1356G>A), V424F (1577G>T), R429K (1593G>A), D435G (1611A>G), E470G (1716A>G), Q491R (1779A>G), L567S (2007T>C), E582K (2051G>A), T616I (2154C>T) | | | | | | | | |
| Codon mutations: | GTA117ATA (656G>A), AAA120GAA (665A>G), AAC131ACC (699A>C), GCG172GTG (822C>T), CTA173CGA (825T>G), AGG199AAG (903G>A), GAA208GAG (931A>G), CAG217CAA (958G>A), TTG243TTA (1036G>A), ATG273ATA (1126G>A), ACT275ACG (1132T>G), TTC280TTT (1147C>T), TCC288TCT (1171C>T), CTA299CTG (1204A>G), ACA309ACT (1234A>T), CAT331CAC (1300T>C), GCC338ACC (1319G>A), CAC340CAT (1327C>T), GGA350GAA (1356G>A), GTC424TTC (1577G>T), AGA429AAA (1593G>A), GAC435GGC (1611A>G), GAG470GGG (1716A>G), CTA483TTA (1754C>T), CAA491CGA (1779A>G), ATC529ATA (1894C>A), ATA534ATC (1909A>C), GAG536GAA (1915G>A), CCC550CCA (1957C>A), TTA567TCA (2007T>C), GAG582AAG (2051G>A), ACC616ATC (2154C>T) | | | | | | | | |
| Rap1 | 1 | 42 | 100% | 277 | 96.5% | 42 (100%) | 41 (97.6%) | 0/0/0/0 | 1 |
| Protein mutations: | R29G (1716A>G) | | | | | | | | |
| Codon mutations: | AGA29GGA (1716A>G), CTC41CTT (1754C>T) | | | | | | | | |
| ORF3a | 1 | 46 | 100% | 289 | 96.7% | 46 (100%) | 44 (95.7%) | 0/0/0/0 | 1 |
| Protein mutations: | S14T (3614T>A), F26L (3652T>A) | | | | | | | | |
| Codon mutations: | TCA14ACA (3614T>A), TCC15TCT (3619C>T), TTT26TTA (3652T>A) | | | | | | | | |
| PLRVgp4 | 1 | 718 | 100% | 4695 | 93.4% | 717 (99.7%) | 705 (98.1%) | 1/1/2/2 | 2 |
| Protein mutations: | V10G (3721T>G), M19del (3748_3750delTGC), T83S (3939A>T), V141I (4113G>A), P449L (5038C>T), A457V (5062C>T), H460R (5071A>G 5072C>T), P503S (5199C>T), T526A (5268A>G), E666_R667insX (5689_5690insGA), F683L (5741T>G), T686A (5748A>G), E697K (5781G>A) | | | | | | | | |
| Codon mutations: | GTC10GGC (3721T>G), ATG19del (3748_3750delTGC), CGA20AGA (3748_3750delTGC), GTT32GTG (3788T>G), CGC46CGT (3830C>T), ACC83TCC (3939A>T), ACA113ACG (4031A>G), GAG134GAA (4094G>A), GTA141ATA (4113G>A), GCC158GCT (4166C>T), ACT160ACC (4172T>C), CAC172CAT (4208C>T), TCT174TCG (4214T>G), CGG322CGA (4658G>A), GGT323GGC (4661T>C), GTC347GTG (4733C>G), GGT375GGC (4817T>C), TTC397TTT (4883C>T), CTT398CTC (4886T>C), GCA407GCT (4913A>T), ACG419ACA (4949G>A), GAA424GAG (4964A>G), ACC429ACT (4979C>T), CCA449CTA (5038C>T), GCC457GTC (5062C>T), CAC460CGT (5071A>G 5072C>T), ACG479ACA (5129G>A), GAC490GAT (5162C>T), AAC499AAT (5189C>T), CCA503TCA (5199C>T), CGA508CGC (5216A>C), ACA526GCA (5268A>G), GAA666GAG (5689_5690insGA), GAA666_CGT667insA-A (5689_5690insGA), CGT667C-- (5692_5693delGT), TTT683TTG (5741T>G), ACT686GCT (5748A>G), GAA697AAA (5781G>A) | | | | | | | | |
| PLRVgp5 | 1 | 209 | 100% | 1411 | 99.2% | 208 (99.5%) | 205 (98.1%) | 0/1/0/0 | 1 |
| Protein mutations: | V10G (3721T>G), M19del (3748_3750delTGC), T83S (3939A>T), V141I (4113G>A) | | | | | | | | |
| Codon mutations: | GTC10GGC (3721T>G), ATG19del (3748_3750delTGC), CGA20AGA (3748_3750delTGC), GTT32GTG (3788T>G), CGC46CGT (3830C>T), ACC83TCC (3939A>T), ACA113ACG (4031A>G), GAG134GAA (4094G>A), GTA141ATA (4113G>A), GCC158GCT (4166C>T), ACT160ACC (4172T>C), CAC172CAT (4208C>T), TCT174TCG (4214T>G) | | | | | | | | |
| PLRVgp6 | 1 | 157 | 100% | 1030 | 95.5% | 156 (99.4%) | 149 (94.9%) | 0/1/0/0 | 1 |
| Protein mutations: | S2A (3721T>G), C11del (3748_3750delTGC), F24C (3788T>G), A38V (3830C>T), Q105R (4031A>G), S126N (4094G>A), P150L (4166C>T), L152P (4172T>C) | | | | | | | | |
| Codon mutations: | TCA2GCA (3721T>G), TGC11del (3748_3750delTGC), TTC24TGC (3788T>G), GCC38GTC (3830C>T), ACA74ACT (3939A>T), CAA105CGA (4031A>G), AGT126AAT (4094G>A), AAG132AAA (4113G>A), CCA150CTA (4166C>T), CTT152CCT (4172T>C) | | | | | | | | |
| **Proteins** | | | | | | | | | |
| P0 protein (NP_056746.1) | 1 | 248 | 100% | 1715 | 99.0% | 248 (100%) | 245 (98.8%) | 0/0/0/0 | 1 |
| Protein mutations: | S161N (656G>A), Q164R (665A>G), Q175H (699A>C) | | | | | | | | |
| Codon mutations: | AGT161AAT (656G>A), CAA164CGA (665A>G), CAA175CAC (699A>C), CGC216CGT (822C>T), GCT217GCG (825T>G), AAG243AAA (903G>A) | | | | | | | | |

| | Begin | End | Coverage | Score | Concordance | Matches | Identities | I/D/M/F* | Stop Codons |
|---|---|---|---|---|---|---|---|---|---|
| **NT** | **122** | **5948** | **97.3%** | **11226** | **96.6%** | **5822 (99.9%)** | **5726 (98.2%)** | **2/5** | |
| RNA-dependent RNA polymerase (NP_056748.3) | 1 | 1063 | 100% | 7327 | 98.2% | 1063 (100%) | 1041 (97.9%) | 0/0/0/0 | 1 |
| Protein mutations: | V117I (656G>A), K120E (665A>G), N131T (699A>C), A172V (822C>T), L173R (825T>G), R199K (903G>A), M273I (1126G>A), A338T (1319G>A), G350E (1356G>A), V424F (1577G>T), R429K (1593G>A), D435G (1611A>G), E470G (1716A>G), Q530K (1894C>A), E537K (1915G>A), Q551K (1957C>A), G582E (2051G>A), A722T (2470G>A), A750V (2555C>T), N808S (2729A>G), P848L (2849C>T), I1025V (3379A>G) | | | | | | | | |
| Codon mutations: | GTA117ATA (656G>A), AAA120GAA (665A>G), AAC131ACC (699A>C), GCG172GTG (822C>T), CTA173CGA (825T>G), AGG199AAG (903G>A), GAA208GAG (931A>G), CAG217CAA (958G>A), TTG243TTA (1036G>A), ATG273ATA (1126G>A), ACT275ACG (1132T>G), TTC280TTT (1147C>T), TCC288TCT (1171C>T), CTA299CTG (1204A>G), ACA309ACT (1234A>T), CAT331CAC (1300T>C), GCC338ACC (1319G>A), CAC340CAT (1327C>T), GGA350GAA (1356G>A), GTC424TTC (1577G>T), AGA429AAA (1593G>A), GAC435GGC (1611A>G), GAG470GGG (1716A>G), CTA483TTA (1754C>T), GCA491GCG (1779A>G), CAA530AAA (1894C>A), AGA535CGA (1909A>C), GAA537AAA (1915G>A), CAA551AAA (1957C>A), TTT567TTC (2007T>C), GGA582GAA (2051G>A), CAC616CAT (2154C>T), TCA653TCG (2265A>G), GCG659GCA (2283G>A), ACA669ACG (2313A>G), GTT670GTG (2316T>G), GCA722ACA (2470G>A), GCC750GTC (2555C>T), CTA760CTG (2586A>G), CAC778CAT (2640C>T), AGC781AGT (2649C>T), AAT808AGT (2729A>G), CCA848CTA (2849C>T), GTG850GTA (2856G>A), CAC857CAT (2877C>T), ACT864ACC (2898T>C), TAT875TAC (2931T>C), CGT898CGC (3000T>C), AAC909AAT (3033C>T), TCC910TCT (3036C>T), GCC921GCT (3069C>T), TCC937TCT (3117C>T), GCA955GCG (3171A>G), GAT960GAC (3186T>C), AAC1005AAT (3321C>T), GTG1024GTT (3378G>T), ATT1025GTT (3379A>G), AAC1027AAT (3387C>T) | | | | | | | | |
| P1 protein (NP_056747.1) | 1 | 640 | 100% | 4199 | 97.8% | 640 (100%) | 623 (97.3%) | 0/0/0/0 | 1 |
| Protein mutations: | V117I (656G>A), K120E (665A>G), N131T (699A>C), A172V (822C>T), L173R (825T>G), R199K (903G>A), M273I (1126G>A), A338T (1319G>A), G350E (1356G>A), V424F (1577G>T), R429K (1593G>A), D435G (1611A>G), E470G (1716A>G), Q491R (1779A>G), L567S (2007T>C), E582K (2051G>A), T616I (2154C>T) | | | | | | | | |
| Codon mutations: | GTA117ATA (656G>A), AAA120GAA (665A>G), AAC131ACC (699A>C), GCG172GTG (822C>T), CTA173CGA (825T>G), AGG199AAG (903G>A), GAA208GAG (931A>G), CAG217CAA (958G>A), TTG243TTA (1036G>A), ATG273ATA (1126G>A), ACT275ACG (1132T>G), TTC280TTT (1147C>T), TCC288TCT (1171C>T), CTA299CTG (1204A>G), ACA309ACT (1234A>T), CAT331CAC (1300T>C), GCC338ACC (1319G>A), CAC340CAT (1327C>T), GGA350GAA (1356G>A), GTC424TTC (1577G>T), AGA429AAA (1593G>A), GAC435GGC (1611A>G), GAG470GGG (1716A>G), CTA483TTA (1754C>T), CAA491CGA (1779A>G), ATC529ATA (1894C>A), ATA534ATC (1909A>C), GAG536GAA (1915G>A), CCC550CCA (1957C>A), TTA567TCA (2007T>C), GAG582AAG (2051G>A), ACC616ATC (2154C>T) | | | | | | | | |
| Replication-associated protein (YP_006355442.1) | 1 | 42 | 100% | 277 | 96.5% | 42 (100%) | 41 (97.6%) | 0/0/0/0 | 1 |
| Protein mutations: | R29G (1716A>G) | | | | | | | | |
| Codon mutations: | AGA29GGA (1716A>G), CTC41CTT (1754C>T) | | | | | | | | |
| protein 3a (YP_009179365.2) | 1 | 46 | 100% | 289 | 96.7% | 46 (100%) | 44 (95.7%) | 0/0/0/0 | 1 |
| Protein mutations: | S14T (3614T>A), F26L (3652T>A) | | | | | | | | |
| Codon mutations: | TCA14ACA (3614T>A), TCC15TCT (3619C>T), TTT26TTA (3652T>A) | | | | | | | | |
| CP read-through protein (NP_056751.2) | 1 | 718 | 100% | 4695 | 93.4% | 717 (99.7%) | 705 (98.1%) | 1/1/2/2 | 2 |
| Protein mutations: | V10G (3721T>G), M19del (3748_3750delTGC), T83S (3939A>T), V141I (4113G>A), P449L (5038C>T), A457V (5062C>T), H460R (5071A>G 5072C>T), P503S (5199C>T), T526A (5268A>G), E666_R667insX (5689_5690insGA), F683L (5741T>G), T686A (5748A>G), E697K (5781G>A) | | | | | | | | |
| Codon mutations: | GTC10GGC (3721T>G), ATG19del (3748_3750delTGC), CGA20AGA (3748_3750delTGC), GTT32GTG (3788T>G), CGC46CGT (3830C>T), ACC83TCC (3939A>T), ACA113ACG (4031A>G), GAG134GAA (4094G>A), GTA141ATA (4113G>A), GCC158GCT (4166C>T), ACT160ACC (4172T>C), CAC172CAT (4208C>T), TCT174TCG (4214T>G), CGG322CGA (4658G>A), GGT323GGC (4661T>C), GTC347GTG (4733C>G), GGT375GGC (4817T>C), TTC397TTT (4883C>T), CTT398CTC (4886T>C), GCA407GCT (4913A>T), ACG419ACA (4949G>A), GAA424GAG (4964A>G), ACC429ACT (4979C>T), CCA449CTA (5038C>T), GCC457GTC (5062C>T), CAC460CGT (5071A>G 5072C>T), ACG479ACA (5129G>A), GAC490GAT (5162C>T), AAC499AAT (5189C>T), CCA503TCA (5199C>T), CGA508CGC (5216A>C), ACA526GCA (5268A>G), GAA666GAG (5689_5690insGA), GAA666_CGT667insA-A (5689_5690insGA), CGT667C-- (5692_5693delGT), TTT683TTG (5741T>G), ACT686GCT (5748A>G), GAA697AAA (5781G>A) | | | | | | | | |
| coat protein (NP_056749.1) | 1 | 209 | 100% | 1411 | 99.2% | 208 (99.5%) | 205 (98.1%) | 0/1/0/0 | 1 |
| Protein mutations: | V10G (3721T>G), M19del (3748_3750delTGC), T83S (3939A>T), V141I (4113G>A) | | | | | | | | |
| Codon mutations: | GTC10GGC (3721T>G), ATG19del (3748_3750delTGC), CGA20AGA (3748_3750delTGC), GTT32GTG (3788T>G), CGC46CGT (3830C>T), ACC83TCC (3939A>T), ACA113ACG (4031A>G), GAG134GAA (4094G>A), GTA141ATA (4113G>A), GCC158GCT (4166C>T), ACT160ACC (4172T>C), CAC172CAT (4208C>T), TCT174TCG (4214T>G) | | | | | | | | |
| movement protein (NP_056750.1) | 1 | 157 | 100% | 1030 | 95.5% | 156 (99.4%) | 149 (94.9%) | 0/1/0/0 | 1 |
| Protein mutations: | S2A (3721T>G), C11del (3748_3750delTGC), F24C (3788T>G), A38V (3830C>T), Q105R (4031A>G), S126N (4094G>A), P150L (4166C>T), L152P (4172T>C) | | | | | | | | |
| Codon mutations: | TCA2GCA (3721T>G), TGC11del (3748_3750delTGC), TTC24TGC (3788T>G), GCC38GTC (3830C>T), ACA74ACT (3939A>T), CAA105CGA (4031A>G), AGT126AAT (4094G>A), AAG132AAA (4113G>A), CCA150CTA (4166C>T), CTT152CCT (4172T>C) | | | | | | | | |

*: Inserts / Deletes / Misaligned / Frameshifts

#### Analysis details

This analysis was performed with panviral2.64

### NGS Details (UN8): Duamitovirus soch1

#### Assembly

| | |
|---|---|
| **Coverage Length** | 2255 (2 contig(s)) |
| **Depth Of Coverage** | 25.7 |
| **Number Of Reads** | 508 |
| **Reads Per Million** | 8.49 rpm (after QC) |
| **Ambiguities** | 0 |
| **Assembly Method** | de novo + reference guided assembly |
| **Consensus Caller** | Bcf Tools |

#### Coverage Map

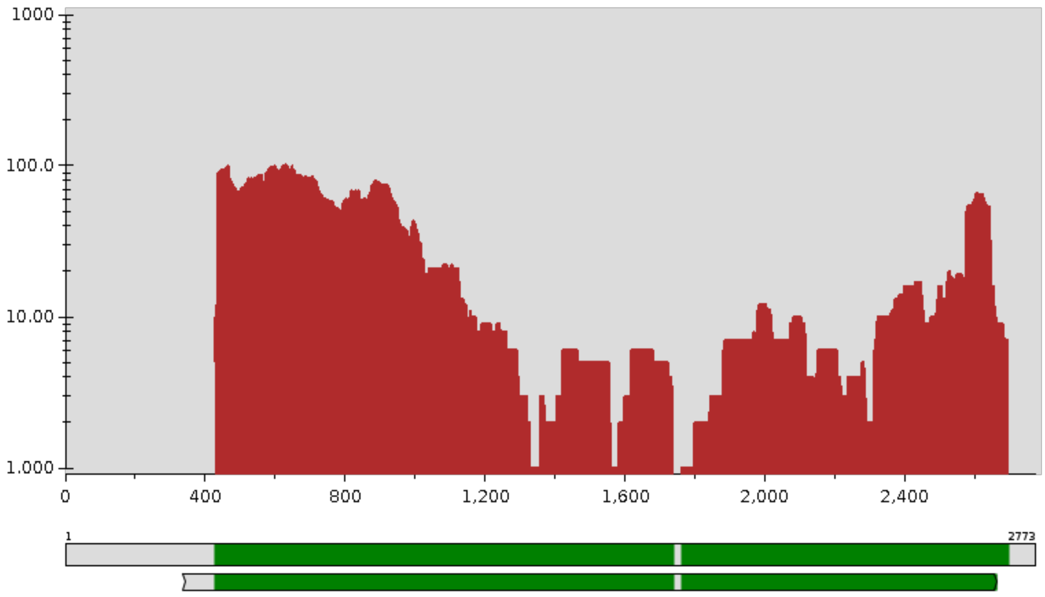

#### Assignment

| | |
|---|---|
| **Type** | Duamitovirus soch1 (Taxonomy ID: 2955838) |
| **Reference Genome** | NC_076524.1 |
| **NT Identity (%)** | 69.0561 |
| **AA Identity (%)** | 69.4708 |
| **Number Of Stop Codons** | 5 |
| **Number Of CDS** | 1 |

#### Alignment

| | |
|---|---|
| **Alignment Score** | 1676.0 (NT) + 3439.0 (AA) = 5115.0 |
| **Concordance (%)** | 54.6692 |

| Alignment Method | Global, seeded, nucleotide + amino acids (AGA) |
|---|---|

#### Genome Region

Sequence starts at position 428 and ends at position 2700 relative to NC_076524.1 reference sequence.

#### Alignment Detailed Statistics

| | Begin | End | Coverage | Score | Concordance | Matches | Identities | I/D/M/F* | Stop Codons |
|---|---|---|---|---|---|---|---|---|---|
| **NT** | **428** | **2700** | **81.3%** | **1676** | **37.6%** | **2246 (99.6%)** | **1551 (68.8%)** | **0/9** | |
| Mutations: | 437A>G, 441A>C, 443C>T, 447C>T, 459C>T, 466C>T, 470C>T, 471G>A, 472C>T, 473T>G, 477T>A, 480A>C, 481C>A, 482A>T, 485A>G, 489A>G, 494T>C, 497T>A, 500T>G, 503A>C, 506T>C, 509C>T, 512T>C, 513G>A, 516T>A, 517C>A, 518T>A, 520A>C, 524G>T, 525A>G, 526C>T, 527T>G, 529C>A, 530C>A, 531T>C, 533A>G, 534C>A, 535G>A, 536C>A, 539G>A, 540A>C, 541A>G, 542G>T, 545G>A, 548C>T, 550A>T, 556T>A, 557C>T, 560A>T, 569C>T, 575G>A, 581C>T, 582G>A, 587A>T, 589G>C, 591T>C, 593A>T, 599A>T, 602C>T, 612T>A, 613C>G, 614C>T, 616T>A, 618C>T, 619G>C, 620A>T, 624G>A, 626A>T, 629A>C, 630C>A, 632A>C, 633C>A, 635T>G, 638G>T, 641G>T, 644T>A, 647G>T, 651C>T, 653C>G, 656A>C, 657A>C, 661C>G, 662G>T, 665T>A, 668C>T, 672C>A, 673G>A, 674T>G, 680C>A, 683C>T, 684A>G, 685A>C, 686A>C, 692G>A, 695T>A, 699G>C, 700C>A, 701A>C, 702A>G, 705C>A, 707T>G, 708A>G, 709G>C, 710G>A, 713A>G, 714G>T, 715A>C, 719A>T, 722C>T, 737T>C, 746C>T, 752A>T, 761G>A, 765G>T, 767T>A, 770G>A, 771G>T, 772T>A, 773T>G, 774T>A, 776G>T, 777A>G, 778A>C, 779A>G, 782T>G, 796G>A, 797T>G, 803A>C, 809T>C, 810G>C, 815C>A, 818A>T, 821T>G, 824C>T, 827A>T, 831C>T, 833T>A, 841G>A, 842G>T, 850T>G, 851A>T, 857G>T, 860G>T, 862A>G, 869T>G, 875G>T, 880T>G, 881C>A, 885C>A, 887T>G, 892A>G, 905A>T, 908C>A, 909T>C, 910T>A, 916G>A, 918G>T, 919G>C, 923T>A, 926A>T, 927A>C, 929G>T, 934T>A, 935T>G, 938G>T, 942T>A, 943C>A, 948C>A, 971C>A, 972C>T, 975A>C, 976A>T, 977T>C, 983T>C, 990A>C, 992G>C, 993T>C, 994C>G, 995A>G, 997A>T, 998C>T, 999G>C, 1000G>A, 1001T>A, 1003T>A, 1004A>T, 1005G>T, 1007C>G, 1010A>G, 1016C>T, 1017A>G, 1018A>T, 1019A>T, 1022T>A, 1025T>C, 1026C>G, 1027A>G, 1030A>C, 1031G>A, 1034T>G, 1035A>G, 1036G>C, 1044T>G, 1046C>T, 1049G>A, 1050T>G, 1051T>A, 1052T>A, 1054C>A, 1055A>C, 1061T>C, 1067T>G, 1073G>A, 1076A>G, 1083C>A, 1088G>T, 1091C>G, 1099G>T, 1104A>G, 1105C>T, 1106A>G, 1115T>G, 1121A>C, 1122T>A, 1124C>T, 1127G>A, 1130C>T, 1133C>G, 1142A>C, 1145C>T, 1148G>T, 1151G>T, 1152T>C, 1154G>T, 1160C>T, 1161A>C, 1164T>A, 1165C>T, 1166T>G, 1169C>T, 1172C>A, 1174T>A, 1175T>C, 1178T>A, 1181A>T, 1184C>T, 1185A>T, 1186G>C, 1187G>T, 1188G>A, 1189C>A, 1191G>A, 1192C>A, 1193A>T, 1196T>C, 1197_1199delGAA, 1205A>T, 1211C>T, 1212T>G, 1213T>A, 1217T>C, 1220C>T, 1223A>T, 1224C>T, 1225A>T, 1226C>A, 1232G>A, 1236A>C, 1237G>A, 1238T>A, 1241C>A, 1247A>C, 1248T>C, 1249G>T, 1250G>T, 1253T>C, 1256A>T, 1258C>G, 1265G>A, 1266A>G, 1268T>G, 1269C>G, 1271T>C, 1274T>C, 1276G>A, 1280G>A, 1283G>T, 1292T>G, 1301G>A, 1302C>T, 1304T>G, 1306G>C, 1308G>C, 1309C>A, 1310A>G, 1312T>C, 1313G>C, 1316A>T, 1319G>A, 1322C>T, 1328A>G, 1332A>C, 1334A>G, 1338G>A, 1340A>C, 1345T>C, 1346A>T, 1349G>A, 1355C>T, 1357A>C, 1358C>T, 1370A>G, 1371C>T, 1372T>G, 1373T>A, 1376C>G, 1377C>T, 1378A>T, 1379C>T, 1382T>C, 1385T>C, 1389G>C, 1391T>A, 1397T>A, 1402C>A, 1409G>A, 1413A>C, 1414A>G, 1415A>C, 1416A>C, 1418A>T, 1419A>C, 1420A>C, 1422A>G, 1424A>G, 1427C>T, 1430G>A, 1431A>G, 1433T>A, 1445G>A, 1446A>C, 1447A>G, 1448A>T, 1451T>G, 1455T>G, 1457G>A, 1458G>C, 1459C>G, 1460T>A, 1464A>C, 1465A>G, 1466G>A, 1467C>T, 1468A>C, 1469A>T, 1470A>C, 1472G>A, 1475C>G, 1476T>A, 1477T>A, 1478T>A, 1479A>G, 1480A>G, 1481A>T, 1486G>C, 1489A>T, 1490T>C, 1502A>C, 1505G>A, 1508A>G, 1526G>A, 1538G>A, 1541T>C, 1545C>T, 1547T>C, 1553T>C, 1554A>T, 1555T>G, 1556G>C, 1559A>G, 1566C>A, 1567C>G, 1571C>A, 1572T>C, 1580C>A, 1589T>C, 1593_1597delAGCTC, 1598T>A, 1607T>C, 1613G>T, 1614A>T, 1617C>T, 1619T>G, 1620G>A, 1622T>C, 1623A>G, 1624A>T, 1626A>C, 1627A>C, 1628A>T, 1632T>A, 1634A>G, 1635A>G, 1636C>T, 1637A>C, 1643A>G, 1647T>A, 1648A>G, 1655A>T, 1661T>C, 1662C>T, 1664G>A, 1667T>A, 1670G>A, 1676T>G, 1679G>A, 1682C>T, 1684T>A, 1686C>T, 1691T>A, 1694T>A, 1698G>T, 1700A>C, 1703T>A, 1706C>T, 1707T>G, 1710T>C, 1712G>T, 1715T>C, 1721C>T, 1727A>G, 1734T>C, 1739A>T, 1761T>A, 1765A>C, 1766C>T, 1767T>A, 1769T>C, 1772A>T, 1777G>C, 1778G>T, 1784C>T, 1787T>C, 1788T>C, 1790A>G, 1819G>C, 1820T>A, 1821A>G, 1823A>G, 1829C>G, 1830A>C, 1831A>G, 1832T>G, 1833G>C, 1840A>G, 1856A>G, 1859C>T, 1860A>G, 1861A>G, 1865C>T, 1867A>C, 1868A>C, 1874G>T, 1875A>T, 1878C>G, 1880A>T, 1883A>G, 1886T>A, 1887T>A, 1889G>T, 1892A>T, 1895T>G, 1898T>G, 1904A>T, 1907C>T, 1919G>C, 1922G>A, 1923A>C, 1940A>G, 1947A>G, 1967C>T, 1968C>G, 1970T>G, 1972G>A, 1973A>G, 1976T>A, 1980T>A, 1981T>C, 1984C>T, 1985T>C, 1991T>C, 2000T>G, 2003A>T, 2006T>G, 2010G>A, 2011C>A, 2013C>A, 2015C>T, 2020T>C, 2030T>C, 2036C>T, 2039A>G, 2052A>C, 2054A>G, 2060G>T, 2063T>A, 2066C>T, 2071A>T, 2074A>G, 2084G>C, 2087C>T, 2088T>C, 2090A>T, 2093C>T, 2102A>G, 2103A>T, 2104G>C, 2105C>A, 2108G>A, 2111T>A, 2117G>A, 2118C>A, 2120T>G, 2121C>T, 2128C>T, 2129T>G, 2133C>A, 2134A>G, 2135T>G, 2138T>C, 2141G>A, 2143C>T, 2146T>G, 2148C>A, 2152C>T, 2158T>C, 2167A>T, 2168C>A, 2173G>T, 2174G>T, 2177G>T, 2186G>A, 2191T>A, 2192G>A, 2195C>T, 2204C>T, 2210G>A, 2222A>G, 2225A>G, 2230A>T, 2231C>T, 2234C>T, 2237T>A, 2239A>G, 2240A>G, 2241G>A, 2243A>G, 2250G>C, 2251T>C, 2252T>C, 2264G>T, 2271C>A, 2273A>T, 2274A>G, 2276G>A, 2280C>A, 2285G>A, 2288C>T, 2298G>T, 2299T>G, 2300T>G, 2303A>G, 2306A>C, 2309A>T, 2313C>T, 2315T>C, 2322C>T, 2326G>T, 2328C>A, 2330A>T, 2336A>G, 2337G>A, 2346T>C, 2348A>T, 2349A>C, 2350A>G, 2355C>G, 2357C>T, 2358A>T, 2359A>C, 2360C>T, 2369T>C, 2372G>T, 2377G>T, 2378T>G, 2382A>T, 2383A>C, 2384C>T, 2385C>T, 2386C>T, 2387A>T, 2390T>A, 2396C>T, 2399T>C, 2400A>G, 2402G>A, 2403A>C, 2404G>A, 2405T>A, 2406C>T, 2408C>G, 2411C>G, 2414C>T, 2415T>G, 2417C>T, 2420T>C, 2423A>T, 2431C>A, 2432A>G, 2435C>A, 2441A>G, 2442A>C, 2444G>A, 2445C>T, 2446T>C, 2447C>G, 2448G>A, 2450A>T, 2451A>G, 2452C>A, 2453T>A, 2456T>C, 2457T>C, 2459T>G, 2460G>A, 2462T>A, 2465A>G, 2466A>G, 2477C>T, 2478C>T, 2489A>G, 2492T>C, 2495C>T, 2504T>A, 2510T>C, 2514A>G, 2518C>T, 2519C>G, 2525C>T, 2528T>A, 2532C>A, 2533A>G, 2536G>A, 2537G>T, 2541T>C, 2543T>G, 2551A>G, 2552C>T, 2555T>C, 2556G>T, 2557T>C, 2561T>C, 2568A>G, 2571A>C, 2572A>G, 2582A>G, 2585G>A, 2597G>A, 2598A>G, 2599A>C, 2600C>T, 2609T>A, 2613T>G, 2615T>A, 2618T>A, 2624A>T, 2626A>T, 2627C>A, 2629A>C, 2631A>T, 2634delA, 2636A>T, 2639A>T, 2640A>G, 2643G>A, 2644A>G, 2647A>T, 2648A>G, 2649G>T, 2650T>A, 2653G>T, 2655T>A, 2681G>A, 2686T>C, 2689T>C, 2690G>C, 2694A>G, 2695T>A | | | | | | | | |

##### CDS

| | Begin | End | Coverage | Score | Concordance | Matches | Identities | I/D/M/F* | Stop Codons |
|---|---|---|---|---|---|---|---|---|---|
| RdRp | 32 | 777 | 95.1% | 3439 | 66.5% | 737 (99.7%) | 512 (69.3%) | 0/2/2/2 | 5 |
| Protein mutations: | I36L (441A>C 443C>T), L38F (447C>T), A44V (466C>T), A46M (471G>A 472C>T 473T>G), L48M (477T>A), T49H (480A>C 481C>A 482A>T), I52V (489A>G), V60I (513G>A), S61K (516T>A 517C>A 518T>A), K62T (520A>C), T64V (525A>G 526C>T 527T>G), T65K (529C>A 530C>A), R67K (534C>A 535G>A 536C>A), K69R (540A>C 541A>G 542G>T), Y72F (550A>T), F74Y (556T>A 557C>T), A83T (582G>A), C85S (589G>C), F94Y (616T>A), R95S (618C>T 619G>C 620A>T), E97N (624G>A 626A>T), E98D (629A>C), P99T (630C>A 632A>C), L100M (633C>A 635T>G), S109C (661C>G 662G>T), R113K (672C>A 673G>A 674T>G), K117A (684A>G 685A>C 686A>C), A122H (699G>C 700C>A 701A>C), I123V (702A>G), R125A (708A>G 709G>C 710G>A), D127S (714G>T 715A>C), E128D (719A>T), A144S (765G>T 767T>A), V146* (771G>T 772T>A 773T>G), L147I (774T>A 776G>T), K148A (777A>G 778A>C 779A>G), S154K (796G>A 797T>G), E159Q (810G>C), P166S (831C>T 833T>A), G169D (841G>A 842G>T), L172R (850T>G 851A>T), E176G (862A>G), I178M (869T>G), F182* (880T>G 881C>A), H184K (885C>A 887T>G), Q186R (892A>G), F192H (909T>C 910T>A), R194Q (916G>A), G195S (918G>T 919G>C), M198L (927A>C 929G>T), I200K (934T>A 935T>G), S203N (942T>A 943C>A), Q205K (948C>A), N214L (975A>C 976A>T 977T>C), S220R (993T>C 994C>G 995A>G), Y221F (997A>T 998C>T), G222Q (999G>C 1000G>A 1001T>A), L223Y (1003T>A 1004A>T), A224S (1005G>T 1007C>G), K228V (1017A>G 1018A>T 1019A>T), Q231G (1026C>G 1027A>G), E232A (1030A>C 1031G>A), S234A (1035A>G 1036G>C), F237V (1044T>G 1046C>T), F239E (1050T>G 1051T>A 1052T>A), P240H (1054C>A 1055A>C), L250I (1083C>A), W255L (1099G>T), T257V (1104A>G 1105C>T 1106A>G), I260M (1115T>G), F263I (1122T>A 1124C>T), M272I (1151G>T), K276Q (1161A>C), S277M (1164T>A 1165C>T 1166T>G), F279L (1172C>A), F280Y (1174T>A 1175T>C), L282F (1181A>T), R284S (1185A>T 1186G>C 1187G>T), A285N (1188G>A 1189C>A), A286N (1191G>A 1192C>A 1193A>T), E288del (1197_1199delGAA), L290F (1205A>T), L293E (1212T>G 1213T>A), E296D (1223A>T), H297L (1224C>T 1225A>T 1226C>A), S301Q (1236A>C 1237G>A 1238T>A), W305L (1248T>C 1249G>T 1250G>T), T308S (1258C>G), S311G (1266A>G 1268T>G), P312A (1269C>G 1271T>C), R314Q (1276G>A), G324A (1306G>C), A325Q (1308G>C 1309C>A 1310A>G), V326A (1312T>C 1313G>C), V335I (1338G>A 1340A>C), V337A (1345T>C 1346A>T), M338I (1349G>A), Y341S (1357A>C 1358C>T), L346* (1371C>T 1372T>G 1373T>A), H348F (1377C>T 1378A>T 1379C>T), D352Q (1389G>C 1391T>A), T356K (1402C>A), K360R (1413A>C 1414A>G 1415A>C), I361L (1416A>C 1418A>T), K362P (1419A>C 1420A>C), T363A (1422A>G 1424A>G), T366A (1431A>G 1433T>A), K371R (1446A>C 1447A>G 1448A>T), L374V (1455T>G 1457G>A), A375R (1458G>C 1459C>G 1460T>A), K377R (1464A>C 1465A>G 1466G>A), Q378S (1467C>T 1468A>C 1469A>T), K379Q (1470A>C 1472G>A), N380K (1475C>G), F381K (1476T>A 1477T>A 1478T>A), K382G (1479A>G 1480A>G 1481A>T), C384S (1486G>C), Y385F (1489A>T 1490T>C), L389F (1502A>C), M401I (1538G>A), L404F (1545C>T 1547T>C), M407C (1554A>T 1555T>G 1556G>C), I408M (1559A>G), P411S (1566C>A 1567C>G), S420del (1593_1597delAGCTC), I427F (1614A>T), V429I (1620G>A 1622T>C), N430V (1623A>G 1624A>T), K431P (1626A>C 1627A>C 1628A>T), L433M (1632T>A 1634A>G), T434V (1635A>G 1636C>T 1637A>C), Y438S (1647T>A 1648A>G), F450Y (1684T>A), H451Y (1686C>T), A455S (1698G>T 1700A>C), S458A (1707T>G), I464M (1727A>G), C476S (1761T>A), N477T (1765A>C 1766C>T), S478T (1767T>A 1769T>C), G481A (1777G>C 1778G>T), S495T (1819G>C 1820T>A), K496E (1821A>G 1823A>G), N499R (1830A>C 1831A>G 1832T>G), E500Q (1833G>C), K502R (1840A>G), N509G (1860A>G 1861A>G), K511T (1867A>C 1868A>C), I514L (1875A>T), P515A (1878C>G 1880A>T), L518I (1887T>A 1889G>T), D521E (1898T>G), I538V (1947A>G), L545V (1968C>G 1970T>G), R546K (1972G>A 1973A>G), L549T (1980T>A 1981T>C), S550F (1984C>T 1985T>C), A559N (2010G>A 2011C>A), L560I (2013C>A 2015C>T), I562T (2020T>C), I568M (2039A>G), Y579F (2071A>T), K580R (2074A>G), A598V (2128C>T 2129T>G), H600R (2133C>A 2134A>G 2135T>G), P603L (2143C>T), V604G (2146T>G), S606L (2152C>T), L608S (2158T>C), Y611L (2167A>T 2168C>A), W613F (2173G>T 2174G>T), W614C (2177G>T), M619K (2191T>A 2192G>A), I629M (2222A>G), Y632F (2230A>T 2231C>T), K635R (2239A>G 2240A>G), E636K (2241G>A 2243A>G), V639P (2250G>C 2251T>C 2252T>C), Q643H (2264G>T), P646T (2271C>A 2273A>T), K647E (2274A>G 2276G>A), L649M (2280C>A), V655W (2298G>T 2299T>G 2300T>G), R660C (2313C>T 2315T>C), R664L (2326G>T), Q665N (2328C>A 2330A>T), E668K (2337G>A), K672R (2349A>C 2350A>G), L674V (2355C>G 2357C>T), N675S (2358A>T 2359A>C 2360C>T), C681L (2377G>T 2378T>G), N683S (2382A>T 2383A>C 2384C>T), P684F (2385C>T 2386C>T 2387A>T), D685E (2390T>A), K689E (2400A>G 2402G>A), S690Q (2403A>C 2404G>A 2405T>A), I692M (2411C>G), S694A (2415T>G 2417C>T), T699K (2431C>A 2432A>G), L704S (2445C>T 2446T>C 2447C>G), E705N (2448G>A 2450A>T), T706E (2451A>G 2452C>A 2453T>A), F708L (2457T>C 2459T>G), D709K (2460G>A 2462T>A), I711V (2466A>G), N727D (2514A>G), P728L (2518C>T 2519C>G), Q733R (2532C>A 2533A>G), W734Y (2536G>A 2537G>T), F736L (2541T>C 2543T>G), N739S (2551A>G 2552C>T), V741S (2556G>T 2557T>C), N745D (2568A>G), K746R (2571A>C 2572A>G), N755A (2598A>G 2599A>C 2600C>T), S760A (2613T>G 2615T>A), E763D (2624A>T), Y764L (2626A>T 2627C>A), Q765P (2629A>C), T766S (2631A>T), K769E (2640A>G), E770R (2643G>A 2644A>G), E771V (2647A>T 2648A>G), V772* (2649G>T 2650T>A), G773V (2653G>T), L774I (2655T>A) | | | | | | | | |

| | Begin | End | Coverage | Score | Concordance | Matches | Identities | I/D/M/F* | Stop Codons |
|---|---|---|---|---|---|---|---|---|---|
| **NT** | **428** | **2700** | **81.3%** | **1676** | **37.6%** | **2246 (99.6%)** | **1551 (68.8%)** | **0/9** | |

Codon mutations:

AAA34AAG (437A>G), ATC36CTT (441A>C 443C>T), CTT38TTT (447C>T), CTA42TTA (459C>T), GCC44GTC (466C>T), TCC45TCT (470C>T), GCT46ATG (471G>A 472C>T 473T>G), TTG48ATG (477T>A), ACA49CAT (480A>C 481C>A 482A>T), AAA50AAG (485A>G), ATC52GTC (489A>G), GCT53GCC (494T>C), CTT54CTA (497T>A), GCT55GCG (500T>G), GGA56GGC (503A>C), TGT57TGC (506T>C), GAC58GAT (509C>T), TTT59TTC (512T>C), GTA60ATA (513G>A), TCT61AAA (516T>A 517C>A 518T>A), AAA62ACA (520A>C), GTG63GTT (524G>T), ACT64GTG (525A>G 526C>T 527T>G), ACC65AAA (529C>A 530C>A), TTA66CTG (531T>C 533A>G), CGC67AAA (534C>A 535G>A 536C>A), CGG68CGA (539G>A), AAG69CGT (540A>C 541A>G 542G>T), TCG70TCA (545G>A), GGC71GGT (548C>T), TAT72TTT (550A>T), TTC74TAT (556T>A 557C>T), ACA75ACT (560A>T), TAC78TAT (569C>T), AAG80AAA (575G>A), TGC82TGT (581C>T), GCA83ACA (582G>A), GTA84GTT (587A>T), TGC85TCC (589G>C), TTA86CTT (591T>C 593A>T), CGA88CGT (599A>T), TAC89TAT (602C>T), TCC93AGT (612T>A 613C>G 614C>T), TTT94TAT (616T>A), CGA95TCT (618C>T 619G>C 620A>T), GAA97AAT (624G>A 626A>T), GAA98GAC (629A>C), CCA99ACC (630C>A 632A>C), CTT100ATG (633C>A 635T>G), TCG101TCT (638G>T), GTG102GTT (641G>T), CCT103CCA (644T>A), GTG104GTT (647G>T), CTC106TTG (651C>T 653C>G), ACA107ACC (656A>C), AGA108CGA (657A>C), TCG109TGT (661C>G 662G>T), GGT110GGA (665T>A), ATC111ATT (668C>T), CGT113AAG (672C>A 673G>A 674T>G), ATC115ATA (680C>A), CCC116CCT (683C>T), AAA117GCC (684A>G 685A>C 686A>C), TTG119TTA (692G>A), CGT120CGA (695T>A), GCA122CAC (699G>C 700C>A 701A>C), ATA123GTA (702A>G), CGT124AGG (705C>A 707T>G), AGG125GCA (708A>G 709G>C 710G>A), AAA126AAG (713A>G), GAT127TCT (714G>T 715A>C), GAA128GAT (719A>T), CAC129CAT (722C>T), GTT134GTC (737T>C), TAC137TAT (746C>T), TCA139TCT (752A>T), GGG142GGA (761G>A), GCT144TCA (765G>T 767T>A), AAG145AAA (770G>A), GTT146TAG (771G>T 772T>A 773T>G), TTG147ATT (774T>A 776G>T), AAA148GCG (777A>G 778A>C 779A>G), GTT149GTG (782T>G), AGT154AAG (796G>A 797T>G), GCA156GCC (803A>C), TTT158TTC (809T>C), GAA159CAA (810G>C), TCC160TCA (815C>A), ATA161ATT (818A>T), ACT162ACG (821T>G), ACC163ACT (824C>T), CCA164CCT (827A>T), CCT166TCA (831C>T 833T>A), GGG169GAT (841G>A 842G>T), CTA172CGT (850T>G 851A>T), GTG174GTT (857G>T), CTG175CTT (860G>T), GAA176GGA (862A>G), ATT178ATG (869T>G), ACG180ACT (875G>T), TTC182TGA (880T>G 881C>A), CAT184AAG (885C>A 887T>G), CAA186CGA (892A>G), CTA190CTT (905A>T), CCC191CCA (908C>A), TTT192CAT (909T>C 910T>A), CGA194CAA (916G>A), GGC195TCC (918G>T 919G>C), ATT196ATA (923T>A), CCA197CCT (926A>T), ATG198CTT (927A>C 929G>T), ATT200AAG (934T>A 935T>G), GGG201GGT (938G>T), TCT203AAT (942T>A 943C>A), CAA205AAA (948C>A), CCC212CCA (971C>A), CTA213TTA (972C>T), AAT214CTC (975A>C 976A>T 977T>C), AAT216AAC (983T>C), AGG219CGC (990A>C 992G>C), TCA220CGG (993T>C 994C>G 995A>G), TAC221TTT (997A>T 998C>T), GGT222CAA (999G>C 1000G>A 1001T>A), TTA223TAT (1003T>A 1004A>T), GCC224TCG (1005G>T 1007C>G), GTA225GTG (1010A>G), GAC227GAT (1016C>T), AAA228GTT (1017A>G 1018A>T 1019A>T), GTT229GTA (1022T>A), GAT230GAC (1025T>C), CAA231GGA (1026C>G 1027A>G), GAG232GCA (1030A>C 1031G>A), GCT233GCG (1034T>G), AGT234GCT (1035A>G 1036G>C), TTC237GTT (1044T>G 1046C>T), AAG238AAA (1049G>A), TTT239GAA (1050T>G 1051T>A 1052T>A), CCA240CAC (1054C>A 1055A>C), ATT242ATC (1061T>C), GTT244GTG (1067T>G), TTG246TTA (1073G>A), AAA247AAG (1076A>G), CTA250ATA (1083C>A), GCG251GCT (1088G>T), GCC252GCG (1091C>G), TGG255TTG (1099G>T), ACA257GTG (1104A>G 1105C>T 1106A>G), ATT260ATG (1115T>G), TCA262TCC (1121A>C), TTC263ATT (1122T>A 1124C>T), CAG264CAA (1127G>A), GAC265GAT (1130C>T), GGC266GGG (1133C>G), TCA269TCC (1142A>C), CCC270CCT (1145C>T), GGG271GGT (1148G>T), ATG272ATT (1151G>T), TTG273CTT (1152T>C 1154G>T), TCC275TCT (1160C>T), AAA276CAA (1161A>C), TCT277ATG (1164T>A 1165C>T 1166T>G), GTC278GTT (1169C>T), TTC279TTA (1172C>A), TTT280TAC (1174T>A 1175T>C), CCT281CCA (1178T>A), TTA282TTT (1181A>T), GAC283GAT (1184C>T), AGG284TCT (1185A>T 1186G>C 1187G>T), GCT285AAT (1188G>A 1189C>A), GCA286AAT (1191G>A 1192C>A 1193A>T), ACT287ACC (1196T>C), GAA288del (1197_1199delGAA), TTA290TTT (1205A>T), AAC292AAT (1211C>T), TTA293GAA (1212T>G 1213T>A), TCT294TCC (1217T>C), CTC295CTT (1220C>T), GAA296GAT (1223A>T), CAC297TTA (1224C>T 1225A>T 1226C>A), GAG299GAA (1232G>A), AGT301CAA (1236A>C 1237G>A 1238T>A), GTC302GTA (1241C>A), CCA304CCC (1247A>C), TGG305CTT (1248T>C 1249G>T 1250G>T), TTT306TTC (1253T>C), TCA307TCT (1256A>T), ACC308AGC (1258C>G), TTG310TTA (1265G>A), AGT311GGG (1266A>G 1268T>G), CCT312GCC (1269C>G 1271T>C), TAT313TAC (1274T>C), CGA314CAA (1276G>A), GGG315GGA (1280G>A), GTG316GTT (1283G>T), GTT319GTG (1292T>G), AGG322AGA (1301G>A), CTT323TTG (1302C>T 1304T>G), GGT324GCT (1306G>C), GCA325CAG (1308G>C 1309C>A 1310A>G), GTG326GCC (1312T>C 1313G>C), ATA327ATT (1316A>T), GAG328GAA (1319G>A), GGC329GGT (1322C>T), GGA331GGG (1328A>G), AGA333CGG (1332A>C 1334A>G), GTA335ATC (1338G>A 1340A>C), GTA337GCT (1345T>C 1346A>T), ATG338ATA (1349G>A), AAC340AAT (1355C>T), TAC341TCT (1357A>C 1358C>T), AGA345AGG (1370A>G), CTT346TGA (1371C>T 1372T>G 1373T>A), CTC347CTG (1376C>G), CAC348TTT (1377C>T 1378A>T 1379C>T), CCT349CCC (1382T>C), GTT350GTC (1385T>C), GAT352CAA (1389G>C 1391T>A), GCT354GCA (1397T>A), ACA356AAA (1402C>A), CTG358CTA (1409G>A), AAA360CGC (1413A>C 1414A>G 1415A>C), ATA361CTT (1416A>C 1418A>T), AAG362CCG (1419A>C 1420A>C), ACA363GCG (1422A>G 1424A>G), GAC364GAT (1427C>T), GGG365GGA (1430G>A), ACT366GCA (1431A>G 1433T>A), GAG370GAA (1445G>A), AAA371CGT (1446A>C 1447A>G 1448A>T), CCT372CCG (1451T>G), TTG374GTA (1455T>G 1457G>A), GCT375CGA (1458G>C 1459C>G 1460T>A), AAG377CGA (1464A>C 1465A>G 1466G>A), CAA378TCT (1467C>T 1468A>C 1469A>T), AAG379CAA (1470A>C 1472G>A), AAC380AAG (1475C>G), TTT381AAA (1476T>A 1477T>A 1478T>A), AAA382GGT (1479A>G 1480A>G 1481A>T), TGT384TCT (1486G>C), TAT385TTC (1489A>T 1490T>C), TTA389TTC (1502A>C), AAG390AAA (1505G>A), TCA391TCG (1508A>G), CCG397CCA (1526G>A), ATG401ATA (1538G>A), TAT402TAC (1541T>C), CTT404TTC (1545C>T 1547T>C), TCT406TCC (1553T>C), ATG407TGC (1554A>T 1555T>G 1556G>C), ATA408ATG (1559A>G), CCT411AGT (1566C>A 1567C>G), ACC412ACA (1571C>A), TTG413CTG (1572T>C), TCC415TCA (1580C>A), GTT418GTC (1589T>C), AGC420del (1593_1597delAGCTC), TCT421--A (1593_1597delAGCTC 1598T>A), CTT424CTC (1607T>C), ACG426ACT (1613G>T), ATC427TTC (1614A>T), CTT428TTG (1617C>T 1619T>G), GTT429ATC (1620G>A 1622T>C), AAC430GTC (1623A>G 1624A>T), AAA431CCT (1626A>C 1627A>C 1628A>T), TTA433ATG (1632T>A 1634A>G), ACA434GTC (1635A>G 1636C>T 1637A>C), AGA436AGG (1643A>G), TAT438AGT (1647T>A 1648A>G), ATA440ATT (1655A>T), TTT442TTC (1661T>C), CTG443TTA (1662C>T 1664G>A), ACT444ACA (1667T>A), GGG445GGA (1670G>A), CCT447CCG (1676T>G), TTG448TTA (1679G>A), GGC449GGT (1682C>T), TTC450TAC (1684T>A), CAC451TAC (1686C>T), GGT452GGA (1691T>A), TCT453TCA (1694T>A), GCA455TCC (1698G>T 1700A>C), CTT456CTA (1703T>A), TTC457TTT (1706C>T), TCG458GCG (1707T>G), TTG459CTT (1710T>C 1712G>T), TCT460TCC (1715T>C), CAC462CAT (1721C>T), ATA464ATG (1727A>G), TTA467CTA (1734T>C), GCA468GCT (1739A>T), TGT476AGT (1761T>A), AAC477ACT (1765A>C 1766C>T), TCT478ACC (1767T>A 1769T>C), CCA479CCT (1772A>T), GGG481GCT (1777G>C 1778G>T), TAC483TAT (1784C>T), GCT484GCC (1787T>C), TTA485CTG (1788T>C 1790A>G), AGT495ACA (1819G>C 1820T>A), AAA496GAG (1821A>G 1823A>G), GCC498GCG (1829C>G), AAT499CGG (1830A>C 1831A>G 1832T>G), GAG500CAG (1833G>C), AAG502AGG (1840A>G), AGA507AGG (1856A>G), CTC508CTT (1859C>T), AAT509GGT (1860A>G 1861A>G), GTC510GTT (1865C>T), AAA511ACC (1867A>C 1868A>C), TCG513TCT (1874G>T), ATA514TTA (1875A>T), CCA515GCT (1878C>G 1880A>T), AAA516AAG (1883A>G), TCT517TCA (1886T>A), TTG518ATT (1887T>A 1889G>T), ATA519ATT (1892A>T), TCT520TCG (1895T>G), GAT521GAG (1898T>G), GGA523GGT (1904A>T), ACC524ACT (1907C>T), GCG528GCC (1919G>C), AAG529AAA (1922G>A), AGG530CGG (1923A>C), TCA535TCG (1940A>G), ATT538GTT (1947A>G), TCC544TCT (1967C>T), CTT545GTG (1968C>G 1970T>G), AGA546AAG (1972G>A 1973A>G), GCT547GCA (1976T>A), TTA549ACA (1980T>A 1981T>C), TCT550TTC (1984C>T 1985T>C), CGT552CGC (1991T>C), GTT555GTG (2000T>G), GGA556GGT (2003A>T), CTT557CTG (2006T>G), GCC559AAC (2010G>A 2011C>A), CTC560ATT (2013C>A 2015C>T), ATC562ACC (2020T>C), GAT565GAC (2030T>C), AAC567AAT (2036C>T), ATA568ATG (2039A>G), AGA573CGG (2052A>C 2054A>G), GGG575GGT (2060G>T), GGT576GGA (2063T>A), GCC577GCT (2066C>T), TAC579TTC (2071A>T), AAG580AGG (2074A>G), GCG583GCC (2084G>C), CGC584CGT (2087C>T), TTA585CTT (2088T>C 2090A>T), TTC586TTT (2093C>T), CAA589CAG (2102A>G), AGC590TCA (2103A>T 2104G>C 2105C>A), AAG591AAA (2108G>A), CGT592CGA (2111T>A), GAG594GAA (2117G>A), CGT595AGG (2118C>A 2120T>G), CTA596TTA (2121C>T), GCT598GTG (2128C>T 2129T>G), CAT600AGG (2133C>A 2134A>G 2135T>G), CTT601CTC (2138T>C), AAG602AAA (2141G>A), CCA603CTA (2143C>T), GTT604GGT (2146T>G), CGG605AGG (2148C>A), TCA606TTA (2152C>T), TTA608TCA (2158T>C), TAC611TTA (2167A>T 2168C>A), TGG613TTT (2173G>T 2174G>T), TGG614TGT (2177G>T), AGG617AGA (2186G>A), ATG619AAA (2191T>A 2192G>A), CCC620CCT (2195C>T), CCC623CCT (2204C>T), TTG625TTA (2210G>A), ATA629ATG (2222A>G), GTA630GTG (2225A>G), TAC632TTT (2230A>T 2231C>T), CTC633CTT (2234C>T), CTT634CTA (2237T>A), AAA635AGG (2239A>G 2240A>G), GAA636AAG (2241G>A 2243A>G), GTT639CCC (2250G>C 2251T>C 2252T>C), CAG643CAT (2264G>T), CCA646ACT (2271C>A 2273A>T), AAG647GAA (2274A>G 2276G>A), CTG649ATG (2280C>A), GTG650GTA (2285G>A), TTC651TTT (2288C>T), GTT655TGG (2298G>T 2299T>G 2300T>G), GAA656GAG (2303A>G), ATA657ATC (2306A>C), CTA658CTT (2309A>T), CGT660TGC (2313C>T 2315T>C), CTA663TTA (2322C>T), CGC664CTC (2326G>T), CAA665AAT (2328C>A 2330A>T), GTA667GTG (2336A>G), GAA668AAA (2337G>A), TTA671CTT (2346T>C 2348A>T), AAG672CGG (2349A>C 2350A>G), CTC674GTT (2355C>G 2357C>T), AAC675TCT (2358A>T 2359A>C 2360C>T), CAT678CAC (2369T>C), ACG679ACT (2372G>T), TGT681TTG (2377G>T 2378T>G), AAC683TCT (2382A>T 2383A>C 2384C>T), CCA684TTT (2385C>T 2386C>T 2387A>T), GAT685GAA (2390T>A), TCC687TCT (2396C>T), ATT688ATC (2399T>C), AAG689GAA (2400A>G 2402G>A), AGT690CAA (2403A>C 2404G>A 2405T>A), CTC691TTG (2406C>T 2408C>G), ATC692ATG (2411C>G), GAC693GAT (2414C>T), TCC694GCT (2415T>G 2417C>T), CCT695CCC (2420T>C), ATA696ATT (2423A>T), ACA699AAG (2431C>A 2432A>G), TCC700TCA (2435C>A), AAA702AAG (2441A>G), AGG703CGA (2442A>C 2444G>A), CTC704TCG (2445C>T 2446T>C 2447C>G), GAA705AAT (2448G>A 2450A>T), ACT706GAA (2451A>G 2452C>A 2453T>A), GAT707GAC (2456T>C), TTT708CTG (2457T>C 2459T>G), GAT709AAA (2460G>A 2462T>A), TTA710TTG (2465A>G), ATC711GTC (2466A>G), GGC714GGT (2477C>T), CTA715TTA (2478C>T), AAA718AAG (2489A>G), TGT719TGC (2492T>C), TAC720TAT (2495C>T), GGT723GGA (2504T>A), GGT725GGC (2510T>C), AAT727GAT (2514A>G), CCC728CTG (2518C>T 2519C>G), ACC730ACT (2525C>T), ACT731ACA (2528T>A), CAG733AGG (2532C>A 2533A>G), TGG734TAT (2536G>A 2537G>T), TTT736CTG (2541T>C 2543T>G), AAC739AGT (2551A>G 2552C>T), ACT740ACC (2555T>C), GTT741TCT (2556G>T 2557T>C), ATT742ATC (2561T>C), AAT745GAT (2568A>G), AAA746CGA (2571A>C 2572A>G), TTA749TTG (2582A>G), GGG750GGA (2585G>A), GGG754GGA (2597G>A), AAC755GCT (2598A>G 2599A>C 2600C>T), ATT758ATA (2609T>A), TCT760GCA (2613T>G 2615T>A), CCT761CCA (2618T>A), GAA763GAT (2624A>T), TAC764TTA (2626A>T 2627C>A), CAA765CCA (2629A>C), ACT766TCT (2631A>T), ACA767-CT (2634delA 2636A>T), GGA768GGT (2639A>T), AAA769GAA (2640A>G), GAG770AGG (2643G>A 2644A>G), GAA771GTG (2647A>T 2648A>G), GTG772TAG (2649G>T 2650T>A), GGC773GTC (2653G>T), TTA774ATA (2655T>A)

#### Proteins

| | Begin | End | Coverage | Score | Concordance | Matches | Identities | I/D/M/F* | Stop Codons |
|---|---|---|---|---|---|---|---|---|---|
| RNA-dependent RNA polymerase (YP_010798874.1) | 32 | 777 | 95.1% | 3439 | 66.5% | 737 (99.7%) | 512 (69.3%) | 0/2/2/2 | 5 |

| | Begin | End | Coverage | Score | Concordance | Matches | Identities | I/D/M/F* | Stop Codons |
|---|---|---|---|---|---|---|---|---|---|
| **NT** | **428** | **2700** | **81.3%** | **1676** | **37.6%** | **2246 (99.6%)** | **1551 (68.8%)** | **0/9** | |
| Protein mutations: | I36L (441A>C 443C>T), L38F (447C>T), A44V (466C>T), A46M (471G>A 472C>T 473T>G), L48M (477T>A), T49H (480A>C 481C>A 482A>T), I52V (489A>G), V60I (513G>A), S61K (516T>A 517C>A 518T>A), K62T (520A>C), T64V (525A>G 526C>T 527T>G), T65K (529C>A 530C>A), R67K (534C>A 535G>A 536C>A), K69R (540A>C 541A>G 542G>T), Y72F (550A>T), F74Y (556T>A 557C>T), A83T (582G>A), C85S (589G>C), F94Y (616T>A), R95S (618C>T 619G>C 620A>T), E97N (624G>A 626A>T), E98D (629A>C), P99T (630C>A 632A>C), L100M (633C>A 635T>G), S109C (661C>G 662G>T), R113K (672C>A 673G>A 674T>G), K117A (684A>G 685A>C 686A>C), A122H (699G>C 700C>A 701A>C), I123V (702A>G), R125A (708A>G 709G>C 710G>A), D127S (714G>T 715A>C), E128D (719A>T), A144S (765G>T 767T>A), V146* (771G>T 772T>A 773T>G), L147I (774T>A 776G>T), K148A (777A>G 778A>C 779A>G), S154K (796G>A 797T>G), E159Q (810G>C), P166S (831C>T 833T>A), G169D (841G>A 842G>T), L172R (850T>G 851A>T), E176G (862A>G), I178M (869T>G), F182* (880T>G 881C>A), H184K (885C>A 887T>G), Q186R (892A>G), F192H (909T>C 910T>A), R194Q (916G>A), G195S (918G>T 919G>C), M198L (927A>C 929G>T), I200K (934T>A 935T>G), S203N (942T>A 943C>A), Q205K (948C>A), N214L (975A>C 976A>T 977T>C), S220R (993T>C 994C>G 995A>G), Y221F (997A>T 998C>T), G222Q (999G>C 1000G>A 1001T>A), L223Y (1003T>A 1004A>T), A224S (1005G>T 1007C>G), K228V (1017A>G 1018A>T 1019A>T), Q231G (1026C>G 1027A>G), E232A (1030A>C 1031G>A), S234A (1035A>G 1036G>C), F237V (1044T>G 1046C>T), F239E (1050T>G 1051T>A 1052T>A), P240H (1054C>A 1055A>C), L250I (1083C>A), W255L (1099G>T), T257V (1104A>G 1105C>T 1106A>G), I260M (1115T>G), F263I (1122T>A 1124C>T), M272I (1151G>T), K276Q (1161A>C), S277M (1164T>A 1165C>T 1166T>G), F279L (1172C>A), F280Y (1174T>A 1175T>C), L282F (1181A>T), R284S (1185A>T 1186G>C 1187G>T), A285N (1188G>A 1189C>A), A286N (1191G>A 1192C>A 1193A>T), E288del (1197_1199delGAA), L290F (1205A>T), L293E (1212T>G 1213T>A), E296D (1223A>T), H297L (1224C>T 1225A>T 1226C>A), S301Q (1236A>C 1237G>A 1238T>A), W305L (1248T>C 1249G>T 1250G>T), T308S (1258C>G), S311G (1266A>G 1268T>G), P312A (1269C>G 1271T>C), R314Q (1276G>A), G324A (1306G>C), A325Q (1308G>C 1309C>A 1310A>G), V326A (1312T>C 1313G>C), V335I (1338G>A 1340A>C), V337A (1345T>C 1346A>T), M338I (1349G>A), Y341S (1357A>C 1358C>T), L346* (1371C>T 1372T>G 1373T>A), H348F (1377C>T 1378A>T 1379C>T), D352Q (1389G>C 1391T>A), T356K (1402C>A), K360R (1413A>C 1414A>G 1415A>C), I361L (1416A>C 1418A>T), K362P (1419A>C 1420A>C), T363A (1422A>G 1424A>G), T366A (1431A>G 1433T>A), K371R (1446A>C 1447A>G 1448A>T), L374V (1455T>G 1457G>A), A375R (1458G>C 1459C>G 1460T>A), K377R (1464A>C 1465A>G 1466G>A), Q378S (1467C>T 1468A>C 1469A>T), K379Q (1470A>C 1472G>A), N380K (1475C>G), F381K (1476T>A 1477T>A 1478T>A), K382G (1479A>G 1480A>G 1481A>T), C384S (1486G>C), Y385F (1489A>T 1490T>C), L389F (1502A>C), M401I (1538G>A), L404F (1545C>T 1547T>C), M407C (1554A>T 1555T>G 1556G>C), I408M (1559A>G), P411S (1566C>A 1567C>G), S420del (1593_1597delAGCTC), I427F (1614A>T), V429I (1620G>A 1622T>C), N430V (1623A>G 1624A>T), K431P (1626A>C 1627A>C 1628A>T), L433M (1632T>A 1634A>G), T434V (1635A>G 1636C>T 1637A>C), Y438S (1647T>A 1648A>G), F450Y (1684T>A), H451Y (1686C>T), A455S (1698G>T 1700A>C), S458A (1707T>G), I464M (1727A>G), C476S (1761T>A), N477T (1765A>C 1766C>T), S478T (1767T>A 1769T>C), G481A (1777G>C 1778G>T), S495T (1819G>C 1820T>A), K496E (1821A>G 1823A>G), N499R (1830A>C 1831A>G 1832T>G), E500Q (1833G>C), K502R (1840A>G), N509G (1860A>G 1861A>G), K511T (1867A>C 1868A>C), I514L (1875A>T), P515A (1878C>G 1880A>T), L518I (1887T>A 1889G>T), D521E (1898T>G), I538V (1947A>G), L545V (1968C>G 1970T>G), R546K (1972G>A 1973A>G), L549T (1980T>A 1981T>C), S550F (1984C>T 1985T>C), A559N (2010G>A 2011C>A), L560I (2013C>A 2015C>T), I562T (2020T>C), I568M (2039A>G), Y579F (2071A>T), K580R (2074A>G), A598V (2128C>T 2129T>G), H600R (2133C>A 2134A>G 2135T>G), P603L (2143C>T), V604G (2146T>G), S606L (2152C>T), L608S (2158T>C), Y611L (2167A>T 2168C>A), W613F (2173G>T 2174G>T), W614C (2177G>T), M619K (2191T>A 2192G>A), I629M (2222A>G), Y632F (2230A>T 2231C>T), K635R (2239A>G 2240A>G), E636K (2241G>A 2243A>G), V639P (2250G>C 2251T>C 2252T>C), Q643H (2264G>T), P646T (2271C>A 2273A>T), K647E (2274A>G 2276G>A), L649M (2280C>A), V655W (2298G>T 2299T>G 2300T>G), R660C (2313C>T 2315T>C), R664L (2326G>T), Q665N (2328C>A 2330A>T), E668K (2337G>A), K672R (2349A>C 2350A>G), L674V (2355C>G 2357C>T), N675S (2358A>T 2359A>C 2360C>T), C681L (2377G>T 2378T>G), N683S (2382A>T 2383A>C 2384C>T), P684F (2385C>T 2386C>T 2387A>T), D685E (2390T>A), K689E (2400A>G 2402G>A), S690Q (2403A>C 2404G>A 2405T>A), I692M (2411C>G), S694A (2415T>G 2417C>T), T699K (2431C>A 2432A>G), L704S (2445C>T 2446T>C 2447C>G), E705N (2448G>A 2450A>T), T706E (2451A>G 2452C>A 2453T>A), F708L (2457T>C 2459T>G), D709K (2460G>A 2462T>A), I711V (2466A>G), N727D (2514A>G), P728L (2518C>T 2519C>G), Q733R (2532C>A 2533A>G), W734Y (2536G>A 2537G>T), F736L (2541T>C 2543T>G), N739S (2551A>G 2552C>T), V741S (2556G>T 2557T>C), N745D (2568A>G), K746R (2571A>C 2572A>G), N755A (2598A>G 2599A>C 2600C>T), S760A (2613T>G 2615T>A), E763D (2624A>T), Y764L (2626A>T 2627C>A), Q765P (2629A>C), T766S (2631A>T), K769E (2640A>G), E770R (2643G>A 2644A>G), E771V (2647A>T 2648A>G), V772* (2649G>T 2650T>A), G773V (2653G>T), L774I (2655T>A) | | | | | | | | |

| | Begin | End | Coverage | Score | Concordance | Matches | Identities | I/D/M/F* | Stop Codons |
|---|---|---|---|---|---|---|---|---|---|
| NT | 428 | 2700 | 81.3% | 1676 | 37.6% | 2246 (99.6%) | 1551 (68.8%) | 0/9 | |
| Codon mutations: | AAA34AAG (437A>G), ATC36CTT (441A>C 443C>T), CTT38TTT (447C>T), CTA42TTA (459C>T), GCC44GTC (466C>T), TCC45TCT (470C>T), GCT46ATG (471G>A 472C>T 473T>G), TTG48ATG (477T>A), ACA49CAT (480A>C 481C>A 482A>T), AAA50AAG (485A>G), ATC52GTC (489A>G), GCT53GCC (494T>C), CTT54CTA (497T>A), GCT55GCG (500T>G), GGA56GGC (503A>C), TGT57TGC (506T>C), GAC58GAT (509C>T), TTT59TTC (512T>C), GTA60ATA (513G>A), TCT61AAA (516T>A 517C>A 518T>A), AAA62ACA (520A>C), GTG63GTT (524G>T), ACT64GTG (525A>G 526C>T 527T>G), ACC65AAA (529C>A 530C>A), TTA66CTG (531T>C 533A>G), CGC67AAA (534C>A 535G>A 536C>A), CGG68CGA (539G>A), AAG69CGT (540A>C 541A>G 542G>T), TCG70TCA (545G>A), GGC71GGT (548C>T), TAT72TTT (550A>T), TTC74TAT (556T>A 557C>T), ACA75ACT (560A>T), TAC78TAT (569C>T), AAG80AAA (575G>A), TGC82TGT (581C>T), GCA83ACA (582G>A), GTA84GTT (587A>T), TGC85TCC (589G>C), TTA86CTT (591T>C 593A>T), CGA88CGT (599A>T), TAC89TAT (602C>T), TCC93AGT (612T>A 613C>G 614C>T), TTT94TAT (616T>A), CGA95TCT (618C>T 619G>C 620A>T), GAA97AAT (624G>A 626A>T), GAA98GAC (629A>C), CCA99ACC (630C>A 632A>C), CTT100ATG (633C>A 635T>G), TCG101TCT (638G>T), GTG102GTT (641G>T), CCT103CCA (644T>A), GTG104GTT (647G>T), CTC106TTG (651C>T 653C>G), ACA107ACC (656A>C), AGA108CGA (657A>C), TCG109TGT (661C>G 662G>T), GGT110GGA (665T>A), ATC111ATT (668C>T), CGT113AAG (672C>A 673G>A 674T>G), ATC115ATA (680C>A), CCC116CCT (683C>T), AAA117GCC (684A>G 685A>C 686A>C), TTG119TTA (692G>A), CGT120CGA (695T>A), GCA122CAC (699G>C 700C>A 701A>C), ATA123GTA (702A>G), CGT124AGG (705C>A 707T>G), AGG125GCA (708A>G 709G>C 710G>A), AAA126AAG (713A>G), GAT127TCT (714G>T 715A>C), GAA128GAT (719A>T), CAC129CAT (722C>T), GTT134GTC (737T>C), TAC137TAT (746C>T), TCA139TCT (752A>T), GGG142GGA (761G>A), GCT144TCA (765G>T 767T>A), AAG145AAA (770G>A), GTT146TAG (771G>T 772T>A 773T>G), TTG147ATT (774T>A 776G>T), AAA148GCG (777A>G 778A>C 779A>G), GTT149GTG (782T>G), AGT154AAG (796G>A 797T>G), GCA156GCC (803A>C), TTT158TTC (809T>C), GAA159CAA (810G>C), TCC160TCA (815C>A), ATA161ATT (818A>T), ACT162ACG (821T>G), ACC163ACT (824C>T), CCA164CCT (827A>T), CCT166TCA (831C>T 833T>A), GGG169GAT (841G>A 842G>T), CTA172CGT (850T>G 851A>T), GTG174GTT (857G>T), CTG175CTT (860G>T), GAA176GGA (862A>G), ATT178ATG (869T>G), ACG180ACT (875G>T), TTC182TGA (880T>G 881C>A), CAT184AAG (885C>A 887T>G), CAA186CGA (892A>G), CTA190CTT (905A>T), CCC191CCA (908C>A), TTT192CAT (909T>C 910T>A), CGA194CAA (916G>A), GGC195TCC (918G>T 919G>C), ATT196ATA (923T>A), CCA197CCT (926A>T), ATG198CTT (927A>C 929G>T), ATT200AAG (934T>A 935T>G), GGG201GGT (938G>T), TCT203AAT (942T>A 943C>A), CAA205AAA (948C>A), CCC212CCA (971C>A), CTA213TTA (972C>T), AAT214CTC (975A>C 976A>T 977T>C), AAT216AAC (983T>C), AGG219CGC (990A>C 992G>C), TCA220CGG (993T>C 994C>G 995A>G), TAC221TTT (997A>T 998C>T), GGT222CAA (999G>C 1000G>A 1001T>A), TTA223TAT (1003T>A 1004A>T), GCC224TCG (1005G>T 1007C>G), GTA225GTG (1010A>G), GAC227GAT (1016C>T), AAA228GTT (1017A>G 1018A>T 1019A>T), GTT229GTA (1022T>A), GAT230GAC (1025T>C), CAA231GGA (1026C>G 1027A>G), GAG232GCA (1030A>C 1031G>A), GCT233GCG (1034T>G), AGT234GCT (1035A>G 1036G>C), TTC237GTT (1044T>G 1046C>T), AAG238AAA (1049G>A), TTT239GAA (1050T>G 1051T>A 1052T>A), CCA240CAC (1054C>A 1055A>C), ATT242ATC (1061T>C), GTT244GTG (1067T>G), TTG246TTA (1073G>A), AAA247AAG (1076A>G), CTA250ATA (1083C>A), GCG251GCT (1088G>T), GCC252GCG (1091C>G), TGG255TTG (1099G>T), ACA257GTG (1104A>G 1105C>T 1106A>G), ATT260ATG (1115T>G), TCA262TCC (1121A>C), TTC263ATT (1122T>A 1124C>T), CAG264CAA (1127G>A), GAC265GAT (1130C>T), GGC266GGG (1133C>G), TCA269TCC (1142A>C), CCC270CCT (1145C>T), GGG271GGT (1148G>T), ATG272ATT (1151G>T), TTG273CTT (1152T>C 1154G>T), TCC275TCT (1160C>T), AAA276CAA (1161A>C), TCT277ATG (1164T>A 1165C>T 1166T>G), GTC278GTT (1169C>T), TTC279TTA (1172C>A), TTT280TAC (1174T>A 1175T>C), CCT281CCA (1178T>A), TTA282TTT (1181A>T), GAC283GAT (1184C>T), AGG284TCT (1185A>T 1186G>C 1187G>T), GCT285AAT (1188G>A 1189C>A), GCA286AAT (1191G>A 1192C>A 1193A>T), ACT287ACC (1196T>C), GAA288del (1197_1199delGAA), TTA290TTT (1205A>T), AAC292AAT (1211C>T), TTA293GAA (1212T>G 1213T>A), TCT294TCC (1217T>C), CTC295CTT (1220C>T), GAA296GAT (1223A>T), CAC297TTA (1224C>T 1225A>T 1226C>A), GAG299GAA (1232G>A), AGT301CAA (1236A>C 1237G>A 1238T>A), GTC302GTA (1241C>A), CCA304CCC (1247A>C), TGG305CTT (1248T>C 1249G>T 1250G>T), TTT306TTC (1253T>C), TCA307TCT (1256A>T), ACC308AGC (1258C>G), TTG310TTA (1265G>A), AGT311GGG (1266A>G 1268T>G), CCT312GCC (1269C>G 1271T>C), TAT313TAC (1274T>C), CGA314CAA (1276G>A), GGG315GGA (1280G>A), GTG316GTT (1283G>T), GTT319GTG (1292T>G), AGG322AGA (1301G>A), CTT323TTG (1302C>T 1304T>G), GGT324GCT (1306G>C), GCA325CAG (1308G>C 1309C>A 1310A>G), GTG326GCC (1312T>C 1313G>C), ATA327ATT (1316A>T), GAG328GAA (1319G>A), GGC329GGT (1322C>T), GGA331GGG (1328A>G), AGA333CGG (1332A>C 1334A>G), GTA335ATC (1338G>A 1340A>C), GTA337GCT (1345T>C 1346A>T), ATG338ATA (1349G>A), AAC340AAT (1355C>T), TAC341TCT (1357A>C 1358C>T), AGA345AGG (1370A>G), CTT346TGA (1371C>T 1372T>G 1373T>A), CTC347CTG (1376C>G), CAC348TTT (1377C>T 1378A>T 1379C>T), CCT349CCC (1382T>C), GTT350GTC (1385T>C), GAT352CAA (1389G>C 1391T>A), GCT354GCA (1397T>A), ACA356AAA (1402C>A), CTG358CTA (1409G>A), AAA360CGC (1413A>C 1414A>G 1415A>C), ATA361CTT (1416A>C 1418A>T), AAG362CCG (1419A>C 1420A>C), ACA363GCG (1422A>G 1424A>G), GAC364GAT (1427C>T), GGG365GGA (1430G>A), ACT366GCA (1431A>G 1433T>A), GAG370GAA (1445G>A), AAA371CGT (1446A>C 1447A>G 1448A>T), CCT372CCG (1451T>G), TTG374GTA (1455T>G 1457G>A), GCT375CGA (1458G>C 1459C>G 1460T>A), AAG377CGA (1464A>C 1465A>G 1466G>A), CAA378TCT (1467C>T 1468A>C 1469A>T), AAG379CAA (1470A>C 1472G>A), AAC380AAG (1475C>G), TTT381AAA (1476T>A 1477T>A 1478T>A), AAA382GGT (1479A>G 1480A>G 1481A>T), TGT384TCT (1486G>C), TAT385TTC (1489A>T 1490T>C), TTA389TTC (1502A>C), AAG390AAA (1505G>A), TCA391TCG (1508A>G), CCG397CCA (1526G>A), ATG401ATA (1538G>A), TAT402TAC (1541T>C), CTT404TTC (1545C>T 1547T>C), TCT406TCC (1553T>C), ATG407TGC (1554A>T 1555T>G 1556G>C), ATA408ATG (1559A>G), CCT411AGT (1566C>A 1567C>G), ACC412ACA (1571C>A), TTG413CTG (1572T>C), TCC415TCA (1580C>A), GTT418GTC (1589T>C), AGC420del (1593_1597delAGCTC), TCT421--A (1593_1597delAGCTC 1598T>A), CTT424CTC (1607T>C), ACG426ACT (1613G>T), ATC427TTC (1614A>T), CTT428TTG (1617C>T 1619T>G), GTT429ATC (1620G>A 1622T>C), AAC430GTC (1623A>G 1624A>T), AAA431CCT (1626A>C 1627A>C 1628A>T), TTA433ATG (1632T>A 1634A>G), ACA434GTC (1635A>G 1636C>T 1637A>C), AGA436AGG (1643A>G), TAT438AGT (1647T>A 1648A>G), ATA440ATT (1655A>T), TTT442TTC (1661T>C), CTG443TTA (1662C>T 1664G>A), ACT444ACA (1667T>A), GGG445GGA (1670G>A), CCT447CCG (1676T>G), TTG448TTA (1679G>A), GGC449GGT (1682C>T), TTC450TAC (1684T>A), CAC451TAC (1686C>T), GGT452GGA (1691T>A), TCT453TCA (1694T>A), GCA455TCC (1698G>T 1700A>C), CTT456CTA (1703T>A), TTC457TTT (1706C>T), TCG458GCG (1707T>G), TTG459CTT (1710T>C 1712G>T), TCT460TCC (1715T>C), CAC462CAT (1721C>T), ATA464ATG (1727A>G), TTA467CTA (1734T>C), GCA468GCT (1739A>T), TGT476AGT (1761T>A), AAC477ACT (1765A>C 1766C>T), TCT478ACC (1767T>A 1769T>C), CCA479CCT (1772A>T), GGG481GCT (1777G>C 1778G>T), TAC483TAT (1784C>T), GCT484GCC (1787T>C), TTA485CTG (1788T>C 1790A>G), AGT495ACA (1819G>C 1820T>A), AAA496GAG (1821A>G 1823A>G), GCC498GCG (1829C>G), AAT499CGG (1830A>C 1831A>G 1832T>G), GAG500CAG (1833G>C), AAG502AGG (1840A>G), AGA507AGG (1856A>G), CTC508CTT (1859C>T), AAT509GGT (1860A>G 1861A>G), GTC510GTT (1865C>T), AAA511ACC (1867A>C 1868A>C), TCG513TCT (1874G>T), ATA514TTA (1875A>T), CCA515GCT (1878C>G 1880A>T), AAA516AAG (1883A>G), TCT517TCA (1886T>A), TTG518ATT (1887T>A 1889G>T), ATA519ATT (1892A>T), TCT520TCG (1895T>G), GAT521GAG (1898T>G), GGA523GGT (1904A>T), ACC524ACT (1907C>T), GCG528GCC (1919G>C), AAG529AAA (1922G>A), AGG530CGG (1923A>C), TCA535TCG (1940A>G), ATT538GTT (1947A>G), TCC544TCT (1967C>T), CTT545GTG (1968C>G 1970T>G), AGA546AAG (1972G>A 1973A>G), GCT547GCA (1976T>A), TTA549ACA (1980T>A 1981T>C), TCT550TTC (1984C>T 1985T>C), CGT552CGC (1991T>C), GTT555GTG (2000T>G), GGA556GGT (2003A>T), CTT557CTG (2006T>G), GCC559AAC (2010G>A 2011C>A), CTC560ATT (2013C>A 2015C>T), ATC562ACC (2020T>C), GAT565GAC (2030T>C), AAC567AAT (2036C>T), ATA568ATG (2039A>G), AGA573CGG (2052A>C 2054A>G), GGG575GGT (2060G>T), GGT576GGA (2063T>A), GCC577GCT (2066C>T), TAC579TTC (2071A>T), AAG580AGG (2074A>G), GCG583GCC (2084G>C), CGC584CGT (2087C>T), TTA585CTT (2088T>C 2090A>T), TTC586TTT (2093C>T), CAA589CAG (2102A>G), AGC590TCA (2103A>T 2104G>C 2105C>A), AAG591AAA (2108G>A), CGT592CGA (2111T>A), GAG594GAA (2117G>A), CGT595AGG (2118C>A 2120T>G), CTA596TTA (2121C>T), GCT598GTG (2128C>T 2129T>G), CAT600AGG (2133C>A 2134A>G 2135T>G), CTT601CTC (2138T>C), AAG602AAA (2141G>A), CCA603CTA (2143C>T), GTT604GGT (2146T>G), CGG605AGG (2148C>A), TCA606TTA (2152C>T), TTA608TCA (2158T>C), TAC611TTA (2167A>T 2168C>A), TGG613TTT (2173G>T 2174G>T), TGG614TGT (2177G>T), AGG617AGA (2186G>A), ATG619AAA (2191T>A 2192G>A), CCC620CCT (2195C>T), CCC623CCT (2204C>T), TTG625TTA (2210G>A), ATA629ATG (2222A>G), GTA630GTG (2225A>G), TAC632TTT (2230A>T 2231C>T), CTC633CTT (2234C>T), CTT634CTA (2237T>A), AAA635AGG (2239A>G 2240A>G), GAA636AAG (2241G>A 2243A>G), GTT639CCC (2250G>C 2251T>C 2252T>C), CAG643CAT (2264G>T), CCA646ACT (2271C>A 2273A>T), AAG647GAA (2274A>G 2276G>A), CTG649ATG (2280C>A), GTG650GTA (2285G>A), TTC651TTT (2288C>T), GTT655TGG (2298G>T 2299T>G 2300T>G), GAA656GAG (2303A>G), ATA657ATC (2306A>C), CTA658CTT (2309A>T), CGT660TGC (2313C>T 2315T>C), CTA663TTA (2322C>T), CGC664CTC (2326G>T), CAA665AAT (2328C>A 2330A>T), GTA667GTG (2336A>G), GAA668AAA (2337G>A), TTA671CTT (2346T>C 2348A>T), AAG672CGG (2349A>C 2350A>G), CTC674GTT (2355C>G 2357C>T), AAC675TCT (2358A>T 2359A>C 2360C>T), CAT678CAC (2369T>C), ACG679ACT (2372G>T), TGT681TTG (2377G>T 2378T>G), AAC683TCT (2382A>T 2383A>C 2384C>T), CCA684TTT (2385C>T 2386C>T 2387A>T), GAT685GAA (2390T>A), TCC687TCT (2396C>T), ATT688ATC (2399T>C), AAG689GAA (2400A>G 2402G>A), AGT690CAA (2403A>C 2404G>A 2405T>A), CTC691TTG (2406C>T 2408C>G), ATC692ATG (2411C>G), GAC693GAT (2414C>T), TCC694GCT (2415T>G 2417C>T), CCT695CCC (2420T>C), ATA696ATT (2423A>T), ACA699AAG (2431C>A 2432A>G), TCC700TCA (2435C>A), AAA702AAG (2441A>G), AGG703CGA (2442A>C 2444G>A), CTC704TCG (2445C>T 2446T>C 2447C>G), GAA705AAT (2448G>A 2450A>T), ACT706GAA (2451A>G 2452C>A 2453T>A), GAT707GAC (2456T>C), TTT708CTG (2457T>C 2459T>G), GAT709AAA (2460G>A 2462T>A), TTA710TTG (2465A>G), ATC711GTC (2466A>G), GGC714GGT (2477C>T), CTA715TTA (2478C>T), AAA718AAG (2489A>G), TGT719TGC (2492T>C), TAC720TAT (2495C>T), GGT723GGA (2504T>A), GGT725GGC (2510T>C), AAT727GAT (2514A>G), CCC728CTG (2518C>T 2519C>G), ACC730ACT (2525C>T), ACT731ACA (2528T>A), CAG733AGG (2532C>A 2533A>G), TGG734TAT (2536G>A 2537G>T), TTT736CTG (2541T>C 2543T>G), AAC739AGT (2551A>G 2552C>T), ACT740ACC (2555T>C), GTT741TCT (2556G>T 2557T>C), ATT742ATC (2561T>C), AAT745GAT (2568A>G), AAA746CGA (2571A>C 2572A>G), TTA749TTG (2582A>G), GGG750GGA (2585G>A), GGG754GGA (2597G>A), AAC755GCT (2598A>G 2599A>C 2600C>T), ATT758ATA (2609T>A), TCT760GCA (2613T>G 2615T>A), CCT761CCA (2618T>A), GAA763GAT (2624A>T), TAC764TTA (2626A>T 2627C>A), CAA765CCA (2629A>C), ACT766TCT (2631A>T), ACA767-CT (2634delA 2636A>T), GGA768GGT (2639A>T), AAA769GAA (2640A>G), GAG770AGG (2643G>A 2644A>G), GAA771GTG (2647A>T 2648A>G), GTG772TAG (2649G>T 2650T>A), GGC773GTC (2653G>T), TTA774ATA (2655T>A) | | | | | | | | |

*: Inserts / Deletes / Misaligned / Frameshifts

#### Analysis details

This analysis was performed with panviral2.64

### NGS Details (UN8): Solendovirus venanicotianae

#### Assembly

| | |
|---|---|
| **Coverage Length** | 2332 (4 contig(s)) |
| **Depth Of Coverage** | 21.1 |
| **Number Of Reads** | 430 |
| **Reads Per Million** | 7.19 rpm (after QC) |
| **Ambiguities** | 7 |
| **Assembly Method** | de novo + reference guided assembly |
| **Consensus Caller** | Bcf Tools |

#### Coverage Map

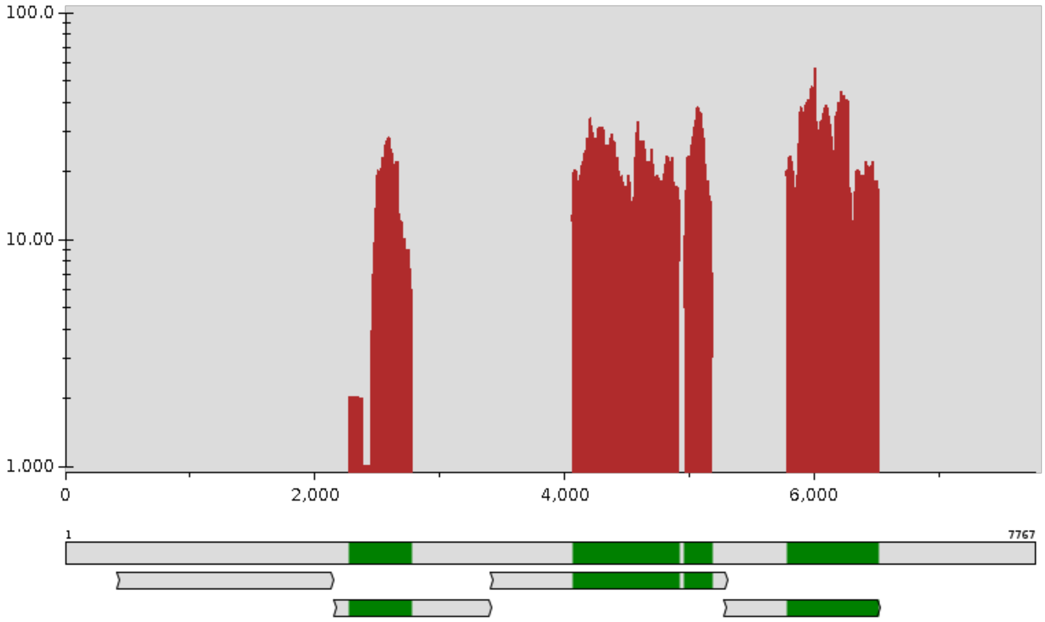

#### Assignment

| | |
|---|---|
| **Type** | Solendovirus venanicotianae (Taxonomy ID: 3048371) |
| **Reference Genome** | NC_003378.1 |
| **NT Identity (%)** | 79.7655 |
| **AA Identity (%)** | 76.7927 |
| **Number Of Stop Codons** | 4 |
| **Number Of CDS** | 4 |

#### Alignment

| | |
|---|---|
| **Alignment Score** | 2691.0 (NT) + 3770.0 (AA) = 6461.0 |
| **Concordance (%)** | 67.8676 |

| Alignment Method | Global, seeded, nucleotide + amino acids (AGA) |
|---|---|

#### Genome Region

Sequence starts at position 2271 and ends at position 6519 relative to NC_003378.1 reference sequence.

#### Alignment Detailed Statistics

| | Begin | End | Coverage | Score | Concordance | Matches | Identities | I/D/M/F* | Stop Codons |
|---|---|---|---|---|---|---|---|---|---|
| **NT** | **2271** | **6519** | **30.0%** | **2691** | **59.3%** | **2303 (98.8%)** | **1837 (78.8%)** | **0/29** | |
| Mutations: | 2274C>T, 2280T>C, 2281G>A, 2283C>A, 2294T>A, 2295T>C, 2299G>A, 2301A>G, 2304T>G, 2306C>A, 2307G>A, 2308A>T, 2313A>C, 2314A>T, 2316T>C, 2328A>G, 2330A>C, 2332A>G, 2333A>T, 2343C>T, 2347G>A, 2350G>A, 2352T>A, 2355C>T, 2368A>G, 2370A>G, 2373T>A, 2376T>G, 2388A>C, 2389G>A, 2390G>A, 2393G>A, 2394T>A, 2397G>A, 2398G>A, 2400C>T, 2402A>G, 2412A>C, 2418C>T, 2422A>T, 2424A>G, 2437G>A, 2443C>T, 2448T>A, 2452A>C, 2454G>A, 2461A>G, 2463C>T, 2472G>A, 2475T>C, 2481C>T, 2482T>C, 2493T>A, 2511A>T, 2535C>A, 2551C>T, 2568A>T, 2571C>A, 2572C>G, 2589C>T, 2602A>G, 2622T>C, 2628G>A, 2637T>C, 2638A>G, 2643A>C, 2646C>T, 2652C>T, 2665C>A, 2668A>G, 2670T>A, 2676A>T, 2687A>G, 2692T>C, 2709G>A, 2717G>R, 2718A>R, 2724G>A, 2725T>C, 2728G>A, 2729C>M, 2730A>M, 2733C>A, 2737A>K, 2743G>A, 2754C>A, 2756G>R, 2759G>S, 2763G>A, 2766C>T, 2767C>T, 2772C>T, 2775A>T, 2776T>C, 4079A>G, 4086A>G, 4091T>A, 4098G>A, 4100T>A, 4103T>A, 4104A>T, 4105A>G, 4124T>C, 4127C>T, 4181A>T, 4190G>A, 4196C>T, 4199A>T, 4202C>T, 4235T>C, 4241T>C, 4250C>T, 4253T>C, 4260T>C, 4265G>A, 4284T>A, 4292C>T, 4298T>C, 4307C>T, 4313G>A, 4325T>C, 4328T>C, 4329T>C, 4331G>A, 4334A>G, 4340G>A, 4343T>A, 4355A>G, 4362G>T, 4364T>A, 4400T>A, 4430A>T, 4436C>T, 4451C>T, 4463T>C, 4470G>C, 4475T>C, 4478T>C, 4484A>T, 4500C>A, 4503C>T, 4508T>C, 4513G>A, 4523T>A, 4530A>T, 4550C>T, 4551G>A, 4552C>T, 4554C>A, 4559T>A, 4568C>T, 4571T>A, 4574T>A, 4581T>C, 4591C>A, 4592C>G, 4598A>T, 4601A>G, 4604T>A, 4605A>T, 4607G>T, 4626T>C, 4643T>C, 4653A>G, 4660T>C, 4665A>C, 4666C>A, 4667T>A, 4679A>G, 4687T>C, 4688C>A, 4691T>C, 4692C>T, 4693T>C, 4694T>A, 4697T>A, 4701A>G, 4703C>A, 4704A>T, 4706T>A, 4709T>C, 4721A>G, 4734C>T, 4740A>T, 4755A>C, 4757G>A, 4769T>A, 4773T>C, 4784C>T, 4785T>C, 4793T>A, 4806C>T, 4811A>T, 4814G>A, 4817T>C, 4828G>A, 4831T>A, 4836G>A, 4839A>G, 4847G>A, 4848G>A, 4850A>C, 4854A>G, 4857A>C, 4859G>A, 4865T>A, 4868G>A, 4869A>T, 4872T>C, 4874G>T, 4877T>C, 4896T>C, 4913C>T, 4920T>A, 4970T>A, 4971C>T, 4973T>A, 4976G>A, 4979T>C, 4982A>G, 4986A>G, 4988T>A, 5011G>A, 5012C>T, 5018C>T, 5021T>C, 5027A>T, 5028T>A, 5030T>A, 5032A>T, 5033T>C, 5034A>T, 5037G>A, 5039A>C, 5040C>T, 5043C>G, 5046G>A, 5047A>T, 5053G>A, 5055G>A, 5076T>C, 5077T>A, 5079G>T, 5094C>T, 5117A>T, 5120T>G, 5123C>T, 5125C>A, 5128G>A, 5135T>A, 5136G>A, 5147C>T, 5150T>C, 5156A>G, 5174T>A, 5781A>C, 5784G>A, 5786A>C, 5787T>A, 5789T>A, 5796T>C, 5797A>T, 5798A>T, 5802T>A, 5805A>C, 5807G>A, 5811C>A, 5812C>A, 5814A>T, 5815T>A, 5816C>T, 5817T>C, 5820C>T, 5821C>A, 5824A>G, 5826T>A, 5831A>C, 5833G>C, 5838C>T, 5844T>A, 5850C>A, 5851C>A, 5853T>A, 5857A>C, 5858C>A, 5865C>T, 5872T>C, 5874G>T, 5877A>T, 5880T>A, 5881T>C, 5882T>A, 5886A>C, 5889T>A, 5892A>T, 5894G>A, 5897C>T, 5898A>C, 5901A>T, 5902T>C, 5907A>T, 5908G>A, 5909C>G, 5910C>A, 5913C>A, 5916C>T, 5919C>T, 5921A>G, 5922C>T, 5925C>T, 5935G>A, 5936A>G, 5940T>C, 5950C>T, 5952G>C, 5953A>G, 5954C>A, 5955A>C, 5959C>G, 5964G>A, 5968G>T, 5970C>A, 5972A>C, 5979T>A, 5991G>A, 5997C>T, 5999T>C, 6000G>A, 6001C>A, 6002A>C, 6009G>A, 6011G>A, 6012G>A, 6014T>A, 6015A>T, 6018T>A, 6020A>G, 6024G>A, 6026C>A, 6027A>T, 6031T>G, 6033C>T, 6035A>T, 6036C>T, 6037G>A, 6040C>A, 6041G>A, 6044T>G, 6045C>T, 6049T>A, 6051G>A, 6054T>C, 6055A>C, 6060A>G, 6067C>T, 6071A>T, 6075A>T, 6084G>A, 6087T>A, 6088A>G, 6090C>T, 6094C>T, 6097G>A, 6099T>A, 6108T>A, 6112C>T, 6114C>A, 6117A>C, 6120G>A, 6123A>T, 6129C>T, 6132A>T, 6139A>G, 6141G>T, 6144A>T, 6145G>A, 6146A>C, 6148G>C, 6150A>G, 6153A>G, 6154G>C, 6155A>C, 6156A>G, 6163G>C, 6164A>C, 6170T>A, 6172A>G, 6174T>G, 6177T>A, 6179C>T, 6181G>A, 6183A>T, 6186C>T, 6192T>C, 6201G>A, 6204A>T, 6211A>C, 6219T>A, 6223C>A, 6225A>T, 6232C>T, 6236C>G, 6240C>T, 6247C>A, 6253C>G, 6254A>G, 6258T>C, 6259G>C, 6264T>G, 6269G>A, 6270C>T, 6271T>C, 6277_6283delTCAAGGG, 6285A>C, 6290C>T, 6298_6315delTATTCAAATTGCAGAGAT, 6318A>C, 6319_6321delAGA, 6325G>A, 6326C>G, 6340C>G, 6341T>C, 6344G>T, 6345A>G, 6352G>A, 6354C>A, 6355T>A, 6357A>G, 6359G>C, 6360T>A, 6361C>T, 6363T>G, 6369A>T, 6375A>G, 6376C>A, 6380C>A, 6381T>A, 6382A>C, 6384A>T, 6386C>T, 6387C>T, 6401G>A, 6402G>A, 6404G>A, 6406delA, 6407A>G, 6408T>A, 6411C>T, 6414C>T, 6417C>G, 6418C>G, 6419C>A, 6420A>T, 6423T>A, 6424C>T, 6426G>A, 6427T>C, 6446T>A, 6451A>G, 6453T>G, 6456C>A, 6462T>C, 6471C>T, 6477C>T, 6480A>G, 6489A>C, 6502A>G, 6504T>A, 6505G>A, 6507T>C | | | | | | | | |

##### CDS

| | Begin | End | Coverage | Score | Concordance | Matches | Identities | I/D/M/F* | Stop Codons |
|---|---|---|---|---|---|---|---|---|---|
| Tvvgp2 | 40 | 209 | 40.5% | 821 | 71.7% | 170 (100%) | 133 (78.2%) | 0/0/5/0 | 1 |
| Protein mutations: | V43I (2281G>A 2283C>A), F47Y (2294T>A 2295T>C), V49M (2299G>A 2301A>G), T51K (2306C>A 2307G>A), I52L (2308A>T), K53N (2313A>C), N54Y (2314A>T 2316T>C), Q59P (2330A>C), K60V (2332A>G 2333A>T), V65I (2347G>A), V66I (2350G>A 2352T>A), I72V (2368A>G 2370A>G), D74E (2376T>G), G79K (2389G>A 2390G>A), G80E (2393G>A 2394T>A), A82T (2398G>A 2400C>T), K83R (2402A>G), R90W (2422A>T 2424A>G), E95K (2437G>A), Q97* (2443C>T), D98E (2448T>A), K100Q (2452A>C 2454G>A), N103D (2461A>G 2463C>T), Q140E (2572C>G), I150V (2602A>G), I162V (2638A>G), L171I (2665C>A), N172E (2668A>G 2670T>A), R174S (2676A>T), K178R (2687A>G), V197I (2743G>A) | | | | | | | | |
| Codon mutations: | GAC40GAT (2274C>T), AGT42AGC (2280T>C), GTC43ATA (2281G>A 2283C>A), TTT47TAC (2294T>A 2295T>C), GTA49ATG (2299G>A 2301A>G), CCT50CCG (2304T>G), ACG51AAA (2306C>A 2307G>A), ATA52TTA (2308A>T), AAA53AAC (2313A>C), AAT54TAC (2314A>T 2316T>C), AGA58AGG (2328A>G), CAA59CCA (2330A>C), AAA60GTA (2332A>G 2333A>T), TAC63TAT (2343C>T), GTA65ATA (2347G>A), GTT66ATA (2350G>A 2352T>A), AGC67AGT (2355C>T), ATA72GTG (2368A>G 2370A>G), ATT73ATA (2373T>A), GAT74GAG (2376T>G), ACA78ACC (2388A>C), GGA79AAA (2389G>A 2390G>A), GGT80GAA (2393G>A 2394T>A), AAG81AAA (2397G>A), GCC82ACT (2398G>A 2400C>T), AAA83AGA (2402A>G), ATA86ATC (2412A>C), AAC88AAT (2418C>T), AGA90TGG (2422A>T 2424A>G), GAA95AAA (2437G>A), CAA97TAA (2443C>T), GAT98GAA (2448T>A), AAG100CAA (2452A>C 2454G>A), AAC103GAT (2461A>G 2463C>T), AAG106AAA (2472G>A), TAT107TAC (2475T>C), CAC109CAT (2481C>T), TTA110CTA (2482T>C), ACT113ACA (2493T>A), GCA119GCT (2511A>T), ACC127ACA (2535C>A), CTA133TTA (2551C>T), ATA138ATT (2568A>T), ATC139ATA (2571C>A), CAA140GAA (2572C>G), AGC145AGT (2589C>T), ATA150GTA (2602A>G), TAT156TAC (2622T>C), AAG158AAA (2628G>A), TTT161TTC (2637T>C), ATA162GTA (2638A>G), ATA163ATC (2643A>C), AGC164AGT (2646C>T), AAC166AAT (2652C>T), CTA171ATA (2665C>A), AAT172GAA (2668A>G 2670T>A), AGA174AGT (2676A>T), AAA178AGA (2687A>G), TTA180CTA (2692T>C), AAG185AAA (2709G>A), GGA188GRR (2717G>R 2718A>R), GAG190GAA (2724G>A), TTA191CTA (2725T>C), GCA192AMM (2728G>A 2729C>M 2730A>M), CCC193CCA (2733C>A), AGT195KGT (2737A>K), GTA197ATA (2743G>A), GCC200GCA (2754C>A), AGA201ARA (2756G>R), TGT202TST (2759G>S), AAG203AAA (2763G>A), AAC204AAT (2766C>T), CTA205TTA (2767C>T), TAC206TAT (2772C>T), GTA207GTT (2775A>T), TTA208CTA (2776T>C) | | | | | | | | |
| Tvvgp3 | 219 | 591 | 56.6% | 2134 | 85.0% | 360 (100%) | 306 (85.0%) | 0/0/0/0 | 2 |
| Protein mutations: | I223M (4079A>G), K226E (4086A>G), D227E (4091T>A), D230K (4098G>A 4100T>A), N231K (4103T>A), K232* (4104A>T 4105A>G), Y292N (4284T>A), D311E (4343T>A), A318S (4362G>T 4364T>A), E354Q (4470G>C), L364I (4500C>A), R368K (4513G>A), D371E (4523T>A), I374L (4530A>T), A381I (4551G>A 4552C>T), H382N (4554C>A), T394K (4591C>A 4592C>G), M399F (4605A>T 4607G>T), I415V (4653A>G), M417T (4660T>C), T419Q (4665A>C 4666C>A 4667T>A), I426T (4687T>C 4688C>A), L428S (4692C>T 4693T>C 4694T>A), D429E (4697T>A), N431E (4701A>G 4703C>A), I432L (4704A>T 4706T>A), I444L (4740A>T), K467N (4811A>T), S473N (4828G>A), F474Y (4831T>A), E476K (4836G>A), K477E (4839A>G), E480N (4848G>A 4850A>C), I482V (4854A>G), K483Q (4857A>C 4859G>A), I487L (4869A>T), Y496H (4896T>C), N526E (4986A>G 4988T>A), C534Y (5011G>A 5012C>T), S540T (5028T>A 5030T>A), Y541F (5032A>T 5033T>C), T542S (5034A>T), E543N (5037G>A 5039A>C), P544S (5040C>T), Q545E (5043C>G), E546I (5046G>A 5047A>T), W548* (5053G>A), E549K (5055G>A), F556H (5076T>C 5077T>A), A557S (5079G>T), T572N (5125C>A), R573K (5128G>A), V576I (5136G>A) | | | | | | | | |
| Codon mutations: | ATA223ATG (4079A>G), AAA226GAA (4086A>G), GAT227GAA (4091T>A), GAT230AAA (4098G>A 4100T>A), AAT231AAA (4103T>A), AAA232TGA (4104A>T 4105A>G), AAT238AAC (4124T>C), AGC239AGT (4127C>T), GGA257GGT (4181A>T), AGG260AGA (4190G>A), GTC262GTT (4196C>T), ATA263ATT (4199A>T), GAC264GAT (4202C>T), TAT275TAC (4235T>C), TAT277TAC (4241T>C), CCC280CCT (4250C>T), AAT281AAC (4253T>C), TTA284CTA (4260T>C), AAG285AAA (4265G>A), TAT292AAT (4284T>A), TAC294TAT (4292C>T), AGT296AGC (4298T>C), GAC299GAT (4307C>T), AAG301AAA (4313G>A), TAT305TAC (4325T>C), CAT306CAC (4328T>C), TTG307CTA (4329T>C 4331G>A), AAA308AAG (4334A>G), GAG310GAA (4340G>A), GAT311GAA (4343T>A), AAA315AAG (4355A>G), GCT318TCA (4362G>T 4364T>A), GTT330GTA (4400T>A), GGA340GGT (4430A>T), TAC342TAT (4436C>T), GAC347GAT (4451C>T), AAT351AAC (4463T>C), GAA354CAA (4470G>C), AAT355AAC (4475T>C), TGT356TGC (4478T>C), GTA358GTT (4484A>T), CTA364ATA (4500C>A), CTA365TTA (4503C>T), TAT366TAC (4508T>C), AGA368AAA (4513G>A), GAT371GAA (4523T>A), ATA374TTA (4530A>T), TTC380TTT (4550C>T), GCA381ATA (4551G>A 4552C>T), CAT382AAT (4554C>A), ATT383ATA (4559T>A), AAC386AAT (4568C>T), TCT387TCA (4571T>A), GGT388GGA (4574T>A), TTA391CTA (4581T>C), ACC394AAG (4591C>A 4592C>G), GCA396GCT (4598A>T), GAA397GAG (4601A>G), ATT398ATA (4604T>A), ATG399TTT (4605A>T 4607G>T), TTA406CTA (4626T>C), GAT411GAC (4643T>C), ATA415GTA (4653A>G), ATG417ACG (4660T>C), ACT419CAA (4665A>C 4666C>A 4667T>A), CAA423CAG (4679A>G), ATC426ACA (4687T>C 4688C>A), AAT427AAC (4691T>C), CTT428TCA (4692C>T 4693T>C 4694T>A), GAT429GAA (4697T>A), AAC431GAA (4701A>G 4703C>A), ATT432TTA (4704A>T 4706T>A), GAT433GAC (4709T>C), AAA437AAG (4721A>G), CTA442TTA (4734C>T), ATA444TTA (4740A>T), AGG449CGA (4755A>C 4757G>A), CCT453CCA (4769T>A), TTA455CTA (4773T>C), AAC458AAT (4784C>T), TTA459CTA (4785T>C), CCT461CCA (4793T>A), CTA466TTA (4806C>T), AAA467AAT (4811A>T), AAG468AAA (4814G>A), GAT469GAC (4817T>C), AGT473AAT (4828G>A), TTT474TAT (4831T>A), GAA476AAA (4836G>A), AAA477GAA (4839A>G), AAG479AAA (4847G>A), GAA480AAC (4848G>A 4850A>C), ATA482GTA (4854A>G), AAG483CAA (4857A>C 4859G>A), ATT485ATA (4865T>A), AAG486AAA (4868G>A), ATA487TTA (4869A>T), TTG488CTT (4872T>C 4874G>T), TGT489TGC (4877T>C), TAC496CAC (4896T>C), AAC501AAT (4913C>T), TTT504AT. (4920T>A), GTT520GTA (4970T>A), CTT521TTA (4971C>T 4973T>A), AAG522AAA (4976G>A), TAT523TAC (4979T>C), AGA524AGG (4982A>G), AAT526GAA (4986A>G 4988T>A), TGC534TAT (5011G>A 5012C>T), TAC536TAT (5018C>T), TAT537TAC (5021T>C), GGA539GGT (5027A>T), TCT540ACA (5028T>A 5030T>A), TAT541TTC (5032A>T 5033T>C), ACA542TCA (5034A>T), GAA543AAC (5037G>A 5039A>C), CCA544TCA (5040C>T), CAA545GAA (5043C>G), GAA546ATA (5046G>A 5047A>T), TGG548TAG (5053G>A), GAA549AAA (5055G>A), TTT556CAT (5076T>C 5077T>A), GCA557TCA (5079G>T), CTA562TTA (5094C>T), ATA569ATT (5117A>T), GTT570GTG (5120T>G), TAC571TAT (5123C>T), ACC572AAC (5125C>A), AGA573AAA (5128G>A), ATT575ATA (5135T>A), GTA576ATA (5136G>A), GAC579GAT (5147C>T), AAT580AAC (5150T>C), CAA582CAG (5156A>G), ACT588ACA (5174T>A) | | | | | | | | |
| Tvvgp4 | 169 | 415 | 58.9% | 815 | 50.7% | 237 (96.3%) | 150 (61.0%) | 0/9/2/2 | 1 |

| | Begin | End | Coverage | Score | Concordance | Matches | Identities | I/D/M/F* | Stop Codons |
|---|---|---|---|---|---|---|---|---|---|
| **NT** | **2271** | **6519** | **30.0%** | **2691** | **59.3%** | **2303 (98.8%)** | **1837 (78.8%)** | **0/29** | |
| Protein mutations: | N171T (5786A>C 5787T>A), F172Y (5789T>A), K175L (5797A>T 5798A>T), N176K (5802T>A), R178K (5807G>A), Q180N (5812C>A 5814A>T), S181I (5815T>A 5816C>T 5817T>C), Q183K (5821C>A), N184E (5824A>G 5826T>A), N186T (5831A>C), E187Q (5833G>C), H192Q (5850C>A), H193K (5851C>A 5853T>A), T195Q (5857A>C 5858C>A), L203Q (5881T>C 5882T>A), N205K (5889T>A), S207N (5894G>A), A208V (5897C>T 5898A>C), K209N (5901A>T), A212R (5908G>A 5909C>G 5910C>A), N216S (5921A>G 5922C>T), D221S (5935G>A 5936A>G), Q226Y (5950C>T 5952G>C), T227D (5953A>G 5954C>A 5955A>C), Q229E (5959C>G), A232S (5968G>T 5970C>A), N233T (5972A>C), M242T (5999T>C 6000G>A), Q243T (6001C>A 6002A>C), R246K (6011G>A 6012G>A), I247N (6014T>A 6015A>T), K249R (6020A>G), T251N (6026C>A 6027A>T), F253V (6031T>G 6033C>T), Y254F (6035A>T 6036C>T), V255I (6037G>A), R256K (6040C>A 6041G>A), F257C (6044T>G 6045C>T), S259T (6049T>A 6051G>A), T261P (6055A>C), Y266F (6071A>T), E267D (6075A>T), I272V (6088A>G 6090C>T), Q274* (6094C>T), V275I (6097G>A 6099T>A), E283D (6123A>T), K289D (6139A>G 6141G>T), E291T (6145G>A 6146A>C), E292Q (6148G>C 6150A>G), E294P (6154G>C 6155A>C 6156A>G), E297P (6163G>C 6164A>C), V299E (6170T>A), N300E (6172A>G 6174T>G), P302L (6179C>T), E303N (6181G>A 6183A>T), I313L (6211A>C), L317I (6223C>A 6225A>T), A321G (6236C>G), L325I (6247C>A), Q327G (6253C>G 6254A>G), A329P (6259G>C), I330M (6264T>G), S332N (6269G>A 6270C>T), Y333H (6271T>C), S335_R336del (6277_6283delTCAAGGG), T339I (6290C>T), Y342_D347del (6298_6315delTATTCAAATTGCAGAGAT), R349del (6319_6321delAGA), A351R (6325G>A 6326C>G), L356A (6340C>G 6341T>C), R357M (6344G>T 6345A>G), V360I (6352G>A 6354C>A), L361M (6355T>A 6357A>G), S362T (6359G>C 6360T>A), K365N (6369A>T), Q368K (6376C>A), P369Q (6380C>A 6381T>A), T370P (6382A>C 6384A>T), T371I (6386C>T 6387C>T), R376K (6401G>A 6402G>A), R377K (6404G>A), P382D (6418C>G 6419C>A 6420A>T), D383E (6423T>A), L391Q (6446T>A), S393G (6451A>G 6453T>G), H394Q (6456C>A), K405N (6489A>C), I410V (6502A>G 6504T>A), V411I (6505G>A 6507T>C) | | | | | | | | |
| Codon mutations: | ATA169ATC (5781A>C), CAG170CAA (5784G>A), AAT171ACA (5786A>C 5787T>A), TTT172TAT (5789T>A), AAT174AAC (5796T>C), AAA175TTA (5797A>T 5798A>T), AAT176AAA (5802T>A), CCA177CCC (5805A>C), AGA178AAA (5807G>A), TCC179TCA (5811C>A), CAA180AAT (5812C>A 5814A>T), TCT181ATC (5815T>A 5816C>T 5817T>C), ACC182ACT (5820C>T), CAA183AAA (5821C>A), AAT184GAA (5824A>G 5826T>A), AAC186ACC (5831A>C), GAA187CAA (5833G>C), GAC188GAT (5838C>T), ATT190ATA (5844T>A), CAC192CAA (5850C>A), CAT193AAA (5851C>A 5853T>A), ACA195CAA (5857A>C 5858C>A), TAC197TAT (5865C>T), TTG200CTT (5872T>C 5874G>T), ATA201ATT (5877A>T), GCT202GCA (5880T>A), TTA203CAA (5881T>C 5882T>A), CCA204CCC (5886A>C), AAT205AAA (5889T>A), ACA206ACT (5892A>T), AGT207AAT (5894G>A), GCA208GTC (5897C>T 5898A>C), AAA209AAT (5901A>T), TTA210CTA (5902T>C), GTA211GTT (5907A>T), GCC212AGA (5908G>A 5909C>G 5910C>A), ACC213ACA (5913C>A), TGC214TGT (5916C>T), TAC215TAT (5919C>T), AAC216AGT (5921A>G 5922C>T), TAC217TAT (5925C>T), GAT221AGT (5935G>A 5936A>G), ACT222ACC (5940T>C), CAG226TAC (5950C>T 5952G>C), ACA227GAC (5953A>G 5954C>A 5955A>C), CAA229GAA (5959C>G), GAG230GAA (5964G>A), GCC232TCA (5968G>T 5970C>A), AAT233ACT (5972A>C), CCT235CCA (5979T>A), AAG239AAA (5991G>A), TTC241TTT (5997C>T), ATG242ACA (5999T>C 6000G>A), CAA243ACA (6001C>A 6002A>C), AAG245AAA (6009G>A), AGG246AAA (6011G>A 6012G>A), ATA247AAT (6014T>A 6015A>T), ACT248ACA (6018T>A), AAA249AGA (6020A>G), GGG250GGA (6024G>A), ACA251AAT (6026C>A 6027A>T), TTC253GTT (6031T>G 6033C>T), TAC254TTT (6035A>T 6036C>T), GTA255ATA (6037G>A), CGA256AAA (6040C>A 6041G>A), TTC257TGT (6044T>G 6045C>T), TCG259ACA (6049T>A 6051G>A), GCT260GCC (6054T>C), ACA261CCA (6055A>C), GCA262GCG (6060A>G), CTA265TTA (6067C>T), TAT266TTT (6071A>T), GAA267GAT (6075A>T), AAG270AAA (6084G>A), CCT271CCA (6087T>A), ATC272GTT (6088A>G 6090C>T), CAA274TAA (6094C>T), GTT275ATA (6097G>A 6099T>A), ATT278ATA (6108T>A), CTC280TTA (6112C>T 6114C>A), ACA281ACC (6117A>C), AGG282AGA (6120G>A), GAA283GAT (6123A>T), ATC285ATT (6129C>T), ATA286ATT (6132A>T), AAG289GAT (6139A>G 6141G>T), ATA290ATT (6144A>T), GAA291ACA (6145G>A 6146A>C), GAA292CAG (6148G>C 6150A>G), CAA293CAG (6153A>G), GAA294CCG (6154G>C 6155A>C 6156A>G), GAA297CCA (6163G>C 6164A>C), GTA299GAA (6170T>A), AAT300GAG (6172A>G 6174T>G), ATT301ATA (6177T>A), CCA302CTA (6179C>T), GAA303AAT (6181G>A 6183A>T), TTC304TTT (6186C>T), GCT306GCC (6192T>C), AGG309AGA (6201G>A), ATA310ATT (6204A>T), ATA313CTA (6211A>C), ACT315ACA (6219T>A), CTA317ATT (6223C>A 6225A>T), CTA320TTA (6232C>T), GCA321GGA (6236C>G), AAC322AAT (6240C>T), CTA325ATA (6247C>A), CAG327GGG (6253C>G 6254A>G), AAT328AAC (6258T>C), GCA329CCA (6259G>C), ATT330ATG (6264T>G), AGC332AAT (6269G>A 6270C>T), TAT333CAT (6271T>C), TCA335_AGG336del (6277_6283delTCAAGGG), GAA337-AC (6277_6283delTCAAGGG 6285A>C), ACA339ATA (6290C>T), TAT342_GAT347del (6298_6315delTATTCAAATTGCAGAGAT), ATA348ATC (6318A>C), AGA349del (6319_6321delAGA), GCA351AGA (6325G>A 6326C>G), CTA356GCA (6340C>G 6341T>C), AGA357ATG (6344G>T 6345A>G), GTC360ATA (6352G>A 6354C>A), TTA361ATG (6355T>A 6357A>G), AGT362ACA (6359G>C 6360T>A), CTT363TTG (6361C>T 6363T>G), AAA365AAT (6369A>T), GAA367GAG (6375A>G), CAA368AAA (6376C>A), CCT369CAA (6380C>A 6381T>A), ACA370CCT (6382A>C 6384A>T), ACC371ATT (6386C>T 6387C>T), AGG376AAA (6401G>A 6402G>A), AGG377AAG (6404G>A), AAT378-GA (6406delA 6407A>G 6408T>A), TTC379TTT (6411C>T), ATC380ATT (6414C>T), TCC381TCG (6417C>G), CCA382GAT (6418C>G 6419C>A 6420A>T), GAT383GAA (6423T>A), CTG384TTA (6424C>T 6426G>A), TTA385CTA (6427T>C), CTA391CAA (6446T>A), AGT393GGG (6451A>G 6453T>G), CAC394CAA (6456C>A), TAT396TAC (6462T>C), CAC399CAT (6471C>T), TGC401TGT (6477C>T), TCA402TCG (6480A>G), AAA405AAC (6489A>C), ATT410GTA (6502A>G 6504T>A), GTT411ATC (6505G>A 6507T>C) | | | | | | | | |

#### Proteins

| | Begin | End | Coverage | Score | Concordance | Matches | Identities | I/D/M/F* | Stop Codons |
|---|---|---|---|---|---|---|---|---|---|
| putative cell-to-cell movement protein (NP_569140.1) | 40 | 209 | 40.5% | 821 | 71.7% | 170 (100%) | 133 (78.2%) | 0/0/5/0 | 1 |
| Protein mutations: | V43I (2281G>A 2283C>A), F47Y (2294T>A 2295T>C), V49M (2299G>A 2301A>G), T51K (2306C>A 2307G>A), I52L (2308A>T), K53N (2313A>C), N54Y (2314A>T 2316T>C), Q59P (2330A>C), K60V (2332A>G 2333A>T), V65I (2347G>A), V66I (2350G>A 2352T>A), I72V (2368A>G 2370A>G), D74E (2376T>G), G79K (2389G>A 2390G>A), G80E (2393G>A 2394T>A), A82T (2398G>A 2400C>T), K83R (2402A>G), R90W (2422A>T 2424A>G), E95K (2437G>A), Q97* (2443C>T), D98E (2448T>A), K100Q (2452A>C 2454G>A), N103D (2461A>G 2463C>T), Q140E (2572C>G), I150V (2602A>G), I162V (2638A>G), L171I (2665C>A), N172E (2668A>G 2670T>A), R174S (2676A>T), K178R (2687A>G), V197I (2743G>A) | | | | | | | | |
| Codon mutations: | GAC40GAT (2274C>T), AGT42AGC (2280T>C), GTC43ATA (2281G>A 2283C>A), TTT47TAC (2294T>A 2295T>C), GTA49ATG (2299G>A 2301A>G), CCT50CCG (2304T>G), ACG51AAA (2306C>A 2307G>A), ATA52TTA (2308A>T), AAA53AAC (2313A>C), AAT54TAC (2314A>T 2316T>C), AGA58AGG (2328A>G), CAA59CCA (2330A>C), AAA60GTA (2332A>G 2333A>T), TAC63TAT (2343C>T), GTA65ATA (2347G>A), GTT66ATA (2350G>A 2352T>A), AGC67AGT (2355C>T), ATA72GTG (2368A>G 2370A>G), ATT73ATA (2373T>A), GAT74GAG (2376T>G), ACA78ACC (2388A>C), GGA79AAA (2389G>A 2390G>A), GGT80GAA (2393G>A 2394T>A), AAG81AAA (2397G>A), GCC82ACT (2398G>A 2400C>T), AAA83AGA (2402A>G), ATA86ATC (2412A>C), AAC88AAT (2418C>T), AGA90TGG (2422A>T 2424A>G), GAA95AAA (2437G>A), CAA97TAA (2443C>T), GAT98GAA (2448T>A), AAG100CAA (2452A>C 2454G>A), AAC103GAT (2461A>G 2463C>T), AAG106AAA (2472G>A), TAT107TAC (2475T>C), CAC109CAT (2481C>T), TTA110CTA (2482T>C), ACT113ACA (2493T>A), GCA119GCT (2511A>T), ACC127ACA (2535C>A), CTA133TTA (2551C>T), ATA138ATT (2568A>T), ATC139ATA (2571C>A), CAA140GAA (2572C>G), AGC145AGT (2589C>T), ATA150GTA (2602A>G), TAT156TAC (2622T>C), AAG158AAA (2628G>A), TTT161TTC (2637T>C), ATA162GTA (2638A>G), ATA163ATC (2643A>C), AGC164AGT (2646C>T), AAC166AAT (2652C>T), CTA171ATA (2665C>A), AAT172GAA (2668A>G 2670T>A), AGA174AGT (2676A>T), AAA178AGA (2687A>G), TTA180CTA (2692T>C), AAG185AAA (2709G>A), GGA188GRR (2717G>R 2718A>R), GAG190GAA (2724G>A), TTA191CTA (2725T>C), GCA192AMM (2728G>A 2729C>M 2730A>M), CCC193CCA (2733C>A), AGT195KGT (2737A>K), GTA197ATA (2743G>A), GCC200GCA (2754C>A), AGA201ARA (2756G>R), TGT202TST (2759G>S), AAG203AAA (2763G>A), AAC204AAT (2766C>T), CTA205TTA (2767C>T), TAC206TAT (2772C>T), GTA207GTT (2775A>T), TTA208CTA (2776T>C) | | | | | | | | |
| polyprotein (NP_569141.1) | 219 | 591 | 56.6% | 2134 | 85.0% | 360 (100%) | 306 (85.0%) | 0/0/0/0 | 2 |
| Protein mutations: | I223M (4079A>G), K226E (4086A>G), D227E (4091T>A), D230K (4098G>A 4100T>A), N231K (4103T>A), K232* (4104A>T 4105A>G), Y292N (4284T>A), D311E (4343T>A), A318S (4362G>T 4364T>A), E354Q (4470G>C), L364I (4500C>A), R368K (4513G>A), D371E (4523T>A), I374L (4530A>T), A381I (4551G>A 4552C>T), H382N (4554C>A), T394K (4591C>A 4592C>G), M399F (4605A>T 4607G>T), I415V (4653A>G), M417T (4660T>C), T419Q (4665A>C 4666C>A 4667T>A), I426T (4687T>C 4688C>A), L428S (4692C>T 4693T>C 4694T>A), D429E (4697T>A), N431E (4701A>G 4703C>A), I432L (4704A>T 4706T>A), I444L (4740A>T), K467N (4811A>T), S473N (4828G>A), F474Y (4831T>A), E476K (4836G>A), K477E (4839A>G), E480N (4848G>A 4850A>C), I482V (4854A>G), K483Q (4857A>C 4859G>A), I487L (4869A>T), Y496H (4896T>C), N526E (4986A>G 4988T>A), C534Y (5011G>A 5012C>T), S540T (5028T>A 5030T>A), Y541F (5032A>T 5033T>C), T542S (5034A>T), E543N (5037G>A 5039A>C), P544S (5040C>T), Q545E (5043C>G), E546I (5046G>A 5047A>T), W548* (5053G>A), E549K (5055G>A), F556H (5076T>C 5077T>A), A557S (5079G>T), T572N (5125C>A), R573K (5128G>A), V576I (5136G>A) | | | | | | | | |
| Codon mutations: | ATA223ATG (4079A>G), AAA226GAA (4086A>G), GAT227GAA (4091T>A), GAT230AAA (4098G>A 4100T>A), AAT231AAA (4103T>A), AAA232TGA (4104A>T 4105A>G), AAT238AAC (4124T>C), AGC239AGT (4127C>T), GGA257GGT (4181A>T), AGG260AGA (4190G>A), GTC262GTT (4196C>T), ATA263ATT (4199A>T), GAC264GAT (4202C>T), TAT275TAC (4235T>C), TAT277TAC (4241T>C), CCC280CCT (4250C>T), AAT281AAC (4253T>C), TTA284CTA (4260T>C), AAG285AAA (4265G>A), TAT292AAT (4284T>A), TAC294TAT (4292C>T), AGT296AGC (4298T>C), GAC299GAT (4307C>T), AAG301AAA (4313G>A), TAT305TAC (4325T>C), CAT306CAC (4328T>C), TTG307CTA (4329T>C 4331G>A), AAA308AAG (4334A>G), GAG310GAA (4340G>A), GAT311GAA (4343T>A), AAA315AAG (4355A>G), GCT318TCA (4362G>T 4364T>A), GTT330GTA (4400T>A), GGA340GGT (4430A>T), TAC342TAT (4436C>T), GAC347GAT (4451C>T), AAT351AAC (4463T>C), GAA354CAA (4470G>C), AAT355AAC (4475T>C), TGT356TGC (4478T>C), GTA358GTT (4484A>T), CTA364ATA (4500C>A), CTA365TTA (4503C>T), TAT366TAC (4508T>C), AGA368AAA (4513G>A), GAT371GAA (4523T>A), ATA374TTA (4530A>T), TTC380TTT (4550C>T), GCA381ATA (4551G>A 4552C>T), CAT382AAT (4554C>A), ATT383ATA (4559T>A), AAC386AAT (4568C>T), TCT387TCA (4571T>A), GGT388GGA (4574T>A), TTA391CTA (4581T>C), ACC394AAG (4591C>A 4592C>G), GCA396GCT (4598A>T), GAA397GAG (4601A>G), ATT398ATA (4604T>A), ATG399TTT (4605A>T 4607G>T), TTA406CTA (4626T>C), GAT411GAC (4643T>C), ATA415GTA (4653A>G), ATG417ACG (4660T>C), ACT419CAA (4665A>C 4666C>A 4667T>A), CAA423CAG (4679A>G), ATC426ACA (4687T>C 4688C>A), AAT427AAC (4691T>C), CTT428TCA (4692C>T 4693T>C 4694T>A), GAT429GAA (4697T>A), AAC431GAA (4701A>G 4703C>A), ATT432TTA (4704A>T 4706T>A), GAT433GAC (4709T>C), AAA437AAG (4721A>G), CTA442TTA (4734C>T), ATA444TTA (4740A>T), AGG449CGA (4755A>C 4757G>A), CCT453CCA (4769T>A), TTA455CTA (4773T>C), AAC458AAT (4784C>T), TTA459CTA (4785T>C), CCT461CCA (4793T>A), CTA466TTA (4806C>T), AAA467AAT (4811A>T), AAG468AAA (4814G>A), GAT469GAC (4817T>C), AGT473AAT (4828G>A), TTT474TAT (4831T>A), GAA476AAA (4836G>A), AAA477GAA (4839A>G), AAG479AAA (4847G>A), GAA480AAC (4848G>A 4850A>C), ATA482GTA (4854A>G), AAG483CAA (4857A>C 4859G>A), ATT485ATA (4865T>A), AAG486AAA (4868G>A), ATA487TTA (4869A>T), TTG488CTT (4872T>C 4874G>T), TGT489TGC (4877T>C), TAC496CAC (4896T>C), AAC501AAT (4913C>T), TTT504AT. (4920T>A), GTT520GTA (4970T>A), CTT521TTA (4971C>T 4973T>A), AAG522AAA (4976G>A), TAT523TAC (4979T>C), AGA524AGG (4982A>G), AAT526GAA (4986A>G 4988T>A), TGC534TAT (5011G>A 5012C>T), TAC536TAT (5018C>T), TAT537TAC (5021T>C), GGA539GGT (5027A>T), TCT540ACA (5028T>A 5030T>A), TAT541TTC (5032A>T 5033T>C), ACA542TCA (5034A>T), GAA543AAC (5037G>A 5039A>C), CCA544TCA (5040C>T), CAA545GAA (5043C>G), GAA546ATA (5046G>A 5047A>T), TGG548TAG (5053G>A), GAA549AAA (5055G>A), TTT556CAT (5076T>C 5077T>A), GCA557TCA (5079G>T), CTA562TTA (5094C>T), ATA569ATT (5117A>T), GTT570GTG (5120T>G), TAC571TAT (5123C>T), ACC572AAC (5125C>A), AGA573AAA (5128G>A), ATT575ATA (5135T>A), GTA576ATA (5136G>A), GAC579GAT (5147C>T), AAT580AAC (5150T>C), CAA582CAG (5156A>G), ACT588ACA (5174T>A) | | | | | | | | |
| putative transactivator factor (NP_569142.1) | 169 | 415 | 58.9% | 815 | 50.7% | 237 (96.3%) | 150 (61.0%) | 0/9/2/2 | 1 |

| | Begin | End | Coverage | Score | Concordance | Matches | Identities | I/D/M/F* | Stop Codons |
|---|---|---|---|---|---|---|---|---|---|
| **NT** | **2271** | **6519** | **30.0%** | **2691** | **59.3%** | **2303 (98.8%)** | **1837 (78.8%)** | **0/29** | |
| Protein mutations: | N171T (5786A>C 5787T>A), F172Y (5789T>A), K175L (5797A>T 5798A>T), N176K (5802T>A), R178K (5807G>A), Q180N (5812C>A 5814A>T), S181I (5815T>A 5816C>T 5817T>C), Q183K (5821C>A), N184E (5824A>G 5826T>A), N186T (5831A>C), E187Q (5833G>C), H192Q (5850C>A), H193K (5851C>A 5853T>A), T195Q (5857A>C 5858C>A), L203Q (5881T>C 5882T>A), N205K (5889T>A), S207N (5894G>A), A208V (5897C>T 5898A>C), K209N (5901A>T), A212R (5908G>A 5909C>G 5910C>A), N216S (5921A>G 5922C>T), D221S (5935G>A 5936A>G), Q226Y (5950C>T 5952G>C), T227D (5953A>G 5954C>A 5955A>C), Q229E (5959C>G), A232S (5968G>T 5970C>A), N233T (5972A>C), M242T (5999T>C 6000G>A), Q243T (6001C>A 6002A>C), R246K (6011G>A 6012G>A), I247N (6014T>A 6015A>T), K249R (6020A>G), T251N (6026C>A 6027A>T), F253V (6031T>G 6033C>T), Y254F (6035A>T 6036C>T), V255I (6037G>A), R256K (6040C>A 6041G>A), F257C (6044T>G 6045C>T), S259T (6049T>A 6051G>A), T261P (6055A>C), Y266F (6071A>T), E267D (6075A>T), I272V (6088A>G 6090C>T), Q274* (6094C>T), V275I (6097G>A 6099T>A), E283D (6123A>T), K289D (6139A>G 6141G>T), E291T (6145G>A 6146A>C), E292Q (6148G>C 6150A>G), E294P (6154G>C 6155A>C 6156A>G), E297P (6163G>C 6164A>C), V299E (6170T>A), N300E (6172A>G 6174T>G), P302L (6179C>T), E303N (6181G>A 6183A>T), I313L (6211A>C), L317I (6223C>A 6225A>T), A321G (6236C>G), L325I (6247C>A), Q327G (6253C>G 6254A>G), A329P (6259G>C), I330M (6264T>G), S332N (6269G>A 6270C>T), Y333H (6271T>C), S335_R336del (6277_6283delTCAAGGG), T339I (6290C>T), Y342_D347del (6298_6315delTATTCAAATTGCAGAGAT), R349del (6319_6321delAGA), A351R (6325G>A 6326C>G), L356A (6340C>G 6341T>C), R357M (6344G>T 6345A>G), V360I (6352G>A 6354C>A), L361M (6355T>A 6357A>G), S362T (6359G>C 6360T>A), K365N (6369A>T), Q368K (6376C>A), P369Q (6380C>A 6381T>A), T370P (6382A>C 6384A>T), T371I (6386C>T 6387C>T), R376K (6401G>A 6402G>A), R377K (6404G>A), P382D (6418C>G 6419C>A 6420A>T), D383E (6423T>A), L391Q (6446T>A), S393G (6451A>G 6453T>G), H394Q (6456C>A), K405N (6489A>C), I410V (6502A>G 6504T>A), V411I (6505G>A 6507T>C) | | | | | | | | |
| Codon mutations: | ATA169ATC (5781A>C), CAG170CAA (5784G>A), AAT171ACA (5786A>C 5787T>A), TTT172TAT (5789T>A), AAT174AAC (5796T>C), AAA175TTA (5797A>T 5798A>T), AAT176AAA (5802T>A), CCA177CCC (5805A>C), AGA178AAA (5807G>A), TCC179TCA (5811C>A), CAA180AAT (5812C>A 5814A>T), TCT181ATC (5815T>A 5816C>T 5817T>C), ACC182ACT (5820C>T), CAA183AAA (5821C>A), AAT184GAA (5824A>G 5826T>A), AAC186ACC (5831A>C), GAA187CAA (5833G>C), GAC188GAT (5838C>T), ATT190ATA (5844T>A), CAC192CAA (5850C>A), CAT193AAA (5851C>A 5853T>A), ACA195CAA (5857A>C 5858C>A), TAC197TAT (5865C>T), TTG200CTT (5872T>C 5874G>T), ATA201ATT (5877A>T), GCT202GCA (5880T>A), TTA203CAA (5881T>C 5882T>A), CCA204CCC (5886A>C), AAT205AAA (5889T>A), ACA206ACT (5892A>T), AGT207AAT (5894G>A), GCA208GTC (5897C>T 5898A>C), AAA209AAT (5901A>T), TTA210CTA (5902T>C), GTA211GTT (5907A>T), GCC212AGA (5908G>A 5909C>G 5910C>A), ACC213ACA (5913C>A), TGC214TGT (5916C>T), TAC215TAT (5919C>T), AAC216AGT (5921A>G 5922C>T), TAC217TAT (5925C>T), GAT221AGT (5935G>A 5936A>G), ACT222ACC (5940T>C), CAG226TAC (5950C>T 5952G>C), ACA227GAC (5953A>G 5954C>A 5955A>C), CAA229GAA (5959C>G), GAG230GAA (5964G>A), GCC232TCA (5968G>T 5970C>A), AAT233ACT (5972A>C), CCT235CCA (5979T>A), AAG239AAA (5991G>A), TTC241TTT (5997C>T), ATG242ACA (5999T>C 6000G>A), CAA243ACA (6001C>A 6002A>C), AAG245AAA (6009G>A), AGG246AAA (6011G>A 6012G>A), ATA247AAT (6014T>A 6015A>T), ACT248ACA (6018T>A), AAA249AGA (6020A>G), GGG250GGA (6024G>A), ACA251AAT (6026C>A 6027A>T), TTC253GTT (6031T>G 6033C>T), TAC254TTT (6035A>T 6036C>T), GTA255ATA (6037G>A), CGA256AAA (6040C>A 6041G>A), TTC257TGT (6044T>G 6045C>T), TCG259ACA (6049T>A 6051G>A), GCT260GCC (6054T>C), ACA261CCA (6055A>C), GCA262GCG (6060A>G), CTA265TTA (6067C>T), TAT266TTT (6071A>T), GAA267GAT (6075A>T), AAG270AAA (6084G>A), CCT271CCA (6087T>A), ATC272GTT (6088A>G 6090C>T), CAA274TAA (6094C>T), GTT275ATA (6097G>A 6099T>A), ATT278ATA (6108T>A), CTC280TTA (6112C>T 6114C>A), ACA281ACC (6117A>C), AGG282AGA (6120G>A), GAA283GAT (6123A>T), ATC285ATT (6129C>T), ATA286ATT (6132A>T), AAG289GAT (6139A>G 6141G>T), ATA290ATT (6144A>T), GAA291ACA (6145G>A 6146A>C), GAA292CAG (6148G>C 6150A>G), CAA293CAG (6153A>G), GAA294CCG (6154G>C 6155A>C 6156A>G), GAA297CCA (6163G>C 6164A>C), GTA299GAA (6170T>A), AAT300GAG (6172A>G 6174T>G), ATT301ATA (6177T>A), CCA302CTA (6179C>T), GAA303AAT (6181G>A 6183A>T), TTC304TTT (6186C>T), GCT306GCC (6192T>C), AGG309AGA (6201G>A), ATA310ATT (6204A>T), ATA313CTA (6211A>C), ACT315ACA (6219T>A), CTA317ATT (6223C>A 6225A>T), CTA320TTA (6232C>T), GCA321GGA (6236C>G), AAC322AAT (6240C>T), CTA325ATA (6247C>A), CAG327GGG (6253C>G 6254A>G), AAT328AAC (6258T>C), GCA329CCA (6259G>C), ATT330ATG (6264T>G), AGC332AAT (6269G>A 6270C>T), TAT333CAT (6271T>C), TCA335_AGG336del (6277_6283delTCAAGGG), GAA337-AC (6277_6283delTCAAGGG 6285A>C), ACA339ATA (6290C>T), TAT342_GAT347del (6298_6315delTATTCAAATTGCAGAGAT), ATA348ATC (6318A>C), AGA349del (6319_6321delAGA), GCA351AGA (6325G>A 6326C>G), CTA356GCA (6340C>G 6341T>C), AGA357ATG (6344G>T 6345A>G), GTC360ATA (6352G>A 6354C>A), TTA361ATG (6355T>A 6357A>G), AGT362ACA (6359G>C 6360T>A), CTT363TTG (6361C>T 6363T>G), AAA365AAT (6369A>T), GAA367GAG (6375A>G), CAA368AAA (6376C>A), CCT369CAA (6380C>A 6381T>A), ACA370CCT (6382A>C 6384A>T), ACC371ATT (6386C>T 6387C>T), AGG376AAA (6401G>A 6402G>A), AGG377AAG (6404G>A), AAT378-GA (6406delA 6407A>G 6408T>A), TTC379TTT (6411C>T), ATC380ATT (6414C>T), TCC381TCG (6417C>G), CCA382GAT (6418C>G 6419C>A 6420A>T), GAT383GAA (6423T>A), CTG384TTA (6424C>T 6426G>A), TTA385CTA (6427T>C), CTA391CAA (6446T>A), AGT393GGG (6451A>G 6453T>G), CAC394CAA (6456C>A), TAT396TAC (6462T>C), CAC399CAT (6471C>T), TGC401TGT (6477C>T), TCA402TCG (6480A>G), AAA405AAC (6489A>C), ATT410GTA (6502A>G 6504T>A), GTT411ATC (6505G>A 6507T>C) | | | | | | | | |

*: Inserts / Deletes / Misaligned / Frameshifts

#### Analysis details

This analysis was performed with panviral2.64

### NGS Details (UN8): Harvey murine sarcoma virus

#### Assembly

| | |
|---|---|
| **Coverage Length** | 279 (1 contig(s)) |
| **Depth Of Coverage** | 3.9 |
| **Number Of Reads** | 8 |
| **Reads Per Million** | 0.13 rpm (after QC) |
| **Ambiguities** | 0 |
| **Assembly Method** | de novo + reference guided assembly |
| **Consensus Caller** | Bcf Tools |

#### Coverage Map

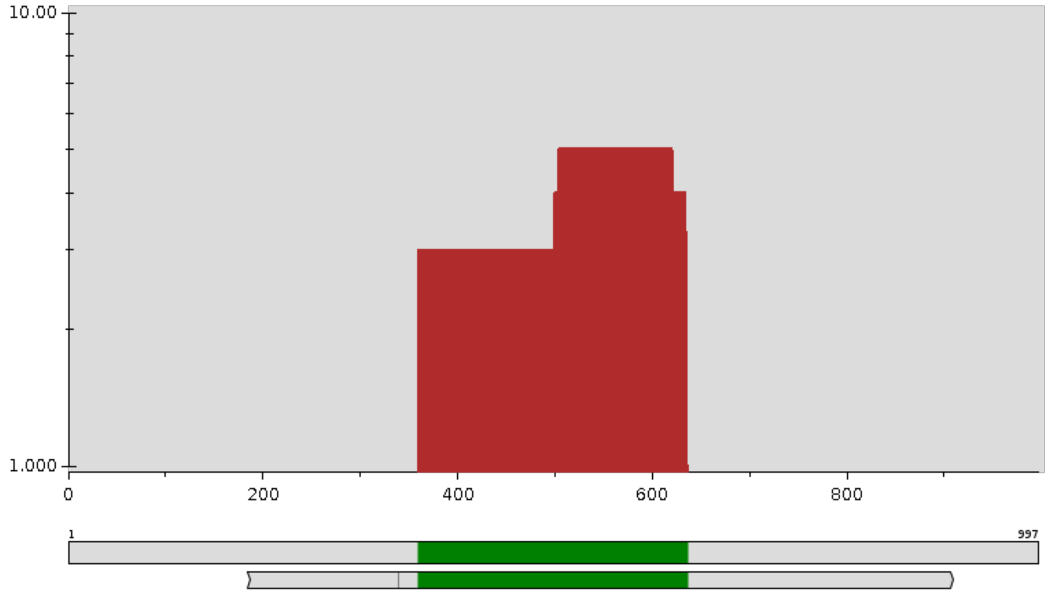

#### Assignment

| | |
|---|---|
| **Type** | Harvey murine sarcoma virus (Taxonomy ID: 11807) |
| **Reference Genome** | NC_038668.1 |
| **NT Identity (%)** | 73.3333 |
| **AA Identity (%)** | 90.0 |
| **Number Of Stop Codons** | 0 |
| **Number Of CDS** | 2 |

#### Alignment

| | |
|---|---|
| **Alignment Score** | 234.0 (NT) + 1076.0 (AA) = 1310.0 |
| **Concordance (%)** | 75.1147 |

| Alignment Method | Global, seeded, nucleotide + amino acids (AGA) |
|---|---|

#### Genome Region

Sequence starts at position 360 and ends at position 638 relative to NC_038668.1 reference sequence.

#### Alignment Detailed Statistics

| | Begin | End | Coverage | Score | Concordance | Matches | Identities | I/D/M/F* | Stop Codons |
|---|---|---|---|---|---|---|---|---|---|
| **NT** | **360** | **638** | **28.0%** | **234** | **44.8%** | **270 (96.8%)** | **198 (71.0%)** | **0/9** | |
| Mutations: | 367G>C, 370C>T, 373T>C, 374A>G, 376A>T, 379C>T, 382G>T, 385A>T, 388G>A, 394C>T, 397G>C, 403C>T, 407C>T, 409G>A, 412C>T, 415G>A, 418C>T, 427G>T, 430C>T, 433G>A, 442C>A, 448A>T, 451G>A, 454C>T, 460C>T, 463G>T, 466A>G, 469G>A, 472A>C, 475C>G, 481T>C, 484G>A, 487G>A, 490G>A, 491_499delTGTTTACTG, 500G>T, 502C>T, 505C>T, 508A>G, 514A>T, 515A>G, 517A>C, 520T>A, 532T>C, 542C>A, 544G>A, 547C>T, 550G>A, 553C>T, 562A>T, 568G>A, 571C>T, 574C>T, 575C>T, 577C>G, 578T>C, 579G>T, 590A>C, 592C>T, 595C>T, 598C>T, 599A>G, 601C>G, 604G>A, 607C>A, 619C>T, 620C>T, 621A>C, 622T>A, 623C>A, 624A>T, 631G>A, 634G>A | | | | | | | | |
| **CDS** | | | | | | | | | |
| D1R95_gp1 | 60 | 152 | 38.4% | 538 | 86.5% | 90 (96.8%) | 81 (87.1%) | 0/3/0/0 | 0 |
| Protein mutations: | R64G (374A>G 376A>T), C103_L105del (491_499delTGTTTACTG), D106Y (500G>T 502C>T), T111A (515A>G 517A>C), C132L (578T>C 579G>T), I136L (590A>C 592C>T), T139A (599A>G 601C>G), H146S (620C>T 621A>C 622T>A), Q147M (623C>A 624A>T) | | | | | | | | |
| Codon mutations: | GTG61GTC (367G>C), GGC62GGT (370C>T), GCT63GCC (373T>C), AGA64GGT (374A>G 376A>T), GGC65GGT (379C>T), GTG66GTT (382G>T), GGA67GGT (385A>T), AAG68AAA (388G>A), GCC70GCT (394C>T), CTG71CTC (397G>C), ATC73ATT (403C>T), CTG75TTA (407C>T 409G>A), ATC76ATT (412C>T), CAG77CAA (415G>A), AAC78AAT (418C>T), GTG81GTT (427G>T), GAC82GAT (430C>T), GAG83GAA (433G>A), CCC86CCA (442C>A), ATA88ATT (448A>T), GAG89GAA (451G>A), GAC90GAT (454C>T), TAC92TAT (460C>T), CGG93CGT (463G>T), AAA94AAG (466A>G), CAG95CAA (469G>A), GTA96GTC (472A>C), GTC97GTG (475C>G), GAT99GAC (481T>C), GGG100GGA (484G>A), GAG101GAA (487G>A), ACG102ACA (490G>A), TGT103_CTG105del (491_499delTGTTTACTG), GAC106TAT (500G>T 502C>T), ATC107ATT (505C>T), TTA108TTG (508A>G), ACA110ACT (514A>T), ACA111GCC (515A>G 517A>C), GGT112GGA (520T>A), TAT116TAC (532T>C), CGG120AGA (542C>A 544G>A), GAC121GAT (547C>T), CAG122CAA (550G>A), TAC123TAT (553C>T), ACA126ACT (562A>T), GAG128GAA (568G>A), GGC129GGT (571C>T), TTC130TTT (574C>T), CTC131TTG (575C>T 577C>G), TGT132CTT (578T>C 579G>T), ATC136CTT (590A>C 592C>T), AAC137AAT (595C>T), AAC138AAT (598C>T), ACC139GCG (599A>G 601C>G), AAG140AAA (604G>A), TCC141TCA (607C>A), ATC145ATT (619C>T), CAT146TCA (620C>T 621A>C 622T>A), CAG147ATG (623C>A 624A>T), AGG149AGA (631G>A), GAG150GAA (634G>A) | | | | | | | | |
| D1R95_gp2 | 8 | 100 | 48.9% | 538 | 86.5% | 90 (96.8%) | 81 (87.1%) | 0/3/0/0 | 0 |
| Protein mutations: | R12G (374A>G 376A>T), C51_L53del (491_499delTGTTTACTG), D54Y (500G>T 502C>T), T59A (515A>G 517A>C), C80L (578T>C 579G>T), I84L (590A>C 592C>T), T87A (599A>G 601C>G), H94S (620C>T 621A>C 622T>A), Q95M (623C>A 624A>T) | | | | | | | | |
| Codon mutations: | GTG9GTC (367G>C), GGC10GGT (370C>T), GCT11GCC (373T>C), AGA12GGT (374A>G 376A>T), GGC13GGT (379C>T), GTG14GTT (382G>T), GGA15GGT (385A>T), AAG16AAA (388G>A), GCC18GCT (394C>T), CTG19CTC (397G>C), ATC21ATT (403C>T), CTG23TTA (407C>T 409G>A), ATC24ATT (412C>T), CAG25CAA (415G>A), AAC26AAT (418C>T), GTG29GTT (427G>T), GAC30GAT (430C>T), GAG31GAA (433G>A), CCC34CCA (442C>A), ATA36ATT (448A>T), GAG37GAA (451G>A), GAC38GAT (454C>T), TAC40TAT (460C>T), CGG41CGT (463G>T), AAA42AAG (466A>G), CAG43CAA (469G>A), GTA44GTC (472A>C), GTC45GTG (475C>G), GAT47GAC (481T>C), GGG48GGA (484G>A), GAG49GAA (487G>A), ACG50ACA (490G>A), TGT51_CTG53del (491_499delTGTTTACTG), GAC54TAT (500G>T 502C>T), ATC55ATT (505C>T), TTA56TTG (508A>G), ACA58ACT (514A>T), ACA59GCC (515A>G 517A>C), GGT60GGA (520T>A), TAT64TAC (532T>C), CGG68AGA (542C>A 544G>A), GAC69GAT (547C>T), CAG70CAA (550G>A), TAC71TAT (553C>T), ACA74ACT (562A>T), GAG76GAA (568G>A), GGC77GGT (571C>T), TTC78TTT (574C>T), CTC79TTG (575C>T 577C>G), TGT80CTT (578T>C 579G>T), ATC84CTT (590A>C 592C>T), AAC85AAT (595C>T), AAC86AAT (598C>T), ACC87GCG (599A>G 601C>G), AAG88AAA (604G>A), TCC89TCA (607C>A), ATC93ATT (619C>T), CAT94TCA (620C>T 621A>C 622T>A), CAG95ATG (623C>A 624A>T), AGG97AGA (631G>A), GAG98GAA (634G>A) | | | | | | | | |
| **Proteins** | | | | | | | | | |
| hypothetical protein (YP_009507788.1) | 60 | 152 | 38.4% | 538 | 86.5% | 90 (96.8%) | 81 (87.1%) | 0/3/0/0 | 0 |
| Protein mutations: | R64G (374A>G 376A>T), C103_L105del (491_499delTGTTTACTG), D106Y (500G>T 502C>T), T111A (515A>G 517A>C), C132L (578T>C 579G>T), I136L (590A>C 592C>T), T139A (599A>G 601C>G), H146S (620C>T 621A>C 622T>A), Q147M (623C>A 624A>T) | | | | | | | | |
| Codon mutations: | GTG61GTC (367G>C), GGC62GGT (370C>T), GCT63GCC (373T>C), AGA64GGT (374A>G 376A>T), GGC65GGT (379C>T), GTG66GTT (382G>T), GGA67GGT (385A>T), AAG68AAA (388G>A), GCC70GCT (394C>T), CTG71CTC (397G>C), ATC73ATT (403C>T), CTG75TTA (407C>T 409G>A), ATC76ATT (412C>T), CAG77CAA (415G>A), AAC78AAT (418C>T), GTG81GTT (427G>T), GAC82GAT (430C>T), GAG83GAA (433G>A), CCC86CCA (442C>A), ATA88ATT (448A>T), GAG89GAA (451G>A), GAC90GAT (454C>T), TAC92TAT (460C>T), CGG93CGT (463G>T), AAA94AAG (466A>G), CAG95CAA (469G>A), GTA96GTC (472A>C), GTC97GTG (475C>G), GAT99GAC (481T>C), GGG100GGA (484G>A), GAG101GAA (487G>A), ACG102ACA (490G>A), TGT103_CTG105del (491_499delTGTTTACTG), GAC106TAT (500G>T 502C>T), ATC107ATT (505C>T), TTA108TTG (508A>G), ACA110ACT (514A>T), ACA111GCC (515A>G 517A>C), GGT112GGA (520T>A), TAT116TAC (532T>C), CGG120AGA (542C>A 544G>A), GAC121GAT (547C>T), CAG122CAA (550G>A), TAC123TAT (553C>T), ACA126ACT (562A>T), GAG128GAA (568G>A), GGC129GGT (571C>T), TTC130TTT (574C>T), CTC131TTG (575C>T 577C>G), TGT132CTT (578T>C 579G>T), ATC136CTT (590A>C 592C>T), AAC137AAT (595C>T), AAC138AAT (598C>T), ACC139GCG (599A>G 601C>G), AAG140AAA (604G>A), TCC141TCA (607C>A), ATC145ATT (619C>T), CAT146TCA (620C>T 621A>C 622T>A), CAG147ATG (623C>A 624A>T), AGG149AGA (631G>A), GAG150GAA (634G>A) | | | | | | | | |
| hypothetical protein (YP_009507789.1) | 8 | 100 | 48.9% | 538 | 86.5% | 90 (96.8%) | 81 (87.1%) | 0/3/0/0 | 0 |
| Protein mutations: | R12G (374A>G 376A>T), C51_L53del (491_499delTGTTTACTG), D54Y (500G>T 502C>T), T59A (515A>G 517A>C), C80L (578T>C 579G>T), I84L (590A>C 592C>T), T87A (599A>G 601C>G), H94S (620C>T 621A>C 622T>A), Q95M (623C>A 624A>T) | | | | | | | | |
| Codon mutations: | GTG9GTC (367G>C), GGC10GGT (370C>T), GCT11GCC (373T>C), AGA12GGT (374A>G 376A>T), GGC13GGT (379C>T), GTG14GTT (382G>T), GGA15GGT (385A>T), AAG16AAA (388G>A), GCC18GCT (394C>T), CTG19CTC (397G>C), ATC21ATT (403C>T), CTG23TTA (407C>T 409G>A), ATC24ATT (412C>T), CAG25CAA (415G>A), AAC26AAT (418C>T), GTG29GTT (427G>T), GAC30GAT (430C>T), GAG31GAA (433G>A), CCC34CCA (442C>A), ATA36ATT (448A>T), GAG37GAA (451G>A), GAC38GAT (454C>T), TAC40TAT (460C>T), CGG41CGT (463G>T), AAA42AAG (466A>G), CAG43CAA (469G>A), GTA44GTC (472A>C), GTC45GTG (475C>G), GAT47GAC (481T>C), GGG48GGA (484G>A), GAG49GAA (487G>A), ACG50ACA (490G>A), TGT51_CTG53del (491_499delTGTTTACTG), GAC54TAT (500G>T 502C>T), ATC55ATT (505C>T), TTA56TTG (508A>G), ACA58ACT (514A>T), ACA59GCC (515A>G 517A>C), GGT60GGA (520T>A), TAT64TAC (532T>C), CGG68AGA (542C>A 544G>A), GAC69GAT (547C>T), CAG70CAA (550G>A), TAC71TAT (553C>T), ACA74ACT (562A>T), GAG76GAA (568G>A), GGC77GGT (571C>T), TTC78TTT (574C>T), CTC79TTG (575C>T 577C>G), TGT80CTT (578T>C 579G>T), ATC84CTT (590A>C 592C>T), AAC85AAT (595C>T), AAC86AAT (598C>T), ACC87GCG (599A>G 601C>G), AAG88AAA (604G>A), TCC89TCA (607C>A), ATC93ATT (619C>T), CAT94TCA (620C>T 621A>C 622T>A), CAG95ATG (623C>A 624A>T), AGG97AGA (631G>A), GAG98GAA (634G>A) | | | | | | | | |

*: Inserts / Deletes / Misaligned / Frameshifts

#### Analysis details

This analysis was performed with panviral2.64

### NGS Details (UN8): Tomato chocolate spot virus (segment RNA2)

#### Assembly

| | |
|---|---|
| **Coverage Length** | 393 (1 contig(s)) |
| **Depth Of Coverage** | 4272.8 |
| **Number Of Reads** | 14918 |
| **Reads Per Million** | 249.32 rpm (after QC) |
| **Ambiguities** | 0 |
| **Assembly Method** | de novo + reference guided assembly |
| **Consensus Caller** | Bcf Tools |

#### Coverage Map

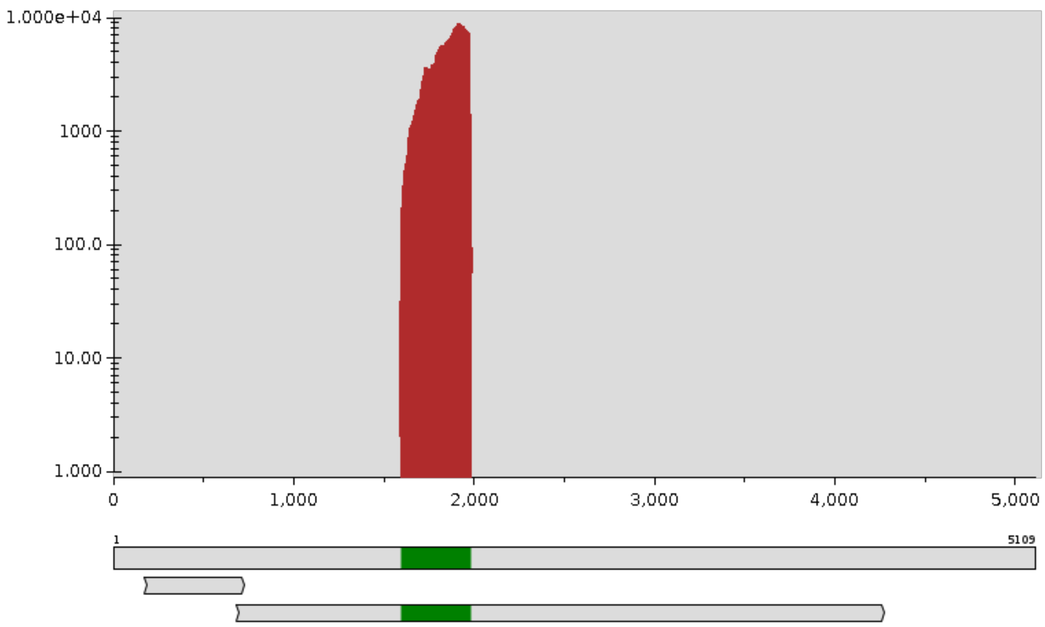

#### Assignment

| | |
|---|---|
| **Type** | Tomato chocolate spot virus (Taxonomy ID: 661101) |
| **Reference Genome** | NC_013076.1 |
| **NT Identity (%)** | 56.2341 |
| **AA Identity (%)** | 51.9084 |
| **Number Of Stop Codons** | 0 |
| **Number Of CDS** | 2 |

#### Alignment

| | |
|---|---|
| **Alignment Score** | 98.0 (NT) + 555.0 (AA) = 653.0 |
| **Concordance (%)** | 37.4427 |

| Alignment Method | Global, seeded, nucleotide + amino acids (AGA) |
|---|---|

#### Genome Region

Sequence starts at position 1593 and ends at position 1985 relative to NC_013076.1 reference sequence.

#### Alignment Detailed Statistics

| | Begin | End | Coverage | Score | Concordance | Matches | Identities | I/D/M/F* | Stop Codons |
|---|---|---|---|---|---|---|---|---|---|
| **NT** | **1593** | **1985** | **7.7%** | **98** | **12.5%** | **393 (100%)** | **221 (56.2%)** | **0/0** | |
| Mutations: | 1603A>C, 1606T>C, 1610A>G, 1618A>G, 1620G>C, 1622T>A, 1624C>T, 1625A>G, 1627C>T, 1628A>T, 1630G>T, 1633T>C, 1634G>C, 1635T>A, 1636T>G, 1645T>C, 1649C>T, 1650T>A, 1651G>T, 1657A>C, 1658G>A, 1659A>C, 1660C>T, 1661T>G, 1662C>A, 1663C>T, 1668C>A, 1669A>T, 1670C>T, 1671A>C, 1672C>T, 1673A>T, 1678C>T, 1690T>G, 1696T>A, 1697G>A, 1698A>C, 1699G>C, 1700C>T, 1702C>G, 1703A>C, 1704C>A, 1705G>T, 1708T>A, 1709G>T, 1711G>A, 1712C>A, 1713A>G, 1714A>G, 1717G>A, 1718G>A, 1721C>G, 1723A>T, 1725A>T, 1726C>T, 1727A>G, 1729T>G, 1730C>T, 1734T>C, 1736G>A, 1737A>C, 1738T>C, 1739G>C, 1740A>T, 1741G>C, 1742G>C, 1743C>T, 1744T>C, 1745T>C, 1746C>G, 1747A>C, 1749G>A, 1750A>G, 1753C>G, 1755A>G, 1756T>A, 1757G>T, 1758T>G, 1759G>T, 1760A>C, 1761T>A, 1762C>T, 1763G>C, 1768G>C, 1771A>G, 1774T>C, 1775G>A, 1777A>T, 1778T>G, 1779C>A, 1780A>G, 1782A>G, 1783G>A, 1784A>C, 1786G>A, 1789G>C, 1790C>T, 1791A>C, 1792G>C, 1798T>C, 1801T>A, 1802T>A, 1805A>C, 1806A>T, 1807G>A, 1811C>T, 1816C>T, 1817A>G, 1818C>A, 1823G>A, 1825C>T, 1832G>C, 1834T>A, 1835G>C, 1837T>G, 1838A>G, 1840T>G, 1843T>C, 1844G>A, 1845C>A, 1846T>G, 1848C>A, 1861C>A, 1864G>T, 1867A>G, 1870A>T, 1871A>T, 1872A>C, 1873T>C, 1876C>A, 1877T>G, 1878C>A, 1879T>G, 1883T>A, 1885A>T, 1887T>G, 1888G>C, 1889G>A, 1897T>C, 1898G>T, 1899A>C, 1901A>T, 1903G>A, 1904T>A, 1906G>T, 1909T>C, 1910C>A, 1911A>G, 1912T>G, 1915G>A, 1918A>G, 1921C>G, 1924T>A, 1926C>A, 1927C>A, 1928C>T, 1930A>G, 1936C>T, 1939A>G, 1945T>C, 1948C>T, 1952G>T, 1953T>A, 1954G>C, 1959T>G, 1960A>G, 1961A>T, 1962T>A, 1963G>T, 1970T>G, 1971T>A, 1972T>G | | | | | | | | |

##### CDS

| | Begin | End | Coverage | Score | Concordance | Matches | Identities | I/D/M/F* | Stop Codons |
|---|---|---|---|---|---|---|---|---|---|
| ToChV_s2_gp2 | 304 | 434 | 10.9% | 555 | 56.7% | 131 (100%) | 68 (51.9%) | 0/0/0/0 | 0 |
| Protein mutations: | K309E (1610A>G), G312A (1620G>C), F313I (1622T>A 1624C>T), N314D (1625A>G 1627C>T), M315F (1628A>T 1630G>T), V317Q (1634G>C 1635T>A 1636T>G), L322Y (1649C>T 1650T>A 1651G>T), D325T (1658G>A 1659A>C 1660C>T), S326D (1661T>G 1662C>A 1663C>T), T328N (1668C>A 1669A>T), H329S (1670C>T 1671A>C 1672C>T), I330F (1673A>T), D335E (1690T>G), E338T (1697G>A 1698A>C 1699G>C), R339W (1700C>T 1702C>G), T340H (1703A>C 1704C>A 1705G>T), A342S (1709G>T 1711G>A), Q343R (1712C>A 1713A>G 1714A>G), V345I (1718G>A), Q346D (1721C>G 1723A>T), Y347F (1725A>T 1726C>T), I348V (1727A>G 1729T>G), F350S (1734T>C), D351T (1736G>A 1737A>C 1738T>C), E352L (1739G>C 1740A>T 1741G>C), A353L (1742G>C 1743C>T 1744T>C), S354R (1745T>C 1746C>G 1747A>C), R355Q (1749G>A 1750A>G), N357R (1755A>G 1756T>A), V358C (1757G>T 1758T>G 1759G>T), I359H (1760A>C 1761T>A 1762C>T), V360L (1763G>C), V364I (1775G>A 1777A>T), S365E (1778T>G 1779C>A 1780A>G), K366R (1782A>G 1783G>A), E368D (1789G>C), Q369S (1790C>T 1791A>C 1792G>C), Y373N (1802T>A), K374L (1805A>C 1806A>T 1807G>A), H376Y (1811C>T), T378D (1817A>G 1818C>A), D380N (1823G>A 1825C>T), V383L (1832G>C 1834T>A), D384Q (1835G>C 1837T>G), I385V (1838A>G 1840T>G), A387K (1844G>A 1845C>A 1846T>G), A388E (1848C>A), K393N (1864G>T), N396S (1871A>T 1872A>C 1873T>C), S398E (1877T>G 1878C>A 1879T>G), S400T (1883T>A 1885A>T), L401C (1887T>G 1888G>C), D402N (1889G>A), D405S (1898G>T 1899A>C), M406L (1901A>T 1903G>A), S407T (1904T>A 1906G>T), H409R (1910C>A 1911A>G 1912T>G), A414E (1926C>A 1927C>A), V423Y (1952G>T 1953T>A 1954G>C), I425R (1959T>G 1960A>G), M426Y (1961A>T 1962T>A 1963G>T), F429E (1970T>G 1971T>A 1972T>G) | | | | | | | | |
| Codon mutations: | CCA306CCC (1603A>C), GTT307GTC (1606T>C), AAG309GAG (1610A>G), CAA311CAG (1618A>G), GGC312GCC (1620G>C), TTC313ATT (1622T>A 1624C>T), AAC314GAT (1625A>G 1627C>T), ATG315TTT (1628A>T 1630G>T), GGT316GGC (1633T>C), GTT317CAG (1634G>C 1635T>A 1636T>G), GAT320GAC (1645T>C), CTG322TAT (1649C>T 1650T>A 1651G>T), ATA324ATC (1657A>C), GAC325ACT (1658G>A 1659A>C 1660C>T), TCC326GAT (1661T>G 1662C>A 1663C>T), ACA328AAT (1668C>A 1669A>T), CAC329TCT (1670C>T 1671A>C 1672C>T), ATT330TTT (1673A>T), GAC331GAT (1678C>T), GAT335GAG (1690T>G), ACT337ACA (1696T>A), GAG338ACC (1697G>A 1698A>C 1699G>C), CGC339TGG (1700C>T 1702C>G), ACG340CAT (1703A>C 1704C>A 1705G>T), ATT341ATA (1708T>A), GCG342TCA (1709G>T 1711G>A), CAA343AGG (1712C>A 1713A>G 1714A>G), CAG344CAA (1717G>A), GTT345ATT (1718G>A), CAA346GAT (1721C>G 1723A>T), TAC347TTT (1725A>T 1726C>T), ATT348GTG (1727A>G 1729T>G), CTG349TTG (1730C>T), TTT350TCT (1734T>C), GAT351ACC (1736G>A 1737A>C 1738T>C), GAG352CTC (1739G>C 1740A>T 1741G>C), GCT353CTC (1742G>C 1743C>T 1744T>C), TCA354CGC (1745T>C 1746C>G 1747A>C), CGA355CAG (1749G>A 1750A>G), GGC356GGG (1753C>G), AAT357AGA (1755A>G 1756T>A), GTG358TGT (1757G>T 1758T>G 1759G>T), ATC359CAT (1760A>C 1761T>A 1762C>T), GTG360CTG (1763G>C), CCG361CCC (1768G>C), AAA362AAG (1771A>G), CAT363CAC (1774T>C), GTA364ATT (1775G>A 1777A>T), TCA365GAG (1778T>G 1779C>A 1780A>G), AAG366AGA (1782A>G 1783G>A), AGG367CGA (1784A>C 1786G>A), GAG368GAC (1789G>C), CAG369TCC (1790C>T 1791A>C 1792G>C), AAT371AAC (1798T>C), ATT372ATA (1801T>A), TAC373AAC (1802T>A), AAG374CTA (1805A>C 1806A>T 1807G>A), CAC376TAC (1811C>T), ATC377ATT (1816C>T), ACT378GAT (1817A>G 1818C>A), GAC380AAT (1823G>A 1825C>T), GTT383CTA (1832G>C 1834T>A), GAT384CAG (1835G>C 1837T>G), ATT385GTG (1838A>G 1840T>G), CTT386CTC (1843T>C), GCT387AAG (1844G>A 1845C>A 1846T>G), GCA388GAA (1848C>A), ACC392ACA (1861C>A), AAG393AAT (1864G>T), GTA394GTG (1867A>G), GGA395GGT (1870A>T), AAT396TCC (1871A>T 1872A>C 1873T>C), CTC397CTA (1876C>A), TCT398GAG (1877T>G 1878C>A 1879T>G), TCA400ACT (1883T>A 1885A>T), TTG401TGC (1887T>G 1888G>C), GAT402AAT (1889G>A), TAT404TAC (1897T>C), GAT405TCT (1898G>T 1899A>C), ATG406TTA (1901A>T 1903G>A), TCG407ACT (1904T>A 1906G>T), CTT408CTC (1909T>C), CAT409AGG (1910C>A 1911A>G 1912T>G), GAG410GAA (1915G>A), AGA411AGG (1918A>G), GTC412GTG (1921C>G), ATT413ATA (1924T>A), GCC414GAA (1926C>A 1927C>A), CTA415TTG (1928C>T 1930A>G), CAC417CAT (1936C>T), CAA418CAG (1939A>G), GAT420GAC (1945T>C), CAC421CAT (1948C>T), GTG423TAC (1952G>T 1953T>A 1954G>C), ATA425AGG (1959T>G 1960A>G), ATG426TAT (1961A>T 1962T>A 1963G>T), TTT429GAG (1970T>G 1971T>A 1972T>G) | | | | | | | | |

##### Proteins

| | Begin | End | Coverage | Score | Concordance | Matches | Identities | I/D/M/F* | Stop Codons |
|---|---|---|---|---|---|---|---|---|---|
| polyprotein (YP_003097231.1) | 304 | 434 | 10.9% | 555 | 56.7% | 131 (100%) | 68 (51.9%) | 0/0/0/0 | 0 |
| Protein mutations: | K309E (1610A>G), G312A (1620G>C), F313I (1622T>A 1624C>T), N314D (1625A>G 1627C>T), M315F (1628A>T 1630G>T), V317Q (1634G>C 1635T>A 1636T>G), L322Y (1649C>T 1650T>A 1651G>T), D325T (1658G>A 1659A>C 1660C>T), S326D (1661T>G 1662C>A 1663C>T), T328N (1668C>A 1669A>T), H329S (1670C>T 1671A>C 1672C>T), I330F (1673A>T), D335E (1690T>G), E338T (1697G>A 1698A>C 1699G>C), R339W (1700C>T 1702C>G), T340H (1703A>C 1704C>A 1705G>T), A342S (1709G>T 1711G>A), Q343R (1712C>A 1713A>G 1714A>G), V345I (1718G>A), Q346D (1721C>G 1723A>T), Y347F (1725A>T 1726C>T), I348V (1727A>G 1729T>G), F350S (1734T>C), D351T (1736G>A 1737A>C 1738T>C), E352L (1739G>C 1740A>T 1741G>C), A353L (1742G>C 1743C>T 1744T>C), S354R (1745T>C 1746C>G 1747A>C), R355Q (1749G>A 1750A>G), N357R (1755A>G 1756T>A), V358C (1757G>T 1758T>G 1759G>T), I359H (1760A>C 1761T>A 1762C>T), V360L (1763G>C), V364I (1775G>A 1777A>T), S365E (1778T>G 1779C>A 1780A>G), K366R (1782A>G 1783G>A), E368D (1789G>C), Q369S (1790C>T 1791A>C 1792G>C), Y373N (1802T>A), K374L (1805A>C 1806A>T 1807G>A), H376Y (1811C>T), T378D (1817A>G 1818C>A), D380N (1823G>A 1825C>T), V383L (1832G>C 1834T>A), D384Q (1835G>C 1837T>G), I385V (1838A>G 1840T>G), A387K (1844G>A 1845C>A 1846T>G), A388E (1848C>A), K393N (1864G>T), N396S (1871A>T 1872A>C 1873T>C), S398E (1877T>G 1878C>A 1879T>G), S400T (1883T>A 1885A>T), L401C (1887T>G 1888G>C), D402N (1889G>A), D405S (1898G>T 1899A>C), M406L (1901A>T 1903G>A), S407T (1904T>A 1906G>T), H409R (1910C>A 1911A>G 1912T>G), A414E (1926C>A 1927C>A), V423Y (1952G>T 1953T>A 1954G>C), I425R (1959T>G 1960A>G), M426Y (1961A>T 1962T>A 1963G>T), F429E (1970T>G 1971T>A 1972T>G) | | | | | | | | |
| Codon mutations: | CCA306CCC (1603A>C), GTT307GTC (1606T>C), AAG309GAG (1610A>G), CAA311CAG (1618A>G), GGC312GCC (1620G>C), TTC313ATT (1622T>A 1624C>T), AAC314GAT (1625A>G 1627C>T), ATG315TTT (1628A>T 1630G>T), GGT316GGC (1633T>C), GTT317CAG (1634G>C 1635T>A 1636T>G), GAT320GAC (1645T>C), CTG322TAT (1649C>T 1650T>A 1651G>T), ATA324ATC (1657A>C), GAC325ACT (1658G>A 1659A>C 1660C>T), TCC326GAT (1661T>G 1662C>A 1663C>T), ACA328AAT (1668C>A 1669A>T), CAC329TCT (1670C>T 1671A>C 1672C>T), ATT330TTT (1673A>T), GAC331GAT (1678C>T), GAT335GAG (1690T>G), ACT337ACA (1696T>A), GAG338ACC (1697G>A 1698A>C 1699G>C), CGC339TGG (1700C>T 1702C>G), ACG340CAT (1703A>C 1704C>A 1705G>T), ATT341ATA (1708T>A), GCG342TCA (1709G>T 1711G>A), CAA343AGG (1712C>A 1713A>G 1714A>G), CAG344CAA (1717G>A), GTT345ATT (1718G>A), CAA346GAT (1721C>G 1723A>T), TAC347TTT (1725A>T 1726C>T), ATT348GTG (1727A>G 1729T>G), CTG349TTG (1730C>T), TTT350TCT (1734T>C), GAT351ACC (1736G>A 1737A>C 1738T>C), GAG352CTC (1739G>C 1740A>T 1741G>C), GCT353CTC (1742G>C 1743C>T 1744T>C), TCA354CGC (1745T>C 1746C>G 1747A>C), CGA355CAG (1749G>A 1750A>G), GGC356GGG (1753C>G), AAT357AGA (1755A>G 1756T>A), GTG358TGT (1757G>T 1758T>G 1759G>T), ATC359CAT (1760A>C 1761T>A 1762C>T), GTG360CTG (1763G>C), CCG361CCC (1768G>C), AAA362AAG (1771A>G), CAT363CAC (1774T>C), GTA364ATT (1775G>A 1777A>T), TCA365GAG (1778T>G 1779C>A 1780A>G), AAG366AGA (1782A>G 1783G>A), AGG367CGA (1784A>C 1786G>A), GAG368GAC (1789G>C), CAG369TCC (1790C>T 1791A>C 1792G>C), AAT371AAC (1798T>C), ATT372ATA (1801T>A), TAC373AAC (1802T>A), AAG374CTA (1805A>C 1806A>T 1807G>A), CAC376TAC (1811C>T), ATC377ATT (1816C>T), ACT378GAT (1817A>G 1818C>A), GAC380AAT (1823G>A 1825C>T), GTT383CTA (1832G>C 1834T>A), GAT384CAG (1835G>C 1837T>G), ATT385GTG (1838A>G 1840T>G), CTT386CTC (1843T>C), GCT387AAG (1844G>A 1845C>A 1846T>G), GCA388GAA (1848C>A), ACC392ACA (1861C>A), AAG393AAT (1864G>T), GTA394GTG (1867A>G), GGA395GGT (1870A>T), AAT396TCC (1871A>T 1872A>C 1873T>C), CTC397CTA (1876C>A), TCT398GAG (1877T>G 1878C>A 1879T>G), TCA400ACT (1883T>A 1885A>T), TTG401TGC (1887T>G 1888G>C), GAT402AAT (1889G>A), TAT404TAC (1897T>C), GAT405TCT (1898G>T 1899A>C), ATG406TTA (1901A>T 1903G>A), TCG407ACT (1904T>A 1906G>T), CTT408CTC (1909T>C), CAT409AGG (1910C>A 1911A>G 1912T>G), GAG410GAA (1915G>A), AGA411AGG (1918A>G), GTC412GTG (1921C>G), ATT413ATA (1924T>A), GCC414GAA (1926C>A 1927C>A), CTA415TTG (1928C>T 1930A>G), CAC417CAT (1936C>T), CAA418CAG (1939A>G), GAT420GAC (1945T>C), CAC421CAT (1948C>T), GTG423TAC (1952G>T 1953T>A 1954G>C), ATA425AGG (1959T>G 1960A>G), ATG426TAT (1961A>T 1962T>A 1963G>T), TTT429GAG (1970T>G 1971T>A 1972T>G) | | | | | | | | |

*: Inserts / Deletes / Misaligned / Frameshifts

#### Analysis details

This analysis was performed with panviral2.64

### NGS Details (UN8): Tomato chocolate spot virus (segment RNA 1)

#### Assembly

| | |
|---|---|
| **Coverage Length** | 277 (1 contig(s)) |
| **Depth Of Coverage** | 15605.2 |
| **Number Of Reads** | 36615 |
| **Reads Per Million** | 611.93 rpm (after QC) |
| **Ambiguities** | 0 |
| **Assembly Method** | de novo + reference guided assembly |
| **Consensus Caller** | Bcf Tools |

#### Coverage Map

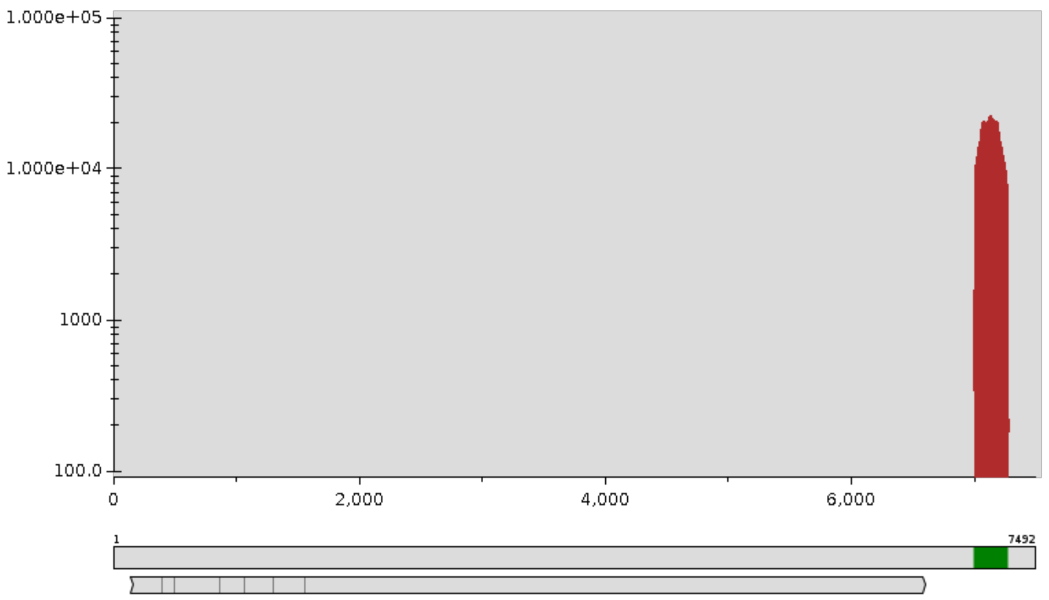

#### Assignment

| | |
|---|---|
| **Type** | Tomato chocolate spot virus (Taxonomy ID: 661101) |
| **Reference Genome** | NC_013075.1 |
| **NT Identity (%)** | 82.3741 |
| **AA Identity (%)** | 0.0 |
| **Number Of Stop Codons** | 0 |
| **Number Of CDS** | 1 |

#### Alignment

| | |
|---|---|
| **Alignment Score** | 352.0 (NT) + 0.0 (AA) = 352.0 |
| **Concordance (%)** | 63.5379 |

| Alignment Method | Global, seeded, nucleotide + amino acids (AGA) |
|---|---|

#### Genome Region

Sequence starts at position 6999 and ends at position 7275 relative to NC_013075.1 reference sequence.

#### Alignment Detailed Statistics

| | Begin | End | Coverage | Score | Concordance | Matches | Identities | I/D/M/F* | Stop Codons |
|---|---|---|---|---|---|---|---|---|---|
| **NT** | **6999** | **7275** | **3.7%** | **352** | **63.5%** | **277 (99.6%)** | **229 (82.4%)** | **1/0** | |
| Mutations: | 7012T>G, 7018C>T, 7019A>G, 7020A>G, 7020_7021insT, 7023C>A, 7042G>T, 7044A>T, 7047G>A, 7048C>G, 7066G>A, 7082G>A, 7083T>C, 7098C>T, 7099T>C, 7104T>A, 7107G>A, 7116C>G, 7121G>A, 7127A>C, 7140C>T, 7151A>T, 7152T>C, 7183C>T, 7184T>C, 7185G>T, 7186T>G, 7187T>C, 7188G>A, 7196C>T, 7197A>G, 7198A>C, 7199C>A, 7200A>G, 7201G>A, 7207A>T, 7208A>T, 7209T>G, 7210C>T, 7212T>A, 7219T>A, 7224T>C, 7242A>T, 7255A>T, 7257G>A, 7258G>C, 7260T>C, 7262C>G, 7264T>C | | | | | | | | |

**CDS**

**Proteins**

*: Inserts / Deletes / Misaligned / Frameshifts

#### Analysis details

This analysis was performed with panviral2.64

### NGS Details (UN8): Tomato chocolate spot virus (segment RNA2)

#### Assembly

| | |
|---|---|
| **Coverage Length** | 1253 (2 contig(s)) |
| **Depth Of Coverage** | 5116.5 |
| **Number Of Reads** | 50187 |
| **Reads Per Million** | 838.75 rpm (after QC) |
| **Ambiguities** | 0 |
| **Assembly Method** | de novo + reference guided assembly |
| **Consensus Caller** | Bcf Tools |

#### Coverage Map

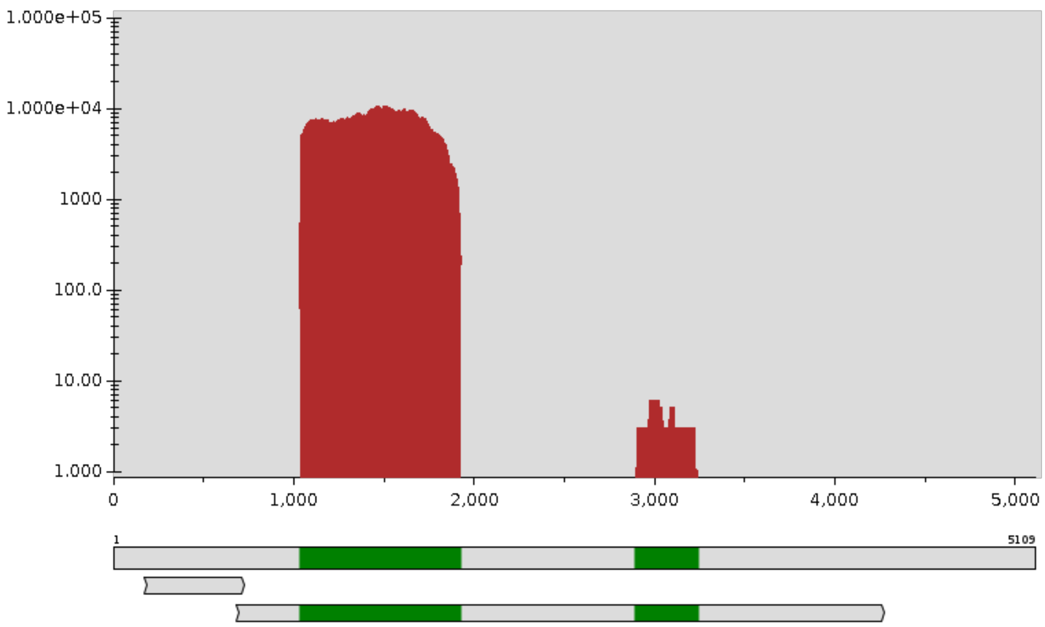

#### Assignment

| | |
|---|---|
| **Type** | Tomato chocolate spot virus (Taxonomy ID: 661101) |
| **Reference Genome** | NC_013076.1 |
| **NT Identity (%)** | 65.2105 |
| **AA Identity (%)** | 70.4762 |
| **Number Of Stop Codons** | 0 |
| **Number Of CDS** | 2 |

#### Alignment

| | |
|---|---|
| **Alignment Score** | 763.0 (NT) + 2195.0 (AA) = 2958.0 |
| **Concordance (%)** | 54.3051 |

| | |
|---|---|
| **Alignment Method** | Global, seeded, nucleotide + amino acids (AGA) |

#### Genome Region

Sequence starts at position 1033 and ends at position 3248 relative to NC_013076.1 reference sequence.

#### Alignment Detailed Statistics

| | Begin | End | Coverage | Score | Concordance | Matches | Identities | I/D/M/F* | Stop Codons |
|---|---|---|---|---|---|---|---|---|---|
| **NT** | **1033** | **3248** | **24.5%** | **763** | **30.4%** | **1253 (99.5%)** | **821 (65.2%)** | **6/0** | |
| Mutations: | 1042G>C, 1054C>A, 1056T>C, 1057C>T, 1058A>G, 1059G>A, 1063G>T, 1066G>T, 1069T>A, 1072A>G, 1075G>A, 1079T>A, 1081C>G, 1084G>A, 1086C>T, 1087A>C, 1088C>T, 1090A>T, 1091G>T, 1093T>A, 1095A>G, 1096T>G, 1099T>A, 1102A>T, 1105A>G, 1108C>T, 1111G>C, 1112C>T, 1114C>G, 1124A>G, 1125G>A, 1126T>A, 1129C>T, 1138C>T, 1142C>G, 1144C>T, 1153A>G, 1156A>C, 1162C>T, 1181G>C, 1182A>T, 1183G>T, 1186T>C, 1192T>G, 1195A>T, 1197A>T, 1198G>A, 1201A>T, 1204T>A, 1210C>T, 1213G>T, 1216A>G, 1219G>A, 1222A>T, 1228A>G, 1231A>G, 1238G>A, 1239A>G, 1240T>C, 1241A>C, 1246G>A, 1252C>T, 1255A>T, 1258C>T, 1259T>A, 1261T>G, 1262C>G, 1263A>C, 1267G>A, 1273T>A, 1277T>A, 1278C>G, 1285C>A, 1291A>G, 1294A>T, 1295T>C, 1297G>C, 1300C>T, 1306T>A, 1309A>G, 1312T>C, 1315C>T, 1316C>T, 1317A>C, 1318A>T, 1327G>A, 1333T>A, 1336C>A, 1351A>T, 1354A>C, 1360T>G, 1366G>A, 1367C>G, 1369A>G, 1372T>G, 1373T>G, 1375C>G, 1376C>T, 1377A>C, 1378G>T, 1379A>C, 1380T>A, 1381G>A, 1385T>C, 1387G>A, 1394C>A, 1396T>A, 1397G>C, 1405T>A, 1408A>T, 1410A>C, 1411G>A, 1412A>C, 1413G>T, 1414G>T, 1423T>C, 1426A>T, 1428G>A, 1429A>C, 1433G>A, 1438A>T, 1440T>C, 1441G>C, 1442A>C, 1443G>A, 1444T>G, 1448A>T, 1449G>C, 1450C>T, 1453C>A, 1454A>T, 1456G>A, 1458C>T, 1459T>G, 1463A>G, 1464G>C, 1465C>G, 1468T>A, 1471C>G, 1474G>T, 1477A>C, 1483G>A, 1489C>A, 1493G>A, 1495G>C, 1501T>C, 1503G>A, 1505C>A, 1513A>T, 1516C>G, 1522A>G, 1525A>G, 1531C>T, 1532G>T, 1534G>T, 1535C>T, 1537T>A, 1539A>G, 1540C>G, 1541A>G, 1547C>T, 1548G>C, 1551T>C, 1552A>G, 1555_1556insGTAGGT, 1556A>G, 1559C>A, 1572C>G, 1573T>G, 1574C>G, 1575A>T, 1576C>A, 1582A>T, 1585C>T, 1586A>T, 1588T>C, 1589G>A, 1590G>A, 1592G>T, 1593G>T, 1595T>C, 1603A>C, 1606T>C, 1610A>G, 1618A>G, 1620G>C, 1622T>A, 1624C>T, 1625A>G, 1627C>T, 1628A>T, 1630G>T, 1633T>C, 1634G>C, 1635T>A, 1636T>G, 1645T>C, 1649C>T, 1650T>A, 1651G>T, 1657A>C, 1658G>A, 1659A>C, 1660C>T, 1661T>G, 1662C>A, 1663C>T, 1668C>A, 1669A>T, 1670C>T, 1671A>C, 1672C>T, 1673A>T, 1678C>T, 1690T>G, 1696T>A, 1697G>A, 1698A>C, 1699G>C, 1700C>T, 1702C>G, 1703A>C, 1704C>A, 1705G>T, 1708T>A, 1709G>T, 1711G>A, 1712C>A, 1713A>G, 1714A>G, 1717G>A, 1718G>A, 1721C>G, 1723A>T, 1725A>T, 1726C>T, 1727A>G, 1729T>G, 1730C>T, 1734T>C, 1736G>A, 1737A>C, 1738T>C, 1739G>C, 1740A>T, 1741G>C, 1742G>C, 1743C>T, 1744T>C, 1745T>C, 1746C>G, 1747A>C, 1749G>A, 1750A>G, 1753C>G, 1755A>G, 1756T>A, 1757G>T, 1758T>G, 1759G>T, 1760A>C, 1761T>A, 1762C>T, 1763G>C, 1768G>C, 1771A>G, 1774T>C, 1775G>A, 1777A>T, 1778T>G, 1779C>A, 1780A>G, 1782A>G, 1783G>A, 1784A>C, 1786G>A, 1789G>C, 1790C>T, 1791A>C, 1792G>C, 1798T>C, 1801T>A, 1802T>A, 1805A>C, 1806A>T, 1807G>A, 1811C>T, 1816C>T, 1817A>G, 1818C>A, 1823G>A, 1825C>T, 1832G>C, 1834T>A, 1835G>C, 1837T>G, 1838A>G, 1840T>G, 1843T>C, 1844G>A, 1845C>A, 1846T>G, 1848C>A, 1861C>A, 1864G>T, 1867A>G, 1870A>T, 1871A>T, 1872A>C, 1873T>C, 1876C>A, 1877T>G, 1878C>A, 1879T>G, 1883T>A, 1885A>T, 1887T>G, 1888G>C, 1889G>A, 1897T>C, 1898G>T, 1899A>C, 1901A>T, 1903G>A, 1904T>A, 1906G>T, 1909T>C, 1910C>A, 1911A>G, 1912T>G, 1915G>A, 1918A>G, 2890G>A, 2894C>A, 2895A>C, 2899A>C, 2901A>T, 2902T>C, 2905C>T, 2906C>A, 2907C>G, 2908G>T, 2911G>C, 2917C>T, 2920C>A, 2926A>G, 2929T>G, 2930T>C, 2932G>T, 2933T>C, 2935A>G, 2938T>C, 2941T>C, 2948G>A, 2950A>T, 2951A>G, 2952C>T, 2953C>A, 2962A>C, 2968C>A, 2975C>T, 2977T>G, 2980G>A, 2983C>T, 2984A>G, 2986A>C, 2992G>C, 2995T>A, 2998G>A, 3001T>A, 3004C>T, 3010G>A, 3020G>A, 3022A>C, 3023C>T, 3025G>A, 3028A>G, 3034C>A, 3035T>C, 3037A>G, 3043T>A, 3046T>C, 3047G>A, 3049A>T, 3052G>A, 3053C>A, 3055G>A, 3072C>G, 3073T>A, 3076G>C, 3079G>C, 3085C>T, 3091G>A, 3092A>G, 3094T>G, 3098A>G, 3109G>T, 3112C>G, 3115C>T, 3118C>T, 3121A>T, 3124A>G, 3128T>C, 3130G>T, 3133C>T, 3139A>G, 3140C>T, 3142C>G, 3145G>C, 3148A>G, 3152A>G, 3153A>C, 3154C>A, 3158A>G, 3160A>G, 3166A>C, 3168A>C, 3169T>A, 3172A>G, 3178T>C, 3181T>A, 3184T>G, 3193T>C, 3199A>G, 3205C>T, 3209C>T, 3211T>G, 3214G>A, 3220C>G, 3223C>A, 3225A>G, 3231G>A, 3232C>A | | | | | | | | |

##### CDS

| | | | | | | | | | |
|---|---|---|---|---|---|---|---|---|---|
| ToChV_s2_gp2 | 117 | 855 | 34.9% | 2195 | 74.0% | 418 (99.5%) | 296 (70.5%) | 2/0/0/0 | 0 |
| Protein mutations: | I124T (1056T>C 1057C>T), S125D (1058A>G 1059G>A), S132T (1079T>A 1081C>G), S134F (1086C>T 1087A>C), L135F (1088C>T 1090A>T), A136S (1091G>T 1093T>A), D137G (1095A>G 1096T>G), S147E (1124A>G 1125G>A 1126T>A), L153V (1142C>G 1144C>T), E166L (1181G>C 1182A>T 1183G>T), Q171L (1197A>T 1198G>A), D185S (1238G>A 1239A>G 1240T>C), T186P (1241A>C), S192T (1259T>A 1261T>G), Q193A (1262C>G 1263A>C), Q211S (1316C>T 1317A>C 1318A>T), Q228E (1367C>G 1369A>G), S229R (1372T>G), S230A (1373T>G 1375C>G), Q231S (1376C>T 1377A>C 1378G>T), M232Q (1379A>C 1380T>A 1381G>A), A238P (1397G>C), E242A (1410A>C 1411G>A), R243L (1412A>C 1413G>T 1414G>T), R248H (1428G>A 1429A>C), D250N (1433G>A), L252P (1440T>C 1441G>C), S253Q (1442A>C 1443G>A 1444T>G), M257L (1454A>T 1456G>A), T258M (1458C>T 1459T>G), S260A (1463A>G 1464G>C 1465C>G), V270I (1493G>A 1495G>C), R273K (1503G>A), V283F (1532G>T 1534G>T), P284S (1535C>T 1537T>A), N285R (1539A>G 1540C>G), T286A (1541A>G), R288S (1547C>T 1548G>C), L289S (1551T>C 1552A>G), T290_M291insVG (1555_1556insGTAGGT), M291V (1556A>G), A296G (1572C>G 1573T>G), H297V (1574C>G 1575A>T 1576C>A), E299D (1582A>T), T301S (1586A>T 1588T>C), G302N (1589G>A 1590G>A), G303L (1592G>T 1593G>T), K309E (1610A>G), G312A (1620G>C), F313I (1622T>A 1624C>T), N314D (1625A>G 1627C>T), M315F (1628A>T 1630G>T), V317Q (1634G>C 1635T>A 1636T>G), L322Y (1649C>T 1650T>A 1651G>T), D325T (1658G>A 1659A>C 1660C>T), S326D (1661T>G 1662C>A 1663C>T), T328N (1668C>A 1669A>T), H329S (1670C>T 1671A>C 1672C>T), I330F (1673A>T), D335E (1690T>G), E338T (1697G>A 1698A>C 1699G>C), R339W (1700C>T 1702C>G), T340H (1703A>C 1704C>A 1705G>T), A342S (1709G>T 1711G>A), Q343R (1712C>A 1713A>G 1714A>G), V345I (1718G>A), Q346D (1721C>G 1723A>T), Y347F (1725A>T 1726C>T), I348V (1727A>G 1729T>G), F350S (1734T>C), D351T (1736G>A 1737A>C 1738T>C), E352L (1739G>C 1740A>T 1741G>C), A353L (1742G>C 1743C>T 1744T>C), S354R (1745T>C 1746C>G 1747A>C), R355Q (1749G>A 1750A>G), N357R (1755A>G 1756T>A), V358C (1757G>T 1758T>G 1759G>T), I359H (1760A>C 1761T>A 1762C>T), V360L (1763G>C), V364I (1775G>A 1777A>T), S365E (1778T>G 1779C>A 1780A>G), K366R (1782A>G 1783G>A), E368D (1789G>C), Q369S (1790C>T 1791A>C 1792G>C), Y373N (1802T>A), K374L (1805A>C 1806A>T 1807G>A), H376Y (1811C>T), T378D (1817A>G 1818C>A), D380N (1823G>A 1825C>T), V383L (1832G>C 1834T>A), D384Q (1835G>C 1837T>G), I385V (1838A>G 1840T>G), A387K (1844G>A 1845C>A 1846T>G), A388E (1848C>A), K393N (1864G>T), N396S (1871A>T 1872A>C 1873T>C), S398E (1877T>G 1878C>A 1879T>G), S400T (1883T>A 1885A>T), L401C (1887T>G 1888G>C), D402N (1889G>A), D405S (1898G>T 1899A>C), M406L (1901A>T 1903G>A), S407T (1904T>A 1906G>T), H409R (1910C>A 1911A>G 1912T>G), Q737T (2894C>A 2895A>C), Y739F (2901A>T 2902T>C), P741S (2906C>A 2907C>G 2908G>T), G755S (2948G>A 2950A>T), T756V (2951A>G 2952C>T 2953C>A), I767V (2984A>G 2986A>C), V779I (3020G>A 3022A>C), V788I (3047G>A 3049A>T), T796R (3072C>G 3073T>A), I803V (3092A>G 3094T>G), I805V (3098A>G), N823A (3152A>G 3153A>C 3154C>A), K825E (3158A>G 3160A>G), N828T (3168A>C 3169T>A), F833L (3184T>G), K847R (3225A>G), R849Q (3231G>A 3232C>A) | | | | | | | | |

| | Begin | End | Coverage | Score | Concordance | Matches | Identities | I/D/M/F* | Stop Codons |
|---|---|---|---|---|---|---|---|---|---|
| **NT** | **1033** | **3248** | **24.5%** | **763** | **30.4%** | **1253 (99.5%)** | **821 (65.2%)** | **6/0** | |
| Codon mutations: | TCG119TCC (1042G>C), CCC123CCA (1054C>A), ATC124ACT (1056T>C 1057C>T), AGC125GAC (1058A>G 1059G>A), CCG126CCT (1063G>T), GTG127GTT (1066G>T), ATT128ATA (1069T>A), CAA129CAG (1072A>G), GAG130GAA (1075G>A), TCC132ACG (1079T>A 1081C>G), TTG133TTA (1084G>A), TCA134TTC (1086C>T 1087A>C), CTA135TTT (1088C>T 1090A>T), GCT136TCA (1091G>T 1093T>A), GAT137GGG (1095A>G 1096T>G), GGT138GGA (1099T>A), GCA139GCT (1102A>T), CAA140CAG (1105A>G), GCC141GCT (1108C>T), GTG142GTC (1111G>C), CTC143TTG (1112C>T 1114C>G), AGT147GAA (1124A>G 1125G>A 1126T>A), ATC148ATT (1129C>T), CCC151CCT (1138C>T), CTC153GTT (1142C>G 1144C>T), GGA156GGG (1153A>G), CCA157CCC (1156A>C), TTC159TTT (1162C>T), GAG166CTT (1181G>C 1182A>T 1183G>T), AAT167AAC (1186T>C), CGT169CGG (1192T>G), GCA170GCT (1195A>T), CAG171CTA (1197A>T 1198G>A), ATA172ATT (1201A>T), CCT173CCA (1204T>A), TCC175TCT (1210C>T), GTG176GTT (1213G>T), GTA177GTG (1216A>G), GCG178GCA (1219G>A), ATA179ATT (1222A>T), AAA181AAG (1228A>G), CAA182CAG (1231A>G), GAT185AGC (1238G>A 1239A>G 1240T>C), ACA186CCA (1241A>C), AGG187AGA (1246G>A), GCC189GCT (1252C>T), GTA190GTT (1255A>T), TAC191TAT (1258C>T), TCT192ACG (1259T>A 1261T>G), CAA193GCA (1262C>G 1263A>C), CAG194CAA (1267G>A), ACT196ACA (1273T>A), TCT198AGT (1277T>A 1278C>G), GCC200GCA (1285C>A), GAA202GAG (1291A>G), GCA203GCT (1294A>T), TTG204CTC (1295T>C 1297G>C), AAC205AAT (1300C>T), CCT207CCA (1306T>A), CAA208CAG (1309A>G), TTT209TTC (1312T>C), TTC210TTT (1315C>T), CAA211TCT (1316C>T 1317A>C 1318A>T), CAG214CAA (1327G>A), GCT216GCA (1333T>A), GCC217GCA (1336C>A), TCA222TCT (1351A>T), GTA223GTC (1354A>C), TCT225TCG (1360T>G), GGG227GGA (1366G>A), CAA228GAG (1367C>G 1369A>G), AGT229AGG (1372T>G), TCC230GCG (1373T>G 1375C>G), CAG231TCT (1376C>T 1377A>C 1378G>T), ATG232CAA (1379A>C 1380T>A 1381G>A), TTG234CTA (1385T>C 1387G>A), CGT237AGA (1394C>A 1396T>A), GCT238CCT (1397G>C), GTT240GTA (1405T>A), GGA241GGT (1408A>T), GAG242GCA (1410A>C 1411G>A), AGG243CTT (1412A>C 1413G>T 1414G>T), AAT246AAC (1423T>C), ACA247ACT (1426A>T), CGA248CAC (1428G>A 1429A>C), GAT250AAT (1433G>A), GTA251GTT (1438A>T), CTG252CCC (1440T>C 1441G>C), AGT253CAG (1442A>C 1443G>A 1444T>G), AGC255TCT (1448A>T 1449G>C 1450C>T), TCC256TCA (1453C>A), ATG257TTA (1454A>T 1456G>A), ACT258ATG (1458C>T 1459T>G), AGC260GCG (1463A>G 1464G>C 1465C>G), TCT261TCA (1468T>A), TCC262TCG (1471C>G), CTG263CTT (1474G>T), CGA264CGC (1477A>C), CCG266CCA (1483G>A), CTC268CTA (1489C>A), GTG270ATC (1493G>A 1495G>C), AGT272AGC (1501T>C), AGG273AAG (1503G>A), CGG274AGG (1505C>A), CCA276CCT (1513A>T), TCC277TCG (1516C>G), GAA279GAG (1522A>G), CTA280CTG (1525A>G), GAC282GAT (1531C>T), GTG283TTT (1532G>T 1534G>T), CCT284TCA (1535C>T 1537T>A), AAC285AGG (1539A>G 1540C>G), ACC286GCC (1541A>G), CGA288TCA (1547C>T 1548G>C), TTA289TCG (1551T>C 1552A>G), ACA290_ATG291insGTAGGT (1555_1556insGTAGGT), ATG291GTG (1556A>G), CGA292AGA (1559C>A), GCT296GGG (1572C>G 1573T>G), CAC297GTA (1574C>G 1575A>T 1576C>A), GAA299GAT (1582A>T), GAC300GAT (1585C>T), ACT301TCC (1586A>T 1588T>C), GGT302AAT (1589G>A 1590G>A), GGA303TTA (1592G>T 1593G>T), TTG304CTG (1595T>C), CCA306CCC (1603A>C), GTT307GTC (1606T>C), AAG309GAG (1610A>G), CAA311CAG (1618A>G), GGC312GCC (1620G>C), TTC313ATT (1622T>A 1624C>T), AAC314GAT (1625A>G 1627C>T), ATG315TTT (1628A>T 1630G>T), GGT316GGC (1633T>C), GTT317CAG (1634G>C 1635T>A 1636T>G), GAT320GAC (1645T>C), CTG322TAT (1649C>T 1650T>A 1651G>T), ATA324ATC (1657A>C), GAC325ACT (1658G>A 1659A>C 1660C>T), TCC326GAT (1661T>G 1662C>A 1663C>T), ACA328AAT (1668C>A 1669A>T), CAC329TCT (1670C>T 1671A>C 1672C>T), ATT330TTT (1673A>T), GAC331GAT (1678C>T), GAT335GAG (1690T>G), ACT337ACA (1696T>A), GAG338ACC (1697G>A 1698A>C 1699G>C), CGC339TGG (1700C>T 1702C>G), ACG340CAT (1703A>C 1704C>A 1705G>T), ATT341ATA (1708T>A), GCG342TCA (1709G>T 1711G>A), CAA343AGG (1712C>A 1713A>G 1714A>G), CAG344CAA (1717G>A), GTT345ATT (1718G>A), CAA346GAT (1721C>G 1723A>T), TAC347TTT (1725A>T 1726C>T), ATT348GTG (1727A>G 1729T>G), CTG349TTG (1730C>T), TTT350TCT (1734T>C), GAT351ACC (1736G>A 1737A>C 1738T>C), GAG352CTC (1739G>C 1740A>T 1741G>C), GCT353CTC (1742G>C 1743C>T 1744T>C), TCA354CGC (1745T>C 1746C>G 1747A>C), CGA355CAG (1749G>A 1750A>G), GGC356GGG (1753C>G), AAT357AGA (1755A>G 1756T>A), GTG358TGT (1757G>T 1758T>G 1759G>T), ATC359CAT (1760A>C 1761T>A 1762C>T), GTG360CTG (1763G>C), CCG361CCC (1768G>C), AAA362AAG (1771A>G), CAT363CAC (1774T>C), GTA364ATT (1775G>A 1777A>T), TCA365GAG (1778T>G 1779C>A 1780A>G), AAG366AGA (1782A>G 1783G>A), AGG367CGA (1784A>C 1786G>A), GAG368GAC (1789G>C), CAG369TCC (1790C>T 1791A>C 1792G>C), AAT371AAC (1798T>C), ATT372ATA (1801T>A), TAC373AAC (1802T>A), AAG374CTA (1805A>C 1806A>T 1807G>A), CAC376TAC (1811C>T), ATC377ATT (1816C>T), ACT378GAT (1817A>G 1818C>A), GAC380AAT (1823G>A 1825C>T), GTT383CTA (1832G>C 1834T>A), GAT384CAG (1835G>C 1837T>G), ATT385GTG (1838A>G 1840T>G), CTT386CTC (1843T>C), GCT387AAG (1844G>A 1845C>A 1846T>G), GCA388GAA (1848C>A), ACC392ACA (1861C>A), AAG393AAT (1864G>T), GTA394GTG (1867A>G), GGA395GGT (1870A>T), AAT396TCC (1871A>T 1872A>C 1873T>C), CTC397CTA (1876C>A), TCT398GAG (1877T>G 1878C>A 1879T>G), TCA400ACT (1883T>A 1885A>T), TTG401TGC (1887T>G 1888G>C), GAT402AAT (1889G>A), TAT404TAC (1897T>C), GAT405TCT (1898G>T 1899A>C), ATG406TTA (1901A>T 1903G>A), TCG407ACT (1904T>A 1906G>T), CTT408CTC (1909T>C), CAT409AGG (1910C>A 1911A>G 1912T>G), GAG410GAA (1915G>A), AGA411AGG (1918A>G), CCG735..A (2890G>A), CAA737ACA (2894C>A 2895A>C), ACA738ACC (2899A>C), TAT739TTC (2901A>T 2902T>C), TTC740TTT (2905C>T), CCG741AGT (2906C>A 2907C>G 2908G>T), CGG742CGC (2911G>C), CCC744CCT (2917C>T), ACC745ACA (2920C>A), CAA747CAG (2926A>G), GTT748GTG (2929T>G), TTG749CTT (2930T>C 2932G>T), TTA750CTG (2933T>C 2935A>G), CAT751CAC (2938T>C), TAT752TAC (2941T>C), GGA755AGT (2948G>A 2950A>T), ACC756GTA (2951A>G 2952C>T 2953C>A), GCA759GCC (2962A>C), GGC761GGA (2968C>A), CTT764TTG (2975C>T 2977T>G), GTG765GTA (2980G>A), AGC766AGT (2983C>T), ATA767GTC (2984A>G 2986A>C), TCG769TCC (2992G>C), CCT770CCA (2995T>A), TCG771TCA (2998G>A), GGT772GGA (3001T>A), ATC773ATT (3004C>T), GAG775GAA (3010G>A), GTA779ATC (3020G>A 3022A>C), CTG780TTA (3023C>T 3025G>A), CAA781CAG (3028A>G), TCC783TCA (3034C>A), TTA784CTG (3035T>C 3037A>G), GGT786GGA (3043T>A), AAT787AAC (3046T>C), GTA788ATT (3047G>A 3049A>T), GCG789GCA (3052G>A), CGG790AGA (3053C>A 3055G>A), ACT796AGA (3072C>G 3073T>A), GGG797GGC (3076G>C), ACG798ACC (3079G>C), GTC800GTT (3085C>T), GAG802GAA (3091G>A), ATT803GTG (3092A>G 3094T>G), ATT805GTT (3098A>G), ACG808ACT (3109G>T), CTC809CTG (3112C>G), TTC810TTT (3115C>T), CAC811CAT (3118C>T), TCA812TCT (3121A>T), GGA813GGG (3124A>G), TTG815CTT (3128T>C 3130G>T), GCC816GCT (3133C>T), GGA818GGG (3139A>G), CTC819TTG (3140C>T 3142C>G), GGG820GGC (3145G>C), ACA821ACG (3148A>G), AAC823GCA (3152A>G 3153A>C 3154C>A), AAA825GAG (3158A>G 3160A>G), TCA827TCC (3166A>C), AAT828ACA (3168A>C 3169T>A), GCA829GCG (3172A>G), GAT831GAC (3178T>C), ATT832ATA (3181T>A), TTT833TTG (3184T>G), CCT836CCC (3193T>C), GTA838GTG (3199A>G), TGC840TGT (3205C>T), CTT842TTG (3209C>T 3211T>G), GAG843GAA (3214G>A), GGC845GGG (3220C>G), CGC846CGA (3223C>A), AAG847AGG (3225A>G), CGC849CAA (3231G>A 3232C>A) | | | | | | | | |

#### Proteins

| | Begin | End | Coverage | Score | Concordance | Matches | Identities | I/D/M/F* | Stop Codons |
|---|---|---|---|---|---|---|---|---|---|
| polyprotein (YP_003097231.1) | 117 | 855 | 34.9% | 2195 | 74.0% | 418 (99.5%) | 296 (70.5%) | 2/0/0/0 | 0 |
| Protein mutations: | I124T (1056T>C 1057C>T), S125D (1058A>G 1059G>A), S132T (1079T>A 1081C>G), S134F (1086C>T 1087A>C), L135F (1088C>T 1090A>T), A136S (1091G>T 1093T>A), D137G (1095A>G 1096T>G), S147E (1124A>G 1125G>A 1126T>A), L153V (1142C>G 1144C>T), E166L (1181G>C 1182A>T 1183G>T), Q171L (1197A>T 1198G>A), D185S (1238G>A 1239A>G 1240T>C), T186P (1241A>C), S192T (1259T>A 1261T>G), Q193A (1262C>G 1263A>C), Q211S (1316C>T 1317A>C 1318A>T), Q228E (1367C>G 1369A>G), S229R (1372T>G), S230A (1373T>G 1375C>G), Q231S (1376C>T 1377A>C 1378G>T), M232Q (1379A>C 1380T>A 1381G>A), A238P (1397G>C), E242A (1410A>C 1411G>A), R243L (1412A>C 1413G>T 1414G>T), R248H (1428G>A 1429A>C), D250N (1433G>A), L252P (1440T>C 1441G>C), S253Q (1442A>C 1443G>A 1444T>G), M257L (1454A>T 1456G>A), T258M (1458C>T 1459T>G), S260A (1463A>G 1464G>C 1465C>G), V270I (1493G>A 1495G>C), R273K (1503G>A), V283F (1532G>T 1534G>T), P284S (1535C>T 1537T>A), N285R (1539A>G 1540C>G), T286A (1541A>G), R288S (1547C>T 1548G>C), L289S (1551T>C 1552A>G), T290_M291insVG (1555_1556insGTAGGT), M291V (1556A>G), A296G (1572C>G 1573T>G), H297V (1574C>G 1575A>T 1576C>A), E299D (1582A>T), T301S (1586A>T 1588T>C), G302N (1589G>A 1590G>A), G303L (1592G>T 1593G>T), K309E (1610A>G), G312A (1620G>C), F313I (1622T>A 1624C>T), N314D (1625A>G 1627C>T), M315F (1628A>T 1630G>T), V317Q (1634G>C 1635T>A 1636T>G), L322Y (1649C>T 1650T>A 1651G>T), D325T (1658G>A 1659A>C 1660C>T), S326D (1661T>G 1662C>A 1663C>T), T328N (1668C>A 1669A>T), H329S (1670C>T 1671A>C 1672C>T), I330F (1673A>T), D335E (1690T>G), E338T (1697G>A 1698A>C 1699G>C), R339W (1700C>T 1702C>G), T340H (1703A>C 1704C>A 1705G>T), A342S (1709G>T 1711G>A), Q343R (1712C>A 1713A>G 1714A>G), V345I (1718G>A), Q346D (1721C>G 1723A>T), Y347F (1725A>T 1726C>T), I348V (1727A>G 1729T>G), F350S (1734T>C), D351T (1736G>A 1737A>C 1738T>C), E352L (1739G>C 1740A>T 1741G>C), A353L (1742G>C 1743C>T 1744T>C), S354R (1745T>C 1746C>G 1747A>C), R355Q (1749G>A 1750A>G), N357R (1755A>G 1756T>A), V358C (1757G>T 1758T>G 1759G>T), I359H (1760A>C 1761T>A 1762C>T), V360L (1763G>C), V364I (1775G>A 1777A>T), S365E (1778T>G 1779C>A 1780A>G), K366R (1782A>G 1783G>A), E368D (1789G>C), Q369S (1790C>T 1791A>C 1792G>C), Y373N (1802T>A), K374L (1805A>C 1806A>T 1807G>A), H376Y (1811C>T), T378D (1817A>G 1818C>A), D380N (1823G>A 1825C>T), V383L (1832G>C 1834T>A), D384Q (1835G>C 1837T>G), I385V (1838A>G 1840T>G), A387K (1844G>A 1845C>A 1846T>G), A388E (1848C>A), K393N (1864G>T), N396S (1871A>T 1872A>C 1873T>C), S398E (1877T>G 1878C>A 1879T>G), S400T (1883T>A 1885A>T), L401C (1887T>G 1888G>C), D402N (1889G>A), D405S (1898G>T 1899A>C), M406L (1901A>T 1903G>A), S407T (1904T>A 1906G>T), H409R (1910C>A 1911A>G 1912T>G), Q737T (2894C>A 2895A>C), Y739F (2901A>T 2902T>C), P741S (2906C>A 2907C>G 2908G>T), G755S (2948G>A 2950A>T), T756V (2951A>G 2952C>T 2953C>A), I767V (2984A>G 2986A>C), V779I (3020G>A 3022A>C), V788I (3047G>A 3049A>T), T796R (3072C>G 3073T>A), I803V (3092A>G 3094T>G), I805V (3098A>G), N823A (3152A>G 3153A>C 3154C>A), K825E (3158A>G 3160A>G), N828T (3168A>C 3169T>A), F833L (3184T>G), K847R (3225A>G), R849Q (3231G>A 3232C>A) | | | | | | | | |

| | Begin | End | Coverage | Score | Concordance | Matches | Identities | I/D/M/F* | Stop Codons |
|---|---|---|---|---|---|---|---|---|---|
| **NT** | **1033** | **3248** | **24.5%** | **763** | **30.4%** | **1253 (99.5%)** | **821 (65.2%)** | **6/0** | |

Codon mutations:

TCG119TCC (1042G>C), CCC123CCA (1054C>A), ATC124ACT (1056T>C 1057C>T), AGC125GAC (1058A>G 1059G>A), CCG126CCT (1063G>T), GTG127GTT (1066G>T), ATT128ATA (1069T>A), CAA129CAG (1072A>G), GAG130GAA (1075G>A), TCC132ACG (1079T>A 1081C>G), TTG133TTA (1084G>A), TCA134TTC (1086C>T 1087A>C), CTA135TTT (1088C>T 1090A>T), GCT136TCA (1091G>T 1093T>A), GAT137GGG (1095A>G 1096T>G), GGT138GGA (1099T>A), GCA139GCT (1102A>T), CAA140CAG (1105A>G), GCC141GCT (1108C>T), GTG142GTC (1111G>C), CTC143TTG (1112C>T 1114C>G), AGT147GAA (1124A>G 1125G>A 1126T>A), ATC148ATT (1129C>T), CCC151CCT (1138C>T), CTC153GTT (1142C>G 1144C>T), GGA156GGG (1153A>G), CCA157CCC (1156A>C), TTC159TTT (1162C>T), GAG166CTT (1181G>C 1182A>T 1183G>T), AAT167AAC (1186T>C), CGT169CGG (1192T>G), GCA170GCT (1195A>T), CAG171CTA (1197A>T 1198G>A), ATA172ATT (1201A>T), CCT173CCA (1204T>A), TCC175TCT (1210C>T), GTG176GTT (1213G>T), GTA177GTG (1216A>G), GCG178GCA (1219G>A), ATA179ATT (1222A>T), AAA181AAG (1228A>G), CAA182CAG (1231A>G), GAT185AGC (1238G>A 1239A>G 1240T>C), ACA186CCA (1241A>C), AGG187AGA (1246G>A), GCC189GCT (1252C>T), GTA190GTT (1255A>T), TAC191TAT (1258C>T), TCT192ACG (1259T>A 1261T>G), CAA193GCA (1262C>G 1263A>C), CAG194CAA (1267G>A), ACT196ACA (1273T>A), TCT198AGT (1277T>A 1278C>G), GCC200GCA (1285C>A), GAA202GAG (1291A>G), GCA203GCT (1294A>T), TTG204CTC (1295T>C 1297G>C), AAC205AAT (1300C>T), CCT207CCA (1306T>A), CAA208CAG (1309A>G), TTT209TTC (1312T>C), TTC210TTT (1315C>T), CAA211TCT (1316C>T 1317A>C 1318A>T), CAG214CAA (1327G>A), GCT216GCA (1333T>A), GCC217GCA (1336C>A), TCA222TCT (1351A>T), GTA223GTC (1354A>C), TCT225TCG (1360T>G), GGG227GGA (1366G>A), CAA228GAG (1367C>G 1369A>G), AGT229AGG (1372T>G), TCC230GCG (1373T>G 1375C>G), CAG231TCT (1376C>T 1377A>C 1378G>T), ATG232CAA (1379A>C 1380T>A 1381G>A), TTG234CTA (1385T>C 1387G>A), CGT237AGA (1394C>A 1396T>A), GCT238CCT (1397G>C), GTT240GTA (1405T>A), GGA241GGT (1408A>T), GAG242GCA (1410A>C 1411G>A), AGG243CTT (1412A>C 1413G>T 1414G>T), AAT246AAC (1423T>C), ACA247ACT (1426A>T), CGA248CAC (1428G>A 1429A>C), GAT250AAT (1433G>A), GTA251GTT (1438A>T), CTG252CCC (1440T>C 1441G>C), AGT253CAG (1442A>C 1443G>A 1444T>G), AGC255TCT (1448A>T 1449G>C 1450C>T), TCC256TCA (1453C>A), ATG257TTA (1454A>T 1456G>A), ACT258ATG (1458C>T 1459T>G), AGC260GCG (1463A>G 1464G>C 1465C>G), TCT261TCA (1468T>A), TCC262TCG (1471C>G), CTG263CTT (1474G>T), CGA264CGC (1477A>C), CCG266CCA (1483G>A), CTC268CTA (1489C>A), GTG270ATC (1493G>A 1495G>C), AGT272AGC (1501T>C), AGG273AAG (1503G>A), CGG274AGG (1505C>A), CCA276CCT (1513A>T), TCC277TCG (1516C>G), GAA279GAG (1522A>G), CTA280CTG (1525A>G), GAC282GAT (1531C>T), GTG283TTT (1532G>T 1534G>T), CCT284TCA (1535C>T 1537T>A), AAC285AGG (1539A>G 1540C>G), ACC286GCC (1541A>G), CGA288TCA (1547C>T 1548G>C), TTA289TCG (1551T>C 1552A>G), ACA290_ATG291insGTAGGT (1555_1556insGTAGGT), ATG291GTG (1556A>G), CGA292AGA (1559C>A), GCT296GGG (1572C>G 1573T>G), CAC297GTA (1574C>G 1575A>T 1576C>A), GAA299GAT (1582A>T), GAC300GAT (1585C>T), ACT301TCC (1586A>T 1588T>C), GGT302AAT (1589G>A 1590G>A), GGA303TTA (1592G>T 1593G>T), TTG304CTG (1595T>C), CCA306CCC (1603A>C), GTT307GTC (1606T>C), AAG309GAG (1610A>G), CAA311CAG (1618A>G), GGC312GCC (1620G>C), TTC313ATT (1622T>A 1624C>T), AAC314GAT (1625A>G 1627C>T), ATG315TTT (1628A>T 1630G>T), GGT316GGC (1633T>C), GTT317CAG (1634G>C 1635T>A 1636T>G), GAT320GAC (1645T>C), CTG322TAT (1649C>T 1650T>A 1651G>T), ATA324ATC (1657A>C), GAC325ACT (1658G>A 1659A>C 1660C>T), TCC326GAT (1661T>G 1662C>A 1663C>T), ACA328AAT (1668C>A 1669A>T), CAC329TCT (1670C>T 1671A>C 1672C>T), ATT330TTT (1673A>T), GAC331GAT (1678C>T), GAT335GAG (1690T>G), ACT337ACA (1696T>A), GAG338ACC (1697G>A 1698A>C 1699G>C), CGC339TGG (1700C>T 1702C>G), ACG340CAT (1703A>C 1704C>A 1705G>T), ATT341ATA (1708T>A), GCG342TCA (1709G>T 1711G>A), CAA343AGG (1712C>A 1713A>G 1714A>G), CAG344CAA (1717G>A), GTT345ATT (1718G>A), CAA346GAT (1721C>G 1723A>T), TAC347TTT (1725A>T 1726C>T), ATT348GTG (1727A>G 1729T>G), CTG349TTG (1730C>T), TTT350TCT (1734T>C), GAT351ACC (1736G>A 1737A>C 1738T>C), GAG352CTC (1739G>C 1740A>T 1741G>C), GCT353CTC (1742G>C 1743C>T 1744T>C), TCA354CGC (1745T>C 1746C>G 1747A>C), CGA355CAG (1749G>A 1750A>G), GGC356GGG (1753C>G), AAT357AGA (1755A>G 1756T>A), GTG358TGT (1757G>T 1758T>G 1759G>T), ATC359CAT (1760A>C 1761T>A 1762C>T), GTG360CTG (1763G>C), CCG361CCC (1768G>C), AAA362AAG (1771A>G), CAT363CAC (1774T>C), GTA364ATT (1775G>A 1777A>T), TCA365GAG (1778T>G 1779C>A 1780A>G), AAG366AGA (1782A>G 1783G>A), AGG367CGA (1784A>C 1786G>A), GAG368GAC (1789G>C), CAG369TCC (1790C>T 1791A>C 1792G>C), AAT371AAC (1798T>C), ATT372ATA (1801T>A), TAC373AAC (1802T>A), AAG374CTA (1805A>C 1806A>T 1807G>A), CAC376TAC (1811C>T), ATC377ATT (1816C>T), ACT378GAT (1817A>G 1818C>A), GAC380AAT (1823G>A 1825C>T), GTT383CTA (1832G>C 1834T>A), GAT384CAG (1835G>C 1837T>G), ATT385GTG (1838A>G 1840T>G), CTT386CTC (1843T>C), GCT387AAG (1844G>A 1845C>A 1846T>G), GCA388GAA (1848C>A), ACC392ACA (1861C>A), AAG393AAT (1864G>T), GTA394GTG (1867A>G), GGA395GGT (1870A>T), AAT396TCC (1871A>T 1872A>C 1873T>C), CTC397CTA (1876C>A), TCT398GAG (1877T>G 1878C>A 1879T>G), TCA400ACT (1883T>A 1885A>T), TTG401TGC (1887T>G 1888G>C), GAT402AAT (1889G>A), TAT404TAC (1897T>C), GAT405TCT (1898G>T 1899A>C), ATG406TTA (1901A>T 1903G>A), TCG407ACT (1904T>A 1906G>T), CTT408CTC (1909T>C), CAT409AGG (1910C>A 1911A>G 1912T>G), GAG410GAA (1915G>A), AGA411AGG (1918A>G), CCG735..A (2890G>A), CAA737ACA (2894C>A 2895A>C), ACA738ACC (2899A>C), TAT739TTC (2901A>T 2902T>C), TTC740TTT (2905C>T), CCG741AGT (2906C>A 2907C>G 2908G>T), CGG742CGC (2911G>C), CCC744CCT (2917C>T), ACC745ACA (2920C>A), CAA747CAG (2926A>G), GTT748GTG (2929T>G), TTG749CTT (2930T>C 2932G>T), TTA750CTG (2933T>C 2935A>G), CAT751CAC (2938T>C), TAT752TAC (2941T>C), GGA755AGT (2948G>A 2950A>T), ACC756GTA (2951A>G 2952C>T 2953C>A), GCA759GCC (2962A>C), GGC761GGA (2968C>A), CTT764TTG (2975C>T 2977T>G), GTG765GTA (2980G>A), AGC766AGT (2983C>T), ATA767GTC (2984A>G 2986A>C), TCG769TCC (2992G>C), CCT770CCA (2995T>A), TCG771TCA (2998G>A), GGT772GGA (3001T>A), ATC773ATT (3004C>T), GAG775GAA (3010G>A), GTA779ATC (3020G>A 3022A>C), CTG780TTA (3023C>T 3025G>A), CAA781CAG (3028A>G), TCC783TCA (3034C>A), TTA784CTG (3035T>C 3037A>G), GGT786GGA (3043T>A), AAT787AAC (3046T>C), GTA788ATT (3047G>A 3049A>T), GCG789GCA (3052G>A), CGG790AGA (3053C>A 3055G>A), ACT796AGA (3072C>G 3073T>A), GGG797GGC (3076G>C), ACG798ACC (3079G>C), GTC800GTT (3085C>T), GAG802GAA (3091G>A), ATT803GTG (3092A>G 3094T>G), ATT805GTT (3098A>G), ACG808ACT (3109G>T), CTC809CTG (3112C>G), TTC810TTT (3115C>T), CAC811CAT (3118C>T), TCA812TCT (3121A>T), GGA813GGG (3124A>G), TTG815CTT (3128T>C 3130G>T), GCC816GCT (3133C>T), GGA818GGG (3139A>G), CTC819TTG (3140C>T 3142C>G), GGG820GGC (3145G>C), ACA821ACG (3148A>G), AAC823GCA (3152A>G 3153A>C 3154C>A), AAA825GAG (3158A>G 3160A>G), TCA827TCC (3166A>C), AAT828ACA (3168A>C 3169T>A), GCA829GCG (3172A>G), GAT831GAC (3178T>C), ATT832ATA (3181T>A), TTT833TTG (3184T>G), CCT836CCC (3193T>C), GTA838GTG (3199A>G), TGC840TGT (3205C>T), CTT842TTG (3209C>T 3211T>G), GAG843GAA (3214G>A), GGC845GGG (3220C>G), CGC846CGA (3223C>A), AAG847AGG (3225A>G), CGC849CAA (3231G>A 3232C>A)

*: Inserts / Deletes / Misaligned / Frameshifts

#### Analysis details

This analysis was performed with panviral2.64

### NGS Details (UN8): Lausannevirus

#### Assembly

| | |
|---|---|
| **Coverage Length** | 137 (1 contig(s)) |
| **Depth Of Coverage** | 23002.5 |
| **Number Of Reads** | 34961 |
| **Reads Per Million** | 584.29 rpm (after QC) |
| **Ambiguities** | 0 |
| **Assembly Method** | de novo + reference guided assembly |
| **Consensus Caller** | Bcf Tools |

#### Coverage Map

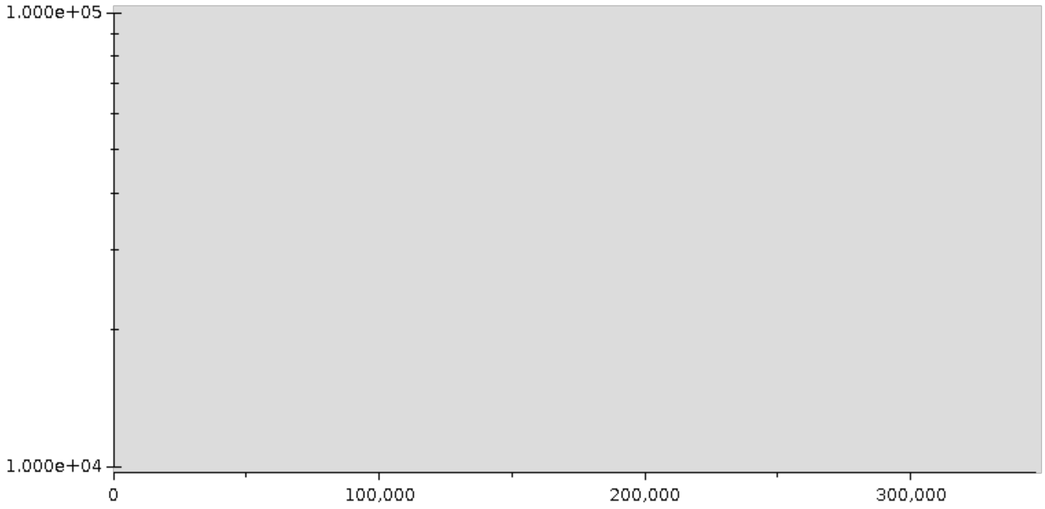

#### Assignment

| | |
|---|---|
| **Type** | Lausannevirus (Taxonomy ID: 999883) |
| **Reference Genome** | NC_015326.1 |
| **NT Identity (%)** | 80.292 |
| **AA Identity (%)** | 93.4783 |
| **Number Of Stop Codons** | 0 |
| **Number Of CDS** | 444 |

#### Alignment

| | |
|---|---|
| **Alignment Score** | 166.0 (NT) + 291.0 (AA) = 457.0 |
| **Concordance (%)** | 79.2028 |
| **Alignment Method** | Local, heuristic, nucleotide (BLASTN) |

#### Genome Region

Sequence starts at position 194953 and ends at position 195089 relative to NC_015326.1 reference sequence.

#### Alignment Detailed Statistics

| | Begin | End | Coverage | Score | Concordance | Matches | Identities | I/D/M/F* | Stop Codons |
|---|---|---|---|---|---|---|---|---|---|
| **NT** | **194953** | **195089** | **0.1%** | **166** | **60.6%** | **137 (100%)** | **110 (80.3%)** | **0/0** | |

Mutations: 194967A>G, 194970G>A, 194973A>T, 194976A>G, 194979T>G, 194981T>G, 194982C>G, 194991G>C, 194993G>A, 195000A>T, 195006G>A, 195012A>C, 195014G>A, 195015T>C, 195017G>T, 195018T>C, 195021T>C, 195030A>G, 195042T>C, 195045G>A, 195048T>C, 195051A>G, 195054T>C, 195057T>A, 195058T>G, 195059G>C, 195069C>G

*: Inserts / Deletes / Misaligned / Frameshifts

#### Analysis details

This analysis was performed with panviral2.64

### NGS Details (UN8): Yellowstone lake phycodnavirus 1

#### Assembly

| | |
|---|---|
| **Coverage Length** | 173 (1 contig(s)) |
| **Depth Of Coverage** | 15316.6 |
| **Number Of Reads** | 29336 |
| **Reads Per Million** | 490.28 rpm (after QC) |
| **Ambiguities** | 0 |
| **Assembly Method** | de novo + reference guided assembly |
| **Consensus Caller** | Bcf Tools |

#### Coverage Map

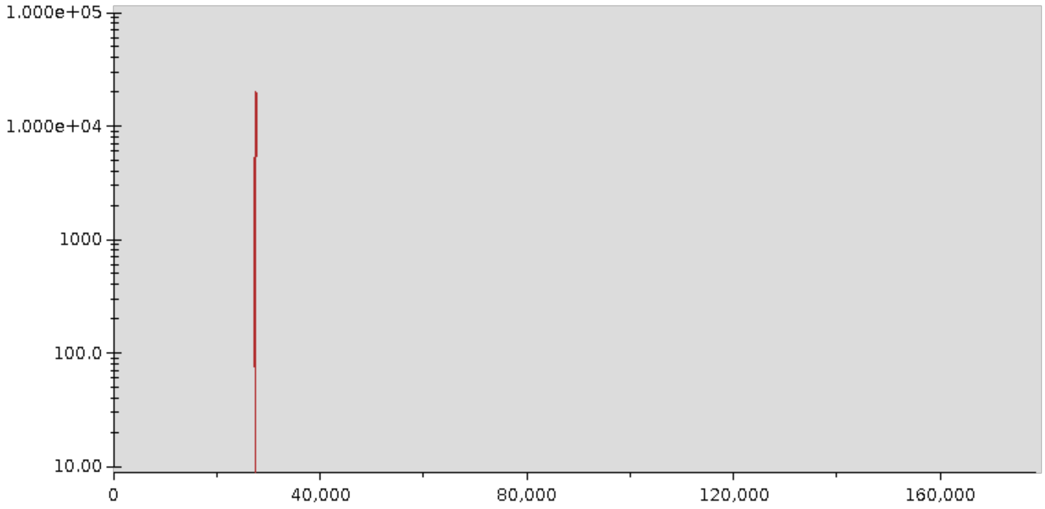

#### Assignment

| | |
|---|---|
| **Type** | Yellowstone lake phycodnavirus 1 (Taxonomy ID: 1586713) |
| **Reference Genome** | NC_028112.1 |
| **NT Identity (%)** | 82.0809 |
| **AA Identity (%)** | 93.1034 |
| **Number Of Stop Codons** | 0 |
| **Number Of CDS** | 248 |

#### Alignment

| | |
|---|---|
| **Alignment Score** | 222.0 (NT) + 350.0 (AA) = 572.0 |
| **Concordance (%)** | 80.5634 |
| **Alignment Method** | Local, heuristic, nucleotide (BLASTN) |

#### Genome Region

Sequence starts at position 27469 and ends at position 27641 relative to NC_028112.1 reference sequence.

#### Alignment Detailed Statistics

| | Begin | End | Coverage | Score | Concordance | Matches | Identities | I/D/M/F* | Stop Codons |
|---|---|---|---|---|---|---|---|---|---|
| **NT** | **27469** | **27641** | **0.1%** | **222** | **64.2%** | **173 (100%)** | **142 (82.1%)** | **0/0** | |

Mutations: 27481T>G, 27486G>A, 27492A>G, 27510A>G, 27513T>C, 27516T>A, 27529C>A, 27531T>G, 27534A>G, 27537T>A, 27540T>C, 27543T>A, 27552A>G, 27558G>A, 27563A>G, 27564T>C, 27567T>C, 27573A>T, 27576A>G, 27582T>C, 27586G>A, 27591A>G, 27597A>G, 27600C>A, 27606T>G, 27609T>C, 27612A>T, 27616T>C, 27618G>C, 27624T>C, 27627C>T

*: Inserts / Deletes / Misaligned / Frameshifts

#### Analysis details

This analysis was performed with panviral2.64

### NGS Details (UN8): Euproctis digramma nucleopolyhedrovirus

#### Assembly

| | |
|---|---|
| **Coverage Length** | 151 (1 contig(s)) |
| **Depth Of Coverage** | 15809.8 |
| **Number Of Reads** | 26156 |
| **Reads Per Million** | 437.13 rpm (after QC) |
| **Ambiguities** | 0 |
| **Assembly Method** | de novo + reference guided assembly |
| **Consensus Caller** | Bcf Tools |

#### Coverage Map

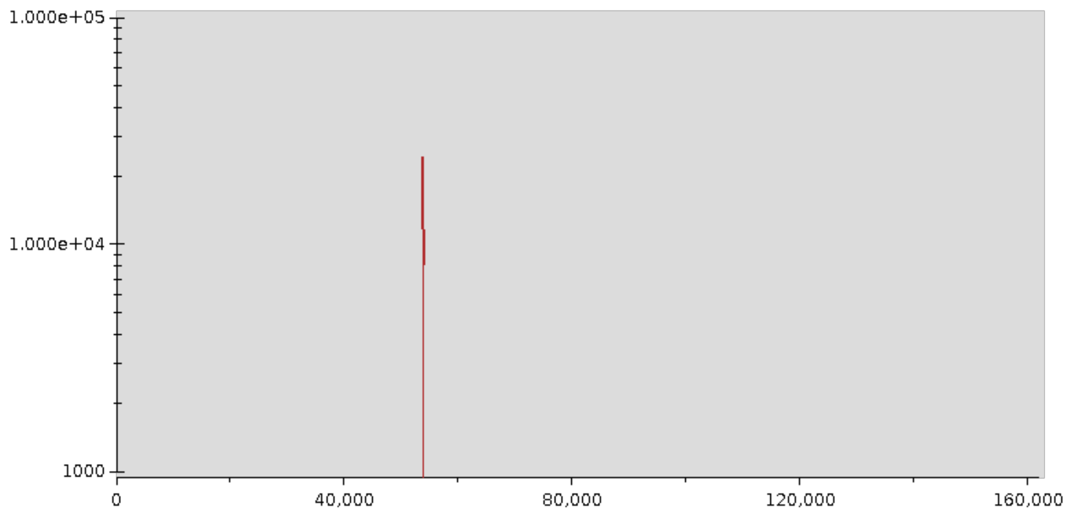

#### Assignment

| | |
|---|---|
| **Type** | Euproctis digramma nucleopolyhedrovirus (Taxonomy ID: 307466) |
| **Reference Genome** | NC_076728.1 |
| **NT Identity (%)** | 79.4702 |
| **AA Identity (%)** | 92.0 |
| **Number Of Stop Codons** | 1 |
| **Number Of CDS** | 149 |

#### Alignment

| | |
|---|---|
| **Alignment Score** | 178.0 (NT) + 311.0 (AA) = 489.0 |
| **Concordance (%)** | 77.619 |
| **Alignment Method** | Local, heuristic, nucleotide (BLASTN) |

#### Genome Region

Sequence starts at position 53764 and ends at position 53914 relative to NC_076728.1 reference sequence.

#### Alignment Detailed Statistics

| | Begin | End | Coverage | Score | Concordance | Matches | Identities | I/D/M/F* | Stop Codons |
|---|---|---|---|---|---|---|---|---|---|
| NT | 53764 | 53914 | 0.1% | 178 | 58.9% | 151 (100%) | 120 (79.5%) | 0/0 | |

Mutations: 53777C>A, 53780C>G, 53782A>G, 53783T>A, 53785T>G, 53786C>A, 53804G>A, 53807T>C, 53810T>C, 53813T>C, 53816A>G, 53819A>G, 53825A>G, 53828G>A, 53829C>G, 53830T>C, 53831A>T, 53833T>G, 53834C>A, 53840C>G, 53841G>C, 53842A>C, 53843G>A, 53855T>C, 53858G>T, 53861G>T, 53865T>A, 53875G>T, 53882A>G, 53885G>T, 53900T>C

*: Inserts / Deletes / Misaligned / Frameshifts

#### Analysis details

This analysis was performed with panviral2.64

#### NGS Details (UN8): Betabaculovirus clanastomosis

##### Assembly

| | |
|---|---|
| **Coverage Length** | 134 (1 contig(s)) |
| **Depth Of Coverage** | 13042.5 |
| **Number Of Reads** | 22579 |
| **Reads Per Million** | 377.35 rpm (after QC) |
| **Ambiguities** | 0 |
| **Assembly Method** | de novo + reference guided assembly |
| **Consensus Caller** | Bcf Tools |

##### Coverage Map

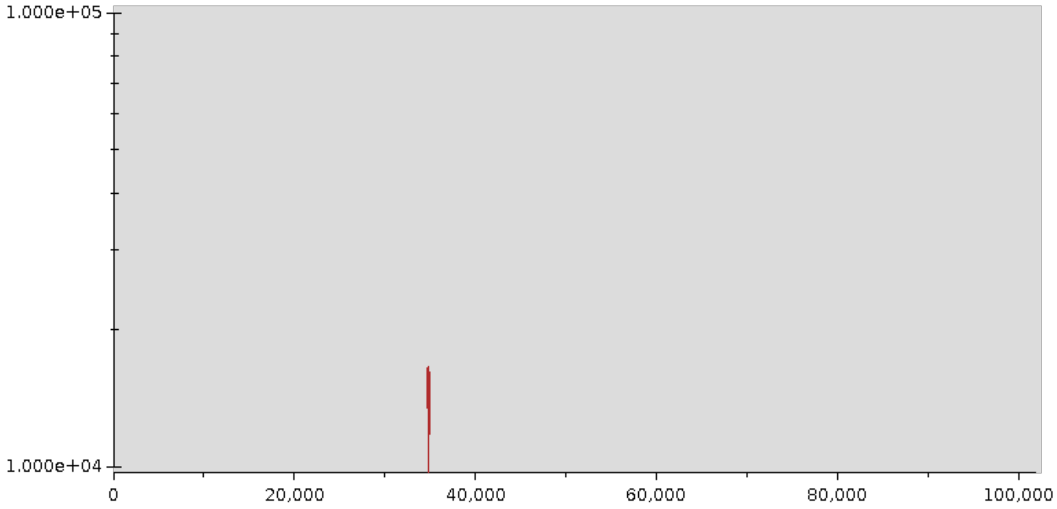

##### Assignment

| | |
|---|---|
| **Type** | Betabaculovirus clanastomosis (Taxonomy ID: 3047763) |
| **Reference Genome** | NC_022646.1 |
| **NT Identity (%)** | 83.5821 |
| **AA Identity (%)** | 86.6667 |
| **Number Of Stop Codons** | 0 |
| **Number Of CDS** | 122 |

##### Alignment

| | |
|---|---|
| **Alignment Score** | 180.0 (NT) + 289.0 (AA) = 469.0 |
| **Concordance (%)** | 80.5842 |
| **Alignment Method** | Local, heuristic, nucleotide (BLASTN) |

##### Genome Region

Sequence starts at position 34740 and ends at position 34873 relative to NC_022646.1 reference sequence.

#### Alignment Detailed Statistics

| | Begin | End | Coverage | Score | Concordance | Matches | Identities | I/D/M/F* | Stop Codons |
|---|---|---|---|---|---|---|---|---|---|
| **NT** | **34740** | **34873** | **0.1%** | **180** | **67.2%** | **134 (100%)** | **112 (83.6%)** | **0/0** | |

Mutations: 34753T>A, 34761A>G, 34764T>G, 34768G>A, 34773G>A, 34779T>G, 34784C>G, 34785T>A, 34801A>G, 34804G>A, 34809A>C, 34810T>A, 34814C>G, 34819C>G, 34820T>C, 34825G>C, 34828C>A, 34831T>C, 34837T>A, 34840G>A, 34851G>A, 34852T>C

*: Inserts / Deletes / Misaligned / Frameshifts

#### Analysis details

This analysis was performed with panviral2.64

### NGS Details (UN8): Lausannevirus

#### Assembly

| | |
|---|---|
| **Coverage Length** | 146 (1 contig(s)) |
| **Depth Of Coverage** | 7006.5 |
| **Number Of Reads** | 11975 |
| **Reads Per Million** | 200.13 rpm (after QC) |
| **Ambiguities** | 0 |
| **Assembly Method** | de novo + reference guided assembly |
| **Consensus Caller** | Bcf Tools |

#### Coverage Map

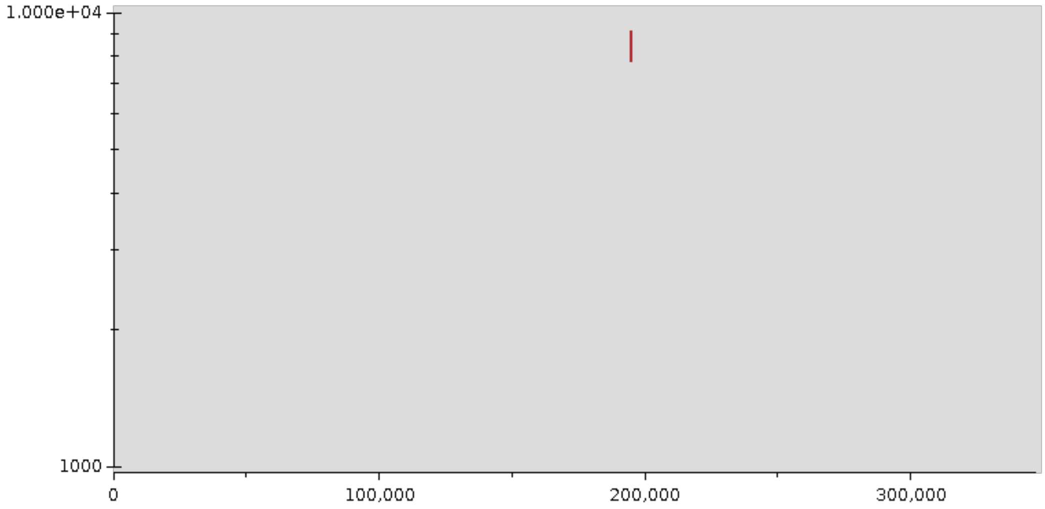

#### Assignment

| | |
|---|---|
| **Type** | Lausannevirus (Taxonomy ID: 999883) |
| **Reference Genome** | NC_015326.1 |
| **NT Identity (%)** | 84.2466 |
| **AA Identity (%)** | 100.0 |
| **Number Of Stop Codons** | 1 |
| **Number Of CDS** | 444 |

#### Alignment

| | |
|---|---|
| **Alignment Score** | 200.0 (NT) + 309.0 (AA) = 509.0 |
| **Concordance (%)** | 84.6922 |
| **Alignment Method** | Local, heuristic, nucleotide (BLASTN) |

#### Genome Region

Sequence starts at position 194911 and ends at position 195056 relative to NC_015326.1 reference sequence.

#### Alignment Detailed Statistics

| | Begin | End | Coverage | Score | Concordance | Matches | Identities | I/D/M/F* | Stop Codons |
|---|---|---|---|---|---|---|---|---|---|
| **NT** | **194911** | **195056** | **0.1%** | **200** | **68.5%** | **146 (100%)** | **123 (84.2%)** | **0/0** | |

Mutations: 194925T>C, 194931A>C, 194937G>A, 194943A>G, 194946G>C, 194949C>T, 194955T>C, 194967A>G, 194970G>A, 194979T>C, 194982C>T, 194991G>C, 194993G>A, 195003T>A, 195006G>A, 195009G>A, 195012A>C, 195015T>C, 195017G>T, 195021T>C, 195030A>G, 195039T>C, 195042T>C

*: Inserts / Deletes / Misaligned / Frameshifts

#### Analysis details

This analysis was performed with panviral2.64

#### NGS Details (UN8): Yellowstone lake phycodnavirus 1

##### Assembly

| | |
|---|---|
| **Coverage Length** | 153 (1 contig(s)) |
| **Depth Of Coverage** | 6804.1 |
| **Number Of Reads** | 11161 |
| **Reads Per Million** | 186.53 rpm (after QC) |
| **Ambiguities** | 0 |
| **Assembly Method** | de novo + reference guided assembly |
| **Consensus Caller** | Bcf Tools |

##### Coverage Map

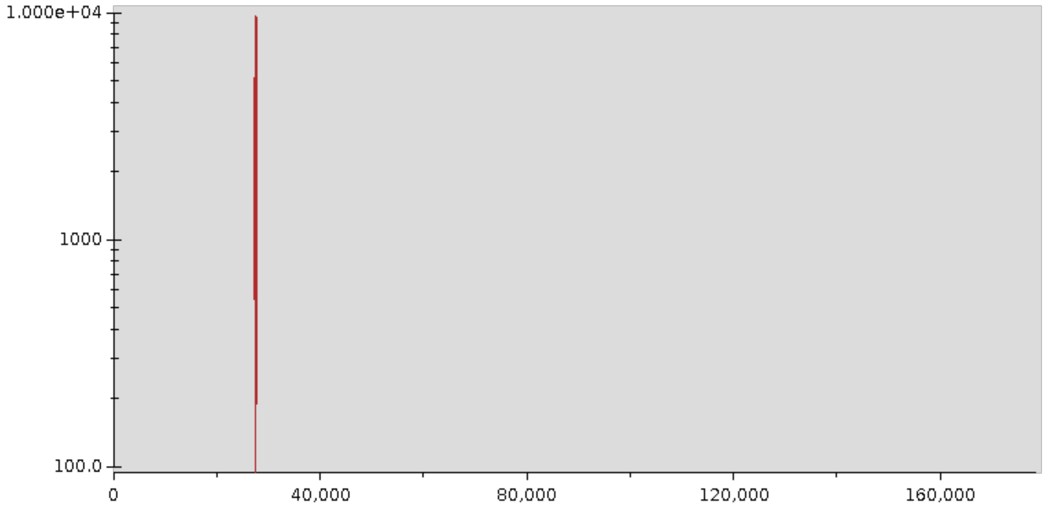

##### Assignment

| | |
|---|---|
| **Type** | Yellowstone lake phycodnavirus 1 (Taxonomy ID: 1586713) |
| **Reference Genome** | NC_028112.1 |
| **NT Identity (%)** | 83.0065 |
| **AA Identity (%)** | 90.1961 |
| **Number Of Stop Codons** | 0 |
| **Number Of CDS** | 248 |

##### Alignment

| | |
|---|---|
| **Alignment Score** | 202.0 (NT) + 312.0 (AA) = 514.0 |
| **Concordance (%)** | 80.8176 |
| **Alignment Method** | Local, heuristic, nucleotide (BLASTN) |

##### Genome Region

Sequence starts at position 27462 and ends at position 27614 relative to NC_028112.1 reference sequence.

#### Alignment Detailed Statistics

| | Begin | End | Coverage | Score | Concordance | Matches | Identities | I/D/M/F* | Stop Codons |
|---|---|---|---|---|---|---|---|---|---|
| **NT** | **27462** | **27614** | **0.1%** | **202** | **66.0%** | **153 (100%)** | **127 (83.0%)** | **0/0** | |

Mutations: 27474T>C, 27476C>A, 27477A>C, 27481T>G, 27486G>A, 27492A>G, 27507G>A, 27510A>G, 27513T>C, 27529C>A, 27531T>G, 27534A>G, 27537T>C, 27540T>C, 27552A>G, 27563A>G, 27564T>C, 27567T>C, 27570C>T, 27573A>T, 27576A>G, 27582T>C, 27586G>A, 27591A>G, 27597A>G, 27600C>A

*: Inserts / Deletes / Misaligned / Frameshifts

#### Analysis details

This analysis was performed with panviral2.64

### NGS Details (UN8): Lausannevirus

#### Assembly

| | |
|---|---|
| **Coverage Length** | 151 (1 contig(s)) |
| **Depth Of Coverage** | 3607.8 |
| **Number Of Reads** | 6209 |
| **Reads Per Million** | 103.77 rpm (after QC) |
| **Ambiguities** | 0 |
| **Assembly Method** | de novo + reference guided assembly |
| **Consensus Caller** | Bcf Tools |

#### Coverage Map

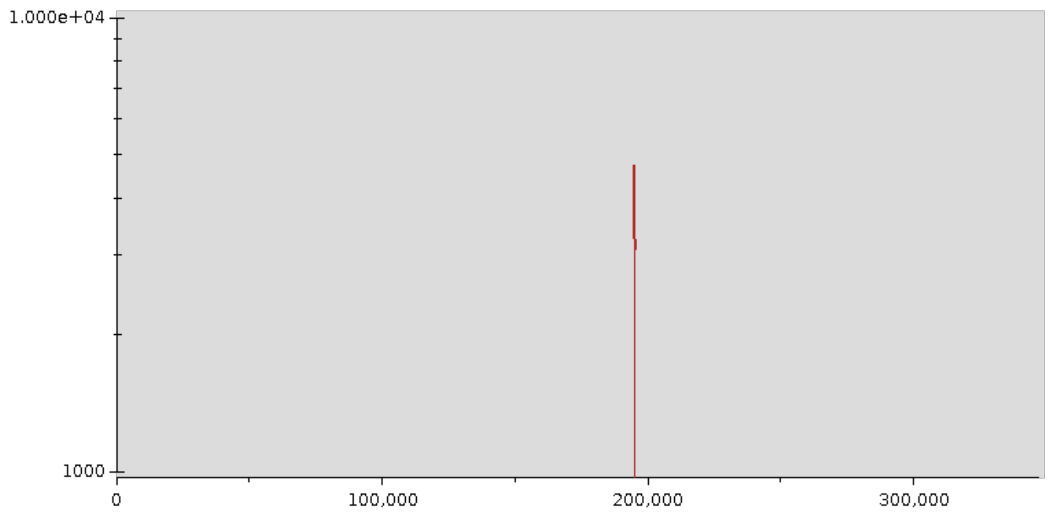

#### Assignment

| | |
|---|---|
| **Type** | Lausannevirus (Taxonomy ID: 999883) |
| **Reference Genome** | NC_015326.1 |
| **NT Identity (%)** | 82.1192 |
| **AA Identity (%)** | 94.1176 |
| **Number Of Stop Codons** | 0 |
| **Number Of CDS** | 444 |

#### Alignment

| | |
|---|---|
| **Alignment Score** | 194.0 (NT) + 316.0 (AA) = 510.0 |
| **Concordance (%)** | 80.9524 |
| **Alignment Method** | Local, heuristic, nucleotide (BLASTN) |

#### Genome Region

Sequence starts at position 194954 and ends at position 195104 relative to NC_015326.1 reference sequence.

#### Alignment Detailed Statistics

| | Begin | End | Coverage | Score | Concordance | Matches | Identities | I/D/M/F* | Stop Codons |
|---|---|---|---|---|---|---|---|---|---|
| **NT** | **194954** | **195104** | **0.1%** | **194** | **64.2%** | **151 (100%)** | **124 (82.1%)** | **0/0** | |

Mutations: 194967A>G, 194970G>T, 194973A>C, 194979T>C, 194982C>T, 194991G>C, 195000A>C, 195006G>A, 195012A>C, 195015T>C, 195017G>T, 195018T>C, 195021T>C, 195030A>G, 195042T>C, 195048T>C, 195054T>C, 195057T>A, 195058T>G, 195059G>C, 195063G>C, 195066G>A, 195069C>G, 195075G>A, 195078G>A, 195081C>A, 195093T>C

*: Inserts / Deletes / Misaligned / Frameshifts

#### Analysis details

This analysis was performed with panviral2.64

### NGS Details (UN8): Synechococcus phage S-CRM01

#### Assembly

| | |
|---|---|
| **Coverage Length** | 151 (1 contig(s)) |
| **Depth Of Coverage** | 3685.1 |
| **Number Of Reads** | 6209 |
| **Reads Per Million** | 103.77 rpm (after QC) |
| **Ambiguities** | 0 |
| **Assembly Method** | de novo + reference guided assembly |
| **Consensus Caller** | Bcf Tools |

#### Coverage Map

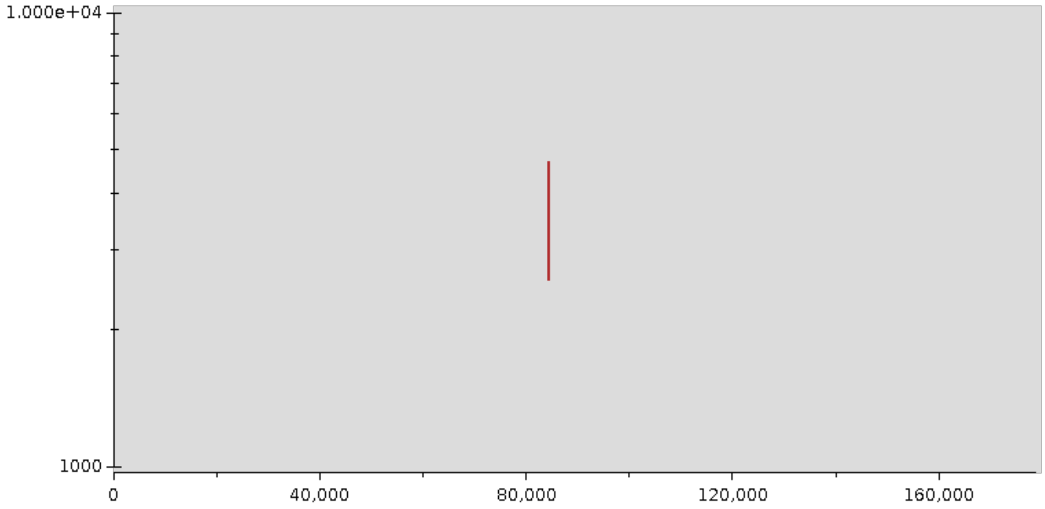

#### Assignment

| | |
|---|---|
| **Type** | Synechococcus phage S-CRM01 (Taxonomy ID: 1026955) |
| **Reference Genome** | NC_015569.1 |
| **NT Identity (%)** | 78.8079 |
| **AA Identity (%)** | 92.0 |
| **Number Of Stop Codons** | 0 |
| **Number Of CDS** | 297 |

#### Alignment

| | |
|---|---|
| **Alignment Score** | 174.0 (NT) + 311.0 (AA) = 485.0 |
| **Concordance (%)** | 75.5452 |
| **Alignment Method** | Local, heuristic, nucleotide (BLASTN) |

#### Genome Region

Sequence starts at position 84284 and ends at position 84434 relative to NC_015569.1 reference sequence.

#### Alignment Detailed Statistics

| | Begin | End | Coverage | Score | Concordance | Matches | Identities | I/D/M/F* | Stop Codons |
|---|---|---|---|---|---|---|---|---|---|
| NT | 84284 | 84434 | 0.1% | 174 | 57.6% | 151 (100%) | 119 (78.8%) | 0/0 | |

Mutations: 84299A>T, 84301A>C, 84302G>A, 84305T>A, 84306T>G, 84307G>C, 84308T>A, 84311G>T, 84312A>T, 84323A>G, 84329T>C, 84331G>T, 84335T>C, 84337G>A, 84339G>C, 84340A>T, 84347G>T, 84350A>C, 84352G>A, 84353T>G, 84365T>A, 84369G>C, 84370A>T, 84377A>G, 84379C>A, 84380T>G, 84383A>G, 84398A>C, 84401T>G, 84404C>T, 84416G>T, 84419A>G

*: Inserts / Deletes / Misaligned / Frameshifts

#### Analysis details

This analysis was performed with panviral2.64

### NGS Details (UN8): Marseillevirus marseillevirus

#### Assembly

| | |
|---|---|
| **Coverage Length** | 126 (1 contig(s)) |
| **Depth Of Coverage** | 3706.7 |
| **Number Of Reads** | 5547 |
| **Reads Per Million** | 92.70 rpm (after QC) |
| **Ambiguities** | 0 |
| **Assembly Method** | de novo + reference guided assembly |
| **Consensus Caller** | Bcf Tools |

#### Coverage Map

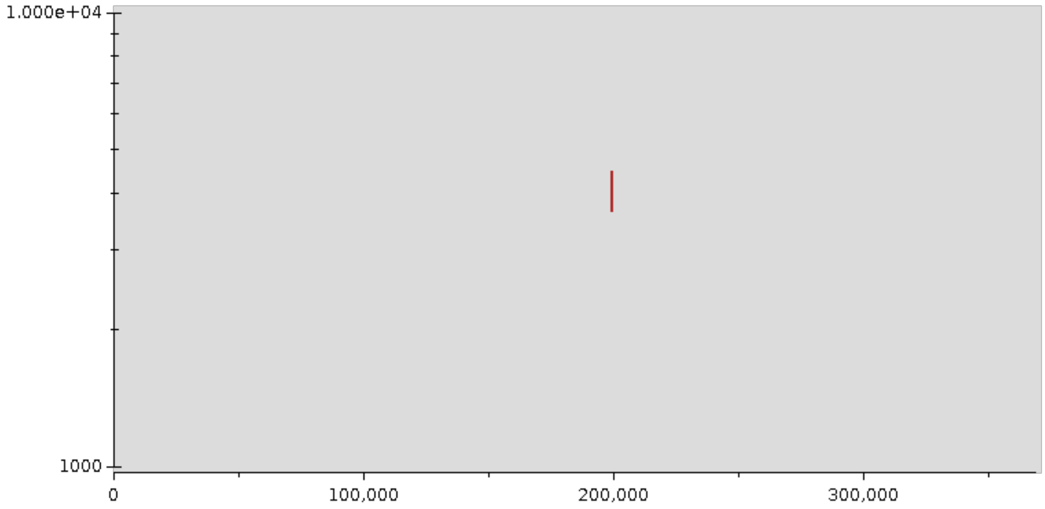

#### Assignment

| | |
|---|---|
| **Type** | Marseillevirus marseillevirus (Taxonomy ID: 694581) |
| **Reference Genome** | NC_013756.1 |
| **NT Identity (%)** | 85.7143 |
| **AA Identity (%)** | 97.5 |
| **Number Of Stop Codons** | 1 |
| **Number Of CDS** | 428 |

#### Alignment

| | |
|---|---|
| **Alignment Score** | 180.0 (NT) + 261.0 (AA) = 441.0 |
| **Concordance (%)** | 85.7977 |
| **Alignment Method** | Local, heuristic, nucleotide (BLASTN) |

#### Genome Region

Sequence starts at position 199400 and ends at position 199525 relative to NC_013756.1 reference sequence.

#### Alignment Detailed Statistics

| | Begin | End | Coverage | Score | Concordance | Matches | Identities | I/D/M/F* | Stop Codons |
|---|---|---|---|---|---|---|---|---|---|
| **NT** | **199400** | **199525** | **0.1%** | **180** | **71.4%** | **126 (100%)** | **108 (85.7%)** | **0/0** | |

Mutations: 199411A>G, 199414T>A, 199416T>G, 199417C>G, 199429G>C, 199431G>A, 199435C>G, 199449C>T, 199459A>C, 199464A>G, 199471T>A, 199489T>C, 199492G>T, 199495G>A, 199498G>A, 199501G>C, 199504A>C, 199506G>T

*: Inserts / Deletes / Misaligned / Frameshifts

#### Analysis details

This analysis was performed with panviral2.64

### NGS Details (UN8): Lausannevirus

#### Assembly

| | |
|---|---|
| **Coverage Length** | 125 (1 contig(s)) |
| **Depth Of Coverage** | 3324.6 |
| **Number Of Reads** | 4798 |
| **Reads Per Million** | 80.19 rpm (after QC) |
| **Ambiguities** | 0 |
| **Assembly Method** | de novo + reference guided assembly |
| **Consensus Caller** | Bcf Tools |

#### Coverage Map

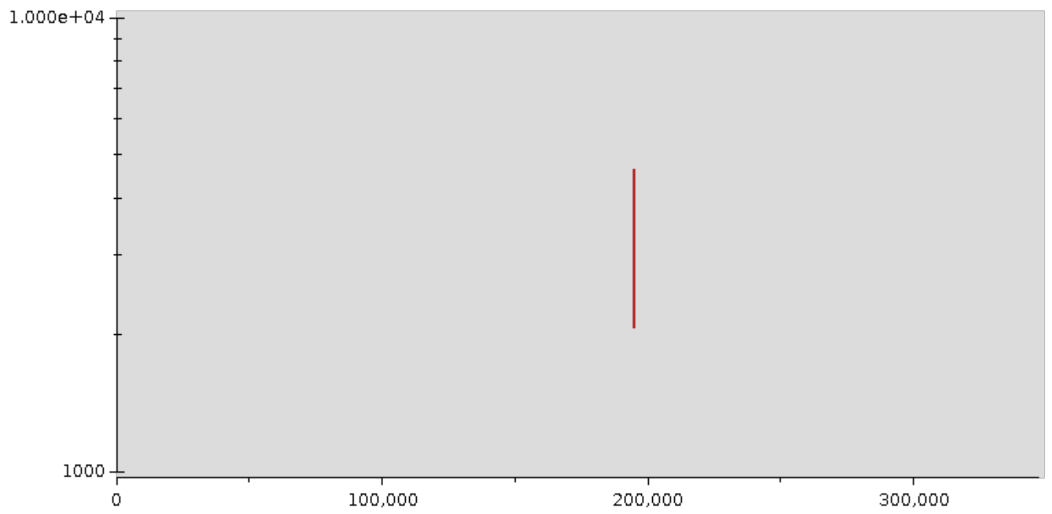

#### Assignment

| | |
|---|---|
| **Type** | Lausannevirus (Taxonomy ID: 999883) |
| **Reference Genome** | NC_015326.1 |
| **NT Identity (%)** | 80.0 |
| **AA Identity (%)** | 95.2381 |
| **Number Of Stop Codons** | 0 |
| **Number Of CDS** | 444 |

#### Alignment

| | |
|---|---|
| **Alignment Score** | 150.0 (NT) + 275.0 (AA) = 425.0 |
| **Concordance (%)** | 79.7373 |
| **Alignment Method** | Local, heuristic, nucleotide (BLASTN) |

#### Genome Region

Sequence starts at position 194935 and ends at position 195059 relative to NC_015326.1 reference sequence.

#### Alignment Detailed Statistics

| | Begin | End | Coverage | Score | Concordance | Matches | Identities | I/D/M/F* | Stop Codons |
|---|---|---|---|---|---|---|---|---|---|
| NT | 194935 | 195059 | 0.1% | 150 | 60.0% | 125 (100%) | 100 (80.0%) | 0/0 | |

Mutations: 194940A>C, 194946G>T, 194952T>C, 194955T>C, 194958A>G, 194967A>G, 194970G>C, 194976A>G, 194979T>G, 194981T>G, 194982C>G, 195009G>T, 195012A>C, 195015T>C, 195017G>T, 195018T>C, 195021T>C, 195030A>T, 195033A>G, 195036G>C, 195039T>C, 195042T>C, 195048T>C, 195054T>C, 195057T>A

*: Inserts / Deletes / Misaligned / Frameshifts

#### Analysis details

This analysis was performed with panviral2.64

### NGS Details (UN8): Betabaculovirus disaccharalis

#### Assembly

| | |
|---|---|
| **Coverage Length** | 125 (1 contig(s)) |
| **Depth Of Coverage** | 1794.9 |
| **Number Of Reads** | 2967 |
| **Reads Per Million** | 49.59 rpm (after QC) |
| **Ambiguities** | 0 |
| **Assembly Method** | de novo + reference guided assembly |
| **Consensus Caller** | Bcf Tools |

#### Coverage Map

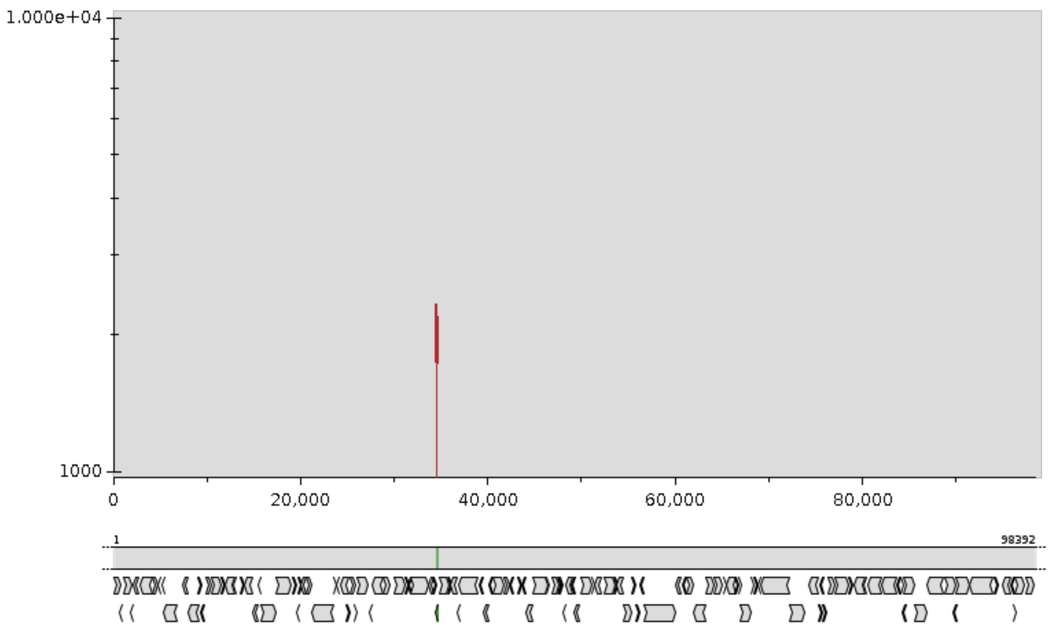

#### Assignment

| | |
|---|---|
| **Type** | Betabaculovirus disaccharalis (Taxonomy ID: 3047795) |
| **Reference Genome** | NC_028491.1 |
| **NT Identity (%)** | 82.4 |
| **AA Identity (%)** | 92.8571 |
| **Number Of Stop Codons** | 0 |
| **Number Of CDS** | 125 |

#### Alignment

| | |
|---|---|
| **Alignment Score** | 162.0 (NT) + 265.0 (AA) = 427.0 |
| **Concordance (%)** | 81.1787 |

| | |
|---|---|
| **Alignment Method** | Local, heuristic, nucleotide (BLASTN) |

#### Genome Region

Sequence starts at position 34471 and ends at position 34595 relative to NC_028491.1 reference sequence.

#### Alignment Detailed Statistics

| | Begin | End | Coverage | Score | Concordance | Matches | Identities | I/D/M/F* | Stop Codons |
|---|---|---|---|---|---|---|---|---|---|
| **NT** | **34471** | **34595** | **0.1%** | **162** | **64.8%** | **125 (100%)** | **103 (82.4%)** | **0/0** | |
| Mutations: | 34482G>T, 34485T>C, 34488A>C, 34490G>A, 34491A>C, 34512G>T, 34513C>G, 34514T>A, 34521T>C, 34524A>G, 34533A>G, 34538A>C, 34545T>C, 34547G>T, 34548C>G, 34549T>C, 34551G>A, 34554T>C, 34569G>A, 34575T>A, 34578T>C, 34580A>G | | | | | | | | |
| **CDS** | | | | | | | | | |
| v-ubq | 61 | 102 | 35.3% | 265 | 88.9% | 42 (100%) | 39 (92.9%) | 0/0/0/0 | 0 |
| Protein mutations: | E76G (34548C>G 34549T>C), S80A (34538A>C) | | | | | | | | |
| Codon mutations: | TTA66CTG (34578T>C 34580A>G), ATA67ATT (34575T>A), GCC69GCT (34569G>A), GAA74GAG (34554T>C), GAC75GAT (34551G>A), GAG76GGC (34548C>G 34549T>C), CGA77AGG (34545T>C 34547G>T), TCA80GCA (34538A>C), GAT81GAC (34533A>G), ATT84ATC (34524A>G), CAA85CAG (34521T>C), AGC88TCA (34512G>T 34513C>G 34514T>A), CGT95CGG (34491A>C), CTT96TTG (34488A>C 34490G>A), AGA97AGG (34485T>C), GGC98GGA (34482G>T) | | | | | | | | |
| **Proteins** | | | | | | | | | |
| ubiquitin-like protein (YP_009182246.1) | 61 | 102 | 35.3% | 265 | 88.9% | 42 (100%) | 39 (92.9%) | 0/0/0/0 | 0 |
| Protein mutations: | E76G (34548C>G 34549T>C), S80A (34538A>C) | | | | | | | | |
| Codon mutations: | TTA66CTG (34578T>C 34580A>G), ATA67ATT (34575T>A), GCC69GCT (34569G>A), GAA74GAG (34554T>C), GAC75GAT (34551G>A), GAG76GGC (34548C>G 34549T>C), CGA77AGG (34545T>C 34547G>T), TCA80GCA (34538A>C), GAT81GAC (34533A>G), ATT84ATC (34524A>G), CAA85CAG (34521T>C), AGC88TCA (34512G>T 34513C>G 34514T>A), CGT95CGG (34491A>C), CTT96TTG (34488A>C 34490G>A), AGA97AGG (34485T>C), GGC98GGA (34482G>T) | | | | | | | | |

*: Inserts / Deletes / Misaligned / Frameshifts

#### Analysis details

This analysis was performed with panviral2.64

### NGS Details (UN8): Lausannevirus

#### Assembly

| | |
|---|---|
| **Coverage Length** | 151 (1 contig(s)) |
| **Depth Of Coverage** | 1791.9 |
| **Number Of Reads** | 2902 |
| **Reads Per Million** | 48.50 rpm (after QC) |
| **Ambiguities** | 0 |
| **Assembly Method** | de novo + reference guided assembly |
| **Consensus Caller** | Bcf Tools |

#### Coverage Map

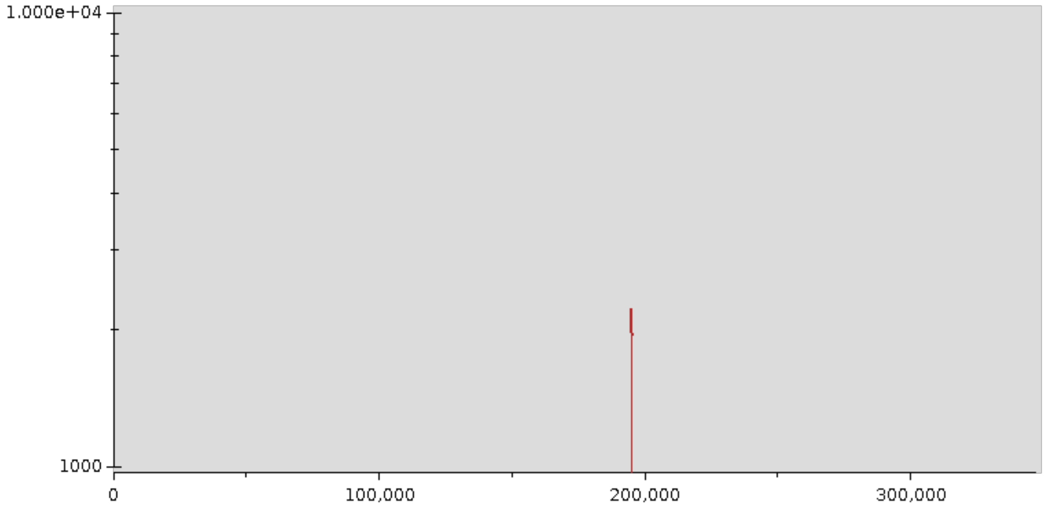

#### Assignment

| | |
|---|---|
| **Type** | Lausannevirus (Taxonomy ID: 999883) |
| **Reference Genome** | NC_015326.1 |
| **NT Identity (%)** | 80.7947 |
| **AA Identity (%)** | 94.1176 |
| **Number Of Stop Codons** | 0 |
| **Number Of CDS** | 444 |

#### Alignment

| | |
|---|---|
| **Alignment Score** | 186.0 (NT) + 316.0 (AA) = 502.0 |
| **Concordance (%)** | 79.6825 |
| **Alignment Method** | Local, heuristic, nucleotide (BLASTN) |

#### Genome Region

Sequence starts at position 194954 and ends at position 195104 relative to NC_015326.1 reference sequence.

#### Alignment Detailed Statistics

| | Begin | End | Coverage | Score | Concordance | Matches | Identities | I/D/M/F* | Stop Codons |
|---|---|---|---|---|---|---|---|---|---|
| **NT** | **194954** | **195104** | **0.1%** | **186** | **61.6%** | **151 (100%)** | **122 (80.8%)** | **0/0** | |

Mutations: 194970G>T, 194973A>T, 194979T>C, 194988T>C, 194991G>A, 195000A>G, 195003T>A, 195006G>A, 195009G>A, 195012A>T, 195014G>A, 195015T>A, 195018T>C, 195021T>C, 195030A>T, 195033A>G, 195036G>C, 195045G>A, 195051A>G, 195054T>C, 195058T>G, 195059G>C, 195060C>T, 195063G>C, 195066G>A, 195069C>A, 195081C>T, 195090C>T, 195093T>C

*: Inserts / Deletes / Misaligned / Frameshifts

#### Analysis details

This analysis was performed with panviral2.64

### NGS Details (UN8): Lausannevirus

#### Assembly

| | |
|---|---|
| **Coverage Length** | 130 (1 contig(s)) |
| **Depth Of Coverage** | 631.3 |
| **Number Of Reads** | 1319 |
| **Reads Per Million** | 22.04 rpm (after QC) |
| **Ambiguities** | 0 |
| **Assembly Method** | de novo + reference guided assembly |
| **Consensus Caller** | Bcf Tools |

#### Coverage Map

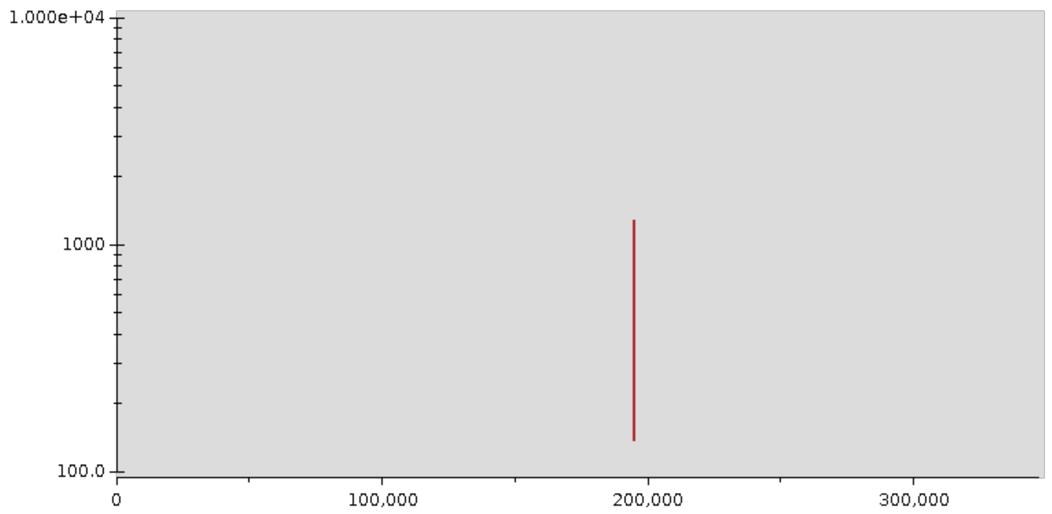

#### Assignment

| | |
|---|---|
| **Type** | Lausannevirus (Taxonomy ID: 999883) |
| **Reference Genome** | NC_015326.1 |
| **NT Identity (%)** | 83.8462 |
| **AA Identity (%)** | 95.3488 |
| **Number Of Stop Codons** | 1 |
| **Number Of CDS** | 444 |

#### Alignment

| | |
|---|---|
| **Alignment Score** | 176.0 (NT) + 264.0 (AA) = 440.0 |
| **Concordance (%)** | 81.1808 |
| **Alignment Method** | Local, heuristic, nucleotide (BLASTN) |

#### Genome Region

Sequence starts at position 194915 and ends at position 195044 relative to NC_015326.1 reference sequence.

#### Alignment Detailed Statistics

| | Begin | End | Coverage | Score | Concordance | Matches | Identities | I/D/M/F* | Stop Codons |
|---|---|---|---|---|---|---|---|---|---|
| **NT** | **194915** | **195044** | **0.1%** | **176** | **67.7%** | **130 (100%)** | **109 (83.8%)** | **0/0** | |

Mutations: 194925T>C, 194931A>C, 194937G>A, 194943A>G, 194946G>C, 194949C>T, 194955T>C, 194967A>T, 194970G>A, 194979T>C, 194982C>T, 194987C>A, 194991G>C, 194993G>A, 195003T>A, 195006G>A, 195009G>A, 195015T>C, 195017G>T, 195021T>C, 195030A>G

*: Inserts / Deletes / Misaligned / Frameshifts

#### Analysis details

This analysis was performed with panviral2.64

### NGS Details (UN8): Cladosporium fulvum T-1 virus

#### Assembly

| | |
|---|---|
| **Coverage Length** | 1429 (6 contig(s)) |
| **Depth Of Coverage** | 75.0 |
| **Number Of Reads** | 1081 |
| **Reads Per Million** | 18.07 rpm (after QC) |
| **Ambiguities** | 0 |
| **Assembly Method** | de novo + reference guided assembly |
| **Consensus Caller** | Bcf Tools |

#### Coverage Map

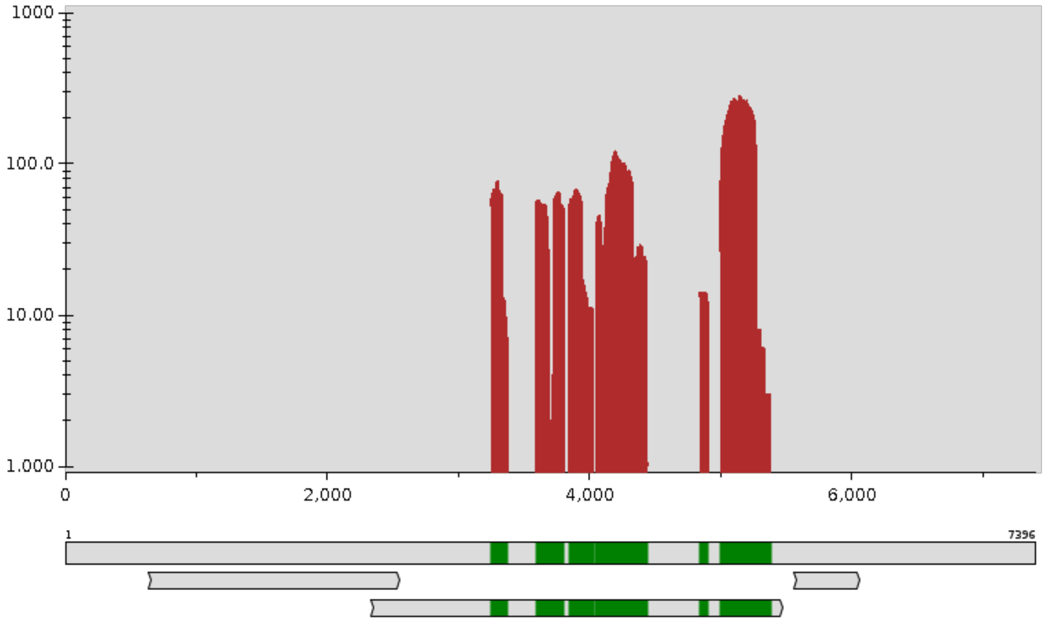

#### Assignment

| | |
|---|---|
| **Type** | Cladosporium fulvum T-1 virus (Taxonomy ID: 2052899) |
| **Reference Genome** | NC_043491.1 |
| **NT Identity (%)** | 53.1825 |
| **AA Identity (%)** | 48.0932 |
| **Number Of Stop Codons** | 0 |
| **Number Of CDS** | 3 |

#### Alignment

| | |
|---|---|
| **Alignment Score** | 138.0 (NT) + 1641.0 (AA) = 1779.0 |
| **Concordance (%)** | 29.2647 |

| **Alignment Method** | Global, seeded, nucleotide + amino acids (AGA) |
|---|---|

#### Genome Region

Sequence starts at position 3246 and ends at position 5386 relative to NC_043491.1 reference sequence.

#### Alignment Detailed Statistics

| | Begin | End | Coverage | Score | Concordance | Matches | Identities | I/D/M/F* | Stop Codons |
|---|---|---|---|---|---|---|---|---|---|
| **NT** | **3246** | **5386** | **19.3%** | **138** | **5.0%** | **1414 (99.0%)** | **752 (52.6%)** | **0/15** | |
| Mutations: | 3247A>G, 3248C>G, 3249C>A, 3250A>G, 3251T>A, 3252G>T, 3253G>A, 3257C>T, 3258A>T, 3259C>T, 3260A>G, 3261A>G, 3262G>C, 3265A>T, 3266A>G, 3269A>T, 3271T>G, 3272C>T, 3273A>T, 3277A>T, 3279G>A, 3280G>C, 3282A>C, 3283A>C, 3284G>C, 3286G>A, 3289T>A, 3290C>A, 3291C>T, 3292A>C, 3294G>C, 3295G>C, 3296G>A, 3297G>T, 3298A>T, 3301C>T, 3303T>C, 3304A>G, 3309A>G, 3310A>T, 3314T>G, 3317G>C, 3318A>C, 3320A>G, 3321A>C, 3325G>A, 3326C>T, 3328A>G, 3329C>A, 3332A>G, 3333C>A, 3334C>G, 3338C>A, 3339G>A, 3343A>G, 3344T>C, 3345G>A, 3347C>T, 3349G>A, 3355G>T, 3356A>T, 3357A>T, 3359C>T, 3361A>G, 3363C>A, 3367A>G, 3370A>T, 3372G>T, 3373G>T, 3376A>T, 3381G>C, 3584C>A, 3586A>G, 3589G>A, 3598A>G, 3601A>T, 3604A>G, 3605T>C, 3606G>C, 3607G>T, 3610A>G, 3613C>T, 3614G>A, 3618T>G, 3619C>T, 3620A>G, 3621G>T, 3622G>A, 3625A>T, 3628A>G, 3631C>T, 3634A>G, 3636T>A, 3637C>A, 3640C>T, 3646C>T, 3647T>C, 3649G>T, 3652C>A, 3658A>T, 3659A>T, 3661G>T, 3664A>G, 3667G>A, 3670C>T, 3673C>T, 3676A>C, 3679C>G, 3682A>T, 3683T>A, 3687G>T, 3688C>T, 3689C>A, 3690A>T, 3692G>A, 3695C>T, 3697T>A, 3698G>A, 3700C>G, 3703C>T, 3704G>A, 3706A>G, 3707A>G, 3708C>T, 3710C>T, 3712T>C, 3714G>A, 3715A>T, 3716G>C, 3717A>C, 3719C>T, 3720T>A, 3721A>T, 3724C>T, 3728G>A, 3729T>A, 3732G>T, 3733C>T, 3736C>G, 3739T>A, 3741C>T, 3742T>C, 3748G>A, 3751C>T, 3757A>T, 3758C>G, 3760G>C, 3761G>A, 3763C>A, 3766C>T, 3768C>G, 3769A>T, 3770A>C, 3771A>G, 3774G>A, 3775A>T, 3776T>A, 3778C>G, 3779C>T, 3781C>G, 3782C>G, 3791A>G, 3792C>T, 3793C>G, 3796G>C, 3799A>T, 3800G>T, 3802T>G, 3803C>A, 3804A>G, 3805A>G, 3806G>A, 3807A>T, 3808T>G, 3838C>G, 3839A>T, 3841G>T, 3842A>G, 3843C>T, 3844A>T, 3845G>A, 3846C>A, 3848C>A, 3849C>T, 3850C>G, 3856A>G, 3859C>T, 3862A>G, 3865C>T, 3866C>T, 3867A>C, 3868C>G, 3869A>C, 3870A>T, 3871G>A, 3874A>G, 3877A>G, 3880C>G, 3886T>C, 3887T>C, 3893T>C, 3894T>A, 3898C>A, 3899A>G, 3901C>T, 3905A>G, 3906C>A, 3908A>G, 3909C>A, 3910A>C, 3913G>A, 3918C>G, 3922C>G, 3925C>T, 3929G>A, 3930C>A, 3931A>G, 3934G>A, 3935A>G, 3936C>T, 3937A>G, 3938C>G, 3940G>A, 3941T>G, 3943A>G, 3946C>T, 3948G>T, 3949A>G, 3952A>T, 3959G>A, 3960A>G, 3964G>C, 3970A>T, 3973C>G, 3975A>C, 3980G>C, 3982A>T, 3983C>A, 3984A>G, 3986T>A, 3987C>G, 3988A>T, 3992C>T, 3994T>A, 3997A>G, 3998C>T, 4000C>G, 4003C>A, 4009C>T, 4010A>T, 4025A>G, 4027G>T, 4029A>G, 4030C>A, 4033T>C, 4045A>T, 4052A>T, 4057G>T, 4058A>G, 4059T>A, 4060G>T, 4061C>T, 4063T>G, 4064A>C, 4065C>T, 4066A>G, 4068G>A, 4069A>G, 4072A>G, 4075C>G, 4081C>G, 4085A>G, 4086A>T, 4091G>T, 4092G>C, 4093A>C, 4094A>G, 4096A>G, 4100C>T, 4101A>G, 4102G>C, 4103A>G, 4104C>A, 4105C>A, 4106G>A, 4111G>T, 4115A>C, 4119G>A, 4120A>G, 4123C>G, 4129A>C, 4130C>G, 4131A>C, 4132G>T, 4133T>A, 4134G>T, 4135C>T, 4141A>T, 4143C>A, 4144C>A, 4149C>T, 4150G>A, 4153T>G, 4154C>A, 4155G>A, 4156A>G, 4159A>G, 4162C>T, 4165T>G, 4166G>C, 4167G>T, 4168T>A, 4169A>C, 4170G>C, 4171C>T, 4172A>T, 4173A>T, 4174G>T, 4177A>G, 4180C>A, 4184_4189delATCGAG, 4192C>T, 4195C>T, 4201T>G, 4204T>C, 4206T>A, 4207G>A, 4210A>T, 4211A>G, 4213A>G, 4216C>A, 4218C>G, 4219A>C, 4220T>G, 4221G>T, 4222T>A, 4223C>T, 4225A>G, 4226A>G, 4227C>T, 4228A>G, 4231G>A, 4233C>A, 4234A>T, 4235C>G, 4240T>A, 4243G>T, 4244_4249delAAAAGA, 4252C>T, 4255A>G, 4263A>T, 4265T>G, 4267T>A, 4268T>A, 4269C>G, 4271C>A, 4277A>T, 4279G>A, 4281C>A, 4282C>T, 4283A>G, 4284C>A, 4285A>T, 4288G>T, 4291A>G, 4296A>G, 4297C>G, 4301G>T, 4302A>C, 4303C>T, 4304A>G, 4305T>C, 4306C>A, 4312C>A, 4319C>A, 4321T>G, 4322C>G, 4324A>G, 4327C>A, 4328A>T, 4330T>G, 4331G>C, 4333T>C, 4334G>C, 4335C>A, 4336C>T, 4337G>T, 4338C>G, 4339C>T, 4340A>C, 4342G>T, 4346C>A, 4347A>C, 4348T>A, 4354A>G, 4360C>T, 4361G>C, 4364G>T, 4365A>T, 4366G>A, 4369C>A, 4370C>A, 4373_4375delCCG, 4381A>C, 4382A>G, 4383C>T, 4384G>A, 4385A>G, 4387T>A, 4388C>A, 4389T>A, 4390T>G, 4391T>A, 4393A>G, 4397C>A, 4399C>T, 4400A>G, 4401A>T, 4403A>G, 4404A>C, 4406C>A, 4407T>A, 4408C>T, 4411G>T, 4419C>A, 4420G>A, 4423G>T, 4424A>C, 4425C>A, 4426G>A, 4427A>C, 4430G>A, 4433C>T, 4435T>G, 4838A>G, 4840C>G, 4841A>C, 4845A>T, 4846A>T, 4849C>T, 4850G>C, 4852A>T, 4853C>G, 4855C>G, 4865A>G, 4870A>G, 4872C>T, 4873T>A, 4875C>A, 4876A>G, 4878G>A, 4881A>G, 4882C>A, 4883G>C, 4884C>A, 4885G>A, 4886A>C, 4887A>C, 4888A>C, 4889T>G, 4891C>A, 4894T>A, 4896A>T, 4897C>T, 4898C>T, 4901C>G, 4904T>C, 4995C>G, 4996C>G, 5002C>T, 5003A>T, 5004T>C, 5005G>C, 5008A>T, 5009C>T, 5010T>G, 5011A>G, 5014C>G, 5017G>T, 5020C>G, 5027C>T, 5029C>G, 5032A>T, 5035A>G, 5037A>C, 5038T>A, 5041A>G, 5047C>T, 5048A>C, 5055C>T, 5056A>G, 5057T>A, 5058C>G, 5059A>C, 5060G>A, 5061A>C, 5062A>T, 5064T>G, 5065A>C, 5070C>G, 5079A>G, 5080G>A, 5081C>T, 5083C>G, 5085G>C, 5086A>T, 5087T>A, 5088A>G, 5089C>A, 5090C>T, 5092C>G, 5093G>T, 5094T>A, 5095A>T, 5096C>G, 5099G>C, 5100A>G, 5101C>T, 5102A>G, 5103G>A, 5104A>G, 5105T>A, 5107G>T, 5108A>G, 5111C>A, 5115A>T, 5116T>G, 5119C>T, 5123T>G, 5125C>G, 5128G>T, 5130A>T, 5131A>T, 5132G>T, 5133T>C, 5134A>T, 5135T>A, 5137C>T, 5138A>G, 5140T>G, 5141A>T, 5143A>T, 5146C>T, 5147A>C, 5151A>G, 5154A>C, 5156C>G, 5158C>G, 5161C>T, 5164A>C, 5167A>G, 5168A>C, 5169A>G, 5170C>T, 5172A>T, 5178A>G, 5179G>A, 5180A>G, 5182G>C, 5183C>T, 5185C>A, 5186A>C, 5187T>A, 5189G>A, 5190G>A, 5191A>G, 5192A>G, 5193C>A, 5194G>T, 5195A>T, 5197T>G, 5200A>T, 5202T>C, 5205A>G, 5206G>A, 5207C>T, 5208A>T, 5209C>G, 5210A>G, 5212G>T, 5213T>A, 5216T>A, 5217C>G, 5218A>T, 5219A>T, 5226A>T, 5227C>T, 5233A>T, 5234G>C, 5239G>C, 5245G>C, 5248A>G, 5249A>T, 5251G>T, 5254A>G, 5257A>G, 5260G>C, 5262A>T, 5263C>T, 5266G>A, 5269A>T, 5272C>T, 5276C>G, 5278A>C, 5279T>A, 5280A>T, 5281C>G, 5282C>T, 5284A>G, 5285C>A, 5287G>A, 5288C>G, 5289A>C, 5290C>T, 5292A>G, 5294A>G, 5296C>G, 5298A>T, 5299C>G, 5300T>G, 5302C>A, 5303G>T, 5304C>T, 5305A>G, 5306C>A, 5310A>G, 5311C>G, 5313A>G, 5314C>T, 5319T>A, 5321T>G, 5322C>A, 5323A>T, 5324T>C, 5325T>A, 5326A>T, 5332A>C, 5338G>A, 5341G>A, 5342A>T, 5344C>T, 5345G>T, 5346C>G, 5347A>C, 5348C>T, 5349T>A, 5350G>C, 5352A>G, 5353C>T, 5354A>C, 5355A>T, 5356C>G, 5359C>T, 5362A>G, 5366G>T, 5367A>C, 5369A>G, 5374T>A, 5375T>A, 5376C>A, 5379C>T, 5382C>G, 5383G>T, 5384C>T, 5386A>G | | | | | | | | |
| **CDS** | | | | | | | | | |
| homologue_of_retroviral_POL_genes | 308 | 1020 | 45.6% | 1641 | 49.0% | 472 (99.0%) | 227 (47.6%) | 0/5/0/0 | 0 |

| | Begin | End | Coverage | Score | Concordance | Matches | Identities | I/D/M/F* | Stop Codons |
|---|---|---|---|---|---|---|---|---|---|
| **NT** | **3246** | **5386** | **19.3%** | **138** | **5.0%** | **1414 (99.0%)** | **752 (52.6%)** | **0/15** | |
| Protein mutations: | P308E (3248C>G 3249C>A 3250A>G), W309I (3251T>A 3252G>T 3253G>A), H311F (3257C>T 3258A>T 3259C>T), K312G (3260A>G 3261A>G 3262G>C), N314D (3266A>G), I315L (3269A>T 3271T>G), Q316L (3272C>T 3273A>T), G318D (3279G>A 3280G>C), K319T (3282A>C 3283A>C), E320Q (3284G>C 3286G>A), P322I (3290C>A 3291C>T 3292A>C), W323S (3294G>C 3295G>C), G324I (3296G>A 3297G>T 3298A>T), L326P (3303T>C 3304A>G), Q328R (3309A>G 3310A>T), S330A (3314T>G), E331P (3317G>C 3318A>C), K332A (3320A>G 3321A>C), Q335K (3329C>A), T336E (3332A>G 3333C>A 3334C>G), R338K (3338C>A 3339G>A), W340Q (3344T>C 3345G>A), E343D (3355G>T), K344L (3356A>T 3357A>T), A346D (3363C>A), W349F (3372G>T 3373G>T), M421I (3589G>A), E425D (3601A>T), W427P (3605T>C 3606G>C 3607G>T), A430T (3614G>A), F431C (3618T>G 3619C>T), R432V (3620A>G 3621G>T 3622G>A), L437Q (3636T>A 3637C>A), M445F (3659A>T 3661G>T), S453T (3683T>A), C454F (3687G>T 3688C>T), Q455M (3689C>A 3690A>T), D456N (3692G>A), V458M (3698G>A 3700C>G), E460K (3704G>A 3706A>G), T461V (3707A>G 3708C>T), L462F (3710C>T 3712T>C), R463N (3714G>A 3715A>T), D464P (3716G>C 3717A>C), L465Y (3719C>T 3720T>A 3721A>T), V468K (3728G>A 3729T>A), C469F (3732G>T 3733C>T), A472V (3741C>T 3742T>C), M474I (3748G>A), L478V (3758C>G 3760G>C), V479I (3761G>A 3763C>A), T481S (3768C>G 3769A>T), K482R (3770A>C 3771A>G), G483D (3774G>A 3775A>T), S484T (3776T>A 3778C>G), Q486E (3782C>G), T489V (3791A>G 3792C>T 3793C>G), K490N (3796G>C), Q491H (3799A>T), V492L (3800G>T 3802T>G), Q493R (3803C>A 3804A>G 3805A>G), D494M (3806G>A 3807A>T 3808T>G), K505Y (3839A>T 3841G>T), T506V (3842A>G 3843C>T 3844A>T), A507K (3845G>A 3846C>A), P508M (3848C>A 3849C>T 3850C>G), H514S (3866C>T 3867A>C 3868C>G), K515L (3869A>C 3870A>T 3871G>A), F523H (3893T>C 3894T>A), I525V (3899A>G 3901C>T), T527E (3905A>G 3906C>A), T528D (3908A>G 3909C>A 3910A>C), T531R (3918C>G), I532M (3922C>G), A535K (3929G>A 3930C>A 3931A>G), T537V (3935A>G 3936C>T 3937A>G), Q538E (3938C>G 3940G>A), S539A (3941T>G 3943A>G), R541M (3948G>T 3949A>G), E542D (3952A>T), E545R (3959G>A 3960A>G), K550T (3975A>C), V552L (3980G>C 3982A>T), Q553R (3983C>A 3984A>G), N562Y (4010A>T), K567D (4025A>G 4027G>T), D568G (4029A>G 4030C>A), M576L (4052A>T), M578D (4058A>G 4059T>A 4060G>T), T580L (4064A>C 4065C>T 4066A>G), R581K (4068G>A 4069A>G), D583E (4075C>G), N585K (4081C>G), K587V (4085A>G 4086A>T), G589S (4091G>T 4092G>C 4093A>C), K590E (4094A>G 4096A>G), Q592C (4100C>T 4101A>G 4102G>C), T593E (4103A>G 4104C>A 4105C>A), E594K (4106G>A), K597Q (4115A>C), R598K (4119G>A 4120A>G), E601D (4129A>C), Q602A (4130C>G 4131A>C 4132G>T), C603I (4133T>A 4134G>T 4135C>T), A606E (4143C>A 4144C>A), T608I (4149C>T 4150G>A), R610K (4154C>A 4155G>A 4156A>G), D613E (4165T>G), G614L (4166G>C 4167G>T 4168T>A), S615P (4169A>C 4170G>C 4171C>T), K616F (4172A>T 4173A>T 4174G>T), I620_E621del (4184_4189delATCGAG), M627K (4206T>A 4207G>A), I629V (4211A>G 4213A>G), A631G (4218C>G 4219A>C), C632V (4220T>G 4221G>T 4222T>A), T634V (4226A>G 4227C>T 4228A>G), T636N (4233C>A 4234A>T), H637D (4235C>G), D638E (4240T>A), K640_R641del (4244_4249delAAAAGA), Y646F (4263A>T), Y647E (4265T>G 4267T>A), M651L (4277A>T 4279G>A), T652N (4281C>A 4282C>T), T653D (4283A>G 4284C>A 4285A>T), N657R (4296A>G 4297C>G), D659S (4301G>T 4302A>C 4303C>T), I660A (4304A>G 4305T>C 4306C>A), D662E (4312C>A), L665M (4319C>A 4321T>G), L666V (4322C>G 4324A>G), I668L (4328A>T 4330T>G), V669L (4331G>C 4333T>C), A670H (4334G>C 4335C>A 4336C>T), A671C (4337G>T 4338C>G 4339C>T), M672L (4340A>C 4342G>T), H674T (4346C>A 4347A>C 4348T>A), V679L (4361G>C), E680L (4364G>T 4365A>T 4366G>A), P682T (4370C>A), P683del (4373_4375delCCG), L685F (4381A>C), T686V (4382A>G 4383C>T 4384G>A), I687V (4385A>G 4387T>A), L688K (4388C>A 4389T>A 4390T>G), S689T (4391T>A 4393A>G), H691N (4397C>A 4399C>T), K692V (4400A>G 4401A>T), N693A (4403A>G 4404A>C), L694N (4406C>A 4407T>A 4408C>T), T698K (4419C>A 4420G>A), T700Q (4424A>C 4425C>A 4426G>A), K701Q (4427A>C), E702K (4430G>A), I838V (4838A>G 4840C>G), K839Q (4841A>C), K840I (4845A>T 4846A>T), V842L (4850G>C 4852A>T), H843E (4853C>G 4855C>G), N847D (4865A>G), A849V (4872C>T 4873T>A), A850E (4875C>A 4876A>G), R851Q (4878G>A), H852R (4881A>G 4882C>A), A853Q (4883G>C 4884C>A 4885G>A), K854P (4886A>C 4887A>C 4888A>C), Y855E (4889T>G 4891C>A), H857L (4896A>T 4897C>T), Q859E (4901C>G), F860L (4904T>C), A890G (4995C>G 4996C>G), M893S (5003A>T 5004T>C 5005G>C), L895W (5009C>T 5010T>G 5011A>G), M897I (5017G>T), Y904S (5037A>C 5038T>A), I908L (5048A>C), A910V (5055C>T 5056A>G), E912T (5060G>A 5061A>C 5062A>T), I913S (5064T>G 5065A>C), T915S (5070C>G), Q918R (5079A>G 5080G>A), G920A (5085G>C 5086A>T), Y921R (5087T>A 5088A>G 5089C>A), V923Y (5093G>T 5094T>A 5095A>T), L924V (5096C>G), D925R (5099G>C 5100A>G 5101C>T), R926E (5102A>G 5103G>A 5104A>G), L927I (5105T>A 5107G>T), I928V (5108A>G), Y930L (5115A>T 5116T>G), F933V (5123T>G 5125C>G), E935V (5130A>T 5131A>T), V936S (5132G>T 5133T>C 5134A>T), F937I (5135T>A 5137C>T), I938V (5138A>G 5140T>G), T939S (5141A>T 5143A>T), D942G (5151A>G), K943T (5154A>C), L944V (5156C>G 5158C>G), N948R (5168A>C 5169A>G 5170C>T), Y949F (5172A>T), K951R (5178A>G 5179G>A), T952A (5180A>G 5182G>C), M954Q (5186A>C 5187T>A), G955K (5189G>A 5190G>A 5191A>G), T956D (5192A>G 5193C>A 5194G>T), I957L (5195A>T 5197T>G), I959T (5202T>C), K960R (5205A>G 5206G>A), H961L (5207C>T 5208A>T 5209C>G), K962D (5210A>G 5212G>T), L963M (5213T>A), T965S (5219A>T), Y967F (5226A>T 5227C>T), E970Q (5234G>C), T975S (5249A>T 5251G>T), N979I (5262A>T 5263C>T), Q984D (5276C>G 5278A>C), Y985M (5279T>A 5280A>T 5281C>G), H988A (5288C>G 5289A>C 5290C>T), Y989C (5292A>G), I990V (5294A>G 5296C>G), N991M (5298A>T 5299C>G), Y992E (5300T>G 5302C>A), A993L (5303G>T 5304C>T 5305A>G), Q994K (5306C>A), D995G (5310A>G 5311C>G), N996S (5313A>G 5314C>T), V998D (5319T>A), S999D (5321T>G 5322C>A 5323A>T), L1000H (5324T>C 5325T>A 5326A>T), I1006F (5342A>T 5344C>T), A1007C (5345G>T 5346C>G 5347A>C), L1008Y (5348C>T 5349T>A 5350G>C), N1009S (5352A>G 5353C>T), N1010L (5354A>C 5355A>T 5356C>G), E1014S (5366G>T 5367A>C), T1015A (5369A>G), S1017K (5375T>A 5376C>A), T1018M (5379C>T), T1019S (5382C>G 5383G>T), P1020S (5384C>T 5386A>G) | | | | | | | | |

| | Begin | End | Coverage | Score | Concordance | Matches | Identities | I/D/M/F* | Stop Codons |
|---|---|---|---|---|---|---|---|---|---|
| **NT** | **3246** | **5386** | **19.3%** | **138** | **5.0%** | **1414 (99.0%)** | **752 (52.6%)** | **0/15** | |
| Codon mutations: | CAA307.AG (3247A>G), CCA308GAG (3248C>G 3249C>A 3250A>G), TGG309ATA (3251T>A 3252G>T 3253G>A), CAC311TTT (3257C>T 3258A>T 3259C>T), AAG312GGC (3260A>G 3261A>G 3262G>C), ATA313ATT (3265A>T), AAC314GAC (3266A>G), ATT315TTG (3269A>T 3271T>G), CAG316TTG (3272C>T 3273A>T), CCA317CCT (3277A>T), GGG318GAC (3279G>A 3280G>C), AAA319ACC (3282A>C 3283A>C), GAG320CAA (3284G>C 3286G>A), CCT321CCA (3289T>A), CCA322ATC (3290C>A 3291C>T 3292A>C), TGG323TCC (3294G>C 3295G>C), GGA324ATT (3296G>A 3297G>T 3298A>T), CCC325CCT (3301C>T), CTA326CCG (3303T>C 3304A>G), CAA328CGT (3309A>G 3310A>T), TCT330GCT (3314T>G), GAG331CCG (3317G>C 3318A>C), AAA332GCA (3320A>G 3321A>C), GAG333GAA (3325G>A), CTA334TTG (3326C>T 3328A>G), CAG335AAG (3329C>A), ACC336GAG (3332A>G 3333C>A 3334C>G), CGA338AAA (3338C>A 3339G>A), GAA339GAG (3343A>G), TGG340CAG (3344T>C 3345G>A), CTG341TTA (3347C>T 3349G>A), GAG343GAT (3355G>T), AAG344TTG (3356A>T 3357A>T), CTA345TTG (3359C>T 3361A>G), GCC346GAC (3363C>A), AAA347AAG (3367A>G), GGA348GGT (3370A>T), TGG349TTT (3372G>T 3373G>T), ATA350ATT (3376A>T), CGA352CC. (3381G>C), CGA420AGG (3584C>A 3586A>G), ATG421ATA (3589G>A), GGA424GGG (3598A>G), GAA425GAT (3601A>T), GAA426GAG (3604A>G), TGG427CCT (3605T>C 3606G>C 3607G>T), AAA428AAG (3610A>G), ACC429ACT (3613C>T), GCT430ACT (3614G>A), TTC431TGT (3618T>G 3619C>T), AGG432GTA (3620A>G 3621G>T 3622G>A), ACA433ACT (3625A>T), AGA434AGG (3628A>G), TAC435TAT (3631C>T), GGA436GGG (3634A>G), CTC437CAA (3636T>A 3637C>A), TAC438TAT (3640C>T), TTC440TTT (3646C>T), TTG441CTT (3647T>C 3649G>T), GTC442GTA (3652C>A), CCA444CCT (3658A>T), ATG445TTT (3659A>T 3661G>T), GGA446GGG (3664A>G), TTG447TTA (3667G>A), ACC448ACT (3670C>T), AAC449AAT (3673C>T), GCA450GCC (3676A>C), CCC451CCG (3679C>G), GCA452GCT (3682A>T), TCC453ACC (3683T>A), TGC454TTT (3687G>T 3688C>T), CAG455ATG (3689C>A 3690A>T), GAC456AAC (3692G>A), CTT457TTA (3695C>T 3697T>A), GTC458ATG (3698G>A 3700C>G), AAC459AAT (3703C>T), GAA460AAG (3704G>A 3706A>G), ACA461GTA (3707A>G 3708C>T), CTT462TTC (3710C>T 3712T>C), AGA463AAT (3714G>A 3715A>T), GAC464CCC (3716G>C 3717A>C), CTA465TAT (3719C>T 3720T>A 3721A>T), CTC466CTT (3724C>T), GTG468AAG (3728G>A 3729T>A), TGC469TTT (3732G>T 3733C>T), GTC470GTG (3736C>G), GTT471GTA (3739T>A), GCT472GTC (3741C>T 3742T>C), ATG474ATA (3748G>A), GAC475GAT (3751C>T), ATA477ATT (3757A>T), CTG478GTC (3758C>G 3760G>C), GTC479ATA (3761G>A 3763C>A), TAC480TAT (3766C>T), ACA481AGT (3768C>G 3769A>T), AAA482CGA (3770A>C 3771A>G), GGA483GAT (3774G>A 3775A>T), TCC484ACG (3776T>A 3778C>G), CTC485TTG (3779C>T 3781C>G), CAG486GAG (3782C>G), ACC489GTG (3791A>G 3792C>T 3793C>G), AAG490AAC (3796G>C), CAA491CAT (3799A>T), GTT492TTG (3800G>T 3802T>G), CAA493AGG (3803C>A 3804A>G 3805A>G), GAT494ATG (3806G>A 3807A>T 3808T>G), TTC504..G (3838C>G), AAG505TAT (3839A>T 3841G>T), ACA506GTT (3842A>G 3843C>T 3844A>T), GCA507AAA (3845G>A 3846C>A), CCC508ATG (3848C>A 3849C>T 3850C>G), AAA510AAG (3856A>G), TGC511TGT (3859C>T), GAA512GAG (3862A>G), TTC513TTT (3865C>T), CAC514TCG (3866C>T 3867A>C 3868C>G), AAG515CTA (3869A>C 3870A>T 3871G>A), AAA516AAG (3874A>G), GAA517GAG (3877A>G), GTC518GTG (3880C>G), TTT520TTC (3886T>C), TTA521CTA (3887T>C), TTT523CAT (3893T>C 3894T>A), ATC524ATA (3898C>A), ATC525GTT (3899A>G 3901C>T), ACA527GAA (3905A>G 3906C>A), ACA528GAC (3908A>G 3909C>A 3910A>C), GGG529GGA (3913G>A), ACG531AGG (3918C>G), ATC532ATG (3922C>G), GAC533GAT (3925C>T), GCA535AAG (3929G>A 3930C>A 3931A>G), AAG536AAA (3934G>A), ACA537GTG (3935A>G 3936C>T 3937A>G), CAG538GAA (3938C>G 3940G>A), TCA539GCG (3941T>G 3943A>G), ATC540ATT (3946C>T), AGA541ATG (3948G>T 3949A>G), GAA542GAT (3952A>T), GAA545AGA (3959G>A 3960A>G), CCG546CCC (3964G>C), ACA548ACT (3970A>T), GTC549GTG (3973C>G), AAG550ACG (3975A>C), GTA552CTT (3980G>C 3982A>T), CAG553AGG (3983C>A 3984A>G), TCA554AGT (3986T>A 3987C>G 3988A>T), CTT556TTA (3992C>T 3994T>A), GGA557GGG (3997A>G), CTC558TTG (3998C>T 4000C>G), GCC559GCA (4003C>A), TAC561TAT (4009C>T), AAC562TAC (4010A>T), AAG567GAT (4025A>G 4027G>T), GAC568GGA (4029A>G 4030C>A), TAT569TAC (4033T>C), GCA573.CT (4045A>T), ATG576TTG (4052A>T), ACG577ACT (4057G>T), ATG578GAT (4058A>G 4059T>A 4060G>T), CTT579TTG (4061C>T 4063T>G), ACA580CTG (4064A>C 4065C>T 4066A>G), AGA581AAG (4068G>A 4069A>G), AAA582AAG (4072A>G), GAC583GAG (4075C>G), AAC585AAG (4081C>G), AAA587GTA (4085A>G 4086A>T), GGA589TCC (4091G>T 4092G>C 4093A>C), AAA590GAG (4094A>G 4096A>G), CAG592TGC (4100C>T 4101A>G 4102G>C), ACC593GAA (4103A>G 4104C>A 4105C>A), GAA594AAA (4106G>A), GCG595GCT (4111G>T), AAA597CAA (4115A>C), AGA598AAG (4119G>A 4120A>G), CTC599CTG (4123C>G), GAA601GAC (4129A>C), CAG602GCT (4130C>G 4131A>C 4132G>T), TGC603ATT (4133T>A 4134G>T 4135C>T), TCA605TCT (4141A>T), GCC606GAA (4143C>A 4144C>A), ACG608ATA (4149C>T 4150G>A), CTT609CTG (4153T>G), CGA610AAG (4154C>A 4155G>A 4156A>G), CTA611CTG (4159A>G), TTC612TTT (4162C>T), GAT613GAG (4165T>G), GGT614CTA (4166G>C 4167G>T 4168T>A), AGC615CCT (4169A>C 4170G>C 4171C>T), AAG616TTT (4172A>T 4173A>T 4174G>T), GAA617GAG (4177A>G), GTC618GTA (4180C>A), ATC620_GAG621del (4184_4189delATCGAG), ACC622ACT (4192C>T), GAC623GAT (4195C>T), TCT625TCG (4201T>G), GAT626GAC (4204T>C), ATG627AAA (4206T>A 4207G>A), GCA628GCT (4210A>T), ATA629GTG (4211A>G 4213A>G), GGC630GGA (4216C>A), GCA631GGC (4218C>G 4219A>C), TGT632GTA (4220T>G 4221G>T 4222T>A), CTA633TTG (4223C>T 4225A>G), ACA634GTG (4226A>G 4227C>T 4228A>G), CAG635CAA (4231G>A), ACA636AAT (4233C>A 4234A>T), CAC637GAC (4235C>G), GAT638GAA (4240T>A), GGG639GGT (4243G>T), AAA640_AGA641del (4244_4249delAAAAGA), CAC642CAT (4252C>T), CCA643CCG (4255A>G), TAT646TTT (4263A>T), TAT647GAA (4265T>G 4267T>A), TCC648AGC (4268T>A 4269C>G), CGG649AGG (4271C>A), ATG651TTA (4277A>T 4279G>A), ACC652AAT (4281C>A 4282C>T), ACA653GAT (4283A>G 4284C>A 4285A>T), GCG654GCT (4288G>T), GAA655GAG (4291A>G), AAC657AGG (4296A>G 4297C>G), GAC659TCT (4301G>T 4302A>C 4303C>T), ATC660GCA (4304A>G 4305T>C 4306C>A), GAC662GAA (4312C>A), CTT665ATG (4319C>A 4321T>G), CTA666GTG (4322C>G 4324A>G), GCC667GCA (4327C>A), ATT668TTG (4328A>T 4330T>G), GTT669CTC (4331G>C 4333T>C), GCC670CAT (4334G>C 4335C>A 4336C>T), GCC671TGT (4337G>T 4338C>G 4339C>T), ATG672CTT (4340A>C 4342G>T), CAT674ACA (4346C>A 4347A>C 4348T>A), AGA676AGG (4354A>G), TAC678TAT (4360C>T), GTC679CTC (4361G>C), GAG680TTA (4364G>T 4365A>T 4366G>A), GGC681GGA (4369C>A), CCA682ACA (4370C>A), CCG683del (4373_4375delCCG), TTA685TTC (4381A>C), ACG686GTA (4382A>G 4383C>T 4384G>A), ATT687GTA (4385A>G 4387T>A), CTT688AAG (4388C>A 4389T>A 4390T>G), TCA689ACG (4391T>A 4393A>G), CAC691AAT (4397C>A 4399C>T), AAG692GTG (4400A>G 4401A>T), AAT693GCT (4403A>G 4404A>C), CTC694AAT (4406C>A 4407T>A 4408C>T), ACG695ACT (4411G>T), ACG698AAA (4419C>A 4420G>A), ACG699ACT (4423G>T), ACG700CAA (4424A>C 4425C>A 4426G>A), AAG701CAG (4427A>C), GAA702AAA (4430G>A), CTT703TTG (4433C>T 4435T>G), ATC838GTG (4838A>G 4840C>G), AAG839CAG (4841A>C), AAA840ATT (4845A>T 4846A>T), TGC841TGT (4849C>T), GTA842CTT (4850G>C 4852A>T), CAC843GAG (4853C>G 4855C>G), AAC847GAC (4865A>G), AAA848AAG (4870A>G), GCT849GTA (4872C>T 4873T>A), GCA850GAG (4875C>A 4876A>G), CGG851CAG (4878G>A), CAC852CGA (4881A>G 4882C>A), GCG853CAA (4883G>C 4884C>A 4885G>A), AAA854CCC (4886A>C 4887A>C 4888A>C), TAC855GAA (4889T>G 4891C>A), GGT856GGA (4894T>A), CAC857CTT (4896A>T 4897C>T), CTA858TTA (4898C>T), CAG859GAG (4901C>G), TTC860CTC (4904T>C), GCC890GGG (4995C>G 4996C>G), GAC892GAT (5002C>T), ATG893TCC (5003A>T 5004T>C 5005G>C), ATA894ATT (5008A>T), CTA895TGG (5009C>T 5010T>G 5011A>G), GTC896GTG (5014C>G), ATG897ATT (5017G>T), GTC898GTG (5020C>G), CTC901TTG (5027C>T 5029C>G), ACA902ACT (5032A>T), AAA903AAG (5035A>G), TAT904TCA (5037A>C 5038T>A), GCA905GCG (5041A>G), TTC907TTT (5047C>T), ATT908CTT (5048A>C), GCA910GTG (5055C>T 5056A>G), TCA911AGC (5057T>A 5058C>G 5059A>C), GAA912ACT (5060G>A 5061A>C 5062A>T), ATA913AGC (5064T>G 5065A>C), ACT915AGT (5070C>G), CAG918CGA (5079A>G 5080G>A), CTC919TTG (5081C>T 5083C>G), GGA920GCT (5085G>C 5086A>T), TAC921AGA (5087T>A 5088A>G 5089C>A), CTC922TTG (5090C>T 5092C>G), GTA923TAT (5093G>T 5094T>A 5095A>T), CTG924GTG (5096C>G), GAC925CGT (5099G>C 5100A>G 5101C>T), AGA926GAG (5102A>G 5103G>A 5104A>G), TTG927ATT (5105T>A 5107G>T), ATC928GTC (5108A>G), CGA929AGA (5111C>A), TAT930TTG (5115A>T 5116T>G), CAC931CAT (5119C>T), TTC933GTG (5123T>G 5125C>G), CCG934CCT (5128G>T), GAA935GTT (5130A>T 5131A>T), GTA936TCT (5132G>T 5133T>C 5134A>T), TTC937ATT (5135T>A 5137C>T), ATT938GTG (5138A>G 5140T>G), ACA939TCT (5141A>T 5143A>T), GAC940GAT (5146C>T), AGA941CGA (5147A>C), GAC942GGC (5151A>G), AAG943ACG (5154A>C), CTC944GTG (5156C>G 5158C>G), TTC945TTT (5161C>T), ACA946ACC (5164A>C), TCA947TCG (5167A>G), AAC948CGT (5168A>C 5169A>G 5170C>T), TAC949TTC (5172A>T), AAG951AGA (5178A>G 5179G>A), ACG952GCC (5180A>G 5182G>C), CTC953TTA (5183C>T 5185C>A), ATG954CAG (5186A>C 5187T>A), GGA955AAG (5189G>A 5190G>A 5191A>G), ACG956GAT (5192A>G 5193C>A 5194G>T), ATT957TTG (5195A>T 5197T>G), GGA958GGT (5200A>T), ATC959ACC (5202T>C), AAG960AGA (5205A>G 5206G>A), CAC961TTG (5207C>T 5208A>T 5209C>G), AAG962GAT (5210A>G 5212G>T), TTG963ATG (5213T>A), TCA964AGT (5216T>A 5217C>G 5218A>T), ACA965TCA (5219A>T), TAC967TTT (5226A>T 5227C>T), CCA969CCT (5233A>T), GAG970CAG (5234G>C), ACG971ACC (5239G>C), GGG973GGC (5245G>C), CAA974CAG (5248A>G), ACG975TCT (5249A>T 5251G>T), GAA976GAG (5254A>G), AGA977AGG (5257A>G), ACG978ACC (5260G>C), AAC979ATT (5262A>T 5263C>T), CAG980CAA (5266G>A), ACA981ACT (5269A>T), CTC982CTT (5272C>T), CAA984GAC (5276C>G 5278A>C), TAC985ATG (5279T>A 5280A>T 5281C>G), CTA986TTG (5282C>T 5284A>G), CGG987AGA (5285C>A 5287G>A), CAC988GCT (5288C>G 5289A>C 5290C>T), TAC989TGC (5292A>G), ATC990GTG (5294A>G 5296C>G), AAC991ATG (5298A>T 5299C>G), TAC992GAA (5300T>G 5302C>A), GCA993TTG (5303G>T 5304C>T 5305A>G), CAA994AAA (5306C>A), GAC995GGG (5310A>G 5311C>G), AAC996AGT (5313A>G 5314C>T), GTT998GAT (5319T>A), TCA999GAT (5321T>G 5322C>A 5323A>T), TTA1000CAT (5324T>C 5325T>A 5326A>T), CCA1002CCC (5332A>C), GCG1004GCA (5338G>A), CAG1005CAA (5341G>A), ATC1006TTT (5342A>T 5344C>T), GCA1007TGC (5345G>T 5346C>G 5347A>C), CTG1008TAC (5348C>T 5349T>A 5350G>C), AAC1009AGT (5352A>G 5353C>T), AAC1010CTG (5354A>C 5355A>T 5356C>G), CAC1011CAT (5359C>T), AAA1012AAG (5362A>G), GAG1014TCG (5366G>T 5367A>C), ACA1015GCA (5369A>G), ACT1016ACA (5374T>A), TCG1017AAG (5375T>A 5376C>A), ACG1018ATG (5379C>T), ACG1019AGT (5382C>G 5383G>T), CCA1020TCG (5384C>T 5386A>G) | | | | | | | | |
| **Proteins** | | | | | | | | | |
| Reverse Transcriptase (YP_009666308.1) | 308 | 1020 | 45.6% | 1641 | 49.0% | 472 (99.0%) | 227 (47.6%) | 0/5/0/0 | 0 |

| | Begin | End | Coverage | Score | Concordance | Matches | Identities | I/D/M/F* | Stop Codons |
|---|---|---|---|---|---|---|---|---|---|
| **NT** | **3246** | **5386** | **19.3%** | **138** | **5.0%** | **1414 (99.0%)** | **752 (52.6%)** | **0/15** | |
| Protein mutations: | P308E (3248C>G 3249C>A 3250A>G), W309I (3251T>A 3252G>T 3253G>A), H311F (3257C>T 3258A>T 3259C>T), K312G (3260A>G 3261A>G 3262G>C), N314D (3266A>G), I315L (3269A>T 3271T>G), Q316L (3272C>T 3273A>T), G318D (3279G>A 3280G>C), K319T (3282A>C 3283A>C), E320Q (3284G>C 3286G>A), P322I (3290C>A 3291C>T 3292A>C), W323S (3294G>C 3295G>C), G324I (3296G>A 3297G>T 3298A>T), L326P (3303T>C 3304A>G), Q328R (3309A>G 3310A>T), S330A (3314T>G), E331P (3317G>C 3318A>C), K332A (3320A>G 3321A>C), Q335K (3329C>A), T336E (3332A>G 3333C>A 3334C>G), R338K (3338C>A 3339G>A), W340Q (3344T>C 3345G>A), E343D (3355G>T), K344L (3356A>T 3357A>T), A346D (3363C>A), W349F (3372G>T 3373G>T), M421I (3589G>A), E425D (3601A>T), W427P (3605T>C 3606G>C 3607G>T), A430T (3614G>A), F431C (3618T>G 3619C>T), R432V (3620A>G 3621G>T 3622G>A), L437Q (3636T>A 3637C>A), M445F (3659A>T 3661G>T), S453T (3683T>A), C454F (3687G>T 3688C>T), Q455M (3689C>A 3690A>T), D456N (3692G>A), V458M (3698G>A 3700C>G), E460K (3704G>A 3706A>G), T461V (3707A>G 3708C>T), L462F (3710C>T 3712T>C), R463N (3714G>A 3715A>T), D464P (3716G>C 3717A>C), L465Y (3719C>T 3720T>A 3721A>T), V468K (3728G>A 3729T>A), C469F (3732G>T 3733C>T), A472V (3741C>T 3742T>C), M474I (3748G>A), L478V (3758C>G 3760G>C), V479I (3761G>A 3763C>A), T481S (3768C>G 3769A>T), K482R (3770A>C 3771A>G), G483D (3774G>A 3775A>T), S484T (3776T>A 3778C>G), Q486E (3782C>G), T489V (3791A>G 3792C>T 3793C>G), K490N (3796G>C), Q491H (3799A>T), V492L (3800G>T 3802T>G), Q493R (3803C>A 3804A>G 3805A>G), D494M (3806G>A 3807A>T 3808T>G), K505Y (3839A>T 3841G>T), T506V (3842A>G 3843C>T 3844A>T), A507K (3845G>A 3846C>A), P508M (3848C>A 3849C>T 3850C>G), H514S (3866C>T 3867A>C 3868C>G), K515L (3869A>C 3870A>T 3871G>A), F523H (3893T>C 3894T>A), I525V (3899A>G 3901C>T), T527E (3905A>G 3906C>A), T528D (3908A>G 3909C>A 3910A>C), T531R (3918C>G), I532M (3922C>G), A535K (3929G>A 3930C>A 3931A>G), T537V (3935A>G 3936C>T 3937A>G), Q538E (3938C>G 3940G>A), S539A (3941T>G 3943A>G), R541M (3948G>T 3949A>G), E542D (3952A>T), E545R (3959G>A 3960A>G), K550T (3975A>C), V552L (3980G>C 3982A>T), Q553R (3983C>A 3984A>G), N562Y (4010A>T), K567D (4025A>G 4027G>T), D568G (4029A>G 4030C>A), M576L (4052A>T), M578D (4058A>G 4059T>A 4060G>T), T580L (4064A>C 4065C>T 4066A>G), R581K (4068G>A 4069A>G), D583E (4075C>G), N585K (4081C>G), K587V (4085A>G 4086A>T), G589S (4091G>T 4092G>C 4093A>C), K590E (4094A>G 4096A>G), Q592C (4100C>T 4101A>G 4102G>C), T593E (4103A>G 4104C>A 4105C>A), E594K (4106G>A), K597Q (4115A>C), R598K (4119G>A 4120A>G), E601D (4129A>C), Q602A (4130C>G 4131A>C 4132G>T), C603I (4133T>A 4134G>T 4135C>T), A606E (4143C>A 4144C>A), T608I (4149C>T 4150G>A), R610K (4154C>A 4155G>A 4156A>G), D613E (4165T>G), G614L (4166G>C 4167G>T 4168T>A), S615P (4169A>C 4170G>C 4171C>T), K616F (4172A>T 4173A>T 4174G>T), I620_E621del (4184_4189delATCGAG), M627K (4206T>A 4207G>A), I629V (4211A>G 4213A>G), A631G (4218C>G 4219A>C), C632V (4220T>G 4221G>T 4222T>A), T634V (4226A>G 4227C>T 4228A>G), T636N (4233C>A 4234A>T), H637D (4235C>G), D638E (4240T>A), K640_R641del (4244_4249delAAAAGA), Y646F (4263A>T), Y647E (4265T>G 4267T>A), M651L (4277A>T 4279G>A), T652N (4281C>A 4282C>T), T653D (4283A>G 4284C>A 4285A>T), N657R (4296A>G 4297C>G), D659S (4301G>T 4302A>C 4303C>T), I660A (4304A>G 4305T>C 4306C>A), D662E (4312C>A), L665M (4319C>A 4321T>G), L666V (4322C>G 4324A>G), I668L (4328A>T 4330T>G), V669L (4331G>C 4333T>C), A670H (4334G>C 4335C>A 4336C>T), A671C (4337G>T 4338C>G 4339C>T), M672L (4340A>C 4342G>T), H674T (4346C>A 4347A>C 4348T>A), V679L (4361G>C), E680L (4364G>T 4365A>T 4366G>A), P682T (4370C>A), P683del (4373_4375delCCG), L685F (4381A>C), T686V (4382A>G 4383C>T 4384G>A), I687V (4385A>G 4387T>A), L688K (4388C>A 4389T>A 4390T>G), S689T (4391T>A 4393A>G), H691N (4397C>A 4399C>T), K692V (4400A>G 4401A>T), N693A (4403A>G 4404A>C), L694N (4406C>A 4407T>A 4408C>T), T698K (4419C>A 4420G>A), T700Q (4424A>C 4425C>A 4426G>A), K701Q (4427A>C), E702K (4430G>A), I838V (4838A>G 4840C>G), K839Q (4841A>C), K840I (4845A>T 4846A>T), V842L (4850G>C 4852A>T), H843E (4853C>G 4855C>G), N847D (4865A>G), A849V (4872C>T 4873T>A), A850E (4875C>A 4876A>G), R851Q (4878G>A), H852R (4881A>G 4882C>A), A853Q (4883G>C 4884C>A 4885G>A), K854P (4886A>C 4887A>C 4888A>C), Y855E (4889T>G 4891C>A), H857L (4896A>T 4897C>T), Q859E (4901C>G), F860L (4904T>C), A890G (4995C>G 4996C>G), M893S (5003A>T 5004T>C 5005G>C), L895W (5009C>T 5010T>G 5011A>G), M897I (5017G>T), Y904S (5037A>C 5038T>A), I908L (5048A>C), A910V (5055C>T 5056A>G), E912T (5060G>A 5061A>C 5062A>T), I913S (5064T>G 5065A>C), T915S (5070C>G), Q918R (5079A>G 5080G>A), G920A (5085G>C 5086A>T), Y921R (5087T>A 5088A>G 5089C>A), V923Y (5093G>T 5094T>A 5095A>T), L924V (5096C>G), D925R (5099G>C 5100A>G 5101C>T), R926E (5102A>G 5103G>A 5104A>G), L927I (5105T>A 5107G>T), I928V (5108A>G), Y930L (5115A>T 5116T>G), F933V (5123T>G 5125C>G), E935V (5130A>T 5131A>T), V936S (5132G>T 5133T>C 5134A>T), F937I (5135T>A 5137C>T), I938V (5138A>G 5140T>G), T939S (5141A>T 5143A>T), D942G (5151A>G), K943T (5154A>C), L944V (5156C>G 5158C>G), N948R (5168A>C 5169A>G 5170C>T), Y949F (5172A>T), K951R (5178A>G 5179G>A), T952A (5180A>G 5182G>C), M954Q (5186A>C 5187T>A), G955K (5189G>A 5190G>A 5191A>G), T956D (5192A>G 5193C>A 5194G>T), I957L (5195A>T 5197T>G), I959T (5202T>C), K960R (5205A>G 5206G>A), H961L (5207C>T 5208A>T 5209C>G), K962D (5210A>G 5212G>T), L963M (5213T>A), T965S (5219A>T), Y967F (5226A>T 5227C>T), E970Q (5234G>C), T975S (5249A>T 5251G>T), N979I (5262A>T 5263C>T), Q984D (5276C>G 5278A>C), Y985M (5279T>A 5280A>T 5281C>G), H988A (5288C>G 5289A>C 5290C>T), Y989C (5292A>G), I990V (5294A>G 5296C>G), N991M (5298A>T 5299C>G), Y992E (5300T>G 5302C>A), A993L (5303G>T 5304C>T 5305A>G), Q994K (5306C>A), D995G (5310A>G 5311C>G), N996S (5313A>G 5314C>T), V998D (5319T>A), S999D (5321T>G 5322C>A 5323A>T), L1000H (5324T>C 5325T>A 5326A>T), I1006F (5342A>T 5344C>T), A1007C (5345G>T 5346C>G 5347A>C), L1008Y (5348C>T 5349T>A 5350G>C), N1009S (5352A>G 5353C>T), N1010L (5354A>C 5355A>T 5356C>G), E1014S (5366G>T 5367A>C), T1015A (5369A>G), S1017K (5375T>A 5376C>A), T1018M (5379C>T), T1019S (5382C>G 5383G>T), P1020S (5384C>T 5386A>G) | | | | | | | | |

| | Begin | End | Coverage | Score | Concordance | Matches | Identities | I/D/M/F* | Stop Codons |
|---|---|---|---|---|---|---|---|---|---|
| NT | 3246 | 5386 | 19.3% | 138 | 5.0% | 1414 (99.0%) | 752 (52.6%) | 0/15 | |
| Codon mutations: | CAA307.AG (3247A>G), CCA308GAG (3248C>G 3249C>A 3250A>G), TGG309ATA (3251T>A 3252G>T 3253G>A), CAC311TTT (3257C>T 3258A>T 3259C>T), AAG312GGC (3260A>G 3261A>G 3262G>C), ATA313ATT (3265A>T), AAC314GAC (3266A>G), ATT315TTG (3269A>T 3271T>G), CAG316TTG (3272C>T 3273A>T), CCA317CCT (3277A>T), GGG318GAC (3279G>A 3280G>C), AAA319ACC (3282A>C 3283A>C), GAG320CAA (3284G>C 3286G>A), CCT321CCA (3289T>A), CCA322ATC (3290C>A 3291C>T 3292A>C), TGG323TCC (3294G>C 3295G>C), GGA324ATT (3296G>A 3297G>T 3298A>T), CCC325CCT (3301C>T), CTA326CCG (3303T>C 3304A>G), CAA328CGT (3309A>G 3310A>T), TCT330GCT (3314T>G), GAG331CCG (3317G>C 3318A>C), AAA332GCA (3320A>G 3321A>C), GAG333GAA (3325G>A), CTA334TTG (3326C>T 3328A>G), CAG335AAG (3329C>A), ACC336GAG (3332A>G 3333C>A 3334C>G), CGA338AAA (3338C>A 3339G>A), GAA339GAG (3343A>G), TGG340CAG (3344T>C 3345G>A), CTG341TTA (3347C>T 3349G>A), GAG343GAT (3355G>T), AAG344TTG (3356A>T 3357A>T), CTA345TTG (3359C>T 3361A>G), GCC346GAC (3363C>A), AAA347AAG (3367A>G), GGA348GGT (3370A>T), TGG349TTT (3372G>T 3373G>T), ATA350ATT (3376A>T), CGA352CC. (3381G>C), CGA420AGG (3584C>A 3586A>G), ATG421ATA (3589G>A), GGA424GGG (3598A>G), GAA425GAT (3601A>T), GAA426GAG (3604A>G), TGG427CCT (3605T>C 3606G>C 3607G>T), AAA428AAG (3610A>G), ACC429ACT (3613C>T), GCT430ACT (3614G>A), TTC431TGT (3618T>G 3619C>T), AGG432GTA (3620A>G 3621G>T 3622G>A), ACA433ACT (3625A>T), AGA434AGG (3628A>G), TAC435TAT (3631C>T), GGA436GGG (3634A>G), CTC437CAA (3636T>A 3637C>A), TAC438TAT (3640C>T), TTC440TTT (3646C>T), TTG441CTT (3647T>C 3649G>T), GTC442GTA (3652C>A), CCA444CCT (3658A>T), ATG445TTT (3659A>T 3661G>T), GGA446GGG (3664A>G), TTG447TTA (3667G>A), ACC448ACT (3670C>T), AAC449AAT (3673C>T), GCA450GCC (3676A>C), CCC451CCG (3679C>G), GCA452GCT (3682A>T), TCC453ACC (3683T>A), TGC454TTT (3687G>T 3688C>T), CAG455ATG (3689C>A 3690A>T), GAC456AAC (3692G>A), CTT457TTA (3695C>T 3697T>A), GTC458ATG (3698G>A 3700C>G), AAC459AAT (3703C>T), GAA460AAG (3704G>A 3706A>G), ACA461GTA (3707A>G 3708C>T), CTT462TTC (3710C>T 3712T>C), AGA463AAT (3714G>A 3715A>T), GAC464CCC (3716G>C 3717A>C), CTA465TAT (3719C>T 3720T>A 3721A>T), CTC466CTT (3724C>T), GTG468AAG (3728G>A 3729T>A), TGC469TTT (3732G>T 3733C>T), GTC470GTG (3736C>G), GTT471GTA (3739T>A), GCT472GTC (3741C>T 3742T>C), ATG474ATA (3748G>A), GAC475GAT (3751C>T), ATA477ATT (3757A>T), CTG478GTC (3758C>G 3760G>C), GTC479ATA (3761G>A 3763C>A), TAC480TAT (3766C>T), ACA481AGT (3768C>G 3769A>T), AAA482CGA (3770A>C 3771A>G), GGA483GAT (3774G>A 3775A>T), TCC484ACG (3776T>A 3778C>G), CTC485TTG (3779C>T 3781C>G), CAG486GAG (3782C>G), ACC489GTG (3791A>G 3792C>T 3793C>G), AAG490AAC (3796G>C), CAA491CAT (3799A>T), GTT492TTG (3800G>T 3802T>G), CAA493AGG (3803C>A 3804A>G 3805A>G), GAT494ATG (3806G>A 3807A>T 3808T>G), TTC504..G (3838C>G), AAG505TAT (3839A>T 3841G>T), ACA506GTT (3842A>G 3843C>T 3844A>T), GCA507AAA (3845G>A 3846C>A), CCC508ATG (3848C>A 3849C>T 3850C>G), AAA510AAG (3856A>G), TGC511TGT (3859C>T), GAA512GAG (3862A>G), TTC513TTT (3865C>T), CAC514TCG (3866C>T 3867A>C 3868C>G), AAG515CTA (3869A>C 3870A>T 3871G>A), AAA516AAG (3874A>G), GAA517GAG (3877A>G), GTC518GTG (3880C>G), TTT520TTC (3886T>C), TTA521CTA (3887T>C), TTT523CAT (3893T>C 3894T>A), ATC524ATA (3898C>A), ATC525GTT (3899A>G 3901C>T), ACA527GAA (3905A>G 3906C>A), ACA528GAC (3908A>G 3909C>A 3910A>C), GGG529GGA (3913G>A), ACG531AGG (3918C>G), ATC532ATG (3922C>G), GAC533GAT (3925C>T), GCA535AAG (3929G>A 3930C>A 3931A>G), AAG536AAA (3934G>A), ACA537GTG (3935A>G 3936C>T 3937A>G), CAG538GAA (3938C>G 3940G>A), TCA539GCG (3941T>G 3943A>G), ATC540ATT (3946C>T), AGA541ATG (3948G>T 3949A>G), GAA542GAT (3952A>T), GAA545AGA (3959G>A 3960A>G), CCG546CCC (3964G>C), ACA548ACT (3970A>T), GTC549GTG (3973C>G), AAG550ACG (3975A>C), GTA552CTT (3980G>C 3982A>T), CAG553AGG (3983C>A 3984A>G), TCA554AGT (3986T>A 3987C>G 3988A>T), CTT556TTA (3992C>T 3994T>A), GGA557GGG (3997A>G), CTC558TTG (3998C>T 4000C>G), GCC559GCA (4003C>A), TAC561TAT (4009C>T), AAC562TAC (4010A>T), AAG567GAT (4025A>G 4027G>T), GAC568GGA (4029A>G 4030C>A), TAT569TAC (4033T>C), GCA573.CT (4045A>T), ATG576TTG (4052A>T), ACG577ACT (4057G>T), ATG578GAT (4058A>G 4059T>A 4060G>T), CTT579TTG (4061C>T 4063T>G), ACA580CTG (4064A>C 4065C>T 4066A>G), AGA581AAG (4068G>A 4069A>G), AAA582AAG (4072A>G), GAC583GAG (4075C>G), AAC585AAG (4081C>G), AAA587GTA (4085A>G 4086A>T), GGA589TCC (4091G>T 4092G>C 4093A>C), AAA590GAG (4094A>G 4096A>G), CAG592TGC (4100C>T 4101A>G 4102G>C), ACC593GAA (4103A>G 4104C>A 4105C>A), GAA594AAA (4106G>A), GCG595GCT (4111G>T), AAA597CAA (4115A>C), AGA598AAG (4119G>A 4120A>G), CTC599CTG (4123C>G), GAA601GAC (4129A>C), CAG602GCT (4130C>G 4131A>C 4132G>T), TGC603ATT (4133T>A 4134G>T 4135C>T), TCA605TCT (4141A>T), GCC606GAA (4143C>A 4144C>A), ACG608ATA (4149C>T 4150G>A), CTT609CTG (4153T>G), CGA610AAG (4154C>A 4155G>A 4156A>G), CTA611CTG (4159A>G), TTC612TTT (4162C>T), GAT613GAG (4165T>G), GGT614CTA (4166G>C 4167G>T 4168T>A), AGC615CCT (4169A>C 4170G>C 4171C>T), AAG616TTT (4172A>T 4173A>T 4174G>T), GAA617GAG (4177A>G), GTC618GTA (4180C>A), ATC620_GAG621del (4184_4189delATCGAG), ACC622ACT (4192C>T), GAC623GAT (4195C>T), TCT625TCG (4201T>G), GAT626GAC (4204T>C), ATG627AAA (4206T>A 4207G>A), GCA628GCT (4210A>T), ATA629GTG (4211A>G 4213A>G), GGC630GGA (4216C>A), GCA631GGC (4218C>G 4219A>C), TGT632GTA (4220T>G 4221G>T 4222T>A), CTA633TTG (4223C>T 4225A>G), ACA634GTG (4226A>G 4227C>T 4228A>G), CAG635CAA (4231G>A), ACA636AAT (4233C>A 4234A>T), CAC637GAC (4235C>G), GAT638GAA (4240T>A), GGG639GGT (4243G>T), AAA640_AGA641del (4244_4249delAAAAGA), CAC642CAT (4252C>T), CCA643CCG (4255A>G), TAT646TTT (4263A>T), TAT647GAA (4265T>G 4267T>A), TCC648AGC (4268T>A 4269C>G), CGG649AGG (4271C>A), ATG651TTA (4277A>T 4279G>A), ACC652AAT (4281C>A 4282C>T), ACA653GAT (4283A>G 4284C>A 4285A>T), GCG654GCT (4288G>T), GAA655GAG (4291A>G), AAC657AGG (4296A>G 4297C>G), GAC659TCT (4301G>T 4302A>C 4303C>T), ATC660GCA (4304A>G 4305T>C 4306C>A), GAC662GAA (4312C>A), CTT665ATG (4319C>A 4321T>G), CTA666GTG (4322C>G 4324A>G), GCC667GCA (4327C>A), ATT668TTG (4328A>T 4330T>G), GTT669CTC (4331G>C 4333T>C), GCC670CAT (4334G>C 4335C>A 4336C>T), GCC671TGT (4337G>T 4338C>G 4339C>T), ATG672CTT (4340A>C 4342G>T), CAT674ACA (4346C>A 4347A>C 4348T>A), AGA676AGG (4354A>G), TAC678TAT (4360C>T), GTC679CTC (4361G>C), GAG680TTA (4364G>T 4365A>T 4366G>A), GGC681GGA (4369C>A), CCA682ACA (4370C>A), CCG683del (4373_4375delCCG), TTA685TTC (4381A>C), ACG686GTA (4382A>G 4383C>T 4384G>A), ATT687GTA (4385A>G 4387T>A), CTT688AAG (4388C>A 4389T>A 4390T>G), TCA689ACG (4391T>A 4393A>G), CAC691AAT (4397C>A 4399C>T), AAG692GTG (4400A>G 4401A>T), AAT693GCT (4403A>G 4404A>C), CTC694AAT (4406C>A 4407T>A 4408C>T), ACG695ACT (4411G>T), ACG698AAA (4419C>A 4420G>A), ACG699ACT (4423G>T), ACG700CAA (4424A>C 4425C>A 4426G>A), AAG701CAG (4427A>C), GAA702AAA (4430G>A), CTT703TTG (4433C>T 4435T>G), ATC838GTG (4838A>G 4840C>G), AAG839CAG (4841A>C), AAA840ATT (4845A>T 4846A>T), TGC841TGT (4849C>T), GTA842CTT (4850G>C 4852A>T), CAC843GAG (4853C>G 4855C>G), AAC847GAC (4865A>G), AAA848AAG (4870A>G), GCT849GTA (4872C>T 4873T>A), GCA850GAG (4875C>A 4876A>G), CGG851CAG (4878G>A), CAC852CGA (4881A>G 4882C>A), GCG853CAA (4883G>C 4884C>A 4885G>A), AAA854CCC (4886A>C 4887A>C 4888A>C), TAC855GAA (4889T>G 4891C>A), GGT856GGA (4894T>A), CAC857CTT (4896A>T 4897C>T), CTA858TTA (4898C>T), CAG859GAG (4901C>G), TTC860CTC (4904T>C), GCC890GGG (4995C>G 4996C>G), GAC892GAT (5002C>T), ATG893TCC (5003A>T 5004T>C 5005G>C), ATA894ATT (5008A>T), CTA895TGG (5009C>T 5010T>G 5011A>G), GTC896GTG (5014C>G), ATG897ATT (5017G>T), GTC898GTG (5020C>G), CTC901TTG (5027C>T 5029C>G), ACA902ACT (5032A>T), AAA903AAG (5035A>G), TAT904TCA (5037A>C 5038T>A), GCA905GCG (5041A>G), TTC907TTT (5047C>T), ATT908CTT (5048A>C), GCA910GTG (5055C>T 5056A>G), TCA911AGC (5057T>A 5058C>G 5059A>C), GAA912ACT (5060G>A 5061A>C 5062A>T), ATA913AGC (5064T>G 5065A>C), ACT915AGT (5070C>G), CAG918CGA (5079A>G 5080G>A), CTC919TTG (5081C>T 5083C>G), GGA920GCT (5085G>C 5086A>T), TAC921AGA (5087T>A 5088A>G 5089C>A), CTC922TTG (5090C>T 5092C>G), GTA923TAT (5093G>T 5094T>A 5095A>T), CTG924GTG (5096C>G), GAC925CGT (5099G>C 5100A>G 5101C>T), AGA926GAG (5102A>G 5103G>A 5104A>G), TTG927ATT (5105T>A 5107G>T), ATC928GTC (5108A>G), CGA929AGA (5111C>A), TAT930TTG (5115A>T 5116T>G), CAC931CAT (5119C>T), TTC933GTG (5123T>G 5125C>G), CCG934CCT (5128G>T), GAA935GTT (5130A>T 5131A>T), GTA936TCT (5132G>T 5133T>C 5134A>T), TTC937ATT (5135T>A 5137C>T), ATT938GTG (5138A>G 5140T>G), ACA939TCT (5141A>T 5143A>T), GAC940GAT (5146C>T), AGA941CGA (5147A>C), GAC942GGC (5151A>G), AAG943ACG (5154A>C), CTC944GTG (5156C>G 5158C>G), TTC945TTT (5161C>T), ACA946ACC (5164A>C), TCA947TCG (5167A>G), AAC948CGT (5168A>C 5169A>G 5170C>T), TAC949TTC (5172A>T), AAG951AGA (5178A>G 5179G>A), ACG952GCC (5180A>G 5182G>C), CTC953TTA (5183C>T 5185C>A), ATG954CAG (5186A>C 5187T>A), GGA955AAG (5189G>A 5190G>A 5191A>G), ACG956GAT (5192A>G 5193C>A 5194G>T), ATT957TTG (5195A>T 5197T>G), GGA958GGT (5200A>T), ATC959ACC (5202T>C), AAG960AGA (5205A>G 5206G>A), CAC961TTG (5207C>T 5208A>T 5209C>G), AAG962GAT (5210A>G 5212G>T), TTG963ATG (5213T>A), TCA964AGT (5216T>A 5217C>G 5218A>T), ACA965TCA (5219A>T), TAC967TTT (5226A>T 5227C>T), CCA969CCT (5233A>T), GAG970CAG (5234G>C), ACG971ACC (5239G>C), GGG973GGC (5245G>C), CAA974CAG (5248A>G), ACG975TCT (5249A>T 5251G>T), GAA976GAG (5254A>G), AGA977AGG (5257A>G), ACG978ACC (5260G>C), AAC979ATT (5262A>T 5263C>T), CAG980CAA (5266G>A), ACA981ACT (5269A>T), CTC982CTT (5272C>T), CAA984GAC (5276C>G 5278A>C), TAC985ATG (5279T>A 5280A>T 5281C>G), CTA986TTG (5282C>T 5284A>G), CGG987AGA (5285C>A 5287G>A), CAC988GCT (5288C>G 5289A>C 5290C>T), TAC989TGC (5292A>G), ATC990GTG (5294A>G 5296C>G), AAC991ATG (5298A>T 5299C>G), TAC992GAA (5300T>G 5302C>A), GCA993TTG (5303G>T 5304C>T 5305A>G), CAA994AAA (5306C>A), GAC995GGG (5310A>G 5311C>G), AAC996AGT (5313A>G 5314C>T), GTT998GAT (5319T>A), TCA999GAT (5321T>G 5322C>A 5323A>T), TTA1000CAT (5324T>C 5325T>A 5326A>T), CCA1002CCC (5332A>C), GCG1004GCA (5338G>A), CAG1005CAA (5341G>A), ATC1006TTT (5342A>T 5344C>T), GCA1007TGC (5345G>T 5346C>G 5347A>C), CTG1008TAC (5348C>T 5349T>A 5350G>C), AAC1009AGT (5352A>G 5353C>T), AAC1010CTG (5354A>C 5355A>T 5356C>G), CAC1011CAT (5359C>T), AAA1012AAG (5362A>G), GAG1014TCG (5366G>T 5367A>C), ACA1015GCA (5369A>G), ACT1016ACA (5374T>A), TCG1017AAG (5375T>A 5376C>A), ACG1018ATG (5379C>T), ACG1019AGT (5382C>G 5383G>T), CCA1020TCG (5384C>T 5386A>G) | | | | | | | | |

*: Inserts / Deletes / Misaligned / Frameshifts

#### Analysis details

This analysis was performed with panviral2.64

#### NGS Details (UN8): Errantivirus

##### Assembly

| | |
|---|---|
| **Coverage Length** | 698 (1 contig(s)) |
| **Depth Of Coverage** | 139.9 |
| **Number Of Reads** | 891 |
| **Reads Per Million** | 14.89 rpm (after QC) |
| **Ambiguities** | 0 |
| **Assembly Method** | de novo + reference guided assembly |
| **Consensus Caller** | Bcf Tools |

##### Coverage Map

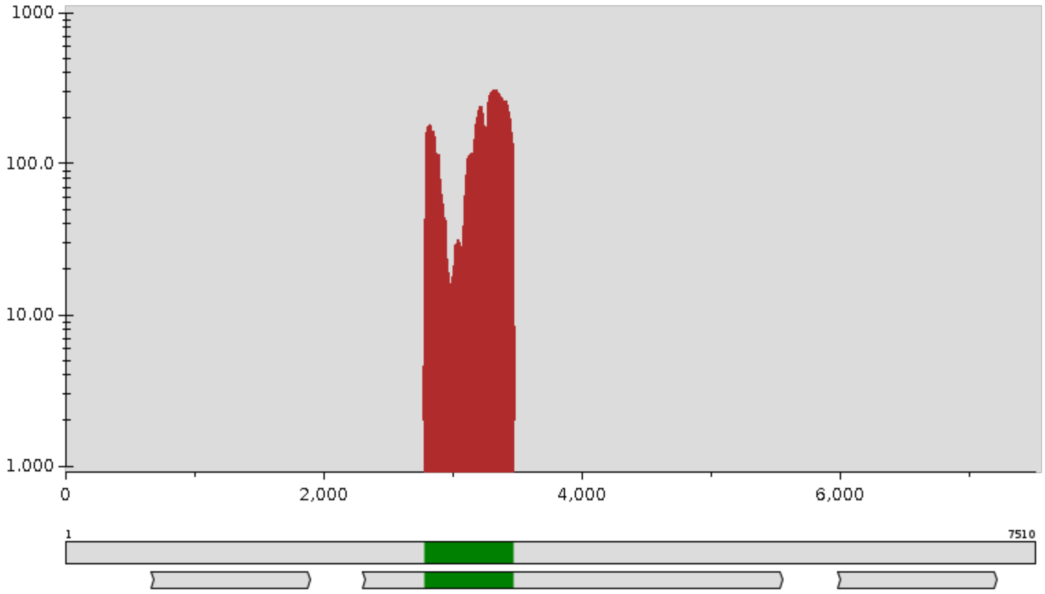

##### Assignment

| | |
|---|---|
| **Type** | Errantivirus (Taxonomy ID: 186666) |
| **Reference Genome** | NC_038512.1 |
| **NT Identity (%)** | 53.1479 |
| **AA Identity (%)** | 40.7895 |
| **Number Of Stop Codons** | 0 |
| **Number Of CDS** | 3 |

##### Alignment

| | |
|---|---|
| **Alignment Score** | 44.0 (NT) + 709.0 (AA) = 753.0 |
| **Concordance (%)** | 26.0734 |

| Alignment Method | Global, seeded, nucleotide + amino acids (AGA) |
|---|---|

#### Genome Region

Sequence starts at position 2779 and ends at position 3476 relative to NC_038512.1 reference sequence.

#### Alignment Detailed Statistics

| | Begin | End | Coverage | Score | Concordance | Matches | Identities | I/D/M/F* | Stop Codons |
|---|---|---|---|---|---|---|---|---|---|
| **NT** | **2779** | **3476** | **9.3%** | **44** | **3.3%** | **683 (97.9%)** | **363 (52.0%)** | **0/15** | |
| Mutations: | 2786A>T, 2791G>A, 2792G>A, 2794C>G, 2795C>A, 2797A>G, 2801A>T, 2806A>C, 2809A>G, 2812A>T, 2813T>A, 2814C>G, 2815A>T, 2817A>T, 2818C>G, 2821T>A, 2822G>C, 2824A>T, 2828A>G, 2831T>G, 2836C>T, 2839A>G, 2841G>T, 2843G>T, 2849C>A, 2850C>A, 2851C>A, 2856A>G, 2858_2860delATC, 2863C>T, 2865C>G, 2870_2872delGGG, 2874A>T, 2875A>G, 2878A>C, 2880A>T, 2884G>T, 2885_2893delCGTCTCGTA, 2894G>A, 2899C>T, 2901T>A, 2902C>T, 2905T>C, 2906A>C, 2915G>A, 2918A>G, 2919A>T, 2927G>A, 2929T>G, 2930G>A, 2935A>G, 2938C>T, 2941G>C, 2942A>T, 2944A>G, 2947A>C, 2948A>C, 2949A>G, 2953A>T, 2954A>G, 2955G>A, 2960G>T, 2963C>T, 2969A>C, 2972T>C, 2975G>C, 2976G>A, 2977T>G, 2978A>G, 2980G>A, 2981T>G, 2982G>C, 2984C>T, 2985A>C, 2986A>C, 2987T>G, 2989C>G, 2992C>T, 2993A>T, 2995C>T, 2997C>A, 2998C>G, 2999T>A, 3001A>C, 3003A>G, 3004T>C, 3008G>A, 3009C>G, 3010A>G, 3011A>T, 3012G>C, 3013T>G, 3016G>C, 3018T>A, 3019T>C, 3020T>C, 3022T>C, 3026G>T, 3029G>A, 3031G>A, 3032A>G, 3034G>T, 3035G>C, 3036A>G, 3037C>G, 3038C>G, 3039C>A, 3040T>A, 3041C>T, 3042A>C, 3047A>G, 3049A>C, 3050T>C, 3055A>G, 3059G>A, 3061G>C, 3065A>C, 3066A>G, 3067C>G, 3068G>A, 3069T>C, 3071G>C, 3072A>G, 3073A>T, 3074C>T, 3076C>T, 3082T>C, 3084T>A, 3085T>C, 3088A>G, 3092C>G, 3094T>G, 3095C>G, 3096G>T, 3097A>C, 3101C>T, 3104A>T, 3106G>C, 3109A>G, 3110T>C, 3112A>T, 3114A>C, 3115A>T, 3118C>T, 3119T>G, 3121A>C, 3124A>T, 3125T>G, 3128A>G, 3134C>A, 3135A>T, 3136A>G, 3137A>G, 3138G>A, 3139A>G, 3140G>T, 3142T>G, 3146G>A, 3148C>T, 3149A>C, 3151T>G, 3154C>G, 3155C>T, 3157A>T, 3159G>A, 3161G>C, 3162G>C, 3163T>C, 3164C>T, 3165T>A, 3167C>T, 3168A>T, 3170A>G, 3173A>C, 3175C>G, 3176A>T, 3179T>G, 3180G>T, 3181T>G, 3182C>A, 3184C>T, 3187C>G, 3189A>T, 3190C>T, 3191C>A, 3193T>A, 3196C>T, 3202T>C, 3203A>T, 3205T>G, 3208C>T, 3212A>T, 3213G>C, 3214T>C, 3216C>A, 3217T>G, 3218T>A, 3219C>G, 3220C>T, 3221C>G, 3222T>A, 3223A>G, 3224C>A, 3225A>G, 3226G>T, 3229A>C, 3232C>T, 3233C>G, 3236G>A, 3237A>G, 3238G>A, 3239A>C, 3245G>A, 3246A>G, 3247A>G, 3248C>A, 3249G>T, 3250A>T, 3253T>G, 3256C>G, 3259A>G, 3263C>T, 3265T>G, 3272A>G, 3273G>A, 3274T>A, 3277C>G, 3280C>A, 3281A>T, 3283A>T, 3284A>G, 3285T>C, 3286T>C, 3287C>A, 3289A>G, 3290A>T, 3292G>C, 3293G>T, 3294A>C, 3295C>A, 3300C>G, 3301C>T, 3304A>G, 3307C>T, 3309T>G, 3311A>C, 3312A>T, 3313G>A, 3315T>A, 3316C>G, 3317G>T, 3318A>C, 3320A>G, 3321C>T, 3322T>A, 3323G>T, 3327A>T, 3331T>A, 3337C>T, 3338A>G, 3340C>G, 3341A>G, 3343A>G, 3345G>C, 3346C>T, 3348G>A, 3352C>T, 3355T>C, 3358C>T, 3360A>T, 3362C>G, 3363C>T, 3365A>G, 3367C>T, 3370T>G, 3372A>C, 3373T>C, 3376G>A, 3377A>G, 3379T>G, 3380T>G, 3382C>T, 3385T>G, 3386A>G, 3388T>G, 3389C>A, 3390A>G, 3392A>G, 3396A>G, 3397T>G, 3398C>G, 3399T>C, 3400G>C, 3402T>G, 3403T>G, 3406A>T, 3408A>C, 3410A>T, 3413C>G, 3414C>T, 3415T>G, 3417A>C, 3418G>T, 3421A>G, 3424A>T, 3425A>C, 3426A>G, 3427A>T, 3428C>A, 3429A>G, 3430A>T, 3433T>C, 3434T>C, 3436A>T, 3440C>T, 3442T>G, 3443C>G, 3444T>C, 3445C>A, 3457A>G, 3459A>G, 3463C>T, 3464A>G | | | | | | | | |

##### CDS

| | Begin | End | Coverage | Score | Concordance | Matches | Identities | I/D/M/F* | Stop Codons |
|---|---|---|---|---|---|---|---|---|---|
| D1R33_gp2 | 159 | 391 | 21.5% | 709 | 45.0% | 228 (97.9%) | 93 (39.9%) | 0/5/0/0 | 0 |
| Protein mutations: | M161L (2786A>T), D163K (2792G>A 2794C>G), Q164K (2795C>A 2797A>G), I166F (2801A>T), D171V (2817A>T 2818C>G), A173P (2822G>C 2824A>T), S175G (2828A>G), S176A (2831T>G), I178M (2839A>G), W179L (2841G>T), V180F (2843G>T), P182K (2849C>A 2850C>A 2851C>A), K184R (2856A>G), I185del (2858_2860delATC), A187G (2865C>G), G189del (2870_2872delGGG), K190M (2874A>T 2875A>G), Q191H (2878A>C), K192M (2880A>T), W193C (2884G>T), R194_V196del (2885_2893delCGTCTCGTA), V197I (2894G>A), F199Y (2901T>A 2902C>T), K201Q (2906A>C), E204K (2915G>A), K205V (2918A>G 2919A>T), D208K (2927G>A 2929T>G), D209N (2930G>A), I213L (2942A>T 2944A>G), N215R (2948A>C 2949A>G), S217D (2954A>G 2955G>A), V219L (2960G>T), L220F (2963C>T), K222Q (2969A>C), G224Q (2975G>C 2976G>A 2977T>G), K225E (2978A>G 2980G>A), C226A (2981T>G 2982G>C), Q227S (2984C>T 2985A>C 2986A>C), Y228E (2987T>G 2989C>G), T230S (2993A>T 2995C>T), T231K (2997C>A 2998C>G), L232I (2999T>A 3001A>C), D233G (3003A>G 3004T>C), A235R (3008G>A 3009C>G 3010A>G), F238Y (3018T>A 3019T>C), Y239H (3020T>C 3022T>C), V241L (3026G>T), E242K (3029G>A 3031G>A), M243V (3032A>G 3034G>T), D244R (3035G>C 3036A>G 3037C>G), P245E (3038C>G 3039C>A 3040T>A), Q246S (3041C>T 3042A>C), I248V (3047A>G 3049A>C), S249P (3050T>C), A252T (3059G>A 3061G>C), N254R (3065A>C 3066A>G 3067C>G), V255T (3068G>A 3069T>C), E256R (3071G>C 3072A>G 3073A>T), H257Y (3074C>T 3076C>T), F260Y (3084T>A 3085T>C), L263V (3092C>G 3094T>G), R264V (3095C>G 3096G>T 3097A>C), P266S (3101C>T), M267F (3104A>T 3106G>C), K270T (3114A>C 3115A>T), S272A (3119T>G 3121A>C), S274A (3125T>G), T275A (3128A>G), Q277M (3134C>A 3135A>T 3136A>G), R278E (3137A>G 3138G>A 3139A>G), V279L (3140G>T 3142T>G), D281N (3146G>A 3148C>T), N282Q (3149A>C 3151T>G), L284F (3155C>T 3157A>T), R285K (3159G>A), G286P (3161G>C 3162G>C 3163T>C), L287Y (3164C>T 3165T>A), Q288L (3167C>T 3168A>T), N289D (3170A>G), N290Q (3173A>C 3175C>G), I291F (3176A>T), C292V (3179T>G 3180G>T 3181T>G), L293I (3182C>A 3184C>T), Y295F (3189A>T 3190C>T), L296I (3191C>A 3193T>A), I300L (3203A>T 3205T>G), T304K (3216C>A 3217T>G), L306E (3221C>G 3222T>A 3223A>G), Q307S (3224C>A 3225A>G 3226G>T), E308D (3229A>C), L310V (3233C>G), E311R (3236G>A 3237A>G 3238G>A), N312H (3239A>C), E314R (3245G>A 3246A>G 3247A>G), R315I (3248C>A 3249G>T 3250A>T), F317L (3256C>G), S323E (3272A>G 3273G>A 3274T>A), N324K (3277C>G), F325L (3280C>A), K326Y (3281A>T 3283A>T), I327A (3284A>G 3285T>C 3286T>C), Q328K (3287C>A 3289A>G), M329F (3290A>T 3292G>C), D330S (3293G>T 3294A>C 3295C>A), S332C (3300C>G 3301C>T), L335W (3309T>G), K336L (3311A>C 3312A>T 3313G>A), L337Q (3315T>A 3316C>G), E338S (3317G>T 3318A>C), T339V (3320A>G 3321C>T 3322T>A), A340S (3323G>T), Y341F (3327A>T), I345V (3338A>G 3340C>G), I346V (3341A>G 3343A>G), S347T (3345G>C 3346C>T), R348K (3348G>A), K352M (3360A>T), P353V (3362C>G 3363C>T), N354D (3365A>G 3367C>T), D356A (3372A>C 3373T>C), I358V (3377A>G 3379T>G), S359A (3380T>G 3382C>T), I361V (3386A>G 3388T>G), Q362R (3389C>A 3390A>G), K363E (3392A>G), Y364W (3396A>G 3397T>G), L365A (3398C>G 3399T>C 3400G>C), I366R (3402T>G 3403T>G), K368T (3408A>C), T369S (3410A>T), P370V (3413C>G 3414C>T 3415T>G), K371T (3417A>C 3418G>T), K374R (3425A>C 3426A>G 3427A>T), Q375S (3428C>A 3429A>G 3430A>T), L380A (3443C>G 3444T>C 3445C>A), K385R (3459A>G), I387V (3464A>G) | | | | | | | | |
| Codon mutations: | ATG161TTG (2786A>T), TTG162TTA (2791G>A), GAC163AAG (2792G>A 2794C>G), CAA164AAG (2795C>A 2797A>G), ATT166TTT (2801A>T), ATA167ATC (2806A>C), AGA168AGG (2809A>G), CCA169CCT (2812A>T), TCA170AGT (2813T>A 2814C>G 2815A>T), GAC171GTG (2817A>T 2818C>G), TCT172TCA (2821T>A), GCA173CCT (2822G>C 2824A>T), AGC175GGC (2828A>G), TCA176GCA (2831T>G), CCC177CCT (2836C>T), ATA178ATG (2839A>G), TGG179TTG (2841G>T), GTT180TTT (2843G>T), CCC182AAA (2849C>A 2850C>A 2851C>A), AAA184AGA (2856A>G), ATC185del (2858_2860delATC), GAC186GAT (2863C>T), GCT187GGT (2865C>G), GGG189del (2870_2872delGGG), AAA190ATG (2874A>T 2875A>G), CAA191CAC (2878A>C), AAG192ATG (2880A>T), TGG193TGT (2884G>T), CGT194_GTA196del (2885_2893delCGTCTCGTA), GTT197ATT (2894G>A), GAC198GAT (2899C>T), TTC199TAT (2901T>A 2902C>T), CGT200CGC (2905T>C), AAG201CAG (2906A>C), GAG204AAG (2915G>A), AAG205GTG (2918A>G 2919A>T), GAT208AAG (2927G>A 2929T>G), GAC209AAC (2930G>A), AAA210AAG (2935A>G), TAC211TAT (2938C>T), CCG212CCC (2941G>C), ATA213TTG (2942A>T 2944A>G), CCA214CCC (2947A>C), AAC215CGC (2948A>C 2949A>G), ATA216ATT (2953A>T), AGT217GAT (2954A>G 2955G>A), GTA219TTA (2960G>T), CTT220TTT (2963C>T), AAG222CAG (2969A>C), TTA223CTA (2972T>C), GGT224CAG (2975G>C 2976G>A 2977T>G), AAG225GAA (2978A>G 2980G>A), TGC226GCC (2981T>G 2982G>C), CAA227TCC (2984C>T 2985A>C 2986A>C), TAC228GAG (2987T>G 2989C>G), TTC229TTT (2992C>T), ACC230TCT (2993A>T 2995C>T), ACC231AAG (2997C>A 2998C>G), TTA232ATC (2999T>A 3001A>C), GAT233GGC (3003A>G 3004T>C), GCA235AGG (3008G>A 3009C>G 3010A>G), AGT236TCG (3011A>T 3012G>C 3013T>G), GGG237GGC (3016G>C), TTT238TAC (3018T>A 3019T>C), TAT239CAC (3020T>C 3022T>C), GTG241TTG (3026G>T), GAG242AAA (3029G>A 3031G>A), ATG243GTT (3032A>G 3034G>T), GAC244CGG (3035G>C 3036A>G 3037C>G), CCT245GAA (3038C>G 3039C>A 3040T>A), CAA246TCA (3041C>T 3042A>C), ATA248GTC (3047A>G 3049A>C), TCG249CCG (3050T>C), AAA250AAG (3055A>G), GCG252ACC (3059G>A 3061G>C), AAC254CGG (3065A>C 3066A>G 3067C>G), GTA255ACA (3068G>A 3069T>C), GAA256CGT (3071G>C 3072A>G 3073A>T), CAC257TAT (3074C>T 3076C>T), CAT259CAC (3082T>C), TTT260TAC (3084T>A 3085T>C), GAA261GAG (3088A>G), CTT263GTG (3092C>G 3094T>G), CGA264GTC (3095C>G 3096G>T 3097A>C), CCT266TCT (3101C>T), ATG267TTC (3104A>T 3106G>C), GGA268GGG (3109A>G), TTA269CTT (3110T>C 3112A>T), AAA270ACT (3114A>C 3115A>T), AAC271AAT (3118C>T), TCA272GCC (3119T>G 3121A>C), CCA273CCT (3124A>T), TCT274GCT (3125T>G), ACT275GCT (3128A>G), CAA277ATG (3134C>A 3135A>T 3136A>G), AGA278GAG (3137A>G 3138G>A 3139A>G), GTT279TTG (3140G>T 3142T>G), GAC281AAT (3146G>A 3148C>T), AAT282CAG (3149A>C 3151T>G), GTC283GTG (3154C>G), CTA284TTT (3155C>T 3157A>T), AGA285AAA (3159G>A), GGT286CCC (3161G>C 3162G>C 3163T>C), CTC287TAC (3164C>T 3165T>A), CAA288TTA (3167C>T 3168A>T), AAT289GAT (3170A>G), AAC290CAG (3173A>C 3175C>G), ATC291TTC (3176A>T), TGT292GTG (3179T>G 3180G>T 3181T>G), CTC293ATT (3182C>A 3184C>T), GTC294GTG (3187C>G), TAC295TTT (3189A>T 3190C>T), CTT296ATA (3191C>A 3193T>A), GAC297GAT (3196C>T), ATT299ATC (3202T>C), ATT300TTG (3203A>T 3205T>G), GTC301GTT (3208C>T), AGT303TCC (3212A>T 3213G>C 3214T>C), ACT304AAG (3216C>A 3217T>G), TCC305AGT (3218T>A 3219C>G 3220C>T), CTA306GAG (3221C>G 3222T>A 3223A>G), CAG307AGT (3224C>A 3225A>G 3226G>T), GAA308GAC (3229A>C), CAC309CAT (3232C>T), CTG310GTG (3233C>G), GAG311AGA (3236G>A 3237A>G 3238G>A), AAC312CAC (3239A>C), GAA314AGG (3245G>A 3246A>G 3247A>G), CGA315ATT (3248C>A 3249G>T 3250A>T), GTT316GTG (3253T>G), TTC317TTG (3256C>G), CAA318CAG (3259A>G), CTT320TTG (3263C>T 3265T>G), AGT323GAA (3272A>G 3273G>A 3274T>A), AAC324AAG (3277C>G), TTC325TTA (3280C>A), AAA326TAT (3281A>T 3283A>T), ATT327GCC (3284A>G 3285T>C 3286T>C), CAA328AAG (3287C>A 3289A>G), ATG329TTC (3290A>T 3292G>C), GAC330TCA (3293G>T 3294A>C 3295C>A), TCC332TGT (3300C>G 3301C>T), GAA333GAG (3304A>G), TTC334TTT (3307C>T), TTG335TGG (3309T>G), AAG336CTA (3311A>C 3312A>T 3313G>A), CTC337CAG (3315T>A 3316C>G), GAA338TCA (3317G>T 3318A>C), ACT339GTA (3320A>G 3321C>T 3322T>A), GCT340TCT (3323G>T), TAT341TTT (3327A>T), CTT342CTA (3331T>A), CAC344CAT (3337C>T), ATC345GTG (3338A>G 3340C>G), ATA346GTG (3341A>G 3343A>G), AGC347ACT (3345G>C 3346C>T), AGG348AAG (3348G>A), GAC349GAT (3352C>T), GGT350GGC (3355T>C), ATC351ATT (3358C>T), AAG352ATG (3360A>T), CCT353GTT (3362C>G 3363C>T), AAC354GAT (3365A>G 3367C>T), CCT355CCG (3370T>G), GAT356GCC (3372A>C 3373T>C), AAG357AAA (3376G>A), ATT358GTG (3377A>G 3379T>G), TCC359GCT (3380T>G 3382C>T), GCT360GCG (3385T>G), ATT361GTG (3386A>G 3388T>G), CAA362AGA (3389C>A 3390A>G), AAA363GAA (3392A>G), TAT364TGG (3396A>G 3397T>G), CTG365GCC (3398C>G 3399T>C 3400G>C), ATT366AGG (3402T>G 3403T>G), CCA367CCT (3406A>T), AAG368ACG (3408A>C), ACC369TCC (3410A>T), CCT370GTG (3413C>G 3414C>T 3415T>G), AAG371ACT (3417A>C 3418G>T), GAA372GAG (3421A>G), ATA373ATT (3424A>T), AAA374CGT (3425A>C 3426A>G 3427A>T), CAA375AGT (3428C>A 3429A>G 3430A>T), TTT376TTC (3433T>C), TTA377CTT (3434T>C 3436A>T), CTT379TTG (3440C>T 3442T>G), CTC380GCA (3443C>G 3444T>C 3445C>A), CGA384CGG (3457A>G), AAA385AGA (3459A>G), TTC386TTT (3463C>T), ATT387GTT (3464A>G) | | | | | | | | |

##### Proteins

| | Begin | End | Coverage | Score | Concordance | Matches | Identities | I/D/M/F* | Stop Codons |
|---|---|---|---|---|---|---|---|---|---|
| ORF B (YP_009507248.1) | 159 | 391 | 21.5% | 709 | 45.0% | 228 (97.9%) | 93 (39.9%) | 0/5/0/0 | 0 |

| | Begin | End | Coverage | Score | Concordance | Matches | Identities | I/D/M/F* | Stop Codons |
|---|---|---|---|---|---|---|---|---|---|
| **NT** | **2779** | **3476** | **9.3%** | **44** | **3.3%** | **683 (97.9%)** | **363 (52.0%)** | **0/15** | |
| Protein mutations: | M161L (2786A>T), D163K (2792G>A 2794C>G), Q164K (2795C>A 2797A>G), I166F (2801A>T), D171V (2817A>T 2818C>G), A173P (2822G>C 2824A>T), S175G (2828A>G), S176A (2831T>G), I178M (2839A>G), W179L (2841G>T), V180F (2843G>T), P182K (2849C>A 2850C>A 2851C>A), K184R (2856A>G), I185del (2858_2860delATC), A187G (2865C>G), G189del (2870_2872delGGG), K190M (2874A>T 2875A>G), Q191H (2878A>C), K192M (2880A>T), W193C (2884G>T), R194_V196del (2885_2893delCGTCTCGTA), V197I (2894G>A), F199Y (2901T>A 2902C>T), K201Q (2906A>C), E204K (2915G>A), K205V (2918A>G 2919A>T), D208K (2927G>A 2929T>G), D209N (2930G>A), I213L (2942A>T 2944A>G), N215R (2948A>C 2949A>G), S217D (2954A>G 2955G>A), V219L (2960G>T), L220F (2963C>T), K222Q (2969A>C), G224Q (2975G>C 2976G>A 2977T>G), K225E (2978A>G 2980G>A), C226A (2981T>G 2982G>C), Q227S (2984C>T 2985A>C 2986A>C), Y228E (2987T>G 2989C>G), T230S (2993A>T 2995C>T), T231K (2997C>A 2998C>G), L232I (2999T>A 3001A>C), D233G (3003A>G 3004T>C), A235R (3008G>A 3009C>G 3010A>G), F238Y (3018T>A 3019T>C), Y239H (3020T>C 3022T>C), V241L (3026G>T), E242K (3029G>A 3031G>A), M243V (3032A>G 3034G>T), D244R (3035G>C 3036A>G 3037C>G), P245E (3038C>G 3039C>A 3040T>A), Q246S (3041C>T 3042A>C), I248V (3047A>G 3049A>C), S249P (3050T>C), A252T (3059G>A 3061G>C), N254R (3065A>C 3066A>G 3067C>G), V255T (3068G>A 3069T>C), E256R (3071G>C 3072A>G 3073A>T), H257Y (3074C>T 3076C>T), F260Y (3084T>A 3085T>C), L263V (3092C>G 3094T>G), R264V (3095C>G 3096G>T 3097A>C), P266S (3101C>T), M267F (3104A>T 3106G>C), K270T (3114A>C 3115A>T), S272A (3119T>G 3121A>C), S274A (3125T>G), T275A (3128A>G), Q277M (3134C>A 3135A>T 3136A>G), R278E (3137A>G 3138G>A 3139A>G), V279L (3140G>T 3142T>G), D281N (3146G>A 3148C>T), N282Q (3149A>C 3151T>G), L284F (3155C>T 3157A>T), R285K (3159G>A), G286P (3161G>C 3162G>C 3163T>C), L287Y (3164C>T 3165T>A), Q288L (3167C>T 3168A>T), N289D (3170A>G), N290Q (3173A>C 3175C>G), I291F (3176A>T), C292V (3179T>G 3180G>T 3181T>G), L293I (3182C>A 3184C>T), Y295F (3189A>T 3190C>T), L296I (3191C>A 3193T>A), I300L (3203A>T 3205T>G), T304K (3216C>A 3217T>G), L306E (3221C>G 3222T>A 3223A>G), Q307S (3224C>A 3225A>G 3226G>T), E308D (3229A>C), L310V (3233C>G), E311R (3236G>A 3237A>G 3238G>A), N312H (3239A>C), E314R (3245G>A 3246A>G 3247A>G), R315I (3248C>A 3249G>T 3250A>T), F317L (3256C>G), S323E (3272A>G 3273G>A 3274T>A), N324K (3277C>G), F325L (3280C>A), K326Y (3281A>T 3283A>T), I327A (3284A>G 3285T>C 3286T>C), Q328K (3287C>A 3289A>G), M329F (3290A>T 3292G>C), D330S (3293G>T 3294A>C 3295C>A), S332C (3300C>G 3301C>T), L335W (3309T>G), K336L (3311A>C 3312A>T 3313G>A), L337Q (3315T>A 3316C>G), E338S (3317G>T 3318A>C), T339V (3320A>G 3321C>T 3322T>A), A340S (3323G>T), Y341F (3327A>T), I345V (3338A>G 3340C>G), I346V (3341A>G 3343A>G), S347T (3345G>C 3346C>T), R348K (3348G>A), K352M (3360A>T), P353V (3362C>G 3363C>T), N354D (3365A>G 3367C>T), D356A (3372A>C 3373T>C), I358V (3377A>G 3379T>G), S359A (3380T>G 3382C>T), I361V (3386A>G 3388T>G), Q362R (3389C>A 3390A>G), K363E (3392A>G), Y364W (3396A>G 3397T>G), L365A (3398C>G 3399T>C 3400G>C), I366R (3402T>G 3403T>G), K368T (3408A>C), T369S (3410A>T), P370V (3413C>G 3414C>T 3415T>G), K371T (3417A>C 3418G>T), K374R (3425A>C 3426A>G 3427A>T), Q375S (3428C>A 3429A>G 3430A>T), L380A (3443C>G 3444T>C 3445C>A), K385R (3459A>G), I387V (3464A>G) | | | | | | | | |
| Codon mutations: | ATG161TTG (2786A>T), TTG162TTA (2791G>A), GAC163AAG (2792G>A 2794C>G), CAA164AAG (2795C>A 2797A>G), ATT166TTT (2801A>T), ATA167ATC (2806A>C), AGA168AGG (2809A>G), CCA169CCT (2812A>T), TCA170AGT (2813T>A 2814C>G 2815A>T), GAC171GTG (2817A>T 2818C>G), TCT172TCA (2821T>A), GCA173CCT (2822G>C 2824A>T), AGC175GGC (2828A>G), TCA176GCA (2831T>G), CCC177CCT (2836C>T), ATA178ATG (2839A>G), TGG179TTG (2841G>T), GTT180TTT (2843G>T), CCC182AAA (2849C>A 2850C>A 2851C>A), AAA184AGA (2856A>G), ATC185del (2858_2860delATC), GAC186GAT (2863C>T), GCT187GGT (2865C>G), GGG189del (2870_2872delGGG), AAA190ATG (2874A>T 2875A>G), CAA191CAC (2878A>C), AAG192ATG (2880A>T), TGG193TGT (2884G>T), CGT194_GTA196del (2885_2893delCGTCTCGTA), GTT197ATT (2894G>A), GAC198GAT (2899C>T), TTC199TAT (2901T>A 2902C>T), CGT200CGC (2905T>C), AAG201CAG (2906A>C), GAG204AAG (2915G>A), AAG205GTG (2918A>G 2919A>T), GAT208AAG (2927G>A 2929T>G), GAC209AAC (2930G>A), AAA210AAG (2935A>G), TAC211TAT (2938C>T), CCG212CCC (2941G>C), ATA213TTG (2942A>T 2944A>G), CCA214CCC (2947A>C), AAC215CGC (2948A>C 2949A>G), ATA216ATT (2953A>T), AGT217GAT (2954A>G 2955G>A), GTA219TTA (2960G>T), CTT220TTT (2963C>T), AAG222CAG (2969A>C), TTA223CTA (2972T>C), GGT224CAG (2975G>C 2976G>A 2977T>G), AAG225GAA (2978A>G 2980G>A), TGC226GCC (2981T>G 2982G>C), CAA227TCC (2984C>T 2985A>C 2986A>C), TAC228GAG (2987T>G 2989C>G), TTC229TTT (2992C>T), ACC230TCT (2993A>T 2995C>T), ACC231AAG (2997C>A 2998C>G), TTA232ATC (2999T>A 3001A>C), GAT233GGC (3003A>G 3004T>C), GCA235AGG (3008G>A 3009C>G 3010A>G), AGT236TCG (3011A>T 3012G>C 3013T>G), GGG237GGC (3016G>C), TTT238TAC (3018T>A 3019T>C), TAT239CAC (3020T>C 3022T>C), GTG241TTG (3026G>T), GAG242AAA (3029G>A 3031G>A), ATG243GTT (3032A>G 3034G>T), GAC244CGG (3035G>C 3036A>G 3037C>G), CCT245GAA (3038C>G 3039C>A 3040T>A), CAA246TCA (3041C>T 3042A>C), ATA248GTC (3047A>G 3049A>C), TCG249CCG (3050T>C), AAA250AAG (3055A>G), GCG252ACC (3059G>A 3061G>C), AAC254CGG (3065A>C 3066A>G 3067C>G), GTA255ACA (3068G>A 3069T>C), GAA256CGT (3071G>C 3072A>G 3073A>T), CAC257TAT (3074C>T 3076C>T), CAT259CAC (3082T>C), TTT260TAC (3084T>A 3085T>C), GAA261GAG (3088A>G), CTT263GTG (3092C>G 3094T>G), CGA264GTC (3095C>G 3096G>T 3097A>C), CCT266TCT (3101C>T), ATG267TTC (3104A>T 3106G>C), GGA268GGG (3109A>G), TTA269CTT (3110T>C 3112A>T), AAA270ACT (3114A>C 3115A>T), AAC271AAT (3118C>T), TCA272GCC (3119T>G 3121A>C), CCA273CCT (3124A>T), TCT274GCT (3125T>G), ACT275GCT (3128A>G), CAA277ATG (3134C>A 3135A>T 3136A>G), AGA278GAG (3137A>G 3138G>A 3139A>G), GTT279TTG (3140G>T 3142T>G), GAC281AAT (3146G>A 3148C>T), AAT282CAG (3149A>C 3151T>G), GTC283GTG (3154C>G), CTA284TTT (3155C>T 3157A>T), AGA285AAA (3159G>A), GGT286CCC (3161G>C 3162G>C 3163T>C), CTC287TAC (3164C>T 3165T>A), CAA288TTA (3167C>T 3168A>T), AAT289GAT (3170A>G), AAC290CAG (3173A>C 3175C>G), ATC291TTC (3176A>T), TGT292GTG (3179T>G 3180G>T 3181T>G), CTC293ATT (3182C>A 3184C>T), GTC294GTG (3187C>G), TAC295TTT (3189A>T 3190C>T), CTT296ATA (3191C>A 3193T>A), GAC297GAT (3196C>T), ATT299ATC (3202T>C), ATT300TTG (3203A>T 3205T>G), GTC301GTT (3208C>T), AGT303TCC (3212A>T 3213G>C 3214T>C), ACT304AAG (3216C>A 3217T>G), TCC305AGT (3218T>A 3219C>G 3220C>T), CTA306GAG (3221C>G 3222T>A 3223A>G), CAG307AGT (3224C>A 3225A>G 3226G>T), GAA308GAC (3229A>C), CAC309CAT (3232C>T), CTG310GTG (3233C>G), GAG311AGA (3236G>A 3237A>G 3238G>A), AAC312CAC (3239A>C), GAA314AGG (3245G>A 3246A>G 3247A>G), CGA315ATT (3248C>A 3249G>T 3250A>T), GTT316GTG (3253T>G), TTC317TTG (3256C>G), CAA318CAG (3259A>G), CTT320TTG (3263C>T 3265T>G), AGT323GAA (3272A>G 3273G>A 3274T>A), AAC324AAG (3277C>G), TTC325TTA (3280C>A), AAA326TAT (3281A>T 3283A>T), ATT327GCC (3284A>G 3285T>C 3286T>C), CAA328AAG (3287C>A 3289A>G), ATG329TTC (3290A>T 3292G>C), GAC330TCA (3293G>T 3294A>C 3295C>A), TCC332TGT (3300C>G 3301C>T), GAA333GAG (3304A>G), TTC334TTT (3307C>T), TTG335TGG (3309T>G), AAG336CTA (3311A>C 3312A>T 3313G>A), CTC337CAG (3315T>A 3316C>G), GAA338TCA (3317G>T 3318A>C), ACT339GTA (3320A>G 3321C>T 3322T>A), GCT340TCT (3323G>T), TAT341TTT (3327A>T), CTT342CTA (3331T>A), CAC344CAT (3337C>T), ATC345GTG (3338A>G 3340C>G), ATA346GTG (3341A>G 3343A>G), AGC347ACT (3345G>C 3346C>T), AGG348AAG (3348G>A), GAC349GAT (3352C>T), GGT350GGC (3355T>C), ATC351ATT (3358C>T), AAG352ATG (3360A>T), CCT353GTT (3362C>G 3363C>T), AAC354GAT (3365A>G 3367C>T), CCT355CCG (3370T>G), GAT356GCC (3372A>C 3373T>C), AAG357AAA (3376G>A), ATT358GTG (3377A>G 3379T>G), TCC359GCT (3380T>G 3382C>T), GCT360GCG (3385T>G), ATT361GTG (3386A>G 3388T>G), CAA362AGA (3389C>A 3390A>G), AAA363GAA (3392A>G), TAT364TGG (3396A>G 3397T>G), CTG365GCC (3398C>G 3399T>C 3400G>C), ATT366AGG (3402T>G 3403T>G), CCA367CCT (3406A>T), AAG368ACG (3408A>C), ACC369TCC (3410A>T), CCT370GTG (3413C>G 3414C>T 3415T>G), AAG371ACT (3417A>C 3418G>T), GAA372GAG (3421A>G), ATA373ATT (3424A>T), AAA374CGT (3425A>C 3426A>G 3427A>T), CAA375AGT (3428C>A 3429A>G 3430A>T), TTT376TTC (3433T>C), TTA377CTT (3434T>C 3436A>T), CTT379TTG (3440C>T 3442T>G), CTC380GCA (3443C>G 3444T>C 3445C>A), CGA384CGG (3457A>G), AAA385AGA (3459A>G), TTC386TTT (3463C>T), ATT387GTT (3464A>G) | | | | | | | | |

*: Inserts / Deletes / Misaligned / Frameshifts

#### Analysis details

This analysis was performed with panviral2.64

### NGS Details (UN8): Human gammaherpesvirus 8

#### Assembly

| | |
|---|---|
| **Coverage Length** | 135 (1 contig(s)) |
| **Depth Of Coverage** | 228.3 |
| **Number Of Reads** | 368 |
| **Reads Per Million** | 6.15 rpm (after QC) |
| **Ambiguities** | 0 |
| **Assembly Method** | de novo + reference guided assembly |
| **Consensus Caller** | Bcf Tools |

#### Coverage Map

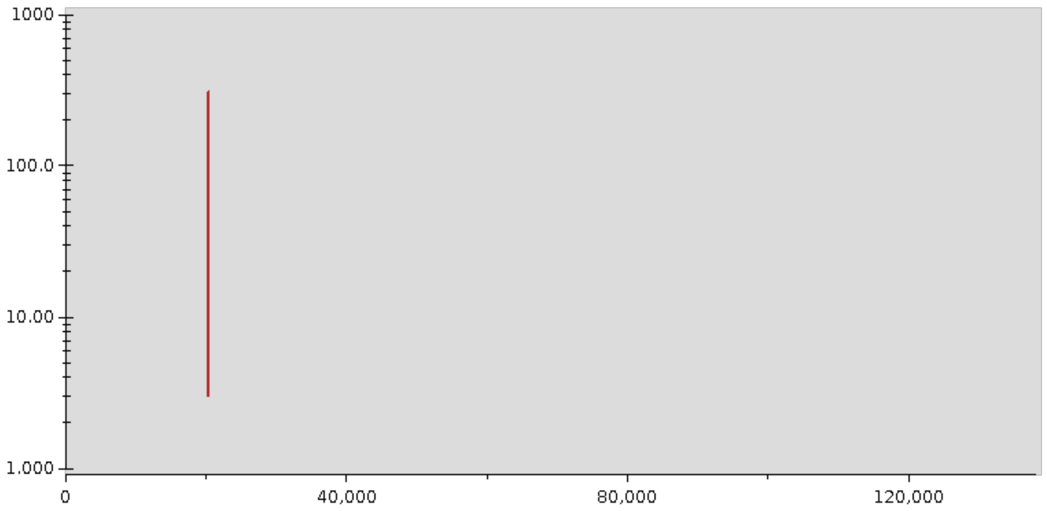

#### Assignment

| | |
|---|---|
| **Type** | Human gammaherpesvirus 8 (Taxonomy ID: 37296) |
| **Subtype** | Could not assign |
| **Reference Genome** | NC_009333.1 |
| **NT Identity (%)** | 83.7037 |
| **AA Identity (%)** | 84.4444 |
| **Number Of Stop Codons** | 0 |
| **Number Of CDS** | 86 |

#### Alignment

| | |
|---|---|
| **Alignment Score** | 182.0 (NT) + 287.0 (AA) = 469.0 |
| **Concordance (%)** | 78.2972 |
| **Alignment Method** | Local, heuristic, nucleotide (BLASTN) |

#### Genome Region

Sequence starts at position 20273 and ends at position 20407 relative to NC_009333.1 reference sequence.

#### Alignment Detailed Statistics

| | Begin | End | Coverage | Score | Concordance | Matches | Identities | I/D/M/F* | Stop Codons |
|---|---|---|---|---|---|---|---|---|---|
| **NT** | **20273** | **20407** | **0.1%** | **182** | **67.4%** | **135 (100%)** | **113 (83.7%)** | **0/0** | |

Mutations: 20293A>T, 20296A>C, 20311G>A, 20314T>G, 20315C>G, 20317C>A, 20320C>T, 20322T>G, 20331G>T, 20341A>C, 20347A>C, 20350G>A, 20352C>T, 20353A>G, 20369C>G, 20370A>C, 20371T>A, 20376A>T, 20383G>A, 20386C>T, 20389A>G, 20392C>T

*: Inserts / Deletes / Misaligned / Frameshifts

#### Analysis details

This analysis was performed with panviral2.64

### NGS Details (UN8): Epiphyllum badnavirus 1

#### Assembly

| | |
|---|---|
| **Coverage Length** | 256 (1 contig(s)) |
| **Depth Of Coverage** | 148.9 |
| **Number Of Reads** | 360 |
| **Reads Per Million** | 6.02 rpm (after QC) |
| **Ambiguities** | 0 |
| **Assembly Method** | de novo + reference guided assembly |
| **Consensus Caller** | Bcf Tools |

#### Coverage Map

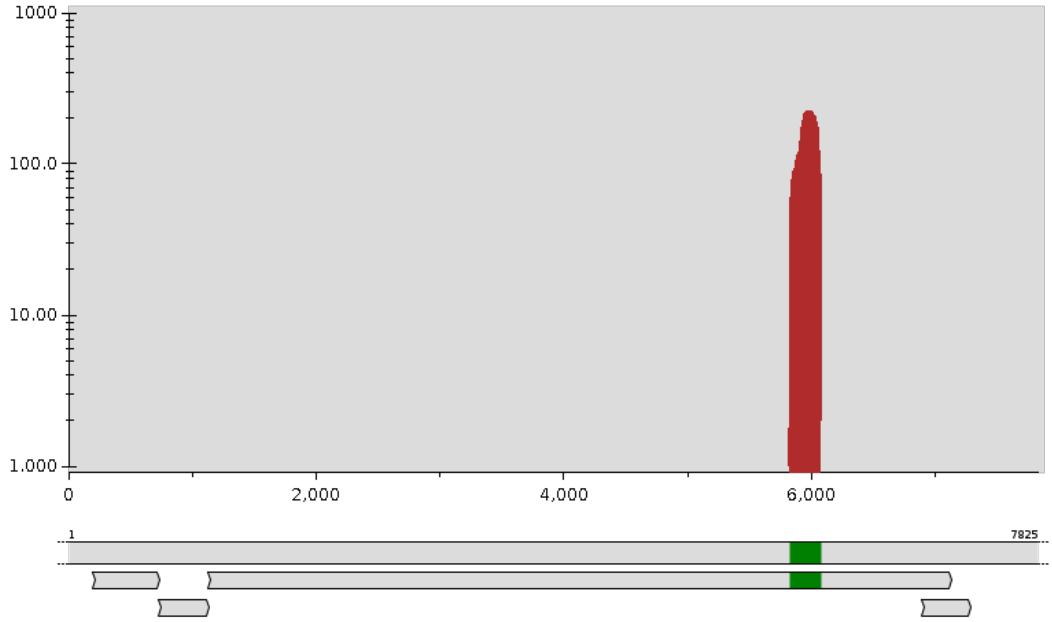

#### Assignment

| | |
|---|---|
| **Type** | Epiphyllum badnavirus 1 (Taxonomy ID: 2518008) |
| **Reference Genome** | NC_076247.1 |
| **NT Identity (%)** | 57.529 |
| **AA Identity (%)** | 56.9767 |
| **Number Of Stop Codons** | 0 |
| **Number Of CDS** | 4 |

#### Alignment

| | |
|---|---|
| **Alignment Score** | 72.0 (NT) + 324.0 (AA) = 396.0 |
| **Concordance (%)** | 34.7673 |

| Alignment Method | Global, seeded, nucleotide + amino acids (AGA) |
|---|---|

#### Genome Region

Sequence starts at position 5823 and ends at position 6078 relative to NC_076247.1 reference sequence.

#### Alignment Detailed Statistics

| | Begin | End | Coverage | Score | Concordance | Matches | Identities | I/D/M/F* | Stop Codons |
|---|---|---|---|---|---|---|---|---|---|
| **NT** | **5823** | **6078** | **3.3%** | **72** | **14.1%** | **256 (98.8%)** | **149 (57.5%)** | **3/0** | |
| Mutations: | 5832T>G, 5835C>T, 5839G>T, 5840T>G, 5847C>A, 5851T>A, 5853C>A, 5859G>T, 5860A>C, 5861A>G, 5862G>T, 5863A>T, 5864G>C, 5865C>G, 5868A>T, 5870T>A, 5874C>T, 5878A>T, 5880T>G, 5882T>A, 5884A>G, 5886G>C, 5887G>A, 5888A>G, 5889C>G, 5892A>G, 5893G>A, 5894A>G, 5895A>T, 5896T>G, 5897C>A, 5898C>T, 5905T>A, 5906G>A, 5910C>A, 5913C>G, 5918T>G, 5922C>A, 5923C>A, 5924C>G, 5925A>G, 5926G>T, 5933T>A, 5936A>T, 5937C>T, 5940A>G, 5942G>T, 5943G>T, 5944T>C, 5946A>T, 5949T>C, 5953C>T, 5955A>C, 5958C>T, 5961A>T, 5964G>A, 5966A>C, 5967G>T, 5970T>C, 5973C>T, 5976T>G, 5982G>C, 5985C>T, 5986C>A, 5987A>T, 5988A>G, 5989C>G, 5990G>A, 5991A>T, 5992A>C, 5993A>T, 5994G>T, 5998G>A, 6000C>T, 6002A>G, 6003C>G, 6004T>A, 6005G>T, 6006T>A, 6011G>A, 6012A>G, 6012_6013insCCT, 6013G>T, 6018G>T, 6019T>G, 6020C>A, 6021A>C, 6022G>A, 6023G>T, 6024C>G, 6026T>A, 6030A>G, 6032C>T, 6033A>G, 6036T>G, 6038A>T, 6039C>T, 6054G>A, 6059T>A, 6060C>T, 6061A>T, 6062G>C, 6063T>G, 6065A>G, 6066C>A, 6067A>G, 6068C>A, 6069T>C | | | | | | | | |
| **CDS** | | | | | | | | | |
| QKM20_gp3 | 1566 | 1650 | 4.2% | 324 | 51.3% | 85 (98.8%) | 49 (57.0%) | 1/0/0/0 | 0 |
| Protein mutations: | N1568K (5832T>G), V1571C (5839G>T 5840T>G), F1575I (5851T>A 5853C>A), K1578R (5860A>C 5861A>G 5862G>T), F1581Y (5870T>A), I1584L (5878A>T 5880T>G), M1585K (5882T>A), M1586V (5884A>G 5886G>C), D1587R (5887G>A 5888A>G 5889C>G), E1589S (5893G>A 5894A>G 5895A>T), S1590D (5896T>G 5897C>A 5898C>T), W1593K (5905T>A 5906G>A), L1597R (5918T>G), P1599R (5923C>A 5924C>G 5925A>G), D1600Y (5926G>T), L1602H (5933T>A), Y1603F (5936A>T 5937C>T), W1605F (5942G>T 5943G>T), P1609S (5953C>T 5955A>C), K1613T (5966A>C 5967G>T), Q1620M (5986C>A 5987A>T 5988A>G), R1621D (5989C>G 5990G>A 5991A>T), K1622L (5992A>C 5993A>T 5994G>T), D1624N (5998G>A 6000C>T), N1625R (6002A>G 6003C>G), C1626I (6004T>A 6005G>T 6006T>A), R1628K (6011G>A 6012A>G), R1628_D1629insP (6012_6013insCCT), D1629Y (6013G>T), S1631D (6019T>G 6020C>A 6021A>C), G1632M (6022G>A 6023G>T 6024C>G), F1633Y (6026T>A), A1635V (6032C>T 6033A>G), Y1637F (6038A>T 6039C>T), F1644Y (6059T>A 6060C>T), N1646R (6065A>G 6066C>A), T1647D (6067A>G 6068C>A 6069T>C) | | | | | | | | |
| Codon mutations: | AAT1568AAG (5832T>G), GCC1569GCT (5835C>T), GTC1571TGC (5839G>T 5840T>G), TCC1573TCA (5847C>A), TTC1575ATA (5851T>A 5853C>A), CTG1577CTT (5859G>T), AAG1578CGT (5860A>C 5861A>G 5862G>T), AGC1579TCG (5863A>T 5864G>C 5865C>G), GGA1580GGT (5868A>T), TTC1581TAC (5870T>A), CAC1582CAT (5874C>T), ATT1584TTG (5878A>T 5880T>G), ATG1585AAG (5882T>A), ATG1586GTC (5884A>G 5886G>C), GAC1587AGG (5887G>A 5888A>G 5889C>G), GAA1588GAG (5892A>G), GAA1589AGT (5893G>A 5894A>G 5895A>T), TCC1590GAT (5896T>G 5897C>A 5898C>T), TGG1593AAG (5905T>A 5906G>A), ACC1594ACA (5910C>A), GCC1595GCG (5913C>G), CTG1597CGG (5918T>G), ACC1598ACA (5922C>A), CCA1599AGG (5923C>A 5924C>G 5925A>G), GAT1600TAT (5926G>T), CTT1602CAT (5933T>A), TAC1603TTT (5936A>T 5937C>T), GAA1604GAG (5940A>G), TGG1605TTT (5942G>T 5943G>T), TTA1606CTT (5944T>C 5946A>T), GTT1607GTC (5949T>C), CCA1609TCC (5953C>T 5955A>C), TTC1610TTT (5958C>T), GGA1611GGT (5961A>T), TTG1612TTA (5964G>A), AAG1613ACT (5966A>C 5967G>T), AAT1614AAC (5970T>C), GCC1615GCT (5973C>T), CCT1616CCG (5976T>G), GTG1618GTC (5982G>C), TTC1619TTT (5985C>T), CAA1620ATG (5986C>A 5987A>T 5988A>G), CGA1621GAT (5989C>G 5990G>A 5991A>T), AAG1622CTT (5992A>C 5993A>T 5994G>T), GAC1624AAT (5998G>A 6000C>T), AAC1625AGG (6002A>G 6003C>G), TGT1626ATA (6004T>A 6005G>T 6006T>A), AGA1628AAG (6011G>A 6012A>G), AGA1628_GAC1629insCCT (6012_6013insCCT), GAC1629TAC (6013G>T), CTG1630CTT (6018G>T), TCA1631GAC (6019T>G 6020C>A 6021A>C), GGC1632ATG (6022G>A 6023G>T 6024C>G), TTT1633TAT (6026T>A), GTA1634GTG (6030A>G), GCA1635GTG (6032C>T 6033A>G), GTT1636GTG (6036T>G), TAC1637TTT (6038A>T 6039C>T), CTG1642CTA (6054G>A), TTC1644TAT (6059T>A 6060C>T), AGT1645TCG (6061A>T 6062G>C 6063T>G), AAC1646AGA (6065A>G 6066C>A), ACT1647GAC (6067A>G 6068C>A 6069T>C) | | | | | | | | |
| **Proteins** | | | | | | | | | |
| polyprotein (YP_010797894.1) | 1566 | 1650 | 4.2% | 324 | 51.3% | 85 (98.8%) | 49 (57.0%) | 1/0/0/0 | 0 |
| Protein mutations: | N1568K (5832T>G), V1571C (5839G>T 5840T>G), F1575I (5851T>A 5853C>A), K1578R (5860A>C 5861A>G 5862G>T), F1581Y (5870T>A), I1584L (5878A>T 5880T>G), M1585K (5882T>A), M1586V (5884A>G 5886G>C), D1587R (5887G>A 5888A>G 5889C>G), E1589S (5893G>A 5894A>G 5895A>T), S1590D (5896T>G 5897C>A 5898C>T), W1593K (5905T>A 5906G>A), L1597R (5918T>G), P1599R (5923C>A 5924C>G 5925A>G), D1600Y (5926G>T), L1602H (5933T>A), Y1603F (5936A>T 5937C>T), W1605F (5942G>T 5943G>T), P1609S (5953C>T 5955A>C), K1613T (5966A>C 5967G>T), Q1620M (5986C>A 5987A>T 5988A>G), R1621D (5989C>G 5990G>A 5991A>T), K1622L (5992A>C 5993A>T 5994G>T), D1624N (5998G>A 6000C>T), N1625R (6002A>G 6003C>G), C1626I (6004T>A 6005G>T 6006T>A), R1628K (6011G>A 6012A>G), R1628_D1629insP (6012_6013insCCT), D1629Y (6013G>T), S1631D (6019T>G 6020C>A 6021A>C), G1632M (6022G>A 6023G>T 6024C>G), F1633Y (6026T>A), A1635V (6032C>T 6033A>G), Y1637F (6038A>T 6039C>T), F1644Y (6059T>A 6060C>T), N1646R (6065A>G 6066C>A), T1647D (6067A>G 6068C>A 6069T>C) | | | | | | | | |
| Codon mutations: | AAT1568AAG (5832T>G), GCC1569GCT (5835C>T), GTC1571TGC (5839G>T 5840T>G), TCC1573TCA (5847C>A), TTC1575ATA (5851T>A 5853C>A), CTG1577CTT (5859G>T), AAG1578CGT (5860A>C 5861A>G 5862G>T), AGC1579TCG (5863A>T 5864G>C 5865C>G), GGA1580GGT (5868A>T), TTC1581TAC (5870T>A), CAC1582CAT (5874C>T), ATT1584TTG (5878A>T 5880T>G), ATG1585AAG (5882T>A), ATG1586GTC (5884A>G 5886G>C), GAC1587AGG (5887G>A 5888A>G 5889C>G), GAA1588GAG (5892A>G), GAA1589AGT (5893G>A 5894A>G 5895A>T), TCC1590GAT (5896T>G 5897C>A 5898C>T), TGG1593AAG (5905T>A 5906G>A), ACC1594ACA (5910C>A), GCC1595GCG (5913C>G), CTG1597CGG (5918T>G), ACC1598ACA (5922C>A), CCA1599AGG (5923C>A 5924C>G 5925A>G), GAT1600TAT (5926G>T), CTT1602CAT (5933T>A), TAC1603TTT (5936A>T 5937C>T), GAA1604GAG (5940A>G), TGG1605TTT (5942G>T 5943G>T), TTA1606CTT (5944T>C 5946A>T), GTT1607GTC (5949T>C), CCA1609TCC (5953C>T 5955A>C), TTC1610TTT (5958C>T), GGA1611GGT (5961A>T), TTG1612TTA (5964G>A), AAG1613ACT (5966A>C 5967G>T), AAT1614AAC (5970T>C), GCC1615GCT (5973C>T), CCT1616CCG (5976T>G), GTG1618GTC (5982G>C), TTC1619TTT (5985C>T), CAA1620ATG (5986C>A 5987A>T 5988A>G), CGA1621GAT (5989C>G 5990G>A 5991A>T), AAG1622CTT (5992A>C 5993A>T 5994G>T), GAC1624AAT (5998G>A 6000C>T), AAC1625AGG (6002A>G 6003C>G), TGT1626ATA (6004T>A 6005G>T 6006T>A), AGA1628AAG (6011G>A 6012A>G), AGA1628_GAC1629insCCT (6012_6013insCCT), GAC1629TAC (6013G>T), CTG1630CTT (6018G>T), TCA1631GAC (6019T>G 6020C>A 6021A>C), GGC1632ATG (6022G>A 6023G>T 6024C>G), TTT1633TAT (6026T>A), GTA1634GTG (6030A>G), GCA1635GTG (6032C>T 6033A>G), GTT1636GTG (6036T>G), TAC1637TTT (6038A>T 6039C>T), CTG1642CTA (6054G>A), TTC1644TAT (6059T>A 6060C>T), AGT1645TCG (6061A>T 6062G>C 6063T>G), AAC1646AGA (6065A>G 6066C>A), ACT1647GAC (6067A>G 6068C>A 6069T>C) | | | | | | | | |

*: Inserts / Deletes / Misaligned / Frameshifts

#### Analysis details

This analysis was performed with panviral2.64

#### NGS Details (UN8): Cassava brown streak virus

##### Assembly

| | |
|---|---|
| **Coverage Length** | 527 (1 contig(s)) |
| **Depth Of Coverage** | 70.8 |
| **Number Of Reads** | 308 |
| **Reads Per Million** | 5.15 rpm (after QC) |
| **Ambiguities** | 0 |
| **Assembly Method** | de novo + reference guided assembly |
| **Consensus Caller** | Bcf Tools |

##### Coverage Map

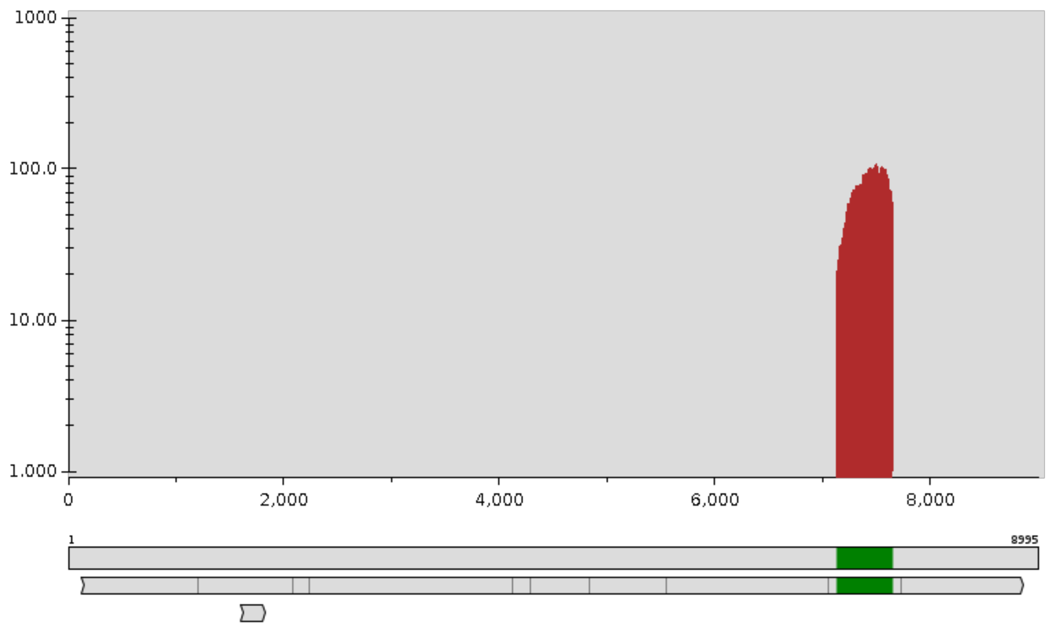

##### Assignment

| | |
|---|---|
| **Type** | Cassava brown streak virus (Taxonomy ID: 137758) |
| **Reference Genome** | NC_012698.2 |
| **NT Identity (%)** | 60.9108 |
| **AA Identity (%)** | 57.9545 |
| **Number Of Stop Codons** | 0 |
| **Number Of CDS** | 2 |

##### Alignment

| | |
|---|---|
| **Alignment Score** | 230.0 (NT) + 742.0 (AA) = 972.0 |
| **Concordance (%)** | 42.7253 |

| Alignment Method | Global, seeded, nucleotide + amino acids (AGA) |
|---|---|

#### Genome Region

Sequence starts at position 7124 and ends at position 7650 relative to NC_012698.2 reference sequence.

#### Alignment Detailed Statistics

| | Begin | End | Coverage | Score | Concordance | Matches | Identities | I/D/M/F* | Stop Codons |
|---|---|---|---|---|---|---|---|---|---|
| **NT** | **7124** | **7650** | **5.9%** | **230** | **21.8%** | **527 (100%)** | **321 (60.9%)** | **0/0** | |
| Mutations: | 7130A>G, 7133A>G, 7134A>G, 7136T>G, 7142C>T, 7145A>T, 7148A>C, 7151G>A, 7157A>T, 7158C>A, 7163A>G, 7167A>G, 7175A>T, 7177A>G, 7178A>G, 7179C>G, 7180A>C, 7184A>T, 7185T>C, 7190T>C, 7192C>A, 7193C>G, 7194A>T, 7202A>C, 7203A>T, 7205C>T, 7206A>C, 7207T>A, 7208C>G, 7211T>C, 7213G>T, 7214G>T, 7217A>G, 7218G>C, 7229A>T, 7234C>T, 7235A>T, 7241A>G, 7242A>G, 7243C>A, 7244A>G, 7245G>C, 7246T>C, 7250G>A, 7253A>T, 7256T>A, 7257A>T, 7258T>C, 7270T>C, 7271A>T, 7272C>A, 7273G>A, 7274T>A, 7275G>A, 7277A>T, 7280G>T, 7281T>G, 7283T>C, 7284G>A, 7286G>A, 7287T>G, 7288T>A, 7290A>G, 7292T>G, 7293G>A, 7294G>A, 7295A>T, 7298G>A, 7304T>G, 7307C>A, 7310A>T, 7313A>G, 7319A>T, 7320A>T, 7323T>C, 7325G>C, 7331T>C, 7332G>A, 7334T>C, 7337T>C, 7340C>T, 7343T>G, 7355A>G, 7358A>T, 7361C>T, 7364T>C, 7374A>T, 7377G>C, 7380G>A, 7381G>A, 7385T>C, 7388A>T, 7390T>A, 7391A>T, 7397A>T, 7398T>C, 7400A>T, 7401T>A, 7403T>C, 7406G>C, 7409G>A, 7410G>T, 7413G>A, 7414A>T, 7416C>G, 7418G>T, 7422C>G, 7424A>G, 7425A>G, 7432T>C, 7433G>A, 7436T>A, 7437A>T, 7438G>A, 7442T>C, 7443C>A, 7445C>G, 7448C>T, 7449G>A, 7451G>C, 7455G>T, 7457T>A, 7458T>C, 7462T>C, 7463A>T, 7464A>C, 7465A>T, 7467A>G, 7468A>G, 7469A>G, 7470G>C, 7471T>C, 7472T>A, 7473G>A, 7474G>A, 7476G>A, 7477A>C, 7481T>G, 7484T>G, 7487A>G, 7489T>C, 7490C>T, 7494A>G, 7495A>T, 7496G>T, 7499T>A, 7500G>A, 7501T>A, 7502G>A, 7503C>A, 7504T>C, 7505A>G, 7506A>C, 7507G>T, 7508A>G, 7511T>A, 7512G>A, 7517T>A, 7520A>G, 7521A>C, 7522T>C, 7523G>A, 7524C>G, 7526A>T, 7527C>A, 7529C>G, 7535A>C, 7539T>A, 7540C>A, 7541A>T, 7544T>C, 7547G>A, 7553C>T, 7556A>G, 7559T>A, 7562C>T, 7568A>T, 7570T>A, 7572A>G, 7573A>G, 7576G>A, 7577G>T, 7578A>G, 7579G>A, 7580A>C, 7581A>C, 7586A>T, 7588T>A, 7589T>C, 7599A>C, 7600T>C, 7601G>C, 7602G>A, 7603C>A, 7604C>G, 7607G>A, 7610G>A, 7616T>C, 7618T>A, 7622A>T, 7628T>C, 7631A>C, 7632T>G, 7633T>G, 7635C>A, 7636G>A | | | | | | | | |

##### CDS

| | Begin | End | Coverage | Score | Concordance | Matches | Identities | I/D/M/F* | Stop Codons |
|---|---|---|---|---|---|---|---|---|---|
| CBSV_gp1 | 2334 | 2509 | 6.0% | 742 | 60.2% | 176 (100%) | 102 (58.0%) | 0/0/0/0 | 0 |
| Protein mutations: | I2337V (7134A>G 7136T>G), Q2345K (7158C>A), K2348E (7167A>G), K2351R (7177A>G 7178A>G), Q2352A (7179C>G 7180A>C), F2354L (7185T>C), P2356Q (7192C>A 7193C>G), T2357S (7194A>T), I2360F (7203A>T 7205C>T), I2361Q (7206A>C 7207T>A 7208C>G), R2363L (7213G>T 7214G>T), V2365L (7218G>C), P2370L (7234C>T 7235A>T), T2373E (7242A>G 7243C>A 7244A>G), V2374P (7245G>C 7246T>C), E2376D (7253A>T), I2378S (7257A>T 7258T>C), V2382A (7270T>C 7271A>T), R2383K (7272C>A 7273G>A 7274T>A), V2384I (7275G>A 7277A>T), S2386A (7281T>G 7283T>C), E2387K (7284G>A 7286G>A), L2388E (7287T>G 7288T>A), I2389V (7290A>G 7292T>G), G2390N (7293G>A 7294G>A 7295A>T), S2399C (7320A>T), D2403N (7332G>A 7334T>C), N2406K (7343T>G), M2417L (7374A>T), E2418Q (7377G>C), G2419K (7380G>A 7381G>A), L2422H (7390T>A 7391A>T), Y2426N (7401T>A 7403T>C), K2427N (7406G>C), V2429L (7410G>T), E2430M (7413G>A 7414A>T), P2431A (7416C>G 7418G>T), Q2433E (7422C>G 7424A>G), N2434D (7425A>G), M2436T (7432T>C 7433G>A), S2438Y (7437A>T 7438G>A), L2440M (7443C>A 7445C>G), V2442I (7449G>A 7451G>C), A2444S (7455G>T 7457T>A), F2445L (7458T>C), V2446A (7462T>C 7463A>T), N2447L (7464A>C 7465A>T), K2448G (7467A>G 7468A>G 7469A>G), V2449P (7470G>C 7471T>C 7472T>A), G2450N (7473G>A 7474G>A), D2451T (7476G>A 7477A>C), D2452E (7481T>G), I2454M (7487A>G), I2455T (7489T>C 7490C>T), K2457V (7494A>G 7495A>T 7496G>T), V2459K (7500G>A 7501T>A 7502G>A), L2460T (7503C>A 7504T>C 7505A>G), R2461L (7506A>C 7507G>T 7508A>G), E2463K (7512G>A), M2466P (7521A>C 7522T>C 7523G>A), P2467A (7524C>G 7526A>T), S2472N (7539T>A 7540C>A 7541A>T), L2482H (7570T>A), N2483G (7572A>G 7573A>G), W2484Y (7576G>A 7577G>T), R2485D (7578A>G 7579G>A 7580A>C), K2486Q (7581A>C), F2488Y (7588T>A 7589T>C), M2492P (7599A>C 7600T>C 7601G>C), A2493K (7602G>A 7603C>A 7604C>G), M2498K (7618T>A), F2503G (7632T>G 7633T>G), R2504K (7635C>A 7636G>A) | | | | | | | | |
| Codon mutations: | GCA2335GCG (7130A>G), CCA2336CCG (7133A>G), ATT2337GTG (7134A>G 7136T>G), TTC2339TTT (7142C>T), GTA2340GTT (7145A>T), ACA2341ACC (7148A>C), GGG2342GGA (7151G>A), GCA2344GCT (7157A>T), CAA2345AAA (7158C>A), AAA2346AAG (7163A>G), AAG2348GAG (7167A>G), GTA2350GTT (7175A>T), AAA2351AGG (7177A>G 7178A>G), CAA2352GCA (7179C>G 7180A>C), ATA2353ATT (7184A>T), TTT2354CTT (7185T>C), GGT2355GGC (7190T>C), CCC2356CAG (7192C>A 7193C>G), ACT2357TCT (7194A>T), CCA2359CCC (7202A>C), ATC2360TTT (7203A>T 7205C>T), ATC2361CAG (7206A>C 7207T>A 7208C>G), TCT2362TCC (7211T>C), CGG2363CTT (7213G>T 7214G>T), AAA2364AAG (7217A>G), GTT2365CTT (7218G>C), CCA2368CCT (7229A>T), CCA2370CTT (7234C>T 7235A>T), GGA2372GGG (7241A>G), ACA2373GAG (7242A>G 7243C>A 7244A>G), GTT2374CCT (7245G>C 7246T>C), GAG2375GAA (7250G>A), GAA2376GAT (7253A>T), ATT2377ATA (7256T>A), ATC2378TCC (7257A>T 7258T>C), GTA2382GCT (7270T>C 7271A>T), CGT2383AAA (7272C>A 7273G>A 7274T>A), GTA2384ATT (7275G>A 7277A>T), GCG2385GCT (7280G>T), TCT2386GCC (7281T>G 7283T>C), GAG2387AAA (7284G>A 7286G>A), TTG2388GAG (7287T>G 7288T>A), ATT2389GTG (7290A>G 7292T>G), GGA2390AAT (7293G>A 7294G>A 7295A>T), GGG2391GGA (7298G>A), GTT2393GTG (7304T>G), CTC2394CTA (7307C>A), GTA2395GTT (7310A>T), GAA2396GAG (7313A>G), ACA2398ACT (7319A>T), AGT2399TGT (7320A>T), TTG2400CTC (7323T>C 7325G>C), TTT2402TTC (7331T>C), GAT2403AAC (7332G>A 7334T>C), GCT2404GCC (7337T>C), CTC2405CTT (7340C>T), AAT2406AAG (7343T>G), GGA2410GGG (7355A>G), CCA2411CCT (7358A>T), TAC2412TAT (7361C>T), ATT2413ATC (7364T>C), ATG2417TTG (7374A>T), GAA2418CAA (7377G>C), GGG2419AAG (7380G>A 7381G>A), ATT2420ATC (7385T>C), GGA2421GGT (7388A>T), CTA2422CAT (7390T>A 7391A>T), GGA2424GGT (7397A>T), TTA2425CTT (7398T>C 7400A>T), TAT2426AAC (7401T>A 7403T>C), AAG2427AAC (7406G>C), TTG2428TTA (7409G>A), GTG2429TTG (7410G>T), GAG2430ATG (7413G>A 7414A>T), CCG2431GCT (7416C>G 7418G>T), CAA2433GAG (7422C>G 7424A>G), AAT2434GAT (7425A>G), ATG2436ACA (7432T>C 7433G>A), GCT2437GCA (7436T>A), AGT2438TAT (7437A>T 7438G>A), GCT2439GCC (7442T>C), CTC2440ATG (7443C>A 7445C>G), TGC2441TGT (7448C>T), GTG2442ATC (7449G>A 7451G>C), GCT2444TCA (7455G>T 7457T>A), TTT2445CTT (7458T>C), GTA2446GCT (7462T>C 7463A>T), AAT2447CTT (7464A>C 7465A>T), AAA2448GGG (7467A>G 7468A>G 7469A>G), GTT2449CCA (7470G>C 7471T>C 7472T>A), GGT2450AAT (7473G>A 7474G>A), GAT2451ACT (7476G>A 7477A>C), GAT2452GAG (7481T>G), CCT2453CCG (7484T>G), ATA2454ATG (7487A>G), ATC2455ACT (7489T>C 7490C>T), AAG2457GTT (7494A>G 7495A>T 7496G>T), GGT2458GGA (7499T>A), GTG2459AAA (7500G>A 7501T>A 7502G>A), CTA2460ACG (7503C>A 7504T>C 7505A>G), AGA2461CTG (7506A>C 7507G>T 7508A>G), GGT2462GGA (7511T>A), GAG2463AAG (7512G>A), ATT2464ATA (7517T>A), GTA2465GTG (7520A>G), ATG2466CCA (7521A>C 7522T>C 7523G>A), CCA2467GCT (7524C>G 7526A>T), CGC2468AGG (7527C>A 7529C>G), CCA2470CCC (7535A>C), TCA2472AAT (7539T>A 7540C>A 7541A>T), TTT2473TTC (7544T>C), GGG2474GGA (7547G>A), GAC2476GAT (7553C>T), CCA2477CCG (7556A>G), ATT2478ATA (7559T>A), TTC2479TTT (7562C>T), CCA2481CCT (7568A>T), CTT2482CAT (7570T>A), AAC2483GGC (7572A>G 7573A>G), TGG2484TAT (7576G>A 7577G>T), AGA2485GAC (7578A>G 7579G>A 7580A>C), AAG2486CAG (7581A>C), ACA2487ACT (7586A>T), TTT2488TAC (7588T>A 7589T>C), ATG2492CCC (7599A>C 7600T>C 7601G>C), GCC2493AAG (7602G>A 7603C>A 7604C>G), GAG2494GAA (7607G>A), GAG2495GAA (7610G>A), AAT2497AAC (7616T>C), ATG2498AAG (7618T>A), ATA2499ATT (7622A>T), CAT2501CAC (7628T>C), CGA2502CGC (7631A>C), TTT2503GGT (7632T>G 7633T>G), CGA2504AAA (7635C>A 7636G>A) | | | | | | | | |

##### Proteins

| | Begin | End | Coverage | Score | Concordance | Matches | Identities | I/D/M/F* | Stop Codons |
|---|---|---|---|---|---|---|---|---|---|
| polyprotein (YP_007027011.1) | 2334 | 2509 | 6.0% | 742 | 60.2% | 176 (100%) | 102 (58.0%) | 0/0/0/0 | 0 |
| Protein mutations: | I2337V (7134A>G 7136T>G), Q2345K (7158C>A), K2348E (7167A>G), K2351R (7177A>G 7178A>G), Q2352A (7179C>G 7180A>C), F2354L (7185T>C), P2356Q (7192C>A 7193C>G), T2357S (7194A>T), I2360F (7203A>T 7205C>T), I2361Q (7206A>C 7207T>A 7208C>G), R2363L (7213G>T 7214G>T), V2365L (7218G>C), P2370L (7234C>T 7235A>T), T2373E (7242A>G 7243C>A 7244A>G), V2374P (7245G>C 7246T>C), E2376D (7253A>T), I2378S (7257A>T 7258T>C), V2382A (7270T>C 7271A>T), R2383K (7272C>A 7273G>A 7274T>A), V2384I (7275G>A 7277A>T), S2386A (7281T>G 7283T>C), E2387K (7284G>A 7286G>A), L2388E (7287T>G 7288T>A), I2389V (7290A>G 7292T>G), G2390N (7293G>A 7294G>A 7295A>T), S2399C (7320A>T), D2403N (7332G>A 7334T>C), N2406K (7343T>G), M2417L (7374A>T), E2418Q (7377G>C), G2419K (7380G>A 7381G>A), L2422H (7390T>A 7391A>T), Y2426N (7401T>A 7403T>C), K2427N (7406G>C), V2429L (7410G>T), E2430M (7413G>A 7414A>T), P2431A (7416C>G 7418G>T), Q2433E (7422C>G 7424A>G), N2434D (7425A>G), M2436T (7432T>C 7433G>A), S2438Y (7437A>T 7438G>A), L2440M (7443C>A 7445C>G), V2442I (7449G>A 7451G>C), A2444S (7455G>T 7457T>A), F2445L (7458T>C), V2446A (7462T>C 7463A>T), N2447L (7464A>C 7465A>T), K2448G (7467A>G 7468A>G 7469A>G), V2449P (7470G>C 7471T>C 7472T>A), G2450N (7473G>A 7474G>A), D2451T (7476G>A 7477A>C), D2452E (7481T>G), I2454M (7487A>G), I2455T (7489T>C 7490C>T), K2457V (7494A>G 7495A>T 7496G>T), V2459K (7500G>A 7501T>A 7502G>A), L2460T (7503C>A 7504T>C 7505A>G), R2461L (7506A>C 7507G>T 7508A>G), E2463K (7512G>A), M2466P (7521A>C 7522T>C 7523G>A), P2467A (7524C>G 7526A>T), S2472N (7539T>A 7540C>A 7541A>T), L2482H (7570T>A), N2483G (7572A>G 7573A>G), W2484Y (7576G>A 7577G>T), R2485D (7578A>G 7579G>A 7580A>C), K2486Q (7581A>C), F2488Y (7588T>A 7589T>C), M2492P (7599A>C 7600T>C 7601G>C), A2493K (7602G>A 7603C>A 7604C>G), M2498K (7618T>A), F2503G (7632T>G 7633T>G), R2504K (7635C>A 7636G>A) | | | | | | | | |

| | Begin | End | Coverage | Score | Concordance | Matches | Identities | I/D/M/F* | Stop Codons |
|---|---|---|---|---|---|---|---|---|---|
| **NT** | **7124** | **7650** | **5.9%** | **230** | **21.8%** | **527 (100%)** | **321 (60.9%)** | **0/0** | |
| Codon mutations: | GCA2335GCG (7130A>G), CCA2336CCG (7133A>G), ATT2337GTG (7134A>G 7136T>G), TTC2339TTT (7142C>T), GTA2340GTT (7145A>T), ACA2341ACC (7148A>C), GGG2342GGA (7151G>A), GCA2344GCT (7157A>T), CAA2345AAA (7158C>A), AAA2346AAG (7163A>G), AAG2348GAG (7167A>G), GTA2350GTT (7175A>T), AAA2351AGG (7177A>G 7178A>G), CAA2352GCA (7179C>G 7180A>C), ATA2353ATT (7184A>T), TTT2354CTT (7185T>C), GGT2355GGC (7190T>C), CCC2356CAG (7192C>A 7193C>G), ACT2357TCT (7194A>T), CCA2359CCC (7202A>C), ATC2360TTT (7203A>T 7205C>T), ATC2361CAG (7206A>C 7207T>A 7208C>G), TCT2362TCC (7211T>C), CGG2363CTT (7213G>T 7214G>T), AAA2364AAG (7217A>G), GTT2365CTT (7218G>C), CCA2368CCT (7229A>T), CCA2370CTT (7234C>T 7235A>T), GGA2372GGG (7241A>G), ACA2373GAG (7242A>G 7243C>A 7244A>G), GTT2374CCT (7245G>C 7246T>C), GAG2375GAA (7250G>A), GAA2376GAT (7253A>T), ATT2377ATA (7256T>A), ATC2378TCC (7257A>T 7258T>C), GTA2382GCT (7270T>C 7271A>T), CGT2383AAA (7272C>A 7273G>A 7274T>A), GTA2384ATT (7275G>A 7277A>T), GCG2385GCT (7280G>T), TCT2386GCC (7281T>G 7283T>C), GAG2387AAA (7284G>A 7286G>A), TTG2388GAG (7287T>G 7288T>A), ATT2389GTG (7290A>G 7292T>G), GGA2390AAT (7293G>A 7294G>A 7295A>T), GGG2391GGA (7298G>A), GTT2393GTG (7304T>G), CTC2394CTA (7307C>A), GTA2395GTT (7310A>T), GAA2396GAG (7313A>G), ACA2398ACT (7319A>T), AGT2399TGT (7320A>T), TTG2400CTC (7323T>C 7325G>C), TTT2402TTC (7331T>C), GAT2403AAC (7332G>A 7334T>C), GCT2404GCC (7337T>C), CTC2405CTT (7340C>T), AAT2406AAG (7343T>G), GGA2410GGG (7355A>G), CCA2411CCT (7358A>T), TAC2412TAT (7361C>T), ATT2413ATC (7364T>C), ATG2417TTG (7374A>T), GAA2418CAA (7377G>C), GGG2419AAG (7380G>A 7381G>A), ATT2420ATC (7385T>C), GGA2421GGT (7388A>T), CTA2422CAT (7390T>A 7391A>T), GGA2424GGT (7397A>T), TTA2425CTT (7398T>C 7400A>T), TAT2426AAC (7401T>A 7403T>C), AAG2427AAC (7406G>C), TTG2428TTA (7409G>A), GTG2429TTG (7410G>T), GAG2430ATG (7413G>A 7414A>T), CCG2431GCT (7416C>G 7418G>T), CAA2433GAG (7422C>G 7424A>G), AAT2434GAT (7425A>G), ATG2436ACA (7432T>C 7433G>A), GCT2437GCA (7436T>A), AGT2438TAT (7437A>T 7438G>A), GCT2439GCC (7442T>C), CTC2440ATG (7443C>A 7445C>G), TGC2441TGT (7448C>T), GTG2442ATC (7449G>A 7451G>C), GCT2444TCA (7455G>T 7457T>A), TTT2445CTT (7458T>C), GTA2446GCT (7462T>C 7463A>T), AAT2447CTT (7464A>C 7465A>T), AAA2448GGG (7467A>G 7468A>G 7469A>G), GTT2449CCA (7470G>C 7471T>C 7472T>A), GGT2450AAT (7473G>A 7474G>A), GAT2451ACT (7476G>A 7477A>C), GAT2452GAG (7481T>G), CCT2453CCG (7484T>G), ATA2454ATG (7487A>G), ATC2455ACT (7489T>C 7490C>T), AAG2457GTT (7494A>G 7495A>T 7496G>T), GGT2458GGA (7499T>A), GTG2459AAA (7500G>A 7501T>A 7502G>A), CTA2460ACG (7503C>A 7504T>C 7505A>G), AGA2461CTG (7506A>C 7507G>T 7508A>G), GGT2462GGA (7511T>A), GAG2463AAG (7512G>A), ATT2464ATA (7517T>A), GTA2465GTG (7520A>G), ATG2466CCA (7521A>C 7522T>C 7523G>A), CCA2467GCT (7524C>G 7526A>T), CGC2468AGG (7527C>A 7529C>G), CCA2470CCC (7535A>C), TCA2472AAT (7539T>A 7540C>A 7541A>T), TTT2473TTC (7544T>C), GGG2474GGA (7547G>A), GAC2476GAT (7553C>T), CCA2477CCG (7556A>G), ATT2478ATA (7559T>A), TTC2479TTT (7562C>T), CCA2481CCT (7568A>T), CTT2482CAT (7570T>A), AAC2483GGC (7572A>G 7573A>G), TGG2484TAT (7576G>A 7577G>T), AGA2485GAC (7578A>G 7579G>A 7580A>C), AAG2486CAG (7581A>C), ACA2487ACT (7586A>T), TTT2488TAC (7588T>A 7589T>C), ATG2492CCC (7599A>C 7600T>C 7601G>C), GCC2493AAG (7602G>A 7603C>A 7604C>G), GAG2494GAA (7607G>A), GAG2495GAA (7610G>A), AAT2497AAC (7616T>C), ATG2498AAG (7618T>A), ATA2499ATT (7622A>T), CAT2501CAC (7628T>C), CGA2502CGC (7631A>C), TTT2503GGT (7632T>G 7633T>G), CGA2504AAA (7635C>A 7636G>A) | | | | | | | | |
| HAM1-like protein (YP_007032446.1) | 26 | 201 | 77.9% | 742 | 60.2% | 176 (100%) | 102 (58.0%) | 0/0/0/0 | 0 |
| Protein mutations: | I29V (7134A>G 7136T>G), Q37K (7158C>A), K40E (7167A>G), K43R (7177A>G 7178A>G), Q44A (7179C>G 7180A>C), F46L (7185T>C), P48Q (7192C>A 7193C>G), T49S (7194A>T), I52F (7203A>T 7205C>T), I53Q (7206A>C 7207T>A 7208C>G), R55L (7213G>T 7214G>T), V57L (7218G>C), P62L (7234C>T 7235A>T), T65E (7242A>G 7243C>A 7244A>G), V66P (7245G>C 7246T>C), E68D (7253A>T), I70S (7257A>T 7258T>C), V74A (7270T>C 7271A>T), R75K (7272C>A 7273G>A 7274T>A), V76I (7275G>A 7277A>T), S78A (7281T>G 7283T>C), E79K (7284G>A 7286G>A), L80E (7287T>G 7288T>A), I81V (7290A>G 7292T>G), G82N (7293G>A 7294G>A 7295A>T), S91C (7320A>T), D95N (7332G>A 7334T>C), N98K (7343T>G), M109L (7374A>T), E110Q (7377G>C), G111K (7380G>A 7381G>A), L114H (7390T>A 7391A>T), Y118N (7401T>A 7403T>C), K119N (7406G>C), V121L (7410G>T), E122M (7413G>A 7414A>T), P123A (7416C>G 7418G>T), Q125E (7422C>G 7424A>G), N126D (7425A>G), M128T (7432T>C 7433G>A), S130Y (7437A>T 7438G>A), L132M (7443C>A 7445C>G), V134I (7449G>A 7451G>C), A136S (7455G>T 7457T>A), F137L (7458T>C), V138A (7462T>C 7463A>T), N139L (7464A>C 7465A>T), K140G (7467A>G 7468A>G 7469A>G), V141P (7470G>C 7471T>C 7472T>A), G142N (7473G>A 7474G>A), D143T (7476G>A 7477A>C), D144E (7481T>G), I146M (7487A>G), I147T (7489T>C 7490C>T), K149V (7494A>G 7495A>T 7496G>T), V151K (7500G>A 7501T>A 7502G>A), L152T (7503C>A 7504T>C 7505A>G), R153L (7506A>C 7507G>T 7508A>G), E155K (7512G>A), M158P (7521A>C 7522T>C 7523G>A), P159A (7524C>G 7526A>T), S164N (7539T>A 7540C>A 7541A>T), L174H (7570T>A), N175G (7572A>G 7573A>G), W176Y (7576G>A 7577G>T), R177D (7578A>G 7579G>A 7580A>C), K178Q (7581A>C), F180Y (7588T>A 7589T>C), M184P (7599A>C 7600T>C 7601G>C), A185K (7602G>A 7603C>A 7604C>G), M190K (7618T>A), F195G (7632T>G 7633T>G), R196K (7635C>A 7636G>A) | | | | | | | | |
| Codon mutations: | GCA27GCG (7130A>G), CCA28CCG (7133A>G), ATT29GTG (7134A>G 7136T>G), TTC31TTT (7142C>T), GTA32GTT (7145A>T), ACA33ACC (7148A>C), GGG34GGA (7151G>A), GCA36GCT (7157A>T), CAA37AAA (7158C>A), AAA38AAG (7163A>G), AAG40GAG (7167A>G), GTA42GTT (7175A>T), AAA43AGG (7177A>G 7178A>G), CAA44GCA (7179C>G 7180A>C), ATA45ATT (7184A>T), TTT46CTT (7185T>C), GGT47GGC (7190T>C), CCC48CAG (7192C>A 7193C>G), ACT49TCT (7194A>T), CCA51CCC (7202A>C), ATC52TTT (7203A>T 7205C>T), ATC53CAG (7206A>C 7207T>A 7208C>G), TCT54TCC (7211T>C), CGG55CTT (7213G>T 7214G>T), AAA56AAG (7217A>G), GTT57CTT (7218G>C), CCA60CCT (7229A>T), CCA62CTT (7234C>T 7235A>T), GGA64GGG (7241A>G), ACA65GAG (7242A>G 7243C>A 7244A>G), GTT66CCT (7245G>C 7246T>C), GAG67GAA (7250G>A), GAA68GAT (7253A>T), ATT69ATA (7256T>A), ATC70TCC (7257A>T 7258T>C), GTA74GCT (7270T>C 7271A>T), CGT75AAA (7272C>A 7273G>A 7274T>A), GTA76ATT (7275G>A 7277A>T), GCG77GCT (7280G>T), TCT78GCC (7281T>G 7283T>C), GAG79AAA (7284G>A 7286G>A), TTG80GAG (7287T>G 7288T>A), ATT81GTG (7290A>G 7292T>G), GGA82AAT (7293G>A 7294G>A 7295A>T), GGG83GGA (7298G>A), GTT85GTG (7304T>G), CTC86CTA (7307C>A), GTA87GTT (7310A>T), GAA88GAG (7313A>G), ACA90ACT (7319A>T), AGT91TGT (7320A>T), TTG92CTC (7323T>C 7325G>C), TTT94TTC (7331T>C), GAT95AAC (7332G>A 7334T>C), GCT96GCC (7337T>C), CTC97CTT (7340C>T), AAT98AAG (7343T>G), GGA102GGG (7355A>G), CCA103CCT (7358A>T), TAC104TAT (7361C>T), ATT105ATC (7364T>C), ATG109TTG (7374A>T), GAA110CAA (7377G>C), GGG111AAG (7380G>A 7381G>A), ATT112ATC (7385T>C), GGA113GGT (7388A>T), CTA114CAT (7390T>A 7391A>T), GGA116GGT (7397A>T), TTA117CTT (7398T>C 7400A>T), TAT118AAC (7401T>A 7403T>C), AAG119AAC (7406G>C), TTG120TTA (7409G>A), GTG121TTG (7410G>T), GAG122ATG (7413G>A 7414A>T), CCG123GCT (7416C>G 7418G>T), CAA125GAG (7422C>G 7424A>G), AAT126GAT (7425A>G), ATG128ACA (7432T>C 7433G>A), GCT129GCA (7436T>A), AGT130TAT (7437A>T 7438G>A), GCT131GCC (7442T>C), CTC132ATG (7443C>A 7445C>G), TGC133TGT (7448C>T), GTG134ATC (7449G>A 7451G>C), GCT136TCA (7455G>T 7457T>A), TTT137CTT (7458T>C), GTA138GCT (7462T>C 7463A>T), AAT139CTT (7464A>C 7465A>T), AAA140GGG (7467A>G 7468A>G 7469A>G), GTT141CCA (7470G>C 7471T>C 7472T>A), GGT142AAT (7473G>A 7474G>A), GAT143ACT (7476G>A 7477A>C), GAT144GAG (7481T>G), CCT145CCG (7484T>G), ATA146ATG (7487A>G), ATC147ACT (7489T>C 7490C>T), AAG149GTT (7494A>G 7495A>T 7496G>T), GGT150GGA (7499T>A), GTG151AAA (7500G>A 7501T>A 7502G>A), CTA152ACG (7503C>A 7504T>C 7505A>G), AGA153CTG (7506A>C 7507G>T 7508A>G), GGT154GGA (7511T>A), GAG155AAG (7512G>A), ATT156ATA (7517T>A), GTA157GTG (7520A>G), ATG158CCA (7521A>C 7522T>C 7523G>A), CCA159GCT (7524C>G 7526A>T), CGC160AGG (7527C>A 7529C>G), CCA162CCC (7535A>C), TCA164AAT (7539T>A 7540C>A 7541A>T), TTT165TTC (7544T>C), GGG166GGA (7547G>A), GAC168GAT (7553C>T), CCA169CCG (7556A>G), ATT170ATA (7559T>A), TTC171TTT (7562C>T), CCA173CCT (7568A>T), CTT174CAT (7570T>A), AAC175GGC (7572A>G 7573A>G), TGG176TAT (7576G>A 7577G>T), AGA177GAC (7578A>G 7579G>A 7580A>C), AAG178CAG (7581A>C), ACA179ACT (7586A>T), TTT180TAC (7588T>A 7589T>C), ATG184CCC (7599A>C 7600T>C 7601G>C), GCC185AAG (7602G>A 7603C>A 7604C>G), GAG186GAA (7607G>A), GAG187GAA (7610G>A), AAT189AAC (7616T>C), ATG190AAG (7618T>A), ATA191ATT (7622A>T), CAT193CAC (7628T>C), CGA194CGC (7631A>C), TTT195GGT (7632T>G 7633T>G), CGA196AAA (7635C>A 7636G>A) | | | | | | | | |

*: Inserts / Deletes / Misaligned / Frameshifts

#### Analysis details

This analysis was performed with panviral2.64

### NGS Details (UN8): Epiphyllum badnavirus 1

#### Assembly

| | |
|---|---|
| **Coverage Length** | 275 (1 contig(s)) |
| **Depth Of Coverage** | 50.7 |
| **Number Of Reads** | 137 |
| **Reads Per Million** | 2.29 rpm (after QC) |
| **Ambiguities** | 0 |
| **Assembly Method** | de novo + reference guided assembly |
| **Consensus Caller** | Bcf Tools |

#### Coverage Map

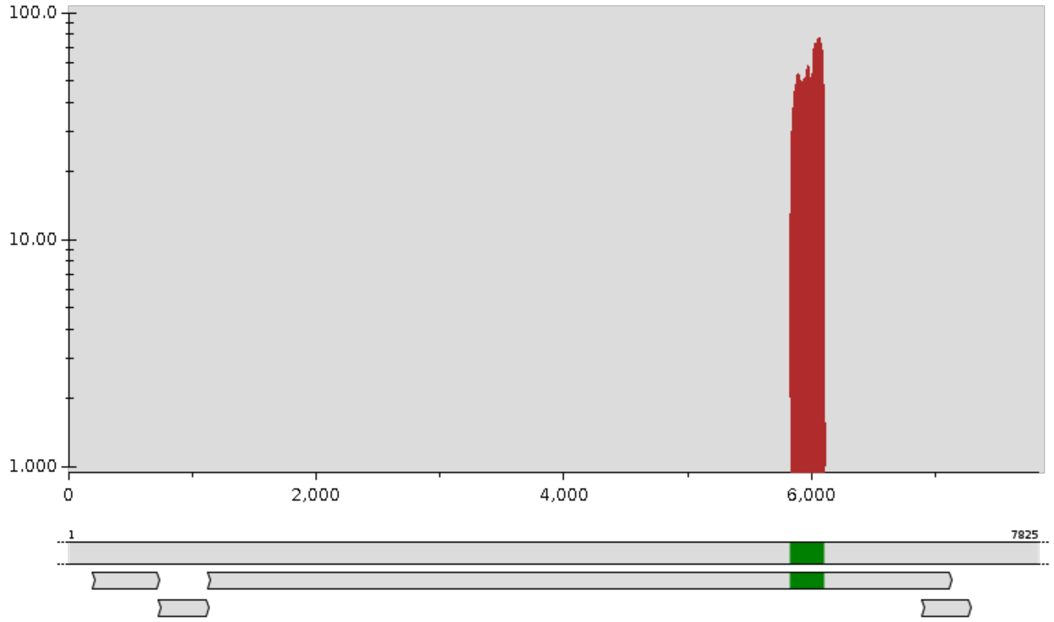

#### Assignment

| | |
|---|---|
| **Type** | Epiphyllum badnavirus 1 (Taxonomy ID: 2518008) |
| **Reference Genome** | NC_076247.1 |
| **NT Identity (%)** | 57.554 |
| **AA Identity (%)** | 52.6882 |
| **Number Of Stop Codons** | 0 |
| **Number Of CDS** | 4 |

#### Alignment

| | |
|---|---|
| **Alignment Score** | 78.0 (NT) + 337.0 (AA) = 415.0 |
| **Concordance (%)** | 34.2127 |

| Alignment Method | Global, seeded, nucleotide + amino acids (AGA) |
|---|---|

#### Genome Region

Sequence starts at position 5829 and ends at position 6103 relative to NC_076247.1 reference sequence.

#### Alignment Detailed Statistics

| | Begin | End | Coverage | Score | Concordance | Matches | Identities | I/D/M/F* | Stop Codons |
|---|---|---|---|---|---|---|---|---|---|
| **NT** | **5829** | **6103** | **3.5%** | **78** | **14.2%** | **275 (98.9%)** | **160 (57.6%)** | **3/0** | |
| Mutations: | 5832T>A, 5835C>T, 5837G>C, 5841C>G, 5844C>T, 5847C>T, 5851T>A, 5853C>T, 5856T>A, 5857C>T, 5861A>G, 5863A>T, 5864G>C, 5868A>T, 5870T>A, 5871C>T, 5874C>T, 5878A>T, 5880T>C, 5882T>G, 5886G>C, 5887G>A, 5888A>G, 5889C>G, 5891A>C, 5895A>G, 5896T>G, 5897C>A, 5898C>T, 5901C>A, 5905T>A, 5906G>A, 5910C>T, 5916C>T, 5918T>G, 5924C>A, 5925A>T, 5926G>T, 5931A>C, 5933T>A, 5937C>T, 5940A>G, 5942G>T, 5943G>T, 5944T>C, 5949T>G, 5953C>T, 5958C>T, 5961A>G, 5962T>C, 5964G>T, 5966A>C, 5967G>T, 5976T>A, 5979T>A, 5981T>C, 5982G>T, 5983T>C, 5985C>T, 5986C>A, 5987A>T, 5988A>G, 5989C>G, 5990G>A, 5991A>C, 5992A>T, 5993A>T, 5998G>A, 5999A>C, 6000C>A, 6001A>T, 6002A>G, 6003C>G, 6004T>G, 6005G>T, 6006T>A, 6009C>T, 6012A>G, 6013G>C, 6014A>C, 6015_6016insTAT, 6016C>A, 6018G>T, 6019T>G, 6020C>A, 6021A>C, 6022G>T, 6023G>C, 6027T>C, 6030A>G, 6032C>T, 6033A>T, 6038A>T, 6039C>T, 6042T>C, 6048C>T, 6051C>A, 6054G>T, 6057C>G, 6059T>A, 6060C>T, 6061A>T, 6062G>C, 6063T>A, 6064A>C, 6065A>G, 6066C>G, 6068C>G, 6070G>A, 6071A>G, 6078C>G, 6083C>T, 6094A>C, 6095A>G, 6096G>T, 6097T>A | | | | | | | | |

##### CDS

| | Begin | End | Coverage | Score | Concordance | Matches | Identities | I/D/M/F* | Stop Codons |
|---|---|---|---|---|---|---|---|---|---|
| QKM20_gp3 | 1568 | 1659 | 4.6% | 337 | 49.3% | 92 (98.9%) | 49 (52.7%) | 1/0/0/0 | 0 |
| Protein mutations: | N1568K (5832T>A), S1570T (5837G>C), F1575I (5851T>A 5853C>T), D1576E (5856T>A), K1578R (5861A>G), F1581Y (5870T>A 5871C>T), I1584F (5878A>T 5880T>C), M1585R (5882T>G), M1586I (5886G>C), D1587R (5887G>A 5888A>G 5889C>G), E1588A (5891A>C), S1590D (5896T>G 5897C>A 5898C>T), W1593K (5905T>A 5906G>A), L1597R (5918T>G), P1599H (5924C>A 5925A>T), D1600Y (5926G>T), L1602H (5933T>A), W1605F (5942G>T 5943G>T), P1609S (5953C>T), K1613T (5966A>C 5967G>T), V1618A (5981T>C 5982G>T), F1619L (5983T>C 5985C>T), Q1620M (5986C>A 5987A>T 5988A>G), R1621D (5989C>G 5990G>A 5991A>C), K1622L (5992A>T 5993A>T), D1624T (5998G>A 5999A>C 6000C>A), N1625W (6001A>T 6002A>G 6003C>G), C1626V (6004T>G 6005G>T 6006T>A), D1629P (6013G>C 6014A>C), D1629_L1630insY (6015_6016insTAT), L1630I (6016C>A 6018G>T), S1631D (6019T>G 6020C>A 6021A>C), G1632S (6022G>T 6023G>C), A1635V (6032C>T 6033A>T), Y1637F (6038A>T 6039C>T), F1644Y (6059T>A 6060C>T), N1646R (6064A>C 6065A>G 6066C>G), T1647S (6068C>G), E1648R (6070G>A 6071A>G), D1650E (6078C>G), A1652V (6083C>T), K1656R (6094A>C 6095A>G 6096G>T), S1657T (6097T>A) | | | | | | | | |
| Codon mutations: | AAT1568AAA (5832T>A), GCC1569GCT (5835C>T), AGT1570ACT (5837G>C), GTC1571GTG (5841C>G), TTC1572TTT (5844C>T), TCC1573TCT (5847C>T), TTC1575ATT (5851T>A 5853C>T), GAT1576GAA (5856T>A), CTG1577TTG (5857C>T), AAG1578AGG (5861A>G), AGC1579TCC (5863A>T 5864G>C), GGA1580GGT (5868A>T), TTC1581TAT (5870T>A 5871C>T), CAC1582CAT (5874C>T), ATT1584TTC (5878A>T 5880T>C), ATG1585AGG (5882T>G), ATG1586ATC (5886G>C), GAC1587AGG (5887G>A 5888A>G 5889C>G), GAA1588GCA (5891A>C), GAA1589GAG (5895A>G), TCC1590GAT (5896T>G 5897C>A 5898C>T), ATC1591ATA (5901C>A), TGG1593AAG (5905T>A 5906G>A), ACC1594ACT (5910C>T), TTC1596TTT (5916C>T), CTG1597CGG (5918T>G), CCA1599CAT (5924C>A 5925A>T), GAT1600TAT (5926G>T), GGA1601GGC (5931A>C), CTT1602CAT (5933T>A), TAC1603TAT (5937C>T), GAA1604GAG (5940A>G), TGG1605TTT (5942G>T 5943G>T), TTA1606CTA (5944T>C), GTT1607GTG (5949T>G), CCA1609TCA (5953C>T), TTC1610TTT (5958C>T), GGA1611GGG (5961A>G), TTG1612CTT (5962T>C 5964G>T), AAG1613ACT (5966A>C 5967G>T), CCT1616CCA (5976T>A), GCT1617GCA (5979T>A), GTG1618GCT (5981T>C 5982G>T), TTC1619CTT (5983T>C 5985C>T), CAA1620ATG (5986C>A 5987A>T 5988A>G), CGA1621GAC (5989C>G 5990G>A 5991A>C), AAG1622TTG (5992A>T 5993A>T), GAC1624ACA (5998G>A 5999A>C 6000C>A), AAC1625TGG (6001A>T 6002A>G 6003C>G), TGT1626GTA (6004T>G 6005G>T 6006T>A), TTC1627TTT (6009C>T), AGA1628AGG (6012A>G), GAC1629CCC (6013G>C 6014A>C), GAC1629_CTG1630insTAT (6015_6016insTAT), CTG1630ATT (6016C>A 6018G>T), TCA1631GAC (6019T>G 6020C>A 6021A>C), GGC1632TCC (6022G>T 6023G>C), TTT1633TTC (6027T>C), GTA1634GTG (6030A>G), GCA1635GTT (6032C>T 6033A>T), TAC1637TTT (6038A>T 6039C>T), ATT1638ATC (6042T>C), GAC1640GAT (6048C>T), ATC1641ATA (6051C>A), CTG1642CTT (6054G>T), GTC1643GTG (6057C>G), TTC1644TAT (6059T>A 6060C>T), AGT1645TCA (6061A>T 6062G>C 6063T>A), AAC1646CGG (6064A>C 6065A>G 6066C>G), ACT1647AGT (6068C>G), GAA1648AGA (6070G>A 6071A>G), GAC1650GAG (6078C>G), GCA1652GTA (6083C>T), AAG1656CGT (6094A>C 6095A>G 6096G>T), TCC1657ACC (6097T>A) | | | | | | | | |

##### Proteins

| | Begin | End | Coverage | Score | Concordance | Matches | Identities | I/D/M/F* | Stop Codons |
|---|---|---|---|---|---|---|---|---|---|
| polyprotein (YP_010797894.1) | 1568 | 1659 | 4.6% | 337 | 49.3% | 92 (98.9%) | 49 (52.7%) | 1/0/0/0 | 0 |
| Protein mutations: | N1568K (5832T>A), S1570T (5837G>C), F1575I (5851T>A 5853C>T), D1576E (5856T>A), K1578R (5861A>G), F1581Y (5870T>A 5871C>T), I1584F (5878A>T 5880T>C), M1585R (5882T>G), M1586I (5886G>C), D1587R (5887G>A 5888A>G 5889C>G), E1588A (5891A>C), S1590D (5896T>G 5897C>A 5898C>T), W1593K (5905T>A 5906G>A), L1597R (5918T>G), P1599H (5924C>A 5925A>T), D1600Y (5926G>T), L1602H (5933T>A), W1605F (5942G>T 5943G>T), P1609S (5953C>T), K1613T (5966A>C 5967G>T), V1618A (5981T>C 5982G>T), F1619L (5983T>C 5985C>T), Q1620M (5986C>A 5987A>T 5988A>G), R1621D (5989C>G 5990G>A 5991A>C), K1622L (5992A>T 5993A>T), D1624T (5998G>A 5999A>C 6000C>A), N1625W (6001A>T 6002A>G 6003C>G), C1626V (6004T>G 6005G>T 6006T>A), D1629P (6013G>C 6014A>C), D1629_L1630insY (6015_6016insTAT), L1630I (6016C>A 6018G>T), S1631D (6019T>G 6020C>A 6021A>C), G1632S (6022G>T 6023G>C), A1635V (6032C>T 6033A>T), Y1637F (6038A>T 6039C>T), F1644Y (6059T>A 6060C>T), N1646R (6064A>C 6065A>G 6066C>G), T1647S (6068C>G), E1648R (6070G>A 6071A>G), D1650E (6078C>G), A1652V (6083C>T), K1656R (6094A>C 6095A>G 6096G>T), S1657T (6097T>A) | | | | | | | | |
| Codon mutations: | AAT1568AAA (5832T>A), GCC1569GCT (5835C>T), AGT1570ACT (5837G>C), GTC1571GTG (5841C>G), TTC1572TTT (5844C>T), TCC1573TCT (5847C>T), TTC1575ATT (5851T>A 5853C>T), GAT1576GAA (5856T>A), CTG1577TTG (5857C>T), AAG1578AGG (5861A>G), AGC1579TCC (5863A>T 5864G>C), GGA1580GGT (5868A>T), TTC1581TAT (5870T>A 5871C>T), CAC1582CAT (5874C>T), ATT1584TTC (5878A>T 5880T>C), ATG1585AGG (5882T>G), ATG1586ATC (5886G>C), GAC1587AGG (5887G>A 5888A>G 5889C>G), GAA1588GCA (5891A>C), GAA1589GAG (5895A>G), TCC1590GAT (5896T>G 5897C>A 5898C>T), ATC1591ATA (5901C>A), TGG1593AAG (5905T>A 5906G>A), ACC1594ACT (5910C>T), TTC1596TTT (5916C>T), CTG1597CGG (5918T>G), CCA1599CAT (5924C>A 5925A>T), GAT1600TAT (5926G>T), GGA1601GGC (5931A>C), CTT1602CAT (5933T>A), TAC1603TAT (5937C>T), GAA1604GAG (5940A>G), TGG1605TTT (5942G>T 5943G>T), TTA1606CTA (5944T>C), GTT1607GTG (5949T>G), CCA1609TCA (5953C>T), TTC1610TTT (5958C>T), GGA1611GGG (5961A>G), TTG1612CTT (5962T>C 5964G>T), AAG1613ACT (5966A>C 5967G>T), CCT1616CCA (5976T>A), GCT1617GCA (5979T>A), GTG1618GCT (5981T>C 5982G>T), TTC1619CTT (5983T>C 5985C>T), CAA1620ATG (5986C>A 5987A>T 5988A>G), CGA1621GAC (5989C>G 5990G>A 5991A>C), AAG1622TTG (5992A>T 5993A>T), GAC1624ACA (5998G>A 5999A>C 6000C>A), AAC1625TGG (6001A>T 6002A>G 6003C>G), TGT1626GTA (6004T>G 6005G>T 6006T>A), TTC1627TTT (6009C>T), AGA1628AGG (6012A>G), GAC1629CCC (6013G>C 6014A>C), GAC1629_CTG1630insTAT (6015_6016insTAT), CTG1630ATT (6016C>A 6018G>T), TCA1631GAC (6019T>G 6020C>A 6021A>C), GGC1632TCC (6022G>T 6023G>C), TTT1633TTC (6027T>C), GTA1634GTG (6030A>G), GCA1635GTT (6032C>T 6033A>T), TAC1637TTT (6038A>T 6039C>T), ATT1638ATC (6042T>C), GAC1640GAT (6048C>T), ATC1641ATA (6051C>A), CTG1642CTT (6054G>T), GTC1643GTG (6057C>G), TTC1644TAT (6059T>A 6060C>T), AGT1645TCA (6061A>T 6062G>C 6063T>A), AAC1646CGG (6064A>C 6065A>G 6066C>G), ACT1647AGT (6068C>G), GAA1648AGA (6070G>A 6071A>G), GAC1650GAG (6078C>G), GCA1652GTA (6083C>T), AAG1656CGT (6094A>C 6095A>G 6096G>T), TCC1657ACC (6097T>A) | | | | | | | | |

*: Inserts / Deletes / Misaligned / Frameshifts

#### Analysis details

This analysis was performed with panviral2.64

### NGS Details (UN8): Errantivirus

#### Assembly

| | |
|---|---|
| **Coverage Length** | 716 (1 contig(s)) |
| **Depth Of Coverage** | 22.2 |
| **Number Of Reads** | 133 |
| **Reads Per Million** | 2.22 rpm (after QC) |
| **Ambiguities** | 0 |
| **Assembly Method** | de novo + reference guided assembly |
| **Consensus Caller** | Bcf Tools |

#### Coverage Map

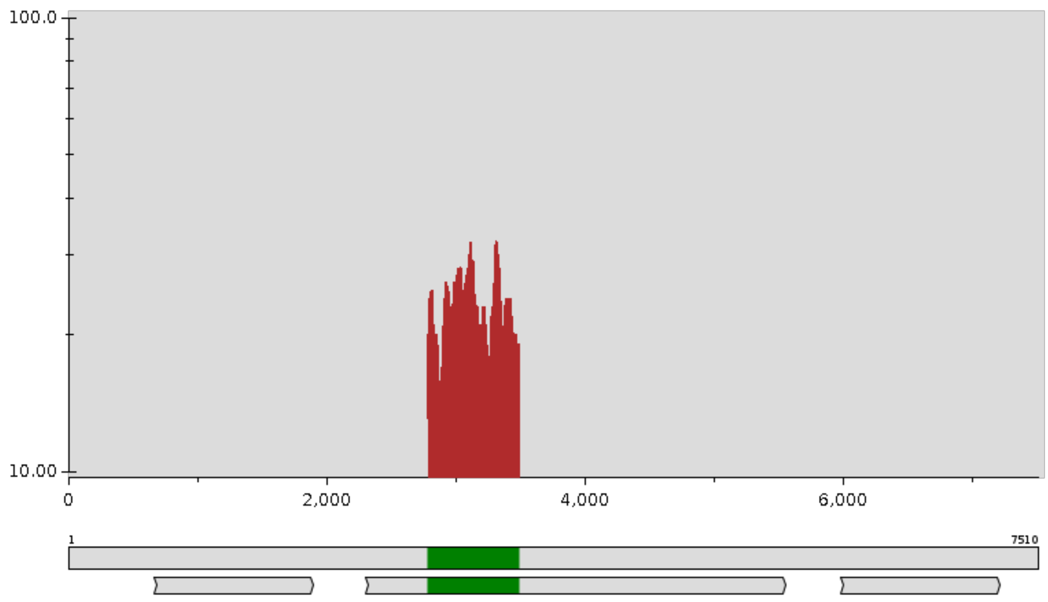

#### Assignment

| | |
|---|---|
| **Type** | Errantivirus (Taxonomy ID: 186666) |
| **Reference Genome** | NC_038512.1 |
| **NT Identity (%)** | 52.3605 |
| **AA Identity (%)** | 46.1538 |
| **Number Of Stop Codons** | 1 |
| **Number Of CDS** | 3 |

#### Alignment

| | |
|---|---|
| **Alignment Score** | 22.0 (NT) + 632.0 (AA) = 654.0 |
| **Concordance (%)** | 23.0769 |

| Alignment Method | Global, seeded, nucleotide + amino acids (AGA) |
|---|---|

#### Genome Region

Sequence starts at position 2783 and ends at position 3498 relative to NC_038512.1 reference sequence.

#### Alignment Detailed Statistics

| | Begin | End | Coverage | Score | Concordance | Matches | Identities | I/D/M/F* | Stop Codons |
|---|---|---|---|---|---|---|---|---|---|
| **NT** | **2783** | **3498** | **9.5%** | **22** | **1.6%** | **699 (97.6%)** | **366 (51.1%)** | **0/17** | |
| Mutations: | 2783A>G, 2785A>T, 2789T>C, 2791G>T, 2794C>T, 2800A>C, 2803T>C, 2806A>C, 2807A>C, 2808G>A, 2812A>C, 2813T>A, 2814C>G, 2815A>C, 2816G>T, 2817A>G, 2819T>A, 2820C>A, 2821T>C, 2822G>C, 2824A>T, 2826G>A, 2827G>T, 2828A>G, 2829G>C, 2832C>T, 2836C>T, 2837A>G, 2840T>G, 2841G>T, 2842G>C, 2843G>C, 2845T>C, 2849C>G, 2850C>G, 2851C>A, 2857A>G, 2858_2860delATC, 2863C>T, 2864G>A, 2865C>G, 2869T>A, 2870_2881delGGGAAACAAAAG, 2885C>A, 2887T>G, 2888C>T, 2890C>A, 2891G>T, 2892T>G, 2893A>T, 2896T>A, 2901T>A, 2902C>T, 2903C>A, 2905T>G, 2906A>G, 2907A>C, 2908G>T, 2914C>T, 2915G>A, 2918A>T, 2919A>T, 2927G>A, 2929T>A, 2932C>T, 2937A>T, 2938C>T, 2939C>A, 2941G>A, 2944A>T, 2947A>C, 2949A>T, 2953A>T, 2954A>G, 2955G>A, 2956T>G, 2959C>G, 2960G>T, 2965T>G, 2968C>T, 2969A>G, 2971G>A, 2974A>G, 2977T>A, 2978A>G, 2979A>G, 2980G>T, 2982G>C, 2986A>G, 2987T>A, 2988A>T, 2997C>A, 2998C>G, 2999T>C, 3001A>T, 3005T>C, 3008G>A, 3009C>G, 3011A>T, 3012G>C, 3013T>A, 3016G>T, 3018T>A, 3020T>C, 3025G>A, 3026G>A, 3028G>T, 3029G>A, 3030A>G, 3035G>A, 3036A>C, 3037C>A, 3038C>A, 3039C>A, 3040T>G, 3041C>G, 3043A>G, 3047A>G, 3049A>C, 3050T>C, 3051C>A, 3058C>T, 3059G>A, 3061G>A, 3066A>G, 3067C>G, 3068G>A, 3069T>C, 3070A>T, 3071G>C, 3073A>T, 3074C>T, 3075A>C, 3076C>T, 3079G>C, 3082T>C, 3084T>A, 3088A>G, 3090T>A, 3094T>A, 3095C>G, 3096G>T, 3103T>C, 3104A>T, 3106G>T, 3109A>C, 3110T>C, 3112A>C, 3114A>C, 3115A>T, 3118C>T, 3119T>G, 3124A>T, 3127T>A, 3129C>G, 3137A>G, 3139A>G, 3140G>C, 3142T>G, 3146G>A, 3148C>T, 3149A>C, 3154C>T, 3155C>T, 3157A>T, 3159G>A, 3160A>G, 3162G>A, 3163T>A, 3165T>A, 3168A>T, 3169A>T, 3171A>G, 3172T>G, 3175C>G, 3176A>T, 3179T>A, 3180G>T, 3182C>T, 3184C>G, 3187C>T, 3189A>T, 3190C>T, 3191_3192delCT, 3196C>T, 3203A>C, 3205T>G, 3206G>A, 3211T>C, 3214T>C, 3216C>A, 3217T>G, 3218T>A, 3219C>G, 3220C>T, 3224C>G, 3226G>A, 3229A>T, 3233C>T, 3236G>C, 3238G>A, 3239A>C, 3245G>A, 3246A>G, 3247A>G, 3248C>G, 3249G>T, 3250A>T, 3251G>A, 3252T>C, 3256C>T, 3257C>A, 3259A>C, 3260A>T, 3261G>T, 3263C>T, 3265T>G, 3267G>T, 3268A>C, 3269G>C, 3272A>C, 3273G>A, 3276A>G, 3277C>A, 3278T>C, 3280C>G, 3281A>T, 3282A>T, 3283A>G, 3286T>C, 3287C>A, 3289A>T, 3291T>A, 3293G>T, 3294A>C, 3295C>A, 3300C>G, 3301C>T, 3302G>T, 3303A>C, 3304A>T, 3307C>T, 3308T>G, 3309T>G, 3310G>A, 3311A>G, 3312A>C, 3313G>A, 3314C>T, 3315T>C, 3316C>A, 3317G>A, 3318A>G, 3319A>G, 3320A>G, 3321C>T, 3324C>A, 3325T>A, 3328T>C, 3333G>A, 3338A>T, 3340C>T, 3344A>T, 3345G>C, 3346C>T, 3347A>G, 3348G>C, 3349G>T, 3352C>G, 3355T>A, 3356A>G, 3358C>G, 3359A>G, 3360A>C, 3361G>T, 3362C>A, 3364T>A, 3365A>G, 3367C>T, 3371G>A, 3373T>G, 3380T>G, 3382C>A, 3385T>A, 3386A>G, 3388T>A, 3389C>T, 3392A>G, 3396A>G, 3397T>G, 3399T>C, 3400G>A, 3406A>T, 3407A>C, 3409G>T, 3412C>T, 3413C>A, 3414C>T, 3415T>A, 3421A>G, 3422A>T, 3424A>G, 3426A>G, 3427A>G, 3428C>G, 3429A>G, 3434T>C, 3436A>T, 3439C>G, 3440C>T, 3442T>G, 3443C>A, 3444T>C, 3445C>A, 3454C>T, 3455C>A, 3457A>G, 3459A>G, 3460A>G, 3467C>A, 3468C>G, 3470G>A, 3471A>G, 3472T>A, 3474T>A, 3477C>G, 3478A>G, 3479C>A, 3480G>T, 3481A>C, 3482C>A, 3484C>T, 3486C>G, 3487A>T, 3493C>T, 3498C>A | | | | | | | | |

##### CDS

| | Begin | End | Coverage | Score | Concordance | Matches | Identities | I/D/M/F* | Stop Codons |
|---|---|---|---|---|---|---|---|---|---|
| D1R33_gp2 | 160 | 398 | 22.0% | 632 | 39.3% | 234 (97.9%) | 108 (45.2%) | 0/5/1/1 | 1 |
| Protein mutations: | K160D (2783A>G 2785A>T), R168Q (2807A>C 2808G>A), D171C (2816G>T 2817A>G), S172N (2819T>A 2820C>A 2821T>C), A173P (2822G>C 2824A>T), W174Y (2826G>A 2827G>T), S175A (2828A>G 2829G>C), S176L (2832C>T), I178V (2837A>G), W179V (2840T>G 2841G>T 2842G>C), V180L (2843G>C 2845T>C), P182G (2849C>G 2850C>G 2851C>A), I185del (2858_2860delATC), A187S (2864G>A 2865C>G), G189_K192del (2870_2881delGGGAAACAAAAG), V196C (2891G>T 2892T>G 2893A>T), F199Y (2901T>A 2902C>T), K201A (2906A>G 2907A>C 2908G>T), E204K (2915G>A), K205L (2918A>T 2919A>T), D208K (2927G>A 2929T>A), Y211F (2937A>T 2938C>T), P212T (2939C>A 2941G>A), N215I (2949A>T), S217E (2954A>G 2955G>A 2956T>G), D218E (2959C>G), V219L (2960G>T), K222E (2969A>G 2971G>A), K225G (2978A>G 2979A>G 2980G>T), C226S (2982G>C), Y228I (2987T>A 2988A>T), T231K (2997C>A 2998C>G), A235R (3008G>A 3009C>G), F238Y (3018T>A), Y239H (3020T>C), V241I (3026G>A 3028G>T), E242R (3029G>A 3030A>G), D244T (3035G>A 3036A>C 3037C>A), P245K (3038C>A 3039C>A 3040T>G), Q246E (3041C>G 3043A>G), I248V (3047A>G 3049A>C), S249Q (3050T>C 3051C>A), A252T (3059G>A 3061G>A), N254R (3066A>G 3067C>G), V255T (3068G>A 3069T>C 3070A>T), E256H (3071G>C 3073A>T), H257S (3074C>T 3075A>C 3076C>T), F260Y (3084T>A), F262Y (3090T>A), R264V (3095C>G 3096G>T), M267F (3104A>T 3106G>T), K270T (3114A>C 3115A>T), S272A (3119T>G), T275S (3129C>G), R278G (3137A>G 3139A>G), V279L (3140G>C 3142T>G), D281N (3146G>A 3148C>T), N282H (3149A>C), L284F (3155C>T 3157A>T), R285K (3159G>A 3160A>G), G286E (3162G>A 3163T>A), L287H (3165T>A), Q288L (3168A>T 3169A>T), N289R (3171A>G 3172T>G), N290K (3175C>G), I291F (3176A>T), C292I (3179T>A 3180G>T), Y295F (3189A>T 3190C>T), I300L (3203A>C 3205T>G), V301I (3206G>A), T304K (3216C>A 3217T>G), Q307E (3224C>G 3226G>A), E308D (3229A>T), E311Q (3236G>C 3238G>A), N312H (3239A>C), E314R (3245G>A 3246A>G 3247A>G), R315V (3248C>G 3249G>T 3250A>T), V316T (3251G>A 3252T>C), Q318N (3257C>A 3259A>C), R319L (3260A>T 3261G>T), R321I (3267G>T 3268A>C), E322Q (3269G>C), S323H (3272A>C 3273G>A), N324R (3276A>G 3277C>A), F325L (3278T>C 3280C>G), K326L (3281A>T 3282A>T 3283A>G), Q328N (3287C>A 3289A>T), M329K (3291T>A), D330S (3293G>T 3294A>C 3295C>A), S332C (3300C>G 3301C>T), E333S (3302G>T 3303A>C 3304A>T), L335G (3308T>G 3309T>G 3310G>A), K336A (3311A>G 3312A>C 3313G>A), L337S (3314C>T 3315T>C 3316C>A), E338R (3317G>A 3318A>G 3319A>G), T339V (3320A>G 3321C>T), A340E (3324C>A 3325T>A), G343D (3333G>A), I345F (3338A>T 3340C>T), R348A (3347A>G 3348G>C 3349G>T), D349E (3352C>G), I351V (3356A>G 3358C>G), K352A (3359A>G 3360A>C 3361G>T), P353T (3362C>A 3364T>A), N354D (3365A>G 3367C>T), D356K (3371G>A 3373T>G), S359A (3380T>G 3382C>A), I361V (3386A>G 3388T>A), Q362* (3389C>T), K363E (3392A>G), Y364W (3396A>G 3397T>G), L365P (3399T>C 3400G>A), K368H (3407A>C 3409G>T), P370I (3413C>A 3414C>T 3415T>A), I373L (3422A>T 3424A>G), K374R (3426A>G 3427A>G), Q375G (3428C>G 3429A>G), L380T (3443C>A 3444T>C 3445C>A), K385R (3459A>G 3460A>G), P388R (3467C>A 3468C>G), D389R (3470G>A 3471A>G 3472T>A), F390Y (3474T>A), A391G (3477C>G 3478A>G), R392I (3479C>A 3480G>T 3481A>C), L393I (3482C>A 3484C>T), T394S (3486C>G 3487A>T) | | | | | | | | |
| Codon mutations: | AAA160GAT (2783A>G 2785A>T), TTG162CTT (2789T>C 2791G>T), GAC163GAT (2794C>T), GGA165GGC (2800A>C), ATT166ATC (2803T>C), ATA167ATC (2806A>C), AGA168CAA (2807A>C 2808G>A), CCA169CCC (2812A>C), TCA170AGC (2813T>A 2814C>G 2815A>C), GAC171TGC (2816G>T 2817A>G), TCT172AAC (2819T>A 2820C>A 2821T>C), GCA173CCT (2822G>C 2824A>T), TGG174TAT (2826G>A 2827G>T), AGC175GCC (2828A>G 2829G>C), TCA176TTA (2832C>T), CCC177CCT (2836C>T), ATA178GTA (2837A>G), TGG179GTC (2840T>G 2841G>T 2842G>C), GTT180CTC (2843G>C 2845T>C), CCC182GGA (2849C>G 2850C>G 2851C>A), AAA184AAG (2857A>G), ATC185del (2858_2860delATC), GAC186GAT (2863C>T), GCT187AGT (2864G>A 2865C>G), TCT188TCA (2869T>A), GGG189_AAG192del (2870_2881delGGGAAACAAAAG), CGT194AGG (2885C>A 2887T>G), CTC195TTA (2888C>T 2890C>A), GTA196TGT (2891G>T 2892T>G 2893A>T), GTT197GTA (2896T>A), TTC199TAT (2901T>A 2902C>T), CGT200AGG (2903C>A 2905T>G), AAG201GCT (2906A>G 2907A>C 2908G>T), AAC203AAT (2914C>T), GAG204AAG (2915G>A), AAG205TTG (2918A>T 2919A>T), GAT208AAA (2927G>A 2929T>A), GAC209GAT (2932C>T), TAC211TTT (2937A>T 2938C>T), CCG212ACA (2939C>A 2941G>A), ATA213ATT (2944A>T), CCA214CCC (2947A>C), AAC215ATC (2949A>T), ATA216ATT (2953A>T), AGT217GAG (2954A>G 2955G>A 2956T>G), GAC218GAG (2959C>G), GTA219TTA (2960G>T), CTT220CTG (2965T>G), GAC221GAT (2968C>T), AAG222GAA (2969A>G 2971G>A), TTA223TTG (2974A>G), GGT224GGA (2977T>A), AAG225GGT (2978A>G 2979A>G 2980G>T), TGC226TCC (2982G>C), CAA227CAG (2986A>G), TAC228ATC (2987T>A 2988A>T), ACC231AAG (2997C>A 2998C>G), TTA232CTT (2999T>C 3001A>T), TTG234CTG (3005T>C), GCA235AGA (3008G>A 3009C>G), AGT236TCA (3011A>T 3012G>C 3013T>A), GGG237GGT (3016G>T), TTT238TAT (3018T>A), TAT239CAT (3020T>C), CAG240CAA (3025G>A), GTG241ATT (3026G>A 3028G>T), GAG242AGG (3029G>A 3030A>G), GAC244ACA (3035G>A 3036A>C 3037C>A), CCT245AAG (3038C>A 3039C>A 3040T>G), CAA246GAG (3041C>G 3043A>G), ATA248GTC (3047A>G 3049A>C), TCG249CAG (3050T>C 3051C>A), ACC251ACT (3058C>T), GCG252ACA (3059G>A 3061G>A), AAC254AGG (3066A>G 3067C>G), GTA255ACT (3068G>A 3069T>C 3070A>T), GAA256CAT (3071G>C 3073A>T), CAC257TCT (3074C>T 3075A>C 3076C>T), GGG258GGC (3079G>C), CAT259CAC (3082T>C), TTT260TAT (3084T>A), GAA261GAG (3088A>G), TTC262TAC (3090T>A), CTT263CTA (3094T>A), CGA264GTA (3095C>G 3096G>T), CCT266CCC (3103T>C), ATG267TTT (3104A>T 3106G>T), GGA268GGC (3109A>C), TTA269CTC (3110T>C 3112A>C), AAA270ACT (3114A>C 3115A>T), AAC271AAT (3118C>T), TCA272GCA (3119T>G), CCA273CCT (3124A>T), TCT274TCA (3127T>A), ACT275AGT (3129C>G), AGA278GGG (3137A>G 3139A>G), GTT279CTG (3140G>C 3142T>G), GAC281AAT (3146G>A 3148C>T), AAT282CAT (3149A>C), GTC283GTT (3154C>T), CTA284TTT (3155C>T 3157A>T), AGA285AAG (3159G>A 3160A>G), GGT286GAA (3162G>A 3163T>A), CTC287CAC (3165T>A), CAA288CTT (3168A>T 3169A>T), AAT289AGG (3171A>G 3172T>G), AAC290AAG (3175C>G), ATC291TTC (3176A>T), TGT292ATT (3179T>A 3180G>T), CTC293TTG (3182C>T 3184C>G), GTC294GTT (3187C>T), TAC295TTT (3189A>T 3190C>T), CTT296--T (3191_3192delCT), GAC297GAT (3196C>T), ATT300CTG (3203A>C 3205T>G), GTC301ATC (3206G>A), TAT302TAC (3211T>C), AGT303AGC (3214T>C), ACT304AAG (3216C>A 3217T>G), TCC305AGT (3218T>A 3219C>G 3220C>T), CAG307GAA (3224C>G 3226G>A), GAA308GAT (3229A>T), CTG310TTG (3233C>T), GAG311CAA (3236G>C 3238G>A), AAC312CAC (3239A>C), GAA314AGG (3245G>A 3246A>G 3247A>G), CGA315GTT (3248C>G 3249G>T 3250A>T), GTT316ACT (3251G>A 3252T>C), TTC317TTT (3256C>T), CAA318AAC (3257C>A 3259A>C), AGA319TTA (3260A>T 3261G>T), CTT320TTG (3263C>T 3265T>G), AGA321ATC (3267G>T 3268A>C), GAA322CAA (3269G>C), AGT323CAT (3272A>C 3273G>A), AAC324AGA (3276A>G 3277C>A), TTC325CTG (3278T>C 3280C>G), AAA326TTG (3281A>T 3282A>T 3283A>G), ATT327ATC (3286T>C), CAA328AAT (3287C>A 3289A>T), ATG329AAG (3291T>A), GAC330TCA (3293G>T 3294A>C 3295C>A), TCC332TGT (3300C>G 3301C>T), GAA333TCT (3302G>T 3303A>C 3304A>T), TTC334TTT (3307C>T), TTG335GGA (3308T>G 3309T>G 3310G>A), AAG336GCA (3311A>G 3312A>C 3313G>A), CTC337TCA (3314C>T 3315T>C 3316C>A), GAA338AGG (3317G>A 3318A>G 3319A>G), ACT339GTT (3320A>G 3321C>T), GCT340GAA (3324C>A 3325T>A), TAT341TAC (3328T>C), GGT343GAT (3333G>A), ATC345TTT (3338A>T 3340C>T), AGC347TCT (3344A>T 3345G>C 3346C>T), AGG348GCT (3347A>G 3348G>C 3349G>T), GAC349GAG (3352C>G), GGT350GGA (3355T>A), ATC351GTG (3356A>G 3358C>G), AAG352GCT (3359A>G 3360A>C 3361G>T), CCT353ACA (3362C>A 3364T>A), AAC354GAT (3365A>G 3367C>T), GAT356AAG (3371G>A 3373T>G), TCC359GCA (3380T>G 3382C>A), GCT360GCA (3385T>A), ATT361GTA (3386A>G 3388T>A), CAA362TAA (3389C>T), AAA363GAA (3392A>G), TAT364TGG (3396A>G 3397T>G), CTG365CCA (3399T>C 3400G>A), CCA367CCT (3406A>T), AAG368CAT (3407A>C 3409G>T), ACC369ACT (3412C>T), CCT370ATA (3413C>A 3414C>T 3415T>A), GAA372GAG (3421A>G), ATA373TTG (3422A>T 3424A>G), AAA374AGG (3426A>G 3427A>G), CAA375GGA (3428C>G 3429A>G), TTA377CTT (3434T>C 3436A>T), GGC378GGG (3439C>G), CTT379TTG (3440C>T 3442T>G), CTC380ACA (3443C>A 3444T>C 3445C>A), TAC383TAT (3454C>T), CGA384AGG (3455C>A 3457A>G), AAA385AGG (3459A>G 3460A>G), CCA388AGA (3467C>A 3468C>G), GAT389AGA (3470G>A 3471A>G 3472T>A), TTT390TAT (3474T>A), GCA391GGG (3477C>G 3478A>G), CGA392ATC (3479C>A 3480G>T 3481A>C), CTC393ATT (3482C>A 3484C>T), ACA394AGT (3486C>G 3487A>T), CCC396CCT (3493C>T), ACA398AA. (3498C>A) | | | | | | | | |

##### Proteins

| | Begin | End | Coverage | Score | Concordance | Matches | Identities | I/D/M/F* | Stop Codons |
|---|---|---|---|---|---|---|---|---|---|
| ORF B (YP_009507248.1) | 160 | 398 | 22.0% | 632 | 39.3% | 234 (97.9%) | 108 (45.2%) | 0/5/1/1 | 1 |

| | Begin | End | Coverage | Score | Concordance | Matches | Identities | I/D/M/F* | Stop Codons |
|---|---|---|---|---|---|---|---|---|---|
| **NT** | **2783** | **3498** | **9.5%** | **22** | **1.6%** | **699 (97.6%)** | **366 (51.1%)** | **0/17** | |
| Protein mutations: | K160D (2783A>G 2785A>T), R168Q (2807A>C 2808G>A), D171C (2816G>T 2817A>G), S172N (2819T>A 2820C>A 2821T>C), A173P (2822G>C 2824A>T), W174Y (2826G>A 2827G>T), S175A (2828A>G 2829G>C), S176L (2832C>T), I178V (2837A>G), W179V (2840T>G 2841G>T 2842G>C), V180L (2843G>C 2845T>C), P182G (2849C>G 2850C>G 2851C>A), I185del (2858_2860delATC), A187S (2864G>A 2865C>G), G189_K192del (2870_2881delGGGAAACAAAAG), V196C (2891G>T 2892T>G 2893A>T), F199Y (2901T>A 2902C>T), K201A (2906A>G 2907A>C 2908G>T), E204K (2915G>A), K205L (2918A>T 2919A>T), D208K (2927G>A 2929T>A), Y211F (2937A>T 2938C>T), P212T (2939C>A 2941G>A), N215I (2949A>T), S217E (2954A>G 2955G>A 2956T>G), D218E (2959C>G), V219L (2960G>T), K222E (2969A>G 2971G>A), K225G (2978A>G 2979A>G 2980G>T), C226S (2982G>C), Y228I (2987T>A 2988A>T), T231K (2997C>A 2998C>G), A235R (3008G>A 3009C>G), F238Y (3018T>A), Y239H (3020T>C), V241I (3026G>A 3028G>T), E242R (3029G>A 3030A>G), D244T (3035G>A 3036A>C 3037C>A), P245K (3038C>A 3039C>A 3040T>G), Q246E (3041C>G 3043A>G), I248V (3047A>G 3049A>C), S249Q (3050T>C 3051C>A), A252T (3059G>A 3061G>A), N254R (3066A>G 3067C>G), V255T (3068G>A 3069T>C 3070A>T), E256H (3071G>C 3073A>T), H257S (3074C>T 3075A>C 3076C>T), F260Y (3084T>A), F262Y (3090T>A), R264V (3095C>G 3096G>T), M267F (3104A>T 3106G>T), K270T (3114A>C 3115A>T), S272A (3119T>G), T275S (3129C>G), R278G (3137A>G 3139A>G), V279L (3140G>C 3142T>G), D281N (3146G>A 3148C>T), N282H (3149A>C), L284F (3155C>T 3157A>T), R285K (3159G>A 3160A>G), G286E (3162G>A 3163T>A), L287H (3165T>A), Q288L (3168A>T 3169A>T), N289R (3171A>G 3172T>G), N290K (3175C>G), I291F (3176A>T), C292I (3179T>A 3180G>T), Y295F (3189A>T 3190C>T), I300L (3203A>C 3205T>G), V301I (3206G>A), T304K (3216C>A 3217T>G), Q307E (3224C>G 3226G>A), E308D (3229A>T), E311Q (3236G>C 3238G>A), N312H (3239A>C), E314R (3245G>A 3246A>G 3247A>G), R315V (3248C>G 3249G>T 3250A>T), V316T (3251G>A 3252T>C), Q318N (3257C>A 3259A>C), R319L (3260A>T 3261G>T), R321I (3267G>T 3268A>C), E322Q (3269G>C), S323H (3272A>C 3273G>A), N324R (3276A>G 3277C>A), F325L (3278T>C 3280C>G), K326L (3281A>T 3282A>T 3283A>G), Q328N (3287C>A 3289A>T), M329K (3291T>A), D330S (3293G>T 3294A>C 3295C>A), S332C (3300C>G 3301C>T), E333S (3302G>T 3303A>C 3304A>T), L335G (3308T>G 3309T>G 3310G>A), K336A (3311A>G 3312A>C 3313G>A), L337S (3314C>T 3315T>C 3316C>A), E338R (3317G>A 3318A>G 3319A>G), T339V (3320A>G 3321C>T), A340E (3324C>A 3325T>A), G343D (3333G>A), I345F (3338A>T 3340C>T), R348A (3347A>G 3348G>C 3349G>T), D349E (3352C>G), I351V (3356A>G 3358C>G), K352A (3359A>G 3360A>C 3361G>T), P353T (3362C>A 3364T>A), N354D (3365A>G 3367C>T), D356K (3371G>A 3373T>G), S359A (3380T>G 3382C>A), I361V (3386A>G 3388T>A), Q362* (3389C>T), K363E (3392A>G), Y364W (3396A>G 3397T>G), L365P (3399T>C 3400G>A), K368H (3407A>C 3409G>T), P370I (3413C>A 3414C>T 3415T>A), I373L (3422A>T 3424A>G), K374R (3426A>G 3427A>G), Q375G (3428C>G 3429A>G), L380T (3443C>A 3444T>C 3445C>A), K385R (3459A>G 3460A>G), P388R (3467C>A 3468C>G), D389R (3470G>A 3471A>G 3472T>A), F390Y (3474T>A), A391G (3477C>G 3478A>G), R392I (3479C>A 3480G>T 3481A>C), L393I (3482C>A 3484C>T), T394S (3486C>G 3487A>T) | | | | | | | | |
| Codon mutations: | AAA160GAT (2783A>G 2785A>T), TTG162CTT (2789T>C 2791G>T), GAC163GAT (2794C>T), GGA165GGC (2800A>C), ATT166ATC (2803T>C), ATA167ATC (2806A>C), AGA168CAA (2807A>C 2808G>A), CCA169CCC (2812A>C), TCA170AGC (2813T>A 2814C>G 2815A>C), GAC171TGC (2816G>T 2817A>G), TCT172AAC (2819T>A 2820C>A 2821T>C), GCA173CCT (2822G>C 2824A>T), TGG174TAT (2826G>A 2827G>T), AGC175GCC (2828A>G 2829G>C), TCA176TTA (2832C>T), CCC177CCT (2836C>T), ATA178GTA (2837A>G), TGG179GTC (2840T>G 2841G>T 2842G>C), GTT180CTC (2843G>C 2845T>C), CCC182GGA (2849C>G 2850C>G 2851C>A), AAA184AAG (2857A>G), ATC185del (2858_2860delATC), GAC186GAT (2863C>T), GCT187AGT (2864G>A 2865C>G), TCT188TCA (2869T>A), GGG189_AAG192del (2870_2881delGGGAAACAAAAG), CGT194AGG (2885C>A 2887T>G), CTC195TTA (2888C>T 2890C>A), GTA196TGT (2891G>T 2892T>G 2893A>T), GTT197GTA (2896T>A), TTC199TAT (2901T>A 2902C>T), CGT200AGG (2903C>A 2905T>G), AAG201GCT (2906A>G 2907A>C 2908G>T), AAC203AAT (2914C>T), GAG204AAG (2915G>A), AAG205TTG (2918A>T 2919A>T), GAT208AAA (2927G>A 2929T>A), GAC209GAT (2932C>T), TAC211TTT (2937A>T 2938C>T), CCG212ACA (2939C>A 2941G>A), ATA213ATT (2944A>T), CCA214CCC (2947A>C), AAC215ATC (2949A>T), ATA216ATT (2953A>T), AGT217GAG (2954A>G 2955G>A 2956T>G), GAC218GAG (2959C>G), GTA219TTA (2960G>T), CTT220CTG (2965T>G), GAC221GAT (2968C>T), AAG222GAA (2969A>G 2971G>A), TTA223TTG (2974A>G), GGT224GGA (2977T>A), AAG225GGT (2978A>G 2979A>G 2980G>T), TGC226TCC (2982G>C), CAA227CAG (2986A>G), TAC228ATC (2987T>A 2988A>T), ACC231AAG (2997C>A 2998C>G), TTA232CTT (2999T>C 3001A>T), TTG234CTG (3005T>C), GCA235AGA (3008G>A 3009C>G), AGT236TCA (3011A>T 3012G>C 3013T>A), GGG237GGT (3016G>T), TTT238TAT (3018T>A), TAT239CAT (3020T>C), CAG240CAA (3025G>A), GTG241ATT (3026G>A 3028G>T), GAG242AGG (3029G>A 3030A>G), GAC244ACA (3035G>A 3036A>C 3037C>A), CCT245AAG (3038C>A 3039C>A 3040T>G), CAA246GAG (3041C>G 3043A>G), ATA248GTC (3047A>G 3049A>C), TCG249CAG (3050T>C 3051C>A), ACC251ACT (3058C>T), GCG252ACA (3059G>A 3061G>A), AAC254AGG (3066A>G 3067C>G), GTA255ACT (3068G>A 3069T>C 3070A>T), GAA256CAT (3071G>C 3073A>T), CAC257TCT (3074C>T 3075A>C 3076C>T), GGG258GGC (3079G>C), CAT259CAC (3082T>C), TTT260TAT (3084T>A), GAA261GAG (3088A>G), TTC262TAC (3090T>A), CTT263CTA (3094T>A), CGA264GTA (3095C>G 3096G>T), CCT266CCC (3103T>C), ATG267TTT (3104A>T 3106G>T), GGA268GGC (3109A>C), TTA269CTC (3110T>C 3112A>C), AAA270ACT (3114A>C 3115A>T), AAC271AAT (3118C>T), TCA272GCA (3119T>G), CCA273CCT (3124A>T), TCT274TCA (3127T>A), ACT275AGT (3129C>G), AGA278GGG (3137A>G 3139A>G), GTT279CTG (3140G>C 3142T>G), GAC281AAT (3146G>A 3148C>T), AAT282CAT (3149A>C), GTC283GTT (3154C>T), CTA284TTT (3155C>T 3157A>T), AGA285AAG (3159G>A 3160A>G), GGT286GAA (3162G>A 3163T>A), CTC287CAC (3165T>A), CAA288CTT (3168A>T 3169A>T), AAT289AGG (3171A>G 3172T>G), AAC290AAG (3175C>G), ATC291TTC (3176A>T), TGT292ATT (3179T>A 3180G>T), CTC293TTG (3182C>T 3184C>G), GTC294GTT (3187C>T), TAC295TTT (3189A>T 3190C>T), CTT296--T (3191_3192delCT), GAC297GAT (3196C>T), ATT300CTG (3203A>C 3205T>G), GTC301ATC (3206G>A), TAT302TAC (3211T>C), AGT303AGC (3214T>C), ACT304AAG (3216C>A 3217T>G), TCC305AGT (3218T>A 3219C>G 3220C>T), CAG307GAA (3224C>G 3226G>A), GAA308GAT (3229A>T), CTG310TTG (3233C>T), GAG311CAA (3236G>C 3238G>A), AAC312CAC (3239A>C), GAA314AGG (3245G>A 3246A>G 3247A>G), CGA315GTT (3248C>G 3249G>T 3250A>T), GTT316ACT (3251G>A 3252T>C), TTC317TTT (3256C>T), CAA318AAC (3257C>A 3259A>C), AGA319TTA (3260A>T 3261G>T), CTT320TTG (3263C>T 3265T>G), AGA321ATC (3267G>T 3268A>C), GAA322CAA (3269G>C), AGT323CAT (3272A>C 3273G>A), AAC324AGA (3276A>G 3277C>A), TTC325CTG (3278T>C 3280C>G), AAA326TTG (3281A>T 3282A>T 3283A>G), ATT327ATC (3286T>C), CAA328AAT (3287C>A 3289A>T), ATG329AAG (3291T>A), GAC330TCA (3293G>T 3294A>C 3295C>A), TCC332TGT (3300C>G 3301C>T), GAA333TCT (3302G>T 3303A>C 3304A>T), TTC334TTT (3307C>T), TTG335GGA (3308T>G 3309T>G 3310G>A), AAG336GCA (3311A>G 3312A>C 3313G>A), CTC337TCA (3314C>T 3315T>C 3316C>A), GAA338AGG (3317G>A 3318A>G 3319A>G), ACT339GTT (3320A>G 3321C>T), GCT340GAA (3324C>A 3325T>A), TAT341TAC (3328T>C), GGT343GAT (3333G>A), ATC345TTT (3338A>T 3340C>T), AGC347TCT (3344A>T 3345G>C 3346C>T), AGG348GCT (3347A>G 3348G>C 3349G>T), GAC349GAG (3352C>G), GGT350GGA (3355T>A), ATC351GTG (3356A>G 3358C>G), AAG352GCT (3359A>G 3360A>C 3361G>T), CCT353ACA (3362C>A 3364T>A), AAC354GAT (3365A>G 3367C>T), GAT356AAG (3371G>A 3373T>G), TCC359GCA (3380T>G 3382C>A), GCT360GCA (3385T>A), ATT361GTA (3386A>G 3388T>A), CAA362TAA (3389C>T), AAA363GAA (3392A>G), TAT364TGG (3396A>G 3397T>G), CTG365CCA (3399T>C 3400G>A), CCA367CCT (3406A>T), AAG368CAT (3407A>C 3409G>T), ACC369ACT (3412C>T), CCT370ATA (3413C>A 3414C>T 3415T>A), GAA372GAG (3421A>G), ATA373TTG (3422A>T 3424A>G), AAA374AGG (3426A>G 3427A>G), CAA375GGA (3428C>G 3429A>G), TTA377CTT (3434T>C 3436A>T), GGC378GGG (3439C>G), CTT379TTG (3440C>T 3442T>G), CTC380ACA (3443C>A 3444T>C 3445C>A), TAC383TAT (3454C>T), CGA384AGG (3455C>A 3457A>G), AAA385AGG (3459A>G 3460A>G), CCA388AGA (3467C>A 3468C>G), GAT389AGA (3470G>A 3471A>G 3472T>A), TTT390TAT (3474T>A), GCA391GGG (3477C>G 3478A>G), CGA392ATC (3479C>A 3480G>T 3481A>C), CTC393ATT (3482C>A 3484C>T), ACA394AGT (3486C>G 3487A>T), CCC396CCT (3493C>T), ACA398AA. (3498C>A) | | | | | | | | |

*: Inserts / Deletes / Misaligned / Frameshifts

#### Analysis details

This analysis was performed with panviral2.64

### NGS Details (UN8): Dioscovirus dioscoreae

#### Assembly

| | |
|---|---|
| **Coverage Length** | 1064 (2 contig(s)) |
| **Depth Of Coverage** | 10.3 |
| **Number Of Reads** | 87 |
| **Reads Per Million** | 1.45 rpm (after QC) |
| **Ambiguities** | 0 |
| **Assembly Method** | de novo + reference guided assembly |
| **Consensus Caller** | Bcf Tools |

#### Coverage Map

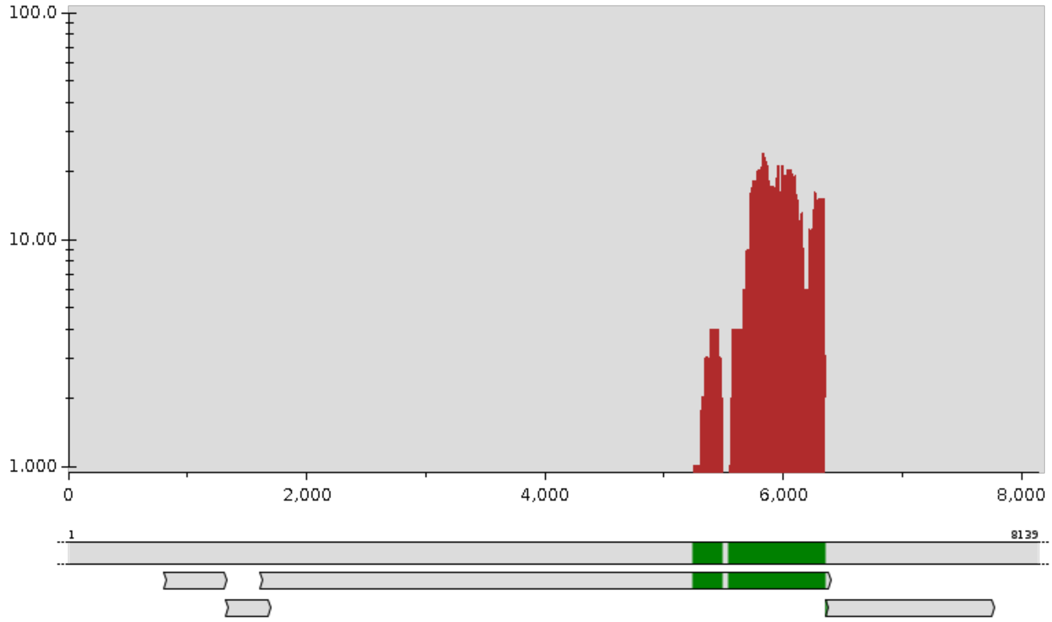

#### Assignment

| | |
|---|---|
| **Type** | Dioscovirus dioscoreae (Taxonomy ID: 3052184) |
| **Reference Genome** | NC_040712.1 |
| **NT Identity (%)** | 61.7121 |
| **AA Identity (%)** | 49.8592 |
| **Number Of Stop Codons** | 1 |
| **Number Of CDS** | 4 |

#### Alignment

| | |
|---|---|
| **Alignment Score** | 461.0 (NT) + 1175.0 (AA) = 1636.0 |
| **Concordance (%)** | 37.5933 |

| Alignment Method | Global, seeded, nucleotide + amino acids (AGA) |
|---|---|

#### Genome Region

Sequence starts at position 5241 and ends at position 6355 relative to NC_040712.1 reference sequence.

#### Alignment Detailed Statistics

| | Begin | End | Coverage | Score | Concordance | Matches | Identities | I/D/M/F* | Stop Codons |
|---|---|---|---|---|---|---|---|---|---|
| **NT** | **5241** | **6355** | **13.1%** | **461** | **22.2%** | **1057 (98.8%)** | **656 (61.3%)** | **6/7** | |
| Mutations: | 5248G>C, 5250C>A, 5252T>A, 5255G>A, 5259T>A, 5260A>T, 5261C>A, 5270G>A, 5271C>A, 5276A>T, 5280G>A, 5288A>T, 5289T>A, 5290A>G, 5291C>A, 5293A>C, 5294A>G, 5297C>T, 5298C>G, 5299A>C, 5303C>T, 5305C>A, 5307C>T, 5309C>A, 5313A>G, 5314G>A, 5317T>A, 5318T>A, 5319G>A, 5320A>G, 5321C>T, 5322T>G, 5325C>T, 5327C>A, 5328C>A, 5330A>T, 5331C>A, 5332T>A, 5335A>G, 5339T>G, 5340A>T, 5345C>T, 5348G>A, 5350T>A, 5351T>A, 5352G>T, 5354C>A, 5356A>T, 5360C>T, 5370C>T, 5371T>G, 5372A>T, 5375G>A, 5379G>C, 5380G>T, 5381A>C, 5383T>A, 5384C>T, 5385C>T, 5386A>G, 5387C>G, 5390G>A, 5391A>G, 5395T>A, 5396G>A, 5400G>C, 5406C>G, 5408G>C, 5412delA, 5424G>A, 5429T>C, 5430G>A, 5431T>C, 5432C>A, 5435C>T, 5438A>T, 5439C>G, 5444T>A, 5457A>C, 5461T>C, 5462A>C, 5471C>T, 5474A>T, 5475C>T, 5477C>A, 5482A>C, 5483T>A, 5486A>G, 5549T>C, 5553A>G, 5558C>T, 5561T>C, 5564T>C, 5565T>C, 5567G>A, 5568A>G, 5573T>C, 5574T>A, 5575C>G, 5577G>A, 5578A>G, 5580A>G, 5581G>A, 5585A>G, 5589C>G, 5595G>C, 5598C>G, 5607T>C, 5608T>A, 5610C>A, 5611A>C, 5612G>A, 5613T>A, 5615C>T, 5618C>T, 5619C>A, 5620A>G, 5621A>G, 5622G>T, 5626G>T, 5627T>C, 5628A>G, 5629A>T, 5630G>A, 5631G>A, 5632A>G, 5633A>T, 5634G>A, 5639G>A, 5640C>A, 5642T>A, 5645T>A, 5646T>A, 5648G>A, 5649T>A, 5651A>C, 5656C>A, 5657A>G, 5661T>A, 5664A>G, 5667A>T, 5670G>T, 5672A>T, 5673G>A, 5674T>A, 5676G>A, 5679A>C, 5685G>A, 5687A>C, 5691C>T, 5693T>A, 5696C>A, 5697C>G, 5700G>A, 5701A>C, 5702A>T, 5703A>T, 5711A>T, 5714A>G, 5717A>G, 5718G>A, 5720T>A, 5721C>A, 5726C>A, 5739T>G, 5740T>C, 5741A>T, 5748A>G, 5756A>T, 5757T>A, 5759T>G, 5762T>A, 5763_5765delGAA, 5768C>T, 5769C>A, 5777A>T, 5779C>A, 5781T>A, 5782T>C, 5786G>A, 5788G>A, 5790C>T, 5795G>A, 5798G>A, 5807G>A, 5808A>T, 5811T>G, 5813A>G, 5816C>T, 5822C>T, 5828T>C, 5830A>T, 5834C>A, 5836C>A, 5843G>A, 5844A>G, 5846T>A, 5850T>A, 5851A>T, 5853A>G, 5855T>A, 5856A>G, 5859A>C, 5860T>C, 5864T>A, 5868A>G, 5869A>C, 5870T>A, 5873G>A, 5874T>A, 5875G>T, 5876T>A, 5877T>G, 5878C>G, 5879C>A, 5882C>T, 5889G>C, 5893G>A, 5895A>C, 5897A>C, 5903T>C, 5904T>A, 5906T>A, 5907C>G, 5910G>A, 5912C>T, 5913T>A, 5914G>T, 5915G>T, 5918G>A, 5919A>C, 5924T>A, 5925A>C, 5926G>A, 5928A>C, 5933C>T, 5940G>A, 5941T>A, 5945T>A, 5947C>A, 5948C>A, 5951A>C, 5952T>A, 5957C>A, 5958C>G, 5959C>A, 5960T>C, 5963G>A, 5965T>G, 5966T>C, 5967T>A, 5973A>C, 5974A>T, 5976C>G, 5977C>A, 5979C>T, 5980A>C, 5982A>G, 5983G>A, 5988A>C, 5990T>A, 5999G>A, 6005T>C, 6009T>A, 6010C>G, 6011T>C, 6014A>C, 6015G>C, 6026G>A, 6028G>C, 6029A>T, 6030A>G, 6032T>A, 6039T>G, 6040G>C, 6041G>A, 6047C>A, 6048T>A, 6049C>A, 6050T>C, 6051C>A, 6054C>T, 6056A>C, 6057G>A, 6058A>G, 6060T>C, 6062A>T, 6063C>A, 6064C>A, 6065T>A, 6066T>A, 6067C>A, 6069A>G, 6070C>A, 6082C>G, 6083A>T, 6084C>G, 6085G>C, 6086G>T, 6089T>C, 6090T>C, 6091G>A, 6092T>A, 6100G>A, 6101C>A, 6104T>C, 6107_6108insGAAAAG, 6108T>G, 6109C>G, 6111G>A, 6112C>A, 6116C>G, 6117T>A, 6118C>G, 6120G>A, 6121C>G, 6122A>T, 6130C>T, 6131T>C, 6135A>G, 6138A>T, 6140G>A, 6144T>A, 6145G>T, 6146T>A, 6154T>G, 6156T>C, 6159G>A, 6163A>G, 6164G>T, 6167T>C, 6171A>T, 6175T>A, 6177T>A, 6179A>T, 6181A>T, 6182T>A, 6183G>A, 6185G>C, 6190A>G, 6191A>T, 6195T>A, 6197T>A, 6198A>G, 6199C>T, 6200C>A, 6201C>G, 6203T>A, 6206G>A, 6212C>T, 6215T>C, 6216A>G, 6217T>A, 6218T>A, 6221A>T, 6224T>A, 6230T>A, 6234T>C, 6237A>C, 6239C>A, 6243C>A, 6245T>A, 6246A>G, 6248T>C, 6249A>C, 6251T>G, 6257A>T, 6258T>A, 6259T>G, 6260G>T, 6262A>G, 6265T>A, 6268A>G, 6269T>A, 6270C>A, 6272T>A, 6276G>C, 6284C>T, 6285T>A, 6286G>T, 6287T>A, 6290T>C, 6291T>A, 6292T>C, 6293A>C, 6297A>T, 6299A>T, 6300G>A, 6301G>T, 6304T>A, 6305A>C, 6306G>C, 6307G>C, 6309_6311delTTA, 6313C>A, 6318A>G, 6319A>T, 6320A>T, 6321A>T, 6323A>C, 6327C>T, 6332C>A, 6333C>A, 6335G>A, 6338T>A | | | | | | | | |

**CDS**

| | Begin | End | Coverage | Score | Concordance | Matches | Identities | I/D/M/F* | Stop Codons |
|---|---|---|---|---|---|---|---|---|---|
| EXK67_gp3 | 1211 | 1582 | 22.2% | 1175 | 47.7% | 353 (98.9%) | 177 (49.6%) | 2/2/1/1 | 1 |
| Protein mutations: | S1213T (5248G>C), M1215I (5255G>A), Y1217I (5259T>A 5260A>T 5261C>A), D1224N (5280G>A), Y1227R (5289T>A 5290A>G 5291C>A), K1228T (5293A>C 5294A>G), Q1230A (5298C>G 5299A>C), T1232K (5305C>A), S1235D (5313A>G 5314G>A), I1236K (5317T>A 5318T>A), D1237S (5319G>A 5320A>G 5321C>T), Y1238D (5322T>G), L1240I (5328C>A 5330A>T), L1241N (5331C>A 5332T>A), K1242R (5335A>G), I1243M (5339T>G), K1244* (5340A>T), I1247K (5350T>A 5351T>A), V1248L (5352G>T 5354C>A), Y1249F (5356A>T), L1254C (5370C>T 5371T>G 5372A>T), G1257L (5379G>C 5380G>T 5381A>C), F1258Y (5383T>A 5384C>T), H1259W (5385C>T 5386A>G 5387C>G), I1261V (5391A>G), M1262K (5395T>A 5396G>A), D1264H (5400G>C), Q1266D (5406C>G 5408G>C), A1272T (5424G>A), V1274T (5430G>A 5431T>C 5432C>A), Q1277E (5439C>G), I1283L (5457A>C), V1284A (5461T>C 5462A>C), N1291T (5482A>C 5483T>A), I1315V (5553A>G), I1320V (5568A>G), E1323R (5577G>A 5578A>G), S1324D (5580A>G 5581G>A), I1325M (5585A>G), Q1327E (5589C>G), V1329L (5595G>C), Q1330E (5598C>G), L1333Q (5607T>C 5608T>A), Q1334T (5610C>A 5611A>C 5612G>A), F1335I (5613T>A 5615C>T), Q1337R (5619C>A 5620A>G 5621A>G), V1338L (5622G>T), C1339F (5626G>T 5627T>C), K1340V (5628A>G 5629A>T 5630G>A), E1341S (5631G>A 5632A>G 5633A>T), E1342K (5634G>A), L1344I (5640C>A 5642T>A), L1346I (5646T>A 5648G>A), S1347T (5649T>A 5651A>C), T1349K (5656C>A 5657A>G), L1351M (5661T>A), K1352E (5664A>G), I1353L (5667A>T), G1354C (5670G>T 5672A>T), V1355K (5673G>A 5674T>A), A1356T (5676G>A), N1357H (5679A>C), E1359N (5685G>A 5687A>C), L1363V (5697C>G), E1364T (5700G>A 5701A>C 5702A>T), I1365L (5703A>T), E1367D (5711A>T), V1370I (5718G>A 5720T>A), Q1371K (5721C>A), L1377A (5739T>G 5740T>C 5741A>T), I1380V (5748A>G), E1382D (5756A>T), F1383M (5757T>A 5759T>G), E1385del (5763_5765delGAA), Q1387K (5769C>A), E1389D (5777A>T), T1390K (5779C>A), L1391T (5781T>A 5782T>C), G1393D (5788G>A), I1400L (5808A>T), L1401V (5811T>G 5813A>G), Y1407F (5830A>T), P1409Q (5836C>A), S1412G (5844A>G 5846T>A), Y1414I (5850T>A 5851A>T), T1415A (5853A>G 5855T>A), R1416G (5856A>G), I1417P (5859A>C 5860T>C), F1418L (5864T>A), N1420A (5868A>G 5869A>C 5870T>A), C1422I (5874T>A 5875G>T 5876T>A), S1423G (5877T>G 5878C>G 5879C>A), E1427Q (5889G>C), R1428K (5893G>A), K1429H (5895A>C 5897A>C), S1432T (5904T>A 5906T>A), Q1433E (5907C>G), D1434N (5910G>A 5912C>T), W1435I (5913T>A 5914G>T 5915G>T), M1437L (5919A>C), R1439Q (5925A>C 5926G>A), K1440Q (5928A>C), V1444K (5940G>A 5941T>A), T1446K (5947C>A 5948C>A), K1447N (5951A>C), L1448I (5952T>A), P1450D (5958C>G 5959C>A 5960T>C), I1452S (5965T>G 5966T>C), L1453I (5967T>A), K1455L (5973A>C 5974A>T), P1456E (5976C>G 5977C>A), Q1457S (5979C>T 5980A>C), S1458D (5982A>G 5983G>A), I1460L (5988A>C 5990T>A), E1469Q (6015G>C), G1473A (6028G>C 6029A>T), I1474V (6030A>G 6032T>A), W1477A (6039T>G 6040G>C 6041G>A), S1480N (6048T>A 6049C>A 6050T>C), Q1481K (6051C>A), Q1482Y (6054C>T 6056A>C), D1483S (6057G>A 6058A>G), S1484P (6060T>C 6062A>T), P1485K (6063C>A 6064C>A 6065T>A), S1486K (6066T>A 6067C>A), T1487E (6069A>G 6070C>A), S1491C (6082C>G 6083A>T), R1492A (6084C>G 6085G>C 6086G>T), C1494Q (6090T>C 6091G>A 6092T>A), S1497K (6100G>A 6101C>A), K1499_S1500insEK (6107_6108insGAAAAG), S1500G (6108T>G 6109C>G), A1501N (6111G>A 6112C>A), I1502M (6116C>G), A1504S (6120G>A 6121C>G 6122A>T), A1507V (6130C>T 6131T>C), I1509V (6135A>G), M1510L (6138A>T 6140G>A), C1512I (6144T>A 6145G>T 6146T>A), V1515G (6154T>G), D1517N (6159G>A), K1518S (6163A>G 6164G>T), I1521L (6171A>T), F1522Y (6175T>A), L1523I (6177T>A 6179A>T), Y1524L (6181A>T 6182T>A), E1525N (6183G>A 6185G>C), K1527S (6190A>G 6191A>T), F1529I (6195T>A 6197T>A), T1530V (6198A>G 6199C>T 6200C>A), L1531V (6201C>G 6203T>A), I1536E (6216A>G 6217T>A 6218T>A), N1540K (6230T>A), Y1542H (6234T>C), N1543Q (6237A>C 6239C>A), L1545I (6243C>A 6245T>A), N1546D (6246A>G 6248T>C), N1547Q (6249A>C 6251T>G), K1549N (6257A>T), L1550S (6258T>A 6259T>G 6260G>T), N1551S (6262A>G), I1552K (6265T>A), N1553R (6268A>G 6269T>A), V1556L (6276G>C), C1559I (6285T>A 6286G>T 6287T>A), L1561T (6291T>A 6292T>C 6293A>C), T1563S (6297A>T 6299A>T), G1564I (6300G>A 6301G>T), L1565Y (6304T>A 6305A>C), G1566P (6306G>C 6307G>C), L1567del (6309_6311delTTA), T1568N (6313C>A), K1570V (6318A>G 6319A>T 6320A>T), I1571F (6321A>T 6323A>C), H1573Y (6327C>T), Q1575K (6333C>A 6335G>A) | | | | | | | | |

| | Begin | End | Coverage | Score | Concordance | Matches | Identities | I/D/M/F* | Stop Codons |
|---|---|---|---|---|---|---|---|---|---|
| **NT** | **5241** | **6355** | **13.1%** | **461** | **22.2%** | **1057 (98.8%)** | **656 (61.3%)** | **6/7** | |
| Codon mutations: | AGT1213ACT (5248G>C), CGT1214AGA (5250C>A 5252T>A), ATG1215ATA (5255G>A), TAC1217ATA (5259T>A 5260A>T 5261C>A), AAG1220AAA (5270G>A), CGA1221AGA (5271C>A), CTA1222CTT (5276A>T), GAT1224AAT (5280G>A), ACA1226ACT (5288A>T), TAC1227AGA (5289T>A 5290A>G 5291C>A), AAA1228ACG (5293A>C 5294A>G), GAC1229GAT (5297C>T), CAA1230GCA (5298C>G 5299A>C), TAC1231TAT (5303C>T), ACA1232AAA (5305C>A), CTC1233TTA (5307C>T 5309C>A), AGT1235GAT (5313A>G 5314G>A), ATT1236AAA (5317T>A 5318T>A), GAC1237AGT (5319G>A 5320A>G 5321C>T), TAT1238GAT (5322T>G), CTC1239TTA (5325C>T 5327C>A), CTA1240ATT (5328C>A 5330A>T), CTT1241AAT (5331C>A 5332T>A), AAA1242AGA (5335A>G), ATT1243ATG (5339T>G), AAA1244TAA (5340A>T), GAC1245GAT (5345C>T), AAG1246AAA (5348G>A), ATT1247AAA (5350T>A 5351T>A), GTC1248TTA (5352G>T 5354C>A), TAC1249TTC (5356A>T), AGC1250AGT (5360C>T), CTA1254TGT (5370C>T 5371T>G 5372A>T), AAG1255AAA (5375G>A), GGA1257CTC (5379G>C 5380G>T 5381A>C), TTC1258TAT (5383T>A 5384C>T), CAC1259TGG (5385C>T 5386A>G 5387C>G), CAG1260CAA (5390G>A), ATT1261GTT (5391A>G), ATG1262AAA (5395T>A 5396G>A), GAT1264CAT (5400G>C), CAG1266GAC (5406C>G 5408G>C), ATA1268-TA (5412delA), GCA1272ACA (5424G>A), TTT1273TTC (5429T>C), GTC1274ACA (5430G>A 5431T>C 5432C>A), TGC1275TGT (5435C>T), CCA1276CCT (5438A>T), CAG1277GAG (5439C>G), GGT1278GGA (5444T>A), ATA1283CTA (5457A>C), GTA1284GCC (5461T>C 5462A>C), TTC1287TTT (5471C>T), GGA1288GGT (5474A>T), CTC1289TTA (5475C>T 5477C>A), AAT1291ACA (5482A>C 5483T>A), GCA1292GCG (5486A>G), GTT1313.TC (5549T>C), ATA1315GTA (5553A>G), GAC1316GAT (5558C>T), GAT1317GAC (5561T>C), ATT1318ATC (5564T>C), TTG1319CTA (5565T>C 5567G>A), ATA1320GTA (5568A>G), TTT1321TTC (5573T>C), TCT1322AGT (5574T>A 5575C>G), GAA1323AGA (5577G>A 5578A>G), AGT1324GAT (5580A>G 5581G>A), ATA1325ATG (5585A>G), CAA1327GAA (5589C>G), GTA1329CTA (5595G>C), CAA1330GAA (5598C>G), TTA1333CAA (5607T>C 5608T>A), CAG1334ACA (5610C>A 5611A>C 5612G>A), TTC1335ATT (5613T>A 5615C>T), TTC1336TTT (5618C>T), CAA1337AGG (5619C>A 5620A>G 5621A>G), GTA1338TTA (5622G>T), TGT1339TTC (5626G>T 5627T>C), AAG1340GTA (5628A>G 5629A>T 5630G>A), GAA1341AGT (5631G>A 5632A>G 5633A>T), GAA1342AAA (5634G>A), GGG1343GGA (5639G>A), CTT1344ATA (5640C>A 5642T>A), ATT1345ATA (5645T>A), TTG1346ATA (5646T>A 5648G>A), TCA1347ACC (5649T>A 5651A>C), ACA1349AAG (5656C>A 5657A>G), TTG1351ATG (5661T>A), AAA1352GAA (5664A>G), ATA1353TTA (5667A>T), GGA1354TGT (5670G>T 5672A>T), GTA1355AAA (5673G>A 5674T>A), GCA1356ACA (5676G>A), AAT1357CAT (5679A>C), GAA1359AAC (5685G>A 5687A>C), CTT1361TTA (5691C>T 5693T>A), GGC1362GGA (5696C>A), CTA1363GTA (5697C>G), GAA1364ACT (5700G>A 5701A>C 5702A>T), ATA1365TTA (5703A>T), GAA1367GAT (5711A>T), GGA1368GGG (5714A>G), AAA1369AAG (5717A>G), GTT1370ATA (5718G>A 5720T>A), CAA1371AAA (5721C>A), CTC1372CTA (5726C>A), TTA1377GCT (5739T>G 5740T>C 5741A>T), ATA1380GTA (5748A>G), GAA1382GAT (5756A>T), TTT1383ATG (5757T>A 5759T>G), CCT1384CCA (5762T>A), GAA1385del (5763_5765delGAA), GAC1386GAT (5768C>T), CAA1387AAA (5769C>A), GAA1389GAT (5777A>T), ACA1390AAA (5779C>A), TTA1391ACA (5781T>A 5782T>C), AAG1392AAA (5786G>A), GGT1393GAT (5788G>A), CTA1394TTA (5790C>T), CAG1395CAA (5795G>A), AAG1396AAA (5798G>A), GGG1399GGA (5807G>A), ATA1400TTA (5808A>T), TTA1401GTG (5811T>G 5813A>G), AAC1402AAT (5816C>T), GCC1404GCT (5822C>T), AAT1406AAC (5828T>C), TAT1407TTT (5830A>T), ATC1408ATA (5834C>A), CCA1409CAA (5836C>A), TTG1411TTA (5843G>A), AGT1412GGA (5844A>G 5846T>A), TAT1414ATT (5850T>A 5851A>T), ACT1415GCA (5853A>G 5855T>A), AGA1416GGA (5856A>G), ATA1417CCA (5859A>C 5860T>C), TTT1418TTA (5864T>A), AAT1420GCA (5868A>G 5869A>C 5870T>A), AAG1421AAA (5873G>A), TGT1422ATA (5874T>A 5875G>T 5876T>A), TCC1423GGA (5877T>G 5878C>G 5879C>A), AGC1424AGT (5882C>T), GAA1427CAA (5889G>C), AGA1428AAA (5893G>A), AAA1429CAC (5895A>C 5897A>C), AAT1431AAC (5903T>C), TCT1432ACA (5904T>A 5906T>A), CAA1433GAA (5907C>G), GAC1434AAT (5910G>A 5912C>T), TGG1435ATT (5913T>A 5914G>T 5915G>T), AAG1436AAA (5918G>A), ATG1437CTG (5919A>C), GTT1438GTA (5924T>A), AGA1439CAA (5925A>C 5926G>A), AAA1440CAA (5928A>C), ATC1441ATT (5933C>T), GTA1444AAA (5940G>A 5941T>A), ATT1445ATA (5945T>A), ACC1446AAA (5947C>A 5948C>A), AAA1447AAC (5951A>C), TTA1448ATA (5952T>A), CCC1449CCA (5957C>A), CCT1450GAC (5958C>G 5959C>A 5960T>C), TTG1451TTA (5963G>A), ATT1452AGC (5965T>G 5966T>C), TTA1453ATA (5967T>A), AAA1455CTA (5973A>C 5974A>T), CCA1456GAA (5976C>G 5977C>A), CAA1457TCA (5979C>T 5980A>C), AGT1458GAT (5982A>G 5983G>A), ATT1460CTA (5988A>C 5990T>A), GAG1463GAA (5999G>A), GAT1465GAC (6005T>C), TCT1467AGC (6009T>A 6010C>G 6011T>C), CTA1468CTC (6014A>C), GAA1469CAA (6015G>C), GGG1472GGA (6026G>A), GGA1473GCT (6028G>C 6029A>T), ATT1474GTA (6030A>G 6032T>A), TGG1477GCA (6039T>G 6040G>C 6041G>A), CCC1479CCA (6047C>A), TCT1480AAC (6048T>A 6049C>A 6050T>C), CAA1481AAA (6051C>A), CAA1482TAC (6054C>T 6056A>C), GAT1483AGT (6057G>A 6058A>G), TCA1484CCT (6060T>C 6062A>T), CCT1485AAA (6063C>A 6064C>A 6065T>A), TCA1486AAA (6066T>A 6067C>A), ACA1487GAA (6069A>G 6070C>A), TCA1491TGT (6082C>G 6083A>T), CGG1492GCT (6084C>G 6085G>C 6086G>T), TAT1493TAC (6089T>C), TGT1494CAA (6090T>C 6091G>A 6092T>A), AGC1497AAA (6100G>A 6101C>A), TAT1498TAC (6104T>C), AAA1499_TCT1500insGAAAAG (6107_6108insGAAAAG), TCT1500GGT (6108T>G 6109C>G), GCC1501AAC (6111G>A 6112C>A), ATC1502ATG (6116C>G), TCT1503AGT (6117T>A 6118C>G), GCA1504AGT (6120G>A 6121C>G 6122A>T), GCT1507GTC (6130C>T 6131T>C), ATA1509GTA (6135A>G), ATG1510TTA (6138A>T 6140G>A), TGT1512ATA (6144T>A 6145G>T 6146T>A), GTA1515GGA (6154T>G), TTA1516CTA (6156T>C), GAT1517AAT (6159G>A), AAG1518AGT (6163A>G 6164G>T), TTT1519TTC (6167T>C), ATA1521TTA (6171A>T), TTT1522TAT (6175T>A), TTA1523ATT (6177T>A 6179A>T), TAT1524TTA (6181A>T 6182T>A), GAG1525AAC (6183G>A 6185G>C), AAA1527AGT (6190A>G 6191A>T), TTT1529ATA (6195T>A 6197T>A), ACC1530GTA (6198A>G 6199C>T 6200C>A), CTT1531GTA (6201C>G 6203T>A), AGG1532AGA (6206G>A), GAC1534GAT (6212C>T), TGT1535TGC (6215T>C), ATT1536GAA (6216A>G 6217T>A 6218T>A), GCA1537GCT (6221A>T), ATT1538ATA (6224T>A), AAT1540AAA (6230T>A), TAT1542CAT (6234T>C), AAC1543CAA (6237A>C 6239C>A), CTT1545ATA (6243C>A 6245T>A), AAT1546GAC (6246A>G 6248T>C), AAT1547CAG (6249A>C 6251T>G), AAA1549AAT (6257A>T), TTG1550AGT (6258T>A 6259T>G 6260G>T), AAT1551AGT (6262A>G), ATA1552AAA (6265T>A), AAT1553AGA (6268A>G 6269T>A), CGT1554AGA (6270C>A 6272T>A), GTA1556CTA (6276G>C), TTC1558TTT (6284C>T), TGT1559ATA (6285T>A 6286G>T 6287T>A), GAT1560GAC (6290T>C), TTA1561ACC (6291T>A 6292T>C 6293A>C), ACA1563TCT (6297A>T 6299A>T), GGA1564ATA (6300G>A 6301G>T), TTA1565TAC (6304T>A 6305A>C), GGT1566CCT (6306G>C 6307G>C), TTA1567del (6309_6311delTTA), ACT1568AAT (6313C>A), AAA1570GTT (6318A>G 6319A>T 6320A>T), ATA1571TTC (6321A>T 6323A>C), CAT1573TAT (6327C>T), ATC1574ATA (6332C>A), CAG1575AAA (6333C>A 6335G>A), GGT1576GGA (6338T>A) | | | | | | | | |

#### Proteins

| | | | | | | | | | |
|---|---|---|---|---|---|---|---|---|---|
| ORF3 (YP_009553219.1) | 1211 | 1582 | 22.2% | 1175 | 47.7% | 353 (98.9%) | 177 (49.6%) | 2/2/1/1 | 1 |
| Protein mutations: | S1213T (5248G>C), M1215I (5255G>A), Y1217I (5259T>A 5260A>T 5261C>A), D1224N (5280G>A), Y1227R (5289T>A 5290A>G 5291C>A), K1228T (5293A>C 5294A>G), Q1230A (5298C>G 5299A>C), T1232K (5305C>A), S1235D (5313A>G 5314G>A), I1236K (5317T>A 5318T>A), D1237S (5319G>A 5320A>G 5321C>T), Y1238D (5322T>G), L1240I (5328C>A 5330A>T), L1241N (5331C>A 5332T>A), K1242R (5335A>G), I1243M (5339T>G), K1244* (5340A>T), I1247K (5350T>A 5351T>A), V1248L (5352G>T 5354C>A), Y1249F (5356A>T), L1254C (5370C>T 5371T>G 5372A>T), G1257L (5379G>C 5380G>T 5381A>C), F1258Y (5383T>A 5384C>T), H1259W (5385C>T 5386A>G 5387C>G), I1261V (5391A>G), M1262K (5395T>A 5396G>A), D1264H (5400G>C), Q1266D (5406C>G 5408G>C), A1272T (5424G>A), V1274T (5430G>A 5431T>C 5432C>A), Q1277E (5439C>G), I1283L (5457A>C), V1284A (5461T>C 5462A>C), N1291T (5482A>C 5483T>A), I1315V (5553A>G), I1320V (5568A>G), E1323R (5577G>A 5578A>G), S1324D (5580A>G 5581G>A), I1325M (5585A>G), Q1327E (5589C>G), V1329L (5595G>C), Q1330E (5598C>G), L1333Q (5607T>C 5608T>A), Q1334T (5610C>A 5611A>C 5612G>A), F1335I (5613T>A 5615C>T), Q1337R (5619C>A 5620A>G 5621A>G), V1338L (5622G>T), C1339F (5626G>T 5627T>C), K1340V (5628A>G 5629A>T 5630G>A), E1341S (5631G>A 5632A>G 5633A>T), E1342K (5634G>A), L1344I (5640C>A 5642T>A), L1346I (5646T>A 5648G>A), S1347T (5649T>A 5651A>C), T1349K (5656C>A 5657A>G), L1351M (5661T>A), K1352E (5664A>G), I1353L (5667A>T), G1354C (5670G>T 5672A>T), V1355K (5673G>A 5674T>A), A1356T (5676G>A), N1357H (5679A>C), E1359N (5685G>A 5687A>C), L1363V (5697C>G), E1364T (5700G>A 5701A>C 5702A>T), I1365L (5703A>T), E1367D (5711A>T), V1370I (5718G>A 5720T>A), Q1371K (5721C>A), L1377A (5739T>G 5740T>C 5741A>T), I1380V (5748A>G), E1382D (5756A>T), F1383M (5757T>A 5759T>G), E1385del (5763_5765delGAA), Q1387K (5769C>A), E1389D (5777A>T), T1390K (5779C>A), L1391T (5781T>A 5782T>C), G1393D (5788G>A), I1400L (5808A>T), L1401V (5811T>G 5813A>G), Y1407F (5830A>T), P1409Q (5836C>A), S1412G (5844A>G 5846T>A), Y1414I (5850T>A 5851A>T), T1415A (5853A>G 5855T>A), R1416G (5856A>G), I1417P (5859A>C 5860T>C), F1418L (5864T>A), N1420A (5868A>G 5869A>C 5870T>A), C1422I (5874T>A 5875G>T 5876T>A), S1423G (5877T>G 5878C>G 5879C>A), E1427Q (5889G>C), R1428K (5893G>A), K1429H (5895A>C 5897A>C), S1432T (5904T>A 5906T>A), Q1433E (5907C>G), D1434N (5910G>A 5912C>T), W1435I (5913T>A 5914G>T 5915G>T), M1437L (5919A>C), R1439Q (5925A>C 5926G>A), K1440Q (5928A>C), V1444K (5940G>A 5941T>A), T1446K (5947C>A 5948C>A), K1447N (5951A>C), L1448I (5952T>A), P1450D (5958C>G 5959C>A 5960T>C), I1452S (5965T>G 5966T>C), L1453I (5967T>A), K1455L (5973A>C 5974A>T), P1456E (5976C>G 5977C>A), Q1457S (5979C>T 5980A>C), S1458D (5982A>G 5983G>A), I1460L (5988A>C 5990T>A), E1469Q (6015G>C), G1473A (6028G>C 6029A>T), I1474V (6030A>G 6032T>A), W1477A (6039T>G 6040G>C 6041G>A), S1480N (6048T>A 6049C>A 6050T>C), Q1481K (6051C>A), Q1482Y (6054C>T 6056A>C), D1483S (6057G>A 6058A>G), S1484P (6060T>C 6062A>T), P1485K (6063C>A 6064C>A 6065T>A), S1486K (6066T>A 6067C>A), T1487E (6069A>G 6070C>A), S1491C (6082C>G 6083A>T), R1492A (6084C>G 6085G>C 6086G>T), C1494Q (6090T>C 6091G>A 6092T>A), S1497K (6100G>A 6101C>A), K1499_S1500insEK (6107_6108insGAAAAG), S1500G (6108T>G 6109C>G), A1501N (6111G>A 6112C>A), I1502M (6116C>G), A1504S (6120G>A 6121C>G 6122A>T), A1507V (6130C>T 6131T>C), I1509V (6135A>G), M1510L (6138A>T 6140G>A), C1512I (6144T>A 6145G>T 6146T>A), V1515G (6154T>G), D1517N (6159G>A), K1518S (6163A>G 6164G>T), I1521L (6171A>T), F1522Y (6175T>A), L1523I (6177T>A 6179A>T), Y1524L (6181A>T 6182T>A), E1525N (6183G>A 6185G>C), K1527S (6190A>G 6191A>T), F1529I (6195T>A 6197T>A), T1530V (6198A>G 6199C>T 6200C>A), L1531V (6201C>G 6203T>A), I1536E (6216A>G 6217T>A 6218T>A), N1540K (6230T>A), Y1542H (6234T>C), N1543Q (6237A>C 6239C>A), L1545I (6243C>A 6245T>A), N1546D (6246A>G 6248T>C), N1547Q (6249A>C 6251T>G), K1549N (6257A>T), L1550S (6258T>A 6259T>G 6260G>T), N1551S (6262A>G), I1552K (6265T>A), N1553R (6268A>G 6269T>A), V1556L (6276G>C), C1559I (6285T>A 6286G>T 6287T>A), L1561T (6291T>A 6292T>C 6293A>C), T1563S (6297A>T 6299A>T), G1564I (6300G>A 6301G>T), L1565Y (6304T>A 6305A>C), G1566P (6306G>C 6307G>C), L1567del (6309_6311delTTA), T1568N (6313C>A), K1570V (6318A>G 6319A>T 6320A>T), I1571F (6321A>T 6323A>C), H1573Y (6327C>T), Q1575K (6333C>A 6335G>A) | | | | | | | | |

| | Begin | End | Coverage | Score | Concordance | Matches | Identities | I/D/M/F* | Stop Codons |
|---|---|---|---|---|---|---|---|---|---|
| **NT** | **5241** | **6355** | **13.1%** | **461** | **22.2%** | **1057 (98.8%)** | **656 (61.3%)** | **6/7** | |
| Codon mutations: | AGT1213ACT (5248G>C), CGT1214AGA (5250C>A 5252T>A), ATG1215ATA (5255G>A), TAC1217ATA (5259T>A 5260A>T 5261C>A), AAG1220AAA (5270G>A), CGA1221AGA (5271C>A), CTA1222CTT (5276A>T), GAT1224AAT (5280G>A), ACA1226ACT (5288A>T), TAC1227AGA (5289T>A 5290A>G 5291C>A), AAA1228ACG (5293A>C 5294A>G), GAC1229GAT (5297C>T), CAA1230GCA (5298C>G 5299A>C), TAC1231TAT (5303C>T), ACA1232AAA (5305C>A), CTC1233TTA (5307C>T 5309C>A), AGT1235GAT (5313A>G 5314G>A), ATT1236AAA (5317T>A 5318T>A), GAC1237AGT (5319G>A 5320A>G 5321C>T), TAT1238GAT (5322T>G), CTC1239TTA (5325C>T 5327C>A), CTA1240ATT (5328C>A 5330A>T), CTT1241AAT (5331C>A 5332T>A), AAA1242AGA (5335A>G), ATT1243ATG (5339T>G), AAA1244TAA (5340A>T), GAC1245GAT (5345C>T), AAG1246AAA (5348G>A), ATT1247AAA (5350T>A 5351T>A), GTC1248TTA (5352G>T 5354C>A), TAC1249TTC (5356A>T), AGC1250AGT (5360C>T), CTA1254TGT (5370C>T 5371T>G 5372A>T), AAG1255AAA (5375G>A), GGA1257CTC (5379G>C 5380G>T 5381A>C), TTC1258TAT (5383T>A 5384C>T), CAC1259TGG (5385C>T 5386A>G 5387C>G), CAG1260CAA (5390G>A), ATT1261GTT (5391A>G), ATG1262AAA (5395T>A 5396G>A), GAT1264CAT (5400G>C), CAG1266GAC (5406C>G 5408G>C), ATA1268-TA (5412delA), GCA1272ACA (5424G>A), TTT1273TTC (5429T>C), GTC1274ACA (5430G>A 5431T>C 5432C>A), TGC1275TGT (5435C>T), CCA1276CCT (5438A>T), CAG1277GAG (5439C>G), GGT1278GGA (5444T>A), ATA1283CTA (5457A>C), GTA1284GCC (5461T>C 5462A>C), TTC1287TTT (5471C>T), GGA1288GGT (5474A>T), CTC1289TTA (5475C>T 5477C>A), AAT1291ACA (5482A>C 5483T>A), GCA1292GCG (5486A>G), GTT1313.TC (5549T>C), ATA1315GTA (5553A>G), GAC1316GAT (5558C>T), GAT1317GAC (5561T>C), ATT1318ATC (5564T>C), TTG1319CTA (5565T>C 5567G>A), ATA1320GTA (5568A>G), TTT1321TTC (5573T>C), TCT1322AGT (5574T>A 5575C>G), GAA1323AGA (5577G>A 5578A>G), AGT1324GAT (5580A>G 5581G>A), ATA1325ATG (5585A>G), CAA1327GAA (5589C>G), GTA1329CTA (5595G>C), CAA1330GAA (5598C>G), TTA1333CAA (5607T>C 5608T>A), CAG1334ACA (5610C>A 5611A>C 5612G>A), TTC1335ATT (5613T>A 5615C>T), TTC1336TTT (5618C>T), CAA1337AGG (5619C>A 5620A>G 5621A>G), GTA1338TTA (5622G>T), TGT1339TTC (5626G>T 5627T>C), AAG1340GTA (5628A>G 5629A>T 5630G>A), GAA1341AGT (5631G>A 5632A>G 5633A>T), GAA1342AAA (5634G>A), GGG1343GGA (5639G>A), CTT1344ATA (5640C>A 5642T>A), ATT1345ATA (5645T>A), TTG1346ATA (5646T>A 5648G>A), TCA1347ACC (5649T>A 5651A>C), ACA1349AAG (5656C>A 5657A>G), TTG1351ATG (5661T>A), AAA1352GAA (5664A>G), ATA1353TTA (5667A>T), GGA1354TGT (5670G>T 5672A>T), GTA1355AAA (5673G>A 5674T>A), GCA1356ACA (5676G>A), AAT1357CAT (5679A>C), GAA1359AAC (5685G>A 5687A>C), CTT1361TTA (5691C>T 5693T>A), GGC1362GGA (5696C>A), CTA1363GTA (5697C>G), GAA1364ACT (5700G>A 5701A>C 5702A>T), ATA1365TTA (5703A>T), GAA1367GAT (5711A>T), GGA1368GGG (5714A>G), AAA1369AAG (5717A>G), GTT1370ATA (5718G>A 5720T>A), CAA1371AAA (5721C>A), CTC1372CTA (5726C>A), TTA1377GCT (5739T>G 5740T>C 5741A>T), ATA1380GTA (5748A>G), GAA1382GAT (5756A>T), TTT1383ATG (5757T>A 5759T>G), CCT1384CCA (5762T>A), GAA1385del (5763_5765delGAA), GAC1386GAT (5768C>T), CAA1387AAA (5769C>A), GAA1389GAT (5777A>T), ACA1390AAA (5779C>A), TTA1391ACA (5781T>A 5782T>C), AAG1392AAA (5786G>A), GGT1393GAT (5788G>A), CTA1394TTA (5790C>T), CAG1395CAA (5795G>A), AAG1396AAA (5798G>A), GGG1399GGA (5807G>A), ATA1400TTA (5808A>T), TTA1401GTG (5811T>G 5813A>G), AAC1402AAT (5816C>T), GCC1404GCT (5822C>T), AAT1406AAC (5828T>C), TAT1407TTT (5830A>T), ATC1408ATA (5834C>A), CCA1409CAA (5836C>A), TTG1411TTA (5843G>A), AGT1412GGA (5844A>G 5846T>A), TAT1414ATT (5850T>A 5851A>T), ACT1415GCA (5853A>G 5855T>A), AGA1416GGA (5856A>G), ATA1417CCA (5859A>C 5860T>C), TTT1418TTA (5864T>A), AAT1420GCA (5868A>G 5869A>C 5870T>A), AAG1421AAA (5873G>A), TGT1422ATA (5874T>A 5875G>T 5876T>A), TCC1423GGA (5877T>G 5878C>G 5879C>A), AGC1424AGT (5882C>T), GAA1427CAA (5889G>C), AGA1428AAA (5893G>A), AAA1429CAC (5895A>C 5897A>C), AAT1431AAC (5903T>C), TCT1432ACA (5904T>A 5906T>A), CAA1433GAA (5907C>G), GAC1434AAT (5910G>A 5912C>T), TGG1435ATT (5913T>A 5914G>T 5915G>T), AAG1436AAA (5918G>A), ATG1437CTG (5919A>C), GTT1438GTA (5924T>A), AGA1439CAA (5925A>C 5926G>A), AAA1440CAA (5928A>C), ATC1441ATT (5933C>T), GTA1444AAA (5940G>A 5941T>A), ATT1445ATA (5945T>A), ACC1446AAA (5947C>A 5948C>A), AAA1447AAC (5951A>C), TTA1448ATA (5952T>A), CCC1449CCA (5957C>A), CCT1450GAC (5958C>G 5959C>A 5960T>C), TTG1451TTA (5963G>A), ATT1452AGC (5965T>G 5966T>C), TTA1453ATA (5967T>A), AAA1455CTA (5973A>C 5974A>T), CCA1456GAA (5976C>G 5977C>A), CAA1457TCA (5979C>T 5980A>C), AGT1458GAT (5982A>G 5983G>A), ATT1460CTA (5988A>C 5990T>A), GAG1463GAA (5999G>A), GAT1465GAC (6005T>C), TCT1467AGC (6009T>A 6010C>G 6011T>C), CTA1468CTC (6014A>C), GAA1469CAA (6015G>C), GGG1472GGA (6026G>A), GGA1473GCT (6028G>C 6029A>T), ATT1474GTA (6030A>G 6032T>A), TGG1477GCA (6039T>G 6040G>C 6041G>A), CCC1479CCA (6047C>A), TCT1480AAC (6048T>A 6049C>A 6050T>C), CAA1481AAA (6051C>A), CAA1482TAC (6054C>T 6056A>C), GAT1483AGT (6057G>A 6058A>G), TCA1484CCT (6060T>C 6062A>T), CCT1485AAA (6063C>A 6064C>A 6065T>A), TCA1486AAA (6066T>A 6067C>A), ACA1487GAA (6069A>G 6070C>A), TCA1491TGT (6082C>G 6083A>T), CGG1492GCT (6084C>G 6085G>C 6086G>T), TAT1493TAC (6089T>C), TGT1494CAA (6090T>C 6091G>A 6092T>A), AGC1497AAA (6100G>A 6101C>A), TAT1498TAC (6104T>C), AAA1499_TCT1500insGAAAAG (6107_6108insGAAAAG), TCT1500GGT (6108T>G 6109C>G), GCC1501AAC (6111G>A 6112C>A), ATC1502ATG (6116C>G), TCT1503AGT (6117T>A 6118C>G), GCA1504AGT (6120G>A 6121C>G 6122A>T), GCT1507GTC (6130C>T 6131T>C), ATA1509GTA (6135A>G), ATG1510TTA (6138A>T 6140G>A), TGT1512ATA (6144T>A 6145G>T 6146T>A), GTA1515GGA (6154T>G), TTA1516CTA (6156T>C), GAT1517AAT (6159G>A), AAG1518AGT (6163A>G 6164G>T), TTT1519TTC (6167T>C), ATA1521TTA (6171A>T), TTT1522TAT (6175T>A), TTA1523ATT (6177T>A 6179A>T), TAT1524TTA (6181A>T 6182T>A), GAG1525AAC (6183G>A 6185G>C), AAA1527AGT (6190A>G 6191A>T), TTT1529ATA (6195T>A 6197T>A), ACC1530GTA (6198A>G 6199C>T 6200C>A), CTT1531GTA (6201C>G 6203T>A), AGG1532AGA (6206G>A), GAC1534GAT (6212C>T), TGT1535TGC (6215T>C), ATT1536GAA (6216A>G 6217T>A 6218T>A), GCA1537GCT (6221A>T), ATT1538ATA (6224T>A), AAT1540AAA (6230T>A), TAT1542CAT (6234T>C), AAC1543CAA (6237A>C 6239C>A), CTT1545ATA (6243C>A 6245T>A), AAT1546GAC (6246A>G 6248T>C), AAT1547CAG (6249A>C 6251T>G), AAA1549AAT (6257A>T), TTG1550AGT (6258T>A 6259T>G 6260G>T), AAT1551AGT (6262A>G), ATA1552AAA (6265T>A), AAT1553AGA (6268A>G 6269T>A), CGT1554AGA (6270C>A 6272T>A), GTA1556CTA (6276G>C), TTC1558TTT (6284C>T), TGT1559ATA (6285T>A 6286G>T 6287T>A), GAT1560GAC (6290T>C), TTA1561ACC (6291T>A 6292T>C 6293A>C), ACA1563TCT (6297A>T 6299A>T), GGA1564ATA (6300G>A 6301G>T), TTA1565TAC (6304T>A 6305A>C), GGT1566CCT (6306G>C 6307G>C), TTA1567del (6309_6311delTTA), ACT1568AAT (6313C>A), AAA1570GTT (6318A>G 6319A>T 6320A>T), ATA1571TTC (6321A>T 6323A>C), CAT1573TAT (6327C>T), ATC1574ATA (6332C>A), CAG1575AAA (6333C>A 6335G>A), GGT1576GGA (6338T>A) | | | | | | | | |

*: Inserts / Deletes / Misaligned / Frameshifts

#### Analysis details

This analysis was performed with panviral2.64

### NGS Details (UN8): Badnavirus occultipomeae

#### Assembly

| | |
|---|---|
| **Coverage Length** | 322 (1 contig(s)) |
| **Depth Of Coverage** | 29.0 |
| **Number Of Reads** | 84 |
| **Reads Per Million** | 1.40 rpm (after QC) |
| **Ambiguities** | 0 |
| **Assembly Method** | de novo + reference guided assembly |
| **Consensus Caller** | Bcf Tools |

#### Coverage Map

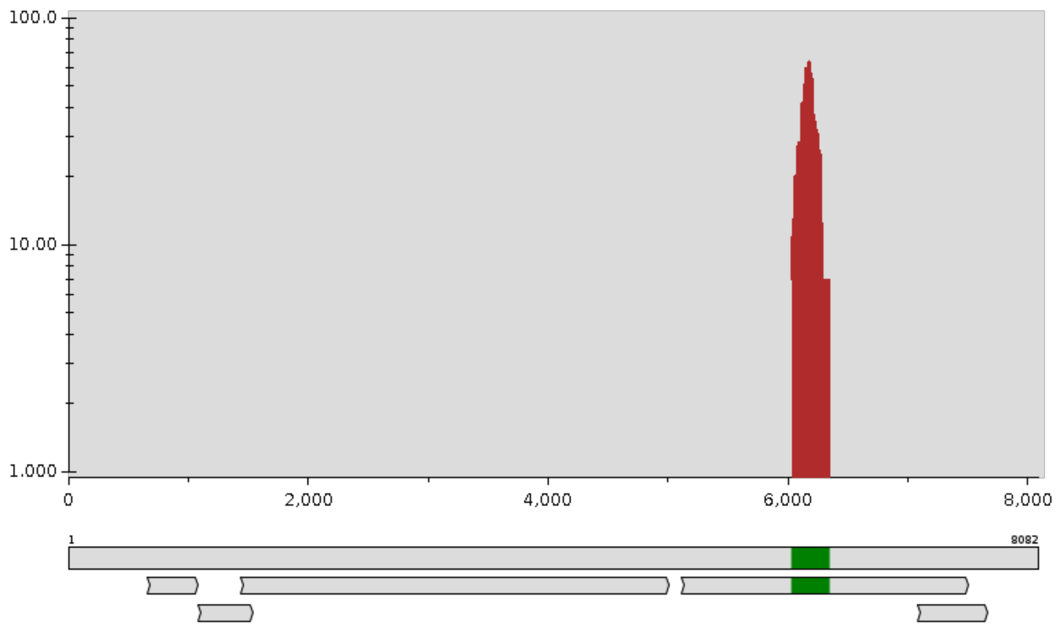

#### Assignment

| | |
|---|---|
| **Type** | Badnavirus occultipomeae (Taxonomy ID: 3048353) |
| **Reference Genome** | NC_015655.1 |
| **NT Identity (%)** | 51.0769 |
| **AA Identity (%)** | 41.2844 |
| **Number Of Stop Codons** | 2 |
| **Number Of CDS** | 5 |

#### Alignment

| | |
|---|---|
| **Alignment Score** | 8.0 (NT) + 331.0 (AA) = 339.0 |
| **Concordance (%)** | 24.2663 |

| Alignment Method | Global, seeded, nucleotide + amino acids (AGA) |
|---|---|

#### Genome Region

Sequence starts at position 6028 and ends at position 6349 relative to NC_015655.1 reference sequence.

#### Alignment Detailed Statistics

| | Begin | End | Coverage | Score | Concordance | Matches | Identities | I/D/M/F* | Stop Codons |
|---|---|---|---|---|---|---|---|---|---|
| **NT** | **6028** | **6349** | **4.0%** | **8** | **1.2%** | **322 (99.1%)** | **166 (51.1%)** | **3/0** | |
| Mutations: | 6029A>G, 6030G>T, 6031A>T, 6032T>G, 6033A>T, 6036C>T, 6042A>G, 6043T>A, 6045T>C, 6051C>T, 6052A>C, 6053A>G, 6054A>T, 6055G>T, 6057A>T, 6060C>G, 6062T>A, 6066T>C, 6069G>A, 6070A>C, 6072A>T, 6075A>G, 6076A>G, 6078G>A, 6079G>A, 6084A>G, 6086A>G, 6087G>T, 6088T>G, 6089C>A, 6090C>T, 6092A>T, 6093A>T, 6097T>A, 6098G>A, 6099G>A, 6101C>T, 6102T>A, 6114T>G, 6115C>A, 6116C>G, 6117A>G, 6118G>T, 6120A>T, 6124C>A, 6125T>A, 6126C>T, 6128A>T, 6129T>C, 6130G>A, 6135T>C, 6136G>C, 6137A>T, 6138A>T, 6139G>T, 6145C>T, 6147C>T, 6153A>T, 6154C>T, 6156G>A, 6158T>C, 6159G>T, 6165A>C, 6168A>G, 6169G>A, 6173A>T, 6174T>G, 6178C>A, 6179A>T, 6181A>G, 6182G>A, 6183A>T, 6184A>T, 6185A>T, 6186G>A, 6190G>A, 6192T>C, 6194A>G, 6195T>G, 6197C>T, 6198A>T, 6201T>C, 6205G>C, 6206G>C, 6207A>T, 6207_6208insTTT, 6208A>C, 6209C>T, 6210A>C, 6211G>A, 6213T>C, 6214G>A, 6216A>G, 6220A>T, 6222C>G, 6224C>T, 6225A>G, 6228A>C, 6230A>T, 6234T>C, 6240C>T, 6241A>G, 6243A>G, 6244T>C, 6246G>A, 6247G>A, 6251T>A, 6255T>C, 6256G>C, 6257A>G, 6260A>G, 6262G>A, 6263A>G, 6264A>G, 6270G>A, 6279T>G, 6282T>C, 6283C>T, 6285C>A, 6286T>C, 6287T>G, 6290A>T, 6291T>G, 6292T>G, 6295G>T, 6296C>T, 6297C>A, 6298C>T, 6300A>G, 6303T>A, 6304G>T, 6307A>T, 6311G>A, 6312A>T, 6315A>G, 6316G>A, 6317G>A, 6319C>T, 6321T>G, 6322A>T, 6323T>A, 6325C>G, 6327A>G, 6329G>A, 6330C>G, 6331C>T, 6332C>T, 6333A>T, 6335C>A, 6336A>C, 6340A>T, 6341T>G, 6342G>T, 6343A>G, 6345A>G, 6346A>T, 6349G>T | | | | | | | | |

##### CDS

| | Begin | End | Coverage | Score | Concordance | Matches | Identities | I/D/M/F* | Stop Codons |
|---|---|---|---|---|---|---|---|---|---|
| SPBVa_gp4 | 305 | 412 | 13.6% | 331 | 43.4% | 108 (99.1%) | 45 (41.3%) | 1/0/0/0 | 2 |
| Protein mutations: | K305S (6029A>G 6030G>T), I306C (6031A>T 6032T>G 6033A>T), F310I (6043T>A 6045T>C), K313R (6052A>C 6053A>G 6054A>T), A314S (6055G>T 6057A>T), F316Y (6062T>A), I319L (6070A>C 6072A>T), M321V (6076A>G 6078G>A), E322K (6079G>A), K324S (6086A>G 6087G>T), S325D (6088T>G 6089C>A 6090C>T), K326I (6092A>T 6093A>T), W328K (6097T>A 6098G>A 6099G>A), T329I (6101C>T 6102T>A), P334R (6115C>A 6116C>G 6117A>G), E335Y (6118G>T 6120A>T), L337N (6124C>A 6125T>A 6126C>T), Y338F (6128A>T 6129T>C), E339K (6130G>A), E341L (6136G>C 6137A>T 6138A>T), V342F (6139G>T), P344S (6145C>T 6147C>T), M348T (6158T>C 6159G>T), A352T (6169G>A), D353V (6173A>T 6174T>G), Q355M (6178C>A 6179A>T), R356D (6181A>G 6182G>A 6183A>T), K357L (6184A>T 6185A>T 6186G>A), D359N (6190G>A 6192T>C), N360R (6194A>G 6195T>G), A361V (6197C>T 6198A>T), G364P (6205G>C 6206G>C 6207A>T), G364_T365insF (6207_6208insTTT), T365L (6208A>C 6209C>T 6210A>C), D366N (6211G>A 6213T>C), A367T (6214G>A 6216A>G), I369L (6220A>T 6222C>G), A370V (6224C>T 6225A>G), Y372F (6230A>T), I376V (6241A>G 6243A>G), V378I (6247G>A), F379Y (6251T>A), E381R (6256G>C 6257A>G), N382S (6260A>G), E383R (6262G>A 6263A>G 6264A>G), D388E (6279T>G), L391R (6286T>C 6287T>G), N392M (6290A>T 6291T>G), L393V (6292T>G), A394L (6295G>T 6296C>T 6297C>A), Q395* (6298C>T 6300A>G), V397L (6304G>T), R398* (6307A>T), R399N (6311G>A 6312A>T), G401K (6316G>A 6317G>A), I403Y (6322A>T 6323T>A), L404V (6325C>G 6327A>G), S405K (6329G>A 6330C>G), P406F (6331C>T 6332C>T 6333A>T), T407N (6335C>A 6336A>C), M409C (6340A>T 6341T>G 6342G>T), K410E (6343A>G 6345A>G), I411F (6346A>T) | | | | | | | | |
| Codon mutations: | AAG305AGT (6029A>G 6030G>T), ATA306TGT (6031A>T 6032T>G 6033A>T), TTC307TTT (6036C>T), AAA309AAG (6042A>G), TTT310ATC (6043T>A 6045T>C), CTC312CTT (6051C>T), AAA313CGT (6052A>C 6053A>G 6054A>T), GCA314TCT (6055G>T 6057A>T), GGC315GGG (6060C>G), TTT316TAT (6062T>A), CAT317CAC (6066T>C), CAG318CAA (6069G>A), ATA319CTT (6070A>C 6072A>T), AGA320AGG (6075A>G), ATG321GTA (6076A>G 6078G>A), GAG322AAG (6079G>A), GAA323GAG (6084A>G), AAG324AGT (6086A>G 6087G>T), TCC325GAT (6088T>G 6089C>A 6090C>T), AAA326ATT (6092A>T 6093A>T), TGG328AAA (6097T>A 6098G>A 6099G>A), ACT329ATA (6101C>T 6102T>A), ACT333ACG (6114T>G), CCA334AGG (6115C>A 6116C>G 6117A>G), GAA335TAT (6118G>T 6120A>T), CTC337AAT (6124C>A 6125T>A 6126C>T), TAT338TTC (6128A>T 6129T>C), GAA339AAA (6130G>A), TTT340TTC (6135T>C), GAA341CTT (6136G>C 6137A>T 6138A>T), GTC342TTC (6139G>T), CCC344TCT (6145C>T 6147C>T), GGA346GGT (6153A>T), CTG347TTA (6154C>T 6156G>A), ATG348ACT (6158T>C 6159G>T), GCA350GCC (6165A>C), CCA351CCG (6168A>G), GCA352ACA (6169G>A), GAT353GTG (6173A>T 6174T>G), CAG355ATG (6178C>A 6179A>T), AGA356GAT (6181A>G 6182G>A 6183A>T), AAG357TTA (6184A>T 6185A>T 6186G>A), GAT359AAC (6190G>A 6192T>C), AAT360AGG (6194A>G 6195T>G), GCA361GTT (6197C>T 6198A>T), TTT362TTC (6201T>C), GGA364CCT (6205G>C 6206G>C 6207A>T), GGA364_ACA365insTTT (6207_6208insTTT), ACA365CTC (6208A>C 6209C>T 6210A>C), GAT366AAC (6211G>A 6213T>C), GCA367ACG (6214G>A 6216A>G), ATC369TTG (6220A>T 6222C>G), GCA370GTG (6224C>T 6225A>G), GTA371GTC (6228A>C), TAT372TTT (6230A>T), ATT373ATC (6234T>C), GAC375GAT (6240C>T), ATA376GTG (6241A>G 6243A>G), TTG377CTA (6244T>C 6246G>A), GTA378ATA (6247G>A), TTC379TAC (6251T>A), TCT380TCC (6255T>C), GAA381CGA (6256G>C 6257A>G), AAT382AGT (6260A>G), GAA383AGG (6262G>A 6263A>G 6264A>G), GAG385GAA (6270G>A), GAT388GAG (6279T>G), CAT389CAC (6282T>C), CTC390TTA (6283C>T 6285C>A), TTA391CGA (6286T>C 6287T>G), AAT392ATG (6290A>T 6291T>G), TTG393GTG (6292T>G), GCC394TTA (6295G>T 6296C>T 6297C>A), CAA395TAG (6298C>T 6300A>G), ATT396ATA (6303T>A), GTG397TTG (6304G>T), AGA398TGA (6307A>T), AGA399AAT (6311G>A 6312A>T), GAA400GAG (6315A>G), GGG401AAG (6316G>A 6317G>A), CTT402TTG (6319C>T 6321T>G), ATT403TAT (6322A>T 6323T>A), CTA404GTG (6325C>G 6327A>G), AGC405AAG (6329G>A 6330C>G), CCA406TTT (6331C>T 6332C>T 6333A>T), ACA407AAC (6335C>A 6336A>C), ATG409TGT (6340A>T 6341T>G 6342G>T), AAA410GAG (6343A>G 6345A>G), ATT411TTT (6346A>T), GGA412T.. (6349G>T) | | | | | | | | |

##### Proteins

| | Begin | End | Coverage | Score | Concordance | Matches | Identities | I/D/M/F* | Stop Codons |
|---|---|---|---|---|---|---|---|---|---|
| RNaseH/reverse transcriptase (YP_004581513.1) | 305 | 412 | 13.6% | 331 | 43.4% | 108 (99.1%) | 45 (41.3%) | 1/0/0/0 | 2 |
| Protein mutations: | K305S (6029A>G 6030G>T), I306C (6031A>T 6032T>G 6033A>T), F310I (6043T>A 6045T>C), K313R (6052A>C 6053A>G 6054A>T), A314S (6055G>T 6057A>T), F316Y (6062T>A), I319L (6070A>C 6072A>T), M321V (6076A>G 6078G>A), E322K (6079G>A), K324S (6086A>G 6087G>T), S325D (6088T>G 6089C>A 6090C>T), K326I (6092A>T 6093A>T), W328K (6097T>A 6098G>A 6099G>A), T329I (6101C>T 6102T>A), P334R (6115C>A 6116C>G 6117A>G), E335Y (6118G>T 6120A>T), L337N (6124C>A 6125T>A 6126C>T), Y338F (6128A>T 6129T>C), E339K (6130G>A), E341L (6136G>C 6137A>T 6138A>T), V342F (6139G>T), P344S (6145C>T 6147C>T), M348T (6158T>C 6159G>T), A352T (6169G>A), D353V (6173A>T 6174T>G), Q355M (6178C>A 6179A>T), R356D (6181A>G 6182G>A 6183A>T), K357L (6184A>T 6185A>T 6186G>A), D359N (6190G>A 6192T>C), N360R (6194A>G 6195T>G), A361V (6197C>T 6198A>T), G364P (6205G>C 6206G>C 6207A>T), G364_T365insF (6207_6208insTTT), T365L (6208A>C 6209C>T 6210A>C), D366N (6211G>A 6213T>C), A367T (6214G>A 6216A>G), I369L (6220A>T 6222C>G), A370V (6224C>T 6225A>G), Y372F (6230A>T), I376V (6241A>G 6243A>G), V378I (6247G>A), F379Y (6251T>A), E381R (6256G>C 6257A>G), N382S (6260A>G), E383R (6262G>A 6263A>G 6264A>G), D388E (6279T>G), L391R (6286T>C 6287T>G), N392M (6290A>T 6291T>G), L393V (6292T>G), A394L (6295G>T 6296C>T 6297C>A), Q395* (6298C>T 6300A>G), V397L (6304G>T), R398* (6307A>T), R399N (6311G>A 6312A>T), G401K (6316G>A 6317G>A), I403Y (6322A>T 6323T>A), L404V (6325C>G 6327A>G), S405K (6329G>A 6330C>G), P406F (6331C>T 6332C>T 6333A>T), T407N (6335C>A 6336A>C), M409C (6340A>T 6341T>G 6342G>T), K410E (6343A>G 6345A>G), I411F (6346A>T) | | | | | | | | |
| Codon mutations: | AAG305AGT (6029A>G 6030G>T), ATA306TGT (6031A>T 6032T>G 6033A>T), TTC307TTT (6036C>T), AAA309AAG (6042A>G), TTT310ATC (6043T>A 6045T>C), CTC312CTT (6051C>T), AAA313CGT (6052A>C 6053A>G 6054A>T), GCA314TCT (6055G>T 6057A>T), GGC315GGG (6060C>G), TTT316TAT (6062T>A), CAT317CAC (6066T>C), CAG318CAA (6069G>A), ATA319CTT (6070A>C 6072A>T), AGA320AGG (6075A>G), ATG321GTA (6076A>G 6078G>A), GAG322AAG (6079G>A), GAA323GAG (6084A>G), AAG324AGT (6086A>G 6087G>T), TCC325GAT (6088T>G 6089C>A 6090C>T), AAA326ATT (6092A>T 6093A>T), TGG328AAA (6097T>A 6098G>A 6099G>A), ACT329ATA (6101C>T 6102T>A), ACT333ACG (6114T>G), CCA334AGG (6115C>A 6116C>G 6117A>G), GAA335TAT (6118G>T 6120A>T), CTC337AAT (6124C>A 6125T>A 6126C>T), TAT338TTC (6128A>T 6129T>C), GAA339AAA (6130G>A), TTT340TTC (6135T>C), GAA341CTT (6136G>C 6137A>T 6138A>T), GTC342TTC (6139G>T), CCC344TCT (6145C>T 6147C>T), GGA346GGT (6153A>T), CTG347TTA (6154C>T 6156G>A), ATG348ACT (6158T>C 6159G>T), GCA350GCC (6165A>C), CCA351CCG (6168A>G), GCA352ACA (6169G>A), GAT353GTG (6173A>T 6174T>G), CAG355ATG (6178C>A 6179A>T), AGA356GAT (6181A>G 6182G>A 6183A>T), AAG357TTA (6184A>T 6185A>T 6186G>A), GAT359AAC (6190G>A 6192T>C), AAT360AGG (6194A>G 6195T>G), GCA361GTT (6197C>T 6198A>T), TTT362TTC (6201T>C), GGA364CCT (6205G>C 6206G>C 6207A>T), GGA364_ACA365insTTT (6207_6208insTTT), ACA365CTC (6208A>C 6209C>T 6210A>C), GAT366AAC (6211G>A 6213T>C), GCA367ACG (6214G>A 6216A>G), ATC369TTG (6220A>T 6222C>G), GCA370GTG (6224C>T 6225A>G), GTA371GTC (6228A>C), TAT372TTT (6230A>T), ATT373ATC (6234T>C), GAC375GAT (6240C>T), ATA376GTG (6241A>G 6243A>G), TTG377CTA (6244T>C 6246G>A), GTA378ATA (6247G>A), TTC379TAC (6251T>A), TCT380TCC (6255T>C), GAA381CGA (6256G>C 6257A>G), AAT382AGT (6260A>G), GAA383AGG (6262G>A 6263A>G 6264A>G), GAG385GAA (6270G>A), GAT388GAG (6279T>G), CAT389CAC (6282T>C), CTC390TTA (6283C>T 6285C>A), TTA391CGA (6286T>C 6287T>G), AAT392ATG (6290A>T 6291T>G), TTG393GTG (6292T>G), GCC394TTA (6295G>T 6296C>T 6297C>A), CAA395TAG (6298C>T 6300A>G), ATT396ATA (6303T>A), GTG397TTG (6304G>T), AGA398TGA (6307A>T), AGA399AAT (6311G>A 6312A>T), GAA400GAG (6315A>G), GGG401AAG (6316G>A 6317G>A), CTT402TTG (6319C>T 6321T>G), ATT403TAT (6322A>T 6323T>A), CTA404GTG (6325C>G 6327A>G), AGC405AAG (6329G>A 6330C>G), CCA406TTT (6331C>T 6332C>T 6333A>T), ACA407AAC (6335C>A 6336A>C), ATG409TGT (6340A>T 6341T>G 6342G>T), AAA410GAG (6343A>G 6345A>G), ATT411TTT (6346A>T), GGA412T.. (6349G>T) | | | | | | | | |

*: Inserts / Deletes / Misaligned / Frameshifts

#### Analysis details

This analysis was performed with panviral2.64

### NGS Details (UN8): Errantivirus

#### Assembly

| | |
|---|---|
| **Coverage Length** | 380 (1 contig(s)) |
| **Depth Of Coverage** | 22.1 |
| **Number Of Reads** | 73 |
| **Reads Per Million** | 1.22 rpm (after QC) |
| **Ambiguities** | 0 |
| **Assembly Method** | de novo + reference guided assembly |
| **Consensus Caller** | Bcf Tools |

#### Coverage Map

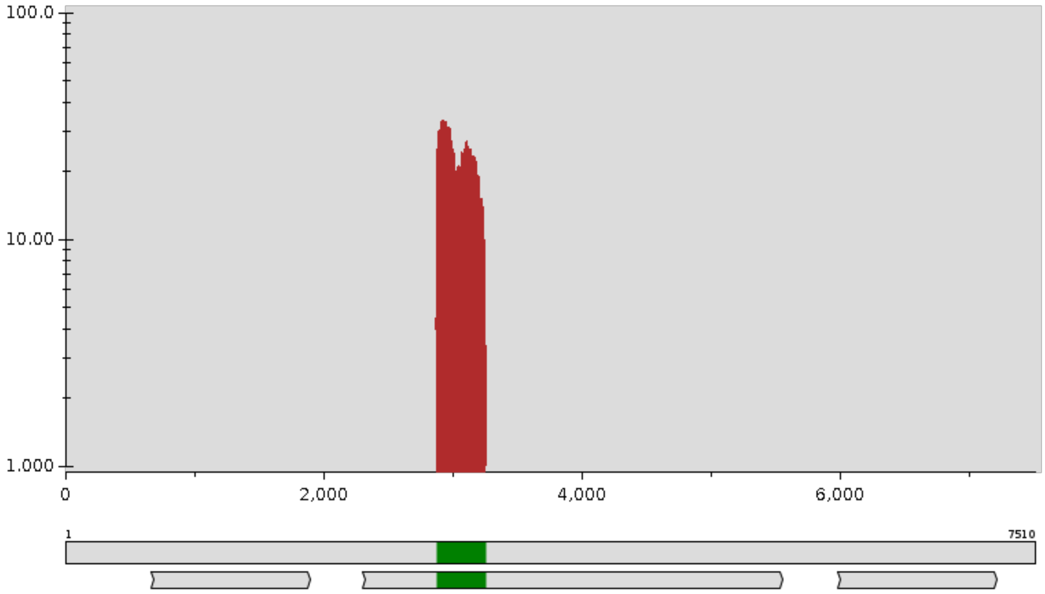

#### Assignment

| | |
|---|---|
| **Type** | Errantivirus (Taxonomy ID: 186666) |
| **Reference Genome** | NC_038512.1 |
| **NT Identity (%)** | 54.5932 |
| **AA Identity (%)** | 45.3125 |
| **Number Of Stop Codons** | 1 |
| **Number Of CDS** | 3 |

#### Alignment

| | |
|---|---|
| **Alignment Score** | 62.0 (NT) + 310.0 (AA) = 372.0 |
| **Concordance (%)** | 24.1505 |

| Alignment Method | Global, seeded, nucleotide + amino acids (AGA) |
|---|---|

#### Genome Region

Sequence starts at position 2876 and ends at position 3255 relative to NC_038512.1 reference sequence.

#### Alignment Detailed Statistics

| | Begin | End | Coverage | Score | Concordance | Matches | Identities | I/D/M/F* | Stop Codons |
|---|---|---|---|---|---|---|---|---|---|
| **NT** | **2876** | **3255** | **5.1%** | **62** | **8.2%** | **380 (99.7%)** | **208 (54.6%)** | **1/0** | |
| Mutations: | 2885C>A, 2887T>G, 2888C>T, 2890C>G, 2891G>T, 2892T>G, 2893A>T, 2896T>A, 2901T>A, 2902C>T, 2903C>A, 2905T>G, 2906A>G, 2908G>T, 2909T>C, 2911G>T, 2914C>T, 2915G>A, 2918A>C, 2920G>C, 2924A>G, 2926C>T, 2927G>A, 2929T>G, 2930G>A, 2937A>T, 2941G>T, 2947A>T, 2948A>C, 2949A>T, 2950C>A, 2951A>G, 2954A>G, 2955G>A, 2956T>A, 2959C>T, 2960G>T, 2962A>G, 2963C>T, 2965T>G, 2968C>T, 2969A>G, 2971G>A, 2972T>C, 2974A>T, 2977T>G, 2978A>G, 2979A>G, 2980G>T, 2982G>C, 2983C>A, 2984C>G, 2987T>A, 2988A>T, 2989C>T, 2992C>T, 2993A>T, 2997C>A, 2998C>A, 2999T>A, 3004T>C, 3005T>C, 3007G>T, 3008G>A, 3009C>G, 3010A>G, 3011A>G, 3012G>C, 3014G>A, 3018T>A, 3019T>C, 3020T>C, 3026G>T, 3029G>A, 3030A>G, 3031G>A, 3036A>C, 3038C>A, 3039C>A, 3040T>A, 3041C>G, 3046T>C, 3050T>C, 3052G>A, 3058C>A, 3061G>A, 3064T>C, 3066A>G, 3067C>A, 3068G>A, 3069T>C, 3070A>T, 3071G>C, 3073A>C, 3074C>T, 3075A>C, 3076C>A, 3090T>A, 3091C>T, 3092C>T, 3094T>G, 3095C>G, 3096G>T, 3103T>A, 3104A>T, 3106G>T, 3110T>C, 3113A>T, 3114A>C, 3115A>T, 3118C>T, 3119T>G, 3125T>G, 3127T>A, 3130T>C, 3137A>G, 3139A>G, 3140G>T, 3142T>G, 3146G>A, 3148C>T, 3150A>C, 3154C>G, 3155C>T, 3157A>T, 3158A>T, 3159G>A, 3161G>A, 3162G>A, 3163T>G, 3164C>T, 3166C>T, 3167C>T, 3168A>T, 3169A>G, 3171A>G, 3172T>G, 3175C>G, 3176A>T, 3177T>C, 3178C>A, 3179T>G, 3180G>T, 3182C>T, 3184C>A, 3187C>T, 3189A>T, 3190C>T, 3191C>T, 3193_3194insT, 3196C>T, 3203A>C, 3205T>G, 3206G>A, 3209T>G, 3215A>C, 3216C>A, 3217T>G, 3220C>T, 3221C>A, 3223A>T, 3224C>G, 3226G>A, 3229A>C, 3232C>T, 3235G>A, 3236G>T, 3237A>T, 3239A>C, 3245G>A, 3246A>G, 3248C>A, 3249G>A | | | | | | | | |

##### CDS

| | Begin | End | Coverage | Score | Concordance | Matches | Identities | I/D/M/F* | Stop Codons |
|---|---|---|---|---|---|---|---|---|---|
| D1R33_gp2 | 191 | 317 | 11.7% | 310 | 34.5% | 127 (99.2%) | 58 (45.3%) | 1/0/1/1 | 1 |
| Protein mutations: | V196C (2891G>T 2892T>G 2893A>T), F199Y (2901T>A 2902C>T), K201D (2906A>G 2908G>T), E204K (2915G>A), K205H (2918A>C 2920G>C), I207V (2924A>G 2926C>T), D208K (2927G>A 2929T>G), D209N (2930G>A), Y211F (2937A>T), N215L (2948A>C 2949A>T 2950C>A), I216V (2951A>G), S217E (2954A>G 2955G>A 2956T>A), V219L (2960G>T 2962A>G), K222E (2969A>G 2971G>A), K225G (2978A>G 2979A>G 2980G>T), C226S (2982G>C 2983C>A), Q227E (2984C>G), Y228I (2987T>A 2988A>T 2989C>T), T230S (2993A>T), T231K (2997C>A 2998C>A), L232I (2999T>A), A235R (3008G>A 3009C>G 3010A>G), S236A (3011A>G 3012G>C), G237R (3014G>A), F238Y (3018T>A 3019T>C), Y239H (3020T>C), V241L (3026G>T), E242R (3029G>A 3030A>G 3031G>A), D244A (3036A>C), P245K (3038C>A 3039C>A 3040T>A), Q246E (3041C>G), S249P (3050T>C 3052G>A), N254R (3066A>G 3067C>A), V255T (3068G>A 3069T>C 3070A>T), E256H (3071G>C 3073A>C), H257S (3074C>T 3075A>C 3076C>A), F262Y (3090T>A 3091C>T), R264V (3095C>G 3096G>T), M267F (3104A>T 3106G>T), K270S (3113A>T 3114A>C 3115A>T), S272A (3119T>G), S274A (3125T>G 3127T>A), R278G (3137A>G 3139A>G), V279L (3140G>T 3142T>G), D281N (3146G>A 3148C>T), N282T (3150A>C), L284F (3155C>T 3157A>T), R285* (3158A>T 3159G>A), G286K (3161G>A 3162G>A 3163T>G), L287F (3164C>T 3166C>T), Q288L (3167C>T 3168A>T 3169A>G), N289R (3171A>G 3172T>G), N290K (3175C>G), I291S (3176A>T 3177T>C 3178C>A), C292V (3179T>G 3180G>T), Y295F (3189A>T 3190C>T), L296F (3191C>T), L296_D297insX (3193_3194insT), I300L (3203A>C 3205T>G), V301I (3206G>A), Y302D (3209T>G), T304Q (3215A>C 3216C>A 3217T>G), L306I (3221C>A 3223A>T), Q307E (3224C>G 3226G>A), E308D (3229A>C), E311L (3236G>T 3237A>T), N312H (3239A>C), E314R (3245G>A 3246A>G), R315K (3248C>A 3249G>A) | | | | | | | | |
| Codon mutations: | CGT194AGG (2885C>A 2887T>G), CTC195TTG (2888C>T 2890C>G), GTA196TGT (2891G>T 2892T>G 2893A>T), GTT197GTA (2896T>A), TTC199TAT (2901T>A 2902C>T), CGT200AGG (2903C>A 2905T>G), AAG201GAT (2906A>G 2908G>T), TTG202CTT (2909T>C 2911G>T), AAC203AAT (2914C>T), GAG204AAG (2915G>A), AAG205CAC (2918A>C 2920G>C), ATC207GTT (2924A>G 2926C>T), GAT208AAG (2927G>A 2929T>G), GAC209AAC (2930G>A), TAC211TTC (2937A>T), CCG212CCT (2941G>T), CCA214CCT (2947A>T), AAC215CTA (2948A>C 2949A>T 2950C>A), ATA216GTA (2951A>G), AGT217GAA (2954A>G 2955G>A 2956T>A), GAC218GAT (2959C>T), GTA219TTG (2960G>T 2962A>G), CTT220TTG (2963C>T 2965T>G), GAC221GAT (2968C>T), AAG222GAA (2969A>G 2971G>A), TTA223CTT (2972T>C 2974A>T), GGT224GGG (2977T>G), AAG225GGT (2978A>G 2979A>G 2980G>T), TGC226TCA (2982G>C 2983C>A), CAA227GAA (2984C>G), TAC228ATT (2987T>A 2988A>T 2989C>T), TTC229TTT (2992C>T), ACC230TCC (2993A>T), ACC231AAA (2997C>A 2998C>A), TTA232ATA (2999T>A), GAT233GAC (3004T>C), TTG234CTT (3005T>C 3007G>T), GCA235AGG (3008G>A 3009C>G 3010A>G), AGT236GCT (3011A>G 3012G>C), GGG237AGG (3014G>A), TTT238TAC (3018T>A 3019T>C), TAT239CAT (3020T>C), GTG241TTG (3026G>T), GAG242AGA (3029G>A 3030A>G 3031G>A), GAC244GCC (3036A>C), CCT245AAA (3038C>A 3039C>A 3040T>A), CAA246GAA (3041C>G), GAT247GAC (3046T>C), TCG249CCA (3050T>C 3052G>A), ACC251ACA (3058C>A), GCG252GCA (3061G>A), TTT253TTC (3064T>C), AAC254AGA (3066A>G 3067C>A), GTA255ACT (3068G>A 3069T>C 3070A>T), GAA256CAC (3071G>C 3073A>C), CAC257TCA (3074C>T 3075A>C 3076C>A), TTC262TAT (3090T>A 3091C>T), CTT263TTG (3092C>T 3094T>G), CGA264GTA (3095C>G 3096G>T), CCT266CCA (3103T>A), ATG267TTT (3104A>T 3106G>T), TTA269CTA (3110T>C), AAA270TCT (3113A>T 3114A>C 3115A>T), AAC271AAT (3118C>T), TCA272GCA (3119T>G), TCT274GCA (3125T>G 3127T>A), ACT275ACC (3130T>C), AGA278GGG (3137A>G 3139A>G), GTT279TTG (3140G>T 3142T>G), GAC281AAT (3146G>A 3148C>T), AAT282ACT (3150A>C), GTC283GTG (3154C>G), CTA284TTT (3155C>T 3157A>T), AGA285TAA (3158A>T 3159G>A), GGT286AAG (3161G>A 3162G>A 3163T>G), CTC287TTT (3164C>T 3166C>T), CAA288TTG (3167C>T 3168A>T 3169A>G), AAT289AGG (3171A>G 3172T>G), AAC290AAG (3175C>G), ATC291TCA (3176A>T 3177T>C 3178C>A), TGT292GTT (3179T>G 3180G>T), CTC293TTA (3182C>T 3184C>A), GTC294GTT (3187C>T), TAC295TTT (3189A>T 3190C>T), CTT296TTT (3191C>T), CTT296_GAC297insT-- (3193_3194insT), GAC297GAT (3196C>T), ATT300CTG (3203A>C 3205T>G), GTC301ATC (3206G>A), TAT302GAT (3209T>G), ACT304CAG (3215A>C 3216C>A 3217T>G), TCC305TCT (3220C>T), CTA306ATT (3221C>A 3223A>T), CAG307GAA (3224C>G 3226G>A), GAA308GAC (3229A>C), CAC309CAT (3232C>T), CTG310CTA (3235G>A), GAG311TTG (3236G>T 3237A>T), AAC312CAC (3239A>C), GAA314AGA (3245G>A 3246A>G), CGA315AAA (3248C>A 3249G>A) | | | | | | | | |

##### Proteins

| | Begin | End | Coverage | Score | Concordance | Matches | Identities | I/D/M/F* | Stop Codons |
|---|---|---|---|---|---|---|---|---|---|
| ORF B (YP_009507248.1) | 191 | 317 | 11.7% | 310 | 34.5% | 127 (99.2%) | 58 (45.3%) | 1/0/1/1 | 1 |
| Protein mutations: | V196C (2891G>T 2892T>G 2893A>T), F199Y (2901T>A 2902C>T), K201D (2906A>G 2908G>T), E204K (2915G>A), K205H (2918A>C 2920G>C), I207V (2924A>G 2926C>T), D208K (2927G>A 2929T>G), D209N (2930G>A), Y211F (2937A>T), N215L (2948A>C 2949A>T 2950C>A), I216V (2951A>G), S217E (2954A>G 2955G>A 2956T>A), V219L (2960G>T 2962A>G), K222E (2969A>G 2971G>A), K225G (2978A>G 2979A>G 2980G>T), C226S (2982G>C 2983C>A), Q227E (2984C>G), Y228I (2987T>A 2988A>T 2989C>T), T230S (2993A>T), T231K (2997C>A 2998C>A), L232I (2999T>A), A235R (3008G>A 3009C>G 3010A>G), S236A (3011A>G 3012G>C), G237R (3014G>A), F238Y (3018T>A 3019T>C), Y239H (3020T>C), V241L (3026G>T), E242R (3029G>A 3030A>G 3031G>A), D244A (3036A>C), P245K (3038C>A 3039C>A 3040T>A), Q246E (3041C>G), S249P (3050T>C 3052G>A), N254R (3066A>G 3067C>A), V255T (3068G>A 3069T>C 3070A>T), E256H (3071G>C 3073A>C), H257S (3074C>T 3075A>C 3076C>A), F262Y (3090T>A 3091C>T), R264V (3095C>G 3096G>T), M267F (3104A>T 3106G>T), K270S (3113A>T 3114A>C 3115A>T), S272A (3119T>G), S274A (3125T>G 3127T>A), R278G (3137A>G 3139A>G), V279L (3140G>T 3142T>G), D281N (3146G>A 3148C>T), N282T (3150A>C), L284F (3155C>T 3157A>T), R285* (3158A>T 3159G>A), G286K (3161G>A 3162G>A 3163T>G), L287F (3164C>T 3166C>T), Q288L (3167C>T 3168A>T 3169A>G), N289R (3171A>G 3172T>G), N290K (3175C>G), I291S (3176A>T 3177T>C 3178C>A), C292V (3179T>G 3180G>T), Y295F (3189A>T 3190C>T), L296F (3191C>T), L296_D297insX (3193_3194insT), I300L (3203A>C 3205T>G), V301I (3206G>A), Y302D (3209T>G), T304Q (3215A>C 3216C>A 3217T>G), L306I (3221C>A 3223A>T), Q307E (3224C>G 3226G>A), E308D (3229A>C), E311L (3236G>T 3237A>T), N312H (3239A>C), E314R (3245G>A 3246A>G), R315K (3248C>A 3249G>A) | | | | | | | | |
| Codon mutations: | CGT194AGG (2885C>A 2887T>G), CTC195TTG (2888C>T 2890C>G), GTA196TGT (2891G>T 2892T>G 2893A>T), GTT197GTA (2896T>A), TTC199TAT (2901T>A 2902C>T), CGT200AGG (2903C>A 2905T>G), AAG201GAT (2906A>G 2908G>T), TTG202CTT (2909T>C 2911G>T), AAC203AAT (2914C>T), GAG204AAG (2915G>A), AAG205CAC (2918A>C 2920G>C), ATC207GTT (2924A>G 2926C>T), GAT208AAG (2927G>A 2929T>G), GAC209AAC (2930G>A), TAC211TTC (2937A>T), CCG212CCT (2941G>T), CCA214CCT (2947A>T), AAC215CTA (2948A>C 2949A>T 2950C>A), ATA216GTA (2951A>G), AGT217GAA (2954A>G 2955G>A 2956T>A), GAC218GAT (2959C>T), GTA219TTG (2960G>T 2962A>G), CTT220TTG (2963C>T 2965T>G), GAC221GAT (2968C>T), AAG222GAA (2969A>G 2971G>A), TTA223CTT (2972T>C 2974A>T), GGT224GGG (2977T>G), AAG225GGT (2978A>G 2979A>G 2980G>T), TGC226TCA (2982G>C 2983C>A), CAA227GAA (2984C>G), TAC228ATT (2987T>A 2988A>T 2989C>T), TTC229TTT (2992C>T), ACC230TCC (2993A>T), ACC231AAA (2997C>A 2998C>A), TTA232ATA (2999T>A), GAT233GAC (3004T>C), TTG234CTT (3005T>C 3007G>T), GCA235AGG (3008G>A 3009C>G 3010A>G), AGT236GCT (3011A>G 3012G>C), GGG237AGG (3014G>A), TTT238TAC (3018T>A 3019T>C), TAT239CAT (3020T>C), GTG241TTG (3026G>T), GAG242AGA (3029G>A 3030A>G 3031G>A), GAC244GCC (3036A>C), CCT245AAA (3038C>A 3039C>A 3040T>A), CAA246GAA (3041C>G), GAT247GAC (3046T>C), TCG249CCA (3050T>C 3052G>A), ACC251ACA (3058C>A), GCG252GCA (3061G>A), TTT253TTC (3064T>C), AAC254AGA (3066A>G 3067C>A), GTA255ACT (3068G>A 3069T>C 3070A>T), GAA256CAC (3071G>C 3073A>C), CAC257TCA (3074C>T 3075A>C 3076C>A), TTC262TAT (3090T>A 3091C>T), CTT263TTG (3092C>T 3094T>G), CGA264GTA (3095C>G 3096G>T), CCT266CCA (3103T>A), ATG267TTT (3104A>T 3106G>T), TTA269CTA (3110T>C), AAA270TCT (3113A>T 3114A>C 3115A>T), AAC271AAT (3118C>T), TCA272GCA (3119T>G), TCT274GCA (3125T>G 3127T>A), ACT275ACC (3130T>C), AGA278GGG (3137A>G 3139A>G), GTT279TTG (3140G>T 3142T>G), GAC281AAT (3146G>A 3148C>T), AAT282ACT (3150A>C), GTC283GTG (3154C>G), CTA284TTT (3155C>T 3157A>T), AGA285TAA (3158A>T 3159G>A), GGT286AAG (3161G>A 3162G>A 3163T>G), CTC287TTT (3164C>T 3166C>T), CAA288TTG (3167C>T 3168A>T 3169A>G), AAT289AGG (3171A>G 3172T>G), AAC290AAG (3175C>G), ATC291TCA (3176A>T 3177T>C 3178C>A), TGT292GTT (3179T>G 3180G>T), CTC293TTA (3182C>T 3184C>A), GTC294GTT (3187C>T), TAC295TTT (3189A>T 3190C>T), CTT296TTT (3191C>T), CTT296_GAC297insT-- (3193_3194insT), GAC297GAT (3196C>T), ATT300CTG (3203A>C 3205T>G), GTC301ATC (3206G>A), TAT302GAT (3209T>G), ACT304CAG (3215A>C 3216C>A 3217T>G), TCC305TCT (3220C>T), CTA306ATT (3221C>A 3223A>T), CAG307GAA (3224C>G 3226G>A), GAA308GAC (3229A>C), CAC309CAT (3232C>T), CTG310CTA (3235G>A), GAG311TTG (3236G>T 3237A>T), AAC312CAC (3239A>C), GAA314AGA (3245G>A 3246A>G), CGA315AAA (3248C>A 3249G>A) | | | | | | | | |

*: Inserts / Deletes / Misaligned / Frameshifts

#### Analysis details

This analysis was performed with panviral2.64

### NGS Details (UN8): Errantivirus

#### Assembly

| | |
|---|---|
| **Coverage Length** | 353 (1 contig(s)) |
| **Depth Of Coverage** | 20.6 |
| **Number Of Reads** | 64 |
| **Reads Per Million** | 1.07 rpm (after QC) |
| **Ambiguities** | 0 |
| **Assembly Method** | de novo + reference guided assembly |
| **Consensus Caller** | Bcf Tools |

#### Coverage Map

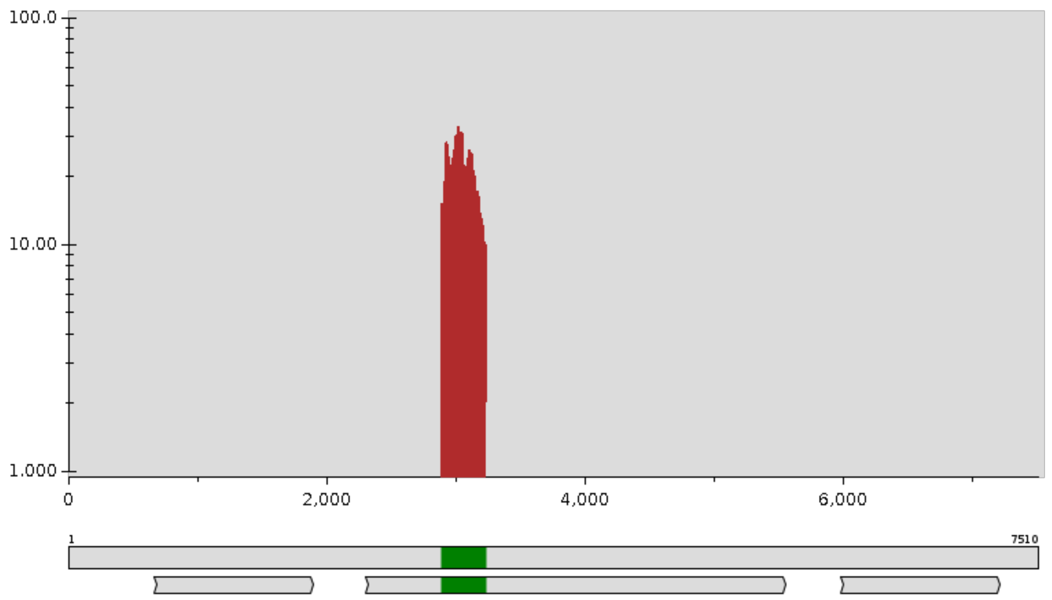

#### Assignment

| | |
|---|---|
| **Type** | Errantivirus (Taxonomy ID: 186666) |
| **Reference Genome** | NC_038512.1 |
| **NT Identity (%)** | 56.9405 |
| **AA Identity (%)** | 47.4576 |
| **Number Of Stop Codons** | 0 |
| **Number Of CDS** | 3 |

#### Alignment

| | |
|---|---|
| **Alignment Score** | 98.0 (NT) + 431.0 (AA) = 529.0 |
| **Concordance (%)** | 34.7113 |

| **Alignment Method** | Global, seeded, nucleotide + amino acids (AGA) |
|---|---|

#### Genome Region

Sequence starts at position 2887 and ends at position 3239 relative to NC_038512.1 reference sequence.

#### Alignment Detailed Statistics

| | Begin | End | Coverage | Score | Concordance | Matches | Identities | I/D/M/F* | Stop Codons |
|---|---|---|---|---|---|---|---|---|---|
| **NT** | **2887** | **3239** | **4.7%** | **98** | **13.9%** | **353 (100%)** | **201 (56.9%)** | **0/0** | |
| Mutations: | 2893A>T, 2896T>A, 2901T>A, 2902C>T, 2903C>A, 2905T>G, 2906A>G, 2908G>C, 2911G>A, 2914C>T, 2915G>A, 2918A>T, 2919A>C, 2920G>C, 2927G>A, 2929T>G, 2930G>A, 2935A>G, 2937A>T, 2938C>T, 2941G>A, 2944A>T, 2947A>T, 2949A>T, 2950C>A, 2951A>G, 2953A>G, 2954A>G, 2955G>A, 2956T>G, 2960G>C, 2963C>T, 2965T>G, 2969A>G, 2971G>A, 2977T>A, 2978A>G, 2979A>G, 2982G>C, 2984C>A, 2987T>A, 2988A>T, 2989C>A, 2993A>T, 2995C>T, 2997C>A, 2998C>A, 2999T>A, 3004T>C, 3007G>A, 3008G>A, 3009C>G, 3011A>T, 3012G>C, 3016G>A, 3018T>A, 3019T>C, 3020T>C, 3025G>A, 3026G>C, 3028G>T, 3029G>A, 3030A>G, 3031G>A, 3036A>C, 3038C>A, 3040T>A, 3041C>G, 3043A>G, 3047A>G, 3049A>G, 3050T>C, 3052G>A, 3058C>T, 3061G>C, 3066A>G, 3067C>A, 3068G>A, 3069T>C, 3070A>T, 3071G>C, 3073A>C, 3074C>T, 3075A>C, 3076C>T, 3088A>G, 3090T>A, 3092C>T, 3094T>A, 3095C>G, 3096G>T, 3097A>C, 3103T>C, 3104A>T, 3106G>T, 3109A>T, 3110T>C, 3112A>G, 3113A>T, 3114A>C, 3118C>T, 3119T>G, 3121A>C, 3125T>G, 3127T>C, 3130T>A, 3137A>G, 3140G>T, 3142T>A, 3146G>A, 3148C>T, 3149A>T, 3150A>C, 3154C>G, 3155C>T, 3157A>C, 3158A>C, 3159G>A, 3161G>C, 3162G>A, 3163T>A, 3164C>T, 3166C>T, 3168A>T, 3171A>G, 3172T>G, 3175C>G, 3176A>C, 3177T>A, 3178C>T, 3179T>A, 3180G>T, 3181T>G, 3182C>T, 3184C>G, 3187C>A, 3189A>T, 3190C>T, 3191C>T, 3196C>T, 3202T>A, 3203A>C, 3206G>A, 3208C>A, 3214T>C, 3216C>A, 3217T>A, 3218T>A, 3221C>A, 3224C>G, 3226G>A, 3229A>T | | | | | | | | |

##### CDS

| | Begin | End | Coverage | Score | Concordance | Matches | Identities | I/D/M/F* | Stop Codons |
|---|---|---|---|---|---|---|---|---|---|
| D1R33_gp2 | 195 | 312 | 10.9% | 431 | 51.7% | 118 (100%) | 56 (47.5%) | 0/0/0/0 | 0 |
| Protein mutations: | F199Y (2901T>A 2902C>T), K201D (2906A>G 2908G>C), E204K (2915G>A), K205S (2918A>T 2919A>C 2920G>C), D208K (2927G>A 2929T>G), D209N (2930G>A), Y211F (2937A>T 2938C>T), N215I (2949A>T 2950C>A), I216V (2951A>G 2953A>G), S217E (2954A>G 2955G>A 2956T>G), V219L (2960G>C), K222E (2969A>G 2971G>A), K225G (2978A>G 2979A>G), C226S (2982G>C), Q227K (2984C>A), Y228I (2987T>A 2988A>T 2989C>A), T230S (2993A>T 2995C>T), T231K (2997C>A 2998C>A), L232I (2999T>A), A235R (3008G>A 3009C>G), F238Y (3018T>A 3019T>C), Y239H (3020T>C), V241L (3026G>C 3028G>T), E242R (3029G>A 3030A>G 3031G>A), D244A (3036A>C), P245T (3038C>A 3040T>A), Q246E (3041C>G 3043A>G), I248V (3047A>G 3049A>G), S249P (3050T>C 3052G>A), N254R (3066A>G 3067C>A), V255T (3068G>A 3069T>C 3070A>T), E256H (3071G>C 3073A>C), H257S (3074C>T 3075A>C 3076C>T), F262Y (3090T>A), R264V (3095C>G 3096G>T 3097A>C), M267F (3104A>T 3106G>T), K270S (3113A>T 3114A>C), S272A (3119T>G 3121A>C), S274A (3125T>G 3127T>C), R278G (3137A>G), V279L (3140G>T 3142T>A), D281N (3146G>A 3148C>T), N282S (3149A>T 3150A>C), L284F (3155C>T 3157A>C), R285Q (3158A>C 3159G>A), G286Q (3161G>C 3162G>A 3163T>A), L287F (3164C>T 3166C>T), Q288L (3168A>T), N289R (3171A>G 3172T>G), N290K (3175C>G), I291H (3176A>C 3177T>A 3178C>T), C292M (3179T>A 3180G>T 3181T>G), Y295F (3189A>T 3190C>T), L296F (3191C>T), I300L (3203A>C), V301I (3206G>A 3208C>A), T304K (3216C>A 3217T>A), S305T (3218T>A), L306I (3221C>A), Q307E (3224C>G 3226G>A), E308D (3229A>T) | | | | | | | | |
| Codon mutations: | GTA196GTT (2893A>T), GTT197GTA (2896T>A), TTC199TAT (2901T>A 2902C>T), CGT200AGG (2903C>A 2905T>G), AAG201GAC (2906A>G 2908G>C), TTG202TTA (2911G>A), AAC203AAT (2914C>T), GAG204AAG (2915G>A), AAG205TCC (2918A>T 2919A>C 2920G>C), GAT208AAG (2927G>A 2929T>G), GAC209AAC (2930G>A), AAA210AAG (2935A>G), TAC211TTT (2937A>T 2938C>T), CCG212CCA (2941G>A), ATA213ATT (2944A>T), CCA214CCT (2947A>T), AAC215ATA (2949A>T 2950C>A), ATA216GTG (2951A>G 2953A>G), AGT217GAG (2954A>G 2955G>A 2956T>G), GTA219CTA (2960G>C), CTT220TTG (2963C>T 2965T>G), AAG222GAA (2969A>G 2971G>A), GGT224GGA (2977T>A), AAG225GGG (2978A>G 2979A>G), TGC226TCC (2982G>C), CAA227AAA (2984C>A), TAC228ATA (2987T>A 2988A>T 2989C>A), ACC230TCT (2993A>T 2995C>T), ACC231AAA (2997C>A 2998C>A), TTA232ATA (2999T>A), GAT233GAC (3004T>C), TTG234TTA (3007G>A), GCA235AGA (3008G>A 3009C>G), AGT236TCT (3011A>T 3012G>C), GGG237GGA (3016G>A), TTT238TAC (3018T>A 3019T>C), TAT239CAT (3020T>C), CAG240CAA (3025G>A), GTG241CTT (3026G>C 3028G>T), GAG242AGA (3029G>A 3030A>G 3031G>A), GAC244GCC (3036A>C), CCT245ACA (3038C>A 3040T>A), CAA246GAG (3041C>G 3043A>G), ATA248GTG (3047A>G 3049A>G), TCG249CCA (3050T>C 3052G>A), ACC251ACT (3058C>T), GCG252GCC (3061G>C), AAC254AGA (3066A>G 3067C>A), GTA255ACT (3068G>A 3069T>C 3070A>T), GAA256CAC (3071G>C 3073A>C), CAC257TCT (3074C>T 3075A>C 3076C>T), GAA261GAG (3088A>G), TTC262TAC (3090T>A), CTT263TTA (3092C>T 3094T>A), CGA264GTC (3095C>G 3096G>T 3097A>C), CCT266CCC (3103T>C), ATG267TTT (3104A>T 3106G>T), GGA268GGT (3109A>T), TTA269CTG (3110T>C 3112A>G), AAA270TCA (3113A>T 3114A>C), AAC271AAT (3118C>T), TCA272GCC (3119T>G 3121A>C), TCT274GCC (3125T>G 3127T>C), ACT275ACA (3130T>A), AGA278GGA (3137A>G), GTT279TTA (3140G>T 3142T>A), GAC281AAT (3146G>A 3148C>T), AAT282TCT (3149A>T 3150A>C), GTC283GTG (3154C>G), CTA284TTC (3155C>T 3157A>C), AGA285CAA (3158A>C 3159G>A), GGT286CAA (3161G>C 3162G>A 3163T>A), CTC287TTT (3164C>T 3166C>T), CAA288CTA (3168A>T), AAT289AGG (3171A>G 3172T>G), AAC290AAG (3175C>G), ATC291CAT (3176A>C 3177T>A 3178C>T), TGT292ATG (3179T>A 3180G>T 3181T>G), CTC293TTG (3182C>T 3184C>G), GTC294GTA (3187C>A), TAC295TTT (3189A>T 3190C>T), CTT296TTT (3191C>T), GAC297GAT (3196C>T), ATT299ATA (3202T>A), ATT300CTT (3203A>C), GTC301ATA (3206G>A 3208C>A), AGT303AGC (3214T>C), ACT304AAA (3216C>A 3217T>A), TCC305ACC (3218T>A), CTA306ATA (3221C>A), CAG307GAA (3224C>G 3226G>A), GAA308GAT (3229A>T) | | | | | | | | |

##### Proteins

| | Begin | End | Coverage | Score | Concordance | Matches | Identities | I/D/M/F* | Stop Codons |
|---|---|---|---|---|---|---|---|---|---|
| ORF B (YP_009507248.1) | 195 | 312 | 10.9% | 431 | 51.7% | 118 (100%) | 56 (47.5%) | 0/0/0/0 | 0 |
| Protein mutations: | F199Y (2901T>A 2902C>T), K201D (2906A>G 2908G>C), E204K (2915G>A), K205S (2918A>T 2919A>C 2920G>C), D208K (2927G>A 2929T>G), D209N (2930G>A), Y211F (2937A>T 2938C>T), N215I (2949A>T 2950C>A), I216V (2951A>G 2953A>G), S217E (2954A>G 2955G>A 2956T>G), V219L (2960G>C), K222E (2969A>G 2971G>A), K225G (2978A>G 2979A>G), C226S (2982G>C), Q227K (2984C>A), Y228I (2987T>A 2988A>T 2989C>A), T230S (2993A>T 2995C>T), T231K (2997C>A 2998C>A), L232I (2999T>A), A235R (3008G>A 3009C>G), F238Y (3018T>A 3019T>C), Y239H (3020T>C), V241L (3026G>C 3028G>T), E242R (3029G>A 3030A>G 3031G>A), D244A (3036A>C), P245T (3038C>A 3040T>A), Q246E (3041C>G 3043A>G), I248V (3047A>G 3049A>G), S249P (3050T>C 3052G>A), N254R (3066A>G 3067C>A), V255T (3068G>A 3069T>C 3070A>T), E256H (3071G>C 3073A>C), H257S (3074C>T 3075A>C 3076C>T), F262Y (3090T>A), R264V (3095C>G 3096G>T 3097A>C), M267F (3104A>T 3106G>T), K270S (3113A>T 3114A>C), S272A (3119T>G 3121A>C), S274A (3125T>G 3127T>C), R278G (3137A>G), V279L (3140G>T 3142T>A), D281N (3146G>A 3148C>T), N282S (3149A>T 3150A>C), L284F (3155C>T 3157A>C), R285Q (3158A>C 3159G>A), G286Q (3161G>C 3162G>A 3163T>A), L287F (3164C>T 3166C>T), Q288L (3168A>T), N289R (3171A>G 3172T>G), N290K (3175C>G), I291H (3176A>C 3177T>A 3178C>T), C292M (3179T>A 3180G>T 3181T>G), Y295F (3189A>T 3190C>T), L296F (3191C>T), I300L (3203A>C), V301I (3206G>A 3208C>A), T304K (3216C>A 3217T>A), S305T (3218T>A), L306I (3221C>A), Q307E (3224C>G 3226G>A), E308D (3229A>T) | | | | | | | | |
| Codon mutations: | GTA196GTT (2893A>T), GTT197GTA (2896T>A), TTC199TAT (2901T>A 2902C>T), CGT200AGG (2903C>A 2905T>G), AAG201GAC (2906A>G 2908G>C), TTG202TTA (2911G>A), AAC203AAT (2914C>T), GAG204AAG (2915G>A), AAG205TCC (2918A>T 2919A>C 2920G>C), GAT208AAG (2927G>A 2929T>G), GAC209AAC (2930G>A), AAA210AAG (2935A>G), TAC211TTT (2937A>T 2938C>T), CCG212CCA (2941G>A), ATA213ATT (2944A>T), CCA214CCT (2947A>T), AAC215ATA (2949A>T 2950C>A), ATA216GTG (2951A>G 2953A>G), AGT217GAG (2954A>G 2955G>A 2956T>G), GTA219CTA (2960G>C), CTT220TTG (2963C>T 2965T>G), AAG222GAA (2969A>G 2971G>A), GGT224GGA (2977T>A), AAG225GGG (2978A>G 2979A>G), TGC226TCC (2982G>C), CAA227AAA (2984C>A), TAC228ATA (2987T>A 2988A>T 2989C>A), ACC230TCT (2993A>T 2995C>T), ACC231AAA (2997C>A 2998C>A), TTA232ATA (2999T>A), GAT233GAC (3004T>C), TTG234TTA (3007G>A), GCA235AGA (3008G>A 3009C>G), AGT236TCT (3011A>T 3012G>C), GGG237GGA (3016G>A), TTT238TAC (3018T>A 3019T>C), TAT239CAT (3020T>C), CAG240CAA (3025G>A), GTG241CTT (3026G>C 3028G>T), GAG242AGA (3029G>A 3030A>G 3031G>A), GAC244GCC (3036A>C), CCT245ACA (3038C>A 3040T>A), CAA246GAG (3041C>G 3043A>G), ATA248GTG (3047A>G 3049A>G), TCG249CCA (3050T>C 3052G>A), ACC251ACT (3058C>T), GCG252GCC (3061G>C), AAC254AGA (3066A>G 3067C>A), GTA255ACT (3068G>A 3069T>C 3070A>T), GAA256CAC (3071G>C 3073A>C), CAC257TCT (3074C>T 3075A>C 3076C>T), GAA261GAG (3088A>G), TTC262TAC (3090T>A), CTT263TTA (3092C>T 3094T>A), CGA264GTC (3095C>G 3096G>T 3097A>C), CCT266CCC (3103T>C), ATG267TTT (3104A>T 3106G>T), GGA268GGT (3109A>T), TTA269CTG (3110T>C 3112A>G), AAA270TCA (3113A>T 3114A>C), AAC271AAT (3118C>T), TCA272GCC (3119T>G 3121A>C), TCT274GCC (3125T>G 3127T>C), ACT275ACA (3130T>A), AGA278GGA (3137A>G), GTT279TTA (3140G>T 3142T>A), GAC281AAT (3146G>A 3148C>T), AAT282TCT (3149A>T 3150A>C), GTC283GTG (3154C>G), CTA284TTC (3155C>T 3157A>C), AGA285CAA (3158A>C 3159G>A), GGT286CAA (3161G>C 3162G>A 3163T>A), CTC287TTT (3164C>T 3166C>T), CAA288CTA (3168A>T), AAT289AGG (3171A>G 3172T>G), AAC290AAG (3175C>G), ATC291CAT (3176A>C 3177T>A 3178C>T), TGT292ATG (3179T>A 3180G>T 3181T>G), CTC293TTG (3182C>T 3184C>G), GTC294GTA (3187C>A), TAC295TTT (3189A>T 3190C>T), CTT296TTT (3191C>T), GAC297GAT (3196C>T), ATT299ATA (3202T>A), ATT300CTT (3203A>C), GTC301ATA (3206G>A 3208C>A), AGT303AGC (3214T>C), ACT304AAA (3216C>A 3217T>A), TCC305ACC (3218T>A), CTA306ATA (3221C>A), CAG307GAA (3224C>G 3226G>A), GAA308GAT (3229A>T) | | | | | | | | |

*: Inserts / Deletes / Misaligned / Frameshifts

#### Analysis details

This analysis was performed with panviral2.64

### NGS Details (UN8): Errantivirus

#### Assembly

| | |
|---|---|
| **Coverage Length** | 372 (1 contig(s)) |
| **Depth Of Coverage** | 18.4 |
| **Number Of Reads** | 59 |
| **Reads Per Million** | 0.99 rpm (after QC) |
| **Ambiguities** | 0 |
| **Assembly Method** | de novo + reference guided assembly |
| **Consensus Caller** | Bcf Tools |

#### Coverage Map

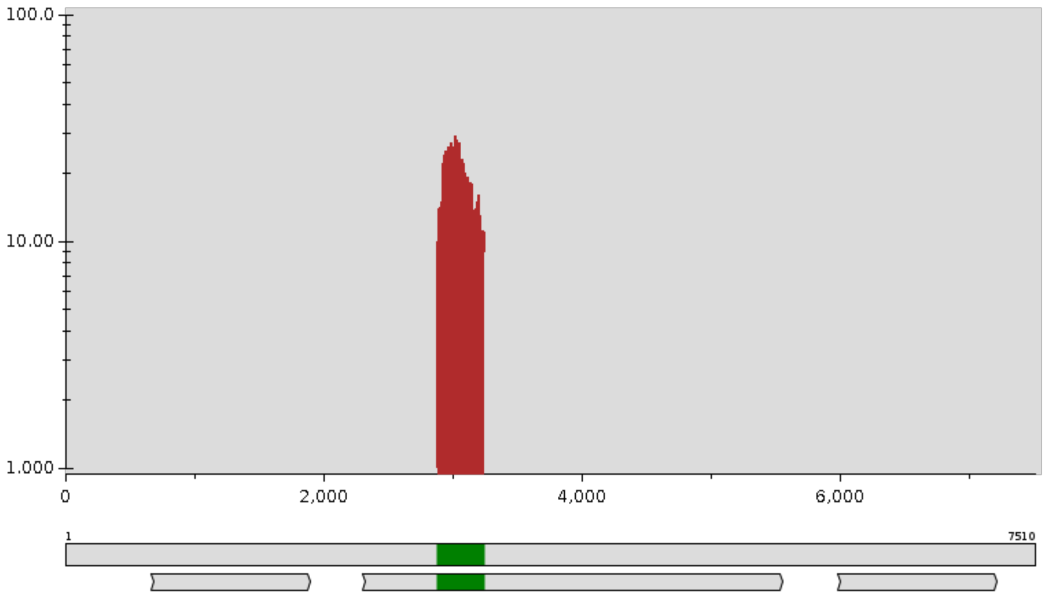

#### Assignment

| | |
|---|---|
| **Type** | Errantivirus (Taxonomy ID: 186666) |
| **Reference Genome** | NC_038512.1 |
| **NT Identity (%)** | 53.0667 |
| **AA Identity (%)** | 42.4 |
| **Number Of Stop Codons** | 0 |
| **Number Of CDS** | 3 |

#### Alignment

| | |
|---|---|
| **Alignment Score** | 40.0 (NT) + 399.0 (AA) = 439.0 |
| **Concordance (%)** | 27.2502 |

| Alignment Method | Global, seeded, nucleotide + amino acids (AGA) |
|---|---|

#### Genome Region

Sequence starts at position 2878 and ends at position 3249 relative to NC_038512.1 reference sequence.

#### Alignment Detailed Statistics

| | Begin | End | Coverage | Score | Concordance | Matches | Identities | I/D/M/F* | Stop Codons |
|---|---|---|---|---|---|---|---|---|---|
| **NT** | **2878** | **3249** | **5.0%** | **40** | **5.4%** | **372 (99.2%)** | **199 (53.1%)** | **3/0** | |
| Mutations: | 2885C>A, 2887T>G, 2888C>T, 2890C>G, 2891G>T, 2892T>G, 2893A>T, 2896T>A, 2901T>A, 2902C>T, 2903C>A, 2905T>G, 2906A>G, 2908G>T, 2909T>C, 2911G>T, 2914C>T, 2915G>A, 2918A>C, 2920G>C, 2924A>G, 2926C>T, 2927G>A, 2929T>G, 2930G>A, 2937A>T, 2939C>T, 2941G>T, 2947A>T, 2948A>C, 2949A>T, 2950C>A, 2951A>G, 2954A>G, 2955G>A, 2956T>A, 2959C>T, 2960G>C, 2963C>T, 2965T>G, 2968C>T, 2969A>G, 2971G>A, 2972T>C, 2974A>T, 2977T>G, 2978A>G, 2979A>G, 2980G>T, 2982G>C, 2983C>A, 2984C>G, 2985A>T, 2987T>A, 2988A>T, 2991T>G, 2992C>T, 2993A>T, 2994C>T, 2997C>A, 2998C>A, 2999T>A, 3004T>C, 3005T>C, 3007G>T, 3008G>A, 3009C>G, 3010A>G, 3011A>G, 3012G>C, 3018T>A, 3019T>C, 3020T>C, 3026G>T, 3029G>A, 3030A>G, 3031G>A, 3036A>C, 3038C>A, 3039C>A, 3040T>A, 3041C>G, 3046T>C, 3050T>C, 3052G>A, 3058C>A, 3061G>A, 3064T>G, 3066A>G, 3067C>A, 3068G>A, 3069T>C, 3070A>T, 3071G>C, 3073A>C, 3074C>T, 3075A>T, 3076C>A, 3088A>G, 3090T>A, 3091C>T, 3092C>T, 3094T>G, 3095C>G, 3096G>T, 3103T>A, 3104A>T, 3106G>T, 3113A>T, 3114A>C, 3115A>T, 3118C>T, 3119T>G, 3125T>A, 3127T>A, 3130T>C, 3137A>G, 3139A>G, 3140G>T, 3142T>G, 3146G>A, 3148C>T, 3150A>C, 3154C>G, 3155C>T, 3157A>T, 3158A>C, 3159G>A, 3160_3161insAAG, 3161G>T, 3162G>T, 3164C>T, 3166C>G, 3167C>A, 3168A>G, 3169A>G, 3172T>G, 3173A>C, 3174A>C, 3175C>T, 3176A>G, 3178C>T, 3180G>T, 3181T>A, 3182C>G, 3184C>T, 3185G>T, 3187C>T, 3189A>T, 3190C>T, 3191C>T, 3194G>T, 3196C>T, 3203A>C, 3205T>G, 3206G>A, 3215A>C, 3216C>A, 3217T>G, 3220C>T, 3221C>A, 3223A>T, 3224C>G, 3226G>A, 3229A>C, 3232C>T, 3235G>A, 3236G>T, 3237A>T, 3239A>C, 3245G>A, 3246A>G, 3248C>A, 3249G>A | | | | | | | | |
| **CDS** | | | | | | | | | |
| D1R33_gp2 | 192 | 315 | 11.4% | 399 | 45.1% | 124 (99.2%) | 53 (42.4%) | 1/0/0/0 | 0 |
| Protein mutations: | V196C (2891G>T 2892T>G 2893A>T), F199Y (2901T>A 2902C>T), K201D (2906A>G 2908G>T), E204K (2915G>A), K205H (2918A>C 2920G>C), I207V (2924A>G 2926C>T), D208K (2927G>A 2929T>G), D209N (2930G>A), Y211F (2937A>T), P212S (2939C>T 2941G>T), N215L (2948A>C 2949A>T 2950C>A), I216V (2951A>G), S217E (2954A>G 2955G>A 2956T>A), V219L (2960G>C), K222E (2969A>G 2971G>A), K225G (2978A>G 2979A>G 2980G>T), C226S (2982G>C 2983C>A), Q227V (2984C>G 2985A>T), Y228I (2987T>A 2988A>T), F229C (2991T>G 2992C>T), T230F (2993A>T 2994C>T), T231K (2997C>A 2998C>A), L232I (2999T>A), A235R (3008G>A 3009C>G 3010A>G), S236A (3011A>G 3012G>C), F238Y (3018T>A 3019T>C), Y239H (3020T>C), V241L (3026G>T), E242R (3029G>A 3030A>G 3031G>A), D244A (3036A>C), P245K (3038C>A 3039C>A 3040T>A), Q246E (3041C>G), S249P (3050T>C 3052G>A), F253L (3064T>G), N254R (3066A>G 3067C>A), V255T (3068G>A 3069T>C 3070A>T), E256H (3071G>C 3073A>C), H257L (3074C>T 3075A>T 3076C>A), F262Y (3090T>A 3091C>T), R264V (3095C>G 3096G>T), M267F (3104A>T 3106G>T), K270S (3113A>T 3114A>C 3115A>T), S272A (3119T>G), S274T (3125T>A 3127T>A), R278G (3137A>G 3139A>G), V279L (3140G>T 3142T>G), D281N (3146G>A 3148C>T), N282T (3150A>C), L284F (3155C>T 3157A>T), R285Q (3158A>C 3159G>A), R285_G286insK (3160_3161insAAG), G286F (3161G>T 3162G>T), Q288R (3167C>A 3168A>G 3169A>G), N289K (3172T>G), N290P (3173A>C 3174A>C 3175C>T), I291V (3176A>G 3178C>T), C292L (3180G>T 3181T>A), L293V (3182C>G 3184C>T), V294F (3185G>T 3187C>T), Y295F (3189A>T 3190C>T), L296F (3191C>T), D297Y (3194G>T 3196C>T), I300L (3203A>C 3205T>G), V301I (3206G>A), T304Q (3215A>C 3216C>A 3217T>G), L306I (3221C>A 3223A>T), Q307E (3224C>G 3226G>A), E308D (3229A>C), E311L (3236G>T 3237A>T), N312H (3239A>C), E314R (3245G>A 3246A>G) | | | | | | | | |
| Codon mutations: | CGT194AGG (2885C>A 2887T>G), CTC195TTG (2888C>T 2890C>G), GTA196TGT (2891G>T 2892T>G 2893A>T), GTT197GTA (2896T>A), TTC199TAT (2901T>A 2902C>T), CGT200AGG (2903C>A 2905T>G), AAG201GAT (2906A>G 2908G>T), TTG202CTT (2909T>C 2911G>T), AAC203AAT (2914C>T), GAG204AAG (2915G>A), AAG205CAC (2918A>C 2920G>C), ATC207GTT (2924A>G 2926C>T), GAT208AAG (2927G>A 2929T>G), GAC209AAC (2930G>A), TAC211TTC (2937A>T), CCG212TCT (2939C>T 2941G>T), CCA214CCT (2947A>T), AAC215CTA (2948A>C 2949A>T 2950C>A), ATA216GTA (2951A>G), AGT217GAA (2954A>G 2955G>A 2956T>A), GAC218GAT (2959C>T), GTA219CTA (2960G>C), CTT220TTG (2963C>T 2965T>G), GAC221GAT (2968C>T), AAG222GAA (2969A>G 2971G>A), TTA223CTT (2972T>C 2974A>T), GGT224GGG (2977T>G), AAG225GGT (2978A>G 2979A>G 2980G>T), TGC226TCA (2982G>C 2983C>A), CAA227GTA (2984C>G 2985A>T), TAC228ATC (2987T>A 2988A>T), TTC229TGT (2991T>G 2992C>T), ACC230TTC (2993A>T 2994C>T), ACC231AAA (2997C>A 2998C>A), TTA232ATA (2999T>A), GAT233GAC (3004T>C), TTG234CTT (3005T>C 3007G>T), GCA235AGG (3008G>A 3009C>G 3010A>G), AGT236GCT (3011A>G 3012G>C), TTT238TAC (3018T>A 3019T>C), TAT239CAT (3020T>C), GTG241TTG (3026G>T), GAG242AGA (3029G>A 3030A>G 3031G>A), GAC244GCC (3036A>C), CCT245AAA (3038C>A 3039C>A 3040T>A), CAA246GAA (3041C>G), GAT247GAC (3046T>C), TCG249CCA (3050T>C 3052G>A), ACC251ACA (3058C>A), GCG252GCA (3061G>A), TTT253TTG (3064T>G), AAC254AGA (3066A>G 3067C>A), GTA255ACT (3068G>A 3069T>C 3070A>T), GAA256CAC (3071G>C 3073A>C), CAC257TTA (3074C>T 3075A>T 3076C>A), GAA261GAG (3088A>G), TTC262TAT (3090T>A 3091C>T), CTT263TTG (3092C>T 3094T>G), CGA264GTA (3095C>G 3096G>T), CCT266CCA (3103T>A), ATG267TTT (3104A>T 3106G>T), AAA270TCT (3113A>T 3114A>C 3115A>T), AAC271AAT (3118C>T), TCA272GCA (3119T>G), TCT274ACA (3125T>A 3127T>A), ACT275ACC (3130T>C), AGA278GGG (3137A>G 3139A>G), GTT279TTG (3140G>T 3142T>G), GAC281AAT (3146G>A 3148C>T), AAT282ACT (3150A>C), GTC283GTG (3154C>G), CTA284TTT (3155C>T 3157A>T), AGA285CAA (3158A>C 3159G>A), AGA285_GGT286insAAG (3160_3161insAAG), GGT286TTT (3161G>T 3162G>T), CTC287TTG (3164C>T 3166C>G), CAA288AGG (3167C>A 3168A>G 3169A>G), AAT289AAG (3172T>G), AAC290CCT (3173A>C 3174A>C 3175C>T), ATC291GTT (3176A>G 3178C>T), TGT292TTA (3180G>T 3181T>A), CTC293GTT (3182C>G 3184C>T), GTC294TTT (3185G>T 3187C>T), TAC295TTT (3189A>T 3190C>T), CTT296TTT (3191C>T), GAC297TAT (3194G>T 3196C>T), ATT300CTG (3203A>C 3205T>G), GTC301ATC (3206G>A), ACT304CAG (3215A>C 3216C>A 3217T>G), TCC305TCT (3220C>T), CTA306ATT (3221C>A 3223A>T), CAG307GAA (3224C>G 3226G>A), GAA308GAC (3229A>C), CAC309CAT (3232C>T), CTG310CTA (3235G>A), GAG311TTG (3236G>T 3237A>T), AAC312CAC (3239A>C), GAA314AGA (3245G>A 3246A>G), CGA315AA. (3248C>A 3249G>A) | | | | | | | | |
| **Proteins** | | | | | | | | | |
| ORF B (YP_009507248.1) | 192 | 315 | 11.4% | 399 | 45.1% | 124 (99.2%) | 53 (42.4%) | 1/0/0/0 | 0 |
| Protein mutations: | V196C (2891G>T 2892T>G 2893A>T), F199Y (2901T>A 2902C>T), K201D (2906A>G 2908G>T), E204K (2915G>A), K205H (2918A>C 2920G>C), I207V (2924A>G 2926C>T), D208K (2927G>A 2929T>G), D209N (2930G>A), Y211F (2937A>T), P212S (2939C>T 2941G>T), N215L (2948A>C 2949A>T 2950C>A), I216V (2951A>G), S217E (2954A>G 2955G>A 2956T>A), V219L (2960G>C), K222E (2969A>G 2971G>A), K225G (2978A>G 2979A>G 2980G>T), C226S (2982G>C 2983C>A), Q227V (2984C>G 2985A>T), Y228I (2987T>A 2988A>T), F229C (2991T>G 2992C>T), T230F (2993A>T 2994C>T), T231K (2997C>A 2998C>A), L232I (2999T>A), A235R (3008G>A 3009C>G 3010A>G), S236A (3011A>G 3012G>C), F238Y (3018T>A 3019T>C), Y239H (3020T>C), V241L (3026G>T), E242R (3029G>A 3030A>G 3031G>A), D244A (3036A>C), P245K (3038C>A 3039C>A 3040T>A), Q246E (3041C>G), S249P (3050T>C 3052G>A), F253L (3064T>G), N254R (3066A>G 3067C>A), V255T (3068G>A 3069T>C 3070A>T), E256H (3071G>C 3073A>C), H257L (3074C>T 3075A>T 3076C>A), F262Y (3090T>A 3091C>T), R264V (3095C>G 3096G>T), M267F (3104A>T 3106G>T), K270S (3113A>T 3114A>C 3115A>T), S272A (3119T>G), S274T (3125T>A 3127T>A), R278G (3137A>G 3139A>G), V279L (3140G>T 3142T>G), D281N (3146G>A 3148C>T), N282T (3150A>C), L284F (3155C>T 3157A>T), R285Q (3158A>C 3159G>A), R285_G286insK (3160_3161insAAG), G286F (3161G>T 3162G>T), Q288R (3167C>A 3168A>G 3169A>G), N289K (3172T>G), N290P (3173A>C 3174A>C 3175C>T), I291V (3176A>G 3178C>T), C292L (3180G>T 3181T>A), L293V (3182C>G 3184C>T), V294F (3185G>T 3187C>T), Y295F (3189A>T 3190C>T), L296F (3191C>T), D297Y (3194G>T 3196C>T), I300L (3203A>C 3205T>G), V301I (3206G>A), T304Q (3215A>C 3216C>A 3217T>G), L306I (3221C>A 3223A>T), Q307E (3224C>G 3226G>A), E308D (3229A>C), E311L (3236G>T 3237A>T), N312H (3239A>C), E314R (3245G>A 3246A>G) | | | | | | | | |
| Codon mutations: | CGT194AGG (2885C>A 2887T>G), CTC195TTG (2888C>T 2890C>G), GTA196TGT (2891G>T 2892T>G 2893A>T), GTT197GTA (2896T>A), TTC199TAT (2901T>A 2902C>T), CGT200AGG (2903C>A 2905T>G), AAG201GAT (2906A>G 2908G>T), TTG202CTT (2909T>C 2911G>T), AAC203AAT (2914C>T), GAG204AAG (2915G>A), AAG205CAC (2918A>C 2920G>C), ATC207GTT (2924A>G 2926C>T), GAT208AAG (2927G>A 2929T>G), GAC209AAC (2930G>A), TAC211TTC (2937A>T), CCG212TCT (2939C>T 2941G>T), CCA214CCT (2947A>T), AAC215CTA (2948A>C 2949A>T 2950C>A), ATA216GTA (2951A>G), AGT217GAA (2954A>G 2955G>A 2956T>A), GAC218GAT (2959C>T), GTA219CTA (2960G>C), CTT220TTG (2963C>T 2965T>G), GAC221GAT (2968C>T), AAG222GAA (2969A>G 2971G>A), TTA223CTT (2972T>C 2974A>T), GGT224GGG (2977T>G), AAG225GGT (2978A>G 2979A>G 2980G>T), TGC226TCA (2982G>C 2983C>A), CAA227GTA (2984C>G 2985A>T), TAC228ATC (2987T>A 2988A>T), TTC229TGT (2991T>G 2992C>T), ACC230TTC (2993A>T 2994C>T), ACC231AAA (2997C>A 2998C>A), TTA232ATA (2999T>A), GAT233GAC (3004T>C), TTG234CTT (3005T>C 3007G>T), GCA235AGG (3008G>A 3009C>G 3010A>G), AGT236GCT (3011A>G 3012G>C), TTT238TAC (3018T>A 3019T>C), TAT239CAT (3020T>C), GTG241TTG (3026G>T), GAG242AGA (3029G>A 3030A>G 3031G>A), GAC244GCC (3036A>C), CCT245AAA (3038C>A 3039C>A 3040T>A), CAA246GAA (3041C>G), GAT247GAC (3046T>C), TCG249CCA (3050T>C 3052G>A), ACC251ACA (3058C>A), GCG252GCA (3061G>A), TTT253TTG (3064T>G), AAC254AGA (3066A>G 3067C>A), GTA255ACT (3068G>A 3069T>C 3070A>T), GAA256CAC (3071G>C 3073A>C), CAC257TTA (3074C>T 3075A>T 3076C>A), GAA261GAG (3088A>G), TTC262TAT (3090T>A 3091C>T), CTT263TTG (3092C>T 3094T>G), CGA264GTA (3095C>G 3096G>T), CCT266CCA (3103T>A), ATG267TTT (3104A>T 3106G>T), AAA270TCT (3113A>T 3114A>C 3115A>T), AAC271AAT (3118C>T), TCA272GCA (3119T>G), TCT274ACA (3125T>A 3127T>A), ACT275ACC (3130T>C), AGA278GGG (3137A>G 3139A>G), GTT279TTG (3140G>T 3142T>G), GAC281AAT (3146G>A 3148C>T), AAT282ACT (3150A>C), GTC283GTG (3154C>G), CTA284TTT (3155C>T 3157A>T), AGA285CAA (3158A>C 3159G>A), AGA285_GGT286insAAG (3160_3161insAAG), GGT286TTT (3161G>T 3162G>T), CTC287TTG (3164C>T 3166C>G), CAA288AGG (3167C>A 3168A>G 3169A>G), AAT289AAG (3172T>G), AAC290CCT (3173A>C 3174A>C 3175C>T), ATC291GTT (3176A>G 3178C>T), TGT292TTA (3180G>T 3181T>A), CTC293GTT (3182C>G 3184C>T), GTC294TTT (3185G>T 3187C>T), TAC295TTT (3189A>T 3190C>T), CTT296TTT (3191C>T), GAC297TAT (3194G>T 3196C>T), ATT300CTG (3203A>C 3205T>G), GTC301ATC (3206G>A), ACT304CAG (3215A>C 3216C>A 3217T>G), TCC305TCT (3220C>T), CTA306ATT (3221C>A 3223A>T), CAG307GAA (3224C>G 3226G>A), GAA308GAC (3229A>C), CAC309CAT (3232C>T), CTG310CTA (3235G>A), GAG311TTG (3236G>T 3237A>T), AAC312CAC (3239A>C), GAA314AGA (3245G>A 3246A>G), CGA315AA. (3248C>A 3249G>A) | | | | | | | | |

*: Inserts / Deletes / Misaligned / Frameshifts

#### Analysis details

This analysis was performed with panviral2.64

### NGS Details (UN8): Badnavirus maculacommelinae

#### Assembly

| | |
|---|---|
| **Coverage Length** | 909 (1 contig(s)) |
| **Depth Of Coverage** | 7.8 |
| **Number Of Reads** | 55 |
| **Reads Per Million** | 0.92 rpm (after QC) |
| **Ambiguities** | 0 |
| **Assembly Method** | de novo + reference guided assembly |
| **Consensus Caller** | Bcf Tools |

#### Coverage Map

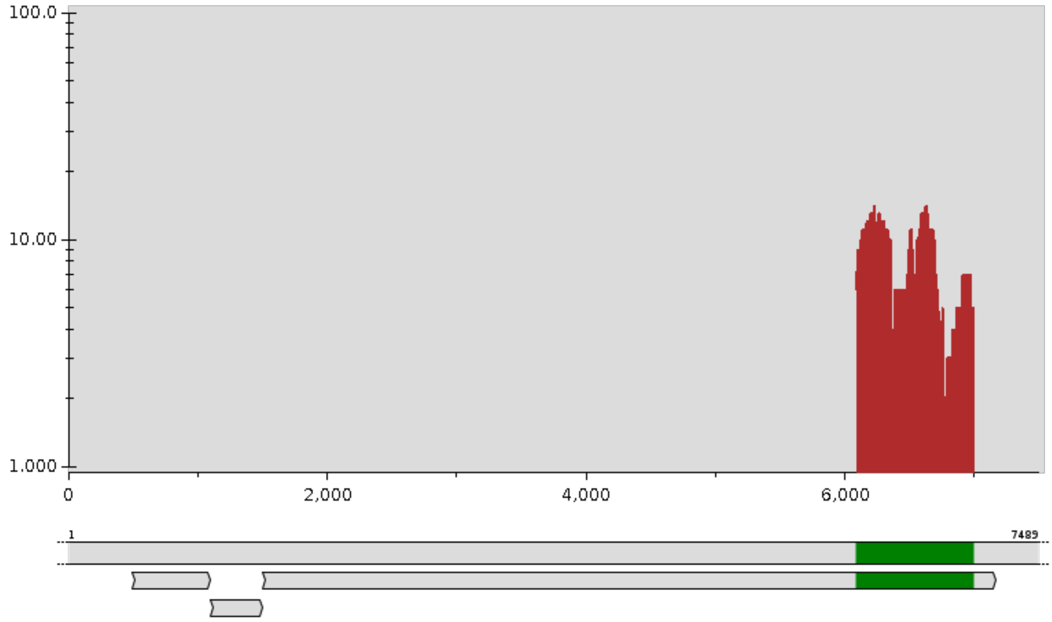

#### Assignment

| | |
|---|---|
| **Type** | Badnavirus maculacommelinae (Taxonomy ID: 3047769) |
| **Reference Genome** | NC_001343.1 |
| **NT Identity (%)** | 54.6858 |
| **AA Identity (%)** | 47.8548 |
| **Number Of Stop Codons** | 3 |
| **Number Of CDS** | 3 |

#### Alignment

| | |
|---|---|
| **Alignment Score** | 123.0 (NT) + 750.0 (AA) = 873.0 |
| **Concordance (%)** | 24.9451 |

| Alignment Method | Global, seeded, nucleotide + amino acids (AGA) |
|---|---|

#### Genome Region

Sequence starts at position 6090 and ends at position 6998 relative to NC_001343.1 reference sequence.

#### Alignment Detailed Statistics

| | Begin | End | Coverage | Score | Concordance | Matches | Identities | I/D/M/F* | Stop Codons |
|---|---|---|---|---|---|---|---|---|---|
| **NT** | **6090** | **6998** | **12.1%** | **123** | **7.0%** | **901 (98.5%)** | **496 (54.2%)** | **6/8** | |
| Mutations: | 6091T>A, 6094A>T, 6102C>T, 6104T>A, 6111C>T, 6119T>A, 6120C>T, 6122C>G, 6125A>G, 6127A>C, 6128T>A, 6131G>A, 6134C>A, 6135G>C, 6143T>C, 6149G>A, 6161C>T, 6162G>A, 6164C>T, 6167C>T, 6168A>G, 6169A>G, 6172G>A, 6173G>T, 6174A>T, 6175C>A, 6176A>C, 6177G>A, 6181A>G, 6182G>A, 6185C>T, 6186A>G, 6188A>C, 6189G>C, 6190C>T, 6191T>A, 6194T>A, 6197C>T, 6198A>G, 6200T>A, 6206T>C, 6209A>T, 6212T>A, 6215T>A, 6218C>T, 6219T>A, 6220C>A, 6221A>T, 6222G>A, 6225A>G, 6226C>A, 6227A>C, 6228G>A, 6229C>T, 6231G>A, 6232A>G, 6233A>G, 6234C>G, 6236A>G, 6240T>C, 6241C>T, 6242G>A, 6243C>G, 6244A>G, 6245G>T, 6251C>A, 6252T>C, 6254T>G, 6258A>G, 6260G>A, 6261C>T, 6263G>T, 6264C>A, 6265A>G, 6266G>A, 6269T>A, 6271G>T, 6272C>T, 6273A>G, 6274A>T, 6275G>A, 6276G>A, 6277A>G, 6278A>T, 6281T>C, 6284G>T, 6285T>A, 6290C>A, 6291C>G, 6293T>A, 6297C>A, 6298C>A, 6299T>A, 6301C>A, 6302C>G, 6311G>A, 6312A>T, 6315G>T, 6319C>A, 6320T>A, 6321C>A, 6323G>C, 6324G>C, 6326G>T, 6329C>A, 6330G>A, 6340G>A, 6342G>A, 6343C>T, 6345T>A, 6346C>T, 6347T>A, 6348C>T, 6350C>A, 6354T>G, 6355G>A, 6356T>A, 6357A>T, 6358C>G, 6362A>G, 6365C>A, 6368G>A, 6369C>T, 6371T>A, 6374G>A, 6378_6380delCAC, 6383C>A, 6384A>T, 6385T>A, 6386T>G, 6387T>C, 6388C>T, 6394T>G, 6395A>G, 6397G>C, 6399G>T, 6401C>G, 6402T>A, 6403T>G, 6404C>A, 6406C>G, 6407A>C, 6408G>C, 6411G>A, 6414A>T, 6415A>T, 6417C>G, 6418T>A, 6420G>A, 6421C>A, 6422A>C, 6426C>A, 6427C>A, 6428T>A, 6432delG, 6433G>T, 6436T>C, 6437G>A, 6439G>A, 6442G>T, 6443C>T, 6444T>C, 6445G>T, 6446G>T, 6447T>A, 6449G>A, 6453A>T, 6455C>A, 6456C>A, 6458C>A, 6459T>A, 6460C>A, 6461A>T, 6475A>T, 6479T>A, 6480C>A, 6482G>A, 6485T>C, 6486A>T, 6488C>A, 6491C>A, 6495T>A, 6497G>A, 6500G>A, 6501C>G, 6502A>G, 6510A>T, 6511G>A, 6512A>C, 6513C>T, 6514A>C, 6515A>T, 6520T>C, 6521G>A, 6523C>G, 6525C>A, 6526C>G, 6527A>T, 6529C>A, 6534G>C, 6536C>A, 6540A>T, 6541G>A, 6542A>T, 6543A>T, 6545G>T, 6549C>A, 6550C>A, 6551A>T, 6555A>G, 6556C>A, 6557A>T, 6558T>A, 6559G>T, 6560G>C, 6563G>A, 6564A>T, 6566G>A, 6569A>C, 6570A>C, 6571G>A, 6573C>A, 6578A>C, 6584A>T, 6587G>A, 6590G>T, 6596T>C, 6597C>A, 6602T>A, 6606C>T, 6608T>G, 6609C>G, 6610A>C, 6611G>T, 6612T>A, 6614A>T, 6617A>T, 6619C>T, 6620T>A, 6621A>G, 6624G>T, 6625A>C, 6626T>C, 6627T>G, 6628C>A, 6629A>T, 6631T>A, 6632C>T, 6633A>T, 6635C>A, 6644G>A, 6647G>A, 6653T>G, 6654T>A, 6656T>C, 6657A>T, 6658T>C, 6659G>C, 6662T>A, 6665C>A, 6674C>A, 6675G>A, 6677C>T, 6679G>T, 6680C>G, 6683A>G, 6684T>G, 6685G>C, 6686G>C, 6690A>C, 6691T>C, 6692G>A, 6693T>A, 6694C>G, 6695A>T, 6698G>A, 6699C>T, 6702G>A, 6703A>G, 6705C>G, 6706C>A, 6707A>T, 6709G>A, 6712G>A, 6713C>T, 6716C>A, 6721G>A, 6722A>G, 6725T>A, 6730C>G, 6731C>T, 6735G>C, 6736C>A, 6737T>A, 6740T>C, 6743A>T, 6744T>A, 6745C>A, 6748T>A, 6752T>A, 6753C>G, 6754C>A, 6757T>A, 6758_6759insGGAAAT, 6760A>T, 6761A>G, 6762T>A, 6763C>G, 6764A>T, 6766C>G, 6767C>T, 6770C>A, 6771G>A, 6774G>T, 6775C>T, 6776A>T, 6779G>A, 6782T>A, 6784A>T, 6785G>T, 6788G>A, 6790C>T, 6791A>T, 6793T>A, 6794C>T, 6795delC, 6800C>T, 6801C>T, 6803G>A, 6804G>A, 6808A>G, 6809A>T, 6812C>T, 6814A>G, 6816A>C, 6818T>A, 6821T>C, 6822T>A, 6823A>T, 6825C>A, 6827T>A, 6828G>A, 6830T>C, 6836G>A, 6839G>A, 6840C>A, 6842C>A, 6843A>C, 6845A>G, 6846A>G, 6848T>G, 6849C>A, 6851C>A, 6852T>A, 6869T>A, 6870A>G, 6872C>A, 6879T>C, 6881C>T, 6882A>C, 6884C>A, 6889C>T, 6890G>A, 6893C>T, 6899T>A, 6901A>C, 6902G>T, 6903C>A, 6904C>G, 6905G>T, 6906T>A, 6907C>G, 6912G>A, 6913T>G, 6914T>A, 6915A>C, 6921T>C, 6923A>T, 6925C>A, 6926A>T, 6930T>C, 6931C>T, 6936T>A, 6937T>C, 6938C>A, 6939T>A, 6942A>T, 6944A>T, 6945G>A, 6946G>C, 6948C>T, 6949T>A, 6950T>C, 6952G>A, 6953A>G, 6954_6956delATC, 6958C>A, 6959A>T, 6960G>A, 6964C>T, 6965A>T, 6968C>T, 6974C>T, 6978G>A, 6980T>A, 6986G>A, 6987C>A, 6989T>A, 6993G>A, 6994G>A | | | | | | | | |

##### CDS

| | Begin | End | Coverage | Score | Concordance | Matches | Identities | I/D/M/F* | Stop Codons |
|---|---|---|---|---|---|---|---|---|---|
| ComYMVgp3 | 1529 | 1831 | 16.1% | 750 | 35.5% | 301 (98.7%) | 145 (47.5%) | 2/2/2/2 | 3 |
| Protein mutations: | L1529H (6091T>A), Y1530F (6094A>T), P1536S (6111C>T), N1541T (6127A>C 6128T>A), A1544P (6135G>C), V1553I (6162G>A 6164C>T), K1555G (6168A>G 6169A>G), G1556D (6172G>A 6173G>T), T1557Y (6174A>T 6175C>A 6176A>C), E1558K (6177G>A), K1559R (6181A>G 6182G>A), I1561V (6186A>G 6188A>C), A1562L (6189G>C 6190C>T 6191T>A), I1565V (6198A>G 6200T>A), S1572N (6219T>A 6220C>A 6221A>T), E1573K (6222G>A), T1574D (6225A>G 6226C>A 6227A>C), A1575I (6228G>A 6229C>T), E1576R (6231G>A 6232A>G 6233A>G), Q1577E (6234C>G 6236A>G), S1579L (6240T>C 6241C>T 6242G>A), Q1580G (6243C>G 6244A>G 6245G>T), Y1583Q (6252T>C 6254T>G), M1585V (6258A>G 6260G>A), L1586F (6261C>T 6263G>T), Q1587R (6264C>A 6265A>G 6266G>A), C1589F (6271G>T 6272C>T), K1590V (6273A>G 6274A>T 6275G>A), E1591S (6276G>A 6277A>G 6278A>T), L1594I (6285T>A), L1596V (6291C>G 6293T>A), P1598K (6297C>A 6298C>A 6299T>A), T1599K (6301C>A 6302C>G), I1603L (6312A>T), G1604C (6315G>T), T1605K (6319C>A 6320T>A), P1606T (6321C>A 6323G>C), E1607H (6324G>C 6326G>T), D1609N (6330G>A), G1612E (6340G>A), A1613I (6342G>A 6343C>T), S1614I (6345T>A 6346C>T 6347T>A), C1617E (6354T>G 6355G>A 6356T>A), T1618* (6357A>T 6358C>G), H1625del (6378_6380delCAC), I1627* (6384A>T 6385T>A 6386T>G), S1628L (6387T>C 6388C>T), I1630R (6394T>G 6395A>G), C1631S (6397G>C), D1632* (6399G>T 6401C>G), F1633R (6402T>A 6403T>G 6404C>A), S1634C (6406C>G 6407A>C), D1635H (6408G>C), E1636K (6411G>A), K1637L (6414A>T 6415A>T), L1638E (6417C>G 6418T>A), A1639N (6420G>A 6421C>A 6422A>C), P1641K (6426C>A 6427C>A 6428T>A), M1644T (6436T>C 6437G>A), R1645K (6439G>A), S1646I (6442G>T 6443C>T), W1647L (6444T>C 6445G>T 6446G>T), L1648I (6447T>A 6449G>A), I1650L (6453A>T 6455C>A), L1651I (6456C>A 6458C>A), S1652N (6459T>A 6460C>A 6461A>T), Y1657F (6475A>T), Q1659K (6480C>A 6482G>A), I1661L (6486A>T 6488C>A), L1664I (6495T>A 6497G>A), Q1666G (6501C>G 6502A>G), R1669Y (6510A>T 6511G>A 6512A>C), Q1670S (6513C>T 6514A>C 6515A>T), M1672T (6520T>C 6521G>A), A1673G (6523C>G), P1674S (6525C>A 6526C>G 6527A>T), T1675K (6529C>A), D1677Q (6534G>C 6536C>A), R1679Y (6540A>T 6541G>A 6542A>T), M1680F (6543A>T 6545G>T), P1682N (6549C>A 6550C>A 6551A>T), T1684D (6555A>G 6556C>A 6557A>T), W1685I (6558T>A 6559G>T 6560G>C), M1687L (6564A>T 6566G>A), R1689Q (6570A>C 6571G>A), Q1690K (6573C>A), E1693D (6584A>T), L1698I (6597C>A), Q1702A (6609C>G 6610A>C 6611G>T), L1703I (6612T>A 6614A>T), P1705L (6619C>T 6620T>A), K1706E (6621A>G), D1707S (6624G>T 6625A>C 6626T>C), S1708D (6627T>G 6628C>A 6629A>T), F1709Y (6631T>A 6632C>T), I1710L (6633A>T 6635C>A), C1717S (6654T>A 6656T>C), M1718S (6657A>T 6658T>C 6659G>C), V1724I (6675G>A 6677C>T), C1725L (6679G>T 6680C>G), W1727A (6684T>G 6685G>C 6686G>C), M1729P (6690A>C 6691T>C 6692G>A), H1732Y (6699C>T), D1733S (6702G>A 6703A>G), P1734D (6705C>G 6706C>A 6707A>T), R1735K (6709G>A), S1736N (6712G>A 6713C>T), R1739K (6721G>A 6722A>G), A1742G (6730C>G 6731C>T), A1744Q (6735G>C 6736C>A 6737T>A), S1747K (6744T>A 6745C>A), F1748Y (6748T>A), N1749K (6752T>A), P1750E (6753C>G 6754C>A), I1751K (6757T>A), I1751_K1752insGN (6758_6759insGGAAAT), K1752M (6760A>T 6761A>G), T1754S (6766C>G 6767C>T), D1756N (6771G>A), A1757F (6774G>T 6775C>T 6776A>T), Q1760L (6784A>T 6785G>T), A1762V (6790C>T 6791A>T), I1763N (6793T>A 6794C>T), D1767N (6804G>A), K1768S (6808A>G 6809A>T), K1770R (6814A>G), I1771L (6816A>C 6818T>A), Y1773I (6822T>A 6823A>T), L1774I (6825C>A 6827T>A), D1775N (6828G>A 6830T>C), L1779I (6840C>A 6842C>A), I1780L (6843A>C 6845A>G), I1781V (6846A>G 6848T>G), S1783T (6852T>A), I1789V (6870A>G 6872C>A), Y1792H (6879T>C 6881C>T), N1793Q (6882A>C 6884C>A), T1795I (6889C>T 6890G>A), N1798K (6899T>A), K1799T (6901A>C 6902G>T), P1800S (6903C>A 6904C>G 6905G>T), V1803R (6912G>A 6913T>G 6914T>A), T1807N (6925C>A 6926A>T), S1809L (6930T>C 6931C>T), F1811T (6936T>A 6937T>C 6938C>A), L1812I (6939T>A), T1813S (6942A>T 6944A>T), G1814T (6945G>A 6946G>C), L1815Y (6948C>T 6949T>A 6950T>C), G1816E (6952G>A 6953A>G), I1817del (6954_6956delATC), T1818N (6958C>A 6959A>T), V1819I (6960G>A), T1820I (6964C>T 6965A>T), D1825K (6978G>A 6980T>A), H1828K (6987C>A 6989T>A), G1830N (6993G>A 6994G>A) | | | | | | | | |

| | Begin | End | Coverage | Score | Concordance | Matches | Identities | I/D/M/F* | Stop Codons |
|---|---|---|---|---|---|---|---|---|---|
| **NT** | **6090** | **6998** | **12.1%** | **123** | **7.0%** | **901 (98.5%)** | **496 (54.2%)** | **6/8** | |
| Codon mutations: | CTT1529CAT (6091T>A), TAT1530TTT (6094A>T), CTT1533TTA (6102C>T 6104T>A), CCA1536TCA (6111C>T), GGT1538GGA (6119T>A), CTC1539TTG (6120C>T 6122C>G), AAA1540AAG (6125A>G), AAT1541ACA (6127A>C 6128T>A), GCG1542GCA (6131G>A), CCC1543CCA (6134C>A), GCT1544CCT (6135G>C), TTT1546TTC (6143T>C), AGG1548AGA (6149G>A), AAC1552AAT (6161C>T), GTC1553ATT (6162G>A 6164C>T), TTC1554TTT (6167C>T), AAA1555GGA (6168A>G 6169A>G), GGG1556GAT (6172G>A 6173G>T), ACA1557TAC (6174A>T 6175C>A 6176A>C), GAA1558AAA (6177G>A), AAG1559AGA (6181A>G 6182G>A), TTC1560TTT (6185C>T), ATA1561GTC (6186A>G 6188A>C), GCT1562CTA (6189G>C 6190C>T 6191T>A), GTT1563GTA (6194T>A), TAC1564TAT (6197C>T), ATT1565GTA (6198A>G 6200T>A), GAT1567GAC (6206T>C), ATA1568ATT (6209A>T), CTT1569CTA (6212T>A), GTT1570GTA (6215T>A), TTC1571TTT (6218C>T), TCA1572AAT (6219T>A 6220C>A 6221A>T), GAA1573AAA (6222G>A), ACA1574GAC (6225A>G 6226C>A 6227A>C), GCA1575ATA (6228G>A 6229C>T), GAA1576AGG (6231G>A 6232A>G 6233A>G), CAA1577GAG (6234C>G 6236A>G), TCG1579CTA (6240T>C 6241C>T 6242G>A), CAG1580GGT (6243C>G 6244A>G 6245G>T), CTC1582CTA (6251C>A), TAT1583CAG (6252T>C 6254T>G), ATG1585GTA (6258A>G 6260G>A), CTG1586TTT (6261C>T 6263G>T), CAG1587AGA (6264C>A 6265A>G 6266G>A), CTT1588CTA (6269T>A), TGC1589TTT (6271G>T 6272C>T), AAG1590GTA (6273A>G 6274A>T 6275G>A), GAA1591AGT (6276G>A 6277A>G 6278A>T), AAT1592AAC (6281T>C), GGG1593GGT (6284G>T), TTA1594ATA (6285T>A), ATC1595ATA (6290C>A), CTT1596GTA (6291C>G 6293T>A), CCT1598AAA (6297C>A 6298C>A 6299T>A), ACC1599AAG (6301C>A 6302C>G), AAG1602AAA (6311G>A), ATA1603TTA (6312A>T), GGT1604TGT (6315G>T), ACT1605AAA (6319C>A 6320T>A), CCG1606ACC (6321C>A 6323G>C), GAG1607CAT (6324G>C 6326G>T), ATC1608ATA (6329C>A), GAT1609AAT (6330G>A), GGG1612GAG (6340G>A), GCT1613ATT (6342G>A 6343C>T), TCT1614ATA (6345T>A 6346C>T 6347T>A), CTC1615TTA (6348C>T 6350C>A), TGT1617GAA (6354T>G 6355G>A 6356T>A), ACA1618TGA (6357A>T 6358C>G), AAA1619AAG (6362A>G), ATC1620ATA (6365C>A), AAG1621AAA (6368G>A), CTT1622TTA (6369C>T 6371T>A), CAG1623CAA (6374G>A), CAC1625del (6378_6380delCAC), ATC1626ATA (6383C>A), ATT1627TAG (6384A>T 6385T>A 6386T>G), TCA1628CTA (6387T>C 6388C>T), ATA1630AGG (6394T>G 6395A>G), TGT1631TCT (6397G>C), GAC1632TAG (6399G>T 6401C>G), TTC1633AGA (6402T>A 6403T>G 6404C>A), TCA1634TGC (6406C>G 6407A>C), GAT1635CAT (6408G>C), GAA1636AAA (6411G>A), AAA1637TTA (6414A>T 6415A>T), CTA1638GAA (6417C>G 6418T>A), GCA1639AAC (6420G>A 6421C>A 6422A>C), CCT1641AAA (6426C>A 6427C>A 6428T>A), GGT1643-TT (6432delG 6433G>T), ATG1644ACA (6436T>C 6437G>A), AGA1645AAA (6439G>A), AGC1646ATT (6442G>T 6443C>T), TGG1647CTT (6444T>C 6445G>T 6446G>T), TTG1648ATA (6447T>A 6449G>A), ATC1650TTA (6453A>T 6455C>A), CTC1651ATA (6456C>A 6458C>A), TCA1652AAT (6459T>A 6460C>A 6461A>T), TAT1657TTT (6475A>T), ATT1658ATA (6479T>A), CAG1659AAA (6480C>A 6482G>A), GAT1660GAC (6485T>C), ATC1661TTA (6486A>T 6488C>A), GGC1662GGA (6491C>A), TTG1664ATA (6495T>A 6497G>A), GTG1665GTA (6500G>A), CAA1666GGA (6501C>G 6502A>G), AGA1669TAC (6510A>T 6511G>A 6512A>C), CAA1670TCT (6513C>T 6514A>C 6515A>T), ATG1672ACA (6520T>C 6521G>A), GCA1673GGA (6523C>G), CCA1674AGT (6525C>A 6526C>G 6527A>T), ACA1675AAA (6529C>A), GAC1677CAA (6534G>C 6536C>A), AGA1679TAT (6540A>T 6541G>A 6542A>T), ATG1680TTT (6543A>T 6545G>T), CCA1682AAT (6549C>A 6550C>A 6551A>T), ACA1684GAT (6555A>G 6556C>A 6557A>T), TGG1685ATC (6558T>A 6559G>T 6560G>C), AAG1686AAA (6563G>A), ATG1687TTA (6564A>T 6566G>A), GTA1688GTC (6569A>C), AGA1689CAA (6570A>C 6571G>A), CAG1690AAG (6573C>A), ATA1691ATC (6578A>C), GAA1693GAT (6584A>T), AAG1694AAA (6587G>A), GTG1695GTT (6590G>T), AAT1697AAC (6596T>C), CTC1698ATC (6597C>A), CCT1699CCA (6602T>A), CTT1701TTG (6606C>T 6608T>G), CAG1702GCT (6609C>G 6610A>C 6611G>T), TTA1703ATT (6612T>A 6614A>T), CCA1704CCT (6617A>T), CCT1705CTA (6619C>T 6620T>A), AAA1706GAA (6621A>G), GAT1707TCC (6624G>T 6625A>C 6626T>C), TCA1708GAT (6627T>G 6628C>A 6629A>T), TTC1709TAT (6631T>A 6632C>T), ATC1710TTA (6633A>T 6635C>A), GAG1713GAA (6644G>A), ACG1714ACA (6647G>A), GGT1716GGG (6653T>G), TGT1717AGC (6654T>A 6656T>C), ATG1718TCC (6657A>T 6658T>C 6659G>C), ACT1719ACA (6662T>A), GGC1720GGA (6665C>A), GCC1723GCA (6674C>A), GTC1724ATT (6675G>A 6677C>T), TGC1725TTG (6679G>T 6680C>G), AAA1726AAG (6683A>G), TGG1727GCC (6684T>G 6685G>C 6686G>C), ATG1729CCA (6690A>C 6691T>C 6692G>A), TCA1730AGT (6693T>A 6694C>G 6695A>T), AAG1731AAA (6698G>A), CAT1732TAT (6699C>T), GAT1733AGT (6702G>A 6703A>G), CCA1734GAT (6705C>G 6706C>A 6707A>T), AGA1735AAA (6709G>A), AGC1736AAT (6712G>A 6713C>T), ACC1737ACA (6716C>A), AGA1739AAG (6721G>A 6722A>G), ATT1740ATA (6725T>A), GCC1742GGT (6730C>G 6731C>T), GCT1744CAA (6735G>C 6736C>A 6737T>A), AGT1745AGC (6740T>C), GGA1746GGT (6743A>T), TCA1747AAA (6744T>A 6745C>A), TTC1748TAC (6748T>A), AAT1749AAA (6752T>A), CCA1750GAA (6753C>G 6754C>A), ATA1751AAA (6757T>A), ATA1751_AAA1752insGGAAAT (6758_6759insGGAAAT), AAA1752ATG (6760A>T 6761A>G), TCA1753AGT (6762T>A 6763C>G 6764A>T), ACC1754AGT (6766C>G 6767C>T), ATC1755ATA (6770C>A), GAT1756AAT (6771G>A), GCA1757TTT (6774G>T 6775C>T 6776A>T), GAG1758GAA (6779G>A), ATT1759ATA (6782T>A), CAG1760CTT (6784A>T 6785G>T), GCG1761GCA (6788G>A), GCA1762GTT (6790C>T 6791A>T), ATC1763AAT (6793T>A 6794C>T), CAT1764-AT (6795delC), GGC1765GGT (6800C>T), CTG1766TTA (6801C>T 6803G>A), GAT1767AAT (6804G>A), AAA1768AGT (6808A>G 6809A>T), TTC1769TTT (6812C>T), AAA1770AGA (6814A>G), ATT1771CTA (6816A>C 6818T>A), TAT1772TAC (6821T>C), TAT1773ATT (6822T>A 6823A>T), CTT1774ATA (6825C>A 6827T>A), GAT1775AAC (6828G>A 6830T>C), AAG1777AAA (6836G>A), GAG1778GAA (6839G>A), CTC1779ATA (6840C>A 6842C>A), ATA1780CTG (6843A>C 6845A>G), ATT1781GTG (6846A>G 6848T>G), CGC1782AGA (6849C>A 6851C>A), TCA1783ACA (6852T>A), ATT1788ATA (6869T>A), ATC1789GTA (6870A>G 6872C>A), TAC1792CAT (6879T>C 6881C>T), AAC1793CAA (6882A>C 6884C>A), ACG1795ATA (6889C>T 6890G>A), AAC1796AAT (6893C>T), AAT1798AAA (6899T>A), AAG1799ACT (6901A>C 6902G>T), CCG1800AGT (6903C>A 6904C>G 6905G>T), TCT1801AGT (6906T>A 6907C>G), GTT1803AGA (6912G>A 6913T>G 6914T>A), AGA1804CGA (6915A>C), TTA1806CTT (6921T>C 6923A>T), ACA1807AAT (6925C>A 6926A>T), TCA1809CTA (6930T>C 6931C>T), TTC1811ACA (6936T>A 6937T>C 6938C>A), TTA1812ATA (6939T>A), ACA1813TCT (6942A>T 6944A>T), GGT1814ACT (6945G>A 6946G>C), CTT1815TAC (6948C>T 6949T>A 6950T>C), GGA1816GAG (6952G>A 6953A>G), ATC1817del (6954_6956delATC), ACA1818AAT (6958C>A 6959A>T), GTT1819ATT (6960G>A), ACA1820ATT (6964C>T 6965A>T), TTC1821TTT (6968C>T), CAC1823CAT (6974C>T), GAT1825AAA (6978G>A 6980T>A), AAG1827AAA (6986G>A), CAT1828AAA (6987C>A 6989T>A), GGC1830AAC (6993G>A 6994G>A) | | | | | | | | |

#### Proteins

| | Begin | End | Coverage | Score | Concordance | Matches | Identities | I/D/M/F* | Stop Codons |
|---|---|---|---|---|---|---|---|---|---|
| polyprotein (NP_039820.1) | 1529 | 1831 | 16.1% | 750 | 35.5% | 301 (98.7%) | 145 (47.5%) | 2/2/2/2 | 3 |
| Protein mutations: | L1529H (6091T>A), Y1530F (6094A>T), P1536S (6111C>T), N1541T (6127A>C 6128T>A), A1544P (6135G>C), V1553I (6162G>A 6164C>T), K1555G (6168A>G 6169A>G), G1556D (6172G>A 6173G>T), T1557Y (6174A>T 6175C>A 6176A>C), E1558K (6177G>A), K1559R (6181A>G 6182G>A), I1561V (6186A>G 6188A>C), A1562L (6189G>C 6190C>T 6191T>A), I1565V (6198A>G 6200T>A), S1572N (6219T>A 6220C>A 6221A>T), E1573K (6222G>A), T1574D (6225A>G 6226C>A 6227A>C), A1575I (6228G>A 6229C>T), E1576R (6231G>A 6232A>G 6233A>G), Q1577E (6234C>G 6236A>G), S1579L (6240T>C 6241C>T 6242G>A), Q1580G (6243C>G 6244A>G 6245G>T), Y1583Q (6252T>C 6254T>G), M1585V (6258A>G 6260G>A), L1586F (6261C>T 6263G>T), Q1587R (6264C>A 6265A>G 6266G>A), C1589F (6271G>T 6272C>T), K1590V (6273A>G 6274A>T 6275G>A), E1591S (6276G>A 6277A>G 6278A>T), L1594I (6285T>A), L1596V (6291C>G 6293T>A), P1598K (6297C>A 6298C>A 6299T>A), T1599K (6301C>A 6302C>G), I1603L (6312A>T), G1604C (6315G>T), T1605K (6319C>A 6320T>A), P1606T (6321C>A 6323G>C), E1607H (6324G>C 6326G>T), D1609N (6330G>A), G1612E (6340G>A), A1613I (6342G>A 6343C>T), S1614I (6345T>A 6346C>T 6347T>A), C1617E (6354T>G 6355G>A 6356T>A), T1618* (6357A>T 6358C>G), H1625del (6378_6380delCAC), I1627* (6384A>T 6385T>A 6386T>G), S1628L (6387T>C 6388C>T), I1630R (6394T>G 6395A>G), C1631S (6397G>C), D1632* (6399G>T 6401C>G), F1633R (6402T>A 6403T>G 6404C>A), S1634C (6406C>G 6407A>C), D1635H (6408G>C), E1636K (6411G>A), K1637L (6414A>T 6415A>T), L1638E (6417C>G 6418T>A), A1639N (6420G>A 6421C>A 6422A>C), P1641K (6426C>A 6427C>A 6428T>A), M1644T (6436T>C 6437G>A), R1645K (6439G>A), S1646I (6442G>T 6443C>T), W1647L (6444T>C 6445G>T 6446G>T), L1648I (6447T>A 6449G>A), I1650L (6453A>T 6455C>A), L1651I (6456C>A 6458C>A), S1652N (6459T>A 6460C>A 6461A>T), Y1657F (6475A>T), Q1659K (6480C>A 6482G>A), I1661L (6486A>T 6488C>A), L1664I (6495T>A 6497G>A), Q1666G (6501C>G 6502A>G), R1669Y (6510A>T 6511G>A 6512A>C), Q1670S (6513C>T 6514A>C 6515A>T), M1672T (6520T>C 6521G>A), A1673G (6523C>G), P1674S (6525C>A 6526C>G 6527A>T), T1675K (6529C>A), D1677Q (6534G>C 6536C>A), R1679Y (6540A>T 6541G>A 6542A>T), M1680F (6543A>T 6545G>T), P1682N (6549C>A 6550C>A 6551A>T), T1684D (6555A>G 6556C>A 6557A>T), W1685I (6558T>A 6559G>T 6560G>C), M1687L (6564A>T 6566G>A), R1689Q (6570A>C 6571G>A), Q1690K (6573C>A), E1693D (6584A>T), L1698I (6597C>A), Q1702A (6609C>G 6610A>C 6611G>T), L1703I (6612T>A 6614A>T), P1705L (6619C>T 6620T>A), K1706E (6621A>G), D1707S (6624G>T 6625A>C 6626T>C), S1708D (6627T>G 6628C>A 6629A>T), F1709Y (6631T>A 6632C>T), I1710L (6633A>T 6635C>A), C1717S (6654T>A 6656T>C), M1718S (6657A>T 6658T>C 6659G>C), V1724I (6675G>A 6677C>T), C1725L (6679G>T 6680C>G), W1727A (6684T>G 6685G>C 6686G>C), M1729P (6690A>C 6691T>C 6692G>A), H1732Y (6699C>T), D1733S (6702G>A 6703A>G), P1734D (6705C>G 6706C>A 6707A>T), R1735K (6709G>A), S1736N (6712G>A 6713C>T), R1739K (6721G>A 6722A>G), A1742G (6730C>G 6731C>T), A1744Q (6735G>C 6736C>A 6737T>A), S1747K (6744T>A 6745C>A), F1748Y (6748T>A), N1749K (6752T>A), P1750E (6753C>G 6754C>A), I1751K (6757T>A), I1751_K1752insGN (6758_6759insGGAAAT), K1752M (6760A>T 6761A>G), T1754S (6766C>G 6767C>T), D1756N (6771G>A), A1757F (6774G>T 6775C>T 6776A>T), Q1760L (6784A>T 6785G>T), A1762V (6790C>T 6791A>T), I1763N (6793T>A 6794C>T), D1767N (6804G>A), K1768S (6808A>G 6809A>T), K1770R (6814A>G), I1771L (6816A>C 6818T>A), Y1773I (6822T>A 6823A>T), L1774I (6825C>A 6827T>A), D1775N (6828G>A 6830T>C), L1779I (6840C>A 6842C>A), I1780L (6843A>C 6845A>G), I1781V (6846A>G 6848T>G), S1783T (6852T>A), I1789V (6870A>G 6872C>A), Y1792H (6879T>C 6881C>T), N1793Q (6882A>C 6884C>A), T1795I (6889C>T 6890G>A), N1798K (6899T>A), K1799T (6901A>C 6902G>T), P1800S (6903C>A 6904C>G 6905G>T), V1803R (6912G>A 6913T>G 6914T>A), T1807N (6925C>A 6926A>T), S1809L (6930T>C 6931C>T), F1811T (6936T>A 6937T>C 6938C>A), L1812I (6939T>A), T1813S (6942A>T 6944A>T), G1814T (6945G>A 6946G>C), L1815Y (6948C>T 6949T>A 6950T>C), G1816E (6952G>A 6953A>G), I1817del (6954_6956delATC), T1818N (6958C>A 6959A>T), V1819I (6960G>A), T1820I (6964C>T 6965A>T), D1825K (6978G>A 6980T>A), H1828K (6987C>A 6989T>A), G1830N (6993G>A 6994G>A) | | | | | | | | |

| | Begin | End | Coverage | Score | Concordance | Matches | Identities | I/D/M/F* | Stop Codons |
|---|---|---|---|---|---|---|---|---|---|
| **NT** | **6090** | **6998** | **12.1%** | **123** | **7.0%** | **901 (98.5%)** | **496 (54.2%)** | **6/8** | |
| Codon mutations: | CTT1529CAT (6091T>A), TAT1530TTT (6094A>T), CTT1533TTA (6102C>T 6104T>A), CCA1536TCA (6111C>T), GGT1538GGA (6119T>A), CTC1539TTG (6120C>T 6122C>G), AAA1540AAG (6125A>G), AAT1541ACA (6127A>C 6128T>A), GCG1542GCA (6131G>A), CCC1543CCA (6134C>A), GCT1544CCT (6135G>C), TTT1546TTC (6143T>C), AGG1548AGA (6149G>A), AAC1552AAT (6161C>T), GTC1553ATT (6162G>A 6164C>T), TTC1554TTT (6167C>T), AAA1555GGA (6168A>G 6169A>G), GGG1556GAT (6172G>A 6173G>T), ACA1557TAC (6174A>T 6175C>A 6176A>C), GAA1558AAA (6177G>A), AAG1559AGA (6181A>G 6182G>A), TTC1560TTT (6185C>T), ATA1561GTC (6186A>G 6188A>C), GCT1562CTA (6189G>C 6190C>T 6191T>A), GTT1563GTA (6194T>A), TAC1564TAT (6197C>T), ATT1565GTA (6198A>G 6200T>A), GAT1567GAC (6206T>C), ATA1568ATT (6209A>T), CTT1569CTA (6212T>A), GTT1570GTA (6215T>A), TTC1571TTT (6218C>T), TCA1572AAT (6219T>A 6220C>A 6221A>T), GAA1573AAA (6222G>A), ACA1574GAC (6225A>G 6226C>A 6227A>C), GCA1575ATA (6228G>A 6229C>T), GAA1576AGG (6231G>A 6232A>G 6233A>G), CAA1577GAG (6234C>G 6236A>G), TCG1579CTA (6240T>C 6241C>T 6242G>A), CAG1580GGT (6243C>G 6244A>G 6245G>T), CTC1582CTA (6251C>A), TAT1583CAG (6252T>C 6254T>G), ATG1585GTA (6258A>G 6260G>A), CTG1586TTT (6261C>T 6263G>T), CAG1587AGA (6264C>A 6265A>G 6266G>A), CTT1588CTA (6269T>A), TGC1589TTT (6271G>T 6272C>T), AAG1590GTA (6273A>G 6274A>T 6275G>A), GAA1591AGT (6276G>A 6277A>G 6278A>T), AAT1592AAC (6281T>C), GGG1593GGT (6284G>T), TTA1594ATA (6285T>A), ATC1595ATA (6290C>A), CTT1596GTA (6291C>G 6293T>A), CCT1598AAA (6297C>A 6298C>A 6299T>A), ACC1599AAG (6301C>A 6302C>G), AAG1602AAA (6311G>A), ATA1603TTA (6312A>T), GGT1604TGT (6315G>T), ACT1605AAA (6319C>A 6320T>A), CCG1606ACC (6321C>A 6323G>C), GAG1607CAT (6324G>C 6326G>T), ATC1608ATA (6329C>A), GAT1609AAT (6330G>A), GGG1612GAG (6340G>A), GCT1613ATT (6342G>A 6343C>T), TCT1614ATA (6345T>A 6346C>T 6347T>A), CTC1615TTA (6348C>T 6350C>A), TGT1617GAA (6354T>G 6355G>A 6356T>A), ACA1618TGA (6357A>T 6358C>G), AAA1619AAG (6362A>G), ATC1620ATA (6365C>A), AAG1621AAA (6368G>A), CTT1622TTA (6369C>T 6371T>A), CAG1623CAA (6374G>A), CAC1625del (6378_6380delCAC), ATC1626ATA (6383C>A), ATT1627TAG (6384A>T 6385T>A 6386T>G), TCA1628CTA (6387T>C 6388C>T), ATA1630AGG (6394T>G 6395A>G), TGT1631TCT (6397G>C), GAC1632TAG (6399G>T 6401C>G), TTC1633AGA (6402T>A 6403T>G 6404C>A), TCA1634TGC (6406C>G 6407A>C), GAT1635CAT (6408G>C), GAA1636AAA (6411G>A), AAA1637TTA (6414A>T 6415A>T), CTA1638GAA (6417C>G 6418T>A), GCA1639AAC (6420G>A 6421C>A 6422A>C), CCT1641AAA (6426C>A 6427C>A 6428T>A), GGT1643-TT (6432delG 6433G>T), ATG1644ACA (6436T>C 6437G>A), AGA1645AAA (6439G>A), AGC1646ATT (6442G>T 6443C>T), TGG1647CTT (6444T>C 6445G>T 6446G>T), TTG1648ATA (6447T>A 6449G>A), ATC1650TTA (6453A>T 6455C>A), CTC1651ATA (6456C>A 6458C>A), TCA1652AAT (6459T>A 6460C>A 6461A>T), TAT1657TTT (6475A>T), ATT1658ATA (6479T>A), CAG1659AAA (6480C>A 6482G>A), GAT1660GAC (6485T>C), ATC1661TTA (6486A>T 6488C>A), GGC1662GGA (6491C>A), TTG1664ATA (6495T>A 6497G>A), GTG1665GTA (6500G>A), CAA1666GGA (6501C>G 6502A>G), AGA1669TAC (6510A>T 6511G>A 6512A>C), CAA1670TCT (6513C>T 6514A>C 6515A>T), ATG1672ACA (6520T>C 6521G>A), GCA1673GGA (6523C>G), CCA1674AGT (6525C>A 6526C>G 6527A>T), ACA1675AAA (6529C>A), GAC1677CAA (6534G>C 6536C>A), AGA1679TAT (6540A>T 6541G>A 6542A>T), ATG1680TTT (6543A>T 6545G>T), CCA1682AAT (6549C>A 6550C>A 6551A>T), ACA1684GAT (6555A>G 6556C>A 6557A>T), TGG1685ATC (6558T>A 6559G>T 6560G>C), AAG1686AAA (6563G>A), ATG1687TTA (6564A>T 6566G>A), GTA1688GTC (6569A>C), AGA1689CAA (6570A>C 6571G>A), CAG1690AAG (6573C>A), ATA1691ATC (6578A>C), GAA1693GAT (6584A>T), AAG1694AAA (6587G>A), GTG1695GTT (6590G>T), AAT1697AAC (6596T>C), CTC1698ATC (6597C>A), CCT1699CCA (6602T>A), CTT1701TTG (6606C>T 6608T>G), CAG1702GCT (6609C>G 6610A>C 6611G>T), TTA1703ATT (6612T>A 6614A>T), CCA1704CCT (6617A>T), CCT1705CTA (6619C>T 6620T>A), AAA1706GAA (6621A>G), GAT1707TCC (6624G>T 6625A>C 6626T>C), TCA1708GAT (6627T>G 6628C>A 6629A>T), TTC1709TAT (6631T>A 6632C>T), ATC1710TTA (6633A>T 6635C>A), GAG1713GAA (6644G>A), ACG1714ACA (6647G>A), GGT1716GGG (6653T>G), TGT1717AGC (6654T>A 6656T>C), ATG1718TCC (6657A>T 6658T>C 6659G>C), ACT1719ACA (6662T>A), GGC1720GGA (6665C>A), GCC1723GCA (6674C>A), GTC1724ATT (6675G>A 6677C>T), TGC1725TTG (6679G>T 6680C>G), AAA1726AAG (6683A>G), TGG1727GCC (6684T>G 6685G>C 6686G>C), ATG1729CCA (6690A>C 6691T>C 6692G>A), TCA1730AGT (6693T>A 6694C>G 6695A>T), AAG1731AAA (6698G>A), CAT1732TAT (6699C>T), GAT1733AGT (6702G>A 6703A>G), CCA1734GAT (6705C>G 6706C>A 6707A>T), AGA1735AAA (6709G>A), AGC1736AAT (6712G>A 6713C>T), ACC1737ACA (6716C>A), AGA1739AAG (6721G>A 6722A>G), ATT1740ATA (6725T>A), GCC1742GGT (6730C>G 6731C>T), GCT1744CAA (6735G>C 6736C>A 6737T>A), AGT1745AGC (6740T>C), GGA1746GGT (6743A>T), TCA1747AAA (6744T>A 6745C>A), TTC1748TAC (6748T>A), AAT1749AAA (6752T>A), CCA1750GAA (6753C>G 6754C>A), ATA1751AAA (6757T>A), ATA1751_AAA1752insGGAAAT (6758_6759insGGAAAT), AAA1752ATG (6760A>T 6761A>G), TCA1753AGT (6762T>A 6763C>G 6764A>T), ACC1754AGT (6766C>G 6767C>T), ATC1755ATA (6770C>A), GAT1756AAT (6771G>A), GCA1757TTT (6774G>T 6775C>T 6776A>T), GAG1758GAA (6779G>A), ATT1759ATA (6782T>A), CAG1760CTT (6784A>T 6785G>T), GCG1761GCA (6788G>A), GCA1762GTT (6790C>T 6791A>T), ATC1763AAT (6793T>A 6794C>T), CAT1764-AT (6795delC), GGC1765GGT (6800C>T), CTG1766TTA (6801C>T 6803G>A), GAT1767AAT (6804G>A), AAA1768AGT (6808A>G 6809A>T), TTC1769TTT (6812C>T), AAA1770AGA (6814A>G), ATT1771CTA (6816A>C 6818T>A), TAT1772TAC (6821T>C), TAT1773ATT (6822T>A 6823A>T), CTT1774ATA (6825C>A 6827T>A), GAT1775AAC (6828G>A 6830T>C), AAG1777AAA (6836G>A), GAG1778GAA (6839G>A), CTC1779ATA (6840C>A 6842C>A), ATA1780CTG (6843A>C 6845A>G), ATT1781GTG (6846A>G 6848T>G), CGC1782AGA (6849C>A 6851C>A), TCA1783ACA (6852T>A), ATT1788ATA (6869T>A), ATC1789GTA (6870A>G 6872C>A), TAC1792CAT (6879T>C 6881C>T), AAC1793CAA (6882A>C 6884C>A), ACG1795ATA (6889C>T 6890G>A), AAC1796AAT (6893C>T), AAT1798AAA (6899T>A), AAG1799ACT (6901A>C 6902G>T), CCG1800AGT (6903C>A 6904C>G 6905G>T), TCT1801AGT (6906T>A 6907C>G), GTT1803AGA (6912G>A 6913T>G 6914T>A), AGA1804CGA (6915A>C), TTA1806CTT (6921T>C 6923A>T), ACA1807AAT (6925C>A 6926A>T), TCA1809CTA (6930T>C 6931C>T), TTC1811ACA (6936T>A 6937T>C 6938C>A), TTA1812ATA (6939T>A), ACA1813TCT (6942A>T 6944A>T), GGT1814ACT (6945G>A 6946G>C), CTT1815TAC (6948C>T 6949T>A 6950T>C), GGA1816GAG (6952G>A 6953A>G), ATC1817del (6954_6956delATC), ACA1818AAT (6958C>A 6959A>T), GTT1819ATT (6960G>A), ACA1820ATT (6964C>T 6965A>T), TTC1821TTT (6968C>T), CAC1823CAT (6974C>T), GAT1825AAA (6978G>A 6980T>A), AAG1827AAA (6986G>A), CAT1828AAA (6987C>A 6989T>A), GGC1830AAC (6993G>A 6994G>A) | | | | | | | | |

*: Inserts / Deletes / Misaligned / Frameshifts

#### Analysis details

This analysis was performed with panviral2.64

### NGS Details (UN8): Epiphyllum badnavirus 1

#### Assembly

| | |
|---|---|
| **Coverage Length** | 405 (1 contig(s)) |
| **Depth Of Coverage** | 16.6 |
| **Number Of Reads** | 55 |
| **Reads Per Million** | 0.92 rpm (after QC) |
| **Ambiguities** | 0 |
| **Assembly Method** | de novo + reference guided assembly |
| **Consensus Caller** | Bcf Tools |

#### Coverage Map

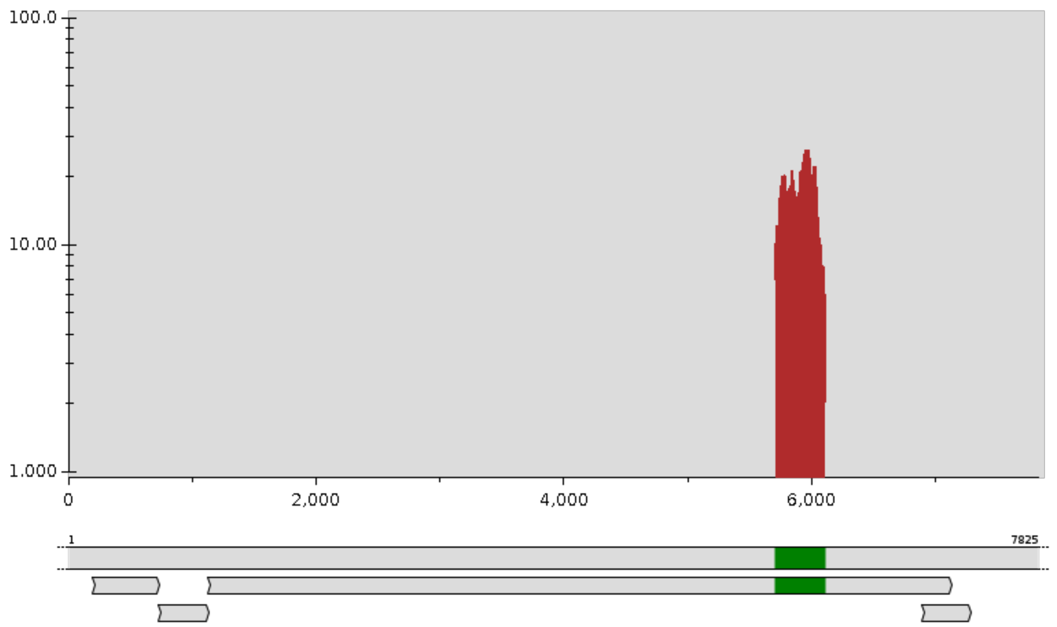

#### Assignment

| | |
|---|---|
| **Type** | Epiphyllum badnavirus 1 (Taxonomy ID: 2518008) |
| **Reference Genome** | NC_076247.1 |
| **NT Identity (%)** | 52.3573 |
| **AA Identity (%)** | 41.4815 |
| **Number Of Stop Codons** | 1 |
| **Number Of CDS** | 4 |

#### Alignment

| | |
|---|---|
| **Alignment Score** | 9.0 (NT) + 220.0 (AA) = 229.0 |
| **Concordance (%)** | 15.6984 |

| Alignment Method | Global, seeded, nucleotide + amino acids (AGA) |
|---|---|

#### Genome Region

Sequence starts at position 5706 and ends at position 6110 relative to NC_076247.1 reference sequence.

#### Alignment Detailed Statistics

| | Begin | End | Coverage | Score | Concordance | Matches | Identities | I/D/M/F* | Stop Codons |
|---|---|---|---|---|---|---|---|---|---|
| **NT** | **5706** | **6110** | **5.2%** | **9** | **1.2%** | **400 (98.0%)** | **211 (51.7%)** | **3/5** | |
| Mutations: | 5706T>C, 5707G>T, 5708C>T, 5709A>G, 5710A>T, 5711C>T, 5714G>T, 5715A>T, 5716G>A, 5718A>G, 5719G>A, 5725A>G, 5727A>C, 5730A>G, 5732A>G, 5733G>C, 5734G>A, 5735A>T, 5736G>A, 5737A>T, 5739A>G, 5742G>A, 5743G>T, 5744T>G, 5745C>T, 5746T>A, 5748C>T, 5749A>G, 5756A>G, 5758A>C, 5759C>A, 5760C>G, 5767G>A, 5769C>G, 5770A>G, 5771A>T, 5772C>A, 5776T>G, 5778C>G, 5781A>G, 5782G>A, 5787G>T, 5791T>C, 5792C>T, 5793T>C, 5794C>A, 5796A>G, 5800G>C, 5806A>G, 5808C>T, 5809_5810delAC, 5811A>G, 5812A>T, 5816T>A, 5817A>C, 5821_5823delAAG, 5824G>C, 5826C>A, 5828G>A, 5829T>G, 5830A>G, 5831A>G, 5835C>T, 5836A>G, 5837G>T, 5838T>C, 5841C>G, 5844C>T, 5847C>T, 5851T>A, 5853C>T, 5857C>G, 5861A>G, 5862G>T, 5863A>T, 5864G>C, 5868A>C, 5870T>A, 5871C>T, 5872C>A, 5874C>T, 5878A>T, 5880T>G, 5882T>G, 5886G>T, 5887G>A, 5889C>G, 5890G>A, 5891A>C, 5892A>T, 5894A>T, 5895A>T, 5896T>G, 5897C>A, 5898C>T, 5901C>T, 5905T>A, 5906G>A, 5908A>G, 5909C>A, 5910C>T, 5913C>T, 5916C>T, 5918T>G, 5919G>A, 5922C>T, 5924C>G, 5925A>C, 5926G>T, 5931A>C, 5933T>A, 5934T>C, 5938G>A, 5940A>G, 5942G>T, 5943G>C, 5944T>C, 5946A>T, 5949T>G, 5953C>T, 5955A>T, 5958C>T, 5961A>G, 5962T>C, 5964G>T, 5966A>C, 5967G>T, 5975C>T, 5976T>C, 5981T>C, 5982G>C, 5985C>T, 5986C>A, 5987A>T, 5989C>G, 5990G>A, 5991A>T, 5992A>C, 5993A>G, 5998G>A, 5999A>C, 6000C>T, 6001A>C, 6002A>G, 6003C>A, 6004T>G, 6005G>T, 6006T>A, 6009C>T, 6012A>G, 6012_6013insCCT, 6013G>T, 6015C>T, 6016C>A, 6018G>T, 6019T>G, 6020C>A, 6021A>T, 6022G>T, 6023G>C, 6030A>G, 6031G>A, 6032C>T, 6033A>T, 6036T>G, 6038A>T, 6051C>T, 6052C>T, 6054G>A, 6057C>A, 6059T>A, 6061A>T, 6062G>C, 6063T>G, 6064A>T, 6065A>G, 6066C>G, 6068C>G, 6070G>A, 6071A>G, 6075T>G, 6078C>G, 6079C>A, 6087G>A, 6090C>T, 6091C>T, 6095A>G, 6097T>G, 6098C>T, 6099C>T, 6100A>G, 6105G>C | | | | | | | | |

##### CDS

| | Begin | End | Coverage | Score | Concordance | Matches | Identities | I/D/M/F* | Stop Codons |
|---|---|---|---|---|---|---|---|---|---|
| QKM20_gp3 | 1527 | 1661 | 6.7% | 220 | 23.0% | 134 (98.5%) | 56 (41.2%) | 1/1/1/1 | 1 |
| Protein mutations: | A1527L (5707G>T 5708C>T 5709A>G), T1528F (5710A>T 5711C>T), G1529V (5714G>T 5715A>T), E1530K (5716G>A 5718A>G), E1531K (5719G>A), K1533D (5725A>G 5727A>C), K1535S (5732A>G 5733G>C), E1536I (5734G>A 5735A>T 5736G>A), R1537W (5737A>T 5739A>G), M1538I (5742G>A), V1539C (5743G>T 5744T>G 5745C>T), F1540I (5746T>A 5748C>T), N1541D (5749A>G), K1543R (5756A>G), T1544Q (5758A>C 5759C>A 5760C>G), D1547K (5767G>A 5769C>G), N1548V (5770A>G 5771A>T 5772C>A), F1550V (5776T>G 5778C>G), D1552N (5782G>A), Q1553H (5787G>T), S1555L (5791T>C 5792C>T 5793T>C), L1556M (5794C>A 5796A>G), G1558R (5800G>C), N1560D (5806A>G 5808C>T), I1562F (5812A>T), I1563N (5816T>A 5817A>C), K1565del (5821_5823delAAG), V1566L (5824G>C 5826C>A), C1567* (5828G>A 5829T>G), N1568G (5830A>G 5831A>G), S1570V (5836A>G 5837G>T 5838T>C), F1575I (5851T>A 5853C>T), L1577V (5857C>G), K1578S (5861A>G 5862G>T), F1581Y (5870T>A 5871C>T), H1582N (5872C>A 5874C>T), I1584L (5878A>T 5880T>G), M1585R (5882T>G), M1586I (5886G>T), D1587K (5887G>A 5889C>G), E1588T (5890G>A 5891A>C 5892A>T), E1589V (5894A>T 5895A>T), S1590D (5896T>G 5897C>A 5898C>T), W1593K (5905T>A 5906G>A), T1594D (5908A>G 5909C>A 5910C>T), L1597R (5918T>G 5919G>A), P1599R (5924C>G 5925A>C), D1600Y (5926G>T), L1602H (5933T>A 5934T>C), E1604K (5938G>A 5940A>G), W1605F (5942G>T 5943G>C), P1609S (5953C>T 5955A>T), K1613T (5966A>C 5967G>T), P1616L (5975C>T 5976T>C), V1618A (5981T>C 5982G>C), Q1620I (5986C>A 5987A>T), R1621D (5989C>G 5990G>A 5991A>T), K1622R (5992A>C 5993A>G), D1624T (5998G>A 5999A>C 6000C>T), N1625R (6001A>C 6002A>G 6003C>A), C1626V (6004T>G 6005G>T 6006T>A), R1628_D1629insP (6012_6013insCCT), D1629Y (6013G>T 6015C>T), L1630I (6016C>A 6018G>T), S1631D (6019T>G 6020C>A 6021A>T), G1632S (6022G>T 6023G>C), A1635I (6031G>A 6032C>T 6033A>T), Y1637F (6038A>T), F1644Y (6059T>A), N1646W (6064A>T 6065A>G 6066C>G), T1647S (6068C>G), E1648R (6070G>A 6071A>G), D1649E (6075T>G), D1650E (6078C>G), H1651N (6079C>A), K1656R (6095A>G), S1657V (6097T>G 6098C>T 6099C>T), M1658V (6100A>G) | | | | | | | | |
| Codon mutations: | CCT1526..C (5706T>C), GCA1527TTG (5707G>T 5708C>T 5709A>G), ACT1528TTT (5710A>T 5711C>T), GGA1529GTT (5714G>T 5715A>T), GAA1530AAG (5716G>A 5718A>G), GAG1531AAG (5719G>A), AAA1533GAC (5725A>G 5727A>C), GGA1534GGG (5730A>G), AAG1535AGC (5732A>G 5733G>C), GAG1536ATA (5734G>A 5735A>T 5736G>A), AGA1537TGG (5737A>T 5739A>G), ATG1538ATA (5742G>A), GTC1539TGT (5743G>T 5744T>G 5745C>T), TTC1540ATT (5746T>A 5748C>T), AAC1541GAC (5749A>G), AAA1543AGA (5756A>G), ACC1544CAG (5758A>C 5759C>A 5760C>G), GAC1547AAG (5767G>A 5769C>G), AAC1548GTA (5770A>G 5771A>T 5772C>A), TTC1550GTG (5776T>G 5778C>G), AAA1551AAG (5781A>G), GAT1552AAT (5782G>A), CAG1553CAT (5787G>T), TCT1555CTC (5791T>C 5792C>T 5793T>C), CTA1556ATG (5794C>A 5796A>G), GGC1558CGC (5800G>C), AAC1560GAT (5806A>G 5808C>T), ACA1561--G (5809_5810delAC 5811A>G), ATC1562TTC (5812A>T), ATA1563AAC (5816T>A 5817A>C), AAG1565del (5821_5823delAAG), GTC1566CTA (5824G>C 5826C>A), TGT1567TAG (5828G>A 5829T>G), AAT1568GGT (5830A>G 5831A>G), GCC1569GCT (5835C>T), AGT1570GTC (5836A>G 5837G>T 5838T>C), GTC1571GTG (5841C>G), TTC1572TTT (5844C>T), TCC1573TCT (5847C>T), TTC1575ATT (5851T>A 5853C>T), CTG1577GTG (5857C>G), AAG1578AGT (5861A>G 5862G>T), AGC1579TCC (5863A>T 5864G>C), GGA1580GGC (5868A>C), TTC1581TAT (5870T>A 5871C>T), CAC1582AAT (5872C>A 5874C>T), ATT1584TTG (5878A>T 5880T>G), ATG1585AGG (5882T>G), ATG1586ATT (5886G>T), GAC1587AAG (5887G>A 5889C>G), GAA1588ACT (5890G>A 5891A>C 5892A>T), GAA1589GTT (5894A>T 5895A>T), TCC1590GAT (5896T>G 5897C>A 5898C>T), ATC1591ATT (5901C>T), TGG1593AAG (5905T>A 5906G>A), ACC1594GAT (5908A>G 5909C>A 5910C>T), GCC1595GCT (5913C>T), TTC1596TTT (5916C>T), CTG1597CGA (5918T>G 5919G>A), ACC1598ACT (5922C>T), CCA1599CGC (5924C>G 5925A>C), GAT1600TAT (5926G>T), GGA1601GGC (5931A>C), CTT1602CAC (5933T>A 5934T>C), GAA1604AAG (5938G>A 5940A>G), TGG1605TTC (5942G>T 5943G>C), TTA1606CTT (5944T>C 5946A>T), GTT1607GTG (5949T>G), CCA1609TCT (5953C>T 5955A>T), TTC1610TTT (5958C>T), GGA1611GGG (5961A>G), TTG1612CTT (5962T>C 5964G>T), AAG1613ACT (5966A>C 5967G>T), CCT1616CTC (5975C>T 5976T>C), GTG1618GCC (5981T>C 5982G>C), TTC1619TTT (5985C>T), CAA1620ATA (5986C>A 5987A>T), CGA1621GAT (5989C>G 5990G>A 5991A>T), AAG1622CGG (5992A>C 5993A>G), GAC1624ACT (5998G>A 5999A>C 6000C>T), AAC1625CGA (6001A>C 6002A>G 6003C>A), TGT1626GTA (6004T>G 6005G>T 6006T>A), TTC1627TTT (6009C>T), AGA1628AGG (6012A>G), AGA1628_GAC1629insCCT (6012_6013insCCT), GAC1629TAT (6013G>T 6015C>T), CTG1630ATT (6016C>A 6018G>T), TCA1631GAT (6019T>G 6020C>A 6021A>T), GGC1632TCC (6022G>T 6023G>C), GTA1634GTG (6030A>G), GCA1635ATT (6031G>A 6032C>T 6033A>T), GTT1636GTG (6036T>G), TAC1637TTC (6038A>T), ATC1641ATT (6051C>T), CTG1642TTA (6052C>T 6054G>A), GTC1643GTA (6057C>A), TTC1644TAC (6059T>A), AGT1645TCG (6061A>T 6062G>C 6063T>G), AAC1646TGG (6064A>T 6065A>G 6066C>G), ACT1647AGT (6068C>G), GAA1648AGA (6070G>A 6071A>G), GAT1649GAG (6075T>G), GAC1650GAG (6078C>G), CAT1651AAT (6079C>A), CAG1653CAA (6087G>A), CAC1654CAT (6090C>T), CTG1655TTG (6091C>T), AAG1656AGG (6095A>G), TCC1657GTT (6097T>G 6098C>T 6099C>T), ATG1658GTG (6100A>G), CTG1659CTC (6105G>C) | | | | | | | | |

##### Proteins

| | Begin | End | Coverage | Score | Concordance | Matches | Identities | I/D/M/F* | Stop Codons |
|---|---|---|---|---|---|---|---|---|---|
| polyprotein (YP_010797894.1) | 1527 | 1661 | 6.7% | 220 | 23.0% | 134 (98.5%) | 56 (41.2%) | 1/1/1/1 | 1 |
| Protein mutations: | A1527L (5707G>T 5708C>T 5709A>G), T1528F (5710A>T 5711C>T), G1529V (5714G>T 5715A>T), E1530K (5716G>A 5718A>G), E1531K (5719G>A), K1533D (5725A>G 5727A>C), K1535S (5732A>G 5733G>C), E1536I (5734G>A 5735A>T 5736G>A), R1537W (5737A>T 5739A>G), M1538I (5742G>A), V1539C (5743G>T 5744T>G 5745C>T), F1540I (5746T>A 5748C>T), N1541D (5749A>G), K1543R (5756A>G), T1544Q (5758A>C 5759C>A 5760C>G), D1547K (5767G>A 5769C>G), N1548V (5770A>G 5771A>T 5772C>A), F1550V (5776T>G 5778C>G), D1552N (5782G>A), Q1553H (5787G>T), S1555L (5791T>C 5792C>T 5793T>C), L1556M (5794C>A 5796A>G), G1558R (5800G>C), N1560D (5806A>G 5808C>T), I1562F (5812A>T), I1563N (5816T>A 5817A>C), K1565del (5821_5823delAAG), V1566L (5824G>C 5826C>A), C1567* (5828G>A 5829T>G), N1568G (5830A>G 5831A>G), S1570V (5836A>G 5837G>T 5838T>C), F1575I (5851T>A 5853C>T), L1577V (5857C>G), K1578S (5861A>G 5862G>T), F1581Y (5870T>A 5871C>T), H1582N (5872C>A 5874C>T), I1584L (5878A>T 5880T>G), M1585R (5882T>G), M1586I (5886G>T), D1587K (5887G>A 5889C>G), E1588T (5890G>A 5891A>C 5892A>T), E1589V (5894A>T 5895A>T), S1590D (5896T>G 5897C>A 5898C>T), W1593K (5905T>A 5906G>A), T1594D (5908A>G 5909C>A 5910C>T), L1597R (5918T>G 5919G>A), P1599R (5924C>G 5925A>C), D1600Y (5926G>T), L1602H (5933T>A 5934T>C), E1604K (5938G>A 5940A>G), W1605F (5942G>T 5943G>C), P1609S (5953C>T 5955A>T), K1613T (5966A>C 5967G>T), P1616L (5975C>T 5976T>C), V1618A (5981T>C 5982G>C), Q1620I (5986C>A 5987A>T), R1621D (5989C>G 5990G>A 5991A>T), K1622R (5992A>C 5993A>G), D1624T (5998G>A 5999A>C 6000C>T), N1625R (6001A>C 6002A>G 6003C>A), C1626V (6004T>G 6005G>T 6006T>A), R1628_D1629insP (6012_6013insCCT), D1629Y (6013G>T 6015C>T), L1630I (6016C>A 6018G>T), S1631D (6019T>G 6020C>A 6021A>T), G1632S (6022G>T 6023G>C), A1635I (6031G>A 6032C>T 6033A>T), Y1637F (6038A>T), F1644Y (6059T>A), N1646W (6064A>T 6065A>G 6066C>G), T1647S (6068C>G), E1648R (6070G>A 6071A>G), D1649E (6075T>G), D1650E (6078C>G), H1651N (6079C>A), K1656R (6095A>G), S1657V (6097T>G 6098C>T 6099C>T), M1658V (6100A>G) | | | | | | | | |

| | Begin | End | Coverage | Score | Concordance | Matches | Identities | I/D/M/F* | Stop Codons |
|---|---|---|---|---|---|---|---|---|---|
| **NT** | **5706** | **6110** | **5.2%** | **9** | **1.2%** | **400 (98.0%)** | **211 (51.7%)** | **3/5** | |
| Codon mutations: | CCT1526..C (5706T>C), GCA1527TTG (5707G>T 5708C>T 5709A>G), ACT1528TTT (5710A>T 5711C>T), GGA1529GTT (5714G>T 5715A>T), GAA1530AAG (5716G>A 5718A>G), GAG1531AAG (5719G>A), AAA1533GAC (5725A>G 5727A>C), GGA1534GGG (5730A>G), AAG1535AGC (5732A>G 5733G>C), GAG1536ATA (5734G>A 5735A>T 5736G>A), AGA1537TGG (5737A>T 5739A>G), ATG1538ATA (5742G>A), GTC1539TGT (5743G>T 5744T>G 5745C>T), TTC1540ATT (5746T>A 5748C>T), AAC1541GAC (5749A>G), AAA1543AGA (5756A>G), ACC1544CAG (5758A>C 5759C>A 5760C>G), GAC1547AAG (5767G>A 5769C>G), AAC1548GTA (5770A>G 5771A>T 5772C>A), TTC1550GTG (5776T>G 5778C>G), AAA1551AAG (5781A>G), GAT1552AAT (5782G>A), CAG1553CAT (5787G>T), TCT1555CTC (5791T>C 5792C>T 5793T>C), CTA1556ATG (5794C>A 5796A>G), GGC1558CGC (5800G>C), AAC1560GAT (5806A>G 5808C>T), ACA1561--G (5809_5810delAC 5811A>G), ATC1562TTC (5812A>T), ATA1563AAC (5816T>A 5817A>C), AAG1565del (5821_5823delAAG), GTC1566CTA (5824G>C 5826C>A), TGT1567TAG (5828G>A 5829T>G), AAT1568GGT (5830A>G 5831A>G), GCC1569GCT (5835C>T), AGT1570GTC (5836A>G 5837G>T 5838T>C), GTC1571GTG (5841C>G), TTC1572TTT (5844C>T), TCC1573TCT (5847C>T), TTC1575ATT (5851T>A 5853C>T), CTG1577GTG (5857C>G), AAG1578AGT (5861A>G 5862G>T), AGC1579TCC (5863A>T 5864G>C), GGA1580GGC (5868A>C), TTC1581TAT (5870T>A 5871C>T), CAC1582AAT (5872C>A 5874C>T), ATT1584TTG (5878A>T 5880T>G), ATG1585AGG (5882T>G), ATG1586ATT (5886G>T), GAC1587AAG (5887G>A 5889C>G), GAA1588ACT (5890G>A 5891A>C 5892A>T), GAA1589GTT (5894A>T 5895A>T), TCC1590GAT (5896T>G 5897C>A 5898C>T), ATC1591ATT (5901C>T), TGG1593AAG (5905T>A 5906G>A), ACC1594GAT (5908A>G 5909C>A 5910C>T), GCC1595GCT (5913C>T), TTC1596TTT (5916C>T), CTG1597CGA (5918T>G 5919G>A), ACC1598ACT (5922C>T), CCA1599CGC (5924C>G 5925A>C), GAT1600TAT (5926G>T), GGA1601GGC (5931A>C), CTT1602CAC (5933T>A 5934T>C), GAA1604AAG (5938G>A 5940A>G), TGG1605TTC (5942G>T 5943G>C), TTA1606CTT (5944T>C 5946A>T), GTT1607GTG (5949T>G), CCA1609TCT (5953C>T 5955A>T), TTC1610TTT (5958C>T), GGA1611GGG (5961A>G), TTG1612CTT (5962T>C 5964G>T), AAG1613ACT (5966A>C 5967G>T), CCT1616CTC (5975C>T 5976T>C), GTG1618GCC (5981T>C 5982G>C), TTC1619TTT (5985C>T), CAA1620ATA (5986C>A 5987A>T), CGA1621GAT (5989C>G 5990G>A 5991A>T), AAG1622CGG (5992A>C 5993A>G), GAC1624ACT (5998G>A 5999A>C 6000C>T), AAC1625CGA (6001A>C 6002A>G 6003C>A), TGT1626GTA (6004T>G 6005G>T 6006T>A), TTC1627TTT (6009C>T), AGA1628AGG (6012A>G), AGA1628_GAC1629insCCT (6012_6013insCCT), GAC1629TAT (6013G>T 6015C>T), CTG1630ATT (6016C>A 6018G>T), TCA1631GAT (6019T>G 6020C>A 6021A>T), GGC1632TCC (6022G>T 6023G>C), GTA1634GTG (6030A>G), GCA1635ATT (6031G>A 6032C>T 6033A>T), GTT1636GTG (6036T>G), TAC1637TTC (6038A>T), ATC1641ATT (6051C>T), CTG1642TTA (6052C>T 6054G>A), GTC1643GTA (6057C>A), TTC1644TAC (6059T>A), AGT1645TCG (6061A>T 6062G>C 6063T>G), AAC1646TGG (6064A>T 6065A>G 6066C>G), ACT1647AGT (6068C>G), GAA1648AGA (6070G>A 6071A>G), GAT1649GAG (6075T>G), GAC1650GAG (6078C>G), CAT1651AAT (6079C>A), CAG1653CAA (6087G>A), CAC1654CAT (6090C>T), CTG1655TTG (6091C>T), AAG1656AGG (6095A>G), TCC1657GTT (6097T>G 6098C>T 6099C>T), ATG1658GTG (6100A>G), CTG1659CTC (6105G>C) | | | | | | | | |

*: Inserts / Deletes / Misaligned / Frameshifts

#### Analysis details

This analysis was performed with panviral2.64

#### NGS Details (UN8): Petuvirus venapetuniae

##### Assembly

| | |
|---|---|
| **Coverage Length** | 822 (1 contig(s)) |
| **Depth Of Coverage** | 7.5 |
| **Number Of Reads** | 50 |
| **Reads Per Million** | 0.84 rpm (after QC) |
| **Ambiguities** | 0 |
| **Assembly Method** | de novo + reference guided assembly |
| **Consensus Caller** | Bcf Tools |

##### Coverage Map

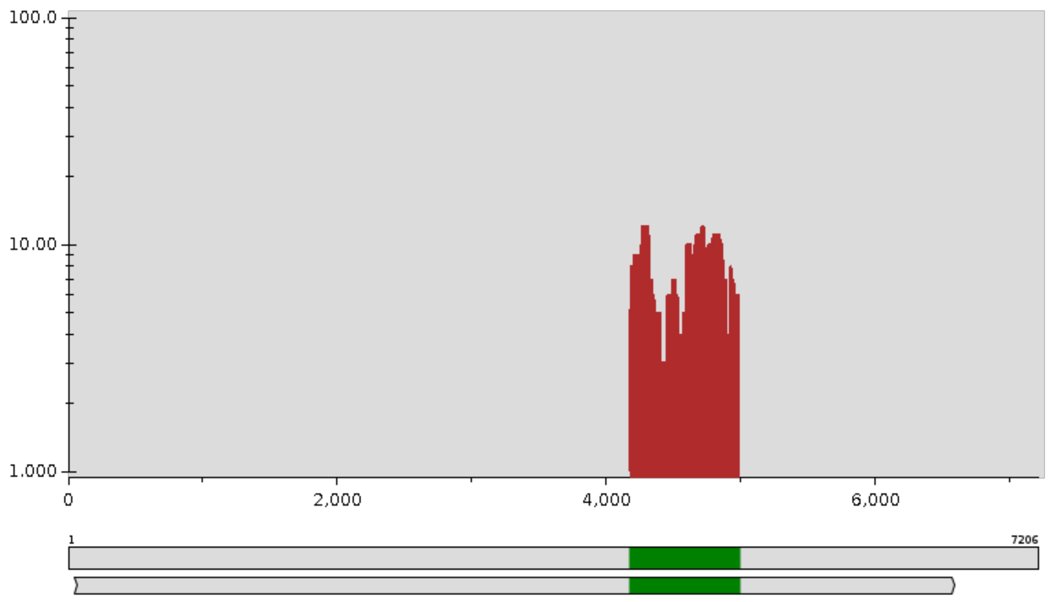

##### Assignment

| | |
|---|---|
| **Type** | Petuvirus venapetuniae (Taxonomy ID: 3048265) |
| **Reference Genome** | NC_001839.2 |
| **NT Identity (%)** | 51.7032 |
| **AA Identity (%)** | 43.0657 |
| **Number Of Stop Codons** | 1 |
| **Number Of CDS** | 1 |

##### Alignment

| | |
|---|---|
| **Alignment Score** | 56.0 (NT) + 808.0 (AA) = 864.0 |
| **Concordance (%)** | 24.2356 |

| **Alignment Method** | Global, seeded, nucleotide + amino acids (AGA) |
|---|---|

#### Genome Region

Sequence starts at position 4174 and ends at position 4995 relative to NC_001839.2 reference sequence.

#### Alignment Detailed Statistics

| | Begin | End | Coverage | Score | Concordance | Matches | Identities | I/D/M/F* | Stop Codons |
|---|---|---|---|---|---|---|---|---|---|
| **NT** | **4174** | **4995** | **11.4%** | **56** | **3.4%** | **822 (100%)** | **425 (51.7%)** | **0/0** | |
| Mutations: | 4180T>A, 4183A>G, 4185A>T, 4186A>T, 4187G>C, 4193A>G, 4196C>G, 4197A>G, 4198T>C, 4199C>T, 4200A>T, 4201A>T, 4203C>T, 4205C>T, 4208A>T, 4211G>A, 4214A>C, 4217C>G, 4219A>C, 4220T>C, 4221T>A, 4222C>T, 4223T>C, 4224G>C, 4225G>A, 4232T>C, 4234C>A, 4235A>T, 4239C>T, 4240A>T, 4241T>A, 4242C>A, 4244C>G, 4245C>G, 4247G>A, 4248T>A, 4250A>C, 4251G>T, 4252C>G, 4253A>T, 4255T>A, 4259G>A, 4263T>A, 4264G>T, 4265T>C, 4268T>C, 4269G>A, 4270A>C, 4271G>T, 4275C>T, 4276A>T, 4277A>G, 4278C>G, 4280A>T, 4281T>A, 4282T>G, 4284G>A, 4286T>A, 4287C>A, 4289A>C, 4292C>T, 4293G>A, 4294A>G, 4298C>T, 4300C>T, 4302G>A, 4304T>A, 4307A>T, 4309A>C, 4310A>T, 4314G>A, 4315C>G, 4316A>C, 4320G>T, 4321A>C, 4322A>C, 4328T>C, 4334C>T, 4337C>T, 4340G>T, 4341A>G, 4342G>T, 4343G>T, 4344T>G, 4346A>T, 4349G>A, 4350C>A, 4354T>A, 4355G>A, 4356A>T, 4358A>C, 4360G>A, 4361A>G, 4362A>T, 4363A>G, 4364A>T, 4365T>C, 4366T>C, 4370G>A, 4371T>C, 4373A>T, 4376C>T, 4379A>C, 4386C>A, 4391A>T, 4394A>G, 4397C>T, 4398C>A, 4399A>C, 4401T>G, 4402T>C, 4403C>T, 4404C>T, 4406C>A, 4407C>A, 4410G>T, 4411A>G, 4412T>G, 4413G>A, 4414A>T, 4415C>T, 4417A>G, 4418G>A, 4420T>A, 4421T>C, 4424C>T, 4430A>T, 4433T>C, 4436G>A, 4437C>A, 4438T>A, 4439C>G, 4440A>G, 4441C>A, 4442A>T, 4445C>T, 4446T>C, 4448C>A, 4449T>A, 4450C>A, 4451T>A, 4452C>A, 4453A>G, 4454C>A, 4455T>A, 4456T>C, 4457A>C, 4459C>T, 4460C>T, 4469G>C, 4470C>A, 4472C>T, 4474T>A, 4475T>C, 4476T>A, 4477C>G, 4478A>T, 4481G>A, 4488C>A, 4490T>G, 4493G>A, 4499A>G, 4509T>A, 4511A>T, 4512G>C, 4513G>A, 4514A>G, 4517C>T, 4518C>T, 4519A>C, 4521C>G, 4522C>A, 4523T>A, 4526T>A, 4529G>C, 4531G>A, 4533C>T, 4534C>A, 4535A>G, 4538A>G, 4539A>T, 4541A>C, 4543G>C, 4544G>T, 4548T>A, 4549G>A, 4550C>T, 4551A>G, 4556T>G, 4557G>T, 4558A>T, 4560A>G, 4562A>T, 4565T>A, 4566T>C, 4567T>A, 4569C>G, 4577G>C, 4580C>T, 4585C>T, 4586A>T, 4589C>T, 4592T>G, 4600C>A, 4601A>T, 4607T>C, 4610T>A, 4611T>G, 4612T>A, 4613G>A, 4616C>T, 4623G>A, 4624C>T, 4625C>T, 4630T>A, 4631C>T, 4632A>G, 4634A>T, 4638T>A, 4640C>T, 4641C>A, 4643G>T, 4645C>A, 4646C>T, 4647A>T, 4649C>G, 4650C>A, 4651T>G, 4652T>C, 4653T>C, 4654T>A, 4656T>A, 4657C>T, 4658A>G, 4659G>T, 4661T>A, 4662T>A, 4664G>A, 4667C>T, 4670C>T, 4677G>A, 4680A>G, 4682C>T, 4683C>T, 4685T>A, 4686C>A, 4688T>C, 4691T>C, 4697A>G, 4698A>G, 4699C>A, 4700A>C, 4701C>A, 4703G>C, 4706A>T, 4707G>T, 4708A>C, 4712C>T, 4713A>T, 4715A>G, 4719T>C, 4720T>A, 4721G>C, 4724T>A, 4727C>T, 4728C>A, 4729A>T, 4730G>T, 4733C>A, 4734A>T, 4736A>T, 4738G>A, 4739T>C, 4740T>A, 4741T>C, 4742G>C, 4745T>C, 4750A>G, 4752T>A, 4753T>A, 4757A>T, 4758G>T, 4760C>G, 4761A>G, 4763G>A, 4764T>G, 4767T>A, 4768C>G, 4769G>C, 4770G>A, 4772A>T, 4781G>T, 4783T>A, 4784C>A, 4788G>T, 4789C>T, 4795A>C, 4796T>A, 4803C>A, 4804A>G, 4808C>T, 4811A>G, 4814A>T, 4815A>C, 4816T>A, 4817G>T, 4823T>A, 4824G>T, 4825C>A, 4826T>C, 4827G>C, 4829T>A, 4832A>T, 4833A>T, 4837T>G, 4838T>C, 4839T>A, 4840C>G, 4844T>A, 4845G>A, 4846C>T, 4848G>T, 4850T>C, 4851C>A, 4852A>G, 4853C>A, 4854A>G, 4855T>C, 4857T>A, 4858C>T, 4860T>G, 4861T>A, 4863G>T, 4864A>T, 4865A>T, 4866C>T, 4867T>C, 4868C>T, 4869C>A, 4871A>T, 4874G>A, 4880T>C, 4881G>A, 4884A>G, 4885C>T, 4886A>T, 4888A>T, 4890C>A, 4891T>C, 4892C>T, 4893T>G, 4894C>A, 4895G>C, 4896G>A, 4897T>A, 4900A>C, 4901G>C, 4904G>A, 4905A>C, 4907A>G, 4911C>A, 4912A>G, 4913A>G, 4917C>T, 4919T>G, 4922G>A, 4924T>G, 4926G>C, 4931T>C, 4935A>G, 4938A>G, 4939G>C, 4940A>T, 4943T>C, 4946T>C, 4952A>T, 4953G>A, 4960C>G, 4961T>A, 4962G>C, 4964G>A, 4965C>G, 4966A>T, 4968A>T, 4969T>G, 4971T>A, 4972C>A, 4973C>A, 4974C>T, 4977C>T, 4979T>G, 4980A>T, 4981G>T, 4985T>C, 4987T>G, 4989C>T, 4994G>A | | | | | | | | |

##### CDS

| | Begin | End | Coverage | Score | Concordance | Matches | Identities | I/D/M/F* | Stop Codons |
|---|---|---|---|---|---|---|---|---|---|
| PVCVgp1 | 1376 | 1649 | 12.6% | 808 | 41.9% | 274 (100%) | 118 (43.1%) | 0/0/0/0 | 1 |
| Protein mutations: | F1377Y (4180T>A), K1378R (4183A>G), K1379F (4185A>T 4186A>T 4187G>C), N1382K (4196C>G), I1383A (4197A>G 4198T>C 4199C>T), N1384F (4200A>T 4201A>T), P1385S (4203C>T 4205C>T), S1389R (4217C>G), H1390P (4219A>C 4220T>C), S1391I (4221T>A 4222C>T 4223T>C), G1392Q (4224G>C 4225G>A), P1395H (4234C>A 4235A>T), H1397L (4239C>T 4240A>T 4241T>A), L1398M (4242C>A 4244C>G), Q1399E (4245C>G 4247G>A), L1400I (4248T>A 4250A>C), A1401C (4251G>T 4252C>G 4253A>T), I1402K (4255T>A), C1405I (4263T>A 4264G>T 4265T>C), E1407T (4269G>A 4270A>C 4271G>T), Q1409L (4275C>T 4276A>T 4277A>G), Q1410D (4278C>G 4280A>T), F1411S (4281T>A 4282T>G), D1412K (4284G>A 4286T>A), L1413I (4287C>A 4289A>C), E1415R (4293G>A 4294A>G), S1417F (4300C>T), D1418K (4302G>A 4304T>A), Q1420P (4309A>C 4310A>T), A1422S (4314G>A 4315C>G 4316A>C), E1424S (4320G>T 4321A>C 4322A>C), K1430N (4340G>T), R1431V (4341A>G 4342G>T 4343G>T), S1432A (4344T>G 4346A>T), Q1434K (4350C>A), V1435E (4354T>A 4355G>A), R1436C (4356A>T 4358A>C), G1437E (4360G>A 4361A>G), K1438C (4362A>T 4363A>G 4364A>T), L1439P (4365T>C 4366T>C), Q1446K (4386C>A), H1450T (4398C>A 4399A>C), F1451A (4401T>G 4402T>C 4403C>T), Q1453K (4407C>A), D1454W (4410G>T 4411A>G 4412T>G), D1455I (4413G>A 4414A>T 4415C>T), K1456R (4417A>G 4418G>A), F1457Y (4420T>A 4421T>C), L1463K (4437C>A 4438T>A 4439C>G), T1464D (4440A>G 4441C>A 4442A>T), F1466L (4446T>C 4448C>A), S1467K (4449T>A 4450C>A 4451T>A), H1468R (4452C>A 4453A>G 4454C>A), L1469T (4455T>A 4456T>C 4457A>C), S1470F (4459C>T 4460C>T), K1473N (4469G>C), L1474I (4470C>A 4472C>T), F1475Y (4474T>A 4475T>C), L1480M (4488C>A 4490T>G), L1487I (4509T>A 4511A>T), G1488Q (4512G>C 4513G>A 4514A>G), H1490S (4518C>T 4519A>C), P1491E (4521C>G 4522C>A 4523T>A), N1492K (4526T>A), E1493D (4529G>C), R1494K (4531G>A), P1495* (4533C>T 4534C>A 4535A>G), T1497S (4539A>T 4541A>C), G1498A (4543G>C 4544G>T), C1500N (4548T>A 4549G>A 4550C>T), I1501V (4551A>G), D1503F (4557G>T 4558A>T), R1504G (4560A>G 4562A>T), H1505Q (4565T>A), F1506H (4566T>C 4567T>A), Q1507E (4569C>G), K1509N (4577G>C), P1512L (4585C>T 4586A>T), T1517N (4600C>A 4601A>T), L1521E (4611T>G 4612T>A 4613G>A), A1525I (4623G>A 4624C>T 4625C>T), I1527N (4630T>A 4631C>T), K1528D (4632A>G 4634A>T), F1530I (4638T>A 4640C>T), Q1531N (4641C>A 4643G>T), P1532H (4645C>A 4646C>T), I1533L (4647A>T 4649C>G), L1534S (4650C>A 4651T>G 4652T>C), F1535H (4653T>C 4654T>A), S1536M (4656T>A 4657C>T 4658A>G), A1537S (4659G>T 4661T>A), L1538I (4662T>A 4664G>A), D1543N (4677G>A), I1544V (4680A>G 4682C>T), L1546I (4686C>A 4688T>C), T1550D (4698A>G 4699C>A 4700A>C), L1551I (4701C>A 4703G>C), E1552D (4706A>T), D1553S (4707G>T 4708A>C), I1555L (4713A>T 4715A>G), L1557H (4719T>C 4720T>A 4721G>C), Q1560I (4728C>A 4729A>T 4730G>T), F1561L (4733C>A), I1562F (4734A>T 4736A>T), S1563N (4738G>A 4739T>C), L1564T (4740T>A 4741T>C 4742G>C), K1567R (4750A>G), F1568N (4752T>A 4753T>A), V1570L (4758G>T 4760C>G), M1571V (4761A>G 4763G>A), L1572V (4764T>G), A1574T (4770G>A 4772A>T), M1577I (4781G>T), I1578K (4783T>A 4784C>A), A1580F (4788G>T 4789C>T), N1582T (4795A>C 4796T>A), Q1585R (4803C>A 4804A>G), M1589H (4815A>C 4816T>A 4817G>T), F1591L (4823T>A), A1592Y (4824G>T 4825C>A 4826T>C), D1593Q (4827G>C 4829T>A), T1595S (4833A>T), F1596C (4837T>G 4838T>C), S1597R (4839T>A 4840C>G), A1599I (4845G>A 4846C>T), G1600C (4848G>T 4850T>C), H1601R (4851C>A 4852A>G 4853C>A), I1602A (4854A>G 4855T>C), S1603I (4857T>A 4858C>T), L1604E (4860T>G 4861T>A), E1605F (4863G>T 4864A>T 4865A>T), L1606S (4866C>T 4867T>C 4868C>T), Q1607N (4869C>A 4871A>T), D1611N (4881G>A), T1612V (4884A>G 4885C>T 4886A>T), N1613I (4888A>T), L1614T (4890C>A 4891T>C 4892C>T), S1615D (4893T>G 4894C>A 4895G>C), V1616K (4896G>A 4897T>A), K1617T (4900A>C 4901G>C), I1619L (4905A>C 4907A>G), Q1621R (4911C>A 4912A>G 4913A>G), I1625S (4924T>G), V1626L (4926G>C), I1629V (4935A>G), R1630A (4938A>G 4939G>C 4940A>T), E1635K (4953G>A), T1637R (4960C>G 4961T>A), E1638Q (4962G>C 4964G>A), H1639V (4965C>G 4966A>T), I1640C (4968A>T 4969T>G), S1641K (4971T>A 4972C>A 4973C>A), P1642S (4974C>T), S1644F (4980A>T 4981G>T), M1646R (4987T>G) | | | | | | | | |

| | Begin | End | Coverage | Score | Concordance | Matches | Identities | I/D/M/F* | Stop Codons |
|---|---|---|---|---|---|---|---|---|---|
| **NT** | **4174** | **4995** | **11.4%** | **56** | **3.4%** | **822 (100%)** | **425 (51.7%)** | **0/0** | |
| Codon mutations: | TTT1377TAT (4180T>A), AAA1378AGA (4183A>G), AAG1379TTC (4185A>T 4186A>T 4187G>C), GAA1381GAG (4193A>G), AAC1382AAG (4196C>G), ATC1383GCT (4197A>G 4198T>C 4199C>T), AAT1384TTT (4200A>T 4201A>T), CCC1385TCT (4203C>T 4205C>T), ACA1386ACT (4208A>T), AAG1387AAA (4211G>A), GCA1388GCC (4214A>C), AGC1389AGG (4217C>G), CAT1390CCC (4219A>C 4220T>C), TCT1391ATC (4221T>A 4222C>T 4223T>C), GGA1392CAA (4224G>C 4225G>A), AAT1394AAC (4232T>C), CCA1395CAT (4234C>A 4235A>T), CAT1397TTA (4239C>T 4240A>T 4241T>A), CTC1398ATG (4242C>A 4244C>G), CAG1399GAA (4245C>G 4247G>A), TTA1400ATC (4248T>A 4250A>C), GCA1401TGT (4251G>T 4252C>G 4253A>T), ATA1402AAA (4255T>A), AAG1403AAA (4259G>A), TGT1405ATC (4263T>A 4264G>T 4265T>C), GAT1406GAC (4268T>C), GAG1407ACT (4269G>A 4270A>C 4271G>T), CAA1409TTG (4275C>T 4276A>T 4277A>G), CAA1410GAT (4278C>G 4280A>T), TTT1411AGT (4281T>A 4282T>G), GAT1412AAA (4284G>A 4286T>A), CTA1413ATC (4287C>A 4289A>C), ATC1414ATT (4292C>T), GAA1415AGA (4293G>A 4294A>G), CCC1416CCT (4298C>T), TCT1417TTT (4300C>T), GAT1418AAA (4302G>A 4304T>A), TCA1419TCT (4307A>T), CAA1420CCT (4309A>C 4310A>T), GCA1422AGC (4314G>A 4315C>G 4316A>C), GAA1424TCC (4320G>T 4321A>C 4322A>C), TTT1426TTC (4328T>C), GTC1428GTT (4334C>T), AAC1429AAT (4337C>T), AAG1430AAT (4340G>T), AGG1431GTT (4341A>G 4342G>T 4343G>T), TCA1432GCT (4344T>G 4346A>T), GAG1433GAA (4349G>A), CAA1434AAA (4350C>A), GTG1435GAA (4354T>A 4355G>A), AGA1436TGC (4356A>T 4358A>C), GGA1437GAG (4360G>A 4361A>G), AAA1438TGT (4362A>T 4363A>G 4364A>T), TTA1439CCA (4365T>C 4366T>C), AGG1440AGA (4370G>A), TTA1441CTT (4371T>C 4373A>T), GTC1442GTT (4376C>T), ATA1443ATC (4379A>C), CAA1446AAA (4386C>A), CCA1447CCT (4391A>T), TTA1448TTG (4394A>G), AAC1449AAT (4397C>T), CAT1450ACT (4398C>A 4399A>C), TTC1451GCT (4401T>G 4402T>C 4403C>T), CTC1452TTA (4404C>T 4406C>A), CAA1453AAA (4407C>A), GAT1454TGG (4410G>T 4411A>G 4412T>G), GAC1455ATT (4413G>A 4414A>T 4415C>T), AAG1456AGA (4417A>G 4418G>A), TTT1457TAC (4420T>A 4421T>C), CCC1458CCT (4424C>T), CCA1460CCT (4430A>T), AAT1461AAC (4433T>C), AAG1462AAA (4436G>A), CTC1463AAG (4437C>A 4438T>A 4439C>G), ACA1464GAT (4440A>G 4441C>A 4442A>T), CTC1465CTT (4445C>T), TTC1466CTA (4446T>C 4448C>A), TCT1467AAA (4449T>A 4450C>A 4451T>A), CAC1468AGA (4452C>A 4453A>G 4454C>A), TTA1469ACC (4455T>A 4456T>C 4457A>C), TCC1470TTT (4459C>T 4460C>T), AAG1473AAC (4469G>C), CTC1474ATT (4470C>A 4472C>T), TTT1475TAC (4474T>A 4475T>C), TCA1476AGT (4476T>A 4477C>G 4478A>T), AAG1477AAA (4481G>A), CTT1480ATG (4488C>A 4490T>G), AAG1481AAA (4493G>A), GGA1483GGG (4499A>G), TTA1487ATT (4509T>A 4511A>T), GGA1488CAG (4512G>C 4513G>A 4514A>G), ATC1489ATT (4517C>T), CAT1490TCT (4518C>T 4519A>C), CCT1491GAA (4521C>G 4522C>A 4523T>A), AAT1492AAA (4526T>A), GAG1493GAC (4529G>C), AGA1494AAA (4531G>A), CCA1495TAG (4533C>T 4534C>A 4535A>G), AAA1496AAG (4538A>G), ACA1497TCC (4539A>T 4541A>C), GGG1498GCT (4543G>C 4544G>T), TGC1500AAT (4548T>A 4549G>A 4550C>T), ATT1501GTT (4551A>G), CCT1502CCG (4556T>G), GAT1503TTT (4557G>T 4558A>T), AGA1504GGT (4560A>G 4562A>T), CAT1505CAA (4565T>A), TTT1506CAT (4566T>C 4567T>A), CAA1507GAA (4569C>G), AAG1509AAC (4577G>C), GTC1510GTT (4580C>T), CCA1512CTT (4585C>T 4586A>T), TTC1513TTT (4589C>T), GGT1514GGG (4592T>G), ACA1517AAT (4600C>A 4601A>T), CCT1519CCC (4607T>C), TCT1520TCA (4610T>A), TTG1521GAA (4611T>G 4612T>A 4613G>A), TTC1522TTT (4616C>T), GCC1525ATT (4623G>A 4624C>T 4625C>T), ATC1527AAT (4630T>A 4631C>T), AAA1528GAT (4632A>G 4634A>T), TTC1530ATT (4638T>A 4640C>T), CAG1531AAT (4641C>A 4643G>T), CCC1532CAT (4645C>A 4646C>T), ATC1533TTG (4647A>T 4649C>G), CTT1534AGC (4650C>A 4651T>G 4652T>C), TTT1535CAT (4653T>C 4654T>A), TCA1536ATG (4656T>A 4657C>T 4658A>G), GCT1537TCA (4659G>T 4661T>A), TTG1538ATA (4662T>A 4664G>A), GTC1539GTT (4667C>T), TAC1540TAT (4670C>T), GAT1543AAT (4677G>A), ATC1544GTT (4680A>G 4682C>T), CTT1545TTA (4683C>T 4685T>A), CTT1546ATC (4686C>A 4688T>C), TTT1547TTC (4691T>C), GAA1549GAG (4697A>G), ACA1550GAC (4698A>G 4699C>A 4700A>C), CTG1551ATC (4701C>A 4703G>C), GAA1552GAT (4706A>T), GAT1553TCT (4707G>T 4708A>C), CAC1554CAT (4712C>T), ATA1555TTG (4713A>T 4715A>G), TTG1557CAC (4719T>C 4720T>A 4721G>C), CTT1558CTA (4724T>A), AAC1559AAT (4727C>T), CAG1560ATT (4728C>A 4729A>T 4730G>T), TTC1561TTA (4733C>A), ATA1562TTT (4734A>T 4736A>T), AGT1563AAC (4738G>A 4739T>C), TTG1564ACC (4740T>A 4741T>C 4742G>C), GTT1565GTC (4745T>C), AAA1567AGA (4750A>G), TTT1568AAT (4752T>A 4753T>A), GGA1569GGT (4757A>T), GTC1570TTG (4758G>T 4760C>G), ATG1571GTA (4761A>G 4763G>A), TTA1572GTA (4764T>G), TCG1573AGC (4767T>A 4768C>G 4769G>C), GCA1574ACT (4770G>A 4772A>T), ATG1577ATT (4781G>T), ATC1578AAA (4783T>A 4784C>A), GCT1580TTT (4788G>T 4789C>T), AAT1582ACA (4795A>C 4796T>A), CAA1585AGA (4803C>A 4804A>G), TTC1586TTT (4808C>T), TTA1587TTG (4811A>G), GGA1588GGT (4814A>T), ATG1589CAT (4815A>C 4816T>A 4817G>T), TTT1591TTA (4823T>A), GCT1592TAC (4824G>T 4825C>A 4826T>C), GAT1593CAA (4827G>C 4829T>A), GGA1594GGT (4832A>T), ACT1595TCT (4833A>T), TTT1596TGC (4837T>G 4838T>C), TCA1597AGA (4839T>A 4840C>G), CCT1598CCA (4844T>A), GCT1599ATT (4845G>A 4846C>T), GGT1600TGC (4848G>T 4850T>C), CAC1601AGA (4851C>A 4852A>G 4853C>A), ATC1602GCC (4854A>G 4855T>C), TCT1603ATT (4857T>A 4858C>T), TTG1604GAG (4860T>G 4861T>A), GAA1605TTT (4863G>T 4864A>T 4865A>T), CTC1606TCT (4866C>T 4867T>C 4868C>T), CAA1607AAT (4869C>A 4871A>T), AAG1608AAA (4874G>A), CCT1610CCC (4880T>C), GAT1611AAT (4881G>A), ACA1612GTT (4884A>G 4885C>T 4886A>T), AAT1613ATT (4888A>T), CTC1614ACT (4890C>A 4891T>C 4892C>T), TCG1615GAC (4893T>G 4894C>A 4895G>C), GTA1616AAA (4896G>A 4897T>A), AAG1617ACC (4900A>C 4901G>C), CAG1618CAA (4904G>A), ATA1619CTG (4905A>C 4907A>G), CAA1621AGG (4911C>A 4912A>G 4913A>G), CTT1623TTG (4917C>T 4919T>G), GGG1624GGA (4922G>A), ATT1625AGT (4924T>G), GTT1626CTT (4926G>C), AAT1627AAC (4931T>C), ATC1629GTC (4935A>G), AGA1630GCT (4938A>G 4939G>C 4940A>T), GAT1631GAC (4943T>C), TTT1632TTC (4946T>C), CCA1634CCT (4952A>T), GAG1635AAG (4953G>A), ACT1637AGA (4960C>G 4961T>A), GAG1638CAA (4962G>C 4964G>A), CAT1639GTT (4965C>G 4966A>T), ATT1640TGT (4968A>T 4969T>G), TCC1641AAA (4971T>A 4972C>A 4973C>A), CCT1642TCT (4974C>T), CTT1643TTG (4977C>T 4979T>G), AGT1644TTT (4980A>T 4981G>T), GAT1645GAC (4985T>C), ATG1646AGG (4987T>G), CTA1647TTA (4989C>T), AAG1648AAA (4994G>A) | | | | | | | | |

#### Proteins

| | Begin | End | Coverage | Score | Concordance | Matches | Identities | I/D/M/F* | Stop Codons |
|---|---|---|---|---|---|---|---|---|---|
| ORF I polyprotein (NP_127504.1) | 1376 | 1649 | 12.6% | 808 | 41.9% | 274 (100%) | 118 (43.1%) | 0/0/0/0 | 1 |
| Protein mutations: | F1377Y (4180T>A), K1378R (4183A>G), K1379F (4185A>T 4186A>T 4187G>C), N1382K (4196C>G), I1383A (4197A>G 4198T>C 4199C>T), N1384F (4200A>T 4201A>T), P1385S (4203C>T 4205C>T), S1389R (4217C>G), H1390P (4219A>C 4220T>C), S1391I (4221T>A 4222C>T 4223T>C), G1392Q (4224G>C 4225G>A), P1395H (4234C>A 4235A>T), H1397L (4239C>T 4240A>T 4241T>A), L1398M (4242C>A 4244C>G), Q1399E (4245C>G 4247G>A), L1400I (4248T>A 4250A>C), A1401C (4251G>T 4252C>G 4253A>T), I1402K (4255T>A), C1405I (4263T>A 4264G>T 4265T>C), E1407T (4269G>A 4270A>C 4271G>T), Q1409L (4275C>T 4276A>T 4277A>G), Q1410D (4278C>G 4280A>T), F1411S (4281T>A 4282T>G), D1412K (4284G>A 4286T>A), L1413I (4287C>A 4289A>C), E1415R (4293G>A 4294A>G), S1417F (4300C>T), D1418K (4302G>A 4304T>A), Q1420P (4309A>C 4310A>T), A1422S (4314G>A 4315C>G 4316A>C), E1424S (4320G>T 4321A>C 4322A>C), K1430N (4340G>T), R1431V (4341A>G 4342G>T 4343G>T), S1432A (4344T>G 4346A>T), Q1434K (4350C>A), V1435E (4354T>A 4355G>A), R1436C (4356A>T 4358A>C), G1437E (4360G>A 4361A>G), K1438C (4362A>T 4363A>G 4364A>T), L1439P (4365T>C 4366T>C), Q1446K (4386C>A), H1450T (4398C>A 4399A>C), F1451A (4401T>G 4402T>C 4403C>T), Q1453K (4407C>A), D1454W (4410G>T 4411A>G 4412T>G), D1455I (4413G>A 4414A>T 4415C>T), K1456R (4417A>G 4418G>A), F1457Y (4420T>A 4421T>C), L1463K (4437C>A 4438T>A 4439C>G), T1464D (4440A>G 4441C>A 4442A>T), F1466L (4446T>C 4448C>A), S1467K (4449T>A 4450C>A 4451T>A), H1468R (4452C>A 4453A>G 4454C>A), L1469T (4455T>A 4456T>C 4457A>C), S1470F (4459C>T 4460C>T), K1473N (4469G>C), L1474I (4470C>A 4472C>T), F1475Y (4474T>A 4475T>C), L1480M (4488C>A 4490T>G), L1487I (4509T>A 4511A>T), G1488Q (4512G>C 4513G>A 4514A>G), H1490S (4518C>T 4519A>C), P1491E (4521C>G 4522C>A 4523T>A), N1492K (4526T>A), E1493D (4529G>C), R1494K (4531G>A), P1495* (4533C>T 4534C>A 4535A>G), T1497S (4539A>T 4541A>C), G1498A (4543G>C 4544G>T), C1500N (4548T>A 4549G>A 4550C>T), I1501V (4551A>G), D1503F (4557G>T 4558A>T), R1504G (4560A>G 4562A>T), H1505Q (4565T>A), F1506H (4566T>C 4567T>A), Q1507E (4569C>G), K1509N (4577G>C), P1512L (4585C>T 4586A>T), T1517N (4600C>A 4601A>T), L1521E (4611T>G 4612T>A 4613G>A), A1525I (4623G>A 4624C>T 4625C>T), I1527N (4630T>A 4631C>T), K1528D (4632A>G 4634A>T), F1530I (4638T>A 4640C>T), Q1531N (4641C>A 4643G>T), P1532H (4645C>A 4646C>T), I1533L (4647A>T 4649C>G), L1534S (4650C>A 4651T>G 4652T>C), F1535H (4653T>C 4654T>A), S1536M (4656T>A 4657C>T 4658A>G), A1537S (4659G>T 4661T>A), L1538I (4662T>A 4664G>A), D1543N (4677G>A), I1544V (4680A>G 4682C>T), L1546I (4686C>A 4688T>C), T1550D (4698A>G 4699C>A 4700A>C), L1551I (4701C>A 4703G>C), E1552D (4706A>T), D1553S (4707G>T 4708A>C), I1555L (4713A>T 4715A>G), L1557H (4719T>C 4720T>A 4721G>C), Q1560I (4728C>A 4729A>T 4730G>T), F1561L (4733C>A), I1562F (4734A>T 4736A>T), S1563N (4738G>A 4739T>C), L1564T (4740T>A 4741T>C 4742G>C), K1567R (4750A>G), F1568N (4752T>A 4753T>A), V1570L (4758G>T 4760C>G), M1571V (4761A>G 4763G>A), L1572V (4764T>G), A1574T (4770G>A 4772A>T), M1577I (4781G>T), I1578K (4783T>A 4784C>A), A1580F (4788G>T 4789C>T), N1582T (4795A>C 4796T>A), Q1585R (4803C>A 4804A>G), M1589H (4815A>C 4816T>A 4817G>T), F1591L (4823T>A), A1592Y (4824G>T 4825C>A 4826T>C), D1593Q (4827G>C 4829T>A), T1595S (4833A>T), F1596C (4837T>G 4838T>C), S1597R (4839T>A 4840C>G), A1599I (4845G>A 4846C>T), G1600C (4848G>T 4850T>C), H1601R (4851C>A 4852A>G 4853C>A), I1602A (4854A>G 4855T>C), S1603I (4857T>A 4858C>T), L1604E (4860T>G 4861T>A), E1605F (4863G>T 4864A>T 4865A>T), L1606S (4866C>T 4867T>C 4868C>T), Q1607N (4869C>A 4871A>T), D1611N (4881G>A), T1612V (4884A>G 4885C>T 4886A>T), N1613I (4888A>T), L1614T (4890C>A 4891T>C 4892C>T), S1615D (4893T>G 4894C>A 4895G>C), V1616K (4896G>A 4897T>A), K1617T (4900A>C 4901G>C), I1619L (4905A>C 4907A>G), Q1621R (4911C>A 4912A>G 4913A>G), I1625S (4924T>G), V1626L (4926G>C), I1629V (4935A>G), R1630A (4938A>G 4939G>C 4940A>T), E1635K (4953G>A), T1637R (4960C>G 4961T>A), E1638Q (4962G>C 4964G>A), H1639V (4965C>G 4966A>T), I1640C (4968A>T 4969T>G), S1641K (4971T>A 4972C>A 4973C>A), P1642S (4974C>T), S1644F (4980A>T 4981G>T), M1646R (4987T>G) | | | | | | | | |

| | Begin | End | Coverage | Score | Concordance | Matches | Identities | I/D/M/F* | Stop Codons |
|---|---|---|---|---|---|---|---|---|---|
| **NT** | **4174** | **4995** | **11.4%** | **56** | **3.4%** | **822 (100%)** | **425 (51.7%)** | **0/0** | |

Codon mutations: TTT1377TAT (4180T>A), AAA1378AGA (4183A>G), AAG1379TTC (4185A>T 4186A>T 4187G>C), GAA1381GAG (4193A>G), AAC1382AAG (4196C>G), ATC1383GCT (4197A>G 4198T>C 4199C>T), AAT1384TTT (4200A>T 4201A>T), CCC1385TCT (4203C>T 4205C>T), ACA1386ACT (4208A>T), AAG1387AAA (4211G>A), GCA1388GCC (4214A>C), AGC1389AGG (4217C>G), CAT1390CCC (4219A>C 4220T>C), TCT1391ATC (4221T>A 4222C>T 4223T>C), GGA1392CAA (4224G>C 4225G>A), AAT1394AAC (4232T>C), CCA1395CAT (4234C>A 4235A>T), CAT1397TTA (4239C>T 4240A>T 4241T>A), CTC1398ATG (4242C>A 4244C>G), CAG1399GAA (4245C>G 4247G>A), TTA1400ATC (4248T>A 4250A>C), GCA1401TGT (4251G>T 4252C>G 4253A>T), ATA1402AAA (4255T>A), AAG1403AAA (4259G>A), TGT1405ATC (4263T>A 4264G>T 4265T>C), GAT1406GAC (4268T>C), GAG1407ACT (4269G>A 4270A>C 4271G>T), CAA1409TTG (4275C>T 4276A>T 4277A>G), CAA1410GAT (4278C>G 4280A>T), TTT1411AGT (4281T>A 4282T>G), GAT1412AAA (4284G>A 4286T>A), CTA1413ATC (4287C>A 4289A>C), ATC1414ATT (4292C>T), GAA1415AGA (4293G>A 4294A>G), CCC1416CCT (4298C>T), TCT1417TTT (4300C>T), GAT1418AAA (4302G>A 4304T>A), TCA1419TCT (4307A>T), CAA1420CCT (4309A>C 4310A>T), GCA1422AGC (4314G>A 4315C>G 4316A>C), GAA1424TCC (4320G>T 4321A>C 4322A>C), TTT1426TTC (4328T>C), GTC1428GTT (4334C>T), AAC1429AAT (4337C>T), AAG1430AAT (4340G>T), AGG1431GTT (4341A>G 4342G>T 4343G>T), TCA1432GCT (4344T>G 4346A>T), GAG1433GAA (4349G>A), CAA1434AAA (4350C>A), GTG1435GAA (4354T>A 4355G>A), AGA1436TGC (4356A>T 4358A>C), GGA1437GAG (4360G>A 4361A>G), AAA1438TGT (4362A>T 4363A>G 4364A>T), TTA1439CCA (4365T>C 4366T>C), AGG1440AGA (4370G>A), TTA1441CTT (4371T>C 4373A>T), GTC1442GTT (4376C>T), ATA1443ATC (4379A>C), CAA1446AAA (4386C>A), CCA1447CCT (4391A>T), TTA1448TTG (4394A>G), AAC1449AAT (4397C>T), CAT1450ACT (4398C>A 4399A>C), TTC1451GCT (4401T>G 4402T>C 4403C>T), CTC1452TTA (4404C>T 4406C>A), CAA1453AAA (4407C>A), GAT1454TGG (4410G>T 4411A>G 4412T>G), GAC1455ATT (4413G>A 4414A>T 4415C>T), AAG1456AGA (4417A>G 4418G>A), TTT1457TAC (4420T>A 4421T>C), CCC1458CCT (4424C>T), CCA1460CCT (4430A>T), AAT1461AAC (4433T>C), AAG1462AAA (4436G>A), CTC1463AAG (4437C>A 4438T>A 4439C>G), ACA1464GAT (4440A>G 4441C>A 4442A>T), CTC1465CTT (4445C>T), TTC1466CTA (4446T>C 4448C>A), TCT1467AAA (4449T>A 4450C>A 4451T>A), CAC1468AGA (4452C>A 4453A>G 4454C>A), TTA1469ACC (4455T>A 4456T>C 4457A>C), TCC1470TTT (4459C>T 4460C>T), AAG1473AAC (4469G>C), CTC1474ATT (4470C>A 4472C>T), TTT1475TAC (4474T>A 4475T>C), TCA1476AGT (4476T>A 4477C>G 4478A>T), AAG1477AAA (4481G>A), CTT1480ATG (4488C>A 4490T>G), AAG1481AAA (4493G>A), GGA1483GGG (4499A>G), TTA1487ATT (4509T>A 4511A>T), GGA1488CAG (4512G>C 4513G>A 4514A>G), ATC1489ATT (4517C>T), CAT1490TCT (4518C>T 4519A>C), CCT1491GAA (4521C>G 4522C>A 4523T>A), AAT1492AAA (4526T>A), GAG1493GAC (4529G>C), AGA1494AAA (4531G>A), CCA1495TAG (4533C>T 4534C>A 4535A>G), AAA1496AAG (4538A>G), ACA1497TCC (4539A>T 4541A>C), GGG1498GCT (4543G>C 4544G>T), TGC1500AAT (4548T>A 4549G>A 4550C>T), ATT1501GTT (4551A>G), CCT1502CCG (4556T>G), GAT1503TTT (4557G>T 4558A>T), AGA1504GGT (4560A>G 4562A>T), CAT1505CAA (4565T>A), TTT1506CAT (4566T>C 4567T>A), CAA1507GAA (4569C>G), AAG1509AAC (4577G>C), GTC1510GTT (4580C>T), CCA1512CTT (4585C>T 4586A>T), TTC1513TTT (4589C>T), GGT1514GGG (4592T>G), ACA1517AAT (4600C>A 4601A>T), CCT1519CCC (4607T>C), TCT1520TCA (4610T>A), TTG1521GAA (4611T>G 4612T>A 4613G>A), TTC1522TTT (4616C>T), GCC1525ATT (4623G>A 4624C>T 4625C>T), ATC1527AAT (4630T>A 4631C>T), AAA1528GAT (4632A>G 4634A>T), TTC1530ATT (4638T>A 4640C>T), CAG1531AAT (4641C>A 4643G>T), CCC1532CAT (4645C>A 4646C>T), ATC1533TTG (4647A>T 4649C>G), CTT1534AGC (4650C>A 4651T>G 4652T>C), TTT1535CAT (4653T>C 4654T>A), TCA1536ATG (4656T>A 4657C>T 4658A>G), GCT1537TCA (4659G>T 4661T>A), TTG1538ATA (4662T>A 4664G>A), GTC1539GTT (4667C>T), TAC1540TAT (4670C>T), GAT1543AAT (4677G>A), ATC1544GTT (4680A>G 4682C>T), CTT1545TTA (4683C>T 4685T>A), CTT1546ATC (4686C>A 4688T>C), TTT1547TTC (4691T>C), GAA1549GAG (4697A>G), ACA1550GAC (4698A>G 4699C>A 4700A>C), CTG1551ATC (4701C>A 4703G>C), GAA1552GAT (4706A>T), GAT1553TCT (4707G>T 4708A>C), CAC1554CAT (4712C>T), ATA1555TTG (4713A>T 4715A>G), TTG1557CAC (4719T>C 4720T>A 4721G>C), CTT1558CTA (4724T>A), AAC1559AAT (4727C>T), CAG1560ATT (4728C>A 4729A>T 4730G>T), TTC1561TTA (4733C>A), ATA1562TTT (4734A>T 4736A>T), AGT1563AAC (4738G>A 4739T>C), TTG1564ACC (4740T>A 4741T>C 4742G>C), GTT1565GTC (4745T>C), AAA1567AGA (4750A>G), TTT1568AAT (4752T>A 4753T>A), GGA1569GGT (4757A>T), GTC1570TTG (4758G>T 4760C>G), ATG1571GTA (4761A>G 4763G>A), TTA1572GTA (4764T>G), TCG1573AGC (4767T>A 4768C>G 4769G>C), GCA1574ACT (4770G>A 4772A>T), ATG1577ATT (4781G>T), ATC1578AAA (4783T>A 4784C>A), GCT1580TTT (4788G>T 4789C>T), AAT1582ACA (4795A>C 4796T>A), CAA1585AGA (4803C>A 4804A>G), TTC1586TTT (4808C>T), TTA1587TTG (4811A>G), GGA1588GGT (4814A>T), ATG1589CAT (4815A>C 4816T>A 4817G>T), TTT1591TTA (4823T>A), GCT1592TAC (4824G>T 4825C>A 4826T>C), GAT1593CAA (4827G>C 4829T>A), GGA1594GGT (4832A>T), ACT1595TCT (4833A>T), TTT1596TGC (4837T>G 4838T>C), TCA1597AGA (4839T>A 4840C>G), CCT1598CCA (4844T>A), GCT1599ATT (4845G>A 4846C>T), GGT1600TGC (4848G>T 4850T>C), CAC1601AGA (4851C>A 4852A>G 4853C>A), ATC1602GCC (4854A>G 4855T>C), TCT1603ATT (4857T>A 4858C>T), TTG1604GAG (4860T>G 4861T>A), GAA1605TTT (4863G>T 4864A>T 4865A>T), CTC1606TCT (4866C>T 4867T>C 4868C>T), CAA1607AAT (4869C>A 4871A>T), AAG1608AAA (4874G>A), CCT1610CCC (4880T>C), GAT1611AAT (4881G>A), ACA1612GTT (4884A>G 4885C>T 4886A>T), AAT1613ATT (4888A>T), CTC1614ACT (4890C>A 4891T>C 4892C>T), TCG1615GAC (4893T>G 4894C>A 4895G>C), GTA1616AAA (4896G>A 4897T>A), AAG1617ACC (4900A>C 4901G>C), CAG1618CAA (4904G>A), ATA1619CTG (4905A>C 4907A>G), CAA1621AGG (4911C>A 4912A>G 4913A>G), CTT1623TTG (4917C>T 4919T>G), GGG1624GGA (4922G>A), ATT1625AGT (4924T>G), GTT1626CTT (4926G>C), AAT1627AAC (4931T>C), ATC1629GTC (4935A>G), AGA1630GCT (4938A>G 4939G>C 4940A>T), GAT1631GAC (4943T>C), TTT1632TTC (4946T>C), CCA1634CCT (4952A>T), GAG1635AAG (4953G>A), ACT1637AGA (4960C>G 4961T>A), GAG1638CAA (4962G>C 4964G>A), CAT1639GTT (4965C>G 4966A>T), ATT1640TGT (4968A>T 4969T>G), TCC1641AAA (4971T>A 4972C>A 4973C>A), CCT1642TCT (4974C>T), CTT1643TTG (4977C>T 4979T>G), AGT1644TTT (4980A>T 4981G>T), GAT1645GAC (4985T>C), ATG1646AGG (4987T>G), CTA1647TTA (4989C>T), AAG1648AAA (4994G>A)

*: Inserts / Deletes / Misaligned / Frameshifts

#### Analysis details

This analysis was performed with panviral2.64

### NGS Details (UN8): Badnavirus occultipomeae

#### Assembly

| | |
|---|---|
| **Coverage Length** | 366 (1 contig(s)) |
| **Depth Of Coverage** | 11.3 |
| **Number Of Reads** | 40 |
| **Reads Per Million** | 0.67 rpm (after QC) |
| **Ambiguities** | 0 |
| **Assembly Method** | de novo + reference guided assembly |
| **Consensus Caller** | Bcf Tools |

#### Coverage Map

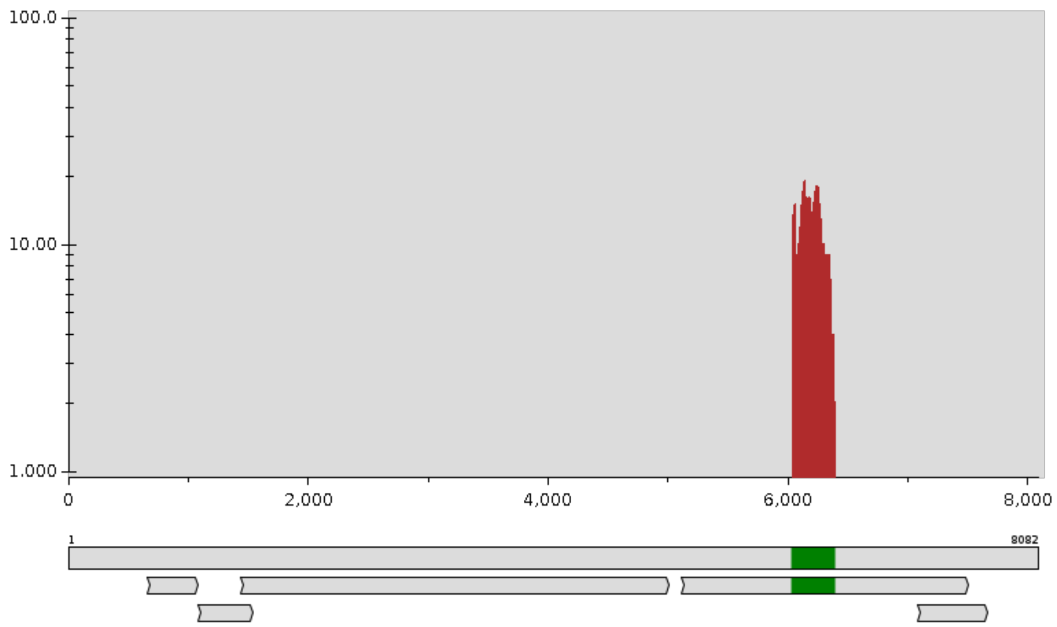

#### Assignment

| | |
|---|---|
| **Type** | Badnavirus occultipomeae (Taxonomy ID: 3048353) |
| **Reference Genome** | NC_015655.1 |
| **NT Identity (%)** | 55.0136 |
| **AA Identity (%)** | 40.6504 |
| **Number Of Stop Codons** | 2 |
| **Number Of CDS** | 5 |

#### Alignment

| | |
|---|---|
| **Alignment Score** | 68.0 (NT) + 320.0 (AA) = 388.0 |
| **Concordance (%)** | 24.5725 |

| Alignment Method | Global, seeded, nucleotide + amino acids (AGA) |
|---|---|

#### Genome Region

Sequence starts at position 6033 and ends at position 6398 relative to NC_015655.1 reference sequence.

#### Alignment Detailed Statistics

| | Begin | End | Coverage | Score | Concordance | Matches | Identities | I/D/M/F* | Stop Codons |
|---|---|---|---|---|---|---|---|---|---|
| **NT** | **6033** | **6398** | **4.5%** | **68** | **9.3%** | **366 (99.2%)** | **203 (55.0%)** | **3/0** | |
| Mutations: | 6033A>T, 6042A>G, 6043T>A, 6045T>A, 6048T>C, 6051C>A, 6053A>G, 6055G>T, 6060C>T, 6062T>A, 6063T>C, 6067C>T, 6070A>T, 6072A>G, 6074G>A, 6075A>G, 6076A>G, 6080A>G, 6081G>A, 6085A>T, 6087G>T, 6088T>G, 6089C>A, 6090C>T, 6092A>T, 6093A>C, 6095C>T, 6096A>G, 6097T>A, 6098G>A, 6099G>A, 6101C>T, 6102T>A, 6103G>A, 6105T>C, 6109T>C, 6114T>C, 6116C>G, 6117A>T, 6118G>T, 6120A>T, 6123G>C, 6125T>A, 6126C>T, 6128A>T, 6130G>A, 6135T>C, 6136G>C, 6137A>T, 6138A>G, 6141C>T, 6145C>T, 6154C>T, 6158T>C, 6159G>T, 6162T>C, 6167C>T, 6168A>T, 6173A>C, 6178C>A, 6179A>T, 6181A>G, 6182G>A, 6183A>T, 6184A>C, 6185A>T, 6186G>T, 6190G>A, 6193A>T, 6194A>G, 6195T>A, 6196G>C, 6197C>T, 6198A>G, 6201T>C, 6203G>A, 6205G>C, 6206G>C, 6207_6208insTGC, 6210A>G, 6214G>A, 6215C>T, 6216A>G, 6219T>C, 6222C>A, 6223G>A, 6224C>T, 6225A>T, 6230A>T, 6243A>T, 6249A>C, 6251T>A, 6255T>C, 6256G>A, 6258A>G, 6264A>G, 6275A>C, 6276G>C, 6277G>A, 6282T>C, 6285C>T, 6286T>A, 6287T>G, 6289A>T, 6290A>T, 6292T>G, 6294G>T, 6295G>C, 6296C>T, 6298C>A, 6302T>C, 6304G>C, 6306G>T, 6308G>A, 6309A>G, 6310A>G, 6311G>A, 6312A>T, 6313G>A, 6314A>G, 6315A>G, 6317G>A, 6319C>T, 6321T>G, 6322A>T, 6323T>A, 6325C>G, 6326T>C, 6329G>A, 6330C>A, 6331C>T, 6332C>T, 6333A>C, 6334A>T, 6339A>G, 6340A>T, 6341T>G, 6342G>T, 6343A>G, 6345A>T, 6346A>T, 6349G>T, 6351A>G, 6352G>A, 6354C>T, 6355A>G, 6356A>C, 6358G>T, 6359A>C, 6365A>T, 6366T>A, 6376A>C, 6377T>A, 6378A>T, 6380A>T, 6381G>T, 6384C>A, 6385C>T, 6386A>C, 6387A>T, 6391A>G, 6394A>G, 6395A>G, 6396G>T | | | | | | | | |

##### CDS

| | Begin | End | Coverage | Score | Concordance | Matches | Identities | I/D/M/F* | Stop Codons |
|---|---|---|---|---|---|---|---|---|---|
| SPBVa_gp4 | 307 | 428 | 15.3% | 320 | 37.3% | 122 (99.2%) | 50 (40.7%) | 1/0/0/0 | 2 |
| Protein mutations: | F310I (6043T>A 6045T>A), K313R (6053A>G), A314S (6055G>T), F316Y (6062T>A 6063T>C), Q318* (6067C>T), I319L (6070A>T 6072A>G), R320K (6074G>A 6075A>G), M321V (6076A>G), E322G (6080A>G 6081G>A), K324Y (6085A>T 6087G>T), S325D (6088T>G 6089C>A 6090C>T), K326I (6092A>T 6093A>C), P327L (6095C>T 6096A>G), W328K (6097T>A 6098G>A 6099G>A), T329I (6101C>T 6102T>A), A330T (6103G>A 6105T>C), W332R (6109T>C), P334R (6116C>G 6117A>T), E335Y (6118G>T 6120A>T), L337H (6125T>A 6126C>T), Y338F (6128A>T), E339K (6130G>A), E341L (6136G>C 6137A>T 6138A>G), P344S (6145C>T), M348T (6158T>C 6159G>T), P351L (6167C>T 6168A>T), D353A (6173A>C), Q355M (6178C>A 6179A>T), R356D (6181A>G 6182G>A 6183A>T), K357L (6184A>C 6185A>T 6186G>T), D359N (6190G>A), N360* (6193A>T 6194A>G 6195T>A), A361L (6196G>C 6197C>T 6198A>G), R363K (6203G>A), G364P (6205G>C 6206G>C), G364_T365insC (6207_6208insTGC), A367M (6214G>A 6215C>T 6216A>G), A370I (6223G>A 6224C>T 6225A>T), Y372F (6230A>T), F379Y (6251T>A), E381K (6256G>A 6258A>G), E387A (6275A>C 6276G>C), D388N (6277G>A), L391R (6286T>A 6287T>G), N392F (6289A>T 6290A>T), L393V (6292T>G 6294G>T), A394L (6295G>C 6296C>T), Q395K (6298C>A), I396T (6302T>C), V397L (6304G>C 6306G>T), R398K (6308G>A 6309A>G), R399D (6310A>G 6311G>A 6312A>T), E400R (6313G>A 6314A>G 6315A>G), G401E (6317G>A), I403Y (6322A>T 6323T>A), L404A (6325C>G 6326T>C), S405K (6329G>A 6330C>A), P406F (6331C>T 6332C>T 6333A>C), T407S (6334A>T), M409C (6340A>T 6341T>G 6342G>T), K410D (6343A>G 6345A>T), I411F (6346A>T), G412W (6349G>T 6351A>G), V413I (6352G>A 6354C>T), K414A (6355A>G 6356A>C), E415S (6358G>T 6359A>C), D417V (6365A>T 6366T>A), I421H (6376A>C 6377T>A 6378A>T), K422I (6380A>T 6381G>T), Q424S (6385C>T 6386A>C 6387A>T), N426D (6391A>G), K427G (6394A>G 6395A>G 6396G>T) | | | | | | | | |
| Codon mutations: | ATA306..T (6033A>T), AAA309AAG (6042A>G), TTT310ATA (6043T>A 6045T>A), GAT311GAC (6048T>C), CTC312CTA (6051C>A), AAA313AGA (6053A>G), GCA314TCA (6055G>T), GGC315GGT (6060C>T), TTT316TAC (6062T>A 6063T>C), CAG318TAG (6067C>T), ATA319TTG (6070A>T 6072A>G), AGA320AAG (6074G>A 6075A>G), ATG321GTG (6076A>G), GAG322GGA (6080A>G 6081G>A), AAG324TAT (6085A>T 6087G>T), TCC325GAT (6088T>G 6089C>A 6090C>T), AAA326ATC (6092A>T 6093A>C), CCA327CTG (6095C>T 6096A>G), TGG328AAA (6097T>A 6098G>A 6099G>A), ACT329ATA (6101C>T 6102T>A), GCT330ACC (6103G>A 6105T>C), TGG332CGG (6109T>C), ACT333ACC (6114T>C), CCA334CGT (6116C>G 6117A>T), GAA335TAT (6118G>T 6120A>T), GGG336GGC (6123G>C), CTC337CAT (6125T>A 6126C>T), TAT338TTT (6128A>T), GAA339AAA (6130G>A), TTT340TTC (6135T>C), GAA341CTG (6136G>C 6137A>T 6138A>G), GTC342GTT (6141C>T), CCC344TCC (6145C>T), CTG347TTG (6154C>T), ATG348ACT (6158T>C 6159G>T), AAT349AAC (6162T>C), CCA351CTT (6167C>T 6168A>T), GAT353GCT (6173A>C), CAG355ATG (6178C>A 6179A>T), AGA356GAT (6181A>G 6182G>A 6183A>T), AAG357CTT (6184A>C 6185A>T 6186G>T), GAT359AAT (6190G>A), AAT360TGA (6193A>T 6194A>G 6195T>A), GCA361CTG (6196G>C 6197C>T 6198A>G), TTT362TTC (6201T>C), AGG363AAG (6203G>A), GGA364CCA (6205G>C 6206G>C), GGA364_ACA365insTGC (6207_6208insTGC), ACA365ACG (6210A>G), GCA367ATG (6214G>A 6215C>T 6216A>G), TTT368TTC (6219T>C), ATC369ATA (6222C>A), GCA370ATT (6223G>A 6224C>T 6225A>T), TAT372TTT (6230A>T), ATA376ATT (6243A>T), GTA378GTC (6249A>C), TTC379TAC (6251T>A), TCT380TCC (6255T>C), GAA381AAG (6256G>A 6258A>G), GAA383GAG (6264A>G), GAG387GCC (6275A>C 6276G>C), GAT388AAT (6277G>A), CAT389CAC (6282T>C), CTC390CTT (6285C>T), TTA391AGA (6286T>A 6287T>G), AAT392TTT (6289A>T 6290A>T), TTG393GTT (6292T>G 6294G>T), GCC394CTC (6295G>C 6296C>T), CAA395AAA (6298C>A), ATT396ACT (6302T>C), GTG397CTT (6304G>C 6306G>T), AGA398AAG (6308G>A 6309A>G), AGA399GAT (6310A>G 6311G>A 6312A>T), GAA400AGG (6313G>A 6314A>G 6315A>G), GGG401GAG (6317G>A), CTT402TTG (6319C>T 6321T>G), ATT403TAT (6322A>T 6323T>A), CTA404GCA (6325C>G 6326T>C), AGC405AAA (6329G>A 6330C>A), CCA406TTC (6331C>T 6332C>T 6333A>C), ACA407TCA (6334A>T), AAA408AAG (6339A>G), ATG409TGT (6340A>T 6341T>G 6342G>T), AAA410GAT (6343A>G 6345A>T), ATT411TTT (6346A>T), GGA412TGG (6349G>T 6351A>G), GTC413ATT (6352G>A 6354C>T), AAA414GCA (6355A>G 6356A>C), GAA415TCA (6358G>T 6359A>C), GAT417GTA (6365A>T 6366T>A), ATA421CAT (6376A>C 6377T>A 6378A>T), AAG422ATT (6380A>T 6381G>T), ATC423ATA (6384C>A), CAA424TCT (6385C>T 6386A>C 6387A>T), AAT426GAT (6391A>G), AAG427GGT (6394A>G 6395A>G 6396G>T) | | | | | | | | |

##### Proteins

| | Begin | End | Coverage | Score | Concordance | Matches | Identities | I/D/M/F* | Stop Codons |
|---|---|---|---|---|---|---|---|---|---|
| RNaseH/reverse transcriptase (YP_004581513.1) | 307 | 428 | 15.3% | 320 | 37.3% | 122 (99.2%) | 50 (40.7%) | 1/0/0/0 | 2 |
| Protein mutations: | F310I (6043T>A 6045T>A), K313R (6053A>G), A314S (6055G>T), F316Y (6062T>A 6063T>C), Q318* (6067C>T), I319L (6070A>T 6072A>G), R320K (6074G>A 6075A>G), M321V (6076A>G), E322G (6080A>G 6081G>A), K324Y (6085A>T 6087G>T), S325D (6088T>G 6089C>A 6090C>T), K326I (6092A>T 6093A>C), P327L (6095C>T 6096A>G), W328K (6097T>A 6098G>A 6099G>A), T329I (6101C>T 6102T>A), A330T (6103G>A 6105T>C), W332R (6109T>C), P334R (6116C>G 6117A>T), E335Y (6118G>T 6120A>T), L337H (6125T>A 6126C>T), Y338F (6128A>T), E339K (6130G>A), E341L (6136G>C 6137A>T 6138A>G), P344S (6145C>T), M348T (6158T>C 6159G>T), P351L (6167C>T 6168A>T), D353A (6173A>C), Q355M (6178C>A 6179A>T), R356D (6181A>G 6182G>A 6183A>T), K357L (6184A>C 6185A>T 6186G>T), D359N (6190G>A), N360* (6193A>T 6194A>G 6195T>A), A361L (6196G>C 6197C>T 6198A>G), R363K (6203G>A), G364P (6205G>C 6206G>C), G364_T365insC (6207_6208insTGC), A367M (6214G>A 6215C>T 6216A>G), A370I (6223G>A 6224C>T 6225A>T), Y372F (6230A>T), F379Y (6251T>A), E381K (6256G>A 6258A>G), E387A (6275A>C 6276G>C), D388N (6277G>A), L391R (6286T>A 6287T>G), N392F (6289A>T 6290A>T), L393V (6292T>G 6294G>T), A394L (6295G>C 6296C>T), Q395K (6298C>A), I396T (6302T>C), V397L (6304G>C 6306G>T), R398K (6308G>A 6309A>G), R399D (6310A>G 6311G>A 6312A>T), E400R (6313G>A 6314A>G 6315A>G), G401E (6317G>A), I403Y (6322A>T 6323T>A), L404A (6325C>G 6326T>C), S405K (6329G>A 6330C>A), P406F (6331C>T 6332C>T 6333A>C), T407S (6334A>T), M409C (6340A>T 6341T>G 6342G>T), K410D (6343A>G 6345A>T), I411F (6346A>T), G412W (6349G>T 6351A>G), V413I (6352G>A 6354C>T), K414A (6355A>G 6356A>C), E415S (6358G>T 6359A>C), D417V (6365A>T 6366T>A), I421H (6376A>C 6377T>A 6378A>T), K422I (6380A>T 6381G>T), Q424S (6385C>T 6386A>C 6387A>T), N426D (6391A>G), K427G (6394A>G 6395A>G 6396G>T) | | | | | | | | |
| Codon mutations: | ATA306..T (6033A>T), AAA309AAG (6042A>G), TTT310ATA (6043T>A 6045T>A), GAT311GAC (6048T>C), CTC312CTA (6051C>A), AAA313AGA (6053A>G), GCA314TCA (6055G>T), GGC315GGT (6060C>T), TTT316TAC (6062T>A 6063T>C), CAG318TAG (6067C>T), ATA319TTG (6070A>T 6072A>G), AGA320AAG (6074G>A 6075A>G), ATG321GTG (6076A>G), GAG322GGA (6080A>G 6081G>A), AAG324TAT (6085A>T 6087G>T), TCC325GAT (6088T>G 6089C>A 6090C>T), AAA326ATC (6092A>T 6093A>C), CCA327CTG (6095C>T 6096A>G), TGG328AAA (6097T>A 6098G>A 6099G>A), ACT329ATA (6101C>T 6102T>A), GCT330ACC (6103G>A 6105T>C), TGG332CGG (6109T>C), ACT333ACC (6114T>C), CCA334CGT (6116C>G 6117A>T), GAA335TAT (6118G>T 6120A>T), GGG336GGC (6123G>C), CTC337CAT (6125T>A 6126C>T), TAT338TTT (6128A>T), GAA339AAA (6130G>A), TTT340TTC (6135T>C), GAA341CTG (6136G>C 6137A>T 6138A>G), GTC342GTT (6141C>T), CCC344TCC (6145C>T), CTG347TTG (6154C>T), ATG348ACT (6158T>C 6159G>T), AAT349AAC (6162T>C), CCA351CTT (6167C>T 6168A>T), GAT353GCT (6173A>C), CAG355ATG (6178C>A 6179A>T), AGA356GAT (6181A>G 6182G>A 6183A>T), AAG357CTT (6184A>C 6185A>T 6186G>T), GAT359AAT (6190G>A), AAT360TGA (6193A>T 6194A>G 6195T>A), GCA361CTG (6196G>C 6197C>T 6198A>G), TTT362TTC (6201T>C), AGG363AAG (6203G>A), GGA364CCA (6205G>C 6206G>C), GGA364_ACA365insTGC (6207_6208insTGC), ACA365ACG (6210A>G), GCA367ATG (6214G>A 6215C>T 6216A>G), TTT368TTC (6219T>C), ATC369ATA (6222C>A), GCA370ATT (6223G>A 6224C>T 6225A>T), TAT372TTT (6230A>T), ATA376ATT (6243A>T), GTA378GTC (6249A>C), TTC379TAC (6251T>A), TCT380TCC (6255T>C), GAA381AAG (6256G>A 6258A>G), GAA383GAG (6264A>G), GAG387GCC (6275A>C 6276G>C), GAT388AAT (6277G>A), CAT389CAC (6282T>C), CTC390CTT (6285C>T), TTA391AGA (6286T>A 6287T>G), AAT392TTT (6289A>T 6290A>T), TTG393GTT (6292T>G 6294G>T), GCC394CTC (6295G>C 6296C>T), CAA395AAA (6298C>A), ATT396ACT (6302T>C), GTG397CTT (6304G>C 6306G>T), AGA398AAG (6308G>A 6309A>G), AGA399GAT (6310A>G 6311G>A 6312A>T), GAA400AGG (6313G>A 6314A>G 6315A>G), GGG401GAG (6317G>A), CTT402TTG (6319C>T 6321T>G), ATT403TAT (6322A>T 6323T>A), CTA404GCA (6325C>G 6326T>C), AGC405AAA (6329G>A 6330C>A), CCA406TTC (6331C>T 6332C>T 6333A>C), ACA407TCA (6334A>T), AAA408AAG (6339A>G), ATG409TGT (6340A>T 6341T>G 6342G>T), AAA410GAT (6343A>G 6345A>T), ATT411TTT (6346A>T), GGA412TGG (6349G>T 6351A>G), GTC413ATT (6352G>A 6354C>T), AAA414GCA (6355A>G 6356A>C), GAA415TCA (6358G>T 6359A>C), GAT417GTA (6365A>T 6366T>A), ATA421CAT (6376A>C 6377T>A 6378A>T), AAG422ATT (6380A>T 6381G>T), ATC423ATA (6384C>A), CAA424TCT (6385C>T 6386A>C 6387A>T), AAT426GAT (6391A>G), AAG427GGT (6394A>G 6395A>G 6396G>T) | | | | | | | | |

*: Inserts / Deletes / Misaligned / Frameshifts

#### Analysis details

This analysis was performed with panviral2.64

### NGS Details (UN8): Caulimovirus tessellobrassicae

#### Assembly

| | |
|---|---|
| **Coverage Length** | 725 (2 contig(s)) |
| **Depth Of Coverage** | 6.6 |
| **Number Of Reads** | 39 |
| **Reads Per Million** | 0.65 rpm (after QC) |
| **Ambiguities** | 0 |
| **Assembly Method** | de novo + reference guided assembly |
| **Consensus Caller** | Bcf Tools |

#### Coverage Map

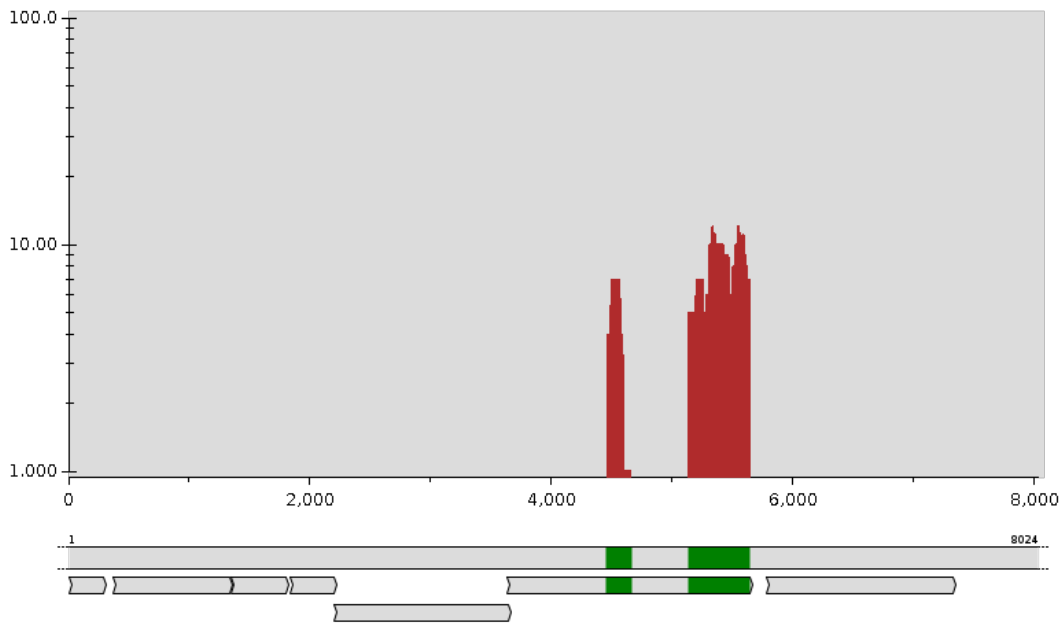

#### Assignment

| | |
|---|---|
| **Type** | Caulimovirus tessellobrassicae (Taxonomy ID: 3047738) |
| **Reference Genome** | NC_001497.2 |
| **NT Identity (%)** | 56.6761 |
| **AA Identity (%)** | 46.383 |
| **Number Of Stop Codons** | 0 |
| **Number Of CDS** | 7 |

#### Alignment

| | |
|---|---|
| **Alignment Score** | 149.0 (NT) + 792.0 (AA) = 941.0 |
| **Concordance (%)** | 31.5772 |

| Alignment Method | Global, seeded, nucleotide + amino acids (AGA) |
|---|---|

#### Genome Region

Sequence starts at position 4456 and ends at position 5647 relative to NC_001497.2 reference sequence.

#### Alignment Detailed Statistics

| | Begin | End | Coverage | Score | Concordance | Matches | Identities | I/D/M/F* | Stop Codons |
|---|---|---|---|---|---|---|---|---|---|
| **NT** | **4456** | **5647** | **9.0%** | **149** | **10.9%** | **704 (97.1%)** | **399 (55.0%)** | **0/21** | |
| Mutations: | 4465A>T, 4467T>C, 4469A>G, 4473C>T, 4476T>A, 4481T>G, 4482G>C, 4483G>A, 4484C>G, 4485A>T, 4488A>C, 4491C>G, 4495T>A, 4500C>A, 4502A>G, 4503C>A, 4506T>C, 4507G>C, 4509A>T, 4512C>G, 4515G>A, 4516A>C, 4518G>A, 4519C>A, 4520G>A, 4522A>C, 4527A>T, 4531A>G, 4532A>C, 4533A>T, 4535G>A, 4536T>A, 4542A>C, 4543G>A, 4545C>A, 4548C>T, 4551C>T, 4556C>A, 4557T>G, 4558A>C, 4560G>T, 4563C>T, 4567G>A, 4568C>A, 4570A>T, 4571C>G, 4575A>T, 4576G>T, 4577G>T, 4578A>T, 4581T>C, 4583C>G, 4584C>A, 4588A>T, 4589A>T, 4591C>A, 4596C>T, 4603G>A, 4606G>A, 4608G>T, 4609T>C, 4611A>T, 4612C>A, 4613T>C, 4616C>A, 4617A>T, 4620C>A, 4621A>G, 4622T>C, 4623T>A, 4624C>A, 4625G>A, 4627G>A, 4632G>A, 4634A>T, 4635G>T, 4636A>T, 4637T>A, 4641C>T, 4644T>A, 4645T>A, 4646C>A, 4647C>A, 4650C>T, 4653C>T, 4660T>A, 4661C>G, 5132T>A, 5133C>G, 5139G>T, 5141C>T, 5143C>T, 5145G>A, 5149G>A, 5150C>G, 5151C>G, 5152A>C, 5153A>T, 5154G>T, 5155C>A, 5158A>C, 5159A>G, 5161G>A, 5164A>C, 5165A>C, 5167G>A, 5168T>A, 5169T>C, 5171C>G, 5173T>A, 5174G>T, 5176A>G, 5185A>G, 5187A>T, 5190G>A, 5191G>C, 5196C>T, 5197C>A, 5198T>G, 5199C>A, 5201A>G, 5202C>T, 5203A>G, 5205G>T, 5210A>G, 5211G>C, 5212G>C, 5214G>A, 5217G>A, 5218A>C, 5221A>G, 5223T>A, 5224C>T, 5225T>G, 5226G>T, 5227C>T, 5228A>C, 5229A>C, 5230G>A, 5231G>A, 5235T>A, 5238T>A, 5239C>A, 5240C>A, 5241A>G, 5245C>A, 5246A>G, 5247T>A, 5249A>T, 5253C>G, 5254T>G, 5255T>A, 5262G>A, 5265G>C, 5268G>C, 5269C>T, 5271G>A, 5272A>G, 5274C>T, 5275A>T, 5277C>A, 5278G>C, 5280G>A, 5283C>A, 5289A>T, 5295C>T, 5296G>T, 5299T>C, 5301C>T, 5309G>C, 5310T>A, 5311A>T, 5313G>A, 5317A>C, 5320G>A, 5323_5325delATC, 5326A>G, 5328A>C, 5329A>C, 5334C>T, 5338_5355delGGTACTAATACTGAGTTA, 5361C>T, 5364A>G, 5367C>T, 5368G>A, 5371T>A, 5372C>G, 5376A>T, 5378G>C, 5379C>A, 5385A>T, 5387C>A, 5388T>A, 5389G>A, 5391A>C, 5396A>C, 5397G>T, 5400T>A, 5404C>T, 5405A>C, 5406C>A, 5408G>C, 5409C>A, 5415C>A, 5421G>A, 5422A>T, 5423C>T, 5430G>C, 5431G>A, 5434A>G, 5438A>G, 5439T>A, 5440A>G, 5441C>G, 5442T>C, 5446A>C, 5447A>G, 5448G>A, 5455A>T, 5456G>C, 5458A>G, 5459T>C, 5462A>T, 5463T>C, 5467A>C, 5468C>T, 5469T>A, 5472T>A, 5473G>A, 5474T>A, 5475T>G, 5478T>A, 5481T>C, 5482C>A, 5484G>C, 5487T>C, 5490G>A, 5491A>T, 5496T>C, 5502T>A, 5505T>G, 5506T>G, 5508C>A, 5509A>T, 5510A>C, 5511G>A, 5512A>G, 5517C>G, 5518G>A, 5520T>A, 5521A>T, 5522A>T, 5524C>A, 5525T>A, 5526C>T, 5529T>A, 5530T>C, 5531A>T, 5532C>T, 5533A>C, 5534A>C, 5536G>T, 5537G>C, 5538A>T, 5541T>A, 5542T>C, 5544G>T, 5545A>C, 5548C>T, 5549T>A, 5551G>A, 5557A>T, 5558A>T, 5559C>A, 5560A>C, 5561T>A, 5562C>T, 5573C>T, 5576G>T, 5578C>T, 5580T>A, 5581A>T, 5582G>C, 5583C>T, 5584C>T, 5585A>T, 5586C>T, 5590T>A, 5592A>C, 5595T>C, 5596G>A, 5598T>A, 5599G>A, 5605C>A, 5607C>T, 5616A>T, 5617A>T, 5619C>T, 5622C>T, 5625C>T, 5626C>G, 5627A>T, 5628C>T, 5634G>A, 5637C>T, 5641C>T, 5643T>A | | | | | | | | |

##### CDS

| | Begin | End | Coverage | Score | Concordance | Matches | Identities | I/D/M/F* | Stop Codons |
|---|---|---|---|---|---|---|---|---|---|
| CaMVgp6 | 275 | 672 | 35.6% | 792 | 48.5% | 235 (97.1%) | 109 (45.0%) | 0/7/0/0 | 0 |
| Protein mutations: | S278C (4465A>T 4467T>C), K279R (4469A>G), M283S (4481T>G 4482G>C), A284S (4483G>A 4484C>G 4485A>T), L288M (4495T>A), N290R (4502A>G 4503C>A), E292H (4507G>C 4509A>T), K295Q (4516A>C 4518G>A), R296K (4519C>A 4520G>A), K300A (4531A>G 4532A>C 4533A>T), R301Q (4535G>A 4536T>A), V304I (4543G>A 4545C>A), A308E (4556C>A 4557T>G), M309L (4558A>C 4560G>T), A312N (4567G>A 4568C>A), T313C (4570A>T 4571C>G), G315F (4576G>T 4577G>T 4578A>T), A317G (4583C>G 4584C>A), N319F (4588A>T 4589A>T), L320I (4591C>A), D324N (4603G>A), E325N (4606G>A 4608G>T), L327T (4612C>A 4613T>C), T328N (4616C>A 4617A>T), I330A (4621A>G 4622T>C 4623T>A), R331K (4624C>A 4625G>A), G332R (4627G>A), K334I (4634A>T 4635G>T), I335Y (4636A>T 4637T>A), S338K (4645T>A 4646C>A 4647C>A), I500K (5132T>A 5133C>G), K502N (5139G>T), P503L (5141C>T), A506R (5149G>A 5150C>G 5151C>G), K507L (5152A>C 5153A>T 5154G>T), L508I (5155C>A), K509R (5158A>C 5159A>G), E510K (5161G>A), N511P (5164A>C 5165A>C), V512N (5167G>A 5168T>A 5169T>C), P513R (5171C>G), W514M (5173T>A 5174G>T), R515G (5176A>G), K518D (5185A>G 5187A>T), D520H (5191G>C), L522R (5197C>A 5198T>G 5199C>A), Y523C (5201A>G 5202C>T), M524V (5203A>G 5205G>T), K526S (5210A>G 5211G>C), V527L (5212G>C 5214G>A), K529Q (5218A>C), N530E (5221A>G 5223T>A), L531C (5224C>T 5225T>G 5226G>T), Q532S (5227C>T 5228A>C 5229A>C), G533K (5230G>A 5231G>A), F534L (5235T>A), P536K (5239C>A 5240C>A 5241A>G), H538R (5245C>A 5246A>G 5247T>A), H539L (5249A>T), L541E (5254T>G 5255T>A), E544D (5265G>C), K545N (5268G>C), I547V (5272A>G 5274C>T), I548L (5275A>T 5277C>A), E549Q (5278G>C 5280G>A), D555Y (5296G>T), Y556H (5299T>C 5301C>T), G559A (5309G>C 5310T>A), M560L (5311A>T 5313G>A), K562Q (5317A>C), A563T (5320G>A), I564del (5323_5325delATC), K565D (5326A>G 5328A>C), I566L (5329A>C), G569_L574del (5338_5355delGGTACTAATACTGAGTTA), A579T (5368G>A), S582T (5378G>C 5379C>A), K584N (5385A>T), A585E (5387C>A 5388T>A), A586T (5389G>A 5391A>C), K588T (5396A>C 5397G>T), N589K (5400T>A), H591S (5404C>T 5405A>C 5406C>A), S592T (5408G>C 5409C>A), D594E (5415C>A), T597L (5422A>T 5423C>T), V600I (5431G>A), I601V (5434A>G), N602R (5438A>G 5439T>A), T603G (5440A>G 5441C>G 5442T>C), K605R (5446A>C 5447A>G 5448G>A), I609A (5458A>G 5459T>C), Y610F (5462A>T 5463T>C), T612L (5467A>C 5468C>T 5469T>A), V614K (5473G>A 5474T>A 5475T>G), H615Q (5478T>A), L617I (5482C>A 5484G>C), T620S (5491A>T), H624Q (5505T>G), F625V (5506T>G 5508C>A), K626S (5509A>T 5510A>C 5511G>A), S627G (5512A>G), F628L (5517C>G), V629I (5518G>A 5520T>A), N630F (5521A>T 5522A>T), L631N (5524C>A 5525T>A 5526C>T), N632K (5529T>A), Y633L (5530T>C 5531A>T 5532C>T), K634P (5533A>C 5534A>C), G635S (5536G>T 5537G>C 5538A>T), D636E (5541T>A), S637P (5542T>C 5544G>T), K638Q (5545A>C), L639Y (5548C>T 5549T>A), G640R (5551G>A), N642L (5557A>T 5558A>T 5559C>A), I643H (5560A>C 5561T>A 5562C>T), A647V (5573C>T), W648L (5576G>T), H651F (5584C>T 5585A>T 5586C>T), S653T (5590T>A 5592A>C), D655K (5596G>A 5598T>A), V656I (5599G>A), H658N (5605C>A 5607C>T), T662S (5617A>T 5619C>T), H665V (5626C>G 5627A>T 5628C>T) | | | | | | | | |
| Codon mutations: | AGT278TGC (4465A>T 4467T>C), AAA279AGA (4469A>G), AGC280AGT (4473C>T), CCT281CCA (4476T>A), ATG283AGC (4481T>G 4482G>C), GCA284AGT (4483G>A 4484C>G 4485A>T), CCA285CCC (4488A>C), GCC286GCG (4491C>G), TTG288ATG (4495T>A), GTC289GTA (4500C>A), AAC290AGA (4502A>G 4503C>A), AAT291AAC (4506T>C), GAA292CAT (4507G>C 4509A>T), GCC293GCG (4512C>G), GAG294GAA (4515G>A), AAG295CAA (4516A>C 4518G>A), CGA296AAA (4519C>A 4520G>A), AGA297CGA (4522A>C), GGA298GGT (4527A>T), AAA300GCT (4531A>G 4532A>C 4533A>T), CGT301CAA (4535G>A 4536T>A), GTA303GTC (4542A>C), GTC304ATA (4543G>A 4545C>A), AAC305AAT (4548C>T), TAC306TAT (4551C>T), GCT308GAG (4556C>A 4557T>G), ATG309CTT (4558A>C 4560G>T), AAC310AAT (4563C>T), GCT312AAT (4567G>A 4568C>A), ACT313TGT (4570A>T 4571C>G), GTA314GTT (4575A>T), GGA315TTT (4576G>T 4577G>T 4578A>T), GAT316GAC (4581T>C), GCC317GGA (4583C>G 4584C>A), AAT319TTT (4588A>T 4589A>T), CTT320ATT (4591C>A), CCC321CCT (4596C>T), GAC324AAC (4603G>A), GAG325AAT (4606G>A 4608G>T), TTA326CTT (4609T>C 4611A>T), CTT327ACT (4612C>A 4613T>C), ACA328AAT (4616C>A 4617A>T), CTC329CTA (4620C>A), ATT330GCA (4621A>G 4622T>C 4623T>A), CGA331AAA (4624C>A 4625G>A), GGA332AGA (4627G>A), AAG333AAA (4632G>A), AAG334ATT (4634A>T 4635G>T), ATC335TAC (4636A>T 4637T>A), TTC336TTT (4641C>T), TCT337TCA (4644T>A), TCC338AAA (4645T>A 4646C>A 4647C>A), TTC339TTT (4650C>T), GAC340GAT (4653C>T), TCA343AG. (4660T>A 4661C>G), ATC500AAG (5132T>A 5133C>G), AAG502AAT (5139G>T), CCT503CTT (5141C>T), CTG504TTA (5143C>T 5145G>A), GCC506AGG (5149G>A 5150C>G 5151C>G), AAG507CTT (5152A>C 5153A>T 5154G>T), CTT508ATT (5155C>A), AAA509CGA (5158A>C 5159A>G), GAA510AAA (5161G>A), AAC511CCC (5164A>C 5165A>C), GTT512AAC (5167G>A 5168T>A 5169T>C), CCA513CGA (5171C>G), TGG514ATG (5173T>A 5174G>T), AGA515GGA (5176A>G), AAA518GAT (5185A>G 5187A>T), GAG519GAA (5190G>A), GAT520CAT (5191G>C), ACC521ACT (5196C>T), CTC522AGA (5197C>A 5198T>G 5199C>A), TAC523TGT (5201A>G 5202C>T), ATG524GTT (5203A>G 5205G>T), AAG526AGC (5210A>G 5211G>C), GTG527CTA (5212G>C 5214G>A), AAG528AAA (5217G>A), AAA529CAA (5218A>C), AAT530GAA (5221A>G 5223T>A), CTG531TGT (5224C>T 5225T>G 5226G>T), CAA532TCC (5227C>T 5228A>C 5229A>C), GGA533AAA (5230G>A 5231G>A), TTT534TTA (5235T>A), CCT535CCA (5238T>A), CCA536AAG (5239C>A 5240C>A 5241A>G), CAT538AGA (5245C>A 5246A>G 5247T>A), CAT539CTT (5249A>T), CCC540CCG (5253C>G), TTA541GAA (5254T>G 5255T>A), GAG543GAA (5262G>A), GAG544GAC (5265G>C), AAG545AAC (5268G>C), CTG546TTA (5269C>T 5271G>A), ATC547GTT (5272A>G 5274C>T), ATC548TTA (5275A>T 5277C>A), GAG549CAA (5278G>C 5280G>A), ACC550ACA (5283C>A), GCA552GCT (5289A>T), GAC554GAT (5295C>T), GAC555TAC (5296G>T), TAC556CAT (5299T>C 5301C>T), GGT559GCA (5309G>C 5310T>A), ATG560TTA (5311A>T 5313G>A), AAA562CAA (5317A>C), GCT563ACT (5320G>A), ATC564del (5323_5325delATC), AAA565GAC (5326A>G 5328A>C), ATT566CTT (5329A>C), AAC567AAT (5334C>T), GGT569_TTA574del (5338_5355delGGTACTAATACTGAGTTA), TGC576TGT (5361C>T), AGA577AGG (5364A>G), TAC578TAT (5367C>T), GCA579ACA (5368G>A), TCT580AGT (5371T>A 5372C>G), GGA581GGT (5376A>T), AGC582ACA (5378G>C 5379C>A), AAA584AAT (5385A>T), GCT585GAA (5387C>A 5388T>A), GCA586ACC (5389G>A 5391A>C), AAG588ACT (5396A>C 5397G>T), AAT589AAA (5400T>A), CAC591TCA (5404C>T 5405A>C 5406C>A), AGC592ACA (5408G>C 5409C>A), GAC594GAA (5415C>A), GAG596GAA (5421G>A), ACA597TTA (5422A>T 5423C>T), GCG599GCC (5430G>C), GTA600ATA (5431G>A), ATA601GTA (5434A>G), AAT602AGA (5438A>G 5439T>A), ACT603GGC (5440A>G 5441C>G 5442T>C), AAG605CGA (5446A>C 5447A>G 5448G>A), AGT608TCT (5455A>T 5456G>C), ATT609GCT (5458A>G 5459T>C), TAT610TTC (5462A>T 5463T>C), ACT612CTA (5467A>C 5468C>T 5469T>A), CCT613CCA (5472T>A), GTT614AAG (5473G>A 5474T>A 5475T>G), CAT615CAA (5478T>A), TTT616TTC (5481T>C), CTG617ATC (5482C>A 5484G>C), ATT618ATC (5487T>C), AGG619AGA (5490G>A), ACA620TCA (5491A>T), GAT621GAC (5496T>C), ACT623ACA (5502T>A), CAT624CAG (5505T>G), TTC625GTA (5506T>G 5508C>A), AAG626TCA (5509A>T 5510A>C 5511G>A), AGT627GGT (5512A>G), TTC628TTG (5517C>G), GTT629ATA (5518G>A 5520T>A), AAT630TTT (5521A>T 5522A>T), CTC631AAT (5524C>A 5525T>A 5526C>T), AAT632AAA (5529T>A), TAC633CTT (5530T>C 5531A>T 5532C>T), AAA634CCA (5533A>C 5534A>C), GGA635TCT (5536G>T 5537G>C 5538A>T), GAT636GAA (5541T>A), TCG637CCT (5542T>C 5544G>T), AAA638CAA (5545A>C), CTT639TAT (5548C>T 5549T>A), GGA640AGA (5551G>A), AAC642TTA (5557A>T 5558A>T 5559C>A), ATC643CAT (5560A>C 5561T>A 5562C>T), GCA647GTA (5573C>T), TGG648TTG (5576G>T), CTT649TTA (5578C>T 5580T>A), AGC650TCT (5581A>T 5582G>C 5583C>T), CAC651TTT (5584C>T 5585A>T 5586C>T), TCA653ACC (5590T>A 5592A>C), TTT654TTC (5595T>C), GAT655AAA (5596G>A 5598T>A), GTT656ATT (5599G>A), CAC658AAT (5605C>A 5607C>T), GGA661GGT (5616A>T), ACC662TCT (5617A>T 5619C>T), GAC663GAT (5622C>T), AAC664AAT (5625C>T), CAC665GTT (5626C>G 5627A>T 5628C>T), GCG667GCA (5634G>A), GAC668GAT (5637C>T), CTT670TTA (5641C>T 5643T>A) | | | | | | | | |

##### Proteins

| | Begin | End | Coverage | Score | Concordance | Matches | Identities | I/D/M/F* | Stop Codons |
|---|---|---|---|---|---|---|---|---|---|
| reverse transcriptase (NP_056728.1) | 275 | 672 | 35.6% | 792 | 48.5% | 235 (97.1%) | 109 (45.0%) | 0/7/0/0 | 0 |

| | Begin | End | Coverage | Score | Concordance | Matches | Identities | I/D/M/F* | Stop Codons |
|---|---|---|---|---|---|---|---|---|---|
| NT | 4456 | 5647 | 9.0% | 149 | 10.9% | 704 (97.1%) | 399 (55.0%) | 0/21 | |
| Protein mutations: | S278C (4465A>T 4467T>C), K279R (4469A>G), M283S (4481T>G 4482G>C), A284S (4483G>A 4484C>G 4485A>T), L288M (4495T>A), N290R (4502A>G 4503C>A), E292H (4507G>C 4509A>T), K295Q (4516A>C 4518G>A), R296K (4519C>A 4520G>A), K300A (4531A>G 4532A>C 4533A>T), R301Q (4535G>A 4536T>A), V304I (4543G>A 4545C>A), A308E (4556C>A 4557T>G), M309L (4558A>C 4560G>T), A312N (4567G>A 4568C>A), T313C (4570A>T 4571C>G), G315F (4576G>T 4577G>T 4578A>T), A317G (4583C>G 4584C>A), N319F (4588A>T 4589A>T), L320I (4591C>A), D324N (4603G>A), E325N (4606G>A 4608G>T), L327T (4612C>A 4613T>C), T328N (4616C>A 4617A>T), I330A (4621A>G 4622T>C 4623T>A), R331K (4624C>A 4625G>A), G332R (4627G>A), K334I (4634A>T 4635G>T), I335Y (4636A>T 4637T>A), S338K (4645T>A 4646C>A 4647C>A), I500K (5132T>A 5133C>G), K502N (5139G>T), P503L (5141C>T), A506R (5149G>A 5150C>G 5151C>G), K507L (5152A>C 5153A>T 5154G>T), L508I (5155C>A), K509R (5158A>C 5159A>G), E510K (5161G>A), N511P (5164A>C 5165A>C), V512N (5167G>A 5168T>A 5169T>C), P513R (5171C>G), W514M (5173T>A 5174G>T), R515G (5176A>G), K518D (5185A>G 5187A>T), D520H (5191G>C), L522R (5197C>A 5198T>G 5199C>A), Y523C (5201A>G 5202C>T), M524V (5203A>G 5205G>T), K526S (5210A>G 5211G>C), V527L (5212G>C 5214G>A), K529Q (5218A>C), N530E (5221A>G 5223T>A), L531C (5224C>T 5225T>G 5226G>T), Q532S (5227C>T 5228A>C 5229A>C), G533K (5230G>A 5231G>A), F534L (5235T>A), P536K (5239C>A 5240C>A 5241A>G), H538R (5245C>A 5246A>G 5247T>A), H539L (5249A>T), L541E (5254T>G 5255T>A), E544D (5265G>C), K545N (5268G>C), I547V (5272A>G 5274C>T), I548L (5275A>T 5277C>A), E549Q (5278G>C 5280G>A), D555Y (5296G>T), Y556H (5299T>C 5301C>T), G559A (5309G>C 5310T>A), M560L (5311A>T 5313G>A), K562Q (5317A>C), A563T (5320G>A), I564del (5323_5325delATC), K565D (5326A>G 5328A>C), I566L (5329A>C), G569_L574del (5338_5355delGGTACTAATACTGAGTTA), A579T (5368G>A), S582T (5378G>C 5379C>A), K584N (5385A>T), A585E (5387C>A 5388T>A), A586T (5389G>A 5391A>C), K588T (5396A>C 5397G>T), N589K (5400T>A), H591S (5404C>T 5405A>C 5406C>A), S592T (5408G>C 5409C>A), D594E (5415C>A), T597L (5422A>T 5423C>T), V600I (5431G>A), I601V (5434A>G), N602R (5438A>G 5439T>A), T603G (5440A>G 5441C>G 5442T>C), K605R (5446A>C 5447A>G 5448G>A), I609A (5458A>G 5459T>C), Y610F (5462A>T 5463T>C), T612L (5467A>C 5468C>T 5469T>A), V614K (5473G>A 5474T>A 5475T>G), H615Q (5478T>A), L617I (5482C>A 5484G>C), T620S (5491A>T), H624Q (5505T>G), F625V (5506T>G 5508C>A), K626S (5509A>T 5510A>C 5511G>A), S627G (5512A>G), F628L (5517C>G), V629I (5518G>A 5520T>A), N630F (5521A>T 5522A>T), L631N (5524C>A 5525T>A 5526C>T), N632K (5529T>A), Y633L (5530T>C 5531A>T 5532C>T), K634P (5533A>C 5534A>C), G635S (5536G>T 5537G>C 5538A>T), D636E (5541T>A), S637P (5542T>C 5544G>T), K638Q (5545A>C), L639Y (5548C>T 5549T>A), G640R (5551G>A), N642L (5557A>T 5558A>T 5559C>A), I643H (5560A>C 5561T>A 5562C>T), A647V (5573C>T), W648L (5576G>T), H651F (5584C>T 5585A>T 5586C>T), S653T (5590T>A 5592A>C), D655K (5596G>A 5598T>A), V656I (5599G>A), H658N (5605C>A 5607C>T), T662S (5617A>T 5619C>T), H665V (5626C>G 5627A>T 5628C>T) | | | | | | | | |
| Codon mutations: | AGT278TGC (4465A>T 4467T>C), AAA279AGA (4469A>G), AGC280AGT (4473C>T), CCT281CCA (4476T>A), ATG283AGC (4481T>G 4482G>C), GCA284AGT (4483G>A 4484C>G 4485A>T), CCA285CCC (4488A>C), GCC286GCG (4491C>G), TTG288ATG (4495T>A), GTC289GTA (4500C>A), AAC290AGA (4502A>G 4503C>A), AAT291AAC (4506T>C), GAA292CAT (4507G>C 4509A>T), GCC293GCG (4512C>G), GAG294GAA (4515G>A), AAG295CAA (4516A>C 4518G>A), CGA296AAA (4519C>A 4520G>A), AGA297CGA (4522A>C), GGA298GGT (4527A>T), AAA300GCT (4531A>G 4532A>C 4533A>T), CGT301CAA (4535G>A 4536T>A), GTA303GTC (4542A>C), GTC304ATA (4543G>A 4545C>A), AAC305AAT (4548C>T), TAC306TAT (4551C>T), GCT308GAG (4556C>A 4557T>G), ATG309CTT (4558A>C 4560G>T), AAC310AAT (4563C>T), GCT312AAT (4567G>A 4568C>A), ACT313TGT (4570A>T 4571C>G), GTA314GTT (4575A>T), GGA315TTT (4576G>T 4577G>T 4578A>T), GAT316GAC (4581T>C), GCC317GGA (4583C>G 4584C>A), AAT319TTT (4588A>T 4589A>T), CTT320ATT (4591C>A), CCC321CCT (4596C>T), GAC324AAC (4603G>A), GAG325AAT (4606G>A 4608G>T), TTA326CTT (4609T>C 4611A>T), CTT327ACT (4612C>A 4613T>C), ACA328AAT (4616C>A 4617A>T), CTC329CTA (4620C>A), ATT330GCA (4621A>G 4622T>C 4623T>A), CGA331AAA (4624C>A 4625G>A), GGA332AGA (4627G>A), AAG333AAA (4632G>A), AAG334ATT (4634A>T 4635G>T), ATC335TAC (4636A>T 4637T>A), TTC336TTT (4641C>T), TCT337TCA (4644T>A), TCC338AAA (4645T>A 4646C>A 4647C>A), TTC339TTT (4650C>T), GAC340GAT (4653C>T), TCA343AG. (4660T>A 4661C>G), ATC500AAG (5132T>A 5133C>G), AAG502AAT (5139G>T), CCT503CTT (5141C>T), CTG504TTA (5143C>T 5145G>A), GCC506AGG (5149G>A 5150C>G 5151C>G), AAG507CTT (5152A>C 5153A>T 5154G>T), CTT508ATT (5155C>A), AAA509CGA (5158A>C 5159A>G), GAA510AAA (5161G>A), AAC511CCC (5164A>C 5165A>C), GTT512AAC (5167G>A 5168T>A 5169T>C), CCA513CGA (5171C>G), TGG514ATG (5173T>A 5174G>T), AGA515GGA (5176A>G), AAA518GAT (5185A>G 5187A>T), GAG519GAA (5190G>A), GAT520CAT (5191G>C), ACC521ACT (5196C>T), CTC522AGA (5197C>A 5198T>G 5199C>A), TAC523TGT (5201A>G 5202C>T), ATG524GTT (5203A>G 5205G>T), AAG526AGC (5210A>G 5211G>C), GTG527CTA (5212G>C 5214G>A), AAG528AAA (5217G>A), AAA529CAA (5218A>C), AAT530GAA (5221A>G 5223T>A), CTG531TGT (5224C>T 5225T>G 5226G>T), CAA532TCC (5227C>T 5228A>C 5229A>C), GGA533AAA (5230G>A 5231G>A), TTT534TTA (5235T>A), CCT535CCA (5238T>A), CCA536AAG (5239C>A 5240C>A 5241A>G), CAT538AGA (5245C>A 5246A>G 5247T>A), CAT539CTT (5249A>T), CCC540CCG (5253C>G), TTA541GAA (5254T>G 5255T>A), GAG543GAA (5262G>A), GAG544GAC (5265G>C), AAG545AAC (5268G>C), CTG546TTA (5269C>T 5271G>A), ATC547GTT (5272A>G 5274C>T), ATC548TTA (5275A>T 5277C>A), GAG549CAA (5278G>C 5280G>A), ACC550ACA (5283C>A), GCA552GCT (5289A>T), GAC554GAT (5295C>T), GAC555TAC (5296G>T), TAC556CAT (5299T>C 5301C>T), GGT559GCA (5309G>C 5310T>A), ATG560TTA (5311A>T 5313G>A), AAA562CAA (5317A>C), GCT563ACT (5320G>A), ATC564del (5323_5325delATC), AAA565GAC (5326A>G 5328A>C), ATT566CTT (5329A>C), AAC567AAT (5334C>T), GGT569_TTA574del (5338_5355delGGTACTAATACTGAGTTA), TGC576TGT (5361C>T), AGA577AGG (5364A>G), TAC578TAT (5367C>T), GCA579ACA (5368G>A), TCT580AGT (5371T>A 5372C>G), GGA581GGT (5376A>T), AGC582ACA (5378G>C 5379C>A), AAA584AAT (5385A>T), GCT585GAA (5387C>A 5388T>A), GCA586ACC (5389G>A 5391A>C), AAG588ACT (5396A>C 5397G>T), AAT589AAA (5400T>A), CAC591TCA (5404C>T 5405A>C 5406C>A), AGC592ACA (5408G>C 5409C>A), GAC594GAA (5415C>A), GAG596GAA (5421G>A), ACA597TTA (5422A>T 5423C>T), GCG599GCC (5430G>C), GTA600ATA (5431G>A), ATA601GTA (5434A>G), AAT602AGA (5438A>G 5439T>A), ACT603GGC (5440A>G 5441C>G 5442T>C), AAG605CGA (5446A>C 5447A>G 5448G>A), AGT608TCT (5455A>T 5456G>C), ATT609GCT (5458A>G 5459T>C), TAT610TTC (5462A>T 5463T>C), ACT612CTA (5467A>C 5468C>T 5469T>A), CCT613CCA (5472T>A), GTT614AAG (5473G>A 5474T>A 5475T>G), CAT615CAA (5478T>A), TTT616TTC (5481T>C), CTG617ATC (5482C>A 5484G>C), ATT618ATC (5487T>C), AGG619AGA (5490G>A), ACA620TCA (5491A>T), GAT621GAC (5496T>C), ACT623ACA (5502T>A), CAT624CAG (5505T>G), TTC625GTA (5506T>G 5508C>A), AAG626TCA (5509A>T 5510A>C 5511G>A), AGT627GGT (5512A>G), TTC628TTG (5517C>G), GTT629ATA (5518G>A 5520T>A), AAT630TTT (5521A>T 5522A>T), CTC631AAT (5524C>A 5525T>A 5526C>T), AAT632AAA (5529T>A), TAC633CTT (5530T>C 5531A>T 5532C>T), AAA634CCA (5533A>C 5534A>C), GGA635TCT (5536G>T 5537G>C 5538A>T), GAT636GAA (5541T>A), TCG637CCT (5542T>C 5544G>T), AAA638CAA (5545A>C), CTT639TAT (5548C>T 5549T>A), GGA640AGA (5551G>A), AAC642TTA (5557A>T 5558A>T 5559C>A), ATC643CAT (5560A>C 5561T>A 5562C>T), GCA647GTA (5573C>T), TGG648TTG (5576G>T), CTT649TTA (5578C>T 5580T>A), AGC650TCT (5581A>T 5582G>C 5583C>T), CAC651TTT (5584C>T 5585A>T 5586C>T), TCA653ACC (5590T>A 5592A>C), TTT654TTC (5595T>C), GAT655AAA (5596G>A 5598T>A), GTT656ATT (5599G>A), CAC658AAT (5605C>A 5607C>T), GGA661GGT (5616A>T), ACC662TCT (5617A>T 5619C>T), GAC663GAT (5622C>T), AAC664AAT (5625C>T), CAC665GTT (5626C>G 5627A>T 5628C>T), GCG667GCA (5634G>A), GAC668GAT (5637C>T), CTT670TTA (5641C>T 5643T>A) | | | | | | | | |

*: Inserts / Deletes / Misaligned / Frameshifts

#### Analysis details

This analysis was performed with panviral2.64

### NGS Details (UN8): Caulimovirus venafragariae

#### Assembly

| | |
|---|---|
| **Coverage Length** | 425 (1 contig(s)) |
| **Depth Of Coverage** | 11.4 |
| **Number Of Reads** | 39 |
| **Reads Per Million** | 0.65 rpm (after QC) |
| **Ambiguities** | 0 |
| **Assembly Method** | de novo + reference guided assembly |
| **Consensus Caller** | Bcf Tools |

#### Coverage Map

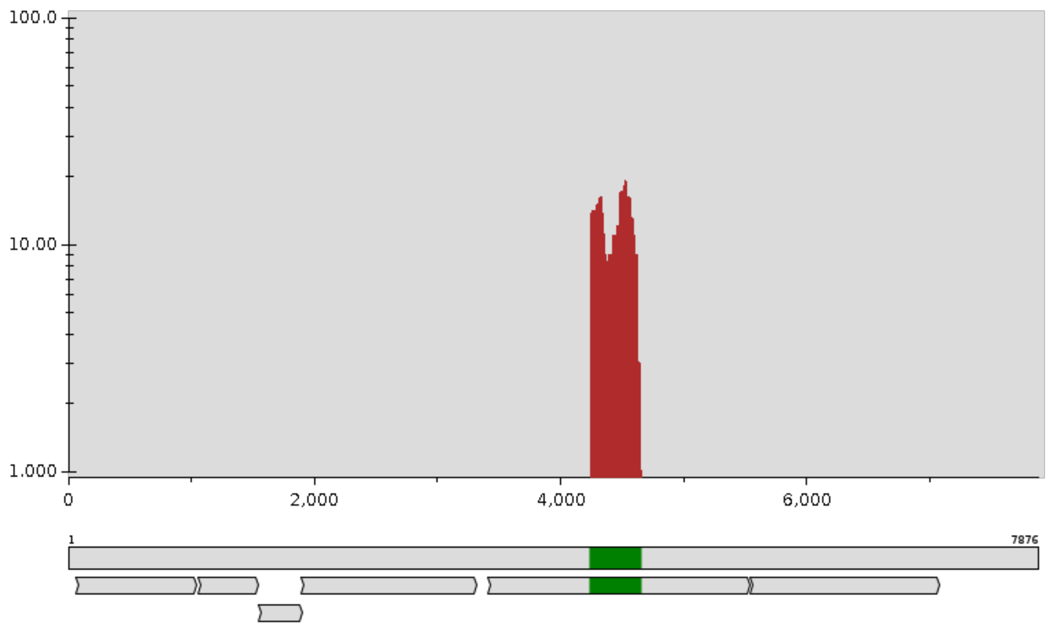

#### Assignment

| | |
|---|---|
| **Type** | Caulimovirus venafragariae (Taxonomy ID: 3048344) |
| **Reference Genome** | NC_001725.1 |
| **NT Identity (%)** | 64.0662 |
| **AA Identity (%)** | 67.6056 |
| **Number Of Stop Codons** | 0 |
| **Number Of CDS** | 6 |

#### Alignment

| | |
|---|---|
| **Alignment Score** | 227.0 (NT) + 590.0 (AA) = 817.0 |
| **Concordance (%)** | 48.2386 |

| **Alignment Method** | Global, seeded, nucleotide + amino acids (AGA) |
|---|---|

#### Genome Region

Sequence starts at position 4236 and ends at position 4660 relative to NC_001725.1 reference sequence.

#### Alignment Detailed Statistics

| | Begin | End | Coverage | Score | Concordance | Matches | Identities | I/D/M/F* | Stop Codons |
|---|---|---|---|---|---|---|---|---|---|
| **NT** | **4236** | **4660** | **5.4%** | **227** | **27.2%** | **423 (99.5%)** | **271 (63.8%)** | **0/2** | |
| Mutations: | 4236A>C, 4242T>A, 4244A>G, 4245G>A, 4246T>G, 4247G>T, 4259A>T, 4260G>A, 4264T>A, 4265C>A, 4266A>T, 4267C>G, 4268C>A, 4269T>G, 4275C>T, 4276C>A, 4277G>A, 4279G>A, 4280A>C, 4281A>T, 4284G>A, 4292C>T, 4299C>T, 4300G>C, 4306C>T, 4309C>T, 4311C>A, 4314G>T, 4315C>A, 4317T>G, 4318G>A, 4319G>A, 4320T>A, 4321A>T, 4323C>A, 4326C>A, 4330_4331delCC, 4332T>C, 4335C>T, 4338G>A, 4341T>C, 4344A>G, 4348T>A, 4349C>G, 4350C>T, 4351T>A, 4352C>G, 4353A>T, 4359C>A, 4371G>A, 4374T>C, 4380C>G, 4381G>A, 4383G>A, 4384A>C, 4385T>A, 4386C>A, 4395C>A, 4401A>T, 4402A>C, 4413T>A, 4423A>G, 4426T>C, 4428A>T, 4432G>A, 4434C>A, 4435C>A, 4438A>T, 4439C>G, 4440A>T, 4441A>G, 4442A>T, 4443G>C, 4444G>T, 4445G>T, 4446A>C, 4449T>C, 4452C>A, 4455C>T, 4456C>T, 4458A>C, 4459C>A, 4461T>C, 4464T>A, 4467C>T, 4470G>A, 4471G>A, 4473G>T, 4474C>A, 4476A>C, 4479G>T, 4480C>A, 4482T>A, 4483C>A, 4485A>T, 4486A>C, 4487G>T, 4488A>T, 4489A>G, 4490T>C, 4491C>A, 4492G>A, 4493G>A, 4497T>A, 4500G>A, 4505T>A, 4506T>C, 4508A>T, 4509C>T, 4512T>A, 4513T>A, 4514C>A, 4515C>A, 4521C>T, 4524T>C, 4528T>A, 4529C>G, 4542G>A, 4543G>A, 4546C>A, 4547G>A, 4548C>A, 4551T>A, 4553C>A, 4554T>A, 4555C>G, 4556C>A, 4558G>A, 4560A>T, 4561A>T, 4563A>T, 4566T>A, 4567C>G, 4568A>C, 4569G>T, 4570C>T, 4575C>A, 4578T>A, 4584C>T, 4585T>A, 4586G>C, 4587T>A, 4590C>A, 4591C>A, 4593A>T, 4614C>G, 4633C>A, 4635A>T, 4642G>C, 4643C>A, 4644T>A, 4647T>C, 4650C>T | | | | | | | | |

##### CDS

| | Begin | End | Coverage | Score | Concordance | Matches | Identities | I/D/M/F* | Stop Codons |
|---|---|---|---|---|---|---|---|---|---|
| ORF_V | 278 | 419 | 20.0% | 590 | 57.0% | 142 (100%) | 96 (67.6%) | 0/0/1/1 | 0 |
| Protein mutations: | K280R (4244A>G 4245G>A), C281V (4246T>G 4247G>T), Q285L (4259A>T 4260G>A), S287N (4264T>A 4265C>A 4266A>T), P288E (4267C>G 4268C>A 4269T>G), R291K (4276C>A 4277G>A), E292T (4279G>A 4280A>C 4281A>T), T296I (4292C>T), E299Q (4300G>C), K303N (4314G>T), L304M (4315C>A 4317T>G), G305K (4318G>A 4319G>A 4320T>A), I306L (4321A>T 4323C>A), E326K (4381G>A 4383G>A), I327Q (4384A>C 4385T>A 4386C>A), K340E (4423A>G), D343K (4432G>A 4434C>A), H344N (4435C>A), T345C (4438A>T 4439C>G 4440A>T), K346V (4441A>G 4442A>T 4443G>C), G347F (4444G>T 4445G>T 4446A>C), L351F (4456C>T 4458A>C), L352I (4459C>A 4461T>C), E356N (4471G>A 4473G>T), Q357N (4474C>A 4476A>C), L359I (4480C>A 4482T>A), Q360N (4483C>A 4485A>T), R361L (4486A>C 4487G>T 4488A>T), I362A (4489A>G 4490T>C 4491C>A), G363K (4492G>A 4493G>A), F367Y (4505T>A 4506T>C), Y368F (4508A>T 4509C>T), S370K (4513T>A 4514C>A 4515C>A), V380I (4543G>A), R381K (4546C>A 4547G>A 4548C>A), A383E (4553C>A 4554T>A), P384E (4555C>G 4556C>A), E385N (4558G>A 4560A>T), T386S (4561A>T 4563A>T), Q388A (4567C>G 4568A>C 4569G>T), C394T (4585T>A 4586G>C 4587T>A), Q396N (4591C>A 4593A>T), Q410N (4633C>A 4635A>T), A413Q (4642G>C 4643C>A 4644T>A) | | | | | | | | |
| Codon mutations: | AAA277..C (4236A>C), ATT279ATA (4242T>A), AAG280AGA (4244A>G 4245G>A), TGT281GTT (4246T>G 4247G>T), CAG285CTA (4259A>T 4260G>A), TCA287AAT (4264T>A 4265C>A 4266A>T), CCT288GAG (4267C>G 4268C>A 4269T>G), GAC290GAT (4275C>T), CGA291AAA (4276C>A 4277G>A), GAA292ACT (4279G>A 4280A>C 4281A>T), GAG293GAA (4284G>A), ACT296ATT (4292C>T), ATC298ATT (4299C>T), GAA299CAA (4300G>C), CTA301TTA (4306C>T), CTC302TTA (4309C>T 4311C>A), AAG303AAT (4314G>T), CTT304ATG (4315C>A 4317T>G), GGT305AAA (4318G>A 4319G>A 4320T>A), ATC306TTA (4321A>T 4323C>A), ATC307ATA (4326C>A), CCT309--C (4330_4331delCC 4332T>C), AGC310AGT (4335C>T), AAG311AAA (4338G>A), AGT312AGC (4341T>C), CCA313CCG (4344A>G), TCC315AGT (4348T>A 4349C>G 4350C>T), TCA316AGT (4351T>A 4352C>G 4353A>T), GCC318GCA (4359C>A), AGG322AGA (4371G>A), AAT323AAC (4374T>C), GCC325GCG (4380C>G), GAG326AAA (4381G>A 4383G>A), ATC327CAA (4384A>C 4385T>A 4386C>A), GGC330GGA (4395C>A), GCA332GCT (4401A>T), AGA333CGA (4402A>C), ATT336ATA (4413T>A), AAG340GAG (4423A>G), TTA341CTT (4426T>C 4428A>T), GAC343AAA (4432G>A 4434C>A), CAT344AAT (4435C>A), ACA345TGT (4438A>T 4439C>G 4440A>T), AAG346GTC (4441A>G 4442A>T 4443G>C), GGA347TTC (4444G>T 4445G>T 4446A>C), GAT348GAC (4449T>C), GGC349GGA (4452C>A), TAC350TAT (4455C>T), CTA351TTC (4456C>T 4458A>C), CTT352ATC (4459C>A 4461T>C), CCT353CCA (4464T>A), AAC354AAT (4467C>T), AAG355AAA (4470G>A), GAG356AAT (4471G>A 4473G>T), CAA357AAC (4474C>A 4476A>C), CTG358CTT (4479G>T), CTT359ATA (4480C>A 4482T>A), CAA360AAT (4483C>A 4485A>T), AGA361CTT (4486A>C 4487G>T 4488A>T), ATC362GCA (4489A>G 4490T>C 4491C>A), GGA363AAA (4492G>A 4493G>A), GGT364GGA (4497T>A), AAG365AAA (4500G>A), TTT367TAC (4505T>A 4506T>C), TAC368TTT (4508A>T 4509C>T), TCT369TCA (4512T>A), TCC370AAA (4513T>A 4514C>A 4515C>A), GAC372GAT (4521C>T), TGT373TGC (4524T>C), TCT375AGT (4528T>A 4529C>G), CAG379CAA (4542G>A), GTA380ATA (4543G>A), CGC381AAA (4546C>A 4547G>A 4548C>A), CTT382CTA (4551T>A), GCT383GAA (4553C>A 4554T>A), CCA384GAA (4555C>G 4556C>A), GAA385AAT (4558G>A 4560A>T), ACA386TCT (4561A>T 4563A>T), ATT387ATA (4566T>A), CAG388GCT (4567C>G 4568A>C 4569G>T), CTA389TTA (4570C>T), ACC390ACA (4575C>A), GCT391GCA (4578T>A), AGC393AGT (4584C>T), TGT394ACA (4585T>A 4586G>C 4587T>A), CCC395CCA (4590C>A), CAA396AAT (4591C>A 4593A>T), GTC403GTG (4614C>G), CAA410AAT (4633C>A 4635A>T), GCT413CAA (4642G>C 4643C>A 4644T>A), ATT414ATC (4647T>C), TTC415TTT (4650C>T) | | | | | | | | |

##### Proteins

| | Begin | End | Coverage | Score | Concordance | Matches | Identities | I/D/M/F* | Stop Codons |
|---|---|---|---|---|---|---|---|---|---|
| hypothetical protein (NP_043933.1) | 278 | 419 | 20.0% | 590 | 57.0% | 142 (100%) | 96 (67.6%) | 0/0/1/1 | 0 |
| Protein mutations: | K280R (4244A>G 4245G>A), C281V (4246T>G 4247G>T), Q285L (4259A>T 4260G>A), S287N (4264T>A 4265C>A 4266A>T), P288E (4267C>G 4268C>A 4269T>G), R291K (4276C>A 4277G>A), E292T (4279G>A 4280A>C 4281A>T), T296I (4292C>T), E299Q (4300G>C), K303N (4314G>T), L304M (4315C>A 4317T>G), G305K (4318G>A 4319G>A 4320T>A), I306L (4321A>T 4323C>A), E326K (4381G>A 4383G>A), I327Q (4384A>C 4385T>A 4386C>A), K340E (4423A>G), D343K (4432G>A 4434C>A), H344N (4435C>A), T345C (4438A>T 4439C>G 4440A>T), K346V (4441A>G 4442A>T 4443G>C), G347F (4444G>T 4445G>T 4446A>C), L351F (4456C>T 4458A>C), L352I (4459C>A 4461T>C), E356N (4471G>A 4473G>T), Q357N (4474C>A 4476A>C), L359I (4480C>A 4482T>A), Q360N (4483C>A 4485A>T), R361L (4486A>C 4487G>T 4488A>T), I362A (4489A>G 4490T>C 4491C>A), G363K (4492G>A 4493G>A), F367Y (4505T>A 4506T>C), Y368F (4508A>T 4509C>T), S370K (4513T>A 4514C>A 4515C>A), V380I (4543G>A), R381K (4546C>A 4547G>A 4548C>A), A383E (4553C>A 4554T>A), P384E (4555C>G 4556C>A), E385N (4558G>A 4560A>T), T386S (4561A>T 4563A>T), Q388A (4567C>G 4568A>C 4569G>T), C394T (4585T>A 4586G>C 4587T>A), Q396N (4591C>A 4593A>T), Q410N (4633C>A 4635A>T), A413Q (4642G>C 4643C>A 4644T>A) | | | | | | | | |
| Codon mutations: | AAA277..C (4236A>C), ATT279ATA (4242T>A), AAG280AGA (4244A>G 4245G>A), TGT281GTT (4246T>G 4247G>T), CAG285CTA (4259A>T 4260G>A), TCA287AAT (4264T>A 4265C>A 4266A>T), CCT288GAG (4267C>G 4268C>A 4269T>G), GAC290GAT (4275C>T), CGA291AAA (4276C>A 4277G>A), GAA292ACT (4279G>A 4280A>C 4281A>T), GAG293GAA (4284G>A), ACT296ATT (4292C>T), ATC298ATT (4299C>T), GAA299CAA (4300G>C), CTA301TTA (4306C>T), CTC302TTA (4309C>T 4311C>A), AAG303AAT (4314G>T), CTT304ATG (4315C>A 4317T>G), GGT305AAA (4318G>A 4319G>A 4320T>A), ATC306TTA (4321A>T 4323C>A), ATC307ATA (4326C>A), CCT309--C (4330_4331delCC 4332T>C), AGC310AGT (4335C>T), AAG311AAA (4338G>A), AGT312AGC (4341T>C), CCA313CCG (4344A>G), TCC315AGT (4348T>A 4349C>G 4350C>T), TCA316AGT (4351T>A 4352C>G 4353A>T), GCC318GCA (4359C>A), AGG322AGA (4371G>A), AAT323AAC (4374T>C), GCC325GCG (4380C>G), GAG326AAA (4381G>A 4383G>A), ATC327CAA (4384A>C 4385T>A 4386C>A), GGC330GGA (4395C>A), GCA332GCT (4401A>T), AGA333CGA (4402A>C), ATT336ATA (4413T>A), AAG340GAG (4423A>G), TTA341CTT (4426T>C 4428A>T), GAC343AAA (4432G>A 4434C>A), CAT344AAT (4435C>A), ACA345TGT (4438A>T 4439C>G 4440A>T), AAG346GTC (4441A>G 4442A>T 4443G>C), GGA347TTC (4444G>T 4445G>T 4446A>C), GAT348GAC (4449T>C), GGC349GGA (4452C>A), TAC350TAT (4455C>T), CTA351TTC (4456C>T 4458A>C), CTT352ATC (4459C>A 4461T>C), CCT353CCA (4464T>A), AAC354AAT (4467C>T), AAG355AAA (4470G>A), GAG356AAT (4471G>A 4473G>T), CAA357AAC (4474C>A 4476A>C), CTG358CTT (4479G>T), CTT359ATA (4480C>A 4482T>A), CAA360AAT (4483C>A 4485A>T), AGA361CTT (4486A>C 4487G>T 4488A>T), ATC362GCA (4489A>G 4490T>C 4491C>A), GGA363AAA (4492G>A 4493G>A), GGT364GGA (4497T>A), AAG365AAA (4500G>A), TTT367TAC (4505T>A 4506T>C), TAC368TTT (4508A>T 4509C>T), TCT369TCA (4512T>A), TCC370AAA (4513T>A 4514C>A 4515C>A), GAC372GAT (4521C>T), TGT373TGC (4524T>C), TCT375AGT (4528T>A 4529C>G), CAG379CAA (4542G>A), GTA380ATA (4543G>A), CGC381AAA (4546C>A 4547G>A 4548C>A), CTT382CTA (4551T>A), GCT383GAA (4553C>A 4554T>A), CCA384GAA (4555C>G 4556C>A), GAA385AAT (4558G>A 4560A>T), ACA386TCT (4561A>T 4563A>T), ATT387ATA (4566T>A), CAG388GCT (4567C>G 4568A>C 4569G>T), CTA389TTA (4570C>T), ACC390ACA (4575C>A), GCT391GCA (4578T>A), AGC393AGT (4584C>T), TGT394ACA (4585T>A 4586G>C 4587T>A), CCC395CCA (4590C>A), CAA396AAT (4591C>A 4593A>T), GTC403GTG (4614C>G), CAA410AAT (4633C>A 4635A>T), GCT413CAA (4642G>C 4643C>A 4644T>A), ATT414ATC (4647T>C), TTC415TTT (4650C>T) | | | | | | | | |

*: Inserts / Deletes / Misaligned / Frameshifts

#### Analysis details

This analysis was performed with panviral2.64

#### NGS Details (UN8): Caulimovirus deformatiolamii

##### Assembly

| | |
|---|---|
| **Coverage Length** | 576 (2 contig(s)) |
| **Depth Of Coverage** | 5.8 |
| **Number Of Reads** | 27 |
| **Reads Per Million** | 0.45 rpm (after QC) |
| **Ambiguities** | 0 |
| **Assembly Method** | de novo + reference guided assembly |
| **Consensus Caller** | Bcf Tools |

##### Coverage Map

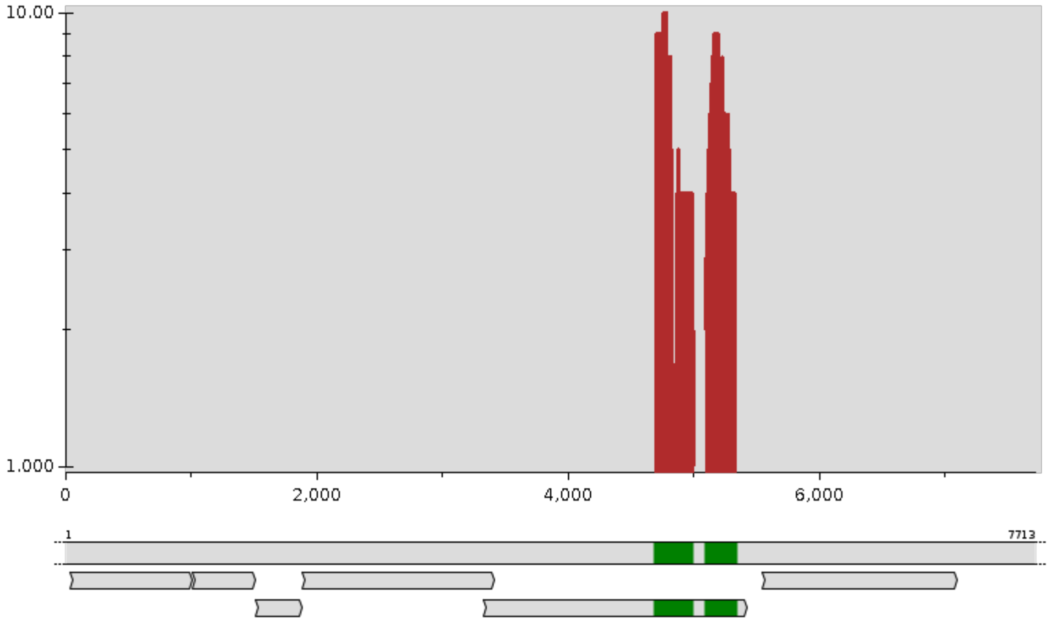

##### Assignment

| | |
|---|---|
| **Type** | Caulimovirus deformatiolamii (Taxonomy ID: 3048173) |
| **Reference Genome** | NC_010737.1 |
| **NT Identity (%)** | 54.4503 |
| **AA Identity (%)** | 45.0262 |
| **Number Of Stop Codons** | 0 |
| **Number Of CDS** | 6 |

##### Alignment

| | |
|---|---|
| **Alignment Score** | 81.0 (NT) + 632.0 (AA) = 713.0 |
| **Concordance (%)** | 28.7964 |

| Alignment Method | Global, seeded, nucleotide + amino acids (AGA) |
|---|---|

#### Genome Region

Sequence starts at position 4686 and ends at position 5347 relative to NC_010737.1 reference sequence.

#### Alignment Detailed Statistics

| | Begin | End | Coverage | Score | Concordance | Matches | Identities | I/D/M/F* | Stop Codons |
|---|---|---|---|---|---|---|---|---|---|
| **NT** | **4686** | **5347** | **7.5%** | **81** | **7.2%** | **570 (98.4%)** | **312 (53.9%)** | **3/6** | |
| Mutations: | 4690G>A, 4691C>T, 4692A>G, 4693C>A, 4696C>T, 4698A>T, 4702A>C, 4704G>A, 4705T>A, 4706G>C, 4707C>A, 4709G>A, 4713T>C, 4715A>G, 4716C>A, 4725C>G, 4727T>A, 4732A>T, 4734T>A, 4735G>T, 4737C>T, 4738G>C, 4743G>A, 4746T>A, 4747C>T, 4749C>T, 4750C>A, 4751G>A, 4752T>A, 4755T>C, 4756C>A, 4757A>T, 4758G>C, 4759A>T, 4760A>G, 4761T>C, 4762C>A, 4763A>G, 4764T>A, 4765A>G, 4766T>C, 4767T>C, 4768C>A, 4774C>T, 4775A>T, 4777A>T, 4778T>C, 4780C>G, 4781A>G, 4785G>A, 4791A>T, 4795A>G, 4800T>C, 4801G>A, 4802A>C, 4803A>T, 4811A>C, 4812A>T, 4816C>T, 4818T>A, 4821A>G, 4827T>C, 4828C>T, 4830T>G, 4833T>A, 4835T>G, 4836T>C, 4837T>C, 4839A>C, 4841C>A, 4847C>T, 4848C>T, 4849A>G, 4850G>C, 4851T>A, 4854T>C, 4856A>T, 4857T>C, 4864C>A, 4867T>G, 4870G>A, 4871C>G, 4873T>C, 4874C>A, 4875T>G, 4876A>G, 4878G>T, 4879C>T, 4881T>C, 4882G>A, 4883C>A, 4884T>A, 4887T>C, 4888T>C, 4890G>C, 4891C>T, 4893G>T, 4896A>T, 4898A>G, 4899G>A, 4902C>T, 4906G>C, 4908A>G, 4908_4909insACC, 4909G>C, 4910A>C, 4917A>C, 4923T>A, 4924T>G, 4925G>A, 4927A>G, 4929G>A, 4932T>A, 4933T>A, 4936G>C, 4937A>T, 4938C>A, 4939A>G, 4941T>G, 4942_4947delGAGTAT, 4951G>C, 4953A>G, 4954G>A, 4962G>A, 4965A>G, 4967G>T, 4968T>G, 4969C>G, 4973C>T, 4974A>G, 4975G>A, 4978T>C, 4980T>C, 4983T>A, 4984A>T, 4985A>G, 4986G>T, 4987C>T, 4990C>G, 4991A>G, 4992T>A, 5088T>A, 5091C>T, 5094C>T, 5097G>A, 5103C>T, 5104T>A, 5105C>G, 5106A>T, 5109T>A, 5111G>C, 5112T>C, 5118A>T, 5119A>G, 5121G>A, 5122G>A, 5128T>A, 5129T>C, 5130A>T, 5132A>G, 5133T>G, 5136T>C, 5137C>T, 5138A>T, 5139T>A, 5140T>A, 5142T>A, 5155A>T, 5157T>A, 5160T>A, 5163A>T, 5164G>A, 5166T>A, 5167A>G, 5169C>A, 5174T>G, 5175T>C, 5178C>A, 5179G>C, 5180C>G, 5181T>A, 5187C>T, 5188A>T, 5191A>G, 5192T>C, 5193T>A, 5195A>T, 5196T>C, 5197C>T, 5200A>C, 5201C>T, 5202T>A, 5205T>A, 5206C>A, 5207T>A, 5208T>G, 5209G>C, 5211G>A, 5214T>C, 5215C>A, 5217G>C, 5220A>C, 5224A>T, 5226A>G, 5229T>C, 5232C>T, 5234A>C, 5236A>C, 5238C>G, 5239T>G, 5241T>A, 5242A>T, 5244C>A, 5245T>G, 5246T>G, 5248T>C, 5250T>A, 5253G>A, 5254A>T, 5255A>T, 5258C>A, 5259C>T, 5262T>A, 5263G>C, 5265G>T, 5266A>T, 5267A>C, 5269G>T, 5270G>C, 5274T>A, 5275T>C, 5276A>C, 5278A>C, 5281C>T, 5283G>T, 5284G>A, 5286A>G, 5290C>T, 5292T>A, 5293G>C, 5294T>A, 5295C>T, 5298G>A, 5304A>G, 5305C>G, 5306A>T, 5309G>T, 5311C>T, 5316C>T, 5317A>T, 5318G>T, 5319A>T, 5322C>T, 5323A>T, 5324G>C, 5331G>A, 5332G>A, 5338C>A, 5340C>T, 5345C>A | | | | | | | | |

##### CDS

| | Begin | End | Coverage | Score | Concordance | Matches | Identities | I/D/M/F* | Stop Codons |
|---|---|---|---|---|---|---|---|---|---|
| LLDV_gp5 | 455 | 675 | 27.5% | 632 | 46.1% | 190 (98.4%) | 86 (44.6%) | 1/2/0/0 | 0 |
| Protein mutations: | A456M (4690G>A 4691C>T 4692A>G), Q457K (4693C>A), L458F (4696C>T 4698A>T), K460Q (4702A>C 4704G>A), C461T (4705T>A 4706G>C 4707C>A), R462K (4709G>A), N464R (4715A>G 4716C>A), L468H (4727T>A), I470L (4732A>T 4734T>A), D471Y (4735G>T 4737C>T), E472Q (4738G>C), H475Y (4747C>T 4749C>T), R476K (4750C>A 4751G>A 4752T>A), Q478I (4756C>A 4757A>T 4758G>C), N479C (4759A>T 4760A>G 4761T>C), H480R (4762C>A 4763A>G 4764T>A), I481A (4765A>G 4766T>C 4767T>C), L482I (4768C>A), H484F (4774C>T 4775A>T), I485S (4777A>T 4778T>C), H486G (4780C>G 4781A>G), K491E (4795A>G), E493T (4801G>A 4802A>C 4803A>T), K496T (4811A>C 4812A>T), I504S (4835T>G 4836T>C), T506N (4841C>A), A508V (4847C>T 4848C>T), S509A (4849A>G 4850G>C 4851T>A), Y511F (4856A>T 4857T>C), Q514K (4864C>A), L515V (4867T>G), A516R (4870G>A 4871C>G), S517Q (4873T>C 4874C>A 4875T>G), M518V (4876A>G 4878G>T), R519C (4879C>T 4881T>C), A520K (4882G>A 4883C>A 4884T>A), Q523Y (4891C>T 4893G>T), E524D (4896A>T), K525R (4898A>G 4899G>A), E528Q (4906G>C 4908A>G), E528_D529insT (4908_4909insACC), D529P (4909G>C 4910A>C), N533K (4923T>A), W534E (4924T>G 4925G>A), K535E (4927A>G 4929G>A), H536Q (4932T>A), S537T (4933T>A), D538L (4936G>C 4937A>T 4938C>A), T539A (4939A>G 4941T>G), E540_Y541del (4942_4947delGAGTAT), E543Q (4951G>C 4953A>G), E544K (4954G>A), S548M (4967G>T 4968T>G), L549V (4969C>G), T550M (4973C>T 4974A>G), D551N (4975G>A), F552L (4978T>C 4980T>C), K554C (4984A>T 4985A>G 4986G>T), H556G (4990C>G 4991A>G 4992T>A), S596T (5111G>C 5112T>C), K598N (5118A>T), K599E (5119A>G 5121G>A), A600T (5122G>A), L602T (5128T>A 5129T>C 5130A>T), N603R (5132A>G 5133T>G), H605L (5137C>T 5138A>T 5139T>A), S606T (5140T>A 5142T>A), I611L (5155A>T 5157T>A), V614I (5164G>A 5166T>A), I615V (5167A>G 5169C>A), V617G (5174T>G 5175T>C), A619R (5179G>C 5180C>G 5181T>A), T622S (5188A>T), I623A (5191A>G 5192T>C 5193T>A), Y624F (5195A>T 5196T>C), T626L (5200A>C 5201C>T 5202T>A), L628K (5206C>A 5207T>A 5208T>G), E629Q (5209G>C 5211G>A), L631I (5215C>A 5217G>C), T634S (5224A>T 5226A>G), K637T (5234A>C), N638Q (5236A>C 5238C>G), F639V (5239T>G 5241T>A), T640S (5242A>T 5244C>A), F641G (5245T>G 5246T>G), F642L (5248T>C 5250T>A), M643I (5253G>A), N644F (5254A>T 5255A>T), T645N (5258C>A 5259C>T), N646K (5262T>A), V647L (5263G>C 5265G>T), K648S (5266A>T 5267A>C), G649S (5269G>T 5270G>C), D650E (5274T>A), Y651P (5275T>C 5276A>C), K652Q (5278A>C), Q653Y (5281C>T 5283G>T), G654R (5284G>A 5286A>G), V657H (5293G>C 5294T>A 5295C>T), Q661V (5305C>G 5306A>T), W662L (5309G>T), R665F (5317A>T 5318G>T 5319A>T), V670I (5332G>A), H672N (5338C>A 5340C>T), T674K (5345C>A) | | | | | | | | |
| Codon mutations: | GCA456ATG (4690G>A 4691C>T 4692A>G), CAA457AAA (4693C>A), CTA458TTT (4696C>T 4698A>T), AAG460CAA (4702A>C 4704G>A), TGC461ACA (4705T>A 4706G>C 4707C>A), AGA462AAA (4709G>A), ATT463ATC (4713T>C), AAC464AGA (4715A>G 4716C>A), GGC467GGG (4725C>G), CTT468CAT (4727T>A), ATT470TTA (4732A>T 4734T>A), GAC471TAT (4735G>T 4737C>T), GAA472CAA (4738G>C), GGG473GGA (4743G>A), ACT474ACA (4746T>A), CAC475TAT (4747C>T 4749C>T), CGT476AAA (4750C>A 4751G>A 4752T>A), CCT477CCC (4755T>C), CAG478ATC (4756C>A 4757A>T 4758G>C), AAT479TGC (4759A>T 4760A>G 4761T>C), CAT480AGA (4762C>A 4763A>G 4764T>A), ATT481GCC (4765A>G 4766T>C 4767T>C), CTC482ATC (4768C>A), CAT484TTT (4774C>T 4775A>T), ATT485TCT (4777A>T 4778T>C), CAT486GGT (4780C>G 4781A>G), AAG487AAA (4785G>A), CCA489CCT (4791A>T), AAA491GAA (4795A>G), ATT492ATC (4800T>C), GAA493ACT (4801G>A 4802A>C 4803A>T), AAA496ACT (4811A>C 4812A>T), CTT498TTA (4816C>T 4818T>A), CAA499CAG (4821A>G), TTT501TTC (4827T>C), CTT502TTG (4828C>T 4830T>G), GGT503GGA (4833T>A), ATT504AGC (4835T>G 4836T>C), TTA505CTC (4837T>C 4839A>C), ACT506AAT (4841C>A), GCC508GTT (4847C>T 4848C>T), AGT509GCA (4849A>G 4850G>C 4851T>A), GAT510GAC (4854T>C), TAT511TTC (4856A>T 4857T>C), CAA514AAA (4864C>A), TTA515GTA (4867T>G), GCA516AGA (4870G>A 4871C>G), TCT517CAG (4873T>C 4874C>A 4875T>G), ATG518GTT (4876A>G 4878G>T), CGT519TGC (4879C>T 4881T>C), GCT520AAA (4882G>A 4883C>A 4884T>A), CCT521CCC (4887T>C), TTG522CTC (4888T>C 4890G>C), CAG523TAT (4891C>T 4893G>T), GAA524GAT (4896A>T), AAG525AGA (4898A>G 4899G>A), CTC526CTT (4902C>T), GAA528CAG (4906G>C 4908A>G), GAA528_GAT529insACC (4908_4909insACC), GAT529CCT (4909G>C 4910A>C), CCA531CCC (4917A>C), AAT533AAA (4923T>A), TGG534GAG (4924T>G 4925G>A), AAG535GAA (4927A>G 4929G>A), CAT536CAA (4932T>A), TCT537ACT (4933T>A), GAC538CTA (4936G>C 4937A>T 4938C>A), ACT539GCG (4939A>G 4941T>G), GAG540_TAT541del (4942_4947delGAGTAT), GAA543CAG (4951G>C 4953A>G), GAA544AAA (4954G>A), AAG546AAA (4962G>A), AAA547AAG (4965A>G), AGT548ATG (4967G>T 4968T>G), CTC549GTC (4969C>G), ACA550ATG (4973C>T 4974A>G), GAC551AAC (4975G>A), TTT552CTC (4978T>C 4980T>C), CCT553CCA (4983T>A), AAG554TGT (4984A>T 4985A>G 4986G>T), CTA555TTA (4987C>T), CAT556GGA (4990C>G 4991A>G 4992T>A), CGT588..A (5088T>A), ATC589ATT (5091C>T), TGC590TGT (5094C>T), AGG591AGA (5097G>A), ACC593ACT (5103C>T), TCA594AGT (5104T>A 5105C>G 5106A>T), GGT595GGA (5109T>A), AGT596ACC (5111G>C 5112T>C), AAA598AAT (5118A>T), AAG599GAA (5119A>G 5121G>A), GCA600ACA (5122G>A), TTA602ACT (5128T>A 5129T>C 5130A>T), AAT603AGG (5132A>G 5133T>G), TAT604TAC (5136T>C), CAT605TTA (5137C>T 5138A>T 5139T>A), TCT606ACA (5140T>A 5142T>A), ATT611TTA (5155A>T 5157T>A), CTT612CTA (5160T>A), GCA613GCT (5163A>T), GTT614ATA (5164G>A 5166T>A), ATC615GTA (5167A>G 5169C>A), GTT617GGC (5174T>G 5175T>C), ATC618ATA (5178C>A), GCT619CGA (5179G>C 5180C>G 5181T>A), TTC621TTT (5187C>T), ACT622TCT (5188A>T), ATT623GCA (5191A>G 5192T>C 5193T>A), TAT624TTC (5195A>T 5196T>C), CTA625TTA (5197C>T), ACT626CTA (5200A>C 5201C>T 5202T>A), CCT627CCA (5205T>A), CTT628AAG (5206C>A 5207T>A 5208T>G), GAG629CAA (5209G>C 5211G>A), TTT630TTC (5214T>C), CTG631ATC (5215C>A 5217G>C), ATA632ATC (5220A>C), ACA634TCG (5224A>T 5226A>G), GAT635GAC (5229T>C), AAC636AAT (5232C>T), AAG637ACG (5234A>C), AAC638CAG (5236A>C 5238C>G), TTT639GTA (5239T>G 5241T>A), ACC640TCA (5242A>T 5244C>A), TTT641GGT (5245T>G 5246T>G), TTT642CTA (5248T>C 5250T>A), ATG643ATA (5253G>A), AAC644TTC (5254A>T 5255A>T), ACC645AAT (5258C>A 5259C>T), AAT646AAA (5262T>A), GTG647CTT (5263G>C 5265G>T), AAA648TCA (5266A>T 5267A>C), GGT649TCT (5269G>T 5270G>C), GAT650GAA (5274T>A), TAT651CCT (5275T>C 5276A>C), AAA652CAA (5278A>C), CAG653TAT (5281C>T 5283G>T), GGA654AGG (5284G>A 5286A>G), CTT656TTA (5290C>T 5292T>A), GTC657CAT (5293G>C 5294T>A 5295C>T), AGG658AGA (5298G>A), CAA660CAG (5304A>G), CAA661GTA (5305C>G 5306A>T), TGG662TTG (5309G>T), CTA663TTA (5311C>T), TCC664TCT (5316C>T), AGA665TTT (5317A>T 5318G>T 5319A>T), TAC666TAT (5322C>T), AGT667TCT (5323A>T 5324G>C), AAG669AAA (5331G>A), GTT670ATT (5332G>A), CAC672AAT (5338C>A 5340C>T), ACA674AAA (5345C>A) | | | | | | | | |

##### Proteins

| | Begin | End | Coverage | Score | Concordance | Matches | Identities | I/D/M/F* | Stop Codons |
|---|---|---|---|---|---|---|---|---|---|
| replicase (YP_001931961.1) | 455 | 675 | 27.5% | 632 | 46.1% | 190 (98.4%) | 86 (44.6%) | 1/2/0/0 | 0 |
| Protein mutations: | A456M (4690G>A 4691C>T 4692A>G), Q457K (4693C>A), L458F (4696C>T 4698A>T), K460Q (4702A>C 4704G>A), C461T (4705T>A 4706G>C 4707C>A), R462K (4709G>A), N464R (4715A>G 4716C>A), L468H (4727T>A), I470L (4732A>T 4734T>A), D471Y (4735G>T 4737C>T), E472Q (4738G>C), H475Y (4747C>T 4749C>T), R476K (4750C>A 4751G>A 4752T>A), Q478I (4756C>A 4757A>T 4758G>C), N479C (4759A>T 4760A>G 4761T>C), H480R (4762C>A 4763A>G 4764T>A), I481A (4765A>G 4766T>C 4767T>C), L482I (4768C>A), H484F (4774C>T 4775A>T), I485S (4777A>T 4778T>C), H486G (4780C>G 4781A>G), K491E (4795A>G), E493T (4801G>A 4802A>C 4803A>T), K496T (4811A>C 4812A>T), I504S (4835T>G 4836T>C), T506N (4841C>A), A508V (4847C>T 4848C>T), S509A (4849A>G 4850G>C 4851T>A), Y511F (4856A>T 4857T>C), Q514K (4864C>A), L515V (4867T>G), A516R (4870G>A 4871C>G), S517Q (4873T>C 4874C>A 4875T>G), M518V (4876A>G 4878G>T), R519C (4879C>T 4881T>C), A520K (4882G>A 4883C>A 4884T>A), Q523Y (4891C>T 4893G>T), E524D (4896A>T), K525R (4898A>G 4899G>A), E528Q (4906G>C 4908A>G), E528_D529insT (4908_4909insACC), D529P (4909G>C 4910A>C), N533K (4923T>A), W534E (4924T>G 4925G>A), K535E (4927A>G 4929G>A), H536Q (4932T>A), S537T (4933T>A), D538L (4936G>C 4937A>T 4938C>A), T539A (4939A>G 4941T>G), E540_Y541del (4942_4947delGAGTAT), E543Q (4951G>C 4953A>G), E544K (4954G>A), S548M (4967G>T 4968T>G), L549V (4969C>G), T550M (4973C>T 4974A>G), D551N (4975G>A), F552L (4978T>C 4980T>C), K554C (4984A>T 4985A>G 4986G>T), H556G (4990C>G 4991A>G 4992T>A), S596T (5111G>C 5112T>C), K598N (5118A>T), K599E (5119A>G 5121G>A), A600T (5122G>A), L602T (5128T>A 5129T>C 5130A>T), N603R (5132A>G 5133T>G), H605L (5137C>T 5138A>T 5139T>A), S606T (5140T>A 5142T>A), I611L (5155A>T 5157T>A), V614I (5164G>A 5166T>A), I615V (5167A>G 5169C>A), V617G (5174T>G 5175T>C), A619R (5179G>C 5180C>G 5181T>A), T622S (5188A>T), I623A (5191A>G 5192T>C 5193T>A), Y624F (5195A>T 5196T>C), T626L (5200A>C 5201C>T 5202T>A), L628K (5206C>A 5207T>A 5208T>G), E629Q (5209G>C 5211G>A), L631I (5215C>A 5217G>C), T634S (5224A>T 5226A>G), K637T (5234A>C), N638Q (5236A>C 5238C>G), F639V (5239T>G 5241T>A), T640S (5242A>T 5244C>A), F641G (5245T>G 5246T>G), F642L (5248T>C 5250T>A), M643I (5253G>A), N644F (5254A>T 5255A>T), T645N (5258C>A 5259C>T), N646K (5262T>A), V647L (5263G>C 5265G>T), K648S (5266A>T 5267A>C), G649S (5269G>T 5270G>C), D650E (5274T>A), Y651P (5275T>C 5276A>C), K652Q (5278A>C), Q653Y (5281C>T 5283G>T), G654R (5284G>A 5286A>G), V657H (5293G>C 5294T>A 5295C>T), Q661V (5305C>G 5306A>T), W662L (5309G>T), R665F (5317A>T 5318G>T 5319A>T), V670I (5332G>A), H672N (5338C>A 5340C>T), T674K (5345C>A) | | | | | | | | |

| | Begin | End | Coverage | Score | Concordance | Matches | Identities | I/D/M/F* | Stop Codons |
|---|---|---|---|---|---|---|---|---|---|
| **NT** | **4686** | **5347** | **7.5%** | **81** | **7.2%** | **570 (98.4%)** | **312 (53.9%)** | **3/6** | |

Codon mutations: GCA456ATG (4690G>A 4691C>T 4692A>G), CAA457AAA (4693C>A), CTA458TTT (4696C>T 4698A>T), AAG460CAA (4702A>C 4704G>A), TGC461ACA (4705T>A 4706G>C 4707C>A), AGA462AAA (4709G>A), ATT463ATC (4713T>C), AAC464AGA (4715A>G 4716C>A), GGC467GGG (4725C>G), CTT468CAT (4727T>A), ATT470TTA (4732A>T 4734T>A), GAC471TAT (4735G>T 4737C>T), GAA472CAA (4738G>C), GGG473GGA (4743G>A), ACT474ACA (4746T>A), CAC475TAT (4747C>T 4749C>T), CGT476AAA (4750C>A 4751G>A 4752T>A), CCT477CCC (4755T>C), CAG478ATC (4756C>A 4757A>T 4758G>C), AAT479TGC (4759A>T 4760A>G 4761T>C), CAT480AGA (4762C>A 4763A>G 4764T>A), ATT481GCC (4765A>G 4766T>C 4767T>C), CTC482ATC (4768C>A), CAT484TTT (4774C>T 4775A>T), ATT485TCT (4777A>T 4778T>C), CAT486GGT (4780C>G 4781A>G), AAG487AAA (4785G>A), CCA489CCT (4791A>T), AAA491GAA (4795A>G), ATT492ATC (4800T>C), GAA493ACT (4801G>A 4802A>C 4803A>T), AAA496ACT (4811A>C 4812A>T), CTT498TTA (4816C>T 4818T>A), CAA499CAG (4821A>G), TTT501TTC (4827T>C), CTT502TTG (4828C>T 4830T>G), GGT503GGA (4833T>A), ATT504AGC (4835T>G 4836T>C), TTA505CTC (4837T>C 4839A>C), ACT506AAT (4841C>A), GCC508GTT (4847C>T 4848C>T), AGT509GCA (4849A>G 4850G>C 4851T>A), GAT510GAC (4854T>C), TAT511TTC (4856A>T 4857T>C), CAA514AAA (4864C>A), TTA515GTA (4867T>G), GCA516AGA (4870G>A 4871C>G), TCT517CAG (4873T>C 4874C>A 4875T>G), ATG518GTT (4876A>G 4878G>T), CGT519TGC (4879C>T 4881T>C), GCT520AAA (4882G>A 4883C>A 4884T>A), CCT521CCC (4887T>C), TTG522CTC (4888T>C 4890G>C), CAG523TAT (4891C>T 4893G>T), GAA524GAT (4896A>T), AAG525AGA (4898A>G 4899G>A), CTC526CTT (4902C>T), GAA528CAG (4906G>C 4908A>G), GAA528_GAT529insACC (4908_4909insACC), GAT529CCT (4909G>C 4910A>C), CCA531CCC (4917A>C), AAT533AAA (4923T>A), TGG534GAG (4924T>G 4925G>A), AAG535GAA (4927A>G 4929G>A), CAT536CAA (4932T>A), TCT537ACT (4933T>A), GAC538CTA (4936G>C 4937A>T 4938C>A), ACT539GCG (4939A>G 4941T>G), GAG540_TAT541del (4942_4947delGAGTAT), GAA543CAG (4951G>C 4953A>G), GAA544AAA (4954G>A), AAG546AAA (4962G>A), AAA547AAG (4965A>G), AGT548ATG (4967G>T 4968T>G), CTC549GTC (4969C>G), ACA550ATG (4973C>T 4974A>G), GAC551AAC (4975G>A), TTT552CTC (4978T>C 4980T>C), CCT553CCA (4983T>A), AAG554TGT (4984A>T 4985A>G 4986G>T), CTA555TTA (4987C>T), CAT556GGA (4990C>G 4991A>G 4992T>A), CGT588..A (5088T>A), ATC589ATT (5091C>T), TGC590TGT (5094C>T), AGG591AGA (5097G>A), ACC593ACT (5103C>T), TCA594AGT (5104T>A 5105C>G 5106A>T), GGT595GGA (5109T>A), AGT596ACC (5111G>C 5112T>C), AAA598AAT (5118A>T), AAG599GAA (5119A>G 5121G>A), GCA600ACA (5122G>A), TTA602ACT (5128T>A 5129T>C 5130A>T), AAT603AGG (5132A>G 5133T>G), TAT604TAC (5136T>C), CAT605TTA (5137C>T 5138A>T 5139T>A), TCT606ACA (5140T>A 5142T>A), ATT611TTA (5155A>T 5157T>A), CTT612CTA (5160T>A), GCA613GCT (5163A>T), GTT614ATA (5164G>A 5166T>A), ATC615GTA (5167A>G 5169C>A), GTT617GGC (5174T>G 5175T>C), ATC618ATA (5178C>A), GCT619CGA (5179G>C 5180C>G 5181T>A), TTC621TTT (5187C>T), ACT622TCT (5188A>T), ATT623GCA (5191A>G 5192T>C 5193T>A), TAT624TTC (5195A>T 5196T>C), CTA625TTA (5197C>T), ACT626CTA (5200A>C 5201C>T 5202T>A), CCT627CCA (5205T>A), CTT628AAG (5206C>A 5207T>A 5208T>G), GAG629CAA (5209G>C 5211G>A), TTT630TTC (5214T>C), CTG631ATC (5215C>A 5217G>C), ATA632ATC (5220A>C), ACA634TCG (5224A>T 5226A>G), GAT635GAC (5229T>C), AAC636AAT (5232C>T), AAG637ACG (5234A>C), AAC638CAG (5236A>C 5238C>G), TTT639GTA (5239T>G 5241T>A), ACC640TCA (5242A>T 5244C>A), TTT641GGT (5245T>G 5246T>G), TTT642CTA (5248T>C 5250T>A), ATG643ATA (5253G>A), AAC644TTC (5254A>T 5255A>T), ACC645AAT (5258C>A 5259C>T), AAT646AAA (5262T>A), GTG647CTT (5263G>C 5265G>T), AAA648TCA (5266A>T 5267A>C), GGT649TCT (5269G>T 5270G>C), GAT650GAA (5274T>A), TAT651CCT (5275T>C 5276A>C), AAA652CAA (5278A>C), CAG653TAT (5281C>T 5283G>T), GGA654AGG (5284G>A 5286A>G), CTT656TTA (5290C>T 5292T>A), GTC657CAT (5293G>C 5294T>A 5295C>T), AGG658AGA (5298G>A), CAA660CAG (5304A>G), CAA661GTA (5305C>G 5306A>T), TGG662TTG (5309G>T), CTA663TTA (5311C>T), TCC664TCT (5316C>T), AGA665TTT (5317A>T 5318G>T 5319A>T), TAC666TAT (5322C>T), AGT667TCT (5323A>T 5324G>C), AAG669AAA (5331G>A), GTT670ATT (5332G>A), CAC672AAT (5338C>A 5340C>T), ACA674AAA (5345C>A)

*: Inserts / Deletes / Misaligned / Frameshifts

#### Analysis details

This analysis was performed with panviral2.64

### NGS Details (UN8): Petuvirus venapetuniae

#### Assembly

| | |
|---|---|
| **Coverage Length** | 551 (1 contig(s)) |
| **Depth Of Coverage** | 5.8 |
| **Number Of Reads** | 25 |
| **Reads Per Million** | 0.42 rpm (after QC) |
| **Ambiguities** | 0 |
| **Assembly Method** | de novo + reference guided assembly |
| **Consensus Caller** | Bcf Tools |

#### Coverage Map

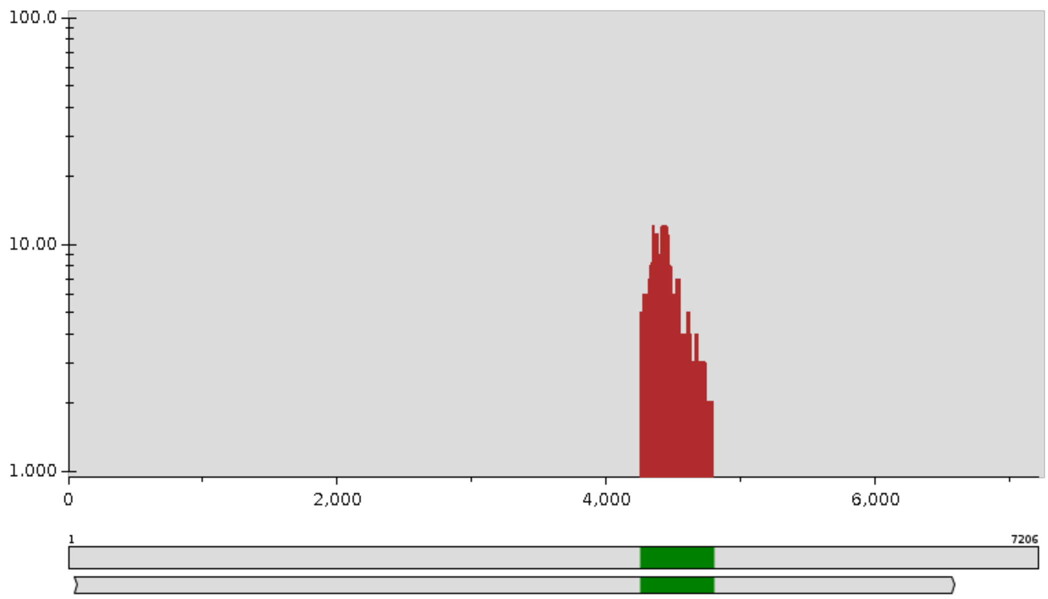

#### Assignment

| | |
|---|---|
| **Type** | Petuvirus venapetuniae (Taxonomy ID: 3048265) |
| **Reference Genome** | NC_001839.2 |
| **NT Identity (%)** | 55.1724 |
| **AA Identity (%)** | 49.4565 |
| **Number Of Stop Codons** | 0 |
| **Number Of CDS** | 1 |

#### Alignment

| | |
|---|---|
| **Alignment Score** | 114.0 (NT) + 660.0 (AA) = 774.0 |
| **Concordance (%)** | 32.3714 |

| **Alignment Method** | Global, seeded, nucleotide + amino acids (AGA) |
|---|---|

#### Genome Region

Sequence starts at position 4248 and ends at position 4798 relative to NC_001839.2 reference sequence.

#### Alignment Detailed Statistics

| | Begin | End | Coverage | Score | Concordance | Matches | Identities | I/D/M/F* | Stop Codons |
|---|---|---|---|---|---|---|---|---|---|
| **NT** | **4248** | **4798** | **7.6%** | **114** | **10.3%** | **551 (100%)** | **304 (55.2%)** | **0/0** | |
| Mutations: | 4250A>T, 4251G>T, 4252C>G, 4253A>T, 4255T>A, 4259G>A, 4263T>A, 4264G>T, 4265T>C, 4266G>A, 4269G>T, 4270A>C, 4271G>T, 4275C>T, 4276A>T, 4278C>G, 4281T>A, 4282T>A, 4283T>C, 4284G>A, 4286T>A, 4287C>A, 4289A>T, 4293G>A, 4294A>G, 4298C>T, 4302G>A, 4304T>A, 4307A>T, 4309A>C, 4310A>T, 4314G>A, 4315C>G, 4316A>C, 4320G>T, 4321A>C, 4325T>C, 4328T>C, 4334C>T, 4337C>T, 4340G>A, 4342G>C, 4343G>T, 4344T>G, 4346A>T, 4349G>A, 4350C>A, 4354T>A, 4355G>A, 4361A>C, 4362A>T, 4363A>C, 4364A>C, 4365T>C, 4366T>C, 4367A>T, 4376C>T, 4379A>T, 4386C>A, 4388A>G, 4391A>T, 4397C>T, 4398C>T, 4399A>C, 4401T>G, 4403C>T, 4404C>T, 4406C>A, 4407C>A, 4410G>T, 4411A>G, 4412T>G, 4413G>A, 4414A>T, 4415C>T, 4417A>G, 4418G>A, 4420T>A, 4421T>C, 4424C>G, 4430A>T, 4436G>A, 4437C>A, 4439C>G, 4440A>G, 4441C>A, 4442A>T, 4445C>T, 4448C>G, 4449T>A, 4450C>A, 4451T>A, 4452C>A, 4453A>G, 4454C>A, 4455T>A, 4456T>C, 4457A>T, 4459C>T, 4460C>T, 4464G>A, 4466A>C, 4469G>T, 4470C>A, 4472C>T, 4474T>A, 4475T>C, 4476T>A, 4477C>G, 4478A>T, 4481G>A, 4488C>A, 4490T>G, 4493G>A, 4499A>T, 4502T>C, 4509T>A, 4511A>T, 4512G>C, 4513G>A, 4517C>T, 4518C>T, 4519A>C, 4520T>G, 4521C>A, 4522C>A, 4523T>A, 4526T>A, 4529G>T, 4531G>A, 4533C>T, 4534C>A, 4535A>T, 4541A>T, 4543G>C, 4544G>T, 4547T>C, 4548T>A, 4549G>A, 4551A>G, 4553T>C, 4556T>C, 4557G>T, 4558A>T, 4559T>C, 4560A>G, 4565T>A, 4567T>A, 4569C>G, 4577G>T, 4580C>T, 4586A>C, 4589C>T, 4592T>A, 4593C>T, 4595T>A, 4600C>A, 4601A>T, 4611T>G, 4612T>A, 4613G>A, 4616C>T, 4622A>C, 4624C>T, 4625C>T, 4630T>A, 4631C>T, 4632A>T, 4633A>C, 4634A>T, 4637C>T, 4640C>T, 4641C>A, 4643G>T, 4645C>A, 4650C>A, 4651T>G, 4652T>C, 4653T>A, 4654T>A, 4655T>C, 4656T>A, 4657C>T, 4658A>C, 4659G>T, 4662T>A, 4664G>T, 4667C>G, 4670C>T, 4680A>G, 4682C>T, 4683C>T, 4685T>G, 4686C>A, 4688T>A, 4697A>G, 4698A>G, 4699C>A, 4700A>T, 4701C>A, 4703G>T, 4706A>T, 4707G>T, 4708A>C, 4712C>T, 4713A>T, 4715A>T, 4719T>C, 4720T>A, 4721G>T, 4722C>T, 4724T>A, 4727C>T, 4728C>A, 4729A>C, 4730G>C, 4734A>T, 4736A>C, 4738G>A, 4739T>A, 4740T>A, 4742G>T, 4745T>C, 4750A>G, 4752T>A, 4753T>A, 4757A>T, 4758G>C, 4760C>T, 4761A>G, 4763G>C, 4764T>A, 4766A>C, 4767T>A, 4768C>G, 4769G>T, 4772A>T, 4775A>G, 4778G>A, 4783T>A, 4784C>A, 4785C>T, 4787T>G, 4788G>T, 4789C>T, 4795A>C, 4796T>A | | | | | | | | |

##### CDS

| | Begin | End | Coverage | Score | Concordance | Matches | Identities | I/D/M/F* | Stop Codons |
|---|---|---|---|---|---|---|---|---|---|
| PVCVgp1 | 1400 | 1583 | 8.4% | 660 | 51.0% | 184 (100%) | 91 (49.5%) | 0/0/0/0 | 0 |
| Protein mutations: | L1400F (4250A>T), A1401C (4251G>T 4252C>G 4253A>T), I1402K (4255T>A), C1405I (4263T>A 4264G>T 4265T>C), D1406N (4266G>A), E1407S (4269G>T 4270A>C 4271G>T), Q1409L (4275C>T 4276A>T), Q1410E (4278C>G), F1411N (4281T>A 4282T>A 4283T>C), D1412K (4284G>A 4286T>A), L1413I (4287C>A 4289A>T), E1415R (4293G>A 4294A>G), D1418K (4302G>A 4304T>A), Q1420P (4309A>C 4310A>T), A1422S (4314G>A 4315C>G 4316A>C), E1424S (4320G>T 4321A>C), R1431T (4342G>C 4343G>T), S1432A (4344T>G 4346A>T), Q1434K (4350C>A), V1435E (4354T>A 4355G>A), K1438S (4362A>T 4363A>C 4364A>C), L1439P (4365T>C 4366T>C 4367A>T), Q1446K (4386C>A 4388A>G), H1450S (4398C>T 4399A>C), F1451V (4401T>G 4403C>T), Q1453K (4407C>A), D1454W (4410G>T 4411A>G 4412T>G), D1455I (4413G>A 4414A>T 4415C>T), K1456R (4417A>G 4418G>A), F1457Y (4420T>A 4421T>C), L1463M (4437C>A 4439C>G), T1464D (4440A>G 4441C>A 4442A>T), F1466L (4448C>G), S1467K (4449T>A 4450C>A 4451T>A), H1468R (4452C>A 4453A>G 4454C>A), L1469T (4455T>A 4456T>C 4457A>T), S1470F (4459C>T 4460C>T), A1472T (4464G>A 4466A>C), K1473N (4469G>T), L1474I (4470C>A 4472C>T), F1475Y (4474T>A 4475T>C), L1480M (4488C>A 4490T>G), L1487I (4509T>A 4511A>T), G1488Q (4512G>C 4513G>A), H1490S (4518C>T 4519A>C 4520T>G), P1491K (4521C>A 4522C>A 4523T>A), N1492K (4526T>A), E1493D (4529G>T), R1494K (4531G>A), P1495Y (4533C>T 4534C>A 4535A>T), G1498A (4543G>C 4544G>T), C1500N (4548T>A 4549G>A), I1501V (4551A>G 4553T>C), D1503F (4557G>T 4558A>T 4559T>C), R1504G (4560A>G), H1505Q (4565T>A), F1506Y (4567T>A), Q1507E (4569C>G), K1509N (4577G>T), T1517N (4600C>A 4601A>T), L1521E (4611T>G 4612T>A 4613G>A), K1524N (4622A>C), A1525V (4624C>T 4625C>T), I1527N (4630T>A 4631C>T), K1528S (4632A>T 4633A>C 4634A>T), Q1531N (4641C>A 4643G>T), P1532H (4645C>A), L1534S (4650C>A 4651T>G 4652T>C), F1535N (4653T>A 4654T>A 4655T>C), S1536I (4656T>A 4657C>T 4658A>C), A1537S (4659G>T), L1538I (4662T>A 4664G>T), I1544V (4680A>G 4682C>T), L1546I (4686C>A 4688T>A), T1550D (4698A>G 4699C>A 4700A>T), L1551I (4701C>A 4703G>T), E1552D (4706A>T), D1553S (4707G>T 4708A>C), I1555F (4713A>T 4715A>T), L1557H (4719T>C 4720T>A 4721G>T), Q1560T (4728C>A 4729A>C 4730G>C), I1562F (4734A>T 4736A>C), S1563K (4738G>A 4739T>A), L1564I (4740T>A 4742G>T), K1567R (4750A>G), F1568N (4752T>A 4753T>A), V1570L (4758G>C 4760C>T), M1571V (4761A>G 4763G>C), L1572I (4764T>A 4766A>C), I1578K (4783T>A 4784C>A), A1580F (4788G>T 4789C>T), N1582T (4795A>C 4796T>A) | | | | | | | | |
| Codon mutations: | TTA1400TTT (4250A>T), GCA1401TGT (4251G>T 4252C>G 4253A>T), ATA1402AAA (4255T>A), AAG1403AAA (4259G>A), TGT1405ATC (4263T>A 4264G>T 4265T>C), GAT1406AAT (4266G>A), GAG1407TCT (4269G>T 4270A>C 4271G>T), CAA1409TTA (4275C>T 4276A>T), CAA1410GAA (4278C>G), TTT1411AAC (4281T>A 4282T>A 4283T>C), GAT1412AAA (4284G>A 4286T>A), CTA1413ATT (4287C>A 4289A>T), GAA1415AGA (4293G>A 4294A>G), CCC1416CCT (4298C>T), GAT1418AAA (4302G>A 4304T>A), TCA1419TCT (4307A>T), CAA1420CCT (4309A>C 4310A>T), GCA1422AGC (4314G>A 4315C>G 4316A>C), GAA1424TCA (4320G>T 4321A>C), GCT1425GCC (4325T>C), TTT1426TTC (4328T>C), GTC1428GTT (4334C>T), AAC1429AAT (4337C>T), AAG1430AAA (4340G>A), AGG1431ACT (4342G>C 4343G>T), TCA1432GCT (4344T>G 4346A>T), GAG1433GAA (4349G>A), CAA1434AAA (4350C>A), GTG1435GAA (4354T>A 4355G>A), GGA1437GGC (4361A>C), AAA1438TCC (4362A>T 4363A>C 4364A>C), TTA1439CCT (4365T>C 4366T>C 4367A>T), GTC1442GTT (4376C>T), ATA1443ATT (4379A>T), CAA1446AAG (4386C>A 4388A>G), CCA1447CCT (4391A>T), AAC1449AAT (4397C>T), CAT1450TCT (4398C>T 4399A>C), TTC1451GTT (4401T>G 4403C>T), CTC1452TTA (4404C>T 4406C>A), CAA1453AAA (4407C>A), GAT1454TGG (4410G>T 4411A>G 4412T>G), GAC1455ATT (4413G>A 4414A>T 4415C>T), AAG1456AGA (4417A>G 4418G>A), TTT1457TAC (4420T>A 4421T>C), CCC1458CCG (4424C>G), CCA1460CCT (4430A>T), AAG1462AAA (4436G>A), CTC1463ATG (4437C>A 4439C>G), ACA1464GAT (4440A>G 4441C>A 4442A>T), CTC1465CTT (4445C>T), TTC1466TTG (4448C>G), TCT1467AAA (4449T>A 4450C>A 4451T>A), CAC1468AGA (4452C>A 4453A>G 4454C>A), TTA1469ACT (4455T>A 4456T>C 4457A>T), TCC1470TTT (4459C>T 4460C>T), GCA1472ACC (4464G>A 4466A>C), AAG1473AAT (4469G>T), CTC1474ATT (4470C>A 4472C>T), TTT1475TAC (4474T>A 4475T>C), TCA1476AGT (4476T>A 4477C>G 4478A>T), AAG1477AAA (4481G>A), CTT1480ATG (4488C>A 4490T>G), AAG1481AAA (4493G>A), GGA1483GGT (4499A>T), TTT1484TTC (4502T>C), TTA1487ATT (4509T>A 4511A>T), GGA1488CAA (4512G>C 4513G>A), ATC1489ATT (4517C>T), CAT1490TCG (4518C>T 4519A>C 4520T>G), CCT1491AAA (4521C>A 4522C>A 4523T>A), AAT1492AAA (4526T>A), GAG1493GAT (4529G>T), AGA1494AAA (4531G>A), CCA1495TAT (4533C>T 4534C>A 4535A>T), ACA1497ACT (4541A>T), GGG1498GCT (4543G>C 4544G>T), TTT1499TTC (4547T>C), TGC1500AAC (4548T>A 4549G>A), ATT1501GTC (4551A>G 4553T>C), CCT1502CCC (4556T>C), GAT1503TTC (4557G>T 4558A>T 4559T>C), AGA1504GGA (4560A>G), CAT1505CAA (4565T>A), TTT1506TAT (4567T>A), CAA1507GAA (4569C>G), AAG1509AAT (4577G>T), GTC1510GTT (4580C>T), CCA1512CCC (4586A>C), TTC1513TTT (4589C>T), GGT1514GGA (4592T>A), CTT1515TTA (4593C>T 4595T>A), ACA1517AAT (4600C>A 4601A>T), TTG1521GAA (4611T>G 4612T>A 4613G>A), TTC1522TTT (4616C>T), AAA1524AAC (4622A>C), GCC1525GTT (4624C>T 4625C>T), ATC1527AAT (4630T>A 4631C>T), AAA1528TCT (4632A>T 4633A>C 4634A>T), ATC1529ATT (4637C>T), TTC1530TTT (4640C>T), CAG1531AAT (4641C>A 4643G>T), CCC1532CAC (4645C>A), CTT1534AGC (4650C>A 4651T>G 4652T>C), TTT1535AAC (4653T>A 4654T>A 4655T>C), TCA1536ATC (4656T>A 4657C>T 4658A>C), GCT1537TCT (4659G>T), TTG1538ATT (4662T>A 4664G>T), GTC1539GTG (4667C>G), TAC1540TAT (4670C>T), ATC1544GTT (4680A>G 4682C>T), CTT1545TTG (4683C>T 4685T>G), CTT1546ATA (4686C>A 4688T>A), GAA1549GAG (4697A>G), ACA1550GAT (4698A>G 4699C>A 4700A>T), CTG1551ATT (4701C>A 4703G>T), GAA1552GAT (4706A>T), GAT1553TCT (4707G>T 4708A>C), CAC1554CAT (4712C>T), ATA1555TTT (4713A>T 4715A>T), TTG1557CAT (4719T>C 4720T>A 4721G>T), CTT1558TTA (4722C>T 4724T>A), AAC1559AAT (4727C>T), CAG1560ACC (4728C>A 4729A>C 4730G>C), ATA1562TTC (4734A>T 4736A>C), AGT1563AAA (4738G>A 4739T>A), TTG1564ATT (4740T>A 4742G>T), GTT1565GTC (4745T>C), AAA1567AGA (4750A>G), TTT1568AAT (4752T>A 4753T>A), GGA1569GGT (4757A>T), GTC1570CTT (4758G>C 4760C>T), ATG1571GTC (4761A>G 4763G>C), TTA1572ATC (4764T>A 4766A>C), TCG1573AGT (4767T>A 4768C>G 4769G>T), GCA1574GCT (4772A>T), AAA1575AAG (4775A>G), AAG1576AAA (4778G>A), ATC1578AAA (4783T>A 4784C>A), CTT1579TTG (4785C>T 4787T>G), GCT1580TTT (4788G>T 4789C>T), AAT1582ACA (4795A>C 4796T>A) | | | | | | | | |

##### Proteins

| | Begin | End | Coverage | Score | Concordance | Matches | Identities | I/D/M/F* | Stop Codons |
|---|---|---|---|---|---|---|---|---|---|
| ORF I polyprotein (NP_127504.1) | 1400 | 1583 | 8.4% | 660 | 51.0% | 184 (100%) | 91 (49.5%) | 0/0/0/0 | 0 |
| Protein mutations: | L1400F (4250A>T), A1401C (4251G>T 4252C>G 4253A>T), I1402K (4255T>A), C1405I (4263T>A 4264G>T 4265T>C), D1406N (4266G>A), E1407S (4269G>T 4270A>C 4271G>T), Q1409L (4275C>T 4276A>T), Q1410E (4278C>G), F1411N (4281T>A 4282T>A 4283T>C), D1412K (4284G>A 4286T>A), L1413I (4287C>A 4289A>T), E1415R (4293G>A 4294A>G), D1418K (4302G>A 4304T>A), Q1420P (4309A>C 4310A>T), A1422S (4314G>A 4315C>G 4316A>C), E1424S (4320G>T 4321A>C), R1431T (4342G>C 4343G>T), S1432A (4344T>G 4346A>T), Q1434K (4350C>A), V1435E (4354T>A 4355G>A), K1438S (4362A>T 4363A>C 4364A>C), L1439P (4365T>C 4366T>C 4367A>T), Q1446K (4386C>A 4388A>G), H1450S (4398C>T 4399A>C), F1451V (4401T>G 4403C>T), Q1453K (4407C>A), D1454W (4410G>T 4411A>G 4412T>G), D1455I (4413G>A 4414A>T 4415C>T), K1456R (4417A>G 4418G>A), F1457Y (4420T>A 4421T>C), L1463M (4437C>A 4439C>G), T1464D (4440A>G 4441C>A 4442A>T), F1466L (4448C>G), S1467K (4449T>A 4450C>A 4451T>A), H1468R (4452C>A 4453A>G 4454C>A), L1469T (4455T>A 4456T>C 4457A>T), S1470F (4459C>T 4460C>T), A1472T (4464G>A 4466A>C), K1473N (4469G>T), L1474I (4470C>A 4472C>T), F1475Y (4474T>A 4475T>C), L1480M (4488C>A 4490T>G), L1487I (4509T>A 4511A>T), G1488Q (4512G>C 4513G>A), H1490S (4518C>T 4519A>C 4520T>G), P1491K (4521C>A 4522C>A 4523T>A), N1492K (4526T>A), E1493D (4529G>T), R1494K (4531G>A), P1495Y (4533C>T 4534C>A 4535A>T), G1498A (4543G>C 4544G>T), C1500N (4548T>A 4549G>A), I1501V (4551A>G 4553T>C), D1503F (4557G>T 4558A>T 4559T>C), R1504G (4560A>G), H1505Q (4565T>A), F1506Y (4567T>A), Q1507E (4569C>G), K1509N (4577G>T), T1517N (4600C>A 4601A>T), L1521E (4611T>G 4612T>A 4613G>A), K1524N (4622A>C), A1525V (4624C>T 4625C>T), I1527N (4630T>A 4631C>T), K1528S (4632A>T 4633A>C 4634A>T), Q1531N (4641C>A 4643G>T), P1532H (4645C>A), L1534S (4650C>A 4651T>G 4652T>C), F1535N (4653T>A 4654T>A 4655T>C), S1536I (4656T>A 4657C>T 4658A>C), A1537S (4659G>T), L1538I (4662T>A 4664G>T), I1544V (4680A>G 4682C>T), L1546I (4686C>A 4688T>A), T1550D (4698A>G 4699C>A 4700A>T), L1551I (4701C>A 4703G>T), E1552D (4706A>T), D1553S (4707G>T 4708A>C), I1555F (4713A>T 4715A>T), L1557H (4719T>C 4720T>A 4721G>T), Q1560T (4728C>A 4729A>C 4730G>C), I1562F (4734A>T 4736A>C), S1563K (4738G>A 4739T>A), L1564I (4740T>A 4742G>T), K1567R (4750A>G), F1568N (4752T>A 4753T>A), V1570L (4758G>C 4760C>T), M1571V (4761A>G 4763G>C), L1572I (4764T>A 4766A>C), I1578K (4783T>A 4784C>A), A1580F (4788G>T 4789C>T), N1582T (4795A>C 4796T>A) | | | | | | | | |

| | Begin | End | Coverage | Score | Concordance | Matches | Identities | I/D/M/F* | Stop Codons |
|---|---|---|---|---|---|---|---|---|---|
| **NT** | **4248** | **4798** | **7.6%** | **114** | **10.3%** | **551 (100%)** | **304 (55.2%)** | **0/0** | |
| Codon mutations: | TTA1400TTT (4250A>T), GCA1401TGT (4251G>T 4252C>G 4253A>T), ATA1402AAA (4255T>A), AAG1403AAA (4259G>A), TGT1405ATC (4263T>A 4264G>T 4265T>C), GAT1406AAT (4266G>A), GAG1407TCT (4269G>T 4270A>C 4271G>T), CAA1409TTA (4275C>T 4276A>T), CAA1410GAA (4278C>G), TTT1411AAC (4281T>A 4282T>A 4283T>C), GAT1412AAA (4284G>A 4286T>A), CTA1413ATT (4287C>A 4289A>T), GAA1415AGA (4293G>A 4294A>G), CCC1416CCT (4298C>T), GAT1418AAA (4302G>A 4304T>A), TCA1419TCT (4307A>T), CAA1420CCT (4309A>C 4310A>T), GCA1422AGC (4314G>A 4315C>G 4316A>C), GAA1424TCA (4320G>T 4321A>C), GCT1425GCC (4325T>C), TTT1426TTC (4328T>C), GTC1428GTT (4334C>T), AAC1429AAT (4337C>T), AAG1430AAA (4340G>A), AGG1431ACT (4342G>C 4343G>T), TCA1432GCT (4344T>G 4346A>T), GAG1433GAA (4349G>A), CAA1434AAA (4350C>A), GTG1435GAA (4354T>A 4355G>A), GGA1437GGC (4361A>C), AAA1438TCC (4362A>T 4363A>C 4364A>C), TTA1439CCT (4365T>C 4366T>C 4367A>T), GTC1442GTT (4376C>T), ATA1443ATT (4379A>T), CAA1446AAG (4386C>A 4388A>G), CCA1447CCT (4391A>T), AAC1449AAT (4397C>T), CAT1450TCT (4398C>T 4399A>C), TTC1451GTT (4401T>G 4403C>T), CTC1452TTA (4404C>T 4406C>A), CAA1453AAA (4407C>A), GAT1454TGG (4410G>T 4411A>G 4412T>G), GAC1455ATT (4413G>A 4414A>T 4415C>T), AAG1456AGA (4417A>G 4418G>A), TTT1457TAC (4420T>A 4421T>C), CCC1458CCG (4424C>G), CCA1460CCT (4430A>T), AAG1462AAA (4436G>A), CTC1463ATG (4437C>A 4439C>G), ACA1464GAT (4440A>G 4441C>A 4442A>T), CTC1465CTT (4445C>T), TTC1466TTG (4448C>G), TCT1467AAA (4449T>A 4450C>A 4451T>A), CAC1468AGA (4452C>A 4453A>G 4454C>A), TTA1469ACT (4455T>A 4456T>C 4457A>T), TCC1470TTT (4459C>T 4460C>T), GCA1472ACC (4464G>A 4466A>C), AAG1473AAT (4469G>T), CTC1474ATT (4470C>A 4472C>T), TTT1475TAC (4474T>A 4475T>C), TCA1476AGT (4476T>A 4477C>G 4478A>T), AAG1477AAA (4481G>A), CTT1480ATG (4488C>A 4490T>G), AAG1481AAA (4493G>A), GGA1483GGT (4499A>T), TTT1484TTC (4502T>C), TTA1487ATT (4509T>A 4511A>T), GGA1488CAA (4512G>C 4513G>A), ATC1489ATT (4517C>T), CAT1490TCG (4518C>T 4519A>C 4520T>G), CCT1491AAA (4521C>A 4522C>A 4523T>A), AAT1492AAA (4526T>A), GAG1493GAT (4529G>T), AGA1494AAA (4531G>A), CCA1495TAT (4533C>T 4534C>A 4535A>T), ACA1497ACT (4541A>T), GGG1498GCT (4543G>C 4544G>T), TTT1499TTC (4547T>C), TGC1500AAC (4548T>A 4549G>A), ATT1501GTC (4551A>G 4553T>C), CCT1502CCC (4556T>C), GAT1503TTC (4557G>T 4558A>T 4559T>C), AGA1504GGA (4560A>G), CAT1505CAA (4565T>A), TTT1506TAT (4567T>A), CAA1507GAA (4569C>G), AAG1509AAT (4577G>T), GTC1510GTT (4580C>T), CCA1512CCC (4586A>C), TTC1513TTT (4589C>T), GGT1514GGA (4592T>A), CTT1515TTA (4593C>T 4595T>A), ACA1517AAT (4600C>A 4601A>T), TTG1521GAA (4611T>G 4612T>A 4613G>A), TTC1522TTT (4616C>T), AAA1524AAC (4622A>C), GCC1525GTT (4624C>T 4625C>T), ATC1527AAT (4630T>A 4631C>T), AAA1528TCT (4632A>T 4633A>C 4634A>T), ATC1529ATT (4637C>T), TTC1530TTT (4640C>T), CAG1531AAT (4641C>A 4643G>T), CCC1532CAC (4645C>A), CTT1534AGC (4650C>A 4651T>G 4652T>C), TTT1535AAC (4653T>A 4654T>A 4655T>C), TCA1536ATC (4656T>A 4657C>T 4658A>C), GCT1537TCT (4659G>T), TTG1538ATT (4662T>A 4664G>T), GTC1539GTG (4667C>G), TAC1540TAT (4670C>T), ATC1544GTT (4680A>G 4682C>T), CTT1545TTG (4683C>T 4685T>G), CTT1546ATA (4686C>A 4688T>A), GAA1549GAG (4697A>G), ACA1550GAT (4698A>G 4699C>A 4700A>T), CTG1551ATT (4701C>A 4703G>T), GAA1552GAT (4706A>T), GAT1553TCT (4707G>T 4708A>C), CAC1554CAT (4712C>T), ATA1555TTT (4713A>T 4715A>T), TTG1557CAT (4719T>C 4720T>A 4721G>T), CTT1558TTA (4722C>T 4724T>A), AAC1559AAT (4727C>T), CAG1560ACC (4728C>A 4729A>C 4730G>C), ATA1562TTC (4734A>T 4736A>C), AGT1563AAA (4738G>A 4739T>A), TTG1564ATT (4740T>A 4742G>T), GTT1565GTC (4745T>C), AAA1567AGA (4750A>G), TTT1568AAT (4752T>A 4753T>A), GGA1569GGT (4757A>T), GTC1570CTT (4758G>C 4760C>T), ATG1571GTC (4761A>G 4763G>C), TTA1572ATC (4764T>A 4766A>C), TCG1573AGT (4767T>A 4768C>G 4769G>T), GCA1574GCT (4772A>T), AAA1575AAG (4775A>G), AAG1576AAA (4778G>A), ATC1578AAA (4783T>A 4784C>A), CTT1579TTG (4785C>T 4787T>G), GCT1580TTT (4788G>T 4789C>T), AAT1582ACA (4795A>C 4796T>A) | | | | | | | | |

*: Inserts / Deletes / Misaligned / Frameshifts

#### Analysis details

This analysis was performed with panviral2.64

### NGS Details (UN8): Errantivirus

#### Assembly

| | |
|---|---|
| **Coverage Length** | 361 (1 contig(s)) |
| **Depth Of Coverage** | 7.6 |
| **Number Of Reads** | 23 |
| **Reads Per Million** | 0.38 rpm (after QC) |
| **Ambiguities** | 0 |
| **Assembly Method** | de novo + reference guided assembly |
| **Consensus Caller** | Bcf Tools |

#### Coverage Map

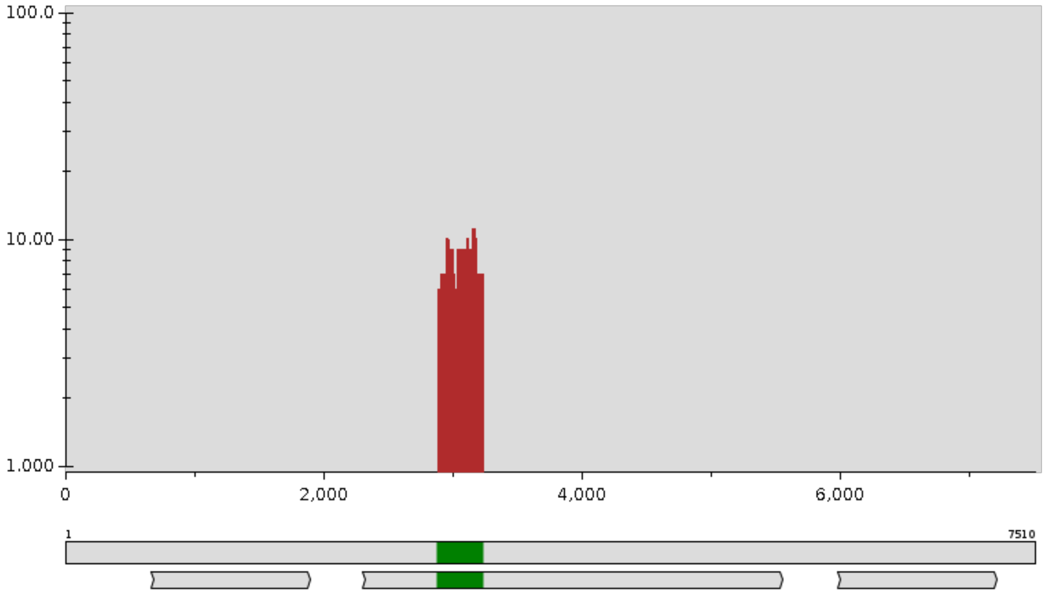

#### Assignment

| | |
|---|---|
| **Type** | Errantivirus (Taxonomy ID: 186666) |
| **Reference Genome** | NC_038512.1 |
| **NT Identity (%)** | 55.4017 |
| **AA Identity (%)** | 47.5 |
| **Number Of Stop Codons** | 1 |
| **Number Of CDS** | 3 |

#### Alignment

| | |
|---|---|
| **Alignment Score** | 78.0 (NT) + 430.0 (AA) = 508.0 |
| **Concordance (%)** | 32.2335 |

| **Alignment Method** | Global, seeded, nucleotide + amino acids (AGA) |
|---|---|

#### Genome Region

Sequence starts at position 2881 and ends at position 3241 relative to NC_038512.1 reference sequence.

#### Alignment Detailed Statistics

| | Begin | End | Coverage | Score | Concordance | Matches | Identities | I/D/M/F* | Stop Codons |
|---|---|---|---|---|---|---|---|---|---|
| **NT** | **2881** | **3241** | **4.8%** | **78** | **10.8%** | **361 (100%)** | **200 (55.4%)** | **0/0** | |
| Mutations: | 2881G>T, 2887T>G, 2888C>T, 2890C>T, 2891G>T, 2892T>G, 2893A>T, 2896T>G, 2899C>T, 2901T>A, 2907A>G, 2908G>C, 2909T>C, 2911G>T, 2915G>A, 2917G>A, 2919A>T, 2920G>C, 2924A>G, 2927G>C, 2928A>C, 2929T>G, 2930G>A, 2938C>T, 2941G>C, 2944A>T, 2947A>C, 2953A>T, 2954A>G, 2955G>A, 2959C>G, 2960G>T, 2962A>G, 2963C>T, 2965T>A, 2968C>T, 2969A>G, 2972T>C, 2975G>C, 2976G>A, 2978A>G, 2979A>G, 2980G>A, 2981T>G, 2982G>C, 2984C>A, 2985A>T, 2987T>A, 2988A>T, 2989C>T, 2992C>T, 2993A>C, 2995C>A, 2997C>A, 2998C>A, 2999T>A, 3001A>T, 3005T>C, 3007G>T, 3008G>T, 3009C>G, 3010A>T, 3011A>T, 3012G>T, 3013T>G, 3016G>A, 3018T>A, 3019T>C, 3020T>C, 3025G>A, 3026G>A, 3028G>T, 3029G>C, 3030A>G, 3031G>T, 3032A>G, 3034G>C, 3035G>C, 3037C>A, 3040T>A, 3041C>A, 3043A>T, 3047A>G, 3049A>C, 3051C>A, 3052G>T, 3058C>T, 3061G>C, 3065A>C, 3066A>G, 3068G>A, 3069T>C, 3070A>T, 3071G>C, 3073A>T, 3074C>T, 3075A>C, 3076C>A, 3079G>C, 3084T>A, 3088A>G, 3091C>T, 3094T>C, 3095C>G, 3096G>T, 3104A>T, 3106G>T, 3110T>C, 3112A>C, 3114A>C, 3115A>C, 3118C>T, 3119T>G, 3127T>C, 3130T>G, 3134C>T, 3136A>G, 3137A>G, 3139A>C, 3141T>C, 3142T>C, 3146G>A, 3148C>T, 3149A>G, 3152G>C, 3154C>T, 3155C>T, 3157A>C, 3160A>G, 3161G>C, 3162G>C, 3163T>A, 3164C>T, 3165T>A, 3168A>T, 3169A>T, 3171A>G, 3172T>A, 3175C>A, 3176A>T, 3178C>T, 3179T>G, 3180G>T, 3181T>C, 3184C>A, 3187C>G, 3189A>T, 3191C>T, 3196C>T, 3203A>C, 3205T>A, 3206G>A, 3215A>C, 3216C>T, 3218T>G, 3219C>A, 3222T>C, 3223A>T, 3226G>A, 3227G>A, 3228A>C | | | | | | | | |

##### CDS

| | Begin | End | Coverage | Score | Concordance | Matches | Identities | I/D/M/F* | Stop Codons |
|---|---|---|---|---|---|---|---|---|---|
| D1R33_gp2 | 193 | 312 | 11.1% | 430 | 50.1% | 120 (100%) | 57 (47.5%) | 0/0/0/0 | 1 |
| Protein mutations: | L195F (2888C>T 2890C>T), V196C (2891G>T 2892T>G 2893A>T), F199Y (2901T>A), K201S (2907A>G 2908G>C), E204K (2915G>A 2917G>A), K205I (2919A>T 2920G>C), I207V (2924A>G), D208P (2927G>C 2928A>C 2929T>G), D209N (2930G>A), S217D (2954A>G 2955G>A), D218E (2959C>G), V219L (2960G>T 2962A>G), K222E (2969A>G), G224H (2975G>C 2976G>A), K225G (2978A>G 2979A>G 2980G>A), C226A (2981T>G 2982G>C), Q227I (2984C>A 2985A>T), Y228I (2987T>A 2988A>T 2989C>T), T230P (2993A>C 2995C>A), T231K (2997C>A 2998C>A), L232I (2999T>A 3001A>T), A235C (3008G>T 3009C>G 3010A>T), S236L (3011A>T 3012G>T 3013T>G), F238Y (3018T>A 3019T>C), Y239H (3020T>C), V241I (3026G>A 3028G>T), E242R (3029G>C 3030A>G 3031G>T), M243V (3032A>G 3034G>C), D244Q (3035G>C 3037C>A), Q246N (3041C>A 3043A>T), I248V (3047A>G 3049A>C), S249Y (3051C>A 3052G>T), N254R (3065A>C 3066A>G), V255T (3068G>A 3069T>C 3070A>T), E256H (3071G>C 3073A>T), H257S (3074C>T 3075A>C 3076C>A), F260Y (3084T>A), R264V (3095C>G 3096G>T), M267F (3104A>T 3106G>T), K270T (3114A>C 3115A>C), S272A (3119T>G), Q277* (3134C>T 3136A>G), R278G (3137A>G 3139A>C), V279A (3141T>C 3142T>C), D281N (3146G>A 3148C>T), N282D (3149A>G), V283L (3152G>C 3154C>T), L284F (3155C>T 3157A>C), G286P (3161G>C 3162G>C 3163T>A), L287Y (3164C>T 3165T>A), Q288L (3168A>T 3169A>T), N289R (3171A>G 3172T>A), N290K (3175C>A), I291F (3176A>T 3178C>T), C292V (3179T>G 3180G>T 3181T>C), Y295F (3189A>T), L296F (3191C>T), I300L (3203A>C 3205T>A), V301I (3206G>A), T304L (3215A>C 3216C>T), S305D (3218T>G 3219C>A), L306P (3222T>C 3223A>T), E308T (3227G>A 3228A>C) | | | | | | | | |
| Codon mutations: | AAG192..T (2881G>T), CGT194CGG (2887T>G), CTC195TTT (2888C>T 2890C>T), GTA196TGT (2891G>T 2892T>G 2893A>T), GTT197GTG (2896T>G), GAC198GAT (2899C>T), TTC199TAC (2901T>A), AAG201AGC (2907A>G 2908G>C), TTG202CTT (2909T>C 2911G>T), GAG204AAA (2915G>A 2917G>A), AAG205ATC (2919A>T 2920G>C), ATC207GTC (2924A>G), GAT208CCG (2927G>C 2928A>C 2929T>G), GAC209AAC (2930G>A), TAC211TAT (2938C>T), CCG212CCC (2941G>C), ATA213ATT (2944A>T), CCA214CCC (2947A>C), ATA216ATT (2953A>T), AGT217GAT (2954A>G 2955G>A), GAC218GAG (2959C>G), GTA219TTG (2960G>T 2962A>G), CTT220TTA (2963C>T 2965T>A), GAC221GAT (2968C>T), AAG222GAG (2969A>G), TTA223CTA (2972T>C), GGT224CAT (2975G>C 2976G>A), AAG225GGA (2978A>G 2979A>G 2980G>A), TGC226GCC (2981T>G 2982G>C), CAA227ATA (2984C>A 2985A>T), TAC228ATT (2987T>A 2988A>T 2989C>T), TTC229TTT (2992C>T), ACC230CCA (2993A>C 2995C>A), ACC231AAA (2997C>A 2998C>A), TTA232ATT (2999T>A 3001A>T), TTG234CTT (3005T>C 3007G>T), GCA235TGT (3008G>T 3009C>G 3010A>T), AGT236TTG (3011A>T 3012G>T 3013T>G), GGG237GGA (3016G>A), TTT238TAC (3018T>A 3019T>C), TAT239CAT (3020T>C), CAG240CAA (3025G>A), GTG241ATT (3026G>A 3028G>T), GAG242CGT (3029G>C 3030A>G 3031G>T), ATG243GTC (3032A>G 3034G>C), GAC244CAA (3035G>C 3037C>A), CCT245CCA (3040T>A), CAA246AAT (3041C>A 3043A>T), ATA248GTC (3047A>G 3049A>C), TCG249TAT (3051C>A 3052G>T), ACC251ACT (3058C>T), GCG252GCC (3061G>C), AAC254CGC (3065A>C 3066A>G), GTA255ACT (3068G>A 3069T>C 3070A>T), GAA256CAT (3071G>C 3073A>T), CAC257TCA (3074C>T 3075A>C 3076C>A), GGG258GGC (3079G>C), TTT260TAT (3084T>A), GAA261GAG (3088A>G), TTC262TTT (3091C>T), CTT263CTC (3094T>C), CGA264GTA (3095C>G 3096G>T), ATG267TTT (3104A>T 3106G>T), TTA269CTC (3110T>C 3112A>C), AAA270ACC (3114A>C 3115A>C), AAC271AAT (3118C>T), TCA272GCA (3119T>G), TCT274TCC (3127T>C), ACT275ACG (3130T>G), CAA277TAG (3134C>T 3136A>G), AGA278GGC (3137A>G 3139A>C), GTT279GCC (3141T>C 3142T>C), GAC281AAT (3146G>A 3148C>T), AAT282GAT (3149A>G), GTC283CTT (3152G>C 3154C>T), CTA284TTC (3155C>T 3157A>C), AGA285AGG (3160A>G), GGT286CCA (3161G>C 3162G>C 3163T>A), CTC287TAC (3164C>T 3165T>A), CAA288CTT (3168A>T 3169A>T), AAT289AGA (3171A>G 3172T>A), AAC290AAA (3175C>A), ATC291TTT (3176A>T 3178C>T), TGT292GTC (3179T>G 3180G>T 3181T>C), CTC293CTA (3184C>A), GTC294GTG (3187C>G), TAC295TTC (3189A>T), CTT296TTT (3191C>T), GAC297GAT (3196C>T), ATT300CTA (3203A>C 3205T>A), GTC301ATC (3206G>A), ACT304CTT (3215A>C 3216C>T), TCC305GAC (3218T>G 3219C>A), CTA306CCT (3222T>C 3223A>T), CAG307CAA (3226G>A), GAA308ACA (3227G>A 3228A>C) | | | | | | | | |

##### Proteins

| | Begin | End | Coverage | Score | Concordance | Matches | Identities | I/D/M/F* | Stop Codons |
|---|---|---|---|---|---|---|---|---|---|
| ORF B (YP_009507248.1) | 193 | 312 | 11.1% | 430 | 50.1% | 120 (100%) | 57 (47.5%) | 0/0/0/0 | 1 |
| Protein mutations: | L195F (2888C>T 2890C>T), V196C (2891G>T 2892T>G 2893A>T), F199Y (2901T>A), K201S (2907A>G 2908G>C), E204K (2915G>A 2917G>A), K205I (2919A>T 2920G>C), I207V (2924A>G), D208P (2927G>C 2928A>C 2929T>G), D209N (2930G>A), S217D (2954A>G 2955G>A), D218E (2959C>G), V219L (2960G>T 2962A>G), K222E (2969A>G), G224H (2975G>C 2976G>A), K225G (2978A>G 2979A>G 2980G>A), C226A (2981T>G 2982G>C), Q227I (2984C>A 2985A>T), Y228I (2987T>A 2988A>T 2989C>T), T230P (2993A>C 2995C>A), T231K (2997C>A 2998C>A), L232I (2999T>A 3001A>T), A235C (3008G>T 3009C>G 3010A>T), S236L (3011A>T 3012G>T 3013T>G), F238Y (3018T>A 3019T>C), Y239H (3020T>C), V241I (3026G>A 3028G>T), E242R (3029G>C 3030A>G 3031G>T), M243V (3032A>G 3034G>C), D244Q (3035G>C 3037C>A), Q246N (3041C>A 3043A>T), I248V (3047A>G 3049A>C), S249Y (3051C>A 3052G>T), N254R (3065A>C 3066A>G), V255T (3068G>A 3069T>C 3070A>T), E256H (3071G>C 3073A>T), H257S (3074C>T 3075A>C 3076C>A), F260Y (3084T>A), R264V (3095C>G 3096G>T), M267F (3104A>T 3106G>T), K270T (3114A>C 3115A>C), S272A (3119T>G), Q277* (3134C>T 3136A>G), R278G (3137A>G 3139A>C), V279A (3141T>C 3142T>C), D281N (3146G>A 3148C>T), N282D (3149A>G), V283L (3152G>C 3154C>T), L284F (3155C>T 3157A>C), G286P (3161G>C 3162G>C 3163T>A), L287Y (3164C>T 3165T>A), Q288L (3168A>T 3169A>T), N289R (3171A>G 3172T>A), N290K (3175C>A), I291F (3176A>T 3178C>T), C292V (3179T>G 3180G>T 3181T>C), Y295F (3189A>T), L296F (3191C>T), I300L (3203A>C 3205T>A), V301I (3206G>A), T304L (3215A>C 3216C>T), S305D (3218T>G 3219C>A), L306P (3222T>C 3223A>T), E308T (3227G>A 3228A>C) | | | | | | | | |
| Codon mutations: | AAG192..T (2881G>T), CGT194CGG (2887T>G), CTC195TTT (2888C>T 2890C>T), GTA196TGT (2891G>T 2892T>G 2893A>T), GTT197GTG (2896T>G), GAC198GAT (2899C>T), TTC199TAC (2901T>A), AAG201AGC (2907A>G 2908G>C), TTG202CTT (2909T>C 2911G>T), GAG204AAA (2915G>A 2917G>A), AAG205ATC (2919A>T 2920G>C), ATC207GTC (2924A>G), GAT208CCG (2927G>C 2928A>C 2929T>G), GAC209AAC (2930G>A), TAC211TAT (2938C>T), CCG212CCC (2941G>C), ATA213ATT (2944A>T), CCA214CCC (2947A>C), ATA216ATT (2953A>T), AGT217GAT (2954A>G 2955G>A), GAC218GAG (2959C>G), GTA219TTG (2960G>T 2962A>G), CTT220TTA (2963C>T 2965T>A), GAC221GAT (2968C>T), AAG222GAG (2969A>G), TTA223CTA (2972T>C), GGT224CAT (2975G>C 2976G>A), AAG225GGA (2978A>G 2979A>G 2980G>A), TGC226GCC (2981T>G 2982G>C), CAA227ATA (2984C>A 2985A>T), TAC228ATT (2987T>A 2988A>T 2989C>T), TTC229TTT (2992C>T), ACC230CCA (2993A>C 2995C>A), ACC231AAA (2997C>A 2998C>A), TTA232ATT (2999T>A 3001A>T), TTG234CTT (3005T>C 3007G>T), GCA235TGT (3008G>T 3009C>G 3010A>T), AGT236TTG (3011A>T 3012G>T 3013T>G), GGG237GGA (3016G>A), TTT238TAC (3018T>A 3019T>C), TAT239CAT (3020T>C), CAG240CAA (3025G>A), GTG241ATT (3026G>A 3028G>T), GAG242CGT (3029G>C 3030A>G 3031G>T), ATG243GTC (3032A>G 3034G>C), GAC244CAA (3035G>C 3037C>A), CCT245CCA (3040T>A), CAA246AAT (3041C>A 3043A>T), ATA248GTC (3047A>G 3049A>C), TCG249TAT (3051C>A 3052G>T), ACC251ACT (3058C>T), GCG252GCC (3061G>C), AAC254CGC (3065A>C 3066A>G), GTA255ACT (3068G>A 3069T>C 3070A>T), GAA256CAT (3071G>C 3073A>T), CAC257TCA (3074C>T 3075A>C 3076C>A), GGG258GGC (3079G>C), TTT260TAT (3084T>A), GAA261GAG (3088A>G), TTC262TTT (3091C>T), CTT263CTC (3094T>C), CGA264GTA (3095C>G 3096G>T), ATG267TTT (3104A>T 3106G>T), TTA269CTC (3110T>C 3112A>C), AAA270ACC (3114A>C 3115A>C), AAC271AAT (3118C>T), TCA272GCA (3119T>G), TCT274TCC (3127T>C), ACT275ACG (3130T>G), CAA277TAG (3134C>T 3136A>G), AGA278GGC (3137A>G 3139A>C), GTT279GCC (3141T>C 3142T>C), GAC281AAT (3146G>A 3148C>T), AAT282GAT (3149A>G), GTC283CTT (3152G>C 3154C>T), CTA284TTC (3155C>T 3157A>C), AGA285AGG (3160A>G), GGT286CCA (3161G>C 3162G>C 3163T>A), CTC287TAC (3164C>T 3165T>A), CAA288CTT (3168A>T 3169A>T), AAT289AGA (3171A>G 3172T>A), AAC290AAA (3175C>A), ATC291TTT (3176A>T 3178C>T), TGT292GTC (3179T>G 3180G>T 3181T>C), CTC293CTA (3184C>A), GTC294GTG (3187C>G), TAC295TTC (3189A>T), CTT296TTT (3191C>T), GAC297GAT (3196C>T), ATT300CTA (3203A>C 3205T>A), GTC301ATC (3206G>A), ACT304CTT (3215A>C 3216C>T), TCC305GAC (3218T>G 3219C>A), CTA306CCT (3222T>C 3223A>T), CAG307CAA (3226G>A), GAA308ACA (3227G>A 3228A>C) | | | | | | | | |

*: Inserts / Deletes / Misaligned / Frameshifts

#### Analysis details

This analysis was performed with panviral2.64

### NGS Details (UN8): Badnavirus maculaucubae

#### Assembly

| | |
|---|---|
| **Coverage Length** | 340 (1 contig(s)) |
| **Depth Of Coverage** | 8.4 |
| **Number Of Reads** | 20 |
| **Reads Per Million** | 0.33 rpm (after QC) |
| **Ambiguities** | 0 |
| **Assembly Method** | de novo + reference guided assembly |
| **Consensus Caller** | Bcf Tools |

#### Coverage Map

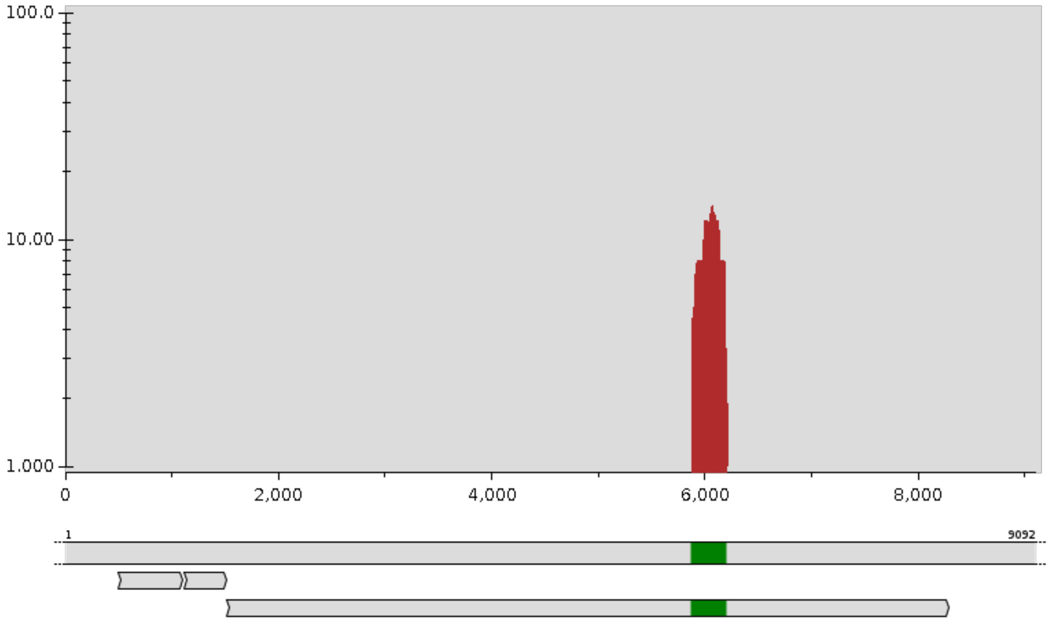

#### Assignment

| | |
|---|---|
| **Type** | Badnavirus maculaucubae (Taxonomy ID: 3051986) |
| **Reference Genome** | NC_076606.1 |
| **NT Identity (%)** | 57.2327 |
| **AA Identity (%)** | 57.5472 |
| **Number Of Stop Codons** | 0 |
| **Number Of CDS** | 3 |

#### Alignment

| | |
|---|---|
| **Alignment Score** | 52.0 (NT) + 297.0 (AA) = 349.0 |
| **Concordance (%)** | 30.7435 |

| Alignment Method | Global, seeded, nucleotide + amino acids (AGA) |
|---|---|

#### Genome Region

Sequence starts at position 5868 and ends at position 6207 relative to NC_076606.1 reference sequence.

#### Alignment Detailed Statistics

| | Begin | End | Coverage | Score | Concordance | Matches | Identities | I/D/M/F* | Stop Codons |
|---|---|---|---|---|---|---|---|---|---|
| **NT** | **5868** | **6207** | **3.7%** | **52** | **8.7%** | **318 (93.5%)** | **182 (53.5%)** | **0/22** | |
| Mutations: | 5868C>A, 5871T>A, 5872A>G, 5873T>A, 5874T>A, 5875C>T, 5877G>A, 5878C>T, 5880G>A, 5884C>T, 5886T>A, 5889T>A, 5892T>A, 5895C>A, 5896C>A, 5899C>A, 5900C>G, 5901C>A, 5902A>T, 5903G>C, 5904T>A, 5906A>C, 5907G>A, 5908A>T, 5909G>C, 5910C>T, 5916T>C, 5920A>T, 5922G>A, 5928T>A, 5934T>G, 5937C>A, 5941T>A, 5944G>A, 5945G>T, 5949T>G, 5950delA, 5951C>A, 5955T>A, 5957A>T, 5958T>C, 5959_5979delCCAGAGACGGGAAAGGAGAAG, 5981A>G, 5985C>A, 5990A>C, 5991A>C, 5992C>A, 5994T>A, 6000C>A, 6001T>A, 6003T>A, 6009C>T, 6012A>G, 6014T>G, 6015G>A, 6018G>C, 6031C>A, 6032A>G, 6033T>A, 6035A>T, 6036A>G, 6040C>G, 6041A>C, 6042A>T, 6045C>T, 6047G>A, 6048C>A, 6049C>T, 6051T>A, 6054C>A, 6056G>A, 6057G>C, 6059T>A, 6060C>A, 6063C>T, 6064T>G, 6065C>A, 6066C>A, 6067A>C, 6069T>A, 6070A>C, 6075A>T, 6076G>A, 6077C>G, 6078C>A, 6081T>A, 6082G>C, 6083G>A, 6085A>G, 6086A>G, 6087T>A, 6088G>A, 6089C>A, 6092A>G, 6094G>A, 6096C>T, 6102C>T, 6111T>C, 6113T>G, 6114G>T, 6120T>A, 6123T>G, 6125T>A, 6127C>T, 6128A>G, 6129C>G, 6132G>A, 6135G>A, 6136G>A, 6137C>A, 6142G>C, 6144A>C, 6145G>C, 6146A>C, 6147A>T, 6148T>G, 6149C>A, 6150A>T, 6153C>T, 6156T>A, 6157C>G, 6158C>A, 6165G>A, 6166G>A, 6168T>A, 6171C>T, 6173T>C, 6174A>T, 6175A>T, 6176C>G, 6183T>A, 6188T>A, 6189A>T, 6191A>T, 6192C>T, 6195A>G, 6199C>T, 6203T>C | | | | | | | | |

##### CDS

| | Begin | End | Coverage | Score | Concordance | Matches | Identities | I/D/M/F* | Stop Codons |
|---|---|---|---|---|---|---|---|---|---|
| QKP79_gp3 | 1454 | 1566 | 5.0% | 297 | 42.0% | 106 (93.8%) | 61 (54.0%) | 0/7/1/1 | 0 |
| Protein mutations: | N1454K (5871T>A), I1455E (5872A>G 5873T>A 5874T>A), P1464R (5899C>A 5900C>G 5901C>A), K1466T (5906A>C 5907G>A), T1471S (5920A>T 5922G>A), I1475M (5934T>G), S1478T (5941T>A), G1479I (5944G>A 5945G>T), D1483V (5957A>T 5958T>C), P1484_K1490del (5959_5979delCCAGAGACGGGAAAGGAGAAG), K1491R (5981A>G), E1494A (5990A>C 5991A>C), F1498I (6001T>A 6003T>A), M1502R (6014T>G 6015G>A), H1508R (6031C>A 6032A>G 6033T>A), K1509M (6035A>T 6036A>G), Q1511A (6040C>G 6041A>C 6042A>T), S1513K (6047G>A 6048C>A), G1516D (6056G>A 6057G>C), I1517K (6059T>A 6060C>A), S1519E (6064T>G 6065C>A 6066C>A), I1520L (6067A>C 6069T>A), I1521L (6070A>C), K1522N (6075A>T), A1523R (6076G>A 6077C>G 6078C>A), G1525Q (6082G>C 6083G>A), N1526G (6085A>G 6086A>G 6087T>A), A1527K (6088G>A 6089C>A), K1528R (6092A>G), V1529I (6094G>A 6096C>T), L1535C (6113T>G 6114G>T), F1539Y (6125T>A), H1540W (6127C>T 6128A>G 6129C>G), A1543K (6136G>A 6137C>A), E1545H (6142G>C 6144A>C), E1546P (6145G>C 6146A>C 6147A>T), S1547D (6148T>G 6149C>A 6150A>T), P1550E (6157C>G 6158C>A), A1553T (6166G>A 6168T>A), I1555T (6173T>C 6174A>T), T1556C (6175A>T 6176C>G), D1558E (6183T>A), L1560H (6188T>A 6189A>T), Y1561F (6191A>T 6192C>T), V1565A (6203T>C) | | | | | | | | |
| Codon mutations: | ATC1453..A (5868C>A), AAT1454AAA (5871T>A), ATT1455GAA (5872A>G 5873T>A 5874T>A), CTG1456TTA (5875C>T 5877G>A), CTG1457TTA (5878C>T 5880G>A), CTT1459TTA (5884C>T 5886T>A), GGT1460GGA (5889T>A), GTT1461GTA (5892T>A), ATC1462ATA (5895C>A), CGA1463AGA (5896C>A), CCC1464AGA (5899C>A 5900C>G 5901C>A), AGT1465TCA (5902A>T 5903G>C 5904T>A), AAG1466ACA (5906A>C 5907G>A), AGC1467TCT (5908A>T 5909G>C 5910C>T), CAT1469CAC (5916T>C), ACG1471TCA (5920A>T 5922G>A), GCT1473GCA (5928T>A), ATT1475ATG (5934T>G), GTC1476GTA (5937C>A), TCC1478ACC (5941T>A), GGA1479ATA (5944G>A 5945G>T), ACT1480ACG (5949T>G), ACA1481-AA (5950delA 5951C>A), ATT1482ATA (5955T>A), GAT1483GTC (5957A>T 5958T>C), CCA1484_AAG1490del (5959_5979delCCAGAGACGGGAAAGGAGAAG), AAA1491AGA (5981A>G), GGC1492GGA (5985C>A), GAA1494GCC (5990A>C 5991A>C), CGT1495AGA (5992C>A 5994T>A), GTC1497GTA (6000C>A), TTT1498ATA (6001T>A 6003T>A), TAC1500TAT (6009C>T), AAA1501AAG (6012A>G), ATG1502AGA (6014T>G 6015G>A), CTG1503CTC (6018G>C), CAT1508AGA (6031C>A 6032A>G 6033T>A), AAA1509ATG (6035A>T 6036A>G), CAA1511GCT (6040C>G 6041A>C 6042A>T), TAC1512TAT (6045C>T), AGC1513AAA (6047G>A 6048C>A), CTT1514TTA (6049C>T 6051T>A), CCC1515CCA (6054C>A), GGG1516GAC (6056G>A 6057G>C), ATC1517AAA (6059T>A 6060C>A), AAC1518AAT (6063C>T), TCC1519GAA (6064T>G 6065C>A 6066C>A), ATT1520CTA (6067A>C 6069T>A), ATA1521CTA (6070A>C), AAA1522AAT (6075A>T), GCC1523AGA (6076G>A 6077C>G 6078C>A), ATT1524ATA (6081T>A), GGA1525CAA (6082G>C 6083G>A), AAT1526GGA (6085A>G 6086A>G 6087T>A), GCA1527AAA (6088G>A 6089C>A), AAA1528AGA (6092A>G), GTC1529ATT (6094G>A 6096C>T), AGC1531AGT (6102C>T), GAT1534GAC (6111T>C), TTG1535TGT (6113T>G 6114G>T), TCT1537TCA (6120T>A), GGT1538GGG (6123T>G), TTT1539TAT (6125T>A), CAC1540TGG (6127C>T 6128A>G 6129C>G), CAG1541CAA (6132G>A), GTG1542GTA (6135G>A), GCA1543AAA (6136G>A 6137C>A), GAA1545CAC (6142G>C 6144A>C), GAA1546CCT (6145G>C 6146A>C 6147A>T), TCA1547GAT (6148T>G 6149C>A 6150A>T), AGC1548AGT (6153C>T), ATT1549ATA (6156T>A), CCA1550GAA (6157C>G 6158C>A), ACG1552ACA (6165G>A), GCT1553ACA (6166G>A 6168T>A), TTC1554TTT (6171C>T), ATA1555ACT (6173T>C 6174A>T), ACT1556TGT (6175A>T 6176C>G), GAT1558GAA (6183T>A), CTA1560CAT (6188T>A 6189A>T), TAC1561TTT (6191A>T 6192C>T), GAA1562GAG (6195A>G), CTA1564TTA (6199C>T), GTA1565GCA (6203T>C) | | | | | | | | |

##### Proteins

| | Begin | End | Coverage | Score | Concordance | Matches | Identities | I/D/M/F* | Stop Codons |
|---|---|---|---|---|---|---|---|---|---|
| hypothetical protein (YP_010799265.1) | 1454 | 1566 | 5.0% | 297 | 42.0% | 106 (93.8%) | 61 (54.0%) | 0/7/1/1 | 0 |
| Protein mutations: | N1454K (5871T>A), I1455E (5872A>G 5873T>A 5874T>A), P1464R (5899C>A 5900C>G 5901C>A), K1466T (5906A>C 5907G>A), T1471S (5920A>T 5922G>A), I1475M (5934T>G), S1478T (5941T>A), G1479I (5944G>A 5945G>T), D1483V (5957A>T 5958T>C), P1484_K1490del (5959_5979delCCAGAGACGGGAAAGGAGAAG), K1491R (5981A>G), E1494A (5990A>C 5991A>C), F1498I (6001T>A 6003T>A), M1502R (6014T>G 6015G>A), H1508R (6031C>A 6032A>G 6033T>A), K1509M (6035A>T 6036A>G), Q1511A (6040C>G 6041A>C 6042A>T), S1513K (6047G>A 6048C>A), G1516D (6056G>A 6057G>C), I1517K (6059T>A 6060C>A), S1519E (6064T>G 6065C>A 6066C>A), I1520L (6067A>C 6069T>A), I1521L (6070A>C), K1522N (6075A>T), A1523R (6076G>A 6077C>G 6078C>A), G1525Q (6082G>C 6083G>A), N1526G (6085A>G 6086A>G 6087T>A), A1527K (6088G>A 6089C>A), K1528R (6092A>G), V1529I (6094G>A 6096C>T), L1535C (6113T>G 6114G>T), F1539Y (6125T>A), H1540W (6127C>T 6128A>G 6129C>G), A1543K (6136G>A 6137C>A), E1545H (6142G>C 6144A>C), E1546P (6145G>C 6146A>C 6147A>T), S1547D (6148T>G 6149C>A 6150A>T), P1550E (6157C>G 6158C>A), A1553T (6166G>A 6168T>A), I1555T (6173T>C 6174A>T), T1556C (6175A>T 6176C>G), D1558E (6183T>A), L1560H (6188T>A 6189A>T), Y1561F (6191A>T 6192C>T), V1565A (6203T>C) | | | | | | | | |
| Codon mutations: | ATC1453..A (5868C>A), AAT1454AAA (5871T>A), ATT1455GAA (5872A>G 5873T>A 5874T>A), CTG1456TTA (5875C>T 5877G>A), CTG1457TTA (5878C>T 5880G>A), CTT1459TTA (5884C>T 5886T>A), GGT1460GGA (5889T>A), GTT1461GTA (5892T>A), ATC1462ATA (5895C>A), CGA1463AGA (5896C>A), CCC1464AGA (5899C>A 5900C>G 5901C>A), AGT1465TCA (5902A>T 5903G>C 5904T>A), AAG1466ACA (5906A>C 5907G>A), AGC1467TCT (5908A>T 5909G>C 5910C>T), CAT1469CAC (5916T>C), ACG1471TCA (5920A>T 5922G>A), GCT1473GCA (5928T>A), ATT1475ATG (5934T>G), GTC1476GTA (5937C>A), TCC1478ACC (5941T>A), GGA1479ATA (5944G>A 5945G>T), ACT1480ACG (5949T>G), ACA1481-AA (5950delA 5951C>A), ATT1482ATA (5955T>A), GAT1483GTC (5957A>T 5958T>C), CCA1484_AAG1490del (5959_5979delCCAGAGACGGGAAAGGAGAAG), AAA1491AGA (5981A>G), GGC1492GGA (5985C>A), GAA1494GCC (5990A>C 5991A>C), CGT1495AGA (5992C>A 5994T>A), GTC1497GTA (6000C>A), TTT1498ATA (6001T>A 6003T>A), TAC1500TAT (6009C>T), AAA1501AAG (6012A>G), ATG1502AGA (6014T>G 6015G>A), CTG1503CTC (6018G>C), CAT1508AGA (6031C>A 6032A>G 6033T>A), AAA1509ATG (6035A>T 6036A>G), CAA1511GCT (6040C>G 6041A>C 6042A>T), TAC1512TAT (6045C>T), AGC1513AAA (6047G>A 6048C>A), CTT1514TTA (6049C>T 6051T>A), CCC1515CCA (6054C>A), GGG1516GAC (6056G>A 6057G>C), ATC1517AAA (6059T>A 6060C>A), AAC1518AAT (6063C>T), TCC1519GAA (6064T>G 6065C>A 6066C>A), ATT1520CTA (6067A>C 6069T>A), ATA1521CTA (6070A>C), AAA1522AAT (6075A>T), GCC1523AGA (6076G>A 6077C>G 6078C>A), ATT1524ATA (6081T>A), GGA1525CAA (6082G>C 6083G>A), AAT1526GGA (6085A>G 6086A>G 6087T>A), GCA1527AAA (6088G>A 6089C>A), AAA1528AGA (6092A>G), GTC1529ATT (6094G>A 6096C>T), AGC1531AGT (6102C>T), GAT1534GAC (6111T>C), TTG1535TGT (6113T>G 6114G>T), TCT1537TCA (6120T>A), GGT1538GGG (6123T>G), TTT1539TAT (6125T>A), CAC1540TGG (6127C>T 6128A>G 6129C>G), CAG1541CAA (6132G>A), GTG1542GTA (6135G>A), GCA1543AAA (6136G>A 6137C>A), GAA1545CAC (6142G>C 6144A>C), GAA1546CCT (6145G>C 6146A>C 6147A>T), TCA1547GAT (6148T>G 6149C>A 6150A>T), AGC1548AGT (6153C>T), ATT1549ATA (6156T>A), CCA1550GAA (6157C>G 6158C>A), ACG1552ACA (6165G>A), GCT1553ACA (6166G>A 6168T>A), TTC1554TTT (6171C>T), ATA1555ACT (6173T>C 6174A>T), ACT1556TGT (6175A>T 6176C>G), GAT1558GAA (6183T>A), CTA1560CAT (6188T>A 6189A>T), TAC1561TTT (6191A>T 6192C>T), GAA1562GAG (6195A>G), CTA1564TTA (6199C>T), GTA1565GCA (6203T>C) | | | | | | | | |

*: Inserts / Deletes / Misaligned / Frameshifts

#### Analysis details

This analysis was performed with panviral2.64

### NGS Details (UN8): Caulimovirus maculatractylodei

#### Assembly

| | |
|---|---|
| **Coverage Length** | 509 (1 contig(s)) |
| **Depth Of Coverage** | 4.7 |
| **Number Of Reads** | 18 |
| **Reads Per Million** | 0.30 rpm (after QC) |
| **Ambiguities** | 0 |
| **Assembly Method** | de novo + reference guided assembly |
| **Consensus Caller** | Bcf Tools |

#### Coverage Map

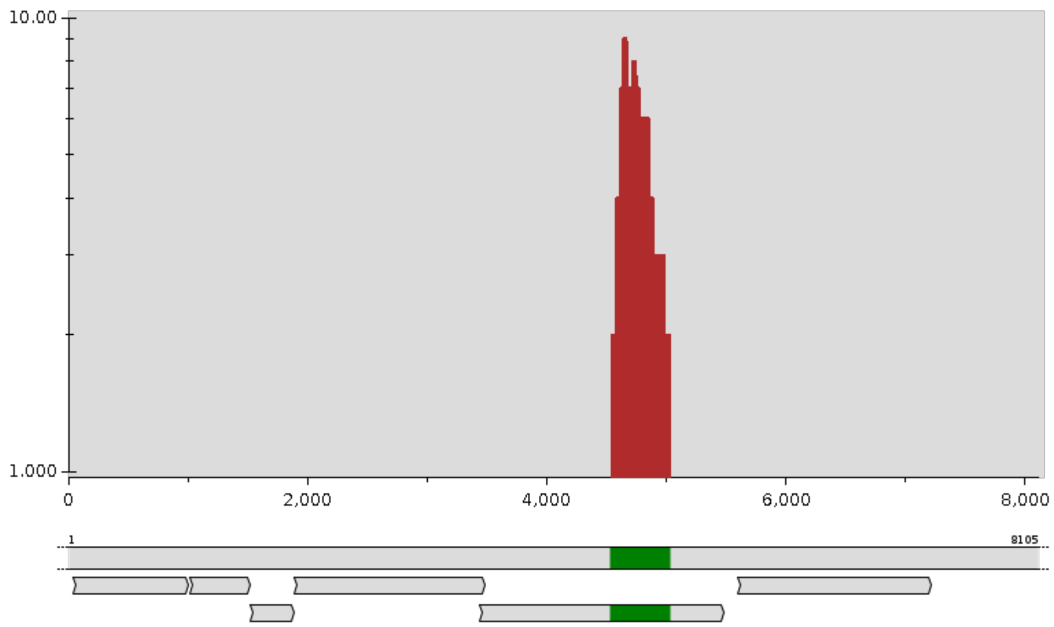

#### Assignment

| | |
|---|---|
| **Type** | Caulimovirus maculatractylodei (Taxonomy ID: 3047382) |
| **Reference Genome** | NC_027924.1 |
| **NT Identity (%)** | 56.66 |
| **AA Identity (%)** | 47.0238 |
| **Number Of Stop Codons** | 0 |
| **Number Of CDS** | 6 |

#### Alignment

| | |
|---|---|
| **Alignment Score** | 110.0 (NT) + 569.0 (AA) = 679.0 |
| **Concordance (%)** | 31.0329 |

| Alignment Method | Global, seeded, nucleotide + amino acids (AGA) |
|---|---|

#### Genome Region

Sequence starts at position 4531 and ends at position 5039 relative to NC_027924.1 reference sequence.

#### Alignment Detailed Statistics

| | Begin | End | Coverage | Score | Concordance | Matches | Identities | I/D/M/F* | Stop Codons |
|---|---|---|---|---|---|---|---|---|---|
| **NT** | **4531** | **5039** | **6.3%** | **110** | **11.2%** | **503 (98.8%)** | **285 (56.0%)** | **0/6** | |
| Mutations: | 4533G>T, 4538C>A, 4539C>T, 4540T>G, 4541G>T, 4542T>C, 4545T>C, 4546A>T, 4547C>T, 4548G>T, 4551T>A, 4554T>A, 4558C>G, 4560G>A, 4566G>T, 4569A>T, 4570G>A, 4572C>G, 4575A>T, 4581A>G, 4584C>T, 4588C>A, 4590G>T, 4592C>T, 4593T>C, 4596G>A, 4599T>A, 4600A>G, 4601T>A, 4605C>T, 4610G>A, 4611A>T, 4612C>A, 4613A>T, 4614C>T, 4620C>T, 4621A>T, 4622A>C, 4623T>A, 4624G>A, 4625C>T, 4626G>T, 4629C>T, 4630_4632delAGG, 4633G>A, 4635G>T, 4636T>C, 4639G>A, 4640A>G, 4641A>C, 4642G>C, 4644A>T, 4649G>C, 4651T>A, 4652G>T, 4656T>C, 4659T>C, 4660G>A, 4669A>G, 4671C>T, 4672C>T, 4674C>A, 4675G>A, 4677C>A, 4680C>T, 4681A>T, 4682G>C, 4683C>A, 4684A>G, 4687A>G, 4690G>A, 4691A>T, 4692A>T, 4693A>G, 4694C>A, 4695G>T, 4696G>T, 4697A>C, 4698A>T, 4701C>T, 4702A>T, 4703G>T, 4704G>C, 4711G>C, 4715T>A, 4720A>T, 4723C>T, 4725A>C, 4726G>A, 4729C>A, 4730A>C, 4731G>T, 4732T>A, 4733G>T, 4739A>T, 4741T>A, 4742T>A, 4743G>C, 4746A>C, 4747A>T, 4749C>T, 4750A>G, 4753C>A, 4756T>A, 4757C>G, 4758C>T, 4759A>G, 4760A>C, 4761G>A, 4768G>A, 4769C>T, 4771C>A, 4773T>G, 4774T>C, 4776G>T, 4779C>T, 4780A>C, 4785A>T, 4786A>C, 4791C>A, 4793A>G, 4794T>A, 4797T>C, 4800A>T, 4803C>A, 4804T>C, 4805T>A, 4806A>T, 4809A>T, 4810A>T, 4812C>A, 4813G>T, 4821C>G, 4822T>A, 4825C>T, 4827C>T, 4834C>A, 4835A>T, 4836G>T, 4837A>T, 4838A>G, 4839T>C, 4840C>A, 4841A>G, 4842T>A, 4844T>C, 4846C>A, 4848G>T, 4852C>T, 4853A>T, 4855A>G, 4856T>C, 4858C>G, 4863G>A, 4869A>T, 4874G>C, 4876T>A, 4878A>C, 4879G>A, 4880A>C, 4881A>T, 4884T>C, 4887G>A, 4889A>C, 4890G>A, 4905C>T, 4908C>T, 4911G>A, 4913T>G, 4919C>A, 4923C>T, 4925C>T, 4927T>G, 4934A>T, 4941T>C, 4944G>A, 4945T>G, 4947A>C, 4948G>A, 4949C>G, 4950T>G, 4953G>C, 4956A>T, 4957A>T, 4959G>C, 4965T>C, 4968T>G, 4969C>T, 4970A>T, 4971A>T, 4973C>A, 4974T>C, 4976A>G, 4977A>G, 4980T>C, 4983G>A, 4986A>G, 4990_4992delGTG, 4993A>C, 4995T>C, 4996T>C, 4997G>A, 4998G>A, 4999T>C, 5007T>C, 5008G>A, 5010T>G, 5011A>G, 5012G>A, 5014G>C, 5016T>C, 5020A>G, 5021G>A, 5023C>G, 5024T>C, 5026A>G, 5028G>A, 5031C>T, 5034G>T, 5037C>T | | | | | | | | |

##### CDS

| | Begin | End | Coverage | Score | Concordance | Matches | Identities | I/D/M/F* | Stop Codons |
|---|---|---|---|---|---|---|---|---|---|
| APL28_gp5 | 364 | 533 | 25.0% | 569 | 47.0% | 168 (98.8%) | 79 (46.5%) | 0/2/0/0 | 0 |
| Protein mutations: | T366N (4538C>A 4539C>T), C367V (4540T>G 4541G>T 4542T>C), T369F (4546A>T 4547C>T 4548G>T), H371Q (4554T>A), Q373E (4558C>G 4560G>A), K375N (4566G>T), V377M (4570G>A 4572C>G), Q383N (4588C>A 4590G>T), A384V (4592C>T 4593T>C), I387E (4600A>G 4601T>A), R390N (4610G>A 4611A>T), H391I (4612C>A 4613A>T 4614C>T), N394S (4621A>T 4622A>C 4623T>A), A395I (4624G>A 4625C>T 4626G>T), R397del (4630_4632delAGG), E398N (4633G>A 4635G>T), F399L (4636T>C), E400S (4639G>A 4640A>G 4641A>C), E401H (4642G>C 4644A>T), C403S (4649G>C), C404I (4651T>A 4652G>T), V407I (4660G>A), I410V (4669A>G 4671C>T), V412I (4675G>A 4677C>A), K415E (4684A>G), N416D (4687A>G), E417I (4690G>A 4691A>T 4692A>T), T418D (4693A>G 4694C>A 4695G>T), E419S (4696G>T 4697A>C 4698A>T), R421F (4702A>T 4703G>T 4704G>C), V424L (4711G>C), I425N (4715T>A), I427F (4720A>T), L428F (4723C>T 4725A>C), E429K (4726G>A), Q430T (4729C>A 4730A>C 4731G>T), C431I (4732T>A 4733G>T), N433I (4739A>T), L434N (4741T>A 4742T>A 4743G>C), I436F (4747A>T 4749C>T), I437V (4750A>G), L438I (4753C>A), K440A (4759A>G 4760A>C 4761G>A), A443I (4768G>A 4769C>T), H444K (4771C>A 4773T>G), K447Q (4780A>C), K449Q (4786A>C), N451R (4793A>G 4794T>A), L455H (4804T>C 4805T>A 4806A>T), E456D (4809A>T), I457L (4810A>T 4812C>A), D458Y (4813G>T), S461T (4822T>A), H462Y (4825C>T 4827C>T), Q465I (4834C>A 4835A>T 4836G>T), N466C (4837A>T 4838A>G 4839T>C), H467R (4840C>A 4841A>G 4842T>A), I468T (4844T>C), L469I (4846C>A 4848G>T), H471F (4852C>T 4853A>T), I472A (4855A>G 4856T>C), H473D (4858C>G), R478T (4874G>C), L479I (4876T>A 4878A>C), E480T (4879G>A 4880A>C 4881A>T), K483T (4889A>C 4890G>A), I491S (4913T>G), T493N (4919C>A), A495V (4925C>T), S496A (4927T>G), Y498F (4934A>T), L502V (4945T>G 4947A>C), A503R (4948G>A 4949C>G 4950T>G), Q504H (4953G>C), R506C (4957A>T 4959G>C), F509L (4968T>G), Q510F (4969C>T 4970A>T 4971A>T), A511D (4973C>A 4974T>C), K512R (4976A>G 4977A>G), V517del (4990_4992delGTG), T518P (4993A>C 4995T>C), W519Q (4996T>C 4997G>A 4998G>A), S520P (4999T>C), D523K (5008G>A 5010T>G), S524D (5011A>G 5012G>A), D525H (5014G>C 5016T>C), S527D (5020A>G 5021G>A), L528A (5023C>G 5024T>C), M529V (5026A>G 5028G>A), K531N (5034G>T) | | | | | | | | |
| Codon mutations: | GCG364GCT (4533G>T), ACC366AAT (4538C>A 4539C>T), TGT367GTC (4540T>G 4541G>T 4542T>C), CCT368CCC (4545T>C), ACG369TTT (4546A>T 4547C>T 4548G>T), GGT370GGA (4551T>A), CAT371CAA (4554T>A), CAG373GAA (4558C>G 4560G>A), AAG375AAT (4566G>T), GTA376GTT (4569A>T), GTC377ATG (4570G>A 4572C>G), CCA378CCT (4575A>T), GGA380GGG (4581A>G), CTC381CTT (4584C>T), CAG383AAT (4588C>A 4590G>T), GCT384GTC (4592C>T 4593T>C), CCG385CCA (4596G>A), TCT386TCA (4599T>A), ATA387GAA (4600A>G 4601T>A), TTC388TTT (4605C>T), AGA390AAT (4610G>A 4611A>T), CAC391ATT (4612C>A 4613A>T 4614C>T), AAC393AAT (4620C>T), AAT394TCA (4621A>T 4622A>C 4623T>A), GCG395ATT (4624G>A 4625C>T 4626G>T), TTC396TTT (4629C>T), AGG397del (4630_4632delAGG), GAG398AAT (4633G>A 4635G>T), TTT399CTT (4636T>C), GAA400AGC (4639G>A 4640A>G 4641A>C), GAA401CAT (4642G>C 4644A>T), TGT403TCT (4649G>C), TGT404ATT (4651T>A 4652G>T), GTT405GTC (4656T>C), TAT406TAC (4659T>C), GTT407ATT (4660G>A), ATC410GTT (4669A>G 4671C>T), CTC411TTA (4672C>T 4674C>A), GTC412ATA (4675G>A 4677C>A), TTC413TTT (4680C>T), AGC414TCA (4681A>T 4682G>C 4683C>A), AAA415GAA (4684A>G), AAT416GAT (4687A>G), GAA417ATT (4690G>A 4691A>T 4692A>T), ACG418GAT (4693A>G 4694C>A 4695G>T), GAA419TCT (4696G>T 4697A>C 4698A>T), CAC420CAT (4701C>T), AGG421TTC (4702A>T 4703G>T 4704G>C), GTC424CTC (4711G>C), ATT425AAT (4715T>A), ATC427TTC (4720A>T), CTA428TTC (4723C>T 4725A>C), GAA429AAA (4726G>A), CAG430ACT (4729C>A 4730A>C 4731G>T), TGT431ATT (4732T>A 4733G>T), AAC433ATC (4739A>T), TTG434AAC (4741T>A 4742T>A 4743G>C), GGA435GGC (4746A>C), ATC436TTT (4747A>T 4749C>T), ATC437GTC (4750A>G), CTT438ATT (4753C>A), TCC439AGT (4756T>A 4757C>G 4758C>T), AAG440GCA (4759A>G 4760A>C 4761G>A), GCA443ATA (4768G>A 4769C>T), CAT444AAG (4771C>A 4773T>G), TTG445CTT (4774T>C 4776G>T), TTC446TTT (4779C>T), AAA447CAA (4780A>C), ACA448ACT (4785A>T), AAG449CAG (4786A>C), ATC450ATA (4791C>A), AAT451AGA (4793A>G 4794T>A), TTT452TTC (4797T>C), CTA453CTT (4800A>T), GGC454GGA (4803C>A), TTA455CAT (4804T>C 4805T>A 4806A>T), GAA456GAT (4809A>T), ATC457TTA (4810A>T 4812C>A), GAT458TAT (4813G>T), GGC460GGG (4821C>G), TCT461ACT (4822T>A), CAC462TAT (4825C>T 4827C>T), CAG465ATT (4834C>A 4835A>T 4836G>T), AAT466TGC (4837A>T 4838A>G 4839T>C), CAT467AGA (4840C>A 4841A>G 4842T>A), ATT468ACT (4844T>C), CTG469ATT (4846C>A 4848G>T), CAT471TTT (4852C>T 4853A>T), ATT472GCT (4855A>G 4856T>C), CAT473GAT (4858C>G), AAG474AAA (4863G>A), CCA476CCT (4869A>T), AGA478ACA (4874G>C), TTA479ATC (4876T>A 4878A>C), GAA480ACT (4879G>A 4880A>C 4881A>T), GAT481GAC (4884T>C), AAG482AAA (4887G>A), AAG483ACA (4889A>C 4890G>A), TTC488TTT (4905C>T), CTC489CTT (4908C>T), GGG490GGA (4911G>A), ATC491AGC (4913T>G), ACT493AAT (4919C>A), TAC494TAT (4923C>T), GCT495GTT (4925C>T), TCT496GCT (4927T>G), TAC498TTC (4934A>T), CCT500CCC (4941T>C), AAG501AAA (4944G>A), TTA502GTC (4945T>G 4947A>C), GCT503AGG (4948G>A 4949C>G 4950T>G), CAG504CAC (4953G>C), ATA505ATT (4956A>T), AGG506TGC (4957A>T 4959G>C), CCT508CCC (4965T>C), TTT509TTG (4968T>G), CAA510TTT (4969C>T 4970A>T 4971A>T), GCT511GAC (4973C>A 4974T>C), AAA512AGG (4976A>G 4977A>G), CTT513CTC (4980T>C), AAG514AAA (4983G>A), AAA515AAG (4986A>G), GTG517del (4990_4992delGTG), ACT518CCC (4993A>C 4995T>C), TGG519CAA (4996T>C 4997G>A 4998G>A), TCT520CCT (4999T>C), ACT522ACC (5007T>C), GAT523AAG (5008G>A 5010T>G), AGT524GAT (5011A>G 5012G>A), GAT525CAC (5014G>C 5016T>C), AGT527GAT (5020A>G 5021G>A), CTT528GCT (5023C>G 5024T>C), ATG529GTA (5026A>G 5028G>A), ATC530ATT (5031C>T), AAG531AAT (5034G>T), ATC532ATT (5037C>T) | | | | | | | | |

##### Proteins

| | Begin | End | Coverage | Score | Concordance | Matches | Identities | I/D/M/F* | Stop Codons |
|---|---|---|---|---|---|---|---|---|---|
| ORF5 (YP_009165750.1) | 364 | 533 | 25.0% | 569 | 47.0% | 168 (98.8%) | 79 (46.5%) | 0/2/0/0 | 0 |
| Protein mutations: | T366N (4538C>A 4539C>T), C367V (4540T>G 4541G>T 4542T>C), T369F (4546A>T 4547C>T 4548G>T), H371Q (4554T>A), Q373E (4558C>G 4560G>A), K375N (4566G>T), V377M (4570G>A 4572C>G), Q383N (4588C>A 4590G>T), A384V (4592C>T 4593T>C), I387E (4600A>G 4601T>A), R390N (4610G>A 4611A>T), H391I (4612C>A 4613A>T 4614C>T), N394S (4621A>T 4622A>C 4623T>A), A395I (4624G>A 4625C>T 4626G>T), R397del (4630_4632delAGG), E398N (4633G>A 4635G>T), F399L (4636T>C), E400S (4639G>A 4640A>G 4641A>C), E401H (4642G>C 4644A>T), C403S (4649G>C), C404I (4651T>A 4652G>T), V407I (4660G>A), I410V (4669A>G 4671C>T), V412I (4675G>A 4677C>A), K415E (4684A>G), N416D (4687A>G), E417I (4690G>A 4691A>T 4692A>T), T418D (4693A>G 4694C>A 4695G>T), E419S (4696G>T 4697A>C 4698A>T), R421F (4702A>T 4703G>T 4704G>C), V424L (4711G>C), I425N (4715T>A), I427F (4720A>T), L428F (4723C>T 4725A>C), E429K (4726G>A), Q430T (4729C>A 4730A>C 4731G>T), C431I (4732T>A 4733G>T), N433I (4739A>T), L434N (4741T>A 4742T>A 4743G>C), I436F (4747A>T 4749C>T), I437V (4750A>G), L438I (4753C>A), K440A (4759A>G 4760A>C 4761G>A), A443I (4768G>A 4769C>T), H444K (4771C>A 4773T>G), K447Q (4780A>C), K449Q (4786A>C), N451R (4793A>G 4794T>A), L455H (4804T>C 4805T>A 4806A>T), E456D (4809A>T), I457L (4810A>T 4812C>A), D458Y (4813G>T), S461T (4822T>A), H462Y (4825C>T 4827C>T), Q465I (4834C>A 4835A>T 4836G>T), N466C (4837A>T 4838A>G 4839T>C), H467R (4840C>A 4841A>G 4842T>A), I468T (4844T>C), L469I (4846C>A 4848G>T), H471F (4852C>T 4853A>T), I472A (4855A>G 4856T>C), H473D (4858C>G), R478T (4874G>C), L479I (4876T>A 4878A>C), E480T (4879G>A 4880A>C 4881A>T), K483T (4889A>C 4890G>A), I491S (4913T>G), T493N (4919C>A), A495V (4925C>T), S496A (4927T>G), Y498F (4934A>T), L502V (4945T>G 4947A>C), A503R (4948G>A 4949C>G 4950T>G), Q504H (4953G>C), R506C (4957A>T 4959G>C), F509L (4968T>G), Q510F (4969C>T 4970A>T 4971A>T), A511D (4973C>A 4974T>C), K512R (4976A>G 4977A>G), V517del (4990_4992delGTG), T518P (4993A>C 4995T>C), W519Q (4996T>C 4997G>A 4998G>A), S520P (4999T>C), D523K (5008G>A 5010T>G), S524D (5011A>G 5012G>A), D525H (5014G>C 5016T>C), S527D (5020A>G 5021G>A), L528A (5023C>G 5024T>C), M529V (5026A>G 5028G>A), K531N (5034G>T) | | | | | | | | |

| | Begin | End | Coverage | Score | Concordance | Matches | Identities | I/D/M/F* | Stop Codons |
|---|---|---|---|---|---|---|---|---|---|
| **NT** | **4531** | **5039** | **6.3%** | **110** | **11.2%** | **503 (98.8%)** | **285 (56.0%)** | **0/6** | |
| Codon mutations: | GCG364GCT (4533G>T), ACC366AAT (4538C>A 4539C>T), TGT367GTC (4540T>G 4541G>T 4542T>C), CCT368CCC (4545T>C), ACG369TTT (4546A>T 4547C>T 4548G>T), GGT370GGA (4551T>A), CAT371CAA (4554T>A), CAG373GAA (4558C>G 4560G>A), AAG375AAT (4566G>T), GTA376GTT (4569A>T), GTC377ATG (4570G>A 4572C>G), CCA378CCT (4575A>T), GGA380GGG (4581A>G), CTC381CTT (4584C>T), CAG383AAT (4588C>A 4590G>T), GCT384GTC (4592C>T 4593T>C), CCG385CCA (4596G>A), TCT386TCA (4599T>A), ATA387GAA (4600A>G 4601T>A), TTC388TTT (4605C>T), AGA390AAT (4610G>A 4611A>T), CAC391ATT (4612C>A 4613A>T 4614C>T), AAC393AAT (4620C>T), AAT394TCA (4621A>T 4622A>C 4623T>A), GCG395ATT (4624G>A 4625C>T 4626G>T), TTC396TTT (4629C>T), AGG397del (4630_4632delAGG), GAG398AAT (4633G>A 4635G>T), TTT399CTT (4636T>C), GAA400AGC (4639G>A 4640A>G 4641A>C), GAA401CAT (4642G>C 4644A>T), TGT403TCT (4649G>C), TGT404ATT (4651T>A 4652G>T), GTT405GTC (4656T>C), TAT406TAC (4659T>C), GTT407ATT (4660G>A), ATC410GTT (4669A>G 4671C>T), CTC411TTA (4672C>T 4674C>A), GTC412ATA (4675G>A 4677C>A), TTC413TTT (4680C>T), AGC414TCA (4681A>T 4682G>C 4683C>A), AAA415GAA (4684A>G), AAT416GAT (4687A>G), GAA417ATT (4690G>A 4691A>T 4692A>T), ACG418GAT (4693A>G 4694C>A 4695G>T), GAA419TCT (4696G>T 4697A>C 4698A>T), CAC420CAT (4701C>T), AGG421TTC (4702A>T 4703G>T 4704G>C), GTC424CTC (4711G>C), ATT425AAT (4715T>A), ATC427TTC (4720A>T), CTA428TTC (4723C>T 4725A>C), GAA429AAA (4726G>A), CAG430ACT (4729C>A 4730A>C 4731G>T), TGT431ATT (4732T>A 4733G>T), AAC433ATC (4739A>T), TTG434AAC (4741T>A 4742T>A 4743G>C), GGA435GGC (4746A>C), ATC436TTT (4747A>T 4749C>T), ATC437GTC (4750A>G), CTT438ATT (4753C>A), TCC439AGT (4756T>A 4757C>G 4758C>T), AAG440GCA (4759A>G 4760A>C 4761G>A), GCA443ATA (4768G>A 4769C>T), CAT444AAG (4771C>A 4773T>G), TTG445CTT (4774T>C 4776G>T), TTC446TTT (4779C>T), AAA447CAA (4780A>C), ACA448ACT (4785A>T), AAG449CAG (4786A>C), ATC450ATA (4791C>A), AAT451AGA (4793A>G 4794T>A), TTT452TTC (4797T>C), CTA453CTT (4800A>T), GGC454GGA (4803C>A), TTA455CAT (4804T>C 4805T>A 4806A>T), GAA456GAT (4809A>T), ATC457TTA (4810A>T 4812C>A), GAT458TAT (4813G>T), GGC460GGG (4821C>G), TCT461ACT (4822T>A), CAC462TAT (4825C>T 4827C>T), CAG465ATT (4834C>A 4835A>T 4836G>T), AAT466TGC (4837A>T 4838A>G 4839T>C), CAT467AGA (4840C>A 4841A>G 4842T>A), ATT468ACT (4844T>C), CTG469ATT (4846C>A 4848G>T), CAT471TTT (4852C>T 4853A>T), ATT472GCT (4855A>G 4856T>C), CAT473GAT (4858C>G), AAG474AAA (4863G>A), CCA476CCT (4869A>T), AGA478ACA (4874G>C), TTA479ATC (4876T>A 4878A>C), GAA480ACT (4879G>A 4880A>C 4881A>T), GAT481GAC (4884T>C), AAG482AAA (4887G>A), AAG483ACA (4889A>C 4890G>A), TTC488TTT (4905C>T), CTC489CTT (4908C>T), GGG490GGA (4911G>A), ATC491AGC (4913T>G), ACT493AAT (4919C>A), TAC494TAT (4923C>T), GCT495GTT (4925C>T), TCT496GCT (4927T>G), TAC498TTC (4934A>T), CCT500CCC (4941T>C), AAG501AAA (4944G>A), TTA502GTC (4945T>G 4947A>C), GCT503AGG (4948G>A 4949C>G 4950T>G), CAG504CAC (4953G>C), ATA505ATT (4956A>T), AGG506TGC (4957A>T 4959G>C), CCT508CCC (4965T>C), TTT509TTG (4968T>G), CAA510TTT (4969C>T 4970A>T 4971A>T), GCT511GAC (4973C>A 4974T>C), AAA512AGG (4976A>G 4977A>G), CTT513CTC (4980T>C), AAG514AAA (4983G>A), AAA515AAG (4986A>G), GTG517del (4990_4992delGTG), ACT518CCC (4993A>C 4995T>C), TGG519CAA (4996T>C 4997G>A 4998G>A), TCT520CCT (4999T>C), ACT522ACC (5007T>C), GAT523AAG (5008G>A 5010T>G), AGT524GAT (5011A>G 5012G>A), GAT525CAC (5014G>C 5016T>C), AGT527GAT (5020A>G 5021G>A), CTT528GCT (5023C>G 5024T>C), ATG529GTA (5026A>G 5028G>A), ATC530ATT (5031C>T), AAG531AAT (5034G>T), ATC532ATT (5037C>T) | | | | | | | | |

*: Inserts / Deletes / Misaligned / Frameshifts

#### Analysis details

This analysis was performed with panviral2.64

#### NGS Details (UN8): Cavemovirus venamanihotis

##### Assembly

| | |
|---|---|
| Coverage Length | 379 (1 contig(s)) |
| Depth Of Coverage | 4.8 |
| Number Of Reads | 15 |
| Reads Per Million | 0.25 rpm (after QC) |
| Ambiguities | 0 |
| Assembly Method | de novo + reference guided assembly |
| Consensus Caller | Bcf Tools |

##### Coverage Map

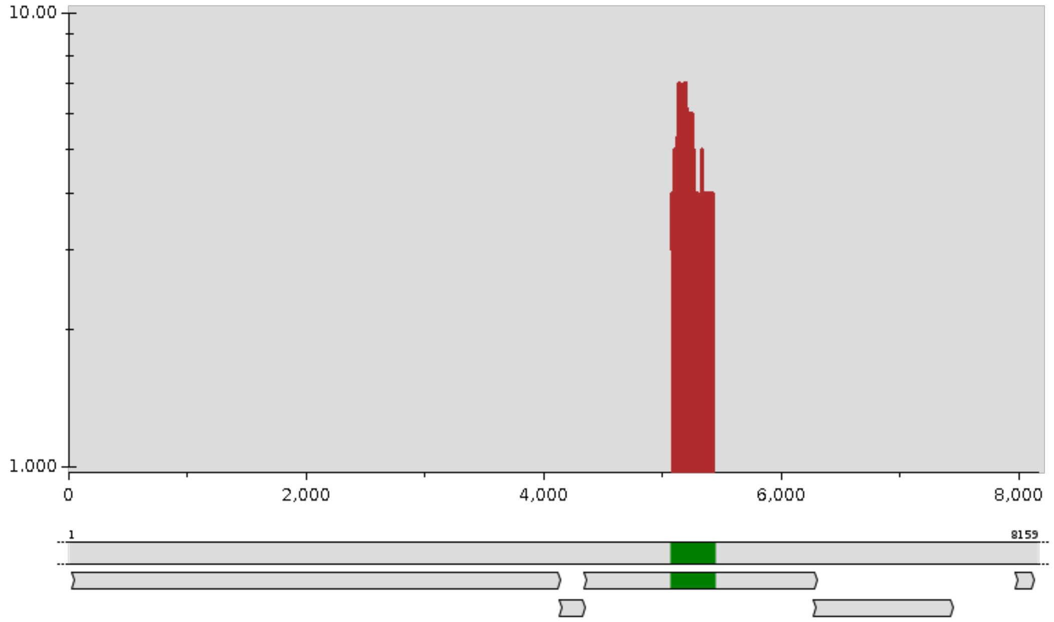

##### Assignment

| | |
|---|---|
| Type | Cavemovirus venamanihotis (Taxonomy ID: 3047736) |
| Reference Genome | NC_001648.1 |
| NT Identity (%) | 60.1583 |
| AA Identity (%) | 50.0 |
| Number Of Stop Codons | 2 |
| Number Of CDS | 5 |

##### Alignment

| | |
|---|---|
| Alignment Score | 154.0 (NT) + 452.0 (AA) = 606.0 |
| Concordance (%) | 36.2657 |

| **Alignment Method** | Global, seeded, nucleotide + amino acids (AGA) |
|---|---|

#### Genome Region

Sequence starts at position 5072 and ends at position 5450 relative to NC_001648.1 reference sequence.

#### Alignment Detailed Statistics

| | Begin | End | Coverage | Score | Concordance | Matches | Identities | I/D/M/F* | Stop Codons |
|---|---|---|---|---|---|---|---|---|---|
| **NT** | **5072** | **5450** | **4.6%** | **154** | **20.3%** | **379 (100%)** | **228 (60.2%)** | **0/0** | |
| Mutations: | 5075T>G, 5076A>T, 5079C>A, 5081A>T, 5088A>T, 5089T>A, 5090A>T, 5093A>C, 5099A>T, 5100C>A, 5101A>C, 5102T>G, 5103T>G, 5109C>A, 5112A>G, 5115A>C, 5118G>A, 5119G>C, 5121A>G, 5122A>T, 5123A>T, 5124A>C, 5127A>C, 5130A>T, 5132G>A, 5135A>T, 5139G>A, 5141T>C, 5148G>T, 5149A>C, 5160A>T, 5161A>T, 5162A>T, 5163G>T, 5164C>T, 5165T>A, 5168A>G, 5169G>T, 5170T>G, 5172G>A, 5174A>T, 5176A>G, 5177A>G, 5186C>T, 5196G>A, 5197A>G, 5198T>G, 5199A>G, 5200C>A, 5201A>C, 5205A>T, 5208C>T, 5209A>C, 5214A>T, 5215G>T, 5216T>A, 5217A>T, 5218T>A, 5219A>T, 5220C>G, 5222G>T, 5227G>A, 5228A>C, 5229T>A, 5230A>T, 5231C>T, 5233A>T, 5235A>T, 5236G>C, 5237T>A, 5248G>T, 5249T>A, 5260T>A, 5263A>G, 5264T>G, 5267T>A, 5270A>T, 5271A>T, 5274T>A, 5276A>T, 5277G>T, 5278A>T, 5279A>T, 5280G>A, 5285T>G, 5286A>C, 5287G>A, 5290A>C, 5291A>T, 5292A>T, 5294A>T, 5295T>A, 5296A>G, 5297T>A, 5300A>T, 5302C>T, 5308C>A, 5309A>T, 5315G>T, 5316C>T, 5317A>T, 5318G>T, 5322T>C, 5323A>C, 5324T>A, 5328C>G, 5330G>A, 5336A>C, 5337G>T, 5339A>T, 5340T>A, 5342A>G, 5345A>T, 5349G>T, 5351A>T, 5353A>T, 5354T>G, 5355C>A, 5357T>A, 5360C>T, 5361T>G, 5362C>T, 5363A>C, 5366A>T, 5367A>T, 5368G>C, 5370A>G, 5371T>A, 5372T>A, 5375C>T, 5378A>G, 5379C>A, 5382T>A, 5384T>C, 5388G>A, 5391A>G, 5392G>A, 5393A>T, 5396A>T, 5401G>A, 5402A>T, 5410A>T, 5411T>A, 5420T>C, 5426G>T, 5438T>C, 5441A>T, 5444A>G, 5445G>A, 5447C>T, 5449T>A | | | | | | | | |

##### CDS

| | Begin | End | Coverage | Score | Concordance | Matches | Identities | I/D/M/F* | Stop Codons |
|---|---|---|---|---|---|---|---|---|---|
| CsVMVgp3 | 244 | 369 | 19.3% | 452 | 49.0% | 126 (100%) | 63 (50.0%) | 0/0/0/0 | 2 |
| Protein mutations: | S245C (5076A>T), P246T (5079C>A 5081A>T), I249Y (5088A>T 5089T>A 5090A>T), K252N (5099A>T), H253T (5100C>A 5101A>C 5102T>G), S254A (5103T>G), Q256K (5109C>A), K257E (5112A>G), G259T (5118G>A 5119G>C), K260V (5121A>G 5122A>T 5123A>T), T261P (5124A>C), M263L (5130A>T 5132G>A), D266N (5139G>A 5141T>C), D269S (5148G>T 5149A>C), K273F (5160A>T 5161A>T 5162A>T), A274L (5163G>T 5164C>T 5165T>A), V276* (5169G>T 5170T>G), V277I (5172G>A 5174A>T), K278R (5176A>G 5177A>G), D285R (5196G>A 5197A>G 5198T>G), T286D (5199A>G 5200C>A 5201A>C), I288L (5205A>T), H289S (5208C>T 5209A>C), S291L (5214A>T 5215G>T 5216T>A), I292Y (5217A>T 5218T>A 5219A>T), Q293D (5220C>G 5222G>T), R295N (5227G>A 5228A>C), Y296I (5229T>A 5230A>T 5231C>T), Y297F (5233A>T), C302L (5248G>T 5249T>A), F306Y (5260T>A), Y307W (5263A>G 5264T>G), H308Q (5267T>A), K310* (5271A>T), L311I (5274T>A 5276A>T), E312F (5277G>T 5278A>T 5279A>T), E313K (5280G>A), D314E (5285T>G), S315H (5286A>C 5287G>A), K316T (5290A>C 5291A>T), K317Y (5292A>T 5294A>T), Y318R (5295T>A 5296A>G 5297T>A), A320V (5302C>T), T322N (5308C>A 5309A>T), Q325F (5316C>T 5317A>T 5318G>T), Y327P (5322T>C 5323A>C 5324T>A), Q329E (5328C>G 5330G>A), K331N (5336A>C), V332F (5337G>T 5339A>T), L333M (5340T>A 5342A>G), G336C (5349G>T 5351A>T), Y337L (5353A>T 5354T>G), H338K (5355C>A 5357T>A), S340V (5361T>G 5362C>T 5363A>C), I343E (5370A>G 5371T>A 5372T>A), Q346K (5379C>A), F347I (5382T>A 5384T>C), D349N (5388G>A), R350D (5391A>G 5392G>A 5393A>T), R353N (5401G>A 5402A>T), Y356L (5410A>T 5411T>A), V368I (5445G>A 5447C>T), F369Y (5449T>A) | | | | | | | | |
| Codon mutations: | TCT244TCG (5075T>G), AGC245TGC (5076A>T), CCA246ACT (5079C>A 5081A>T), ATA249TAT (5088A>T 5089T>A 5090A>T), GTA250GTC (5093A>C), AAA252AAT (5099A>T), CAT253ACG (5100C>A 5101A>C 5102T>G), TCA254GCA (5103T>G), CAA256AAA (5109C>A), AAA257GAA (5112A>G), AGA258CGA (5115A>C), GGA259ACA (5118G>A 5119G>C), AAA260GTT (5121A>G 5122A>T 5123A>T), ACT261CCT (5124A>C), AGA262CGA (5127A>C), ATG263TTA (5130A>T 5132G>A), GTA264GTT (5135A>T), GAT266AAC (5139G>A 5141T>C), GAT269TCT (5148G>T 5149A>C), AAA273TTT (5160A>T 5161A>T 5162A>T), GCT274TTA (5163G>T 5164C>T 5165T>A), AAA275AAG (5168A>G), GTA276TGA (5169G>T 5170T>G), GTA277ATT (5172G>A 5174A>T), AAA278AGG (5176A>G 5177A>G), ATC281ATT (5186C>T), GAT285AGG (5196G>A 5197A>G 5198T>G), ACA286GAC (5199A>G 5200C>A 5201A>C), ATA288TTA (5205A>T), CAT289TCT (5208C>T 5209A>C), AGT291TTA (5214A>T 5215G>T 5216T>A), ATA292TAT (5217A>T 5218T>A 5219A>T), CAG293GAT (5220C>G 5222G>T), AGA295AAC (5227G>A 5228A>C), TAC296ATT (5229T>A 5230A>T 5231C>T), TAT297TTT (5233A>T), AGT298TCA (5235A>T 5236G>C 5237T>A), TGT302TTA (5248G>T 5249T>A), TTT306TAT (5260T>A), TAT307TGG (5263A>G 5264T>G), CAT308CAA (5267T>A), ATA309ATT (5270A>T), AAG310TAG (5271A>T), TTA311ATT (5274T>A 5276A>T), GAA312TTT (5277G>T 5278A>T 5279A>T), GAA313AAA (5280G>A), GAT314GAG (5285T>G), AGT315CAT (5286A>C 5287G>A), AAA316ACT (5290A>C 5291A>T), AAA317TAT (5292A>T 5294A>T), TAT318AGA (5295T>A 5296A>G 5297T>A), ACA319ACT (5300A>T), GCT320GTT (5302C>T), ACA322AAT (5308C>A 5309A>T), CCG324CCT (5315G>T), CAG325TTT (5316C>T 5317A>T 5318G>T), TAT327CCA (5322T>C 5323A>C 5324T>A), CAG329GAA (5328C>G 5330G>A), AAA331AAC (5336A>C), GTA332TTT (5337G>T 5339A>T), TTA333ATG (5340T>A 5342A>G), CCA334CCT (5345A>T), GGA336TGT (5349G>T 5351A>T), TAT337TTG (5353A>T 5354T>G), CAT338AAA (5355C>A 5357T>A), AAC339AAT (5360C>T), TCA340GTC (5361T>G 5362C>T 5363A>C), CCA341CCT (5366A>T), AGT342TCT (5367A>T 5368G>C), ATT343GAA (5370A>G 5371T>A 5372T>A), TTC344TTT (5375C>T), CAA345CAG (5378A>G), CAA346AAA (5379C>A), TTT347ATC (5382T>A 5384T>C), GAT349AAT (5388G>A), AGA350GAT (5391A>G 5392G>A 5393A>T), ATA351ATT (5396A>T), AGA353AAT (5401G>A 5402A>T), TAT356TTA (5410A>T 5411T>A), ATT359ATC (5420T>C), GTG361GTT (5426G>T), GAT365GAC (5438T>C), ATA366ATT (5441A>T), TTA367TTG (5444A>G), GTC368ATT (5445G>A 5447C>T), TTT369TAT (5449T>A) | | | | | | | | |

##### Proteins

| | Begin | End | Coverage | Score | Concordance | Matches | Identities | I/D/M/F* | Stop Codons |
|---|---|---|---|---|---|---|---|---|---|
| aspartic protease/reverse transcriptase (NP_056848.1) | 244 | 369 | 19.3% | 452 | 49.0% | 126 (100%) | 63 (50.0%) | 0/0/0/0 | 2 |
| Protein mutations: | S245C (5076A>T), P246T (5079C>A 5081A>T), I249Y (5088A>T 5089T>A 5090A>T), K252N (5099A>T), H253T (5100C>A 5101A>C 5102T>G), S254A (5103T>G), Q256K (5109C>A), K257E (5112A>G), G259T (5118G>A 5119G>C), K260V (5121A>G 5122A>T 5123A>T), T261P (5124A>C), M263L (5130A>T 5132G>A), D266N (5139G>A 5141T>C), D269S (5148G>T 5149A>C), K273F (5160A>T 5161A>T 5162A>T), A274L (5163G>T 5164C>T 5165T>A), V276* (5169G>T 5170T>G), V277I (5172G>A 5174A>T), K278R (5176A>G 5177A>G), D285R (5196G>A 5197A>G 5198T>G), T286D (5199A>G 5200C>A 5201A>C), I288L (5205A>T), H289S (5208C>T 5209A>C), S291L (5214A>T 5215G>T 5216T>A), I292Y (5217A>T 5218T>A 5219A>T), Q293D (5220C>G 5222G>T), R295N (5227G>A 5228A>C), Y296I (5229T>A 5230A>T 5231C>T), Y297F (5233A>T), C302L (5248G>T 5249T>A), F306Y (5260T>A), Y307W (5263A>G 5264T>G), H308Q (5267T>A), K310* (5271A>T), L311I (5274T>A 5276A>T), E312F (5277G>T 5278A>T 5279A>T), E313K (5280G>A), D314E (5285T>G), S315H (5286A>C 5287G>A), K316T (5290A>C 5291A>T), K317Y (5292A>T 5294A>T), Y318R (5295T>A 5296A>G 5297T>A), A320V (5302C>T), T322N (5308C>A 5309A>T), Q325F (5316C>T 5317A>T 5318G>T), Y327P (5322T>C 5323A>C 5324T>A), Q329E (5328C>G 5330G>A), K331N (5336A>C), V332F (5337G>T 5339A>T), L333M (5340T>A 5342A>G), G336C (5349G>T 5351A>T), Y337L (5353A>T 5354T>G), H338K (5355C>A 5357T>A), S340V (5361T>G 5362C>T 5363A>C), I343E (5370A>G 5371T>A 5372T>A), Q346K (5379C>A), F347I (5382T>A 5384T>C), D349N (5388G>A), R350D (5391A>G 5392G>A 5393A>T), R353N (5401G>A 5402A>T), Y356L (5410A>T 5411T>A), V368I (5445G>A 5447C>T), F369Y (5449T>A) | | | | | | | | |
| Codon mutations: | TCT244TCG (5075T>G), AGC245TGC (5076A>T), CCA246ACT (5079C>A 5081A>T), ATA249TAT (5088A>T 5089T>A 5090A>T), GTA250GTC (5093A>C), AAA252AAT (5099A>T), CAT253ACG (5100C>A 5101A>C 5102T>G), TCA254GCA (5103T>G), CAA256AAA (5109C>A), AAA257GAA (5112A>G), AGA258CGA (5115A>C), GGA259ACA (5118G>A 5119G>C), AAA260GTT (5121A>G 5122A>T 5123A>T), ACT261CCT (5124A>C), AGA262CGA (5127A>C), ATG263TTA (5130A>T 5132G>A), GTA264GTT (5135A>T), GAT266AAC (5139G>A 5141T>C), GAT269TCT (5148G>T 5149A>C), AAA273TTT (5160A>T 5161A>T 5162A>T), GCT274TTA (5163G>T 5164C>T 5165T>A), AAA275AAG (5168A>G), GTA276TGA (5169G>T 5170T>G), GTA277ATT (5172G>A 5174A>T), AAA278AGG (5176A>G 5177A>G), ATC281ATT (5186C>T), GAT285AGG (5196G>A 5197A>G 5198T>G), ACA286GAC (5199A>G 5200C>A 5201A>C), ATA288TTA (5205A>T), CAT289TCT (5208C>T 5209A>C), AGT291TTA (5214A>T 5215G>T 5216T>A), ATA292TAT (5217A>T 5218T>A 5219A>T), CAG293GAT (5220C>G 5222G>T), AGA295AAC (5227G>A 5228A>C), TAC296ATT (5229T>A 5230A>T 5231C>T), TAT297TTT (5233A>T), AGT298TCA (5235A>T 5236G>C 5237T>A), TGT302TTA (5248G>T 5249T>A), TTT306TAT (5260T>A), TAT307TGG (5263A>G 5264T>G), CAT308CAA (5267T>A), ATA309ATT (5270A>T), AAG310TAG (5271A>T), TTA311ATT (5274T>A 5276A>T), GAA312TTT (5277G>T 5278A>T 5279A>T), GAA313AAA (5280G>A), GAT314GAG (5285T>G), AGT315CAT (5286A>C 5287G>A), AAA316ACT (5290A>C 5291A>T), AAA317TAT (5292A>T 5294A>T), TAT318AGA (5295T>A 5296A>G 5297T>A), ACA319ACT (5300A>T), GCT320GTT (5302C>T), ACA322AAT (5308C>A 5309A>T), CCG324CCT (5315G>T), CAG325TTT (5316C>T 5317A>T 5318G>T), TAT327CCA (5322T>C 5323A>C 5324T>A), CAG329GAA (5328C>G 5330G>A), AAA331AAC (5336A>C), GTA332TTT (5337G>T 5339A>T), TTA333ATG (5340T>A 5342A>G), CCA334CCT (5345A>T), GGA336TGT (5349G>T 5351A>T), TAT337TTG (5353A>T 5354T>G), CAT338AAA (5355C>A 5357T>A), AAC339AAT (5360C>T), TCA340GTC (5361T>G 5362C>T 5363A>C), CCA341CCT (5366A>T), AGT342TCT (5367A>T 5368G>C), ATT343GAA (5370A>G 5371T>A 5372T>A), TTC344TTT (5375C>T), CAA345CAG (5378A>G), CAA346AAA (5379C>A), TTT347ATC (5382T>A 5384T>C), GAT349AAT (5388G>A), AGA350GAT (5391A>G 5392G>A 5393A>T), ATA351ATT (5396A>T), AGA353AAT (5401G>A 5402A>T), TAT356TTA (5410A>T 5411T>A), ATT359ATC (5420T>C), GTG361GTT (5426G>T), GAT365GAC (5438T>C), ATA366ATT (5441A>T), TTA367TTG (5444A>G), GTC368ATT (5445G>A 5447C>T), TTT369TAT (5449T>A) | | | | | | | | |

*: Inserts / Deletes / Misaligned / Frameshifts

#### Analysis details

This analysis was performed with panviral2.64

### NGS Details (UN8): Errantivirus

#### Assembly

| | |
|---|---|
| **Coverage Length** | 314 (1 contig(s)) |
| **Depth Of Coverage** | 5.4 |
| **Number Of Reads** | 14 |
| **Reads Per Million** | 0.23 rpm (after QC) |
| **Ambiguities** | 0 |
| **Assembly Method** | de novo + reference guided assembly |
| **Consensus Caller** | Bcf Tools |

#### Coverage Map

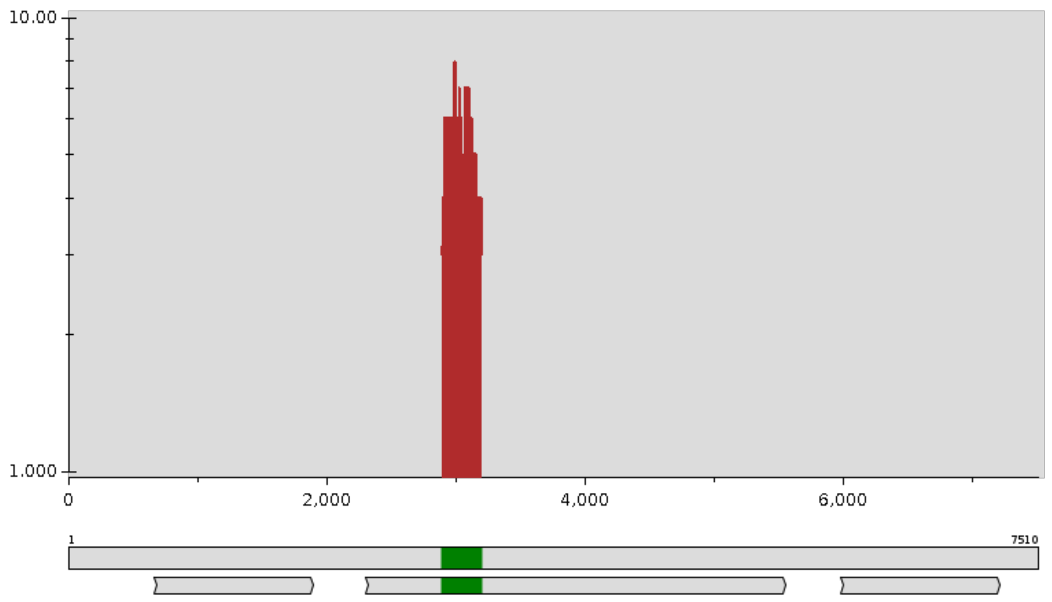

#### Assignment

| | |
|---|---|
| **Type** | Errantivirus (Taxonomy ID: 186666) |
| **Reference Genome** | NC_038512.1 |
| **NT Identity (%)** | 56.3694 |
| **AA Identity (%)** | 48.5714 |
| **Number Of Stop Codons** | 1 |
| **Number Of CDS** | 3 |

#### Alignment

| | |
|---|---|
| **Alignment Score** | 80.0 (NT) + 360.0 (AA) = 440.0 |
| **Concordance (%)** | 32.1637 |

| Alignment Method | Global, seeded, nucleotide + amino acids (AGA) |
|---|---|

#### Genome Region

Sequence starts at position 2893 and ends at position 3206 relative to NC_038512.1 reference sequence.

#### Alignment Detailed Statistics

| | Begin | End | Coverage | Score | Concordance | Matches | Identities | I/D/M/F* | Stop Codons |
|---|---|---|---|---|---|---|---|---|---|
| **NT** | **2893** | **3206** | **4.2%** | **80** | **12.7%** | **314 (100%)** | **177 (56.4%)** | **0/0** | |
| Mutations: | 2893A>T, 2899C>T, 2901T>A, 2902C>T, 2905T>C, 2906A>G, 2914C>T, 2916A>C, 2917G>A, 2920G>A, 2923T>A, 2927G>A, 2929T>A, 2932C>T, 2937A>T, 2938C>T, 2939C>T, 2941G>T, 2944A>T, 2948A>G, 2949A>T, 2950C>A, 2953A>T, 2954A>G, 2956T>A, 2959C>A, 2960G>T, 2962A>G, 2963C>T, 2964T>A, 2965T>G, 2968C>G, 2969A>G, 2971G>A, 2972T>C, 2974A>G, 2975G>T, 2976G>T, 2978A>G, 2979A>G, 2981T>G, 2982G>C, 2983C>T, 2984C>A, 2989C>T, 2993A>T, 2995C>T, 2997C>A, 2998C>A, 3004T>C, 3007G>A, 3008G>C, 3009C>G, 3010A>C, 3011A>C, 3012G>A, 3013T>A, 3016G>T, 3018T>A, 3025G>A, 3028G>T, 3029G>C, 3035G>A, 3037C>A, 3038C>G, 3039C>A, 3040T>G, 3041C>G, 3043A>T, 3047A>G, 3049A>G, 3050T>G, 3051C>A, 3058C>T, 3061G>T, 3065A>C, 3068G>A, 3069T>C, 3070A>T, 3071G>C, 3073A>T, 3076C>T, 3088A>G, 3091C>T, 3092C>T, 3094T>G, 3095C>G, 3096G>T, 3097A>T, 3103T>C, 3104A>T, 3106G>T, 3109A>C, 3114A>C, 3115A>C, 3118C>T, 3119T>G, 3121A>T, 3122C>T, 3130T>C, 3135A>T, 3137A>T, 3138G>C, 3140G>T, 3142T>A, 3146G>A, 3148C>T, 3149A>G, 3151T>G, 3154C>A, 3155C>T, 3157A>T, 3158A>C, 3159G>A, 3160A>G, 3161G>C, 3162G>T, 3165T>A, 3167C>T, 3168A>T, 3169A>G, 3171A>G, 3172T>A, 3175C>A, 3176A>T, 3178C>T, 3179T>A, 3180G>T, 3182C>T, 3184C>G, 3187C>G, 3188T>C, 3189A>T, 3190C>T, 3191C>T, 3196C>T, 3203A>C | | | | | | | | |

##### CDS

| | Begin | End | Coverage | Score | Concordance | Matches | Identities | I/D/M/F* | Stop Codons |
|---|---|---|---|---|---|---|---|---|---|
| D1R33_gp2 | 197 | 301 | 9.7% | 360 | 48.0% | 105 (100%) | 51 (48.6%) | 0/0/0/0 | 1 |
| Protein mutations: | F199Y (2901T>A 2902C>T), K201E (2906A>G), E204A (2916A>C 2917G>A), D208K (2927G>A 2929T>A), Y211F (2937A>T 2938C>T), P212S (2939C>T 2941G>T), N215V (2948A>G 2949A>T 2950C>A), S217G (2954A>G 2956T>A), D218E (2959C>A), V219L (2960G>T 2962A>G), L220* (2963C>T 2964T>A 2965T>G), D221E (2968C>G), K222E (2969A>G 2971G>A), G224F (2975G>T 2976G>T), K225G (2978A>G 2979A>G), C226A (2981T>G 2982G>C 2983C>T), Q227K (2984C>A), T230S (2993A>T 2995C>T), T231K (2997C>A 2998C>A), A235R (3008G>C 3009C>G 3010A>C), S236Q (3011A>C 3012G>A 3013T>A), F238Y (3018T>A), E242Q (3029G>C), D244K (3035G>A 3037C>A), P245E (3038C>G 3039C>A 3040T>G), Q246D (3041C>G 3043A>T), I248V (3047A>G 3049A>G), S249E (3050T>G 3051C>A), N254H (3065A>C), V255T (3068G>A 3069T>C 3070A>T), E256H (3071G>C 3073A>T), R264V (3095C>G 3096G>T 3097A>T), M267F (3104A>T 3106G>T), K270T (3114A>C 3115A>C), S272A (3119T>G 3121A>T), P273S (3122C>T), Q277L (3135A>T), R278S (3137A>T 3138G>C), V279L (3140G>T 3142T>A), D281N (3146G>A 3148C>T), N282E (3149A>G 3151T>G), L284F (3155C>T 3157A>T), R285Q (3158A>C 3159G>A 3160A>G), G286L (3161G>C 3162G>T), L287H (3165T>A), Q288L (3167C>T 3168A>T 3169A>G), N289R (3171A>G 3172T>A), N290K (3175C>A), I291F (3176A>T 3178C>T), C292I (3179T>A 3180G>T), Y295L (3188T>C 3189A>T 3190C>T), L296F (3191C>T), I300L (3203A>C) | | | | | | | | |
| Codon mutations: | GTA196..T (2893A>T), GAC198GAT (2899C>T), TTC199TAT (2901T>A 2902C>T), CGT200CGC (2905T>C), AAG201GAG (2906A>G), AAC203AAT (2914C>T), GAG204GCA (2916A>C 2917G>A), AAG205AAA (2920G>A), ACT206ACA (2923T>A), GAT208AAA (2927G>A 2929T>A), GAC209GAT (2932C>T), TAC211TTT (2937A>T 2938C>T), CCG212TCT (2939C>T 2941G>T), ATA213ATT (2944A>T), AAC215GTA (2948A>G 2949A>T 2950C>A), ATA216ATT (2953A>T), AGT217GGA (2954A>G 2956T>A), GAC218GAA (2959C>A), GTA219TTG (2960G>T 2962A>G), CTT220TAG (2963C>T 2964T>A 2965T>G), GAC221GAG (2968C>G), AAG222GAA (2969A>G 2971G>A), TTA223CTG (2972T>C 2974A>G), GGT224TTT (2975G>T 2976G>T), AAG225GGG (2978A>G 2979A>G), TGC226GCT (2981T>G 2982G>C 2983C>T), CAA227AAA (2984C>A), TAC228TAT (2989C>T), ACC230TCT (2993A>T 2995C>T), ACC231AAA (2997C>A 2998C>A), GAT233GAC (3004T>C), TTG234TTA (3007G>A), GCA235CGC (3008G>C 3009C>G 3010A>C), AGT236CAA (3011A>C 3012G>A 3013T>A), GGG237GGT (3016G>T), TTT238TAT (3018T>A), CAG240CAA (3025G>A), GTG241GTT (3028G>T), GAG242CAG (3029G>C), GAC244AAA (3035G>A 3037C>A), CCT245GAG (3038C>G 3039C>A 3040T>G), CAA246GAT (3041C>G 3043A>T), ATA248GTG (3047A>G 3049A>G), TCG249GAG (3050T>G 3051C>A), ACC251ACT (3058C>T), GCG252GCT (3061G>T), AAC254CAC (3065A>C), GTA255ACT (3068G>A 3069T>C 3070A>T), GAA256CAT (3071G>C 3073A>T), CAC257CAT (3076C>T), GAA261GAG (3088A>G), TTC262TTT (3091C>T), CTT263TTG (3092C>T 3094T>G), CGA264GTT (3095C>G 3096G>T 3097A>T), CCT266CCC (3103T>C), ATG267TTT (3104A>T 3106G>T), GGA268GGC (3109A>C), AAA270ACC (3114A>C 3115A>C), AAC271AAT (3118C>T), TCA272GCT (3119T>G 3121A>T), CCA273TCA (3122C>T), ACT275ACC (3130T>C), CAA277CTA (3135A>T), AGA278TCA (3137A>T 3138G>C), GTT279TTA (3140G>T 3142T>A), GAC281AAT (3146G>A 3148C>T), AAT282GAG (3149A>G 3151T>G), GTC283GTA (3154C>A), CTA284TTT (3155C>T 3157A>T), AGA285CAG (3158A>C 3159G>A 3160A>G), GGT286CTT (3161G>C 3162G>T), CTC287CAC (3165T>A), CAA288TTG (3167C>T 3168A>T 3169A>G), AAT289AGA (3171A>G 3172T>A), AAC290AAA (3175C>A), ATC291TTT (3176A>T 3178C>T), TGT292ATT (3179T>A 3180G>T), CTC293TTG (3182C>T 3184C>G), GTC294GTG (3187C>G), TAC295CTT (3188T>C 3189A>T 3190C>T), CTT296TTT (3191C>T), GAC297GAT (3196C>T), ATT300CTT (3203A>C) | | | | | | | | |

##### Proteins

| | Begin | End | Coverage | Score | Concordance | Matches | Identities | I/D/M/F* | Stop Codons |
|---|---|---|---|---|---|---|---|---|---|
| ORF B (YP_009507248.1) | 197 | 301 | 9.7% | 360 | 48.0% | 105 (100%) | 51 (48.6%) | 0/0/0/0 | 1 |
| Protein mutations: | F199Y (2901T>A 2902C>T), K201E (2906A>G), E204A (2916A>C 2917G>A), D208K (2927G>A 2929T>A), Y211F (2937A>T 2938C>T), P212S (2939C>T 2941G>T), N215V (2948A>G 2949A>T 2950C>A), S217G (2954A>G 2956T>A), D218E (2959C>A), V219L (2960G>T 2962A>G), L220* (2963C>T 2964T>A 2965T>G), D221E (2968C>G), K222E (2969A>G 2971G>A), G224F (2975G>T 2976G>T), K225G (2978A>G 2979A>G), C226A (2981T>G 2982G>C 2983C>T), Q227K (2984C>A), T230S (2993A>T 2995C>T), T231K (2997C>A 2998C>A), A235R (3008G>C 3009C>G 3010A>C), S236Q (3011A>C 3012G>A 3013T>A), F238Y (3018T>A), E242Q (3029G>C), D244K (3035G>A 3037C>A), P245E (3038C>G 3039C>A 3040T>G), Q246D (3041C>G 3043A>T), I248V (3047A>G 3049A>G), S249E (3050T>G 3051C>A), N254H (3065A>C), V255T (3068G>A 3069T>C 3070A>T), E256H (3071G>C 3073A>T), R264V (3095C>G 3096G>T 3097A>T), M267F (3104A>T 3106G>T), K270T (3114A>C 3115A>C), S272A (3119T>G 3121A>T), P273S (3122C>T), Q277L (3135A>T), R278S (3137A>T 3138G>C), V279L (3140G>T 3142T>A), D281N (3146G>A 3148C>T), N282E (3149A>G 3151T>G), L284F (3155C>T 3157A>T), R285Q (3158A>C 3159G>A 3160A>G), G286L (3161G>C 3162G>T), L287H (3165T>A), Q288L (3167C>T 3168A>T 3169A>G), N289R (3171A>G 3172T>A), N290K (3175C>A), I291F (3176A>T 3178C>T), C292I (3179T>A 3180G>T), Y295L (3188T>C 3189A>T 3190C>T), L296F (3191C>T), I300L (3203A>C) | | | | | | | | |
| Codon mutations: | GTA196..T (2893A>T), GAC198GAT (2899C>T), TTC199TAT (2901T>A 2902C>T), CGT200CGC (2905T>C), AAG201GAG (2906A>G), AAC203AAT (2914C>T), GAG204GCA (2916A>C 2917G>A), AAG205AAA (2920G>A), ACT206ACA (2923T>A), GAT208AAA (2927G>A 2929T>A), GAC209GAT (2932C>T), TAC211TTT (2937A>T 2938C>T), CCG212TCT (2939C>T 2941G>T), ATA213ATT (2944A>T), AAC215GTA (2948A>G 2949A>T 2950C>A), ATA216ATT (2953A>T), AGT217GGA (2954A>G 2956T>A), GAC218GAA (2959C>A), GTA219TTG (2960G>T 2962A>G), CTT220TAG (2963C>T 2964T>A 2965T>G), GAC221GAG (2968C>G), AAG222GAA (2969A>G 2971G>A), TTA223CTG (2972T>C 2974A>G), GGT224TTT (2975G>T 2976G>T), AAG225GGG (2978A>G 2979A>G), TGC226GCT (2981T>G 2982G>C 2983C>T), CAA227AAA (2984C>A), TAC228TAT (2989C>T), ACC230TCT (2993A>T 2995C>T), ACC231AAA (2997C>A 2998C>A), GAT233GAC (3004T>C), TTG234TTA (3007G>A), GCA235CGC (3008G>C 3009C>G 3010A>C), AGT236CAA (3011A>C 3012G>A 3013T>A), GGG237GGT (3016G>T), TTT238TAT (3018T>A), CAG240CAA (3025G>A), GTG241GTT (3028G>T), GAG242CAG (3029G>C), GAC244AAA (3035G>A 3037C>A), CCT245GAG (3038C>G 3039C>A 3040T>G), CAA246GAT (3041C>G 3043A>T), ATA248GTG (3047A>G 3049A>G), TCG249GAG (3050T>G 3051C>A), ACC251ACT (3058C>T), GCG252GCT (3061G>T), AAC254CAC (3065A>C), GTA255ACT (3068G>A 3069T>C 3070A>T), GAA256CAT (3071G>C 3073A>T), CAC257CAT (3076C>T), GAA261GAG (3088A>G), TTC262TTT (3091C>T), CTT263TTG (3092C>T 3094T>G), CGA264GTT (3095C>G 3096G>T 3097A>T), CCT266CCC (3103T>C), ATG267TTT (3104A>T 3106G>T), GGA268GGC (3109A>C), AAA270ACC (3114A>C 3115A>C), AAC271AAT (3118C>T), TCA272GCT (3119T>G 3121A>T), CCA273TCA (3122C>T), ACT275ACC (3130T>C), CAA277CTA (3135A>T), AGA278TCA (3137A>T 3138G>C), GTT279TTA (3140G>T 3142T>A), GAC281AAT (3146G>A 3148C>T), AAT282GAG (3149A>G 3151T>G), GTC283GTA (3154C>A), CTA284TTT (3155C>T 3157A>T), AGA285CAG (3158A>C 3159G>A 3160A>G), GGT286CTT (3161G>C 3162G>T), CTC287CAC (3165T>A), CAA288TTG (3167C>T 3168A>T 3169A>G), AAT289AGA (3171A>G 3172T>A), AAC290AAA (3175C>A), ATC291TTT (3176A>T 3178C>T), TGT292ATT (3179T>A 3180G>T), CTC293TTG (3182C>T 3184C>G), GTC294GTG (3187C>G), TAC295CTT (3188T>C 3189A>T 3190C>T), CTT296TTT (3191C>T), GAC297GAT (3196C>T), ATT300CTT (3203A>C) | | | | | | | | |

*: Inserts / Deletes / Misaligned / Frameshifts

#### Analysis details

This analysis was performed with panviral2.64

#### NGS Details (UN8): Caulimovirus venafragariae

##### Assembly

| | |
|---|---|
| **Coverage Length** | 254 (1 contig(s)) |
| **Depth Of Coverage** | 6.4 |
| **Number Of Reads** | 14 |
| **Reads Per Million** | 0.23 rpm (after QC) |
| **Ambiguities** | 0 |
| **Assembly Method** | de novo + reference guided assembly |
| **Consensus Caller** | Bcf Tools |

##### Coverage Map

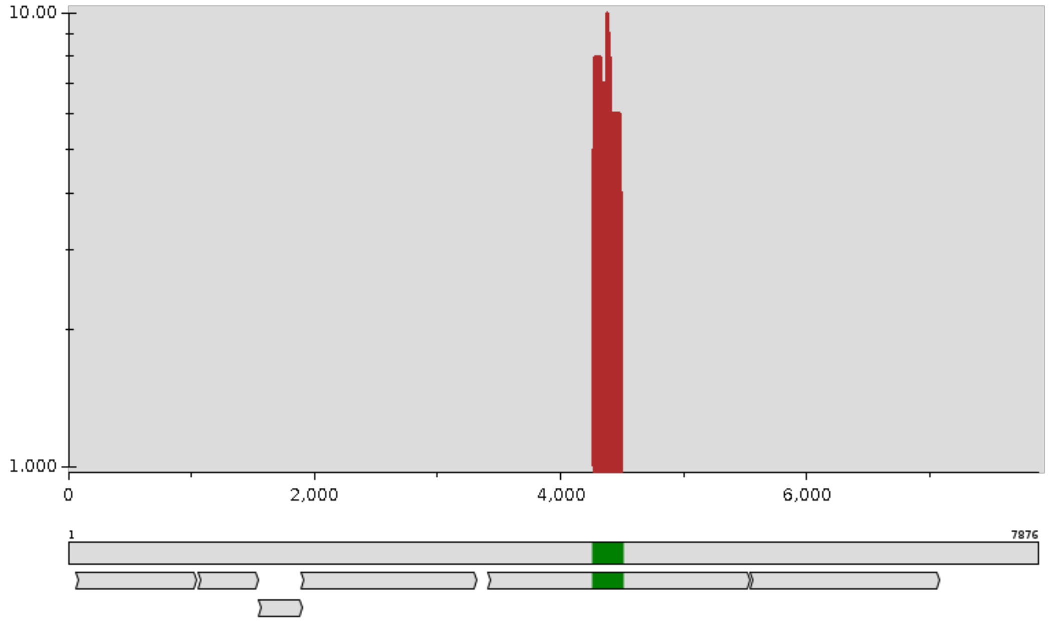

##### Assignment

| | |
|---|---|
| **Type** | Caulimovirus venafragariae (Taxonomy ID: 3048344) |
| **Reference Genome** | NC_001725.1 |
| **NT Identity (%)** | 60.8696 |
| **AA Identity (%)** | 61.9048 |
| **Number Of Stop Codons** | 0 |
| **Number Of CDS** | 6 |

##### Alignment

| | |
|---|---|
| **Alignment Score** | 100.0 (NT) + 212.0 (AA) = 312.0 |
| **Concordance (%)** | 34.6833 |

| Alignment Method | Global, seeded, nucleotide + amino acids (AGA) |
|---|---|

#### Genome Region

Sequence starts at position 4259 and ends at position 4512 relative to NC_001725.1 reference sequence.

#### Alignment Detailed Statistics

| | Begin | End | Coverage | Score | Concordance | Matches | Identities | I/D/M/F* | Stop Codons |
|---|---|---|---|---|---|---|---|---|---|
| **NT** | **4259** | **4512** | **3.2%** | **100** | **20.2%** | **253 (99.6%)** | **154 (60.6%)** | **0/1** | |
| Mutations: | 4262A>C, 4264T>A, 4267C>A, 4268C>A, 4276C>A, 4277G>T, 4284G>T, 4288A>C, 4290A>G, 4292C>T, 4293T>A, 4296G>A, 4299C>A, 4300G>A, 4306C>T, 4309C>T, 4311C>A, 4312A>G, 4314G>T, 4315C>T, 4317T>A, 4320T>A, 4321A>G, 4323C>A, 4326C>A, 4330C>A, 4331C>G, 4332T>A, 4333A>T, 4334G>C, 4335C>A, 4337A>C, 4338G>A, 4339A>T, 4340G>C, 4342C>A, 4343C>G, 4347T>C, 4348T>A, 4349C>G, 4350C>A, 4354C>G, 4359C>A, 4369delA, 4370G>A, 4371G>A, 4374T>C, 4378G>A, 4379C>A, 4383G>A, 4386C>A, 4387A>G, 4388A>T, 4389A>C, 4390C>A, 4395C>A, 4401A>C, 4413T>A, 4416C>T, 4422A>G, 4424A>G, 4425G>A, 4426T>C, 4428A>C, 4434C>T, 4435C>A, 4440A>T, 4442A>G, 4443G>A, 4444G>A, 4445G>T, 4446A>G, 4451G>C, 4452C>T, 4455C>T, 4456C>A, 4457T>A, 4459C>T, 4461T>A, 4464T>A, 4465A>G, 4470G>A, 4471G>A, 4473G>T, 4474C>G, 4479G>A, 4482T>A, 4483C>A, 4485A>T, 4491C>A, 4492G>C, 4493G>A, 4497T>A, 4500G>A, 4502C>G, 4503C>A, 4504T>A, 4508A>T, 4510T>A, 4511C>G | | | | | | | | |

**CDS**

| | Begin | End | Coverage | Score | Concordance | Matches | Identities | I/D/M/F* | Stop Codons |
|---|---|---|---|---|---|---|---|---|---|
| ORF_V | 286 | 369 | 11.8% | 212 | 36.9% | 84 (100%) | 52 (61.9%) | 0/0/1/1 | 0 |
| Protein mutations: | Y286S (4262A>C), S287T (4264T>A), P288N (4267C>A 4268C>A), R291I (4276C>A 4277G>T), E293D (4284G>T), K295Q (4288A>C 4290A>G), T296I (4292C>T 4293T>A), E299K (4300G>A), K303D (4312A>G 4314G>T), I306V (4321A>G 4323C>A), P309R (4330C>A 4331C>G 4332T>A), K311T (4337A>C 4338G>A), P313R (4342C>A 4343C>G), S315R (4348T>A 4349C>G 4350C>A), P317A (4354C>G), A325N (4378G>A 4379C>A), K328V (4387A>G 4388A>T 4389A>C), K340R (4424A>G 4425G>A), H344N (4435C>A), K346R (4442A>G 4443G>A), G347M (4444G>A 4445G>T 4446A>G), G349A (4451G>C 4452C>T), L351K (4456C>A 4457T>A), N354D (4465A>G), E356N (4471G>A 4473G>T), Q357E (4474C>G), Q360N (4483C>A 4485A>T), G363Q (4492G>C 4493G>A), T366R (4502C>G 4503C>A), F367I (4504T>A), Y368F (4508A>T) | | | | | | | | |
| Codon mutations: | TAT286TCT (4262A>C), TCA287ACA (4264T>A), CCT288AAT (4267C>A 4268C>A), CGA291ATA (4276C>A 4277G>T), GAG293GAT (4284G>T), AAA295CAG (4288A>C 4290A>G), ACT296ATA (4292C>T 4293T>A), CAG297CAA (4296G>A), ATC298ATA (4299C>A), GAA299AAA (4300G>A), CTA301TTA (4306C>T), CTC302TTA (4309C>T 4311C>A), AAG303GAT (4312A>G 4314G>T), CTT304TTA (4315C>T 4317T>A), GGT305GGA (4320T>A), ATC306GTA (4321A>G 4323C>A), ATC307ATA (4326C>A), CCT309AGA (4330C>A 4331C>G 4332T>A), AGC310TCA (4333A>T 4334G>C 4335C>A), AAG311ACA (4337A>C 4338G>A), AGT312TCT (4339A>T 4340G>C), CCA313AGA (4342C>A 4343C>G), CAT314CAC (4347T>C), TCC315AGA (4348T>A 4349C>G 4350C>A), CCA317GCA (4354C>G), GCC318GCA (4359C>A), AGG322-AA (4369delA 4370G>A 4371G>A), AAT323AAC (4374T>C), GCC325AAC (4378G>A 4379C>A), GAG326GAA (4383G>A), ATC327ATA (4386C>A), AAA328GTC (4387A>G 4388A>T 4389A>C), CGA329AGA (4390C>A), GGC330GGA (4395C>A), GCA332GCC (4401A>C), ATT336ATA (4413T>A), AAC337AAT (4416C>T), AAA339AAG (4422A>G), AAG340AGA (4424A>G 4425G>A), TTA341CTC (4426T>C 4428A>C), GAC343GAT (4434C>T), CAT344AAT (4435C>A), ACA345ACT (4440A>T), AAG346AGA (4442A>G 4443G>A), GGA347ATG (4444G>A 4445G>T 4446A>G), GGC349GCT (4451G>C 4452C>T), TAC350TAT (4455C>T), CTA351AAA (4456C>A 4457T>A), CTT352TTA (4459C>T 4461T>A), CCT353CCA (4464T>A), AAC354GAC (4465A>G), AAG355AAA (4470G>A), GAG356AAT (4471G>A 4473G>T), CAA357GAA (4474C>G), CTG358CTA (4479G>A), CTT359CTA (4482T>A), CAA360AAT (4483C>A 4485A>T), ATC362ATA (4491C>A), GGA363CAA (4492G>C 4493G>A), GGT364GGA (4497T>A), AAG365AAA (4500G>A), ACC366AGA (4502C>G 4503C>A), TTT367ATT (4504T>A), TAC368TTC (4508A>T), TCT369AGT (4510T>A 4511C>G) | | | | | | | | |

**Proteins**

| | Begin | End | Coverage | Score | Concordance | Matches | Identities | I/D/M/F* | Stop Codons |
|---|---|---|---|---|---|---|---|---|---|
| hypothetical protein (NP_043933.1) | 286 | 369 | 11.8% | 212 | 36.9% | 84 (100%) | 52 (61.9%) | 0/0/1/1 | 0 |
| Protein mutations: | Y286S (4262A>C), S287T (4264T>A), P288N (4267C>A 4268C>A), R291I (4276C>A 4277G>T), E293D (4284G>T), K295Q (4288A>C 4290A>G), T296I (4292C>T 4293T>A), E299K (4300G>A), K303D (4312A>G 4314G>T), I306V (4321A>G 4323C>A), P309R (4330C>A 4331C>G 4332T>A), K311T (4337A>C 4338G>A), P313R (4342C>A 4343C>G), S315R (4348T>A 4349C>G 4350C>A), P317A (4354C>G), A325N (4378G>A 4379C>A), K328V (4387A>G 4388A>T 4389A>C), K340R (4424A>G 4425G>A), H344N (4435C>A), K346R (4442A>G 4443G>A), G347M (4444G>A 4445G>T 4446A>G), G349A (4451G>C 4452C>T), L351K (4456C>A 4457T>A), N354D (4465A>G), E356N (4471G>A 4473G>T), Q357E (4474C>G), Q360N (4483C>A 4485A>T), G363Q (4492G>C 4493G>A), T366R (4502C>G 4503C>A), F367I (4504T>A), Y368F (4508A>T) | | | | | | | | |
| Codon mutations: | TAT286TCT (4262A>C), TCA287ACA (4264T>A), CCT288AAT (4267C>A 4268C>A), CGA291ATA (4276C>A 4277G>T), GAG293GAT (4284G>T), AAA295CAG (4288A>C 4290A>G), ACT296ATA (4292C>T 4293T>A), CAG297CAA (4296G>A), ATC298ATA (4299C>A), GAA299AAA (4300G>A), CTA301TTA (4306C>T), CTC302TTA (4309C>T 4311C>A), AAG303GAT (4312A>G 4314G>T), CTT304TTA (4315C>T 4317T>A), GGT305GGA (4320T>A), ATC306GTA (4321A>G 4323C>A), ATC307ATA (4326C>A), CCT309AGA (4330C>A 4331C>G 4332T>A), AGC310TCA (4333A>T 4334G>C 4335C>A), AAG311ACA (4337A>C 4338G>A), AGT312TCT (4339A>T 4340G>C), CCA313AGA (4342C>A 4343C>G), CAT314CAC (4347T>C), TCC315AGA (4348T>A 4349C>G 4350C>A), CCA317GCA (4354C>G), GCC318GCA (4359C>A), AGG322-AA (4369delA 4370G>A 4371G>A), AAT323AAC (4374T>C), GCC325AAC (4378G>A 4379C>A), GAG326GAA (4383G>A), ATC327ATA (4386C>A), AAA328GTC (4387A>G 4388A>T 4389A>C), CGA329AGA (4390C>A), GGC330GGA (4395C>A), GCA332GCC (4401A>C), ATT336ATA (4413T>A), AAC337AAT (4416C>T), AAA339AAG (4422A>G), AAG340AGA (4424A>G 4425G>A), TTA341CTC (4426T>C 4428A>C), GAC343GAT (4434C>T), CAT344AAT (4435C>A), ACA345ACT (4440A>T), AAG346AGA (4442A>G 4443G>A), GGA347ATG (4444G>A 4445G>T 4446A>G), GGC349GCT (4451G>C 4452C>T), TAC350TAT (4455C>T), CTA351AAA (4456C>A 4457T>A), CTT352TTA (4459C>T 4461T>A), CCT353CCA (4464T>A), AAC354GAC (4465A>G), AAG355AAA (4470G>A), GAG356AAT (4471G>A 4473G>T), CAA357GAA (4474C>G), CTG358CTA (4479G>A), CTT359CTA (4482T>A), CAA360AAT (4483C>A 4485A>T), ATC362ATA (4491C>A), GGA363CAA (4492G>C 4493G>A), GGT364GGA (4497T>A), AAG365AAA (4500G>A), ACC366AGA (4502C>G 4503C>A), TTT367ATT (4504T>A), TAC368TTC (4508A>T), TCT369AGT (4510T>A 4511C>G) | | | | | | | | |

*: Inserts / Deletes / Misaligned / Frameshifts

#### Analysis details

This analysis was performed with panviral2.64

### NGS Details (UN8): Badnavirus maculasmallanthi

#### Assembly

| | |
|---|---|
| **Coverage Length** | 304 (1 contig(s)) |
| **Depth Of Coverage** | 4.7 |
| **Number Of Reads** | 13 |
| **Reads Per Million** | 0.22 rpm (after QC) |
| **Ambiguities** | 0 |
| **Assembly Method** | de novo + reference guided assembly |
| **Consensus Caller** | Bcf Tools |

#### Coverage Map

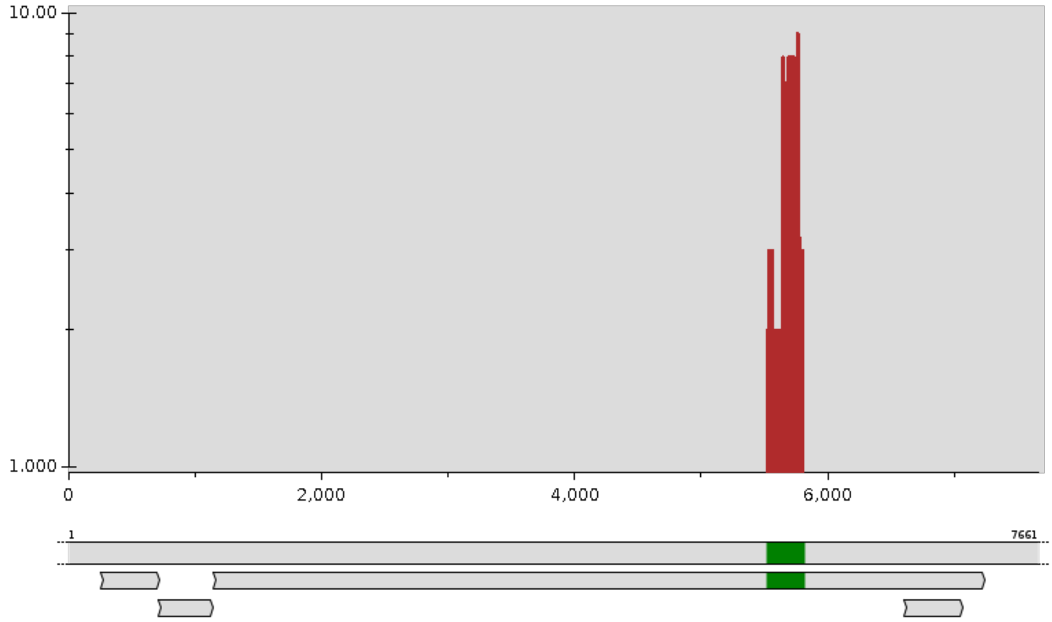

#### Assignment

| | |
|---|---|
| **Type** | Badnavirus maculasmallanthi (Taxonomy ID: 3048453) |
| **Reference Genome** | NC_026472.1 |
| **NT Identity (%)** | 51.1401 |
| **AA Identity (%)** | 41.1765 |
| **Number Of Stop Codons** | 0 |
| **Number Of CDS** | 4 |

#### Alignment

| | |
|---|---|
| **Alignment Score** | 8.0 (NT) + 290.0 (AA) = 298.0 |
| **Concordance (%)** | 22.61 |

| **Alignment Method** | Global, seeded, nucleotide + amino acids (AGA) |
|---|---|

#### Genome Region

Sequence starts at position 5516 and ends at position 5819 relative to NC_026472.1 reference sequence.

#### Alignment Detailed Statistics

| | Begin | End | Coverage | Score | Concordance | Matches | Identities | I/D/M/F* | Stop Codons |
|---|---|---|---|---|---|---|---|---|---|
| **NT** | **5516** | **5819** | **4.0%** | **8** | **1.3%** | **304 (99.0%)** | **157 (51.1%)** | **3/0** | |
| Mutations: | 5517T>C, 5518T>C, 5527G>C, 5529G>T, 5532A>C, 5535C>T, 5536A>G, 5537C>A, 5542C>T, 5544A>T, 5545A>G, 5547A>T, 5548A>C, 5549G>A, 5550A>G, 5551A>C, 5553A>T, 5554G>C, 5555G>A, 5556G>A, 5558A>G, 5559T>G, 5560G>A, 5561C>T, 5562A>G, 5563A>T, 5564A>C, 5565G>C, 5566A>T, 5568A>G, 5574A>T, 5578T>A, 5584C>T, 5588A>G, 5589G>A, 5592T>C, 5595A>G, 5597T>A, 5599C>T, 5604A>G, 5605G>T, 5607T>G, 5608G>A, 5609C>G, 5610C>G, 5613G>T, 5614G>T, 5615A>G, 5616A>G, 5618A>C, 5620G>T, 5621A>C, 5622A>T, 5623T>G, 5624C>A, 5628A>T, 5631C>T, 5632T>A, 5633G>A, 5637G>A, 5638G>A, 5643T>C, 5644T>A, 5645T>A, 5646A>G, 5648T>C, 5649C>T, 5650C>A, 5651C>A, 5653G>T, 5654G>A, 5655A>C, 5657G>A, 5658A>C, 5659T>C, 5660T>A, 5661A>T, 5664C>T, 5667A>G, 5669G>T, 5670G>C, 5674G>A, 5676C>G, 5680C>T, 5685C>T, 5688A>G, 5691C>A, 5693A>C, 5694A>C, 5700A>C, 5702C>T, 5703C>T, 5705C>T, 5706T>A, 5707A>G, 5708T>C, 5709T>A, 5712T>C, 5713C>A, 5714A>T, 5715G>A, 5716A>G, 5717G>A, 5718A>G, 5719A>T, 5720A>T, 5721A>G, 5725G>A, 5727T>C, 5728A>G, 5729A>G, 5731T>G, 5732G>T, 5733C>G, 5736C>T, 5736_5737insCAG, 5737A>C, 5738A>C, 5740G>T, 5743A>T, 5744C>T, 5748A>T, 5749G>T, 5750A>T, 5757C>G, 5758G>A, 5759C>T, 5760C>T, 5761G>A, 5763A>T, 5765A>T, 5766C>T, 5769C>T, 5772C>T, 5779C>T, 5784A>T, 5793C>G, 5795G>C, 5796T>A, 5797G>A, 5799A>G, 5802G>C, 5805T>C, 5808T>C, 5810A>T, 5811A>T, 5814G>A, 5815C>T | | | | | | | | |

##### CDS

| | Begin | End | Coverage | Score | Concordance | Matches | Identities | I/D/M/F* | Stop Codons |
|---|---|---|---|---|---|---|---|---|---|
| UF61_gp3 | 1459 | 1559 | 5.0% | 290 | 40.1% | 101 (99.0%) | 42 (41.2%) | 1/0/0/0 | 0 |
| Protein mutations: | S1459P (5518T>C), G1462R (5527G>C 5529G>T), T1465D (5536A>G 5537C>A), L1467F (5542C>T 5544A>T), K1468D (5545A>G 5547A>T), R1469Q (5548A>C 5549G>A 5550A>G), I1470L (5551A>C 5553A>T), G1471Q (5554G>C 5555G>A 5556G>A), N1472R (5558A>G 5559T>G), A1473M (5560G>A 5561C>T 5562A>G), K1474S (5563A>T 5564A>C 5565G>C), I1475L (5566A>T 5568A>G), F1479I (5578T>A), K1482R (5588A>G 5589G>A), F1485Y (5597T>A), H1486Y (5599C>T), V1488L (5605G>T 5607T>G), A1489R (5608G>A 5609C>G 5610C>G), M1490I (5613G>T), E1491W (5614G>T 5615A>G 5616A>G), E1492A (5618A>C), E1493S (5620G>T 5621A>C 5622A>T), S1494D (5623T>G 5624C>A), W1497K (5632T>A 5633G>A), A1499T (5638G>A), L1501K (5644T>A 5645T>A 5646A>G), I1502T (5648T>C 5649C>T), P1503K (5650C>A 5651C>A), G1504Y (5653G>T 5654G>A 5655A>C), G1505D (5657G>A 5658A>C), L1506H (5659T>C 5660T>A 5661A>T), W1509F (5669G>T 5670G>C), V1511M (5674G>A 5676C>G), P1513S (5680C>T), K1517T (5693A>C 5694A>C), P1520L (5702C>T 5703C>T), A1521V (5705C>T 5706T>A), I1522A (5707A>G 5708T>C 5709T>A), Q1524I (5713C>A 5714A>T 5715G>A), R1525E (5716A>G 5717G>A 5718A>G), K1526L (5719A>T 5720A>T 5721A>G), D1528N (5725G>A 5727T>C), K1529G (5728A>G 5729A>G), C1530V (5731T>G 5732G>T 5733C>G), F1531_K1532insQ (5736_5737insCAG), K1532P (5737A>C 5738A>C), D1533Y (5740G>T), T1534L (5743A>T 5744C>T), E1535D (5748A>T), E1536L (5749G>T 5750A>T), I1538M (5757C>G), A1539I (5758G>A 5759C>T 5760C>T), V1540I (5761G>A 5763A>T), Y1541F (5765A>T 5766C>T), N1550K (5793C>G), S1551T (5795G>C 5796T>A), E1552K (5797G>A 5799A>G), E1556V (5810A>T 5811A>T), H1558Y (5815C>T) | | | | | | | | |
| Codon mutations: | TAT1458.AC (5517T>C), TCC1459CCC (5518T>C), GGG1462CGT (5527G>C 5529G>T), ATA1463ATC (5532A>C), AAC1464AAT (5535C>T), ACC1465GAC (5536A>G 5537C>A), CTA1467TTT (5542C>T 5544A>T), AAA1468GAT (5545A>G 5547A>T), AGA1469CAG (5548A>C 5549G>A 5550A>G), ATA1470CTT (5551A>C 5553A>T), GGG1471CAA (5554G>C 5555G>A 5556G>A), AAT1472AGG (5558A>G 5559T>G), GCA1473ATG (5560G>A 5561C>T 5562A>G), AAG1474TCC (5563A>T 5564A>C 5565G>C), ATA1475TTG (5566A>T 5568A>G), TCA1477TCT (5574A>T), TTT1479ATT (5578T>A), CTG1481TTG (5584C>T), AAG1482AGA (5588A>G 5589G>A), TCT1483TCC (5592T>C), GGA1484GGG (5595A>G), TTT1485TAT (5597T>A), CAT1486TAT (5599C>T), CAA1487CAG (5604A>G), GTT1488TTG (5605G>T 5607T>G), GCC1489AGG (5608G>A 5609C>G 5610C>G), ATG1490ATT (5613G>T), GAA1491TGG (5614G>T 5615A>G 5616A>G), GAA1492GCA (5618A>C), GAA1493TCT (5620G>T 5621A>C 5622A>T), TCC1494GAC (5623T>G 5624C>A), ATA1495ATT (5628A>T), CCC1496CCT (5631C>T), TGG1497AAG (5632T>A 5633G>A), ACG1498ACA (5637G>A), GCC1499ACC (5638G>A), TTT1500TTC (5643T>C), TTA1501AAG (5644T>A 5645T>A 5646A>G), ATC1502ACT (5648T>C 5649C>T), CCA1503AAA (5650C>A 5651C>A), GGA1504TAC (5653G>T 5654G>A 5655A>C), GGA1505GAC (5657G>A 5658A>C), TTA1506CAT (5659T>C 5660T>A 5661A>T), TAC1507TAT (5664C>T), GAA1508GAG (5667A>G), TGG1509TTC (5669G>T 5670G>C), GTC1511ATG (5674G>A 5676C>G), CCA1513TCA (5680C>T), TTC1514TTT (5685C>T), GGA1515GGG (5688A>G), CTC1516CTA (5691C>A), AAA1517ACC (5693A>C 5694A>C), GCA1519GCC (5700A>C), CCC1520CTT (5702C>T 5703C>T), GCT1521GTA (5705C>T 5706T>A), ATT1522GCA (5707A>G 5708T>C 5709T>A), TTT1523TTC (5712T>C), CAG1524ATA (5713C>A 5714A>T 5715G>A), AGA1525GAG (5716A>G 5717G>A 5718A>G), AAA1526TTG (5719A>T 5720A>T 5721A>G), GAT1528AAC (5725G>A 5727T>C), AAA1529GGA (5728A>G 5729A>G), TGC1530GTG (5731T>G 5732G>T 5733C>G), TTC1531TTT (5736C>T), TTC1531_AAA1532insCAG (5736_5737insCAG), AAA1532CCA (5737A>C 5738A>C), GAC1533TAC (5740G>T), ACA1534TTA (5743A>T 5744C>T), GAA1535GAT (5748A>T), GAG1536TTG (5749G>T 5750A>T), ATC1538ATG (5757C>G), GCC1539ATT (5758G>A 5759C>T 5760C>T), GTA1540ATT (5761G>A 5763A>T), TAC1541TTT (5765A>T 5766C>T), ATC1542ATT (5769C>T), GAC1543GAT (5772C>T), CTG1546TTG (5779C>T), GTA1547GTT (5784A>T), AAC1550AAG (5793C>G), AGT1551ACA (5795G>C 5796T>A), GAA1552AAG (5797G>A 5799A>G), GCG1553GCC (5802G>C), GAT1554GAC (5805T>C), CAT1555CAC (5808T>C), GAA1556GTT (5810A>T 5811A>T), AGG1557AGA (5814G>A), CAT1558TAT (5815C>T) | | | | | | | | |

##### Proteins

| | Begin | End | Coverage | Score | Concordance | Matches | Identities | I/D/M/F* | Stop Codons |
|---|---|---|---|---|---|---|---|---|---|
| ORF3 (YP_009121747.1) | 1459 | 1559 | 5.0% | 290 | 40.1% | 101 (99.0%) | 42 (41.2%) | 1/0/0/0 | 0 |
| Protein mutations: | S1459P (5518T>C), G1462R (5527G>C 5529G>T), T1465D (5536A>G 5537C>A), L1467F (5542C>T 5544A>T), K1468D (5545A>G 5547A>T), R1469Q (5548A>C 5549G>A 5550A>G), I1470L (5551A>C 5553A>T), G1471Q (5554G>C 5555G>A 5556G>A), N1472R (5558A>G 5559T>G), A1473M (5560G>A 5561C>T 5562A>G), K1474S (5563A>T 5564A>C 5565G>C), I1475L (5566A>T 5568A>G), F1479I (5578T>A), K1482R (5588A>G 5589G>A), F1485Y (5597T>A), H1486Y (5599C>T), V1488L (5605G>T 5607T>G), A1489R (5608G>A 5609C>G 5610C>G), M1490I (5613G>T), E1491W (5614G>T 5615A>G 5616A>G), E1492A (5618A>C), E1493S (5620G>T 5621A>C 5622A>T), S1494D (5623T>G 5624C>A), W1497K (5632T>A 5633G>A), A1499T (5638G>A), L1501K (5644T>A 5645T>A 5646A>G), I1502T (5648T>C 5649C>T), P1503K (5650C>A 5651C>A), G1504Y (5653G>T 5654G>A 5655A>C), G1505D (5657G>A 5658A>C), L1506H (5659T>C 5660T>A 5661A>T), W1509F (5669G>T 5670G>C), V1511M (5674G>A 5676C>G), P1513S (5680C>T), K1517T (5693A>C 5694A>C), P1520L (5702C>T 5703C>T), A1521V (5705C>T 5706T>A), I1522A (5707A>G 5708T>C 5709T>A), Q1524I (5713C>A 5714A>T 5715G>A), R1525E (5716A>G 5717G>A 5718A>G), K1526L (5719A>T 5720A>T 5721A>G), D1528N (5725G>A 5727T>C), K1529G (5728A>G 5729A>G), C1530V (5731T>G 5732G>T 5733C>G), F1531_K1532insQ (5736_5737insCAG), K1532P (5737A>C 5738A>C), D1533Y (5740G>T), T1534L (5743A>T 5744C>T), E1535D (5748A>T), E1536L (5749G>T 5750A>T), I1538M (5757C>G), A1539I (5758G>A 5759C>T 5760C>T), V1540I (5761G>A 5763A>T), Y1541F (5765A>T 5766C>T), N1550K (5793C>G), S1551T (5795G>C 5796T>A), E1552K (5797G>A 5799A>G), E1556V (5810A>T 5811A>T), H1558Y (5815C>T) | | | | | | | | |
| Codon mutations: | TAT1458.AC (5517T>C), TCC1459CCC (5518T>C), GGG1462CGT (5527G>C 5529G>T), ATA1463ATC (5532A>C), AAC1464AAT (5535C>T), ACC1465GAC (5536A>G 5537C>A), CTA1467TTT (5542C>T 5544A>T), AAA1468GAT (5545A>G 5547A>T), AGA1469CAG (5548A>C 5549G>A 5550A>G), ATA1470CTT (5551A>C 5553A>T), GGG1471CAA (5554G>C 5555G>A 5556G>A), AAT1472AGG (5558A>G 5559T>G), GCA1473ATG (5560G>A 5561C>T 5562A>G), AAG1474TCC (5563A>T 5564A>C 5565G>C), ATA1475TTG (5566A>T 5568A>G), TCA1477TCT (5574A>T), TTT1479ATT (5578T>A), CTG1481TTG (5584C>T), AAG1482AGA (5588A>G 5589G>A), TCT1483TCC (5592T>C), GGA1484GGG (5595A>G), TTT1485TAT (5597T>A), CAT1486TAT (5599C>T), CAA1487CAG (5604A>G), GTT1488TTG (5605G>T 5607T>G), GCC1489AGG (5608G>A 5609C>G 5610C>G), ATG1490ATT (5613G>T), GAA1491TGG (5614G>T 5615A>G 5616A>G), GAA1492GCA (5618A>C), GAA1493TCT (5620G>T 5621A>C 5622A>T), TCC1494GAC (5623T>G 5624C>A), ATA1495ATT (5628A>T), CCC1496CCT (5631C>T), TGG1497AAG (5632T>A 5633G>A), ACG1498ACA (5637G>A), GCC1499ACC (5638G>A), TTT1500TTC (5643T>C), TTA1501AAG (5644T>A 5645T>A 5646A>G), ATC1502ACT (5648T>C 5649C>T), CCA1503AAA (5650C>A 5651C>A), GGA1504TAC (5653G>T 5654G>A 5655A>C), GGA1505GAC (5657G>A 5658A>C), TTA1506CAT (5659T>C 5660T>A 5661A>T), TAC1507TAT (5664C>T), GAA1508GAG (5667A>G), TGG1509TTC (5669G>T 5670G>C), GTC1511ATG (5674G>A 5676C>G), CCA1513TCA (5680C>T), TTC1514TTT (5685C>T), GGA1515GGG (5688A>G), CTC1516CTA (5691C>A), AAA1517ACC (5693A>C 5694A>C), GCA1519GCC (5700A>C), CCC1520CTT (5702C>T 5703C>T), GCT1521GTA (5705C>T 5706T>A), ATT1522GCA (5707A>G 5708T>C 5709T>A), TTT1523TTC (5712T>C), CAG1524ATA (5713C>A 5714A>T 5715G>A), AGA1525GAG (5716A>G 5717G>A 5718A>G), AAA1526TTG (5719A>T 5720A>T 5721A>G), GAT1528AAC (5725G>A 5727T>C), AAA1529GGA (5728A>G 5729A>G), TGC1530GTG (5731T>G 5732G>T 5733C>G), TTC1531TTT (5736C>T), TTC1531_AAA1532insCAG (5736_5737insCAG), AAA1532CCA (5737A>C 5738A>C), GAC1533TAC (5740G>T), ACA1534TTA (5743A>T 5744C>T), GAA1535GAT (5748A>T), GAG1536TTG (5749G>T 5750A>T), ATC1538ATG (5757C>G), GCC1539ATT (5758G>A 5759C>T 5760C>T), GTA1540ATT (5761G>A 5763A>T), TAC1541TTT (5765A>T 5766C>T), ATC1542ATT (5769C>T), GAC1543GAT (5772C>T), CTG1546TTG (5779C>T), GTA1547GTT (5784A>T), AAC1550AAG (5793C>G), AGT1551ACA (5795G>C 5796T>A), GAA1552AAG (5797G>A 5799A>G), GCG1553GCC (5802G>C), GAT1554GAC (5805T>C), CAT1555CAC (5808T>C), GAA1556GTT (5810A>T 5811A>T), AGG1557AGA (5814G>A), CAT1558TAT (5815C>T) | | | | | | | | |

*: Inserts / Deletes / Misaligned / Frameshifts

#### Analysis details

This analysis was performed with panviral2.64

### NGS Details (UN8): Hubei noda-like virus 9 (segment 2)

#### Assembly

| | |
|---|---|
| **Coverage Length** | 328 (2 contig(s)) |
| **Depth Of Coverage** | 2.7 |
| **Number Of Reads** | 9 |
| **Reads Per Million** | 0.15 rpm (after QC) |
| **Ambiguities** | 0 |
| **Assembly Method** | de novo + reference guided assembly |
| **Consensus Caller** | Bcf Tools |

#### Coverage Map

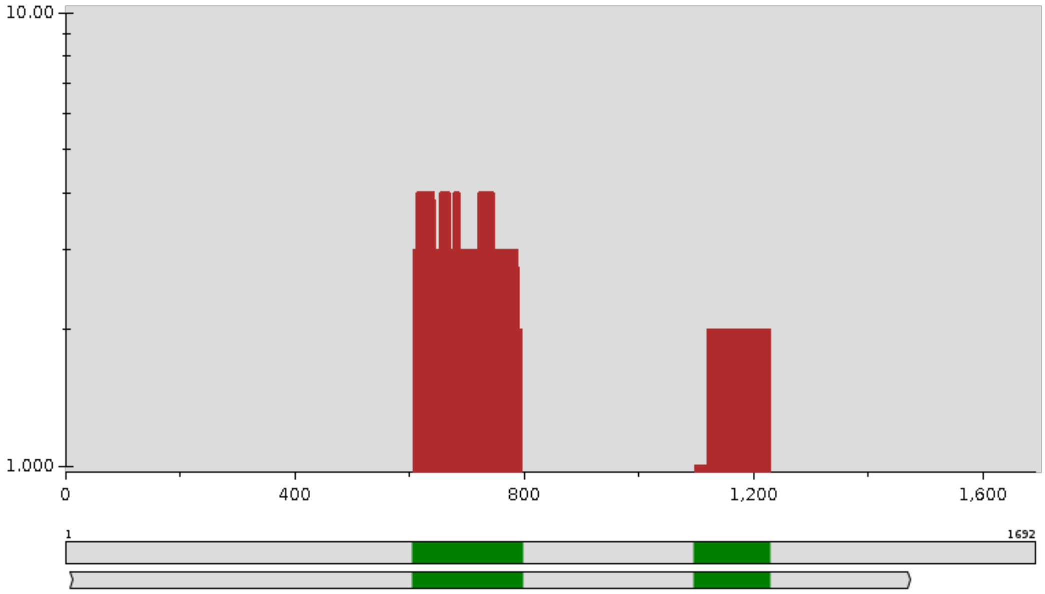

#### Assignment

| | |
|---|---|
| **Type** | Hubei noda-like virus 9 (Taxonomy ID: 1922989) |
| **Reference Genome** | NC_033308.1 |
| **NT Identity (%)** | 66.4634 |
| **AA Identity (%)** | 69.0909 |
| **Number Of Stop Codons** | 0 |
| **Number Of CDS** | 1 |

#### Alignment

| | |
|---|---|
| **Alignment Score** | 216.0 (NT) + 562.0 (AA) = 778.0 |
| **Concordance (%)** | 55.8908 |

| Alignment Method | Global, seeded, nucleotide + amino acids (AGA) |
|---|---|

#### Genome Region

Sequence starts at position 607 and ends at position 1231 relative to NC_033308.1 reference sequence.

#### Alignment Detailed Statistics

| | Begin | End | Coverage | Score | Concordance | Matches | Identities | I/D/M/F* | Stop Codons |
|---|---|---|---|---|---|---|---|---|---|
| **NT** | **607** | **1231** | **19.4%** | **216** | **32.9%** | **328 (100%)** | **218 (66.5%)** | **0/0** | |
| Mutations: | 607G>C, 613T>A, 614C>G, 616C>A, 618T>C, 621A>G, 622G>A, 624A>T, 628T>G, 631T>A, 636A>T, 642A>G, 645T>A, 646T>C, 648A>C, 652A>G, 653C>T, 654T>C, 655G>A, 656C>A, 657A>T, 658C>G, 659C>A, 661G>A, 662G>C, 664A>G, 665A>C, 667T>C, 668T>A, 669C>T, 673A>T, 674A>C, 675T>C, 676C>T, 678T>G, 680T>A, 688A>G, 690C>T, 691C>A, 694A>C, 696A>G, 697A>G, 699C>T, 703G>T, 704C>G, 708T>A, 714A>T, 721T>C, 723G>T, 734C>A, 735C>T, 738T>G, 744C>T, 747G>A, 750T>A, 753T>G, 756A>T, 757G>A, 777C>T, 780T>C, 784T>A, 787G>C, 788A>T, 796A>T, 798T>C, 1098T>A, 1105T>A, 1106C>G, 1107C>T, 1113T>A, 1114G>A, 1115T>C, 1116T>C, 1122T>G, 1126A>T, 1128G>A, 1132C>T, 1134T>A, 1137C>T, 1140G>T, 1146C>T, 1147G>C, 1149G>A, 1152T>A, 1156C>A, 1158C>T, 1167C>T, 1171A>G, 1173T>C, 1174C>T, 1176T>A, 1179A>G, 1188A>C, 1191T>A, 1197T>A, 1200C>G, 1201G>A, 1202T>C, 1203C>A, 1205A>T, 1209T>A, 1210G>C, 1211A>C, 1212T>A, 1219A>T, 1221G>T, 1222C>G, 1223C>T, 1227C>T, 1230C>T | | | | | | | | |
| **CDS** | | | | | | | | | |
| BWW32_s2gp1 | 200 | 408 | 22.5% | 562 | 73.4% | 110 (100%) | 76 (69.1%) | 0/0/0/0 | 0 |
| Protein mutations: | E200Q (607G>C), L203I (616C>A 618T>C), A205T (622G>A 624A>T), S207A (628T>G), S208T (631T>A), L209F (636A>T), T215V (652A>G 653C>T 654T>C), A216N (655G>A 656C>A 657A>T), P217D (658C>G 659C>A), G218T (661G>A 662G>C), N219A (664A>G 665A>C), F220H (667T>C 668T>A 669C>T), N222S (673A>T 674A>C 675T>C), F224Y (680T>A), I227V (688A>G 690C>T), T230A (697A>G 699C>T), A232C (703G>T 704C>G), A242D (734C>A 735C>T), V250I (757G>A), S259T (784T>A), E260L (787G>C 788A>T), I263F (796A>T 798T>C), V369T (1114G>A 1115T>C 1116T>C), M373L (1126A>T 1128G>A), E380Q (1147G>C 1149G>A), L383I (1156C>A 1158C>T), N388D (1171A>G 1173T>C), V398T (1201G>A 1202T>C 1203C>A), Y399F (1205A>T), D401P (1210G>C 1211A>C 1212T>A), T404S (1219A>T 1221G>T), P405V (1222C>G 1223C>T) | | | | | | | | |
| Codon mutations: | GAA200CAA (607G>C), TCT202AGT (613T>A 614C>G), CTT203ATC (616C>A 618T>C), TTA204TTG (621A>G), GCA205ACT (622G>A 624A>T), TCC207GCC (628T>G), TCT208ACT (631T>A), TTA209TTT (636A>T), CCA211CCG (642A>G), GCT212GCA (645T>A), TTA213CTC (646T>C 648A>C), ACT215GTC (652A>G 653C>T 654T>C), GCA216AAT (655G>A 656C>A 657A>T), CCT217GAT (658C>G 659C>A), GGT218ACT (661G>A 662G>C), AAC219GCC (664A>G 665A>C), TTC220CAT (667T>C 668T>A 669C>T), AAT222TCC (673A>T 674A>C 675T>C), CTT223TTG (676C>T 678T>G), TTT224TAT (680T>A), ATC227GTT (688A>G 690C>T), CGA228AGA (691C>A), AGA229CGG (694A>C 696A>G), ACC230GCT (697A>G 699C>T), GCT232TGT (703G>T 704C>G), GGT233GGA (708T>A), ACA235ACT (714A>T), TTG238CTT (721T>C 723G>T), GCC242GAT (734C>A 735C>T), CTT243CTG (738T>G), AAC245AAT (744C>T), CAG246CAA (747G>A), GGT247GGA (750T>A), CGT248CGG (753T>G), GTA249GTT (756A>T), GTT250ATT (757G>A), CCC256CCT (777C>T), TCT257TCC (780T>C), TCT259ACT (784T>A), GAG260CTG (787G>C 788A>T), ATT263TTC (796A>T 798T>C), CGT363.GA (1098T>A), TCC366AGT (1105T>A 1106C>G 1107C>T), GGT368GGA (1113T>A), GTT369ACC (1114G>A 1115T>C 1116T>C), CTT371CTG (1122T>G), ATG373TTA (1126A>T 1128G>A), CTT375TTA (1132C>T 1134T>A), GAC376GAT (1137C>T), CCG377CCT (1140G>T), GCC379GCT (1146C>T), GAG380CAA (1147G>C 1149G>A), CTT381CTA (1152T>A), CTC383ATT (1156C>A 1158C>T), GTC386GTT (1167C>T), AAT388GAC (1171A>G 1173T>C), CTT389TTA (1174C>T 1176T>A), GAA390GAG (1179A>G), CCA393CCC (1188A>C), TCT394TCA (1191T>A), TCT396TCA (1197T>A), TCC397TCG (1200C>G), GTC398ACA (1201G>A 1202T>C 1203C>A), TAT399TTT (1205A>T), TCT400TCA (1209T>A), GAT401CCA (1210G>C 1211A>C 1212T>A), ACG404TCT (1219A>T 1221G>T), CCT405GTT (1222C>G 1223C>T), GGC406GGT (1227C>T), TAC407TAT (1230C>T) | | | | | | | | |
| **Proteins** | | | | | | | | | |
| capsid protein (YP_009337880.1) | 200 | 408 | 22.5% | 562 | 73.4% | 110 (100%) | 76 (69.1%) | 0/0/0/0 | 0 |
| Protein mutations: | E200Q (607G>C), L203I (616C>A 618T>C), A205T (622G>A 624A>T), S207A (628T>G), S208T (631T>A), L209F (636A>T), T215V (652A>G 653C>T 654T>C), A216N (655G>A 656C>A 657A>T), P217D (658C>G 659C>A), G218T (661G>A 662G>C), N219A (664A>G 665A>C), F220H (667T>C 668T>A 669C>T), N222S (673A>T 674A>C 675T>C), F224Y (680T>A), I227V (688A>G 690C>T), T230A (697A>G 699C>T), A232C (703G>T 704C>G), A242D (734C>A 735C>T), V250I (757G>A), S259T (784T>A), E260L (787G>C 788A>T), I263F (796A>T 798T>C), V369T (1114G>A 1115T>C 1116T>C), M373L (1126A>T 1128G>A), E380Q (1147G>C 1149G>A), L383I (1156C>A 1158C>T), N388D (1171A>G 1173T>C), V398T (1201G>A 1202T>C 1203C>A), Y399F (1205A>T), D401P (1210G>C 1211A>C 1212T>A), T404S (1219A>T 1221G>T), P405V (1222C>G 1223C>T) | | | | | | | | |
| Codon mutations: | GAA200CAA (607G>C), TCT202AGT (613T>A 614C>G), CTT203ATC (616C>A 618T>C), TTA204TTG (621A>G), GCA205ACT (622G>A 624A>T), TCC207GCC (628T>G), TCT208ACT (631T>A), TTA209TTT (636A>T), CCA211CCG (642A>G), GCT212GCA (645T>A), TTA213CTC (646T>C 648A>C), ACT215GTC (652A>G 653C>T 654T>C), GCA216AAT (655G>A 656C>A 657A>T), CCT217GAT (658C>G 659C>A), GGT218ACT (661G>A 662G>C), AAC219GCC (664A>G 665A>C), TTC220CAT (667T>C 668T>A 669C>T), AAT222TCC (673A>T 674A>C 675T>C), CTT223TTG (676C>T 678T>G), TTT224TAT (680T>A), ATC227GTT (688A>G 690C>T), CGA228AGA (691C>A), AGA229CGG (694A>C 696A>G), ACC230GCT (697A>G 699C>T), GCT232TGT (703G>T 704C>G), GGT233GGA (708T>A), ACA235ACT (714A>T), TTG238CTT (721T>C 723G>T), GCC242GAT (734C>A 735C>T), CTT243CTG (738T>G), AAC245AAT (744C>T), CAG246CAA (747G>A), GGT247GGA (750T>A), CGT248CGG (753T>G), GTA249GTT (756A>T), GTT250ATT (757G>A), CCC256CCT (777C>T), TCT257TCC (780T>C), TCT259ACT (784T>A), GAG260CTG (787G>C 788A>T), ATT263TTC (796A>T 798T>C), CGT363.GA (1098T>A), TCC366AGT (1105T>A 1106C>G 1107C>T), GGT368GGA (1113T>A), GTT369ACC (1114G>A 1115T>C 1116T>C), CTT371CTG (1122T>G), ATG373TTA (1126A>T 1128G>A), CTT375TTA (1132C>T 1134T>A), GAC376GAT (1137C>T), CCG377CCT (1140G>T), GCC379GCT (1146C>T), GAG380CAA (1147G>C 1149G>A), CTT381CTA (1152T>A), CTC383ATT (1156C>A 1158C>T), GTC386GTT (1167C>T), AAT388GAC (1171A>G 1173T>C), CTT389TTA (1174C>T 1176T>A), GAA390GAG (1179A>G), CCA393CCC (1188A>C), TCT394TCA (1191T>A), TCT396TCA (1197T>A), TCC397TCG (1200C>G), GTC398ACA (1201G>A 1202T>C 1203C>A), TAT399TTT (1205A>T), TCT400TCA (1209T>A), GAT401CCA (1210G>C 1211A>C 1212T>A), ACG404TCT (1219A>T 1221G>T), CCT405GTT (1222C>G 1223C>T), GGC406GGT (1227C>T), TAC407TAT (1230C>T) | | | | | | | | |

*: Inserts / Deletes / Misaligned / Frameshifts

#### Analysis details

This analysis was performed with panviral2.64

### NGS Details (UN8): Hubei noda-like virus 9 (segment 1)

#### Assembly

| | |
|---|---|
| **Coverage Length** | 706 (3 contig(s)) |
| **Depth Of Coverage** | 1.9 |
| **Number Of Reads** | 11 |
| **Reads Per Million** | 0.18 rpm (after QC) |
| **Ambiguities** | 0 |
| **Assembly Method** | de novo + reference guided assembly |
| **Consensus Caller** | Bcf Tools |

#### Coverage Map

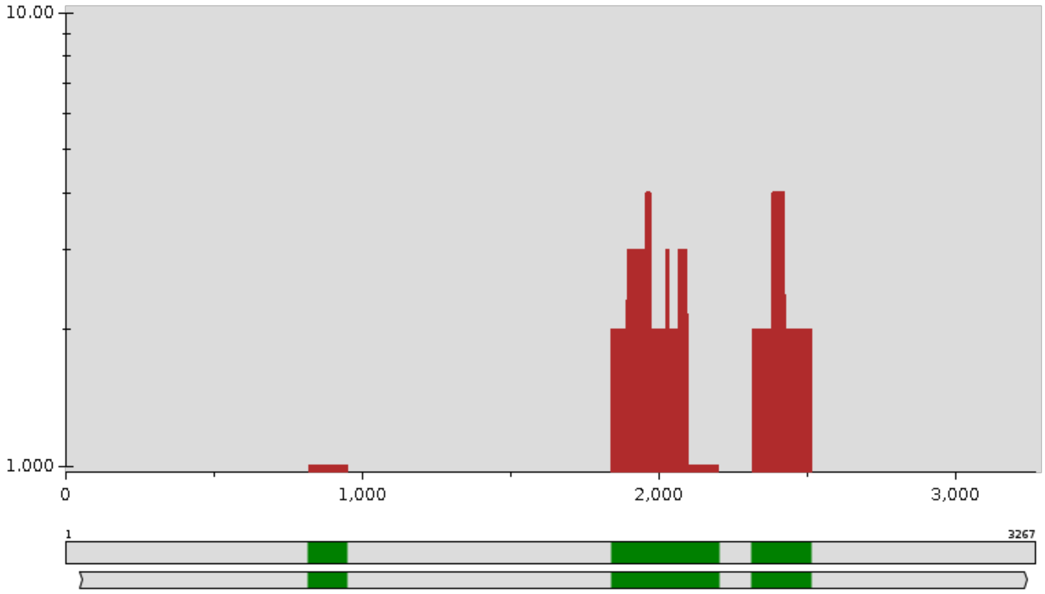

#### Assignment

| | |
|---|---|
| **Type** | Hubei noda-like virus 9 (Taxonomy ID: 1922989) |
| **Reference Genome** | NC_033307.1 |
| **NT Identity (%)** | 66.4306 |
| **AA Identity (%)** | 69.9153 |
| **Number Of Stop Codons** | 0 |
| **Number Of CDS** | 1 |

#### Alignment

| | |
|---|---|
| **Alignment Score** | 464.0 (NT) + 1227.0 (AA) = 1691.0 |
| **Concordance (%)** | 55.479 |

| Alignment Method | Global, seeded, nucleotide + amino acids (AGA) |
|---|---|

#### Genome Region

Sequence starts at position 818 and ends at position 2517 relative to NC_033307.1 reference sequence.

#### Alignment Detailed Statistics

| | Begin | End | Coverage | Score | Concordance | Matches | Identities | I/D/M/F* | Stop Codons |
|---|---|---|---|---|---|---|---|---|---|
| **NT** | **818** | **2517** | **21.6%** | **464** | **32.9%** | **706 (100%)** | **469 (66.4%)** | **0/0** | |
| Mutations: | 824C>G, 833C>T, 836C>T, 839A>C, 840C>A, 842G>A, 843A>G, 848C>T, 866A>G, 878G>C, 880A>G, 881A>G, 885A>G, 886G>A, 887T>A, 896C>T, 902T>C, 908C>T, 909A>G, 910G>C, 917C>T, 918A>T, 920T>A, 921T>C, 922C>A, 923T>A, 924A>G, 927G>A, 928T>C, 932T>C, 933T>A, 934C>A, 935A>G, 936T>A, 937T>A, 942T>A, 943T>A, 944T>C, 1841A>T, 1842C>T, 1847G>T, 1848G>A, 1849C>G, 1852G>A, 1853G>T, 1859T>C, 1862T>A, 1865C>T, 1868G>A, 1871A>C, 1877T>C, 1880A>G, 1885A>G, 1886T>C, 1889C>T, 1894C>G, 1895A>T, 1898C>T, 1901A>G, 1907C>G, 1910A>G, 1913A>T, 1917C>T, 1919G>A, 1920C>G, 1923T>A, 1925T>C, 1927C>T, 1928T>G, 1931A>G, 1932C>T, 1934C>G, 1937G>A, 1941G>A, 1942A>G, 1946T>C, 1949T>C, 1950C>G, 1951A>C, 1952A>T, 1954A>G, 1959G>T, 1960C>G, 1962G>A, 1965C>T, 1970C>T, 1979C>G, 1985C>T, 1988T>C, 1991T>A, 1997A>T, 2007C>T, 2009A>G, 2013C>A, 2014G>A, 2015A>G, 2016T>A, 2021T>A, 2028C>A, 2032T>G, 2033C>T, 2036C>T, 2037C>A, 2039A>G, 2040C>T, 2041G>T, 2046A>T, 2047G>C, 2048T>G, 2050G>A, 2054C>T, 2057A>C, 2060C>T, 2061C>A, 2062T>C, 2063G>T, 2065C>T, 2066A>C, 2067A>T, 2068A>C, 2069C>A, 2070T>A, 2072C>T, 2076T>A, 2083A>G, 2084A>T, 2087C>A, 2090C>T, 2093A>G, 2096A>G, 2097T>A, 2103G>A, 2104T>C, 2108C>T, 2109C>A, 2110G>A, 2111T>A, 2114A>G, 2118A>G, 2119A>T, 2121C>A, 2127T>A, 2128C>G, 2129A>T, 2130G>A, 2132C>A, 2134T>A, 2135T>C, 2144A>T, 2145A>C, 2147A>T, 2153C>T, 2156C>T, 2159T>A, 2171T>C, 2174C>T, 2186C>T, 2189C>T, 2313C>A, 2319A>G, 2320T>A, 2321T>A, 2324T>G, 2326T>C, 2327G>T, 2328G>A, 2329T>A, 2333T>C, 2334T>C, 2339A>C, 2343T>A, 2348C>T, 2357G>T, 2360G>T, 2362C>G, 2363T>G, 2364G>A, 2365A>T, 2366T>G, 2368C>A, 2369C>A, 2372T>G, 2374C>A, 2375G>A, 2376T>C, 2378A>T, 2379A>G, 2381C>T, 2382A>T, 2383A>C, 2388T>C, 2390A>T, 2394C>A, 2396C>G, 2399T>C, 2402C>T, 2405T>A, 2409A>C, 2411C>T, 2412C>G, 2413T>C, 2414T>A, 2415G>C, 2416C>G, 2418T>A, 2420T>A, 2421C>A, 2422T>C, 2423T>A, 2425G>C, 2426T>A, 2429A>C, 2432C>T, 2438C>T, 2443C>A, 2444A>G, 2450A>T, 2456A>C, 2459A>G, 2460T>G, 2472A>G, 2473C>T, 2474A>G, 2477T>C, 2480G>A, 2481A>C, 2482T>A, 2483G>A, 2486A>T, 2489C>T, 2490C>A, 2492T>A, 2495C>G, 2501T>A, 2504A>T, 2506A>T, 2510C>T | | | | | | | | |

##### CDS

| | Begin | End | Coverage | Score | Concordance | Matches | Identities | I/D/M/F* | Stop Codons |
|---|---|---|---|---|---|---|---|---|---|
| BWW32_s1gp1 | 257 | 823 | 22.2% | 1227 | 74.4% | 236 (100%) | 165 (69.9%) | 0/0/0/0 | 0 |
| Protein mutations: | D258E (824C>G), Q264K (840C>A 842G>A), N265D (843A>G), K277R (880A>G 881A>G), S279E (885A>G 886G>A 887T>A), S287A (909A>G 910G>C), I290L (918A>T 920T>A), S291Q (921T>C 922C>A 923T>A), I292V (924A>G), V293T (927G>A 928T>C), S295K (933T>A 934C>A 935A>G), F296N (936T>A 937T>A), F298N (942T>A 943T>A 944T>C), A600R (1848G>A 1849C>G), W601Y (1852G>A 1853G>T), E607D (1871A>C), H612R (1885A>G 1886T>C), S615C (1894C>G 1895A>T), Q624E (1920C>G), S625T (1923T>A 1925T>C), A626V (1927C>T 1928T>G), D631S (1941G>A 1942A>G), Q634A (1950C>G 1951A>C 1952A>T), N635S (1954A>G), A637C (1959G>T 1960C>G), V638I (1962G>A), R655K (2013C>A 2014G>A 2015A>G), Y656N (2016T>A), I661S (2032T>G 2033C>T), Q663K (2037C>A 2039A>G), R664L (2040C>T 2041G>T), S667N (2050G>A), L671T (2061C>A 2062T>C 2063G>T), T672I (2065C>T 2066A>C), N673S (2067A>T 2068A>C 2069C>A), S674T (2070T>A 2072C>T), F676I (2076T>A), K678S (2083A>G 2084A>T), S683T (2097T>A), V685T (2103G>A 2104T>C), R687K (2109C>A 2110G>A 2111T>A), K690V (2118A>G 2119A>T), A694T (2130G>A 2132C>A), F695Y (2134T>A 2135T>C), L755I (2313C>A), I757E (2319A>G 2320T>A 2321T>A), V759A (2326T>C 2327G>T), V760N (2328G>A 2329T>A), S765T (2343T>A), E769D (2357G>T), T771R (2362C>G 2363T>G), D772M (2364G>A 2365A>T 2366T>G), P773Q (2368C>A 2369C>A), A775E (2374C>A 2375G>A), S776P (2376T>C 2378A>T), I777V (2379A>G 2381C>T), N778S (2382A>T 2383A>C), I787L (2409A>C 2411C>T), L788A (2412C>G 2413T>C 2414T>A), A789R (2415G>C 2416C>G), S790T (2418T>A 2420T>A), L791T (2421C>A 2422T>C 2423T>A), S792T (2425G>C 2426T>A), T798K (2443C>A 2444A>G), L804V (2460T>G), T808V (2472A>G 2473C>T 2474A>G), M811Q (2481A>C 2482T>A 2483G>A), D815E (2495C>G), Y819F (2506A>T) | | | | | | | | |
| Codon mutations: | GAC258GAG (824C>G), TAC261TAT (833C>T), TAC262TAT (836C>T), ATA263ATC (839A>C), CAG264AAA (840C>A 842G>A), AAT265GAT (843A>G), AAC266AAT (848C>T), GTA272GTG (866A>G), GGG276GGC (878G>C), AAA277AGG (880A>G 881A>G), AGT279GAA (885A>G 886G>A 887T>A), GTC282GTT (896C>T), GAT284GAC (902T>C), GAC286GAT (908C>T), AGT287GCT (909A>G 910G>C), ACC289ACT (917C>T), ATT290TTA (918A>T 920T>A), TCT291CAA (921T>C 922C>A 923T>A), ATC292GTC (924A>G), GTT293ACT (927G>A 928T>C), GAT294GAC (932T>C), TCA295AAG (933T>A 934C>A 935A>G), TTT296AAT (936T>A 937T>A), TTT298AAC (942T>A 943T>A 944T>C), GCA597GCT (1841A>T), CTA598TTA (1842C>T), TCG599TCT (1847G>T), GCA600AGA (1848G>A 1849C>G), TGG601TAT (1852G>A 1853G>T), TAT603TAC (1859T>C), GCT604GCA (1862T>A), TTC605TTT (1865C>T), AAG606AAA (1868G>A), GAA607GAC (1871A>C), TGT609TGC (1877T>C), TTA610TTG (1880A>G), CAT612CGC (1885A>G 1886T>C), CTC613CTT (1889C>T), TCA615TGT (1894C>G 1895A>T), TAC616TAT (1898C>T), GGA617GGG (1901A>G), GGC619GGG (1907C>G), AAA620AAG (1910A>G), ACA621ACT (1913A>T), CTG623TTA (1917C>T 1919G>A), CAG624GAG (1920C>G), TCT625ACC (1923T>A 1925T>C), GCT626GTG (1927C>T 1928T>G), AAA627AAG (1931A>G), CTC628TTG (1932C>T 1934C>G), CTG629CTA (1937G>A), GAT631AGT (1941G>A 1942A>G), ATT632ATC (1946T>C), TGT633TGC (1949T>C), CAA634GCT (1950C>G 1951A>C 1952A>T), AAT635AGT (1954A>G), GCT637TGT (1959G>T 1960C>G), GTC638ATC (1962G>A), CTA639TTA (1965C>T), ACC640ACT (1970C>T), ACC643ACG (1979C>G), TTC645TTT (1985C>T), GAT646GAC (1988T>C), GGT647GGA (1991T>A), ATA649ATT (1997A>T), CTA653TTG (2007C>T 2009A>G), CGA655AAG (2013C>A 2014G>A 2015A>G), TAT656AAT (2016T>A), GTT657GTA (2021T>A), CGG660AGG (2028C>A), ATC661AGT (2032T>G 2033C>T), TAC662TAT (2036C>T), CAA663AAG (2037C>A 2039A>G), CGA664TTA (2040C>T 2041G>T), AGT666TCG (2046A>T 2047G>C 2048T>G), AGT667AAT (2050G>A), GAC668GAT (2054C>T), CCA669CCC (2057A>C), GCC670GCT (2060C>T), CTG671ACT (2061C>A 2062T>C 2063G>T), ACA672ATC (2065C>T 2066A>C), AAC673TCA (2067A>T 2068A>C 2069C>A), TCC674ACT (2070T>A 2072C>T), TTT676ATT (2076T>A), AAA678AGT (2083A>G 2084A>T), ATC679ATA (2087C>A), TTC680TTT (2090C>T), AAA681AAG (2093A>G), CAA682CAG (2096A>G), TCA683ACA (2097T>A), GTA685ACA (2103G>A 2104T>C), ACC686ACT (2108C>T), CGT687AAA (2109C>A 2110G>A 2111T>A), GAA688GAG (2114A>G), AAA690GTA (2118A>G 2119A>T), CGA691AGA (2121C>A), TCA693AGT (2127T>A 2128C>G 2129A>T), GCC694ACA (2130G>A 2132C>A), TTT695TAC (2134T>A 2135T>C), ACA698ACT (2144A>T), AGA699CGT (2145A>C 2147A>T), GGC701GGT (2153C>T), AGC702AGT (2156C>T), CCT703CCA (2159T>A), GAT707GAC (2171T>C), GGC708GGT (2174C>T), ATC712ATT (2186C>T), AAC713AAT (2189C>T), CTT755ATT (2313C>A), ATT757GAA (2319A>G 2320T>A 2321T>A), GTT758GTG (2324T>G), GTG759GCT (2326T>C 2327G>T), GTT760AAT (2328G>A 2329T>A), GAT761GAC (2333T>C), TTA762CTA (2334T>C), GGA763GGC (2339A>C), TCA765ACA (2343T>A), GTC766GTT (2348C>T), GAG769GAT (2357G>T), GTG770GTT (2360G>T), ACT771AGG (2362C>G 2363T>G), GAT772ATG (2364G>A 2365A>T 2366T>G), CCC773CAA (2368C>A 2369C>A), GGT774GGG (2372T>G), GCG775GAA (2374C>A 2375G>A), TCA776CCT (2376T>C 2378A>T), ATC777GTT (2379A>G 2381C>T), AAC778TCC (2382A>T 2383A>C), TTA780CTT (2388T>C 2390A>T), CGC782AGG (2394C>A 2396C>G), ATT783ATC (2399T>C), TTC784TTT (2402C>T), CCT785CCA (2405T>A), ATC787CTT (2409A>C 2411C>T), CTT788GCA (2412C>G 2413T>C 2414T>A), GCA789CGA (2415G>C 2416C>G), TCT790ACA (2418T>A 2420T>A), CTT791ACA (2421C>A 2422T>C 2423T>A), AGT792ACA (2425G>C 2426T>A), TCA793TCC (2429A>C), CAC794CAT (2432C>T), GAC796GAT (2438C>T), ACA798AAG (2443C>A 2444A>G), ACA800ACT (2450A>T), CCA802CCC (2456A>C), AAA803AAG (2459A>G), TTA804GTA (2460T>G), ACA808GTG (2472A>G 2473C>T 2474A>G), AAT809AAC (2477T>C), CAG810CAA (2480G>A), ATG811CAA (2481A>C 2482T>A 2483G>A), GTA812GTT (2486A>T), ACC813ACT (2489C>T), CGT814AGA (2490C>A 2492T>A), GAC815GAG (2495C>G), GCT817GCA (2501T>A), GCA818GCT (2504A>T), TAT819TTT (2506A>T), AAC820AAT (2510C>T) | | | | | | | | |

##### Proteins

| | Begin | End | Coverage | Score | Concordance | Matches | Identities | I/D/M/F* | Stop Codons |
|---|---|---|---|---|---|---|---|---|---|
| RNA-dependent RNA polymerase (YP_009337879.1) | 257 | 823 | 22.2% | 1227 | 74.4% | 236 (100%) | 165 (69.9%) | 0/0/0/0 | 0 |
| Protein mutations: | D258E (824C>G), Q264K (840C>A 842G>A), N265D (843A>G), K277R (880A>G 881A>G), S279E (885A>G 886G>A 887T>A), S287A (909A>G 910G>C), I290L (918A>T 920T>A), S291Q (921T>C 922C>A 923T>A), I292V (924A>G), V293T (927G>A 928T>C), S295K (933T>A 934C>A 935A>G), F296N (936T>A 937T>A), F298N (942T>A 943T>A 944T>C), A600R (1848G>A 1849C>G), W601Y (1852G>A 1853G>T), E607D (1871A>C), H612R (1885A>G 1886T>C), S615C (1894C>G 1895A>T), Q624E (1920C>G), S625T (1923T>A 1925T>C), A626V (1927C>T 1928T>G), D631S (1941G>A 1942A>G), Q634A (1950C>G 1951A>C 1952A>T), N635S (1954A>G), A637C (1959G>T 1960C>G), V638I (1962G>A), R655K (2013C>A 2014G>A 2015A>G), Y656N (2016T>A), I661S (2032T>G 2033C>T), Q663K (2037C>A 2039A>G), R664L (2040C>T 2041G>T), S667N (2050G>A), L671T (2061C>A 2062T>C 2063G>T), T672I (2065C>T 2066A>C), N673S (2067A>T 2068A>C 2069C>A), S674T (2070T>A 2072C>T), F676I (2076T>A), K678S (2083A>G 2084A>T), S683T (2097T>A), V685T (2103G>A 2104T>C), R687K (2109C>A 2110G>A 2111T>A), K690V (2118A>G 2119A>T), A694T (2130G>A 2132C>A), F695Y (2134T>A 2135T>C), L755I (2313C>A), I757E (2319A>G 2320T>A 2321T>A), V759A (2326T>C 2327G>T), V760N (2328G>A 2329T>A), S765T (2343T>A), E769D (2357G>T), T771R (2362C>G 2363T>G), D772M (2364G>A 2365A>T 2366T>G), P773Q (2368C>A 2369C>A), A775E (2374C>A 2375G>A), S776P (2376T>C 2378A>T), I777V (2379A>G 2381C>T), N778S (2382A>T 2383A>C), I787L (2409A>C 2411C>T), L788A (2412C>G 2413T>C 2414T>A), A789R (2415G>C 2416C>G), S790T (2418T>A 2420T>A), L791T (2421C>A 2422T>C 2423T>A), S792T (2425G>C 2426T>A), T798K (2443C>A 2444A>G), L804V (2460T>G), T808V (2472A>G 2473C>T 2474A>G), M811Q (2481A>C 2482T>A 2483G>A), D815E (2495C>G), Y819F (2506A>T) | | | | | | | | |

| | Begin | End | Coverage | Score | Concordance | Matches | Identities | I/D/M/F* | Stop Codons |
|---|---|---|---|---|---|---|---|---|---|
| **NT** | **818** | **2517** | **21.6%** | **464** | **32.9%** | **706 (100%)** | **469 (66.4%)** | **0/0** | |
| Codon mutations: | GAC258GAG (824C>G), TAC261TAT (833C>T), TAC262TAT (836C>T), ATA263ATC (839A>C), CAG264AAA (840C>A 842G>A), AAT265GAT (843A>G), AAC266AAT (848C>T), GTA272GTG (866A>G), GGG276GGC (878G>C), AAA277AGG (880A>G 881A>G), AGT279GAA (885A>G 886G>A 887T>A), GTC282GTT (896C>T), GAT284GAC (902T>C), GAC286GAT (908C>T), AGT287GCT (909A>G 910G>C), ACC289ACT (917C>T), ATT290TTA (918A>T 920T>A), TCT291CAA (921T>C 922C>A 923T>A), ATC292GTC (924A>G), GTT293ACT (927G>A 928T>C), GAT294GAC (932T>C), TCA295AAG (933T>A 934C>A 935A>G), TTT296AAT (936T>A 937T>A), TTT298AAC (942T>A 943T>A 944T>C), GCA597GCT (1841A>T), CTA598TTA (1842C>T), TCG599TCT (1847G>T), GCA600AGA (1848G>A 1849C>G), TGG601TAT (1852G>A 1853G>T), TAT603TAC (1859T>C), GCT604GCA (1862T>A), TTC605TTT (1865C>T), AAG606AAA (1868G>A), GAA607GAC (1871A>C), TGT609TGC (1877T>C), TTA610TTG (1880A>G), CAT612CGC (1885A>G 1886T>C), CTC613CTT (1889C>T), TCA615TGT (1894C>G 1895A>T), TAC616TAT (1898C>T), GGA617GGG (1901A>G), GGC619GGG (1907C>G), AAA620AAG (1910A>G), ACA621ACT (1913A>T), CTG623TTA (1917C>T 1919G>A), CAG624GAG (1920C>G), TCT625ACC (1923T>A 1925T>C), GCT626GTG (1927C>T 1928T>G), AAA627AAG (1931A>G), CTC628TTG (1932C>T 1934C>G), CTG629CTA (1937G>A), GAT631AGT (1941G>A 1942A>G), ATT632ATC (1946T>C), TGT633TGC (1949T>C), CAA634GCT (1950C>G 1951A>C 1952A>T), AAT635AGT (1954A>G), GCT637TGT (1959G>T 1960C>G), GTC638ATC (1962G>A), CTA639TTA (1965C>T), ACC640ACT (1970C>T), ACC643ACG (1979C>G), TTC645TTT (1985C>T), GAT646GAC (1988T>C), GGT647GGA (1991T>A), ATA649ATT (1997A>T), CTA653TTG (2007C>T 2009A>G), CGA655AAG (2013C>A 2014G>A 2015A>G), TAT656AAT (2016T>A), GTT657GTA (2021T>A), CGG660AGG (2028C>A), ATC661AGT (2032T>G 2033C>T), TAC662TAT (2036C>T), CAA663AAG (2037C>A 2039A>G), CGA664TTA (2040C>T 2041G>T), AGT666TCG (2046A>T 2047G>C 2048T>G), AGT667AAT (2050G>A), GAC668GAT (2054C>T), CCA669CCC (2057A>C), GCC670GCT (2060C>T), CTG671ACT (2061C>A 2062T>C 2063G>T), ACA672ATC (2065C>T 2066A>C), AAC673TCA (2067A>T 2068A>C 2069C>A), TCC674ACT (2070T>A 2072C>T), TTT676ATT (2076T>A), AAA678AGT (2083A>G 2084A>T), ATC679ATA (2087C>A), TTC680TTT (2090C>T), AAA681AAG (2093A>G), CAA682CAG (2096A>G), TCA683ACA (2097T>A), GTA685ACA (2103G>A 2104T>C), ACC686ACT (2108C>T), CGT687AAA (2109C>A 2110G>A 2111T>A), GAA688GAG (2114A>G), AAA690GTA (2118A>G 2119A>T), CGA691AGA (2121C>A), TCA693AGT (2127T>A 2128C>G 2129A>T), GCC694ACA (2130G>A 2132C>A), TTT695TAC (2134T>A 2135T>C), ACA698ACT (2144A>T), AGA699CGT (2145A>C 2147A>T), GGC701GGT (2153C>T), AGC702AGT (2156C>T), CCT703CCA (2159T>A), GAT707GAC (2171T>C), GGC708GGT (2174C>T), ATC712ATT (2186C>T), AAC713AAT (2189C>T), CTT755ATT (2313C>A), ATT757GAA (2319A>G 2320T>A 2321T>A), GTT758GTG (2324T>G), GTG759GCT (2326T>C 2327G>T), GTT760AAT (2328G>A 2329T>A), GAT761GAC (2333T>C), TTA762CTA (2334T>C), GGA763GGC (2339A>C), TCA765ACA (2343T>A), GTC766GTT (2348C>T), GAG769GAT (2357G>T), GTG770GTT (2360G>T), ACT771AGG (2362C>G 2363T>G), GAT772ATG (2364G>A 2365A>T 2366T>G), CCC773CAA (2368C>A 2369C>A), GGT774GGG (2372T>G), GCG775GAA (2374C>A 2375G>A), TCA776CCT (2376T>C 2378A>T), ATC777GTT (2379A>G 2381C>T), AAC778TCC (2382A>T 2383A>C), TTA780CTT (2388T>C 2390A>T), CGC782AGG (2394C>A 2396C>G), ATT783ATC (2399T>C), TTC784TTT (2402C>T), CCT785CCA (2405T>A), ATC787CTT (2409A>C 2411C>T), CTT788GCA (2412C>G 2413T>C 2414T>A), GCA789CGA (2415G>C 2416C>G), TCT790ACA (2418T>A 2420T>A), CTT791ACA (2421C>A 2422T>C 2423T>A), AGT792ACA (2425G>C 2426T>A), TCA793TCC (2429A>C), CAC794CAT (2432C>T), GAC796GAT (2438C>T), ACA798AAG (2443C>A 2444A>G), ACA800ACT (2450A>T), CCA802CCC (2456A>C), AAA803AAG (2459A>G), TTA804GTA (2460T>G), ACA808GTG (2472A>G 2473C>T 2474A>G), AAT809AAC (2477T>C), CAG810CAA (2480G>A), ATG811CAA (2481A>C 2482T>A 2483G>A), GTA812GTT (2486A>T), ACC813ACT (2489C>T), CGT814AGA (2490C>A 2492T>A), GAC815GAG (2495C>G), GCT817GCA (2501T>A), GCA818GCT (2504A>T), TAT819TTT (2506A>T), AAC820AAT (2510C>T) | | | | | | | | |

*: Inserts / Deletes / Misaligned / Frameshifts

#### Analysis details

This analysis was performed with panviral2.64

### NGS Details (UN8): Caulimovirus venafragariae

#### Assembly

| | |
|---|---|
| Coverage Length | 253 (1 contig(s)) |
| Depth Of Coverage | 4.2 |
| Number Of Reads | 9 |
| Reads Per Million | 0.15 rpm (after QC) |
| Ambiguities | 0 |
| Assembly Method | de novo + reference guided assembly |
| Consensus Caller | Bcf Tools |

#### Coverage Map

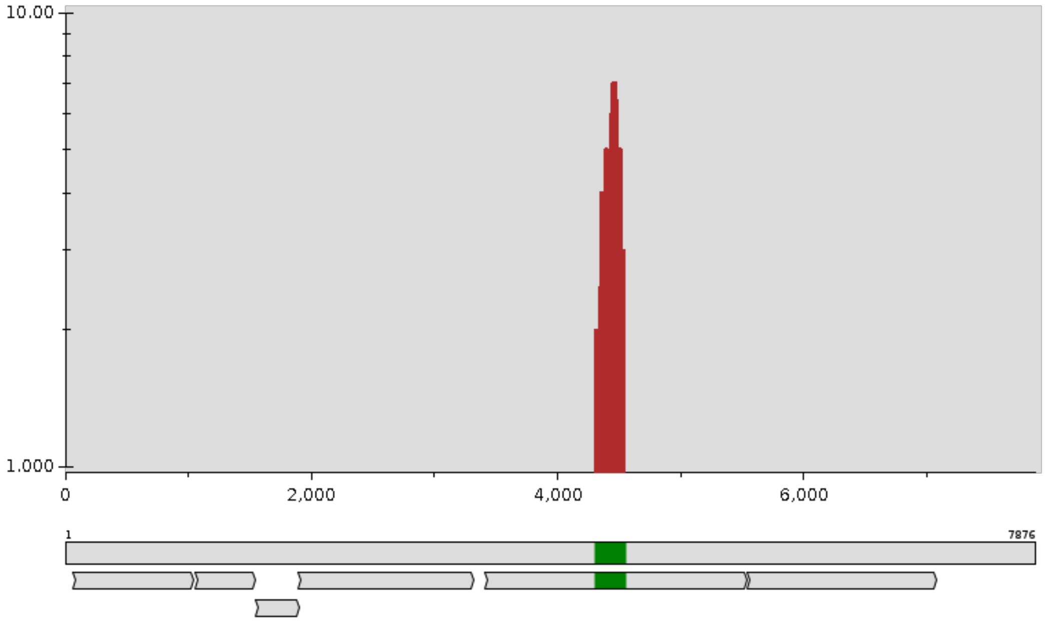

#### Assignment

| | |
|---|---|
| Type | Caulimovirus venafragariae (Taxonomy ID: 3048344) |
| Reference Genome | NC_001725.1 |
| NT Identity (%) | 63.6364 |
| AA Identity (%) | 63.5294 |
| Number Of Stop Codons | 0 |
| Number Of CDS | 6 |

#### Alignment

| | |
|---|---|
| Alignment Score | 138.0 (NT) + 390.0 (AA) = 528.0 |
| Concordance (%) | 48.4404 |

| Alignment Method | Global, seeded, nucleotide + amino acids (AGA) |
|---|---|

#### Genome Region

Sequence starts at position 4300 and ends at position 4552 relative to NC_001725.1 reference sequence.

#### Alignment Detailed Statistics

| | Begin | End | Coverage | Score | Concordance | Matches | Identities | I/D/M/F* | Stop Codons |
|---|---|---|---|---|---|---|---|---|---|
| **NT** | **4300** | **4552** | **3.2%** | **138** | **27.3%** | **253 (100%)** | **161 (63.6%)** | **0/0** | |
| Mutations: | 4300G>A, 4311C>A, 4314G>C, 4315C>T, 4317T>A, 4320T>A, 4321A>G, 4323C>T, 4326C>A, 4330C>A, 4331C>G, 4332T>A, 4333A>T, 4334G>C, 4335C>T, 4337A>C, 4338G>A, 4339A>T, 4340G>C, 4341T>A, 4342C>A, 4343C>A, 4348T>A, 4349C>G, 4350C>A, 4354C>G, 4362C>T, 4371G>A, 4378G>A, 4379C>G, 4383G>A, 4386C>A, 4387A>G, 4388A>T, 4390C>A, 4395C>T, 4402A>C, 4403G>A, 4408G>A, 4413T>A, 4419T>C, 4424A>G, 4425G>A, 4426T>C, 4428A>T, 4434C>T, 4435C>A, 4437T>C, 4440A>T, 4442A>G, 4443G>A, 4444G>A, 4445G>C, 4446A>G, 4447G>T, 4451G>C, 4452C>T, 4455C>T, 4456C>A, 4457T>A, 4458A>G, 4461T>A, 4464T>A, 4465A>G, 4467C>T, 4470G>A, 4471G>A, 4472A>G, 4473G>T, 4474C>G, 4477C>T, 4479G>A, 4480C>A, 4482T>A, 4483C>A, 4485A>T, 4491C>A, 4492G>C, 4493G>A, 4503C>A, 4504T>G, 4506T>A, 4508A>T, 4510T>A, 4511C>A, 4513T>A, 4514C>A, 4515C>A, 4521C>T, 4530T>A, 4535T>A, 4542G>A | | | | | | | | |
| **CDS** | | | | | | | | | |
| ORF_V | 299 | 383 | 12.0% | 390 | 66.3% | 85 (100%) | 54 (63.5%) | 0/0/0/0 | 0 |
| Protein mutations: | E299K (4300G>A), K303N (4314G>C), I306V (4321A>G 4323C>T), P309R (4330C>A 4331C>G 4332T>A), K311T (4337A>C 4338G>A), P313K (4342C>A 4343C>A), S315R (4348T>A 4349C>G 4350C>A), P317A (4354C>G), A325S (4378G>A 4379C>G), K328V (4387A>G 4388A>T), R333Q (4402A>C 4403G>A), V335I (4408G>A), K340R (4424A>G 4425G>A), H344N (4435C>A 4437T>C), K346R (4442A>G 4443G>A), G347T (4444G>A 4445G>C 4446A>G), D348Y (4447G>T), G349A (4451G>C 4452C>T), L351K (4456C>A 4457T>A 4458A>G), N354D (4465A>G 4467C>T), E356S (4471G>A 4472A>G 4473G>T), Q357E (4474C>G), L359I (4480C>A 4482T>A), Q360N (4483C>A 4485A>T), G363Q (4492G>C 4493G>A), F367V (4504T>G 4506T>A), Y368F (4508A>T), S369N (4510T>A 4511C>A), S370K (4513T>A 4514C>A 4515C>A), F377Y (4535T>A) | | | | | | | | |
| Codon mutations: | GAA299AAA (4300G>A), CTC302CTA (4311C>A), AAG303AAC (4314G>C), CTT304TTA (4315C>T 4317T>A), GGT305GGA (4320T>A), ATC306GTT (4321A>G 4323C>T), ATC307ATA (4326C>A), CCT309AGA (4330C>A 4331C>G 4332T>A), AGC310TCT (4333A>T 4334G>C 4335C>T), AAG311ACA (4337A>C 4338G>A), AGT312TCA (4339A>T 4340G>C 4341T>A), CCA313AAA (4342C>A 4343C>A), TCC315AGA (4348T>A 4349C>G 4350C>A), CCA317GCA (4354C>G), TTC319TTT (4362C>T), AGG322AGA (4371G>A), GCC325AGC (4378G>A 4379C>G), GAG326GAA (4383G>A), ATC327ATA (4386C>A), AAA328GTA (4387A>G 4388A>T), CGA329AGA (4390C>A), GGC330GGT (4395C>T), AGA333CAA (4402A>C 4403G>A), GTA335ATA (4408G>A), ATT336ATA (4413T>A), TAT338TAC (4419T>C), AAG340AGA (4424A>G 4425G>A), TTA341CTT (4426T>C 4428A>T), GAC343GAT (4434C>T), CAT344AAC (4435C>A 4437T>C), ACA345ACT (4440A>T), AAG346AGA (4442A>G 4443G>A), GGA347ACG (4444G>A 4445G>C 4446A>G), GAT348TAT (4447G>T), GGC349GCT (4451G>C 4452C>T), TAC350TAT (4455C>T), CTA351AAG (4456C>A 4457T>A 4458A>G), CTT352CTA (4461T>A), CCT353CCA (4464T>A), AAC354GAT (4465A>G 4467C>T), AAG355AAA (4470G>A), GAG356AGT (4471G>A 4472A>G 4473G>T), CAA357GAA (4474C>G), CTG358TTA (4477C>T 4479G>A), CTT359ATA (4480C>A 4482T>A), CAA360AAT (4483C>A 4485A>T), ATC362ATA (4491C>A), GGA363CAA (4492G>C 4493G>A), ACC366ACA (4503C>A), TTT367GTA (4504T>G 4506T>A), TAC368TTC (4508A>T), TCT369AAT (4510T>A 4511C>A), TCC370AAA (4513T>A 4514C>A 4515C>A), GAC372GAT (4521C>T), TCT375TCA (4530T>A), TTT377TAT (4535T>A), CAG379CAA (4542G>A) | | | | | | | | |
| **Proteins** | | | | | | | | | |
| hypothetical protein (NP_043933.1) | 299 | 383 | 12.0% | 390 | 66.3% | 85 (100%) | 54 (63.5%) | 0/0/0/0 | 0 |
| Protein mutations: | E299K (4300G>A), K303N (4314G>C), I306V (4321A>G 4323C>T), P309R (4330C>A 4331C>G 4332T>A), K311T (4337A>C 4338G>A), P313K (4342C>A 4343C>A), S315R (4348T>A 4349C>G 4350C>A), P317A (4354C>G), A325S (4378G>A 4379C>G), K328V (4387A>G 4388A>T), R333Q (4402A>C 4403G>A), V335I (4408G>A), K340R (4424A>G 4425G>A), H344N (4435C>A 4437T>C), K346R (4442A>G 4443G>A), G347T (4444G>A 4445G>C 4446A>G), D348Y (4447G>T), G349A (4451G>C 4452C>T), L351K (4456C>A 4457T>A 4458A>G), N354D (4465A>G 4467C>T), E356S (4471G>A 4472A>G 4473G>T), Q357E (4474C>G), L359I (4480C>A 4482T>A), Q360N (4483C>A 4485A>T), G363Q (4492G>C 4493G>A), F367V (4504T>G 4506T>A), Y368F (4508A>T), S369N (4510T>A 4511C>A), S370K (4513T>A 4514C>A 4515C>A), F377Y (4535T>A) | | | | | | | | |
| Codon mutations: | GAA299AAA (4300G>A), CTC302CTA (4311C>A), AAG303AAC (4314G>C), CTT304TTA (4315C>T 4317T>A), GGT305GGA (4320T>A), ATC306GTT (4321A>G 4323C>T), ATC307ATA (4326C>A), CCT309AGA (4330C>A 4331C>G 4332T>A), AGC310TCT (4333A>T 4334G>C 4335C>T), AAG311ACA (4337A>C 4338G>A), AGT312TCA (4339A>T 4340G>C 4341T>A), CCA313AAA (4342C>A 4343C>A), TCC315AGA (4348T>A 4349C>G 4350C>A), CCA317GCA (4354C>G), TTC319TTT (4362C>T), AGG322AGA (4371G>A), GCC325AGC (4378G>A 4379C>G), GAG326GAA (4383G>A), ATC327ATA (4386C>A), AAA328GTA (4387A>G 4388A>T), CGA329AGA (4390C>A), GGC330GGT (4395C>T), AGA333CAA (4402A>C 4403G>A), GTA335ATA (4408G>A), ATT336ATA (4413T>A), TAT338TAC (4419T>C), AAG340AGA (4424A>G 4425G>A), TTA341CTT (4426T>C 4428A>T), GAC343GAT (4434C>T), CAT344AAC (4435C>A 4437T>C), ACA345ACT (4440A>T), AAG346AGA (4442A>G 4443G>A), GGA347ACG (4444G>A 4445G>C 4446A>G), GAT348TAT (4447G>T), GGC349GCT (4451G>C 4452C>T), TAC350TAT (4455C>T), CTA351AAG (4456C>A 4457T>A 4458A>G), CTT352CTA (4461T>A), CCT353CCA (4464T>A), AAC354GAT (4465A>G 4467C>T), AAG355AAA (4470G>A), GAG356AGT (4471G>A 4472A>G 4473G>T), CAA357GAA (4474C>G), CTG358TTA (4477C>T 4479G>A), CTT359ATA (4480C>A 4482T>A), CAA360AAT (4483C>A 4485A>T), ATC362ATA (4491C>A), GGA363CAA (4492G>C 4493G>A), ACC366ACA (4503C>A), TTT367GTA (4504T>G 4506T>A), TAC368TTC (4508A>T), TCT369AAT (4510T>A 4511C>A), TCC370AAA (4513T>A 4514C>A 4515C>A), GAC372GAT (4521C>T), TCT375TCA (4530T>A), TTT377TAT (4535T>A), CAG379CAA (4542G>A) | | | | | | | | |

*: Inserts / Deletes / Misaligned / Frameshifts

#### Analysis details

This analysis was performed with panviral2.64

### NGS Details (UN8): Cavemovirus venamanihotis

#### Assembly

| | |
|---|---|
| **Coverage Length** | 273 (1 contig(s)) |
| **Depth Of Coverage** | 3.2 |
| **Number Of Reads** | 8 |
| **Reads Per Million** | 0.13 rpm (after QC) |
| **Ambiguities** | 0 |
| **Assembly Method** | de novo + reference guided assembly |
| **Consensus Caller** | Bcf Tools |

#### Coverage Map

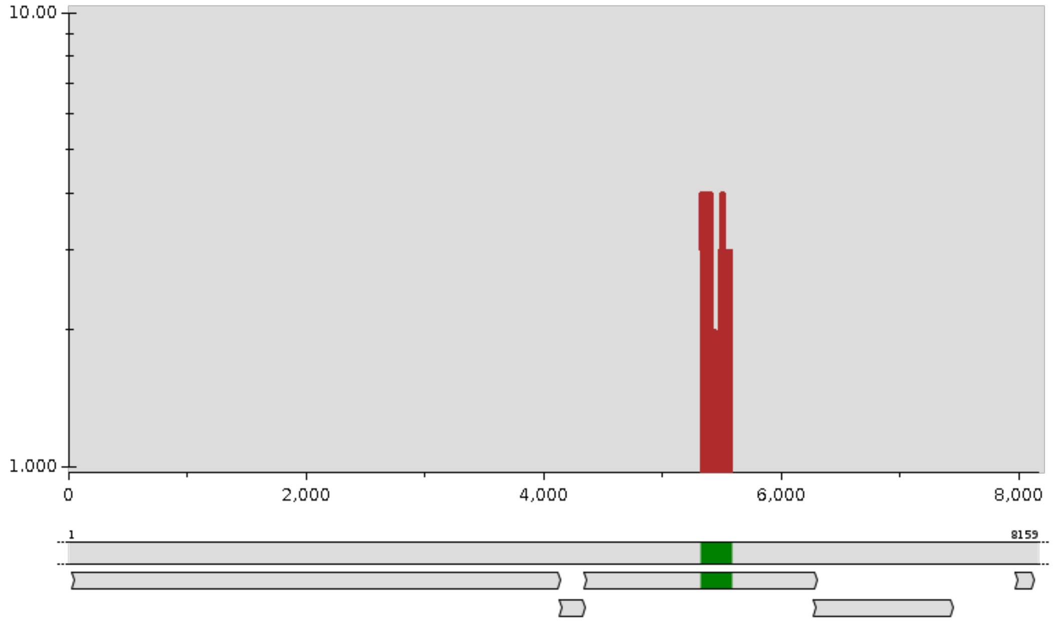

#### Assignment

| | |
|---|---|
| **Type** | Cavemovirus venamanihotis (Taxonomy ID: 3047736) |
| **Reference Genome** | NC_001648.1 |
| **NT Identity (%)** | 60.0733 |
| **AA Identity (%)** | 51.6484 |
| **Number Of Stop Codons** | 0 |
| **Number Of CDS** | 5 |

#### Alignment

| | |
|---|---|
| **Alignment Score** | 110.0 (NT) + 360.0 (AA) = 470.0 |
| **Concordance (%)** | 39.2977 |

| **Alignment Method** | Global, seeded, nucleotide + amino acids (AGA) |
|---|---|

#### Genome Region

Sequence starts at position 5318 and ends at position 5590 relative to NC_001648.1 reference sequence.

#### Alignment Detailed Statistics

| | Begin | End | Coverage | Score | Concordance | Matches | Identities | I/D/M/F* | Stop Codons |
|---|---|---|---|---|---|---|---|---|---|
| **NT** | **5318** | **5590** | **3.3%** | **110** | **20.1%** | **273 (100%)** | **164 (60.1%)** | **0/0** | |
| Mutations: | 5318G>T, 5321A>G, 5322T>C, 5324T>A, 5328C>G, 5330G>A, 5336A>T, 5339A>T, 5340T>A, 5342A>G, 5345A>T, 5351A>C, 5353A>T, 5354T>A, 5355C>A, 5357T>A, 5360C>T, 5361T>G, 5363A>C, 5366A>T, 5367A>T, 5368G>C, 5370A>G, 5371T>A, 5372T>A, 5375C>T, 5379C>A, 5382T>A, 5388G>A, 5391A>G, 5392G>A, 5393A>T, 5396A>T, 5401G>A, 5402A>T, 5410A>T, 5411T>G, 5426G>C, 5428A>G, 5429T>C, 5438T>C, 5444A>G, 5447C>A, 5449T>A, 5452C>T, 5453T>A, 5456G>A, 5458C>T, 5459T>A, 5460A>T, 5466G>A, 5467A>T, 5473A>T, 5474A>T, 5476T>A, 5477A>G, 5481A>C, 5485C>A, 5486A>C, 5488A>T, 5489A>T, 5494G>A, 5495A>G, 5496G>A, 5498T>A, 5501A>T, 5502A>G, 5503C>T, 5504A>G, 5505C>A, 5507T>A, 5508G>C, 5509C>A, 5516A>T, 5519A>G, 5520A>G, 5522T>G, 5525T>A, 5526A>T, 5527G>C, 5531G>A, 5532A>C, 5533A>C, 5539C>T, 5541G>A, 5542A>G, 5543A>T, 5546A>G, 5548G>T, 5550A>C, 5553G>A, 5554A>C, 5556A>G, 5559A>G, 5561A>C, 5562G>C, 5563A>G, 5564T>A, 5567C>T, 5568C>T, 5570T>A, 5574G>C, 5575T>A, 5576C>T, 5577C>A, 5579A>T, 5583G>T, 5584A>G, 5585A>T | | | | | | | | |
| **CDS** | | | | | | | | | |
| CsVMVgp3 | 326 | 416 | 13.9% | 360 | 54.0% | 91 (100%) | 47 (51.6%) | 0/0/0/0 | 0 |
| Protein mutations: | Y327Q (5322T>C 5324T>A), Q329E (5328C>G 5330G>A), K331N (5336A>T), L333M (5340T>A 5342A>G), Y337L (5353A>T 5354T>A), H338K (5355C>A 5357T>A), S340A (5361T>G 5363A>C), I343E (5370A>G 5371T>A 5372T>A), Q346K (5379C>A), F347I (5382T>A), D349N (5388G>A), R350D (5391A>G 5392G>A 5393A>T), R353N (5401G>A 5402A>T), Y356L (5410A>T 5411T>G), Y362C (5428A>G 5429T>C), F369Y (5449T>A), S370L (5452C>T 5453T>A), T372I (5458C>T 5459T>A), I373F (5460A>T), E375M (5466G>A 5467A>T), K377I (5473A>T 5474A>T), I378K (5476T>A 5477A>G), I380L (5481A>C), A381D (5485C>A 5486A>C), K382I (5488A>T 5489A>T), R384K (5494G>A 5495A>G), D385K (5496G>A 5498T>A), T387V (5502A>G 5503C>T 5504A>G), L388I (5505C>A 5507T>A), A389Q (5508G>C 5509C>A), I393V (5520A>G 5522T>G), K397P (5532A>C 5533A>C), T399I (5539C>T), E400S (5541G>A 5542A>G 5543A>T), C402F (5548G>T), K403Q (5550A>C), E404T (5553G>A 5554A>C), K405E (5556A>G), I406V (5559A>G 5561A>C), D407R (5562G>C 5563A>G 5564T>A), V411H (5574G>C 5575T>A 5576C>T), Q412N (5577C>A 5579A>T), E414C (5583G>T 5584A>G 5585A>T) | | | | | | | | |
| Codon mutations: | CAG325..T (5318G>T), GGA326GGG (5321A>G), TAT327CAA (5322T>C 5324T>A), CAG329GAA (5328C>G 5330G>A), AAA331AAT (5336A>T), GTA332GTT (5339A>T), TTA333ATG (5340T>A 5342A>G), CCA334CCT (5345A>T), GGA336GGC (5351A>C), TAT337TTA (5353A>T 5354T>A), CAT338AAA (5355C>A 5357T>A), AAC339AAT (5360C>T), TCA340GCC (5361T>G 5363A>C), CCA341CCT (5366A>T), AGT342TCT (5367A>T 5368G>C), ATT343GAA (5370A>G 5371T>A 5372T>A), TTC344TTT (5375C>T), CAA346AAA (5379C>A), TTT347ATT (5382T>A), GAT349AAT (5388G>A), AGA350GAT (5391A>G 5392G>A 5393A>T), ATA351ATT (5396A>T), AGA353AAT (5401G>A 5402A>T), TAT356TTG (5410A>T 5411T>G), GTG361GTC (5426G>C), TAT362TGC (5428A>G 5429T>C), GAT365GAC (5438T>C), TTA367TTG (5444A>G), GTC368GTA (5447C>A), TTT369TAT (5449T>A), TCT370TTA (5452C>T 5453T>A), AAG371AAA (5456G>A), ACT372ATA (5458C>T 5459T>A), ATT373TTT (5460A>T), GAG375ATG (5466G>A 5467A>T), AAA377ATT (5473A>T 5474A>T), ATA378AAG (5476T>A 5477A>G), ATT380CTT (5481A>C), GCA381GAC (5485C>A 5486A>C), AAA382ATT (5488A>T 5489A>T), AGA384AAG (5494G>A 5495A>G), GAT385AAA (5496G>A 5498T>A), ATA386ATT (5501A>T), ACA387GTG (5502A>G 5503C>T 5504A>G), CTT388ATA (5505C>A 5507T>A), GCA389CAA (5508G>C 5509C>A), GGA391GGT (5516A>T), TTA392TTG (5519A>G), ATT393GTG (5520A>G 5522T>G), ATT394ATA (5525T>A), AGT395TCT (5526A>T 5527G>C), AAG396AAA (5531G>A), AAA397CCA (5532A>C 5533A>C), ACA399ATA (5539C>T), GAA400AGT (5541G>A 5542A>G 5543A>T), TTA401TTG (5546A>G), TGT402TTT (5548G>T), AAA403CAA (5550A>C), GAA404ACA (5553G>A 5554A>C), AAA405GAA (5556A>G), ATA406GTC (5559A>G 5561A>C), GAT407CGA (5562G>C 5563A>G 5564T>A), TTC408TTT (5567C>T), CTT409TTA (5568C>T 5570T>A), GTC411CAT (5574G>C 5575T>A 5576C>T), CAA412AAT (5577C>A 5579A>T), GAA414TGT (5583G>T 5584A>G 5585A>T) | | | | | | | | |
| **Proteins** | | | | | | | | | |
| aspartic protease/reverse transcriptase (NP_056848.1) | 326 | 416 | 13.9% | 360 | 54.0% | 91 (100%) | 47 (51.6%) | 0/0/0/0 | 0 |
| Protein mutations: | Y327Q (5322T>C 5324T>A), Q329E (5328C>G 5330G>A), K331N (5336A>T), L333M (5340T>A 5342A>G), Y337L (5353A>T 5354T>A), H338K (5355C>A 5357T>A), S340A (5361T>G 5363A>C), I343E (5370A>G 5371T>A 5372T>A), Q346K (5379C>A), F347I (5382T>A), D349N (5388G>A), R350D (5391A>G 5392G>A 5393A>T), R353N (5401G>A 5402A>T), Y356L (5410A>T 5411T>G), Y362C (5428A>G 5429T>C), F369Y (5449T>A), S370L (5452C>T 5453T>A), T372I (5458C>T 5459T>A), I373F (5460A>T), E375M (5466G>A 5467A>T), K377I (5473A>T 5474A>T), I378K (5476T>A 5477A>G), I380L (5481A>C), A381D (5485C>A 5486A>C), K382I (5488A>T 5489A>T), R384K (5494G>A 5495A>G), D385K (5496G>A 5498T>A), T387V (5502A>G 5503C>T 5504A>G), L388I (5505C>A 5507T>A), A389Q (5508G>C 5509C>A), I393V (5520A>G 5522T>G), K397P (5532A>C 5533A>C), T399I (5539C>T), E400S (5541G>A 5542A>G 5543A>T), C402F (5548G>T), K403Q (5550A>C), E404T (5553G>A 5554A>C), K405E (5556A>G), I406V (5559A>G 5561A>C), D407R (5562G>C 5563A>G 5564T>A), V411H (5574G>C 5575T>A 5576C>T), Q412N (5577C>A 5579A>T), E414C (5583G>T 5584A>G 5585A>T) | | | | | | | | |
| Codon mutations: | CAG325..T (5318G>T), GGA326GGG (5321A>G), TAT327CAA (5322T>C 5324T>A), CAG329GAA (5328C>G 5330G>A), AAA331AAT (5336A>T), GTA332GTT (5339A>T), TTA333ATG (5340T>A 5342A>G), CCA334CCT (5345A>T), GGA336GGC (5351A>C), TAT337TTA (5353A>T 5354T>A), CAT338AAA (5355C>A 5357T>A), AAC339AAT (5360C>T), TCA340GCC (5361T>G 5363A>C), CCA341CCT (5366A>T), AGT342TCT (5367A>T 5368G>C), ATT343GAA (5370A>G 5371T>A 5372T>A), TTC344TTT (5375C>T), CAA346AAA (5379C>A), TTT347ATT (5382T>A), GAT349AAT (5388G>A), AGA350GAT (5391A>G 5392G>A 5393A>T), ATA351ATT (5396A>T), AGA353AAT (5401G>A 5402A>T), TAT356TTG (5410A>T 5411T>G), GTG361GTC (5426G>C), TAT362TGC (5428A>G 5429T>C), GAT365GAC (5438T>C), TTA367TTG (5444A>G), GTC368GTA (5447C>A), TTT369TAT (5449T>A), TCT370TTA (5452C>T 5453T>A), AAG371AAA (5456G>A), ACT372ATA (5458C>T 5459T>A), ATT373TTT (5460A>T), GAG375ATG (5466G>A 5467A>T), AAA377ATT (5473A>T 5474A>T), ATA378AAG (5476T>A 5477A>G), ATT380CTT (5481A>C), GCA381GAC (5485C>A 5486A>C), AAA382ATT (5488A>T 5489A>T), AGA384AAG (5494G>A 5495A>G), GAT385AAA (5496G>A 5498T>A), ATA386ATT (5501A>T), ACA387GTG (5502A>G 5503C>T 5504A>G), CTT388ATA (5505C>A 5507T>A), GCA389CAA (5508G>C 5509C>A), GGA391GGT (5516A>T), TTA392TTG (5519A>G), ATT393GTG (5520A>G 5522T>G), ATT394ATA (5525T>A), AGT395TCT (5526A>T 5527G>C), AAG396AAA (5531G>A), AAA397CCA (5532A>C 5533A>C), ACA399ATA (5539C>T), GAA400AGT (5541G>A 5542A>G 5543A>T), TTA401TTG (5546A>G), TGT402TTT (5548G>T), AAA403CAA (5550A>C), GAA404ACA (5553G>A 5554A>C), AAA405GAA (5556A>G), ATA406GTC (5559A>G 5561A>C), GAT407CGA (5562G>C 5563A>G 5564T>A), TTC408TTT (5567C>T), CTT409TTA (5568C>T 5570T>A), GTC411CAT (5574G>C 5575T>A 5576C>T), CAA412AAT (5577C>A 5579A>T), GAA414TGT (5583G>T 5584A>G 5585A>T) | | | | | | | | |

*: Inserts / Deletes / Misaligned / Frameshifts

#### Analysis details

This analysis was performed with panviral2.64

### NGS Details (UN8): Alphabaculovirus chofumiferanae

#### Assembly

| | |
|---|---|
| **Coverage Length** | 136 (1 contig(s)) |
| **Depth Of Coverage** | 4.5 |
| **Number Of Reads** | 7 |
| **Reads Per Million** | 0.12 rpm (after QC) |
| **Ambiguities** | 0 |
| **Assembly Method** | de novo + reference guided assembly |
| **Consensus Caller** | Bcf Tools |

#### Coverage Map

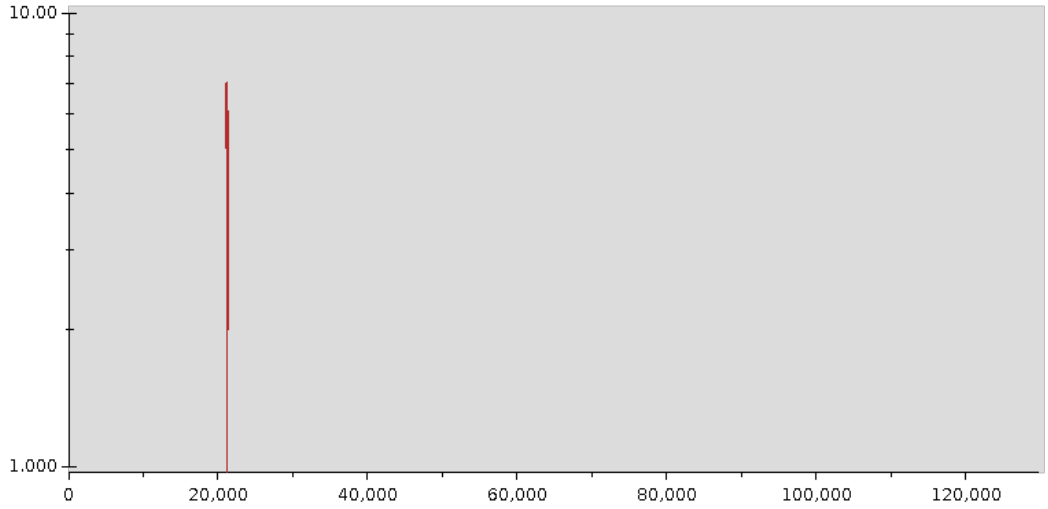

#### Assignment

| | |
|---|---|
| **Type** | Alphabaculovirus chofumiferanae (Taxonomy ID: 3047750) |
| **Reference Genome** | NC_004778.3 |
| **NT Identity (%)** | 78.6765 |
| **AA Identity (%)** | 80.0 |
| **Number Of Stop Codons** | 0 |
| **Number Of CDS** | 145 |

#### Alignment

| | |
|---|---|
| **Alignment Score** | 156.0 (NT) + 309.0 (AA) = 465.0 |
| **Concordance (%)** | 71.9814 |
| **Alignment Method** | Local, heuristic, nucleotide (BLASTN) |

#### Genome Region

Sequence starts at position 21138 and ends at position 21273 relative to NC_004778.3 reference sequence.

#### Alignment Detailed Statistics

| | Begin | End | Coverage | Score | Concordance | Matches | Identities | I/D/M/F* | Stop Codons |
|---|---|---|---|---|---|---|---|---|---|
| **NT** | **21138** | **21273** | **0.1%** | **156** | **57.4%** | **136 (100%)** | **107 (78.7%)** | **0/0** | |

Mutations: 21147C>G, 21153G>A, 21162C>T, 21168T>A, 21189A>C, 21192T>G, 21193T>G, 21194G>C, 21195G>T, 21196T>C, 21197G>C, 21198C>G, 21200T>C, 21201A>G, 21207A>G, 21213C>G, 21214T>G, 21215C>G, 21216G>T, 21219G>A, 21222G>A, 21225C>T, 21231G>C, 21241G>T, 21246G>A, 21254C>G, 21260C>T, 21264A>G, 21271T>C

*: Inserts / Deletes / Misaligned / Frameshifts

#### Analysis details

This analysis was performed with panviral2.64

### NGS Details (UN8): Badnavirus volubetulae

#### Assembly

| | |
|---|---|
| **Coverage Length** | 305 (1 contig(s)) |
| **Depth Of Coverage** | 3.2 |
| **Number Of Reads** | 7 |
| **Reads Per Million** | 0.12 rpm (after QC) |
| **Ambiguities** | 0 |
| **Assembly Method** | de novo + reference guided assembly |
| **Consensus Caller** | Bcf Tools |

#### Coverage Map

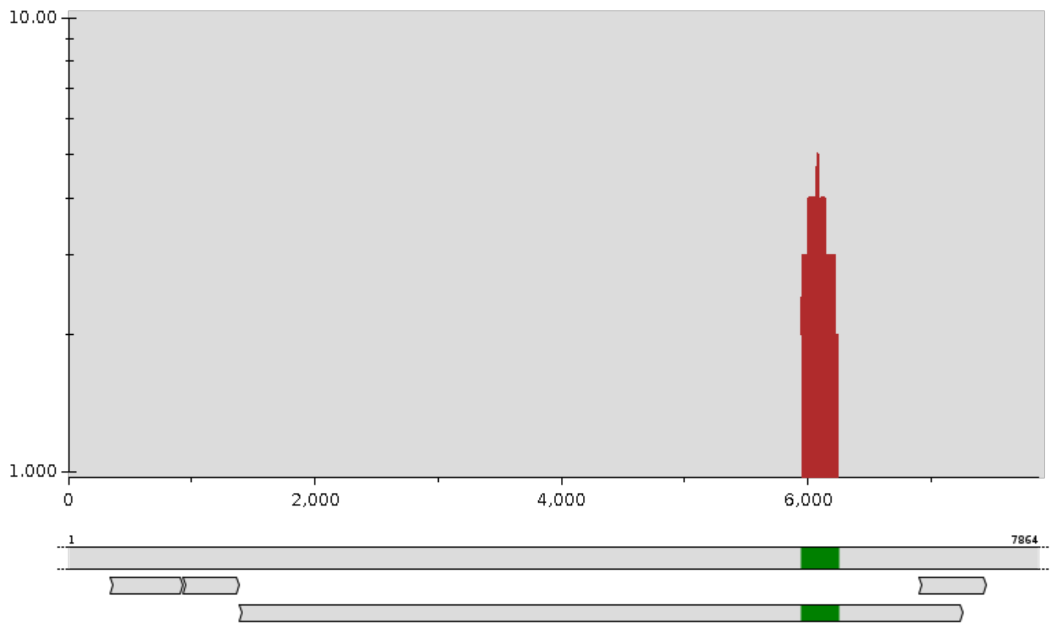

#### Assignment

| | |
|---|---|
| **Type** | Badnavirus volubetulae (Taxonomy ID: 3047683) |
| **Reference Genome** | NC_040635.1 |
| **NT Identity (%)** | 61.0738 |
| **AA Identity (%)** | 47.4747 |
| **Number Of Stop Codons** | 0 |
| **Number Of CDS** | 4 |

#### Alignment

| | |
|---|---|
| **Alignment Score** | 107.0 (NT) + 244.0 (AA) = 351.0 |
| **Concordance (%)** | 30.7895 |

| Alignment Method | Global, seeded, nucleotide + amino acids (AGA) |
|---|---|

#### Genome Region

Sequence starts at position 5945 and ends at position 6249 relative to NC_040635.1 reference sequence.

#### Alignment Detailed Statistics

| | Begin | End | Coverage | Score | Concordance | Matches | Identities | I/D/M/F* | Stop Codons |
|---|---|---|---|---|---|---|---|---|---|
| **NT** | **5945** | **6249** | **3.9%** | **107** | **18.7%** | **298 (97.7%)** | **182 (59.7%)** | **0/7** | |
| Mutations: | 5949C>T, 5956C>T, 5961T>C, 5965C>T, 5973T>G, 5976T>A, 5981A>T, 5982T>G, 5985C>A, 5989G>T, 5992G>A, 5997C>T, 6000G>A, 6009G>T, 6014A>G, 6016T>A, 6017G>T, 6018C>A, 6021T>C, 6023A>G, 6026G>A, 6028A>T, 6029C>A, 6030G>T, 6031G>A, 6034G>A, 6035A>G, 6039C>T, 6042A>T, 6043G>T, 6044C>T, 6045T>A, 6051C>T, 6052A>G, 6054C>A, 6060T>C, 6066G>A, 6069C>A, 6072C>T, 6073T>A, 6074C>G, 6076G>A, 6079A>G, 6082C>A, 6083A>T, 6084T>A, 6087A>G, 6088C>G, 6090G>A, 6092A>G, 6094G>T, 6095A>T, 6099G>A, 6107G>A, 6110A>T, 6111G>A, 6114G>A, 6117G>T, 6118G>A, 6119A>G, 6120G>A, 6121A>C, 6124G>T, 6126C>T, 6127C>G, 6128G>T, 6129C>A, 6130C>A, 6132G>A, 6136G>A, 6137G>A, 6139C>A, 6142G>A, 6144C>A, 6145T>G, 6148_6153delTCACCC, 6155C>G, 6161T>A, 6162G>A, 6165G>T, 6166A>T, 6167T>A, 6168A>T, 6170C>T, 6172T>A, 6174C>A, 6175delT, 6177A>T, 6178G>C, 6180A>T, 6183T>A, 6184G>A, 6190C>T, 6192T>A, 6196G>A, 6197C>T, 6200C>T, 6201T>A, 6202A>T, 6204T>G, 6207C>A, 6208A>G, 6211A>G, 6212G>A, 6213C>A, 6215G>A, 6219T>A, 6220C>A, 6222G>A, 6225G>A, 6231C>T, 6237C>A, 6239T>C, 6240G>T, 6241C>A, 6243G>A, 6246G>A, 6248T>A | | | | | | | | |

##### CDS

| | Begin | End | Coverage | Score | Concordance | Matches | Identities | I/D/M/F* | Stop Codons |
|---|---|---|---|---|---|---|---|---|---|
| BLRaV_gp3 | 1520 | 1620 | 5.2% | 244 | 35.0% | 99 (98.0%) | 47 (46.5%) | 0/2/1/1 | 0 |
| Protein mutations: | P1526S (5965C>T), N1531M (5981A>T 5982T>G), A1534S (5989G>T), V1535I (5992G>A), M1540I (6009G>T), N1542S (6014A>G), C1543I (6016T>A 6017G>T 6018C>A), K1545R (6023A>G), G1546D (6026G>A), T1547Y (6028A>T 6029C>A 6030G>T), E1548K (6031G>A), E1549R (6034G>A 6035A>G), A1552L (6043G>T 6044C>T 6045T>A), I1555V (6052A>G 6054C>A), E1563K (6076G>A), N1564D (6079A>G), H1565I (6082C>A 6083A>T 6084T>A), Q1567E (6088C>G 6090G>A), H1568R (6092A>G), E1569L (6094G>T 6095A>T), R1573Q (6107G>A), K1574I (6110A>T 6111G>A), M1575I (6114G>A), L1576F (6117G>T), E1577R (6118G>A 6119A>G 6120G>A), I1578L (6121A>C), V1579F (6124G>T 6126C>T), R1580V (6127C>G 6128G>T 6129C>A), Q1581K (6130C>A 6132G>A), G1583K (6136G>A 6137G>A), L1584I (6139C>A), V1585I (6142G>A 6144C>A), L1586V (6145T>G), S1587_P1588del (6148_6153delTCACCC), T1589S (6155C>G), M1591K (6161T>A 6162G>A), K1592N (6165G>T), I1593Y (6166A>T 6167T>A 6168A>T), A1594V (6170C>T), C1595R (6172T>A 6174C>A), E1597H (6178G>C 6180A>T), D1599N (6184G>A), A1603I (6196G>A 6197C>T), T1604I (6200C>T 6201T>A), I1605L (6202A>T 6204T>G), N1607D (6208A>G), S1608E (6211A>G 6212G>A 6213C>A), R1609K (6215G>A), Q1611K (6220C>A 6222G>A), V1617A (6239T>C 6240G>T), Q1618K (6241C>A 6243G>A), I1620K (6248T>A) | | | | | | | | |
| Codon mutations: | TAC1520TAT (5949C>T), CTA1523TTA (5956C>T), GTT1524GTC (5961T>C), CCA1526TCA (5965C>T), GGT1528GGG (5973T>G), CTT1529CTA (5976T>A), AAT1531ATG (5981A>T 5982T>G), GCC1532GCA (5985C>A), GCA1534TCA (5989G>T), GTA1535ATA (5992G>A), TTC1536TTT (5997C>T), CAG1537CAA (6000G>A), ATG1540ATT (6009G>T), AAT1542AGT (6014A>G), TGC1543ATA (6016T>A 6017G>T 6018C>A), TTT1544TTC (6021T>C), AAA1545AGA (6023A>G), GGT1546GAT (6026G>A), ACG1547TAT (6028A>T 6029C>A 6030G>T), GAA1548AAA (6031G>A), GAA1549AGA (6034G>A 6035A>G), TTC1550TTT (6039C>T), ATA1551ATT (6042A>T), GCT1552TTA (6043G>T 6044C>T 6045T>A), TAC1554TAT (6051C>T), ATC1555GTA (6052A>G 6054C>A), GAT1557GAC (6060T>C), TTG1559TTA (6066G>A), GTC1560GTA (6069C>A), TTC1561TTT (6072C>T), TCC1562AGC (6073T>A 6074C>G), GAA1563AAA (6076G>A), AAT1564GAT (6079A>G), CAT1565ATA (6082C>A 6083A>T 6084T>A), AAA1566AAG (6087A>G), CAG1567GAA (6088C>G 6090G>A), CAT1568CGT (6092A>G), GAA1569TTA (6094G>T 6095A>T), GGG1570GGA (6099G>A), CGG1573CAG (6107G>A), AAG1574ATA (6110A>T 6111G>A), ATG1575ATA (6114G>A), TTG1576TTT (6117G>T), GAG1577AGA (6118G>A 6119A>G 6120G>A), ATA1578CTA (6121A>C), GTC1579TTT (6124G>T 6126C>T), CGC1580GTA (6127C>G 6128G>T 6129C>A), CAG1581AAA (6130C>A 6132G>A), GGA1583AAA (6136G>A 6137G>A), CTA1584ATA (6139C>A), GTC1585ATA (6142G>A 6144C>A), TTA1586GTA (6145T>G), TCA1587_CCC1588del (6148_6153delTCACCC), ACT1589AGT (6155C>G), ATG1591AAA (6161T>A 6162G>A), AAG1592AAT (6165G>T), ATA1593TAT (6166A>T 6167T>A 6168A>T), GCA1594GTA (6170C>T), TGC1595AGA (6172T>A 6174C>A), TCA1596-CT (6175delT 6177A>T), GAA1597CAT (6178G>C 6180A>T), ATT1598ATA (6183T>A), GAT1599AAT (6184G>A), CTT1601TTA (6190C>T 6192T>A), GCA1603ATA (6196G>A 6197C>T), ACT1604ATA (6200C>T 6201T>A), ATT1605TTG (6202A>T 6204T>G), GGC1606GGA (6207C>A), AAC1607GAC (6208A>G), AGC1608GAA (6211A>G 6212G>A 6213C>A), AGA1609AAA (6215G>A), ATT1610ATA (6219T>A), CAG1611AAA (6220C>A 6222G>A), TTG1612TTA (6225G>A), CCC1614CCT (6231C>T), ATC1616ATA (6237C>A), GTG1617GCT (6239T>C 6240G>T), CAG1618AAA (6241C>A 6243G>A), AAG1619AAA (6246G>A), ATA1620AAA (6248T>A) | | | | | | | | |

##### Proteins

| | Begin | End | Coverage | Score | Concordance | Matches | Identities | I/D/M/F* | Stop Codons |
|---|---|---|---|---|---|---|---|---|---|
| polyprotein (YP_009552737.1) | 1520 | 1620 | 5.2% | 244 | 35.0% | 99 (98.0%) | 47 (46.5%) | 0/2/1/1 | 0 |
| Protein mutations: | P1526S (5965C>T), N1531M (5981A>T 5982T>G), A1534S (5989G>T), V1535I (5992G>A), M1540I (6009G>T), N1542S (6014A>G), C1543I (6016T>A 6017G>T 6018C>A), K1545R (6023A>G), G1546D (6026G>A), T1547Y (6028A>T 6029C>A 6030G>T), E1548K (6031G>A), E1549R (6034G>A 6035A>G), A1552L (6043G>T 6044C>T 6045T>A), I1555V (6052A>G 6054C>A), E1563K (6076G>A), N1564D (6079A>G), H1565I (6082C>A 6083A>T 6084T>A), Q1567E (6088C>G 6090G>A), H1568R (6092A>G), E1569L (6094G>T 6095A>T), R1573Q (6107G>A), K1574I (6110A>T 6111G>A), M1575I (6114G>A), L1576F (6117G>T), E1577R (6118G>A 6119A>G 6120G>A), I1578L (6121A>C), V1579F (6124G>T 6126C>T), R1580V (6127C>G 6128G>T 6129C>A), Q1581K (6130C>A 6132G>A), G1583K (6136G>A 6137G>A), L1584I (6139C>A), V1585I (6142G>A 6144C>A), L1586V (6145T>G), S1587_P1588del (6148_6153delTCACCC), T1589S (6155C>G), M1591K (6161T>A 6162G>A), K1592N (6165G>T), I1593Y (6166A>T 6167T>A 6168A>T), A1594V (6170C>T), C1595R (6172T>A 6174C>A), E1597H (6178G>C 6180A>T), D1599N (6184G>A), A1603I (6196G>A 6197C>T), T1604I (6200C>T 6201T>A), I1605L (6202A>T 6204T>G), N1607D (6208A>G), S1608E (6211A>G 6212G>A 6213C>A), R1609K (6215G>A), Q1611K (6220C>A 6222G>A), V1617A (6239T>C 6240G>T), Q1618K (6241C>A 6243G>A), I1620K (6248T>A) | | | | | | | | |
| Codon mutations: | TAC1520TAT (5949C>T), CTA1523TTA (5956C>T), GTT1524GTC (5961T>C), CCA1526TCA (5965C>T), GGT1528GGG (5973T>G), CTT1529CTA (5976T>A), AAT1531ATG (5981A>T 5982T>G), GCC1532GCA (5985C>A), GCA1534TCA (5989G>T), GTA1535ATA (5992G>A), TTC1536TTT (5997C>T), CAG1537CAA (6000G>A), ATG1540ATT (6009G>T), AAT1542AGT (6014A>G), TGC1543ATA (6016T>A 6017G>T 6018C>A), TTT1544TTC (6021T>C), AAA1545AGA (6023A>G), GGT1546GAT (6026G>A), ACG1547TAT (6028A>T 6029C>A 6030G>T), GAA1548AAA (6031G>A), GAA1549AGA (6034G>A 6035A>G), TTC1550TTT (6039C>T), ATA1551ATT (6042A>T), GCT1552TTA (6043G>T 6044C>T 6045T>A), TAC1554TAT (6051C>T), ATC1555GTA (6052A>G 6054C>A), GAT1557GAC (6060T>C), TTG1559TTA (6066G>A), GTC1560GTA (6069C>A), TTC1561TTT (6072C>T), TCC1562AGC (6073T>A 6074C>G), GAA1563AAA (6076G>A), AAT1564GAT (6079A>G), CAT1565ATA (6082C>A 6083A>T 6084T>A), AAA1566AAG (6087A>G), CAG1567GAA (6088C>G 6090G>A), CAT1568CGT (6092A>G), GAA1569TTA (6094G>T 6095A>T), GGG1570GGA (6099G>A), CGG1573CAG (6107G>A), AAG1574ATA (6110A>T 6111G>A), ATG1575ATA (6114G>A), TTG1576TTT (6117G>T), GAG1577AGA (6118G>A 6119A>G 6120G>A), ATA1578CTA (6121A>C), GTC1579TTT (6124G>T 6126C>T), CGC1580GTA (6127C>G 6128G>T 6129C>A), CAG1581AAA (6130C>A 6132G>A), GGA1583AAA (6136G>A 6137G>A), CTA1584ATA (6139C>A), GTC1585ATA (6142G>A 6144C>A), TTA1586GTA (6145T>G), TCA1587_CCC1588del (6148_6153delTCACCC), ACT1589AGT (6155C>G), ATG1591AAA (6161T>A 6162G>A), AAG1592AAT (6165G>T), ATA1593TAT (6166A>T 6167T>A 6168A>T), GCA1594GTA (6170C>T), TGC1595AGA (6172T>A 6174C>A), TCA1596-CT (6175delT 6177A>T), GAA1597CAT (6178G>C 6180A>T), ATT1598ATA (6183T>A), GAT1599AAT (6184G>A), CTT1601TTA (6190C>T 6192T>A), GCA1603ATA (6196G>A 6197C>T), ACT1604ATA (6200C>T 6201T>A), ATT1605TTG (6202A>T 6204T>G), GGC1606GGA (6207C>A), AAC1607GAC (6208A>G), AGC1608GAA (6211A>G 6212G>A 6213C>A), AGA1609AAA (6215G>A), ATT1610ATA (6219T>A), CAG1611AAA (6220C>A 6222G>A), TTG1612TTA (6225G>A), CCC1614CCT (6231C>T), ATC1616ATA (6237C>A), GTG1617GCT (6239T>C 6240G>T), CAG1618AAA (6241C>A 6243G>A), AAG1619AAA (6246G>A), ATA1620AAA (6248T>A) | | | | | | | | |

*: Inserts / Deletes / Misaligned / Frameshifts

#### Analysis details

This analysis was performed with panviral2.64

### NGS Details (UN8): Epiphyllum badnavirus 1

#### Assembly

| | |
|---|---|
| **Coverage Length** | 307 (1 contig(s)) |
| **Depth Of Coverage** | 2.2 |
| **Number Of Reads** | 6 |
| **Reads Per Million** | 0.10 rpm (after QC) |
| **Ambiguities** | 0 |
| **Assembly Method** | de novo + reference guided assembly |
| **Consensus Caller** | Bcf Tools |

#### Coverage Map

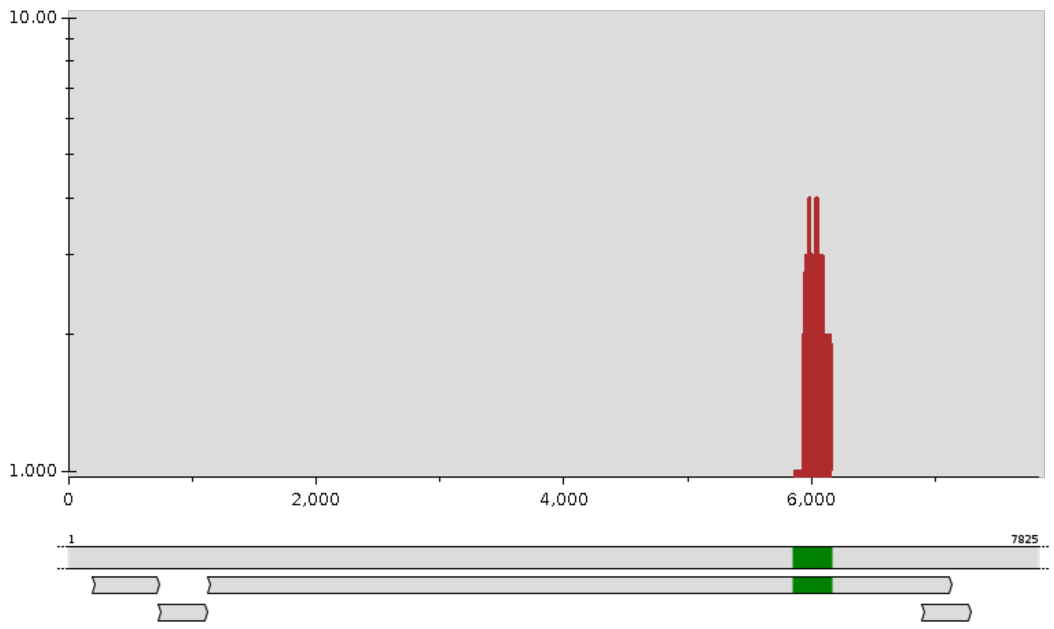

#### Assignment

| | |
|---|---|
| **Type** | Epiphyllum badnavirus 1 (Taxonomy ID: 2518008) |
| **Reference Genome** | NC_076247.1 |
| **NT Identity (%)** | 52.8481 |
| **AA Identity (%)** | 43.8095 |
| **Number Of Stop Codons** | 2 |
| **Number Of CDS** | 4 |

#### Alignment

| | |
|---|---|
| **Alignment Score** | 18.0 (NT) + 299.0 (AA) = 317.0 |
| **Concordance (%)** | 23.6744 |

| Alignment Method | Global, seeded, nucleotide + amino acids (AGA) |
|---|---|

#### Genome Region

Sequence starts at position 5855 and ends at position 6161 relative to NC_076247.1 reference sequence.

#### Alignment Detailed Statistics

| | Begin | End | Coverage | Score | Concordance | Matches | Identities | I/D/M/F* | Stop Codons |
|---|---|---|---|---|---|---|---|---|---|
| **NT** | **5855** | **6161** | **3.9%** | **18** | **2.9%** | **307 (97.2%)** | **167 (52.8%)** | **9/0** | |
| Mutations: | 5857C>T, 5861A>G, 5863A>T, 5864G>C, 5865C>A, 5867G>C, 5870T>A, 5871C>T, 5877G>A, 5880T>G, 5882T>G, 5884A>G, 5887G>A, 5889C>G, 5892A>G, 5893G>A, 5896T>G, 5897C>T, 5901C>A, 5904T>C, 5905T>A, 5906G>A, 5910C>A, 5913C>T, 5916C>T, 5917C>A, 5918T>G, 5919G>A, 5923C>T, 5924C>A, 5926G>T, 5931A>T, 5933T>A, 5936A>T, 5937C>T, 5942G>T, 5943G>T, 5944T>C, 5953C>T, 5958C>T, 5961A>C, 5962T>C, 5964G>A, 5966A>C, 5967G>A, 5973C>A, 5976T>G, 5977G>A, 5979T>A, 5982G>T, 5985C>T, 5986C>A, 5987A>T, 5988A>G, 5989C>G, 5990G>A, 5991A>T, 5992A>T, 5993A>T, 5998G>A, 6000C>T, 6002A>G, 6003C>G, 6004T>G, 6005G>T, 6006T>G, 6011G>A, 6012A>G, 6012_6013insATA, 6013G>T, 6014A>T, 6016C>T, 6018G>A, 6020C>A, 6021A>T, 6023G>T, 6024C>A, 6030A>G, 6031G>A, 6032C>T, 6036T>A, 6038A>T, 6039C>T, 6042T>A, 6043G>A, 6048C>T, 6051C>T, 6052C>T, 6057C>G, 6059T>A, 6060C>T, 6061A>T, 6062G>C, 6063T>A, 6065A>G, 6066C>A, 6067A>T, 6069T>A, 6070G>A, 6075T>G, 6083C>A, 6085C>A, 6087G>T, 6094A>C, 6095A>G, 6097T>C, 6098C>A, 6099C>G, 6100A>G, 6102G>A, 6103C>T, 6105G>A, 6108G>A, 6111_6112insCTT, 6117G>T, 6118A>C, 6121G>C, 6123G>A, 6124G>T, 6125G>T, 6127C>T, 6128T>A, 6129G>T, 6130A>G, 6132_6133insAAG, 6133C>T, 6139C>A, 6140C>A, 6141T>A, 6142A>T, 6143C>G, 6145A>G, 6147G>A, 6148A>T, 6150G>T, 6151A>T, 6152A>G, 6153G>A, 6154A>T, 6157G>A, 6158C>A, 6159T>A, 6160G>T | | | | | | | | |
| **CDS** | | | | | | | | | |
| QKM20_gp3 | 1577 | 1678 | 5.1% | 299 | 40.5% | 102 (97.1%) | 46 (43.8%) | 3/0/0/0 | 2 |
| Protein mutations: | K1578R (5861A>G), G1580A (5867G>C), F1581Y (5870T>A 5871C>T), I1584M (5880T>G), M1585R (5882T>G), M1586V (5884A>G), D1587K (5887G>A 5889C>G), E1589K (5893G>A), S1590V (5896T>G 5897C>T), W1593K (5905T>A 5906G>A), L1597R (5917C>A 5918T>G 5919G>A), P1599* (5923C>T 5924C>A), D1600Y (5926G>T), L1602H (5933T>A), Y1603F (5936A>T 5937C>T), W1605F (5942G>T 5943G>T), P1609S (5953C>T), K1613T (5966A>C 5967G>A), A1617T (5977G>A 5979T>A), Q1620M (5986C>A 5987A>T 5988A>G), R1621D (5989C>G 5990G>A 5991A>T), K1622L (5992A>T 5993A>T), D1624N (5998G>A 6000C>T), N1625R (6002A>G 6003C>G), C1626V (6004T>G 6005G>T 6006T>G), R1628K (6011G>A 6012A>G), R1628_D1629insI (6012_6013insATA), D1629F (6013G>T 6014A>T), S1631Y (6020C>A 6021A>T), G1632V (6023G>T 6024C>A), A1635I (6031G>A 6032C>T), Y1637F (6038A>T 6039C>T), D1639N (6043G>A), F1644Y (6059T>A 6060C>T), N1646R (6065A>G 6066C>A), T1647S (6067A>T 6069T>A), E1648K (6070G>A), D1649E (6075T>G), A1652E (6083C>A), Q1653N (6085C>A 6087G>T), K1656R (6094A>C 6095A>G), S1657Q (6097T>C 6098C>A 6099C>G), M1658V (6100A>G 6102G>A), V1661_C1662insL (6111_6112insCTT), E1663D (6117G>T), E1665Q (6121G>C 6123G>A), G1666L (6124G>T 6125G>T), L1667Y (6127C>T 6128T>A 6129G>T), I1668V (6130A>G), I1668_L1669insK (6132_6133insAAG), L1669F (6133C>T), P1671K (6139C>A 6140C>A 6141T>A), T1672W (6142A>T 6143C>G), K1673E (6145A>G 6147G>A), M1674F (6148A>T 6150G>T), K1675* (6151A>T 6152A>G 6153G>A), I1676L (6154A>T), A1677K (6157G>A 6158C>A 6159T>A) | | | | | | | | |
| Codon mutations: | CTG1577TTG (5857C>T), AAG1578AGG (5861A>G), AGC1579TCA (5863A>T 5864G>C 5865C>A), GGA1580GCA (5867G>C), TTC1581TAT (5870T>A 5871C>T), CAG1583CAA (5877G>A), ATT1584ATG (5880T>G), ATG1585AGG (5882T>G), ATG1586GTG (5884A>G), GAC1587AAG (5887G>A 5889C>G), GAA1588GAG (5892A>G), GAA1589AAA (5893G>A), TCC1590GTC (5896T>G 5897C>T), ATC1591ATA (5901C>A), CCT1592CCC (5904T>C), TGG1593AAG (5905T>A 5906G>A), ACC1594ACA (5910C>A), GCC1595GCT (5913C>T), TTC1596TTT (5916C>T), CTG1597AGA (5917C>A 5918T>G 5919G>A), CCA1599TAA (5923C>T 5924C>A), GAT1600TAT (5926G>T), GGA1601GGT (5931A>T), CTT1602CAT (5933T>A), TAC1603TTT (5936A>T 5937C>T), TGG1605TTT (5942G>T 5943G>T), TTA1606CTA (5944T>C), CCA1609TCA (5953C>T), TTC1610TTT (5958C>T), GGA1611GGC (5961A>C), TTG1612CTA (5962T>C 5964G>A), AAG1613ACA (5966A>C 5967G>A), GCC1615GCA (5973C>A), CCT1616CCG (5976T>G), GCT1617ACA (5977G>A 5979T>A), GTG1618GTT (5982G>T), TTC1619TTT (5985C>T), CAA1620ATG (5986C>A 5987A>T 5988A>G), CGA1621GAT (5989C>G 5990G>A 5991A>T), AAG1622TTG (5992A>T 5993A>T), GAC1624AAT (5998G>A 6000C>T), AAC1625AGG (6002A>G 6003C>G), TGT1626GTG (6004T>G 6005G>T 6006T>G), AGA1628AAG (6011G>A 6012A>G), AGA1628_GAC1629insATA (6012_6013insATA), GAC1629TTC (6013G>T 6014A>T), CTG1630TTA (6016C>T 6018G>A), TCA1631TAT (6020C>A 6021A>T), GGC1632GTA (6023G>T 6024C>A), GTA1634GTG (6030A>G), GCA1635ATA (6031G>A 6032C>T), GTT1636GTA (6036T>A), TAC1637TTT (6038A>T 6039C>T), ATT1638ATA (6042T>A), GAT1639AAT (6043G>A), GAC1640GAT (6048C>T), ATC1641ATT (6051C>T), CTG1642TTG (6052C>T), GTC1643GTG (6057C>G), TTC1644TAT (6059T>A 6060C>T), AGT1645TCA (6061A>T 6062G>C 6063T>A), AAC1646AGA (6065A>G 6066C>A), ACT1647TCA (6067A>T 6069T>A), GAA1648AAA (6070G>A), GAT1649GAG (6075T>G), GCA1652GAA (6083C>A), CAG1653AAT (6085C>A 6087G>T), AAG1656CGG (6094A>C 6095A>G), TCC1657CAG (6097T>C 6098C>A 6099C>G), ATG1658GTA (6100A>G 6102G>A), CTG1659TTA (6103C>T 6105G>A), CAG1660CAA (6108G>A), GTT1661_TGT1662insCTT (6111_6112insCTT), GAG1663GAT (6117G>T), AGA1664CGA (6118A>C), GAG1665CAA (6121G>C 6123G>A), GGG1666TTG (6124G>T 6125G>T), CTG1667TAT (6127C>T 6128T>A 6129G>T), ATC1668GTC (6130A>G), ATC1668_CTC1669insAAG (6132_6133insAAG), CTC1669TTC (6133C>T), CCT1671AAA (6139C>A 6140C>A 6141T>A), ACG1672TGG (6142A>T 6143C>G), AAG1673GAA (6145A>G 6147G>A), ATG1674TTT (6148A>T 6150G>T), AAG1675TGA (6151A>T 6152A>G 6153G>A), ATA1676TTA (6154A>T), GCT1677AAA (6157G>A 6158C>A 6159T>A), GCA1678TC. (6160G>T) | | | | | | | | |
| **Proteins** | | | | | | | | | |
| polyprotein (YP_010797894.1) | 1577 | 1678 | 5.1% | 299 | 40.5% | 102 (97.1%) | 46 (43.8%) | 3/0/0/0 | 2 |
| Protein mutations: | K1578R (5861A>G), G1580A (5867G>C), F1581Y (5870T>A 5871C>T), I1584M (5880T>G), M1585R (5882T>G), M1586V (5884A>G), D1587K (5887G>A 5889C>G), E1589K (5893G>A), S1590V (5896T>G 5897C>T), W1593K (5905T>A 5906G>A), L1597R (5917C>A 5918T>G 5919G>A), P1599* (5923C>T 5924C>A), D1600Y (5926G>T), L1602H (5933T>A), Y1603F (5936A>T 5937C>T), W1605F (5942G>T 5943G>T), P1609S (5953C>T), K1613T (5966A>C 5967G>A), A1617T (5977G>A 5979T>A), Q1620M (5986C>A 5987A>T 5988A>G), R1621D (5989C>G 5990G>A 5991A>T), K1622L (5992A>T 5993A>T), D1624N (5998G>A 6000C>T), N1625R (6002A>G 6003C>G), C1626V (6004T>G 6005G>T 6006T>G), R1628K (6011G>A 6012A>G), R1628_D1629insI (6012_6013insATA), D1629F (6013G>T 6014A>T), S1631Y (6020C>A 6021A>T), G1632V (6023G>T 6024C>A), A1635I (6031G>A 6032C>T), Y1637F (6038A>T 6039C>T), D1639N (6043G>A), F1644Y (6059T>A 6060C>T), N1646R (6065A>G 6066C>A), T1647S (6067A>T 6069T>A), E1648K (6070G>A), D1649E (6075T>G), A1652E (6083C>A), Q1653N (6085C>A 6087G>T), K1656R (6094A>C 6095A>G), S1657Q (6097T>C 6098C>A 6099C>G), M1658V (6100A>G 6102G>A), V1661_C1662insL (6111_6112insCTT), E1663D (6117G>T), E1665Q (6121G>C 6123G>A), G1666L (6124G>T 6125G>T), L1667Y (6127C>T 6128T>A 6129G>T), I1668V (6130A>G), I1668_L1669insK (6132_6133insAAG), L1669F (6133C>T), P1671K (6139C>A 6140C>A 6141T>A), T1672W (6142A>T 6143C>G), K1673E (6145A>G 6147G>A), M1674F (6148A>T 6150G>T), K1675* (6151A>T 6152A>G 6153G>A), I1676L (6154A>T), A1677K (6157G>A 6158C>A 6159T>A) | | | | | | | | |
| Codon mutations: | CTG1577TTG (5857C>T), AAG1578AGG (5861A>G), AGC1579TCA (5863A>T 5864G>C 5865C>A), GGA1580GCA (5867G>C), TTC1581TAT (5870T>A 5871C>T), CAG1583CAA (5877G>A), ATT1584ATG (5880T>G), ATG1585AGG (5882T>G), ATG1586GTG (5884A>G), GAC1587AAG (5887G>A 5889C>G), GAA1588GAG (5892A>G), GAA1589AAA (5893G>A), TCC1590GTC (5896T>G 5897C>T), ATC1591ATA (5901C>A), CCT1592CCC (5904T>C), TGG1593AAG (5905T>A 5906G>A), ACC1594ACA (5910C>A), GCC1595GCT (5913C>T), TTC1596TTT (5916C>T), CTG1597AGA (5917C>A 5918T>G 5919G>A), CCA1599TAA (5923C>T 5924C>A), GAT1600TAT (5926G>T), GGA1601GGT (5931A>T), CTT1602CAT (5933T>A), TAC1603TTT (5936A>T 5937C>T), TGG1605TTT (5942G>T 5943G>T), TTA1606CTA (5944T>C), CCA1609TCA (5953C>T), TTC1610TTT (5958C>T), GGA1611GGC (5961A>C), TTG1612CTA (5962T>C 5964G>A), AAG1613ACA (5966A>C 5967G>A), GCC1615GCA (5973C>A), CCT1616CCG (5976T>G), GCT1617ACA (5977G>A 5979T>A), GTG1618GTT (5982G>T), TTC1619TTT (5985C>T), CAA1620ATG (5986C>A 5987A>T 5988A>G), CGA1621GAT (5989C>G 5990G>A 5991A>T), AAG1622TTG (5992A>T 5993A>T), GAC1624AAT (5998G>A 6000C>T), AAC1625AGG (6002A>G 6003C>G), TGT1626GTG (6004T>G 6005G>T 6006T>G), AGA1628AAG (6011G>A 6012A>G), AGA1628_GAC1629insATA (6012_6013insATA), GAC1629TTC (6013G>T 6014A>T), CTG1630TTA (6016C>T 6018G>A), TCA1631TAT (6020C>A 6021A>T), GGC1632GTA (6023G>T 6024C>A), GTA1634GTG (6030A>G), GCA1635ATA (6031G>A 6032C>T), GTT1636GTA (6036T>A), TAC1637TTT (6038A>T 6039C>T), ATT1638ATA (6042T>A), GAT1639AAT (6043G>A), GAC1640GAT (6048C>T), ATC1641ATT (6051C>T), CTG1642TTG (6052C>T), GTC1643GTG (6057C>G), TTC1644TAT (6059T>A 6060C>T), AGT1645TCA (6061A>T 6062G>C 6063T>A), AAC1646AGA (6065A>G 6066C>A), ACT1647TCA (6067A>T 6069T>A), GAA1648AAA (6070G>A), GAT1649GAG (6075T>G), GCA1652GAA (6083C>A), CAG1653AAT (6085C>A 6087G>T), AAG1656CGG (6094A>C 6095A>G), TCC1657CAG (6097T>C 6098C>A 6099C>G), ATG1658GTA (6100A>G 6102G>A), CTG1659TTA (6103C>T 6105G>A), CAG1660CAA (6108G>A), GTT1661_TGT1662insCTT (6111_6112insCTT), GAG1663GAT (6117G>T), AGA1664CGA (6118A>C), GAG1665CAA (6121G>C 6123G>A), GGG1666TTG (6124G>T 6125G>T), CTG1667TAT (6127C>T 6128T>A 6129G>T), ATC1668GTC (6130A>G), ATC1668_CTC1669insAAG (6132_6133insAAG), CTC1669TTC (6133C>T), CCT1671AAA (6139C>A 6140C>A 6141T>A), ACG1672TGG (6142A>T 6143C>G), AAG1673GAA (6145A>G 6147G>A), ATG1674TTT (6148A>T 6150G>T), AAG1675TGA (6151A>T 6152A>G 6153G>A), ATA1676TTA (6154A>T), GCT1677AAA (6157G>A 6158C>A 6159T>A), GCA1678TC. (6160G>T) | | | | | | | | |

*: Inserts / Deletes / Misaligned / Frameshifts

#### Analysis details

This analysis was performed with panviral2.64

### NGS Details (UN8): Potato virus X

#### Assembly

| | |
|---|---|
| **Coverage Length** | 236 (1 contig(s)) |
| **Depth Of Coverage** | 2.3 |
| **Number Of Reads** | 4 |
| **Reads Per Million** | 0.07 rpm (after QC) |
| **Ambiguities** | 0 |
| **Assembly Method** | de novo + reference guided assembly |
| **Consensus Caller** | Bcf Tools |

#### Coverage Map

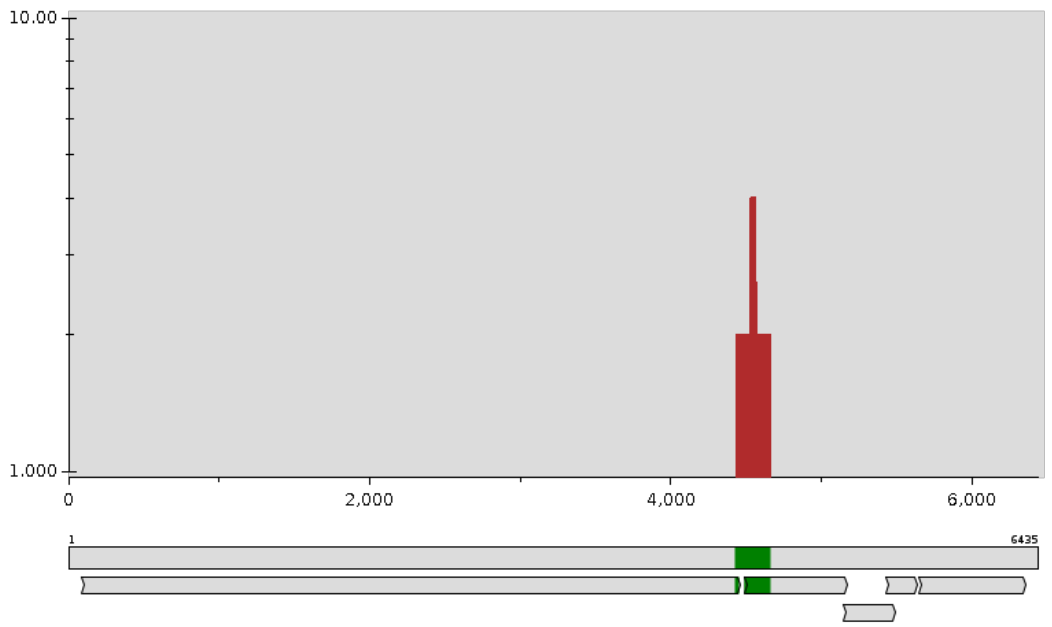

#### Assignment

| | |
|---|---|
| **Type** | Potato virus X (Taxonomy ID: 12183) |
| **Reference Genome** | NC_011620.1 |
| **NT Identity (%)** | 96.6102 |
| **AA Identity (%)** | 95.5882 |
| **Number Of Stop Codons** | 1 |
| **Number Of CDS** | 5 |

#### Alignment

| | |
|---|---|
| **Alignment Score** | 440.0 (NT) + 407.0 (AA) = 847.0 |
| **Concordance (%)** | 94.7427 |

| Alignment Method | Global, seeded, nucleotide + amino acids (AGA) |
|---|---|

#### Genome Region

Sequence starts at position 4430 and ends at position 4665 relative to NC_011620.1 reference sequence.

#### Alignment Detailed Statistics

| | Begin | End | Coverage | Score | Concordance | Matches | Identities | I/D/M/F* | Stop Codons |
|---|---|---|---|---|---|---|---|---|---|
| **NT** | **4430** | **4665** | **3.7%** | **440** | **93.2%** | **236 (100%)** | **228 (96.6%)** | **0/0** | |
| Mutations: | 4439T>A, 4463T>G, 4472T>G, 4502G>T, 4534T>C, 4535C>A, 4584A>C, 4638G>A | | | | | | | | |
| **CDS** | | | | | | | | | |
| PVX_gp1 | 1450 | 1457 | 0.5% | 49 | 84.5% | 8 (100%) | 7 (87.5%) | 0/0/0/0 | 1 |
| Protein mutations: | L1452H (4439T>A) | | | | | | | | |
| Codon mutations: | CTC1452CAC (4439T>A) | | | | | | | | |
| PVX_gp2 | 1 | 60 | 26.4% | 358 | 97.3% | 60 (100%) | 58 (96.7%) | 0/0/0/0 | 0 |
| Protein mutations: | S6I (4502G>T), S17H (4534T>C 4535C>A) | | | | | | | | |
| Codon mutations: | AGT6ATT (4502G>T), TCT17CAT (4534T>C 4535C>A), GCA33GCC (4584A>C), GTG51GTA (4638G>A) | | | | | | | | |
| **Proteins** | | | | | | | | | |
| replicase (YP_002332929.1) | 1450 | 1457 | 0.5% | 49 | 84.5% | 8 (100%) | 7 (87.5%) | 0/0/0/0 | 1 |
| Protein mutations: | L1452H (4439T>A) | | | | | | | | |
| Codon mutations: | CTC1452CAC (4439T>A) | | | | | | | | |
| 25K protein (YP_002332930.1) | 1 | 60 | 26.4% | 358 | 97.3% | 60 (100%) | 58 (96.7%) | 0/0/0/0 | 0 |
| Protein mutations: | S6I (4502G>T), S17H (4534T>C 4535C>A) | | | | | | | | |
| Codon mutations: | AGT6ATT (4502G>T), TCT17CAT (4534T>C 4535C>A), GCA33GCC (4584A>C), GTG51GTA (4638G>A) | | | | | | | | |

*: Inserts / Deletes / Misaligned / Frameshifts

#### Analysis details

This analysis was performed with panviral2.64

### NGS Details (UN8): Petuvirus venapetuniae

#### Assembly

| | |
|---|---|
| **Coverage Length** | 225 (1 contig(s)) |
| **Depth Of Coverage** | 2.4 |
| **Number Of Reads** | 4 |
| **Reads Per Million** | 0.07 rpm (after QC) |
| **Ambiguities** | 0 |
| **Assembly Method** | de novo + reference guided assembly |
| **Consensus Caller** | Bcf Tools |

#### Coverage Map

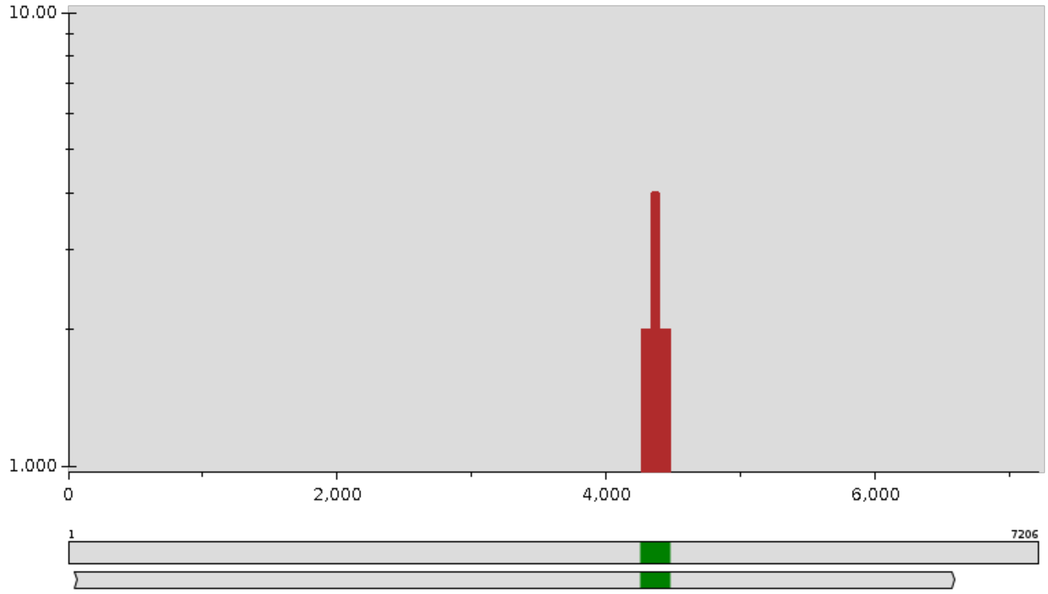

#### Assignment

| | |
|---|---|
| **Type** | Petuvirus venapetuniae (Taxonomy ID: 3048265) |
| **Reference Genome** | NC_001839.2 |
| **NT Identity (%)** | 68.4444 |
| **AA Identity (%)** | 69.3333 |
| **Number Of Stop Codons** | 0 |
| **Number Of CDS** | 1 |

#### Alignment

| | |
|---|---|
| **Alignment Score** | 166.0 (NT) + 387.0 (AA) = 553.0 |
| **Concordance (%)** | 56.1421 |

| Alignment Method | Global, seeded, nucleotide + amino acids (AGA) |
|---|---|

#### Genome Region

Sequence starts at position 4253 and ends at position 4477 relative to NC_001839.2 reference sequence.

#### Alignment Detailed Statistics

| | Begin | End | Coverage | Score | Concordance | Matches | Identities | I/D/M/F* | Stop Codons |
|---|---|---|---|---|---|---|---|---|---|
| **NT** | **4253** | **4477** | **3.1%** | **166** | **36.9%** | **225 (100%)** | **154 (68.4%)** | **0/0** | |
| Mutations: | 4256A>G, 4264G>A, 4269G>A, 4271G>A, 4272C>A, 4275C>T, 4276A>T, 4277A>G, 4278C>A, 4280A>G, 4284G>A, 4285A>G, 4286T>A, 4287C>T, 4292C>T, 4295A>G, 4298C>A, 4307A>C, 4314G>T, 4316A>T, 4320G>A, 4334C>T, 4343G>A, 4344T>G, 4346A>T, 4349G>A, 4354T>C, 4355G>A, 4361A>T, 4365T>C, 4367A>T, 4368A>C, 4370G>A, 4379A>C, 4385T>C, 4388A>G, 4391A>C, 4392T>C, 4394A>C, 4400T>C, 4406C>T, 4407C>T, 4408A>T, 4409A>G, 4413G>A, 4414A>G, 4415C>T, 4418G>A, 4421T>C, 4422C>A, 4424C>T, 4427T>C, 4429C>T, 4436G>T, 4439C>A, 4441C>T, 4442A>T, 4443C>T, 4445C>A, 4450C>T, 4451T>A, 4458T>G, 4459C>T, 4460C>A, 4461A>G, 4466A>C, 4468A>C, 4469G>A, 4470C>T, 4471T>A, 4475T>C | | | | | | | | |

**CDS**

| | Begin | End | Coverage | Score | Concordance | Matches | Identities | I/D/M/F* | Stop Codons |
|---|---|---|---|---|---|---|---|---|---|
| PVCVgp1 | 1402 | 1476 | 3.4% | 387 | 71.3% | 75 (100%) | 52 (69.3%) | 0/0/0/0 | 0 |
| Protein mutations: | I1402M (4256A>G), C1405Y (4264G>A), E1407K (4269G>A 4271G>A), L1408I (4272C>A), Q1409L (4275C>T 4276A>T 4277A>G), Q1410K (4278C>A 4280A>G), D1412R (4284G>A 4285A>G 4286T>A), A1422S (4314G>T 4316A>T), E1424K (4320G>A), S1432A (4344T>G 4346A>T), V1435A (4354T>C 4355G>A), Q1453L (4407C>T 4408A>T 4409A>G), D1455S (4413G>A 4414A>G 4415C>T), P1458T (4422C>A 4424C>T), P1460L (4429C>T), K1462N (4436G>T), T1464I (4441C>T 4442A>T), S1467L (4450C>T 4451T>A), S1470V (4458T>G 4459C>T 4460C>A), K1471E (4461A>G), K1473T (4468A>C 4469G>A), L1474Y (4470C>T 4471T>A) | | | | | | | | |
| Codon mutations: | ATA1402ATG (4256A>G), TGT1405TAT (4264G>A), GAG1407AAA (4269G>A 4271G>A), CTT1408ATT (4272C>A), CAA1409TTG (4275C>T 4276A>T 4277A>G), CAA1410AAG (4278C>A 4280A>G), GAT1412AGA (4284G>A 4285A>G 4286T>A), CTA1413TTA (4287C>T), ATC1414ATT (4292C>T), GAA1415GAG (4295A>G), CCC1416CCA (4298C>A), TCA1419TCC (4307A>C), GCA1422TCT (4314G>T 4316A>T), GAA1424AAA (4320G>A), GTC1428GTT (4334C>T), AGG1431AGA (4343G>A), TCA1432GCT (4344T>G 4346A>T), GAG1433GAA (4349G>A), GTG1435GCA (4354T>C 4355G>A), GGA1437GGT (4361A>T), TTA1439CTT (4365T>C 4367A>T), AGG1440CGA (4368A>C 4370G>A), ATA1443ATC (4379A>C), TAT1445TAC (4385T>C), CAA1446CAG (4388A>G), CCA1447CCC (4391A>C), TTA1448CTC (4392T>C 4394A>C), CAT1450CAC (4400T>C), CTC1452CTT (4406C>T), CAA1453TTG (4407C>T 4408A>T 4409A>G), GAC1455AGT (4413G>A 4414A>G 4415C>T), AAG1456AAA (4418G>A), TTT1457TTC (4421T>C), CCC1458ACT (4422C>A 4424C>T), ATT1459ATC (4427T>C), CCA1460CTA (4429C>T), AAG1462AAT (4436G>T), CTC1463CTA (4439C>A), ACA1464ATT (4441C>T 4442A>T), CTC1465TTA (4443C>T 4445C>A), TCT1467TTA (4450C>T 4451T>A), TCC1470GTA (4458T>G 4459C>T 4460C>A), AAA1471GAA (4461A>G), GCA1472GCC (4466A>C), AAG1473ACA (4468A>C 4469G>A), CTC1474TAC (4470C>T 4471T>A), TTT1475TTC (4475T>C) | | | | | | | | |

**Proteins**

| | Begin | End | Coverage | Score | Concordance | Matches | Identities | I/D/M/F* | Stop Codons |
|---|---|---|---|---|---|---|---|---|---|
| ORF I polyprotein (NP_127504.1) | 1402 | 1476 | 3.4% | 387 | 71.3% | 75 (100%) | 52 (69.3%) | 0/0/0/0 | 0 |
| Protein mutations: | I1402M (4256A>G), C1405Y (4264G>A), E1407K (4269G>A 4271G>A), L1408I (4272C>A), Q1409L (4275C>T 4276A>T 4277A>G), Q1410K (4278C>A 4280A>G), D1412R (4284G>A 4285A>G 4286T>A), A1422S (4314G>T 4316A>T), E1424K (4320G>A), S1432A (4344T>G 4346A>T), V1435A (4354T>C 4355G>A), Q1453L (4407C>T 4408A>T 4409A>G), D1455S (4413G>A 4414A>G 4415C>T), P1458T (4422C>A 4424C>T), P1460L (4429C>T), K1462N (4436G>T), T1464I (4441C>T 4442A>T), S1467L (4450C>T 4451T>A), S1470V (4458T>G 4459C>T 4460C>A), K1471E (4461A>G), K1473T (4468A>C 4469G>A), L1474Y (4470C>T 4471T>A) | | | | | | | | |
| Codon mutations: | ATA1402ATG (4256A>G), TGT1405TAT (4264G>A), GAG1407AAA (4269G>A 4271G>A), CTT1408ATT (4272C>A), CAA1409TTG (4275C>T 4276A>T 4277A>G), CAA1410AAG (4278C>A 4280A>G), GAT1412AGA (4284G>A 4285A>G 4286T>A), CTA1413TTA (4287C>T), ATC1414ATT (4292C>T), GAA1415GAG (4295A>G), CCC1416CCA (4298C>A), TCA1419TCC (4307A>C), GCA1422TCT (4314G>T 4316A>T), GAA1424AAA (4320G>A), GTC1428GTT (4334C>T), AGG1431AGA (4343G>A), TCA1432GCT (4344T>G 4346A>T), GAG1433GAA (4349G>A), GTG1435GCA (4354T>C 4355G>A), GGA1437GGT (4361A>T), TTA1439CTT (4365T>C 4367A>T), AGG1440CGA (4368A>C 4370G>A), ATA1443ATC (4379A>C), TAT1445TAC (4385T>C), CAA1446CAG (4388A>G), CCA1447CCC (4391A>C), TTA1448CTC (4392T>C 4394A>C), CAT1450CAC (4400T>C), CTC1452CTT (4406C>T), CAA1453TTG (4407C>T 4408A>T 4409A>G), GAC1455AGT (4413G>A 4414A>G 4415C>T), AAG1456AAA (4418G>A), TTT1457TTC (4421T>C), CCC1458ACT (4422C>A 4424C>T), ATT1459ATC (4427T>C), CCA1460CTA (4429C>T), AAG1462AAT (4436G>T), CTC1463CTA (4439C>A), ACA1464ATT (4441C>T 4442A>T), CTC1465TTA (4443C>T 4445C>A), TCT1467TTA (4450C>T 4451T>A), TCC1470GTA (4458T>G 4459C>T 4460C>A), AAA1471GAA (4461A>G), GCA1472GCC (4466A>C), AAG1473ACA (4468A>C 4469G>A), CTC1474TAC (4470C>T 4471T>A), TTT1475TTC (4475T>C) | | | | | | | | |

*: Inserts / Deletes / Misaligned / Frameshifts

#### Analysis details

This analysis was performed with panviral2.64

### NGS Details (UN8): Tomato associated geminivirus 1

#### Assembly

| | |
|---|---|
| **Coverage Length** | 236 (1 contig(s)) |
| **Depth Of Coverage** | 1.8 |
| **Number Of Reads** | 4 |
| **Reads Per Million** | 0.07 rpm (after QC) |
| **Ambiguities** | 0 |
| **Assembly Method** | de novo + reference guided assembly |
| **Consensus Caller** | Bcf Tools |

#### Coverage Map

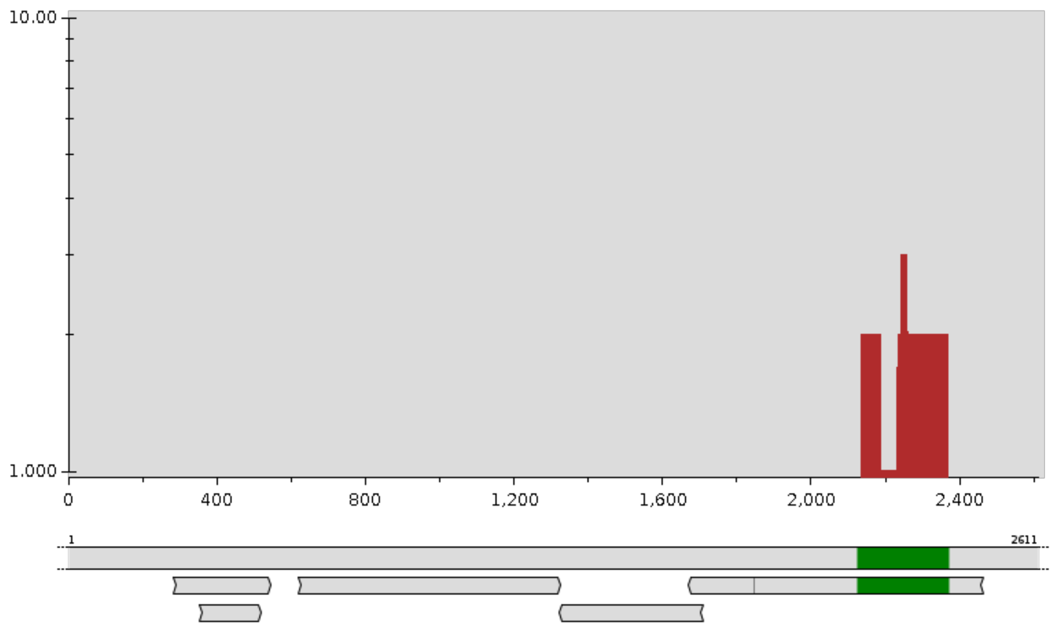

#### Assignment

| | |
|---|---|
| **Type** | Tomato associated geminivirus 1 (Taxonomy ID: 2027645) |
| **Reference Genome** | NC_035676.1 |
| **NT Identity (%)** | 63.7097 |
| **AA Identity (%)** | 59.0361 |
| **Number Of Stop Codons** | 2 |
| **Number Of CDS** | 5 |

#### Alignment

| | |
|---|---|
| **Alignment Score** | 139.0 (NT) + 778.0 (AA) = 917.0 |
| **Concordance (%)** | 54.4537 |

| | |
|---|---|
| **Alignment Method** | Global, seeded, nucleotide + amino acids (AGA) |

#### Genome Region

Sequence starts at position 2135 and ends at position 2370 relative to NC_035676.1 reference sequence.

#### Alignment Detailed Statistics

| | Begin | End | Coverage | Score | Concordance | Matches | Identities | I/D/M/F* | Stop Codons |
|---|---|---|---|---|---|---|---|---|---|
| **NT** | **2135** | **2370** | **9.0%** | **139** | **29.4%** | **236 (95.2%)** | **158 (63.7%)** | **12/0** | |
| Mutations: | 2137G>A, 2139A>T, 2142C>T, 2148G>T, 2151A>G, 2154G>A, 2157G>A, 2160C>G, 2162C>A, 2163A>G, 2166A>G, 2168C>G, 2170G>T, 2171C>T, 2172A>G, 2176C>G, 2177T>A, 2181T>C, 2184G>T, 2190G>A, 2191A>G, 2196T>G, 2199A>G, 2203T>A, 2205A>T, 2210C>T, 2213A>G, 2213_2214insATTTGTTGAGCT, 2218T>A, 2219T>C, 2220A>G, 2223G>C, 2226G>A, 2228T>G, 2229G>A, 2234C>T, 2238G>C, 2239A>T, 2244C>G, 2245T>A, 2250C>T, 2251T>C, 2252G>T, 2253C>A, 2254G>A, 2255T>A, 2256C>A, 2259A>G, 2262C>T, 2263C>T, 2271A>G, 2277T>A, 2278A>C, 2279C>A, 2280C>T, 2283A>G, 2289G>A, 2291T>G, 2292G>A, 2298C>T, 2301T>C, 2313G>T, 2314T>G, 2319C>T, 2322A>G, 2323C>G, 2324A>C, 2326G>A, 2333C>T, 2334G>A, 2337T>A, 2338G>T, 2340G>A, 2341A>G, 2342C>G, 2346A>G, 2354T>G, 2358T>A, 2360A>T | | | | | | | | |

##### CDS

| | Begin | End | Coverage | Score | Concordance | Matches | Identities | I/D/M/F* | Stop Codons |
|---|---|---|---|---|---|---|---|---|---|
| CK833_gp4 | 33 | 111 | 23.7% | 389 | 62.1% | 79 (95.2%) | 49 (59.0%) | 4/0/0/0 | 1 |
| Protein mutations: | L36I (2358T>A 2360A>T), K38Q (2354T>G), V42P (2340G>A 2341A>G 2342C>G), T43N (2337T>A 2338G>T), V45I (2333C>T), A47V (2326G>A), C48A (2322A>G 2323C>G 2324A>C), N51T (2313G>T 2314T>G), I59L (2289G>A 2291T>G), V63C (2277T>A 2278A>C 2279C>A), R68K (2262C>T 2263C>T), Q70H (2256C>A), T71F (2253C>A 2254G>A 2255T>A), Q72R (2250C>T 2251T>C 2252G>T), Q74L (2244C>G 2245T>A), F76* (2238G>C 2239A>T), D78N (2234C>T), T80P (2226G>A 2228T>G), D81E (2223G>C), N83V (2218T>A 2219T>C), G84_S85insSSTN (2213_2214insATTTGTTGAGCT), S85P (2213A>G), G86R (2210C>T), Y88F (2203T>A), I92T (2190G>A 2191A>G), R97S (2176C>G 2177T>A), A99K (2170G>T 2171C>T), D100H (2166A>G 2168C>G), A102S (2160C>G 2162C>A), T110I (2137G>A) | | | | | | | | |
| Codon mutations: | TTA36ATT (2358T>A 2360A>T), AAA38CAA (2354T>G), TAT40TAC (2346A>G), GTC42CCT (2340G>A 2341A>G 2342C>G), ACA43AAT (2337T>A 2338G>T), TAC44TAT (2334G>A), GTT45ATT (2333C>T), GCT47GTT (2326G>A), TGT48GCC (2322A>G 2323C>G 2324A>C), ACG49ACA (2319C>T), AAC51ACA (2313G>T 2314T>G), GGA55GGG (2301T>C), GAG56GAA (2298C>T), CAC58CAT (2292G>A), ATC59CTT (2289G>A 2291T>G), TGT61TGC (2283A>G), CTG62CTA (2280C>T), GTA63TGT (2277T>A 2278A>C 2279C>A), ACT65ACC (2271A>G), AGG68AAA (2262C>T 2263C>T), TTT69TTC (2259A>G), CAG70CAT (2256C>A), ACG71TTT (2253C>A 2254G>A 2255T>A), CAG72AGA (2250C>T 2251T>C 2252G>T), CAG74CTC (2244C>G 2245T>A), TTC76TAG (2238G>C 2239A>T), GAC78AAC (2234C>T), ATC79ATT (2229G>A), ACC80CCT (2226G>A 2228T>G), GAC81GAG (2223G>C), CCT82CCC (2220A>G), AAC83GTC (2218T>A 2219T>C), GGA84_TCG85insAGCTCAACAAAT (2213_2214insATTTGTTGAGCT), TCG85CCG (2213A>G), GGA86AGA (2210C>T), ATT87ATA (2205A>T), TAT88TTT (2203T>A), CAT89CAC (2199A>G), CCA90CCC (2196T>G), ATC92ACT (2190G>A 2191A>G), ATC94ATA (2184G>T), CCA95CCG (2181T>C), AGG97TCG (2176C>G 2177T>A), GAT98GAC (2172A>G), GCA99AAA (2170G>T 2171C>T), GAT100CAC (2166A>G 2168C>G), GTT101GTC (2163A>G), GCG102TCC (2160C>G 2162C>A), GAC103GAT (2157G>A), TAC104TAT (2154G>A), ATT105ATC (2151A>G), TCC106TCA (2148G>T), GGG108GGA (2142C>T), GGT109GGA (2139A>T), ACC110ATC (2137G>A) | | | | | | | | |
| CK833_gp5 | 33 | 111 | 30.2% | 389 | 62.1% | 79 (95.2%) | 49 (59.0%) | 4/0/0/0 | 1 |
| Protein mutations: | L36I (2358T>A 2360A>T), K38Q (2354T>G), V42P (2340G>A 2341A>G 2342C>G), T43N (2337T>A 2338G>T), V45I (2333C>T), A47V (2326G>A), C48A (2322A>G 2323C>G 2324A>C), N51T (2313G>T 2314T>G), I59L (2289G>A 2291T>G), V63C (2277T>A 2278A>C 2279C>A), R68K (2262C>T 2263C>T), Q70H (2256C>A), T71F (2253C>A 2254G>A 2255T>A), Q72R (2250C>T 2251T>C 2252G>T), Q74L (2244C>G 2245T>A), F76* (2238G>C 2239A>T), D78N (2234C>T), T80P (2226G>A 2228T>G), D81E (2223G>C), N83V (2218T>A 2219T>C), G84_S85insSSTN (2213_2214insATTTGTTGAGCT), S85P (2213A>G), G86R (2210C>T), Y88F (2203T>A), I92T (2190G>A 2191A>G), R97S (2176C>G 2177T>A), A99K (2170G>T 2171C>T), D100H (2166A>G 2168C>G), A102S (2160C>G 2162C>A), T110I (2137G>A) | | | | | | | | |
| Codon mutations: | TTA36ATT (2358T>A 2360A>T), AAA38CAA (2354T>G), TAT40TAC (2346A>G), GTC42CCT (2340G>A 2341A>G 2342C>G), ACA43AAT (2337T>A 2338G>T), TAC44TAT (2334G>A), GTT45ATT (2333C>T), GCT47GTT (2326G>A), TGT48GCC (2322A>G 2323C>G 2324A>C), ACG49ACA (2319C>T), AAC51ACA (2313G>T 2314T>G), GGA55GGG (2301T>C), GAG56GAA (2298C>T), CAC58CAT (2292G>A), ATC59CTT (2289G>A 2291T>G), TGT61TGC (2283A>G), CTG62CTA (2280C>T), GTA63TGT (2277T>A 2278A>C 2279C>A), ACT65ACC (2271A>G), AGG68AAA (2262C>T 2263C>T), TTT69TTC (2259A>G), CAG70CAT (2256C>A), ACG71TTT (2253C>A 2254G>A 2255T>A), CAG72AGA (2250C>T 2251T>C 2252G>T), CAG74CTC (2244C>G 2245T>A), TTC76TAG (2238G>C 2239A>T), GAC78AAC (2234C>T), ATC79ATT (2229G>A), ACC80CCT (2226G>A 2228T>G), GAC81GAG (2223G>C), CCT82CCC (2220A>G), AAC83GTC (2218T>A 2219T>C), GGA84_TCG85insAGCTCAACAAAT (2213_2214insATTTGTTGAGCT), TCG85CCG (2213A>G), GGA86AGA (2210C>T), ATT87ATA (2205A>T), TAT88TTT (2203T>A), CAT89CAC (2199A>G), CCA90CCC (2196T>G), ATC92ACT (2190G>A 2191A>G), ATC94ATA (2184G>T), CCA95CCG (2181T>C), AGG97TCG (2176C>G 2177T>A), GAT98GAC (2172A>G), GCA99AAA (2170G>T 2171C>T), GAT100CAC (2166A>G 2168C>G), GTT101GTC (2163A>G), GCG102TCC (2160C>G 2162C>A), GAC103GAT (2157G>A), TAC104TAT (2154G>A), ATT105ATC (2151A>G), TCC106TCA (2148G>T), GGG108GGA (2142C>T), GGT109GGA (2139A>T), ACC110ATC (2137G>A) | | | | | | | | |

##### Proteins

| | Begin | End | Coverage | Score | Concordance | Matches | Identities | I/D/M/F* | Stop Codons |
|---|---|---|---|---|---|---|---|---|---|
| replication-associated protein (YP_009417307.1) | 33 | 111 | 23.7% | 389 | 62.1% | 79 (95.2%) | 49 (59.0%) | 4/0/0/0 | 1 |
| Protein mutations: | L36I (2358T>A 2360A>T), K38Q (2354T>G), V42P (2340G>A 2341A>G 2342C>G), T43N (2337T>A 2338G>T), V45I (2333C>T), A47V (2326G>A), C48A (2322A>G 2323C>G 2324A>C), N51T (2313G>T 2314T>G), I59L (2289G>A 2291T>G), V63C (2277T>A 2278A>C 2279C>A), R68K (2262C>T 2263C>T), Q70H (2256C>A), T71F (2253C>A 2254G>A 2255T>A), Q72R (2250C>T 2251T>C 2252G>T), Q74L (2244C>G 2245T>A), F76* (2238G>C 2239A>T), D78N (2234C>T), T80P (2226G>A 2228T>G), D81E (2223G>C), N83V (2218T>A 2219T>C), G84_S85insSSTN (2213_2214insATTTGTTGAGCT), S85P (2213A>G), G86R (2210C>T), Y88F (2203T>A), I92T (2190G>A 2191A>G), R97S (2176C>G 2177T>A), A99K (2170G>T 2171C>T), D100H (2166A>G 2168C>G), A102S (2160C>G 2162C>A), T110I (2137G>A) | | | | | | | | |
| Codon mutations: | TTA36ATT (2358T>A 2360A>T), AAA38CAA (2354T>G), TAT40TAC (2346A>G), GTC42CCT (2340G>A 2341A>G 2342C>G), ACA43AAT (2337T>A 2338G>T), TAC44TAT (2334G>A), GTT45ATT (2333C>T), GCT47GTT (2326G>A), TGT48GCC (2322A>G 2323C>G 2324A>C), ACG49ACA (2319C>T), AAC51ACA (2313G>T 2314T>G), GGA55GGG (2301T>C), GAG56GAA (2298C>T), CAC58CAT (2292G>A), ATC59CTT (2289G>A 2291T>G), TGT61TGC (2283A>G), CTG62CTA (2280C>T), GTA63TGT (2277T>A 2278A>C 2279C>A), ACT65ACC (2271A>G), AGG68AAA (2262C>T 2263C>T), TTT69TTC (2259A>G), CAG70CAT (2256C>A), ACG71TTT (2253C>A 2254G>A 2255T>A), CAG72AGA (2250C>T 2251T>C 2252G>T), CAG74CTC (2244C>G 2245T>A), TTC76TAG (2238G>C 2239A>T), GAC78AAC (2234C>T), ATC79ATT (2229G>A), ACC80CCT (2226G>A 2228T>G), GAC81GAG (2223G>C), CCT82CCC (2220A>G), AAC83GTC (2218T>A 2219T>C), GGA84_TCG85insAGCTCAACAAAT (2213_2214insATTTGTTGAGCT), TCG85CCG (2213A>G), GGA86AGA (2210C>T), ATT87ATA (2205A>T), TAT88TTT (2203T>A), CAT89CAC (2199A>G), CCA90CCC (2196T>G), ATC92ACT (2190G>A 2191A>G), ATC94ATA (2184G>T), CCA95CCG (2181T>C), AGG97TCG (2176C>G 2177T>A), GAT98GAC (2172A>G), GCA99AAA (2170G>T 2171C>T), GAT100CAC (2166A>G 2168C>G), GTT101GTC (2163A>G), GCG102TCC (2160C>G 2162C>A), GAC103GAT (2157G>A), TAC104TAT (2154G>A), ATT105ATC (2151A>G), TCC106TCA (2148G>T), GGG108GGA (2142C>T), GGT109GGA (2139A>T), ACC110ATC (2137G>A) | | | | | | | | |
| RepA (YP_009417308.1) | 33 | 111 | 30.2% | 389 | 62.1% | 79 (95.2%) | 49 (59.0%) | 4/0/0/0 | 1 |
| Protein mutations: | L36I (2358T>A 2360A>T), K38Q (2354T>G), V42P (2340G>A 2341A>G 2342C>G), T43N (2337T>A 2338G>T), V45I (2333C>T), A47V (2326G>A), C48A (2322A>G 2323C>G 2324A>C), N51T (2313G>T 2314T>G), I59L (2289G>A 2291T>G), V63C (2277T>A 2278A>C 2279C>A), R68K (2262C>T 2263C>T), Q70H (2256C>A), T71F (2253C>A 2254G>A 2255T>A), Q72R (2250C>T 2251T>C 2252G>T), Q74L (2244C>G 2245T>A), F76* (2238G>C 2239A>T), D78N (2234C>T), T80P (2226G>A 2228T>G), D81E (2223G>C), N83V (2218T>A 2219T>C), G84_S85insSSTN (2213_2214insATTTGTTGAGCT), S85P (2213A>G), G86R (2210C>T), Y88F (2203T>A), I92T (2190G>A 2191A>G), R97S (2176C>G 2177T>A), A99K (2170G>T 2171C>T), D100H (2166A>G 2168C>G), A102S (2160C>G 2162C>A), T110I (2137G>A) | | | | | | | | |
| Codon mutations: | TTA36ATT (2358T>A 2360A>T), AAA38CAA (2354T>G), TAT40TAC (2346A>G), GTC42CCT (2340G>A 2341A>G 2342C>G), ACA43AAT (2337T>A 2338G>T), TAC44TAT (2334G>A), GTT45ATT (2333C>T), GCT47GTT (2326G>A), TGT48GCC (2322A>G 2323C>G 2324A>C), ACG49ACA (2319C>T), AAC51ACA (2313G>T 2314T>G), GGA55GGG (2301T>C), GAG56GAA (2298C>T), CAC58CAT (2292G>A), ATC59CTT (2289G>A 2291T>G), TGT61TGC (2283A>G), CTG62CTA (2280C>T), GTA63TGT (2277T>A 2278A>C 2279C>A), ACT65ACC (2271A>G), AGG68AAA (2262C>T 2263C>T), TTT69TTC (2259A>G), CAG70CAT (2256C>A), ACG71TTT (2253C>A 2254G>A 2255T>A), CAG72AGA (2250C>T 2251T>C 2252G>T), CAG74CTC (2244C>G 2245T>A), TTC76TAG (2238G>C 2239A>T), GAC78AAC (2234C>T), ATC79ATT (2229G>A), ACC80CCT (2226G>A 2228T>G), GAC81GAG (2223G>C), CCT82CCC (2220A>G), AAC83GTC (2218T>A 2219T>C), GGA84_TCG85insAGCTCAACAAAT (2213_2214insATTTGTTGAGCT), TCG85CCG (2213A>G), GGA86AGA (2210C>T), ATT87ATA (2205A>T), TAT88TTT (2203T>A), CAT89CAC (2199A>G), CCA90CCC (2196T>G), ATC92ACT (2190G>A 2191A>G), ATC94ATA (2184G>T), CCA95CCG (2181T>C), AGG97TCG (2176C>G 2177T>A), GAT98GAC (2172A>G), GCA99AAA (2170G>T 2171C>T), GAT100CAC (2166A>G 2168C>G), GTT101GTC (2163A>G), GCG102TCC (2160C>G 2162C>A), GAC103GAT (2157G>A), TAC104TAT (2154G>A), ATT105ATC (2151A>G), TCC106TCA (2148G>T), GGG108GGA (2142C>T), GGT109GGA (2139A>T), ACC110ATC (2137G>A) | | | | | | | | |

*: Inserts / Deletes / Misaligned / Frameshifts

#### Analysis details

This analysis was performed with panviral2.64

### NGS Details (UN8): Bracoviriform congregatae (segment Circle 7)

#### Assembly

| | |
|---|---|
| **Coverage Length** | 135 (1 contig(s)) |
| **Depth Of Coverage** | 2.5 |
| **Number Of Reads** | 3 |
| **Reads Per Million** | 0.05 rpm (after QC) |
| **Ambiguities** | 0 |
| **Assembly Method** | de novo + reference guided assembly |
| **Consensus Caller** | Bcf Tools |

#### Coverage Map

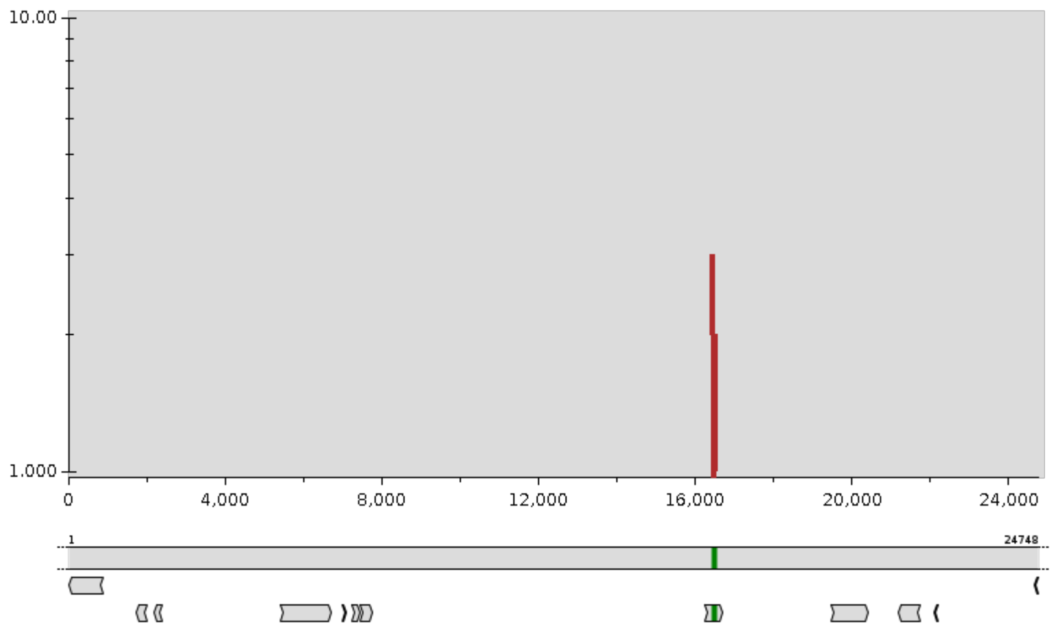

#### Assignment

| | |
|---|---|
| **Type** | Bracoviriform congregatae (Taxonomy ID: 39640) |
| **Reference Genome** | NC_006639.1 |
| **NT Identity (%)** | 75.5556 |
| **AA Identity (%)** | 86.6667 |
| **Number Of Stop Codons** | 0 |
| **Number Of CDS** | 5 |

#### Alignment

| | |
|---|---|
| **Alignment Score** | 138.0 (NT) + 273.0 (AA) = 411.0 |
| **Concordance (%)** | 72.4868 |

| Alignment Method | Global, seeded, nucleotide + amino acids (AGA) |
|---|---|

#### Genome Region

Sequence starts at position 16415 and ends at position 16549 relative to NC_006639.1 reference sequence.

#### Alignment Detailed Statistics

| | Begin | End | Coverage | Score | Concordance | Matches | Identities | I/D/M/F* | Stop Codons |
|---|---|---|---|---|---|---|---|---|---|
| **NT** | **16415** | **16549** | **0.5%** | **138** | **51.1%** | **135 (100%)** | **102 (75.6%)** | **0/0** | |
| Mutations: | 16417A>T, 16418T>C, 16420G>C, 16423A>C, 16426A>G, 16429G>C, 16432C>A, 16435T>C, 16444T>C, 16449G>A, 16450A>G, 16451T>A, 16452C>T, 16454C>T, 16456T>G, 16459A>T, 16462T>C, 16471A>G, 16474A>T, 16486A>C, 16489T>G, 16492C>T, 16501G>C, 16504T>G, 16516A>T, 16526G>A, 16531T>A, 16533G>C, 16534A>C, 16535T>A, 16538G>A, 16543T>C, 16546A>G | | | | | | | | |
| **CDS** | | | | | | | | | |
| Histone | 62 | 106 | 28.8% | 273 | 91.9% | 45 (100%) | 39 (86.7%) | 0/0/0/0 | 0 |
| Protein mutations: | R73K (16449G>A 16450A>G), S74I (16451T>A 16452C>T), V99I (16526G>A), G101A (16533G>C 16534A>C), L102M (16535T>A), V103I (16538G>A) | | | | | | | | |
| Codon mutations: | GGA62GGT (16417A>T), TTG63CTC (16418T>C 16420G>C), GGA64GGC (16423A>C), AAA65AAG (16426A>G), GGG66GGC (16429G>C), GGC67GGA (16432C>A), GCT68GCC (16435T>C), CAT71CAC (16444T>C), AGA73AAG (16449G>A 16450A>G), TCT74ATT (16451T>A 16452C>T), CTT75TTG (16454C>T 16456T>G), CGA76CGT (16459A>T), GAT77GAC (16462T>C), CAA80CAG (16471A>G), GGA81GGT (16474A>T), CCA85CCC (16486A>C), GCT86GCG (16489T>G), ATC87ATT (16492C>T), CTG90CTC (16501G>C), GCT91GCG (16504T>G), GGA95GGT (16516A>T), GTC99ATC (16526G>A), TCT100TCA (16531T>A), GGA101GCC (16533G>C 16534A>C), TTG102ATG (16535T>A), GTC103ATC (16538G>A), TAT104TAC (16543T>C), GAA105GAG (16546A>G) | | | | | | | | |
| **Proteins** | | | | | | | | | |
| hypothetical protein (YP_184795.1) | 62 | 106 | 28.8% | 273 | 91.9% | 45 (100%) | 39 (86.7%) | 0/0/0/0 | 0 |
| Protein mutations: | R73K (16449G>A 16450A>G), S74I (16451T>A 16452C>T), V99I (16526G>A), G101A (16533G>C 16534A>C), L102M (16535T>A), V103I (16538G>A) | | | | | | | | |
| Codon mutations: | GGA62GGT (16417A>T), TTG63CTC (16418T>C 16420G>C), GGA64GGC (16423A>C), AAA65AAG (16426A>G), GGG66GGC (16429G>C), GGC67GGA (16432C>A), GCT68GCC (16435T>C), CAT71CAC (16444T>C), AGA73AAG (16449G>A 16450A>G), TCT74ATT (16451T>A 16452C>T), CTT75TTG (16454C>T 16456T>G), CGA76CGT (16459A>T), GAT77GAC (16462T>C), CAA80CAG (16471A>G), GGA81GGT (16474A>T), CCA85CCC (16486A>C), GCT86GCG (16489T>G), ATC87ATT (16492C>T), CTG90CTC (16501G>C), GCT91GCG (16504T>G), GGA95GGT (16516A>T), GTC99ATC (16526G>A), TCT100TCA (16531T>A), GGA101GCC (16533G>C 16534A>C), TTG102ATG (16535T>A), GTC103ATC (16538G>A), TAT104TAC (16543T>C), GAA105GAG (16546A>G) | | | | | | | | |

*: Inserts / Deletes / Misaligned / Frameshifts

#### Analysis details

This analysis was performed with panviral2.64

### NGS Details (UN8): Tobacco streak virus (segment RNA 1)

#### Assembly

| | |
|---|---|
| **Coverage Length** | 269 (2 contig(s)) |
| **Depth Of Coverage** | 1.0 |
| **Number Of Reads** | 2 |
| **Reads Per Million** | 0.03 rpm (after QC) |
| **Ambiguities** | 0 |
| **Assembly Method** | de novo + reference guided assembly |
| **Consensus Caller** | Bcf Tools |

#### Coverage Map

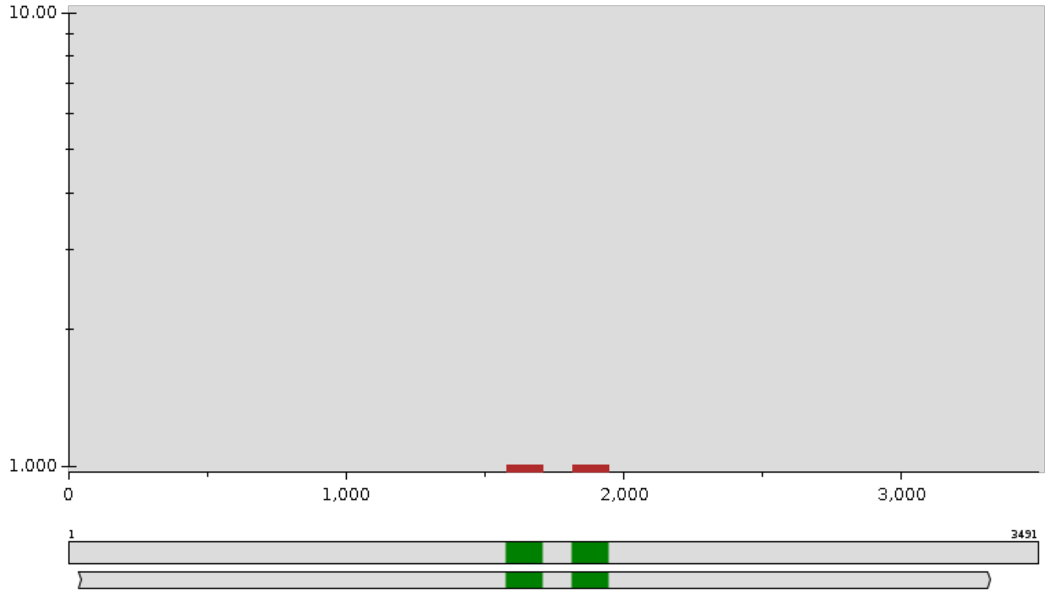

#### Assignment

| | |
|---|---|
| **Type** | Tobacco streak virus (Taxonomy ID: 12317) |
| **Reference Genome** | NC_003844.1 |
| **NT Identity (%)** | 74.3494 |
| **AA Identity (%)** | 80.0 |
| **Number Of Stop Codons** | 0 |
| **Number Of CDS** | 1 |

#### Alignment

| | |
|---|---|
| **Alignment Score** | 262.0 (NT) + 508.0 (AA) = 770.0 |
| **Concordance (%)** | 69.3069 |

| Alignment Method | Global, seeded, nucleotide + amino acids (AGA) |
|---|---|

#### Genome Region

Sequence starts at position 1576 and ends at position 1947 relative to NC_003844.1 reference sequence.

#### Alignment Detailed Statistics

| | Begin | End | Coverage | Score | Concordance | Matches | Identities | I/D/M/F* | Stop Codons |
|---|---|---|---|---|---|---|---|---|---|
| **NT** | **1576** | **1947** | **7.7%** | **262** | **48.7%** | **269 (100%)** | **200 (74.3%)** | **0/0** | |
| Mutations: | 1577A>G, 1579A>G, 1585A>T, 1588G>A, 1590G>A, 1591C>T, 1592C>T, 1595T>C, 1597G>T, 1599G>T, 1603A>G, 1606G>T, 1609T>A, 1612C>T, 1613A>T, 1615G>T, 1618C>G, 1619C>A, 1621A>G, 1623G>A, 1624T>G, 1625G>A, 1626T>C, 1633G>A, 1634C>T, 1636C>G, 1639G>C, 1648C>T, 1651G>T, 1657C>T, 1666T>C, 1669C>T, 1673G>A, 1678T>C, 1681T>C, 1690G>A, 1693G>T, 1694A>G, 1695C>T, 1696T>C, 1699T>C, 1702G>T, 1822T>G, 1825A>G, 1826C>A, 1830C>A, 1831C>T, 1834T>C, 1867A>C, 1870G>C, 1873A>G, 1876T>G, 1879T>C, 1890A>C, 1892A>T, 1894T>C, 1895A>G, 1897C>G, 1899A>G, 1903T>C, 1906T>G, 1912A>G, 1915G>A, 1918T>C, 1930T>C, 1934A>G, 1936T>C, 1938A>G, 1939A>G | | | | | | | | |
| **CDS** | | | | | | | | | |
| TbSVs1gp1 | 514 | 637 | 8.2% | 508 | 86.7% | 90 (100%) | 72 (80.0%) | 0/0/0/0 | 0 |
| Protein mutations: | I514V (1577A>G 1579A>G), S518N (1590G>A 1591C>T), R521L (1599G>T), T526S (1613A>T 1615G>T), S529K (1623G>A 1624T>G), V530T (1625G>A 1626T>C), V546I (1673G>A), T553V (1694A>G 1695C>T 1696T>C), Q597K (1826C>A), A598D (1830C>A 1831C>T), Q618P (1890A>C), T619S (1892A>T 1894T>C), I620V (1895A>G 1897C>G), K621R (1899A>G), I633V (1934A>G 1936T>C), K634R (1938A>G 1939A>G) | | | | | | | | |
| Codon mutations: | ATA514GTG (1577A>G 1579A>G), CCA516CCT (1585A>T), AGG517AGA (1588G>A), AGC518AAT (1590G>A 1591C>T), CTG519TTG (1592C>T), TTG520CTT (1595T>C 1597G>T), CGT521CTT (1599G>T), GAA522GAG (1603A>G), GTG523GTT (1606G>T), ATT524ATA (1609T>A), TAC525TAT (1612C>T), ACG526TCT (1613A>T 1615G>T), GTC527GTG (1618C>G), CGA528AGG (1619C>A 1621A>G), AGT529AAG (1623G>A 1624T>G), GTT530ACT (1625G>A 1626T>C), GGG532GGA (1633G>A), CTC533TTG (1634C>T 1636C>G), ACG534ACC (1639G>C), GAC537GAT (1648C>T), TCG538TCT (1651G>T), TAC540TAT (1657C>T), GTT543GTC (1666T>C), TCC544TCT (1669C>T), GTT546ATT (1673G>A), CCT547CCC (1678T>C), TCT548TCC (1681T>C), AAG551AAA (1690G>A), GCG552GCT (1693G>T), ACT553GTC (1694A>G 1695C>T 1696T>C), CAT554CAC (1699T>C), GTG555GTT (1702G>T), CGT595CGG (1822T>G), CAA596CAG (1825A>G), CAG597AAG (1826C>A), GCC598GAT (1830C>A 1831C>T), TTT599TTC (1834T>C), ATA610ATC (1867A>C), GCG611GCC (1870G>C), AAA612AAG (1873A>G), GCT613GCG (1876T>G), ATT614ATC (1879T>C), CAA618CCA (1890A>C), ACT619TCC (1892A>T 1894T>C), ATC620GTG (1895A>G 1897C>G), AAA621AGA (1899A>G), GAT622GAC (1903T>C), GGT623GGG (1906T>G), TTA625TTG (1912A>G), CCG626CCA (1915G>A), ATT627ATC (1918T>C), TGT631TGC (1930T>C), ATT633GTC (1934A>G 1936T>C), AAA634AGG (1938A>G 1939A>G) | | | | | | | | |
| **Proteins** | | | | | | | | | |
| putative viral replicase (NP_620772.1) | 514 | 637 | 8.2% | 508 | 86.7% | 90 (100%) | 72 (80.0%) | 0/0/0/0 | 0 |
| Protein mutations: | I514V (1577A>G 1579A>G), S518N (1590G>A 1591C>T), R521L (1599G>T), T526S (1613A>T 1615G>T), S529K (1623G>A 1624T>G), V530T (1625G>A 1626T>C), V546I (1673G>A), T553V (1694A>G 1695C>T 1696T>C), Q597K (1826C>A), A598D (1830C>A 1831C>T), Q618P (1890A>C), T619S (1892A>T 1894T>C), I620V (1895A>G 1897C>G), K621R (1899A>G), I633V (1934A>G 1936T>C), K634R (1938A>G 1939A>G) | | | | | | | | |
| Codon mutations: | ATA514GTG (1577A>G 1579A>G), CCA516CCT (1585A>T), AGG517AGA (1588G>A), AGC518AAT (1590G>A 1591C>T), CTG519TTG (1592C>T), TTG520CTT (1595T>C 1597G>T), CGT521CTT (1599G>T), GAA522GAG (1603A>G), GTG523GTT (1606G>T), ATT524ATA (1609T>A), TAC525TAT (1612C>T), ACG526TCT (1613A>T 1615G>T), GTC527GTG (1618C>G), CGA528AGG (1619C>A 1621A>G), AGT529AAG (1623G>A 1624T>G), GTT530ACT (1625G>A 1626T>C), GGG532GGA (1633G>A), CTC533TTG (1634C>T 1636C>G), ACG534ACC (1639G>C), GAC537GAT (1648C>T), TCG538TCT (1651G>T), TAC540TAT (1657C>T), GTT543GTC (1666T>C), TCC544TCT (1669C>T), GTT546ATT (1673G>A), CCT547CCC (1678T>C), TCT548TCC (1681T>C), AAG551AAA (1690G>A), GCG552GCT (1693G>T), ACT553GTC (1694A>G 1695C>T 1696T>C), CAT554CAC (1699T>C), GTG555GTT (1702G>T), CGT595CGG (1822T>G), CAA596CAG (1825A>G), CAG597AAG (1826C>A), GCC598GAT (1830C>A 1831C>T), TTT599TTC (1834T>C), ATA610ATC (1867A>C), GCG611GCC (1870G>C), AAA612AAG (1873A>G), GCT613GCG (1876T>G), ATT614ATC (1879T>C), CAA618CCA (1890A>C), ACT619TCC (1892A>T 1894T>C), ATC620GTG (1895A>G 1897C>G), AAA621AGA (1899A>G), GAT622GAC (1903T>C), GGT623GGG (1906T>G), TTA625TTG (1912A>G), CCG626CCA (1915G>A), ATT627ATC (1918T>C), TGT631TGC (1930T>C), ATT633GTC (1934A>G 1936T>C), AAA634AGG (1938A>G 1939A>G) | | | | | | | | |

*: Inserts / Deletes / Misaligned / Frameshifts

#### Analysis details

This analysis was performed with panviral2.64

### NGS Details (UN8): Nepovirus aeonii (segment RNA2)

#### Assembly

| | |
|---|---|
| **Coverage Length** | 242 (1 contig(s)) |
| **Depth Of Coverage** | 1.1 |
| **Number Of Reads** | 2 |
| **Reads Per Million** | 0.03 rpm (after QC) |
| **Ambiguities** | 0 |
| **Assembly Method** | de novo + reference guided assembly |
| **Consensus Caller** | Bcf Tools |

#### Coverage Map

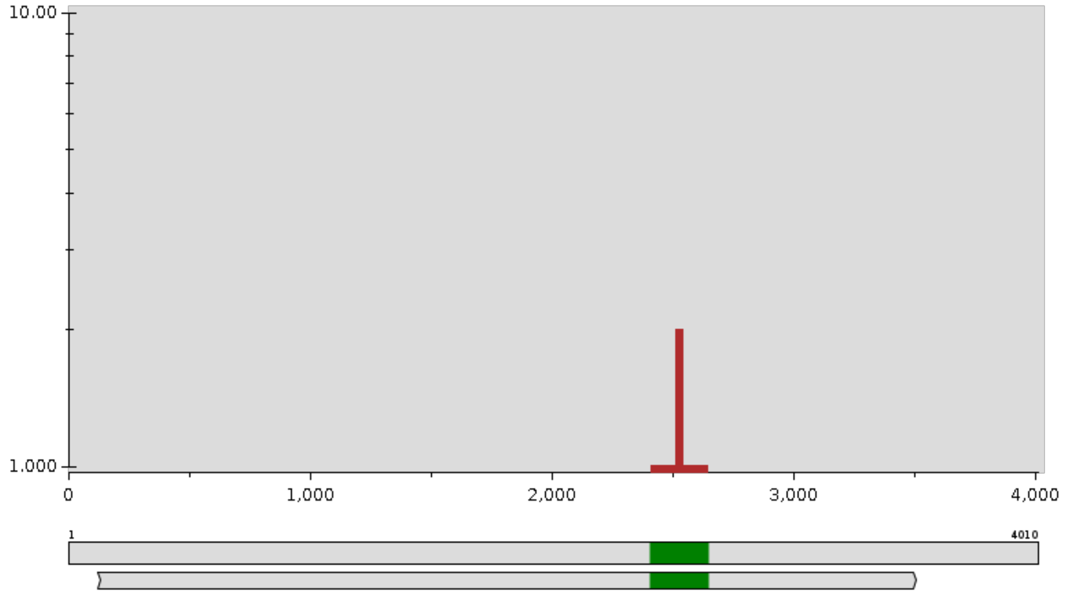

#### Assignment

| | |
|---|---|
| **Type** | Nepovirus aeonii (Taxonomy ID: 3047350) |
| **Reference Genome** | NC_038761.1 |
| **NT Identity (%)** | 69.0083 |
| **AA Identity (%)** | 66.6667 |
| **Number Of Stop Codons** | 0 |
| **Number Of CDS** | 1 |

#### Alignment

| | |
|---|---|
| **Alignment Score** | 184.0 (NT) + 410.0 (AA) = 594.0 |
| **Concordance (%)** | 57.7259 |

| Alignment Method | Global, seeded, nucleotide + amino acids (AGA) |
|---|---|

#### Genome Region

Sequence starts at position 2406 and ends at position 2647 relative to NC_038761.1 reference sequence.

#### Alignment Detailed Statistics

| | Begin | End | Coverage | Score | Concordance | Matches | Identities | I/D/M/F* | Stop Codons |
|---|---|---|---|---|---|---|---|---|---|
| **NT** | **2406** | **2647** | **6.0%** | **184** | **38.0%** | **242 (100%)** | **167 (69.0%)** | **0/0** | |
| Mutations: | 2409T>C, 2412T>C, 2415A>G, 2421G>T, 2422A>C, 2424G>T, 2427G>A, 2428G>T, 2433T>G, 2437A>C, 2439T>G, 2445G>T, 2446G>A, 2447T>C, 2452G>T, 2457T>C, 2460A>G, 2461G>T, 2462A>C, 2463T>A, 2466C>T, 2468C>T, 2469C>T, 2470C>G, 2472G>A, 2474A>C, 2481G>A, 2482T>C, 2483C>A, 2484T>G, 2491C>T, 2493T>G, 2496C>T, 2499C>A, 2502G>A, 2503G>A, 2505T>G, 2514G>T, 2515C>A, 2517G>A, 2519G>C, 2520T>A, 2521G>C, 2522T>A, 2524G>A, 2526G>T, 2529T>C, 2530T>A, 2531C>G, 2532A>T, 2533C>T, 2535C>T, 2536A>G, 2537G>A, 2541T>G, 2542G>A, 2562A>G, 2565G>A, 2571G>A, 2580G>C, 2583T>C, 2592A>G, 2593A>T, 2607T>C, 2608A>G, 2622T>C, 2623A>G, 2629C>T, 2631C>T, 2632A>T, 2634T>A, 2635C>A, 2637A>G, 2643C>T, 2646G>A | | | | | | | | |

##### CDS

| | Begin | End | Coverage | Score | Concordance | Matches | Identities | I/D/M/F* | Stop Codons |
|---|---|---|---|---|---|---|---|---|---|
| D1S54_s2gp1 | 762 | 842 | 7.2% | 410 | 73.7% | 81 (100%) | 54 (66.7%) | 0/0/0/0 | 0 |
| Protein mutations: | M767L (2422A>C 2424G>T), A769S (2428G>T), D770E (2433T>G), N772Q (2437A>C 2439T>G), V775T (2446G>A 2447T>C), V777L (2452G>T), I779M (2460A>G), D780S (2461G>T 2462A>C 2463T>A), T782I (2468C>T 2469C>T), Q783E (2470C>G 2472G>A), Q784P (2474A>C), S787Q (2482T>C 2483C>A 2484T>G), V794M (2503G>A 2505T>G), L798I (2515C>A 2517G>A), S799T (2519G>C 2520T>A), V800H (2521G>C 2522T>A), G801S (2524G>A 2526G>T), L804F (2533C>T 2535C>T), S805D (2536A>G 2537G>A), D807N (2542G>A), M824L (2593A>T), N829D (2608A>G), I834V (2623A>G), L836F (2629C>T 2631C>T), T837S (2632A>T 2634T>A), Q838K (2635C>A 2637A>G) | | | | | | | | |
| Codon mutations: | ACT762ACC (2409T>C), AAT763AAC (2412T>C), CAA764CAG (2415A>G), GTG766GTT (2421G>T), ATG767CTT (2422A>C 2424G>T), GCG768GCA (2427G>A), GCA769TCA (2428G>T), GAT770GAG (2433T>G), AAT772CAG (2437A>C 2439T>G), GTG774GTT (2445G>T), GTA775ACA (2446G>A 2447T>C), GTG777TTG (2452G>T), CAT778CAC (2457T>C), ATA779ATG (2460A>G), GAT780TCA (2461G>T 2462A>C 2463T>A), CGC781CGT (2466C>T), ACC782ATT (2468C>T 2469C>T), CAG783GAA (2470C>G 2472G>A), CAA784CCA (2474A>C), GAG786GAA (2481G>A), TCT787CAG (2482T>C 2483C>A 2484T>G), CTT790TTG (2491C>T 2493T>G), GCC791GCT (2496C>T), CCC792CCA (2499C>A), ACG793ACA (2502G>A), GTT794ATG (2503G>A 2505T>G), CCG797CCT (2514G>T), CTG798ATA (2515C>A 2517G>A), AGT799ACA (2519G>C 2520T>A), GTT800CAT (2521G>C 2522T>A), GGG801AGT (2524G>A 2526G>T), GAT802GAC (2529T>C), TCA803AGT (2530T>A 2531C>G 2532A>T), CTC804TTT (2533C>T 2535C>T), AGT805GAT (2536A>G 2537G>A), GTT806GTG (2541T>G), GAT807AAT (2542G>A), AAA813AAG (2562A>G), GAG814GAA (2565G>A), AAG816AAA (2571G>A), GGG819GGC (2580G>C), ACT820ACC (2583T>C), TTA823TTG (2592A>G), ATG824TTG (2593A>T), TAT828TAC (2607T>C), AAT829GAT (2608A>G), CCT833CCC (2622T>C), ATC834GTC (2623A>G), CTC836TTT (2629C>T 2631C>T), ACT837TCA (2632A>T 2634T>A), CAA838AAG (2635C>A 2637A>G), AGC840AGT (2643C>T), TCG841TCA (2646G>A) | | | | | | | | |

##### Proteins

| | Begin | End | Coverage | Score | Concordance | Matches | Identities | I/D/M/F* | Stop Codons |
|---|---|---|---|---|---|---|---|---|---|
| polyprotein P2 (YP_009507917.1) | 762 | 842 | 7.2% | 410 | 73.7% | 81 (100%) | 54 (66.7%) | 0/0/0/0 | 0 |
| Protein mutations: | M767L (2422A>C 2424G>T), A769S (2428G>T), D770E (2433T>G), N772Q (2437A>C 2439T>G), V775T (2446G>A 2447T>C), V777L (2452G>T), I779M (2460A>G), D780S (2461G>T 2462A>C 2463T>A), T782I (2468C>T 2469C>T), Q783E (2470C>G 2472G>A), Q784P (2474A>C), S787Q (2482T>C 2483C>A 2484T>G), V794M (2503G>A 2505T>G), L798I (2515C>A 2517G>A), S799T (2519G>C 2520T>A), V800H (2521G>C 2522T>A), G801S (2524G>A 2526G>T), L804F (2533C>T 2535C>T), S805D (2536A>G 2537G>A), D807N (2542G>A), M824L (2593A>T), N829D (2608A>G), I834V (2623A>G), L836F (2629C>T 2631C>T), T837S (2632A>T 2634T>A), Q838K (2635C>A 2637A>G) | | | | | | | | |
| Codon mutations: | ACT762ACC (2409T>C), AAT763AAC (2412T>C), CAA764CAG (2415A>G), GTG766GTT (2421G>T), ATG767CTT (2422A>C 2424G>T), GCG768GCA (2427G>A), GCA769TCA (2428G>T), GAT770GAG (2433T>G), AAT772CAG (2437A>C 2439T>G), GTG774GTT (2445G>T), GTA775ACA (2446G>A 2447T>C), GTG777TTG (2452G>T), CAT778CAC (2457T>C), ATA779ATG (2460A>G), GAT780TCA (2461G>T 2462A>C 2463T>A), CGC781CGT (2466C>T), ACC782ATT (2468C>T 2469C>T), CAG783GAA (2470C>G 2472G>A), CAA784CCA (2474A>C), GAG786GAA (2481G>A), TCT787CAG (2482T>C 2483C>A 2484T>G), CTT790TTG (2491C>T 2493T>G), GCC791GCT (2496C>T), CCC792CCA (2499C>A), ACG793ACA (2502G>A), GTT794ATG (2503G>A 2505T>G), CCG797CCT (2514G>T), CTG798ATA (2515C>A 2517G>A), AGT799ACA (2519G>C 2520T>A), GTT800CAT (2521G>C 2522T>A), GGG801AGT (2524G>A 2526G>T), GAT802GAC (2529T>C), TCA803AGT (2530T>A 2531C>G 2532A>T), CTC804TTT (2533C>T 2535C>T), AGT805GAT (2536A>G 2537G>A), GTT806GTG (2541T>G), GAT807AAT (2542G>A), AAA813AAG (2562A>G), GAG814GAA (2565G>A), AAG816AAA (2571G>A), GGG819GGC (2580G>C), ACT820ACC (2583T>C), TTA823TTG (2592A>G), ATG824TTG (2593A>T), TAT828TAC (2607T>C), AAT829GAT (2608A>G), CCT833CCC (2622T>C), ATC834GTC (2623A>G), CTC836TTT (2629C>T 2631C>T), ACT837TCA (2632A>T 2634T>A), CAA838AAG (2635C>A 2637A>G), AGC840AGT (2643C>T), TCG841TCA (2646G>A) | | | | | | | | |

*: Inserts / Deletes / Misaligned / Frameshifts

#### Analysis details

This analysis was performed with panviral2.64

### NGS Details (UN8): Duamitovirus peex1

#### Assembly

| | |
|---|---|
| **Coverage Length** | 135 (1 contig(s)) |
| **Depth Of Coverage** | 1.5 |
| **Number Of Reads** | 2 |
| **Reads Per Million** | 0.03 rpm (after QC) |
| **Ambiguities** | 0 |
| **Assembly Method** | de novo + reference guided assembly |
| **Consensus Caller** | Bcf Tools |

#### Coverage Map

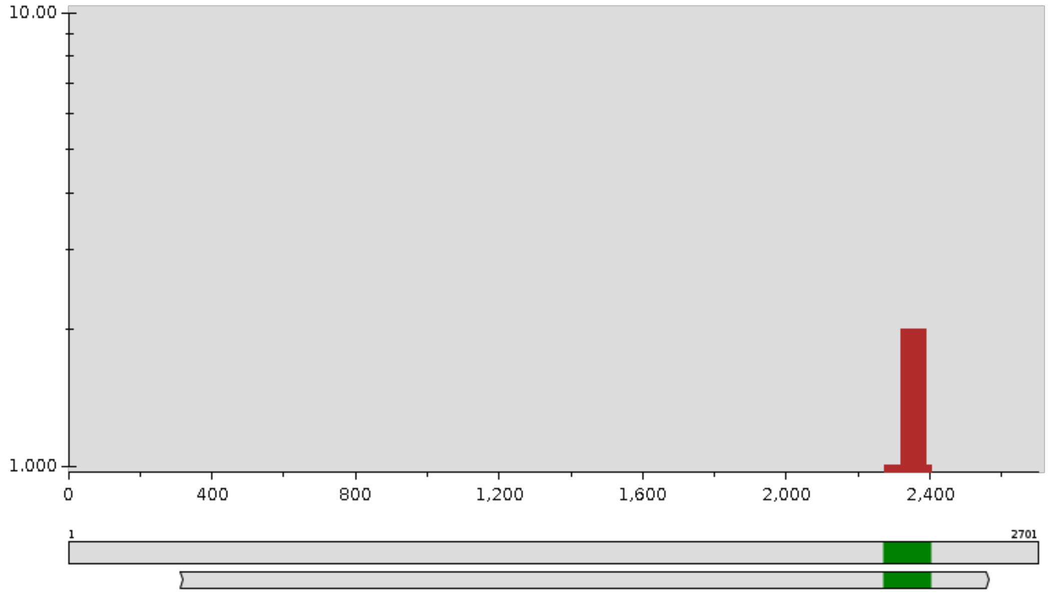

#### Assignment

| | |
|---|---|
| **Type** | Duamitovirus peex1 (Taxonomy ID: 2955799) |
| **Reference Genome** | NC_076525.1 |
| **NT Identity (%)** | 70.3704 |
| **AA Identity (%)** | 73.3333 |
| **Number Of Stop Codons** | 1 |
| **Number Of CDS** | 1 |

#### Alignment

| | |
|---|---|
| **Alignment Score** | 110.0 (NT) + 266.0 (AA) = 376.0 |
| **Concordance (%)** | 58.567 |

| Alignment Method | Global, seeded, nucleotide + amino acids (AGA) |
|---|---|

#### Genome Region

Sequence starts at position 2271 and ends at position 2405 relative to NC_076525.1 reference sequence.

#### Alignment Detailed Statistics

| | Begin | End | Coverage | Score | Concordance | Matches | Identities | I/D/M/F* | Stop Codons |
|---|---|---|---|---|---|---|---|---|---|
| **NT** | **2271** | **2405** | **5.0%** | **110** | **40.7%** | **135 (100%)** | **95 (70.4%)** | **0/0** | |
| Mutations: | 2276T>A, 2280A>G, 2283G>A, 2294T>G, 2295C>A, 2296G>A, 2301C>G, 2303T>C, 2306A>T, 2312T>C, 2315C>T, 2321G>T, 2325A>G, 2326A>C, 2327T>A, 2330C>A, 2331C>G, 2333G>A, 2339G>A, 2345A>T, 2348C>T, 2351A>G, 2352C>T, 2354T>G, 2363G>T, 2373G>A, 2374C>G, 2375G>T, 2377C>A, 2378T>A, 2381T>C, 2384G>A, 2388C>A, 2390T>A, 2393T>A, 2394C>A, 2396G>C, 2397A>G, 2398C>A, 2399T>A | | | | | | | | |
| **CDS** | | | | | | | | | |
| RdRp | 654 | 698 | 6.0% | 266 | 71.5% | 45 (100%) | 33 (73.3%) | 0/0/0/0 | 1 |
| Protein mutations: | H655Q (2276T>A), M657V (2280A>G), E658K (2283G>A), R662K (2295C>A 2296G>A), L664V (2301C>G 2303T>C), N672A (2325A>G 2326A>C 2327T>A), P674A (2331C>G 2333G>A), A688S (2373G>A 2374C>G 2375G>T), T689K (2377C>A 2378T>A), W691* (2384G>A), Q695N (2394C>A 2396G>C), T696E (2397A>G 2398C>A 2399T>A) | | | | | | | | |
| Codon mutations: | CAT655CAA (2276T>A), ATG657GTG (2280A>G), GAA658AAA (2283G>A), CTT661CTG (2294T>G), CGA662AAA (2295C>A 2296G>A), CTT664GTC (2301C>G 2303T>C), TCA665TCT (2306A>T), TAT667TAC (2312T>C), CAC668CAT (2315C>T), GTG670GTT (2321G>T), AAT672GCA (2325A>G 2326A>C 2327T>A), TCC673TCA (2330C>A), CCG674GCA (2331C>G 2333G>A), GTG676GTA (2339G>A), ATA678ATT (2345A>T), GAC679GAT (2348C>T), CAA680CAG (2351A>G), CTT681TTG (2352C>T 2354T>G), GTG684GTT (2363G>T), GCG688AGT (2373G>A 2374C>G 2375G>T), ACT689AAA (2377C>A 2378T>A), TCT690TCC (2381T>C), TGG691TGA (2384G>A), CGT693AGA (2388C>A 2390T>A), ACT694ACA (2393T>A), CAG695AAC (2394C>A 2396G>C), ACT696GAA (2397A>G 2398C>A 2399T>A) | | | | | | | | |
| **Proteins** | | | | | | | | | |
| RNA-dependent RNA polymerase (YP_010798875.1) | 654 | 698 | 6.0% | 266 | 71.5% | 45 (100%) | 33 (73.3%) | 0/0/0/0 | 1 |
| Protein mutations: | H655Q (2276T>A), M657V (2280A>G), E658K (2283G>A), R662K (2295C>A 2296G>A), L664V (2301C>G 2303T>C), N672A (2325A>G 2326A>C 2327T>A), P674A (2331C>G 2333G>A), A688S (2373G>A 2374C>G 2375G>T), T689K (2377C>A 2378T>A), W691* (2384G>A), Q695N (2394C>A 2396G>C), T696E (2397A>G 2398C>A 2399T>A) | | | | | | | | |
| Codon mutations: | CAT655CAA (2276T>A), ATG657GTG (2280A>G), GAA658AAA (2283G>A), CTT661CTG (2294T>G), CGA662AAA (2295C>A 2296G>A), CTT664GTC (2301C>G 2303T>C), TCA665TCT (2306A>T), TAT667TAC (2312T>C), CAC668CAT (2315C>T), GTG670GTT (2321G>T), AAT672GCA (2325A>G 2326A>C 2327T>A), TCC673TCA (2330C>A), CCG674GCA (2331C>G 2333G>A), GTG676GTA (2339G>A), ATA678ATT (2345A>T), GAC679GAT (2348C>T), CAA680CAG (2351A>G), CTT681TTG (2352C>T 2354T>G), GTG684GTT (2363G>T), GCG688AGT (2373G>A 2374C>G 2375G>T), ACT689AAA (2377C>A 2378T>A), TCT690TCC (2381T>C), TGG691TGA (2384G>A), CGT693AGA (2388C>A 2390T>A), ACT694ACA (2393T>A), CAG695AAC (2394C>A 2396G>C), ACT696GAA (2397A>G 2398C>A 2399T>A) | | | | | | | | |

*: Inserts / Deletes / Misaligned / Frameshifts

#### Analysis details

This analysis was performed with panviral2.64

### NGS Details (UN8): Moloney murine leukemia virus

#### Assembly

| | |
|---|---|
| **Coverage Length** | 222 (1 contig(s)) |
| **Depth Of Coverage** | 1.3 |
| **Number Of Reads** | 2 |
| **Reads Per Million** | 0.03 rpm (after QC) |
| **Ambiguities** | 0 |
| **Assembly Method** | de novo + reference guided assembly |
| **Consensus Caller** | Bcf Tools |

#### Coverage Map

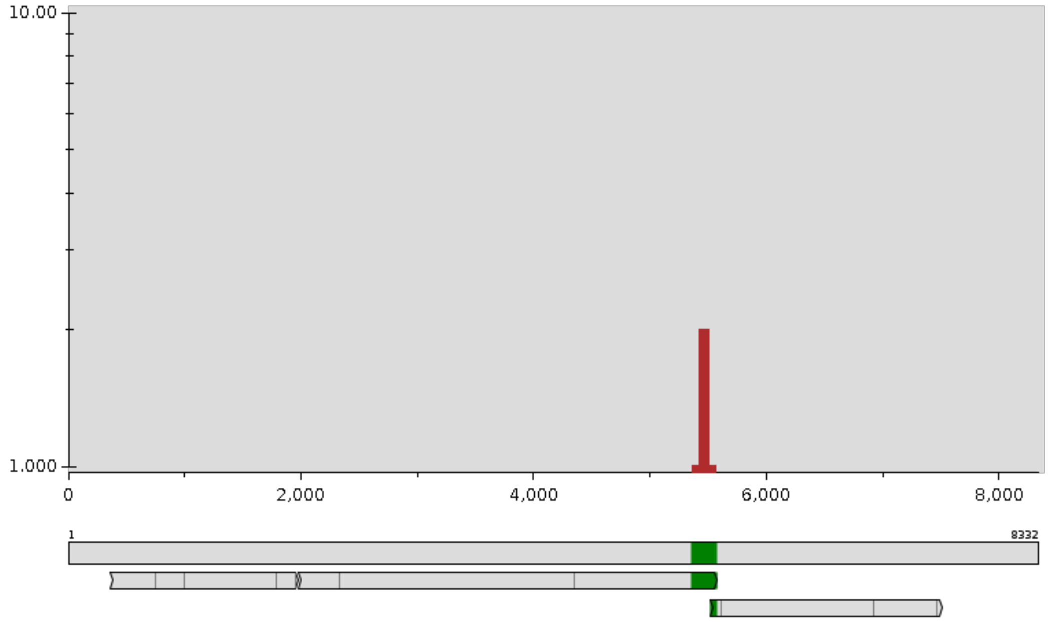

#### Assignment

| | |
|---|---|
| **Type** | Moloney murine leukemia virus (Taxonomy ID: 11801) |
| **Reference Genome** | NC_001501.1 |
| **NT Identity (%)** | 81.2207 |
| **AA Identity (%)** | 78.2609 |
| **Number Of Stop Codons** | 1 |
| **Number Of CDS** | 3 |

#### Alignment

| | |
|---|---|
| **Alignment Score** | 248.0 (NT) + 498.0 (AA) = 746.0 |
| **Concordance (%)** | 72.8516 |

| Alignment Method | Global, seeded, nucleotide + amino acids (AGA) |
|---|---|

#### Genome Region

Sequence starts at position 5355 and ends at position 5576 relative to NC_001501.1 reference sequence.

#### Alignment Detailed Statistics

| | Begin | End | Coverage | Score | Concordance | Matches | Identities | I/D/M/F* | Stop Codons |
|---|---|---|---|---|---|---|---|---|---|
| **NT** | **5355** | **5576** | **2.7%** | **248** | **60.8%** | **213 (95.9%)** | **173 (77.9%)** | **0/9** | |
| Mutations: | 5357A>G, 5373C>T, 5381T>C, 5396T>C, 5402A>C, 5426C>T, 5445G>T, 5447A>T, 5450T>G, 5465C>T, 5471G>A, 5477T>G, 5480C>G, 5481G>A, 5482A>C, 5483C>A, 5484C>A, 5486C>T, 5487_5495delGGGGGTGGA, 5498A>T, 5499T>C, 5501C>A, 5502T>G, 5504T>C, 5505A>G, 5508C>A, 5509T>C, 5510G>A, 5511A>G, 5517C>A, 5518G>A, 5519C>G, 5522T>C, 5525A>G, 5528C>T, 5537C>T, 5546A>G, 5552G>A, 5561C>T, 5563A>G, 5572A>G | | | | | | | | |
| **CDS** | | | | | | | | | |
| gag-pol | 1666 | 1738 | 4.2% | 421 | 86.6% | 70 (95.9%) | 60 (82.2%) | 0/3/0/0 | 1 |
| Protein mutations: | A1696S (5445G>T 5447A>T), D1708T (5481G>A 5482A>C 5483C>A), P1709T (5484C>A 5486C>T), G1710_G1712del (5487_5495delGGGGGTGGA), S1714P (5499T>C 5501C>A), S1715A (5502T>G 5504T>C), R1716G (5505A>G), L1717T (5508C>A 5509T>C 5510G>A), T1718A (5511A>G), R1720K (5517C>A 5518G>A 5519C>G), E1735G (5563A>G) | | | | | | | | |
| Codon mutations: | CGA1666CGG (5357A>G), CTA1672TTA (5373C>T), CCT1674CCC (5381T>C), CCT1679CCC (5396T>C), ACA1681ACC (5402A>C), GCC1689GCT (5426C>T), GCA1696TCT (5445G>T 5447A>T), GCT1697GCG (5450T>G), GCC1702GCT (5465C>T), GTG1704GTA (5471G>A), GCT1706GCG (5477T>G), GCC1707GCG (5480C>G), GAC1708ACA (5481G>A 5482A>C 5483C>A), CCC1709ACT (5484C>A 5486C>T), GGG1710_GGA1712del (5487_5495delGGGGGTGGA), CCA1713CCT (5498A>T), TCC1714CCA (5499T>C 5501C>A), TCT1715GCC (5502T>G 5504T>C), AGA1716GGA (5505A>G), CTG1717ACA (5508C>A 5509T>C 5510G>A), ACA1718GCA (5511A>G), CGC1720AAG (5517C>A 5518G>A 5519C>G), GTT1721GTC (5522T>C), CAA1722CAG (5525A>G), CGC1723CGT (5528C>T), AAC1726AAT (5537C>T), AAA1729AAG (5546A>G), AGG1731AGA (5552G>A), CGC1734CGT (5561C>T), GAG1735GGG (5563A>G), TAA1738TGA (5572A>G) | | | | | | | | |
| env | 1 | 22 | 3.3% | 77 | 54.2% | 22 (100%) | 12 (54.5%) | 0/0/0/0 | 0 |
| Protein mutations: | A2E (5517C>A 5518G>A), R3G (5519C>G), S4P (5522T>C), T5A (5525A>G), L6F (5528C>T), P9S (5537C>T), N12D (5546A>G), V14I (5552G>A), R17W (5561C>T 5563A>G) | | | | | | | | |
| Codon mutations: | GCG2GAA (5517C>A 5518G>A), CGT3GGT (5519C>G), TCA4CCA (5522T>C), ACG5GCG (5525A>G), CTC6TTC (5528C>T), CCC9TCC (5537C>T), AAT12GAT (5546A>G), GTT14ATT (5552G>A), CGA17TGG (5561C>T 5563A>G), CTA20CTG (5572A>G) | | | | | | | | |
| **Proteins** | | | | | | | | | |
| Pr180 (NP_057933.2) | 1666 | 1738 | 4.2% | 421 | 86.6% | 70 (95.9%) | 60 (82.2%) | 0/3/0/0 | 1 |
| Protein mutations: | A1696S (5445G>T 5447A>T), D1708T (5481G>A 5482A>C 5483C>A), P1709T (5484C>A 5486C>T), G1710_G1712del (5487_5495delGGGGGTGGA), S1714P (5499T>C 5501C>A), S1715A (5502T>G 5504T>C), R1716G (5505A>G), L1717T (5508C>A 5509T>C 5510G>A), T1718A (5511A>G), R1720K (5517C>A 5518G>A 5519C>G), E1735G (5563A>G) | | | | | | | | |
| Codon mutations: | CGA1666CGG (5357A>G), CTA1672TTA (5373C>T), CCT1674CCC (5381T>C), CCT1679CCC (5396T>C), ACA1681ACC (5402A>C), GCC1689GCT (5426C>T), GCA1696TCT (5445G>T 5447A>T), GCT1697GCG (5450T>G), GCC1702GCT (5465C>T), GTG1704GTA (5471G>A), GCT1706GCG (5477T>G), GCC1707GCG (5480C>G), GAC1708ACA (5481G>A 5482A>C 5483C>A), CCC1709ACT (5484C>A 5486C>T), GGG1710_GGA1712del (5487_5495delGGGGGTGGA), CCA1713CCT (5498A>T), TCC1714CCA (5499T>C 5501C>A), TCT1715GCC (5502T>G 5504T>C), AGA1716GGA (5505A>G), CTG1717ACA (5508C>A 5509T>C 5510G>A), ACA1718GCA (5511A>G), CGC1720AAG (5517C>A 5518G>A 5519C>G), GTT1721GTC (5522T>C), CAA1722CAG (5525A>G), CGC1723CGT (5528C>T), AAC1726AAT (5537C>T), AAA1729AAG (5546A>G), AGG1731AGA (5552G>A), CGC1734CGT (5561C>T), GAG1735GGG (5563A>G), TAA1738TGA (5572A>G) | | | | | | | | |
| p46 IN (NP_955592.1) | 336 | 407 | 17.7% | 420 | 86.6% | 69 (95.8%) | 59 (81.9%) | 0/3/0/0 | 0 |
| Protein mutations: | A366S (5445G>T 5447A>T), D378T (5481G>A 5482A>C 5483C>A), P379T (5484C>A 5486C>T), G380_G382del (5487_5495delGGGGGTGGA), S384P (5499T>C 5501C>A), S385A (5502T>G 5504T>C), R386G (5505A>G), L387T (5508C>A 5509T>C 5510G>A), T388A (5511A>G), R390K (5517C>A 5518G>A 5519C>G), E405G (5563A>G) | | | | | | | | |
| Codon mutations: | CGA336CGG (5357A>G), CTA342TTA (5373C>T), CCT344CCC (5381T>C), CCT349CCC (5396T>C), ACA351ACC (5402A>C), GCC359GCT (5426C>T), GCA366TCT (5445G>T 5447A>T), GCT367GCG (5450T>G), GCC372GCT (5465C>T), GTG374GTA (5471G>A), GCT376GCG (5477T>G), GCC377GCG (5480C>G), GAC378ACA (5481G>A 5482A>C 5483C>A), CCC379ACT (5484C>A 5486C>T), GGG380_GGA382del (5487_5495delGGGGGTGGA), CCA383CCT (5498A>T), TCC384CCA (5499T>C 5501C>A), TCT385GCC (5502T>G 5504T>C), AGA386GGA (5505A>G), CTG387ACA (5508C>A 5509T>C 5510G>A), ACA388GCA (5511A>G), CGC390AAG (5517C>A 5518G>A 5519C>G), GTT391GTC (5522T>C), CAA392CAG (5525A>G), CGC393CGT (5528C>T), AAC396AAT (5537C>T), AAA399AAG (5546A>G), AGG401AGA (5552G>A), CGC404CGT (5561C>T), GAG405GGG (5563A>G) | | | | | | | | |
| gPr80 (NP_057935.1) | 1 | 22 | 3.3% | 77 | 54.2% | 22 (100%) | 12 (54.5%) | 0/0/0/0 | 0 |
| Protein mutations: | A2E (5517C>A 5518G>A), R3G (5519C>G), S4P (5522T>C), T5A (5525A>G), L6F (5528C>T), P9S (5537C>T), N12D (5546A>G), V14I (5552G>A), R17W (5561C>T 5563A>G) | | | | | | | | |
| Codon mutations: | GCG2GAA (5517C>A 5518G>A), CGT3GGT (5519C>G), TCA4CCA (5522T>C), ACG5GCG (5525A>G), CTC6TTC (5528C>T), CCC9TCC (5537C>T), AAT12GAT (5546A>G), GTT14ATT (5552G>A), CGA17TGG (5561C>T 5563A>G), CTA20CTG (5572A>G) | | | | | | | | |

*: Inserts / Deletes / Misaligned / Frameshifts

#### Analysis details

This analysis was performed with panviral2.64

### NGS Details (UN8): Teseptimavirus EG1

#### Assembly

| | |
|---|---|
| Coverage Length | 209 (1 contig(s)) |
| Depth Of Coverage | 1.2 |
| Number Of Reads | 2 |
| Reads Per Million | 0.03 rpm (after QC) |
| Ambiguities | 0 |
| Assembly Method | de novo + reference guided assembly |
| Consensus Caller | Bcf Tools |

#### Coverage Map

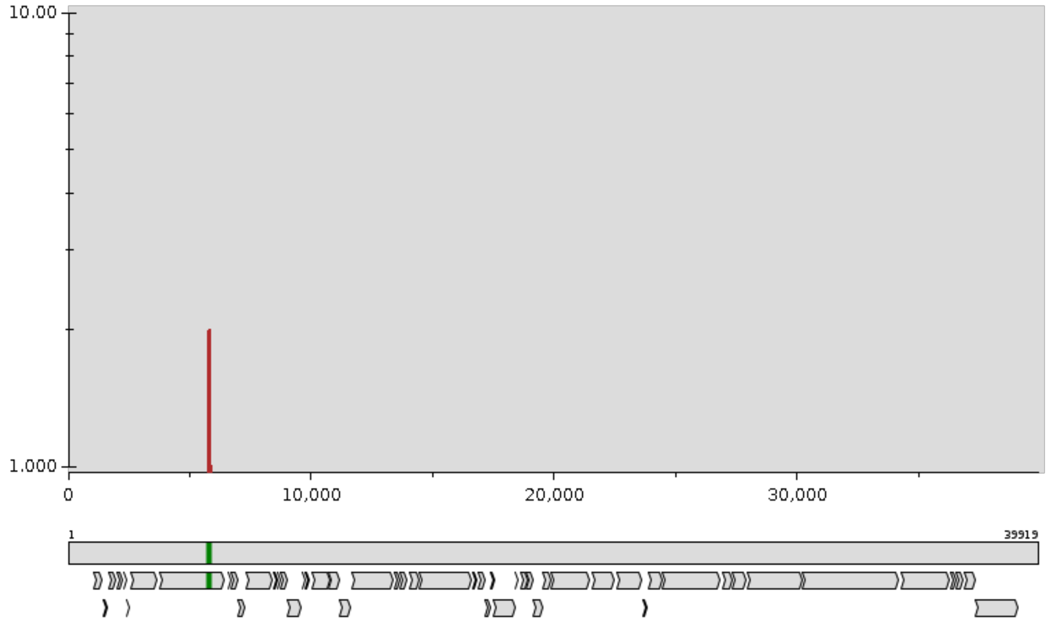

#### Assignment

| | |
|---|---|
| Type | Teseptimavirus EG1 (Taxonomy ID: 2733876) |
| Reference Genome | NC_047895.1 |
| NT Identity (%) | 75.1196 |
| AA Identity (%) | 98.5714 |
| Number Of Stop Codons | 0 |
| Number Of CDS | 51 |

#### Alignment

| | |
|---|---|
| Alignment Score | 210.0 (NT) + 425.0 (AA) = 635.0 |
| Concordance (%) | 75.3262 |

| Alignment Method | Local, heuristic, nucleotide (BLASTN) |
|---|---|

#### Genome Region

Sequence starts at position 5710 and ends at position 5918 relative to NC_047895.1 reference sequence.

#### Alignment Detailed Statistics

| | Begin | End | Coverage | Score | Concordance | Matches | Identities | I/D/M/F* | Stop Codons |
|---|---|---|---|---|---|---|---|---|---|
| **NT** | **5710** | **5918** | **0.5%** | **210** | **50.2%** | **209 (100%)** | **157 (75.1%)** | **0/0** | |
| Mutations: | 5712A>C, 5715T>G, 5718T>C, 5721T>C, 5722T>A, 5723C>G, 5724A>C, 5727C>T, 5733T>A, 5734C>T, 5745C>G, 5751T>C, 5754T>C, 5757G>A, 5760C>A, 5763T>A, 5766G>T, 5775T>G, 5778G>A, 5779C>T, 5791T>A, 5792C>G, 5793A>T, 5796G>T, 5797A>T, 5798G>C, 5799C>G, 5808A>C, 5814T>C, 5820A>T, 5826T>C, 5832C>T, 5838T>G, 5841G>A, 5842T>A, 5843C>G, 5847T>G, 5853G>A, 5854C>T, 5856G>A, 5859G>C, 5862T>A, 5865T>C, 5868G>A, 5871C>T, 5874A>G, 5883G>A, 5886T>G, 5889T>C, 5892G>A, 5895T>C, 5898C>G | | | | | | | | |
| **CDS** | | | | | | | | | |
| HOS48_gp08 | 657 | 726 | 7.9% | 425 | 97.7% | 70 (100%) | 69 (98.6%) | 0/0/0/0 | 0 |
| **Proteins** | | | | | | | | | |
| DNA-dependent RNA polymerase (YP_009795804.1) | 657 | 726 | 7.9% | 425 | 97.7% | 70 (100%) | 69 (98.6%) | 0/0/0/0 | 0 |
| Protein mutations: | *none* | | | | | | | | |
| Codon mutations: | CCA657CCC (5712A>C), GCT658GCG (5715T>G), ATT659ATC (5718T>C), GAT660GAC (5721T>C), TCA661AGC (5722T>A 5723C>G 5724A>C), GGC662GGT (5727C>T), GGT664GGA (5733T>A), CTG665TTG (5734C>T), ACC668ACG (5745C>G), CCT670CCC (5751T>C), AAT671AAC (5754T>C), CAG672CAA (5757G>A), GCC673GCA (5760C>A), GCT674GCA (5763T>A), GGG675GGT (5766G>T), GCT678GCG (5775T>G), AAG679AAA (5778G>A), CTG680TTG (5779C>T), TCA684AGT (5791T>A 5792C>G 5793A>T), GTG685GTT (5796G>T), AGC686TCG (5797A>T 5798G>C 5799C>G), GTA689GTC (5808A>C), GCT691GCC (5814T>C), GTA693GTT (5820A>T), GCT695GCC (5826T>C), AAC697AAT (5832C>T), CTT699CTG (5838T>G), AAG700AAA (5841G>A), TCT701AGT (5842T>A 5843C>G), GCT702GCG (5847T>G), AAG704AAA (5853G>A), CTG705TTA (5854C>T 5856G>A), CTG706CTC (5859G>C), GCT707GCA (5862T>A), GCT708GCC (5865T>C), GAG709GAA (5868G>A), GTC710GTT (5871C>T), AAA711AAG (5874A>G), AAG714AAA (5883G>A), ACT715ACG (5886T>G), GGT716GGC (5889T>C), GAG717GAA (5892G>A), ATT718ATC (5895T>C), CTC719CTG (5898C>G) | | | | | | | | |

*: Inserts / Deletes / Misaligned / Frameshifts

#### Analysis details

This analysis was performed with panviral2.64

### NGS Details (UN8): Duamitovirus peex1

#### Assembly

| | |
|---|---|
| **Coverage Length** | 135 (1 contig(s)) |
| **Depth Of Coverage** | 1.0 |
| **Number Of Reads** | 1 |
| **Reads Per Million** | 0.02 rpm (after QC) |
| **Ambiguities** | 0 |
| **Assembly Method** | de novo + reference guided assembly |
| **Consensus Caller** | Bcf Tools |

#### Coverage Map

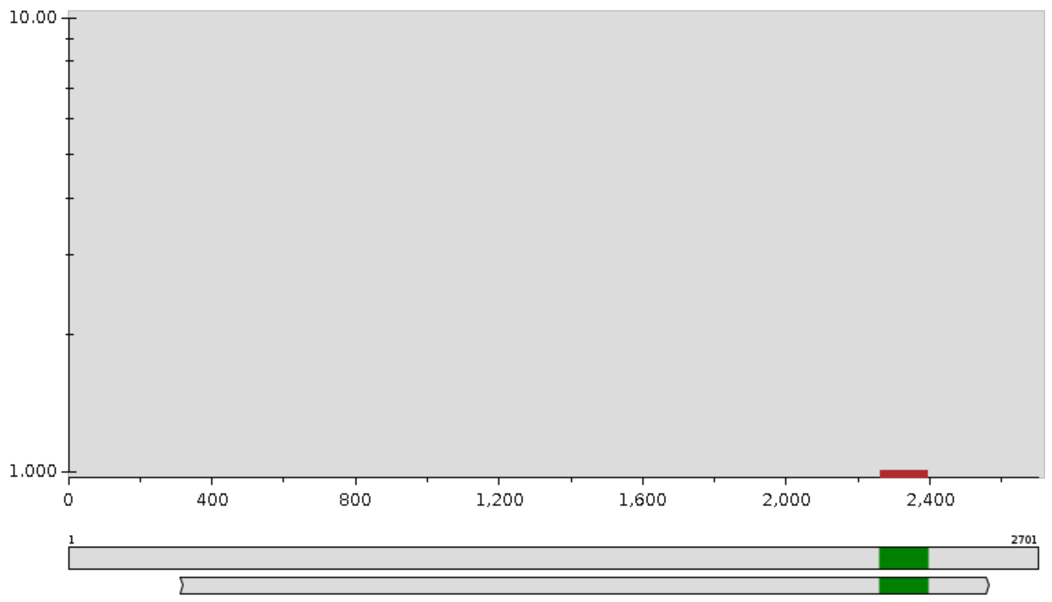

#### Assignment

| | |
|---|---|
| **Type** | Duamitovirus peex1 (Taxonomy ID: 2955799) |
| **Reference Genome** | NC_076525.1 |
| **NT Identity (%)** | 80.0 |
| **AA Identity (%)** | 80.0 |
| **Number Of Stop Codons** | 0 |
| **Number Of CDS** | 1 |

#### Alignment

| | |
|---|---|
| **Alignment Score** | 162.0 (NT) + 300.0 (AA) = 462.0 |
| **Concordance (%)** | 72.3005 |

| Alignment Method | Global, seeded, nucleotide + amino acids (AGA) |
|---|---|

#### Genome Region

Sequence starts at position 2262 and ends at position 2396 relative to NC_076525.1 reference sequence.

#### Alignment Detailed Statistics

| | Begin | End | Coverage | Score | Concordance | Matches | Identities | I/D/M/F* | Stop Codons |
|---|---|---|---|---|---|---|---|---|---|
| **NT** | **2262** | **2396** | **5.0%** | **162** | **60.0%** | **135 (100%)** | **108 (80.0%)** | **0/0** | |
| Mutations: | 2268G>A, 2270A>C, 2272G>T, 2274C>A, 2283G>A, 2292C>T, 2294T>G, 2295C>A, 2296G>C, 2297A>T, 2306A>T, 2315C>T, 2321G>A, 2325A>T, 2326A>T, 2327T>A, 2330C>A, 2333G>A, 2336A>T, 2345A>T, 2348C>T, 2354T>A, 2355A>T, 2357G>T, 2360A>T, 2363G>T, 2375G>C | | | | | | | | |
| **CDS** | | | | | | | | | |
| RdRp | 651 | 695 | 6.0% | 300 | 81.3% | 45 (100%) | 36 (80.0%) | 0/0/0/0 | 0 |
| Protein mutations: | V653I (2268G>A 2270A>C), R654L (2272G>T), H655N (2274C>A), E658K (2283G>A), R662T (2295C>A 2296G>C 2297A>T), N672L (2325A>T 2326A>T 2327T>A), E675D (2336A>T), M682F (2355A>T 2357G>T), E683D (2360A>T) | | | | | | | | |
| Codon mutations: | GTA653ATC (2268G>A 2270A>C), CGT654CTT (2272G>T), CAT655AAT (2274C>A), GAA658AAA (2283G>A), CTT661TTG (2292C>T 2294T>G), CGA662ACT (2295C>A 2296G>C 2297A>T), TCA665TCT (2306A>T), CAC668CAT (2315C>T), GTG670GTA (2321G>A), AAT672TTA (2325A>T 2326A>T 2327T>A), TCC673TCA (2330C>A), CCG674CCA (2333G>A), GAA675GAT (2336A>T), ATA678ATT (2345A>T), GAC679GAT (2348C>T), CTT681CTA (2354T>A), ATG682TTT (2355A>T 2357G>T), GAA683GAT (2360A>T), GTG684GTT (2363G>T), GCG688GCC (2375G>C) | | | | | | | | |
| **Proteins** | | | | | | | | | |
| RNA-dependent RNA polymerase (YP_010798875.1) | 651 | 695 | 6.0% | 300 | 81.3% | 45 (100%) | 36 (80.0%) | 0/0/0/0 | 0 |
| Protein mutations: | V653I (2268G>A 2270A>C), R654L (2272G>T), H655N (2274C>A), E658K (2283G>A), R662T (2295C>A 2296G>C 2297A>T), N672L (2325A>T 2326A>T 2327T>A), E675D (2336A>T), M682F (2355A>T 2357G>T), E683D (2360A>T) | | | | | | | | |
| Codon mutations: | GTA653ATC (2268G>A 2270A>C), CGT654CTT (2272G>T), CAT655AAT (2274C>A), GAA658AAA (2283G>A), CTT661TTG (2292C>T 2294T>G), CGA662ACT (2295C>A 2296G>C 2297A>T), TCA665TCT (2306A>T), CAC668CAT (2315C>T), GTG670GTA (2321G>A), AAT672TTA (2325A>T 2326A>T 2327T>A), TCC673TCA (2330C>A), CCG674CCA (2333G>A), GAA675GAT (2336A>T), ATA678ATT (2345A>T), GAC679GAT (2348C>T), CTT681CTA (2354T>A), ATG682TTT (2355A>T 2357G>T), GAA683GAT (2360A>T), GTG684GTT (2363G>T), GCG688GCC (2375G>C) | | | | | | | | |

*: Inserts / Deletes / Misaligned / Frameshifts

#### Analysis details

This analysis was performed with panviral2.64

### NGS Details (UN8): Cyvirus cyprinidallo3

#### Assembly

| | |
|---|---|
| **Coverage Length** | 90 (1 contig(s)) |
| **Depth Of Coverage** | 1.0 |
| **Number Of Reads** | 1 |
| **Reads Per Million** | 0.02 rpm (after QC) |
| **Ambiguities** | 0 |
| **Assembly Method** | de novo + reference guided assembly |
| **Consensus Caller** | Bcf Tools |

#### Coverage Map

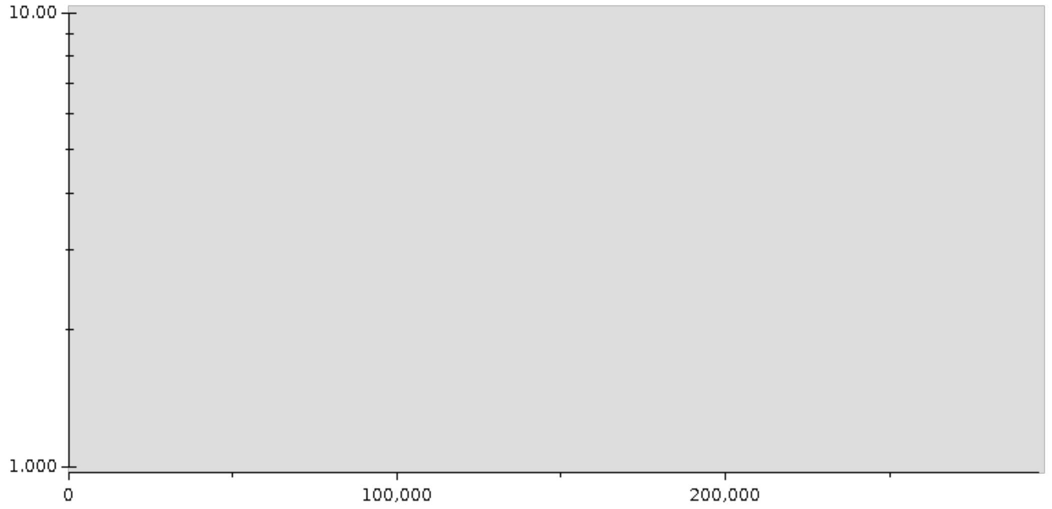

#### Assignment

| | |
|---|---|
| **Type** | Cyvirus cyprinidallo3 (Taxonomy ID: 3050271) |
| **Reference Genome** | NC_009127.1 |
| **NT Identity (%)** | 83.3333 |
| **AA Identity (%)** | 80.0 |
| **Number Of Stop Codons** | 0 |
| **Number Of CDS** | 163 |

#### Alignment

| | |
|---|---|
| **Alignment Score** | 120.0 (NT) + 177.0 (AA) = 297.0 |
| **Concordance (%)** | 74.25 |
| **Alignment Method** | Local, heuristic, nucleotide (BLASTN) |

#### Genome Region

Sequence starts at position 43055 and ends at position 43144 relative to NC_009127.1 reference sequence.

#### Alignment Detailed Statistics

| | Begin | End | Coverage | Score | Concordance | Matches | Identities | I/D/M/F* | Stop Codons |
|---|---|---|---|---|---|---|---|---|---|
| NT | 43055 | 43144 | 0.1% | 120 | 66.7% | 90 (100%) | 75 (83.3%) | 0/0 | |

Mutations: 43062A>G, 43065C>A, 43067G>C, 43068T>A, 43069G>A, 43082T>C, 43088G>A, 43097G>C, 43098T>C, 43100G>T, 43102A>C, 43103G>T, 43118T>C, 43133C>T, 43135C>G

*: Inserts / Deletes / Misaligned / Frameshifts

#### Analysis details

This analysis was performed with panviral2.64

### NGS Details (UN8): Diplodia scrobiculata RNA virus 1

#### Assembly

| | |
|---|---|
| **Coverage Length** | 120 (1 contig(s)) |
| **Depth Of Coverage** | 1.0 |
| **Number Of Reads** | 1 |
| **Reads Per Million** | 0.02 rpm (after QC) |
| **Ambiguities** | 0 |
| **Assembly Method** | de novo + reference guided assembly |
| **Consensus Caller** | Bcf Tools |

#### Coverage Map

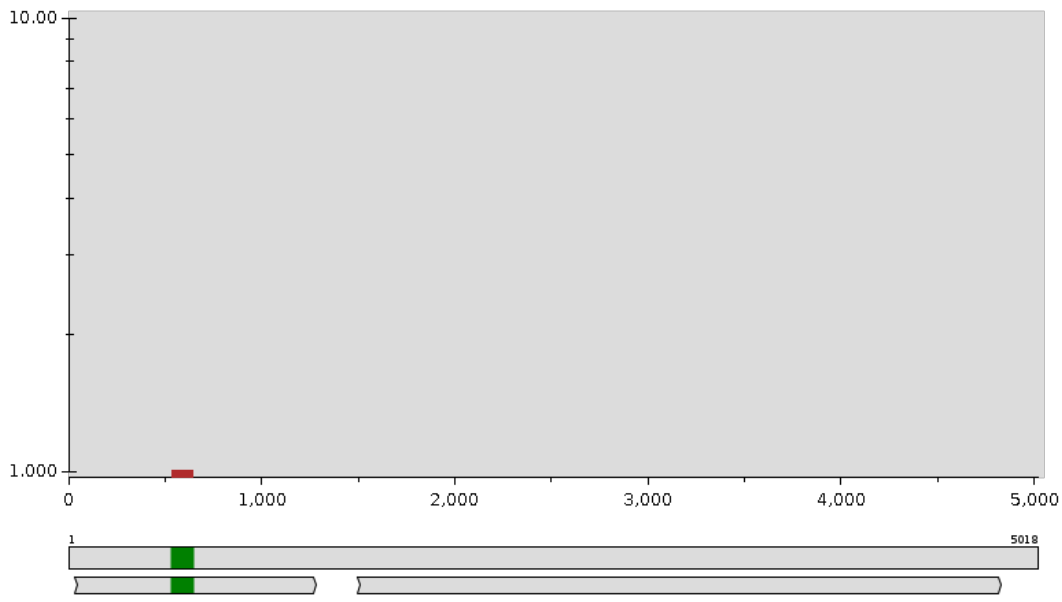

#### Assignment

| | |
|---|---|
| **Type** | Diplodia scrobiculata RNA virus 1 (Taxonomy ID: 459770) |
| **Reference Genome** | NC_013699.1 |
| **NT Identity (%)** | 80.8333 |
| **AA Identity (%)** | 80.0 |
| **Number Of Stop Codons** | 0 |
| **Number Of CDS** | 2 |

#### Alignment

| | |
|---|---|
| **Alignment Score** | 148.0 (NT) + 195.0 (AA) = 343.0 |
| **Concordance (%)** | 72.6695 |

| **Alignment Method** | Global, seeded, nucleotide + amino acids (AGA) |
|---|---|

#### Genome Region

Sequence starts at position 532 and ends at position 651 relative to NC_013699.1 reference sequence.

#### Alignment Detailed Statistics

| | Begin | End | Coverage | Score | Concordance | Matches | Identities | I/D/M/F* | Stop Codons |
|---|---|---|---|---|---|---|---|---|---|
| **NT** | **532** | **651** | **2.4%** | **148** | **61.7%** | **120 (100%)** | **97 (80.8%)** | **0/0** | |
| Mutations: | 536C>G, 560G>A, 564C>G, 566T>G, 581C>T, 585A>C, 587A>C, 590A>T, 603G>T, 608A>G, 616C>G, 617G>C, 621A>G, 623C>G, 624A>C, 625A>G, 626G>C, 628A>G, 629C>T, 632G>T, 641G>A, 644A>C, 648C>T | | | | | | | | |
| **CDS** | | | | | | | | | |
| DSRV-1_gp1 | 169 | 208 | 9.6% | 195 | 79.3% | 40 (100%) | 32 (80.0%) | 0/0/0/0 | 0 |
| Protein mutations: | P179A (564C>G 566T>G), E187D (590A>T), A192S (603G>T), A196G (616C>G 617G>C), I198V (621A>G 623C>G), K199R (624A>C 625A>G 626G>C), N200S (628A>G 629C>T) | | | | | | | | |
| Codon mutations: | CCC169CCG (536C>G), CAG177CAA (560G>A), CCT179GCG (564C>G 566T>G), CTC184CTT (581C>T), AGA186CGC (585A>C 587A>C), GAA187GAT (590A>T), GCG192TCG (603G>T), GAA193GAG (608A>G), GCG196GGC (616C>G 617G>C), ATC198GTG (621A>G 623C>G), AAG199CGC (624A>C 625A>G 626G>C), AAC200AGT (628A>G 629C>T), GTG201GTT (632G>T), CTG204CTA (641G>A), CTA205CTC (644A>C), CTG207TTG (648C>T) | | | | | | | | |
| **Proteins** | | | | | | | | | |
| hypothetical protein (YP_003359177.1) | 169 | 208 | 9.6% | 195 | 79.3% | 40 (100%) | 32 (80.0%) | 0/0/0/0 | 0 |
| Protein mutations: | P179A (564C>G 566T>G), E187D (590A>T), A192S (603G>T), A196G (616C>G 617G>C), I198V (621A>G 623C>G), K199R (624A>C 625A>G 626G>C), N200S (628A>G 629C>T) | | | | | | | | |
| Codon mutations: | CCC169CCG (536C>G), CAG177CAA (560G>A), CCT179GCG (564C>G 566T>G), CTC184CTT (581C>T), AGA186CGC (585A>C 587A>C), GAA187GAT (590A>T), GCG192TCG (603G>T), GAA193GAG (608A>G), GCG196GGC (616C>G 617G>C), ATC198GTG (621A>G 623C>G), AAG199CGC (624A>C 625A>G 626G>C), AAC200AGT (628A>G 629C>T), GTG201GTT (632G>T), CTG204CTA (641G>A), CTA205CTC (644A>C), CTG207TTG (648C>T) | | | | | | | | |

*: Inserts / Deletes / Misaligned / Frameshifts

#### Analysis details

This analysis was performed with panviral2.64

### Analysis of UN9

| | |
|---|---|
| **Technology** | Paired-end short reads |
| **Input Files** | UN9_R1.fq.gz (1.41 GB), UN9_R2.fq.gz (1.45 GB) |
| **Submitted On** | 2023-09-26 13:10:39 UTC |
| **Duration** | 2h 48m 21s |
| **Tool Version** | panviral2.64 |
| **Location** | https://www.genomedetective.com/db/ui/analysis/c136d63b-9925-4a37-b229-014ac43e54d9 |

#### Statistics

| | |
|---|---|
| **Original Read Length** | 20 - 150 |
| **Trimmed Read Length** | 50 - 135 |

| | # Reads | % of Reads |
|---|---|---|
| **Input file** | 44538018 | 100.0% |
| **After QC** | 44242186 | 99.3% |
| **After filtering** | 10664852 | 23.9% |
| **Mapped back to contigs** | 6008674 | 13.5% |

#### Assignments

| Assignment | No. of Reads | Depth of Coverage | Identity | | Genome Coverage | |
|---|---|---|---|---|---|---|
| | | | NT | AA | | |
| Torradovirus lycopersici (2 segments out of 2) | 4175010 | 42283.7 | 91.6% | 97.1% | 99.8% | |
| Torradovirus lycopersici (segment RNA 2) | 2524050 | 60857.9 | 94.8% | 96.5% | 99.6% | 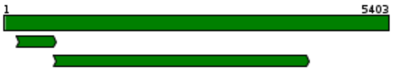 |
| Torradovirus lycopersici (segment RNA 1) | 1650960 | 29487.7 | 89.5% | 97.5% | 100.0% | 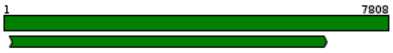 |
| Potato virus Y | 812319 | 11166.5 | 82.6% | 90.0% | 99.9% | 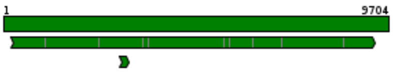 |
| Tomato chocolate spot virus (2 segments out of 2) | 490370 | 17490.7 | 68.2% | 69.6% | 26.5% | |
| Tomato chocolate spot virus (segment RNA2) | 328101 | 19230.5 | 64.0% | 66.1% | 40.8% | 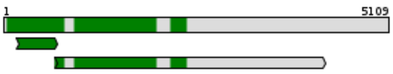 |
| Tomato chocolate spot virus (segment RNA 1) | 162269 | 14600.6 | 75.1% | 76.9% | 16.7% | 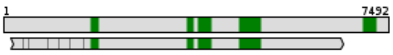 |
| Torradovirus marchitezum (2 segments out of 2) | 401174 | 16110.8 | 63.8% | 63.2% | 23.5% | |

| Assignment | No. of Reads | Depth of Coverage | Identity | | Genome Coverage | |
|---|---|---|---|---|---|---|
| | | | NT | AA | | |
| Torradovirus marchitezum (segment RNA 1) | 77945 | 7049.3 | 65.4% | 66.6% | 17.9% | 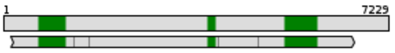 |
| Torradovirus marchitezum (segment RNA 2) | 323229 | 23655.4 | 62.3% | 59.9% | 31.7% | 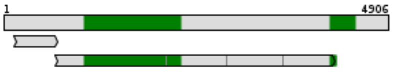 |
| Diachasmimorpha longicaudata entomopoxvirus (segment NC_043455.1) | 7050 | 776.0 | 59.3% | 55.9% | 83.4% | 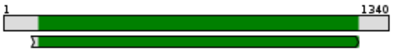 |
| Bracoviriform glomeratae (segment NC_043292.1) | 6038 | 2248.3 | 72.2% | 85.0% | 70.9% | 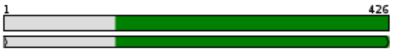 |
| Potato leafroll virus | 4587 | 104.8 | 97.6% | 96.6% | 97.3% | 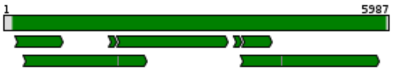 |
| Solendovirus venanicotianae | 233 | 11.9 | 79.9% | 76.3% | 28.5% | 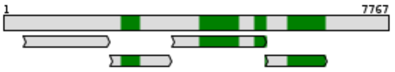 |
| Duamitovirus soch1 | 163 | 11.3 | 69.7% | 67.9% | 62.1% | 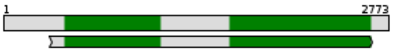 |
| Harvey murine sarcoma virus | 6 | 3.2 | 70.2% | 83.3% | 21.6% | 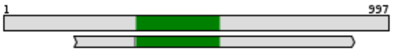 |

#### Discoveries

| Similar to | No. of Reads | Depth of Coverage | Identity | | Genome Coverage | |
|---|---|---|---|---|---|---|
| | | | NT | AA | | |
| Tomato necrotic dwarf virus (segment RNA1) | 65373 | 9641.5 | 71.6% | 77.5% | 11.0% | 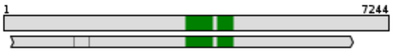 |
| Brazilian marseillevirus | 19918 | 9135.3 | 76.8% | 94.4% | 0.1% | 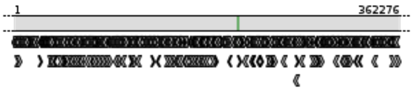 |
| Tokyovirus A1 | 10543 | 6408.0 | 81.1% | 94.4% | 0.0% | 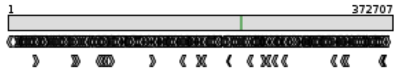 |
| Lausannevirus | 3601 | 2010.8 | 83.6% | 98.0% | 0.0% | 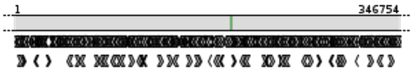 |
| Betabaculovirus disaccharalis | 2310 | 1356.2 | 84.7% | 92.0% | 0.2% | 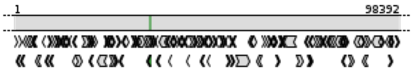 |
| Lausannevirus | 2249 | 390.8 | 78.4% | 91.0% | 0.1% | 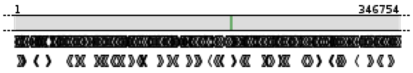 |
| Makelovirus prm1 | 1984 | 253.9 | 76.6% | 87.5% | 0.7% | 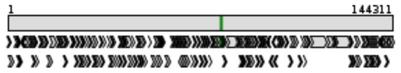 |

| Similar to | No. of Reads | Depth of Coverage | Identity | | Genome Coverage | |
|---|---|---|---|---|---|---|
| | | | NT | AA | | |
| Golden Marseillevirus | 1782 | 1085.8 | 85.8% | 97.4% | 0.0% | 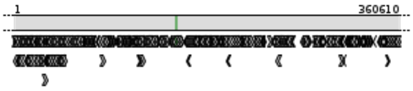 |
| Cladosporium fulvum T-1 virus | 808 | 54.0 | 52.5% | 47.0% | 23.2% | 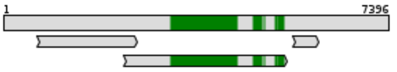 |
| Lausannevirus | 604 | 394.0 | 81.5% | 92.2% | 0.0% | 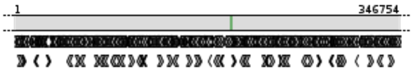 |
| Errantivirus | 402 | 65.5 | 53.6% | 43.9% | 7.6% | 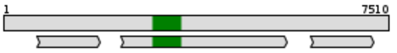 |
| Epiphyllum badnavirus 1 | 324 | 124.8 | 60.3% | 57.6% | 3.5% | 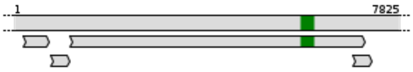 |
| Lowelvirus tuscon4d | 311 | 36.3 | 73.3% | 87.1% | 0.6% | 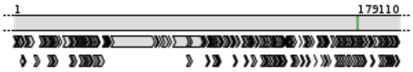 |
| Hibiscus bacilliform virus GD1 | 204 | 73.4 | 59.5% | 51.6% | 3.8% | 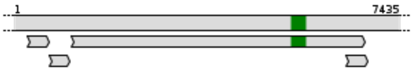 |
| Cassava brown streak virus | 197 | 45.5 | 60.5% | 57.5% | 5.8% | 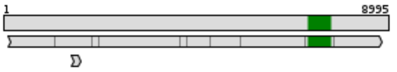 |
| Badnavirus maculaucubae | 137 | 52.7 | 56.1% | 52.6% | 3.1% | 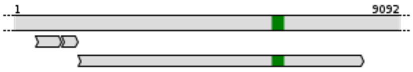 |
| Errantivirus | 111 | 19.5 | 52.9% | 41.2% | 8.0% | 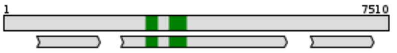 |
| Badnavirus maculaucubae | 81 | 26.2 | 55.8% | 46.5% | 3.6% | 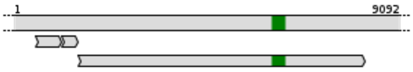 |
| Nodensvirus spm2 | 67 | 39.2 | 80.7% | 86.3% | 0.1% | 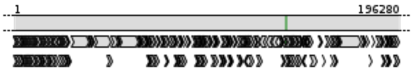 |
| Epiphyllum badnavirus 1 | 66 | 15.2 | 52.2% | 44.4% | 5.4% | 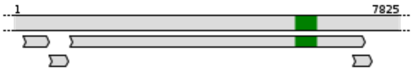 |
| Caulimovirus venafragariae | 60 | 21.2 | 54.2% | 42.1% | 4.1% | 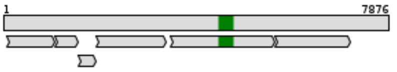 |
| Epiphyllum badnavirus 1 | 55 | 18.6 | 52.9% | 45.1% | 3.9% | 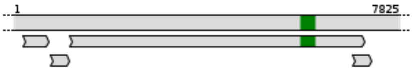 |
| Caulimovirus glycinis | 44 | 13.0 | 57.2% | 43.3% | 5.4% | 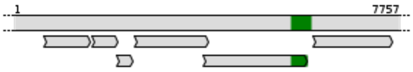 |
| Caulimovirus venafragariae | 41 | 6.4 | 56.2% | 54.5% | 10.1% | 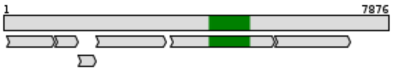 |

| Similar to | No. of Reads | Depth of Coverage | Identity | | Genome Coverage | |
|---|---|---|---|---|---|---|
| | | | NT | AA | | |
| Pinus nigra virus 1 | 41 | 19.8 | 56.3% | 50.0% | 3.4% | 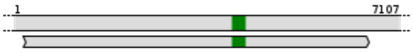 |
| Hibiscus bacilliform virus GD1 | 37 | 9.2 | 52.2% | 43.0% | 6.7% | 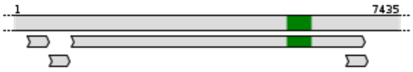 |
| Dioscovirus dioscoreae | 27 | 4.9 | 64.1% | 56.1% | 9.1% | 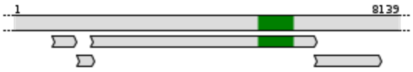 |
| Badnavirus venatheobromae | 25 | 11.2 | 50.6% | 44.8% | 3.5% | 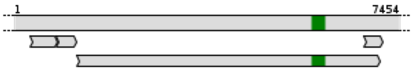 |
| Epiphyllum badnavirus 1 | 25 | 10.2 | 57.0% | 43.0% | 3.3% | 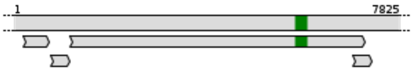 |
| Badnavirus tessellocastaneae | 22 | 6.4 | 57.2% | 47.9% | 5.8% | 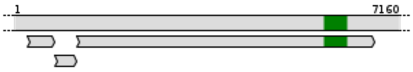 |
| Ruflodivirus deformatiorudbeckiae | 21 | 4.7 | 57.3% | 45.5% | 7.4% | 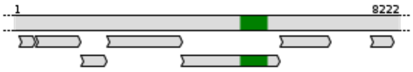 |
| Caulimovirus latensarmoraciae | 21 | 7.3 | 58.8% | 56.2% | 4.6% | 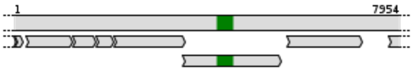 |
| Badnavirus occultipomeae | 18 | 5.4 | 54.9% | 46.2% | 4.3% | 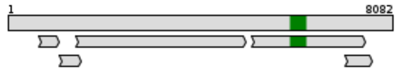 |
| Badnavirus occultipomeae | 15 | 6.7 | 61.5% | 47.2% | 3.2% | 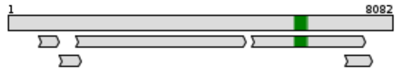 |
| Badnavirus maculaucubae | 15 | 5.3 | 52.7% | 43.2% | 3.8% | 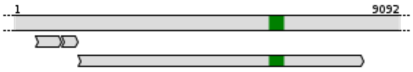 |
| Bracoviriform congregatae (segment Circle 7) | 14 | 9.0 | 78.4% | 91.3% | 0.6% | 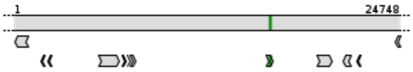 |
| Badnavirus epsiloninflatheobromae | 14 | 5.0 | 59.0% | 49.5% | 4.1% | 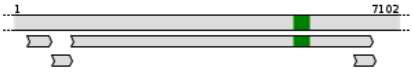 |
| Badnavirus maculasmallanthi | 14 | 4.2 | 58.0% | 45.9% | 4.2% | 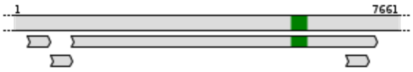 |
| Arhar cryptic virus-I (segment RNA 3) | 14 | 5.6 | 57.7% | 42.6% | 21.7% | 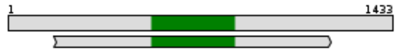 |
| Ichnoviriform fugitivi (2 segments out of 56) | 12 | 3.2 | 60.4% | 50.0% | 0.1% | |
| Ichnoviriform fugitivi (segment B17) | 10 | 5.2 | 61.0% | 50.9% | 4.0% | 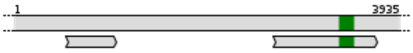 |
| Ichnoviriform fugitivi (segment C16) | 2 | 1.3 | 59.8% | 49.2% | 3.8% | 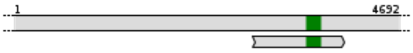 |

| Similar to | No. of Reads | Depth of Coverage | Identity | | Genome Coverage | |
|---|---|---|---|---|---|---|
| | | | NT | AA | | |
| Dioscovirus dioscoreae | 11 | 4.2 | 57.0% | 49.5% | 4.1% | 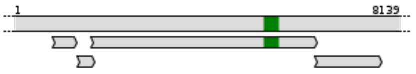 |
| Dioscovirus dioscoreae | 11 | 4.1 | 66.3% | 55.6% | 4.0% | 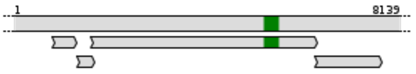 |
| Badnavirus maculasmallanthi | 10 | 4.2 | 53.0% | 48.5% | 3.8% | 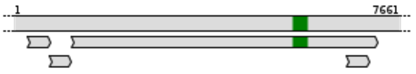 |
| Pinus nigra virus 1 | 10 | 6.0 | 57.7% | 47.4% | 3.4% | 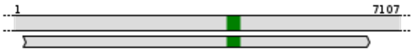 |
| Badnavirus occultipomeae | 9 | 4.2 | 58.7% | 51.8% | 3.1% | 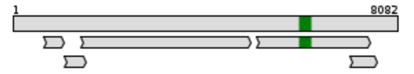 |
| Epiphyllum badnavirus 1 | 9 | 4.0 | 58.0% | 51.9% | 3.1% | 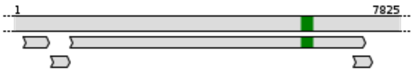 |
| Colombian datura virus | 8 | 2.3 | 75.5% | 79.0% | 4.5% | 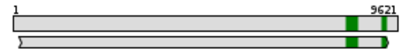 |
| Pinus nigra virus 1 | 8 | 2.7 | 55.3% | 50.4% | 5.0% | 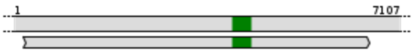 |
| Badnavirus rutilanscamelliae | 7 | 2.6 | 55.3% | 45.1% | 4.1% | 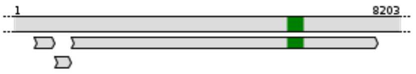 |
| Fig cryptic virus (segment RNA 2) | 6 | 3.1 | 63.4% | 61.8% | 11.6% | 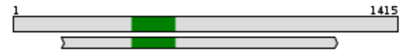 |
| Caulimovirus venafragariae | 6 | 2.8 | 61.3% | 63.0% | 3.1% | 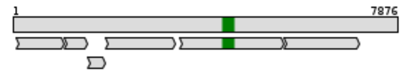 |
| Tomato associated geminivirus 1 | 6 | 2.5 | 67.5% | 62.0% | 10.5% | 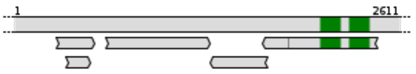 |
| Moloney murine leukemia virus | 2 | 2.0 | 79.1% | 97.7% | 1.6% | 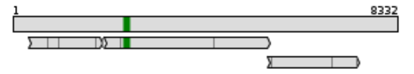 |
| Friend murine leukemia virus | 2 | 2.0 | 78.5% | 97.8% | 1.6% | 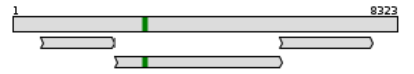 |
| Tokyovirus A1 | 3 | 2.4 | 80.0% | 92.9% | 0.0% | 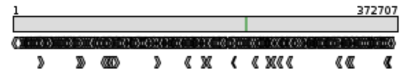 |
| Nepovirus aeonii (segment RNA2) | 2 | 1.1 | 71.2% | 72.3% | 6.2% | 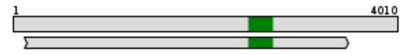 |
| Duamitovirus peex1 | 1 | 1.0 | 71.9% | 66.7% | 5.0% | 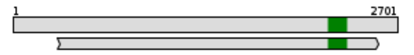 |
| Plantago lanceolata latent virus | 1 | 1.0 | 67.4% | 54.1% | 4.8% | 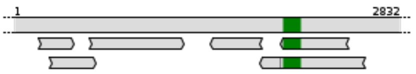 |

| Similar to | No. of Reads | Depth of Coverage | Identity | | Genome Coverage | |
|---|---|---|---|---|---|---|
| | | | NT | AA | | |
| Duamitovirus dapi1 | 1 | 1.0 | 68.7% | 68.2% | 4.8% | 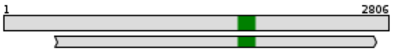  |

### NGS Details (UN9): Torradovirus marchitezum (segment RNA 1)

#### Assembly

| | |
|---|---|
| **Coverage Length** | 1293 (3 contig(s)) |
| **Depth Of Coverage** | 7049.3 |
| **Number Of Reads** | 77945 |
| **Reads Per Million** | 1761.78 rpm (after QC) |
| **Ambiguities** | 0 |
| **Assembly Method** | de novo + reference guided assembly |
| **Consensus Caller** | Bcf Tools |

#### Coverage Map

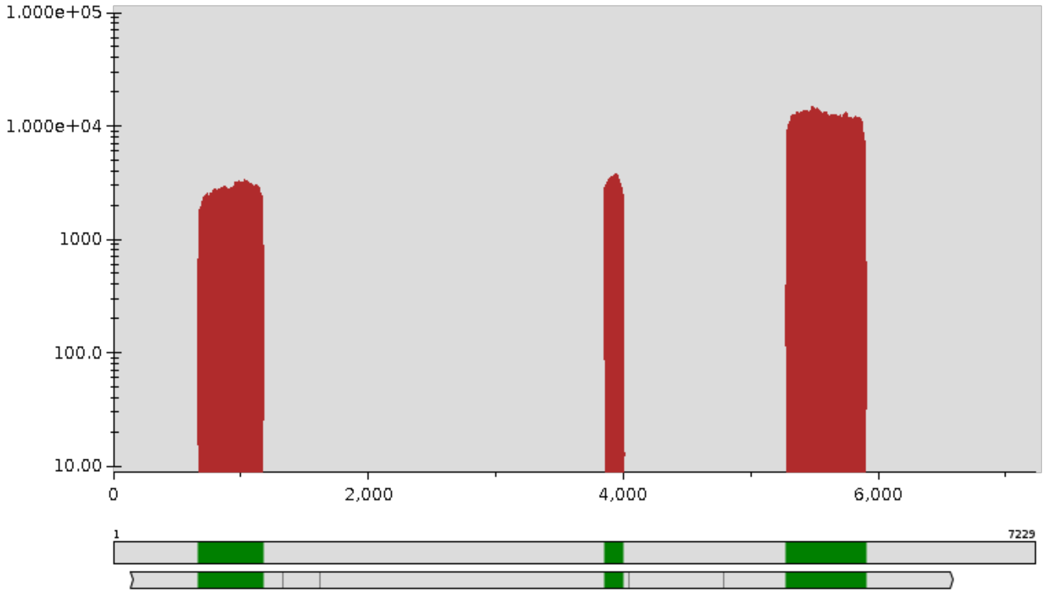

#### Assignment

| | |
|---|---|
| **Type** | Torradovirus marchitezum (Taxonomy ID: 3048376) |
| **Reference Genome** | NC_010987.1 |
| **NT Identity (%)** | 65.4349 |
| **AA Identity (%)** | 66.5899 |
| **Number Of Stop Codons** | 0 |
| **Number Of CDS** | 1 |

#### Alignment

| | |
|---|---|
| **Alignment Score** | 781.0 (NT) + 2123.0 (AA) = 2904.0 |
| **Concordance (%)** | 52.071 |

| | |
|---|---|
| **Alignment Method** | Global, seeded, nucleotide + amino acids (AGA) |

#### Genome Region

Sequence starts at position 666 and ends at position 5905 relative to NC_010987.1 reference sequence.

#### Alignment Detailed Statistics

| | Begin | End | Coverage | Score | Concordance | Matches | Identities | I/D/M/F* | Stop Codons |
|---|---|---|---|---|---|---|---|---|---|
| **NT** | **666** | **5905** | **17.9%** | **781** | **30.4%** | **1290 (99.1%)** | **850 (65.3%)** | **9/3** | |
| Mutations: | 677C>T, 681T>A, 683A>T, 684G>A, 685A>G, 695A>C, 696A>C, 697A>T, 698A>C, 702G>T, 704C>T, 707A>G, 708C>A, 709G>A, 713C>T, 714A>T, 716C>T, 717A>C, 718T>A, 720G>A, 725C>A, 728A>G, 732A>T, 733G>C, 735A>G, 740T>A, 746G>T, 749A>G, 750T>A, 751C>G, 752A>C, 753G>T, 755T>G, 756T>A, 757C>A, 758A>T, 770G>C, 771T>A, 772C>G, 776G>A, 780A>C, 785C>T, 786A>T, 788A>T, 789A>T, 791A>G, 792T>A, 794G>A, 797T>A, 803G>C, 806T>C, 809G>A, 818G>A, 821C>T, 822A>G, 824T>G, 830G>A, 839A>G, 849T>C, 851G>T, 853G>C, 854C>A, 860A>G, 861T>C, 863G>A, 873C>T, 875C>G, 881G>A, 882A>G, 884A>G, 887A>C, 893T>A, 899G>A, 914A>C, 918A>C, 919C>A, 920T>A, 921C>T, 923T>G, 926C>T, 927C>A, 928G>A, 929T>G, 932T>C, 938A>G, 947C>A, 962A>T, 968T>G, 969C>T, 971T>G, 977A>T, 978A>G, 980A>T, 982T>C, 983C>T, 985G>A, 986C>T, 988C>T, 989C>A, 992T>C, 995A>G, 998C>T, 1001A>G, 1004T>C, 1010G>T, 1011T>C, 1016A>T, 1019T>A, 1022A>T, 1031G>A, 1032C>A, 1034G>T, 1046G>C, 1049C>T, 1059C>A, 1060A>C, 1061G>T, 1064G>A, 1068G>T, 1069A>C, 1077T>G, 1078G>C, 1091G>A, 1092C>A, 1094C>G, 1100G>A, 1109C>T, 1113A>C, 1114A>C, 1115G>T, 1121C>T, 1122C>T, 1124T>A, 1126C>T, 1142G>T, 1144G>A, 1145G>A, 1148A>G, 1149G>C, 1151A>G, 1154T>A, 1163C>A, 1169T>C, 3863G>C, 3872T>A, 3875C>T, 3881T>C, 3884C>T, 3890C>T, 3893G>A, 3894A>C, 3899G>T, 3900A>T, 3905G>T, 3907T>C, 3908G>C, 3911C>T, 3914G>A, 3917A>G, 3923T>C, 3926G>T, 3927C>G, 3930T>G, 3932C>A, 3933A>C, 3934C>A, 3935T>G, 3937G>A, 3938C>G, 3939G>A, 3940A>G, 3941A>T, 3942G>A, 3943G>C, 3944A>C, 3945G>C, 3947G>A, 3950A>G, 3953G>A, 3954G>C, 3955A>T, 3962A>G, 3966G>A, 3968C>T, 3969A>T, 3977A>T, 3980T>C, 3981G>C, 3992A>G, 5288T>A, 5294T>G, 5300T>G, 5303A>T, 5309C>T, 5333G>A, 5339G>C, 5346A>T, 5349T>G, 5354T>A, 5357A>C, 5358A>C, 5359T>A, 5364G>C, 5365C>A, 5366C>T, 5370A>C, 5371A>G, 5372G>C, 5375T>G, 5376G>A, 5377T>G, 5378G>T, 5379T>C, 5383C>A, 5386G>T, 5387A>T, 5390A>T, 5391C>G, 5395T>G, 5396A>G, 5399A>T, 5403A>T, 5404A>C, 5409G>A, 5410T>C, 5411T>C, 5417G>T, 5423A>T, 5429A>T, 5435T>A, 5436G>A, 5438C>A, 5440G>C, 5441C>T, 5447T>C, 5456G>A, 5457C>T, 5460A>T, 5462G>T, 5465C>A, 5468A>G, 5471A>C, 5474A>T, 5475A>C, 5476G>A, 5477G>T, 5483T>C, 5484T>A, 5486T>G, 5488T>C, 5489G>C, 5490G>A, 5491A>C, 5492G>A, 5496A>C, 5498A>T, 5501A>T, 5504T>G, 5505A>C, 5507A>C, 5508T>C, 5510G>A, 5513C>A, 5516A>C, 5519T>G, 5521G>C, 5522A>T, 5523A>G, 5524A>G, 5525T>G, 5531T>G, 5534C>T, 5537C>G, 5540T>C, 5546T>C, 5555C>T, 5560T>C, 5564G>A, 5567T>C, 5570T>G, 5573T>C, 5574C>A, 5582G>T, 5583C>G, 5584C>A, 5585T>A, 5586C>G, 5588T>C, 5588_5589insACC, 5589G>T, 5590G>C, 5592G>C, 5597A>G, 5598G>T, 5602C>G, 5603T>A, 5606A>G, 5612T>C, 5613G>A, 5617C>A, 5618G>T, 5619C>G, 5620T>G, 5622_5624delCAC, 5626G>C, 5627T>C, 5629G>C, 5635T>A, 5636G>C, 5642A>T, 5642_5643insCACAGT, 5644G>C, 5645G>A, 5649G>T, 5650A>C, 5652T>G, 5654G>T, 5657C>A, 5658A>C, 5659A>T, 5660C>T, 5663C>A, 5664C>A, 5666C>G, 5669T>A, 5670A>G, 5672A>T, 5674A>C, 5676G>T, 5677G>C, 5681T>C, 5682T>C, 5684A>C, 5685G>T, 5687A>T, 5688A>G, 5691C>A, 5695C>T, 5696A>G, 5700A>G, 5702G>T, 5705G>T, 5706A>T, 5714T>C, 5715G>A, 5716T>C, 5717G>A, 5718G>A, 5720A>T, 5723A>T, 5726A>T, 5727C>G, 5729T>A, 5735A>T, 5740G>C, 5741A>C, 5742A>C, 5744T>G, 5746T>G, 5747T>C, 5750C>A, 5753A>G, 5754C>A, 5757G>A, 5759T>C, 5762T>G, 5763G>A, 5766A>C, 5767G>A, 5769T>G, 5770T>C, 5771G>T, 5772C>T, 5774A>G, 5775C>G, 5777C>G, 5778T>A, 5779A>C, 5780T>A, 5784G>T, 5785A>C, 5786G>T, 5787C>G, 5788A>T, 5789G>C, 5790G>A, 5792A>T, 5798A>T, 5799G>A, 5800A>C, 5801G>T, 5802T>C, 5804A>T, 5805A>G, 5806T>C, 5807T>C, 5808C>G, 5809A>C, 5812A>G, 5813G>A, 5814A>G, 5815A>G, 5816T>A, 5817A>G, 5818C>T, 5819A>C, 5823C>G, 5825A>G, 5828T>A, 5829C>T, 5837C>T, 5843G>A, 5844C>T, 5846G>C, 5847A>T, 5848G>C, 5849T>C, 5850A>C, 5851T>A, 5852G>A, 5853G>C, 5855A>C, 5856A>G, 5859G>T, 5860T>G, 5861G>T, 5862C>G, 5865C>A, 5866A>T, 5867A>G, 5868C>A, 5870G>A, 5871A>C, 5873A>T, 5876T>G, 5880G>C, 5882T>G, 5883A>G, 5885G>A, 5886C>A, 5887T>G, 5891G>A, 5895T>G | | | | | | | | |
| **CDS** | | | | | | | | | |
| ToMarV_RNA1gp1 | 176 | 1922 | 20.1% | 2123 | 70.1% | 431 (99.1%) | 289 (66.4%) | 3/1/0/0 | 0 |
| Protein mutations: | L181I (681T>A 683A>T), D182S (684G>A 685A>G), K186L (696A>C 697A>T 698A>C), A188S (702G>T 704C>T), R190K (708C>A 709G>A), T192S (714A>T 716C>T), M193Q (717A>C 718T>A), V194M (720G>A), M199V (735A>G), V205L (753G>T 755T>G), S206N (756T>A 757C>A 758A>T), M214L (780A>C), T216S (786A>T 788A>T), I217L (789A>T 791A>G), L218I (792T>A 794G>A), I228V (822A>G 824T>G), S238T (853G>C 854C>A), T248A (882A>G 884A>G), T260Q (918A>C 919C>A 920T>A), R263K (927C>A 928G>A 929T>G), I280V (978A>G 980A>T), V281A (982T>C 983C>T), G282D (985G>A 986C>T), T283I (988C>T 989C>A), L298I (1032C>A 1034G>T), Q307T (1059C>A 1060A>C 1061G>T), D310S (1068G>T 1069A>C), C313A (1077T>G 1078G>C), K325P (1113A>C 1114A>C 1115G>T), P328S (1122C>T 1124T>A), T329I (1126C>T), R335K (1144G>A 1145G>A), E337Q (1149G>C 1151A>G), M1252L (3894A>C), M1254L (3900A>T), M1256T (3907T>C 3908G>C), Q1263E (3927C>G), S1264A (3930T>G 3932C>A), T1265Q (3933A>C 3934C>A 3935T>G), G1266E (3937G>A 3938C>G), E1267S (3939G>A 3940A>G 3941A>T), G1268T (3942G>A 3943G>C 3944A>C), E1269Q (3945G>C 3947G>A), E1272L (3954G>C 3955A>T), D1276N (3966G>A 3968C>T), T1277S (3969A>T), E1281Q (3981G>C), N1716K (5288T>A), M1731I (5333G>A), M1736L (5346A>T), S1737A (5349T>G), M1740Q (5358A>C 5359T>A), A1742H (5364G>C 5365C>A 5366C>T), K1744R (5370A>C 5371A>G 5372G>C), N1745K (5375T>G), V1746S (5376G>A 5377T>G 5378G>T), A1748E (5383C>A), G1749V (5386G>T 5387A>T), H1751D (5391C>G), V1752G (5395T>G 5396A>G), N1755S (5403A>T 5404A>C), V1757T (5409G>A 5410T>C 5411T>C), V1766I (5436G>A 5438C>A), G1767A (5440G>C 5441C>T), M1774F (5460A>T 5462G>T), R1779H (5475A>C 5476G>A 5477G>T), C1782R (5484T>A 5486T>G), V1783A (5488T>C 5489G>C), E1784T (5490G>A 5491A>C 5492G>A), I1786L (5496A>C 5498A>T), I1793M (5519T>G), G1794A (5521G>C 5522A>T), N1795G (5523A>G 5524A>G 5525T>G), V1807A (5560T>C), F1810L (5570T>G), Q1812K (5574C>A), P1815E (5583C>G 5584C>A 5585T>A), H1816D (5586C>G 5588T>C), H1816_G1817insT (5588_5589insACC), G1817S (5589G>T 5590G>C), D1818H (5592G>C), V1820L (5598G>T), A1821G (5602C>G 5603T>A), A1825T (5613G>A), S1826Y (5617C>A 5618G>T), L1827G (5619C>G 5620T>G), H1828del (5622_5624delCAC), S1829T (5626G>C 5627T>C), S1830T (5629G>C), M1832N (5635T>A 5636G>C), Q1834H (5642A>T), Q1834_G1835insHS (5642_5643insCACAGT), G1835A (5644G>C 5645G>A), E1837S (5649G>T 5650A>C), L1838V (5652T>G 5654G>T), N1840L (5658A>C 5659A>T 5660C>T), I1844V (5670A>G 5672A>T), H1845P (5674A>C), G1846S (5676G>T 5677G>C), A1849S (5685G>T 5687A>T), K1850E (5688A>G), Q1851K (5691C>A), T1852M (5695C>T 5696A>G), M1854V (5700A>G 5702G>T), M1856L (5706A>T), V1859T (5715G>A 5716T>C 5717G>A), V1860I (5718G>A 5720A>T), H1863E (5727C>G 5729T>A), G1867A (5740G>C 5741A>C), N1868Q (5742A>C 5744T>G), F1869C (5746T>G 5747T>C), Q1872K (5754C>A), D1873N (5757G>A 5759T>C), E1875K (5763G>A), S1876H (5766A>C 5767G>A), L1877A (5769T>G 5770T>C 5771G>T), H1879E (5775C>G 5777C>G), Y1880T (5778T>A 5779A>C 5780T>A), E1882S (5784G>T 5785A>C 5786G>T), Q1883V (5787C>G 5788A>T 5789G>C), G1884S (5790G>A 5792A>T), E1887T (5799G>A 5800A>C 5801G>T), I1889A (5805A>G 5806T>C 5807T>C), Q1890A (5808C>G 5809A>C), K1891R (5812A>G 5813G>A), N1892G (5814A>G 5815A>G 5816T>A), T1893V (5817A>G 5818C>T 5819A>C), L1895V (5823C>G 5825A>G), L1902F (5844C>T 5846G>C), M1904Q (5850A>C 5851T>A 5852G>A), E1905H (5853G>C 5855A>C), N1906D (5856A>G), V1907C (5859G>T 5860T>G 5861G>T), Q1908E (5862C>G), Q1909M (5865C>A 5866A>T 5867A>G), L1910I (5868C>A 5870G>A), I1911L (5871A>C 5873A>T), D1912E (5876T>G), V1914L (5880G>C 5882T>G), K1915E (5883A>G 5885G>A), L1916S (5886C>A 5887T>G), F1919V (5895T>G) | | | | | | | | |

| | Begin | End | Coverage | Score | Concordance | Matches | Identities | I/D/M/F* | Stop Codons |
|---|---|---|---|---|---|---|---|---|---|
| **NT** | **666** | **5905** | **17.9%** | **781** | **30.4%** | **1290 (99.1%)** | **850 (65.3%)** | **9/3** | |

Codon mutations:

TAC179TAT (677C>T), TTA181ATT (681T>A 683A>T), GAT182AGT (684G>A 685A>G), CCA185CCC (695A>C), AAA186CTC (696A>C 697A>T 698A>C), GCC188TCT (702G>T 704C>T), GAA189GAG (707A>G), CGA190AAA (708C>A 709G>A), TTC191TTT (713C>T), ACC192TCT (714A>T 716C>T), ATG193CAG (717A>C 718T>A), GTG194ATG (720G>A), GCC195GCA (725C>A), CAA196CAG (728A>G), AGT198TCT (732A>T 733G>C), ATG199GTG (735A>G), CTT200CTA (740T>A), GGG202GGT (746G>T), GAA203GAG (749A>G), TCA204AGC (750T>A 751C>G 752A>C), GTT205TTG (753G>T 755T>G), TCA206AAT (756T>A 757C>A 758A>T), ACG210ACC (770G>C), TCT211AGT (771T>A 772C>G), CAG212CAA (776G>A), ATG214CTG (780A>C), TCC215TCT (785C>T), ACA216TCT (786A>T 788A>T), ATA217TTG (789A>T 791A>G), TTG218ATA (792T>A 794G>A), GCT219GCA (797T>A), GGG221GGC (803G>C), ATT222ATC (806T>C), TCG223TCA (809G>A), GTG226GTA (818G>A), TTC227TTT (821C>T), ATT228GTG (822A>G 824T>G), GTG230GTA (830G>A), AGA233AGG (839A>G), TTG237CTT (849T>C 851G>T), AGC238ACA (853G>C 854C>A), CCA240CCG (860A>G), TTG241CTA (861T>C 863G>A), CTC245TTG (873C>T 875C>G), TCG247TCA (881G>A), ACA248GCG (882A>G 884A>G), GGA249GGC (887A>C), GGT251GGA (893T>A), CAG253CAA (899G>A), CTA258CTC (914A>C), ACT260CAA (918A>C 919C>A 920T>A), CTT261TTG (921C>T 923T>G), TAC262TAT (926C>T), CGT263AAG (927C>A 928G>A 929T>G), AAT264AAC (932T>C), AAA266AAG (938A>G), ACC269ACA (947C>A), TCA274TCT (962A>T), GTT276GTG (968T>G), CTT277TTG (969C>T 971T>G), ATA279ATT (977A>T), ATA280GTT (978A>G 980A>T), GTC281GCT (982T>C 983C>T), GGC282GAT (985G>A 986C>T), ACC283ATA (988C>T 989C>A), TTT284TTC (992T>C), GGA285GGG (995A>G), TTC286TTT (998C>T), AAA287AAG (1001A>G), AAT288AAC (1004T>C), GTG290GTT (1010G>T), TTG291CTG (1011T>C), TCA292TCT (1016A>T), GCT293GCA (1019T>A), ATA294ATT (1022A>T), ACG297ACA (1031G>A), CTG298ATT (1032C>A 1034G>T), CTG302CTC (1046G>C), TTC303TTT (1049C>T), CAG307ACT (1059C>A 1060A>C 1061G>T), GAG308GAA (1064G>A), GAT310TCT (1068G>T 1069A>C), TGT313GCT (1077T>G 1078G>C), ACG317ACA (1091G>A), CGC318AGG (1092C>A 1094C>G), GAG320GAA (1100G>A), GCC323GCT (1109C>T), AAG325CCT (1113A>C 1114A>C 1115G>T), TTC327TTT (1121C>T), CCT328TCA (1122C>T 1124T>A), ACC329ATC (1126C>T), CTG334CTT (1142G>T), AGG335AAA (1144G>A 1145G>A), GAA336GAG (1148A>G), GAA337CAG (1149G>C 1151A>G), GCT338GCA (1154T>A), ATC341ATA (1163C>A), GCT343GCC (1169T>C), CTG1241CTC (3863G>C), TCT1244TCA (3872T>A), TGC1245TGT (3875C>T), TAT1247TAC (3881T>C), CCC1248CCT (3884C>T), AAC1250AAT (3890C>T), AAG1251AAA (3893G>A), ATG1252CTG (3894A>C), GGG1253GGT (3899G>T), ATG1254TTG (3900A>T), GGG1255GGT (3905G>T), ATG1256ACC (3907T>C 3908G>C), AAC1257AAT (3911C>T), AAG1258AAA (3914G>A), AGA1259AGG (3917A>G), TTT1261TTC (3923T>C), GTG1262GTT (3926G>T), CAA1263GAA (3927C>G), TCC1264GCA (3930T>G 3932C>A), ACT1265CAG (3933A>C 3934C>A 3935T>G), GGC1266GAG (3937G>A 3938C>G), GAA1267AGT (3939G>A 3940A>G 3941A>T), GGA1268ACC (3942G>A 3943G>C 3944A>C), GAG1269CAA (3945G>C 3947G>A), AGA1270AGG (3950A>G), GTG1271GTA (3953G>A), GAA1272CTA (3954G>C 3955A>T), AAA1274AAG (3962A>G), GAC1276AAT (3966G>A 3968C>T), ACT1277TCT (3969A>T), GTA1279GTT (3977A>T), TTT1280TTC (3980T>C), GAA1281CAA (3981G>C), GAA1284GAG (3992A>G), AAT1716AAA (5288T>A), GTT1718GTG (5294T>G), GGT1720GGG (5300T>G), CCA1721CCT (5303A>T), AAC1723AAT (5309C>T), ATG1731ATA (5333G>A), CCG1733CCC (5339G>C), ATG1736TTG (5346A>T), TCA1737GCA (5349T>G), GCT1738GCA (5354T>A), ATA1739ATC (5357A>C), ATG1740CAG (5358A>C 5359T>A), GCC1742CAT (5364G>C 5365C>A 5366C>T), AAG1744CGC (5370A>C 5371A>G 5372G>C), AAT1745AAG (5375T>G), GTG1746AGT (5376G>A 5377T>G 5378G>T), TTG1747CTG (5379T>C), GCA1748GAA (5383C>A), GGA1749GTT (5386G>T 5387A>T), GGA1750GGT (5390A>T), CAT1751GAT (5391C>G), GTA1752GGG (5395T>G 5396A>G), GCA1753GCT (5399A>T), AAT1755TCT (5403A>T 5404A>C), GTT1757ACC (5409G>A 5410T>C 5411T>C), GTG1759GTT (5417G>T), GGA1761GGT (5423A>T), GGA1763GGT (5429A>T), GCT1765GCA (5435T>A), GTC1766ATA (5436G>A 5438C>A), GGC1767GCT (5440G>C 5441C>T), ATT1769ATC (5447T>C), GCG1772GCA (5456G>A), CTG1773TTG (5457C>T), ATG1774TTT (5460A>T 5462G>T), GCC1775GCA (5465C>A), TTA1776TTG (5468A>G), GCA1777GCC (5471A>C), ACA1778ACT (5474A>T), AGG1779CAT (5475A>C 5476G>A 5477G>T), TAT1781TAC (5483T>C), TGT1782AGG (5484T>A 5486T>G), GTG1783GCC (5488T>C 5489G>C), GAG1784ACA (5490G>A 5491A>C 5492G>A), ATA1786CTT (5496A>C 5498A>T), ATA1787ATT (5501A>T), GTT1788GTG (5504T>G), AGA1789CGC (5505A>C 5507A>C), TTG1790CTA (5508T>C 5510G>A), CGC1791CGA (5513C>A), ACA1792ACC (5516A>C), ATT1793ATG (5519T>G), GGA1794GCT (5521G>C 5522A>T), AAT1795GGG (5523A>G 5524A>G 5525T>G), CTT1797CTG (5531T>G), GAC1798GAT (5534C>T), CTC1799CTG (5537C>G), AAT1800AAC (5540T>C), TAT1802TAC (5546T>C), GGC1805GGT (5555C>T), GTG1807GCG (5560T>C), CAG1808CAA (5564G>A), TAT1809TAC (5567T>C), TTT1810TTG (5570T>G), CTT1811CTC (5573T>C), CAA1812AAA (5574C>A), GTG1814GTT (5582G>T), CCT1815GAA (5583C>G 5584C>A 5585T>A), CAT1816GAC (5586C>G 5588T>C), CAT1816_GGA1817insACC (5588_5589insACC), GGA1817TCA (5589G>T 5590G>C), GAC1818CAC (5592G>C), AAA1819AAG (5597A>G), GTG1820TTG (5598G>T), GCT1821GGA (5602C>G 5603T>A), CAA1822CAG (5606A>G), GGT1824GGC (5612T>C), GCA1825ACA (5613G>A), TCG1826TAT (5617C>A 5618G>T), CTC1827GGC (5619C>G 5620T>G), CAC1828del (5622_5624delCAC), AGT1829ACC (5626G>C 5627T>C), AGT1830ACT (5629G>C), ATG1832AAC (5635T>A 5636G>C), CAA1834CAT (5642A>T), CAA1834_GGG1835insCACAGT (5642_5643insCACAGT), GGG1835GCA (5644G>C 5645G>A), GAA1837TCA (5649G>T 5650A>C), TTG1838GTT (5652T>G 5654G>T), GGC1839GGA (5657C>A), AAC1840CTT (5658A>C 5659A>T 5660C>T), CTC1841CTA (5663C>A), CGC1842AGG (5664C>A 5666C>G), ATT1843ATA (5669T>A), ATA1844GTT (5670A>G 5672A>T), CAT1845CCT (5674A>C), GGT1846TCT (5676G>T 5677G>C), GAT1847GAC (5681T>C), TTA1848CTC (5682T>C 5684A>C), GCA1849TCT (5685G>T 5687A>T), AAA1850GAA (5688A>G), CAG1851AAG (5691C>A), ACA1852ATG (5695C>T 5696A>G), ATG1854GTT (5700A>G 5702G>T), CGG1855CGT (5705G>T), ATG1856TTG (5706A>T), TAT1858TAC (5714T>C), GTG1859ACA (5715G>A 5716T>C 5717G>A), GTA1860ATT (5718G>A 5720A>T), GGA1861GGT (5723A>T), CCA1862CCT (5726A>T), CAT1863GAA (5727C>G 5729T>A), GGA1865GGT (5735A>T), GGA1867GCC (5740G>C 5741A>C), AAT1868CAG (5742A>C 5744T>G), TTT1869TGC (5746T>G 5747T>C), TCC1870TCA (5750C>A), ACA1871ACG (5753A>G), CAG1872AAG (5754C>A), GAT1873AAC (5757G>A 5759T>C), CTT1874CTG (5762T>G), GAG1875AAG (5763G>A), AGT1876CAT (5766A>C 5767G>A), TTG1877GCT (5769T>G 5770T>C 5771G>T), CTA1878TTG (5772C>T 5774A>G), CAC1879GAG (5775C>G 5777C>G), TAT1880ACA (5778T>A 5779A>C 5780T>A), GAG1882TCT (5784G>T 5785A>C 5786G>T), CAG1883GTC (5787C>G 5788A>T 5789G>C), GGA1884AGT (5790G>A 5792A>T), GCA1886GCT (5798A>T), GAG1887ACT (5799G>A 5800A>C 5801G>T), TTA1888CTT (5802T>C 5804A>T), ATT1889GCC (5805A>G 5806T>C 5807T>C), CAA1890GCA (5808C>G 5809A>C), AAG1891AGA (5812A>G 5813G>A), AAT1892GGA (5814A>G 5815A>G 5816T>A), ACA1893GTC (5817A>G 5818C>T 5819A>C), CTA1895GTG (5823C>G 5825A>G), ACT1896ACA (5828T>A), CTG1897TTG (5829C>T), TTC1899TTT (5837C>T), GAG1901GAA (5843G>A), CTG1902TTC (5844C>T 5846G>C), AGT1903TCC (5847A>T 5848G>C 5849T>C), ATG1904CAA (5850A>C 5851T>A 5852G>A), GAA1905CAC (5853G>C 5855A>C), AAT1906GAT (5856A>G), GTG1907TGT (5859G>T 5860T>G 5861G>T), CAA1908GAA (5862C>G), CAA1909ATG (5865C>A 5866A>T 5867A>G), CTG1910ATA (5868C>A 5870G>A), ATA1911CTT (5871A>C 5873A>T), GAT1912GAG (5876T>G), GTT1914CTG (5880G>C 5882T>G), AAG1915GAA (5883A>G 5885G>A), CTT1916AGT (5886C>A 5887T>G), CAG1917CAA (5891G>A), TTT1919GTT (5895T>G)

#### Proteins

| | Begin | End | Coverage | Score | Concordance | Matches | Identities | I/D/M/F* | Stop Codons |
|---|---|---|---|---|---|---|---|---|---|
| polyprotein (YP_001976147.1) | 176 | 1922 | 20.1% | 2123 | 70.1% | 431 (99.1%) | 289 (66.4%) | 3/1/0/0 | 0 |

Protein mutations:

L181I (681T>A 683A>T), D182S (684G>A 685A>G), K186L (696A>C 697A>T 698A>C), A188S (702G>T 704C>T), R190K (708C>A 709G>A), T192S (714A>T 716C>T), M193Q (717A>C 718T>A), V194M (720G>A), M199V (735A>G), V205L (753G>T 755T>G), S206N (756T>A 757C>A 758A>T), M214L (780A>C), T216S (786A>T 788A>T), I217L (789A>T 791A>G), L218I (792T>A 794G>A), I228V (822A>G 824T>G), S238T (853G>C 854C>A), T248A (882A>G 884A>G), T260Q (918A>C 919C>A 920T>A), R263K (927C>A 928G>A 929T>G), I280V (978A>G 980A>T), V281A (982T>C 983C>T), G282D (985G>A 986C>T), T283I (988C>T 989C>A), L298I (1032C>A 1034G>T), Q307T (1059C>A 1060A>C 1061G>T), D310S (1068G>T 1069A>C), C313A (1077T>G 1078G>C), K325P (1113A>C 1114A>C 1115G>T), P328S (1122C>T 1124T>A), T329I (1126C>T), R335K (1144G>A 1145G>A), E337Q (1149G>C 1151A>G), M1252L (3894A>C), M1254L (3900A>T), M1256T (3907T>C 3908G>C), Q1263E (3927C>G), S1264A (3930T>G 3932C>A), T1265Q (3933A>C 3934C>A 3935T>G), G1266E (3937G>A 3938C>G), E1267S (3939G>A 3940A>G 3941A>T), G1268T (3942G>A 3943G>C 3944A>C), E1269Q (3945G>C 3947G>A), E1272L (3954G>C 3955A>T), D1276N (3966G>A 3968C>T), T1277S (3969A>T), E1281Q (3981G>C), N1716K (5288T>A), M1731I (5333G>A), M1736L (5346A>T), S1737A (5349T>G), M1740Q (5358A>C 5359T>A), A1742H (5364G>C 5365C>A 5366C>T), K1744R (5370A>C 5371A>G 5372G>C), N1745K (5375T>G), V1746S (5376G>A 5377T>G 5378G>T), A1748E (5383C>A), G1749V (5386G>T 5387A>T), H1751D (5391C>G), V1752G (5395T>G 5396A>G), N1755S (5403A>T 5404A>C), V1757T (5409G>A 5410T>C 5411T>C), V1766I (5436G>A 5438C>A), G1767A (5440G>C 5441C>T), M1774F (5460A>T 5462G>T), R1779H (5475A>C 5476G>A 5477G>T), C1782R (5484T>A 5486T>G), V1783A (5488T>C 5489G>C), E1784T (5490G>A 5491A>C 5492G>A), I1786L (5496A>C 5498A>T), I1793M (5519T>G), G1794A (5521G>C 5522A>T), N1795G (5523A>G 5524A>G 5525T>G), V1807A (5560T>C), F1810L (5570T>G), Q1812K (5574C>A), P1815E (5583C>G 5584C>A 5585T>A), H1816D (5586C>G 5588T>C), H1816_G1817insT (5588_5589insACC), G1817S (5589G>T 5590G>C), D1818H (5592G>C), V1820L (5598G>T), A1821G (5602C>G 5603T>A), A1825T (5613G>A), S1826Y (5617C>A 5618G>T), L1827G (5619C>G 5620T>G), H1828del (5622_5624delCAC), S1829T (5626G>C 5627T>C), S1830T (5629G>C), M1832N (5635T>A 5636G>C), Q1834H (5642A>T), Q1834_G1835insHS (5642_5643insCACAGT), G1835A (5644G>C 5645G>A), E1837S (5649G>T 5650A>C), L1838V (5652T>G 5654G>T), N1840L (5658A>C 5659A>T 5660C>T), I1844V (5670A>G 5672A>T), H1845P (5674A>C), G1846S (5676G>T 5677G>C), A1849S (5685G>T 5687A>T), K1850E (5688A>G), Q1851K (5691C>A), T1852M (5695C>T 5696A>G), M1854V (5700A>G 5702G>T), M1856L (5706A>T), V1859T (5715G>A 5716T>C 5717G>A), V1860I (5718G>A 5720A>T), H1863E (5727C>G 5729T>A), G1867A (5740G>C 5741A>C), N1868Q (5742A>C 5744T>G), F1869C (5746T>G 5747T>C), Q1872K (5754C>A), D1873N (5757G>A 5759T>C), E1875K (5763G>A), S1876H (5766A>C 5767G>A), L1877A (5769T>G 5770T>C 5771G>T), H1879E (5775C>G 5777C>G), Y1880T (5778T>A 5779A>C 5780T>A), E1882S (5784G>T 5785A>C 5786G>T), Q1883V (5787C>G 5788A>T 5789G>C), G1884S (5790G>A 5792A>T), E1887T (5799G>A 5800A>C 5801G>T), I1889A (5805A>G 5806T>C 5807T>C), Q1890A (5808C>G 5809A>C), K1891R (5812A>G 5813G>A), N1892G (5814A>G 5815A>G 5816T>A), T1893V (5817A>G 5818C>T 5819A>C), L1895V (5823C>G 5825A>G), L1902F (5844C>T 5846G>C), M1904Q (5850A>C 5851T>A 5852G>A), E1905H (5853G>C 5855A>C), N1906D (5856A>G), V1907C (5859G>T 5860T>G 5861G>T), Q1908E (5862C>G), Q1909M (5865C>A 5866A>T 5867A>G), L1910I (5868C>A 5870G>A), I1911L (5871A>C 5873A>T), D1912E (5876T>G), V1914L (5880G>C 5882T>G), K1915E (5883A>G 5885G>A), L1916S (5886C>A 5887T>G), F1919V (5895T>G)

| | Begin | End | Coverage | Score | Concordance | Matches | Identities | I/D/M/F* | Stop Codons |
|---|---|---|---|---|---|---|---|---|---|
| **NT** | **666** | **5905** | **17.9%** | **781** | **30.4%** | **1290 (99.1%)** | **850 (65.3%)** | **9/3** | |
| Codon mutations: | TAC179TAT (677C>T), TTA181ATT (681T>A 683A>T), GAT182AGT (684G>A 685A>G), CCA185CCC (695A>C), AAA186CTC (696A>C 697A>T 698A>C), GCC188TCT (702G>T 704C>T), GAA189GAG (707A>G), CGA190AAA (708C>A 709G>A), TTC191TTT (713C>T), ACC192TCT (714A>T 716C>T), ATG193CAG (717A>C 718T>A), GTG194ATG (720G>A), GCC195GCA (725C>A), CAA196CAG (728A>G), AGT198TCT (732A>T 733G>C), ATG199GTG (735A>G), CTT200CTA (740T>A), GGG202GGT (746G>T), GAA203GAG (749A>G), TCA204AGC (750T>A 751C>G 752A>C), GTT205TTG (753G>T 755T>G), TCA206AAT (756T>A 757C>A 758A>T), ACG210ACC (770G>C), TCT211AGT (771T>A 772C>G), CAG212CAA (776G>A), ATG214CTG (780A>C), TCC215TCT (785C>T), ACA216TCT (786A>T 788A>T), ATA217TTG (789A>T 791A>G), TTG218ATA (792T>A 794G>A), GCT219GCA (797T>A), GGG221GGC (803G>C), ATT222ATC (806T>C), TCG223TCA (809G>A), GTG226GTA (818G>A), TTC227TTT (821C>T), ATT228GTG (822A>G 824T>G), GTG230GTA (830G>A), AGA233AGG (839A>G), TTG237CTT (849T>C 851G>T), AGC238ACA (853G>C 854C>A), CCA240CCG (860A>G), TTG241CTA (861T>C 863G>A), CTC245TTG (873C>T 875C>G), TCG247TCA (881G>A), ACA248GCG (882A>G 884A>G), GGA249GGC (887A>C), GGT251GGA (893T>A), CAG253CAA (899G>A), CTA258CTC (914A>C), ACT260CAA (918A>C 919C>A 920T>A), CTT261TTG (921C>T 923T>G), TAC262TAT (926C>T), CGT263AAG (927C>A 928G>A 929T>G), AAT264AAC (932T>C), AAA266AAG (938A>G), ACC269ACA (947C>A), TCA274TCT (962A>T), GTT276GTG (968T>G), CTT277TTG (969C>T 971T>G), ATA279ATT (977A>T), ATA280GTT (978A>G 980A>T), GTC281GCT (982T>C 983C>T), GGC282GAT (985G>A 986C>T), ACC283ATA (988C>T 989C>A), TTT284TTC (992T>C), GGA285GGG (995A>G), TTC286TTT (998C>T), AAA287AAG (1001A>G), AAT288AAC (1004T>C), GTG290GTT (1010G>T), TTG291CTG (1011T>C), TCA292TCT (1016A>T), GCT293GCA (1019T>A), ATA294ATT (1022A>T), ACG297ACA (1031G>A), CTG298ATT (1032C>A 1034G>T), CTG302CTC (1046G>C), TTC303TTT (1049C>T), CAG307ACT (1059C>A 1060A>C 1061G>T), GAG308GAA (1064G>A), GAT310TCT (1068G>T 1069A>C), TGT313GCT (1077T>G 1078G>C), ACG317ACA (1091G>A), CGC318AGG (1092C>A 1094C>G), GAG320GAA (1100G>A), GCC323GCT (1109C>T), AAG325CCT (1113A>C 1114A>C 1115G>T), TTC327TTT (1121C>T), CCT328TCA (1122C>T 1124T>A), ACC329ATC (1126C>T), CTG334CTT (1142G>T), AGG335AAA (1144G>A 1145G>A), GAA336GAG (1148A>G), GAA337CAG (1149G>C 1151A>G), GCT338GCA (1154T>A), ATC341ATA (1163C>A), GCT343GCC (1169T>C), CTG1241CTC (3863G>C), TCT1244TCA (3872T>A), TGC1245TGT (3875C>T), TAT1247TAC (3881T>C), CCC1248CCT (3884C>T), AAC1250AAT (3890C>T), AAG1251AAA (3893G>A), ATG1252CTG (3894A>C), GGG1253GGT (3899G>T), ATG1254TTG (3900A>T), GGG1255GGT (3905G>T), ATG1256ACC (3907T>C 3908G>C), AAC1257AAT (3911C>T), AAG1258AAA (3914G>A), AGA1259AGG (3917A>G), TTT1261TTC (3923T>C), GTG1262GTT (3926G>T), CAA1263GAA (3927C>G), TCC1264GCA (3930T>G 3932C>A), ACT1265CAG (3933A>C 3934C>A 3935T>G), GGC1266GAG (3937G>A 3938C>G), GAA1267AGT (3939G>A 3940A>G 3941A>T), GGA1268ACC (3942G>A 3943G>C 3944A>C), GAG1269CAA (3945G>C 3947G>A), AGA1270AGG (3950A>G), GTG1271GTA (3953G>A), GAA1272CTA (3954G>C 3955A>T), AAA1274AAG (3962A>G), GAC1276AAT (3966G>A 3968C>T), ACT1277TCT (3969A>T), GTA1279GTT (3977A>T), TTT1280TTC (3980T>C), GAA1281CAA (3981G>C), GAA1284GAG (3992A>G), AAT1716AAA (5288T>A), GTT1718GTG (5294T>G), GGT1720GGG (5300T>G), CCA1721CCT (5303A>T), AAC1723AAT (5309C>T), ATG1731ATA (5333G>A), CCG1733CCC (5339G>C), ATG1736TTG (5346A>T), TCA1737GCA (5349T>G), GCT1738GCA (5354T>A), ATA1739ATC (5357A>C), ATG1740CAG (5358A>C 5359T>A), GCC1742CAT (5364G>C 5365C>A 5366C>T), AAG1744CGC (5370A>C 5371A>G 5372G>C), AAT1745AAG (5375T>G), GTG1746AGT (5376G>A 5377T>G 5378G>T), TTG1747CTG (5379T>C), GCA1748GAA (5383C>A), GGA1749GTT (5386G>T 5387A>T), GGA1750GGT (5390A>T), CAT1751GAT (5391C>G), GTA1752GGG (5395T>G 5396A>G), GCA1753GCT (5399A>T), AAT1755TCT (5403A>T 5404A>C), GTT1757ACC (5409G>A 5410T>C 5411T>C), GTG1759GTT (5417G>T), GGA1761GGT (5423A>T), GGA1763GGT (5429A>T), GCT1765GCA (5435T>A), GTC1766ATA (5436G>A 5438C>A), GGC1767GCT (5440G>C 5441C>T), ATT1769ATC (5447T>C), GCG1772GCA (5456G>A), CTG1773TTG (5457C>T), ATG1774TTT (5460A>T 5462G>T), GCC1775GCA (5465C>A), TTA1776TTG (5468A>G), GCA1777GCC (5471A>C), ACA1778ACT (5474A>T), AGG1779CAT (5475A>C 5476G>A 5477G>T), TAT1781TAC (5483T>C), TGT1782AGG (5484T>A 5486T>G), GTG1783GCC (5488T>C 5489G>C), GAG1784ACA (5490G>A 5491A>C 5492G>A), ATA1786CTT (5496A>C 5498A>T), ATA1787ATT (5501A>T), GTT1788GTG (5504T>G), AGA1789CGC (5505A>C 5507A>C), TTG1790CTA (5508T>C 5510G>A), CGC1791CGA (5513C>A), ACA1792ACC (5516A>C), ATT1793ATG (5519T>G), GGA1794GCT (5521G>C 5522A>T), AAT1795GGG (5523A>G 5524A>G 5525T>G), CTT1797CTG (5531T>G), GAC1798GAT (5534C>T), CTC1799CTG (5537C>G), AAT1800AAC (5540T>C), TAT1802TAC (5546T>C), GGC1805GGT (5555C>T), GTG1807GCG (5560T>C), CAG1808CAA (5564G>A), TAT1809TAC (5567T>C), TTT1810TTG (5570T>G), CTT1811CTC (5573T>C), CAA1812AAA (5574C>A), GTG1814GTT (5582G>T), CCT1815GAA (5583C>G 5584C>A 5585T>A), CAT1816GAC (5586C>G 5588T>C), CAT1816_GGA1817insACC (5588_5589insACC), GGA1817TCA (5589G>T 5590G>C), GAC1818CAC (5592G>C), AAA1819AAG (5597A>G), GTG1820TTG (5598G>T), GCT1821GGA (5602C>G 5603T>A), CAA1822CAG (5606A>G), GGT1824GGC (5612T>C), GCA1825ACA (5613G>A), TCG1826TAT (5617C>A 5618G>T), CTC1827GGC (5619C>G 5620T>G), CAC1828del (5622_5624delCAC), AGT1829ACC (5626G>C 5627T>C), AGT1830ACT (5629G>C), ATG1832AAC (5635T>A 5636G>C), CAA1834CAT (5642A>T), CAA1834_GGG1835insCACAGT (5642_5643insCACAGT), GGG1835GCA (5644G>C 5645G>A), GAA1837TCA (5649G>T 5650A>C), TTG1838GTT (5652T>G 5654G>T), GGC1839GGA (5657C>A), AAC1840CTT (5658A>C 5659A>T 5660C>T), CTC1841CTA (5663C>A), CGC1842AGG (5664C>A 5666C>G), ATT1843ATA (5669T>A), ATA1844GTT (5670A>G 5672A>T), CAT1845CCT (5674A>C), GGT1846TCT (5676G>T 5677G>C), GAT1847GAC (5681T>C), TTA1848CTC (5682T>C 5684A>C), GCA1849TCT (5685G>T 5687A>T), AAA1850GAA (5688A>G), CAG1851AAG (5691C>A), ACA1852ATG (5695C>T 5696A>G), ATG1854GTT (5700A>G 5702G>T), CGG1855CGT (5705G>T), ATG1856TTG (5706A>T), TAT1858TAC (5714T>C), GTG1859ACA (5715G>A 5716T>C 5717G>A), GTA1860ATT (5718G>A 5720A>T), GGA1861GGT (5723A>T), CCA1862CCT (5726A>T), CAT1863GAA (5727C>G 5729T>A), GGA1865GGT (5735A>T), GGA1867GCC (5740G>C 5741A>C), AAT1868CAG (5742A>C 5744T>G), TTT1869TGC (5746T>G 5747T>C), TCC1870TCA (5750C>A), ACA1871ACG (5753A>G), CAG1872AAG (5754C>A), GAT1873AAC (5757G>A 5759T>C), CTT1874CTG (5762T>G), GAG1875AAG (5763G>A), AGT1876CAT (5766A>C 5767G>A), TTG1877GCT (5769T>G 5770T>C 5771G>T), CTA1878TTG (5772C>T 5774A>G), CAC1879GAG (5775C>G 5777C>G), TAT1880ACA (5778T>A 5779A>C 5780T>A), GAG1882TCT (5784G>T 5785A>C 5786G>T), CAG1883GTC (5787C>G 5788A>T 5789G>C), GGA1884AGT (5790G>A 5792A>T), GCA1886GCT (5798A>T), GAG1887ACT (5799G>A 5800A>C 5801G>T), TTA1888CTT (5802T>C 5804A>T), ATT1889GCC (5805A>G 5806T>C 5807T>C), CAA1890GCA (5808C>G 5809A>C), AAG1891AGA (5812A>G 5813G>A), AAT1892GGA (5814A>G 5815A>G 5816T>A), ACA1893GTC (5817A>G 5818C>T 5819A>C), CTA1895GTG (5823C>G 5825A>G), ACT1896ACA (5828T>A), CTG1897TTG (5829C>T), TTC1899TTT (5837C>T), GAG1901GAA (5843G>A), CTG1902TTC (5844C>T 5846G>C), AGT1903TCC (5847A>T 5848G>C 5849T>C), ATG1904CAA (5850A>C 5851T>A 5852G>A), GAA1905CAC (5853G>C 5855A>C), AAT1906GAT (5856A>G), GTG1907TGT (5859G>T 5860T>G 5861G>T), CAA1908GAA (5862C>G), CAA1909ATG (5865C>A 5866A>T 5867A>G), CTG1910ATA (5868C>A 5870G>A), ATA1911CTT (5871A>C 5873A>T), GAT1912GAG (5876T>G), GTT1914CTG (5880G>C 5882T>G), AAG1915GAA (5883A>G 5885G>A), CTT1916AGT (5886C>A 5887T>G), CAG1917CAA (5891G>A), TTT1919GTT (5895T>G) | | | | | | | | |

*: Inserts / Deletes / Misaligned / Frameshifts

#### Analysis details

This analysis was performed with panviral2.64

### NGS Details (UN9): Torradovirus marchitezum (segment RNA 2)

#### Assembly

| | |
|---|---|
| **Coverage Length** | 1553 (2 contig(s)) |
| **Depth Of Coverage** | 23655.4 |
| **Number Of Reads** | 323229 |
| **Reads Per Million** | 7305.90 rpm (after QC) |
| **Ambiguities** | 0 |
| **Assembly Method** | de novo + reference guided assembly |
| **Consensus Caller** | Bcf Tools |

#### Coverage Map

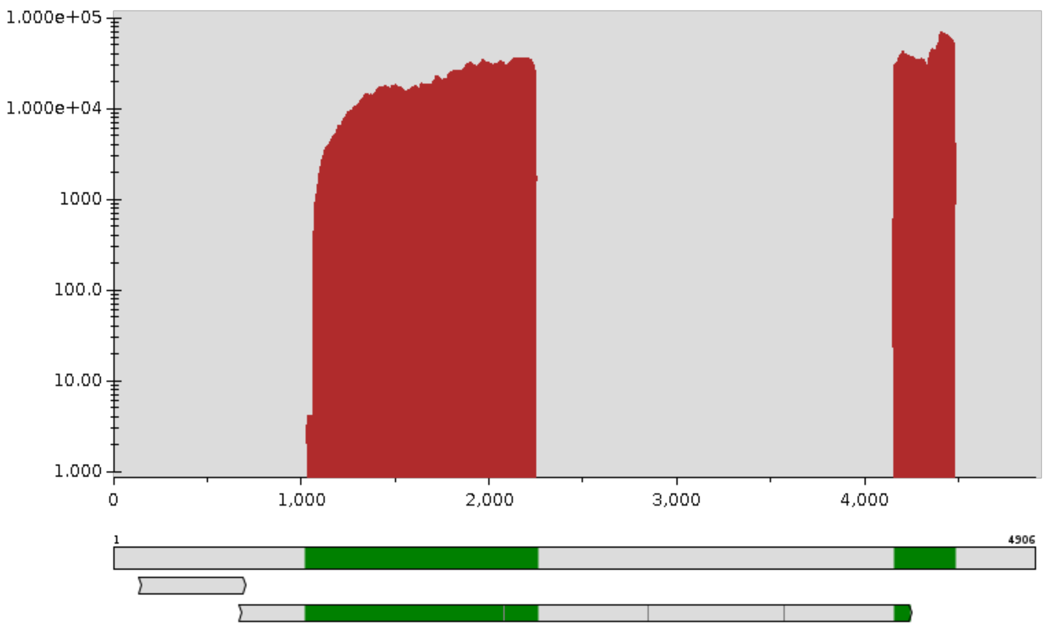

#### Assignment

| | |
|---|---|
| **Type** | Torradovirus marchitezum (Taxonomy ID: 3048376) |
| **Reference Genome** | NC_010988.1 |
| **NT Identity (%)** | 62.3492 |
| **AA Identity (%)** | 59.9099 |
| **Number Of Stop Codons** | 1 |
| **Number Of CDS** | 2 |

#### Alignment

| | |
|---|---|
| **Alignment Score** | 701.0 (NT) + 1905.0 (AA) = 2606.0 |
| **Concordance (%)** | 43.2531 |

| Alignment Method | Global, seeded, nucleotide + amino acids (AGA) |
| --- | --- |

#### Genome Region

Sequence starts at position 1030 and ends at position 4484 relative to NC_010988.1 reference sequence.

#### Alignment Detailed Statistics

| | Begin | End | Coverage | Score | Concordance | Matches | Identities | I/D/M/F* | Stop Codons |
| --- | --- | --- | --- | --- | --- | --- | --- | --- | --- |
| **NT** | **1030** | **4484** | **31.7%** | **701** | **23.0%** | **1541 (97.1%)** | **982 (61.9%)** | **34/12** | |
| Mutations: | 1030T>C, 1039C>A, 1040C>A, 1041T>C, 1042C>T, 1043T>G, 1044C>A, 1045A>T, 1048G>T, 1051A>T, 1057A>G, 1065C>T, 1066A>G, 1067C>T, 1069G>A, 1071C>T, 1075G>T, 1076G>T, 1078C>A, 1080A>G, 1081T>G, 1087G>T, 1093A>T, 1096T>C, 1097C>T, 1099C>G, 1105C>T, 1108C>T, 1109A>G, 1110G>A, 1111C>A, 1114A>T, 1123C>T, 1127C>G, 1129C>T, 1132A>G, 1156C>T, 1159G>A, 1165C>T, 1167A>T, 1168A>T, 1171T>C, 1174C>A, 1175A>C, 1180A>T, 1182A>T, 1183G>A, 1189G>A, 1195C>T, 1198G>T, 1201T>G, 1204C>A, 1207A>T, 1213A>G, 1216A>G, 1222T>A, 1223G>A, 1224A>G, 1226A>C, 1231G>A, 1237A>T, 1244T>A, 1246A>G, 1247C>G, 1248A>C, 1249G>A, 1262T>A, 1263C>G, 1270C>A, 1273A>T, 1276A>G, 1279A>T, 1286C>A, 1288T>G, 1291G>A, 1294A>G, 1297T>C, 1300C>T, 1301C>T, 1302A>C, 1303G>T, 1312G>A, 1318T>A, 1321T>A, 1327T>C, 1330A>T, 1339T>C, 1342T>C, 1351G>A, 1352C>G, 1354A>G, 1357T>G, 1358A>G, 1360T>G, 1361C>T, 1362A>C, 1363A>T, 1364A>C, 1365T>A, 1366G>A, 1369T>C, 1372T>A, 1375A>G, 1378C>T, 1380A>G, 1382G>C, 1384C>T, 1390T>A, 1393A>T, 1395A>C, 1398A>T, 1399G>T, 1402T>C, 1406G>A, 1407G>A, 1411A>T, 1412A>T, 1413G>A, 1414A>C, 1418G>A, 1420C>T, 1423C>T, 1424A>C, 1425T>C, 1426G>C, 1427A>C, 1429T>G, 1432A>G, 1433A>T, 1434G>C, 1435C>T, 1436A>T, 1438T>A, 1439A>T, 1441G>A, 1444C>G, 1447G>T, 1448A>G, 1449G>C, 1450T>G, 1456T>G, 1457T>C, 1459G>T, 1460A>C, 1462A>C, 1463G>A, 1465C>T, 1471T>C, 1474C>A, 1477A>G, 1478G>A, 1480G>C, 1483G>A, 1490C>A, 1492C>G, 1495C>T, 1498G>T, 1501C>G, 1507A>G, 1513G>T, 1515C>A, 1517G>T, 1519A>T, 1520A>T, 1521G>C, 1522T>A, 1522_1523insAGGGCC, 1523A>T, 1531C>G, 1532T>A, 1533T>C, 1535C>G, 1537C>A, 1538A>G, 1540C>T, 1541A>G, 1542C>T, 1543A>G, 1544C>A, 1546T>A, 1549A>G, 1557T>G, 1558T>G, 1559C>G, 1560A>T, 1561T>A, 1565_1567delGAG, 1575G>C, 1576T>C, 1577G>A, 1578G>A, 1579G>T, 1580G>T, 1581G>T, 1583T>C, 1591A>T, 1593C>T, 1594A>C, 1595_1600delCCCAAG, 1603A>G, 1604A>G, 1608G>C, 1609C>A, 1612A>G, 1616T>A, 1618C>T, 1619A>G, 1621C>T, 1622A>T, 1624G>T, 1627T>C, 1628G>C, 1629C>A, 1630C>G, 1633T>C, 1639T>C, 1644T>A, 1645G>T, 1651G>C, 1652G>A, 1653A>C, 1654G>T, 1655T>G, 1656C>A, 1657G>T, 1660C>T, 1661G>A, 1662C>A, 1663A>T, 1664C>T, 1665A>C, 1666C>G, 1667A>T, 1684T>G, 1691G>A, 1692A>C, 1693A>C, 1694C>T, 1696A>G, 1697A>C, 1698C>A, 1699A>T, 1703G>T, 1705T>A, 1706C>A, 1707A>G, 1711G>A, 1712G>A, 1714A>T, 1715C>G, 1717A>T, 1721A>G, 1723C>G, 1724C>T, 1726C>G, 1728T>C, 1731G>C, 1732T>C, 1733G>C, 1734A>T, 1735G>C, 1736G>C, 1737C>T, 1738T>C, 1739A>C, 1740A>G, 1741A>C, 1742A>C, 1743G>A, 1744A>G, 1747A>G, 1749A>G, 1750C>A, 1751G>T, 1752T>G, 1754A>C, 1755T>A, 1757G>C, 1762A>C, 1764G>A, 1765A>G, 1768T>C, 1769G>A, 1773C>A, 1774A>G, 1776A>G, 1777G>A, 1780G>A, 1781C>G, 1783T>C, 1784C>T, 1785T>C, 1786G>C, 1789T>C, 1792T>C, 1795C>A, 1798T>C, 1799A>C, 1800G>T, 1801G>A, 1805C>T, 1810A>T, 1811A>G, 1812C>A, 1813C>T, 1817G>A, 1819C>T, 1822T>C, 1826G>C, 1829G>C, 1831A>G, 1832A>G, 1834A>G, 1837T>C, 1838G>A, 1840A>G, 1842G>A, 1843T>A, 1849T>C, 1850G>A, 1852T>C, 1855C>A, 1859A>G, 1861C>G, 1862C>G, 1863A>G, 1864A>T, 1865G>T, 1866G>C, 1868T>C, 1870G>A, 1871A>G, 1872C>A, 1873T>G, 1876C>T, 1881T>G, 1891T>C, 1892G>T, 1895A>T, 1897G>A, 1898T>A, 1905A>G, 1909G>A, 1912A>G, 1915T>G, 1916G>A, 1918G>A, 1921G>A, 1922C>T, 1924C>G, 1927C>G, 1933A>G, 1939T>C, 1943T>C, 1944C>A, 1945C>A, 1947T>A, 1951T>C, 1953T>G, 1954T>G, 1955C>T, 1957A>T, 1960G>T, 1961C>G, 1963G>T, 1964A>G, 1965C>A, 1966T>G, 1968A>C, 1969C>A, 1975A>C, 1981G>A, 1983A>G, 1984T>A, 1990C>T, 1996T>C, 2002A>C, 2003T>C, 2005A>C, 2008C>A, 2013A>T, 2014A>G, 2017G>C, 2018C>A, 2020T>A, 2023G>C, 2023_2024insAAGGGTGGCAGGGCTAAG, 2026G>A, 2027A>G, 2028T>A, 2032G>A, 2034A>C, 2035G>T, 2037A>G, 2038G>T, 2039C>G, 2040A>C, 2041A>C, 2042G>T, 2044G>A, 2047C>A, 2048_2050delCAA, 2053A>T, 2054A>T, 2055G>T, 2057T>C, 2059T>A, 2060G>C, 2063A>C, 2064G>C, 2065A>T, 2066C>A, 2067A>G, 2070C>T, 2071A>G, 2072C>A, 2078A>G, 2080A>C, 2080_2081insCAG, 2082C>A, 2084A>G, 2086G>A, 2089C>G, 2090A>G, 2091T>G, 2093G>A, 2094C>G, 2095T>C, 2096G>A, 2097A>G, 2098A>G, 2099A>T, 2100G>C, 2101T>C, 2102C>A, 2103A>C, 2104G>C, 2105C>A, 2110A>G, 2111G>A, 2112A>G, 2115G>A, 2116A>G, 2119G>T, 2120A>C, 2121C>T, 2122T>G, 2133T>C, 2134G>A, 2135T>A, 2137A>T, 2140T>A, 2143C>T, 2144A>G, 2146G>A, 2150G>C, 2152_2153insAGC, 2155T>G, 2156C>G, 2157C>G, 2158C>A, 2160C>T, 2164G>C, 2165C>A, 2166C>A, 2167A>G, 2170T>C, 2171A>C, 2173G>A, 2176A>T, 2178A>T, 2182T>A, 2184T>A, 2185T>G, 2187C>T, 2188A>T, 2190A>C, 2191G>T, 2193G>A, 2194A>C, 2195G>C, 2200A>G, 2201G>C, 2203A>G, 2207A>G, 2208C>T, 2209T>C, 2210C>T, 2211A>C, 2217G>A, 2218T>C, 2224T>C, 2225A>T, 2227T>C, 2239A>G, 2240C>G, 2241C>A, 2242A>G, 4180C>T, 4181A>G, 4192T>G, 4195C>T, 4198C>T, 4201T>C, 4204G>A, 4213A>T, 4219T>C, 4222T>A, 4225G>A, 4228A>G, 4229C>A, 4231T>A, 4234T>C, 4237T>C, 4238G>T, 4242T>A, 4243T>C, 4248A>C, 4252T>A, 4279A>T, 4279_4280insT, 4280A>T, 4280_4281insC, 4283A>G, 4286C>T, 4287T>A, 4288T>C, 4291C>A, 4294C>T, 4306G>A, 4309G>A, 4312G>T, 4313C>A, 4313_4314insG, 4315A>G, 4316A>T, 4318T>C, 4319G>T, 4320G>A, 4323T>G, 4324T>A, 4328T>C, 4330G>A, 4333A>G, 4335G>T, 4336A>C, 4338A>G, 4339A>G, 4340G>T, 4341G>C, 4342G>T, 4343A>T, 4345A>T, 4346G>C, 4349A>C, 4364C>T, 4365T>C, 4369A>G, 4372A>G, 4373G>A, 4392T>C, 4393C>T, 4419_4420insG, 4422A>G, 4426G>A, 4430G>A, 4437C>T, 4441T>C, 4449A>T, 4451T>G | | | | | | | | |
| **CDS** | | | | | | | | | |
| ToMarV_RNA2gp2 | 121 | 1192 | 36.7% | 1905 | 63.6% | 434 (96.9%) | 266 (59.4%) | 10/4/0/0 | 1 |
| Protein mutations: | L124T (1040C>A 1041T>C 1042C>T), S125D (1043T>G 1044C>A 1045A>T), T132M (1065C>T 1066A>G), S134F (1071C>T), L135F (1075G>T), A136S (1076G>T 1078C>A), D137G (1080A>G 1081T>G), S147E (1109A>G 1110G>A 1111C>A), L153V (1127C>G 1129C>T), Q166L (1167A>T 1168A>T), Q171L (1182A>T 1183G>A), D185S (1223G>A 1224A>G), T186P (1226A>C), S192T (1244T>A 1246A>G), Q193A (1247C>G 1248A>C 1249G>A), Q211S (1301C>T 1302A>C 1303G>T), Q228E (1352C>G 1354A>G), S229R (1357T>G), T230A (1358A>G 1360T>G), Q231S (1361C>T 1362A>C 1363A>T), M232Q (1364A>C 1365T>A 1366G>A), K237R (1380A>G), A238P (1382G>C 1384C>T), E242A (1395A>C), Q243L (1398A>T 1399G>T), G246N (1406G>A 1407G>A), R248Y (1412A>T 1413G>A 1414A>C), D250N (1418G>A 1420C>T), M252P (1424A>C 1425T>C 1426G>C), N253Q (1427A>C 1429T>G), T256S (1436A>T 1438T>A), M257L (1439A>T 1441G>A), I258M (1444C>G), S260A (1448A>G 1449G>C 1450T>G), V265I (1463G>A 1465C>T), V270I (1478G>A 1480G>C), A282D (1515C>A), V283F (1517G>T 1519A>T), S284_T285insRA (1522_1523insAGGGCC), T285S (1523A>T), L288T (1532T>A 1533T>C), L289V (1535C>G 1537C>A), S290G (1538A>G 1540C>T), T291V (1541A>G 1542C>T 1543A>G), V296G (1557T>G 1558T>G), H297V (1559C>G 1560A>T 1561T>A), E299del (1565_1567delGAG), C302S (1575G>C 1576T>C), G303N (1577G>A 1578G>A 1579G>T), G304L (1580G>T 1581G>T), A308V (1593C>T 1594A>C), P309_K310del (1595_1600delCCCAAG), K312E (1604A>G), G313A (1608G>C 1609C>A), F316I (1616T>A 1618C>T), N317D (1619A>G 1621C>T), M318F (1622A>T 1624G>T), A320Q (1628G>C 1629C>A 1630C>G), L325Y (1644T>A 1645G>T), M327I (1651G>C), E328T (1652G>A 1653A>C 1654G>T), S329D (1655T>G 1656C>A 1657G>T), A331N (1661G>A 1662C>A 1663A>T), H332S (1664C>T 1665A>C 1666C>G), I333F (1667A>T), D338E (1684T>G), E341T (1691G>A 1692A>C 1693A>C), R342W (1694C>T 1696A>G), T343H (1697A>C 1698C>A 1699A>T), A345S (1703G>T 1705T>A), Q346R (1706C>A 1707A>G), V348I (1712G>A 1714A>T), Q349D (1715C>G 1717A>T), I351V (1721A>G 1723C>G), F353S (1728T>C), S354T (1731G>C 1732T>C), E355L (1733G>C 1734A>T 1735G>C), A356L (1736G>C 1737C>T 1738T>C), K357R (1739A>C 1740A>G 1741A>C), R358Q (1742A>C 1743G>A 1744A>G), N360R (1749A>G 1750C>A), V361C (1751G>T 1752T>G), I362H (1754A>C 1755T>A), V363L (1757G>C), R365K (1764G>A 1765A>G), V367I (1769G>A), A368E (1773C>A 1774A>G), K369R (1776A>G 1777G>A), H371D (1781C>G 1783T>C), L372S (1784C>T 1785T>C 1786G>C), R377L (1799A>C 1800G>T 1801G>A), H379Y (1805C>T), T381D (1811A>G 1812C>A 1813C>T), D383N (1817G>A 1819C>T), V386L (1826G>C), E387Q (1829G>C 1831A>G), I388V (1832A>G 1834A>G), E390K (1838G>A 1840A>G), G391E (1842G>A 1843T>A), V394I (1850G>A 1852T>C), I397V (1859A>G 1861C>G), Q398G (1862C>G 1863A>G 1864A>T), G399S (1865G>T 1866G>C), T401E (1871A>G 1872C>A 1873T>G), F404C (1881T>G), A408S (1892G>T), M409L (1895A>T 1897G>A), S410T (1898T>A), K412R (1905A>G), V416I (1916G>A 1918G>A), S425Q (1943T>C 1944C>A 1945C>A), F426Y (1947T>A), I428R (1953T>G 1954T>G), Q429Y (1955C>T 1957A>T), Q431D (1961C>G 1963G>T), T432E (1964A>G 1965C>A 1966T>G), N433T (1968A>C 1969C>A), N438R (1983A>G 1984T>A), K448M (2013A>T 2014A>G), E449D (2017G>C), Q451H (2023G>C), Q451_L452insKGGRAK (2023_2024insAAGGGTGGCAGGGCTAAG), I453D (2027A>G 2028T>A), E455A (2034A>C 2035G>T), E456G (2037A>G 2038G>T), Q457A (2039C>G 2040A>C 2041A>C), V458L (2042G>T 2044G>A), Q460del (2048_2050delCAA), R461S (2053A>T), R462L (2054A>T 2055G>T), S463P (2057T>C 2059T>A), E464Q (2060G>C), R465P (2063A>C 2064G>C 2065A>T), Q466R (2066C>A 2067A>G), A467V (2070C>T 2071A>G), Q468K (2072C>A), T470A (2078A>G 2080A>C), T470_A471insQ (2080_2081insCAG), A471E (2082C>A), R472G (2084A>G 2086G>A), I474G (2090A>G 2091T>G), A475S (2093G>A 2094C>G 2095T>C), E476R (2096G>A 2097A>G 2098A>G), Q478T (2102C>A 2103A>C 2104G>C), P479T (2105C>A), D481S (2111G>A 2112A>G), R482K (2115G>A 2116A>G), T484L (2120A>C 2121C>T 2122T>G), V488A (2133T>C 2134G>A), S489T (2135T>A 2137A>T), T492A (2144A>G 2146G>A), E494Q (2150G>C), E494_D495insS (2152_2153insAGC), D495E (2155T>G), P496G (2156C>G 2157C>G 2158C>A), T497I (2160C>T), K498N (2164G>C), P499K (2165C>A 2166C>A 2167A>G), K501Q (2171A>C 2173G>A), E503V (2178A>T), V505E (2184T>A 2185T>G), A506V (2187C>T 2188A>T), E507A (2190A>C 2191G>T), G508D (2193G>A 2194A>C), A509P (2195G>C), E511Q (2201G>C 2203A>G), T513V (2207A>G 2208C>T 2209T>C), Q514S (2210C>T 2211A>C), G516D (2217G>A 2218T>C), I519F (2225A>T 2227T>C), P524E (2240C>G 2241C>A 2242A>G), N1171D (4181A>G), A1190S (4238G>T), I1191N (4242T>A 4243T>C) | | | | | | | | |

| | Begin | End | Coverage | Score | Concordance | Matches | Identities | I/D/M/F* | Stop Codons |
|---|---|---|---|---|---|---|---|---|---|
| **NT** | **1030** | **4484** | **31.7%** | **701** | **23.0%** | **1541 (97.1%)** | **982 (61.9%)** | **34/12** | |
| Codon mutations: | AGT120..C (1030T>C), CCC123CCA (1039C>A), CTC124ACT (1040C>A 1041T>C 1042C>T), TCA125GAT (1043T>G 1044C>A 1045A>T), CCG126CCT (1048G>T), GTA127GTT (1051A>T), CAA129CAG (1057A>G), ACA132ATG (1065C>T 1066A>G), CTG133TTA (1067C>T 1069G>A), TCC134TTC (1071C>T), TTG135TTT (1075G>T), GCC136TCA (1076G>T 1078C>A), GAT137GGG (1080A>G 1081T>G), GCG139GCT (1087G>T), GCA141GCT (1093A>T), GTT142GTC (1096T>C), CTC143TTG (1097C>T 1099C>G), AGC145AGT (1105C>T), CCC146CCT (1108C>T), AGC147GAA (1109A>G 1110G>A 1111C>A), ATA148ATT (1114A>T), CCC151CCT (1123C>T), CTC153GTT (1127C>G 1129C>T), AAA154AAG (1132A>G), ACC162ACT (1156C>T), CTG163CTA (1159G>A), TGC165TGT (1165C>T), CAA166CTT (1167A>T 1168A>T), AAT167AAC (1171T>C), ACC168ACA (1174C>A), AGG169CGG (1175A>C), GCA170GCT (1180A>T), CAG171CTA (1182A>T 1183G>A), CCG173CCA (1189G>A), TCC175TCT (1195C>T), GTG176GTT (1198G>T), GTT177GTG (1201T>G), GCC178GCA (1204C>A), ATA179ATT (1207A>T), AAA181AAG (1213A>G), CAA182CAG (1216A>G), ATT184ATA (1222T>A), GAC185AGC (1223G>A 1224A>G), ACA186CCA (1226A>C), AGG187AGA (1231G>A), GCA189GCT (1237A>T), TCA192ACG (1244T>A 1246A>G), CAG193GCA (1247C>G 1248A>C 1249G>A), TCT198AGT (1262T>A 1263C>G), GCC200GCA (1270C>A), ATA201ATT (1273A>T), GAA202GAG (1276A>G), GCA203GCT (1279A>T), CGT206AGG (1286C>A 1288T>G), CCG207CCA (1291G>A), CAA208CAG (1294A>G), TTT209TTC (1297T>C), TTC210TTT (1300C>T), CAG211TCT (1301C>T 1302A>C 1303G>T), CAG214CAA (1312G>A), GCT216GCA (1318T>A), GCT217GCA (1321T>A), TAT219TAC (1327T>C), ATA220ATT (1330A>T), GTT223GTC (1339T>C), TAT224TAC (1342T>C), GGG227GGA (1351G>A), CAA228GAG (1352C>G 1354A>G), AGT229AGG (1357T>G), ACT230GCG (1358A>G 1360T>G), CAA231TCT (1361C>T 1362A>C 1363A>T), ATG232CAA (1364A>C 1365T>A 1366G>A), GCT233GCC (1369T>C), CTT234CTA (1372T>A), GAA235GAG (1375A>G), AAC236AAT (1378C>T), AAA237AGA (1380A>G), GCC238CCT (1382G>C 1384C>T), GTT240GTA (1390T>A), GGA241GGT (1393A>T), GAA242GCA (1395A>C), CAG243CTT (1398A>T 1399G>T), CTT244CTC (1402T>C), GGC246AAC (1406G>A 1407G>A), ACA247ACT (1411A>T), AGA248TAC (1412A>T 1413G>A 1414A>C), GAC250AAT (1418G>A 1420C>T), GTC251GTT (1423C>T), ATG252CCC (1424A>C 1425T>C 1426G>C), AAT253CAG (1427A>C 1429T>G), GAA254GAG (1432A>G), AGC255TCT (1433A>T 1434G>C 1435C>T), ACT256TCA (1436A>T 1438T>A), ATG257TTA (1439A>T 1441G>A), ATC258ATG (1444C>G), CGG259CGT (1447G>T), AGT260GCG (1448A>G 1449G>C 1450T>G), TCT262TCG (1456T>G), TTG263CTT (1457T>C 1459G>T), AGA264CGC (1460A>C 1462A>C), GTC265ATT (1463G>A 1465C>T), ACT267ACC (1471T>C), CTC268CTA (1474C>A), AAA269AAG (1477A>G), GTG270ATC (1478G>A 1480G>C), CAG271CAA (1483G>A), CGC274AGG (1490C>A 1492C>G), TTC275TTT (1495C>T), CCG276CCT (1498G>T), TCC277TCG (1501C>G), GAA279GAG (1507A>G), CCG281CCT (1513G>T), GCT282GAT (1515C>A), GTA283TTT (1517G>T 1519A>T), AGT284TCA (1520A>T 1521G>C 1522T>A), AGT284_ACA285insAGGGCC (1522_1523insAGGGCC), ACA285TCA (1523A>T), TCC287TCG (1531C>G), TTA288ACA (1532T>A 1533T>C), CTC289GTA (1535C>G 1537C>A), AGC290GGT (1538A>G 1540C>T), ACA291GTG (1541A>G 1542C>T 1543A>G), CGT292AGA (1544C>A 1546T>A), GAA293GAG (1549A>G), GTT296GGG (1557T>G 1558T>G), CAT297GTA (1559C>G 1560A>T 1561T>A), GAG299del (1565_1567delGAG), TGT302TCC (1575G>C 1576T>C), GGG303AAT (1577G>A 1578G>A 1579G>T), GGA304TTA (1580G>T 1581G>T), TTG305CTG (1583T>C), CCA307CCT (1591A>T), GCA308GTC (1593C>T 1594A>C), CCC309_AAG310del (1595_1600delCCCAAG), AAA311AAG (1603A>G), AAG312GAG (1604A>G), GGC313GCA (1608G>C 1609C>A), CAA314CAG (1612A>G), TTC316ATT (1616T>A 1618C>T), AAC317GAT (1619A>G 1621C>T), ATG318TTT (1622A>T 1624G>T), GGT319GGC (1627T>C), GCC320CAG (1628G>C 1629C>A 1630C>G), ATT321ATC (1633T>C), GAT323GAC (1639T>C), TTG325TAT (1644T>A 1645G>T), ATG327ATC (1651G>C), GAG328ACT (1652G>A 1653A>C 1654G>T), TCG329GAT (1655T>G 1656C>A 1657G>T), TTC330TTT (1660C>T), GCA331AAT (1661G>A 1662C>A 1663A>T), CAC332TCG (1664C>T 1665A>C 1666C>G), ATT333TTT (1667A>T), GAT338GAG (1684T>G), GAA341ACC (1691G>A 1692A>C 1693A>C), CGA342TGG (1694C>T 1696A>G), ACA343CAT (1697A>C 1698C>A 1699A>T), GCT345TCA (1703G>T 1705T>A), CAG346AGG (1706C>A 1707A>G), CAG347CAA (1711G>A), GTA348ATT (1712G>A 1714A>T), CAA349GAT (1715C>G 1717A>T), ATC351GTG (1721A>G 1723C>G), CTC352TTG (1724C>T 1726C>G), TTT353TCT (1728T>C), AGT354ACC (1731G>C 1732T>C), GAG355CTC (1733G>C 1734A>T 1735G>C), GCT356CTC (1736G>C 1737C>T 1738T>C), AAA357CGC (1739A>C 1740A>G 1741A>C), AGA358CAG (1742A>C 1743G>A 1744A>G), GGA359GGG (1747A>G), AAC360AGA (1749A>G 1750C>A), GTT361TGT (1751G>T 1752T>G), ATT362CAT (1754A>C 1755T>A), GTG363CTG (1757G>C), CCA364CCC (1762A>C), AGA365AAG (1764G>A 1765A>G), CAT366CAC (1768T>C), GTT367ATT (1769G>A), GCA368GAG (1773C>A 1774A>G), AAG369AGA (1776A>G 1777G>A), CGG370CGA (1780G>A), CAT371GAC (1781C>G 1783T>C), CTG372TCC (1784C>T 1785T>C 1786G>C), CAT373CAC (1789T>C), AAT374AAC (1792T>C), ATC375ATA (1795C>A), AAT376AAC (1798T>C), AGG377CTA (1799A>C 1800G>T 1801G>A), CAC379TAC (1805C>T), ATA380ATT (1810A>T), ACC381GAT (1811A>G 1812C>A 1813C>T), GAC383AAT (1817G>A 1819C>T), AAT384AAC (1822T>C), GTA386CTA (1826G>C), GAA387CAG (1829G>C 1831A>G), ATA388GTG (1832A>G 1834A>G), CTT389CTC (1837T>C), GAA390AAG (1838G>A 1840A>G), GGT391GAA (1842G>A 1843T>A), GGT393GGC (1849T>C), GTT394ATC (1850G>A 1852T>C), ACC395ACA (1855C>A), ATC397GTG (1859A>G 1861C>G), CAA398GGT (1862C>G 1863A>G 1864A>T), GGC399TCC (1865G>T 1866G>C), TTG400CTA (1868T>C 1870G>A), ACT401GAG (1871A>G 1872C>A 1873T>G), CGC402CGT (1876C>T), TTC404TGC (1881T>G), TAT407TAC (1891T>C), GCT408TCT (1892G>T), ATG409TTA (1895A>T 1897G>A), TCT410ACT (1898T>A), AAG412AGG (1905A>G), GAG413GAA (1909G>A), AGA414AGG (1912A>G), GTT415GTG (1915T>G), GTG416ATA (1916G>A 1918G>A), GAG417GAA (1921G>A), CTC418TTG (1922C>T 1924C>G), GTC419GTG (1927C>G), CAA421CAG (1933A>G), GAT423GAC (1939T>C), TCC425CAA (1943T>C 1944C>A 1945C>A), TTC426TAC (1947T>A), TAT427TAC (1951T>C), ATT428AGG (1953T>G 1954T>G), CAA429TAT (1955C>T 1957A>T), GGG430GGT (1960G>T), CAG431GAT (1961C>G 1963G>T), ACT432GAG (1964A>G 1965C>A 1966T>G), AAC433ACA (1968A>C 1969C>A), CCA435CCC (1975A>C), CCG437CCA (1981G>A), AAT438AGA (1983A>G 1984T>A), GAC440GAT (1990C>T), TAT442TAC (1996T>C), GGA444GGC (2002A>C), TTA445CTC (2003T>C 2005A>C), ACC446ACA (2008C>A), AAA448ATG (2013A>T 2014A>G), GAG449GAC (2017G>C), CGT450AGA (2018C>A 2020T>A), CAG451CAC (2023G>C), CAG451_TTG452insAAGGGTGGCAGGGCTAAG (2023_2024insAAGGGTGGCAGGGCTAAG), TTG452TTA (2026G>A), ATT453GAT (2027A>G 2028T>A), GTG454GTA (2032G>A), GAG455GCT (2034A>C 2035G>T), GAG456GGT (2037A>G 2038G>T), CAA457GCC (2039C>G 2040A>C 2041A>C), GTG458TTA (2042G>T 2044G>A), GCC459GCA (2047C>A), CAA460del (2048_2050delCAA), AGA461AGT (2053A>T), AGG462TTG (2054A>T 2055G>T), TCT463CCA (2057T>C 2059T>A), GAA464CAA (2060G>C), AGA465CCT (2063A>C 2064G>C 2065A>T), CAA466AGA (2066C>A 2067A>G), GCA467GTG (2070C>T 2071A>G), CAG468AAG (2072C>A), ACA470GCC (2078A>G 2080A>C), ACA470_GCA471insCAG (2080_2081insCAG), GCA471GAA (2082C>A), AGG472GGA (2084A>G 2086G>A), GGC473GGG (2089C>G), ATA474GGA (2090A>G 2091T>G), GCT475AGC (2093G>A 2094C>G 2095T>C), GAA476AGG (2096G>A 2097A>G 2098A>G), AGT477TCC (2099A>T 2100G>C 2101T>C), CAG478ACC (2102C>A 2103A>C 2104G>C), CCT479ACT (2105C>A), GAA480GAG (2110A>G), GAC481AGC (2111G>A 2112A>G), AGA482AAG (2115G>A 2116A>G), GTG483GTT (2119G>T), ACT484CTG (2120A>C 2121C>T 2122T>G), GTG488GCA (2133T>C 2134G>A), TCA489ACT (2135T>A 2137A>T), ACT490ACA (2140T>A), ACC491ACT (2143C>T), ACG492GCA (2144A>G 2146G>A), GAA494CAA (2150G>C), GAA494_GAT495insAGC (2152_2153insAGC), GAT495GAG (2155T>G), CCC496GGA (2156C>G 2157C>G 2158C>A), ACC497ATC (2160C>T), AAG498AAC (2164G>C), CCA499AAG (2165C>A 2166C>A 2167A>G), GAT500GAC (2170T>C), AAG501CAA (2171A>C 2173G>A), ATA502ATT (2176A>T), GAA503GTA (2178A>T), ATT504ATA (2182T>A), GTT505GAG (2184T>A 2185T>G), GCA506GTT (2187C>T 2188A>T), GAG507GCT (2190A>C 2191G>T), GGA508GAC (2193G>A 2194A>C), GCA509CCA (2195G>C), GAA510GAG (2200A>G), GAA511CAG (2201G>C 2203A>G), ACT513GTC (2207A>G 2208C>T 2209T>C), CAA514TCA (2210C>T 2211A>C), GGT516GAC (2217G>A 2218T>C), GTT518GTC (2224T>C), ATT519TTC (2225A>T 2227T>C), GGA523GGG (2239A>G), CCA524GAG (2240C>G 2241C>A 2242A>G), GGC1170GGT (4180C>T), AAT1171GAT (4181A>G), ACT1174ACG (4192T>G), CAC1175CAT (4195C>T), ACC1176ACT (4198C>T), TAT1177TAC (4201T>C), CAG1178CAA (4204G>A), TCA1181TCT (4213A>T), TTT1183TTC (4219T>C), TCT1184TCA (4222T>A), GAG1185GAA (4225G>A), CTA1186CTG (4228A>G), CGT1187AGA (4229C>A 4231T>A), GAT1188GAC (4234T>C), TTT1189TTC (4237T>C), GCG1190TCG (4238G>T), ATT1191AAC (4242T>A 4243T>C) | | | | | | | | |

#### Proteins

| | Begin | End | Coverage | Score | Concordance | Matches | Identities | I/D/M/F* | Stop Codons |
|---|---|---|---|---|---|---|---|---|---|
| polyprotein (YP_001976149.1) | 121 | 1192 | 36.7% | 1905 | 63.6% | 434 (96.9%) | 266 (59.4%) | 10/4/0/0 | 1 |
| Protein mutations: | L124T (1040C>A 1041T>C 1042C>T), S125D (1043T>G 1044C>A 1045A>T), T132M (1065C>T 1066A>G), S134F (1071C>T), L135F (1075G>T), A136S (1076G>T 1078C>A), D137G (1080A>G 1081T>G), S147E (1109A>G 1110G>A 1111C>A), L153V (1127C>G 1129C>T), Q166L (1167A>T 1168A>T), Q171L (1182A>T 1183G>A), D185S (1223G>A 1224A>G), T186P (1226A>C), S192T (1244T>A 1246A>G), Q193A (1247C>G 1248A>C 1249G>A), Q211S (1301C>T 1302A>C 1303G>T), Q228E (1352C>G 1354A>G), S229R (1357T>G), T230A (1358A>G 1360T>G), Q231S (1361C>T 1362A>C 1363A>T), M232Q (1364A>C 1365T>A 1366G>A), K237R (1380A>G), A238P (1382G>C 1384C>T), E242A (1395A>C), Q243L (1398A>T 1399G>T), G246N (1406G>A 1407G>A), R248Y (1412A>T 1413G>A 1414A>C), D250N (1418G>A 1420C>T), M252P (1424A>C 1425T>C 1426G>C), N253Q (1427A>C 1429T>G), T256S (1436A>T 1438T>A), M257L (1439A>T 1441G>A), I258M (1444C>G), S260A (1448A>G 1449G>C 1450T>G), V265I (1463G>A 1465C>T), V270I (1478G>A 1480G>C), A282D (1515C>A), V283F (1517G>T 1519A>T), S284_T285insRA (1522_1523insAGGGCC), T285S (1523A>T), L288T (1532T>A 1533T>C), L289V (1535C>G 1537C>A), S290G (1538A>G 1540C>T), T291V (1541A>G 1542C>T 1543A>G), V296G (1557T>G 1558T>G), H297V (1559C>G 1560A>T 1561T>A), E299del (1565_1567delGAG), C302S (1575G>C 1576T>C), G303N (1577G>A 1578G>A 1579G>T), G304L (1580G>T 1581G>T), A308V (1593C>T 1594A>C), P309_K310del (1595_1600delCCCAAG), K312E (1604A>G), G313A (1608G>C 1609C>A), F316I (1616T>A 1618C>T), N317D (1619A>G 1621C>T), M318F (1622A>T 1624G>T), A320Q (1628G>C 1629C>A 1630C>G), L325Y (1644T>A 1645G>T), M327I (1651G>C), E328T (1652G>A 1653A>C 1654G>T), S329D (1655T>G 1656C>A 1657G>T), A331N (1661G>A 1662C>A 1663A>T), H332S (1664C>T 1665A>C 1666C>G), I333F (1667A>T), D338E (1684T>G), E341T (1691G>A 1692A>C 1693A>C), R342W (1694C>T 1696A>G), T343H (1697A>C 1698C>A 1699A>T), A345S (1703G>T 1705T>A), Q346R (1706C>A 1707A>G), V348I (1712G>A 1714A>T), Q349D (1715C>G 1717A>T), I351V (1721A>G 1723C>G), F353S (1728T>C), S354T (1731G>C 1732T>C), E355L (1733G>C 1734A>T 1735G>C), A356L (1736G>C 1737C>T 1738T>C), K357R (1739A>C 1740A>G 1741A>C), R358Q (1742A>C 1743G>A 1744A>G), N360R (1749A>G 1750C>A), V361C (1751G>T 1752T>G), I362H (1754A>C 1755T>A), V363L (1757G>C), R365K (1764G>A 1765A>G), V367I (1769G>A), A368E (1773C>A 1774A>G), K369R (1776A>G 1777G>A), H371D (1781C>G 1783T>C), L372S (1784C>T 1785T>C 1786G>C), R377L (1799A>C 1800G>T 1801G>A), H379Y (1805C>T), T381D (1811A>G 1812C>A 1813C>T), D383N (1817G>A 1819C>T), V386L (1826G>C), E387Q (1829G>C 1831A>G), I388V (1832A>G 1834A>G), E390K (1838G>A 1840A>G), G391E (1842G>A 1843T>A), V394I (1850G>A 1852T>C), I397V (1859A>G 1861C>G), Q398G (1862C>G 1863A>G 1864A>T), G399S (1865G>T 1866G>C), T401E (1871A>G 1872C>A 1873T>G), F404C (1881T>G), A408S (1892G>T), M409L (1895A>T 1897G>A), S410T (1898T>A), K412R (1905A>G), V416I (1916G>A 1918G>A), S425Q (1943T>C 1944C>A 1945C>A), F426Y (1947T>A), I428R (1953T>G 1954T>G), Q429Y (1955C>T 1957A>T), Q431D (1961C>G 1963G>T), T432E (1964A>G 1965C>A 1966T>G), N433T (1968A>C 1969C>A), N438R (1983A>G 1984T>A), K448M (2013A>T 2014A>G), E449D (2017G>C), Q451H (2023G>C), Q451_L452insKGGRAK (2023_2024insAAGGGTGGCAGGGCTAAG), I453D (2027A>G 2028T>A), E455A (2034A>C 2035G>T), E456G (2037A>G 2038G>T), Q457A (2039C>G 2040A>C 2041A>C), V458L (2042G>T 2044G>A), Q460del (2048_2050delCAA), R461S (2053A>T), R462L (2054A>T 2055G>T), S463P (2057T>C 2059T>A), E464Q (2060G>C), R465P (2063A>C 2064G>C 2065A>T), Q466R (2066C>A 2067A>G), A467V (2070C>T 2071A>G), Q468K (2072C>A), T470A (2078A>G 2080A>C), T470_A471insQ (2080_2081insCAG), A471E (2082C>A), R472G (2084A>G 2086G>A), I474G (2090A>G 2091T>G), A475S (2093G>A 2094C>G 2095T>C), E476R (2096G>A 2097A>G 2098A>G), Q478T (2102C>A 2103A>C 2104G>C), P479T (2105C>A), D481S (2111G>A 2112A>G), R482K (2115G>A 2116A>G), T484L (2120A>C 2121C>T 2122T>G), V488A (2133T>C 2134G>A), S489T (2135T>A 2137A>T), T492A (2144A>G 2146G>A), E494Q (2150G>C), E494_D495insS (2152_2153insAGC), D495E (2155T>G), P496G (2156C>G 2157C>G 2158C>A), T497I (2160C>T), K498N (2164G>C), P499K (2165C>A 2166C>A 2167A>G), K501Q (2171A>C 2173G>A), E503V (2178A>T), V505E (2184T>A 2185T>G), A506V (2187C>T 2188A>T), E507A (2190A>C 2191G>T), G508D (2193G>A 2194A>C), A509P (2195G>C), E511Q (2201G>C 2203A>G), T513V (2207A>G 2208C>T 2209T>C), Q514S (2210C>T 2211A>C), G516D (2217G>A 2218T>C), I519F (2225A>T 2227T>C), P524E (2240C>G 2241C>A 2242A>G), N1171D (4181A>G), A1190S (4238G>T), I1191N (4242T>A 4243T>C) | | | | | | | | |

| | Begin | End | Coverage | Score | Concordance | Matches | Identities | I/D/M/F* | Stop Codons |
|---|---|---|---|---|---|---|---|---|---|
| **NT** | **1030** | **4484** | **31.7%** | **701** | **23.0%** | **1541 (97.1%)** | **982 (61.9%)** | **34/12** | |
| Codon mutations: | AGT120..C (1030T>C), CCC123CCA (1039C>A), CTC124ACT (1040C>A 1041T>C 1042C>T), TCA125GAT (1043T>G 1044C>A 1045A>T), CCG126CCT (1048G>T), GTA127GTT (1051A>T), CAA129CAG (1057A>G), ACA132ATG (1065C>T 1066A>G), CTG133TTA (1067C>T 1069G>A), TCC134TTC (1071C>T), TTG135TTT (1075G>T), GCC136TCA (1076G>T 1078C>A), GAT137GGG (1080A>G 1081T>G), GCG139GCT (1087G>T), GCA141GCT (1093A>T), GTT142GTC (1096T>C), CTC143TTG (1097C>T 1099C>G), AGC145AGT (1105C>T), CCC146CCT (1108C>T), AGC147GAA (1109A>G 1110G>A 1111C>A), ATA148ATT (1114A>T), CCC151CCT (1123C>T), CTC153GTT (1127C>G 1129C>T), AAA154AAG (1132A>G), ACC162ACT (1156C>T), CTG163CTA (1159G>A), TGC165TGT (1165C>T), CAA166CTT (1167A>T 1168A>T), AAT167AAC (1171T>C), ACC168ACA (1174C>A), AGG169CGG (1175A>C), GCA170GCT (1180A>T), CAG171CTA (1182A>T 1183G>A), CCG173CCA (1189G>A), TCC175TCT (1195C>T), GTG176GTT (1198G>T), GTT177GTG (1201T>G), GCC178GCA (1204C>A), ATA179ATT (1207A>T), AAA181AAG (1213A>G), CAA182CAG (1216A>G), ATT184ATA (1222T>A), GAC185AGC (1223G>A 1224A>G), ACA186CCA (1226A>C), AGG187AGA (1231G>A), GCA189GCT (1237A>T), TCA192ACG (1244T>A 1246A>G), CAG193GCA (1247C>G 1248A>C 1249G>A), TCT198AGT (1262T>A 1263C>G), GCC200GCA (1270C>A), ATA201ATT (1273A>T), GAA202GAG (1276A>G), GCA203GCT (1279A>T), CGT206AGG (1286C>A 1288T>G), CCG207CCA (1291G>A), CAA208CAG (1294A>G), TTT209TTC (1297T>C), TTC210TTT (1300C>T), CAG211TCT (1301C>T 1302A>C 1303G>T), CAG214CAA (1312G>A), GCT216GCA (1318T>A), GCT217GCA (1321T>A), TAT219TAC (1327T>C), ATA220ATT (1330A>T), GTT223GTC (1339T>C), TAT224TAC (1342T>C), GGG227GGA (1351G>A), CAA228GAG (1352C>G 1354A>G), AGT229AGG (1357T>G), ACT230GCG (1358A>G 1360T>G), CAA231TCT (1361C>T 1362A>C 1363A>T), ATG232CAA (1364A>C 1365T>A 1366G>A), GCT233GCC (1369T>C), CTT234CTA (1372T>A), GAA235GAG (1375A>G), AAC236AAT (1378C>T), AAA237AGA (1380A>G), GCC238CCT (1382G>C 1384C>T), GTT240GTA (1390T>A), GGA241GGT (1393A>T), GAA242GCA (1395A>C), CAG243CTT (1398A>T 1399G>T), CTT244CTC (1402T>C), GGC246AAC (1406G>A 1407G>A), ACA247ACT (1411A>T), AGA248TAC (1412A>T 1413G>A 1414A>C), GAC250AAT (1418G>A 1420C>T), GTC251GTT (1423C>T), ATG252CCC (1424A>C 1425T>C 1426G>C), AAT253CAG (1427A>C 1429T>G), GAA254GAG (1432A>G), AGC255TCT (1433A>T 1434G>C 1435C>T), ACT256TCA (1436A>T 1438T>A), ATG257TTA (1439A>T 1441G>A), ATC258ATG (1444C>G), CGG259CGT (1447G>T), AGT260GCG (1448A>G 1449G>C 1450T>G), TCT262TCG (1456T>G), TTG263CTT (1457T>C 1459G>T), AGA264CGC (1460A>C 1462A>C), GTC265ATT (1463G>A 1465C>T), ACT267ACC (1471T>C), CTC268CTA (1474C>A), AAA269AAG (1477A>G), GTG270ATC (1478G>A 1480G>C), CAG271CAA (1483G>A), CGC274AGG (1490C>A 1492C>G), TTC275TTT (1495C>T), CCG276CCT (1498G>T), TCC277TCG (1501C>G), GAA279GAG (1507A>G), CCG281CCT (1513G>T), GCT282GAT (1515C>A), GTA283TTT (1517G>T 1519A>T), AGT284TCA (1520A>T 1521G>C 1522T>A), AGT284_ACA285insAGGGCC (1522_1523insAGGGCC), ACA285TCA (1523A>T), TCC287TCG (1531C>G), TTA288ACA (1532T>A 1533T>C), CTC289GTA (1535C>G 1537C>A), AGC290GGT (1538A>G 1540C>T), ACA291GTG (1541A>G 1542C>T 1543A>G), CGT292AGA (1544C>A 1546T>A), GAA293GAG (1549A>G), GTT296GGG (1557T>G 1558T>G), CAT297GTA (1559C>G 1560A>T 1561T>A), GAG299del (1565_1567delGAG), TGT302TCC (1575G>C 1576T>C), GGG303AAT (1577G>A 1578G>A 1579G>T), GGA304TTA (1580G>T 1581G>T), TTG305CTG (1583T>C), CCA307CCT (1591A>T), GCA308GTC (1593C>T 1594A>C), CCC309_AAG310del (1595_1600delCCCAAG), AAA311AAG (1603A>G), AAG312GAG (1604A>G), GGC313GCA (1608G>C 1609C>A), CAA314CAG (1612A>G), TTC316ATT (1616T>A 1618C>T), AAC317GAT (1619A>G 1621C>T), ATG318TTT (1622A>T 1624G>T), GGT319GGC (1627T>C), GCC320CAG (1628G>C 1629C>A 1630C>G), ATT321ATC (1633T>C), GAT323GAC (1639T>C), TTG325TAT (1644T>A 1645G>T), ATG327ATC (1651G>C), GAG328ACT (1652G>A 1653A>C 1654G>T), TCG329GAT (1655T>G 1656C>A 1657G>T), TTC330TTT (1660C>T), GCA331AAT (1661G>A 1662C>A 1663A>T), CAC332TCG (1664C>T 1665A>C 1666C>G), ATT333TTT (1667A>T), GAT338GAG (1684T>G), GAA341ACC (1691G>A 1692A>C 1693A>C), CGA342TGG (1694C>T 1696A>G), ACA343CAT (1697A>C 1698C>A 1699A>T), GCT345TCA (1703G>T 1705T>A), CAG346AGG (1706C>A 1707A>G), CAG347CAA (1711G>A), GTA348ATT (1712G>A 1714A>T), CAA349GAT (1715C>G 1717A>T), ATC351GTG (1721A>G 1723C>G), CTC352TTG (1724C>T 1726C>G), TTT353TCT (1728T>C), AGT354ACC (1731G>C 1732T>C), GAG355CTC (1733G>C 1734A>T 1735G>C), GCT356CTC (1736G>C 1737C>T 1738T>C), AAA357CGC (1739A>C 1740A>G 1741A>C), AGA358CAG (1742A>C 1743G>A 1744A>G), GGA359GGG (1747A>G), AAC360AGA (1749A>G 1750C>A), GTT361TGT (1751G>T 1752T>G), ATT362CAT (1754A>C 1755T>A), GTG363CTG (1757G>C), CCA364CCC (1762A>C), AGA365AAG (1764G>A 1765A>G), CAT366CAC (1768T>C), GTT367ATT (1769G>A), GCA368GAG (1773C>A 1774A>G), AAG369AGA (1776A>G 1777G>A), CGG370CGA (1780G>A), CAT371GAC (1781C>G 1783T>C), CTG372TCC (1784C>T 1785T>C 1786G>C), CAT373CAC (1789T>C), AAT374AAC (1792T>C), ATC375ATA (1795C>A), AAT376AAC (1798T>C), AGG377CTA (1799A>C 1800G>T 1801G>A), CAC379TAC (1805C>T), ATA380ATT (1810A>T), ACC381GAT (1811A>G 1812C>A 1813C>T), GAC383AAT (1817G>A 1819C>T), AAT384AAC (1822T>C), GTA386CTA (1826G>C), GAA387CAG (1829G>C 1831A>G), ATA388GTG (1832A>G 1834A>G), CTT389CTC (1837T>C), GAA390AAG (1838G>A 1840A>G), GGT391GAA (1842G>A 1843T>A), GGT393GGC (1849T>C), GTT394ATC (1850G>A 1852T>C), ACC395ACA (1855C>A), ATC397GTG (1859A>G 1861C>G), CAA398GGT (1862C>G 1863A>G 1864A>T), GGC399TCC (1865G>T 1866G>C), TTG400CTA (1868T>C 1870G>A), ACT401GAG (1871A>G 1872C>A 1873T>G), CGC402CGT (1876C>T), TTC404TGC (1881T>G), TAT407TAC (1891T>C), GCT408TCT (1892G>T), ATG409TTA (1895A>T 1897G>A), TCT410ACT (1898T>A), AAG412AGG (1905A>G), GAG413GAA (1909G>A), AGA414AGG (1912A>G), GTT415GTG (1915T>G), GTG416ATA (1916G>A 1918G>A), GAG417GAA (1921G>A), CTC418TTG (1922C>T 1924C>G), GTC419GTG (1927C>G), CAA421CAG (1933A>G), GAT423GAC (1939T>C), TCC425CAA (1943T>C 1944C>A 1945C>A), TTC426TAC (1947T>A), TAT427TAC (1951T>C), ATT428AGG (1953T>G 1954T>G), CAA429TAT (1955C>T 1957A>T), GGG430GGT (1960G>T), CAG431GAT (1961C>G 1963G>T), ACT432GAG (1964A>G 1965C>A 1966T>G), AAC433ACA (1968A>C 1969C>A), CCA435CCC (1975A>C), CCG437CCA (1981G>A), AAT438AGA (1983A>G 1984T>A), GAC440GAT (1990C>T), TAT442TAC (1996T>C), GGA444GGC (2002A>C), TTA445CTC (2003T>C 2005A>C), ACC446ACA (2008C>A), AAA448ATG (2013A>T 2014A>G), GAG449GAC (2017G>C), CGT450AGA (2018C>A 2020T>A), CAG451CAC (2023G>C), CAG451_TTG452insAAGGGTGGCAGGGCTAAG (2023_2024insAAGGGTGGCAGGGCTAAG), TTG452TTA (2026G>A), ATT453GAT (2027A>G 2028T>A), GTG454GTA (2032G>A), GAG455GCT (2034A>C 2035G>T), GAG456GGT (2037A>G 2038G>T), CAA457GCC (2039C>G 2040A>C 2041A>C), GTG458TTA (2042G>T 2044G>A), GCC459GCA (2047C>A), CAA460del (2048_2050delCAA), AGA461AGT (2053A>T), AGG462TTG (2054A>T 2055G>T), TCT463CCA (2057T>C 2059T>A), GAA464CAA (2060G>C), AGA465CCT (2063A>C 2064G>C 2065A>T), CAA466AGA (2066C>A 2067A>G), GCA467GTG (2070C>T 2071A>G), CAG468AAG (2072C>A), ACA470GCC (2078A>G 2080A>C), ACA470_GCA471insCAG (2080_2081insCAG), GCA471GAA (2082C>A), AGG472GGA (2084A>G 2086G>A), GGC473GGG (2089C>G), ATA474GGA (2090A>G 2091T>G), GCT475AGC (2093G>A 2094C>G 2095T>C), GAA476AGG (2096G>A 2097A>G 2098A>G), AGT477TCC (2099A>T 2100G>C 2101T>C), CAG478ACC (2102C>A 2103A>C 2104G>C), CCT479ACT (2105C>A), GAA480GAG (2110A>G), GAC481AGC (2111G>A 2112A>G), AGA482AAG (2115G>A 2116A>G), GTG483GTT (2119G>T), ACT484CTG (2120A>C 2121C>T 2122T>G), GTG488GCA (2133T>C 2134G>A), TCA489ACT (2135T>A 2137A>T), ACT490ACA (2140T>A), ACC491ACT (2143C>T), ACG492GCA (2144A>G 2146G>A), GAA494CAA (2150G>C), GAA494_GAT495insAGC (2152_2153insAGC), GAT495GAG (2155T>G), CCC496GGA (2156C>G 2157C>G 2158C>A), ACC497ATC (2160C>T), AAG498AAC (2164G>C), CCA499AAG (2165C>A 2166C>A 2167A>G), GAT500GAC (2170T>C), AAG501CAA (2171A>C 2173G>A), ATA502ATT (2176A>T), GAA503GTA (2178A>T), ATT504ATA (2182T>A), GTT505GAG (2184T>A 2185T>G), GCA506GTT (2187C>T 2188A>T), GAG507GCT (2190A>C 2191G>T), GGA508GAC (2193G>A 2194A>C), GCA509CCA (2195G>C), GAA510GAG (2200A>G), GAA511CAG (2201G>C 2203A>G), ACT513GTC (2207A>G 2208C>T 2209T>C), CAA514TCA (2210C>T 2211A>C), GGT516GAC (2217G>A 2218T>C), GTT518GTC (2224T>C), ATT519TTC (2225A>T 2227T>C), GGA523GGG (2239A>G), CCA524GAG (2240C>G 2241C>A 2242A>G), GGC1170GGT (4180C>T), AAT1171GAT (4181A>G), ACT1174ACG (4192T>G), CAC1175CAT (4195C>T), ACC1176ACT (4198C>T), TAT1177TAC (4201T>C), CAG1178CAA (4204G>A), TCA1181TCT (4213A>T), TTT1183TTC (4219T>C), TCT1184TCA (4222T>A), GAG1185GAA (4225G>A), CTA1186CTG (4228A>G), CGT1187AGA (4229C>A 4231T>A), GAT1188GAC (4234T>C), TTT1189TTC (4237T>C), GCG1190TCG (4238G>T), ATT1191AAC (4242T>A 4243T>C) | | | | | | | | |
| Coat protein A (YP_001976155.1) | 1 | 60 | 23.6% | 172 | 42.7% | 60 (96.8%) | 26 (41.9%) | 2/0/0/0 | 0 |
| Protein mutations: | T2A (2078A>G 2080A>C), T2_A3insQ (2080_2081insCAG), A3E (2082C>A), R4G (2084A>G 2086G>A), I6G (2090A>G 2091T>G), A7S (2093G>A 2094C>G 2095T>C), E8R (2096G>A 2097A>G 2098A>G), Q10T (2102C>A 2103A>C 2104G>C), P11T (2105C>A), D13S (2111G>A 2112A>G), R14K (2115G>A 2116A>G), T16L (2120A>C 2121C>T 2122T>G), V20A (2133T>C 2134G>A), S21T (2135T>A 2137A>T), T24A (2144A>G 2146G>A), E26Q (2150G>C), E26_D27insS (2152_2153insAGC), D27E (2155T>G), P28G (2156C>G 2157C>G 2158C>A), T29I (2160C>T), K30N (2164G>C), P31K (2165C>A 2166C>A 2167A>G), K33Q (2171A>C 2173G>A), E35V (2178A>T), V37E (2184T>A 2185T>G), A38V (2187C>T 2188A>T), E39A (2190A>C 2191G>T), G40D (2193G>A 2194A>C), A41P (2195G>C), E43Q (2201G>C 2203A>G), T45V (2207A>G 2208C>T 2209T>C), Q46S (2210C>T 2211A>C), G48D (2217G>A 2218T>C), I51F (2225A>T 2227T>C), P56E (2240C>G 2241C>A 2242A>G) | | | | | | | | |
| Codon mutations: | ACA2GCC (2078A>G 2080A>C), ACA2_GCA3insCAG (2080_2081insCAG), GCA3GAA (2082C>A), AGG4GGA (2084A>G 2086G>A), GGC5GGG (2089C>G), ATA6GGA (2090A>G 2091T>G), GCT7AGC (2093G>A 2094C>G 2095T>C), GAA8AGG (2096G>A 2097A>G 2098A>G), AGT9TCC (2099A>T 2100G>C 2101T>C), CAG10ACC (2102C>A 2103A>C 2104G>C), CCT11ACT (2105C>A), GAA12GAG (2110A>G), GAC13AGC (2111G>A 2112A>G), AGA14AAG (2115G>A 2116A>G), GTG15GTT (2119G>T), ACT16CTG (2120A>C 2121C>T 2122T>G), GTG20GCA (2133T>C 2134G>A), TCA21ACT (2135T>A 2137A>T), ACT22ACA (2140T>A), ACC23ACT (2143C>T), ACG24GCA (2144A>G 2146G>A), GAA26CAA (2150G>C), GAA26_GAT27insAGC (2152_2153insAGC), GAT27GAG (2155T>G), CCC28GGA (2156C>G 2157C>G 2158C>A), ACC29ATC (2160C>T), AAG30AAC (2164G>C), CCA31AAG (2165C>A 2166C>A 2167A>G), GAT32GAC (2170T>C), AAG33CAA (2171A>C 2173G>A), ATA34ATT (2176A>T), GAA35GTA (2178A>T), ATT36ATA (2182T>A), GTT37GAG (2184T>A 2185T>G), GCA38GTT (2187C>T 2188A>T), GAG39GCT (2190A>C 2191G>T), GGA40GAC (2193G>A 2194A>C), GCA41CCA (2195G>C), GAA42GAG (2200A>G), GAA43CAG (2201G>C 2203A>G), ACT45GTC (2207A>G 2208C>T 2209T>C), CAA46TCA (2210C>T 2211A>C), GGT48GAC (2217G>A 2218T>C), GTT50GTC (2224T>C), ATT51TTC (2225A>T 2227T>C), GGA55GGG (2239A>G), CCA56GAG (2240C>G 2241C>A 2242A>G) | | | | | | | | |
| Coat protein C (YP_001976157.1) | 197 | 225 | 12.9% | 192 | 90.6% | 29 (100%) | 26 (89.7%) | 0/0/0/0 | 0 |
| Protein mutations: | N205D (4181A>G), A224S (4238G>T), I225N (4242T>A 4243T>C) | | | | | | | | |
| Codon mutations: | GGC204GGT (4180C>T), AAT205GAT (4181A>G), ACT208ACG (4192T>G), CAC209CAT (4195C>T), ACC210ACT (4198C>T), TAT211TAC (4201T>C), CAG212CAA (4204G>A), TCA215TCT (4213A>T), TTT217TTC (4219T>C), TCT218TCA (4222T>A), GAG219GAA (4225G>A), CTA220CTG (4228A>G), CGT221AGA (4229C>A 4231T>A), GAT222GAC (4234T>C), TTT223TTC (4237T>C), GCG224TCG (4238G>T), ATT225AAC (4242T>A 4243T>C) | | | | | | | | |

*: Inserts / Deletes / Misaligned / Frameshifts

#### Analysis details

This analysis was performed with panviral2.64

### NGS Details (UN9): Torradovirus lycopersici (segment RNA 1)

#### Assembly

| | |
|---|---|
| **Coverage Length** | 7808 (1 contig(s)) |
| **Depth Of Coverage** | 29487.7 |
| **Number Of Reads** | 1650960 |
| **Reads Per Million** | 37316.42 rpm (after QC) |
| **Ambiguities** | 0 |
| **Assembly Method** | de novo + reference guided assembly |
| **Consensus Caller** | Bcf Tools |

#### Coverage Map

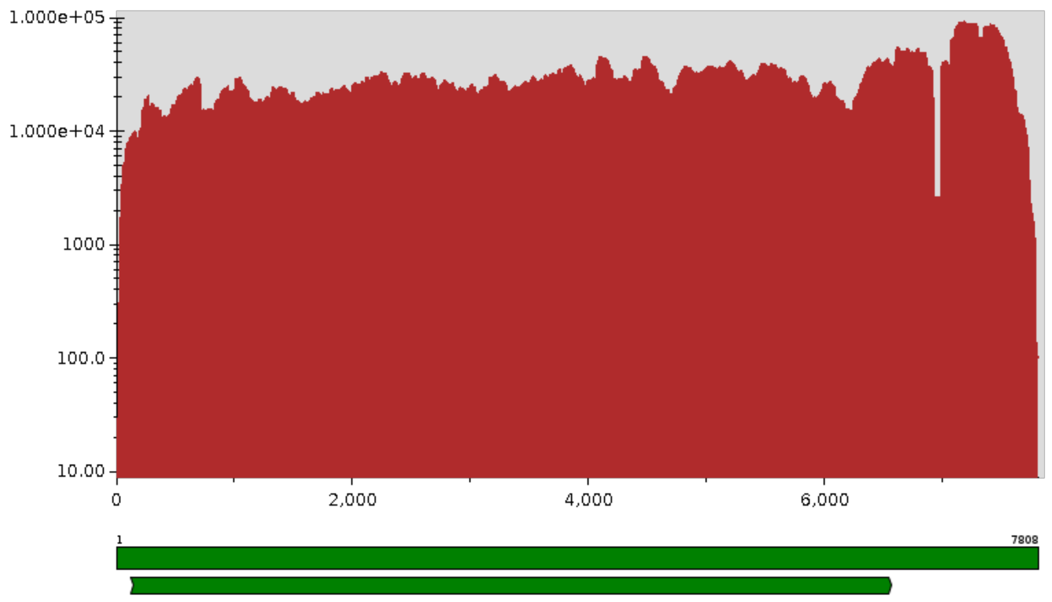

#### Assignment

| | |
|---|---|
| **Type** | Torradovirus lycopersici (Taxonomy ID: 3048378) |
| **Reference Genome** | NC_009013.1 |
| **NT Identity (%)** | 89.4563 |
| **AA Identity (%)** | 97.5452 |
| **Number Of Stop Codons** | 1 |
| **Number Of CDS** | 1 |

#### Alignment

| | |
|---|---|
| **Alignment Score** | 11384.0 (NT) + 14810.0 (AA) = 26194.0 |
| **Concordance (%)** | 88.347 |

| **Alignment Method** | Global, seeded, nucleotide + amino acids (AGA) |
|---|---|

#### Genome Region

Sequence starts at position 1 and ends at position 7808 relative to NC_009013.1 reference sequence.

#### Alignment Detailed Statistics

| | Begin | End | Coverage | Score | Concordance | Matches | Identities | I/D/M/F* | Stop Codons |
|---|---|---|---|---|---|---|---|---|---|
| **NT** | **1** | **7808** | **100%** | **11384** | **78.1%** | **7528 (96.0%)** | **6762 (86.3%)** | **31/280** | |
| Mutations: | 22T>A, 24T>G, 26A>T, 65A>G, 72C>T, 77A>T, 78A>G, 83C>T, 83_84insTACTTTATATTGTTTTGTGCT, 88T>C, 90C>G, 92T>C, 93T>A, 101G>A, 102G>C, 103C>A, 104A>G, 115T>C, 118C>T, 121G>A, 122A>T, 124G>A, 128C>T, 130C>T, 134T>C, 135T>C, 136C>T, 138A>G, 140T>C, 142A>C, 144T>C, 145T>A, 147C>T, 148T>G, 151A>G, 155T>A, 157C>T, 159C>T, 161A>G, 178T>G, 187A>T, 196T>A, 197G>C, 198C>A, 199T>A, 208T>C, 211T>C, 214C>T, 220C>T, 223T>A, 226G>T, 247A>C, 253A>G, 256T>C, 262T>A, 265C>T, 271A>G, 275T>C, 328T>C, 334T>C, 340G>A, 349G>A, 373C>T, 376C>T, 445C>T, 487C>T, 502G>A, 541T>C, 559C>T, 578C>T, 628C>T, 655C>T, 676A>G, 694A>G, 707C>T, 727A>G, 760C>T, 781C>T, 790C>T, 799A>G, 805T>C, 818C>T, 820A>G, 832A>T, 835A>C, 841T>C, 842C>T, 847C>T, 883G>A, 886T>C, 892G>A, 901T>C, 913T>C, 925C>T, 952A>G, 964T>G, 970G>A, 997C>T, 1006T>C, 1033G>A, 1060T>C, 1063C>T, 1064T>C, 1078A>T, 1081C>T, 1090C>T, 1093T>C, 1105A>G, 1108T>C, 1109C>T, 1165C>T, 1180G>A, 1196A>G, 1231G>A, 1240C>T, 1255G>A, 1267T>C, 1276A>G, 1300G>A, 1303A>G, 1319C>T, 1361T>C, 1366C>T, 1408T>C, 1411T>C, 1417T>C, 1423T>C, 1426T>C, 1429C>T, 1430C>T, 1441C>T, 1450T>C, 1451C>T, 1480C>T, 1481A>G, 1489T>C, 1522C>T, 1552C>T, 1555C>T, 1570A>G, 1573T>C, 1582C>T, 1585A>T, 1591C>T, 1594T>C, 1607G>T, 1612G>A, 1618T>G, 1619C>T, 1624C>T, 1630C>T, 1639T>C, 1642G>A, 1645A>G, 1669C>T, 1672G>A, 1681T>A, 1684G>T, 1699T>C, 1705T>C, 1737A>G, 1738A>G, 1750A>T, 1756A>G, 1765T>G, 1781T>C, 1783G>A, 1784C>T, 1789A>G, 1810G>A, 1825C>T, 1897A>G, 1910T>C, 1954T>C, 1960A>G, 1984C>T, 2014A>G, 2056T>C, 2077C>A, 2082T>A, 2092A>G, 2107A>G, 2128T>C, 2138C>T, 2143T>C, 2149T>C, 2155T>C, 2174C>T, 2179A>C, 2182G>A, 2194T>A, 2200T>C, 2215G>A, 2218C>T, 2224C>T, 2236A>G, 2241A>G, 2250G>A, 2251C>T, 2257C>T, 2263G>A, 2275T>C, 2287G>A, 2290C>T, 2314C>T, 2323A>T, 2347C>T, 2350A>G, 2356C>T, 2359G>A, 2362C>T, 2377G>A, 2380T>C, 2392A>G, 2407T>C, 2409G>A, 2443T>C, 2446G>A, 2479G>A, 2485C>T, 2497G>A, 2500T>C, 2503G>A, 2509C>T, 2512T>C, 2542G>A, 2548C>T, 2557C>T, 2620G>A, 2626T>C, 2650G>A, 2665T>C, 2675T>C, 2683A>G, 2692G>A, 2702C>T, 2710A>G, 2719T>C, 2725T>C, 2731C>T, 2734C>T, 2743A>G, 2744T>C, 2752A>G, 2773A>G, 2785C>T, 2788C>T, 2809C>T, 2821G>T, 2827A>G, 2842T>C, 2860C>T, 2866T>A, 2869C>T, 2872A>T, 2893T>A, 2896C>A, 2926G>A, 2929C>T, 2930T>C, 2941A>G, 2944T>C, 2950C>T, 2951C>T, 2980C>T, 2983G>A, 2998A>G, 3007T>C, 3016T>C, 3028T>C, 3031T>C, 3034G>A, 3040T>C, 3058A>G, 3064A>G, 3073G>A, 3079T>C, 3109T>C, 3115G>A, 3145A>G, 3151C>T, 3163T>C, 3169C>T, 3172T>C, 3202T>C, 3241T>C, 3253C>T, 3256G>A, 3265A>G, 3298T>C, 3319C>T, 3406A>C, 3418C>A, 3421T>C, 3463A>G, 3466G>A, 3481C>T, 3484A>T, 3508G>A, 3559G>A, 3568A>G, 3574T>C, 3586T>C, 3591C>T, 3610A>G, 3613T>C, 3625C>T, 3628G>A, 3646C>T, 3661A>G, 3664T>C, 3673G>A, 3676C>T, 3686C>T, 3706T>C, 3716T>C, 3721T>A, 3739A>T, 3751A>G, 3754G>A, 3758G>A, 3766G>A, 3781G>A, 3805C>T, 3808T>C, 3814C>T, 3817G>C, 3824T>C, 3835T>C, 3844T>C, 3857T>C, 3859G>A, 3868C>T, 3871T>C, 3883G>A, 3898T>C, 3907A>T, 3916C>T, 3925A>G, 3928G>A, 3949C>T, 3955G>A, 3961G>A, 3970A>G, 3973T>C, 4034T>C, 4066T>C, 4075A>G, 4084C>T, 4100T>C, 4118C>T, 4129C>T, 4147C>A, 4159G>A, 4162A>T, 4168A>G, 4186A>T, 4195A>G, 4207G>A, 4210T>C, 4228C>T, 4294A>G, 4318C>T, 4324T>C, 4339T>C, 4343C>T, 4348C>T, 4354T>C, 4396T>C, 4408C>T, 4414T>G, 4417T>C, 4420G>A, 4459T>C, 4462C>T, 4465C>T, 4468C>T, 4471T>C, 4474T>C, 4486T>C, 4489T>C, 4504T>C, 4510T>A, 4511C>T, 4514T>C, 4519A>G, 4543T>C, 4555A>G, 4570C>T, 4574C>T, 4576A>G, 4588C>T, 4591T>C, 4612A>G, 4625T>C, 4630T>C, 4648T>C, 4660A>G, 4666G>A, 4672G>C, 4679A>G, 4684A>T, 4687G>A, 4696C>T, 4702G>A, 4705C>T, 4714T>C, 4730C>T, 4735C>T, 4747A>G, 4759C>T, 4768T>C, 4774G>A, 4777C>T, 4783A>G, 4786A>G, 4789C>T, 4813A>G, 4816T>C, 4843G>A, 4898G>A, 4906G>A, 4969G>A, 4988A>G, 4993A>C, 4999T>C, 5012T>C, 5017G>A, 5023C>T, 5044A>C, 5059G>A, 5071A>G, 5074T>C, 5077T>C, 5079T>C, 5086T>C, 5102T>C, 5104G>A, 5119T>C, 5120G>A, 5122T>C, 5123C>T, 5128T>C, 5132T>A, 5146T>C, 5176C>A, 5209C>T, 5244G>A, 5245A>G, 5251A>G, 5287A>G, 5290G>A, 5302T>C, 5332C>T, 5354G>A, 5356G>A, 5365A>T, 5392G>A, 5413C>T, 5419A>C, 5429C>T, 5437T>C, 5464T>C, 5470A>G, 5491T>G, 5492A>T, 5497T>C, 5518T>C, 5533T>C, 5537A>C, 5560A>G, 5575A>T, 5578T>C, 5591G>T, 5603A>G, 5608A>G, 5611A>G, 5617A>T, 5641T>C, 5647T>C, 5656G>T, 5674A>G, 5677G>A, 5686G>A, 5690T>C, 5702G>A, 5704G>A, 5710T>C, 5716G>A, 5719A>C, 5731C>T, 5734T>C, 5737T>C, 5739G>A, 5746T>C, 5750T>C, 5752G>A, 5758C>T, 5773G>A, 5782A>G, 5784T>C, 5786T>C, 5797A>G, 5806C>T, 5836A>T, 5849T>C, 5850T>A, 5851G>A, 5869G>A, 5875A>G, 5878G>C, 5884T>C, 5887C>A, 5890C>T, 5902G>T, 5908C>T, 5909T>C, 5914T>C, 5923T>C, 5927C>T, 5932C>T, 5955A>G, 5965G>A, 5998A>G, 6004C>T, 6016A>G, 6028C>T, 6031A>T, 6033C>T, 6035G>T, 6055T>A, 6067C>A, 6070T>C, 6094T>C, 6100T>C, 6103G>A, 6106G>A, 6109C>T, 6112A>T, 6131T>C, 6139T>C, 6143A>C, 6145T>C, 6154C>T, 6157C>T, 6163C>T, 6166C>T, 6167C>T, 6184A>G, 6187T>C, 6193C>T, 6196A>G, 6202G>A, 6208A>G, 6211A>G, 6217G>A, 6223A>G, 6226G>A, 6230A>G, 6232A>G, 6235G>A, 6244A>T, 6253A>G, 6256C>A, 6265G>A, 6268G>A, 6273G>A, 6274C>T, 6295T>C, 6296G>T, 6298A>G, 6304A>G, 6308T>C, 6315A>T, 6319A>T, 6322G>A, 6323A>G, 6327A>G, 6331T>C, 6344C>T, 6358T>C, 6382C>T, 6385C>T, 6406T>C, 6412C>T, 6414G>A, 6427A>G, 6442C>G, 6445C>T, 6448G>A, 6451T>C, 6457C>A, 6466T>C, 6484A>C, 6490A>G, 6493T>C, 6500A>T, 6517C>T, 6532C>T, 6545C>T, 6571C>T, 6587G>A, 6661T>C, 6670C>T, 6685A>G, 6742A>T, 6766delC, 6786T>C, 6798_6799insA, 6806_6852delTAACGGAGAGTACTGACTTTAACTAGTTGGGAGTCCGGCTCCATTTG, 6854A>T, 6857A>G, 6858C>A, 6859C>T, 6860A>G, 6861A>G, 6864A>T, 6867T>A, 6871_6894delATTGGTTAAAGAGATTTGCACGCC, 6897T>A, 6905T>G, 6907T>A, 6908G>T, 6911T>C, 6912C>G, 6915G>C, 6921T>G, 6923C>G, 6924T>A, 6927G>T, 6928T>G, 6931G>T, 6933_6935delAAG, 6938C>A, 6947A>C, 6948_6951delGGTT, 6957A>T, 6959_6968delTACGGGGAGC, 6975T>A, 6984_6985insG, 6988G>A, 6994G>A, 6997A>T, 6998T>C, 7002A>T, 7003T>A, 7005_7012delATAGTCCT, 7014T>A, 7020_7026delAAGGGTT, 7031T>A, 7033G>A, 7035T>G, 7038G>A, 7043T>C, 7045T>G, 7047_7060delGCCGTGATGAAAGA, 7063A>T, 7069C>T, 7071T>A, 7074C>A, 7077T>C, 7083G>T, 7085A>G, 7086G>T, 7087_7099delGCTGAAACTACAC, 7101T>A, 7108T>A, 7109G>T, 7113T>G, 7117C>T, 7118T>G, 7120A>T, 7123_7125delCCT, 7129_7133delATCAG, 7136C>A, 7137G>C, 7141G>A, 7144_7169delATTCGATAATTTTCCGTAGCTTGCGT, 7173_7177delGCTGC, 7179C>G, 7182G>A, 7183T>C, 7184T>C, 7185A>C, 7189G>A, 7194_7216delAGTGAAGATGCGCCGTACCACGT, 7221C>T, 7224_7270delGGCAATGCCAGTGGTTCAGAGCGGGCCCTCAGAATAGAGGTTAAAAC, 7275_7285delGTGATGGTGTA, 7288T>A, 7294T>G, 7295A>C, 7297A>C, 7300T>G, 7303C>A, 7304_7315delACCCGGGTTGTG, 7320delG, 7325_7326delGT, 7332C>G, 7333G>T, 7336C>T, 7337A>G, 7339_7348delAGGTCCCACC, 7350T>A, 7351A>T, 7353A>T, 7354G>A, 7358A>C, 7360G>A, 7361A>C, 7362A>C, 7364C>T, 7368G>A, 7372A>T, 7375T>G, 7376T>G, 7377T>A, 7379A>T, 7388C>T, 7397G>A, 7398C>A, 7399G>A, 7400A>T, 7404C>T, 7407A>T, 7408G>T, 7423A>T, 7425G>A, 7435T>C, 7436G>A, 7444C>T, 7445A>G, 7457A>G, 7458T>C, 7460G>A, 7498C>T, 7503C>T, 7505A>G, 7511_7512insC, 7513A>C, 7514A>T, 7515A>C, 7522T>C, 7525_7526insG, 7526T>A, 7526_7527insG, 7545T>A, 7554A>G, 7569G>A, 7589T>A, 7600T>G, 7601T>A, 7602T>G, 7603_7604insC, 7612G>A, 7615A>G, 7616A>C, 7617A>T, 7618C>T, 7625A>T, 7628_7631delGCTT, 7637G>T, 7641T>A, 7642T>A, 7644G>T, 7647T>G, 7648G>T, 7649T>C, 7654C>A, 7655T>A, 7657G>C, 7665G>T, 7666T>C, 7667G>T, 7678C>T, 7686T>G, 7687T>A, 7689G>T, 7698G>A, 7713G>A, 7717A>C, 7719T>A, 7739T>A, 7742_7743insC, 7749T>C, 7751T>C, 7752G>A, 7754_7755insG, 7756_7757insT, 7762C>T, 7763A>C, 7764A>T, 7767C>T, 7773G>T, 7775G>A, 7778T>A, 7782A>T, 7783A>C, 7784_7785insT | | | | | | | | |

##### CDS

| | Begin | End | Coverage | Score | Concordance | Matches | Identities | I/D/M/F* | Stop Codons |
|---|---|---|---|---|---|---|---|---|---|
| ToTV_sRNA1gp1 | 1 | 2159 | 100% | 14810 | 98.3% | 2159 (100%) | 2106 (97.5%) | 0/0/0/0 | 1 |
| Protein mutations: | M6L (122A>T 124G>A), P8S (128C>T 130C>T), F10P (134T>C 135T>C 136C>T), N11S (138A>G), S12P (140T>C 142A>C), V13A (144T>C 145T>A), T14M (147C>T 148T>G), C17S (155T>A 157C>T), A18V (159C>T), T19A (161A>G), A31Q (197G>C 198C>A 199T>A), T364A (1196A>G), I459V (1481A>G), A501S (1607G>T), E526D (1684G>T), K544R (1737A>G 1738A>G), F659Y (2082T>A), N712S (2241A>G), S715N (2250G>A 2251C>T), R768K (2409G>A), A1162V (3591C>T), V1218I (3758G>A), P1413S (4343C>T), I1525V (4679A>G), V1598I (4898G>A), T1628A (4988A>G), I1658T (5079T>C), V1672I (5120G>A 5122T>C), S1676T (5132T>A), R1713K (5244G>A 5245A>G), V1750I (5354G>A 5356G>A), T1796S (5492A>T), K1811Q (5537A>C), A1829S (5591G>T), T1833A (5603A>G), V1866I (5702G>A 5704G>A), R1878K (5739G>A), I1893T (5784T>C), L1915Q (5849T>C 5850T>A 5851G>A), N1950S (5955A>G), T1976I (6033C>T), A1977S (6035G>T), I2013L (6143A>C 6145T>C), M2032I (6202G>A), I2042V (6230A>G 6232A>G), S2056N (6273G>A 6274C>T), V2064L (6296G>T 6298A>G), N2070I (6315A>T), I2073V (6323A>G), Q2074R (6327A>G), R2103Q (6414G>A), I2132F (6500A>T), L2147F (6545C>T) | | | | | | | | |

| | Begin | End | Coverage | Score | Concordance | Matches | Identities | I/D/M/F* | Stop Codons |
|---|---|---|---|---|---|---|---|---|---|
| **NT** | **1** | **7808** | **100%** | **11384** | **78.1%** | **7528 (96.0%)** | **6762 (86.3%)** | **31/280** | |
| Codon mutations: | TTT3TTC (115T>C), TCC4TCT (118C>T), AAG5AAA (121G>A), ATG6TTA (122A>T 124G>A), CCC8TCT (128C>T 130C>T), TTC10CCT (134T>C 135T>C 136C>T), AAC11AGC (138A>G), TCA12CCC (140T>C 142A>C), GTT13GCA (144T>C 145T>A), ACT14ATG (147C>T 148T>G), GAA15GAG (151A>G), TGC17AGT (155T>A 157C>T), GCT18GTT (159C>T), ACC19GCC (161A>G), TCT24TCG (178T>G), TCA27TCT (187A>T), ACT30ACA (196T>A), GCT31CAA (197G>C 198C>A 199T>A), AGT34AGC (208T>C), AAT35AAC (211T>C), TTC36TTT (214C>T), CGC38CGT (220C>T), ACT39ACA (223T>A), CTG40CTT (226G>T), TCA47TCC (247A>C), CAA49CAG (253A>G), ATT50ATC (256T>C), TCT52TCA (262T>A), CAC53CAT (265C>T), GAA55GAG (271A>G), TTG57CTG (275T>C), TCT74TCC (328T>C), ACT76ACC (334T>C), TTG78TTA (340G>A), AAG81AAA (349G>A), TTC89TTT (373C>T), ATC90ATT (376C>T), AGC113AGT (445C>T), GAC127GAT (487C>T), GTG132GTA (502G>A), TCT145TCC (541T>C), TTC151TTT (559C>T), CTA158TTA (578C>T), AAC174AAT (628C>T), GAC183GAT (655C>T), CAA190CAG (676A>G), GCA196GCG (694A>G), CTA201TTA (707C>T), TCA207TCG (727A>G), CTC218CTT (760C>T), ACC225ACT (781C>T), TTC228TTT (790C>T), GTA231GTG (799A>G), GGT233GGC (805T>C), CTA238TTG (818C>T 820A>G), CTA242CTT (832A>T), TCA243TCC (835A>C), ATT245ATC (841T>C), CTG246TTG (842C>T), TAC247TAT (847C>T), TTG259TTA (883G>A), TTT260TTC (886T>C), CTG262CTA (892G>A), AAT265AAC (901T>C), TGT269TGC (913T>C), GCC273GCT (925C>T), GTA282GTG (952A>G), GGT286GGG (964T>G), AAG288AAA (970G>A), GCC297GCT (997C>T), TCT300TCC (1006T>C), GAG309GAA (1033G>A), CAT318CAC (1060T>C), CGC319CGT (1063C>T), TTG320CTG (1064T>C), GCA324GCT (1078A>T), AAC325AAT (1081C>T), TTC328TTT (1090C>T), ACT329ACC (1093T>C), CAA333CAG (1105A>G), CAT334CAC (1108T>C), CTA335TTA (1109C>T), GCC353GCT (1165C>T), AGG358AGA (1180G>A), ACC364GCC (1196A>G), CAG375CAA (1231G>A), GGC378GGT (1240C>T), AGG383AGA (1255G>A), TTT387TTC (1267T>C), CAA390CAG (1276A>G), GGG398GGA (1300G>A), AAA399AAG (1303A>G), CTG405TTG (1319C>T), TTG419CTG (1361T>C), AAC420AAT (1366C>T), AAT434AAC (1408T>C), TAT435TAC (1411T>C), CAT437CAC (1417T>C), ACT439ACC (1423T>C), GCT440GCC (1426T>C), ATC441ATT (1429C>T), CTG442TTG (1430C>T), GAC445GAT (1441C>T), GCT448GCC (1450T>C), CTG449TTG (1451C>T), GAC458GAT (1480C>T), ATT459GTT (1481A>G), GAT461GAC (1489T>C), CCC472CCT (1522C>T), GGC482GGT (1552C>T), CGC483CGT (1555C>T), CGA488CGG (1570A>G), TAT489TAC (1573T>C), GCC492GCT (1582C>T), ACA493ACT (1585A>T), AAC495AAT (1591C>T), CTT496CTC (1594T>C), GCT501TCT (1607G>T), CAG502CAA (1612G>A), GGT504GGG (1618T>G), CTA505TTA (1619C>T), ACC506ACT (1624C>T), CCC508CCT (1630C>T), TTT511TTC (1639T>C), GAG512GAA (1642G>A), AGA513AGG (1645A>G), TGC521TGT (1669C>T), AGG522AGA (1672G>A), GGT525GGA (1681T>A), GAG526GAT (1684G>T), GCT531GCC (1699T>C), ACT533ACC (1705T>C), AAA544AGG (1737A>G 1738A>G), GCA548GCT (1750A>T), ACA550ACG (1756A>G), GGT553GGG (1765T>G), TTG559CTA (1781T>C 1783G>A), CTG560TTG (1784C>T), GAA561GAG (1789A>G), AAG568AAA (1810G>A), ATC573ATT (1825C>T), GCA597GCG (1897A>G), TTG602CTG (1910T>C), GTT616GTC (1954T>C), GTA618GTG (1960A>G), TAC626TAT (1984C>T), AAA636AAG (2014A>G), CCT650CCC (2056T>C), GCC657GCA (2077C>A), TTT659TAT (2082T>A), CTA662CTG (2092A>G), AAA667AAG (2107A>G), GTT674GTC (2128T>C), CTG678TTG (2138C>T), TAT679TAC (2143T>C), GGT681GGC (2149T>C), TGT683TGC (2155T>C), CTG690TTG (2174C>T), ATA691ATC (2179A>C), AAG692AAA (2182G>A), GCT696GCA (2194T>A), TAT698TAC (2200T>C), AAG703AAA (2215G>A), AGC704AGT (2218C>T), ATC706ATT (2224C>T), ACA710ACG (2236A>G), AAT712AGT (2241A>G), AGC715AAT (2250G>A 2251C>T), GAC717GAT (2257C>T), CAG719CAA (2263G>A), CCT723CCC (2275T>C), GCG727GCA (2287G>A), CAC728CAT (2290C>T), GAC736GAT (2314C>T), ATA739ATT (2323A>T), CAC747CAT (2347C>T), GTA748GTG (2350A>G), TTC750TTT (2356C>T), TTG751TTA (2359G>A), TAC752TAT (2362C>T), AGG757AGA (2377G>A), TAT758TAC (2380T>C), AGA762AGG (2392A>G), CCT767CCC (2407T>C), AGG768AAG (2409G>A), ATT779ATC (2443T>C), GTG780GTA (2446G>A), CCG791CCA (2479G>A), ATC793ATT (2485C>T), CTG797CTA (2497G>A), ATT798ATC (2500T>C), AAG799AAA (2503G>A), GGC801GGT (2509C>T), CTT802CTC (2512T>C), GGG812GGA (2542G>A), GTC814GTT (2548C>T), TTC817TTT (2557C>T), GAG838GAA (2620G>A), AAT840AAC (2626T>C), AAG848AAA (2650G>A), TTT853TTC (2665T>C), TTG857CTG (2675T>C), GCA859GCG (2683A>G), AAG862AAA (2692G>A), CTG866TTG (2702C>T), AAA868AAG (2710A>G), AGT871AGC (2719T>C), GAT873GAC (2725T>C), TTC875TTT (2731C>T), ATC876ATT (2734C>T), GGA879GGG (2743A>G), TTG880CTG (2744T>C), AAA882AAG (2752A>G), CTA889CTG (2773A>G), GTC893GTT (2785C>T), TAC894TAT (2788C>T), TGC901TGT (2809C>T), GTG905GTT (2821G>T), ACA907ACG (2827A>G), TTT912TTC (2842T>C), GGC918GGT (2860C>T), ACT920ACA (2866T>A), TTC921TTT (2869C>T), TCA922TCT (2872A>T), ATT929ATA (2893T>A), TCC930TCA (2896C>A), CGG940CGA (2926G>A), TTC941TTT (2929C>T), TTG942CTG (2930T>C), CAA945CAG (2941A>G), GAT946GAC (2944T>C), GTC948GTT (2950C>T), CTG949TTG (2951C>T), CCC958CCT (2980C>T), AAG959AAA (2983G>A), AAA964AAG (2998A>G), CCT967CCC (3007T>C), GAT970GAC (3016T>C), TTT974TTC (3028T>C), TCT975TCC (3031T>C), AAG976AAA (3034G>A), ACT978ACC (3040T>C), AGA984AGG (3058A>G), GTA986GTG (3064A>G), AAG989AAA (3073G>A), GCT991GCC (3079T>C), TTT1001TTC (3109T>C), GTG1003GTA (3115G>A), GTA1013GTG (3145A>G), TAC1015TAT (3151C>T), TCT1019TCC (3163T>C), TCC1021TCT (3169C>T), TTT1022TTC (3172T>C), TAT1032TAC (3202T>C), CTT1045CTC (3241T>C), AAC1049AAT (3253C>T), GTG1050GTA (3256G>A), AAA1053AAG (3265A>G), TCT1064TCC (3298T>C), CAC1071CAT (3319C>T), GTA1100GTC (3406A>C), CTC1104CTA (3418C>A), CTT1105CTC (3421T>C), AAA1119AAG (3463A>G), CAG1120CAA (3466G>A), GGC1125GGT (3481C>T), ACA1126ACT (3484A>T), ACG1134ACA (3508G>A), GGG1151GGA (3559G>A), GAA1154GAG (3568A>G), GCT1156GCC (3574T>C), ATT1160ATC (3586T>C), GCT1162GTT (3591C>T), GAA1168GAG (3610A>G), GGT1169GGC (3613T>C), TAC1173TAT (3625C>T), GTG1174GTA (3628G>A), TAC1180TAT (3646C>T), CAA1185CAG (3661A>G), AAT1186AAC (3664T>C), GAG1189GAA (3673G>A), GAC1190GAT (3676C>T), CTA1194TTA (3686C>T), CGT1200CGC (3706T>C), TTG1204CTG (3716T>C), CCT1205CCA (3721T>A), CGA1211CGT (3739A>T), GGA1215GGG (3751A>G), AAG1216AAA (3754G>A), GTC1218ATC (3758G>A), CAG1220CAA (3766G>A), GTG1225GTA (3781G>A), GAC1233GAT (3805C>T), TAT1234TAC (3808T>C), GAC1236GAT (3814C>T), GGG1237GGC (3817G>C), TTG1240CTG (3824T>C), TCT1243TCC (3835T>C), TAT1246TAC (3844T>C), TTG1251CTA (3857T>C 3859G>A), GGC1254GGT (3868C>T), ACT1255ACC (3871T>C), GAG1259GAA (3883G>A), AGT1264AGC (3898T>C), GGA1267GGT (3907A>T), GTC1270GTT (3916C>T), AAA1273AAG (3925A>G), AGG1274AGA (3928G>A), GAC1281GAT (3949C>T), GAG1283GAA (3955G>A), TTG1285TTA (3961G>A), GAA1288GAG (3970A>G), ATT1289ATC (3973T>C), TTG1310CTG (4034T>C), CGT1320CGC (4066T>C), GAA1323GAG (4075A>G), CCC1326CCT (4084C>T), TTG1332CTG (4100T>C), CTG1338TTG (4118C>T), TCC1341TCT (4129C>T), TCC1347TCA (4147C>A), CTG1351CTA (4159G>A), CCA1352CCT (4162A>T), AAA1354AAG (4168A>G), GGA1360GGT (4186A>T), GAA1363GAG (4195A>G), TTG1367TTA (4207G>A), GCT1368GCC (4210T>C), GTC1374GTT (4228C>T), CAA1396CAG (4294A>G), GAC1404GAT (4318C>T), TAT1406TAC (4324T>C), GGT1411GGC (4339T>C), CCA1413TCA (4343C>T), GAC1414GAT (4348C>T), AAT1416AAC (4354T>C), ACT1430ACC (4396T>C), AAC1434AAT (4408C>T), GTT1436GTG (4414T>G), TTT1437TTC (4417T>C), AGG1438AGA (4420G>A), GTT1451GTC (4459T>C), ATC1452ATT (4462C>T), TTC1453TTT (4465C>T), AAC1454AAT (4468C>T), TCT1455TCC (4471T>C), CTT1456CTC (4474T>C), TTT1460TTC (4486T>C), TAT1461TAC (4489T>C), TTT1466TTC (4504T>C), TCT1468TCA (4510T>A), CTA1469TTA (4511C>T), TTG1470CTG (4514T>C), AGA1471AGG (4519A>G), GCT1479GCC (4543T>C), AAA1483AAG (4555A>G), AAC1488AAT (4570C>T), CTA1490TTG (4574C>T 4576A>G), GTC1494GTT (4588C>T), TAT1495TAC (4591T>C), GCA1502GCG (4612A>G), TTG1507CTG (4625T>C), CAT1508CAC (4630T>C), GCT1514GCC (4648T>C), GAA1518GAG (4660A>G), GAG1520GAA (4666G>A), GTG1522GTC (4672G>C), ATT1525GTT (4679A>G), ATA1526ATT (4684A>T), AAG1527AAA (4687G>A), ATC1530ATT (4696C>T), AAG1532AAA (4702G>A), AAC1533AAT (4705C>T), GTT1536GTC (4714T>C), CTA1542TTA (4730C>T), GAC1543GAT (4735C>T), CTA1547CTG (4747A>G), AGC1551AGT (4759C>T), TTT1554TTC (4768T>C), AGG1556AGA (4774G>A), CAC1557CAT (4777C>T), CTA1559CTG (4783A>G), GGA1560GGG (4786A>G), TAC1561TAT (4789C>T), GTA1569GTG (4813A>G), TCT1570TCC (4816T>C), AGG1579AGA (4843G>A), GTT1598ATT (4898G>A), GAG1600GAA (4906G>A), AAG1621AAA (4969G>A), ACC1628GCC (4988A>G), ACA1629ACC (4993A>C), CCT1631CCC (4999T>C), TTG1636CTG (5012T>C), GAG1637GAA (5017G>A), GAC1639GAT (5023C>T), GGA1646GGC (5044A>C), GGG1651GGA (5059G>A), GAA1655GAG (5071A>G), CTT1656CTC (5074T>C), CAT1657CAC (5077T>C), ATT1658ACT (5079T>C), GCT1660GCC (5086T>C), TTG1666CTA (5102T>C 5104G>A), ACT1671ACC (5119T>C), GTT1672ATC (5120G>A 5122T>C), CTG1673TTG (5123C>T), AAT1674AAC (5128T>C), TCT1676ACT (5132T>A), CAT1680CAC (5146T>C), TCC1690TCA (5176C>A), ACC1701ACT (5209C>T), AGA1713AAG (5244G>A 5245A>G), ACA1715ACG (5251A>G), GAA1727GAG (5287A>G), GGG1728GGA (5290G>A), CCT1732CCC (5302T>C), ATC1742ATT (5332C>T), GTG1750ATA (5354G>A 5356G>A), CCA1753CCT (5365A>T), GGG1762GGA (5392G>A), TGC1769TGT (5413C>T), GCA1771GCC (5419A>C), CTG1775TTG (5429C>T), ACT1777ACC (5437T>C), GTT1786GTC (5464T>C), AGA1788AGG (5470A>G), GTT1795GTG (5491T>G), ACA1796TCA (5492A>T), GAT1797GAC (5497T>C), GGT1804GGC (5518T>C), CTT1809CTC (5533T>C), AAA1811CAA (5537A>C), GAA1818GAG (5560A>G), ACA1823ACT (5575A>T), TGT1824TGC (5578T>C), GCC1829TCC (5591G>T), ACA1833GCA (5603A>G), CCA1834CCG (5608A>G), GCA1835GCG (5611A>G), ATA1837ATT (5617A>T), GGT1845GGC (5641T>C), TTT1847TTC (5647T>C), GTG1850GTT (5656G>T), CTA1856CTG (5674A>G), CAG1857CAA (5677G>A), CAG1860CAA (5686G>A), TTG1862CTG (5690T>C), GTG1866ATA (5702G>A 5704G>A), CCT1868CCC (5710T>C), GGG1870GGA (5716G>A), GGA1871GGC (5719A>C), CAC1875CAT (5731C>T), ACT1876ACC (5734T>C), ACT1877ACC (5737T>C), AGA1878AAA (5739G>A), CTT1880CTC (5746T>C), TTG1882CTA (5750T>C 5752G>A), CTC1884CTT (5758C>T), AAG1889AAA (5773G>A), CAA1892CAG (5782A>G), ATT1893ACT (5784T>C), TTA1894CTA (5786T>C), AGA1897AGG (5797A>G), TTC1900TTT (5806C>T), TCA1910TCT (5836A>T), TTG1915CAA (5849T>C 5850T>A 5851G>A), AGG1921AGA (5869G>A), CAA1923CAG (5875A>G), GGG1924GGC (5878G>C), TTT1926TTC (5884T>C), CCC1927CCA (5887C>A), CGC1928CGT (5890C>T), GTG1932GTT (5902G>T), TAC1934TAT (5908C>T), TTG1935CTG (5909T>C), CTT1936CTC (5914T>C), TTT1939TTC (5923T>C), CTG1941TTG (5927C>T), AGC1942AGT (5932C>T), AAC1950AGC (5955A>G), ACG1953ACA (5965G>A), AAA1964AAG (5998A>G), ACC1966ACT (6004C>T), AAA1970AAG (6016A>G), TCC1974TCT (6028C>T), GCA1975GCT (6031A>T), ACA1976ATA (6033C>T), GCT1977TCT (6035G>T), GGT1983GGA (6055T>A), TCC1987TCA (6067C>A), GCT1988GCC (6070T>C), CTT1996CTC (6094T>C), CTT1998CTC (6100T>C), CAG1999CAA (6103G>A), AAG2000AAA (6106G>A), CCC2001CCT (6109C>T), ATA2002ATT (6112A>T), TTG2009CTG (6131T>C), TTT2011TTC (6139T>C), ATT2013CTC (6143A>C 6145T>C), GTC2016GTT (6154C>T), ATC2017ATT (6157C>T), GGC2019GGT (6163C>T), CAC2020CAT (6166C>T), CTG2021TTG (6167C>T), GTA2026GTG (6184A>G), ACT2027ACC (6187T>C), TTC2029TTT (6193C>T), CAA2030CAG (6196A>G), ATG2032ATA (6202G>A), CAA2034CAG (6208A>G), CAA2035CAG (6211A>G), CTG2037CTA (6217G>A), GAA2039GAG (6223A>G), GGG2040GGA (6226G>A), ATA2042GTG (6230A>G 6232A>G), GAG2043GAA (6235G>A), ATA2046ATT (6244A>T), TTA2049TTG (6253A>G), ACC2050ACA (6256C>A), CTG2053CTA (6265G>A), TCG2054TCA (6268G>A), AGC2056AAT (6273G>A 6274C>T), GAT2063GAC (6295T>C), GTA2064TTG (6296G>T 6298A>G), TTA2066TTG (6304A>G), TTG2068CTG (6308T>C), AAC2070ATC (6315A>T), ATA2071ATT (6319A>T), GAG2072GAA (6322G>A), ATC2073GTC (6323A>G), CAG2074CGG (6327A>G), GAT2075GAC (6331T>C), CTA2080TTA (6344C>T), AAT2084AAC (6358T>C), GCC2092GCT (6382C>T), ATC2093ATT (6385C>T), TGT2100TGC (6406T>C), AAC2102AAT (6412C>T), CGA2103CAA (6414G>A), AAA2107AAG (6427A>G), CGC2112CGG (6442C>G), TAC2113TAT (6445C>T), ACG2114ACA (6448G>A), ACT2115ACC (6451T>C), TCC2117TCA (6457C>A), GTT2120GTC (6466T>C), GCA2126GCC (6484A>C), AGA2128AGG (6490A>G), CAT2129CAC (6493T>C), ATT2132TTT (6500A>T), TCC2137TCT (6517C>T), CAC2142CAT (6532C>T), CTT2147TTT (6545C>T), TAC2155TAT (6571C>T) | | | | | | | | |

#### Proteins

| | Begin | End | Coverage | Score | Concordance | Matches | Identities | I/D/M/F* | Stop Codons |
|---|---|---|---|---|---|---|---|---|---|
| polyprotein (YP_001039627.1) | 1 | 2159 | 100% | 14810 | 98.3% | 2159 (100%) | 2106 (97.5%) | 0/0/0/0 | 1 |
| Protein mutations: | M6L (122A>T 124G>A), P8S (128C>T 130C>T), F10P (134T>C 135T>C 136C>T), N11S (138A>G), S12P (140T>C 142A>C), V13A (144T>C 145T>A), T14M (147C>T 148T>G), C17S (155T>A 157C>T), A18V (159C>T), T19A (161A>G), A31Q (197G>C 198C>A 199T>A), T364A (1196A>G), I459V (1481A>G), A501S (1607G>T), E526D (1684G>T), K544R (1737A>G 1738A>G), F659Y (2082T>A), N712S (2241A>G), S715N (2250G>A 2251C>T), R768K (2409G>A), A1162V (3591C>T), V1218I (3758G>A), P1413S (4343C>T), I1525V (4679A>G), V1598I (4898G>A), T1628A (4988A>G), I1658T (5079T>C), V1672I (5120G>A 5122T>C), S1676T (5132T>A), R1713K (5244G>A 5245A>G), V1750I (5354G>A 5356G>A), T1796S (5492A>T), K1811Q (5537A>C), A1829S (5591G>T), T1833A (5603A>G), V1866I (5702G>A 5704G>A), R1878K (5739G>A), I1893T (5784T>C), L1915Q (5849T>C 5850T>A 5851G>A), N1950S (5955A>G), T1976I (6033C>T), A1977S (6035G>T), I2013L (6143A>C 6145T>C), M2032I (6202G>A), I2042V (6230A>G 6232A>G), S2056N (6273G>A 6274C>T), V2064L (6296G>T 6298A>G), N2070I (6315A>T), I2073V (6323A>G), Q2074R (6327A>G), R2103Q (6414G>A), I2132F (6500A>T), L2147F (6545C>T) | | | | | | | | |

| | Begin | End | Coverage | Score | Concordance | Matches | Identities | I/D/M/F* | Stop Codons |
|---|---|---|---|---|---|---|---|---|---|
| **NT** | **1** | **7808** | **100%** | **11384** | **78.1%** | **7528 (96.0%)** | **6762 (86.3%)** | **31/280** | |

Codon mutations:

TTT3TTC (115T>C), TCC4TCT (118C>T), AAG5AAA (121G>A), ATG6TTA (122A>T 124G>A), CCC8TCT (128C>T 130C>T), TTC10CCT (134T>C 135T>C 136C>T), AAC11AGC (138A>G), TCA12CCC (140T>C 142A>C), GTT13GCA (144T>C 145T>A), ACT14ATG (147C>T 148T>G), GAA15GAG (151A>G), TGC17AGT (155T>A 157C>T), GCT18GTT (159C>T), ACC19GCC (161A>G), TCT24TCG (178T>G), TCA27TCT (187A>T), ACT30ACA (196T>A), GCT31CAA (197G>C 198C>A 199T>A), AGT34AGC (208T>C), AAT35AAC (211T>C), TTC36TTT (214C>T), CGC38CGT (220C>T), ACT39ACA (223T>A), CTG40CTT (226G>T), TCA47TCC (247A>C), CAA49CAG (253A>G), ATT50ATC (256T>C), TCT52TCA (262T>A), CAC53CAT (265C>T), GAA55GAG (271A>G), TTG57CTG (275T>C), TCT74TCC (328T>C), ACT76ACC (334T>C), TTG78TTA (340G>A), AAG81AAA (349G>A), TTC89TTT (373C>T), ATC90ATT (376C>T), AGC113AGT (445C>T), GAC127GAT (487C>T), GTG132GTA (502G>A), TCT145TCC (541T>C), TTC151TTT (559C>T), CTA158TTA (578C>T), AAC174AAT (628C>T), GAC183GAT (655C>T), CAA190CAG (676A>G), GCA196GCG (694A>G), CTA201TTA (707C>T), TCA207TCG (727A>G), CTC218CTT (760C>T), ACC225ACT (781C>T), TTC228TTT (790C>T), GTA231GTG (799A>G), GGT233GGC (805T>C), CTA238TTG (818C>T 820A>G), CTA242CTT (832A>T), TCA243TCC (835A>C), ATT245ATC (841T>C), CTG246TTG (842C>T), TAC247TAT (847C>T), TTG259TTA (883G>A), TTT260TTC (886T>C), CTG262CTA (892G>A), AAT265AAC (901T>C), TGT269TGC (913T>C), GCC273GCT (925C>T), GTA282GTG (952A>G), GGT286GGG (964T>G), AAG288AAA (970G>A), GCC297GCT (997C>T), TCT300TCC (1006T>C), GAG309GAA (1033G>A), CAT318CAC (1060T>C), CGC319CGT (1063C>T), TTG320CTG (1064T>C), GCA324GCT (1078A>T), AAC325AAT (1081C>T), TTC328TTT (1090C>T), ACT329ACC (1093T>C), CAA333CAG (1105A>G), CAT334CAC (1108T>C), CTA335TTA (1109C>T), GCC353GCT (1165C>T), AGG358AGA (1180G>A), ACC364GCC (1196A>G), CAG375CAA (1231G>A), GGC378GGT (1240C>T), AGG383AGA (1255G>A), TTT387TTC (1267T>C), CAA390CAG (1276A>G), GGG398GGA (1300G>A), AAA399AAG (1303A>G), CTG405TTG (1319C>T), TTG419CTG (1361T>C), AAC420AAT (1366C>T), AAT434AAC (1408T>C), TAT435TAC (1411T>C), CAT437CAC (1417T>C), ACT439ACC (1423T>C), GCT440GCC (1426T>C), ATC441ATT (1429C>T), CTG442TTG (1430C>T), GAC445GAT (1441C>T), GCT448GCC (1450T>C), CTG449TTG (1451C>T), GAC458GAT (1480C>T), ATT459GTT (1481A>G), GAT461GAC (1489T>C), CCC472CCT (1522C>T), GGC482GGT (1552C>T), CGC483CGT (1555C>T), CGA488CGG (1570A>G), TAT489TAC (1573T>C), GCC492GCT (1582C>T), ACA493ACT (1585A>T), AAC495AAT (1591C>T), CTT496CTC (1594T>C), GCT501TCT (1607G>T), CAG502CAA (1612G>A), GGT504GGG (1618T>G), CTA505TTA (1619C>T), ACC506ACT (1624C>T), CCC508CCT (1630C>T), TTT511TTC (1639T>C), GAG512GAA (1642G>A), AGA513AGG (1645A>G), TGC521TGT (1669C>T), AGG522AGA (1672G>A), GGT525GGA (1681T>A), GAG526GAT (1684G>T), GCT531GCC (1699T>C), ACT533ACC (1705T>C), AAA544AGG (1737A>G 1738A>G), GCA548GCT (1750A>T), ACA550ACG (1756A>G), GGT553GGG (1765T>G), TTG559CTA (1781T>C 1783G>A), CTG560TTG (1784C>T), GAA561GAG (1789A>G), AAG568AAA (1810G>A), ATC573ATT (1825C>T), GCA597GCG (1897A>G), TTG602CTG (1910T>C), GTT616GTC (1954T>C), GTA618GTG (1960A>G), TAC626TAT (1984C>T), AAA636AAG (2014A>G), CCT650CCC (2056T>C), GCC657GCA (2077C>A), TTT659TAT (2082T>A), CTA662CTG (2092A>G), AAA667AAG (2107A>G), GTT674GTC (2128T>C), CTG678TTG (2138C>T), TAT679TAC (2143T>C), GGT681GGC (2149T>C), TGT683TGC (2155T>C), CTG690TTG (2174C>T), ATA691ATC (2179A>C), AAG692AAA (2182G>A), GCT696GCA (2194T>A), TAT698TAC (2200T>C), AAG703AAA (2215G>A), AGC704AGT (2218C>T), ATC706ATT (2224C>T), ACA710ACG (2236A>G), AAT712AGT (2241A>G), AGC715AAT (2250G>A 2251C>T), GAC717GAT (2257C>T), CAG719CAA (2263G>A), CCT723CCC (2275T>C), GCG727GCA (2287G>A), CAC728CAT (2290C>T), GAC736GAT (2314C>T), ATA739ATT (2323A>T), CAC747CAT (2347C>T), GTA748GTG (2350A>G), TTC750TTT (2356C>T), TTG751TTA (2359G>A), TAC752TAT (2362C>T), AGG757AGA (2377G>A), TAT758TAC (2380T>C), AGA762AGG (2392A>G), CCT767CCC (2407T>C), AGG768AAG (2409G>A), ATT779ATC (2443T>C), GTG780GTA (2446G>A), CCG791CCA (2479G>A), ATC793ATT (2485C>T), CTG797CTA (2497G>A), ATT798ATC (2500T>C), AAG799AAA (2503G>A), GGC801GGT (2509C>T), CTT802CTC (2512T>C), GGG812GGA (2542G>A), GTC814GTT (2548C>T), TTC817TTT (2557C>T), GAG838GAA (2620G>A), AAT840AAC (2626T>C), AAG848AAA (2650G>A), TTT853TTC (2665T>C), TTG857CTG (2675T>C), GCA859GCG (2683A>G), AAG862AAA (2692G>A), CTG866TTG (2702C>T), AAA868AAG (2710A>G), AGT871AGC (2719T>C), GAT873GAC (2725T>C), TTC875TTT (2731C>T), ATC876ATT (2734C>T), GGA879GGG (2743A>G), TTG880CTG (2744T>C), AAA882AAG (2752A>G), CTA889CTG (2773A>G), GTC893GTT (2785C>T), TAC894TAT (2788C>T), TGC901TGT (2809C>T), GTG905GTT (2821G>T), ACA907ACG (2827A>G), TTT912TTC (2842T>C), GGC918GGT (2860C>T), ACT920ACA (2866T>A), TTC921TTT (2869C>T), TCA922TCT (2872A>T), ATT929ATA (2893T>A), TCC930TCA (2896C>A), CGG940CGA (2926G>A), TTC941TTT (2929C>T), TTG942CTG (2930T>C), CAA945CAG (2941A>G), GAT946GAC (2944T>C), GTC948GTT (2950C>T), CTG949TTG (2951C>T), CCC958CCT (2980C>T), AAG959AAA (2983G>A), AAA964AAG (2998A>G), CCT967CCC (3007T>C), GAT970GAC (3016T>C), TTT974TTC (3028T>C), TCT975TCC (3031T>C), AAG976AAA (3034G>A), ACT978ACC (3040T>C), AGA984AGG (3058A>G), GTA986GTG (3064A>G), AAG989AAA (3073G>A), GCT991GCC (3079T>C), TTT1001TTC (3109T>C), GTG1003GTA (3115G>A), GTA1013GTG (3145A>G), TAC1015TAT (3151C>T), TCT1019TCC (3163T>C), TCC1021TCT (3169C>T), TTT1022TTC (3172T>C), TAT1032TAC (3202T>C), CTT1045CTC (3241T>C), AAC1049AAT (3253C>T), GTG1050GTA (3256G>A), AAA1053AAG (3265A>G), TCT1064TCC (3298T>C), CAC1071CAT (3319C>T), GTA1100GTC (3406A>C), CTC1104CTA (3418C>A), CTT1105CTC (3421T>C), AAA1119AAG (3463A>G), CAG1120CAA (3466G>A), GGC1125GGT (3481C>T), ACA1126ACT (3484A>T), ACG1134ACA (3508G>A), GGG1151GGA (3559G>A), GAA1154GAG (3568A>G), GCT1156GCC (3574T>C), ATT1160ATC (3586T>C), GCT1162GTT (3591C>T), GAA1168GAG (3610A>G), GGT1169GGC (3613T>C), TAC1173TAT (3625C>T), GTG1174GTA (3628G>A), TAC1180TAT (3646C>T), CAA1185CAG (3661A>G), AAT1186AAC (3664T>C), GAG1189GAA (3673G>A), GAC1190GAT (3676C>T), CTA1194TTA (3686C>T), CGT1200CGC (3706T>C), TTG1204CTG (3716T>C), CCT1205CCA (3721T>A), CGA1211CGT (3739A>T), GGA1215GGG (3751A>G), AAG1216AAA (3754G>A), GTC1218ATC (3758G>A), CAG1220CAA (3766G>A), GTG1225GTA (3781G>A), GAC1233GAT (3805C>T), TAT1234TAC (3808T>C), GAC1236GAT (3814C>T), GGG1237GGC (3817G>C), TTG1240CTG (3824T>C), TCT1243TCC (3835T>C), TAT1246TAC (3844T>C), TTG1251CTA (3857T>C 3859G>A), GGC1254GGT (3868C>T), ACT1255ACC (3871T>C), GAG1259GAA (3883G>A), AGT1264AGC (3898T>C), GGA1267GGT (3907A>T), GTC1270GTT (3916C>T), AAA1273AAG (3925A>G), AGG1274AGA (3928G>A), GAC1281GAT (3949C>T), GAG1283GAA (3955G>A), TTG1285TTA (3961G>A), GAA1288GAG (3970A>G), ATT1289ATC (3973T>C), TTG1310CTG (4034T>C), CGT1320CGC (4066T>C), GAA1323GAG (4075A>G), CCC1326CCT (4084C>T), TTG1332CTG (4100T>C), CTG1338TTG (4118C>T), TCC1341TCT (4129C>T), TCC1347TCA (4147C>A), CTG1351CTA (4159G>A), CCA1352CCT (4162A>T), AAA1354AAG (4168A>G), GGA1360GGT (4186A>T), GAA1363GAG (4195A>G), TTG1367TTA (4207G>A), GCT1368GCC (4210T>C), GTC1374GTT (4228C>T), CAA1396CAG (4294A>G), GAC1404GAT (4318C>T), TAT1406TAC (4324T>C), GGT1411GGC (4339T>C), CCA1413TCA (4343C>T), GAC1414GAT (4348C>T), AAT1416AAC (4354T>C), ACT1430ACC (4396T>C), AAC1434AAT (4408C>T), GTT1436GTG (4414T>G), TTT1437TTC (4417T>C), AGG1438AGA (4420G>A), GTT1451GTC (4459T>C), ATC1452ATT (4462C>T), TTC1453TTT (4465C>T), AAC1454AAT (4468C>T), TCT1455TCC (4471T>C), CTT1456CTC (4474T>C), TTT1460TTC (4486T>C), TAT1461TAC (4489T>C), TTT1466TTC (4504T>C), TCT1468TCA (4510T>A), CTA1469TTA (4511C>T), TTG1470CTG (4514T>C), AGA1471AGG (4519A>G), GCT1479GCC (4543T>C), AAA1483AAG (4555A>G), AAC1488AAT (4570C>T), CTA1490TTG (4574C>T 4576A>G), GTC1494GTT (4588C>T), TAT1495TAC (4591T>C), GCA1502GCG (4612A>G), TTG1507CTG (4625T>C), CAT1508CAC (4630T>C), GCT1514GCC (4648T>C), GAA1518GAG (4660A>G), GAG1520GAA (4666G>A), GTG1522GTC (4672G>C), ATT1525GTT (4679A>G), ATA1526ATT (4684A>T), AAG1527AAA (4687G>A), ATC1530ATT (4696C>T), AAG1532AAA (4702G>A), AAC1533AAT (4705C>T), GTT1536GTC (4714T>C), CTA1542TTA (4730C>T), GAC1543GAT (4735C>T), CTA1547CTG (4747A>G), AGC1551AGT (4759C>T), TTT1554TTC (4768T>C), AGG1556AGA (4774G>A), CAC1557CAT (4777C>T), CTA1559CTG (4783A>G), GGA1560GGG (4786A>G), TAC1561TAT (4789C>T), GTA1569GTG (4813A>G), TCT1570TCC (4816T>C), AGG1579AGA (4843G>A), GTT1598ATT (4898G>A), GAG1600GAA (4906G>A), AAG1621AAA (4969G>A), ACC1628GCC (4988A>G), ACA1629ACC (4993A>C), CCT1631CCC (4999T>C), TTG1636CTG (5012T>C), GAG1637GAA (5017G>A), GAC1639GAT (5023C>T), GGA1646GGC (5044A>C), GGG1651GGA (5059G>A), GAA1655GAG (5071A>G), CTT1656CTC (5074T>C), CAT1657CAC (5077T>C), ATT1658ACT (5079T>C), GCT1660GCC (5086T>C), TTG1666CTA (5102T>C 5104G>A), ACT1671ACC (5119T>C), GTT1672ATC (5120G>A 5122T>C), CTG1673TTG (5123C>T), AAT1674AAC (5128T>C), TCT1676ACT (5132T>A), CAT1680CAC (5146T>C), TCC1690TCA (5176C>A), ACC1701ACT (5209C>T), AGA1713AAG (5244G>A 5245A>G), ACA1715ACG (5251A>G), GAA1727GAG (5287A>G), GGG1728GGA (5290G>A), CCT1732CCC (5302T>C), ATC1742ATT (5332C>T), GTG1750ATA (5354G>A 5356G>A), CCA1753CCT (5365A>T), GGG1762GGA (5392G>A), TGC1769TGT (5413C>T), GCA1771GCC (5419A>C), CTG1775TTG (5429C>T), ACT1777ACC (5437T>C), GTT1786GTC (5464T>C), AGA1788AGG (5470A>G), GTT1795GTG (5491T>G), ACA1796TCA (5492A>T), GAT1797GAC (5497T>C), GGT1804GGC (5518T>C), CTT1809CTC (5533T>C), AAA1811CAA (5537A>C), GAA1818GAG (5560A>G), ACA1823ACT (5575A>T), TGT1824TGC (5578T>C), GCC1829TCC (5591G>T), ACA1833GCA (5603A>G), CCA1834CCG (5608A>G), GCA1835GCG (5611A>G), ATA1837ATT (5617A>T), GGT1845GGC (5641T>C), TTT1847TTC (5647T>C), GTG1850GTT (5656G>T), CTA1856CTG (5674A>G), CAG1857CAA (5677G>A), CAG1860CAA (5686G>A), TTG1862CTG (5690T>C), GTG1866ATA (5702G>A 5704G>A), CCT1868CCC (5710T>C), GGG1870GGA (5716G>A), GGA1871GGC (5719A>C), CAC1875CAT (5731C>T), ACT1876ACC (5734T>C), ACT1877ACC (5737T>C), AGA1878AAA (5739G>A), CTT1880CTC (5746T>C), TTG1882CTA (5750T>C 5752G>A), CTC1884CTT (5758C>T), AAG1889AAA (5773G>A), CAA1892CAG (5782A>G), ATT1893ACT (5784T>C), TTA1894CTA (5786T>C), AGA1897AGG (5797A>G), TTC1900TTT (5806C>T), TCA1910TCT (5836A>T), TTG1915CAA (5849T>C 5850T>A 5851G>A), AGG1921AGA (5869G>A), CAA1923CAG (5875A>G), GGG1924GGC (5878G>C), TTT1926TTC (5884T>C), CCC1927CCA (5887C>A), CGC1928CGT (5890C>T), GTG1932GTT (5902G>T), TAC1934TAT (5908C>T), TTG1935CTG (5909T>C), CTT1936CTC (5914T>C), TTT1939TTC (5923T>C), CTG1941TTG (5927C>T), AGC1942AGT (5932C>T), AAC1950AGC (5955A>G), ACG1953ACA (5965G>A), AAA1964AAG (5998A>G), ACC1966ACT (6004C>T), AAA1970AAG (6016A>G), TCC1974TCT (6028C>T), GCA1975GCT (6031A>T), ACA1976ATA (6033C>T), GCT1977TCT (6035G>T), GGT1983GGA (6055T>A), TCC1987TCA (6067C>A), GCT1988GCC (6070T>C), CTT1996CTC (6094T>C), CTT1998CTC (6100T>C), CAG1999CAA (6103G>A), AAG2000AAA (6106G>A), CCC2001CCT (6109C>T), ATA2002ATT (6112A>T), TTG2009CTG (6131T>C), TTT2011TTC (6139T>C), ATT2013CTC (6143A>C 6145T>C), GTC2016GTT (6154C>T), ATC2017ATT (6157C>T), GGC2019GGT (6163C>T), CAC2020CAT (6166C>T), CTG2021TTG (6167C>T), GTA2026GTG (6184A>G), ACT2027ACC (6187T>C), TTC2029TTT (6193C>T), CAA2030CAG (6196A>G), ATG2032ATA (6202G>A), CAA2034CAG (6208A>G), CAA2035CAG (6211A>G), CTG2037CTA (6217G>A), GAA2039GAG (6223A>G), GGG2040GGA (6226G>A), ATA2042GTG (6230A>G 6232A>G), GAG2043GAA (6235G>A), ATA2046ATT (6244A>T), TTA2049TTG (6253A>G), ACC2050ACA (6256C>A), CTG2053CTA (6265G>A), TCG2054TCA (6268G>A), AGC2056AAT (6273G>A 6274C>T), GAT2063GAC (6295T>C), GTA2064TTG (6296G>T 6298A>G), TTA2066TTG (6304A>G), TTG2068CTG (6308T>C), AAC2070ATC (6315A>T), ATA2071ATT (6319A>T), GAG2072GAA (6322G>A), ATC2073GTC (6323A>G), CAG2074CGG (6327A>G), GAT2075GAC (6331T>C), CTA2080TTA (6344C>T), AAT2084AAC (6358T>C), GCC2092GCT (6382C>T), ATC2093ATT (6385C>T), TGT2100TGC (6406T>C), AAC2102AAT (6412C>T), CGA2103CAA (6414G>A), AAA2107AAG (6427A>G), CGC2112CGG (6442C>G), TAC2113TAT (6445C>T), ACG2114ACA (6448G>A), ACT2115ACC (6451T>C), TCC2117TCA (6457C>A), GTT2120GTC (6466T>C), GCA2126GCC (6484A>C), AGA2128AGG (6490A>G), CAT2129CAC (6493T>C), ATT2132TTT (6500A>T), TCC2137TCT (6517C>T), CAC2142CAT (6532C>T), CTT2147TTT (6545C>T), TAC2155TAT (6571C>T)

*: Inserts / Deletes / Misaligned / Frameshifts

#### Analysis details

This analysis was performed with panviral2.64

### NGS Details (UN9): Torradovirus lycopersici (segment RNA 2)

#### Assembly

| | |
|---|---|
| **Coverage Length** | 5379 (1 contig(s)) |
| **Depth Of Coverage** | 60857.9 |
| **Number Of Reads** | 2524050 |
| **Reads Per Million** | 57050.75 rpm (after QC) |
| **Ambiguities** | 0 |
| **Assembly Method** | de novo + reference guided assembly |
| **Consensus Caller** | Bcf Tools |

#### Coverage Map

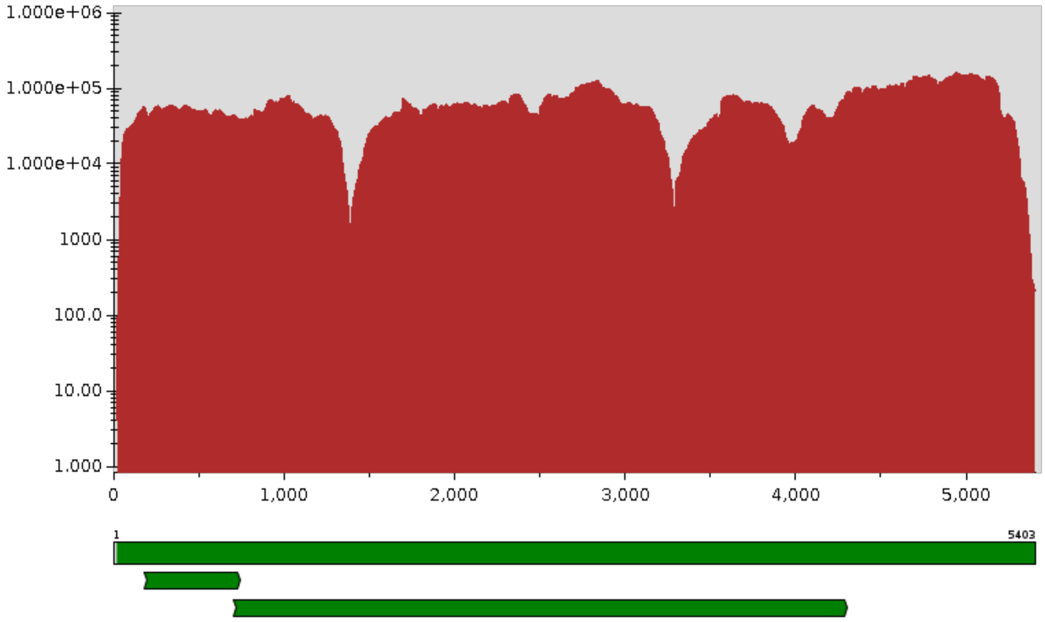

#### Assignment

| | |
|---|---|
| **Type** | Torradovirus lycopersici (Taxonomy ID: 3048378) |
| **Reference Genome** | NC_009032.1 |
| **NT Identity (%)** | 94.8151 |
| **AA Identity (%)** | 96.4672 |
| **Number Of Stop Codons** | 3 |
| **Number Of CDS** | 2 |

#### Alignment

| | |
|---|---|
| **Alignment Score** | 9630.0 (NT) + 9213.0 (AA) = 18843.0 |
| **Concordance (%)** | 93.0748 |

| Alignment Method | Global, seeded, nucleotide + amino acids (AGA) |
|---|---|

#### Genome Region

Sequence starts at position 25 and ends at position 5403 relative to NC_009032.1 reference sequence.

#### Alignment Detailed Statistics

| | Begin | End | Coverage | Score | Concordance | Matches | Identities | I/D/M/F* | Stop Codons |
|---|---|---|---|---|---|---|---|---|---|
| **NT** | **25** | **5403** | **99.6%** | **9630** | **89.5%** | **5379 (99.9%)** | **5102 (94.8%)** | **2/0** | |
| Mutations: | 89G>A, 90C>T, 97C>T, 107G>A, 111C>A, 137T>C, 144A>G, 161A>G, 277C>T, 280T>C, 283C>T, 295T>C, 301A>C, 313C>T, 343T>C, 401T>C, 433C>T, 440C>T, 448C>A, 460T>C, 481C>A, 487A>G, 507G>A, 583A>T, 586A>G, 589A>G, 654T>A, 679G>A, 696T>C, 721C>T, 733A>G, 748A>G, 750A>G, 797T>C, 806C>T, 827A>G, 844A>T, 866A>G, 968A>G, 986G>A, 1004C>T, 1061T>C, 1064T>C, 1085G>A, 1106T>C, 1109C>T, 1139G>A, 1226A>G, 1271T>C, 1277A>G, 1300C>T, 1316C>T, 1337T>C, 1346G>A, 1355G>A, 1361A>T, 1367A>T, 1383A>G, 1384C>T, 1390C>G, 1394A>T, 1412G>T, 1421A>G, 1426A>C, 1436C>T, 1439A>C, 1457C>T, 1460A>G, 1465G>A, 1472C>G, 1475G>C, 1484G>A, 1487C>G, 1490G>A, 1502C>T, 1508G>A, 1547C>T, 1559C>A, 1608T>C, 1619G>A, 1637G>A, 1646T>C, 1651A>G, 1662C>T, 1685T>C, 1694T>C, 1724C>T, 1757A>G, 1794G>A, 1817A>T, 1819T>A, 1820C>G, 1847G>T, 1850G>A, 1859C>T, 1863C>T, 1881C>T, 1892C>T, 1896A>G, 1899G>A, 1900T>C, 1904T>G, 1906C>T, 1914C>T, 1967G>A, 1970A>G, 1994T>C, 2072G>A, 2084G>A, 2108A>G, 2128G>A, 2135G>A, 2138G>T, 2140A>C, 2159T>A, 2181A>G, 2182C>A, 2183C>T, 2189C>T, 2199G>A, 2200T>C, 2205A>G, 2207G>T, 2216C>G, 2225T>A, 2249T>C, 2263A>C, 2270A>G, 2285T>C, 2303A>G, 2312T>C, 2321T>C, 2324G>A, 2375C>T, 2405G>A, 2408A>G, 2414T>C, 2429G>A, 2432G>T, 2436T>G, 2459C>G, 2465T>C, 2468A>G, 2469T>A, 2471C>T, 2501A>C, 2503G>A, 2519A>G, 2570G>A, 2573T>C, 2612C>T, 2630C>T, 2660T>C, 2708A>G, 2720C>T, 2732T>C, 2792C>T, 2819G>A, 2840T>C, 2841C>T, 2885T>C, 2939A>G, 2969C>T, 3003T>C, 3050G>A, 3122T>C, 3128T>C, 3131G>A, 3168C>T, 3215T>C, 3227T>G, 3236C>T, 3239T>G, 3242G>A, 3253A>G, 3265G>A, 3266G>A, 3269T>C, 3270A>T, 3296T>C, 3303A>T, 3305C>T, 3311T>A, 3323T>C, 3324C>T, 3332G>A, 3335C>T, 3354A>C, 3368A>T, 3386A>T, 3395A>C, 3404A>G, 3413C>T, 3428G>A, 3429C>T, 3431T>G, 3437C>T, 3440A>T, 3447T>C, 3461C>T, 3470T>C, 3485C>T, 3491C>T, 3497T>C, 3503A>G, 3508A>G, 3524A>G, 3527C>T, 3533G>A, 3540T>C, 3543T>A, 3560T>A, 3569A>G, 3575G>A, 3581C>T, 3590A>C, 3605A>G, 3626A>G, 3638A>G, 3662A>T, 3674G>A, 3716C>T, 3722C>T, 3743G>A, 3767C>T, 3773C>T, 3800C>T, 3827C>T, 3863C>T, 3869A>T, 3872A>T, 3885A>G, 3893T>C, 3911T>C, 3914T>A, 3920A>T, 3932C>A, 3933A>G, 3941T>G, 3944C>A, 3950T>A, 3960G>A, 3984A>G, 3986G>T, 3990G>A, 3995C>T, 4001T>C, 4013G>C, 4040C>T, 4046C>T, 4059A>G, 4061A>G, 4069A>G, 4100A>T, 4112C>T, 4130C>T, 4157C>T, 4202T>C, 4205T>A, 4208A>T, 4224C>T, 4250C>T, 4457A>G, 4518G>A, 4556A>G, 4590G>A, 4690C>T, 4703G>A, 4749T>A, 4812C>T, 4823A>G, 4974_4975insT, 5089T>C, 5192C>T, 5206C>T, 5218C>T, 5235_5236insT, 5249A>G, 5262A>T | | | | | | | | |
| **CDS** | | | | | | | | | |
| ToTV_sRNA2gp1 | 1 | 188 | 100% | 1189 | 98.3% | 188 (100%) | 185 (98.4%) | 0/0/0/0 | 1 |
| Protein mutations: | R109K (507G>A), L158Q (654T>A), L172P (696T>C) | | | | | | | | |
| Codon mutations: | GGC32GGT (277C>T), TCT33TCC (280T>C), GGC34GGT (283C>T), AGT38AGC (295T>C), CCA40CCC (301A>C), TTC44TTT (313C>T), ATT54ATC (343T>C), TTA74CTA (401T>C), GGC84GGT (433C>T), CTA87TTA (440C>T), GGC89GGA (448C>A), CAT93CAC (460T>C), ACC100ACA (481C>A), AGA102AGG (487A>G), AGG109AAG (507G>A), ACA134ACT (583A>T), CAA135CAG (586A>G), TTA136TTG (589A>G), CTA158CAA (654T>A), TTG166TTA (679G>A), CTA172CCA (696T>C), GGC180GGT (721C>T), GAA184GAG (733A>G) | | | | | | | | |
| ToTV_sRNA2gp2 | 1 | 1199 | 100% | 8024 | 96.9% | 1199 (100%) | 1153 (96.2%) | 0/0/0/0 | 2 |
| Protein mutations: | A7V (721C>T), K11R (733A>G), K16R (748A>G), S17G (750A>G), Y48F (844A>T), A200V (1300C>T), T228V (1383A>G 1384C>T), S230* (1390C>G), Q242P (1426A>C), R255K (1465G>A), I257M (1472C>G), H286Q (1559C>A), Y303H (1608T>C), K317R (1651A>G), V365I (1794G>A), I373K (1819T>A 1820C>G), L382F (1847G>T), H388Y (1863C>T), I399V (1896A>G), V400T (1899G>A 1900T>C), N401K (1904T>G), A402V (1906C>T), R476K (2128G>A), K480T (2140A>C), T494D (2181A>G 2182C>A 2183C>T), V500T (2199G>A 2200T>C), M502V (2205A>G 2207G>T), D508E (2225T>A), E521A (2263A>C), S579A (2436T>G), S590T (2469T>A 2471C>T), R601K (2503G>A), K851R (3253A>G), R855K (3265G>A 3266G>A), T857S (3270A>T), T868S (3303A>T 3305C>T), N936S (3508A>G), S948T (3543T>A), T1062A (3885A>G), T1078A (3933A>G), A1087T (3960G>A), M1095V (3984A>G 3986G>T), D1097N (3990G>A), I1120V (4059A>G 4061A>G), N1123S (4069A>G), H1175Y (4224C>T) | | | | | | | | |
| Codon mutations: | GCT7GTT (721C>T), AAG11AGG (733A>G), AAA16AGA (748A>G), AGT17GGT (750A>G), GTT32GTC (797T>C), GGC35GGT (806C>T), AAA42AAG (827A>G), TAT48TTT (844A>T), CAA55CAG (866A>G), GGA89GGG (968A>G), TTG95TTA (986G>A), TTC101TTT (1004C>T), AGT120AGC (1061T>C), AAT121AAC (1064T>C), CTG128CTA (1085G>A), TTT135TTC (1106T>C), TCC136TCT (1109C>T), CCG146CCA (1139G>A), TCA175TCG (1226A>G), ATT190ATC (1271T>C), TCA192TCG (1277A>G), GCT200GTT (1300C>T), CAC205CAT (1316C>T), GAT212GAC (1337T>C), GAG215GAA (1346G>A), CAG218CAA (1355G>A), ATA220ATT (1361A>T), TCA222TCT (1367A>T), ACC228GTC (1383A>G 1384C>T), TCA230TGA (1390C>G), GCA231GCT (1394A>T), CCG237CCT (1412G>T), GGA240GGG (1421A>G), CAG242CCG (1426A>C), GAC245GAT (1436C>T), ACA246ACC (1439A>C), AGC252AGT (1457C>T), GAA253GAG (1460A>G), AGA255AAA (1465G>A), ATC257ATG (1472C>G), CGG258CGC (1475G>C), TCG261TCA (1484G>A), CTC262CTG (1487C>G), CGG263CGA (1490G>A), CTC267CTT (1502C>T), GTG269GTA (1508G>A), ACC282ACT (1547C>T), CAC286CAA (1559C>A), TAC303CAC (1608T>C), GAG306GAA (1619G>A), TTG312TTA (1637G>A), GTT315GTC (1646T>C), AAG317AGG (1651A>G), CTG321TTG (1662C>T), GAT328GAC (1685T>C), GGT331GGC (1694T>C), CCC341CCT (1724C>T), CAA352CAG (1757A>G), GTT365ATT (1794G>A), ATA372ATT (1817A>T), ATC373AAG (1819T>A 1820C>G), TTG382TTT (1847G>T), GAG383GAA (1850G>A), ACC386ACT (1859C>T), CAC388TAC (1863C>T), CTG394TTG (1881C>T), TAC397TAT (1892C>T), ATT399GTT (1896A>G), GTG400ACG (1899G>A 1900T>C), AAT401AAG (1904T>G), GCC402GTC (1906C>T), CTG405TTG (1914C>T), GAG422GAA (1967G>A), CTA423CTG (1970A>G), TAT431TAC (1994T>C), GAG457GAA (2072G>A), GTG461GTA (2084G>A), AGA469AGG (2108A>G), AGG476AAG (2128G>A), CAG478CAA (2135G>A), GTG479GTT (2138G>T), AAA480ACA (2140A>C), ACT486ACA (2159T>A), ACC494GAT (2181A>G 2182C>A 2183C>T), TTC496TTT (2189C>T), GTT500ACT (2199G>A 2200T>C), ATG502GTT (2205A>G 2207G>T), GGC505GGG (2216C>G), GAT508GAA (2225T>A), GCT516GCC (2249T>C), GAA521GCA (2263A>C), GTA523GTG (2270A>G), GTT528GTC (2285T>C), CAA534CAG (2303A>G), AAT537AAC (2312T>C), GGT540GGC (2321T>C), GCG541GCA (2324G>A), TTC558TTT (2375C>T), AAG568AAA (2405G>A), AAA569AAG (2408A>G), AGT571AGC (2414T>C), GAG576GAA (2429G>A), GTG577GTT (2432G>T), TCA579GCA (2436T>G), CTC586CTG (2459C>G), GGT588GGC (2465T>C), AAA589AAG (2468A>G), TCC590ACT (2469T>A 2471C>T), CTA600CTC (2501A>C), AGG601AAG (2503G>A), GCA606GCG (2519A>G), TCG623TCA (2570G>A), GGT624GGC (2573T>C), AAC637AAT (2612C>T), ATC643ATT (2630C>T), CAT653CAC (2660T>C), ACA669ACG (2708A>G), ATC673ATT (2720C>T), GTT677GTC (2732T>C), CAC697CAT (2792C>T), GAG706GAA (2819G>A), CAT713CAC (2840T>C), CTG714TTG (2841C>T), CGT728CGC (2885T>C), CGA746CGG (2939A>G), TAC756TAT (2969C>T), TTA768CTA (3003T>C), ACG783ACA (3050G>A), ATT807ATC (3122T>C), ATT809ATC (3128T>C), GAG810GAA (3131G>A), CTG823TTG (3168C>T), AAT838AAC (3215T>C), GTT842GTG (3227T>G), AAC845AAT (3236C>T), CTT846CTG (3239T>G), GAG847GAA (3242G>A), AAG851AGG (3253A>G), AGG855AAA (3265G>A 3266G>A), TGT856TGC (3269T>C), ACG857TCG (3270A>T), AAT865AAC (3296T>C), ACC868TCT (3303A>T 3305C>T), GGT870GGA (3311T>A), TCT874TCC (3323T>C), CTA875TTA (3324C>T), CGG877CGA (3332G>A), CCC878CCT (3335C>T), AGG885CGG (3354A>C), ACA889ACT (3368A>T), GTA895GTT (3386A>T), TCA898TCC (3395A>C), CTA901CTG (3404A>G), GTC904GTT (3413C>T), CAG909CAA (3428G>A), CTT910TTG (3429C>T 3431T>G), TGC912TGT (3437C>T), ATA913ATT (3440A>T), TTG916CTG (3447T>C), GGC920GGT (3461C>T), TCT923TCC (3470T>C), TAC928TAT (3485C>T), CAC930CAT (3491C>T), ACT932ACC (3497T>C), GGA934GGG (3503A>G), AAT936AGT (3508A>G), CTA941CTG (3524A>G), TTC942TTT (3527C>T), GAG944GAA (3533G>A), TTG947CTG (3540T>C), TCT948ACT (3543T>A), ATT953ATA (3560T>A), TTA956TTG (3569A>G), AAG958AAA (3575G>A), AAC960AAT (3581C>T), ACA963ACC (3590A>C), GGA968GGG (3605A>G), CAA975CAG (3626A>G), AGA979AGG (3638A>G), GTA987GTT (3662A>T), GTG991GTA (3674G>A), TGC1005TGT (3716C>T), CCC1007CCT (3722C>T), CAG1014CAA (3743G>A), ATC1022ATT (3767C>T), TGC1024TGT (3773C>T), TTC1033TTT (3800C>T), TAC1042TAT (3827C>T), AAC1054AAT (3863C>T), GGA1056GGT (3869A>T), GGA1057GGT (3872A>T), ACC1062GCC (3885A>G), GAT1064GAC (3893T>C), TTT1070TTC (3911T>C), CCT1071CCA (3914T>A), GGA1073GGT (3920A>T), GGC1077GGA (3932C>A), ACC1078GCC (3933A>G), GTT1080GTG (3941T>G), GTC1081GTA (3944C>A), GCT1083GCA (3950T>A), GCA1087ACA (3960G>A), ATG1095GTT (3984A>G 3986G>T), GAC1097AAC (3990G>A), AAC1098AAT (3995C>T), TTT1100TTC (4001T>C), GTG1104GTC (4013G>C), CGC1113CGT (4040C>T), ACC1115ACT (4046C>T), ATA1120GTG (4059A>G 4061A>G), AAT1123AGT (4069A>G), CGA1133CGT (4100A>T), CTC1137CTT (4112C>T), GCC1143GCT (4130C>T), ATC1152ATT (4157C>T), GCT1167GCC (4202T>C), CCT1168CCA (4205T>A), TCA1169TCT (4208A>T), CAT1175TAT (4224C>T), ACC1183ACT (4250C>T) | | | | | | | | |
| **Proteins** | | | | | | | | | |
| Orf1 (YP_001040017.1) | 1 | 188 | 100% | 1189 | 98.3% | 188 (100%) | 185 (98.4%) | 0/0/0/0 | 1 |
| Protein mutations: | R109K (507G>A), L158Q (654T>A), L172P (696T>C) | | | | | | | | |
| Codon mutations: | GGC32GGT (277C>T), TCT33TCC (280T>C), GGC34GGT (283C>T), AGT38AGC (295T>C), CCA40CCC (301A>C), TTC44TTT (313C>T), ATT54ATC (343T>C), TTA74CTA (401T>C), GGC84GGT (433C>T), CTA87TTA (440C>T), GGC89GGA (448C>A), CAT93CAC (460T>C), ACC100ACA (481C>A), AGA102AGG (487A>G), AGG109AAG (507G>A), ACA134ACT (583A>T), CAA135CAG (586A>G), TTA136TTG (589A>G), CTA158CAA (654T>A), TTG166TTA (679G>A), CTA172CCA (696T>C), GGC180GGT (721C>T), GAA184GAG (733A>G) | | | | | | | | |
| polyprotein (YP_001040018.1) | 1 | 1199 | 100% | 8024 | 96.9% | 1199 (100%) | 1153 (96.2%) | 0/0/0/0 | 2 |
| Protein mutations: | A7V (721C>T), K11R (733A>G), K16R (748A>G), S17G (750A>G), Y48F (844A>T), A200V (1300C>T), T228V (1383A>G 1384C>T), S230* (1390C>G), Q242P (1426A>C), R255K (1465G>A), I257M (1472C>G), H286Q (1559C>A), Y303H (1608T>C), K317R (1651A>G), V365I (1794G>A), I373K (1819T>A 1820C>G), L382F (1847G>T), H388Y (1863C>T), I399V (1896A>G), V400T (1899G>A 1900T>C), N401K (1904T>G), A402V (1906C>T), R476K (2128G>A), K480T (2140A>C), T494D (2181A>G 2182C>A 2183C>T), V500T (2199G>A 2200T>C), M502V (2205A>G 2207G>T), D508E (2225T>A), E521A (2263A>C), S579A (2436T>G), S590T (2469T>A 2471C>T), R601K (2503G>A), K851R (3253A>G), R855K (3265G>A 3266G>A), T857S (3270A>T), T868S (3303A>T 3305C>T), N936S (3508A>G), S948T (3543T>A), T1062A (3885A>G), T1078A (3933A>G), A1087T (3960G>A), M1095V (3984A>G 3986G>T), D1097N (3990G>A), I1120V (4059A>G 4061A>G), N1123S (4069A>G), H1175Y (4224C>T) | | | | | | | | |

| | Begin | End | Coverage | Score | Concordance | Matches | Identities | I/D/M/F* | Stop Codons |
|---|---|---|---|---|---|---|---|---|---|
| **NT** | **25** | **5403** | **99.6%** | **9630** | **89.5%** | **5379 (99.9%)** | **5102 (94.8%)** | **2/0** | |
| Codon mutations: | GCT7GTT (721C>T), AAG11AGG (733A>G), AAA16AGA (748A>G), AGT17GGT (750A>G), GTT32GTC (797T>C), GGC35GGT (806C>T), AAA42AAG (827A>G), TAT48TTT (844A>T), CAA55CAG (866A>G), GGA89GGG (968A>G), TTG95TTA (986G>A), TTC101TTT (1004C>T), AGT120AGC (1061T>C), AAT121AAC (1064T>C), CTG128CTA (1085G>A), TTT135TTC (1106T>C), TCC136TCT (1109C>T), CCG146CCA (1139G>A), TCA175TCG (1226A>G), ATT190ATC (1271T>C), TCA192TCG (1277A>G), GCT200GTT (1300C>T), CAC205CAT (1316C>T), GAT212GAC (1337T>C), GAG215GAA (1346G>A), CAG218CAA (1355G>A), ATA220ATT (1361A>T), TCA222TCT (1367A>T), ACC228GTC (1383A>G 1384C>T), TCA230TGA (1390C>G), GCA231GCT (1394A>T), CCG237CCT (1412G>T), GGA240GGG (1421A>G), CAG242CCG (1426A>C), GAC245GAT (1436C>T), ACA246ACC (1439A>C), AGC252AGT (1457C>T), GAA253GAG (1460A>G), AGA255AAA (1465G>A), ATC257ATG (1472C>G), CGG258CGC (1475G>C), TCG261TCA (1484G>A), CTC262CTG (1487C>G), CGG263CGA (1490G>A), CTC267CTT (1502C>T), GTG269GTA (1508G>A), ACC282ACT (1547C>T), CAC286CAA (1559C>A), TAC303CAC (1608T>C), GAG306GAA (1619G>A), TTG312TTA (1637G>A), GTT315GTC (1646T>C), AAG317AGG (1651A>G), CTG321TTG (1662C>T), GAT328GAC (1685T>C), GGT331GGC (1694T>C), CCC341CCT (1724C>T), CAA352CAG (1757A>G), GTT365ATT (1794G>A), ATA372ATT (1817A>T), ATC373AAG (1819T>A 1820C>G), TTG382TTT (1847G>T), GAG383GAA (1850G>A), ACC386ACT (1859C>T), CAC388TAC (1863C>T), CTG394TTG (1881C>T), TAC397TAT (1892C>T), ATT399GTT (1896A>G), GTG400ACG (1899G>A 1900T>C), AAT401AAG (1904T>G), GCC402GTC (1906C>T), CTG405TTG (1914C>T), GAG422GAA (1967G>A), CTA423CTG (1970A>G), TAT431TAC (1994T>C), GAG457GAA (2072G>A), GTG461GTA (2084G>A), AGA469AGG (2108A>G), AGG476AAG (2128G>A), CAG478CAA (2135G>A), GTG479GTT (2138G>T), AAA480ACA (2140A>C), ACT486ACA (2159T>A), ACC494GAT (2181A>G 2182C>A 2183C>T), TTC496TTT (2189C>T), GTT500ACT (2199G>A 2200T>C), ATG502GTT (2205A>G 2207G>T), GGC505GGG (2216C>G), GAT508GAA (2225T>A), GCT516GCC (2249T>C), GAA521GCA (2263A>C), GTA523GTG (2270A>G), GTT528GTC (2285T>C), CAA534CAG (2303A>G), AAT537AAC (2312T>C), GGT540GGC (2321T>C), GCG541GCA (2324G>A), TTC558TTT (2375C>T), AAG568AAA (2405G>A), AAA569AAG (2408A>G), AGT571AGC (2414T>C), GAG576GAA (2429G>A), GTG577GTT (2432G>T), TCA579GCA (2436T>G), CTC586CTG (2459C>G), GGT588GGC (2465T>C), AAA589AAG (2468A>G), TCC590ACT (2469T>A 2471C>T), CTA600CTC (2501A>C), AGG601AAG (2503G>A), GCA606GCG (2519A>G), TCG623TCA (2570G>A), GGT624GGC (2573T>C), AAC637AAT (2612C>T), ATC643ATT (2630C>T), CAT653CAC (2660T>C), ACA669ACG (2708A>G), ATC673ATT (2720C>T), GTT677GTC (2732T>C), CAC697CAT (2792C>T), GAG706GAA (2819G>A), CAT713CAC (2840T>C), CTG714TTG (2841C>T), CGT728CGC (2885T>C), CGA746CGG (2939A>G), TAC756TAT (2969C>T), TTA768CTA (3003T>C), ACG783ACA (3050G>A), ATT807ATC (3122T>C), ATT809ATC (3128T>C), GAG810GAA (3131G>A), CTG823TTG (3168C>T), AAT838AAC (3215T>C), GTT842GTG (3227T>G), AAC845AAT (3236C>T), CTT846CTG (3239T>G), GAG847GAA (3242G>A), AAG851AGG (3253A>G), AGG855AAA (3265G>A 3266G>A), TGT856TGC (3269T>C), ACG857TCG (3270A>T), AAT865AAC (3296T>C), ACC868TCT (3303A>T 3305C>T), GGT870GGA (3311T>A), TCT874TCC (3323T>C), CTA875TTA (3324C>T), CGG877CGA (3332G>A), CCC878CCT (3335C>T), AGG885CGG (3354A>C), ACA889ACT (3368A>T), GTA895GTT (3386A>T), TCA898TCC (3395A>C), CTA901CTG (3404A>G), GTC904GTT (3413C>T), CAG909CAA (3428G>A), CTT910TTG (3429C>T 3431T>G), TGC912TGT (3437C>T), ATA913ATT (3440A>T), TTG916CTG (3447T>C), GGC920GGT (3461C>T), TCT923TCC (3470T>C), TAC928TAT (3485C>T), CAC930CAT (3491C>T), ACT932ACC (3497T>C), GGA934GGG (3503A>G), AAT936AGT (3508A>G), CTA941CTG (3524A>G), TTC942TTT (3527C>T), GAG944GAA (3533G>A), TTG947CTG (3540T>C), TCT948ACT (3543T>A), ATT953ATA (3560T>A), TTA956TTG (3569A>G), AAG958AAA (3575G>A), AAC960AAT (3581C>T), ACA963ACC (3590A>C), GGA968GGG (3605A>G), CAA975CAG (3626A>G), AGA979AGG (3638A>G), GTA987GTT (3662A>T), GTG991GTA (3674G>A), TGC1005TGT (3716C>T), CCC1007CCT (3722C>T), CAG1014CAA (3743G>A), ATC1022ATT (3767C>T), TGC1024TGT (3773C>T), TTC1033TTT (3800C>T), TAC1042TAT (3827C>T), AAC1054AAT (3863C>T), GGA1056GGT (3869A>T), GGA1057GGT (3872A>T), ACC1062GCC (3885A>G), GAT1064GAC (3893T>C), TTT1070TTC (3911T>C), CCT1071CCA (3914T>A), GGA1073GGT (3920A>T), GGC1077GGA (3932C>A), ACC1078GCC (3933A>G), GTT1080GTG (3941T>G), GTC1081GTA (3944C>A), GCT1083GCA (3950T>A), GCA1087ACA (3960G>A), ATG1095GTT (3984A>G 3986G>T), GAC1097AAC (3990G>A), AAC1098AAT (3995C>T), TTT1100TTC (4001T>C), GTG1104GTC (4013G>C), CGC1113CGT (4040C>T), ACC1115ACT (4046C>T), ATA1120GTG (4059A>G 4061A>G), AAT1123AGT (4069A>G), CGA1133CGT (4100A>T), CTC1137CTT (4112C>T), GCC1143GCT (4130C>T), ATC1152ATT (4157C>T), GCT1167GCC (4202T>C), CCT1168CCA (4205T>A), TCA1169TCT (4208A>T), CAT1175TAT (4224C>T), ACC1183ACT (4250C>T) | | | | | | | | |

*: Inserts / Deletes / Misaligned / Frameshifts

#### Analysis details

This analysis was performed with panviral2.64

#### NGS Details (UN9): Potato virus Y

##### Assembly

| | |
|---|---|
| **Coverage Length** | 9700 (1 contig(s)) |
| **Depth Of Coverage** | 11166.5 |
| **Number Of Reads** | 812319 |
| **Reads Per Million** | 18360.73 rpm (after QC) |
| **Ambiguities** | 0 |
| **Assembly Method** | de novo + reference guided assembly |
| **Consensus Caller** | Bcf Tools |

##### Coverage Map

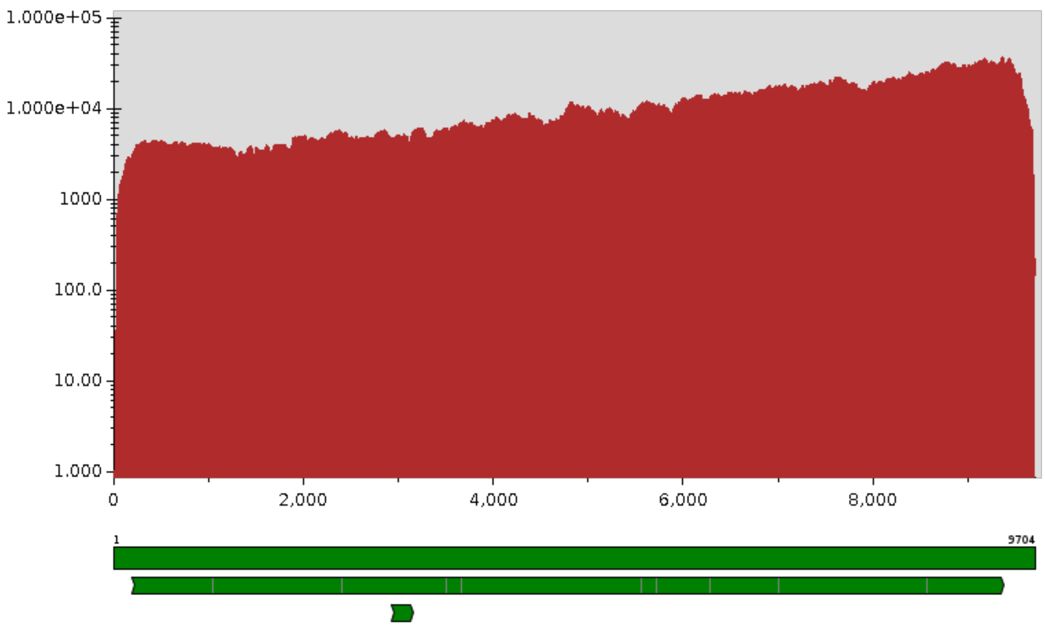

##### Assignment

| | |
|---|---|
| **Type** | Potato virus Y (Taxonomy ID: 12216) |
| **Reference Genome** | NC_001616.1 |
| **NT Identity (%)** | 82.6033 |
| **AA Identity (%)** | 89.9682 |
| **Number Of Stop Codons** | 3 |
| **Number Of CDS** | 2 |

##### Alignment

| | |
|---|---|
| **Alignment Score** | 12522.0 (NT) + 19018.0 (AA) = 31540.0 |
| **Concordance (%)** | 78.0601 |

| Alignment Method | Global, seeded, nucleotide + amino acids (AGA) |
|---|---|

#### Genome Region

Sequence starts at position 5 and ends at position 9704 relative to NC_001616.1 reference sequence.

#### Alignment Detailed Statistics

| | Begin | End | Coverage | Score | Concordance | Matches | Identities | I/D/M/F* | Stop Codons |
|---|---|---|---|---|---|---|---|---|---|
| **NT** | **5** | **9704** | **99.9%** | **12522** | **64.8%** | **9688 (99.7%)** | **8015 (82.5%)** | **15/12** | |
| Mutations: | 5A>G, 7A>G, 12C>G, 13T>G, 14C>G, 21A>C, 22C>T, 23A>T, 32A>T, 42_43insGAAATAT, 45A>T, 47T>A, 50T>G, 53A>G, 54C>T, 55G>T, 56C>T, 58C>T, 60T>A, 60_61insA, 75C>T, 79C>T, 83G>A, 85T>C, 87C>T, 89G>A, 90T>C, 93A>T, 94A>G, 95A>T, 100T>C, 101T>A, 102C>T, 104T>C, 107C>T, 110T>G, 112C>T, 114C>T, 115T>C, 116A>T, 117G>C, 119A>G, 120C>T, 121A>T, 125T>G, 126T>C, 127G>A, 128G>T, 131A>G, 132C>T, 139A>T, 139_140insG, 141C>T, 143C>T, 146C>T, 150C>T, 150_151insT, 154T>C, 156C>A, 158T>A, 160A>C, 164C>T, 168C>A, 169C>G, 171A>C, 172T>C, 173T>A, 174T>G, 177G>C, 178A>G, 179T>C, 180C>T, 181C>A, 182T>G, 183C>T, 193T>C, 194T>C, 196C>A, 202A>G, 208C>T, 209T>C, 210G>T, 211T>G, 220G>A, 221T>A, 223T>G, 226A>G, 235A>T, 241C>T, 247A>T, 251T>C, 255G>T, 256C>T, 258A>G, 262T>C, 263A>G, 265T>G, 267T>C, 273A>T, 277A>T, 280A>G, 283G>T, 286G>A, 287G>A, 289T>C, 293G>A, 294T>C, 301T>C, 304C>T, 305G>A, 306C>T, 309A>G, 311C>T, 316A>T, 317A>G, 318C>T, 319A>G, 322A>G, 331A>T, 339T>A, 340C>A, 342A>G, 347A>G, 348A>C, 349A>G, 350T>C, 355T>A, 356A>G, 357C>T, 364T>C, 365G>A, 366T>C, 367G>A, 368C>T, 369T>C, 373A>G, 376C>T, 384T>G, 388G>A, 391C>T, 394A>G, 400G>A, 406T>C, 408C>T, 409C>T, 412G>A, 415A>T, 416A>G, 417T>A, 424T>C, 427G>A, 433A>G, 436G>C, 437G>A, 440A>G, 441G>A, 443A>C, 444A>G, 445G>A, 448T>A, 454A>G, 456A>C, 460T>C, 461C>A, 469A>G, 475C>G, 481T>A, 502T>C, 505A>T, 511C>T, 517T>A, 520T>A, 523A>T, 531C>T, 535G>A, 536C>T, 539C>T, 540A>G, 541A>G, 549G>A, 550A>G, 556C>T, 559T>C, 562T>C, 568T>C, 571A>G, 576T>C, 580T>C, 585T>C, 586C>G, 593C>T, 594G>A, 595C>T, 597C>G, 599A>G, 601A>G, 602A>C, 603T>C, 604A>G, 613A>G, 616A>G, 619C>G, 622G>A, 625G>A, 628T>C, 631C>T, 635A>G, 637T>C, 644A>G, 646A>G, 655T>C, 661G>A, 663A>C, 664G>C, 667A>G, 679C>T, 682C>A, 683T>C, 685A>G, 691T>C, 699C>G, 703T>A, 706T>A, 719A>G, 721G>A, 722A>G, 724A>G, 725C>T, 727T>G, 730T>A, 731G>T, 734T>C, 735A>C, 736C>T, 737T>C, 738C>G, 739C>T, 744C>T, 745G>C, 749C>T, 751A>C, 754T>C, 765T>A, 770T>C, 773C>A, 774G>A, 775A>G, 776C>A, 790C>T, 801T>A, 813G>C, 814A>G, 816T>A, 818T>C, 823A>G, 825G>A, 826T>A, 833C>A, 835G>A, 841C>T, 851A>G, 853T>C, 859C>T, 860C>T, 862C>T, 863A>G, 865T>C, 867T>C, 869A>G, 871C>T, 872A>T, 874A>G, 876G>A, 879G>A, 886T>C, 898C>T, 903A>G, 906C>A, 907A>C, 909A>C, 910A>C, 912G>A, 913C>T, 922C>A, 931T>A, 938T>C, 939C>T, 940A>T, 943A>G, 952C>T, 967A>G, 973A>G, 976G>C, 980T>A, 982G>C, 991A>G, 994T>C, 999G>A, 1003T>C, 1007C>A, 1008A>G, 1009G>A, 1010A>G, 1012T>G, 1013A>G, 1018A>G, 1019A>C, 1021C>G, 1030C>T, 1039G>A, 1042T>C, 1045C>A, 1048C>A, 1051T>C, 1060G>A, 1063T>A, 1064C>T, 1069C>T, 1083G>A, 1093T>C, 1097T>G, 1099G>A, 1105C>T, 1114A>G, 1117T>G, 1123A>G, 1126T>A, 1129C>T, 1143G>A, 1147A>T, 1150T>A, 1154T>A, 1156G>A, 1165C>T, 1168T>C, 1172C>T, 1174T>A, 1201C>T, 1221G>A, 1229G>A, 1230T>C, 1231T>C, 1233G>A, 1234C>T, 1240G>A, 1246G>A, 1250T>C, 1252G>A, 1255T>C, 1257A>C, 1258A>G, 1267A>G, 1273T>C, 1276G>A, 1278A>G, 1283T>C, 1290C>T, 1291G>A, 1298G>A, 1299A>C, 1300C>T, 1306T>C, 1307A>G, 1309A>G, 1312T>C, 1316A>G, 1318T>A, 1327G>A, 1329T>C, 1333G>T, 1336A>G, 1343C>T, 1345A>G, 1354G>A, 1357G>T, 1360C>T, 1367C>T, 1369C>G, 1373C>A, 1378C>T, 1384G>A, 1385A>G, 1393A>G, 1396C>T, 1405G>T, 1408A>G, 1411G>A, 1415G>T, 1417A>T, 1420G>C, 1429T>C, 1432A>G, 1436G>A, 1438C>T, 1439T>C, 1441A>G, 1444T>C, 1453C>T, 1456G>A, 1468A>G, 1492A>G, 1499T>C, 1501G>A, 1504T>C, 1507G>A, 1510C>T, 1513A>G, 1516A>G, 1519A>T, 1522G>A, 1525C>T, 1528G>A, 1534C>T, 1540T>A, 1546C>T, 1555A>G, 1558T>C, 1561T>C, 1564A>C, 1567T>A, 1573C>T, 1576A>G, 1583T>C, 1588T>A, 1594G>A, 1600C>T, 1609G>A, 1618G>A, 1621C>T, 1624C>T, 1631C>T, 1639A>G, 1651C>T, 1654C>G, 1675T>C, 1693A>G, 1702T>C, 1707A>G, 1718G>A, 1726A>G, 1729C>T, 1732A>C, 1738T>C, 1742A>G, 1744C>T, 1747C>T, 1753T>C, 1758G>A, 1768G>A, 1786C>T, 1789A>G, 1790G>A, 1795C>T, 1805T>C, 1807A>G, 1813G>A, 1819G>A, 1825G>A, 1837C>T, 1840T>C, 1849A>G, 1858C>G, 1864A>G, 1870C>T, 1876T>C, 1882A>G, 1888T>A, 1891T>C, 1906T>C, 1915A>T, 1930T>C, 1936C>T, 1937A>G, 1939T>C, 1945A>G, 1951C>T, 1960A>T, 1978A>G, 1981T>A, 1984C>T, 1987T>C, 1996C>T, 2004T>A, 2020A>G, 2023G>A, 2026T>C, 2029G>T, 2032G>A, 2059T>C, 2068T>G, 2071C>T, 2072G>A, 2074G>A, 2087C>T, 2089G>A, 2101C>T, 2104G>A, 2107G>A, 2116G>A, 2122C>T, 2137C>T, 2158T>C, 2164C>T, 2170A>G, 2182T>C, 2188G>A, 2191C>A, 2208G>A, 2218T>C, 2224C>T, 2227A>T, 2242G>A, 2245C>T, 2254G>A, 2257T>C, 2260C>T, 2266C>G, 2272A>G, 2275G>A, 2293A>G, 2299C>T, 2302G>A, 2305G>A, 2314A>T, 2317T>C, 2326A>G, 2341G>A, 2347A>G, 2350T>C, 2356G>A, 2365T>C, 2371A>G, 2374A>G, 2383T>C, 2386A>T, 2389A>G, 2392T>C, 2395T>C, 2414A>G, 2415A>G, 2416T>G, 2419T>A, 2420A>T, 2440A>T, 2467T>C, 2476T>C, 2482G>A, 2491G>A, 2497G>T, 2503G>A, 2518A>G, 2521T>C, 2523A>G, 2533A>G, 2539G>A, 2545A>G, 2551G>A, 2554T>A, 2558C>T, 2561T>C, 2563G>C, 2572A>G, 2584T>C, 2587C>T, 2593G>C, 2599T>C, 2608T>C, 2620T>C, 2629G>A, 2632G>A, 2634G>A, 2638G>A, 2644T>A, 2653A>G, 2665T>C, 2678C>T, 2683A>G, 2689T>C, 2698A>C, 2707A>G, 2710G>A, 2716A>G, 2719A>G, 2722C>T, 2723G>A, 2725C>T, 2728A>G, 2734G>A, 2737T>A, 2746T>C, 2747G>A, 2749T>A, 2752A>T, 2755T>A, 2761C>T, 2764C>T, 2779T>C, 2785G>T, 2791C>T, 2795C>A, 2806C>T, 2816T>C, 2818G>C, 2822G>A, 2824G>A, 2827G>A, 2836T>C, 2839G>A, 2842T>C, 2863C>A, 2864C>T, 2882C>A, 2885A>C, 2886G>A, 2893C>T, 2895C>T, 2902C>T, 2905A>G, 2911T>C, 2923T>C, 2931A>G, 2940A>G, 2947T>C, 2962T>C, 2971_2972insA, 2989T>C, 2991C>T, 2994A>G, 2999G>A, 3000A>G, 3016delC, 3021G>A, 3024A>T, 3025C>T, 3036C>T, 3039A>G, 3040A>C, 3042A>C, 3043G>A, 3052G>A, 3061A>G, 3082G>A, 3091C>T, 3099C>T, 3100C>A, 3104G>A, 3105T>G, 3106G>A, 3109C>T, 3112A>G, 3118T>C, 3121C>T, 3127A>G, 3132G>A, 3137C>A, 3138G>A, 3139A>G, 3151T>C, 3152T>A, 3154C>T, 3165G>A, 3169A>G, 3199C>T, 3203A>C, 3208T>C, 3214A>G, 3217C>T, 3218T>C, 3223C>A, 3225C>A, 3230G>A, 3235C>T, 3257T>C, 3262C>T, 3265C>T, 3266G>A, 3268A>G, 3271G>A, 3275G>A, 3284G>A, 3301G>A, 3315G>A, 3316A>G, 3318A>C, 3322C>T, 3325G>A, 3328G>A, 3337A>T, 3349G>A, 3355C>T, 3358C>T, 3359A>G, 3367A>G, 3370T>C, 3377T>C, 3379A>G, 3394A>G, 3397C>T, 3400T>G, 3406A>G, 3418C>T, 3421T>A, 3427T>C, 3433G>A, 3436A>T, 3439A>G, 3442C>T, 3445T>C, 3457G>A, 3460T>C, 3463T>A, 3469G>T, 3472G>A, 3496C>T, 3499G>A, 3505C>T, 3513T>G, 3520A>G, 3529A>G, 3532A>G, 3545A>G, 3548G>A, 3551T>C, 3554G>A, 3556C>T, 3583T>C, 3586T>C, 3589C>T, 3595C>T, 3607T>C, 3613T>C, 3616G>A, 3619T>C, 3623C>G, 3628C>T, 3632C>A, 3637T>C, 3638T>C, 3643A>G, 3661A>G, 3670G>A, 3682T>C, 3688G>A, 3697G>A, 3698A>G, 3706T>C, 3709T>C, 3713T>C, 3715G>A, 3717G>A, 3721G>A, 3731C>A, 3732G>A, 3736T>A, 3746C>T, 3748A>G, 3754A>G, 3763T>C, 3778A>G, 3781A>G, 3784C>T, 3796T>C, 3805A>T, 3811C>T, 3814A>G, 3829C>T, 3838T>C, 3844A>G, 3847A>C, 3856T>A, 3862G>T, 3871C>T, 3877C>A, 3883C>T, 3886A>G, 3892A>G, 3895C>T, 3898T>C, 3904A>G, 3905C>A, 3907G>A, 3913T>C, 3928G>A, 3931A>T, 3937G>A, 3955T>C, 3958G>A, 3961C>T, 3964A>G, 3967T>C, 3976A>G, 3979T>C, 3985A>T, 3997A>T, 4000G>A, 4006C>T, 4009T>G, 4012C>T, 4015A>G, 4024T>C, 4030A>G, 4033A>T, 4042G>A, 4052C>T, 4054G>A, 4057T>C, 4063T>A, 4066A>G, 4073A>G, 4078T>C, 4081C>T, 4084T>C, 4096C>T, 4108C>T, 4111A>T, 4117G>A, 4118C>T, 4120A>G, 4126C>T, 4128T>A, 4129C>T, 4132C>T, 4135T>C, 4147A>G, 4148T>C, 4151G>T, 4162C>T, 4166G>A, 4171A>C, 4180G>A, 4190C>T, 4198T>C, 4201C>T, 4210G>T, 4213C>T, 4219T>C, 4220C>T, 4223C>A, 4225G>A, 4231T>G, 4237T>C, 4252A>G, 4253T>C, 4255A>G, 4258A>G, 4270T>C, 4282A>G, 4285G>A, 4291A>G, 4303G>A, 4312C>T, 4319A>G, 4321A>G, 4324G>T, 4327G>A, 4336G>T, 4339T>C, 4348A>T, 4360A>G, 4366T>C, 4369T>A, 4375T>C, 4387T>C, 4395T>A, 4406G>A, 4411T>C, 4438T>C, 4444C>A, 4450C>T, 4456C>T, 4465C>T, 4475A>G, 4477G>T, 4480C>T, 4481A>T, 4483A>G, 4495C>T, 4507G>A, 4510C>T, 4513T>G, 4517C>T, 4528C>T, 4534A>G, 4540C>T, 4544G>T, 4545C>T, 4546G>A, 4548G>A, 4558T>C, 4564A>T, 4570C>T, 4576A>T, 4597G>A, 4600C>T, 4606C>T, 4609G>A, 4615G>A, 4618C>T, 4625T>C, 4627G>T, 4648C>T, 4651T>C, 4660G>A, 4663C>T, 4669T>A, 4679G>A, 4681G>C, 4684T>C, 4687T>G, 4699G>A, 4702A>G, 4708A>G, 4711G>A, 4712C>T, 4738A>G, 4744A>T, 4747A>G, 4748T>C, 4765T>C, 4796G>A, 4798C>T, 4803C>G, 4804T>C, 4807G>A, 4810G>T, 4813T>G, 4816T>C, 4831T>C, 4834C>T, 4840A>T, 4852A>G, 4855C>T, 4867T>C, 4870G>C, 4873T>A, 4882T>C, 4903A>T, 4909G>A, 4915T>C, 4918A>G, 4919T>C, 4921G>A, 4924T>C, 4927C>G, 4933T>C, 4936C>A, 4939G>A, 4942T>C, 4945C>T, 4963A>G, 4972T>C, 4990T>C, 5002A>G, 5023T>C, 5024T>C, 5029C>T, 5032T>C, 5038T>C, 5044A>T, 5053A>T, 5056G>A, 5057A>G, 5060A>G, 5071G>T, 5090C>A, 5095A>G, 5098G>T, 5100C>T, 5104A>G, 5107A>T, 5110T>C, 5113A>G, 5116G>A, 5119A>G, 5120G>T, 5122C>T, 5126A>G, 5137T>C, 5140C>T, 5143A>G, 5146G>A, 5161C>A, 5167A>G, 5173T>G, 5182G>A, 5183G>A, 5188C>T, 5191G>A, 5197A>G, 5200C>T, 5206C>T, 5209A>G, 5218C>T, 5230G>A, 5236C>T, 5239C>T, 5245C>T, 5263A>T, 5269C>T, 5272C>T, 5275C>A, 5278C>T, 5281A>C, 5284A>G, 5287T>C, 5288C>T, 5293A>T, 5294T>C, 5296G>A, 5302G>A, 5311T>A, 5314A>G, 5323A>T, 5347T>C, 5353C>T, 5365C>T, 5386T>C, 5395C>A, 5422A>G, 5425T>C, 5431C>T, 5437G>A, 5452T>C, 5458G>A, 5467T>C, 5470A>G, 5471C>T, 5473A>G, 5479A>G, 5482C>T, 5485A>C, 5488T>C, 5489T>C, 5491G>A, 5497T>C, 5506T>C, 5509G>A, 5512G>A, 5516T>C, 5530T>C, 5533G>A, 5536G>T, 5539G>A, 5548T>G, 5554C>T, 5560T>A, 5561G>A, 5563G>A, 5566G>C, 5569A>G, 5584C>G, 5593G>A, 5596G>A, 5597A>G, 5605C>A, 5617G>T, 5619C>T, 5620T>G, 5629G>A, 5630A>C, 5641C>T, 5650A>T, 5662A>T, 5665A>G, 5689A>G, 5693G>A, 5698G>A, 5713A>G, 5716G>C, 5719A>G, 5743C>A, 5755C>A, 5758T>C, 5764T>C, 5770A>G, 5779C>T, 5782T>C, 5791C>T, 5803C>T, 5836G>A, 5839A>G, 5845A>G, 5848A>G, 5853A>G, 5857T>C, 5860C>T, 5863A>T, 5869T>A, 5875T>C, 5884C>T, 5893C>T, 5894A>G, 5896C>G, 5899C>T, 5908G>A, 5911T>C, 5923G>A, 5938A>G, 5941C>T, 5944T>C, 5947T>C, 5959G>A, 5960C>G, 5961G>C, 5962G>C, 5974A>G, 5986T>A, 5989C>T, 5990A>G, 5998T>C, 6008A>C, 6010A>T, 6019A>T, 6022G>T, 6025A>T, 6030A>C, 6049C>A, 6058G>A, 6061A>G, 6064C>A, 6070T>C, 6072G>A, 6073T>C, 6074A>C, 6079G>A, 6081C>A, 6085A>T, 6091A>T, 6094C>T, 6111G>C, 6112T>C, 6118A>G, 6124G>A, 6127G>A, 6128A>G, 6130T>C, 6133T>C, 6136A>G, 6148C>T, 6157A>G, 6158G>A, 6163T>C, 6169A>G, 6170A>T, 6172A>G, 6178C>T, 6187A>G, 6190T>C, 6196G>A, 6203C>T, 6205C>T, 6208A>G, 6209C>T, 6211A>G, 6217G>A, 6229T>A, 6230G>A, 6238C>T, 6247G>A, 6253A>T, 6257G>A, 6258C>A, 6274T>C, 6282A>G, 6286G>A, 6298C>T, 6301G>A, 6304A>G, 6307C>T, 6322C>T, 6328A>T, 6331T>G, 6334T>C, 6337G>A, 6338C>T, 6358T>C, 6361G>A, 6362G>A, 6367A>T, 6370G>A, 6376C>T, 6379T>G, 6385A>C, 6391A>T, 6394A>G, 6397C>T, 6401G>A, 6405C>T, 6412C>T, 6418A>G, 6421T>C, 6429A>T, 6436T>A, 6445G>A, 6450A>G, 6454C>G, 6460C>T, 6463T>A, 6469C>T, 6472G>A, 6475G>A, 6481T>C, 6485C>A, 6487C>T, 6490T>C, 6496C>T, 6502G>A, 6514T>C, 6517G>A, 6527C>A, 6532C>A, 6541G>A, 6544A>G, 6556C>T, 6559T>C, 6565G>A, 6569T>C, 6571G>A, 6586T>A, 6588C>T, 6598A>G, 6601A>G, 6604T>C, 6607T>C, 6616A>G, 6622C>T, 6631G>A, 6634G>A, 6635T>C, 6640T>A, 6646G>T, 6647A>G, 6652C>T, 6664C>T, 6670T>G, 6677A>G, 6679A>G, 6685C>T, 6691A>T, 6694C>T, 6715A>G, 6716G>A, 6719A>G, 6724A>G, 6734C>T, 6742G>A, 6745G>A, 6754C>T, 6760A>G, 6764A>T, 6766A>G, 6775T>A, 6782C>T, 6787A>T, 6790C>T, 6795C>T, 6796A>G, 6799C>A, 6802C>A, 6808C>T, 6811C>T, 6814C>T, 6823C>T, 6829A>G, 6847C>T, 6853A>C, 6857A>G, 6862G>A, 6871A>G, 6875G>A, 6876T>C, 6877C>A, 6880G>A, 6883T>A, 6889T>G, 6901C>T, 6916C>T, 6919G>A, 6925A>G, 6928T>C, 6934C>A, 6949G>C, 6955C>T, 6958A>G, 6976A>G, 6980C>T, 6985C>T, 6997A>T, 7000G>T, 7001G>A, 7003G>A, 7012T>C, 7015G>A, 7018T>C, 7032T>A, 7039C>T, 7040T>C, 7042G>A, 7048A>G, 7060T>A, 7063C>A, 7078C>T, 7082T>C, 7084A>G, 7087A>G, 7090C>A, 7096T>C, 7099A>G, 7102T>C, 7111G>A, 7117A>G, 7125C>A, 7129A>G, 7132T>C, 7133C>T, 7135G>A, 7141G>A, 7144T>C, 7145G>T, 7147A>G, 7150G>A, 7159A>T, 7160_7165delGAGGCA, 7171C>T, 7189G>T, 7198A>G, 7201C>T, 7205C>T, 7210T>C, 7216T>A, 7219G>A, 7225C>A, 7228G>A, 7240G>A, 7243T>C, 7249A>G, 7252T>G, 7258T>A, 7261T>C, 7264T>A, 7267C>G, 7270G>A, 7273_7274insT, 7274C>G, 7275G>T, 7281A>T, 7284T>C, 7296C>T, 7311C>T, 7314C>T, 7317G>A, 7321T>A, 7322delG, 7327C>T, 7330C>T, 7333C>T, 7334A>C, 7342T>C, 7343G>A, 7344C>A, 7345A>C, 7354T>C, 7363A>G, 7369T>C, 7372C>T, 7378G>T, 7381C>T, 7396A>T, 7399C>T, 7405C>T, 7407G>T, 7408T>G, 7414G>T, 7415T>G, 7420A>G, 7429C>T, 7432T>C, 7441T>C, 7459G>A, 7462A>G, 7468A>T, 7471C>A, 7477A>G, 7480C>T, 7486G>T, 7495T>C, 7498A>G, 7501T>C, 7503T>C, 7504G>A, 7513T>A, 7519C>T, 7522A>G, 7531G>A, 7537G>A, 7540C>T, 7546T>C, 7570G>A, 7573G>A, 7579G>A, 7588T>C, 7595C>T, 7604T>C, 7606G>A, 7615T>A, 7624T>C, 7633C>T, 7636C>T, 7639T>C, 7648T>C, 7666A>G, 7672T>C, 7678A>T, 7684G>A, 7690T>C, 7696T>C, 7702T>G, 7709G>A, 7713A>G, 7714A>G, 7715C>T, 7726T>C, 7735G>A, 7744A>G, 7747C>T, 7762C>T, 7768G>A, 7771T>C, 7780A>G, 7783A>C, 7786T>C, 7793C>T, 7795A>G, 7798C>T, 7804T>C, 7812C>T, 7834A>G, 7837C>T, 7844G>T, 7846G>A, 7855G>A, 7861G>A, 7870A>G, 7873C>T, 7876T>A, 7879G>A, 7882T>A, 7883G>A, 7894A>T, 7897T>C, 7921C>T, 7927G>A, 7936T>C, 7939T>C, 7942T>C, 7966G>A, 7969T>C, 7975T>C, 7987C>T, 7990T>C, 8005T>C, 8032A>G, 8038C>T, 8041C>T, 8043G>A, 8080G>A, 8089G>A, 8095G>A, 8098G>A, 8107C>T, 8110T>C, 8116T>C, 8128A>G, 8146T>C, 8151A>G, 8152C>T, 8164G>A, 8178G>A, 8182G>A, 8191G>A, 8197C>T, 8203C>T, 8208G>A, 8213C>T, 8215G>A, 8218A>G, 8221C>T, 8224G>A, 8239A>G, 8254G>A, 8260T>C, 8263A>G, 8281T>C, 8284A>G, 8294C>G, 8296A>G, 8299G>A, 8302C>T, 8317T>C, 8320C>T, 8326T>A, 8335G>A, 8338C>T, 8349C>T, 8353A>G, 8362C>T, 8365A>G, 8368C>T, 8380C>T, 8383A>G, 8389A>G, 8404T>C, 8408G>T, 8413A>G, 8419G>T, 8434T>C, 8440T>C, 8452G>A, 8458A>G, 8460G>A, 8464A>G, 8467G>A, 8479G>A, 8480G>A, 8482T>A, 8497G>A, 8498C>T, 8500A>G, 8502G>A, 8503A>G, 8506C>T, 8515A>T, 8524C>T, 8527A>C, 8528T>C, 8530A>G, 8539G>C, 8546C>T, 8547T>G, 8552T>A, 8563A>G, 8574C>G, 8585A>G, 8598G>A, 8602C>T, 8603A>T, 8604A>C, 8605C>T, 8620A>G, 8622C>A, 8625A>G, 8626G>A, 8629G>A, 8632C>T, 8638C>T, 8641G>A, 8647C>T, 8648C>T, 8649C>T, 8650G>C, 8653C>T, 8656A>G, 8658G>A, 8664A>C, 8665T>G, 8674T>G, 8679C>T, 8680A>T, 8683C>A, 8692G>A, 8695A>T, 8707G>A, 8708A>C, 8716G>A, 8722C>T, 8725G>A, 8731A>G, 8737A>G, 8743C>T, 8745C>A, 8746A>G, 8752G>A, 8755A>T, 8770C>T, 8771T>C, 8776A>G, 8779T>C, 8782G>A, 8791T>C, 8797A>G, 8800A>G, 8803A>G, 8809T>C, 8812T>C, 8818T>C, 8824G>A, 8827A>G, 8848G>A, 8854T>C, 8856A>G, 8857G>A, 8863G>A, 8865G>A, 8867A>C, 8869G>T, 8872A>G, 8883G>A, 8888A>T, 8929C>T, 8954G>A, 8959C>T, 8962A>T, 8983G>A, 8984A>G, 8986T>C, 8988A>T, 8998G>A, 9004G>A, 9007G>A, 9016C>T, 9037C>A, 9085A>T, 9097C>T, 9131A>G, 9136A>T, 9145G>T, 9149A>G, 9150T>G, 9151G>A, 9152G>A, 9157A>G, 9160G>T, 9163T>C, 9184G>A, 9196A>G, 9214G>A, 9217A>G, 9220G>A, 9244A>T, 9247G>A, 9256C>T, 9260C>T, 9262T>C, 9274G>A, 9280C>T, 9289C>T, 9298A>C, 9304G>A, 9334C>T, 9364C>G, 9377_9378insT, 9387C>T, 9404G>A, 9411C>T, 9412A>G, 9414A>T, 9437C>T, 9454A>G, 9458T>C, 9467G>A, 9470_9471insT, 9482G>A, 9483C>T, 9498T>C, 9516T>G, 9517delA, 9533G>A, 9534G>T, 9548A>T, 9551C>T, 9553_9554delTA, 9556A>T, 9557T>A, 9558A>T, 9574T>A, 9581G>T, 9582T>G, 9583G>C, 9584_9585insG, 9590C>T, 9591T>C, 9607C>T, 9610T>C, 9615delC, 9620A>G, 9648A>G, 9674T>C, 9690C>T | | | | | | | | |

| | Begin | End | Coverage | Score | Concordance | Matches | Identities | I/D/M/F* | Stop Codons |
|---|---|---|---|---|---|---|---|---|---|
| **NT** | **5** | **9704** | **99.9%** | **12522** | **64.8%** | **9688 (99.7%)** | **8015 (82.5%)** | **15/12** | |
| **CDS** | | | | | | | | | |
| PVYgp1 | 1 | 3064 | 100% | 18923 | 89.7% | 3062 (99.9%) | 2771 (90.4%) | 2/2/4/4 | 1 |
| Protein mutations: | Y4Q (194T>C 196C>A), C9L (209T>C 210G>T 211T>G), F13M (221T>A 223T>G), S23P (251T>C), C24F (255G>T 256C>T), E25G (258A>G), I27V (263A>G 265T>G), V28A (267T>C), E30V (273A>T), A35T (287G>A 289T>C), V37T (293G>A 294T>C), A41I (305G>A 306C>T), D42G (309A>G), E44D (316A>T), T45V (317A>G 318C>T 319A>G), L52Q (339T>A 340C>A), K53R (342A>G), K55A (347A>G 348A>C 349A>G), Y56H (350T>C), T58V (356A>G 357C>T), V61T (365G>A 366T>C 367G>A), L62S (368C>T 369T>C), F67C (384T>G), A75V (408C>T 409C>T), M78E (416A>G 417T>A), E85K (437G>A), R86E (440A>G 441G>A), K87R (443A>C 444A>G 445G>A), D88E (448T>A), E91A (456A>C), H93N (461C>A), D111E (517T>A), S116F (531C>T), P118S (536C>T), Q119W (539C>T 540A>G 541A>G), R122K (549G>A 550A>G), M131T (576T>C), V134A (585T>C 586C>G), R137Y (593C>T 594G>A 595C>T), P138R (597C>G), I139V (599A>G 601A>G), I140P (602A>C 603T>C 604A>G), M147I (625G>A), I151V (635A>G 637T>C), I154V (644A>G 646A>G), E160A (663A>C 664G>C), H166Q (682C>A), T172S (699C>G), H174Q (706T>A), K179E (719A>G 721G>A), I180V (722A>G 724A>G), A183S (731G>T), Y184P (734T>C 735A>C 736C>T), S185R (737T>C 738C>G 739C>T), A187V (744C>T 745G>C), R189C (749C>T 751A>C), M194K (765T>A), R197K (773C>A 774G>A 775A>G), M206K (801T>A), G210A (813G>C 814A>G), L211H (816T>A), R214Q (825G>A 826T>A), N223D (851A>G 853T>C), R226C (860C>T 862C>T), T227A (863A>G 865T>C), I228T (867T>C), N229D (869A>G 871C>T), I230L (872A>T 874A>G), R231Q (876G>A), R232K (879G>A), N240S (903A>G), T241N (906C>A 907A>C), K242T (909A>C 910A>C), S243N (912G>A 913C>T), S252L (938T>C 939C>T 940A>T), L266I (980T>A 982G>C), R272K (999G>A), Q275R (1007C>A 1008A>G 1009G>A), S276G (1010A>G 1012T>G), I277V (1013A>G), N279Q (1019A>C 1021C>G), D288E (1048C>A), R300Q (1083G>A), S305A (1097T>G 1099G>A), R320K (1143G>A), L324I (1154T>A 1156G>A), S346N (1221G>A), V349T (1229G>A 1230T>C 1231T>C), S350N (1233G>A 1234C>T), K358T (1257A>C 1258A>G), N365S (1278A>G), A369V (1290C>T 1291G>A), D372T (1298G>A 1299A>C 1300C>T), I375V (1307A>G 1309A>G), N378E (1316A>G 1318T>A), I382T (1329T>C), L397I (1373C>A), I401V (1385A>G), E407D (1405G>T), A411S (1415G>T 1417A>T), V418I (1436G>A 1438C>T), E508G (1707A>G), A512T (1718G>A), I520V (1742A>G 1744C>T), S525N (1758G>A), V536I (1790G>A), I585V (1937A>G 1939T>C), F607Y (2004T>A), I628M (2068T>G), V630I (2072G>A 2074G>A), R675K (2208G>A), D694E (2266C>G), N744G (2414A>G 2415A>G 2416T>G), S746C (2420A>T), K780R (2523A>G), R817K (2634G>A), V847I (2723G>A 2725C>T), A855T (2747G>A 2749T>A), H871N (2795C>A), V880I (2822G>A 2824G>A), P900T (2882C>A), S901H (2885A>C 2886G>A), T904M (2895C>T), N916S (2931A>G), N919S (2940A>G), E929_N930insX (2971_2972insA), T936I (2991C>T), H937R (2994A>G), E939R (2999G>A 3000A>G), R946Q (3021G>A), Y947F (3024A>T 3025C>T), T951I (3036C>T), E952G (3039A>G 3040A>C), K953T (3042A>C 3043G>A), A972V (3099C>T 3100C>A), V974R (3104G>A 3105T>G 3106G>A), S983N (3132G>A), R985K (3137C>A 3138G>A 3139A>G), F990I (3152T>A 3154C>T), C994Y (3165G>A), F1012L (3218T>C), T1014N (3225C>A), V1016I (3230G>A), V1028M (3266G>A 3268A>G), V1031M (3275G>A), A1034T (3284G>A), R1044K (3315G>A 3316A>G), E1045A (3318A>C), M1059V (3359A>G), D1072E (3400T>G), V1110G (3513T>G), M1121V (3545A>G), A1122T (3548G>A), V1124I (3554G>A 3556C>T), L1147V (3623C>G), L1150M (3632C>A), Y1152H (3638T>C), I1172V (3698A>G), S1178N (3717G>A), R1183K (3731C>A 3732G>A), I1297V (4073A>G), F1315Y (4128T>A 4129C>T), A1323S (4151G>T), V1328I (4166G>A), L1347I (4223C>A 4225G>A), I1379V (4319A>G 4321A>G), F1404Y (4395T>A), V1408I (4406G>A), M1431V (4475A>G 4477G>T), T1433S (4481A>T 4483A>G), A1454L (4544G>T 4545C>T 4546G>A), R1455K (4548G>A), V1499I (4679G>A 4681G>C), V1538I (4796G>A 4798C>T), T1540S (4803C>G 4804T>C), S1625G (5057A>G), T1626A (5060A>G), L1636I (5090C>A), A1639V (5100C>T), E1641D (5107A>T), V1646F (5120G>T 5122C>T), I1648V (5126A>G), V1667I (5183G>A), A1793T (5561G>A 5563G>A), I1805V (5597A>G), N1807K (5605C>A), A1812V (5619C>T 5620T>G), I1816L (5630A>C), V1837I (5693G>A), K1890R (5853A>G), I1904V (5894A>G 5896C>G), R1926A (5960C>G 5961G>C 5962G>C), I1936V (5990A>G), E1945D (6019A>T), K1949T (6030A>C), D1955E (6049C>A), M1958I (6058G>A), S1963N (6072G>A 6073T>C), N1964H (6074A>C), T1966N (6081C>A), C1976S (6111G>C 6112T>C), I1982V (6128A>G 6130T>C), V1992I (6158G>A), T1996S (6170A>T 6172A>G), L2007F (6203C>T 6205C>T), V2016I (6230G>A), A2025K (6257G>A 6258C>A), K2033R (6282A>G), A2060T (6362G>A), V2073I (6401G>A), A2074V (6405C>T), Y2082F (6429A>T), Q2089R (6450A>G), H2101N (6485C>A 6487C>T), L2115I (6527C>A), T2135I (6588C>T), Y2151H (6635T>C), I2155V (6647A>G), I2165V (6677A>G 6679A>G), D2178N (6716G>A), N2179D (6719A>G), I2194L (6764A>T 6766A>G), A2204V (6795C>T 6796A>G), H2205Q (6799C>A), N2225D (6857A>G), V2231T (6875G>A 6876T>C 6877C>A), D2250E (6934C>A), V2273I (7001G>A 7003G>A), F2283Y (7032T>A), T2314K (7125C>A), A2321S (7145G>T 7147A>G), E2326_A2327del (7160_7165delGAGGCA), D2344E (7216T>A), D2358E (7258T>A), D2363_R2364insX (7273_7274insT), R2364V (7274C>G 7275G>T), H2366L (7281A>T), L2367S (7284T>C), S2371L (7296C>T), S2376F (7311C>T), T2377I (7314C>T), C2378Y (7317G>A), N2379K (7321T>A), K2384Q (7334A>C), A2387N (7343G>A 7344C>A 7345A>C), S2408M (7407G>T 7408T>G), C2411G (7415T>G), L2440S (7503T>C 7504G>A), D2509N (7709G>A), K2510R (7713A>G 7714A>G), T2543I (7812C>T), V2554L (7844G>T 7846G>A), V2567I (7883G>A), S2620N (8043G>A), N2656S (8151A>G 8152C>T), R2665K (8178G>A), R2675K (8208G>A), P2704A (8294C>G 8296A>G), S2722F (8349C>T), A2742S (8408G>T), M2756I (8452G>A), R2759K (8460G>A), A2766T (8480G>A 8482T>A), R2773K (8502G>A 8503A>G), E2777D (8515A>T), E2785D (8539G>C), L2788C (8546C>T 8547T>G), S2790T (8552T>A), A2797G (8574C>G), I2801V (8585A>G), G2805E (8598G>A), N2807S (8603A>T 8604A>C 8605C>T), P2813Q (8622C>A), E2814G (8625A>G 8626G>A), P2822F (8648C>T 8649C>T 8650G>C), G2825E (8658G>A), D2827A (8664A>C 8665T>G), A2832V (8679C>T 8680A>T), T2854K (8745C>A 8746A>G), E2891G (8856A>G 8857G>A), R2894Q (8865G>A), M2895L (8867A>C 8869G>T), G2900E (8883G>A), T2902S (8888A>T), V2924I (8954G>A), N2934D (8984A>G 8986T>C), E2935V (8988A>T), I2983V (9131A>G), M2989G (9149A>G 9150T>G 9151G>A), G2990S (9152G>A), P3026S (9260C>T 9262T>C), Q3038H (9298A>C) | | | | | | | | |

| | Begin | End | Coverage | Score | Concordance | Matches | Identities | I/D/M/F* | Stop Codons |
|---|---|---|---|---|---|---|---|---|---|
| NT | 5 | 9704 | 99.9% | 12522 | 64.8% | 9688 (99.7%) | 8015 (82.5%) | 15/12 | |

Codon mutations:

ACT3ACC (193T>C), TAC4CAA (194T>C 196C>A), TCA6TCG (202A>G), ATC8ATT (208C>T), TGT9CTG (209T>C 210G>T 211T>G), TCG12TCA (220G>A), TTT13ATG (221T>A 223T>G), GAA14GAG (226A>G), CTA17CTT (235A>T), TAC19TAT (241C>T), CCA21CCT (247A>T), TCT23CCT (251T>C), TGC24TTT (255G>T 256C>T), GAG25GGG (258A>G), CAT26CAC (262T>C), ATT27GTG (263A>G 265T>G), GTG28GCG (267T>C), GAA30GTA (273A>T), CGA31CGT (277A>T), GAA32GAG (280A>G), GTG33GTT (283G>T), CCG34CCA (286G>A), GCT35ACC (287G>A 289T>C), GTT37ACT (293G>A 294T>C), CCT39CCC (301T>C), TTC40TTT (304C>T), GCA41ATA (305G>A 306C>T), GAT42GGT (309A>G), CTG43TTG (311C>T), GAA44GAT (316A>T), ACA45GTG (317A>G 318C>T 319A>G), CAA46CAG (322A>G), GCA49GCT (331A>T), CTC52CAA (339T>A 340C>A), AAG53AGG (342A>G), AAA55GCG (347A>G 348A>C 349A>G), TAT56CAT (350T>C), GCT57GCA (355T>A), ACT58GTT (356A>G 357C>T), CGT60CGC (364T>C), GTG61ACA (365G>A 366T>C 367G>A), CTC62TCC (368C>T 369T>C), AAA63AAG (373A>G), AAC64AAT (376C>T), TTT67TGT (384T>G), ACG68ACA (388G>A), TAC69TAT (391C>T), CGA70CGG (394A>G), AAG72AAA (400G>A), GAT74GAC (406T>C), GCC75GTT (408C>T 409C>T), CAG76CAA (412G>A), ATA77ATT (415A>T), ATG78GAG (416A>G 417T>A), ATT80ATC (424T>C), CAG81CAA (427G>A), AAA83AAG (433A>G), CTG84CTC (436G>C), GAG85AAG (437G>A), AGG86GAG (440A>G 441G>A), AAG87CGA (443A>C 444A>G 445G>A), GAT88GAA (448T>A), GAA90GAG (454A>G), GAA91GCA (456A>C), TAT92TAC (460T>C), CAC93AAC (461C>A), CAA95CAG (469A>G), GCC97GCG (475C>G), CCT99CCA (481T>A), ACT106ACC (502T>C), ATA107ATT (505A>T), GGC109GGT (511C>T), GAT111GAA (517T>A), CCT112CCA (520T>A), CCA113CCT (523A>T), TCT116TTT (531C>T), GAG117GAA (535G>A), CCA118TCA (536C>T), CAA119TGG (539C>T 540A>G 541A>G), AGA122AAG (549G>A 550A>G), ATC124ATT (556C>T), ATT125ATC (559T>C), CAT126CAC (562T>C), ACT128ACC (568T>C), CCA129CCG (571A>G), ATG131ACG (576T>C), CGT132CGC (580T>C), GTC134GCG (585T>C 586C>G), CGC137TAT (593C>T 594G>A 595C>T), CCC138CGC (597C>G), ATA139GTG (599A>G 601A>G), ATA140CCG (602A>C 603T>C 604A>G), ACA143ACG (613A>G), GAA144GAG (616A>G), GGC145GGG (619C>G), CAG146CAA (622G>A), ATG147ATA (625G>A), AAT148AAC (628T>C), CAC149CAT (631C>T), ATT151GTC (635A>G 637T>C), ATA154GTG (644A>G 646A>G), ATT157ATC (655T>C), TCG159TCA (661G>A), GAG160GCC (663A>C 664G>C), AAA161AAG (667A>G), GTC165GTT (679C>T), CAC166CAA (682C>A), TTA167CTG (683T>C 685A>G), AGT169AGC (691T>C), ACC172AGC (699C>G), ACT173ACA (703T>A), CAT174CAA (706T>A), AAG179GAA (719A>G 721G>A), ATA180GTG (722A>G 724A>G), CTT181TTG (725C>T 727T>G), GGT182GGA (730T>A), GCA183TCA (731G>T), TAC184CCT (734T>C 735A>C 736C>T), TCC185CGT (737T>C 738C>G 739C>T), GCG187GTC (744C>T 745G>C), CGA189TGC (749C>T 751A>C), ACT190ACC (754T>C), ATG194AAG (765T>A), TTG196CTG (770T>C), CGA197AAG (773C>A 774G>A 775A>G), CGG198AGG (776C>A), TTC202TTT (790C>T), ATG206AAG (801T>A), GGA210GCG (813G>C 814A>G), CTT211CAT (816T>A), TTG212CTG (818T>C), CAA213CAG (823A>G), CGT214CAA (825G>A 826T>A), CGG217AGA (833C>A 835G>A), GAC219GAT (841C>T), AAT223GAC (851A>G 853T>C), GTC225GTT (859C>T), CGC226TGT (860C>T 862C>T), ACT227GCC (863A>G 865T>C), ATC228ACC (867T>C), AAC229GAT (869A>G 871C>T), ATA230TTG (872A>T 874A>G), CGA231CAA (876G>A), AGG232AAG (879G>A), GAT234GAC (886T>C), ATC238ATT (898C>T), AAC240AGC (903A>G), ACA241AAC (906C>A 907A>C), AAA242ACC (909A>C 910A>C), AGC243AAT (912G>A 913C>T), GGC246GGA (922C>A), GGT249GGA (931T>A), TCA252CTT (938T>C 939C>T 940A>T), GGA253GGG (943A>G), TTC256TTT (952C>T), TCA261TCG (967A>G), GAA263GAG (973A>G), GGG264GGC (976G>C), TTG266ATC (980T>A 982G>C), GCA269GCG (991A>G), CGT270CGC (994T>C), AGA272AAA (999G>A), GTT273GTC (1003T>C), CAG275AGA (1007C>A 1008A>G 1009G>A), AGT276GGG (1010A>G 1012T>G), ATT277GTT (1013A>G), TTA278TTG (1018A>G), AAC279CAG (1019A>C 1021C>G), ATC282ATT (1030C>T), TCG285TCA (1039G>A), AAT286AAC (1042T>C), GCC287GCA (1045C>A), GAC288GAA (1048C>A), AAT289AAC (1051T>C), AAG292AAA (1060G>A), GGT293GGA (1063T>A), CTG294TTG (1064C>T), GAC295GAT (1069C>T), CGA300CAA (1083G>A), TAT303TAC (1093T>C), TCG305GCA (1097T>G 1099G>A), CAC307CAT (1105C>T), GTA310GTG (1114A>G), GCT311GCG (1117T>G), TTA313TTG (1123A>G), CCT314CCA (1126T>A), GTC315GTT (1129C>T), AGG320AAG (1143G>A), GTA321GTT (1147A>T), GCT322GCA (1150T>A), TTG324ATA (1154T>A 1156G>A), CAC327CAT (1165C>T), AGT328AGC (1168T>C), CTT330TTA (1172C>T 1174T>A), ACC339ACT (1201C>T), AGC346AAC (1221G>A), GTT349ACC (1229G>A 1230T>C 1231T>C), AGC350AAT (1233G>A 1234C>T), CTG352CTA (1240G>A), AAG354AAA (1246G>A), TTG356CTA (1250T>C 1252G>A), CAT357CAC (1255T>C), AAA358ACG (1257A>C 1258A>G), AGA361AGG (1267A>G), GGT363GGC (1273T>C), TTG364TTA (1276G>A), AAT365AGT (1278A>G), TTG367CTG (1283T>C), GCG369GTA (1290C>T 1291G>A), GAC372ACT (1298G>A 1299A>C 1300C>T), TTT374TTC (1306T>C), ATA375GTG (1307A>G 1309A>G), CAT376CAC (1312T>C), AAT378GAA (1316A>G 1318T>A), TTG381TTA (1327G>A), ATA382ACA (1329T>C), GCG383GCT (1333G>T), TTA384TTG (1336A>G), CTA387TTG (1343C>T 1345A>G), CCG390CCA (1354G>A), GTG391GTT (1357G>T), GAC392GAT (1360C>T), CTC395TTG (1367C>T 1369C>G), CTT397ATT (1373C>A), TTC398TTT (1378C>T), GAG400GAA (1384G>A), ATA401GTA (1385A>G), AAA403AAG (1393A>G), TCC404TCT (1396C>T), GAG407GAT (1405G>T), AAA408AAG (1408A>G), CAG409CAA (1411G>A), GCA411TCT (1415G>T 1417A>T), CCG412CCC (1420G>C), AAT415AAC (1429T>C), TTA416TTG (1432A>G), GTC418ATT (1436G>A 1438C>T), TTA419CTG (1439T>C 1441A>G), AAT420AAC (1444T>C), TTC423TTT (1453C>T), CTG424CTA (1456G>A), GAA428GAG (1468A>G), GTA436GTG (1492A>G), TTG439CTA (1499T>C 1501G>A), AGT440AGC (1504T>C), TTG441TTA (1507G>A), CTC442CTT (1510C>T), GAA443GAG (1513A>G), TTA444TTG (1516A>G), GCA445GCT (1519A>T), AGG446AGA (1522G>A), TTC447TTT (1525C>T), CAG448CAA (1528G>A), AAC450AAT (1534C>T), ACT452ACA (1540T>A), AAC454AAT (1546C>T), AAA457AAG (1555A>G), GGT458GGC (1558T>C), GAT459GAC (1561T>C), ATA460ATC (1564A>C), TCT461TCA (1567T>A), TTC463TTT (1573C>T), AGA464AGG (1576A>G), TTA467CTA (1583T>C), TCT468TCA (1588T>A), AAG470AAA (1594G>A), AAC472AAT (1600C>T), CTG475CTA (1609G>A), TCG478TCA (1618G>A), TGC479TGT (1621C>T), GAC480GAT (1624C>T), CTG483TTG (1631C>T), AAA485AAG (1639A>G), TTC489TTT (1651C>T), CTC490CTG (1654C>G), CAT497CAC (1675T>C), TCA503TCG (1693A>G), TTT506TTC (1702T>C), GAA508GGA (1707A>G), GCA512ACA (1718G>A), GGA514GGG (1726A>G), TAC515TAT (1729C>T), TCA516TCC (1732A>C), TAT518TAC (1738T>C), ATC520GTT (1742A>G 1744C>T), CGC521CGT (1747C>T), CAT523CAC (1753T>C), AGT525AAT (1758G>A), AGG528AGA (1768G>A), AAC534AAT (1786C>T), TTA535TTG (1789A>G), GTT536ATT (1790G>A), GTC537GTT (1795C>T), TTA541CTG (1805T>C 1807A>G), GAG543GAA (1813G>A), AGG545AGA (1819G>A), AAG547AAA (1825G>A), GAC551GAT (1837C>T), TAT552TAC (1840T>C), CAA555CAG (1849A>G), GTC558GTG (1858C>G), AAA560AAG (1864A>G), TGC562TGT (1870C>T), AGT564AGC (1876T>C), AAA566AAG (1882A>G), GGT568GGA (1888T>A), AAT569AAC (1891T>C), TGT574TGC (1906T>C), ACA577ACT (1915A>T), GGT582GGC (1930T>C), GCC584GCT (1936C>T), ATT585GTC (1937A>G 1939T>C), TCA587TCG (1945A>G), TTC589TTT (1951C>T), CCA592CCT (1960A>T), GTA598GTG (1978A>G), ATT599ATA (1981T>A), GGC600GGT (1984C>T), AAT601AAC (1987T>C), GAC604GAT (1996C>T), TTT607TAT (2004T>A), AAA612AAG (2020A>G), GGG613GGA (2023G>A), GAT614GAC (2026T>C), TCG615TCT (2029G>T), GAG616GAA (2032G>A), TAT625TAC (2059T>C), ATT628ATG (2068T>G), AAC629AAT (2071C>T), GTG630ATA (2072G>A 2074G>A), CTG635TTA (2087C>T 2089G>A), AGC639AGT (2101C>T), GAG640GAA (2104G>A), GAG641GAA (2107G>A), AAG644AAA (2116G>A), TTC646TTT (2122C>T), CGC651CGT (2137C>T), CTT658CTC (2158T>C), ACC660ACT (2164C>T), CCA662CCG (2170A>G), GAT666GAC (2182T>C), GCG668GCA (2188G>A), ACC669ACA (2191C>A), AGA675AAA (2208G>A), TAT678TAC (2218T>C), GAC680GAT (2224C>T), GTA681GTT (2227A>T), TTG686TTA (2242G>A), CCC687CCT (2245C>T), TTG690TTA (2254G>A), GTT691GTC (2257T>C), GAC692GAT (2260C>T), GAC694GAG (2266C>G), CAA696CAG (2272A>G), ACG697ACA (2275G>A), TCA703TCG (2293A>G), GGC705GGT (2299C>T), TCG706TCA (2302G>A), CAG707CAA (2305G>A), GGA710GGT (2314A>T), TAT711TAC (2317T>C), CTA714CTG (2326A>G), GTG719GTA (2341G>A), CAA721CAG (2347A>G), CTT722CTC (2350T>C), TTG724TTA (2356G>A), AAT727AAC (2365T>C), GAA729GAG (2371A>G), TTA730TTG (2374A>G), GAT733GAC (2383T>C), ATA734ATT (2386A>T), AAA735AAG (2389A>G), CAT736CAC (2392T>C), TAT737TAC (2395T>C), AAT744GGG (2414A>G 2415A>G 2416T>G), GCT745GCA (2419T>A), AGC746TGC (2420A>T), ACA752ACT (2440A>T), GTT761GTC (2467T>C), TCT764TCC (2476T>C), TCG766TCA (2482G>A), CTG769CTA (2491G>A), CTG771CTT (2497G>T), TTG773TTA (2503G>A), AGA778AGG (2518A>G), CCT779CCC (2521T>C), AAG780AGG (2523A>G), AGA783AGG (2533A>G), TTG785TTA (2539G>A), TTA787TTG (2545A>G), GAG789GAA (2551G>A), CCT790CCA (2554T>A), CTG792TTG (2558C>T), TTG793CTC (2561T>C 2563G>C), TCA796TCG (2572A>G), CCT800CCC (2584T>C), GGC801GGT (2587C>T), CTG803CTC (2593G>C), GCT805GCC (2599T>C), AAT808AAC (2608T>C), TTT812TTC (2620T>C), GCG815GCA (2629G>A), GTG816GTA (2632G>A), AGG817AAG (2634G>A), TTG818TTA (2638G>A), ATT820ATA (2644T>A), AAA823AAG (2653A>G), GCT827GCC (2665T>C), CTA832TTA (2678C>T), CTA833CTG (2683A>G), GCT835GCC (2689T>C), CTA838CTC (2698A>C), TCA841TCG (2707A>G), GCG842GCA (2710G>A), GAA844GAG (2716A>G), ACA845ACG (2719A>G), CTC846CTT (2722C>T), GTC847ATT (2723G>A 2725C>T), GCA848GCG (2728A>G), AGG850AGA (2734G>A), ATT851ATA (2737T>A), GAT854GAC (2746T>C), GCT855ACA (2747G>A 2749T>A), GCA856GCT (2752A>T), GCT857GCA (2755T>A), GAC859GAT (2761C>T), CTC860CTT (2764C>T), TGT865TGC (2779T>C), GGG867GGT (2785G>T), AAC869AAT (2791C>T), CAT871AAT (2795C>A), TAC874TAT (2806C>T), TTG878CTC (2816T>C 2818G>C), GTG880ATA (2822G>A 2824G>A), TTG881TTA (2827G>A), GTT884GTC (2836T>C), AAG885AAA (2839G>A), AAT886AAC (2842T>C), ACC893ACA (2863C>A), CTA894TTA (2864C>T), CCA900ACA (2882C>A), AGT901CAT (2885A>C 2886G>A), AAC903AAT (2893C>T), ACG904ATG (2895C>T), GTC906GTT (2902C>T), GTA907GTG (2905A>G), ATT909ATC (2911T>C), AAT913AAC (2923T>C), AAT916AGT (2931A>G), AAC919AGC (2940A>G), GCT921GCC (2947T>C), ACT926ACC (2962T>C), GAA929_AAT930insA-- (2971_2972insA), GGT935GGC (2989T>C), ACT936ATT (2991C>T), CAT937CGT (2994A>G), GAG939AGG (2999G>A 3000A>G), TCC944TC- (3016delC), CGG946CAG (3021G>A), TAC947TTT (3024A>T 3025C>T), ACA951ATA (3036C>T), GAA952GGC (3039A>G 3040A>C), AAG953ACA (3042A>C 3043G>A), TTG956TTA (3052G>A), TTA959TTG (3061A>G), GCG966GCA (3082G>A), GGC969GGT (3091C>T), GCC972GTA (3099C>T 3100C>A), GTG974AGA (3104G>A 3105T>G 3106G>A), GTC975GTT (3109C>T), AAA976AAG (3112A>G), ACT978ACC (3118T>C), GCC979GCT (3121C>T), GGA981GGG (3127A>G), AGC983AAC (3132G>A), CGA985AAG (3137C>A 3138G>A 3139A>G), TAT989TAC (3151T>C), TTC990ATT (3152T>A 3154C>T), TGT994TAT (3165G>A), GTA995GTG (3169A>G), ATC1005ATT (3199C>T), AGG1007CGG (3203A>C), CGT1008CGC (3208T>C), CCA1010CCG (3214A>G), ACC1011ACT (3217C>T), TTT1012CTT (3218T>C), GTC1013GTA (3223C>A), ACT1014AAT (3225C>A), GTT1016ATT (3230G>A), AAC1017AAT (3235C>T), TTA1025CTA (3257T>C), ACC1026ACT (3262C>T), AGC1027AGT (3265C>T), GTA1028ATG (3266G>A 3268A>G), GTG1029GTA (3271G>A), GTG1031ATG (3275G>A), GCA1034ACA (3284G>A), CAG1039CAA (3301G>A), AGA1044AAG (3315G>A 3316A>G), GAA1045GCA (3318A>C), ATC1046ATT (3322C>T), GAG1047GAA (3325G>A), TTG1048TTA (3328G>A), ATA1051ATT (3337A>T), GAG1055GAA (3349G>A), GTC1057GTT (3355C>T), TGC1058TGT (3358C>T), ATG1059GTG (3359A>G), CTA1061CTG (3367A>G), TAT1062TAC (3370T>C), TTA1065CTG (3377T>C 3379A>G), GAA1070GAG (3394A>G), CGC1071CGT (3397C>T), GAT1072GAG (3400T>G), ACA1074ACG (3406A>G), TAC1078TAT (3418C>T), ATT1079ATA (3421T>A), TAT1081TAC (3427T>C), AAG1083AAA (3433G>A), TCA1084TCT (3436A>T), GTA1085GTG (3439A>G), AAC1086AAT (3442C>T), CCT1087CCC (3445T>C), CAG1091CAA (3457G>A), TTT1092TTC (3460T>C), GCT1093GCA (3463T>A), GCG1095GCT (3469G>T), CAG1096CAA (3472G>A), CAC1104CAT (3496C>T), CAG1105CAA (3499G>A), TCC1107TCT (3505C>T), GTT1110GGT (3513T>G), AAA1112AAG (3520A>G), GAA1115GAG (3529A>G), CAA1116CAG (3532A>G), ATG1121GTG (3545A>G), GCT1122ACT (3548G>A), TTA1123CTA (3551T>C), GTC1124ATT (3554G>A 3556C>T), AGT1133AGC (3583T>C), GAT1134GAC (3586T>C), TGC1135TGT (3589C>T), TTC1137TTT (3595C>T), AAT1141AAC (3607T>C), TTT1143TTC (3613T>C), AAG1144AAA (3616G>A), GGT1145GGC (3619T>C), CTT1147GTT (3623C>G), TCC1148TCT (3628C>T), CTG1150ATG (3632C>A), GAT1151GAC (3637T>C), TAT1152CAT (3638T>C), GAA1153GAG (3643A>G), TTA1159TTG (3661A>G), GTG1162GTA (3670G>A), TTT1166TTC (3682T>C), GAG1168GAA (3688G>A), GAG1171GAA (3697G>A), ATT1172GTT (3698A>G), GAT1174GAC (3706T>C), TTT1175TTC (3709T>C), TTG1177CTA (3713T>C 3715G>A), AGT1178AAT (3717G>A), GAG1179GAA (3721G>A), CGA1183AAA (3731C>A 3732G>A), ACT1184ACA (3736T>A), CTA1188TTG (3746C>T 3748A>G), ACA1190ACG (3754A>G), AGT1193AGC (3763T>C), CGA1198CGG (3778A>G), CAA1199CAG (3781A>G), ATC1200ATT (3784C>T), CAT1204CAC (3796T>C), CCA1207CCT (3805A>T), TAC1209TAT (3811C>T), AGA1210AGG (3814A>G), TTC1215TTT (3829C>T), TTT1218TTC (3838T>C), AGA1220AGG (3844A>G), GCA1221GCC (3847A>C), GTT1224GTA (3856T>A), GTG1226GTT (3862G>T), GAC1229GAT (3871C>T), GCC1231GCA (3877C>A), AGC1233AGT (3883C>T), GAA1234GAG (3886A>G), CTA1236CTG (3892A>G), GAC1237GAT (3895C>T), TTT1238TTC (3898T>C), GTA1240GTG (3904A>G), CGG1241AGA (3905C>A 3907G>A), GCT1243GCC (3913T>C), AAG1248AAA (3928G>A), TCA1249TCT (3931A>T), GGG1251GGA (3937G>A), AGT1257AGC (3955T>C), GTG1258GTA (3958G>A), GCC1259GCT (3961C>T), GGA1260GGG (3964A>G), TCT1261TCC (3967T>C), TTA1264TTG (3976A>G), ATT1265ATC (3979T>C), CCA1267CCT (3985A>T), CTA1271CTT (3997A>T), GCG1272GCA (4000G>A), AAC1274AAT (4006C>T), GTT1275GTG (4009T>G), TTC1276TTT (4012C>T), AAA1277AAG (4015A>G), TCT1280TCC (4024T>C), GAA1282GAG (4030A>G), CCA1283CCT (4033A>T), AAG1286AAA (4042G>A), CTG1290TTA (4052C>T 4054G>A), CGT1291CGC (4057T>C), CGT1293CGA (4063T>A), GGA1294GGG (4066A>G), ATA1297GTA (4073A>G), TTT1298TTC (4078T>C), GGC1299GGT (4081C>T), TCT1300TCC (4084T>C), TCC1304TCT (4096C>T), AGC1308AGT (4108C>T), GGA1309GGT (4111A>T), GCG1311GCA (4117G>A), CTA1312TTG (4118C>T 4120A>G), TAC1314TAT (4126C>T), TTC1315TAT (4128T>A 4129C>T), GCC1316GCT (4132C>T), AAT1317AAC (4135T>C), CAA1321CAG (4147A>G), TTA1322CTA (4148T>C), GCT1323TCT (4151G>T), AAC1326AAT (4162C>T), GTA1328ATA (4166G>A), ATA1329ATC (4171A>C), GAG1332GAA (4180G>A), CTG1336TTG (4190C>T), CCT1338CCC (4198T>C), TCC1339TCT (4201C>T), GCG1342GCT (4210G>T), TTC1343TTT (4213C>T), AGT1345AGC (4219T>C), CTG1346TTG (4220C>T), CTG1347ATA (4223C>A 4225G>A), GTT1349GTG (4231T>G), CAT1351CAC (4237T>C), GTA1356GTG (4252A>G), TTA1357CTG (4253T>C 4255A>G), AAA1358AAG (4258A>G), ACT1362ACC (4270T>C), AGA1366AGG (4282A>G), GAG1367GAA (4285G>A), GAA1369GAG (4291A>G), CAG1373CAA (4303G>A), GTC1376GTT (4312C>T), ATA1379GTG (4319A>G 4321A>G), GTG1380GTT (4324G>T), GAG1381GAA (4327G>A), CTG1384CTT (4336G>T), TCT1385TCC (4339T>C), TCA1388TCT (4348A>T), GCA1392GCG (4360A>G), GGT1394GGC (4366T>C), TCT1395TCA (4369T>A), ACT1397ACC (4375T>C), GTT1401GTC (4387T>C), TTT1404TAT (4395T>A), GTA1408ATA (4406G>A), CTT1409CTC (4411T>C), GTT1418GTC (4438T>C), ACC1420ACA (4444C>A), GCC1422GCT (4450C>T), CTC1424CTT (4456C>T), GAC1427GAT (4465C>T), ATG1431GTT (4475A>G 4477G>T), GTC1432GTT (4480C>T), ACA1433TCG (4481A>T 4483A>G), GGC1437GGT (4495C>T), AAG1441AAA (4507G>A), CAC1442CAT (4510C>T), GGT1443GGG (4513T>G), CTA1445TTA (4517C>T), GTC1448GTT (4528C>T), AAA1450AAG (4534A>G), ACC1452ACT (4540C>T), GCG1454TTA (4544G>T 4545C>T 4546G>A), AGA1455AAA (4548G>A), TTT1458TTC (4558T>C), GTA1460GTT (4564A>T), ACC1462ACT (4570C>T), ATA1464ATT (4576A>T), TTG1471TTA (4597G>A), GAC1472GAT (4600C>T), GAC1474GAT (4606C>T), GTG1475GTA (4609G>A), GTG1477GTA (4615G>A), GAC1478GAT (4618C>T), TTG1481CTT (4625T>C 4627G>T), GAC1488GAT (4648C>T), ATT1489ATC (4651T>C), AGG1492AGA (4660G>A), AGC1493AGT (4663C>T), GCT1495GCA (4669T>A), GTG1499ATC (4679G>A 4681G>C), AGT1500AGC (4684T>C), GTT1501GTG (4687T>G), GAG1505GAA (4699G>A), AGA1506AGG (4702A>G), CAA1508CAG (4708A>G), AGG1509AGA (4711G>A), CTG1510TTG (4712C>T), AAA1518AAG (4738A>G), GTA1520GTT (4744A>T), GCA1521GCG (4747A>G), TTG1522CTG (4748T>C), ACT1527ACC (4765T>C), GTC1538ATT (4796G>A 4798C>T), ACT1540AGC (4803C>G 4804T>C), GAG1541GAA (4807G>A), GCG1542GCT (4810G>T), GCT1543GCG (4813T>G), CTT1544CTC (4816T>C), TAT1549TAC

| | | | | | | | | | |
|---|---|---|---|---|---|---|---|---|---|
| | (4831T>C), AAC1550AAT (4834C>T), CCA1552CCT (4840A>T), GGA1556GGG (4852A>G), GGC1557GGT (4855C>T), AGT1561AGC (4867T>C), CTG1562CTC (4870G>C), ATT1563ATA (4873T>A), TGT1566TGC (4882T>C), ACA1573ACT (4903A>T), CAG1575CAA (4909G>A), TTT1577TTC (4915T>C), GAA1578GAG (4918A>G), TTG1579CTA (4919T>C 4921G>A), AGT1580AGC (4924T>C), CCC1581CCG (4927C>G), TTT1583TTC (4933T>C), ATC1584ATA (4936C>A), CAG1585CAA (4939G>A), AAT1586AAC (4942T>C), TTC1587TTT (4945C>T), TCA1593TCG (4963A>G), CCT1596CCC (4972T>C), CTT1602CTC (4990T>C), AAA1606AAG (5002A>G), CCT1613CCC (5023T>C), TTG1614CTG (5024T>C), TGC1615TGT (5029C>T), GAT1616GAC (5032T>C), TCT1618TCC (5038T>C), CCA1620CCT (5044A>T), GCA1623GCT (5053A>T), TCG1624TCA (5056G>A), AGT1625GGT (5057A>G), ACT1626GCT (5060A>G), TCG1629TCT (5071G>T), CTT1636ATT (5090C>A), GGA1637GGG (5095A>G), GTG1638GTT (5098G>T), GCC1639GTC (5100C>T), TTA1640TTG (5104A>G), GAA1641GAT (5107A>T), ATT1642ATC (5110T>C), CCA1643CCG (5113A>G), AAG1644AAA (5116G>A), CAA1645CAG (5119A>G), GTC1646TTT (5120G>T 5122C>T), ATT1648GTT (5126A>G), CAT1651CAC (5137T>C), ATC1652ATT (5140C>T), AAA1653AAG (5143A>G), GAG1654GAA (5146G>A), CTC1659CTA (5161C>A), GAA1661GAG (5167A>G), CTT1663CTG (5173T>G), ACG1666ACA (5182G>A), GTT1667ATT (5183G>A), GTC1668GTT (5188C>T), AAG1669AAA (5191G>A), AAA1671AAG (5197A>G), GAC1672GAT (5200C>T), TGC1674TGT (5206C>T), TTA1675TTG (5209A>G), AGC1678AGT (5218C>T), TCG1682TCA (5230G>A), ATC1684ATT (5236C>T), AGC1685AGT (5239C>T), ATC1687ATT (5245C>T), ACA1693ACT (5263A>T), CTC1695CTT (5269C>T), TTC1696TTT (5272C>T), GCC1697GCA (5275C>A), ATC1698ATT (5278C>T), CCA1699CCC (5281A>C), AGA1700AGG (5284A>G), ACT1701ACC (5287T>C), CTA1702TTA (5288C>T), ATA1703ATT (5293A>T), TTG1704CTA (5294T>C 5296G>A), GAG1706GAA (5302G>A), CTT1709CTA (5311T>A), GAA1710GAG (5314A>G), CGA1713CGT (5323A>T), AGT1721AGC (5347T>C), ATC1723ATT (5353C>T), TGC1727TGT (5365C>T), GTT1734GTC (5386T>C), ACC1737ACA (5395C>A), AAA1746AAG (5422A>G), GAT1747GAC (5425T>C), ACC1749ACT (5431C>T), GAG1751GAA (5437G>A), CTT1756CTC (5452T>C), AAG1758AAA (5458G>A), AGT1761AGC (5467T>C), CAA1762CAG (5470A>G), CTA1763TTG (5471C>T 5473A>G), GAA1765GAG (5479A>G), TTC1766TTT (5482C>T), TCA1767TCC (5485A>C), AAT1768AAC (5488T>C), TTG1769CTA (5489T>C 5491G>A), GGT1771GGC (5497T>C), TGT1774TGC (5506T>C), GAG1775GAA (5509G>A), GAG1776GAA (5512G>A), TTA1778CTA (5516T>C), TAT1782TAC (5530T>C), GAG1783GAA (5533G>A), TCG1784TCT (5536G>T), TTG1785TTA (5539G>A), GTT1788GTG (5548T>G), CAC1790CAT (5554C>T), GCT1792GCA (5560T>A), GCG1793ACA (5561G>A 5563G>A), ACG1794ACC (5566G>C), TCA1795TCG (5569A>G), CTC1800CTG (5584C>G), AAG1803AAA (5593G>A), GGG1804GGA (5596G>A), ATT1805GTT (5597A>G), AAC1807AAA (5605C>A), GTG1811GTT (5617G>T), GCT1812GTG (5619C>T 5620T>G), TTG1815TTA (5629G>A), ATC1816CTC (5630A>C), GGC1819GGT (5641C>T), GCA1822GCT (5650A>T), ATA1826ATT (5662A>T), GGA1827GGG (5665A>G), CAA1835CAG (5689A>G), GTT1837ATT (5693G>A), GAG1838GAA (5698G>A), CAA1843CAG (5713A>G), GGG1844GGC (5716G>C), AAA1845AAG (5719A>G), GCC1853GCA (5743C>A), CGC1857CGA (5755C>A), CAT1858CAC (5758T>C), CGT1860CGC (5764T>C), AAA1862AAG (5770A>G), GGC1865GGT (5779C>T), TTT1866TTC (5782T>C), GAC1869GAT (5791C>T), GAC1873GAT (5803C>T), AGG1884AGA (5836G>A), AAA1885AAG (5839A>G), GGA1887GGG (5845A>G), AAA1888AAG (5848A>G), AAA1890AGA (5853A>G), GGT1891GGC (5857T>C), ACC1892ACT (5860C>T), ACA1893ACT (5863A>T), GGT1895GGA (5869T>A), GGT1897GGC (5875T>C), AGC1900AGT (5884C>T), TTC1903TTT (5893C>T), ATC1904GTG (5894A>G 5896C>G), AAC1905AAT (5899C>T), GGG1908GGA (5908G>A), TTT1909TTC (5911T>C), GAG1913GAA (5923G>A), CAA1918CAG (5938A>G), TTC1919TTT (5941C>T), GTT1920GTC (5944T>C), GAT1921GAC (5947T>C), GGG1925GGA (5959G>A), CGG1926GCC (5960C>G 5961G>C 5962G>C), GAA1930GAG (5974A>G), GCT1934GCA (5986T>A), GAC1935GAT (5989C>T), ATT1936GTT (5990A>G), GAT1938GAC (5998T>C), AGA1942CGT (6008A>C 6010A>T), GAA1945GAT (6019A>T), GTG1946GTT (6022G>T), CGA1947CGT (6025A>T), AAA1949ACA (6030A>C), GAC1955GAA (6049C>A), ATG1958ATA (6058G>A), CAA1959CAG (6061A>G), GCC1960GCA (6064C>A), GGT1962GGC (6070T>C), AGT1963AAC (6072G>A 6073T>C), AAC1964CAC (6074A>C), ACG1965ACA (6079G>A), ACC1966AAC (6081C>A), ATA1967ATT (6085A>T), GCA1969GCT (6091A>T), TAC1970TAT (6094C>T), TGT1976TCC (6111G>C 6112T>C), AAA1978AAG (6118A>G), TTG1980TTA (6124G>A), AAG1981AAA (6127G>A), ATT1982GTC (6128A>G 6130T>C), GAT1983GAC (6133T>C), TTA1984TTG (6136A>G), AAC1988AAT (6148C>T), AAA1991AAG (6157A>G), GTT1992ATT (6158G>A), TGT1993TGC (6163T>C), AAA1995AAG (6169A>G), ACA1996TCG (6170A>T 6172A>G), GGC1998GGT (6178C>T), AAA2001AAG (6187A>G), TTT2002TTC (6190T>C), GAG2004GAA (6196G>A), CTC2007TTT (6203C>T 6205C>T), GAA2008GAG (6208A>G), CTA2009TTG (6209C>T 6211A>G), CAG2011CAA (6217G>A), GCT2015GCA (6229T>A), GTA2016ATA (6230G>A), GTC2018GTT (6238C>T), AAG2021AAA (6247G>A), ATA2023ATT (6253A>T), GCA2025AAA (6257G>A 6258C>A), CAT2030CAC (6274T>C), AAA2033AGA (6282A>G), TCG2034TCA (6286G>A), GGC2038GGT (6298C>T), TTG2039TTA (6301G>A), AGA2040AGG (6304A>G), GAC2041GAT (6307C>T), GCC2046GCT (6322C>T), ACA2048ACT (6328A>T), GTT2049GTG (6331T>G), TGT2050TGC (6334T>C), AGG2051AGA (6337G>A), CTG2052TTG (6338C>T), TAT2058TAC (6358T>C), GGG2059GGA (6361G>A), GCA2060ACA (6362G>A), TCA2061TCT (6367A>T), GAG2062GAA (6370G>A), TAC2064TAT (6376C>T), GGT2065GGG (6379T>G), GGA2067GGC (6385A>C), GGA2069GGT (6391A>T), GCA2070GCG (6394A>G), TAC2071TAT (6397C>T), GTA2073ATA (6401G>A), GCG2074GTG (6405C>T), CAC2076CAT (6412C>T), TTA2078TTG (6418A>G), TTT2079TTC (6421T>C), TAC2082TTC (6429A>T), GGT2084GGA (6436T>A), GAG2087GAA (6445G>A), CAA2089CGA (6450A>G), TCC2090TCG (6454C>G), CAC2092CAT (6460C>T), GGT2093GGA (6463T>A), TTC2095TTT (6469C>T), AGG2096AGA (6472G>A), GTG2097GTA (6475G>A), AAT2099AAC (6481T>C), CAC2101AAT (6485C>A 6487C>T), AGT2102AGC (6490T>C), AGC2104AGT (6496C>T), CTG2106CTA (6502G>A), GGT2110GGC (6514T>C), AGG2111AGA (6517G>A), CTC2115ATC (6527C>A), ATC2116ATA (6532C>A), CCG2119CCA (6541G>A), AAA2120AAG (6544A>G), GTC2124GTT (6556C>T), TTT2125TTC (6559T>C), CAG2127CAA (6565G>A), TTG2129CTA (6569T>C 6571G>A), CCT2134CCA (6586T>A), ACA2135ATA (6588C>T), GAA2138GAG (6598A>G), AGA2139AGG (6601A>G), ATT2140ATC (6604T>C), TGT2141TGC (6607T>C), GGA2144GGG (6616A>G), AAC2146AAT (6622C>T), GAG2149GAA (6631G>A), AAG2150AAA (6634G>A), TAT2151CAT (6635T>C), GCT2152GCA (6640T>A), TCG2154TCT (6646G>T), ATC2155GTC (6647A>G), ATC2156ATT (6652C>T), AGC2160AGT (6664C>T), ACT2162ACG (6670T>G), ATA2165GTG (6677A>G 6679A>G), GGC2167GGT (6685C>T), ACA2169ACT (6691A>T), TTC2170TTT (6694C>T), ACA2177ACG (6715A>G), GAT2178AAT (6716G>A), AAT2179GAT (6719A>G), GGA2180GGG (6724A>G), CTA2184TTA (6734C>T), GTG2186GTA (6742G>A), GTG2187GTA (6745G>A), GCC2190GCT (6754C>T), GGA2192GGG (6760A>G), ATA2194TTG (6764A>T 6766A>G), ATT2197ATA (6775T>A), CTG2200TTG (6782C>T), GCA2201GCT (6787A>T), AAC2202AAT (6790C>T), GCA2204GTG (6795C>T 6796A>G), CAC2205CAA (6799C>A), ACC2206ACA (6802C>A), AAC2208AAT (6808C>T), TAC2209TAT (6811C>T), TAC2210TAT (6814C>T), TTC2213TTT (6823C>T), GAA2215GAG (6829A>G), TAC2221TAT (6847C>T), CGA2223CGC (6853A>C), AAT2225GAT (6857A>G), GAG2226GAA (6862G>A), GAA2229GAG (6871A>G), GTC2231ACA (6875G>A 6876T>C 6877C>A), AAG2232AAA (6880G>A), TCT2233TCA (6883T>A), GTT2235GTG (6889T>G), GAC2239GAT (6901C>T), GGC2244GGT (6916C>T), CCG2245CCA (6919G>A), AAA2247AAG (6925A>G), CTT2248CTC (6928T>C), GAC2250GAA (6934C>A), GGG2255GGC (6949G>C), TTC2257TTT (6955C>T), AAA2258AAG (6958A>G), CAA2264CAG (6976A>G), CTA2266TTA (6980C>T), ATC2267ATT (6985C>T), GTA2271GTT (6997A>T), GTG2272GTT (7000G>T), GTG2273ATA (7001G>A 7003G>A), GCT2276GCC (7012T>C), AAG2277AAA (7015G>A), CAT2278CAC (7018T>C), TTT2283TAT (7032T>A), GCC2285GCT (7039C>T), TTG2286CTA (7040T>C 7042G>A), GGA2288GGG (7048A>G), GCT2292GCA (7060T>A), GTC2293GTA (7063C>A), AGC2298AGT (7078C>T), TTA2300CTG (7082T>C 7084A>G), GTA2301GTG (7087A>G), ACC2302ACA (7090C>A), CAT2304CAC (7096T>C), GTA2305GTG (7099A>G), GTT2306GTC (7102T>C), GAG2309GAA (7111G>A), CGA2311CGG (7117A>G), ACA2314AAA (7125C>A), GAA2315GAG (7129A>G), TTT2316TTC (7132T>C), CTG2317TTA (7133C>T 7135G>A), GTG2319GTA (7141G>A), GAT2320GAC (7144T>C), GCA2321TCG (7145G>T 7147A>G), GAG2322GAA (7150G>A), GCA2325GCT (7159A>T), GAG2326_GCA2327del (7160_7165delGAGGCA), TTC2329TTT (7171C>T), GCG2335GCT (7189G>T), AAA2338AAG (7198A>G), AGC2339AGT (7201C>T), CTA2341TTA (7205C>T), AAT2342AAC (7210T>C), GAT2344GAA (7216T>A), GCG2345GCA (7219G>A), ATC2347ATA (7225C>A), AAG2348AAA (7228G>A), AAG2352AAA (7240G>A), TAT2353TAC (7243T>C), AAA2355AAG (7249A>G), CCT2356CCG (7252T>G), GAT2358GAA (7258T>A), GTT2359GTC (7261T>C), GGT2360GGA (7264T>A), GTC2361GTG (7267C>G), GTG2362GTA (7270G>A), GAT2363_CGG2364insT-- (7273_7274insT), CGG2364GTG (7274C>G 7275G>T), CAT2366CTT (7281A>T), TTG2367TCG (7284T>C), TCA2371TTA (7296C>T), TCT2376TTT (7311C>T), ACC2377ATC (7314C>T), TGC2378TAC (7317G>A), AAT2379AAA (7321T>A), GTG2380-TG (7322delG), CAC2381CAT (7327C>T), GGC2382GGT (7330C>T), TTC2383TTT (7333C>T), AAG2384CAG (7334A>C), TGT2386TGC (7342T>C), GCA2387AAC (7343G>A 7344C>A 7345A>C), ACT2390ACC (7354T>C), CAA2393CAG (7363A>G), ATT2395ATC (7369T>C), TTC2396TTT (7372C>T), GCG2398GCT (7378G>T), CTC2399CTT (7381C>T), GCA2404GCT (7396A>T), GTC2405GTT (7399C>T), GCC2407GCT (7405C>T), AGT2408ATG (7407G>T 7408T>G), GGG2410GGT (7414G>T), TGC2411GGC (7415T>G), AAA2412AAG (7420A>G), GAC2415GAT (7429C>T), TAT2416TAC (7432T>C), CAT2419CAC (7441T>C), AAG2425AAA (7459G>A), GAA2426GAG (7462A>G), ATA2428ATT (7468A>T), GTC2429GTA (7471C>A), CAA2431CAG (7477A>G), AGC2432AGT (7480C>T), CTG2434CTT (7486G>T), TAT2437TAC (7495T>C), AAA2438AAG (7498A>G), GGT2439GGC (7501T>C), TTG2440TCA (7503T>C 7504G>A), ATT2443ATA (7513T>A), AAC2445AAT (7519C>T), GGA2446GGG (7522A>G), AAG2449AAA (7531G>A), GAG2451GAA (7537G>A), CTC2452CTT (7540C>T), TGT2454TGC (7546T>C), AAG2462AAA (7570G>A), ACG2463ACA (7573G>A), ACG2465ACA (7579G>A), GCT2468GCC (7588T>C), CTA2471TTA (7595C>T), TTG2474CTA (7604T>C 7606G>A), GGT2477GGA (7615T>A), TGT2480TGC (7624T>C), GAC2483GAT (7633C>T), TTC2484TTT (7636C>T), AAT2485AAC (7639T>C), TTT2488TTC (7648T>C), GAA2494GAG (7666A>G), TGT2496TGC (7672T>C), ACA2498ACT (7678A>T), GGG2500GGA (7684G>A), ACT2502ACC (7690T>C), TTT2504TTC (7696T>C), GGT2506GGG (7702T>G), GAT2509AAT (7709G>A), AAA2510AGG (7713A>G 7714A>G), CTG2511TTG (7715C>T), CGT2514CGC (7726T>C), GAG2517GAA (7735G>A), GTA2520GTG (7744A>G), TAC2521TAT (7747C>T), GGC2526GGT (7762C>T), CAG2528CAA (7768G>A), TTT2529TTC (7771T>C), TCA2532TCG (7780A>G), CTA2533CTC (7783A>C), ACT2534ACC (7786T>C), CTA2537TTG (7793C>T 7795A>G), ATC2538ATT (7798C>T), GCT2540GCC (7804T>C), ACC2543ATC (7812C>T), GAA2550GAG (7834A>G), GAC2551GAT (7837C>T), GTG2554TTA (7844G>T 7846G>A), CAG2557CAA (7855G>A), CTG2559CTA (7861G>A), TTA2562TTG (7870A>G), TAC2563TAT (7873C>T), ACT2564ACA (7876T>A), GAG2565GAA (7879G>A), ATT2566ATA (7882T>A), GTT2567ATT (7883G>A), CCA2570CCT (7894A>T), ATT2571ATC (7897T>C), GTC2579GTT (7921C>T), AAG2581AAA (7927G>A), GGT2584GGC (7936T>C), AAT2585AAC (7939T>C), AAT2586AAC (7942T>C), GTG2594GTA (7966G>A), GAT2595GAC (7969T>C), TCT2597TCC (7975T>C), GTC2601GTT (7987C>T), CTT2602CTC (7990T>C), GCT2607GCC (8005T>C), GAA2616GAG (8032A>G), ATC2618ATT (8038C>T), GAC2619GAT (8041C>T), AGC2620AAC (8043G>A), TTG2632TTA (8080G>A), GTG2635GTA (8089G>A), CCG2637CCA (8095G>A), GAG2638GAA (8098G>A), AGC2641AGT (8107C>T), ATT2642ATC (8110T>C), GAT2644GAC (8116T>C), CAA2648CAG (8128A>G), GGT2654GGC (8146T>C), AAC2656AGT (8151A>G 8152C>T), TCG2660TCA (8164G>A), AGG2665AAG (8178G>A), AAG2666AAA (8182G>A), TTG2669TTA (8191G>A), TTC2671TTT (8197C>T), TCC2673TCT (8203C>T), AGA2675AAA (8208G>A), CTG2677TTA (8213C>T 8215G>A), CTA2678CTG (8218A>G), ATC2679ATT (8221C>T), GAG2680GAA (8224G>A), CCA2685CCG (8239A>G), GAG2690GAA (8254G>A), ATT2692ATC (8260T>C), GTA2693GTG (8263A>G), GAT2699GAC (8281T>C), AGA2700AGG (8284A>G), CCA2704GCG (8294C>G 8296A>G), GAG2705GAA (8299G>A), CAC2706CAT (8302C>T), ATT2711ATC (8317T>C), TGC2712TGT (8320C>T), GCT2714GCA (8326T>A), GAG2717GAA (8335G>A), TCC2718TCT (8338C>T), TCT2722TTT (8349C>T), GAA2723GAG (8353A>G), CAC2726CAT (8362C>T), CAA2727CAG (8365A>G), ATC2728ATT (8368C>T), TAC2732TAT (8380C>T), TCA2733TCG (8383A>G), TTA2735TTG (8389A>G), CCT2740CCC (8404T>C), GCA2742TCA (8408G>T), ACA2743ACG (8413A>G), GCG2745GCT (8419G>T), GCT2750GCC (8434T>C), TAT2752TAC (8440T>C), ATG2756ATA (8452G>A), CTA2758CTG (8458A>G), AGG2759AAG (8460G>A), AAA2760AAG (8464A>G), CTG2761CTA (8467G>A), AGG2765AGA (8479G>A), GCT2766ACA (8480G>A 8482T>A), GAG2771GAA (8497G>A), CTA2772TTG (8498C>T 8500A>G), AGA2773AAG (8502G>A 8503A>G), GCC2774GCT (8506C>T), GAA2777GAT (8515A>T), GTC2780GTT (8524C>T), GCA2781GCC (8527A>C), TTA2782CTG (8528T>C 8530A>G), GAG2785GAC (8539G>C), CTT2788TGT (8546C>T 8547T>G), TCT2790ACT (8552T>A), GTA2793GTG (8563A>G), GCA2797GGA (8574C>G), ATT2801GTT (8585A>G), GGA2805GAA (8598G>A), AGC2806AGT (8602C>T), AAC2807TCT (8603A>T 8604A>C 8605C>T), AAA2812AAG (8620A>G), CCA2813CAA (8622C>A), GAG2814GGA (8625A>G 8626G>A), CAG2815CAA (8629G>A), GGC2816GGT (8632C>T), ATC2818ATT (8638C>T), CAG2819CAA (8641G>A), AAC2821AAT (8647C>T), CCG2822TTC (8648C>T 8649C>T 8650G>C), AAC2823AAT (8653C>T), AAA2824AAG (8656A>G), GGA2825GAA (8658G>A), GAT2827GCG (8664A>C 8665T>G), GTT2830GTG (8674T>G), GCA2832GTT (8679C>T 8680A>T), GGC2833GGA (8683C>A), GGG2836GGA (8692G>A), ACA2837ACT (8695A>T), CCG2841CCA (8707G>A), AGA2842CGA (8708A>C), AAG2844AAA (8716G>A), ATC2846ATT (8722C>T), ACG2847ACA (8725G>A), AAA2849AAG (8731A>G), AGA2851AGG (8737A>G), CCC2853CCT (8743C>T), ACA2854AAG (8745C>A 8746A>G), AAG2856AAA (8752G>A), GGA2857GGT (8755A>T), AAC2862AAT (8770C>T), TTA2863CTA (8771T>C), GAA2864GAG (8776A>G), CAT2865CAC (8779T>C), TTG2866TTA (8782G>A), TAT2869TAC (8791T>C), CCA2871CCG (8797A>G), CAA2872CAG (8800A>G), CAA2873CAG (8803A>G), GAT2875GAC (8809T>C), ATT2876ATC (8812T>C), AAT2878AAC (8818T>C), CGG2880CGA (8824G>A), GCA2881GCG (8827A>G), ACG2888ACA (8848G>A), TAT2890TAC (8854T>C), GAG2891GGA (8856A>G 8857G>A), GTG2893GTA (8863G>A), CGG2894CAG (8865G>A), ATG2895CTT (8867A>C 8869G>T), GCA2896GCG (8872A>G), GGA2900GAA (8883G>A), ACT2902TCT (8888A>T), TGC2915TGT (8929C>T), GTC2924ATC (8954G>A), AAC2925AAT (8959C>T), GGA2926GGT (8962A>T), GGG2933GGA (8983G>A), AAT2934GAC (8984A>G 8986T>C), GAA2935GTA (8988A>T), GAG2938GAA (8998G>A), CCG2940CCA (9004G>A), TTG2941TTA (9007G>A), ATC2944ATT (9016C>T), ACC2951ACA (9037C>A), ATA2967ATT (9085A>T), AAC2971AAT (9097C>T), ATT2983GTT (9131A>G), CGA2984CGT (9136A>T), CGG2987CGT (9145G>T), ATG2989GGA (9149A>G 9150T>G 9151G>A), GGT2990AGT (9152G>A), TTA2991TTG (9157A>G), GCG2992GCT (9160G>T), CGT2993CGC (9163T>C), GAG3000GAA (9184G>A), CGA3004CGG (9196A>G), AGG3010AGA (9214G>A), GAA3011GAG (9217A>G), GCG3012GCA (9220G>A), GCA3020GCT (9244A>T), TTG3021TTA (9247G>A), GCC3024GCT (9256C>T), CCT3026TCC (9260C>T 9262T>C), GGG3030GGA (9274G>A), GAC3032GAT (9280C>T), ATC3035ATT (9289C>T), CAA3038CAC (9298A>C), GAG3040GAA (9304G>A), GTC3050GTT (9334C>T), GTC3060GTG (9364C>G) | | | | | | | | |
| PVYgp2 | 1 | 75 | 100% | 95 | 18.1% | 75 (98.7%) | 54 (71.1%) | 1/0/3/2 | 2 |
| Protein mutations: | R17_K17insX (2971_2972insA), Y23H (2989T>C 2991C>T), R26K (2999G>A 3000A>G), H35Y (3025C>T), K40H (3040A>C 3042A>C), G41S (3043G>A), E44K (3052G>A), I47V (3061A>G), V54I (3082G>A), P57S (3091C>T), P60T (3100C>A), G61E (3104G>A 3105T>G), G62S (3106G>A), Q63* (3109C>T), R64G (3112A>G), C66R (3118T>C), L67F (3121C>T), I69V (3127A>G), A72E (3137C>A 3138G>A), I73V (3139A>G) | | | | | | | | |
| Codon mutations: | TTA2CTA (2923T>C), AAA4AAG (2931A>G), GAA7GAG (2940A>G), TTG10CTG (2947T>C), TTG15CTG (2962T>C), AGA17_AAA17insAA- (2971_2972insA), TAC23CAT (2989T>C 2991C>T), TCA24TCG (2994A>G), AGA26AAG (2999G>A 3000A>G), CAC32-AC (3016delC), TCG33TCA (3021G>A), GTA34GTT (3024A>T), CAT35TAT (3025C>T), CAC38CAT (3036C>T), AGA39AGG (3039A>G), AAA40CAC (3040A>C 3042A>C), GGC41AGC (3043G>A), GAA44AAA (3052G>A), ATA47GTA (3061A>G), GTT54ATT (3082G>A), CCG57TCG (3091C>T), CGC59CGT (3099C>T), CCA60ACA (3100C>A), GGT61GAG (3104G>A 3105T>G), GGT62AGT (3106G>A), CAA63TAA (3109C>T), AGG64GGG (3112A>G), TGC66CGC (3118T>C), CTC67TTC (3121C>T), ATT69GTT (3127A>G), GAG70GAA (3132G>A), GCG72GAA (3137C>A 3138G>A), ATT73GTT (3139A>G) | | | | | | | | |

| | Begin | End | Coverage | Score | Concordance | Matches | Identities | I/D/M/F* | Stop Codons |
|---|---|---|---|---|---|---|---|---|---|
| **NT** | **5** | **9704** | **99.9%** | **12522** | **64.8%** | **9688 (99.7%)** | **8015 (82.5%)** | **15/12** | |
| **Proteins** | | | | | | | | | |
| polyprotein (NP_056759.1) | 1 | 3064 | 100% | 18923 | 89.7% | 3062 (99.9%) | 2771 (90.4%) | 2/2/4/4 | 1 |
| Protein mutations: | Y4Q (194T>C 196C>A), C9L (209T>C 210G>T 211T>G), F13M (221T>A 223T>G), S23P (251T>C), C24F (255G>T 256C>T), E25G (258A>G), I27V (263A>G 265T>G), V28A (267T>C), E30V (273A>T), A35T (287G>A 289T>C), V37T (293G>A 294T>C), A41I (305G>A 306C>T), D42G (309A>G), E44D (316A>T), T45V (317A>G 318C>T 319A>G), L52Q (339T>A 340C>A), K53R (342A>G), K55A (347A>G 348A>C 349A>G), Y56H (350T>C), T58V (356A>G 357C>T), V61T (365G>A 366T>C 367G>A), L62S (368C>T 369T>C), F67C (384T>G), A75V (408C>T 409C>T), M78E (416A>G 417T>A), E85K (437G>A), R86E (440A>G 441G>A), K87R (443A>C 444A>G 445G>A), D88E (448T>A), E91A (456A>C), H93N (461C>A), D111E (517T>A), S116F (531C>T), P118S (536C>T), Q119W (539C>T 540A>G 541A>G), R122K (549G>A 550A>G), M131T (576T>C), V134A (585T>C 586C>G), R137Y (593C>T 594G>A 595C>T), P138R (597C>G), I139V (599A>G 601A>G), I140P (602A>C 603T>C 604A>G), M147I (625G>A), I151V (635A>G 637T>C), I154V (644A>G 646A>G), E160A (663A>C 664G>C), H166Q (682C>A), T172S (699C>G), H174Q (706T>A), K179E (719A>G 721G>A), I180V (722A>G 724A>G), A183S (731G>T), Y184P (734T>C 735A>C 736C>T), S185R (737T>C 738C>G 739C>T), A187V (744C>T 745G>C), R189C (749C>T 751A>C), M194K (765T>A), R197K (773C>A 774G>A 775A>G), M206K (801T>A), G210A (813G>C 814A>G), L211H (816T>A), R214Q (825G>A 826T>A), N223D (851A>G 853T>C), R226C (860C>T 862C>T), T227A (863A>G 865T>C), I228T (867T>C), N229D (869A>G 871C>T), I230L (872A>T 874A>G), R231Q (876G>A), R232K (879G>A), N240S (903A>G), T241N (906C>A 907A>C), K242T (909A>C 910A>C), S243N (912G>A 913C>T), S252L (938T>C 939C>T 940A>T), L266I (980T>A 982G>C), R272K (999G>A), Q275R (1007C>A 1008A>G 1009G>A), S276G (1010A>G 1012T>G), I277V (1013A>G), N279Q (1019A>C 1021C>G), D288E (1048C>A), R300Q (1083G>A), S305A (1097T>G 1099G>A), R320K (1143G>A), L324I (1154T>A 1156G>A), S346N (1221G>A), V349T (1229G>A 1230T>C 1231T>C), S350N (1233G>A 1234C>T), K358T (1257A>C 1258A>G), N365S (1278A>G), A369V (1290C>T 1291G>A), D372T (1298G>A 1299A>C 1300C>T), I375V (1307A>G 1309A>G), N378E (1316A>G 1318T>A), I382T (1329T>C), L397I (1373C>A), I401V (1385A>G), E407D (1405G>T), A411S (1415G>T 1417A>T), V418I (1436G>A 1438C>T), E508G (1707A>G), A512T (1718G>A), I520V (1742A>G 1744C>T), S525N (1758G>A), V536I (1790G>A), I585V (1937A>G 1939T>C), F607Y (2004T>A), I628M (2068T>G), V630I (2072G>A 2074G>A), R675K (2208G>A), D694E (2266C>G), N744G (2414A>G 2415A>G 2416T>G), S746C (2420A>T), K780R (2523A>G), R817K (2634G>A), V847I (2723G>A 2725C>T), A855T (2747G>A 2749T>A), H871N (2795C>A), V880I (2822G>A 2824G>A), P900T (2882C>A), S901H (2885A>C 2886G>A), T904M (2895C>T), N916S (2931A>G), N919S (2940A>G), E929_N930insX (2971_2972insA), T936I (2991C>T), H937R (2994A>G), E939R (2999G>A 3000A>G), R946Q (3021G>A), Y947F (3024A>T 3025C>T), T951I (3036C>T), E952G (3039A>G 3040A>C), K953T (3042A>C 3043G>A), A972V (3099C>T 3100C>A), V974R (3104G>A 3105T>G 3106G>A), S983N (3132G>A), R985K (3137C>A 3138G>A 3139A>G), F990I (3152T>A 3154C>T), C994Y (3165G>A), F1012L (3218T>C), T1014N (3225C>A), V1016I (3230G>A), V1028M (3266G>A 3268A>G), V1031M (3275G>A), A1034T (3284G>A), R1044K (3315G>A 3316A>G), E1045A (3318A>C), M1059V (3359A>G), D1072E (3400T>G), V1110G (3513T>G), M1121V (3545A>G), A1122T (3548G>A), V1124I (3554G>A 3556C>T), L1147V (3623C>G), L1150M (3632C>A), Y1152H (3638T>C), I1172V (3698A>G), S1178N (3717G>A), R1183K (3731C>A 3732G>A), I1297V (4073A>G), F1315Y (4128T>A 4129C>T), A1323S (4151G>T), V1328I (4166G>A), L1347I (4223C>A 4225G>A), I1379V (4319A>G 4321A>G), F1404Y (4395T>A), V1408I (4406G>A), M1431V (4475A>G 4477G>T), T1433S (4481A>T 4483A>G), A1454L (4544G>T 4545C>T 4546G>A), R1455K (4548G>A), V1499I (4679G>A 4681G>C), V1538I (4796G>A 4798C>T), T1540S (4803C>G 4804T>C), S1625G (5057A>G), T1626A (5060A>G), L1636I (5090C>A), A1639V (5100C>T), E1641D (5107A>T), V1646F (5120G>T 5122C>T), I1648V (5126A>G), V1667I (5183G>A), A1793T (5561G>A 5563G>A), I1805V (5597A>G), N1807K (5605C>A), A1812V (5619C>T 5620T>G), I1816L (5630A>C), V1837I (5693G>A), K1890R (5853A>G), I1904V (5894A>G 5896C>G), R1926A (5960C>G 5961G>C 5962G>C), I1936V (5990A>G), E1945D (6019A>T), K1949T (6030A>C), D1955E (6049C>A), M1958I (6058G>A), S1963N (6072G>A 6073T>C), N1964H (6074A>C), T1966N (6081C>A), C1976S (6111G>C 6112T>C), I1982V (6128A>G 6130T>C), V1992I (6158G>A), T1996S (6170A>T 6172A>G), L2007F (6203C>T 6205C>T), V2016I (6230G>A), A2025K (6257G>A 6258C>A), K2033R (6282A>G), A2060T (6362G>A), V2073I (6401G>A), A2074V (6405C>T), Y2082F (6429A>T), Q2089R (6450A>G), H2101N (6485C>A 6487C>T), L2115I (6527C>A), T2135I (6588C>T), Y2151H (6635T>C), I2155V (6647A>G), I2165V (6677A>G 6679A>G), D2178N (6716G>A), N2179D (6719A>G), I2194L (6764A>T 6766A>G), A2204V (6795C>T 6796A>G), H2205Q (6799C>A), N2225D (6857A>G), V2231T (6875G>A 6876T>C 6877C>A), D2250E (6934C>A), V2273I (7001G>A 7003G>A), F2283Y (7032T>A), T2314K (7125C>A), A2321S (7145G>T 7147A>G), E2326_A2327del (7160_7165delGAGGCA), D2344E (7216T>A), D2358E (7258T>A), D2363_R2364insX (7273_7274insT), R2364V (7274C>G 7275G>T), H2366L (7281A>T), L2367S (7284T>C), S2371L (7296C>T), S2376F (7311C>T), T2377I (7314C>T), C2378Y (7317G>A), N2379K (7321T>A), K2384Q (7334A>C), A2387N (7343G>A 7344C>A 7345A>C), S2408M (7407G>T 7408T>G), C2411G (7415T>G), L2440S (7503T>C 7504G>A), D2509N (7709G>A), K2510R (7713A>G 7714A>G), T2543I (7812C>T), V2554L (7844G>T 7846G>A), V2567I (7883G>A), S2620N (8043G>A), N2656S (8151A>G 8152C>T), R2665K (8178G>A), R2675K (8208G>A), P2704A (8294C>G 8296A>G), S2722F (8349C>T), A2742S (8408G>T), M2756I (8452G>A), R2759K (8460G>A), A2766T (8480G>A 8482T>A), R2773K (8502G>A 8503A>G), E2777D (8515A>T), E2785D (8539G>C), L2788C (8546C>T 8547T>G), S2790T (8552T>A), A2797G (8574C>G), I2801V (8585A>G), G2805E (8598G>A), N2807S (8603A>T 8604A>C 8605C>T), P2813Q (8622C>A), E2814G (8625A>G 8626G>A), P2822F (8648C>T 8649C>T 8650G>C), G2825E (8658G>A), D2827A (8664A>C 8665T>G), A2832V (8679C>T 8680A>T), T2854K (8745C>A 8746A>G), E2891G (8856A>G 8857G>A), R2894Q (8865G>A), M2895L (8867A>C 8869G>T), G2900E (8883G>A), T2902S (8888A>T), V2924I (8954G>A), N2934D (8984A>G 8986T>C), E2935V (8988A>T), I2983V (9131A>G), M2989G (9149A>G 9150T>G 9151G>A), G2990S (9152G>A), P3026S (9260C>T 9262T>C), Q3038H (9298A>C) | | | | | | | | |

| | Begin | End | Coverage | Score | Concordance | Matches | Identities | I/D/M/F* | Stop Codons |
|---|---|---|---|---|---|---|---|---|---|
| NT | 5 | 9704 | 99.9% | 12522 | 64.8% | 9688 (99.7%) | 8015 (82.5%) | 15/12 | |

Codon mutations:

ACT3ACC (193T>C), TAC4CAA (194T>C 196C>A), TCA6TCG (202A>G), ATC8ATT (208C>T), TGT9CTG (209T>C 210G>T 211T>G), TCG12TCA (220G>A), TTT13ATG (221T>A 223T>G), GAA14GAG (226A>G), CTA17CTT (235A>T), TAC19TAT (241C>T), CCA21CCT (247A>T), TCT23CCT (251T>C), TGC24TTT (255G>T 256C>T), GAG25GGG (258A>G), CAT26CAC (262T>C), ATT27GTG (263A>G 265T>G), GTG28GCG (267T>C), GAA30GTA (273A>T), CGA31CGT (277A>T), GAA32GAG (280A>G), GTG33GTT (283G>T), CCG34CCA (286G>A), GCT35ACC (287G>A 289T>C), GTT37ACT (293G>A 294T>C), CCT39CCC (301T>C), TTC40TTT (304C>T), GCA41ATA (305G>A 306C>T), GAT42GGT (309A>G), CTG43TTG (311C>T), GAA44GAT (316A>T), ACA45GTG (317A>G 318C>T 319A>G), CAA46CAG (322A>G), GCA49GCT (331A>T), CTC52CAA (339T>A 340C>A), AAG53AGG (342A>G), AAA55GCG (347A>G 348A>C 349A>G), TAT56CAT (350T>C), GCT57GCA (355T>A), ACT58GTT (356A>G 357C>T), CGT60CGC (364T>C), GTG61ACA (365G>A 366T>C 367G>A), CTC62TCC (368C>T 369T>C), AAA63AAG (373A>G), AAC64AAT (376C>T), TTT67TGT (384T>G), ACG68ACA (388G>A), TAC69TAT (391C>T), CGA70CGG (394A>G), AAG72AAA (400G>A), GAT74GAC (406T>C), GCC75GTT (408C>T 409C>T), CAG76CAA (412G>A), ATA77ATT (415A>T), ATG78GAG (416A>G 417T>A), ATT80ATC (424T>C), CAG81CAA (427G>A), AAA83AAG (433A>G), CTG84CTC (436G>C), GAG85AAG (437G>A), AGG86GAG (440A>G 441G>A), AAG87CGA (443A>C 444A>G 445G>A), GAT88GAA (448T>A), GAA90GAG (454A>G), GAA91GCA (456A>C), TAT92TAC (460T>C), CAC93AAC (461C>A), CAA95CAG (469A>G), GCC97GCG (475C>G), CCT99CCA (481T>A), ACT106ACC (502T>C), ATA107ATT (505A>T), GGC109GGT (511C>T), GAT111GAA (517T>A), CCT112CCA (520T>A), CCA113CCT (523A>T), TCT116TTT (531C>T), GAG117GAA (535G>A), CCA118TCA (536C>T), CAA119TGG (539C>T 540A>G 541A>G), AGA122AAG (549G>A 550A>G), ATC124ATT (556C>T), ATT125ATC (559T>C), CAT126CAC (562T>C), ACT128ACC (568T>C), CCA129CCG (571A>G), ATG131ACG (576T>C), CGT132CGC (580T>C), GTC134GCG (585T>C 586C>G), CGC137TAT (593C>T 594G>A 595C>T), CCC138CGC (597C>G), ATA139GTG (599A>G 601A>G), ATA140CCG (602A>C 603T>C 604A>G), ACA143ACG (613A>G), GAA144GAG (616A>G), GGC145GGG (619C>G), CAG146CAA (622G>A), ATG147ATA (625G>A), AAT148AAC (628T>C), CAC149CAT (631C>T), ATT151GTC (635A>G 637T>C), ATA154GTG (644A>G 646A>G), ATT157ATC (655T>C), TCG159TCA (661G>A), GAG160GCC (663A>C 664G>C), AAA161AAG (667A>G), GTC165GTT (679C>T), CAC166CAA (682C>A), TTA167CTG (683T>C 685A>G), AGT169AGC (691T>C), ACC172AGC (699C>G), ACT173ACA (703T>A), CAT174CAA (706T>A), AAG179GAA (719A>G 721G>A), ATA180GTG (722A>G 724A>G), CTT181TTG (725C>T 727T>G), GGT182GGA (730T>A), GCA183TCA (731G>T), TAC184CCT (734T>C 735A>C 736C>T), TCC185CGT (737T>C 738C>G 739C>T), GCG187GTC (744C>T 745G>C), CGA189TGC (749C>T 751A>C), ACT190ACC (754T>C), ATG194AAG (765T>A), TTG196CTG (770T>C), CGA197AAG (773C>A 774G>A 775A>G), CGG198AGG (776C>A), TTC202TTT (790C>T), ATG206AAG (801T>A), GGA210GCG (813G>C 814A>G), CTT211CAT (816T>A), TTG212CTG (818T>C), CAA213CAG (823A>G), CGT214CAA (825G>A 826T>A), CGG217AGA (833C>A 835G>A), GAC219GAT (841C>T), AAT223GAC (851A>G 853T>C), GTC225GTT (859C>T), CGC226TGT (860C>T 862C>T), ACT227GCC (863A>G 865T>C), ATC228ACC (867T>C), AAC229GAT (869A>G 871C>T), ATA230TTG (872A>T 874A>G), CGA231CAA (876G>A), AGG232AAG (879G>A), GAT234GAC (886T>C), ATC238ATT (898C>T), AAC240AGC (903A>G), ACA241AAC (906C>A 907A>C), AAA242ACC (909A>C 910A>C), AGC243AAT (912G>A 913C>T), GGC246GGA (922C>A), GGT249GGA (931T>A), TCA252CTT (938T>C 939C>T 940A>T), GGA253GGG (943A>G), TTC256TTT (952C>T), TCA261TCG (967A>G), GAA263GAG (973A>G), GGG264GGC (976G>C), TTG266ATC (980T>A 982G>C), GCA269GCG (991A>G), CGT270CGC (994T>C), AGA272AAA (999G>A), GTT273GTC (1003T>C), CAG275AGA (1007C>A 1008A>G 1009G>A), AGT276GGG (1010A>G 1012T>G), ATT277GTT (1013A>G), TTA278TTG (1018A>G), AAC279CAG (1019A>C 1021C>G), ATC282ATT (1030C>T), TCG285TCA (1039G>A), AAT286AAC (1042T>C), GCC287GCA (1045C>A), GAC288GAA (1048C>A), AAT289AAC (1051T>C), AAG292AAA (1060G>A), GGT293GGA (1063T>A), CTG294TTG (1064C>T), GAC295GAT (1069C>T), CGA300CAA (1083G>A), TAT303TAC (1093T>C), TCG305GCA (1097T>G 1099G>A), CAC307CAT (1105C>T), GTA310GTG (1114A>G), GCT311GCG (1117T>G), TTA313TTG (1123A>G), CCT314CCA (1126T>A), GTC315GTT (1129C>T), AGG320AAG (1143G>A), GTA321GTT (1147A>T), GCT322GCA (1150T>A), TTG324ATA (1154T>A 1156G>A), CAC327CAT (1165C>T), AGT328AGC (1168T>C), CTT330TTA (1172C>T 1174T>A), ACC339ACT (1201C>T), AGC346AAC (1221G>A), GTT349ACC (1229G>A 1230T>C 1231T>C), AGC350AAT (1233G>A 1234C>T), CTG352CTA (1240G>A), AAG354AAA (1246G>A), TTG356CTA (1250T>C 1252G>A), CAT357CAC (1255T>C), AAA358ACG (1257A>C 1258A>G), AGA361AGG (1267A>G), GGT363GGC (1273T>C), TTG364TTA (1276G>A), AAT365AGT (1278A>G), TTG367CTG (1283T>C), GCG369GTA (1290C>T 1291G>A), GAC372ACT (1298G>A 1299A>C 1300C>T), TTT374TTC (1306T>C), ATA375GTG (1307A>G 1309A>G), CAT376CAC (1312T>C), AAT378GAA (1316A>G 1318T>A), TTG381TTA (1327G>A), ATA382ACA (1329T>C), GCG383GCT (1333G>T), TTA384TTG (1336A>G), CTA387TTG (1343C>T 1345A>G), CCG390CCA (1354G>A), GTG391GTT (1357G>T), GAC392GAT (1360C>T), CTC395TTG (1367C>T 1369C>G), CTT397ATT (1373C>A), TTC398TTT (1378C>T), GAG400GAA (1384G>A), ATA401GTA (1385A>G), AAA403AAG (1393A>G), TCC404TCT (1396C>T), GAG407GAT (1405G>T), AAA408AAG (1408A>G), CAG409CAA (1411G>A), GCA411TCT (1415G>T 1417A>T), CCG412CCC (1420G>C), AAT415AAC (1429T>C), TTA416TTG (1432A>G), GTC418ATT (1436G>A 1438C>T), TTA419CTG (1439T>C 1441A>G), AAT420AAC (1444T>C), TTC423TTT (1453C>T), CTG424CTA (1456G>A), GAA428GAG (1468A>G), GTA436GTG (1492A>G), TTG439CTA (1499T>C 1501G>A), AGT440AGC (1504T>C), TTG441TTA (1507G>A), CTC442CTT (1510C>T), GAA443GAG (1513A>G), TTA444TTG (1516A>G), GCA445GCT (1519A>T), AGG446AGA (1522G>A), TTC447TTT (1525C>T), CAG448CAA (1528G>A), AAC450AAT (1534C>T), ACT452ACA (1540T>A), AAC454AAT (1546C>T), AAA457AAG (1555A>G), GGT458GGC (1558T>C), GAT459GAC (1561T>C), ATA460ATC (1564A>C), TCT461TCA (1567T>A), TTC463TTT (1573C>T), AGA464AGG (1576A>G), TTA467CTA (1583T>C), TCT468TCA (1588T>A), AAG470AAA (1594G>A), AAC472AAT (1600C>T), CTG475CTA (1609G>A), TCG478TCA (1618G>A), TGC479TGT (1621C>T), GAC480GAT (1624C>T), CTG483TTG (1631C>T), AAA485AAG (1639A>G), TTC489TTT (1651C>T), CTC490CTG (1654C>G), CAT497CAC (1675T>C), TCA503TCG (1693A>G), TTT506TTC (1702T>C), GAA508GGA (1707A>G), GCA512ACA (1718G>A), GGA514GGG (1726A>G), TAC515TAT (1729C>T), TCA516TCC (1732A>C), TAT518TAC (1738T>C), ATC520GTT (1742A>G 1744C>T), CGC521CGT (1747C>T), CAT523CAC (1753T>C), AGT525AAT (1758G>A), AGG528AGA (1768G>A), AAC534AAT (1786C>T), TTA535TTG (1789A>G), GTT536ATT (1790G>A), GTC537GTT (1795C>T), TTA541CTG (1805T>C 1807A>G), GAG543GAA (1813G>A), AGG545AGA (1819G>A), AAG547AAA (1825G>A), GAC551GAT (1837C>T), TAT552TAC (1840T>C), CAA555CAG (1849A>G), GTC558GTG (1858C>G), AAA560AAG (1864A>G), TGC562TGT (1870C>T), AGT564AGC (1876T>C), AAA566AAG (1882A>G), GGT568GGA (1888T>A), AAT569AAC (1891T>C), TGT574TGC (1906T>C), ACA577ACT (1915A>T), GGT582GGC (1930T>C), GCC584GCT (1936C>T), ATT585GTC (1937A>G 1939T>C), TCA587TCG (1945A>G), TTC589TTT (1951C>T), CCA592CCT (1960A>T), GTA598GTG (1978A>G), ATT599ATA (1981T>A), GGC600GGT (1984C>T), AAT601AAC (1987T>C), GAC604GAT (1996C>T), TTT607TAT (2004T>A), AAA612AAG (2020A>G), GGG613GGA (2023G>A), GAT614GAC (2026T>C), TCG615TCT (2029G>T), GAG616GAA (2032G>A), TAT625TAC (2059T>C), ATT628ATG (2068T>G), AAC629AAT (2071C>T), GTG630ATA (2072G>A 2074G>A), CTG635TTA (2087C>T 2089G>A), AGC639AGT (2101C>T), GAG640GAA (2104G>A), GAG641GAA (2107G>A), AAG644AAA (2116G>A), TTC646TTT (2122C>T), CGC651CGT (2137C>T), CTT658CTC (2158T>C), ACC660ACT (2164C>T), CCA662CCG (2170A>G), GAT666GAC (2182T>C), GCG668GCA (2188G>A), ACC669ACA (2191C>A), AGA675AAA (2208G>A), TAT678TAC (2218T>C), GAC680GAT (2224C>T), GTA681GTT (2227A>T), TTG686TTA (2242G>A), CCC687CCT (2245C>T), TTG690TTA (2254G>A), GTT691GTC (2257T>C), GAC692GAT (2260C>T), GAC694GAG (2266C>G), CAA696CAG (2272A>G), ACG697ACA (2275G>A), TCA703TCG (2293A>G), GGC705GGT (2299C>T), TCG706TCA (2302G>A), CAG707CAA (2305G>A), GGA710GGT (2314A>T), TAT711TAC (2317T>C), CTA714CTG (2326A>G), GTG719GTA (2341G>A), CAA721CAG (2347A>G), CTT722CTC (2350T>C), TTG724TTA (2356G>A), AAT727AAC (2365T>C), GAA729GAG (2371A>G), TTA730TTG (2374A>G), GAT733GAC (2383T>C), ATA734ATT (2386A>T), AAA735AAG (2389A>G), CAT736CAC (2392T>C), TAT737TAC (2395T>C), AAT744GGG (2414A>G 2415A>G 2416T>G), GCT745GCA (2419T>A), AGC746TGC (2420A>T), ACA752ACT (2440A>T), GTT761GTC (2467T>C), TCT764TCC (2476T>C), TCG766TCA (2482G>A), CTG769CTA (2491G>A), CTG771CTT (2497G>T), TTG773TTA (2503G>A), AGA778AGG (2518A>G), CCT779CCC (2521T>C), AAG780AGG (2523A>G), AGA783AGG (2533A>G), TTG785TTA (2539G>A), TTA787TTG (2545A>G), GAG789GAA (2551G>A), CCT790CCA (2554T>A), CTG792TTG (2558C>T), TTG793CTC (2561T>C 2563G>C), TCA796TCG (2572A>G), CCT800CCC (2584T>C), GGC801GGT (2587C>T), CTG803CTC (2593G>C), GCT805GCC (2599T>C), AAT808AAC (2608T>C), TTT812TTC (2620T>C), GCG815GCA (2629G>A), GTG816GTA (2632G>A), AGG817AAG (2634G>A), TTG818TTA (2638G>A), ATT820ATA (2644T>A), AAA823AAG (2653A>G), GCT827GCC (2665T>C), CTA832TTA (2678C>T), CTA833CTG (2683A>G), GCT835GCC (2689T>C), CTA838CTC (2698A>C), TCA841TCG (2707A>G), GCG842GCA (2710G>A), GAA844GAG (2716A>G), ACA845ACG (2719A>G), CTC846CTT (2722C>T), GTC847ATT (2723G>A 2725C>T), GCA848GCG (2728A>G), AGG850AGA (2734G>A), ATT851ATA (2737T>A), GAT854GAC (2746T>C), GCT855ACA (2747G>A 2749T>A), GCA856GCT (2752A>T), GCT857GCA (2755T>A), GAC859GAT (2761C>T), CTC860CTT (2764C>T), TGT865TGC (2779T>C), GGG867GGT (2785G>T), AAC869AAT (2791C>T), CAT871AAT (2795C>A), TAC874TAT (2806C>T), TTG878CTC (2816T>C 2818G>C), GTG880ATA (2822G>A 2824G>A), TTG881TTA (2827G>A), GTT884GTC (2836T>C), AAG885AAA (2839G>A), AAT886AAC (2842T>C), ACC893ACA (2863C>A), CTA894TTA (2864C>T), CCA900ACA (2882C>A), AGT901CAT (2885A>C 2886G>A), AAC903AAT (2893C>T), ACG904ATG (2895C>T), GTC906GTT (2902C>T), GTA907GTG (2905A>G), ATT909ATC (2911T>C), AAT913AAC (2923T>C), AAT916AGT (2931A>G), AAC919AGC (2940A>G), GCT921GCC (2947T>C), ACT926ACC (2962T>C), GAA929_AAT930insA-- (2971_2972insA), GGT935GGC (2989T>C), ACT936ATT (2991C>T), CAT937CGT (2994A>G), GAG939AGG (2999G>A 3000A>G), TCC944TC- (3016delC), CGG946CAG (3021G>A), TAC947TTT (3024A>T 3025C>T), ACA951ATA (3036C>T), GAA952GGC (3039A>G 3040A>C), AAG953ACA (3042A>C 3043G>A), TTG956TTA (3052G>A), TTA959TTG (3061A>G), GCG966GCA (3082G>A), GGC969GGT (3091C>T), GCC972GTA (3099C>T 3100C>A), GTG974AGA (3104G>A 3105T>G 3106G>A), GTC975GTT (3109C>T), AAA976AAG (3112A>G), ACT978ACC (3118T>C), GCC979GCT (3121C>T), GGA981GGG (3127A>G), AGC983AAC (3132G>A), CGA985AAG (3137C>A 3138G>A 3139A>G), TAT989TAC (3151T>C), TTC990ATT (3152T>A 3154C>T), TGT994TAT (3165G>A), GTA995GTG (3169A>G), ATC1005ATT (3199C>T), AGG1007CGG (3203A>C), CGT1008CGC (3208T>C), CCA1010CCG (3214A>G), ACC1011ACT (3217C>T), TTT1012CTT (3218T>C), GTC1013GTA (3223C>A), ACT1014AAT (3225C>A), GTT1016ATT (3230G>A), AAC1017AAT (3235C>T), TTA1025CTA (3257T>C), ACC1026ACT (3262C>T), AGC1027AGT (3265C>T), GTA1028ATG (3266G>A 3268A>G), GTG1029GTA (3271G>A), GTG1031ATG (3275G>A), GCA1034ACA (3284G>A), CAG1039CAA (3301G>A), AGA1044AAG (3315G>A 3316A>G), GAA1045GCA (3318A>C), ATC1046ATT (3322C>T), GAG1047GAA (3325G>A), TTG1048TTA (3328G>A), ATA1051ATT (3337A>T), GAG1055GAA (3349G>A), GTC1057GTT (3355C>T), TGC1058TGT (3358C>T), ATG1059GTG (3359A>G), CTA1061CTG (3367A>G), TAT1062TAC (3370T>C), TTA1065CTG (3377T>C 3379A>G), GAA1070GAG (3394A>G), CGC1071CGT (3397C>T), GAT1072GAG (3400T>G), ACA1074ACG (3406A>G), TAC1078TAT (3418C>T), ATT1079ATA (3421T>A), TAT1081TAC (3427T>C), AAG1083AAA (3433G>A), TCA1084TCT (3436A>T), GTA1085GTG (3439A>G), AAC1086AAT (3442C>T), CCT1087CCC (3445T>C), CAG1091CAA (3457G>A), TTT1092TTC (3460T>C), GCT1093GCA (3463T>A), GCG1095GCT (3469G>T), CAG1096CAA (3472G>A), CAC1104CAT (3496C>T), CAG1105CAA (3499G>A), TCC1107TCT (3505C>T), GTT1110GGT (3513T>G), AAA1112AAG (3520A>G), GAA1115GAG (3529A>G), CAA1116CAG (3532A>G), ATG1121GTG (3545A>G), GCT1122ACT (3548G>A), TTA1123CTA (3551T>C), GTC1124ATT (3554G>A 3556C>T), AGT1133AGC (3583T>C), GAT1134GAC (3586T>C), TGC1135TGT (3589C>T), TTC1137TTT (3595C>T), AAT1141AAC (3607T>C), TTT1143TTC (3613T>C), AAG1144AAA (3616G>A), GGT1145GGC (3619T>C), CTT1147GTT (3623C>G), TCC1148TCT (3628C>T), CTG1150ATG (3632C>A), GAT1151GAC (3637T>C), TAT1152CAT (3638T>C), GAA1153GAG (3643A>G), TTA1159TTG (3661A>G), GTG1162GTA (3670G>A), TTT1166TTC (3682T>C), GAG1168GAA (3688G>A), GAG1171GAA (3697G>A), ATT1172GTT (3698A>G), GAT1174GAC (3706T>C), TTT1175TTC (3709T>C), TTG1177CTA (3713T>C 3715G>A), AGT1178AAT (3717G>A), GAG1179GAA (3721G>A), CGA1183AAA (3731C>A 3732G>A), ACT1184ACA (3736T>A), CTA1188TTG (3746C>T 3748A>G), ACA1190ACG (3754A>G), AGT1193AGC (3763T>C), CGA1198CGG (3778A>G), CAA1199CAG (3781A>G), ATC1200ATT (3784C>T), CAT1204CAC (3796T>C), CCA1207CCT (3805A>T), TAC1209TAT (3811C>T), AGA1210AGG (3814A>G), TTC1215TTT (3829C>T), TTT1218TTC (3838T>C), AGA1220AGG (3844A>G), GCA1221GCC (3847A>C), GTT1224GTA (3856T>A), GTG1226GTT (3862G>T), GAC1229GAT (3871C>T), GCC1231GCA (3877C>A), AGC1233AGT (3883C>T), GAA1234GAG (3886A>G), CTA1236CTG (3892A>G), GAC1237GAT (3895C>T), TTT1238TTC (3898T>C), GTA1240GTG (3904A>G), CGG1241AGA (3905C>A 3907G>A), GCT1243GCC (3913T>C), AAG1248AAA (3928G>A), TCA1249TCT (3931A>T), GGG1251GGA (3937G>A), AGT1257AGC (3955T>C), GTG1258GTA (3958G>A), GCC1259GCT (3961C>T), GGA1260GGG (3964A>G), TCT1261TCC (3967T>C), TTA1264TTG (3976A>G), ATT1265ATC (3979T>C), CCA1267CCT (3985A>T), CTA1271CTT (3997A>T), GCG1272GCA (4000G>A), AAC1274AAT (4006C>T), GTT1275GTG (4009T>G), TTC1276TTT (4012C>T), AAA1277AAG (4015A>G), TCT1280TCC (4024T>C), GAA1282GAG (4030A>G), CCA1283CCT (4033A>T), AAG1286AAA (4042G>A), CTG1290TTA (4052C>T 4054G>A), CGT1291CGC (4057T>C), CGT1293CGA (4063T>A), GGA1294GGG (4066A>G), ATA1297GTA (4073A>G), TTT1298TTC (4078T>C), GGC1299GGT (4081C>T), TCT1300TCC (4084T>C), TCC1304TCT (4096C>T), AGC1308AGT (4108C>T), GGA1309GGT (4111A>T), GCG1311GCA (4117G>A), CTA1312TTG (4118C>T 4120A>G), TAC1314TAT (4126C>T), TTC1315TAT (4128T>A 4129C>T), GCC1316GCT (4132C>T), AAT1317AAC (4135T>C), CAA1321CAG (4147A>G), TTA1322CTA (4148T>C), GCT1323TCT (4151G>T), AAC1326AAT (4162C>T), GTA1328ATA (4166G>A), ATA1329ATC (4171A>C), GAG1332GAA (4180G>A), CTG1336TTG (4190C>T), CCT1338CCC (4198T>C), TCC1339TCT (4201C>T), GCG1342GCT (4210G>T), TTC1343TTT (4213C>T), AGT1345AGC (4219T>C), CTG1346TTG (4220C>T), CTG1347ATA (4223C>A 4225G>A), GTT1349GTG (4231T>G), CAT1351CAC (4237T>C), GTA1356GTG (4252A>G), TTA1357CTG (4253T>C 4255A>G), AAA1358AAG (4258A>G), ACT1362ACC (4270T>C), AGA1366AGG (4282A>G), GAG1367GAA (4285G>A), GAA1369GAG (4291A>G), CAG1373CAA (4303G>A), GTC1376GTT (4312C>T), ATA1379GTG (4319A>G 4321A>G), GTG1380GTT (4324G>T), GAG1381GAA (4327G>A), CTG1384CTT (4336G>T), TCT1385TCC (4339T>C), TCA1388TCT (4348A>T), GCA1392GCG (4360A>G), GGT1394GGC (4366T>C), TCT1395TCA (4369T>A), ACT1397ACC (4375T>C), GTT1401GTC (4387T>C), TTT1404TAT (4395T>A), GTA1408ATA (4406G>A), CTT1409CTC (4411T>C), GTT1418GTC (4438T>C), ACC1420ACA (4444C>A), GCC1422GCT (4450C>T), CTC1424CTT (4456C>T), GAC1427GAT (4465C>T), ATG1431GTT (4475A>G 4477G>T), GTC1432GTT (4480C>T), ACA1433TCG (4481A>T 4483A>G), GGC1437GGT (4495C>T), AAG1441AAA (4507G>A), CAC1442CAT (4510C>T), GGT1443GGG (4513T>G), CTA1445TTA (4517C>T), GTC1448GTT (4528C>T), AAA1450AAG (4534A>G), ACC1452ACT (4540C>T), GCG1454TTA (4544G>T 4545C>T 4546G>A), AGA1455AAA (4548G>A), TTT1458TTC (4558T>C), GTA1460GTT (4564A>T), ACC1462ACT (4570C>T), ATA1464ATT (4576A>T), TTG1471TTA (4597G>A), GAC1472GAT (4600C>T), GAC1474GAT (4606C>T), GTG1475GTA (4609G>A), GTG1477GTA (4615G>A), GAC1478GAT (4618C>T), TTG1481CTT (4625T>C 4627G>T), GAC1488GAT (4648C>T), ATT1489ATC (4651T>C), AGG1492AGA (4660G>A), AGC1493AGT (4663C>T), GCT1495GCA (4669T>A), GTG1499ATC (4679G>A 4681G>C), AGT1500AGC (4684T>C), GTT1501GTG (4687T>G), GAG1505GAA (4699G>A), AGA1506AGG (4702A>G), CAA1508CAG (4708A>G), AGG1509AGA (4711G>A), CTG1510TTG (4712C>T), AAA1518AAG (4738A>G), GTA1520GTT (4744A>T), GCA1521GCG (4747A>G), TTG1522CTG (4748T>C), ACT1527ACC (4765T>C), GTC1538ATT (4796G>A 4798C>T), ACT1540AGC (4803C>G 4804T>C), GAG1541GAA (4807G>A), GCG1542GCT (4810G>T), GCT1543GCG (4813T>G), CTT1544CTC (4816T>C), TAT1549TAC

| | | | | | | | | | |
|---|---|---|---|---|---|---|---|---|---|
| | (4831T>C), AAC1550AAT (4834C>T), CCA1552CCT (4840A>T), GGA1556GGG (4852A>G), GGC1557GGT (4855C>T), AGT1561AGC (4867T>C), CTG1562CTC (4870G>C), ATT1563ATA (4873T>A), TGT1566TGC (4882T>C), ACA1573ACT (4903A>T), CAG1575CAA (4909G>A), TTT1577TTC (4915T>C), GAA1578GAG (4918A>G), TTG1579CTA (4919T>C 4921G>A), AGT1580AGC (4924T>C), CCC1581CCG (4927C>G), TTT1583TTC (4933T>C), ATC1584ATA (4936C>A), CAG1585CAA (4939G>A), AAT1586AAC (4942T>C), TTC1587TTT (4945C>T), TCA1593TCG (4963A>G), CCT1596CCC (4972T>C), CTT1602CTC (4990T>C), AAA1606AAG (5002A>G), CCT1613CCC (5023T>C), TTG1614CTG (5024T>C), TGC1615TGT (5029C>T), GAT1616GAC (5032T>C), TCT1618TCC (5038T>C), CCA1620CCT (5044A>T), GCA1623GCT (5053A>T), TCG1624TCA (5056G>A), AGT1625GGT (5057A>G), ACT1626GCT (5060A>G), TCG1629TCT (5071G>T), CTT1636ATT (5090C>A), GGA1637GGG (5095A>G), GTG1638GTT (5098G>T), GCC1639GTC (5100C>T), TTA1640TTG (5104A>G), GAA1641GAT (5107A>T), ATT1642ATC (5110T>C), CCA1643CCG (5113A>G), AAG1644AAA (5116G>A), CAA1645CAG (5119A>G), GTC1646TTT (5120G>T 5122C>T), ATT1648GTT (5126A>G), CAT1651CAC (5137T>C), ATC1652ATT (5140C>T), AAA1653AAG (5143A>G), GAG1654GAA (5146G>A), CTC1659CTA (5161C>A), GAA1661GAG (5167A>G), CTT1663CTG (5173T>G), ACG1666ACA (5182G>A), GTT1667ATT (5183G>A), GTC1668GTT (5188C>T), AAG1669AAA (5191G>A), AAA1671AAG (5197A>G), GAC1672GAT (5200C>T), TGC1674TGT (5206C>T), TTA1675TTG (5209A>G), AGC1678AGT (5218C>T), TCG1682TCA (5230G>A), ATC1684ATT (5236C>T), AGC1685AGT (5239C>T), ATC1687ATT (5245C>T), ACA1693ACT (5263A>T), CTC1695CTT (5269C>T), TTC1696TTT (5272C>T), GCC1697GCA (5275C>A), ATC1698ATT (5278C>T), CCA1699CCC (5281A>C), AGA1700AGG (5284A>G), ACT1701ACC (5287T>C), CTA1702TTA (5288C>T), ATA1703ATT (5293A>T), TTG1704CTA (5294T>C 5296G>A), GAG1706GAA (5302G>A), CTT1709CTA (5311T>A), GAA1710GAG (5314A>G), CGA1713CGT (5323A>T), AGT1721AGC (5347T>C), ATC1723ATT (5353C>T), TGC1727TGT (5365C>T), GTT1734GTC (5386T>C), ACC1737ACA (5395C>A), AAA1746AAG (5422A>G), GAT1747GAC (5425T>C), ACC1749ACT (5431C>T), GAG1751GAA (5437G>A), CTT1756CTC (5452T>C), AAG1758AAA (5458G>A), AGT1761AGC (5467T>C), CAA1762CAG (5470A>G), CTA1763TTG (5471C>T 5473A>G), GAA1765GAG (5479A>G), TTC1766TTT (5482C>T), TCA1767TCC (5485A>C), AAT1768AAC (5488T>C), TTG1769CTA (5489T>C 5491G>A), GGT1771GGC (5497T>C), TGT1774TGC (5506T>C), GAG1775GAA (5509G>A), GAG1776GAA (5512G>A), TTA1778CTA (5516T>C), TAT1782TAC (5530T>C), GAG1783GAA (5533G>A), TCG1784TCT (5536G>T), TTG1785TTA (5539G>A), GTT1788GTG (5548T>G), CAC1790CAT (5554C>T), GCT1792GCA (5560T>A), GCG1793ACA (5561G>A 5563G>A), ACG1794ACC (5566G>C), TCA1795TCG (5569A>G), CTC1800CTG (5584C>G), AAG1803AAA (5593G>A), GGG1804GGA (5596G>A), ATT1805GTT (5597A>G), AAC1807AAA (5605C>A), GTG1811GTT (5617G>T), GCT1812GTG (5619C>T 5620T>G), TTG1815TTA (5629G>A), ATC1816CTC (5630A>C), GGC1819GGT (5641C>T), GCA1822GCT (5650A>T), ATA1826ATT (5662A>T), GGA1827GGG (5665A>G), CAA1835CAG (5689A>G), GTT1837ATT (5693G>A), GAG1838GAA (5698G>A), CAA1843CAG (5713A>G), GGG1844GGC (5716G>C), AAA1845AAG (5719A>G), GCC1853GCA (5743C>A), CGC1857CGA (5755C>A), CAT1858CAC (5758T>C), CGT1860CGC (5764T>C), AAA1862AAG (5770A>G), GGC1865GGT (5779C>T), TTT1866TTC (5782T>C), GAC1869GAT (5791C>T), GAC1873GAT (5803C>T), AGG1884AGA (5836G>A), AAA1885AAG (5839A>G), GGA1887GGG (5845A>G), AAA1888AAG (5848A>G), AAA1890AGA (5853A>G), GGT1891GGC (5857T>C), ACC1892ACT (5860C>T), ACA1893ACT (5863A>T), GGT1895GGA (5869T>A), GGT1897GGC (5875T>C), AGC1900AGT (5884C>T), TTC1903TTT (5893C>T), ATC1904GTG (5894A>G 5896C>G), AAC1905AAT (5899C>T), GGG1908GGA (5908G>A), TTT1909TTC (5911T>C), GAG1913GAA (5923G>A), CAA1918CAG (5938A>G), TTC1919TTT (5941C>T), GTT1920GTC (5944T>C), GAT1921GAC (5947T>C), GGG1925GGA (5959G>A), CGG1926GCC (5960C>G 5961G>C 5962G>C), GAA1930GAG (5974A>G), GCT1934GCA (5986T>A), GAC1935GAT (5989C>T), ATT1936GTT (5990A>G), GAT1938GAC (5998T>C), AGA1942CGT (6008A>C 6010A>T), GAA1945GAT (6019A>T), GTG1946GTT (6022G>T), CGA1947CGT (6025A>T), AAA1949ACA (6030A>C), GAC1955GAA (6049C>A), ATG1958ATA (6058G>A), CAA1959CAG (6061A>G), GCC1960GCA (6064C>A), GGT1962GGC (6070T>C), AGT1963AAC (6072G>A 6073T>C), AAC1964CAC (6074A>C), ACG1965ACA (6079G>A), ACC1966AAC (6081C>A), ATA1967ATT (6085A>T), GCA1969GCT (6091A>T), TAC1970TAT (6094C>T), TGT1976TCC (6111G>C 6112T>C), AAA1978AAG (6118A>G), TTG1980TTA (6124G>A), AAG1981AAA (6127G>A), ATT1982GTC (6128A>G 6130T>C), GAT1983GAC (6133T>C), TTA1984TTG (6136A>G), AAC1988AAT (6148C>T), AAA1991AAG (6157A>G), GTT1992ATT (6158G>A), TGT1993TGC (6163T>C), AAA1995AAG (6169A>G), ACA1996TCG (6170A>T 6172A>G), GGC1998GGT (6178C>T), AAA2001AAG (6187A>G), TTT2002TTC (6190T>C), GAG2004GAA (6196G>A), CTC2007TTT (6203C>T 6205C>T), GAA2008GAG (6208A>G), CTA2009TTG (6209C>T 6211A>G), CAG2011CAA (6217G>A), GCT2015GCA (6229T>A), GTA2016ATA (6230G>A), GTC2018GTT (6238C>T), AAG2021AAA (6247G>A), ATA2023ATT (6253A>T), GCA2025AAA (6257G>A 6258C>A), CAT2030CAC (6274T>C), AAA2033AGA (6282A>G), TCG2034TCA (6286G>A), GGC2038GGT (6298C>T), TTG2039TTA (6301G>A), AGA2040AGG (6304A>G), GAC2041GAT (6307C>T), GCC2046GCT (6322C>T), ACA2048ACT (6328A>T), GTT2049GTG (6331T>G), TGT2050TGC (6334T>C), AGG2051AGA (6337G>A), CTG2052TTG (6338C>T), TAT2058TAC (6358T>C), GGG2059GGA (6361G>A), GCA2060ACA (6362G>A), TCA2061TCT (6367A>T), GAG2062GAA (6370G>A), TAC2064TAT (6376C>T), GGT2065GGG (6379T>G), GGA2067GGC (6385A>C), GGA2069GGT (6391A>T), GCA2070GCG (6394A>G), TAC2071TAT (6397C>T), GTA2073ATA (6401G>A), GCG2074GTG (6405C>T), CAC2076CAT (6412C>T), TTA2078TTG (6418A>G), TTT2079TTC (6421T>C), TAC2082TTC (6429A>T), GGT2084GGA (6436T>A), GAG2087GAA (6445G>A), CAA2089CGA (6450A>G), TCC2090TCG (6454C>G), CAC2092CAT (6460C>T), GGT2093GGA (6463T>A), TTC2095TTT (6469C>T), AGG2096AGA (6472G>A), GTG2097GTA (6475G>A), AAT2099AAC (6481T>C), CAC2101AAT (6485C>A 6487C>T), AGT2102AGC (6490T>C), AGC2104AGT (6496C>T), CTG2106CTA (6502G>A), GGT2110GGC (6514T>C), AGG2111AGA (6517G>A), CTC2115ATC (6527C>A), ATC2116ATA (6532C>A), CCG2119CCA (6541G>A), AAA2120AAG (6544A>G), GTC2124GTT (6556C>T), TTT2125TTC (6559T>C), CAG2127CAA (6565G>A), TTG2129CTA (6569T>C 6571G>A), CCT2134CCA (6586T>A), ACA2135ATA (6588C>T), GAA2138GAG (6598A>G), AGA2139AGG (6601A>G), ATT2140ATC (6604T>C), TGT2141TGC (6607T>C), GGA2144GGG (6616A>G), AAC2146AAT (6622C>T), GAG2149GAA (6631G>A), AAG2150AAA (6634G>A), TAT2151CAT (6635T>C), GCT2152GCA (6640T>A), TCG2154TCT (6646G>T), ATC2155GTC (6647A>G), ATC2156ATT (6652C>T), AGC2160AGT (6664C>T), ACT2162ACG (6670T>G), ATA2165GTG (6677A>G 6679A>G), GGC2167GGT (6685C>T), ACA2169ACT (6691A>T), TTC2170TTT (6694C>T), ACA2177ACG (6715A>G), GAT2178AAT (6716G>A), AAT2179GAT (6719A>G), GGA2180GGG (6724A>G), CTA2184TTA (6734C>T), GTG2186GTA (6742G>A), GTG2187GTA (6745G>A), GCC2190GCT (6754C>T), GGA2192GGG (6760A>G), ATA2194TTG (6764A>T 6766A>G), ATT2197ATA (6775T>A), CTG2200TTG (6782C>T), GCA2201GCT (6787A>T), AAC2202AAT (6790C>T), GCA2204GTG (6795C>T 6796A>G), CAC2205CAA (6799C>A), ACC2206ACA (6802C>A), AAC2208AAT (6808C>T), TAC2209TAT (6811C>T), TAC2210TAT (6814C>T), TTC2213TTT (6823C>T), GAA2215GAG (6829A>G), TAC2221TAT (6847C>T), CGA2223CGC (6853A>C), AAT2225GAT (6857A>G), GAG2226GAA (6862G>A), GAA2229GAG (6871A>G), GTC2231ACA (6875G>A 6876T>C 6877C>A), AAG2232AAA (6880G>A), TCT2233TCA (6883T>A), GTT2235GTG (6889T>G), GAC2239GAT (6901C>T), GGC2244GGT (6916C>T), CCG2245CCA (6919G>A), AAA2247AAG (6925A>G), CTT2248CTC (6928T>C), GAC2250GAA (6934C>A), GGG2255GGC (6949G>C), TTC2257TTT (6955C>T), AAA2258AAG (6958A>G), CAA2264CAG (6976A>G), CTA2266TTA (6980C>T), ATC2267ATT (6985C>T), GTA2271GTT (6997A>T), GTG2272GTT (7000G>T), GTG2273ATA (7001G>A 7003G>A), GCT2276GCC (7012T>C), AAG2277AAA (7015G>A), CAT2278CAC (7018T>C), TTT2283TAT (7032T>A), GCC2285GCT (7039C>T), TTG2286CTA (7040T>C 7042G>A), GGA2288GGG (7048A>G), GCT2292GCA (7060T>A), GTC2293GTA (7063C>A), AGC2298AGT (7078C>T), TTA2300CTG (7082T>C 7084A>G), GTA2301GTG (7087A>G), ACC2302ACA (7090C>A), CAT2304CAC (7096T>C), GTA2305GTG (7099A>G), GTT2306GTC (7102T>C), GAG2309GAA (7111G>A), CGA2311CGG (7117A>G), ACA2314AAA (7125C>A), GAA2315GAG (7129A>G), TTT2316TTC (7132T>C), CTG2317TTA (7133C>T 7135G>A), GTG2319GTA (7141G>A), GAT2320GAC (7144T>C), GCA2321TCG (7145G>T 7147A>G), GAG2322GAA (7150G>A), GCA2325GCT (7159A>T), GAG2326_GCA2327del (7160_7165delGAGGCA), TTC2329TTT (7171C>T), GCG2335GCT (7189G>T), AAA2338AAG (7198A>G), AGC2339AGT (7201C>T), CTA2341TTA (7205C>T), AAT2342AAC (7210T>C), GAT2344GAA (7216T>A), GCG2345GCA (7219G>A), ATC2347ATA (7225C>A), AAG2348AAA (7228G>A), AAG2352AAA (7240G>A), TAT2353TAC (7243T>C), AAA2355AAG (7249A>G), CCT2356CCG (7252T>G), GAT2358GAA (7258T>A), GTT2359GTC (7261T>C), GGT2360GGA (7264T>A), GTC2361GTG (7267C>G), GTG2362GTA (7270G>A), GAT2363_CGG2364insT-- (7273_7274insT), CGG2364GTG (7274C>G 7275G>T), CAT2366CTT (7281A>T), TTG2367TCG (7284T>C), TCA2371TTA (7296C>T), TCT2376TTT (7311C>T), ACC2377ATC (7314C>T), TGC2378TAC (7317G>A), AAT2379AAA (7321T>A), GTG2380-TG (7322delG), CAC2381CAT (7327C>T), GGC2382GGT (7330C>T), TTC2383TTT (7333C>T), AAG2384CAG (7334A>C), TGT2386TGC (7342T>C), GCA2387AAC (7343G>A 7344C>A 7345A>C), ACT2390ACC (7354T>C), CAA2393CAG (7363A>G), ATT2395ATC (7369T>C), TTC2396TTT (7372C>T), GCG2398GCT (7378G>T), CTC2399CTT (7381C>T), GCA2404GCT (7396A>T), GTC2405GTT (7399C>T), GCC2407GCT (7405C>T), AGT2408ATG (7407G>T 7408T>G), GGG2410GGT (7414G>T), TGC2411GGC (7415T>G), AAA2412AAG (7420A>G), GAC2415GAT (7429C>T), TAT2416TAC (7432T>C), CAT2419CAC (7441T>C), AAG2425AAA (7459G>A), GAA2426GAG (7462A>G), ATA2428ATT (7468A>T), GTC2429GTA (7471C>A), CAA2431CAG (7477A>G), AGC2432AGT (7480C>T), CTG2434CTT (7486G>T), TAT2437TAC (7495T>C), AAA2438AAG (7498A>G), GGT2439GGC (7501T>C), TTG2440TCA (7503T>C 7504G>A), ATT2443ATA (7513T>A), AAC2445AAT (7519C>T), GGA2446GGG (7522A>G), AAG2449AAA (7531G>A), GAG2451GAA (7537G>A), CTC2452CTT (7540C>T), TGT2454TGC (7546T>C), AAG2462AAA (7570G>A), ACG2463ACA (7573G>A), ACG2465ACA (7579G>A), GCT2468GCC (7588T>C), CTA2471TTA (7595C>T), TTG2474CTA (7604T>C 7606G>A), GGT2477GGA (7615T>A), TGT2480TGC (7624T>C), GAC2483GAT (7633C>T), TTC2484TTT (7636C>T), AAT2485AAC (7639T>C), TTT2488TTC (7648T>C), GAA2494GAG (7666A>G), TGT2496TGC (7672T>C), ACA2498ACT (7678A>T), GGG2500GGA (7684G>A), ACT2502ACC (7690T>C), TTT2504TTC (7696T>C), GGT2506GGG (7702T>G), GAT2509AAT (7709G>A), AAA2510AGG (7713A>G 7714A>G), CTG2511TTG (7715C>T), CGT2514CGC (7726T>C), GAG2517GAA (7735G>A), GTA2520GTG (7744A>G), TAC2521TAT (7747C>T), GGC2526GGT (7762C>T), CAG2528CAA (7768G>A), TTT2529TTC (7771T>C), TCA2532TCG (7780A>G), CTA2533CTC (7783A>C), ACT2534ACC (7786T>C), CTA2537TTG (7793C>T 7795A>G), ATC2538ATT (7798C>T), GCT2540GCC (7804T>C), ACC2543ATC (7812C>T), GAA2550GAG (7834A>G), GAC2551GAT (7837C>T), GTG2554TTA (7844G>T 7846G>A), CAG2557CAA (7855G>A), CTG2559CTA (7861G>A), TTA2562TTG (7870A>G), TAC2563TAT (7873C>T), ACT2564ACA (7876T>A), GAG2565GAA (7879G>A), ATT2566ATA (7882T>A), GTT2567ATT (7883G>A), CCA2570CCT (7894A>T), ATT2571ATC (7897T>C), GTC2579GTT (7921C>T), AAG2581AAA (7927G>A), GGT2584GGC (7936T>C), AAT2585AAC (7939T>C), AAT2586AAC (7942T>C), GTG2594GTA (7966G>A), GAT2595GAC (7969T>C), TCT2597TCC (7975T>C), GTC2601GTT (7987C>T), CTT2602CTC (7990T>C), GCT2607GCC (8005T>C), GAA2616GAG (8032A>G), ATC2618ATT (8038C>T), GAC2619GAT (8041C>T), AGC2620AAC (8043G>A), TTG2632TTA (8080G>A), GTG2635GTA (8089G>A), CCG2637CCA (8095G>A), GAG2638GAA (8098G>A), AGC2641AGT (8107C>T), ATT2642ATC (8110T>C), GAT2644GAC (8116T>C), CAA2648CAG (8128A>G), GGT2654GGC (8146T>C), AAC2656AGT (8151A>G 8152C>T), TCG2660TCA (8164G>A), AGG2665AAG (8178G>A), AAG2666AAA (8182G>A), TTG2669TTA (8191G>A), TTC2671TTT (8197C>T), TCC2673TCT (8203C>T), AGA2675AAA (8208G>A), CTG2677TTA (8213C>T 8215G>A), CTA2678CTG (8218A>G), ATC2679ATT (8221C>T), GAG2680GAA (8224G>A), CCA2685CCG (8239A>G), GAG2690GAA (8254G>A), ATT2692ATC (8260T>C), GTA2693GTG (8263A>G), GAT2699GAC (8281T>C), AGA2700AGG (8284A>G), CCA2704GCG (8294C>G 8296A>G), GAG2705GAA (8299G>A), CAC2706CAT (8302C>T), ATT2711ATC (8317T>C), TGC2712TGT (8320C>T), GCT2714GCA (8326T>A), GAG2717GAA (8335G>A), TCC2718TCT (8338C>T), TCT2722TTT (8349C>T), GAA2723GAG (8353A>G), CAC2726CAT (8362C>T), CAA2727CAG (8365A>G), ATC2728ATT (8368C>T), TAC2732TAT (8380C>T), TCA2733TCG (8383A>G), TTA2735TTG (8389A>G), CCT2740CCC (8404T>C), GCA2742TCA (8408G>T), ACA2743ACG (8413A>G), GCG2745GCT (8419G>T), GCT2750GCC (8434T>C), TAT2752TAC (8440T>C), ATG2756ATA (8452G>A), CTA2758CTG (8458A>G), AGG2759AAG (8460G>A), AAA2760AAG (8464A>G), CTG2761CTA (8467G>A), AGG2765AGA (8479G>A), GCT2766ACA (8480G>A 8482T>A), GAG2771GAA (8497G>A), CTA2772TTG (8498C>T 8500A>G), AGA2773AAG (8502G>A 8503A>G), GCC2774GCT (8506C>T), GAA2777GAT (8515A>T), GTC2780GTT (8524C>T), GCA2781GCC (8527A>C), TTA2782CTG (8528T>C 8530A>G), GAG2785GAC (8539G>C), CTT2788TGT (8546C>T 8547T>G), TCT2790ACT (8552T>A), GTA2793GTG (8563A>G), GCA2797GGA (8574C>G), ATT2801GTT (8585A>G), GGA2805GAA (8598G>A), AGC2806AGT (8602C>T), AAC2807TCT (8603A>T 8604A>C 8605C>T), AAA2812AAG (8620A>G), CCA2813CAA (8622C>A), GAG2814GGA (8625A>G 8626G>A), CAG2815CAA (8629G>A), GGC2816GGT (8632C>T), ATC2818ATT (8638C>T), CAG2819CAA (8641G>A), AAC2821AAT (8647C>T), CCG2822TTC (8648C>T 8649C>T 8650G>C), AAC2823AAT (8653C>T), AAA2824AAG (8656A>G), GGA2825GAA (8658G>A), GAT2827GCG (8664A>C 8665T>G), GTT2830GTG (8674T>G), GCA2832GTT (8679C>T 8680A>T), GGC2833GGA (8683C>A), GGG2836GGA (8692G>A), ACA2837ACT (8695A>T), CCG2841CCA (8707G>A), AGA2842CGA (8708A>C), AAG2844AAA (8716G>A), ATC2846ATT (8722C>T), ACG2847ACA (8725G>A), AAA2849AAG (8731A>G), AGA2851AGG (8737A>G), CCC2853CCT (8743C>T), ACA2854AAG (8745C>A 8746A>G), AAG2856AAA (8752G>A), GGA2857GGT (8755A>T), AAC2862AAT (8770C>T), TTA2863CTA (8771T>C), GAA2864GAG (8776A>G), CAT2865CAC (8779T>C), TTG2866TTA (8782G>A), TAT2869TAC (8791T>C), CCA2871CCG (8797A>G), CAA2872CAG (8800A>G), CAA2873CAG (8803A>G), GAT2875GAC (8809T>C), ATT2876ATC (8812T>C), AAT2878AAC (8818T>C), CGG2880CGA (8824G>A), GCA2881GCG (8827A>G), ACG2888ACA (8848G>A), TAT2890TAC (8854T>C), GAG2891GGA (8856A>G 8857G>A), GTG2893GTA (8863G>A), CGG2894CAG (8865G>A), ATG2895CTT (8867A>C 8869G>T), GCA2896GCG (8872A>G), GGA2900GAA (8883G>A), ACT2902TCT (8888A>T), TGC2915TGT (8929C>T), GTC2924ATC (8954G>A), AAC2925AAT (8959C>T), GGA2926GGT (8962A>T), GGG2933GGA (8983G>A), AAT2934GAC (8984A>G 8986T>C), GAA2935GTA (8988A>T), GAG2938GAA (8998G>A), CCG2940CCA (9004G>A), TTG2941TTA (9007G>A), ATC2944ATT (9016C>T), ACC2951ACA (9037C>A), ATA2967ATT (9085A>T), AAC2971AAT (9097C>T), ATT2983GTT (9131A>G), CGA2984CGT (9136A>T), CGG2987CGT (9145G>T), ATG2989GGA (9149A>G 9150T>G 9151G>A), GGT2990AGT (9152G>A), TTA2991TTG (9157A>G), GCG2992GCT (9160G>T), CGT2993CGC (9163T>C), GAG3000GAA (9184G>A), CGA3004CGG (9196A>G), AGG3010AGA (9214G>A), GAA3011GAG (9217A>G), GCG3012GCA (9220G>A), GCA3020GCT (9244A>T), TTG3021TTA (9247G>A), GCC3024GCT (9256C>T), CCT3026TCC (9260C>T 9262T>C), GGG3030GGA (9274G>A), GAC3032GAT (9280C>T), ATC3035ATT (9289C>T), CAA3038CAC (9298A>C), GAG3040GAA (9304G>A), GTC3050GTT (9334C>T), GTC3060GTG (9364C>G) | | | | | | | | |
| P1 protein (NP_734243.1) | 1 | 284 | 100% | 1394 | 72.7% | 284 (100%) | 203 (71.5%) | 0/0/0/0 | 0 |

| | Begin | End | Coverage | Score | Concordance | Matches | Identities | I/D/M/F* | Stop Codons |
|---|---|---|---|---|---|---|---|---|---|
| **NT** | **5** | **9704** | **99.9%** | **12522** | **64.8%** | **9688 (99.7%)** | **8015 (82.5%)** | **15/12** | |
| Protein mutations: | Y4Q (194T>C 196C>A), C9L (209T>C 210G>T 211T>G), F13M (221T>A 223T>G), S23P (251T>C), C24F (255G>T 256C>T), E25G (258A>G), I27V (263A>G 265T>G), V28A (267T>C), E30V (273A>T), A35T (287G>A 289T>C), V37T (293G>A 294T>C), A41I (305G>A 306C>T), D42G (309A>G), E44D (316A>T), T45V (317A>G 318C>T 319A>G), L52Q (339T>A 340C>A), K53R (342A>G), K55A (347A>G 348A>C 349A>G), Y56H (350T>C), T58V (356A>G 357C>T), V61T (365G>A 366T>C 367G>A), L62S (368C>T 369T>C), F67C (384T>G), A75V (408C>T 409C>T), M78E (416A>G 417T>A), E85K (437G>A), R86E (440A>G 441G>A), K87R (443A>C 444A>G 445G>A), D88E (448T>A), E91A (456A>C), H93N (461C>A), D111E (517T>A), S116F (531C>T), P118S (536C>T), Q119W (539C>T 540A>G 541A>G), R122K (549G>A 550A>G), M131T (576T>C), V134A (585T>C 586C>G), R137Y (593C>T 594G>A 595C>T), P138R (597C>G), I139V (599A>G 601A>G), I140P (602A>C 603T>C 604A>G), M147I (625G>A), I151V (635A>G 637T>C), I154V (644A>G 646A>G), E160A (663A>C 664G>C), H166Q (682C>A), T172S (699C>G), H174Q (706T>A), K179E (719A>G 721G>A), I180V (722A>G 724A>G), A183S (731G>T), Y184P (734T>C 735A>C 736C>T), S185R (737T>C 738C>G 739C>T), A187V (744C>T 745G>C), R189C (749C>T 751A>C), M194K (765T>A), R197K (773C>A 774G>A 775A>G), M206K (801T>A), G210A (813G>C 814A>G), L211H (816T>A), R214Q (825G>A 826T>A), N223D (851A>G 853T>C), R226C (860C>T 862C>T), T227A (863A>G 865T>C), I228T (867T>C), N229D (869A>G 871C>T), I230L (872A>T 874A>G), R231Q (876G>A), R232K (879G>A), N240S (903A>G), T241N (906C>A 907A>C), K242T (909A>C 910A>C), S243N (912G>A 913C>T), S252L (938T>C 939C>T 940A>T), L266I (980T>A 982G>C), R272K (999G>A), Q275R (1007C>A 1008A>G 1009G>A), S276G (1010A>G 1012T>G), I277V (1013A>G), N279Q (1019A>C 1021C>G) | | | | | | | | |
| Codon mutations: | ACT3ACC (193T>C), TAC4CAA (194T>C 196C>A), TCA6TCG (202A>G), ATC8ATT (208C>T), TGT9CTG (209T>C 210G>T 211T>G), TCG12TCA (220G>A), TTT13ATG (221T>A 223T>G), GAA14GAG (226A>G), CTA17CTT (235A>T), TAC19TAT (241C>T), CCA21CCT (247A>T), TCT23CCT (251T>C), TGC24TTT (255G>T 256C>T), GAG25GGG (258A>G), CAT26CAC (262T>C), ATT27GTG (263A>G 265T>G), GTG28GCG (267T>C), GAA30GTA (273A>T), CGA31CGT (277A>T), GAA32GAG (280A>G), GTG33GTT (283G>T), CCG34CCA (286G>A), GCT35ACC (287G>A 289T>C), GTT37ACT (293G>A 294T>C), CCT39CCC (301T>C), TTC40TTT (304C>T), GCA41ATA (305G>A 306C>T), GAT42GGT (309A>G), CTG43TTG (311C>T), GAA44GAT (316A>T), ACA45GTG (317A>G 318C>T 319A>G), CAA46CAG (322A>G), GCA49GCT (331A>T), CTC52CAA (339T>A 340C>A), AAG53AGG (342A>G), AAA55GCG (347A>G 348A>C 349A>G), TAT56CAT (350T>C), GCT57GCA (355T>A), ACT58GTT (356A>G 357C>T), CGT60CGC (364T>C), GTG61ACA (365G>A 366T>C 367G>A), CTC62TCC (368C>T 369T>C), AAA63AAG (373A>G), AAC64AAT (376C>T), TTT67TGT (384T>G), ACG68ACA (388G>A), TAC69TAT (391C>T), CGA70CGG (394A>G), AAG72AAA (400G>A), GAT74GAC (406T>C), GCC75GTT (408C>T 409C>T), CAG76CAA (412G>A), ATA77ATT (415A>T), ATG78GAG (416A>G 417T>A), ATT80ATC (424T>C), CAG81CAA (427G>A), AAA83AAG (433A>G), CTG84CTC (436G>C), GAG85AAG (437G>A), AGG86GAG (440A>G 441G>A), AAG87CGA (443A>C 444A>G 445G>A), GAT88GAA (448T>A), GAA90GAG (454A>G), GAA91GCA (456A>C), TAT92TAC (460T>C), CAC93AAC (461C>A), CAA95CAG (469A>G), GCC97GCG (475C>G), CCT99CCA (481T>A), ACT106ACC (502T>C), ATA107ATT (505A>T), GGC109GGT (511C>T), GAT111GAA (517T>A), CCT112CCA (520T>A), CCA113CCT (523A>T), TCT116TTT (531C>T), GAG117GAA (535G>A), CCA118TCA (536C>T), CAA119TGG (539C>T 540A>G 541A>G), AGA122AAG (549G>A 550A>G), ATC124ATT (556C>T), ATT125ATC (559T>C), CAT126CAC (562T>C), ACT128ACC (568T>C), CCA129CCG (571A>G), ATG131ACG (576T>C), CGT132CGC (580T>C), GTC134GCG (585T>C 586C>G), CGC137TAT (593C>T 594G>A 595C>T), CCC138CGC (597C>G), ATA139GTG (599A>G 601A>G), ATA140CCG (602A>C 603T>C 604A>G), ACA143ACG (613A>G), GAA144GAG (616A>G), GGC145GGG (619C>G), CAG146CAA (622G>A), ATG147ATA (625G>A), AAT148AAC (628T>C), CAC149CAT (631C>T), ATT151GTC (635A>G 637T>C), ATA154GTG (644A>G 646A>G), ATT157ATC (655T>C), TCG159TCA (661G>A), GAG160GCC (663A>C 664G>C), AAA161AAG (667A>G), GTC165GTT (679C>T), CAC166CAA (682C>A), TTA167CTG (683T>C 685A>G), AGT169AGC (691T>C), ACC172AGC (699C>G), ACT173ACA (703T>A), CAT174CAA (706T>A), AAG179GAA (719A>G 721G>A), ATA180GTG (722A>G 724A>G), CTT181TTG (725C>T 727T>G), GGT182GGA (730T>A), GCA183TCA (731G>T), TAC184CCT (734T>C 735A>C 736C>T), TCC185CGT (737T>C 738C>G 739C>T), GCG187GTC (744C>T 745G>C), CGA189TGC (749C>T 751A>C), ACT190ACC (754T>C), ATG194AAG (765T>A), TTG196CTG (770T>C), CGA197AAG (773C>A 774G>A 775A>G), CGG198AGG (776C>A), TTC202TTT (790C>T), ATG206AAG (801T>A), GGA210GCG (813G>C 814A>G), CTT211CAT (816T>A), TTG212CTG (818T>C), CAA213CAG (823A>G), CGT214CAA (825G>A 826T>A), CGG217AGA (833C>A 835G>A), GAC219GAT (841C>T), AAT223GAC (851A>G 853T>C), GTC225GTT (859C>T), CGC226TGT (860C>T 862C>T), ACT227GCC (863A>G 865T>C), ATC228ACC (867T>C), AAC229GAT (869A>G 871C>T), ATA230TTG (872A>T 874A>G), CGA231CAA (876G>A), AGG232AAG (879G>A), GAT234GAC (886T>C), ATC238ATT (898C>T), AAC240AGC (903A>G), ACA241AAC (906C>A 907A>C), AAA242ACC (909A>C 910A>C), AGC243AAT (912G>A 913C>T), GGC246GGA (922C>A), GGT249GGA (931T>A), TCA252CTT (938T>C 939C>T 940A>T), GGA253GGG (943A>G), TTC256TTT (952C>T), TCA261TCG (967A>G), GAA263GAG (973A>G), GGG264GGC (976G>C), TTG266ATC (980T>A 982G>C), GCA269GCG (991A>G), CGT270CGC (994T>C), AGA272AAA (999G>A), GTT273GTC (1003T>C), CAG275AGA (1007C>A 1008A>G 1009G>A), AGT276GGG (1010A>G 1012T>G), ATT277GTT (1013A>G), TTA278TTG (1018A>G), AAC279CAG (1019A>C 1021C>G), ATC282ATT (1030C>T) | | | | | | | | |
| HC-Pro protein (NP_734244.1) | 1 | 456 | 100% | 3144 | 95.7% | 456 (100%) | 425 (93.2%) | 0/0/0/0 | 0 |
| Protein mutations: | D4E (1048C>A), R16Q (1083G>A), S21A (1097T>G 1099G>A), R36K (1143G>A), L40I (1154T>A 1156G>A), S62N (1221G>A), V65T (1229G>A 1230T>C 1231T>C), S66N (1233G>A 1234C>T), K74T (1257A>C 1258A>G), N81S (1278A>G), A85V (1290C>T 1291G>A), D88T (1298G>A 1299A>C 1300C>T), I91V (1307A>G 1309A>G), N94E (1316A>G 1318T>A), I98T (1329T>C), L113I (1373C>A), I117V (1385A>G), E123D (1405G>T), A127S (1415G>T 1417A>T), V134I (1436G>A 1438C>T), E224G (1707A>G), A228T (1718G>A), I236V (1742A>G 1744C>T), S241N (1758G>A), V252I (1790G>A), I301V (1937A>G 1939T>C), F323Y (2004T>A), I344M (2068T>G), V346I (2072G>A 2074G>A), R391K (2208G>A), D410E (2266C>G) | | | | | | | | |
| Codon mutations: | TCG1TCA (1039G>A), AAT2AAC (1042T>C), GCC3GCA (1045C>A), GAC4GAA (1048C>A), AAT5AAC (1051T>C), AAG8AAA (1060G>A), GGT9GGA (1063T>A), CTG10TTG (1064C>T), GAC11GAT (1069C>T), CGA16CAA (1083G>A), TAT19TAC (1093T>C), TCG21GCA (1097T>G 1099G>A), CAC23CAT (1105C>T), GTA26GTG (1114A>G), GCT27GCG (1117T>G), TTA29TTG (1123A>G), CCT30CCA (1126T>A), GTC31GTT (1129C>T), AGG36AAG (1143G>A), GTA37GTT (1147A>T), GCT38GCA (1150T>A), TTG40ATA (1154T>A 1156G>A), CAC43CAT (1165C>T), AGT44AGC (1168T>C), CTT46TTA (1172C>T 1174T>A), ACC55ACT (1201C>T), AGC62AAC (1221G>A), GTT65ACC (1229G>A 1230T>C 1231T>C), AGC66AAT (1233G>A 1234C>T), CTG68CTA (1240G>A), AAG70AAA (1246G>A), TTG72CTA (1250T>C 1252G>A), CAT73CAC (1255T>C), AAA74ACG (1257A>C 1258A>G), AGA77AGG (1267A>G), GGT79GGC (1273T>C), TTG80TTA (1276G>A), AAT81AGT (1278A>G), TTG83CTG (1283T>C), GCG85GTA (1290C>T 1291G>A), GAC88ACT (1298G>A 1299A>C 1300C>T), TTT90TTC (1306T>C), ATA91GTG (1307A>G 1309A>G), CAT92CAC (1312T>C), AAT94GAA (1316A>G 1318T>A), TTG97TTA (1327G>A), ATA98ACA (1329T>C), GCG99GCT (1333G>T), TTA100TTG (1336A>G), CTA103TTG (1343C>T 1345A>G), CCG106CCA (1354G>A), GTG107GTT (1357G>T), GAC108GAT (1360C>T), CTC111TTG (1367C>T 1369C>G), CTT113ATT (1373C>A), TTC114TTT (1378C>T), GAG116GAA (1384G>A), ATA117GTA (1385A>G), AAA119AAG (1393A>G), TCC120TCT (1396C>T), GAG123GAT (1405G>T), AAA124AAG (1408A>G), CAG125CAA (1411G>A), GCA127TCT (1415G>T 1417A>T), CCG128CCC (1420G>C), AAT131AAC (1429T>C), TTA132TTG (1432A>G), GTC134ATT (1436G>A 1438C>T), TTA135CTG (1439T>C 1441A>G), AAT136AAC (1444T>C), TTC139TTT (1453C>T), CTG140CTA (1456G>A), GAA144GAG (1468A>G), GTA152GTG (1492A>G), TTG155CTA (1499T>C 1501G>A), AGT156AGC (1504T>C), TTG157TTA (1507G>A), CTC158CTT (1510C>T), GAA159GAG (1513A>G), TTA160TTG (1516A>G), GCA161GCT (1519A>T), AGG162AGA (1522G>A), TTC163TTT (1525C>T), CAG164CAA (1528G>A), AAC166AAT (1534C>T), ACT168ACA (1540T>A), AAC170AAT (1546C>T), AAA173AAG (1555A>G), GGT174GGC (1558T>C), GAT175GAC (1561T>C), ATA176ATC (1564A>C), TCT177TCA (1567T>A), TTC179TTT (1573C>T), AGA180AGG (1576A>G), TTA183CTA (1583T>C), TCT184TCA (1588T>A), AAG186AAA (1594G>A), AAC188AAT (1600C>T), CTG191CTA (1609G>A), TCG194TCA (1618G>A), TGC195TGT (1621C>T), GAC196GAT (1624C>T), CTG199TTG (1631C>T), AAA201AAG (1639A>G), TTC205TTT (1651C>T), CTC206CTG (1654C>G), CAT213CAC (1675T>C), TCA219TCG (1693A>G), TTT222TTC (1702T>C), GAA224GGA (1707A>G), GCA228ACA (1718G>A), GGA230GGG (1726A>G), TAC231TAT (1729C>T), TCA232TCC (1732A>C), TAT234TAC (1738T>C), ATC236GTT (1742A>G 1744C>T), CGC237CGT (1747C>T), CAT239CAC (1753T>C), AGT241AAT (1758G>A), AGG244AGA (1768G>A), AAC250AAT (1786C>T), TTA251TTG (1789A>G), GTT252ATT (1790G>A), GTC253GTT (1795C>T), TTA257CTG (1805T>C 1807A>G), GAG259GAA (1813G>A), AGG261AGA (1819G>A), AAG263AAA (1825G>A), GAC267GAT (1837C>T), TAT268TAC (1840T>C), CAA271CAG (1849A>G), GTC274GTG (1858C>G), AAA276AAG (1864A>G), TGC278TGT (1870C>T), AGT280AGC (1876T>C), AAA282AAG (1882A>G), GGT284GGA (1888T>A), AAT285AAC (1891T>C), TGT290TGC (1906T>C), ACA293ACT (1915A>T), GGT298GGC (1930T>C), GCC300GCT (1936C>T), ATT301GTC (1937A>G 1939T>C), TCA303TCG (1945A>G), TTC305TTT (1951C>T), CCA308CCT (1960A>T), GTA314GTG (1978A>G), ATT315ATA (1981T>A), GGC316GGT (1984C>T), AAT317AAC (1987T>C), GAC320GAT (1996C>T), TTT323TAT (2004T>A), AAA328AAG (2020A>G), GGG329GGA (2023G>A), GAT330GAC (2026T>C), TCG331TCT (2029G>T), GAG332GAA (2032G>A), TAT341TAC (2059T>C), ATT344ATG (2068T>G), AAC345AAT (2071C>T), GTG346ATA (2072G>A 2074G>A), CTG351TTA (2087C>T 2089G>A), AGC355AGT (2101C>T), GAG356GAA (2104G>A), GAG357GAA (2107G>A), AAG360AAA (2116G>A), TTC362TTT (2122C>T), CGC367CGT (2137C>T), CTT374CTC (2158T>C), ACC376ACT (2164C>T), CCA378CCG (2170A>G), GAT382GAC (2182T>C), GCG384GCA (2188G>A), ACC385ACA (2191C>A), AGA391AAA (2208G>A), TAT394TAC (2218T>C), GAC396GAT (2224C>T), GTA397GTT (2227A>T), TTG402TTA (2242G>A), CCC403CCT (2245C>T), TTG406TTA (2254G>A), GTT407GTC (2257T>C), GAC408GAT (2260C>T), GAC410GAG (2266C>G), CAA412CAG (2272A>G), ACG413ACA (2275G>A), TCA419TCG (2293A>G), GGC421GGT (2299C>T), TCG422TCA (2302G>A), CAG423CAA (2305G>A), GGA426GGT (2314A>T), TAT427TAC (2317T>C), CTA430CTG (2326A>G), GTG435GTA (2341G>A), CAA437CAG (2347A>G), CTT438CTC (2350T>C), TTG440TTA (2356G>A), AAT443AAC (2365T>C), GAA445GAG (2371A>G), TTA446TTG (2374A>G), GAT449GAC (2383T>C), ATA450ATT (2386A>T), AAA451AAG (2389A>G), CAT452CAC (2392T>C), TAT453TAC (2395T>C) | | | | | | | | |
| P3 protein (NP_734251.1) | 1 | 365 | 100% | 1924 | 79.6% | 365 (99.7%) | 327 (89.3%) | 1/0/2/2 | 0 |
| Protein mutations: | N4G (2414A>G 2415A>G 2416T>G), S6C (2420A>T), K40R (2523A>G), R77K (2634G>A), V107I (2723G>A 2725C>T), A115T (2747G>A 2749T>A), H131N (2795C>A), V140I (2822G>A 2824G>A), P160T (2882C>A), S161H (2885A>C 2886G>A), T164M (2895C>T), N176S (2931A>G), N179S (2940A>G), E189_N190insX (2971_2972insA), T196I (2991C>T), H197R (2994A>G), E199R (2999G>A 3000A>G), R206Q (3021G>A), Y207F (3024A>T 3025C>T), T211I (3036C>T), E212G (3039A>G 3040A>C), K213T (3042A>C 3043G>A), A232V (3099C>T 3100C>A), V234R (3104G>A 3105T>G 3106G>A), S243N (3132G>A), R245K (3137C>A 3138G>A 3139A>G), F250I (3152T>A 3154C>T), C254Y (3165G>A), F272L (3218T>C), T274N (3225C>A), V276I (3230G>A), V288M (3266G>A 3268A>G), V291M (3275G>A), A294T (3284G>A), R304K (3315G>A 3316A>G), E305A (3318A>C), M319V (3359A>G), D332E (3400T>G) | | | | | | | | |

| | Begin | End | Coverage | Score | Concordance | Matches | Identities | I/D/M/F* | Stop Codons |
|---|---|---|---|---|---|---|---|---|---|
| **NT** | **5** | **9704** | **99.9%** | **12522** | **64.8%** | **9688 (99.7%)** | **8015 (82.5%)** | **15/12** | |
| Codon mutations: | AAT4GGG (2414A>G 2415A>G 2416T>G), GCT5GCA (2419T>A), AGC6TGC (2420A>T), ACA12ACT (2440A>T), GTT21GTC (2467T>C), TCT24TCC (2476T>C), TCG26TCA (2482G>A), CTG29CTA (2491G>A), CTG31CTT (2497G>T), TTG33TTA (2503G>A), AGA38AGG (2518A>G), CCT39CCC (2521T>C), AAG40AGG (2523A>G), AGA43AGG (2533A>G), TTG45TTA (2539G>A), TTA47TTG (2545A>G), GAG49GAA (2551G>A), CCT50CCA (2554T>A), CTG52TTG (2558C>T), TTG53CTC (2561T>C 2563G>C), TCA56TCG (2572A>G), CCT60CCC (2584T>C), GGC61GGT (2587C>T), CTG63CTC (2593G>C), GCT65GCC (2599T>C), AAT68AAC (2608T>C), TTT72TTC (2620T>C), GCG75GCA (2629G>A), GTG76GTA (2632G>A), AGG77AAG (2634G>A), TTG78TTA (2638G>A), ATT80ATA (2644T>A), AAA83AAG (2653A>G), GCT87GCC (2665T>C), CTA92TTA (2678C>T), CTA93CTG (2683A>G), GCT95GCC (2689T>C), CTA98CTC (2698A>C), TCA101TCG (2707A>G), GCG102GCA (2710G>A), GAA104GAG (2716A>G), ACA105ACG (2719A>G), CTC106CTT (2722C>T), GTC107ATT (2723G>A 2725C>T), GCA108GCG (2728A>G), AGG110AGA (2734G>A), ATT111ATA (2737T>A), GAT114GAC (2746T>C), GCT115ACA (2747G>A 2749T>A), GCA116GCT (2752A>T), GCT117GCA (2755T>A), GAC119GAT (2761C>T), CTC120CTT (2764C>T), TGT125TGC (2779T>C), GGG127GGT (2785G>T), AAC129AAT (2791C>T), CAT131AAT (2795C>A), TAC134TAT (2806C>T), TTG138CTC (2816T>C 2818G>C), GTG140ATA (2822G>A 2824G>A), TTG141TTA (2827G>A), GTT144GTC (2836T>C), AAG145AAA (2839G>A), AAT146AAC (2842T>C), ACC153ACA (2863C>A), CTA154TTA (2864C>T), CCA160ACA (2882C>A), AGT161CAT (2885A>C 2886G>A), AAC163AAT (2893C>T), ACG164ATG (2895C>T), GTC166GTT (2902C>T), GTA167GTG (2905A>G), ATT169ATC (2911T>C), AAT173AAC (2923T>C), AAT176AGT (2931A>G), AAC179AGC (2940A>G), GCT181GCC (2947T>C), ACT186ACC (2962T>C), GAA189_AAT190insA-- (2971_2972insA), GGT195GGC (2989T>C), ACT196ATT (2991C>T), CAT197CGT (2994A>G), GAG199AGG (2999G>A 3000A>G), TCC204TC- (3016delC), CGG206CAG (3021G>A), TAC207TTT (3024A>T 3025C>T), ACA211ATA (3036C>T), GAA212GGC (3039A>G 3040A>C), AAG213ACA (3042A>C 3043G>A), TTG216TTA (3052G>A), TTA219TTG (3061A>G), GCG226GCA (3082G>A), GGC229GGT (3091C>T), GCC232GTA (3099C>T 3100C>A), GTG234AGA (3104G>A 3105T>G 3106G>A), GTC235GTT (3109C>T), AAA236AAG (3112A>G), ACT238ACC (3118T>C), GCC239GCT (3121C>T), GGA241GGG (3127A>G), AGC243AAC (3132G>A), CGA245AAG (3137C>A 3138G>A 3139A>G), TAT249TAC (3151T>C), TTC250ATT (3152T>A 3154C>T), TGT254TAT (3165G>A), GTA255GTG (3169A>G), ATC265ATT (3199C>T), AGG267CGG (3203A>C), CGT268CGC (3208T>C), CCA270CCG (3214A>G), ACC271ACT (3217C>T), TTT272CTT (3218T>C), GTC273GTA (3223C>A), ACT274AAT (3225C>A), GTT276ATT (3230G>A), AAC277AAT (3235C>T), TTA285CTA (3257T>C), ACC286ACT (3262C>T), AGC287AGT (3265C>T), GTA288ATG (3266G>A 3268A>G), GTG289GTA (3271G>A), GTG291ATG (3275G>A), GCA294ACA (3284G>A), CAG299CAA (3301G>A), AGA304AAG (3315G>A 3316A>G), GAA305GCA (3318A>C), ATC306ATT (3322C>T), GAG307GAA (3325G>A), TTG308TTA (3328G>A), ATA311ATT (3337A>T), GAG315GAA (3349G>A), GTC317GTT (3355C>T), TGC318TGT (3358C>T), ATG319GTG (3359A>G), CTA321CTG (3367A>G), TAT322TAC (3370T>C), TTA325CTG (3377T>C 3379A>G), GAA330GAG (3394A>G), CGC331CGT (3397C>T), GAT332GAG (3400T>G), ACA334ACG (3406A>G), TAC338TAT (3418C>T), ATT339ATA (3421T>A), TAT341TAC (3427T>C), AAG343AAA (3433G>A), TCA344TCT (3436A>T), GTA345GTG (3439A>G), AAC346AAT (3442C>T), CCT347CCC (3445T>C), CAG351CAA (3457G>A), TTT352TTC (3460T>C), GCT353GCA (3463T>A), GCG355GCT (3469G>T), CAG356CAA (3472G>A), CAC364CAT (3496C>T), CAG365CAA (3499G>A) | | | | | | | | |
| 6K1 protein (NP_734245.1) | 1 | 52 | 100% | 305 | 90.5% | 52 (100%) | 45 (86.5%) | 0/0/0/0 | 0 |
| Protein mutations: | V5G (3513T>G), M16V (3545A>G), A17T (3548G>A), V19I (3554G>A 3556C>T), L42V (3623C>G), L45M (3632C>A), Y47H (3638T>C) | | | | | | | | |
| Codon mutations: | TCC2TCT (3505C>T), GTT5GGT (3513T>G), AAA7AAG (3520A>G), GAA10GAG (3529A>G), CAA11CAG (3532A>G), ATG16GTG (3545A>G), GCT17ACT (3548G>A), TTA18CTA (3551T>C), GTC19ATT (3554G>A 3556C>T), AGT28AGC (3583T>C), GAT29GAC (3586T>C), TGC30TGT (3589C>T), TTC32TTT (3595C>T), AAT36AAC (3607T>C), TTT38TTC (3613T>C), AAG39AAA (3616G>A), GGT40GGC (3619T>C), CTT42GTT (3623C>G), TCC43TCT (3628C>T), CTG45ATG (3632C>A), GAT46GAC (3637T>C), TAT47CAT (3638T>C), GAA48GAG (3643A>G) | | | | | | | | |
| CI protein (NP_734246.1) | 1 | 634 | 100% | 4190 | 97.9% | 634 (100%) | 608 (95.9%) | 0/0/0/0 | 0 |
| Protein mutations: | I15V (3698A>G), S21N (3717G>A), R26K (3731C>A 3732G>A), I140V (4073A>G), F158Y (4128T>A 4129C>T), A166S (4151G>T), V171I (4166G>A), L190I (4223C>A 4225G>A), I222V (4319A>G 4321A>G), F247Y (4395T>A), V251I (4406G>A), M274V (4475A>G 4477G>T), T276S (4481A>T 4483A>G), A297L (4544G>T 4545C>T 4546G>A), R298K (4548G>A), V342I (4679G>A 4681G>C), V381I (4796G>A 4798C>T), T383S (4803C>G 4804T>C), S468G (5057A>G), T469A (5060A>G), L479I (5090C>A), A482V (5100C>T), E484D (5107A>T), V489F (5120G>T 5122C>T), I491V (5126A>G), V510I (5183G>A) | | | | | | | | |
| Codon mutations: | TTA2TTG (3661A>G), GTG5GTA (3670G>A), TTT9TTC (3682T>C), GAG11GAA (3688G>A), GAG14GAA (3697G>A), ATT15GTT (3698A>G), GAT17GAC (3706T>C), TTT18TTC (3709T>C), TTG20CTA (3713T>C 3715G>A), AGT21AAT (3717G>A), GAG22GAA (3721G>A), CGA26AAA (3731C>A 3732G>A), ACT27ACA (3736T>A), CTA31TTG (3746C>T 3748A>G), ACA33ACG (3754A>G), AGT36AGC (3763T>C), CGA41CGG (3778A>G), CAA42CAG (3781A>G), ATC43ATT (3784C>T), CAT47CAC (3796T>C), CCA50CCT (3805A>T), TAC52TAT (3811C>T), AGA53AGG (3814A>G), TTC58TTT (3829C>T), TTT61TTC (3838T>C), AGA63AGG (3844A>G), GCA64GCC (3847A>C), GTT67GTA (3856T>A), GTG69GTT (3862G>T), GAC72GAT (3871C>T), GCC74GCA (3877C>A), AGC76AGT (3883C>T), GAA77GAG (3886A>G), CTA79CTG (3892A>G), GAC80GAT (3895C>T), TTT81TTC (3898T>C), GTA83GTG (3904A>G), CGG84AGA (3905C>A 3907G>A), GCT86GCC (3913T>C), AAG91AAA (3928G>A), TCA92TCT (3931A>T), GGG94GGA (3937G>A), AGT100AGC (3955T>C), GTG101GTA (3958G>A), GCC102GCT (3961C>T), GGA103GGG (3964A>G), TCT104TCC (3967T>C), TTA107TTG (3976A>G), ATT108ATC (3979T>C), CCA110CCT (3985A>T), CTA114CTT (3997A>T), GCG115GCA (4000G>A), AAC117AAT (4006C>T), GTT118GTG (4009T>G), TTC119TTT (4012C>T), AAA120AAG (4015A>G), TCT123TCC (4024T>C), GAA125GAG (4030A>G), CCA126CCT (4033A>T), AAG129AAA (4042G>A), CTG133TTA (4052C>T 4054G>A), CGT134CGC (4057T>C), CGT136CGA (4063T>A), GGA137GGG (4066A>G), ATA140GTA (4073A>G), TTT141TTC (4078T>C), GGC142GGT (4081C>T), TCT143TCC (4084T>C), TCC147TCT (4096C>T), AGC151AGT (4108C>T), GGA152GGT (4111A>T), GCG154GCA (4117G>A), CTA155TTG (4118C>T 4120A>G), TAC157TAT (4126C>T), TTC158TAT (4128T>A 4129C>T), GCC159GCT (4132C>T), AAT160AAC (4135T>C), CAA164CAG (4147A>G), TTA165CTA (4148T>C), GCT166TCT (4151G>T), AAC169AAT (4162C>T), GTA171ATA (4166G>A), ATA172ATC (4171A>C), GAG175GAA (4180G>A), CTG179TTG (4190C>T), CCT181CCC (4198T>C), TCC182TCT (4201C>T), GCG185GCT (4210G>T), TTC186TTT (4213C>T), AGT188AGC (4219T>C), CTG189TTG (4220C>T), CTG190ATA (4223C>A 4225G>A), GTT192GTG (4231T>G), CAT194CAC (4237T>C), GTA199GTG (4252A>G), TTA200CTG (4253T>C 4255A>G), AAA201AAG (4258A>G), ACT205ACC (4270T>C), AGA209AGG (4282A>G), GAG210GAA (4285G>A), GAA212GAG (4291A>G), CAG216CAA (4303G>A), GTC219GTT (4312C>T), ATA222GTG (4319A>G 4321A>G), GTG223GTT (4324G>T), GAG224GAA (4327G>A), CTG227CTT (4336G>T), TCT228TCC (4339T>C), TCA231TCT (4348A>T), GCA235GCG (4360A>G), GGT237GGC (4366T>C), TCT238TCA (4369T>A), ACT240ACC (4375T>C), GTT244GTC (4387T>C), TTT247TAT (4395T>A), GTA251ATA (4406G>A), CTT252CTC (4411T>C), GTT261GTC (4438T>C), ACC263ACA (4444C>A), GCC265GCT (4450C>T), CTC267CTT (4456C>T), GAC270GAT (4465C>T), ATG274GTT (4475A>G 4477G>T), GTC275GTT (4480C>T), ACA276TCG (4481A>T 4483A>G), GGC280GGT (4495C>T), AAG284AAA (4507G>A), CAC285CAT (4510C>T), GGT286GGG (4513T>G), CTA288TTA (4517C>T), GTC291GTT (4528C>T), AAA293AAG (4534A>G), ACC295ACT (4540C>T), GCG297TTA (4544G>T 4545C>T 4546G>A), AGA298AAA (4548G>A), TTT301TTC (4558T>C), GTA303GTT (4564A>T), ACC305ACT (4570C>T), ATA307ATT (4576A>T), TTG314TTA (4597G>A), GAC315GAT (4600C>T), GAC317GAT (4606C>T), GTG318GTA (4609G>A), GTG320GTA (4615G>A), GAC321GAT (4618C>T), TTG324CTT (4625T>C 4627G>T), GAC331GAT (4648C>T), ATT332ATC (4651T>C), AGG335AGA (4660G>A), AGC336AGT (4663C>T), GCT338GCA (4669T>A), GTG342ATC (4679G>A 4681G>C), AGT343AGC (4684T>C), GTT344GTG (4687T>G), GAG348GAA (4699G>A), AGA349AGG (4702A>G), CAA351CAG (4708A>G), AGG352AGA (4711G>A), CTG353TTG (4712C>T), AAA361AAG (4738A>G), GTA363GTT (4744A>T), GCA364GCG (4747A>G), TTG365CTG (4748T>C), ACT370ACC (4765T>C), GTC381ATT (4796G>A 4798C>T), ACT383AGC (4803C>G 4804T>C), GAG384GAA (4807G>A), GCG385GCT (4810G>T), GCT386GCG (4813T>G), CTT387CTC (4816T>C), TAT392TAC (4831T>C), AAC393AAT (4834C>T), CCA395CCT (4840A>T), GGA399GGG (4852A>G), GGC400GGT (4855C>T), AGT404AGC (4867T>C), CTG405CTC (4870G>C), ATT406ATA (4873T>A), TGT409TGC (4882T>C), ACA416ACT (4903A>T), CAG418CAA (4909G>A), TTT420TTC (4915T>C), GAA421GAG (4918A>G), TTG422CTA (4919T>C 4921G>A), AGT423AGC (4924T>C), CCC424CCG (4927C>G), TTT426TTC (4933T>C), ATC427ATA (4936C>A), CAG428CAA (4939G>A), AAT429AAC (4942T>C), TTC430TTT (4945C>T), TCA436TCG (4963A>G), CCT439CCC (4972T>C), CTT445CTC (4990T>C), AAA449AAG (5002A>G), CCT456CCC (5023T>C), TTG457CTG (5024T>C), TGC458TGT (5029C>T), GAT459GAC (5032T>C), TCT461TCC (5038T>C), CCA463CCT (5044A>T), GCA466GCT (5053A>T), TCG467TCA (5056G>A), AGT468GGT (5057A>G), ACT469GCT (5060A>G), TCG472TCT (5071G>T), CTT479ATT (5090C>A), GGA480GGG (5095A>G), GTG481GTT (5098G>T), GCC482GTC (5100C>T), TTA483TTG (5104A>G), GAA484GAT (5107A>T), ATT485ATC (5110T>C), CCA486CCG (5113A>G), AAG487AAA (5116G>A), CAA488CAG (5119A>G), GTC489TTT (5120G>T 5122C>T), ATT491GTT (5126A>G), CAT494CAC (5137T>C), ATC495ATT (5140C>T), AAA496AAG (5143A>G), GAG497GAA (5146G>A), CTC502CTA (5161C>A), GAA504GAG (5167A>G), CTT506CTG (5173T>G), ACG509ACA (5182G>A), GTT510ATT (5183G>A), GTC511GTT (5188C>T), AAG512AAA (5191G>A), AAA514AAG (5197A>G), GAC515GAT (5200C>T), TGC517TGT (5206C>T), TTA518TTG (5209A>G), AGC521AGT (5218C>T), TCG525TCA (5230G>A), ATC527ATT (5236C>T), AGC528AGT (5239C>T), ATC530ATT (5245C>T), ACA536ACT (5263A>T), CTC538CTT (5269C>T), TTC539TTT (5272C>T), GCC540GCA (5275C>A), ATC541ATT (5278C>T), CCA542CCC (5281A>C), AGA543AGG (5284A>G), ACT544ACC (5287T>C), CTA545TTA (5288C>T), ATA546ATT (5293A>T), TTG547CTA (5294T>C 5296G>A), GAG549GAA (5302G>A), CTT552CTA (5311T>A), GAA553GAG (5314A>G), CGA556CGT (5323A>T), AGT564AGC (5347T>C), ATC566ATT (5353C>T), TGC570TGT (5365C>T), GTT577GTC (5386T>C), ACC580ACA (5395C>A), AAA589AAG (5422A>G), GAT590GAC (5425T>C), ACC592ACT (5431C>T), GAG594GAA (5437G>A), CTT599CTC (5452T>C), AAG601AAA (5458G>A), AGT604AGC (5467T>C), CAA605CAG (5470A>G), CTA606TTG (5471C>T 5473A>G), GAA608GAG (5479A>G), TTC609TTT (5482C>T), TCA610TCC (5485A>C), AAT611AAC (5488T>C), TTG612CTA (5489T>C 5491G>A), GGT614GGC (5497T>C), TGT617TGC (5506T>C), GAG618GAA (5509G>A), GAG619GAA (5512G>A), TTA621CTA (5516T>C), TAT625TAC (5530T>C), GAG626GAA (5533G>A), TCG627TCT (5536G>T), TTG628TTA (5539G>A), GTT631GTG (5548T>G), CAC633CAT (5554C>T) | | | | | | | | |
| 6K2 protein (NP_734247.1) | 1 | 52 | 100% | 303 | 93.5% | 52 (100%) | 46 (88.5%) | 0/0/0/0 | 0 |
| Protein mutations: | A2T (5561G>A 5563G>A), I14V (5597A>G), N16K (5605C>A), A21V (5619C>T 5620T>G), I25L (5630A>C), V46I (5693G>A) | | | | | | | | |
| Codon mutations: | GCT1GCA (5560T>A), GCG2ACA (5561G>A 5563G>A), ACG3ACC (5566G>C), TCA4TCG (5569A>G), CTC9CTG (5584C>G), AAG12AAA (5593G>A), GGG13GGA (5596G>A), ATT14GTT (5597A>G), AAC16AAA (5605C>A), GTG20GTT (5617G>T), GCT21GTG (5619C>T 5620T>G), TTG24TTA (5629G>A), ATC25CTC (5630A>C), GGC28GGT (5641C>T), GCA31GCT (5650A>T), ATA35ATT (5662A>T), GGA36GGG (5665A>G), CAA44CAG (5689A>G), GTT46ATT (5693G>A), GAG47GAA (5698G>A), CAA52CAG (5713A>G) | | | | | | | | |
| NIa-VPg protein (NP_734252.1) | 1 | 188 | 100% | 1227 | 93.3% | 188 (100%) | 170 (90.4%) | 0/0/0/0 | 0 |
| Protein mutations: | K47R (5853A>G), I61V (5894A>G 5896C>G), R83A (5960C>G 5961G>C 5962G>C), I93V (5990A>G), E102D (6019A>T), K106T (6030A>C), D112E (6049C>A), M115I (6058G>A), S120N (6072G>A 6073T>C), N121H (6074A>C), T123N (6081C>A), C133S (6111G>C 6112T>C), I139V (6128A>G 6130T>C), V149I (6158G>A), T153S (6170A>T 6172A>G), L164F (6203C>T 6205C>T), V173I (6230G>A), A182K (6257G>A 6258C>A) | | | | | | | | |

| | Begin | End | Coverage | Score | Concordance | Matches | Identities | I/D/M/F* | Stop Codons |
|---|---|---|---|---|---|---|---|---|---|
| **NT** | **5** | **9704** | **99.9%** | **12522** | **64.8%** | **9688 (99.7%)** | **8015 (82.5%)** | **15/12** | |
| Codon mutations: | GGG1GGC (5716G>C), AAA2AAG (5719A>G), GCC10GCA (5743C>A), CGC14CGA (5755C>A), CAT15CAC (5758T>C), CGT17CGC (5764T>C), AAA19AAG (5770A>G), GGC22GGT (5779C>T), TTT23TTC (5782T>C), GAC26GAT (5791C>T), GAC30GAT (5803C>T), AGG41AGA (5836G>A), AAA42AAG (5839A>G), GGA44GGG (5845A>G), AAA45AAG (5848A>G), AAA47AGA (5853A>G), GGT48GGC (5857T>C), ACC49ACT (5860C>T), ACA50ACT (5863A>T), GGT52GGA (5869T>A), GGT54GGC (5875T>C), AGC57AGT (5884C>T), TTC60TTT (5893C>T), ATC61GTG (5894A>G 5896C>G), AAC62AAT (5899C>T), GGG65GGA (5908G>A), TTT66TTC (5911T>C), GAG70GAA (5923G>A), CAA75CAG (5938A>G), TTC76TTT (5941C>T), GTT77GTC (5944T>C), GAT78GAC (5947T>C), GGG82GGA (5959G>A), CGG83GCC (5960C>G 5961G>C 5962G>C), GAA87GAG (5974A>G), GCT91GCA (5986T>A), GAC92GAT (5989C>T), ATT93GTT (5990A>G), GAT95GAC (5998T>C), AGA99CGT (6008A>C 6010A>T), GAA102GAT (6019A>T), GTG103GTT (6022G>T), CGA104CGT (6025A>T), AAA106ACA (6030A>C), GAC112GAA (6049C>A), ATG115ATA (6058G>A), CAA116CAG (6061A>G), GCC117GCA (6064C>A), GGT119GGC (6070T>C), AGT120AAC (6072G>A 6073T>C), AAC121CAC (6074A>C), ACG122ACA (6079G>A), ACC123AAC (6081C>A), ATA124ATT (6085A>T), GCA126GCT (6091A>T), TAC127TAT (6094C>T), TGT133TCC (6111G>C 6112T>C), AAA135AAG (6118A>G), TTG137TTA (6124G>A), AAG138AAA (6127G>A), ATT139GTC (6128A>G 6130T>C), GAT140GAC (6133T>C), TTA141TTG (6136A>G), AAC145AAT (6148C>T), AAA148AAG (6157A>G), GTT149ATT (6158G>A), TGT150TGC (6163T>C), AAA152AAG (6169A>G), ACA153TCG (6170A>T 6172A>G), GGC155GGT (6178C>T), AAA158AAG (6187A>G), TTT159TTC (6190T>C), GAG161GAA (6196G>A), CTC164TTT (6203C>T 6205C>T), GAA165GAG (6208A>G), CTA166TTG (6209C>T 6211A>G), CAG168CAA (6217G>A), GCT172GCA (6229T>A), GTA173ATA (6230G>A), GTC175GTT (6238C>T), AAG178AAA (6247G>A), ATA180ATT (6253A>T), GCA182AAA (6257G>A 6258C>A), CAT187CAC (6274T>C) | | | | | | | | |
| NIa-Pro protein (NP_734248.1) | 1 | 244 | 100% | 1654 | 93.7% | 244 (100%) | 223 (91.4%) | 0/0/0/0 | 0 |
| Protein mutations: | K2R (6282A>G), A29T (6362G>A), V42I (6401G>A), A43V (6405C>T), Y51F (6429A>T), Q58R (6450A>G), H70N (6485C>A 6487C>T), L84I (6527C>A), T104I (6588C>T), Y120H (6635T>C), I124V (6647A>G), I134V (6677A>G 6679A>G), D147N (6716G>A), N148D (6719A>G), I163L (6764A>T 6766A>G), A173V (6795C>T 6796A>G), H174Q (6799C>A), N194D (6857A>G), V200T (6875G>A 6876T>C 6877C>A), D219E (6934C>A), V242I (7001G>A 7003G>A) | | | | | | | | |
| Codon mutations: | AAA2AGA (6282A>G), TCG3TCA (6286G>A), GGC7GGT (6298C>T), TTG8TTA (6301G>A), AGA9AGG (6304A>G), GAC10GAT (6307C>T), GCC15GCT (6322C>T), ACA17ACT (6328A>T), GTT18GTG (6331T>G), TGT19TGC (6334T>C), AGG20AGA (6337G>A), CTG21TTG (6338C>T), TAT27TAC (6358T>C), GGG28GGA (6361G>A), GCA29ACA (6362G>A), TCA30TCT (6367A>T), GAG31GAA (6370G>A), TAC33TAT (6376C>T), GGT34GGG (6379T>G), GGA36GGC (6385A>C), GGA38GGT (6391A>T), GCA39GCG (6394A>G), TAC40TAT (6397C>T), GTA42ATA (6401G>A), GCG43GTG (6405C>T), CAC45CAT (6412C>T), TTA47TTG (6418A>G), TTT48TTC (6421T>C), TAC51TTC (6429A>T), GGT53GGA (6436T>A), GAG56GAA (6445G>A), CAA58CGA (6450A>G), TCC59TCG (6454C>G), CAC61CAT (6460C>T), GGT62GGA (6463T>A), TTC64TTT (6469C>T), AGG65AGA (6472G>A), GTG66GTA (6475G>A), AAT68AAC (6481T>C), CAC70AAT (6485C>A 6487C>T), AGT71AGC (6490T>C), AGC73AGT (6496C>T), CTG75CTA (6502G>A), GGT79GGC (6514T>C), AGG80AGA (6517G>A), CTC84ATC (6527C>A), ATC85ATA (6532C>A), CCG88CCA (6541G>A), AAA89AAG (6544A>G), GTC93GTT (6556C>T), TTT94TTC (6559T>C), CAG96CAA (6565G>A), TTG98CTA (6569T>C 6571G>A), CCT103CCA (6586T>A), ACA104ATA (6588C>T), GAA107GAG (6598A>G), AGA108AGG (6601A>G), ATT109ATC (6604T>C), TGT110TGC (6607T>C), GGA113GGG (6616A>G), AAC115AAT (6622C>T), GAG118GAA (6631G>A), AAG119AAA (6634G>A), TAT120CAT (6635T>C), GCT121GCA (6640T>A), TCG123TCT (6646G>T), ATC124GTC (6647A>G), ATC125ATT (6652C>T), AGC129AGT (6664C>T), ACT131ACG (6670T>G), ATA134GTG (6677A>G 6679A>G), GGC136GGT (6685C>T), ACA138ACT (6691A>T), TTC139TTT (6694C>T), ACA146ACG (6715A>G), GAT147AAT (6716G>A), AAT148GAT (6719A>G), GGA149GGG (6724A>G), CTA153TTA (6734C>T), GTG155GTA (6742G>A), GTG156GTA (6745G>A), GCC159GCT (6754C>T), GGA161GGG (6760A>G), ATA163TTG (6764A>T 6766A>G), ATT166ATA (6775T>A), CTG169TTG (6782C>T), GCA170GCT (6787A>T), AAC171AAT (6790C>T), GCA173GTG (6795C>T 6796A>G), CAC174CAA (6799C>A), ACC175ACA (6802C>A), AAC177AAT (6808C>T), TAC178TAT (6811C>T), TAC179TAT (6814C>T), TTC182TTT (6823C>T), GAA184GAG (6829A>G), TAC190TAT (6847C>T), CGA192CGC (6853A>C), AAT194GAT (6857A>G), GAG195GAA (6862G>A), GAA198GAG (6871A>G), GTC200ACA (6875G>A 6876T>C 6877C>A), AAG201AAA (6880G>A), TCT202TCA (6883T>A), GTT204GTG (6889T>G), GAC208GAT (6901C>T), GGC213GGT (6916C>T), CCG214CCA (6919G>A), AAA216AAG (6925A>G), CTT217CTC (6928T>C), GAC219GAA (6934C>A), GGG224GGC (6949G>C), TTC226TTT (6955C>T), AAA227AAG (6958A>G), CAA233CAG (6976A>G), CTA235TTA (6980C>T), ATC236ATT (6985C>T), GTA240GTT (6997A>T), GTG241GTT (7000G>T), GTG242ATA (7001G>A 7003G>A) | | | | | | | | |
| NIb protein (NP_734249.1) | 1 | 521 | 100% | 3129 | 86.1% | 519 (99.4%) | 480 (92.0%) | 1/2/2/2 | 0 |
| Protein mutations: | F8Y (7032T>A), T39K (7125C>A), A46S (7145G>T 7147A>G), E51_A52del (7160_7165delGAGGCA), D69E (7216T>A), D83E (7258T>A), D88_R89insX (7273_7274insT), R89V (7274C>G 7275G>T), H91L (7281A>T), L92S (7284T>C), S96L (7296C>T), S101F (7311C>T), T102I (7314C>T), C103Y (7317G>A), N104K (7321T>A), K109Q (7334A>C), A112N (7343G>A 7344C>A 7345A>C), S133M (7407G>T 7408T>G), C136G (7415T>G), L165S (7503T>C 7504G>A), D234N (7709G>A), K235R (7713A>G 7714A>G), T268I (7812C>T), V279L (7844G>T 7846G>A), V292I (7883G>A), S345N (8043G>A), N381S (8151A>G 8152C>T), R390K (8178G>A), R400K (8208G>A), P429A (8294C>G 8296A>G), S447F (8349C>T), A467S (8408G>T), M481I (8452G>A), R484K (8460G>A), A491T (8480G>A 8482T>A), R498K (8502G>A 8503A>G), E502D (8515A>T), E510D (8539G>C), L513C (8546C>T 8547T>G), S515T (8552T>A) | | | | | | | | |
| Codon mutations: | GCT1GCC (7012T>C), AAG2AAA (7015G>A), CAT3CAC (7018T>C), TTT8TAT (7032T>A), GCC10GCT (7039C>T), TTG11CTA (7040T>C 7042G>A), GGA13GGG (7048A>G), GCT17GCA (7060T>A), GTC18GTA (7063C>A), AGC23AGT (7078C>T), TTA25CTG (7082T>C 7084A>G), GTA26GTG (7087A>G), ACC27ACA (7090C>A), CAT29CAC (7096T>C), GTA30GTG (7099A>G), GTT31GTC (7102T>C), GAG34GAA (7111G>A), CGA36CGG (7117A>G), ACA39AAA (7125C>A), GAA40GAG (7129A>G), TTT41TTC (7132T>C), CTG42TTA (7133C>T 7135G>A), GTG44GTA (7141G>A), GAT45GAC (7144T>C), GCA46TCG (7145G>T 7147A>G), GAG47GAA (7150G>A), GCA50GCT (7159A>T), GAG51_GCA52del (7160_7165delGAGGCA), TTC54TTT (7171C>T), GCG60GCT (7189G>T), AAA63AAG (7198A>G), AGC64AGT (7201C>T), CTA66TTA (7205C>T), AAT67AAC (7210T>C), GAT69GAA (7216T>A), GCG70GCA (7219G>A), ATC72ATA (7225C>A), AAG73AAA (7228G>A), AAG77AAA (7240G>A), TAT78TAC (7243T>C), AAA80AAG (7249A>G), CCT81CCG (7252T>G), GAT83GAA (7258T>A), GTT84GTC (7261T>C), GGT85GGA (7264T>A), GTC86GTG (7267C>G), GTG87GTA (7270G>A), GAT88_CGG89insT-- (7273_7274insT), CGG89GTG (7274C>G 7275G>T), CAT91CTT (7281A>T), TTG92TCG (7284T>C), TCA96TTA (7296C>T), TCT101TTT (7311C>T), ACC102ATC (7314C>T), TGC103TAC (7317G>A), AAT104AAA (7321T>A), GTG105-TG (7322delG), CAC106CAT (7327C>T), GGC107GGT (7330C>T), TTC108TTT (7333C>T), AAG109CAG (7334A>C), TGT111TGC (7342T>C), GCA112AAC (7343G>A 7344C>A 7345A>C), ACT115ACC (7354T>C), CAA118CAG (7363A>G), ATT120ATC (7369T>C), TTC121TTT (7372C>T), GCG123GCT (7378G>T), CTC124CTT (7381C>T), GCA129GCT (7396A>T), GTC130GTT (7399C>T), GCC132GCT (7405C>T), AGT133ATG (7407G>T 7408T>G), GGG135GGT (7414G>T), TGC136GGC (7415T>G), AAA137AAG (7420A>G), GAC140GAT (7429C>T), TAT141TAC (7432T>C), CAT144CAC (7441T>C), AAG150AAA (7459G>A), GAA151GAG (7462A>G), ATA153ATT (7468A>T), GTC154GTA (7471C>A), CAA156CAG (7477A>G), AGC157AGT (7480C>T), CTG159CTT (7486G>T), TAT162TAC (7495T>C), AAA163AAG (7498A>G), GGT164GGC (7501T>C), TTG165TCA (7503T>C 7504G>A), ATT168ATA (7513T>A), AAC170AAT (7519C>T), GGA171GGG (7522A>G), AAG174AAA (7531G>A), GAG176GAA (7537G>A), CTC177CTT (7540C>T), TGT179TGC (7546T>C), AAG187AAA (7570G>A), ACG188ACA (7573G>A), ACG190ACA (7579G>A), GCT193GCC (7588T>C), CTA196TTA (7595C>T), TTG199CTA (7604T>C 7606G>A), GGT202GGA (7615T>A), TGT205TGC (7624T>C), GAC208GAT (7633C>T), TTC209TTT (7636C>T), AAT210AAC (7639T>C), TTT213TTC (7648T>C), GAA219GAG (7666A>G), TGT221TGC (7672T>C), ACA223ACT (7678A>T), GGG225GGA (7684G>A), ACT227ACC (7690T>C), TTT229TTC (7696T>C), GGT231GGG (7702T>G), GAT234AAT (7709G>A), AAA235AGG (7713A>G 7714A>G), CTG236TTG (7715C>T), CGT239CGC (7726T>C), GAG242GAA (7735G>A), GTA245GTG (7744A>G), TAC246TAT (7747C>T), GGC251GGT (7762C>T), CAG253CAA (7768G>A), TTT254TTC (7771T>C), TCA257TCG (7780A>G), CTA258CTC (7783A>C), ACT259ACC (7786T>C), CTA262TTG (7793C>T 7795A>G), ATC263ATT (7798C>T), GCT265GCC (7804T>C), ACC268ATC (7812C>T), GAA275GAG (7834A>G), GAC276GAT (7837C>T), GTG279TTA (7844G>T 7846G>A), CAG282CAA (7855G>A), CTG284CTA (7861G>A), TTA287TTG (7870A>G), TAC288TAT (7873C>T), ACT289ACA (7876T>A), GAG290GAA (7879G>A), ATT291ATA (7882T>A), GTT292ATT (7883G>A), CCA295CCT (7894A>T), ATT296ATC (7897T>C), GTC304GTT (7921C>T), AAG306AAA (7927G>A), GGT309GGC (7936T>C), AAT310AAC (7939T>C), AAT311AAC (7942T>C), GTG319GTA (7966G>A), GAT320GAC (7969T>C), TCT322TCC (7975T>C), GTC326GTT (7987C>T), CTT327CTC (7990T>C), GCT332GCC (8005T>C), GAA341GAG (8032A>G), ATC343ATT (8038C>T), GAC344GAT (8041C>T), AGC345AAC (8043G>A), TTG357TTA (8080G>A), GTG360GTA (8089G>A), CCG362CCA (8095G>A), GAG363GAA (8098G>A), AGC366AGT (8107C>T), ATT367ATC (8110T>C), GAT369GAC (8116T>C), CAA373CAG (8128A>G), GGT379GGC (8146T>C), AAC381AGT (8151A>G 8152C>T), TCG385TCA (8164G>A), AGG390AAG (8178G>A), AAG391AAA (8182G>A), TTG394TTA (8191G>A), TTC396TTT (8197C>T), TCC398TCT (8203C>T), AGA400AAA (8208G>A), CTG402TTA (8213C>T 8215G>A), CTA403CTG (8218A>G), ATC404ATT (8221C>T), GAG405GAA (8224G>A), CCA410CCG (8239A>G), GAG415GAA (8254G>A), ATT417ATC (8260T>C), GTA418GTG (8263A>G), GAT424GAC (8281T>C), AGA425AGG (8284A>G), CCA429GCG (8294C>G 8296A>G), GAG430GAA (8299G>A), CAC431CAT (8302C>T), ATT436ATC (8317T>C), TGC437TGT (8320C>T), GCT439GCA (8326T>A), GAG442GAA (8335G>A), TCC443TCT (8338C>T), TCT447TTT (8349C>T), GAA448GAG (8353A>G), CAC451CAT (8362C>T), CAA452CAG (8365A>G), ATC453ATT (8368C>T), TAC457TAT (8380C>T), TCA458TCG (8383A>G), TTA460TTG (8389A>G), CCT465CCC (8404T>C), GCA467TCA (8408G>T), ACA468ACG (8413A>G), GCG470GCT (8419G>T), GCT475GCC (8434T>C), TAT477TAC (8440T>C), ATG481ATA (8452G>A), CTA483CTG (8458A>G), AGG484AAG (8460G>A), AAA485AAG (8464A>G), CTG486CTA (8467G>A), AGG490AGA (8479G>A), GCT491ACA (8480G>A 8482T>A), GAG496GAA (8497G>A), CTA497TTG (8498C>T 8500A>G), AGA498AAG (8502G>A 8503A>G), GCC499GCT (8506C>T), GAA502GAT (8515A>T), GTC505GTT (8524C>T), GCA506GCC (8527A>C), TTA507CTG (8528T>C 8530A>G), GAG510GAC (8539G>C), CTT513TGT (8546C>T 8547T>G), TCT515ACT (8552T>A), GTA518GTG (8563A>G) | | | | | | | | |
| coat protein (NP_734250.1) | 1 | 267 | 100% | 1652 | 90.6% | 267 (100%) | 243 (91.0%) | 0/0/0/0 | 0 |
| Protein mutations: | A1G (8574C>G), I5V (8585A>G), G9E (8598G>A), N11S (8603A>T 8604A>C 8605C>T), P17Q (8622C>A), E18G (8625A>G 8626G>A), P26F (8648C>T 8649C>T 8650G>C), G29E (8658G>A), D31A (8664A>C 8665T>G), A36V (8679C>T 8680A>T), T58K (8745C>A 8746A>G), E95G (8856A>G 8857G>A), R98Q (8865G>A), M99L (8867A>C 8869G>T), G104E (8883G>A), T106S (8888A>T), V128I (8954G>A), N138D (8984A>G 8986T>C), E139V (8988A>T), I187V (9131A>G), M193G (9149A>G 9150T>G 9151G>A), G194S (9152G>A), P230S (9260C>T 9262T>C), Q242H (9298A>C) | | | | | | | | |
| Codon mutations: | GCA1GGA (8574C>G), ATT5GTT (8585A>G), GGA9GAA (8598G>A), AGC10AGT (8602C>T), AAC11TCT (8603A>T 8604A>C 8605C>T), AAA16AAG (8620A>G), CCA17CAA (8622C>A), GAG18GGA (8625A>G 8626G>A), CAG19CAA (8629G>A), GGC20GGT (8632C>T), ATC22ATT (8638C>T), CAG23CAA (8641G>A), AAC25AAT (8647C>T), CCG26TTC (8648C>T 8649C>T 8650G>C), AAC27AAT (8653C>T), AAA28AAG (8656A>G), GGA29GAA (8658G>A), GAT31GCG (8664A>C 8665T>G), GTT34GTG (8674T>G), GCA36GTT (8679C>T 8680A>T), GGC37GGA (8683C>A), GGG40GGA (8692G>A), ACA41ACT (8695A>T), CCG45CCA (8707G>A), AGA46CGA (8708A>C), AAG48AAA (8716G>A), ATC50ATT (8722C>T), ACG51ACA (8725G>A), AAA53AAG (8731A>G), AGA55AGG (8737A>G), CCC57CCT (8743C>T), ACA58AAG (8745C>A 8746A>G), AAG60AAA (8752G>A), GGA61GGT (8755A>T), AAC66AAT (8770C>T), TTA67CTA (8771T>C), GAA68GAG (8776A>G), CAT69CAC (8779T>C), TTG70TTA (8782G>A), TAT73TAC (8791T>C), CCA75CCG (8797A>G), CAA76CAG (8800A>G), CAA77CAG (8803A>G), GAT79GAC (8809T>C), ATT80ATC (8812T>C), AAT82AAC (8818T>C), CGG84CGA (8824G>A), GCA85GCG (8827A>G), ACG92ACA (8848G>A), TAT94TAC (8854T>C), GAG95GGA (8856A>G 8857G>A), GTG97GTA (8863G>A), CGG98CAG (8865G>A), ATG99CTT (8867A>C 8869G>T), GCA100GCG (8872A>G), GGA104GAA (8883G>A), ACT106TCT (8888A>T), TGC119TGT (8929C>T), GTC128ATC (8954G>A), AAC129AAT (8959C>T), GGA130GGT (8962A>T), GGG137GGA (8983G>A), AAT138GAC (8984A>G 8986T>C), GAA139GTA (8988A>T), GAG142GAA (8998G>A), CCG144CCA (9004G>A), TTG145TTA (9007G>A), ATC148ATT (9016C>T), ACC155ACA (9037C>A), ATA171ATT (9085A>T), AAC175AAT (9097C>T), ATT187GTT (9131A>G), CGA188CGT (9136A>T), CGG191CGT (9145G>T), ATG193GGA (9149A>G 9150T>G 9151G>A), GGT194AGT (9152G>A), TTA195TTG (9157A>G), GCG196GCT (9160G>T), CGT197CGC (9163T>C), GAG204GAA (9184G>A), CGA208CGG (9196A>G), AGG214AGA (9214G>A), GAA215GAG (9217A>G), GCG216GCA (9220G>A), GCA224GCT (9244A>T), TTG225TTA (9247G>A), GCC228GCT (9256C>T), CCT230TCC (9260C>T 9262T>C), GGG234GGA (9274G>A), GAC236GAT (9280C>T), ATC239ATT (9289C>T), CAA242CAC (9298A>C), GAG244GAA (9304G>A), GTC254GTT (9334C>T), GTC264GTG (9364C>G) | | | | | | | | |

| | Begin | End | Coverage | Score | Concordance | Matches | Identities | I/D/M/F* | Stop Codons |
|---|---|---|---|---|---|---|---|---|---|
| **NT** | **5** | **9704** | **99.9%** | **12522** | **64.8%** | **9688 (99.7%)** | **8015 (82.5%)** | **15/12** | |
| PIPO (YP_006393460.1) | 1 | 75 | 100% | 95 | 18.1% | 75 (98.7%) | 54 (71.1%) | 1/0/3/2 | 2 |
| Protein mutations: | R17_K17insX (2971_2972insA), Y23H (2989T>C 2991C>T), R26K (2999G>A 3000A>G), H35Y (3025C>T), K40H (3040A>C 3042A>C), G41S (3043G>A), E44K (3052G>A), I47V (3061A>G), V54I (3082G>A), P57S (3091C>T), P60T (3100C>A), G61E (3104G>A 3105T>G), G62S (3106G>A), Q63* (3109C>T), R64G (3112A>G), C66R (3118T>C), L67F (3121C>T), I69V (3127A>G), A72E (3137C>A 3138G>A), I73V (3139A>G) | | | | | | | | |
| Codon mutations: | TTA2CTA (2923T>C), AAA4AAG (2931A>G), GAA7GAG (2940A>G), TTG10CTG (2947T>C), TTG15CTG (2962T>C), AGA17_AAA17insAA- (2971_2972insA), TAC23CAT (2989T>C 2991C>T), TCA24TCG (2994A>G), AGA26AAG (2999G>A 3000A>G), CAC32-AC (3016delC), TCG33TCA (3021G>A), GTA34GTT (3024A>T), CAT35TAT (3025C>T), CAC38CAT (3036C>T), AGA39AGG (3039A>G), AAA40CAC (3040A>C 3042A>C), GGC41AGC (3043G>A), GAA44AAA (3052G>A), ATA47GTA (3061A>G), GTT54ATT (3082G>A), CCG57TCG (3091C>T), CGC59CGT (3099C>T), CCA60ACA (3100C>A), GGT61GAG (3104G>A 3105T>G), GGT62AGT (3106G>A), CAA63TAA (3109C>T), AGG64GGG (3112A>G), TGC66CGC (3118T>C), CTC67TTC (3121C>T), ATT69GTT (3127A>G), GAG70GAA (3132G>A), GCG72GAA (3137C>A 3138G>A), ATT73GTT (3139A>G) | | | | | | | | |

*: Inserts / Deletes / Misaligned / Frameshifts

#### Analysis details

This analysis was performed with panviral2.64

### NGS Details (UN9): Tomato chocolate spot virus (segment RNA 1)

#### Assembly

| | |
|---|---|
| **Coverage Length** | 1254 (5 contig(s)) |
| **Depth Of Coverage** | 14600.6 |
| **Number Of Reads** | 162269 |
| **Reads Per Million** | 3667.74 rpm (after QC) |
| **Ambiguities** | 0 |
| **Assembly Method** | de novo + reference guided assembly |
| **Consensus Caller** | Bcf Tools |

#### Coverage Map

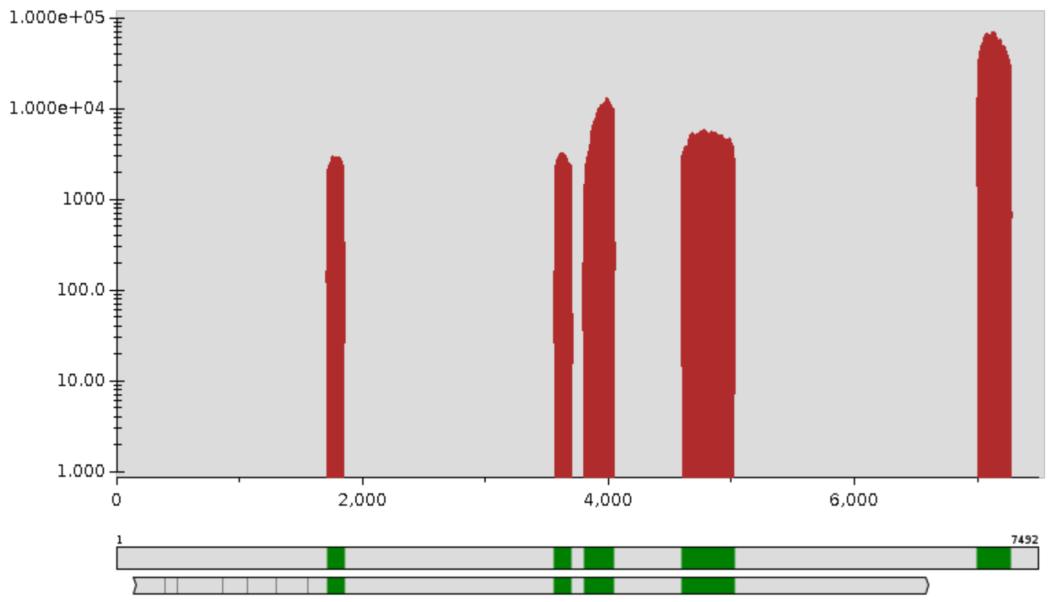

#### Assignment

| | |
|---|---|
| **Type** | Tomato chocolate spot virus (Taxonomy ID: 661101) |
| **Reference Genome** | NC_013075.1 |
| **NT Identity (%)** | 75.1394 |
| **AA Identity (%)** | 76.9231 |
| **Number Of Stop Codons** | 0 |
| **Number Of CDS** | 1 |

#### Alignment

| | |
|---|---|
| **Alignment Score** | 1254.0 (NT) + 1858.0 (AA) = 3112.0 |
| **Concordance (%)** | 64.5509 |

| Alignment Method | Global, seeded, nucleotide + amino acids (AGA) |
|---|---|

#### Genome Region

Sequence starts at position 1710 and ends at position 7275 relative to NC_013075.1 reference sequence.

#### Alignment Detailed Statistics

| | Begin | End | Coverage | Score | Concordance | Matches | Identities | I/D/M/F* | Stop Codons |
|---|---|---|---|---|---|---|---|---|---|
| **NT** | **1710** | **7275** | **16.7%** | **1254** | **50.0%** | **1254 (99.9%)** | **943 (75.1%)** | **1/0** | |
| Mutations: | 1725T>A, 1728A>G, 1729G>A, 1734C>A, 1737T>A, 1739G>C, 1740C>A, 1748A>G, 1749A>C, 1755T>C, 1758T>C, 1759G>A, 1763A>G, 1764G>A, 1771C>A, 1773A>G, 1776A>T, 1785C>A, 1792C>A, 1794T>A, 1796G>A, 1797T>C, 1798T>C, 1800G>C, 1801T>A, 1806C>T, 1807A>T, 1808G>C, 1809C>T, 1812T>C, 1821C>A, 1824C>T, 1833A>C, 1839T>C, 1840C>A, 1845T>C, 3573A>G, 3575G>A, 3579C>T, 3581A>C, 3582G>A, 3594T>A, 3600A>T, 3601C>G, 3602C>A, 3603T>A, 3607G>C, 3612A>T, 3615A>T, 3618A>T, 3622A>C, 3624A>G, 3628A>T, 3630C>T, 3633C>G, 3636T>A, 3639C>T, 3642G>A, 3645G>A, 3648T>G, 3651G>T, 3654C>A, 3663A>T, 3675G>C, 3676G>A, 3678C>T, 3679A>G, 3680T>A, 3808G>A, 3810C>T, 3813A>G, 3819T>A, 3828C>T, 3831T>G, 3834T>A, 3840T>A, 3842G>T, 3843G>T, 3851A>T, 3852C>T, 3857G>C, 3858T>A, 3864G>A, 3865C>T, 3873T>G, 3888C>T, 3898A>T, 3903T>A, 3904A>C, 3906G>T, 3910C>T, 3911T>C, 3912C>T, 3921A>G, 3924G>A, 3934A>G, 3937A>C, 3938C>A, 3939T>A, 3940G>C, 3941G>C, 3942T>A, 3943G>A, 3944T>A, 3945G>C, 3946G>T, 3947G>C, 3949G>C, 3951C>G, 3952C>A, 3954C>G, 3958G>C, 3959A>T, 3960G>T, 3965A>T, 3966G>A, 3967C>A, 3969A>G, 3970T>A, 3971C>A, 3972A>C, 3981G>C, 3987A>T, 3990G>A, 3996G>A, 4005T>C, 4006G>T, 4007T>C, 4008T>G, 4011A>G, 4014G>A, 4017A>G, 4018A>C, 4024G>A, 4025T>C, 4026C>A, 4027C>T, 4028A>T, 4032G>T, 4038G>C, 4039T>G, 4611C>T, 4612C>G, 4614G>C, 4615C>T, 4617T>A, 4620C>T, 4621G>A, 4626T>A, 4635A>G, 4644C>T, 4650T>G, 4656T>G, 4659C>A, 4660T>A, 4666T>C, 4668G>T, 4671G>A, 4677T>C, 4678T>G, 4679C>A, 4680A>G, 4683G>A, 4686C>T, 4687A>G, 4688A>C, 4689C>A, 4692T>A, 4701A>G, 4704G>A, 4707T>A, 4708A>C, 4709T>A, 4710G>A, 4713A>G, 4716T>C, 4720G>A, 4722A>T, 4725T>C, 4728A>G, 4731C>T, 4734T>C, 4735T>C, 4737G>T, 4740C>T, 4744T>A, 4745C>A, 4746A>C, 4750G>A, 4755G>C, 4761G>T, 4764G>C, 4765C>A, 4767G>A, 4770T>C, 4785A>G, 4786C>T, 4791T>G, 4795C>T, 4806G>C, 4813A>C, 4815A>C, 4822C>T, 4827C>T, 4833T>A, 4834A>C, 4836G>A, 4839A>C, 4842G>C, 4845T>C, 4854A>T, 4860A>T, 4861A>G, 4863A>G, 4881A>C, 4882C>A, 4891C>G, 4892C>T, 4893C>T, 4896T>G, 4902T>A, 4906G>A, 4907C>T, 4908A>C, 4909C>A, 4910A>C, 4911G>T, 4914A>C, 4918A>T, 4919T>A, 4920G>T, 4929T>G, 4935T>G, 4938C>T, 4939A>G, 4950C>T, 4953T>C, 4956T>G, 4962T>A, 4965A>G, 4968T>C, 4970C>T, 4971A>T, 4977G>C, 4979A>C, 4984G>A, 4985T>C, 4986G>A, 4988C>A, 4989A>C, 4990G>A, 4995G>C, 4999G>A, 5005C>A, 5007G>A, 5009A>T, 5010T>G, 5011G>C, 5013A>G, 5016T>A, 5019A>C, 7007T>C, 7012T>G, 7018C>T, 7019A>G, 7020A>G, 7020_7021insT, 7023C>A, 7042G>T, 7044A>T, 7047G>A, 7048C>G, 7066G>A, 7082G>A, 7083T>C, 7098C>T, 7099T>C, 7104T>A, 7107G>A, 7116C>G, 7121G>A, 7127A>C, 7140C>T, 7151A>T, 7152T>C, 7177A>C, 7183C>T, 7184T>C, 7185G>T, 7186T>G, 7187T>C, 7188G>A, 7196C>T, 7197A>G, 7198A>C, 7199C>A, 7200A>G, 7201G>A, 7207A>T, 7208A>T, 7209T>G, 7210C>T, 7212T>A, 7219T>A, 7224T>C, 7242A>T, 7255A>T, 7257G>A, 7258G>C, 7260T>C, 7262C>G, 7264T>C | | | | | | | | |

##### CDS

| | Begin | End | Coverage | Score | Concordance | Matches | Identities | I/D/M/F* | Stop Codons |
|---|---|---|---|---|---|---|---|---|---|
| ToChSV_s1_gp1 | 525 | 1631 | 15.1% | 1858 | 78.7% | 325 (100%) | 250 (76.9%) | 0/0/0/0 | 0 |
| Protein mutations: | D531N (1729G>A), G534A (1739G>C 1740C>A), K537S (1748A>G 1749A>C), V541I (1759G>A), E542G (1763A>G 1764G>A), G553D (1796G>A 1797T>C), S555T (1801T>A), D561E (1821C>A), Q568K (1840C>A), R1146K (3575G>A), Q1148P (3581A>C 3582G>A), P1155E (3601C>G 3602C>A 3603T>A), E1157Q (3607G>C), I1162L (3622A>C 3624A>G), T1164S (3628A>T 3630C>T), N1166K (3636T>A), E1171D (3651G>T), V1180I (3676G>A 3678C>T), M1181E (3679A>G 3680T>A), D1224N (3808G>A 3810C>T), W1235F (3842G>T 3843G>T), Y1238F (3851A>T 3852C>T), G1240A (3857G>C 3858T>A), M1254L (3898A>T), M1256L (3904A>C 3906G>T), L1258S (3910C>T 3911T>C 3912C>T), T1266A (3934A>G), T1267Q (3937A>C 3938C>A 3939T>A), G1268P (3940G>C 3941G>C 3942T>A), V1269N (3943G>A 3944T>A 3945G>C), G1270S (3946G>T 3947G>C), D1271Q (3949G>C 3951C>G), E1274L (3958G>C 3959A>T 3960G>T), K1276I (3965A>T 3966G>A), S1278N (3970T>A 3971C>A 3972A>C), E1283D (3987A>T), V1290S (4006G>T 4007T>C 4008T>G), K1294Q (4018A>C), V1296T (4024G>A 4025T>C 4026C>A), H1297F (4027C>T 4028A>T), F1301V (4039T>G), Q1492D (4612C>G 4614G>C), V1495I (4621G>A), L1508M (4660T>A), S1514E (4678T>G 4679C>A 4680A>G), N1517A (4687A>G 4688A>C 4689C>A), D1523E (4707T>A), M1524Q (4708A>C 4709T>A 4710G>A), V1528I (4720G>A 4722A>T), S1536N (4744T>A 4745C>A 4746A>C), D1538N (4750G>A), E1541D (4761G>T), Q1543K (4765C>A 4767G>A), H1565Q (4833T>A), M1566L (4834A>C 4836G>A), I1575V (4861A>G 4863A>G), P1585V (4891C>G 4892C>T 4893C>T), A1590I (4906G>A 4907C>T 4908A>C), Q1591T (4909C>A 4910A>C 4911G>T), M1594Y (4918A>T 4919T>A 4920G>T), I1601V (4939A>G), H1606Q (4956T>G), A1611V (4970C>T 4971A>T), E1613D (4977G>C), K1614T (4979A>C), V1616T (4984G>A 4985T>C 4986G>A), A1617D (4988C>A 4989A>C), D1618N (4990G>A), E1621K (4999G>A), L1623I (5005C>A 5007G>A), N1624M (5009A>T 5010T>G), E1625Q (5011G>C 5013A>G) | | | | | | | | |
| Codon mutations: | ACT529ACA (1725T>A), GAA530GAG (1728A>G), GAC531AAC (1729G>A), CCC532CCA (1734C>A), ACT533ACA (1737T>A), GGC534GCA (1739G>C 1740C>A), AAA537AGC (1748A>G 1749A>C), AAT539AAC (1755T>C), ATT540ATC (1758T>C), GTT541ATT (1759G>A), GAG542GGA (1763A>G 1764G>A), CGA545AGG (1771C>A 1773A>G), CCA546CCT (1776A>T), ATC549ATA (1785C>A), CGT552AGA (1792C>A 1794T>A), GGT553GAC (1796G>A 1797T>C), TTG554CTC (1798T>C 1800G>C), TCT555ACT (1801T>A), TAC556TAT (1806C>T), AGC557TCT (1807A>T 1808G>C 1809C>T), GAT558GAC (1812T>C), GAC561GAA (1821C>A), TAC562TAT (1824C>T), GCA565GCC (1833A>C), TGT567TGC (1839T>C), CAG568AAG (1840C>A), GTT569GTC (1845T>C), AGA1145AGG (3573A>G), AGA1146AAA (3575G>A), AGC1147AGT (3579C>T), CAG1148CCA (3581A>C 3582G>A), GCT1152GCA (3594T>A), GGA1154GGT (3600A>T), CCT1155GAA (3601C>G 3602C>A 3603T>A), GAA1157CAA (3607G>C), ACA1158ACT (3612A>T), GCA1159GCT (3615A>T), CCA1160CCT (3618A>T), ATA1162CTG (3622A>C 3624A>G), ACC1164TCT (3628A>T 3630C>T), GTC1165GTG (3633C>G), AAT1166AAA (3636T>A), GAC1167GAT (3639C>T), AAG1168AAA (3642G>A), AGG1169AGA (3645G>A), GGT1170GGG (3648T>G), GAG1171GAT (3651G>T), GGC1172GGA (3654C>A), CCA1175CCT (3663A>T), GGG1179GGC (3675G>C), GTC1180ATT (3676G>A 3678C>T), ATG1181GAG (3679A>G 3680T>A), GAC1224AAT (3808G>A 3810C>T), GAA1225GAG (3813A>G), GTT1227GTA (3819T>A), AAC1230AAT (3828C>T), GGT1231GGG (3831T>G), ATT1232ATA (3834T>A), GGT1234GGA (3840T>A), TGG1235TTT (3842G>T 3843G>T), TAC1238TTT (3851A>T 3852C>T), GGT1240GCA (3857G>C 3858T>A), GAG1242GAA (3864G>A), CTG1243TTG (3865C>T), ACT1245ACG (3873T>G), CCC1250CCT (3888C>T), ATG1254TTG (3898A>T), GGT1255GGA (3903T>A), ATG1256CTT (3904A>C 3906G>T), CTC1258TCT (3910C>T 3911T>C 3912C>T), AGA1261AGG (3921A>G), GAG1262GAA (3924G>A), ACC1266GCC (3934A>G), ACT1267CAA (3937A>C 3938C>A 3939T>A), GGT1268CCA (3940G>C 3941G>C 3942T>A), GTG1269AAC (3943G>A 3944T>A 3945G>C), GGG1270TCG (3946G>T 3947G>C), GAC1271CAG (3949G>C 3951C>G), CGC1272AGG (3952C>A 3954C>G), GAG1274CTT (3958G>C 3959A>T 3960G>T), AAG1276ATA (3965A>T 3966G>A), CGA1277AGG (3967C>A 3969A>G), TCA1278AAC (3970T>A 3971C>A 3972A>C), GTG1281GTC (3981G>C), GAA1283GAT (3987A>T), GCG1284GCA (3990G>A), GAG1286GAA (3996G>A), GAT1289GAC (4005T>C), GTT1290TCG (4006G>T 4007T>C 4008T>G), CAA1291CAG (4011A>G), GTG1292GTA (4014G>A), AGA1293AGG (4017A>G), AAA1294CAA (4018A>C), GTC1296ACA (4024G>A 4025T>C 4026C>A), CAT1297TTT (4027C>T 4028A>T), GTG1298GTT (4032G>T), CTG1300CTC (4038G>C), TTC1301GTC (4039T>G), AAC1491AAT (4611C>T), CAG1492GAC (4612C>G 4614G>C), CTT1493TTA (4615C>T 4617T>A), TTC1494TTT (4620C>T), GTT1495ATT (4621G>A), GCT1496GCA (4626T>A), GGA1499GGG (4635A>G), AAC1502AAT (4644C>T), GTT1504GTG (4650T>G), GTT1506GTG (4656T>G), CCC1507CCA (4659C>A), TTG1508ATG (4660T>A), TTG1510CTT (4666T>C 4668G>T), CAG1511CAA (4671G>A), TAT1513TAC (4677T>C), TCA1514GAG (4678T>G 4679C>A 4680A>G), TTG1515TTA (4683G>A), CCC1516CCT (4686C>T), AAC1517GCA (4687A>G 4688A>C 4689C>A), ATT1518ATA (4692T>A), GAA1521GAG (4701A>G), CTG1522CTA (4704G>A), GAT1523GAA (4707T>A), ATG1524CAA (4708A>C 4709T>A 4710G>A), GTA1525GTG (4713A>G), AAT1526AAC (4716T>C), GTA1528ATT (4720G>A 4722A>T), ATT1529ATC (4725T>C), AAA1530AAG (4728A>G), AAC1531AAT (4731C>T), GGT1532GGC (4734T>C), TTG1533CTT (4735T>C 4737G>T), GAC1534GAT (4740C>T), TCA1536AAC (4744T>A 4745C>A 4746A>C), GAT1538AAT (4750G>A), GTG1539GTC (4755G>C), GAG1541GAT (4761G>T), GTG1542GTC (4764G>C), CAG1543AAA (4765C>A 4767G>A), TTT1544TTC (4770T>C), GAA1549GAG (4785A>G), CTG1550TTG (4786C>T), ACT1551ACG (4791T>G), CTG1553TTG (4795C>T), GGG1556GGC (4806G>C), AGA1559CGC (4813A>C 4815A>C), CTG1562TTG (4822C>T), GGC1563GGT (4827C>T), CAT1565CAA (4833T>A), ATG1566CTA (4834A>C 4836G>A), GCA1567GCC (4839A>C), CCG1568CCC (4842G>C), CTT1569CTC (4845T>C), GTA1572GTT (4854A>T), ATA1574ATT (4860A>T), ATA1575GTG (4861A>G 4863A>G), ATA1581ATC (4881A>C), CGA1582AGA (4882C>A), CCC1585GTT (4891C>G 4892C>T 4893C>T), GGT1586GGG (4896T>G), CCT1588CCA (4902T>A), GCA1590ATC (4906G>A 4907C>T 4908A>C), CAG1591ACT (4909C>A 4910A>C 4911G>T), GCA1592GCC (4914A>C), ATG1594TAT (4918A>T 4919T>A 4920G>T), GTT1597GTG (4929T>G), ACT1599ACG (4935T>G), GGC1600GGT (4938C>T), ATA1601GTA (4939A>G), GCC1604GCT (4950C>T), TTT1605TTC (4953T>C), CAT1606CAG (4956T>G), GGT1608GGA (4962T>A), CAA1609CAG (4965A>G), GTT1610GTC (4968T>C), GCA1611GTT (4970C>T 4971A>T), GAG1613GAC (4977G>C), AAA1614ACA (4979A>C), GTG1616ACA (4984G>A 4985T>C 4986G>A), GCA1617GAC (4988C>A 4989A>C), GAT1618AAT (4990G>A), GTG1619GTC (4995G>C), GAA1621AAA (4999G>A), CTG1623ATA (5005C>A 5007G>A), AAT1624ATG (5009A>T 5010T>G), GAA1625CAG (5011G>C 5013A>G), CGT1626CGA (5016T>A), GGA1627GGC (5019A>C) | | | | | | | | |

##### Proteins

| | Begin | End | Coverage | Score | Concordance | Matches | Identities | I/D/M/F* | Stop Codons |
|---|---|---|---|---|---|---|---|---|---|
| polyprotein (YP_003097229.1) | 525 | 1631 | 15.1% | 1858 | 78.7% | 325 (100%) | 250 (76.9%) | 0/0/0/0 | 0 |
| Protein mutations: | D531N (1729G>A), G534A (1739G>C 1740C>A), K537S (1748A>G 1749A>C), V541I (1759G>A), E542G (1763A>G 1764G>A), G553D (1796G>A 1797T>C), S555T (1801T>A), D561E (1821C>A), Q568K (1840C>A), R1146K (3575G>A), Q1148P (3581A>C 3582G>A), P1155E (3601C>G 3602C>A 3603T>A), E1157Q (3607G>C), I1162L (3622A>C 3624A>G), T1164S (3628A>T 3630C>T), N1166K (3636T>A), E1171D (3651G>T), V1180I (3676G>A 3678C>T), M1181E (3679A>G 3680T>A), D1224N (3808G>A 3810C>T), W1235F (3842G>T 3843G>T), Y1238F (3851A>T 3852C>T), G1240A (3857G>C 3858T>A), M1254L (3898A>T), M1256L (3904A>C 3906G>T), L1258S (3910C>T 3911T>C 3912C>T), T1266A (3934A>G), T1267Q (3937A>C 3938C>A 3939T>A), G1268P (3940G>C 3941G>C 3942T>A), V1269N (3943G>A 3944T>A 3945G>C), G1270S (3946G>T 3947G>C), D1271Q (3949G>C 3951C>G), E1274L (3958G>C 3959A>T 3960G>T), K1276I (3965A>T 3966G>A), S1278N (3970T>A 3971C>A 3972A>C), E1283D (3987A>T), V1290S (4006G>T 4007T>C 4008T>G), K1294Q (4018A>C), V1296T (4024G>A 4025T>C 4026C>A), H1297F (4027C>T 4028A>T), F1301V (4039T>G), Q1492D (4612C>G 4614G>C), V1495I (4621G>A), L1508M (4660T>A), S1514E (4678T>G 4679C>A 4680A>G), N1517A (4687A>G 4688A>C 4689C>A), D1523E (4707T>A), M1524Q (4708A>C 4709T>A 4710G>A), V1528I (4720G>A 4722A>T), S1536N (4744T>A 4745C>A 4746A>C), D1538N (4750G>A), E1541D (4761G>T), Q1543K (4765C>A 4767G>A), H1565Q (4833T>A), M1566L (4834A>C 4836G>A), I1575V (4861A>G 4863A>G), P1585V (4891C>G 4892C>T 4893C>T), A1590I (4906G>A 4907C>T 4908A>C), Q1591T (4909C>A 4910A>C 4911G>T), M1594Y (4918A>T 4919T>A 4920G>T), I1601V (4939A>G), H1606Q (4956T>G), A1611V (4970C>T 4971A>T), E1613D (4977G>C), K1614T (4979A>C), V1616T (4984G>A 4985T>C 4986G>A), A1617D (4988C>A 4989A>C), D1618N (4990G>A), E1621K (4999G>A), L1623I (5005C>A 5007G>A), N1624M (5009A>T 5010T>G), E1625Q (5011G>C 5013A>G) | | | | | | | | |

| | Begin | End | Coverage | Score | Concordance | Matches | Identities | I/D/M/F* | Stop Codons |
|---|---|---|---|---|---|---|---|---|---|
| NT | 1710 | 7275 | 16.7% | 1254 | 50.0% | 1254 (99.9%) | 943 (75.1%) | 1/0 | |
| Codon mutations: | ACT529ACA (1725T>A), GAA530GAG (1728A>G), GAC531AAC (1729G>A), CCC532CCA (1734C>A), ACT533ACA (1737T>A), GGC534GCA (1739G>C 1740C>A), AAA537AGC (1748A>G 1749A>C), AAT539AAC (1755T>C), ATT540ATC (1758T>C), GTT541ATT (1759G>A), GAG542GGA (1763A>G 1764G>A), CGA545AGG (1771C>A 1773A>G), CCA546CCT (1776A>T), ATC549ATA (1785C>A), CGT552AGA (1792C>A 1794T>A), GGT553GAC (1796G>A 1797T>C), TTG554CTC (1798T>C 1800G>C), TCT555ACT (1801T>A), TAC556TAT (1806C>T), AGC557TCT (1807A>T 1808G>C 1809C>T), GAT558GAC (1812T>C), GAC561GAA (1821C>A), TAC562TAT (1824C>T), GCA565GCC (1833A>C), TGT567TGC (1839T>C), CAG568AAG (1840C>A), GTT569GTC (1845T>C), AGA1145AGG (3573A>G), AGA1146AAA (3575G>A), AGC1147AGT (3579C>T), CAG1148CCA (3581A>C 3582G>A), GCT1152GCA (3594T>A), GGA1154GGT (3600A>T), CCT1155GAA (3601C>G 3602C>A 3603T>A), GAA1157CAA (3607G>C), ACA1158ACT (3612A>T), GCA1159GCT (3615A>T), CCA1160CCT (3618A>T), ATA1162CTG (3622A>C 3624A>G), ACC1164TCT (3628A>T 3630C>T), GTC1165GTG (3633C>G), AAT1166AAA (3636T>A), GAC1167GAT (3639C>T), AAG1168AAA (3642G>A), AGG1169AGA (3645G>A), GGT1170GGG (3648T>G), GAG1171GAT (3651G>T), GGC1172GGA (3654C>A), CCA1175CCT (3663A>T), GGG1179GGC (3675G>C), GTC1180ATT (3676G>A 3678C>T), ATG1181GAG (3679A>G 3680T>A), GAC1224AAT (3808G>A 3810C>T), GAA1225GAG (3813A>G), GTT1227GTA (3819T>A), AAC1230AAT (3828C>T), GGT1231GGG (3831T>G), ATT1232ATA (3834T>A), GGT1234GGA (3840T>A), TGG1235TTT (3842G>T 3843G>T), TAC1238TTT (3851A>T 3852C>T), GGT1240GCA (3857G>C 3858T>A), GAG1242GAA (3864G>A), CTG1243TTG (3865C>T), ACT1245ACG (3873T>G), CCC1250CCT (3888C>T), ATG1254TTG (3898A>T), GGT1255GGA (3903T>A), ATG1256CTT (3904A>C 3906G>T), CTC1258TCT (3910C>T 3911T>C 3912C>T), AGA1261AGG (3921A>G), GAG1262GAA (3924G>A), ACC1266GCC (3934A>G), ACT1267CAA (3937A>C 3938C>A 3939T>A), GGT1268CCA (3940G>C 3941G>C 3942T>A), GTG1269AAC (3943G>A 3944T>A 3945G>C), GGG1270TCG (3946G>T 3947G>C), GAC1271CAG (3949G>C 3951C>G), CGC1272AGG (3952C>A 3954C>G), GAG1274CTT (3958G>C 3959A>T 3960G>T), AAG1276ATA (3965A>T 3966G>A), CGA1277AGG (3967C>A 3969A>G), TCA1278AAC (3970T>A 3971C>A 3972A>C), GTG1281GTC (3981G>C), GAA1283GAT (3987A>T), GCG1284GCA (3990G>A), GAG1286GAA (3996G>A), GAT1289GAC (4005T>C), GTT1290TCG (4006G>T 4007T>C 4008T>G), CAA1291CAG (4011A>G), GTG1292GTA (4014G>A), AGA1293AGG (4017A>G), AAA1294CAA (4018A>C), GTC1296ACA (4024G>A 4025T>C 4026C>A), CAT1297TTT (4027C>T 4028A>T), GTG1298GTT (4032G>T), CTG1300CTC (4038G>C), TTC1301GTC (4039T>G), AAC1491AAT (4611C>T), CAG1492GAC (4612C>G 4614G>C), CTT1493TTA (4615C>T 4617T>A), TTC1494TTT (4620C>T), GTT1495ATT (4621G>A), GCT1496GCA (4626T>A), GGA1499GGG (4635A>G), AAC1502AAT (4644C>T), GTT1504GTG (4650T>G), GTT1506GTG (4656T>G), CCC1507CCA (4659C>A), TTG1508ATG (4660T>A), TTG1510CTT (4666T>C 4668G>T), CAG1511CAA (4671G>A), TAT1513TAC (4677T>C), TCA1514GAG (4678T>G 4679C>A 4680A>G), TTG1515TTA (4683G>A), CCC1516CCT (4686C>T), AAC1517GCA (4687A>G 4688A>C 4689C>A), ATT1518ATA (4692T>A), GAA1521GAG (4701A>G), CTG1522CTA (4704G>A), GAT1523GAA (4707T>A), ATG1524CAA (4708A>C 4709T>A 4710G>A), GTA1525GTG (4713A>G), AAT1526AAC (4716T>C), GTA1528ATT (4720G>A 4722A>T), ATT1529ATC (4725T>C), AAA1530AAG (4728A>G), AAC1531AAT (4731C>T), GGT1532GGC (4734T>C), TTG1533CTT (4735T>C 4737G>T), GAC1534GAT (4740C>T), TCA1536AAC (4744T>A 4745C>A 4746A>C), GAT1538AAT (4750G>A), GTG1539GTC (4755G>C), GAG1541GAT (4761G>T), GTG1542GTC (4764G>C), CAG1543AAA (4765C>A 4767G>A), TTT1544TTC (4770T>C), GAA1549GAG (4785A>G), CTG1550TTG (4786C>T), ACT1551ACG (4791T>G), CTG1553TTG (4795C>T), GGG1556GGC (4806G>C), AGA1559CGC (4813A>C 4815A>C), CTG1562TTG (4822C>T), GGC1563GGT (4827C>T), CAT1565CAA (4833T>A), ATG1566CTA (4834A>C 4836G>A), GCA1567GCC (4839A>C), CCG1568CCC (4842G>C), CTT1569CTC (4845T>C), GTA1572GTT (4854A>T), ATA1574ATT (4860A>T), ATA1575GTG (4861A>G 4863A>G), ATA1581ATC (4881A>C), CGA1582AGA (4882C>A), CCC1585GTT (4891C>G 4892C>T 4893C>T), GGT1586GGG (4896T>G), CCT1588CCA (4902T>A), GCA1590ATC (4906G>A 4907C>T 4908A>C), CAG1591ACT (4909C>A 4910A>C 4911G>T), GCA1592GCC (4914A>C), ATG1594TAT (4918A>T 4919T>A 4920G>T), GTT1597GTG (4929T>G), ACT1599ACG (4935T>G), GGC1600GGT (4938C>T), ATA1601GTA (4939A>G), GCC1604GCT (4950C>T), TTT1605TTC (4953T>C), CAT1606CAG (4956T>G), GGT1608GGA (4962T>A), CAA1609CAG (4965A>G), GTT1610GTC (4968T>C), GCA1611GTT (4970C>T 4971A>T), GAG1613GAC (4977G>C), AAA1614ACA (4979A>C), GTG1616ACA (4984G>A 4985T>C 4986G>A), GCA1617GAC (4988C>A 4989A>C), GAT1618AAT (4990G>A), GTG1619GTC (4995G>C), GAA1621AAA (4999G>A), CTG1623ATA (5005C>A 5007G>A), AAT1624ATG (5009A>T 5010T>G), GAA1625CAG (5011G>C 5013A>G), CGT1626CGA (5016T>A), GGA1627GGC (5019A>C) | | | | | | | | |

*: Inserts / Deletes / Misaligned / Frameshifts

#### Analysis details

This analysis was performed with panviral2.64

### NGS Details (UN9): Tomato chocolate spot virus (segment RNA2)

#### Assembly

| | |
|---|---|
| **Coverage Length** | 2083 (3 contig(s)) |
| **Depth Of Coverage** | 19230.5 |
| **Number Of Reads** | 328101 |
| **Reads Per Million** | 7416.02 rpm (after QC) |
| **Ambiguities** | 0 |
| **Assembly Method** | de novo + reference guided assembly |
| **Consensus Caller** | Bcf Tools |

#### Coverage Map

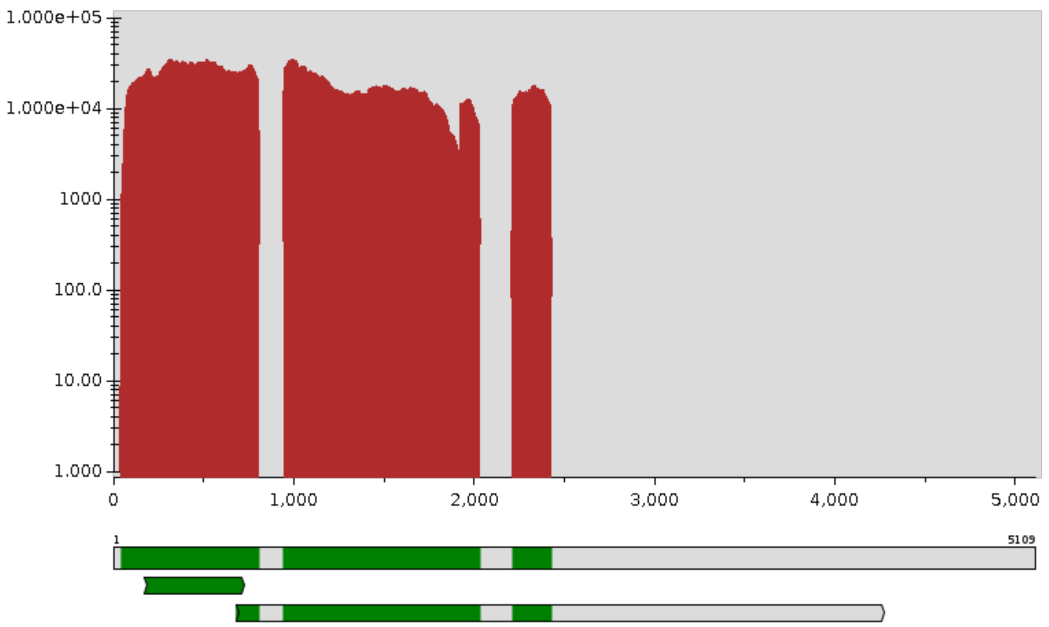

#### Assignment

| | |
|---|---|
| **Type** | Tomato chocolate spot virus (Taxonomy ID: 661101) |
| **Reference Genome** | NC_013076.1 |
| **NT Identity (%)** | 63.9923 |
| **AA Identity (%)** | 66.0633 |
| **Number Of Stop Codons** | 1 |
| **Number Of CDS** | 2 |

#### Alignment

| | |
|---|---|
| **Alignment Score** | 1099.0 (NT) + 3139.0 (AA) = 4238.0 |
| **Concordance (%)** | 49.4978 |

| Alignment Method | Global, seeded, nucleotide + amino acids (AGA) |
|---|---|

#### Genome Region

Sequence starts at position 41 and ends at position 2430 relative to NC_013076.1 reference sequence.

#### Alignment Detailed Statistics

| | Begin | End | Coverage | Score | Concordance | Matches | Identities | I/D/M/F* | Stop Codons |
|---|---|---|---|---|---|---|---|---|---|
| **NT** | **41** | **2430** | **40.8%** | **1099** | **27.0%** | **2063 (98.8%)** | **1324 (63.4%)** | **6/20** | |
| Mutations: | 51A>T, 53C>T, 57T>C, 59T>A, 65A>C, 66A>G, 68C>A, 71G>C, 74T>C, 75T>G, 76T>A, 77A>C, 81G>C, 82A>C, 84A>C, 85T>A, 86C>A, 88T>A, 90G>T, 91T>A, 93T>C, 94T>C, 95G>A, 97G>T, 98T>A, 101T>C, 102delT, 106T>C, 107C>A, 110T>C, 111T>C, 112T>A, 114C>T, 116T>A, 117C>T, 118T>C, 120T>A, 123G>C, 124A>C, 125G>T, 127T>C, 130T>A, 132A>C, 134C>A, 135C>A, 138C>A, 139A>C, 140C>T, 141T>A, 149T>A, 151T>A, 152_158delGATAATC, 160C>T, 161A>C, 164C>A, 165C>T, 168G>T, 169T>C, 177T>C, 180T>C, 183T>C, 184G>T, 185G>C, 186A>T, 188G>A, 198A>T, 199G>C, 200C>A, 201A>G, 202G>T, 203A>T, 211A>C, 215C>A, 216T>A, 219C>T, 223C>A, 225A>T, 231C>A, 240C>A, 243T>G, 247A>G, 249T>G, 252C>T, 253T>A, 254C>G, 255G>T, 258G>T, 261C>T, 262G>A, 263T>C, 267A>T, 268T>G, 269C>T, 270A>T, 273T>A, 274G>A, 276A>T, 277A>G, 288C>T, 291C>G, 294T>G, 297A>T, 303T>C, 307G>A, 309G>C, 310A>T, 311T>G, 312T>C, 315C>G, 321A>T, 331G>A, 336A>T, 339G>C, 340C>A, 342A>C, 343A>T, 345T>C, 346G>A, 348T>C, 351G>C, 357G>A, 363A>T, 366A>T, 369A>G, 372G>A, 373T>C, 378T>A, 381G>T, 384C>T, 390T>C, 405T>A, 408T>G, 412C>G, 413C>A, 414T>C, 417T>C, 421A>G, 422A>C, 423C>T, 426G>C, 429G>A, 430T>C, 432A>C, 435T>A, 439C>T, 441C>G, 442T>A, 444G>A, 447G>T, 449A>G, 450T>G, 453T>A, 456T>A, 457G>A, 459T>C, 460C>A, 461G>A, 462T>A, 474C>A, 475A>C, 477G>C, 480A>G, 481G>C, 486A>T, 490C>G, 491A>C, 492G>A, 501T>A, 504A>T, 508_516delGCTCAACAA, 517A>T, 519T>A, 523A>G, 524A>C, 525T>G, 526T>C, 527C>A, 528T>A, 529G>T, 530T>C, 531A>C, 535G>T, 536A>C, 538A>C, 543G>T, 544A>C, 548C>A, 549A>G, 550A>C, 551C>T, 552C>T, 553G>T, 554A>C, 555A>T, 556A>C, 557C>T, 560C>T, 561T>A, 562T>A, 563G>A, 566A>G, 567G>T, 568T>C, 570C>T, 571A>G, 576G>A, 581T>A, 582T>C, 585T>A, 586C>G, 587A>C, 588T>A, 589C>T, 591C>A, 594C>G, 595G>A, 599A>C, 600G>T, 601C>T, 603T>G, 605A>C, 606G>T, 607C>G, 608G>T, 612A>T, 614G>A, 615A>G, 621A>G, 622A>T, 627A>C, 628G>C, 630A>G, 632A>G, 633A>C, 639A>T, 645A>C, 646T>A, 647C>T, 652T>C, 654G>T, 655C>A, 660T>G, 663G>A, 664C>A, 667T>C, 668C>A, 669C>G, 670A>G, 671A>C, 672T>A, 675G>A, 678T>A, 685A>G, 687T>A, 691A>G, 697A>T, 699A>G, 702T>C, 706T>A, 712G>A, 716G>A, 717G>T, 725A>G, 729G>A, 730G>T, 733G>T, 735A>G, 736G>C, 740_742delGAG, 745T>C, 752C>A, 753G>A, 754G>A, 755C>T, 760C>T, 761A>T, 763A>T, 769T>A, 775G>T, 778T>A, 781T>G, 786C>A, 787T>A, 790A>G, 802G>C, 958T>A, 967T>C, 970G>A, 976T>C, 979T>C, 980C>A, 982T>G, 985C>T, 994C>G, 997T>A, 1012G>A, 1019C>G, 1020G>C, 1021T>A, 1033T>C, 1034T>C, 1039T>C, 1042G>C, 1054C>A, 1056T>C, 1057C>T, 1058A>G, 1059G>A, 1060C>T, 1063G>T, 1066G>T, 1069T>A, 1072A>G, 1075G>A, 1079T>A, 1081C>G, 1084G>A, 1086C>T, 1087A>C, 1088C>T, 1090A>T, 1091G>T, 1093T>A, 1095A>G, 1096T>G, 1099T>A, 1102A>T, 1105A>G, 1108C>T, 1111G>C, 1112C>T, 1114C>G, 1124A>G, 1125G>A, 1126T>A, 1129C>T, 1138C>T, 1142C>G, 1144C>T, 1153A>G, 1156A>C, 1162C>T, 1181G>C, 1182A>T, 1183G>T, 1186T>C, 1192T>G, 1195A>T, 1197A>T, 1198G>A, 1201A>T, 1204T>A, 1210C>T, 1213G>T, 1216A>G, 1219G>A, 1222A>T, 1228A>G, 1231A>G, 1238G>A, 1239A>G, 1240T>C, 1241A>C, 1246G>A, 1252C>T, 1255A>T, 1258C>T, 1259T>A, 1261T>G, 1262C>G, 1263A>C, 1267G>A, 1273T>A, 1277T>A, 1278C>G, 1285C>A, 1291A>G, 1294A>T, 1295T>C, 1297G>C, 1300C>T, 1306T>A, 1309A>G, 1312T>C, 1315C>T, 1316C>T, 1317A>C, 1318A>T, 1327G>A, 1333T>A, 1336C>A, 1351A>T, 1354A>C, 1360T>G, 1366G>A, 1367C>G, 1369A>G, 1372T>G, 1373T>G, 1375C>G, 1376C>T, 1377A>C, 1378G>T, 1379A>C, 1380T>A, 1381G>A, 1385T>C, 1387G>A, 1394C>A, 1396T>A, 1397G>C, 1405T>A, 1408A>T, 1410A>C, 1411G>A, 1412A>C, 1413G>T, 1414G>T, 1423T>C, 1426A>T, 1427C>T, 1428G>A, 1429A>C, 1433G>A, 1438A>T, 1440T>C, 1441G>C, 1442A>C, 1443G>A, 1444T>G, 1448A>T, 1449G>C, 1450C>T, 1453C>A, 1454A>T, 1456G>A, 1458C>T, 1459T>G, 1463A>G, 1464G>C, 1465C>G, 1468T>A, 1471C>G, 1474G>T, 1477A>C, 1483G>A, 1489C>A, 1493G>A, 1495G>C, 1501T>C, 1503G>A, 1505C>A, 1513A>T, 1516C>G, 1522A>G, 1525A>G, 1531C>T, 1532G>T, 1534G>T, 1535C>T, 1537T>A, 1539A>G, 1540C>G, 1541A>G, 1547C>T, 1548G>C, 1551T>C, 1552A>G, 1555_1556insGTAGGT, 1556A>G, 1559C>A, 1572C>G, 1573T>G, 1574C>G, 1575A>T, 1576C>A, 1582A>T, 1585C>T, 1586A>T, 1588T>C, 1589G>A, 1590G>A, 1592G>T, 1593G>T, 1595T>C, 1603A>T, 1606T>C, 1610A>G, 1618A>G, 1620G>C, 1622T>A, 1624C>T, 1625A>G, 1627C>T, 1628A>T, 1630G>T, 1633T>C, 1634G>C, 1635T>A, 1636T>G, 1645T>C, 1649C>T, 1650T>A, 1651G>T, 1657A>C, 1658G>A, 1659A>C, 1660C>T, 1661T>G, 1662C>A, 1663C>T, 1668C>A, 1669A>T, 1670C>T, 1671A>C, 1672C>G, 1673A>T, 1678C>T, 1690T>G, 1696T>A, 1697G>A, 1698A>C, 1699G>C, 1700C>T, 1702C>G, 1703A>C, 1704C>A, 1705G>T, 1708T>A, 1709G>T, 1711G>A, 1712C>A, 1713A>G, 1714A>G, 1717G>A, 1718G>A, 1721C>G, 1723A>T, 1725A>T, 1726C>T, 1727A>G, 1729T>G, 1730C>T, 1734T>C, 1736G>A, 1737A>C, 1738T>C, 1739G>C, 1740A>T, 1741G>C, 1742G>C, 1743C>T, 1744T>C, 1745T>C, 1746C>G, 1747A>C, 1749G>A, 1750A>G, 1753C>G, 1755A>G, 1756T>A, 1757G>T, 1758T>G, 1759G>T, 1760A>C, 1761T>A, 1762C>T, 1763G>C, 1768G>C, 1771A>G, 1774T>C, 1775G>A, 1777A>T, 1778T>G, 1779C>A, 1780A>G, 1782A>G, 1783G>A, 1784A>C, 1786G>A, 1789G>C, 1790C>T, 1791A>C, 1792G>C, 1798T>C, 1801T>A, 1802T>A, 1805A>C, 1806A>T, 1807G>A, 1811C>T, 1816C>T, 1817A>G, 1818C>A, 1823G>A, 1825C>T, 1832G>C, 1834T>A, 1835G>C, 1837T>G, 1838A>G, 1840T>G, 1843T>C, 1844G>A, 1845C>A, 1846T>G, 1848C>A, 1861C>A, 1864G>T, 1867A>G, 1870A>T, 1871A>T, 1872A>C, 1873T>C, 1876C>A, 1877T>G, 1878C>A, 1879T>G, 1883T>A, 1885A>T, 1887T>G, 1888G>C, 1889G>A, 1897T>C, 1898G>T, 1899A>C, 1901A>T, 1903G>A, 1904T>A, 1906G>T, 1909T>C, 1910C>A, 1911A>G, 1912T>G, 1915G>A, 1918A>G, 1921C>G, 1924T>A, 1926C>A, 1927C>G, 1933G>C, 1937C>A, 1938A>T, 1939A>G, 1942A>G, 1945T>C, 1948C>T, 1952G>T, 1954G>T, 1957C>T, 1959T>G, 1960A>G, 1961A>T, 1962T>A, 1963G>T, 1964G>T, 1966T>G, 1972T>A, 1981C>G, 1982C>A, 1984A>G, 1988T>G, 1989C>A, 1990A>C, 1996T>C, 1997T>G, 1998G>T, 1999T>G, 2005C>T, 2011C>T, 2015A>C, 2017T>G, 2019G>C, 2020T>A, 2023G>T, 2218C>G, 2221A>T, 2224G>A, 2225C>G, 2228A>C, 2229T>C, 2230T>G, 2235A>G, 2236T>G, 2237G>A, 2239T>A, 2243A>G, 2244C>A, 2245T>G, 2249G>A, 2250G>A, 2254T>C, 2255G>A, 2258A>T, 2260T>C, 2266C>T, 2269C>T, 2272T>G, 2273T>G, 2274C>A, 2275A>G, 2278C>A, 2282G>A, 2283A>T, 2284C>G, 2285T>G, 2286C>G, 2287A>T, 2288A>G, 2290C>T, 2291C>G, 2293A>C, 2294G>C, 2296C>G, 2297A>T, 2300G>A, 2303T>C, 2305A>G, 2309A>T, 2317A>G, 2322T>C, 2332A>T, 2335C>T, 2341A>G, 2356T>G, 2357G>A, 2359G>T, 2365G>A, 2368C>T, 2369G>C, 2375G>A, 2386A>C, 2392G>T, 2395C>T, 2401A>G, 2404C>T, 2405A>C, 2407G>T, 2408T>G, 2411T>C, 2413G>A, 2414G>T, 2416A>C, 2419A>C | | | | | | | | |

##### CDS

| | | | | | | | | | |
|---|---|---|---|---|---|---|---|---|---|
| ToChV_s2_gp1 | 1 | 185 | 100% | 827 | 69.1% | 182 (98.4%) | 117 (63.2%) | 0/3/0/0 | 1 |
| Protein mutations: | G5S (184G>T 185G>C 186A>T), R6K (188G>A), A10Q (199G>C 200C>A 201A>G), E11L (202G>T 203A>T), K14Q (211A>C), A15E (215C>A 216T>A), Q18N (223C>A 225A>T), N24K (243T>G), I26V (247A>G 249T>G), V31T (262G>A 263T>C), S33V (268T>G 269C>T 270A>T), V35I (274G>A 276A>T), I36V (277A>G), V46I (307G>A 309G>C), I47C (310A>T 311T>G 312T>C), V54I (331G>A), L57I (340C>A 342A>C), T58S (343A>T 345T>C), V59I (346G>A 348T>C), P81D (412C>G 413C>A 414T>C), N84A (421A>G 422A>C 423C>T), L91I (442T>A 444G>A), H93R (449A>G 450T>G), V96I (457G>A 459T>C), R97K (460C>A 461G>A 462T>A), V104L (481G>C), Q107A (490C>G 491A>C 492G>A), A113_Q115del (508_516delGCTCAACAA), T116S (517A>T 519T>A), N118A (523A>G 524A>C 525T>G), S119Q (526T>C 527C>A 528T>A), V120S (529G>T 530T>C 531A>C), D122S (535G>T 536A>C), K123Q (538A>C), K125Q (544A>C), T126K (548C>A 549A>G), T127L (550A>C 551C>T 552C>T), E128S (553G>T 554A>C 555A>T), T129L (556A>C 557C>T), A130V (560C>T 561T>A), W131K (562T>A 563G>A), K132S (566A>G 567G>T), F133L (568T>C 570C>T), K134E (571A>G), I137N (581T>A 582T>C), H139A (586C>G 587A>C 588T>A), N141K (594C>G), A142T (595G>A), E143A (599A>C 600G>T), E145A (605A>C 606G>T), R146V (607C>G 608G>T), R148K (614G>A 615A>G), I151L (622A>T), E153Q (628G>C 630A>G), K154S (632A>G 633A>C), S159I (646T>A 647C>T), Q162K (655C>A), L165I (664C>A), S166Q (667T>C 668C>A 669C>G), N167A (670A>G 671A>C 672T>A), N172E (685A>G 687T>A), I174V (691A>G), T176S (697A>T 699A>G), S179T (706T>A), D181N (712G>A), G182D (716G>A 717G>T) | | | | | | | | |
| Codon mutations: | TCT2TCC (177T>C), TTT3TTC (180T>C), ATT4ATC (183T>C), GGA5TCT (184G>T 185G>C 186A>T), AGG6AAG (188G>A), ACA9ACT (198A>T), GCA10CAG (199G>C 200C>A 201A>G), GAG11TTG (202G>T 203A>T), AAA14CAA (211A>C), GCT15GAA (215C>A 216T>A), TTC16TTT (219C>T), CAA18AAT (223C>A 225A>T), GTC20GTA (231C>A), TCC23TCA (240C>A), AAT24AAG (243T>G), ATT26GTG (247A>G 249T>G), TGC27TGT (252C>T), TCG28AGT (253T>A 254C>G 255G>T), GTG29GTT (258G>T), GAC30GAT (261C>T), GTA31ACA (262G>A 263T>C), GGA32GGT (267A>T), TCA33GTT (268T>G 269C>T 270A>T), GGT34GGA (273T>A), GTA35ATT (274G>A 276A>T), ATT36GTT (277A>G), AAC39AAT (288C>T), CCC40CCG (291C>G), ACT41ACG (294T>G), CTA42CTT (297A>T), TTT44TTC (303T>C), GTG46ATC (307G>A 309G>C), ATT47TGC (310A>T 311T>G 312T>C), CCC48CCG (315C>G), ACA50ACT (321A>T), GTT54ATT (331G>A), TCA55TCT (336A>T), GTG56GTC (339G>C), CTA57ATC (340C>A 342A>C), ACT58TCC (343A>T 345T>C), GTT59ATC (346G>A 348T>C), TCG60TCC (351G>C), GAG62GAA (357G>A), TCA64TCT (363A>T), ACA65ACT (366A>T), CCA66CCG (369A>G), CAG67CAA (372G>A), TTG68CTG (373T>C), GTT69GTA (378T>A), CCG70CCT (381G>T), GGC71GGT (384C>T), TAT73TAC (390T>C), GGT78GGA (405T>A), ACT79ACG (408T>G), CCT81GAC (412C>G 413C>A 414T>C), GTT82GTC (417T>C), AAC84GCT (421A>G 422A>C 423C>T), GTG85GTC (426G>C), AAG86AAA (429G>A), TTA87CTC (430T>C 432A>C), TCT88TCA (435T>A), CTC90TTG (439C>T 441C>G), TTG91ATA (442T>A 444G>A), GTG92GTT (447G>T), CAT93CGG (449A>G 450T>G), CGT94CGA (453T>A), TCT95TCA (456T>A), GTT96ATC (457G>A 459T>C), CGT97AAA (460C>A 461G>A 462T>A), ACC101ACA (474C>A), AGG102CGC (475A>C 477G>C), AAA103AAG (480A>G), GTT104CTT (481G>C), CTA105CTT (486A>T), CAG107GCA (490C>G 491A>C 492G>A), GTT110GTA (501T>A), TCA111TCT (504A>T), GCT113_CAA115del (508_516delGCTCAACAA), ACT116TCA (517A>T 519T>A), AAT118GCG (523A>G 524A>C 525T>G), TCT119CAA (526T>C 527C>A 528T>A), GTA120TCC (529G>T 530T>C 531A>C), GAC122TCC (535G>T 536A>C), AAA123CAA (538A>C), GGG124GGT (543G>T), AAA125CAA (544A>C), ACA126AAG (548C>A 549A>G), ACC127CTT (550A>C 551C>T 552C>T), GAA128TCT (553G>T 554A>C 555A>T), ACA129CTA (556A>C 557C>T), GCT130GTA (560C>T 561T>A), TGG131AAG (562T>A 563G>A), AAG132AGT (566A>G 567G>T), TTC133CTT (568T>C 570C>T), AAA134GAA (571A>G), GAG135GAA (576G>A), ATT137AAC (581T>A 582T>C), GCT138GCA (585T>A), CAT139GCA (586C>G 587A>C 588T>A), CTC140TTA (589C>T 591C>A), AAC141AAG (594C>G), GCT142ACT (595G>A), GAG143GCT (599A>C 600G>T), CTT144TTG (601C>T 603T>G), GAG145GCT (605A>C 606G>T), CGA146GTA (607C>G 608G>T), GCA147GCT (612A>T), AGA148AAG (614G>A 615A>G), GAA150GAG (621A>G), ATA151TTA (622A>T), GCA152GCC (627A>C), GAA153CAG (628G>C 630A>G), AAA154AGC (632A>G 633A>C), TCA156TCT (639A>T), ATA158ATC (645A>C), TCC159ATC (646T>A 647C>T), TTG161CTT (652T>C 654G>T), CAA162AAA (655C>A), CTT163CTG (660T>G), CAG164CAA (663G>A), CTC165ATC (664C>A), TCC166CAG (667T>C 668C>A 669C>G), AAT167GCA (670A>G 671A>C 672T>A), CAG168CAA (675G>A), CCT169CCA (678T>A), AAT172GAA (685A>G 687T>A), ATC174GTC (691A>G), ACA176TCG (697A>T 699A>G), GGT177GGC (702T>C), TCA179ACA (706T>A), GAT181AAT (712G>A), GGG182GAT (716G>A 717G>T), TAA185TGA (725A>G) | | | | | | | | |
| ToChV_s2_gp2 | 1 | 582 | 40.1% | 2312 | 69.6% | 479 (99.4%) | 321 (66.6%) | 2/1/0/0 | 0 |

| | Begin | End | Coverage | Score | Concordance | Matches | Identities | I/D/M/F* | Stop Codons |
|---|---|---|---|---|---|---|---|---|---|
| **NT** | **41** | **2430** | **40.8%** | **1099** | **27.0%** | **2063 (98.8%)** | **1324 (63.4%)** | **6/20** | |
| Protein mutations: | M1K (687T>A), I2M (691A>G), Q5R (699A>G), V6A (702T>C), G11I (716G>A 717G>T), N14D (725A>G), R15N (729G>A 730G>T), Q17R (735A>G 736G>C), E19del (740_742delGAG), R23K (752C>A 753G>A 754G>A), T26S (761A>T 763A>T), A34E (786C>A 787T>A), L99M (980C>A 982T>G), R112A (1019C>G 1020G>C 1021T>A), I124T (1056T>C 1057C>T), S125D (1058A>G 1059G>A 1060C>T), S132T (1079T>A 1081C>G), S134F (1086C>T 1087A>C), L135F (1088C>T 1090A>T), A136S (1091G>T 1093T>A), D137G (1095A>G 1096T>G), S147E (1124A>G 1125G>A 1126T>A), L153V (1142C>G 1144C>T), E166L (1181G>C 1182A>T 1183G>T), Q171L (1197A>T 1198G>A), D185S (1238G>A 1239A>G 1240T>C), T186P (1241A>C), S192T (1259T>A 1261T>G), Q193A (1262C>G 1263A>C), Q211S (1316C>T 1317A>C 1318A>T), Q228E (1367C>G 1369A>G), S229R (1372T>G), S230A (1373T>G 1375C>G), Q231S (1376C>T 1377A>C 1378G>T), M232Q (1379A>C 1380T>A 1381G>A), A238P (1397G>C), E242A (1410A>C 1411G>A), R243L (1412A>C 1413G>T 1414G>T), R248Y (1427C>T 1428G>A 1429A>C), D250N (1433G>A), L252P (1440T>C 1441G>C), S253Q (1442A>C 1443G>A 1444T>G), M257L (1454A>T 1456G>A), T258M (1458C>T 1459T>G), S260A (1463A>G 1464G>C 1465C>G), V270I (1493G>A 1495G>C), R273K (1503G>A), V283F (1532G>T 1534G>T), P284S (1535C>T 1537T>A), N285R (1539A>G 1540C>G), T286A (1541A>G), R288S (1547C>T 1548G>C), L289S (1551T>C 1552A>G), T290_M291insVG (1555_1556insGTAGGT), M291V (1556A>G), A296G (1572C>G 1573T>G), H297V (1574C>G 1575A>T 1576C>A), E299D (1582A>T), T301S (1586A>T 1588T>C), G302N (1589G>A 1590G>A), G303L (1592G>T 1593G>T), K309E (1610A>G), G312A (1620G>C), F313I (1622T>A 1624C>T), N314D (1625A>G 1627C>T), M315F (1628A>T 1630G>T), V317Q (1634G>C 1635T>A 1636T>G), L322Y (1649C>T 1650T>A 1651G>T), D325T (1658G>A 1659A>C 1660C>T), S326D (1661T>G 1662C>A 1663C>T), T328N (1668C>A 1669A>T), H329S (1670C>T 1671A>C 1672C>G), I330F (1673A>T), D335E (1690T>G), E338T (1697G>A 1698A>C 1699G>C), R339W (1700C>T 1702C>G), T340H (1703A>C 1704C>A 1705G>T), A342S (1709G>T 1711G>A), Q343R (1712C>A 1713A>G 1714A>G), V345I (1718G>A), Q346D (1721C>G 1723A>T), Y347F (1725A>T 1726C>T), I348V (1727A>G 1729T>G), F350S (1734T>C), D351T (1736G>A 1737A>C 1738T>C), E352L (1739G>C 1740A>T 1741G>C), A353L (1742G>C 1743C>T 1744T>C), S354R (1745T>C 1746C>G 1747A>C), R355Q (1749G>A 1750A>G), N357R (1755A>G 1756T>A), V358C (1757G>T 1758T>G 1759G>T), I359H (1760A>C 1761T>A 1762C>T), V360L (1763G>C), V364I (1775G>A 1777A>T), S365E (1778T>G 1779C>A 1780A>G), K366R (1782A>G 1783G>A), E368D (1789G>C), Q369S (1790C>T 1791A>C 1792G>C), Y373N (1802T>A), K374L (1805A>C 1806A>T 1807G>A), H376Y (1811C>T), T378D (1817A>G 1818C>A), D380N (1823G>A 1825C>T), V383L (1832G>C 1834T>A), D384Q (1835G>C 1837T>G), I385V (1838A>G 1840T>G), A387K (1844G>A 1845C>A 1846T>G), A388E (1848C>A), K393N (1864G>T), N396S (1871A>T 1872A>C 1873T>C), S398E (1877T>G 1878C>A 1879T>G), S400T (1883T>A 1885A>T), L401C (1887T>G 1888G>C), D402N (1889G>A), D405S (1898G>T 1899A>C), M406L (1901A>T 1903G>A), S407T (1904T>A 1906G>T), H409R (1910C>A 1911A>G 1912T>G), A414E (1926C>A 1927C>G), Q418M (1937C>A 1938A>T 1939A>G), V423F (1952G>T 1954G>T), I425R (1959T>G 1960A>G), M426Y (1961A>T 1962T>A 1963G>T), G427W (1964G>T 1966T>G), F429L (1972T>A), L433M (1982C>A 1984A>G), S435D (1988T>G 1989C>A 1990A>C), C438V (1997T>G 1998G>T 1999T>G), I444L (2015A>C 2017T>G), G445A (2019G>C 2020T>A), E446D (2023G>T), H514D (2225C>G), I515P (2228A>C 2229T>C 2230T>G), D517G (2235A>G 2236T>G), D518K (2237G>A 2239T>A), T520E (2243A>G 2244C>A 2245T>G), G522N (2249G>A 2250G>A), V524I (2255G>A), I525F (2258A>T 2260T>C), S530E (2273T>G 2274C>A 2275A>G), D531E (2278C>A), D533M (2282G>A 2283A>T 2284C>G), S534G (2285T>G 2286C>G 2287A>T), N535D (2288A>G 2290C>T), P536A (2291C>G 2293A>C), A537P (2294G>C 2296C>G), I538L (2297A>T), D539N (2300G>A), M542L (2309A>T), V546A (2322T>C), V558I (2357G>A 2359G>T), V562L (2369G>C), E564K (2375G>A), K567N (2386A>C), M574L (2405A>C 2407G>T), S575A (2408T>G), A577S (2414G>T 2416A>C) | | | | | | | | |
| Codon mutations: | ATG1AAG (687T>A), ATA2ATG (691A>G), TCA4TCT (697A>T), CAG5CGG (699A>G), GTT6GCT (702T>C), GGT7GGA (706T>A), AAG9AAA (712G>A), GGC11ATC (716G>A 717G>T), AAC14GAC (725A>G), AGG15AAT (729G>A 730G>T), GCG16GCT (733G>T), CAG17CGC (735A>G 736G>C), GAG19del (740_742delGAG), GCT20GCC (745T>C), CGG23AAA (752C>A 753G>A 754G>A), CTA24TTA (755C>T), ATC25ATT (760C>T), ACA26TCT (761A>T 763A>T), ATT28ATA (769T>A), GCG30GCT (775G>T), GCT31GCA (778T>A), GTT32GTG (781T>G), GCT34GAA (786C>A 787T>A), GGA35GGG (790A>G), CTG39CTC (802G>C), CTT91CTA (958T>A), CAT94CAC (967T>C), TTG95TTA (970G>A), GGT97GGC (976T>C), GTT98GTC (979T>C), CTT99ATG (980C>A 982T>G), TTC100TTT (985C>T), GTC103GTG (994C>G), CCT104CCA (997T>A), GGG109GGA (1012G>A), CGT112GCA (1019C>G 1020G>C 1021T>A), AGT116AGC (1033T>C), TTG117CTG (1034T>C), TGT118TGC (1039T>C), TCG119TCC (1042G>C), CCC123CCA (1054C>A), ATC124ACT (1056T>C 1057C>T), AGC125GAT (1058A>G 1059G>A 1060C>T), CCG126CCT (1063G>T), GTG127GTT (1066G>T), ATT128ATA (1069T>A), CAA129CAG (1072A>G), GAG130GAA (1075G>A), TCC132ACG (1079T>A 1081C>G), TTG133TTA (1084G>A), TCA134TTC (1086C>T 1087A>C), CTA135TTT (1088C>T 1090A>T), GCT136TCA (1091G>T 1093T>A), GAT137GGG (1095A>G 1096T>G), GGT138GGA (1099T>A), GCA139GCT (1102A>T), CAA140CAG (1105A>G), GCC141GCT (1108C>T), GTG142GTC (1111G>C), CTC143TTG (1112C>T 1114C>G), AGT147GAA (1124A>G 1125G>A 1126T>A), ATC148ATT (1129C>T), CCC151CCT (1138C>T), CTC153GTT (1142C>G 1144C>T), GGA156GGG (1153A>G), CCA157CCC (1156A>C), TTC159TTT (1162C>T), GAG166CTT (1181G>C 1182A>T 1183G>T), AAT167AAC (1186T>C), CGT169CGG (1192T>G), GCA170GCT (1195A>T), CAG171CTA (1197A>T 1198G>A), ATA172ATT (1201A>T), CCT173CCA (1204T>A), TCC175TCT (1210C>T), GTG176GTT (1213G>T), GTA177GTG (1216A>G), GCG178GCA (1219G>A), ATA179ATT (1222A>T), AAA181AAG (1228A>G), CAA182CAG (1231A>G), GAT185AGC (1238G>A 1239A>G 1240T>C), ACA186CCA (1241A>C), AGG187AGA (1246G>A), GCC189GCT (1252C>T), GTA190GTT (1255A>T), TAC191TAT (1258C>T), TCT192ACG (1259T>A 1261T>G), CAA193GCA (1262C>G 1263A>C), CAG194CAA (1267G>A), ACT196ACA (1273T>A), TCT198AGT (1277T>A 1278C>G), GCC200GCA (1285C>A), GAA202GAG (1291A>G), GCA203GCT (1294A>T), TTG204CTC (1295T>C 1297G>C), AAC205AAT (1300C>T), CCT207CCA (1306T>A), CAA208CAG (1309A>G), TTT209TTC (1312T>C), TTC210TTT (1315C>T), CAA211TCT (1316C>T 1317A>C 1318A>T), CAG214CAA (1327G>A), GCT216GCA (1333T>A), GCC217GCA (1336C>A), TCA222TCT (1351A>T), GTA223GTC (1354A>C), TCT225TCG (1360T>G), GGG227GGA (1366G>A), CAA228GAG (1367C>G 1369A>G), AGT229AGG (1372T>G), TCC230GCG (1373T>G 1375C>G), CAG231TCT (1376C>T 1377A>C 1378G>T), ATG232CAA (1379A>C 1380T>A 1381G>A), TTG234CTA (1385T>C 1387G>A), CGT237AGA (1394C>A 1396T>A), GCT238CCT (1397G>C), GTT240GTA (1405T>A), GGA241GGT (1408A>T), GAG242GCA (1410A>C 1411G>A), AGG243CTT (1412A>C 1413G>T 1414G>T), AAT246AAC (1423T>C), ACA247ACT (1426A>T), CGA248TAC (1427C>T 1428G>A 1429A>C), GAT250AAT (1433G>A), GTA251GTT (1438A>T), CTG252CCC (1440T>C 1441G>C), AGT253CAG (1442A>C 1443G>A 1444T>G), AGC255TCT (1448A>T 1449G>C 1450C>T), TCC256TCA (1453C>A), ATG257TTA (1454A>T 1456G>A), ACT258ATG (1458C>T 1459T>G), AGC260GCG (1463A>G 1464G>C 1465C>G), TCT261TCA (1468T>A), TCC262TCG (1471C>G), CTG263CTT (1474G>T), CGA264CGC (1477A>C), CCG266CCA (1483G>A), CTC268CTA (1489C>A), GTG270ATC (1493G>A 1495G>C), AGT272AGC (1501T>C), AGG273AAG (1503G>A), CGG274AGG (1505C>A), CCA276CCT (1513A>T), TCC277TCG (1516C>G), GAA279GAG (1522A>G), CTA280CTG (1525A>G), GAC282GAT (1531C>T), GTG283TTT (1532G>T 1534G>T), CCT284TCA (1535C>T 1537T>A), AAC285AGG (1539A>G 1540C>G), ACC286GCC (1541A>G), CGA288TCA (1547C>T 1548G>C), TTA289TCG (1551T>C 1552A>G), ACA290_ATG291insGTAGGT (1555_1556insGTAGGT), ATG291GTG (1556A>G), CGA292AGA (1559C>A), GCT296GGG (1572C>G 1573T>G), CAC297GTA (1574C>G 1575A>T 1576C>A), GAA299GAT (1582A>T), GAC300GAT (1585C>T), ACT301TCC (1586A>T 1588T>C), GGT302AAT (1589G>A 1590G>A), GGA303TTA (1592G>T 1593G>T), TTG304CTG (1595T>C), CCA306CCT (1603A>T), GTT307GTC (1606T>C), AAG309GAG (1610A>G), CAA311CAG (1618A>G), GGC312GCC (1620G>C), TTC313ATT (1622T>A 1624C>T), AAC314GAT (1625A>G 1627C>T), ATG315TTT (1628A>T 1630G>T), GGT316GGC (1633T>C), GTT317CAG (1634G>C 1635T>A 1636T>G), GAT320GAC (1645T>C), CTG322TAT (1649C>T 1650T>A 1651G>T), ATA324ATC (1657A>C), GAC325ACT (1658G>A 1659A>C 1660C>T), TCC326GAT (1661T>G 1662C>A 1663C>T), ACA328AAT (1668C>A 1669A>T), CAC329TCG (1670C>T 1671A>C 1672C>G), ATT330TTT (1673A>T), GAC331GAT (1678C>T), GAT335GAG (1690T>G), ACT337ACA (1696T>A), GAG338ACC (1697G>A 1698A>C 1699G>C), CGC339TGG (1700C>T 1702C>G), ACG340CAT (1703A>C 1704C>A 1705G>T), ATT341ATA (1708T>A), GCG342TCA (1709G>T 1711G>A), CAA343AGG (1712C>A 1713A>G 1714A>G), CAG344CAA (1717G>A), GTT345ATT (1718G>A), CAA346GAT (1721C>G 1723A>T), TAC347TTT (1725A>T 1726C>T), ATT348GTG (1727A>G 1729T>G), CTG349TTG (1730C>T), TTT350TCT (1734T>C), GAT351ACC (1736G>A 1737A>C 1738T>C), GAG352CTC (1739G>C 1740A>T 1741G>C), GCT353CTC (1742G>C 1743C>T 1744T>C), TCA354CGC (1745T>C 1746C>G 1747A>C), CGA355CAG (1749G>A 1750A>G), GGC356GGG (1753C>G), AAT357AGA (1755A>G 1756T>A), GTG358TGT (1757G>T 1758T>G 1759G>T), ATC359CAT (1760A>C 1761T>A 1762C>T), GTG360CTG (1763G>C), CCG361CCC (1768G>C), AAA362AAG (1771A>G), CAT363CAC (1774T>C), GTA364ATT (1775G>A 1777A>T), TCA365GAG (1778T>G 1779C>A 1780A>G), AAG366AGA (1782A>G 1783G>A), AGG367CGA (1784A>C 1786G>A), GAG368GAC (1789G>C), CAG369TCC (1790C>T 1791A>C 1792G>C), AAT371AAC (1798T>C), ATT372ATA (1801T>A), TAC373AAC (1802T>A), AAG374CTA (1805A>C 1806A>T 1807G>A), CAC376TAC (1811C>T), ATC377ATT (1816C>T), ACT378GAT (1817A>G 1818C>A), GAC380AAT (1823G>A 1825C>T), GTT383CTA (1832G>C 1834T>A), GAT384CAG (1835G>C 1837T>G), ATT385GTG (1838A>G 1840T>G), CTT386CTC (1843T>C), GCT387AAG (1844G>A 1845C>A 1846T>G), GCA388GAA (1848C>A), ACC392ACA (1861C>A), AAG393AAT (1864G>T), GTA394GTG (1867A>G), GGA395GGT (1870A>T), AAT396TCC (1871A>T 1872A>C 1873T>C), CTC397CTA (1876C>A), TCT398GAG (1877T>G 1878C>A 1879T>G), TCA400ACT (1883T>A 1885A>T), TTG401TGC (1887T>G 1888G>C), GAT402AAT (1889G>A), TAT404TAC (1897T>C), GAT405TCT (1898G>T 1899A>C), ATG406TTA (1901A>T 1903G>A), TCG407ACT (1904T>A 1906G>T), CTT408CTC (1909T>C), CAT409AGG (1910C>A 1911A>G 1912T>G), GAG410GAA (1915G>A), AGA411AGG (1918A>G), GTC412GTG (1921C>G), ATT413ATA (1924T>A), GCC414GAG (1926C>A 1927C>G), GTG416GTC (1933G>C), CAA418ATG (1937C>A 1938A>T 1939A>G), AGA419AGG (1942A>G), GAT420GAC (1945T>C), CAC421CAT (1948C>T), GTG423TTT (1952G>T 1954G>T), TAC424TAT (1957C>T), ATA425AGG (1959T>G 1960A>G), ATG426TAT (1961A>T 1962T>A 1963G>T), GGT427TGG (1964G>T 1966T>G), TTT429TTA (1972T>A), CCC432CCG (1981C>G), CTA433ATG (1982C>A 1984A>G), TCA435GAC (1988T>G 1989C>A 1990A>C), GAT437GAC (1996T>C), TGT438GTG (1997T>G 1998G>T 1999T>G), GAC440GAT (2005C>T), CTC442CTT (2011C>T), ATT444CTG (2015A>C 2017T>G), GGT445GCA (2019G>C 2020T>A), GAG446GAT (2023G>T), ACC511ACG (2218C>G), GCA512GCT (2221A>T), GAG513GAA (2224G>A), CAT514GAT (2225C>G), ATT515CCG (2228A>C 2229T>C 2230T>G), GAT517GGG (2235A>G 2236T>G), GAT518AAA (2237G>A 2239T>A), ACT520GAG (2243A>G 2244C>A 2245T>G), GGT522AAT (2249G>A 2250G>A), GAT523GAC (2254T>C), GTT524ATT (2255G>A), ATT525TTC (2258A>T 2260T>C), GAC527GAT (2266C>T), TTC528TTT (2269C>T), GGT529GGG (2272T>G), TCA530GAG (2273T>G 2274C>A 2275A>G), GAC531GAA (2278C>A), GAC533ATG (2282G>A 2283A>T 2284C>G), TCA534GGT (2285T>G 2286C>G 2287A>T), AAC535GAT (2288A>G 2290C>T), CCA536GCC (2291C>G 2293A>C), GCC537CCG (2294G>C 2296C>G), ATA538TTA (2297A>T), GAT539AAT (2300G>A), TTA540CTG (2303T>C 2305A>G), ATG542TTG (2309A>T), CAA544CAG (2317A>G), GTA546GCA (2322T>C), GCA549GCT (2332A>T), AGC550AGT (2335C>T), GAA552GAG (2341A>G), GGT557GGG (2356T>G), GTG558ATT (2357G>A 2359G>T), GAG560GAA (2365G>A), TTC561TTT (2368C>T), GTG562CTG (2369G>C), GAA564AAA (2375G>A), AAA567AAC (2386A>C), GTG569GTT (2392G>T), GCC570GCT (2395C>T), CAA572CAG (2401A>G), GTC573GTT (2404C>T), ATG574CTT (2405A>C 2407G>T), TCA575GCA (2408T>G), TTG576CTA (2411T>C 2413G>A), GCA577TCC (2414G>T 2416A>C), CTA578CTC (2419A>C) | | | | | | | | |

#### Proteins

| | Begin | End | Coverage | Score | Concordance | Matches | Identities | I/D/M/F* | Stop Codons |
|---|---|---|---|---|---|---|---|---|---|
| hypothetical protein (YP_003097230.1) | 1 | 185 | 100% | 827 | 69.1% | 182 (98.4%) | 117 (63.2%) | 0/3/0/0 | 1 |
| Protein mutations: | G5S (184G>T 185G>C 186A>T), R6K (188G>A), A10Q (199G>C 200C>A 201A>G), E11L (202G>T 203A>T), K14Q (211A>C), A15E (215C>A 216T>A), Q18N (223C>A 225A>T), N24K (243T>G), I26V (247A>G 249T>G), V31T (262G>A 263T>C), S33V (268T>G 269C>T 270A>T), V35I (274G>A 276A>T), I36V (277A>G), V46I (307G>A 309G>C), I47C (310A>T 311T>G 312T>C), V54I (331G>A), L57I (340C>A 342A>C), T58S (343A>T 345T>C), V59I (346G>A 348T>C), P81D (412C>G 413C>A 414T>C), N84A (421A>G 422A>C 423C>T), L91I (442T>A 444G>A), H93R (449A>G 450T>G), V96I (457G>A 459T>C), R97K (460C>A 461G>A 462T>A), V104L (481G>C), Q107A (490C>G 491A>C 492G>A), A113_Q115del (508_516delGCTCAACAA), T116S (517A>T 519T>A), N118A (523A>G 524A>C 525T>G), S119Q (526T>C 527C>A 528T>A), V120S (529G>T 530T>C 531A>C), D122S (535G>T 536A>C), K123Q (538A>C), K125Q (544A>C), T126K (548C>A 549A>G), T127L (550A>C 551C>T 552C>T), E128S (553G>T 554A>C 555A>T), T129L (556A>C 557C>T), A130V (560C>T 561T>A), W131K (562T>A 563G>A), K132S (566A>G 567G>T), F133L (568T>C 570C>T), K134E (571A>G), I137N (581T>A 582T>C), H139A (586C>G 587A>C 588T>A), N141K (594C>G), A142T (595G>A), E143A (599A>C 600G>T), E145A (605A>C 606G>T), R146V (607C>G 608G>T), R148K (614G>A 615A>G), I151L (622A>T), E153Q (628G>C 630A>G), K154S (632A>G 633A>C), S159I (646T>A 647C>T), Q162K (655C>A), L165I (664C>A), S166Q (667T>C 668C>A 669C>G), N167A (670A>G 671A>C 672T>A), N172E (685A>G 687T>A), I174V (691A>G), T176S (697A>T 699A>G), S179T (706T>A), D181N (712G>A), G182D (716G>A 717G>T) | | | | | | | | |

| | Begin | End | Coverage | Score | Concordance | Matches | Identities | I/D/M/F* | Stop Codons |
|---|---|---|---|---|---|---|---|---|---|
| **NT** | **41** | **2430** | **40.8%** | **1099** | **27.0%** | **2063 (98.8%)** | **1324 (63.4%)** | **6/20** | |
| Codon mutations: | TCT2TCC (177T>C), TTT3TTC (180T>C), ATT4ATC (183T>C), GGA5TCT (184G>T 185G>C 186A>T), AGG6AAG (188G>A), ACA9ACT (198A>T), GCA10CAG (199G>C 200C>A 201A>G), GAG11TTG (202G>T 203A>T), AAA14CAA (211A>C), GCT15GAA (215C>A 216T>A), TTC16TTT (219C>T), CAA18AAT (223C>A 225A>T), GTC20GTA (231C>A), TCC23TCA (240C>A), AAT24AAG (243T>G), ATT26GTG (247A>G 249T>G), TGC27TGT (252C>T), TCG28AGT (253T>A 254C>G 255G>T), GTG29GTT (258G>T), GAC30GAT (261C>T), GTA31ACA (262G>A 263T>C), GGA32GGT (267A>T), TCA33GTT (268T>G 269C>T 270A>T), GGT34GGA (273T>A), GTA35ATT (274G>A 276A>T), ATT36GTT (277A>G), AAC39AAT (288C>T), CCC40CCG (291C>G), ACT41ACG (294T>G), CTA42CTT (297A>T), TTT44TTC (303T>C), GTG46ATC (307G>A 309G>C), ATT47TGC (310A>T 311T>G 312T>C), CCC48CCG (315C>G), ACA50ACT (321A>T), GTT54ATT (331G>A), TCA55TCT (336A>T), GTG56GTC (339G>C), CTA57ATC (340C>A 342A>C), ACT58TCC (343A>T 345T>C), GTT59ATC (346G>A 348T>C), TCG60TCC (351G>C), GAG62GAA (357G>A), TCA64TCT (363A>T), ACA65ACT (366A>T), CCA66CCG (369A>G), CAG67CAA (372G>A), TTG68CTG (373T>C), GTT69GTA (378T>A), CCG70CCT (381G>T), GGC71GGT (384C>T), TAT73TAC (390T>C), GGT78GGA (405T>A), ACT79ACG (408T>G), CCT81GAC (412C>G 413C>A 414T>C), GTT82GTC (417T>C), AAC84GCT (421A>G 422A>C 423C>T), GTG85GTC (426G>C), AAG86AAA (429G>A), TTA87CTC (430T>C 432A>C), TCT88TCA (435T>A), CTC90TTG (439C>T 441C>G), TTG91ATA (442T>A 444G>A), GTG92GTT (447G>T), CAT93CGG (449A>G 450T>G), CGT94CGA (453T>A), TCT95TCA (456T>A), GTT96ATC (457G>A 459T>C), CGT97AAA (460C>A 461G>A 462T>A), ACC101ACA (474C>A), AGG102CGC (475A>C 477G>C), AAA103AAG (480A>G), GTT104CTT (481G>C), CTA105CTT (486A>T), CAG107GCA (490C>G 491A>C 492G>A), GTT110GTA (501T>A), TCA111TCT (504A>T), GCT113_CAA115del (508_516delGCTCAACAA), ACT116TCA (517A>T 519T>A), AAT118GCG (523A>G 524A>C 525T>G), TCT119CAA (526T>C 527C>A 528T>A), GTA120TCC (529G>T 530T>C 531A>C), GAC122TCC (535G>T 536A>C), AAA123CAA (538A>C), GGG124GGT (543G>T), AAA125CAA (544A>C), ACA126AAG (548C>A 549A>G), ACC127CTT (550A>C 551C>T 552C>T), GAA128TCT (553G>T 554A>C 555A>T), ACA129CTA (556A>C 557C>T), GCT130GTA (560C>T 561T>A), TGG131AAG (562T>A 563G>A), AAG132AGT (566A>G 567G>T), TTC133CTT (568T>C 570C>T), AAA134GAA (571A>G), GAG135GAA (576G>A), ATT137AAC (581T>A 582T>C), GCT138GCA (585T>A), CAT139GCA (586C>G 587A>C 588T>A), CTC140TTA (589C>T 591C>A), AAC141AAG (594C>G), GCT142ACT (595G>A), GAG143GCT (599A>C 600G>T), CTT144TTG (601C>T 603T>G), GAG145GCT (605A>C 606G>T), CGA146GTA (607C>G 608G>T), GCA147GCT (612A>T), AGA148AAG (614G>A 615A>G), GAA150GAG (621A>G), ATA151TTA (622A>T), GCA152GCC (627A>C), GAA153CAG (628G>C 630A>G), AAA154AGC (632A>G 633A>C), TCA156TCT (639A>T), ATA158ATC (645A>C), TCC159ATC (646T>A 647C>T), TTG161CTT (652T>C 654G>T), CAA162AAA (655C>A), CTT163CTG (660T>G), CAG164CAA (663G>A), CTC165ATC (664C>A), TCC166CAG (667T>C 668C>A 669C>G), AAT167GCA (670A>G 671A>C 672T>A), CAG168CAA (675G>A), CCT169CCA (678T>A), AAT172GAA (685A>G 687T>A), ATC174GTC (691A>G), ACA176TCG (697A>T 699A>G), GGT177GGC (702T>C), TCA179ACA (706T>A), GAT181AAT (712G>A), GGG182GAT (716G>A 717G>T), TAA185TGA (725A>G) | | | | | | | | |
| polyprotein (YP_003097231.1) | 1 | 582 | 40.1% | 2312 | 69.6% | 479 (99.4%) | 321 (66.6%) | 2/1/0/0 | 0 |
| Protein mutations: | M1K (687T>A), I2M (691A>G), Q5R (699A>G), V6A (702T>C), G11I (716G>A 717G>T), N14D (725A>G), R15N (729G>A 730G>T), Q17R (735A>G 736G>C), E19del (740_742delGAG), R23K (752C>A 753G>A 754G>A), T26S (761A>T 763A>T), A34E (786C>A 787T>A), L99M (980C>A 982T>G), R112A (1019C>G 1020G>C 1021T>A), I124T (1056T>C 1057C>T), S125D (1058A>G 1059G>A 1060C>T), S132T (1079T>A 1081C>G), S134F (1086C>T 1087A>C), L135F (1088C>T 1090A>T), A136S (1091G>T 1093T>A), D137G (1095A>G 1096T>G), S147E (1124A>G 1125G>A 1126T>A), L153V (1142C>G 1144C>T), E166L (1181G>C 1182A>T 1183G>T), Q171L (1197A>T 1198G>A), D185S (1238G>A 1239A>G 1240T>C), T186P (1241A>C), S192T (1259T>A 1261T>G), Q193A (1262C>G 1263A>C), Q211S (1316C>T 1317A>C 1318A>T), Q228E (1367C>G 1369A>G), S229R (1372T>G), S230A (1373T>G 1375C>G), Q231S (1376C>T 1377A>C 1378G>T), M232Q (1379A>C 1380T>A 1381G>A), A238P (1397G>C), E242A (1410A>C 1411G>A), R243L (1412A>C 1413G>T 1414G>T), R248Y (1427C>T 1428G>A 1429A>C), D250N (1433G>A), L252P (1440T>C 1441G>C), S253Q (1442A>C 1443G>A 1444T>G), M257L (1454A>T 1456G>A), T258M (1458C>T 1459T>G), S260A (1463A>G 1464G>C 1465C>G), V270I (1493G>A 1495G>C), R273K (1503G>A), V283F (1532G>T 1534G>T), P284S (1535C>T 1537T>A), N285R (1539A>G 1540C>G), T286A (1541A>G), R288S (1547C>T 1548G>C), L289S (1551T>C 1552A>G), T290_M291insVG (1555_1556insGTAGGT), M291V (1556A>G), A296G (1572C>G 1573T>G), H297V (1574C>G 1575A>T 1576C>A), E299D (1582A>T), T301S (1586A>T 1588T>C), G302N (1589G>A 1590G>A), G303L (1592G>T 1593G>T), K309E (1610A>G), G312A (1620G>C), F313I (1622T>A 1624C>T), N314D (1625A>G 1627C>T), M315F (1628A>T 1630G>T), V317Q (1634G>C 1635T>A 1636T>G), L322Y (1649C>T 1650T>A 1651G>T), D325T (1658G>A 1659A>C 1660C>T), S326D (1661T>G 1662C>A 1663C>T), T328N (1668C>A 1669A>T), H329S (1670C>T 1671A>C 1672C>G), I330F (1673A>T), D335E (1690T>G), E338T (1697G>A 1698A>C 1699G>C), R339W (1700C>T 1702C>G), T340H (1703A>C 1704C>A 1705G>T), A342S (1709G>T 1711G>A), Q343R (1712C>A 1713A>G 1714A>G), V345I (1718G>A), Q346D (1721C>G 1723A>T), Y347F (1725A>T 1726C>T), I348V (1727A>G 1729T>G), F350S (1734T>C), D351T (1736G>A 1737A>C 1738T>C), E352L (1739G>C 1740A>T 1741G>C), A353L (1742G>C 1743C>T 1744T>C), S354R (1745T>C 1746C>G 1747A>C), R355Q (1749G>A 1750A>G), N357R (1755A>G 1756T>A), V358C (1757G>T 1758T>G 1759G>T), I359H (1760A>C 1761T>A 1762C>T), V360L (1763G>C), V364I (1775G>A 1777A>T), S365E (1778T>G 1779C>A 1780A>G), K366R (1782A>G 1783G>A), E368D (1789G>C), Q369S (1790C>T 1791A>C 1792G>C), Y373N (1802T>A), K374L (1805A>C 1806A>T 1807G>A), H376Y (1811C>T), T378D (1817A>G 1818C>A), D380N (1823G>A 1825C>T), V383L (1832G>C 1834T>A), D384Q (1835G>C 1837T>G), I385V (1838A>G 1840T>G), A387K (1844G>A 1845C>A 1846T>G), A388E (1848C>A), K393N (1864G>T), N396S (1871A>T 1872A>C 1873T>C), S398E (1877T>G 1878C>A 1879T>G), S400T (1883T>A 1885A>T), L401C (1887T>G 1888G>C), D402N (1889G>A), D405S (1898G>T 1899A>C), M406L (1901A>T 1903G>A), S407T (1904T>A 1906G>T), H409R (1910C>A 1911A>G 1912T>G), A414E (1926C>A 1927C>G), Q418M (1937C>A 1938A>T 1939A>G), V423F (1952G>T 1954G>T), I425R (1959T>G 1960A>G), M426Y (1961A>T 1962T>A 1963G>T), G427W (1964G>T 1966T>G), F429L (1972T>A), L433M (1982C>A 1984A>G), S435D (1988T>G 1989C>A 1990A>C), C438V (1997T>G 1998G>T 1999T>G), I444L (2015A>C 2017T>G), G445A (2019G>C 2020T>A), E446D (2023G>T), H514D (2225C>G), I515P (2228A>C 2229T>C 2230T>G), D517G (2235A>G 2236T>G), D518K (2237G>A 2239T>A), T520E (2243A>G 2244C>A 2245T>G), G522N (2249G>A 2250G>A), V524I (2255G>A), I525F (2258A>T 2260T>C), S530E (2273T>G 2274C>A 2275A>G), D531E (2278C>A), D533M (2282G>A 2283A>T 2284C>G), S534G (2285T>G 2286C>G 2287A>T), N535D (2288A>G 2290C>T), P536A (2291C>G 2293A>C), A537P (2294G>C 2296C>G), I538L (2297A>T), D539N (2300G>A), M542L (2309A>T), V546A (2322T>C), V558I (2357G>A 2359G>T), V562L (2369G>C), E564K (2375G>A), K567N (2386A>C), M574L (2405A>C 2407G>T), S575A (2408T>G), A577S (2414G>T 2416A>C) | | | | | | | | |
| Codon mutations: | ATG1AAG (687T>A), ATA2ATG (691A>G), TCA4TCT (697A>T), CAG5CGG (699A>G), GTT6GCT (702T>C), GGT7GGA (706T>A), AAG9AAA (712G>A), GGC11ATC (716G>A 717G>T), AAC14GAC (725A>G), AGG15AAT (729G>A 730G>T), GCG16GCT (733G>T), CAG17CGC (735A>G 736G>C), GAG19del (740_742delGAG), GCT20GCC (745T>C), CGG23AAA (752C>A 753G>A 754G>A), CTA24TTA (755C>T), ATC25ATT (760C>T), ACA26TCT (761A>T 763A>T), ATT28ATA (769T>A), GCG30GCT (775G>T), GCT31GCA (778T>A), GTT32GTG (781T>G), GCT34GAA (786C>A 787T>A), GGA35GGG (790A>G), CTG39CTC (802G>C), CTT91CTA (958T>A), CAT94CAC (967T>C), TTG95TTA (970G>A), GGT97GGC (976T>C), GTT98GTC (979T>C), CTT99ATG (980C>A 982T>G), TTC100TTT (985C>T), GTC103GTG (994C>G), CCT104CCA (997T>A), GGG109GGA (1012G>A), CGT112GCA (1019C>G 1020G>C 1021T>A), AGT116AGC (1033T>C), TTG117CTG (1034T>C), TGT118TGC (1039T>C), TCG119TCC (1042G>C), CCC123CCA (1054C>A), ATC124ACT (1056T>C 1057C>T), AGC125GAT (1058A>G 1059G>A 1060C>T), CCG126CCT (1063G>T), GTG127GTT (1066G>T), ATT128ATA (1069T>A), CAA129CAG (1072A>G), GAG130GAA (1075G>A), TCC132ACG (1079T>A 1081C>G), TTG133TTA (1084G>A), TCA134TTC (1086C>T 1087A>C), CTA135TTT (1088C>T 1090A>T), GCT136TCA (1091G>T 1093T>A), GAT137GGG (1095A>G 1096T>G), GGT138GGA (1099T>A), GCA139GCT (1102A>T), CAA140CAG (1105A>G), GCC141GCT (1108C>T), GTG142GTC (1111G>C), CTC143TTG (1112C>T 1114C>G), AGT147GAA (1124A>G 1125G>A 1126T>A), ATC148ATT (1129C>T), CCC151CCT (1138C>T), CTC153GTT (1142C>G 1144C>T), GGA156GGG (1153A>G), CCA157CCC (1156A>C), TTC159TTT (1162C>T), GAG166CTT (1181G>C 1182A>T 1183G>T), AAT167AAC (1186T>C), CGT169CGG (1192T>G), GCA170GCT (1195A>T), CAG171CTA (1197A>T 1198G>A), ATA172ATT (1201A>T), CCT173CCA (1204T>A), TCC175TCT (1210C>T), GTG176GTT (1213G>T), GTA177GTG (1216A>G), GCG178GCA (1219G>A), ATA179ATT (1222A>T), AAA181AAG (1228A>G), CAA182CAG (1231A>G), GAT185AGC (1238G>A 1239A>G 1240T>C), ACA186CCA (1241A>C), AGG187AGA (1246G>A), GCC189GCT (1252C>T), GTA190GTT (1255A>T), TAC191TAT (1258C>T), TCT192ACG (1259T>A 1261T>G), CAA193GCA (1262C>G 1263A>C), CAG194CAA (1267G>A), ACT196ACA (1273T>A), TCT198AGT (1277T>A 1278C>G), GCC200GCA (1285C>A), GAA202GAG (1291A>G), GCA203GCT (1294A>T), TTG204CTC (1295T>C 1297G>C), AAC205AAT (1300C>T), CCT207CCA (1306T>A), CAA208CAG (1309A>G), TTT209TTC (1312T>C), TTC210TTT (1315C>T), CAA211TCT (1316C>T 1317A>C 1318A>T), CAG214CAA (1327G>A), GCT216GCA (1333T>A), GCC217GCA (1336C>A), TCA222TCT (1351A>T), GTA223GTC (1354A>C), TCT225TCG (1360T>G), GGG227GGA (1366G>A), CAA228GAG (1367C>G 1369A>G), AGT229AGG (1372T>G), TCC230GCG (1373T>G 1375C>G), CAG231TCT (1376C>T 1377A>C 1378G>T), ATG232CAA (1379A>C 1380T>A 1381G>A), TTG234CTA (1385T>C 1387G>A), CGT237AGA (1394C>A 1396T>A), GCT238CCT (1397G>C), GTT240GTA (1405T>A), GGA241GGT (1408A>T), GAG242GCA (1410A>C 1411G>A), AGG243CTT (1412A>C 1413G>T 1414G>T), AAT246AAC (1423T>C), ACA247ACT (1426A>T), CGA248TAC (1427C>T 1428G>A 1429A>C), GAT250AAT (1433G>A), GTA251GTT (1438A>T), CTG252CCC (1440T>C 1441G>C), AGT253CAG (1442A>C 1443G>A 1444T>G), AGC255TCT (1448A>T 1449G>C 1450C>T), TCC256TCA (1453C>A), ATG257TTA (1454A>T 1456G>A), ACT258ATG (1458C>T 1459T>G), AGC260GCG (1463A>G 1464G>C 1465C>G), TCT261TCA (1468T>A), TCC262TCG (1471C>G), CTG263CTT (1474G>T), CGA264CGC (1477A>C), CCG266CCA (1483G>A), CTC268CTA (1489C>A), GTG270ATC (1493G>A 1495G>C), AGT272AGC (1501T>C), AGG273AAG (1503G>A), CGG274AGG (1505C>A), CCA276CCT (1513A>T), TCC277TCG (1516C>G), GAA279GAG (1522A>G), CTA280CTG (1525A>G), GAC282GAT (1531C>T), GTG283TTT (1532G>T 1534G>T), CCT284TCA (1535C>T 1537T>A), AAC285AGG (1539A>G 1540C>G), ACC286GCC (1541A>G), CGA288TCA (1547C>T 1548G>C), TTA289TCG (1551T>C 1552A>G), ACA290_ATG291insGTAGGT (1555_1556insGTAGGT), ATG291GTG (1556A>G), CGA292AGA (1559C>A), GCT296GGG (1572C>G 1573T>G), CAC297GTA (1574C>G 1575A>T 1576C>A), GAA299GAT (1582A>T), GAC300GAT (1585C>T), ACT301TCC (1586A>T 1588T>C), GGT302AAT (1589G>A 1590G>A), GGA303TTA (1592G>T 1593G>T), TTG304CTG (1595T>C), CCA306CCT (1603A>T), GTT307GTC (1606T>C), AAG309GAG (1610A>G), CAA311CAG (1618A>G), GGC312GCC (1620G>C), TTC313ATT (1622T>A 1624C>T), AAC314GAT (1625A>G 1627C>T), ATG315TTT (1628A>T 1630G>T), GGT316GGC (1633T>C), GTT317CAG (1634G>C 1635T>A 1636T>G), GAT320GAC (1645T>C), CTG322TAT (1649C>T 1650T>A 1651G>T), ATA324ATC (1657A>C), GAC325ACT (1658G>A 1659A>C 1660C>T), TCC326GAT (1661T>G 1662C>A 1663C>T), ACA328AAT (1668C>A 1669A>T), CAC329TCG (1670C>T 1671A>C 1672C>G), ATT330TTT (1673A>T), GAC331GAT (1678C>T), GAT335GAG (1690T>G), ACT337ACA (1696T>A), GAG338ACC (1697G>A 1698A>C 1699G>C), CGC339TGG (1700C>T 1702C>G), ACG340CAT (1703A>C 1704C>A 1705G>T), ATT341ATA (1708T>A), GCG342TCA (1709G>T 1711G>A), CAA343AGG (1712C>A 1713A>G 1714A>G), CAG344CAA (1717G>A), GTT345ATT (1718G>A), CAA346GAT (1721C>G 1723A>T), TAC347TTT (1725A>T 1726C>T), ATT348GTG (1727A>G 1729T>G), CTG349TTG (1730C>T), TTT350TCT (1734T>C), GAT351ACC (1736G>A 1737A>C 1738T>C), GAG352CTC (1739G>C 1740A>T 1741G>C), GCT353CTC (1742G>C 1743C>T 1744T>C), TCA354CGC (1745T>C 1746C>G 1747A>C), CGA355CAG (1749G>A 1750A>G), GGC356GGG (1753C>G), AAT357AGA (1755A>G 1756T>A), GTG358TGT (1757G>T 1758T>G 1759G>T), ATC359CAT (1760A>C 1761T>A 1762C>T), GTG360CTG (1763G>C), CCG361CCC (1768G>C), AAA362AAG (1771A>G), CAT363CAC (1774T>C), GTA364ATT (1775G>A 1777A>T), TCA365GAG (1778T>G 1779C>A 1780A>G), AAG366AGA (1782A>G 1783G>A), AGG367CGA (1784A>C 1786G>A), GAG368GAC (1789G>C), CAG369TCC (1790C>T 1791A>C 1792G>C), AAT371AAC (1798T>C), ATT372ATA (1801T>A), TAC373AAC (1802T>A), AAG374CTA (1805A>C 1806A>T 1807G>A), CAC376TAC (1811C>T), ATC377ATT (1816C>T), ACT378GAT (1817A>G 1818C>A), GAC380AAT (1823G>A 1825C>T), GTT383CTA (1832G>C 1834T>A), GAT384CAG (1835G>C 1837T>G), ATT385GTG (1838A>G 1840T>G), CTT386CTC (1843T>C), GCT387AAG (1844G>A 1845C>A 1846T>G), GCA388GAA (1848C>A), ACC392ACA (1861C>A), AAG393AAT (1864G>T), GTA394GTG (1867A>G), GGA395GGT (1870A>T), AAT396TCC (1871A>T 1872A>C 1873T>C), CTC397CTA (1876C>A), TCT398GAG (1877T>G 1878C>A 1879T>G), TCA400ACT (1883T>A 1885A>T), TTG401TGC (1887T>G 1888G>C), GAT402AAT (1889G>A), TAT404TAC (1897T>C), GAT405TCT (1898G>T 1899A>C), ATG406TTA (1901A>T 1903G>A), TCG407ACT (1904T>A 1906G>T), CTT408CTC (1909T>C), CAT409AGG (1910C>A 1911A>G 1912T>G), GAG410GAA (1915G>A), AGA411AGG (1918A>G), GTC412GTG (1921C>G), ATT413ATA (1924T>A), GCC414GAG (1926C>A 1927C>G), GTG416GTC (1933G>C), CAA418ATG (1937C>A 1938A>T 1939A>G), AGA419AGG (1942A>G), GAT420GAC (1945T>C), CAC421CAT (1948C>T), GTG423TTT (1952G>T 1954G>T), TAC424TAT (1957C>T), ATA425AGG (1959T>G 1960A>G), ATG426TAT (1961A>T 1962T>A 1963G>T), GGT427TGG (1964G>T 1966T>G), TTT429TTA (1972T>A), CCC432CCG (1981C>G), CTA433ATG (1982C>A 1984A>G), TCA435GAC (1988T>G 1989C>A 1990A>C), GAT437GAC (1996T>C), TGT438GTG (1997T>G 1998G>T 1999T>G), GAC440GAT (2005C>T), CTC442CTT (2011C>T), ATT444CTG (2015A>C 2017T>G), GGT445GCA (2019G>C 2020T>A), GAG446GAT (2023G>T), ACC511ACG (2218C>G), GCA512GCT (2221A>T), GAG513GAA (2224G>A), CAT514GAT (2225C>G), ATT515CCG (2228A>C 2229T>C 2230T>G), GAT517GGG (2235A>G 2236T>G), GAT518AAA (2237G>A 2239T>A), ACT520GAG (2243A>G 2244C>A 2245T>G), GGT522AAT (2249G>A 2250G>A), GAT523GAC (2254T>C), GTT524ATT (2255G>A), ATT525TTC (2258A>T 2260T>C), GAC527GAT (2266C>T), TTC528TTT (2269C>T), GGT529GGG (2272T>G), TCA530GAG (2273T>G 2274C>A 2275A>G), GAC531GAA (2278C>A), GAC533ATG (2282G>A 2283A>T 2284C>G), TCA534GGT (2285T>G 2286C>G 2287A>T), AAC535GAT (2288A>G 2290C>T), CCA536GCC (2291C>G 2293A>C), GCC537CCG (2294G>C 2296C>G), ATA538TTA (2297A>T), GAT539AAT (2300G>A), TTA540CTG (2303T>C 2305A>G), ATG542TTG (2309A>T), CAA544CAG (2317A>G), GTA546GCA (2322T>C), GCA549GCT (2332A>T), AGC550AGT (2335C>T), GAA552GAG (2341A>G), GGT557GGG (2356T>G), GTG558ATT (2357G>A 2359G>T), GAG560GAA (2365G>A), TTC561TTT (2368C>T), GTG562CTG (2369G>C), GAA564AAA (2375G>A), AAA567AAC (2386A>C), GTG569GTT (2392G>T), GCC570GCT (2395C>T), CAA572CAG (2401A>G), GTC573GTT (2404C>T), ATG574CTT (2405A>C 2407G>T), TCA575GCA (2408T>G), TTG576CTA (2411T>C 2413G>A), GCA577TCC (2414G>T 2416A>C), CTA578CTC (2419A>C) | | | | | | | | |

*: Inserts / Deletes / Misaligned / Frameshifts

#### Analysis details

This analysis was performed with panviral2.64

### NGS Details (UN9): Diachasmimorpha longicaudata entomopoxvirus (segment NC_043455.1)

#### Assembly

| | |
|---|---|
| **Coverage Length** | 1117 (1 contig(s)) |
| **Depth Of Coverage** | 776.0 |
| **Number Of Reads** | 7050 |
| **Reads Per Million** | 159.35 rpm (after QC) |
| **Ambiguities** | 0 |
| **Assembly Method** | de novo + reference guided assembly |
| **Consensus Caller** | Bcf Tools |

#### Coverage Map

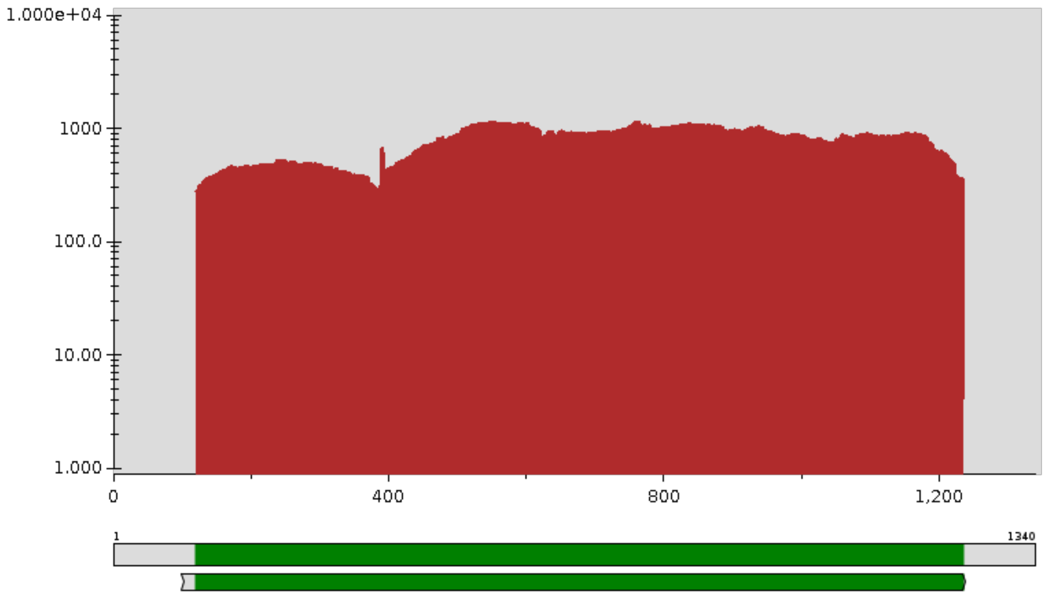

#### Assignment

| | |
|---|---|
| **Type** | Diachasmimorpha longicaudata entomopoxvirus (Taxonomy ID: 109981) |
| **Reference Genome** | NC_043455.1 |
| **NT Identity (%)** | 59.2659 |
| **AA Identity (%)** | 55.914 |
| **Number Of Stop Codons** | 0 |
| **Number Of CDS** | 1 |

#### Alignment

| | |
|---|---|
| **Alignment Score** | 396.0 (NT) + 1471.0 (AA) = 1867.0 |
| **Concordance (%)** | 39.9188 |

| Alignment Method | Global, seeded, nucleotide + amino acids (AGA) |
|---|---|

#### Genome Region

Sequence starts at position 121 and ends at position 1237 relative to NC_043455.1 reference sequence.

#### Alignment Detailed Statistics

| | Begin | End | Coverage | Score | Concordance | Matches | Identities | I/D/M/F* | Stop Codons |
|---|---|---|---|---|---|---|---|---|---|
| **NT** | **121** | **1237** | **83.4%** | **396** | **17.9%** | **1114 (99.5%)** | **662 (59.1%)** | **3/3** | |
| Mutations: | 121A>T, 126A>C, 127C>T, 128C>G, 130T>G, 136C>G, 137C>A, 139C>T, 142G>A, 143A>G, 145A>T, 146A>G, 148T>C, 149A>C, 151C>T, 154A>T, 156A>G, 157G>A, 160T>G, 163A>T, 166T>C, 167T>C, 168C>A, 169T>G, 171G>A, 172T>C, 175A>G, 178T>C, 187T>A, 190A>T, 191A>G, 193T>C, 203A>C, 204A>G, 208C>T, 209A>G, 211T>A, 214T>G, 217A>T, 218T>A, 219G>T, 223T>A, 226T>G, 230A>C, 231A>G, 232A>T, 238G>C, 241A>T, 243T>C, 244C>T, 247A>G, 256G>C, 259A>T, 262A>C, 271A>C, 272G>T, 274A>C, 276C>T, 277A>G, 278T>A, 279A>T, 283A>T, 284A>C, 286C>T, 288G>C, 289T>C, 294T>G, 295A>C, 298G>A, 299C>G, 300A>T, 301A>T, 302A>G, 304A>G, 310T>C, 311T>A, 312C>A, 313T>A, 320A>G, 322T>G, 323A>G, 325C>A, 331T>A, 337C>A, 341A>T, 343C>T, 346A>T, 352T>G, 355A>G, 356T>C, 358G>T, 361T>A, 362C>G, 363T>C, 364C>T, 368G>A, 370T>G, 371C>G, 374C>A, 375G>A, 376T>G, 379A>G, 380T>A, 382G>T, 383C>T, 384A>T, 385A>G, 386A>G, 394G>T, 395A>G, 397T>C, 403A>T, 404_406delTAT, 407A>G, 408A>G, 410T>G, 412T>G, 415A>G, 416T>G, 417G>T, 421A>T, 423T>C, 424T>C, 428A>G, 430T>A, 436C>T, 439A>C, 443A>G, 446A>C, 447A>G, 448G>T, 451A>G, 452A>G, 453G>A, 454C>T, 458G>C, 459A>G, 460A>T, 462C>T, 463T>C, 466A>T, 467A>C, 469G>A, 471A>G, 472A>T, 472_473insGGT, 474C>T, 475C>T, 478G>T, 481T>G, 482T>G, 484G>T, 485A>G, 487T>C, 490A>T, 499A>C, 502A>T, 503A>G, 505G>A, 506A>T, 511T>C, 512C>A, 514A>G, 515C>T, 517C>G, 518A>C, 519C>G, 521C>A, 523A>G, 524A>C, 526A>G, 527A>T, 528G>C, 530A>C, 532C>T, 533G>C, 534A>G, 535T>C, 536A>C, 538C>T, 539A>G, 541A>C, 542G>C, 543C>A, 544A>C, 547T>C, 550A>G, 553A>G, 554G>T, 559A>T, 560A>T, 562A>G, 574T>C, 577G>A, 581T>C, 583G>C, 584A>T, 585T>C, 586A>T, 587G>A, 588A>G, 589T>A, 590A>G, 591A>G, 595T>C, 596T>A, 597T>A, 602A>C, 604A>G, 607A>T, 608C>T, 610A>T, 616T>A, 620G>C, 622A>G, 623T>C, 628T>G, 629G>C, 630A>C, 632A>C, 633G>C, 634T>A, 635C>A, 637T>A, 638G>A, 646T>G, 647A>G, 648T>G, 650T>G, 652A>T, 655A>C, 658A>T, 661A>C, 664C>T, 665G>A, 667T>G, 671T>C, 674A>G, 675G>A, 678T>C, 680A>C, 682C>T, 683A>G, 685T>G, 687C>T, 688A>T, 689T>A, 692C>A, 693A>G, 694G>A, 695G>A, 696T>A, 697C>G, 700T>C, 705G>A, 706A>C, 707G>A, 709T>G, 715A>G, 717A>G, 718A>G, 724G>T, 727A>G, 730A>G, 731A>C, 732A>G, 735C>A, 739A>G, 740T>C, 742G>C, 745A>T, 748C>T, 754C>T, 759G>A, 765A>T, 770A>G, 772A>C, 778A>C, 779A>G, 781G>T, 784A>G, 785A>G, 787T>A, 790T>A, 792T>G, 793T>G, 796A>G, 797G>C, 798C>T, 799A>T, 810T>G, 823C>G, 824C>A, 825A>C, 826C>T, 829A>G, 830A>G, 831G>C, 833C>A, 838T>C, 841A>G, 843C>G, 844A>T, 845C>G, 847A>C, 850A>C, 854T>G, 855G>T, 859T>C, 862A>T, 863C>A, 865T>G, 866A>C, 869C>A, 871A>G, 878G>T, 879T>G, 880A>G, 885T>C, 886G>T, 889A>T, 890T>A, 891G>A, 892T>G, 893T>A, 895A>G, 896A>C, 897C>G, 898A>C, 900A>G, 901T>C, 902A>C, 904A>T, 905A>G, 908T>C, 909T>A, 915C>T, 919A>T, 920A>G, 921G>C, 922C>A, 924T>C, 925C>T, 934T>C, 938T>G, 939C>A, 943A>G, 944C>A, 946A>C, 947G>A, 948A>C, 949A>T, 950C>A, 952T>A, 956T>A, 958T>C, 966A>G, 967A>G, 970A>G, 973T>C, 974A>C, 977G>T, 978A>C, 982T>C, 983A>T, 984A>C, 986A>T, 989A>C, 991A>T, 992A>G, 994A>G, 998C>A, 1000T>C, 1001T>A, 1007A>G, 1012A>C, 1013T>C, 1015A>T, 1018A>T, 1019A>C, 1021A>T, 1024C>T, 1033T>A, 1036A>G, 1040A>G, 1042A>C, 1045T>C, 1046T>C, 1048A>T, 1051T>C, 1054C>T, 1057T>C, 1063T>C, 1064T>C, 1070C>A, 1071A>C, 1073A>C, 1075T>A, 1076A>C, 1077G>C, 1087T>C, 1088A>C, 1090A>G, 1103A>C, 1105A>T, 1111T>A, 1112A>C, 1120C>A, 1121C>A, 1123A>G, 1126A>G, 1129G>T, 1130A>G, 1141T>C, 1145A>G, 1147A>C, 1150T>C, 1151G>A, 1153A>G, 1156C>T, 1159A>T, 1160T>G, 1161T>A, 1163C>A, 1164C>G, 1165A>G, 1167A>T, 1172G>T, 1173C>T, 1177A>C, 1178T>A, 1181G>C, 1183A>G, 1185C>A, 1186G>A, 1192T>C, 1195T>C, 1196A>G, 1197C>T, 1199A>G, 1200A>T, 1204A>C, 1207T>G, 1210A>G, 1211A>C, 1213G>C, 1218A>C, 1219G>C, 1223A>G, 1227T>C, 1232T>C, 1234G>C, 1235A>C, 1237A>G | | | | | | | | |

##### CDS

| | Begin | End | Coverage | Score | Concordance | Matches | Identities | I/D/M/F* | Stop Codons |
|---|---|---|---|---|---|---|---|---|---|
| FLA14_p101 | 8 | 379 | 97.9% | 1471 | 59.6% | 371 (99.5%) | 208 (55.8%) | 1/1/0/0 | 0 |
| Protein mutations: | D9A (126A>C 127C>T), H10E (128C>G 130T>G), L13I (137C>A 139C>T), K15D (143A>G 145A>T), N16D (146A>G 148T>C), I17L (149A>C 151C>T), K19R (156A>G 157G>A), S23Q (167T>C 168C>A 169T>G), C24Y (171G>A 172T>C), T31A (191A>G 193T>C), K35R (203A>C 204A>G), I37V (209A>G 211T>A), F38L (214T>G), C40I (218T>A 219G>T), K44R (230A>C 231A>G 232A>T), V48A (243T>C 244C>T), A58S (272G>T 274A>C), T59M (276C>T 277A>G), Y60I (278T>A 279A>T), I62L (284A>C 286C>T), S63T (288G>C 289T>C), L65C (294T>G 295A>C), Q67V (299C>G 300A>T 301A>T), I68V (302A>G 304A>G), S71K (311T>A 312C>A 313T>A), N74E (320A>G 322T>G), I75V (323A>G 325C>A), T81S (341A>T 343C>T), L88A (362C>G 363T>C 364C>T), A90T (368G>A 370T>G), Q91E (371C>G), R92K (374C>A 375G>A 376T>G), L94I (380T>A 382G>T), Q95L (383C>T 384A>T 385A>G), T96A (386A>G), N99D (395A>G 397T>C), Y102del (404_406delTAT), N103G (407A>G 408A>G), F104V (410T>G 412T>G), C106V (416T>G 417G>T), Q107H (421A>T), V108A (423T>C 424T>C), I110V (428A>G 430T>A), I115V (443A>G), K116R (446A>C 447A>G 448G>T), S118D (452A>G 453G>A 454C>T), E120R (458G>C 459A>G 460A>T), T121I (462C>T 463T>C), K123Q (467A>C 469G>A), K124S (471A>G 472A>T), K124_A125insG (472_473insGGT), A125V (474C>T 475C>T), Q126H (478G>T), L128V (482T>G 484G>T), I129V (485A>G 487T>C), M135V (503A>G 505G>A), I136F (506A>T), L138M (512C>A 514A>G), T140R (518A>C 519C>G), K142Q (524A>C 526A>G), I144L (530A>C 532C>T), D145R (533G>C 534A>G 535T>C), T146P (536A>C 538C>T), K147D (539A>G 541A>C), A148H (542G>C 543C>A 544A>C), I151M (553A>G), V152F (554G>T), I154L (560A>T 562A>G), I162S (584A>T 585T>C 586A>T), D163R (587G>A 588A>G 589T>A), N164G (590A>G 591A>G), L166K (596T>A 597T>A), K168Q (602A>C 604A>G), Q170Y (608C>T 610A>T), E174Q (620G>C 622A>G), F175L (623T>C), F176L (628T>G), E177P (629G>C 630A>C), S178P (632A>C 633G>C 634T>A), H179K (635C>A 637T>A), V180I (638G>A), I183G (647A>G 648T>G), L184V (650T>G 652A>T), L185F (655A>C), V189M (665G>A 667T>G), S191P (671T>C), R192E (674A>G 675G>A), V193A (678T>C), I194L (680A>C 682C>T), N195E (683A>G 685T>G), T196I (687C>T 688A>T), S197T (689T>A), Q198R (692C>A 693A>G 694G>A), V199K (695G>A 696T>A 697C>G), R202N (705G>A 706A>C), D203K (707G>A 709T>G), K206R (717A>G 718A>G), N211R (731A>C 732A>G), A212D (735C>A), R220K (759G>A), Y222F (765A>T), I224V (770A>G 772A>C), K227D (779A>G 781G>T), N229E (785A>G 787T>A), D230E (790T>A), F231W (792T>G 793T>G), A233L (797G>C 798C>T 799A>T), F237C (810T>G), D241E (823C>G), H242T (824C>A 825A>C 826C>T), S244A (830A>G 831G>C), L245I (833C>A), T248S (843C>G 844A>T), L249V (845C>G 847A>C), C252V (854T>G 855G>T), Q257K (869C>A 871A>G), V260W (878G>T 879T>G 880A>G), M262T (885T>C 886G>T), E263D (889A>T), C264K (890T>A 891G>A 892T>G), L265M (893T>A 895A>G), T266R (896A>C 897C>G 898A>C), N267S (900A>G 901T>C), N269D (905A>G), F270H (908T>C 909T>A), A272V (915C>T), S274A (920A>G 921G>C 922C>A), I275T (924T>C 925C>T), S280D (938T>G 939C>A), Q282N (944C>A 946A>C), E283T (947G>A 948A>C 949A>T), F286I (956T>A 958T>C), K289R (966A>G 967A>G), D293S (977G>T 978A>C), K295S (983A>T 984A>C), T296S (986A>T), I298V (992A>G 994A>G), L300I (998C>A 1000T>C), S301T (1001T>A), N303D (1007A>G), I314V (1040A>G 1042A>C), H324T (1070C>A 1071A>C), N325Q (1073A>C 1075T>A), R326P (1076A>C 1077G>C), I330L (1088A>C 1090A>G), I344V (1130A>G), I349V (1145A>G 1147A>C), E351K (1151G>A 1153A>G), E353D (1159A>T), L354E (1160T>G 1161T>A), P355R (1163C>A 1164C>G 1165A>G), K356M (1167A>T), A358F (1172G>T 1173C>T), E359D (1177A>C), L360I (1178T>A), E361Q (1181G>C 1183A>G), T362K (1185C>A 1186G>A), T366V (1196A>G 1197C>T), K367V (1199A>G 1200A>T), D369E (1207T>G), M371L (1211A>C 1213G>C), E373A (1218A>C 1219G>C), I375V (1223A>G), V376A (1227T>C), I379L (1235A>C 1237A>G) | | | | | | | | |

| | Begin | End | Coverage | Score | Concordance | Matches | Identities | I/D/M/F* | Stop Codons |
|---|---|---|---|---|---|---|---|---|---|
| **NT** | **121** | **1237** | **83.4%** | **396** | **17.9%** | **1114 (99.5%)** | **662 (59.1%)** | **3/3** | |
| Codon mutations: | GAA7..T (121A>T), GAC9GCT (126A>C 127C>T), CAT10GAG (128C>G 130T>G), GGC12GGG (136C>G), CTC13ATT (137C>A 139C>T), AAG14AAA (142G>A), AAA15GAT (143A>G 145A>T), AAT16GAC (146A>G 148T>C), ATC17CTT (149A>C 151C>T), CTA18CTT (154A>T), AAG19AGA (156A>G 157G>A), GGT20GGG (160T>G), ATA21ATT (163A>T), TAT22TAC (166T>C), TCT23CAG (167T>C 168C>A 169T>G), TGT24TAC (171G>A 172T>C), GGA25GGG (175A>G), TTT26TTC (178T>C), CCT29CCA (187T>A), TCA30TCT (190A>T), ACT31GCC (191A>G 193T>C), AAA35CGA (203A>C 204A>G), GCC36GCT (208C>T), ATT37GTA (209A>G 211T>A), TTT38TTG (214T>G), CCA39CCT (217A>T), TGT40ATT (218T>A 219G>T), ATT41ATA (223T>A), TCT42TCG (226T>G), AAA44CGT (230A>C 231A>G 232A>T), GTG46GTC (238G>C), ATA47ATT (241A>T), GTC48GCT (243T>C 244C>T), CAA49CAG (247A>G), TCG52TCC (256G>C), GGA53GGT (259A>T), ACA54ACC (262A>C), ACA57ACC (271A>C), GCA58TCC (272G>T 274A>C), ACA59ATG (276C>T 277A>G), TAT60ATT (278T>A 279A>T), GCA61GCT (283A>T), ATC62CTT (284A>C 286C>T), AGT63ACC (288G>C 289T>C), TTA65TGC (294T>G 295A>C), CAG66CAA (298G>A), CAA67GTT (299C>G 300A>T 301A>T), ATA68GTG (302A>G 304A>G), ACT70ACC (310T>C), TCT71AAA (311T>A 312C>A 313T>A), AAT74GAG (320A>G 322T>G), ATC75GTA (323A>G 325C>A), GCT77GCA (331T>A), ATC79ATA (337C>A), ACC81TCT (341A>T 343C>T), CCA82CCT (346A>T), CGT84CGG (352T>G), GAA85GAG (355A>G), TTG86CTT (356T>C 358G>T), GCT87GCA (361T>A), CTC88GCT (362C>G 363T>C 364C>T), GCT90ACG (368G>A 370T>G), CAA91GAA (371C>G), CGT92AAG (374C>A 375G>A 376T>G), GTA93GTG (379A>G), TTG94ATT (380T>A 382G>T), CAA95TTG (383C>T 384A>T 385A>G), ACA96GCA (386A>G), GGG98GGT (394G>T), AAT99GAC (395A>G 397T>C), CTA101CTT (403A>T), TAT102del (404_406delTAT), AAT103GGT (407A>G 408A>G), TTT104GTG (410T>G 412T>G), AAA105AAG (415A>G), TGT106GTT (416T>G 417G>T), CAA107CAT (421A>T), GTT108GCC (423T>C 424T>C), ATT110GTA (428A>G 430T>A), GGC112GGT (436C>T), ACA113ACC (439A>C), ATC115GTC (443A>G), AAG116CGT (446A>C 447A>G 448G>T), GAA117GAG (451A>G), AGC118GAT (452A>G 453G>A 454C>T), GAA120CGT (458G>C 459A>G 460A>T), ACT121ATC (462C>T 463T>C), CTA122CTT (466A>T), AAG123CAA (467A>C 469G>A), AAA124AGT (471A>G 472A>T), AAA124_GCC125insGGT (472_473insGGT), GCC125GTT (474C>T 475C>T), CAG126CAT (478G>T), GTT127GTG (481T>G), TTG128GTT (482T>G 484G>T), ATT129GTC (485A>G 487T>C), GGA130GGT (490A>T), GGA133GGC (499A>C), CGA134CGT (502A>T), ATG135GTA (503A>G 505G>A), ATT136TTT (506A>T), GAT137GAC (511T>C), CTA138ATG (512C>A 514A>G), CTC139TTG (515C>T 517C>G), ACC140CGC (518A>C 519C>G), CGA141AGG (521C>A 523A>G), AAA142CAG (524A>C 526A>G), AGT143TCT (527A>T 528G>C), ATC144CTT (530A>C 532C>T), GAT145CGC (533G>C 534A>G 535T>C), ACC146CCT (536A>C 538C>T), AAA147GAC (539A>G 541A>C), GCA148CAC (542G>C 543C>A 544A>C), ATT149ATC (547T>C), AAA150AAG (550A>G), ATA151ATG (553A>G), GTT152TTT (554G>T), GTA153GTT (559A>T), ATA154TTG (560A>T 562A>G), GAT158GAC (574T>C), GAG159GAA (577G>A), TTG161CTC (581T>C 583G>C), ATA162TCT (584A>T 585T>C 586A>T), GAT163AGA (587G>A 588A>G 589T>A), AAT164GGT (590A>G 591A>G), TTT165TTC (595T>C), TTG166AAG (596T>A 597T>A), AAA168CAG (602A>C 604A>G), ATA169ATT (607A>T), CAA170TAT (608C>T 610A>T), ATT172ATA (616T>A), GAA174CAG (620G>C 622A>G), TTT175CTT (623T>C), TTT176TTG (628T>G), GAA177CCA (629G>C 630A>C), AGT178CCA (632A>C 633G>C 634T>A), CAT179AAA (635C>A 637T>A), GTC180ATC (638G>A), GTT182GTG (646T>G), ATT183GGT (647A>G 648T>G), TTA184GTT (650T>G 652A>T), TTA185TTC (655A>C), TCA186TCT (658A>T), GCA187GCC (661A>C), ACC188ACT (664C>T), GTT189ATG (665G>A 667T>G), TCA191CCA (671T>C), AGG192GAG (674A>G 675G>A), GTT193GCT (678T>C), ATC194CTT (680A>C 682C>T), AAT195GAG (683A>G 685T>G), ACA196ATT (687C>T 688A>T), TCT197ACT (689T>A), CAG198AGA (692C>A 693A>G 694G>A), GTC199AAG (695G>A 696T>A 697C>G), TTT200TTC (700T>C), AGA202AAC (705G>A 706A>C), GAT203AAG (707G>A 709T>G), GTA205GTG (715A>G), AAA206AGG (717A>G 718A>G), CTG208CTT (724G>T), GTA209GTG (727A>G), AAA210AAG (730A>G), AAT211CGT (731A>C 732A>G), GCT212GAT (735C>A), GAA213GAG (739A>G), TTG214CTC (740T>C 742G>C), ACA215ACT (745A>T), CTC216CTT (748C>T), GGC218GGT (754C>T), AGG220AAG (759G>A), TAT222TTT (765A>T), ATA224GTC (770A>G 772A>C), GTA226GTC (778A>C), AAG227GAT (779A>G 781G>T), AAA228AAG (784A>G), AAT229GAA (785A>G 787T>A), GAT230GAA (790T>A), TTT231TGG (792T>G 793T>G), AAA232AAG (796A>G), GCA233CTT (797G>C 798C>T 799A>T), TTT237TGT (810T>G), GAC241GAG (823C>G), CAC242ACT (824C>A 825A>C 826C>T), TTA243TTG (829A>G), AGC244GCC (830A>G 831G>C), CTC245ATC (833C>A), ACT246ACC (838T>C), CAA247CAG (841A>G), ACA248AGT (843C>G 844A>T), CTA249GTC (845C>G 847A>C), ATA250ATC (850A>C), TGT252GTT (854T>G 855G>T), AAT253AAC (859T>C), ACA254ACT (862A>T), CGT255AGG (863C>A 865T>G), AGA256CGA (866A>C), CAA257AAG (869C>A 871A>G), GTA260TGG (878G>T 879T>G 880A>G), ATG262ACT (885T>C 886G>T), GAA263GAT (889A>T), TGT264AAG (890T>A 891G>A 892T>G), TTA265ATG (893T>A 895A>G), ACA266CGC (896A>C 897C>G 898A>C), AAT267AGC (900A>G 901T>C), AGA268CGT (902A>C 904A>T), AAT269GAT (905A>G), TTC270CAC (908T>C 909T>A), GCA272GTA (915C>T), TCA273TCT (919A>T), AGC274GCA (920A>G 921G>C 922C>A), ATC275ACT (924T>C 925C>T), GAT278GAC (934T>C), TCC280GAC (938T>G 939C>A), CAA281CAG (943A>G), CAA282AAC (944C>A 946A>C), GAA283ACT (947G>A 948A>C 949A>T), CGT284AGA (950C>A 952T>A), TTT286ATC (956T>A 958T>C), AAA289AGG (966A>G 967A>G), GAA290GAG (970A>G), TTT291TTC (973T>C), AGA292CGA (974A>C), GAT293TCT (977G>T 978A>C), GGT294GGC (982T>C), AAA295TCA (983A>T 984A>C), ACT296TCT (986A>T), AGA297CGT (989A>C 991A>T), ATA298GTG (992A>G 994A>G), CTT300ATC (998C>A 1000T>C), TCA301ACA (1001T>A), AAT303GAT (1007A>G), CTA304CTC (1012A>C), TTA305CTT (1013T>C 1015A>T), GCA306GCT (1018A>T), AGA307CGT (1019A>C 1021A>T), GGC308GGT (1024C>T), GTT311GTA (1033T>A), CAA312CAG (1036A>G), ATA314GTC (1040A>G 1042A>C), TCT315TCC (1045T>C), TTA316CTT (1046T>C 1048A>T), GTT317GTC (1051T>C), ATC318ATT (1054C>T), AAT319AAC (1057T>C), GAT321GAC (1063T>C), TTG322CTG (1064T>C), CAT324ACT (1070C>A 1071A>C), AAT325CAA (1073A>C 1075T>A), AGA326CCA (1076A>C 1077G>C), TAT329TAC (1087T>C), ATA330CTG (1088A>C 1090A>G), AGA335CGT (1103A>C 1105A>T), GGT337GGA (1111T>A), AGA338CGA (1112A>C), GGC340GGA (1120C>A), CGA341AGG (1121C>A 1123A>G), AAA342AAG (1126A>G), GGG343GGT (1129G>T), ATT344GTT (1130A>G), AAT347AAC (1141T>C), ATA349GTC (1145A>G 1147A>C), ACT350ACC (1150T>C), GAA351AAG (1151G>A 1153A>G), GAC352GAT (1156C>T), GAA353GAT (1159A>T), TTA354GAA (1160T>G 1161T>A), CCA355AGG (1163C>A 1164C>G 1165A>G), AAG356ATG (1167A>T), GCT358TTT (1172G>T 1173C>T), GAA359GAC (1177A>C), TTA360ATA (1178T>A), GAA361CAG (1181G>C 1183A>G), ACG362AAA (1185C>A 1186G>A), TAT364TAC (1192T>C), AAT365AAC (1195T>C), ACC366GTC (1196A>G 1197C>T), AAA367GTA (1199A>G 1200A>T), ATA368ATC (1204A>C), GAT369GAG (1207T>G), GAA370GAG (1210A>G), ATG371CTC (1211A>C 1213G>C), GAG373GCC (1218A>C 1219G>C), ATT375GTT (1223A>G), GTT376GCT (1227T>C), TTG378CTC (1232T>C 1234G>C), ATA379CTG (1235A>C 1237A>G) | | | | | | | | |

#### Proteins

| | Begin | End | Coverage | Score | Concordance | Matches | Identities | I/D/M/F* | Stop Codons |
|---|---|---|---|---|---|---|---|---|---|
| hypothetical protein (YP_009666148.1) | 8 | 379 | 97.9% | 1471 | 59.6% | 371 (99.5%) | 208 (55.8%) | 1/1/0/0 | 0 |
| Protein mutations: | D9A (126A>C 127C>T), H10E (128C>G 130T>G), L13I (137C>A 139C>T), K15D (143A>G 145A>T), N16D (146A>G 148T>C), I17L (149A>C 151C>T), K19R (156A>G 157G>A), S23Q (167T>C 168C>A 169T>G), C24Y (171G>A 172T>C), T31A (191A>G 193T>C), K35R (203A>C 204A>G), I37V (209A>G 211T>A), F38L (214T>G), C40I (218T>A 219G>T), K44R (230A>C 231A>G 232A>T), V48A (243T>C 244C>T), A58S (272G>T 274A>C), T59M (276C>T 277A>G), Y60I (278T>A 279A>T), I62L (284A>C 286C>T), S63T (288G>C 289T>C), L65C (294T>G 295A>C), Q67V (299C>G 300A>T 301A>T), I68V (302A>G 304A>G), S71K (311T>A 312C>A 313T>A), N74E (320A>G 322T>G), I75V (323A>G 325C>A), T81S (341A>T 343C>T), L88A (362C>G 363T>C 364C>T), A90T (368G>A 370T>G), Q91E (371C>G), R92K (374C>A 375G>A 376T>G), L94I (380T>A 382G>T), Q95L (383C>T 384A>T 385A>G), T96A (386A>G), N99D (395A>G 397T>C), Y102del (404_406delTAT), N103G (407A>G 408A>G), F104V (410T>G 412T>G), C106V (416T>G 417G>T), Q107H (421A>T), V108A (423T>C 424T>C), I110V (428A>G 430T>A), I115V (443A>G), K116R (446A>C 447A>G 448G>T), S118D (452A>G 453G>A 454C>T), E120R (458G>C 459A>G 460A>T), T121I (462C>T 463T>C), K123Q (467A>C 469G>A), K124S (471A>G 472A>T), K124_A125insG (472_473insGGT), A125V (474C>T 475C>T), Q126H (478G>T), L128V (482T>G 484G>T), I129V (485A>G 487T>C), M135V (503A>G 505G>A), I136F (506A>T), L138M (512C>A 514A>G), T140R (518A>C 519C>G), K142Q (524A>C 526A>G), I144L (530A>C 532C>T), D145R (533G>C 534A>G 535T>C), T146P (536A>C 538C>T), K147D (539A>G 541A>C), A148H (542G>C 543C>A 544A>C), I151M (553A>G), V152F (554G>T), I154L (560A>T 562A>G), I162S (584A>T 585T>C 586A>T), D163R (587G>A 588A>G 589T>A), N164G (590A>G 591A>G), L166K (596T>A 597T>A), K168Q (602A>C 604A>G), Q170Y (608C>T 610A>T), E174Q (620G>C 622A>G), F175L (623T>C), F176L (628T>G), E177P (629G>C 630A>C), S178P (632A>C 633G>C 634T>A), H179K (635C>A 637T>A), V180I (638G>A), I183G (647A>G 648T>G), L184V (650T>G 652A>T), L185F (655A>C), V189M (665G>A 667T>G), S191P (671T>C), R192E (674A>G 675G>A), V193A (678T>C), I194L (680A>C 682C>T), N195E (683A>G 685T>G), T196I (687C>T 688A>T), S197T (689T>A), Q198R (692C>A 693A>G 694G>A), V199K (695G>A 696T>A 697C>G), R202N (705G>A 706A>C), D203K (707G>A 709T>G), K206R (717A>G 718A>G), N211R (731A>C 732A>G), A212D (735C>A), R220K (759G>A), Y222F (765A>T), I224V (770A>G 772A>C), K227D (779A>G 781G>T), N229E (785A>G 787T>A), D230E (790T>A), F231W (792T>G 793T>G), A233L (797G>C 798C>T 799A>T), F237C (810T>G), D241E (823C>G), H242T (824C>A 825A>C 826C>T), S244A (830A>G 831G>C), L245I (833C>A), T248S (843C>G 844A>T), L249V (845C>G 847A>C), C252V (854T>G 855G>T), Q257K (869C>A 871A>G), V260W (878G>T 879T>G 880A>G), M262T (885T>C 886G>T), E263D (889A>T), C264K (890T>A 891G>A 892T>G), L265M (893T>A 895A>G), T266R (896A>C 897C>G 898A>C), N267S (900A>G 901T>C), N269D (905A>G), F270H (908T>C 909T>A), A272V (915C>T), S274A (920A>G 921G>C 922C>A), I275T (924T>C 925C>T), S280D (938T>G 939C>A), Q282N (944C>A 946A>C), E283T (947G>A 948A>C 949A>T), F286I (956T>A 958T>C), K289R (966A>G 967A>G), D293S (977G>T 978A>C), K295S (983A>T 984A>C), T296S (986A>T), I298V (992A>G 994A>G), L300I (998C>A 1000T>C), S301T (1001T>A), N303D (1007A>G), I314V (1040A>G 1042A>C), H324T (1070C>A 1071A>C), N325Q (1073A>C 1075T>A), R326P (1076A>C 1077G>C), I330L (1088A>C 1090A>G), I344V (1130A>G), I349V (1145A>G 1147A>C), E351K (1151G>A 1153A>G), E353D (1159A>T), L354E (1160T>G 1161T>A), P355R (1163C>A 1164C>G 1165A>G), K356M (1167A>T), A358F (1172G>T 1173C>T), E359D (1177A>C), L360I (1178T>A), E361Q (1181G>C 1183A>G), T362K (1185C>A 1186G>A), T366V (1196A>G 1197C>T), K367V (1199A>G 1200A>T), D369E (1207T>G), M371L (1211A>C 1213G>C), E373A (1218A>C 1219G>C), I375V (1223A>G), V376A (1227T>C), I379L (1235A>C 1237A>G) | | | | | | | | |

| | Begin | End | Coverage | Score | Concordance | Matches | Identities | I/D/M/F* | Stop Codons |
|---|---|---|---|---|---|---|---|---|---|
| **NT** | **121** | **1237** | **83.4%** | **396** | **17.9%** | **1114 (99.5%)** | **662 (59.1%)** | **3/3** | |
| Codon mutations: | GAA7..T (121A>T), GAC9GCT (126A>C 127C>T), CAT10GAG (128C>G 130T>G), GGC12GGG (136C>G), CTC13ATT (137C>A 139C>T), AAG14AAA (142G>A), AAA15GAT (143A>G 145A>T), AAT16GAC (146A>G 148T>C), ATC17CTT (149A>C 151C>T), CTA18CTT (154A>T), AAG19AGA (156A>G 157G>A), GGT20GGG (160T>G), ATA21ATT (163A>T), TAT22TAC (166T>C), TCT23CAG (167T>C 168C>A 169T>G), TGT24TAC (171G>A 172T>C), GGA25GGG (175A>G), TTT26TTC (178T>C), CCT29CCA (187T>A), TCA30TCT (190A>T), ACT31GCC (191A>G 193T>C), AAA35CGA (203A>C 204A>G), GCC36GCT (208C>T), ATT37GTA (209A>G 211T>A), TTT38TTG (214T>G), CCA39CCT (217A>T), TGT40ATT (218T>A 219G>T), ATT41ATA (223T>A), TCT42TCG (226T>G), AAA44CGT (230A>C 231A>G 232A>T), GTG46GTC (238G>C), ATA47ATT (241A>T), GTC48GCT (243T>C 244C>T), CAA49CAG (247A>G), TCG52TCC (256G>C), GGA53GGT (259A>T), ACA54ACC (262A>C), ACA57ACC (271A>C), GCA58TCC (272G>T 274A>C), ACA59ATG (276C>T 277A>G), TAT60ATT (278T>A 279A>T), GCA61GCT (283A>T), ATC62CTT (284A>C 286C>T), AGT63ACC (288G>C 289T>C), TTA65TGC (294T>G 295A>C), CAG66CAA (298G>A), CAA67GTT (299C>G 300A>T 301A>T), ATA68GTG (302A>G 304A>G), ACT70ACC (310T>C), TCT71AAA (311T>A 312C>A 313T>A), AAT74GAG (320A>G 322T>G), ATC75GTA (323A>G 325C>A), GCT77GCA (331T>A), ATC79ATA (337C>A), ACC81TCT (341A>T 343C>T), CCA82CCT (346A>T), CGT84CGG (352T>G), GAA85GAG (355A>G), TTG86CTT (356T>C 358G>T), GCT87GCA (361T>A), CTC88GCT (362C>G 363T>C 364C>T), GCT90ACG (368G>A 370T>G), CAA91GAA (371C>G), CGT92AAG (374C>A 375G>A 376T>G), GTA93GTG (379A>G), TTG94ATT (380T>A 382G>T), CAA95TTG (383C>T 384A>T 385A>G), ACA96GCA (386A>G), GGG98GGT (394G>T), AAT99GAC (395A>G 397T>C), CTA101CTT (403A>T), TAT102del (404_406delTAT), AAT103GGT (407A>G 408A>G), TTT104GTG (410T>G 412T>G), AAA105AAG (415A>G), TGT106GTT (416T>G 417G>T), CAA107CAT (421A>T), GTT108GCC (423T>C 424T>C), ATT110GTA (428A>G 430T>A), GGC112GGT (436C>T), ACA113ACC (439A>C), ATC115GTC (443A>G), AAG116CGT (446A>C 447A>G 448G>T), GAA117GAG (451A>G), AGC118GAT (452A>G 453G>A 454C>T), GAA120CGT (458G>C 459A>G 460A>T), ACT121ATC (462C>T 463T>C), CTA122CTT (466A>T), AAG123CAA (467A>C 469G>A), AAA124AGT (471A>G 472A>T), AAA124_GCC125insGGT (472_473insGGT), GCC125GTT (474C>T 475C>T), CAG126CAT (478G>T), GTT127GTG (481T>G), TTG128GTT (482T>G 484G>T), ATT129GTC (485A>G 487T>C), GGA130GGT (490A>T), GGA133GGC (499A>C), CGA134CGT (502A>T), ATG135GTA (503A>G 505G>A), ATT136TTT (506A>T), GAT137GAC (511T>C), CTA138ATG (512C>A 514A>G), CTC139TTG (515C>T 517C>G), ACC140CGC (518A>C 519C>G), CGA141AGG (521C>A 523A>G), AAA142CAG (524A>C 526A>G), AGT143TCT (527A>T 528G>C), ATC144CTT (530A>C 532C>T), GAT145CGC (533G>C 534A>G 535T>C), ACC146CCT (536A>C 538C>T), AAA147GAC (539A>G 541A>C), GCA148CAC (542G>C 543C>A 544A>C), ATT149ATC (547T>C), AAA150AAG (550A>G), ATA151ATG (553A>G), GTT152TTT (554G>T), GTA153GTT (559A>T), ATA154TTG (560A>T 562A>G), GAT158GAC (574T>C), GAG159GAA (577G>A), TTG161CTC (581T>C 583G>C), ATA162TCT (584A>T 585T>C 586A>T), GAT163AGA (587G>A 588A>G 589T>A), AAT164GGT (590A>G 591A>G), TTT165TTC (595T>C), TTG166AAG (596T>A 597T>A), AAA168CAG (602A>C 604A>G), ATA169ATT (607A>T), CAA170TAT (608C>T 610A>T), ATT172ATA (616T>A), GAA174CAG (620G>C 622A>G), TTT175CTT (623T>C), TTT176TTG (628T>G), GAA177CCA (629G>C 630A>C), AGT178CCA (632A>C 633G>C 634T>A), CAT179AAA (635C>A 637T>A), GTC180ATC (638G>A), GTT182GTG (646T>G), ATT183GGT (647A>G 648T>G), TTA184GTT (650T>G 652A>T), TTA185TTC (655A>C), TCA186TCT (658A>T), GCA187GCC (661A>C), ACC188ACT (664C>T), GTT189ATG (665G>A 667T>G), TCA191CCA (671T>C), AGG192GAG (674A>G 675G>A), GTT193GCT (678T>C), ATC194CTT (680A>C 682C>T), AAT195GAG (683A>G 685T>G), ACA196ATT (687C>T 688A>T), TCT197ACT (689T>A), CAG198AGA (692C>A 693A>G 694G>A), GTC199AAG (695G>A 696T>A 697C>G), TTT200TTC (700T>C), AGA202AAC (705G>A 706A>C), GAT203AAG (707G>A 709T>G), GTA205GTG (715A>G), AAA206AGG (717A>G 718A>G), CTG208CTT (724G>T), GTA209GTG (727A>G), AAA210AAG (730A>G), AAT211CGT (731A>C 732A>G), GCT212GAT (735C>A), GAA213GAG (739A>G), TTG214CTC (740T>C 742G>C), ACA215ACT (745A>T), CTC216CTT (748C>T), GGC218GGT (754C>T), AGG220AAG (759G>A), TAT222TTT (765A>T), ATA224GTC (770A>G 772A>C), GTA226GTC (778A>C), AAG227GAT (779A>G 781G>T), AAA228AAG (784A>G), AAT229GAA (785A>G 787T>A), GAT230GAA (790T>A), TTT231TGG (792T>G 793T>G), AAA232AAG (796A>G), GCA233CTT (797G>C 798C>T 799A>T), TTT237TGT (810T>G), GAC241GAG (823C>G), CAC242ACT (824C>A 825A>C 826C>T), TTA243TTG (829A>G), AGC244GCC (830A>G 831G>C), CTC245ATC (833C>A), ACT246ACC (838T>C), CAA247CAG (841A>G), ACA248AGT (843C>G 844A>T), CTA249GTC (845C>G 847A>C), ATA250ATC (850A>C), TGT252GTT (854T>G 855G>T), AAT253AAC (859T>C), ACA254ACT (862A>T), CGT255AGG (863C>A 865T>G), AGA256CGA (866A>C), CAA257AAG (869C>A 871A>G), GTA260TGG (878G>T 879T>G 880A>G), ATG262ACT (885T>C 886G>T), GAA263GAT (889A>T), TGT264AAG (890T>A 891G>A 892T>G), TTA265ATG (893T>A 895A>G), ACA266CGC (896A>C 897C>G 898A>C), AAT267AGC (900A>G 901T>C), AGA268CGT (902A>C 904A>T), AAT269GAT (905A>G), TTC270CAC (908T>C 909T>A), GCA272GTA (915C>T), TCA273TCT (919A>T), AGC274GCA (920A>G 921G>C 922C>A), ATC275ACT (924T>C 925C>T), GAT278GAC (934T>C), TCC280GAC (938T>G 939C>A), CAA281CAG (943A>G), CAA282AAC (944C>A 946A>C), GAA283ACT (947G>A 948A>C 949A>T), CGT284AGA (950C>A 952T>A), TTT286ATC (956T>A 958T>C), AAA289AGG (966A>G 967A>G), GAA290GAG (970A>G), TTT291TTC (973T>C), AGA292CGA (974A>C), GAT293TCT (977G>T 978A>C), GGT294GGC (982T>C), AAA295TCA (983A>T 984A>C), ACT296TCT (986A>T), AGA297CGT (989A>C 991A>T), ATA298GTG (992A>G 994A>G), CTT300ATC (998C>A 1000T>C), TCA301ACA (1001T>A), AAT303GAT (1007A>G), CTA304CTC (1012A>C), TTA305CTT (1013T>C 1015A>T), GCA306GCT (1018A>T), AGA307CGT (1019A>C 1021A>T), GGC308GGT (1024C>T), GTT311GTA (1033T>A), CAA312CAG (1036A>G), ATA314GTC (1040A>G 1042A>C), TCT315TCC (1045T>C), TTA316CTT (1046T>C 1048A>T), GTT317GTC (1051T>C), ATC318ATT (1054C>T), AAT319AAC (1057T>C), GAT321GAC (1063T>C), TTG322CTG (1064T>C), CAT324ACT (1070C>A 1071A>C), AAT325CAA (1073A>C 1075T>A), AGA326CCA (1076A>C 1077G>C), TAT329TAC (1087T>C), ATA330CTG (1088A>C 1090A>G), AGA335CGT (1103A>C 1105A>T), GGT337GGA (1111T>A), AGA338CGA (1112A>C), GGC340GGA (1120C>A), CGA341AGG (1121C>A 1123A>G), AAA342AAG (1126A>G), GGG343GGT (1129G>T), ATT344GTT (1130A>G), AAT347AAC (1141T>C), ATA349GTC (1145A>G 1147A>C), ACT350ACC (1150T>C), GAA351AAG (1151G>A 1153A>G), GAC352GAT (1156C>T), GAA353GAT (1159A>T), TTA354GAA (1160T>G 1161T>A), CCA355AGG (1163C>A 1164C>G 1165A>G), AAG356ATG (1167A>T), GCT358TTT (1172G>T 1173C>T), GAA359GAC (1177A>C), TTA360ATA (1178T>A), GAA361CAG (1181G>C 1183A>G), ACG362AAA (1185C>A 1186G>A), TAT364TAC (1192T>C), AAT365AAC (1195T>C), ACC366GTC (1196A>G 1197C>T), AAA367GTA (1199A>G 1200A>T), ATA368ATC (1204A>C), GAT369GAG (1207T>G), GAA370GAG (1210A>G), ATG371CTC (1211A>C 1213G>C), GAG373GCC (1218A>C 1219G>C), ATT375GTT (1223A>G), GTT376GCT (1227T>C), TTG378CTC (1232T>C 1234G>C), ATA379CTG (1235A>C 1237A>G) | | | | | | | | |

*: Inserts / Deletes / Misaligned / Frameshifts

#### Analysis details

This analysis was performed with panviral2.64

### NGS Details (UN9): Bracoviriform glomeratae (segment NC_043292.1)

#### Assembly

| | |
|---|---|
| **Coverage Length** | 302 (1 contig(s)) |
| **Depth Of Coverage** | 2248.3 |
| **Number Of Reads** | 6038 |
| **Reads Per Million** | 136.48 rpm (after QC) |
| **Ambiguities** | 0 |
| **Assembly Method** | de novo + reference guided assembly |
| **Consensus Caller** | Bcf Tools |

#### Coverage Map

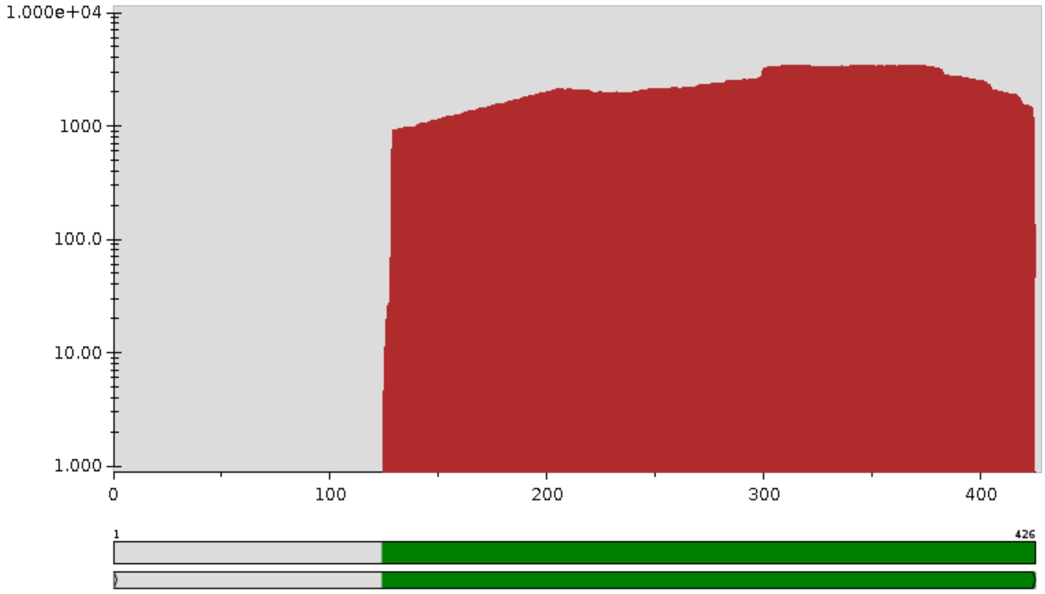

#### Assignment

| | |
|---|---|
| **Type** | Bracoviriform glomeratae (Taxonomy ID: 257816) |
| **Reference Genome** | NC_043292.1 |
| **NT Identity (%)** | 72.1854 |
| **AA Identity (%)** | 85.0 |
| **Number Of Stop Codons** | 1 |
| **Number Of CDS** | 1 |

#### Alignment

| | |
|---|---|
| **Alignment Score** | 268.0 (NT) + 549.0 (AA) = 817.0 |
| **Concordance (%)** | 65.834 |

| Alignment Method | Global, seeded, nucleotide + amino acids (AGA) |
|---|---|

#### Genome Region

Sequence starts at position 125 and ends at position 426 relative to NC_043292.1 reference sequence.

#### Alignment Detailed Statistics

| | Begin | End | Coverage | Score | Concordance | Matches | Identities | I/D/M/F* | Stop Codons |
|---|---|---|---|---|---|---|---|---|---|
| **NT** | **125** | **426** | **70.9%** | **268** | **44.4%** | **302 (100%)** | **218 (72.2%)** | **0/0** | |
| Mutations: | 126G>A, 129C>A, 132A>G, 136A>G, 137C>G, 138G>C, 150A>C, 153A>G, 155C>G, 156G>A, 159C>A, 166C>A, 168T>G, 169T>C, 171T>C, 172C>A, 174C>G, 177A>G, 183T>A, 184C>A, 189T>C, 198A>G, 201A>C, 207C>T, 214A>G, 219C>T, 222T>C, 223C>A, 225T>G, 228G>T, 235C>A, 237T>G, 243A>T, 246C>G, 255C>T, 261A>G, 264A>G, 265G>A, 273A>G, 278T>C, 279T>C, 284A>G, 285T>A, 288T>G, 289T>C, 300C>T, 301C>T, 303C>G, 309C>T, 312T>G, 315C>T, 322G>T, 329T>C, 333T>C, 336C>T, 339A>G, 345A>C, 347A>G, 348A>G, 349C>A, 351T>A, 360C>T, 363C>T, 366C>T, 375C>T, 378C>T, 384A>C, 385T>C, 387G>T, 390A>G, 391C>A, 392A>G, 393T>A, 394A>C, 399C>A, 400C>A, 402T>G, 404T>C, 405G>T, 406A>C, 408C>T, 414T>A, 416A>T, 420A>G | | | | | | | | |

##### CDS

| | Begin | End | Coverage | Score | Concordance | Matches | Identities | I/D/M/F* | Stop Codons |
|---|---|---|---|---|---|---|---|---|---|
| FK954_p501 | 43 | 142 | 70.4% | 549 | 85.6% | 100 (100%) | 85 (85.0%) | 0/0/0/0 | 1 |
| Protein mutations: | T46G (136A>G 137C>G 138G>C), A52G (155C>G 156G>A), Y57H (169T>C 171T>C), T72A (214A>G), V89I (265G>A), I93T (278T>C 279T>C), D95G (284A>G 285T>A), A108S (322G>T), I110T (329T>C), K116R (347A>G 348A>G), H131R (391C>A 392A>G 393T>A), K132Q (394A>C), M135T (404T>C 405G>T), I136L (406A>C 408C>T), Y139F (416A>T) | | | | | | | | |
| Codon mutations: | TTG42.TA (126G>A), GGC43GGA (129C>A), AAA44AAG (132A>G), ACG46GGC (136A>G 137C>G 138G>C), GGA50GGC (150A>C), AAA51AAG (153A>G), GCG52GGA (155C>G 156G>A), GGC53GGA (159C>A), CGT56AGG (166C>A 168T>G), TAT57CAC (169T>C 171T>C), CGC58AGG (172C>A 174C>G), AAA59AAG (177A>G), CTT61CTA (183T>A), CGA62AGA (184C>A), GAT63GAC (189T>C), CAA66CAG (198A>G), GGA67GGC (201A>C), ACC69ACT (207C>T), ACT72GCT (214A>G), ATC73ATT (219C>T), CGT74CGC (222T>C), CGT75AGG (223C>A 225T>G), CTG76CTT (228G>T), CGT79AGG (235C>A 237T>G), GGA81GGT (243A>T), GTC82GTG (246C>G), ATC85ATT (255C>T), GGA87GGG (261A>G), TTA88TTG (264A>G), GTC89ATC (265G>A), GAA91GAG (273A>G), ATT93ACC (278T>C 279T>C), GAT95GGA (284A>G 285T>A), GTT96GTG (288T>G), TTG97CTG (289T>C), TTC100TTT (300C>T), CTC101TTG (301C>T 303C>G), AAC103AAT (309C>T), GTT104GTG (312T>G), ATC105ATT (315C>T), GCT108TCT (322G>T), ATC110ACC (329T>C), TAT111TAC (333T>C), ACC112ACT (336C>T), GAA113GAG (339A>G), GCA115GCC (345A>C), AAA116AGG (347A>G 348A>G), CGT117AGA (349C>A 351T>A), GTC120GTT (360C>T), ACC121ACT (363C>T), GCC122GCT (366C>T), GTC125GTT (375C>T), GTC126GTT (378C>T), GCA128GCC (384A>C), TTG129CTT (385T>C 387G>T), AAA130AAG (390A>G), CAT131AGA (391C>A 392A>G 393T>A), AAA132CAA (394A>C), GGC133GGA (399C>A), CGT134AGG (400C>A 402T>G), ATG135ACT (404T>C 405G>T), ATC136CTT (406A>C 408C>T), GGT138GGA (414T>A), TAT139TTT (416A>T), GGA140GGG (420A>G) | | | | | | | | |

##### Proteins

| | Begin | End | Coverage | Score | Concordance | Matches | Identities | I/D/M/F* | Stop Codons |
|---|---|---|---|---|---|---|---|---|---|
| putative histone 4 (YP_009665791.1) | 43 | 142 | 70.4% | 549 | 85.6% | 100 (100%) | 85 (85.0%) | 0/0/0/0 | 1 |
| Protein mutations: | T46G (136A>G 137C>G 138G>C), A52G (155C>G 156G>A), Y57H (169T>C 171T>C), T72A (214A>G), V89I (265G>A), I93T (278T>C 279T>C), D95G (284A>G 285T>A), A108S (322G>T), I110T (329T>C), K116R (347A>G 348A>G), H131R (391C>A 392A>G 393T>A), K132Q (394A>C), M135T (404T>C 405G>T), I136L (406A>C 408C>T), Y139F (416A>T) | | | | | | | | |
| Codon mutations: | TTG42.TA (126G>A), GGC43GGA (129C>A), AAA44AAG (132A>G), ACG46GGC (136A>G 137C>G 138G>C), GGA50GGC (150A>C), AAA51AAG (153A>G), GCG52GGA (155C>G 156G>A), GGC53GGA (159C>A), CGT56AGG (166C>A 168T>G), TAT57CAC (169T>C 171T>C), CGC58AGG (172C>A 174C>G), AAA59AAG (177A>G), CTT61CTA (183T>A), CGA62AGA (184C>A), GAT63GAC (189T>C), CAA66CAG (198A>G), GGA67GGC (201A>C), ACC69ACT (207C>T), ACT72GCT (214A>G), ATC73ATT (219C>T), CGT74CGC (222T>C), CGT75AGG (223C>A 225T>G), CTG76CTT (228G>T), CGT79AGG (235C>A 237T>G), GGA81GGT (243A>T), GTC82GTG (246C>G), ATC85ATT (255C>T), GGA87GGG (261A>G), TTA88TTG (264A>G), GTC89ATC (265G>A), GAA91GAG (273A>G), ATT93ACC (278T>C 279T>C), GAT95GGA (284A>G 285T>A), GTT96GTG (288T>G), TTG97CTG (289T>C), TTC100TTT (300C>T), CTC101TTG (301C>T 303C>G), AAC103AAT (309C>T), GTT104GTG (312T>G), ATC105ATT (315C>T), GCT108TCT (322G>T), ATC110ACC (329T>C), TAT111TAC (333T>C), ACC112ACT (336C>T), GAA113GAG (339A>G), GCA115GCC (345A>C), AAA116AGG (347A>G 348A>G), CGT117AGA (349C>A 351T>A), GTC120GTT (360C>T), ACC121ACT (363C>T), GCC122GCT (366C>T), GTC125GTT (375C>T), GTC126GTT (378C>T), GCA128GCC (384A>C), TTG129CTT (385T>C 387G>T), AAA130AAG (390A>G), CAT131AGA (391C>A 392A>G 393T>A), AAA132CAA (394A>C), GGC133GGA (399C>A), CGT134AGG (400C>A 402T>G), ATG135ACT (404T>C 405G>T), ATC136CTT (406A>C 408C>T), GGT138GGA (414T>A), TAT139TTT (416A>T), GGA140GGG (420A>G) | | | | | | | | |

*: Inserts / Deletes / Misaligned / Frameshifts

#### Analysis details

This analysis was performed with panviral2.64

### NGS Details (UN9): Potato leafroll virus

#### Assembly

| | |
|---|---|
| **Coverage Length** | 5825 (1 contig(s)) |
| **Depth Of Coverage** | 104.8 |
| **Number Of Reads** | 4587 |
| **Reads Per Million** | 103.68 rpm (after QC) |
| **Ambiguities** | 0 |
| **Assembly Method** | read mapping against reference + variant calling |
| **Consensus Caller** | Bcf Tools |

#### Coverage Map

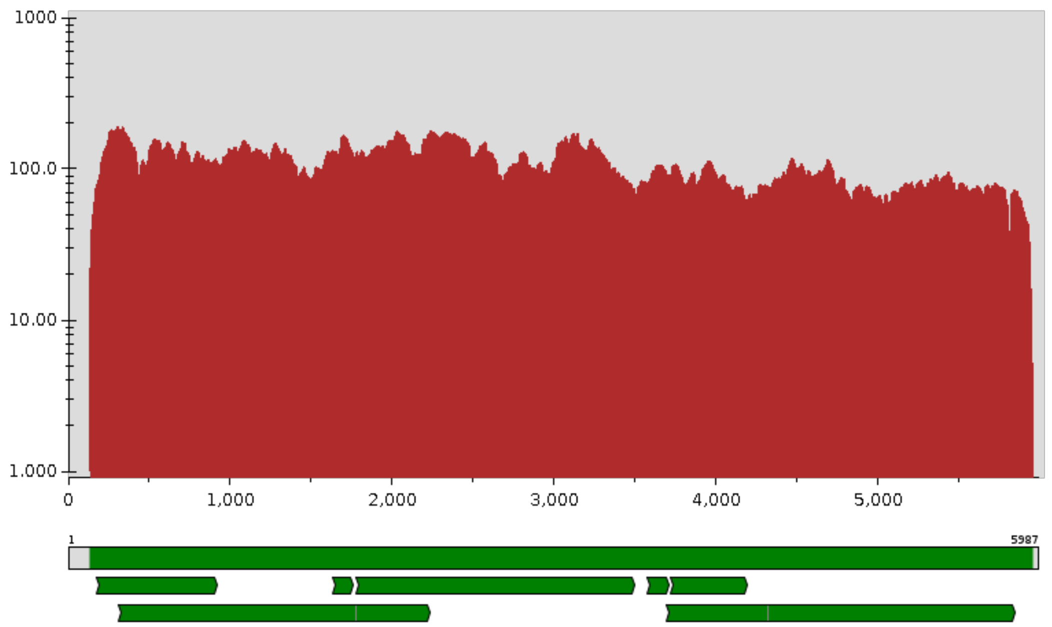

#### Assignment

| | |
|---|---|
| **Type** | Potato leafroll virus (Taxonomy ID: 12045) |
| **Reference Genome** | NC_001747.1 |
| **NT Identity (%)** | 97.6481 |
| **AA Identity (%)** | 96.6091 |
| **Number Of Stop Codons** | 8 |
| **Number Of CDS** | 8 |

#### Alignment

| | |
|---|---|
| **Alignment Score** | 11084.0 (NT) + 20290.0 (AA) = 31374.0 |
| **Concordance (%)** | 95.6095 |

| Alignment Method | Global, seeded, nucleotide + amino acids (AGA) |
|---|---|

#### Genome Region

Sequence starts at position 135 and ends at position 5959 relative to NC_001747.1 reference sequence.

#### Alignment Detailed Statistics

| | Begin | End | Coverage | Score | Concordance | Matches | Identities | I/D/M/F* | Stop Codons |
|---|---|---|---|---|---|---|---|---|---|
| **NT** | **135** | **5959** | **97.3%** | **11084** | **95.3%** | **5823 (99.9%)** | **5688 (97.6%)** | **2/2** | |
| Mutations: | 191A>C, 221T>C, 228C>T, 269T>C, 276G>A, 300C>T, 343C>T, 462T>A, 476_477insTT, 479_480delTT, 492T>C, 525C>T, 561C>T, 600T>C, 665A>G, 697C>A, 699A>C, 822C>T, 825T>G, 922G>A, 931A>T, 958G>A, 1054C>T, 1132T>G, 1147C>T, 1171C>T, 1183T>C, 1204A>G, 1319G>A, 1356G>A, 1381T>C, 1411T>G, 1434A>G, 1465C>T, 1474G>A, 1534G>A, 1577G>T, 1593G>A, 1611A>G, 1618C>T, 1648T>C, 1662G>A, 1701A>G, 1716A>G, 1755T>C, 1757A>G, 1774T>C, 1814G>T, 1830C>T, 1894C>A, 1924G>A, 2007T>C, 2021A>G, 2047G>A, 2064A>G, 2079C>T, 2154C>T, 2187C>T, 2265A>G, 2283G>A, 2316T>G, 2400C>T, 2433T>C, 2470G>A, 2550G>A, 2574C>T, 2580A>G, 2586A>G, 2673C>T, 2704C>T, 2729A>G, 2850A>G, 2856G>A, 2869G>A, 2877C>T, 2898T>C, 2931T>C, 3000T>C, 3033C>T, 3036C>T, 3054T>C, 3096C>T, 3105C>T, 3117C>T, 3195C>T, 3321C>T, 3379A>G, 3420A>G, 3437A>G, 3441C>T, 3619C>T, 3652T>A, 3748T>G, 3750C>A, 3777G>T, 3788T>A, 3939A>T, 4013C>T, 4031A>G, 4034C>T, 4132A>T, 4151T>C, 4214T>G, 4265T>C, 4280C>T, 4379G>A, 4385C>T, 4421A>G, 4430G>A, 4433C>T, 4511T>C, 4520G>T, 4521T>G, 4661T>C, 4700C>T, 4733C>G, 4736A>G, 4817T>C, 4835C>T, 4883C>T, 4886T>C, 4949G>A, 4964A>G, 4979C>T, 5062C>T, 5072C>T, 5095G>A, 5129G>A, 5162C>T, 5375T>G, 5385G>A, 5432T>C, 5493G>A, 5517A>G, 5527C>T, 5530T>C, 5925T>C | | | | | | | | |
| **CDS** | | | | | | | | | |
| PLRVgp1 | 1 | 248 | 100% | 1428 | 82.4% | 248 (99.6%) | 241 (96.8%) | 1/0/2/2 | 1 |
| Protein mutations: | Q6P (191A>C), F16S (221T>C), L32P (269T>C), H57Y (343C>T), G101_L102insX (476_477insTT), Q164R (665A>G), Q175N (697C>A 699A>C) | | | | | | | | |
| Codon mutations: | CAG6CCG (191A>C), TTT16TCT (221T>C), CTC18CTT (228C>T), CTT32CCT (269T>C), CTG34CTA (276G>A), GGC42GGT (300C>T), CAT57TAT (343C>T), GCT96GCA (462T>A), GGC101GGT (476_477insTT), GGC101_CTT102insT-C (476_477insTT), CTT102C-- (479_480delTT), ATT106ATC (492T>C), TAC117TAT (525C>T), TAC129TAT (561C>T), CAT142CAC (600T>C), CAA164CGA (665A>G), CAA175AAC (697C>A 699A>C), CGC216CGT (822C>T), GCT217GCG (825T>G) | | | | | | | | |
| PLRVgp2 | 1 | 1063 | 100% | 7004 | 93.9% | 1063 (99.9%) | 1029 (96.7%) | 1/0/2/2 | 1 |
| Protein mutations: | L52Q (462T>A), W56_A56insV (476_477insTT), L61S (492T>C), T72I (525C>T), T84I (561C>T), M97T (600T>C), K119E (665A>G), N130T (699A>C), A171V (822C>T), L172R (825T>G), E207D (931A>T), A337T (1319G>A), G349E (1356G>A), E375G (1434A>G), V423F (1577G>T), R428K (1593G>A), D434G (1611A>G), R451K (1662G>A), Q464R (1701A>G), E469G (1716A>G), L482P (1755T>C), T483A (1757A>G), W489R (1774T>C), R502L (1814G>T), Q529K (1894C>A), E539K (1924G>A), N571S (2021A>G), E580K (2047G>A), A721T (2470G>A), N807S (2729A>G), A854T (2869G>A), I1024V (3379A>G), E1043G (3437A>G) | | | | | | | | |
| Codon mutations: | TTC12TTT (343C>T), CTA52CAA (462T>A), TGG56_GCC56insGTT (476_477insTT), TTA57--A (479_480delTT), TTA61TCA (492T>C), ACA72ATA (525C>T), ACA84ATA (561C>T), ATG97ACG (600T>C), AAA119GAA (665A>G), ATC129ATA (697C>A), AAC130ACC (699A>C), GCG171GTG (822C>T), CTA172CGA (825T>G), AGG204AGA (922G>A), GAA207GAT (931A>T), CAG216CAA (958G>A), GCC248GCT (1054C>T), ACT274ACG (1132T>G), TTC279TTT (1147C>T), TCC287TCT (1171C>T), GGT291GGC (1183T>C), CTA298CTG (1204A>G), GCC337ACC (1319G>A), GGA349GAA (1356G>A), TTT357TTC (1381T>C), CTT367CTG (1411T>G), GAG375GGG (1434A>G), CCC385CCT (1465C>T), CCG388CCA (1474G>A), TCG408TCA (1534G>A), GTC423TTC (1577G>T), AGA428AAA (1593G>A), GAC434GGC (1611A>G), AAC436AAT (1618C>T), GAT446GAC (1648T>C), AGA451AAA (1662G>A), CAA464CGA (1701A>G), GAG469GGG (1716A>G), CTA482CCA (1755T>C), ACA483GCA (1757A>G), AAT488AAC (1774T>C), TGG489CGG (1774T>C), CGA502CTA (1814G>T), CGC507CGT (1830C>T), CAA529AAA (1894C>A), GAG539AAG (1924G>A), TTT566TTC (2007T>C), AAC571AGC (2021A>G), GAA580AAA (2047G>A), CCA585CCG (2064A>G), TGC590TGT (2079C>T), CAC615CAT (2154C>T), GGC626GGT (2187C>T), TCA652TCG (2265A>G), GCG658GCA (2283G>A), GTT669GTG (2316T>G), ATC697ATT (2400C>T), GCT708GCC (2433T>C), GCA721ACA (2470G>A), TCG747TCA (2550G>A), AGC755AGT (2574C>T), GAA757GAG (2580A>G), CTA759CTG (2586A>G), CGC788CGT (2673C>T), CTG799TTG (2704C>T), AAT807AGT (2729A>G), CCA847CCG (2850A>G), GTG849GTA (2856G>A), GCA854ACA (2869G>A), CAC856CAT (2877C>T), ACT863ACC (2898T>C), TAT874TAC (2931T>C), CGT897CGC (3000T>C), AAC908AAT (3033C>T), TCC909TCT (3036C>T), GAT915GAC (3054T>C), AGC929AGT (3096C>T), TAC932TAT (3105C>T), TCC936TCT (3117C>T), CTC962CTT (3195C>T), AAC1004AAT (3321C>T), ATT1024GTT (3379A>G), GAA1037GAG (3420A>G), GAG1043GGG (3437A>G), CTC1044CTT (3441C>T) | | | | | | | | |
| PLRVgp3 | 1 | 640 | 100% | 3875 | 90.3% | 640 (99.8%) | 609 (95.0%) | 1/0/2/2 | 1 |
| Protein mutations: | L52Q (462T>A), W56_A56insV (476_477insTT), L61S (492T>C), T72I (525C>T), T84I (561C>T), M97T (600T>C), K119E (665A>G), N130T (699A>C), A171V (822C>T), L172R (825T>G), E207D (931A>T), A337T (1319G>A), G349E (1356G>A), E375G (1434A>G), V423F (1577G>T), R428K (1593G>A), D434G (1611A>G), R451K (1662G>A), Q464R (1701A>G), E469G (1716A>G), L482P (1755T>C), T483A (1757A>G), D502Y (1814G>T), A507V (1830C>T), L566S (2007T>C), T571A (2021A>G), Q585R (2064A>G), A590V (2079C>T), T615I (2154C>T), A626V (2187C>T) | | | | | | | | |
| Codon mutations: | TTC12TTT (343C>T), CTA52CAA (462T>A), TGG56_GCC56insGTT (476_477insTT), TTA57--A (479_480delTT), TTA61TCA (492T>C), ACA72ATA (525C>T), ACA84ATA (561C>T), ATG97ACG (600T>C), AAA119GAA (665A>G), ATC129ATA (697C>A), AAC130ACC (699A>C), GCG171GTG (822C>T), CTA172CGA (825T>G), AGG204AGA (922G>A), GAA207GAT (931A>T), CAG216CAA (958G>A), GCC248GCT (1054C>T), ACT274ACG (1132T>G), TTC279TTT (1147C>T), TCC287TCT (1171C>T), GGT291GGC (1183T>C), CTA298CTG (1204A>G), GCC337ACC (1319G>A), GGA349GAA (1356G>A), TTT357TTC (1381T>C), CTT367CTG (1411T>G), GAG375GGG (1434A>G), CCC385CCT (1465C>T), CCG388CCA (1474G>A), TCG408TCA (1534G>A), GTC423TTC (1577G>T), AGA428AAA (1593G>A), GAC434GGC (1611A>G), AAC436AAT (1618C>T), GAT446GAC (1648T>C), AGA451AAA (1662G>A), CAA464CGA (1701A>G), GAG469GGG (1716A>G), CTA482CCA (1755T>C), ACA483GCA (1757A>G), AAT488AAC (1774T>C), GAC502TAC (1814G>T), GCT507GTT (1830C>T), ATC528ATA (1894C>A), GTG538GTA (1924G>A), TTA566TCA (2007T>C), ACG571GCG (2021A>G), AAG579AAA (2047G>A), CAG585CGG (2064A>G), GCG590GTG (2079C>T), ACC615ATC (2154C>T), GCT626GTT (2187C>T) | | | | | | | | |
| Rap1 | 1 | 42 | 100% | 257 | 89.5% | 42 (100%) | 37 (88.1%) | 0/0/0/0 | 0 |
| Protein mutations: | I6T (1648T>C), E11K (1662G>A), K24E (1701A>G), R29G (1716A>G), *42Q (1755T>C 1757A>G) | | | | | | | | |
| Codon mutations: | ATT6ACT (1648T>C), GAG11AAG (1662G>A), AAA24GAA (1701A>G), AGA29GGA (1716A>G), TAA42CAG (1755T>C 1757A>G) | | | | | | | | |
| ORF3a | 1 | 46 | 100% | 291 | 97.3% | 46 (100%) | 45 (97.8%) | 0/0/0/0 | 1 |
| Protein mutations: | F26L (3652T>A) | | | | | | | | |
| Codon mutations: | TCC15TCT (3619C>T), TTT26TTA (3652T>A) | | | | | | | | |
| PLRVgp4 | 1 | 718 | 100% | 4966 | 98.6% | 718 (100%) | 704 (98.1%) | 0/0/0/0 | 2 |
| Protein mutations: | M19R (3748T>G), A29S (3777G>T), T83S (3939A>T), Y147F (4132A>T), K276N (4520G>T), F277V (4521T>G), A457V (5062C>T), G468E (5095G>A), N561K (5375T>G), E565K (5385G>A), V601I (5493G>A), K609E (5517A>G), S612L (5527C>T), L613P (5530T>C) | | | | | | | | |
| Codon mutations: | ATG19AGG (3748T>G), CGA20AGA (3750C>A), GCT29TCT (3777G>T), GTT32GTA (3788T>A), ACC83TCC (3939A>T), TAC107TAT (4013C>T), ACA113ACG (4031A>G), AGC114AGT (4034C>T), TAC147TTC (4132A>T), ATT153ATC (4151T>C), TCT174TCG (4214T>G), GAT191GAC (4265T>C), TTC196TTT (4280C>T), CAG229CAA (4379G>A), CAC231CAT (4385C>T), CTA243CTG (4421A>G), CAG246CAA (4430G>A), GCC247GCT (4433C>T), AAT273AAC (4511T>C), AAG276AAT (4520G>T), TTT277GTT (4521T>G), GGT323GGC (4661T>C), TCC336TCT (4700C>T), GTC347GTG (4733C>G), AAA348AAG (4736A>G), GGT375GGC (4817T>C), GAC381GAT (4835C>T), TTC397TTT (4883C>T), CTT398CTC (4886T>C), ACG419ACA (4949G>A), GAA424GAG (4964A>G), ACC429ACT (4979C>T), GCC457GTC (5062C>T), CAC460CAT (5072C>T), GGA468GAA (5095G>A), ACG479ACA (5129G>A), GAC490GAT (5162C>T), AAT561AAG (5375T>G), GAA565AAA (5385G>A), ACT580ACC (5432T>C), GTC601ATC (5493G>A), AAG609GAG (5517A>G), TCA612TTA (5527C>T), CTT613CCT (5530T>C) | | | | | | | | |
| PLRVgp5 | 1 | 209 | 100% | 1417 | 98.7% | 209 (100%) | 205 (98.1%) | 0/0/0/0 | 1 |
| Protein mutations: | M19R (3748T>G), A29S (3777G>T), T83S (3939A>T), Y147F (4132A>T) | | | | | | | | |
| Codon mutations: | ATG19AGG (3748T>G), CGA20AGA (3750C>A), GCT29TCT (3777G>T), GTT32GTA (3788T>A), ACC83TCC (3939A>T), TAC107TAT (4013C>T), ACA113ACG (4031A>G), AGC114AGT (4034C>T), TAC147TTC (4132A>T), ATT153ATC (4151T>C), TCT174TCG (4214T>G), GAT191GAC (4265T>C), TTC196TTT (4280C>T) | | | | | | | | |
| PLRVgp6 | 1 | 157 | 100% | 1052 | 95.5% | 157 (100%) | 150 (95.5%) | 0/0/0/0 | 1 |
| Protein mutations: | C11G (3748T>G 3750C>A), F24Y (3788T>A), T99I (4013C>T), Q105R (4031A>G), A106V (4034C>T), T139S (4132A>T), L145S (4151T>C) | | | | | | | | |
| Codon mutations: | TGC11GGA (3748T>G 3750C>A), GCG20GCT (3777G>T), TTC24TAC (3788T>A), ACA74ACT (3939A>T), ACC99ATC (4013C>T), CAA105CGA (4031A>G), GCA106GTA (4034C>T), ACG139TCG (4132A>T), TTA145TCA (4151T>C) | | | | | | | | |

| | Begin | End | Coverage | Score | Concordance | Matches | Identities | I/D/M/F* | Stop Codons |
|---|---|---|---|---|---|---|---|---|---|
| **NT** | **135** | **5959** | **97.3%** | **11084** | **95.3%** | **5823 (99.9%)** | **5688 (97.6%)** | **2/2** | |
| **Proteins** | | | | | | | | | |
| P0 protein (NP_056746.1) | 1 | 248 | 100% | 1428 | 82.4% | 248 (99.6%) | 241 (96.8%) | 1/0/2/2 | 1 |
| Protein mutations: | Q6P (191A>C), F16S (221T>C), L32P (269T>C), H57Y (343C>T), G101_L102insX (476_477insTT), Q164R (665A>G), Q175N (697C>A 699A>C) | | | | | | | | |
| Codon mutations: | CAG6CCG (191A>C), TTT16TCT (221T>C), CTC18CTT (228C>T), CTT32CCT (269T>C), CTG34CTA (276G>A), GGC42GGT (300C>T), CAT57TAT (343C>T), GCT96GCA (462T>A), GGC101GGT (476_477insTT), GGC101_CTT102insT-C (476_477insTT), CTT102C-- (479_480delTT), ATT106ATC (492T>C), TAC117TAT (525C>T), TAC129TAT (561C>T), CAT142CAC (600T>C), CAA164CGA (665A>G), CAA175AAC (697C>A 699A>C), CGC216CGT (822C>T), GCT217GCG (825T>G) | | | | | | | | |
| RNA-dependent RNA polymerase (NP_056748.3) | 1 | 1063 | 100% | 7004 | 93.9% | 1063 (99.9%) | 1029 (96.7%) | 1/0/2/2 | 1 |
| Protein mutations: | L52Q (462T>A), W56_A56insV (476_477insTT), L61S (492T>C), T72I (525C>T), T84I (561C>T), M97T (600T>C), K119E (665A>G), N130T (699A>C), A171V (822C>T), L172R (825T>G), E207D (931A>T), A337T (1319G>A), G349E (1356G>A), E375G (1434A>G), V423F (1577G>T), R428K (1593G>A), D434G (1611A>G), R451K (1662G>A), Q464R (1701A>G), E469G (1716A>G), L482P (1755T>C), T483A (1757A>G), W489R (1774T>C), R502L (1814G>T), Q529K (1894C>A), E539K (1924G>A), N571S (2021A>G), E580K (2047G>A), A721T (2470G>A), N807S (2729A>G), A854T (2869G>A), I1024V (3379A>G), E1043G (3437A>G) | | | | | | | | |
| Codon mutations: | TTC12TTT (343C>T), CTA52CAA (462T>A), TGG56_GCC56insGTT (476_477insTT), TTA57--A (479_480delTT), TTA61TCA (492T>C), ACA72ATA (525C>T), ACA84ATA (561C>T), ATG97ACG (600T>C), AAA119GAA (665A>G), ATC129ATA (697C>A), AAC130ACC (699A>C), GCG171GTG (822C>T), CTA172CGA (825T>G), AGG204AGA (922G>A), GAA207GAT (931A>T), CAG216CAA (958G>A), GCC248GCT (1054C>T), ACT274ACG (1132T>G), TTC279TTT (1147C>T), TCC287TCT (1171C>T), GGT291GGC (1183T>C), CTA298CTG (1204A>G), GCC337ACC (1319G>A), GGA349GAA (1356G>A), TTT357TTC (1381T>C), CTT367CTG (1411T>G), GAG375GGG (1434A>G), CCC385CCT (1465C>T), CCG388CCA (1474G>A), TCG408TCA (1534G>A), GTC423TTC (1577G>T), AGA428AAA (1593G>A), GAC434GGC (1611A>G), AAC436AAT (1618C>T), GAT446GAC (1648T>C), AGA451AAA (1662G>A), CAA464CGA (1701A>G), GAG469GGG (1716A>G), CTA482CCA (1755T>C), ACA483GCA (1757A>G), AAT488AAC (1774T>C), TGG489CGG (1774T>C), CGA502CTA (1814G>T), CGC507CGT (1830C>T), CAA529AAA (1894C>A), GAG539AAG (1924G>A), TTT566TTC (2007T>C), AAC571AGC (2021A>G), GAA580AAA (2047G>A), CCA585CCG (2064A>G), TGC590TGT (2079C>T), CAC615CAT (2154C>T), GGC626GGT (2187C>T), TCA652TCG (2265A>G), GCG658GCA (2283G>A), GTT669GTG (2316T>G), ATC697ATT (2400C>T), GCT708GCC (2433T>C), GCA721ACA (2470G>A), TCG747TCA (2550G>A), AGC755AGT (2574C>T), GAA757GAG (2580A>G), CTA759CTG (2586A>G), CGC788CGT (2673C>T), CTG799TTG (2704C>T), AAT807AGT (2729A>G), CCA847CCG (2850A>G), GTG849GTA (2856G>A), GCA854ACA (2869G>A), CAC856CAT (2877C>T), ACT863ACC (2898T>C), TAT874TAC (2931T>C), CGT897CGC (3000T>C), AAC908AAT (3033C>T), TCC909TCT (3036C>T), GAT915GAC (3054T>C), AGC929AGT (3096C>T), TAC932TAT (3105C>T), TCC936TCT (3117C>T), CTC962CTT (3195C>T), AAC1004AAT (3321C>T), ATT1024GTT (3379A>G), GAA1037GAG (3420A>G), GAG1043GGG (3437A>G), CTC1044CTT (3441C>T) | | | | | | | | |
| P1 protein (NP_056747.1) | 1 | 640 | 100% | 3875 | 90.3% | 640 (99.8%) | 609 (95.0%) | 1/0/2/2 | 1 |
| Protein mutations: | L52Q (462T>A), W56_A56insV (476_477insTT), L61S (492T>C), T72I (525C>T), T84I (561C>T), M97T (600T>C), K119E (665A>G), N130T (699A>C), A171V (822C>T), L172R (825T>G), E207D (931A>T), A337T (1319G>A), G349E (1356G>A), E375G (1434A>G), V423F (1577G>T), R428K (1593G>A), D434G (1611A>G), R451K (1662G>A), Q464R (1701A>G), E469G (1716A>G), L482P (1755T>C), T483A (1757A>G), D502Y (1814G>T), A507V (1830C>T), L566S (2007T>C), T571A (2021A>G), Q585R (2064A>G), A590V (2079C>T), T615I (2154C>T), A626V (2187C>T) | | | | | | | | |
| Codon mutations: | TTC12TTT (343C>T), CTA52CAA (462T>A), TGG56_GCC56insGTT (476_477insTT), TTA57--A (479_480delTT), TTA61TCA (492T>C), ACA72ATA (525C>T), ACA84ATA (561C>T), ATG97ACG (600T>C), AAA119GAA (665A>G), ATC129ATA (697C>A), AAC130ACC (699A>C), GCG171GTG (822C>T), CTA172CGA (825T>G), AGG204AGA (922G>A), GAA207GAT (931A>T), CAG216CAA (958G>A), GCC248GCT (1054C>T), ACT274ACG (1132T>G), TTC279TTT (1147C>T), TCC287TCT (1171C>T), GGT291GGC (1183T>C), CTA298CTG (1204A>G), GCC337ACC (1319G>A), GGA349GAA (1356G>A), TTT357TTC (1381T>C), CTT367CTG (1411T>G), GAG375GGG (1434A>G), CCC385CCT (1465C>T), CCG388CCA (1474G>A), TCG408TCA (1534G>A), GTC423TTC (1577G>T), AGA428AAA (1593G>A), GAC434GGC (1611A>G), AAC436AAT (1618C>T), GAT446GAC (1648T>C), AGA451AAA (1662G>A), CAA464CGA (1701A>G), GAG469GGG (1716A>G), CTA482CCA (1755T>C), ACA483GCA (1757A>G), AAT488AAC (1774T>C), GAC502TAC (1814G>T), GCT507GTT (1830C>T), ATC528ATA (1894C>A), GTG538GTA (1924G>A), TTA566TCA (2007T>C), ACG571GCG (2021A>G), AAG579AAA (2047G>A), CAG585CGG (2064A>G), GCG590GTG (2079C>T), ACC615ATC (2154C>T), GCT626GTT (2187C>T) | | | | | | | | |
| Replication-associated protein (YP_006355442.1) | 1 | 42 | 100% | 257 | 89.5% | 42 (100%) | 37 (88.1%) | 0/0/0/0 | 0 |
| Protein mutations: | I6T (1648T>C), E11K (1662G>A), K24E (1701A>G), R29G (1716A>G), *42Q (1755T>C 1757A>G) | | | | | | | | |
| Codon mutations: | ATT6ACT (1648T>C), GAG11AAG (1662G>A), AAA24GAA (1701A>G), AGA29GGA (1716A>G), TAA42CAG (1755T>C 1757A>G) | | | | | | | | |
| protein 3a (YP_009179365.2) | 1 | 46 | 100% | 291 | 97.3% | 46 (100%) | 45 (97.8%) | 0/0/0/0 | 1 |
| Protein mutations: | F26L (3652T>A) | | | | | | | | |
| Codon mutations: | TCC15TCT (3619C>T), TTT26TTA (3652T>A) | | | | | | | | |
| CP read-through protein (NP_056751.2) | 1 | 718 | 100% | 4966 | 98.6% | 718 (100%) | 704 (98.1%) | 0/0/0/0 | 2 |
| Protein mutations: | M19R (3748T>G), A29S (3777G>T), T83S (3939A>T), Y147F (4132A>T), K276N (4520G>T), F277V (4521T>G), A457V (5062C>T), G468E (5095G>A), N561K (5375T>G), E565K (5385G>A), V601I (5493G>A), K609E (5517A>G), S612L (5527C>T), L613P (5530T>C) | | | | | | | | |
| Codon mutations: | ATG19AGG (3748T>G), CGA20AGA (3750C>A), GCT29TCT (3777G>T), GTT32GTA (3788T>A), ACC83TCC (3939A>T), TAC107TAT (4013C>T), ACA113ACG (4031A>G), AGC114AGT (4034C>T), TAC147TTC (4132A>T), ATT153ATC (4151T>C), TCT174TCG (4214T>G), GAT191GAC (4265T>C), TTC196TTT (4280C>T), CAG229CAA (4379G>A), CAC231CAT (4385C>T), CTA243CTG (4421A>G), CAG246CAA (4430G>A), GCC247GCT (4433C>T), AAT273AAC (4511T>C), AAG276AAT (4520G>T), TTT277GTT (4521T>G), GGT323GGC (4661T>C), TCC336TCT (4700C>T), GTC347GTG (4733C>G), AAA348AAG (4736A>G), GGT375GGC (4817T>C), GAC381GAT (4835C>T), TTC397TTT (4883C>T), CTT398CTC (4886T>C), ACG419ACA (4949G>A), GAA424GAG (4964A>G), ACC429ACT (4979C>T), GCC457GTC (5062C>T), CAC460CAT (5072C>T), GGA468GAA (5095G>A), ACG479ACA (5129G>A), GAC490GAT (5162C>T), AAT561AAG (5375T>G), GAA565AAA (5385G>A), ACT580ACC (5432T>C), GTC601ATC (5493G>A), AAG609GAG (5517A>G), TCA612TTA (5527C>T), CTT613CCT (5530T>C) | | | | | | | | |
| coat protein (NP_056749.1) | 1 | 209 | 100% | 1417 | 98.7% | 209 (100%) | 205 (98.1%) | 0/0/0/0 | 1 |
| Protein mutations: | M19R (3748T>G), A29S (3777G>T), T83S (3939A>T), Y147F (4132A>T) | | | | | | | | |
| Codon mutations: | ATG19AGG (3748T>G), CGA20AGA (3750C>A), GCT29TCT (3777G>T), GTT32GTA (3788T>A), ACC83TCC (3939A>T), TAC107TAT (4013C>T), ACA113ACG (4031A>G), AGC114AGT (4034C>T), TAC147TTC (4132A>T), ATT153ATC (4151T>C), TCT174TCG (4214T>G), GAT191GAC (4265T>C), TTC196TTT (4280C>T) | | | | | | | | |
| movement protein (NP_056750.1) | 1 | 157 | 100% | 1052 | 95.5% | 157 (100%) | 150 (95.5%) | 0/0/0/0 | 1 |
| Protein mutations: | C11G (3748T>G 3750C>A), F24Y (3788T>A), T99I (4013C>T), Q105R (4031A>G), A106V (4034C>T), T139S (4132A>T), L145S (4151T>C) | | | | | | | | |
| Codon mutations: | TGC11GGA (3748T>G 3750C>A), GCG20GCT (3777G>T), TTC24TAC (3788T>A), ACA74ACT (3939A>T), ACC99ATC (4013C>T), CAA105CGA (4031A>G), GCA106GTA (4034C>T), ACG139TCG (4132A>T), TTA145TCA (4151T>C) | | | | | | | | |

*: Inserts / Deletes / Misaligned / Frameshifts

#### Analysis details

This analysis was performed with panviral2.64

### NGS Details (UN9): Solendovirus venanicotianae

#### Assembly

| | |
|---|---|
| **Coverage Length** | 2217 (4 contig(s)) |
| **Depth Of Coverage** | 11.9 |
| **Number Of Reads** | 233 |
| **Reads Per Million** | 5.27 rpm (after QC) |
| **Ambiguities** | 0 |
| **Assembly Method** | de novo + reference guided assembly |
| **Consensus Caller** | Bcf Tools |

#### Coverage Map

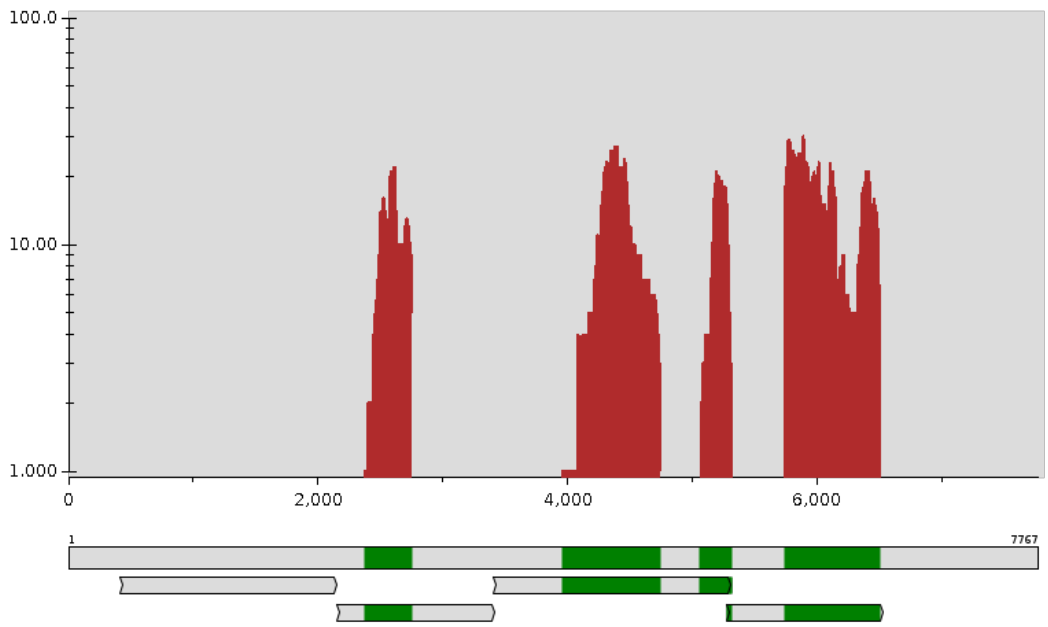

#### Assignment

| | |
|---|---|
| **Type** | Solendovirus venanicotianae (Taxonomy ID: 3048371) |
| **Reference Genome** | NC_003378.1 |
| **NT Identity (%)** | 79.8918 |
| **AA Identity (%)** | 76.2599 |
| **Number Of Stop Codons** | 5 |
| **Number Of CDS** | 4 |

#### Alignment

| | |
|---|---|
| **Alignment Score** | 2644.0 (NT) + 3921.0 (AA) = 6565.0 |
| **Concordance (%)** | 68.8715 |

| **Alignment Method** | Global, seeded, nucleotide + amino acids (AGA) |
|---|---|

#### Genome Region

Sequence starts at position 2369 and ends at position 6509 relative to NC_003378.1 reference sequence.

#### Alignment Detailed Statistics

| | Begin | End | Coverage | Score | Concordance | Matches | Identities | I/D/M/F* | Stop Codons |
|---|---|---|---|---|---|---|---|---|---|
| **NT** | **2369** | **6509** | **28.5%** | **2644** | **59.6%** | **2217 (99.9%)** | **1772 (79.9%)** | **1/0** | |
| Mutations: | 2373T>A, 2376T>C, 2379C>T, 2388A>T, 2389G>A, 2390G>A, 2394T>A, 2397G>A, 2403A>T, 2410A>C, 2412A>T, 2417A>C, 2418C>T, 2420A>G, 2423G>A, 2425A>T, 2428A>G, 2433T>C, 2434A>C, 2443C>A, 2454G>A, 2461A>G, 2463C>A, 2469T>A, 2472G>A, 2481C>T, 2482T>C, 2493T>C, 2500T>C, 2501T>A, 2505A>T, 2535C>A, 2538T>G, 2545A>C, 2551C>T, 2565A>G, 2569A>G, 2571C>A, 2572C>G, 2589C>T, 2602A>G, 2613T>C, 2622T>C, 2628G>A, 2637T>C, 2638A>G, 2643A>C, 2646C>T, 2647G>C, 2649C>A, 2665C>T, 2668A>T, 2669A>C, 2670T>A, 2674A>G, 2675G>A, 2685T>A, 2688A>T, 2691A>T, 2700A>T, 2709G>A, 2712G>A, 2715A>G, 2722G>A, 2724G>A, 2725T>C, 2730A>T, 2733C>A, 2734G>T, 2738G>C, 2743G>A, 2748T>C, 2754C>A, 3957A>G, 3962T>A, 3983T>C, 3986A>G, 3990A>C, 3993G>A, 4013A>C, 4016G>A, 4019T>C, 4022T>C, 4037A>G, 4040T>A, 4045A>G, 4056A>G, 4059G>A, 4061T>C, 4065A>G, 4066C>A, 4076A>G, 4086A>G, 4088A>T, 4091T>A, 4100T>A, 4103T>A, 4104A>G, 4105A>G, 4115A>G, 4116G>A, 4127C>T, 4181A>T, 4187C>T, 4190G>A, 4196C>T, 4199A>T, 4212T>C, 4235T>C, 4241T>C, 4250C>T, 4265G>A, 4286T>C, 4292C>T, 4307C>T, 4313G>A, 4325T>C, 4328T>C, 4329T>C, 4331G>A, 4334A>G, 4340G>A, 4343T>A, 4353A>C, 4355A>G, 4362G>A, 4364T>A, 4388T>C, 4391A>G, 4393G>T, 4400T>A, 4403A>G, 4404C>T, 4411G>A, 4417A>G, 4421T>C, 4436C>T, 4451C>T, 4467C>T, 4470G>C, 4471A>C, 4475T>C, 4478T>C, 4482G>A, 4484A>T, 4497A>G, 4500C>A, 4503C>T, 4508T>C, 4513G>A, 4517A>T, 4518C>A, 4523T>A, 4530A>T, 4540T>C, 4545A>C, 4550C>T, 4551G>A, 4552C>T, 4554C>A, 4559T>A, 4560A>G, 4568C>A, 4571T>A, 4574T>A, 4581T>C, 4591C>A, 4592C>G, 4598A>T, 4601A>G, 4604T>A, 4605A>T, 4607G>T, 4626T>C, 4643T>C, 4653A>G, 4660T>C, 4665A>C, 4666C>A, 4667T>A, 4679A>G, 4687T>C, 4688C>A, 4691T>C, 4692C>T, 4693T>C, 4694T>A, 4697T>A, 4701A>G, 4703C>A, 4704A>T, 4706T>A, 4709T>C, 4721A>G, 4734C>T, 4740A>T, 5077T>A, 5078T>G, 5079G>T, 5080C>T, 5081A>G, 5094C>T, 5097T>C, 5102A>C, 5117A>T, 5120T>G, 5125C>A, 5126C>T, 5128G>A, 5132T>C, 5135T>A, 5136G>A, 5147C>T, 5156A>G, 5174T>A, 5180A>G, 5192A>G, 5195C>T, 5211C>A, 5217C>T, 5219G>A, 5237C>T, 5253G>A, 5256A>G, 5261T>A, 5263A>G, 5273G>A, 5280G>A, 5283G>A, 5286G>T, 5291T>C, 5295T>G, 5301C>T, 5310C>A, 5313A>G, 5745T>C, 5754T>A, 5755A>C, 5756C>A, 5757C>A, 5760T>C, 5763A>T, 5766C>A, 5781A>C, 5784G>A, 5786A>C, 5796T>A, 5797A>C, 5798A>T, 5799A>C, 5801A>C, 5802T>G, 5811C>T, 5812C>A, 5813A>G, 5814A>C, 5815T>A, 5820C>A, 5821C>G, 5826T>A, 5831A>C, 5838C>T, 5844T>A, 5847C>T, 5850C>A, 5851C>A, 5853T>A, 5854C>T, 5857A>C, 5858C>A, 5872T>C, 5874G>A, 5877A>T, 5880T>A, 5881T>C, 5882T>A, 5889T>A, 5892A>T, 5894G>A, 5897C>T, 5898A>C, 5901A>T, 5907A>T, 5908G>A, 5909C>G, 5910C>A, 5913C>A, 5916C>T, 5921A>G, 5932C>T, 5935G>A, 5940T>C, 5950C>T, 5952G>T, 5953A>G, 5954C>A, 5955A>C, 5959C>G, 5964G>A, 5968G>T, 5970C>A, 5971A>G, 5972A>C, 5979T>A, 5982G>A, 5983T>C, 5988C>T, 5991G>A, 5997C>T, 5999T>C, 6000G>A, 6001C>A, 6002A>C, 6009G>A, 6012G>A, 6015A>T, 6018T>A, 6024G>C, 6026C>A, 6027A>C, 6033C>G, 6036C>T, 6037G>A, 6040C>A, 6041G>A, 6042A>G, 6045C>T, 6049T>A, 6050C>T, 6051G>A, 6053C>T, 6054T>A, 6055A>C, 6059C>T, 6067C>T, 6071A>T, 6078G>A, 6078_6079insA, 6081A>T, 6084G>A, 6086C>T, 6087T>A, 6088A>G, 6090C>G, 6102C>T, 6108T>A, 6110G>A, 6114C>A, 6118A>C, 6120G>A, 6123A>T, 6129C>T, 6132A>T, 6135A>C, 6139A>G, 6141G>A, 6144A>T, 6154G>A, 6163G>A, 6164A>C, 6170T>A, 6172A>G, 6174T>G, 6176T>G, 6177T>A, 6180A>T, 6182A>T, 6183A>T, 6186C>T, 6201G>A, 6204A>T, 6211A>T, 6219T>A, 6222A>C, 6223C>A, 6225A>T, 6240C>T, 6243C>T, 6247C>A, 6253C>G, 6255G>A, 6259G>C, 6260C>T, 6264T>G, 6269G>T, 6273T>C, 6276T>C, 6277T>A, 6282G>A, 6285A>C, 6288G>A, 6290C>T, 6308G>C, 6309C>A, 6311G>A, 6315T>G, 6320G>T, 6321A>C, 6325G>A, 6326C>G, 6340C>G, 6341T>C, 6344G>T, 6345A>G, 6352G>A, 6354C>A, 6355T>A, 6357A>G, 6359G>C, 6360T>A, 6361C>T, 6363T>G, 6369A>T, 6375A>G, 6376C>A, 6380C>A, 6381T>A, 6382A>C, 6384A>T, 6386C>T, 6387C>T, 6401G>A, 6402G>A, 6404G>A, 6406A>G, 6407A>G, 6408T>A, 6411C>T, 6414C>T, 6417C>G, 6418C>G, 6419C>A, 6420A>T, 6423T>A, 6424C>T, 6426G>A, 6427T>C, 6446T>A, 6451A>G, 6453T>G, 6456C>A, 6462T>C, 6471C>T, 6477C>T, 6480A>G, 6489A>C | | | | | | | | |
| **CDS** | | | | | | | | | |
| Tvvgp2 | 73 | 201 | 30.7% | 701 | 81.3% | 129 (100%) | 100 (77.5%) | 0/0/0/0 | 1 |
| Protein mutations: | G79K (2389G>A 2390G>A), K83N (2403A>T), I86L (2410A>C 2412A>T), N88T (2417A>C 2418C>T), K89R (2420A>G), R90K (2423G>A), I91L (2425A>T), I92V (2428A>G), K94Q (2434A>C), Q97K (2443C>A), N103E (2461A>G 2463C>A), L116Q (2500T>C 2501T>A), I131L (2545A>C), I139V (2569A>G 2571C>A), Q140E (2572C>G), I150V (2602A>G), I162V (2638A>G), A165P (2647G>C 2649C>A), N172S (2668A>T 2669A>C 2670T>A), R174E (2674A>G 2675G>A), D177E (2685T>A), K178N (2688A>T), L182F (2700A>T), M186I (2712G>A), E190K (2722G>A 2724G>A), G194* (2734G>T), S195T (2738G>C), V197I (2743G>A) | | | | | | | | |
| Codon mutations: | ATT73ATA (2373T>A), GAT74GAC (2376T>C), TGC75TGT (2379C>T), ACA78ACT (2388A>T), GGA79AAA (2389G>A 2390G>A), GGT80GGA (2394T>A), AAG81AAA (2397G>A), AAA83AAT (2403A>T), ATA86CTT (2410A>C 2412A>T), AAC88ACT (2417A>C 2418C>T), AAA89AGA (2420A>G), AGA90AAA (2423G>A), ATA91TTA (2425A>T), ATA92GTA (2428A>G), AAT93AAC (2433T>C), AAA94CAA (2434A>C), CAA97AAA (2443C>A), AAG100AAA (2454G>A), AAC103GAA (2461A>G 2463C>A), ATT105ATA (2469T>A), AAG106AAA (2472G>A), CAC109CAT (2481C>T), TTA110CTA (2482T>C), ACT113ACC (2493T>C), TTA116CAA (2500T>C 2501T>A), ATA117ATT (2505A>T), ACC127ACA (2535C>A), CCT128CCG (2538T>G), ATA131CTA (2545A>C), CTA133TTA (2551C>T), AGA137AGG (2565A>G), ATC139GTA (2569A>G 2571C>A), CAA140GAA (2572C>G), AGC145AGT (2589C>T), ATA150GTA (2602A>G), AAT153AAC (2613T>C), TAT156TAC (2622T>C), AAG158AAA (2628G>A), TTT161TTC (2637T>C), ATA162GTA (2638A>G), ATA163ATC (2643A>C), AGC164AGT (2646C>T), GCC165CCA (2647G>C 2649C>A), CTA171TTA (2665C>T), AAT172TCA (2668A>T 2669A>C 2670T>A), AGA174GAA (2674A>G 2675G>A), GAT177GAA (2685T>A), AAA178AAT (2688A>T), TCA179TCT (2691A>T), TTA182TTT (2700A>T), AAG185AAA (2709G>A), ATG186ATA (2712G>A), TCA187TCG (2715A>G), GAG190AAA (2722G>A 2724G>A), TTA191CTA (2725T>C), GCA192GCT (2730A>T), CCC193CCA (2733C>A), GGA194TGA (2734G>T), AGT195ACT (2738G>C), GTA197ATA (2743G>A), TTT198TTC (2748T>C), GCC200GCA (2754C>A) | | | | | | | | |
| Tvvgp3 | 183 | 636 | 55.0% | 2017 | 83.4% | 350 (100%) | 288 (82.3%) | 0/0/0/0 | 3 |
| Protein mutations: | N183D (3957A>G), K194Q (3990A>C), V195I (3993G>A), E201D (4013A>C), K212R (4045A>G), N216D (4056A>G), D217N (4059G>A 4061T>C), T219E (4065A>G 4066C>A), K226D (4086A>G 4088A>T), D227E (4091T>A), D230E (4100T>A), N231K (4103T>A), K232G (4104A>G 4105A>G), E236K (4116G>A), D311E (4343T>A), K315Q (4353A>C 4355A>G), A318T (4362G>A 4364T>A), W328L (4393G>T), P332S (4404C>T), G334E (4411G>A), K336R (4417A>G), E354P (4470G>C 4471A>C), V358I (4482G>A 4484A>T), I363V (4497A>G), L364I (4500C>A), R368K (4513G>A), Q370K (4518C>A), D371E (4523T>A), I374L (4530A>T), L377S (4540T>C), K379Q (4545A>C), A381I (4551G>A 4552C>T), H382N (4554C>A), I384V (4560A>G), N386K (4568C>A), T394K (4591C>A 4592C>G), M399F (4605A>T 4607G>T), I415V (4653A>G), M417T (4660T>C), T419Q (4665A>C 4666C>A 4667T>A), I426T (4687T>C 4688C>A), L428S (4692C>T 4693T>C 4694T>A), D429E (4697T>A), N431E (4701A>G 4703C>A), I432L (4704A>T 4706T>A), I444L (4740A>T), F556* (5077T>A 5078T>G), A557L (5079G>T 5080C>T 5081A>G), T572N (5125C>A 5126C>T), R573K (5128G>A), V576I (5136G>A), E615K (5253G>A), I616V (5256A>G), N618S (5263A>G), V624I (5280G>A), A625T (5283G>A), D626Y (5286G>T), S629A (5295T>G), Q631* (5301C>T), P634T (5310C>A), N635D (5313A>G) | | | | | | | | |
| Codon mutations: | AAC183GAC (3957A>G), CCT184CCA (3962T>A), CAT191CAC (3983T>C), AAA192AAG (3986A>G), AAA194CAA (3990A>C), GTA195ATA (3993G>A), GAA201GAC (4013A>C), GAG202GAA (4016G>A), AAT203AAC (4019T>C), AGT204AGC (4022T>C), AAA209AAG (4037A>G), CCT210CCA (4040T>A), AAA212AGA (4045A>G), AAT216GAT (4056A>G), GAT217AAC (4059G>A 4061T>C), ACA219GAA (4065A>G 4066C>A), AAA222AAG (4076A>G), AAA226GAT (4086A>G 4088A>T), GAT227GAA (4091T>A), GAT230GAA (4100T>A), AAT231AAA (4103T>A), AAA232GGA (4104A>G 4105A>G), CAA235CAG (4115A>G), GAA236AAA (4116G>A), AGC239AGT (4127C>T), GGA257GGT (4181A>T), AGC259AGT (4187C>T), AGG260AGA (4190G>A), GTC262GTT (4196C>T), ATA263ATT (4199A>T), TTA268CTA (4212T>C), TAT275TAC (4235T>C), TAT277TAC (4241T>C), CCC280CCT (4250C>T), AAG285AAA (4265G>A), TAT292TAC (4286T>C), TAC294TAT (4292C>T), GAC299GAT (4307C>T), AAG301AAA (4313G>A), TAT305TAC (4325T>C), CAT306CAC (4328T>C), TTG307CTA (4329T>C 4331G>A), AAA308AAG (4334A>G), GAG310GAA (4340G>A), GAT311GAA (4343T>A), AAA315CAG (4353A>C 4355A>G), GCT318ACA (4362G>A 4364T>A), TAT326TAC (4388T>C), GAA327GAG (4391A>G), TGG328TTG (4393G>T), GTT330GTA (4400T>A), TTA331TTG (4403A>G), CCA332TCA (4404C>T), GGA334GAA (4411G>A), AAA336AGA (4417A>G), AAT337AAC (4421T>C), TAC342TAT (4436C>T), GAC347GAT (4451C>T), CTA353TTA (4467C>T), GAA354CCA (4470G>C 4471A>C), AAT355AAC (4475T>C), TGT356TGC (4478T>C), GTA358ATT (4482G>A 4484A>T), ATA363GTA (4497A>G), CTA364ATA (4500C>A), CTA365TTA (4503C>T), TAT366TAC (4508T>C), AGA368AAA (4513G>A), ACA369ACT (4517A>T), CAA370AAA (4518C>A), GAT371GAA (4523T>A), ATA374TTA (4530A>T), TTA377TCA (4540T>C), AAA379CAA (4545A>C), TTC380TTT (4550C>T), GCA381ATA (4551G>A 4552C>T), CAT382AAT (4554C>A), ATT383ATA (4559T>A), ATA384GTA (4560A>G), AAC386AAA (4568C>A), TCT387TCA (4571T>A), GGT388GGA (4574T>A), TTA391CTA (4581T>C), ACC394AAG (4591C>A 4592C>G), GCA396GCT (4598A>T), GAA397GAG (4601A>G), ATT398ATA (4604T>A), ATG399TTT (4605A>T 4607G>T), TTA406CTA (4626T>C), GAT411GAC (4643T>C), ATA415GTA (4653A>G), ATG417ACG (4660T>C), ACT419CAA (4665A>C 4666C>A 4667T>A), CAA423CAG (4679A>G), ATC426ACA (4687T>C 4688C>A), AAT427AAC (4691T>C), CTT428TCA (4692C>T 4693T>C 4694T>A), GAT429GAA (4697T>A), AAC431GAA (4701A>G 4703C>A), ATT432TTA (4704A>T 4706T>A), GAT433GAC (4709T>C), AAA437AAG (4721A>G), CTA442TTA (4734C>T), ATA444TTA (4740A>T), TTT556TAG (5077T>A 5078T>G), GCA557TTG (5079G>T 5080C>T 5081A>G), CTA562TTA (5094C>T), TTA563CTA (5097T>C), GCA564GCC (5102A>C), ATA569ATT (5117A>T), GTT570GTG (5120T>G), ACC572AAT (5125C>A 5126C>T), AGA573AAA (5128G>A), TTT574TTC (5132T>C), ATT575ATA (5135T>A), GTA576ATA (5136G>A), GAC579GAT (5147C>T), CAA582CAG (5156A>G), ACT588ACA (5174T>A), AAA590AAG (5180A>G), TCA594TCG (5192A>G), GTC595GTT (5195C>T), CGG601AGG (5211C>A), CTG603TTA (5217C>T 5219G>A), AAC609AAT (5237C>T), GAA615AAA (5253G>A), ATA616GTA (5256A>G), ATT617ATA (5261T>A), AAT618AGT (5263A>G), AAG621AAA (5273G>A), GTT624ATT (5280G>A), GCA625ACA (5283G>A), GAT626TAT (5286G>T), TAT627TAC (5291T>C), TCA629GCA (5295T>G), CAA631TAA (5301C>T), CCA634ACA (5310C>A), AAC635GAC (5313A>G) | | | | | | | | |
| Tvvgp4 | 1 | 412 | 65.6% | 1203 | 63.0% | 274 (99.6%) | 187 (68.0%) | 1/0/1/1 | 1 |
| Protein mutations: | Q4H (5286G>T), I6T (5291T>C), Y7* (5295T>G), T161Q (5755A>C 5756C>A 5757C>A), D164E (5766C>A), N171T (5786A>C), N174K (5796T>A), K175L (5797A>C 5798A>T 5799A>C), N176T (5801A>C 5802T>G), Q180S (5812C>A 5813A>G 5814A>C), S181T (5815T>A), Q183E (5821C>G), N184K (5826T>A), N186T (5831A>C), H192Q (5850C>A), H193K (5851C>A 5853T>A), T195Q (5857A>C 5858C>A), L203Q (5881T>C 5882T>A), N205K (5889T>A), S207N (5894G>A), A208V (5897C>T 5898A>C), K209N (5901A>T), A212R (5908G>A 5909C>G 5910C>A), N216S (5921A>G), D221N (5935G>A), Q226Y (5950C>T 5952G>T), T227D (5953A>G 5954C>A 5955A>C), Q229E (5959C>G), A232S (5968G>T 5970C>A), N233A (5971A>G 5972A>C), M242T (5999T>C 6000G>A), Q243T (6001C>A 6002A>C), T251N (6026C>A 6027A>C), F253L (6033C>G), V255I (6037G>A), R256K (6040C>A 6041G>A 6042A>G), S259I (6049T>A 6050C>T 6051G>A), A260V (6053C>T 6054T>A), T261P (6055A>C), A262V (6059C>T), Y266F (6071A>T), E268_I269insX (6078_6079insA), P271L (6086C>T 6087T>A), I272V (6088A>G 6090C>G), G279E (6110G>A), E283D (6123A>T), K289E (6139A>G 6141G>A), E294K (6154G>A), E297T (6163G>A 6164A>C), V299E (6170T>A), N300E (6172A>G 6174T>G), I301R (6176T>G 6177T>A), E303V (6182A>T 6183A>T), I313L (6211A>T), L317I (6223C>A 6225A>T), L325I (6247C>A), Q327E (6253C>G 6255G>A), A329L (6259G>C 6260C>T), I330M (6264T>G), S332I (6269G>T), S335T (6277T>A), E337D (6285A>C), T339I (6290C>T), C345S (6308G>C 6309C>A), R346K (6311G>A), D347E (6315T>G), R349I (6320G>T 6321A>C), A351R (6325G>A 6326C>G), L356A (6340C>G 6341T>C), R357M (6344G>T 6345A>G), V360I (6352G>A 6354C>A), L361M (6355T>A 6357A>G), S362T (6359G>C 6360T>A), K365N (6369A>T), Q368K (6376C>A), P369Q (6380C>A 6381T>A), T370P (6382A>C 6384A>T), T371I (6386C>T 6387C>T), R376K (6401G>A 6402G>A), R377K (6404G>A), N378G (6406A>G 6407A>G 6408T>A), P382D (6418C>G 6419C>A 6420A>T), D383E (6423T>A), L391Q (6446T>A), S393G (6451A>G 6453T>G), H394Q (6456C>A), K405N (6489A>C) | | | | | | | | |

| | Begin | End | Coverage | Score | Concordance | Matches | Identities | I/D/M/F* | Stop Codons |
|---|---|---|---|---|---|---|---|---|---|
| **NT** | **2369** | **6509** | **28.5%** | **2644** | **59.6%** | **2217 (99.9%)** | **1772 (79.9%)** | **1/0** | |
| Codon mutations: | TTG2TTA (5280G>A), TTG3TTA (5283G>A), CAG4CAT (5286G>T), ATT6ACT (5291T>C), TAT7TAG (5295T>G), GAC9GAT (5301C>T), ACC12ACA (5310C>A), CAA13CAG (5313A>G), CAT157CAC (5745T>C), CGT160CGA (5754T>A), ACC161CAA (5755A>C 5756C>A 5757C>A), TAT162TAC (5760T>C), ATA163ATT (5763A>T), GAC164GAA (5766C>A), ATA169ATC (5781A>C), CAG170CAA (5784G>A), AAT171ACT (5786A>C), AAT174AAA (5796T>A), AAA175CTC (5797A>C 5798A>T 5799A>C), AAT176ACG (5801A>C 5802T>G), TCC179TCT (5811C>T), CAA180AGC (5812C>A 5813A>G 5814A>C), TCT181ACT (5815T>A), ACC182ACA (5820C>A), CAA183GAA (5821C>G), AAT184AAA (5826T>A), AAC186ACC (5831A>C), GAC188GAT (5838C>T), ATT190ATA (5844T>A), ACC191ACT (5847C>T), CAC192CAA (5850C>A), CAT193AAA (5851C>A 5853T>A), CTA194TTA (5854C>T), ACA195CAA (5857A>C 5858C>A), TTG200CTA (5872T>C 5874G>A), ATA201ATT (5877A>T), GCT202GCA (5880T>A), TTA203CAA (5881T>C 5882T>A), AAT205AAA (5889T>A), ACA206ACT (5892A>T), AGT207AAT (5894G>A), GCA208GTC (5897C>T 5898A>C), AAA209AAT (5901A>T), GTA211GTT (5907A>T), GCC212AGA (5908G>A 5909C>G 5910C>A), ACC213ACA (5913C>A), TGC214TGT (5916C>T), AAC216AGC (5921A>G), CTA220TTA (5932C>T), GAT221AAT (5935G>A), ACT222ACC (5940T>C), CAG226TAT (5950C>T 5952G>T), ACA227GAC (5953A>G 5954C>A 5955A>C), CAA229GAA (5959C>G), GAG230GAA (5964G>A), GCC232TCA (5968G>T 5970C>A), AAT233GCT (5971A>G 5972A>C), CCT235CCA (5979T>A), GAG236GAA (5982G>A), TTA237CTA (5983T>C), TAC238TAT (5988C>T), AAG239AAA (5991G>A), TTC241TTT (5997C>T), ATG242ACA (5999T>C 6000G>A), CAA243ACA (6001C>A 6002A>C), AAG245AAA (6009G>A), AGG246AGA (6012G>A), ATA247ATT (6015A>T), ACT248ACA (6018T>A), GGG250GGC (6024G>C), ACA251AAC (6026C>A 6027A>C), TTC253TTG (6033C>G), TAC254TAT (6036C>T), GTA255ATA (6037G>A), CGA256AAG (6040C>A 6041G>A 6042A>G), TTC257TTT (6045C>T), TCG259ATA (6049T>A 6050C>T 6051G>A), GCT260GTA (6053C>T 6054T>A), ACA261CCA (6055A>C), GCA262GTA (6059C>T), CTA265TTA (6067C>T), TAT266TTT (6071A>T), GAG268GAA (6078G>A), GAG268_ATA269insA-- (6078_6079insA), ATA269ATT (6081A>T), AAG270AAA (6084G>A), CCT271CTA (6086C>T 6087T>A), ATC272GTG (6088A>G 6090C>G), ATC276ATT (6102C>T), ATT278ATA (6108T>A), GGA279GAA (6110G>A), CTC280CTA (6114C>A), AGG282CGA (6118A>C 6120G>A), GAA283GAT (6123A>T), ATC285ATT (6129C>T), ATA286ATT (6132A>T), CCA287CCC (6135A>C), AAG289GAA (6139A>G 6141G>A), ATA290ATT (6144A>T), GAA294AAA (6154G>A), GAA297ACA (6163G>A 6164A>C), GTA299GAA (6170T>A), AAT300GAG (6172A>G 6174T>G), ATT301AGA (6176T>G 6177T>A), CCA302CCT (6180A>T), GAA303GTT (6182A>T 6183A>T), TTC304TTT (6186C>T), AGG309AGA (6201G>A), ATA310ATT (6204A>T), ATA313TTA (6211A>T), ACT315ACA (6219T>A), ATA316ATC (6222A>C), CTA317ATT (6223C>A 6225A>T), AAC322AAT (6240C>T), AAC323AAT (6243C>T), CTA325ATA (6247C>A), CAG327GAA (6253C>G 6255G>A), GCA329CTA (6259G>C 6260C>T), ATT330ATG (6264T>G), AGC332ATC (6269G>T), TAT333TAC (6273T>C), TAT334TAC (6276T>C), TCA335ACA (6277T>A), AGG336AGA (6282G>A), GAA337GAC (6285A>C), CAG338CAA (6288G>A), ACA339ATA (6290C>T), TGC345TCA (6308G>C 6309C>A), AGA346AAA (6311G>A), GAT347GAG (6315T>G), AGA349ATC (6320G>T 6321A>C), GCA351AGA (6325G>A 6326C>G), CTA356GCA (6340C>G 6341T>C), AGA357ATG (6344G>T 6345A>G), GTC360ATA (6352G>A 6354C>A), TTA361ATG (6355T>A 6357A>G), AGT362ACA (6359G>C 6360T>A), CTT363TTG (6361C>T 6363T>G), AAA365AAT (6369A>T), GAA367GAG (6375A>G), CAA368AAA (6376C>A), CCT369CAA (6380C>A 6381T>A), ACA370CCT (6382A>C 6384A>T), ACC371ATT (6386C>T 6387C>T), AGG376AAA (6401G>A 6402G>A), AGG377AAG (6404G>A), AAT378GGA (6406A>G 6407A>G 6408T>A), TTC379TTT (6411C>T), ATC380ATT (6414C>T), TCC381TCG (6417C>G), CCA382GAT (6418C>G 6419C>A 6420A>T), GAT383GAA (6423T>A), CTG384TTA (6424C>T 6426G>A), TTA385CTA (6427T>C), CTA391CAA (6446T>A), AGT393GGG (6451A>G 6453T>G), CAC394CAA (6456C>A), TAT396TAC (6462T>C), CAC399CAT (6471C>T), TGC401TGT (6477C>T), TCA402TCG (6480A>G), AAA405AAC (6489A>C) | | | | | | | | |

#### Proteins

| | Begin | End | Coverage | Score | Concordance | Matches | Identities | I/D/M/F* | Stop Codons |
|---|---|---|---|---|---|---|---|---|---|
| putative cell-to-cell movement protein (NP_569140.1) | 73 | 201 | 30.7% | 701 | 81.3% | 129 (100%) | 100 (77.5%) | 0/0/0/0 | 1 |
| Protein mutations: | G79K (2389G>A 2390G>A), K83N (2403A>T), I86L (2410A>C 2412A>T), N88T (2417A>C 2418C>T), K89R (2420A>G), R90K (2423G>A), I91L (2425A>T), I92V (2428A>G), K94Q (2434A>C), Q97K (2443C>A), N103E (2461A>G 2463C>A), L116Q (2500T>C 2501T>A), I131L (2545A>C), I139V (2569A>G 2571C>A), Q140E (2572C>G), I150V (2602A>G), I162V (2638A>G), A165P (2647G>C 2649C>A), N172S (2668A>T 2669A>C 2670T>A), R174E (2674A>G 2675G>A), D177E (2685T>A), K178N (2688A>T), L182F (2700A>T), M186I (2712G>A), E190K (2722G>A 2724G>A), G194* (2734G>T), S195T (2738G>C), V197I (2743G>A) | | | | | | | | |
| Codon mutations: | ATT73ATA (2373T>A), GAT74GAC (2376T>C), TGC75TGT (2379C>T), ACA78ACT (2388A>T), GGA79AAA (2389G>A 2390G>A), GGT80GGA (2394T>A), AAG81AAA (2397G>A), AAA83AAT (2403A>T), ATA86CTT (2410A>C 2412A>T), AAC88ACT (2417A>C 2418C>T), AAA89AGA (2420A>G), AGA90AAA (2423G>A), ATA91TTA (2425A>T), ATA92GTA (2428A>G), AAT93AAC (2433T>C), AAA94CAA (2434A>C), CAA97AAA (2443C>A), AAG100AAA (2454G>A), AAC103GAA (2461A>G 2463C>A), ATT105ATA (2469T>A), AAG106AAA (2472G>A), CAC109CAT (2481C>T), TTA110CTA (2482T>C), ACT113ACC (2493T>C), TTA116CAA (2500T>C 2501T>A), ATA117ATT (2505A>T), ACC127ACA (2535C>A), CCT128CCG (2538T>G), ATA131CTA (2545A>C), CTA133TTA (2551C>T), AGA137AGG (2565A>G), ATC139GTA (2569A>G 2571C>A), CAA140GAA (2572C>G), AGC145AGT (2589C>T), ATA150GTA (2602A>G), AAT153AAC (2613T>C), TAT156TAC (2622T>C), AAG158AAA (2628G>A), TTT161TTC (2637T>C), ATA162GTA (2638A>G), ATA163ATC (2643A>C), AGC164AGT (2646C>T), GCC165CCA (2647G>C 2649C>A), CTA171TTA (2665C>T), AAT172TCA (2668A>T 2669A>C 2670T>A), AGA174GAA (2674A>G 2675G>A), GAT177GAA (2685T>A), AAA178AAT (2688A>T), TCA179TCT (2691A>T), TTA182TTT (2700A>T), AAG185AAA (2709G>A), ATG186ATA (2712G>A), TCA187TCG (2715A>G), GAG190AAA (2722G>A 2724G>A), TTA191CTA (2725T>C), GCA192GCT (2730A>T), CCC193CCA (2733C>A), GGA194TGA (2734G>T), AGT195ACT (2738G>C), GTA197ATA (2743G>A), TTT198TTC (2748T>C), GCC200GCA (2754C>A) | | | | | | | | |
| polyprotein (NP_569141.1) | 183 | 636 | 55.0% | 2017 | 83.4% | 350 (100%) | 288 (82.3%) | 0/0/0/0 | 3 |
| Protein mutations: | N183D (3957A>G), K194Q (3990A>C), V195I (3993G>A), E201D (4013A>C), K212R (4045A>G), N216D (4056A>G), D217N (4059G>A 4061T>C), T219E (4065A>G 4066C>A), K226D (4086A>G 4088A>T), D227E (4091T>A), D230E (4100T>A), N231K (4103T>A), K232G (4104A>G 4105A>G), E236K (4116G>A), D311E (4343T>A), K315Q (4353A>C 4355A>G), A318T (4362G>A 4364T>A), W328L (4393G>T), P332S (4404C>T), G334E (4411G>A), K336R (4417A>G), E354P (4470G>C 4471A>C), V358I (4482G>A 4484A>T), I363V (4497A>G), L364I (4500C>A), R368K (4513G>A), Q370K (4518C>A), D371E (4523T>A), I374L (4530A>T), L377S (4540T>C), K379Q (4545A>C), A381I (4551G>A 4552C>T), H382N (4554C>A), I384V (4560A>G), N386K (4568C>A), T394K (4591C>A 4592C>G), M399F (4605A>T 4607G>T), I415V (4653A>G), M417T (4660T>C), T419Q (4665A>C 4666C>A 4667T>A), I426T (4687T>C 4688C>A), L428S (4692C>T 4693T>C 4694T>A), D429E (4697T>A), N431E (4701A>G 4703C>A), I432L (4704A>T 4706T>A), I444L (4740A>T), F556* (5077T>A 5078T>G), A557L (5079G>T 5080C>T 5081A>G), T572N (5125C>A 5126C>T), R573K (5128G>A), V576I (5136G>A), E615K (5253G>A), I616V (5256A>G), N618S (5263A>G), V624I (5280G>A), A625T (5283G>A), D626Y (5286G>T), S629A (5295T>G), Q631* (5301C>T), P634T (5310C>A), N635D (5313A>G) | | | | | | | | |
| Codon mutations: | AAC183GAC (3957A>G), CCT184CCA (3962T>A), CAT191CAC (3983T>C), AAA192AAG (3986A>G), AAA194CAA (3990A>C), GTA195ATA (3993G>A), GAA201GAC (4013A>C), GAG202GAA (4016G>A), AAT203AAC (4019T>C), AGT204AGC (4022T>C), AAA209AAG (4037A>G), CCT210CCA (4040T>A), AAA212AGA (4045A>G), AAT216GAT (4056A>G), GAT217AAC (4059G>A 4061T>C), ACA219GAA (4065A>G 4066C>A), AAA222AAG (4076A>G), AAA226GAT (4086A>G 4088A>T), GAT227GAA (4091T>A), GAT230GAA (4100T>A), AAT231AAA (4103T>A), AAA232GGA (4104A>G 4105A>G), CAA235CAG (4115A>G), GAA236AAA (4116G>A), AGC239AGT (4127C>T), GGA257GGT (4181A>T), AGC259AGT (4187C>T), AGG260AGA (4190G>A), GTC262GTT (4196C>T), ATA263ATT (4199A>T), TTA268CTA (4212T>C), TAT275TAC (4235T>C), TAT277TAC (4241T>C), CCC280CCT (4250C>T), AAG285AAA (4265G>A), TAT292TAC (4286T>C), TAC294TAT (4292C>T), GAC299GAT (4307C>T), AAG301AAA (4313G>A), TAT305TAC (4325T>C), CAT306CAC (4328T>C), TTG307CTA (4329T>C 4331G>A), AAA308AAG (4334A>G), GAG310GAA (4340G>A), GAT311GAA (4343T>A), AAA315CAG (4353A>C 4355A>G), GCT318ACA (4362G>A 4364T>A), TAT326TAC (4388T>C), GAA327GAG (4391A>G), TGG328TTG (4393G>T), GTT330GTA (4400T>A), TTA331TTG (4403A>G), CCA332TCA (4404C>T), GGA334GAA (4411G>A), AAA336AGA (4417A>G), AAT337AAC (4421T>C), TAC342TAT (4436C>T), GAC347GAT (4451C>T), CTA353TTA (4467C>T), GAA354CCA (4470G>C 4471A>C), AAT355AAC (4475T>C), TGT356TGC (4478T>C), GTA358ATT (4482G>A 4484A>T), ATA363GTA (4497A>G), CTA364ATA (4500C>A), CTA365TTA (4503C>T), TAT366TAC (4508T>C), AGA368AAA (4513G>A), ACA369ACT (4517A>T), CAA370AAA (4518C>A), GAT371GAA (4523T>A), ATA374TTA (4530A>T), TTA377TCA (4540T>C), AAA379CAA (4545A>C), TTC380TTT (4550C>T), GCA381ATA (4551G>A 4552C>T), CAT382AAT (4554C>A), ATT383ATA (4559T>A), ATA384GTA (4560A>G), AAC386AAA (4568C>A), TCT387TCA (4571T>A), GGT388GGA (4574T>A), TTA391CTA (4581T>C), ACC394AAG (4591C>A 4592C>G), GCA396GCT (4598A>T), GAA397GAG (4601A>G), ATT398ATA (4604T>A), ATG399TTT (4605A>T 4607G>T), TTA406CTA (4626T>C), GAT411GAC (4643T>C), ATA415GTA (4653A>G), ATG417ACG (4660T>C), ACT419CAA (4665A>C 4666C>A 4667T>A), CAA423CAG (4679A>G), ATC426ACA (4687T>C 4688C>A), AAT427AAC (4691T>C), CTT428TCA (4692C>T 4693T>C 4694T>A), GAT429GAA (4697T>A), AAC431GAA (4701A>G 4703C>A), ATT432TTA (4704A>T 4706T>A), GAT433GAC (4709T>C), AAA437AAG (4721A>G), CTA442TTA (4734C>T), ATA444TTA (4740A>T), TTT556TAG (5077T>A 5078T>G), GCA557TTG (5079G>T 5080C>T 5081A>G), CTA562TTA (5094C>T), TTA563CTA (5097T>C), GCA564GCC (5102A>C), ATA569ATT (5117A>T), GTT570GTG (5120T>G), ACC572AAT (5125C>A 5126C>T), AGA573AAA (5128G>A), TTT574TTC (5132T>C), ATT575ATA (5135T>A), GTA576ATA (5136G>A), GAC579GAT (5147C>T), CAA582CAG (5156A>G), ACT588ACA (5174T>A), AAA590AAG (5180A>G), TCA594TCG (5192A>G), GTC595GTT (5195C>T), CGG601AGG (5211C>A), CTG603TTA (5217C>T 5219G>A), AAC609AAT (5237C>T), GAA615AAA (5253G>A), ATA616GTA (5256A>G), ATT617ATA (5261T>A), AAT618AGT (5263A>G), AAG621AAA (5273G>A), GTT624ATT (5280G>A), GCA625ACA (5283G>A), GAT626TAT (5286G>T), TAT627TAC (5291T>C), TCA629GCA (5295T>G), CAA631TAA (5301C>T), CCA634ACA (5310C>A), AAC635GAC (5313A>G) | | | | | | | | |
| putative transactivator factor (NP_569142.1) | 1 | 412 | 65.6% | 1203 | 63.0% | 274 (99.6%) | 187 (68.0%) | 1/0/1/1 | 1 |
| Protein mutations: | Q4H (5286G>T), I6T (5291T>C), Y7* (5295T>G), T161Q (5755A>C 5756C>A 5757C>A), D164E (5766C>A), N171T (5786A>C), N174K (5796T>A), K175L (5797A>C 5798A>T 5799A>C), N176T (5801A>C 5802T>G), Q180S (5812C>A 5813A>G 5814A>C), S181T (5815T>A), Q183E (5821C>G), N184K (5826T>A), N186T (5831A>C), H192Q (5850C>A), H193K (5851C>A 5853T>A), T195Q (5857A>C 5858C>A), L203Q (5881T>C 5882T>A), N205K (5889T>A), S207N (5894G>A), A208V (5897C>T 5898A>C), K209N (5901A>T), A212R (5908G>A 5909C>G 5910C>A), N216S (5921A>G), D221N (5935G>A), Q226Y (5950C>T 5952G>T), T227D (5953A>G 5954C>A 5955A>C), Q229E (5959C>G), A232S (5968G>T 5970C>A), N233A (5971A>G 5972A>C), M242T (5999T>C 6000G>A), Q243T (6001C>A 6002A>C), T251N (6026C>A 6027A>C), F253L (6033C>G), V255I (6037G>A), R256K (6040C>A 6041G>A 6042A>G), S259I (6049T>A 6050C>T 6051G>A), A260V (6053C>T 6054T>A), T261P (6055A>C), A262V (6059C>T), Y266F (6071A>T), E268_I269insX (6078_6079insA), P271L (6086C>T 6087T>A), I272V (6088A>G 6090C>G), G279E (6110G>A), E283D (6123A>T), K289E (6139A>G 6141G>A), E294K (6154G>A), E297T (6163G>A 6164A>C), V299E (6170T>A), N300E (6172A>G 6174T>G), I301R (6176T>G 6177T>A), E303V (6182A>T 6183A>T), I313L (6211A>T), L317I (6223C>A 6225A>T), L325I (6247C>A), Q327E (6253C>G 6255G>A), A329L (6259G>C 6260C>T), I330M (6264T>G), S332I (6269G>T), S335T (6277T>A), E337D (6285A>C), T339I (6290C>T), C345S (6308G>C 6309C>A), R346K (6311G>A), D347E (6315T>G), R349I (6320G>T 6321A>C), A351R (6325G>A 6326C>G), L356A (6340C>G 6341T>C), R357M (6344G>T 6345A>G), V360I (6352G>A 6354C>A), L361M (6355T>A 6357A>G), S362T (6359G>C 6360T>A), K365N (6369A>T), Q368K (6376C>A), P369Q (6380C>A 6381T>A), T370P (6382A>C 6384A>T), T371I (6386C>T 6387C>T), R376K (6401G>A 6402G>A), R377K (6404G>A), N378G (6406A>G 6407A>G 6408T>A), P382D (6418C>G 6419C>A 6420A>T), D383E (6423T>A), L391Q (6446T>A), S393G (6451A>G 6453T>G), H394Q (6456C>A), K405N (6489A>C) | | | | | | | | |

| | Begin | End | Coverage | Score | Concordance | Matches | Identities | I/D/M/F* | Stop Codons |
|---|---|---|---|---|---|---|---|---|---|
| **NT** | **2369** | **6509** | **28.5%** | **2644** | **59.6%** | **2217 (99.9%)** | **1772 (79.9%)** | **1/0** | |

Codon mutations: TTG2TTA (5280G>A), TTG3TTA (5283G>A), CAG4CAT (5286G>T), ATT6ACT (5291T>C), TAT7TAG (5295T>G), GAC9GAT (5301C>T), ACC12ACA (5310C>A), CAA13CAG (5313A>G), CAT157CAC (5745T>C), CGT160CGA (5754T>A), ACC161CAA (5755A>C 5756C>A 5757C>A), TAT162TAC (5760T>C), ATA163ATT (5763A>T), GAC164GAA (5766C>A), ATA169ATC (5781A>C), CAG170CAA (5784G>A), AAT171ACT (5786A>C), AAT174AAA (5796T>A), AAA175CTC (5797A>C 5798A>T 5799A>C), AAT176ACG (5801A>C 5802T>G), TCC179TCT (5811C>T), CAA180AGC (5812C>A 5813A>G 5814A>C), TCT181ACT (5815T>A), ACC182ACA (5820C>A), CAA183GAA (5821C>G), AAT184AAA (5826T>A), AAC186ACC (5831A>C), GAC188GAT (5838C>T), ATT190ATA (5844T>A), ACC191ACT (5847C>T), CAC192CAA (5850C>A), CAT193AAA (5851C>A 5853T>A), CTA194TTA (5854C>T), ACA195CAA (5857A>C 5858C>A), TTG200CTA (5872T>C 5874G>A), ATA201ATT (5877A>T), GCT202GCA (5880T>A), TTA203CAA (5881T>C 5882T>A), AAT205AAA (5889T>A), ACA206ACT (5892A>T), AGT207AAT (5894G>A), GCA208GTC (5897C>T 5898A>C), AAA209AAT (5901A>T), GTA211GTT (5907A>T), GCC212AGA (5908G>A 5909C>G 5910C>A), ACC213ACA (5913C>A), TGC214TGT (5916C>T), AAC216AGC (5921A>G), CTA220TTA (5932C>T), GAT221AAT (5935G>A), ACT222ACC (5940T>C), CAG226TAT (5950C>T 5952G>T), ACA227GAC (5953A>G 5954C>A 5955A>C), CAA229GAA (5959C>G), GAG230GAA (5964G>A), GCC232TCA (5968G>T 5970C>A), AAT233GCT (5971A>G 5972A>C), CCT235CCA (5979T>A), GAG236GAA (5982G>A), TTA237CTA (5983T>C), TAC238TAT (5988C>T), AAG239AAA (5991G>A), TTC241TTT (5997C>T), ATG242ACA (5999T>C 6000G>A), CAA243ACA (6001C>A 6002A>C), AAG245AAA (6009G>A), AGG246AGA (6012G>A), ATA247ATT (6015A>T), ACT248ACA (6018T>A), GGG250GGC (6024G>C), ACA251AAC (6026C>A 6027A>C), TTC253TTG (6033C>G), TAC254TAT (6036C>T), GTA255ATA (6037G>A), CGA256AAG (6040C>A 6041G>A 6042A>G), TTC257TTT (6045C>T), TCG259ATA (6049T>A 6050C>T 6051G>A), GCT260GTA (6053C>T 6054T>A), ACA261CCA (6055A>C), GCA262GTA (6059C>T), CTA265TTA (6067C>T), TAT266TTT (6071A>T), GAG268GAA (6078G>A), GAG268_ATA269insA-- (6078_6079insA), ATA269ATT (6081A>T), AAG270AAA (6084G>A), CCT271CTA (6086C>T 6087T>A), ATC272GTG (6088A>G 6090C>G), ATC276ATT (6102C>T), ATT278ATA (6108T>A), GGA279GAA (6110G>A), CTC280CTA (6114C>A), AGG282CGA (6118A>C 6120G>A), GAA283GAT (6123A>T), ATC285ATT (6129C>T), ATA286ATT (6132A>T), CCA287CCC (6135A>C), AAG289GAA (6139A>G 6141G>A), ATA290ATT (6144A>T), GAA294AAA (6154G>A), GAA297ACA (6163G>A 6164A>C), GTA299GAA (6170T>A), AAT300GAG (6172A>G 6174T>G), ATT301AGA (6176T>G 6177T>A), CCA302CCT (6180A>T), GAA303GTT (6182A>T 6183A>T), TTC304TTT (6186C>T), AGG309AGA (6201G>A), ATA310ATT (6204A>T), ATA313TTA (6211A>T), ACT315ACA (6219T>A), ATA316ATC (6222A>C), CTA317ATT (6223C>A 6225A>T), AAC322AAT (6240C>T), AAC323AAT (6243C>T), CTA325ATA (6247C>A), CAG327GAA (6253C>G 6255G>A), GCA329CTA (6259G>C 6260C>T), ATT330ATG (6264T>G), AGC332ATC (6269G>T), TAT333TAC (6273T>C), TAT334TAC (6276T>C), TCA335ACA (6277T>A), AGG336AGA (6282G>A), GAA337GAC (6285A>C), CAG338CAA (6288G>A), ACA339ATA (6290C>T), TGC345TCA (6308G>C 6309C>A), AGA346AAA (6311G>A), GAT347GAG (6315T>G), AGA349ATC (6320G>T 6321A>C), GCA351AGA (6325G>A 6326C>G), CTA356GCA (6340C>G 6341T>C), AGA357ATG (6344G>T 6345A>G), GTC360ATA (6352G>A 6354C>A), TTA361ATG (6355T>A 6357A>G), AGT362ACA (6359G>C 6360T>A), CTT363TTG (6361C>T 6363T>G), AAA365AAT (6369A>T), GAA367GAG (6375A>G), CAA368AAA (6376C>A), CCT369CAA (6380C>A 6381T>A), ACA370CCT (6382A>C 6384A>T), ACC371ATT (6386C>T 6387C>T), AGG376AAA (6401G>A 6402G>A), AGG377AAG (6404G>A), AAT378GGA (6406A>G 6407A>G 6408T>A), TTC379TTT (6411C>T), ATC380ATT (6414C>T), TCC381TCG (6417C>G), CCA382GAT (6418C>G 6419C>A 6420A>T), GAT383GAA (6423T>A), CTG384TTA (6424C>T 6426G>A), TTA385CTA (6427T>C), CTA391CAA (6446T>A), AGT393GGG (6451A>G 6453T>G), CAC394CAA (6456C>A), TAT396TAC (6462T>C), CAC399CAT (6471C>T), TGC401TGT (6477C>T), TCA402TCG (6480A>G), AAA405AAC (6489A>C)

*: Inserts / Deletes / Misaligned / Frameshifts

#### Analysis details

This analysis was performed with panviral2.64

### NGS Details (UN9): Duamitovirus soch1

#### Assembly

| | |
|---|---|
| **Coverage Length** | 1721 (2 contig(s)) |
| **Depth Of Coverage** | 11.3 |
| **Number Of Reads** | 163 |
| **Reads Per Million** | 3.68 rpm (after QC) |
| **Ambiguities** | 0 |
| **Assembly Method** | de novo + reference guided assembly |
| **Consensus Caller** | Bcf Tools |

#### Coverage Map

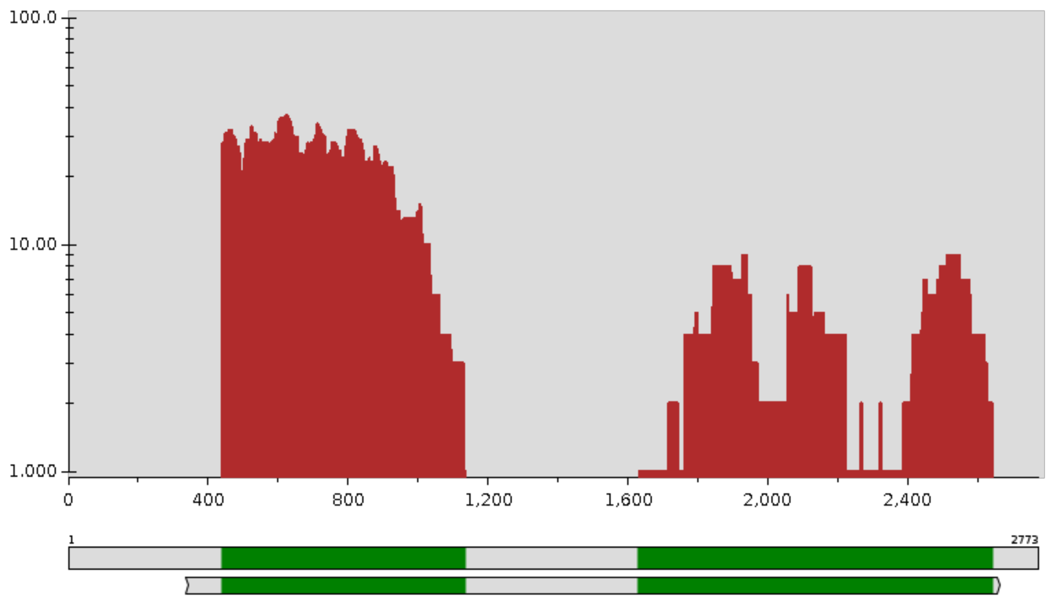

#### Assignment

| | |
|---|---|
| **Type** | Duamitovirus soch1 (Taxonomy ID: 2955838) |
| **Reference Genome** | NC_076524.1 |
| **NT Identity (%)** | 69.7269 |
| **AA Identity (%)** | 67.9443 |
| **Number Of Stop Codons** | 7 |
| **Number Of CDS** | 1 |

#### Alignment

| | |
|---|---|
| **Alignment Score** | 1358.0 (NT) + 2762.0 (AA) = 4120.0 |
| **Concordance (%)** | 55.014 |

| Alignment Method | Global, seeded, nucleotide + amino acids (AGA) |
|---|---|

#### Genome Region

Sequence starts at position 437 and ends at position 2647 relative to NC_076524.1 reference sequence.

#### Alignment Detailed Statistics

| | Begin | End | Coverage | Score | Concordance | Matches | Identities | I/D/M/F* | Stop Codons |
|---|---|---|---|---|---|---|---|---|---|
| **NT** | **437** | **2647** | **62.1%** | **1358** | **39.5%** | **1721 (100%)** | **1200 (69.7%)** | **0/0** | |
| Mutations: | 437A>G, 441A>C, 443C>T, 447C>T, 459C>T, 466C>T, 470C>T, 471G>A, 472C>T, 473T>G, 477T>A, 480A>C, 481C>A, 482A>T, 485A>G, 489A>G, 494T>C, 497T>A, 500T>G, 503A>C, 506T>C, 509C>T, 512T>C, 513G>A, 516T>A, 517C>A, 518T>A, 520A>C, 524G>T, 525A>G, 526C>T, 527T>G, 529C>A, 530C>A, 531T>C, 533A>G, 534C>A, 535G>A, 536C>A, 539G>A, 540A>C, 541A>G, 542G>T, 545G>A, 548C>T, 550A>T, 556T>A, 557C>T, 560A>T, 569C>T, 575G>A, 581C>T, 582G>A, 587A>T, 589G>C, 591T>C, 593A>T, 599A>T, 602C>T, 612T>A, 613C>G, 614C>T, 616T>A, 618C>T, 619G>C, 620A>T, 624G>A, 626A>T, 629A>C, 630C>A, 632A>C, 633C>A, 635T>G, 638G>T, 641G>T, 644T>A, 647G>T, 651C>T, 653C>G, 656A>C, 657A>C, 661C>G, 662G>T, 665T>A, 668C>T, 672C>A, 673G>A, 674T>G, 680C>A, 683C>T, 684A>G, 685A>C, 686A>C, 692G>A, 695T>A, 699G>C, 700C>A, 701A>C, 702A>G, 705C>A, 707T>G, 708A>G, 709G>C, 710G>A, 713A>G, 714G>T, 715A>C, 719A>T, 722C>T, 737T>C, 746C>T, 752A>T, 761G>A, 765G>T, 767T>A, 770G>A, 771G>T, 772T>A, 773T>G, 774T>A, 776G>T, 777A>G, 778A>C, 779A>G, 782T>G, 796G>A, 797T>G, 803A>C, 809T>C, 810G>C, 815C>A, 818A>T, 821T>G, 824C>T, 827A>T, 831C>T, 833T>A, 841G>A, 842G>T, 850T>G, 851A>T, 857G>T, 860G>T, 862A>G, 869T>G, 875G>T, 880T>G, 881C>A, 885C>A, 887T>G, 892A>G, 905A>T, 908C>A, 909T>C, 910T>A, 916G>A, 918G>T, 919G>C, 923T>A, 926A>T, 927A>C, 929G>T, 934T>A, 935T>G, 938G>T, 942T>A, 943C>A, 948C>A, 971C>A, 972C>T, 975A>C, 976A>T, 977T>C, 983T>C, 990A>C, 992G>C, 993T>C, 994C>G, 995A>G, 997A>T, 998C>T, 999G>C, 1000G>A, 1001T>A, 1003T>A, 1004A>T, 1005G>T, 1007C>G, 1010A>G, 1016C>T, 1017A>G, 1018A>T, 1019A>T, 1022T>A, 1025T>C, 1026C>G, 1027A>G, 1030A>C, 1031G>A, 1034T>G, 1035A>G, 1036G>C, 1044T>G, 1046C>T, 1049G>A, 1050T>G, 1051T>A, 1052T>A, 1054C>A, 1055A>C, 1061T>C, 1067T>G, 1073G>A, 1076A>G, 1083C>A, 1088G>T, 1091C>G, 1099G>T, 1104A>G, 1105C>T, 1106A>G, 1115T>G, 1121A>C, 1122T>A, 1124C>T, 1127G>A, 1130C>T, 1629C>T, 1632T>A, 1634A>C, 1644G>A, 1664G>A, 1668G>A, 1670G>A, 1671C>A, 1673G>A, 1676T>A, 1679G>A, 1682C>T, 1686C>T, 1691T>A, 1696G>A, 1698G>T, 1703T>C, 1706C>T, 1707T>G, 1709G>T, 1710T>C, 1712G>A, 1724T>C, 1738C>G, 1742G>A, 1744G>A, 1745C>G, 1759G>A, 1760A>C, 1762G>A, 1766C>A, 1767T>A, 1769T>C, 1772A>C, 1775T>C, 1776G>A, 1777G>A, 1778G>A, 1781T>C, 1783A>C, 1784C>T, 1787T>C, 1790A>G, 1817T>C, 1819G>A, 1820T>G, 1823A>G, 1833G>C, 1840A>G, 1856A>G, 1859C>T, 1860A>G, 1861A>G, 1865C>T, 1867A>C, 1868A>C, 1874G>T, 1875A>T, 1878C>G, 1880A>T, 1886T>A, 1887T>A, 1889G>A, 1895T>C, 1898T>G, 1904A>T, 1907C>T, 1919G>C, 1922G>A, 1923A>T, 1925G>A, 1943G>T, 1947A>G, 1960C>T, 1967C>T, 1968C>G, 1970T>G, 1972G>A, 1973A>G, 1976T>A, 1980T>A, 1981T>C, 1984C>T, 1985T>G, 1991T>C, 1999T>G, 2003A>T, 2006T>A, 2010G>A, 2011C>A, 2013C>A, 2015C>T, 2020T>C, 2030T>C, 2036C>T, 2039A>G, 2049C>T, 2051A>G, 2052A>T, 2054A>G, 2063T>G, 2069G>A, 2071A>T, 2074A>G, 2075G>A, 2078A>T, 2082G>T, 2084G>T, 2087C>T, 2090A>G, 2091T>A, 2093C>G, 2094A>T, 2099C>T, 2103A>T, 2104G>C, 2105C>T, 2108G>A, 2117G>A, 2118C>A, 2120T>A, 2123A>G, 2129T>G, 2133C>G, 2134A>C, 2135T>A, 2136C>A, 2137T>G, 2138T>C, 2141G>A, 2145G>C, 2146T>A, 2147T>C, 2148C>A, 2156C>T, 2157T>C, 2160C>T, 2166T>C, 2167A>T, 2173G>A, 2174G>T, 2183A>G, 2186G>A, 2187G>T, 2188G>T, 2191T>A, 2192G>A, 2195C>T, 2204C>T, 2206A>T, 2213A>G, 2216G>A, 2223G>A, 2226A>G, 2227G>A, 2231C>T, 2236T>G, 2237T>G, 2240A>G, 2248A>C, 2250G>C, 2251T>C, 2259T>A, 2263A>G, 2264G>A, 2265A>C, 2271C>A, 2273A>T, 2274A>G, 2276G>A, 2280C>A, 2285G>A, 2288C>T, 2298G>T, 2299T>G, 2300T>G, 2303A>G, 2306A>C, 2309A>T, 2313C>T, 2315T>C, 2319G>A, 2322C>T, 2325C>G, 2326G>T, 2328C>A, 2330A>T, 2336A>T, 2337G>A, 2346T>C, 2348A>T, 2349A>C, 2350A>G, 2355C>G, 2357C>T, 2358A>T, 2359A>C, 2360C>T, 2362G>A, 2369T>C, 2372G>T, 2377G>T, 2378T>G, 2382A>T, 2383A>C, 2384C>T, 2385C>T, 2386C>T, 2387A>T, 2390T>A, 2396C>T, 2399T>C, 2400A>G, 2402G>A, 2404G>A, 2405T>A, 2406C>T, 2408C>G, 2411C>G, 2414C>T, 2415T>G, 2417C>T, 2420T>C, 2423A>T, 2425G>A, 2430A>T, 2431C>A, 2432A>G, 2435C>A, 2441A>G, 2442A>C, 2444G>A, 2445C>T, 2446T>C, 2447C>G, 2448G>A, 2450A>T, 2451A>G, 2452C>A, 2453T>A, 2456T>C, 2457T>C, 2459T>G, 2462T>A, 2465A>G, 2466A>G, 2477C>T, 2478C>T, 2489A>G, 2492T>C, 2495C>T, 2504T>A, 2510T>C, 2514A>G, 2518C>T, 2519C>G, 2524C>T, 2525C>T, 2528T>A, 2532C>A, 2533A>G, 2536G>A, 2537G>T, 2543T>G, 2551A>G, 2552C>T, 2555T>C, 2556G>T, 2557T>C, 2561T>C, 2568A>G, 2571A>C, 2572A>G, 2582A>T, 2585G>A, 2597G>A, 2598A>G, 2599A>C, 2600C>T, 2601G>A, 2609T>A, 2613T>G, 2615T>A, 2618T>A, 2622G>T, 2624A>T, 2626A>T, 2627C>A, 2629A>C, 2631A>T, 2634A>C, 2635C>T, 2638G>T, 2639A>G | | | | | | | | |
| **CDS** | | | | | | | | | |
| RdRp | 35 | 771 | 73.9% | 2762 | 67.9% | 574 (100%) | 390 (67.9%) | 0/0/0/0 | 7 |
| Protein mutations: | I36L (441A>C 443C>T), L38F (447C>T), A44V (466C>T), A46M (471G>A 472C>T 473T>G), L48M (477T>A), T49H (480A>C 481C>A 482A>T), I52V (489A>G), V60I (513G>A), S61K (516T>A 517C>A 518T>A), K62T (520A>C), T64V (525A>G 526C>T 527T>G), T65K (529C>A 530C>A), R67K (534C>A 535G>A 536C>A), K69R (540A>C 541A>G 542G>T), Y72F (550A>T), F74Y (556T>A 557C>T), A83T (582G>A), C85S (589G>C), F94Y (616T>A), R95S (618C>T 619G>C 620A>T), E97N (624G>A 626A>T), E98D (629A>C), P99T (630C>A 632A>C), L100M (633C>A 635T>G), S109C (661C>G 662G>T), R113K (672C>A 673G>A 674T>G), K117A (684A>G 685A>C 686A>C), A122H (699G>C 700C>A 701A>C), I123V (702A>G), R125A (708A>G 709G>C 710G>A), D127S (714G>T 715A>C), E128D (719A>T), A144S (765G>T 767T>A), V146* (771G>T 772T>A 773T>G), L147I (774T>A 776G>T), K148A (777A>G 778A>C 779A>G), S154K (796G>A 797T>G), E159Q (810G>C), P166S (831C>T 833T>A), G169D (841G>A 842G>T), L172R (850T>G 851A>T), E176G (862A>G), I178M (869T>G), F182* (880T>G 881C>A), H184K (885C>A 887T>G), Q186R (892A>G), F192H (909T>C 910T>A), R194Q (916G>A), G195S (918G>T 919G>C), M198L (927A>C 929G>T), I200K (934T>A 935T>G), S203N (942T>A 943C>A), Q205K (948C>A), N214L (975A>C 976A>T 977T>C), S220R (993T>C 994C>G 995A>G), Y221F (997A>T 998C>T), G222Q (999G>C 1000G>A 1001T>A), L223Y (1003T>A 1004A>T), A224S (1005G>T 1007C>G), K228V (1017A>G 1018A>T 1019A>T), Q231G (1026C>G 1027A>G), E232A (1030A>C 1031G>A), S234A (1035A>G 1036G>C), F237V (1044T>G 1046C>T), F239E (1050T>G 1051T>A 1052T>A), P240H (1054C>A 1055A>C), L250I (1083C>A), W255L (1099G>T), T257V (1104A>G 1105C>T 1106A>G), I260M (1115T>G), F263I (1122T>A 1124C>T), P432S (1629C>T), L433I (1632T>A 1634A>C), V437M (1644G>A), G445R (1668G>A 1670G>A), Q446K (1671C>A 1673G>A), H451Y (1686C>T), W454* (1696G>A), A455S (1698G>T), S458A (1707T>G 1709G>T), A468G (1738C>G), S470K (1744G>A 1745C>G), G475D (1759G>A 1760A>C), C476Y (1762G>A), N477K (1766C>A), S478T (1767T>A 1769T>C), G481K (1776G>A 1777G>A 1778G>A), Y483S (1783A>C 1784C>T), S495K (1819G>A 1820T>G), E500Q (1833G>C), K502R (1840A>G), N509G (1860A>G 1861A>G), K511T (1867A>C 1868A>C), I514L (1875A>T), P515A (1878C>G 1880A>T), L518I (1887T>A 1889G>A), D521E (1898T>G), R530* (1923A>T 1925G>A), M536I (1943G>T), I538V (1947A>G), P542L (1960C>T), L545V (1968C>G 1970T>G), R546K (1972G>A 1973A>G), L549T (1980T>A 1981T>C), S550L (1984C>T 1985T>G), V555G (1999T>G), A559N (2010G>A 2011C>A), L560I (2013C>A 2015C>T), I562T (2020T>C), I568M (2039A>G), Q572* (2049C>T 2051A>G), R573W (2052A>T 2054A>G), Y579F (2071A>T), K580R (2074A>G 2075G>A), A583S (2082G>T 2084G>T), F586M (2091T>A 2093C>G), T587S (2094A>T), H600A (2133C>G 2134A>C 2135T>A), L601S (2136C>A 2137T>G 2138T>C), V604H (2145G>C 2146T>A 2147T>C), Y611L (2166T>C 2167A>T), W613Y (2173G>A 2174G>T), G618F (2187G>T 2188G>T), M619K (2191T>A 2192G>A), Y624F (2206A>T), V630I (2223G>A), S631D (2226A>G 2227G>A), L634R (2236T>G 2237T>G), K638T (2248A>C), V639P (2250G>C 2251T>C), L642I (2259T>A), Q643R (2263A>G 2264G>A), I644L (2265A>C), P646T (2271C>A 2273A>T), K647E (2274A>G 2276G>A), L649M (2280C>A), V655W (2298G>T 2299T>G 2300T>G), R660C (2313C>T 2315T>C), V662I (2319G>A), R664V (2325C>G 2326G>T), Q665N (2328C>A 2330A>T), E668K (2337G>A), K672R (2349A>C 2350A>G), L674V (2355C>G 2357C>T), N675S (2358A>T 2359A>C 2360C>T), W676* (2362G>A), C681L (2377G>T 2378T>G), N683S (2382A>T 2383A>C 2384C>T), P684F (2385C>T 2386C>T 2387A>T), D685E (2390T>A), K689E (2400A>G 2402G>A), S690K (2404G>A 2405T>A), I692M (2411C>G), S694A (2415T>G 2417C>T), C697Y (2425G>A), T699* (2430A>T 2431C>A 2432A>G), L704S (2445C>T 2446T>C 2447C>G), E705N (2448G>A 2450A>T), T706E (2451A>G 2452C>A 2453T>A), F708L (2457T>C 2459T>G), D709E (2462T>A), I711V (2466A>G), N727D (2514A>G), P728L (2518C>T 2519C>G), T730I (2524C>T 2525C>T), Q733R (2532C>A 2533A>G), W734Y (2536G>A 2537G>T), F736L (2543T>G), N739S (2551A>G 2552C>T), V741S (2556G>T 2557T>C), N745D (2568A>G), K746R (2571A>C 2572A>G), L749F (2582A>T), N755A (2598A>G 2599A>C 2600C>T), D756N (2601G>A), S760A (2613T>G 2615T>A), E763Y (2622G>T 2624A>T), Y764L (2626A>T 2627C>A), Q765P (2629A>C), T766S (2631A>T), T767L (2634A>C 2635C>T), G768V (2638G>T 2639A>G) | | | | | | | | |

| | Begin | End | Coverage | Score | Concordance | Matches | Identities | I/D/M/F* | Stop Codons |
|---|---|---|---|---|---|---|---|---|---|
| **NT** | **437** | **2647** | **62.1%** | **1358** | **39.5%** | **1721 (100%)** | **1200 (69.7%)** | **0/0** | |
| Codon mutations: | AAA34..G (437A>G), ATC36CTT (441A>C 443C>T), CTT38TTT (447C>T), CTA42TTA (459C>T), GCC44GTC (466C>T), TCC45TCT (470C>T), GCT46ATG (471G>A 472C>T 473T>G), TTG48ATG (477T>A), ACA49CAT (480A>C 481C>A 482A>T), AAA50AAG (485A>G), ATC52GTC (489A>G), GCT53GCC (494T>C), CTT54CTA (497T>A), GCT55GCG (500T>G), GGA56GGC (503A>C), TGT57TGC (506T>C), GAC58GAT (509C>T), TTT59TTC (512T>C), GTA60ATA (513G>A), TCT61AAA (516T>A 517C>A 518T>A), AAA62ACA (520A>C), GTG63GTT (524G>T), ACT64GTG (525A>G 526C>T 527T>G), ACC65AAA (529C>A 530C>A), TTA66CTG (531T>C 533A>G), CGC67AAA (534C>A 535G>A 536C>A), CGG68CGA (539G>A), AAG69CGT (540A>C 541A>G 542G>T), TCG70TCA (545G>A), GGC71GGT (548C>T), TAT72TTT (550A>T), TTC74TAT (556T>A 557C>T), ACA75ACT (560A>T), TAC78TAT (569C>T), AAG80AAA (575G>A), TGC82TGT (581C>T), GCA83ACA (582G>A), GTA84GTT (587A>T), TGC85TCC (589G>C), TTA86CTT (591T>C 593A>T), CGA88CGT (599A>T), TAC89TAT (602C>T), TCC93AGT (612T>A 613C>G 614C>T), TTT94TAT (616T>A), CGA95TCT (618C>T 619G>C 620A>T), GAA97AAT (624G>A 626A>T), GAA98GAC (629A>C), CCA99ACC (630C>A 632A>C), CTT100ATG (633C>A 635T>G), TCG101TCT (638G>T), GTG102GTT (641G>T), CCT103CCA (644T>A), GTG104GTT (647G>T), CTC106TTG (651C>T 653C>G), ACA107ACC (656A>C), AGA108CGA (657A>C), TCG109TGT (661C>G 662G>T), GGT110GGA (665T>A), ATC111ATT (668C>T), CGT113AAG (672C>A 673G>A 674T>G), ATC115ATA (680C>A), CCC116CCT (683C>T), AAA117GCC (684A>G 685A>C 686A>C), TTG119TTA (692G>A), CGT120CGA (695T>A), GCA122CAC (699G>C 700C>A 701A>C), ATA123GTA (702A>G), CGT124AGG (705C>A 707T>G), AGG125GCA (708A>G 709G>C 710G>A), AAA126AAG (713A>G), GAT127TCT (714G>T 715A>C), GAA128GAT (719A>T), CAC129CAT (722C>T), GTT134GTC (737T>C), TAC137TAT (746C>T), TCA139TCT (752A>T), GGG142GGA (761G>A), GCT144TCA (765G>T 767T>A), AAG145AAA (770G>A), GTT146TAG (771G>T 772T>A 773T>G), TTG147ATT (774T>A 776G>T), AAA148GCG (777A>G 778A>C 779A>G), GTT149GTG (782T>G), AGT154AAG (796G>A 797T>G), GCA156GCC (803A>C), TTT158TTC (809T>C), GAA159CAA (810G>C), TCC160TCA (815C>A), ATA161ATT (818A>T), ACT162ACG (821T>G), ACC163ACT (824C>T), CCA164CCT (827A>T), CCT166TCA (831C>T 833T>A), GGG169GAT (841G>A 842G>T), CTA172CGT (850T>G 851A>T), GTG174GTT (857G>T), CTG175CTT (860G>T), GAA176GGA (862A>G), ATT178ATG (869T>G), ACG180ACT (875G>T), TTC182TGA (880T>G 881C>A), CAT184AAG (885C>A 887T>G), CAA186CGA (892A>G), CTA190CTT (905A>T), CCC191CCA (908C>A), TTT192CAT (909T>C 910T>A), CGA194CAA (916G>A), GGC195TCC (918G>T 919G>C), ATT196ATA (923T>A), CCA197CCT (926A>T), ATG198CTT (927A>C 929G>T), ATT200AAG (934T>A 935T>G), GGG201GGT (938G>T), TCT203AAT (942T>A 943C>A), CAA205AAA (948C>A), CCC212CCA (971C>A), CTA213TTA (972C>T), AAT214CTC (975A>C 976A>T 977T>C), AAT216AAC (983T>C), AGG219CGC (990A>C 992G>C), TCA220CGG (993T>C 994C>G 995A>G), TAC221TTT (997A>T 998C>T), GGT222CAA (999G>C 1000G>A 1001T>A), TTA223TAT (1003T>A 1004A>T), GCC224TCG (1005G>T 1007C>G), GTA225GTG (1010A>G), GAC227GAT (1016C>T), AAA228GTT (1017A>G 1018A>T 1019A>T), GTT229GTA (1022T>A), GAT230GAC (1025T>C), CAA231GGA (1026C>G 1027A>G), GAG232GCA (1030A>C 1031G>A), GCT233GCG (1034T>G), AGT234GCT (1035A>G 1036G>C), TTC237GTT (1044T>G 1046C>T), AAG238AAA (1049G>A), TTT239GAA (1050T>G 1051T>A 1052T>A), CCA240CAC (1054C>A 1055A>C), ATT242ATC (1061T>C), GTT244GTG (1067T>G), TTG246TTA (1073G>A), AAA247AAG (1076A>G), CTA250ATA (1083C>A), GCG251GCT (1088G>T), GCC252GCG (1091C>G), TGG255TTG (1099G>T), ACA257GTG (1104A>G 1105C>T 1106A>G), ATT260ATG (1115T>G), TCA262TCC (1121A>C), TTC263ATT (1122T>A 1124C>T), CAG264CAA (1127G>A), GAC265GAT (1130C>T), CCG432TCG (1629C>T), TTA433ATC (1632T>A 1634A>C), GTG437ATG (1644G>A), CTG443CTA (1664G>A), GGG445AGA (1668G>A 1670G>A), CAG446AAA (1671C>A 1673G>A), CCT447CCA (1676T>A), TTG448TTA (1679G>A), GGC449GGT (1682C>T), CAC451TAC (1686C>T), GGT452GGA (1691T>A), TGG454TAG (1696G>A), GCA455TCA (1698G>T), CTT456CTC (1703T>C), TTC457TTT (1706C>T), TCG458GCT (1707T>G 1709G>T), TTG459CTA (1710T>C 1712G>A), TAT463TAC (1724T>C), GCA468GGA (1738C>G), GCG469GCA (1742G>A), AGC470AAG (1744G>A 1745C>G), GGA475GAC (1759G>A 1760A>C), TGT476TAT (1762G>A), AAC477AAA (1766C>A), TCT478ACC (1767T>A 1769T>C), CCA479CCC (1772A>C), TTT480TTC (1775T>C), GGG481AAA (1776G>A 1777G>A 1778G>A), GAT482GAC (1781T>C), TAC483TCT (1783A>C 1784C>T), GCT484GCC (1787T>C), TTA485TTG (1790A>G), GAT494GAC (1817T>C), AGT495AAG (1819G>A 1820T>G), AAA496AAG (1823A>G), GAG500CAG (1833G>C), AAG502AGG (1840A>G), AGA507AGG (1856A>G), CTC508CTT (1859C>T), AAT509GGT (1860A>G 1861A>G), GTC510GTT (1865C>T), AAA511ACC (1867A>C 1868A>C), TCG513TCT (1874G>T), ATA514TTA (1875A>T), CCA515GCT (1878C>G 1880A>T), TCT517TCA (1886T>A), TTG518ATA (1887T>A 1889G>A), TCT520TCC (1895T>C), GAT521GAG (1898T>G), GGA523GGT (1904A>T), ACC524ACT (1907C>T), GCG528GCC (1919G>C), AAG529AAA (1922G>A), AGG530TGA (1923A>T 1925G>A), ATG536ATT (1943G>T), ATT538GTT (1947A>G), CCA542CTA (1960C>T), TCC544TCT (1967C>T), CTT545GTG (1968C>G 1970T>G), AGA546AAG (1972G>A 1973A>G), GCT547GCA (1976T>A), TTA549ACA (1980T>A 1981T>C), TCT550TTG (1984C>T 1985T>G), CGT552CGC (1991T>C), GTT555GGT (1999T>G), GGA556GGT (2003A>T), CTT557CTA (2006T>A), GCC559AAC (2010G>A 2011C>A), CTC560ATT (2013C>A 2015C>T), ATC562ACC (2020T>C), GAT565GAC (2030T>C), AAC567AAT (2036C>T), ATA568ATG (2039A>G), CAA572TAG (2049C>T 2051A>G), AGA573TGG (2052A>T 2054A>G), GGT576GGG (2063T>G), GGG578GGA (2069G>A), TAC579TTC (2071A>T), AAG580AGA (2074A>G 2075G>A), GTA581GTT (2078A>T), GCG583TCT (2082G>T 2084G>T), CGC584CGT (2087C>T), TTA585TTG (2090A>G), TTC586ATG (2091T>A 2093C>G), ACT587TCT (2094A>T), ACC588ACT (2099C>T), AGC590TCT (2103A>T 2104G>C 2105C>T), AAG591AAA (2108G>A), GAG594GAA (2117G>A), CGT595AGA (2118C>A 2120T>A), CTA596CTG (2123A>G), GCT598GCG (2129T>G), CAT600GCA (2133C>G 2134A>C 2135T>A), CTT601AGC (2136C>A 2137T>G 2138T>C), AAG602AAA (2141G>A), GTT604CAC (2145G>C 2146T>A 2147T>C), CGG605AGG (2148C>A), CAC607CAT (2156C>T), TTA608CTA (2157T>C), CTA609TTA (2160C>T), TAC611CTC (2166T>C 2167A>T), TGG613TAT (2173G>A 2174G>T), GGA616GGG (2183A>G), AGG617AGA (2186G>A), GGT618TTT (2187G>T 2188G>T), ATG619AAA (2191T>A 2192G>A), CCC620CCT (2195C>T), CCC623CCT (2204C>T), TAT624TTT (2206A>T), AAA626AAG (2213A>G), GGG627GGA (2216G>A), GTA630ATA (2223G>A), AGT631GAT (2226A>G 2227G>A), TAC632TAT (2231C>T), CTT634CGG (2236T>G 2237T>G), AAA635AAG (2240A>G), AAA638ACA (2248A>C), GTT639CCT (2250G>C 2251T>C), TTA642ATA (2259T>A), CAG643CGA (2263A>G 2264G>A), ATC644CTC (2265A>C), CCA646ACT (2271C>A 2273A>T), AAG647GAA (2274A>G 2276G>A), CTG649ATG (2280C>A), GTG650GTA (2285G>A), TTC651TTT (2288C>T), GTT655TGG (2298G>T 2299T>G 2300T>G), GAA656GAG (2303A>G), ATA657ATC (2306A>C), CTA658CTT (2309A>T), CGT660TGC (2313C>T 2315T>C), GTT662ATT (2319G>A), CTA663TTA (2322C>T), CGC664GTC (2325C>G 2326G>T), CAA665AAT (2328C>A 2330A>T), GTA667GTT (2336A>T), GAA668AAA (2337G>A), TTA671CTT (2346T>C 2348A>T), AAG672CGG (2349A>C 2350A>G), CTC674GTT (2355C>G 2357C>T), AAC675TCT (2358A>T 2359A>C 2360C>T), TGG676TAG (2362G>A), CAT678CAC (2369T>C), ACG679ACT (2372G>T), TGT681TTG (2377G>T 2378T>G), AAC683TCT (2382A>T 2383A>C 2384C>T), CCA684TTT (2385C>T 2386C>T 2387A>T), GAT685GAA (2390T>A), TCC687TCT (2396C>T), ATT688ATC (2399T>C), AAG689GAA (2400A>G 2402G>A), AGT690AAA (2404G>A 2405T>A), CTC691TTG (2406C>T 2408C>G), ATC692ATG (2411C>G), GAC693GAT (2414C>T), TCC694GCT (2415T>G 2417C>T), CCT695CCC (2420T>C), ATA696ATT (2423A>T), TGC697TAC (2425G>A), ACA699TAG (2430A>T 2431C>A 2432A>G), TCC700TCA (2435C>A), AAA702AAG (2441A>G), AGG703CGA (2442A>C 2444G>A), CTC704TCG (2445C>T 2446T>C 2447C>G), GAA705AAT (2448G>A 2450A>T), ACT706GAA (2451A>G 2452C>A 2453T>A), GAT707GAC (2456T>C), TTT708CTG (2457T>C 2459T>G), GAT709GAA (2462T>A), TTA710TTG (2465A>G), ATC711GTC (2466A>G), GGC714GGT (2477C>T), CTA715TTA (2478C>T), AAA718AAG (2489A>G), TGT719TGC (2492T>C), TAC720TAT (2495C>T), GGT723GGA (2504T>A), GGT725GGC (2510T>C), AAT727GAT (2514A>G), CCC728CTG (2518C>T 2519C>G), ACC730ATT (2524C>T 2525C>T), ACT731ACA (2528T>A), CAG733AGG (2532C>A 2533A>G), TGG734TAT (2536G>A 2537G>T), TTT736TTG (2543T>G), AAC739AGT (2551A>G 2552C>T), ACT740ACC (2555T>C), GTT741TCT (2556G>T 2557T>C), ATT742ATC (2561T>C), AAT745GAT (2568A>G), AAA746CGA (2571A>C 2572A>G), TTA749TTT (2582A>T), GGG750GGA (2585G>A), GGG754GGA (2597G>A), AAC755GCT (2598A>G 2599A>C 2600C>T), GAT756AAT (2601G>A), ATT758ATA (2609T>A), TCT760GCA (2613T>G 2615T>A), CCT761CCA (2618T>A), GAA763TAT (2622G>T 2624A>T), TAC764TTA (2626A>T 2627C>A), CAA765CCA (2629A>C), ACT766TCT (2631A>T), ACA767CTA (2634A>C 2635C>T), GGA768GTG (2638G>T 2639A>G) | | | | | | | | |

#### Proteins

| | Begin | End | Coverage | Score | Concordance | Matches | Identities | I/D/M/F* | Stop Codons |
|---|---|---|---|---|---|---|---|---|---|
| RNA-dependent RNA polymerase (YP_010798874.1) | 35 | 771 | 73.9% | 2762 | 67.9% | 574 (100%) | 390 (67.9%) | 0/0/0/0 | 7 |
| Protein mutations: | I36L (441A>C 443C>T), L38F (447C>T), A44V (466C>T), A46M (471G>A 472C>T 473T>G), L48M (477T>A), T49H (480A>C 481C>A 482A>T), I52V (489A>G), V60I (513G>A), S61K (516T>A 517C>A 518T>A), K62T (520A>C), T64V (525A>G 526C>T 527T>G), T65K (529C>A 530C>A), R67K (534C>A 535G>A 536C>A), K69R (540A>C 541A>G 542G>T), Y72F (550A>T), F74Y (556T>A 557C>T), A83T (582G>A), C85S (589G>C), F94Y (616T>A), R95S (618C>T 619G>C 620A>T), E97N (624G>A 626A>T), E98D (629A>C), P99T (630C>A 632A>C), L100M (633C>A 635T>G), S109C (661C>G 662G>T), R113K (672C>A 673G>A 674T>G), K117A (684A>G 685A>C 686A>C), A122H (699G>C 700C>A 701A>C), I123V (702A>G), R125A (708A>G 709G>C 710G>A), D127S (714G>T 715A>C), E128D (719A>T), A144S (765G>T 767T>A), V146* (771G>T 772T>A 773T>G), L147I (774T>A 776G>T), K148A (777A>G 778A>C 779A>G), S154K (796G>A 797T>G), E159Q (810G>C), P166S (831C>T 833T>A), G169D (841G>A 842G>T), L172R (850T>G 851A>T), E176G (862A>G), I178M (869T>G), F182* (880T>G 881C>A), H184K (885C>A 887T>G), Q186R (892A>G), F192H (909T>C 910T>A), R194Q (916G>A), G195S (918G>T 919G>C), M198L (927A>C 929G>T), I200K (934T>A 935T>G), S203N (942T>A 943C>A), Q205K (948C>A), N214L (975A>C 976A>T 977T>C), S220R (993T>C 994C>G 995A>G), Y221F (997A>T 998C>T), G222Q (999G>C 1000G>A 1001T>A), L223Y (1003T>A 1004A>T), A224S (1005G>T 1007C>G), K228V (1017A>G 1018A>T 1019A>T), Q231G (1026C>G 1027A>G), E232A (1030A>C 1031G>A), S234A (1035A>G 1036G>C), F237V (1044T>G 1046C>T), F239E (1050T>G 1051T>A 1052T>A), P240H (1054C>A 1055A>C), L250I (1083C>A), W255L (1099G>T), T257V (1104A>G 1105C>T 1106A>G), I260M (1115T>G), F263I (1122T>A 1124C>T), P432S (1629C>T), L433I (1632T>A 1634A>C), V437M (1644G>A), G445R (1668G>A 1670G>A), Q446K (1671C>A 1673G>A), H451Y (1686C>T), W454* (1696G>A), A455S (1698G>T), S458A (1707T>G 1709G>T), A468G (1738C>G), S470K (1744G>A 1745C>G), G475D (1759G>A 1760A>C), C476Y (1762G>A), N477K (1766C>A), S478T (1767T>A 1769T>C), G481K (1776G>A 1777G>A 1778G>A), Y483S (1783A>C 1784C>T), S495K (1819G>A 1820T>G), E500Q (1833G>C), K502R (1840A>G), N509G (1860A>G 1861A>G), K511T (1867A>C 1868A>C), I514L (1875A>T), P515A (1878C>G 1880A>T), L518I (1887T>A 1889G>A), D521E (1898T>G), R530* (1923A>T 1925G>A), M536I (1943G>T), I538V (1947A>G), P542L (1960C>T), L545V (1968C>G 1970T>G), R546K (1972G>A 1973A>G), L549T (1980T>A 1981T>C), S550L (1984C>T 1985T>G), V555G (1999T>G), A559N (2010G>A 2011C>A), L560I (2013C>A 2015C>T), I562T (2020T>C), I568M (2039A>G), Q572* (2049C>T 2051A>G), R573W (2052A>T 2054A>G), Y579F (2071A>T), K580R (2074A>G 2075G>A), A583S (2082G>T 2084G>T), F586M (2091T>A 2093C>G), T587S (2094A>T), H600A (2133C>G 2134A>C 2135T>A), L601S (2136C>A 2137T>G 2138T>C), V604H (2145G>C 2146T>A 2147T>C), Y611L (2166T>C 2167A>T), W613Y (2173G>A 2174G>T), G618F (2187G>T 2188G>T), M619K (2191T>A 2192G>A), Y624F (2206A>T), V630I (2223G>A), S631D (2226A>G 2227G>A), L634R (2236T>G 2237T>G), K638T (2248A>C), V639P (2250G>C 2251T>C), L642I (2259T>A), Q643R (2263A>G 2264G>A), I644L (2265A>C), P646T (2271C>A 2273A>T), K647E (2274A>G 2276G>A), L649M (2280C>A), V655W (2298G>T 2299T>G 2300T>G), R660C (2313C>T 2315T>C), V662I (2319G>A), R664V (2325C>G 2326G>T), Q665N (2328C>A 2330A>T), E668K (2337G>A), K672R (2349A>C 2350A>G), L674V (2355C>G 2357C>T), N675S (2358A>T 2359A>C 2360C>T), W676* (2362G>A), C681L (2377G>T 2378T>G), N683S (2382A>T 2383A>C 2384C>T), P684F (2385C>T 2386C>T 2387A>T), D685E (2390T>A), K689E (2400A>G 2402G>A), S690K (2404G>A 2405T>A), I692M (2411C>G), S694A (2415T>G 2417C>T), C697Y (2425G>A), T699* (2430A>T 2431C>A 2432A>G), L704S (2445C>T 2446T>C 2447C>G), E705N (2448G>A 2450A>T), T706E (2451A>G 2452C>A 2453T>A), F708L (2457T>C 2459T>G), D709E (2462T>A), I711V (2466A>G), N727D (2514A>G), P728L (2518C>T 2519C>G), T730I (2524C>T 2525C>T), Q733R (2532C>A 2533A>G), W734Y (2536G>A 2537G>T), F736L (2543T>G), N739S (2551A>G 2552C>T), V741S (2556G>T 2557T>C), N745D (2568A>G), K746R (2571A>C 2572A>G), L749F (2582A>T), N755A (2598A>G 2599A>C 2600C>T), D756N (2601G>A), S760A (2613T>G 2615T>A), E763Y (2622G>T 2624A>T), Y764L (2626A>T 2627C>A), Q765P (2629A>C), T766S (2631A>T), T767L (2634A>C 2635C>T), G768V (2638G>T 2639A>G) | | | | | | | | |

| | Begin | End | Coverage | Score | Concordance | Matches | Identities | I/D/M/F* | Stop Codons |
|---|---|---|---|---|---|---|---|---|---|
| **NT** | **437** | **2647** | **62.1%** | **1358** | **39.5%** | **1721 (100%)** | **1200 (69.7%)** | **0/0** | |
| Codon mutations: | AAA34..G (437A>G), ATC36CTT (441A>C 443C>T), CTT38TTT (447C>T), CTA42TTA (459C>T), GCC44GTC (466C>T), TCC45TCT (470C>T), GCT46ATG (471G>A 472C>T 473T>G), TTG48ATG (477T>A), ACA49CAT (480A>C 481C>A 482A>T), AAA50AAG (485A>G), ATC52GTC (489A>G), GCT53GCC (494T>C), CTT54CTA (497T>A), GCT55GCG (500T>G), GGA56GGC (503A>C), TGT57TGC (506T>C), GAC58GAT (509C>T), TTT59TTC (512T>C), GTA60ATA (513G>A), TCT61AAA (516T>A 517C>A 518T>A), AAA62ACA (520A>C), GTG63GTT (524G>T), ACT64GTG (525A>G 526C>T 527T>G), ACC65AAA (529C>A 530C>A), TTA66CTG (531T>C 533A>G), CGC67AAA (534C>A 535G>A 536C>A), CGG68CGA (539G>A), AAG69CGT (540A>C 541A>G 542G>T), TCG70TCA (545G>A), GGC71GGT (548C>T), TAT72TTT (550A>T), TTC74TAT (556T>A 557C>T), ACA75ACT (560A>T), TAC78TAT (569C>T), AAG80AAA (575G>A), TGC82TGT (581C>T), GCA83ACA (582G>A), GTA84GTT (587A>T), TGC85TCC (589G>C), TTA86CTT (591T>C 593A>T), CGA88CGT (599A>T), TAC89TAT (602C>T), TCC93AGT (612T>A 613C>G 614C>T), TTT94TAT (616T>A), CGA95TCT (618C>T 619G>C 620A>T), GAA97AAT (624G>A 626A>T), GAA98GAC (629A>C), CCA99ACC (630C>A 632A>C), CTT100ATG (633C>A 635T>G), TCG101TCT (638G>T), GTG102GTT (641G>T), CCT103CCA (644T>A), GTG104GTT (647G>T), CTC106TTG (651C>T 653C>G), ACA107ACC (656A>C), AGA108CGA (657A>C), TCG109TGT (661C>G 662G>T), GGT110GGA (665T>A), ATC111ATT (668C>T), CGT113AAG (672C>A 673G>A 674T>G), ATC115ATA (680C>A), CCC116CCT (683C>T), AAA117GCC (684A>G 685A>C 686A>C), TTG119TTA (692G>A), CGT120CGA (695T>A), GCA122CAC (699G>C 700C>A 701A>C), ATA123GTA (702A>G), CGT124AGG (705C>A 707T>G), AGG125GCA (708A>G 709G>C 710G>A), AAA126AAG (713A>G), GAT127TCT (714G>T 715A>C), GAA128GAT (719A>T), CAC129CAT (722C>T), GTT134GTC (737T>C), TAC137TAT (746C>T), TCA139TCT (752A>T), GGG142GGA (761G>A), GCT144TCA (765G>T 767T>A), AAG145AAA (770G>A), GTT146TAG (771G>T 772T>A 773T>G), TTG147ATT (774T>A 776G>T), AAA148GCG (777A>G 778A>C 779A>G), GTT149GTG (782T>G), AGT154AAG (796G>A 797T>G), GCA156GCC (803A>C), TTT158TTC (809T>C), GAA159CAA (810G>C), TCC160TCA (815C>A), ATA161ATT (818A>T), ACT162ACG (821T>G), ACC163ACT (824C>T), CCA164CCT (827A>T), CCT166TCA (831C>T 833T>A), GGG169GAT (841G>A 842G>T), CTA172CGT (850T>G 851A>T), GTG174GTT (857G>T), CTG175CTT (860G>T), GAA176GGA (862A>G), ATT178ATG (869T>G), ACG180ACT (875G>T), TTC182TGA (880T>G 881C>A), CAT184AAG (885C>A 887T>G), CAA186CGA (892A>G), CTA190CTT (905A>T), CCC191CCA (908C>A), TTT192CAT (909T>C 910T>A), CGA194CAA (916G>A), GGC195TCC (918G>T 919G>C), ATT196ATA (923T>A), CCA197CCT (926A>T), ATG198CTT (927A>C 929G>T), ATT200AAG (934T>A 935T>G), GGG201GGT (938G>T), TCT203AAT (942T>A 943C>A), CAA205AAA (948C>A), CCC212CCA (971C>A), CTA213TTA (972C>T), AAT214CTC (975A>C 976A>T 977T>C), AAT216AAC (983T>C), AGG219CGC (990A>C 992G>C), TCA220CGG (993T>C 994C>G 995A>G), TAC221TTT (997A>T 998C>T), GGT222CAA (999G>C 1000G>A 1001T>A), TTA223TAT (1003T>A 1004A>T), GCC224TCG (1005G>T 1007C>G), GTA225GTG (1010A>G), GAC227GAT (1016C>T), AAA228GTT (1017A>G 1018A>T 1019A>T), GTT229GTA (1022T>A), GAT230GAC (1025T>C), CAA231GGA (1026C>G 1027A>G), GAG232GCA (1030A>C 1031G>A), GCT233GCG (1034T>G), AGT234GCT (1035A>G 1036G>C), TTC237GTT (1044T>G 1046C>T), AAG238AAA (1049G>A), TTT239GAA (1050T>G 1051T>A 1052T>A), CCA240CAC (1054C>A 1055A>C), ATT242ATC (1061T>C), GTT244GTG (1067T>G), TTG246TTA (1073G>A), AAA247AAG (1076A>G), CTA250ATA (1083C>A), GCG251GCT (1088G>T), GCC252GCG (1091C>G), TGG255TTG (1099G>T), ACA257GTG (1104A>G 1105C>T 1106A>G), ATT260ATG (1115T>G), TCA262TCC (1121A>C), TTC263ATT (1122T>A 1124C>T), CAG264CAA (1127G>A), GAC265GAT (1130C>T), CCG432TCG (1629C>T), TTA433ATC (1632T>A 1634A>C), GTG437ATG (1644G>A), CTG443CTA (1664G>A), GGG445AGA (1668G>A 1670G>A), CAG446AAA (1671C>A 1673G>A), CCT447CCA (1676T>A), TTG448TTA (1679G>A), GGC449GGT (1682C>T), CAC451TAC (1686C>T), GGT452GGA (1691T>A), TGG454TAG (1696G>A), GCA455TCA (1698G>T), CTT456CTC (1703T>C), TTC457TTT (1706C>T), TCG458GCT (1707T>G 1709G>T), TTG459CTA (1710T>C 1712G>A), TAT463TAC (1724T>C), GCA468GGA (1738C>G), GCG469GCA (1742G>A), AGC470AAG (1744G>A 1745C>G), GGA475GAC (1759G>A 1760A>C), TGT476TAT (1762G>A), AAC477AAA (1766C>A), TCT478ACC (1767T>A 1769T>C), CCA479CCC (1772A>C), TTT480TTC (1775T>C), GGG481AAA (1776G>A 1777G>A 1778G>A), GAT482GAC (1781T>C), TAC483TCT (1783A>C 1784C>T), GCT484GCC (1787T>C), TTA485TTG (1790A>G), GAT494GAC (1817T>C), AGT495AAG (1819G>A 1820T>G), AAA496AAG (1823A>G), GAG500CAG (1833G>C), AAG502AGG (1840A>G), AGA507AGG (1856A>G), CTC508CTT (1859C>T), AAT509GGT (1860A>G 1861A>G), GTC510GTT (1865C>T), AAA511ACC (1867A>C 1868A>C), TCG513TCT (1874G>T), ATA514TTA (1875A>T), CCA515GCT (1878C>G 1880A>T), TCT517TCA (1886T>A), TTG518ATA (1887T>A 1889G>A), TCT520TCC (1895T>C), GAT521GAG (1898T>G), GGA523GGT (1904A>T), ACC524ACT (1907C>T), GCG528GCC (1919G>C), AAG529AAA (1922G>A), AGG530TGA (1923A>T 1925G>A), ATG536ATT (1943G>T), ATT538GTT (1947A>G), CCA542CTA (1960C>T), TCC544TCT (1967C>T), CTT545GTG (1968C>G 1970T>G), AGA546AAG (1972G>A 1973A>G), GCT547GCA (1976T>A), TTA549ACA (1980T>A 1981T>C), TCT550TTG (1984C>T 1985T>G), CGT552CGC (1991T>C), GTT555GGT (1999T>G), GGA556GGT (2003A>T), CTT557CTA (2006T>A), GCC559AAC (2010G>A 2011C>A), CTC560ATT (2013C>A 2015C>T), ATC562ACC (2020T>C), GAT565GAC (2030T>C), AAC567AAT (2036C>T), ATA568ATG (2039A>G), CAA572TAG (2049C>T 2051A>G), AGA573TGG (2052A>T 2054A>G), GGT576GGG (2063T>G), GGG578GGA (2069G>A), TAC579TTC (2071A>T), AAG580AGA (2074A>G 2075G>A), GTA581GTT (2078A>T), GCG583TCT (2082G>T 2084G>T), CGC584CGT (2087C>T), TTA585TTG (2090A>G), TTC586ATG (2091T>A 2093C>G), ACT587TCT (2094A>T), ACC588ACT (2099C>T), AGC590TCT (2103A>T 2104G>C 2105C>T), AAG591AAA (2108G>A), GAG594GAA (2117G>A), CGT595AGA (2118C>A 2120T>A), CTA596CTG (2123A>G), GCT598GCG (2129T>G), CAT600GCA (2133C>G 2134A>C 2135T>A), CTT601AGC (2136C>A 2137T>G 2138T>C), AAG602AAA (2141G>A), GTT604CAC (2145G>C 2146T>A 2147T>C), CGG605AGG (2148C>A), CAC607CAT (2156C>T), TTA608CTA (2157T>C), CTA609TTA (2160C>T), TAC611CTC (2166T>C 2167A>T), TGG613TAT (2173G>A 2174G>T), GGA616GGG (2183A>G), AGG617AGA (2186G>A), GGT618TTT (2187G>T 2188G>T), ATG619AAA (2191T>A 2192G>A), CCC620CCT (2195C>T), CCC623CCT (2204C>T), TAT624TTT (2206A>T), AAA626AAG (2213A>G), GGG627GGA (2216G>A), GTA630ATA (2223G>A), AGT631GAT (2226A>G 2227G>A), TAC632TAT (2231C>T), CTT634CGG (2236T>G 2237T>G), AAA635AAG (2240A>G), AAA638ACA (2248A>C), GTT639CCT (2250G>C 2251T>C), TTA642ATA (2259T>A), CAG643CGA (2263A>G 2264G>A), ATC644CTC (2265A>C), CCA646ACT (2271C>A 2273A>T), AAG647GAA (2274A>G 2276G>A), CTG649ATG (2280C>A), GTG650GTA (2285G>A), TTC651TTT (2288C>T), GTT655TGG (2298G>T 2299T>G 2300T>G), GAA656GAG (2303A>G), ATA657ATC (2306A>C), CTA658CTT (2309A>T), CGT660TGC (2313C>T 2315T>C), GTT662ATT (2319G>A), CTA663TTA (2322C>T), CGC664GTC (2325C>G 2326G>T), CAA665AAT (2328C>A 2330A>T), GTA667GTT (2336A>T), GAA668AAA (2337G>A), TTA671CTT (2346T>C 2348A>T), AAG672CGG (2349A>C 2350A>G), CTC674GTT (2355C>G 2357C>T), AAC675TCT (2358A>T 2359A>C 2360C>T), TGG676TAG (2362G>A), CAT678CAC (2369T>C), ACG679ACT (2372G>T), TGT681TTG (2377G>T 2378T>G), AAC683TCT (2382A>T 2383A>C 2384C>T), CCA684TTT (2385C>T 2386C>T 2387A>T), GAT685GAA (2390T>A), TCC687TCT (2396C>T), ATT688ATC (2399T>C), AAG689GAA (2400A>G 2402G>A), AGT690AAA (2404G>A 2405T>A), CTC691TTG (2406C>T 2408C>G), ATC692ATG (2411C>G), GAC693GAT (2414C>T), TCC694GCT (2415T>G 2417C>T), CCT695CCC (2420T>C), ATA696ATT (2423A>T), TGC697TAC (2425G>A), ACA699TAG (2430A>T 2431C>A 2432A>G), TCC700TCA (2435C>A), AAA702AAG (2441A>G), AGG703CGA (2442A>C 2444G>A), CTC704TCG (2445C>T 2446T>C 2447C>G), GAA705AAT (2448G>A 2450A>T), ACT706GAA (2451A>G 2452C>A 2453T>A), GAT707GAC (2456T>C), TTT708CTG (2457T>C 2459T>G), GAT709GAA (2462T>A), TTA710TTG (2465A>G), ATC711GTC (2466A>G), GGC714GGT (2477C>T), CTA715TTA (2478C>T), AAA718AAG (2489A>G), TGT719TGC (2492T>C), TAC720TAT (2495C>T), GGT723GGA (2504T>A), GGT725GGC (2510T>C), AAT727GAT (2514A>G), CCC728CTG (2518C>T 2519C>G), ACC730ATT (2524C>T 2525C>T), ACT731ACA (2528T>A), CAG733AGG (2532C>A 2533A>G), TGG734TAT (2536G>A 2537G>T), TTT736TTG (2543T>G), AAC739AGT (2551A>G 2552C>T), ACT740ACC (2555T>C), GTT741TCT (2556G>T 2557T>C), ATT742ATC (2561T>C), AAT745GAT (2568A>G), AAA746CGA (2571A>C 2572A>G), TTA749TTT (2582A>T), GGG750GGA (2585G>A), GGG754GGA (2597G>A), AAC755GCT (2598A>G 2599A>C 2600C>T), GAT756AAT (2601G>A), ATT758ATA (2609T>A), TCT760GCA (2613T>G 2615T>A), CCT761CCA (2618T>A), GAA763TAT (2622G>T 2624A>T), TAC764TTA (2626A>T 2627C>A), CAA765CCA (2629A>C), ACT766TCT (2631A>T), ACA767CTA (2634A>C 2635C>T), GGA768GTG (2638G>T 2639A>G) | | | | | | | | |

*: Inserts / Deletes / Misaligned / Frameshifts

#### Analysis details

This analysis was performed with panviral2.64

### NGS Details (UN9): Harvey murine sarcoma virus

#### Assembly

| | |
|---|---|
| **Coverage Length** | 215 (1 contig(s)) |
| **Depth Of Coverage** | 3.2 |
| **Number Of Reads** | 6 |
| **Reads Per Million** | 0.14 rpm (after QC) |
| **Ambiguities** | 0 |
| **Assembly Method** | de novo + reference guided assembly |
| **Consensus Caller** | Bcf Tools |

#### Coverage Map

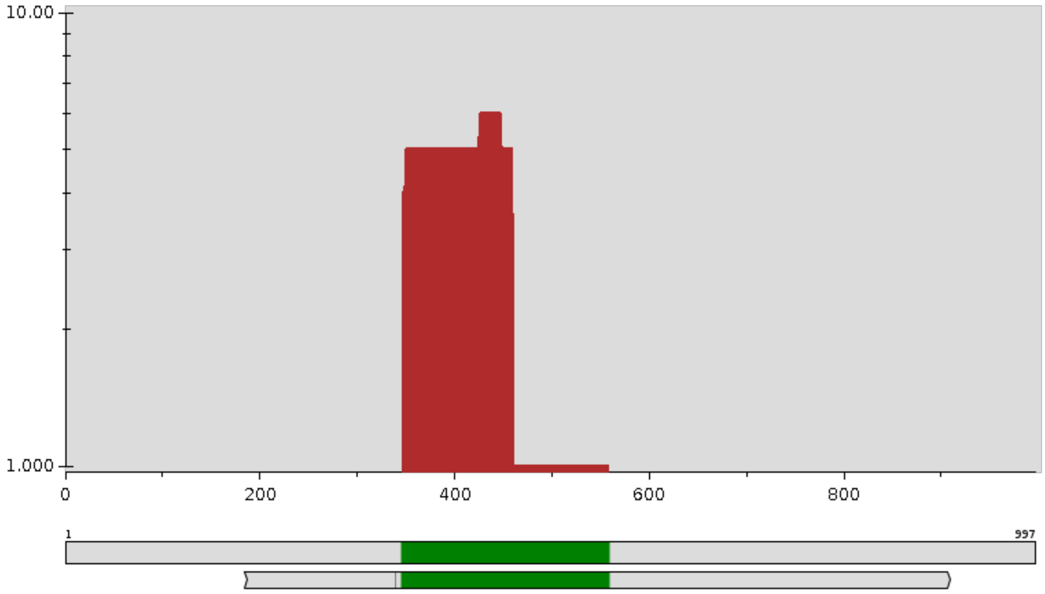

#### Assignment

| | |
|---|---|
| **Type** | Harvey murine sarcoma virus (Taxonomy ID: 11807) |
| **Reference Genome** | NC_038668.1 |
| **NT Identity (%)** | 70.2326 |
| **AA Identity (%)** | 83.3333 |
| **Number Of Stop Codons** | 0 |
| **Number Of CDS** | 2 |

#### Alignment

| | |
|---|---|
| **Alignment Score** | 174.0 (NT) + 798.0 (AA) = 972.0 |
| **Concordance (%)** | 70.1299 |

| Alignment Method | Global, seeded, nucleotide + amino acids (AGA) |
|---|---|

#### Genome Region

Sequence starts at position 346 and ends at position 560 relative to NC_038668.1 reference sequence.

#### Alignment Detailed Statistics

| | Begin | End | Coverage | Score | Concordance | Matches | Identities | I/D/M/F* | Stop Codons |
|---|---|---|---|---|---|---|---|---|---|
| **NT** | **346** | **560** | **21.6%** | **174** | **40.5%** | **215 (100%)** | **151 (70.2%)** | **0/0** | |
| Mutations: | 346A>G, 355G>A, 358T>C, 361G>C, 364G>C, 367G>C, 372C>G, 374A>G, 376A>T, 379C>T, 382G>C, 389A>T, 390G>C, 391T>C, 394C>T, 397G>C, 400C>G, 403C>A, 406G>A, 407C>T, 409G>C, 412C>A, 415G>A, 416A>T, 417A>C, 418C>G, 424T>C, 427G>C, 430C>T, 439T>C, 442C>T, 448A>C, 451G>A, 457C>A, 463G>C, 469G>A, 470G>T, 471T>G, 472A>C, 475C>G, 478T>A, 481T>C, 483G>A, 484G>A, 487G>A, 488A>G, 489C>T, 490G>T, 491T>G, 492G>C, 493T>A, 503A>G, 511C>T, 515A>G, 517A>C, 520T>C, 529G>A, 532T>C, 533A>T, 534G>C, 542C>A, 547C>G, 550G>A, 559C>G | | | | | | | | |
| **CDS** | | | | | | | | | |
| D1R95_gp1 | 55 | 126 | 29.8% | 399 | 81.8% | 72 (100%) | 60 (83.3%) | 0/0/0/0 | 0 |
| Protein mutations: | A63G (372C>G), R64G (374A>G 376A>T), L75F (407C>T 409G>C), N78S (416A>T 417A>C 418C>G), V96C (470G>T 471T>G 472A>C), G100E (483G>A 484G>A), T102V (488A>G 489C>T 490G>T), C103A (491T>G 492G>C 493T>A), I107V (503A>G), T111A (515A>G 517A>C), D121E (547C>G) | | | | | | | | |
| Codon mutations: | ACA54..G (346A>G), AAG57AAA (355G>A), CTT58CTC (358T>C), GTG59GTC (361G>C), GTG60GTC (364G>C), GTG61GTC (367G>C), GCT63GGT (372C>G), AGA64GGT (374A>G 376A>T), GGC65GGT (379C>T), GTG66GTC (382G>C), AGT69TCC (389A>T 390G>C 391T>C), GCC70GCT (394C>T), CTG71CTC (397G>C), ACC72ACG (400C>G), ATC73ATA (403C>A), CAG74CAA (406G>A), CTG75TTC (407C>T 409G>C), ATC76ATA (412C>A), CAG77CAA (415G>A), AAC78TCG (416A>T 417A>C 418C>G), TTT80TTC (424T>C), GTG81GTC (427G>C), GAC82GAT (430C>T), GAT85GAC (439T>C), CCC86CCT (442C>T), ATA88ATC (448A>C), GAG89GAA (451G>A), TCC91TCA (457C>A), CGG93CGC (463G>C), CAG95CAA (469G>A), GTA96TGC (470G>T 471T>G 472A>C), GTC97GTG (475C>G), ATT98ATA (478T>A), GAT99GAC (481T>C), GGG100GAA (483G>A 484G>A), GAG101GAA (487G>A), ACG102GTT (488A>G 489C>T 490G>T), TGT103GCA (491T>G 492G>C 493T>A), ATC107GTC (503A>G), GAC109GAT (511C>T), ACA111GCC (515A>G 517A>C), GGT112GGC (520T>C), GAG115GAA (529G>A), TAT116TAC (532T>C), AGT117TCT (533A>T 534G>C), CGG120AGG (542C>A), GAC121GAG (547C>G), CAG122CAA (550G>A), CGC125CGG (559C>G) | | | | | | | | |
| D1R95_gp2 | 3 | 74 | 37.9% | 399 | 81.8% | 72 (100%) | 60 (83.3%) | 0/0/0/0 | 0 |
| Protein mutations: | A11G (372C>G), R12G (374A>G 376A>T), L23F (407C>T 409G>C), N26S (416A>T 417A>C 418C>G), V44C (470G>T 471T>G 472A>C), G48E (483G>A 484G>A), T50V (488A>G 489C>T 490G>T), C51A (491T>G 492G>C 493T>A), I55V (503A>G), T59A (515A>G 517A>C), D69E (547C>G) | | | | | | | | |
| Codon mutations: | ACA2..G (346A>G), AAG5AAA (355G>A), CTT6CTC (358T>C), GTG7GTC (361G>C), GTG8GTC (364G>C), GTG9GTC (367G>C), GCT11GGT (372C>G), AGA12GGT (374A>G 376A>T), GGC13GGT (379C>T), GTG14GTC (382G>C), AGT17TCC (389A>T 390G>C 391T>C), GCC18GCT (394C>T), CTG19CTC (397G>C), ACC20ACG (400C>G), ATC21ATA (403C>A), CAG22CAA (406G>A), CTG23TTC (407C>T 409G>C), ATC24ATA (412C>A), CAG25CAA (415G>A), AAC26TCG (416A>T 417A>C 418C>G), TTT28TTC (424T>C), GTG29GTC (427G>C), GAC30GAT (430C>T), GAT33GAC (439T>C), CCC34CCT (442C>T), ATA36ATC (448A>C), GAG37GAA (451G>A), TCC39TCA (457C>A), CGG41CGC (463G>C), CAG43CAA (469G>A), GTA44TGC (470G>T 471T>G 472A>C), GTC45GTG (475C>G), ATT46ATA (478T>A), GAT47GAC (481T>C), GGG48GAA (483G>A 484G>A), GAG49GAA (487G>A), ACG50GTT (488A>G 489C>T 490G>T), TGT51GCA (491T>G 492G>C 493T>A), ATC55GTC (503A>G), GAC57GAT (511C>T), ACA59GCC (515A>G 517A>C), GGT60GGC (520T>C), GAG63GAA (529G>A), TAT64TAC (532T>C), AGT65TCT (533A>T 534G>C), CGG68AGG (542C>A), GAC69GAG (547C>G), CAG70CAA (550G>A), CGC73CGG (559C>G) | | | | | | | | |
| **Proteins** | | | | | | | | | |
| hypothetical protein (YP_009507788.1) | 55 | 126 | 29.8% | 399 | 81.8% | 72 (100%) | 60 (83.3%) | 0/0/0/0 | 0 |
| Protein mutations: | A63G (372C>G), R64G (374A>G 376A>T), L75F (407C>T 409G>C), N78S (416A>T 417A>C 418C>G), V96C (470G>T 471T>G 472A>C), G100E (483G>A 484G>A), T102V (488A>G 489C>T 490G>T), C103A (491T>G 492G>C 493T>A), I107V (503A>G), T111A (515A>G 517A>C), D121E (547C>G) | | | | | | | | |
| Codon mutations: | ACA54..G (346A>G), AAG57AAA (355G>A), CTT58CTC (358T>C), GTG59GTC (361G>C), GTG60GTC (364G>C), GTG61GTC (367G>C), GCT63GGT (372C>G), AGA64GGT (374A>G 376A>T), GGC65GGT (379C>T), GTG66GTC (382G>C), AGT69TCC (389A>T 390G>C 391T>C), GCC70GCT (394C>T), CTG71CTC (397G>C), ACC72ACG (400C>G), ATC73ATA (403C>A), CAG74CAA (406G>A), CTG75TTC (407C>T 409G>C), ATC76ATA (412C>A), CAG77CAA (415G>A), AAC78TCG (416A>T 417A>C 418C>G), TTT80TTC (424T>C), GTG81GTC (427G>C), GAC82GAT (430C>T), GAT85GAC (439T>C), CCC86CCT (442C>T), ATA88ATC (448A>C), GAG89GAA (451G>A), TCC91TCA (457C>A), CGG93CGC (463G>C), CAG95CAA (469G>A), GTA96TGC (470G>T 471T>G 472A>C), GTC97GTG (475C>G), ATT98ATA (478T>A), GAT99GAC (481T>C), GGG100GAA (483G>A 484G>A), GAG101GAA (487G>A), ACG102GTT (488A>G 489C>T 490G>T), TGT103GCA (491T>G 492G>C 493T>A), ATC107GTC (503A>G), GAC109GAT (511C>T), ACA111GCC (515A>G 517A>C), GGT112GGC (520T>C), GAG115GAA (529G>A), TAT116TAC (532T>C), AGT117TCT (533A>T 534G>C), CGG120AGG (542C>A), GAC121GAG (547C>G), CAG122CAA (550G>A), CGC125CGG (559C>G) | | | | | | | | |
| hypothetical protein (YP_009507789.1) | 3 | 74 | 37.9% | 399 | 81.8% | 72 (100%) | 60 (83.3%) | 0/0/0/0 | 0 |
| Protein mutations: | A11G (372C>G), R12G (374A>G 376A>T), L23F (407C>T 409G>C), N26S (416A>T 417A>C 418C>G), V44C (470G>T 471T>G 472A>C), G48E (483G>A 484G>A), T50V (488A>G 489C>T 490G>T), C51A (491T>G 492G>C 493T>A), I55V (503A>G), T59A (515A>G 517A>C), D69E (547C>G) | | | | | | | | |
| Codon mutations: | ACA2..G (346A>G), AAG5AAA (355G>A), CTT6CTC (358T>C), GTG7GTC (361G>C), GTG8GTC (364G>C), GTG9GTC (367G>C), GCT11GGT (372C>G), AGA12GGT (374A>G 376A>T), GGC13GGT (379C>T), GTG14GTC (382G>C), AGT17TCC (389A>T 390G>C 391T>C), GCC18GCT (394C>T), CTG19CTC (397G>C), ACC20ACG (400C>G), ATC21ATA (403C>A), CAG22CAA (406G>A), CTG23TTC (407C>T 409G>C), ATC24ATA (412C>A), CAG25CAA (415G>A), AAC26TCG (416A>T 417A>C 418C>G), TTT28TTC (424T>C), GTG29GTC (427G>C), GAC30GAT (430C>T), GAT33GAC (439T>C), CCC34CCT (442C>T), ATA36ATC (448A>C), GAG37GAA (451G>A), TCC39TCA (457C>A), CGG41CGC (463G>C), CAG43CAA (469G>A), GTA44TGC (470G>T 471T>G 472A>C), GTC45GTG (475C>G), ATT46ATA (478T>A), GAT47GAC (481T>C), GGG48GAA (483G>A 484G>A), GAG49GAA (487G>A), ACG50GTT (488A>G 489C>T 490G>T), TGT51GCA (491T>G 492G>C 493T>A), ATC55GTC (503A>G), GAC57GAT (511C>T), ACA59GCC (515A>G 517A>C), GGT60GGC (520T>C), GAG63GAA (529G>A), TAT64TAC (532T>C), AGT65TCT (533A>T 534G>C), CGG68AGG (542C>A), GAC69GAG (547C>G), CAG70CAA (550G>A), CGC73CGG (559C>G) | | | | | | | | |

*: Inserts / Deletes / Misaligned / Frameshifts

#### Analysis details

This analysis was performed with panviral2.64

### NGS Details (UN9): Tomato necrotic dwarf virus (segment RNA1)

#### Assembly

| | |
|---|---|
| **Coverage Length** | 799 (2 contig(s)) |
| **Depth Of Coverage** | 9641.5 |
| **Number Of Reads** | 65373 |
| **Reads Per Million** | 1477.62 rpm (after QC) |
| **Ambiguities** | 0 |
| **Assembly Method** | de novo + reference guided assembly |
| **Consensus Caller** | Bcf Tools |

#### Coverage Map

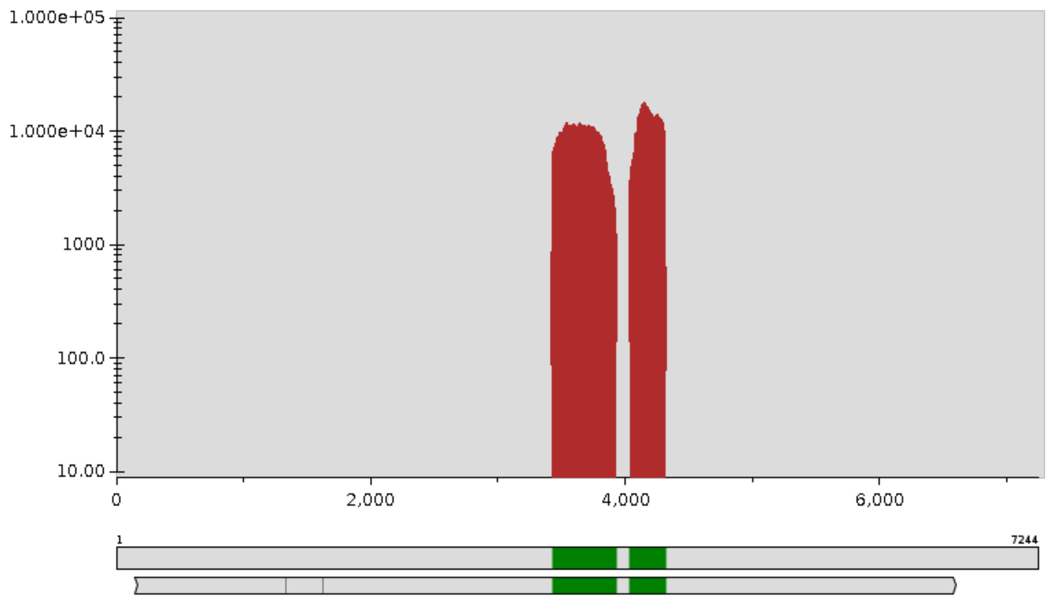

#### Assignment

| | |
|---|---|
| **Type** | Tomato necrotic dwarf virus (Taxonomy ID: 1481465) |
| **Reference Genome** | NC_027926.1 |
| **NT Identity (%)** | 71.5895 |
| **AA Identity (%)** | 77.5281 |
| **Number Of Stop Codons** | 0 |
| **Number Of CDS** | 1 |

#### Alignment

| | |
|---|---|
| **Alignment Score** | 690.0 (NT) + 1456.0 (AA) = 2146.0 |
| **Concordance (%)** | 62.9326 |

| Alignment Method | Global, seeded, nucleotide + amino acids (AGA) |
|---|---|

#### Genome Region

Sequence starts at position 3424 and ends at position 4321 relative to NC_027926.1 reference sequence.

#### Alignment Detailed Statistics

| | Begin | End | Coverage | Score | Concordance | Matches | Identities | I/D/M/F* | Stop Codons |
|---|---|---|---|---|---|---|---|---|---|
| **NT** | **3424** | **4321** | **11.0%** | **690** | **43.2%** | **799 (100%)** | **572 (71.6%)** | **0/0** | |
| Mutations: | 3433G>A, 3440C>T, 3444A>T, 3453A>C, 3459C>A, 3460A>C, 3462T>G, 3463C>T, 3465C>G, 3468A>G, 3469G>C, 3470C>A, 3472A>C, 3474T>G, 3477G>A, 3478A>T, 3479G>C, 3480T>C, 3482G>A, 3483A>G, 3486A>G, 3487A>G, 3488C>A, 3489C>A, 3491C>G, 3492T>A, 3493C>T, 3495T>C, 3501A>G, 3507C>A, 3510A>G, 3513C>G, 3514T>G, 3516C>T, 3519T>C, 3520C>T, 3522T>G, 3531T>A, 3534T>G, 3535C>T, 3536A>T, 3537A>G, 3541T>A, 3546C>T, 3549A>T, 3555A>G, 3558C>G, 3561T>C, 3567T>A, 3570A>C, 3573C>T, 3575G>A, 3577A>C, 3578A>G, 3579G>C, 3584A>C, 3585A>C, 3586T>C, 3597A>T, 3603T>G, 3604C>G, 3605C>A, 3606T>C, 3610G>C, 3612A>G, 3615C>T, 3618T>C, 3621C>A, 3624A>C, 3627C>T, 3630C>A, 3631A>T, 3633C>T, 3635T>C, 3639T>A, 3646C>A, 3654G>A, 3657A>G, 3660C>T, 3666A>C, 3675A>T, 3681G>C, 3682A>G, 3683T>A, 3687A>G, 3690C>T, 3699T>A, 3703C>T, 3704A>C, 3705T>C, 3706G>A, 3707G>A, 3708T>C, 3714T>G, 3717T>G, 3718G>C, 3719A>C, 3720T>A, 3726T>A, 3729A>G, 3730T>C, 3733T>G, 3735C>T, 3741A>G, 3742A>G, 3743C>G, 3747C>T, 3751A>G, 3752G>C, 3753T>A, 3756A>C, 3757C>A, 3759G>T, 3762G>T, 3765A>T, 3766A>G, 3767C>T, 3768A>G, 3769A>C, 3771G>T, 3774A>G, 3776A>T, 3777C>G, 3780A>G, 3781A>C, 3782A>G, 3795C>G, 3801A>T, 3802C>G, 3803A>C, 3804G>A, 3805G>A, 3806A>T, 3807G>C, 3811G>A, 3822G>A, 3828G>C, 3837T>A, 3845G>T, 3854A>T, 3860G>C, 3861C>A, 3868C>T, 3870C>G, 3876A>G, 3879C>A, 3888C>T, 3894C>T, 3901A>T, 3906C>A, 3907A>C, 3909G>T, 3912A>T, 3913A>T, 3914T>C, 3915G>T, 3921G>A, 3922C>A, 4047C>A, 4050A>C, 4053A>G, 4059C>G, 4062G>A, 4068G>T, 4071C>G, 4072C>T, 4074T>G, 4077T>A, 4080T>G, 4083A>C, 4089T>A, 4092T>C, 4104A>G, 4107A>C, 4108A>C, 4110A>C, 4111T>C, 4113A>G, 4116T>C, 4119G>A, 4122A>C, 4137C>T, 4149C>G, 4150A>C, 4161T>C, 4173C>T, 4176C>T, 4177A>T, 4178C>G, 4179C>T, 4188A>T, 4189T>G, 4190T>C, 4197T>C, 4200C>A, 4201A>C, 4206A>C, 4209T>C, 4215T>A, 4218A>T, 4221A>C, 4222G>A, 4230C>G, 4233T>C, 4234T>C, 4235C>A, 4243T>A, 4245T>A, 4246C>G, 4248T>G, 4251T>G, 4254C>T, 4256A>G, 4257C>T, 4259A>G, 4263C>T, 4267G>T, 4269T>A, 4279A>G, 4280C>A, 4281T>G, 4284G>A, 4287T>C, 4293T>C, 4302A>G, 4305T>C, 4311C>T | | | | | | | | |

##### CDS

| | Begin | End | Coverage | Score | Concordance | Matches | Identities | I/D/M/F* | Stop Codons |
|---|---|---|---|---|---|---|---|---|---|
| APL33_sRA11gp1 | 1092 | 1391 | 12.4% | 1456 | 79.4% | 267 (100%) | 207 (77.5%) | 0/0/0/0 | 0 |
| Protein mutations: | V1095I (3433G>A), A1097V (3440C>T), D1103E (3459C>A), I1104L (3460A>C 3462T>G), A1107H (3469G>C 3470C>A), I1108L (3472A>C 3474T>G), R1111K (3482G>A 3483A>G), T1113E (3487A>G 3488C>A 3489C>A), T1114R (3491C>G 3492T>A), L1115F (3493C>T 3495T>C), N1119K (3507C>A), S1122A (3514T>G 3516C>T), Q1129L (3535C>T 3536A>T 3537A>G), L1131M (3541T>A), R1142K (3575G>A), K1143R (3577A>C 3578A>G 3579G>C), Q1145P (3584A>C 3585A>C), P1152D (3604C>G 3605C>A 3606T>C), E1154Q (3610G>C 3612A>G), T1161S (3631A>T 3633C>T), V1162A (3635T>C), N1163K (3639T>A), M1178E (3682A>G 3683T>A), H1185S (3703C>T 3704A>C 3705T>C), G1186N (3706G>A 3707G>A 3708T>C), D1188E (3714T>G), D1189E (3717T>G), D1190P (3718G>C 3719A>C 3720T>A), S1195A (3733T>G 3735C>T), T1198G (3742A>G 3743C>G), S1201A (3751A>G 3752G>C 3753T>A), L1203I (3757C>A 3759G>T), T1206V (3766A>G 3767C>T 3768A>G), M1207L (3769A>C 3771G>T), N1209M (3776A>T 3777C>G), K1211R (3781A>C 3782A>G), Q1218A (3802C>G 3803A>C 3804G>A), E1219I (3805G>A 3806A>T 3807G>C), D1221N (3811G>A), C1232F (3845G>T), Y1235F (3854A>T), G1237A (3860G>C 3861C>A), M1251L (3901A>T), M1253L (3907A>C 3909G>T), M1255S (3913A>T 3914T>C 3915G>T), E1306D (4068G>T), E1311D (4083A>C), T1343C (4177A>T 4178C>G 4179C>T), L1347A (4189T>G 4190T>C), I1351L (4201A>C), D1358N (4222G>A), S1362H (4234T>C 4235C>A), S1365T (4243T>A 4245T>A), L1366V (4246C>G 4248T>G), N1369S (4256A>G 4257C>T), N1370S (4259A>G), A1373S (4267G>T 4269T>A), T1377E (4279A>G 4280C>A 4281T>G) | | | | | | | | |
| Codon mutations: | GTT1095ATT (3433G>A), GCG1097GTG (3440C>T), GCA1098GCT (3444A>T), CGA1101CGC (3453A>C), GAC1103GAA (3459C>A), ATT1104CTG (3460A>C 3462T>G), CTC1105TTG (3463C>T 3465C>G), AAA1106AAG (3468A>G), GCC1107CAC (3469G>C 3470C>A), ATT1108CTG (3472A>C 3474T>G), AGG1109AGA (3477G>A), AGT1110TCC (3478A>T 3479G>C 3480T>C), AGA1111AAG (3482G>A 3483A>G), GAA1112GAG (3486A>G), ACC1113GAA (3487A>G 3488C>A 3489C>A), ACT1114AGA (3491C>G 3492T>A), CTT1115TTC (3493C>T 3495T>C), GAA1117GAG (3501A>G), AAC1119AAA (3507C>A), CAA1120CAG (3510A>G), GTC1121GTG (3513C>G), TCC1122GCT (3514T>G 3516C>T), TTT1123TTC (3519T>C), CTT1124TTG (3520C>T 3522T>G), GTT1127GTA (3531T>A), CCT1128CCG (3534T>G), CAA1129TTG (3535C>T 3536A>T 3537A>G), TTG1131ATG (3541T>A), GCC1132GCT (3546C>T), GCA1133GCT (3549A>T), GTA1135GTG (3555A>G), CCC1136CCG (3558C>G), CAT1137CAC (3561T>C), ACT1139ACA (3567T>A), ACA1140ACC (3570A>C), CTC1141CTT (3573C>T), AGA1142AAA (3575G>A), AAG1143CGC (3577A>C 3578A>G 3579G>C), CAA1145CCC (3584A>C 3585A>C), TTG1146CTG (3586T>C), GCA1149GCT (3597A>T), GGT1151GGG (3603T>G), CCT1152GAC (3604C>G 3605C>A 3606T>C), GAA1154CAG (3610G>C 3612A>G), ACC1155ACT (3615C>T), GCT1156GCC (3618T>C), CCC1157CCA (3621C>A), TCA1158TCC (3624A>C), ATC1159ATT (3627C>T), CTC1160CTA (3630C>A), ACC1161TCT (3631A>T 3633C>T), GTC1162GCC (3635T>C), AAT1163AAA (3639T>A), CGG1166AGG (3646C>A), GAG1168GAA (3654G>A), GGA1169GGG (3657A>G), TTC1170TTT (3660C>T), CCA1172CCC (3666A>C), GCA1175GCT (3675A>T), GTG1177GTC (3681G>C), ATG1178GAG (3682A>G 3683T>A), AAA1179AAG (3687A>G), TAC1180TAT (3690C>T), ACT1183ACA (3699T>A), CAT1185TCC (3703C>T 3704A>C 3705T>C), GGT1186AAC (3706G>A 3707G>A 3708T>C), GAT1188GAG (3714T>G), GAT1189GAG (3717T>G), GAT1190CCA (3718G>C 3719A>C 3720T>A), GCT1192GCA (3726T>A), AAA1193AAG (3729A>G), TTG1194CTG (3730T>C), TCC1195GCT (3733T>G 3735C>T), GAA1197GAG (3741A>G), ACT1198GGT (3742A>G 3743C>G), CTC1199CTT (3747C>T), AGT1201GCA (3751A>G 3752G>C 3753T>A), TCA1202TCC (3756A>C), CTG1203ATT (3757C>A 3759G>T), CTG1204CTT (3762G>T), CCA1205CCT (3765A>T), ACA1206GTG (3766A>G 3767C>T 3768A>G), ATG1207CTT (3769A>C 3771G>T), AAA1208AAG (3774A>G), AAC1209ATG (3776A>T 3777C>G), CAA1210CAG (3780A>G), AAA1211CGA (3781A>C 3782A>G), GGC1215GGG (3795C>G), CCA1217CCT (3801A>T), CAG1218GCA (3802C>G 3803A>C 3804G>A), GAG1219ATC (3805G>A 3806A>T 3807G>C), GAT1221AAT (3811G>A), GTG1224GTA (3822G>A), CTG1226CTC (3828G>C), ATT1229ATA (3837T>A), TGT1232TTT (3845G>T), TAT1235TTT (3854A>T), GGC1237GCA (3860G>C 3861C>A), CTC1240TTG (3868C>T 3870C>G), ACA1242ACG (3876A>G), TCC1243TCA (3879C>A), TAC1246TAT (3888C>T), TTC1248TTT (3894C>T), ATG1251TTG (3901A>T), GGC1252GGA (3906C>A), ATG1253CTT (3907A>C 3909G>T), GGA1254GGT (3912A>T), ATG1255TCT (3913A>T 3914T>C 3915G>T), AAG1257AAA (3921G>A), CGG1258AGG (3922C>A), ACC1299ACA (4047C>A), ACA1300ACC (4050A>C), CAA1301CAG (4053A>G), GCC1303GCG (4059C>G), AAG1304AAA (4062G>A), GAG1306GAT (4068G>T), CGC1307CGG (4071C>G), CTT1308TTG (4072C>T 4074T>G), CCT1309CCA (4077T>A), CTT1310CTG (4080T>G), GAA1311GAC (4083A>C), ATT1313ATA (4089T>A), TAT1314TAC (4092T>C), AAA1318AAG (4104A>G), ACA1319ACC (4107A>C), AGA1320CGC (4108A>C 4110A>C), TTA1321CTG (4111T>C 4113A>G), TTT1322TTC (4116T>C), GAG1323GAA (4119G>A), ATA1324ATC (4122A>C), TAC1329TAT (4137C>T), GTC1333GTG (4149C>G), AGG1334CGG (4150A>C), TTT1337TTC (4161T>C), TCC1341TCT (4173C>T), GCC1342GCT (4176C>T), ACC1343TGT (4177A>T 4178C>G 4179C>T), GCA1346GCT (4188A>T), TTA1347GCA (4189T>G 4190T>C), AAT1349AAC (4197T>C), GTC1350GTA (4200C>A), ATC1351CTC (4201A>C), CCA1352CCC (4206A>C), TGT1353TGC (4209T>C), GTT1355GTA (4215T>A), GGA1356GGT (4218A>T), ATA1357ATC (4221A>C), GAT1358AAT (4222G>A), ACC1360ACG (4230C>G), AGT1361AGC (4233T>C), TCT1362CAT (4234T>C 4235C>A), TCT1365ACA (4243T>A 4245T>A), CTT1366GTG (4246C>G 4248T>G), CTT1367CTG (4251T>G), GCC1368GCT (4254C>T), AAC1369AGT (4256A>G 4257C>T), AAC1370AGC (4259A>G), TTC1371TTT (4263C>T), GCT1373TCA (4267G>T 4269T>A), ACT1377GAG (4279A>G 4280C>A 4281T>G), GGG1378GGA (4284G>A), TTT1379TTC (4287T>C), GCT1381GCC (4293T>C), TCA1384TCG (4302A>G), AGT1385AGC (4305T>C), GAC1387GAT (4311C>T) | | | | | | | | |

##### Proteins

| | Begin | End | Coverage | Score | Concordance | Matches | Identities | I/D/M/F* | Stop Codons |
|---|---|---|---|---|---|---|---|---|---|
| polyprotein (YP_009165993.1) | 1092 | 1391 | 12.4% | 1456 | 79.4% | 267 (100%) | 207 (77.5%) | 0/0/0/0 | 0 |
| Protein mutations: | V1095I (3433G>A), A1097V (3440C>T), D1103E (3459C>A), I1104L (3460A>C 3462T>G), A1107H (3469G>C 3470C>A), I1108L (3472A>C 3474T>G), R1111K (3482G>A 3483A>G), T1113E (3487A>G 3488C>A 3489C>A), T1114R (3491C>G 3492T>A), L1115F (3493C>T 3495T>C), N1119K (3507C>A), S1122A (3514T>G 3516C>T), Q1129L (3535C>T 3536A>T 3537A>G), L1131M (3541T>A), R1142K (3575G>A), K1143R (3577A>C 3578A>G 3579G>C), Q1145P (3584A>C 3585A>C), P1152D (3604C>G 3605C>A 3606T>C), E1154Q (3610G>C 3612A>G), T1161S (3631A>T 3633C>T), V1162A (3635T>C), N1163K (3639T>A), M1178E (3682A>G 3683T>A), H1185S (3703C>T 3704A>C 3705T>C), G1186N (3706G>A 3707G>A 3708T>C), D1188E (3714T>G), D1189E (3717T>G), D1190P (3718G>C 3719A>C 3720T>A), S1195A (3733T>G 3735C>T), T1198G (3742A>G 3743C>G), S1201A (3751A>G 3752G>C 3753T>A), L1203I (3757C>A 3759G>T), T1206V (3766A>G 3767C>T 3768A>G), M1207L (3769A>C 3771G>T), N1209M (3776A>T 3777C>G), K1211R (3781A>C 3782A>G), Q1218A (3802C>G 3803A>C 3804G>A), E1219I (3805G>A 3806A>T 3807G>C), D1221N (3811G>A), C1232F (3845G>T), Y1235F (3854A>T), G1237A (3860G>C 3861C>A), M1251L (3901A>T), M1253L (3907A>C 3909G>T), M1255S (3913A>T 3914T>C 3915G>T), E1306D (4068G>T), E1311D (4083A>C), T1343C (4177A>T 4178C>G 4179C>T), L1347A (4189T>G 4190T>C), I1351L (4201A>C), D1358N (4222G>A), S1362H (4234T>C 4235C>A), S1365T (4243T>A 4245T>A), L1366V (4246C>G 4248T>G), N1369S (4256A>G 4257C>T), N1370S (4259A>G), A1373S (4267G>T 4269T>A), T1377E (4279A>G 4280C>A 4281T>G) | | | | | | | | |

| | Begin | End | Coverage | Score | Concordance | Matches | Identities | I/D/M/F* | Stop Codons |
|---|---|---|---|---|---|---|---|---|---|
| **NT** | **3424** | **4321** | **11.0%** | **690** | **43.2%** | **799 (100%)** | **572 (71.6%)** | **0/0** | |
| Codon mutations: | GTT1095ATT (3433G>A), GCG1097GTG (3440C>T), GCA1098GCT (3444A>T), CGA1101CGC (3453A>C), GAC1103GAA (3459C>A), ATT1104CTG (3460A>C 3462T>G), CTC1105TTG (3463C>T 3465C>G), AAA1106AAG (3468A>G), GCC1107CAC (3469G>C 3470C>A), ATT1108CTG (3472A>C 3474T>G), AGG1109AGA (3477G>A), AGT1110TCC (3478A>T 3479G>C 3480T>C), AGA1111AAG (3482G>A 3483A>G), GAA1112GAG (3486A>G), ACC1113GAA (3487A>G 3488C>A 3489C>A), ACT1114AGA (3491C>G 3492T>A), CTT1115TTC (3493C>T 3495T>C), GAA1117GAG (3501A>G), AAC1119AAA (3507C>A), CAA1120CAG (3510A>G), GTC1121GTG (3513C>G), TCC1122GCT (3514T>G 3516C>T), TTT1123TTC (3519T>C), CTT1124TTG (3520C>T 3522T>G), GTT1127GTA (3531T>A), CCT1128CCG (3534T>G), CAA1129TTG (3535C>T 3536A>T 3537A>G), TTG1131ATG (3541T>A), GCC1132GCT (3546C>T), GCA1133GCT (3549A>T), GTA1135GTG (3555A>G), CCC1136CCG (3558C>G), CAT1137CAC (3561T>C), ACT1139ACA (3567T>A), ACA1140ACC (3570A>C), CTC1141CTT (3573C>T), AGA1142AAA (3575G>A), AAG1143CGC (3577A>C 3578A>G 3579G>C), CAA1145CCC (3584A>C 3585A>C), TTG1146CTG (3586T>C), GCA1149GCT (3597A>T), GGT1151GGG (3603T>G), CCT1152GAC (3604C>G 3605C>A 3606T>C), GAA1154CAG (3610G>C 3612A>G), ACC1155ACT (3615C>T), GCT1156GCC (3618T>C), CCC1157CCA (3621C>A), TCA1158TCC (3624A>C), ATC1159ATT (3627C>T), CTC1160CTA (3630C>A), ACC1161TCT (3631A>T 3633C>T), GTC1162GCC (3635T>C), AAT1163AAA (3639T>A), CGG1166AGG (3646C>A), GAG1168GAA (3654G>A), GGA1169GGG (3657A>G), TTC1170TTT (3660C>T), CCA1172CCC (3666A>C), GCA1175GCT (3675A>T), GTG1177GTC (3681G>C), ATG1178GAG (3682A>G 3683T>A), AAA1179AAG (3687A>G), TAC1180TAT (3690C>T), ACT1183ACA (3699T>A), CAT1185TCC (3703C>T 3704A>C 3705T>C), GGT1186AAC (3706G>A 3707G>A 3708T>C), GAT1188GAG (3714T>G), GAT1189GAG (3717T>G), GAT1190CCA (3718G>C 3719A>C 3720T>A), GCT1192GCA (3726T>A), AAA1193AAG (3729A>G), TTG1194CTG (3730T>C), TCC1195GCT (3733T>G 3735C>T), GAA1197GAG (3741A>G), ACT1198GGT (3742A>G 3743C>G), CTC1199CTT (3747C>T), AGT1201GCA (3751A>G 3752G>C 3753T>A), TCA1202TCC (3756A>C), CTG1203ATT (3757C>A 3759G>T), CTG1204CTT (3762G>T), CCA1205CCT (3765A>T), ACA1206GTG (3766A>G 3767C>T 3768A>G), ATG1207CTT (3769A>C 3771G>T), AAA1208AAG (3774A>G), AAC1209ATG (3776A>T 3777C>G), CAA1210CAG (3780A>G), AAA1211CGA (3781A>C 3782A>G), GGC1215GGG (3795C>G), CCA1217CCT (3801A>T), CAG1218GCA (3802C>G 3803A>C 3804G>A), GAG1219ATC (3805G>A 3806A>T 3807G>C), GAT1221AAT (3811G>A), GTG1224GTA (3822G>A), CTG1226CTC (3828G>C), ATT1229ATA (3837T>A), TGT1232TTT (3845G>T), TAT1235TTT (3854A>T), GGC1237GCA (3860G>C 3861C>A), CTC1240TTG (3868C>T 3870C>G), ACA1242ACG (3876A>G), TCC1243TCA (3879C>A), TAC1246TAT (3888C>T), TTC1248TTT (3894C>T), ATG1251TTG (3901A>T), GGC1252GGA (3906C>A), ATG1253CTT (3907A>C 3909G>T), GGA1254GGT (3912A>T), ATG1255TCT (3913A>T 3914T>C 3915G>T), AAG1257AAA (3921G>A), CGG1258AGG (3922C>A), ACC1299ACA (4047C>A), ACA1300ACC (4050A>C), CAA1301CAG (4053A>G), GCC1303GCG (4059C>G), AAG1304AAA (4062G>A), GAG1306GAT (4068G>T), CGC1307CGG (4071C>G), CTT1308TTG (4072C>T 4074T>G), CCT1309CCA (4077T>A), CTT1310CTG (4080T>G), GAA1311GAC (4083A>C), ATT1313ATA (4089T>A), TAT1314TAC (4092T>C), AAA1318AAG (4104A>G), ACA1319ACC (4107A>C), AGA1320CGC (4108A>C 4110A>C), TTA1321CTG (4111T>C 4113A>G), TTT1322TTC (4116T>C), GAG1323GAA (4119G>A), ATA1324ATC (4122A>C), TAC1329TAT (4137C>T), GTC1333GTG (4149C>G), AGG1334CGG (4150A>C), TTT1337TTC (4161T>C), TCC1341TCT (4173C>T), GCC1342GCT (4176C>T), ACC1343TGT (4177A>T 4178C>G 4179C>T), GCA1346GCT (4188A>T), TTA1347GCA (4189T>G 4190T>C), AAT1349AAC (4197T>C), GTC1350GTA (4200C>A), ATC1351CTC (4201A>C), CCA1352CCC (4206A>C), TGT1353TGC (4209T>C), GTT1355GTA (4215T>A), GGA1356GGT (4218A>T), ATA1357ATC (4221A>C), GAT1358AAT (4222G>A), ACC1360ACG (4230C>G), AGT1361AGC (4233T>C), TCT1362CAT (4234T>C 4235C>A), TCT1365ACA (4243T>A 4245T>A), CTT1366GTG (4246C>G 4248T>G), CTT1367CTG (4251T>G), GCC1368GCT (4254C>T), AAC1369AGT (4256A>G 4257C>T), AAC1370AGC (4259A>G), TTC1371TTT (4263C>T), GCT1373TCA (4267G>T 4269T>A), ACT1377GAG (4279A>G 4280C>A 4281T>G), GGG1378GGA (4284G>A), TTT1379TTC (4287T>C), GCT1381GCC (4293T>C), TCA1384TCG (4302A>G), AGT1385AGC (4305T>C), GAC1387GAT (4311C>T) | | | | | | | | |

*: Inserts / Deletes / Misaligned / Frameshifts

#### Analysis details

This analysis was performed with panviral2.64

### NGS Details (UN9): Brazilian marseillevirus

#### Assembly

| | |
|---|---|
| **Coverage Length** | 224 (1 contig(s)) |
| **Depth Of Coverage** | 9135.3 |
| **Number Of Reads** | 19918 |
| **Reads Per Million** | 450.20 rpm (after QC) |
| **Ambiguities** | 0 |
| **Assembly Method** | de novo + reference guided assembly |
| **Consensus Caller** | Bcf Tools |

#### Coverage Map

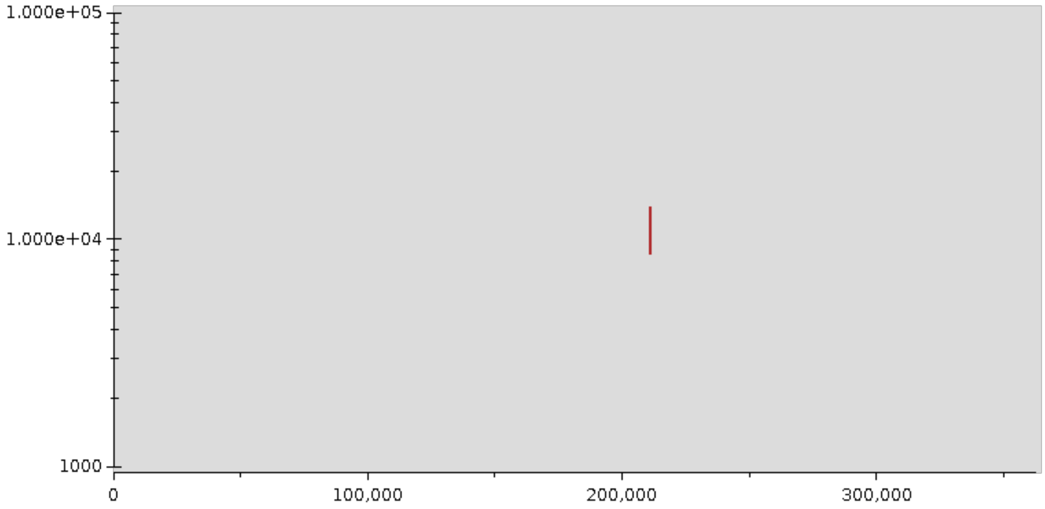

#### Assignment

| | |
|---|---|
| **Type** | Brazilian marseillevirus (Taxonomy ID: 1813599) |
| **Reference Genome** | NC_029692.1 |
| **NT Identity (%)** | 76.7857 |
| **AA Identity (%)** | 94.4444 |
| **Number Of Stop Codons** | 0 |
| **Number Of CDS** | 491 |

#### Alignment

| | |
|---|---|
| **Alignment Score** | 240.0 (NT) + 428.0 (AA) = 668.0 |
| **Concordance (%)** | 74.4705 |
| **Alignment Method** | Local, heuristic, nucleotide (BLASTN) |

#### Genome Region

Sequence starts at position 210884 and ends at position 211107 relative to NC_029692.1 reference sequence.

#### Alignment Detailed Statistics

| | Begin | End | Coverage | Score | Concordance | Matches | Identities | I/D/M/F* | Stop Codons |
|---|---|---|---|---|---|---|---|---|---|
| **NT** | **210884** | **211107** | **0.1%** | **240** | **53.6%** | **224 (100%)** | **172 (76.8%)** | **0/0** | |
| Mutations: | 210884A>G, 210888T>C, 210889T>C, 210890T>A, 210892A>C, 210896A>G, 210899T>A, 210901T>G, 210902T>G, 210914A>C, 210920G>T, 210926T>C, 210929T>C, 210935G>A, 210938A>G, 210941A>G, 210953C>G, 210955T>G, 210956T>G, 210965C>G, 210974G>A, 210983G>T, 210986A>C, 210989G>C, 210991G>T, 210992T>C, 210995T>C, 211001A>T, 211004A>T, 211007A>G, 211009C>T, 211010T>C, 211013T>C, 211025G>A, 211031T>A, 211032T>G, 211033G>C, 211037C>A, 211040G>A, 211043T>G, 211046G>A, 211058G>A, 211064A>C, 211070G>A, 211073A>G, 211076G>A, 211079C>A, 211085T>G, 211088C>A, 211094A>T, 211097C>T, 211100T>G | | | | | | | | |

*: Inserts / Deletes / Misaligned / Frameshifts

#### Analysis details

This analysis was performed with panviral2.64

### NGS Details (UN9): Tokyovirus A1

#### Assembly

| | |
|---|---|
| **Coverage Length** | 106 (1 contig(s)) |
| **Depth Of Coverage** | 6408.0 |
| **Number Of Reads** | 10543 |
| **Reads Per Million** | 238.30 rpm (after QC) |
| **Ambiguities** | 0 |
| **Assembly Method** | de novo + reference guided assembly |
| **Consensus Caller** | Bcf Tools |

#### Coverage Map

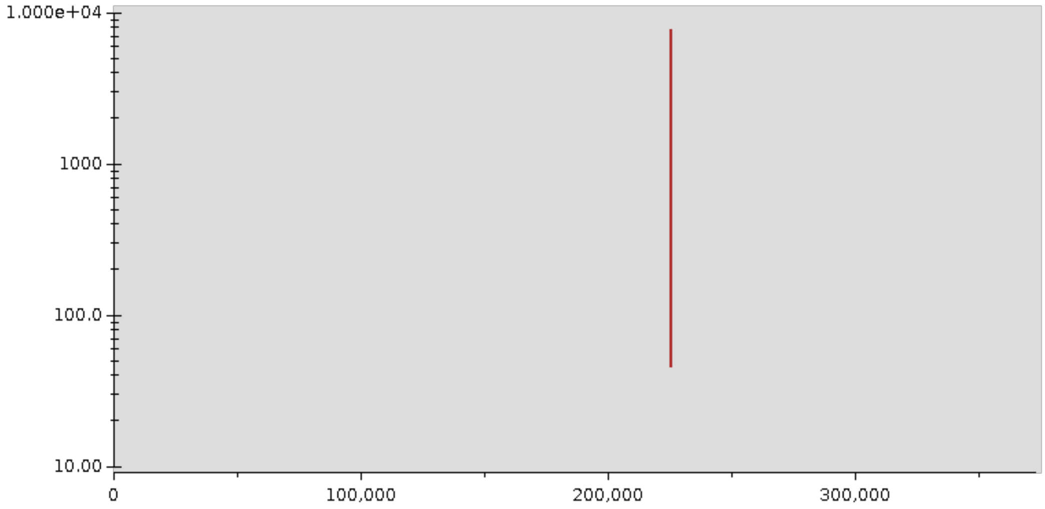

#### Assignment

| | |
|---|---|
| **Type** | Tokyovirus A1 (Taxonomy ID: 1826170) |
| **Reference Genome** | NC_030230.1 |
| **NT Identity (%)** | 81.1321 |
| **AA Identity (%)** | 94.4444 |
| **Number Of Stop Codons** | 0 |
| **Number Of CDS** | 470 |

#### Alignment

| | |
|---|---|
| **Alignment Score** | 132.0 (NT) + 232.0 (AA) = 364.0 |
| **Concordance (%)** | 81.7978 |
| **Alignment Method** | Local, heuristic, nucleotide (BLASTN) |

#### Genome Region

Sequence starts at position 225497 and ends at position 225602 relative to NC_030230.1 reference sequence.

#### Alignment Detailed Statistics

| | Begin | End | Coverage | Score | Concordance | Matches | Identities | I/D/M/F* | Stop Codons |
|---|---|---|---|---|---|---|---|---|---|
| **NT** | **225497** | **225602** | **0.1%** | **132** | **62.3%** | **106 (100%)** | **86 (81.1%)** | **0/0** | |

Mutations: 225513G>C, 225519G>T, 225522T>C, 225525T>C, 225528T>C, 225533C>T, 225540A>G, 225543G>C, 225546C>A, 225548A>G, 225549A>G, 225552T>G, 225554T>G, 225555C>G, 225561T>C, 225564T>A, 225573C>A, 225585A>C, 225587G>A, 225588T>C

*: Inserts / Deletes / Misaligned / Frameshifts

#### Analysis details

This analysis was performed with panviral2.64

### NGS Details (UN9): Lausannevirus

#### Assembly

| | |
|---|---|
| **Coverage Length** | 152 (1 contig(s)) |
| **Depth Of Coverage** | 2010.8 |
| **Number Of Reads** | 3601 |
| **Reads Per Million** | 81.39 rpm (after QC) |
| **Ambiguities** | 1 |
| **Assembly Method** | de novo + reference guided assembly |
| **Consensus Caller** | Bcf Tools |

#### Coverage Map

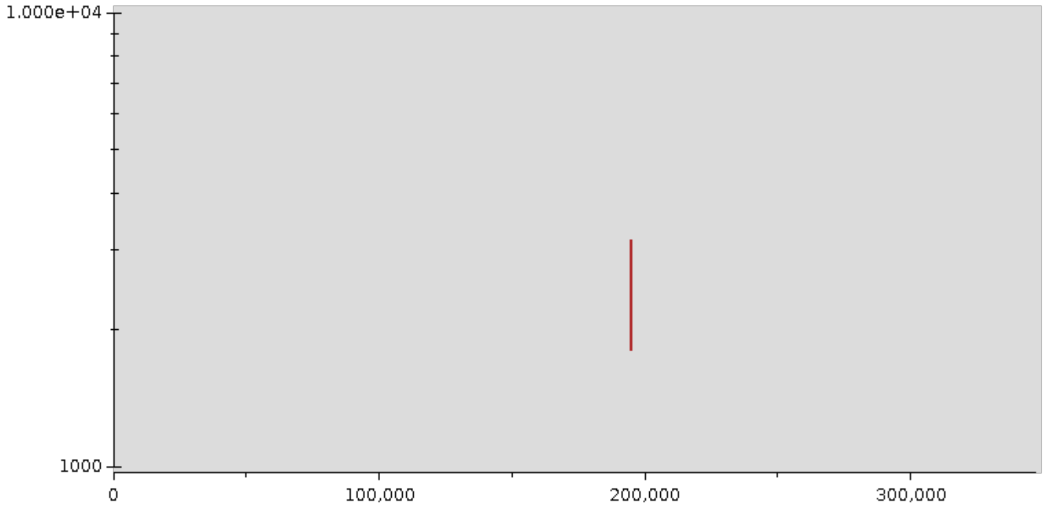

#### Assignment

| | |
|---|---|
| **Type** | Lausannevirus (Taxonomy ID: 999883) |
| **Reference Genome** | NC_015326.1 |
| **NT Identity (%)** | 83.5526 |
| **AA Identity (%)** | 97.9592 |
| **Number Of Stop Codons** | 1 |
| **Number Of CDS** | 444 |

#### Alignment

| | |
|---|---|
| **Alignment Score** | 206.0 (NT) + 294.0 (AA) = 500.0 |
| **Concordance (%)** | 80.0 |
| **Alignment Method** | Local, heuristic, nucleotide (BLASTN) |

#### Genome Region

Sequence starts at position 194911 and ends at position 195062 relative to NC_015326.1 reference sequence.

#### Alignment Detailed Statistics

| | Begin | End | Coverage | Score | Concordance | Matches | Identities | I/D/M/F* | Stop Codons |
|---|---|---|---|---|---|---|---|---|---|
| **NT** | **194911** | **195062** | **0.1%** | **206** | **67.8%** | **152 (100%)** | **127 (83.6%)** | **0/0** | |

Mutations: 194925T>C, 194931A>C, 194937G>A, 194943A>G, 194946G>C, 194949C>T, 194955T>C, 194967A>G, 194970G>A, 194979T>C, 194982C>T, 194990C>M, 194991G>C, 194993G>A, 195003T>A, 195006G>A, 195009G>A, 195012A>C, 195015T>C, 195017G>T, 195021T>C, 195030A>G, 195039T>C, 195042T>C, 195048T>C

*: Inserts / Deletes / Misaligned / Frameshifts

#### Analysis details

This analysis was performed with panviral2.64

### NGS Details (UN9): Betabaculovirus disaccharalis

#### Assembly

| | |
|---|---|
| **Coverage Length** | 150 (1 contig(s)) |
| **Depth Of Coverage** | 1356.2 |
| **Number Of Reads** | 2310 |
| **Reads Per Million** | 52.21 rpm (after QC) |
| **Ambiguities** | 0 |
| **Assembly Method** | de novo + reference guided assembly |
| **Consensus Caller** | Bcf Tools |

#### Coverage Map

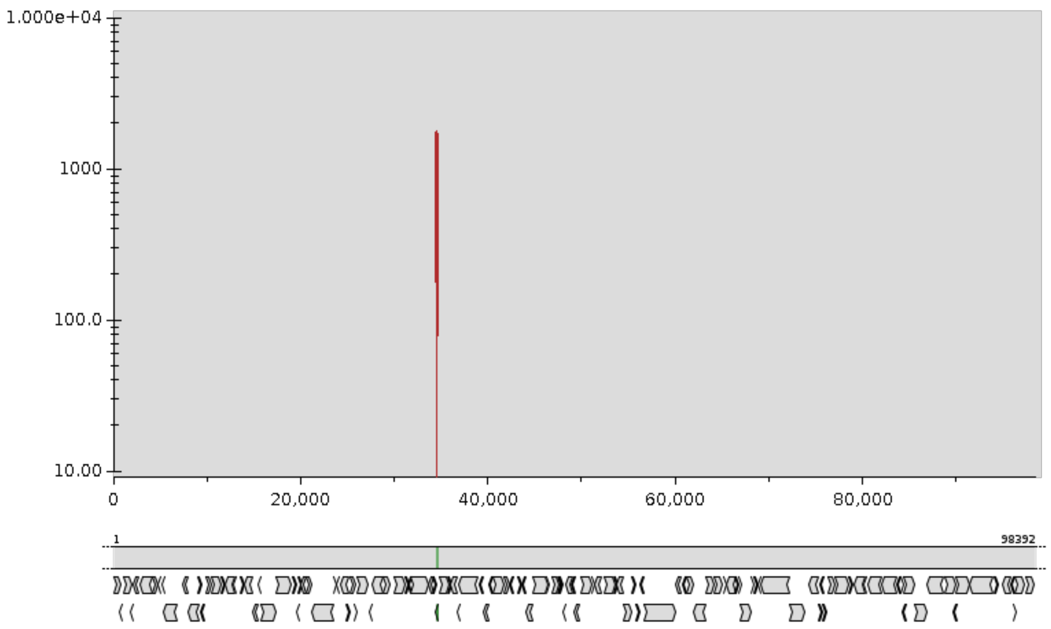

#### Assignment

| | |
|---|---|
| **Type** | Betabaculovirus disaccharalis (Taxonomy ID: 3047795) |
| **Reference Genome** | NC_028491.1 |
| **NT Identity (%)** | 84.6667 |
| **AA Identity (%)** | 92.0 |
| **Number Of Stop Codons** | 0 |
| **Number Of CDS** | 125 |

#### Alignment

| | |
|---|---|
| **Alignment Score** | 208.0 (NT) + 314.0 (AA) = 522.0 |
| **Concordance (%)** | 83.3866 |

| Alignment Method | Local, heuristic, nucleotide (BLASTN) |
|---|---|

#### Genome Region

Sequence starts at position 34474 and ends at position 34623 relative to NC_028491.1 reference sequence.

#### Alignment Detailed Statistics

| | Begin | End | Coverage | Score | Concordance | Matches | Identities | I/D/M/F* | Stop Codons |
|---|---|---|---|---|---|---|---|---|---|
| **NT** | **34474** | **34623** | **0.2%** | **208** | **69.3%** | **150 (100%)** | **127 (84.7%)** | **0/0** | |
| Mutations: | 34485T>C, 34488A>C, 34490G>A, 34513C>G, 34514T>A, 34521T>C, 34524A>G, 34533A>G, 34538A>C, 34542T>A, 34545T>C, 34547G>T, 34549T>C, 34554T>C, 34569G>A, 34575T>G, 34578T>C, 34587T>C, 34593C>T, 34596C>G, 34599C>A, 34601C>T, 34602T>A | | | | | | | | |
| **CDS** | | | | | | | | | |
| v-ubq | 52 | 101 | 42.0% | 314 | 91.3% | 50 (100%) | 46 (92.0%) | 0/0/0/0 | 0 |
| Protein mutations: | V59I (34599C>A 34601C>T), E76G (34549T>C), S80A (34538A>C) | | | | | | | | |
| Codon mutations: | GGA58GGT (34602T>A), GTG59ATT (34599C>A 34601C>T), CCG60CCC (34596C>G), CCG61CCA (34593C>T), CAA63CAG (34587T>C), TTA66TTG (34578T>C), ATA67ATC (34575T>G), GCC69GCT (34569G>A), GAA74GAG (34554T>C), GAG76GGG (34549T>C), CGA77AGG (34545T>C 34547G>T), ACA78ACT (34542T>A), TCA80GCA (34538A>C), GAT81GAC (34533A>G), ATT84ATC (34524A>G), CAA85CAG (34521T>C), AGC88TCC (34513C>G 34514T>A), CTT96TTG (34488A>C 34490G>A), AGA97AGG (34485T>C) | | | | | | | | |
| **Proteins** | | | | | | | | | |
| ubiquitin-like protein (YP_009182246.1) | 52 | 101 | 42.0% | 314 | 91.3% | 50 (100%) | 46 (92.0%) | 0/0/0/0 | 0 |
| Protein mutations: | V59I (34599C>A 34601C>T), E76G (34549T>C), S80A (34538A>C) | | | | | | | | |
| Codon mutations: | GGA58GGT (34602T>A), GTG59ATT (34599C>A 34601C>T), CCG60CCC (34596C>G), CCG61CCA (34593C>T), CAA63CAG (34587T>C), TTA66TTG (34578T>C), ATA67ATC (34575T>G), GCC69GCT (34569G>A), GAA74GAG (34554T>C), GAG76GGG (34549T>C), CGA77AGG (34545T>C 34547G>T), ACA78ACT (34542T>A), TCA80GCA (34538A>C), GAT81GAC (34533A>G), ATT84ATC (34524A>G), CAA85CAG (34521T>C), AGC88TCC (34513C>G 34514T>A), CTT96TTG (34488A>C 34490G>A), AGA97AGG (34485T>C) | | | | | | | | |

*: Inserts / Deletes / Misaligned / Frameshifts

#### Analysis details

This analysis was performed with panviral2.64

### NGS Details (UN9): Lausannevirus

#### Assembly

| | |
|---|---|
| **Coverage Length** | 208 (1 contig(s)) |
| **Depth Of Coverage** | 390.8 |
| **Number Of Reads** | 2249 |
| **Reads Per Million** | 50.83 rpm (after QC) |
| **Ambiguities** | 0 |
| **Assembly Method** | de novo + reference guided assembly |
| **Consensus Caller** | Bcf Tools |

#### Coverage Map

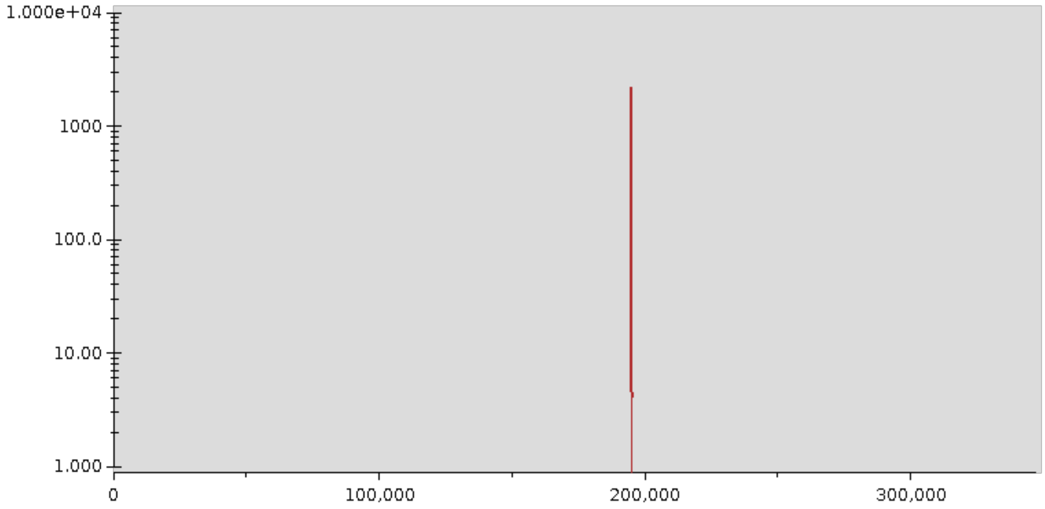

#### Assignment

| | |
|---|---|
| **Type** | Lausannevirus (Taxonomy ID: 999883) |
| **Reference Genome** | NC_015326.1 |
| **NT Identity (%)** | 78.3654 |
| **AA Identity (%)** | 91.0448 |
| **Number Of Stop Codons** | 1 |
| **Number Of CDS** | 444 |

#### Alignment

| | |
|---|---|
| **Alignment Score** | 236.0 (NT) + 399.0 (AA) = 635.0 |
| **Concordance (%)** | 75.8662 |
| **Alignment Method** | Local, heuristic, nucleotide (BLASTN) |

#### Genome Region

Sequence starts at position 194910 and ends at position 195117 relative to NC_015326.1 reference sequence.

#### Alignment Detailed Statistics

| | Begin | End | Coverage | Score | Concordance | Matches | Identities | I/D/M/F* | Stop Codons |
|---|---|---|---|---|---|---|---|---|---|
| **NT** | **194910** | **195117** | **0.1%** | **236** | **56.7%** | **208 (100%)** | **163 (78.4%)** | **0/0** | |

Mutations: 194922G>C, 194928C>G, 194930A>G, 194931A>G, 194934A>C, 194936G>A, 194943A>G, 194952T>C, 194955T>C, 194961A>G, 194967A>G, 194970G>A, 194972C>A, 194973A>G, 194976A>G, 194982C>G, 194985A>G, 194994C>T, 195000A>T, 195003T>G, 195017G>T, 195021T>C, 195024G>A, 195027T>G, 195030A>T, 195033A>G, 195036G>A, 195042T>C, 195054T>C, 195057T>G, 195058T>G, 195059G>C, 195075G>A, 195077T>A, 195081C>A, 195084A>G, 195085C>G, 195086T>C, 195090C>G, 195093T>G, 195102A>G, 195104C>T, 195105C>G, 195111T>G, 195114T>A

*: Inserts / Deletes / Misaligned / Frameshifts

#### Analysis details

This analysis was performed with panviral2.64

### NGS Details (UN9): Makelovirus prm1

#### Assembly

| | |
|---|---|
| **Coverage Length** | 1005 (1 contig(s)) |
| **Depth Of Coverage** | 253.9 |
| **Number Of Reads** | 1984 |
| **Reads Per Million** | 44.84 rpm (after QC) |
| **Ambiguities** | 0 |
| **Assembly Method** | de novo + reference guided assembly |
| **Consensus Caller** | Bcf Tools |

#### Coverage Map

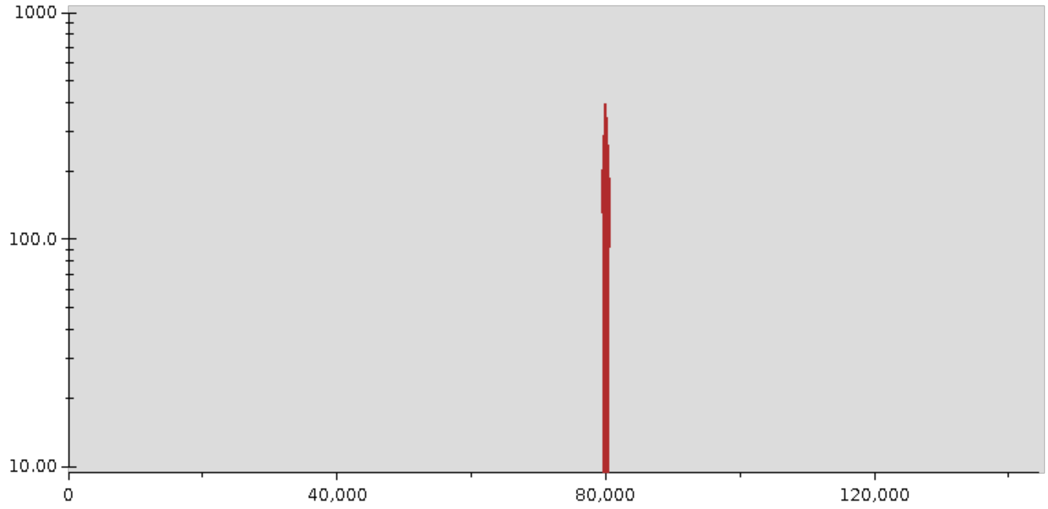

#### Assignment

| | |
|---|---|
| **Type** | Makelovirus prm1 (Taxonomy ID: 2956181) |
| **Reference Genome** | NC_055761.1 |
| **NT Identity (%)** | 76.6169 |
| **AA Identity (%)** | 87.4627 |
| **Number Of Stop Codons** | 0 |
| **Number Of CDS** | 190 |

#### Alignment

| | |
|---|---|
| **Alignment Score** | 1070.0 (NT) + 2242.0 (AA) = 3312.0 |
| **Concordance (%)** | 74.2601 |
| **Alignment Method** | Local, heuristic, nucleotide (BLASTN) |

#### Genome Region

Sequence starts at position 79512 and ends at position 80516 relative to NC_055761.1 reference sequence.

#### Alignment Detailed Statistics

| | Begin | End | Coverage | Score | Concordance | Matches | Identities | I/D/M/F* | Stop Codons |
|---|---|---|---|---|---|---|---|---|---|
| **NT** | **79512** | **80516** | **0.7%** | **1070** | **53.2%** | **1005 (100%)** | **770 (76.6%)** | **0/0** | |
| Mutations: | 79519A>G, 79520A>T, 79522A>G, 79523A>C, 79529C>T, 79530G>A, 79532G>C, 79536G>A, 79541G>T, 79542T>A, 79543C>G, 79544T>C, 79547A>T, 79548A>G, 79550C>A, 79563G>A, 79565C>T, 79568T>A, 79574C>T, 79580G>T, 79581C>T, 79592A>T, 79595T>C, 79596C>T, 79598G>A, 79602G>A, 79604A>G, 79610C>T, 79611G>T, 79612T>C, 79613C>T, 79614T>G, 79615G>T, 79616C>A, 79619C>T, 79624C>T, 79628G>C, 79632G>A, 79643C>T, 79646T>A, 79649G>A, 79655C>T, 79658C>T, 79664C>T, 79673A>T, 79676C>T, 79677G>T, 79679T>A, 79682T>G, 79688G>A, 79689A>C, 79691G>T, 79694T>C, 79703C>T, 79712T>C, 79718T>C, 79719G>A, 79722G>A, 79724A>T, 79728T>A, 79733C>T, 79734A>G, 79735A>C, 79736C>A, 79739A>T, 79742T>A, 79746C>T, 79748T>A, 79751C>T, 79754C>T, 79760C>A, 79763T>C, 79772T>G, 79779C>G, 79791C>T, 79793C>A, 79799T>C, 79811C>T, 79812C>G, 79817G>A, 79818G>A, 79820C>T, 79823A>T, 79824T>C, 79826T>A, 79833C>T, 79835G>A, 79836A>C, 79838C>T, 79842A>G, 79844C>A, 79845T>G, 79846T>C, 79847C>T, 79850C>T, 79871A>G, 79874C>T, 79875T>A, 79876C>G, 79879A>T, 79880T>C, 79895T>A, 79898C>T, 79904C>T, 79905T>G, 79906G>C, 79907C>T, 79916C>T, 79919T>A, 79922A>T, 79931T>A, 79937A>T, 79938T>A, 79940T>C, 79946C>T, 79950C>T, 79953G>A, 79958T>C, 79961T>A, 79962T>A, 79973T>A, 79974T>A, 79975C>G, 79976G>T, 79979C>T, 79985C>T, 79994C>T, 79997C>A, 80000C>A, 80018C>T, 80025C>A, 80030C>A, 80039A>T, 80042A>G, 80054G>T, 80063C>A, 80066C>T, 80069T>C, 80073C>T, 80075T>A, 80078T>C, 80090C>A, 80096T>C, 80099T>C, 80105G>A, 80114A>T, 80120C>T, 80126T>C, 80127C>T, 80132T>A, 80139T>A, 80140C>G, 80141G>T, 80142C>T, 80145G>A, 80147T>C, 80148C>A, 80150T>G, 80159C>A, 80162G>A, 80165C>T, 80168G>A, 80171C>T, 80172C>G, 80173A>C, 80174G>T, 80177C>T, 80178T>G, 80180T>A, 80188A>G, 80189G>A, 80201A>G, 80207G>A, 80210C>T, 80213C>T, 80216C>T, 80222G>A, 80225T>C, 80237C>T, 80240C>T, 80243T>C, 80246C>A, 80247C>T, 80261C>T, 80265T>A, 80266C>G, 80267C>T, 80279C>T, 80286C>T, 80288G>A, 80291T>C, 80300T>A, 80306A>T, 80315T>A, 80318C>A, 80333T>C, 80336A>T, 80337C>T, 80339T>A, 80343G>A, 80346T>A, 80347C>G, 80351C>T, 80357A>T, 80366G>A, 80369C>T, 80378C>T, 80387G>A, 80390C>T, 80391A>G, 80393C>A, 80394A>G, 80396G>T, 80399T>C, 80400G>A, 80409A>C, 80410A>G, 80411A>T, 80414C>A, 80415C>A, 80417C>T, 80433G>A, 80435T>C, 80436C>A, 80438A>C, 80447C>T, 80453G>T, 80456G>T, 80462G>A, 80465A>T, 80471C>T, 80474G>A, 80486C>T, 80498G>A, 80504G>A | | | | | | | | |

*: Inserts / Deletes / Misaligned / Frameshifts

#### Analysis details

This analysis was performed with panviral2.64

### NGS Details (UN9): Golden Marseillevirus

#### Assembly

| | |
|---|---|
| **Coverage Length** | 120 (1 contig(s)) |
| **Depth Of Coverage** | 1085.8 |
| **Number Of Reads** | 1782 |
| **Reads Per Million** | 40.28 rpm (after QC) |
| **Ambiguities** | 0 |
| **Assembly Method** | de novo + reference guided assembly |
| **Consensus Caller** | Bcf Tools |

#### Coverage Map

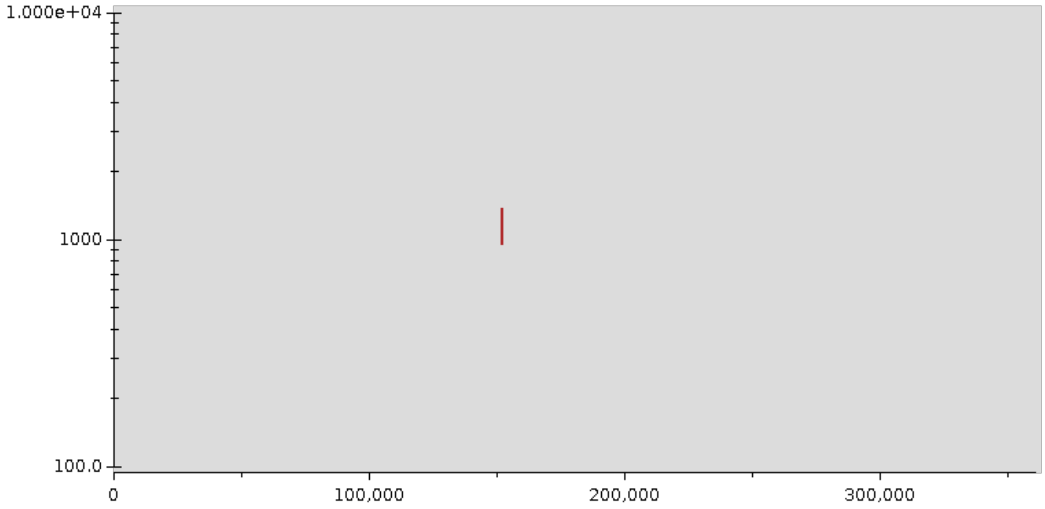

#### Assignment

| | |
|---|---|
| **Type** | Golden Marseillevirus (Taxonomy ID: 1720526) |
| **Reference Genome** | NC_031465.1 |
| **NT Identity (%)** | 85.8333 |
| **AA Identity (%)** | 97.4359 |
| **Number Of Stop Codons** | 1 |
| **Number Of CDS** | 296 |

#### Alignment

| | |
|---|---|
| **Alignment Score** | 172.0 (NT) + 235.0 (AA) = 407.0 |
| **Concordance (%)** | 84.7917 |
| **Alignment Method** | Local, heuristic, nucleotide (BLASTN) |

#### Genome Region

Sequence starts at position 151878 and ends at position 151997 relative to NC_031465.1 reference sequence.

#### Alignment Detailed Statistics

| | Begin | End | Coverage | Score | Concordance | Matches | Identities | I/D/M/F* | Stop Codons |
|---|---|---|---|---|---|---|---|---|---|
| NT | 151878 | 151997 | 0.1% | 172 | 71.7% | 120 (100%) | 103 (85.8%) | 0/0 | |

Mutations: 151889T>G, 151891T>G, 151907A>G, 151910A>G, 151916T>C, 151931A>G, 151939A>G, 151940A>T, 151945T>G, 151946C>G, 151955T>A, 151958G>C, 151959C>T, 151960T>G, 151964C>T, 151970G>A, 151976A>C

*: Inserts / Deletes / Misaligned / Frameshifts

#### Analysis details

This analysis was performed with panviral2.64

### NGS Details (UN9): Cladosporium fulvum T-1 virus

#### Assembly

| | |
|---|---|
| **Coverage Length** | 1715 (5 contig(s)) |
| **Depth Of Coverage** | 54.0 |
| **Number Of Reads** | 808 |
| **Reads Per Million** | 18.26 rpm (after QC) |
| **Ambiguities** | 0 |
| **Assembly Method** | de novo + reference guided assembly |
| **Consensus Caller** | Bcf Tools |

#### Coverage Map

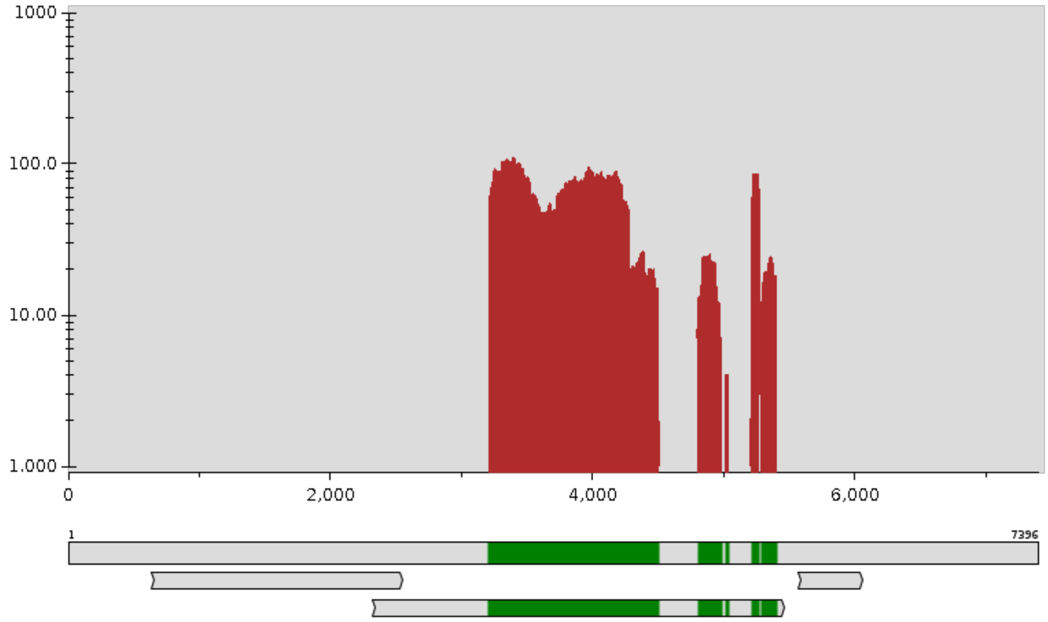

#### Assignment

| | |
|---|---|
| **Type** | Cladosporium fulvum T-1 virus (Taxonomy ID: 2052899) |
| **Reference Genome** | NC_043491.1 |
| **NT Identity (%)** | 52.5117 |
| **AA Identity (%)** | 47.028 |
| **Number Of Stop Codons** | 1 |
| **Number Of CDS** | 3 |

#### Alignment

| | |
|---|---|
| **Alignment Score** | 115.0 (NT) + 1929.0 (AA) = 2044.0 |
| **Concordance (%)** | 28.0769 |

| | |
|---|---|
| **Alignment Method** | Global, seeded, nucleotide + amino acids (AGA) |

#### Genome Region

Sequence starts at position 3205 and ends at position 5406 relative to NC_043491.1 reference sequence.

#### Alignment Detailed Statistics

| | Begin | End | Coverage | Score | Concordance | Matches | Identities | I/D/M/F* | Stop Codons |
|---|---|---|---|---|---|---|---|---|---|
| **NT** | **3205** | **5406** | **23.2%** | **115** | **3.4%** | **1697 (98.1%)** | **899 (52.0%)** | **15/18** | |
| Mutations: | 3214G>T, 3216A>T, 3217A>G, 3219G>T, 3220A>G, 3221G>C, 3222G>C, 3223C>A, 3224A>C, 3225A>C, 3226G>T, 3229C>G, 3230G>T, 3231C>T, 3232C>G, 3233T>C, 3234T>C, 3236C>A, 3237C>A, 3238T>G, 3243A>T, 3244C>G, 3246A>C, 3251T>A, 3253_3254insAGGGATATT, 3261A>G, 3262G>A, 3265A>T, 3266A>G, 3268C>G, 3269A>T, 3271T>G, 3272C>A, 3273A>T, 3280G>T, 3281A>T, 3282A>C, 3283A>T, 3284G>A, 3285A>C, 3289T>G, 3292A>T, 3293T>G, 3294G>C, 3295G>A, 3296G>C, 3297G>A, 3299C>G, 3301C>T, 3303T>C, 3304A>T, 3307T>C, 3308C>A, 3309A>G, 3314T>G, 3317G>C, 3318A>C, 3319G>T, 3326C>T, 3328A>G, 3329C>G, 3330A>C, 3331G>T, 3332A>G, 3333C>A, 3334C>A, 3337A>G, 3338C>A, 3340A>G, 3341G>A, 3343A>G, 3344T>C, 3345G>A, 3352G>C, 3355G>A, 3356A>T, 3357A>T, 3361A>T, 3363C>A, 3364C>T, 3365A>T, 3366A>C, 3371T>C, 3372G>T, 3376A>T, 3378G>A, 3379A>G, 3381G>C, 3382A>T, 3385C>T, 3387C>A, 3388C>G, 3389T>G, 3391A>T, 3392A>C, 3393G>C, 3394T>C, 3395G>T, 3396C>A, 3397A>T, 3400A>T, 3401A>G, 3403T>A, 3406A>T, 3407T>G, 3408G>T, 3409C>T, 3410A>T, 3412G>A, 3416G>C, 3417T>A, 3418T>G, 3419C>A, 3420C>A, 3425G>C, 3426C>A, 3427A>G, 3428A>G, 3430C>T, 3434A>T, 3435A>C, 3437C>T, 3440C>A, 3442A>G, 3443C>A, 3445C>G, 3446G>T, 3447T>G, 3448A>C, 3449C>G, 3450A>T, 3451A>G, 3454C>T, 3457C>T, 3460A>G, 3461A>G, 3462A>C, 3464T>C, 3470G>A, 3473A>G, 3475C>A, 3478G>T, 3479A>G, 3481C>G, 3488C>A, 3489G>A, 3490A>G, 3493T>C, 3496G>A, 3497C>G, 3499A>T, 3502C>T, 3503A>T, 3504A>T, 3505C>G, 3506A>G, 3508C>G, 3509G>C, 3511A>G, 3514A>T, 3515G>C, 3516C>T, 3517A>G, 3518C>A, 3519A>T, 3520A>G, 3526A>G, 3529A>G, 3531C>G, 3533G>A, 3534G>A, 3536T>G, 3538A>G, 3539G>T, 3540A>G, 3546A>T, 3550G>T, 3554A>C, 3556T>A, 3559C>T, 3562A>T, 3563C>A, 3565A>G, 3566G>T, 3567A>C, 3568C>A, 3570C>G, 3573T>A, 3574C>T, 3576A>G, 3577T>G, 3578G>C, 3579C>A, 3580T>G, 3581A>G, 3583C>T, 3584C>A, 3586A>G, 3589G>A, 3598A>G, 3601A>T, 3604A>G, 3605T>C, 3606G>C, 3607G>T, 3610A>G, 3613C>T, 3614G>A, 3618T>G, 3619C>T, 3620A>G, 3621G>T, 3622G>A, 3625A>T, 3628A>G, 3631C>T, 3634A>G, 3635C>T, 3636T>C, 3637C>A, 3640C>T, 3647T>C, 3649G>T, 3652C>A, 3658A>G, 3659A>T, 3661G>T, 3664A>G, 3670C>T, 3673C>T, 3676A>C, 3679C>G, 3682A>T, 3683T>A, 3687G>T, 3688C>T, 3689C>T, 3690A>G, 3691G>C, 3692G>A, 3694C>T, 3695C>T, 3697T>A, 3698G>A, 3700C>G, 3703C>T, 3706A>C, 3707A>G, 3708C>T, 3712T>G, 3713A>T, 3714G>T, 3715A>T, 3719C>T, 3720T>A, 3721A>T, 3724C>T, 3727C>T, 3729T>A, 3730G>T, 3732G>T, 3733C>T, 3736C>A, 3739T>A, 3741C>T, 3742T>C, 3745C>T, 3746A>C, 3748G>A, 3751C>T, 3757A>T, 3758C>G, 3760G>C, 3761G>A, 3763C>A, 3766C>T, 3768C>G, 3769A>T, 3770A>C, 3773_3775delGGA, 3776T>A, 3778C>G, 3779C>T, 3781C>G, 3782C>G, 3784G>T, 3791A>G, 3792C>T, 3793C>T, 3795A>G, 3796G>C, 3799A>T, 3800G>T, 3802T>G, 3803C>A, 3804A>G, 3805A>T, 3806G>T, 3807A>T, 3808T>G, 3811G>A, 3814C>A, 3815G>T, 3816A>C, 3817A>T, 3820A>T, 3821C>T, 3823C>A, 3825C>G, 3827A>C, 3829G>A, 3831C>A, 3833G>A, 3834G>C, 3835A>T, 3836T>C, 3839A>T, 3841G>T, 3842A>G, 3843C>T, 3844A>T, 3845G>A, 3846C>A, 3848C>A, 3849C>T, 3850C>G, 3856A>G, 3859C>T, 3862A>G, 3865C>T, 3866C>G, 3867A>C, 3869A>C, 3871G>A, 3877A>G, 3878G>A, 3880C>A, 3886T>C, 3887T>C, 3892C>G, 3893T>C, 3894T>A, 3896A>C, 3898C>A, 3899A>G, 3901C>T, 3905A>G, 3906C>A, 3909C>A, 3910A>C, 3911G>C, 3912G>A, 3913G>A, 3914A>G, 3918C>G, 3922C>G, 3925C>T, 3929G>A, 3930C>A, 3931A>G, 3934G>A, 3935A>G, 3936C>T, 3937A>G, 3940G>A, 3941T>G, 3943A>C, 3946C>T, 3947A>G, 3948G>T, 3949A>T, 3952A>T, 3957C>A, 3960A>C, 3964G>C, 3967G>A, 3970A>T, 3973C>G, 3980G>C, 3982A>G, 3983C>A, 3984A>G, 3988A>T, 3992C>T, 3994T>G, 3997A>G, 4000C>A, 4003C>T, 4010A>T, 4012C>T, 4025A>G, 4026A>C, 4027G>T, 4029A>G, 4030C>A, 4033T>C, 4041C>A, 4048A>T, 4049C>G, 4052A>C, 4057G>T, 4058A>G, 4059T>A, 4060G>T, 4061C>T, 4063T>G, 4064A>C, 4065C>T, 4066A>G, 4068G>A, 4069A>G, 4072A>G, 4075C>A, 4081C>G, 4085A>G, 4086A>T, 4091G>T, 4092G>C, 4093A>G, 4094A>G, 4095A>T, 4097G>A, 4098A>G, 4100C>T, 4101A>G, 4102G>C, 4103A>G, 4104C>A, 4105C>A, 4111G>T, 4114C>T, 4115A>C, 4117A>G, 4120A>T, 4121C>T, 4123C>G, 4130C>G, 4131A>C, 4132G>T, 4133T>A, 4134G>T, 4135C>T, 4138T>A, 4143C>A, 4144C>A, 4149C>T, 4150G>A, 4153T>G, 4154C>A, 4155G>A, 4156A>G, 4159A>G, 4159_4160insCCAAAT, 4162C>T, 4165T>G, 4166G>C, 4167G>T, 4168T>A, 4169A>C, 4170G>C, 4171C>T, 4172A>T, 4173A>T, 4174G>T, 4177A>G, 4180C>A, 4184_4189delATCGAG, 4192C>T, 4195C>T, 4201T>G, 4204T>C, 4206T>A, 4207G>A, 4210A>T, 4211A>G, 4213A>G, 4216C>A, 4218C>G, 4219A>C, 4220T>G, 4221G>T, 4222T>A, 4223C>T, 4225A>G, 4226A>G, 4227C>T, 4228A>G, 4231G>A, 4233C>A, 4234A>T, 4235C>G, 4240T>A, 4243G>T, 4244_4249delAAAAGA, 4252C>T, 4255A>G, 4263A>T, 4265T>G, 4267T>G, 4268T>A, 4269C>G, 4271C>A, 4277A>T, 4281C>A, 4282C>T, 4283A>C, 4285A>G, 4286G>A, 4294G>A, 4296A>G, 4297C>A, 4300C>T, 4301G>T, 4302A>C, 4303C>T, 4304A>G, 4305T>C, 4306C>A, 4312C>A, 4319C>A, 4321T>G, 4322C>G, 4324A>G, 4327C>A, 4328A>G, 4330T>G, 4331G>C, 4333T>C, 4334G>C, 4335C>A, 4336C>T, 4337G>T, 4338C>G, 4339C>T, 4340A>C, 4342G>T, 4346C>A, 4347A>C, 4348T>A, 4354A>G, 4360C>T, 4361G>C, 4364G>T, 4365A>T, 4366G>A, 4369C>A, 4370C>A, 4373_4375delCCG, 4381A>T, 4382A>G, 4383C>T, 4384G>A, 4385A>G, 4387T>A, 4388C>A, 4389T>A, 4390T>G, 4391T>A, 4393A>G, 4397C>A, 4399C>T, 4400A>G, 4401A>T, 4403A>G, 4404A>C, 4406C>A, 4407T>A, 4408C>T, 4411G>T, 4419C>A, 4420G>A, 4423G>T, 4424A>C, 4425C>A, 4426G>A, 4427A>C, 4430G>A, 4433C>T, 4435T>G, 4437C>G, 4438C>T, 4439C>T, 4440G>T, 4441A>G, 4443G>A, 4444A>G, 4445C>T, 4447A>G, 4450C>G, 4451C>A, 4453C>A, 4457T>C, 4458C>A, 4459G>A, 4463C>A, 4465G>A, 4468T>A, 4470G>C, 4471G>A, 4472C>G, 4474G>A, 4477C>T, 4478A>G, 4480G>C, 4483C>T, 4485A>T, 4486A>G, 4487A>T, 4488T>G, 4489C>G, 4491A>T, 4492A>T, 4493T>C, 4495C>T, 4497C>A, 4498T>A, 4797G>T, 4800T>A, 4803C>A, 4804A>T, 4806T>G, 4807C>G, 4812A>G, 4813G>C, 4817A>C, 4819A>G, 4820C>G, 4821T>A, 4822T>G, 4823A>G, 4825G>T, 4826G>A, 4828G>A, 4829C>G, 4830T>A, 4838A>G, 4840C>G, 4841A>C, 4843G>A, 4845A>C, 4846A>T, 4849C>T, 4850G>C, 4852A>T, 4853C>G, 4854A>T, 4855C>G, 4865A>G, 4870A>G, 4872C>T, 4873T>A, 4875C>A, 4876A>G, 4878G>A, 4881A>G, 4882C>A, 4883G>C, 4884C>A, 4885G>A, 4886A>C, 4887A>C, 4888A>C, 4889T>G, 4890A>G, 4891C>A, 4894T>A, 4896A>T, 4897C>T, 4898C>T, 4901C>G, 4904T>C, 4905T>C, 4906C>A, 4907A>T, 4908G>T, 4909G>A, 4910A>C, 4911C>A, 4912A>T, 4913C>G, 4914C>T, 4916C>G, 4918A>C, 4919A>G, 4920C>A, 4922A>C, 4923A>G, 4924A>T, 4933C>G, 4934G>A, 4935A>G, 4936G>T, 4937G>A, 4939T>C, 4954T>A, 4957G>C, 4958A>T, 4959A>G, 4960A>T, 4961C>T, 4963C>G, 4966G>A, 4968G>A, 4972A>C, 4973A>G, 4975G>T, 5009C>T, 5010T>G, 5011A>G, 5014C>G, 5017G>T, 5020C>G, 5026A>G, 5027C>T, 5029C>G, 5032A>T, 5035A>G, 5037A>C, 5207C>T, 5208A>T, 5209C>G, 5212G>T, 5216T>A, 5217C>G, 5218A>T, 5226A>T, 5227C>T, 5233A>T, 5234G>C, 5239G>C, 5245G>C, 5248A>G, 5249A>T, 5251G>A, 5254A>G, 5257A>G, 5260G>T, 5262A>T, 5263C>T, 5269A>T, 5270C>T, 5278A>C, 5284A>T, 5285C>A, 5292A>T, 5294A>G, 5296C>G, 5298A>G, 5301A>C, 5302C>T, 5303G>C, 5304C>A, 5305A>T, 5309G>A, 5311C>A, 5312A>G, 5314C>T, 5319T>C, 5321T>A, 5322C>A, 5323A>G, 5326A>G, 5329G>A, 5330C>G, 5331C>A, 5332A>T, 5333A>G, 5338G>C, 5341G>A, 5342A>T, 5345G>T, 5348C>T, 5349T>A, 5350G>T, 5353C>T, 5354A>C, 5355A>T, 5356C>A, 5359C>G, 5360A>C, 5361A>G, 5362A>G, 5363T>A, 5364C>G, 5365A>T, 5368G>A, 5369A>G, 5371A>C, 5375T>G, 5376C>G, 5377G>C, 5379C>A, 5390A>G, 5391T>A, 5394G>T, 5396A>G, 5398T>C, 5399T>G, 5400T>C, 5401G>A, 5403C>G, 5404A>C, 5405A>C, 5406G>A | | | | | | | | |

**CDS**

| | | | | | | | | | |
|---|---|---|---|---|---|---|---|---|---|
| homologue_of_retroviral_POL_genes | 294 | 1027 | 54.8% | 1929 | 48.4% | 567 (98.1%) | 269 (46.5%) | 5/6/0/0 | 1 |

| | Begin | End | Coverage | Score | Concordance | Matches | Identities | I/D/M/F* | Stop Codons |
|---|---|---|---|---|---|---|---|---|---|
| **NT** | **3205** | **5406** | **23.2%** | **115** | **3.4%** | **1697 (98.1%)** | **899 (52.0%)** | **15/18** | |
| Protein mutations: | E296D (3214G>T), E297V (3216A>T 3217A>G), R298M (3219G>T 3220A>G), G299P (3221G>C 3222G>C 3223C>A), K300P (3224A>C 3225A>C 3226G>T), D301E (3229C>G), A302L (3230G>T 3231C>T 3232C>G), L303P (3233T>C 3234T>C), P304K (3236C>A 3237C>A 3238T>G), H306L (3243A>T 3244C>G), Q307P (3246A>C), W309R (3251T>A), W309_D310insRDI (3253_3254insAGGGATATT), K312R (3261A>G 3262G>A), N314E (3266A>G 3268C>G), I315L (3269A>T 3271T>G), Q316M (3272C>A 3273A>T), K319S (3281A>T 3282A>C 3283A>T), E320T (3284G>A 3285A>C), W323A (3293T>G 3294G>C 3295G>A), G324Q (3296G>C 3297G>A), P325A (3299C>G 3301C>T), L326P (3303T>C 3304A>T), Q328R (3308C>A 3309A>G), S330A (3314T>G), E331P (3317G>C 3318A>C 3319G>T), Q335A (3329C>G 3330A>C 3331G>T), T336E (3332A>G 3333C>A 3334C>A), E339K (3341G>A 3343A>G), W340Q (3344T>C 3345G>A), K342N (3352G>C), K344L (3356A>T 3357A>T), A346D (3363C>A 3364C>T), K347S (3365A>T 3366A>C), W349L (3371T>C 3372G>T), R351Q (3378G>A 3379A>G), R352P (3381G>C 3382A>T), T354K (3387C>A 3388C>G), S355A (3389T>G 3391A>T), S356P (3392A>C 3393G>C 3394T>C), A357Y (3395G>T 3396C>A 3397A>T), T359A (3401A>G 3403T>A), C361V (3407T>G 3408G>T 3409C>T), M362L (3410A>T 3412G>A), V364Q (3416G>C 3417T>A 3418T>G), P365K (3419C>A 3420C>A), A367Q (3425G>C 3426C>A 3427A>G), N368D (3428A>G 3430C>T), K370S (3434A>T 3435A>C), L373M (3443C>A 3445C>G), V374C (3446G>T 3447T>G 3448A>C), Q375V (3449C>G 3450A>T 3451A>G), K379A (3461A>G 3462A>C), E382K (3470G>A), I383V (3473A>G 3475C>A), I385V (3479A>G 3481C>G), R388K (3488C>A 3489G>A 3490A>G), L391V (3497C>G 3499A>T), N393L (3503A>T 3504A>T 3505C>G), I394V (3506A>G 3508C>G), E395Q (3509G>C 3511A>G), E396D (3514A>T), A397L (3515G>C 3516C>T 3517A>G), Q398M (3518C>A 3519A>T 3520A>G), T402S (3531C>G), G403K (3533G>A 3534G>A), S404A (3536T>G 3538A>G), D405C (3539G>T 3540A>G), Y407F (3546A>T), I410L (3554A>C 3556T>A), D414S (3566G>T 3567A>C 3568C>A), A415G (3570C>G), F416Y (3573T>A 3574C>T), Y417W (3576A>G 3577T>G), A418Q (3578G>C 3579C>A 3580T>G), I419V (3581A>G 3583C>T), M421I (3589G>A), E425D (3601A>T), W427P (3605T>C 3606G>C 3607G>T), A430T (3614G>A), F431C (3618T>G 3619C>T), R432V (3620A>G 3621G>T 3622G>A), L437S (3635C>T 3636T>C 3637C>A), M445F (3659A>T 3661G>T), S453T (3683T>A), C454F (3687G>T 3688C>T), Q455C (3689C>T 3690A>G 3691G>C), D456N (3692G>A 3694C>T), V458M (3698G>A 3700C>G), E460D (3706A>C), T461V (3707A>G 3708C>T), R463F (3713A>T 3714G>T 3715A>T), L465Y (3719C>T 3720T>A 3721A>T), V468D (3729T>A 3730G>T), C469F (3732G>T 3733C>T), A472V (3741C>T 3742T>C), M474L (3746A>C 3748G>A), L478V (3758C>G 3760G>C), V479I (3761G>A 3763C>A), T481S (3768C>G 3769A>T), K482Q (3770A>C), G483del (3773_3775delGGA), S484T (3776T>A 3778C>G), Q486D (3782C>G 3784G>T), T489V (3791A>G 3792C>T 3793C>T), K490S (3795A>G 3796G>C), Q491H (3799A>T), V492L (3800G>T 3802T>G), Q493S (3803C>A 3804A>G 3805A>T), D494L (3806G>T 3807A>T 3808T>G), F496L (3814C>A), E497S (3815G>T 3816A>C 3817A>T), T500R (3825C>G), K501Q (3827A>C 3829G>A), S502Y (3831C>A), G503T (3833G>A 3834G>C 3835A>T), F504L (3836T>C), K505Y (3839A>T 3841G>T), T506V (3842A>G 3843C>T 3844A>T), A507K (3845G>A 3846C>A), P508M (3848C>A 3849C>T 3850C>G), H514A (3866C>G 3867A>C), K515Q (3869A>C 3871G>A), V518I (3878G>A 3880C>A), F523H (3893T>C 3894T>A), I524L (3896A>C 3898C>A), I525V (3899A>G 3901C>T), T527E (3905A>G 3906C>A), T528N (3909C>A 3910A>C), G529Q (3911G>C 3912G>A 3913G>A), I530V (3914A>G), T531R (3918C>G), I532M (3922C>G), A535K (3929G>A 3930C>A 3931A>G), T537V (3935A>G 3936C>T 3937A>G), S539A (3941T>G 3943A>C), R541V (3947A>G 3948G>T 3949A>T), E542D (3952A>T), P544Q (3957C>A), E545A (3960A>C), V552L (3980G>C 3982A>G), Q553R (3983C>A 3984A>G), N562Y (4010A>T 4012C>T), K567A (4025A>G 4026A>C 4027G>T), D568G (4029A>G 4030C>A), T572K (4041C>A), P575A (4049C>G), M576L (4052A>C), M578D (4058A>G 4059T>A 4060G>T), T580L (4064A>C 4065C>T 4066A>G), R581K (4068G>A 4069A>G), D583E (4075C>A), N585K (4081C>G), K587V (4085A>G 4086A>T), G589S (4091G>T 4092G>C 4093A>G), K590V (4094A>G 4095A>T), E591R (4097G>A 4098A>G), Q592C (4100C>T 4101A>G 4102G>C), T593E (4103A>G 4104C>A 4105C>A), K597Q (4115A>C 4117A>G), R598S (4120A>T), Q602A (4130C>G 4131A>C 4132G>T), C603I (4133T>A 4134G>T 4135C>T), A606E (4143C>A 4144C>A), T608I (4149C>T 4150G>A), R610K (4154C>A 4155G>A 4156A>G), L611_F612insPN (4159_4160insCCAAAT), D613E (4165T>G), G614L (4166G>C 4167G>T 4168T>A), S615P (4169A>C 4170G>C 4171C>T), K616F (4172A>T 4173A>T 4174G>T), I620_E621del (4184_4189delATCGAG), M627K (4206T>A 4207G>A), I629V (4211A>G 4213A>G), A631G (4218C>G 4219A>C), C632V (4220T>G 4221G>T 4222T>A), T634V (4226A>G 4227C>T 4228A>G), T636N (4233C>A 4234A>T), H637D (4235C>G), D638E (4240T>A), K640_R641del (4244_4249delAAAAGA), Y646F (4263A>T), Y647E (4265T>G 4267T>G), M651L (4277A>T), T652N (4281C>A 4282C>T), T653P (4283A>C 4285A>G), A654T (4286G>A), N657R (4296A>G 4297C>A), D659S (4301G>T 4302A>C 4303C>T), I660A (4304A>G 4305T>C 4306C>A), D662E (4312C>A), L665M (4319C>A 4321T>G), L666V (4322C>G 4324A>G), I668V (4328A>G 4330T>G), V669L (4331G>C 4333T>C), A670H (4334G>C 4335C>A 4336C>T), A671C (4337G>T 4338C>G 4339C>T), M672L (4340A>C 4342G>T), H674T (4346C>A 4347A>C 4348T>A), V679L (4361G>C), E680L (4364G>T 4365A>T 4366G>A), P682T (4370C>A), P683del (4373_4375delCCG), L685F (4381A>T), T686V (4382A>G 4383C>T 4384G>A), I687V (4385A>G 4387T>A), L688K (4388C>A 4389T>A 4390T>G), S689T (4391T>A 4393A>G), H691N (4397C>A 4399C>T), K692V (4400A>G 4401A>T), N693A (4403A>G 4404A>C), L694N (4406C>A 4407T>A 4408C>T), T698K (4419C>A 4420G>A), T700Q (4424A>C 4425C>A 4426G>A), K701Q (4427A>C), E702K (4430G>A), T704S (4437C>G 4438C>T), R705L (4439C>T 4440G>T 4441A>G), R706K (4443G>A 4444A>G), Q707* (4445C>T 4447A>G), S711Q (4457T>C 4458C>A 4459G>A), L713I (4463C>A 4465G>A), G715A (4470G>C 4471G>A), Q716E (4472C>G 4474G>A), K718D (4478A>G 4480G>C), E720V (4485A>T 4486A>G), I721W (4487A>T 4488T>G 4489C>G), K722I (4491A>T 4492A>T), Y723H (4493T>C 4495C>T), T724K (4497C>A 4498T>A), F825Y (4800T>A), S826Y (4803C>A 4804A>T), F827W (4806T>G 4807C>G), Q829R (4812A>G 4813G>C), L832E (4820C>G 4821T>A 4822T>G), K833D (4823A>G 4825G>T), V834I (4826G>A 4828G>A), L835E (4829C>G 4830T>A), I838V (4838A>G 4840C>G), K839Q (4841A>C 4843G>A), K840T (4845A>C 4846A>T), V842L (4850G>C 4852A>T), H843V (4853C>G 4854A>T 4855C>G), N847D (4865A>G), A849V (4872C>T 4873T>A), A850E (4875C>A 4876A>G), R851Q (4878G>A), H852R (4881A>G 4882C>A), A853Q (4883G>C 4884C>A 4885G>A), K854P (4886A>C 4887A>C 4888A>C), Y855G (4889T>G 4890A>G 4891C>A), H857L (4896A>T 4897C>T), Q859E (4901C>G), F860P (4904T>C 4905T>C 4906C>A), R861L (4907A>T 4908G>T 4909G>A), T862H (4910A>C 4911C>A 4912A>T), P863V (4913C>G 4914C>T), P864A (4916C>G 4918A>C), T865E (4919A>G 4920C>A), K866R (4922A>C 4923A>G 4924A>T), D869E (4933C>G), E870S (4934G>A 4935A>G 4936G>T), V871I (4937G>A 4939T>C), K878C (4958A>T 4959A>G 4960A>T), R881K (4968G>A), K883D (4973A>G 4975G>T), L895W (5009C>T 5010T>G 5011A>G), M897I (5017G>T), H961L (5207C>T 5208A>T 5209C>G), K962N (5212G>T), Y967F (5226A>T 5227C>T), E970Q (5234G>C), T975S (5249A>T 5251G>A), N979I (5262A>T 5263C>T), Y989F (5292A>T), I990V (5294A>G 5296C>G), N991S (5298A>G), Y992S (5301A>C 5302C>T), A993H (5303G>C 5304C>A 5305A>T), D995K (5309G>A 5311C>A), N996D (5312A>G 5314C>T), V998A (5319T>C), S999K (5321T>A 5322C>A 5323A>G), P1002D (5330C>G 5331C>A 5332A>T), M1003V (5333A>G), I1006F (5342A>T), A1007S (5345G>T), L1008Y (5348C>T 5349T>A 5350G>T), N1010L (5354A>C 5355A>T 5356C>A), H1011Q (5359C>G), K1012R (5360A>C 5361A>G 5362A>G), T1015A (5369A>G 5371A>C), S1017G (5375T>G 5376C>G 5377G>C), T1018K (5379C>A), M1022E (5390A>G 5391T>A), R1023L (5394G>T), T1024A (5396A>G 5398T>C), L1025A (5399T>G 5400T>C 5401G>A), A1026G (5403C>G 5404A>C) | | | | | | | | |

| | Begin | End | Coverage | Score | Concordance | Matches | Identities | I/D/M/F* | Stop Codons |
|---|---|---|---|---|---|---|---|---|---|
| **NT** | **3205** | **5406** | **23.2%** | **115** | **3.4%** | **1697 (98.1%)** | **899 (52.0%)** | **15/18** | |

Codon mutations:

GAG296GAT (3214G>T), GAA297GTG (3216A>T 3217A>G), AGA298ATG (3219G>T 3220A>G), GGC299CCA (3221G>C 3222G>C 3223C>A), AAG300CCT (3224A>C 3225A>C 3226G>T), GAC301GAG (3229C>G), GCC302TTG (3230G>T 3231C>T 3232C>G), TTA303CCA (3233T>C 3234T>C), CCT304AAG (3236C>A 3237C>A 3238T>G), CAC306CTG (3243A>T 3244C>G), CAA307CCA (3246A>C), TGG309AGG (3251T>A), TGG309_GAT310insAGGGATATT (3253_3254insAGGGATATT), AAG312AGA (3261A>G 3262G>A), ATA313ATT (3265A>T), AAC314GAG (3266A>G 3268C>G), ATT315TTG (3269A>T 3271T>G), CAG316ATG (3272C>A 3273A>T), GGG318GGT (3280G>T), AAA319TCT (3281A>T 3282A>C 3283A>T), GAG320ACG (3284G>A 3285A>C), CCT321CCG (3289T>G), CCA322CCT (3292A>T), TGG323GCA (3293T>G 3294G>C 3295G>A), GGA324CAA (3296G>C 3297G>A), CCC325GCT (3299C>G 3301C>T), CTA326CCT (3303T>C 3304A>T), TAT327TAC (3307T>C), CAA328AGA (3308C>A 3309A>G), TCT330GCT (3314T>G), GAG331CCT (3317G>C 3318A>C 3319G>T), CTA334TTG (3326C>T 3328A>G), CAG335GCT (3329C>G 3330A>C 3331G>T), ACC336GAA (3332A>G 3333C>A 3334C>A), CTA337CTG (3337A>G), CGA338AGG (3338C>A 3340A>G), GAA339AAG (3341G>A 3343A>G), TGG340CAG (3344T>C 3345G>A), AAG342AAC (3352G>C), GAG343GAA (3355G>A), AAG344TTG (3356A>T 3357A>T), CTA345CTT (3361A>T), GCC346GAT (3363C>A 3364C>T), AAA347TCA (3365A>T 3366A>C), TGG349CTG (3371T>C 3372G>T), ATA350ATT (3376A>T), CGA351CAG (3378G>A 3379A>G), CGA352CCT (3381G>C 3382A>T), TCC353TCT (3385C>T), ACC354AAG (3387C>A 3388C>G), TCA355GCT (3389T>G 3391A>T), AGT356CCC (3392A>C 3393G>C 3394T>C), GCA357TAT (3395G>T 3396C>A 3397A>T), GGA358GGT (3400A>T), ACT359GCA (3401A>G 3403T>A), CCA360CCT (3406A>T), TGC361GTT (3407T>G 3408G>T 3409C>T), ATG362TTA (3410A>T 3412G>A), GTT364CAG (3416G>C 3417T>A 3418T>G), CCA365AAA (3419C>A 3420C>A), GCA367CAG (3425G>C 3426C>A 3427A>G), AAC368GAT (3428A>G 3430C>T), AAA370TCA (3434A>T 3435A>C), CTA371TTA (3437C>T), CGA372AGG (3440C>A 3442A>G), CTC373ATG (3443C>A 3445C>G), GTA374TGC (3446G>T 3447T>G 3448A>C), CAA375GTG (3449C>G 3450A>T 3451A>G), GAC376GAT (3454C>T), TAC377TAT (3457C>T), CGA378CGG (3460A>G), AAG379GCG (3461A>G 3462A>C), TTG380CTG (3464T>C), GAG382AAG (3470G>A), ATC383GTA (3473A>G 3475C>A), ACG384ACT (3478G>T), ATC385GTG (3479A>G 3481C>G), CGA388AAG (3488C>A 3489G>A 3490A>G), TAT389TAC (3493T>C), CCG390CCA (3496G>A), CTA391GTT (3497C>G 3499A>T), CCC392CCT (3502C>T), AAC393TTG (3503A>T 3504A>T 3505C>G), ATC394GTG (3506A>G 3508C>G), GAA395CAG (3509G>C 3511A>G), GAA396GAT (3514A>T), GCA397CTG (3515G>C 3516C>T 3517A>G), CAA398ATG (3518C>A 3519A>T 3520A>G), AGA400AGG (3526A>G), TTA401TTG (3529A>G), ACC402AGC (3531C>G), GGA403AAA (3533G>A 3534G>A), TCA404GCG (3536T>G 3538A>G), GAC405TGC (3539G>T 3540A>G), TAC407TTC (3546A>T), ACG408ACT (3550G>T), ATT410CTA (3554A>C 3556T>A), GAC411GAT (3559C>T), CTA412CTT (3562A>T), CGA413AGG (3563C>A 3565A>G), GAC414TCA (3566G>T 3567A>C 3568C>A), GCC415GGC (3570C>G), TTC416TAT (3573T>A 3574C>T), TAT417TGG (3576A>G 3577T>G), GCT418CAG (3578G>C 3579C>A 3580T>G), ATC419GTT (3581A>G 3583C>T), CGA420AGG (3584C>A 3586A>G), ATG421ATA (3589G>A), GGA424GGG (3598A>G), GAA425GAT (3601A>T), GAA426GAG (3604A>G), TGG427CCT (3605T>C 3606G>C 3607G>T), AAA428AAG (3610A>G), ACC429ACT (3613C>T), GCT430ACT (3614G>A), TTC431TGT (3618T>G 3619C>T), AGG432GTA (3620A>G 3621G>T 3622G>A), ACA433ACT (3625A>T), AGA434AGG (3628A>G), TAC435TAT (3631C>T), GGA436GGG (3634A>G), CTC437TCA (3635C>T 3636T>C 3637C>A), TAC438TAT (3640C>T), TTG441CTT (3647T>C 3649G>T), GTC442GTA (3652C>A), CCA444CCG (3658A>G), ATG445TTT (3659A>T 3661G>T), GGA446GGG (3664A>G), ACC448ACT (3670C>T), AAC449AAT (3673C>T), GCA450GCC (3676A>C), CCC451CCG (3679C>G), GCA452GCT (3682A>T), TCC453ACC (3683T>A), TGC454TTT (3687G>T 3688C>T), CAG455TGC (3689C>T 3690A>G 3691G>C), GAC456AAT (3692G>A 3694C>T), CTT457TTA (3695C>T 3697T>A), GTC458ATG (3698G>A 3700C>G), AAC459AAT (3703C>T), GAA460GAC (3706A>C), ACA461GTA (3707A>G 3708C>T), CTT462CTG (3712T>G), AGA463TTT (3713A>T 3714G>T 3715A>T), CTA465TAT (3719C>T 3720T>A 3721A>T), CTC466CTT (3724C>T), GAC467GAT (3727C>T), GTG468GAT (3729T>A 3730G>T), TGC469TTT (3732G>T 3733C>T), GTC470GTA (3736C>A), GTT471GTA (3739T>A), GCT472GTC (3741C>T 3742T>C), TAC473TAT (3745C>T), ATG474CTA (3746A>C 3748G>A), GAC475GAT (3751C>T), ATA477ATT (3757A>T), CTG478GTC (3758C>G 3760G>C), GTC479ATA (3761G>A 3763C>A), TAC480TAT (3766C>T), ACA481AGT (3768C>G 3769A>T), AAA482CAA (3770A>C), GGA483del (3773_3775delGGA), TCC484ACG (3776T>A 3778C>G), CTC485TTG (3779C>T 3781C>G), CAG486GAT (3782C>G 3784G>T), ACC489GTT (3791A>G 3792C>T 3793C>T), AAG490AGC (3795A>G 3796G>C), CAA491CAT (3799A>T), GTT492TTG (3800G>T 3802T>G), CAA493AGT (3803C>A 3804A>G 3805A>T), GAT494TTG (3806G>T 3807A>T 3808T>G), GTG495GTA (3811G>A), TTC496TTA (3814C>A), GAA497TCT (3815G>T 3816A>C 3817A>T), CGA498CGT (3820A>T), CTC499TTA (3821C>T 3823C>A), ACG500AGG (3825C>G), AAG501CAA (3827A>C 3829G>A), TCC502TAC (3831C>A), GGA503ACT (3833G>A 3834G>C 3835A>T), TTC504CTC (3836T>C), AAG505TAT (3839A>T 3841G>T), ACA506GTT (3842A>G 3843C>T 3844A>T), GCA507AAA (3845G>A 3846C>A), CCC508ATG (3848C>A 3849C>T 3850C>G), AAA510AAG (3856A>G), TGC511TGT (3859C>T), GAA512GAG (3862A>G), TTC513TTT (3865C>T), CAC514GCC (3866C>G 3867A>C), AAG515CAA (3869A>C 3871G>A), GAA517GAG (3877A>G), GTC518ATA (3878G>A 3880C>A), TTT520TTC (3886T>C), TTA521CTA (3887T>C), GGC522GGG (3892C>G), TTT523CAT (3893T>C 3894T>A), ATC524CTA (3896A>C 3898C>A), ATC525GTT (3899A>G 3901C>T), ACA527GAA (3905A>G 3906C>A), ACA528AAC (3909C>A 3910A>C), GGG529CAA (3911G>C 3912G>A 3913G>A), ATA530GTA (3914A>G), ACG531AGG (3918C>G), ATC532ATG (3922C>G), GAC533GAT (3925C>T), GCA535AAG (3929G>A 3930C>A 3931A>G), AAG536AAA (3934G>A), ACA537GTG (3935A>G 3936C>T 3937A>G), CAG538CAA (3940G>A), TCA539GCC (3941T>G 3943A>C), ATC540ATT (3946C>T), AGA541GTT (3947A>G 3948G>T 3949A>T), GAA542GAT (3952A>T), CCA544CAA (3957C>A), GAA545GCA (3960A>C), CCG546CCC (3964G>C), AAG547AAA (3967G>A), ACA548ACT (3970A>T), GTC549GTG (3973C>G), GTA552CTG (3980G>C 3982A>G), CAG553AGG (3983C>A 3984A>G), TCA554TCT (3988A>T), CTT556TTG (3992C>T 3994T>G), GGA557GGG (3997A>G), CTC558CTA (4000C>A), GCC559GCT (4003C>T), AAC562TAT (4010A>T 4012C>T), AAG567GCT (4025A>G 4026A>C 4027G>T), GAC568GGA (4029A>G 4030C>A), TAT569TAC (4033T>C), ACA572AAA (4041C>A), GCA574GCT (4048A>T), CCA575GCA (4049C>G), ATG576CTG (4052A>C), ACG577ACT (4057G>T), ATG578GAT (4058A>G 4059T>A 4060G>T), CTT579TTG (4061C>T 4063T>G), ACA580CTG (4064A>C 4065C>T 4066A>G), AGA581AAG (4068G>A 4069A>G), AAA582AAG (4072A>G), GAC583GAA (4075C>A), AAC585AAG (4081C>G), AAA587GTA (4085A>G 4086A>T), GGA589TCG (4091G>T 4092G>C 4093A>G), AAA590GTA (4094A>G 4095A>T), GAA591AGA (4097G>A 4098A>G), CAG592TGC (4100C>T 4101A>G 4102G>C), ACC593GAA (4103A>G 4104C>A 4105C>A), GCG595GCT (4111G>T), TTC596TTT (4114C>T), AAA597CAG (4115A>C 4117A>G), AGA598AGT (4120A>T), CTC599TTG (4121C>T 4123C>G), CAG602GCT (4130C>G 4131A>C 4132G>T), TGC603ATT (4133T>A 4134G>T 4135C>T), GCT604GCA (4138T>A), GCC606GAA (4143C>A 4144C>A), ACG608ATA (4149C>T 4150G>A), CTT609CTG (4153T>G), CGA610AAG (4154C>A 4155G>A 4156A>G), CTA611CTG (4159A>G), CTA611_TTC612insCCAAAT (4159_4160insCCAAAT), TTC612TTT (4162C>T), GAT613GAG (4165T>G), GGT614CTA (4166G>C 4167G>T 4168T>A), AGC615CCT (4169A>C 4170G>C 4171C>T), AAG616TTT (4172A>T 4173A>T 4174G>T), GAA617GAG (4177A>G), GTC618GTA (4180C>A), ATC620_GAG621del (4184_4189delATCGAG), ACC622ACT (4192C>T), GAC623GAT (4195C>T), TCT625TCG (4201T>G), GAT626GAC (4204T>C), ATG627AAA (4206T>A 4207G>A), GCA628GCT (4210A>T), ATA629GTG (4211A>G 4213A>G), GGC630GGA (4216C>A), GCA631GGC (4218C>G 4219A>C), TGT632GTA (4220T>G 4221G>T 4222T>A), CTA633TTG (4223C>T 4225A>G), ACA634GTG (4226A>G 4227C>T 4228A>G), CAG635CAA (4231G>A), ACA636AAT (4233C>A 4234A>T), CAC637GAC (4235C>G), GAT638GAA (4240T>A), GGG639GGT (4243G>T), AAA640_AGA641del (4244_4249delAAAAGA), CAC642CAT (4252C>T), CCA643CCG (4255A>G), TAT646TTT (4263A>T), TAT647GAG (4265T>G 4267T>G), TCC648AGC (4268T>A 4269C>G), CGG649AGG (4271C>A), ATG651TTG (4277A>T), ACC652AAT (4281C>A 4282C>T), ACA653CCG (4283A>C 4285A>G), GCG654ACG (4286G>A), CAG656CAA (4294G>A), AAC657AGA (4296A>G 4297C>A), TAC658TAT (4300C>T), GAC659TCT (4301G>T 4302A>C 4303C>T), ATC660GCA (4304A>G 4305T>C 4306C>A), GAC662GAA (4312C>A), CTT665ATG (4319C>A 4321T>G), CTA666GTG (4322C>G 4324A>G), GCC667GCA (4327C>A), ATT668GTG (4328A>G 4330T>G), GTT669CTC (4331G>C 4333T>C), GCC670CAT (4334G>C 4335C>A 4336C>T), GCC671TGT (4337G>T 4338C>G 4339C>T), ATG672CTT (4340A>C 4342G>T), CAT674ACA (4346C>A 4347A>C 4348T>A), AGA676AGG (4354A>G), TAC678TAT (4360C>T), GTC679CTC (4361G>C), GAG680TTA (4364G>T 4365A>T 4366G>A), GGC681GGA (4369C>A), CCA682ACA (4370C>A), CCG683del (4373_4375delCCG), TTA685TTT (4381A>T), ACG686GTA (4382A>G 4383C>T 4384G>A), ATT687GTA (4385A>G 4387T>A), CTT688AAG (4388C>A 4389T>A 4390T>G), TCA689ACG (4391T>A 4393A>G), CAC691AAT (4397C>A 4399C>T), AAG692GTG (4400A>G 4401A>T), AAT693GCT (4403A>G 4404A>C), CTC694AAT (4406C>A 4407T>A 4408C>T), ACG695ACT (4411G>T), ACG698AAA (4419C>A 4420G>A), ACG699ACT (4423G>T), ACG700CAA (4424A>C 4425C>A 4426G>A), AAG701CAG (4427A>C), GAA702AAA (4430G>A), CTT703TTG (4433C>T 4435T>G), ACC704AGT (4437C>G 4438C>T), CGA705TTG (4439C>T 4440G>T 4441A>G), AGA706AAG (4443G>A 4444A>G), CAA707TAG (4445C>T 4447A>G), GCC708GCG (4450C>G), CGC709AGA (4451C>A 4453C>A), TCG711CAA (4457T>C 4458C>A 4459G>A), CTG713ATA (4463C>A 4465G>A), CTT714CTA (4468T>A), GGG715GCA (4470G>C 4471G>A), CAG716GAA (4472C>G 4474G>A), TAC717TAT (4477C>T), AAG718GAC (4478A>G 4480G>C), TTC719TTT (4483C>T), GAA720GTG (4485A>T 4486A>G), ATC721TGG (4487A>T 4488T>G 4489C>G), AAA722ATT (4491A>T 4492A>T), TAC723CAT (4493T>C 4495C>T), ACT724AAA (4497C>A 4498T>A), AGC824.TC (4797G>T), TTC825TAC (4800T>A), TCA826TAT (4803C>A 4804A>T), TTC827TGG (4806T>G 4807C>G), CAG829CGC (4812A>G 4813G>C), AGA831CGG (4817A>C 4819A>G), CTT832GAG (4820C>G 4821T>A 4822T>G), AAG833GAT (4823A>G 4825G>T), GTG834ATA (4826G>A 4828G>A), CTA835GAA (4829C>G 4830T>A), ATC838GTG (4838A>G 4840C>G), AAG839CAA (4841A>C 4843G>A), AAA840ACT (4845A>C 4846A>T), TGC841TGT (4849C>T), GTA842CTT (4850G>C 4852A>T), CAC843GTG (4853C>G 4854A>T 4855C>G), AAC847GAC (4865A>G), AAA848AAG (4870A>G), GCT849GTA (4872C>T 4873T>A), GCA850GAG (4875C>A 4876A>G), CGG851CAG (4878G>A), CAC852CGA (4881A>G 4882C>A), GCG853CAA (4883G>C 4884C>A 4885G>A), AAA854CCC (4886A>C 4887A>C 4888A>C), TAC855GGA (4889T>G 4890A>G 4891C>A), GGT856GGA (4894T>A), CAC857CTT (4896A>T 4897C>T), CTA858TTA (4898C>T), CAG859GAG (4901C>G), TTC860CCA (4904T>C 4905T>C 4906C>A), AGG861TTA (4907A>T 4908G>T 4909G>A), ACA862CAT (4910A>C 4911C>A 4912A>T), CCA863GTA (4913C>G 4914C>T), CCA864GCC (4916C>G 4918A>C), ACG865GAG (4919A>G 4920C>A), AAA866CGT (4922A>C 4923A>G 4924A>T), GAC869GAG (4933C>G), GAG870AGT (4934G>A 4935A>G 4936G>T), GTT871ATC (4937G>A 4939T>C), ATT876ATA (4954T>A), ACG877ACC (4957G>C), AAA878TGT (4958A>T 4959A>G 4960A>T), CTC879TTG (4961C>T 4963C>G), CCG880CCA (4966G>A), AGG881AAG (4968G>A), TCA882TCC (4972A>C), AAG883GAT (4973A>G 4975G>T), CTA895TGG (5009C>T 5010T>G 5011A>G), GTC896GTG (5014C>G), ATG897ATT (5017G>T), GTC898GTG (5020C>G), AGA900AGG (5026A>G), CTC901TTG (5027C>T 5029C>G), ACA902ACT (5032A>T), AAA903AAG (5035A>G), TAT904TC. (5037A>C), CAC961TTG (5207C>T 5208A>T 5209C>G), AAG962AAT (5212G>T), TCA964AGT (5216T>A 5217C>G 5218A>T), TAC967TTT (5226A>T 5227C>T), CCA969CCT (5233A>T), GAG970CAG (5234G>C), ACG971ACC (5239G>C), GGG973GGC (5245G>C), CAA974CAG (5248A>G), ACG975TCA (5249A>T 5251G>A), GAA976GAG (5254A>G), AGA977AGG (5257A>G), ACG978ACT (5260G>T), AAC979ATT (5262A>T 5263C>T), ACA981ACT (5269A>T), CTC982T.. (5270C>T), CAA984..C (5278A>C), CTA986CTT (5284A>T), CGG987AGG (5285C>A), TAC989TTC (5292A>T), ATC990GTG (5294A>G 5296C>G), AAC991AGC (5298A>G), TAC992TCT (5301A>C 5302C>T), GCA993CAT (5303G>C 5304C>A 5305A>T), GAC995AAA (5309G>A 5311C>A), AAC996GAT (5312A>G 5314C>T), GTT998GCT (5319T>C), TCA999AAG (5321T>A 5322C>A 5323A>G), TTA1000TTG (5326A>G), CTG1001CTA (5329G>A), CCA1002GAT (5330C>G 5331C>A 5332A>T), ATG1003GTG (5333A>G), GCG1004GCC (5338G>C), CAG1005CAA (5341G>A), ATC1006TTC (5342A>T), GCA1007TCA (5345G>T), CTG1008TAT (5348C>T 5349T>A 5350G>T), AAC1009AAT (5353C>T), AAC1010CTA (5354A>C 5355A>T 5356C>A), CAC1011CAG (5359C>G), AAA1012CGG (5360A>C 5361A>G 5362A>G), TCA1013AGT (5363T>A 5364C>G 5365A>T), GAG1014GAA (5368G>A), ACA1015GCC (5369A>G 5371A>C), TCG1017GGC (5375T>G 5376C>G 5377G>C), ACG1018AAG (5379C>A), ATG1022GAG (5390A>G 5391T>A), CGA1023CTA (5394G>T), ACT1024GCC (5396A>G 5398T>C), TTG1025GCA (5399T>G 5400T>C 5401G>A), GCA1026GGC (5403C>G 5404A>C), AGG1027CA. (5405A>C 5406G>A)

#### Proteins

| | Begin | End | Coverage | Score | Concordance | Matches | Identities | I/D/M/F* | Stop Codons |
|---|---|---|---|---|---|---|---|---|---|
| **NT** | **3205** | **5406** | **23.2%** | **115** | **3.4%** | **1697 (98.1%)** | **899 (52.0%)** | **15/18** | |
| Reverse Transcriptase (YP_009666308.1) | 294 | 1027 | 54.8% | 1929 | 48.4% | 567 (98.1%) | 269 (46.5%) | 5/6/0/0 | 1 |
| Protein mutations: | E296D (3214G>T), E297V (3216A>T 3217A>G), R298M (3219G>T 3220A>G), G299P (3221G>C 3222G>C 3223C>A), K300P (3224A>C 3225A>C 3226G>T), D301E (3229C>G), A302L (3230G>T 3231C>T 3232C>G), L303P (3233T>C 3234T>C), P304K (3236C>A 3237C>A 3238T>G), H306L (3243A>T 3244C>G), Q307P (3246A>C), W309R (3251T>A), W309_D310insRDI (3253_3254insAGGGATATT), K312R (3261A>G 3262G>A), N314E (3266A>G 3268C>G), I315L (3269A>T 3271T>G), Q316M (3272C>A 3273A>T), K319S (3281A>T 3282A>C 3283A>T), E320T (3284G>A 3285A>C), W323A (3293T>G 3294G>C 3295G>A), G324Q (3296G>C 3297G>A), P325A (3299C>G 3301C>T), L326P (3303T>C 3304A>T), Q328R (3308C>A 3309A>G), S330A (3314T>G), E331P (3317G>C 3318A>C 3319G>T), Q335A (3329C>G 3330A>C 3331G>T), T336E (3332A>G 3333C>A 3334C>A), E339K (3341G>A 3343A>G), W340Q (3344T>C 3345G>A), K342N (3352G>C), K344L (3356A>T 3357A>T), A346D (3363C>A 3364C>T), K347S (3365A>T 3366A>C), W349L (3371T>C 3372G>T), R351Q (3378G>A 3379A>G), R352P (3381G>C 3382A>T), T354K (3387C>A 3388C>G), S355A (3389T>G 3391A>T), S356P (3392A>C 3393G>C 3394T>C), A357Y (3395G>T 3396C>A 3397A>T), T359A (3401A>G 3403T>A), C361V (3407T>G 3408G>T 3409C>T), M362L (3410A>T 3412G>A), V364Q (3416G>C 3417T>A 3418T>G), P365K (3419C>A 3420C>A), A367Q (3425G>C 3426C>A 3427A>G), N368D (3428A>G 3430C>T), K370S (3434A>T 3435A>C), L373M (3443C>A 3445C>G), V374C (3446G>T 3447T>G 3448A>C), Q375V (3449C>G 3450A>T 3451A>G), K379A (3461A>G 3462A>C), E382K (3470G>A), I383V (3473A>G 3475C>A), I385V (3479A>G 3481C>G), R388K (3488C>A 3489G>A 3490A>G), L391V (3497C>G 3499A>T), N393L (3503A>T 3504A>T 3505C>G), I394V (3506A>G 3508C>G), E395Q (3509G>C 3511A>G), E396D (3514A>T), A397L (3515G>C 3516C>T 3517A>G), Q398M (3518C>A 3519A>T 3520A>G), T402S (3531C>G), G403K (3533G>A 3534G>A), S404A (3536T>G 3538A>G), D405C (3539G>T 3540A>G), Y407F (3546A>T), I410L (3554A>C 3556T>A), D414S (3566G>T 3567A>C 3568C>A), A415G (3570C>G), F416Y (3573T>A 3574C>T), Y417W (3576A>G 3577T>G), A418Q (3578G>C 3579C>A 3580T>G), I419V (3581A>G 3583C>T), M421I (3589G>A), E425D (3601A>T), W427P (3605T>C 3606G>C 3607G>T), A430T (3614G>A), F431C (3618T>G 3619C>T), R432V (3620A>G 3621G>T 3622G>A), L437S (3635C>T 3636T>C 3637C>A), M445F (3659A>T 3661G>T), S453T (3683T>A), C454F (3687G>T 3688C>T), Q455C (3689C>T 3690A>G 3691G>C), D456N (3692G>A 3694C>T), V458M (3698G>A 3700C>G), E460D (3706A>C), T461V (3707A>G 3708C>T), R463F (3713A>T 3714G>T 3715A>T), L465Y (3719C>T 3720T>A 3721A>T), V468D (3729T>A 3730G>T), C469F (3732G>T 3733C>T), A472V (3741C>T 3742T>C), M474L (3746A>C 3748G>A), L478V (3758C>G 3760G>C), V479I (3761G>A 3763C>A), T481S (3768C>G 3769A>T), K482Q (3770A>C), G483del (3773_3775delGGA), S484T (3776T>A 3778C>G), Q486D (3782C>G 3784G>T), T489V (3791A>G 3792C>T 3793C>T), K490S (3795A>G 3796G>C), Q491H (3799A>T), V492L (3800G>T 3802T>G), Q493S (3803C>A 3804A>G 3805A>T), D494L (3806G>T 3807A>T 3808T>G), F496L (3814C>A), E497S (3815G>T 3816A>C 3817A>T), T500R (3825C>G), K501Q (3827A>C 3829G>A), S502Y (3831C>A), G503T (3833G>A 3834G>C 3835A>T), F504L (3836T>C), K505Y (3839A>T 3841G>T), T506V (3842A>G 3843C>T 3844A>T), A507K (3845G>A 3846C>A), P508M (3848C>A 3849C>T 3850C>G), H514A (3866C>G 3867A>C), K515Q (3869A>C 3871G>A), V518I (3878G>A 3880C>A), F523H (3893T>C 3894T>A), I524L (3896A>C 3898C>A), I525V (3899A>G 3901C>T), T527E (3905A>G 3906C>A), T528N (3909C>A 3910A>C), G529Q (3911G>C 3912G>A 3913G>A), I530V (3914A>G), T531R (3918C>G), I532M (3922C>G), A535K (3929G>A 3930C>A 3931A>G), T537V (3935A>G 3936C>T 3937A>G), S539A (3941T>G 3943A>C), R541V (3947A>G 3948G>T 3949A>T), E542D (3952A>T), P544Q (3957C>A), E545A (3960A>C), V552L (3980G>C 3982A>G), Q553R (3983C>A 3984A>G), N562Y (4010A>T 4012C>T), K567A (4025A>G 4026A>C 4027G>T), D568G (4029A>G 4030C>A), T572K (4041C>A), P575A (4049C>G), M576L (4052A>C), M578D (4058A>G 4059T>A 4060G>T), T580L (4064A>C 4065C>T 4066A>G), R581K (4068G>A 4069A>G), D583E (4075C>A), N585K (4081C>G), K587V (4085A>G 4086A>T), G589S (4091G>T 4092G>C 4093A>G), K590V (4094A>G 4095A>T), E591R (4097G>A 4098A>G), Q592C (4100C>T 4101A>G 4102G>C), T593E (4103A>G 4104C>A 4105C>A), K597Q (4115A>C 4117A>G), R598S (4120A>T), Q602A (4130C>G 4131A>C 4132G>T), C603I (4133T>A 4134G>T 4135C>T), A606E (4143C>A 4144C>A), T608I (4149C>T 4150G>A), R610K (4154C>A 4155G>A 4156A>G), L611_F612insPN (4159_4160insCCAAAT), D613E (4165T>G), G614L (4166G>C 4167G>T 4168T>A), S615P (4169A>C 4170G>C 4171C>T), K616F (4172A>T 4173A>T 4174G>T), I620_E621del (4184_4189delATCGAG), M627K (4206T>A 4207G>A), I629V (4211A>G 4213A>G), A631G (4218C>G 4219A>C), C632V (4220T>G 4221G>T 4222T>A), T634V (4226A>G 4227C>T 4228A>G), T636N (4233C>A 4234A>T), H637D (4235C>G), D638E (4240T>A), K640_R641del (4244_4249delAAAAGA), Y646F (4263A>T), Y647E (4265T>G 4267T>G), M651L (4277A>T), T652N (4281C>A 4282C>T), T653P (4283A>C 4285A>G), A654T (4286G>A), N657R (4296A>G 4297C>A), D659S (4301G>T 4302A>C 4303C>T), I660A (4304A>G 4305T>C 4306C>A), D662E (4312C>A), L665M (4319C>A 4321T>G), L666V (4322C>G 4324A>G), I668V (4328A>G 4330T>G), V669L (4331G>C 4333T>C), A670H (4334G>C 4335C>A 4336C>T), A671C (4337G>T 4338C>G 4339C>T), M672L (4340A>C 4342G>T), H674T (4346C>A 4347A>C 4348T>A), V679L (4361G>C), E680L (4364G>T 4365A>T 4366G>A), P682T (4370C>A), P683del (4373_4375delCCG), L685F (4381A>T), T686V (4382A>G 4383C>T 4384G>A), I687V (4385A>G 4387T>A), L688K (4388C>A 4389T>A 4390T>G), S689T (4391T>A 4393A>G), H691N (4397C>A 4399C>T), K692V (4400A>G 4401A>T), N693A (4403A>G 4404A>C), L694N (4406C>A 4407T>A 4408C>T), T698K (4419C>A 4420G>A), T700Q (4424A>C 4425C>A 4426G>A), K701Q (4427A>C), E702K (4430G>A), T704S (4437C>G 4438C>T), R705L (4439C>T 4440G>T 4441A>G), R706K (4443G>A 4444A>G), Q707* (4445C>T 4447A>G), S711Q (4457T>C 4458C>A 4459G>A), L713I (4463C>A 4465G>A), G715A (4470G>C 4471G>A), Q716E (4472C>G 4474G>A), K718D (4478A>G 4480G>C), E720V (4485A>T 4486A>G), I721W (4487A>T 4488T>G 4489C>G), K722I (4491A>T 4492A>T), Y723H (4493T>C 4495C>T), T724K (4497C>A 4498T>A), F825Y (4800T>A), S826Y (4803C>A 4804A>T), F827W (4806T>G 4807C>G), Q829R (4812A>G 4813G>C), L832E (4820C>G 4821T>A 4822T>G), K833D (4823A>G 4825G>T), V834I (4826G>A 4828G>A), L835E (4829C>G 4830T>A), I838V (4838A>G 4840C>G), K839Q (4841A>C 4843G>A), K840T (4845A>C 4846A>T), V842L (4850G>C 4852A>T), H843V (4853C>G 4854A>T 4855C>G), N847D (4865A>G), A849V (4872C>T 4873T>A), A850E (4875C>A 4876A>G), R851Q (4878G>A), H852R (4881A>G 4882C>A), A853Q (4883G>C 4884C>A 4885G>A), K854P (4886A>C 4887A>C 4888A>C), Y855G (4889T>G 4890A>G 4891C>A), H857L (4896A>T 4897C>T), Q859E (4901C>G), F860P (4904T>C 4905T>C 4906C>A), R861L (4907A>T 4908G>T 4909G>A), T862H (4910A>C 4911C>A 4912A>T), P863V (4913C>G 4914C>T), P864A (4916C>G 4918A>C), T865E (4919A>G 4920C>A), K866R (4922A>C 4923A>G 4924A>T), D869E (4933C>G), E870S (4934G>A 4935A>G 4936G>T), V871I (4937G>A 4939T>C), K878C (4958A>T 4959A>G 4960A>T), R881K (4968G>A), K883D (4973A>G 4975G>T), L895W (5009C>T 5010T>G 5011A>G), M897I (5017G>T), H961L (5207C>T 5208A>T 5209C>G), K962N (5212G>T), Y967F (5226A>T 5227C>T), E970Q (5234G>C), T975S (5249A>T 5251G>A), N979I (5262A>T 5263C>T), Y989F (5292A>T), I990V (5294A>G 5296C>G), N991S (5298A>G), Y992S (5301A>C 5302C>T), A993H (5303G>C 5304C>A 5305A>T), D995K (5309G>A 5311C>A), N996D (5312A>G 5314C>T), V998A (5319T>C), S999K (5321T>A 5322C>A 5323A>G), P1002D (5330C>G 5331C>A 5332A>T), M1003V (5333A>G), I1006F (5342A>T), A1007S (5345G>T), L1008Y (5348C>T 5349T>A 5350G>T), N1010L (5354A>C 5355A>T 5356C>A), H1011Q (5359C>G), K1012R (5360A>C 5361A>G 5362A>G), T1015A (5369A>G 5371A>C), S1017G (5375T>G 5376C>G 5377G>C), T1018K (5379C>A), M1022E (5390A>G 5391T>A), R1023L (5394G>T), T1024A (5396A>G 5398T>C), L1025A (5399T>G 5400T>C 5401G>A), A1026G (5403C>G 5404A>C) | | | | | | | | |

| | Begin | End | Coverage | Score | Concordance | Matches | Identities | I/D/M/F* | Stop Codons |
|---|---|---|---|---|---|---|---|---|---|
| **NT** | **3205** | **5406** | **23.2%** | **115** | **3.4%** | **1697 (98.1%)** | **899 (52.0%)** | **15/18** | |

Codon mutations: GAG296GAT (3214G>T), GAA297GTG (3216A>T 3217A>G), AGA298ATG (3219G>T 3220A>G), GGC299CCA (3221G>C 3222G>C 3223C>A), AAG300CCT (3224A>C 3225A>C 3226G>T), GAC301GAG (3229C>G), GCC302TTG (3230G>T 3231C>T 3232C>G), TTA303CCA (3233T>C 3234T>C), CCT304AAG (3236C>A 3237C>A 3238T>G), CAC306CTG (3243A>T 3244C>G), CAA307CCA (3246A>C), TGG309AGG (3251T>A), TGG309_GAT310insAGGGATATT (3253_3254insAGGGATATT), AAG312AGA (3261A>G 3262G>A), ATA313ATT (3265A>T), AAC314GAG (3266A>G 3268C>G), ATT315TTG (3269A>T 3271T>G), CAG316ATG (3272C>A 3273A>T), GGG318GGT (3280G>T), AAA319TCT (3281A>T 3282A>C 3283A>T), GAG320ACG (3284G>A 3285A>C), CCT321CCG (3289T>G), CCA322CCT (3292A>T), TGG323GCA (3293T>G 3294G>C 3295G>A), GGA324CAA (3296G>C 3297G>A), CCC325GCT (3299C>G 3301C>T), CTA326CCT (3303T>C 3304A>T), TAT327TAC (3307T>C), CAA328AGA (3308C>A 3309A>G), TCT330GCT (3314T>G), GAG331CCT (3317G>C 3318A>C 3319G>T), CTA334TTG (3326C>T 3328A>G), CAG335GCT (3329C>G 3330A>C 3331G>T), ACC336GAA (3332A>G 3333C>A 3334C>A), CTA337CTG (3337A>G), CGA338AGG (3338C>A 3340A>G), GAA339AAG (3341G>A 3343A>G), TGG340CAG (3344T>C 3345G>A), AAG342AAC (3352G>C), GAG343GAA (3355G>A), AAG344TTG (3356A>T 3357A>T), CTA345CTT (3361A>T), GCC346GAT (3363C>A 3364C>T), AAA347TCA (3365A>T 3366A>C), TGG349CTG (3371T>C 3372G>T), ATA350ATT (3376A>T), CGA351CAG (3378G>A 3379A>G), CGA352CCT (3381G>C 3382A>T), TCC353TCT (3385C>T), ACC354AAG (3387C>A 3388C>G), TCA355GCT (3389T>G 3391A>T), AGT356CCC (3392A>C 3393G>C 3394T>C), GCA357TAT (3395G>T 3396C>A 3397A>T), GGA358GGT (3400A>T), ACT359GCA (3401A>G 3403T>A), CCA360CCT (3406A>T), TGC361GTT (3407T>G 3408G>T 3409C>T), ATG362TTA (3410A>T 3412G>A), GTT364CAG (3416G>C 3417T>A 3418T>G), CCA365AAA (3419C>A 3420C>A), GCA367CAG (3425G>C 3426C>A 3427A>G), AAC368GAT (3428A>G 3430C>T), AAA370TCA (3434A>T 3435A>C), CTA371TTA (3437C>T), CGA372AGG (3440C>A 3442A>G), CTC373ATG (3443C>A 3445C>G), GTA374TGC (3446G>T 3447T>G 3448A>C), CAA375GTG (3449C>G 3450A>T 3451A>G), GAC376GAT (3454C>T), TAC377TAT (3457C>T), CGA378CGG (3460A>G), AAG379GCG (3461A>G 3462A>C), TTG380CTG (3464T>C), GAG382AAG (3470G>A), ATC383GTA (3473A>G 3475C>A), ACG384ACT (3478G>T), ATC385GTG (3479A>G 3481C>G), CGA388AAG (3488C>A 3489G>A 3490A>G), TAT389TAC (3493T>C), CCG390CCA (3496G>A), CTA391GTT (3497C>G 3499A>T), CCC392CCT (3502C>T), AAC393TTG (3503A>T 3504A>T 3505C>G), ATC394GTG (3506A>G 3508C>G), GAA395CAG (3509G>C 3511A>G), GAA396GAT (3514A>T), GCA397CTG (3515G>C 3516C>T 3517A>G), CAA398ATG (3518C>A 3519A>T 3520A>G), AGA400AGG (3526A>G), TTA401TTG (3529A>G), ACC402AGC (3531C>G), GGA403AAA (3533G>A 3534G>A), TCA404GCG (3536T>G 3538A>G), GAC405TGC (3539G>T 3540A>G), TAC407TTC (3546A>T), ACG408ACT (3550G>T), ATT410CTA (3554A>C 3556T>A), GAC411GAT (3559C>T), CTA412CTT (3562A>T), CGA413AGG (3563C>A 3565A>G), GAC414TCA (3566G>T 3567A>C 3568C>A), GCC415GGC (3570C>G), TTC416TAT (3573T>A 3574C>T), TAT417TGG (3576A>G 3577T>G), GCT418CAG (3578G>C 3579C>A 3580T>G), ATC419GTT (3581A>G 3583C>T), CGA420AGG (3584C>A 3586A>G), ATG421ATA (3589G>A), GGA424GGG (3598A>G), GAA425GAT (3601A>T), GAA426GAG (3604A>G), TGG427CCT (3605T>C 3606G>C 3607G>T), AAA428AAG (3610A>G), ACC429ACT (3613C>T), GCT430ACT (3614G>A), TTC431TGT (3618T>G 3619C>T), AGG432GTA (3620A>G 3621G>T 3622G>A), ACA433ACT (3625A>T), AGA434AGG (3628A>G), TAC435TAT (3631C>T), GGA436GGG (3634A>G), CTC437TCA (3635C>T 3636T>C 3637C>A), TAC438TAT (3640C>T), TTG441CTT (3647T>C 3649G>T), GTC442GTA (3652C>A), CCA444CCG (3658A>G), ATG445TTT (3659A>T 3661G>T), GGA446GGG (3664A>G), ACC448ACT (3670C>T), AAC449AAT (3673C>T), GCA450GCC (3676A>C), CCC451CCG (3679C>G), GCA452GCT (3682A>T), TCC453ACC (3683T>A), TGC454TTT (3687G>T 3688C>T), CAG455TGC (3689C>T 3690A>G 3691G>C), GAC456AAT (3692G>A 3694C>T), CTT457TTA (3695C>T 3697T>A), GTC458ATG (3698G>A 3700C>G), AAC459AAT (3703C>T), GAA460GAC (3706A>C), ACA461GTA (3707A>G 3708C>T), CTT462CTG (3712T>G), AGA463TTT (3713A>T 3714G>T 3715A>T), CTA465TAT (3719C>T 3720T>A 3721A>T), CTC466CTT (3724C>T), GAC467GAT (3727C>T), GTG468GAT (3729T>A 3730G>T), TGC469TTT (3732G>T 3733C>T), GTC470GTA (3736C>A), GTT471GTA (3739T>A), GCT472GTC (3741C>T 3742T>C), TAC473TAT (3745C>T), ATG474CTA (3746A>C 3748G>A), GAC475GAT (3751C>T), ATA477ATT (3757A>T), CTG478GTC (3758C>G 3760G>C), GTC479ATA (3761G>A 3763C>A), TAC480TAT (3766C>T), ACA481AGT (3768C>G 3769A>T), AAA482CAA (3770A>C), GGA483del (3773_3775delGGA), TCC484ACG (3776T>A 3778C>G), CTC485TTG (3779C>T 3781C>G), CAG486GAT (3782C>G 3784G>T), ACC489GTT (3791A>G 3792C>T 3793C>T), AAG490AGC (3795A>G 3796G>C), CAA491CAT (3799A>T), GTT492TTG (3800G>T 3802T>G), CAA493AGT (3803C>A 3804A>G 3805A>T), GAT494TTG (3806G>T 3807A>T 3808T>G), GTG495GTA (3811G>A), TTC496TTA (3814C>A), GAA497TCT (3815G>T 3816A>C 3817A>T), CGA498CGT (3820A>T), CTC499TTA (3821C>T 3823C>A), ACG500AGG (3825C>G), AAG501CAA (3827A>C 3829G>A), TCC502TAC (3831C>A), GGA503ACT (3833G>A 3834G>C 3835A>T), TTC504CTC (3836T>C), AAG505TAT (3839A>T 3841G>T), ACA506GTT (3842A>G 3843C>T 3844A>T), GCA507AAA (3845G>A 3846C>A), CCC508ATG (3848C>A 3849C>T 3850C>G), AAA510AAG (3856A>G), TGC511TGT (3859C>T), GAA512GAG (3862A>G), TTC513TTT (3865C>T), CAC514GCC (3866C>G 3867A>C), AAG515CAA (3869A>C 3871G>A), GAA517GAG (3877A>G), GTC518ATA (3878G>A 3880C>A), TTT520TTC (3886T>C), TTA521CTA (3887T>C), GGC522GGG (3892C>G), TTT523CAT (3893T>C 3894T>A), ATC524CTA (3896A>C 3898C>A), ATC525GTT (3899A>G 3901C>T), ACA527GAA (3905A>G 3906C>A), ACA528AAC (3909C>A 3910A>C), GGG529CAA (3911G>C 3912G>A 3913G>A), ATA530GTA (3914A>G), ACG531AGG (3918C>G), ATC532ATG (3922C>G), GAC533GAT (3925C>T), GCA535AAG (3929G>A 3930C>A 3931A>G), AAG536AAA (3934G>A), ACA537GTG (3935A>G 3936C>T 3937A>G), CAG538CAA (3940G>A), TCA539GCC (3941T>G 3943A>C), ATC540ATT (3946C>T), AGA541GTT (3947A>G 3948G>T 3949A>T), GAA542GAT (3952A>T), CCA544CAA (3957C>A), GAA545GCA (3960A>C), CCG546CCC (3964G>C), AAG547AAA (3967G>A), ACA548ACT (3970A>T), GTC549GTG (3973C>G), GTA552CTG (3980G>C 3982A>G), CAG553AGG (3983C>A 3984A>G), TCA554TCT (3988A>T), CTT556TTG (3992C>T 3994T>G), GGA557GGG (3997A>G), CTC558CTA (4000C>A), GCC559GCT (4003C>T), AAC562TAT (4010A>T 4012C>T), AAG567GCT (4025A>G 4026A>C 4027G>T), GAC568GGA (4029A>G 4030C>A), TAT569TAC (4033T>C), ACA572AAA (4041C>A), GCA574GCT (4048A>T), CCA575GCA (4049C>G), ATG576CTG (4052A>C), ACG577ACT (4057G>T), ATG578GAT (4058A>G 4059T>A 4060G>T), CTT579TTG (4061C>T 4063T>G), ACA580CTG (4064A>C 4065C>T 4066A>G), AGA581AAG (4068G>A 4069A>G), AAA582AAG (4072A>G), GAC583GAA (4075C>A), AAC585AAG (4081C>G), AAA587GTA (4085A>G 4086A>T), GGA589TCG (4091G>T 4092G>C 4093A>G), AAA590GTA (4094A>G 4095A>T), GAA591AGA (4097G>A 4098A>G), CAG592TGC (4100C>T 4101A>G 4102G>C), ACC593GAA (4103A>G 4104C>A 4105C>A), GCG595GCT (4111G>T), TTC596TTT (4114C>T), AAA597CAG (4115A>C 4117A>G), AGA598AGT (4120A>T), CTC599TTG (4121C>T 4123C>G), CAG602GCT (4130C>G 4131A>C 4132G>T), TGC603ATT (4133T>A 4134G>T 4135C>T), GCT604GCA (4138T>A), GCC606GAA (4143C>A 4144C>A), ACG608ATA (4149C>T 4150G>A), CTT609CTG (4153T>G), CGA610AAG (4154C>A 4155G>A 4156A>G), CTA611CTG (4159A>G), CTA611_TTC612insCCAAAT (4159_4160insCCAAAT), TTC612TTT (4162C>T), GAT613GAG (4165T>G), GGT614CTA (4166G>C 4167G>T 4168T>A), AGC615CCT (4169A>C 4170G>C 4171C>T), AAG616TTT (4172A>T 4173A>T 4174G>T), GAA617GAG (4177A>G), GTC618GTA (4180C>A), ATC620_GAG621del (4184_4189delATCGAG), ACC622ACT (4192C>T), GAC623GAT (4195C>T), TCT625TCG (4201T>G), GAT626GAC (4204T>C), ATG627AAA (4206T>A 4207G>A), GCA628GCT (4210A>T), ATA629GTG (4211A>G 4213A>G), GGC630GGA (4216C>A), GCA631GGC (4218C>G 4219A>C), TGT632GTA (4220T>G 4221G>T 4222T>A), CTA633TTG (4223C>T 4225A>G), ACA634GTG (4226A>G 4227C>T 4228A>G), CAG635CAA (4231G>A), ACA636AAT (4233C>A 4234A>T), CAC637GAC (4235C>G), GAT638GAA (4240T>A), GGG639GGT (4243G>T), AAA640_AGA641del (4244_4249delAAAAGA), CAC642CAT (4252C>T), CCA643CCG (4255A>G), TAT646TTT (4263A>T), TAT647GAG (4265T>G 4267T>G), TCC648AGC (4268T>A 4269C>G), CGG649AGG (4271C>A), ATG651TTG (4277A>T), ACC652AAT (4281C>A 4282C>T), ACA653CCG (4283A>C 4285A>G), GCG654ACG (4286G>A), CAG656CAA (4294G>A), AAC657AGA (4296A>G 4297C>A), TAC658TAT (4300C>T), GAC659TCT (4301G>T 4302A>C 4303C>T), ATC660GCA (4304A>G 4305T>C 4306C>A), GAC662GAA (4312C>A), CTT665ATG (4319C>A 4321T>G), CTA666GTG (4322C>G 4324A>G), GCC667GCA (4327C>A), ATT668GTG (4328A>G 4330T>G), GTT669CTC (4331G>C 4333T>C), GCC670CAT (4334G>C 4335C>A 4336C>T), GCC671TGT (4337G>T 4338C>G 4339C>T), ATG672CTT (4340A>C 4342G>T), CAT674ACA (4346C>A 4347A>C 4348T>A), AGA676AGG (4354A>G), TAC678TAT (4360C>T), GTC679CTC (4361G>C), GAG680TTA (4364G>T 4365A>T 4366G>A), GGC681GGA (4369C>A), CCA682ACA (4370C>A), CCG683del (4373_4375delCCG), TTA685TTT (4381A>T), ACG686GTA (4382A>G 4383C>T 4384G>A), ATT687GTA (4385A>G 4387T>A), CTT688AAG (4388C>A 4389T>A 4390T>G), TCA689ACG (4391T>A 4393A>G), CAC691AAT (4397C>A 4399C>T), AAG692GTG (4400A>G 4401A>T), AAT693GCT (4403A>G 4404A>C), CTC694AAT (4406C>A 4407T>A 4408C>T), ACG695ACT (4411G>T), ACG698AAA (4419C>A 4420G>A), ACG699ACT (4423G>T), ACG700CAA (4424A>C 4425C>A 4426G>A), AAG701CAG (4427A>C), GAA702AAA (4430G>A), CTT703TTG (4433C>T 4435T>G), ACC704AGT (4437C>G 4438C>T), CGA705TTG (4439C>T 4440G>T 4441A>G), AGA706AAG (4443G>A 4444A>G), CAA707TAG (4445C>T 4447A>G), GCC708GCG (4450C>G), CGC709AGA (4451C>A 4453C>A), TCG711CAA (4457T>C 4458C>A 4459G>A), CTG713ATA (4463C>A 4465G>A), CTT714CTA (4468T>A), GGG715GCA (4470G>C 4471G>A), CAG716GAA (4472C>G 4474G>A), TAC717TAT (4477C>T), AAG718GAC (4478A>G 4480G>C), TTC719TTT (4483C>T), GAA720GTG (4485A>T 4486A>G), ATC721TGG (4487A>T 4488T>G 4489C>G), AAA722ATT (4491A>T 4492A>T), TAC723CAT (4493T>C 4495C>T), ACT724AAA (4497C>A 4498T>A), AGC824.TC (4797G>T), TTC825TAC (4800T>A), TCA826TAT (4803C>A 4804A>T), TTC827TGG (4806T>G 4807C>G), CAG829CGC (4812A>G 4813G>C), AGA831CGG (4817A>C 4819A>G), CTT832GAG (4820C>G 4821T>A 4822T>G), AAG833GAT (4823A>G 4825G>T), GTG834ATA (4826G>A 4828G>A), CTA835GAA (4829C>G 4830T>A), ATC838GTG (4838A>G 4840C>G), AAG839CAA (4841A>C 4843G>A), AAA840ACT (4845A>C 4846A>T), TGC841TGT (4849C>T), GTA842CTT (4850G>C 4852A>T), CAC843GTG (4853C>G 4854A>T 4855C>G), AAC847GAC (4865A>G), AAA848AAG (4870A>G), GCT849GTA (4872C>T 4873T>A), GCA850GAG (4875C>A 4876A>G), CGG851CAG (4878G>A), CAC852CGA (4881A>G 4882C>A), GCG853CAA (4883G>C 4884C>A 4885G>A), AAA854CCC (4886A>C 4887A>C 4888A>C), TAC855GGA (4889T>G 4890A>G 4891C>A), GGT856GGA (4894T>A), CAC857CTT (4896A>T 4897C>T), CTA858TTA (4898C>T), CAG859GAG (4901C>G), TTC860CCA (4904T>C 4905T>C 4906C>A), AGG861TTA (4907A>T 4908G>T 4909G>A), ACA862CAT (4910A>C 4911C>A 4912A>T), CCA863GTA (4913C>G 4914C>T), CCA864GCC (4916C>G 4918A>C), ACG865GAG (4919A>G 4920C>A), AAA866CGT (4922A>C 4923A>G 4924A>T), GAC869GAG (4933C>G), GAG870AGT (4934G>A 4935A>G 4936G>T), GTT871ATC (4937G>A 4939T>C), ATT876ATA (4954T>A), ACG877ACC (4957G>C), AAA878TGT (4958A>T 4959A>G 4960A>T), CTC879TTG (4961C>T 4963C>G), CCG880CCA (4966G>A), AGG881AAG (4968G>A), TCA882TCC (4972A>C), AAG883GAT (4973A>G 4975G>T), CTA895TGG (5009C>T 5010T>G 5011A>G), GTC896GTG (5014C>G), ATG897ATT (5017G>T), GTC898GTG (5020C>G), AGA900AGG (5026A>G), CTC901TTG (5027C>T 5029C>G), ACA902ACT (5032A>T), AAA903AAG (5035A>G), TAT904TC. (5037A>C), CAC961TTG (5207C>T 5208A>T 5209C>G), AAG962AAT (5212G>T), TCA964AGT (5216T>A 5217C>G 5218A>T), TAC967TTT (5226A>T 5227C>T), CCA969CCT (5233A>T), GAG970CAG (5234G>C), ACG971ACC (5239G>C), GGG973GGC (5245G>C), CAA974CAG (5248A>G), ACG975TCA (5249A>T 5251G>A), GAA976GAG (5254A>G), AGA977AGG (5257A>G), ACG978ACT (5260G>T), AAC979ATT (5262A>T 5263C>T), ACA981ACT (5269A>T), CTC982T.. (5270C>T), CAA984..C (5278A>C), CTA986CTT (5284A>T), CGG987AGG (5285C>A), TAC989TTC (5292A>T), ATC990GTG (5294A>G 5296C>G), AAC991AGC (5298A>G), TAC992TCT (5301A>C 5302C>T), GCA993CAT (5303G>C 5304C>A 5305A>T), GAC995AAA (5309G>A 5311C>A), AAC996GAT (5312A>G 5314C>T), GTT998GCT (5319T>C), TCA999AAG (5321T>A 5322C>A 5323A>G), TTA1000TTG (5326A>G), CTG1001CTA (5329G>A), CCA1002GAT (5330C>G 5331C>A 5332A>T), ATG1003GTG (5333A>G), GCG1004GCC (5338G>C), CAG1005CAA (5341G>A), ATC1006TTC (5342A>T), GCA1007TCA (5345G>T), CTG1008TAT (5348C>T 5349T>A 5350G>T), AAC1009AAT (5353C>T), AAC1010CTA (5354A>C 5355A>T 5356C>A), CAC1011CAG (5359C>G), AAA1012CGG (5360A>C 5361A>G 5362A>G), TCA1013AGT (5363T>A 5364C>G 5365A>T), GAG1014GAA (5368G>A), ACA1015GCC (5369A>G 5371A>C), TCG1017GGC (5375T>G 5376C>G 5377G>C), ACG1018AAG (5379C>A), ATG1022GAG (5390A>G 5391T>A), CGA1023CTA (5394G>T), ACT1024GCC (5396A>G 5398T>C), TTG1025GCA (5399T>G 5400T>C 5401G>A), GCA1026GGC (5403C>G 5404A>C), AGG1027CA. (5405A>C 5406G>A)

*: Inserts / Deletes / Misaligned / Frameshifts

#### Analysis details

This analysis was performed with panviral2.64

### NGS Details (UN9): Lausannevirus

#### Assembly

| | |
|---|---|
| **Coverage Length** | 151 (1 contig(s)) |
| **Depth Of Coverage** | 394.0 |
| **Number Of Reads** | 604 |
| **Reads Per Million** | 13.65 rpm (after QC) |
| **Ambiguities** | 0 |
| **Assembly Method** | de novo + reference guided assembly |
| **Consensus Caller** | Bcf Tools |

#### Coverage Map

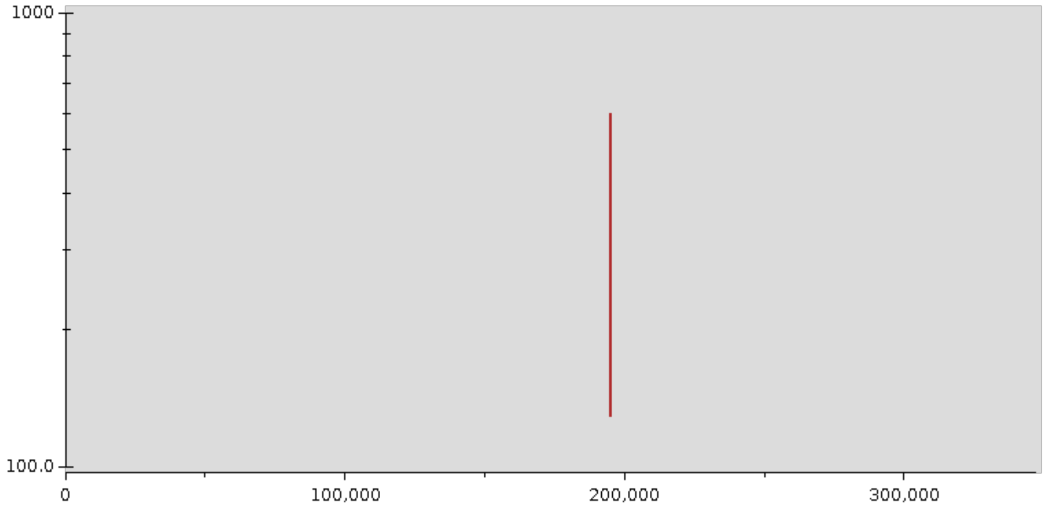

#### Assignment

| | |
|---|---|
| **Type** | Lausannevirus (Taxonomy ID: 999883) |
| **Reference Genome** | NC_015326.1 |
| **NT Identity (%)** | 81.457 |
| **AA Identity (%)** | 92.1569 |
| **Number Of Stop Codons** | 1 |
| **Number Of CDS** | 444 |

#### Alignment

| | |
|---|---|
| **Alignment Score** | 190.0 (NT) + 308.0 (AA) = 498.0 |
| **Concordance (%)** | 78.3019 |
| **Alignment Method** | Local, heuristic, nucleotide (BLASTN) |

#### Genome Region

Sequence starts at position 194933 and ends at position 195083 relative to NC_015326.1 reference sequence.

#### Alignment Detailed Statistics

| | Begin | End | Coverage | Score | Concordance | Matches | Identities | I/D/M/F* | Stop Codons |
|---|---|---|---|---|---|---|---|---|---|
| **NT** | **194933** | **195083** | **0.1%** | **190** | **62.9%** | **151 (100%)** | **123 (81.5%)** | **0/0** | |

Mutations: 194943A>G, 194946G>C, 194949C>T, 194955T>C, 194967A>G, 194970G>A, 194979T>C, 194982C>T, 194990C>A, 194991G>C, 194993G>A, 195003T>A, 195006G>A, 195009G>A, 195015T>C, 195017G>T, 195021T>C, 195030A>G, 195039T>C, 195042T>C, 195043T>G, 195048T>C, 195054T>C, 195057T>G, 195058T>G, 195059G>T, 195063G>T, 195069C>T

*: Inserts / Deletes / Misaligned / Frameshifts

#### Analysis details

This analysis was performed with panviral2.64

### NGS Details (UN9): Errantivirus

#### Assembly

| | |
|---|---|
| **Coverage Length** | 572 (1 contig(s)) |
| **Depth Of Coverage** | 65.5 |
| **Number Of Reads** | 402 |
| **Reads Per Million** | 9.09 rpm (after QC) |
| **Ambiguities** | 0 |
| **Assembly Method** | de novo + reference guided assembly |
| **Consensus Caller** | Bcf Tools |

#### Coverage Map

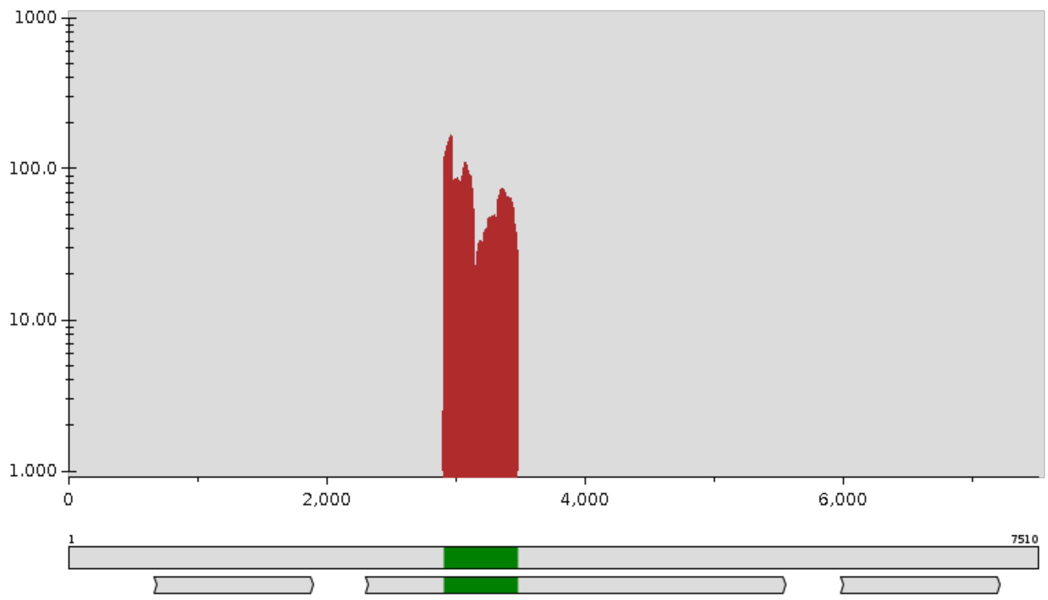

#### Assignment

| | |
|---|---|
| **Type** | Errantivirus (Taxonomy ID: 186666) |
| **Reference Genome** | NC_038512.1 |
| **NT Identity (%)** | 53.5587 |
| **AA Identity (%)** | 43.8503 |
| **Number Of Stop Codons** | 0 |
| **Number Of CDS** | 3 |

#### Alignment

| | |
|---|---|
| **Alignment Score** | 61.0 (NT) + 478.0 (AA) = 539.0 |
| **Concordance (%)** | 23.8813 |

| Alignment Method | Global, seeded, nucleotide + amino acids (AGA) |
|---|---|

#### Genome Region

Sequence starts at position 2909 and ends at position 3480 relative to NC_038512.1 reference sequence.

#### Alignment Detailed Statistics

| | Begin | End | Coverage | Score | Concordance | Matches | Identities | I/D/M/F* | Stop Codons |
|---|---|---|---|---|---|---|---|---|---|
| **NT** | **2909** | **3480** | **7.6%** | **61** | **5.5%** | **562 (98.3%)** | **301 (52.6%)** | **0/10** | |

Mutations:

2919A>T, 2920G>C, 2924A>G, 2926C>G, 2927G>A, 2928A>G, 2929T>G, 2932C>T, 2934A>G, 2935A>G, 2937A>T, 2941G>A, 2944A>C, 2949A>C, 2950C>T, 2953A>T, 2954A>G, 2955G>A, 2959C>G, 2960G>C, 2963C>T, 2968C>T, 2969A>G, 2972T>G, 2975G>A, 2976G>A, 2978A>G, 2979A>G, 2980G>C, 2981T>G, 2982G>C, 2983C>T, 2984C>G, 2985A>T, 2986A>T, 2989C>T, 2993A>T, 2995C>A, 2997C>A, 2998C>G, 2999T>C, 3001A>T, 3008G>T, 3009C>T, 3011A>T, 3012G>C, 3016G>A, 3018T>A, 3019T>C, 3020T>G, 3025G>A, 3026G>A, 3028G>T, 3029G>C, 3030A>G, 3032A>G, 3034G>T, 3035G>A, 3037C>G, 3038C>G, 3039C>T, 3040T>G, 3041C>G, 3042A>C, 3050T>C, 3052G>C, 3058C>T, 3061G>T, 3066A>G, 3067C>G, 3068G>A, 3069T>C, 3071G>C, 3073A>C, 3074C>G, 3076C>T, 3079G>T, 3084T>A, 3085T>C, 3091C>T, 3092C>T, 3094T>A, 3095C>G, 3096G>T, 3097A>C, 3104A>T, 3106G>T, 3109A>G, 3112A>G, 3113A>T, 3114A>C, 3115A>T, 3119T>G, 3121A>G, 3122C>T, 3124A>G, 3127T>C, 3130T>A, 3135A>G, 3137_3146delAGAGTTATGG, 3150A>G, 3151T>G, 3154C>T, 3155C>T, 3157A>C, 3159G>A, 3160A>G, 3161G>T, 3162G>C, 3164C>T, 3168A>T, 3169A>C, 3170A>G, 3172T>C, 3174A>T, 3175C>G, 3176A>T, 3178C>T, 3179T>G, 3180G>T, 3181T>G, 3182C>G, 3184C>A, 3189A>T, 3191C>A, 3196C>T, 3200A>G, 3202T>A, 3203A>T, 3205T>G, 3206G>A, 3208C>A, 3212A>T, 3213G>C, 3215A>C, 3216C>G, 3217T>A, 3219C>G, 3220C>T, 3221C>A, 3222T>A, 3223A>G, 3224C>G, 3232C>T, 3233C>G, 3234T>A, 3238G>A, 3239A>C, 3241C>T, 3242C>T, 3245G>A, 3246A>G, 3247A>G, 3248C>A, 3249G>T, 3253T>A, 3254T>C, 3256C>G, 3259A>G, 3261G>C, 3263C>T, 3265T>A, 3268A>G, 3271A>T, 3272A>G, 3273G>A, 3274T>G, 3277C>G, 3280C>G, 3281A>T, 3283A>T, 3284A>G, 3285T>C, 3286T>C, 3287C>A, 3289A>G, 3290A>T, 3292G>T, 3295C>T, 3300C>G, 3307C>T, 3309T>G, 3311A>C, 3312A>T, 3314C>A, 3315T>A, 3316C>T, 3320A>G, 3321C>T, 3322T>G, 3325T>C, 3327A>T, 3328T>C, 3334T>G, 3337C>T, 3340C>A, 3341A>G, 3343A>T, 3344A>T, 3345G>C, 3346C>G, 3347A>G, 3349G>T, 3352C>T, 3355T>A, 3362C>G, 3363C>T, 3364T>G, 3365A>G, 3370T>G, 3371G>A, 3373T>G, 3376G>A, 3378T>C, 3380T>G, 3381C>A, 3382C>G, 3385T>G, 3386A>G, 3388T>C, 3389C>A, 3390A>G, 3391A>G, 3394A>T, 3396A>G, 3397T>G, 3399T>C, 3400G>A, 3402T>G, 3403T>G, 3406A>T, 3407A>T, 3408A>T, 3410A>T, 3412C>T, 3415T>G, 3417A>C, 3418G>T, 3421A>T, 3424A>T, 3426A>G, 3428C>A, 3429A>G, 3430A>C, 3436A>G, 3439C>A, 3440C>T, 3442T>G, 3443C>G, 3444T>C, 3445C>A, 3451T>C, 3454C>T, 3455C>A, 3457A>G, 3459A>G, 3464A>G, 3466T>A, 3467C>G, 3468C>A, 3471A>G, 3476G>T

##### CDS

| | Begin | End | Coverage | Score | Concordance | Matches | Identities | I/D/M/F* | Stop Codons |
|---|---|---|---|---|---|---|---|---|---|
| D1R33_gp2 | 202 | 392 | 17.5% | 478 | 37.1% | 187 (98.4%) | 82 (43.2%) | 0/3/1/1 | 0 |

Protein mutations:

K205I (2919A>T 2920G>C), I207V (2924A>G 2926C>G), D208R (2927G>A 2928A>G 2929T>G), K210R (2934A>G 2935A>G), Y211F (2937A>T), N215T (2949A>C 2950C>T), S217D (2954A>G 2955G>A), D218E (2959C>G), V219L (2960G>C), L220F (2963C>T), K222E (2969A>G), L223V (2972T>G), G224N (2975G>A 2976G>A), K225G (2978A>G 2979A>G 2980G>C), C226A (2981T>G 2982G>C 2983C>T), Q227V (2984C>G 2985A>T 2986A>T), T230S (2993A>T 2995C>A), T231K (2997C>A 2998C>G), A235L (3008G>T 3009C>T), F238Y (3018T>A 3019T>C), Y239D (3020T>G), V241I (3026G>A 3028G>T), E242R (3029G>C 3030A>G), M243V (3032A>G 3034G>T), D244K (3035G>A 3037C>G), P245V (3038C>G 3039C>T 3040T>G), Q246A (3041C>G 3042A>C), S249P (3050T>C 3052G>C), N254R (3066A>G 3067C>G), V255T (3068G>A 3069T>C), E256H (3071G>C 3073A>C), H257D (3074C>G 3076C>T), F260Y (3084T>A 3085T>C), R264V (3095C>G 3096G>T 3097A>C), M267F (3104A>T 3106G>T), K270S (3113A>T 3114A>C 3115A>T), S272A (3119T>G 3121A>G), P273S (3122C>T 3124A>G), Q277R (3135A>G), R278_M280del (3137_3146delAGAGTTATGG), N282R (3150A>G 3151T>G), L284F (3155C>T 3157A>C), R285K (3159G>A 3160A>G), G286S (3161G>T 3162G>C), L287F (3164C>T), Q288L (3168A>T 3169A>C), N289D (3170A>G 3172T>C), N290M (3174A>T 3175C>G), I291F (3176A>T 3178C>T), C292V (3179T>G 3180G>T 3181T>G), L293V (3182C>G 3184C>A), Y295F (3189A>T), L296I (3191C>A), I299V (3200A>G 3202T>A), I300L (3203A>T 3205T>G), V301I (3206G>A 3208C>A), T304R (3215A>C 3216C>G 3217T>A), S305C (3219C>G 3220C>T), L306K (3221C>A 3222T>A 3223A>G), Q307E (3224C>G), L310E (3233C>G 3234T>A), N312H (3239A>C 3241C>T), E314R (3245G>A 3246A>G 3247A>G), R315I (3248C>A 3249G>T), F317L (3254T>C 3256C>G), R319T (3261G>C), E322D (3271A>T), S323E (3272A>G 3273G>A 3274T>G), N324K (3277C>G), F325L (3280C>G), K326Y (3281A>T 3283A>T), I327A (3284A>G 3285T>C 3286T>C), Q328K (3287C>A 3289A>G), M329F (3290A>T 3292G>T), S332C (3300C>G), L335W (3309T>G), K336L (3311A>C 3312A>T), L337N (3314C>A 3315T>A 3316C>T), T339V (3320A>G 3321C>T 3322T>G), Y341F (3327A>T 3328T>C), I346V (3341A>G 3343A>T), R348G (3347A>G 3349G>T), P353V (3362C>G 3363C>T 3364T>G), N354D (3365A>G), D356K (3371G>A 3373T>G), I358T (3378T>C), S359E (3380T>G 3381C>A 3382C>G), I361V (3386A>G 3388T>C), Q362R (3389C>A 3390A>G 3391A>G), K363N (3394A>T), Y364W (3396A>G 3397T>G), L365P (3399T>C 3400G>A), I366R (3402T>G 3403T>G), K368L (3407A>T 3408A>T), T369S (3410A>T 3412C>T), K371T (3417A>C 3418G>T), E372D (3421A>T), K374R (3426A>G), Q375S (3428C>A 3429A>G 3430A>C), L380A (3443C>G 3444T>C 3445C>A), K385R (3459A>G), I387V (3464A>G 3466T>A), P388E (3467C>G 3468C>A), D389G (3471A>G), A391S (3476G>T)

Codon mutations:

AAG205ATC (2919A>T 2920G>C), ATC207GTG (2924A>G 2926C>G), GAT208AGG (2927G>A 2928A>G 2929T>G), GAC209GAT (2932C>T), AAA210AGG (2934A>G 2935A>G), TAC211TTC (2937A>T), CCG212CCA (2941G>A), ATA213ATC (2944A>C), AAC215ACT (2949A>C 2950C>T), ATA216ATT (2953A>T), AGT217GAT (2954A>G 2955G>A), GAC218GAG (2959C>G), GTA219CTA (2960G>C), CTT220TTT (2963C>T), GAC221GAT (2968C>T), AAG222GAG (2969A>G), TTA223GTA (2972T>G), GGT224AAT (2975G>A 2976G>A), AAG225GGC (2978A>G 2979A>G 2980G>C), TGC226GCT (2981T>G 2982G>C 2983C>T), CAA227GTT (2984C>G 2985A>T 2986A>T), TAC228TAT (2989C>T), ACC230TCA (2993A>T 2995C>A), ACC231AAG (2997C>A 2998C>G), TTA232CTT (2999T>C 3001A>T), GCA235TTA (3008G>T 3009C>T), AGT236TCT (3011A>T 3012G>C), GGG237GGA (3016G>A), TTT238TAC (3018T>A 3019T>C), TAT239GAT (3020T>G), CAG240CAA (3025G>A), GTG241ATT (3026G>A 3028G>T), GAG242CGG (3029G>C 3030A>G), ATG243GTT (3032A>G 3034G>T), GAC244AAG (3035G>A 3037C>G), CCT245GTG (3038C>G 3039C>T 3040T>G), CAA246GCA (3041C>G 3042A>C), TCG249CCC (3050T>C 3052G>C), ACC251ACT (3058C>T), GCG252GCT (3061G>T), AAC254AGG (3066A>G 3067C>G), GTA255ACA (3068G>A 3069T>C), GAA256CAC (3071G>C 3073A>C), CAC257GAT (3074C>G 3076C>T), GGG258GGT (3079G>T), TTT260TAC (3084T>A 3085T>C), TTC262TTT (3091C>T), CTT263TTA (3092C>T 3094T>A), CGA264GTC (3095C>G 3096G>T 3097A>C), ATG267TTT (3104A>T 3106G>T), GGA268GGG (3109A>G), TTA269TTG (3112A>G), AAA270TCT (3113A>T 3114A>C 3115A>T), TCA272GCG (3119T>G 3121A>G), CCA273TCG (3122C>T 3124A>G), TCT274TCC (3127T>C), ACT275ACA (3130T>A), CAA277CGA (3135A>G), AGA278_ATG280del (3137_3146delAGAGTTATGG), GAC281-AC (3137_3146delAGAGTTATGG), AAT282AGG (3150A>G 3151T>G), GTC283GTT (3154C>T), CTA284TTC (3155C>T 3157A>C), AGA285AAG (3159G>A 3160A>G), GGT286TCT (3161G>T 3162G>C), CTC287TTC (3164C>T), CAA288CTC (3168A>T 3169A>C), AAT289GAC (3170A>G 3172T>C), AAC290ATG (3174A>T 3175C>G), ATC291TTT (3176A>T 3178C>T), TGT292GTG (3179T>G 3180G>T 3181T>G), CTC293GTA (3182C>G 3184C>A), TAC295TTC (3189A>T), CTT296ATT (3191C>A), GAC297GAT (3196C>T), ATT299GTA (3200A>G 3202T>A), ATT300TTG (3203A>T 3205T>G), GTC301ATA (3206G>A 3208C>A), AGT303TCT (3212A>T 3213G>C), ACT304CGA (3215A>C 3216C>G 3217T>A), TCC305TGT (3219C>G 3220C>T), CTA306AAG (3221C>A 3222T>A 3223A>G), CAG307GAG (3224C>G), CAC309CAT (3232C>T), CTG310GAG (3233C>G 3234T>A), GAG311GAA (3238G>A), AAC312CAT (3239A>C 3241C>T), CTG313TTG (3242C>T), GAA314AGG (3245G>A 3246A>G 3247A>G), CGA315ATA (3248C>A 3249G>T), GTT316GTA (3253T>A), TTC317CTG (3254T>C 3256C>G), CAA318CAG (3259A>G), AGA319ACA (3261G>C), CTT320TTA (3263C>T 3265T>A), AGA321AGG (3268A>G), GAA322GAT (3271A>T), AGT323GAG (3272A>G 3273G>A 3274T>G), AAC324AAG (3277C>G), TTC325TTG (3280C>G), AAA326TAT (3281A>T 3283A>T), ATT327GCC (3284A>G 3285T>C 3286T>C), CAA328AAG (3287C>A 3289A>G), ATG329TTT (3290A>T 3292G>T), GAC330GAT (3295C>T), TCC332TGC (3300C>G), TTC334TTT (3307C>T), TTG335TGG (3309T>G), AAG336CTG (3311A>C 3312A>T), CTC337AAT (3314C>A 3315T>A 3316C>T), ACT339GTG (3320A>G 3321C>T 3322T>G), GCT340GCC (3325T>C), TAT341TTC (3327A>T 3328T>C), GGT343GGG (3334T>G), CAC344CAT (3337C>T), ATC345ATA (3340C>A), ATA346GTT (3341A>G 3343A>T), AGC347TCG (3344A>T 3345G>C 3346C>G), AGG348GGT (3347A>G 3349G>T), GAC349GAT (3352C>T), GGT350GGA (3355T>A), CCT353GTG (3362C>G 3363C>T 3364T>G), AAC354GAC (3365A>G), CCT355CCG (3370T>G), GAT356AAG (3371G>A 3373T>G), AAG357AAA (3376G>A), ATT358ACT (3378T>C), TCC359GAG (3380T>G 3381C>A 3382C>G), GCT360GCG (3385T>G), ATT361GTC (3386A>G 3388T>C), CAA362AGG (3389C>A 3390A>G 3391A>G), AAA363AAT (3394A>T), TAT364TGG (3396A>G 3397T>G), CTG365CCA (3399T>C 3400G>A), ATT366AGG (3402T>G 3403T>G), CCA367CCT (3406A>T), AAG368TTG (3407A>T 3408A>T), ACC369TCT (3410A>T 3412C>T), CCT370CCG (3415T>G), AAG371ACT (3417A>C 3418G>T), GAA372GAT (3421A>T), ATA373ATT (3424A>T), AAA374AGA (3426A>G), CAA375AGC (3428C>A 3429A>G 3430A>C), TTA377TTG (3436A>G), GGC378GGA (3439C>A), CTT379TTG (3440C>T 3442T>G), CTC380GCA (3443C>G 3444T>C 3445C>A), TAT382TAC (3451T>C), TAC383TAT (3454C>T), CGA384AGG (3455C>A 3457A>G), AAA385AGA (3459A>G), ATT387GTA (3464A>G 3466T>A), CCA388GAA (3467C>G 3468C>A), GAT389GGT (3471A>G), GCA391TCA (3476G>T)

##### Proteins

| | Begin | End | Coverage | Score | Concordance | Matches | Identities | I/D/M/F* | Stop Codons |
|---|---|---|---|---|---|---|---|---|---|
| ORF B (YP_009507248.1) | 202 | 392 | 17.5% | 478 | 37.1% | 187 (98.4%) | 82 (43.2%) | 0/3/1/1 | 0 |

Protein mutations:

K205I (2919A>T 2920G>C), I207V (2924A>G 2926C>G), D208R (2927G>A 2928A>G 2929T>G), K210R (2934A>G 2935A>G), Y211F (2937A>T), N215T (2949A>C 2950C>T), S217D (2954A>G 2955G>A), D218E (2959C>G), V219L (2960G>C), L220F (2963C>T), K222E (2969A>G), L223V (2972T>G), G224N (2975G>A 2976G>A), K225G (2978A>G 2979A>G 2980G>C), C226A (2981T>G 2982G>C 2983C>T), Q227V (2984C>G 2985A>T 2986A>T), T230S (2993A>T 2995C>A), T231K (2997C>A 2998C>G), A235L (3008G>T 3009C>T), F238Y (3018T>A 3019T>C), Y239D (3020T>G), V241I (3026G>A 3028G>T), E242R (3029G>C 3030A>G), M243V (3032A>G 3034G>T), D244K (3035G>A 3037C>G), P245V (3038C>G 3039C>T 3040T>G), Q246A (3041C>G 3042A>C), S249P (3050T>C 3052G>C), N254R (3066A>G 3067C>G), V255T (3068G>A 3069T>C), E256H (3071G>C 3073A>C), H257D (3074C>G 3076C>T), F260Y (3084T>A 3085T>C), R264V (3095C>G 3096G>T 3097A>C), M267F (3104A>T 3106G>T), K270S (3113A>T 3114A>C 3115A>T), S272A (3119T>G 3121A>G), P273S (3122C>T 3124A>G), Q277R (3135A>G), R278_M280del (3137_3146delAGAGTTATGG), N282R (3150A>G 3151T>G), L284F (3155C>T 3157A>C), R285K (3159G>A 3160A>G), G286S (3161G>T 3162G>C), L287F (3164C>T), Q288L (3168A>T 3169A>C), N289D (3170A>G 3172T>C), N290M (3174A>T 3175C>G), I291F (3176A>T 3178C>T), C292V (3179T>G 3180G>T 3181T>G), L293V (3182C>G 3184C>A), Y295F (3189A>T), L296I (3191C>A), I299V (3200A>G 3202T>A), I300L (3203A>T 3205T>G), V301I (3206G>A 3208C>A), T304R (3215A>C 3216C>G 3217T>A), S305C (3219C>G 3220C>T), L306K (3221C>A 3222T>A 3223A>G), Q307E (3224C>G), L310E (3233C>G 3234T>A), N312H (3239A>C 3241C>T), E314R (3245G>A 3246A>G 3247A>G), R315I (3248C>A 3249G>T), F317L (3254T>C 3256C>G), R319T (3261G>C), E322D (3271A>T), S323E (3272A>G 3273G>A 3274T>G), N324K (3277C>G), F325L (3280C>G), K326Y (3281A>T 3283A>T), I327A (3284A>G 3285T>C 3286T>C), Q328K (3287C>A 3289A>G), M329F (3290A>T 3292G>T), S332C (3300C>G), L335W (3309T>G), K336L (3311A>C 3312A>T), L337N (3314C>A 3315T>A 3316C>T), T339V (3320A>G 3321C>T 3322T>G), Y341F (3327A>T 3328T>C), I346V (3341A>G 3343A>T), R348G (3347A>G 3349G>T), P353V (3362C>G 3363C>T 3364T>G), N354D (3365A>G), D356K (3371G>A 3373T>G), I358T (3378T>C), S359E (3380T>G 3381C>A 3382C>G), I361V (3386A>G 3388T>C), Q362R (3389C>A 3390A>G 3391A>G), K363N (3394A>T), Y364W (3396A>G 3397T>G), L365P (3399T>C 3400G>A), I366R (3402T>G 3403T>G), K368L (3407A>T 3408A>T), T369S (3410A>T 3412C>T), K371T (3417A>C 3418G>T), E372D (3421A>T), K374R (3426A>G), Q375S (3428C>A 3429A>G 3430A>C), L380A (3443C>G 3444T>C 3445C>A), K385R (3459A>G), I387V (3464A>G 3466T>A), P388E (3467C>G 3468C>A), D389G (3471A>G), A391S (3476G>T)

| | Begin | End | Coverage | Score | Concordance | Matches | Identities | I/D/M/F* | Stop Codons |
|---|---|---|---|---|---|---|---|---|---|
| **NT** | **2909** | **3480** | **7.6%** | **61** | **5.5%** | **562 (98.3%)** | **301 (52.6%)** | **0/10** | |
| Codon mutations: | AAG205ATC (2919A>T 2920G>C), ATC207GTG (2924A>G 2926C>G), GAT208AGG (2927G>A 2928A>G 2929T>G), GAC209GAT (2932C>T), AAA210AGG (2934A>G 2935A>G), TAC211TTC (2937A>T), CCG212CCA (2941G>A), ATA213ATC (2944A>C), AAC215ACT (2949A>C 2950C>T), ATA216ATT (2953A>T), AGT217GAT (2954A>G 2955G>A), GAC218GAG (2959C>G), GTA219CTA (2960G>C), CTT220TTT (2963C>T), GAC221GAT (2968C>T), AAG222GAG (2969A>G), TTA223GTA (2972T>G), GGT224AAT (2975G>A 2976G>A), AAG225GGC (2978A>G 2979A>G 2980G>C), TGC226GCT (2981T>G 2982G>C 2983C>T), CAA227GTT (2984C>G 2985A>T 2986A>T), TAC228TAT (2989C>T), ACC230TCA (2993A>T 2995C>A), ACC231AAG (2997C>A 2998C>G), TTA232CTT (2999T>C 3001A>T), GCA235TTA (3008G>T 3009C>T), AGT236TCT (3011A>T 3012G>C), GGG237GGA (3016G>A), TTT238TAC (3018T>A 3019T>C), TAT239GAT (3020T>G), CAG240CAA (3025G>A), GTG241ATT (3026G>A 3028G>T), GAG242CGG (3029G>C 3030A>G), ATG243GTT (3032A>G 3034G>T), GAC244AAG (3035G>A 3037C>G), CCT245GTG (3038C>G 3039C>T 3040T>G), CAA246GCA (3041C>G 3042A>C), TCG249CCC (3050T>C 3052G>C), ACC251ACT (3058C>T), GCG252GCT (3061G>T), AAC254AGG (3066A>G 3067C>G), GTA255ACA (3068G>A 3069T>C), GAA256CAC (3071G>C 3073A>C), CAC257GAT (3074C>G 3076C>T), GGG258GGT (3079G>T), TTT260TAC (3084T>A 3085T>C), TTC262TTT (3091C>T), CTT263TTA (3092C>T 3094T>A), CGA264GTC (3095C>G 3096G>T 3097A>C), ATG267TTT (3104A>T 3106G>T), GGA268GGG (3109A>G), TTA269TTG (3112A>G), AAA270TCT (3113A>T 3114A>C 3115A>T), TCA272GCG (3119T>G 3121A>G), CCA273TCG (3122C>T 3124A>G), TCT274TCC (3127T>C), ACT275ACA (3130T>A), CAA277CGA (3135A>G), AGA278_ATG280del (3137_3146delAGAGTTATGG), GAC281-AC (3137_3146delAGAGTTATGG), AAT282AGG (3150A>G 3151T>G), GTC283GTT (3154C>T), CTA284TTC (3155C>T 3157A>C), AGA285AAG (3159G>A 3160A>G), GGT286TCT (3161G>T 3162G>C), CTC287TTC (3164C>T), CAA288CTC (3168A>T 3169A>C), AAT289GAC (3170A>G 3172T>C), AAC290ATG (3174A>T 3175C>G), ATC291TTT (3176A>T 3178C>T), TGT292GTG (3179T>G 3180G>T 3181T>G), CTC293GTA (3182C>G 3184C>A), TAC295TTC (3189A>T), CTT296ATT (3191C>A), GAC297GAT (3196C>T), ATT299GTA (3200A>G 3202T>A), ATT300TTG (3203A>T 3205T>G), GTC301ATA (3206G>A 3208C>A), AGT303TCT (3212A>T 3213G>C), ACT304CGA (3215A>C 3216C>G 3217T>A), TCC305TGT (3219C>G 3220C>T), CTA306AAG (3221C>A 3222T>A 3223A>G), CAG307GAG (3224C>G), CAC309CAT (3232C>T), CTG310GAG (3233C>G 3234T>A), GAG311GAA (3238G>A), AAC312CAT (3239A>C 3241C>T), CTG313TTG (3242C>T), GAA314AGG (3245G>A 3246A>G 3247A>G), CGA315ATA (3248C>A 3249G>T), GTT316GTA (3253T>A), TTC317CTG (3254T>C 3256C>G), CAA318CAG (3259A>G), AGA319ACA (3261G>C), CTT320TTA (3263C>T 3265T>A), AGA321AGG (3268A>G), GAA322GAT (3271A>T), AGT323GAG (3272A>G 3273G>A 3274T>G), AAC324AAG (3277C>G), TTC325TTG (3280C>G), AAA326TAT (3281A>T 3283A>T), ATT327GCC (3284A>G 3285T>C 3286T>C), CAA328AAG (3287C>A 3289A>G), ATG329TTT (3290A>T 3292G>T), GAC330GAT (3295C>T), TCC332TGC (3300C>G), TTC334TTT (3307C>T), TTG335TGG (3309T>G), AAG336CTG (3311A>C 3312A>T), CTC337AAT (3314C>A 3315T>A 3316C>T), ACT339GTG (3320A>G 3321C>T 3322T>G), GCT340GCC (3325T>C), TAT341TTC (3327A>T 3328T>C), GGT343GGG (3334T>G), CAC344CAT (3337C>T), ATC345ATA (3340C>A), ATA346GTT (3341A>G 3343A>T), AGC347TCG (3344A>T 3345G>C 3346C>G), AGG348GGT (3347A>G 3349G>T), GAC349GAT (3352C>T), GGT350GGA (3355T>A), CCT353GTG (3362C>G 3363C>T 3364T>G), AAC354GAC (3365A>G), CCT355CCG (3370T>G), GAT356AAG (3371G>A 3373T>G), AAG357AAA (3376G>A), ATT358ACT (3378T>C), TCC359GAG (3380T>G 3381C>A 3382C>G), GCT360GCG (3385T>G), ATT361GTC (3386A>G 3388T>C), CAA362AGG (3389C>A 3390A>G 3391A>G), AAA363AAT (3394A>T), TAT364TGG (3396A>G 3397T>G), CTG365CCA (3399T>C 3400G>A), ATT366AGG (3402T>G 3403T>G), CCA367CCT (3406A>T), AAG368TTG (3407A>T 3408A>T), ACC369TCT (3410A>T 3412C>T), CCT370CCG (3415T>G), AAG371ACT (3417A>C 3418G>T), GAA372GAT (3421A>T), ATA373ATT (3424A>T), AAA374AGA (3426A>G), CAA375AGC (3428C>A 3429A>G 3430A>C), TTA377TTG (3436A>G), GGC378GGA (3439C>A), CTT379TTG (3440C>T 3442T>G), CTC380GCA (3443C>G 3444T>C 3445C>A), TAT382TAC (3451T>C), TAC383TAT (3454C>T), CGA384AGG (3455C>A 3457A>G), AAA385AGA (3459A>G), ATT387GTA (3464A>G 3466T>A), CCA388GAA (3467C>G 3468C>A), GAT389GGT (3471A>G), GCA391TCA (3476G>T) | | | | | | | | |

*: Inserts / Deletes / Misaligned / Frameshifts

#### Analysis details

This analysis was performed with panviral2.64

### NGS Details (UN9): Epiphyllum badnavirus 1

#### Assembly

| | |
|---|---|
| **Coverage Length** | 274 (1 contig(s)) |
| **Depth Of Coverage** | 124.8 |
| **Number Of Reads** | 324 |
| **Reads Per Million** | 7.32 rpm (after QC) |
| **Ambiguities** | 0 |
| **Assembly Method** | de novo + reference guided assembly |
| **Consensus Caller** | Bcf Tools |

#### Coverage Map

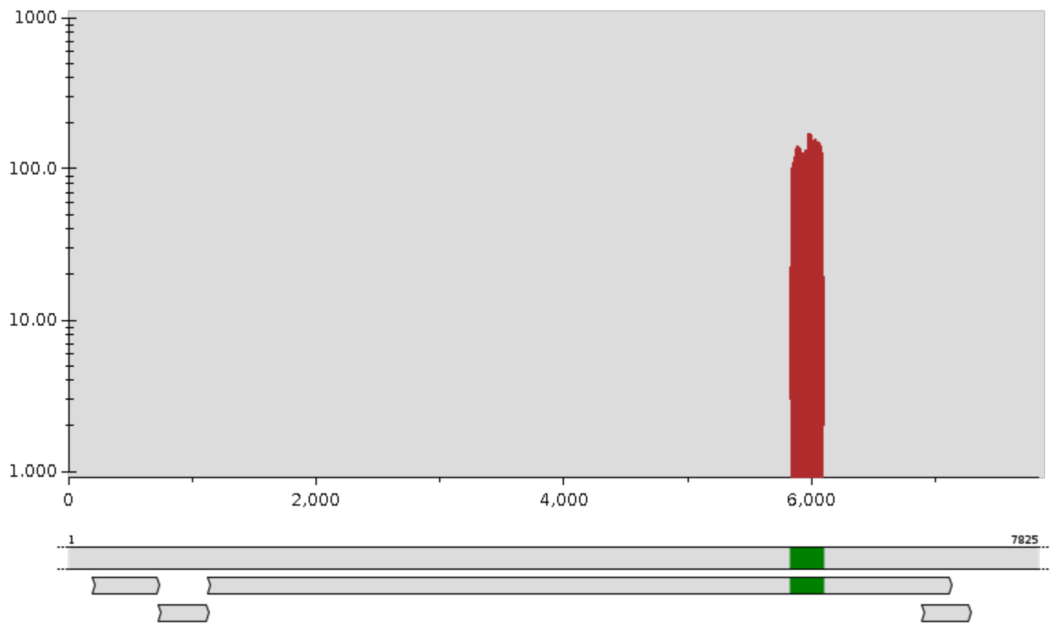

#### Assignment

| | |
|---|---|
| **Type** | Epiphyllum badnavirus 1 (Taxonomy ID: 2518008) |
| **Reference Genome** | NC_076247.1 |
| **NT Identity (%)** | 60.2888 |
| **AA Identity (%)** | 57.6087 |
| **Number Of Stop Codons** | 0 |
| **Number Of CDS** | 4 |

#### Alignment

| | |
|---|---|
| **Alignment Score** | 108.0 (NT) + 362.0 (AA) = 470.0 |
| **Concordance (%)** | 39.0041 |

| Alignment Method | Global, seeded, nucleotide + amino acids (AGA) |
|---|---|

#### Genome Region

Sequence starts at position 5828 and ends at position 6101 relative to NC_076247.1 reference sequence.

#### Alignment Detailed Statistics

| | Begin | End | Coverage | Score | Concordance | Matches | Identities | I/D/M/F* | Stop Codons |
|---|---|---|---|---|---|---|---|---|---|
| **NT** | **5828** | **6101** | **3.5%** | **108** | **19.7%** | **274 (98.9%)** | **167 (60.3%)** | **3/0** | |
| Mutations: | 5837G>C, 5841C>G, 5847C>T, 5851T>A, 5853C>T, 5856T>C, 5857C>T, 5861A>G, 5862G>A, 5863A>T, 5864G>C, 5865C>G, 5868A>T, 5870T>A, 5871C>T, 5874C>T, 5878A>T, 5880T>G, 5882T>G, 5883G>A, 5884A>G, 5886G>T, 5887G>C, 5888A>G, 5889C>G, 5892A>G, 5893G>T, 5894A>C, 5896T>G, 5897C>A, 5898C>T, 5904T>G, 5905T>A, 5906G>A, 5910C>T, 5913C>G, 5917C>A, 5918T>G, 5924C>G, 5925A>C, 5926G>T, 5931A>T, 5933T>A, 5937C>T, 5940A>G, 5942G>T, 5943G>T, 5944T>G, 5946A>G, 5953C>T, 5955A>T, 5958C>T, 5961A>G, 5966A>C, 5967G>T, 5981T>C, 5982G>T, 5985C>T, 5986C>A, 5987A>T, 5988A>G, 5989C>G, 5990G>A, 5991A>G, 5992A>T, 5993A>T, 5998G>A, 6001A>C, 6002A>G, 6003C>A, 6004T>G, 6005G>T, 6006T>G, 6011G>A, 6013G>C, 6014A>C, 6015_6016insTAT, 6018G>T, 6019T>G, 6020C>A, 6021A>T, 6022G>C, 6023G>A, 6024C>G, 6030A>G, 6031G>A, 6032C>T, 6033A>T, 6036T>G, 6038A>T, 6042T>A, 6048C>T, 6052C>T, 6057C>T, 6061A>T, 6062G>C, 6063T>C, 6066C>G, 6068C>G, 6072A>G, 6073G>A, 6074A>G, 6078C>T, 6083C>T, 6084A>G, 6085C>A, 6090C>T, 6091C>T | | | | | | | | |

##### CDS

| | Begin | End | Coverage | Score | Concordance | Matches | Identities | I/D/M/F* | Stop Codons |
|---|---|---|---|---|---|---|---|---|---|
| QKM20_gp3 | 1568 | 1658 | 4.5% | 362 | 53.2% | 91 (98.9%) | 53 (57.6%) | 1/0/0/0 | 0 |
| Protein mutations: | S1570T (5837G>C), F1575I (5851T>A 5853C>T), K1578R (5861A>G 5862G>A), F1581Y (5870T>A 5871C>T), I1584L (5878A>T 5880T>G), M1585R (5882T>G 5883G>A), M1586V (5884A>G 5886G>T), D1587R (5887G>C 5888A>G 5889C>G), E1589S (5893G>T 5894A>C), S1590D (5896T>G 5897C>A 5898C>T), W1593K (5905T>A 5906G>A), L1597R (5917C>A 5918T>G), P1599R (5924C>G 5925A>C), D1600Y (5926G>T), L1602H (5933T>A), W1605F (5942G>T 5943G>T), L1606V (5944T>G 5946A>G), P1609S (5953C>T 5955A>T), K1613T (5966A>C 5967G>T), V1618A (5981T>C 5982G>T), Q1620M (5986C>A 5987A>T 5988A>G), R1621E (5989C>G 5990G>A 5991A>G), K1622L (5992A>T 5993A>T), D1624N (5998G>A), N1625R (6001A>C 6002A>G 6003C>A), C1626V (6004T>G 6005G>T 6006T>G), R1628K (6011G>A), D1629P (6013G>C 6014A>C), D1629_L1630insY (6015_6016insTAT), S1631D (6019T>G 6020C>A 6021A>T), G1632Q (6022G>C 6023G>A 6024C>G), A1635I (6031G>A 6032C>T 6033A>T), Y1637F (6038A>T), N1646K (6066C>G), T1647S (6068C>G), D1649S (6073G>A 6074A>G), A1652V (6083C>T 6084A>G), Q1653K (6085C>A) | | | | | | | | |
| Codon mutations: | AGT1570ACT (5837G>C), GTC1571GTG (5841C>G), TCC1573TCT (5847C>T), TTC1575ATT (5851T>A 5853C>T), GAT1576GAC (5856T>C), CTG1577TTG (5857C>T), AAG1578AGA (5861A>G 5862G>A), AGC1579TCG (5863A>T 5864G>C 5865C>G), GGA1580GGT (5868A>T), TTC1581TAT (5870T>A 5871C>T), CAC1582CAT (5874C>T), ATT1584TTG (5878A>T 5880T>G), ATG1585AGA (5882T>G 5883G>A), ATG1586GTT (5884A>G 5886G>T), GAC1587CGG (5887G>C 5888A>G 5889C>G), GAA1588GAG (5892A>G), GAA1589TCA (5893G>T 5894A>C), TCC1590GAT (5896T>G 5897C>A 5898C>T), CCT1592CCG (5904T>G), TGG1593AAG (5905T>A 5906G>A), ACC1594ACT (5910C>T), GCC1595GCG (5913C>G), CTG1597AGG (5917C>A 5918T>G), CCA1599CGC (5924C>G 5925A>C), GAT1600TAT (5926G>T), GGA1601GGT (5931A>T), CTT1602CAT (5933T>A), TAC1603TAT (5937C>T), GAA1604GAG (5940A>G), TGG1605TTT (5942G>T 5943G>T), TTA1606GTG (5944T>G 5946A>G), CCA1609TCT (5953C>T 5955A>T), TTC1610TTT (5958C>T), GGA1611GGG (5961A>G), AAG1613ACT (5966A>C 5967G>T), GTG1618GCT (5981T>C 5982G>T), TTC1619TTT (5985C>T), CAA1620ATG (5986C>A 5987A>T 5988A>G), CGA1621GAG (5989C>G 5990G>A 5991A>G), AAG1622TTG (5992A>T 5993A>T), GAC1624AAC (5998G>A), AAC1625CGA (6001A>C 6002A>G 6003C>A), TGT1626GTG (6004T>G 6005G>T 6006T>G), AGA1628AAA (6011G>A), GAC1629CCC (6013G>C 6014A>C), GAC1629_CTG1630insTAT (6015_6016insTAT), CTG1630CTT (6018G>T), TCA1631GAT (6019T>G 6020C>A 6021A>T), GGC1632CAG (6022G>C 6023G>A 6024C>G), GTA1634GTG (6030A>G), GCA1635ATT (6031G>A 6032C>T 6033A>T), GTT1636GTG (6036T>G), TAC1637TTC (6038A>T), ATT1638ATA (6042T>A), GAC1640GAT (6048C>T), CTG1642TTG (6052C>T), GTC1643GTT (6057C>T), AGT1645TCC (6061A>T 6062G>C 6063T>C), AAC1646AAG (6066C>G), ACT1647AGT (6068C>G), GAA1648GAG (6072A>G), GAT1649AGT (6073G>A 6074A>G), GAC1650GAT (6078C>T), GCA1652GTG (6083C>T 6084A>G), CAG1653AAG (6085C>A), CAC1654CAT (6090C>T), CTG1655TTG (6091C>T) | | | | | | | | |

##### Proteins

| | Begin | End | Coverage | Score | Concordance | Matches | Identities | I/D/M/F* | Stop Codons |
|---|---|---|---|---|---|---|---|---|---|
| polyprotein (YP_010797894.1) | 1568 | 1658 | 4.5% | 362 | 53.2% | 91 (98.9%) | 53 (57.6%) | 1/0/0/0 | 0 |
| Protein mutations: | S1570T (5837G>C), F1575I (5851T>A 5853C>T), K1578R (5861A>G 5862G>A), F1581Y (5870T>A 5871C>T), I1584L (5878A>T 5880T>G), M1585R (5882T>G 5883G>A), M1586V (5884A>G 5886G>T), D1587R (5887G>C 5888A>G 5889C>G), E1589S (5893G>T 5894A>C), S1590D (5896T>G 5897C>A 5898C>T), W1593K (5905T>A 5906G>A), L1597R (5917C>A 5918T>G), P1599R (5924C>G 5925A>C), D1600Y (5926G>T), L1602H (5933T>A), W1605F (5942G>T 5943G>T), L1606V (5944T>G 5946A>G), P1609S (5953C>T 5955A>T), K1613T (5966A>C 5967G>T), V1618A (5981T>C 5982G>T), Q1620M (5986C>A 5987A>T 5988A>G), R1621E (5989C>G 5990G>A 5991A>G), K1622L (5992A>T 5993A>T), D1624N (5998G>A), N1625R (6001A>C 6002A>G 6003C>A), C1626V (6004T>G 6005G>T 6006T>G), R1628K (6011G>A), D1629P (6013G>C 6014A>C), D1629_L1630insY (6015_6016insTAT), S1631D (6019T>G 6020C>A 6021A>T), G1632Q (6022G>C 6023G>A 6024C>G), A1635I (6031G>A 6032C>T 6033A>T), Y1637F (6038A>T), N1646K (6066C>G), T1647S (6068C>G), D1649S (6073G>A 6074A>G), A1652V (6083C>T 6084A>G), Q1653K (6085C>A) | | | | | | | | |
| Codon mutations: | AGT1570ACT (5837G>C), GTC1571GTG (5841C>G), TCC1573TCT (5847C>T), TTC1575ATT (5851T>A 5853C>T), GAT1576GAC (5856T>C), CTG1577TTG (5857C>T), AAG1578AGA (5861A>G 5862G>A), AGC1579TCG (5863A>T 5864G>C 5865C>G), GGA1580GGT (5868A>T), TTC1581TAT (5870T>A 5871C>T), CAC1582CAT (5874C>T), ATT1584TTG (5878A>T 5880T>G), ATG1585AGA (5882T>G 5883G>A), ATG1586GTT (5884A>G 5886G>T), GAC1587CGG (5887G>C 5888A>G 5889C>G), GAA1588GAG (5892A>G), GAA1589TCA (5893G>T 5894A>C), TCC1590GAT (5896T>G 5897C>A 5898C>T), CCT1592CCG (5904T>G), TGG1593AAG (5905T>A 5906G>A), ACC1594ACT (5910C>T), GCC1595GCG (5913C>G), CTG1597AGG (5917C>A 5918T>G), CCA1599CGC (5924C>G 5925A>C), GAT1600TAT (5926G>T), GGA1601GGT (5931A>T), CTT1602CAT (5933T>A), TAC1603TAT (5937C>T), GAA1604GAG (5940A>G), TGG1605TTT (5942G>T 5943G>T), TTA1606GTG (5944T>G 5946A>G), CCA1609TCT (5953C>T 5955A>T), TTC1610TTT (5958C>T), GGA1611GGG (5961A>G), AAG1613ACT (5966A>C 5967G>T), GTG1618GCT (5981T>C 5982G>T), TTC1619TTT (5985C>T), CAA1620ATG (5986C>A 5987A>T 5988A>G), CGA1621GAG (5989C>G 5990G>A 5991A>G), AAG1622TTG (5992A>T 5993A>T), GAC1624AAC (5998G>A), AAC1625CGA (6001A>C 6002A>G 6003C>A), TGT1626GTG (6004T>G 6005G>T 6006T>G), AGA1628AAA (6011G>A), GAC1629CCC (6013G>C 6014A>C), GAC1629_CTG1630insTAT (6015_6016insTAT), CTG1630CTT (6018G>T), TCA1631GAT (6019T>G 6020C>A 6021A>T), GGC1632CAG (6022G>C 6023G>A 6024C>G), GTA1634GTG (6030A>G), GCA1635ATT (6031G>A 6032C>T 6033A>T), GTT1636GTG (6036T>G), TAC1637TTC (6038A>T), ATT1638ATA (6042T>A), GAC1640GAT (6048C>T), CTG1642TTG (6052C>T), GTC1643GTT (6057C>T), AGT1645TCC (6061A>T 6062G>C 6063T>C), AAC1646AAG (6066C>G), ACT1647AGT (6068C>G), GAA1648GAG (6072A>G), GAT1649AGT (6073G>A 6074A>G), GAC1650GAT (6078C>T), GCA1652GTG (6083C>T 6084A>G), CAG1653AAG (6085C>A), CAC1654CAT (6090C>T), CTG1655TTG (6091C>T) | | | | | | | | |

*: Inserts / Deletes / Misaligned / Frameshifts

#### Analysis details

This analysis was performed with panviral2.64

### NGS Details (UN9): Lowelvirus tuscon4d

#### Assembly

| | |
|---|---|
| **Coverage Length** | 1030 (1 contig(s)) |
| **Depth Of Coverage** | 36.3 |
| **Number Of Reads** | 311 |
| **Reads Per Million** | 7.03 rpm (after QC) |
| **Ambiguities** | 0 |
| **Assembly Method** | de novo + reference guided assembly |
| **Consensus Caller** | Bcf Tools |

#### Coverage Map

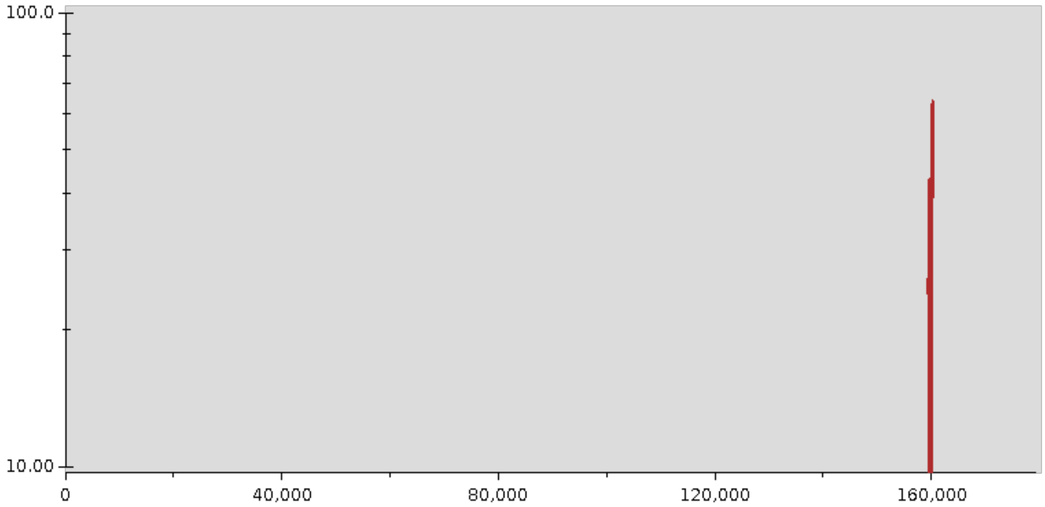

#### Assignment

| | |
|---|---|
| **Type** | Lowelvirus tuscon4d (Taxonomy ID: 2956131) |
| **Reference Genome** | NC_026923.1 |
| **NT Identity (%)** | 73.301 |
| **AA Identity (%)** | 87.0588 |
| **Number Of Stop Codons** | 1 |
| **Number Of CDS** | 218 |

#### Alignment

| | |
|---|---|
| **Alignment Score** | 960.0 (NT) + 2288.0 (AA) = 3248.0 |
| **Concordance (%)** | 71.2593 |
| **Alignment Method** | Local, heuristic, nucleotide (BLASTN) |

#### Genome Region

Sequence starts at position 159375 and ends at position 160404 relative to NC_026923.1 reference sequence.

#### Alignment Detailed Statistics

| | Begin | End | Coverage | Score | Concordance | Matches | Identities | I/D/M/F* | Stop Codons |
|---|---|---|---|---|---|---|---|---|---|
| **NT** | **159375** | **160404** | **0.6%** | **960** | **46.6%** | **1030 (100%)** | **755 (73.3%)** | **0/0** | |
| Mutations: | 159385G>A, 159387C>T, 159388C>A, 159400C>T, 159402T>A, 159403A>C, 159404A>G, 159405A>G, 159406C>A, 159408A>G, 159411T>C, 159414C>T, 159417T>C, 159420C>T, 159426G>A, 159429T>C, 159435T>C, 159438A>T, 159441C>A, 159442C>T, 159444A>G, 159447T>C, 159448C>T, 159453C>T, 159454A>T, 159455C>G, 159459T>C, 159463A>T, 159465G>C, 159468C>T, 159469A>C, 159471T>A, 159474C>G, 159481C>T, 159483T>C, 159486T>A, 159489C>T, 159492G>A, 159495T>C, 159499A>G, 159501C>A, 159504G>T, 159505A>T, 159506G>C, 159507T>A, 159513C>T, 159519C>T, 159522T>A, 159523C>T, 159525T>G, 159528T>C, 159534C>T, 159537C>T, 159538C>T, 159540T>G, 159546T>C, 159547G>T, 159548C>G, 159549T>C, 159555T>C, 159556C>T, 159558C>A, 159561G>T, 159564A>C, 159567T>G, 159570C>T, 159573A>T, 159576G>T, 159583G>A, 159586G>A, 159587C>G, 159589A>T, 159591G>A, 159593G>C, 159594T>A, 159600T>G, 159601C>T, 159603T>G, 159604C>T, 159606T>G, 159607C>T, 159612T>A, 159624G>A, 159625T>G, 159627T>A, 159633C>A, 159636C>T, 159639C>T, 159640C>A, 159641A>C, 159642A>T, 159645C>T, 159651C>T, 159655C>T, 159657T>G, 159660A>G, 159663G>T, 159664A>C, 159666T>G, 159670T>A, 159672C>T, 159687C>T, 159690T>A, 159693A>T, 159696C>T, 159698C>G, 159699T>C, 159702C>A, 159711T>C, 159715C>T, 159720G>T, 159726T>C, 159729A>G, 159730A>C, 159738T>A, 159739C>T, 159740T>C, 159741A>T, 159745G>C, 159746G>A, 159747T>A, 159748A>T, 159750C>G, 159751C>A, 159753T>A, 159759C>T, 159765T>A, 159768T>C, 159771G>A, 159783T>A, 159798C>T, 159799A>T, 159800G>C, 159801C>T, 159807C>T, 159810T>G, 159813C>T, 159822G>A, 159832T>G, 159833C>G, 159834A>T, 159855C>T, 159858G>T, 159867A>T, 159870C>A, 159874A>C, 159880C>A, 159882G>C, 159883T>C, 159885G>A, 159888C>T, 159889C>T, 159894G>A, 159897T>G, 159900C>T, 159903C>T, 159906C>T, 159913C>T, 159915C>G, 159924C>T, 159942T>C, 159946A>G, 159949C>T, 159951A>G, 159954T>C, 159956G>C, 159957A>T, 159960A>T, 159961C>T, 159964T>C, 159966G>A, 159969T>C, 159972A>C, 159975C>T, 159987T>C, 159993G>A, 159996C>T, 159999A>T, 160002G>A, 160011T>C, 160014A>T, 160017A>T, 160018C>G, 160019A>G, 160020A>T, 160026C>T, 160032T>C, 160033A>C, 160034A>G, 160035G>T, 160038G>T, 160042G>A, 160044G>C, 160047T>A, 160055A>C, 160056A>C, 160062G>A, 160071T>A, 160080T>C, 160083A>T, 160092C>T, 160098G>C, 160101G>A, 160107C>T, 160110T>G, 160116A>T, 160122T>C, 160125C>T, 160128G>A, 160131G>T, 160135C>T, 160137T>A, 160143T>C, 160146C>T, 160152G>A, 160158G>A, 160161T>A, 160164T>A, 160166T>C, 160167G>C, 160171C>T, 160178C>T, 160179A>G, 160180T>A, 160181C>G, 160182C>T, 160183T>G, 160186A>C, 160191C>A, 160192A>G, 160194T>A, 160195A>G, 160197T>C, 160203T>A, 160207T>C, 160209C>G, 160215T>A, 160221T>C, 160224C>T, 160227T>C, 160233G>T, 160234A>T, 160235G>C, 160242G>A, 160246A>C, 160248A>C, 160254A>G, 160275C>T, 160287G>A, 160290C>T, 160293C>T, 160296A>C, 160299G>A, 160302T>C, 160308A>T, 160309C>A, 160314T>C, 160326A>G, 160327C>G, 160330G>C, 160331C>A, 160332T>A, 160338A>G, 160341A>T, 160351T>C, 160356C>A, 160368A>G, 160371A>T, 160377T>A, 160378A>C, 160380A>T, 160383T>A, 160389A>T, 160392G>T, 160394G>A | | | | | | | | |

*: Inserts / Deletes / Misaligned / Frameshifts

#### Analysis details

This analysis was performed with panviral2.64

### NGS Details (UN9): Hibiscus bacilliform virus GD1

#### Assembly

| | |
|---|---|
| **Coverage Length** | 281 (1 contig(s)) |
| **Depth Of Coverage** | 73.4 |
| **Number Of Reads** | 204 |
| **Reads Per Million** | 4.61 rpm (after QC) |
| **Ambiguities** | 0 |
| **Assembly Method** | de novo + reference guided assembly |
| **Consensus Caller** | Bcf Tools |

#### Coverage Map

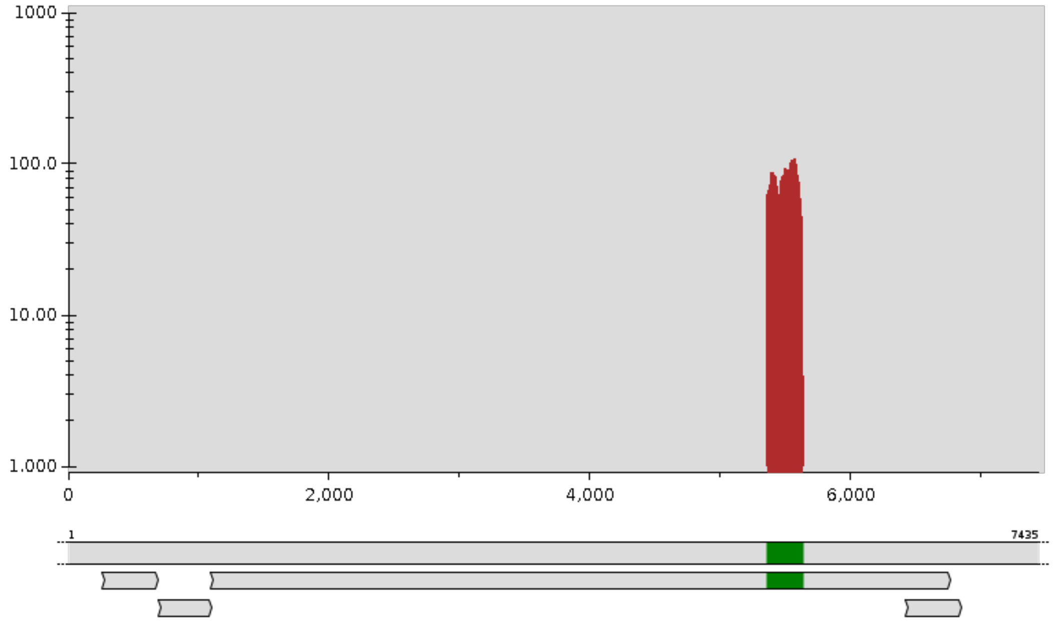

#### Assignment

| | |
|---|---|
| **Type** | Hibiscus bacilliform virus GD1 (Taxonomy ID: 1459800) |
| **Reference Genome** | NC_023485.1 |
| **NT Identity (%)** | 59.507 |
| **AA Identity (%)** | 51.5789 |
| **Number Of Stop Codons** | 0 |
| **Number Of CDS** | 4 |

#### Alignment

| | |
|---|---|
| **Alignment Score** | 102.0 (NT) + 360.0 (AA) = 462.0 |
| **Concordance (%)** | 37.6835 |

| Alignment Method | Global, seeded, nucleotide + amino acids (AGA) |
|---|---|

#### Genome Region

Sequence starts at position 5357 and ends at position 5637 relative to NC_023485.1 reference sequence.

#### Alignment Detailed Statistics

| | Begin | End | Coverage | Score | Concordance | Matches | Identities | I/D/M/F* | Stop Codons |
|---|---|---|---|---|---|---|---|---|---|
| **NT** | **5357** | **5637** | **3.8%** | **102** | **18.1%** | **281 (98.9%)** | **169 (59.5%)** | **3/0** | |
| Mutations: | 5364A>G, 5366C>G, 5368A>T, 5372A>T, 5376T>A, 5382C>T, 5384C>G, 5386A>G, 5387G>A, 5388A>T, 5389G>T, 5390T>G, 5393A>T, 5395T>A, 5396T>C, 5403G>T, 5406G>A, 5407C>G, 5408T>A, 5409A>G, 5411G>T, 5414C>G, 5415C>G, 5416C>A, 5418G>T, 5419A>C, 5420G>A, 5421T>G, 5422C>A, 5427A>C, 5428A>C, 5430T>A, 5431G>A, 5438T>A, 5441C>T, 5445G>A, 5446T>C, 5447T>A, 5449C>G, 5450A>C, 5451G>T, 5456A>T, 5458T>A, 5459G>T, 5467G>T, 5468G>C, 5469C>G, 5471A>G, 5478C>T, 5480A>T, 5486A>G, 5491A>C, 5492A>C, 5495C>T, 5501A>T, 5504A>T, 5506T>C, 5510T>C, 5511C>A, 5512A>T, 5514C>G, 5515G>A, 5516A>G, 5517A>C, 5518A>T, 5523G>A, 5525C>A, 5526A>C, 5527A>G, 5528C>A, 5531C>G, 5534C>T, 5537A>G, 5538G>T, 5539G>C, 5541A>T, 5542C>A, 5543_5544insTTG, 5546A>T, 5547G>C, 5548C>A, 5549C>G, 5553A>G, 5555C>G, 5556G>A, 5557C>T, 5558T>C, 5561A>G, 5563A>T, 5564C>T, 5567T>A, 5576A>C, 5577C>T, 5579C>G, 5582A>T, 5584T>A, 5585T>C, 5589C>A, 5592T>A, 5593C>G, 5598G>A, 5599A>G, 5600G>T, 5603A>T, 5607A>G, 5609C>G, 5612G>A, 5616A>T, 5618A>G, 5619C>A, 5622G>A, 5624C>T, 5625A>G | | | | | | | | |

##### CDS

| | Begin | End | Coverage | Score | Concordance | Matches | Identities | I/D/M/F* | Stop Codons |
|---|---|---|---|---|---|---|---|---|---|
| BU35_gp3 | 1423 | 1516 | 5.0% | 360 | 51.6% | 94 (98.9%) | 49 (51.6%) | 1/0/0/0 | 0 |
| Protein mutations: | I1425V (5364A>G 5366C>G), Y1426F (5368A>T), F1429I (5376T>A), K1432R (5386A>G 5387G>A), S1433L (5388A>T 5389G>T 5390T>G), F1435Y (5395T>A 5396T>C), V1438L (5403G>T), A1439R (5406G>A 5407C>G 5408T>A), M1440V (5409A>G 5411G>T), H1441Q (5414C>G), P1442E (5415C>G 5416C>A), E1443S (5418G>T 5419A>C 5420G>A), S1444D (5421T>G 5422C>A), K1446P (5427A>C 5428A>C), W1447K (5430T>A 5431G>A), V1452T (5445G>A 5446T>C 5447T>A), P1453R (5449C>G 5450A>C), D1454Y (5451G>T), L1456H (5458T>A 5459G>T), W1459F (5467G>T 5468G>C), L1460V (5469C>G 5471A>G), P1463S (5478C>T 5480A>T), K1467T (5491A>C 5492A>C), V1472A (5506T>C), Q1474M (5511C>A 5512A>T), R1475E (5514C>G 5515G>A 5516A>G), K1476L (5517A>C 5518A>T), D1478K (5523G>A 5525C>A), N1479R (5526A>C 5527A>G 5528C>A), G1483S (5538G>T 5539G>C), T1484Y (5541A>T 5542C>A), T1484_E1485insL (5543_5544insTTG), E1485D (5546A>T), A1486Q (5547G>C 5548C>A 5549C>G), I1488V (5553A>G 5555C>G), A1489I (5556G>A 5557C>T 5558T>C), Y1491F (5563A>T 5564C>T), F1498Y (5584T>A 5585T>C), Q1500K (5589C>A), E1503S (5598G>A 5599A>G 5600G>T), E1504D (5603A>T), I1506V (5607A>G 5609C>G), I1509L (5616A>T 5618A>G), V1511I (5622G>A 5624C>T), M1512V (5625A>G) | | | | | | | | |
| Codon mutations: | ATC1425GTG (5364A>G 5366C>G), TAC1426TTC (5368A>T), TCA1427TCT (5372A>T), TTT1429ATT (5376T>A), CTC1431TTG (5382C>T 5384C>G), AAG1432AGA (5386A>G 5387G>A), AGT1433TTG (5388A>T 5389G>T 5390T>G), GGA1434GGT (5393A>T), TTT1435TAC (5395T>A 5396T>C), GTG1438TTG (5403G>T), GCT1439AGA (5406G>A 5407C>G 5408T>A), ATG1440GTT (5409A>G 5411G>T), CAC1441CAG (5414C>G), CCG1442GAG (5415C>G 5416C>A), GAG1443TCA (5418G>T 5419A>C 5420G>A), TCT1444GAT (5421T>G 5422C>A), AAG1446CCG (5427A>C 5428A>C), TGG1447AAG (5430T>A 5431G>A), GCT1449GCA (5438T>A), TTC1450TTT (5441C>T), GTT1452ACA (5445G>A 5446T>C 5447T>A), CCA1453CGC (5449C>G 5450A>C), GAT1454TAT (5451G>T), GGA1455GGT (5456A>T), CTG1456CAT (5458T>A 5459G>T), TGG1459TTC (5467G>T 5468G>C), CTA1460GTG (5469C>G 5471A>G), CCA1463TCT (5478C>T 5480A>T), GGA1465GGG (5486A>G), AAA1467ACC (5491A>C 5492A>C), AAC1468AAT (5495C>T), CCA1470CCT (5501A>T), GCA1471GCT (5504A>T), GTA1472GCA (5506T>C), TTT1473TTC (5510T>C), CAG1474ATG (5511C>A 5512A>T), CGA1475GAG (5514C>G 5515G>A 5516A>G), AAG1476CTG (5517A>C 5518A>T), GAC1478AAA (5523G>A 5525C>A), AAC1479CGA (5526A>C 5527A>G 5528C>A), GTC1480GTG (5531C>G), TTC1481TTT (5534C>T), AAA1482AAG (5537A>G), GGT1483TCT (5538G>T 5539G>C), ACT1484TAT (5541A>T 5542C>A), ACT1484_GAA1485insTTG (5543_5544insTTG), GAA1485GAT (5546A>T), GCC1486CAG (5547G>C 5548C>A 5549C>G), ATC1488GTG (5553A>G 5555C>G), GCT1489ATC (5556G>A 5557C>T 5558T>C), GTA1490GTG (5561A>G), TAC1491TTT (5563A>T 5564C>T), ATT1492ATA (5567T>A), ATA1495ATC (5576A>C), CTC1496TTG (5577C>T 5579C>G), GTA1497GTT (5582A>T), TTT1498TAC (5584T>A 5585T>C), CAG1500AAG (5589C>A), TCT1501AGT (5592T>A 5593C>G), GAG1503AGT (5598G>A 5599A>G 5600G>T), GAA1504GAT (5603A>T), ATC1506GTG (5607A>G 5609C>G), AAG1507AAA (5612G>A), ATA1509TTG (5616A>T 5618A>G), CGG1510AGG (5619C>A), GTC1511ATT (5622G>A 5624C>T), ATG1512GTG (5625A>G) | | | | | | | | |

##### Proteins

| | Begin | End | Coverage | Score | Concordance | Matches | Identities | I/D/M/F* | Stop Codons |
|---|---|---|---|---|---|---|---|---|---|
| polyprotein (YP_009002585.1) | 1423 | 1516 | 5.0% | 360 | 51.6% | 94 (98.9%) | 49 (51.6%) | 1/0/0/0 | 0 |
| Protein mutations: | I1425V (5364A>G 5366C>G), Y1426F (5368A>T), F1429I (5376T>A), K1432R (5386A>G 5387G>A), S1433L (5388A>T 5389G>T 5390T>G), F1435Y (5395T>A 5396T>C), V1438L (5403G>T), A1439R (5406G>A 5407C>G 5408T>A), M1440V (5409A>G 5411G>T), H1441Q (5414C>G), P1442E (5415C>G 5416C>A), E1443S (5418G>T 5419A>C 5420G>A), S1444D (5421T>G 5422C>A), K1446P (5427A>C 5428A>C), W1447K (5430T>A 5431G>A), V1452T (5445G>A 5446T>C 5447T>A), P1453R (5449C>G 5450A>C), D1454Y (5451G>T), L1456H (5458T>A 5459G>T), W1459F (5467G>T 5468G>C), L1460V (5469C>G 5471A>G), P1463S (5478C>T 5480A>T), K1467T (5491A>C 5492A>C), V1472A (5506T>C), Q1474M (5511C>A 5512A>T), R1475E (5514C>G 5515G>A 5516A>G), K1476L (5517A>C 5518A>T), D1478K (5523G>A 5525C>A), N1479R (5526A>C 5527A>G 5528C>A), G1483S (5538G>T 5539G>C), T1484Y (5541A>T 5542C>A), T1484_E1485insL (5543_5544insTTG), E1485D (5546A>T), A1486Q (5547G>C 5548C>A 5549C>G), I1488V (5553A>G 5555C>G), A1489I (5556G>A 5557C>T 5558T>C), Y1491F (5563A>T 5564C>T), F1498Y (5584T>A 5585T>C), Q1500K (5589C>A), E1503S (5598G>A 5599A>G 5600G>T), E1504D (5603A>T), I1506V (5607A>G 5609C>G), I1509L (5616A>T 5618A>G), V1511I (5622G>A 5624C>T), M1512V (5625A>G) | | | | | | | | |
| Codon mutations: | ATC1425GTG (5364A>G 5366C>G), TAC1426TTC (5368A>T), TCA1427TCT (5372A>T), TTT1429ATT (5376T>A), CTC1431TTG (5382C>T 5384C>G), AAG1432AGA (5386A>G 5387G>A), AGT1433TTG (5388A>T 5389G>T 5390T>G), GGA1434GGT (5393A>T), TTT1435TAC (5395T>A 5396T>C), GTG1438TTG (5403G>T), GCT1439AGA (5406G>A 5407C>G 5408T>A), ATG1440GTT (5409A>G 5411G>T), CAC1441CAG (5414C>G), CCG1442GAG (5415C>G 5416C>A), GAG1443TCA (5418G>T 5419A>C 5420G>A), TCT1444GAT (5421T>G 5422C>A), AAG1446CCG (5427A>C 5428A>C), TGG1447AAG (5430T>A 5431G>A), GCT1449GCA (5438T>A), TTC1450TTT (5441C>T), GTT1452ACA (5445G>A 5446T>C 5447T>A), CCA1453CGC (5449C>G 5450A>C), GAT1454TAT (5451G>T), GGA1455GGT (5456A>T), CTG1456CAT (5458T>A 5459G>T), TGG1459TTC (5467G>T 5468G>C), CTA1460GTG (5469C>G 5471A>G), CCA1463TCT (5478C>T 5480A>T), GGA1465GGG (5486A>G), AAA1467ACC (5491A>C 5492A>C), AAC1468AAT (5495C>T), CCA1470CCT (5501A>T), GCA1471GCT (5504A>T), GTA1472GCA (5506T>C), TTT1473TTC (5510T>C), CAG1474ATG (5511C>A 5512A>T), CGA1475GAG (5514C>G 5515G>A 5516A>G), AAG1476CTG (5517A>C 5518A>T), GAC1478AAA (5523G>A 5525C>A), AAC1479CGA (5526A>C 5527A>G 5528C>A), GTC1480GTG (5531C>G), TTC1481TTT (5534C>T), AAA1482AAG (5537A>G), GGT1483TCT (5538G>T 5539G>C), ACT1484TAT (5541A>T 5542C>A), ACT1484_GAA1485insTTG (5543_5544insTTG), GAA1485GAT (5546A>T), GCC1486CAG (5547G>C 5548C>A 5549C>G), ATC1488GTG (5553A>G 5555C>G), GCT1489ATC (5556G>A 5557C>T 5558T>C), GTA1490GTG (5561A>G), TAC1491TTT (5563A>T 5564C>T), ATT1492ATA (5567T>A), ATA1495ATC (5576A>C), CTC1496TTG (5577C>T 5579C>G), GTA1497GTT (5582A>T), TTT1498TAC (5584T>A 5585T>C), CAG1500AAG (5589C>A), TCT1501AGT (5592T>A 5593C>G), GAG1503AGT (5598G>A 5599A>G 5600G>T), GAA1504GAT (5603A>T), ATC1506GTG (5607A>G 5609C>G), AAG1507AAA (5612G>A), ATA1509TTG (5616A>T 5618A>G), CGG1510AGG (5619C>A), GTC1511ATT (5622G>A 5624C>T), ATG1512GTG (5625A>G) | | | | | | | | |

*: Inserts / Deletes / Misaligned / Frameshifts

#### Analysis details

This analysis was performed with panviral2.64

### NGS Details (UN9): Cassava brown streak virus

#### Assembly

| | |
|---|---|
| **Coverage Length** | 522 (1 contig(s)) |
| **Depth Of Coverage** | 45.5 |
| **Number Of Reads** | 197 |
| **Reads Per Million** | 4.45 rpm (after QC) |
| **Ambiguities** | 0 |
| **Assembly Method** | de novo + reference guided assembly |
| **Consensus Caller** | Bcf Tools |

#### Coverage Map

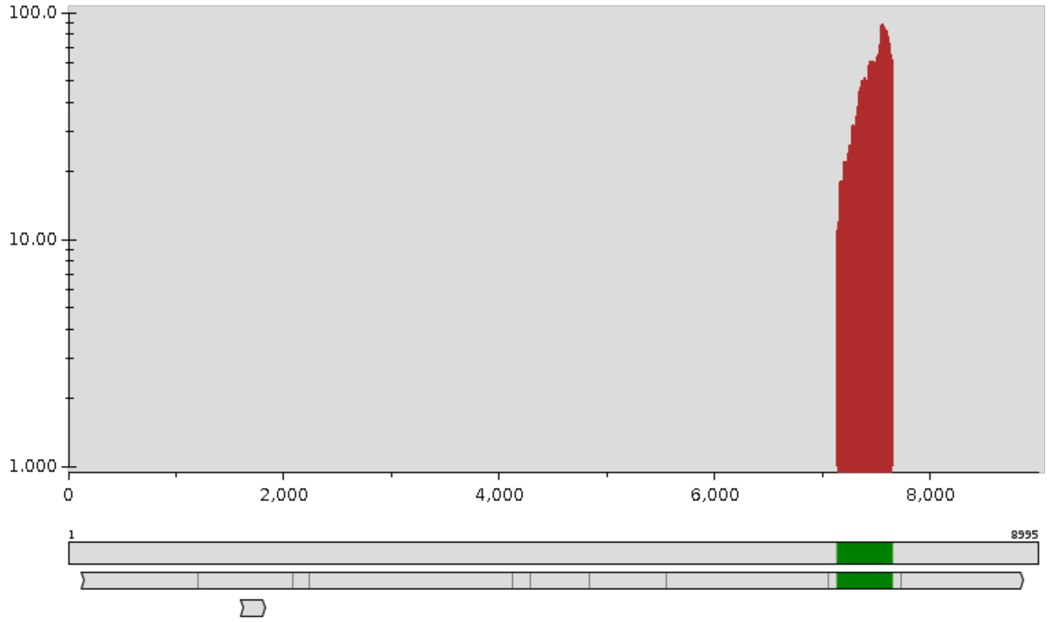

#### Assignment

| | |
|---|---|
| **Type** | Cassava brown streak virus (Taxonomy ID: 137758) |
| **Reference Genome** | NC_012698.2 |
| **NT Identity (%)** | 60.5364 |
| **AA Identity (%)** | 57.4713 |
| **Number Of Stop Codons** | 0 |
| **Number Of CDS** | 2 |

#### Alignment

| | |
|---|---|
| **Alignment Score** | 220.0 (NT) + 732.0 (AA) = 952.0 |
| **Concordance (%)** | 42.2173 |

| Alignment Method | Global, seeded, nucleotide + amino acids (AGA) |
|---|---|

#### Genome Region

Sequence starts at position 7129 and ends at position 7650 relative to NC_012698.2 reference sequence.

#### Alignment Detailed Statistics

| | Begin | End | Coverage | Score | Concordance | Matches | Identities | I/D/M/F* | Stop Codons |
|---|---|---|---|---|---|---|---|---|---|
| **NT** | **7129** | **7650** | **5.8%** | **220** | **21.1%** | **522 (100%)** | **316 (60.5%)** | **0/0** | |
| Mutations: | 7130A>G, 7133A>G, 7134A>G, 7136T>G, 7142C>T, 7145A>T, 7148A>C, 7151G>A, 7157A>T, 7158C>A, 7163A>G, 7167A>G, 7175A>T, 7177A>G, 7178A>G, 7179C>G, 7180A>C, 7184A>T, 7185T>C, 7190T>C, 7192C>A, 7193C>G, 7194A>T, 7202A>C, 7203A>T, 7205C>T, 7206A>C, 7207T>A, 7208C>G, 7211T>C, 7213G>T, 7214G>T, 7217A>G, 7218G>C, 7229A>T, 7234C>T, 7235A>T, 7241A>G, 7242A>G, 7243C>A, 7244A>G, 7245G>C, 7246T>C, 7250G>A, 7253A>T, 7256T>A, 7257A>T, 7258T>C, 7270T>C, 7271A>T, 7272C>A, 7273G>A, 7274T>A, 7275G>A, 7277A>T, 7280G>T, 7281T>G, 7283T>C, 7284G>A, 7286G>A, 7287T>G, 7288T>A, 7290A>G, 7292T>G, 7293G>A, 7294G>A, 7295A>T, 7298G>A, 7304T>G, 7307C>A, 7310A>T, 7313A>G, 7319A>T, 7320A>T, 7323T>C, 7325G>C, 7331T>C, 7332G>A, 7334T>C, 7337T>C, 7340C>T, 7343T>G, 7355A>G, 7358A>T, 7361C>T, 7364T>C, 7374A>T, 7377G>C, 7380G>A, 7381G>A, 7385T>C, 7388A>T, 7390T>A, 7391A>T, 7397A>T, 7398T>C, 7400A>T, 7401T>A, 7403T>C, 7406G>C, 7409G>A, 7410G>T, 7413G>A, 7414A>T, 7416C>G, 7418G>T, 7422C>G, 7424A>G, 7425A>G, 7432T>C, 7433G>A, 7436T>A, 7437A>T, 7438G>A, 7442T>C, 7443C>A, 7445C>G, 7448C>T, 7449G>A, 7451G>C, 7455G>T, 7457T>A, 7458T>C, 7462T>C, 7463A>T, 7464A>C, 7465A>T, 7467A>G, 7468A>G, 7469A>G, 7470G>C, 7471T>C, 7472T>A, 7473G>A, 7474G>A, 7476G>A, 7477A>C, 7481T>G, 7484T>G, 7487A>G, 7489T>C, 7490C>T, 7494A>G, 7495A>T, 7496G>T, 7499T>A, 7500G>A, 7501T>A, 7502G>A, 7503C>A, 7504T>C, 7505A>G, 7506A>C, 7507G>T, 7508A>G, 7511T>A, 7512G>A, 7517T>A, 7520A>G, 7521A>C, 7522T>C, 7523G>A, 7524C>G, 7526A>T, 7527C>A, 7529C>G, 7535A>C, 7539T>A, 7540C>A, 7541A>T, 7544T>C, 7547G>A, 7553C>T, 7556A>G, 7559T>A, 7562C>T, 7568A>T, 7570T>A, 7572A>G, 7573A>G, 7576G>A, 7577G>T, 7578A>G, 7579G>A, 7580A>C, 7581A>C, 7586A>T, 7588T>A, 7589T>C, 7599A>C, 7600T>C, 7601G>C, 7602G>A, 7603C>A, 7604C>G, 7607G>A, 7610G>A, 7616T>C, 7618T>A, 7622A>T, 7628T>C, 7631A>C, 7632T>G, 7633T>G, 7635C>A, 7636G>A | | | | | | | | |
| **CDS** | | | | | | | | | |
| CBSV_gp1 | 2336 | 2509 | 6.0% | 732 | 60.0% | 174 (100%) | 100 (57.5%) | 0/0/0/0 | 0 |
| Protein mutations: | I2337V (7134A>G 7136T>G), Q2345K (7158C>A), K2348E (7167A>G), K2351R (7177A>G 7178A>G), Q2352A (7179C>G 7180A>C), F2354L (7185T>C), P2356Q (7192C>A 7193C>G), T2357S (7194A>T), I2360F (7203A>T 7205C>T), I2361Q (7206A>C 7207T>A 7208C>G), R2363L (7213G>T 7214G>T), V2365L (7218G>C), P2370L (7234C>T 7235A>T), T2373E (7242A>G 7243C>A 7244A>G), V2374P (7245G>C 7246T>C), E2376D (7253A>T), I2378S (7257A>T 7258T>C), V2382A (7270T>C 7271A>T), R2383K (7272C>A 7273G>A 7274T>A), V2384I (7275G>A 7277A>T), S2386A (7281T>G 7283T>C), E2387K (7284G>A 7286G>A), L2388E (7287T>G 7288T>A), I2389V (7290A>G 7292T>G), G2390N (7293G>A 7294G>A 7295A>T), S2399C (7320A>T), D2403N (7332G>A 7334T>C), N2406K (7343T>G), M2417L (7374A>T), E2418Q (7377G>C), G2419K (7380G>A 7381G>A), L2422H (7390T>A 7391A>T), Y2426N (7401T>A 7403T>C), K2427N (7406G>C), V2429L (7410G>T), E2430M (7413G>A 7414A>T), P2431A (7416C>G 7418G>T), Q2433E (7422C>G 7424A>G), N2434D (7425A>G), M2436T (7432T>C 7433G>A), S2438Y (7437A>T 7438G>A), L2440M (7443C>A 7445C>G), V2442I (7449G>A 7451G>C), A2444S (7455G>T 7457T>A), F2445L (7458T>C), V2446A (7462T>C 7463A>T), N2447L (7464A>C 7465A>T), K2448G (7467A>G 7468A>G 7469A>G), V2449P (7470G>C 7471T>C 7472T>A), G2450N (7473G>A 7474G>A), D2451T (7476G>A 7477A>C), D2452E (7481T>G), I2454M (7487A>G), I2455T (7489T>C 7490C>T), K2457V (7494A>G 7495A>T 7496G>T), V2459K (7500G>A 7501T>A 7502G>A), L2460T (7503C>A 7504T>C 7505A>G), R2461L (7506A>C 7507G>T 7508A>G), E2463K (7512G>A), M2466P (7521A>C 7522T>C 7523G>A), P2467A (7524C>G 7526A>T), S2472N (7539T>A 7540C>A 7541A>T), L2482H (7570T>A), N2483G (7572A>G 7573A>G), W2484Y (7576G>A 7577G>T), R2485D (7578A>G 7579G>A 7580A>C), K2486Q (7581A>C), F2488Y (7588T>A 7589T>C), M2492P (7599A>C 7600T>C 7601G>C), A2493K (7602G>A 7603C>A 7604C>G), M2498K (7618T>A), F2503G (7632T>G 7633T>G), R2504K (7635C>A 7636G>A) | | | | | | | | |
| Codon mutations: | GCA2335.CG (7130A>G), CCA2336CCG (7133A>G), ATT2337GTG (7134A>G 7136T>G), TTC2339TTT (7142C>T), GTA2340GTT (7145A>T), ACA2341ACC (7148A>C), GGG2342GGA (7151G>A), GCA2344GCT (7157A>T), CAA2345AAA (7158C>A), AAA2346AAG (7163A>G), AAG2348GAG (7167A>G), GTA2350GTT (7175A>T), AAA2351AGG (7177A>G 7178A>G), CAA2352GCA (7179C>G 7180A>C), ATA2353ATT (7184A>T), TTT2354CTT (7185T>C), GGT2355GGC (7190T>C), CCC2356CAG (7192C>A 7193C>G), ACT2357TCT (7194A>T), CCA2359CCC (7202A>C), ATC2360TTT (7203A>T 7205C>T), ATC2361CAG (7206A>C 7207T>A 7208C>G), TCT2362TCC (7211T>C), CGG2363CTT (7213G>T 7214G>T), AAA2364AAG (7217A>G), GTT2365CTT (7218G>C), CCA2368CCT (7229A>T), CCA2370CTT (7234C>T 7235A>T), GGA2372GGG (7241A>G), ACA2373GAG (7242A>G 7243C>A 7244A>G), GTT2374CCT (7245G>C 7246T>C), GAG2375GAA (7250G>A), GAA2376GAT (7253A>T), ATT2377ATA (7256T>A), ATC2378TCC (7257A>T 7258T>C), GTA2382GCT (7270T>C 7271A>T), CGT2383AAA (7272C>A 7273G>A 7274T>A), GTA2384ATT (7275G>A 7277A>T), GCG2385GCT (7280G>T), TCT2386GCC (7281T>G 7283T>C), GAG2387AAA (7284G>A 7286G>A), TTG2388GAG (7287T>G 7288T>A), ATT2389GTG (7290A>G 7292T>G), GGA2390AAT (7293G>A 7294G>A 7295A>T), GGG2391GGA (7298G>A), GTT2393GTG (7304T>G), CTC2394CTA (7307C>A), GTA2395GTT (7310A>T), GAA2396GAG (7313A>G), ACA2398ACT (7319A>T), AGT2399TGT (7320A>T), TTG2400CTC (7323T>C 7325G>C), TTT2402TTC (7331T>C), GAT2403AAC (7332G>A 7334T>C), GCT2404GCC (7337T>C), CTC2405CTT (7340C>T), AAT2406AAG (7343T>G), GGA2410GGG (7355A>G), CCA2411CCT (7358A>T), TAC2412TAT (7361C>T), ATT2413ATC (7364T>C), ATG2417TTG (7374A>T), GAA2418CAA (7377G>C), GGG2419AAG (7380G>A 7381G>A), ATT2420ATC (7385T>C), GGA2421GGT (7388A>T), CTA2422CAT (7390T>A 7391A>T), GGA2424GGT (7397A>T), TTA2425CTT (7398T>C 7400A>T), TAT2426AAC (7401T>A 7403T>C), AAG2427AAC (7406G>C), TTG2428TTA (7409G>A), GTG2429TTG (7410G>T), GAG2430ATG (7413G>A 7414A>T), CCG2431GCT (7416C>G 7418G>T), CAA2433GAG (7422C>G 7424A>G), AAT2434GAT (7425A>G), ATG2436ACA (7432T>C 7433G>A), GCT2437GCA (7436T>A), AGT2438TAT (7437A>T 7438G>A), GCT2439GCC (7442T>C), CTC2440ATG (7443C>A 7445C>G), TGC2441TGT (7448C>T), GTG2442ATC (7449G>A 7451G>C), GCT2444TCA (7455G>T 7457T>A), TTT2445CTT (7458T>C), GTA2446GCT (7462T>C 7463A>T), AAT2447CTT (7464A>C 7465A>T), AAA2448GGG (7467A>G 7468A>G 7469A>G), GTT2449CCA (7470G>C 7471T>C 7472T>A), GGT2450AAT (7473G>A 7474G>A), GAT2451ACT (7476G>A 7477A>C), GAT2452GAG (7481T>G), CCT2453CCG (7484T>G), ATA2454ATG (7487A>G), ATC2455ACT (7489T>C 7490C>T), AAG2457GTT (7494A>G 7495A>T 7496G>T), GGT2458GGA (7499T>A), GTG2459AAA (7500G>A 7501T>A 7502G>A), CTA2460ACG (7503C>A 7504T>C 7505A>G), AGA2461CTG (7506A>C 7507G>T 7508A>G), GGT2462GGA (7511T>A), GAG2463AAG (7512G>A), ATT2464ATA (7517T>A), GTA2465GTG (7520A>G), ATG2466CCA (7521A>C 7522T>C 7523G>A), CCA2467GCT (7524C>G 7526A>T), CGC2468AGG (7527C>A 7529C>G), CCA2470CCC (7535A>C), TCA2472AAT (7539T>A 7540C>A 7541A>T), TTT2473TTC (7544T>C), GGG2474GGA (7547G>A), GAC2476GAT (7553C>T), CCA2477CCG (7556A>G), ATT2478ATA (7559T>A), TTC2479TTT (7562C>T), CCA2481CCT (7568A>T), CTT2482CAT (7570T>A), AAC2483GGC (7572A>G 7573A>G), TGG2484TAT (7576G>A 7577G>T), AGA2485GAC (7578A>G 7579G>A 7580A>C), AAG2486CAG (7581A>C), ACA2487ACT (7586A>T), TTT2488TAC (7588T>A 7589T>C), ATG2492CCC (7599A>C 7600T>C 7601G>C), GCC2493AAG (7602G>A 7603C>A 7604C>G), GAG2494GAA (7607G>A), GAG2495GAA (7610G>A), AAT2497AAC (7616T>C), ATG2498AAG (7618T>A), ATA2499ATT (7622A>T), CAT2501CAC (7628T>C), CGA2502CGC (7631A>C), TTT2503GGT (7632T>G 7633T>G), CGA2504AAA (7635C>A 7636G>A) | | | | | | | | |
| **Proteins** | | | | | | | | | |
| polyprotein (YP_007027011.1) | 2336 | 2509 | 6.0% | 732 | 60.0% | 174 (100%) | 100 (57.5%) | 0/0/0/0 | 0 |
| Protein mutations: | I2337V (7134A>G 7136T>G), Q2345K (7158C>A), K2348E (7167A>G), K2351R (7177A>G 7178A>G), Q2352A (7179C>G 7180A>C), F2354L (7185T>C), P2356Q (7192C>A 7193C>G), T2357S (7194A>T), I2360F (7203A>T 7205C>T), I2361Q (7206A>C 7207T>A 7208C>G), R2363L (7213G>T 7214G>T), V2365L (7218G>C), P2370L (7234C>T 7235A>T), T2373E (7242A>G 7243C>A 7244A>G), V2374P (7245G>C 7246T>C), E2376D (7253A>T), I2378S (7257A>T 7258T>C), V2382A (7270T>C 7271A>T), R2383K (7272C>A 7273G>A 7274T>A), V2384I (7275G>A 7277A>T), S2386A (7281T>G 7283T>C), E2387K (7284G>A 7286G>A), L2388E (7287T>G 7288T>A), I2389V (7290A>G 7292T>G), G2390N (7293G>A 7294G>A 7295A>T), S2399C (7320A>T), D2403N (7332G>A 7334T>C), N2406K (7343T>G), M2417L (7374A>T), E2418Q (7377G>C), G2419K (7380G>A 7381G>A), L2422H (7390T>A 7391A>T), Y2426N (7401T>A 7403T>C), K2427N (7406G>C), V2429L (7410G>T), E2430M (7413G>A 7414A>T), P2431A (7416C>G 7418G>T), Q2433E (7422C>G 7424A>G), N2434D (7425A>G), M2436T (7432T>C 7433G>A), S2438Y (7437A>T 7438G>A), L2440M (7443C>A 7445C>G), V2442I (7449G>A 7451G>C), A2444S (7455G>T 7457T>A), F2445L (7458T>C), V2446A (7462T>C 7463A>T), N2447L (7464A>C 7465A>T), K2448G (7467A>G 7468A>G 7469A>G), V2449P (7470G>C 7471T>C 7472T>A), G2450N (7473G>A 7474G>A), D2451T (7476G>A 7477A>C), D2452E (7481T>G), I2454M (7487A>G), I2455T (7489T>C 7490C>T), K2457V (7494A>G 7495A>T 7496G>T), V2459K (7500G>A 7501T>A 7502G>A), L2460T (7503C>A 7504T>C 7505A>G), R2461L (7506A>C 7507G>T 7508A>G), E2463K (7512G>A), M2466P (7521A>C 7522T>C 7523G>A), P2467A (7524C>G 7526A>T), S2472N (7539T>A 7540C>A 7541A>T), L2482H (7570T>A), N2483G (7572A>G 7573A>G), W2484Y (7576G>A 7577G>T), R2485D (7578A>G 7579G>A 7580A>C), K2486Q (7581A>C), F2488Y (7588T>A 7589T>C), M2492P (7599A>C 7600T>C 7601G>C), A2493K (7602G>A 7603C>A 7604C>G), M2498K (7618T>A), F2503G (7632T>G 7633T>G), R2504K (7635C>A 7636G>A) | | | | | | | | |

| | Begin | End | Coverage | Score | Concordance | Matches | Identities | I/D/M/F* | Stop Codons |
|---|---|---|---|---|---|---|---|---|---|
| **NT** | **7129** | **7650** | **5.8%** | **220** | **21.1%** | **522 (100%)** | **316 (60.5%)** | **0/0** | |
| Codon mutations: | GCA2335.CG (7130A>G), CCA2336CCG (7133A>G), ATT2337GTG (7134A>G 7136T>G), TTC2339TTT (7142C>T), GTA2340GTT (7145A>T), ACA2341ACC (7148A>C), GGG2342GGA (7151G>A), GCA2344GCT (7157A>T), CAA2345AAA (7158C>A), AAA2346AAG (7163A>G), AAG2348GAG (7167A>G), GTA2350GTT (7175A>T), AAA2351AGG (7177A>G 7178A>G), CAA2352GCA (7179C>G 7180A>C), ATA2353ATT (7184A>T), TTT2354CTT (7185T>C), GGT2355GGC (7190T>C), CCC2356CAG (7192C>A 7193C>G), ACT2357TCT (7194A>T), CCA2359CCC (7202A>C), ATC2360TTT (7203A>T 7205C>T), ATC2361CAG (7206A>C 7207T>A 7208C>G), TCT2362TCC (7211T>C), CGG2363CTT (7213G>T 7214G>T), AAA2364AAG (7217A>G), GTT2365CTT (7218G>C), CCA2368CCT (7229A>T), CCA2370CTT (7234C>T 7235A>T), GGA2372GGG (7241A>G), ACA2373GAG (7242A>G 7243C>A 7244A>G), GTT2374CCT (7245G>C 7246T>C), GAG2375GAA (7250G>A), GAA2376GAT (7253A>T), ATT2377ATA (7256T>A), ATC2378TCC (7257A>T 7258T>C), GTA2382GCT (7270T>C 7271A>T), CGT2383AAA (7272C>A 7273G>A 7274T>A), GTA2384ATT (7275G>A 7277A>T), GCG2385GCT (7280G>T), TCT2386GCC (7281T>G 7283T>C), GAG2387AAA (7284G>A 7286G>A), TTG2388GAG (7287T>G 7288T>A), ATT2389GTG (7290A>G 7292T>G), GGA2390AAT (7293G>A 7294G>A 7295A>T), GGG2391GGA (7298G>A), GTT2393GTG (7304T>G), CTC2394CTA (7307C>A), GTA2395GTT (7310A>T), GAA2396GAG (7313A>G), ACA2398ACT (7319A>T), AGT2399TGT (7320A>T), TTG2400CTC (7323T>C 7325G>C), TTT2402TTC (7331T>C), GAT2403AAC (7332G>A 7334T>C), GCT2404GCC (7337T>C), CTC2405CTT (7340C>T), AAT2406AAG (7343T>G), GGA2410GGG (7355A>G), CCA2411CCT (7358A>T), TAC2412TAT (7361C>T), ATT2413ATC (7364T>C), ATG2417TTG (7374A>T), GAA2418CAA (7377G>C), GGG2419AAG (7380G>A 7381G>A), ATT2420ATC (7385T>C), GGA2421GGT (7388A>T), CTA2422CAT (7390T>A 7391A>T), GGA2424GGT (7397A>T), TTA2425CTT (7398T>C 7400A>T), TAT2426AAC (7401T>A 7403T>C), AAG2427AAC (7406G>C), TTG2428TTA (7409G>A), GTG2429TTG (7410G>T), GAG2430ATG (7413G>A 7414A>T), CCG2431GCT (7416C>G 7418G>T), CAA2433GAG (7422C>G 7424A>G), AAT2434GAT (7425A>G), ATG2436ACA (7432T>C 7433G>A), GCT2437GCA (7436T>A), AGT2438TAT (7437A>T 7438G>A), GCT2439GCC (7442T>C), CTC2440ATG (7443C>A 7445C>G), TGC2441TGT (7448C>T), GTG2442ATC (7449G>A 7451G>C), GCT2444TCA (7455G>T 7457T>A), TTT2445CTT (7458T>C), GTA2446GCT (7462T>C 7463A>T), AAT2447CTT (7464A>C 7465A>T), AAA2448GGG (7467A>G 7468A>G 7469A>G), GTT2449CCA (7470G>C 7471T>C 7472T>A), GGT2450AAT (7473G>A 7474G>A), GAT2451ACT (7476G>A 7477A>C), GAT2452GAG (7481T>G), CCT2453CCG (7484T>G), ATA2454ATG (7487A>G), ATC2455ACT (7489T>C 7490C>T), AAG2457GTT (7494A>G 7495A>T 7496G>T), GGT2458GGA (7499T>A), GTG2459AAA (7500G>A 7501T>A 7502G>A), CTA2460ACG (7503C>A 7504T>C 7505A>G), AGA2461CTG (7506A>C 7507G>T 7508A>G), GGT2462GGA (7511T>A), GAG2463AAG (7512G>A), ATT2464ATA (7517T>A), GTA2465GTG (7520A>G), ATG2466CCA (7521A>C 7522T>C 7523G>A), CCA2467GCT (7524C>G 7526A>T), CGC2468AGG (7527C>A 7529C>G), CCA2470CCC (7535A>C), TCA2472AAT (7539T>A 7540C>A 7541A>T), TTT2473TTC (7544T>C), GGG2474GGA (7547G>A), GAC2476GAT (7553C>T), CCA2477CCG (7556A>G), ATT2478ATA (7559T>A), TTC2479TTT (7562C>T), CCA2481CCT (7568A>T), CTT2482CAT (7570T>A), AAC2483GGC (7572A>G 7573A>G), TGG2484TAT (7576G>A 7577G>T), AGA2485GAC (7578A>G 7579G>A 7580A>C), AAG2486CAG (7581A>C), ACA2487ACT (7586A>T), TTT2488TAC (7588T>A 7589T>C), ATG2492CCC (7599A>C 7600T>C 7601G>C), GCC2493AAG (7602G>A 7603C>A 7604C>G), GAG2494GAA (7607G>A), GAG2495GAA (7610G>A), AAT2497AAC (7616T>C), ATG2498AAG (7618T>A), ATA2499ATT (7622A>T), CAT2501CAC (7628T>C), CGA2502CGC (7631A>C), TTT2503GGT (7632T>G 7633T>G), CGA2504AAA (7635C>A 7636G>A) | | | | | | | | |
| HAM1-like protein (YP_007032446.1) | 28 | 201 | 77.0% | 732 | 60.0% | 174 (100%) | 100 (57.5%) | 0/0/0/0 | 0 |
| Protein mutations: | I29V (7134A>G 7136T>G), Q37K (7158C>A), K40E (7167A>G), K43R (7177A>G 7178A>G), Q44A (7179C>G 7180A>C), F46L (7185T>C), P48Q (7192C>A 7193C>G), T49S (7194A>T), I52F (7203A>T 7205C>T), I53Q (7206A>C 7207T>A 7208C>G), R55L (7213G>T 7214G>T), V57L (7218G>C), P62L (7234C>T 7235A>T), T65E (7242A>G 7243C>A 7244A>G), V66P (7245G>C 7246T>C), E68D (7253A>T), I70S (7257A>T 7258T>C), V74A (7270T>C 7271A>T), R75K (7272C>A 7273G>A 7274T>A), V76I (7275G>A 7277A>T), S78A (7281T>G 7283T>C), E79K (7284G>A 7286G>A), L80E (7287T>G 7288T>A), I81V (7290A>G 7292T>G), G82N (7293G>A 7294G>A 7295A>T), S91C (7320A>T), D95N (7332G>A 7334T>C), N98K (7343T>G), M109L (7374A>T), E110Q (7377G>C), G111K (7380G>A 7381G>A), L114H (7390T>A 7391A>T), Y118N (7401T>A 7403T>C), K119N (7406G>C), V121L (7410G>T), E122M (7413G>A 7414A>T), P123A (7416C>G 7418G>T), Q125E (7422C>G 7424A>G), N126D (7425A>G), M128T (7432T>C 7433G>A), S130Y (7437A>T 7438G>A), L132M (7443C>A 7445C>G), V134I (7449G>A 7451G>C), A136S (7455G>T 7457T>A), F137L (7458T>C), V138A (7462T>C 7463A>T), N139L (7464A>C 7465A>T), K140G (7467A>G 7468A>G 7469A>G), V141P (7470G>C 7471T>C 7472T>A), G142N (7473G>A 7474G>A), D143T (7476G>A 7477A>C), D144E (7481T>G), I146M (7487A>G), I147T (7489T>C 7490C>T), K149V (7494A>G 7495A>T 7496G>T), V151K (7500G>A 7501T>A 7502G>A), L152T (7503C>A 7504T>C 7505A>G), R153L (7506A>C 7507G>T 7508A>G), E155K (7512G>A), M158P (7521A>C 7522T>C 7523G>A), P159A (7524C>G 7526A>T), S164N (7539T>A 7540C>A 7541A>T), L174H (7570T>A), N175G (7572A>G 7573A>G), W176Y (7576G>A 7577G>T), R177D (7578A>G 7579G>A 7580A>C), K178Q (7581A>C), F180Y (7588T>A 7589T>C), M184P (7599A>C 7600T>C 7601G>C), A185K (7602G>A 7603C>A 7604C>G), M190K (7618T>A), F195G (7632T>G 7633T>G), R196K (7635C>A 7636G>A) | | | | | | | | |
| Codon mutations: | GCA27.CG (7130A>G), CCA28CCG (7133A>G), ATT29GTG (7134A>G 7136T>G), TTC31TTT (7142C>T), GTA32GTT (7145A>T), ACA33ACC (7148A>C), GGG34GGA (7151G>A), GCA36GCT (7157A>T), CAA37AAA (7158C>A), AAA38AAG (7163A>G), AAG40GAG (7167A>G), GTA42GTT (7175A>T), AAA43AGG (7177A>G 7178A>G), CAA44GCA (7179C>G 7180A>C), ATA45ATT (7184A>T), TTT46CTT (7185T>C), GGT47GGC (7190T>C), CCC48CAG (7192C>A 7193C>G), ACT49TCT (7194A>T), CCA51CCC (7202A>C), ATC52TTT (7203A>T 7205C>T), ATC53CAG (7206A>C 7207T>A 7208C>G), TCT54TCC (7211T>C), CGG55CTT (7213G>T 7214G>T), AAA56AAG (7217A>G), GTT57CTT (7218G>C), CCA60CCT (7229A>T), CCA62CTT (7234C>T 7235A>T), GGA64GGG (7241A>G), ACA65GAG (7242A>G 7243C>A 7244A>G), GTT66CCT (7245G>C 7246T>C), GAG67GAA (7250G>A), GAA68GAT (7253A>T), ATT69ATA (7256T>A), ATC70TCC (7257A>T 7258T>C), GTA74GCT (7270T>C 7271A>T), CGT75AAA (7272C>A 7273G>A 7274T>A), GTA76ATT (7275G>A 7277A>T), GCG77GCT (7280G>T), TCT78GCC (7281T>G 7283T>C), GAG79AAA (7284G>A 7286G>A), TTG80GAG (7287T>G 7288T>A), ATT81GTG (7290A>G 7292T>G), GGA82AAT (7293G>A 7294G>A 7295A>T), GGG83GGA (7298G>A), GTT85GTG (7304T>G), CTC86CTA (7307C>A), GTA87GTT (7310A>T), GAA88GAG (7313A>G), ACA90ACT (7319A>T), AGT91TGT (7320A>T), TTG92CTC (7323T>C 7325G>C), TTT94TTC (7331T>C), GAT95AAC (7332G>A 7334T>C), GCT96GCC (7337T>C), CTC97CTT (7340C>T), AAT98AAG (7343T>G), GGA102GGG (7355A>G), CCA103CCT (7358A>T), TAC104TAT (7361C>T), ATT105ATC (7364T>C), ATG109TTG (7374A>T), GAA110CAA (7377G>C), GGG111AAG (7380G>A 7381G>A), ATT112ATC (7385T>C), GGA113GGT (7388A>T), CTA114CAT (7390T>A 7391A>T), GGA116GGT (7397A>T), TTA117CTT (7398T>C 7400A>T), TAT118AAC (7401T>A 7403T>C), AAG119AAC (7406G>C), TTG120TTA (7409G>A), GTG121TTG (7410G>T), GAG122ATG (7413G>A 7414A>T), CCG123GCT (7416C>G 7418G>T), CAA125GAG (7422C>G 7424A>G), AAT126GAT (7425A>G), ATG128ACA (7432T>C 7433G>A), GCT129GCA (7436T>A), AGT130TAT (7437A>T 7438G>A), GCT131GCC (7442T>C), CTC132ATG (7443C>A 7445C>G), TGC133TGT (7448C>T), GTG134ATC (7449G>A 7451G>C), GCT136TCA (7455G>T 7457T>A), TTT137CTT (7458T>C), GTA138GCT (7462T>C 7463A>T), AAT139CTT (7464A>C 7465A>T), AAA140GGG (7467A>G 7468A>G 7469A>G), GTT141CCA (7470G>C 7471T>C 7472T>A), GGT142AAT (7473G>A 7474G>A), GAT143ACT (7476G>A 7477A>C), GAT144GAG (7481T>G), CCT145CCG (7484T>G), ATA146ATG (7487A>G), ATC147ACT (7489T>C 7490C>T), AAG149GTT (7494A>G 7495A>T 7496G>T), GGT150GGA (7499T>A), GTG151AAA (7500G>A 7501T>A 7502G>A), CTA152ACG (7503C>A 7504T>C 7505A>G), AGA153CTG (7506A>C 7507G>T 7508A>G), GGT154GGA (7511T>A), GAG155AAG (7512G>A), ATT156ATA (7517T>A), GTA157GTG (7520A>G), ATG158CCA (7521A>C 7522T>C 7523G>A), CCA159GCT (7524C>G 7526A>T), CGC160AGG (7527C>A 7529C>G), CCA162CCC (7535A>C), TCA164AAT (7539T>A 7540C>A 7541A>T), TTT165TTC (7544T>C), GGG166GGA (7547G>A), GAC168GAT (7553C>T), CCA169CCG (7556A>G), ATT170ATA (7559T>A), TTC171TTT (7562C>T), CCA173CCT (7568A>T), CTT174CAT (7570T>A), AAC175GGC (7572A>G 7573A>G), TGG176TAT (7576G>A 7577G>T), AGA177GAC (7578A>G 7579G>A 7580A>C), AAG178CAG (7581A>C), ACA179ACT (7586A>T), TTT180TAC (7588T>A 7589T>C), ATG184CCC (7599A>C 7600T>C 7601G>C), GCC185AAG (7602G>A 7603C>A 7604C>G), GAG186GAA (7607G>A), GAG187GAA (7610G>A), AAT189AAC (7616T>C), ATG190AAG (7618T>A), ATA191ATT (7622A>T), CAT193CAC (7628T>C), CGA194CGC (7631A>C), TTT195GGT (7632T>G 7633T>G), CGA196AAA (7635C>A 7636G>A) | | | | | | | | |

*: Inserts / Deletes / Misaligned / Frameshifts

#### Analysis details

This analysis was performed with panviral2.64

### NGS Details (UN9): Badnavirus maculaucubae

#### Assembly

| | |
|---|---|
| **Coverage Length** | 282 (1 contig(s)) |
| **Depth Of Coverage** | 52.7 |
| **Number Of Reads** | 137 |
| **Reads Per Million** | 3.10 rpm (after QC) |
| **Ambiguities** | 0 |
| **Assembly Method** | de novo + reference guided assembly |
| **Consensus Caller** | Bcf Tools |

#### Coverage Map

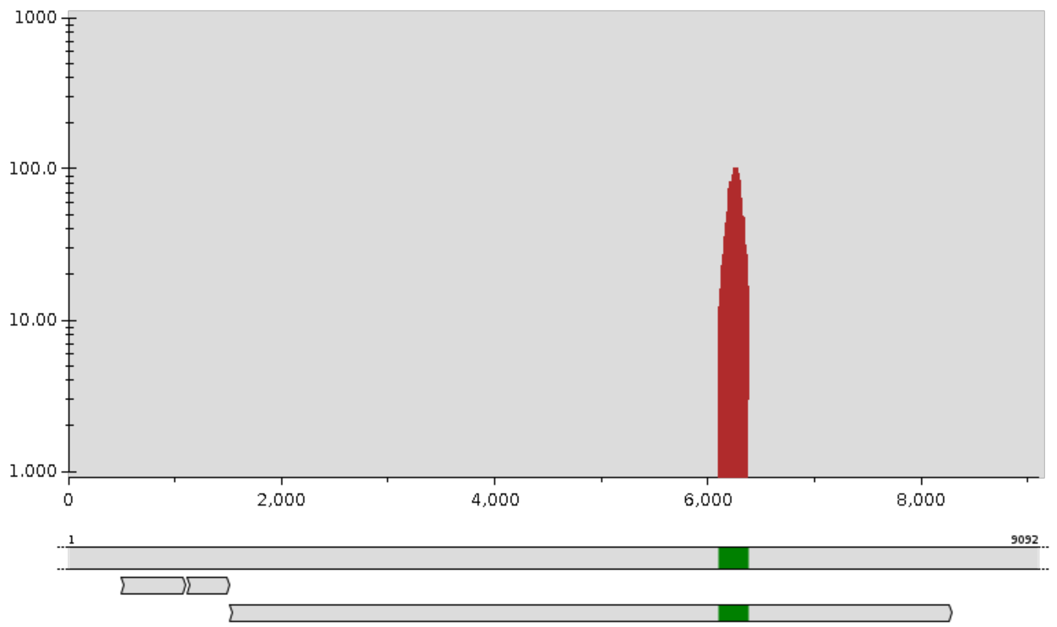

#### Assignment

| | |
|---|---|
| **Type** | Badnavirus maculaucubae (Taxonomy ID: 3051986) |
| **Reference Genome** | NC_076606.1 |
| **NT Identity (%)** | 56.1404 |
| **AA Identity (%)** | 52.6316 |
| **Number Of Stop Codons** | 0 |
| **Number Of CDS** | 3 |

#### Alignment

| | |
|---|---|
| **Alignment Score** | 64.0 (NT) + 355.0 (AA) = 419.0 |
| **Concordance (%)** | 33.6276 |

| Alignment Method | Global, seeded, nucleotide + amino acids (AGA) |
|---|---|

#### Genome Region

Sequence starts at position 6095 and ends at position 6376 relative to NC_076606.1 reference sequence.

#### Alignment Detailed Statistics

| | Begin | End | Coverage | Score | Concordance | Matches | Identities | I/D/M/F* | Stop Codons |
|---|---|---|---|---|---|---|---|---|---|
| **NT** | **6095** | **6376** | **3.1%** | **64** | **11.3%** | **282 (98.9%)** | **160 (56.1%)** | **3/0** | |
| Mutations: | 6095T>G, 6100A>T, 6101G>C, 6102C>A, 6105A>G, 6106T>A, 6108T>A, 6112T>C, 6114G>T, 6115A>C, 6116A>G, 6120T>A, 6125T>A, 6133G>T, 6136G>A, 6137C>A, 6138A>G, 6139A>G, 6141G>C, 6142G>A, 6144A>G, 6147A>G, 6148T>A, 6149C>G, 6150A>T, 6151A>G, 6152G>A, 6159A>C, 6160T>A, 6161G>A, 6165G>T, 6168T>G, 6172A>C, 6173T>G, 6177T>A, 6178C>A, 6179C>G, 6180A>G, 6181G>T, 6186A>G, 6187C>A, 6188T>A, 6189A>T, 6191A>T, 6195A>G, 6197G>T, 6198G>T, 6201A>T, 6204A>C, 6208C>T, 6210A>C, 6217C>T, 6221A>G, 6222G>T, 6228A>T, 6231T>G, 6234A>T, 6237G>A, 6240C>T, 6241C>A, 6242A>T, 6243A>G, 6244A>G, 6245G>A, 6246G>T, 6247A>C, 6249A>T, 6253G>A, 6255C>T, 6258C>G, 6259T>A, 6260G>T, 6261T>A, 6266G>A, 6268G>C, 6269G>C, 6271A>T, 6272C>A, 6273A>C, 6274G>C, 6275A>T, 6276A>T, 6279T>C, 6279_6280insATG, 6281T>A, 6282C>T, 6283A>G, 6285C>G, 6287C>T, 6288T>G, 6291T>G, 6293A>T, 6294C>T, 6303T>C, 6306T>C, 6312A>C, 6314T>A, 6315C>T, 6318T>G, 6320A>G, 6322A>G, 6324T>C, 6325C>G, 6326T>A, 6327T>A, 6330A>G, 6333G>T, 6337G>A, 6338A>T, 6342G>A, 6346C>T, 6348C>G, 6350A>G, 6354A>T, 6355A>G, 6358C>T, 6361G>C, 6363A>G, 6364A>G, 6366A>T, 6368G>T, 6369C>G, 6373A>G | | | | | | | | |

##### CDS

| | Begin | End | Coverage | Score | Concordance | Matches | Identities | I/D/M/F* | Stop Codons |
|---|---|---|---|---|---|---|---|---|---|
| QKP79_gp3 | 1530 | 1623 | 4.2% | 355 | 51.1% | 94 (98.9%) | 50 (52.6%) | 1/0/0/0 | 0 |
| Protein mutations: | F1533I (6106T>A 6108T>A), K1536R (6115A>C 6116A>G), F1539Y (6125T>A), V1542L (6133G>T), A1543K (6136G>A 6137C>A 6138A>G), M1544V (6139A>G 6141G>C), E1545K (6142G>A 6144A>G), S1548D (6151A>G 6152G>A), W1551K (6160T>A 6161G>A), I1555R (6172A>C 6173T>G), P1557R (6178C>A 6179C>G 6180A>G), D1558Y (6181G>T), L1560N (6187C>A 6188T>A 6189A>T), Y1561F (6191A>T), W1563F (6197G>T 6198G>T), P1567S (6208C>T 6210A>C), K1571S (6221A>G 6222G>T), Q1578M (6241C>A 6242A>T 6243A>G), R1579D (6244A>G 6245G>A 6246G>T), K1580H (6247A>C 6249A>T), D1582N (6253G>A 6255C>T), N1583K (6258C>G), C1584I (6259T>A 6260G>T 6261T>A), R1586K (6266G>A), G1587P (6268G>C 6269G>C), T1588Y (6271A>T 6272C>A 6273A>C), E1589L (6274G>C 6275A>T 6276A>T), D1590_F1591insM (6279_6280insATG), F1591Y (6281T>A 6282C>T), I1592V (6283A>G 6285C>G), A1593V (6287C>T 6288T>G), Y1595F (6293A>T 6294C>T), F1602Y (6314T>A 6315C>T), E1604G (6320A>G), N1605D (6322A>G 6324T>C), L1606E (6325C>G 6326T>A 6327T>A), E1608D (6333G>T), E1610M (6337G>A 6338A>T), K1614R (6350A>G), M1616V (6355A>G), E1618Q (6361G>C 6363A>G), I1619V (6364A>G 6366A>T), C1620L (6368G>T 6369C>G), K1622E (6373A>G) | | | | | | | | |
| Codon mutations: | GTC1529.GC (6095T>G), AGC1531TCA (6100A>T 6101G>C 6102C>A), AAA1532AAG (6105A>G), TTT1533ATA (6106T>A 6108T>A), TTG1535CTT (6112T>C 6114G>T), AAA1536CGA (6115A>C 6116A>G), TCT1537TCA (6120T>A), TTT1539TAT (6125T>A), GTG1542TTG (6133G>T), GCA1543AAG (6136G>A 6137C>A 6138A>G), ATG1544GTC (6139A>G 6141G>C), GAA1545AAG (6142G>A 6144A>G), GAA1546GAG (6147A>G), TCA1547AGT (6148T>A 6149C>G 6150A>T), AGC1548GAC (6151A>G 6152G>A), CCA1550CCC (6159A>C), TGG1551AAG (6160T>A 6161G>A), ACG1552ACT (6165G>T), GCT1553GCG (6168T>G), ATA1555CGA (6172A>C 6173T>G), ACT1556ACA (6177T>A), CCA1557AGG (6178C>A 6179C>G 6180A>G), GAT1558TAT (6181G>T), GGA1559GGG (6186A>G), CTA1560AAT (6187C>A 6188T>A 6189A>T), TAC1561TTC (6191A>T), GAA1562GAG (6195A>G), TGG1563TTT (6197G>T 6198G>T), CTA1564CTT (6201A>T), GTA1565GTC (6204A>C), CCA1567TCC (6208C>T 6210A>C), CTA1570TTA (6217C>T), AAG1571AGT (6221A>G 6222G>T), GCA1573GCT (6228A>T), CCT1574CCG (6231T>G), GCA1575GCT (6234A>T), GTG1576GTA (6237G>A), TTC1577TTT (6240C>T), CAA1578ATG (6241C>A 6242A>T 6243A>G), AGG1579GAT (6244A>G 6245G>A 6246G>T), AAA1580CAT (6247A>C 6249A>T), GAC1582AAT (6253G>A 6255C>T), AAC1583AAG (6258C>G), TGT1584ATA (6259T>A 6260G>T 6261T>A), AGG1586AAG (6266G>A), GGT1587CCT (6268G>C 6269G>C), ACA1588TAC (6271A>T 6272C>A 6273A>C), GAA1589CTT (6274G>C 6275A>T 6276A>T), GAT1590GAC (6279T>C), GAT1590_TTC1591insATG (6279_6280insATG), TTC1591TAT (6281T>A 6282C>T), ATC1592GTG (6283A>G 6285C>G), GCT1593GTG (6287C>T 6288T>G), GTT1594GTG (6291T>G), TAC1595TTT (6293A>T 6294C>T), GAT1598GAC (6303T>C), ATT1599ATC (6306T>C), GTA1601GTC (6312A>C), TTC1602TAT (6314T>A 6315C>T), TCT1603TCG (6318T>G), GAA1604GGA (6320A>G), AAT1605GAC (6322A>G 6324T>C), CTT1606GAA (6325C>G 6326T>A 6327T>A), CAA1607CAG (6330A>G), GAG1608GAT (6333G>T), GAG1610ATG (6337G>A 6338A>T), GAG1611GAA (6342G>A), CTC1613TTG (6346C>T 6348C>G), AAA1614AGA (6350A>G), ATA1615ATT (6354A>T), ATG1616GTG (6355A>G), CTG1617TTG (6358C>T), GAA1618CAG (6361G>C 6363A>G), ATA1619GTT (6364A>G 6366A>T), TGC1620TTG (6368G>T 6369C>G), AAG1622GAG (6373A>G) | | | | | | | | |

##### Proteins

| | Begin | End | Coverage | Score | Concordance | Matches | Identities | I/D/M/F* | Stop Codons |
|---|---|---|---|---|---|---|---|---|---|
| hypothetical protein (YP_010799265.1) | 1530 | 1623 | 4.2% | 355 | 51.1% | 94 (98.9%) | 50 (52.6%) | 1/0/0/0 | 0 |
| Protein mutations: | F1533I (6106T>A 6108T>A), K1536R (6115A>C 6116A>G), F1539Y (6125T>A), V1542L (6133G>T), A1543K (6136G>A 6137C>A 6138A>G), M1544V (6139A>G 6141G>C), E1545K (6142G>A 6144A>G), S1548D (6151A>G 6152G>A), W1551K (6160T>A 6161G>A), I1555R (6172A>C 6173T>G), P1557R (6178C>A 6179C>G 6180A>G), D1558Y (6181G>T), L1560N (6187C>A 6188T>A 6189A>T), Y1561F (6191A>T), W1563F (6197G>T 6198G>T), P1567S (6208C>T 6210A>C), K1571S (6221A>G 6222G>T), Q1578M (6241C>A 6242A>T 6243A>G), R1579D (6244A>G 6245G>A 6246G>T), K1580H (6247A>C 6249A>T), D1582N (6253G>A 6255C>T), N1583K (6258C>G), C1584I (6259T>A 6260G>T 6261T>A), R1586K (6266G>A), G1587P (6268G>C 6269G>C), T1588Y (6271A>T 6272C>A 6273A>C), E1589L (6274G>C 6275A>T 6276A>T), D1590_F1591insM (6279_6280insATG), F1591Y (6281T>A 6282C>T), I1592V (6283A>G 6285C>G), A1593V (6287C>T 6288T>G), Y1595F (6293A>T 6294C>T), F1602Y (6314T>A 6315C>T), E1604G (6320A>G), N1605D (6322A>G 6324T>C), L1606E (6325C>G 6326T>A 6327T>A), E1608D (6333G>T), E1610M (6337G>A 6338A>T), K1614R (6350A>G), M1616V (6355A>G), E1618Q (6361G>C 6363A>G), I1619V (6364A>G 6366A>T), C1620L (6368G>T 6369C>G), K1622E (6373A>G) | | | | | | | | |
| Codon mutations: | GTC1529.GC (6095T>G), AGC1531TCA (6100A>T 6101G>C 6102C>A), AAA1532AAG (6105A>G), TTT1533ATA (6106T>A 6108T>A), TTG1535CTT (6112T>C 6114G>T), AAA1536CGA (6115A>C 6116A>G), TCT1537TCA (6120T>A), TTT1539TAT (6125T>A), GTG1542TTG (6133G>T), GCA1543AAG (6136G>A 6137C>A 6138A>G), ATG1544GTC (6139A>G 6141G>C), GAA1545AAG (6142G>A 6144A>G), GAA1546GAG (6147A>G), TCA1547AGT (6148T>A 6149C>G 6150A>T), AGC1548GAC (6151A>G 6152G>A), CCA1550CCC (6159A>C), TGG1551AAG (6160T>A 6161G>A), ACG1552ACT (6165G>T), GCT1553GCG (6168T>G), ATA1555CGA (6172A>C 6173T>G), ACT1556ACA (6177T>A), CCA1557AGG (6178C>A 6179C>G 6180A>G), GAT1558TAT (6181G>T), GGA1559GGG (6186A>G), CTA1560AAT (6187C>A 6188T>A 6189A>T), TAC1561TTC (6191A>T), GAA1562GAG (6195A>G), TGG1563TTT (6197G>T 6198G>T), CTA1564CTT (6201A>T), GTA1565GTC (6204A>C), CCA1567TCC (6208C>T 6210A>C), CTA1570TTA (6217C>T), AAG1571AGT (6221A>G 6222G>T), GCA1573GCT (6228A>T), CCT1574CCG (6231T>G), GCA1575GCT (6234A>T), GTG1576GTA (6237G>A), TTC1577TTT (6240C>T), CAA1578ATG (6241C>A 6242A>T 6243A>G), AGG1579GAT (6244A>G 6245G>A 6246G>T), AAA1580CAT (6247A>C 6249A>T), GAC1582AAT (6253G>A 6255C>T), AAC1583AAG (6258C>G), TGT1584ATA (6259T>A 6260G>T 6261T>A), AGG1586AAG (6266G>A), GGT1587CCT (6268G>C 6269G>C), ACA1588TAC (6271A>T 6272C>A 6273A>C), GAA1589CTT (6274G>C 6275A>T 6276A>T), GAT1590GAC (6279T>C), GAT1590_TTC1591insATG (6279_6280insATG), TTC1591TAT (6281T>A 6282C>T), ATC1592GTG (6283A>G 6285C>G), GCT1593GTG (6287C>T 6288T>G), GTT1594GTG (6291T>G), TAC1595TTT (6293A>T 6294C>T), GAT1598GAC (6303T>C), ATT1599ATC (6306T>C), GTA1601GTC (6312A>C), TTC1602TAT (6314T>A 6315C>T), TCT1603TCG (6318T>G), GAA1604GGA (6320A>G), AAT1605GAC (6322A>G 6324T>C), CTT1606GAA (6325C>G 6326T>A 6327T>A), CAA1607CAG (6330A>G), GAG1608GAT (6333G>T), GAG1610ATG (6337G>A 6338A>T), GAG1611GAA (6342G>A), CTC1613TTG (6346C>T 6348C>G), AAA1614AGA (6350A>G), ATA1615ATT (6354A>T), ATG1616GTG (6355A>G), CTG1617TTG (6358C>T), GAA1618CAG (6361G>C 6363A>G), ATA1619GTT (6364A>G 6366A>T), TGC1620TTG (6368G>T 6369C>G), AAG1622GAG (6373A>G) | | | | | | | | |

*: Inserts / Deletes / Misaligned / Frameshifts

#### Analysis details

This analysis was performed with panviral2.64

### NGS Details (UN9): Errantivirus

#### Assembly

| | |
|---|---|
| **Coverage Length** | 600 (2 contig(s)) |
| **Depth Of Coverage** | 19.5 |
| **Number Of Reads** | 111 |
| **Reads Per Million** | 2.51 rpm (after QC) |
| **Ambiguities** | 0 |
| **Assembly Method** | de novo + reference guided assembly |
| **Consensus Caller** | Bcf Tools |

#### Coverage Map

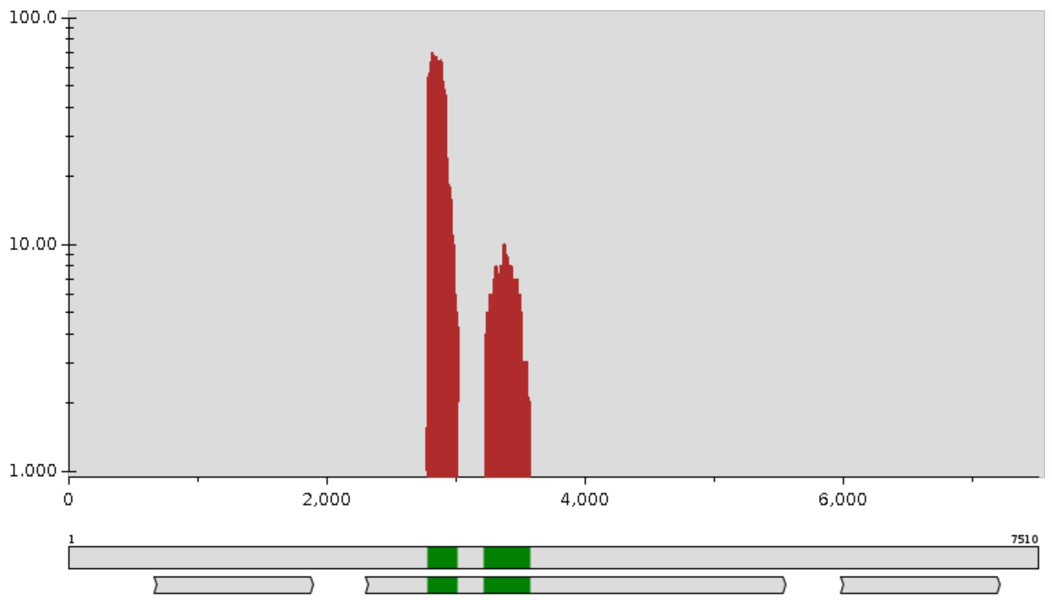

#### Assignment

| | |
|---|---|
| **Type** | Errantivirus (Taxonomy ID: 186666) |
| **Reference Genome** | NC_038512.1 |
| **NT Identity (%)** | 52.921 |
| **AA Identity (%)** | 41.2371 |
| **Number Of Stop Codons** | 0 |
| **Number Of CDS** | 3 |

#### Alignment

| | |
|---|---|
| **Alignment Score** | 14.0 (NT) + 544.0 (AA) = 558.0 |
| **Concordance (%)** | 23.25 |

| Alignment Method | Global, seeded, nucleotide + amino acids (AGA) |
|---|---|

#### Genome Region

Sequence starts at position 2779 and ends at position 3581 relative to NC_038512.1 reference sequence.

#### Alignment Detailed Statistics

| | Begin | End | Coverage | Score | Concordance | Matches | Identities | I/D/M/F* | Stop Codons |
|---|---|---|---|---|---|---|---|---|---|
| **NT** | **2779** | **3581** | **8.0%** | **14** | **1.3%** | **579 (96.0%)** | **308 (51.1%)** | **3/21** | |
| Mutations: | 2786A>T, 2794C>G, 2795C>A, 2797A>G, 2800A>T, 2801A>T, 2809A>G, 2812A>T, 2813T>A, 2814C>G, 2815A>T, 2816G>A, 2818C>G, 2821T>C, 2822G>C, 2828A>G, 2830C>T, 2831T>G, 2832C>T, 2833A>G, 2836C>A, 2837A>G, 2841G>T, 2843G>T, 2849C>A, 2850C>G, 2851C>A, 2858_2860delATC, 2863C>T, 2865C>G, 2866T>C, 2869T>G, 2870_2881delGGGAAACAAAAG, 2883G>T, 2885C>A, 2887T>A, 2888C>A, 2890C>G, 2891G>T, 2892T>G, 2893A>T, 2894G>A, 2896T>A, 2901T>A, 2905T>G, 2908G>A, 2909T>C, 2911G>T, 2913A>G, 2914C>T, 2915G>A, 2918A>G, 2919A>T, 2920G>C, 2923T>C, 2926C>T, 2927G>A, 2929T>G, 2930G>A, 2932C>T, 2934A>G, 2935A>G, 2937A>T, 2944A>T, 2947A>C, 2949A>T, 2950C>G, 2953A>T, 2954A>G, 2955G>A, 2959C>G, 2960G>C, 2963C>T, 2968C>T, 2969A>G, 2974A>G, 2975G>C, 2976G>A, 2977T>A, 2978A>G, 2979A>G, 2981T>G, 2982G>C, 2983C>T, 2984C>A, 2985A>G, 2988A>T, 2993A>T, 2995C>T, 2997C>A, 2998C>A, 2999T>A, 3005T>C, 3007G>T, 3008G>A, 3009C>G, 3010A>T, 3011A>T, 3012G>C, 3013T>G, 3016G>C, 3018T>A, 3019T>C, 3020T>C, 3229A>T, 3233C>T, 3236G>T, 3237A>T, 3239A>C, 3242C>T, 3248C>G, 3249G>A, 3257C>A, 3258A>G, 3259A>G, 3261G>C, 3265T>G, 3269G>C, 3272A>C, 3273G>A, 3274T>A, 3275A>C, 3277C>T, 3280C>A, 3281A>T, 3282A>T, 3283A>G, 3284A>G, 3285T>C, 3286T>A, 3287C>A, 3290A>C, 3291T>A, 3292G>A, 3293G>T, 3294A>C, 3298G>A, 3300C>G, 3301C>T, 3302G>A, 3303A>T, 3304A>G, 3307C>T, 3308T>G, 3309T>G, 3310G>A, 3312A>T, 3313G>T, 3314C>T, 3315T>C, 3316C>T, 3317G>A, 3320A>G, 3321C>T, 3322T>G, 3324C>A, 3325T>G, 3329C>T, 3331T>A, 3334T>C, 3337C>T, 3338A>T, 3344A>T, 3345G>C, 3346C>T, 3347A>T, 3349G>T, 3352C>T, 3356A>G, 3358C>T, 3359A>T, 3360A>C, 3361G>T, 3362C>A, 3365A>G, 3367C>T, 3371G>A, 3372A>G, 3373T>G, 3376G>A, 3380T>C, 3381C>A, 3382C>A, 3384C>T, 3386A>G, 3388T>G, 3391A>G, 3392A>C, 3394A>C, 3396A>G, 3397T>G, 3399T>C, 3400G>T, 3402T>C, 3406A>T, 3408A>G, 3409G>C, 3411C>A, 3414C>T, 3415T>C, 3418G>A, 3419G>C, 3422A>C, 3424A>C, 3426A>G, 3428C>G, 3429A>G, 3430A>T, 3433T>C, 3442T>A, 3443C>G, 3444T>C, 3445C>A, 3448T>G, 3455C>A, 3457A>G, 3467C>A, 3468C>A, 3471A>G, 3472T>G, 3474T>A, 3477C>G, 3478A>T, 3479C>A, 3480G>T, 3481A>T, 3482C>A, 3484C>T, 3486C>G, 3487A>T, 3489A>G, 3490A>G, 3493C>A, 3499A>G, 3500C>G, 3502G>T, 3504G>T, 3505C>G, 3506T>C, 3508A>G, 3514A>G, 3516G>A, 3517T>C, 3519G>A, 3520T>C, 3521A>T, 3522A>T, 3523A>C, 3524G>A, 3527_3532delACTCTT, 3533A>T, 3535T>G, 3536C>T, 3538C>T, 3542T>A, 3543A>T, 3546T>C, 3547A>T, 3549A>C, 3550_3551insCAA, 3551G>C, 3557G>C, 3559A>T, 3560C>A, 3564G>T, 3565T>G, 3568A>G, 3569A>G, 3570C>T, 3571T>A, 3572C>G, 3573T>C, 3574G>A, 3580C>A, 3581A>T | | | | | | | | |

##### CDS

| | Begin | End | Coverage | Score | Concordance | Matches | Identities | I/D/M/F* | Stop Codons |
|---|---|---|---|---|---|---|---|---|---|
| D1R33_gp2 | 159 | 426 | 18.4% | 544 | 41.1% | 193 (96.0%) | 80 (39.8%) | 1/7/0/0 | 0 |
| Protein mutations: | M161L (2786A>T), D163E (2794C>G), Q164K (2795C>A 2797A>G), I166F (2801A>T), D171K (2816G>A 2818C>G), A173P (2822G>C), S175G (2828A>G 2830C>T), S176V (2831T>G 2832C>T 2833A>G), I178V (2837A>G), W179L (2841G>T), V180F (2843G>T), P182R (2849C>A 2850C>G 2851C>A), I185del (2858_2860delATC), A187G (2865C>G 2866T>C), G189_K192del (2870_2881delGGGAAACAAAAG), W193L (2883G>T), L195M (2888C>A 2890C>G), V196C (2891G>T 2892T>G 2893A>T), V197I (2894G>A 2896T>A), F199Y (2901T>A), N203S (2913A>G 2914C>T), E204K (2915G>A), K205V (2918A>G 2919A>T 2920G>C), D208K (2927G>A 2929T>G), D209N (2930G>A 2932C>T), K210R (2934A>G 2935A>G), Y211F (2937A>T), N215M (2949A>T 2950C>G), S217D (2954A>G 2955G>A), D218E (2959C>G), V219L (2960G>C), L220F (2963C>T), K222E (2969A>G), G224Q (2975G>C 2976G>A 2977T>A), K225G (2978A>G 2979A>G), C226A (2981T>G 2982G>C 2983C>T), Q227R (2984C>A 2985A>G), Y228F (2988A>T), T230S (2993A>T 2995C>T), T231K (2997C>A 2998C>A), L232I (2999T>A), A235S (3008G>A 3009C>G 3010A>T), F238Y (3018T>A 3019T>C), E308D (3229A>T), E311L (3236G>T 3237A>T), N312H (3239A>C), R315E (3248C>G 3249G>A), Q318R (3257C>A 3258A>G 3259A>G), R319T (3261G>C), E322Q (3269G>C), S323Q (3272A>C 3273G>A 3274T>A), N324H (3275A>C 3277C>T), F325L (3280C>A), K326L (3281A>T 3282A>T 3283A>G), I327A (3284A>G 3285T>C 3286T>A), Q328K (3287C>A), M329Q (3290A>C 3291T>A 3292G>A), D330S (3293G>T 3294A>C), S332C (3300C>G 3301C>T), E333M (3302G>A 3303A>T 3304A>G), L335G (3308T>G 3309T>G 3310G>A), K336I (3312A>T 3313G>T), L337S (3314C>T 3315T>C 3316C>T), E338K (3317G>A), T339V (3320A>G 3321C>T 3322T>G), A340E (3324C>A 3325T>G), I345F (3338A>T), R348C (3347A>T 3349G>T), I351V (3356A>G 3358C>T), K352S (3359A>T 3360A>C 3361G>T), P353T (3362C>A), N354D (3365A>G 3367C>T), D356R (3371G>A 3372A>G 3373T>G), S359Q (3380T>C 3381C>A 3382C>A), A360V (3384C>T), I361V (3386A>G 3388T>G), K363H (3392A>C 3394A>C), Y364W (3396A>G 3397T>G), L365P (3399T>C 3400G>T), I366T (3402T>C), K368S (3408A>G 3409G>C), T369N (3411C>A), P370L (3414C>T 3415T>C), E372Q (3419G>C), I373L (3422A>C 3424A>C), K374R (3426A>G), Q375G (3428C>G 3429A>G 3430A>T), L380A (3443C>G 3444T>C 3445C>A), P388K (3467C>A 3468C>A), D389G (3471A>G 3472T>G), F390Y (3474T>A), A391G (3477C>G 3478A>T), R392I (3479C>A 3480G>T 3481A>T), L393I (3482C>A 3484C>T), T394S (3486C>G 3487A>T), K395R (3489A>G 3490A>G), Q399D (3500C>G 3502G>T), C400L (3504G>T 3505C>G), G404D (3516G>A 3517T>C), S405N (3519G>A 3520T>C), K406F (3521A>T 3522A>T 3523A>C), V407I (3524G>A), T408_L409del (3527_3532delACTCTT), S410W (3533A>T 3535T>G), P411S (3536C>T 3538C>T), Y413I (3542T>A 3543A>T), V414A (3546T>C 3547A>T), N415T (3549A>C), N415_A416insQ (3550_3551insCAA), A416P (3551G>C), E418H (3557G>C 3559A>T), H419N (3560C>A), C420L (3564G>T 3565T>G), T422V (3569A>G 3570C>T 3571T>A), L423A (3572C>G 3573T>C 3574G>A) | | | | | | | | |
| Codon mutations: | ATG161TTG (2786A>T), GAC163GAG (2794C>G), CAA164AAG (2795C>A 2797A>G), GGA165GGT (2800A>T), ATT166TTT (2801A>T), AGA168AGG (2809A>G), CCA169CCT (2812A>T), TCA170AGT (2813T>A 2814C>G 2815A>T), GAC171AAG (2816G>A 2818C>G), TCT172TCC (2821T>C), GCA173CCA (2822G>C), AGC175GGT (2828A>G 2830C>T), TCA176GTG (2831T>G 2832C>T 2833A>G), CCC177CCA (2836C>A), ATA178GTA (2837A>G), TGG179TTG (2841G>T), GTT180TTT (2843G>T), CCC182AGA (2849C>A 2850C>G 2851C>A), ATC185del (2858_2860delATC), GAC186GAT (2863C>T), GCT187GGC (2865C>G 2866T>C), TCT188TCG (2869T>G), GGG189_AAG192del (2870_2881delGGGAAACAAAAG), TGG193TTG (2883G>T), CGT194AGA (2885C>A 2887T>A), CTC195ATG (2888C>A 2890C>G), GTA196TGT (2891G>T 2892T>G 2893A>T), GTT197ATA (2894G>A 2896T>A), TTC199TAC (2901T>A), CGT200CGG (2905T>G), AAG201AAA (2908G>A), TTG202CTT (2909T>C 2911G>T), AAC203AGT (2913A>G 2914C>T), GAG204AAG (2915G>A), AAG205GTC (2918A>G 2919A>T 2920G>C), ACT206ACC (2923T>C), ATC207ATT (2926C>T), GAT208AAG (2927G>A 2929T>G), GAC209AAT (2930G>A 2932C>T), AAA210AGG (2934A>G 2935A>G), TAC211TTC (2937A>T), ATA213ATT (2944A>T), CCA214CCC (2947A>C), AAC215ATG (2949A>T 2950C>G), ATA216ATT (2953A>T), AGT217GAT (2954A>G 2955G>A), GAC218GAG (2959C>G), GTA219CTA (2960G>C), CTT220TTT (2963C>T), GAC221GAT (2968C>T), AAG222GAG (2969A>G), TTA223TTG (2974A>G), GGT224CAA (2975G>C 2976G>A 2977T>A), AAG225GGG (2978A>G 2979A>G), TGC226GCT (2981T>G 2982G>C 2983C>T), CAA227AGA (2984C>A 2985A>G), TAC228TTC (2988A>T), ACC230TCT (2993A>T 2995C>T), ACC231AAA (2997C>A 2998C>A), TTA232ATA (2999T>A), TTG234CTT (3005T>C 3007G>T), GCA235AGT (3008G>A 3009C>G 3010A>T), AGT236TCG (3011A>T 3012G>C 3013T>G), GGG237GGC (3016G>C), TTT238TAC (3018T>A 3019T>C), TAT239CA. (3020T>C), GAA308GAT (3229A>T), CTG310TTG (3233C>T), GAG311TTG (3236G>T 3237A>T), AAC312CAC (3239A>C), CTG313TTG (3242C>T), CGA315GAA (3248C>G 3249G>A), CAA318AGG (3257C>A 3258A>G 3259A>G), AGA319ACA (3261G>C), CTT320CTG (3265T>G), GAA322CAA (3269G>C), AGT323CAA (3272A>C 3273G>A 3274T>A), AAC324CAT (3275A>C 3277C>T), TTC325TTA (3280C>A), AAA326TTG (3281A>T 3282A>T 3283A>G), ATT327GCA (3284A>G 3285T>C 3286T>A), CAA328AAA (3287C>A), ATG329CAA (3290A>C 3291T>A 3292G>A), GAC330TCC (3293G>T 3294A>C), AAG331AAA (3298G>A), TCC332TGT (3300C>G 3301C>T), GAA333ATG (3302G>A 3303A>T 3304A>G), TTC334TTT (3307C>T), TTG335GGA (3308T>G 3309T>G 3310G>A), AAG336ATT (3312A>T 3313G>T), CTC337TCT (3314C>T 3315T>C 3316C>T), GAA338AAA (3317G>A), ACT339GTG (3320A>G 3321C>T 3322T>G), GCT340GAG (3324C>A 3325T>G), CTT342TTA (3329C>T 3331T>A), GGT343GGC (3334T>C), CAC344CAT (3337C>T), ATC345TTC (3338A>T), AGC347TCT (3344A>T 3345G>C 3346C>T), AGG348TGT (3347A>T 3349G>T), GAC349GAT (3352C>T), ATC351GTT (3356A>G 3358C>T), AAG352TCT (3359A>T 3360A>C 3361G>T), CCT353ACT (3362C>A), AAC354GAT (3365A>G 3367C>T), GAT356AGG (3371G>A 3372A>G 3373T>G), AAG357AAA (3376G>A), TCC359CAA (3380T>C 3381C>A 3382C>A), GCT360GTT (3384C>T), ATT361GTG (3386A>G 3388T>G), CAA362CAG (3391A>G), AAA363CAC (3392A>C 3394A>C), TAT364TGG (3396A>G 3397T>G), CTG365CCT (3399T>C 3400G>T), ATT366ACT (3402T>C), CCA367CCT (3406A>T), AAG368AGC (3408A>G 3409G>C), ACC369AAC (3411C>A), CCT370CTC (3414C>T 3415T>C), AAG371AAA (3418G>A), GAA372CAA (3419G>C), ATA373CTC (3422A>C 3424A>C), AAA374AGA (3426A>G), CAA375GGT (3428C>G 3429A>G 3430A>T), TTT376TTC (3433T>C), CTT379CTA (3442T>A), CTC380GCA (3443C>G 3444T>C 3445C>A), GGT381GGG (3448T>G), CGA384AGG (3455C>A 3457A>G), CCA388AAA (3467C>A 3468C>A), GAT389GGG (3471A>G 3472T>G), TTT390TAT (3474T>A), GCA391GGT (3477C>G 3478A>T), CGA392ATT (3479C>A 3480G>T 3481A>T), CTC393ATT (3482C>A 3484C>T), ACA394AGT (3486C>G 3487A>T), AAA395AGG (3489A>G 3490A>G), CCC396CCA (3493C>A), ACA398ACG (3499A>G), CAG399GAT (3500C>G 3502G>T), TGC400TTG (3504G>T 3505C>G), TTA401CTG (3506T>C 3508A>G), AAA403AAG (3514A>G), GGT404GAC (3516G>A 3517T>C), AGT405AAC (3519G>A 3520T>C), AAA406TTC (3521A>T 3522A>T 3523A>C), GTA407ATA (3524G>A), ACT408_CTT409del (3527_3532delACTCTT), AGT410TGG (3533A>T 3535T>G), CCC411TCT (3536C>T 3538C>T), TAT413ATT (3542T>A 3543A>T), GTA414GCT (3546T>C 3547A>T), AAT415ACT (3549A>C), AAT415_GCT416insCAA (3550_3551insCAA), GCT416CCT (3551G>C), GAA418CAT (3557G>C 3559A>T), CAC419AAC (3560C>A), TGT420TTG (3564G>T 3565T>G), AAA421AAG (3568A>G), ACT422GTA (3569A>G 3570C>T 3571T>A), CTG423GCA (3572C>G 3573T>C 3574G>A), ACC425ACA (3580C>A), AAC426T.. (3581A>T) | | | | | | | | |

##### Proteins

| | Begin | End | Coverage | Score | Concordance | Matches | Identities | I/D/M/F* | Stop Codons |
|---|---|---|---|---|---|---|---|---|---|
| ORF B (YP_009507248.1) | 159 | 426 | 18.4% | 544 | 41.1% | 193 (96.0%) | 80 (39.8%) | 1/7/0/0 | 0 |

| | Begin | End | Coverage | Score | Concordance | Matches | Identities | I/D/M/F* | Stop Codons |
|---|---|---|---|---|---|---|---|---|---|
| **NT** | **2779** | **3581** | **8.0%** | **14** | **1.3%** | **579 (96.0%)** | **308 (51.1%)** | **3/21** | |
| Protein mutations: | M161L (2786A>T), D163E (2794C>G), Q164K (2795C>A 2797A>G), I166F (2801A>T), D171K (2816G>A 2818C>G), A173P (2822G>C), S175G (2828A>G 2830C>T), S176V (2831T>G 2832C>T 2833A>G), I178V (2837A>G), W179L (2841G>T), V180F (2843G>T), P182R (2849C>A 2850C>G 2851C>A), I185del (2858_2860delATC), A187G (2865C>G 2866T>C), G189_K192del (2870_2881delGGGAAACAAAAG), W193L (2883G>T), L195M (2888C>A 2890C>G), V196C (2891G>T 2892T>G 2893A>T), V197I (2894G>A 2896T>A), F199Y (2901T>A), N203S (2913A>G 2914C>T), E204K (2915G>A), K205V (2918A>G 2919A>T 2920G>C), D208K (2927G>A 2929T>G), D209N (2930G>A 2932C>T), K210R (2934A>G 2935A>G), Y211F (2937A>T), N215M (2949A>T 2950C>G), S217D (2954A>G 2955G>A), D218E (2959C>G), V219L (2960G>C), L220F (2963C>T), K222E (2969A>G), G224Q (2975G>C 2976G>A 2977T>A), K225G (2978A>G 2979A>G), C226A (2981T>G 2982G>C 2983C>T), Q227R (2984C>A 2985A>G), Y228F (2988A>T), T230S (2993A>T 2995C>T), T231K (2997C>A 2998C>A), L232I (2999T>A), A235S (3008G>A 3009C>G 3010A>T), F238Y (3018T>A 3019T>C), E308D (3229A>T), E311L (3236G>T 3237A>T), N312H (3239A>C), R315E (3248C>G 3249G>A), Q318R (3257C>A 3258A>G 3259A>G), R319T (3261G>C), E322Q (3269G>C), S323Q (3272A>C 3273G>A 3274T>A), N324H (3275A>C 3277C>T), F325L (3280C>A), K326L (3281A>T 3282A>T 3283A>G), I327A (3284A>G 3285T>C 3286T>A), Q328K (3287C>A), M329Q (3290A>C 3291T>A 3292G>A), D330S (3293G>T 3294A>C), S332C (3300C>G 3301C>T), E333M (3302G>A 3303A>T 3304A>G), L335G (3308T>G 3309T>G 3310G>A), K336I (3312A>T 3313G>T), L337S (3314C>T 3315T>C 3316C>T), E338K (3317G>A), T339V (3320A>G 3321C>T 3322T>G), A340E (3324C>A 3325T>G), I345F (3338A>T), R348C (3347A>T 3349G>T), I351V (3356A>G 3358C>T), K352S (3359A>T 3360A>C 3361G>T), P353T (3362C>A), N354D (3365A>G 3367C>T), D356R (3371G>A 3372A>G 3373T>G), S359Q (3380T>C 3381C>A 3382C>A), A360V (3384C>T), I361V (3386A>G 3388T>G), K363H (3392A>C 3394A>C), Y364W (3396A>G 3397T>G), L365P (3399T>C 3400G>T), I366T (3402T>C), K368S (3408A>G 3409G>C), T369N (3411C>A), P370L (3414C>T 3415T>C), E372Q (3419G>C), I373L (3422A>C 3424A>C), K374R (3426A>G), Q375G (3428C>G 3429A>G 3430A>T), L380A (3443C>G 3444T>C 3445C>A), P388K (3467C>A 3468C>A), D389G (3471A>G 3472T>G), F390Y (3474T>A), A391G (3477C>G 3478A>T), R392I (3479C>A 3480G>T 3481A>T), L393I (3482C>A 3484C>T), T394S (3486C>G 3487A>T), K395R (3489A>G 3490A>G), Q399D (3500C>G 3502G>T), C400L (3504G>T 3505C>G), G404D (3516G>A 3517T>C), S405N (3519G>A 3520T>C), K406F (3521A>T 3522A>T 3523A>C), V407I (3524G>A), T408_L409del (3527_3532delACTCTT), S410W (3533A>T 3535T>G), P411S (3536C>T 3538C>T), Y413I (3542T>A 3543A>T), V414A (3546T>C 3547A>T), N415T (3549A>C), N415_A416insQ (3550_3551insCAA), A416P (3551G>C), E418H (3557G>C 3559A>T), H419N (3560C>A), C420L (3564G>T 3565T>G), T422V (3569A>G 3570C>T 3571T>A), L423A (3572C>G 3573T>C 3574G>A) | | | | | | | | |
| Codon mutations: | ATG161TTG (2786A>T), GAC163GAG (2794C>G), CAA164AAG (2795C>A 2797A>G), GGA165GGT (2800A>T), ATT166TTT (2801A>T), AGA168AGG (2809A>G), CCA169CCT (2812A>T), TCA170AGT (2813T>A 2814C>G 2815A>T), GAC171AAG (2816G>A 2818C>G), TCT172TCC (2821T>C), GCA173CCA (2822G>C), AGC175GGT (2828A>G 2830C>T), TCA176GTG (2831T>G 2832C>T 2833A>G), CCC177CCA (2836C>A), ATA178GTA (2837A>G), TGG179TTG (2841G>T), GTT180TTT (2843G>T), CCC182AGA (2849C>A 2850C>G 2851C>A), ATC185del (2858_2860delATC), GAC186GAT (2863C>T), GCT187GGC (2865C>G 2866T>C), TCT188TCG (2869T>G), GGG189_AAG192del (2870_2881delGGGAAACAAAAG), TGG193TTG (2883G>T), CGT194AGA (2885C>A 2887T>A), CTC195ATG (2888C>A 2890C>G), GTA196TGT (2891G>T 2892T>G 2893A>T), GTT197ATA (2894G>A 2896T>A), TTC199TAC (2901T>A), CGT200CGG (2905T>G), AAG201AAA (2908G>A), TTG202CTT (2909T>C 2911G>T), AAC203AGT (2913A>G 2914C>T), GAG204AAG (2915G>A), AAG205GTC (2918A>G 2919A>T 2920G>C), ACT206ACC (2923T>C), ATC207ATT (2926C>T), GAT208AAG (2927G>A 2929T>G), GAC209AAT (2930G>A 2932C>T), AAA210AGG (2934A>G 2935A>G), TAC211TTC (2937A>T), ATA213ATT (2944A>T), CCA214CCC (2947A>C), AAC215ATG (2949A>T 2950C>G), ATA216ATT (2953A>T), AGT217GAT (2954A>G 2955G>A), GAC218GAG (2959C>G), GTA219CTA (2960G>C), CTT220TTT (2963C>T), GAC221GAT (2968C>T), AAG222GAG (2969A>G), TTA223TTG (2974A>G), GGT224CAA (2975G>C 2976G>A 2977T>A), AAG225GGG (2978A>G 2979A>G), TGC226GCT (2981T>G 2982G>C 2983C>T), CAA227AGA (2984C>A 2985A>G), TAC228TTC (2988A>T), ACC230TCT (2993A>T 2995C>T), ACC231AAA (2997C>A 2998C>A), TTA232ATA (2999T>A), TTG234CTT (3005T>C 3007G>T), GCA235AGT (3008G>A 3009C>G 3010A>T), AGT236TCG (3011A>T 3012G>C 3013T>G), GGG237GGC (3016G>C), TTT238TAC (3018T>A 3019T>C), TAT239CA. (3020T>C), GAA308GAT (3229A>T), CTG310TTG (3233C>T), GAG311TTG (3236G>T 3237A>T), AAC312CAC (3239A>C), CTG313TTG (3242C>T), CGA315GAA (3248C>G 3249G>A), CAA318AGG (3257C>A 3258A>G 3259A>G), AGA319ACA (3261G>C), CTT320CTG (3265T>G), GAA322CAA (3269G>C), AGT323CAA (3272A>C 3273G>A 3274T>A), AAC324CAT (3275A>C 3277C>T), TTC325TTA (3280C>A), AAA326TTG (3281A>T 3282A>T 3283A>G), ATT327GCA (3284A>G 3285T>C 3286T>A), CAA328AAA (3287C>A), ATG329CAA (3290A>C 3291T>A 3292G>A), GAC330TCC (3293G>T 3294A>C), AAG331AAA (3298G>A), TCC332TGT (3300C>G 3301C>T), GAA333ATG (3302G>A 3303A>T 3304A>G), TTC334TTT (3307C>T), TTG335GGA (3308T>G 3309T>G 3310G>A), AAG336ATT (3312A>T 3313G>T), CTC337TCT (3314C>T 3315T>C 3316C>T), GAA338AAA (3317G>A), ACT339GTG (3320A>G 3321C>T 3322T>G), GCT340GAG (3324C>A 3325T>G), CTT342TTA (3329C>T 3331T>A), GGT343GGC (3334T>C), CAC344CAT (3337C>T), ATC345TTC (3338A>T), AGC347TCT (3344A>T 3345G>C 3346C>T), AGG348TGT (3347A>T 3349G>T), GAC349GAT (3352C>T), ATC351GTT (3356A>G 3358C>T), AAG352TCT (3359A>T 3360A>C 3361G>T), CCT353ACT (3362C>A), AAC354GAT (3365A>G 3367C>T), GAT356AGG (3371G>A 3372A>G 3373T>G), AAG357AAA (3376G>A), TCC359CAA (3380T>C 3381C>A 3382C>A), GCT360GTT (3384C>T), ATT361GTG (3386A>G 3388T>G), CAA362CAG (3391A>G), AAA363CAC (3392A>C 3394A>C), TAT364TGG (3396A>G 3397T>G), CTG365CCT (3399T>C 3400G>T), ATT366ACT (3402T>C), CCA367CCT (3406A>T), AAG368AGC (3408A>G 3409G>C), ACC369AAC (3411C>A), CCT370CTC (3414C>T 3415T>C), AAG371AAA (3418G>A), GAA372CAA (3419G>C), ATA373CTC (3422A>C 3424A>C), AAA374AGA (3426A>G), CAA375GGT (3428C>G 3429A>G 3430A>T), TTT376TTC (3433T>C), CTT379CTA (3442T>A), CTC380GCA (3443C>G 3444T>C 3445C>A), GGT381GGG (3448T>G), CGA384AGG (3455C>A 3457A>G), CCA388AAA (3467C>A 3468C>A), GAT389GGG (3471A>G 3472T>G), TTT390TAT (3474T>A), GCA391GGT (3477C>G 3478A>T), CGA392ATT (3479C>A 3480G>T 3481A>T), CTC393ATT (3482C>A 3484C>T), ACA394AGT (3486C>G 3487A>T), AAA395AGG (3489A>G 3490A>G), CCC396CCA (3493C>A), ACA398ACG (3499A>G), CAG399GAT (3500C>G 3502G>T), TGC400TTG (3504G>T 3505C>G), TTA401CTG (3506T>C 3508A>G), AAA403AAG (3514A>G), GGT404GAC (3516G>A 3517T>C), AGT405AAC (3519G>A 3520T>C), AAA406TTC (3521A>T 3522A>T 3523A>C), GTA407ATA (3524G>A), ACT408_CTT409del (3527_3532delACTCTT), AGT410TGG (3533A>T 3535T>G), CCC411TCT (3536C>T 3538C>T), TAT413ATT (3542T>A 3543A>T), GTA414GCT (3546T>C 3547A>T), AAT415ACT (3549A>C), AAT415_GCT416insCAA (3550_3551insCAA), GCT416CCT (3551G>C), GAA418CAT (3557G>C 3559A>T), CAC419AAC (3560C>A), TGT420TTG (3564G>T 3565T>G), AAA421AAG (3568A>G), ACT422GTA (3569A>G 3570C>T 3571T>A), CTG423GCA (3572C>G 3573T>C 3574G>A), ACC425ACA (3580C>A), AAC426T.. (3581A>T) | | | | | | | | |

*: Inserts / Deletes / Misaligned / Frameshifts

#### Analysis details

This analysis was performed with panviral2.64

### NGS Details (UN9): Badnavirus maculaucubae

#### Assembly

| | |
|---|---|
| **Coverage Length** | 329 (1 contig(s)) |
| **Depth Of Coverage** | 26.2 |
| **Number Of Reads** | 81 |
| **Reads Per Million** | 1.83 rpm (after QC) |
| **Ambiguities** | 0 |
| **Assembly Method** | de novo + reference guided assembly |
| **Consensus Caller** | Bcf Tools |

#### Coverage Map

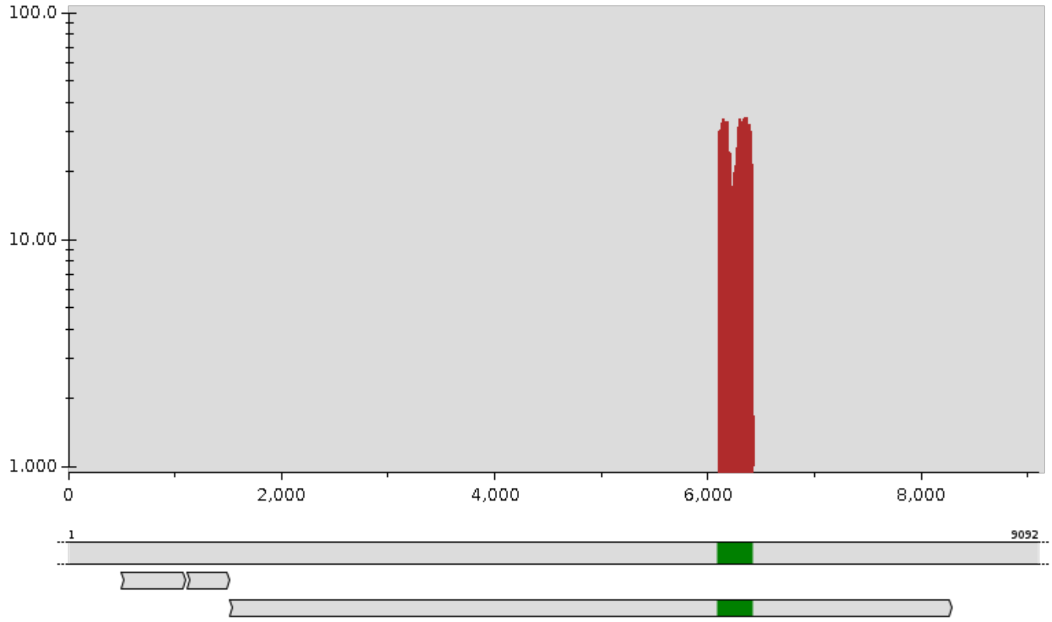

#### Assignment

| | |
|---|---|
| **Type** | Badnavirus maculaucubae (Taxonomy ID: 3051986) |
| **Reference Genome** | NC_076606.1 |
| **NT Identity (%)** | 55.7522 |
| **AA Identity (%)** | 46.4912 |
| **Number Of Stop Codons** | 1 |
| **Number Of CDS** | 3 |

#### Alignment

| | |
|---|---|
| **Alignment Score** | 61.0 (NT) + 239.0 (AA) = 300.0 |
| **Concordance (%)** | 24.1119 |

| Alignment Method | Global, seeded, nucleotide + amino acids (AGA) |
|---|---|

#### Genome Region

Sequence starts at position 6094 and ends at position 6422 relative to NC_076606.1 reference sequence.

#### Alignment Detailed Statistics

| | Begin | End | Coverage | Score | Concordance | Matches | Identities | I/D/M/F* | Stop Codons |
|---|---|---|---|---|---|---|---|---|---|
| **NT** | **6094** | **6422** | **3.6%** | **61** | **9.5%** | **327 (95.9%)** | **189 (55.4%)** | **12/2** | |
| Mutations: | 6102C>T, 6105A>G, 6106T>A, 6112T>C, 6114G>T, 6115A>C, 6116A>G, 6120T>C, 6122G>A, 6125T>A, 6132G>A, 6133G>T, 6136G>A, 6137C>G, 6138A>G, 6139A>G, 6141G>A, 6142G>A, 6147A>G, 6149C>G, 6150A>T, 6151_6152delAG, 6153C>G, 6156T>C, 6159A>G, 6160T>A, 6161G>A, 6165G>T, 6171C>T, 6173T>G, 6179C>A, 6181G>T, 6184G>A, 6186A>T, 6188T>A, 6189A>C, 6191A>T, 6192C>T, 6197G>T, 6198G>T, 6199C>T, 6201A>G, 6204A>T, 6207G>A, 6208C>T, 6210A>T, 6216T>A, 6221A>C, 6222G>A, 6228A>T, 6231T>C, 6235G>A, 6236T>C, 6237G>T, 6240C>T, 6241C>A, 6242A>T, 6243A>G, 6244A>G, 6245G>A, 6246G>C, 6247A>T, 6248A>T, 6253G>A, 6257A>G, 6258C>G, 6259T>G, 6260G>T, 6261T>A, 6266G>A, 6268G>C, 6269G>C, 6270T>G, 6271A>T, 6272C>A, 6273A>T, 6274G>T, 6275A>T, 6276A>G, 6279T>C, 6279_6280insATG, 6282C>T, 6283A>G, 6285C>A, 6287C>T, 6288T>G, 6291T>C, 6292T>C, 6293A>T, 6297T>A, 6303T>C, 6306T>A, 6307C>T, 6312A>T, 6314T>A, 6319G>C, 6320A>G, 6321A>G, 6325C>G, 6326T>C, 6327T>C, 6328C>A, 6330A>G, 6333G>A, 6336C>T, 6342G>T, 6345T>C, 6346C>T, 6348C>G, 6350A>G, 6351A>G, 6354A>T, 6355A>G, 6358C>T, 6361G>C, 6365T>C, 6366A>T, 6371G>A, 6372G>A, 6373A>C, 6374A>T, 6375G>T, 6376A>T, 6377A>G, 6378T>A, 6382C>A, 6383T>C, 6384A>C, 6386T>A, 6387_6388insAATTGTTCG, 6390C>A, 6393C>T, 6394C>T, 6397A>G, 6398C>T, 6402A>T, 6403A>G, 6408A>T, 6409A>T, 6411A>T, 6413C>G | | | | | | | | |

##### CDS

| | Begin | End | Coverage | Score | Concordance | Matches | Identities | I/D/M/F* | Stop Codons |
|---|---|---|---|---|---|---|---|---|---|
| QKP79_gp3 | 1529 | 1638 | 4.9% | 239 | 30.4% | 110 (96.5%) | 53 (46.5%) | 4/0/1/1 | 1 |
| Protein mutations: | F1533I (6106T>A), K1536R (6115A>C 6116A>G), G1538D (6122G>A), F1539Y (6125T>A), V1542L (6133G>T), A1543R (6136G>A 6137C>G 6138A>G), M1544V (6139A>G 6141G>A), E1545K (6142G>A), S1547C (6149C>G 6150A>T), W1551K (6160T>A 6161G>A), I1555R (6173T>G), P1557Q (6179C>A), D1558Y (6181G>T), G1559S (6184G>A 6186A>T), L1560H (6188T>A 6189A>C), Y1561F (6191A>T 6192C>T), W1563F (6197G>T 6198G>T), M1566I (6207G>A), P1567S (6208C>T 6210A>T), K1571T (6221A>C 6222G>A), V1576T (6235G>A 6236T>C 6237G>T), Q1578M (6241C>A 6242A>T 6243A>G), R1579D (6244A>G 6245G>A 6246G>C), K1580L (6247A>T 6248A>T), D1582N (6253G>A), N1583R (6257A>G 6258C>G), C1584V (6259T>G 6260G>T 6261T>A), R1586K (6266G>A), G1587P (6268G>C 6269G>C 6270T>G), T1588Y (6271A>T 6272C>A 6273A>T), E1589L (6274G>T 6275A>T 6276A>G), D1590_F1591insM (6279_6280insATG), I1592V (6283A>G 6285C>A), A1593V (6287C>T 6288T>G), Y1595L (6292T>C 6293A>T), F1602Y (6314T>A), E1604R (6319G>C 6320A>G 6321A>G), L1606A (6325C>G 6326T>C 6327T>C), Q1607K (6328C>A 6330A>G), E1611D (6342G>T), K1614R (6350A>G 6351A>G), M1616V (6355A>G), E1618Q (6361G>C), I1619T (6365T>C 6366A>T), R1621K (6371G>A 6372G>A), K1622L (6373A>C 6374A>T 6375G>T), N1623* (6376A>T 6377A>G 6378T>A), L1625T (6382C>A 6383T>C 6384A>C), V1626D (6386T>A), V1626_L1627insNCS (6387_6388insAATTGTTCG), P1629S (6394C>T), T1630V (6397A>G 6398C>T), K1631N (6402A>T), M1632V (6403A>G), K1633N (6408A>T), I1634F (6409A>T 6411A>T), A1635G (6413C>G) | | | | | | | | |
| Codon mutations: | AGC1531AGT (6102C>T), AAA1532AAG (6105A>G), TTT1533ATT (6106T>A), TTG1535CTT (6112T>C 6114G>T), AAA1536CGA (6115A>C 6116A>G), TCT1537TCC (6120T>C), GGT1538GAT (6122G>A), TTT1539TAT (6125T>A), CAG1541CAA (6132G>A), GTG1542TTG (6133G>T), GCA1543AGG (6136G>A 6137C>G 6138A>G), ATG1544GTA (6139A>G 6141G>A), GAA1545AAA (6142G>A), GAA1546GAG (6147A>G), TCA1547TGT (6149C>G 6150A>T), AGC1548--G (6151_6152delAG 6153C>G), ATT1549ATC (6156T>C), CCA1550CCG (6159A>G), TGG1551AAG (6160T>A 6161G>A), ACG1552ACT (6165G>T), TTC1554TTT (6171C>T), ATA1555AGA (6173T>G), CCA1557CAA (6179C>A), GAT1558TAT (6181G>T), GGA1559AGT (6184G>A 6186A>T), CTA1560CAC (6188T>A 6189A>C), TAC1561TTT (6191A>T 6192C>T), TGG1563TTT (6197G>T 6198G>T), CTA1564TTG (6199C>T 6201A>G), GTA1565GTT (6204A>T), ATG1566ATA (6207G>A), CCA1567TCT (6208C>T 6210A>T), GGT1569GGA (6216T>A), AAG1571ACA (6221A>C 6222G>A), GCA1573GCT (6228A>T), CCT1574CCC (6231T>C), GTG1576ACT (6235G>A 6236T>C 6237G>T), TTC1577TTT (6240C>T), CAA1578ATG (6241C>A 6242A>T 6243A>G), AGG1579GAC (6244A>G 6245G>A 6246G>C), AAA1580TTA (6247A>T 6248A>T), GAC1582AAC (6253G>A), AAC1583AGG (6257A>G 6258C>G), TGT1584GTA (6259T>G 6260G>T 6261T>A), AGG1586AAG (6266G>A), GGT1587CCG (6268G>C 6269G>C 6270T>G), ACA1588TAT (6271A>T 6272C>A 6273A>T), GAA1589TTG (6274G>T 6275A>T 6276A>G), GAT1590GAC (6279T>C), GAT1590_TTC1591insATG (6279_6280insATG), TTC1591TTT (6282C>T), ATC1592GTA (6283A>G 6285C>A), GCT1593GTG (6287C>T 6288T>G), GTT1594GTC (6291T>C), TAC1595CTC (6292T>C 6293A>T), ATT1596ATA (6297T>A), GAT1598GAC (6303T>C), ATT1599ATA (6306T>A), CTG1600TTG (6307C>T), GTA1601GTT (6312A>T), TTC1602TAC (6314T>A), GAA1604CGG (6319G>C 6320A>G 6321A>G), CTT1606GCC (6325C>G 6326T>C 6327T>C), CAA1607AAG (6328C>A 6330A>G), GAG1608GAA (6333G>A), CAC1609CAT (6336C>T), GAG1611GAT (6342G>T), CAT1612CAC (6345T>C), CTC1613TTG (6346C>T 6348C>G), AAA1614AGG (6350A>G 6351A>G), ATA1615ATT (6354A>T), ATG1616GTG (6355A>G), CTG1617TTG (6358C>T), GAA1618CAA (6361G>C), ATA1619ACT (6365T>C 6366A>T), AGG1621AAA (6371G>A 6372G>A), AAG1622CTT (6373A>C 6374A>T 6375G>T), AAT1623TGA (6376A>T 6377A>G 6378T>A), CTA1625ACC (6382C>A 6383T>C 6384A>C), GTC1626GAC (6386T>A), GTC1626_CTC1627insAATTGTTCG (6387_6388insAATTGTTCG), CTC1627CTA (6390C>A), AGC1628AGT (6393C>T), CCA1629TCA (6394C>T), ACA1630GTA (6397A>G 6398C>T), AAA1631AAT (6402A>T), ATG1632GTG (6403A>G), AAA1633AAT (6408A>T), ATA1634TTT (6409A>T 6411A>T), GCT1635GGT (6413C>G) | | | | | | | | |

##### Proteins

| | Begin | End | Coverage | Score | Concordance | Matches | Identities | I/D/M/F* | Stop Codons |
|---|---|---|---|---|---|---|---|---|---|
| hypothetical protein (YP_010799265.1) | 1529 | 1638 | 4.9% | 239 | 30.4% | 110 (96.5%) | 53 (46.5%) | 4/0/1/1 | 1 |
| Protein mutations: | F1533I (6106T>A), K1536R (6115A>C 6116A>G), G1538D (6122G>A), F1539Y (6125T>A), V1542L (6133G>T), A1543R (6136G>A 6137C>G 6138A>G), M1544V (6139A>G 6141G>A), E1545K (6142G>A), S1547C (6149C>G 6150A>T), W1551K (6160T>A 6161G>A), I1555R (6173T>G), P1557Q (6179C>A), D1558Y (6181G>T), G1559S (6184G>A 6186A>T), L1560H (6188T>A 6189A>C), Y1561F (6191A>T 6192C>T), W1563F (6197G>T 6198G>T), M1566I (6207G>A), P1567S (6208C>T 6210A>T), K1571T (6221A>C 6222G>A), V1576T (6235G>A 6236T>C 6237G>T), Q1578M (6241C>A 6242A>T 6243A>G), R1579D (6244A>G 6245G>A 6246G>C), K1580L (6247A>T 6248A>T), D1582N (6253G>A), N1583R (6257A>G 6258C>G), C1584V (6259T>G 6260G>T 6261T>A), R1586K (6266G>A), G1587P (6268G>C 6269G>C 6270T>G), T1588Y (6271A>T 6272C>A 6273A>T), E1589L (6274G>T 6275A>T 6276A>G), D1590_F1591insM (6279_6280insATG), I1592V (6283A>G 6285C>A), A1593V (6287C>T 6288T>G), Y1595L (6292T>C 6293A>T), F1602Y (6314T>A), E1604R (6319G>C 6320A>G 6321A>G), L1606A (6325C>G 6326T>C 6327T>C), Q1607K (6328C>A 6330A>G), E1611D (6342G>T), K1614R (6350A>G 6351A>G), M1616V (6355A>G), E1618Q (6361G>C), I1619T (6365T>C 6366A>T), R1621K (6371G>A 6372G>A), K1622L (6373A>C 6374A>T 6375G>T), N1623* (6376A>T 6377A>G 6378T>A), L1625T (6382C>A 6383T>C 6384A>C), V1626D (6386T>A), V1626_L1627insNCS (6387_6388insAATTGTTCG), P1629S (6394C>T), T1630V (6397A>G 6398C>T), K1631N (6402A>T), M1632V (6403A>G), K1633N (6408A>T), I1634F (6409A>T 6411A>T), A1635G (6413C>G) | | | | | | | | |
| Codon mutations: | AGC1531AGT (6102C>T), AAA1532AAG (6105A>G), TTT1533ATT (6106T>A), TTG1535CTT (6112T>C 6114G>T), AAA1536CGA (6115A>C 6116A>G), TCT1537TCC (6120T>C), GGT1538GAT (6122G>A), TTT1539TAT (6125T>A), CAG1541CAA (6132G>A), GTG1542TTG (6133G>T), GCA1543AGG (6136G>A 6137C>G 6138A>G), ATG1544GTA (6139A>G 6141G>A), GAA1545AAA (6142G>A), GAA1546GAG (6147A>G), TCA1547TGT (6149C>G 6150A>T), AGC1548--G (6151_6152delAG 6153C>G), ATT1549ATC (6156T>C), CCA1550CCG (6159A>G), TGG1551AAG (6160T>A 6161G>A), ACG1552ACT (6165G>T), TTC1554TTT (6171C>T), ATA1555AGA (6173T>G), CCA1557CAA (6179C>A), GAT1558TAT (6181G>T), GGA1559AGT (6184G>A 6186A>T), CTA1560CAC (6188T>A 6189A>C), TAC1561TTT (6191A>T 6192C>T), TGG1563TTT (6197G>T 6198G>T), CTA1564TTG (6199C>T 6201A>G), GTA1565GTT (6204A>T), ATG1566ATA (6207G>A), CCA1567TCT (6208C>T 6210A>T), GGT1569GGA (6216T>A), AAG1571ACA (6221A>C 6222G>A), GCA1573GCT (6228A>T), CCT1574CCC (6231T>C), GTG1576ACT (6235G>A 6236T>C 6237G>T), TTC1577TTT (6240C>T), CAA1578ATG (6241C>A 6242A>T 6243A>G), AGG1579GAC (6244A>G 6245G>A 6246G>C), AAA1580TTA (6247A>T 6248A>T), GAC1582AAC (6253G>A), AAC1583AGG (6257A>G 6258C>G), TGT1584GTA (6259T>G 6260G>T 6261T>A), AGG1586AAG (6266G>A), GGT1587CCG (6268G>C 6269G>C 6270T>G), ACA1588TAT (6271A>T 6272C>A 6273A>T), GAA1589TTG (6274G>T 6275A>T 6276A>G), GAT1590GAC (6279T>C), GAT1590_TTC1591insATG (6279_6280insATG), TTC1591TTT (6282C>T), ATC1592GTA (6283A>G 6285C>A), GCT1593GTG (6287C>T 6288T>G), GTT1594GTC (6291T>C), TAC1595CTC (6292T>C 6293A>T), ATT1596ATA (6297T>A), GAT1598GAC (6303T>C), ATT1599ATA (6306T>A), CTG1600TTG (6307C>T), GTA1601GTT (6312A>T), TTC1602TAC (6314T>A), GAA1604CGG (6319G>C 6320A>G 6321A>G), CTT1606GCC (6325C>G 6326T>C 6327T>C), CAA1607AAG (6328C>A 6330A>G), GAG1608GAA (6333G>A), CAC1609CAT (6336C>T), GAG1611GAT (6342G>T), CAT1612CAC (6345T>C), CTC1613TTG (6346C>T 6348C>G), AAA1614AGG (6350A>G 6351A>G), ATA1615ATT (6354A>T), ATG1616GTG (6355A>G), CTG1617TTG (6358C>T), GAA1618CAA (6361G>C), ATA1619ACT (6365T>C 6366A>T), AGG1621AAA (6371G>A 6372G>A), AAG1622CTT (6373A>C 6374A>T 6375G>T), AAT1623TGA (6376A>T 6377A>G 6378T>A), CTA1625ACC (6382C>A 6383T>C 6384A>C), GTC1626GAC (6386T>A), GTC1626_CTC1627insAATTGTTCG (6387_6388insAATTGTTCG), CTC1627CTA (6390C>A), AGC1628AGT (6393C>T), CCA1629TCA (6394C>T), ACA1630GTA (6397A>G 6398C>T), AAA1631AAT (6402A>T), ATG1632GTG (6403A>G), AAA1633AAT (6408A>T), ATA1634TTT (6409A>T 6411A>T), GCT1635GGT (6413C>G) | | | | | | | | |

*: Inserts / Deletes / Misaligned / Frameshifts

#### Analysis details

This analysis was performed with panviral2.64

#### NGS Details (UN9): Nodensvirus spm2

##### Assembly

| | |
|---|---|
| **Coverage Length** | 163 (1 contig(s)) |
| **Depth Of Coverage** | 39.2 |
| **Number Of Reads** | 67 |
| **Reads Per Million** | 1.51 rpm (after QC) |
| **Ambiguities** | 0 |
| **Assembly Method** | de novo + reference guided assembly |
| **Consensus Caller** | Bcf Tools |

##### Coverage Map

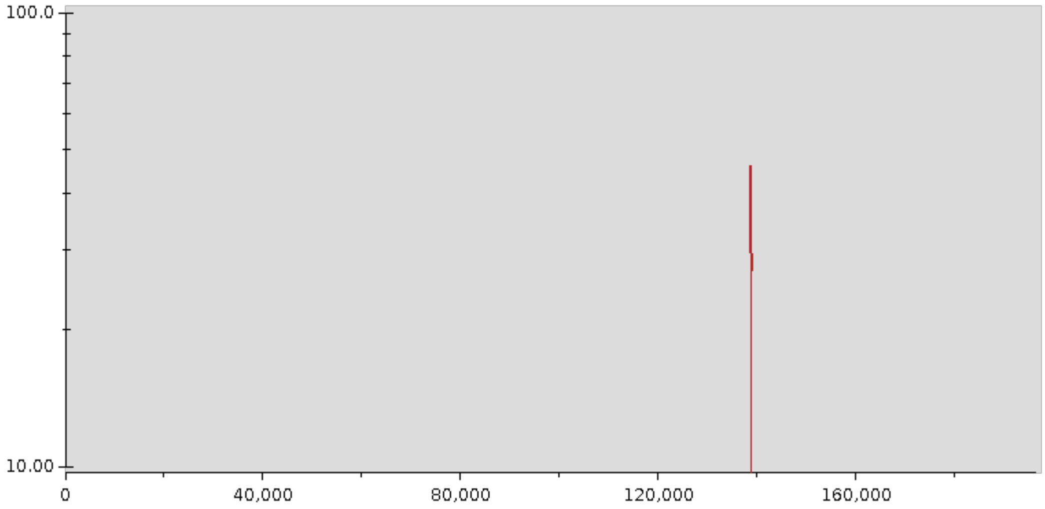

##### Assignment

| | |
|---|---|
| **Type** | Nodensvirus spm2 (Taxonomy ID: 2734126) |
| **Reference Genome** | NC_006820.1 |
| **NT Identity (%)** | 80.7453 |
| **AA Identity (%)** | 86.2745 |
| **Number Of Stop Codons** | 1 |
| **Number Of CDS** | 243 |

##### Alignment

| | |
|---|---|
| **Alignment Score** | 187.0 (NT) + 209.0 (AA) = 396.0 |
| **Concordance (%)** | 72.7903 |
| **Alignment Method** | Local, heuristic, nucleotide (BLASTN) |

##### Genome Region

Sequence starts at position 138728 and ends at position 138890 relative to NC_006820.1 reference sequence.

#### Alignment Detailed Statistics

| | Begin | End | Coverage | Score | Concordance | Matches | Identities | I/D/M/F* | Stop Codons |
|---|---|---|---|---|---|---|---|---|---|
| **NT** | **138728** | **138890** | **0.1%** | **187** | **60.1%** | **161 (98.8%)** | **130 (79.8%)** | **0/2** | |

Mutations: 138734T>C, 138740A>G, 138755C>T, 138756G>A, 138761C>T, 138773G>A, 138776C>T, 138778T>C, 138783_138784delCT, 138785G>A, 138788T>C, 138795C>A, 138797G>T, 138803C>T, 138812T>G, 138813C>G, 138816G>C, 138817T>A, 138824A>G, 138830C>T, 138836C>T, 138840G>A, 138842T>A, 138851A>G, 138854A>G, 138857A>T, 138858T>C, 138860G>A, 138864A>C, 138866A>T, 138869C>A, 138878G>T

*: Inserts / Deletes / Misaligned / Frameshifts

#### Analysis details

This analysis was performed with panviral2.64

### NGS Details (UN9): Epiphyllum badnavirus 1

#### Assembly

| | |
|---|---|
| **Coverage Length** | 426 (1 contig(s)) |
| **Depth Of Coverage** | 15.2 |
| **Number Of Reads** | 66 |
| **Reads Per Million** | 1.49 rpm (after QC) |
| **Ambiguities** | 0 |
| **Assembly Method** | de novo + reference guided assembly |
| **Consensus Caller** | Bcf Tools |

#### Coverage Map

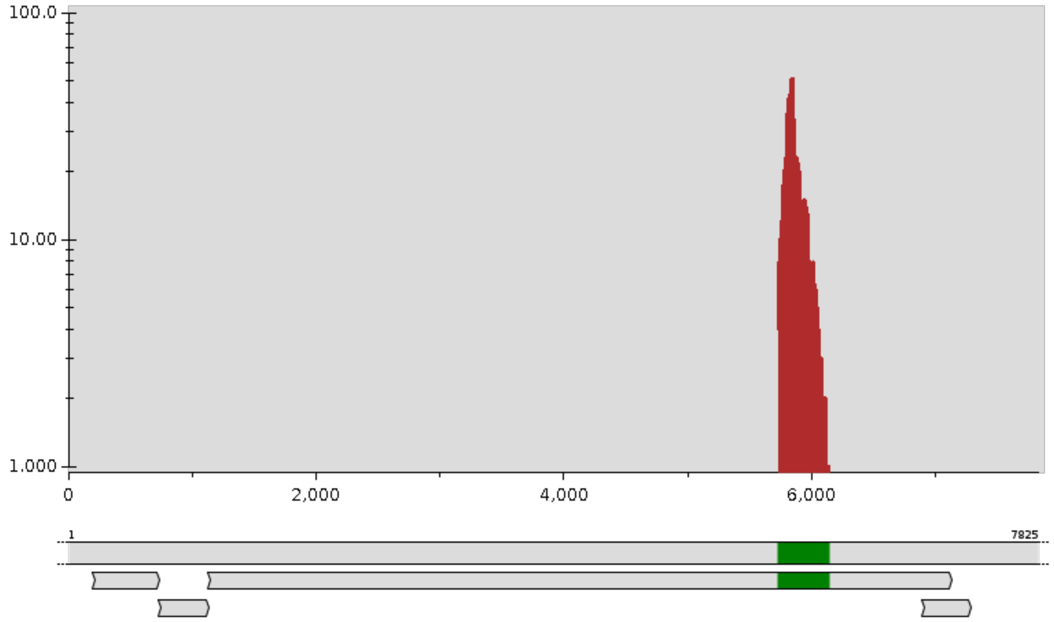

#### Assignment

| | |
|---|---|
| **Type** | Epiphyllum badnavirus 1 (Taxonomy ID: 2518008) |
| **Reference Genome** | NC_076247.1 |
| **NT Identity (%)** | 52.2042 |
| **AA Identity (%)** | 44.4444 |
| **Number Of Stop Codons** | 0 |
| **Number Of CDS** | 4 |

#### Alignment

| | |
|---|---|
| **Alignment Score** | 25.0 (NT) + 292.0 (AA) = 317.0 |
| **Concordance (%)** | 18.4607 |

| Alignment Method | Global, seeded, nucleotide + amino acids (AGA) |
|---|---|

#### Genome Region

Sequence starts at position 5724 and ends at position 6149 relative to NC_076247.1 reference sequence.

#### Alignment Detailed Statistics

| | Begin | End | Coverage | Score | Concordance | Matches | Identities | I/D/M/F* | Stop Codons |
|---|---|---|---|---|---|---|---|---|---|
| **NT** | **5724** | **6149** | **5.4%** | **25** | **2.9%** | **426 (98.8%)** | **225 (52.2%)** | **5/0** | |
| Mutations: | 5724G>A, 5725A>G, 5727A>T, 5728G>A, 5731A>T, 5732A>C, 5734G>T, 5735A>T, 5739A>G, 5743G>T, 5744T>G, 5745C>T, 5746T>A, 5748C>A, 5749A>G, 5754C>T, 5755A>C, 5757A>G, 5758A>C, 5759C>A, 5760C>A, 5761T>C, 5763G>C, 5766C>T, 5767G>A, 5769C>G, 5770A>G, 5771A>T, 5772C>T, 5775G>T, 5776T>G, 5778C>T, 5782G>A, 5785C>A, 5791T>C, 5793T>A, 5794C>A, 5799T>C, 5800G>A, 5802C>T, 5806A>G, 5808C>T, 5809A>G, 5810C>A, 5811A>T, 5814C>G, 5815A>T, 5817A>T, 5818C>G, 5820G>C, 5821A>C, 5823G>A, 5824G>T, 5826C>A, 5827T>C, 5828G>A, 5829T>A, 5830A>G, 5831A>G, 5835C>T, 5839G>T, 5840T>G, 5847C>A, 5851T>A, 5853C>A, 5859G>C, 5860A>T, 5862G>T, 5863A>T, 5864G>C, 5868A>T, 5870T>A, 5871C>T, 5874C>T, 5878A>T, 5880T>G, 5882T>A, 5884A>G, 5887G>A, 5888A>G, 5889C>G, 5893G>C, 5894A>G, 5895A>T, 5896T>G, 5897C>A, 5898C>T, 5901C>T, 5905T>A, 5906G>A, 5907G>A, 5910C>G, 5913C>T, 5916C>T, 5918T>G, 5922C>A, 5923C>A, 5924C>G, 5926G>T, 5931A>G, 5933T>A, 5936A>T, 5937C>T, 5942G>T, 5943G>T, 5944T>C, 5946A>T, 5953C>T, 5955A>C, 5958C>T, 5961A>T, 5964G>A, 5966A>C, 5967G>T, 5973C>T, 5976T>G, 5977G>A, 5982G>A, 5985C>T, 5986C>A, 5987A>T, 5988A>G, 5989C>G, 5990G>A, 5991A>C, 5992A>C, 5993A>T, 5994G>C, 5998G>A, 6000C>T, 6002A>G, 6003C>G, 6003_6004insGTG, 6008T>C, 6009C>A, 6012A>C, 6012_6013insCC, 6013G>T, 6018G>T, 6019T>G, 6020C>A, 6021A>T, 6023G>T, 6024C>G, 6026T>A, 6030A>G, 6032C>T, 6033A>G, 6036T>G, 6038A>T, 6042T>C, 6048C>T, 6051C>T, 6059T>A, 6060C>T, 6061A>T, 6062G>C, 6063T>C, 6064A>C, 6065A>T, 6066C>A, 6067A>G, 6068C>G, 6069T>C, 6073G>C, 6075T>G, 6078C>T, 6081T>C, 6082G>A, 6083C>T, 6084A>G, 6085C>G, 6091C>T, 6095A>G, 6096G>A, 6097T>A, 6099C>G, 6102G>A, 6103C>T, 6108G>A, 6113G>T, 6114T>G, 6115G>A, 6116A>G, 6118A>G, 6119G>A, 6123G>A, 6124G>C, 6127C>T, 6130A>T, 6133C>G, 6134T>C, 6136T>A, 6137C>A, 6138T>A, 6139C>T, 6140C>A, 6142A>G, 6143C>A, 6144G>C, 6147G>A, 6148A>T, 6149T>G | | | | | | | | |

##### CDS

| | Begin | End | Coverage | Score | Concordance | Matches | Identities | I/D/M/F* | Stop Codons |
|---|---|---|---|---|---|---|---|---|---|
| QKM20_gp3 | 1533 | 1674 | 7.1% | 292 | 28.7% | 142 (98.6%) | 64 (44.4%) | 2/0/1/1 | 0 |
| Protein mutations: | K1533D (5725A>G 5727A>T), G1534R (5728G>A), K1535S (5731A>T 5732A>C), E1536L (5734G>T 5735A>T), V1539C (5743G>T 5744T>G 5745C>T), F1540I (5746T>A 5748C>A), N1541D (5749A>G), K1543Q (5755A>C 5757A>G), T1544Q (5758A>C 5759C>A 5760C>A), D1547K (5767G>A 5769C>G), N1548V (5770A>G 5771A>T 5772C>T), F1550V (5776T>G 5778C>T), D1552N (5782G>A), Q1553K (5785C>A), S1555P (5791T>C 5793T>A), L1556I (5794C>A), G1558S (5800G>A 5802C>T), N1560D (5806A>G 5808C>T), T1561D (5809A>G 5810C>A 5811A>T), I1562M (5814C>G), I1563F (5815A>T 5817A>T), Q1564D (5818C>G 5820G>C), K1565Q (5821A>C 5823G>A), V1566L (5824G>T 5826C>A), C1567Q (5827T>C 5828G>A 5829T>A), N1568G (5830A>G 5831A>G), V1571C (5839G>T 5840T>G), F1575I (5851T>A 5853C>A), K1578Y (5860A>T 5862G>T), F1581Y (5870T>A 5871C>T), I1584L (5878A>T 5880T>G), M1585K (5882T>A), M1586V (5884A>G), D1587R (5887G>A 5888A>G 5889C>G), E1589R (5893G>C 5894A>G 5895A>T), S1590D (5896T>G 5897C>A 5898C>T), W1593K (5905T>A 5906G>A 5907G>A), L1597R (5918T>G), P1599R (5923C>A 5924C>G), D1600Y (5926G>T), L1602H (5933T>A), Y1603F (5936A>T 5937C>T), W1605F (5942G>T 5943G>T), P1609S (5953C>T 5955A>C), K1613T (5966A>C 5967G>T), A1617T (5977G>A), Q1620M (5986C>A 5987A>T 5988A>G), R1621D (5989C>G 5990G>A 5991A>C), K1622L (5992A>C 5993A>T 5994G>C), D1624N (5998G>A 6000C>T), N1625R (6002A>G 6003C>G), N1625_C1626insV (6003_6004insGTG), F1627S (6008T>C 6009C>A), R1628S (6012A>C), R1628_D1629insX (6012_6013insCC), D1629Y (6013G>T), S1631D (6019T>G 6020C>A 6021A>T), G1632V (6023G>T 6024C>G), F1633Y (6026T>A), A1635V (6032C>T 6033A>G), Y1637F (6038A>T), F1644Y (6059T>A 6060C>T), N1646L (6064A>C 6065A>T 6066C>A), T1647G (6067A>G 6068C>G 6069T>C), D1649Q (6073G>C 6075T>G), A1652M (6082G>A 6083C>T 6084A>G), Q1653E (6085C>G), K1656R (6095A>G 6096G>A), S1657T (6097T>A 6099C>G), M1658I (6102G>A), C1662L (6113G>T 6114T>G), E1663R (6115G>A 6116A>G), R1664E (6118A>G 6119G>A), G1666R (6124G>C), I1668F (6130A>T), L1669A (6133C>G 6134T>C), S1670K (6136T>A 6137C>A 6138T>A), P1671Y (6139C>T 6140C>A), T1672D (6142A>G 6143C>A 6144G>C) | | | | | | | | |
| Codon mutations: | AAG1532..A (5724G>A), AAA1533GAT (5725A>G 5727A>T), GGA1534AGA (5728G>A), AAG1535TCG (5731A>T 5732A>C), GAG1536TTG (5734G>T 5735A>T), AGA1537AGG (5739A>G), GTC1539TGT (5743G>T 5744T>G 5745C>T), TTC1540ATA (5746T>A 5748C>A), AAC1541GAC (5749A>G), TAC1542TAT (5754C>T), AAA1543CAG (5755A>C 5757A>G), ACC1544CAA (5758A>C 5759C>A 5760C>A), TTG1545CTC (5761T>C 5763G>C), AAC1546AAT (5766C>T), GAC1547AAG (5767G>A 5769C>G), AAC1548GTT (5770A>G 5771A>T 5772C>T), ACG1549ACT (5775G>T), TTC1550GTT (5776T>G 5778C>T), GAT1552AAT (5782G>A), CAG1553AAG (5785C>A), TCT1555CCA (5791T>C 5793T>A), CTA1556ATA (5794C>A), CCT1557CCC (5799T>C), GGC1558AGT (5800G>A 5802C>T), AAC1560GAT (5806A>G 5808C>T), ACA1561GAT (5809A>G 5810C>A 5811A>T), ATC1562ATG (5814C>G), ATA1563TTT (5815A>T 5817A>T), CAG1564GAC (5818C>G 5820G>C), AAG1565CAA (5821A>C 5823G>A), GTC1566TTA (5824G>T 5826C>A), TGT1567CAA (5827T>C 5828G>A 5829T>A), AAT1568GGT (5830A>G 5831A>G), GCC1569GCT (5835C>T), GTC1571TGC (5839G>T 5840T>G), TCC1573TCA (5847C>A), TTC1575ATA (5851T>A 5853C>A), CTG1577CTC (5859G>C), AAG1578TAT (5860A>T 5862G>T), AGC1579TCC (5863A>T 5864G>C), GGA1580GGT (5868A>T), TTC1581TAT (5870T>A 5871C>T), CAC1582CAT (5874C>T), ATT1584TTG (5878A>T 5880T>G), ATG1585AAG (5882T>A), ATG1586GTG (5884A>G), GAC1587AGG (5887G>A 5888A>G 5889C>G), GAA1589CGT (5893G>C 5894A>G 5895A>T), TCC1590GAT (5896T>G 5897C>A 5898C>T), ATC1591ATT (5901C>T), TGG1593AAA (5905T>A 5906G>A 5907G>A), ACC1594ACG (5910C>G), GCC1595GCT (5913C>T), TTC1596TTT (5916C>T), CTG1597CGG (5918T>G), ACC1598ACA (5922C>A), CCA1599AGA (5923C>A 5924C>G), GAT1600TAT (5926G>T), GGA1601GGG (5931A>G), CTT1602CAT (5933T>A), TAC1603TTT (5936A>T 5937C>T), TGG1605TTT (5942G>T 5943G>T), TTA1606CTT (5944T>C 5946A>T), CCA1609TCC (5953C>T 5955A>C), TTC1610TTT (5958C>T), GGA1611GGT (5961A>T), TTG1612TTA (5964G>A), AAG1613ACT (5966A>C 5967G>T), GCC1615GCT (5973C>T), CCT1616CCG (5976T>G), GCT1617ACT (5977G>A), GTG1618GTA (5982G>A), TTC1619TTT (5985C>T), CAA1620ATG (5986C>A 5987A>T 5988A>G), CGA1621GAC (5989C>G 5990G>A 5991A>C), AAG1622CTC (5992A>C 5993A>T 5994G>C), GAC1624AAT (5998G>A 6000C>T), AAC1625AGG (6002A>G 6003C>G), AAC1625_TGT1626insGTG (6003_6004insGTG), TTC1627TCA (6008T>C 6009C>A), AGA1628AGC (6012A>C), AGA1628_GAC1629insCC- (6012_6013insCC), GAC1629TAC (6013G>T), CTG1630CTT (6018G>T), TCA1631GAT (6019T>G 6020C>A 6021A>T), GGC1632GTG (6023G>T 6024C>G), TTT1633TAT (6026T>A), GTA1634GTG (6030A>G), GCA1635GTG (6032C>T 6033A>G), GTT1636GTG (6036T>G), TAC1637TTC (6038A>T), ATT1638ATC (6042T>C), GAC1640GAT (6048C>T), ATC1641ATT (6051C>T), TTC1644TAT (6059T>A 6060C>T), AGT1645TCC (6061A>T 6062G>C 6063T>C), AAC1646CTA (6064A>C 6065A>T 6066C>A), ACT1647GGC (6067A>G 6068C>G 6069T>C), GAT1649CAG (6073G>C 6075T>G), GAC1650GAT (6078C>T), CAT1651CAC (6081T>C), GCA1652ATG (6082G>A 6083C>T 6084A>G), CAG1653GAG (6085C>G), CTG1655TTG (6091C>T), AAG1656AGA (6095A>G 6096G>A), TCC1657ACG (6097T>A 6099C>G), ATG1658ATA (6102G>A), CTG1659TTG (6103C>T), CAG1660CAA (6108G>A), TGT1662TTG (6113G>T 6114T>G), GAG1663AGG (6115G>A 6116A>G), AGA1664GAA (6118A>G 6119G>A), GAG1665GAA (6123G>A), GGG1666CGG (6124G>C), CTG1667TTG (6127C>T), ATC1668TTC (6130A>T), CTC1669GCC (6133C>G 6134T>C), TCT1670AAA (6136T>A 6137C>A 6138T>A), CCT1671TAT (6139C>T 6140C>A), ACG1672GAC (6142A>G 6143C>A 6144G>C), AAG1673AAA (6147G>A), ATG1674TG. (6148A>T 6149T>G) | | | | | | | | |

##### Proteins

| | Begin | End | Coverage | Score | Concordance | Matches | Identities | I/D/M/F* | Stop Codons |
|---|---|---|---|---|---|---|---|---|---|
| polyprotein (YP_010797894.1) | 1533 | 1674 | 7.1% | 292 | 28.7% | 142 (98.6%) | 64 (44.4%) | 2/0/1/1 | 0 |
| Protein mutations: | K1533D (5725A>G 5727A>T), G1534R (5728G>A), K1535S (5731A>T 5732A>C), E1536L (5734G>T 5735A>T), V1539C (5743G>T 5744T>G 5745C>T), F1540I (5746T>A 5748C>A), N1541D (5749A>G), K1543Q (5755A>C 5757A>G), T1544Q (5758A>C 5759C>A 5760C>A), D1547K (5767G>A 5769C>G), N1548V (5770A>G 5771A>T 5772C>T), F1550V (5776T>G 5778C>T), D1552N (5782G>A), Q1553K (5785C>A), S1555P (5791T>C 5793T>A), L1556I (5794C>A), G1558S (5800G>A 5802C>T), N1560D (5806A>G 5808C>T), T1561D (5809A>G 5810C>A 5811A>T), I1562M (5814C>G), I1563F (5815A>T 5817A>T), Q1564D (5818C>G 5820G>C), K1565Q (5821A>C 5823G>A), V1566L (5824G>T 5826C>A), C1567Q (5827T>C 5828G>A 5829T>A), N1568G (5830A>G 5831A>G), V1571C (5839G>T 5840T>G), F1575I (5851T>A 5853C>A), K1578Y (5860A>T 5862G>T), F1581Y (5870T>A 5871C>T), I1584L (5878A>T 5880T>G), M1585K (5882T>A), M1586V (5884A>G), D1587R (5887G>A 5888A>G 5889C>G), E1589R (5893G>C 5894A>G 5895A>T), S1590D (5896T>G 5897C>A 5898C>T), W1593K (5905T>A 5906G>A 5907G>A), L1597R (5918T>G), P1599R (5923C>A 5924C>G), D1600Y (5926G>T), L1602H (5933T>A), Y1603F (5936A>T 5937C>T), W1605F (5942G>T 5943G>T), P1609S (5953C>T 5955A>C), K1613T (5966A>C 5967G>T), A1617T (5977G>A), Q1620M (5986C>A 5987A>T 5988A>G), R1621D (5989C>G 5990G>A 5991A>C), K1622L (5992A>C 5993A>T 5994G>C), D1624N (5998G>A 6000C>T), N1625R (6002A>G 6003C>G), N1625_C1626insV (6003_6004insGTG), F1627S (6008T>C 6009C>A), R1628S (6012A>C), R1628_D1629insX (6012_6013insCC), D1629Y (6013G>T), S1631D (6019T>G 6020C>A 6021A>T), G1632V (6023G>T 6024C>G), F1633Y (6026T>A), A1635V (6032C>T 6033A>G), Y1637F (6038A>T), F1644Y (6059T>A 6060C>T), N1646L (6064A>C 6065A>T 6066C>A), T1647G (6067A>G 6068C>G 6069T>C), D1649Q (6073G>C 6075T>G), A1652M (6082G>A 6083C>T 6084A>G), Q1653E (6085C>G), K1656R (6095A>G 6096G>A), S1657T (6097T>A 6099C>G), M1658I (6102G>A), C1662L (6113G>T 6114T>G), E1663R (6115G>A 6116A>G), R1664E (6118A>G 6119G>A), G1666R (6124G>C), I1668F (6130A>T), L1669A (6133C>G 6134T>C), S1670K (6136T>A 6137C>A 6138T>A), P1671Y (6139C>T 6140C>A), T1672D (6142A>G 6143C>A 6144G>C) | | | | | | | | |

| | Begin | End | Coverage | Score | Concordance | Matches | Identities | I/D/M/F* | Stop Codons |
|---|---|---|---|---|---|---|---|---|---|
| **NT** | **5724** | **6149** | **5.4%** | **25** | **2.9%** | **426 (98.8%)** | **225 (52.2%)** | **5/0** | |
| Codon mutations: | AAG1532..A (5724G>A), AAA1533GAT (5725A>G 5727A>T), GGA1534AGA (5728G>A), AAG1535TCG (5731A>T 5732A>C), GAG1536TTG (5734G>T 5735A>T), AGA1537AGG (5739A>G), GTC1539TGT (5743G>T 5744T>G 5745C>T), TTC1540ATA (5746T>A 5748C>A), AAC1541GAC (5749A>G), TAC1542TAT (5754C>T), AAA1543CAG (5755A>C 5757A>G), ACC1544CAA (5758A>C 5759C>A 5760C>A), TTG1545CTC (5761T>C 5763G>C), AAC1546AAT (5766C>T), GAC1547AAG (5767G>A 5769C>G), AAC1548GTT (5770A>G 5771A>T 5772C>T), ACG1549ACT (5775G>T), TTC1550GTT (5776T>G 5778C>T), GAT1552AAT (5782G>A), CAG1553AAG (5785C>A), TCT1555CCA (5791T>C 5793T>A), CTA1556ATA (5794C>A), CCT1557CCC (5799T>C), GGC1558AGT (5800G>A 5802C>T), AAC1560GAT (5806A>G 5808C>T), ACA1561GAT (5809A>G 5810C>A 5811A>T), ATC1562ATG (5814C>G), ATA1563TTT (5815A>T 5817A>T), CAG1564GAC (5818C>G 5820G>C), AAG1565CAA (5821A>C 5823G>A), GTC1566TTA (5824G>T 5826C>A), TGT1567CAA (5827T>C 5828G>A 5829T>A), AAT1568GGT (5830A>G 5831A>G), GCC1569GCT (5835C>T), GTC1571TGC (5839G>T 5840T>G), TCC1573TCA (5847C>A), TTC1575ATA (5851T>A 5853C>A), CTG1577CTC (5859G>C), AAG1578TAT (5860A>T 5862G>T), AGC1579TCC (5863A>T 5864G>C), GGA1580GGT (5868A>T), TTC1581TAT (5870T>A 5871C>T), CAC1582CAT (5874C>T), ATT1584TTG (5878A>T 5880T>G), ATG1585AAG (5882T>A), ATG1586GTG (5884A>G), GAC1587AGG (5887G>A 5888A>G 5889C>G), GAA1589CGT (5893G>C 5894A>G 5895A>T), TCC1590GAT (5896T>G 5897C>A 5898C>T), ATC1591ATT (5901C>T), TGG1593AAA (5905T>A 5906G>A 5907G>A), ACC1594ACG (5910C>G), GCC1595GCT (5913C>T), TTC1596TTT (5916C>T), CTG1597CGG (5918T>G), ACC1598ACA (5922C>A), CCA1599AGA (5923C>A 5924C>G), GAT1600TAT (5926G>T), GGA1601GGG (5931A>G), CTT1602CAT (5933T>A), TAC1603TTT (5936A>T 5937C>T), TGG1605TTT (5942G>T 5943G>T), TTA1606CTT (5944T>C 5946A>T), CCA1609TCC (5953C>T 5955A>C), TTC1610TTT (5958C>T), GGA1611GGT (5961A>T), TTG1612TTA (5964G>A), AAG1613ACT (5966A>C 5967G>T), GCC1615GCT (5973C>T), CCT1616CCG (5976T>G), GCT1617ACT (5977G>A), GTG1618GTA (5982G>A), TTC1619TTT (5985C>T), CAA1620ATG (5986C>A 5987A>T 5988A>G), CGA1621GAC (5989C>G 5990G>A 5991A>C), AAG1622CTC (5992A>C 5993A>T 5994G>C), GAC1624AAT (5998G>A 6000C>T), AAC1625AGG (6002A>G 6003C>G), AAC1625_TGT1626insGTG (6003_6004insGTG), TTC1627TCA (6008T>C 6009C>A), AGA1628AGC (6012A>C), AGA1628_GAC1629insCC- (6012_6013insCC), GAC1629TAC (6013G>T), CTG1630CTT (6018G>T), TCA1631GAT (6019T>G 6020C>A 6021A>T), GGC1632GTG (6023G>T 6024C>G), TTT1633TAT (6026T>A), GTA1634GTG (6030A>G), GCA1635GTG (6032C>T 6033A>G), GTT1636GTG (6036T>G), TAC1637TTC (6038A>T), ATT1638ATC (6042T>C), GAC1640GAT (6048C>T), ATC1641ATT (6051C>T), TTC1644TAT (6059T>A 6060C>T), AGT1645TCC (6061A>T 6062G>C 6063T>C), AAC1646CTA (6064A>C 6065A>T 6066C>A), ACT1647GGC (6067A>G 6068C>G 6069T>C), GAT1649CAG (6073G>C 6075T>G), GAC1650GAT (6078C>T), CAT1651CAC (6081T>C), GCA1652ATG (6082G>A 6083C>T 6084A>G), CAG1653GAG (6085C>G), CTG1655TTG (6091C>T), AAG1656AGA (6095A>G 6096G>A), TCC1657ACG (6097T>A 6099C>G), ATG1658ATA (6102G>A), CTG1659TTG (6103C>T), CAG1660CAA (6108G>A), TGT1662TTG (6113G>T 6114T>G), GAG1663AGG (6115G>A 6116A>G), AGA1664GAA (6118A>G 6119G>A), GAG1665GAA (6123G>A), GGG1666CGG (6124G>C), CTG1667TTG (6127C>T), ATC1668TTC (6130A>T), CTC1669GCC (6133C>G 6134T>C), TCT1670AAA (6136T>A 6137C>A 6138T>A), CCT1671TAT (6139C>T 6140C>A), ACG1672GAC (6142A>G 6143C>A 6144G>C), AAG1673AAA (6147G>A), ATG1674TG. (6148A>T 6149T>G) | | | | | | | | |

*: Inserts / Deletes / Misaligned / Frameshifts

#### Analysis details

This analysis was performed with panviral2.64

### NGS Details (UN9): Caulimovirus venafragariae

#### Assembly

| | |
|---|---|
| **Coverage Length** | 321 (1 contig(s)) |
| **Depth Of Coverage** | 21.2 |
| **Number Of Reads** | 60 |
| **Reads Per Million** | 1.36 rpm (after QC) |
| **Ambiguities** | 0 |
| **Assembly Method** | de novo + reference guided assembly |
| **Consensus Caller** | Bcf Tools |

#### Coverage Map

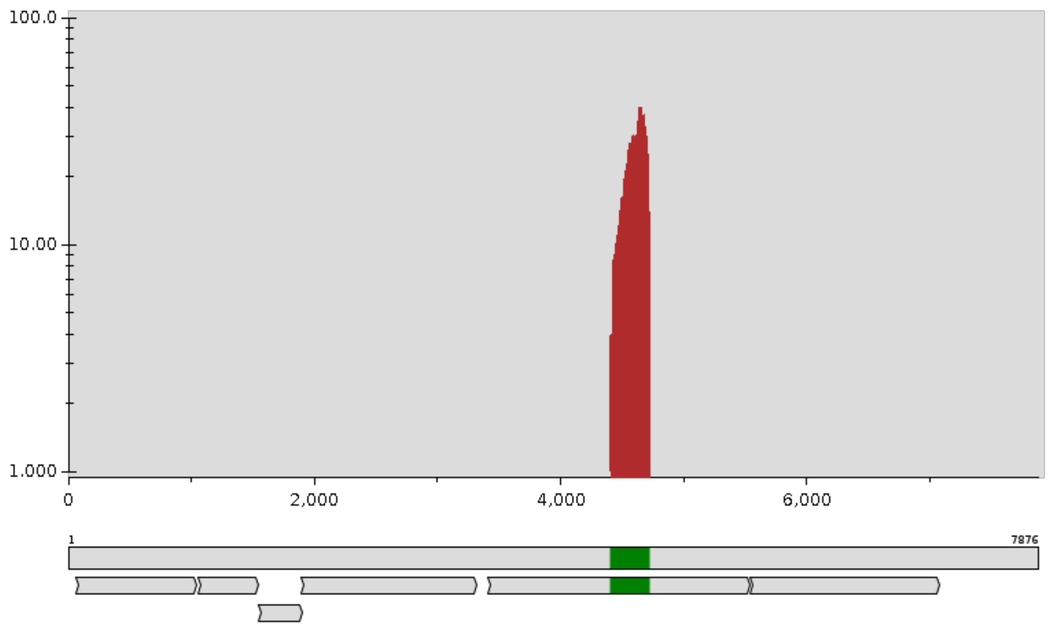

#### Assignment

| | |
|---|---|
| **Type** | Caulimovirus venafragariae (Taxonomy ID: 3048344) |
| **Reference Genome** | NC_001725.1 |
| **NT Identity (%)** | 54.2056 |
| **AA Identity (%)** | 42.0561 |
| **Number Of Stop Codons** | 0 |
| **Number Of CDS** | 6 |

#### Alignment

| | |
|---|---|
| **Alignment Score** | 54.0 (NT) + 321.0 (AA) = 375.0 |
| **Concordance (%)** | 26.0236 |

| Alignment Method | Global, seeded, nucleotide + amino acids (AGA) |
|---|---|

#### Genome Region

Sequence starts at position 4405 and ends at position 4725 relative to NC_001725.1 reference sequence.

#### Alignment Detailed Statistics

| | Begin | End | Coverage | Score | Concordance | Matches | Identities | I/D/M/F* | Stop Codons |
|---|---|---|---|---|---|---|---|---|---|
| **NT** | **4405** | **4725** | **4.1%** | **54** | **8.4%** | **321 (100%)** | **174 (54.2%)** | **0/0** | |
| Mutations: | 4410A>C, 4413T>G, 4414A>G, 4419T>C, 4425G>A, 4426T>C, 4428A>T, 4434C>T, 4435C>G, 4436A>C, 4440A>C, 4441A>C, 4444G>A, 4445G>A, 4446A>G, 4449T>C, 4450G>C, 4451G>A, 4457T>C, 4459C>G, 4461T>G, 4464T>C, 4465A>T, 4466A>T, 4469A>T, 4470G>T, 4471G>A, 4473G>A, 4474C>A, 4477C>A, 4483C>G, 4485A>C, 4489A>T, 4491C>G, 4493G>C, 4494A>T, 4497T>G, 4498A>C, 4500G>A, 4501A>G, 4502C>A, 4503C>T, 4505T>A, 4509C>T, 4511C>G, 4512T>C, 4514C>T, 4518T>G, 4521C>T, 4522T>G, 4524T>G, 4525A>T, 4527A>C, 4535T>A, 4537T>A, 4538G>A, 4539G>T, 4542G>A, 4543G>A, 4545A>C, 4546C>A, 4547G>C, 4548C>A, 4549C>A, 4554T>A, 4560A>G, 4561A>G, 4562C>A, 4563A>C, 4564A>C, 4565T>A, 4566T>A, 4567C>G, 4570C>A, 4571T>A, 4572A>G, 4576G>A, 4583G>C, 4584C>A, 4587T>C, 4590C>T, 4591C>T, 4593A>T, 4596A>G, 4597C>A, 4598A>C, 4599C>A, 4602C>T, 4604A>C, 4607G>T, 4608G>C, 4609T>A, 4610T>A, 4611A>G, 4612G>C, 4613T>G, 4614C>G, 4626A>G, 4630A>T, 4631A>G, 4632A>C, 4633C>A, 4635A>T, 4638C>A, 4641T>G, 4642G>A, 4644T>A, 4646T>C, 4647T>A, 4650C>T, 4656G>A, 4657C>T, 4658A>G, 4663G>A, 4664A>T, 4665C>G, 4667A>C, 4668A>C, 4670G>T, 4671T>A, 4672C>T, 4675A>C, 4676G>A, 4678A>G, 4684T>G, 4685A>T, 4686T>A, 4687C>G, 4688C>A, 4690C>G, 4692G>T, 4696T>G, 4697G>T, 4700C>A, 4701T>G, 4704A>C, 4706A>T, 4707T>C, 4708G>A, 4710T>G, 4714G>A, 4716C>T, 4717A>T, 4719C>T, 4720A>T, 4721T>C, 4722C>A | | | | | | | | |
| **CDS** | | | | | | | | | |
| ORF_V | 334 | 440 | 15.1% | 321 | 40.2% | 107 (100%) | 45 (42.1%) | 0/0/0/0 | 0 |
| Protein mutations: | I336M (4413T>G), N337D (4414A>G), H344A (4435C>G 4436A>C), K346Q (4441A>C), G347K (4444G>A 4445G>A 4446A>G), G349H (4450G>C 4451G>A), L351P (4457T>C), L352V (4459C>G 4461T>G), N354F (4465A>T 4466A>T), K355I (4469A>T 4470G>T), E356K (4471G>A 4473G>A), Q357K (4474C>A), L358M (4477C>A), Q360D (4483C>G 4485A>C), I362L (4489A>T 4491C>G), G363A (4493G>C 4494A>T), K365Q (4498A>C 4500G>A), T366D (4501A>G 4502C>A 4503C>T), F367Y (4505T>A), S369C (4511C>G 4512T>C), S370F (4514C>T), F371L (4518T>G), C373G (4522T>G 4524T>G), K374Y (4525A>T 4527A>C), F377Y (4535T>A), W378N (4537T>A 4538G>A 4539G>T), V380I (4543G>A 4545A>C), R381T (4546C>A 4547G>C 4548C>A), L382I (4549C>A), T386D (4561A>G 4562C>A 4563A>C), I387Q (4564A>C 4565T>A 4566T>A), Q388E (4567C>G), L389K (4570C>A 4571T>A 4572A>G), A391T (4576G>A), S393T (4583G>C 4584C>A), Q396Y (4591C>T 4593A>T), H398T (4597C>A 4598A>C 4599C>A), E400A (4604A>C), W401F (4607G>T 4608G>C), L402K (4609T>A 4610T>A 4611A>G), V403R (4612G>C 4613T>G 4614C>G), K409C (4630A>T 4631A>G 4632A>C), Q410N (4633C>A 4635A>T), A413T (4642G>A 4644T>A), I414T (4646T>C 4647T>A), H418C (4657C>T 4658A>G), D420M (4663G>A 4664A>T 4665C>G), E421A (4667A>C 4668A>C), S422I (4670G>T 4671T>A), L423F (4672C>T), S424H (4675A>C 4676G>A), N425D (4678A>G), Y427V (4684T>G 4685A>T 4686T>A), P428E (4687C>G 4688C>A), Q429D (4690C>G 4692G>T), C431V (4696T>G 4697G>T), A432E (4700C>A 4701T>G), Y434F (4706A>T 4707T>C), V435M (4708G>A 4710T>G), D437N (4714G>A 4716C>T), I438F (4717A>T 4719C>T), I439S (4720A>T 4721T>C 4722C>A) | | | | | | | | |
| Codon mutations: | GTA335GTC (4410A>C), ATT336ATG (4413T>G), AAC337GAC (4414A>G), TAT338TAC (4419T>C), AAG340AAA (4425G>A), TTA341CTT (4426T>C 4428A>T), GAC343GAT (4434C>T), CAT344GCT (4435C>G 4436A>C), ACA345ACC (4440A>C), AAG346CAG (4441A>C), GGA347AAG (4444G>A 4445G>A 4446A>G), GAT348GAC (4449T>C), GGC349CAC (4450G>C 4451G>A), CTA351CCA (4457T>C), CTT352GTG (4459C>G 4461T>G), CCT353CCC (4464T>C), AAC354TTC (4465A>T 4466A>T), AAG355ATT (4469A>T 4470G>T), GAG356AAA (4471G>A 4473G>A), CAA357AAA (4474C>A), CTG358ATG (4477C>A), CAA360GAC (4483C>G 4485A>C), ATC362TTG (4489A>T 4491C>G), GGA363GCT (4493G>C 4494A>T), GGT364GGG (4497T>G), AAG365CAA (4498A>C 4500G>A), ACC366GAT (4501A>G 4502C>A 4503C>T), TTT367TAT (4505T>A), TAC368TAT (4509C>T), TCT369TGC (4511C>G 4512T>C), TCC370TTC (4514C>T), TTT371TTG (4518T>G), GAC372GAT (4521C>T), TGT373GGG (4522T>G 4524T>G), AAA374TAC (4525A>T 4527A>C), TTT377TAT (4535T>A), TGG378AAT (4537T>A 4538G>A 4539G>T), CAG379CAA (4542G>A), GTA380ATC (4543G>A 4545A>C), CGC381ACA (4546C>A 4547G>C 4548C>A), CTT382ATT (4549C>A), GCT383GCA (4554T>A), GAA385GAG (4560A>G), ACA386GAC (4561A>G 4562C>A 4563A>C), ATT387CAA (4564A>C 4565T>A 4566T>A), CAG388GAG (4567C>G), CTA389AAG (4570C>A 4571T>A 4572A>G), GCT391ACT (4576G>A), AGC393ACA (4583G>C 4584C>A), TGT394TGC (4587T>C), CCC395CCT (4590C>T), CAA396TAT (4591C>T 4593A>T), GGA397GGG (4596A>G), CAC398ACA (4597C>A 4598A>C 4599C>A), TAC399TAT (4602C>T), GAA400GCA (4604A>C), TGG401TTC (4607G>T 4608G>C), TTA402AAG (4609T>A 4610T>A 4611A>G), GTC403CGG (4612G>C 4613T>G 4614C>G), GGA407GGG (4626A>G), AAA409TGC (4630A>T 4631A>G 4632A>C), CAA410AAT (4633C>A 4635A>T), GCC411GCA (4638C>A), CCT412CCG (4641T>G), GCT413ACA (4642G>A 4644T>A), ATT414ACA (4646T>C 4647T>A), TTC415TTT (4650C>T), AGG417AGA (4656G>A), CAC418TGC (4657C>T 4658A>G), GAC420ATG (4663G>A 4664A>T 4665C>G), GAA421GCC (4667A>C 4668A>C), AGT422ATA (4670G>T 4671T>A), CTC423TTC (4672C>T), AGC424CAC (4675A>C 4676G>A), AAC425GAC (4678A>G), TAT427GTA (4684T>G 4685A>T 4686T>A), CCA428GAA (4687C>G 4688C>A), CAG429GAT (4690C>G 4692G>T), TGT431GTT (4696T>G 4697G>T), GCT432GAG (4700C>A 4701T>G), GTA433GTC (4704A>C), TAT434TTC (4706A>T 4707T>C), GTT435ATG (4708G>A 4710T>G), GAC437AAT (4714G>A 4716C>T), ATC438TTT (4717A>T 4719C>T), ATC439TCA (4720A>T 4721T>C 4722C>A) | | | | | | | | |
| **Proteins** | | | | | | | | | |
| hypothetical protein (NP_043933.1) | 334 | 440 | 15.1% | 321 | 40.2% | 107 (100%) | 45 (42.1%) | 0/0/0/0 | 0 |
| Protein mutations: | I336M (4413T>G), N337D (4414A>G), H344A (4435C>G 4436A>C), K346Q (4441A>C), G347K (4444G>A 4445G>A 4446A>G), G349H (4450G>C 4451G>A), L351P (4457T>C), L352V (4459C>G 4461T>G), N354F (4465A>T 4466A>T), K355I (4469A>T 4470G>T), E356K (4471G>A 4473G>A), Q357K (4474C>A), L358M (4477C>A), Q360D (4483C>G 4485A>C), I362L (4489A>T 4491C>G), G363A (4493G>C 4494A>T), K365Q (4498A>C 4500G>A), T366D (4501A>G 4502C>A 4503C>T), F367Y (4505T>A), S369C (4511C>G 4512T>C), S370F (4514C>T), F371L (4518T>G), C373G (4522T>G 4524T>G), K374Y (4525A>T 4527A>C), F377Y (4535T>A), W378N (4537T>A 4538G>A 4539G>T), V380I (4543G>A 4545A>C), R381T (4546C>A 4547G>C 4548C>A), L382I (4549C>A), T386D (4561A>G 4562C>A 4563A>C), I387Q (4564A>C 4565T>A 4566T>A), Q388E (4567C>G), L389K (4570C>A 4571T>A 4572A>G), A391T (4576G>A), S393T (4583G>C 4584C>A), Q396Y (4591C>T 4593A>T), H398T (4597C>A 4598A>C 4599C>A), E400A (4604A>C), W401F (4607G>T 4608G>C), L402K (4609T>A 4610T>A 4611A>G), V403R (4612G>C 4613T>G 4614C>G), K409C (4630A>T 4631A>G 4632A>C), Q410N (4633C>A 4635A>T), A413T (4642G>A 4644T>A), I414T (4646T>C 4647T>A), H418C (4657C>T 4658A>G), D420M (4663G>A 4664A>T 4665C>G), E421A (4667A>C 4668A>C), S422I (4670G>T 4671T>A), L423F (4672C>T), S424H (4675A>C 4676G>A), N425D (4678A>G), Y427V (4684T>G 4685A>T 4686T>A), P428E (4687C>G 4688C>A), Q429D (4690C>G 4692G>T), C431V (4696T>G 4697G>T), A432E (4700C>A 4701T>G), Y434F (4706A>T 4707T>C), V435M (4708G>A 4710T>G), D437N (4714G>A 4716C>T), I438F (4717A>T 4719C>T), I439S (4720A>T 4721T>C 4722C>A) | | | | | | | | |
| Codon mutations: | GTA335GTC (4410A>C), ATT336ATG (4413T>G), AAC337GAC (4414A>G), TAT338TAC (4419T>C), AAG340AAA (4425G>A), TTA341CTT (4426T>C 4428A>T), GAC343GAT (4434C>T), CAT344GCT (4435C>G 4436A>C), ACA345ACC (4440A>C), AAG346CAG (4441A>C), GGA347AAG (4444G>A 4445G>A 4446A>G), GAT348GAC (4449T>C), GGC349CAC (4450G>C 4451G>A), CTA351CCA (4457T>C), CTT352GTG (4459C>G 4461T>G), CCT353CCC (4464T>C), AAC354TTC (4465A>T 4466A>T), AAG355ATT (4469A>T 4470G>T), GAG356AAA (4471G>A 4473G>A), CAA357AAA (4474C>A), CTG358ATG (4477C>A), CAA360GAC (4483C>G 4485A>C), ATC362TTG (4489A>T 4491C>G), GGA363GCT (4493G>C 4494A>T), GGT364GGG (4497T>G), AAG365CAA (4498A>C 4500G>A), ACC366GAT (4501A>G 4502C>A 4503C>T), TTT367TAT (4505T>A), TAC368TAT (4509C>T), TCT369TGC (4511C>G 4512T>C), TCC370TTC (4514C>T), TTT371TTG (4518T>G), GAC372GAT (4521C>T), TGT373GGG (4522T>G 4524T>G), AAA374TAC (4525A>T 4527A>C), TTT377TAT (4535T>A), TGG378AAT (4537T>A 4538G>A 4539G>T), CAG379CAA (4542G>A), GTA380ATC (4543G>A 4545A>C), CGC381ACA (4546C>A 4547G>C 4548C>A), CTT382ATT (4549C>A), GCT383GCA (4554T>A), GAA385GAG (4560A>G), ACA386GAC (4561A>G 4562C>A 4563A>C), ATT387CAA (4564A>C 4565T>A 4566T>A), CAG388GAG (4567C>G), CTA389AAG (4570C>A 4571T>A 4572A>G), GCT391ACT (4576G>A), AGC393ACA (4583G>C 4584C>A), TGT394TGC (4587T>C), CCC395CCT (4590C>T), CAA396TAT (4591C>T 4593A>T), GGA397GGG (4596A>G), CAC398ACA (4597C>A 4598A>C 4599C>A), TAC399TAT (4602C>T), GAA400GCA (4604A>C), TGG401TTC (4607G>T 4608G>C), TTA402AAG (4609T>A 4610T>A 4611A>G), GTC403CGG (4612G>C 4613T>G 4614C>G), GGA407GGG (4626A>G), AAA409TGC (4630A>T 4631A>G 4632A>C), CAA410AAT (4633C>A 4635A>T), GCC411GCA (4638C>A), CCT412CCG (4641T>G), GCT413ACA (4642G>A 4644T>A), ATT414ACA (4646T>C 4647T>A), TTC415TTT (4650C>T), AGG417AGA (4656G>A), CAC418TGC (4657C>T 4658A>G), GAC420ATG (4663G>A 4664A>T 4665C>G), GAA421GCC (4667A>C 4668A>C), AGT422ATA (4670G>T 4671T>A), CTC423TTC (4672C>T), AGC424CAC (4675A>C 4676G>A), AAC425GAC (4678A>G), TAT427GTA (4684T>G 4685A>T 4686T>A), CCA428GAA (4687C>G 4688C>A), CAG429GAT (4690C>G 4692G>T), TGT431GTT (4696T>G 4697G>T), GCT432GAG (4700C>A 4701T>G), GTA433GTC (4704A>C), TAT434TTC (4706A>T 4707T>C), GTT435ATG (4708G>A 4710T>G), GAC437AAT (4714G>A 4716C>T), ATC438TTT (4717A>T 4719C>T), ATC439TCA (4720A>T 4721T>C 4722C>A) | | | | | | | | |

*: Inserts / Deletes / Misaligned / Frameshifts

#### Analysis details

This analysis was performed with panviral2.64

### NGS Details (UN9): Epiphyllum badnavirus 1

#### Assembly

| | |
|---|---|
| **Coverage Length** | 306 (1 contig(s)) |
| **Depth Of Coverage** | 18.6 |
| **Number Of Reads** | 55 |
| **Reads Per Million** | 1.24 rpm (after QC) |
| **Ambiguities** | 0 |
| **Assembly Method** | de novo + reference guided assembly |
| **Consensus Caller** | Bcf Tools |

#### Coverage Map

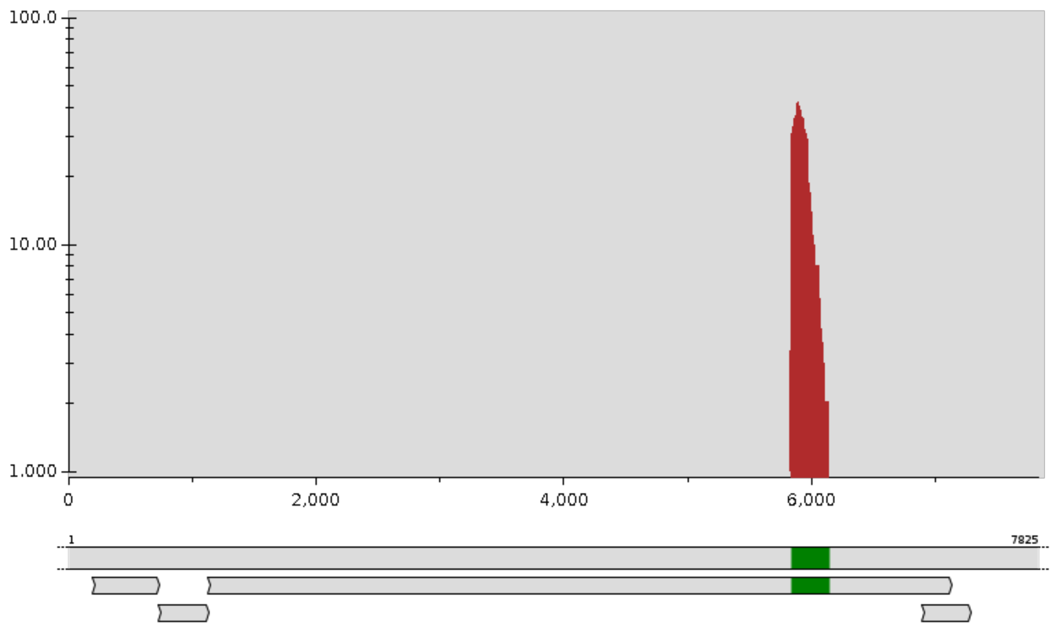

#### Assignment

| | |
|---|---|
| **Type** | Epiphyllum badnavirus 1 (Taxonomy ID: 2518008) |
| **Reference Genome** | NC_076247.1 |
| **NT Identity (%)** | 52.9412 |
| **AA Identity (%)** | 45.098 |
| **Number Of Stop Codons** | 0 |
| **Number Of CDS** | 4 |

#### Alignment

| | |
|---|---|
| **Alignment Score** | 18.0 (NT) + 312.0 (AA) = 330.0 |
| **Concordance (%)** | 25.1524 |

| **Alignment Method** | Global, seeded, nucleotide + amino acids (AGA) |
|---|---|

#### Genome Region

Sequence starts at position 5835 and ends at position 6140 relative to NC_076247.1 reference sequence.

#### Alignment Detailed Statistics

| | Begin | End | Coverage | Score | Concordance | Matches | Identities | I/D/M/F* | Stop Codons |
|---|---|---|---|---|---|---|---|---|---|
| **NT** | **5835** | **6140** | **3.9%** | **18** | **3.0%** | **303 (98.1%)** | **162 (52.4%)** | **3/3** | |
| Mutations: | 5835C>G, 5839G>T, 5847C>A, 5851T>A, 5853C>A, 5859G>T, 5860A>C, 5861A>G, 5862G>T, 5863A>T, 5864G>C, 5865C>T, 5868A>G, 5870T>A, 5871C>T, 5878A>T, 5880T>G, 5882T>A, 5883G>A, 5884A>G, 5886G>T, 5887G>A, 5888A>G, 5889C>G, 5893G>A, 5894A>G, 5895A>T, 5896T>G, 5897C>A, 5898C>T, 5901C>T, 5904T>C, 5905T>A, 5906G>A, 5910C>A, 5911G>A, 5913C>A, 5918T>G, 5922C>A, 5923C>A, 5924C>G, 5925A>G, 5926G>T, 5933T>A, 5934T>C, 5936A>T, 5937C>T, 5938G>A, 5942G>T, 5943G>T, 5944T>C, 5946A>C, 5949T>C, 5953C>T, 5955A>C, 5958C>T, 5961A>T, 5964G>A, 5966A>C, 5967G>T, 5970T>C, 5971G>A, 5973C>T, 5982G>C, 5985C>T, 5986C>A, 5987A>T, 5988A>G, 5989C>G, 5990G>A, 5991A>T, 5992A>C, 5993A>T, 5994G>T, 5997G>A, 5998G>A, 6000C>T, 6002A>G, 6003C>G, 6004T>A, 6005G>T, 6006T>A, 6011G>A, 6012A>G, 6013G>C, 6014A>C, 6015_6016insCAT, 6018G>C, 6019T>G, 6020C>A, 6021A>T, 6022G>A, 6023G>T, 6024C>G, 6026T>A, 6030A>G, 6032C>T, 6033A>G, 6036T>G, 6038A>T, 6039C>T, 6059T>A, 6060C>T, 6061A>T, 6062G>C, 6063T>A, 6065A>G, 6066C>A, 6067A>G, 6068C>A, 6069T>C, 6070G>A, 6072A>G, 6073G>C, 6074A>G, 6075T>A, 6078C>G, 6081T>C, 6082G>A, 6083C>T, 6085C>G, 6088C>A, 6095A>G, 6096G>A, 6097T>A, 6099C>T, 6100A>G, 6113G>T, 6114T>G, 6115_6117delGAG, 6123G>A, 6125G>A, 6126G>A, 6128T>A, 6130A>T, 6132C>A, 6133C>T, 6134T>A, 6135C>T, 6136T>G, 6138T>A, 6139C>A, 6140C>A | | | | | | | | |

##### CDS

| | Begin | End | Coverage | Score | Concordance | Matches | Identities | I/D/M/F* | Stop Codons |
|---|---|---|---|---|---|---|---|---|---|
| QKM20_gp3 | 1570 | 1671 | 5.1% | 312 | 42.5% | 101 (98.1%) | 46 (44.7%) | 1/1/0/0 | 0 |
| Protein mutations: | V1571F (5839G>T), F1575I (5851T>A 5853C>A), K1578R (5860A>C 5861A>G 5862G>T), F1581Y (5870T>A 5871C>T), I1584L (5878A>T 5880T>G), M1585K (5882T>A 5883G>A), M1586V (5884A>G 5886G>T), D1587R (5887G>A 5888A>G 5889C>G), E1589S (5893G>A 5894A>G 5895A>T), S1590D (5896T>G 5897C>A 5898C>T), W1593K (5905T>A 5906G>A), A1595T (5911G>A 5913C>A), L1597R (5918T>G), P1599R (5923C>A 5924C>G 5925A>G), D1600Y (5926G>T), L1602H (5933T>A 5934T>C), Y1603F (5936A>T 5937C>T), E1604K (5938G>A), W1605F (5942G>T 5943G>T), P1609S (5953C>T 5955A>C), K1613T (5966A>C 5967G>T), A1615T (5971G>A 5973C>T), Q1620M (5986C>A 5987A>T 5988A>G), R1621D (5989C>G 5990G>A 5991A>T), K1622L (5992A>C 5993A>T 5994G>T), M1623I (5997G>A), D1624N (5998G>A 6000C>T), N1625R (6002A>G 6003C>G), C1626I (6004T>A 6005G>T 6006T>A), R1628K (6011G>A 6012A>G), D1629P (6013G>C 6014A>C), D1629_L1630insH (6015_6016insCAT), S1631D (6019T>G 6020C>A 6021A>T), G1632M (6022G>A 6023G>T 6024C>G), F1633Y (6026T>A), A1635V (6032C>T 6033A>G), Y1637F (6038A>T 6039C>T), F1644Y (6059T>A 6060C>T), N1646R (6065A>G 6066C>A), T1647D (6067A>G 6068C>A 6069T>C), E1648K (6070G>A 6072A>G), D1649R (6073G>C 6074A>G 6075T>A), D1650E (6078C>G), A1652I (6082G>A 6083C>T), Q1653E (6085C>G), H1654N (6088C>A), K1656R (6095A>G 6096G>A), S1657T (6097T>A 6099C>T), M1658V (6100A>G), C1662L (6113G>T 6114T>G), E1663del (6115_6117delGAG), G1666E (6125G>A 6126G>A), L1667Q (6128T>A), I1668L (6130A>T 6132C>A), L1669Y (6133C>T 6134T>A 6135C>T), S1670A (6136T>G 6138T>A) | | | | | | | | |
| Codon mutations: | GCC1569..G (5835C>G), GTC1571TTC (5839G>T), TCC1573TCA (5847C>A), TTC1575ATA (5851T>A 5853C>A), CTG1577CTT (5859G>T), AAG1578CGT (5860A>C 5861A>G 5862G>T), AGC1579TCT (5863A>T 5864G>C 5865C>T), GGA1580GGG (5868A>G), TTC1581TAT (5870T>A 5871C>T), ATT1584TTG (5878A>T 5880T>G), ATG1585AAA (5882T>A 5883G>A), ATG1586GTT (5884A>G 5886G>T), GAC1587AGG (5887G>A 5888A>G 5889C>G), GAA1589AGT (5893G>A 5894A>G 5895A>T), TCC1590GAT (5896T>G 5897C>A 5898C>T), ATC1591ATT (5901C>T), CCT1592CCC (5904T>C), TGG1593AAG (5905T>A 5906G>A), ACC1594ACA (5910C>A), GCC1595ACA (5911G>A 5913C>A), CTG1597CGG (5918T>G), ACC1598ACA (5922C>A), CCA1599AGG (5923C>A 5924C>G 5925A>G), GAT1600TAT (5926G>T), CTT1602CAC (5933T>A 5934T>C), TAC1603TTT (5936A>T 5937C>T), GAA1604AAA (5938G>A), TGG1605TTT (5942G>T 5943G>T), TTA1606CTC (5944T>C 5946A>C), GTT1607GTC (5949T>C), CCA1609TCC (5953C>T 5955A>C), TTC1610TTT (5958C>T), GGA1611GGT (5961A>T), TTG1612TTA (5964G>A), AAG1613ACT (5966A>C 5967G>T), AAT1614AAC (5970T>C), GCC1615ACT (5971G>A 5973C>T), GTG1618GTC (5982G>C), TTC1619TTT (5985C>T), CAA1620ATG (5986C>A 5987A>T 5988A>G), CGA1621GAT (5989C>G 5990G>A 5991A>T), AAG1622CTT (5992A>C 5993A>T 5994G>T), ATG1623ATA (5997G>A), GAC1624AAT (5998G>A 6000C>T), AAC1625AGG (6002A>G 6003C>G), TGT1626ATA (6004T>A 6005G>T 6006T>A), AGA1628AAG (6011G>A 6012A>G), GAC1629CCC (6013G>C 6014A>C), GAC1629_CTG1630insCAT (6015_6016insCAT), CTG1630CTC (6018G>C), TCA1631GAT (6019T>G 6020C>A 6021A>T), GGC1632ATG (6022G>A 6023G>T 6024C>G), TTT1633TAT (6026T>A), GTA1634GTG (6030A>G), GCA1635GTG (6032C>T 6033A>G), GTT1636GTG (6036T>G), TAC1637TTT (6038A>T 6039C>T), TTC1644TAT (6059T>A 6060C>T), AGT1645TCA (6061A>T 6062G>C 6063T>A), AAC1646AGA (6065A>G 6066C>A), ACT1647GAC (6067A>G 6068C>A 6069T>C), GAA1648AAG (6070G>A 6072A>G), GAT1649CGA (6073G>C 6074A>G 6075T>A), GAC1650GAG (6078C>G), CAT1651CAC (6081T>C), GCA1652ATA (6082G>A 6083C>T), CAG1653GAG (6085C>G), CAC1654AAC (6088C>A), AAG1656AGA (6095A>G 6096G>A), TCC1657ACT (6097T>A 6099C>T), ATG1658GTG (6100A>G), TGT1662TTG (6113G>T 6114T>G), GAG1663del (6115_6117delGAG), GAG1665GAA (6123G>A), GGG1666GAA (6125G>A 6126G>A), CTG1667CAG (6128T>A), ATC1668TTA (6130A>T 6132C>A), CTC1669TAT (6133C>T 6134T>A 6135C>T), TCT1670GCA (6136T>G 6138T>A), CCT1671AA. (6139C>A 6140C>A) | | | | | | | | |

##### Proteins

| | Begin | End | Coverage | Score | Concordance | Matches | Identities | I/D/M/F* | Stop Codons |
|---|---|---|---|---|---|---|---|---|---|
| polyprotein (YP_010797894.1) | 1570 | 1671 | 5.1% | 312 | 42.5% | 101 (98.1%) | 46 (44.7%) | 1/1/0/0 | 0 |
| Protein mutations: | V1571F (5839G>T), F1575I (5851T>A 5853C>A), K1578R (5860A>C 5861A>G 5862G>T), F1581Y (5870T>A 5871C>T), I1584L (5878A>T 5880T>G), M1585K (5882T>A 5883G>A), M1586V (5884A>G 5886G>T), D1587R (5887G>A 5888A>G 5889C>G), E1589S (5893G>A 5894A>G 5895A>T), S1590D (5896T>G 5897C>A 5898C>T), W1593K (5905T>A 5906G>A), A1595T (5911G>A 5913C>A), L1597R (5918T>G), P1599R (5923C>A 5924C>G 5925A>G), D1600Y (5926G>T), L1602H (5933T>A 5934T>C), Y1603F (5936A>T 5937C>T), E1604K (5938G>A), W1605F (5942G>T 5943G>T), P1609S (5953C>T 5955A>C), K1613T (5966A>C 5967G>T), A1615T (5971G>A 5973C>T), Q1620M (5986C>A 5987A>T 5988A>G), R1621D (5989C>G 5990G>A 5991A>T), K1622L (5992A>C 5993A>T 5994G>T), M1623I (5997G>A), D1624N (5998G>A 6000C>T), N1625R (6002A>G 6003C>G), C1626I (6004T>A 6005G>T 6006T>A), R1628K (6011G>A 6012A>G), D1629P (6013G>C 6014A>C), D1629_L1630insH (6015_6016insCAT), S1631D (6019T>G 6020C>A 6021A>T), G1632M (6022G>A 6023G>T 6024C>G), F1633Y (6026T>A), A1635V (6032C>T 6033A>G), Y1637F (6038A>T 6039C>T), F1644Y (6059T>A 6060C>T), N1646R (6065A>G 6066C>A), T1647D (6067A>G 6068C>A 6069T>C), E1648K (6070G>A 6072A>G), D1649R (6073G>C 6074A>G 6075T>A), D1650E (6078C>G), A1652I (6082G>A 6083C>T), Q1653E (6085C>G), H1654N (6088C>A), K1656R (6095A>G 6096G>A), S1657T (6097T>A 6099C>T), M1658V (6100A>G), C1662L (6113G>T 6114T>G), E1663del (6115_6117delGAG), G1666E (6125G>A 6126G>A), L1667Q (6128T>A), I1668L (6130A>T 6132C>A), L1669Y (6133C>T 6134T>A 6135C>T), S1670A (6136T>G 6138T>A) | | | | | | | | |
| Codon mutations: | GCC1569..G (5835C>G), GTC1571TTC (5839G>T), TCC1573TCA (5847C>A), TTC1575ATA (5851T>A 5853C>A), CTG1577CTT (5859G>T), AAG1578CGT (5860A>C 5861A>G 5862G>T), AGC1579TCT (5863A>T 5864G>C 5865C>T), GGA1580GGG (5868A>G), TTC1581TAT (5870T>A 5871C>T), ATT1584TTG (5878A>T 5880T>G), ATG1585AAA (5882T>A 5883G>A), ATG1586GTT (5884A>G 5886G>T), GAC1587AGG (5887G>A 5888A>G 5889C>G), GAA1589AGT (5893G>A 5894A>G 5895A>T), TCC1590GAT (5896T>G 5897C>A 5898C>T), ATC1591ATT (5901C>T), CCT1592CCC (5904T>C), TGG1593AAG (5905T>A 5906G>A), ACC1594ACA (5910C>A), GCC1595ACA (5911G>A 5913C>A), CTG1597CGG (5918T>G), ACC1598ACA (5922C>A), CCA1599AGG (5923C>A 5924C>G 5925A>G), GAT1600TAT (5926G>T), CTT1602CAC (5933T>A 5934T>C), TAC1603TTT (5936A>T 5937C>T), GAA1604AAA (5938G>A), TGG1605TTT (5942G>T 5943G>T), TTA1606CTC (5944T>C 5946A>C), GTT1607GTC (5949T>C), CCA1609TCC (5953C>T 5955A>C), TTC1610TTT (5958C>T), GGA1611GGT (5961A>T), TTG1612TTA (5964G>A), AAG1613ACT (5966A>C 5967G>T), AAT1614AAC (5970T>C), GCC1615ACT (5971G>A 5973C>T), GTG1618GTC (5982G>C), TTC1619TTT (5985C>T), CAA1620ATG (5986C>A 5987A>T 5988A>G), CGA1621GAT (5989C>G 5990G>A 5991A>T), AAG1622CTT (5992A>C 5993A>T 5994G>T), ATG1623ATA (5997G>A), GAC1624AAT (5998G>A 6000C>T), AAC1625AGG (6002A>G 6003C>G), TGT1626ATA (6004T>A 6005G>T 6006T>A), AGA1628AAG (6011G>A 6012A>G), GAC1629CCC (6013G>C 6014A>C), GAC1629_CTG1630insCAT (6015_6016insCAT), CTG1630CTC (6018G>C), TCA1631GAT (6019T>G 6020C>A 6021A>T), GGC1632ATG (6022G>A 6023G>T 6024C>G), TTT1633TAT (6026T>A), GTA1634GTG (6030A>G), GCA1635GTG (6032C>T 6033A>G), GTT1636GTG (6036T>G), TAC1637TTT (6038A>T 6039C>T), TTC1644TAT (6059T>A 6060C>T), AGT1645TCA (6061A>T 6062G>C 6063T>A), AAC1646AGA (6065A>G 6066C>A), ACT1647GAC (6067A>G 6068C>A 6069T>C), GAA1648AAG (6070G>A 6072A>G), GAT1649CGA (6073G>C 6074A>G 6075T>A), GAC1650GAG (6078C>G), CAT1651CAC (6081T>C), GCA1652ATA (6082G>A 6083C>T), CAG1653GAG (6085C>G), CAC1654AAC (6088C>A), AAG1656AGA (6095A>G 6096G>A), TCC1657ACT (6097T>A 6099C>T), ATG1658GTG (6100A>G), TGT1662TTG (6113G>T 6114T>G), GAG1663del (6115_6117delGAG), GAG1665GAA (6123G>A), GGG1666GAA (6125G>A 6126G>A), CTG1667CAG (6128T>A), ATC1668TTA (6130A>T 6132C>A), CTC1669TAT (6133C>T 6134T>A 6135C>T), TCT1670GCA (6136T>G 6138T>A), CCT1671AA. (6139C>A 6140C>A) | | | | | | | | |

*: Inserts / Deletes / Misaligned / Frameshifts

#### Analysis details

This analysis was performed with panviral2.64

### NGS Details (UN9): Caulimovirus glycinis

#### Assembly

| | |
|---|---|
| **Coverage Length** | 418 (1 contig(s)) |
| **Depth Of Coverage** | 13.0 |
| **Number Of Reads** | 44 |
| **Reads Per Million** | 0.99 rpm (after QC) |
| **Ambiguities** | 0 |
| **Assembly Method** | de novo + reference guided assembly |
| **Consensus Caller** | Bcf Tools |

#### Coverage Map

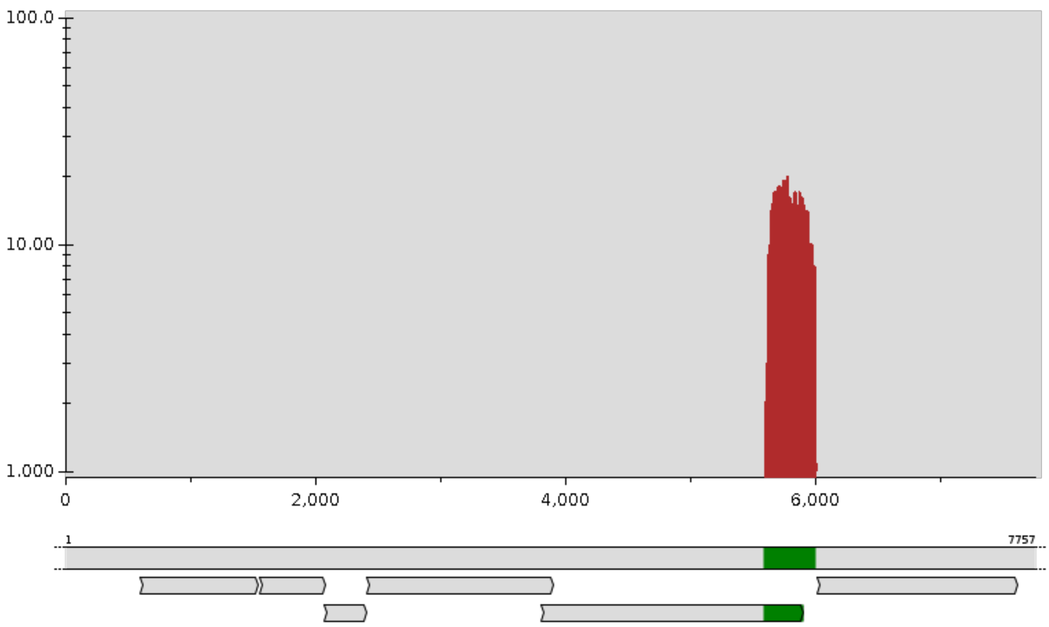

#### Assignment

| | |
|---|---|
| **Type** | Caulimovirus glycinis (Taxonomy ID: 3048324) |
| **Reference Genome** | NC_018505.1 |
| **NT Identity (%)** | 57.2127 |
| **AA Identity (%)** | 43.2692 |
| **Number Of Stop Codons** | 2 |
| **Number Of CDS** | 6 |

#### Alignment

| | |
|---|---|
| **Alignment Score** | 73.0 (NT) + 321.0 (AA) = 394.0 |
| **Concordance (%)** | 26.041 |

| Alignment Method | Global, seeded, nucleotide + amino acids (AGA) |
|---|---|

#### Genome Region

Sequence starts at position 5592 and ends at position 6009 relative to NC_018505.1 reference sequence.

#### Alignment Detailed Statistics

| | Begin | End | Coverage | Score | Concordance | Matches | Identities | I/D/M/F* | Stop Codons |
|---|---|---|---|---|---|---|---|---|---|
| **NT** | **5592** | **6009** | **5.4%** | **73** | **9.4%** | **405 (96.0%)** | **234 (55.5%)** | **4/13** | |
| Mutations: | 5595T>A, 5599A>T, 5611A>T, 5612T>A, 5613C>G, 5617T>A, 5618T>A, 5620A>C, 5623C>T, 5626A>T, 5628C>A, 5630G>A, 5632T>A, 5636T>A, 5637T>C, 5638A>T, 5640A>G, 5641C>G, 5644T>C, 5645C>T, 5646A>T, 5647T>A, 5648T>A, 5650C>A, 5659G>A, 5663A>T, 5665T>A, 5668G>A, 5672G>A, 5674T>A, 5675A>G, 5677G>A, 5679A>G, 5680T>A, 5681A>G, 5682C>G, 5683A>C, 5686T>A, 5687A>C, 5688A>G, 5689G>A, 5692A>G, 5696A>T, 5700G>C, 5701T>A, 5703A>T, 5704T>C, 5705C>T, 5707T>A, 5708A>C, 5709C>T, 5713T>A, 5714G>A, 5715T>A, 5716A>G, 5717A>C, 5723T>A, 5725A>C, 5728A>C, 5729C>A, 5732A>T, 5734A>G, 5737T>C, 5742A>C, 5744A>C, 5746T>G, 5747T>G, 5749T>A, 5750A>T, 5752C>A, 5753C>G, 5754A>G, 5756T>C, 5758T>A, 5759C>A, 5761T>A, 5762A>T, 5763A>T, 5764A>C, 5766T>A, 5770C>A, 5774A>C, 5775A>C, 5777G>T, 5778G>C, 5782T>A, 5783T>C, 5784A>C, 5786A>C, 5788G>A, 5789C>T, 5791G>T, 5792G>A, 5794T>G, 5801G>C, 5802T>A, 5803A>T, 5809G>A, 5812A>G, 5813A>G, 5815G>A, 5817G>T, 5818G>A, 5821T>A, 5824A>T, 5825A>T, 5826G>T, 5827A>T, 5831G>A, 5832T>C, 5836T>C, 5837G>A, 5840G>A, 5842C>T, 5846C>A, 5849T>A, 5851G>T, 5852T>A, 5853C>A, 5858G>T, 5859A>C, 5860C>T, 5861A>G, 5863A>T, 5881T>C, 5882C>T, 5885A>T, 5887C>A, 5888C>A, 5890T>G, 5891G>A, 5893A>C, 5894T>G, 5899T>A, 5903T>G, 5906T>C, 5907T>A, 5908T>A, 5909C>T, 5911G>A, 5912G>A, 5913T>A, 5915T>G, 5915_5916insAAG, 5917C>T, 5918A>C, 5924T>A, 5925A>G, 5928G>A, 5931T>A, 5935C>T, 5940C>A, 5941T>A, 5944G>A, 5945_5956delAACCTTAACCGG, 5958T>G, 5959C>T, 5961A>T, 5962T>C, 5966G>T, 5969T>A, 5971A>G, 5972C>A, 5976_5977insA, 5980G>A, 5981G>T, 5984C>A, 5985G>A, 5987T>A, 5988T>A, 5989delT, 5993C>A, 5997C>T, 5998T>C, 6002G>T | | | | | | | | |

##### CDS

| | Begin | End | Coverage | Score | Concordance | Matches | Identities | I/D/M/F* | Stop Codons |
|---|---|---|---|---|---|---|---|---|---|
| C186_gp5 | 597 | 700 | 14.9% | 321 | 43.4% | 104 (100%) | 45 (43.3%) | 0/0/0/0 | 2 |
| Protein mutations: | L597* (5595T>A), S605T (5618T>A 5620A>C), K607N (5626A>T), A608E (5628C>A), A609T (5630G>A 5632T>A), L611T (5636T>A 5637T>C 5638A>T), N612R (5640A>G 5641C>G), H614L (5645C>T 5646A>T 5647T>A), S615T (5648T>A 5650C>A), I620L (5663A>T 5665T>A), V623I (5672G>A 5674T>A), M624V (5675A>G 5677G>A), N625R (5679A>G 5680T>A), T626G (5681A>G 5682C>G 5683A>C), K628R (5687A>C 5688A>G 5689G>A), T631S (5696A>T), G632A (5700G>C 5701T>A), Y633F (5703A>T 5704T>C), T635L (5708A>C 5709C>T), V637K (5714G>A 5715T>A 5716A>G), K638Q (5717A>C), L640I (5723T>A 5725A>C), T643S (5732A>T 5734A>G), K646T (5742A>C), N647Q (5744A>C 5746T>G), F648V (5747T>G 5749T>A), T649S (5750A>T 5752C>A), H650G (5753C>G 5754A>G), F651L (5756T>C 5758T>A), L652I (5759C>A 5761T>A), K653F (5762A>T 5763A>T 5764A>C), I654N (5766T>A), N655K (5770C>A), K657P (5774A>C 5775A>C), G658S (5777G>T 5778G>C), D659E (5782T>A), Y660P (5783T>C 5784A>C), K661Q (5786A>C 5788G>A), Q662Y (5789C>T 5791G>T), G663R (5792G>A 5794T>G), V666H (5801G>C 5802T>A 5803A>T), W668* (5809G>A), M670V (5813A>G 5815G>A), W671L (5817G>T 5818G>A), F672L (5821T>A), R674F (5825A>T 5826G>T 5827A>T), V676T (5831G>A 5832T>C), E678K (5837G>A), V679I (5840G>A 5842C>T), H681N (5846C>A), L682I (5849T>A 5851G>T), S683K (5852T>A 5853C>A), D685S (5858G>T 5859A>C 5860C>T), K686D (5861A>G 5863A>T), T694S (5885A>T 5887C>A), E696N (5891G>A 5893A>C), F697V (5894T>G), H698Q (5899T>A), *700G (5903T>G) | | | | | | | | |
| Codon mutations: | TTA597TAA (5595T>A), ATA598ATT (5599A>T), ACA602ACT (5611A>T), TCT603AGT (5612T>A 5613C>G), GGT604GGA (5617T>A), TCA605ACC (5618T>A 5620A>C), TTC606TTT (5623C>T), AAA607AAT (5626A>T), GCA608GAA (5628C>A), GCT609ACA (5630G>A 5632T>A), TTA611ACT (5636T>A 5637T>C 5638A>T), AAC612AGG (5640A>G 5641C>G), TAT613TAC (5644T>C), CAT614TTA (5645C>T 5646A>T 5647T>A), TCC615ACA (5648T>A 5650C>A), AAG618AAA (5659G>A), ATT620TTA (5663A>T 5665T>A), CTG621CTA (5668G>A), GTT623ATA (5672G>A 5674T>A), ATG624GTA (5675A>G 5677G>A), AAT625AGA (5679A>G 5680T>A), ACA626GGC (5681A>G 5682C>G 5683A>C), ATT627ATA (5686T>A), AAG628CGA (5687A>C 5688A>G 5689G>A), AAA629AAG (5692A>G), ACT631TCT (5696A>T), GGT632GCA (5700G>C 5701T>A), TAT633TTC (5703A>T 5704T>C), CTT634TTA (5705C>T 5707T>A), ACA635CTA (5708A>C 5709C>T), CCT636CCA (5713T>A), GTA637AAG (5714G>A 5715T>A 5716A>G), AAA638CAA (5717A>C), TTA640ATC (5723T>A 5725A>C), ATA641ATC (5728A>C), CGA642AGA (5729C>A), ACA643TCG (5732A>T 5734A>G), GAT644GAC (5737T>C), AAG646ACG (5742A>C), AAT647CAG (5744A>C 5746T>G), TTT648GTA (5747T>G 5749T>A), ACC649TCA (5750A>T 5752C>A), CAT650GGT (5753C>G 5754A>G), TTT651CTA (5756T>C 5758T>A), CTT652ATA (5759C>A 5761T>A), AAA653TTC (5762A>T 5763A>T 5764A>C), ATT654AAT (5766T>A), AAC655AAA (5770C>A), AAA657CCA (5774A>C 5775A>C), GGT658TCT (5777G>T 5778G>C), GAT659GAA (5782T>A), TAT660CCT (5783T>C 5784A>C), AAG661CAA (5786A>C 5788G>A), CAG662TAT (5789C>T 5791G>T), GGT663AGG (5792G>A 5794T>G), GTA666CAT (5801G>C 5802T>A 5803A>T), TGG668TGA (5809G>A), CAA669CAG (5812A>G), ATG670GTA (5813A>G 5815G>A), TGG671TTA (5817G>T 5818G>A), TTT672TTA (5821T>A), TCA673TCT (5824A>T), AGA674TTT (5825A>T 5826G>T 5827A>T), GTT676ACT (5831G>A 5832T>C), TTT677TTC (5836T>C), GAA678AAA (5837G>A), GTC679ATT (5840G>A 5842C>T), CAT681AAT (5846C>A), TTG682ATT (5849T>A 5851G>T), TCA683AAA (5852T>A 5853C>A), GAC685TCT (5858G>T 5859A>C 5860C>T), AAA686GAT (5861A>G 5863A>T), TTT692TTC (5881T>C), CTA693TTA (5882C>T), ACC694TCA (5885A>T 5887C>A), CGT695AGG (5888C>A 5890T>G), GAA696AAC (5891G>A 5893A>C), TTT697GTT (5894T>G), CAT698CAA (5899T>A), TGA700GGA (5903T>G) | | | | | | | | |

##### Proteins

| | Begin | End | Coverage | Score | Concordance | Matches | Identities | I/D/M/F* | Stop Codons |
|---|---|---|---|---|---|---|---|---|---|
| reverse transcriptase (YP_006607892.1) | 597 | 700 | 14.9% | 321 | 43.4% | 104 (100%) | 45 (43.3%) | 0/0/0/0 | 2 |
| Protein mutations: | L597* (5595T>A), S605T (5618T>A 5620A>C), K607N (5626A>T), A608E (5628C>A), A609T (5630G>A 5632T>A), L611T (5636T>A 5637T>C 5638A>T), N612R (5640A>G 5641C>G), H614L (5645C>T 5646A>T 5647T>A), S615T (5648T>A 5650C>A), I620L (5663A>T 5665T>A), V623I (5672G>A 5674T>A), M624V (5675A>G 5677G>A), N625R (5679A>G 5680T>A), T626G (5681A>G 5682C>G 5683A>C), K628R (5687A>C 5688A>G 5689G>A), T631S (5696A>T), G632A (5700G>C 5701T>A), Y633F (5703A>T 5704T>C), T635L (5708A>C 5709C>T), V637K (5714G>A 5715T>A 5716A>G), K638Q (5717A>C), L640I (5723T>A 5725A>C), T643S (5732A>T 5734A>G), K646T (5742A>C), N647Q (5744A>C 5746T>G), F648V (5747T>G 5749T>A), T649S (5750A>T 5752C>A), H650G (5753C>G 5754A>G), F651L (5756T>C 5758T>A), L652I (5759C>A 5761T>A), K653F (5762A>T 5763A>T 5764A>C), I654N (5766T>A), N655K (5770C>A), K657P (5774A>C 5775A>C), G658S (5777G>T 5778G>C), D659E (5782T>A), Y660P (5783T>C 5784A>C), K661Q (5786A>C 5788G>A), Q662Y (5789C>T 5791G>T), G663R (5792G>A 5794T>G), V666H (5801G>C 5802T>A 5803A>T), W668* (5809G>A), M670V (5813A>G 5815G>A), W671L (5817G>T 5818G>A), F672L (5821T>A), R674F (5825A>T 5826G>T 5827A>T), V676T (5831G>A 5832T>C), E678K (5837G>A), V679I (5840G>A 5842C>T), H681N (5846C>A), L682I (5849T>A 5851G>T), S683K (5852T>A 5853C>A), D685S (5858G>T 5859A>C 5860C>T), K686D (5861A>G 5863A>T), T694S (5885A>T 5887C>A), E696N (5891G>A 5893A>C), F697V (5894T>G), H698Q (5899T>A), *700G (5903T>G) | | | | | | | | |
| Codon mutations: | TTA597TAA (5595T>A), ATA598ATT (5599A>T), ACA602ACT (5611A>T), TCT603AGT (5612T>A 5613C>G), GGT604GGA (5617T>A), TCA605ACC (5618T>A 5620A>C), TTC606TTT (5623C>T), AAA607AAT (5626A>T), GCA608GAA (5628C>A), GCT609ACA (5630G>A 5632T>A), TTA611ACT (5636T>A 5637T>C 5638A>T), AAC612AGG (5640A>G 5641C>G), TAT613TAC (5644T>C), CAT614TTA (5645C>T 5646A>T 5647T>A), TCC615ACA (5648T>A 5650C>A), AAG618AAA (5659G>A), ATT620TTA (5663A>T 5665T>A), CTG621CTA (5668G>A), GTT623ATA (5672G>A 5674T>A), ATG624GTA (5675A>G 5677G>A), AAT625AGA (5679A>G 5680T>A), ACA626GGC (5681A>G 5682C>G 5683A>C), ATT627ATA (5686T>A), AAG628CGA (5687A>C 5688A>G 5689G>A), AAA629AAG (5692A>G), ACT631TCT (5696A>T), GGT632GCA (5700G>C 5701T>A), TAT633TTC (5703A>T 5704T>C), CTT634TTA (5705C>T 5707T>A), ACA635CTA (5708A>C 5709C>T), CCT636CCA (5713T>A), GTA637AAG (5714G>A 5715T>A 5716A>G), AAA638CAA (5717A>C), TTA640ATC (5723T>A 5725A>C), ATA641ATC (5728A>C), CGA642AGA (5729C>A), ACA643TCG (5732A>T 5734A>G), GAT644GAC (5737T>C), AAG646ACG (5742A>C), AAT647CAG (5744A>C 5746T>G), TTT648GTA (5747T>G 5749T>A), ACC649TCA (5750A>T 5752C>A), CAT650GGT (5753C>G 5754A>G), TTT651CTA (5756T>C 5758T>A), CTT652ATA (5759C>A 5761T>A), AAA653TTC (5762A>T 5763A>T 5764A>C), ATT654AAT (5766T>A), AAC655AAA (5770C>A), AAA657CCA (5774A>C 5775A>C), GGT658TCT (5777G>T 5778G>C), GAT659GAA (5782T>A), TAT660CCT (5783T>C 5784A>C), AAG661CAA (5786A>C 5788G>A), CAG662TAT (5789C>T 5791G>T), GGT663AGG (5792G>A 5794T>G), GTA666CAT (5801G>C 5802T>A 5803A>T), TGG668TGA (5809G>A), CAA669CAG (5812A>G), ATG670GTA (5813A>G 5815G>A), TGG671TTA (5817G>T 5818G>A), TTT672TTA (5821T>A), TCA673TCT (5824A>T), AGA674TTT (5825A>T 5826G>T 5827A>T), GTT676ACT (5831G>A 5832T>C), TTT677TTC (5836T>C), GAA678AAA (5837G>A), GTC679ATT (5840G>A 5842C>T), CAT681AAT (5846C>A), TTG682ATT (5849T>A 5851G>T), TCA683AAA (5852T>A 5853C>A), GAC685TCT (5858G>T 5859A>C 5860C>T), AAA686GAT (5861A>G 5863A>T), TTT692TTC (5881T>C), CTA693TTA (5882C>T), ACC694TCA (5885A>T 5887C>A), CGT695AGG (5888C>A 5890T>G), GAA696AAC (5891G>A 5893A>C), TTT697GTT (5894T>G), CAT698CAA (5899T>A), TGA700GGA (5903T>G) | | | | | | | | |

*: Inserts / Deletes / Misaligned / Frameshifts

#### Analysis details

This analysis was performed with panviral2.64

### NGS Details (UN9): Caulimovirus venafragariae

#### Assembly

| | |
|---|---|
| **Coverage Length** | 798 (2 contig(s)) |
| **Depth Of Coverage** | 6.4 |
| **Number Of Reads** | 41 |
| **Reads Per Million** | 0.93 rpm (after QC) |
| **Ambiguities** | 0 |
| **Assembly Method** | de novo + reference guided assembly |
| **Consensus Caller** | Bcf Tools |

#### Coverage Map

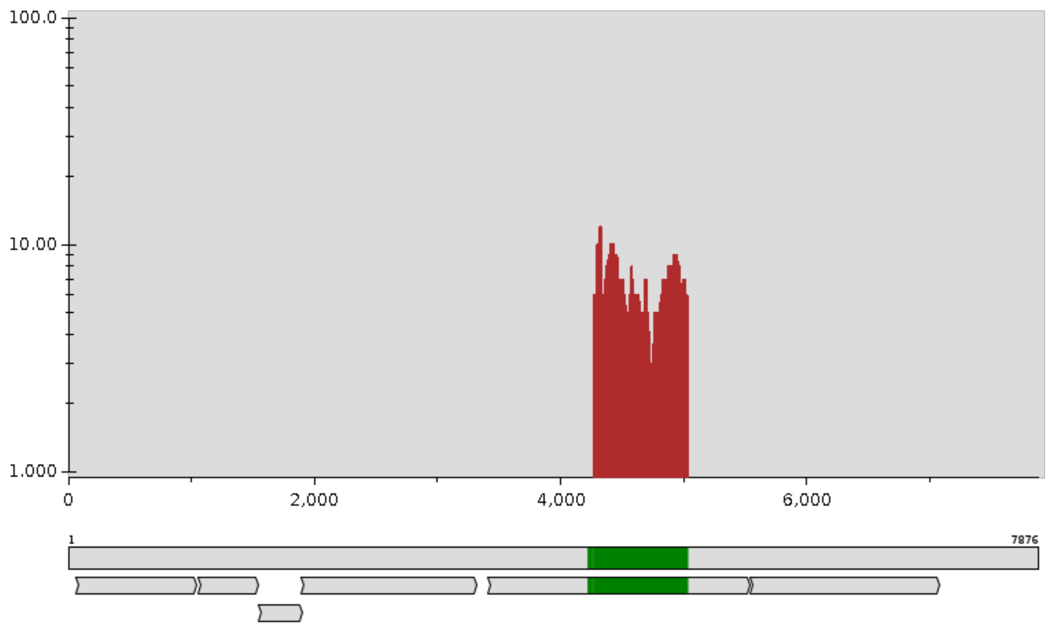

#### Assignment

| | |
|---|---|
| **Type** | Caulimovirus venafragariae (Taxonomy ID: 3048344) |
| **Reference Genome** | NC_001725.1 |
| **NT Identity (%)** | 56.2025 |
| **AA Identity (%)** | 54.4747 |
| **Number Of Stop Codons** | 1 |
| **Number Of CDS** | 6 |

#### Alignment

| | |
|---|---|
| **Alignment Score** | 170.0 (NT) + 883.0 (AA) = 1053.0 |
| **Concordance (%)** | 32.9062 |

| Alignment Method | Global, seeded, nucleotide + amino acids (AGA) |
|---|---|

#### Genome Region

Sequence starts at position 4219 and ends at position 5042 relative to NC_001725.1 reference sequence.

#### Alignment Detailed Statistics

| | Begin | End | Coverage | Score | Concordance | Matches | Identities | I/D/M/F* | Stop Codons |
|---|---|---|---|---|---|---|---|---|---|
| **NT** | **4219** | **5042** | **10.1%** | **170** | **10.9%** | **790 (99.0%)** | **444 (55.6%)** | **0/8** | |
| Mutations: | 4219T>C, 4222T>A, 4223T>A, 4227C>T, 4228A>C, 4230T>C, 4232A>G, 4235A>T, 4236A>C, 4239T>A, 4242T>A, 4244A>G, 4245G>A, 4246T>G, 4247G>T, 4254A>T, 4259A>T, 4260G>A, 4263T>C, 4264T>A, 4265C>A, 4266A>T, 4267C>G, 4268C>A, 4269T>A, 4275C>T, 4276C>A, 4277G>A, 4279G>A, 4280A>C, 4281A>T, 4282G>A, 4284G>A, 4287T>C, 4289A>G, 4292C>T, 4299C>T, 4300G>C, 4308A>T, 4314G>A, 4315C>T, 4317T>A, 4318G>A, 4319G>A, 4320T>A, 4321A>T, 4322T>A, 4323C>A, 4326C>A, 4330C>G, 4331C>A, 4332T>A, 4335C>T, 4348T>A, 4349C>G, 4351T>A, 4352C>G, 4353A>T, 4356A>T, 4359C>A, 4362C>T, 4366G>A, 4371G>A, 4380C>G, 4383G>A, 4384A>C, 4385T>A, 4386C>A, 4390C>A, 4395C>A, 4398A>G, 4401A>T, 4402A>C, 4410A>T, 4413T>A, 4423A>G, 4426T>C, 4428A>T, 4432G>A, 4434C>A, 4435C>A, 4437T>C, 4438A>T, 4439C>G, 4440A>T, 4442A>T, 4443G>T, 4444G>T, 4445G>T, 4446A>T, 4449T>C, 4452C>A, 4456C>T, 4458A>T, 4459C>A, 4461T>C, 4467C>T, 4470G>A, 4471G>A, 4472A>C, 4473G>T, 4474C>A, 4476A>T, 4479G>T, 4480C>A, 4482T>A, 4483C>A, 4485A>T, 4486A>C, 4487G>T, 4489A>G, 4492G>A, 4493G>A, 4496G>T, 4497T>A, 4500G>A, 4505T>A, 4506T>C, 4508A>T, 4509C>T, 4512T>C, 4513T>A, 4514C>A, 4515C>A, 4521C>T, 4528T>A, 4529C>G, 4543G>A, 4546C>A, 4548C>A, 4551T>C, 4552G>T, 4554T>A, 4555C>G, 4556C>A, 4557A>G, 4561A>T, 4568A>C, 4569G>T, 4575C>A, 4578T>A, 4585T>A, 4586G>C, 4590C>T, 4591C>A, 4593A>T, 4599C>T, 4602C>T, 4614C>A, 4618C>T, 4620T>A, 4633C>A, 4635A>T, 4638C>A, 4641T>A, 4642G>C, 4643C>A, 4644T>A, 4650C>T, 4656G>A, 4657C>A, 4659C>A, 4663G>T, 4665C>T, 4666G>A, 4670G>T, 4671T>A, 4672C>T, 4674C>T, 4675_4679delAGCAA, 4680C>T, 4681A>C, 4684T>A, 4686T>A, 4687C>A, 4688C>A, 4689A>G, 4690C>A, 4692G>A, 4697G>T, 4698T>G, 4699G>A, 4700C>T, 4701T>A, 4707T>C, 4708G>A, 4710T>A, 4719C>T, 4720A>T, 4722C>G, 4725C>A, 4727T>G, 4729A>T, 4730G>C, 4731C>A, 4733A>C, 4737T>A, 4738G>T, 4740A>T, 4741G>C, 4743A>G, 4750T>C, 4752A>T, 4753G>A, 4754G>C, 4759G>C, 4761C>T, 4765A>G, 4766T>A, 4767T>A, 4768G>T, 4771C>T, 4772T>C, 4776C>A, 4777A>C, 4778G>T, 4779A>C, 4782C>T, 4783A>G, 4784A>T, 4785A>C, 4786G>A, 4787C>A, 4788C>T, 4789C>A, 4790T>A, 4791A>T, 4793G>A, 4794A>C, 4800G>A, 4801C>T, 4803T>A, 4806C>T, 4807A>C, 4809A>G, 4812G>A, 4818C>A, 4819C>G, 4821G>A, 4822T>A, 4824G>A, 4825T>A, 4826G>A, 4827C>T, 4832C>A, 4833A>T, 4835C>A, 4836T>A, 4839C>A, 4840A>G, 4842T>A, 4851T>A, 4852C>A, 4854A>G, 4855G>A, 4857A>T, 4860A>T, 4861G>T, 4862A>C, 4865G>A, 4867G>A, 4868G>A, 4869A>T, 4870A>G, 4871A>G, 4872T>A, 4873C>A, 4875C>A, 4876A>G, 4879G>T, 4881C>A, 4884G>A, 4886G>C, 4890T>C, 4893C>A, 4894G>T, 4895G>C, 4896A>C, 4897C>A, 4898T>A, 4900C>A, 4902C>G, 4903T>A, 4906G>T, 4908T>A, 4910C>A, 4911T>G, 4917A>T, 4920T>C, 4921C>A, 4923A>G, 4924C>T, 4926T>A, 4928G>T, 4929T>A, 4930G>A, 4931A>C, 4932T>C, 4934G>A, 4935G>A, 4936A>C, 4938T>A, 4940C>A, 4941C>A, 4942T>A, 4954T>C, 4956A>T, 4959C>A, 4960C>T, 4961T>G, 4963C>T, 4968T>C, 4972A>G, 4973T>C, 4974C>A, 4975T>G, 4976C>G, 4977C>A, 4979C>A, 4980T>A, 4982A>T, 4983T>C, 4984T>A, 4986T>A, 4990A>G, 4993A>T, 5001C>G, 5002C>A, 5003T>A, 5004T>G, 5007G>A, 5008T>A, 5009C>A, 5010A>T, 5012C>T, 5014T>C, 5016A>T, 5019G>A, 5020_5022delGTT, 5028T>C, 5030A>T, 5031G>C, 5033A>G, 5035G>A, 5037G>A, 5038A>T, 5039T>C, 5042C>A | | | | | | | | |

##### CDS

| | Begin | End | Coverage | Score | Concordance | Matches | Identities | I/D/M/F* | Stop Codons |
|---|---|---|---|---|---|---|---|---|---|
| ORF_V | 287 | 546 | 36.5% | 883 | 47.1% | 257 (99.2%) | 140 (54.1%) | 0/2/1/1 | 1 |
| Protein mutations: | S287N (4264T>A 4265C>A 4266A>T), P288E (4267C>G 4268C>A 4269T>A), R291K (4276C>A 4277G>A), E292T (4279G>A 4280A>C 4281A>T), E293K (4282G>A 4284G>A), K295R (4289A>G), T296I (4292C>T), E299Q (4300G>C), G305K (4318G>A 4319G>A 4320T>A), I306* (4321A>T 4322T>A 4323C>A), P309E (4330C>G 4331C>A 4332T>A), V321I (4366G>A), I327Q (4384A>C 4385T>A 4386C>A), K340E (4423A>G), D343K (4432G>A 4434C>A), H344N (4435C>A 4437T>C), T345C (4438A>T 4439C>G 4440A>T), K346I (4442A>T 4443G>T), G347F (4444G>T 4445G>T 4446A>T), L351F (4456C>T 4458A>T), L352I (4459C>A 4461T>C), E356T (4471G>A 4472A>C 4473G>T), Q357N (4474C>A 4476A>T), L359I (4480C>A 4482T>A), Q360N (4483C>A 4485A>T), R361L (4486A>C 4487G>T), I362V (4489A>G), G363K (4492G>A 4493G>A), G364V (4496G>T 4497T>A), F367Y (4505T>A 4506T>C), Y368F (4508A>T 4509C>T), S370K (4513T>A 4514C>A 4515C>A), V380I (4543G>A), A383S (4552G>T 4554T>A), P384E (4555C>G 4556C>A 4557A>G), T386S (4561A>T), Q388P (4568A>C 4569G>T), C394T (4585T>A 4586G>C), Q396N (4591C>A 4593A>T), P405S (4618C>T 4620T>A), Q410N (4633C>A 4635A>T), A413Q (4642G>C 4643C>A 4644T>A), H418K (4657C>A 4659C>A), D420Y (4663G>T 4665C>T), E421K (4666G>A), S422I (4670G>T 4671T>A), L423F (4672C>T 4674C>T), S424del (4675_4679delAGCAA), M426L (4681A>C), Y427K (4684T>A 4686T>A), P428K (4687C>A 4688C>A 4689A>G), Q429K (4690C>A 4692G>A), C431L (4697G>T 4698T>G), A432I (4699G>A 4700C>T 4701T>A), V435I (4708G>A 4710T>A), I439L (4720A>T 4722C>G), F441C (4727T>G), K443T (4733A>C), E445Y (4738G>T 4740A>T), E446Q (4741G>C 4743A>G), G450T (4753G>A 4754G>C), V452L (4759G>C 4761C>T), I454E (4765A>G 4766T>A 4767T>A), V455F (4768G>T), L456S (4771C>T 4772T>C), N457K (4776C>A), R458L (4777A>C 4778G>T 4779A>C), K460V (4783A>G 4784A>T 4785A>C), A461N (4786G>A 4787C>A 4788C>T), L462N (4789C>A 4790T>A 4791A>T), G463D (4793G>A 4794A>C), K468Q (4807A>C 4809A>G), Q472E (4819C>G 4821G>A), L473I (4822T>A 4824G>A), C474N (4825T>A 4826G>A 4827C>T), T476N (4832C>A 4833A>T), T477K (4835C>A 4836T>A), N479E (4840A>G 4842T>A), L483M (4852C>A 4854A>G), V484I (4855G>A 4857A>T), E486S (4861G>T 4862A>C), R487K (4865G>A), G488N (4867G>A 4868G>A 4869A>T), N489G (4870A>G 4871A>G 4872T>A), L490I (4873C>A 4875C>A), K491E (4876A>G), V492L (4879G>T 4881C>A), S494T (4886G>C), G497S (4894G>T 4895G>C 4896A>C), L498K (4897C>A 4898T>A), H499K (4900C>A 4902C>G), L500I (4903T>A), V501L (4906G>T 4908T>A), A502E (4910C>A 4911T>G), Q506K (4921C>A 4923A>G), S508I (4928G>T 4929T>A), D509T (4930G>A 4931A>C 4932T>C), R510K (4934G>A 4935G>A), N511Q (4936A>C 4938T>A), A512E (4940C>A 4941C>A), L513I (4942T>A), L519C (4960C>T 4961T>G), I523A (4972A>G 4973T>C 4974C>A), S524G (4975T>G 4976C>G 4977C>A), A525E (4979C>A 4980T>A), Y526F (4982A>T 4983T>C), F527I (4984T>A 4986T>A), K529E (4990A>G), I530L (4993A>T), N532K (5001C>G), L533K (5002C>A 5003T>A 5004T>G), S535N (5008T>A 5009C>A 5010A>T), P536L (5012C>T), V539del (5020_5022delGTT), K542I (5030A>T 5031G>C), K543R (5033A>G), E544K (5035G>A 5037G>A), I545S (5038A>T 5039T>C) | | | | | | | | |
| Codon mutations: | TTA272C.. (4219T>C), TTA273AA. (4222T>A 4223T>A), GAC274..T (4227C>T), ACT275C.C (4228A>C 4230T>C), AAC276.G. (4232A>G), AAA277.TC (4235A>T 4236A>C), ATT278..A (4239T>A), ATT279..A (4242T>A), AAG280.GA (4244A>G 4245G>A), TGT281GT. (4246T>G 4247G>T), CCA283..T (4254A>T), CAG285.TA (4259A>T 4260G>A), TAT286..C (4263T>C), TCA287AAT (4264T>A 4265C>A 4266A>T), CCT288GAA (4267C>G 4268C>A 4269T>A), GAC290GAT (4275C>T), CGA291AAA (4276C>A 4277G>A), GAA292ACT (4279G>A 4280A>C 4281A>T), GAG293AAA (4282G>A 4284G>A), TTT294TTC (4287T>C), AAA295AGA (4289A>G), ACT296ATT (4292C>T), ATC298ATT (4299C>T), GAA299CAA (4300G>C), CTA301CTT (4308A>T), AAG303AAA (4314G>A), CTT304TTA (4315C>T 4317T>A), GGT305AAA (4318G>A 4319G>A 4320T>A), ATC306TAA (4321A>T 4322T>A 4323C>A), ATC307ATA (4326C>A), CCT309GAA (4330C>G 4331C>A 4332T>A), AGC310AGT (4335C>T), TCC315AGC (4348T>A 4349C>G), TCA316AGT (4351T>A 4352C>G 4353A>T), CCA317CCT (4356A>T), GCC318GCA (4359C>A), TTC319TTT (4362C>T), GTA321ATA (4366G>A), AGG322AGA (4371G>A), GCC325GCG (4380C>G), GAG326GAA (4383G>A), ATC327CAA (4384A>C 4385T>A 4386C>A), CGA329AGA (4390C>A), GGC330GGA (4395C>A), AAA331AAG (4398A>G), GCA332GCT (4401A>T), AGA333CGA (4402A>C), GTA335GTT (4410A>T), ATT336ATA (4413T>A), AAG340GAG (4423A>G), TTA341CTT (4426T>C 4428A>T), GAC343AAA (4432G>A 4434C>A), CAT344AAC (4435C>A 4437T>C), ACA345TGT (4438A>T 4439C>G 4440A>T), AAG346ATT (4442A>T 4443G>T), GGA347TTT (4444G>T 4445G>T 4446A>T), GAT348GAC (4449T>C), GGC349GGA (4452C>A), CTA351TTT (4456C>T 4458A>T), CTT352ATC (4459C>A 4461T>C), AAC354AAT (4467C>T), AAG355AAA (4470G>A), GAG356ACT (4471G>A 4472A>C 4473G>T), CAA357AAT (4474C>A 4476A>T), CTG358CTT (4479G>T), CTT359ATA (4480C>A 4482T>A), CAA360AAT (4483C>A 4485A>T), AGA361CTA (4486A>C 4487G>T), ATC362GTC (4489A>G), GGA363AAA (4492G>A 4493G>A), GGT364GTA (4496G>T 4497T>A), AAG365AAA (4500G>A), TTT367TAC (4505T>A 4506T>C), TAC368TTT (4508A>T 4509C>T), TCT369TCC (4512T>C), TCC370AAA (4513T>A 4514C>A 4515C>A), GAC372GAT (4521C>T), TCT375AGT (4528T>A 4529C>G), GTA380ATA (4543G>A), CGC381AGA (4546C>A 4548C>A), CTT382CTC (4551T>C), GCT383TCA (4552G>T 4554T>A), CCA384GAG (4555C>G 4556C>A 4557A>G), ACA386TCA (4561A>T), CAG388CCT (4568A>C 4569G>T), ACC390ACA (4575C>A), GCT391GCA (4578T>A), TGT394ACT (4585T>A 4586G>C), CCC395CCT (4590C>T), CAA396AAT (4591C>A 4593A>T), CAC398CAT (4599C>T), TAC399TAT (4602C>T), GTC403GTA (4614C>A), CCT405TCA (4618C>T 4620T>A), CAA410AAT (4633C>A 4635A>T), GCC411GCA (4638C>A), CCT412CCA (4641T>A), GCT413CAA (4642G>C 4643C>A 4644T>A), TTC415TTT (4650C>T), AGG417AGA (4656G>A), CAC418AAA (4657C>A 4659C>A), GAC420TAT (4663G>T 4665C>T), GAA421AAA (4666G>A), AGT422ATA (4670G>T 4671T>A), CTC423TTT (4672C>T 4674C>T), AGC424del (4675_4679delAGCAA), AAC425--T (4675_4679delAGCAA 4680C>T), ATG426CTG (4681A>C), TAT427AAA (4684T>A 4686T>A), CCA428AAG (4687C>A 4688C>A 4689A>G), CAG429AAA (4690C>A 4692G>A), TGT431TTG (4697G>T 4698T>G), GCT432ATA (4699G>A 4700C>T 4701T>A), TAT434TAC (4707T>C), GTT435ATA (4708G>A 4710T>A), ATC438ATT (4719C>T), ATC439TTG (4720A>T 4722C>G), GTC440GTA (4725C>A), TTC441TGC (4727T>G), AGC442TCA (4729A>T 4730G>C 4731C>A), AAA443ACA (4733A>C), ACT444ACA (4737T>A), GAA445TAT (4738G>T 4740A>T), GAA446CAG (4741G>C 4743A>G), TTA449CTT (4750T>C 4752A>T), GGA450ACA (4753G>A 4754G>C), GTC452CTT (4759G>C 4761C>T), ATT454GAA (4765A>G 4766T>A 4767T>A), GTT455TTT (4768G>T), CTA456TCA (4771C>T 4772T>C), AAC457AAA (4776C>A), AGA458CTC (4777A>C 4778G>T 4779A>C), TGC459TGT (4782C>T), AAA460GTC (4783A>G 4784A>T 4785A>C), GCC461AAT (4786G>A 4787C>A 4788C>T), CTA462AAT (4789C>A 4790T>A 4791A>T), GGA463GAC (4793G>A 4794A>C), GTG465GTA (4800G>A), CTT466TTA (4801C>T 4803T>A), AGC467AGT (4806C>T), AAA468CAG (4807A>C 4809A>G), AAG469AAA (4812G>A), GCC471GCA (4818C>A), CAG472GAA (4819C>G 4821G>A), TTG473ATA (4822T>A 4824G>A), TGC474AAT (4825T>A 4826G>A 4827C>T), ACA476AAT (4832C>A 4833A>T), ACT477AAA (4835C>A 4836T>A), ATC478ATA (4839C>A), AAT479GAA (4840A>G 4842T>A), GGT482GGA (4851T>A), CTA483ATG (4852C>A 4854A>G), GTA484ATT (4855G>A 4857A>T), ATA485ATT (4860A>T), GAA486TCA (4861G>T 4862A>C), AGA487AAA (4865G>A), GGA488AAT (4867G>A 4868G>A 4869A>T), AAT489GGA (4870A>G 4871A>G 4872T>A), CTC490ATA (4873C>A 4875C>A), AAA491GAA (4876A>G), GTC492TTA (4879G>T 4881C>A), CAG493CAA (4884G>A), AGT494ACT (4886G>C), CAT495CAC (4890T>C), ATC496ATA (4893C>A), GGA497TCC (4894G>T 4895G>C 4896A>C), CTA498AAA (4897C>A 4898T>A), CAC499AAG (4900C>A 4902C>G), TTA500ATA (4903T>A), GTT501TTA (4906G>T 4908T>A), GCT502GAG (4910C>A 4911T>G), CCA504CCT (4917A>T), GAT505GAC (4920T>C), CAA506AAG (4921C>A 4923A>G), CTT507TTA (4924C>T 4926T>A), AGT508ATA (4928G>T 4929T>A), GAT509ACC (4930G>A 4931A>C 4932T>C), AGG510AAA (4934G>A 4935G>A), AAT511CAA (4936A>C 4938T>A), GCC512GAA (4940C>A 4941C>A), TTA513ATA (4942T>A), TTA517CTT (4954T>C 4956A>T), GGC518GGA (4959C>A), CTC519TGC (4960C>T 4961T>G), CTA520TTA (4963C>T), AAT521AAC (4968T>C), ATC523GCA (4972A>G 4973T>C 4974C>A), TCC524GGA (4975T>G 4976C>G 4977C>A), GCT525GAA (4979C>A 4980T>A), TAT526TTC (4982A>T 4983T>C), TTT527ATA (4984T>A 4986T>A), AAA529GAA (4990A>G), ATA530TTA (4993A>T), AAC532AAG (5001C>G), CTT533AAG (5002C>A 5003T>A 5004T>G), AGG534AGA (5007G>A), TCA535AAT (5008T>A 5009C>A 5010A>T), CCT536CTT (5012C>T), TTA537CTT (5014T>C 5016A>T), CAG538CAA (5019G>A), GTT539del (5020_5022delGTT), CTT541CTC (5028T>C), AAG542ATC (5030A>T 5031G>C), AAA543AGA (5033A>G), GAG544AAA (5035G>A 5037G>A), ATA545TCA (5038A>T 5039T>C), ACT546AA. (5042C>A) | | | | | | | | |

##### Proteins

| | Begin | End | Coverage | Score | Concordance | Matches | Identities | I/D/M/F* | Stop Codons |
|---|---|---|---|---|---|---|---|---|---|
| hypothetical protein (NP_043933.1) | 287 | 546 | 36.5% | 883 | 47.1% | 257 (99.2%) | 140 (54.1%) | 0/2/1/1 | 1 |

| | Begin | End | Coverage | Score | Concordance | Matches | Identities | I/D/M/F* | Stop Codons |
|---|---|---|---|---|---|---|---|---|---|
| **NT** | **4219** | **5042** | **10.1%** | **170** | **10.9%** | **790 (99.0%)** | **444 (55.6%)** | **0/8** | |
| Protein mutations: | S287N (4264T>A 4265C>A 4266A>T), P288E (4267C>G 4268C>A 4269T>A), R291K (4276C>A 4277G>A), E292T (4279G>A 4280A>C 4281A>T), E293K (4282G>A 4284G>A), K295R (4289A>G), T296I (4292C>T), E299Q (4300G>C), G305K (4318G>A 4319G>A 4320T>A), I306* (4321A>T 4322T>A 4323C>A), P309E (4330C>G 4331C>A 4332T>A), V321I (4366G>A), I327Q (4384A>C 4385T>A 4386C>A), K340E (4423A>G), D343K (4432G>A 4434C>A), H344N (4435C>A 4437T>C), T345C (4438A>T 4439C>G 4440A>T), K346I (4442A>T 4443G>T), G347F (4444G>T 4445G>T 4446A>T), L351F (4456C>T 4458A>T), L352I (4459C>A 4461T>C), E356T (4471G>A 4472A>C 4473G>T), Q357N (4474C>A 4476A>T), L359I (4480C>A 4482T>A), Q360N (4483C>A 4485A>T), R361L (4486A>C 4487G>T), I362V (4489A>G), G363K (4492G>A 4493G>A), G364V (4496G>T 4497T>A), F367Y (4505T>A 4506T>C), Y368F (4508A>T 4509C>T), S370K (4513T>A 4514C>A 4515C>A), V380I (4543G>A), A383S (4552G>T 4554T>A), P384E (4555C>G 4556C>A 4557A>G), T386S (4561A>T), Q388P (4568A>C 4569G>T), C394T (4585T>A 4586G>C), Q396N (4591C>A 4593A>T), P405S (4618C>T 4620T>A), Q410N (4633C>A 4635A>T), A413Q (4642G>C 4643C>A 4644T>A), H418K (4657C>A 4659C>A), D420Y (4663G>T 4665C>T), E421K (4666G>A), S422I (4670G>T 4671T>A), L423F (4672C>T 4674C>T), S424del (4675_4679delAGCAA), M426L (4681A>C), Y427K (4684T>A 4686T>A), P428K (4687C>A 4688C>A 4689A>G), Q429K (4690C>A 4692G>A), C431L (4697G>T 4698T>G), A432I (4699G>A 4700C>T 4701T>A), V435I (4708G>A 4710T>A), I439L (4720A>T 4722C>G), F441C (4727T>G), K443T (4733A>C), E445Y (4738G>T 4740A>T), E446Q (4741G>C 4743A>G), G450T (4753G>A 4754G>C), V452L (4759G>C 4761C>T), I454E (4765A>G 4766T>A 4767T>A), V455F (4768G>T), L456S (4771C>T 4772T>C), N457K (4776C>A), R458L (4777A>C 4778G>T 4779A>C), K460V (4783A>G 4784A>T 4785A>C), A461N (4786G>A 4787C>A 4788C>T), L462N (4789C>A 4790T>A 4791A>T), G463D (4793G>A 4794A>C), K468Q (4807A>C 4809A>G), Q472E (4819C>G 4821G>A), L473I (4822T>A 4824G>A), C474N (4825T>A 4826G>A 4827C>T), T476N (4832C>A 4833A>T), T477K (4835C>A 4836T>A), N479E (4840A>G 4842T>A), L483M (4852C>A 4854A>G), V484I (4855G>A 4857A>T), E486S (4861G>T 4862A>C), R487K (4865G>A), G488N (4867G>A 4868G>A 4869A>T), N489G (4870A>G 4871A>G 4872T>A), L490I (4873C>A 4875C>A), K491E (4876A>G), V492L (4879G>T 4881C>A), S494T (4886G>C), G497S (4894G>T 4895G>C 4896A>C), L498K (4897C>A 4898T>A), H499K (4900C>A 4902C>G), L500I (4903T>A), V501L (4906G>T 4908T>A), A502E (4910C>A 4911T>G), Q506K (4921C>A 4923A>G), S508I (4928G>T 4929T>A), D509T (4930G>A 4931A>C 4932T>C), R510K (4934G>A 4935G>A), N511Q (4936A>C 4938T>A), A512E (4940C>A 4941C>A), L513I (4942T>A), L519C (4960C>T 4961T>G), I523A (4972A>G 4973T>C 4974C>A), S524G (4975T>G 4976C>G 4977C>A), A525E (4979C>A 4980T>A), Y526F (4982A>T 4983T>C), F527I (4984T>A 4986T>A), K529E (4990A>G), I530L (4993A>T), N532K (5001C>G), L533K (5002C>A 5003T>A 5004T>G), S535N (5008T>A 5009C>A 5010A>T), P536L (5012C>T), V539del (5020_5022delGTT), K542I (5030A>T 5031G>C), K543R (5033A>G), E544K (5035G>A 5037G>A), I545S (5038A>T 5039T>C) | | | | | | | | |
| Codon mutations: | TTA272C.. (4219T>C), TTA273AA. (4222T>A 4223T>A), GAC274..T (4227C>T), ACT275C.C (4228A>C 4230T>C), AAC276.G. (4232A>G), AAA277.TC (4235A>T 4236A>C), ATT278..A (4239T>A), ATT279..A (4242T>A), AAG280.GA (4244A>G 4245G>A), TGT281GT. (4246T>G 4247G>T), CCA283..T (4254A>T), CAG285.TA (4259A>T 4260G>A), TAT286..C (4263T>C), TCA287AAT (4264T>A 4265C>A 4266A>T), CCT288GAA (4267C>G 4268C>A 4269T>A), GAC290GAT (4275C>T), CGA291AAA (4276C>A 4277G>A), GAA292ACT (4279G>A 4280A>C 4281A>T), GAG293AAA (4282G>A 4284G>A), TTT294TTC (4287T>C), AAA295AGA (4289A>G), ACT296ATT (4292C>T), ATC298ATT (4299C>T), GAA299CAA (4300G>C), CTA301CTT (4308A>T), AAG303AAA (4314G>A), CTT304TTA (4315C>T 4317T>A), GGT305AAA (4318G>A 4319G>A 4320T>A), ATC306TAA (4321A>T 4322T>A 4323C>A), ATC307ATA (4326C>A), CCT309GAA (4330C>G 4331C>A 4332T>A), AGC310AGT (4335C>T), TCC315AGC (4348T>A 4349C>G), TCA316AGT (4351T>A 4352C>G 4353A>T), CCA317CCT (4356A>T), GCC318GCA (4359C>A), TTC319TTT (4362C>T), GTA321ATA (4366G>A), AGG322AGA (4371G>A), GCC325GCG (4380C>G), GAG326GAA (4383G>A), ATC327CAA (4384A>C 4385T>A 4386C>A), CGA329AGA (4390C>A), GGC330GGA (4395C>A), AAA331AAG (4398A>G), GCA332GCT (4401A>T), AGA333CGA (4402A>C), GTA335GTT (4410A>T), ATT336ATA (4413T>A), AAG340GAG (4423A>G), TTA341CTT (4426T>C 4428A>T), GAC343AAA (4432G>A 4434C>A), CAT344AAC (4435C>A 4437T>C), ACA345TGT (4438A>T 4439C>G 4440A>T), AAG346ATT (4442A>T 4443G>T), GGA347TTT (4444G>T 4445G>T 4446A>T), GAT348GAC (4449T>C), GGC349GGA (4452C>A), CTA351TTT (4456C>T 4458A>T), CTT352ATC (4459C>A 4461T>C), AAC354AAT (4467C>T), AAG355AAA (4470G>A), GAG356ACT (4471G>A 4472A>C 4473G>T), CAA357AAT (4474C>A 4476A>T), CTG358CTT (4479G>T), CTT359ATA (4480C>A 4482T>A), CAA360AAT (4483C>A 4485A>T), AGA361CTA (4486A>C 4487G>T), ATC362GTC (4489A>G), GGA363AAA (4492G>A 4493G>A), GGT364GTA (4496G>T 4497T>A), AAG365AAA (4500G>A), TTT367TAC (4505T>A 4506T>C), TAC368TTT (4508A>T 4509C>T), TCT369TCC (4512T>C), TCC370AAA (4513T>A 4514C>A 4515C>A), GAC372GAT (4521C>T), TCT375AGT (4528T>A 4529C>G), GTA380ATA (4543G>A), CGC381AGA (4546C>A 4548C>A), CTT382CTC (4551T>C), GCT383TCA (4552G>T 4554T>A), CCA384GAG (4555C>G 4556C>A 4557A>G), ACA386TCA (4561A>T), CAG388CCT (4568A>C 4569G>T), ACC390ACA (4575C>A), GCT391GCA (4578T>A), TGT394ACT (4585T>A 4586G>C), CCC395CCT (4590C>T), CAA396AAT (4591C>A 4593A>T), CAC398CAT (4599C>T), TAC399TAT (4602C>T), GTC403GTA (4614C>A), CCT405TCA (4618C>T 4620T>A), CAA410AAT (4633C>A 4635A>T), GCC411GCA (4638C>A), CCT412CCA (4641T>A), GCT413CAA (4642G>C 4643C>A 4644T>A), TTC415TTT (4650C>T), AGG417AGA (4656G>A), CAC418AAA (4657C>A 4659C>A), GAC420TAT (4663G>T 4665C>T), GAA421AAA (4666G>A), AGT422ATA (4670G>T 4671T>A), CTC423TTT (4672C>T 4674C>T), AGC424del (4675_4679delAGCAA), AAC425--T (4675_4679delAGCAA 4680C>T), ATG426CTG (4681A>C), TAT427AAA (4684T>A 4686T>A), CCA428AAG (4687C>A 4688C>A 4689A>G), CAG429AAA (4690C>A 4692G>A), TGT431TTG (4697G>T 4698T>G), GCT432ATA (4699G>A 4700C>T 4701T>A), TAT434TAC (4707T>C), GTT435ATA (4708G>A 4710T>A), ATC438ATT (4719C>T), ATC439TTG (4720A>T 4722C>G), GTC440GTA (4725C>A), TTC441TGC (4727T>G), AGC442TCA (4729A>T 4730G>C 4731C>A), AAA443ACA (4733A>C), ACT444ACA (4737T>A), GAA445TAT (4738G>T 4740A>T), GAA446CAG (4741G>C 4743A>G), TTA449CTT (4750T>C 4752A>T), GGA450ACA (4753G>A 4754G>C), GTC452CTT (4759G>C 4761C>T), ATT454GAA (4765A>G 4766T>A 4767T>A), GTT455TTT (4768G>T), CTA456TCA (4771C>T 4772T>C), AAC457AAA (4776C>A), AGA458CTC (4777A>C 4778G>T 4779A>C), TGC459TGT (4782C>T), AAA460GTC (4783A>G 4784A>T 4785A>C), GCC461AAT (4786G>A 4787C>A 4788C>T), CTA462AAT (4789C>A 4790T>A 4791A>T), GGA463GAC (4793G>A 4794A>C), GTG465GTA (4800G>A), CTT466TTA (4801C>T 4803T>A), AGC467AGT (4806C>T), AAA468CAG (4807A>C 4809A>G), AAG469AAA (4812G>A), GCC471GCA (4818C>A), CAG472GAA (4819C>G 4821G>A), TTG473ATA (4822T>A 4824G>A), TGC474AAT (4825T>A 4826G>A 4827C>T), ACA476AAT (4832C>A 4833A>T), ACT477AAA (4835C>A 4836T>A), ATC478ATA (4839C>A), AAT479GAA (4840A>G 4842T>A), GGT482GGA (4851T>A), CTA483ATG (4852C>A 4854A>G), GTA484ATT (4855G>A 4857A>T), ATA485ATT (4860A>T), GAA486TCA (4861G>T 4862A>C), AGA487AAA (4865G>A), GGA488AAT (4867G>A 4868G>A 4869A>T), AAT489GGA (4870A>G 4871A>G 4872T>A), CTC490ATA (4873C>A 4875C>A), AAA491GAA (4876A>G), GTC492TTA (4879G>T 4881C>A), CAG493CAA (4884G>A), AGT494ACT (4886G>C), CAT495CAC (4890T>C), ATC496ATA (4893C>A), GGA497TCC (4894G>T 4895G>C 4896A>C), CTA498AAA (4897C>A 4898T>A), CAC499AAG (4900C>A 4902C>G), TTA500ATA (4903T>A), GTT501TTA (4906G>T 4908T>A), GCT502GAG (4910C>A 4911T>G), CCA504CCT (4917A>T), GAT505GAC (4920T>C), CAA506AAG (4921C>A 4923A>G), CTT507TTA (4924C>T 4926T>A), AGT508ATA (4928G>T 4929T>A), GAT509ACC (4930G>A 4931A>C 4932T>C), AGG510AAA (4934G>A 4935G>A), AAT511CAA (4936A>C 4938T>A), GCC512GAA (4940C>A 4941C>A), TTA513ATA (4942T>A), TTA517CTT (4954T>C 4956A>T), GGC518GGA (4959C>A), CTC519TGC (4960C>T 4961T>G), CTA520TTA (4963C>T), AAT521AAC (4968T>C), ATC523GCA (4972A>G 4973T>C 4974C>A), TCC524GGA (4975T>G 4976C>G 4977C>A), GCT525GAA (4979C>A 4980T>A), TAT526TTC (4982A>T 4983T>C), TTT527ATA (4984T>A 4986T>A), AAA529GAA (4990A>G), ATA530TTA (4993A>T), AAC532AAG (5001C>G), CTT533AAG (5002C>A 5003T>A 5004T>G), AGG534AGA (5007G>A), TCA535AAT (5008T>A 5009C>A 5010A>T), CCT536CTT (5012C>T), TTA537CTT (5014T>C 5016A>T), CAG538CAA (5019G>A), GTT539del (5020_5022delGTT), CTT541CTC (5028T>C), AAG542ATC (5030A>T 5031G>C), AAA543AGA (5033A>G), GAG544AAA (5035G>A 5037G>A), ATA545TCA (5038A>T 5039T>C), ACT546AA. (5042C>A) | | | | | | | | |

*: Inserts / Deletes / Misaligned / Frameshifts

#### Analysis details

This analysis was performed with panviral2.64

### NGS Details (UN9): Pinus nigra virus 1

#### Assembly

| | |
|---|---|
| **Coverage Length** | 241 (1 contig(s)) |
| **Depth Of Coverage** | 19.8 |
| **Number Of Reads** | 41 |
| **Reads Per Million** | 0.93 rpm (after QC) |
| **Ambiguities** | 0 |
| **Assembly Method** | de novo + reference guided assembly |
| **Consensus Caller** | Bcf Tools |

#### Coverage Map

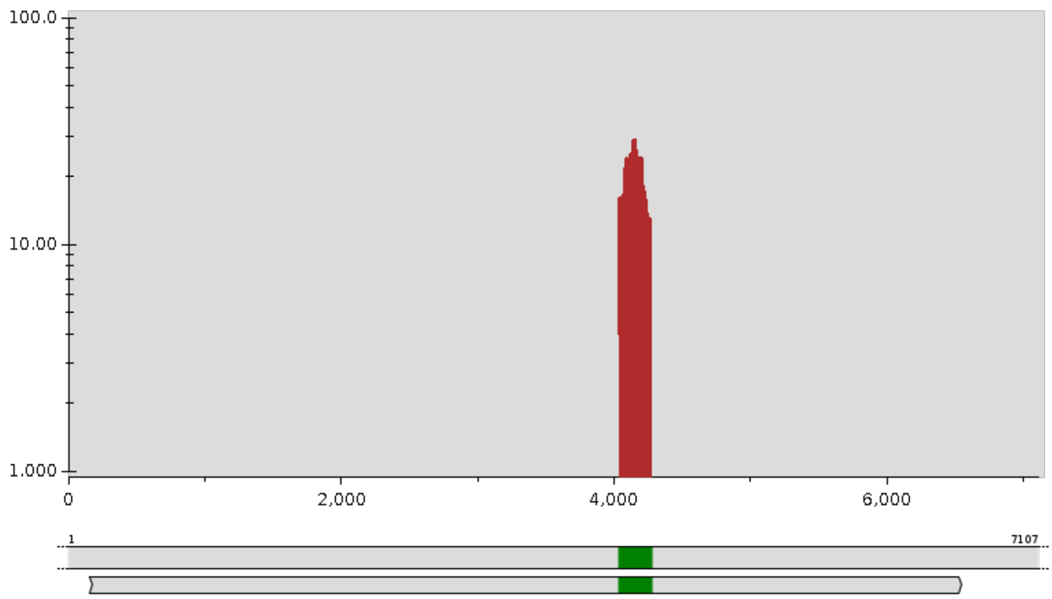

#### Assignment

| | |
|---|---|
| **Type** | Pinus nigra virus 1 (Taxonomy ID: 2267679) |
| **Reference Genome** | NC_040841.1 |
| **NT Identity (%)** | 56.2753 |
| **AA Identity (%)** | 50.0 |
| **Number Of Stop Codons** | 0 |
| **Number Of CDS** | 1 |

#### Alignment

| | |
|---|---|
| **Alignment Score** | 59.0 (NT) + 311.0 (AA) = 370.0 |
| **Concordance (%)** | 33.6364 |

| Alignment Method | Global, seeded, nucleotide + amino acids (AGA) |
|---|---|

#### Genome Region

Sequence starts at position 4036 and ends at position 4276 relative to NC_040841.1 reference sequence.

#### Alignment Detailed Statistics

| | Begin | End | Coverage | Score | Concordance | Matches | Identities | I/D/M/F* | Stop Codons |
|---|---|---|---|---|---|---|---|---|---|
| **NT** | **4036** | **4276** | **3.4%** | **59** | **12.2%** | **241 (97.6%)** | **139 (56.3%)** | **6/0** | |
| Mutations: | 4040G>A, 4045T>C, 4046T>C, 4051G>A, 4054A>T, 4056A>G, 4057A>T, 4058A>G, 4060T>C, 4068A>C, 4073A>G, 4074G>C, 4075A>C, 4078C>T, 4090C>A, 4091A>C, 4092T>G, 4094T>A, 4095G>C, 4098C>A, 4099A>T, 4100G>T, 4102A>T, 4105A>G, 4106T>C, 4107T>A, 4108C>T, 4112C>G, 4114G>A, 4116G>T, 4117G>C, 4118A>C, 4119A>T, 4120A>T, 4123T>A, 4124G>A, 4126T>G, 4129A>T, 4132T>C, 4135T>A, 4136T>C, 4137T>A, 4138G>C, 4140A>C, 4141G>T, 4147G>T, 4150A>C, 4154T>A, 4155T>C, 4156C>T, 4159T>C, 4163A>G, 4164G>C, 4165A>C, 4167G>C, 4168G>T, 4172G>A, 4175T>G, 4177T>C, 4178A>C, 4180C>T, 4183T>C, 4184G>A, 4185C>G, 4186T>A, 4187A>C, 4188A>C, 4192_4193insCTCAGA, 4193G>A, 4195T>A, 4198T>C, 4201A>C, 4202G>T, 4204T>G, 4205A>G, 4206C>T, 4207A>T, 4209A>T, 4211A>T, 4213A>T, 4214G>A, 4225A>G, 4228T>C, 4229C>T, 4237A>T, 4238G>A, 4239A>C, 4240T>C, 4241G>C, 4243A>T, 4247A>C, 4249T>G, 4252T>C, 4253T>C, 4254T>A, 4255G>A, 4258G>C, 4261C>T, 4262T>C, 4264A>C, 4265G>C, 4267A>G, 4270T>A | | | | | | | | |

##### CDS

| | Begin | End | Coverage | Score | Concordance | Matches | Identities | I/D/M/F* | Stop Codons |
|---|---|---|---|---|---|---|---|---|---|
| EXL67_gp1 | 1295 | 1374 | 3.8% | 311 | 50.0% | 80 (97.6%) | 41 (50.0%) | 2/0/0/0 | 0 |
| Protein mutations: | G1296R (4040G>A), Y1298H (4046T>C), Q1301R (4056A>G 4057A>T), I1302V (4058A>G 4060T>C), E1305A (4068A>C), R1307A (4073A>G 4074G>C 4075A>C), F1312L (4090C>A), I1313R (4091A>C 4092T>G), C1314T (4094T>A 4095G>C), P1315H (4098C>A 4099A>T), A1316S (4100G>T 4102A>T), F1318H (4106T>C 4107T>A 4108C>T), Q1320E (4112C>G 4114G>A), W1321F (4116G>T 4117G>C), K1322L (4118A>C 4119A>T 4120A>T), V1324M (4124G>A 4126T>G), L1328H (4136T>C 4137T>A 4138G>C), K1329T (4140A>C 4141G>T), F1334T (4154T>A 4155T>C 4156C>T), R1337A (4163A>G 4164G>C 4165A>C), R1338T (4167G>C 4168G>T), D1340N (4172G>A), Y1341D (4175T>G 4177T>C), I1342L (4178A>C 4180C>T), A1344R (4184G>A 4185C>G 4186T>A), K1345P (4187A>C 4188A>C), Y1346_D1347insLR (4192_4193insCTCAGA), D1347K (4193G>A 4195T>A), V1350L (4202G>T 4204T>G), T1351V (4205A>G 4206C>T 4207A>T), Y1352F (4209A>T), I1353F (4211A>T 4213A>T), D1354N (4214G>A), H1359Y (4229C>T), D1362T (4238G>A 4239A>C 4240T>C), V1363L (4241G>C 4243A>T), N1365Q (4247A>C 4249T>G), L1367Q (4253T>C 4254T>A 4255G>A), K1368N (4258G>C), E1371Q (4265G>C 4267A>G) | | | | | | | | |
| Codon mutations: | GGG1296AGG (4040G>A), TAT1297TAC (4045T>C), TAT1298CAT (4046T>C), CAG1299CAA (4051G>A), ATA1300ATT (4054A>T), CAA1301CGT (4056A>G 4057A>T), ATT1302GTC (4058A>G 4060T>C), GAA1305GCA (4068A>C), AGA1307GCC (4073A>G 4074G>C 4075A>C), CAC1308CAT (4078C>T), TTC1312TTA (4090C>A), ATA1313CGA (4091A>C 4092T>G), TGT1314ACT (4094T>A 4095G>C), CCA1315CAT (4098C>A 4099A>T), GCA1316TCT (4100G>T 4102A>T), GGA1317GGG (4105A>G), TTC1318CAT (4106T>C 4107T>A 4108C>T), CAG1320GAA (4112C>G 4114G>A), TGG1321TTC (4116G>T 4117G>C), AAA1322CTT (4118A>C 4119A>T 4120A>T), GTT1323GTA (4123T>A), GTT1324ATG (4124G>A 4126T>G), CCA1325CCT (4129A>T), TTT1326TTC (4132T>C), GGT1327GGA (4135T>A), TTG1328CAC (4136T>C 4137T>A 4138G>C), AAG1329ACT (4140A>C 4141G>T), GCG1331GCT (4147G>T), CCA1332CCC (4150A>C), TTC1334ACT (4154T>A 4155T>C 4156C>T), TTT1335TTC (4159T>C), AGA1337GCC (4163A>G 4164G>C 4165A>C), AGG1338ACT (4167G>C 4168G>T), GAT1340AAT (4172G>A), TAT1341GAC (4175T>G 4177T>C), ATC1342CTT (4178A>C 4180C>T), TTT1343TTC (4183T>C), GCT1344AGA (4184G>A 4185C>G 4186T>A), AAA1345CCA (4187A>C 4188A>C), TAT1346_GAT1347insCTCAGA (4192_4193insCTCAGA), GAT1347AAA (4193G>A 4195T>A), TTT1348TTC (4198T>C), ATA1349ATC (4201A>C), GTT1350TTG (4202G>T 4204T>G), ACA1351GTT (4205A>G 4206C>T 4207A>T), TAC1352TTC (4209A>T), ATA1353TTT (4211A>T 4213A>T), GAT1354AAT (4214G>A), TTA1357TTG (4225A>G), ATT1358ATC (4228T>C), CAT1359TAT (4229C>T), CCA1361CCT (4237A>T), GAT1362ACC (4238G>A 4239A>C 4240T>C), GTA1363CTT (4241G>C 4243A>T), AAT1365CAG (4247A>C 4249T>G), CAT1366CAC (4252T>C), TTG1367CAA (4253T>C 4254T>A 4255G>A), AAG1368AAC (4258G>C), CAC1369CAT (4261C>T), TTA1370CTC (4262T>C 4264A>C), GAA1371CAG (4265G>C 4267A>G), ATT1372ATA (4270T>A) | | | | | | | | |

##### Proteins

| | Begin | End | Coverage | Score | Concordance | Matches | Identities | I/D/M/F* | Stop Codons |
|---|---|---|---|---|---|---|---|---|---|
| polyprotein (YP_009553669.1) | 1295 | 1374 | 3.8% | 311 | 50.0% | 80 (97.6%) | 41 (50.0%) | 2/0/0/0 | 0 |
| Protein mutations: | G1296R (4040G>A), Y1298H (4046T>C), Q1301R (4056A>G 4057A>T), I1302V (4058A>G 4060T>C), E1305A (4068A>C), R1307A (4073A>G 4074G>C 4075A>C), F1312L (4090C>A), I1313R (4091A>C 4092T>G), C1314T (4094T>A 4095G>C), P1315H (4098C>A 4099A>T), A1316S (4100G>T 4102A>T), F1318H (4106T>C 4107T>A 4108C>T), Q1320E (4112C>G 4114G>A), W1321F (4116G>T 4117G>C), K1322L (4118A>C 4119A>T 4120A>T), V1324M (4124G>A 4126T>G), L1328H (4136T>C 4137T>A 4138G>C), K1329T (4140A>C 4141G>T), F1334T (4154T>A 4155T>C 4156C>T), R1337A (4163A>G 4164G>C 4165A>C), R1338T (4167G>C 4168G>T), D1340N (4172G>A), Y1341D (4175T>G 4177T>C), I1342L (4178A>C 4180C>T), A1344R (4184G>A 4185C>G 4186T>A), K1345P (4187A>C 4188A>C), Y1346_D1347insLR (4192_4193insCTCAGA), D1347K (4193G>A 4195T>A), V1350L (4202G>T 4204T>G), T1351V (4205A>G 4206C>T 4207A>T), Y1352F (4209A>T), I1353F (4211A>T 4213A>T), D1354N (4214G>A), H1359Y (4229C>T), D1362T (4238G>A 4239A>C 4240T>C), V1363L (4241G>C 4243A>T), N1365Q (4247A>C 4249T>G), L1367Q (4253T>C 4254T>A 4255G>A), K1368N (4258G>C), E1371Q (4265G>C 4267A>G) | | | | | | | | |
| Codon mutations: | GGG1296AGG (4040G>A), TAT1297TAC (4045T>C), TAT1298CAT (4046T>C), CAG1299CAA (4051G>A), ATA1300ATT (4054A>T), CAA1301CGT (4056A>G 4057A>T), ATT1302GTC (4058A>G 4060T>C), GAA1305GCA (4068A>C), AGA1307GCC (4073A>G 4074G>C 4075A>C), CAC1308CAT (4078C>T), TTC1312TTA (4090C>A), ATA1313CGA (4091A>C 4092T>G), TGT1314ACT (4094T>A 4095G>C), CCA1315CAT (4098C>A 4099A>T), GCA1316TCT (4100G>T 4102A>T), GGA1317GGG (4105A>G), TTC1318CAT (4106T>C 4107T>A 4108C>T), CAG1320GAA (4112C>G 4114G>A), TGG1321TTC (4116G>T 4117G>C), AAA1322CTT (4118A>C 4119A>T 4120A>T), GTT1323GTA (4123T>A), GTT1324ATG (4124G>A 4126T>G), CCA1325CCT (4129A>T), TTT1326TTC (4132T>C), GGT1327GGA (4135T>A), TTG1328CAC (4136T>C 4137T>A 4138G>C), AAG1329ACT (4140A>C 4141G>T), GCG1331GCT (4147G>T), CCA1332CCC (4150A>C), TTC1334ACT (4154T>A 4155T>C 4156C>T), TTT1335TTC (4159T>C), AGA1337GCC (4163A>G 4164G>C 4165A>C), AGG1338ACT (4167G>C 4168G>T), GAT1340AAT (4172G>A), TAT1341GAC (4175T>G 4177T>C), ATC1342CTT (4178A>C 4180C>T), TTT1343TTC (4183T>C), GCT1344AGA (4184G>A 4185C>G 4186T>A), AAA1345CCA (4187A>C 4188A>C), TAT1346_GAT1347insCTCAGA (4192_4193insCTCAGA), GAT1347AAA (4193G>A 4195T>A), TTT1348TTC (4198T>C), ATA1349ATC (4201A>C), GTT1350TTG (4202G>T 4204T>G), ACA1351GTT (4205A>G 4206C>T 4207A>T), TAC1352TTC (4209A>T), ATA1353TTT (4211A>T 4213A>T), GAT1354AAT (4214G>A), TTA1357TTG (4225A>G), ATT1358ATC (4228T>C), CAT1359TAT (4229C>T), CCA1361CCT (4237A>T), GAT1362ACC (4238G>A 4239A>C 4240T>C), GTA1363CTT (4241G>C 4243A>T), AAT1365CAG (4247A>C 4249T>G), CAT1366CAC (4252T>C), TTG1367CAA (4253T>C 4254T>A 4255G>A), AAG1368AAC (4258G>C), CAC1369CAT (4261C>T), TTA1370CTC (4262T>C 4264A>C), GAA1371CAG (4265G>C 4267A>G), ATT1372ATA (4270T>A) | | | | | | | | |

*: Inserts / Deletes / Misaligned / Frameshifts

#### Analysis details

This analysis was performed with panviral2.64

### NGS Details (UN9): Hibiscus bacilliform virus GD1

#### Assembly

| | |
|---|---|
| **Coverage Length** | 501 (1 contig(s)) |
| **Depth Of Coverage** | 9.2 |
| **Number Of Reads** | 37 |
| **Reads Per Million** | 0.84 rpm (after QC) |
| **Ambiguities** | 0 |
| **Assembly Method** | de novo + reference guided assembly |
| **Consensus Caller** | Bcf Tools |

#### Coverage Map

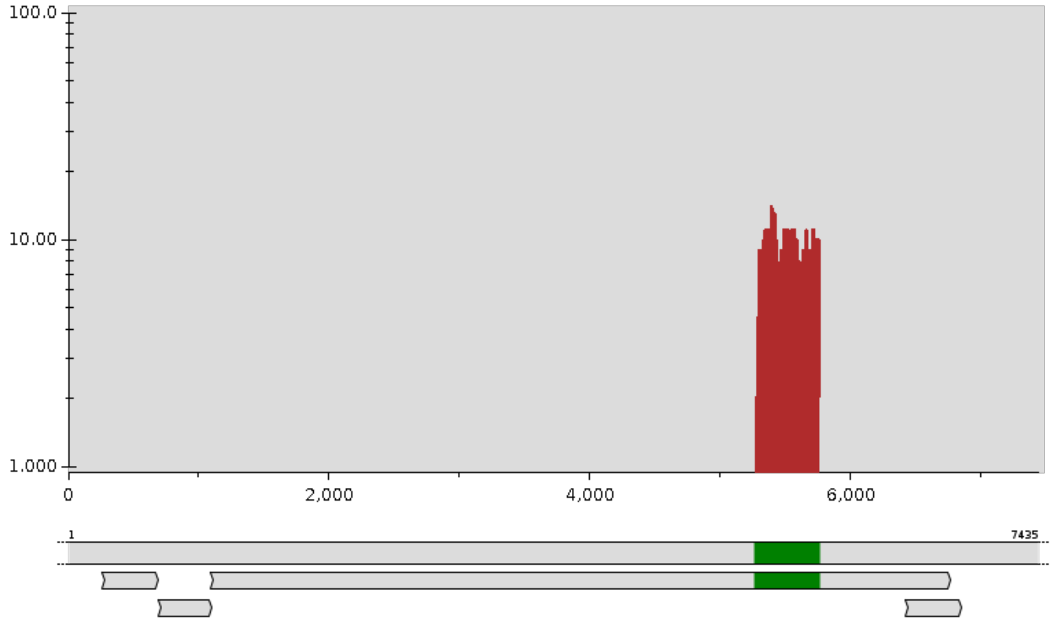

#### Assignment

| | |
|---|---|
| **Type** | Hibiscus bacilliform virus GD1 (Taxonomy ID: 1459800) |
| **Reference Genome** | NC_023485.1 |
| **NT Identity (%)** | 52.2358 |
| **AA Identity (%)** | 43.0303 |
| **Number Of Stop Codons** | 1 |
| **Number Of CDS** | 4 |

#### Alignment

| | |
|---|---|
| **Alignment Score** | 8.0 (NT) + 194.0 (AA) = 202.0 |
| **Concordance (%)** | 13.0457 |

| **Alignment Method** | Global, seeded, nucleotide + amino acids (AGA) |
|---|---|

#### Genome Region

Sequence starts at position 5261 and ends at position 5761 relative to NC_023485.1 reference sequence.

#### Alignment Detailed Statistics

| | Begin | End | Coverage | Score | Concordance | Matches | Identities | I/D/M/F* | Stop Codons |
|---|---|---|---|---|---|---|---|---|---|
| **NT** | **5261** | **5761** | **6.7%** | **8** | **0.8%** | **490 (97.4%)** | **257 (51.1%)** | **2/11** | |
| Mutations: | 5267G>A, 5268G>T, 5269T>G, 5270G>T, 5271T>A, 5281A>G, 5282A>G, 5283A>T, 5284G>A, 5292G>A, 5294T>G, 5295C>G, 5296T>C, 5297C>A, 5300T>A, 5301G>A, 5302A>T, 5303G>C, 5306A>G, 5307G>A, 5310C>A, 5312G>A, 5316A>C, 5317G>T, 5318T>A, 5319C>T, 5324A>T, 5325G>C, 5327C>T, 5330C>T, 5331A>G, 5333C>T, 5335C>A, 5336C>T, 5337A>C, 5339T>A, 5340A>T, 5342G>C, 5345G>T, 5346A>C, 5347G>A, 5348G>A, 5349G>T, 5351T>A, 5352_5353delGG, 5355C>A, 5356A>G, 5357T>G, 5362A>C, 5363G>T, 5364A>T, 5365T>A, 5367T>A, 5368A>T, 5369C>T, 5372A>C, 5375G>A, 5376T>A, 5384C>G, 5386A>G, 5387G>A, 5388A>T, 5389G>C, 5390T>A, 5393A>T, 5395T>A, 5403G>C, 5406G>A, 5407C>G, 5408T>A, 5411G>T, 5412C>A, 5414C>G, 5415C>A, 5417G>T, 5420G>A, 5422C>A, 5427A>T, 5428A>C, 5429G>A, 5430T>A, 5431G>A, 5435T>A, 5438T>C, 5441C>T, 5445G>A, 5446T>C, 5447T>C, 5448C>A, 5449C>G, 5450A>G, 5451G>T, 5456A>G, 5458T>A, 5459G>C, 5461A>T, 5467G>T, 5468G>T, 5469C>T, 5471A>G, 5474T>G, 5478C>T, 5480A>T, 5487C>T, 5489T>G, 5491A>C, 5492A>T, 5495C>T, 5498C>A, 5501A>T, 5503C>T, 5506T>A, 5511C>A, 5512A>T, 5514C>G, 5515G>A, 5516A>C, 5517A>C, 5518A>T, 5519G>C, 5523G>A, 5525C>T, 5527A>G, 5528C>G, 5531C>A, 5533T>C, 5537A>G, 5538G>C, 5539G>C, 5540T>A, 5541A>T, 5542C>A, 5543T>C, 5544G>T, 5545A>T, 5548C>A, 5549_5550insCA, 5553A>T, 5555C>A, 5556G>A, 5557C>T, 5561A>G, 5563A>T, 5573C>T, 5576A>T, 5577C>T, 5579C>G, 5582A>C, 5584T>A, 5585T>C, 5588C>T, 5591G>A, 5592T>G, 5593C>A, 5599A>C, 5600G>C, 5603A>G, 5607A>G, 5608T>A, 5609C>A, 5610A>C, 5612G>T, 5615C>T, 5616A>T, 5618A>G, 5619C>A, 5621G>A, 5624C>T, 5625A>G, 5627G>A, 5630A>G, 5631G>A, 5633A>G, 5635A>C, 5637T>C, 5638G>T, 5639C>T, 5640C>A, 5642G>A, 5645G>A, 5646A>C, 5647A>C, 5648T>A, 5649G>A, 5650G>A, 5655G>T, 5656T>A, 5658C>G, 5659T>C, 5662G>A, 5663C>G, 5664C>T, 5665C>T, 5666A>T, 5667A>T, 5673A>T, 5674T>G, 5675G>T, 5676A>G, 5679A>T, 5681A>T, 5682G>T, 5683C>G, 5684C>G, 5685C>T, 5686A>T, 5688A>G, 5690G>C, 5692A>T, 5693G>A, 5696G>T, 5697G>T, 5698A>C, 5699A>C, 5703C>T, 5705G>A, 5709G>C, 5710C>A, 5711T>C, 5714T>G, 5715C>G, 5718G>T, 5719A>C, 5721G>C, 5722C>G, 5725G>A, 5726C>T, 5727C>G, 5729T>A, 5735G>A, 5736C>G, 5738G>T, 5739C>G, 5741G>C, 5744C>A, 5745_5753delCATATCATA | | | | | | | | |

##### CDS

| | Begin | End | Coverage | Score | Concordance | Matches | Identities | I/D/M/F* | Stop Codons |
|---|---|---|---|---|---|---|---|---|---|
| BU35_gp3 | 1391 | 1557 | 8.8% | 194 | 17.2% | 164 (97.6%) | 71 (42.3%) | 1/3/2/2 | 1 |
| Protein mutations: | M1392I (5267G>A), V1393C (5268G>T 5269T>G 5270G>T), F1394I (5271T>A), K1397R (5281A>G 5282A>G), R1398* (5283A>T 5284G>A), D1401K (5292G>A 5294T>G), L1402A (5295C>G 5296T>C 5297C>A), E1404I (5301G>A 5302A>T 5303G>C), D1406N (5307G>A), Q1407K (5310C>A 5312G>A), S1409L (5316A>C 5317G>T 5318T>A), G1412R (5325G>C 5327C>T), N1414D (5331A>G 5333C>T), T1415N (5335C>A 5336C>T), I1416L (5337A>C 5339T>A), M1417F (5340A>T 5342G>C), K1418N (5345G>T), R1419Q (5346A>C 5347G>A 5348G>A), V1420L (5349G>T 5351T>A), H1422R (5355C>A 5356A>G 5357T>G), K1424T (5362A>C 5363G>T), I1425Y (5364A>T 5365T>A), Y1426I (5367T>A 5368A>T 5369C>T), F1429I (5376T>A), K1432R (5386A>G 5387G>A), F1435Y (5395T>A), V1438L (5403G>C), A1439R (5406G>A 5407C>G 5408T>A), M1440I (5411G>T), H1441K (5412C>A 5414C>G), P1442T (5415C>A 5417G>T), S1444Y (5422C>A), K1446S (5427A>T 5428A>C 5429G>A), W1447K (5430T>A 5431G>A), V1452T (5445G>A 5446T>C 5447T>C), P1453R (5448C>A 5449C>G 5450A>G), D1454Y (5451G>T), L1456H (5458T>A 5459G>C), Y1457F (5461A>T), W1459F (5467G>T 5468G>T), P1463S (5478C>T 5480A>T), K1467T (5491A>C 5492A>T), A1471V (5503C>T), V1472E (5506T>A), Q1474M (5511C>A 5512A>T), R1475D (5514C>G 5515G>A 5516A>C), K1476L (5517A>C 5518A>T 5519G>C), D1478N (5523G>A 5525C>T), N1479R (5527A>G 5528C>G), F1481S (5533T>C), G1483P (5538G>C 5539G>C 5540T>A), T1484Y (5541A>T 5542C>A 5543T>C), E1485L (5544G>T 5545A>T), A1486D (5548C>A), A1486_F1487insX (5549_5550insCA), I1488L (5553A>T 5555C>A), A1489I (5556G>A 5557C>T), Y1491F (5563A>T), F1498Y (5584T>A 5585T>C), S1501D (5592T>G 5593C>A), E1503A (5599A>C 5600G>C), I1506E (5607A>G 5608T>A 5609C>A), K1507H (5610A>C 5612G>T), I1509L (5616A>T 5618A>G), M1512V (5625A>G 5627G>A), E1514K (5631G>A 5633A>G), K1515T (5635A>C), C1516L (5637T>C 5638G>T 5639C>T), N1519P (5646A>C 5647A>C 5648T>A), G1520K (5649G>A 5650G>A), V1522Y (5655G>T 5656T>A), L1523A (5658C>G 5659T>C), S1524K (5662G>A 5663C>G), P1525F (5664C>T 5665C>T 5666A>T), T1526S (5667A>T), M1528C (5673A>T 5674T>G 5675G>T), K1529E (5676A>G), I1530F (5679A>T 5681A>T), A1531W (5682G>T 5683C>G 5684C>G), Q1532L (5685C>T 5686A>T), R1533G (5688A>G 5690G>C), K1534I (5692A>T 5693G>A), E1536S (5697G>T 5698A>C 5699A>C), A1540H (5709G>C 5710C>A 5711T>C), I1541M (5714T>G), L1542V (5715C>G), E1543S (5718G>T 5719A>C), A1544R (5721G>C 5722C>G), G1545D (5725G>A 5726C>T), R1546G (5727C>G 5729T>A), L1549V (5736C>G 5738G>T), Q1550D (5739C>G 5741G>C), H1552_I1554del (5745_5753delCATATCATA) | | | | | | | | |
| Codon mutations: | ATG1392ATA (5267G>A), GTG1393TGT (5268G>T 5269T>G 5270G>T), TTC1394ATC (5271T>A), AAA1397AGG (5281A>G 5282A>G), AGG1398TAG (5283A>T 5284G>A), GAT1401AAG (5292G>A 5294T>G), CTC1402GCA (5295C>G 5296T>C 5297C>A), ACT1403ACA (5300T>A), GAG1404ATC (5301G>A 5302A>T 5303G>C), AAA1405AAG (5306A>G), GAT1406AAT (5307G>A), CAG1407AAA (5310C>A 5312G>A), AGT1409CTA (5316A>C 5317G>T 5318T>A), CTG1410TTG (5319C>T), CCA1411CCT (5324A>T), GGC1412CGT (5325G>C 5327C>T), ATC1413ATT (5330C>T), AAC1414GAT (5331A>G 5333C>T), ACC1415AAT (5335C>A 5336C>T), ATT1416CTA (5337A>C 5339T>A), ATG1417TTC (5340A>T 5342G>C), AAG1418AAT (5345G>T), AGG1419CAA (5346A>C 5347G>A 5348G>A), GTT1420TTA (5349G>T 5351T>A), GGC1421--C (5352_5353delGG), CAT1422AGG (5355C>A 5356A>G 5357T>G), AAG1424ACT (5362A>C 5363G>T), ATC1425TAC (5364A>T 5365T>A), TAC1426ATT (5367T>A 5368A>T 5369C>T), TCA1427TCC (5372A>C), AAG1428AAA (5375G>A), TTT1429ATT (5376T>A), CTC1431CTG (5384C>G), AAG1432AGA (5386A>G 5387G>A), AGT1433TCA (5388A>T 5389G>C 5390T>A), GGA1434GGT (5393A>T), TTT1435TAT (5395T>A), GTG1438CTG (5403G>C), GCT1439AGA (5406G>A 5407C>G 5408T>A), ATG1440ATT (5411G>T), CAC1441AAG (5412C>A 5414C>G), CCG1442ACT (5415C>A 5417G>T), GAG1443GAA (5420G>A), TCT1444TAT (5422C>A), AAG1446TCA (5427A>T 5428A>C 5429G>A), TGG1447AAG (5430T>A 5431G>A), ACT1448ACA (5435T>A), GCT1449GCC (5438T>C), TTC1450TTT (5441C>T), GTT1452ACC (5445G>A 5446T>C 5447T>C), CCA1453AGG (5448C>A 5449C>G 5450A>G), GAT1454TAT (5451G>T), GGA1455GGG (5456A>G), CTG1456CAC (5458T>A 5459G>C), TAT1457TTT (5461A>T), TGG1459TTT (5467G>T 5468G>T), CTA1460TTG (5469C>T 5471A>G), GTT1461GTG (5474T>G), CCA1463TCT (5478C>T 5480A>T), CTT1466TTG (5487C>T 5489T>G), AAA1467ACT (5491A>C 5492A>T), AAC1468AAT (5495C>T), GCC1469GCA (5498C>A), CCA1470CCT (5501A>T), GCA1471GTA (5503C>T), GTA1472GAA (5506T>A), CAG1474ATG (5511C>A 5512A>T), CGA1475GAC (5514C>G 5515G>A 5516A>C), AAG1476CTC (5517A>C 5518A>T 5519G>C), GAC1478AAT (5523G>A 5525C>T), AAC1479AGG (5527A>G 5528C>G), GTC1480GTA (5531C>A), TTC1481TCC (5533T>C), AAA1482AAG (5537A>G), GGT1483CCA (5538G>C 5539G>C 5540T>A), ACT1484TAC (5541A>T 5542C>A 5543T>C), GAA1485TTA (5544G>T 5545A>T), GCC1486GAC (5548C>A), GCC1486_TTT1487insCA- (5549_5550insCA), ATC1488TTA (5553A>T 5555C>A), GCT1489ATT (5556G>A 5557C>T), GTA1490GTG (5561A>G), TAC1491TTC (5563A>T), GAC1494GAT (5573C>T), ATA1495ATT (5576A>T), CTC1496TTG (5577C>T 5579C>G), GTA1497GTC (5582A>C), TTT1498TAC (5584T>A 5585T>C), TCC1499TCT (5588C>T), CAG1500CAA (5591G>A), TCT1501GAT (5592T>G 5593C>A), GAG1503GCC (5599A>C 5600G>C), GAA1504GAG (5603A>G), ATC1506GAA (5607A>G 5608T>A 5609C>A), AAG1507CAT (5610A>C 5612G>T), CAC1508CAT (5615C>T), ATA1509TTG (5616A>T 5618A>G), CGG1510AGA (5619C>A 5621G>A), GTC1511GTT (5624C>T), ATG1512GTA (5625A>G 5627G>A), TTA1513TTG (5630A>G), GAA1514AAG (5631G>A 5633A>G), AAA1515ACA (5635A>C), TGC1516CTT (5637T>C 5638G>T 5639C>T), CGG1517AGA (5640C>A 5642G>A), GAG1518GAA (5645G>A), AAT1519CCA (5646A>C 5647A>C 5648T>A), GGG1520AAG (5649G>A 5650G>A), GTT1522TAT (5655G>T 5656T>A), CTT1523GCT (5658C>G 5659T>C), AGC1524AAG (5662G>A 5663C>G), CCA1525TTT (5664C>T 5665C>T 5666A>T), ACA1526TCA (5667A>T), ATG1528TGT (5673A>T 5674T>G 5675G>T), AAG1529GAG (5676A>G), ATA1530TTT (5679A>T 5681A>T), GCC1531TGG (5682G>T 5683C>G 5684C>G), CAG1532TTG (5685C>T 5686A>T), AGG1533GGC (5688A>G 5690G>C), AAG1534ATA (5692A>T 5693G>A), GTG1535GTT (5696G>T), GAA1536TCC (5697G>T 5698A>C 5699A>C), CTG1538TTA (5703C>T 5705G>A), GCT1540CAC (5709G>C 5710C>A 5711T>C), ATT1541ATG (5714T>G), CTG1542GTG (5715C>G), GAA1543TCA (5718G>T 5719A>C), GCA1544CGA (5721G>C 5722C>G), GGC1545GAT (5725G>A 5726C>T), CGT1546GGA (5727C>G 5729T>A), CAG1548CAA (5735G>A), CTG1549GTT (5736C>G 5738G>T), CAG1550GAC (5739C>G 5741G>C), CCC1551CCA (5744C>A), CAT1552_ATA1554del (5745_5753delCATATCATA) | | | | | | | | |

##### Proteins

| | Begin | End | Coverage | Score | Concordance | Matches | Identities | I/D/M/F* | Stop Codons |
|---|---|---|---|---|---|---|---|---|---|
| polyprotein (YP_009002585.1) | 1391 | 1557 | 8.8% | 194 | 17.2% | 164 (97.6%) | 71 (42.3%) | 1/3/2/2 | 1 |
| Protein mutations: | M1392I (5267G>A), V1393C (5268G>T 5269T>G 5270G>T), F1394I (5271T>A), K1397R (5281A>G 5282A>G), R1398* (5283A>T 5284G>A), D1401K (5292G>A 5294T>G), L1402A (5295C>G 5296T>C 5297C>A), E1404I (5301G>A 5302A>T 5303G>C), D1406N (5307G>A), Q1407K (5310C>A 5312G>A), S1409L (5316A>C 5317G>T 5318T>A), G1412R (5325G>C 5327C>T), N1414D (5331A>G 5333C>T), T1415N (5335C>A 5336C>T), I1416L (5337A>C 5339T>A), M1417F (5340A>T 5342G>C), K1418N (5345G>T), R1419Q (5346A>C 5347G>A 5348G>A), V1420L (5349G>T 5351T>A), H1422R (5355C>A 5356A>G 5357T>G), K1424T (5362A>C 5363G>T), I1425Y (5364A>T 5365T>A), Y1426I (5367T>A 5368A>T 5369C>T), F1429I (5376T>A), K1432R (5386A>G 5387G>A), F1435Y (5395T>A), V1438L (5403G>C), A1439R (5406G>A 5407C>G 5408T>A), M1440I (5411G>T), H1441K (5412C>A 5414C>G), P1442T (5415C>A 5417G>T), S1444Y (5422C>A), K1446S (5427A>T 5428A>C 5429G>A), W1447K (5430T>A 5431G>A), V1452T (5445G>A 5446T>C 5447T>C), P1453R (5448C>A 5449C>G 5450A>G), D1454Y (5451G>T), L1456H (5458T>A 5459G>C), Y1457F (5461A>T), W1459F (5467G>T 5468G>T), P1463S (5478C>T 5480A>T), K1467T (5491A>C 5492A>T), A1471V (5503C>T), V1472E (5506T>A), Q1474M (5511C>A 5512A>T), R1475D (5514C>G 5515G>A 5516A>C), K1476L (5517A>C 5518A>T 5519G>C), D1478N (5523G>A 5525C>T), N1479R (5527A>G 5528C>G), F1481S (5533T>C), G1483P (5538G>C 5539G>C 5540T>A), T1484Y (5541A>T 5542C>A 5543T>C), E1485L (5544G>T 5545A>T), A1486D (5548C>A), A1486_F1487insX (5549_5550insCA), I1488L (5553A>T 5555C>A), A1489I (5556G>A 5557C>T), Y1491F (5563A>T), F1498Y (5584T>A 5585T>C), S1501D (5592T>G 5593C>A), E1503A (5599A>C 5600G>C), I1506E (5607A>G 5608T>A 5609C>A), K1507H (5610A>C 5612G>T), I1509L (5616A>T 5618A>G), M1512V (5625A>G 5627G>A), E1514K (5631G>A 5633A>G), K1515T (5635A>C), C1516L (5637T>C 5638G>T 5639C>T), N1519P (5646A>C 5647A>C 5648T>A), G1520K (5649G>A 5650G>A), V1522Y (5655G>T 5656T>A), L1523A (5658C>G 5659T>C), S1524K (5662G>A 5663C>G), P1525F (5664C>T 5665C>T 5666A>T), T1526S (5667A>T), M1528C (5673A>T 5674T>G 5675G>T), K1529E (5676A>G), I1530F (5679A>T 5681A>T), A1531W (5682G>T 5683C>G 5684C>G), Q1532L (5685C>T 5686A>T), R1533G (5688A>G 5690G>C), K1534I (5692A>T 5693G>A), E1536S (5697G>T 5698A>C 5699A>C), A1540H (5709G>C 5710C>A 5711T>C), I1541M (5714T>G), L1542V (5715C>G), E1543S (5718G>T 5719A>C), A1544R (5721G>C 5722C>G), G1545D (5725G>A 5726C>T), R1546G (5727C>G 5729T>A), L1549V (5736C>G 5738G>T), Q1550D (5739C>G 5741G>C), H1552_I1554del (5745_5753delCATATCATA) | | | | | | | | |

| | Begin | End | Coverage | Score | Concordance | Matches | Identities | I/D/M/F* | Stop Codons |
|---|---|---|---|---|---|---|---|---|---|
| **NT** | **5261** | **5761** | **6.7%** | **8** | **0.8%** | **490 (97.4%)** | **257 (51.1%)** | **2/11** | |
| Codon mutations: | ATG1392ATA (5267G>A), GTG1393TGT (5268G>T 5269T>G 5270G>T), TTC1394ATC (5271T>A), AAA1397AGG (5281A>G 5282A>G), AGG1398TAG (5283A>T 5284G>A), GAT1401AAG (5292G>A 5294T>G), CTC1402GCA (5295C>G 5296T>C 5297C>A), ACT1403ACA (5300T>A), GAG1404ATC (5301G>A 5302A>T 5303G>C), AAA1405AAG (5306A>G), GAT1406AAT (5307G>A), CAG1407AAA (5310C>A 5312G>A), AGT1409CTA (5316A>C 5317G>T 5318T>A), CTG1410TTG (5319C>T), CCA1411CCT (5324A>T), GGC1412CGT (5325G>C 5327C>T), ATC1413ATT (5330C>T), AAC1414GAT (5331A>G 5333C>T), ACC1415AAT (5335C>A 5336C>T), ATT1416CTA (5337A>C 5339T>A), ATG1417TTC (5340A>T 5342G>C), AAG1418AAT (5345G>T), AGG1419CAA (5346A>C 5347G>A 5348G>A), GTT1420TTA (5349G>T 5351T>A), GGC1421--C (5352_5353delGG), CAT1422AGG (5355C>A 5356A>G 5357T>G), AAG1424ACT (5362A>C 5363G>T), ATC1425TAC (5364A>T 5365T>A), TAC1426ATT (5367T>A 5368A>T 5369C>T), TCA1427TCC (5372A>C), AAG1428AAA (5375G>A), TTT1429ATT (5376T>A), CTC1431CTG (5384C>G), AAG1432AGA (5386A>G 5387G>A), AGT1433TCA (5388A>T 5389G>C 5390T>A), GGA1434GGT (5393A>T), TTT1435TAT (5395T>A), GTG1438CTG (5403G>C), GCT1439AGA (5406G>A 5407C>G 5408T>A), ATG1440ATT (5411G>T), CAC1441AAG (5412C>A 5414C>G), CCG1442ACT (5415C>A 5417G>T), GAG1443GAA (5420G>A), TCT1444TAT (5422C>A), AAG1446TCA (5427A>T 5428A>C 5429G>A), TGG1447AAG (5430T>A 5431G>A), ACT1448ACA (5435T>A), GCT1449GCC (5438T>C), TTC1450TTT (5441C>T), GTT1452ACC (5445G>A 5446T>C 5447T>C), CCA1453AGG (5448C>A 5449C>G 5450A>G), GAT1454TAT (5451G>T), GGA1455GGG (5456A>G), CTG1456CAC (5458T>A 5459G>C), TAT1457TTT (5461A>T), TGG1459TTT (5467G>T 5468G>T), CTA1460TTG (5469C>T 5471A>G), GTT1461GTG (5474T>G), CCA1463TCT (5478C>T 5480A>T), CTT1466TTG (5487C>T 5489T>G), AAA1467ACT (5491A>C 5492A>T), AAC1468AAT (5495C>T), GCC1469GCA (5498C>A), CCA1470CCT (5501A>T), GCA1471GTA (5503C>T), GTA1472GAA (5506T>A), CAG1474ATG (5511C>A 5512A>T), CGA1475GAC (5514C>G 5515G>A 5516A>C), AAG1476CTC (5517A>C 5518A>T 5519G>C), GAC1478AAT (5523G>A 5525C>T), AAC1479AGG (5527A>G 5528C>G), GTC1480GTA (5531C>A), TTC1481TCC (5533T>C), AAA1482AAG (5537A>G), GGT1483CCA (5538G>C 5539G>C 5540T>A), ACT1484TAC (5541A>T 5542C>A 5543T>C), GAA1485TTA (5544G>T 5545A>T), GCC1486GAC (5548C>A), GCC1486_TTT1487insCA- (5549_5550insCA), ATC1488TTA (5553A>T 5555C>A), GCT1489ATT (5556G>A 5557C>T), GTA1490GTG (5561A>G), TAC1491TTC (5563A>T), GAC1494GAT (5573C>T), ATA1495ATT (5576A>T), CTC1496TTG (5577C>T 5579C>G), GTA1497GTC (5582A>C), TTT1498TAC (5584T>A 5585T>C), TCC1499TCT (5588C>T), CAG1500CAA (5591G>A), TCT1501GAT (5592T>G 5593C>A), GAG1503GCC (5599A>C 5600G>C), GAA1504GAG (5603A>G), ATC1506GAA (5607A>G 5608T>A 5609C>A), AAG1507CAT (5610A>C 5612G>T), CAC1508CAT (5615C>T), ATA1509TTG (5616A>T 5618A>G), CGG1510AGA (5619C>A 5621G>A), GTC1511GTT (5624C>T), ATG1512GTA (5625A>G 5627G>A), TTA1513TTG (5630A>G), GAA1514AAG (5631G>A 5633A>G), AAA1515ACA (5635A>C), TGC1516CTT (5637T>C 5638G>T 5639C>T), CGG1517AGA (5640C>A 5642G>A), GAG1518GAA (5645G>A), AAT1519CCA (5646A>C 5647A>C 5648T>A), GGG1520AAG (5649G>A 5650G>A), GTT1522TAT (5655G>T 5656T>A), CTT1523GCT (5658C>G 5659T>C), AGC1524AAG (5662G>A 5663C>G), CCA1525TTT (5664C>T 5665C>T 5666A>T), ACA1526TCA (5667A>T), ATG1528TGT (5673A>T 5674T>G 5675G>T), AAG1529GAG (5676A>G), ATA1530TTT (5679A>T 5681A>T), GCC1531TGG (5682G>T 5683C>G 5684C>G), CAG1532TTG (5685C>T 5686A>T), AGG1533GGC (5688A>G 5690G>C), AAG1534ATA (5692A>T 5693G>A), GTG1535GTT (5696G>T), GAA1536TCC (5697G>T 5698A>C 5699A>C), CTG1538TTA (5703C>T 5705G>A), GCT1540CAC (5709G>C 5710C>A 5711T>C), ATT1541ATG (5714T>G), CTG1542GTG (5715C>G), GAA1543TCA (5718G>T 5719A>C), GCA1544CGA (5721G>C 5722C>G), GGC1545GAT (5725G>A 5726C>T), CGT1546GGA (5727C>G 5729T>A), CAG1548CAA (5735G>A), CTG1549GTT (5736C>G 5738G>T), CAG1550GAC (5739C>G 5741G>C), CCC1551CCA (5744C>A), CAT1552_ATA1554del (5745_5753delCATATCATA) | | | | | | | | |

*: Inserts / Deletes / Misaligned / Frameshifts

#### Analysis details

This analysis was performed with panviral2.64

### NGS Details (UN9): Dioscovirus dioscoreae

#### Assembly

| | |
|---|---|
| **Coverage Length** | 740 (1 contig(s)) |
| **Depth Of Coverage** | 4.9 |
| **Number Of Reads** | 27 |
| **Reads Per Million** | 0.61 rpm (after QC) |
| **Ambiguities** | 0 |
| **Assembly Method** | de novo + reference guided assembly |
| **Consensus Caller** | Bcf Tools |

#### Coverage Map

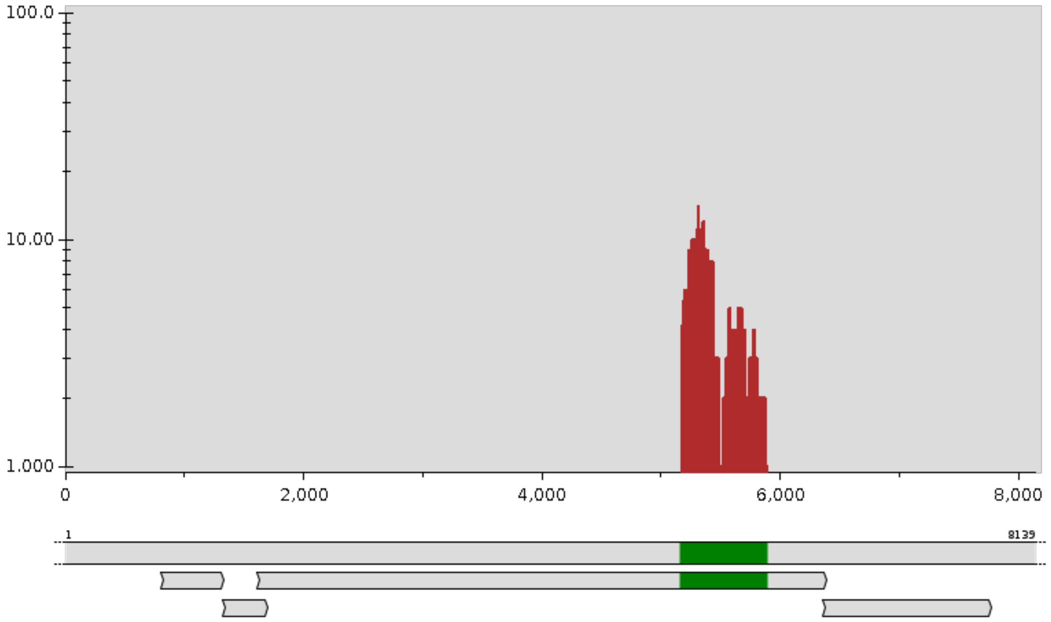

#### Assignment

| | |
|---|---|
| **Type** | Dioscovirus dioscoreae (Taxonomy ID: 3052184) |
| **Reference Genome** | NC_040712.1 |
| **NT Identity (%)** | 64.1304 |
| **AA Identity (%)** | 56.0976 |
| **Number Of Stop Codons** | 0 |
| **Number Of CDS** | 4 |

#### Alignment

| | |
|---|---|
| **Alignment Score** | 394.0 (NT) + 914.0 (AA) = 1308.0 |
| **Concordance (%)** | 43.484 |

| | |
|---|---|
| **Alignment Method** | Global, seeded, nucleotide + amino acids (AGA) |

#### Genome Region

Sequence starts at position 5162 and ends at position 5901 relative to NC_040712.1 reference sequence.

#### Alignment Detailed Statistics

| | Begin | End | Coverage | Score | Concordance | Matches | Identities | I/D/M/F* | Stop Codons |
|---|---|---|---|---|---|---|---|---|---|
| **NT** | **5162** | **5901** | **9.1%** | **394** | **27.2%** | **736 (99.5%)** | **472 (63.8%)** | **0/4** | |
| Mutations: | 5168G>A, 5171T>A, 5174C>A, 5178C>A, 5179C>G, 5180T>A, 5183C>A, 5184C>A, 5185A>C, 5187A>T, 5188G>C, 5190C>A, 5192C>A, 5195T>C, 5198G>A, 5199A>T, 5207T>A, 5213T>G, 5216C>A, 5219C>A, 5220delA, 5222A>C, 5226T>A, 5227C>A, 5228T>C, 5232C>A, 5233A>T, 5237A>C, 5243T>A, 5247A>G, 5248G>C, 5249T>C, 5250C>A, 5252T>A, 5259T>A, 5260A>T, 5261C>A, 5271C>A, 5276A>C, 5288A>T, 5289T>A, 5290A>G, 5291C>A, 5293A>T, 5294A>G, 5297C>T, 5298C>G, 5299A>C, 5300A>T, 5303C>T, 5305C>A, 5307C>T, 5309C>A, 5313A>G, 5314G>A, 5315T>C, 5317T>A, 5318T>A, 5319G>A, 5321C>T, 5322T>G, 5324T>A, 5327C>A, 5331C>A, 5332T>A, 5335A>G, 5339T>A, 5340A>C, 5344A>G, 5345C>A, 5348G>A, 5350T>G, 5351T>A, 5352G>A, 5354C>T, 5356A>T, 5360C>T, 5369T>C, 5370C>T, 5371T>G, 5372A>T, 5375G>A, 5381A>G, 5383T>A, 5384C>T, 5385C>T, 5386A>G, 5387C>G, 5390G>A, 5391A>G, 5393T>A, 5395T>A, 5396G>A, 5400G>C, 5402T>C, 5405A>T, 5406C>G, 5408G>T, 5424G>A, 5430G>A, 5431T>C, 5432C>T, 5435C>T, 5439C>G, 5441G>A, 5444T>A, 5457A>T, 5461T>C, 5466C>T, 5471C>T, 5475C>T, 5477C>G, 5480A>G, 5482A>C, 5483T>A, 5489T>A, 5490T>C, 5492A>T, 5493G>A, 5502C>A, 5507A>G, 5513T>C, 5519A>T, 5523A>G, 5524A>G, 5526A>G, 5532T>A, 5533C>A, 5535G>A, 5538T>A, 5543C>A, 5544T>C, 5545G>T, 5546T>A, 5549T>C, 5553A>G, 5558C>T, 5561T>C, 5564T>C, 5565T>C, 5567G>A, 5568A>G, 5573T>C, 5574T>A, 5575C>G, 5577G>A, 5578A>G, 5580A>G, 5581G>A, 5585A>G, 5589C>G, 5595G>C, 5598C>G, 5607T>C, 5608T>A, 5610C>A, 5611A>C, 5612G>A, 5613T>A, 5615C>T, 5618C>T, 5619C>A, 5620A>G, 5621A>G, 5622G>T, 5626G>T, 5627T>C, 5628A>G, 5629A>T, 5630G>A, 5631G>A, 5632A>G, 5633A>T, 5634G>A, 5639G>A, 5640C>A, 5642T>A, 5645T>A, 5646T>A, 5648G>A, 5649T>A, 5651A>C, 5656C>A, 5657A>G, 5661T>A, 5664A>G, 5667A>T, 5670G>T, 5672A>T, 5673G>A, 5674T>A, 5676G>A, 5679A>C, 5685G>A, 5687A>C, 5691C>T, 5693T>A, 5696C>A, 5697C>G, 5700G>A, 5701A>C, 5702A>T, 5703A>T, 5711A>T, 5714A>G, 5717A>G, 5718G>A, 5720T>A, 5721C>A, 5726C>A, 5739T>G, 5740T>C, 5741A>T, 5748A>G, 5756A>T, 5757T>A, 5759T>G, 5762T>A, 5763_5765delGAA, 5768C>T, 5769C>A, 5777A>T, 5779C>A, 5781T>A, 5782T>C, 5786G>A, 5788G>A, 5790C>T, 5795G>A, 5798G>A, 5807G>A, 5808A>T, 5811T>G, 5813A>G, 5816C>T, 5822C>T, 5828T>C, 5830A>T, 5834C>A, 5836C>A, 5843G>A, 5844A>G, 5846T>A, 5850T>A, 5851A>T, 5853A>G, 5855T>A, 5856A>G, 5859A>C, 5860T>C, 5864T>A, 5868A>G, 5869A>C, 5870T>A, 5873G>A, 5874T>A, 5875G>T, 5876T>A, 5877T>G, 5878C>G, 5879C>A, 5882C>T, 5889G>C, 5893G>A, 5895A>C, 5897A>C | | | | | | | | |

##### CDS

| | Begin | End | Coverage | Score | Concordance | Matches | Identities | I/D/M/F* | Stop Codons |
|---|---|---|---|---|---|---|---|---|---|
| EXK67_gp3 | 1185 | 1431 | 15.5% | 914 | 53.3% | 246 (99.6%) | 138 (55.9%) | 0/1/1/1 | 0 |
| Protein mutations: | P1190R (5178C>A 5179C>G 5180T>A), Q1192T (5184C>A 5185A>C), T1197S (5199A>T), I1201M (5213T>G), N1203K (5219C>A), S1206N (5226T>A 5227C>A 5228T>C), Q1208I (5232C>A 5233A>T), S1213A (5247A>G 5248G>C 5249T>C), Y1217I (5259T>A 5260A>T 5261C>A), Y1227R (5289T>A 5290A>G 5291C>A), K1228M (5293A>T 5294A>G), Q1230A (5298C>G 5299A>C 5300A>T), T1232K (5305C>A), S1235D (5313A>G 5314G>A 5315T>C), I1236K (5317T>A 5318T>A), D1237N (5319G>A 5321C>T), Y1238E (5322T>G 5324T>A), L1241N (5331C>A 5332T>A), K1242R (5335A>G), K1244Q (5340A>C), D1245G (5344A>G 5345C>A), I1247R (5350T>G 5351T>A), V1248I (5352G>A 5354C>T), Y1249F (5356A>T), L1254C (5370C>T 5371T>G 5372A>T), F1258Y (5383T>A 5384C>T), H1259W (5385C>T 5386A>G 5387C>G), I1261V (5391A>G 5393T>A), M1262K (5395T>A 5396G>A), D1264H (5400G>C 5402T>C), Q1266D (5406C>G 5408G>T), A1272T (5424G>A), V1274T (5430G>A 5431T>C 5432C>T), Q1277E (5439C>G 5441G>A), I1283L (5457A>T), V1284A (5461T>C), P1286S (5466C>T), N1291T (5482A>C 5483T>A), S1294P (5490T>C 5492A>T), V1295I (5493G>A), K1305G (5523A>G 5524A>G), K1306E (5526A>G), S1308K (5532T>A 5533C>A), E1309K (5535G>A), F1310I (5538T>A), C1312L (5544T>C 5545G>T 5546T>A), I1315V (5553A>G), I1320V (5568A>G), E1323R (5577G>A 5578A>G), S1324D (5580A>G 5581G>A), I1325M (5585A>G), Q1327E (5589C>G), V1329L (5595G>C), Q1330E (5598C>G), L1333Q (5607T>C 5608T>A), Q1334T (5610C>A 5611A>C 5612G>A), F1335I (5613T>A 5615C>T), Q1337R (5619C>A 5620A>G 5621A>G), V1338L (5622G>T), C1339F (5626G>T 5627T>C), K1340V (5628A>G 5629A>T 5630G>A), E1341S (5631G>A 5632A>G 5633A>T), E1342K (5634G>A), L1344I (5640C>A 5642T>A), L1346I (5646T>A 5648G>A), S1347T (5649T>A 5651A>C), T1349K (5656C>A 5657A>G), L1351M (5661T>A), K1352E (5664A>G), I1353L (5667A>T), G1354C (5670G>T 5672A>T), V1355K (5673G>A 5674T>A), A1356T (5676G>A), N1357H (5679A>C), E1359N (5685G>A 5687A>C), L1363V (5697C>G), E1364T (5700G>A 5701A>C 5702A>T), I1365L (5703A>T), E1367D (5711A>T), V1370I (5718G>A 5720T>A), Q1371K (5721C>A), L1377A (5739T>G 5740T>C 5741A>T), I1380V (5748A>G), E1382D (5756A>T), F1383M (5757T>A 5759T>G), E1385del (5763_5765delGAA), Q1387K (5769C>A), E1389D (5777A>T), T1390K (5779C>A), L1391T (5781T>A 5782T>C), G1393D (5788G>A), I1400L (5808A>T), L1401V (5811T>G 5813A>G), Y1407F (5830A>T), P1409Q (5836C>A), S1412G (5844A>G 5846T>A), Y1414I (5850T>A 5851A>T), T1415A (5853A>G 5855T>A), R1416G (5856A>G), I1417P (5859A>C 5860T>C), F1418L (5864T>A), N1420A (5868A>G 5869A>C 5870T>A), C1422I (5874T>A 5875G>T 5876T>A), S1423G (5877T>G 5878C>G 5879C>A), E1427Q (5889G>C), R1428K (5893G>A), K1429H (5895A>C 5897A>C) | | | | | | | | |
| Codon mutations: | AAG1186AAA (5168G>A), GTT1187GTA (5171T>A), ATC1188ATA (5174C>A), CCT1190AGA (5178C>A 5179C>G 5180T>A), TCC1191TCA (5183C>A), CAA1192ACA (5184C>A 5185A>C), AGT1193TCT (5187A>T 5188G>C), CGC1194AGA (5190C>A 5192C>A), CAT1195CAC (5195T>C), AGG1196AGA (5198G>A), ACA1197TCA (5199A>T), GCT1199GCA (5207T>A), ATT1201ATG (5213T>G), GTC1202GTA (5216C>A), AAC1203AAA (5219C>A), AAA1204-AC (5220delA 5222A>C), TCT1206AAC (5226T>A 5227C>A 5228T>C), CAA1208ATA (5232C>A 5233A>T), GTA1209GTC (5237A>C), GGT1211GGA (5243T>A), AGT1213GCC (5247A>G 5248G>C 5249T>C), CGT1214AGA (5250C>A 5252T>A), TAC1217ATA (5259T>A 5260A>T 5261C>A), CGA1221AGA (5271C>A), CTA1222CTC (5276A>C), ACA1226ACT (5288A>T), TAC1227AGA (5289T>A 5290A>G 5291C>A), AAA1228ATG (5293A>T 5294A>G), GAC1229GAT (5297C>T), CAA1230GCT (5298C>G 5299A>C 5300A>T), TAC1231TAT (5303C>T), ACA1232AAA (5305C>A), CTC1233TTA (5307C>T 5309C>A), AGT1235GAC (5313A>G 5314G>A 5315T>C), ATT1236AAA (5317T>A 5318T>A), GAC1237AAT (5319G>A 5321C>T), TAT1238GAA (5322T>G 5324T>A), CTC1239CTA (5327C>A), CTT1241AAT (5331C>A 5332T>A), AAA1242AGA (5335A>G), ATT1243ATA (5339T>A), AAA1244CAA (5340A>C), GAC1245GGA (5344A>G 5345C>A), AAG1246AAA (5348G>A), ATT1247AGA (5350T>G 5351T>A), GTC1248ATT (5352G>A 5354C>T), TAC1249TTC (5356A>T), AGC1250AGT (5360C>T), GAT1253GAC (5369T>C), CTA1254TGT (5370C>T 5371T>G 5372A>T), AAG1255AAA (5375G>A), GGA1257GGG (5381A>G), TTC1258TAT (5383T>A 5384C>T), CAC1259TGG (5385C>T 5386A>G 5387C>G), CAG1260CAA (5390G>A), ATT1261GTA (5391A>G 5393T>A), ATG1262AAA (5395T>A 5396G>A), GAT1264CAC (5400G>C 5402T>C), CCA1265CCT (5405A>T), CAG1266GAT (5406C>G 5408G>T), GCA1272ACA (5424G>A), GTC1274ACT (5430G>A 5431T>C 5432C>T), TGC1275TGT (5435C>T), CAG1277GAA (5439C>G 5441G>A), GGT1278GGA (5444T>A), ATA1283TTA (5457A>T), GTA1284GCA (5461T>C), CCA1286TCA (5466C>T), TTC1287TTT (5471C>T), CTC1289TTG (5475C>T 5477C>G), AAA1290AAG (5480A>G), AAT1291ACA (5482A>C 5483T>A), CCT1293CCA (5489T>A), TCA1294CCT (5490T>C 5492A>T), GTA1295ATA (5493G>A), CGA1298AGA (5502C>A), AAA1299AAG (5507A>G), GAT1301GAC (5513T>C), ATA1303ATT (5519A>T), AAA1305GGA (5523A>G 5524A>G), AAA1306GAA (5526A>G), TCA1308AAA (5532T>A 5533C>A), GAA1309AAA (5535G>A), TTT1310ATT (5538T>A), GTC1311GTA (5543C>A), TGT1312CTA (5544T>C 5545G>T 5546T>A), GTT1313GTC (5549T>C), ATA1315GTA (5553A>G), GAC1316GAT (5558C>T), GAT1317GAC (5561T>C), ATT1318ATC (5564T>C), TTG1319CTA (5565T>C 5567G>A), ATA1320GTA (5568A>G), TTT1321TTC (5573T>C), TCT1322AGT (5574T>A 5575C>G), GAA1323AGA (5577G>A 5578A>G), AGT1324GAT (5580A>G 5581G>A), ATA1325ATG (5585A>G), CAA1327GAA (5589C>G), GTA1329CTA (5595G>C), CAA1330GAA (5598C>G), TTA1333CAA (5607T>C 5608T>A), CAG1334ACA (5610C>A 5611A>C 5612G>A), TTC1335ATT (5613T>A 5615C>T), TTC1336TTT (5618C>T), CAA1337AGG (5619C>A 5620A>G 5621A>G), GTA1338TTA (5622G>T), TGT1339TTC (5626G>T 5627T>C), AAG1340GTA (5628A>G 5629A>T 5630G>A), GAA1341AGT (5631G>A 5632A>G 5633A>T), GAA1342AAA (5634G>A), GGG1343GGA (5639G>A), CTT1344ATA (5640C>A 5642T>A), ATT1345ATA (5645T>A), TTG1346ATA (5646T>A 5648G>A), TCA1347ACC (5649T>A 5651A>C), ACA1349AAG (5656C>A 5657A>G), TTG1351ATG (5661T>A), AAA1352GAA (5664A>G), ATA1353TTA (5667A>T), GGA1354TGT (5670G>T 5672A>T), GTA1355AAA (5673G>A 5674T>A), GCA1356ACA (5676G>A), AAT1357CAT (5679A>C), GAA1359AAC (5685G>A 5687A>C), CTT1361TTA (5691C>T 5693T>A), GGC1362GGA (5696C>A), CTA1363GTA (5697C>G), GAA1364ACT (5700G>A 5701A>C 5702A>T), ATA1365TTA (5703A>T), GAA1367GAT (5711A>T), GGA1368GGG (5714A>G), AAA1369AAG (5717A>G), GTT1370ATA (5718G>A 5720T>A), CAA1371AAA (5721C>A), CTC1372CTA (5726C>A), TTA1377GCT (5739T>G 5740T>C 5741A>T), ATA1380GTA (5748A>G), GAA1382GAT (5756A>T), TTT1383ATG (5757T>A 5759T>G), CCT1384CCA (5762T>A), GAA1385del (5763_5765delGAA), GAC1386GAT (5768C>T), CAA1387AAA (5769C>A), GAA1389GAT (5777A>T), ACA1390AAA (5779C>A), TTA1391ACA (5781T>A 5782T>C), AAG1392AAA (5786G>A), GGT1393GAT (5788G>A), CTA1394TTA (5790C>T), CAG1395CAA (5795G>A), AAG1396AAA (5798G>A), GGG1399GGA (5807G>A), ATA1400TTA (5808A>T), TTA1401GTG (5811T>G 5813A>G), AAC1402AAT (5816C>T), GCC1404GCT (5822C>T), AAT1406AAC (5828T>C), TAT1407TTT (5830A>T), ATC1408ATA (5834C>A), CCA1409CAA (5836C>A), TTG1411TTA (5843G>A), AGT1412GGA (5844A>G 5846T>A), TAT1414ATT (5850T>A 5851A>T), ACT1415GCA (5853A>G 5855T>A), AGA1416GGA (5856A>G), ATA1417CCA (5859A>C 5860T>C), TTT1418TTA (5864T>A), AAT1420GCA (5868A>G 5869A>C 5870T>A), AAG1421AAA (5873G>A), TGT1422ATA (5874T>A 5875G>T 5876T>A), TCC1423GGA (5877T>G 5878C>G 5879C>A), AGC1424AGT (5882C>T), GAA1427CAA (5889G>C), AGA1428AAA (5893G>A), AAA1429CAC (5895A>C 5897A>C) | | | | | | | | |

##### Proteins

| | Begin | End | Coverage | Score | Concordance | Matches | Identities | I/D/M/F* | Stop Codons |
|---|---|---|---|---|---|---|---|---|---|
| ORF3 (YP_009553219.1) | 1185 | 1431 | 15.5% | 914 | 53.3% | 246 (99.6%) | 138 (55.9%) | 0/1/1/1 | 0 |
| Protein mutations: | P1190R (5178C>A 5179C>G 5180T>A), Q1192T (5184C>A 5185A>C), T1197S (5199A>T), I1201M (5213T>G), N1203K (5219C>A), S1206N (5226T>A 5227C>A 5228T>C), Q1208I (5232C>A 5233A>T), S1213A (5247A>G 5248G>C 5249T>C), Y1217I (5259T>A 5260A>T 5261C>A), Y1227R (5289T>A 5290A>G 5291C>A), K1228M (5293A>T 5294A>G), Q1230A (5298C>G 5299A>C 5300A>T), T1232K (5305C>A), S1235D (5313A>G 5314G>A 5315T>C), I1236K (5317T>A 5318T>A), D1237N (5319G>A 5321C>T), Y1238E (5322T>G 5324T>A), L1241N (5331C>A 5332T>A), K1242R (5335A>G), K1244Q (5340A>C), D1245G (5344A>G 5345C>A), I1247R (5350T>G 5351T>A), V1248I (5352G>A 5354C>T), Y1249F (5356A>T), L1254C (5370C>T 5371T>G 5372A>T), F1258Y (5383T>A 5384C>T), H1259W (5385C>T 5386A>G 5387C>G), I1261V (5391A>G 5393T>A), M1262K (5395T>A 5396G>A), D1264H (5400G>C 5402T>C), Q1266D (5406C>G 5408G>T), A1272T (5424G>A), V1274T (5430G>A 5431T>C 5432C>T), Q1277E (5439C>G 5441G>A), I1283L (5457A>T), V1284A (5461T>C), P1286S (5466C>T), N1291T (5482A>C 5483T>A), S1294P (5490T>C 5492A>T), V1295I (5493G>A), K1305G (5523A>G 5524A>G), K1306E (5526A>G), S1308K (5532T>A 5533C>A), E1309K (5535G>A), F1310I (5538T>A), C1312L (5544T>C 5545G>T 5546T>A), I1315V (5553A>G), I1320V (5568A>G), E1323R (5577G>A 5578A>G), S1324D (5580A>G 5581G>A), I1325M (5585A>G), Q1327E (5589C>G), V1329L (5595G>C), Q1330E (5598C>G), L1333Q (5607T>C 5608T>A), Q1334T (5610C>A 5611A>C 5612G>A), F1335I (5613T>A 5615C>T), Q1337R (5619C>A 5620A>G 5621A>G), V1338L (5622G>T), C1339F (5626G>T 5627T>C), K1340V (5628A>G 5629A>T 5630G>A), E1341S (5631G>A 5632A>G 5633A>T), E1342K (5634G>A), L1344I (5640C>A 5642T>A), L1346I (5646T>A 5648G>A), S1347T (5649T>A 5651A>C), T1349K (5656C>A 5657A>G), L1351M (5661T>A), K1352E (5664A>G), I1353L (5667A>T), G1354C (5670G>T 5672A>T), V1355K (5673G>A 5674T>A), A1356T (5676G>A), N1357H (5679A>C), E1359N (5685G>A 5687A>C), L1363V (5697C>G), E1364T (5700G>A 5701A>C 5702A>T), I1365L (5703A>T), E1367D (5711A>T), V1370I (5718G>A 5720T>A), Q1371K (5721C>A), L1377A (5739T>G 5740T>C 5741A>T), I1380V (5748A>G), E1382D (5756A>T), F1383M (5757T>A 5759T>G), E1385del (5763_5765delGAA), Q1387K (5769C>A), E1389D (5777A>T), T1390K (5779C>A), L1391T (5781T>A 5782T>C), G1393D (5788G>A), I1400L (5808A>T), L1401V (5811T>G 5813A>G), Y1407F (5830A>T), P1409Q (5836C>A), S1412G (5844A>G 5846T>A), Y1414I (5850T>A 5851A>T), T1415A (5853A>G 5855T>A), R1416G (5856A>G), I1417P (5859A>C 5860T>C), F1418L (5864T>A), N1420A (5868A>G 5869A>C 5870T>A), C1422I (5874T>A 5875G>T 5876T>A), S1423G (5877T>G 5878C>G 5879C>A), E1427Q (5889G>C), R1428K (5893G>A), K1429H (5895A>C 5897A>C) | | | | | | | | |

| | Begin | End | Coverage | Score | Concordance | Matches | Identities | I/D/M/F* | Stop Codons |
|---|---|---|---|---|---|---|---|---|---|
| **NT** | **5162** | **5901** | **9.1%** | **394** | **27.2%** | **736 (99.5%)** | **472 (63.8%)** | **0/4** | |
| Codon mutations: | AAG1186AAA (5168G>A), GTT1187GTA (5171T>A), ATC1188ATA (5174C>A), CCT1190AGA (5178C>A 5179C>G 5180T>A), TCC1191TCA (5183C>A), CAA1192ACA (5184C>A 5185A>C), AGT1193TCT (5187A>T 5188G>C), CGC1194AGA (5190C>A 5192C>A), CAT1195CAC (5195T>C), AGG1196AGA (5198G>A), ACA1197TCA (5199A>T), GCT1199GCA (5207T>A), ATT1201ATG (5213T>G), GTC1202GTA (5216C>A), AAC1203AAA (5219C>A), AAA1204-AC (5220delA 5222A>C), TCT1206AAC (5226T>A 5227C>A 5228T>C), CAA1208ATA (5232C>A 5233A>T), GTA1209GTC (5237A>C), GGT1211GGA (5243T>A), AGT1213GCC (5247A>G 5248G>C 5249T>C), CGT1214AGA (5250C>A 5252T>A), TAC1217ATA (5259T>A 5260A>T 5261C>A), CGA1221AGA (5271C>A), CTA1222CTC (5276A>C), ACA1226ACT (5288A>T), TAC1227AGA (5289T>A 5290A>G 5291C>A), AAA1228ATG (5293A>T 5294A>G), GAC1229GAT (5297C>T), CAA1230GCT (5298C>G 5299A>C 5300A>T), TAC1231TAT (5303C>T), ACA1232AAA (5305C>A), CTC1233TTA (5307C>T 5309C>A), AGT1235GAC (5313A>G 5314G>A 5315T>C), ATT1236AAA (5317T>A 5318T>A), GAC1237AAT (5319G>A 5321C>T), TAT1238GAA (5322T>G 5324T>A), CTC1239CTA (5327C>A), CTT1241AAT (5331C>A 5332T>A), AAA1242AGA (5335A>G), ATT1243ATA (5339T>A), AAA1244CAA (5340A>C), GAC1245GGA (5344A>G 5345C>A), AAG1246AAA (5348G>A), ATT1247AGA (5350T>G 5351T>A), GTC1248ATT (5352G>A 5354C>T), TAC1249TTC (5356A>T), AGC1250AGT (5360C>T), GAT1253GAC (5369T>C), CTA1254TGT (5370C>T 5371T>G 5372A>T), AAG1255AAA (5375G>A), GGA1257GGG (5381A>G), TTC1258TAT (5383T>A 5384C>T), CAC1259TGG (5385C>T 5386A>G 5387C>G), CAG1260CAA (5390G>A), ATT1261GTA (5391A>G 5393T>A), ATG1262AAA (5395T>A 5396G>A), GAT1264CAC (5400G>C 5402T>C), CCA1265CCT (5405A>T), CAG1266GAT (5406C>G 5408G>T), GCA1272ACA (5424G>A), GTC1274ACT (5430G>A 5431T>C 5432C>T), TGC1275TGT (5435C>T), CAG1277GAA (5439C>G 5441G>A), GGT1278GGA (5444T>A), ATA1283TTA (5457A>T), GTA1284GCA (5461T>C), CCA1286TCA (5466C>T), TTC1287TTT (5471C>T), CTC1289TTG (5475C>T 5477C>G), AAA1290AAG (5480A>G), AAT1291ACA (5482A>C 5483T>A), CCT1293CCA (5489T>A), TCA1294CCT (5490T>C 5492A>T), GTA1295ATA (5493G>A), CGA1298AGA (5502C>A), AAA1299AAG (5507A>G), GAT1301GAC (5513T>C), ATA1303ATT (5519A>T), AAA1305GGA (5523A>G 5524A>G), AAA1306GAA (5526A>G), TCA1308AAA (5532T>A 5533C>A), GAA1309AAA (5535G>A), TTT1310ATT (5538T>A), GTC1311GTA (5543C>A), TGT1312CTA (5544T>C 5545G>T 5546T>A), GTT1313GTC (5549T>C), ATA1315GTA (5553A>G), GAC1316GAT (5558C>T), GAT1317GAC (5561T>C), ATT1318ATC (5564T>C), TTG1319CTA (5565T>C 5567G>A), ATA1320GTA (5568A>G), TTT1321TTC (5573T>C), TCT1322AGT (5574T>A 5575C>G), GAA1323AGA (5577G>A 5578A>G), AGT1324GAT (5580A>G 5581G>A), ATA1325ATG (5585A>G), CAA1327GAA (5589C>G), GTA1329CTA (5595G>C), CAA1330GAA (5598C>G), TTA1333CAA (5607T>C 5608T>A), CAG1334ACA (5610C>A 5611A>C 5612G>A), TTC1335ATT (5613T>A 5615C>T), TTC1336TTT (5618C>T), CAA1337AGG (5619C>A 5620A>G 5621A>G), GTA1338TTA (5622G>T), TGT1339TTC (5626G>T 5627T>C), AAG1340GTA (5628A>G 5629A>T 5630G>A), GAA1341AGT (5631G>A 5632A>G 5633A>T), GAA1342AAA (5634G>A), GGG1343GGA (5639G>A), CTT1344ATA (5640C>A 5642T>A), ATT1345ATA (5645T>A), TTG1346ATA (5646T>A 5648G>A), TCA1347ACC (5649T>A 5651A>C), ACA1349AAG (5656C>A 5657A>G), TTG1351ATG (5661T>A), AAA1352GAA (5664A>G), ATA1353TTA (5667A>T), GGA1354TGT (5670G>T 5672A>T), GTA1355AAA (5673G>A 5674T>A), GCA1356ACA (5676G>A), AAT1357CAT (5679A>C), GAA1359AAC (5685G>A 5687A>C), CTT1361TTA (5691C>T 5693T>A), GGC1362GGA (5696C>A), CTA1363GTA (5697C>G), GAA1364ACT (5700G>A 5701A>C 5702A>T), ATA1365TTA (5703A>T), GAA1367GAT (5711A>T), GGA1368GGG (5714A>G), AAA1369AAG (5717A>G), GTT1370ATA (5718G>A 5720T>A), CAA1371AAA (5721C>A), CTC1372CTA (5726C>A), TTA1377GCT (5739T>G 5740T>C 5741A>T), ATA1380GTA (5748A>G), GAA1382GAT (5756A>T), TTT1383ATG (5757T>A 5759T>G), CCT1384CCA (5762T>A), GAA1385del (5763_5765delGAA), GAC1386GAT (5768C>T), CAA1387AAA (5769C>A), GAA1389GAT (5777A>T), ACA1390AAA (5779C>A), TTA1391ACA (5781T>A 5782T>C), AAG1392AAA (5786G>A), GGT1393GAT (5788G>A), CTA1394TTA (5790C>T), CAG1395CAA (5795G>A), AAG1396AAA (5798G>A), GGG1399GGA (5807G>A), ATA1400TTA (5808A>T), TTA1401GTG (5811T>G 5813A>G), AAC1402AAT (5816C>T), GCC1404GCT (5822C>T), AAT1406AAC (5828T>C), TAT1407TTT (5830A>T), ATC1408ATA (5834C>A), CCA1409CAA (5836C>A), TTG1411TTA (5843G>A), AGT1412GGA (5844A>G 5846T>A), TAT1414ATT (5850T>A 5851A>T), ACT1415GCA (5853A>G 5855T>A), AGA1416GGA (5856A>G), ATA1417CCA (5859A>C 5860T>C), TTT1418TTA (5864T>A), AAT1420GCA (5868A>G 5869A>C 5870T>A), AAG1421AAA (5873G>A), TGT1422ATA (5874T>A 5875G>T 5876T>A), TCC1423GGA (5877T>G 5878C>G 5879C>A), AGC1424AGT (5882C>T), GAA1427CAA (5889G>C), AGA1428AAA (5893G>A), AAA1429CAC (5895A>C 5897A>C) | | | | | | | | |

*: Inserts / Deletes / Misaligned / Frameshifts

#### Analysis details

This analysis was performed with panviral2.64

### NGS Details (UN9): Badnavirus venatheobromae

#### Assembly

| | |
|---|---|
| **Coverage Length** | 258 (1 contig(s)) |
| **Depth Of Coverage** | 11.2 |
| **Number Of Reads** | 25 |
| **Reads Per Million** | 0.57 rpm (after QC) |
| **Ambiguities** | 0 |
| **Assembly Method** | de novo + reference guided assembly |
| **Consensus Caller** | Bcf Tools |

#### Coverage Map

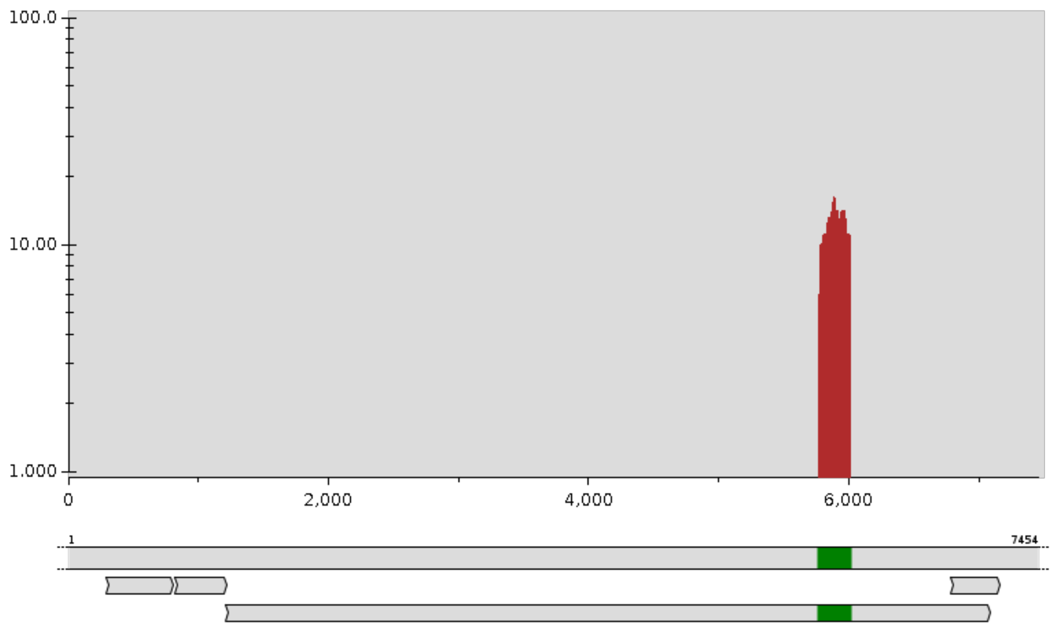

#### Assignment

| | |
|---|---|
| **Type** | Badnavirus venatheobromae (Taxonomy ID: 3047715) |
| **Reference Genome** | NC_033739.1 |
| **NT Identity (%)** | 50.5747 |
| **AA Identity (%)** | 44.8276 |
| **Number Of Stop Codons** | 0 |
| **Number Of CDS** | 4 |

#### Alignment

| | |
|---|---|
| **Alignment Score** | 0.0 (NT) + 287.0 (AA) = 287.0 |
| **Concordance (%)** | 25.4658 |

| Alignment Method | Global, seeded, nucleotide + amino acids (AGA) |
|---|---|

#### Genome Region

Sequence starts at position 5764 and ends at position 6021 relative to NC_033739.1 reference sequence.

#### Alignment Detailed Statistics

| | Begin | End | Coverage | Score | Concordance | Matches | Identities | I/D/M/F* | Stop Codons |
|---|---|---|---|---|---|---|---|---|---|
| **NT** | **5764** | **6021** | **3.5%** | **0** | **0%** | **258 (98.9%)** | **132 (50.6%)** | **3/0** | |
| Mutations: | 5765T>C, 5768A>G, 5770G>T, 5771G>C, 5772C>A, 5773T>A, 5774A>G, 5777C>G, 5786C>T, 5789A>C, 5790T>C, 5792A>T, 5794A>C, 5795G>A, 5807C>T, 5809T>C, 5813T>C, 5817A>G, 5818G>C, 5819A>C, 5820A>T, 5821A>T, 5822G>C, 5826G>A, 5828C>T, 5829A>C, 5831C>G, 5832T>G, 5833G>T, 5834C>T, 5837T>C, 5838A>G, 5841G>C, 5842G>C, 5843A>T, 5843_5844insTTT, 5844A>C, 5845C>T, 5846A>G, 5847G>A, 5848A>G, 5850G>A, 5852A>G, 5856A>G, 5858A>T, 5859G>T, 5860C>T, 5861A>G, 5864T>A, 5866A>T, 5868A>T, 5879C>A, 5882A>T, 5885C>T, 5887T>A, 5889T>A, 5890C>G, 5891A>T, 5892C>T, 5893A>T, 5897T>A, 5899A>T, 5900G>A, 5901G>A, 5902A>G, 5903A>T, 5906A>T, 5912C>T, 5914G>A, 5915T>G, 5918T>C, 5919C>T, 5922C>A, 5923A>G, 5924A>G, 5925A>G, 5926G>T, 5927G>T, 5928T>G, 5933C>T, 5934C>G, 5938T>C, 5939A>T, 5940T>C, 5941G>T, 5943C>A, 5946A>T, 5947A>G, 5948G>T, 5952G>T, 5953G>C, 5954G>A, 5955T>C, 5957A>C, 5958G>T, 5959T>A, 5960C>T, 5961C>G, 5962T>C, 5963G>T, 5965G>A, 5966T>G, 5968A>G, 5970G>T, 5971A>C, 5972T>A, 5976A>T, 5977T>G, 5978G>T, 5979G>T, 5981A>C, 5982A>T, 5984A>T, 5986C>G, 5987A>C, 5988G>C, 5989T>A, 5990C>G, 5997A>G, 6004T>A, 6006C>T, 6008T>A, 6011A>T, 6012G>C, 6013C>A, 6017T>A, 6021G>A | | | | | | | | |

##### CDS

| | Begin | End | Coverage | Score | Concordance | Matches | Identities | I/D/M/F* | Stop Codons |
|---|---|---|---|---|---|---|---|---|---|
| B1U04_gp3 | 1519 | 1604 | 4.4% | 287 | 45.6% | 86 (98.9%) | 39 (44.8%) | 1/0/0/0 | 0 |
| Protein mutations: | W1520F (5770G>T 5771G>C), L1521K (5772C>A 5773T>A 5774A>G), K1528T (5794A>C 5795G>A), I1533T (5809T>C), R1536A (5817A>G 5818G>C 5819A>C), K1537F (5820A>T 5821A>T 5822G>C), D1539N (5826G>A 5828C>T), N1540Q (5829A>C 5831C>G), C1541V (5832T>G 5833G>T 5834C>T), K1543E (5838A>G), G1544P (5841G>C 5842G>C 5843A>T), G1544_T1545insF (5843_5844insTTT), T1545L (5844A>C 5845C>T 5846A>G), E1546R (5847G>A 5848A>G), E1547K (5850G>A 5852A>G), I1549V (5856A>G 5858A>T), A1550L (5859G>T 5860C>T 5861A>G), Y1552F (5866A>T), I1553F (5868A>T), F1559Y (5887T>A), Q1561L (5892C>T 5893A>T), E1563V (5899A>T 5900G>A), E1564S (5901G>A 5902A>G 5903A>T), E1565D (5906A>T), R1568Q (5914G>A 5915T>G), Q1571R (5922C>A 5923A>G 5924A>G), R1572V (5925A>G 5926G>T 5927G>T), F1573V (5928T>G), Q1575E (5934C>G), I1576T (5938T>C 5939A>T), C1577L (5940T>C 5941G>T), Q1578K (5943C>A), K1579C (5946A>T 5947A>G 5948G>T), G1581S (5952G>T 5953G>C 5954G>A), V1583Y (5958G>T 5959T>A 5960C>T), L1584A (5961C>G 5962T>C 5963G>T), S1585K (5965G>A 5966T>G), K1586R (5968A>G), D1587S (5970G>T 5971A>C 5972T>A), M1589C (5976A>T 5977T>G 5978G>T), A1590S (5979G>T 5981A>C), I1591F (5982A>T 5984A>T), A1592G (5986C>G 5987A>C), V1593Q (5988G>C 5989T>A 5990C>G), I1596V (5997A>G), F1598Y (6004T>A), A1601H (6012G>C 6013C>A) | | | | | | | | |
| Codon mutations: | TAT1518.AC (5765T>C), GAA1519GAG (5768A>G), TGG1520TTC (5770G>T 5771G>C), CTA1521AAG (5772C>A 5773T>A 5774A>G), GTC1522GTG (5777C>G), TTC1525TTT (5786C>T), GGA1526GGC (5789A>C), TTA1527CTT (5790T>C 5792A>T), AAG1528ACA (5794A>C 5795G>A), GCC1532GCT (5807C>T), ATT1533ACT (5809T>C), TTT1534TTC (5813T>C), AGA1536GCC (5817A>G 5818G>C 5819A>C), AAG1537TTC (5820A>T 5821A>T 5822G>C), GAC1539AAT (5826G>A 5828C>T), AAC1540CAG (5829A>C 5831C>G), TGC1541GTT (5832T>G 5833G>T 5834C>T), TTT1542TTC (5837T>C), AAA1543GAA (5838A>G), GGA1544CCT (5841G>C 5842G>C 5843A>T), GGA1544_ACA1545insTTT (5843_5844insTTT), ACA1545CTG (5844A>C 5845C>T 5846A>G), GAA1546AGA (5847G>A 5848A>G), GAA1547AAG (5850G>A 5852A>G), ATA1549GTT (5856A>G 5858A>T), GCA1550TTG (5859G>T 5860C>T 5861A>G), GTT1551GTA (5864T>A), TAT1552TTT (5866A>T), ATT1553TTT (5868A>T), ATC1556ATA (5879C>A), CTA1557CTT (5882A>T), GTC1558GTT (5885C>T), TTC1559TAC (5887T>A), TCA1560AGT (5889T>A 5890C>G 5891A>T), CAG1561TTG (5892C>T 5893A>T), ACT1562ACA (5897T>A), GAG1563GTA (5899A>T 5900G>A), GAA1564AGT (5901G>A 5902A>G 5903A>T), GAA1565GAT (5906A>T), GTC1567GTT (5912C>T), CGT1568CAG (5914G>A 5915T>G), CAT1569CAC (5918T>C), CTA1570TTA (5919C>T), CAA1571AGG (5922C>A 5923A>G 5924A>G), AGG1572GTT (5925A>G 5926G>T 5927G>T), TTC1573GTC (5928T>G), TTC1574TTT (5933C>T), CAG1575GAG (5934C>G), ATA1576ACT (5938T>C 5939A>T), TGC1577CTC (5940T>C 5941G>T), CAG1578AAG (5943C>A), AAG1579TGT (5946A>T 5947A>G 5948G>T), GGG1581TCA (5952G>T 5953G>C 5954G>A), TTA1582CTC (5955T>C 5957A>C), GTC1583TAT (5958G>T 5959T>A 5960C>T), CTG1584GCT (5961C>G 5962T>C 5963G>T), AGT1585AAG (5965G>A 5966T>G), AAG1586AGG (5968A>G), GAT1587TCA (5970G>T 5971A>C 5972T>A), ATG1589TGT (5976A>T 5977T>G 5978G>T), GCA1590TCC (5979G>T 5981A>C), ATA1591TTT (5982A>T 5984A>T), GCA1592GGC (5986C>G 5987A>C), GTC1593CAG (5988G>C 5989T>A 5990C>G), ATT1596GTT (5997A>G), TTC1598TAC (6004T>A), CTT1599TTA (6006C>T 6008T>A), GGA1600GGT (6011A>T), GCC1601CAC (6012G>C 6013C>A), ATT1602ATA (6017T>A), GGT1604A.. (6021G>A) | | | | | | | | |

##### Proteins

| | Begin | End | Coverage | Score | Concordance | Matches | Identities | I/D/M/F* | Stop Codons |
|---|---|---|---|---|---|---|---|---|---|
| polyprotein (YP_009345075.1) | 1519 | 1604 | 4.4% | 287 | 45.6% | 86 (98.9%) | 39 (44.8%) | 1/0/0/0 | 0 |
| Protein mutations: | W1520F (5770G>T 5771G>C), L1521K (5772C>A 5773T>A 5774A>G), K1528T (5794A>C 5795G>A), I1533T (5809T>C), R1536A (5817A>G 5818G>C 5819A>C), K1537F (5820A>T 5821A>T 5822G>C), D1539N (5826G>A 5828C>T), N1540Q (5829A>C 5831C>G), C1541V (5832T>G 5833G>T 5834C>T), K1543E (5838A>G), G1544P (5841G>C 5842G>C 5843A>T), G1544_T1545insF (5843_5844insTTT), T1545L (5844A>C 5845C>T 5846A>G), E1546R (5847G>A 5848A>G), E1547K (5850G>A 5852A>G), I1549V (5856A>G 5858A>T), A1550L (5859G>T 5860C>T 5861A>G), Y1552F (5866A>T), I1553F (5868A>T), F1559Y (5887T>A), Q1561L (5892C>T 5893A>T), E1563V (5899A>T 5900G>A), E1564S (5901G>A 5902A>G 5903A>T), E1565D (5906A>T), R1568Q (5914G>A 5915T>G), Q1571R (5922C>A 5923A>G 5924A>G), R1572V (5925A>G 5926G>T 5927G>T), F1573V (5928T>G), Q1575E (5934C>G), I1576T (5938T>C 5939A>T), C1577L (5940T>C 5941G>T), Q1578K (5943C>A), K1579C (5946A>T 5947A>G 5948G>T), G1581S (5952G>T 5953G>C 5954G>A), V1583Y (5958G>T 5959T>A 5960C>T), L1584A (5961C>G 5962T>C 5963G>T), S1585K (5965G>A 5966T>G), K1586R (5968A>G), D1587S (5970G>T 5971A>C 5972T>A), M1589C (5976A>T 5977T>G 5978G>T), A1590S (5979G>T 5981A>C), I1591F (5982A>T 5984A>T), A1592G (5986C>G 5987A>C), V1593Q (5988G>C 5989T>A 5990C>G), I1596V (5997A>G), F1598Y (6004T>A), A1601H (6012G>C 6013C>A) | | | | | | | | |
| Codon mutations: | TAT1518.AC (5765T>C), GAA1519GAG (5768A>G), TGG1520TTC (5770G>T 5771G>C), CTA1521AAG (5772C>A 5773T>A 5774A>G), GTC1522GTG (5777C>G), TTC1525TTT (5786C>T), GGA1526GGC (5789A>C), TTA1527CTT (5790T>C 5792A>T), AAG1528ACA (5794A>C 5795G>A), GCC1532GCT (5807C>T), ATT1533ACT (5809T>C), TTT1534TTC (5813T>C), AGA1536GCC (5817A>G 5818G>C 5819A>C), AAG1537TTC (5820A>T 5821A>T 5822G>C), GAC1539AAT (5826G>A 5828C>T), AAC1540CAG (5829A>C 5831C>G), TGC1541GTT (5832T>G 5833G>T 5834C>T), TTT1542TTC (5837T>C), AAA1543GAA (5838A>G), GGA1544CCT (5841G>C 5842G>C 5843A>T), GGA1544_ACA1545insTTT (5843_5844insTTT), ACA1545CTG (5844A>C 5845C>T 5846A>G), GAA1546AGA (5847G>A 5848A>G), GAA1547AAG (5850G>A 5852A>G), ATA1549GTT (5856A>G 5858A>T), GCA1550TTG (5859G>T 5860C>T 5861A>G), GTT1551GTA (5864T>A), TAT1552TTT (5866A>T), ATT1553TTT (5868A>T), ATC1556ATA (5879C>A), CTA1557CTT (5882A>T), GTC1558GTT (5885C>T), TTC1559TAC (5887T>A), TCA1560AGT (5889T>A 5890C>G 5891A>T), CAG1561TTG (5892C>T 5893A>T), ACT1562ACA (5897T>A), GAG1563GTA (5899A>T 5900G>A), GAA1564AGT (5901G>A 5902A>G 5903A>T), GAA1565GAT (5906A>T), GTC1567GTT (5912C>T), CGT1568CAG (5914G>A 5915T>G), CAT1569CAC (5918T>C), CTA1570TTA (5919C>T), CAA1571AGG (5922C>A 5923A>G 5924A>G), AGG1572GTT (5925A>G 5926G>T 5927G>T), TTC1573GTC (5928T>G), TTC1574TTT (5933C>T), CAG1575GAG (5934C>G), ATA1576ACT (5938T>C 5939A>T), TGC1577CTC (5940T>C 5941G>T), CAG1578AAG (5943C>A), AAG1579TGT (5946A>T 5947A>G 5948G>T), GGG1581TCA (5952G>T 5953G>C 5954G>A), TTA1582CTC (5955T>C 5957A>C), GTC1583TAT (5958G>T 5959T>A 5960C>T), CTG1584GCT (5961C>G 5962T>C 5963G>T), AGT1585AAG (5965G>A 5966T>G), AAG1586AGG (5968A>G), GAT1587TCA (5970G>T 5971A>C 5972T>A), ATG1589TGT (5976A>T 5977T>G 5978G>T), GCA1590TCC (5979G>T 5981A>C), ATA1591TTT (5982A>T 5984A>T), GCA1592GGC (5986C>G 5987A>C), GTC1593CAG (5988G>C 5989T>A 5990C>G), ATT1596GTT (5997A>G), TTC1598TAC (6004T>A), CTT1599TTA (6006C>T 6008T>A), GGA1600GGT (6011A>T), GCC1601CAC (6012G>C 6013C>A), ATT1602ATA (6017T>A), GGT1604A.. (6021G>A) | | | | | | | | |

*: Inserts / Deletes / Misaligned / Frameshifts

#### Analysis details

This analysis was performed with panviral2.64

### NGS Details (UN9): Epiphyllum badnavirus 1

#### Assembly

| | |
|---|---|
| **Coverage Length** | 258 (1 contig(s)) |
| **Depth Of Coverage** | 10.2 |
| **Number Of Reads** | 25 |
| **Reads Per Million** | 0.57 rpm (after QC) |
| **Ambiguities** | 0 |
| **Assembly Method** | de novo + reference guided assembly |
| **Consensus Caller** | Bcf Tools |

#### Coverage Map

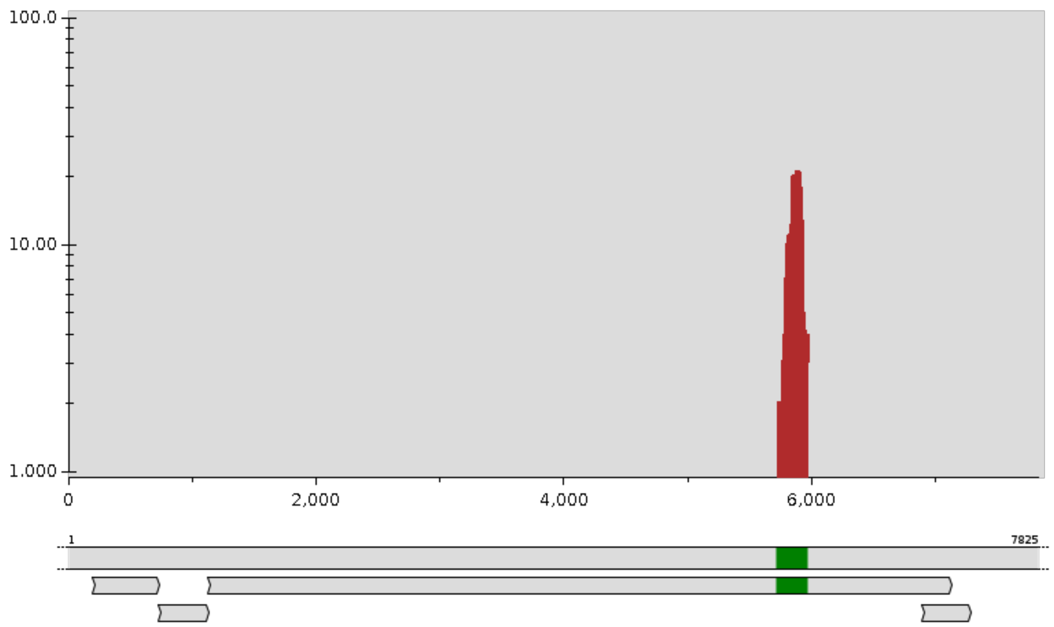

#### Assignment

| | |
|---|---|
| **Type** | Epiphyllum badnavirus 1 (Taxonomy ID: 2518008) |
| **Reference Genome** | NC_076247.1 |
| **NT Identity (%)** | 56.9767 |
| **AA Identity (%)** | 43.0233 |
| **Number Of Stop Codons** | 0 |
| **Number Of CDS** | 4 |

#### Alignment

| | |
|---|---|
| **Alignment Score** | 72.0 (NT) + 240.0 (AA) = 312.0 |
| **Concordance (%)** | 27.932 |

| Alignment Method | Global, seeded, nucleotide + amino acids (AGA) |
|---|---|

#### Genome Region

Sequence starts at position 5717 and ends at position 5974 relative to NC_076247.1 reference sequence.

#### Alignment Detailed Statistics

| | Begin | End | Coverage | Score | Concordance | Matches | Identities | I/D/M/F* | Stop Codons |
|---|---|---|---|---|---|---|---|---|---|
| **NT** | **5717** | **5974** | **3.3%** | **72** | **14.0%** | **258 (100%)** | **147 (57.0%)** | **0/0** | |
| Mutations: | 5719G>A, 5725A>G, 5727A>T, 5730A>C, 5732A>C, 5734G>T, 5735A>T, 5737A>C, 5739A>C, 5743G>T, 5744T>G, 5746T>A, 5748C>T, 5749A>G, 5751C>T, 5754C>T, 5755A>T, 5756A>G, 5757A>G, 5758A>C, 5759C>A, 5760C>G, 5766C>T, 5767G>A, 5769C>G, 5770A>G, 5771A>T, 5772C>G, 5773A>C, 5776T>G, 5778C>G, 5781A>G, 5782G>A, 5784T>C, 5786A>G, 5787G>T, 5789A>G, 5790C>T, 5791T>C, 5794C>A, 5796A>G, 5800G>C, 5802C>T, 5806A>G, 5808C>T, 5809A>G, 5810C>A, 5811A>C, 5812A>C, 5814C>G, 5815A>T, 5817A>T, 5818C>G, 5820G>T, 5821A>C, 5824G>C, 5827T>C, 5828G>A, 5829T>G, 5830A>G, 5837G>C, 5841C>G, 5844C>T, 5847C>T, 5851T>A, 5853C>T, 5856T>C, 5857C>T, 5861A>G, 5863A>T, 5864G>C, 5868A>T, 5870T>A, 5871C>T, 5878A>T, 5880T>G, 5882T>G, 5886G>C, 5887G>A, 5888A>G, 5889C>G, 5891A>T, 5895A>T, 5896T>G, 5897C>A, 5898C>T, 5901C>A, 5905T>A, 5906G>A, 5910C>T, 5916C>T, 5918T>G, 5924C>G, 5926G>T, 5931A>C, 5933T>A, 5934T>C, 5936A>T, 5937C>T, 5940A>G, 5942G>T, 5943G>T, 5949T>A, 5953C>T, 5955A>T, 5958C>T, 5961A>G, 5962T>C, 5964G>T, 5966A>C, 5967G>T | | | | | | | | |
| **CDS** | | | | | | | | | |
| QKM20_gp3 | 1531 | 1616 | 4.3% | 240 | 38.8% | 86 (100%) | 37 (43.0%) | 0/0/0/0 | 0 |
| Protein mutations: | E1531K (5719G>A), K1533D (5725A>G 5727A>T), K1535T (5732A>C), E1536L (5734G>T 5735A>T), V1539C (5743G>T 5744T>G), F1540I (5746T>A 5748C>T), N1541D (5749A>G 5751C>T), K1543W (5755A>T 5756A>G 5757A>G), T1544Q (5758A>C 5759C>A 5760C>G), D1547K (5767G>A 5769C>G), N1548V (5770A>G 5771A>T 5772C>G), T1549P (5773A>C), F1550V (5776T>G 5778C>G), D1552N (5782G>A 5784T>C), Q1553R (5786A>G 5787G>T), Y1554C (5789A>G 5790C>T), S1555P (5791T>C), L1556M (5794C>A 5796A>G), G1558R (5800G>C 5802C>T), N1560D (5806A>G 5808C>T), T1561D (5809A>G 5810C>A 5811A>C), I1562L (5812A>C 5814C>G), I1563F (5815A>T 5817A>T), Q1564D (5818C>G 5820G>T), K1565Q (5821A>C), V1566L (5824G>C), C1567Q (5827T>C 5828G>A 5829T>G), N1568D (5830A>G), S1570T (5837G>C), F1575I (5851T>A 5853C>T), K1578R (5861A>G), F1581Y (5870T>A 5871C>T), I1584L (5878A>T 5880T>G), M1585R (5882T>G), M1586I (5886G>C), D1587R (5887G>A 5888A>G 5889C>G), E1588V (5891A>T), E1589D (5895A>T), S1590D (5896T>G 5897C>A 5898C>T), W1593K (5905T>A 5906G>A), L1597R (5918T>G), P1599R (5924C>G), D1600Y (5926G>T), L1602H (5933T>A 5934T>C), Y1603F (5936A>T 5937C>T), W1605F (5942G>T 5943G>T), P1609S (5953C>T 5955A>T), K1613T (5966A>C 5967G>T) | | | | | | | | |
| Codon mutations: | GAG1531AAG (5719G>A), AAA1533GAT (5725A>G 5727A>T), GGA1534GGC (5730A>C), AAG1535ACG (5732A>C), GAG1536TTG (5734G>T 5735A>T), AGA1537CGC (5737A>C 5739A>C), GTC1539TGC (5743G>T 5744T>G), TTC1540ATT (5746T>A 5748C>T), AAC1541GAT (5749A>G 5751C>T), TAC1542TAT (5754C>T), AAA1543TGG (5755A>T 5756A>G 5757A>G), ACC1544CAG (5758A>C 5759C>A 5760C>G), AAC1546AAT (5766C>T), GAC1547AAG (5767G>A 5769C>G), AAC1548GTG (5770A>G 5771A>T 5772C>G), ACG1549CCG (5773A>C), TTC1550GTG (5776T>G 5778C>G), AAA1551AAG (5781A>G), GAT1552AAC (5782G>A 5784T>C), CAG1553CGT (5786A>G 5787G>T), TAC1554TGT (5789A>G 5790C>T), TCT1555CCT (5791T>C), CTA1556ATG (5794C>A 5796A>G), GGC1558CGT (5800G>C 5802C>T), AAC1560GAT (5806A>G 5808C>T), ACA1561GAC (5809A>G 5810C>A 5811A>C), ATC1562CTG (5812A>C 5814C>G), ATA1563TTT (5815A>T 5817A>T), CAG1564GAT (5818C>G 5820G>T), AAG1565CAG (5821A>C), GTC1566CTC (5824G>C), TGT1567CAG (5827T>C 5828G>A 5829T>G), AAT1568GAT (5830A>G), AGT1570ACT (5837G>C), GTC1571GTG (5841C>G), TTC1572TTT (5844C>T), TCC1573TCT (5847C>T), TTC1575ATT (5851T>A 5853C>T), GAT1576GAC (5856T>C), CTG1577TTG (5857C>T), AAG1578AGG (5861A>G), AGC1579TCC (5863A>T 5864G>C), GGA1580GGT (5868A>T), TTC1581TAT (5870T>A 5871C>T), ATT1584TTG (5878A>T 5880T>G), ATG1585AGG (5882T>G), ATG1586ATC (5886G>C), GAC1587AGG (5887G>A 5888A>G 5889C>G), GAA1588GTA (5891A>T), GAA1589GAT (5895A>T), TCC1590GAT (5896T>G 5897C>A 5898C>T), ATC1591ATA (5901C>A), TGG1593AAG (5905T>A 5906G>A), ACC1594ACT (5910C>T), TTC1596TTT (5916C>T), CTG1597CGG (5918T>G), CCA1599CGA (5924C>G), GAT1600TAT (5926G>T), GGA1601GGC (5931A>C), CTT1602CAC (5933T>A 5934T>C), TAC1603TTT (5936A>T 5937C>T), GAA1604GAG (5940A>G), TGG1605TTT (5942G>T 5943G>T), GTT1607GTA (5949T>A), CCA1609TCT (5953C>T 5955A>T), TTC1610TTT (5958C>T), GGA1611GGG (5961A>G), TTG1612CTT (5962T>C 5964G>T), AAG1613ACT (5966A>C 5967G>T) | | | | | | | | |
| **Proteins** | | | | | | | | | |
| polyprotein (YP_010797894.1) | 1531 | 1616 | 4.3% | 240 | 38.8% | 86 (100%) | 37 (43.0%) | 0/0/0/0 | 0 |
| Protein mutations: | E1531K (5719G>A), K1533D (5725A>G 5727A>T), K1535T (5732A>C), E1536L (5734G>T 5735A>T), V1539C (5743G>T 5744T>G), F1540I (5746T>A 5748C>T), N1541D (5749A>G 5751C>T), K1543W (5755A>T 5756A>G 5757A>G), T1544Q (5758A>C 5759C>A 5760C>G), D1547K (5767G>A 5769C>G), N1548V (5770A>G 5771A>T 5772C>G), T1549P (5773A>C), F1550V (5776T>G 5778C>G), D1552N (5782G>A 5784T>C), Q1553R (5786A>G 5787G>T), Y1554C (5789A>G 5790C>T), S1555P (5791T>C), L1556M (5794C>A 5796A>G), G1558R (5800G>C 5802C>T), N1560D (5806A>G 5808C>T), T1561D (5809A>G 5810C>A 5811A>C), I1562L (5812A>C 5814C>G), I1563F (5815A>T 5817A>T), Q1564D (5818C>G 5820G>T), K1565Q (5821A>C), V1566L (5824G>C), C1567Q (5827T>C 5828G>A 5829T>G), N1568D (5830A>G), S1570T (5837G>C), F1575I (5851T>A 5853C>T), K1578R (5861A>G), F1581Y (5870T>A 5871C>T), I1584L (5878A>T 5880T>G), M1585R (5882T>G), M1586I (5886G>C), D1587R (5887G>A 5888A>G 5889C>G), E1588V (5891A>T), E1589D (5895A>T), S1590D (5896T>G 5897C>A 5898C>T), W1593K (5905T>A 5906G>A), L1597R (5918T>G), P1599R (5924C>G), D1600Y (5926G>T), L1602H (5933T>A 5934T>C), Y1603F (5936A>T 5937C>T), W1605F (5942G>T 5943G>T), P1609S (5953C>T 5955A>T), K1613T (5966A>C 5967G>T) | | | | | | | | |
| Codon mutations: | GAG1531AAG (5719G>A), AAA1533GAT (5725A>G 5727A>T), GGA1534GGC (5730A>C), AAG1535ACG (5732A>C), GAG1536TTG (5734G>T 5735A>T), AGA1537CGC (5737A>C 5739A>C), GTC1539TGC (5743G>T 5744T>G), TTC1540ATT (5746T>A 5748C>T), AAC1541GAT (5749A>G 5751C>T), TAC1542TAT (5754C>T), AAA1543TGG (5755A>T 5756A>G 5757A>G), ACC1544CAG (5758A>C 5759C>A 5760C>G), AAC1546AAT (5766C>T), GAC1547AAG (5767G>A 5769C>G), AAC1548GTG (5770A>G 5771A>T 5772C>G), ACG1549CCG (5773A>C), TTC1550GTG (5776T>G 5778C>G), AAA1551AAG (5781A>G), GAT1552AAC (5782G>A 5784T>C), CAG1553CGT (5786A>G 5787G>T), TAC1554TGT (5789A>G 5790C>T), TCT1555CCT (5791T>C), CTA1556ATG (5794C>A 5796A>G), GGC1558CGT (5800G>C 5802C>T), AAC1560GAT (5806A>G 5808C>T), ACA1561GAC (5809A>G 5810C>A 5811A>C), ATC1562CTG (5812A>C 5814C>G), ATA1563TTT (5815A>T 5817A>T), CAG1564GAT (5818C>G 5820G>T), AAG1565CAG (5821A>C), GTC1566CTC (5824G>C), TGT1567CAG (5827T>C 5828G>A 5829T>G), AAT1568GAT (5830A>G), AGT1570ACT (5837G>C), GTC1571GTG (5841C>G), TTC1572TTT (5844C>T), TCC1573TCT (5847C>T), TTC1575ATT (5851T>A 5853C>T), GAT1576GAC (5856T>C), CTG1577TTG (5857C>T), AAG1578AGG (5861A>G), AGC1579TCC (5863A>T 5864G>C), GGA1580GGT (5868A>T), TTC1581TAT (5870T>A 5871C>T), ATT1584TTG (5878A>T 5880T>G), ATG1585AGG (5882T>G), ATG1586ATC (5886G>C), GAC1587AGG (5887G>A 5888A>G 5889C>G), GAA1588GTA (5891A>T), GAA1589GAT (5895A>T), TCC1590GAT (5896T>G 5897C>A 5898C>T), ATC1591ATA (5901C>A), TGG1593AAG (5905T>A 5906G>A), ACC1594ACT (5910C>T), TTC1596TTT (5916C>T), CTG1597CGG (5918T>G), CCA1599CGA (5924C>G), GAT1600TAT (5926G>T), GGA1601GGC (5931A>C), CTT1602CAC (5933T>A 5934T>C), TAC1603TTT (5936A>T 5937C>T), GAA1604GAG (5940A>G), TGG1605TTT (5942G>T 5943G>T), GTT1607GTA (5949T>A), CCA1609TCT (5953C>T 5955A>T), TTC1610TTT (5958C>T), GGA1611GGG (5961A>G), TTG1612CTT (5962T>C 5964G>T), AAG1613ACT (5966A>C 5967G>T) | | | | | | | | |

*: Inserts / Deletes / Misaligned / Frameshifts

#### Analysis details

This analysis was performed with panviral2.64

### NGS Details (UN9): Badnavirus tessellocastaneae

#### Assembly

| | |
|---|---|
| **Coverage Length** | 415 (1 contig(s)) |
| **Depth Of Coverage** | 6.4 |
| **Number Of Reads** | 22 |
| **Reads Per Million** | 0.50 rpm (after QC) |
| **Ambiguities** | 0 |
| **Assembly Method** | de novo + reference guided assembly |
| **Consensus Caller** | Bcf Tools |

#### Coverage Map

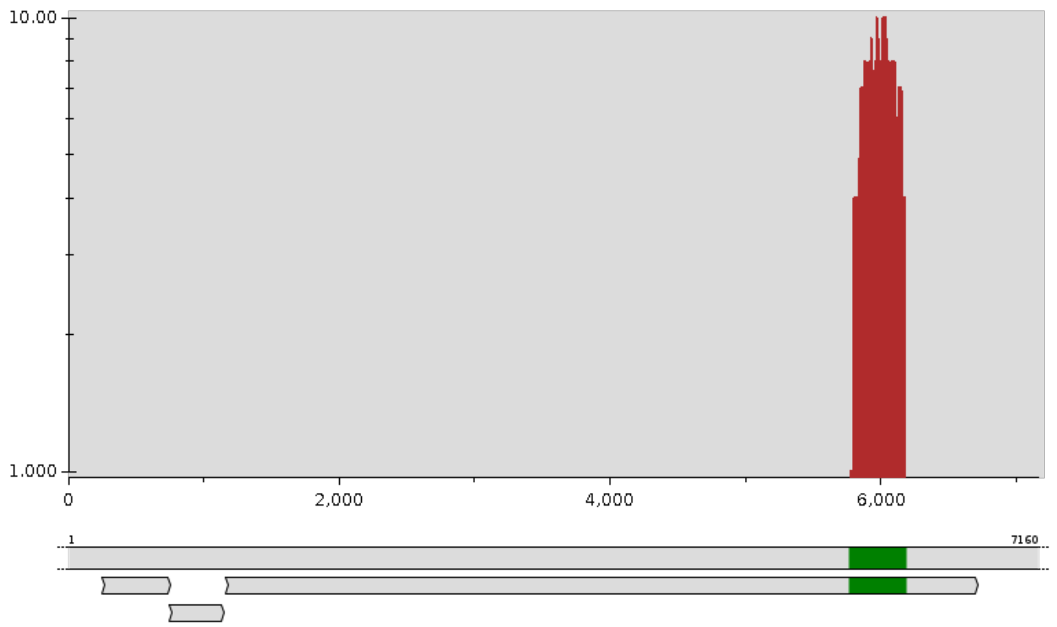

#### Assignment

| | |
|---|---|
| **Type** | Badnavirus tessellocastaneae (Taxonomy ID: 3051987) |
| **Reference Genome** | NC_076885.1 |
| **NT Identity (%)** | 57.177 |
| **AA Identity (%)** | 47.8571 |
| **Number Of Stop Codons** | 0 |
| **Number Of CDS** | 3 |

#### Alignment

| | |
|---|---|
| **Alignment Score** | 105.0 (NT) + 490.0 (AA) = 595.0 |
| **Concordance (%)** | 33.1661 |

| | |
|---|---|
| **Alignment Method** | Global, seeded, nucleotide + amino acids (AGA) |

#### Genome Region

Sequence starts at position 5774 and ends at position 6188 relative to NC_076885.1 reference sequence.

#### Alignment Detailed Statistics

| | Begin | End | Coverage | Score | Concordance | Matches | Identities | I/D/M/F* | Stop Codons |
|---|---|---|---|---|---|---|---|---|---|
| **NT** | **5774** | **6188** | **5.8%** | **105** | **12.9%** | **412 (97.9%)** | **239 (56.8%)** | **6/3** | |
| Mutations: | 5774C>A, 5777C>A, 5779G>C, 5780C>A, 5782C>A, 5789C>T, 5792G>A, 5793A>G, 5794A>C, 5800C>A, 5802C>T, 5807G>T, 5808G>C, 5810A>G, 5811G>A, 5812C>T, 5815C>T, 5816A>C, 5822C>A, 5837T>A, 5840A>C, 5842G>C, 5843G>C, 5854T>A, 5856G>C, 5857A>T, 5862G>T, 5863T>A, 5867T>G, 5868G>C, 5869G>A, 5870G>A, 5871C>G, 5875G>A, 5876T>A, 5877T>A, 5882G>T, 5883G>A, 5884A>C, 5885G>A, 5886C>G, 5887C>T, 5888A>C, 5889A>C, 5890G>T, 5895G>A, 5896T>A, 5902G>A, 5905G>A, 5906G>A, 5908C>T, 5917T>C, 5918G>C, 5919C>A, 5921T>A, 5922C>G, 5923A>T, 5926A>T, 5929G>A, 5931T>A, 5933C>A, 5934C>A, 5935C>A, 5936A>G, 5937C>A, 5938_5939insAAGGGT, 5939C>A, 5940T>A, 5943A>T, 5944A>G, 5947C>T, 5949C>G, 5950C>T, 5953C>A, 5956T>C, 5958C>T, 5959A>C, 5963A>G, 5965C>A, 5967A>T, 5968T>A, 5971G>T, 5972G>A, 5974C>A, 5977G>A, 5978G>T, 5980A>T, 5981A>G, 5982C>G, 5983C>A, 5984A>C, 5986G>A, 5987G>A, 5989A>T, 5990T>A, 5991C>G, 5995G>C, 5999A>T, 6001C>A, 6005T>A, 6006A>T, 6007C>T, 6011G>A, 6013T>C, 6014C>A, 6016G>A, 6018A>G, 6019G>T, 6021C>A, 6026A>G, 6027C>T, 6034G>A, 6040C>T, 6044C>G, 6049A>T, 6052C>A, 6053A>G, 6056G>A, 6057C>A, 6061C>T, 6064C>T, 6065G>C, 6067C>A, 6071C>A, 6072A>T, 6074C>G, 6075T>A, 6076T>C, 6079C>G, 6085G>T, 6086C>A, 6087C>G, 6088C>T, 6089T>A, 6090C>G, 6091A>T, 6093G>A, 6095G>A, 6096T>G, 6097G>A, 6098C>A, 6104A>C, 6106C>A, 6109C>T, 6113T>A, 6114G>T, 6115T>A, 6119T>A, 6120A>C, 6121T>C, 6124A>T, 6125A>T, 6127G>T, 6128G>A, 6129G>T, 6131A>T, 6132C>A, 6134G>C, 6135G>C, 6136G>T, 6137_6139delGTC, 6140G>A, 6142A>T, 6146A>G, 6147A>T, 6148A>T, 6155C>T, 6160C>A, 6167G>A, 6170A>G, 6172C>T, 6175C>T, 6176C>A, 6178G>T, 6187C>T | | | | | | | | |

##### CDS

| | Begin | End | Coverage | Score | Concordance | Matches | Identities | I/D/M/F* | Stop Codons |
|---|---|---|---|---|---|---|---|---|---|
| ORF3 | 1538 | 1676 | 7.5% | 490 | 49.1% | 138 (97.9%) | 67 (47.5%) | 2/1/0/0 | 0 |
| Protein mutations: | Q1538K (5774C>A), Q1539N (5777C>A 5779G>C), L1540I (5780C>A 5782C>A), E1544S (5792G>A 5793A>G 5794A>C), P1547L (5802C>T), G1549S (5807G>T 5808G>C), S1550D (5810A>G 5811G>A 5812C>T), I1552L (5816A>C), L1554I (5822C>A), C1559S (5837T>A), M1560L (5840A>C 5842G>C), E1561Q (5843G>C), G1565A (5856G>C 5857A>T), C1567L (5862G>T 5863T>A), W1569A (5867T>G 5868G>C 5869G>A), A1570R (5870G>A 5871C>G), F1572N (5876T>A 5877T>A), G1574Y (5882G>T 5883G>A 5884A>C), A1575S (5885G>A 5886C>G 5887C>T), K1576P (5888A>C 5889A>C 5890G>T), S1578K (5895G>A 5896T>A), V1582I (5906G>A 5908C>T), A1586Q (5918G>C 5919C>A), F1590Y (5931T>A), P1591K (5933C>A 5934C>A 5935C>A), T1592E (5936A>G 5937C>A), T1592_L1593insKG (5938_5939insAAGGGT), L1593N (5939C>A 5940T>A), K1594M (5943A>T 5944A>G), T1596S (5949C>G 5950C>T), A1599V (5958C>T 5959A>C), I1601V (5963A>G 5965C>A), Y1602L (5967A>T 5968T>A), V1604I (5972G>A 5974C>A), M1605I (5977G>A), E1606Y (5978G>T 5980A>T), T1607G (5981A>G 5982C>G 5983C>A), M1608L (5984A>C 5986G>A), E1609N (5987G>A 5989A>T), L1611F (5995G>C), I1613L (5999A>T 6001C>A), Y1615I (6005T>A 6006A>T 6007C>T), D1617N (6011G>A 6013T>C), Q1618K (6014C>A 6016G>A), K1619S (6018A>G 6019G>T), A1620E (6021C>A), T1622V (6026A>G 6027C>T), Q1628E (6044C>G), I1631V (6053A>G), A1632K (6056G>A 6057C>A), D1635Q (6065G>C 6067C>A), Q1637I (6071C>A 6072A>T), L1638D (6074C>G 6075T>A 6076T>C), H1639Q (6079C>G), K1641N (6085G>T), P1642S (6086C>A 6087C>G 6088C>T), R1644K (6093G>A), V1645R (6095G>A 6096T>G 6097G>A), I1648L (6104A>C 6106C>A), C1651I (6113T>A 6114G>T 6115T>A), Y1653T (6119T>A 6120A>C 6121T>C), T1655S (6125A>T 6127G>T), G1656I (6128G>A 6129G>T), T1657Y (6131A>T 6132C>A), G1658P (6134G>C 6135G>C 6136G>T), V1659del (6137_6139delGTC), E1660N (6140G>A 6142A>T), K1662V (6146A>G 6147A>T 6148A>T), H1665Y (6155C>T), E1669K (6167G>A), N1670D (6170A>G 6172C>T), Q1672N (6176C>A 6178G>T) | | | | | | | | |
| Codon mutations: | CAA1538AAA (5774C>A), CAG1539AAC (5777C>A 5779G>C), CTC1540ATA (5780C>A 5782C>A), CTA1543TTA (5789C>T), GAA1544AGC (5792G>A 5793A>G 5794A>C), CCC1546CCA (5800C>A), CCA1547CTA (5802C>T), GGA1549TCA (5807G>T 5808G>C), AGC1550GAT (5810A>G 5811G>A 5812C>T), TAC1551TAT (5815C>T), ATA1552CTA (5816A>C), CTA1554ATA (5822C>A), TGC1559AGC (5837T>A), ATG1560CTC (5840A>C 5842G>C), GAA1561CAA (5843G>C), GGT1564GGA (5854T>A), GGA1565GCT (5856G>C 5857A>T), TGT1567TTA (5862G>T 5863T>A), TGG1569GCA (5867T>G 5868G>C 5869G>A), GCA1570AGA (5870G>A 5871C>G), CCG1571CCA (5875G>A), TTC1572AAC (5876T>A 5877T>A), GGA1574TAC (5882G>T 5883G>A 5884A>C), GCC1575AGT (5885G>A 5886C>G 5887C>T), AAG1576CCT (5888A>C 5889A>C 5890G>T), AGT1578AAA (5895G>A 5896T>A), GAG1580GAA (5902G>A), AAG1581AAA (5905G>A), GTC1582ATT (5906G>A 5908C>T), TAT1585TAC (5917T>C), GCA1586CAA (5918G>C 5919C>A), TCA1587AGT (5921T>A 5922C>G 5923A>T), GGA1588GGT (5926A>T), AAG1589AAA (5929G>A), TTC1590TAC (5931T>A), CCC1591AAA (5933C>A 5934C>A 5935C>A), ACA1592GAA (5936A>G 5937C>A), ACA1592_CTC1593insAAGGGT (5938_5939insAAGGGT), CTC1593AAC (5939C>A 5940T>A), AAA1594ATG (5943A>T 5944A>G), AGC1595AGT (5947C>T), ACC1596AGT (5949C>G 5950C>T), ATC1597ATA (5953C>A), GAT1598GAC (5956T>C), GCA1599GTC (5958C>T 5959A>C), ATC1601GTA (5963A>G 5965C>A), TAT1602TTA (5967A>T 5968T>A), GCG1603GCT (5971G>T), GTC1604ATA (5972G>A 5974C>A), ATG1605ATA (5977G>A), GAA1606TAT (5978G>T 5980A>T), ACC1607GGA (5981A>G 5982C>G 5983C>A), ATG1608CTA (5984A>C 5986G>A), GAA1609AAT (5987G>A 5989A>T), TCT1610AGT (5990T>A 5991C>G), TTG1611TTC (5995G>C), ATC1613TTA (5999A>T 6001C>A), TAC1615ATT (6005T>A 6006A>T 6007C>T), GAT1617AAC (6011G>A 6013T>C), CAG1618AAA (6014C>A 6016G>A), AAG1619AGT (6018A>G 6019G>T), GCA1620GAA (6021C>A), ACA1622GTA (6026A>G 6027C>T), AGG1624AGA (6034G>A), GAC1626GAT (6040C>T), CAA1628GAA (6044C>G), GCA1629GCT (6049A>T), ATC1630ATA (6052C>A), ATA1631GTA (6053A>G), GCA1632AAA (6056G>A 6057C>A), TTC1633TTT (6061C>T), CAC1634CAT (6064C>T), GAC1635CAA (6065G>C 6067C>A), CAA1637ATA (6071C>A 6072A>T), CTT1638GAC (6074C>G 6075T>A 6076T>C), CAC1639CAG (6079C>G), AAG1641AAT (6085G>T), CCC1642AGT (6086C>A 6087C>G 6088C>T), TCA1643AGT (6089T>A 6090C>G 6091A>T), AGA1644AAA (6093G>A), GTG1645AGA (6095G>A 6096T>G 6097G>A), CGA1646AGA (6098C>A), ATC1648CTA (6104A>C 6106C>A), AAC1649AAT (6109C>T), TGT1651ATA (6113T>A 6114G>T 6115T>A), TAT1653ACC (6119T>A 6120A>C 6121T>C), ATA1654ATT (6124A>T), ACG1655TCT (6125A>T 6127G>T), GGA1656ATA (6128G>A 6129G>T), ACC1657TAC (6131A>T 6132C>A), GGG1658CCT (6134G>C 6135G>C 6136G>T), GTC1659del (6137_6139delGTC), GAA1660AAT (6140G>A 6142A>T), AAA1662GTT (6146A>G 6147A>T 6148A>T), CAT1665TAT (6155C>T), ATC1666ATA (6160C>A), GAA1669AAA (6167G>A), AAC1670GAT (6170A>G 6172C>T), AAC1671AAT (6175C>T), CAG1672AAT (6176C>A 6178G>T), GAC1675GAT (6187C>T) | | | | | | | | |

##### Proteins

| | Begin | End | Coverage | Score | Concordance | Matches | Identities | I/D/M/F* | Stop Codons |
|---|---|---|---|---|---|---|---|---|---|
| polyprotein (YP_010800602.1) | 1538 | 1676 | 7.5% | 490 | 49.1% | 138 (97.9%) | 67 (47.5%) | 2/1/0/0 | 0 |
| Protein mutations: | Q1538K (5774C>A), Q1539N (5777C>A 5779G>C), L1540I (5780C>A 5782C>A), E1544S (5792G>A 5793A>G 5794A>C), P1547L (5802C>T), G1549S (5807G>T 5808G>C), S1550D (5810A>G 5811G>A 5812C>T), I1552L (5816A>C), L1554I (5822C>A), C1559S (5837T>A), M1560L (5840A>C 5842G>C), E1561Q (5843G>C), G1565A (5856G>C 5857A>T), C1567L (5862G>T 5863T>A), W1569A (5867T>G 5868G>C 5869G>A), A1570R (5870G>A 5871C>G), F1572N (5876T>A 5877T>A), G1574Y (5882G>T 5883G>A 5884A>C), A1575S (5885G>A 5886C>G 5887C>T), K1576P (5888A>C 5889A>C 5890G>T), S1578K (5895G>A 5896T>A), V1582I (5906G>A 5908C>T), A1586Q (5918G>C 5919C>A), F1590Y (5931T>A), P1591K (5933C>A 5934C>A 5935C>A), T1592E (5936A>G 5937C>A), T1592_L1593insKG (5938_5939insAAGGGT), L1593N (5939C>A 5940T>A), K1594M (5943A>T 5944A>G), T1596S (5949C>G 5950C>T), A1599V (5958C>T 5959A>C), I1601V (5963A>G 5965C>A), Y1602L (5967A>T 5968T>A), V1604I (5972G>A 5974C>A), M1605I (5977G>A), E1606Y (5978G>T 5980A>T), T1607G (5981A>G 5982C>G 5983C>A), M1608L (5984A>C 5986G>A), E1609N (5987G>A 5989A>T), L1611F (5995G>C), I1613L (5999A>T 6001C>A), Y1615I (6005T>A 6006A>T 6007C>T), D1617N (6011G>A 6013T>C), Q1618K (6014C>A 6016G>A), K1619S (6018A>G 6019G>T), A1620E (6021C>A), T1622V (6026A>G 6027C>T), Q1628E (6044C>G), I1631V (6053A>G), A1632K (6056G>A 6057C>A), D1635Q (6065G>C 6067C>A), Q1637I (6071C>A 6072A>T), L1638D (6074C>G 6075T>A 6076T>C), H1639Q (6079C>G), K1641N (6085G>T), P1642S (6086C>A 6087C>G 6088C>T), R1644K (6093G>A), V1645R (6095G>A 6096T>G 6097G>A), I1648L (6104A>C 6106C>A), C1651I (6113T>A 6114G>T 6115T>A), Y1653T (6119T>A 6120A>C 6121T>C), T1655S (6125A>T 6127G>T), G1656I (6128G>A 6129G>T), T1657Y (6131A>T 6132C>A), G1658P (6134G>C 6135G>C 6136G>T), V1659del (6137_6139delGTC), E1660N (6140G>A 6142A>T), K1662V (6146A>G 6147A>T 6148A>T), H1665Y (6155C>T), E1669K (6167G>A), N1670D (6170A>G 6172C>T), Q1672N (6176C>A 6178G>T) | | | | | | | | |
| Codon mutations: | CAA1538AAA (5774C>A), CAG1539AAC (5777C>A 5779G>C), CTC1540ATA (5780C>A 5782C>A), CTA1543TTA (5789C>T), GAA1544AGC (5792G>A 5793A>G 5794A>C), CCC1546CCA (5800C>A), CCA1547CTA (5802C>T), GGA1549TCA (5807G>T 5808G>C), AGC1550GAT (5810A>G 5811G>A 5812C>T), TAC1551TAT (5815C>T), ATA1552CTA (5816A>C), CTA1554ATA (5822C>A), TGC1559AGC (5837T>A), ATG1560CTC (5840A>C 5842G>C), GAA1561CAA (5843G>C), GGT1564GGA (5854T>A), GGA1565GCT (5856G>C 5857A>T), TGT1567TTA (5862G>T 5863T>A), TGG1569GCA (5867T>G 5868G>C 5869G>A), GCA1570AGA (5870G>A 5871C>G), CCG1571CCA (5875G>A), TTC1572AAC (5876T>A 5877T>A), GGA1574TAC (5882G>T 5883G>A 5884A>C), GCC1575AGT (5885G>A 5886C>G 5887C>T), AAG1576CCT (5888A>C 5889A>C 5890G>T), AGT1578AAA (5895G>A 5896T>A), GAG1580GAA (5902G>A), AAG1581AAA (5905G>A), GTC1582ATT (5906G>A 5908C>T), TAT1585TAC (5917T>C), GCA1586CAA (5918G>C 5919C>A), TCA1587AGT (5921T>A 5922C>G 5923A>T), GGA1588GGT (5926A>T), AAG1589AAA (5929G>A), TTC1590TAC (5931T>A), CCC1591AAA (5933C>A 5934C>A 5935C>A), ACA1592GAA (5936A>G 5937C>A), ACA1592_CTC1593insAAGGGT (5938_5939insAAGGGT), CTC1593AAC (5939C>A 5940T>A), AAA1594ATG (5943A>T 5944A>G), AGC1595AGT (5947C>T), ACC1596AGT (5949C>G 5950C>T), ATC1597ATA (5953C>A), GAT1598GAC (5956T>C), GCA1599GTC (5958C>T 5959A>C), ATC1601GTA (5963A>G 5965C>A), TAT1602TTA (5967A>T 5968T>A), GCG1603GCT (5971G>T), GTC1604ATA (5972G>A 5974C>A), ATG1605ATA (5977G>A), GAA1606TAT (5978G>T 5980A>T), ACC1607GGA (5981A>G 5982C>G 5983C>A), ATG1608CTA (5984A>C 5986G>A), GAA1609AAT (5987G>A 5989A>T), TCT1610AGT (5990T>A 5991C>G), TTG1611TTC (5995G>C), ATC1613TTA (5999A>T 6001C>A), TAC1615ATT (6005T>A 6006A>T 6007C>T), GAT1617AAC (6011G>A 6013T>C), CAG1618AAA (6014C>A 6016G>A), AAG1619AGT (6018A>G 6019G>T), GCA1620GAA (6021C>A), ACA1622GTA (6026A>G 6027C>T), AGG1624AGA (6034G>A), GAC1626GAT (6040C>T), CAA1628GAA (6044C>G), GCA1629GCT (6049A>T), ATC1630ATA (6052C>A), ATA1631GTA (6053A>G), GCA1632AAA (6056G>A 6057C>A), TTC1633TTT (6061C>T), CAC1634CAT (6064C>T), GAC1635CAA (6065G>C 6067C>A), CAA1637ATA (6071C>A 6072A>T), CTT1638GAC (6074C>G 6075T>A 6076T>C), CAC1639CAG (6079C>G), AAG1641AAT (6085G>T), CCC1642AGT (6086C>A 6087C>G 6088C>T), TCA1643AGT (6089T>A 6090C>G 6091A>T), AGA1644AAA (6093G>A), GTG1645AGA (6095G>A 6096T>G 6097G>A), CGA1646AGA (6098C>A), ATC1648CTA (6104A>C 6106C>A), AAC1649AAT (6109C>T), TGT1651ATA (6113T>A 6114G>T 6115T>A), TAT1653ACC (6119T>A 6120A>C 6121T>C), ATA1654ATT (6124A>T), ACG1655TCT (6125A>T 6127G>T), GGA1656ATA (6128G>A 6129G>T), ACC1657TAC (6131A>T 6132C>A), GGG1658CCT (6134G>C 6135G>C 6136G>T), GTC1659del (6137_6139delGTC), GAA1660AAT (6140G>A 6142A>T), AAA1662GTT (6146A>G 6147A>T 6148A>T), CAT1665TAT (6155C>T), ATC1666ATA (6160C>A), GAA1669AAA (6167G>A), AAC1670GAT (6170A>G 6172C>T), AAC1671AAT (6175C>T), CAG1672AAT (6176C>A 6178G>T), GAC1675GAT (6187C>T) | | | | | | | | |

*: Inserts / Deletes / Misaligned / Frameshifts

#### Analysis details

This analysis was performed with panviral2.64

### NGS Details (UN9): Ruflodivirus deformatiorudbeckiae

#### Assembly

| | |
|---|---|
| **Coverage Length** | 605 (1 contig(s)) |
| **Depth Of Coverage** | 4.7 |
| **Number Of Reads** | 21 |
| **Reads Per Million** | 0.47 rpm (after QC) |
| **Ambiguities** | 0 |
| **Assembly Method** | de novo + reference guided assembly |
| **Consensus Caller** | Bcf Tools |

#### Coverage Map

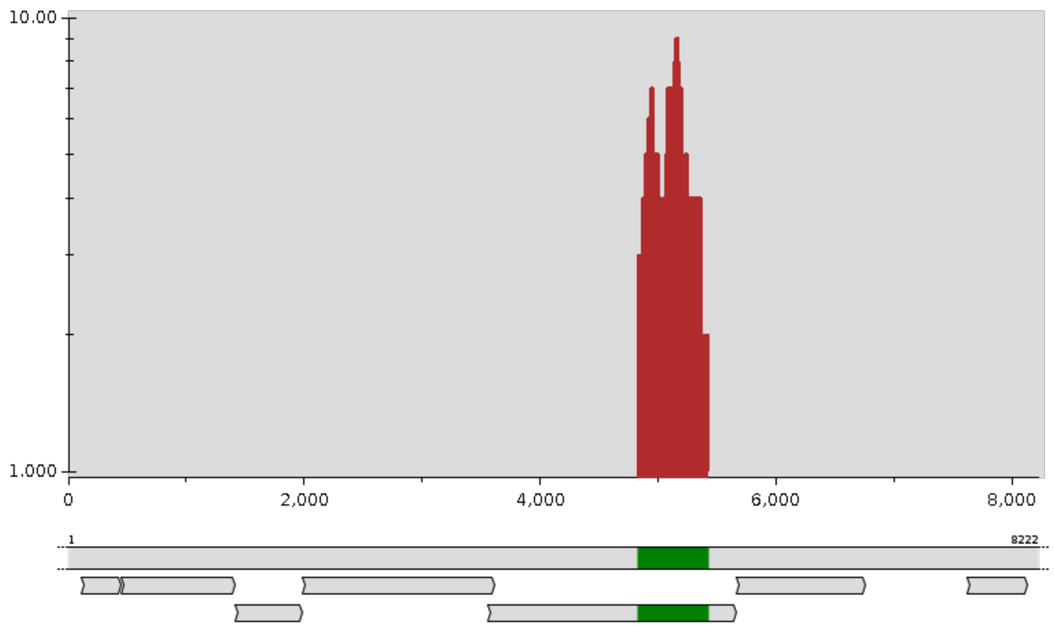

#### Assignment

| | |
|---|---|
| **Type** | Ruflodivirus deformatiorudbeckiae (Taxonomy ID: 3048307) |
| **Reference Genome** | NC_011920.1 |
| **NT Identity (%)** | 57.3356 |
| **AA Identity (%)** | 45.4545 |
| **Number Of Stop Codons** | 0 |
| **Number Of CDS** | 7 |

#### Alignment

| | |
|---|---|
| **Alignment Score** | 144.0 (NT) + 646.0 (AA) = 790.0 |
| **Concordance (%)** | 31.6887 |

| Alignment Method | Global, seeded, nucleotide + amino acids (AGA) |
|---|---|

#### Genome Region

Sequence starts at position 4826 and ends at position 5430 relative to NC_011920.1 reference sequence.

#### Alignment Detailed Statistics

| | Begin | End | Coverage | Score | Concordance | Matches | Identities | I/D/M/F* | Stop Codons |
|---|---|---|---|---|---|---|---|---|---|
| **NT** | **4826** | **5430** | **7.4%** | **144** | **12.5%** | **593 (98.0%)** | **340 (56.2%)** | **0/12** | |
| Mutations: | 4835G>A, 4843C>T, 4847C>T, 4849T>A, 4850G>A, 4852C>A, 4854T>G, 4855C>T, 4858A>C, 4861A>C, 4862A>T, 4865C>T, 4866T>A, 4868G>C, 4870T>A, 4878A>T, 4879T>A, 4880G>A, 4881A>C, 4888A>T, 4889G>A, 4892A>C, 4893C>A, 4894T>A, 4895G>T, 4897G>T, 4898C>T, 4899T>C, 4901A>C, 4902G>A, 4905A>T, 4906G>A, 4909C>T, 4910A>T, 4911T>G, 4912T>C, 4917C>A, 4921C>A, 4924A>T, 4925A>G, 4927C>G, 4928C>T, 4930T>A, 4931T>A, 4932C>G, 4933T>C, 4934A>C, 4936A>G, 4939G>A, 4942A>G, 4945T>A, 4948G>C, 4949G>A, 4951T>A, 4952G>A, 4953C>A, 4959C>A, 4961A>G, 4966C>T, 4967A>G, 4969T>A, 4971A>T, 4975T>C, 4978T>A, 4979T>A, 4981T>G, 4983C>T, 4984T>A, 4987C>T, 4990C>A, 4993T>A, 4994G>A, 4995G>A, 4996A>T, 4998A>G, 4999A>C, 5006C>T, 5008T>A, 5012T>A, 5020C>A, 5021C>T, 5022T>C, 5023T>A, 5024G>A, 5026G>A, 5029T>G, 5032A>C, 5033A>T, 5034A>G, 5035A>T, 5036C>G, 5037T>A, 5038T>A, 5045A>G, 5046G>A, 5049G>A, 5051A>C, 5053T>A, 5054C>A, 5055C>A, 5057G>A, 5058A>C, 5059T>A, 5063A>C, 5066T>G, 5067C>A, 5069C>A, 5071A>C, 5087A>T, 5088T>G, 5089A>T, 5090C>T, 5094C>A, 5095G>T, 5098T>C, 5102G>A, 5103A>G, 5105C>G, 5109A>T, 5110C>T, 5115G>C, 5117A>G, 5119A>C, 5122T>A, 5125C>A, 5126G>A, 5129T>A, 5130G>A, 5132C>A, 5137A>T, 5138C>T, 5139C>T, 5141C>T, 5143T>A, 5146G>A, 5149A>G, 5150A>C, 5151A>T, 5152A>T, 5153T>A, 5155G>T, 5156A>C, 5157A>G, 5158G>A, 5162G>T, 5163A>C, 5164T>C, 5166C>A, 5167T>C, 5168G>C, 5169T>G, 5171T>A, 5172G>T, 5175A>G, 5181A>C, 5182T>A, 5186T>G, 5187C>A, 5188C>A, 5189G>C, 5194T>C, 5195C>A, 5197G>A, 5199A>G, 5200C>T, 5203G>T, 5204G>C, 5208A>G, 5209A>C, 5210T>C, 5212T>A, 5215G>A, 5216A>C, 5217G>A, 5219A>G, 5221T>A, 5222C>T, 5223T>G, 5225A>T, 5226A>C, 5227A>C, 5228G>A, 5233C>A, 5239A>G, 5240T>C, 5243C>A, 5244A>G, 5245C>A, 5247A>T, 5251T>A, 5252T>G, 5253T>A, 5257G>A, 5260T>A, 5263A>C, 5264T>A, 5269G>A, 5270A>G, 5272C>T, 5273A>T, 5275C>A, 5276G>C, 5281T>A, 5287A>T, 5291C>G, 5294G>T, 5296A>C, 5299C>T, 5304C>G, 5305T>A, 5307G>C, 5308T>A, 5309G>T, 5311C>A, 5312C>T, 5315A>C, 5317G>A, 5318_5326delTCCCGAGGG, 5333A>C, 5334A>T, 5335C>T, 5336T>A, 5337T>A, 5338A>T, 5342_5344delCGA, 5345C>A, 5347A>T, 5351C>A, 5353T>G, 5362C>T, 5365A>T, 5366T>A, 5371C>T, 5374A>T, 5375C>G, 5376C>A, 5377C>A, 5378G>A, 5380T>C, 5385T>C, 5386C>T, 5389C>A, 5392T>C, 5393C>T, 5394A>C, 5395C>A, 5397G>C, 5398C>A, 5401C>T, 5411G>T, 5413G>A, 5416T>A, 5420G>A, 5423A>G, 5424A>T | | | | | | | | |

##### CDS

| | Begin | End | Coverage | Score | Concordance | Matches | Identities | I/D/M/F* | Stop Codons |
|---|---|---|---|---|---|---|---|---|---|
| RuFDV_gp5 | 424 | 625 | 28.8% | 646 | 48.1% | 198 (98.0%) | 90 (44.6%) | 0/4/0/0 | 0 |
| Protein mutations: | V427I (4835G>A), V432I (4850G>A 4852C>A), F433C (4854T>G 4855C>T), K435N (4861A>C), T436S (4862A>T), L437Y (4865C>T 4866T>A), D438Q (4868G>C 4870T>A), Y441L (4878A>T 4879T>A), D442T (4880G>A 4881A>C), E445K (4889G>A), T446Q (4892A>C 4893C>A 4894T>A), V447F (4895G>T 4897G>T), L448S (4898C>T 4899T>C), R449Q (4901A>C 4902G>A), K450I (4905A>T 4906G>A), I452C (4910A>T 4911T>G 4912T>C), T454N (4917C>A), I457V (4925A>G 4927C>G), K460Q (4934A>C 4936A>G), E464D (4948G>C), V465I (4949G>A 4951T>A), A466N (4952G>A 4953C>A), T468N (4959C>A), K469E (4961A>G), N471E (4967A>G 4969T>A), Y472F (4971A>T), F475M (4979T>A 4981T>G), T476I (4983C>T 4984T>A), N479K (4993T>A), G480N (4994G>A 4995G>A 4996A>T), E481G (4998A>G 4999A>C), S486T (5012T>A), L489S (5021C>T 5022T>C 5023T>A), E490K (5024G>A 5026G>A), N491K (5029T>G), K493C (5033A>T 5034A>G 5035A>T), L494E (5036C>G 5037T>A 5038T>A), S497D (5045A>G 5046G>A), R498K (5049G>A), I499L (5051A>C 5053T>A), P500N (5054C>A 5055C>A), D501T (5057G>A 5058A>C 5059T>A), K503Q (5063A>C), S504E (5066T>G 5067C>A), L505I (5069C>A 5071A>C), I511C (5087A>T 5088T>G 5089A>T), T513N (5094C>A 5095G>T), D516S (5102G>A 5103A>G), Q517E (5105C>G), Y518F (5109A>T 5110C>T), R520P (5115G>C), K521D (5117A>G 5119A>C), E524K (5126G>A), W525K (5129T>A 5130G>A), K527N (5137A>T), P528F (5138C>T 5139C>T), K532L (5150A>C 5151A>T 5152A>T), L533I (5153T>A 5155G>T), K534R (5156A>C 5157A>G 5158G>A), D536S (5162G>T 5163A>C 5164T>C), T537N (5166C>A 5167T>C), V538R (5168G>C 5169T>G), W539M (5171T>A 5172G>T), E540G (5175A>G), N542T (5181A>C 5182T>A), S544E (5186T>G 5187C>A 5188C>A), D545H (5189G>C), Q547K (5195C>A 5197G>A), Y548C (5199A>G 5200C>T), E550Q (5204G>C), K551S (5208A>G 5209A>C), F552L (5210T>C 5212T>A), R554Q (5216A>C 5217G>A), N555E (5219A>G 5221T>A), L556C (5222C>T 5223T>G), K557S (5225A>T 5226A>C 5227A>C), E558K (5228G>A), F559L (5233C>A), H563R (5243C>A 5244A>G 5245C>A), H564L (5247A>T), L566E (5252T>G 5253T>A), D568E (5260T>A), E569D (5263A>C), Y570N (5264T>A), I572V (5270A>G 5272C>T), I573L (5273A>T 5275C>A), E574Q (5276G>C), H579D (5291C>G), E580Y (5294G>T 5296A>C), A583G (5304C>G 5305T>A), G584A (5307G>C 5308T>A), V585L (5309G>T 5311C>A), K587Q (5315A>C 5317G>A), S588_G590del (5318_5326delTCCCGAGGG), N593L (5333A>C 5334A>T 5335C>T), L594N (5336T>A 5337T>A 5338A>T), R596del (5342_5344delCGA), L597I (5345C>A 5347A>T), S604T (5366T>A), K606N (5374A>T), P607E (5375C>G 5376C>A 5377C>A), A608T (5378G>A 5380T>C), I610T (5385T>C 5386C>T), N611K (5389C>A), H613S (5393C>T 5394A>C 5395C>A), S614T (5397G>C 5398C>A), V619L (5411G>T 5413G>A), V622I (5420G>A), K623V (5423A>G 5424A>T) | | | | | | | | |
| Codon mutations: | GTA427ATA (4835G>A), GAC429GAT (4843C>T), CTT431TTA (4847C>T 4849T>A), GTC432ATA (4850G>A 4852C>A), TTC433TGT (4854T>G 4855C>T), TCA434TCC (4858A>C), AAA435AAC (4861A>C), ACA436TCA (4862A>T), CTT437TAT (4865C>T 4866T>A), GAT438CAA (4868G>C 4870T>A), TAT441TTA (4878A>T 4879T>A), GAT442ACT (4880G>A 4881A>C), CTA444CTT (4888A>T), GAA445AAA (4889G>A), ACT446CAA (4892A>C 4893C>A 4894T>A), GTG447TTT (4895G>T 4897G>T), CTT448TCT (4898C>T 4899T>C), AGA449CAA (4901A>C 4902G>A), AAG450ATA (4905A>T 4906G>A), TGC451TGT (4909C>T), ATT452TGC (4910A>T 4911T>G 4912T>C), ACT454AAT (4917C>A), GGC455GGA (4921C>A), ATA456ATT (4924A>T), ATC457GTG (4925A>G 4927C>G), CTT458TTA (4928C>T 4930T>A), TCT459AGC (4931T>A 4932C>G 4933T>C), AAA460CAG (4934A>C 4936A>G), AAG461AAA (4939G>A), AAA462AAG (4942A>G), GCT463GCA (4945T>A), GAG464GAC (4948G>C), GTT465ATA (4949G>A 4951T>A), GCT466AAT (4952G>A 4953C>A), ACT468AAT (4959C>A), AAA469GAA (4961A>G), ATC470ATT (4966C>T), AAT471GAA (4967A>G 4969T>A), TAT472TTT (4971A>T), CTT473CTC (4975T>C), GGT474GGA (4978T>A), TTT475ATG (4979T>A 4981T>G), ACT476ATA (4983C>T 4984T>A), ATC477ATT (4987C>T), TCC478TCA (4990C>A), AAT479AAA (4993T>A), GGA480AAT (4994G>A 4995G>A 4996A>T), GAA481GGC (4998A>G 4999A>C), CTT484TTA (5006C>T 5008T>A), TCT486ACT (5012T>A), ATC488ATA (5020C>A), CTT489TCA (5021C>T 5022T>C 5023T>A), GAG490AAA (5024G>A 5026G>A), AAT491AAG (5029T>G), ATA492ATC (5032A>C), AAA493TGT (5033A>T 5034A>G 5035A>T), CTT494GAA (5036C>G 5037T>A 5038T>A), AGT497GAT (5045A>G 5046G>A), AGA498AAA (5049G>A), ATT499CTA (5051A>C 5053T>A), CCT500AAT (5054C>A 5055C>A), GAT501ACA (5057G>A 5058A>C 5059T>A), AAA503CAA (5063A>C), TCA504GAA (5066T>G 5067C>A), CTA505ATC (5069C>A 5071A>C), ATA511TGT (5087A>T 5088T>G 5089A>T), CTA512TTA (5090C>T), ACG513AAT (5094C>A 5095G>T), TAT514TAC (5098T>C), GAT516AGT (5102G>A 5103A>G), CAA517GAA (5105C>G), TAC518TTT (5109A>T 5110C>T), CGA520CCA (5115G>C), AAA521GAC (5117A>G 5119A>C), CTT522CTA (5122T>A), GCC523GCA (5125C>A), GAA524AAA (5126G>A), TGG525AAG (5129T>A 5130G>A), CGA526AGA (5132C>A), AAA527AAT (5137A>T), CCT528TTT (5138C>T 5139C>T), CTT529TTA (5141C>T 5143T>A), CAG530CAA (5146G>A), AGA531AGG (5149A>G), AAA532CTT (5150A>C 5151A>T 5152A>T), TTG533ATT (5153T>A 5155G>T), AAG534CGA (5156A>C 5157A>G 5158G>A), GAT536TCC (5162G>T 5163A>C 5164T>C), ACT537AAC (5166C>A 5167T>C), GTA538CGA (5168G>C 5169T>G), TGG539ATG (5171T>A 5172G>T), GAA540GGA (5175A>G), AAT542ACA (5181A>C 5182T>A), TCC544GAA (5186T>G 5187C>A 5188C>A), GAT545CAT (5189G>C), ACT546ACC (5194T>C), CAG547AAA (5195C>A 5197G>A), TAC548TGT (5199A>G 5200C>T), GTG549GTT (5203G>T), GAA550CAA (5204G>C), AAA551AGC (5208A>G 5209A>C), TTT552CTA (5210T>C 5212T>A), AAG553AAA (5215G>A), AGA554CAA (5216A>C 5217G>A), AAT555GAA (5219A>G 5221T>A), CTT556TGT (5222C>T 5223T>G), AAA557TCC (5225A>T 5226A>C 5227A>C), GAA558AAA (5228G>A), TTC559TTA (5233C>A), AAA561AAG (5239A>G), TTA562CTA (5240T>C), CAC563AGA (5243C>A 5244A>G 5245C>A), CAT564CTT (5247A>T), CCT565CCA (5251T>A), TTA566GAA (5252T>G 5253T>A), CCG567CCA (5257G>A), GAT568GAA (5260T>A), GAA569GAC (5263A>C), TAC570AAC (5264T>A), TTG571TTA (5269G>A), ATC572GTT (5270A>G 5272C>T), ATC573TTA (5273A>T 5275C>A), GAA574CAA (5276G>C), ACT575ACA (5281T>A), GCA577GCT (5287A>T), CAT579GAT (5291C>G), GAA580TAC (5294G>T 5296A>C), CAC581CAT (5299C>T), GCT583GGA (5304C>G 5305T>A), GGT584GCA (5307G>C 5308T>A), GTC585TTA (5309G>T 5311C>A), CTA586TTA (5312C>T), AAG587CAA (5315A>C 5317G>A), TCC588_GGG590del (5318_5326delTCCCGAGGG), AAC593CTT (5333A>C 5334A>T 5335C>T), TTA594AAT (5336T>A 5337T>A 5338A>T), CGA596del (5342_5344delCGA), CTA597ATT (5345C>A 5347A>T), CGT599AGG (5351C>A 5353T>G), AGC602AGT (5362C>T), GGA603GGT (5365A>T), TCA604ACA (5366T>A), TTC605TTT (5371C>T), AAA606AAT (5374A>T), CCC607GAA (5375C>G 5376C>A 5377C>A), GCT608ACC (5378G>A 5380T>C), ATC610ACT (5385T>C 5386C>T), AAC611AAA (5389C>A), TAT612TAC (5392T>C), CAC613TCA (5393C>T 5394A>C 5395C>A), AGC614ACA (5397G>C 5398C>A), AAC615AAT (5401C>T), GTG619TTA (5411G>T 5413G>A), CTT620CTA (5416T>A), GTA622ATA (5420G>A), AAA623GTA (5423A>G 5424A>T) | | | | | | | | |

##### Proteins

| | Begin | End | Coverage | Score | Concordance | Matches | Identities | I/D/M/F* | Stop Codons |
|---|---|---|---|---|---|---|---|---|---|
| putative enzymatic polyprotein (YP_002519387.1) | 424 | 625 | 28.8% | 646 | 48.1% | 198 (98.0%) | 90 (44.6%) | 0/4/0/0 | 0 |
| Protein mutations: | V427I (4835G>A), V432I (4850G>A 4852C>A), F433C (4854T>G 4855C>T), K435N (4861A>C), T436S (4862A>T), L437Y (4865C>T 4866T>A), D438Q (4868G>C 4870T>A), Y441L (4878A>T 4879T>A), D442T (4880G>A 4881A>C), E445K (4889G>A), T446Q (4892A>C 4893C>A 4894T>A), V447F (4895G>T 4897G>T), L448S (4898C>T 4899T>C), R449Q (4901A>C 4902G>A), K450I (4905A>T 4906G>A), I452C (4910A>T 4911T>G 4912T>C), T454N (4917C>A), I457V (4925A>G 4927C>G), K460Q (4934A>C 4936A>G), E464D (4948G>C), V465I (4949G>A 4951T>A), A466N (4952G>A 4953C>A), T468N (4959C>A), K469E (4961A>G), N471E (4967A>G 4969T>A), Y472F (4971A>T), F475M (4979T>A 4981T>G), T476I (4983C>T 4984T>A), N479K (4993T>A), G480N (4994G>A 4995G>A 4996A>T), E481G (4998A>G 4999A>C), S486T (5012T>A), L489S (5021C>T 5022T>C 5023T>A), E490K (5024G>A 5026G>A), N491K (5029T>G), K493C (5033A>T 5034A>G 5035A>T), L494E (5036C>G 5037T>A 5038T>A), S497D (5045A>G 5046G>A), R498K (5049G>A), I499L (5051A>C 5053T>A), P500N (5054C>A 5055C>A), D501T (5057G>A 5058A>C 5059T>A), K503Q (5063A>C), S504E (5066T>G 5067C>A), L505I (5069C>A 5071A>C), I511C (5087A>T 5088T>G 5089A>T), T513N (5094C>A 5095G>T), D516S (5102G>A 5103A>G), Q517E (5105C>G), Y518F (5109A>T 5110C>T), R520P (5115G>C), K521D (5117A>G 5119A>C), E524K (5126G>A), W525K (5129T>A 5130G>A), K527N (5137A>T), P528F (5138C>T 5139C>T), K532L (5150A>C 5151A>T 5152A>T), L533I (5153T>A 5155G>T), K534R (5156A>C 5157A>G 5158G>A), D536S (5162G>T 5163A>C 5164T>C), T537N (5166C>A 5167T>C), V538R (5168G>C 5169T>G), W539M (5171T>A 5172G>T), E540G (5175A>G), N542T (5181A>C 5182T>A), S544E (5186T>G 5187C>A 5188C>A), D545H (5189G>C), Q547K (5195C>A 5197G>A), Y548C (5199A>G 5200C>T), E550Q (5204G>C), K551S (5208A>G 5209A>C), F552L (5210T>C 5212T>A), R554Q (5216A>C 5217G>A), N555E (5219A>G 5221T>A), L556C (5222C>T 5223T>G), K557S (5225A>T 5226A>C 5227A>C), E558K (5228G>A), F559L (5233C>A), H563R (5243C>A 5244A>G 5245C>A), H564L (5247A>T), L566E (5252T>G 5253T>A), D568E (5260T>A), E569D (5263A>C), Y570N (5264T>A), I572V (5270A>G 5272C>T), I573L (5273A>T 5275C>A), E574Q (5276G>C), H579D (5291C>G), E580Y (5294G>T 5296A>C), A583G (5304C>G 5305T>A), G584A (5307G>C 5308T>A), V585L (5309G>T 5311C>A), K587Q (5315A>C 5317G>A), S588_G590del (5318_5326delTCCCGAGGG), N593L (5333A>C 5334A>T 5335C>T), L594N (5336T>A 5337T>A 5338A>T), R596del (5342_5344delCGA), L597I (5345C>A 5347A>T), S604T (5366T>A), K606N (5374A>T), P607E (5375C>G 5376C>A 5377C>A), A608T (5378G>A 5380T>C), I610T (5385T>C 5386C>T), N611K (5389C>A), H613S (5393C>T 5394A>C 5395C>A), S614T (5397G>C 5398C>A), V619L (5411G>T 5413G>A), V622I (5420G>A), K623V (5423A>G 5424A>T) | | | | | | | | |

| | Begin | End | Coverage | Score | Concordance | Matches | Identities | I/D/M/F* | Stop Codons |
|---|---|---|---|---|---|---|---|---|---|
| NT | 4826 | 5430 | 7.4% | 144 | 12.5% | 593 (98.0%) | 340 (56.2%) | 0/12 | |
| Codon mutations: | GTA427ATA (4835G>A), GAC429GAT (4843C>T), CTT431TTA (4847C>T 4849T>A), GTC432ATA (4850G>A 4852C>A), TTC433TGT (4854T>G 4855C>T), TCA434TCC (4858A>C), AAA435AAC (4861A>C), ACA436TCA (4862A>T), CTT437TAT (4865C>T 4866T>A), GAT438CAA (4868G>C 4870T>A), TAT441TTA (4878A>T 4879T>A), GAT442ACT (4880G>A 4881A>C), CTA444CTT (4888A>T), GAA445AAA (4889G>A), ACT446CAA (4892A>C 4893C>A 4894T>A), GTG447TTT (4895G>T 4897G>T), CTT448TCT (4898C>T 4899T>C), AGA449CAA (4901A>C 4902G>A), AAG450ATA (4905A>T 4906G>A), TGC451TGT (4909C>T), ATT452TGC (4910A>T 4911T>G 4912T>C), ACT454AAT (4917C>A), GGC455GGA (4921C>A), ATA456ATT (4924A>T), ATC457GTG (4925A>G 4927C>G), CTT458TTA (4928C>T 4930T>A), TCT459AGC (4931T>A 4932C>G 4933T>C), AAA460CAG (4934A>C 4936A>G), AAG461AAA (4939G>A), AAA462AAG (4942A>G), GCT463GCA (4945T>A), GAG464GAC (4948G>C), GTT465ATA (4949G>A 4951T>A), GCT466AAT (4952G>A 4953C>A), ACT468AAT (4959C>A), AAA469GAA (4961A>G), ATC470ATT (4966C>T), AAT471GAA (4967A>G 4969T>A), TAT472TTT (4971A>T), CTT473CTC (4975T>C), GGT474GGA (4978T>A), TTT475ATG (4979T>A 4981T>G), ACT476ATA (4983C>T 4984T>A), ATC477ATT (4987C>T), TCC478TCA (4990C>A), AAT479AAA (4993T>A), GGA480AAT (4994G>A 4995G>A 4996A>T), GAA481GGC (4998A>G 4999A>C), CTT484TTA (5006C>T 5008T>A), TCT486ACT (5012T>A), ATC488ATA (5020C>A), CTT489TCA (5021C>T 5022T>C 5023T>A), GAG490AAA (5024G>A 5026G>A), AAT491AAG (5029T>G), ATA492ATC (5032A>C), AAA493TGT (5033A>T 5034A>G 5035A>T), CTT494GAA (5036C>G 5037T>A 5038T>A), AGT497GAT (5045A>G 5046G>A), AGA498AAA (5049G>A), ATT499CTA (5051A>C 5053T>A), CCT500AAT (5054C>A 5055C>A), GAT501ACA (5057G>A 5058A>C 5059T>A), AAA503CAA (5063A>C), TCA504GAA (5066T>G 5067C>A), CTA505ATC (5069C>A 5071A>C), ATA511TGT (5087A>T 5088T>G 5089A>T), CTA512TTA (5090C>T), ACG513AAT (5094C>A 5095G>T), TAT514TAC (5098T>C), GAT516AGT (5102G>A 5103A>G), CAA517GAA (5105C>G), TAC518TTT (5109A>T 5110C>T), CGA520CCA (5115G>C), AAA521GAC (5117A>G 5119A>C), CTT522CTA (5122T>A), GCC523GCA (5125C>A), GAA524AAA (5126G>A), TGG525AAG (5129T>A 5130G>A), CGA526AGA (5132C>A), AAA527AAT (5137A>T), CCT528TTT (5138C>T 5139C>T), CTT529TTA (5141C>T 5143T>A), CAG530CAA (5146G>A), AGA531AGG (5149A>G), AAA532CTT (5150A>C 5151A>T 5152A>T), TTG533ATT (5153T>A 5155G>T), AAG534CGA (5156A>C 5157A>G 5158G>A), GAT536TCC (5162G>T 5163A>C 5164T>C), ACT537AAC (5166C>A 5167T>C), GTA538CGA (5168G>C 5169T>G), TGG539ATG (5171T>A 5172G>T), GAA540GGA (5175A>G), AAT542ACA (5181A>C 5182T>A), TCC544GAA (5186T>G 5187C>A 5188C>A), GAT545CAT (5189G>C), ACT546ACC (5194T>C), CAG547AAA (5195C>A 5197G>A), TAC548TGT (5199A>G 5200C>T), GTG549GTT (5203G>T), GAA550CAA (5204G>C), AAA551AGC (5208A>G 5209A>C), TTT552CTA (5210T>C 5212T>A), AAG553AAA (5215G>A), AGA554CAA (5216A>C 5217G>A), AAT555GAA (5219A>G 5221T>A), CTT556TGT (5222C>T 5223T>G), AAA557TCC (5225A>T 5226A>C 5227A>C), GAA558AAA (5228G>A), TTC559TTA (5233C>A), AAA561AAG (5239A>G), TTA562CTA (5240T>C), CAC563AGA (5243C>A 5244A>G 5245C>A), CAT564CTT (5247A>T), CCT565CCA (5251T>A), TTA566GAA (5252T>G 5253T>A), CCG567CCA (5257G>A), GAT568GAA (5260T>A), GAA569GAC (5263A>C), TAC570AAC (5264T>A), TTG571TTA (5269G>A), ATC572GTT (5270A>G 5272C>T), ATC573TTA (5273A>T 5275C>A), GAA574CAA (5276G>C), ACT575ACA (5281T>A), GCA577GCT (5287A>T), CAT579GAT (5291C>G), GAA580TAC (5294G>T 5296A>C), CAC581CAT (5299C>T), GCT583GGA (5304C>G 5305T>A), GGT584GCA (5307G>C 5308T>A), GTC585TTA (5309G>T 5311C>A), CTA586TTA (5312C>T), AAG587CAA (5315A>C 5317G>A), TCC588_GGG590del (5318_5326delTCCCGAGGG), AAC593CTT (5333A>C 5334A>T 5335C>T), TTA594AAT (5336T>A 5337T>A 5338A>T), CGA596del (5342_5344delCGA), CTA597ATT (5345C>A 5347A>T), CGT599AGG (5351C>A 5353T>G), AGC602AGT (5362C>T), GGA603GGT (5365A>T), TCA604ACA (5366T>A), TTC605TTT (5371C>T), AAA606AAT (5374A>T), CCC607GAA (5375C>G 5376C>A 5377C>A), GCT608ACC (5378G>A 5380T>C), ATC610ACT (5385T>C 5386C>T), AAC611AAA (5389C>A), TAT612TAC (5392T>C), CAC613TCA (5393C>T 5394A>C 5395C>A), AGC614ACA (5397G>C 5398C>A), AAC615AAT (5401C>T), GTG619TTA (5411G>T 5413G>A), CTT620CTA (5416T>A), GTA622ATA (5420G>A), AAA623GTA (5423A>G 5424A>T) | | | | | | | | |

*: Inserts / Deletes / Misaligned / Frameshifts

#### Analysis details

This analysis was performed with panviral2.64

#### NGS Details (UN9): Caulimovirus latensarmoraciae

##### Assembly

| | |
|---|---|
| **Coverage Length** | 364 (1 contig(s)) |
| **Depth Of Coverage** | 7.3 |
| **Number Of Reads** | 21 |
| **Reads Per Million** | 0.47 rpm (after QC) |
| **Ambiguities** | 0 |
| **Assembly Method** | de novo + reference guided assembly |
| **Consensus Caller** | Bcf Tools |

##### Coverage Map

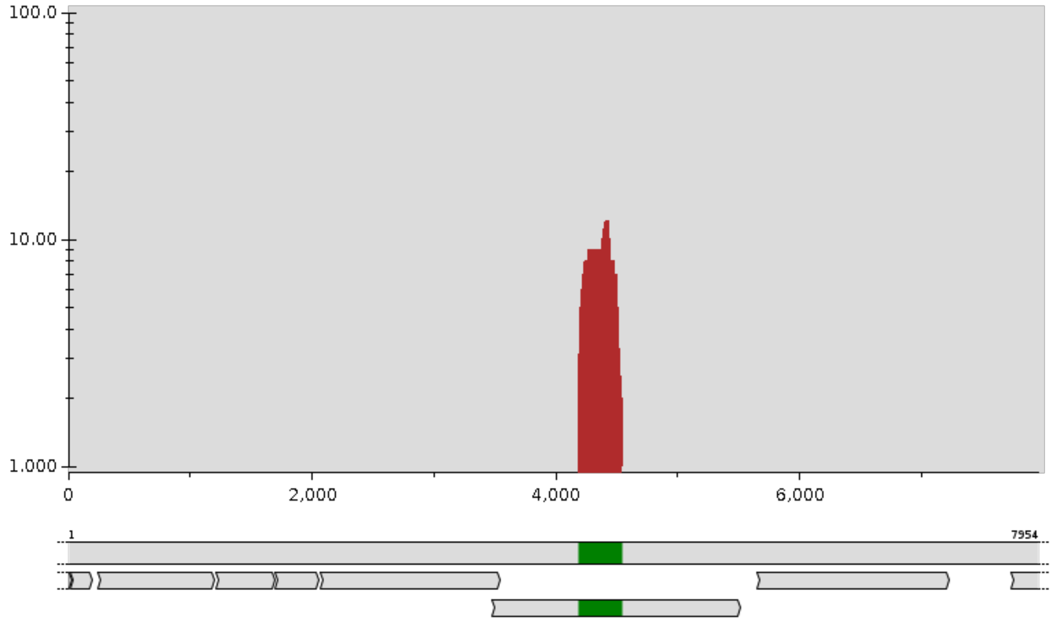

##### Assignment

| | |
|---|---|
| **Type** | Caulimovirus latensarmoraciae (Taxonomy ID: 3047955) |
| **Reference Genome** | NC_018858.1 |
| **NT Identity (%)** | 58.7912 |
| **AA Identity (%)** | 56.1983 |
| **Number Of Stop Codons** | 0 |
| **Number Of CDS** | 8 |

##### Alignment

| | |
|---|---|
| **Alignment Score** | 128.0 (NT) + 500.0 (AA) = 628.0 |
| **Concordance (%)** | 41.5344 |

| | |
|---|---|
| **Alignment Method** | Global, seeded, nucleotide + amino acids (AGA) |

#### Genome Region

Sequence starts at position 4182 and ends at position 4545 relative to NC_018858.1 reference sequence.

#### Alignment Detailed Statistics

| | Begin | End | Coverage | Score | Concordance | Matches | Identities | I/D/M/F* | Stop Codons |
|---|---|---|---|---|---|---|---|---|---|
| **NT** | **4182** | **4545** | **4.6%** | **128** | **17.6%** | **364 (100%)** | **214 (58.8%)** | **0/0** | |
| Mutations: | 4182A>G, 4186C>A, 4189C>T, 4190A>T, 4192C>A, 4194A>C, 4195A>T, 4199T>A, 4200C>A, 4201G>A, 4209C>G, 4210G>T, 4212A>T, 4213G>C, 4214G>A, 4216T>A, 4219C>A, 4221A>G, 4222G>A, 4231C>T, 4235A>C, 4236A>T, 4240T>C, 4242G>A, 4243C>T, 4244C>G, 4245C>A, 4247A>C, 4248T>A, 4249G>A, 4252C>T, 4253C>A, 4254G>A, 4256G>A, 4257A>C, 4258A>T, 4265G>A, 4267G>A, 4269A>T, 4270G>T, 4273A>G, 4286C>T, 4288G>A, 4289G>A, 4291C>T, 4292C>A, 4294C>G, 4297G>A, 4298G>T, 4303C>A, 4306G>A, 4308C>A, 4309C>T, 4318C>T, 4326T>G, 4327G>T, 4328G>A, 4329C>G, 4330A>C, 4333A>C, 4336G>A, 4339C>T, 4340T>A, 4342A>G, 4345C>A, 4347A>G, 4348C>A, 4352G>C, 4354A>T, 4356C>G, 4357T>C, 4360G>A, 4361A>C, 4364C>A, 4365G>A, 4366C>A, 4367A>C, 4369A>T, 4375G>A, 4376A>G, 4377A>C, 4378G>T, 4381T>A, 4387G>A, 4388G>A, 4390C>A, 4396C>T, 4401C>A, 4402A>G, 4403A>C, 4405G>T, 4409G>A, 4411T>A, 4412G>A, 4413C>A, 4414G>T, 4415A>T, 4416C>G, 4420A>C, 4421G>T, 4422G>T, 4428C>G, 4432C>T, 4433A>T, 4434A>T, 4435T>C, 4436C>A, 4438T>C, 4441C>A, 4444C>T, 4447G>A, 4448G>A, 4451G>A, 4453A>C, 4456C>T, 4457C>A, 4459T>A, 4461C>A, 4462G>T, 4465C>T, 4466A>G, 4467T>C, 4468C>A, 4470G>A, 4477G>A, 4479A>C, 4480A>C, 4481A>T, 4482T>A, 4486C>T, 4489T>A, 4490T>A, 4491C>A, 4492C>A, 4498C>T, 4505T>A, 4506C>G, 4507A>T, 4513C>T, 4520G>A, 4522T>A, 4524G>A, 4525G>A, 4528T>A, 4531C>A, 4535G>A, 4537A>T, 4540A>T, 4542A>T, 4544T>G | | | | | | | | |

##### CDS

| | Begin | End | Coverage | Score | Concordance | Matches | Identities | I/D/M/F* | Stop Codons |
|---|---|---|---|---|---|---|---|---|---|
| D306_gp7 | 236 | 356 | 17.8% | 500 | 62.8% | 121 (100%) | 68 (56.2%) | 0/0/0/0 | 0 |
| Protein mutations: | I238L (4190A>T 4192C>A), K239T (4194A>C 4195A>T), S241K (4199T>A 4200C>A 4201G>A), T244S (4209C>G 4210G>T), K245I (4212A>T 4213G>C), V246I (4214G>A 4216T>A), K248R (4221A>G 4222G>A), K253L (4235A>C 4236A>T), S255N (4242G>A 4243C>T), P256E (4244C>G 4245C>A), M257Q (4247A>C 4248T>A 4249G>A), R259K (4253C>A 4254G>A), E260T (4256G>A 4257A>C 4258A>T), E263K (4265G>A 4267G>A), K264I (4269A>T 4270G>T), D271N (4289G>A 4291C>T), L272M (4292C>A 4294C>G), V274L (4298G>T), P277H (4308C>A 4309C>T), M283S (4326T>G 4327G>T), A284S (4328G>A 4329C>G 4330A>C), L288M (4340T>A 4342A>G), N290R (4347A>G 4348C>A), E292H (4352G>C 4354A>T), A293G (4356C>G 4357T>C), K295Q (4361A>C), R296K (4364C>A 4365G>A 4366C>A), K300A (4376A>G 4377A>C 4378G>T), V304I (4388G>A 4390C>A), A308E (4401C>A 4402A>G), M309L (4403A>C 4405G>T), D311K (4409G>A 4411T>A), A312N (4412G>A 4413C>A 4414G>T), T313C (4415A>T 4416C>G), G315F (4421G>T 4422G>T), A317G (4428C>G), N319F (4433A>T 4434A>T 4435T>C), L320I (4436C>A 4438T>C), D324N (4448G>A), E325N (4451G>A 4453A>C), L327I (4457C>A 4459T>A), T328N (4461C>A 4462G>T), I330A (4466A>G 4467T>C 4468C>A), R331K (4470G>A), K334T (4479A>C 4480A>C), I335Y (4481A>T 4482T>A), S338K (4490T>A 4491C>A 4492C>A), V348I (4520G>A 4522T>A), R349K (4524G>A 4525G>A), D351E (4531C>A), E353N (4535G>A 4537A>T), K355I (4542A>T) | | | | | | | | |
| Codon mutations: | CAA235.GA (4182A>G), GCC236GCA (4186C>A), AGC237AGT (4189C>T), ATC238TTA (4190A>T 4192C>A), AAA239ACT (4194A>C 4195A>T), TCG241AAA (4199T>A 4200C>A 4201G>A), ACG244AGT (4209C>G 4210G>T), AAG245ATC (4212A>T 4213G>C), GTT246ATA (4214G>A 4216T>A), ATC247ATA (4219C>A), AAG248AGA (4221A>G 4222G>A), CCC251CCT (4231C>T), AAA253CTA (4235A>C 4236A>T), TAT254TAC (4240T>C), AGC255AAT (4242G>A 4243C>T), CCA256GAA (4244C>G 4245C>A), ATG257CAA (4247A>C 4248T>A 4249G>A), GAC258GAT (4252C>T), CGA259AAA (4253C>A 4254G>A), GAA260ACT (4256G>A 4257A>C 4258A>T), GAG263AAA (4265G>A 4267G>A), AAG264ATT (4269A>T 4270G>T), CAA265CAG (4273A>G), CTG270TTA (4286C>T 4288G>A), GAC271AAT (4289G>A 4291C>T), CTC272ATG (4292C>A 4294C>G), AAG273AAA (4297G>A), GTA274TTA (4298G>T), ATC275ATA (4303C>A), AGG276AGA (4306G>A), CCC277CAT (4308C>A 4309C>T), AGC280AGT (4318C>T), ATG283AGT (4326T>G 4327G>T), GCA284AGC (4328G>A 4329C>G 4330A>C), CCA285CCC (4333A>C), GCG286GCA (4336G>A), TTC287TTT (4339C>T), TTA288ATG (4340T>A 4342A>G), GTC289GTA (4345C>A), AAC290AGA (4347A>G 4348C>A), GAA292CAT (4352G>C 4354A>T), GCT293GGC (4356C>G 4357T>C), GAG294GAA (4360G>A), AAA295CAA (4361A>C), CGC296AAA (4364C>A 4365G>A 4366C>A), AGA297CGT (4367A>C 4369A>T), AAG299AAA (4375G>A), AAG300GCT (4376A>G 4377A>C 4378G>T), CGT301CGA (4381T>A), GTG303GTA (4387G>A), GTC304ATA (4388G>A 4390C>A), TAC306TAT (4396C>T), GCA308GAG (4401C>A 4402A>G), ATG309CTT (4403A>C 4405G>T), GAT311AAA (4409G>A 4411T>A), GCG312AAT (4412G>A 4413C>A 4414G>T), ACT313TGT (4415A>T 4416C>G), GTA314GTC (4420A>C), GGC315TTC (4421G>T 4422G>T), GCA317GGA (4428C>G), TAC318TAT (4432C>T), AAT319TTC (4433A>T 4434A>T 4435T>C), CTT320ATC (4436C>A 4438T>C), CCC321CCA (4441C>A), AAC322AAT (4444C>T), AAG323AAA (4447G>A), GAT324AAT (4448G>A), GAA325AAC (4451G>A 4453A>C), CTC326CTT (4456C>T), CTT327ATA (4457C>A 4459T>A), ACG328AAT (4461C>A 4462G>T), CTC329CTT (4465C>T), ATC330GCA (4466A>G 4467T>C 4468C>A), AGA331AAA (4470G>A), AAG333AAA (4477G>A), AAA334ACC (4479A>C 4480A>C), ATC335TAC (4481A>T 4482T>A), TTC336TTT (4486C>T), TCT337TCA (4489T>A), TCC338AAA (4490T>A 4491C>A 4492C>A), GAC340GAT (4498C>T), TCA343AGT (4505T>A 4506C>G 4507A>T), TTC345TTT (4513C>T), GTT348ATA (4520G>A 4522T>A), AGG349AAA (4524G>A 4525G>A), CTT350CTA (4528T>A), GAC351GAA (4531C>A), GAA353AAT (4535G>A 4537A>T), TCA354TCT (4540A>T), AAA355ATA (4542A>T), TCC356GC. (4544T>G) | | | | | | | | |

##### Proteins

| | Begin | End | Coverage | Score | Concordance | Matches | Identities | I/D/M/F* | Stop Codons |
|---|---|---|---|---|---|---|---|---|---|
| polyprotein (YP_006907834.1) | 236 | 356 | 17.8% | 500 | 62.8% | 121 (100%) | 68 (56.2%) | 0/0/0/0 | 0 |
| Protein mutations: | I238L (4190A>T 4192C>A), K239T (4194A>C 4195A>T), S241K (4199T>A 4200C>A 4201G>A), T244S (4209C>G 4210G>T), K245I (4212A>T 4213G>C), V246I (4214G>A 4216T>A), K248R (4221A>G 4222G>A), K253L (4235A>C 4236A>T), S255N (4242G>A 4243C>T), P256E (4244C>G 4245C>A), M257Q (4247A>C 4248T>A 4249G>A), R259K (4253C>A 4254G>A), E260T (4256G>A 4257A>C 4258A>T), E263K (4265G>A 4267G>A), K264I (4269A>T 4270G>T), D271N (4289G>A 4291C>T), L272M (4292C>A 4294C>G), V274L (4298G>T), P277H (4308C>A 4309C>T), M283S (4326T>G 4327G>T), A284S (4328G>A 4329C>G 4330A>C), L288M (4340T>A 4342A>G), N290R (4347A>G 4348C>A), E292H (4352G>C 4354A>T), A293G (4356C>G 4357T>C), K295Q (4361A>C), R296K (4364C>A 4365G>A 4366C>A), K300A (4376A>G 4377A>C 4378G>T), V304I (4388G>A 4390C>A), A308E (4401C>A 4402A>G), M309L (4403A>C 4405G>T), D311K (4409G>A 4411T>A), A312N (4412G>A 4413C>A 4414G>T), T313C (4415A>T 4416C>G), G315F (4421G>T 4422G>T), A317G (4428C>G), N319F (4433A>T 4434A>T 4435T>C), L320I (4436C>A 4438T>C), D324N (4448G>A), E325N (4451G>A 4453A>C), L327I (4457C>A 4459T>A), T328N (4461C>A 4462G>T), I330A (4466A>G 4467T>C 4468C>A), R331K (4470G>A), K334T (4479A>C 4480A>C), I335Y (4481A>T 4482T>A), S338K (4490T>A 4491C>A 4492C>A), V348I (4520G>A 4522T>A), R349K (4524G>A 4525G>A), D351E (4531C>A), E353N (4535G>A 4537A>T), K355I (4542A>T) | | | | | | | | |
| Codon mutations: | CAA235.GA (4182A>G), GCC236GCA (4186C>A), AGC237AGT (4189C>T), ATC238TTA (4190A>T 4192C>A), AAA239ACT (4194A>C 4195A>T), TCG241AAA (4199T>A 4200C>A 4201G>A), ACG244AGT (4209C>G 4210G>T), AAG245ATC (4212A>T 4213G>C), GTT246ATA (4214G>A 4216T>A), ATC247ATA (4219C>A), AAG248AGA (4221A>G 4222G>A), CCC251CCT (4231C>T), AAA253CTA (4235A>C 4236A>T), TAT254TAC (4240T>C), AGC255AAT (4242G>A 4243C>T), CCA256GAA (4244C>G 4245C>A), ATG257CAA (4247A>C 4248T>A 4249G>A), GAC258GAT (4252C>T), CGA259AAA (4253C>A 4254G>A), GAA260ACT (4256G>A 4257A>C 4258A>T), GAG263AAA (4265G>A 4267G>A), AAG264ATT (4269A>T 4270G>T), CAA265CAG (4273A>G), CTG270TTA (4286C>T 4288G>A), GAC271AAT (4289G>A 4291C>T), CTC272ATG (4292C>A 4294C>G), AAG273AAA (4297G>A), GTA274TTA (4298G>T), ATC275ATA (4303C>A), AGG276AGA (4306G>A), CCC277CAT (4308C>A 4309C>T), AGC280AGT (4318C>T), ATG283AGT (4326T>G 4327G>T), GCA284AGC (4328G>A 4329C>G 4330A>C), CCA285CCC (4333A>C), GCG286GCA (4336G>A), TTC287TTT (4339C>T), TTA288ATG (4340T>A 4342A>G), GTC289GTA (4345C>A), AAC290AGA (4347A>G 4348C>A), GAA292CAT (4352G>C 4354A>T), GCT293GGC (4356C>G 4357T>C), GAG294GAA (4360G>A), AAA295CAA (4361A>C), CGC296AAA (4364C>A 4365G>A 4366C>A), AGA297CGT (4367A>C 4369A>T), AAG299AAA (4375G>A), AAG300GCT (4376A>G 4377A>C 4378G>T), CGT301CGA (4381T>A), GTG303GTA (4387G>A), GTC304ATA (4388G>A 4390C>A), TAC306TAT (4396C>T), GCA308GAG (4401C>A 4402A>G), ATG309CTT (4403A>C 4405G>T), GAT311AAA (4409G>A 4411T>A), GCG312AAT (4412G>A 4413C>A 4414G>T), ACT313TGT (4415A>T 4416C>G), GTA314GTC (4420A>C), GGC315TTC (4421G>T 4422G>T), GCA317GGA (4428C>G), TAC318TAT (4432C>T), AAT319TTC (4433A>T 4434A>T 4435T>C), CTT320ATC (4436C>A 4438T>C), CCC321CCA (4441C>A), AAC322AAT (4444C>T), AAG323AAA (4447G>A), GAT324AAT (4448G>A), GAA325AAC (4451G>A 4453A>C), CTC326CTT (4456C>T), CTT327ATA (4457C>A 4459T>A), ACG328AAT (4461C>A 4462G>T), CTC329CTT (4465C>T), ATC330GCA (4466A>G 4467T>C 4468C>A), AGA331AAA (4470G>A), AAG333AAA (4477G>A), AAA334ACC (4479A>C 4480A>C), ATC335TAC (4481A>T 4482T>A), TTC336TTT (4486C>T), TCT337TCA (4489T>A), TCC338AAA (4490T>A 4491C>A 4492C>A), GAC340GAT (4498C>T), TCA343AGT (4505T>A 4506C>G 4507A>T), TTC345TTT (4513C>T), GTT348ATA (4520G>A 4522T>A), AGG349AAA (4524G>A 4525G>A), CTT350CTA (4528T>A), GAC351GAA (4531C>A), GAA353AAT (4535G>A 4537A>T), TCA354TCT (4540A>T), AAA355ATA (4542A>T), TCC356GC. (4544T>G) | | | | | | | | |

*: Inserts / Deletes / Misaligned / Frameshifts

#### Analysis details

This analysis was performed with panviral2.64

### NGS Details (UN9): Badnavirus occultipomeae

#### Assembly

| | |
|---|---|
| **Coverage Length** | 347 (1 contig(s)) |
| **Depth Of Coverage** | 5.4 |
| **Number Of Reads** | 18 |
| **Reads Per Million** | 0.41 rpm (after QC) |
| **Ambiguities** | 0 |
| **Assembly Method** | de novo + reference guided assembly |
| **Consensus Caller** | Bcf Tools |

#### Coverage Map

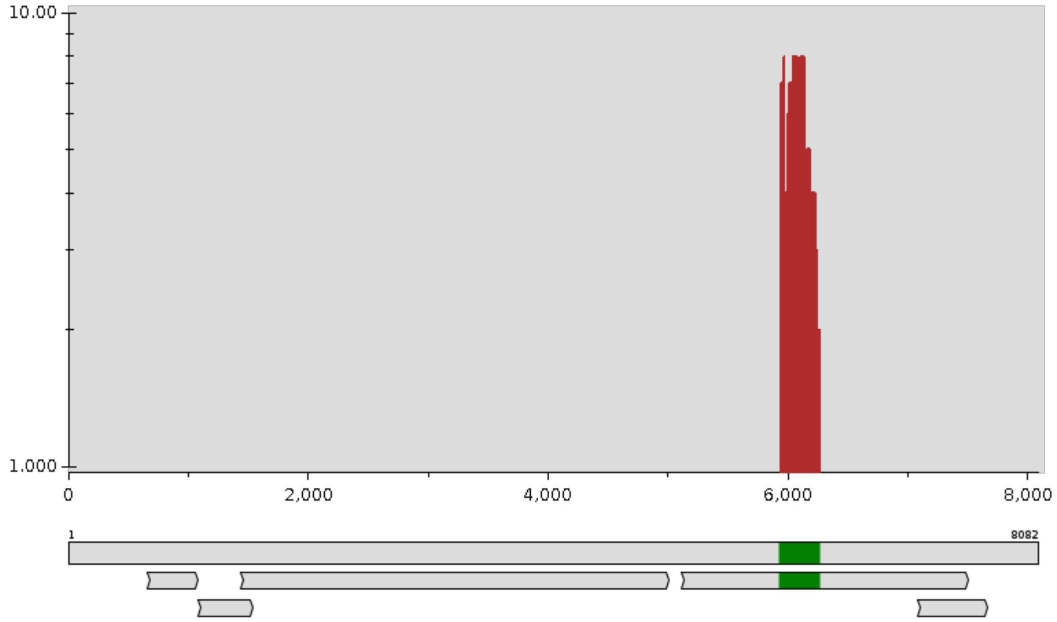

#### Assignment

| | |
|---|---|
| **Type** | Badnavirus occultipomeae (Taxonomy ID: 3048353) |
| **Reference Genome** | NC_015655.1 |
| **NT Identity (%)** | 54.8571 |
| **AA Identity (%)** | 46.1538 |
| **Number Of Stop Codons** | 0 |
| **Number Of CDS** | 5 |

#### Alignment

| | |
|---|---|
| **Alignment Score** | 62.0 (NT) + 380.0 (AA) = 442.0 |
| **Concordance (%)** | 29.4079 |

| **Alignment Method** | Global, seeded, nucleotide + amino acids (AGA) |
|---|---|

#### Genome Region

Sequence starts at position 5926 and ends at position 6272 relative to NC_015655.1 reference sequence.

#### Alignment Detailed Statistics

| | Begin | End | Coverage | Score | Concordance | Matches | Identities | I/D/M/F* | Stop Codons |
|---|---|---|---|---|---|---|---|---|---|
| **NT** | **5926** | **6272** | **4.3%** | **62** | **8.9%** | **347 (99.1%)** | **192 (54.9%)** | **3/0** | |
| Mutations: | 5927A>T, 5928A>T, 5929A>C, 5935G>T, 5936T>G, 5938T>A, 5939A>T, 5940C>T, 5941A>G, 5943C>T, 5946C>T, 5947A>C, 5948A>G, 5949G>T, 5951G>A, 5952C>A, 5953C>T, 5955C>G, 5959G>A, 5961C>A, 5962A>G, 5963A>T, 5964T>C, 5967A>C, 5968G>A, 5969A>T, 5970A>C, 5973A>G, 5974G>A, 5976T>C, 5977C>A, 5985C>A, 5986C>A, 5988A>T, 5991G>T, 5992G>A, 5998A>G, 6002C>A, 6004A>T, 6006T>G, 6007G>T, 6009C>T, 6010A>G, 6011G>A, 6013A>C, 6014G>A, 6016A>T, 6018A>G, 6019T>C, 6020C>A, 6021T>A, 6024A>G, 6025A>G, 6026A>G, 6027A>T, 6029A>G, 6030G>T, 6031A>C, 6032T>A, 6033A>C, 6039A>C, 6042A>G, 6043T>A, 6048T>C, 6051C>T, 6052A>C, 6053A>G, 6054A>T, 6055G>T, 6056C>T, 6057A>C, 6060C>T, 6062T>A, 6069G>A, 6072A>G, 6075A>G, 6076A>G, 6078G>A, 6079G>A, 6080A>G, 6085A>T, 6086A>G, 6087G>T, 6088T>G, 6089C>A, 6090C>T, 6091A>G, 6092A>T, 6093A>T, 6096A>C, 6097T>A, 6098G>A, 6099G>A, 6102T>G, 6103G>A, 6109T>C, 6114T>A, 6115C>A, 6116C>G, 6117A>G, 6118G>T, 6120A>C, 6125T>A, 6128A>T, 6137A>T, 6138A>G, 6141C>G, 6145C>T, 6147C>T, 6153A>G, 6154C>T, 6156G>A, 6168A>G, 6171A>G, 6173A>T, 6174T>G, 6177T>C, 6178C>A, 6179A>T, 6182G>A, 6183A>T, 6184A>C, 6185A>T, 6186G>A, 6190G>A, 6194A>G, 6197C>T, 6198A>G, 6201T>C, 6203G>A, 6205G>C, 6206G>C, 6207A>T, 6207_6208insTAT, 6208A>C, 6209C>T, 6210A>T, 6214G>A, 6220A>G, 6222C>G, 6224C>T, 6225A>G, 6228A>C, 6230A>T, 6231T>C, 6234T>A, 6246G>A, 6247G>A, 6249A>C, 6251T>A, 6256G>C, 6257A>G, 6258A>G, 6259A>G, 6260A>G, 6270G>A | | | | | | | | |

##### CDS

| | Begin | End | Coverage | Score | Concordance | Matches | Identities | I/D/M/F* | Stop Codons |
|---|---|---|---|---|---|---|---|---|---|
| SPBVa_gp4 | 271 | 386 | 14.6% | 380 | 46.1% | 116 (99.1%) | 54 (46.2%) | 1/0/0/0 | 0 |
| Protein mutations: | Q271L (5927A>T 5928A>T), V274C (5935G>T 5936T>G), Y275I (5938T>A 5939A>T 5940C>T), N276D (5941A>G 5943C>T), K278R (5947A>C 5948A>G 5949G>T), R279Q (5951G>A 5952C>A), D282K (5959G>A 5961C>A), N283V (5962A>G 5963A>T 5964T>C), E285I (5968G>A 5969A>T 5970A>C), D287N (5974G>A 5976T>C), Q288K (5977C>A), L291I (5986C>A 5988A>T), G293R (5992G>A), N295D (5998A>G), T296N (6002C>A), I297L (6004A>T 6006T>G), V298F (6007G>T 6009C>T), S299D (6010A>G 6011G>A), R300Q (6013A>C 6014G>A), I301L (6016A>T 6018A>G), S302Q (6019T>C 6020C>A 6021T>A), K304G (6025A>G 6026A>G 6027A>T), K305S (6029A>G 6030G>T), I306H (6031A>C 6032T>A 6033A>C), F310I (6043T>A), K313R (6052A>C 6053A>G 6054A>T), A314F (6055G>T 6056C>T 6057A>C), F316Y (6062T>A), I319M (6072A>G), M321V (6076A>G 6078G>A), E322R (6079G>A 6080A>G), K324C (6085A>T 6086A>G 6087G>T), S325D (6088T>G 6089C>A 6090C>T), K326V (6091A>G 6092A>T 6093A>T), W328K (6097T>A 6098G>A 6099G>A), A330T (6103G>A), W332R (6109T>C), P334R (6115C>A 6116C>G 6117A>G), E335Y (6118G>T 6120A>C), L337H (6125T>A), Y338F (6128A>T), E341V (6137A>T 6138A>G), P344S (6145C>T 6147C>T), D353V (6173A>T 6174T>G), Q355M (6178C>A 6179A>T), R356N (6182G>A 6183A>T), K357L (6184A>C 6185A>T 6186G>A), D359N (6190G>A), N360S (6194A>G), A361V (6197C>T 6198A>G), R363K (6203G>A), G364P (6205G>C 6206G>C 6207A>T), G364_T365insY (6207_6208insTAT), T365L (6208A>C 6209C>T 6210A>T), A367T (6214G>A), I369V (6220A>G 6222C>G), A370V (6224C>T 6225A>G), Y372F (6230A>T 6231T>C), V378I (6247G>A 6249A>C), F379Y (6251T>A), E381R (6256G>C 6257A>G 6258A>G), N382G (6259A>G 6260A>G) | | | | | | | | |
| Codon mutations: | CAA271CTT (5927A>T 5928A>T), AGG272CGG (5929A>C), GTT274TGT (5935G>T 5936T>G), TAC275ATT (5938T>A 5939A>T 5940C>T), AAC276GAT (5941A>G 5943C>T), TAC277TAT (5946C>T), AAG278CGT (5947A>C 5948A>G 5949G>T), CGC279CAA (5951G>A 5952C>A), CTC280TTG (5953C>T 5955C>G), GAC282AAA (5959G>A 5961C>A), AAT283GTC (5962A>G 5963A>T 5964T>C), ACA284ACC (5967A>C), GAA285ATC (5968G>A 5969A>T 5970A>C), AAA286AAG (5973A>G), GAT287AAC (5974G>A 5976T>C), CAG288AAG (5977C>A), TCC290TCA (5985C>A), CTA291ATT (5986C>A 5988A>T), CCG292CCT (5991G>T), GGA293AGA (5992G>A), AAC295GAC (5998A>G), ACT296AAT (6002C>A), ATT297TTG (6004A>T 6006T>G), GTC298TTT (6007G>T 6009C>T), AGT299GAT (6010A>G 6011G>A), AGA300CAA (6013A>C 6014G>A), ATA301TTG (6016A>T 6018A>G), TCT302CAA (6019T>C 6020C>A 6021T>A), GGA303GGG (6024A>G), AAA304GGT (6025A>G 6026A>G 6027A>T), AAG305AGT (6029A>G 6030G>T), ATA306CAC (6031A>C 6032T>A 6033A>C), TCA308TCC (6039A>C), AAA309AAG (6042A>G), TTT310ATT (6043T>A), GAT311GAC (6048T>C), CTC312CTT (6051C>T), AAA313CGT (6052A>C 6053A>G 6054A>T), GCA314TTC (6055G>T 6056C>T 6057A>C), GGC315GGT (6060C>T), TTT316TAT (6062T>A), CAG318CAA (6069G>A), ATA319ATG (6072A>G), AGA320AGG (6075A>G), ATG321GTA (6076A>G 6078G>A), GAG322AGG (6079G>A 6080A>G), AAG324TGT (6085A>T 6086A>G 6087G>T), TCC325GAT (6088T>G 6089C>A 6090C>T), AAA326GTT (6091A>G 6092A>T 6093A>T), CCA327CCC (6096A>C), TGG328AAA (6097T>A 6098G>A 6099G>A), ACT329ACG (6102T>G), GCT330ACT (6103G>A), TGG332CGG (6109T>C), ACT333ACA (6114T>A), CCA334AGG (6115C>A 6116C>G 6117A>G), GAA335TAC (6118G>T 6120A>C), CTC337CAC (6125T>A), TAT338TTT (6128A>T), GAA341GTG (6137A>T 6138A>G), GTC342GTG (6141C>G), CCC344TCT (6145C>T 6147C>T), GGA346GGG (6153A>G), CTG347TTA (6154C>T 6156G>A), CCA351CCG (6168A>G), GCA352GCG (6171A>G), GAT353GTG (6173A>T 6174T>G), TTT354TTC (6177T>C), CAG355ATG (6178C>A 6179A>T), AGA356AAT (6182G>A 6183A>T), AAG357CTA (6184A>C 6185A>T 6186G>A), GAT359AAT (6190G>A), AAT360AGT (6194A>G), GCA361GTG (6197C>T 6198A>G), TTT362TTC (6201T>C), AGG363AAG (6203G>A), GGA364CCT (6205G>C 6206G>C 6207A>T), GGA364_ACA365insTAT (6207_6208insTAT), ACA365CTT (6208A>C 6209C>T 6210A>T), GCA367ACA (6214G>A), ATC369GTG (6220A>G 6222C>G), GCA370GTG (6224C>T 6225A>G), GTA371GTC (6228A>C), TAT372TTC (6230A>T 6231T>C), ATT373ATA (6234T>A), TTG377TTA (6246G>A), GTA378ATC (6247G>A 6249A>C), TTC379TAC (6251T>A), GAA381CGG (6256G>C 6257A>G 6258A>G), AAT382GGT (6259A>G 6260A>G), GAG385GAA (6270G>A) | | | | | | | | |

##### Proteins

| | Begin | End | Coverage | Score | Concordance | Matches | Identities | I/D/M/F* | Stop Codons |
|---|---|---|---|---|---|---|---|---|---|
| RNaseH/reverse transcriptase (YP_004581513.1) | 271 | 386 | 14.6% | 380 | 46.1% | 116 (99.1%) | 54 (46.2%) | 1/0/0/0 | 0 |
| Protein mutations: | Q271L (5927A>T 5928A>T), V274C (5935G>T 5936T>G), Y275I (5938T>A 5939A>T 5940C>T), N276D (5941A>G 5943C>T), K278R (5947A>C 5948A>G 5949G>T), R279Q (5951G>A 5952C>A), D282K (5959G>A 5961C>A), N283V (5962A>G 5963A>T 5964T>C), E285I (5968G>A 5969A>T 5970A>C), D287N (5974G>A 5976T>C), Q288K (5977C>A), L291I (5986C>A 5988A>T), G293R (5992G>A), N295D (5998A>G), T296N (6002C>A), I297L (6004A>T 6006T>G), V298F (6007G>T 6009C>T), S299D (6010A>G 6011G>A), R300Q (6013A>C 6014G>A), I301L (6016A>T 6018A>G), S302Q (6019T>C 6020C>A 6021T>A), K304G (6025A>G 6026A>G 6027A>T), K305S (6029A>G 6030G>T), I306H (6031A>C 6032T>A 6033A>C), F310I (6043T>A), K313R (6052A>C 6053A>G 6054A>T), A314F (6055G>T 6056C>T 6057A>C), F316Y (6062T>A), I319M (6072A>G), M321V (6076A>G 6078G>A), E322R (6079G>A 6080A>G), K324C (6085A>T 6086A>G 6087G>T), S325D (6088T>G 6089C>A 6090C>T), K326V (6091A>G 6092A>T 6093A>T), W328K (6097T>A 6098G>A 6099G>A), A330T (6103G>A), W332R (6109T>C), P334R (6115C>A 6116C>G 6117A>G), E335Y (6118G>T 6120A>C), L337H (6125T>A), Y338F (6128A>T), E341V (6137A>T 6138A>G), P344S (6145C>T 6147C>T), D353V (6173A>T 6174T>G), Q355M (6178C>A 6179A>T), R356N (6182G>A 6183A>T), K357L (6184A>C 6185A>T 6186G>A), D359N (6190G>A), N360S (6194A>G), A361V (6197C>T 6198A>G), R363K (6203G>A), G364P (6205G>C 6206G>C 6207A>T), G364_T365insY (6207_6208insTAT), T365L (6208A>C 6209C>T 6210A>T), A367T (6214G>A), I369V (6220A>G 6222C>G), A370V (6224C>T 6225A>G), Y372F (6230A>T 6231T>C), V378I (6247G>A 6249A>C), F379Y (6251T>A), E381R (6256G>C 6257A>G 6258A>G), N382G (6259A>G 6260A>G) | | | | | | | | |
| Codon mutations: | CAA271CTT (5927A>T 5928A>T), AGG272CGG (5929A>C), GTT274TGT (5935G>T 5936T>G), TAC275ATT (5938T>A 5939A>T 5940C>T), AAC276GAT (5941A>G 5943C>T), TAC277TAT (5946C>T), AAG278CGT (5947A>C 5948A>G 5949G>T), CGC279CAA (5951G>A 5952C>A), CTC280TTG (5953C>T 5955C>G), GAC282AAA (5959G>A 5961C>A), AAT283GTC (5962A>G 5963A>T 5964T>C), ACA284ACC (5967A>C), GAA285ATC (5968G>A 5969A>T 5970A>C), AAA286AAG (5973A>G), GAT287AAC (5974G>A 5976T>C), CAG288AAG (5977C>A), TCC290TCA (5985C>A), CTA291ATT (5986C>A 5988A>T), CCG292CCT (5991G>T), GGA293AGA (5992G>A), AAC295GAC (5998A>G), ACT296AAT (6002C>A), ATT297TTG (6004A>T 6006T>G), GTC298TTT (6007G>T 6009C>T), AGT299GAT (6010A>G 6011G>A), AGA300CAA (6013A>C 6014G>A), ATA301TTG (6016A>T 6018A>G), TCT302CAA (6019T>C 6020C>A 6021T>A), GGA303GGG (6024A>G), AAA304GGT (6025A>G 6026A>G 6027A>T), AAG305AGT (6029A>G 6030G>T), ATA306CAC (6031A>C 6032T>A 6033A>C), TCA308TCC (6039A>C), AAA309AAG (6042A>G), TTT310ATT (6043T>A), GAT311GAC (6048T>C), CTC312CTT (6051C>T), AAA313CGT (6052A>C 6053A>G 6054A>T), GCA314TTC (6055G>T 6056C>T 6057A>C), GGC315GGT (6060C>T), TTT316TAT (6062T>A), CAG318CAA (6069G>A), ATA319ATG (6072A>G), AGA320AGG (6075A>G), ATG321GTA (6076A>G 6078G>A), GAG322AGG (6079G>A 6080A>G), AAG324TGT (6085A>T 6086A>G 6087G>T), TCC325GAT (6088T>G 6089C>A 6090C>T), AAA326GTT (6091A>G 6092A>T 6093A>T), CCA327CCC (6096A>C), TGG328AAA (6097T>A 6098G>A 6099G>A), ACT329ACG (6102T>G), GCT330ACT (6103G>A), TGG332CGG (6109T>C), ACT333ACA (6114T>A), CCA334AGG (6115C>A 6116C>G 6117A>G), GAA335TAC (6118G>T 6120A>C), CTC337CAC (6125T>A), TAT338TTT (6128A>T), GAA341GTG (6137A>T 6138A>G), GTC342GTG (6141C>G), CCC344TCT (6145C>T 6147C>T), GGA346GGG (6153A>G), CTG347TTA (6154C>T 6156G>A), CCA351CCG (6168A>G), GCA352GCG (6171A>G), GAT353GTG (6173A>T 6174T>G), TTT354TTC (6177T>C), CAG355ATG (6178C>A 6179A>T), AGA356AAT (6182G>A 6183A>T), AAG357CTA (6184A>C 6185A>T 6186G>A), GAT359AAT (6190G>A), AAT360AGT (6194A>G), GCA361GTG (6197C>T 6198A>G), TTT362TTC (6201T>C), AGG363AAG (6203G>A), GGA364CCT (6205G>C 6206G>C 6207A>T), GGA364_ACA365insTAT (6207_6208insTAT), ACA365CTT (6208A>C 6209C>T 6210A>T), GCA367ACA (6214G>A), ATC369GTG (6220A>G 6222C>G), GCA370GTG (6224C>T 6225A>G), GTA371GTC (6228A>C), TAT372TTC (6230A>T 6231T>C), ATT373ATA (6234T>A), TTG377TTA (6246G>A), GTA378ATC (6247G>A 6249A>C), TTC379TAC (6251T>A), GAA381CGG (6256G>C 6257A>G 6258A>G), AAT382GGT (6259A>G 6260A>G), GAG385GAA (6270G>A) | | | | | | | | |

*: Inserts / Deletes / Misaligned / Frameshifts

#### Analysis details

This analysis was performed with panviral2.64

### NGS Details (UN9): Badnavirus occultipomeae

#### Assembly

| | |
|---|---|
| **Coverage Length** | 257 (1 contig(s)) |
| **Depth Of Coverage** | 6.7 |
| **Number Of Reads** | 15 |
| **Reads Per Million** | 0.34 rpm (after QC) |
| **Ambiguities** | 0 |
| **Assembly Method** | de novo + reference guided assembly |
| **Consensus Caller** | Bcf Tools |

#### Coverage Map

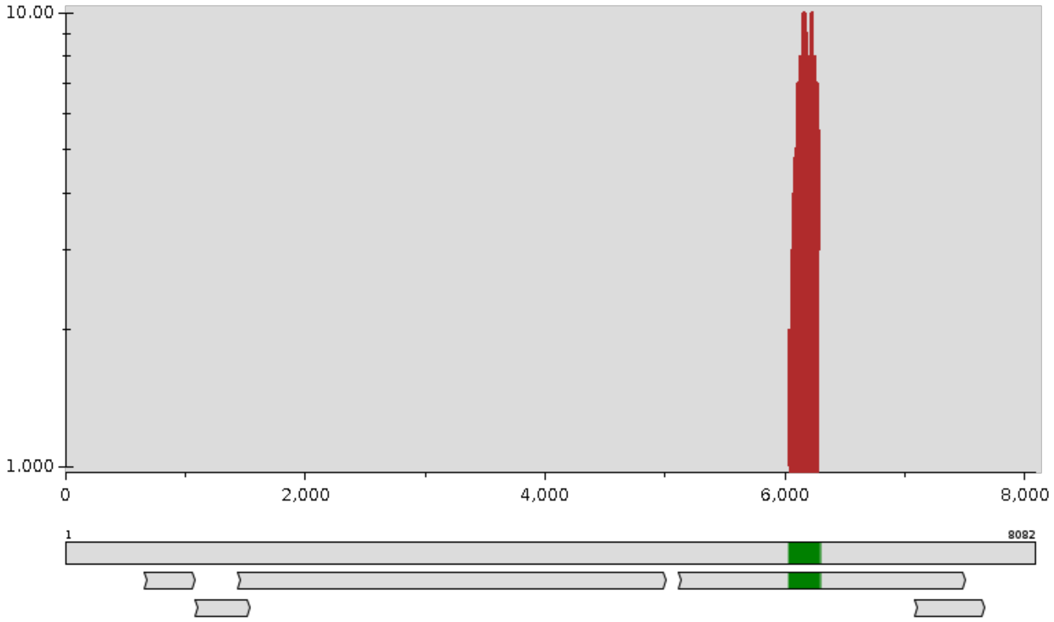

#### Assignment

| | |
|---|---|
| **Type** | Badnavirus occultipomeae (Taxonomy ID: 3048353) |
| **Reference Genome** | NC_015655.1 |
| **NT Identity (%)** | 61.4504 |
| **AA Identity (%)** | 47.191 |
| **Number Of Stop Codons** | 0 |
| **Number Of CDS** | 5 |

#### Alignment

| | |
|---|---|
| **Alignment Score** | 107.0 (NT) + 177.0 (AA) = 284.0 |
| **Concordance (%)** | 27.6539 |

| Alignment Method | Global, seeded, nucleotide + amino acids (AGA) |
|---|---|

#### Genome Region

Sequence starts at position 6031 and ends at position 6289 relative to NC_015655.1 reference sequence.

#### Alignment Detailed Statistics

| | Begin | End | Coverage | Score | Concordance | Matches | Identities | I/D/M/F* | Stop Codons |
|---|---|---|---|---|---|---|---|---|---|
| **NT** | **6031** | **6289** | **3.2%** | **107** | **20.8%** | **257 (98.1%)** | **161 (61.5%)** | **5/0** | |
| Mutations: | 6031A>G, 6033A>G, 6043T>A, 6051C>A, 6053A>G, 6055G>T, 6057A>T, 6060C>T, 6062T>A, 6063T>C, 6070A>T, 6074G>A, 6075A>G, 6077T>A, 6078G>T, 6079G>A, 6082G>C, 6083A>C, 6088T>G, 6089C>A, 6090C>T, 6091A>G, 6092A>T, 6093A>G, 6096A>T, 6097T>A, 6109T>A, 6111G>A, 6116C>G, 6117A>G, 6118G>T, 6120A>T, 6125T>A, 6129T>C, 6135T>C, 6136G>T, 6137A>T, 6138A>G, 6141C>G, 6145C>T, 6147C>G, 6148T>A, 6153A>G, 6154C>T, 6158T>C, 6159G>C, 6165A>G, 6173A>C, 6174T>C, 6177T>C, 6178C>A, 6179A>T, 6183A>T, 6184A>T, 6185A>T, 6190G>A, 6193A>G, 6194A>G, 6195T>G, 6197C>T, 6198A>G, 6203G>A, 6205G>C, 6206G>C, 6207_6208insTTT, 6208A>T, 6209C>T, 6210A>G, 6214G>T, 6216A>G, 6220A>G, 6222C>G, 6223G>A, 6224C>T, 6225A>T, 6227T>G, 6228A>G, 6230A>T, 6243A>T, 6244T>C, 6246G>T, 6247G>A, 6249A>T, 6251T>A, 6252C>T, 6255T>A, 6256G>A, 6258_6259insAG, 6260A>G, 6262G>A, 6264A>G, 6275A>C, 6276G>A, 6277G>C, 6279T>C, 6285C>A, 6286T>A, 6289A>T | | | | | | | | |
| **CDS** | | | | | | | | | |
| SPBVa_gp4 | 306 | 392 | 10.9% | 177 | 27.7% | 87 (97.8%) | 42 (47.2%) | 2/0/2/1 | 0 |
| Protein mutations: | I306V (6031A>G 6033A>G), F310I (6043T>A), K313R (6053A>G), A314S (6055G>T 6057A>T), F316Y (6062T>A 6063T>C), I319L (6070A>T), R320K (6074G>A 6075A>G), M321N (6077T>A 6078G>T), E322K (6079G>A), E323P (6082G>C 6083A>C), S325D (6088T>G 6089C>A 6090C>T), K326V (6091A>G 6092A>T 6093A>G), W328R (6097T>A), W332R (6109T>A 6111G>A), P334R (6116C>G 6117A>G), E335Y (6118G>T 6120A>T), L337H (6125T>A), E341L (6136G>T 6137A>T 6138A>G), P344S (6145C>T 6147C>G), F345I (6148T>A), M348T (6158T>C 6159G>C), D353A (6173A>C 6174T>C), Q355M (6178C>A 6179A>T), R356S (6183A>T), K357L (6184A>T 6185A>T), D359N (6190G>A), N360G (6193A>G 6194A>G 6195T>G), A361V (6197C>T 6198A>G), R363K (6203G>A), G364P (6205G>C 6206G>C), G364_T365insF (6207_6208insTTT), T365L (6208A>T 6209C>T 6210A>G), A367S (6214G>T 6216A>G), I369V (6220A>G 6222C>G), A370I (6223G>A 6224C>T 6225A>T), V371G (6227T>G 6228A>G), Y372F (6230A>T), V378I (6247G>A 6249A>T), F379Y (6251T>A 6252C>T), E381K (6256G>A), E381_N382insX (6258_6259insAG), N382S (6260A>G), E383K (6262G>A 6264A>G), E387A (6275A>C 6276G>A), D388H (6277G>C 6279T>C) | | | | | | | | |
| Codon mutations: | ATA306GTG (6031A>G 6033A>G), TTT310ATT (6043T>A), CTC312CTA (6051C>A), AAA313AGA (6053A>G), GCA314TCT (6055G>T 6057A>T), GGC315GGT (6060C>T), TTT316TAC (6062T>A 6063T>C), ATA319TTA (6070A>T), AGA320AAG (6074G>A 6075A>G), ATG321AAT (6077T>A 6078G>T), GAG322AAG (6079G>A), GAA323CCA (6082G>C 6083A>C), TCC325GAT (6088T>G 6089C>A 6090C>T), AAA326GTG (6091A>G 6092A>T 6093A>G), CCA327CCT (6096A>T), TGG328AGG (6097T>A), TGG332AGA (6109T>A 6111G>A), CCA334CGG (6116C>G 6117A>G), GAA335TAT (6118G>T 6120A>T), CTC337CAC (6125T>A), TAT338TAC (6129T>C), TTT340TTC (6135T>C), GAA341TTG (6136G>T 6137A>T 6138A>G), GTC342GTG (6141C>G), CCC344TCG (6145C>T 6147C>G), TTT345ATT (6148T>A), GGA346GGG (6153A>G), CTG347TTG (6154C>T), ATG348ACC (6158T>C 6159G>C), GCA350GCG (6165A>G), GAT353GCC (6173A>C 6174T>C), TTT354TTC (6177T>C), CAG355ATG (6178C>A 6179A>T), AGA356AGT (6183A>T), AAG357TTG (6184A>T 6185A>T), GAT359AAT (6190G>A), AAT360GGG (6193A>G 6194A>G 6195T>G), GCA361GTG (6197C>T 6198A>G), AGG363AAG (6203G>A), GGA364CCA (6205G>C 6206G>C), GGA364_ACA365insTTT (6207_6208insTTT), ACA365TTG (6208A>T 6209C>T 6210A>G), GCA367TCG (6214G>T 6216A>G), ATC369GTG (6220A>G 6222C>G), GCA370ATT (6223G>A 6224C>T 6225A>T), GTA371GGG (6227T>G 6228A>G), TAT372TTT (6230A>T), ATA376ATT (6243A>T), TTG377CTT (6244T>C 6246G>T), GTA378ATT (6247G>A 6249A>T), TTC379TAT (6251T>A 6252C>T), TCT380TCA (6255T>A), GAA381AAA (6256G>A), GAA381_AAT382insAG- (6258_6259insAG), AAT382AGT (6260A>G), GAA383AAG (6262G>A 6264A>G), GAG387GCA (6275A>C 6276G>A), GAT388CAC (6277G>C 6279T>C), CTC390CTA (6285C>A), TTA391A.. (6286T>A), AAT392T.. (6289A>T) | | | | | | | | |
| **Proteins** | | | | | | | | | |
| RNaseH/reverse transcriptase (YP_004581513.1) | 306 | 392 | 10.9% | 177 | 27.7% | 87 (97.8%) | 42 (47.2%) | 2/0/2/1 | 0 |
| Protein mutations: | I306V (6031A>G 6033A>G), F310I (6043T>A), K313R (6053A>G), A314S (6055G>T 6057A>T), F316Y (6062T>A 6063T>C), I319L (6070A>T), R320K (6074G>A 6075A>G), M321N (6077T>A 6078G>T), E322K (6079G>A), E323P (6082G>C 6083A>C), S325D (6088T>G 6089C>A 6090C>T), K326V (6091A>G 6092A>T 6093A>G), W328R (6097T>A), W332R (6109T>A 6111G>A), P334R (6116C>G 6117A>G), E335Y (6118G>T 6120A>T), L337H (6125T>A), E341L (6136G>T 6137A>T 6138A>G), P344S (6145C>T 6147C>G), F345I (6148T>A), M348T (6158T>C 6159G>C), D353A (6173A>C 6174T>C), Q355M (6178C>A 6179A>T), R356S (6183A>T), K357L (6184A>T 6185A>T), D359N (6190G>A), N360G (6193A>G 6194A>G 6195T>G), A361V (6197C>T 6198A>G), R363K (6203G>A), G364P (6205G>C 6206G>C), G364_T365insF (6207_6208insTTT), T365L (6208A>T 6209C>T 6210A>G), A367S (6214G>T 6216A>G), I369V (6220A>G 6222C>G), A370I (6223G>A 6224C>T 6225A>T), V371G (6227T>G 6228A>G), Y372F (6230A>T), V378I (6247G>A 6249A>T), F379Y (6251T>A 6252C>T), E381K (6256G>A), E381_N382insX (6258_6259insAG), N382S (6260A>G), E383K (6262G>A 6264A>G), E387A (6275A>C 6276G>A), D388H (6277G>C 6279T>C) | | | | | | | | |
| Codon mutations: | ATA306GTG (6031A>G 6033A>G), TTT310ATT (6043T>A), CTC312CTA (6051C>A), AAA313AGA (6053A>G), GCA314TCT (6055G>T 6057A>T), GGC315GGT (6060C>T), TTT316TAC (6062T>A 6063T>C), ATA319TTA (6070A>T), AGA320AAG (6074G>A 6075A>G), ATG321AAT (6077T>A 6078G>T), GAG322AAG (6079G>A), GAA323CCA (6082G>C 6083A>C), TCC325GAT (6088T>G 6089C>A 6090C>T), AAA326GTG (6091A>G 6092A>T 6093A>G), CCA327CCT (6096A>T), TGG328AGG (6097T>A), TGG332AGA (6109T>A 6111G>A), CCA334CGG (6116C>G 6117A>G), GAA335TAT (6118G>T 6120A>T), CTC337CAC (6125T>A), TAT338TAC (6129T>C), TTT340TTC (6135T>C), GAA341TTG (6136G>T 6137A>T 6138A>G), GTC342GTG (6141C>G), CCC344TCG (6145C>T 6147C>G), TTT345ATT (6148T>A), GGA346GGG (6153A>G), CTG347TTG (6154C>T), ATG348ACC (6158T>C 6159G>C), GCA350GCG (6165A>G), GAT353GCC (6173A>C 6174T>C), TTT354TTC (6177T>C), CAG355ATG (6178C>A 6179A>T), AGA356AGT (6183A>T), AAG357TTG (6184A>T 6185A>T), GAT359AAT (6190G>A), AAT360GGG (6193A>G 6194A>G 6195T>G), GCA361GTG (6197C>T 6198A>G), AGG363AAG (6203G>A), GGA364CCA (6205G>C 6206G>C), GGA364_ACA365insTTT (6207_6208insTTT), ACA365TTG (6208A>T 6209C>T 6210A>G), GCA367TCG (6214G>T 6216A>G), ATC369GTG (6220A>G 6222C>G), GCA370ATT (6223G>A 6224C>T 6225A>T), GTA371GGG (6227T>G 6228A>G), TAT372TTT (6230A>T), ATA376ATT (6243A>T), TTG377CTT (6244T>C 6246G>T), GTA378ATT (6247G>A 6249A>T), TTC379TAT (6251T>A 6252C>T), TCT380TCA (6255T>A), GAA381AAA (6256G>A), GAA381_AAT382insAG- (6258_6259insAG), AAT382AGT (6260A>G), GAA383AAG (6262G>A 6264A>G), GAG387GCA (6275A>C 6276G>A), GAT388CAC (6277G>C 6279T>C), CTC390CTA (6285C>A), TTA391A.. (6286T>A), AAT392T.. (6289A>T) | | | | | | | | |

*: Inserts / Deletes / Misaligned / Frameshifts

#### Analysis details

This analysis was performed with panviral2.64

### NGS Details (UN9): Badnavirus maculaucubae

#### Assembly

| | |
|---|---|
| Coverage Length | 350 (1 contig(s)) |
| Depth Of Coverage | 5.3 |
| Number Of Reads | 15 |
| Reads Per Million | 0.34 rpm (after QC) |
| Ambiguities | 0 |
| Assembly Method | de novo + reference guided assembly |
| Consensus Caller | Bcf Tools |

#### Coverage Map

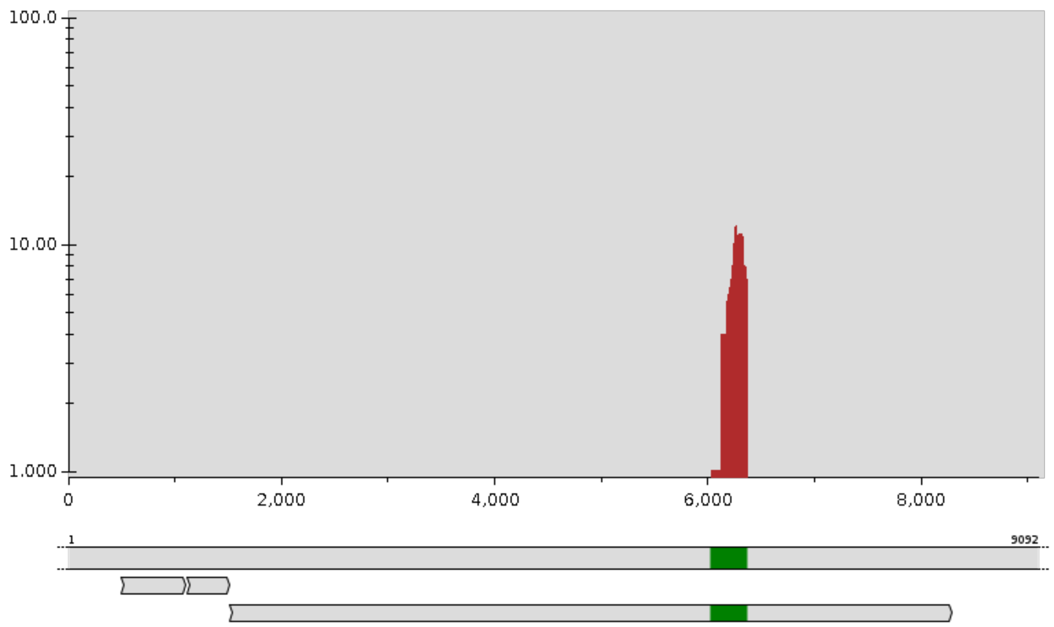

#### Assignment

| | |
|---|---|
| Type | Badnavirus maculaucubae (Taxonomy ID: 3051986) |
| Reference Genome | NC_076606.1 |
| NT Identity (%) | 52.6912 |
| AA Identity (%) | 43.2203 |
| Number Of Stop Codons | 1 |
| Number Of CDS | 3 |

#### Alignment

| | |
|---|---|
| Alignment Score | 32.0 (NT) + 342.0 (AA) = 374.0 |
| Concordance (%) | 24.3648 |

| Alignment Method | Global, seeded, nucleotide + amino acids (AGA) |
|---|---|

#### Genome Region

Sequence starts at position 6024 and ends at position 6373 relative to NC_076606.1 reference sequence.

#### Alignment Detailed Statistics

| | Begin | End | Coverage | Score | Concordance | Matches | Identities | I/D/M/F* | Stop Codons |
|---|---|---|---|---|---|---|---|---|---|
| **NT** | **6024** | **6373** | **3.8%** | **32** | **4.6%** | **350 (99.2%)** | **186 (52.7%)** | **3/0** | |
| Mutations: | 6024T>G, 6025A>G, 6026A>T, 6027T>C, 6030T>G, 6031C>G, 6032A>T, 6033T>G, 6036A>G, 6037G>A, 6040C>A, 6041A>G, 6046A>T, 6047G>T, 6048C>G, 6049C>T, 6051T>G, 6055G>A, 6056G>A, 6060C>T, 6061A>G, 6063C>T, 6064T>G, 6065C>A, 6067A>C, 6069T>A, 6070A>T, 6072A>T, 6073A>G, 6075A>T, 6076G>C, 6077C>A, 6078C>A, 6079A>C, 6082G>T, 6083G>A, 6085A>G, 6087T>A, 6092A>G, 6093A>T, 6094G>T, 6095T>A, 6096C>T, 6100A>T, 6101G>C, 6102C>A, 6105A>G, 6106T>A, 6108T>C, 6112T>C, 6114G>T, 6115A>C, 6117A>T, 6123T>A, 6125T>A, 6132G>A, 6133G>C, 6135G>T, 6136G>A, 6137C>A, 6138A>G, 6139A>G, 6142G>A, 6144A>G, 6147A>G, 6148T>A, 6149C>G, 6150A>C, 6151A>G, 6152G>A, 6153C>T, 6154A>G, 6158C>T, 6159A>G, 6160T>A, 6161G>A, 6164C>T, 6167C>A, 6171C>T, 6172A>C, 6173T>G, 6174A>G, 6176C>T, 6177T>G, 6178C>A, 6179C>G, 6180A>G, 6181G>T, 6186A>T, 6188T>A, 6189A>T, 6191A>T, 6192C>T, 6197G>T, 6198G>C, 6201A>T, 6204A>C, 6208C>T, 6210A>T, 6216T>G, 6217C>T, 6219A>G, 6221A>C, 6222G>T, 6228A>C, 6231T>G, 6234A>T, 6239T>G, 6240C>T, 6241C>A, 6242A>T, 6243A>G, 6244A>G, 6245G>A, 6246G>T, 6247A>T, 6248A>T, 6249A>G, 6252G>A, 6253G>A, 6257A>G, 6258C>G, 6259T>G, 6260G>T, 6264C>T, 6266G>A, 6268G>C, 6269G>C, 6271A>T, 6272C>T, 6273A>C, 6274G>C, 6275A>T, 6276A>T, 6279_6280insATG, 6285C>A, 6287C>T, 6288T>G, 6291T>G, 6293A>T, 6294C>T, 6304A>G, 6306T>G, 6310G>A, 6314T>A, 6318T>C, 6319G>C, 6320A>G, 6322A>G, 6325C>A, 6326T>G, 6327T>G, 6328C>G, 6336C>T, 6345T>C, 6346C>T, 6348C>A, 6350A>G, 6355A>G, 6361G>C, 6362A>G, 6363A>G, 6365T>C, 6368G>T, 6369C>G | | | | | | | | |

##### CDS

| | Begin | End | Coverage | Score | Concordance | Matches | Identities | I/D/M/F* | Stop Codons |
|---|---|---|---|---|---|---|---|---|---|
| QKP79_gp3 | 1506 | 1622 | 5.2% | 342 | 40.3% | 117 (99.2%) | 51 (43.2%) | 1/0/0/0 | 1 |
| Protein mutations: | N1506V (6025A>G 6026A>T 6027T>C), H1508V (6031C>G 6032A>T 6033T>G), D1510N (6037G>A), Q1511R (6040C>A 6041A>G), S1513L (6046A>T 6047G>T 6048C>G), G1516K (6055G>A 6056G>A), N1518D (6061A>G 6063C>T), S1519D (6064T>G 6065C>A), I1520L (6067A>C 6069T>A), I1521F (6070A>T 6072A>T), K1522D (6073A>G 6075A>T), A1523Q (6076G>C 6077C>A 6078C>A), I1524L (6079A>C), G1525* (6082G>T 6083G>A), N1526E (6085A>G 6087T>A), K1528S (6092A>G 6093A>T), V1529Y (6094G>T 6095T>A 6096C>T), F1533I (6106T>A 6108T>C), K1536H (6115A>C 6117A>T), F1539Y (6125T>A), V1542L (6133G>C 6135G>T), A1543K (6136G>A 6137C>A 6138A>G), M1544V (6139A>G), E1545K (6142G>A 6144A>G), S1548D (6151A>G 6152G>A 6153C>T), I1549V (6154A>G), P1550L (6158C>T 6159A>G), W1551K (6160T>A 6161G>A), T1552M (6164C>T), A1553D (6167C>A), I1555R (6172A>C 6173T>G 6174A>G), T1556M (6176C>T 6177T>G), P1557R (6178C>A 6179C>G 6180A>G), D1558Y (6181G>T), L1560H (6188T>A 6189A>T), Y1561F (6191A>T 6192C>T), W1563F (6197G>T 6198G>C), P1567S (6208C>T 6210A>T), K1571T (6221A>C 6222G>T), F1577C (6239T>G 6240C>T), Q1578M (6241C>A 6242A>T 6243A>G), R1579D (6244A>G 6245G>A 6246G>T), K1580L (6247A>T 6248A>T 6249A>G), M1581I (6252G>A), D1582N (6253G>A), N1583R (6257A>G 6258C>G), C1584V (6259T>G 6260G>T), R1586K (6266G>A), G1587P (6268G>C 6269G>C), T1588F (6271A>T 6272C>T 6273A>C), E1589L (6274G>C 6275A>T 6276A>T), D1590_F1591insM (6279_6280insATG), A1593V (6287C>T 6288T>G), Y1595F (6293A>T 6294C>T), I1599V (6304A>G 6306T>G), V1601I (6310G>A), F1602Y (6314T>A), E1604R (6319G>C 6320A>G), N1605D (6322A>G), L1606R (6325C>A 6326T>G 6327T>G), Q1607E (6328C>G), K1614R (6350A>G), M1616V (6355A>G), E1618R (6361G>C 6362A>G 6363A>G), I1619T (6365T>C), C1620L (6368G>T 6369C>G) | | | | | | | | |
| Codon mutations: | GAT1505..G (6024T>G), AAT1506GTC (6025A>G 6026A>T 6027T>C), ACT1507ACG (6030T>G), CAT1508GTG (6031C>G 6032A>T 6033T>G), AAA1509AAG (6036A>G), GAT1510AAT (6037G>A), CAA1511AGA (6040C>A 6041A>G), AGC1513TTG (6046A>T 6047G>T 6048C>G), CTT1514TTG (6049C>T 6051T>G), GGG1516AAG (6055G>A 6056G>A), ATC1517ATT (6060C>T), AAC1518GAT (6061A>G 6063C>T), TCC1519GAC (6064T>G 6065C>A), ATT1520CTA (6067A>C 6069T>A), ATA1521TTT (6070A>T 6072A>T), AAA1522GAT (6073A>G 6075A>T), GCC1523CAA (6076G>C 6077C>A 6078C>A), ATT1524CTT (6079A>C), GGA1525TAA (6082G>T 6083G>A), AAT1526GAA (6085A>G 6087T>A), AAA1528AGT (6092A>G 6093A>T), GTC1529TAT (6094G>T 6095T>A 6096C>T), AGC1531TCA (6100A>T 6101G>C 6102C>A), AAA1532AAG (6105A>G), TTT1533ATC (6106T>A 6108T>C), TTG1535CTT (6112T>C 6114G>T), AAA1536CAT (6115A>C 6117A>T), GGT1538GGA (6123T>A), TTT1539TAT (6125T>A), CAG1541CAA (6132G>A), GTG1542CTT (6133G>C 6135G>T), GCA1543AAG (6136G>A 6137C>A 6138A>G), ATG1544GTG (6139A>G), GAA1545AAG (6142G>A 6144A>G), GAA1546GAG (6147A>G), TCA1547AGC (6148T>A 6149C>G 6150A>C), AGC1548GAT (6151A>G 6152G>A 6153C>T), ATT1549GTT (6154A>G), CCA1550CTG (6158C>T 6159A>G), TGG1551AAG (6160T>A 6161G>A), ACG1552ATG (6164C>T), GCT1553GAT (6167C>A), TTC1554TTT (6171C>T), ATA1555CGG (6172A>C 6173T>G 6174A>G), ACT1556ATG (6176C>T 6177T>G), CCA1557AGG (6178C>A 6179C>G 6180A>G), GAT1558TAT (6181G>T), GGA1559GGT (6186A>T), CTA1560CAT (6188T>A 6189A>T), TAC1561TTT (6191A>T 6192C>T), TGG1563TTC (6197G>T 6198G>C), CTA1564CTT (6201A>T), GTA1565GTC (6204A>C), CCA1567TCT (6208C>T 6210A>T), GGT1569GGG (6216T>G), CTA1570TTG (6217C>T 6219A>G), AAG1571ACT (6221A>C 6222G>T), GCA1573GCC (6228A>C), CCT1574CCG (6231T>G), GCA1575GCT (6234A>T), TTC1577TGT (6239T>G 6240C>T), CAA1578ATG (6241C>A 6242A>T 6243A>G), AGG1579GAT (6244A>G 6245G>A 6246G>T), AAA1580TTG (6247A>T 6248A>T 6249A>G), ATG1581ATA (6252G>A), GAC1582AAC (6253G>A), AAC1583AGG (6257A>G 6258C>G), TGT1584GTT (6259T>G 6260G>T), TTC1585TTT (6264C>T), AGG1586AAG (6266G>A), GGT1587CCT (6268G>C 6269G>C), ACA1588TTC (6271A>T 6272C>T 6273A>C), GAA1589CTT (6274G>C 6275A>T 6276A>T), GAT1590_TTC1591insATG (6279_6280insATG), ATC1592ATA (6285C>A), GCT1593GTG (6287C>T 6288T>G), GTT1594GTG (6291T>G), TAC1595TTT (6293A>T 6294C>T), ATT1599GTG (6304A>G 6306T>G), GTA1601ATA (6310G>A), TTC1602TAC (6314T>A), TCT1603TCC (6318T>C), GAA1604CGA (6319G>C 6320A>G), AAT1605GAT (6322A>G), CTT1606AGG (6325C>A 6326T>G 6327T>G), CAA1607GAA (6328C>G), CAC1609CAT (6336C>T), CAT1612CAC (6345T>C), CTC1613TTA (6346C>T 6348C>A), AAA1614AGA (6350A>G), ATG1616GTG (6355A>G), GAA1618CGG (6361G>C 6362A>G 6363A>G), ATA1619ACA (6365T>C), TGC1620TTG (6368G>T 6369C>G) | | | | | | | | |

##### Proteins

| | Begin | End | Coverage | Score | Concordance | Matches | Identities | I/D/M/F* | Stop Codons |
|---|---|---|---|---|---|---|---|---|---|
| hypothetical protein (YP_010799265.1) | 1506 | 1622 | 5.2% | 342 | 40.3% | 117 (99.2%) | 51 (43.2%) | 1/0/0/0 | 1 |
| Protein mutations: | N1506V (6025A>G 6026A>T 6027T>C), H1508V (6031C>G 6032A>T 6033T>G), D1510N (6037G>A), Q1511R (6040C>A 6041A>G), S1513L (6046A>T 6047G>T 6048C>G), G1516K (6055G>A 6056G>A), N1518D (6061A>G 6063C>T), S1519D (6064T>G 6065C>A), I1520L (6067A>C 6069T>A), I1521F (6070A>T 6072A>T), K1522D (6073A>G 6075A>T), A1523Q (6076G>C 6077C>A 6078C>A), I1524L (6079A>C), G1525* (6082G>T 6083G>A), N1526E (6085A>G 6087T>A), K1528S (6092A>G 6093A>T), V1529Y (6094G>T 6095T>A 6096C>T), F1533I (6106T>A 6108T>C), K1536H (6115A>C 6117A>T), F1539Y (6125T>A), V1542L (6133G>C 6135G>T), A1543K (6136G>A 6137C>A 6138A>G), M1544V (6139A>G), E1545K (6142G>A 6144A>G), S1548D (6151A>G 6152G>A 6153C>T), I1549V (6154A>G), P1550L (6158C>T 6159A>G), W1551K (6160T>A 6161G>A), T1552M (6164C>T), A1553D (6167C>A), I1555R (6172A>C 6173T>G 6174A>G), T1556M (6176C>T 6177T>G), P1557R (6178C>A 6179C>G 6180A>G), D1558Y (6181G>T), L1560H (6188T>A 6189A>T), Y1561F (6191A>T 6192C>T), W1563F (6197G>T 6198G>C), P1567S (6208C>T 6210A>T), K1571T (6221A>C 6222G>T), F1577C (6239T>G 6240C>T), Q1578M (6241C>A 6242A>T 6243A>G), R1579D (6244A>G 6245G>A 6246G>T), K1580L (6247A>T 6248A>T 6249A>G), M1581I (6252G>A), D1582N (6253G>A), N1583R (6257A>G 6258C>G), C1584V (6259T>G 6260G>T), R1586K (6266G>A), G1587P (6268G>C 6269G>C), T1588F (6271A>T 6272C>T 6273A>C), E1589L (6274G>C 6275A>T 6276A>T), D1590_F1591insM (6279_6280insATG), A1593V (6287C>T 6288T>G), Y1595F (6293A>T 6294C>T), I1599V (6304A>G 6306T>G), V1601I (6310G>A), F1602Y (6314T>A), E1604R (6319G>C 6320A>G), N1605D (6322A>G), L1606R (6325C>A 6326T>G 6327T>G), Q1607E (6328C>G), K1614R (6350A>G), M1616V (6355A>G), E1618R (6361G>C 6362A>G 6363A>G), I1619T (6365T>C), C1620L (6368G>T 6369C>G) | | | | | | | | |
| Codon mutations: | GAT1505..G (6024T>G), AAT1506GTC (6025A>G 6026A>T 6027T>C), ACT1507ACG (6030T>G), CAT1508GTG (6031C>G 6032A>T 6033T>G), AAA1509AAG (6036A>G), GAT1510AAT (6037G>A), CAA1511AGA (6040C>A 6041A>G), AGC1513TTG (6046A>T 6047G>T 6048C>G), CTT1514TTG (6049C>T 6051T>G), GGG1516AAG (6055G>A 6056G>A), ATC1517ATT (6060C>T), AAC1518GAT (6061A>G 6063C>T), TCC1519GAC (6064T>G 6065C>A), ATT1520CTA (6067A>C 6069T>A), ATA1521TTT (6070A>T 6072A>T), AAA1522GAT (6073A>G 6075A>T), GCC1523CAA (6076G>C 6077C>A 6078C>A), ATT1524CTT (6079A>C), GGA1525TAA (6082G>T 6083G>A), AAT1526GAA (6085A>G 6087T>A), AAA1528AGT (6092A>G 6093A>T), GTC1529TAT (6094G>T 6095T>A 6096C>T), AGC1531TCA (6100A>T 6101G>C 6102C>A), AAA1532AAG (6105A>G), TTT1533ATC (6106T>A 6108T>C), TTG1535CTT (6112T>C 6114G>T), AAA1536CAT (6115A>C 6117A>T), GGT1538GGA (6123T>A), TTT1539TAT (6125T>A), CAG1541CAA (6132G>A), GTG1542CTT (6133G>C 6135G>T), GCA1543AAG (6136G>A 6137C>A 6138A>G), ATG1544GTG (6139A>G), GAA1545AAG (6142G>A 6144A>G), GAA1546GAG (6147A>G), TCA1547AGC (6148T>A 6149C>G 6150A>C), AGC1548GAT (6151A>G 6152G>A 6153C>T), ATT1549GTT (6154A>G), CCA1550CTG (6158C>T 6159A>G), TGG1551AAG (6160T>A 6161G>A), ACG1552ATG (6164C>T), GCT1553GAT (6167C>A), TTC1554TTT (6171C>T), ATA1555CGG (6172A>C 6173T>G 6174A>G), ACT1556ATG (6176C>T 6177T>G), CCA1557AGG (6178C>A 6179C>G 6180A>G), GAT1558TAT (6181G>T), GGA1559GGT (6186A>T), CTA1560CAT (6188T>A 6189A>T), TAC1561TTT (6191A>T 6192C>T), TGG1563TTC (6197G>T 6198G>C), CTA1564CTT (6201A>T), GTA1565GTC (6204A>C), CCA1567TCT (6208C>T 6210A>T), GGT1569GGG (6216T>G), CTA1570TTG (6217C>T 6219A>G), AAG1571ACT (6221A>C 6222G>T), GCA1573GCC (6228A>C), CCT1574CCG (6231T>G), GCA1575GCT (6234A>T), TTC1577TGT (6239T>G 6240C>T), CAA1578ATG (6241C>A 6242A>T 6243A>G), AGG1579GAT (6244A>G 6245G>A 6246G>T), AAA1580TTG (6247A>T 6248A>T 6249A>G), ATG1581ATA (6252G>A), GAC1582AAC (6253G>A), AAC1583AGG (6257A>G 6258C>G), TGT1584GTT (6259T>G 6260G>T), TTC1585TTT (6264C>T), AGG1586AAG (6266G>A), GGT1587CCT (6268G>C 6269G>C), ACA1588TTC (6271A>T 6272C>T 6273A>C), GAA1589CTT (6274G>C 6275A>T 6276A>T), GAT1590_TTC1591insATG (6279_6280insATG), ATC1592ATA (6285C>A), GCT1593GTG (6287C>T 6288T>G), GTT1594GTG (6291T>G), TAC1595TTT (6293A>T 6294C>T), ATT1599GTG (6304A>G 6306T>G), GTA1601ATA (6310G>A), TTC1602TAC (6314T>A), TCT1603TCC (6318T>C), GAA1604CGA (6319G>C 6320A>G), AAT1605GAT (6322A>G), CTT1606AGG (6325C>A 6326T>G 6327T>G), CAA1607GAA (6328C>G), CAC1609CAT (6336C>T), CAT1612CAC (6345T>C), CTC1613TTA (6346C>T 6348C>A), AAA1614AGA (6350A>G), ATG1616GTG (6355A>G), GAA1618CGG (6361G>C 6362A>G 6363A>G), ATA1619ACA (6365T>C), TGC1620TTG (6368G>T 6369C>G) | | | | | | | | |

*: Inserts / Deletes / Misaligned / Frameshifts

#### Analysis details

This analysis was performed with panviral2.64

### NGS Details (UN9): Bracoviriform congregatae (segment Circle 7)

#### Assembly

| | |
|---|---|
| **Coverage Length** | 139 (1 contig(s)) |
| **Depth Of Coverage** | 9.0 |
| **Number Of Reads** | 14 |
| **Reads Per Million** | 0.32 rpm (after QC) |
| **Ambiguities** | 0 |
| **Assembly Method** | de novo + reference guided assembly |
| **Consensus Caller** | Bcf Tools |

#### Coverage Map

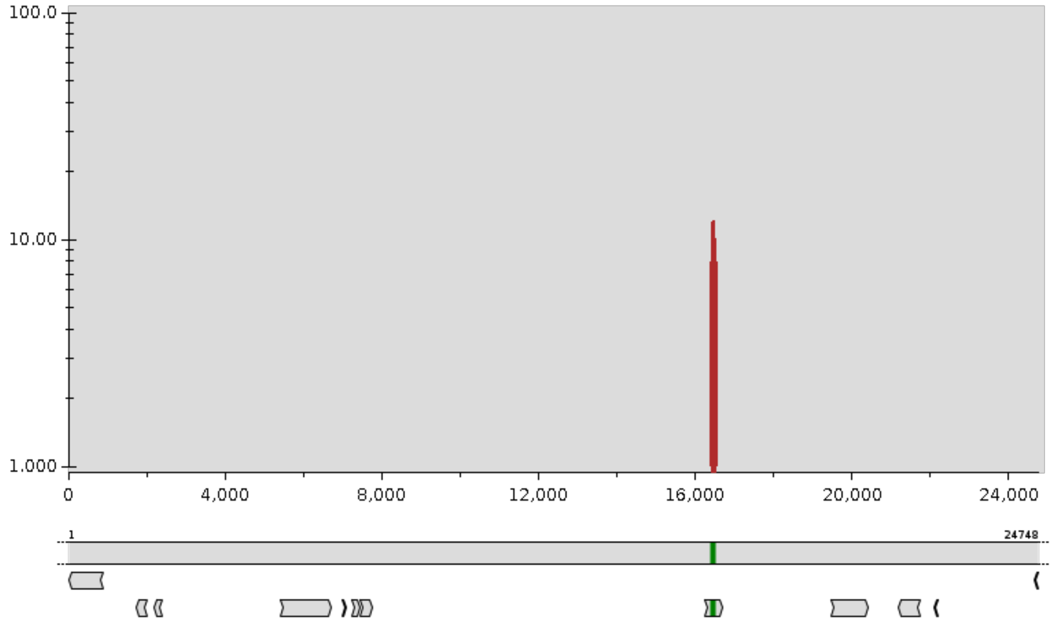

#### Assignment

| | |
|---|---|
| **Type** | Bracoviriform congregatae (Taxonomy ID: 39640) |
| **Reference Genome** | NC_006639.1 |
| **NT Identity (%)** | 78.4173 |
| **AA Identity (%)** | 91.3043 |
| **Number Of Stop Codons** | 0 |
| **Number Of CDS** | 5 |

#### Alignment

| | |
|---|---|
| **Alignment Score** | 158.0 (NT) + 282.0 (AA) = 440.0 |
| **Concordance (%)** | 76.7888 |

| Alignment Method | Global, seeded, nucleotide + amino acids (AGA) |
|---|---|

#### Genome Region

Sequence starts at position 16401 and ends at position 16539 relative to NC_006639.1 reference sequence.

#### Alignment Detailed Statistics

| | Begin | End | Coverage | Score | Concordance | Matches | Identities | I/D/M/F* | Stop Codons |
|---|---|---|---|---|---|---|---|---|---|
| **NT** | **16401** | **16539** | **0.6%** | **158** | **56.8%** | **139 (100%)** | **109 (78.4%)** | **0/0** | |
| Mutations: | 16402A>T, 16408A>T, 16411G>A, 16414A>G, 16423A>C, 16426A>G, 16429G>C, 16435T>G, 16439C>A, 16441T>A, 16444T>C, 16449G>A, 16450A>G, 16451T>A, 16452C>T, 16453T>C, 16454C>T, 16456T>G, 16462T>C, 16471A>G, 16474A>T, 16493C>A, 16495T>A, 16501G>A, 16504T>A, 16507T>A, 16508C>A, 16510T>A, 16516A>T, 16526G>A | | | | | | | | |
| **CDS** | | | | | | | | | |
| Histone | 58 | 103 | 29.5% | 282 | 92.5% | 46 (100%) | 42 (91.3%) | 0/0/0/0 | 0 |
| Protein mutations: | R73K (16449G>A 16450A>G), S74I (16451T>A 16452C>T 16453T>C), V99I (16526G>A) | | | | | | | | |
| Codon mutations: | GTA57.TT (16402A>T), GGA59GGT (16408A>T), GGG60GGA (16411G>A), AAA61AAG (16414A>G), GGA64GGC (16423A>C), AAA65AAG (16426A>G), GGG66GGC (16429G>C), GCT68GCG (16435T>G), CGT70AGA (16439C>A 16441T>A), CAT71CAC (16444T>C), AGA73AAG (16449G>A 16450A>G), TCT74ATC (16451T>A 16452C>T 16453T>C), CTT75TTG (16454C>T 16456T>G), GAT77GAC (16462T>C), CAA80CAG (16471A>G), GGA81GGT (16474A>T), CGT88AGA (16493C>A 16495T>A), CTG90CTA (16501G>A), GCT91GCA (16504T>A), CGT92CGA (16507T>A), CGT93AGA (16508C>A 16510T>A), GGA95GGT (16516A>T), GTC99ATC (16526G>A) | | | | | | | | |
| **Proteins** | | | | | | | | | |
| hypothetical protein (YP_184795.1) | 58 | 103 | 29.5% | 282 | 92.5% | 46 (100%) | 42 (91.3%) | 0/0/0/0 | 0 |
| Protein mutations: | R73K (16449G>A 16450A>G), S74I (16451T>A 16452C>T 16453T>C), V99I (16526G>A) | | | | | | | | |
| Codon mutations: | GTA57.TT (16402A>T), GGA59GGT (16408A>T), GGG60GGA (16411G>A), AAA61AAG (16414A>G), GGA64GGC (16423A>C), AAA65AAG (16426A>G), GGG66GGC (16429G>C), GCT68GCG (16435T>G), CGT70AGA (16439C>A 16441T>A), CAT71CAC (16444T>C), AGA73AAG (16449G>A 16450A>G), TCT74ATC (16451T>A 16452C>T 16453T>C), CTT75TTG (16454C>T 16456T>G), GAT77GAC (16462T>C), CAA80CAG (16471A>G), GGA81GGT (16474A>T), CGT88AGA (16493C>A 16495T>A), CTG90CTA (16501G>A), GCT91GCA (16504T>A), CGT92CGA (16507T>A), CGT93AGA (16508C>A 16510T>A), GGA95GGT (16516A>T), GTC99ATC (16526G>A) | | | | | | | | |

*: Inserts / Deletes / Misaligned / Frameshifts

#### Analysis details

This analysis was performed with panviral2.64

### NGS Details (UN9): Badnavirus epsiloninflatheobromae

#### Assembly

| | |
|---|---|
| Coverage Length | 290 (1 contig(s)) |
| Depth Of Coverage | 5.0 |
| Number Of Reads | 14 |
| Reads Per Million | 0.32 rpm (after QC) |
| Ambiguities | 0 |
| Assembly Method | de novo + reference guided assembly |
| Consensus Caller | Bcf Tools |

#### Coverage Map

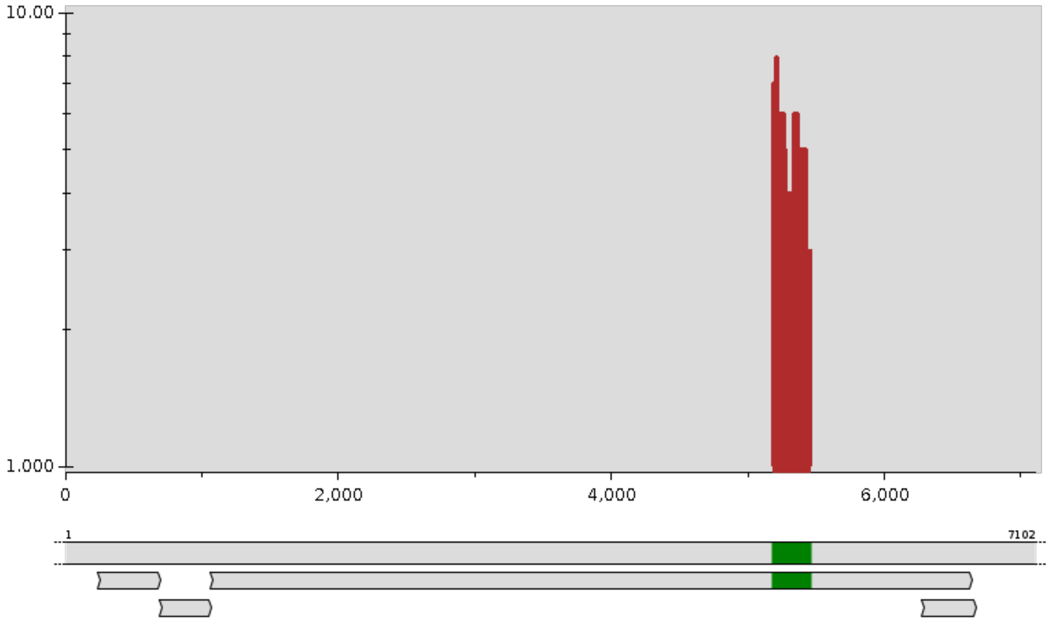

#### Assignment

| | |
|---|---|
| Type | Badnavirus epsiloninflatheobromae (Taxonomy ID: 3047711) |
| Reference Genome | NC_043535.1 |
| NT Identity (%) | 59.0444 |
| AA Identity (%) | 49.4845 |
| Number Of Stop Codons | 1 |
| Number Of CDS | 4 |

#### Alignment

| | |
|---|---|
| Alignment Score | 100.0 (NT) + 347.0 (AA) = 447.0 |
| Concordance (%) | 34.2005 |

| Alignment Method | Global, seeded, nucleotide + amino acids (AGA) |
|---|---|

#### Genome Region

Sequence starts at position 5176 and ends at position 5465 relative to NC_043535.1 reference sequence.

#### Alignment Detailed Statistics

| | Begin | End | Coverage | Score | Concordance | Matches | Identities | I/D/M/F* | Stop Codons |
|---|---|---|---|---|---|---|---|---|---|
| **NT** | **5176** | **5465** | **4.1%** | **100** | **17.2%** | **290 (99.0%)** | **173 (59.0%)** | **3/0** | |
| Mutations: | 5179A>T, 5182C>A, 5183A>T, 5186G>A, 5187A>C, 5192C>A, 5198A>C, 5201A>G, 5202A>T, 5203T>A, 5209G>C, 5213A>G, 5214T>G, 5216C>G, 5220T>C, 5222G>C, 5225A>G, 5227G>A, 5228T>G, 5231A>C, 5233T>A, 5234T>C, 5235C>T, 5237T>C, 5241A>G, 5243C>G, 5244G>C, 5245C>G, 5246A>C, 5249G>A, 5250C>G, 5251A>C, 5252T>A, 5257A>G, 5259T>G, 5260C>A, 5261A>C, 5262A>G, 5263T>A, 5264C>G, 5267A>G, 5268T>A, 5269G>A, 5273C>G, 5274G>A, 5278T>A, 5280T>G, 5281G>T, 5283G>A, 5285A>G, 5286C>A, 5287C>G, 5288T>A, 5289C>T, 5291A>T, 5292G>A, 5294A>G, 5295C>G, 5296T>C, 5303A>G, 5307C>T, 5312T>G, 5318A>T, 5325C>T, 5327C>A, 5329A>C, 5330A>C, 5336C>A, 5339T>C, 5341C>T, 5344T>C, 5345C>T, 5349C>T, 5350A>G, 5351A>C, 5353G>C, 5355A>C, 5356A>T, 5361G>A, 5363T>C, 5364C>G, 5366T>G, 5367T>A, 5368G>T, 5369T>C, 5371T>A, 5372_5373insCAT, 5373G>C, 5375A>C, 5376G>T, 5377G>A, 5378A>C, 5379A>T, 5380C>T, 5381A>G, 5384A>T, 5385G>T, 5390T>C, 5393T>G, 5395C>T, 5396T>A, 5402C>T, 5403A>T, 5405T>G, 5415A>G, 5420T>C, 5422T>A, 5423C>T, 5429T>C, 5432A>C, 5433G>T, 5434A>C, 5439C>G, 5444T>C, 5446A>T, 5450A>G, 5454C>T, 5456G>A | | | | | | | | |

**CDS**

| | Begin | End | Coverage | Score | Concordance | Matches | Identities | I/D/M/F* | Stop Codons |
|---|---|---|---|---|---|---|---|---|---|
| FLA70_gp3 | 1373 | 1468 | 5.2% | 347 | 47.3% | 96 (99.0%) | 48 (49.5%) | 1/0/0/0 | 1 |
| Protein mutations: | H1373L (5179A>T), A1374D (5182C>A 5183A>T), I1376L (5187A>C), I1381Y (5202A>T 5203T>A), S1383T (5209G>C), F1385V (5214T>G 5216C>G), S1389K (5227G>A 5228T>G), F1391Y (5233T>A 5234T>C), H1392Y (5235C>T 5237T>C), I1394V (5241A>G 5243C>G), A1395R (5244G>C 5245C>G 5246A>C), M1396I (5249G>A), H1397A (5250C>G 5251A>C 5252T>A), E1399G (5257A>G), S1400D (5259T>G 5260C>A 5261A>C), I1401E (5262A>G 5263T>A 5264C>G), W1403K (5268T>A 5269G>A), A1405T (5274G>A), F1406Y (5278T>A), W1407V (5280T>G 5281G>T), V1408M (5283G>A 5285A>G), P1409R (5286C>A 5287C>G 5288T>A), Q1410Y (5289C>T 5291A>T), G1411R (5292G>A 5294A>G), L1412A (5295C>G 5296T>C), K1423T (5329A>C 5330A>C), A1427V (5341C>T), I1428T (5344T>C 5345C>T), Q1430C (5349C>T 5350A>G 5351A>C), R1431T (5353G>C), K1432L (5355A>C 5356A>T), D1434N (5361G>A 5363T>C), H1435E (5364C>G 5366T>G), C1436I (5367T>A 5368G>T 5369T>C), F1437Y (5371T>A), F1437_A1438insH (5372_5373insCAT), A1438P (5373G>C 5375A>C), G1439Y (5376G>T 5377G>A 5378A>C), T1440L (5379A>T 5380C>T 5381A>G), E1441D (5384A>T), E1442* (5385G>T), I1444M (5393T>G), A1445V (5395C>T 5396T>A), I1448L (5403A>T 5405T>G), I1452V (5415A>G), F1454Y (5422T>A 5423C>T), E1458S (5433G>T 5434A>C), Q1460E (5439C>G), K1462M (5446A>T) | | | | | | | | |
| Codon mutations: | CAT1373CTT (5179A>T), GCA1374GAT (5182C>A 5183A>T), AGG1375AGA (5186G>A), ATT1376CTT (5187A>C), GGC1377GGA (5192C>A), GCA1379GCC (5198A>C), AAA1380AAG (5201A>G), ATC1381TAC (5202A>T 5203T>A), AGC1383ACC (5209G>C), AAA1384AAG (5213A>G), TTC1385GTG (5214T>G 5216C>G), TTG1387CTC (5220T>C 5222G>C), AAA1388AAG (5225A>G), AGT1389AAG (5227G>A 5228T>G), GGA1390GGC (5231A>C), TTT1391TAC (5233T>A 5234T>C), CAT1392TAC (5235C>T 5237T>C), ATC1394GTG (5241A>G 5243C>G), GCA1395CGC (5244G>C 5245C>G 5246A>C), ATG1396ATA (5249G>A), CAT1397GCA (5250C>G 5251A>C 5252T>A), GAG1399GGG (5257A>G), TCA1400GAC (5259T>G 5260C>A 5261A>C), ATC1401GAG (5262A>G 5263T>A 5264C>G), CCA1402CCG (5267A>G), TGG1403AAG (5268T>A 5269G>A), ACC1404ACG (5273C>G), GCG1405ACG (5274G>A), TTT1406TAT (5278T>A), TGG1407GTG (5280T>G 5281G>T), GTA1408ATG (5283G>A 5285A>G), CCT1409AGA (5286C>A 5287C>G 5288T>A), CAA1410TAT (5289C>T 5291A>T), GGA1411AGG (5292G>A 5294A>G), CTT1412GCT (5295C>G 5296T>C), GAA1414GAG (5303A>G), CTG1416TTG (5307C>T), GTT1417GTG (5312T>G), CCA1419CCT (5318A>T), CTC1422TTA (5325C>T 5327C>A), AAA1423ACC (5329A>C 5330A>C), GCC1425GCA (5336C>A), CCT1426CCC (5339T>C), GCC1427GTC (5341C>T), ATC1428ACT (5344T>C 5345C>T), CAA1430TGC (5349C>T 5350A>G 5351A>C), AGA1431ACA (5353G>C), AAA1432CTA (5355A>C 5356A>T), GAT1434AAC (5361G>A 5363T>C), CAT1435GAG (5364C>G 5366T>G), TGT1436ATC (5367T>A 5368G>T 5369T>C), TTT1437TAT (5371T>A), TTT1437_GCA1438insCAT (5372_5373insCAT), GCA1438CCC (5373G>C 5375A>C), GGA1439TAC (5376G>T 5377G>A 5378A>C), ACA1440TTG (5379A>T 5380C>T 5381A>G), GAA1441GAT (5384A>T), GAG1442TAG (5385G>T), TTT1443TTC (5390T>C), ATT1444ATG (5393T>G), GCT1445GTA (5395C>T 5396T>A), TAC1447TAT (5402C>T), ATT1448TTG (5403A>T 5405T>G), ATC1452GTC (5415A>G), ATT1453ATC (5420T>C), TTC1454TAT (5422T>A 5423C>T), AAT1456AAC (5429T>C), ACA1457ACC (5432A>C), GAG1458TCG (5433G>T 5434A>C), CAG1460GAG (5439C>G), CAT1461CAC (5444T>C), AAG1462ATG (5446A>T), GAA1463GAG (5450A>G), CTG1465TTA (5454C>T 5456G>A) | | | | | | | | |

**Proteins**

| | Begin | End | Coverage | Score | Concordance | Matches | Identities | I/D/M/F* | Stop Codons |
|---|---|---|---|---|---|---|---|---|---|
| ORF3 polyprotein (YP_009666830.1) | 1373 | 1468 | 5.2% | 347 | 47.3% | 96 (99.0%) | 48 (49.5%) | 1/0/0/0 | 1 |
| Protein mutations: | H1373L (5179A>T), A1374D (5182C>A 5183A>T), I1376L (5187A>C), I1381Y (5202A>T 5203T>A), S1383T (5209G>C), F1385V (5214T>G 5216C>G), S1389K (5227G>A 5228T>G), F1391Y (5233T>A 5234T>C), H1392Y (5235C>T 5237T>C), I1394V (5241A>G 5243C>G), A1395R (5244G>C 5245C>G 5246A>C), M1396I (5249G>A), H1397A (5250C>G 5251A>C 5252T>A), E1399G (5257A>G), S1400D (5259T>G 5260C>A 5261A>C), I1401E (5262A>G 5263T>A 5264C>G), W1403K (5268T>A 5269G>A), A1405T (5274G>A), F1406Y (5278T>A), W1407V (5280T>G 5281G>T), V1408M (5283G>A 5285A>G), P1409R (5286C>A 5287C>G 5288T>A), Q1410Y (5289C>T 5291A>T), G1411R (5292G>A 5294A>G), L1412A (5295C>G 5296T>C), K1423T (5329A>C 5330A>C), A1427V (5341C>T), I1428T (5344T>C 5345C>T), Q1430C (5349C>T 5350A>G 5351A>C), R1431T (5353G>C), K1432L (5355A>C 5356A>T), D1434N (5361G>A 5363T>C), H1435E (5364C>G 5366T>G), C1436I (5367T>A 5368G>T 5369T>C), F1437Y (5371T>A), F1437_A1438insH (5372_5373insCAT), A1438P (5373G>C 5375A>C), G1439Y (5376G>T 5377G>A 5378A>C), T1440L (5379A>T 5380C>T 5381A>G), E1441D (5384A>T), E1442* (5385G>T), I1444M (5393T>G), A1445V (5395C>T 5396T>A), I1448L (5403A>T 5405T>G), I1452V (5415A>G), F1454Y (5422T>A 5423C>T), E1458S (5433G>T 5434A>C), Q1460E (5439C>G), K1462M (5446A>T) | | | | | | | | |
| Codon mutations: | CAT1373CTT (5179A>T), GCA1374GAT (5182C>A 5183A>T), AGG1375AGA (5186G>A), ATT1376CTT (5187A>C), GGC1377GGA (5192C>A), GCA1379GCC (5198A>C), AAA1380AAG (5201A>G), ATC1381TAC (5202A>T 5203T>A), AGC1383ACC (5209G>C), AAA1384AAG (5213A>G), TTC1385GTG (5214T>G 5216C>G), TTG1387CTC (5220T>C 5222G>C), AAA1388AAG (5225A>G), AGT1389AAG (5227G>A 5228T>G), GGA1390GGC (5231A>C), TTT1391TAC (5233T>A 5234T>C), CAT1392TAC (5235C>T 5237T>C), ATC1394GTG (5241A>G 5243C>G), GCA1395CGC (5244G>C 5245C>G 5246A>C), ATG1396ATA (5249G>A), CAT1397GCA (5250C>G 5251A>C 5252T>A), GAG1399GGG (5257A>G), TCA1400GAC (5259T>G 5260C>A 5261A>C), ATC1401GAG (5262A>G 5263T>A 5264C>G), CCA1402CCG (5267A>G), TGG1403AAG (5268T>A 5269G>A), ACC1404ACG (5273C>G), GCG1405ACG (5274G>A), TTT1406TAT (5278T>A), TGG1407GTG (5280T>G 5281G>T), GTA1408ATG (5283G>A 5285A>G), CCT1409AGA (5286C>A 5287C>G 5288T>A), CAA1410TAT (5289C>T 5291A>T), GGA1411AGG (5292G>A 5294A>G), CTT1412GCT (5295C>G 5296T>C), GAA1414GAG (5303A>G), CTG1416TTG (5307C>T), GTT1417GTG (5312T>G), CCA1419CCT (5318A>T), CTC1422TTA (5325C>T 5327C>A), AAA1423ACC (5329A>C 5330A>C), GCC1425GCA (5336C>A), CCT1426CCC (5339T>C), GCC1427GTC (5341C>T), ATC1428ACT (5344T>C 5345C>T), CAA1430TGC (5349C>T 5350A>G 5351A>C), AGA1431ACA (5353G>C), AAA1432CTA (5355A>C 5356A>T), GAT1434AAC (5361G>A 5363T>C), CAT1435GAG (5364C>G 5366T>G), TGT1436ATC (5367T>A 5368G>T 5369T>C), TTT1437TAT (5371T>A), TTT1437_GCA1438insCAT (5372_5373insCAT), GCA1438CCC (5373G>C 5375A>C), GGA1439TAC (5376G>T 5377G>A 5378A>C), ACA1440TTG (5379A>T 5380C>T 5381A>G), GAA1441GAT (5384A>T), GAG1442TAG (5385G>T), TTT1443TTC (5390T>C), ATT1444ATG (5393T>G), GCT1445GTA (5395C>T 5396T>A), TAC1447TAT (5402C>T), ATT1448TTG (5403A>T 5405T>G), ATC1452GTC (5415A>G), ATT1453ATC (5420T>C), TTC1454TAT (5422T>A 5423C>T), AAT1456AAC (5429T>C), ACA1457ACC (5432A>C), GAG1458TCG (5433G>T 5434A>C), CAG1460GAG (5439C>G), CAT1461CAC (5444T>C), AAG1462ATG (5446A>T), GAA1463GAG (5450A>G), CTG1465TTA (5454C>T 5456G>A) | | | | | | | | |

*: Inserts / Deletes / Misaligned / Frameshifts

#### Analysis details

This analysis was performed with panviral2.64

### NGS Details (UN9): Badnavirus maculasmallanthi

#### Assembly

| | |
|---|---|
| **Coverage Length** | 323 (1 contig(s)) |
| **Depth Of Coverage** | 4.2 |
| **Number Of Reads** | 14 |
| **Reads Per Million** | 0.32 rpm (after QC) |
| **Ambiguities** | 0 |
| **Assembly Method** | de novo + reference guided assembly |
| **Consensus Caller** | Bcf Tools |

#### Coverage Map

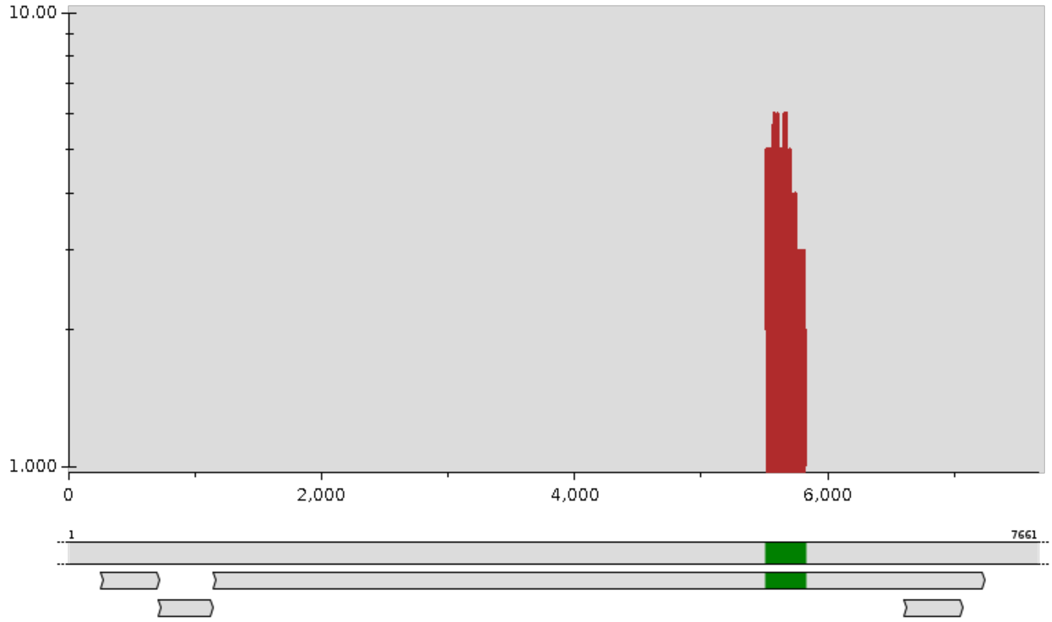

#### Assignment

| | |
|---|---|
| **Type** | Badnavirus maculasmallanthi (Taxonomy ID: 3048453) |
| **Reference Genome** | NC_026472.1 |
| **NT Identity (%)** | 57.9755 |
| **AA Identity (%)** | 45.8716 |
| **Number Of Stop Codons** | 0 |
| **Number Of CDS** | 4 |

#### Alignment

| | |
|---|---|
| **Alignment Score** | 98.0 (NT) + 335.0 (AA) = 433.0 |
| **Concordance (%)** | 30.8845 |

| Alignment Method | Global, seeded, nucleotide + amino acids (AGA) |
|---|---|

#### Genome Region

Sequence starts at position 5509 and ends at position 5831 relative to NC_026472.1 reference sequence.

#### Alignment Detailed Statistics

| | Begin | End | Coverage | Score | Concordance | Matches | Identities | I/D/M/F* | Stop Codons |
|---|---|---|---|---|---|---|---|---|---|
| **NT** | **5509** | **5831** | **4.2%** | **98** | **15.2%** | **323 (99.1%)** | **189 (58.0%)** | **3/0** | |
| Mutations: | 5509G>A, 5512C>A, 5518T>C, 5521C>A, 5526C>G, 5527G>A, 5532A>T, 5533A>G, 5535C>T, 5536A>G, 5537C>A, 5538C>T, 5542C>T, 5544A>C, 5545A>G, 5547A>T, 5549G>A, 5551A>C, 5553A>T, 5554G>A, 5555G>A, 5556G>A, 5557A>G, 5558A>G, 5564A>G, 5565G>T, 5566A>T, 5567T>A, 5568A>C, 5574A>T, 5578T>A, 5586G>T, 5587A>C, 5588A>G, 5589G>T, 5592T>C, 5597T>A, 5600A>T, 5601T>C, 5605G>T, 5607T>G, 5608G>A, 5609C>G, 5610C>G, 5611A>G, 5614G>A, 5615A>G, 5616A>G, 5619A>G, 5620G>T, 5621A>G, 5622A>T, 5623T>G, 5624C>A, 5628A>T, 5631C>A, 5632T>A, 5633G>A, 5638G>A, 5640C>T, 5644T>A, 5645T>G, 5648T>C, 5651C>G, 5653G>T, 5654G>A, 5655A>T, 5658A>T, 5659T>C, 5660T>A, 5661A>T, 5663A>T, 5664C>T, 5669G>T, 5670G>T, 5671C>T, 5673T>G, 5676C>T, 5680C>T, 5682A>T, 5685C>T, 5686G>A, 5691C>A, 5692A>G, 5693A>C, 5700A>T, 5706T>A, 5708T>C, 5713C>A, 5714A>T, 5716A>G, 5717G>A, 5718A>C, 5719A>T, 5720A>T, 5721A>G, 5725G>A, 5727T>C, 5730A>G, 5731T>G, 5732G>T, 5733C>T, 5739_5740insCCG, 5740G>T, 5743A>T, 5744C>T, 5748A>T, 5749G>A, 5750A>T, 5755A>G, 5757C>G, 5759C>T, 5760C>G, 5763A>C, 5765A>T, 5769C>A, 5772C>T, 5778C>A, 5779C>T, 5784A>T, 5787C>T, 5791A>C, 5793C>A, 5801C>A, 5802G>A, 5805T>G, 5809G>A, 5810A>T, 5811A>G, 5812A>G, 5814G>T, 5820A>G, 5822A>G, 5827A>G, 5830C>T | | | | | | | | |

##### CDS

| | Begin | End | Coverage | Score | Concordance | Matches | Identities | I/D/M/F* | Stop Codons |
|---|---|---|---|---|---|---|---|---|---|
| UF61_gp3 | 1456 | 1563 | 5.3% | 335 | 44.1% | 108 (99.1%) | 50 (45.9%) | 1/0/0/0 | 0 |
| Protein mutations: | D1456N (5509G>A), Q1457K (5512C>A), S1459P (5518T>C), L1460I (5521C>A), G1462R (5527G>A), N1464D (5533A>G 5535C>T), T1465D (5536A>G 5537C>A 5538C>T), L1467F (5542C>T 5544A>C), K1468D (5545A>G 5547A>T), R1469K (5549G>A), I1470L (5551A>C 5553A>T), G1471K (5554G>A 5555G>A 5556G>A), N1472G (5557A>G 5558A>G), K1474S (5564A>G 5565G>T), I1475Y (5566A>T 5567T>A 5568A>C), F1479I (5578T>A), K1482R (5587A>C 5588A>G 5589G>T), F1485Y (5597T>A), H1486L (5600A>T 5601T>C), V1488L (5605G>T 5607T>G), A1489R (5608G>A 5609C>G 5610C>G), M1490V (5611A>G), E1491R (5614G>A 5615A>G 5616A>G), E1493C (5620G>T 5621A>G 5622A>T), S1494D (5623T>G 5624C>A), W1497K (5632T>A 5633G>A), A1499T (5638G>A 5640C>T), L1501R (5644T>A 5645T>G), I1502T (5648T>C), P1503R (5651C>G), G1504Y (5653G>T 5654G>A 5655A>T), L1506H (5659T>C 5660T>A 5661A>T), Y1507F (5663A>T 5664C>T), W1509F (5669G>T 5670G>T), P1513S (5680C>T 5682A>T), G1515R (5686G>A), K1517A (5692A>G 5693A>C), I1522T (5708T>C), Q1524M (5713C>A 5714A>T), R1525D (5716A>G 5717G>A 5718A>C), K1526L (5719A>T 5720A>T 5721A>G), D1528N (5725G>A 5727T>C), C1530V (5731T>G 5732G>T 5733C>T), K1532_D1533insP (5739_5740insCCG), D1533Y (5740G>T), T1534L (5743A>T 5744C>T), E1535D (5748A>T), E1536M (5749G>A 5750A>T), I1538V (5755A>G 5757C>G), A1539V (5759C>T 5760C>G), Y1541F (5765A>T), N1550Q (5791A>C 5793C>A), A1553E (5801C>A 5802G>A), D1554E (5805T>G), E1556M (5809G>A 5810A>T 5811A>G), R1557G (5812A>G 5814G>T), K1560R (5822A>G), M1562V (5827A>G) | | | | | | | | |
| Codon mutations: | GAT1456AAT (5509G>A), CAG1457AAG (5512C>A), TCC1459CCC (5518T>C), CTT1460ATT (5521C>A), CCC1461CCG (5526C>G), GGG1462AGG (5527G>A), ATA1463ATT (5532A>T), AAC1464GAT (5533A>G 5535C>T), ACC1465GAT (5536A>G 5537C>A 5538C>T), CTA1467TTC (5542C>T 5544A>C), AAA1468GAT (5545A>G 5547A>T), AGA1469AAA (5549G>A), ATA1470CTT (5551A>C 5553A>T), GGG1471AAA (5554G>A 5555G>A 5556G>A), AAT1472GGT (5557A>G 5558A>G), AAG1474AGT (5564A>G 5565G>T), ATA1475TAC (5566A>T 5567T>A 5568A>C), TCA1477TCT (5574A>T), TTT1479ATT (5578T>A), CTG1481CTT (5586G>T), AAG1482CGT (5587A>C 5588A>G 5589G>T), TCT1483TCC (5592T>C), TTT1485TAT (5597T>A), CAT1486CTC (5600A>T 5601T>C), GTT1488TTG (5605G>T 5607T>G), GCC1489AGG (5608G>A 5609C>G 5610C>G), ATG1490GTG (5611A>G), GAA1491AGG (5614G>A 5615A>G 5616A>G), GAA1492GAG (5619A>G), GAA1493TGT (5620G>T 5621A>G 5622A>T), TCC1494GAC (5623T>G 5624C>A), ATA1495ATT (5628A>T), CCC1496CCA (5631C>A), TGG1497AAG (5632T>A 5633G>A), GCC1499ACT (5638G>A 5640C>T), TTA1501AGA (5644T>A 5645T>G), ATC1502ACC (5648T>C), CCA1503CGA (5651C>G), GGA1504TAT (5653G>T 5654G>A 5655A>T), GGA1505GGT (5658A>T), TTA1506CAT (5659T>C 5660T>A 5661A>T), TAC1507TTT (5663A>T 5664C>T), TGG1509TTT (5669G>T 5670G>T), CTT1510TTG (5671C>T 5673T>G), GTC1511GTT (5676C>T), CCA1513TCT (5680C>T 5682A>T), TTC1514TTT (5685C>T), GGA1515AGA (5686G>A), CTC1516CTA (5691C>A), AAA1517GCA (5692A>G 5693A>C), GCA1519GCT (5700A>T), GCT1521GCA (5706T>A), ATT1522ACT (5708T>C), CAG1524ATG (5713C>A 5714A>T), AGA1525GAC (5716A>G 5717G>A 5718A>C), AAA1526TTG (5719A>T 5720A>T 5721A>G), GAT1528AAC (5725G>A 5727T>C), AAA1529AAG (5730A>G), TGC1530GTT (5731T>G 5732G>T 5733C>T), AAA1532_GAC1533insCCG (5739_5740insCCG), GAC1533TAC (5740G>T), ACA1534TTA (5743A>T 5744C>T), GAA1535GAT (5748A>T), GAG1536ATG (5749G>A 5750A>T), ATC1538GTG (5755A>G 5757C>G), GCC1539GTG (5759C>T 5760C>G), GTA1540GTC (5763A>C), TAC1541TTC (5765A>T), ATC1542ATA (5769C>A), GAC1543GAT (5772C>T), ATC1545ATA (5778C>A), CTG1546TTG (5779C>T), GTA1547GTT (5784A>T), TAC1548TAT (5787C>T), AAC1550CAA (5791A>C 5793C>A), GCG1553GAA (5801C>A 5802G>A), GAT1554GAG (5805T>G), GAA1556ATG (5809G>A 5810A>T 5811A>G), AGG1557GGT (5812A>G 5814G>T), TTA1559TTG (5820A>G), AAG1560AGG (5822A>G), ATG1562GTG (5827A>G), CTG1563TT. (5830C>T) | | | | | | | | |

##### Proteins

| | Begin | End | Coverage | Score | Concordance | Matches | Identities | I/D/M/F* | Stop Codons |
|---|---|---|---|---|---|---|---|---|---|
| ORF3 (YP_009121747.1) | 1456 | 1563 | 5.3% | 335 | 44.1% | 108 (99.1%) | 50 (45.9%) | 1/0/0/0 | 0 |
| Protein mutations: | D1456N (5509G>A), Q1457K (5512C>A), S1459P (5518T>C), L1460I (5521C>A), G1462R (5527G>A), N1464D (5533A>G 5535C>T), T1465D (5536A>G 5537C>A 5538C>T), L1467F (5542C>T 5544A>C), K1468D (5545A>G 5547A>T), R1469K (5549G>A), I1470L (5551A>C 5553A>T), G1471K (5554G>A 5555G>A 5556G>A), N1472G (5557A>G 5558A>G), K1474S (5564A>G 5565G>T), I1475Y (5566A>T 5567T>A 5568A>C), F1479I (5578T>A), K1482R (5587A>C 5588A>G 5589G>T), F1485Y (5597T>A), H1486L (5600A>T 5601T>C), V1488L (5605G>T 5607T>G), A1489R (5608G>A 5609C>G 5610C>G), M1490V (5611A>G), E1491R (5614G>A 5615A>G 5616A>G), E1493C (5620G>T 5621A>G 5622A>T), S1494D (5623T>G 5624C>A), W1497K (5632T>A 5633G>A), A1499T (5638G>A 5640C>T), L1501R (5644T>A 5645T>G), I1502T (5648T>C), P1503R (5651C>G), G1504Y (5653G>T 5654G>A 5655A>T), L1506H (5659T>C 5660T>A 5661A>T), Y1507F (5663A>T 5664C>T), W1509F (5669G>T 5670G>T), P1513S (5680C>T 5682A>T), G1515R (5686G>A), K1517A (5692A>G 5693A>C), I1522T (5708T>C), Q1524M (5713C>A 5714A>T), R1525D (5716A>G 5717G>A 5718A>C), K1526L (5719A>T 5720A>T 5721A>G), D1528N (5725G>A 5727T>C), C1530V (5731T>G 5732G>T 5733C>T), K1532_D1533insP (5739_5740insCCG), D1533Y (5740G>T), T1534L (5743A>T 5744C>T), E1535D (5748A>T), E1536M (5749G>A 5750A>T), I1538V (5755A>G 5757C>G), A1539V (5759C>T 5760C>G), Y1541F (5765A>T), N1550Q (5791A>C 5793C>A), A1553E (5801C>A 5802G>A), D1554E (5805T>G), E1556M (5809G>A 5810A>T 5811A>G), R1557G (5812A>G 5814G>T), K1560R (5822A>G), M1562V (5827A>G) | | | | | | | | |
| Codon mutations: | GAT1456AAT (5509G>A), CAG1457AAG (5512C>A), TCC1459CCC (5518T>C), CTT1460ATT (5521C>A), CCC1461CCG (5526C>G), GGG1462AGG (5527G>A), ATA1463ATT (5532A>T), AAC1464GAT (5533A>G 5535C>T), ACC1465GAT (5536A>G 5537C>A 5538C>T), CTA1467TTC (5542C>T 5544A>C), AAA1468GAT (5545A>G 5547A>T), AGA1469AAA (5549G>A), ATA1470CTT (5551A>C 5553A>T), GGG1471AAA (5554G>A 5555G>A 5556G>A), AAT1472GGT (5557A>G 5558A>G), AAG1474AGT (5564A>G 5565G>T), ATA1475TAC (5566A>T 5567T>A 5568A>C), TCA1477TCT (5574A>T), TTT1479ATT (5578T>A), CTG1481CTT (5586G>T), AAG1482CGT (5587A>C 5588A>G 5589G>T), TCT1483TCC (5592T>C), TTT1485TAT (5597T>A), CAT1486CTC (5600A>T 5601T>C), GTT1488TTG (5605G>T 5607T>G), GCC1489AGG (5608G>A 5609C>G 5610C>G), ATG1490GTG (5611A>G), GAA1491AGG (5614G>A 5615A>G 5616A>G), GAA1492GAG (5619A>G), GAA1493TGT (5620G>T 5621A>G 5622A>T), TCC1494GAC (5623T>G 5624C>A), ATA1495ATT (5628A>T), CCC1496CCA (5631C>A), TGG1497AAG (5632T>A 5633G>A), GCC1499ACT (5638G>A 5640C>T), TTA1501AGA (5644T>A 5645T>G), ATC1502ACC (5648T>C), CCA1503CGA (5651C>G), GGA1504TAT (5653G>T 5654G>A 5655A>T), GGA1505GGT (5658A>T), TTA1506CAT (5659T>C 5660T>A 5661A>T), TAC1507TTT (5663A>T 5664C>T), TGG1509TTT (5669G>T 5670G>T), CTT1510TTG (5671C>T 5673T>G), GTC1511GTT (5676C>T), CCA1513TCT (5680C>T 5682A>T), TTC1514TTT (5685C>T), GGA1515AGA (5686G>A), CTC1516CTA (5691C>A), AAA1517GCA (5692A>G 5693A>C), GCA1519GCT (5700A>T), GCT1521GCA (5706T>A), ATT1522ACT (5708T>C), CAG1524ATG (5713C>A 5714A>T), AGA1525GAC (5716A>G 5717G>A 5718A>C), AAA1526TTG (5719A>T 5720A>T 5721A>G), GAT1528AAC (5725G>A 5727T>C), AAA1529AAG (5730A>G), TGC1530GTT (5731T>G 5732G>T 5733C>T), AAA1532_GAC1533insCCG (5739_5740insCCG), GAC1533TAC (5740G>T), ACA1534TTA (5743A>T 5744C>T), GAA1535GAT (5748A>T), GAG1536ATG (5749G>A 5750A>T), ATC1538GTG (5755A>G 5757C>G), GCC1539GTG (5759C>T 5760C>G), GTA1540GTC (5763A>C), TAC1541TTC (5765A>T), ATC1542ATA (5769C>A), GAC1543GAT (5772C>T), ATC1545ATA (5778C>A), CTG1546TTG (5779C>T), GTA1547GTT (5784A>T), TAC1548TAT (5787C>T), AAC1550CAA (5791A>C 5793C>A), GCG1553GAA (5801C>A 5802G>A), GAT1554GAG (5805T>G), GAA1556ATG (5809G>A 5810A>T 5811A>G), AGG1557GGT (5812A>G 5814G>T), TTA1559TTG (5820A>G), AAG1560AGG (5822A>G), ATG1562GTG (5827A>G), CTG1563TT. (5830C>T) | | | | | | | | |

*: Inserts / Deletes / Misaligned / Frameshifts

#### Analysis details

This analysis was performed with panviral2.64

### NGS Details (UN9): Arhar cryptic virus-I (segment RNA 3)

#### Assembly

| | |
|---|---|
| **Coverage Length** | 311 (1 contig(s)) |
| **Depth Of Coverage** | 5.6 |
| **Number Of Reads** | 14 |
| **Reads Per Million** | 0.32 rpm (after QC) |
| **Ambiguities** | 0 |
| **Assembly Method** | de novo + reference guided assembly |
| **Consensus Caller** | Bcf Tools |

#### Coverage Map

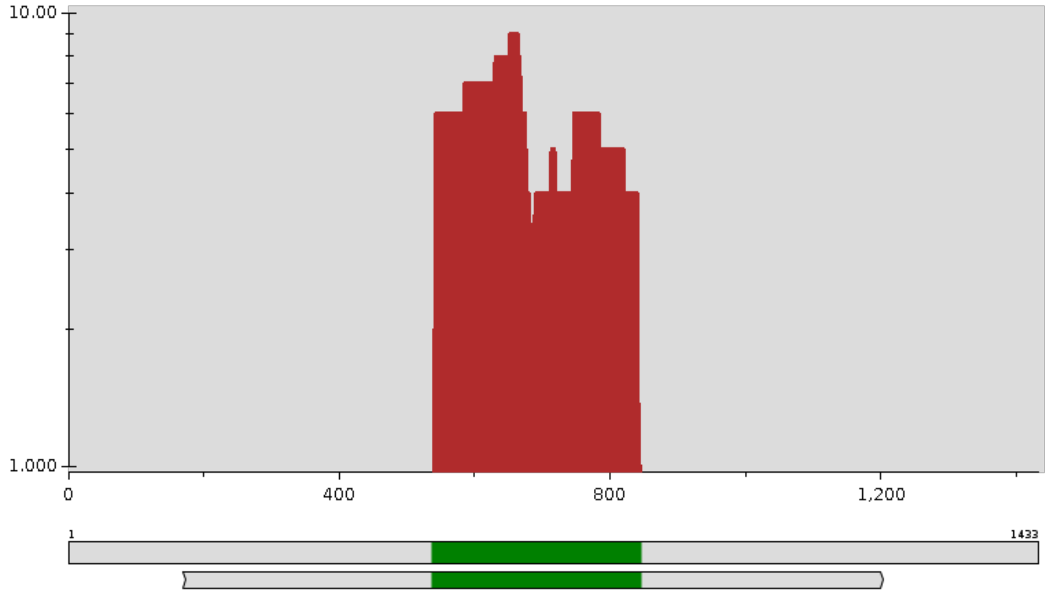

#### Assignment

| | |
|---|---|
| **Type** | Arhar cryptic virus-I (Taxonomy ID: 1585924) |
| **Reference Genome** | NC_024010.1 |
| **NT Identity (%)** | 57.7049 |
| **AA Identity (%)** | 42.5743 |
| **Number Of Stop Codons** | 0 |
| **Number Of CDS** | 1 |

#### Alignment

| | |
|---|---|
| **Alignment Score** | 79.0 (NT) + 309.0 (AA) = 388.0 |
| **Concordance (%)** | 30.4553 |

| Alignment Method | Global, seeded, nucleotide + amino acids (AGA) |
|---|---|

#### Genome Region

Sequence starts at position 538 and ends at position 848 relative to NC_024010.1 reference sequence.

#### Alignment Detailed Statistics

| | Begin | End | Coverage | Score | Concordance | Matches | Identities | I/D/M/F* | Stop Codons |
|---|---|---|---|---|---|---|---|---|---|
| **NT** | **538** | **848** | **21.7%** | **79** | **13.3%** | **305 (98.1%)** | **176 (56.6%)** | **0/6** | |
| Mutations: | 543G>A, 544G>A, 549A>C, 553G>A, 557A>T, 559C>A, 560T>A, 561_566delGAAATT, 570A>G, 572G>T, 576A>G, 578A>T, 581T>G, 582A>G, 583C>A, 585T>C, 586T>A, 589T>C, 591C>A, 592C>T, 593G>T, 594C>A, 595A>T, 599T>G, 601T>C, 603A>G, 608T>G, 617T>A, 620T>A, 623T>C, 629T>G, 635T>G, 637G>A, 638C>T, 640G>A, 642C>A, 645A>C, 648A>C, 649A>G, 650A>T, 651T>G, 655A>T, 656C>T, 660C>A, 661G>C, 662C>T, 663C>T, 666G>T, 667G>C, 668A>C, 670C>A, 674A>G, 677C>T, 678G>A, 680C>A, 681A>G, 682C>A, 684G>C, 686C>T, 690G>C, 691T>G, 692C>A, 696A>G, 698C>A, 699A>G, 700A>T, 704T>C, 706T>C, 707C>T, 711A>T, 713A>G, 716A>C, 717T>G, 718T>C, 719A>C, 722G>T, 723C>T, 728T>A, 729C>A, 730T>A, 731C>G, 734A>G, 737A>C, 738C>T, 739A>C, 740A>T, 741G>A, 742A>G, 744C>A, 750T>G, 751C>T, 756C>A, 758G>T, 761C>T, 762C>G, 764G>A, 765A>T, 766A>T, 767T>G, 768C>A, 769T>C, 772A>T, 778A>T, 779C>T, 781T>A, 782C>G, 784T>C, 788G>T, 789A>G, 791A>G, 797C>A, 803A>C, 806A>G, 809T>A, 810G>A, 811T>C, 812G>T, 813C>G, 815G>A, 816C>A, 824A>T, 825C>T, 827C>T, 828A>T, 830A>C, 831G>A, 832C>G, 834G>C, 835C>A, 836T>G | | | | | | | | |

##### CDS

| | Begin | End | Coverage | Score | Concordance | Matches | Identities | I/D/M/F* | Stop Codons |
|---|---|---|---|---|---|---|---|---|---|
| CP | 124 | 226 | 29.9% | 309 | 45.2% | 101 (98.1%) | 43 (41.7%) | 0/2/0/0 | 0 |
| Protein mutations: | G125N (543G>A 544G>A), M127L (549A>C), R128H (553G>A), Q129H (557A>T), T130K (559C>A 560T>A), E131_I132del (561_566delGAAATT), T134A (570A>G 572G>T), I136V (576A>G 578A>T), T138D (582A>G 583C>A), F139H (585T>C 586T>A), V140A (589T>C), P141I (591C>A 592C>T 593G>T), H142I (594C>A 595A>T), I143M (599T>G), V144A (601T>C), N145D (603A>G), R156H (637G>A 638C>T), S157N (640G>A), L158I (642C>A), N159H (645A>C), K160R (648A>C 649A>G 650A>T), S161A (651T>G), Y162F (655A>T 656C>T), R164T (660C>A 661G>C 662C>T), H165Y (663C>T), G166S (666G>T 667G>C 668A>C), T167N (670C>A), A170T (678G>A 680C>A), T171D (681A>G 682C>A), A172P (684G>C 686C>T), V174R (690G>C 691T>G 692C>A), T176A (696A>G 698C>A), K177V (699A>G 700A>T), I179T (706T>C 707C>T), I181L (711A>T 713A>G), L183A (717T>G 718T>C 719A>C), L185F (723C>T), L187K (729C>A 730T>A 731C>G), Q190S (738C>T 739A>C 740A>T), E191R (741G>A 742A>G), L192I (744C>A), S194V (750T>G 751C>T), Q196N (756C>A 758G>T), Q198E (762C>G 764G>A), N199L (765A>T 766A>T 767T>G), L200T (768C>A 769T>C), E201V (772A>T), N203I (778A>T 779C>T), V204E (781T>A 782C>G), I205T (784T>C), M206I (788G>T), I207V (789A>G 791A>G), V214T (810G>A 811T>C 812G>T), Q215E (813C>G 815G>A), H216N (816C>A), R219C (825C>T 827C>T), T220S (828A>T 830A>C), A221S (831G>A 832C>G), A222Q (834G>C 835C>A 836T>G) | | | | | | | | |
| Codon mutations: | GGC125AAC (543G>A 544G>A), ATG127CTG (549A>C), CGC128CAC (553G>A), CAA129CAT (557A>T), ACT130AAA (559C>A 560T>A), GAA131_ATT132del (561_566delGAAATT), ACG134GCT (570A>G 572G>T), ATA136GTT (576A>G 578A>T), GCT137GCG (581T>G), ACT138GAT (582A>G 583C>A), TTC139CAC (585T>C 586T>A), GTT140GCT (589T>C), CCG141ATT (591C>A 592C>T 593G>T), CAC142ATC (594C>A 595A>T), ATT143ATG (599T>G), GTT144GCT (601T>C), AAT145GAT (603A>G), GTT146GTG (608T>G), TCT149TCA (617T>A), GCT150GCA (620T>A), CTT151CTC (623T>C), GCT153GCG (629T>G), CTT155CTG (635T>G), CGC156CAT (637G>A 638C>T), AGC157AAC (640G>A), CTT158ATT (642C>A), AAC159CAC (645A>C), AAA160CGT (648A>C 649A>G 650A>T), TCC161GCC (651T>G), TAC162TTT (655A>T 656C>T), CGC164ACT (660C>A 661G>C 662C>T), CAT165TAT (663C>T), GGA166TCC (666G>T 667G>C 668A>C), ACC167AAC (670C>A), AGA168AGG (674A>G), TAC169TAT (677C>T), GCC170ACA (678G>A 680C>A), ACT171GAT (681A>G 682C>A), GCC172CCT (684G>C 686C>T), GTC174CGA (690G>C 691T>G 692C>A), ACC176GCA (696A>G 698C>A), AAG177GTG (699A>G 700A>T), GAT178GAC (704T>C), ATC179ACT (706T>C 707C>T), ATA181TTG (711A>T 713A>G), CCA182CCC (716A>C), TTA183GCC (717T>G 718T>C 719A>C), CCG184CCT (722G>T), CTT185TTT (723C>T), GCT186GCA (728T>A), CTC187AAG (729C>A 730T>A 731C>G), GCA188GCG (734A>G), ATA189ATC (737A>C), CAA190TCT (738C>T 739A>C 740A>T), GAA191AGA (741G>A 742A>G), CTT192ATT (744C>A), TCC194GTC (750T>G 751C>T), CAG196AAT (756C>A 758G>T), ACC197ACT (761C>T), CAG198GAA (762C>G 764G>A), AAT199TTG (765A>T 766A>T 767T>G), CTT200ACT (768C>A 769T>C), GAG201GTG (772A>T), AAC203ATT (778A>T 779C>T), GTC204GAG (781T>A 782C>G), ATT205ACT (784T>C), ATG206ATT (788G>T), ATA207GTG (789A>G 791A>G), ACC209ACA (797C>A), CCA211CCC (803A>C), GAA212GAG (806A>G), GGT213GGA (809T>A), GTG214ACT (810G>A 811T>C 812G>T), CAG215GAA (813C>G 815G>A), CAT216AAT (816C>A), GGA218GGT (824A>T), CGC219TGT (825C>T 827C>T), ACA220TCC (828A>T 830A>C), GCC221AGC (831G>A 832C>G), GCT222CAG (834G>C 835C>A 836T>G) | | | | | | | | |

##### Proteins

| | Begin | End | Coverage | Score | Concordance | Matches | Identities | I/D/M/F* | Stop Codons |
|---|---|---|---|---|---|---|---|---|---|
| Coat Protein (YP_009026398.1) | 124 | 226 | 29.9% | 309 | 45.2% | 101 (98.1%) | 43 (41.7%) | 0/2/0/0 | 0 |
| Protein mutations: | G125N (543G>A 544G>A), M127L (549A>C), R128H (553G>A), Q129H (557A>T), T130K (559C>A 560T>A), E131_I132del (561_566delGAAATT), T134A (570A>G 572G>T), I136V (576A>G 578A>T), T138D (582A>G 583C>A), F139H (585T>C 586T>A), V140A (589T>C), P141I (591C>A 592C>T 593G>T), H142I (594C>A 595A>T), I143M (599T>G), V144A (601T>C), N145D (603A>G), R156H (637G>A 638C>T), S157N (640G>A), L158I (642C>A), N159H (645A>C), K160R (648A>C 649A>G 650A>T), S161A (651T>G), Y162F (655A>T 656C>T), R164T (660C>A 661G>C 662C>T), H165Y (663C>T), G166S (666G>T 667G>C 668A>C), T167N (670C>A), A170T (678G>A 680C>A), T171D (681A>G 682C>A), A172P (684G>C 686C>T), V174R (690G>C 691T>G 692C>A), T176A (696A>G 698C>A), K177V (699A>G 700A>T), I179T (706T>C 707C>T), I181L (711A>T 713A>G), L183A (717T>G 718T>C 719A>C), L185F (723C>T), L187K (729C>A 730T>A 731C>G), Q190S (738C>T 739A>C 740A>T), E191R (741G>A 742A>G), L192I (744C>A), S194V (750T>G 751C>T), Q196N (756C>A 758G>T), Q198E (762C>G 764G>A), N199L (765A>T 766A>T 767T>G), L200T (768C>A 769T>C), E201V (772A>T), N203I (778A>T 779C>T), V204E (781T>A 782C>G), I205T (784T>C), M206I (788G>T), I207V (789A>G 791A>G), V214T (810G>A 811T>C 812G>T), Q215E (813C>G 815G>A), H216N (816C>A), R219C (825C>T 827C>T), T220S (828A>T 830A>C), A221S (831G>A 832C>G), A222Q (834G>C 835C>A 836T>G) | | | | | | | | |
| Codon mutations: | GGC125AAC (543G>A 544G>A), ATG127CTG (549A>C), CGC128CAC (553G>A), CAA129CAT (557A>T), ACT130AAA (559C>A 560T>A), GAA131_ATT132del (561_566delGAAATT), ACG134GCT (570A>G 572G>T), ATA136GTT (576A>G 578A>T), GCT137GCG (581T>G), ACT138GAT (582A>G 583C>A), TTC139CAC (585T>C 586T>A), GTT140GCT (589T>C), CCG141ATT (591C>A 592C>T 593G>T), CAC142ATC (594C>A 595A>T), ATT143ATG (599T>G), GTT144GCT (601T>C), AAT145GAT (603A>G), GTT146GTG (608T>G), TCT149TCA (617T>A), GCT150GCA (620T>A), CTT151CTC (623T>C), GCT153GCG (629T>G), CTT155CTG (635T>G), CGC156CAT (637G>A 638C>T), AGC157AAC (640G>A), CTT158ATT (642C>A), AAC159CAC (645A>C), AAA160CGT (648A>C 649A>G 650A>T), TCC161GCC (651T>G), TAC162TTT (655A>T 656C>T), CGC164ACT (660C>A 661G>C 662C>T), CAT165TAT (663C>T), GGA166TCC (666G>T 667G>C 668A>C), ACC167AAC (670C>A), AGA168AGG (674A>G), TAC169TAT (677C>T), GCC170ACA (678G>A 680C>A), ACT171GAT (681A>G 682C>A), GCC172CCT (684G>C 686C>T), GTC174CGA (690G>C 691T>G 692C>A), ACC176GCA (696A>G 698C>A), AAG177GTG (699A>G 700A>T), GAT178GAC (704T>C), ATC179ACT (706T>C 707C>T), ATA181TTG (711A>T 713A>G), CCA182CCC (716A>C), TTA183GCC (717T>G 718T>C 719A>C), CCG184CCT (722G>T), CTT185TTT (723C>T), GCT186GCA (728T>A), CTC187AAG (729C>A 730T>A 731C>G), GCA188GCG (734A>G), ATA189ATC (737A>C), CAA190TCT (738C>T 739A>C 740A>T), GAA191AGA (741G>A 742A>G), CTT192ATT (744C>A), TCC194GTC (750T>G 751C>T), CAG196AAT (756C>A 758G>T), ACC197ACT (761C>T), CAG198GAA (762C>G 764G>A), AAT199TTG (765A>T 766A>T 767T>G), CTT200ACT (768C>A 769T>C), GAG201GTG (772A>T), AAC203ATT (778A>T 779C>T), GTC204GAG (781T>A 782C>G), ATT205ACT (784T>C), ATG206ATT (788G>T), ATA207GTG (789A>G 791A>G), ACC209ACA (797C>A), CCA211CCC (803A>C), GAA212GAG (806A>G), GGT213GGA (809T>A), GTG214ACT (810G>A 811T>C 812G>T), CAG215GAA (813C>G 815G>A), CAT216AAT (816C>A), GGA218GGT (824A>T), CGC219TGT (825C>T 827C>T), ACA220TCC (828A>T 830A>C), GCC221AGC (831G>A 832C>G), GCT222CAG (834G>C 835C>A 836T>G) | | | | | | | | |

*: Inserts / Deletes / Misaligned / Frameshifts

#### Analysis details

This analysis was performed with panviral2.64

### NGS Details (UN9): Dioscovirus dioscoreae

#### Assembly

| | |
|---|---|
| **Coverage Length** | 332 (1 contig(s)) |
| **Depth Of Coverage** | 4.2 |
| **Number Of Reads** | 11 |
| **Reads Per Million** | 0.25 rpm (after QC) |
| **Ambiguities** | 0 |
| **Assembly Method** | de novo + reference guided assembly |
| **Consensus Caller** | Bcf Tools |

#### Coverage Map

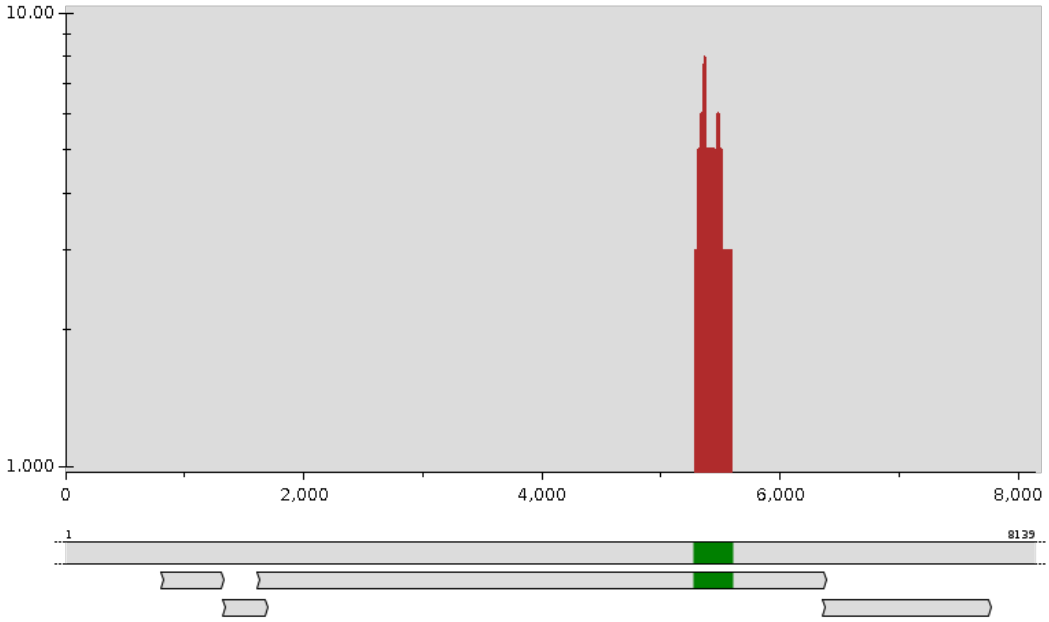

#### Assignment

| | |
|---|---|
| **Type** | Dioscovirus dioscoreae (Taxonomy ID: 3052184) |
| **Reference Genome** | NC_040712.1 |
| **NT Identity (%)** | 57.0122 |
| **AA Identity (%)** | 49.5413 |
| **Number Of Stop Codons** | 0 |
| **Number Of CDS** | 4 |

#### Alignment

| | |
|---|---|
| **Alignment Score** | 70.0 (NT) + 258.0 (AA) = 328.0 |
| **Concordance (%)** | 25.4658 |

| **Alignment Method** | Global, seeded, nucleotide + amino acids (AGA) |
|---|---|

#### Genome Region

Sequence starts at position 5274 and ends at position 5605 relative to NC_040712.1 reference sequence.

#### Alignment Detailed Statistics

<table>
<tr><th></th><th>Begin</th><th>End</th><th>Coverage</th><th>Score</th><th>Concordance</th><th>Matches</th><th>Identities</th><th>I/D/M/F*</th><th>Stop Codons</th></tr>
<tr><td>NT</td><td>5274</td><td>5605</td><td>4.1%</td><td>70</td><td>11.0%</td><td>325 (97.0%)</td><td>187 (55.8%)</td><td>3/7</td><td></td></tr>
<tr><td>Mutations:</td><td colspan="9">5274C>T, 5276A>G, 5280G>C, 5282T>C, 5284A>T, 5288A>C, 5289T>A, 5290A>T, 5298C>A, 5303C>T, 5304A>C, 5306A>T, 5307C>A, 5312A>T, 5314G>T, 5321C>A, 5322T>G, 5324T>A, 5327C>G, 5328C>T, 5330A>G, 5331C>G, 5332T>A, 5334A>G, 5337A>C, 5340A>G, 5341A>G, 5342A>G, 5345C>T, 5346A>G, 5347A>C, 5348G>A, 5349A>C, 5350T>A, 5351T>A, 5352G>C, 5353T>G, 5354C>G, 5356A>T, 5357C>T, 5358A>T, 5359G>C, 5363A>G, 5364T>G, 5372A>C, 5374A>G, 5378A>T, 5381A>G, 5383T>A, 5387C>T, 5393T>C, 5395T>G, 5401A>C, 5402T>C, 5403C>A, 5407A>C, 5408G>A, 5410G>T, 5414A>C, 5415G>C, 5417A>T, 5418T>A, 5419G>A, 5420G>A, 5423A>T, 5426A>C, 5430G>A, 5431T>G, 5432C>G, 5433T>A, 5434G>A, 5435C>A, 5437C>A, 5438A>T, 5439C>T, 5440A>C, 5441G>T, 5444T>C, 5447T>C, 5449T>A, 5451_5457delGAATGGA, 5458T>C, 5462A>G, 5468A>T, 5471C>T, 5477C>A, 5479A>C, 5480A>T, 5485C>A, 5486A>C, 5492A>T, 5493G>A, 5494T>G, 5495A>T, 5498C>T, 5501A>G, 5502C>A, 5504A>T, 5505A>T, 5506A>T, 5511G>A, 5514A>C, 5515A>G, 5516T>A, 5522C>T, 5523A>C, 5528A>G, 5529T>C, 5531T>C, 5532T>C, 5533C>T, 5534_5535insCGG, 5535G>A, 5536A>G, 5541G>A, 5545G>T, 5546T>G, 5549T>G, 5551A>T, 5553A>T, 5555A>T, 5558C>T, 5564T>C, 5565T>C, 5567G>C, 5570A>T, 5574T>A, 5575C>G, 5577G>A, 5580A>T, 5583A>C, 5585A>G, 5586C>G, 5589C>G, 5591A>T, 5597A>G, 5598C>G, 5599A>T, 5600A>T, 5603T>C</td></tr>
<tr><td colspan="10">CDS</td></tr>
<tr><td>EXK67_gp3</td><td>1222</td><td>1332</td><td>6.9%</td><td>258</td><td>32.9%</td><td>108 (97.3%)</td><td>54 (48.6%)</td><td>1/2/1/1</td><td>0</td></tr>
<tr><td>Protein mutations:</td><td colspan="9">D1224H (5280G>C 5282T>C), N1225I (5284A>T), Y1227I (5289T>A 5290A>T), Q1230K (5298C>A), T1232P (5304A>C 5306A>T), L1233I (5307C>A), S1235I (5314G>T), D1237E (5321C>A), Y1238E (5322T>G 5324T>A), L1241D (5331C>G 5332T>A), K1242E (5334A>G), I1243L (5337A>C), K1244G (5340A>G 5341A>G 5342A>G), K1246A (5346A>G 5347A>C 5348G>A), I1247Q (5349A>C 5350T>A 5351T>A), V1248R (5352G>C 5353T>G 5354C>G), Y1249F (5356A>T 5357C>T), F1252V (5364T>G), K1255R (5374A>G), F1258Y (5383T>A), M1262R (5395T>G), D1264A (5401A>C 5402T>C), P1265T (5403C>A), Q1266P (5407A>C 5408G>A), S1267I (5410G>T), E1269H (5415G>C 5417A>T), W1270K (5418T>A 5419G>A 5420G>A), V1274R (5430G>A 5431T>G 5432C>G), C1275K (5433T>A 5434G>A 5435C>A), P1276H (5437C>A 5438A>T), Q1277S (5439C>T 5440A>C 5441G>T), F1280Y (5449T>A), E1281_W1282del (5451_5457delGAATGGA), K1290T (5479A>C 5480A>T), A1292D (5485C>A 5486A>C), V1295S (5493G>A 5494T>G 5495A>T), R1298S (5502C>A 5504A>T), K1299L (5505A>T 5506A>T), D1301N (5511G>A), N1302R (5514A>C 5515A>G 5516T>A), K1305Q (5523A>C), Y1307H (5529T>C 5531T>C), S1308L (5532T>C 5533C>T), S1308_E1309insR (5534_5535insCGG), E1309R (5535G>A 5536A>G), V1311I (5541G>A), C1312L (5545G>T 5546T>G), Y1314F (5551A>T), I1315F (5553A>T 5555A>T), E1323K (5577G>A), S1324C (5580A>T), I1325L (5583A>C 5585A>G), Q1326E (5586C>G), Q1327D (5589C>G 5591A>T), Q1330V (5598C>G 5599A>T 5600A>T)</td></tr>
<tr><td>Codon mutations:</td><td colspan="9">CTA1222TTG (5274C>T 5276A>G), GAT1224CAC (5280G>C 5282T>C), AAT1225ATT (5284A>T), ACA1226ACC (5288A>C), TAC1227ATC (5289T>A 5290A>T), CAA1230AAA (5298C>A), TAC1231TAT (5303C>T), ACA1232CCT (5304A>C 5306A>T), CTC1233ATC (5307C>A), CCA1234CCT (5312A>T), AGT1235ATT (5314G>T), GAC1237GAA (5321C>A), TAT1238GAA (5322T>G 5324T>A), CTC1239CTG (5327C>G), CTA1240TTG (5328C>T 5330A>G), CTT1241GAT (5331C>G 5332T>A), AAA1242GAA (5334A>G), ATT1243CTT (5337A>C), AAA1244GGG (5340A>G 5341A>G 5342A>G), GAC1245GAT (5345C>T), AAG1246GCA (5346A>G 5347A>C 5348G>A), ATT1247CAA (5349A>C 5350T>A 5351T>A), GTC1248CGG (5352G>C 5353T>G 5354C>G), TAC1249TTT (5356A>T 5357C>T), AGC1250TCC (5358A>T 5359G>C), AAA1251AAG (5363A>G), TTT1252GTT (5364T>G), CTA1254CTC (5372A>C), AAG1255AGG (5374A>G), TCA1256TCT (5378A>T), GGA1257GGG (5381A>G), TTC1258TAC (5383T>A), CAC1259CAT (5387C>T), ATT1261ATC (5393T>C), ATG1262AGG (5395T>G), GAT1264GCC (5401A>C 5402T>C), CCA1265ACA (5403C>A), CAG1266CCA (5407A>C 5408G>A), AGT1267ATT (5410G>T), ATA1268ATC (5414A>C), GAA1269CAT (5415G>C 5417A>T), TGG1270AAA (5418T>A 5419G>A 5420G>A), ACA1271ACT (5423A>T), GCA1272GCC (5426A>C), GTC1274AGG (5430G>A 5431T>G 5432C>G), TGC1275AAA (5433T>A 5434G>A 5435C>A), CCA1276CAT (5437C>A 5438A>T), CAG1277TCT (5439C>T 5440A>C 5441G>T), GGT1278GGC (5444T>C), CAT1279CAC (5447T>C), TTT1280TAT (5449T>A), GAA1281_TGG1282del (5451_5457delGAATGGA), ATA1283-CA (5451_5457delGAATGGA 5458T>C), GTA1284GTG (5462A>G), CCA1286CCT (5468A>T), TTC1287TTT (5471C>T), CTC1289CTA (5477C>A), AAA1290ACT (5479A>C 5480A>T), GCA1292GAC (5485C>A 5486A>C), TCA1294TCT (5492A>T), GTA1295AGT (5493G>A 5494T>G 5495A>T), TTC1296TTT (5498C>T), CAA1297CAG (5501A>G), CGA1298AGT (5502C>A 5504A>T), AAA1299TTA (5505A>T 5506A>T), GAT1301AAT (5511G>A), AAT1302CGA (5514A>C 5515A>G 5516T>A), TTC1304TTT (5522C>T), AAA1305CAA (5523A>C), AAA1306AAG (5528A>G), TAT1307CAC (5529T>C 5531T>C), TCA1308CTA (5532T>C 5533C>T), TCA1308_GAA1309insCGG (5534_5535insCGG), GAA1309AGA (5535G>A 5536A>G), GTC1311ATC (5541G>A), TGT1312TTG (5545G>T 5546T>G), GTT1313GTG (5549T>G), TAC1314TTC (5551A>T), ATA1315TTT (5553A>T 5555A>T), GAC1316GAT (5558C>T), ATT1318ATC (5564T>C), TTG1319CTC (5565T>C 5567G>C), ATA1320ATT (5570A>T), TCT1322AGT (5574T>A 5575C>G), GAA1323AAA (5577G>A), AGT1324TGT (5580A>T), ATA1325CTG (5583A>C 5585A>G), CAA1326GAA (5586C>G), CAA1327GAT (5589C>G 5591A>T), GTA1329GTG (5597A>G), CAA1330GTT (5598C>G 5599A>T 5600A>T), CAT1331CAC (5603T>C)</td></tr>
<tr><td colspan="10">Proteins</td></tr>
<tr><td>ORF3 (YP_009553219.1)</td><td>1222</td><td>1332</td><td>6.9%</td><td>258</td><td>32.9%</td><td>108 (97.3%)</td><td>54 (48.6%)</td><td>1/2/1/1</td><td>0</td></tr>
<tr><td>Protein mutations:</td><td colspan="9">D1224H (5280G>C 5282T>C), N1225I (5284A>T), Y1227I (5289T>A 5290A>T), Q1230K (5298C>A), T1232P (5304A>C 5306A>T), L1233I (5307C>A), S1235I (5314G>T), D1237E (5321C>A), Y1238E (5322T>G 5324T>A), L1241D (5331C>G 5332T>A), K1242E (5334A>G), I1243L (5337A>C), K1244G (5340A>G 5341A>G 5342A>G), K1246A (5346A>G 5347A>C 5348G>A), I1247Q (5349A>C 5350T>A 5351T>A), V1248R (5352G>C 5353T>G 5354C>G), Y1249F (5356A>T 5357C>T), F1252V (5364T>G), K1255R (5374A>G), F1258Y (5383T>A), M1262R (5395T>G), D1264A (5401A>C 5402T>C), P1265T (5403C>A), Q1266P (5407A>C 5408G>A), S1267I (5410G>T), E1269H (5415G>C 5417A>T), W1270K (5418T>A 5419G>A 5420G>A), V1274R (5430G>A 5431T>G 5432C>G), C1275K (5433T>A 5434G>A 5435C>A), P1276H (5437C>A 5438A>T), Q1277S (5439C>T 5440A>C 5441G>T), F1280Y (5449T>A), E1281_W1282del (5451_5457delGAATGGA), K1290T (5479A>C 5480A>T), A1292D (5485C>A 5486A>C), V1295S (5493G>A 5494T>G 5495A>T), R1298S (5502C>A 5504A>T), K1299L (5505A>T 5506A>T), D1301N (5511G>A), N1302R (5514A>C 5515A>G 5516T>A), K1305Q (5523A>C), Y1307H (5529T>C 5531T>C), S1308L (5532T>C 5533C>T), S1308_E1309insR (5534_5535insCGG), E1309R (5535G>A 5536A>G), V1311I (5541G>A), C1312L (5545G>T 5546T>G), Y1314F (5551A>T), I1315F (5553A>T 5555A>T), E1323K (5577G>A), S1324C (5580A>T), I1325L (5583A>C 5585A>G), Q1326E (5586C>G), Q1327D (5589C>G 5591A>T), Q1330V (5598C>G 5599A>T 5600A>T)</td></tr>
<tr><td>Codon mutations:</td><td colspan="9">CTA1222TTG (5274C>T 5276A>G), GAT1224CAC (5280G>C 5282T>C), AAT1225ATT (5284A>T), ACA1226ACC (5288A>C), TAC1227ATC (5289T>A 5290A>T), CAA1230AAA (5298C>A), TAC1231TAT (5303C>T), ACA1232CCT (5304A>C 5306A>T), CTC1233ATC (5307C>A), CCA1234CCT (5312A>T), AGT1235ATT (5314G>T), GAC1237GAA (5321C>A), TAT1238GAA (5322T>G 5324T>A), CTC1239CTG (5327C>G), CTA1240TTG (5328C>T 5330A>G), CTT1241GAT (5331C>G 5332T>A), AAA1242GAA (5334A>G), ATT1243CTT (5337A>C), AAA1244GGG (5340A>G 5341A>G 5342A>G), GAC1245GAT (5345C>T), AAG1246GCA (5346A>G 5347A>C 5348G>A), ATT1247CAA (5349A>C 5350T>A 5351T>A), GTC1248CGG (5352G>C 5353T>G 5354C>G), TAC1249TTT (5356A>T 5357C>T), AGC1250TCC (5358A>T 5359G>C), AAA1251AAG (5363A>G), TTT1252GTT (5364T>G), CTA1254CTC (5372A>C), AAG1255AGG (5374A>G), TCA1256TCT (5378A>T), GGA1257GGG (5381A>G), TTC1258TAC (5383T>A), CAC1259CAT (5387C>T), ATT1261ATC (5393T>C), ATG1262AGG (5395T>G), GAT1264GCC (5401A>C 5402T>C), CCA1265ACA (5403C>A), CAG1266CCA (5407A>C 5408G>A), AGT1267ATT (5410G>T), ATA1268ATC (5414A>C), GAA1269CAT (5415G>C 5417A>T), TGG1270AAA (5418T>A 5419G>A 5420G>A), ACA1271ACT (5423A>T), GCA1272GCC (5426A>C), GTC1274AGG (5430G>A 5431T>G 5432C>G), TGC1275AAA (5433T>A 5434G>A 5435C>A), CCA1276CAT (5437C>A 5438A>T), CAG1277TCT (5439C>T 5440A>C 5441G>T), GGT1278GGC (5444T>C), CAT1279CAC (5447T>C), TTT1280TAT (5449T>A), GAA1281_TGG1282del (5451_5457delGAATGGA), ATA1283-CA (5451_5457delGAATGGA 5458T>C), GTA1284GTG (5462A>G), CCA1286CCT (5468A>T), TTC1287TTT (5471C>T), CTC1289CTA (5477C>A), AAA1290ACT (5479A>C 5480A>T), GCA1292GAC (5485C>A 5486A>C), TCA1294TCT (5492A>T), GTA1295AGT (5493G>A 5494T>G 5495A>T), TTC1296TTT (5498C>T), CAA1297CAG (5501A>G), CGA1298AGT (5502C>A 5504A>T), AAA1299TTA (5505A>T 5506A>T), GAT1301AAT (5511G>A), AAT1302CGA (5514A>C 5515A>G 5516T>A), TTC1304TTT (5522C>T), AAA1305CAA (5523A>C), AAA1306AAG (5528A>G), TAT1307CAC (5529T>C 5531T>C), TCA1308CTA (5532T>C 5533C>T), TCA1308_GAA1309insCGG (5534_5535insCGG), GAA1309AGA (5535G>A 5536A>G), GTC1311ATC (5541G>A), TGT1312TTG (5545G>T 5546T>G), GTT1313GTG (5549T>G), TAC1314TTC (5551A>T), ATA1315TTT (5553A>T 5555A>T), GAC1316GAT (5558C>T), ATT1318ATC (5564T>C), TTG1319CTC (5565T>C 5567G>C), ATA1320ATT (5570A>T), TCT1322AGT (5574T>A 5575C>G), GAA1323AAA (5577G>A), AGT1324TGT (5580A>T), ATA1325CTG (5583A>C 5585A>G), CAA1326GAA (5586C>G), CAA1327GAT (5589C>G 5591A>T), GTA1329GTG (5597A>G), CAA1330GTT (5598C>G 5599A>T 5600A>T), CAT1331CAC (5603T>C)</td></tr>
</table>

*: Inserts / Deletes / Misaligned / Frameshifts

#### Analysis details

This analysis was performed with panviral2.64

### NGS Details (UN9): Dioscovirus dioscoreae

#### Assembly

| | |
|---|---|
| **Coverage Length** | 323 (1 contig(s)) |
| **Depth Of Coverage** | 4.1 |
| **Number Of Reads** | 11 |
| **Reads Per Million** | 0.25 rpm (after QC) |
| **Ambiguities** | 0 |
| **Assembly Method** | de novo + reference guided assembly |
| **Consensus Caller** | Bcf Tools |

#### Coverage Map

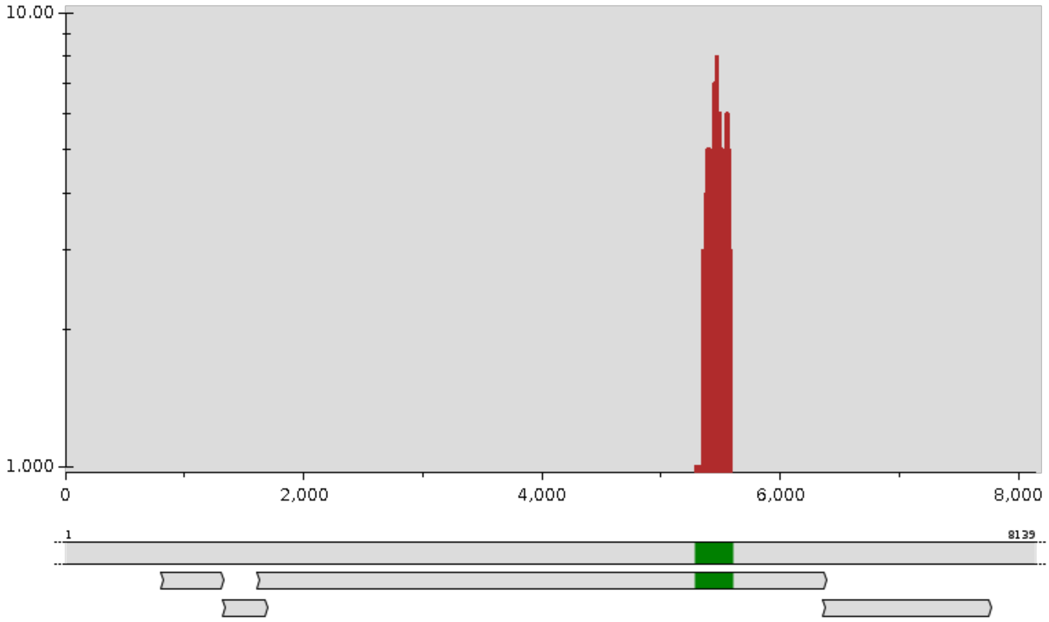

#### Assignment

| | |
|---|---|
| **Type** | Dioscovirus dioscoreae (Taxonomy ID: 3052184) |
| **Reference Genome** | NC_040712.1 |
| **NT Identity (%)** | 66.2539 |
| **AA Identity (%)** | 55.5556 |
| **Number Of Stop Codons** | 1 |
| **Number Of CDS** | 4 |

#### Alignment

| | |
|---|---|
| **Alignment Score** | 210.0 (NT) + 482.0 (AA) = 692.0 |
| **Concordance (%)** | 48.089 |

| **Alignment Method** | Global, seeded, nucleotide + amino acids (AGA) |
|---|---|

#### Genome Region

Sequence starts at position 5283 and ends at position 5605 relative to NC_040712.1 reference sequence.

#### Alignment Detailed Statistics

| | Begin | End | Coverage | Score | Concordance | Matches | Identities | I/D/M/F* | Stop Codons |
|---|---|---|---|---|---|---|---|---|---|
| **NT** | **5283** | **5605** | **4.0%** | **210** | **32.5%** | **323 (100%)** | **214 (66.3%)** | **0/0** | |
| Mutations: | 5288A>T, 5289T>A, 5290A>G, 5291C>A, 5293A>C, 5297C>T, 5298C>G, 5299A>C, 5303C>T, 5305C>A, 5307C>T, 5309C>A, 5310C>T, 5313A>G, 5314G>A, 5317T>A, 5318T>A, 5322T>A, 5324T>A, 5325C>T, 5327C>G, 5328C>A, 5330A>T, 5331C>T, 5332T>G, 5335A>G, 5340A>C, 5344A>G, 5345C>T, 5348G>A, 5350T>A, 5351T>A, 5352G>A, 5356A>T, 5357C>T, 5360C>T, 5361A>G, 5370C>T, 5371T>G, 5372A>T, 5375G>A, 5378A>G, 5383T>A, 5385C>T, 5387C>G, 5390G>A, 5391A>G, 5393T>A, 5395T>A, 5396G>A, 5400G>C, 5402T>C, 5406C>G, 5408G>C, 5429T>C, 5430G>A, 5431T>C, 5434G>A, 5435C>T, 5437C>T, 5438A>T, 5439C>G, 5444T>A, 5457A>C, 5461T>C, 5462A>C, 5471C>T, 5474A>T, 5475C>T, 5477C>A, 5482A>C, 5483T>A, 5486A>G, 5492A>T, 5493G>A, 5498C>T, 5502C>A, 5511G>A, 5514A>G, 5519A>G, 5523A>G, 5524A>G, 5525A>C, 5526A>G, 5528A>C, 5532T>A, 5533C>A, 5535G>A, 5537A>C, 5543C>T, 5545G>T, 5546T>A, 5549T>C, 5553A>G, 5558C>T, 5567G>A, 5568A>G, 5574T>A, 5575C>G, 5576T>C, 5577G>A, 5580A>G, 5581G>A, 5589C>G, 5594T>C, 5595G>C, 5598C>G, 5599A>G, 5604T>C | | | | | | | | |

##### CDS

| | Begin | End | Coverage | Score | Concordance | Matches | Identities | I/D/M/F* | Stop Codons |
|---|---|---|---|---|---|---|---|---|---|
| EXK67_gp3 | 1225 | 1332 | 6.8% | 482 | 60.5% | 108 (100%) | 60 (55.6%) | 0/0/0/0 | 1 |
| Protein mutations: | Y1227R (5289T>A 5290A>G 5291C>A), K1228T (5293A>C), Q1230A (5298C>G 5299A>C), T1232K (5305C>A), P1234S (5310C>T), S1235D (5313A>G 5314G>A), I1236K (5317T>A 5318T>A), Y1238K (5322T>A 5324T>A), L1240I (5328C>A 5330A>T), L1241C (5331C>T 5332T>G), K1242R (5335A>G), K1244Q (5340A>C), D1245G (5344A>G 5345C>T), I1247K (5350T>A 5351T>A), V1248I (5352G>A), Y1249F (5356A>T 5357C>T), K1251E (5361A>G), L1254C (5370C>T 5371T>G 5372A>T), F1258Y (5383T>A), H1259* (5385C>T 5387C>G), I1261V (5391A>G 5393T>A), M1262K (5395T>A 5396G>A), D1264H (5400G>C 5402T>C), Q1266D (5406C>G 5408G>C), V1274T (5430G>A 5431T>C), C1275Y (5434G>A 5435C>T), P1276L (5437C>T 5438A>T), Q1277E (5439C>G), I1283L (5457A>C), V1284A (5461T>C 5462A>C), N1291T (5482A>C 5483T>A), V1295I (5493G>A), D1301N (5511G>A), N1302D (5514A>G), I1303M (5519A>G), K1305G (5523A>G 5524A>G 5525A>C), K1306D (5526A>G 5528A>C), S1308K (5532T>A 5533C>A), E1309N (5535G>A 5537A>C), C1312L (5545G>T 5546T>A), I1315V (5553A>G), I1320V (5568A>G), E1323K (5577G>A), S1324D (5580A>G 5581G>A), Q1327E (5589C>G), V1329L (5595G>C), Q1330G (5598C>G 5599A>G) | | | | | | | | |
| Codon mutations: | ACA1226ACT (5288A>T), TAC1227AGA (5289T>A 5290A>G 5291C>A), AAA1228ACA (5293A>C), GAC1229GAT (5297C>T), CAA1230GCA (5298C>G 5299A>C), TAC1231TAT (5303C>T), ACA1232AAA (5305C>A), CTC1233TTA (5307C>T 5309C>A), CCA1234TCA (5310C>T), AGT1235GAT (5313A>G 5314G>A), ATT1236AAA (5317T>A 5318T>A), TAT1238AAA (5322T>A 5324T>A), CTC1239TTG (5325C>T 5327C>G), CTA1240ATT (5328C>A 5330A>T), CTT1241TGT (5331C>T 5332T>G), AAA1242AGA (5335A>G), AAA1244CAA (5340A>C), GAC1245GGT (5344A>G 5345C>T), AAG1246AAA (5348G>A), ATT1247AAA (5350T>A 5351T>A), GTC1248ATC (5352G>A), TAC1249TTT (5356A>T 5357C>T), AGC1250AGT (5360C>T), AAA1251GAA (5361A>G), CTA1254TGT (5370C>T 5371T>G 5372A>T), AAG1255AAA (5375G>A), TCA1256TCG (5378A>G), TTC1258TAC (5383T>A), CAC1259TAG (5385C>T 5387C>G), CAG1260CAA (5390G>A), ATT1261GTA (5391A>G 5393T>A), ATG1262AAA (5395T>A 5396G>A), GAT1264CAC (5400G>C 5402T>C), CAG1266GAC (5406C>G 5408G>C), TTT1273TTC (5429T>C), GTC1274ACC (5430G>A 5431T>C), TGC1275TAT (5434G>A 5435C>T), CCA1276CTT (5437C>T 5438A>T), CAG1277GAG (5439C>G), GGT1278GGA (5444T>A), ATA1283CTA (5457A>C), GTA1284GCC (5461T>C 5462A>C), TTC1287TTT (5471C>T), GGA1288GGT (5474A>T), CTC1289TTA (5475C>T 5477C>A), AAT1291ACA (5482A>C 5483T>A), GCA1292GCG (5486A>G), TCA1294TCT (5492A>T), GTA1295ATA (5493G>A), TTC1296TTT (5498C>T), CGA1298AGA (5502C>A), GAT1301AAT (5511G>A), AAT1302GAT (5514A>G), ATA1303ATG (5519A>G), AAA1305GGC (5523A>G 5524A>G 5525A>C), AAA1306GAC (5526A>G 5528A>C), TCA1308AAA (5532T>A 5533C>A), GAA1309AAC (5535G>A 5537A>C), GTC1311GTT (5543C>T), TGT1312TTA (5545G>T 5546T>A), GTT1313GTC (5549T>C), ATA1315GTA (5553A>G), GAC1316GAT (5558C>T), TTG1319TTA (5567G>A), ATA1320GTA (5568A>G), TCT1322AGC (5574T>A 5575C>G 5576T>C), GAA1323AAA (5577G>A), AGT1324GAT (5580A>G 5581G>A), CAA1327GAA (5589C>G), CAT1328CAC (5594T>C), GTA1329CTA (5595G>C), CAA1330GGA (5598C>G 5599A>G), TTA1332CT. (5604T>C) | | | | | | | | |

##### Proteins

| | Begin | End | Coverage | Score | Concordance | Matches | Identities | I/D/M/F* | Stop Codons |
|---|---|---|---|---|---|---|---|---|---|
| ORF3 (YP_009553219.1) | 1225 | 1332 | 6.8% | 482 | 60.5% | 108 (100%) | 60 (55.6%) | 0/0/0/0 | 1 |
| Protein mutations: | Y1227R (5289T>A 5290A>G 5291C>A), K1228T (5293A>C), Q1230A (5298C>G 5299A>C), T1232K (5305C>A), P1234S (5310C>T), S1235D (5313A>G 5314G>A), I1236K (5317T>A 5318T>A), Y1238K (5322T>A 5324T>A), L1240I (5328C>A 5330A>T), L1241C (5331C>T 5332T>G), K1242R (5335A>G), K1244Q (5340A>C), D1245G (5344A>G 5345C>T), I1247K (5350T>A 5351T>A), V1248I (5352G>A), Y1249F (5356A>T 5357C>T), K1251E (5361A>G), L1254C (5370C>T 5371T>G 5372A>T), F1258Y (5383T>A), H1259* (5385C>T 5387C>G), I1261V (5391A>G 5393T>A), M1262K (5395T>A 5396G>A), D1264H (5400G>C 5402T>C), Q1266D (5406C>G 5408G>C), V1274T (5430G>A 5431T>C), C1275Y (5434G>A 5435C>T), P1276L (5437C>T 5438A>T), Q1277E (5439C>G), I1283L (5457A>C), V1284A (5461T>C 5462A>C), N1291T (5482A>C 5483T>A), V1295I (5493G>A), D1301N (5511G>A), N1302D (5514A>G), I1303M (5519A>G), K1305G (5523A>G 5524A>G 5525A>C), K1306D (5526A>G 5528A>C), S1308K (5532T>A 5533C>A), E1309N (5535G>A 5537A>C), C1312L (5545G>T 5546T>A), I1315V (5553A>G), I1320V (5568A>G), E1323K (5577G>A), S1324D (5580A>G 5581G>A), Q1327E (5589C>G), V1329L (5595G>C), Q1330G (5598C>G 5599A>G) | | | | | | | | |
| Codon mutations: | ACA1226ACT (5288A>T), TAC1227AGA (5289T>A 5290A>G 5291C>A), AAA1228ACA (5293A>C), GAC1229GAT (5297C>T), CAA1230GCA (5298C>G 5299A>C), TAC1231TAT (5303C>T), ACA1232AAA (5305C>A), CTC1233TTA (5307C>T 5309C>A), CCA1234TCA (5310C>T), AGT1235GAT (5313A>G 5314G>A), ATT1236AAA (5317T>A 5318T>A), TAT1238AAA (5322T>A 5324T>A), CTC1239TTG (5325C>T 5327C>G), CTA1240ATT (5328C>A 5330A>T), CTT1241TGT (5331C>T 5332T>G), AAA1242AGA (5335A>G), AAA1244CAA (5340A>C), GAC1245GGT (5344A>G 5345C>T), AAG1246AAA (5348G>A), ATT1247AAA (5350T>A 5351T>A), GTC1248ATC (5352G>A), TAC1249TTT (5356A>T 5357C>T), AGC1250AGT (5360C>T), AAA1251GAA (5361A>G), CTA1254TGT (5370C>T 5371T>G 5372A>T), AAG1255AAA (5375G>A), TCA1256TCG (5378A>G), TTC1258TAC (5383T>A), CAC1259TAG (5385C>T 5387C>G), CAG1260CAA (5390G>A), ATT1261GTA (5391A>G 5393T>A), ATG1262AAA (5395T>A 5396G>A), GAT1264CAC (5400G>C 5402T>C), CAG1266GAC (5406C>G 5408G>C), TTT1273TTC (5429T>C), GTC1274ACC (5430G>A 5431T>C), TGC1275TAT (5434G>A 5435C>T), CCA1276CTT (5437C>T 5438A>T), CAG1277GAG (5439C>G), GGT1278GGA (5444T>A), ATA1283CTA (5457A>C), GTA1284GCC (5461T>C 5462A>C), TTC1287TTT (5471C>T), GGA1288GGT (5474A>T), CTC1289TTA (5475C>T 5477C>A), AAT1291ACA (5482A>C 5483T>A), GCA1292GCG (5486A>G), TCA1294TCT (5492A>T), GTA1295ATA (5493G>A), TTC1296TTT (5498C>T), CGA1298AGA (5502C>A), GAT1301AAT (5511G>A), AAT1302GAT (5514A>G), ATA1303ATG (5519A>G), AAA1305GGC (5523A>G 5524A>G 5525A>C), AAA1306GAC (5526A>G 5528A>C), TCA1308AAA (5532T>A 5533C>A), GAA1309AAC (5535G>A 5537A>C), GTC1311GTT (5543C>T), TGT1312TTA (5545G>T 5546T>A), GTT1313GTC (5549T>C), ATA1315GTA (5553A>G), GAC1316GAT (5558C>T), TTG1319TTA (5567G>A), ATA1320GTA (5568A>G), TCT1322AGC (5574T>A 5575C>G 5576T>C), GAA1323AAA (5577G>A), AGT1324GAT (5580A>G 5581G>A), CAA1327GAA (5589C>G), CAT1328CAC (5594T>C), GTA1329CTA (5595G>C), CAA1330GGA (5598C>G 5599A>G), TTA1332CT. (5604T>C) | | | | | | | | |

*: Inserts / Deletes / Misaligned / Frameshifts

#### Analysis details

This analysis was performed with panviral2.64

### NGS Details (UN9): Ichnoviriform fugitivi (segment C16)

#### Assembly

| | |
|---|---|
| **Coverage Length** | 179 (1 contig(s)) |
| **Depth Of Coverage** | 1.3 |
| **Number Of Reads** | 2 |
| **Reads Per Million** | 0.05 rpm (after QC) |
| **Ambiguities** | 0 |
| **Assembly Method** | de novo + reference guided assembly |
| **Consensus Caller** | Bcf Tools |

#### Coverage Map

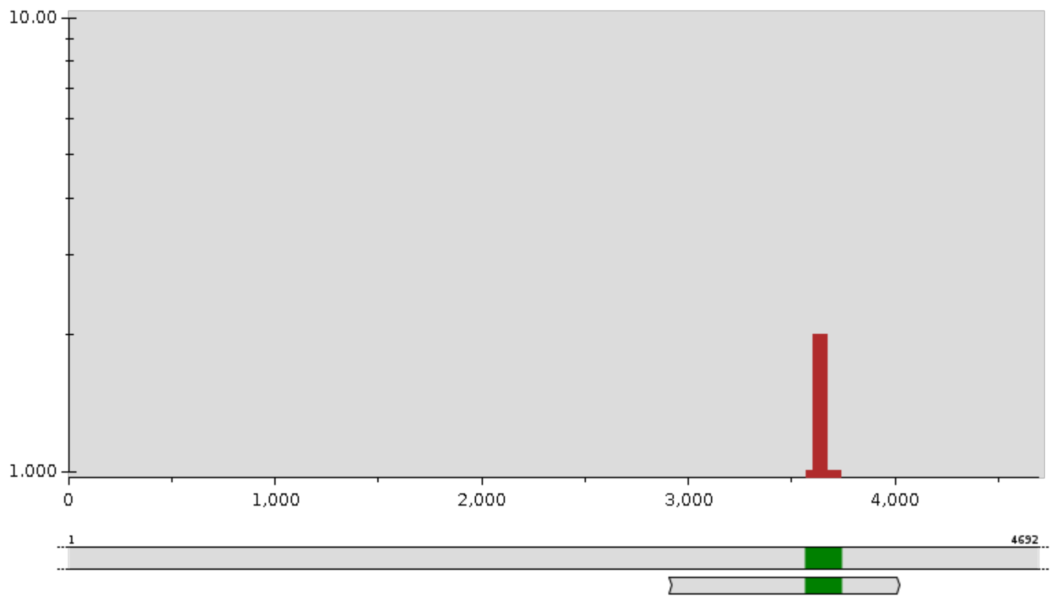

#### Assignment

| | |
|---|---|
| **Type** | Ichnoviriform fugitivi (Taxonomy ID: 265522) |
| **Reference Genome** | NC_008946.1 |
| **NT Identity (%)** | 59.7765 |
| **AA Identity (%)** | 49.1525 |
| **Number Of Stop Codons** | 0 |
| **Number Of CDS** | 1 |

#### Alignment

| | |
|---|---|
| **Alignment Score** | 70.0 (NT) + 268.0 (AA) = 338.0 |
| **Concordance (%)** | 41.8835 |

| | |
|---|---|
| **Alignment Method** | Global, seeded, nucleotide + amino acids (AGA) |

#### Genome Region

Sequence starts at position 3567 and ends at position 3745 relative to NC_008946.1 reference sequence.

#### Alignment Detailed Statistics

| | Begin | End | Coverage | Score | Concordance | Matches | Identities | I/D/M/F* | Stop Codons |
|---|---|---|---|---|---|---|---|---|---|
| **NT** | **3567** | **3745** | **3.8%** | **70** | **19.6%** | **179 (100%)** | **107 (59.8%)** | **0/0** | |
| Mutations: | 3567G>C, 3570G>A, 3572A>G, 3576T>G, 3577G>A, 3581A>G, 3586T>A, 3591A>T, 3592A>T, 3595A>G, 3598G>T, 3601C>T, 3604A>T, 3605A>C, 3606C>G, 3607G>T, 3608A>G, 3610C>T, 3614C>A, 3621T>C, 3622T>C, 3624A>T, 3625T>C, 3626G>C, 3628A>C, 3634C>T, 3637A>T, 3638C>T, 3646G>A, 3647A>G, 3648C>A, 3650A>G, 3652G>T, 3653G>C, 3657G>A, 3658C>G, 3659A>C, 3660G>A, 3661G>T, 3664C>T, 3665G>T, 3666G>C, 3667A>T, 3668A>C, 3670C>T, 3673C>T, 3674G>A, 3676G>T, 3677A>C, 3679G>T, 3680G>C, 3683C>T, 3684A>T, 3689T>A, 3690C>T, 3691A>T, 3692G>A, 3694C>T, 3698T>G, 3699C>A, 3703G>A, 3706C>T, 3710G>A, 3712T>C, 3716T>C, 3718G>C, 3724C>T, 3731C>G, 3732A>T, 3733C>T, 3734A>G, 3743C>G | | | | | | | | |

##### CDS

| | Begin | End | Coverage | Score | Concordance | Matches | Identities | I/D/M/F* | Stop Codons |
|---|---|---|---|---|---|---|---|---|---|
| HfIV_sC16gp1 | 221 | 279 | 15.9% | 268 | 58.6% | 59 (100%) | 29 (49.2%) | 0/0/0/0 | 0 |
| Protein mutations: | G221E (3570G>A), N222D (3572A>G), M223R (3576T>G 3577G>A), N225D (3581A>G), E228V (3591A>T 3592A>T), T233R (3605A>C 3606C>G 3607G>T), I234V (3608A>G 3610C>T), Q236K (3614C>A), I238T (3621T>C 3622T>C), Y239F (3624A>T 3625T>C), E240H (3626G>C 3628A>C), P244S (3638C>T), T247D (3647A>G 3648C>A), M248V (3650A>G 3652G>T), E249Q (3653G>C), S250K (3657G>A 3658C>G), R251H (3659A>C 3660G>A 3661G>T), G253S (3665G>T 3666G>C 3667A>T), I254L (3668A>C 3670C>T), V256I (3674G>A 3676G>T), M257L (3677A>C 3679G>T), A258P (3680G>C), Q259L (3683C>T 3684A>T), S261I (3689T>A 3690C>T 3691A>T), V262I (3692G>A 3694C>T), S264E (3698T>G 3699C>A), V268I (3710G>A 3712T>C), H275V (3731C>G 3732A>T 3733C>T), I276V (3734A>G), L279V (3743C>G) | | | | | | | | |
| Codon mutations: | GGA220.CA (3567G>C), GGG221GAG (3570G>A), AAT222GAT (3572A>G), ATG223AGA (3576T>G 3577G>A), AAT225GAT (3581A>G), CCT226CCA (3586T>A), GAA228GTT (3591A>T 3592A>T), AAA229AAG (3595A>G), GTG230GTT (3598G>T), TTC231TTT (3601C>T), CCA232CCT (3604A>T), ACG233CGT (3605A>C 3606C>G 3607G>T), ATC234GTT (3608A>G 3610C>T), CAG236AAG (3614C>A), ATT238ACC (3621T>C 3622T>C), TAT239TTC (3624A>T 3625T>C), GAA240CAC (3626G>C 3628A>C), TAC242TAT (3634C>T), GGA243GGT (3637A>T), CCT244TCT (3638C>T), GGG246GGA (3646G>A), ACT247GAT (3647A>G 3648C>A), ATG248GTT (3650A>G 3652G>T), GAG249CAG (3653G>C), AGC250AAG (3657G>A 3658C>G), AGG251CAT (3659A>C 3660G>A 3661G>T), GAC252GAT (3664C>T), GGA253TCT (3665G>T 3666G>C 3667A>T), ATC254CTT (3668A>C 3670C>T), TGC255TGT (3673C>T), GTG256ATT (3674G>A 3676G>T), ATG257CTT (3677A>C 3679G>T), GCA258CCA (3680G>C), CAG259TTG (3683C>T 3684A>T), TCA261ATT (3689T>A 3690C>T 3691A>T), GTC262ATT (3692G>A 3694C>T), TCG264GAG (3698T>G 3699C>A), AAG265AAA (3703G>A), ATC266ATT (3706C>T), GTT268ATC (3710G>A 3712T>C), TTG270CTC (3716T>C 3718G>C), TTC272TTT (3724C>T), CAC275GTT (3731C>G 3732A>T 3733C>T), ATT276GTT (3734A>G), CTT279GTT (3743C>G) | | | | | | | | |

##### Proteins

| | Begin | End | Coverage | Score | Concordance | Matches | Identities | I/D/M/F* | Stop Codons |
|---|---|---|---|---|---|---|---|---|---|
| viral inexin-like protein (YP_001031223.1) | 221 | 279 | 15.9% | 268 | 58.6% | 59 (100%) | 29 (49.2%) | 0/0/0/0 | 0 |
| Protein mutations: | G221E (3570G>A), N222D (3572A>G), M223R (3576T>G 3577G>A), N225D (3581A>G), E228V (3591A>T 3592A>T), T233R (3605A>C 3606C>G 3607G>T), I234V (3608A>G 3610C>T), Q236K (3614C>A), I238T (3621T>C 3622T>C), Y239F (3624A>T 3625T>C), E240H (3626G>C 3628A>C), P244S (3638C>T), T247D (3647A>G 3648C>A), M248V (3650A>G 3652G>T), E249Q (3653G>C), S250K (3657G>A 3658C>G), R251H (3659A>C 3660G>A 3661G>T), G253S (3665G>T 3666G>C 3667A>T), I254L (3668A>C 3670C>T), V256I (3674G>A 3676G>T), M257L (3677A>C 3679G>T), A258P (3680G>C), Q259L (3683C>T 3684A>T), S261I (3689T>A 3690C>T 3691A>T), V262I (3692G>A 3694C>T), S264E (3698T>G 3699C>A), V268I (3710G>A 3712T>C), H275V (3731C>G 3732A>T 3733C>T), I276V (3734A>G), L279V (3743C>G) | | | | | | | | |
| Codon mutations: | GGA220.CA (3567G>C), GGG221GAG (3570G>A), AAT222GAT (3572A>G), ATG223AGA (3576T>G 3577G>A), AAT225GAT (3581A>G), CCT226CCA (3586T>A), GAA228GTT (3591A>T 3592A>T), AAA229AAG (3595A>G), GTG230GTT (3598G>T), TTC231TTT (3601C>T), CCA232CCT (3604A>T), ACG233CGT (3605A>C 3606C>G 3607G>T), ATC234GTT (3608A>G 3610C>T), CAG236AAG (3614C>A), ATT238ACC (3621T>C 3622T>C), TAT239TTC (3624A>T 3625T>C), GAA240CAC (3626G>C 3628A>C), TAC242TAT (3634C>T), GGA243GGT (3637A>T), CCT244TCT (3638C>T), GGG246GGA (3646G>A), ACT247GAT (3647A>G 3648C>A), ATG248GTT (3650A>G 3652G>T), GAG249CAG (3653G>C), AGC250AAG (3657G>A 3658C>G), AGG251CAT (3659A>C 3660G>A 3661G>T), GAC252GAT (3664C>T), GGA253TCT (3665G>T 3666G>C 3667A>T), ATC254CTT (3668A>C 3670C>T), TGC255TGT (3673C>T), GTG256ATT (3674G>A 3676G>T), ATG257CTT (3677A>C 3679G>T), GCA258CCA (3680G>C), CAG259TTG (3683C>T 3684A>T), TCA261ATT (3689T>A 3690C>T 3691A>T), GTC262ATT (3692G>A 3694C>T), TCG264GAG (3698T>G 3699C>A), AAG265AAA (3703G>A), ATC266ATT (3706C>T), GTT268ATC (3710G>A 3712T>C), TTG270CTC (3716T>C 3718G>C), TTC272TTT (3724C>T), CAC275GTT (3731C>G 3732A>T 3733C>T), ATT276GTT (3734A>G), CTT279GTT (3743C>G) | | | | | | | | |

*: Inserts / Deletes / Misaligned / Frameshifts

#### Analysis details

This analysis was performed with panviral2.64

### NGS Details (UN9): Ichnoviriform fugitivi (segment B17)

#### Assembly

| | |
|---|---|
| **Coverage Length** | 159 (1 contig(s)) |
| **Depth Of Coverage** | 5.2 |
| **Number Of Reads** | 10 |
| **Reads Per Million** | 0.23 rpm (after QC) |
| **Ambiguities** | 0 |
| **Assembly Method** | de novo + reference guided assembly |
| **Consensus Caller** | Bcf Tools |

#### Coverage Map

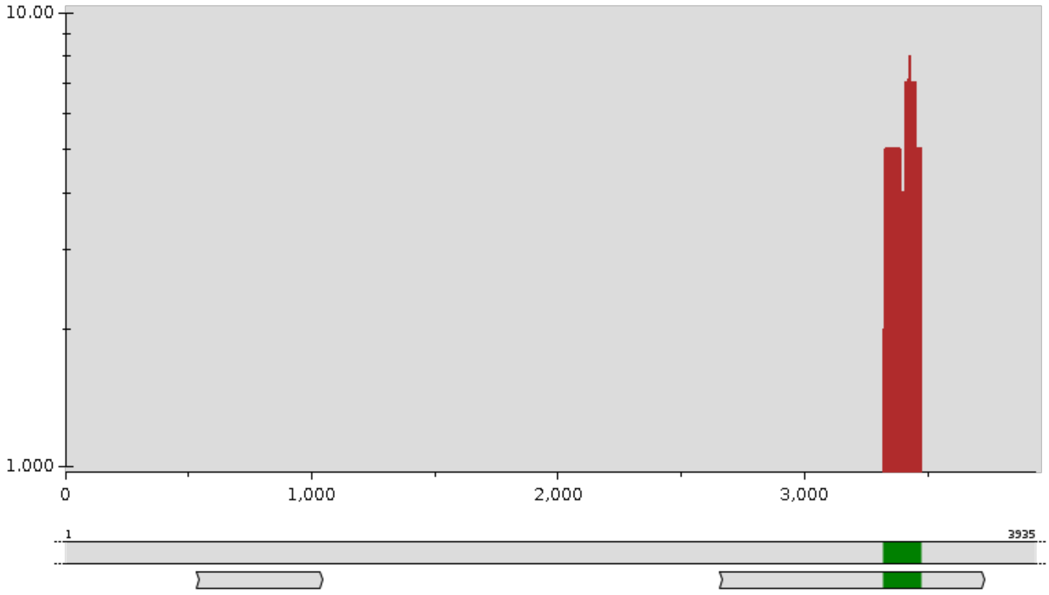

#### Assignment

| | |
|---|---|
| **Type** | Ichnoviriform fugitivi (Taxonomy ID: 265522) |
| **Reference Genome** | NC_008953.1 |
| **NT Identity (%)** | 61.0063 |
| **AA Identity (%)** | 50.9434 |
| **Number Of Stop Codons** | 0 |
| **Number Of CDS** | 2 |

#### Alignment

| | |
|---|---|
| **Alignment Score** | 70.0 (NT) + 258.0 (AA) = 328.0 |
| **Concordance (%)** | 45.7462 |

| Alignment Method | Global, seeded, nucleotide + amino acids (AGA) |
|---|---|

#### Genome Region

Sequence starts at position 3318 and ends at position 3476 relative to NC_008953.1 reference sequence.

#### Alignment Detailed Statistics

| | Begin | End | Coverage | Score | Concordance | Matches | Identities | I/D/M/F* | Stop Codons |
|---|---|---|---|---|---|---|---|---|---|
| **NT** | **3318** | **3476** | **4.0%** | **70** | **22.0%** | **159 (100%)** | **97 (61.0%)** | **0/0** | |
| Mutations: | 3326T>A, 3329C>T, 3330G>C, 3336C>A, 3337A>T, 3339G>A, 3341A>C, 3342G>T, 3348C>G, 3354A>T, 3356C>A, 3360C>G, 3363T>C, 3364C>A, 3365G>A, 3366A>G, 3371C>A, 3372T>C, 3376C>A, 3378C>T, 3379A>C, 3381A>T, 3384C>T, 3388C>T, 3390T>C, 3393G>T, 3396C>T, 3397T>G, 3398C>A, 3399C>T, 3400C>G, 3402G>T, 3403G>C, 3407A>T, 3408C>T, 3411C>T, 3414G>T, 3415G>A, 3416G>A, 3417A>C, 3418C>T, 3419T>A, 3423C>T, 3424A>T, 3426C>A, 3429C>G, 3432G>A, 3433G>T, 3434A>T, 3439G>A, 3441G>T, 3442G>A, 3444C>T, 3447C>T, 3450G>A, 3456C>T, 3463T>G, 3464T>C, 3465C>T, 3466C>T, 3472T>C, 3474C>T | | | | | | | | |
| **CDS** | | | | | | | | | |
| HfIV_sB17gp2 | 222 | 274 | 14.8% | 258 | 59.3% | 53 (100%) | 27 (50.9%) | 0/0/0/0 | 0 |
| Protein mutations: | M224K (3326T>A), T225I (3329C>T 3330G>C), M228L (3337A>T 3339G>A), K229T (3341A>C 3342G>T), T234K (3356C>A), I235M (3360C>G), R237K (3364C>A 3365G>A 3366A>G), T239N (3371C>A 3372T>C), H241N (3376C>A 3378C>T), K242H (3379A>C 3381A>T), P245S (3388C>T 3390T>C), S248D (3397T>G 3398C>A 3399C>T), L249V (3400C>G 3402G>T), E250Q (3403G>C), N251I (3407A>T 3408C>T), E253D (3414G>T), G254N (3415G>A 3416G>A 3417A>C), L255Y (3418C>T 3419T>A), I257L (3424A>T 3426C>A), E260L (3433G>T 3434A>T), V262I (3439G>A 3441G>T), V263I (3442G>A 3444C>T), F270A (3463T>G 3464T>C 3465C>T), L271F (3466C>T), F273L (3472T>C 3474C>T) | | | | | | | | |
| Codon mutations: | ATG224AAG (3326T>A), ACG225ATC (3329C>T 3330G>C), CCC227CCA (3336C>A), ATG228TTA (3337A>T 3339G>A), AAG229ACT (3341A>C 3342G>T), GTC231GTG (3348C>G), CCA233CCT (3354A>T), ACA234AAA (3356C>A), ATC235ATG (3360C>G), ACT236ACC (3363T>C), CGA237AAG (3364C>A 3365G>A 3366A>G), ACT239AAC (3371C>A 3372T>C), CAC241AAT (3376C>A 3378C>T), AAA242CAT (3379A>C 3381A>T), TAC243TAT (3384C>T), CCT245TCC (3388C>T 3390T>C), TCG246TCT (3393G>T), GGC247GGT (3396C>T), TCC248GAT (3397T>G 3398C>A 3399C>T), CTG249GTT (3400C>G 3402G>T), GAA250CAA (3403G>C), AAC251ATT (3407A>T 3408C>T), TAC252TAT (3411C>T), GAG253GAT (3414G>T), GGA254AAC (3415G>A 3416G>A 3417A>C), CTC255TAC (3418C>T 3419T>A), TGC256TGT (3423C>T), ATC257TTA (3424A>T 3426C>A), CTC258CTG (3429C>G), CCG259CCA (3432G>A), GAG260TTG (3433G>T 3434A>T), GTG262ATT (3439G>A 3441G>T), GTC263ATT (3442G>A 3444C>T), AAC264AAT (3447C>T), GAG265GAA (3450G>A), ATC267ATT (3456C>T), TTC270GCT (3463T>G 3464T>C 3465C>T), CTC271TTC (3466C>T), TTC273CTT (3472T>C 3474C>T) | | | | | | | | |
| **Proteins** | | | | | | | | | |
| viral inexin-like protein (YP_001031233.1) | 222 | 274 | 14.8% | 258 | 59.3% | 53 (100%) | 27 (50.9%) | 0/0/0/0 | 0 |
| Protein mutations: | M224K (3326T>A), T225I (3329C>T 3330G>C), M228L (3337A>T 3339G>A), K229T (3341A>C 3342G>T), T234K (3356C>A), I235M (3360C>G), R237K (3364C>A 3365G>A 3366A>G), T239N (3371C>A 3372T>C), H241N (3376C>A 3378C>T), K242H (3379A>C 3381A>T), P245S (3388C>T 3390T>C), S248D (3397T>G 3398C>A 3399C>T), L249V (3400C>G 3402G>T), E250Q (3403G>C), N251I (3407A>T 3408C>T), E253D (3414G>T), G254N (3415G>A 3416G>A 3417A>C), L255Y (3418C>T 3419T>A), I257L (3424A>T 3426C>A), E260L (3433G>T 3434A>T), V262I (3439G>A 3441G>T), V263I (3442G>A 3444C>T), F270A (3463T>G 3464T>C 3465C>T), L271F (3466C>T), F273L (3472T>C 3474C>T) | | | | | | | | |
| Codon mutations: | ATG224AAG (3326T>A), ACG225ATC (3329C>T 3330G>C), CCC227CCA (3336C>A), ATG228TTA (3337A>T 3339G>A), AAG229ACT (3341A>C 3342G>T), GTC231GTG (3348C>G), CCA233CCT (3354A>T), ACA234AAA (3356C>A), ATC235ATG (3360C>G), ACT236ACC (3363T>C), CGA237AAG (3364C>A 3365G>A 3366A>G), ACT239AAC (3371C>A 3372T>C), CAC241AAT (3376C>A 3378C>T), AAA242CAT (3379A>C 3381A>T), TAC243TAT (3384C>T), CCT245TCC (3388C>T 3390T>C), TCG246TCT (3393G>T), GGC247GGT (3396C>T), TCC248GAT (3397T>G 3398C>A 3399C>T), CTG249GTT (3400C>G 3402G>T), GAA250CAA (3403G>C), AAC251ATT (3407A>T 3408C>T), TAC252TAT (3411C>T), GAG253GAT (3414G>T), GGA254AAC (3415G>A 3416G>A 3417A>C), CTC255TAC (3418C>T 3419T>A), TGC256TGT (3423C>T), ATC257TTA (3424A>T 3426C>A), CTC258CTG (3429C>G), CCG259CCA (3432G>A), GAG260TTG (3433G>T 3434A>T), GTG262ATT (3439G>A 3441G>T), GTC263ATT (3442G>A 3444C>T), AAC264AAT (3447C>T), GAG265GAA (3450G>A), ATC267ATT (3456C>T), TTC270GCT (3463T>G 3464T>C 3465C>T), CTC271TTC (3466C>T), TTC273CTT (3472T>C 3474C>T) | | | | | | | | |

*: Inserts / Deletes / Misaligned / Frameshifts

#### Analysis details

This analysis was performed with panviral2.64

### NGS Details (UN9): Badnavirus maculasmallanthi

#### Assembly

| | |
|---|---|
| **Coverage Length** | 293 (1 contig(s)) |
| **Depth Of Coverage** | 4.2 |
| **Number Of Reads** | 10 |
| **Reads Per Million** | 0.23 rpm (after QC) |
| **Ambiguities** | 0 |
| **Assembly Method** | de novo + reference guided assembly |
| **Consensus Caller** | Bcf Tools |

#### Coverage Map

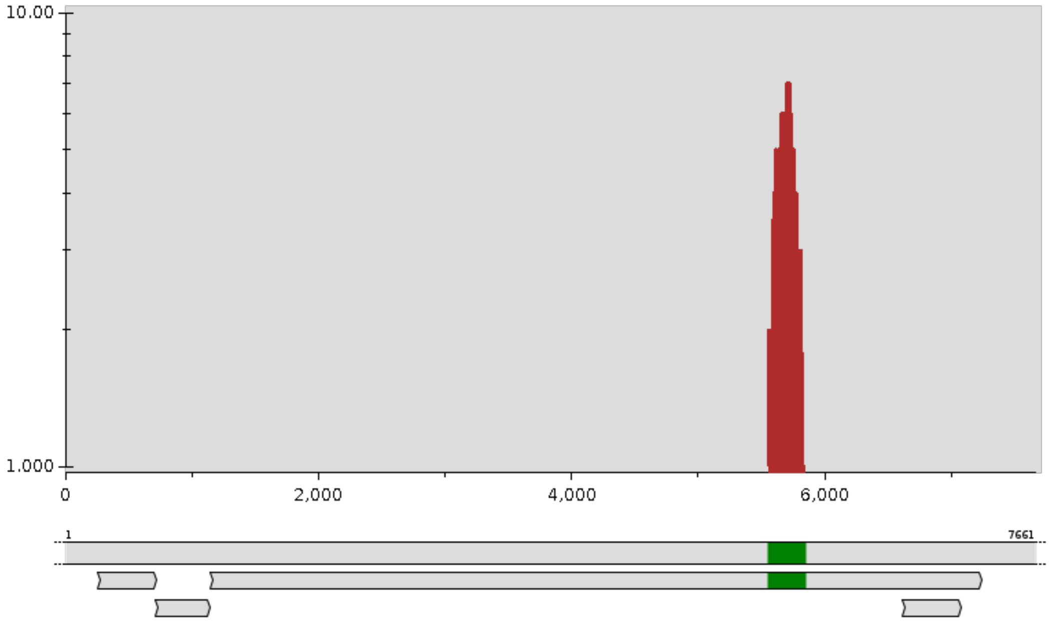

#### Assignment

| | |
|---|---|
| **Type** | Badnavirus maculasmallanthi (Taxonomy ID: 3048453) |
| **Reference Genome** | NC_026472.1 |
| **NT Identity (%)** | 52.9801 |
| **AA Identity (%)** | 48.5149 |
| **Number Of Stop Codons** | 0 |
| **Number Of CDS** | 4 |

#### Alignment

| | |
|---|---|
| **Alignment Score** | 27.0 (NT) + 343.0 (AA) = 370.0 |
| **Concordance (%)** | 28.8387 |

| Alignment Method | Global, seeded, nucleotide + amino acids (AGA) |
|---|---|

#### Genome Region

Sequence starts at position 5557 and ends at position 5849 relative to NC_026472.1 reference sequence.

#### Alignment Detailed Statistics

| | Begin | End | Coverage | Score | Concordance | Matches | Identities | I/D/M/F* | Stop Codons |
|---|---|---|---|---|---|---|---|---|---|
| **NT** | **5557** | **5849** | **3.8%** | **27** | **4.6%** | **293 (97.0%)** | **160 (53.0%)** | **9/0** | |
| Mutations: | 5558A>G, 5562A>G, 5565G>A, 5566A>T, 5567T>A, 5568A>C, 5571C>T, 5574A>T, 5578T>C, 5580T>G, 5583T>C, 5586G>T, 5588A>G, 5592T>G, 5595A>G, 5597T>A, 5605G>A, 5608G>A, 5609C>G, 5610C>A, 5614G>C, 5616A>T, 5620G>C, 5621A>C, 5622A>C, 5623T>G, 5624C>A, 5628A>T, 5632T>A, 5633G>A, 5634G>A, 5637G>T, 5644T>A, 5645T>G, 5648T>C, 5649C>T, 5651C>A, 5652A>C, 5653G>C, 5654G>A, 5655A>T, 5658A>T, 5659T>C, 5660T>A, 5661A>T, 5664C>T, 5669G>T, 5670G>C, 5671C>T, 5673T>G, 5682A>T, 5685C>T, 5688A>C, 5693A>C, 5697T>C, 5700A>T, 5701C>T, 5703C>A, 5704G>T, 5706T>G, 5708T>C, 5709T>C, 5712T>C, 5715G>A, 5718A>C, 5719A>C, 5720A>T, 5721A>C, 5725G>A, 5727T>C, 5730A>G, 5731T>A, 5732G>T, 5736C>T, 5737A>C, 5739A>C, 5739_5740insTCT, 5740G>T, 5743A>C, 5744C>T, 5745A>C, 5746G>A, 5747A>G, 5749G>A, 5754T>C, 5757C>T, 5758G>T, 5759C>T, 5760C>G, 5765A>T, 5766C>T, 5767A>T, 5772C>T, 5775C>T, 5781G>C, 5782G>A, 5784A>C, 5787C>T, 5788T>A, 5789C>G, 5790C>T, 5791A>T, 5792A>C, 5793C>T, 5794A>G, 5795G>A, 5797G>T, 5798A>G, 5799A>G, 5802G>A, 5804A>G, 5805T>G, 5809G>T, 5810A>T, 5812A>T, 5813G>C, 5814G>T, 5821A>G, 5823G>A, 5823_5824insGAGGCT, 5824A>T, 5826T>C, 5828T>A, 5830C>G, 5832G>A, 5833G>C, 5834G>T, 5835T>G, 5837A>G, 5844A>C, 5845G>A, 5846A>C, 5847A>T, 5848A>T, 5849A>T | | | | | | | | |

##### CDS

| | Begin | End | Coverage | Score | Concordance | Matches | Identities | I/D/M/F* | Stop Codons |
|---|---|---|---|---|---|---|---|---|---|
| UF61_gp3 | 1472 | 1569 | 4.8% | 343 | 48.7% | 98 (97.0%) | 49 (48.5%) | 3/0/0/0 | 0 |
| Protein mutations: | N1472S (5558A>G), I1475Y (5566A>T 5567T>A 5568A>C), F1479L (5578T>C 5580T>G), K1482R (5588A>G), F1485Y (5597T>A), V1488I (5605G>A), A1489R (5608G>A 5609C>G 5610C>A), E1491H (5614G>C 5616A>T), E1493P (5620G>C 5621A>C 5622A>C), S1494D (5623T>G 5624C>A), W1497K (5632T>A 5633G>A 5634G>A), L1501R (5644T>A 5645T>G), I1502T (5648T>C 5649C>T), P1503H (5651C>A 5652A>C), G1504H (5653G>C 5654G>A 5655A>T), L1506H (5659T>C 5660T>A 5661A>T), W1509F (5669G>T 5670G>C), K1517T (5693A>C), P1520S (5701C>T 5703C>A), A1521S (5704G>T 5706T>G), I1522T (5708T>C 5709T>C), R1525S (5718A>C), K1526L (5719A>C 5720A>T 5721A>C), D1528N (5725G>A 5727T>C), C1530I (5731T>A 5732G>T), K1532H (5737A>C 5739A>C), K1532_D1533insS (5739_5740insTCT), D1533Y (5740G>T), T1534L (5743A>C 5744C>T 5745A>C), E1535R (5746G>A 5747A>G), E1536K (5749G>A), A1539L (5758G>T 5759C>T 5760C>G), Y1541F (5765A>T 5766C>T), I1542F (5767A>T), V1547I (5782G>A 5784A>C), N1550S (5791A>T 5792A>C 5793C>T), S1551D (5794A>G 5795G>A), E1552W (5797G>T 5798A>G 5799A>G), D1554G (5804A>G 5805T>G), E1556L (5809G>T 5810A>T), R1557S (5812A>T 5813G>C 5814G>T), K1560E (5821A>G 5823G>A), K1560_I1561insEA (5823_5824insGAGGCT), I1561F (5824A>T 5826T>C), M1562K (5828T>A), L1563V (5830C>G 5832G>A), G1564L (5833G>C 5834G>T 5835T>G), K1565R (5837A>G), Q1567H (5844A>C), E1568T (5845G>A 5846A>C 5847A>T) | | | | | | | | |
| Codon mutations: | AAT1472AGT (5558A>G), GCA1473GCG (5562A>G), AAG1474AAA (5565G>A), ATA1475TAC (5566A>T 5567T>A 5568A>C), TTC1476TTT (5571C>T), TCA1477TCT (5574A>T), TTT1479CTG (5578T>C 5580T>G), GAT1480GAC (5583T>C), CTG1481CTT (5586G>T), AAG1482AGG (5588A>G), TCT1483TCG (5592T>G), GGA1484GGG (5595A>G), TTT1485TAT (5597T>A), GTT1488ATT (5605G>A), GCC1489AGA (5608G>A 5609C>G 5610C>A), GAA1491CAT (5614G>C 5616A>T), GAA1493CCC (5620G>C 5621A>C 5622A>C), TCC1494GAC (5623T>G 5624C>A), ATA1495ATT (5628A>T), TGG1497AAA (5632T>A 5633G>A 5634G>A), ACG1498ACT (5637G>T), TTA1501AGA (5644T>A 5645T>G), ATC1502ACT (5648T>C 5649C>T), CCA1503CAC (5651C>A 5652A>C), GGA1504CAT (5653G>C 5654G>A 5655A>T), GGA1505GGT (5658A>T), TTA1506CAT (5659T>C 5660T>A 5661A>T), TAC1507TAT (5664C>T), TGG1509TTC (5669G>T 5670G>C), CTT1510TTG (5671C>T 5673T>G), CCA1513CCT (5682A>T), TTC1514TTT (5685C>T), GGA1515GGC (5688A>C), AAA1517ACA (5693A>C), AAT1518AAC (5697T>C), GCA1519GCT (5700A>T), CCC1520TCA (5701C>T 5703C>A), GCT1521TCG (5704G>T 5706T>G), ATT1522ACC (5708T>C 5709T>C), TTT1523TTC (5712T>C), CAG1524CAA (5715G>A), AGA1525AGC (5718A>C), AAA1526CTC (5719A>C 5720A>T 5721A>C), GAT1528AAC (5725G>A 5727T>C), AAA1529AAG (5730A>G), TGC1530ATC (5731T>A 5732G>T), TTC1531TTT (5736C>T), AAA1532CAC (5737A>C 5739A>C), AAA1532_GAC1533insTCT (5739_5740insTCT), GAC1533TAC (5740G>T), ACA1534CTC (5743A>C 5744C>T 5745A>C), GAA1535AGA (5746G>A 5747A>G), GAG1536AAG (5749G>A), TTT1537TTC (5754T>C), ATC1538ATT (5757C>T), GCC1539TTG (5758G>T 5759C>T 5760C>G), TAC1541TTT (5765A>T 5766C>T), ATC1542TTC (5767A>T), GAC1543GAT (5772C>T), GAC1544GAT (5775C>T), CTG1546CTC (5781G>C), GTA1547ATC (5782G>A 5784A>C), TAC1548TAT (5787C>T), TCC1549AGT (5788T>A 5789C>G 5790C>T), AAC1550TCT (5791A>T 5792A>C 5793C>T), AGT1551GAT (5794A>G 5795G>A), GAA1552TGG (5797G>T 5798A>G 5799A>G), GCG1553GCA (5802G>A), GAT1554GGG (5804A>G 5805T>G), GAA1556TTA (5809G>T 5810A>T), AGG1557TCT (5812A>T 5813G>C 5814G>T), AAG1560GAA (5821A>G 5823G>A), AAG1560_ATT1561insGAGGCT (5823_5824insGAGGCT), ATT1561TTC (5824A>T 5826T>C), ATG1562AAG (5828T>A), CTG1563GTA (5830C>G 5832G>A), GGT1564CTG (5833G>C 5834G>T 5835T>G), AAA1565AGA (5837A>G), CAA1567CAC (5844A>C), GAA1568ACT (5845G>A 5846A>C 5847A>T), AAT1569TT. (5848A>T 5849A>T) | | | | | | | | |

##### Proteins

| | Begin | End | Coverage | Score | Concordance | Matches | Identities | I/D/M/F* | Stop Codons |
|---|---|---|---|---|---|---|---|---|---|
| ORF3 (YP_009121747.1) | 1472 | 1569 | 4.8% | 343 | 48.7% | 98 (97.0%) | 49 (48.5%) | 3/0/0/0 | 0 |
| Protein mutations: | N1472S (5558A>G), I1475Y (5566A>T 5567T>A 5568A>C), F1479L (5578T>C 5580T>G), K1482R (5588A>G), F1485Y (5597T>A), V1488I (5605G>A), A1489R (5608G>A 5609C>G 5610C>A), E1491H (5614G>C 5616A>T), E1493P (5620G>C 5621A>C 5622A>C), S1494D (5623T>G 5624C>A), W1497K (5632T>A 5633G>A 5634G>A), L1501R (5644T>A 5645T>G), I1502T (5648T>C 5649C>T), P1503H (5651C>A 5652A>C), G1504H (5653G>C 5654G>A 5655A>T), L1506H (5659T>C 5660T>A 5661A>T), W1509F (5669G>T 5670G>C), K1517T (5693A>C), P1520S (5701C>T 5703C>A), A1521S (5704G>T 5706T>G), I1522T (5708T>C 5709T>C), R1525S (5718A>C), K1526L (5719A>C 5720A>T 5721A>C), D1528N (5725G>A 5727T>C), C1530I (5731T>A 5732G>T), K1532H (5737A>C 5739A>C), K1532_D1533insS (5739_5740insTCT), D1533Y (5740G>T), T1534L (5743A>C 5744C>T 5745A>C), E1535R (5746G>A 5747A>G), E1536K (5749G>A), A1539L (5758G>T 5759C>T 5760C>G), Y1541F (5765A>T 5766C>T), I1542F (5767A>T), V1547I (5782G>A 5784A>C), N1550S (5791A>T 5792A>C 5793C>T), S1551D (5794A>G 5795G>A), E1552W (5797G>T 5798A>G 5799A>G), D1554G (5804A>G 5805T>G), E1556L (5809G>T 5810A>T), R1557S (5812A>T 5813G>C 5814G>T), K1560E (5821A>G 5823G>A), K1560_I1561insEA (5823_5824insGAGGCT), I1561F (5824A>T 5826T>C), M1562K (5828T>A), L1563V (5830C>G 5832G>A), G1564L (5833G>C 5834G>T 5835T>G), K1565R (5837A>G), Q1567H (5844A>C), E1568T (5845G>A 5846A>C 5847A>T) | | | | | | | | |
| Codon mutations: | AAT1472AGT (5558A>G), GCA1473GCG (5562A>G), AAG1474AAA (5565G>A), ATA1475TAC (5566A>T 5567T>A 5568A>C), TTC1476TTT (5571C>T), TCA1477TCT (5574A>T), TTT1479CTG (5578T>C 5580T>G), GAT1480GAC (5583T>C), CTG1481CTT (5586G>T), AAG1482AGG (5588A>G), TCT1483TCG (5592T>G), GGA1484GGG (5595A>G), TTT1485TAT (5597T>A), GTT1488ATT (5605G>A), GCC1489AGA (5608G>A 5609C>G 5610C>A), GAA1491CAT (5614G>C 5616A>T), GAA1493CCC (5620G>C 5621A>C 5622A>C), TCC1494GAC (5623T>G 5624C>A), ATA1495ATT (5628A>T), TGG1497AAA (5632T>A 5633G>A 5634G>A), ACG1498ACT (5637G>T), TTA1501AGA (5644T>A 5645T>G), ATC1502ACT (5648T>C 5649C>T), CCA1503CAC (5651C>A 5652A>C), GGA1504CAT (5653G>C 5654G>A 5655A>T), GGA1505GGT (5658A>T), TTA1506CAT (5659T>C 5660T>A 5661A>T), TAC1507TAT (5664C>T), TGG1509TTC (5669G>T 5670G>C), CTT1510TTG (5671C>T 5673T>G), CCA1513CCT (5682A>T), TTC1514TTT (5685C>T), GGA1515GGC (5688A>C), AAA1517ACA (5693A>C), AAT1518AAC (5697T>C), GCA1519GCT (5700A>T), CCC1520TCA (5701C>T 5703C>A), GCT1521TCG (5704G>T 5706T>G), ATT1522ACC (5708T>C 5709T>C), TTT1523TTC (5712T>C), CAG1524CAA (5715G>A), AGA1525AGC (5718A>C), AAA1526CTC (5719A>C 5720A>T 5721A>C), GAT1528AAC (5725G>A 5727T>C), AAA1529AAG (5730A>G), TGC1530ATC (5731T>A 5732G>T), TTC1531TTT (5736C>T), AAA1532CAC (5737A>C 5739A>C), AAA1532_GAC1533insTCT (5739_5740insTCT), GAC1533TAC (5740G>T), ACA1534CTC (5743A>C 5744C>T 5745A>C), GAA1535AGA (5746G>A 5747A>G), GAG1536AAG (5749G>A), TTT1537TTC (5754T>C), ATC1538ATT (5757C>T), GCC1539TTG (5758G>T 5759C>T 5760C>G), TAC1541TTT (5765A>T 5766C>T), ATC1542TTC (5767A>T), GAC1543GAT (5772C>T), GAC1544GAT (5775C>T), CTG1546CTC (5781G>C), GTA1547ATC (5782G>A 5784A>C), TAC1548TAT (5787C>T), TCC1549AGT (5788T>A 5789C>G 5790C>T), AAC1550TCT (5791A>T 5792A>C 5793C>T), AGT1551GAT (5794A>G 5795G>A), GAA1552TGG (5797G>T 5798A>G 5799A>G), GCG1553GCA (5802G>A), GAT1554GGG (5804A>G 5805T>G), GAA1556TTA (5809G>T 5810A>T), AGG1557TCT (5812A>T 5813G>C 5814G>T), AAG1560GAA (5821A>G 5823G>A), AAG1560_ATT1561insGAGGCT (5823_5824insGAGGCT), ATT1561TTC (5824A>T 5826T>C), ATG1562AAG (5828T>A), CTG1563GTA (5830C>G 5832G>A), GGT1564CTG (5833G>C 5834G>T 5835T>G), AAA1565AGA (5837A>G), CAA1567CAC (5844A>C), GAA1568ACT (5845G>A 5846A>C 5847A>T), AAT1569TT. (5848A>T 5849A>T) | | | | | | | | |

*: Inserts / Deletes / Misaligned / Frameshifts

#### Analysis details

This analysis was performed with panviral2.64

### NGS Details (UN9): Pinus nigra virus 1

#### Assembly

| | |
|---|---|
| **Coverage Length** | 239 (1 contig(s)) |
| **Depth Of Coverage** | 6.0 |
| **Number Of Reads** | 10 |
| **Reads Per Million** | 0.23 rpm (after QC) |
| **Ambiguities** | 0 |
| **Assembly Method** | de novo + reference guided assembly |
| **Consensus Caller** | Bcf Tools |

#### Coverage Map

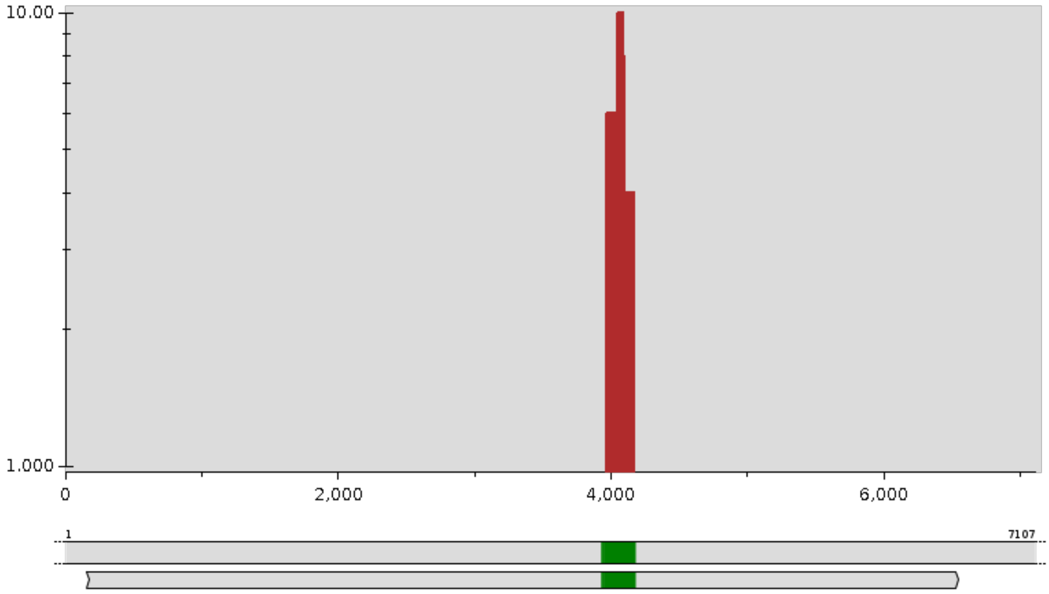

#### Assignment

| | |
|---|---|
| **Type** | Pinus nigra virus 1 (Taxonomy ID: 2267679) |
| **Reference Genome** | NC_040841.1 |
| **NT Identity (%)** | 57.7406 |
| **AA Identity (%)** | 47.4359 |
| **Number Of Stop Codons** | 0 |
| **Number Of CDS** | 1 |

#### Alignment

| | |
|---|---|
| **Alignment Score** | 74.0 (NT) + 260.0 (AA) = 334.0 |
| **Concordance (%)** | 31.8702 |

| Alignment Method | Global, seeded, nucleotide + amino acids (AGA) |
|---|---|

#### Genome Region

Sequence starts at position 3935 and ends at position 4180 relative to NC_040841.1 reference sequence.

#### Alignment Detailed Statistics

| | Begin | End | Coverage | Score | Concordance | Matches | Identities | I/D/M/F* | Stop Codons |
|---|---|---|---|---|---|---|---|---|---|
| **NT** | **3935** | **4180** | **3.4%** | **74** | **15.5%** | **239 (100%)** | **138 (57.7%)** | **0/0** | |
| Mutations: | 3935C>T, 3937T>A, 3941G>T, 3943C>T, 3944A>G, 3946C>T, 3947A>T, 3948C>T, 3949T>A, 3950C>A, 3952G>A, 3953A>T, 3954A>G, 3955T>G, 3956T>A, 3960A>G, 3961T>A, 3967A>G, 3968T>A, 3970G>T, 3973A>T, 3975G>A, 3976A>T, 3977C>A, 3978C>A, 3980G>A, 3981A>T, 3982A>G, 3984T>A, 3985G>T, 3986A>C, 3989A>T, 3992C>A, 3996A>G, 3999T>C, 4001C>T, 4002A>T, 4003G>T, 4004C>A, 4006T>A, 4008G>C, 4009T>C, 4012G>T, 4013G>A, 4015G>T, 4017T>A, 4021C>T, 4024G>A, 4030C>T, 4036G>A, 4042G>T, 4044A>T, 4045T>C, 4047A>G, 4048T>G, 4051G>A, 4054A>T, 4061C>T, 4062A>C, 4064C>A, 4065C>A, 4067G>A, 4074G>A, 4076C>T, 4078C>T, 4084A>T, 4092T>A, 4093A>C, 4094T>G, 4095G>T, 4096T>C, 4099A>C, 4100G>T, 4101C>T, 4102A>C, 4106T>C, 4107T>A, 4108C>A, 4112C>G, 4114G>A, 4120A>T, 4124G>A, 4126T>G, 4129A>C, 4135T>A, 4138G>A, 4141G>A, 4147G>C, 4150A>T, 4151G>T, 4153C>T, 4154T>G, 4155T>A, 4156C>A, 4164G>A, 4165A>C, 4166A>G, 4167G>T, 4168G>T, 4172G>A, 4176A>C | | | | | | | | |

##### CDS

| | Begin | End | Coverage | Score | Concordance | Matches | Identities | I/D/M/F* | Stop Codons |
|---|---|---|---|---|---|---|---|---|---|
| EXL67_gp1 | 1265 | 1342 | 3.7% | 260 | 42.5% | 78 (100%) | 37 (47.4%) | 0/0/1/0 | 0 |
| Protein mutations: | T1265L (3947A>T 3948C>T 3949T>A), N1267W (3953A>T 3954A>G 3955T>G), F1268I (3956T>A), N1269R (3960A>G 3961T>A), L1272I (3968T>A 3970G>T), R1274N (3975G>A 3976A>T), P1275K (3977C>A 3978C>A), E1276M (3980G>A 3981A>T 3982A>G), V1277D (3984T>A 3985G>T), I1278L (3986A>C), M1279L (3989A>T), Q1280K (3992C>A), K1281R (3996A>G), I1282T (3999T>C), Q1283F (4001C>T 4002A>T 4003G>T), H1284K (4004C>A 4006T>A), S1285T (4008G>C 4009T>C), K1286N (4012G>T), V1287I (4013G>A 4015G>T), F1288Y (4017T>A), Y1297F (4044A>T 4045T>C), Y1298W (4047A>G 4048T>G), Q1303S (4061C>T 4062A>C), P1304K (4064C>A 4065C>A), E1305K (4067G>A), R1307K (4074G>A), H1308Y (4076C>T 4078C>T), I1313N (4092T>A 4093A>C), C1314V (4094T>G 4095G>T 4096T>C), A1316F (4100G>T 4101C>T 4102A>C), F1318Q (4106T>C 4107T>A 4108C>A), Q1320E (4112C>G 4114G>A), K1322N (4120A>T), V1324M (4124G>A 4126T>G), A1333S (4151G>T 4153C>T), F1334E (4154T>G 4155T>A 4156C>A), R1337N (4164G>A 4165A>C), R1338V (4166A>G 4167G>T 4168G>T), D1340N (4172G>A), Y1341S (4176A>C) | | | | | | | | |
| Codon mutations: | CTT1261T.A (3935C>T 3937T>A), GCC1263T.T (3941G>T 3943C>T), ATC1264G.T (3944A>G 3946C>T), ACT1265TTA (3947A>T 3948C>T 3949T>A), CAG1266A.A (3950C>A 3952G>A), AAT1267TGG (3953A>T 3954A>G 3955T>G), TTT1268ATT (3956T>A), AAT1269AGA (3960A>G 3961T>A), CCA1271CCG (3967A>G), TTG1272ATT (3968T>A 3970G>T), CCA1273CCT (3973A>T), AGA1274AAT (3975G>A 3976A>T), CCA1275AAA (3977C>A 3978C>A), GAA1276ATG (3980G>A 3981A>T 3982A>G), GTG1277GAT (3984T>A 3985G>T), ATT1278CTT (3986A>C), ATG1279TTG (3989A>T), CAA1280AAA (3992C>A), AAA1281AGA (3996A>G), ATT1282ACT (3999T>C), CAG1283TTT (4001C>T 4002A>T 4003G>T), CAT1284AAA (4004C>A 4006T>A), AGT1285ACC (4008G>C 4009T>C), AAG1286AAT (4012G>T), GTG1287ATT (4013G>A 4015G>T), TTC1288TAC (4017T>A), AGC1289AGT (4021C>T), AAG1290AAA (4024G>A), GAC1292GAT (4030C>T), AAG1294AAA (4036G>A), GGG1296GGT (4042G>T), TAT1297TTC (4044A>T 4045T>C), TAT1298TGG (4047A>G 4048T>G), CAG1299CAA (4051G>A), ATA1300ATT (4054A>T), CAG1303TCG (4061C>T 4062A>C), CCA1304AAA (4064C>A 4065C>A), GAA1305AAA (4067G>A), AGA1307AAA (4074G>A), CAC1308TAT (4076C>T 4078C>T), ACA1310ACT (4084A>T), ATA1313AAC (4092T>A 4093A>C), TGT1314GTC (4094T>G 4095G>T 4096T>C), CCA1315CCC (4099A>C), GCA1316TTC (4100G>T 4101C>T 4102A>C), TTC1318CAA (4106T>C 4107T>A 4108C>A), CAG1320GAA (4112C>G 4114G>A), AAA1322AAT (4120A>T), GTT1324ATG (4124G>A 4126T>G), CCA1325CCC (4129A>C), GGT1327GGA (4135T>A), TTG1328TTA (4138G>A), AAG1329AAA (4141G>A), GCG1331GCC (4147G>C), CCA1332CCT (4150A>T), GCC1333TCT (4151G>T 4153C>T), TTC1334GAA (4154T>G 4155T>A 4156C>A), AGA1337AAC (4164G>A 4165A>C), AGG1338GTT (4166A>G 4167G>T 4168G>T), GAT1340AAT (4172G>A), TAT1341TCT (4176A>C) | | | | | | | | |

##### Proteins

| | Begin | End | Coverage | Score | Concordance | Matches | Identities | I/D/M/F* | Stop Codons |
|---|---|---|---|---|---|---|---|---|---|
| polyprotein (YP_009553669.1) | 1265 | 1342 | 3.7% | 260 | 42.5% | 78 (100%) | 37 (47.4%) | 0/0/1/0 | 0 |
| Protein mutations: | T1265L (3947A>T 3948C>T 3949T>A), N1267W (3953A>T 3954A>G 3955T>G), F1268I (3956T>A), N1269R (3960A>G 3961T>A), L1272I (3968T>A 3970G>T), R1274N (3975G>A 3976A>T), P1275K (3977C>A 3978C>A), E1276M (3980G>A 3981A>T 3982A>G), V1277D (3984T>A 3985G>T), I1278L (3986A>C), M1279L (3989A>T), Q1280K (3992C>A), K1281R (3996A>G), I1282T (3999T>C), Q1283F (4001C>T 4002A>T 4003G>T), H1284K (4004C>A 4006T>A), S1285T (4008G>C 4009T>C), K1286N (4012G>T), V1287I (4013G>A 4015G>T), F1288Y (4017T>A), Y1297F (4044A>T 4045T>C), Y1298W (4047A>G 4048T>G), Q1303S (4061C>T 4062A>C), P1304K (4064C>A 4065C>A), E1305K (4067G>A), R1307K (4074G>A), H1308Y (4076C>T 4078C>T), I1313N (4092T>A 4093A>C), C1314V (4094T>G 4095G>T 4096T>C), A1316F (4100G>T 4101C>T 4102A>C), F1318Q (4106T>C 4107T>A 4108C>A), Q1320E (4112C>G 4114G>A), K1322N (4120A>T), V1324M (4124G>A 4126T>G), A1333S (4151G>T 4153C>T), F1334E (4154T>G 4155T>A 4156C>A), R1337N (4164G>A 4165A>C), R1338V (4166A>G 4167G>T 4168G>T), D1340N (4172G>A), Y1341S (4176A>C) | | | | | | | | |
| Codon mutations: | CTT1261T.A (3935C>T 3937T>A), GCC1263T.T (3941G>T 3943C>T), ATC1264G.T (3944A>G 3946C>T), ACT1265TTA (3947A>T 3948C>T 3949T>A), CAG1266A.A (3950C>A 3952G>A), AAT1267TGG (3953A>T 3954A>G 3955T>G), TTT1268ATT (3956T>A), AAT1269AGA (3960A>G 3961T>A), CCA1271CCG (3967A>G), TTG1272ATT (3968T>A 3970G>T), CCA1273CCT (3973A>T), AGA1274AAT (3975G>A 3976A>T), CCA1275AAA (3977C>A 3978C>A), GAA1276ATG (3980G>A 3981A>T 3982A>G), GTG1277GAT (3984T>A 3985G>T), ATT1278CTT (3986A>C), ATG1279TTG (3989A>T), CAA1280AAA (3992C>A), AAA1281AGA (3996A>G), ATT1282ACT (3999T>C), CAG1283TTT (4001C>T 4002A>T 4003G>T), CAT1284AAA (4004C>A 4006T>A), AGT1285ACC (4008G>C 4009T>C), AAG1286AAT (4012G>T), GTG1287ATT (4013G>A 4015G>T), TTC1288TAC (4017T>A), AGC1289AGT (4021C>T), AAG1290AAA (4024G>A), GAC1292GAT (4030C>T), AAG1294AAA (4036G>A), GGG1296GGT (4042G>T), TAT1297TTC (4044A>T 4045T>C), TAT1298TGG (4047A>G 4048T>G), CAG1299CAA (4051G>A), ATA1300ATT (4054A>T), CAG1303TCG (4061C>T 4062A>C), CCA1304AAA (4064C>A 4065C>A), GAA1305AAA (4067G>A), AGA1307AAA (4074G>A), CAC1308TAT (4076C>T 4078C>T), ACA1310ACT (4084A>T), ATA1313AAC (4092T>A 4093A>C), TGT1314GTC (4094T>G 4095G>T 4096T>C), CCA1315CCC (4099A>C), GCA1316TTC (4100G>T 4101C>T 4102A>C), TTC1318CAA (4106T>C 4107T>A 4108C>A), CAG1320GAA (4112C>G 4114G>A), AAA1322AAT (4120A>T), GTT1324ATG (4124G>A 4126T>G), CCA1325CCC (4129A>C), GGT1327GGA (4135T>A), TTG1328TTA (4138G>A), AAG1329AAA (4141G>A), GCG1331GCC (4147G>C), CCA1332CCT (4150A>T), GCC1333TCT (4151G>T 4153C>T), TTC1334GAA (4154T>G 4155T>A 4156C>A), AGA1337AAC (4164G>A 4165A>C), AGG1338GTT (4166A>G 4167G>T 4168G>T), GAT1340AAT (4172G>A), TAT1341TCT (4176A>C) | | | | | | | | |

*: Inserts / Deletes / Misaligned / Frameshifts

#### Analysis details

This analysis was performed with panviral2.64

### NGS Details (UN9): Badnavirus occultipomeae

#### Assembly

| | |
|---|---|
| **Coverage Length** | 251 (1 contig(s)) |
| **Depth Of Coverage** | 4.2 |
| **Number Of Reads** | 9 |
| **Reads Per Million** | 0.20 rpm (after QC) |
| **Ambiguities** | 0 |
| **Assembly Method** | de novo + reference guided assembly |
| **Consensus Caller** | Bcf Tools |

#### Coverage Map

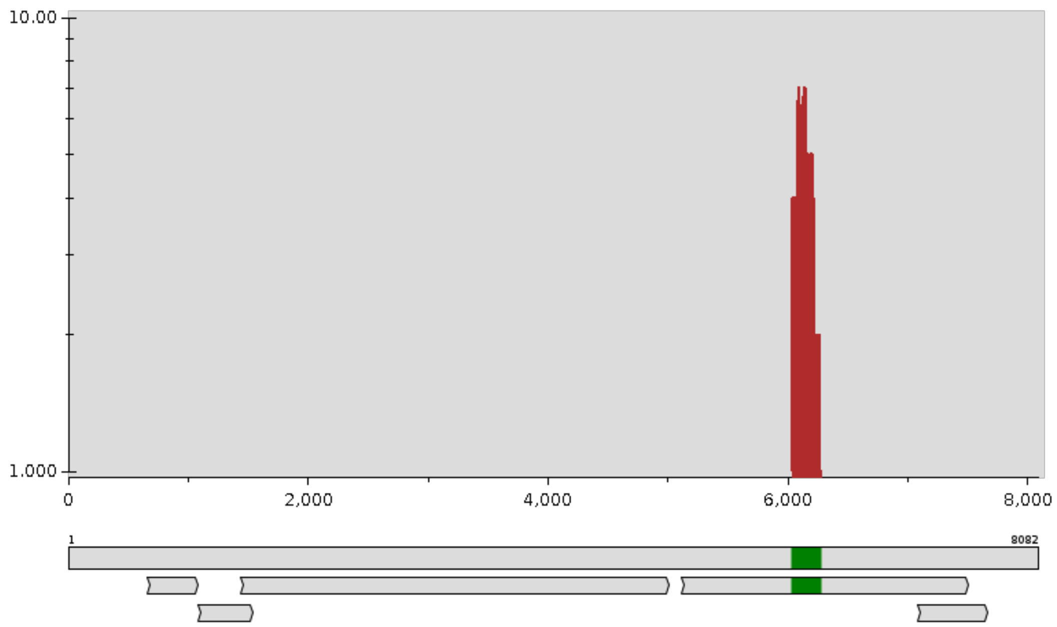

#### Assignment

| | |
|---|---|
| **Type** | Badnavirus occultipomeae (Taxonomy ID: 3048353) |
| **Reference Genome** | NC_015655.1 |
| **NT Identity (%)** | 58.6614 |
| **AA Identity (%)** | 51.7647 |
| **Number Of Stop Codons** | 0 |
| **Number Of CDS** | 5 |

#### Alignment

| | |
|---|---|
| **Alignment Score** | 82.0 (NT) + 327.0 (AA) = 409.0 |
| **Concordance (%)** | 36.8468 |

| **Alignment Method** | Global, seeded, nucleotide + amino acids (AGA) |
|---|---|

#### Genome Region

Sequence starts at position 6031 and ends at position 6281 relative to NC_015655.1 reference sequence.

#### Alignment Detailed Statistics

| | Begin | End | Coverage | Score | Concordance | Matches | Identities | I/D/M/F* | Stop Codons |
|---|---|---|---|---|---|---|---|---|---|
| **NT** | **6031** | **6281** | **3.1%** | **82** | **16.3%** | **251 (98.8%)** | **149 (58.7%)** | **3/0** | |
| Mutations: | 6033A>T, 6039A>T, 6042A>G, 6043T>A, 6049C>T, 6051C>T, 6052A>T, 6053A>G, 6054A>G, 6055G>T, 6056C>T, 6057A>C, 6060C>G, 6062T>A, 6069G>A, 6070A>G, 6072A>G, 6075A>G, 6076A>G, 6078G>A, 6079G>A, 6080A>G, 6085A>T, 6086A>G, 6087G>T, 6088T>G, 6089C>A, 6090C>T, 6092A>T, 6093A>T, 6096A>C, 6097T>A, 6098G>A, 6099G>A, 6102T>G, 6108T>C, 6115C>A, 6116C>G, 6117A>G, 6118G>T, 6120A>T, 6125T>A, 6130G>A, 6137A>T, 6138A>T, 6141C>G, 6145C>T, 6147C>T, 6154C>T, 6158T>C, 6159G>C, 6168A>G, 6171A>C, 6172G>A, 6173A>T, 6174T>C, 6177T>C, 6178C>A, 6179A>T, 6181A>G, 6182G>A, 6183A>C, 6184A>C, 6185A>T, 6186G>T, 6190G>A, 6194A>G, 6195T>G, 6197C>T, 6198A>C, 6201T>C, 6203G>A, 6204G>A, 6205G>C, 6206G>C, 6207A>C, 6207_6208insTAC, 6208A>C, 6209C>T, 6210A>T, 6213T>C, 6214G>T, 6216A>T, 6220A>G, 6222C>T, 6224C>T, 6225A>G, 6228A>G, 6230A>T, 6231T>C, 6234T>C, 6243A>T, 6247G>A, 6249A>C, 6251T>A, 6256G>C, 6258A>T, 6260A>G, 6264A>G, 6267A>G, 6275A>T, 6278A>G, 6279T>C | | | | | | | | |

##### CDS

| | Begin | End | Coverage | Score | Concordance | Matches | Identities | I/D/M/F* | Stop Codons |
|---|---|---|---|---|---|---|---|---|---|
| SPBVa_gp4 | 306 | 389 | 10.6% | 327 | 52.5% | 84 (98.8%) | 44 (51.8%) | 1/0/0/0 | 0 |
| Protein mutations: | F310I (6043T>A), L312F (6049C>T 6051C>T), K313W (6052A>T 6053A>G 6054A>G), A314F (6055G>T 6056C>T 6057A>C), F316Y (6062T>A), I319V (6070A>G 6072A>G), M321V (6076A>G 6078G>A), E322R (6079G>A 6080A>G), K324C (6085A>T 6086A>G 6087G>T), S325D (6088T>G 6089C>A 6090C>T), K326I (6092A>T 6093A>T), W328K (6097T>A 6098G>A 6099G>A), P334R (6115C>A 6116C>G 6117A>G), E335Y (6118G>T 6120A>T), L337H (6125T>A), E339K (6130G>A), E341V (6137A>T 6138A>T), P344S (6145C>T 6147C>T), M348T (6158T>C 6159G>C), D353I (6172G>A 6173A>T 6174T>C), Q355M (6178C>A 6179A>T), R356D (6181A>G 6182G>A 6183A>C), K357L (6184A>C 6185A>T 6186G>T), D359N (6190G>A), N360R (6194A>G 6195T>G), A361V (6197C>T 6198A>C), R363K (6203G>A 6204G>A), G364P (6205G>C 6206G>C 6207A>C), G364_T365insY (6207_6208insTAC), T365L (6208A>C 6209C>T 6210A>T), A367S (6214G>T 6216A>T), I369V (6220A>G 6222C>T), A370V (6224C>T 6225A>G), Y372F (6230A>T 6231T>C), V378I (6247G>A 6249A>C), F379Y (6251T>A), E381H (6256G>C 6258A>T), N382S (6260A>G), E387V (6275A>T), D388G (6278A>G 6279T>C) | | | | | | | | |
| Codon mutations: | ATA306ATT (6033A>T), TCA308TCT (6039A>T), AAA309AAG (6042A>G), TTT310ATT (6043T>A), CTC312TTT (6049C>T 6051C>T), AAA313TGG (6052A>T 6053A>G 6054A>G), GCA314TTC (6055G>T 6056C>T 6057A>C), GGC315GGG (6060C>G), TTT316TAT (6062T>A), CAG318CAA (6069G>A), ATA319GTG (6070A>G 6072A>G), AGA320AGG (6075A>G), ATG321GTA (6076A>G 6078G>A), GAG322AGG (6079G>A 6080A>G), AAG324TGT (6085A>T 6086A>G 6087G>T), TCC325GAT (6088T>G 6089C>A 6090C>T), AAA326ATT (6092A>T 6093A>T), CCA327CCC (6096A>C), TGG328AAA (6097T>A 6098G>A 6099G>A), ACT329ACG (6102T>G), TTT331TTC (6108T>C), CCA334AGG (6115C>A 6116C>G 6117A>G), GAA335TAT (6118G>T 6120A>T), CTC337CAC (6125T>A), GAA339AAA (6130G>A), GAA341GTT (6137A>T 6138A>T), GTC342GTG (6141C>G), CCC344TCT (6145C>T 6147C>T), CTG347TTG (6154C>T), ATG348ACC (6158T>C 6159G>C), CCA351CCG (6168A>G), GCA352GCC (6171A>C), GAT353ATC (6172G>A 6173A>T 6174T>C), TTT354TTC (6177T>C), CAG355ATG (6178C>A 6179A>T), AGA356GAC (6181A>G 6182G>A 6183A>C), AAG357CTT (6184A>C 6185A>T 6186G>T), GAT359AAT (6190G>A), AAT360AGG (6194A>G 6195T>G), GCA361GTC (6197C>T 6198A>C), TTT362TTC (6201T>C), AGG363AAA (6203G>A 6204G>A), GGA364CCC (6205G>C 6206G>C 6207A>C), GGA364_ACA365insTAC (6207_6208insTAC), ACA365CTT (6208A>C 6209C>T 6210A>T), GAT366GAC (6213T>C), GCA367TCT (6214G>T 6216A>T), ATC369GTT (6220A>G 6222C>T), GCA370GTG (6224C>T 6225A>G), GTA371GTG (6228A>G), TAT372TTC (6230A>T 6231T>C), ATT373ATC (6234T>C), ATA376ATT (6243A>T), GTA378ATC (6247G>A 6249A>C), TTC379TAC (6251T>A), GAA381CAT (6256G>C 6258A>T), AAT382AGT (6260A>G), GAA383GAG (6264A>G), GAA384GAG (6267A>G), GAG387GTG (6275A>T), GAT388GGC (6278A>G 6279T>C) | | | | | | | | |

##### Proteins

| | Begin | End | Coverage | Score | Concordance | Matches | Identities | I/D/M/F* | Stop Codons |
|---|---|---|---|---|---|---|---|---|---|
| RNaseH/reverse transcriptase (YP_004581513.1) | 306 | 389 | 10.6% | 327 | 52.5% | 84 (98.8%) | 44 (51.8%) | 1/0/0/0 | 0 |
| Protein mutations: | F310I (6043T>A), L312F (6049C>T 6051C>T), K313W (6052A>T 6053A>G 6054A>G), A314F (6055G>T 6056C>T 6057A>C), F316Y (6062T>A), I319V (6070A>G 6072A>G), M321V (6076A>G 6078G>A), E322R (6079G>A 6080A>G), K324C (6085A>T 6086A>G 6087G>T), S325D (6088T>G 6089C>A 6090C>T), K326I (6092A>T 6093A>T), W328K (6097T>A 6098G>A 6099G>A), P334R (6115C>A 6116C>G 6117A>G), E335Y (6118G>T 6120A>T), L337H (6125T>A), E339K (6130G>A), E341V (6137A>T 6138A>T), P344S (6145C>T 6147C>T), M348T (6158T>C 6159G>C), D353I (6172G>A 6173A>T 6174T>C), Q355M (6178C>A 6179A>T), R356D (6181A>G 6182G>A 6183A>C), K357L (6184A>C 6185A>T 6186G>T), D359N (6190G>A), N360R (6194A>G 6195T>G), A361V (6197C>T 6198A>C), R363K (6203G>A 6204G>A), G364P (6205G>C 6206G>C 6207A>C), G364_T365insY (6207_6208insTAC), T365L (6208A>C 6209C>T 6210A>T), A367S (6214G>T 6216A>T), I369V (6220A>G 6222C>T), A370V (6224C>T 6225A>G), Y372F (6230A>T 6231T>C), V378I (6247G>A 6249A>C), F379Y (6251T>A), E381H (6256G>C 6258A>T), N382S (6260A>G), E387V (6275A>T), D388G (6278A>G 6279T>C) | | | | | | | | |
| Codon mutations: | ATA306ATT (6033A>T), TCA308TCT (6039A>T), AAA309AAG (6042A>G), TTT310ATT (6043T>A), CTC312TTT (6049C>T 6051C>T), AAA313TGG (6052A>T 6053A>G 6054A>G), GCA314TTC (6055G>T 6056C>T 6057A>C), GGC315GGG (6060C>G), TTT316TAT (6062T>A), CAG318CAA (6069G>A), ATA319GTG (6070A>G 6072A>G), AGA320AGG (6075A>G), ATG321GTA (6076A>G 6078G>A), GAG322AGG (6079G>A 6080A>G), AAG324TGT (6085A>T 6086A>G 6087G>T), TCC325GAT (6088T>G 6089C>A 6090C>T), AAA326ATT (6092A>T 6093A>T), CCA327CCC (6096A>C), TGG328AAA (6097T>A 6098G>A 6099G>A), ACT329ACG (6102T>G), TTT331TTC (6108T>C), CCA334AGG (6115C>A 6116C>G 6117A>G), GAA335TAT (6118G>T 6120A>T), CTC337CAC (6125T>A), GAA339AAA (6130G>A), GAA341GTT (6137A>T 6138A>T), GTC342GTG (6141C>G), CCC344TCT (6145C>T 6147C>T), CTG347TTG (6154C>T), ATG348ACC (6158T>C 6159G>C), CCA351CCG (6168A>G), GCA352GCC (6171A>C), GAT353ATC (6172G>A 6173A>T 6174T>C), TTT354TTC (6177T>C), CAG355ATG (6178C>A 6179A>T), AGA356GAC (6181A>G 6182G>A 6183A>C), AAG357CTT (6184A>C 6185A>T 6186G>T), GAT359AAT (6190G>A), AAT360AGG (6194A>G 6195T>G), GCA361GTC (6197C>T 6198A>C), TTT362TTC (6201T>C), AGG363AAA (6203G>A 6204G>A), GGA364CCC (6205G>C 6206G>C 6207A>C), GGA364_ACA365insTAC (6207_6208insTAC), ACA365CTT (6208A>C 6209C>T 6210A>T), GAT366GAC (6213T>C), GCA367TCT (6214G>T 6216A>T), ATC369GTT (6220A>G 6222C>T), GCA370GTG (6224C>T 6225A>G), GTA371GTG (6228A>G), TAT372TTC (6230A>T 6231T>C), ATT373ATC (6234T>C), ATA376ATT (6243A>T), GTA378ATC (6247G>A 6249A>C), TTC379TAC (6251T>A), GAA381CAT (6256G>C 6258A>T), AAT382AGT (6260A>G), GAA383GAG (6264A>G), GAA384GAG (6267A>G), GAG387GTG (6275A>T), GAT388GGC (6278A>G 6279T>C) | | | | | | | | |

*: Inserts / Deletes / Misaligned / Frameshifts

#### Analysis details

This analysis was performed with panviral2.64

### NGS Details (UN9): Epiphyllum badnavirus 1

#### Assembly

| | |
|---|---|
| **Coverage Length** | 240 (1 contig(s)) |
| **Depth Of Coverage** | 4.0 |
| **Number Of Reads** | 9 |
| **Reads Per Million** | 0.20 rpm (after QC) |
| **Ambiguities** | 0 |
| **Assembly Method** | de novo + reference guided assembly |
| **Consensus Caller** | Bcf Tools |

#### Coverage Map

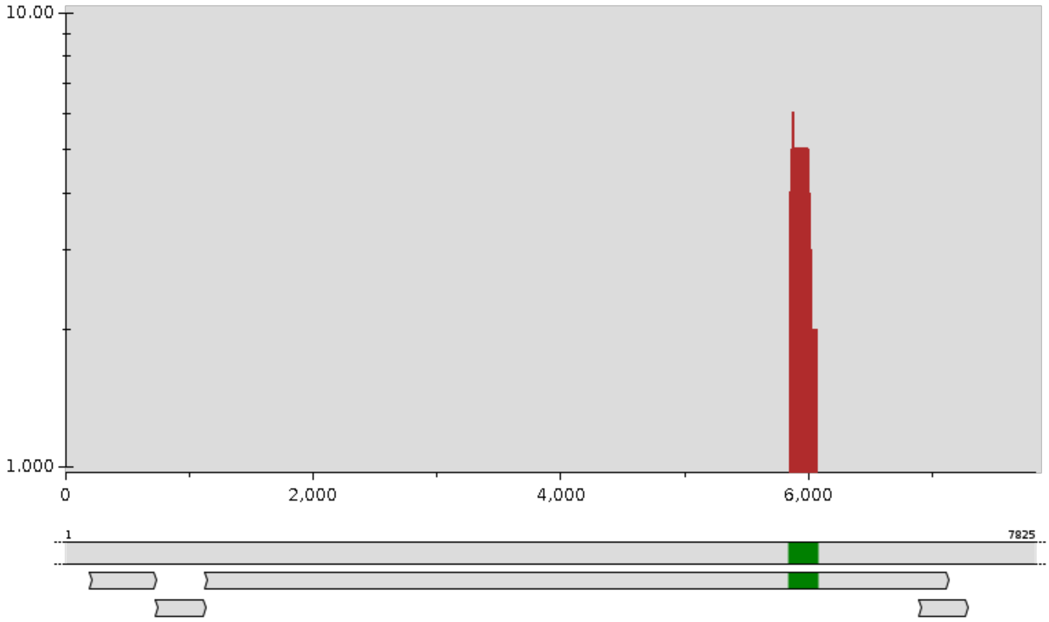

#### Assignment

| | |
|---|---|
| **Type** | Epiphyllum badnavirus 1 (Taxonomy ID: 2518008) |
| **Reference Genome** | NC_076247.1 |
| **NT Identity (%)** | 58.0247 |
| **AA Identity (%)** | 51.8519 |
| **Number Of Stop Codons** | 0 |
| **Number Of CDS** | 4 |

#### Alignment

| | |
|---|---|
| **Alignment Score** | 72.0 (NT) + 288.0 (AA) = 360.0 |
| **Concordance (%)** | 33.9943 |

| Alignment Method | Global, seeded, nucleotide + amino acids (AGA) |
|---|---|

#### Genome Region

Sequence starts at position 5838 and ends at position 6077 relative to NC_076247.1 reference sequence.

#### Alignment Detailed Statistics

| | Begin | End | Coverage | Score | Concordance | Matches | Identities | I/D/M/F* | Stop Codons |
|---|---|---|---|---|---|---|---|---|---|
| **NT** | **5838** | **6077** | **3.1%** | **72** | **15.0%** | **240 (98.8%)** | **141 (58.0%)** | **3/0** | |
| Mutations: | 5838T>G, 5841C>G, 5851T>A, 5853C>A, 5857C>T, 5861A>G, 5863A>T, 5864G>C, 5865C>G, 5868A>G, 5870T>A, 5871C>T, 5874C>T, 5877G>A, 5878A>C, 5880T>G, 5882T>G, 5886G>T, 5887G>A, 5888A>G, 5889C>G, 5892A>G, 5895A>C, 5896T>G, 5897C>A, 5905T>A, 5906G>A, 5910C>T, 5912C>T, 5916C>T, 5917C>A, 5918T>G, 5922C>T, 5924C>G, 5925A>T, 5926G>T, 5931A>C, 5933T>A, 5934T>C, 5940A>G, 5942G>T, 5943G>C, 5949T>C, 5953C>T, 5955A>T, 5958C>T, 5966A>C, 5967G>T, 5976T>C, 5977G>A, 5979T>A, 5980G>A, 5981T>C, 5982G>A, 5985C>T, 5986C>A, 5987A>T, 5988A>G, 5989C>G, 5990G>A, 5991A>T, 5992A>T, 5993A>T, 5998G>A, 6001A>C, 6002A>G, 6003C>T, 6004T>G, 6005G>T, 6006T>A, 6009C>T, 6012_6013insTCT, 6013G>T, 6019T>G, 6020C>A, 6021A>C, 6022G>A, 6023G>T, 6030A>T, 6031G>A, 6032C>T, 6033A>T, 6036T>A, 6038A>T, 6042T>A, 6048C>T, 6051C>T, 6055G>A, 6059T>A, 6061A>T, 6062G>C, 6063T>C, 6065A>G, 6066C>G, 6068C>G, 6069T>C, 6070G>A, 6071A>G, 6075T>G, 6076G>C | | | | | | | | |

**CDS**

| | Begin | End | Coverage | Score | Concordance | Matches | Identities | I/D/M/F* | Stop Codons |
|---|---|---|---|---|---|---|---|---|---|
| QKM20_gp3 | 1571 | 1650 | 4.0% | 288 | 48.6% | 80 (98.8%) | 42 (51.9%) | 1/0/0/0 | 0 |
| Protein mutations: | F1575I (5851T>A 5853C>A), K1578R (5861A>G), F1581Y (5870T>A 5871C>T), I1584L (5878A>C 5880T>G), M1585R (5882T>G), M1586I (5886G>T), D1587R (5887G>A 5888A>G 5889C>G), E1589D (5895A>C), S1590D (5896T>G 5897C>A), W1593K (5905T>A 5906G>A), A1595V (5912C>T), L1597R (5917C>A 5918T>G), P1599R (5924C>G 5925A>T), D1600Y (5926G>T), L1602H (5933T>A 5934T>C), W1605F (5942G>T 5943G>C), P1609S (5953C>T 5955A>T), K1613T (5966A>C 5967G>T), A1617T (5977G>A 5979T>A), V1618T (5980G>A 5981T>C 5982G>A), Q1620M (5986C>A 5987A>T 5988A>G), R1621D (5989C>G 5990G>A 5991A>T), K1622L (5992A>T 5993A>T), D1624N (5998G>A), N1625R (6001A>C 6002A>G 6003C>T), C1626V (6004T>G 6005G>T 6006T>A), R1628_D1629insS (6012_6013insTCT), D1629Y (6013G>T), S1631D (6019T>G 6020C>A 6021A>C), G1632I (6022G>A 6023G>T), A1635I (6031G>A 6032C>T 6033A>T), Y1637F (6038A>T), V1643I (6055G>A), F1644Y (6059T>A), N1646R (6065A>G 6066C>G), T1647S (6068C>G 6069T>C), E1648R (6070G>A 6071A>G), D1649E (6075T>G) | | | | | | | | |
| Codon mutations: | AGT1570..G (5838T>G), GTC1571GTG (5841C>G), TTC1575ATA (5851T>A 5853C>A), CTG1577TTG (5857C>T), AAG1578AGG (5861A>G), AGC1579TCG (5863A>T 5864G>C 5865C>G), GGA1580GGG (5868A>G), TTC1581TAT (5870T>A 5871C>T), CAC1582CAT (5874C>T), CAG1583CAA (5877G>A), ATT1584CTG (5878A>C 5880T>G), ATG1585AGG (5882T>G), ATG1586ATT (5886G>T), GAC1587AGG (5887G>A 5888A>G 5889C>G), GAA1588GAG (5892A>G), GAA1589GAC (5895A>C), TCC1590GAC (5896T>G 5897C>A), TGG1593AAG (5905T>A 5906G>A), ACC1594ACT (5910C>T), GCC1595GTC (5912C>T), TTC1596TTT (5916C>T), CTG1597AGG (5917C>A 5918T>G), ACC1598ACT (5922C>T), CCA1599CGT (5924C>G 5925A>T), GAT1600TAT (5926G>T), GGA1601GGC (5931A>C), CTT1602CAC (5933T>A 5934T>C), GAA1604GAG (5940A>G), TGG1605TTC (5942G>T 5943G>C), GTT1607GTC (5949T>C), CCA1609TCT (5953C>T 5955A>T), TTC1610TTT (5958C>T), AAG1613ACT (5966A>C 5967G>T), CCT1616CCC (5976T>C), GCT1617ACA (5977G>A 5979T>A), GTG1618ACA (5980G>A 5981T>C 5982G>A), TTC1619TTT (5985C>T), CAA1620ATG (5986C>A 5987A>T 5988A>G), CGA1621GAT (5989C>G 5990G>A 5991A>T), AAG1622TTG (5992A>T 5993A>T), GAC1624AAC (5998G>A), AAC1625CGT (6001A>C 6002A>G 6003C>T), TGT1626GTA (6004T>G 6005G>T 6006T>A), TTC1627TTT (6009C>T), AGA1628_GAC1629insTCT (6012_6013insTCT), GAC1629TAC (6013G>T), TCA1631GAC (6019T>G 6020C>A 6021A>C), GGC1632ATC (6022G>A 6023G>T), GTA1634GTT (6030A>T), GCA1635ATT (6031G>A 6032C>T 6033A>T), GTT1636GTA (6036T>A), TAC1637TTC (6038A>T), ATT1638ATA (6042T>A), GAC1640GAT (6048C>T), ATC1641ATT (6051C>T), GTC1643ATC (6055G>A), TTC1644TAC (6059T>A), AGT1645TCC (6061A>T 6062G>C 6063T>C), AAC1646AGG (6065A>G 6066C>G), ACT1647AGC (6068C>G 6069T>C), GAA1648AGA (6070G>A 6071A>G), GAT1649GAG (6075T>G), GAC1650CA. (6076G>C) | | | | | | | | |

**Proteins**

| | Begin | End | Coverage | Score | Concordance | Matches | Identities | I/D/M/F* | Stop Codons |
|---|---|---|---|---|---|---|---|---|---|
| polyprotein (YP_010797894.1) | 1571 | 1650 | 4.0% | 288 | 48.6% | 80 (98.8%) | 42 (51.9%) | 1/0/0/0 | 0 |
| Protein mutations: | F1575I (5851T>A 5853C>A), K1578R (5861A>G), F1581Y (5870T>A 5871C>T), I1584L (5878A>C 5880T>G), M1585R (5882T>G), M1586I (5886G>T), D1587R (5887G>A 5888A>G 5889C>G), E1589D (5895A>C), S1590D (5896T>G 5897C>A), W1593K (5905T>A 5906G>A), A1595V (5912C>T), L1597R (5917C>A 5918T>G), P1599R (5924C>G 5925A>T), D1600Y (5926G>T), L1602H (5933T>A 5934T>C), W1605F (5942G>T 5943G>C), P1609S (5953C>T 5955A>T), K1613T (5966A>C 5967G>T), A1617T (5977G>A 5979T>A), V1618T (5980G>A 5981T>C 5982G>A), Q1620M (5986C>A 5987A>T 5988A>G), R1621D (5989C>G 5990G>A 5991A>T), K1622L (5992A>T 5993A>T), D1624N (5998G>A), N1625R (6001A>C 6002A>G 6003C>T), C1626V (6004T>G 6005G>T 6006T>A), R1628_D1629insS (6012_6013insTCT), D1629Y (6013G>T), S1631D (6019T>G 6020C>A 6021A>C), G1632I (6022G>A 6023G>T), A1635I (6031G>A 6032C>T 6033A>T), Y1637F (6038A>T), V1643I (6055G>A), F1644Y (6059T>A), N1646R (6065A>G 6066C>G), T1647S (6068C>G 6069T>C), E1648R (6070G>A 6071A>G), D1649E (6075T>G) | | | | | | | | |
| Codon mutations: | AGT1570..G (5838T>G), GTC1571GTG (5841C>G), TTC1575ATA (5851T>A 5853C>A), CTG1577TTG (5857C>T), AAG1578AGG (5861A>G), AGC1579TCG (5863A>T 5864G>C 5865C>G), GGA1580GGG (5868A>G), TTC1581TAT (5870T>A 5871C>T), CAC1582CAT (5874C>T), CAG1583CAA (5877G>A), ATT1584CTG (5878A>C 5880T>G), ATG1585AGG (5882T>G), ATG1586ATT (5886G>T), GAC1587AGG (5887G>A 5888A>G 5889C>G), GAA1588GAG (5892A>G), GAA1589GAC (5895A>C), TCC1590GAC (5896T>G 5897C>A), TGG1593AAG (5905T>A 5906G>A), ACC1594ACT (5910C>T), GCC1595GTC (5912C>T), TTC1596TTT (5916C>T), CTG1597AGG (5917C>A 5918T>G), ACC1598ACT (5922C>T), CCA1599CGT (5924C>G 5925A>T), GAT1600TAT (5926G>T), GGA1601GGC (5931A>C), CTT1602CAC (5933T>A 5934T>C), GAA1604GAG (5940A>G), TGG1605TTC (5942G>T 5943G>C), GTT1607GTC (5949T>C), CCA1609TCT (5953C>T 5955A>T), TTC1610TTT (5958C>T), AAG1613ACT (5966A>C 5967G>T), CCT1616CCC (5976T>C), GCT1617ACA (5977G>A 5979T>A), GTG1618ACA (5980G>A 5981T>C 5982G>A), TTC1619TTT (5985C>T), CAA1620ATG (5986C>A 5987A>T 5988A>G), CGA1621GAT (5989C>G 5990G>A 5991A>T), AAG1622TTG (5992A>T 5993A>T), GAC1624AAC (5998G>A), AAC1625CGT (6001A>C 6002A>G 6003C>T), TGT1626GTA (6004T>G 6005G>T 6006T>A), TTC1627TTT (6009C>T), AGA1628_GAC1629insTCT (6012_6013insTCT), GAC1629TAC (6013G>T), TCA1631GAC (6019T>G 6020C>A 6021A>C), GGC1632ATC (6022G>A 6023G>T), GTA1634GTT (6030A>T), GCA1635ATT (6031G>A 6032C>T 6033A>T), GTT1636GTA (6036T>A), TAC1637TTC (6038A>T), ATT1638ATA (6042T>A), GAC1640GAT (6048C>T), ATC1641ATT (6051C>T), GTC1643ATC (6055G>A), TTC1644TAC (6059T>A), AGT1645TCC (6061A>T 6062G>C 6063T>C), AAC1646AGG (6065A>G 6066C>G), ACT1647AGC (6068C>G 6069T>C), GAA1648AGA (6070G>A 6071A>G), GAT1649GAG (6075T>G), GAC1650CA. (6076G>C) | | | | | | | | |

*: Inserts / Deletes / Misaligned / Frameshifts

#### Analysis details

This analysis was performed with panviral2.64

### NGS Details (UN9): Colombian datura virus

#### Assembly

| | |
|---|---|
| **Coverage Length** | 429 (2 contig(s)) |
| **Depth Of Coverage** | 2.3 |
| **Number Of Reads** | 8 |
| **Reads Per Million** | 0.18 rpm (after QC) |
| **Ambiguities** | 1 |
| **Assembly Method** | de novo + reference guided assembly |
| **Consensus Caller** | Bcf Tools |

#### Coverage Map

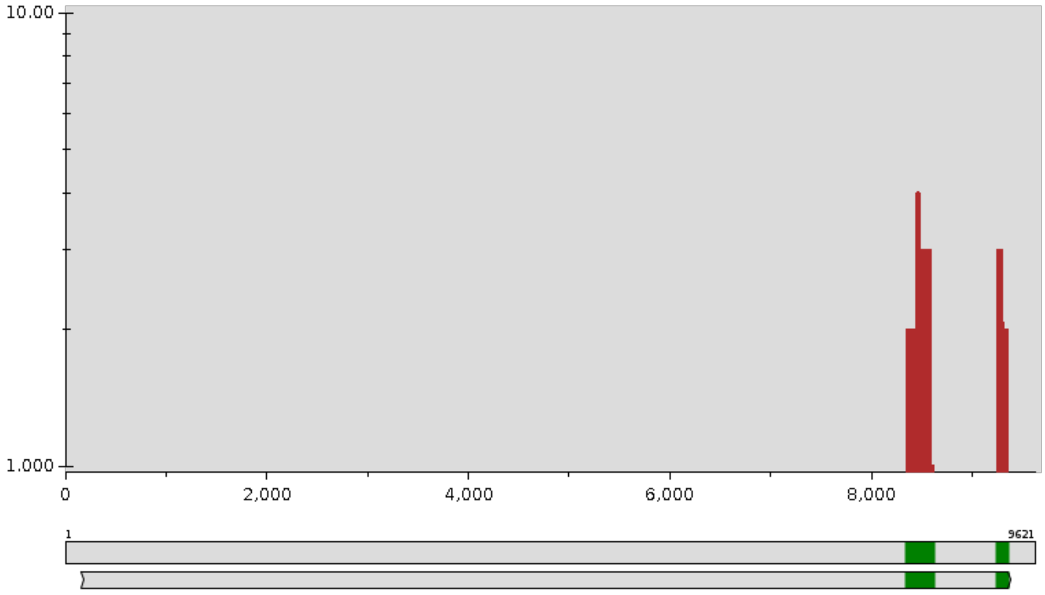

#### Assignment

| | |
|---|---|
| **Type** | Colombian datura virus (Taxonomy ID: 91613) |
| **Reference Genome** | NC_020072.1 |
| **NT Identity (%)** | 75.5245 |
| **AA Identity (%)** | 79.021 |
| **Number Of Stop Codons** | 0 |
| **Number Of CDS** | 1 |

#### Alignment

| | |
|---|---|
| **Alignment Score** | 440.0 (NT) + 731.0 (AA) = 1171.0 |
| **Concordance (%)** | 66.0462 |

| Alignment Method | Global, seeded, nucleotide + amino acids (AGA) |
|---|---|

#### Genome Region

Sequence starts at position 8339 and ends at position 9368 relative to NC_020072.1 reference sequence.

#### Alignment Detailed Statistics

| | Begin | End | Coverage | Score | Concordance | Matches | Identities | I/D/M/F* | Stop Codons |
|---|---|---|---|---|---|---|---|---|---|
| **NT** | **8339** | **9368** | **4.5%** | **440** | **51.3%** | **429 (100%)** | **324 (75.5%)** | **0/0** | |
| Mutations: | 8341A>G, 8342C>T, 8347T>A, 8349A>T, 8353G>A, 8357A>C, 8369C>T, 8371T>G, 8383G>A, 8395C>T, 8411G>C, 8416G>A, 8418G>A, 8422T>A, 8428C>T, 8431A>G, 8434T>C, 8437A>G, 8440T>G, 8443T>A, 8444C>T, 8455A>G, 8461T>A, 8464T>C, 8465G>A, 8467T>C, 8473C>A, 8476T>G, 8479A>T, 8480T>A, 8481T>A, 8482G>A, 8485A>G, 8488G>A, 8489C>T, 8491T>G, 8499C>T, 8500A>T, 8503G>T, 8506C>T, 8507A>G, 8515T>A, 8518T>G, 8520A>C, 8524T>C, 8525G>T, 8526A>T, 8527T>G, 8528A>G, 8529G>A, 8530C>T, 8531T>G, 8533G>A, 8535C>A, 8537T>C, 8538C>A, 8539T>A, 8542A>G, 8543G>C, 8545T>C, 8548A>G, 8549T>A, 8550A>T, 8551T>A, 8552C>T, 8560T>A, 8563T>C, 8565C>A, 8578T>A, 8581T>C, 8587A>C, 8591G>A, 8596A>T, 8599G>A, 8601A>G, 8602C>T, 8606T>G, 8607C>G, 8624A>C, 8625G>T, 8626T>C, 8628T>C, 9244T>A, 9256A>T, 9261A>C, 9262C>A, 9265A>G, 9268G>C, 9271G>A, 9272T>C, 9274A>C, 9283G>C, 9307A>G, 9316C>A, 9325C>T, 9332C>A, 9333A>C, 9334A>T, 9339T>W, 9340T>G, 9346A>G, 9347G>A, 9348G>A, 9349T>C, 9364T>A | | | | | | | | |
| **CDS** | | | | | | | | | |
| G357_gp1 | 2729 | 3072 | 4.6% | 731 | 78.8% | 143 (100%) | 113 (79.0%) | 0/0/1/0 | 0 |
| Protein mutations: | K2732M (8349A>T), E2753Q (8411G>C), R2755K (8418G>A), A2771T (8465G>A 8467T>C), E2775D (8479A>T), L2776K (8480T>A 8481T>A 8482G>A), A2782V (8499C>T 8500A>T), I2785V (8507A>G), D2788E (8518T>G), Y2789S (8520A>C), D2791L (8525G>T 8526A>T 8527T>G), S2792D (8528A>G 8529G>A 8530C>T), L2793V (8531T>G 8533G>A), A2794E (8535C>A), S2795Q (8537T>C 8538C>A 8539T>A), D2797H (8543G>C 8545T>C), I2798M (8548A>G), Y2799I (8549T>A 8550A>T 8551T>A), L2800F (8552C>T), T2804K (8565C>A), A2813T (8591G>A), N2816S (8601A>G 8602C>T), S2818G (8606T>G 8607C>G), S2824L (8624A>C 8625G>T 8626T>C), V2825A (8628T>C), N3036T (9261A>C 9262C>A), Q3060T (9332C>A 9333A>C 9334A>T), G3065N (9347G>A 9348G>A 9349T>C) | | | | | | | | |
| Codon mutations: | GAA2729GAG (8341A>G), CTG2730TTG (8342C>T), ACT2731ACA (8347T>A), AAG2732ATG (8349A>T), AGG2733AGA (8353G>A), AGA2735CGA (8357A>C), CTT2739TTG (8369C>T 8371T>G), GAG2743GAA (8383G>A), TAC2747TAT (8395C>T), GAA2753CAA (8411G>C), GGG2754GGA (8416G>A), AGA2755AAA (8418G>A), GCT2756GCA (8422T>A), TAC2758TAT (8428C>T), CTA2759CTG (8431A>G), GCT2760GCC (8434T>C), GAA2761GAG (8437A>G), ACT2762ACG (8440T>G), GCT2763GCA (8443T>A), CTG2764TTG (8444C>T), TTA2767TTG (8455A>G), CTT2769CTA (8461T>A), GAT2770GAC (8464T>C), GCT2771ACC (8465G>A 8467T>C), CCC2773CCA (8473C>A), ACT2774ACG (8476T>G), GAA2775GAT (8479A>T), TTG2776AAA (8480T>A 8481T>A 8482G>A), GAA2777GAG (8485A>G), TTG2778TTA (8488G>A), CTT2779TTG (8489C>T 8491T>G), GCA2782GTT (8499C>T 8500A>T), GGG2783GGT (8503G>T), TGC2784TGT (8506C>T), ATC2785GTC (8507A>G), GGT2787GGA (8515T>A), GAT2788GAG (8518T>G), TAT2789TCT (8520A>C), GAT2790GAC (8524T>C), GAT2791TTG (8525G>T 8526A>T 8527T>G), AGC2792GAT (8528A>G 8529G>A 8530C>T), TTG2793GTA (8531T>G 8533G>A), GCA2794GAA (8535C>A), TCT2795CAA (8537T>C 8538C>A 8539T>A), GAA2796GAG (8542A>G), GAT2797CAC (8543G>C 8545T>C), ATA2798ATG (8548A>G), TAT2799ATA (8549T>A 8550A>T 8551T>A), CTC2800TTC (8552C>T), GCT2802GCA (8560T>A), GAT2803GAC (8563T>C), ACA2804AAA (8565C>A), GGT2808GGA (8578T>A), AGT2809AGC (8581T>C), ACA2811ACC (8587A>C), GCC2813ACC (8591G>A), GGA2814GGT (8596A>T), AAG2815AAA (8599G>A), AAC2816AGT (8601A>G 8602C>T), TCA2818GGA (8606T>G 8607C>G), AGT2824CTC (8624A>C 8625G>T 8626T>C), GTT2825GCT (8628T>C), GCT3030GCA (9244T>A), CGA3034CGT (9256A>T), AAC3036ACA (9261A>C 9262C>A), ACA3037ACG (9265A>G), ACG3038ACC (9268G>C), AAG3039AAA (9271G>A), TTA3040CTC (9272T>C 9274A>C), CTG3043CTC (9283G>C), GAA3051GAG (9307A>G), ACC3054ACA (9316C>A), CAC3057CAT (9325C>T), CAA3060ACT (9332C>A 9333A>C 9334A>T), GTT3062GWG (9339T>W 9340T>G), AGA3064AGG (9346A>G), GGT3065AAC (9347G>A 9348G>A 9349T>C), CTT3070CTA (9364T>A) | | | | | | | | |
| **Proteins** | | | | | | | | | |
| polyprotein (YP_007346986.1) | 2729 | 3072 | 4.6% | 731 | 78.8% | 143 (100%) | 113 (79.0%) | 0/0/1/0 | 0 |
| Protein mutations: | K2732M (8349A>T), E2753Q (8411G>C), R2755K (8418G>A), A2771T (8465G>A 8467T>C), E2775D (8479A>T), L2776K (8480T>A 8481T>A 8482G>A), A2782V (8499C>T 8500A>T), I2785V (8507A>G), D2788E (8518T>G), Y2789S (8520A>C), D2791L (8525G>T 8526A>T 8527T>G), S2792D (8528A>G 8529G>A 8530C>T), L2793V (8531T>G 8533G>A), A2794E (8535C>A), S2795Q (8537T>C 8538C>A 8539T>A), D2797H (8543G>C 8545T>C), I2798M (8548A>G), Y2799I (8549T>A 8550A>T 8551T>A), L2800F (8552C>T), T2804K (8565C>A), A2813T (8591G>A), N2816S (8601A>G 8602C>T), S2818G (8606T>G 8607C>G), S2824L (8624A>C 8625G>T 8626T>C), V2825A (8628T>C), N3036T (9261A>C 9262C>A), Q3060T (9332C>A 9333A>C 9334A>T), G3065N (9347G>A 9348G>A 9349T>C) | | | | | | | | |
| Codon mutations: | GAA2729GAG (8341A>G), CTG2730TTG (8342C>T), ACT2731ACA (8347T>A), AAG2732ATG (8349A>T), AGG2733AGA (8353G>A), AGA2735CGA (8357A>C), CTT2739TTG (8369C>T 8371T>G), GAG2743GAA (8383G>A), TAC2747TAT (8395C>T), GAA2753CAA (8411G>C), GGG2754GGA (8416G>A), AGA2755AAA (8418G>A), GCT2756GCA (8422T>A), TAC2758TAT (8428C>T), CTA2759CTG (8431A>G), GCT2760GCC (8434T>C), GAA2761GAG (8437A>G), ACT2762ACG (8440T>G), GCT2763GCA (8443T>A), CTG2764TTG (8444C>T), TTA2767TTG (8455A>G), CTT2769CTA (8461T>A), GAT2770GAC (8464T>C), GCT2771ACC (8465G>A 8467T>C), CCC2773CCA (8473C>A), ACT2774ACG (8476T>G), GAA2775GAT (8479A>T), TTG2776AAA (8480T>A 8481T>A 8482G>A), GAA2777GAG (8485A>G), TTG2778TTA (8488G>A), CTT2779TTG (8489C>T 8491T>G), GCA2782GTT (8499C>T 8500A>T), GGG2783GGT (8503G>T), TGC2784TGT (8506C>T), ATC2785GTC (8507A>G), GGT2787GGA (8515T>A), GAT2788GAG (8518T>G), TAT2789TCT (8520A>C), GAT2790GAC (8524T>C), GAT2791TTG (8525G>T 8526A>T 8527T>G), AGC2792GAT (8528A>G 8529G>A 8530C>T), TTG2793GTA (8531T>G 8533G>A), GCA2794GAA (8535C>A), TCT2795CAA (8537T>C 8538C>A 8539T>A), GAA2796GAG (8542A>G), GAT2797CAC (8543G>C 8545T>C), ATA2798ATG (8548A>G), TAT2799ATA (8549T>A 8550A>T 8551T>A), CTC2800TTC (8552C>T), GCT2802GCA (8560T>A), GAT2803GAC (8563T>C), ACA2804AAA (8565C>A), GGT2808GGA (8578T>A), AGT2809AGC (8581T>C), ACA2811ACC (8587A>C), GCC2813ACC (8591G>A), GGA2814GGT (8596A>T), AAG2815AAA (8599G>A), AAC2816AGT (8601A>G 8602C>T), TCA2818GGA (8606T>G 8607C>G), AGT2824CTC (8624A>C 8625G>T 8626T>C), GTT2825GCT (8628T>C), GCT3030GCA (9244T>A), CGA3034CGT (9256A>T), AAC3036ACA (9261A>C 9262C>A), ACA3037ACG (9265A>G), ACG3038ACC (9268G>C), AAG3039AAA (9271G>A), TTA3040CTC (9272T>C 9274A>C), CTG3043CTC (9283G>C), GAA3051GAG (9307A>G), ACC3054ACA (9316C>A), CAC3057CAT (9325C>T), CAA3060ACT (9332C>A 9333A>C 9334A>T), GTT3062GWG (9339T>W 9340T>G), AGA3064AGG (9346A>G), GGT3065AAC (9347G>A 9348G>A 9349T>C), CTT3070CTA (9364T>A) | | | | | | | | |

*: Inserts / Deletes / Misaligned / Frameshifts

#### Analysis details

This analysis was performed with panviral2.64

### NGS Details (UN9): Pinus nigra virus 1

#### Assembly

| | |
|---|---|
| **Coverage Length** | 356 (1 contig(s)) |
| **Depth Of Coverage** | 2.7 |
| **Number Of Reads** | 8 |
| **Reads Per Million** | 0.18 rpm (after QC) |
| **Ambiguities** | 0 |
| **Assembly Method** | de novo + reference guided assembly |
| **Consensus Caller** | Bcf Tools |

#### Coverage Map

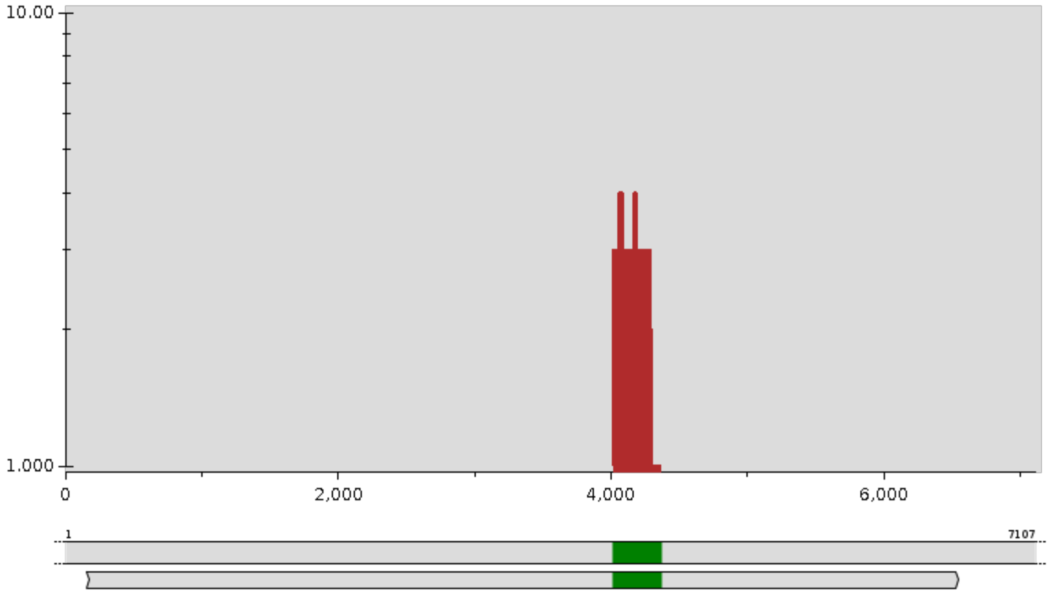

#### Assignment

| | |
|---|---|
| **Type** | Pinus nigra virus 1 (Taxonomy ID: 2267679) |
| **Reference Genome** | NC_040841.1 |
| **NT Identity (%)** | 55.3073 |
| **AA Identity (%)** | 50.4202 |
| **Number Of Stop Codons** | 1 |
| **Number Of CDS** | 1 |

#### Alignment

| | |
|---|---|
| **Alignment Score** | 60.0 (NT) + 317.0 (AA) = 377.0 |
| **Concordance (%)** | 27.1175 |

| Alignment Method | Global, seeded, nucleotide + amino acids (AGA) |
|---|---|

#### Genome Region

Sequence starts at position 4017 and ends at position 4372 relative to NC_040841.1 reference sequence.

#### Alignment Detailed Statistics

| | Begin | End | Coverage | Score | Concordance | Matches | Identities | I/D/M/F* | Stop Codons |
|---|---|---|---|---|---|---|---|---|---|
| **NT** | **4017** | **4372** | **5.0%** | **60** | **8.6%** | **355 (98.9%)** | **198 (55.2%)** | **3/1** | |
| Mutations: | 4018C>T, 4021C>T, 4024G>A, 4036G>A, 4042G>C, 4044A>T, 4047A>G, 4048T>G, 4051G>A, 4054A>C, 4061C>T, 4062A>T, 4063G>C, 4064C>G, 4065C>A, 4067G>A, 4069A>G, 4074G>A, 4076C>T, 4078C>T, 4084A>T, 4092T>A, 4093A>T, 4094T>G, 4095G>T, 4096T>C, 4099A>G, 4100G>T, 4101C>T, 4102A>T, 4106T>C, 4107T>A, 4108C>A, 4112C>G, 4114G>A, 4120A>T, 4122T>C, 4124G>A, 4126T>G, 4135T>A, 4136T>C, 4138G>A, 4141G>A, 4144T>C, 4147G>C, 4150A>T, 4151G>T, 4152C>T, 4154T>G, 4155T>A, 4156C>G, 4159T>C, 4164G>A, 4165A>C, 4167G>T, 4168G>C, 4171G>A, 4172G>A, 4175T>A, 4180C>T, 4183T>C, 4184delG, 4185C>A, 4186T>A, 4187A>C, 4188A>C, 4193G>C, 4194A>G, 4195T>C, 4196T>A, 4197T>A, 4201A>T, 4201_4202insTCT, 4202G>A, 4204T>C, 4205A>G, 4206C>T, 4210C>T, 4213A>T, 4220A>G, 4222T>A, 4223T>C, 4228T>A, 4229C>T, 4230A>T, 4231T>C, 4232A>T, 4233G>C, 4235C>A, 4236C>A, 4237A>G, 4240T>C, 4241G>A, 4244C>G, 4246G>T, 4247A>T, 4248A>C, 4252T>C, 4255G>T, 4258G>A, 4261C>T, 4264A>G, 4265G>A, 4267A>C, 4269T>C, 4270T>C, 4273T>C, 4274C>T, 4276T>C, 4277G>A, 4280G>A, 4281A>T, 4282A>C, 4283G>A, 4285C>T, 4289A>C, 4291A>C, 4292C>A, 4297A>T, 4298A>C, 4300T>C, 4303T>C, 4304T>G, 4306G>T, 4307T>A, 4308C>G, 4309A>T, 4311A>C, 4312A>T, 4314G>A, 4315A>T, 4318G>A, 4322T>A, 4323C>A, 4324C>A, 4325T>A, 4327A>T, 4330T>C, 4333G>A, 4334G>A, 4335A>C, 4336T>A, 4339T>A, 4342T>C, 4343G>A, 4344A>G, 4345T>A, 4349T>A, 4351G>T, 4354A>G, 4355A>C, 4356T>A, 4358A>T, 4361G>T, 4363T>G, 4364G>T, 4365C>A, 4367A>C, 4369T>A | | | | | | | | |

##### CDS

| | | | | | | | | | |
|---|---|---|---|---|---|---|---|---|---|
| EXL67_gp1 | 1289 | 1406 | 5.5% | 317 | 36.3% | 118 (99.2%) | 60 (50.4%) | 1/0/1/1 | 1 |
| Protein mutations: | Y1297F (4044A>T), Y1298W (4047A>G 4048T>G), Q1303F (4061C>T 4062A>T 4063G>C), P1304E (4064C>G 4065C>A), E1305K (4067G>A 4069A>G), R1307K (4074G>A), H1308Y (4076C>T 4078C>T), I1313N (4092T>A 4093A>T), C1314V (4094T>G 4095G>T 4096T>C), A1316F (4100G>T 4101C>T 4102A>T), F1318Q (4106T>C 4107T>A 4108C>A), Q1320E (4112C>G 4114G>A), K1322N (4120A>T), V1323A (4122T>C), V1324M (4124G>A 4126T>G), A1333F (4151G>T 4152C>T), F1334E (4154T>G 4155T>A 4156C>G), R1337N (4164G>A 4165A>C), R1338I (4167G>T 4168G>C), M1339I (4171G>A), D1340N (4172G>A), Y1341N (4175T>A), K1345P (4187A>C 4188A>C), D1347R (4193G>C 4194A>G 4195T>C), F1348N (4196T>A 4197T>A), I1349_V1350insS (4201_4202insTCT), V1350I (4202G>A 4204T>C), T1351V (4205A>G 4206C>T), I1356V (4220A>G 4222T>A), H1359F (4229C>T 4230A>T 4231T>C), P1361K (4235C>A 4236C>A 4237A>G), V1363I (4241G>A), Q1364D (4244C>G 4246G>T), N1365S (4247A>T 4248A>C), L1367F (4255G>T), E1371N (4265G>A 4267A>C), I1372T (4269T>C 4270T>C), L1374F (4274C>T 4276T>C), E1375K (4277G>A), E1376I (4280G>A 4281A>T 4282A>C), V1377I (4283G>A 4285C>T), K1379H (4289A>C 4291A>C), H1380N (4292C>A), I1382L (4298A>C 4300T>C), L1384V (4304T>G 4306G>T), E1386A (4311A>C 4312A>T), R1387N (4314G>A 4315A>T), S1390K (4322T>A 4323C>A 4324C>A), L1391I (4325T>A 4327A>T), D1394T (4334G>A 4335A>C 4336T>A), N1395K (4339T>A), D1397R (4343G>A 4344A>G 4345T>A), L1399I (4349T>A 4351G>T), I1401H (4355A>C 4356T>A), N1402Y (4358A>T), V1403L (4361G>T 4363T>G), A1404* (4364G>T 4365C>A), N1405Q (4367A>C 4369T>A) | | | | | | | | |
| Codon mutations: | TTC1288.TT (4018C>T), AGC1289AGT (4021C>T), AAG1290AAA (4024G>A), AAG1294AAA (4036G>A), GGG1296GGC (4042G>C), TAT1297TTT (4044A>T), TAT1298TGG (4047A>G 4048T>G), CAG1299CAA (4051G>A), ATA1300ATC (4054A>C), CAG1303TTC (4061C>T 4062A>T 4063G>C), CCA1304GAA (4064C>G 4065C>A), GAA1305AAG (4067G>A 4069A>G), AGA1307AAA (4074G>A), CAC1308TAT (4076C>T 4078C>T), ACA1310ACT (4084A>T), ATA1313AAT (4092T>A 4093A>T), TGT1314GTC (4094T>G 4095G>T 4096T>C), CCA1315CCG (4099A>G), GCA1316TTT (4100G>T 4101C>T 4102A>T), TTC1318CAA (4106T>C 4107T>A 4108C>A), CAG1320GAA (4112C>G 4114G>A), AAA1322AAT (4120A>T), GTT1323GCT (4122T>C), GTT1324ATG (4124G>A 4126T>G), GGT1327GGA (4135T>A), TTG1328CTA (4136T>C 4138G>A), AAG1329AAA (4141G>A), AAT1330AAC (4144T>C), GCG1331GCC (4147G>C), CCA1332CCT (4150A>T), GCC1333TTC (4151G>T 4152C>T), TTC1334GAG (4154T>G 4155T>A 4156C>G), TTT1335TTC (4159T>C), AGA1337AAC (4164G>A 4165A>C), AGG1338ATC (4167G>T 4168G>C), ATG1339ATA (4171G>A), GAT1340AAT (4172G>A), TAT1341AAT (4175T>A), ATC1342ATT (4180C>T), TTT1343TTC (4183T>C), GCT1344-AA (4184delG 4185C>A 4186T>A), AAA1345CCA (4187A>C 4188A>C), GAT1347CGC (4193G>C 4194A>G 4195T>C), TTT1348AAT (4196T>A 4197T>A), ATA1349ATT (4201A>T), ATA1349_GTT1350insTCT (4201_4202insTCT), GTT1350ATC (4202G>A 4204T>C), ACA1351GTA (4205A>G 4206C>T), TAC1352TAT (4210C>T), ATA1353ATT (4213A>T), ATT1356GTA (4220A>G 4222T>A), TTA1357CTA (4223T>C), ATT1358ATA (4228T>A), CAT1359TTC (4229C>T 4230A>T 4231T>C), AGT1360TCT (4232A>T 4233G>C), CCA1361AAG (4235C>A 4236C>A 4237A>G), GAT1362GAC (4240T>C), GTA1363ATA (4241G>A), CAG1364GAT (4244C>G 4246G>T), AAT1365TCT (4247A>T 4248A>C), CAT1366CAC (4252T>C), TTG1367TTT (4255G>T), AAG1368AAA (4258G>A), CAC1369CAT (4261C>T), TTA1370TTG (4264A>G), GAA1371AAC (4265G>A 4267A>C), ATT1372ACC (4269T>C 4270T>C), TTT1373TTC (4273T>C), CTT1374TTC (4274C>T 4276T>C), GAA1375AAA (4277G>A), GAA1376ATC (4280G>A 4281A>T 4282A>C), GTC1377ATT (4283G>A 4285C>T), AAA1379CAC (4289A>C 4291A>C), CAT1380AAT (4292C>A), GGA1381GGT (4297A>T), ATT1382CTC (4298A>C 4300T>C), GTT1383GTC (4303T>C), TTG1384GTT (4304T>G 4306G>T), TCA1385AGT (4307T>A 4308C>G 4309A>T), GAA1386GCT (4311A>C 4312A>T), AGA1387AAT (4314G>A 4315A>T), AAG1388AAA (4318G>A), TCC1390AAA (4322T>A 4323C>A 4324C>A), TTA1391ATT (4325T>A 4327A>T), TTT1392TTC (4330T>C), CAG1393CAA (4333G>A), GAT1394ACA (4334G>A 4335A>C 4336T>A), AAT1395AAA (4339T>A), ATT1396ATC (4342T>C), GAT1397AGA (4343G>A 4344A>G 4345T>A), TTG1399ATT (4349T>A 4351G>T), GGA1400GGG (4354A>G), ATT1401CAT (4355A>C 4356T>A), AAT1402TAT (4358A>T), GTT1403TTG (4361G>T 4363T>G), GCA1404TAA (4364G>T 4365C>A), AAT1405CAA (4367A>C 4369T>A) | | | | | | | | |

##### Proteins

| | | | | | | | | | |
|---|---|---|---|---|---|---|---|---|---|
| polyprotein (YP_009553669.1) | 1289 | 1406 | 5.5% | 317 | 36.3% | 118 (99.2%) | 60 (50.4%) | 1/0/1/1 | 1 |
| Protein mutations: | Y1297F (4044A>T), Y1298W (4047A>G 4048T>G), Q1303F (4061C>T 4062A>T 4063G>C), P1304E (4064C>G 4065C>A), E1305K (4067G>A 4069A>G), R1307K (4074G>A), H1308Y (4076C>T 4078C>T), I1313N (4092T>A 4093A>T), C1314V (4094T>G 4095G>T 4096T>C), A1316F (4100G>T 4101C>T 4102A>T), F1318Q (4106T>C 4107T>A 4108C>A), Q1320E (4112C>G 4114G>A), K1322N (4120A>T), V1323A (4122T>C), V1324M (4124G>A 4126T>G), A1333F (4151G>T 4152C>T), F1334E (4154T>G 4155T>A 4156C>G), R1337N (4164G>A 4165A>C), R1338I (4167G>T 4168G>C), M1339I (4171G>A), D1340N (4172G>A), Y1341N (4175T>A), K1345P (4187A>C 4188A>C), D1347R (4193G>C 4194A>G 4195T>C), F1348N (4196T>A 4197T>A), I1349_V1350insS (4201_4202insTCT), V1350I (4202G>A 4204T>C), T1351V (4205A>G 4206C>T), I1356V (4220A>G 4222T>A), H1359F (4229C>T 4230A>T 4231T>C), P1361K (4235C>A 4236C>A 4237A>G), V1363I (4241G>A), Q1364D (4244C>G 4246G>T), N1365S (4247A>T 4248A>C), L1367F (4255G>T), E1371N (4265G>A 4267A>C), I1372T (4269T>C 4270T>C), L1374F (4274C>T 4276T>C), E1375K (4277G>A), E1376I (4280G>A 4281A>T 4282A>C), V1377I (4283G>A 4285C>T), K1379H (4289A>C 4291A>C), H1380N (4292C>A), I1382L (4298A>C 4300T>C), L1384V (4304T>G 4306G>T), E1386A (4311A>C 4312A>T), R1387N (4314G>A 4315A>T), S1390K (4322T>A 4323C>A 4324C>A), L1391I (4325T>A 4327A>T), D1394T (4334G>A 4335A>C 4336T>A), N1395K (4339T>A), D1397R (4343G>A 4344A>G 4345T>A), L1399I (4349T>A 4351G>T), I1401H (4355A>C 4356T>A), N1402Y (4358A>T), V1403L (4361G>T 4363T>G), A1404* (4364G>T 4365C>A), N1405Q (4367A>C 4369T>A) | | | | | | | | |
| Codon mutations: | TTC1288.TT (4018C>T), AGC1289AGT (4021C>T), AAG1290AAA (4024G>A), AAG1294AAA (4036G>A), GGG1296GGC (4042G>C), TAT1297TTT (4044A>T), TAT1298TGG (4047A>G 4048T>G), CAG1299CAA (4051G>A), ATA1300ATC (4054A>C), CAG1303TTC (4061C>T 4062A>T 4063G>C), CCA1304GAA (4064C>G 4065C>A), GAA1305AAG (4067G>A 4069A>G), AGA1307AAA (4074G>A), CAC1308TAT (4076C>T 4078C>T), ACA1310ACT (4084A>T), ATA1313AAT (4092T>A 4093A>T), TGT1314GTC (4094T>G 4095G>T 4096T>C), CCA1315CCG (4099A>G), GCA1316TTT (4100G>T 4101C>T 4102A>T), TTC1318CAA (4106T>C 4107T>A 4108C>A), CAG1320GAA (4112C>G 4114G>A), AAA1322AAT (4120A>T), GTT1323GCT (4122T>C), GTT1324ATG (4124G>A 4126T>G), GGT1327GGA (4135T>A), TTG1328CTA (4136T>C 4138G>A), AAG1329AAA (4141G>A), AAT1330AAC (4144T>C), GCG1331GCC (4147G>C), CCA1332CCT (4150A>T), GCC1333TTC (4151G>T 4152C>T), TTC1334GAG (4154T>G 4155T>A 4156C>G), TTT1335TTC (4159T>C), AGA1337AAC (4164G>A 4165A>C), AGG1338ATC (4167G>T 4168G>C), ATG1339ATA (4171G>A), GAT1340AAT (4172G>A), TAT1341AAT (4175T>A), ATC1342ATT (4180C>T), TTT1343TTC (4183T>C), GCT1344-AA (4184delG 4185C>A 4186T>A), AAA1345CCA (4187A>C 4188A>C), GAT1347CGC (4193G>C 4194A>G 4195T>C), TTT1348AAT (4196T>A 4197T>A), ATA1349ATT (4201A>T), ATA1349_GTT1350insTCT (4201_4202insTCT), GTT1350ATC (4202G>A 4204T>C), ACA1351GTA (4205A>G 4206C>T), TAC1352TAT (4210C>T), ATA1353ATT (4213A>T), ATT1356GTA (4220A>G 4222T>A), TTA1357CTA (4223T>C), ATT1358ATA (4228T>A), CAT1359TTC (4229C>T 4230A>T 4231T>C), AGT1360TCT (4232A>T 4233G>C), CCA1361AAG (4235C>A 4236C>A 4237A>G), GAT1362GAC (4240T>C), GTA1363ATA (4241G>A), CAG1364GAT (4244C>G 4246G>T), AAT1365TCT (4247A>T 4248A>C), CAT1366CAC (4252T>C), TTG1367TTT (4255G>T), AAG1368AAA (4258G>A), CAC1369CAT (4261C>T), TTA1370TTG (4264A>G), GAA1371AAC (4265G>A 4267A>C), ATT1372ACC (4269T>C 4270T>C), TTT1373TTC (4273T>C), CTT1374TTC (4274C>T 4276T>C), GAA1375AAA (4277G>A), GAA1376ATC (4280G>A 4281A>T 4282A>C), GTC1377ATT (4283G>A 4285C>T), AAA1379CAC (4289A>C 4291A>C), CAT1380AAT (4292C>A), GGA1381GGT (4297A>T), ATT1382CTC (4298A>C 4300T>C), GTT1383GTC (4303T>C), TTG1384GTT (4304T>G 4306G>T), TCA1385AGT (4307T>A 4308C>G 4309A>T), GAA1386GCT (4311A>C 4312A>T), AGA1387AAT (4314G>A 4315A>T), AAG1388AAA (4318G>A), TCC1390AAA (4322T>A 4323C>A 4324C>A), TTA1391ATT (4325T>A 4327A>T), TTT1392TTC (4330T>C), CAG1393CAA (4333G>A), GAT1394ACA (4334G>A 4335A>C 4336T>A), AAT1395AAA (4339T>A), ATT1396ATC (4342T>C), GAT1397AGA (4343G>A 4344A>G 4345T>A), TTG1399ATT (4349T>A 4351G>T), GGA1400GGG (4354A>G), ATT1401CAT (4355A>C 4356T>A), AAT1402TAT (4358A>T), GTT1403TTG (4361G>T 4363T>G), GCA1404TAA (4364G>T 4365C>A), AAT1405CAA (4367A>C 4369T>A) | | | | | | | | |

*: Inserts / Deletes / Misaligned / Frameshifts

#### Analysis details

This analysis was performed with panviral2.64

### NGS Details (UN9): Badnavirus rutilanscamelliae

#### Assembly

| | |
|---|---|
| **Coverage Length** | 335 (1 contig(s)) |
| **Depth Of Coverage** | 2.6 |
| **Number Of Reads** | 7 |
| **Reads Per Million** | 0.16 rpm (after QC) |
| **Ambiguities** | 0 |
| **Assembly Method** | de novo + reference guided assembly |
| **Consensus Caller** | Bcf Tools |

#### Coverage Map

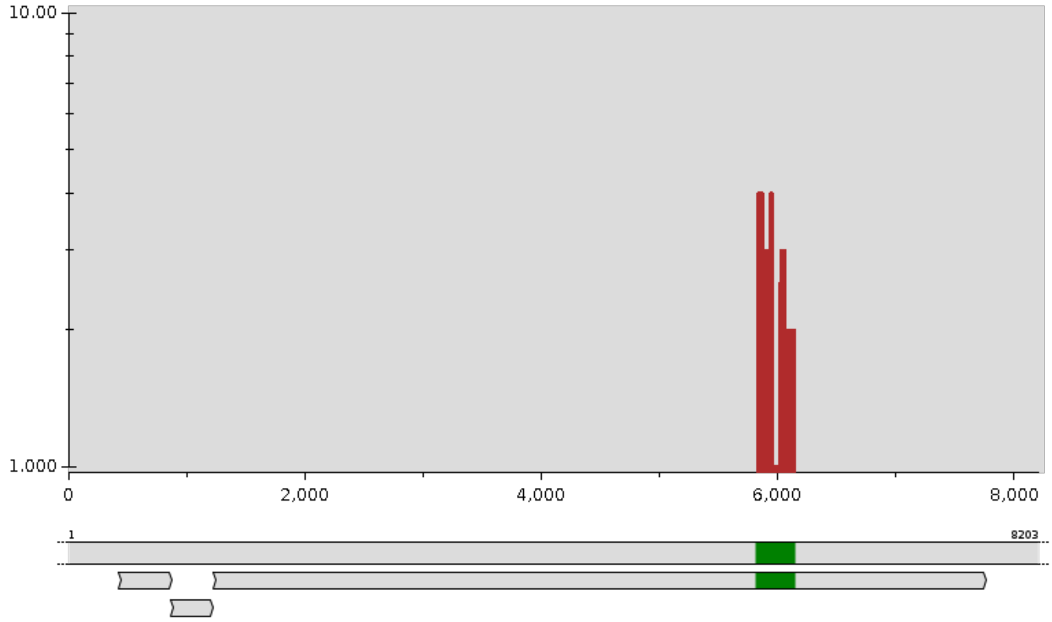

#### Assignment

| | |
|---|---|
| **Type** | Badnavirus rutilanscamelliae (Taxonomy ID: 3047719) |
| **Reference Genome** | NC_055598.1 |
| **NT Identity (%)** | 55.3254 |
| **AA Identity (%)** | 45.1327 |
| **Number Of Stop Codons** | 0 |
| **Number Of CDS** | 3 |

#### Alignment

| | |
|---|---|
| **Alignment Score** | 66.0 (NT) + 368.0 (AA) = 434.0 |
| **Concordance (%)** | 28.972 |

| **Alignment Method** | Global, seeded, nucleotide + amino acids (AGA) |
|---|---|

#### Genome Region

Sequence starts at position 5822 and ends at position 6156 relative to NC_055598.1 reference sequence.

#### Alignment Detailed Statistics

| | Begin | End | Coverage | Score | Concordance | Matches | Identities | I/D/M/F* | Stop Codons |
|---|---|---|---|---|---|---|---|---|---|
| **NT** | **5822** | **6156** | **4.1%** | **66** | **9.9%** | **335 (99.1%)** | **187 (55.3%)** | **3/0** | |
| Mutations: | 5822G>A, 5824A>G, 5828A>G, 5830A>T, 5834A>T, 5835A>C, 5836G>A, 5837G>A, 5838A>T, 5842G>A, 5846G>T, 5847T>G, 5848C>T, 5849T>A, 5850G>T, 5851C>T, 5852A>G, 5854C>T, 5857C>T, 5859A>G, 5860G>A, 5861A>C, 5862G>A, 5864C>T, 5866T>G, 5870G>A, 5872C>G, 5873A>G, 5874A>T, 5875C>G, 5879C>G, 5880A>T, 5881C>A, 5884A>G, 5885G>A, 5888C>A, 5894A>C, 5895G>C, 5899T>C, 5902A>C, 5903G>C, 5905G>T, 5908T>C, 5911C>T, 5913C>A, 5915A>T, 5917T>A, 5918G>T, 5920G>T, 5922C>A, 5923G>T, 5924A>C, 5925G>A, 5927A>T, 5930G>C, 5931G>A, 5933C>G, 5934A>G, 5935T>A, 5936A>G, 5937G>C, 5938C>A, 5939A>G, 5940A>C, 5941A>C, 5944C>A, 5946A>T, 5947T>C, 5950G>T, 5954T>A, 5956C>T, 5959C>T, 5964A>G, 5965G>A, 5967C>T, 5968A>C, 5971G>A, 5973T>A, 5980A>G, 5981G>T, 5983T>G, 5984G>A, 5985C>A, 5986T>G, 5989G>T, 5990G>C, 5996C>T, 5997A>C, 5998A>T, 5999C>G, 6001C>T, 6005G>C, 6006A>C, 6007A>T, 6008T>A, 6009G>A, 6013C>A, 6019T>C, 6020T>C, 6025T>C, 6026C>A, 6027C>G, 6028A>G, 6029G>T, 6030T>A, 6031G>T, 6035T>C, 6036T>A, 6037A>T, 6043A>G, 6045G>T, 6046G>T, 6049G>T, 6056C>T, 6058A>G, 6065C>T, 6067T>G, 6069A>C, 6070G>T, 6079A>G, 6080G>A, 6084T>C, 6089C>A, 6090A>T, 6091A>G, 6092A>G, 6093G>A, 6094G>A, 6095A>T, 6096A>T, 6097G>A, 6101G>A, 6103C>T, 6104C>G, 6106A>G, 6107A>G, 6109C>G, 6113A>C, 6118A>G, 6121_6122insTTG, 6123C>A, 6125G>T, 6126G>C, 6131A>G, 6133T>G, 6134G>A, 6135C>T, 6139G>A, 6141A>T | | | | | | | | |

##### CDS

| | Begin | End | Coverage | Score | Concordance | Matches | Identities | I/D/M/F* | Stop Codons |
|---|---|---|---|---|---|---|---|---|---|
| KM754_gp3 | 1534 | 1645 | 5.1% | 368 | 44.2% | 112 (99.1%) | 51 (45.1%) | 1/0/0/0 | 0 |
| Protein mutations: | E1534K (5822G>A 5824A>G), K1536D (5828A>G 5830A>T), K1538S (5834A>T 5835A>C 5836G>A), E1539M (5837G>A 5838A>T), V1542C (5846G>T 5847T>G 5848C>T), C1543I (5849T>A 5850G>T 5851C>T), N1544D (5852A>G 5854C>T), K1546R (5859A>G 5860G>A), R1547Q (5861A>C 5862G>A), D1550K (5870G>A 5872C>G), N1551V (5873A>G 5874A>T 5875C>G), H1553V (5879C>G 5880A>T 5881C>A), D1555N (5885G>A), Q1556K (5888C>A), S1558P (5894A>C 5895G>C), G1561R (5903G>C 5905G>T), T1564N (5913C>A), I1565L (5915A>T 5917T>A), V1566F (5918G>T 5920G>T), A1567D (5922C>A 5923G>T), R1568Q (5924A>C 5925G>A), I1569L (5927A>T), G1570Q (5930G>C 5931G>A), H1571G (5933C>G 5934A>G 5935T>A), S1572A (5936A>G 5937G>C 5938C>A), K1573A (5939A>G 5940A>C 5941A>C), Y1575F (5946A>T 5947T>C), F1578I (5954T>A 5956C>T), K1581R (5964A>G 5965G>A), S1582F (5967C>T 5968A>C), F1584Y (5973T>A), V1587L (5981G>T 5983T>G), A1588K (5984G>A 5985C>A 5986T>G), M1589I (5989G>T), D1590H (5990G>C), Q1592S (5996C>T 5997A>C 5998A>T), H1593D (5999C>G 6001C>T), E1595P (6005G>C 6006A>C 6007A>T), W1596K (6008T>A 6009G>A), W1600R (6020T>C), P1602R (6026C>A 6027C>G 6028A>G), V1603Y (6029G>T 6030T>A 6031G>T), L1605H (6035T>C 6036T>A 6037A>T), W1608F (6045G>T 6046G>T), P1612S (6056C>T 6058A>G), K1616T (6069A>C 6070G>T), A1620T (6080G>A), V1621A (6084T>C), Q1623M (6089C>A 6090A>T 6091A>G), R1624E (6092A>G 6093G>A 6094G>A), K1625L (6095A>T 6096A>T 6097G>A), D1627N (6101G>A 6103C>T), R1628G (6104C>G 6106A>G), I1629V (6107A>G 6109C>G), N1631H (6113A>C), Y1633_A1634insL (6121_6122insTTG), A1634D (6123C>A), G1635S (6125G>T 6126G>C), I1637V (6131A>G 6133T>G), A1638I (6134G>A 6135C>T), Y1640F (6141A>T) | | | | | | | | |
| Codon mutations: | GAA1534AAG (5822G>A 5824A>G), AAA1536GAT (5828A>G 5830A>T), AAG1538TCA (5834A>T 5835A>C 5836G>A), GAG1539ATG (5837G>A 5838A>T), AGG1540AGA (5842G>A), GTC1542TGT (5846G>T 5847T>G 5848C>T), TGC1543ATT (5849T>A 5850G>T 5851C>T), AAC1544GAT (5852A>G 5854C>T), TAC1545TAT (5857C>T), AAG1546AGA (5859A>G 5860G>A), AGG1547CAG (5861A>C 5862G>A), CTT1548TTG (5864C>T 5866T>G), GAC1550AAG (5870G>A 5872C>G), AAC1551GTG (5873A>G 5874A>T 5875C>G), CAC1553GTA (5879C>G 5880A>T 5881C>A), AAA1554AAG (5884A>G), GAC1555AAC (5885G>A), CAA1556AAA (5888C>A), AGC1558CCC (5894A>C 5895G>C), CTT1559CTC (5899T>C), CCA1560CCC (5902A>C), GGG1561CGT (5903G>C 5905G>T), ATT1562ATC (5908T>C), GAC1563GAT (5911C>T), ACT1564AAT (5913C>A), ATT1565TTA (5915A>T 5917T>A), GTG1566TTT (5918G>T 5920G>T), GCG1567GAT (5922C>A 5923G>T), AGG1568CAG (5924A>C 5925G>A), ATA1569TTA (5927A>T), GGG1570CAG (5930G>C 5931G>A), CAT1571GGA (5933C>G 5934A>G 5935T>A), AGC1572GCA (5936A>G 5937G>C 5938C>A), AAA1573GCC (5939A>G 5940A>C 5941A>C), ATC1574ATA (5944C>A), TAT1575TTC (5946A>T 5947T>C), TCG1576TCT (5950G>T), TTC1578ATT (5954T>A 5956C>T), GAC1579GAT (5959C>T), AAG1581AGA (5964A>G 5965G>A), TCA1582TTC (5967C>T 5968A>C), GGG1583GGA (5971G>A), TTT1584TAT (5973T>A), CAA1586CAG (5980A>G), GTT1587TTG (5981G>T 5983T>G), GCT1588AAG (5984G>A 5985C>A 5986T>G), ATG1589ATT (5989G>T), GAT1590CAT (5990G>C), CAA1592TCT (5996C>T 5997A>C 5998A>T), CAC1593GAT (5999C>G 6001C>T), GAA1595CCT (6005G>C 6006A>C 6007A>T), TGG1596AAG (6008T>A 6009G>A), ACC1597ACA (6013C>A), TTT1599TTC (6019T>C), TGG1600CGG (6020T>C), ACT1601ACC (6025T>C), CCA1602AGG (6026C>A 6027C>G 6028A>G), GTG1603TAT (6029G>T 6030T>A 6031G>T), TTA1605CAT (6035T>C 6036T>A 6037A>T), GAA1607GAG (6043A>G), TGG1608TTT (6045G>T 6046G>T), CTG1609CTT (6049G>T), CCA1612TCG (6056C>T 6058A>G), CTT1615TTG (6065C>T 6067T>G), AAG1616ACT (6069A>C 6070G>T), CCA1619CCG (6079A>G), GCA1620ACA (6080G>A), GTA1621GCA (6084T>C), CAA1623ATG (6089C>A 6090A>T 6091A>G), AGG1624GAA (6092A>G 6093G>A 6094G>A), AAG1625TTA (6095A>T 6096A>T 6097G>A), GAC1627AAT (6101G>A 6103C>T), CGA1628GGG (6104C>G 6106A>G), ATC1629GTG (6107A>G 6109C>G), AAT1631CAT (6113A>C), CCA1632CCG (6118A>G), TAC1633_GCT1634insTTG (6121_6122insTTG), GCT1634GAT (6123C>A), GGA1635TCA (6125G>T 6126G>C), ATT1637GTG (6131A>G 6133T>G), GCT1638ATT (6134G>A 6135C>T), GTG1639GTA (6139G>A), TAT1640TTT (6141A>T) | | | | | | | | |

##### Proteins

| | Begin | End | Coverage | Score | Concordance | Matches | Identities | I/D/M/F* | Stop Codons |
|---|---|---|---|---|---|---|---|---|---|
| polyprotein (YP_010087856.1) | 1534 | 1645 | 5.1% | 368 | 44.2% | 112 (99.1%) | 51 (45.1%) | 1/0/0/0 | 0 |
| Protein mutations: | E1534K (5822G>A 5824A>G), K1536D (5828A>G 5830A>T), K1538S (5834A>T 5835A>C 5836G>A), E1539M (5837G>A 5838A>T), V1542C (5846G>T 5847T>G 5848C>T), C1543I (5849T>A 5850G>T 5851C>T), N1544D (5852A>G 5854C>T), K1546R (5859A>G 5860G>A), R1547Q (5861A>C 5862G>A), D1550K (5870G>A 5872C>G), N1551V (5873A>G 5874A>T 5875C>G), H1553V (5879C>G 5880A>T 5881C>A), D1555N (5885G>A), Q1556K (5888C>A), S1558P (5894A>C 5895G>C), G1561R (5903G>C 5905G>T), T1564N (5913C>A), I1565L (5915A>T 5917T>A), V1566F (5918G>T 5920G>T), A1567D (5922C>A 5923G>T), R1568Q (5924A>C 5925G>A), I1569L (5927A>T), G1570Q (5930G>C 5931G>A), H1571G (5933C>G 5934A>G 5935T>A), S1572A (5936A>G 5937G>C 5938C>A), K1573A (5939A>G 5940A>C 5941A>C), Y1575F (5946A>T 5947T>C), F1578I (5954T>A 5956C>T), K1581R (5964A>G 5965G>A), S1582F (5967C>T 5968A>C), F1584Y (5973T>A), V1587L (5981G>T 5983T>G), A1588K (5984G>A 5985C>A 5986T>G), M1589I (5989G>T), D1590H (5990G>C), Q1592S (5996C>T 5997A>C 5998A>T), H1593D (5999C>G 6001C>T), E1595P (6005G>C 6006A>C 6007A>T), W1596K (6008T>A 6009G>A), W1600R (6020T>C), P1602R (6026C>A 6027C>G 6028A>G), V1603Y (6029G>T 6030T>A 6031G>T), L1605H (6035T>C 6036T>A 6037A>T), W1608F (6045G>T 6046G>T), P1612S (6056C>T 6058A>G), K1616T (6069A>C 6070G>T), A1620T (6080G>A), V1621A (6084T>C), Q1623M (6089C>A 6090A>T 6091A>G), R1624E (6092A>G 6093G>A 6094G>A), K1625L (6095A>T 6096A>T 6097G>A), D1627N (6101G>A 6103C>T), R1628G (6104C>G 6106A>G), I1629V (6107A>G 6109C>G), N1631H (6113A>C), Y1633_A1634insL (6121_6122insTTG), A1634D (6123C>A), G1635S (6125G>T 6126G>C), I1637V (6131A>G 6133T>G), A1638I (6134G>A 6135C>T), Y1640F (6141A>T) | | | | | | | | |
| Codon mutations: | GAA1534AAG (5822G>A 5824A>G), AAA1536GAT (5828A>G 5830A>T), AAG1538TCA (5834A>T 5835A>C 5836G>A), GAG1539ATG (5837G>A 5838A>T), AGG1540AGA (5842G>A), GTC1542TGT (5846G>T 5847T>G 5848C>T), TGC1543ATT (5849T>A 5850G>T 5851C>T), AAC1544GAT (5852A>G 5854C>T), TAC1545TAT (5857C>T), AAG1546AGA (5859A>G 5860G>A), AGG1547CAG (5861A>C 5862G>A), CTT1548TTG (5864C>T 5866T>G), GAC1550AAG (5870G>A 5872C>G), AAC1551GTG (5873A>G 5874A>T 5875C>G), CAC1553GTA (5879C>G 5880A>T 5881C>A), AAA1554AAG (5884A>G), GAC1555AAC (5885G>A), CAA1556AAA (5888C>A), AGC1558CCC (5894A>C 5895G>C), CTT1559CTC (5899T>C), CCA1560CCC (5902A>C), GGG1561CGT (5903G>C 5905G>T), ATT1562ATC (5908T>C), GAC1563GAT (5911C>T), ACT1564AAT (5913C>A), ATT1565TTA (5915A>T 5917T>A), GTG1566TTT (5918G>T 5920G>T), GCG1567GAT (5922C>A 5923G>T), AGG1568CAG (5924A>C 5925G>A), ATA1569TTA (5927A>T), GGG1570CAG (5930G>C 5931G>A), CAT1571GGA (5933C>G 5934A>G 5935T>A), AGC1572GCA (5936A>G 5937G>C 5938C>A), AAA1573GCC (5939A>G 5940A>C 5941A>C), ATC1574ATA (5944C>A), TAT1575TTC (5946A>T 5947T>C), TCG1576TCT (5950G>T), TTC1578ATT (5954T>A 5956C>T), GAC1579GAT (5959C>T), AAG1581AGA (5964A>G 5965G>A), TCA1582TTC (5967C>T 5968A>C), GGG1583GGA (5971G>A), TTT1584TAT (5973T>A), CAA1586CAG (5980A>G), GTT1587TTG (5981G>T 5983T>G), GCT1588AAG (5984G>A 5985C>A 5986T>G), ATG1589ATT (5989G>T), GAT1590CAT (5990G>C), CAA1592TCT (5996C>T 5997A>C 5998A>T), CAC1593GAT (5999C>G 6001C>T), GAA1595CCT (6005G>C 6006A>C 6007A>T), TGG1596AAG (6008T>A 6009G>A), ACC1597ACA (6013C>A), TTT1599TTC (6019T>C), TGG1600CGG (6020T>C), ACT1601ACC (6025T>C), CCA1602AGG (6026C>A 6027C>G 6028A>G), GTG1603TAT (6029G>T 6030T>A 6031G>T), TTA1605CAT (6035T>C 6036T>A 6037A>T), GAA1607GAG (6043A>G), TGG1608TTT (6045G>T 6046G>T), CTG1609CTT (6049G>T), CCA1612TCG (6056C>T 6058A>G), CTT1615TTG (6065C>T 6067T>G), AAG1616ACT (6069A>C 6070G>T), CCA1619CCG (6079A>G), GCA1620ACA (6080G>A), GTA1621GCA (6084T>C), CAA1623ATG (6089C>A 6090A>T 6091A>G), AGG1624GAA (6092A>G 6093G>A 6094G>A), AAG1625TTA (6095A>T 6096A>T 6097G>A), GAC1627AAT (6101G>A 6103C>T), CGA1628GGG (6104C>G 6106A>G), ATC1629GTG (6107A>G 6109C>G), AAT1631CAT (6113A>C), CCA1632CCG (6118A>G), TAC1633_GCT1634insTTG (6121_6122insTTG), GCT1634GAT (6123C>A), GGA1635TCA (6125G>T 6126G>C), ATT1637GTG (6131A>G 6133T>G), GCT1638ATT (6134G>A 6135C>T), GTG1639GTA (6139G>A), TAT1640TTT (6141A>T) | | | | | | | | |

*: Inserts / Deletes / Misaligned / Frameshifts

#### Analysis details

This analysis was performed with panviral2.64

### NGS Details (UN9): Fig cryptic virus (segment RNA 2)

#### Assembly

| | |
|---|---|
| **Coverage Length** | 164 (1 contig(s)) |
| **Depth Of Coverage** | 3.1 |
| **Number Of Reads** | 6 |
| **Reads Per Million** | 0.14 rpm (after QC) |
| **Ambiguities** | 0 |
| **Assembly Method** | de novo + reference guided assembly |
| **Consensus Caller** | Bcf Tools |

#### Coverage Map

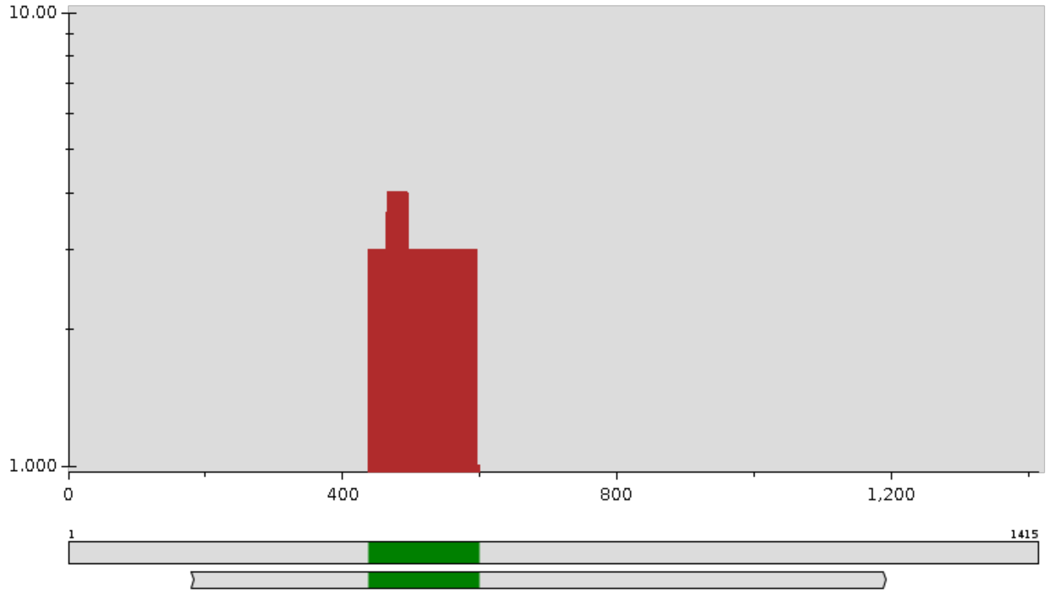

#### Assignment

| | |
|---|---|
| **Type** | Fig cryptic virus (Taxonomy ID: 882768) |
| **Reference Genome** | NC_015495.1 |
| **NT Identity (%)** | 63.4146 |
| **AA Identity (%)** | 61.8182 |
| **Number Of Stop Codons** | 0 |
| **Number Of CDS** | 1 |

#### Alignment

| | |
|---|---|
| **Alignment Score** | 88.0 (NT) + 226.0 (AA) = 314.0 |
| **Concordance (%)** | 43.6718 |

| Alignment Method | Global, seeded, nucleotide + amino acids (AGA) |
|---|---|

#### Genome Region

Sequence starts at position 438 and ends at position 601 relative to NC_015495.1 reference sequence.

#### Alignment Detailed Statistics

| | Begin | End | Coverage | Score | Concordance | Matches | Identities | I/D/M/F* | Stop Codons |
|---|---|---|---|---|---|---|---|---|---|
| **NT** | **438** | **601** | **11.6%** | **88** | **26.8%** | **164 (100%)** | **104 (63.4%)** | **0/0** | |
| Mutations: | 438A>T, 441T>C, 444T>C, 447C>T, 450A>G, 456A>C, 457T>A, 460G>A, 468A>G, 469T>C, 471C>T, 475C>T, 476A>C, 477C>T, 479A>C, 480C>A, 481C>A, 483A>C, 486G>C, 487C>A, 489T>G, 493T>A, 495G>T, 498A>C, 504C>T, 505A>C, 507A>T, 516A>T, 517A>T, 519T>G, 520T>A, 521C>G, 522T>C, 529A>C, 538C>T, 540A>G, 541A>G, 542G>T, 547C>A, 549T>C, 555T>G, 557T>C, 558G>T, 561T>C, 562G>T, 563A>C, 564A>T, 567T>A, 571A>G, 574T>C, 575G>C, 576G>A, 580C>A, 581G>T, 582T>C, 585T>C, 588C>T, 590A>G, 591C>T, 592G>A | | | | | | | | |

##### CDS

| | Begin | End | Coverage | Score | Concordance | Matches | Identities | I/D/M/F* | Stop Codons |
|---|---|---|---|---|---|---|---|---|---|
| CP | 87 | 141 | 16.3% | 226 | 55.3% | 55 (100%) | 34 (61.8%) | 0/0/0/0 | 0 |
| Protein mutations: | S93T (457T>A), V94I (460G>A), C97R (469T>C 471C>T), H99S (475C>T 476A>C 477C>T), H100P (479A>C 480C>A), P101T (481C>A 483A>C), S105T (493T>A 495G>T), T109P (505A>C 507A>T), I113L (517A>T 519T>G), M117L (529A>C), P120S (538C>T 540A>G), S121V (541A>G 542G>T), L123I (547C>A 549T>C), M126T (557T>C 558G>T), E128S (562G>T 563A>C 564A>T), S131G (571A>G), W132P (574T>C 575G>C 576G>A), R134I (580C>A 581G>T 582T>C), D137G (590A>G 591C>T), G138R (592G>A) | | | | | | | | |
| Codon mutations: | CAA86..T (438A>T), CTT87CTC (441T>C), TAT88TAC (444T>C), AAC89AAT (447C>T), AGA90AGG (450A>G), CGA92CGC (456A>C), TCT93ACT (457T>A), GTT94ATT (460G>A), GAA96GAG (468A>G), TGC97CGT (469T>C 471C>T), CAC99TCT (475C>T 476A>C 477C>T), CAC100CCA (479A>C 480C>A), CCA101ACC (481C>A 483A>C), ACG102ACC (486G>C), CGT103AGG (487C>A 489T>G), TCG105ACT (493T>A 495G>T), CGA106CGC (498A>C), AAC108AAT (504C>T), ACA109CCT (505A>C 507A>T), CCA112CCT (516A>T), ATT113TTG (517A>T 519T>G), TCT114AGC (520T>A 521C>G 522T>C), ATG117CTG (529A>C), CCA120TCG (538C>T 540A>G), AGC121GTC (541A>G 542G>T), CTT123ATC (547C>A 549T>C), TCT125TCG (555T>G), ATG126ACT (557T>C 558G>T), CTT127CTC (561T>C), GAA128TCT (562G>T 563A>C 564A>T), TCT129TCA (567T>A), AGC131GGC (571A>G), TGG132CCA (574T>C 575G>C 576G>A), CGT134ATC (580C>A 581G>T 582T>C), ATT135ATC (585T>C), AAC136AAT (588C>T), GAC137GGT (590A>G 591C>T), GGA138AGA (592G>A) | | | | | | | | |

##### Proteins

| | Begin | End | Coverage | Score | Concordance | Matches | Identities | I/D/M/F* | Stop Codons |
|---|---|---|---|---|---|---|---|---|---|
| putative capsid protein (YP_004429259.1) | 87 | 141 | 16.3% | 226 | 55.3% | 55 (100%) | 34 (61.8%) | 0/0/0/0 | 0 |
| Protein mutations: | S93T (457T>A), V94I (460G>A), C97R (469T>C 471C>T), H99S (475C>T 476A>C 477C>T), H100P (479A>C 480C>A), P101T (481C>A 483A>C), S105T (493T>A 495G>T), T109P (505A>C 507A>T), I113L (517A>T 519T>G), M117L (529A>C), P120S (538C>T 540A>G), S121V (541A>G 542G>T), L123I (547C>A 549T>C), M126T (557T>C 558G>T), E128S (562G>T 563A>C 564A>T), S131G (571A>G), W132P (574T>C 575G>C 576G>A), R134I (580C>A 581G>T 582T>C), D137G (590A>G 591C>T), G138R (592G>A) | | | | | | | | |
| Codon mutations: | CAA86..T (438A>T), CTT87CTC (441T>C), TAT88TAC (444T>C), AAC89AAT (447C>T), AGA90AGG (450A>G), CGA92CGC (456A>C), TCT93ACT (457T>A), GTT94ATT (460G>A), GAA96GAG (468A>G), TGC97CGT (469T>C 471C>T), CAC99TCT (475C>T 476A>C 477C>T), CAC100CCA (479A>C 480C>A), CCA101ACC (481C>A 483A>C), ACG102ACC (486G>C), CGT103AGG (487C>A 489T>G), TCG105ACT (493T>A 495G>T), CGA106CGC (498A>C), AAC108AAT (504C>T), ACA109CCT (505A>C 507A>T), CCA112CCT (516A>T), ATT113TTG (517A>T 519T>G), TCT114AGC (520T>A 521C>G 522T>C), ATG117CTG (529A>C), CCA120TCG (538C>T 540A>G), AGC121GTC (541A>G 542G>T), CTT123ATC (547C>A 549T>C), TCT125TCG (555T>G), ATG126ACT (557T>C 558G>T), CTT127CTC (561T>C), GAA128TCT (562G>T 563A>C 564A>T), TCT129TCA (567T>A), AGC131GGC (571A>G), TGG132CCA (574T>C 575G>C 576G>A), CGT134ATC (580C>A 581G>T 582T>C), ATT135ATC (585T>C), AAC136AAT (588C>T), GAC137GGT (590A>G 591C>T), GGA138AGA (592G>A) | | | | | | | | |

*: Inserts / Deletes / Misaligned / Frameshifts

#### Analysis details

This analysis was performed with panviral2.64

### NGS Details (UN9): Caulimovirus venafragariae

#### Assembly

| | |
|---|---|
| **Coverage Length** | 243 (1 contig(s)) |
| **Depth Of Coverage** | 2.8 |
| **Number Of Reads** | 6 |
| **Reads Per Million** | 0.14 rpm (after QC) |
| **Ambiguities** | 0 |
| **Assembly Method** | de novo + reference guided assembly |
| **Consensus Caller** | Bcf Tools |

#### Coverage Map

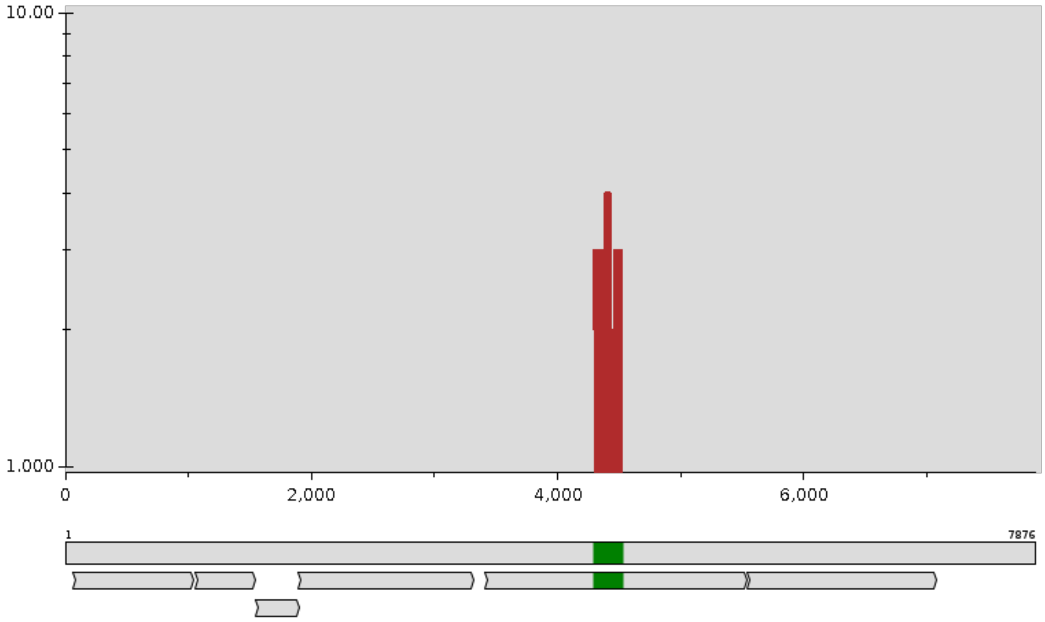

#### Assignment

| | |
|---|---|
| **Type** | Caulimovirus venafragariae (Taxonomy ID: 3048344) |
| **Reference Genome** | NC_001725.1 |
| **NT Identity (%)** | 61.3169 |
| **AA Identity (%)** | 62.963 |
| **Number Of Stop Codons** | 0 |
| **Number Of CDS** | 6 |

#### Alignment

| | |
|---|---|
| **Alignment Score** | 110.0 (NT) + 359.0 (AA) = 469.0 |
| **Concordance (%)** | 45.1396 |

| Alignment Method | Global, seeded, nucleotide + amino acids (AGA) |
|---|---|

#### Genome Region

Sequence starts at position 4294 and ends at position 4536 relative to NC_001725.1 reference sequence.

#### Alignment Detailed Statistics

| | Begin | End | Coverage | Score | Concordance | Matches | Identities | I/D/M/F* | Stop Codons |
|---|---|---|---|---|---|---|---|---|---|
| **NT** | **4294** | **4536** | **3.1%** | **110** | **22.6%** | **243 (100%)** | **149 (61.3%)** | **0/0** | |
| Mutations: | 4296G>A, 4299C>T, 4300G>A, 4311C>A, 4314G>C, 4315C>T, 4317T>A, 4320T>A, 4321A>G, 4323C>T, 4326C>A, 4330C>A, 4331C>G, 4332T>A, 4333A>T, 4334G>C, 4335C>T, 4337A>C, 4338G>A, 4339A>T, 4340G>C, 4341T>A, 4342C>A, 4343C>A, 4348T>A, 4349C>G, 4350C>A, 4354C>G, 4362C>T, 4371G>A, 4378G>A, 4379C>G, 4383G>A, 4386C>A, 4387A>G, 4388A>T, 4390C>A, 4395C>T, 4402A>C, 4403G>A, 4408G>A, 4413T>A, 4419T>C, 4424A>G, 4425G>A, 4426T>C, 4428A>T, 4434C>T, 4435C>A, 4437T>C, 4440A>T, 4442A>G, 4443G>A, 4444G>A, 4445G>C, 4446A>G, 4447G>T, 4451G>C, 4452C>T, 4455C>T, 4456C>A, 4457T>A, 4458A>G, 4461T>A, 4464T>A, 4465A>G, 4467C>T, 4470G>A, 4471G>A, 4472A>G, 4473G>T, 4474C>G, 4477C>T, 4479G>A, 4480C>A, 4482T>A, 4483C>A, 4485A>T, 4491C>A, 4492G>C, 4493G>A, 4503C>A, 4504T>G, 4506T>A, 4508A>T, 4510T>A, 4511C>A, 4513T>A, 4514C>A, 4515C>A, 4521C>T, 4530T>A, 4533A>G, 4535T>A | | | | | | | | |

##### CDS

| | Begin | End | Coverage | Score | Concordance | Matches | Identities | I/D/M/F* | Stop Codons |
|---|---|---|---|---|---|---|---|---|---|
| ORF_V | 297 | 377 | 11.4% | 359 | 64.9% | 81 (100%) | 51 (63.0%) | 0/0/0/0 | 0 |
| Protein mutations: | E299K (4300G>A), K303N (4314G>C), I306V (4321A>G 4323C>T), P309R (4330C>A 4331C>G 4332T>A), K311T (4337A>C 4338G>A), P313K (4342C>A 4343C>A), S315R (4348T>A 4349C>G 4350C>A), P317A (4354C>G), A325S (4378G>A 4379C>G), K328V (4387A>G 4388A>T), R333Q (4402A>C 4403G>A), V335I (4408G>A), K340R (4424A>G 4425G>A), H344N (4435C>A 4437T>C), K346R (4442A>G 4443G>A), G347T (4444G>A 4445G>C 4446A>G), D348Y (4447G>T), G349A (4451G>C 4452C>T), L351K (4456C>A 4457T>A 4458A>G), N354D (4465A>G 4467C>T), E356S (4471G>A 4472A>G 4473G>T), Q357E (4474C>G), L359I (4480C>A 4482T>A), Q360N (4483C>A 4485A>T), G363Q (4492G>C 4493G>A), F367V (4504T>G 4506T>A), Y368F (4508A>T), S369N (4510T>A 4511C>A), S370K (4513T>A 4514C>A 4515C>A), F377Y (4535T>A) | | | | | | | | |
| Codon mutations: | CAG297CAA (4296G>A), ATC298ATT (4299C>T), GAA299AAA (4300G>A), CTC302CTA (4311C>A), AAG303AAC (4314G>C), CTT304TTA (4315C>T 4317T>A), GGT305GGA (4320T>A), ATC306GTT (4321A>G 4323C>T), ATC307ATA (4326C>A), CCT309AGA (4330C>A 4331C>G 4332T>A), AGC310TCT (4333A>T 4334G>C 4335C>T), AAG311ACA (4337A>C 4338G>A), AGT312TCA (4339A>T 4340G>C 4341T>A), CCA313AAA (4342C>A 4343C>A), TCC315AGA (4348T>A 4349C>G 4350C>A), CCA317GCA (4354C>G), TTC319TTT (4362C>T), AGG322AGA (4371G>A), GCC325AGC (4378G>A 4379C>G), GAG326GAA (4383G>A), ATC327ATA (4386C>A), AAA328GTA (4387A>G 4388A>T), CGA329AGA (4390C>A), GGC330GGT (4395C>T), AGA333CAA (4402A>C 4403G>A), GTA335ATA (4408G>A), ATT336ATA (4413T>A), TAT338TAC (4419T>C), AAG340AGA (4424A>G 4425G>A), TTA341CTT (4426T>C 4428A>T), GAC343GAT (4434C>T), CAT344AAC (4435C>A 4437T>C), ACA345ACT (4440A>T), AAG346AGA (4442A>G 4443G>A), GGA347ACG (4444G>A 4445G>C 4446A>G), GAT348TAT (4447G>T), GGC349GCT (4451G>C 4452C>T), TAC350TAT (4455C>T), CTA351AAG (4456C>A 4457T>A 4458A>G), CTT352CTA (4461T>A), CCT353CCA (4464T>A), AAC354GAT (4465A>G 4467C>T), AAG355AAA (4470G>A), GAG356AGT (4471G>A 4472A>G 4473G>T), CAA357GAA (4474C>G), CTG358TTA (4477C>T 4479G>A), CTT359ATA (4480C>A 4482T>A), CAA360AAT (4483C>A 4485A>T), ATC362ATA (4491C>A), GGA363CAA (4492G>C 4493G>A), ACC366ACA (4503C>A), TTT367GTA (4504T>G 4506T>A), TAC368TTC (4508A>T), TCT369AAT (4510T>A 4511C>A), TCC370AAA (4513T>A 4514C>A 4515C>A), GAC372GAT (4521C>T), TCT375TCA (4530T>A), GGA376GGG (4533A>G), TTT377TAT (4535T>A) | | | | | | | | |

##### Proteins

| | Begin | End | Coverage | Score | Concordance | Matches | Identities | I/D/M/F* | Stop Codons |
|---|---|---|---|---|---|---|---|---|---|
| hypothetical protein (NP_043933.1) | 297 | 377 | 11.4% | 359 | 64.9% | 81 (100%) | 51 (63.0%) | 0/0/0/0 | 0 |
| Protein mutations: | E299K (4300G>A), K303N (4314G>C), I306V (4321A>G 4323C>T), P309R (4330C>A 4331C>G 4332T>A), K311T (4337A>C 4338G>A), P313K (4342C>A 4343C>A), S315R (4348T>A 4349C>G 4350C>A), P317A (4354C>G), A325S (4378G>A 4379C>G), K328V (4387A>G 4388A>T), R333Q (4402A>C 4403G>A), V335I (4408G>A), K340R (4424A>G 4425G>A), H344N (4435C>A 4437T>C), K346R (4442A>G 4443G>A), G347T (4444G>A 4445G>C 4446A>G), D348Y (4447G>T), G349A (4451G>C 4452C>T), L351K (4456C>A 4457T>A 4458A>G), N354D (4465A>G 4467C>T), E356S (4471G>A 4472A>G 4473G>T), Q357E (4474C>G), L359I (4480C>A 4482T>A), Q360N (4483C>A 4485A>T), G363Q (4492G>C 4493G>A), F367V (4504T>G 4506T>A), Y368F (4508A>T), S369N (4510T>A 4511C>A), S370K (4513T>A 4514C>A 4515C>A), F377Y (4535T>A) | | | | | | | | |
| Codon mutations: | CAG297CAA (4296G>A), ATC298ATT (4299C>T), GAA299AAA (4300G>A), CTC302CTA (4311C>A), AAG303AAC (4314G>C), CTT304TTA (4315C>T 4317T>A), GGT305GGA (4320T>A), ATC306GTT (4321A>G 4323C>T), ATC307ATA (4326C>A), CCT309AGA (4330C>A 4331C>G 4332T>A), AGC310TCT (4333A>T 4334G>C 4335C>T), AAG311ACA (4337A>C 4338G>A), AGT312TCA (4339A>T 4340G>C 4341T>A), CCA313AAA (4342C>A 4343C>A), TCC315AGA (4348T>A 4349C>G 4350C>A), CCA317GCA (4354C>G), TTC319TTT (4362C>T), AGG322AGA (4371G>A), GCC325AGC (4378G>A 4379C>G), GAG326GAA (4383G>A), ATC327ATA (4386C>A), AAA328GTA (4387A>G 4388A>T), CGA329AGA (4390C>A), GGC330GGT (4395C>T), AGA333CAA (4402A>C 4403G>A), GTA335ATA (4408G>A), ATT336ATA (4413T>A), TAT338TAC (4419T>C), AAG340AGA (4424A>G 4425G>A), TTA341CTT (4426T>C 4428A>T), GAC343GAT (4434C>T), CAT344AAC (4435C>A 4437T>C), ACA345ACT (4440A>T), AAG346AGA (4442A>G 4443G>A), GGA347ACG (4444G>A 4445G>C 4446A>G), GAT348TAT (4447G>T), GGC349GCT (4451G>C 4452C>T), TAC350TAT (4455C>T), CTA351AAG (4456C>A 4457T>A 4458A>G), CTT352CTA (4461T>A), CCT353CCA (4464T>A), AAC354GAT (4465A>G 4467C>T), AAG355AAA (4470G>A), GAG356AGT (4471G>A 4472A>G 4473G>T), CAA357GAA (4474C>G), CTG358TTA (4477C>T 4479G>A), CTT359ATA (4480C>A 4482T>A), CAA360AAT (4483C>A 4485A>T), ATC362ATA (4491C>A), GGA363CAA (4492G>C 4493G>A), ACC366ACA (4503C>A), TTT367GTA (4504T>G 4506T>A), TAC368TTC (4508A>T), TCT369AAT (4510T>A 4511C>A), TCC370AAA (4513T>A 4514C>A 4515C>A), GAC372GAT (4521C>T), TCT375TCA (4530T>A), GGA376GGG (4533A>G), TTT377TAT (4535T>A) | | | | | | | | |

*: Inserts / Deletes / Misaligned / Frameshifts

#### Analysis details

This analysis was performed with panviral2.64

### NGS Details (UN9): Tomato associated geminivirus 1

#### Assembly

| | |
|---|---|
| **Coverage Length** | 274 (2 contig(s)) |
| **Depth Of Coverage** | 2.5 |
| **Number Of Reads** | 6 |
| **Reads Per Million** | 0.14 rpm (after QC) |
| **Ambiguities** | 0 |
| **Assembly Method** | de novo + reference guided assembly |
| **Consensus Caller** | Bcf Tools |

#### Coverage Map

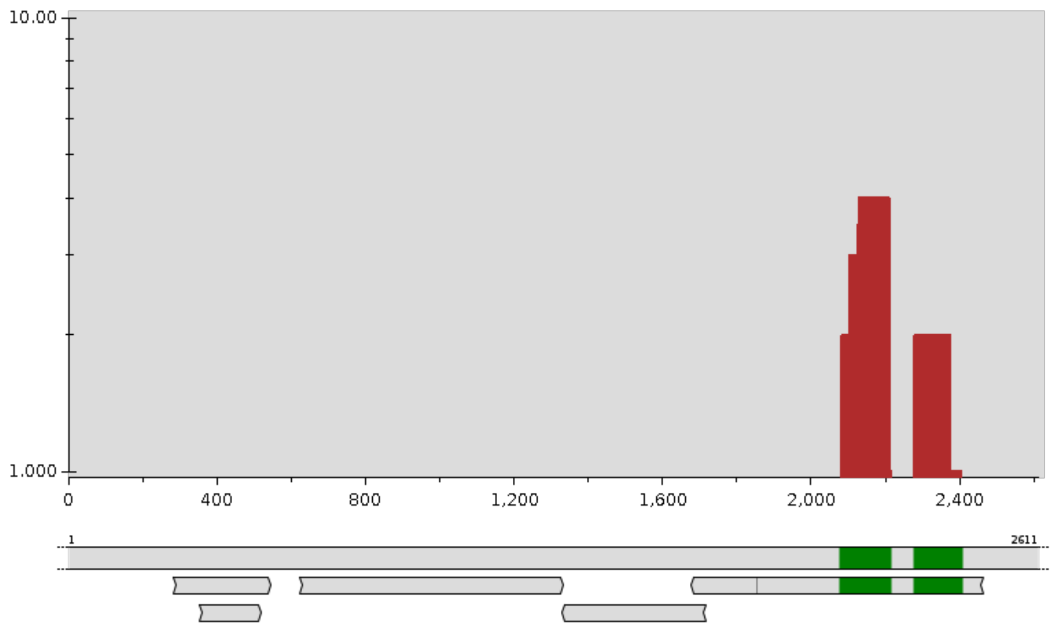

#### Assignment

| | |
|---|---|
| **Type** | Tomato associated geminivirus 1 (Taxonomy ID: 2027645) |
| **Reference Genome** | NC_035676.1 |
| **NT Identity (%)** | 67.5182 |
| **AA Identity (%)** | 61.9565 |
| **Number Of Stop Codons** | 0 |
| **Number Of CDS** | 5 |

#### Alignment

| | |
|---|---|
| **Alignment Score** | 192.0 (NT) + 976.0 (AA) = 1168.0 |
| **Concordance (%)** | 60.5181 |

| **Alignment Method** | Global, seeded, nucleotide + amino acids (AGA) |
|---|---|

#### Genome Region

Sequence starts at position 2078 and ends at position 2408 relative to NC_035676.1 reference sequence.

#### Alignment Detailed Statistics

| | Begin | End | Coverage | Score | Concordance | Matches | Identities | I/D/M/F* | Stop Codons |
|---|---|---|---|---|---|---|---|---|---|
| **NT** | **2078** | **2408** | **10.5%** | **192** | **35.0%** | **274 (100%)** | **185 (67.5%)** | **0/0** | |
| Mutations: | 2082T>C, 2086T>G, 2090T>G, 2091C>T, 2094C>A, 2099T>C, 2102T>G, 2104C>T, 2105G>T, 2106G>T, 2108A>T, 2110G>T, 2111C>T, 2112T>C, 2113C>T, 2117G>A, 2118A>T, 2120T>C, 2121A>T, 2124T>A, 2127C>T, 2136G>T, 2137G>A, 2139A>T, 2142C>T, 2148G>T, 2151A>G, 2154G>A, 2157G>A, 2160C>G, 2162C>A, 2163A>G, 2166A>G, 2168C>G, 2170G>T, 2171C>T, 2172A>G, 2176C>G, 2177T>A, 2181T>C, 2184G>T, 2190G>A, 2191A>G, 2196T>G, 2199A>G, 2203T>A, 2205A>T, 2210C>T, 2280C>T, 2283A>G, 2289G>A, 2291T>G, 2292G>A, 2298C>T, 2301T>C, 2313G>T, 2314T>G, 2319C>T, 2322A>G, 2323C>G, 2324A>C, 2326G>A, 2333C>T, 2334G>A, 2337T>A, 2338G>T, 2340G>A, 2341A>G, 2342C>G, 2346A>G, 2354T>G, 2358T>A, 2360A>T, 2363G>T, 2364A>T, 2365A>T, 2366A>T, 2367A>G, 2369A>G, 2371T>A, 2373A>G, 2376G>T, 2379A>G, 2381T>C, 2385A>T, 2390G>C, 2391A>G, 2400A>G, 2406T>G | | | | | | | | |
| **CDS** | | | | | | | | | |
| CK833_gp4 | 20 | 130 | 27.6% | 488 | 68.6% | 92 (100%) | 57 (62.0%) | 0/0/0/0 | 0 |
| Protein mutations: | P26A (2390G>C), I29V (2379A>G 2381T>C), Y32F (2371T>A), F33L (2367A>G 2369A>G), F34K (2364A>T 2365A>T 2366A>T), Q35K (2363G>T), L36I (2358T>A 2360A>T), K38Q (2354T>G), V42P (2340G>A 2341A>G 2342C>G), T43N (2337T>A 2338G>T), V45I (2333C>T), A47V (2326G>A), C48A (2322A>G 2323C>G 2324A>C), N51T (2313G>T 2314T>G), I59L (2289G>A 2291T>G), G86R (2210C>T), Y88F (2203T>A), I92T (2190G>A 2191A>G), R97S (2176C>G 2177T>A), A99K (2170G>T 2171C>T), D100H (2166A>G 2168C>G), A102S (2160C>G 2162C>A), T110I (2136G>T 2137G>A), R114S (2124T>A), I116V (2118A>T 2120T>C), L117F (2117G>A), R118K (2112T>C 2113C>T), A119N (2110G>T 2111C>T), S120T (2106G>T 2108A>T), R121K (2104C>T 2105G>T), S123G (2099T>C), K126Q (2090T>G), N127T (2086T>G) | | | | | | | | |
| Codon mutations: | CCA20CCC (2406T>G), TGT22TGC (2400A>G), ATT25ATC (2391A>G), CCA26GCA (2390G>C), ATT27ATA (2385A>T), ATT29GTC (2379A>G 2381T>C), ATC30ATA (2376G>T), GAT31GAC (2373A>G), TAC32TTC (2371T>A), TTT33CTC (2367A>G 2369A>G), TTT34AAA (2364A>T 2365A>T 2366A>T), CAA35AAA (2363G>T), TTA36ATT (2358T>A 2360A>T), AAA38CAA (2354T>G), TAT40TAC (2346A>G), GTC42CCT (2340G>A 2341A>G 2342C>G), ACA43AAT (2337T>A 2338G>T), TAC44TAT (2334G>A), GTT45ATT (2333C>T), GCT47GTT (2326G>A), TGT48GCC (2322A>G 2323C>G 2324A>C), ACG49ACA (2319C>T), AAC51ACA (2313G>T 2314T>G), GGA55GGG (2301T>C), GAG56GAA (2298C>T), CAC58CAT (2292G>A), ATC59CTT (2289G>A 2291T>G), TGT61TGC (2283A>G), CTG62CTA (2280C>T), GGA86AGA (2210C>T), ATT87ATA (2205A>T), TAT88TTT (2203T>A), CAT89CAC (2199A>G), CCA90CCC (2196T>G), ATC92ACT (2190G>A 2191A>G), ATC94ATA (2184G>T), CCA95CCG (2181T>C), AGG97TCG (2176C>G 2177T>A), GAT98GAC (2172A>G), GCA99AAA (2170G>T 2171C>T), GAT100CAC (2166A>G 2168C>G), GTT101GTC (2163A>G), GCG102TCC (2160C>G 2162C>A), GAC103GAT (2157G>A), TAC104TAT (2154G>A), ATT105ATC (2151A>G), TCC106TCA (2148G>T), GGG108GGA (2142C>T), GGT109GGA (2139A>T), ACC110ATA (2136G>T 2137G>A), GAG113GAA (2127C>T), AGA114AGT (2124T>A), GGT115GGA (2121A>T), ATT116GTA (2118A>T 2120T>C), CTT117TTT (2117G>A), AGA118AAG (2112T>C 2113C>T), GCT119AAT (2110G>T 2111C>T), TCC120ACA (2106G>T 2108A>T), CGA121AAA (2104C>T 2105G>T), AGA122CGA (2102T>G), AGT123GGT (2099T>C), CCG124CCT (2094C>A), AAG125AAA (2091C>T), AAG126CAG (2090T>G), AAT127ACT (2086T>G), AGA128AGG (2082T>C) | | | | | | | | |
| CK833_gp5 | 20 | 130 | 35.1% | 488 | 68.6% | 92 (100%) | 57 (62.0%) | 0/0/0/0 | 0 |
| Protein mutations: | P26A (2390G>C), I29V (2379A>G 2381T>C), Y32F (2371T>A), F33L (2367A>G 2369A>G), F34K (2364A>T 2365A>T 2366A>T), Q35K (2363G>T), L36I (2358T>A 2360A>T), K38Q (2354T>G), V42P (2340G>A 2341A>G 2342C>G), T43N (2337T>A 2338G>T), V45I (2333C>T), A47V (2326G>A), C48A (2322A>G 2323C>G 2324A>C), N51T (2313G>T 2314T>G), I59L (2289G>A 2291T>G), G86R (2210C>T), Y88F (2203T>A), I92T (2190G>A 2191A>G), R97S (2176C>G 2177T>A), A99K (2170G>T 2171C>T), D100H (2166A>G 2168C>G), A102S (2160C>G 2162C>A), T110I (2136G>T 2137G>A), R114S (2124T>A), I116V (2118A>T 2120T>C), L117F (2117G>A), R118K (2112T>C 2113C>T), A119N (2110G>T 2111C>T), S120T (2106G>T 2108A>T), R121K (2104C>T 2105G>T), S123G (2099T>C), K126Q (2090T>G), N127T (2086T>G) | | | | | | | | |
| Codon mutations: | CCA20CCC (2406T>G), TGT22TGC (2400A>G), ATT25ATC (2391A>G), CCA26GCA (2390G>C), ATT27ATA (2385A>T), ATT29GTC (2379A>G 2381T>C), ATC30ATA (2376G>T), GAT31GAC (2373A>G), TAC32TTC (2371T>A), TTT33CTC (2367A>G 2369A>G), TTT34AAA (2364A>T 2365A>T 2366A>T), CAA35AAA (2363G>T), TTA36ATT (2358T>A 2360A>T), AAA38CAA (2354T>G), TAT40TAC (2346A>G), GTC42CCT (2340G>A 2341A>G 2342C>G), ACA43AAT (2337T>A 2338G>T), TAC44TAT (2334G>A), GTT45ATT (2333C>T), GCT47GTT (2326G>A), TGT48GCC (2322A>G 2323C>G 2324A>C), ACG49ACA (2319C>T), AAC51ACA (2313G>T 2314T>G), GGA55GGG (2301T>C), GAG56GAA (2298C>T), CAC58CAT (2292G>A), ATC59CTT (2289G>A 2291T>G), TGT61TGC (2283A>G), CTG62CTA (2280C>T), GGA86AGA (2210C>T), ATT87ATA (2205A>T), TAT88TTT (2203T>A), CAT89CAC (2199A>G), CCA90CCC (2196T>G), ATC92ACT (2190G>A 2191A>G), ATC94ATA (2184G>T), CCA95CCG (2181T>C), AGG97TCG (2176C>G 2177T>A), GAT98GAC (2172A>G), GCA99AAA (2170G>T 2171C>T), GAT100CAC (2166A>G 2168C>G), GTT101GTC (2163A>G), GCG102TCC (2160C>G 2162C>A), GAC103GAT (2157G>A), TAC104TAT (2154G>A), ATT105ATC (2151A>G), TCC106TCA (2148G>T), GGG108GGA (2142C>T), GGT109GGA (2139A>T), ACC110ATA (2136G>T 2137G>A), GAG113GAA (2127C>T), AGA114AGT (2124T>A), GGT115GGA (2121A>T), ATT116GTA (2118A>T 2120T>C), CTT117TTT (2117G>A), AGA118AAG (2112T>C 2113C>T), GCT119AAT (2110G>T 2111C>T), TCC120ACA (2106G>T 2108A>T), CGA121AAA (2104C>T 2105G>T), AGA122CGA (2102T>G), AGT123GGT (2099T>C), CCG124CCT (2094C>A), AAG125AAA (2091C>T), AAG126CAG (2090T>G), AAT127ACT (2086T>G), AGA128AGG (2082T>C) | | | | | | | | |
| **Proteins** | | | | | | | | | |
| replication-associated protein (YP_009417307.1) | 20 | 130 | 27.6% | 488 | 68.6% | 92 (100%) | 57 (62.0%) | 0/0/0/0 | 0 |
| Protein mutations: | P26A (2390G>C), I29V (2379A>G 2381T>C), Y32F (2371T>A), F33L (2367A>G 2369A>G), F34K (2364A>T 2365A>T 2366A>T), Q35K (2363G>T), L36I (2358T>A 2360A>T), K38Q (2354T>G), V42P (2340G>A 2341A>G 2342C>G), T43N (2337T>A 2338G>T), V45I (2333C>T), A47V (2326G>A), C48A (2322A>G 2323C>G 2324A>C), N51T (2313G>T 2314T>G), I59L (2289G>A 2291T>G), G86R (2210C>T), Y88F (2203T>A), I92T (2190G>A 2191A>G), R97S (2176C>G 2177T>A), A99K (2170G>T 2171C>T), D100H (2166A>G 2168C>G), A102S (2160C>G 2162C>A), T110I (2136G>T 2137G>A), R114S (2124T>A), I116V (2118A>T 2120T>C), L117F (2117G>A), R118K (2112T>C 2113C>T), A119N (2110G>T 2111C>T), S120T (2106G>T 2108A>T), R121K (2104C>T 2105G>T), S123G (2099T>C), K126Q (2090T>G), N127T (2086T>G) | | | | | | | | |
| Codon mutations: | CCA20CCC (2406T>G), TGT22TGC (2400A>G), ATT25ATC (2391A>G), CCA26GCA (2390G>C), ATT27ATA (2385A>T), ATT29GTC (2379A>G 2381T>C), ATC30ATA (2376G>T), GAT31GAC (2373A>G), TAC32TTC (2371T>A), TTT33CTC (2367A>G 2369A>G), TTT34AAA (2364A>T 2365A>T 2366A>T), CAA35AAA (2363G>T), TTA36ATT (2358T>A 2360A>T), AAA38CAA (2354T>G), TAT40TAC (2346A>G), GTC42CCT (2340G>A 2341A>G 2342C>G), ACA43AAT (2337T>A 2338G>T), TAC44TAT (2334G>A), GTT45ATT (2333C>T), GCT47GTT (2326G>A), TGT48GCC (2322A>G 2323C>G 2324A>C), ACG49ACA (2319C>T), AAC51ACA (2313G>T 2314T>G), GGA55GGG (2301T>C), GAG56GAA (2298C>T), CAC58CAT (2292G>A), ATC59CTT (2289G>A 2291T>G), TGT61TGC (2283A>G), CTG62CTA (2280C>T), GGA86AGA (2210C>T), ATT87ATA (2205A>T), TAT88TTT (2203T>A), CAT89CAC (2199A>G), CCA90CCC (2196T>G), ATC92ACT (2190G>A 2191A>G), ATC94ATA (2184G>T), CCA95CCG (2181T>C), AGG97TCG (2176C>G 2177T>A), GAT98GAC (2172A>G), GCA99AAA (2170G>T 2171C>T), GAT100CAC (2166A>G 2168C>G), GTT101GTC (2163A>G), GCG102TCC (2160C>G 2162C>A), GAC103GAT (2157G>A), TAC104TAT (2154G>A), ATT105ATC (2151A>G), TCC106TCA (2148G>T), GGG108GGA (2142C>T), GGT109GGA (2139A>T), ACC110ATA (2136G>T 2137G>A), GAG113GAA (2127C>T), AGA114AGT (2124T>A), GGT115GGA (2121A>T), ATT116GTA (2118A>T 2120T>C), CTT117TTT (2117G>A), AGA118AAG (2112T>C 2113C>T), GCT119AAT (2110G>T 2111C>T), TCC120ACA (2106G>T 2108A>T), CGA121AAA (2104C>T 2105G>T), AGA122CGA (2102T>G), AGT123GGT (2099T>C), CCG124CCT (2094C>A), AAG125AAA (2091C>T), AAG126CAG (2090T>G), AAT127ACT (2086T>G), AGA128AGG (2082T>C) | | | | | | | | |
| RepA (YP_009417308.1) | 20 | 130 | 35.1% | 488 | 68.6% | 92 (100%) | 57 (62.0%) | 0/0/0/0 | 0 |
| Protein mutations: | P26A (2390G>C), I29V (2379A>G 2381T>C), Y32F (2371T>A), F33L (2367A>G 2369A>G), F34K (2364A>T 2365A>T 2366A>T), Q35K (2363G>T), L36I (2358T>A 2360A>T), K38Q (2354T>G), V42P (2340G>A 2341A>G 2342C>G), T43N (2337T>A 2338G>T), V45I (2333C>T), A47V (2326G>A), C48A (2322A>G 2323C>G 2324A>C), N51T (2313G>T 2314T>G), I59L (2289G>A 2291T>G), G86R (2210C>T), Y88F (2203T>A), I92T (2190G>A 2191A>G), R97S (2176C>G 2177T>A), A99K (2170G>T 2171C>T), D100H (2166A>G 2168C>G), A102S (2160C>G 2162C>A), T110I (2136G>T 2137G>A), R114S (2124T>A), I116V (2118A>T 2120T>C), L117F (2117G>A), R118K (2112T>C 2113C>T), A119N (2110G>T 2111C>T), S120T (2106G>T 2108A>T), R121K (2104C>T 2105G>T), S123G (2099T>C), K126Q (2090T>G), N127T (2086T>G) | | | | | | | | |

| | Begin | End | Coverage | Score | Concordance | Matches | Identities | I/D/M/F* | Stop Codons |
|---|---|---|---|---|---|---|---|---|---|
| NT | 2078 | 2408 | 10.5% | 192 | 35.0% | 274 (100%) | 185 (67.5%) | 0/0 | |
| Codon mutations: | CCA20CCC (2406T>G), TGT22TGC (2400A>G), ATT25ATC (2391A>G), CCA26GCA (2390G>C), ATT27ATA (2385A>T), ATT29GTC (2379A>G 2381T>C), ATC30ATA (2376G>T), GAT31GAC (2373A>G), TAC32TTC (2371T>A), TTT33CTC (2367A>G 2369A>G), TTT34AAA (2364A>T 2365A>T 2366A>T), CAA35AAA (2363G>T), TTA36ATT (2358T>A 2360A>T), AAA38CAA (2354T>G), TAT40TAC (2346A>G), GTC42CCT (2340G>A 2341A>G 2342C>G), ACA43AAT (2337T>A 2338G>T), TAC44TAT (2334G>A), GTT45ATT (2333C>T), GCT47GTT (2326G>A), TGT48GCC (2322A>G 2323C>G 2324A>C), ACG49ACA (2319C>T), AAC51ACA (2313G>T 2314T>G), GGA55GGG (2301T>C), GAG56GAA (2298C>T), CAC58CAT (2292G>A), ATC59CTT (2289G>A 2291T>G), TGT61TGC (2283A>G), CTG62CTA (2280C>T), GGA86AGA (2210C>T), ATT87ATA (2205A>T), TAT88TTT (2203T>A), CAT89CAC (2199A>G), CCA90CCC (2196T>G), ATC92ACT (2190G>A 2191A>G), ATC94ATA (2184G>T), CCA95CCG (2181T>C), AGG97TCG (2176C>G 2177T>A), GAT98GAC (2172A>G), GCA99AAA (2170G>T 2171C>T), GAT100CAC (2166A>G 2168C>G), GTT101GTC (2163A>G), GCG102TCC (2160C>G 2162C>A), GAC103GAT (2157G>A), TAC104TAT (2154G>A), ATT105ATC (2151A>G), TCC106TCA (2148G>T), GGG108GGA (2142C>T), GGT109GGA (2139A>T), ACC110ATA (2136G>T 2137G>A), GAG113GAA (2127C>T), AGA114AGT (2124T>A), GGT115GGA (2121A>T), ATT116GTA (2118A>T 2120T>C), CTT117TTT (2117G>A), AGA118AAG (2112T>C 2113C>T), GCT119AAT (2110G>T 2111C>T), TCC120ACA (2106G>T 2108A>T), CGA121AAA (2104C>T 2105G>T), AGA122CGA (2102T>G), AGT123GGT (2099T>C), CCG124CCT (2094C>A), AAG125AAA (2091C>T), AAG126CAG (2090T>G), AAT127ACT (2086T>G), AGA128AGG (2082T>C) | | | | | | | | |

*: Inserts / Deletes / Misaligned / Frameshifts

#### Analysis details

This analysis was performed with panviral2.64

### NGS Details (UN9): Tokyovirus A1

#### Assembly

| | |
|---|---|
| **Coverage Length** | 125 (1 contig(s)) |
| **Depth Of Coverage** | 2.4 |
| **Number Of Reads** | 3 |
| **Reads Per Million** | 0.07 rpm (after QC) |
| **Ambiguities** | 0 |
| **Assembly Method** | de novo + reference guided assembly |
| **Consensus Caller** | Bcf Tools |

#### Coverage Map

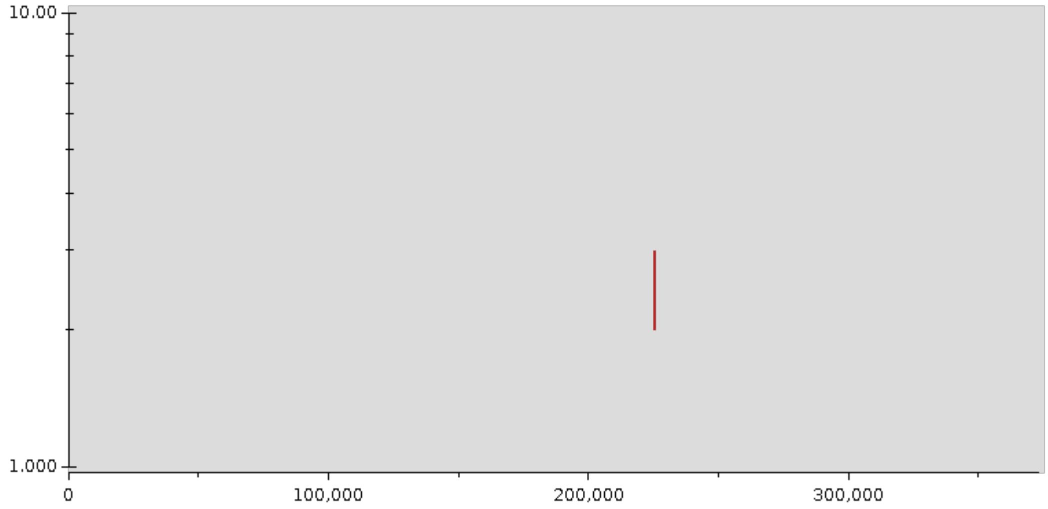

#### Assignment

| | |
|---|---|
| **Type** | Tokyovirus A1 (Taxonomy ID: 1826170) |
| **Reference Genome** | NC_030230.1 |
| **NT Identity (%)** | 80.0 |
| **AA Identity (%)** | 92.8571 |
| **Number Of Stop Codons** | 0 |
| **Number Of CDS** | 470 |

#### Alignment

| | |
|---|---|
| **Alignment Score** | 150.0 (NT) + 273.0 (AA) = 423.0 |
| **Concordance (%)** | 80.2657 |
| **Alignment Method** | Local, heuristic, nucleotide (BLASTN) |

#### Genome Region

Sequence starts at position 225496 and ends at position 225620 relative to NC_030230.1 reference sequence.

#### Alignment Detailed Statistics

| | Begin | End | Coverage | Score | Concordance | Matches | Identities | I/D/M/F* | Stop Codons |
|---|---|---|---|---|---|---|---|---|---|
| NT | 225496 | 225620 | 0.1% | 150 | 60.0% | 125 (100%) | 100 (80.0%) | 0/0 | |

Mutations: 225501C>G, 225507A>C, 225509G>A, 225519G>A, 225525T>C, 225528T>C, 225533C>T, 225534A>G, 225540A>G, 225543G>A, 225545C>A, 225548A>G, 225555C>G, 225558A>G, 225564T>G, 225573C>G, 225579G>A, 225585A>G, 225590G>T, 225594T>C, 225597G>A, 225600G>A, 225603C>T, 225606A>G, 225609A>G

*: Inserts / Deletes / Misaligned / Frameshifts

#### Analysis details

This analysis was performed with panviral2.64

### NGS Details (UN9): Nepovirus aeonii (segment RNA2)

#### Assembly

| | |
|---|---|
| **Coverage Length** | 250 (1 contig(s)) |
| **Depth Of Coverage** | 1.1 |
| **Number Of Reads** | 2 |
| **Reads Per Million** | 0.05 rpm (after QC) |
| **Ambiguities** | 0 |
| **Assembly Method** | de novo + reference guided assembly |
| **Consensus Caller** | Bcf Tools |

#### Coverage Map

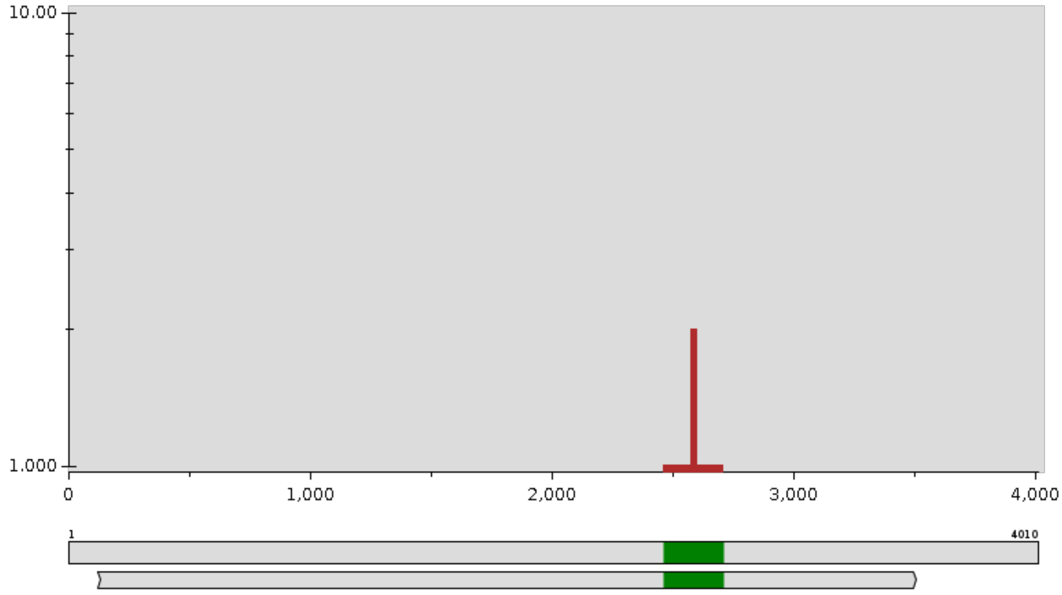

#### Assignment

| | |
|---|---|
| **Type** | Nepovirus aeonii (Taxonomy ID: 3047350) |
| **Reference Genome** | NC_038761.1 |
| **NT Identity (%)** | 71.2 |
| **AA Identity (%)** | 72.2892 |
| **Number Of Stop Codons** | 0 |
| **Number Of CDS** | 1 |

#### Alignment

| | |
|---|---|
| **Alignment Score** | 212.0 (NT) + 425.0 (AA) = 637.0 |
| **Concordance (%)** | 61.6054 |

| Alignment Method | Global, seeded, nucleotide + amino acids (AGA) |
|---|---|

#### Genome Region

Sequence starts at position 2462 and ends at position 2711 relative to NC_038761.1 reference sequence.

#### Alignment Detailed Statistics

| | Begin | End | Coverage | Score | Concordance | Matches | Identities | I/D/M/F* | Stop Codons |
|---|---|---|---|---|---|---|---|---|---|
| **NT** | **2462** | **2711** | **6.2%** | **212** | **42.4%** | **250 (100%)** | **178 (71.2%)** | **0/0** | |
| Mutations: | 2470C>G, 2472G>A, 2474A>C, 2481G>A, 2482T>C, 2483C>A, 2484T>G, 2491C>T, 2493T>G, 2496C>T, 2499C>A, 2502G>A, 2503G>A, 2505T>G, 2514G>T, 2515C>A, 2517G>A, 2519G>C, 2520T>A, 2521G>C, 2522T>A, 2524G>A, 2526G>T, 2529T>C, 2530T>A, 2531C>G, 2532A>T, 2533C>T, 2535C>T, 2536A>G, 2537G>A, 2541T>G, 2542G>A, 2562A>G, 2565G>A, 2571G>A, 2580G>C, 2583T>C, 2592A>G, 2593A>T, 2607T>C, 2608A>G, 2622T>C, 2623A>G, 2629C>T, 2631C>T, 2632A>T, 2634T>A, 2635C>A, 2637A>G, 2643C>T, 2646G>A, 2649T>A, 2652T>G, 2658T>C, 2661T>A, 2664T>A, 2666T>C, 2667T>A, 2671G>T, 2673A>T, 2674C>A, 2676T>A, 2682A>T, 2685A>G, 2687T>G, 2688T>G, 2695T>A, 2696C>G, 2697T>A, 2698A>C, 2703A>G | | | | | | | | |

##### CDS

| | | | | | | | | | |
|---|---|---|---|---|---|---|---|---|---|
| D1S54_s2gp1 | 781 | 863 | 7.4% | 425 | 77.4% | 83 (100%) | 60 (72.3%) | 0/0/0/0 | 0 |
| Protein mutations: | Q783E (2470C>G 2472G>A), Q784P (2474A>C), S787Q (2482T>C 2483C>A 2484T>G), V794M (2503G>A 2505T>G), L798I (2515C>A 2517G>A), S799T (2519G>C 2520T>A), V800H (2521G>C 2522T>A), G801S (2524G>A 2526G>T), L804F (2533C>T 2535C>T), S805D (2536A>G 2537G>A), D807N (2542G>A), M824L (2593A>T), N829D (2608A>G), I834V (2623A>G), L836F (2629C>T 2631C>T), T837S (2632A>T 2634T>A), Q838K (2635C>A 2637A>G), V848A (2666T>C 2667T>A), A850S (2671G>T 2673A>T), F855W (2687T>G 2688T>G), S858R (2695T>A 2696C>G 2697T>A), I859L (2698A>C) | | | | | | | | |
| Codon mutations: | CAG783GAA (2470C>G 2472G>A), CAA784CCA (2474A>C), GAG786GAA (2481G>A), TCT787CAG (2482T>C 2483C>A 2484T>G), CTT790TTG (2491C>T 2493T>G), GCC791GCT (2496C>T), CCC792CCA (2499C>A), ACG793ACA (2502G>A), GTT794ATG (2503G>A 2505T>G), CCG797CCT (2514G>T), CTG798ATA (2515C>A 2517G>A), AGT799ACA (2519G>C 2520T>A), GTT800CAT (2521G>C 2522T>A), GGG801AGT (2524G>A 2526G>T), GAT802GAC (2529T>C), TCA803AGT (2530T>A 2531C>G 2532A>T), CTC804TTT (2533C>T 2535C>T), AGT805GAT (2536A>G 2537G>A), GTT806GTG (2541T>G), GAT807AAT (2542G>A), AAA813AAG (2562A>G), GAG814GAA (2565G>A), AAG816AAA (2571G>A), GGG819GGC (2580G>C), ACT820ACC (2583T>C), TTA823TTG (2592A>G), ATG824TTG (2593A>T), TAT828TAC (2607T>C), AAT829GAT (2608A>G), CCT833CCC (2622T>C), ATC834GTC (2623A>G), CTC836TTT (2629C>T 2631C>T), ACT837TCA (2632A>T 2634T>A), CAA838AAG (2635C>A 2637A>G), AGC840AGT (2643C>T), TCG841TCA (2646G>A), ATT842ATA (2649T>A), TCT843TCG (2652T>G), CCT845CCC (2658T>C), CGT846CGA (2661T>A), GCT847GCA (2664T>A), GTT848GCA (2666T>C 2667T>A), GCA850TCT (2671G>T 2673A>T), CGT851AGA (2674C>A 2676T>A), ATA853ATT (2682A>T), GGA854GGG (2685A>G), TTT855TGG (2687T>G 2688T>G), TCT858AGA (2695T>A 2696C>G 2697T>A), ATC859CTC (2698A>C), AAA860AAG (2703A>G) | | | | | | | | |

##### Proteins

| | | | | | | | | | |
|---|---|---|---|---|---|---|---|---|---|
| polyprotein P2 (YP_009507917.1) | 781 | 863 | 7.4% | 425 | 77.4% | 83 (100%) | 60 (72.3%) | 0/0/0/0 | 0 |
| Protein mutations: | Q783E (2470C>G 2472G>A), Q784P (2474A>C), S787Q (2482T>C 2483C>A 2484T>G), V794M (2503G>A 2505T>G), L798I (2515C>A 2517G>A), S799T (2519G>C 2520T>A), V800H (2521G>C 2522T>A), G801S (2524G>A 2526G>T), L804F (2533C>T 2535C>T), S805D (2536A>G 2537G>A), D807N (2542G>A), M824L (2593A>T), N829D (2608A>G), I834V (2623A>G), L836F (2629C>T 2631C>T), T837S (2632A>T 2634T>A), Q838K (2635C>A 2637A>G), V848A (2666T>C 2667T>A), A850S (2671G>T 2673A>T), F855W (2687T>G 2688T>G), S858R (2695T>A 2696C>G 2697T>A), I859L (2698A>C) | | | | | | | | |
| Codon mutations: | CAG783GAA (2470C>G 2472G>A), CAA784CCA (2474A>C), GAG786GAA (2481G>A), TCT787CAG (2482T>C 2483C>A 2484T>G), CTT790TTG (2491C>T 2493T>G), GCC791GCT (2496C>T), CCC792CCA (2499C>A), ACG793ACA (2502G>A), GTT794ATG (2503G>A 2505T>G), CCG797CCT (2514G>T), CTG798ATA (2515C>A 2517G>A), AGT799ACA (2519G>C 2520T>A), GTT800CAT (2521G>C 2522T>A), GGG801AGT (2524G>A 2526G>T), GAT802GAC (2529T>C), TCA803AGT (2530T>A 2531C>G 2532A>T), CTC804TTT (2533C>T 2535C>T), AGT805GAT (2536A>G 2537G>A), GTT806GTG (2541T>G), GAT807AAT (2542G>A), AAA813AAG (2562A>G), GAG814GAA (2565G>A), AAG816AAA (2571G>A), GGG819GGC (2580G>C), ACT820ACC (2583T>C), TTA823TTG (2592A>G), ATG824TTG (2593A>T), TAT828TAC (2607T>C), AAT829GAT (2608A>G), CCT833CCC (2622T>C), ATC834GTC (2623A>G), CTC836TTT (2629C>T 2631C>T), ACT837TCA (2632A>T 2634T>A), CAA838AAG (2635C>A 2637A>G), AGC840AGT (2643C>T), TCG841TCA (2646G>A), ATT842ATA (2649T>A), TCT843TCG (2652T>G), CCT845CCC (2658T>C), CGT846CGA (2661T>A), GCT847GCA (2664T>A), GTT848GCA (2666T>C 2667T>A), GCA850TCT (2671G>T 2673A>T), CGT851AGA (2674C>A 2676T>A), ATA853ATT (2682A>T), GGA854GGG (2685A>G), TTT855TGG (2687T>G 2688T>G), TCT858AGA (2695T>A 2696C>G 2697T>A), ATC859CTC (2698A>C), AAA860AAG (2703A>G) | | | | | | | | |

*: Inserts / Deletes / Misaligned / Frameshifts

#### Analysis details

This analysis was performed with panviral2.64

### NGS Details (UN9): Moloney murine leukemia virus

#### Assembly

| | |
|---|---|
| **Coverage Length** | 134 (1 contig(s)) |
| **Depth Of Coverage** | 2.0 |
| **Number Of Reads** | 2 |
| **Reads Per Million** | 0.05 rpm (after QC) |
| **Ambiguities** | 0 |
| **Assembly Method** | de novo + reference guided assembly |
| **Consensus Caller** | Bcf Tools |

#### Coverage Map

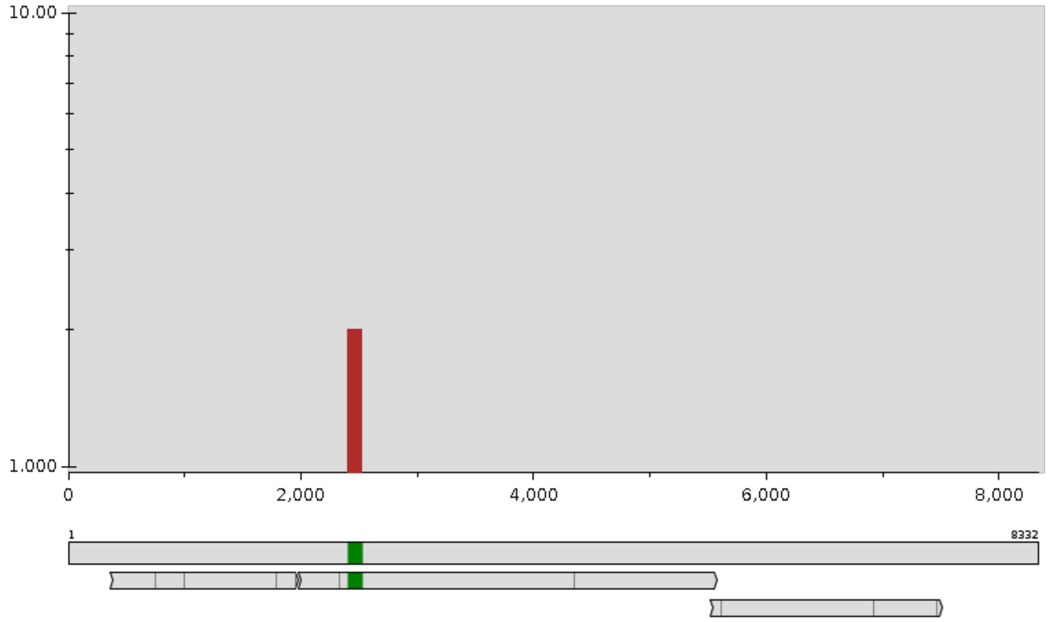

#### Assignment

| | |
|---|---|
| **Type** | Moloney murine leukemia virus (Taxonomy ID: 11801) |
| **Reference Genome** | NC_001501.1 |
| **NT Identity (%)** | 79.1045 |
| **AA Identity (%)** | 97.7273 |
| **Number Of Stop Codons** | 0 |
| **Number Of CDS** | 3 |

#### Alignment

| | |
|---|---|
| **Alignment Score** | 156.0 (NT) + 299.0 (AA) = 455.0 |
| **Concordance (%)** | 79.4066 |

| Alignment Method | Global, seeded, nucleotide + amino acids (AGA) |
|---|---|

#### Genome Region

Sequence starts at position 2398 and ends at position 2531 relative to NC_001501.1 reference sequence.

#### Alignment Detailed Statistics

| | Begin | End | Coverage | Score | Concordance | Matches | Identities | I/D/M/F* | Stop Codons |
|---|---|---|---|---|---|---|---|---|---|
| **NT** | **2398** | **2531** | **1.6%** | **156** | **58.2%** | **134 (100%)** | **106 (79.1%)** | **0/0** | |
| Mutations: | 2401C>G, 2405A>C, 2412T>A, 2413C>G, 2414T>C, 2423T>G, 2429C>G, 2444G>C, 2453A>C, 2459A>G, 2462T>G, 2468A>G, 2471T>G, 2474T>G, 2480C>T, 2483A>T, 2486T>G, 2495A>G, 2499T>A, 2500C>G, 2501T>C, 2507C>G, 2511T>A, 2512C>G, 2516A>T, 2522A>G, 2525C>T, 2528C>G | | | | | | | | |
| **CDS** | | | | | | | | | |
| gag-pol | 681 | 724 | 2.5% | 299 | 95.5% | 44 (100%) | 43 (97.7%) | 0/0/0/0 | 0 |
| Protein mutations: | S681C (2401C>G) | | | | | | | | |
| Codon mutations: | TCC681TGC (2401C>G), ACA682ACC (2405A>C), TCT685AGC (2412T>A 2413C>G 2414T>C), CCT688CCG (2423T>G), GCC690GCG (2429C>G), GGG695GGC (2444G>C), GGA698GGC (2453A>C), GCA700GCG (2459A>G), GTT701GTG (2462T>G), CAA703CAG (2468A>G), GCT704GCG (2471T>G), CCT705CCG (2474T>G), ATC707ATT (2480C>T), ATA708ATT (2483A>T), CCT709CCG (2486T>G), GCA712GCG (2495A>G), TCT714AGC (2499T>A 2500C>G 2501T>C), CCC716CCG (2507C>G), TCC718AGC (2511T>A 2512C>G), ATA719ATT (2516A>T), CAA721CAG (2522A>G), TAC722TAT (2525C>T), CCC723CCG (2528C>G) | | | | | | | | |
| **Proteins** | | | | | | | | | |
| Pr180 (NP_057933.2) | 681 | 724 | 2.5% | 299 | 95.5% | 44 (100%) | 43 (97.7%) | 0/0/0/0 | 0 |
| Protein mutations: | S681C (2401C>G) | | | | | | | | |
| Codon mutations: | TCC681TGC (2401C>G), ACA682ACC (2405A>C), TCT685AGC (2412T>A 2413C>G 2414T>C), CCT688CCG (2423T>G), GCC690GCG (2429C>G), GGG695GGC (2444G>C), GGA698GGC (2453A>C), GCA700GCG (2459A>G), GTT701GTG (2462T>G), CAA703CAG (2468A>G), GCT704GCG (2471T>G), CCT705CCG (2474T>G), ATC707ATT (2480C>T), ATA708ATT (2483A>T), CCT709CCG (2486T>G), GCA712GCG (2495A>G), TCT714AGC (2499T>A 2500C>G 2501T>C), CCC716CCG (2507C>G), TCC718AGC (2511T>A 2512C>G), ATA719ATT (2516A>T), CAA721CAG (2522A>G), TAC722TAT (2525C>T), CCC723CCG (2528C>G) | | | | | | | | |
| p80 RT (NP_955591.1) | 22 | 65 | 6.6% | 299 | 95.5% | 44 (100%) | 43 (97.7%) | 0/0/0/0 | 0 |
| Protein mutations: | S22C (2401C>G) | | | | | | | | |
| Codon mutations: | TCC22TGC (2401C>G), ACA23ACC (2405A>C), TCT26AGC (2412T>A 2413C>G 2414T>C), CCT29CCG (2423T>G), GCC31GCG (2429C>G), GGG36GGC (2444G>C), GGA39GGC (2453A>C), GCA41GCG (2459A>G), GTT42GTG (2462T>G), CAA44CAG (2468A>G), GCT45GCG (2471T>G), CCT46CCG (2474T>G), ATC48ATT (2480C>T), ATA49ATT (2483A>T), CCT50CCG (2486T>G), GCA53GCG (2495A>G), TCT55AGC (2499T>A 2500C>G 2501T>C), CCC57CCG (2507C>G), TCC59AGC (2511T>A 2512C>G), ATA60ATT (2516A>T), CAA62CAG (2522A>G), TAC63TAT (2525C>T), CCC64CCG (2528C>G) | | | | | | | | |

*: Inserts / Deletes / Misaligned / Frameshifts

#### Analysis details

This analysis was performed with panviral2.64

### NGS Details (UN9): Friend murine leukemia virus

#### Assembly

| | |
|---|---|
| **Coverage Length** | 135 (1 contig(s)) |
| **Depth Of Coverage** | 2.0 |
| **Number Of Reads** | 2 |
| **Reads Per Million** | 0.05 rpm (after QC) |
| **Ambiguities** | 0 |
| **Assembly Method** | de novo + reference guided assembly |
| **Consensus Caller** | Bcf Tools |

#### Coverage Map

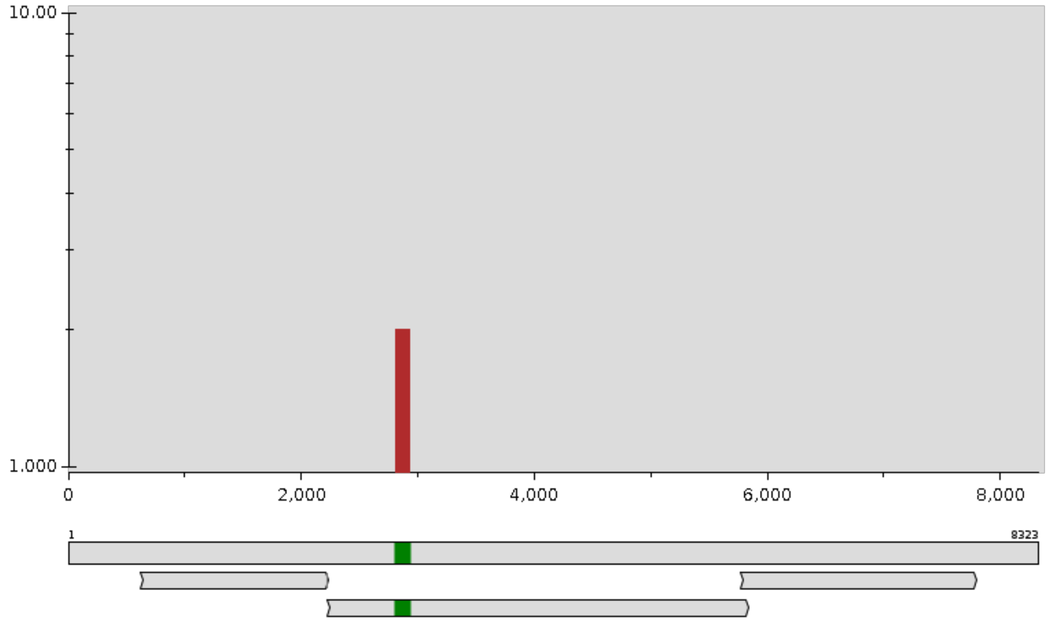

#### Assignment

| | |
|---|---|
| **Type** | Friend murine leukemia virus (Taxonomy ID: 11795) |
| **Reference Genome** | NC_001362.1 |
| **NT Identity (%)** | 78.5185 |
| **AA Identity (%)** | 97.7778 |
| **Number Of Stop Codons** | 0 |
| **Number Of CDS** | 3 |

#### Alignment

| | |
|---|---|
| **Alignment Score** | 154.0 (NT) + 343.0 (AA) = 497.0 |
| **Concordance (%)** | 81.0767 |

| Alignment Method | Global, seeded, nucleotide + amino acids (AGA) |
|---|---|

#### Genome Region

Sequence starts at position 2805 and ends at position 2939 relative to NC_001362.1 reference sequence.

#### Alignment Detailed Statistics

| | Begin | End | Coverage | Score | Concordance | Matches | Identities | I/D/M/F* | Stop Codons |
|---|---|---|---|---|---|---|---|---|---|
| **NT** | **2805** | **2939** | **1.6%** | **154** | **57.0%** | **135 (100%)** | **106 (78.5%)** | **0/0** | |
| Mutations: | 2806A>C, 2808A>C, 2814G>C, 2817C>T, 2820G>A, 2823C>G, 2826C>T, 2829A>T, 2833A>C, 2835A>C, 2850A>C, 2859A>G, 2862C>G, 2869T>A, 2870C>G, 2874C>G, 2883G>C, 2886C>G, 2892A>G, 2895C>G, 2898T>G, 2901G>A, 2910G>C, 2913T>C, 2916T>C, 2923A>C, 2925G>C, 2928T>G, 2931C>G | | | | | | | | |
| **CDS** | | | | | | | | | |
| pol | 196 | 240 | 3.7% | 343 | 97.7% | 45 (100%) | 44 (97.8%) | 0/0/0/0 | 0 |
| **Proteins** | | | | | | | | | |
| RNA-dependent DNA polymerase (NP_040333.1) | 196 | 240 | 3.7% | 343 | 97.7% | 45 (100%) | 44 (97.8%) | 0/0/0/0 | 0 |
| Protein mutations: | *none* | | | | | | | | |
| Codon mutations: | AGA196CGC (2806A>C 2808A>C), GGG198GGC (2814G>C), ATC199ATT (2817C>T), AAG200AAA (2820G>A), CCC201CCG (2823C>G), CAC202CAT (2826C>T), ATA203ATT (2829A>T), AGA205CGC (2833A>C 2835A>C), GGA210GGC (2850A>C), GTA213GTG (2859A>G), CCC214CCG (2862C>G), TCC217AGC (2869T>A 2870C>G), CCC218CCG (2874C>G), ACG221ACC (2883G>C), CCC222CCG (2886C>G), CTA224CTG (2892A>G), CCC225CCG (2895C>G), GTT226GTG (2898T>G), AAG227AAA (2901G>A), GGG230GGC (2910G>C), ACT231ACC (2913T>C), AAT232AAC (2916T>C), AGG235CGC (2923A>C 2925G>C), CCT236CCG (2928T>G), GTC237GTG (2931C>G) | | | | | | | | |

*: Inserts / Deletes / Misaligned / Frameshifts

#### Analysis details

This analysis was performed with panviral2.64

### NGS Details (UN9): Duamitovirus peex1

#### Assembly

| | |
|---|---|
| **Coverage Length** | 135 (1 contig(s)) |
| **Depth Of Coverage** | 1.0 |
| **Number Of Reads** | 1 |
| **Reads Per Million** | 0.02 rpm (after QC) |
| **Ambiguities** | 0 |
| **Assembly Method** | de novo + reference guided assembly |
| **Consensus Caller** | Bcf Tools |

#### Coverage Map

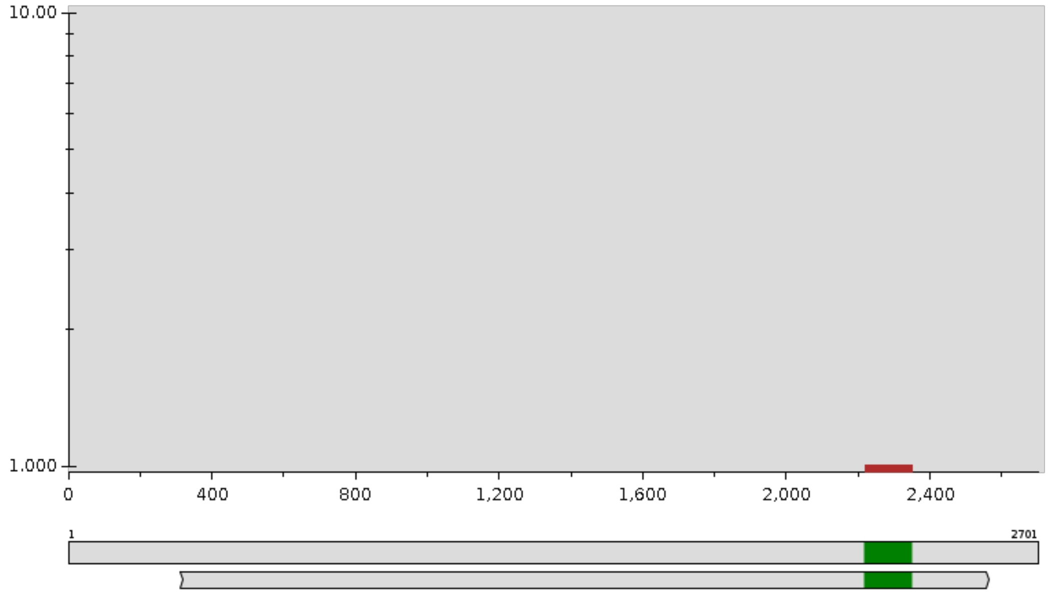

#### Assignment

| | |
|---|---|
| **Type** | Duamitovirus peex1 (Taxonomy ID: 2955799) |
| **Reference Genome** | NC_076525.1 |
| **NT Identity (%)** | 71.8519 |
| **AA Identity (%)** | 66.6667 |
| **Number Of Stop Codons** | 0 |
| **Number Of CDS** | 1 |

#### Alignment

| | |
|---|---|
| **Alignment Score** | 118.0 (NT) + 249.0 (AA) = 367.0 |
| **Concordance (%)** | 59.3851 |

| Alignment Method | Global, seeded, nucleotide + amino acids (AGA) |
|---|---|

#### Genome Region

Sequence starts at position 2218 and ends at position 2352 relative to NC_076525.1 reference sequence.

#### Alignment Detailed Statistics

| | Begin | End | Coverage | Score | Concordance | Matches | Identities | I/D/M/F* | Stop Codons |
|---|---|---|---|---|---|---|---|---|---|
| **NT** | **2218** | **2352** | **5.0%** | **118** | **43.7%** | **135 (100%)** | **97 (71.9%)** | **0/0** | |
| Mutations: | 2222A>T, 2223C>G, 2225A>G, 2226C>T, 2234A>T, 2243A>G, 2244C>A, 2245A>G, 2246T>A, 2249G>A, 2255A>T, 2268G>T, 2272G>T, 2273T>C, 2274C>A, 2275A>G, 2276T>A, 2280A>G, 2283G>A, 2294T>G, 2295C>A, 2296G>A, 2301C>G, 2303T>C, 2306A>T, 2312T>C, 2315C>T, 2321G>T, 2322G>T, 2325A>G, 2326A>C, 2327T>A, 2330C>A, 2331C>G, 2333G>A, 2339G>A, 2345A>T, 2348C>T | | | | | | | | |
| **CDS** | | | | | | | | | |
| RdRp | 637 | 681 | 6.0% | 249 | 68.6% | 45 (100%) | 30 (66.7%) | 0/0/0/0 | 0 |
| Protein mutations: | K637N (2222A>T), Q638E (2223C>G 2225A>G), L641F (2234A>T), H645R (2244C>A 2245A>G 2246T>A), V653L (2268G>T), R654L (2272G>T 2273T>C), H655R (2274C>A 2275A>G 2276T>A), M657V (2280A>G), E658K (2283G>A), R662K (2295C>A 2296G>A), L664V (2301C>G 2303T>C), A671S (2322G>T), N672A (2325A>G 2326A>C 2327T>A), P674A (2331C>G 2333G>A) | | | | | | | | |
| Codon mutations: | AAA637AAT (2222A>T), CAA638GAG (2223C>G 2225A>G), CTG639TTG (2226C>T), TTA641TTT (2234A>T), GAA644GAG (2243A>G), CAT645AGA (2244C>A 2245A>G 2246T>A), GAG646GAA (2249G>A), CTA648CTT (2255A>T), GTA653TTA (2268G>T), CGT654CTC (2272G>T 2273T>C), CAT655AGA (2274C>A 2275A>G 2276T>A), ATG657GTG (2280A>G), GAA658AAA (2283G>A), CTT661CTG (2294T>G), CGA662AAA (2295C>A 2296G>A), CTT664GTC (2301C>G 2303T>C), TCA665TCT (2306A>T), TAT667TAC (2312T>C), CAC668CAT (2315C>T), GTG670GTT (2321G>T), GCA671TCA (2322G>T), AAT672GCA (2325A>G 2326A>C 2327T>A), TCC673TCA (2330C>A), CCG674GCA (2331C>G 2333G>A), GTG676GTA (2339G>A), ATA678ATT (2345A>T), GAC679GAT (2348C>T) | | | | | | | | |
| **Proteins** | | | | | | | | | |
| RNA-dependent RNA polymerase (YP_010798875.1) | 637 | 681 | 6.0% | 249 | 68.6% | 45 (100%) | 30 (66.7%) | 0/0/0/0 | 0 |
| Protein mutations: | K637N (2222A>T), Q638E (2223C>G 2225A>G), L641F (2234A>T), H645R (2244C>A 2245A>G 2246T>A), V653L (2268G>T), R654L (2272G>T 2273T>C), H655R (2274C>A 2275A>G 2276T>A), M657V (2280A>G), E658K (2283G>A), R662K (2295C>A 2296G>A), L664V (2301C>G 2303T>C), A671S (2322G>T), N672A (2325A>G 2326A>C 2327T>A), P674A (2331C>G 2333G>A) | | | | | | | | |
| Codon mutations: | AAA637AAT (2222A>T), CAA638GAG (2223C>G 2225A>G), CTG639TTG (2226C>T), TTA641TTT (2234A>T), GAA644GAG (2243A>G), CAT645AGA (2244C>A 2245A>G 2246T>A), GAG646GAA (2249G>A), CTA648CTT (2255A>T), GTA653TTA (2268G>T), CGT654CTC (2272G>T 2273T>C), CAT655AGA (2274C>A 2275A>G 2276T>A), ATG657GTG (2280A>G), GAA658AAA (2283G>A), CTT661CTG (2294T>G), CGA662AAA (2295C>A 2296G>A), CTT664GTC (2301C>G 2303T>C), TCA665TCT (2306A>T), TAT667TAC (2312T>C), CAC668CAT (2315C>T), GTG670GTT (2321G>T), GCA671TCA (2322G>T), AAT672GCA (2325A>G 2326A>C 2327T>A), TCC673TCA (2330C>A), CCG674GCA (2331C>G 2333G>A), GTG676GTA (2339G>A), ATA678ATT (2345A>T), GAC679GAT (2348C>T) | | | | | | | | |

*: Inserts / Deletes / Misaligned / Frameshifts

#### Analysis details

This analysis was performed with panviral2.64

### NGS Details (UN9): Plantago lanceolata latent virus

#### Assembly

| | |
|---|---|
| **Coverage Length** | 135 (1 contig(s)) |
| **Depth Of Coverage** | 1.0 |
| **Number Of Reads** | 1 |
| **Reads Per Million** | 0.02 rpm (after QC) |
| **Ambiguities** | 0 |
| **Assembly Method** | de novo + reference guided assembly |
| **Consensus Caller** | Bcf Tools |

#### Coverage Map

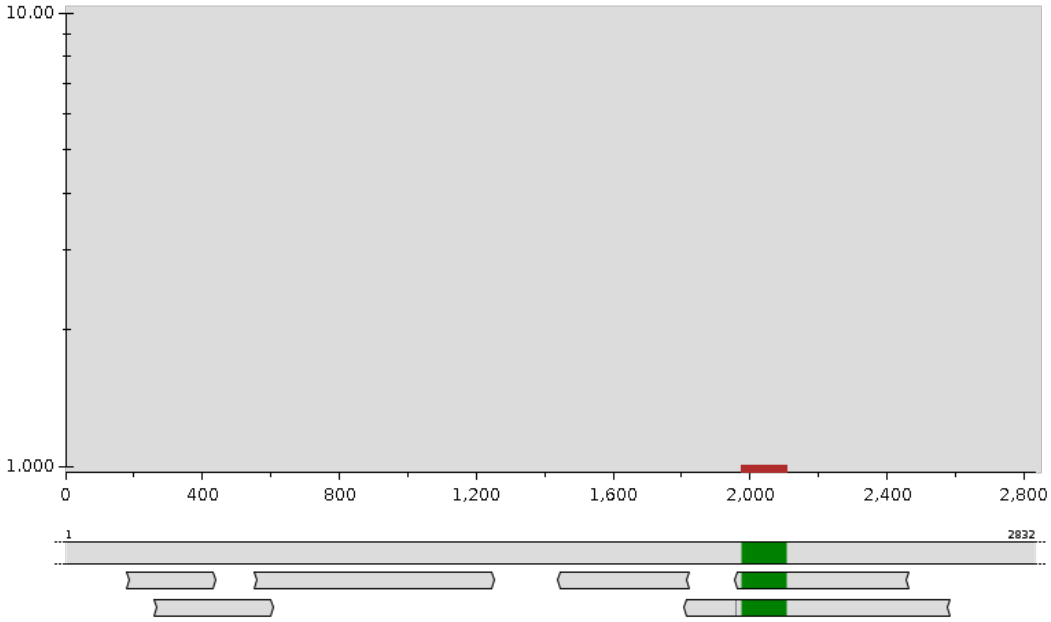

#### Assignment

| | |
|---|---|
| **Type** | Plantago lanceolata latent virus (Taxonomy ID: 1830242) |
| **Reference Genome** | NC_038477.1 |
| **NT Identity (%)** | 67.4074 |
| **AA Identity (%)** | 54.0741 |
| **Number Of Stop Codons** | 4 |
| **Number Of CDS** | 6 |

#### Alignment

| | |
|---|---|
| **Alignment Score** | 94.0 (NT) + 589.0 (AA) = 683.0 |
| **Concordance (%)** | 51.4695 |

| Alignment Method | Global, seeded, nucleotide + amino acids (AGA) |
|---|---|

#### Genome Region

Sequence starts at position 1975 and ends at position 2109 relative to NC_038477.1 reference sequence.

#### Alignment Detailed Statistics

| | Begin | End | Coverage | Score | Concordance | Matches | Identities | I/D/M/F* | Stop Codons |
|---|---|---|---|---|---|---|---|---|---|
| **NT** | **1975** | **2109** | **4.8%** | **94** | **34.8%** | **135 (100%)** | **91 (67.4%)** | **0/0** | |
| Mutations: | 1976T>C, 1979G>T, 1980G>A, 1985T>G, 1987G>C, 1994G>T, 1995G>T, 1996G>C, 1997C>T, 2000A>G, 2005A>C, 2008T>G, 2010G>A, 2011G>T, 2012G>A, 2016G>T, 2017C>T, 2018C>T, 2019C>A, 2021G>C, 2022T>C, 2024T>C, 2027G>T, 2032C>G, 2042A>G, 2045G>T, 2046G>T, 2057T>C, 2058C>T, 2059T>G, 2060G>T, 2061G>T, 2062A>T, 2063A>G, 2075C>T, 2078C>A, 2084G>T, 2086G>A, 2089A>G, 2090C>T, 2102T>G, 2103G>A, 2106T>A, 2108G>C | | | | | | | | |
| **CDS** | | | | | | | | | |
| D1Q99_gp4 | 160 | 204 | 13.4% | 233 | 63.7% | 45 (100%) | 28 (62.2%) | 0/0/0/0 | 1 |
| Protein mutations: | Y160F (2106T>A), T161I (2102T>G 2103G>A), R167* (2084G>T 2086G>A), S175K (2060G>T 2061G>T 2062A>T), R176Q (2057T>C 2058C>T 2059T>G), P180Q (2045G>T 2046G>T), E185Q (2032C>G), Y188W (2021G>C 2022T>C), R189I (2018C>T 2019C>A), A190N (2016G>T 2017C>T), P192I (2010G>A 2011G>T), N193H (2008T>G), L194V (2005A>C), P197E (1994G>T 1995G>T 1996G>C), Q200D (1985T>G 1987G>C), A202V (1979G>T 1980G>A) | | | | | | | | |
| Codon mutations: | CCC159.CG (2108G>C), TAC160TTC (2106T>A), ACA161ATC (2102T>G 2103G>A), CAG165CAA (2090C>T), TTA166CTA (2089A>G), CGC167TGA (2084G>T 2086G>A), CTG169CTT (2078C>A), GAG170GAA (2075C>T), AAT174AAC (2063A>G), TCC175AAA (2060G>T 2061G>T 2062A>T), AGA176CAG (2057T>C 2058C>T 2059T>G), CCC180CAA (2045G>T 2046G>T), CCT181CCC (2042A>G), GAA185CAA (2032C>G), CCC186CCA (2027G>T), AGA187AGG (2024T>C), TAC188TGG (2021G>C 2022T>C), AGG189ATA (2018C>T 2019C>A), GCT190AAT (2016G>T 2017C>T), TTC191TTT (2012G>A), CCT192ATT (2010G>A 2011G>T), AAT193CAT (2008T>G), TTA194GTA (2005A>C), CCT195CCC (2000A>G), GAG196GAA (1997C>T), CCC197GAA (1994G>T 1995G>T 1996G>C), CAA200GAC (1985T>G 1987G>C), GCC202GTA (1979G>T 1980G>A), GAA203GAG (1976T>C) | | | | | | | | |
| D1Q99_gp5 | 160 | 204 | 17.4% | 233 | 63.7% | 45 (100%) | 28 (62.2%) | 0/0/0/0 | 1 |
| Protein mutations: | Y160F (2106T>A), T161I (2102T>G 2103G>A), R167* (2084G>T 2086G>A), S175K (2060G>T 2061G>T 2062A>T), R176Q (2057T>C 2058C>T 2059T>G), P180Q (2045G>T 2046G>T), E185Q (2032C>G), Y188W (2021G>C 2022T>C), R189I (2018C>T 2019C>A), A190N (2016G>T 2017C>T), P192I (2010G>A 2011G>T), N193H (2008T>G), L194V (2005A>C), P197E (1994G>T 1995G>T 1996G>C), Q200D (1985T>G 1987G>C), A202V (1979G>T 1980G>A) | | | | | | | | |
| Codon mutations: | CCC159.CG (2108G>C), TAC160TTC (2106T>A), ACA161ATC (2102T>G 2103G>A), CAG165CAA (2090C>T), TTA166CTA (2089A>G), CGC167TGA (2084G>T 2086G>A), CTG169CTT (2078C>A), GAG170GAA (2075C>T), AAT174AAC (2063A>G), TCC175AAA (2060G>T 2061G>T 2062A>T), AGA176CAG (2057T>C 2058C>T 2059T>G), CCC180CAA (2045G>T 2046G>T), CCT181CCC (2042A>G), GAA185CAA (2032C>G), CCC186CCA (2027G>T), AGA187AGG (2024T>C), TAC188TGG (2021G>C 2022T>C), AGG189ATA (2018C>T 2019C>A), GCT190AAT (2016G>T 2017C>T), TTC191TTT (2012G>A), CCT192ATT (2010G>A 2011G>T), AAT193CAT (2008T>G), TTA194GTA (2005A>C), CCT195CCC (2000A>G), GAG196GAA (1997C>T), CCC197GAA (1994G>T 1995G>T 1996G>C), CAA200GAC (1985T>G 1987G>C), GCC202GTA (1979G>T 1980G>A), GAA203GAG (1976T>C) | | | | | | | | |
| D1Q99_gp6 | 120 | 164 | 26.5% | 123 | 34.5% | 45 (100%) | 17 (37.8%) | 0/0/0/0 | 2 |
| Protein mutations: | P120R (2108G>C), T121S (2106T>A), H122S (2102T>G 2103G>A), S126N (2089A>G 2090C>T), A128E (2084G>T), W130L (2078C>A), S131N (2075C>T), I135T (2062A>T 2063A>G), P136N (2059T>G 2060G>T 2061G>T), D137S (2057T>C 2058C>T), P141N (2045G>T 2046G>T), L142P (2042A>G), M145I (2032C>G), P147Q (2027G>T), D148G (2024T>C), T149G (2021G>C 2022T>C), G150* (2017C>T 2018C>T 2019C>A), L151I (2016G>T), S152L (2011G>T 2012G>A), L153F (2008T>G 2010G>A), I154M (2005A>C), L156P (2000A>G), S157K (1996G>C 1997C>T), P158K (1994G>T 1995G>T), N160K (1987G>C), N161T (1985T>G), P163* (1979G>T 1980G>A), K164R (1976T>C) | | | | | | | | |
| Codon mutations: | CCT120CGT (2108G>C), ACA121TCA (2106T>A), CAT122TCT (2102T>G 2103G>A), AGT126AAC (2089A>G 2090C>T), TAC127TAT (2086G>A), GCA128GAA (2084G>T), TGG130TTG (2078C>A), AGT131AAT (2075C>T), ATT135ACA (2062A>T 2063A>G), CCA136AAC (2059T>G 2060G>T 2061G>T), GAT137AGT (2057T>C 2058C>T), CCC141AAC (2045G>T 2046G>T), CTG142CCG (2042A>G), ATG145ATC (2032C>G), CCA147CAA (2027G>T), GAT148GGT (2024T>C), ACA149GGA (2021G>C 2022T>C), GGG150TAA (2017C>T 2018C>T 2019C>A), CTT151ATT (2016G>T), TCC152TTA (2011G>T 2012G>A), CTA153TTC (2008T>G 2010G>A), ATT154ATG (2005A>C), CTG156CCG (2000A>G), AGC157AAG (1996G>C 1997C>T), CCA158AAA (1994G>T 1995G>T), AAC160AAG (1987G>C), AAT161ACT (1985T>G), CCG163TAG (1979G>T 1980G>A), AAA164AGA (1976T>C) | | | | | | | | |
| **Proteins** | | | | | | | | | |
| replication-associated protein (YP_009506572.1) | 160 | 204 | 13.4% | 233 | 63.7% | 45 (100%) | 28 (62.2%) | 0/0/0/0 | 1 |
| Protein mutations: | Y160F (2106T>A), T161I (2102T>G 2103G>A), R167* (2084G>T 2086G>A), S175K (2060G>T 2061G>T 2062A>T), R176Q (2057T>C 2058C>T 2059T>G), P180Q (2045G>T 2046G>T), E185Q (2032C>G), Y188W (2021G>C 2022T>C), R189I (2018C>T 2019C>A), A190N (2016G>T 2017C>T), P192I (2010G>A 2011G>T), N193H (2008T>G), L194V (2005A>C), P197E (1994G>T 1995G>T 1996G>C), Q200D (1985T>G 1987G>C), A202V (1979G>T 1980G>A) | | | | | | | | |
| Codon mutations: | CCC159.CG (2108G>C), TAC160TTC (2106T>A), ACA161ATC (2102T>G 2103G>A), CAG165CAA (2090C>T), TTA166CTA (2089A>G), CGC167TGA (2084G>T 2086G>A), CTG169CTT (2078C>A), GAG170GAA (2075C>T), AAT174AAC (2063A>G), TCC175AAA (2060G>T 2061G>T 2062A>T), AGA176CAG (2057T>C 2058C>T 2059T>G), CCC180CAA (2045G>T 2046G>T), CCT181CCC (2042A>G), GAA185CAA (2032C>G), CCC186CCA (2027G>T), AGA187AGG (2024T>C), TAC188TGG (2021G>C 2022T>C), AGG189ATA (2018C>T 2019C>A), GCT190AAT (2016G>T 2017C>T), TTC191TTT (2012G>A), CCT192ATT (2010G>A 2011G>T), AAT193CAT (2008T>G), TTA194GTA (2005A>C), CCT195CCC (2000A>G), GAG196GAA (1997C>T), CCC197GAA (1994G>T 1995G>T 1996G>C), CAA200GAC (1985T>G 1987G>C), GCC202GTA (1979G>T 1980G>A), GAA203GAG (1976T>C) | | | | | | | | |
| RepA (YP_009506573.1) | 160 | 204 | 17.4% | 233 | 63.7% | 45 (100%) | 28 (62.2%) | 0/0/0/0 | 1 |
| Protein mutations: | Y160F (2106T>A), T161I (2102T>G 2103G>A), R167* (2084G>T 2086G>A), S175K (2060G>T 2061G>T 2062A>T), R176Q (2057T>C 2058C>T 2059T>G), P180Q (2045G>T 2046G>T), E185Q (2032C>G), Y188W (2021G>C 2022T>C), R189I (2018C>T 2019C>A), A190N (2016G>T 2017C>T), P192I (2010G>A 2011G>T), N193H (2008T>G), L194V (2005A>C), P197E (1994G>T 1995G>T 1996G>C), Q200D (1985T>G 1987G>C), A202V (1979G>T 1980G>A) | | | | | | | | |
| Codon mutations: | CCC159.CG (2108G>C), TAC160TTC (2106T>A), ACA161ATC (2102T>G 2103G>A), CAG165CAA (2090C>T), TTA166CTA (2089A>G), CGC167TGA (2084G>T 2086G>A), CTG169CTT (2078C>A), GAG170GAA (2075C>T), AAT174AAC (2063A>G), TCC175AAA (2060G>T 2061G>T 2062A>T), AGA176CAG (2057T>C 2058C>T 2059T>G), CCC180CAA (2045G>T 2046G>T), CCT181CCC (2042A>G), GAA185CAA (2032C>G), CCC186CCA (2027G>T), AGA187AGG (2024T>C), TAC188TGG (2021G>C 2022T>C), AGG189ATA (2018C>T 2019C>A), GCT190AAT (2016G>T 2017C>T), TTC191TTT (2012G>A), CCT192ATT (2010G>A 2011G>T), AAT193CAT (2008T>G), TTA194GTA (2005A>C), CCT195CCC (2000A>G), GAG196GAA (1997C>T), CCC197GAA (1994G>T 1995G>T 1996G>C), CAA200GAC (1985T>G 1987G>C), GCC202GTA (1979G>T 1980G>A), GAA203GAG (1976T>C) | | | | | | | | |
| C3 (YP_009506574.1) | 120 | 164 | 26.5% | 123 | 34.5% | 45 (100%) | 17 (37.8%) | 0/0/0/0 | 2 |
| Protein mutations: | P120R (2108G>C), T121S (2106T>A), H122S (2102T>G 2103G>A), S126N (2089A>G 2090C>T), A128E (2084G>T), W130L (2078C>A), S131N (2075C>T), I135T (2062A>T 2063A>G), P136N (2059T>G 2060G>T 2061G>T), D137S (2057T>C 2058C>T), P141N (2045G>T 2046G>T), L142P (2042A>G), M145I (2032C>G), P147Q (2027G>T), D148G (2024T>C), T149G (2021G>C 2022T>C), G150* (2017C>T 2018C>T 2019C>A), L151I (2016G>T), S152L (2011G>T 2012G>A), L153F (2008T>G 2010G>A), I154M (2005A>C), L156P (2000A>G), S157K (1996G>C 1997C>T), P158K (1994G>T 1995G>T), N160K (1987G>C), N161T (1985T>G), P163* (1979G>T 1980G>A), K164R (1976T>C) | | | | | | | | |
| Codon mutations: | CCT120CGT (2108G>C), ACA121TCA (2106T>A), CAT122TCT (2102T>G 2103G>A), AGT126AAC (2089A>G 2090C>T), TAC127TAT (2086G>A), GCA128GAA (2084G>T), TGG130TTG (2078C>A), AGT131AAT (2075C>T), ATT135ACA (2062A>T 2063A>G), CCA136AAC (2059T>G 2060G>T 2061G>T), GAT137AGT (2057T>C 2058C>T), CCC141AAC (2045G>T 2046G>T), CTG142CCG (2042A>G), ATG145ATC (2032C>G), CCA147CAA (2027G>T), GAT148GGT (2024T>C), ACA149GGA (2021G>C 2022T>C), GGG150TAA (2017C>T 2018C>T 2019C>A), CTT151ATT (2016G>T), TCC152TTA (2011G>T 2012G>A), CTA153TTC (2008T>G 2010G>A), ATT154ATG (2005A>C), CTG156CCG (2000A>G), AGC157AAG (1996G>C 1997C>T), CCA158AAA (1994G>T 1995G>T), AAC160AAG (1987G>C), AAT161ACT (1985T>G), CCG163TAG (1979G>T 1980G>A), AAA164AGA (1976T>C) | | | | | | | | |

*: Inserts / Deletes / Misaligned / Frameshifts

#### Analysis details

This analysis was performed with panviral2.64

### NGS Details (UN9): Duamitovirus dapi1

#### Assembly

| | |
|---|---|
| **Coverage Length** | 134 (1 contig(s)) |
| **Depth Of Coverage** | 1.0 |
| **Number Of Reads** | 1 |
| **Reads Per Million** | 0.02 rpm (after QC) |
| **Ambiguities** | 0 |
| **Assembly Method** | de novo + reference guided assembly |
| **Consensus Caller** | Bcf Tools |

#### Coverage Map

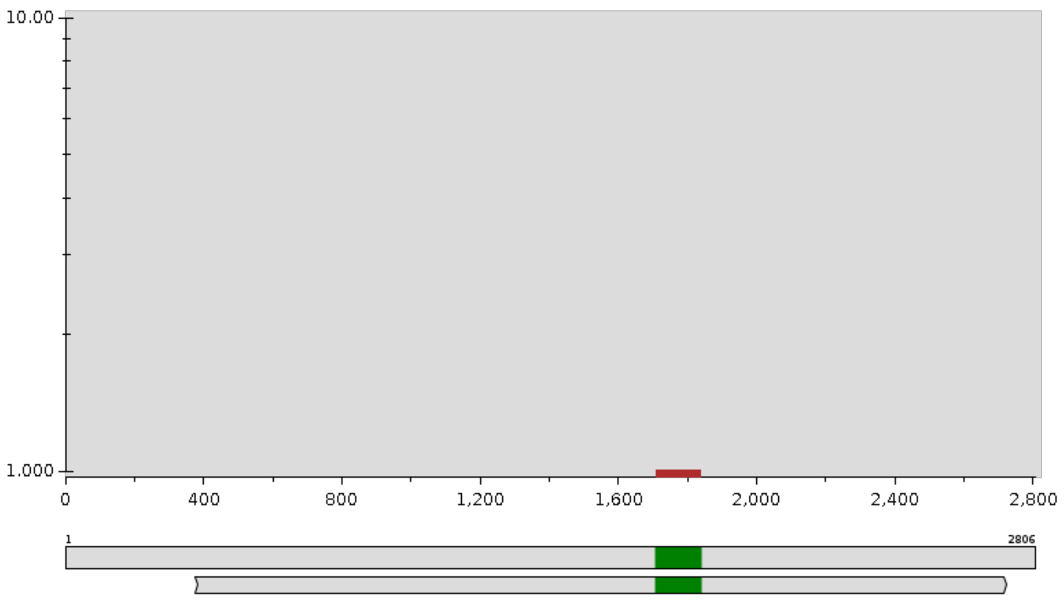

#### Assignment

| | |
|---|---|
| **Type** | Duamitovirus dapi1 (Taxonomy ID: 2955778) |
| **Reference Genome** | NC_076528.1 |
| **NT Identity (%)** | 68.6567 |
| **AA Identity (%)** | 68.1818 |
| **Number Of Stop Codons** | 0 |
| **Number Of CDS** | 1 |

#### Alignment

| | |
|---|---|
| **Alignment Score** | 100.0 (NT) + 249.0 (AA) = 349.0 |
| **Concordance (%)** | 57.0261 |

| **Alignment Method** | Global, seeded, nucleotide + amino acids (AGA) |
|---|---|

#### Genome Region

Sequence starts at position 1709 and ends at position 1842 relative to NC_076528.1 reference sequence.

#### Alignment Detailed Statistics

| | Begin | End | Coverage | Score | Concordance | Matches | Identities | I/D/M/F* | Stop Codons |
|---|---|---|---|---|---|---|---|---|---|
| **NT** | **1709** | **1842** | **4.8%** | **100** | **37.3%** | **134 (100%)** | **92 (68.7%)** | **0/0** | |
| Mutations: | 1710C>T, 1713A>G, 1715G>A, 1725T>C, 1728G>C, 1734C>T, 1736A>T, 1737C>G, 1738T>A, 1739C>G, 1743C>T, 1749G>T, 1750T>C, 1751C>T, 1755C>T, 1756G>A, 1763C>A, 1774G>C, 1776A>T, 1783C>A, 1785G>A, 1788G>T, 1789G>A, 1791C>T, 1792T>G, 1793G>A, 1794G>A, 1800A>G, 1804C>G, 1805C>G, 1806T>C, 1809T>G, 1810C>A, 1811A>C, 1816T>C, 1819A>T, 1820A>G, 1821G>C, 1827C>T, 1830T>C, 1831C>A, 1839T>C | | | | | | | | |
| **CDS** | | | | | | | | | |
| RdRp | 446 | 489 | 5.6% | 249 | 70.3% | 44 (100%) | 30 (68.2%) | 0/0/0/0 | 0 |
| Protein mutations: | R447K (1715G>A), Y454L (1736A>T 1737C>G), S459L (1750T>C 1751C>T), A461T (1756G>A), T463K (1763C>A), V467L (1774G>C 1776A>T), L470I (1783C>A 1785G>A), A472T (1789G>A 1791C>T), W473E (1792T>G 1793G>A 1794G>A), P477G (1804C>G 1805C>G 1806T>C), Q479T (1810C>A 1811A>C), F481L (1816T>C), K482C (1819A>T 1820A>G 1821G>C), L486I (1831C>A) | | | | | | | | |
| Codon mutations: | TTC445.TT (1710C>T), ACA446ACG (1713A>G), AGG447AAG (1715G>A), CCT450CCC (1725T>C), CTG451CTC (1728G>C), TTC453TTT (1734C>T), TAC454TTG (1736A>T 1737C>G), TCT455AGT (1738T>A 1739C>G), TCC456TCT (1743C>T), CCG458CCT (1749G>T), TCA459CTA (1750T>C 1751C>T), TTC460TTT (1755C>T), GCA461ACA (1756G>A), ACA463AAA (1763C>A), GTA467CTT (1774G>C 1776A>T), CTG470ATA (1783C>A 1785G>A), GCG471GCT (1788G>T), GCC472ACT (1789G>A 1791C>T), TGG473GAA (1792T>G 1793G>A 1794G>A), GTA475GTG (1800A>G), CCT477GGC (1804C>G 1805C>G 1806T>C), ACT478ACG (1809T>G), CAG479ACG (1810C>A 1811A>C), TTT481CTT (1816T>C), AAG482TGC (1819A>T 1820A>G 1821G>C), TAC484TAT (1827C>T), GCT485GCC (1830T>C), CTT486ATT (1831C>A), GGT488GGC (1839T>C) | | | | | | | | |
| **Proteins** | | | | | | | | | |
| RNA-dependent RNA polymerase (YP_010798878.1) | 446 | 489 | 5.6% | 249 | 70.3% | 44 (100%) | 30 (68.2%) | 0/0/0/0 | 0 |
| Protein mutations: | R447K (1715G>A), Y454L (1736A>T 1737C>G), S459L (1750T>C 1751C>T), A461T (1756G>A), T463K (1763C>A), V467L (1774G>C 1776A>T), L470I (1783C>A 1785G>A), A472T (1789G>A 1791C>T), W473E (1792T>G 1793G>A 1794G>A), P477G (1804C>G 1805C>G 1806T>C), Q479T (1810C>A 1811A>C), F481L (1816T>C), K482C (1819A>T 1820A>G 1821G>C), L486I (1831C>A) | | | | | | | | |
| Codon mutations: | TTC445.TT (1710C>T), ACA446ACG (1713A>G), AGG447AAG (1715G>A), CCT450CCC (1725T>C), CTG451CTC (1728G>C), TTC453TTT (1734C>T), TAC454TTG (1736A>T 1737C>G), TCT455AGT (1738T>A 1739C>G), TCC456TCT (1743C>T), CCG458CCT (1749G>T), TCA459CTA (1750T>C 1751C>T), TTC460TTT (1755C>T), GCA461ACA (1756G>A), ACA463AAA (1763C>A), GTA467CTT (1774G>C 1776A>T), CTG470ATA (1783C>A 1785G>A), GCG471GCT (1788G>T), GCC472ACT (1789G>A 1791C>T), TGG473GAA (1792T>G 1793G>A 1794G>A), GTA475GTG (1800A>G), CCT477GGC (1804C>G 1805C>G 1806T>C), ACT478ACG (1809T>G), CAG479ACG (1810C>A 1811A>C), TTT481CTT (1816T>C), AAG482TGC (1819A>T 1820A>G 1821G>C), TAC484TAT (1827C>T), GCT485GCC (1830T>C), CTT486ATT (1831C>A), GGT488GGC (1839T>C) | | | | | | | | |

*: Inserts / Deletes / Misaligned / Frameshifts

#### Analysis details

This analysis was performed with panviral2.64

### Analysis of UN18_val

| | |
|---|---|
| **Technology** | Paired-end short reads |
| **Input Files** | UN18_val_R1.fq.gz (1.71 GB), UN18_val_R2.fq.gz (1.73 GB) |
| **Submitted On** | 2023-09-26 06:46:26 UTC |
| **Duration** | 3h 31m 58s |
| **Tool Version** | panviral2.64 |
| **Location** | https://www.genomedetective.com/db/ui/analysis/01461f33-a0b9-4228-b7b0-a45adba24853 |

#### Statistics

| | |
|---|---|
| **Original Read Length** | 20 - 150 |
| **Trimmed Read Length** | 50 - 135 |

| | **# Reads** | **% of Reads** |
|---|---|---|
| **Input file** | 53669874 | 100.0% |
| **After QC** | 53280916 | 99.3% |
| **After filtering** | 2405890 | 4.5% |
| **Mapped back to contigs** | 232586 | 0.4% |

#### Assignments

| **Assignment** | **No. of Reads** | **Depth of Coverage** | **Identity** | | **Genome Coverage** | |
|---|---|---|---|---|---|---|
| | | | **NT** | **AA** | | |
| Torradovirus lycopersici (2 segments out of 2) | 51427 | 582.9 | 86.8% | 91.7% | 81.0% | |
| Torradovirus lycopersici (segment RNA 2) | 38768 | 1183.7 | 83.8% | 86.2% | 72.7% | 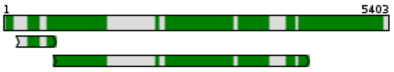  |
| Torradovirus lycopersici (segment RNA 1) | 12659 | 234.6 | 88.5% | 94.1% | 86.7% | 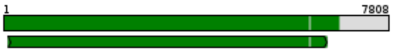  |
| Torradovirus marchitezum (2 segments out of 2) | 31334 | 381.8 | 65.1% | 68.6% | 85.5% | |
| Torradovirus marchitezum (segment RNA 2) | 26259 | 709.3 | 64.3% | 68.0% | 94.7% | 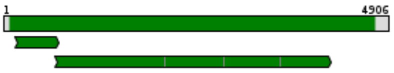  |
| Torradovirus marchitezum (segment RNA 1) | 5075 | 116.5 | 65.8% | 69.0% | 79.3% | 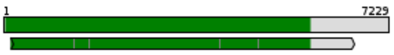  |
| Tomato necrotic dwarf virus (segment RNA1) | 21581 | 468.2 | 64.6% | 66.1% | 84.2% | 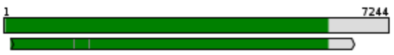  |
| Diachasmimorpha longicaudata entomopoxvirus (2 segments out of 13) | 18634 | 1786.1 | 61.3% | 59.4% | N/A | |

| Assignment | No. of Reads | Depth of Coverage | Identity | | Genome Coverage | |
|---|---|---|---|---|---|---|
| | | | NT | AA | | |
| Diachasmimorpha longicaudata entomopoxvirus | 581 | 361.0 | 76.9% | 90.9% | 0.1% | 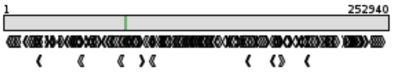 |
| Diachasmimorpha longicaudata entomopoxvirus (segment NC_043455.1) | 18053 | 1953.9 | 59.5% | 55.8% | 84.9% | 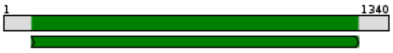 |
| Bracoviriform glomeratae (segment NC_043292.1) | 11330 | 4199.5 | 72.8% | 84.1% | 75.8% | 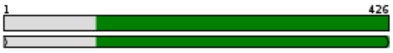 |
| Potato leafroll virus | 4303 | 98.0 | 98.1% | 97.3% | 97.4% | 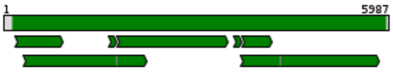 |
| Duamitovirus soch1 | 853 | 45.1 | 68.9% | 68.4% | 82.0% | 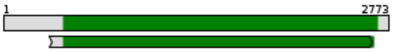 |
| Solendovirus venanicotianae | 218 | 15.1 | 78.0% | 74.0% | 21.4% | 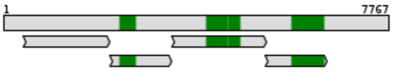 |
| Potato virus Y | 45 | 3.1 | 85.2% | 91.1% | 19.3% | 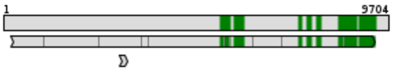 |
| Duamitovirus peex1 | 32 | 5.3 | 69.7% | 67.5% | 26.1% | 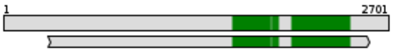 |
| Harvey murine sarcoma virus | 6 | 2.2 | 73.4% | 85.0% | 22.3% | 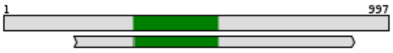 |

#### Discoveries

| Similar to | No. of Reads | Depth of Coverage | Identity | | Genome Coverage | |
|---|---|---|---|---|---|---|
| | | | NT | AA | | |
| Brazilian marseillevirus | 31766 | 13067.1 | 81.7% | 94.7% | 0.1% | 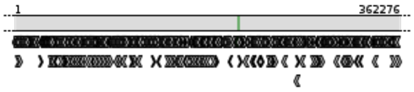 |
| Tomato chocolate spot virus (3 segments out of 2) | 23700 | 1253.2 | 72.1% | 71.7% | 18.2% | |
| Tomato chocolate spot virus (segment RNA2) | 14899 | 1610.5 | 67.0% | 74.8% | 21.8% | 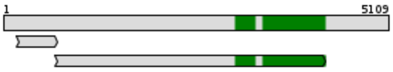 |
| Tomato chocolate spot virus (segment RNA 1) | 6736 | 1872.0 | 78.3% | 74.4% | 5.4% | 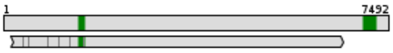 |
| Tomato chocolate spot virus (segment RNA 1) | 2065 | 416.7 | 76.2% | 13.6% | 10.3% | 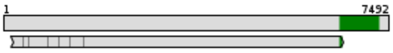 |
| Tokyovirus A1 | 12764 | 4800.1 | 76.8% | 90.7% | 0.0% | 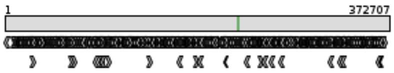 |
| Leucotheavirus sp4 | 6064 | 880.3 | 75.9% | 85.4% | 0.5% | 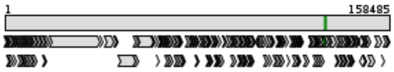 |

| Similar to | No. of Reads | Depth of Coverage | Identity | | Genome Coverage | |
|---|---|---|---|---|---|---|
| | | | NT | AA | | |
| Noumeavirus | 4184 | 2766.6 | 81.7% | 92.2% | 0.0% | 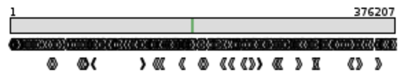  |
| Betabaculovirus disaccharalis | 2638 | 1580.3 | 82.4% | 92.9% | 0.1% | 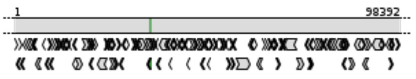  |
| Lausannevirus | 2565 | 1524.8 | 83.3% | 100.0% | 0.0% | 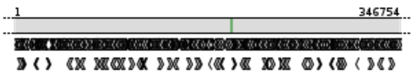  |
| Brazilian marseillevirus | 2433 | 558.0 | 81.6% | 95.9% | 0.1% | 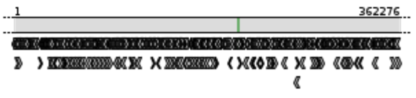  |
| Betabaculovirus disaccharalis | 2060 | 1390.5 | 83.7% | 88.4% | 0.1% | 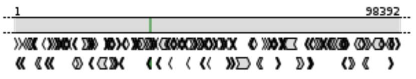  |
| Bracoviriform congregatae (segment Circle 7) | 1498 | 965.9 | 77.3% | 91.2% | 0.7% | 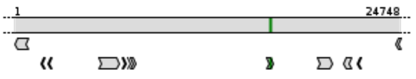  |
| Tunisvirus fontaine2 | 895 | 465.1 | 80.0% | 91.4% | 0.0% | 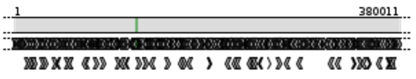  |
| Lowelvirus tuscon4d | 634 | 106.0 | 76.8% | 88.2% | 0.3% | 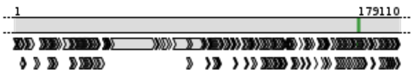  |
| Cassava brown streak virus | 450 | 103.8 | 60.5% | 57.5% | 5.8% | 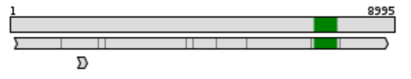  |
| Human gammaherpesvirus 8 (subtype: Could not assign) | 294 | 183.0 | 83.2% | 84.6% | 0.1% | 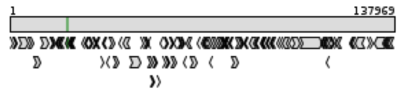  |
| Cladosporium fulvum T-1 virus | 179 | 15.0 | 53.1% | 42.6% | 16.3% | 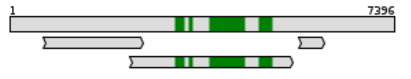  |
| Errantivirus | 149 | 16.0 | 56.1% | 44.8% | 9.6% | 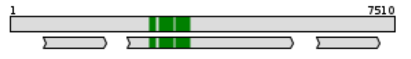  |
| Marseillevirus marseillevirus | 75 | 52.2 | 81.7% | 97.4% | 0.0% | 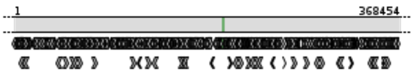  |
| Badnavirus occultipomeae | 66 | 25.5 | 58.8% | 49.5% | 3.2% | 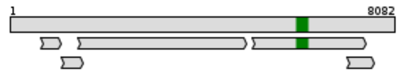  |
| Caulimovirus venafragariae | 48 | 9.9 | 58.3% | 54.9% | 7.2% | 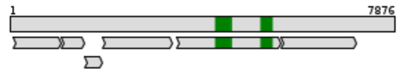  |
| Badnavirus occultipomeae | 38 | 14.4 | 60.5% | 51.8% | 3.1% | 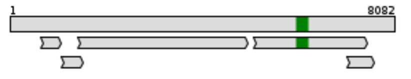  |

| Similar to | No. of Reads | Depth of Coverage | Identity | | Genome Coverage | |
|---|---|---|---|---|---|---|
| | | | NT | AA | | |
| Brazilian marseillevirus | 33 | 15.2 | 78.5% | 91.2% | 0.0% | 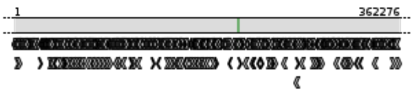 |
| Rahariannevirus rahariannе | 31 | 2.4 | 69.8% | 65.3% | 3.5% | 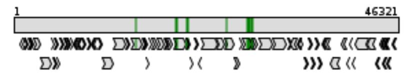 |
| Caulimovirus venafragariae | 30 | 4.8 | 58.1% | 52.9% | 10.6% | 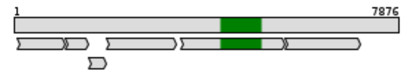 |
| Badnavirus maculapiperis | 30 | 13.7 | 57.9% | 53.2% | 3.1% | 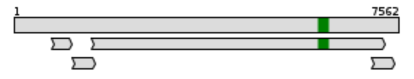 |
| Epiphyllum badnavirus 1 | 30 | 12.9 | 57.1% | 49.4% | 3.1% | 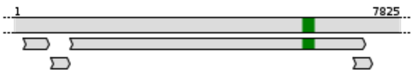 |
| Badnavirus epsiloninflatheobromae | 23 | 3.9 | 57.4% | 48.6% | 8.7% | 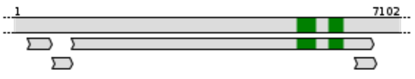 |
| Dioscovirus dioscoreae | 23 | 3.2 | 61.8% | 52.6% | 11.2% | 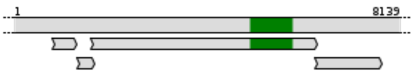 |
| Caulimovirus minutangelicae | 17 | 6.0 | 54.9% | 46.5% | 4.7% | 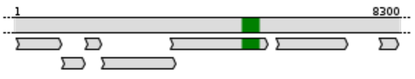 |
| Caulimovirus venafragariae | 17 | 3.8 | 56.1% | 51.2% | 7.7% | 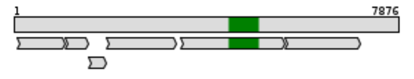 |
| Badnavirus maculasmallanthi | 13 | 5.8 | 56.3% | 48.9% | 3.7% | 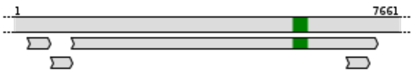 |
| Pinus nigra virus 1 | 13 | 3.8 | 58.0% | 48.3% | 6.1% | 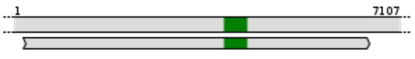 |
| Hibiscus bacilliform virus GD1 | 12 | 6.1 | 56.3% | 47.5% | 3.2% | 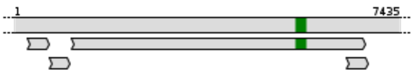 |
| Caulimovirus tessellomirabilis | 11 | 3.1 | 58.2% | 47.7% | 5.9% | 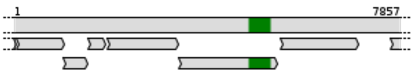 |
| Badnavirus tessellocastaneae | 6 | 2.8 | 55.9% | 54.5% | 3.7% | 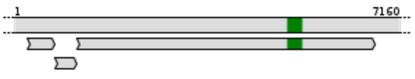 |
| Betabaculovirus erellonis | 6 | 5.9 | 82.4% | 93.3% | 0.1% | 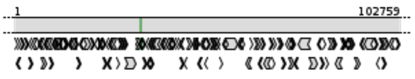 |
| Gihfavirus pelohabitans | 4 | 2.3 | 97.4% | 0.0% | 3.5% | 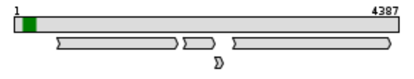 |
| Duamitovirus dapi1 | 4 | 3.8 | 69.4% | 66.7% | 4.8% | 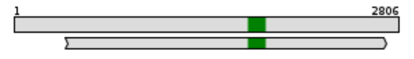 |

| Similar to | No. of Reads | Depth of Coverage | Identity | | Genome Coverage | |
|---|---|---|---|---|---|---|
| | | | NT | AA | | |
| Fusarium graminearum mycotymovirus 1 | 4 | 2.0 | 69.5% | 76.7% | 3.4% | 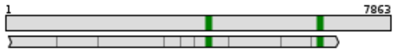 |
| Tokyovirus A1 | 4 | 2.9 | 77.0% | 93.3% | 0.0% | 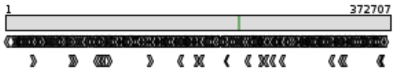 |
| Botryotinia fuckeliana partitivirus 1 (segment RNA 1) | 4 | 3.6 | 68.1% | 71.1% | 7.5% | 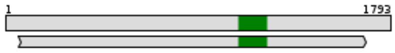 |
| Gammaflexivirus PaGFV-1 | 2 | 2.0 | 80.0% | 86.7% | 1.4% | 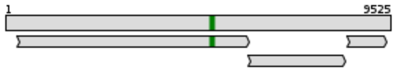 |
| Sugarcane chlorotic streak virus | 2 | 1.9 | 62.8% | 55.8% | 4.4% | 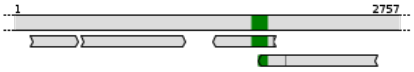 |
| Seussvirus seuss | 2 | 1.1 | 76.3% | 79.5% | 0.3% | 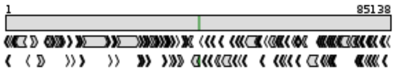 |
| Fusarium poae narnavirus 2 | 2 | 1.0 | 74.8% | 74.4% | 12.0% | 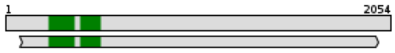 |

### NGS Details (UN18_val): Diachasmimorpha longicaudata entomopoxvirus

#### Assembly

| | |
|---|---|
| **Coverage Length** | 134 (1 contig(s)) |
| **Depth Of Coverage** | 361.0 |
| **Number Of Reads** | 581 |
| **Reads Per Million** | 10.90 rpm (after QC) |
| **Ambiguities** | 0 |
| **Assembly Method** | de novo + reference guided assembly |
| **Consensus Caller** | Bcf Tools |

#### Coverage Map

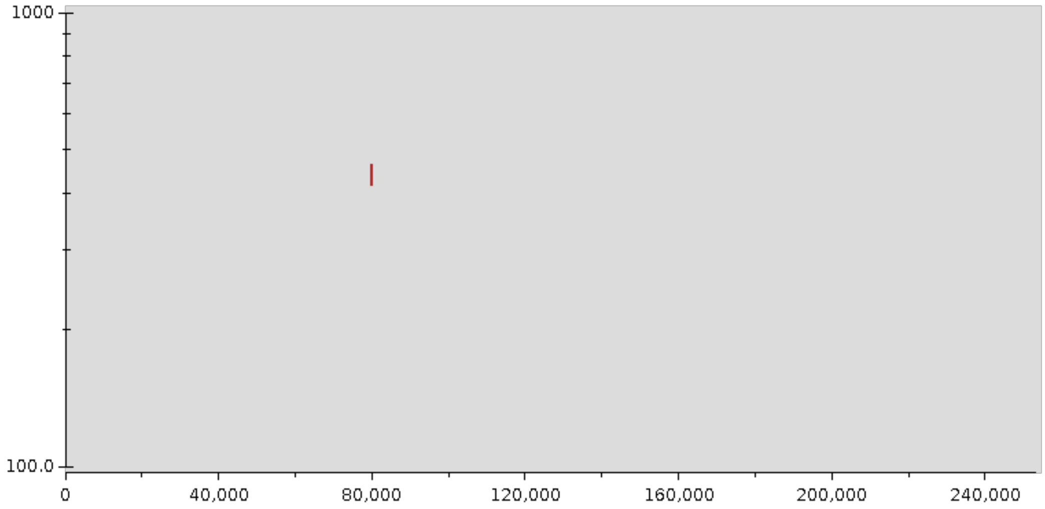

#### Assignment

| | |
|---|---|
| **Type** | Diachasmimorpha longicaudata entomopoxvirus (Taxonomy ID: 109981) |
| **Reference Genome** | NC_076102.1 |
| **NT Identity (%)** | 76.8657 |
| **AA Identity (%)** | 90.9091 |
| **Number Of Stop Codons** | 0 |
| **Number Of CDS** | 193 |

#### Alignment

| | |
|---|---|
| **Alignment Score** | 144.0 (NT) + 293.0 (AA) = 437.0 |
| **Concordance (%)** | 74.0678 |
| **Alignment Method** | Local, heuristic, nucleotide (BLASTN) |

#### Genome Region

Sequence starts at position 79871 and ends at position 80004 relative to NC_076102.1 reference sequence.

#### Alignment Detailed Statistics

| | Begin | End | Coverage | Score | Concordance | Matches | Identities | I/D/M/F* | Stop Codons |
|---|---|---|---|---|---|---|---|---|---|
| NT | 79871 | 80004 | 0.1% | 144 | 53.7% | 134 (100%) | 103 (76.9%) | 0/0 | |

Mutations: 79886G>A, 79894T>C, 79895C>A, 79898T>C, 79904G>A, 79910T>A, 79912T>G, 79917C>G, 79918T>A, 79919T>G, 79921T>G, 79928A>G, 79934T>A, 79937A>G, 79940A>G, 79941T>A, 79942T>G, 79943T>C, 79949A>G, 79952A>G, 79954G>T, 79958C>A, 79960A>G, 79961A>G, 79964G>A, 79970G>A, 79973A>C, 79976T>C, 79982T>C, 79984T>C, 79985T>C

*: Inserts / Deletes / Misaligned / Frameshifts

#### Analysis details

This analysis was performed with panviral2.64

### NGS Details (UN18_val): Diachasmimorpha longicaudata entomopoxvirus (segment NC_043455.1)

#### Assembly

| | |
|---|---|
| **Coverage Length** | 1138 (1 contig(s)) |
| **Depth Of Coverage** | 1953.9 |
| **Number Of Reads** | 18053 |
| **Reads Per Million** | 338.83 rpm (after QC) |
| **Ambiguities** | 0 |
| **Assembly Method** | de novo + reference guided assembly |
| **Consensus Caller** | Bcf Tools |

#### Coverage Map

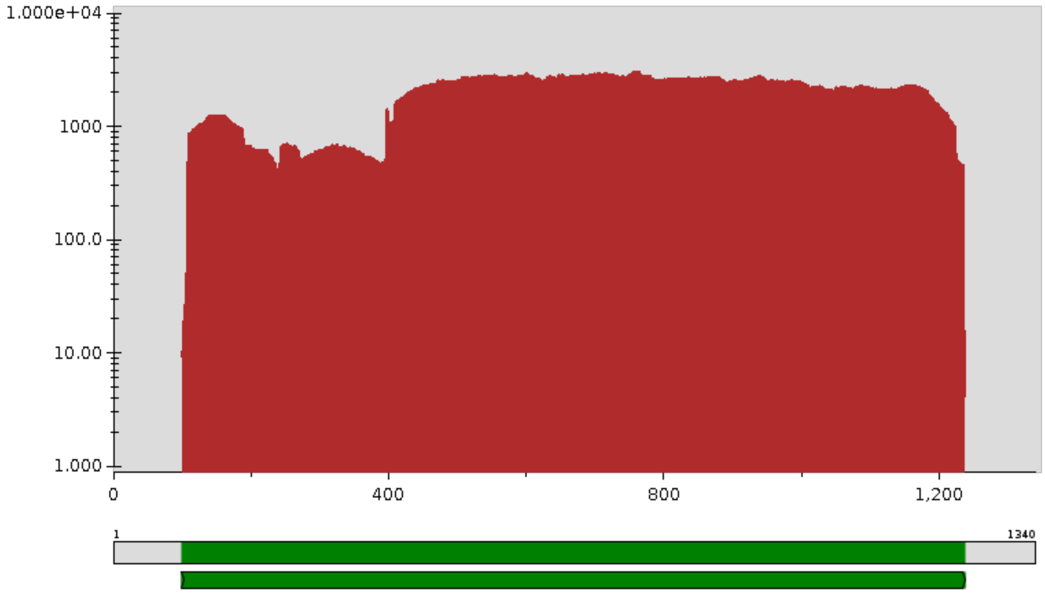

#### Assignment

| | |
|---|---|
| **Type** | Diachasmimorpha longicaudata entomopoxvirus (Taxonomy ID: 109981) |
| **Reference Genome** | NC_043455.1 |
| **NT Identity (%)** | 59.4903 |
| **AA Identity (%)** | 55.7895 |
| **Number Of Stop Codons** | 0 |
| **Number Of CDS** | 1 |

#### Alignment

| | |
|---|---|
| **Alignment Score** | 414.0 (NT) + 1484.0 (AA) = 1898.0 |
| **Concordance (%)** | 39.9159 |

| Alignment Method | Global, seeded, nucleotide + amino acids (AGA) |
|---|---|

#### Genome Region

Sequence starts at position 101 and ends at position 1238 relative to NC_043455.1 reference sequence.

#### Alignment Detailed Statistics

| | Begin | End | Coverage | Score | Concordance | Matches | Identities | I/D/M/F* | Stop Codons |
|---|---|---|---|---|---|---|---|---|---|
| **NT** | **101** | **1238** | **84.9%** | **414** | **18.3%** | **1135 (99.5%)** | **677 (59.3%)** | **3/3** | |
| Mutations: | 110A>G, 112G>T, 117C>A, 119G>A, 120A>G, 121A>T, 127C>T, 128C>A, 129A>C, 130T>C, 136C>T, 137C>T, 139C>G, 140A>C, 142G>A, 143A>G, 148T>C, 149A>C, 151C>T, 154A>G, 156A>G, 157G>A, 160T>C, 163A>C, 167T>G, 169T>C, 171G>A, 175A>T, 181A>G, 187T>A, 190A>T, 191A>G, 193T>C, 199A>G, 204A>G, 207C>G, 208C>T, 211T>A, 212T>G, 214T>G, 217A>T, 219G>T, 221A>T, 222T>G, 223T>C, 224T>A, 225C>A, 226T>G, 230A>C, 231A>T, 232A>T, 235C>T, 238G>C, 241A>T, 242G>C, 243T>A, 244C>T, 247A>G, 256G>T, 262A>C, 268A>G, 271A>C, 274A>C, 276C>T, 277A>G, 278T>A, 279A>T, 283A>T, 284A>C, 286C>T, 288G>C, 289T>C, 294T>G, 295A>C, 298G>A, 299C>G, 300A>T, 301A>T, 302A>G, 304A>G, 310T>C, 311T>A, 312C>A, 313T>A, 320A>G, 322T>G, 323A>G, 325C>T, 335A>G, 337C>A, 341A>G, 343C>T, 346A>T, 352T>G, 355A>G, 356T>C, 358G>T, 361T>A, 362C>G, 363T>C, 364C>T, 368G>A, 370T>G, 371C>G, 374C>A, 375G>A, 376T>G, 379A>G, 380T>A, 382G>T, 383C>T, 384A>T, 385A>G, 386A>G, 394G>A, 395A>G, 397T>C, 403A>G, 404_406delTAT, 407A>G, 408A>G, 409T>C, 410T>G, 412T>G, 415A>G, 416T>G, 417G>T, 421A>T, 423T>C, 428A>G, 433A>G, 436C>T, 439A>C, 443A>G, 446A>C, 447A>G, 448G>T, 451A>G, 452A>G, 453G>A, 454C>T, 458G>C, 459A>G, 460A>T, 462C>T, 463T>C, 466A>T, 467A>C, 471A>G, 472A>T, 472_473insGGT, 474C>T, 475C>T, 478G>T, 481T>A, 482T>G, 485A>G, 490A>T, 496T>A, 499A>C, 502A>T, 503A>G, 505G>C, 506A>T, 511T>C, 512C>A, 514A>G, 517C>G, 518A>C, 519C>G, 521C>A, 523A>G, 524A>C, 526A>G, 527A>T, 528G>C, 530A>C, 532C>T, 533G>C, 534A>G, 535T>C, 536A>C, 538C>T, 539A>G, 541A>C, 542G>C, 543C>A, 544A>C, 547T>C, 550A>G, 553A>G, 554G>T, 559A>T, 560A>T, 562A>G, 574T>C, 577G>A, 581T>C, 583G>C, 584A>T, 585T>C, 586A>T, 587G>A, 588A>G, 589T>A, 590A>G, 591A>G, 595T>C, 596T>A, 597T>A, 602A>C, 604A>G, 607A>T, 608C>T, 610A>T, 616T>A, 620G>C, 622A>G, 623T>C, 628T>G, 629G>C, 630A>C, 632A>C, 633G>C, 634T>A, 635C>A, 637T>A, 638G>A, 646T>G, 647A>G, 648T>G, 650T>G, 652A>T, 655A>C, 658A>T, 661A>C, 664C>T, 665G>A, 667T>G, 671T>C, 674A>G, 675G>A, 678T>C, 680A>C, 682C>T, 683A>G, 685T>G, 687C>T, 688A>T, 689T>A, 692C>A, 693A>G, 694G>A, 695G>A, 696T>A, 697C>G, 700T>C, 705G>A, 706A>C, 707G>A, 709T>A, 712C>T, 715A>G, 717A>G, 718A>G, 724G>T, 727A>G, 730A>G, 731A>C, 732A>G, 735C>A, 739A>G, 740T>C, 742G>C, 745A>T, 748C>T, 757T>C, 759G>A, 765A>T, 770A>G, 772A>C, 778A>T, 779A>G, 781G>C, 784A>G, 785A>G, 787T>A, 790T>G, 792T>G, 793T>G, 796A>G, 797G>C, 798C>T, 799A>T, 805A>T, 810T>G, 811T>C, 820T>C, 823C>G, 824C>A, 825A>C, 830A>G, 831G>C, 833C>A, 838T>C, 841A>G, 843C>G, 844A>T, 845C>G, 847A>C, 850A>C, 854T>G, 855G>T, 856T>G, 859T>C, 862A>C, 863C>A, 865T>G, 866A>C, 868A>T, 869C>A, 878G>T, 879T>G, 880A>G, 885T>C, 886G>T, 889A>T, 890T>A, 891G>A, 892T>G, 893T>A, 895A>G, 896A>C, 897C>G, 898A>C, 900A>G, 901T>C, 902A>C, 904A>T, 905A>G, 908T>C, 909T>A, 915C>T, 919A>T, 920A>G, 921G>C, 922C>A, 924T>C, 925C>T, 934T>C, 938T>G, 939C>A, 943A>G, 944C>A, 946A>C, 947G>A, 948A>C, 949A>T, 950C>A, 952T>A, 956T>A, 958T>C, 965A>C, 966A>G, 970A>G, 973T>C, 974A>C, 977G>T, 978A>C, 982T>G, 983A>T, 984A>C, 986A>T, 989A>C, 991A>T, 992A>G, 994A>G, 997T>C, 998C>A, 1000T>C, 1001T>A, 1007A>G, 1012A>C, 1013T>C, 1015A>T, 1018A>T, 1019A>C, 1021A>T, 1024C>T, 1033T>A, 1036A>G, 1040A>G, 1042A>C, 1045T>C, 1046T>C, 1048A>T, 1051T>C, 1054C>T, 1057T>C, 1063T>C, 1064T>C, 1070C>A, 1071A>C, 1073A>C, 1075T>A, 1076A>C, 1077G>C, 1087T>C, 1088A>C, 1090A>G, 1103A>C, 1105A>T, 1111T>A, 1112A>C, 1120C>A, 1121C>A, 1123A>G, 1126A>G, 1129G>T, 1130A>G, 1141T>C, 1145A>G, 1147A>C, 1150T>C, 1151G>A, 1153A>G, 1156C>T, 1159A>T, 1160T>G, 1161T>A, 1163C>A, 1164C>G, 1165A>G, 1167A>T, 1172G>T, 1173C>T, 1177A>C, 1178T>A, 1181G>C, 1183A>G, 1185C>A, 1186G>A, 1192T>C, 1195T>C, 1196A>G, 1197C>T, 1199A>G, 1200A>T, 1204A>C, 1207T>G, 1210A>G, 1211A>C, 1218A>C, 1219G>C, 1223A>G, 1227T>C, 1232T>C, 1234G>C, 1235A>C, 1237A>G | | | | | | | | |
| **CDS** | | | | | | | | | |
| FLA14_p101 | 1 | 380 | 100% | 1484 | 59.2% | 379 (99.5%) | 212 (55.6%) | 1/1/0/0 | 0 |
| Protein mutations: | M4V (110A>G 112G>T), A6D (117C>A), E7S (119G>A 120A>G 121A>T), H10T (128C>A 129A>C 130T>C), K14Q (140A>C 142G>A), K15E (143A>G), I17L (149A>C 151C>T), K19R (156A>G 157G>A), S23A (167T>G 169T>C), C24Y (171G>A), T31A (191A>G 193T>C), K35R (204A>G), A36G (207C>G 208C>T), F38V (212T>G 214T>G), C40F (219G>T), I41C (221A>T 222T>G 223T>C), S42K (224T>A 225C>A 226T>G), K44L (230A>C 231A>T 232A>T), V48H (242G>C 243T>A 244C>T), T59M (276C>T 277A>G), Y60I (278T>A 279A>T), I62L (284A>C 286C>T), S63T (288G>C 289T>C), L65C (294T>G 295A>C), Q67V (299C>G 300A>T 301A>T), I68V (302A>G 304A>G), S71K (311T>A 312C>A 313T>A), N74E (320A>G 322T>G), I75V (323A>G 325C>T), I79V (335A>G 337C>A), T81A (341A>G 343C>T), L88A (362C>G 363T>C 364C>T), A90T (368G>A 370T>G), Q91E (371C>G), R92K (374C>A 375G>A 376T>G), L94I (380T>A 382G>T), Q95L (383C>T 384A>T 385A>G), T96A (386A>G), N99D (395A>G 397T>C), Y102del (404_406delTAT), N103G (407A>G 408A>G 409T>C), F104V (410T>G 412T>G), C106V (416T>G 417G>T), Q107H (421A>T), V108A (423T>C), I110V (428A>G), I115V (443A>G), K116R (446A>C 447A>G 448G>T), S118D (452A>G 453G>A 454C>T), E120R (458G>C 459A>G 460A>T), T121I (462C>T 463T>C), K123Q (467A>C), K124S (471A>G 472A>T), K124_A125insG (472_473insGGT), A125V (474C>T 475C>T), Q126H (478G>T), L128V (482T>G), I129V (485A>G), M135V (503A>G 505G>C), I136F (506A>T), L138M (512C>A 514A>G), T140R (518A>C 519C>G), K142Q (524A>C 526A>G), I144L (530A>C 532C>T), D145R (533G>C 534A>G 535T>C), T146P (536A>C 538C>T), K147D (539A>G 541A>C), A148H (542G>C 543C>A 544A>C), I151M (553A>G), V152F (554G>T), I154L (560A>T 562A>G), I162S (584A>T 585T>C 586A>T), D163R (587G>A 588A>G 589T>A), N164G (590A>G 591A>G), L166K (596T>A 597T>A), K168Q (602A>C 604A>G), Q170Y (608C>T 610A>T), E174Q (620G>C 622A>G), F175L (623T>C), F176L (628T>G), E177P (629G>C 630A>C), S178P (632A>C 633G>C 634T>A), H179K (635C>A 637T>A), V180I (638G>A), I183G (647A>G 648T>G), L184V (650T>G 652A>T), L185F (655A>C), V189M (665G>A 667T>G), S191P (671T>C), R192E (674A>G 675G>A), V193A (678T>C), I194L (680A>C 682C>T), N195E (683A>G 685T>G), T196I (687C>T 688A>T), S197T (689T>A), Q198R (692C>A 693A>G 694G>A), V199K (695G>A 696T>A 697C>G), R202N (705G>A 706A>C), D203K (707G>A 709T>A), K206R (717A>G 718A>G), N211R (731A>C 732A>G), A212D (735C>A), R220K (759G>A), Y222F (765A>T), I224V (770A>G 772A>C), K227D (779A>G 781G>C), N229E (785A>G 787T>A), D230E (790T>G), F231W (792T>G 793T>G), A233L (797G>C 798C>T 799A>T), F237C (810T>G 811T>C), D241E (823C>G), H242T (824C>A 825A>C), S244A (830A>G 831G>C), L245I (833C>A), T248S (843C>G 844A>T), L249V (845C>G 847A>C), C252V (854T>G 855G>T 856T>G), Q257K (869C>A), V260W (878G>T 879T>G 880A>G), M262T (885T>C 886G>T), E263D (889A>T), C264K (890T>A 891G>A 892T>G), L265M (893T>A 895A>G), T266R (896A>C 897C>G 898A>C), N267S (900A>G 901T>C), N269D (905A>G), F270H (908T>C 909T>A), A272V (915C>T), S274A (920A>G 921G>C 922C>A), I275T (924T>C 925C>T), S280D (938T>G 939C>A), Q282N (944C>A 946A>C), E283T (947G>A 948A>C 949A>T), F286I (956T>A 958T>C), K289R (965A>C 966A>G), D293S (977G>T 978A>C), K295S (983A>T 984A>C), T296S (986A>T), I298V (992A>G 994A>G), L300I (998C>A 1000T>C), S301T (1001T>A), N303D (1007A>G), I314V (1040A>G 1042A>C), H324T (1070C>A 1071A>C), N325Q (1073A>C 1075T>A), R326P (1076A>C 1077G>C), I330L (1088A>C 1090A>G), I344V (1130A>G), I349V (1145A>G 1147A>C), E351K (1151G>A 1153A>G), E353D (1159A>T), L354E (1160T>G 1161T>A), P355R (1163C>A 1164C>G 1165A>G), K356M (1167A>T), A358F (1172G>T 1173C>T), E359D (1177A>C), L360I (1178T>A), E361Q (1181G>C 1183A>G), T362K (1185C>A 1186G>A), T366V (1196A>G 1197C>T), K367V (1199A>G 1200A>T), D369E (1207T>G), M371L (1211A>C), E373A (1218A>C 1219G>C), I375V (1223A>G), V376A (1227T>C), I379L (1235A>C 1237A>G) | | | | | | | | |

| | Begin | End | Coverage | Score | Concordance | Matches | Identities | I/D/M/F* | Stop Codons |
|---|---|---|---|---|---|---|---|---|---|
| **NT** | **101** | **1238** | **84.9%** | **414** | **18.3%** | **1135 (99.5%)** | **677 (59.3%)** | **3/3** | |
| Codon mutations: | ATG4GTT (110A>G 112G>T), GCT6GAT (117C>A), GAA7AGT (119G>A 120A>G 121A>T), GAC9GAT (127C>T), CAT10ACC (128C>A 129A>C 130T>C), GGC12GGT (136C>T), CTC13TTG (137C>T 139C>G), AAG14CAA (140A>C 142G>A), AAA15GAA (143A>G), AAT16AAC (148T>C), ATC17CTT (149A>C 151C>T), CTA18CTG (154A>G), AAG19AGA (156A>G 157G>A), GGT20GGC (160T>C), ATA21ATC (163A>C), TCT23GCC (167T>G 169T>C), TGT24TAT (171G>A), GGA25GGT (175A>T), GAA27GAG (181A>G), CCT29CCA (187T>A), TCA30TCT (190A>T), ACT31GCC (191A>G 193T>C), CAA33CAG (199A>G), AAA35AGA (204A>G), GCC36GGT (207C>G 208C>T), ATT37ATA (211T>A), TTT38GTG (212T>G 214T>G), CCA39CCT (217A>T), TGT40TTT (219G>T), ATT41TGC (221A>T 222T>G 223T>C), TCT42AAG (224T>A 225C>A 226T>G), AAA44CTT (230A>C 231A>T 232A>T), GAC45GAT (235C>T), GTG46GTC (238G>C), ATA47ATT (241A>T), GTC48CAT (242G>C 243T>A 244C>T), CAA49CAG (247A>G), TCG52TCT (256G>T), ACA54ACC (262A>C), AAA56AAG (268A>G), ACA57ACC (271A>C), GCA58GCC (274A>C), ACA59ATG (276C>T 277A>G), TAT60ATT (278T>A 279A>T), GCA61GCT (283A>T), ATC62CTT (284A>C 286C>T), AGT63ACC (288G>C 289T>C), TTA65TGC (294T>G 295A>C), CAG66CAA (298G>A), CAA67GTT (299C>G 300A>T 301A>T), ATA68GTG (302A>G 304A>G), ACT70ACC (310T>C), TCT71AAA (311T>A 312C>A 313T>A), AAT74GAG (320A>G 322T>G), ATC75GTT (323A>G 325C>T), ATC79GTA (335A>G 337C>A), ACC81GCT (341A>G 343C>T), CCA82CCT (346A>T), CGT84CGG (352T>G), GAA85GAG (355A>G), TTG86CTT (356T>C 358G>T), GCT87GCA (361T>A), CTC88GCT (362C>G 363T>C 364C>T), GCT90ACG (368G>A 370T>G), CAA91GAA (371C>G), CGT92AAG (374C>A 375G>A 376T>G), GTA93GTG (379A>G), TTG94ATT (380T>A 382G>T), CAA95TTG (383C>T 384A>T 385A>G), ACA96GCA (386A>G), GGG98GGA (394G>A), AAT99GAC (395A>G 397T>C), CTA101CTG (403A>G), TAT102del (404_406delTAT), AAT103GGC (407A>G 408A>G 409T>C), TTT104GTG (410T>G 412T>G), AAA105AAG (415A>G), TGT106GTT (416T>G 417G>T), CAA107CAT (421A>T), GTT108GCT (423T>C), ATT110GTT (428A>G), GGA111GGG (433A>G), GGC112GGT (436C>T), ACA113ACC (439A>C), ATC115GTC (443A>G), AAG116CGT (446A>C 447A>G 448G>T), GAA117GAG (451A>G), AGC118GAT (452A>G 453G>A 454C>T), GAA120CGT (458G>C 459A>G 460A>T), ACT121ATC (462C>T 463T>C), CTA122CTT (466A>T), AAG123CAG (467A>C), AAA124AGT (471A>G 472A>T), AAA124_GCC125insGGT (472_473insGGT), GCC125GTT (474C>T 475C>T), CAG126CAT (478G>T), GTT127GTA (481T>A), TTG128GTG (482T>G), ATT129GTT (485A>G), GGA130GGT (490A>T), CCT132CCA (496T>A), GGA133GGC (499A>C), CGA134CGT (502A>T), ATG135GTC (503A>G 505G>C), ATT136TTT (506A>T), GAT137GAC (511T>C), CTA138ATG (512C>A 514A>G), CTC139CTG (517C>G), ACC140CGC (518A>C 519C>G), CGA141AGG (521C>A 523A>G), AAA142CAG (524A>C 526A>G), AGT143TCT (527A>T 528G>C), ATC144CTT (530A>C 532C>T), GAT145CGC (533G>C 534A>G 535T>C), ACC146CCT (536A>C 538C>T), AAA147GAC (539A>G 541A>C), GCA148CAC (542G>C 543C>A 544A>C), ATT149ATC (547T>C), AAA150AAG (550A>G), ATA151ATG (553A>G), GTT152TTT (554G>T), GTA153GTT (559A>T), ATA154TTG (560A>T 562A>G), GAT158GAC (574T>C), GAG159GAA (577G>A), TTG161CTC (581T>C 583G>C), ATA162TCT (584A>T 585T>C 586A>T), GAT163AGA (587G>A 588A>G 589T>A), AAT164GGT (590A>G 591A>G), TTT165TTC (595T>C), TTG166AAG (596T>A 597T>A), AAA168CAG (602A>C 604A>G), ATA169ATT (607A>T), CAA170TAT (608C>T 610A>T), ATT172ATA (616T>A), GAA174CAG (620G>C 622A>G), TTT175CTT (623T>C), TTT176TTG (628T>G), GAA177CCA (629G>C 630A>C), AGT178CCA (632A>C 633G>C 634T>A), CAT179AAA (635C>A 637T>A), GTC180ATC (638G>A), GTT182GTG (646T>G), ATT183GGT (647A>G 648T>G), TTA184GTT (650T>G 652A>T), TTA185TTC (655A>C), TCA186TCT (658A>T), GCA187GCC (661A>C), ACC188ACT (664C>T), GTT189ATG (665G>A 667T>G), TCA191CCA (671T>C), AGG192GAG (674A>G 675G>A), GTT193GCT (678T>C), ATC194CTT (680A>C 682C>T), AAT195GAG (683A>G 685T>G), ACA196ATT (687C>T 688A>T), TCT197ACT (689T>A), CAG198AGA (692C>A 693A>G 694G>A), GTC199AAG (695G>A 696T>A 697C>G), TTT200TTC (700T>C), AGA202AAC (705G>A 706A>C), GAT203AAA (707G>A 709T>A), CCC204CCT (712C>T), GTA205GTG (715A>G), AAA206AGG (717A>G 718A>G), CTG208CTT (724G>T), GTA209GTG (727A>G), AAA210AAG (730A>G), AAT211CGT (731A>C 732A>G), GCT212GAT (735C>A), GAA213GAG (739A>G), TTG214CTC (740T>C 742G>C), ACA215ACT (745A>T), CTC216CTT (748C>T), ATT219ATC (757T>C), AGG220AAG (759G>A), TAT222TTT (765A>T), ATA224GTC (770A>G 772A>C), GTA226GTT (778A>T), AAG227GAC (779A>G 781G>C), AAA228AAG (784A>G), AAT229GAA (785A>G 787T>A), GAT230GAG (790T>G), TTT231TGG (792T>G 793T>G), AAA232AAG (796A>G), GCA233CTT (797G>C 798C>T 799A>T), ACA235ACT (805A>T), TTT237TGC (810T>G 811T>C), TAT240TAC (820T>C), GAC241GAG (823C>G), CAC242ACC (824C>A 825A>C), AGC244GCC (830A>G 831G>C), CTC245ATC (833C>A), ACT246ACC (838T>C), CAA247CAG (841A>G), ACA248AGT (843C>G 844A>T), CTA249GTC (845C>G 847A>C), ATA250ATC (850A>C), TGT252GTG (854T>G 855G>T 856T>G), AAT253AAC (859T>C), ACA254ACC (862A>C), CGT255AGG (863C>A 865T>G), AGA256CGT (866A>C 868A>T), CAA257AAA (869C>A), GTA260TGG (878G>T 879T>G 880A>G), ATG262ACT (885T>C 886G>T), GAA263GAT (889A>T), TGT264AAG (890T>A 891G>A 892T>G), TTA265ATG (893T>A 895A>G), ACA266CGC (896A>C 897C>G 898A>C), AAT267AGC (900A>G 901T>C), AGA268CGT (902A>C 904A>T), AAT269GAT (905A>G), TTC270CAC (908T>C 909T>A), GCA272GTA (915C>T), TCA273TCT (919A>T), AGC274GCA (920A>G 921G>C 922C>A), ATC275ACT (924T>C 925C>T), GAT278GAC (934T>C), TCC280GAC (938T>G 939C>A), CAA281CAG (943A>G), CAA282AAC (944C>A 946A>C), GAA283ACT (947G>A 948A>C 949A>T), CGT284AGA (950C>A 952T>A), TTT286ATC (956T>A 958T>C), AAA289CGA (965A>C 966A>G), GAA290GAG (970A>G), TTT291TTC (973T>C), AGA292CGA (974A>C), GAT293TCT (977G>T 978A>C), GGT294GGG (982T>G), AAA295TCA (983A>T 984A>C), ACT296TCT (986A>T), AGA297CGT (989A>C 991A>T), ATA298GTG (992A>G 994A>G), CTT299CTC (997T>C), CTT300ATC (998C>A 1000T>C), TCA301ACA (1001T>A), AAT303GAT (1007A>G), CTA304CTC (1012A>C), TTA305CTT (1013T>C 1015A>T), GCA306GCT (1018A>T), AGA307CGT (1019A>C 1021A>T), GGC308GGT (1024C>T), GTT311GTA (1033T>A), CAA312CAG (1036A>G), ATA314GTC (1040A>G 1042A>C), TCT315TCC (1045T>C), TTA316CTT (1046T>C 1048A>T), GTT317GTC (1051T>C), ATC318ATT (1054C>T), AAT319AAC (1057T>C), GAT321GAC (1063T>C), TTG322CTG (1064T>C), CAT324ACT (1070C>A 1071A>C), AAT325CAA (1073A>C 1075T>A), AGA326CCA (1076A>C 1077G>C), TAT329TAC (1087T>C), ATA330CTG (1088A>C 1090A>G), AGA335CGT (1103A>C 1105A>T), GGT337GGA (1111T>A), AGA338CGA (1112A>C), GGC340GGA (1120C>A), CGA341AGG (1121C>A 1123A>G), AAA342AAG (1126A>G), GGG343GGT (1129G>T), ATT344GTT (1130A>G), AAT347AAC (1141T>C), ATA349GTC (1145A>G 1147A>C), ACT350ACC (1150T>C), GAA351AAG (1151G>A 1153A>G), GAC352GAT (1156C>T), GAA353GAT (1159A>T), TTA354GAA (1160T>G 1161T>A), CCA355AGG (1163C>A 1164C>G 1165A>G), AAG356ATG (1167A>T), GCT358TTT (1172G>T 1173C>T), GAA359GAC (1177A>C), TTA360ATA (1178T>A), GAA361CAG (1181G>C 1183A>G), ACG362AAA (1185C>A 1186G>A), TAT364TAC (1192T>C), AAT365AAC (1195T>C), ACC366GTC (1196A>G 1197C>T), AAA367GTA (1199A>G 1200A>T), ATA368ATC (1204A>C), GAT369GAG (1207T>G), GAA370GAG (1210A>G), ATG371CTG (1211A>C), GAG373GCC (1218A>C 1219G>C), ATT375GTT (1223A>G), GTT376GCT (1227T>C), TTG378CTC (1232T>C 1234G>C), ATA379CTG (1235A>C 1237A>G) | | | | | | | | |

#### Proteins

| | Begin | End | Coverage | Score | Concordance | Matches | Identities | I/D/M/F* | Stop Codons |
|---|---|---|---|---|---|---|---|---|---|
| hypothetical protein (YP_009666148.1) | 1 | 380 | 100% | 1484 | 59.2% | 379 (99.5%) | 212 (55.6%) | 1/1/0/0 | 0 |
| Protein mutations: | M4V (110A>G 112G>T), A6D (117C>A), E7S (119G>A 120A>G 121A>T), H10T (128C>A 129A>C 130T>C), K14Q (140A>C 142G>A), K15E (143A>G), I17L (149A>C 151C>T), K19R (156A>G 157G>A), S23A (167T>G 169T>C), C24Y (171G>A), T31A (191A>G 193T>C), K35R (204A>G), A36G (207C>G 208C>T), F38V (212T>G 214T>G), C40F (219G>T), I41C (221A>T 222T>G 223T>C), S42K (224T>A 225C>A 226T>G), K44L (230A>C 231A>T 232A>T), V48H (242G>C 243T>A 244C>T), T59M (276C>T 277A>G), Y60I (278T>A 279A>T), I62L (284A>C 286C>T), S63T (288G>C 289T>C), L65C (294T>G 295A>C), Q67V (299C>G 300A>T 301A>T), I68V (302A>G 304A>G), S71K (311T>A 312C>A 313T>A), N74E (320A>G 322T>G), I75V (323A>G 325C>T), I79V (335A>G 337C>A), T81A (341A>G 343C>T), L88A (362C>G 363T>C 364C>T), A90T (368G>A 370T>G), Q91E (371C>G), R92K (374C>A 375G>A 376T>G), L94I (380T>A 382G>T), Q95L (383C>T 384A>T 385A>G), T96A (386A>G), N99D (395A>G 397T>C), Y102del (404_406delTAT), N103G (407A>G 408A>G 409T>C), F104V (410T>G 412T>G), C106V (416T>G 417G>T), Q107H (421A>T), V108A (423T>C), I110V (428A>G), I115V (443A>G), K116R (446A>C 447A>G 448G>T), S118D (452A>G 453G>A 454C>T), E120R (458G>C 459A>G 460A>T), T121I (462C>T 463T>C), K123Q (467A>C), K124S (471A>G 472A>T), K124_A125insG (472_473insGGT), A125V (474C>T 475C>T), Q126H (478G>T), L128V (482T>G), I129V (485A>G), M135V (503A>G 505G>C), I136F (506A>T), L138M (512C>A 514A>G), T140R (518A>C 519C>G), K142Q (524A>C 526A>G), I144L (530A>C 532C>T), D145R (533G>C 534A>G 535T>C), T146P (536A>C 538C>T), K147D (539A>G 541A>C), A148H (542G>C 543C>A 544A>C), I151M (553A>G), V152F (554G>T), I154L (560A>T 562A>G), I162S (584A>T 585T>C 586A>T), D163R (587G>A 588A>G 589T>A), N164G (590A>G 591A>G), L166K (596T>A 597T>A), K168Q (602A>C 604A>G), Q170Y (608C>T 610A>T), E174Q (620G>C 622A>G), F175L (623T>C), F176L (628T>G), E177P (629G>C 630A>C), S178P (632A>C 633G>C 634T>A), H179K (635C>A 637T>A), V180I (638G>A), I183G (647A>G 648T>G), L184V (650T>G 652A>T), L185F (655A>C), V189M (665G>A 667T>G), S191P (671T>C), R192E (674A>G 675G>A), V193A (678T>C), I194L (680A>C 682C>T), N195E (683A>G 685T>G), T196I (687C>T 688A>T), S197T (689T>A), Q198R (692C>A 693A>G 694G>A), V199K (695G>A 696T>A 697C>G), R202N (705G>A 706A>C), D203K (707G>A 709T>A), K206R (717A>G 718A>G), N211R (731A>C 732A>G), A212D (735C>A), R220K (759G>A), Y222F (765A>T), I224V (770A>G 772A>C), K227D (779A>G 781G>C), N229E (785A>G 787T>A), D230E (790T>G), F231W (792T>G 793T>G), A233L (797G>C 798C>T 799A>T), F237C (810T>G 811T>C), D241E (823C>G), H242T (824C>A 825A>C), S244A (830A>G 831G>C), L245I (833C>A), T248S (843C>G 844A>T), L249V (845C>G 847A>C), C252V (854T>G 855G>T 856T>G), Q257K (869C>A), V260W (878G>T 879T>G 880A>G), M262T (885T>C 886G>T), E263D (889A>T), C264K (890T>A 891G>A 892T>G), L265M (893T>A 895A>G), T266R (896A>C 897C>G 898A>C), N267S (900A>G 901T>C), N269D (905A>G), F270H (908T>C 909T>A), A272V (915C>T), S274A (920A>G 921G>C 922C>A), I275T (924T>C 925C>T), S280D (938T>G 939C>A), Q282N (944C>A 946A>C), E283T (947G>A 948A>C 949A>T), F286I (956T>A 958T>C), K289R (965A>C 966A>G), D293S (977G>T 978A>C), K295S (983A>T 984A>C), T296S (986A>T), I298V (992A>G 994A>G), L300I (998C>A 1000T>C), S301T (1001T>A), N303D (1007A>G), I314V (1040A>G 1042A>C), H324T (1070C>A 1071A>C), N325Q (1073A>C 1075T>A), R326P (1076A>C 1077G>C), I330L (1088A>C 1090A>G), I344V (1130A>G), I349V (1145A>G 1147A>C), E351K (1151G>A 1153A>G), E353D (1159A>T), L354E (1160T>G 1161T>A), P355R (1163C>A 1164C>G 1165A>G), K356M (1167A>T), A358F (1172G>T 1173C>T), E359D (1177A>C), L360I (1178T>A), E361Q (1181G>C 1183A>G), T362K (1185C>A 1186G>A), T366V (1196A>G 1197C>T), K367V (1199A>G 1200A>T), D369E (1207T>G), M371L (1211A>C), E373A (1218A>C 1219G>C), I375V (1223A>G), V376A (1227T>C), I379L (1235A>C 1237A>G) | | | | | | | | |

| | Begin | End | Coverage | Score | Concordance | Matches | Identities | I/D/M/F* | Stop Codons |
|---|---|---|---|---|---|---|---|---|---|
| **NT** | **101** | **1238** | **84.9%** | **414** | **18.3%** | **1135 (99.5%)** | **677 (59.3%)** | **3/3** | |
| Codon mutations: | ATG4GTT (110A>G 112G>T), GCT6GAT (117C>A), GAA7AGT (119G>A 120A>G 121A>T), GAC9GAT (127C>T), CAT10ACC (128C>A 129A>C 130T>C), GGC12GGT (136C>T), CTC13TTG (137C>T 139C>G), AAG14CAA (140A>C 142G>A), AAA15GAA (143A>G), AAT16AAC (148T>C), ATC17CTT (149A>C 151C>T), CTA18CTG (154A>G), AAG19AGA (156A>G 157G>A), GGT20GGC (160T>C), ATA21ATC (163A>C), TCT23GCC (167T>G 169T>C), TGT24TAT (171G>A), GGA25GGT (175A>T), GAA27GAG (181A>G), CCT29CCA (187T>A), TCA30TCT (190A>T), ACT31GCC (191A>G 193T>C), CAA33CAG (199A>G), AAA35AGA (204A>G), GCC36GGT (207C>G 208C>T), ATT37ATA (211T>A), TTT38GTG (212T>G 214T>G), CCA39CCT (217A>T), TGT40TTT (219G>T), ATT41TGC (221A>T 222T>G 223T>C), TCT42AAG (224T>A 225C>A 226T>G), AAA44CTT (230A>C 231A>T 232A>T), GAC45GAT (235C>T), GTG46GTC (238G>C), ATA47ATT (241A>T), GTC48CAT (242G>C 243T>A 244C>T), CAA49CAG (247A>G), TCG52TCT (256G>T), ACA54ACC (262A>C), AAA56AAG (268A>G), ACA57ACC (271A>C), GCA58GCC (274A>C), ACA59ATG (276C>T 277A>G), TAT60ATT (278T>A 279A>T), GCA61GCT (283A>T), ATC62CTT (284A>C 286C>T), AGT63ACC (288G>C 289T>C), TTA65TGC (294T>G 295A>C), CAG66CAA (298G>A), CAA67GTT (299C>G 300A>T 301A>T), ATA68GTG (302A>G 304A>G), ACT70ACC (310T>C), TCT71AAA (311T>A 312C>A 313T>A), AAT74GAG (320A>G 322T>G), ATC75GTT (323A>G 325C>T), ATC79GTA (335A>G 337C>A), ACC81GCT (341A>G 343C>T), CCA82CCT (346A>T), CGT84CGG (352T>G), GAA85GAG (355A>G), TTG86CTT (356T>C 358G>T), GCT87GCA (361T>A), CTC88GCT (362C>G 363T>C 364C>T), GCT90ACG (368G>A 370T>G), CAA91GAA (371C>G), CGT92AAG (374C>A 375G>A 376T>G), GTA93GTG (379A>G), TTG94ATT (380T>A 382G>T), CAA95TTG (383C>T 384A>T 385A>G), ACA96GCA (386A>G), GGG98GGA (394G>A), AAT99GAC (395A>G 397T>C), CTA101CTG (403A>G), TAT102del (404_406delTAT), AAT103GGC (407A>G 408A>G 409T>C), TTT104GTG (410T>G 412T>G), AAA105AAG (415A>G), TGT106GTT (416T>G 417G>T), CAA107CAT (421A>T), GTT108GCT (423T>C), ATT110GTT (428A>G), GGA111GGG (433A>G), GGC112GGT (436C>T), ACA113ACC (439A>C), ATC115GTC (443A>G), AAG116CGT (446A>C 447A>G 448G>T), GAA117GAG (451A>G), AGC118GAT (452A>G 453G>A 454C>T), GAA120CGT (458G>C 459A>G 460A>T), ACT121ATC (462C>T 463T>C), CTA122CTT (466A>T), AAG123CAG (467A>C), AAA124AGT (471A>G 472A>T), AAA124_GCC125insGGT (472_473insGGT), GCC125GTT (474C>T 475C>T), CAG126CAT (478G>T), GTT127GTA (481T>A), TTG128GTG (482T>G), ATT129GTT (485A>G), GGA130GGT (490A>T), CCT132CCA (496T>A), GGA133GGC (499A>C), CGA134CGT (502A>T), ATG135GTC (503A>G 505G>C), ATT136TTT (506A>T), GAT137GAC (511T>C), CTA138ATG (512C>A 514A>G), CTC139CTG (517C>G), ACC140CGC (518A>C 519C>G), CGA141AGG (521C>A 523A>G), AAA142CAG (524A>C 526A>G), AGT143TCT (527A>T 528G>C), ATC144CTT (530A>C 532C>T), GAT145CGC (533G>C 534A>G 535T>C), ACC146CCT (536A>C 538C>T), AAA147GAC (539A>G 541A>C), GCA148CAC (542G>C 543C>A 544A>C), ATT149ATC (547T>C), AAA150AAG (550A>G), ATA151ATG (553A>G), GTT152TTT (554G>T), GTA153GTT (559A>T), ATA154TTG (560A>T 562A>G), GAT158GAC (574T>C), GAG159GAA (577G>A), TTG161CTC (581T>C 583G>C), ATA162TCT (584A>T 585T>C 586A>T), GAT163AGA (587G>A 588A>G 589T>A), AAT164GGT (590A>G 591A>G), TTT165TTC (595T>C), TTG166AAG (596T>A 597T>A), AAA168CAG (602A>C 604A>G), ATA169ATT (607A>T), CAA170TAT (608C>T 610A>T), ATT172ATA (616T>A), GAA174CAG (620G>C 622A>G), TTT175CTT (623T>C), TTT176TTG (628T>G), GAA177CCA (629G>C 630A>C), AGT178CCA (632A>C 633G>C 634T>A), CAT179AAA (635C>A 637T>A), GTC180ATC (638G>A), GTT182GTG (646T>G), ATT183GGT (647A>G 648T>G), TTA184GTT (650T>G 652A>T), TTA185TTC (655A>C), TCA186TCT (658A>T), GCA187GCC (661A>C), ACC188ACT (664C>T), GTT189ATG (665G>A 667T>G), TCA191CCA (671T>C), AGG192GAG (674A>G 675G>A), GTT193GCT (678T>C), ATC194CTT (680A>C 682C>T), AAT195GAG (683A>G 685T>G), ACA196ATT (687C>T 688A>T), TCT197ACT (689T>A), CAG198AGA (692C>A 693A>G 694G>A), GTC199AAG (695G>A 696T>A 697C>G), TTT200TTC (700T>C), AGA202AAC (705G>A 706A>C), GAT203AAA (707G>A 709T>A), CCC204CCT (712C>T), GTA205GTG (715A>G), AAA206AGG (717A>G 718A>G), CTG208CTT (724G>T), GTA209GTG (727A>G), AAA210AAG (730A>G), AAT211CGT (731A>C 732A>G), GCT212GAT (735C>A), GAA213GAG (739A>G), TTG214CTC (740T>C 742G>C), ACA215ACT (745A>T), CTC216CTT (748C>T), ATT219ATC (757T>C), AGG220AAG (759G>A), TAT222TTT (765A>T), ATA224GTC (770A>G 772A>C), GTA226GTT (778A>T), AAG227GAC (779A>G 781G>C), AAA228AAG (784A>G), AAT229GAA (785A>G 787T>A), GAT230GAG (790T>G), TTT231TGG (792T>G 793T>G), AAA232AAG (796A>G), GCA233CTT (797G>C 798C>T 799A>T), ACA235ACT (805A>T), TTT237TGC (810T>G 811T>C), TAT240TAC (820T>C), GAC241GAG (823C>G), CAC242ACC (824C>A 825A>C), AGC244GCC (830A>G 831G>C), CTC245ATC (833C>A), ACT246ACC (838T>C), CAA247CAG (841A>G), ACA248AGT (843C>G 844A>T), CTA249GTC (845C>G 847A>C), ATA250ATC (850A>C), TGT252GTG (854T>G 855G>T 856T>G), AAT253AAC (859T>C), ACA254ACC (862A>C), CGT255AGG (863C>A 865T>G), AGA256CGT (866A>C 868A>T), CAA257AAA (869C>A), GTA260TGG (878G>T 879T>G 880A>G), ATG262ACT (885T>C 886G>T), GAA263GAT (889A>T), TGT264AAG (890T>A 891G>A 892T>G), TTA265ATG (893T>A 895A>G), ACA266CGC (896A>C 897C>G 898A>C), AAT267AGC (900A>G 901T>C), AGA268CGT (902A>C 904A>T), AAT269GAT (905A>G), TTC270CAC (908T>C 909T>A), GCA272GTA (915C>T), TCA273TCT (919A>T), AGC274GCA (920A>G 921G>C 922C>A), ATC275ACT (924T>C 925C>T), GAT278GAC (934T>C), TCC280GAC (938T>G 939C>A), CAA281CAG (943A>G), CAA282AAC (944C>A 946A>C), GAA283ACT (947G>A 948A>C 949A>T), CGT284AGA (950C>A 952T>A), TTT286ATC (956T>A 958T>C), AAA289CGA (965A>C 966A>G), GAA290GAG (970A>G), TTT291TTC (973T>C), AGA292CGA (974A>C), GAT293TCT (977G>T 978A>C), GGT294GGG (982T>G), AAA295TCA (983A>T 984A>C), ACT296TCT (986A>T), AGA297CGT (989A>C 991A>T), ATA298GTG (992A>G 994A>G), CTT299CTC (997T>C), CTT300ATC (998C>A 1000T>C), TCA301ACA (1001T>A), AAT303GAT (1007A>G), CTA304CTC (1012A>C), TTA305CTT (1013T>C 1015A>T), GCA306GCT (1018A>T), AGA307CGT (1019A>C 1021A>T), GGC308GGT (1024C>T), GTT311GTA (1033T>A), CAA312CAG (1036A>G), ATA314GTC (1040A>G 1042A>C), TCT315TCC (1045T>C), TTA316CTT (1046T>C 1048A>T), GTT317GTC (1051T>C), ATC318ATT (1054C>T), AAT319AAC (1057T>C), GAT321GAC (1063T>C), TTG322CTG (1064T>C), CAT324ACT (1070C>A 1071A>C), AAT325CAA (1073A>C 1075T>A), AGA326CCA (1076A>C 1077G>C), TAT329TAC (1087T>C), ATA330CTG (1088A>C 1090A>G), AGA335CGT (1103A>C 1105A>T), GGT337GGA (1111T>A), AGA338CGA (1112A>C), GGC340GGA (1120C>A), CGA341AGG (1121C>A 1123A>G), AAA342AAG (1126A>G), GGG343GGT (1129G>T), ATT344GTT (1130A>G), AAT347AAC (1141T>C), ATA349GTC (1145A>G 1147A>C), ACT350ACC (1150T>C), GAA351AAG (1151G>A 1153A>G), GAC352GAT (1156C>T), GAA353GAT (1159A>T), TTA354GAA (1160T>G 1161T>A), CCA355AGG (1163C>A 1164C>G 1165A>G), AAG356ATG (1167A>T), GCT358TTT (1172G>T 1173C>T), GAA359GAC (1177A>C), TTA360ATA (1178T>A), GAA361CAG (1181G>C 1183A>G), ACG362AAA (1185C>A 1186G>A), TAT364TAC (1192T>C), AAT365AAC (1195T>C), ACC366GTC (1196A>G 1197C>T), AAA367GTA (1199A>G 1200A>T), ATA368ATC (1204A>C), GAT369GAG (1207T>G), GAA370GAG (1210A>G), ATG371CTG (1211A>C), GAG373GCC (1218A>C 1219G>C), ATT375GTT (1223A>G), GTT376GCT (1227T>C), TTG378CTC (1232T>C 1234G>C), ATA379CTG (1235A>C 1237A>G) | | | | | | | | |

*: Inserts / Deletes / Misaligned / Frameshifts

#### Analysis details

This analysis was performed with panviral2.64

### NGS Details (UN18_val): Torradovirus lycopersici (segment RNA 1)

#### Assembly

| | |
|---|---|
| **Coverage Length** | 6773 (2 contig(s)) |
| **Depth Of Coverage** | 234.6 |
| **Number Of Reads** | 12659 |
| **Reads Per Million** | 237.59 rpm (after QC) |
| **Ambiguities** | 0 |
| **Assembly Method** | de novo + reference guided assembly |
| **Consensus Caller** | Bcf Tools |

#### Coverage Map

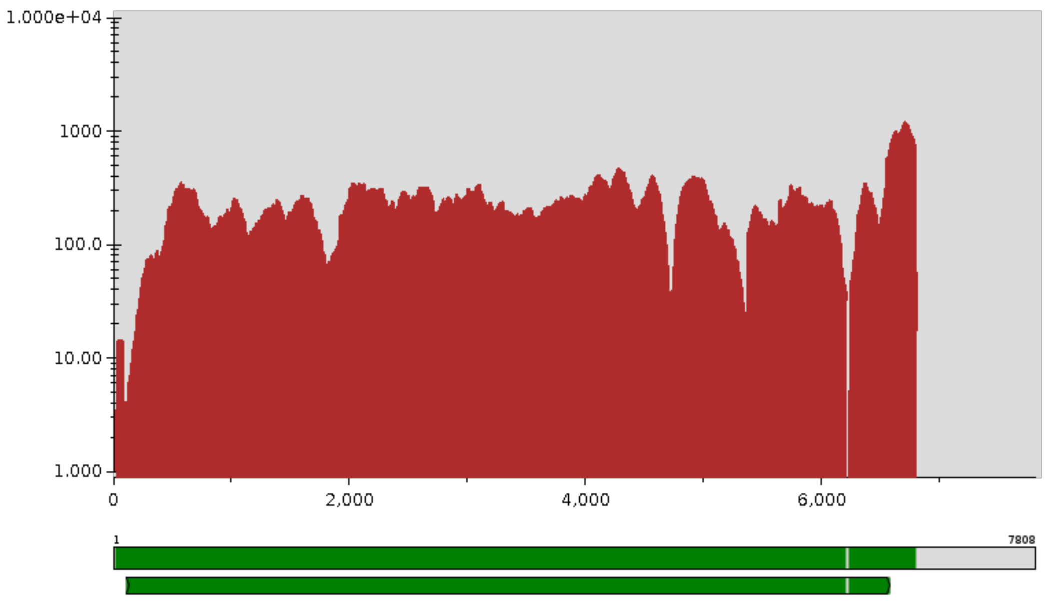

#### Assignment

| | |
|---|---|
| **Type** | Torradovirus lycopersici (Taxonomy ID: 3048378) |
| **Reference Genome** | NC_009013.1 |
| **NT Identity (%)** | 88.5183 |
| **AA Identity (%)** | 94.1067 |
| **Number Of Stop Codons** | 1 |
| **Number Of CDS** | 1 |

#### Alignment

| | |
|---|---|
| **Alignment Score** | 10416.0 (NT) + 14340.0 (AA) = 24756.0 |
| **Concordance (%)** | 86.6321 |

| **Alignment Method** | Global, seeded, nucleotide + amino acids (AGA) |
|---|---|

#### Genome Region

Sequence starts at position 21 and ends at position 6806 relative to NC_009013.1 reference sequence.

#### Alignment Detailed Statistics

| | Begin | End | Coverage | Score | Concordance | Matches | Identities | I/D/M/F* | Stop Codons |
|---|---|---|---|---|---|---|---|---|---|
| **NT** | **21** | **6806** | **86.7%** | **10416** | **77.0%** | **6772 (99.9%)** | **5998 (88.5%)** | **4/1** | |
| Mutations: | 22T>G, 69C>T, 72C>T, 77A>T, 83C>T, 83_84insT, 88T>C, 90C>G, 93T>A, 101G>A, 102G>C, 115T>C, 122A>T, 128C>T, 130C>T, 134T>C, 135T>C, 138A>G, 140T>C, 142A>C, 144T>C, 145T>A, 146A>G, 147C>T, 148T>G, 155T>A, 157C>T, 159C>T, 161A>G, 178T>A, 187A>T, 190G>A, 196T>A, 197G>C, 198C>A, 199T>A, 220C>T, 223T>A, 226G>T, 241G>A, 247A>T, 259T>C, 262T>A, 265C>T, 271A>G, 280G>A, 289T>G, 292A>T, 301T>A, 302T>G, 307C>T, 310T>C, 311A>T, 313C>T, 315G>A, 322T>C, 324A>G, 325C>T, 328T>A, 331T>C, 334T>A, 340G>A, 343A>G, 349G>A, 358T>A, 359T>C, 361A>T, 364T>A, 367A>G, 370C>T, 376C>T, 382T>A, 385C>T, 388T>C, 394C>T, 397A>C, 409A>G, 412A>C, 427T>G, 433G>A, 439T>C, 443A>G, 451A>G, 454T>C, 473A>G, 485G>A, 487C>T, 490A>G, 493T>C, 496G>A, 502G>T, 541T>C, 559C>T, 578C>T, 586A>T, 599A>T, 604C>T, 606T>C, 615A>G, 618C>T, 619A>T, 655C>T, 661A>G, 676A>G, 694A>G, 707C>T, 718A>G, 719T>G, 748G>T, 751A>T, 760C>T, 778T>A, 781C>T, 790C>T, 799A>T, 805T>C, 806A>C, 820A>C, 821T>A, 832A>T, 835A>C, 841T>C, 842C>T, 847C>T, 883G>A, 886T>C, 892G>A, 901T>C, 913T>C, 925C>T, 938C>T, 952A>G, 955C>T, 964T>G, 970G>A, 973T>C, 977G>A, 997C>T, 1001T>C, 1006T>C, 1018C>T, 1019A>G, 1020C>T, 1033G>A, 1060T>C, 1063C>T, 1081C>T, 1093T>C, 1096C>A, 1109C>T, 1111A>G, 1129A>G, 1135T>A, 1165C>A, 1180G>A, 1186G>A, 1195A>T, 1196A>T, 1198C>T, 1204T>G, 1207T>G, 1231G>A, 1240C>T, 1255G>A, 1267T>C, 1300G>A, 1303A>G, 1319C>T, 1324C>T, 1342C>T, 1357A>C, 1361T>C, 1390A>T, 1408T>C, 1426T>A, 1429C>T, 1441C>T, 1451C>T, 1462G>T, 1480C>T, 1481A>G, 1489T>C, 1495A>T, 1522C>A, 1552C>T, 1553C>A, 1555C>G, 1568C>A, 1570A>G, 1573T>C, 1582C>T, 1585A>C, 1591C>T, 1607G>T, 1612G>A, 1618T>G, 1619C>T, 1624C>A, 1630C>T, 1643A>C, 1645A>G, 1669C>T, 1681T>A, 1684G>A, 1692C>T, 1699T>G, 1706T>G, 1714T>G, 1737A>G, 1738A>G, 1739A>C, 1750A>T, 1755C>T, 1756A>C, 1771C>T, 1781T>C, 1783G>A, 1789A>G, 1810G>A, 1813T>G, 1825C>A, 1834A>G, 1839A>T, 1844G>A, 1849A>C, 1850T>A, 1858G>A, 1867G>A, 1877T>A, 1879A>T, 1885G>A, 1889A>G, 1894A>T, 1906T>A, 1907A>C, 1908T>C, 1909G>A, 1911T>G, 1927C>A, 1942G>A, 1960A>G, 1966A>G, 1984C>T, 1990G>A, 1993G>A, 2009T>C, 2023T>C, 2056T>C, 2062C>T, 2077C>A, 2082T>A, 2092A>G, 2120A>C, 2128T>C, 2130A>C, 2138C>T, 2141T>C, 2143T>A, 2146A>G, 2149T>C, 2155T>C, 2174C>T, 2179A>C, 2182G>A, 2185T>C, 2194T>A, 2200T>C, 2212T>C, 2224C>T, 2227C>T, 2240A>T, 2241A>G, 2250G>A, 2251C>A, 2257C>T, 2263G>A, 2287G>A, 2290C>T, 2299A>T, 2314C>T, 2318C>T, 2323A>T, 2347C>T, 2356C>T, 2359G>A, 2365A>T, 2375A>C, 2377G>T, 2380T>C, 2387C>A, 2392A>G, 2407T>C, 2409G>A, 2443T>C, 2446G>A, 2479G>A, 2485C>T, 2494T>G, 2497G>A, 2500T>C, 2503G>A, 2506T>C, 2509C>T, 2512T>C, 2542G>A, 2548C>T, 2557C>T, 2620G>A, 2626T>C, 2641G>A, 2650G>A, 2662C>T, 2692G>A, 2695T>A, 2710A>G, 2716A>C, 2726G>A, 2731C>T, 2734C>T, 2743A>G, 2744T>C, 2765T>C, 2767G>C, 2768A>C, 2773A>G, 2788C>T, 2809C>T, 2821G>A, 2827A>G, 2842T>C, 2848C>T, 2852C>A, 2854T>A, 2860C>G, 2866T>A, 2869C>T, 2881T>C, 2882G>T, 2926G>T, 2929C>T, 2930T>C, 2941A>G, 2947T>A, 2950C>T, 2974T>C, 2980C>G, 2998A>G, 3031T>C, 3034G>A, 3040T>C, 3049A>C, 3064A>T, 3079T>C, 3083C>A, 3085A>G, 3101T>C, 3103G>T, 3109T>C, 3136T>G, 3146A>T, 3151C>T, 3163T>C, 3169C>T, 3172T>C, 3202T>C, 3241T>C, 3253C>T, 3256G>A, 3262C>T, 3283T>C, 3289T>C, 3298T>C, 3307A>G, 3358A>T, 3376T>G, 3382T>A, 3400A>C, 3406A>G, 3413C>G, 3418C>A, 3421T>C, 3431C>A, 3445G>T, 3463A>G, 3466G>A, 3481C>T, 3484A>T, 3508G>T, 3530C>A, 3532A>G, 3534G>A, 3553T>A, 3559G>A, 3574T>C, 3583C>A, 3586T>C, 3589A>G, 3591C>T, 3593T>C, 3594C>A, 3610A>G, 3625C>T, 3628G>A, 3629G>T, 3646C>T, 3658A>C, 3661A>G, 3664T>C, 3676C>T, 3688A>G, 3691C>T, 3693A>T, 3701C>T, 3706T>C, 3710G>T, 3713A>G, 3716T>C, 3739A>T, 3754G>A, 3758G>A, 3766G>A, 3778G>T, 3781G>A, 3808T>C, 3814C>T, 3817G>A, 3824T>C, 3868C>T, 3871T>C, 3883G>A, 3895A>C, 3907A>T, 3911C>A, 3925A>G, 3928G>A, 3949C>T, 3955G>A, 3961G>A, 3970A>G, 3973T>C, 4034T>C, 4055C>A, 4066T>C, 4084C>T, 4093C>T, 4118C>T, 4129C>T, 4147C>A, 4157C>T, 4159G>A, 4162A>T, 4189T>A, 4192T>G, 4195A>G, 4207G>A, 4210T>C, 4225A>T, 4228C>T, 4234T>C, 4252C>T, 4282C>T, 4288C>A, 4294A>G, 4309T>G, 4318C>T, 4339T>C, 4348C>T, 4354T>C, 4358A>C, 4360A>G, 4396T>C, 4414T>G, 4417T>C, 4426G>T, 4429A>G, 4441A>T, 4459T>C, 4468C>T, 4471T>C, 4474T>C, 4501A>T, 4510T>A, 4514T>C, 4537T>A, 4570C>T, 4576A>T, 4588C>T, 4591T>C, 4625T>C, 4636T>C, 4648T>A, 4651T>A, 4660A>G, 4666G>A, 4678T>C, 4684A>T, 4687G>A, 4696C>T, 4699C>T, 4702G>A, 4705C>T, 4714T>C, 4716G>A, 4719G>A, 4735C>T, 4736T>C, 4747A>G, 4753T>C, 4756A>C, 4759C>T, 4765T>C, 4768T>C, 4774G>A, 4780T>C, 4783A>G, 4786A>G, 4789C>T, 4802C>T, 4813A>G, 4816T>C, 4834C>T, 4843G>A, 4897A>C, 4898G>A, 4903A>G, 4958A>C, 4960C>T, 4969G>A, 4975G>A, 4981C>T, 4999T>C, 5005T>C, 5012T>C, 5017G>A, 5023C>T, 5030G>A, 5041A>T, 5044A>C, 5059G>T, 5074T>C, 5077T>C, 5079T>C, 5083A>T, 5090G>T, 5102T>C, 5113C>T, 5119T>C, 5120G>A, 5122T>C, 5123C>T, 5131T>C, 5132T>A, 5143G>T, 5146T>C, 5167T>C, 5176C>T, 5200G>A, 5206C>T, 5209C>A, 5218T>G, 5269C>A, 5278G>A, 5290G>T, 5293T>C, 5302T>A, 5326C>T, 5347C>A, 5353G>A, 5357A>T, 5360A>C, 5369G>A, 5371G>T, 5386C>G, 5392G>A, 5399A>G, 5407A>T, 5410G>T, 5413C>T, 5419A>C, 5429C>T, 5437T>C, 5459G>A, 5462G>A, 5486G>A, 5491T>G, 5492A>T, 5498C>T, 5500C>G, 5515A>T, 5531C>T, 5533T>G, 5537A>C, 5558G>C, 5578T>C, 5582A>G, 5591G>T, 5602G>A, 5604C>T, 5608A>G, 5611A>T, 5613A>T, 5617A>T, 5626A>T, 5638T>G, 5641T>C, 5656G>A, 5674A>T, 5677G>T, 5686G>A, 5689C>A, 5690T>C, 5710T>A, 5711T>C, 5716G>T, 5719A>C, 5731C>T, 5734T>C, 5737T>C, 5739G>A, 5746T>C, 5774A>G, 5779T>C, 5786T>C, 5797A>G, 5804T>A, 5805T>C, 5806C>T, 5826A>T, 5836A>T, 5849T>C, 5850T>A, 5851G>A, 5869G>A, 5878G>T, 5881T>C, 5884T>C, 5887C>T, 5890C>T, 5894A>C, 5907A>T, 5914T>C, 5923T>C, 5927C>T, 5942C>T, 5944C>T, 5955A>G, 5965G>A, 5974T>C, 5975A>C, 5980G>A, 5986T>C, 5998A>G, 6002A>G, 6003C>T, 6004C>T, 6009G>A, 6016A>G, 6019A>G, 6028C>T, 6031A>T, 6033C>T, 6035G>T, 6055T>A, 6067C>A, 6070T>C, 6079C>G, 6089C>T, 6092C>T, 6100T>G, 6103G>A, 6106G>A, 6112A>G, 6121A>T, 6130G>T, 6139T>C, 6145T>C, 6154C>T, 6163C>G, 6166C>T, 6184A>G, 6185A>G, 6186C>A, 6187T>C, 6189A>G, 6193C>T, 6196A>G, 6199C>T, 6202G>C, 6208A>G, 6211A>G, 6214T>G, 6215C>G, 6230A>G, 6232A>G, 6235G>A, 6239G>A, 6240T>C, 6253A>G, 6256C>A, 6262A>T, 6263C>T, 6265G>A, 6268G>T, 6274C>G, 6291T>A, 6292T>A, 6295T>C, 6296G>C, 6298A>G, 6301T>A, 6304A>G, 6308T>C, 6317A>G, 6319A>T, 6322G>A, 6323A>G, 6344C>T, 6358T>C, 6371G>A, 6382C>T, 6385C>T, 6404T>C, 6405G>A, 6406T>C, 6412C>T, 6414G>T, 6415A>G, 6436C>T, 6437T>A, 6445C>T, 6448G>A, 6457C>A, 6458T>A, 6461C>T, 6481T>A, 6485C>A, 6490A>C, 6493T>C, 6496G>A, 6499A>T, 6500A>G, 6501T>A, 6517C>A, 6532C>T, 6545C>T, 6548T>C, 6556A>C, 6562A>T, 6566G>A, 6567G>A, 6568C>T, 6571C>T, 6574A>G, 6587G>A, 6591T>A, 6592A>T, 6593A>T, 6595T>C, 6602G>A, 6603C>T, 6647_6648insGTT, 6669G>A, 6670C>T, 6675C>T, 6685A>G, 6692A>C, 6715C>T, 6716G>A, 6721A>T, 6727T>C, 6742A>T, 6752T>C, 6759G>A, 6766delC, 6776C>A, 6786T>C, 6787G>A | | | | | | | | |

##### CDS

| | Begin | End | Coverage | Score | Concordance | Matches | Identities | I/D/M/F* | Stop Codons |
|---|---|---|---|---|---|---|---|---|---|
| ToTV_sRNA1gp1 | 1 | 2159 | 99.8% | 14340 | 95.3% | 2155 (100%) | 2028 (94.1%) | 0/0/0/0 | 1 |
| Protein mutations: | M6L (122A>T), P8S (128C>T 130C>T), F10P (134T>C 135T>C), N11S (138A>G), S12P (140T>C 142A>C), V13A (144T>C 145T>A), T14V (146A>G 147C>T 148T>G), C17S (155T>A 157C>T), A18V (159C>T), T19A (161A>G), A31Q (197G>C 198C>A 199T>A), S66A (302T>G), T69S (311A>T 313C>T), S70N (315G>A), N73S (324A>G 325C>T), S113G (443A>G), N123D (473A>G), D127N (485G>A 487C>T), T165S (599A>T), I167T (606T>C), N170S (615A>G), T171I (618C>T 619A>T), S205A (719T>G), S239T (821T>A), V291I (977G>A), T305V (1019A>G 1020C>T), T364S (1196A>T 1198C>T), D367E (1207T>G), I459V (1481A>G), A501S (1607G>T), T529I (1692C>T), S534A (1706T>G), K544R (1737A>G 1738A>G), T550I (1755C>T 1756A>C), Y578F (1839A>T), A580T (1844G>A), S582T (1850T>A), S591T (1877T>A 1879A>T), I595V (1889A>G), D600E (1906T>A), M601P (1907A>C 1908T>C 1909G>A), L602W (1911T>G), F659Y (2082T>A), N675T (2130A>C), Y679Q (2141T>C 2143T>A), N712C (2240A>T 2241A>G), S715K (2250G>A 2251C>A), R761S (2387C>A), R768K (2409G>A), G874S (2726G>A), A926S (2882G>T), Q993K (3083C>A 3085A>G), T1014S (3146A>T), Q1103E (3413C>G), R1143K (3534G>A), A1162V (3591C>T), S1163Q (3593T>C 3594C>A), A1175S (3629G>T), Y1196F (3693A>T), A1202S (3710G>T), I1203V (3713A>G), V1218I (3758G>A), I1362M (4192T>G), S1537N (4716G>A), S1538N (4719G>A), V1598I (4898G>A), N1618H (4958A>C 4960C>T), V1642M (5030G>A), I1658T (5079T>C), A1662S (5090G>T), V1672I (5120G>A 5122T>C), S1676T (5132T>A), T1751S (5357A>T), K1752Q (5360A>C), V1755I (5369G>A 5371G>T), I1765V (5399A>G), V1785I (5459G>A), V1786I (5462G>A), D1794N (5486G>A), T1796S (5492A>T), K1811Q (5537A>C), E1818Q (5558G>C), I1826V (5582A>G), A1829S (5591G>T), T1833I (5604C>T), Y1836F (5613A>T), Q1857H (5677G>T), Y1869H (5711T>C), R1878K (5739G>A), I1890V (5774A>G), F1900T (5804T>A 5805T>C 5806C>T), Y1907F (5826A>T), L1915Q (5849T>C 5850T>A 5851G>A), N1930H (5894A>C), Y1934F (5907A>T), H1946Y (5942C>T 5944C>T), N1950S (5955A>G), T1966V (6002A>G 6003C>T 6004C>T), G1968E (6009G>A), T1976I (6033C>T), A1977S (6035G>T), L1996F (6092C>T), I2002M (6112A>G), E2008D (6130G>T), T2027D (6185A>G 6186C>A 6187T>C), H2028R (6189A>G), M2032I (6202G>C), D2036E (6214T>G), I2042V (6230A>G 6232A>G), V2045T (6239G>A 6240T>C), S2056R (6274C>G), V2062E (6291T>A 6292T>A), V2064L (6296G>C 6298A>G), I2071V (6317A>G 6319A>T), I2073V (6323A>G), A2089T (6371G>A), C2100H (6404T>C 6405G>A 6406T>C), R2103L (6414G>T 6415A>G), C2111S (6437T>A), S2118T (6458T>A), L2119F (6461C>T), H2127N (6485C>A), R2128S (6490A>C), I2132D (6500A>G 6501T>A), L2147F (6545C>T), G2154N (6566G>A 6567G>A 6568C>T) | | | | | | | | |

| | Begin | End | Coverage | Score | Concordance | Matches | Identities | I/D/M/F* | Stop Codons |
|---|---|---|---|---|---|---|---|---|---|
| **NT** | **21** | **6806** | **86.7%** | **10416** | **77.0%** | **6772 (99.9%)** | **5998 (88.5%)** | **4/1** | |

Codon mutations: TTT3TTC (115T>C), ATG6TTG (122A>T), CCC8TCT (128C>T 130C>T), TTC10CCC (134T>C 135T>C), AAC11AGC (138A>G), TCA12CCC (140T>C 142A>C), GTT13GCA (144T>C 145T>A), ACT14GTG (146A>G 147C>T 148T>G), TGC17AGT (155T>A 157C>T), GCT18GTT (159C>T), ACC19GCC (161A>G), TCT24TCA (178T>A), TCA27TCT (187A>T), GAG28GAA (190G>A), ACT30ACA (196T>A), GCT31CAA (197G>C 198C>A 199T>A), CGC38CGT (220C>T), ACT39ACA (223T>A), CTG40CTT (226G>T), AAG45AAA (241G>A), TCA47TCT (247A>T), TCT51TCC (259T>C), TCT52TCA (262T>A), CAC53CAT (265C>T), GAA55GAG (271A>G), AAG58AAA (280G>A), GTT61GTG (289T>G), ACA62ACT (292A>T), GCT65GCA (301T>A), TCT66GCT (302T>G), TCC67TCT (307C>T), TTT68TTC (310T>C), ACC69TCT (311A>T 313C>T), AGC70AAC (315G>A), TGT72TGC (322T>C), AAC73AGT (324A>G 325C>T), TCT74TCA (328T>A), GTT75GTC (331T>C), ACT76ACA (334T>A), TTG78TTA (340G>A), TTA79TTG (343A>G), AAG81AAA (349G>A), ACT84ACA (358T>A), TTA85CTT (359T>C 361A>T), ATT86ATA (364T>A), GAA87GAG (367A>G), CCC88CCT (370C>T), ATC90ATT (376C>T), GCT92GCA (382T>A), TAC93TAT (385C>T), TCT94TCC (388T>C), GTC96GTT (394C>T), GCA97GCC (397A>C), AAA101AAG (409A>G), TCA102TCC (412A>C), GTT107GTG (427T>G), AAG109AAA (433G>A), GTT111GTC (439T>C), AGC113GGC (443A>G), AAA115AAG (451A>G), GAT116GAC (454T>C), AAC123GAC (473A>G), GAC127AAT (485G>A 487C>T), AAA128AAG (490A>G), TCT129TCC (493T>C), GAG130GAA (496G>A), GTG132GTT (502G>T), TCT145TCC (541T>C), TTC151TTT (559C>T), CTA158TTA (578C>T), GGA160GGT (586A>T), ACT165TCT (599A>T), CAC166CAT (604C>T), ATA167ACA (606T>C), AAT170AGT (615A>G), ACA171ATT (618C>T 619A>T), GAC183GAT (655C>T), TTA185TTG (661A>G), CAA190CAG (676A>G), GCA196GCG (694A>G), CTA201TTA (707C>T), GAA204GAG (718A>G), TCC205GCC (719T>G), GTG214GTT (748G>T), GTA215GTT (751A>T), CTC218CTT (760C>T), TCT224TCA (778T>A), ACC225ACT (781C>T), TTC228TTT (790C>T), GTA231GTT (799A>T), GGT233GGC (805T>C), AGA234CGA (806A>C), CTA238CTC (820A>C), TCC239ACC (821T>A), CTA242CTT (832A>T), TCA243TCC (835A>C), ATT245ATC (841T>C), CTG246TTG (842C>T), TAC247TAT (847C>T), TTG259TTA (883G>A), TTT260TTC (886T>C), CTG262CTA (892G>A), AAT265AAC (901T>C), TGT269TGC (913T>C), GCC273GCT (925C>T), CTG278TTG (938C>T), GTA282GTG (952A>G), GAC283GAT (955C>T), GGT286GGG (964T>G), AAG288AAA (970G>A), AAT289AAC (973T>C), GTT291ATT (977G>A), GCC297GCT (997C>T), TTG299CTG (1001T>C), TCT300TCC (1006T>C), TTC304TTT (1018C>T), ACG305GTG (1019A>G 1020C>T), GAG309GAA (1033G>A), CAT318CAC (1060T>C), CGC319CGT (1063C>T), AAC325AAT (1081C>T), ACT329ACC (1093T>C), ATC330ATA (1096C>A), CTA335TTG (1109C>T 1111A>G), AAA341AAG (1129A>G), TCT343TCA (1135T>A), GCC353GCA (1165C>A), AGG358AGA (1180G>A), ACG360ACA (1186G>A), ATA363ATT (1195A>T), ACC364TCT (1196A>T 1198C>T), CTT366CTG (1204T>G), GAT367GAG (1207T>G), CAG375CAA (1231G>A), GGC378GGT (1240C>T), AGG383AGA (1255G>A), TTT387TTC (1267T>C), GGG398GGA (1300G>A), AAA399AAG (1303A>G), CTG405TTG (1319C>T), TTC406TTT (1324C>T), GAC412GAT (1342C>T), CCA417CCC (1357A>C), TTG419CTG (1361T>C), CGA428CGT (1390A>T), AAT434AAC (1408T>C), GCT440GCA (1426T>A), ATC441ATT (1429C>T), GAC445GAT (1441C>T), CTG449TTG (1451C>T), GGG452GGT (1462G>T), GAC458GAT (1480C>T), ATT459GTT (1481A>G), GAT461GAC (1489T>C), ATA463ATT (1495A>T), CCC472CCA (1522C>A), GGC482GGT (1552C>T), CGC483AGG (1553C>A 1555C>G), CGA488AGG (1568C>A 1570A>G), TAT489TAC (1573T>C), GCC492GCT (1582C>T), ACA493ACC (1585A>C), AAC495AAT (1591C>T), GCT501TCT (1607G>T), CAG502CAA (1612G>A), GGT504GGG (1618T>G), CTA505TTA (1619C>T), ACC506ACA (1624C>A), CCC508CCT (1630C>T), AGA513CGG (1643A>C 1645A>G), TGC521TGT (1669C>T), GGT525GGA (1681T>A), GAG526GAA (1684G>A), ACT529ATT (1692C>T), GCT531GCG (1699T>G), TCT534GCT (1706T>G), CTT536CTG (1714T>G), AAA544AGG (1737A>G 1738A>G), AGA545CGA (1739A>C), GCA548GCT (1750A>T), ACA550ATC (1755C>T 1756A>C), AGC555AGT (1771C>T), TTG559CTA (1781T>C 1783G>A), GAA561GAG (1789A>G), AAG568AAA (1810G>A), GTT569GTG (1813T>G), ATC573ATA (1825C>A), AAA576AAG (1834A>G), TAT578TTT (1839A>T), GCC580ACC (1844G>A), ACA581ACC (1849A>C), TCA582ACA (1850T>A), AAG584AAA (1858G>A), CAG587CAA (1867G>A), TCA591ACT (1877T>A 1879A>T), GAG593GAA (1885G>A), ATA595GTA (1889A>G), ATA596ATT (1894A>T), GAT600GAA (1906T>A), ATG601CCA (1907A>C 1908T>C 1909G>A), TTG602TGG (1911T>G), ACC607ACA (1927C>A), AAG612AAA (1942G>A), GTA618GTG (1960A>G), GAA620GAG (1966A>G), TAC626TAT (1984C>T), CAG628CAA (1990G>A), GAG629GAA (1993G>A), TTG635CTG (2009T>C), TAT639TAC (2023T>C), CCT650CCC (2056T>C), GAC652GAT (2062C>T), GCC657GCA (2077C>A), TTT659TAT (2082T>A), CTA662CTG (2092A>G), AGG672CGG (2120A>C), GTT674GTC (2128T>C), AAT675ACT (2130A>C), CTG678TTG (2138C>T), TAT679CAA (2141T>C 2143T>A), CAA680CAG (2146A>G), GGT681GGC (2149T>C), TGT683TGC (2155T>C), CTG690TTG (2174C>T), ATA691ATC (2179A>C), AAG692AAA (2182G>A), AAT693AAC (2185T>C), GCT696GCA (2194T>A), TAT698TAC (2200T>C), TTT702TTC (2212T>C), ATC706ATT (2224C>T), TAC707TAT (2227C>T), AAT712TGT (2240A>T 2241A>G), AGC715AAA (2250G>A 2251C>A), GAC717GAT (2257C>T), CAG719CAA (2263G>A), GCG727GCA (2287G>A), CAC728CAT (2290C>T), ATA731ATT (2299A>T), GAC736GAT (2314C>T), CTG738TTG (2318C>T), ATA739ATT (2323A>T), CAC747CAT (2347C>T), TTC750TTT (2356C>T), TTG751TTA (2359G>A), ACA753ACT (2365A>T), AGG757CGT (2375A>C 2377G>T), TAT758TAC (2380T>C), CGC761AGC (2387C>A), AGA762AGG (2392A>G), CCT767CCC (2407T>C), AGG768AAG (2409G>A), ATT779ATC (2443T>C), GTG780GTA (2446G>A), CCG791CCA (2479G>A), ATC793ATT (2485C>T), GTT796GTG (2494T>G), CTG797CTA (2497G>A), ATT798ATC (2500T>C), AAG799AAA (2503G>A), GCT800GCC (2506T>C), GGC801GGT (2509C>T), CTT802CTC (2512T>C), GGG812GGA (2542G>A), GTC814GTT (2548C>T), TTC817TTT (2557C>T), GAG838GAA (2620G>A), AAT840AAC (2626T>C), CGG845CGA (2641G>A), AAG848AAA (2650G>A), ATC852ATT (2662C>T), AAG862AAA (2692G>A), GGT863GGA (2695T>A), AAA868AAG (2710A>G), GCA870GCC (2716A>C), GGC874AGC (2726G>A), TTC875TTT (2731C>T), ATC876ATT (2734C>T), GGA879GGG (2743A>G), TTG880CTG (2744T>C), TTG887CTC (2765T>C 2767G>C), AGA888CGA (2768A>C), CTA889CTG (2773A>G), TAC894TAT (2788C>T), TGC901TGT (2809C>T), GTG905GTA (2821G>A), ACA907ACG (2827A>G), TTT912TTC (2842T>C), AGC914AGT (2848C>T), CGT916AGA (2852C>A 2854T>A), GGC918GGG (2860C>G), ACT920ACA (2866T>A), TTC921TTT (2869C>T), CAT925CAC (2881T>C), GCC926TCC (2882G>T), CGG940CGT (2926G>T), TTC941TTT (2929C>T), TTG942CTG (2930T>C), CAA945CAG (2941A>G), ATT947ATA (2947T>A), GTC948GTT (2950C>T), GAT956GAC (2974T>C), CCC958CCG (2980C>G), AAA964AAG (2998A>G), TCT975TCC (3031T>C), AAG976AAA (3034G>A), ACT978ACC (3040T>C), GTA981GTC (3049A>C), GTA986GTT (3064A>T), GCT991GCC (3079T>C), CAA993AAG (3083C>A 3085A>G), TTG999CTT (3101T>C 3103G>T), TTT1001TTC (3109T>C), CTT1010CTG (3136T>G), ACG1014TCG (3146A>T), TAC1015TAT (3151C>T), TCT1019TCC (3163T>C), TCC1021TCT (3169C>T), TTT1022TTC (3172T>C), TAT1032TAC (3202T>C), CTT1045CTC (3241T>C), AAC1049AAT (3253C>T), GTG1050GTA (3256G>A), GAC1052GAT (3262C>T), GGT1059GGC (3283T>C), CAT1061CAC (3289T>C), TCT1064TCC (3298T>C), GAA1067GAG (3307A>G), CGA1084CGT (3358A>T), CTT1090CTG (3376T>G), ACT1092ACA (3382T>A), GCA1098GCC (3400A>C), GTA1100GTG (3406A>G), CAG1103GAG (3413C>G), CTC1104CTA (3418C>A), CTT1105CTC (3421T>C), CGA1109AGA (3431C>A), GCG1113GCT (3445G>T), AAA1119AAG (3463A>G), CAG1120CAA (3466G>A), GGC1125GGT (3481C>T), ACA1126ACT (3484A>T), ACG1134ACT (3508G>T), CGA1142AGG (3530C>A 3532A>G), AGG1143AAG (3534G>A), GCT1149GCA (3553T>A), GGG1151GGA (3559G>A), GCT1156GCC (3574T>C), ATC1159ATA (3583C>A), ATT1160ATC (3586T>C), TCA1161TCG (3589A>G), GCT1162GTT (3591C>T), TCA1163CAA (3593T>C 3594C>A), GAA1168GAG (3610A>G), TAC1173TAT (3625C>T), GTG1174GTA (3628G>A), GCT1175TCT (3629G>T), TAC1180TAT (3646C>T), GCA1184GCC (3658A>C), CAA1185CAG (3661A>G), AAT1186AAC (3664T>C), GAC1190GAT (3676C>T), CTA1194CTG (3688A>G), GCC1195GCT (3691C>T), TAT1196TTT (3693A>T), CTA1199TTA (3701C>T), CGT1200CGC (3706T>C), GCA1202TCA (3710G>T), ATT1203GTT (3713A>G), TTG1204CTG (3716T>C), CGA1211CGT (3739A>T), AAG1216AAA (3754G>A), GTC1218ATC (3758G>A), CAG1220CAA (3766G>A), GTG1224GTT (3778G>T), GTG1225GTA (3781G>A), TAT1234TAC (3808T>C), GAC1236GAT (3814C>T), GGG1237GGA (3817G>A), TTG1240CTG (3824T>C), GGC1254GGT (3868C>T), ACT1255ACC (3871T>C), GAG1259GAA (3883G>A), CCA1263CCC (3895A>C), GGA1267GGT (3907A>T), CGA1269AGA (3911C>A), AAA1273AAG (3925A>G), AGG1274AGA (3928G>A), GAC1281GAT (3949C>T), GAG1283GAA (3955G>A), TTG1285TTA (3961G>A), GAA1288GAG (3970A>G), ATT1289ATC (3973T>C), TTG1310CTG (4034T>C), CGG1317AGG (4055C>A), CGT1320CGC (4066T>C), CCC1326CCT (4084C>T), TAC1329TAT (4093C>T), CTG1338TTG (4118C>T), TCC1341TCT (4129C>T), TCC1347TCA (4147C>A), CTG1351TTA (4157C>T 4159G>A), CCA1352CCT (4162A>T), GGT1361GGA (4189T>A), ATT1362ATG (4192T>G), GAA1363GAG (4195A>G), TTG1367TTA (4207G>A), GCT1368GCC (4210T>C), GCA1373GCT (4225A>T), GTC1374GTT (4228C>T), GAT1376GAC (4234T>C), GAC1382GAT (4252C>T), ATC1392ATT (4282C>T), GCC1394GCA (4288C>A), CAA1396CAG (4294A>G), CTT1401CTG (4309T>G), GAC1404GAT (4318C>T), GGT1411GGC (4339T>C), GAC1414GAT (4348C>T), AAT1416AAC (4354T>C), AGA1418CGG (4358A>C 4360A>G), ACT1430ACC (4396T>C), GTT1436GTG (4414T>G), TTT1437TTC (4417T>C), GTG1440GTT (4426G>T), GGA1441GGG (4429A>G), TCA1445TCT (4441A>T), GTT1451GTC (4459T>C), AAC1454AAT (4468C>T), TCT1455TCC (4471T>C), CTT1456CTC (4474T>C), GCA1465GCT (4501A>T), TCT1468TCA (4510T>A), TTG1470CTG (4514T>C), GCT1477GCA (4537T>A), AAC1488AAT (4570C>T), CTA1490CTT (4576A>T), GTC1494GTT (4588C>T), TAT1495TAC (4591T>C), TTG1507CTG (4625T>C), TAT1510TAC (4636T>C), GCT1514GCA (4648T>A), ATT1515ATA (4651T>A), GAA1518GAG (4660A>G), GAG1520GAA (4666G>A), GTT1524GTC (4678T>C), ATA1526ATT (4684A>T), AAG1527AAA (4687G>A), ATC1530ATT (4696C>T), GAC1531GAT (4699C>T), AAG1532AAA (4702G>A), AAC1533AAT (4705C>T), GTT1536GTC (4714T>C), AGC1537AAC (4716G>A), AGT1538AAT (4719G>A), GAC1543GAT (4735C>T), TTG1544CTG (4736T>C), CTA1547CTG (4747A>G), TTT1549TTC (4753T>C), CTA1550CTC (4756A>C), AGC1551AGT (4759C>T), GGT1553GGC (4765T>C), TTT1554TTC (4768T>C), AGG1556AGA (4774G>A), CGT1558CGC (4780T>C), CTA1559CTG (4783A>G), GGA1560GGG (4786A>G), TAC1561TAT (4789C>T), CTG1566TTG (4802C>T), GTA1569GTG (4813A>G), TCT1570TCC (4816T>C), TAC1576TAT (4834C>T), AGG1579AGA (4843G>A), GGA1597GGC (4897A>C), GTT1598ATT (4898G>A), AGA1599AGG (4903A>G), AAC1618CAT (4958A>C 4960C>T), AAG1621AAA (4969G>A), CGG1623CGA (4975G>A), TTC1625TTT (4981C>T), CCT1631CCC (4999T>C), TAT1633TAC (5005T>C), TTG1636CTG (5012T>C), GAG1637GAA (5017G>A), GAC1639GAT (5023C>T), GTG1642ATG (5030G>A), ACA1645ACT (5041A>T), GGA1646GGC (5044A>C), GGG1651GGT (5059G>T), CTT1656CTC (5074T>C), CAT1657CAC (5077T>C), ATT1658ACT (5079T>C), GCA1659GCT (5083A>T), GCT1662TCT (5090G>T), TTG1666CTG (5102T>C), GGC1669GGT (5113C>T), ACT1671ACC (5119T>C), GTT1672ATC (5120G>A 5122T>C), CTG1673TTG (5123C>T), TTT1675TTC (5131T>C), TCT1676ACT (5132T>A), GTG1679GTT (5143G>T), CAT1680CAC (5146T>C), AGT1687AGC (5167T>C), TCC1690TCT (5176C>T), CAG1698CAA (5200G>A), GAC1700GAT (5206C>T), ACC1701ACA (5209C>A), GTT1704GTG (5218T>G), ACC1721ACA (5269C>A), AGG1724AGA (5278G>A), GGG1728GGT (5290G>T), CTT1729CTC (5293T>C), CCT1732CCA (5302T>A), GAC1740GAT (5326C>T), ACC1747ACA (5347C>A), GGG1749GGA (5353G>A), ACC1751TCC (5357A>T), AAA1752CAA (5360A>C), GTG1755ATT (5369G>A 5371G>T), GGC1760GGG (5386C>G), GGG1762GGA (5392G>A), ATT1765GTT (5399A>G), CCA1767CCT (5407A>T), GTG1768GTT (5410G>T), TGC1769TGT (5413C>T), GCA1771GCC (5419A>C), CTG1775TTG (5429C>T), ACT1777ACC (5437T>C), GTT1785ATT (5459G>A), GTT1786ATT (5462G>A), GAT1794AAT (5486G>A), GTT1795GTG (5491T>G), ACA1796TCA (5492A>T), CTC1798TTG (5498C>T 5500C>G), GGA1803GGT (5515A>T), CTT1809TTG (5531C>T 5533T>G), AAA1811CAA (5537A>C), GAA1818CAA (5558G>C), TGT1824TGC (5578T>C), ATT1826GTT (5582A>G), GCC1829TCC (5591G>T), GAG1832GAA (5602G>A), ACA1833ATA (5604C>T), CCA1834CCG (5608A>G), GCA1835GCT (5611A>T), TAC1836TTC (5613A>T), ATA1837ATT (5617A>T), GGA1840GGT (5626A>T), CTT1844CTG (5638T>G), GGT1845GGC (5641T>C), GTG1850GTA (5656G>A), CTA1856CTT (5674A>T), CAG1857CAT (5677G>T), CAG1860CAA (5686G>A), CGC1861CGA (5689C>A), TTG1862CTG (5690T>C), CCT1868CCA (5710T>A), TAT1869CAT (5711T>C), GGG1870GGT (5716G>T), GGA1871GGC (5719A>C), CAC1875CAT (5731C>T), ACT1876ACC (5734T>C), ACT1877ACC (5737T>C), AGA1878AAA (5739G>A), CTT1880CTC (5746T>C), ATA1890GTA (5774A>G), TAT1891TAC (5779T>C), TTA1894CTA (5786T>C), AGA1897AGG (5797A>G), TTC1900ACT (5804T>A 5805T>C 5806C>T), TAT1907TTT (5826A>T), TCA1910TCT (5836A>T), TTG1915CAA (5849T>C 5850T>A 5851G>A), AGG1921AGA (5869G>A), GGG1924GGT (5878G>T), TTT1925TTC (5881T>C), TTT1926TTC (5884T>C), CCC1927CCT (5887C>T), CGC1928CGT (5890C>T), AAT1930CAT (5894A>C), TAC1934TTC (5907A>T), CTT1936CTC (5914T>C), TTT1939TTC (5923T>C), CTG1941TTG (5927C>T), CAC1946TAT (5942C>T 5944C>T), AAC1950AGC (5955A>G), ACG1953ACA (5965G>A), TTT1956TTC (5974T>C), AGA1957CGA (5975A>C), AAG1958AAA (5980G>A), TTT1960TTC (5986T>C), AAA1964AAG (5998A>G), ACC1966GTT (6002A>G 6003C>T 6004C>T), GGG1968GAG (6009G>A), AAA1970AAG (6016A>G), GAA1971GAG (6019A>G), TCC1974TCT (6028C>T), GCA1975GCT (6031A>T), ACA1976ATA (6033C>T), GCT1977TCT (6035G>T), GGT1983GGA (6055T>A), TCC1987TCA (6067C>A), GCT1988GCC (6070T>C), CTC1991CTG (6079C>G), CTA1995TTA (6089C>T), CTT1996TTT (6092C>T), CTT1998CTG (6100T>G), CAG1999CAA (6103G>A), AAG2000AAA (6106G>A), ATA2002ATG (6112A>G), TCA2005TCT (6121A>T), GAG2008GAT (6130G>T), TTT2011TTC (6139T>C), ATT2013ATC (6145T>C), GTC2016GTT (6154C>T), GGC2019GGG (6163C>G), CAC2020CAT (6166C>T), GTA2026GTG (6184A>G), ACT2027GAC (6185A>G 6186C>A 6187T>C), CAC2028CGC (6189A>G), TTC2029TTT (6193C>T), CAA2030CAG (6196A>G), CGC2031CGT (6199C>T), ATG2032ATC (6202G>C), CAA2034CAG (6208A>G), CAA2035CAG (6211A>G), GAT2036GAG (6214T>G), CTG2037GT. (6215C>G), ATA2042GTG (6230A>G 6232A>G), GAG2043GAA (6235G>A), GTG2045ACG (6239G>A 6240T>C), TTA2049TTG (6253A>G), ACC2050ACA (6256C>A), ACA2052ACT (6262A>T), CTG2053TTA (6263C>T 6265G>A), TCG2054TCT (6268G>T), AGC2056AGG (6274C>G), GTT2062GAA (6291T>A 6292T>A), GAT2063GAC (6295T>C), GTA2064CTG (6296G>C 6298A>G), CCT2065CCA (6301T>A), TTA2066TTG (6304A>G), TTG2068CTG (6308T>C), ATA2071GTT (6317A>G 6319A>T), GAG2072GAA (6322G>A), ATC2073GTC (6323A>G), CTA2080TTA (6344C>T), AAT2084AAC (6358T>C), GCA2089ACA (6371G>A), GCC2092GCT (6382C>T), ATC2093ATT (6385C>T), TGT2100CAC (6404T>C 6405G>A 6406T>C), AAC2102AAT (6412C>T), CGA2103CTG (6414G>T 6415A>G), TTC2110TTT (6436C>T), TGT2111AGT (6437T>A), TAC2113TAT (6445C>T), ACG2114ACA (6448G>A), TCC2117TCA (6457C>A), TCT2118ACT (6458T>A), CTT2119TTT (6461C>T), CTT2125CTA (6481T>A), CAT2127AAT (6485C>A), AGA2128AGC (6490A>C), CAT2129CAC (6493T>C), CAG2130CAA (6496G>A), ACA2131ACT (6499A>T), ATT2132GAT (6500A>G 6501T>A), TCC2137TCA (6517C>A), CAC2142CAT (6532C>T), CTT2147TTT (6545C>T), TTG2148CTG (6548T>C), CCA2150CCC (6556A>C), CTA2152CTT (6562A>T), GGC2154AAT (6566G>A 6567G>A 6568C>T), TAC2155TAT (6571C>T), AAA2156AAG (6574A>G)

#### Proteins

| | Begin | End | Coverage | Score | Concordance | Matches | Identities | I/D/M/F* | Stop Codons |
|---|---|---|---|---|---|---|---|---|---|
| polyprotein (YP_001039627.1) | 1 | 2159 | 99.8% | 14340 | 95.3% | 2155 (100%) | 2028 (94.1%) | 0/0/0/0 | 1 |

| | Begin | End | Coverage | Score | Concordance | Matches | Identities | I/D/M/F* | Stop Codons |
|---|---|---|---|---|---|---|---|---|---|
| **NT** | **21** | **6806** | **86.7%** | **10416** | **77.0%** | **6772 (99.9%)** | **5998 (88.5%)** | **4/1** | |

| | |
|---|---|
| Protein mutations: | M6L (122A>T), P8S (128C>T 130C>T), F10P (134T>C 135T>C), N11S (138A>G), S12P (140T>C 142A>C), V13A (144T>C 145T>A), T14V (146A>G 147C>T 148T>G), C17S (155T>A 157C>T), A18V (159C>T), T19A (161A>G), A31Q (197G>C 198C>A 199T>A), S66A (302T>G), T69S (311A>T 313C>T), S70N (315G>A), N73S (324A>G 325C>T), S113G (443A>G), N123D (473A>G), D127N (485G>A 487C>T), T165S (599A>T), I167T (606T>C), N170S (615A>G), T171I (618C>T 619A>T), S205A (719T>G), S239T (821T>A), V291I (977G>A), T305V (1019A>G 1020C>T), T364S (1196A>T 1198C>T), D367E (1207T>G), I459V (1481A>G), A501S (1607G>T), T529I (1692C>T), S534A (1706T>G), K544R (1737A>G 1738A>G), T550I (1755C>T 1756A>C), Y578F (1839A>T), A580T (1844G>A), S582T (1850T>A), S591T (1877T>A 1879A>T), I595V (1889A>G), D600E (1906T>A), M601P (1907A>C 1908T>C 1909G>A), L602W (1911T>G), F659Y (2082T>A), N675T (2130A>C), Y679Q (2141T>C 2143T>A), N712C (2240A>T 2241A>G), S715K (2250G>A 2251C>A), R761S (2387C>A), R768K (2409G>A), G874S (2726G>A), A926S (2882G>T), Q993K (3083C>A 3085A>G), T1014S (3146A>T), Q1103E (3413C>G), R1143K (3534G>A), A1162V (3591C>T), S1163Q (3593T>C 3594C>A), A1175S (3629G>T), Y1196F (3693A>T), A1202S (3710G>T), I1203V (3713A>G), V1218I (3758G>A), I1362M (4192T>G), S1537N (4716G>A), S1538N (4719G>A), V1598I (4898G>A), N1618H (4958A>C 4960C>T), V1642M (5030G>A), I1658T (5079T>C), A1662S (5090G>T), V1672I (5120G>A 5122T>C), S1676T (5132T>A), T1751S (5357A>T), K1752Q (5360A>C), V1755I (5369G>A 5371G>T), I1765V (5399A>G), V1785I (5459G>A), V1786I (5462G>A), D1794N (5486G>A), T1796S (5492A>T), K1811Q (5537A>C), E1818Q (5558G>C), I1826V (5582A>G), A1829S (5591G>T), T1833I (5604C>T), Y1836F (5613A>T), Q1857H (5677G>T), Y1869H (5711T>C), R1878K (5739G>A), I1890V (5774A>G), F1900T (5804T>A 5805T>C 5806C>T), Y1907F (5826A>T), L1915Q (5849T>C 5850T>A 5851G>A), N1930H (5894A>C), Y1934F (5907A>T), H1946Y (5942C>T 5944C>T), N1950S (5955A>G), T1966V (6002A>G 6003C>T 6004C>T), G1968E (6009G>A), T1976I (6033C>T), A1977S (6035G>T), L1996F (6092C>T), I2002M (6112A>G), E2008D (6130G>T), T2027D (6185A>G 6186C>A 6187T>C), H2028R (6189A>G), M2032I (6202G>C), D2036E (6214T>G), I2042V (6230A>G 6232A>G), V2045T (6239G>A 6240T>C), S2056R (6274C>G), V2062E (6291T>A 6292T>A), V2064L (6296G>C 6298A>G), I2071V (6317A>G 6319A>T), I2073V (6323A>G), A2089T (6371G>A), C2100H (6404T>C 6405G>A 6406T>C), R2103L (6414G>T 6415A>G), C2111S (6437T>A), S2118T (6458T>A), L2119F (6461C>T), H2127N (6485C>A), R2128S (6490A>C), I2132D (6500A>G 6501T>A), L2147F (6545C>T), G2154N (6566G>A 6567G>A 6568C>T) |

| | Begin | End | Coverage | Score | Concordance | Matches | Identities | I/D/M/F* | Stop Codons |
|---|---|---|---|---|---|---|---|---|---|
| **NT** | **21** | **6806** | **86.7%** | **10416** | **77.0%** | **6772 (99.9%)** | **5998 (88.5%)** | **4/1** | |
| Codon mutations: | TTT3TTC (115T>C), ATG6TTG (122A>T), CCC8TCT (128C>T 130C>T), TTC10CCC (134T>C 135T>C), AAC11AGC (138A>G), TCA12CCC (140T>C 142A>C), GTT13GCA (144T>C 145T>A), ACT14GTG (146A>G 147C>T 148T>G), TGC17AGT (155T>A 157C>T), GCT18GTT (159C>T), ACC19GCC (161A>G), TCT24TCA (178T>A), TCA27TCT (187A>T), GAG28GAA (190G>A), ACT30ACA (196T>A), GCT31CAA (197G>C 198C>A 199T>A), CGC38CGT (220C>T), ACT39ACA (223T>A), CTG40CTT (226G>T), AAG45AAA (241G>A), TCA47TCT (247A>T), TCT51TCC (259T>C), TCT52TCA (262T>A), CAC53CAT (265C>T), GAA55GAG (271A>G), AAG58AAA (280G>A), GTT61GTG (289T>G), ACA62ACT (292A>T), GCT65GCA (301T>A), TCT66GCT (302T>G), TCC67TCT (307C>T), TTT68TTC (310T>C), ACC69TCT (311A>T 313C>T), AGC70AAC (315G>A), TGT72TGC (322T>C), AAC73AGT (324A>G 325C>T), TCT74TCA (328T>A), GTT75GTC (331T>C), ACT76ACA (334T>A), TTG78TTA (340G>A), TTA79TTG (343A>G), AAG81AAA (349G>A), ACT84ACA (358T>A), TTA85CTT (359T>C 361A>T), ATT86ATA (364T>A), GAA87GAG (367A>G), CCC88CCT (370C>T), ATC90ATT (376C>T), GCT92GCA (382T>A), TAC93TAT (385C>T), TCT94TCC (388T>C), GTC96GTT (394C>T), GCA97GCC (397A>C), AAA101AAG (409A>G), TCA102TCC (412A>C), GTT107GTG (427T>G), AAG109AAA (433G>A), GTT111GTC (439T>C), AGC113GGC (443A>G), AAA115AAG (451A>G), GAT116GAC (454T>C), AAC123GAC (473A>G), GAC127AAT (485G>A 487C>T), AAA128AAG (490A>G), TCT129TCC (493T>C), GAG130GAA (496G>A), GTG132GTT (502G>T), TCT145TCC (541T>C), TTC151TTT (559C>T), CTA158TTA (578C>T), GGA160GGT (586A>T), ACT165TCT (599A>T), CAC166CAT (604C>T), ATA167ACA (606T>C), AAT170AGT (615A>G), ACA171ATT (618C>T 619A>T), GAC183GAT (655C>T), TTA185TTG (661A>G), CAA190CAG (676A>G), GCA196GCG (694A>G), CTA201TTA (707C>T), GAA204GAG (718A>G), TCC205GCC (719T>G), GTG214GTT (748G>T), GTA215GTT (751A>T), CTC218CTT (760C>T), TCT224TCA (778T>A), ACC225ACT (781C>T), TTC228TTT (790C>T), GTA231GTT (799A>T), GGT233GGC (805T>C), AGA234CGA (806A>C), CTA238CTC (820A>C), TCC239ACC (821T>A), CTA242CTT (832A>T), TCA243TCC (835A>C), ATT245ATC (841T>C), CTG246TTG (842C>T), TAC247TAT (847C>T), TTG259TTA (883G>A), TTT260TTC (886T>C), CTG262CTA (892G>A), AAT265AAC (901T>C), TGT269TGC (913T>C), GCC273GCT (925C>T), CTG278TTG (938C>T), GTA282GTG (952A>G), GAC283GAT (955C>T), GGT286GGG (964T>G), AAG288AAA (970G>A), AAT289AAC (973T>C), GTT291ATT (977G>A), GCC297GCT (997C>T), TTG299CTG (1001T>C), TCT300TCC (1006T>C), TTC304TTT (1018C>T), ACG305GTG (1019A>G 1020C>T), GAG309GAA (1033G>A), CAT318CAC (1060T>C), CGC319CGT (1063C>T), AAC325AAT (1081C>T), ACT329ACC (1093T>C), ATC330ATA (1096C>A), CTA335TTG (1109C>T 1111A>G), AAA341AAG (1129A>G), TCT343TCA (1135T>A), GCC353GCA (1165C>A), AGG358AGA (1180G>A), ACG360ACA (1186G>A), ATA363ATT (1195A>T), ACC364TCT (1196A>T 1198C>T), CTT366CTG (1204T>G), GAT367GAG (1207T>G), CAG375CAA (1231G>A), GGC378GGT (1240C>T), AGG383AGA (1255G>A), TTT387TTC (1267T>C), GGG398GGA (1300G>A), AAA399AAG (1303A>G), CTG405TTG (1319C>T), TTC406TTT (1324C>T), GAC412GAT (1342C>T), CCA417CCC (1357A>C), TTG419CTG (1361T>C), CGA428CGT (1390A>T), AAT434AAC (1408T>C), GCT440GCA (1426T>A), ATC441ATT (1429C>T), GAC445GAT (1441C>T), CTG449TTG (1451C>T), GGG452GGT (1462G>T), GAC458GAT (1480C>T), ATT459GTT (1481A>G), GAT461GAC (1489T>C), ATA463ATT (1495A>T), CCC472CCA (1522C>A), GGC482GGT (1552C>T), CGC483AGG (1553C>A 1555C>G), CGA488AGG (1568C>A 1570A>G), TAT489TAC (1573T>C), GCC492GCT (1582C>T), ACA493ACC (1585A>C), AAC495AAT (1591C>T), GCT501TCT (1607G>T), CAG502CAA (1612G>A), GGT504GGG (1618T>G), CTA505TTA (1619C>T), ACC506ACA (1624C>A), CCC508CCT (1630C>T), AGA513CGG (1643A>C 1645A>G), TGC521TGT (1669C>T), GGT525GGA (1681T>A), GAG526GAA (1684G>A), ACT529ATT (1692C>T), GCT531GCG (1699T>G), TCT534GCT (1706T>G), CTT536CTG (1714T>G), AAA544AGG (1737A>G 1738A>G), AGA545CGA (1739A>C), GCA548GCT (1750A>T), ACA550ATC (1755C>T 1756A>C), AGC555AGT (1771C>T), TTG559CTA (1781T>C 1783G>A), GAA561GAG (1789A>G), AAG568AAA (1810G>A), GTT569GTG (1813T>G), ATC573ATA (1825C>A), AAA576AAG (1834A>G), TAT578TTT (1839A>T), GCC580ACC (1844G>A), ACA581ACC (1849A>C), TCA582ACA (1850T>A), AAG584AAA (1858G>A), CAG587CAA (1867G>A), TCA591ACT (1877T>A 1879A>T), GAG593GAA (1885G>A), ATA595GTA (1889A>G), ATA596ATT (1894A>T), GAT600GAA (1906T>A), ATG601CCA (1907A>C 1908T>C 1909G>A), TTG602TGG (1911T>G), ACC607ACA (1927C>A), AAG612AAA (1942G>A), GTA618GTG (1960A>G), GAA620GAG (1966A>G), TAC626TAT (1984C>T), CAG628CAA (1990G>A), GAG629GAA (1993G>A), TTG635CTG (2009T>C), TAT639TAC (2023T>C), CCT650CCC (2056T>C), GAC652GAT (2062C>T), GCC657GCA (2077C>A), TTT659TAT (2082T>A), CTA662CTG (2092A>G), AGG672CGG (2120A>C), GTT674GTC (2128T>C), AAT675ACT (2130A>C), CTG678TTG (2138C>T), TAT679CAA (2141T>C 2143T>A), CAA680CAG (2146A>G), GGT681GGC (2149T>C), TGT683TGC (2155T>C), CTG690TTG (2174C>T), ATA691ATC (2179A>C), AAG692AAA (2182G>A), AAT693AAC (2185T>C), GCT696GCA (2194T>A), TAT698TAC (2200T>C), TTT702TTC (2212T>C), ATC706ATT (2224C>T), TAC707TAT (2227C>T), AAT712TGT (2240A>T 2241A>G), AGC715AAA (2250G>A 2251C>A), GAC717GAT (2257C>T), CAG719CAA (2263G>A), GCG727GCA (2287G>A), CAC728CAT (2290C>T), ATA731ATT (2299A>T), GAC736GAT (2314C>T), CTG738TTG (2318C>T), ATA739ATT (2323A>T), CAC747CAT (2347C>T), TTC750TTT (2356C>T), TTG751TTA (2359G>A), ACA753ACT (2365A>T), AGG757CGT (2375A>C 2377G>T), TAT758TAC (2380T>C), CGC761AGC (2387C>A), AGA762AGG (2392A>G), CCT767CCC (2407T>C), AGG768AAG (2409G>A), ATT779ATC (2443T>C), GTG780GTA (2446G>A), CCG791CCA (2479G>A), ATC793ATT (2485C>T), GTT796GTG (2494T>G), CTG797CTA (2497G>A), ATT798ATC (2500T>C), AAG799AAA (2503G>A), GCT800GCC (2506T>C), GGC801GGT (2509C>T), CTT802CTC (2512T>C), GGG812GGA (2542G>A), GTC814GTT (2548C>T), TTC817TTT (2557C>T), GAG838GAA (2620G>A), AAT840AAC (2626T>C), CGG845CGA (2641G>A), AAG848AAA (2650G>A), ATC852ATT (2662C>T), AAG862AAA (2692G>A), GGT863GGA (2695T>A), AAA868AAG (2710A>G), GCA870GCC (2716A>C), GGC874AGC (2726G>A), TTC875TTT (2731C>T), ATC876ATT (2734C>T), GGA879GGG (2743A>G), TTG880CTG (2744T>C), TTG887CTC (2765T>C 2767G>C), AGA888CGA (2768A>C), CTA889CTG (2773A>G), TAC894TAT (2788C>T), TGC901TGT (2809C>T), GTG905GTA (2821G>A), ACA907ACG (2827A>G), TTT912TTC (2842T>C), AGC914AGT (2848C>T), CGT916AGA (2852C>A 2854T>A), GGC918GGG (2860C>G), ACT920ACA (2866T>A), TTC921TTT (2869C>T), CAT925CAC (2881T>C), GCC926TCC (2882G>T), CGG940CGT (2926G>T), TTC941TTT (2929C>T), TTG942CTG (2930T>C), CAA945CAG (2941A>G), ATT947ATA (2947T>A), GTC948GTT (2950C>T), GAT956GAC (2974T>C), CCC958CCG (2980C>G), AAA964AAG (2998A>G), TCT975TCC (3031T>C), AAG976AAA (3034G>A), ACT978ACC (3040T>C), GTA981GTC (3049A>C), GTA986GTT (3064A>T), GCT991GCC (3079T>C), CAA993AAG (3083C>A 3085A>G), TTG999CTT (3101T>C 3103G>T), TTT1001TTC (3109T>C), CTT1010CTG (3136T>G), ACG1014TCG (3146A>T), TAC1015TAT (3151C>T), TCT1019TCC (3163T>C), TCC1021TCT (3169C>T), TTT1022TTC (3172T>C), TAT1032TAC (3202T>C), CTT1045CTC (3241T>C), AAC1049AAT (3253C>T), GTG1050GTA (3256G>A), GAC1052GAT (3262C>T), GGT1059GGC (3283T>C), CAT1061CAC (3289T>C), TCT1064TCC (3298T>C), GAA1067GAG (3307A>G), CGA1084CGT (3358A>T), CTT1090CTG (3376T>G), ACT1092ACA (3382T>A), GCA1098GCC (3400A>C), GTA1100GTG (3406A>G), CAG1103GAG (3413C>G), CTC1104CTA (3418C>A), CTT1105CTC (3421T>C), CGA1109AGA (3431C>A), GCG1113GCT (3445G>T), AAA1119AAG (3463A>G), CAG1120CAA (3466G>A), GGC1125GGT (3481C>T), ACA1126ACT (3484A>T), ACG1134ACT (3508G>T), CGA1142AGG (3530C>A 3532A>G), AGG1143AAG (3534G>A), GCT1149GCA (3553T>A), GGG1151GGA (3559G>A), GCT1156GCC (3574T>C), ATC1159ATA (3583C>A), ATT1160ATC (3586T>C), TCA1161TCG (3589A>G), GCT1162GTT (3591C>T), TCA1163CAA (3593T>C 3594C>A), GAA1168GAG (3610A>G), TAC1173TAT (3625C>T), GTG1174GTA (3628G>A), GCT1175TCT (3629G>T), TAC1180TAT (3646C>T), GCA1184GCC (3658A>C), CAA1185CAG (3661A>G), AAT1186AAC (3664T>C), GAC1190GAT (3676C>T), CTA1194CTG (3688A>G), GCC1195GCT (3691C>T), TAT1196TTT (3693A>T), CTA1199TTA (3701C>T), CGT1200CGC (3706T>C), GCA1202TCA (3710G>T), ATT1203GTT (3713A>G), TTG1204CTG (3716T>C), CGA1211CGT (3739A>T), AAG1216AAA (3754G>A), GTC1218ATC (3758G>A), CAG1220CAA (3766G>A), GTG1224GTT (3778G>T), GTG1225GTA (3781G>A), TAT1234TAC (3808T>C), GAC1236GAT (3814C>T), GGG1237GGA (3817G>A), TTG1240CTG (3824T>C), GGC1254GGT (3868C>T), ACT1255ACC (3871T>C), GAG1259GAA (3883G>A), CCA1263CCC (3895A>C), GGA1267GGT (3907A>T), CGA1269AGA (3911C>A), AAA1273AAG (3925A>G), AGG1274AGA (3928G>A), GAC1281GAT (3949C>T), GAG1283GAA (3955G>A), TTG1285TTA (3961G>A), GAA1288GAG (3970A>G), ATT1289ATC (3973T>C), TTG1310CTG (4034T>C), CGG1317AGG (4055C>A), CGT1320CGC (4066T>C), CCC1326CCT (4084C>T), TAC1329TAT (4093C>T), CTG1338TTG (4118C>T), TCC1341TCT (4129C>T), TCC1347TCA (4147C>A), CTG1351TTA (4157C>T 4159G>A), CCA1352CCT (4162A>T), GGT1361GGA (4189T>A), ATT1362ATG (4192T>G), GAA1363GAG (4195A>G), TTG1367TTA (4207G>A), GCT1368GCC (4210T>C), GCA1373GCT (4225A>T), GTC1374GTT (4228C>T), GAT1376GAC (4234T>C), GAC1382GAT (4252C>T), ATC1392ATT (4282C>T), GCC1394GCA (4288C>A), CAA1396CAG (4294A>G), CTT1401CTG (4309T>G), GAC1404GAT (4318C>T), GGT1411GGC (4339T>C), GAC1414GAT (4348C>T), AAT1416AAC (4354T>C), AGA1418CGG (4358A>C 4360A>G), ACT1430ACC (4396T>C), GTT1436GTG (4414T>G), TTT1437TTC (4417T>C), GTG1440GTT (4426G>T), GGA1441GGG (4429A>G), TCA1445TCT (4441A>T), GTT1451GTC (4459T>C), AAC1454AAT (4468C>T), TCT1455TCC (4471T>C), CTT1456CTC (4474T>C), GCA1465GCT (4501A>T), TCT1468TCA (4510T>A), TTG1470CTG (4514T>C), GCT1477GCA (4537T>A), AAC1488AAT (4570C>T), CTA1490CTT (4576A>T), GTC1494GTT (4588C>T), TAT1495TAC (4591T>C), TTG1507CTG (4625T>C), TAT1510TAC (4636T>C), GCT1514GCA (4648T>A), ATT1515ATA (4651T>A), GAA1518GAG (4660A>G), GAG1520GAA (4666G>A), GTT1524GTC (4678T>C), ATA1526ATT (4684A>T), AAG1527AAA (4687G>A), ATC1530ATT (4696C>T), GAC1531GAT (4699C>T), AAG1532AAA (4702G>A), AAC1533AAT (4705C>T), GTT1536GTC (4714T>C), AGC1537AAC (4716G>A), AGT1538AAT (4719G>A), GAC1543GAT (4735C>T), TTG1544CTG (4736T>C), CTA1547CTG (4747A>G), TTT1549TTC (4753T>C), CTA1550CTC (4756A>C), AGC1551AGT (4759C>T), GGT1553GGC (4765T>C), TTT1554TTC (4768T>C), AGG1556AGA (4774G>A), CGT1558CGC (4780T>C), CTA1559CTG (4783A>G), GGA1560GGG (4786A>G), TAC1561TAT (4789C>T), CTG1566TTG (4802C>T), GTA1569GTG (4813A>G), TCT1570TCC (4816T>C), TAC1576TAT (4834C>T), AGG1579AGA (4843G>A), GGA1597GGC (4897A>C), GTT1598ATT (4898G>A), AGA1599AGG (4903A>G), AAC1618CAT (4958A>C 4960C>T), AAG1621AAA (4969G>A), CGG1623CGA (4975G>A), TTC1625TTT (4981C>T), CCT1631CCC (4999T>C), TAT1633TAC (5005T>C), TTG1636CTG (5012T>C), GAG1637GAA (5017G>A), GAC1639GAT (5023C>T), GTG1642ATG (5030G>A), ACA1645ACT (5041A>T), GGA1646GGC (5044A>C), GGG1651GGT (5059G>T), CTT1656CTC (5074T>C), CAT1657CAC (5077T>C), ATT1658ACT (5079T>C), GCA1659GCT (5083A>T), GCT1662TCT (5090G>T), TTG1666CTG (5102T>C), GGC1669GGT (5113C>T), ACT1671ACC (5119T>C), GTT1672ATC (5120G>A 5122T>C), CTG1673TTG (5123C>T), TTT1675TTC (5131T>C), TCT1676ACT (5132T>A), GTG1679GTT (5143G>T), CAT1680CAC (5146T>C), AGT1687AGC (5167T>C), TCC1690TCT (5176C>T), CAG1698CAA (5200G>A), GAC1700GAT (5206C>T), ACC1701ACA (5209C>A), GTT1704GTG (5218T>G), ACC1721ACA (5269C>A), AGG1724AGA (5278G>A), GGG1728GGT (5290G>T), CTT1729CTC (5293T>C), CCT1732CCA (5302T>A), GAC1740GAT (5326C>T), ACC1747ACA (5347C>A), GGG1749GGA (5353G>A), ACC1751TCC (5357A>T), AAA1752CAA (5360A>C), GTG1755ATT (5369G>A 5371G>T), GGC1760GGG (5386C>G), GGG1762GGA (5392G>A), ATT1765GTT (5399A>G), CCA1767CCT (5407A>T), GTG1768GTT (5410G>T), TGC1769TGT (5413C>T), GCA1771GCC (5419A>C), CTG1775TTG (5429C>T), ACT1777ACC (5437T>C), GTT1785ATT (5459G>A), GTT1786ATT (5462G>A), GAT1794AAT (5486G>A), GTT1795GTG (5491T>G), ACA1796TCA (5492A>T), CTC1798TTG (5498C>T 5500C>G), GGA1803GGT (5515A>T), CTT1809TTG (5531C>T 5533T>G), AAA1811CAA (5537A>C), GAA1818CAA (5558G>C), TGT1824TGC (5578T>C), ATT1826GTT (5582A>G), GCC1829TCC (5591G>T), GAG1832GAA (5602G>A), ACA1833ATA (5604C>T), CCA1834CCG (5608A>G), GCA1835GCT (5611A>T), TAC1836TTC (5613A>T), ATA1837ATT (5617A>T), GGA1840GGT (5626A>T), CTT1844CTG (5638T>G), GGT1845GGC (5641T>C), GTG1850GTA (5656G>A), CTA1856CTT (5674A>T), CAG1857CAT (5677G>T), CAG1860CAA (5686G>A), CGC1861CGA (5689C>A), TTG1862CTG (5690T>C), CCT1868CCA (5710T>A), TAT1869CAT (5711T>C), GGG1870GGT (5716G>T), GGA1871GGC (5719A>C), CAC1875CAT (5731C>T), ACT1876ACC (5734T>C), ACT1877ACC (5737T>C), AGA1878AAA (5739G>A), CTT1880CTC (5746T>C), ATA1890GTA (5774A>G), TAT1891TAC (5779T>C), TTA1894CTA (5786T>C), AGA1897AGG (5797A>G), TTC1900ACT (5804T>A 5805T>C 5806C>T), TAT1907TTT (5826A>T), TCA1910TCT (5836A>T), TTG1915CAA (5849T>C 5850T>A 5851G>A), AGG1921AGA (5869G>A), GGG1924GGT (5878G>T), TTT1925TTC (5881T>C), TTT1926TTC (5884T>C), CCC1927CCT (5887C>T), CGC1928CGT (5890C>T), AAT1930CAT (5894A>C), TAC1934TTC (5907A>T), CTT1936CTC (5914T>C), TTT1939TTC (5923T>C), CTG1941TTG (5927C>T), CAC1946TAT (5942C>T 5944C>T), AAC1950AGC (5955A>G), ACG1953ACA (5965G>A), TTT1956TTC (5974T>C), AGA1957CGA (5975A>C), AAG1958AAA (5980G>A), TTT1960TTC (5986T>C), AAA1964AAG (5998A>G), ACC1966GTT (6002A>G 6003C>T 6004C>T), GGG1968GAG (6009G>A), AAA1970AAG (6016A>G), GAA1971GAG (6019A>G), TCC1974TCT (6028C>T), GCA1975GCT (6031A>T), ACA1976ATA (6033C>T), GCT1977TCT (6035G>T), GGT1983GGA (6055T>A), TCC1987TCA (6067C>A), GCT1988GCC (6070T>C), CTC1991CTG (6079C>G), CTA1995TTA (6089C>T), CTT1996TTT (6092C>T), CTT1998CTG (6100T>G), CAG1999CAA (6103G>A), AAG2000AAA (6106G>A), ATA2002ATG (6112A>G), TCA2005TCT (6121A>T), GAG2008GAT (6130G>T), TTT2011TTC (6139T>C), ATT2013ATC (6145T>C), GTC2016GTT (6154C>T), GGC2019GGG (6163C>G), CAC2020CAT (6166C>T), GTA2026GTG (6184A>G), ACT2027GAC (6185A>G 6186C>A 6187T>C), CAC2028CGC (6189A>G), TTC2029TTT (6193C>T), CAA2030CAG (6196A>G), CGC2031CGT (6199C>T), ATG2032ATC (6202G>C), CAA2034CAG (6208A>G), CAA2035CAG (6211A>G), GAT2036GAG (6214T>G), CTG2037GT. (6215C>G), ATA2042GTG (6230A>G 6232A>G), GAG2043GAA (6235G>A), GTG2045ACG (6239G>A 6240T>C), TTA2049TTG (6253A>G), ACC2050ACA (6256C>A), ACA2052ACT (6262A>T), CTG2053TTA (6263C>T 6265G>A), TCG2054TCT (6268G>T), AGC2056AGG (6274C>G), GTT2062GAA (6291T>A 6292T>A), GAT2063GAC (6295T>C), GTA2064CTG (6296G>C 6298A>G), CCT2065CCA (6301T>A), TTA2066TTG (6304A>G), TTG2068CTG (6308T>C), ATA2071GTT (6317A>G 6319A>T), GAG2072GAA (6322G>A), ATC2073GTC (6323A>G), CTA2080TTA (6344C>T), AAT2084AAC (6358T>C), GCA2089ACA (6371G>A), GCC2092GCT (6382C>T), ATC2093ATT (6385C>T), TGT2100CAC (6404T>C 6405G>A 6406T>C), AAC2102AAT (6412C>T), CGA2103CTG (6414G>T 6415A>G), TTC2110TTT (6436C>T), TGT2111AGT (6437T>A), TAC2113TAT (6445C>T), ACG2114ACA (6448G>A), TCC2117TCA (6457C>A), TCT2118ACT (6458T>A), CTT2119TTT (6461C>T), CTT2125CTA (6481T>A), CAT2127AAT (6485C>A), AGA2128AGC (6490A>C), CAT2129CAC (6493T>C), CAG2130CAA (6496G>A), ACA2131ACT (6499A>T), ATT2132GAT (6500A>G 6501T>A), TCC2137TCA (6517C>A), CAC2142CAT (6532C>T), CTT2147TTT (6545C>T), TTG2148CTG (6548T>C), CCA2150CCC (6556A>C), CTA2152CTT (6562A>T), GGC2154AAT (6566G>A 6567G>A 6568C>T), TAC2155TAT (6571C>T), AAA2156AAG (6574A>G) | | | | | | | | |

*: Inserts / Deletes / Misaligned / Frameshifts

#### Analysis details

This analysis was performed with panviral2.64

### NGS Details (UN18_val): Torradovirus lycopersici (segment RNA 2)

#### Assembly

| | |
|---|---|
| **Coverage Length** | 3928 (8 contig(s)) |
| **Depth Of Coverage** | 1183.7 |
| **Number Of Reads** | 38768 |
| **Reads Per Million** | 727.62 rpm (after QC) |
| **Ambiguities** | 0 |
| **Assembly Method** | de novo + reference guided assembly |
| **Consensus Caller** | Bcf Tools |

#### Coverage Map

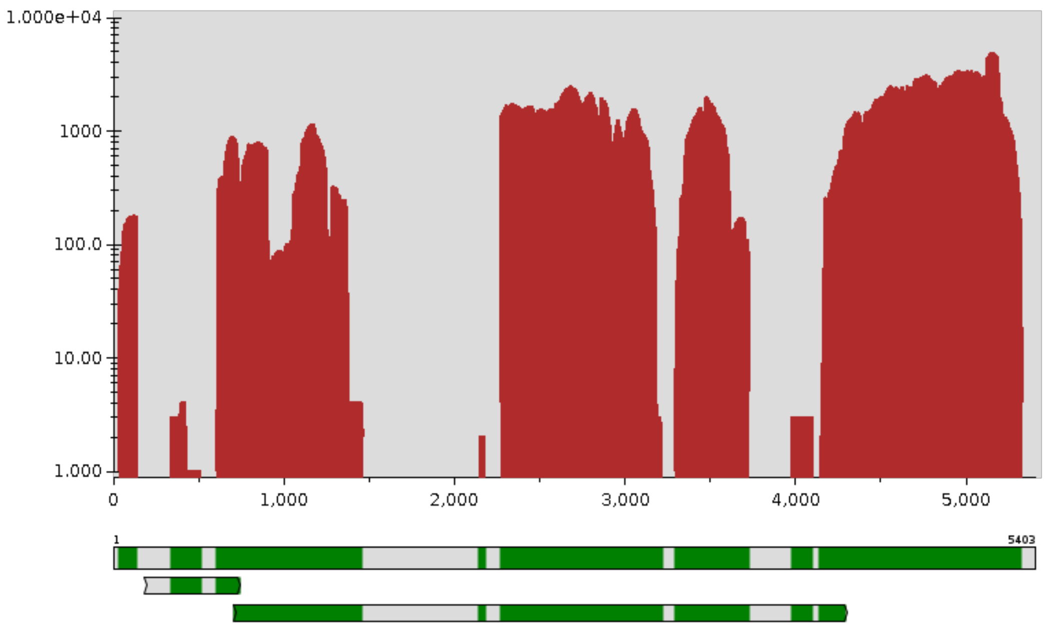

#### Assignment

| | |
|---|---|
| **Type** | Torradovirus lycopersici (Taxonomy ID: 3048378) |
| **Reference Genome** | NC_009032.1 |
| **NT Identity (%)** | 83.834 |
| **AA Identity (%)** | 86.1702 |
| **Number Of Stop Codons** | 4 |
| **Number Of CDS** | 2 |

#### Alignment

| | |
|---|---|
| **Alignment Score** | 5298.0 (NT) + 5645.0 (AA) = 10943.0 |
| **Concordance (%)** | 76.9442 |

| Alignment Method | Global, seeded, nucleotide + amino acids (AGA) |
|---|---|

#### Genome Region

Sequence starts at position 30 and ends at position 5329 relative to NC_009032.1 reference sequence.

#### Alignment Detailed Statistics

| | Begin | End | Coverage | Score | Concordance | Matches | Identities | I/D/M/F* | Stop Codons |
|---|---|---|---|---|---|---|---|---|---|
| **NT** | **30** | **5329** | **72.7%** | **5298** | **67.5%** | **3927 (99.9%)** | **3293 (83.8%)** | **1/1** | |
| Mutations: | 52C>T, 58T>C, 60G>A, 75T>C, 77T>G, 78C>T, 80A>T, 90C>T, 96C>T, 104A>G, 105G>T, 107G>A, 110C>T, 111C>A, 112G>A, 117C>T, 121A>C, 137T>C, 139C>T, 144A>G, 340A>T, 350C>A, 352A>C, 355T>C, 356G>A, 370C>T, 372A>C, 376C>T, 388T>A, 400T>C, 401T>C, 403A>G, 406G>A, 418T>G, 424A>C, 430T>A, 432G>C, 433C>T, 436T>C, 439G>A, 442A>C, 445T>A, 448C>A, 451A>G, 460T>G, 461A>C, 472T>A, 484C>G, 487A>G, 496A>T, 502T>A, 505G>T, 507G>A, 508G>A, 509A>G, 511T>A, 514T>C, 515G>A, 520C>A, 604C>A, 607G>A, 610C>A, 613A>G, 618G>C, 619C>T, 622A>G, 626C>G, 628C>A, 631A>T, 632G>A, 637A>G, 644C>G, 645T>C, 646T>C, 649A>G, 654T>A, 658C>T, 664A>C, 665T>A, 670A>G, 671T>C, 673A>T, 677T>C, 679G>A, 683T>A, 685G>C, 686T>C, 687C>A, 688C>G, 689A>G, 690A>C, 691C>A, 696T>C, 703T>C, 706T>A, 718T>G, 731G>A, 733A>G, 750A>G, 751G>C, 755A>C, 758T>A, 761A>C, 788A>T, 797T>G, 806C>G, 818G>C, 821T>G, 827A>G, 828C>T, 833C>A, 844A>T, 848G>C, 854C>G, 860C>A, 863C>T, 866A>G, 869G>A, 884A>T, 887G>T, 911T>G, 929T>A, 932A>G, 935T>C, 956G>C, 986G>A, 995T>C, 1010T>G, 1031A>G, 1034T>G, 1040A>C, 1052A>G, 1061T>C, 1073C>T, 1083C>A, 1095G>A, 1097T>G, 1100G>A, 1109C>T, 1115G>A, 1118A>T, 1130A>G, 1139G>T, 1142C>A, 1149C>T, 1151A>G, 1163A>G, 1169C>G, 1178C>T, 1187A>T, 1205T>A, 1220T>A, 1226A>T, 1235C>A, 1247A>G, 1262T>A, 1277A>G, 1313T>C, 1316C>T, 1317C>A, 1319A>G, 1322T>A, 1325A>G, 1328T>C, 1331C>T, 1334G>T, 1343G>A, 1355G>A, 1361A>T, 1364A>T, 1367A>T, 1370A>C, 1376T>G, 1383A>G, 1385C>G, 1388C>T, 1390C>A, 1394A>C, 1395A>C, 1397T>A, 1400T>G, 1403A>T, 1406G>A, 1407C>T, 1409T>A, 1412G>T, 1418G>A, 1421A>T, 1426A>T, 1427G>T, 1433C>G, 1434G>A, 1439A>T, 1451G>T, 1454A>C, 1455A>C, 1456G>A, 1457C>G, 1460A>G, 1461G>T, 2140A>G, 2142A>G, 2144C>G, 2159T>A, 2162T>A, 2164T>G, 2167A>G, 2169G>A, 2170A>G, 2171A>C, 2173T>G, 2174A>G, 2175C>T, 2177A>C, 2178T>A, 2183C>T, 2271A>T, 2272G>C, 2273G>A, 2276C>T, 2277A>G, 2279T>C, 2282A>T, 2285T>C, 2286A>T, 2288G>C, 2294A>C, 2301C>G, 2303A>G, 2311A>T, 2312T>G, 2313A>G, 2315T>A, 2318A>T, 2320G>C, 2321T>C, 2322G>C, 2324G>C, 2326C>T, 2327A>T, 2328G>C, 2330G>A, 2331C>T, 2339A>G, 2345A>G, 2357G>T, 2363T>C, 2366C>G, 2367T>G, 2368C>A, 2369C>G, 2375C>T, 2376A>C, 2381T>C, 2384A>C, 2387T>C, 2393G>A, 2397G>A, 2398C>A, 2399A>C, 2409A>T, 2410G>C, 2411C>T, 2412A>G, 2413G>C, 2414T>C, 2415T>A, 2416C>A, 2417T>C, 2420T>G, 2424G>C, 2425C>A, 2426T>A, 2430G>C, 2432G>A, 2436T>G, 2438A>T, 2444G>C, 2445C>T, 2447C>A, 2453G>A, 2456T>A, 2459C>G, 2462T>C, 2463G>A, 2469T>A, 2470C>A, 2471C>G, 2474G>A, 2477G>A, 2479T>C, 2480G>T, 2483A>C, 2489A>C, 2492G>A, 2495A>G, 2501A>G, 2503G>A, 2507T>C, 2516C>T, 2521T>C, 2522A>C, 2523A>C, 2525T>G, 2528G>T, 2530A>T, 2537C>A, 2540C>T, 2546T>C, 2549T>A, 2552C>T, 2555C>T, 2561T>G, 2565A>G, 2567T>A, 2570G>A, 2573T>A, 2582C>A, 2583C>T, 2588T>A, 2597G>A, 2605T>C, 2609C>A, 2613A>C, 2615A>T, 2617A>C, 2618G>C, 2627C>T, 2628A>G, 2630C>G, 2633C>T, 2639C>A, 2640T>A, 2643C>G, 2645G>C, 2660T>C, 2661A>C, 2662G>A, 2663G>T, 2672A>T, 2678T>A, 2681A>G, 2685A>T, 2686G>C, 2687T>A, 2693G>T, 2696A>T, 2700T>G, 2701G>T, 2702T>A, 2708A>T, 2711T>G, 2712A>C, 2713C>A, 2714G>A, 2717T>C, 2720C>A, 2725A>C, 2726A>C, 2729C>T, 2730G>A, 2735A>T, 2736C>T, 2738T>G, 2747C>T, 2748T>G, 2750G>T, 2753G>C, 2755C>T, 2756T>G, 2792C>T, 2795C>T, 2798G>T, 2801T>C, 2819G>A, 2820A>T, 2828T>C, 2835A>G, 2837C>T, 2841C>T, 2843G>A, 2846G>T, 2852A>G, 2861G>C, 2864C>T, 2870C>A, 2879T>A, 2882T>G, 2885T>G, 2886T>C, 2894A>T, 2900A>T, 2909T>A, 2915A>G, 2921C>G, 2936A>T, 2939A>G, 2957A>T, 2958C>T, 2966C>T, 2972T>C, 2978A>G, 2981G>A, 2982G>A, 2987T>A, 2989G>C, 2996T>C, 2999T>A, 3003T>C, 3011C>T, 3012A>G, 3014C>A, 3020T>C, 3023G>T, 3032A>T, 3044C>T, 3047T>C, 3050G>A, 3062C>A, 3063T>C, 3077A>T, 3080A>T, 3083C>A, 3089C>T, 3101T>A, 3107T>A, 3113G>T, 3122T>A, 3126A>G, 3128T>C, 3131G>A, 3137T>C, 3143C>T, 3146T>C, 3152T>A, 3156T>C, 3158G>T, 3167A>T, 3176A>T, 3179G>A, 3182C>A, 3187G>A, 3197C>A, 3206C>T, 3210T>C, 3221G>C, 3293A>C, 3299T>G, 3303A>T, 3305C>T, 3308T>A, 3311T>G, 3323T>A, 3324C>T, 3329G>A, 3335C>A, 3350C>T, 3354A>C, 3357T>C, 3359G>A, 3368A>G, 3371C>G, 3372T>A, 3386A>T, 3398A>T, 3404A>G, 3410G>T, 3413C>T, 3419A>C, 3422A>G, 3428G>A, 3437C>A, 3443A>G, 3449G>A, 3452C>G, 3455T>A, 3458G>C, 3461C>A, 3473T>G, 3479C>T, 3482T>C, 3485C>T, 3488A>T, 3491C>T, 3503A>G, 3508A>G, 3512T>C, 3515G>T, 3521C>T, 3522C>T, 3524A>G, 3527C>T, 3533G>A, 3543T>A, 3551C>T, 3563G>A, 3567T>C, 3569A>G, 3572C>A, 3575G>A, 3581C>T, 3590A>C, 3617A>C, 3623A>T, 3624C>G, 3625A>C, 3627G>C, 3629G>T, 3630C>G, 3631C>G, 3635C>T, 3638A>G, 3641C>A, 3650A>T, 3656G>A, 3662A>T, 3674G>T, 3689C>T, 3698A>C, 3702T>G, 3708T>C, 3716C>T, 3722C>G, 3728C>T, 3983T>A, 3984A>G, 3986G>T, 3988G>C, 3990G>A, 3991A>C, 3995C>T, 3998C>T, 4016T>A, 4025G>A, 4031T>C, 4035A>C, 4037A>G, 4040C>G, 4046C>T, 4049A>G, 4052C>A, 4059A>G, 4061A>T, 4067A>T, 4069A>G, 4070T>G, 4072C>G, 4076A>T, 4077A>C, 4079A>T, 4082A>T, 4094A>G, 4103A>G, 4106A>T, 4139G>A, 4142A>T, 4148T>C, 4149C>A, 4151G>T, 4158T>A, 4163C>T, 4172A>G, 4181C>T, 4187G>A, 4188C>G, 4196T>C, 4208A>G, 4211A>T, 4223G>A, 4224C>T, 4227T>C, 4229G>T, 4250C>T, 4256T>A, 4262T>C, 4271T>C, 4280A>G, 4284A>G, 4286T>C, 4289T>C, 4300T>C, 4354G>C, 4355T>A, 4381A>G, 4382A>T, 4383C>G, 4388A>T, 4393A>G, 4424C>T, 4429T>C, 4457A>G, 4468T>C, 4496T>A, 4506A>T, 4525T>C, 4605T>C, 4610delT, 4633T>A, 4634G>A, 4643G>A, 4648G>A, 4650T>C, 4651G>A, 4659A>G, 4675A>T, 4812C>T, 4813C>T, 4860T>C, 4915G>A, 5020C>T, 5079C>T, 5089T>C, 5109A>G, 5123C>T, 5206C>T, 5213A>G, 5214C>T, 5222C>T, 5224G>A, 5225C>A, 5226T>G, 5231A>T, 5233G>T, 5235T>A, 5236G>A, 5239_5240insA, 5242T>G, 5249A>G, 5258A>G, 5259T>A, 5262A>T, 5282T>C, 5313A>T, 5315T>C, 5325G>A, 5327A>C | | | | | | | | |

##### CDS

| | Begin | End | Coverage | Score | Concordance | Matches | Identities | I/D/M/F* | Stop Codons |
|---|---|---|---|---|---|---|---|---|---|
| ToTV_sRNA2gp1 | 52 | 188 | 58.0% | 561 | 81.1% | 109 (100%) | 86 (78.9%) | 0/0/0/0 | 1 |
| Protein mutations: | L57I (350C>A 352A>C), V59I (356G>A), N64T (372A>C), G84A (432G>C 433C>T), H93Q (460T>G), Q108H (505G>T), R109K (507G>A 508G>A), I110V (509A>G 511T>A), V112I (515G>A), R146P (618G>C 619C>T), L149V (626C>G 628C>A), K150N (631A>T), E151K (632G>A), L155A (644C>G 645T>C 646T>C), L158Q (654T>A), S162T (665T>A), L168I (683T>A 685G>C), S169Q (686T>C 687C>A 688C>G), N170A (689A>G 690A>C 691C>A), L172P (696T>C), D175E (706T>A), S179R (718T>G), E184K (731G>A 733A>G) | | | | | | | | |
| Codon mutations: | GCA53GCT (340A>T), CTA57ATC (350C>A 352A>C), TCT58TCC (355T>C), GTT59ATT (356G>A), AAC63AAT (370C>T), AAT64ACT (372A>C), AGC65AGT (376C>T), GTT69GTA (388T>A), TAT73TAC (400T>C), TTA74CTG (401T>C 403A>G), CTG75CTA (406G>A), ACT79ACG (418T>G), CCA81CCC (424A>C), ACT83ACA (430T>A), GGC84GCT (432G>C 433C>T), GTT85GTC (436T>C), AAG86AAA (439G>A), CTA87CTC (442A>C), TCT88TCA (445T>A), GGC89GGA (448C>A), TTA90TTG (451A>G), CAT93CAG (460T>G), AGA94CGA (461A>C), CGT97CGA (472T>A), ACC101ACG (484C>G), AGA102AGG (487A>G), CTA105CTT (496A>T), GCT107GCA (502T>A), CAG108CAT (505G>T), AGG109AAA (507G>A 508G>A), ATT110GTA (509A>G 511T>A), TCT111TCC (514T>C), GTA112ATA (515G>A), TCC113TCA (520C>A), GAC141..A (604C>A), GAG142GAA (607G>A), GCC143GCA (610C>A), AAA144AAG (613A>G), CGC146CCT (618G>C 619C>T), TTA147TTG (622A>G), CTC149GTA (626C>G 628C>A), AAA150AAT (631A>T), GAG151AAG (632G>A), AAA152AAG (637A>G), CTT155GCC (644C>G 645T>C 646T>C), GAA156GAG (649A>G), CTA158CAA (654T>A), TCC159TCT (658C>T), ATA161ATC (664A>C), TCC162ACC (665T>A), AAA163AAG (670A>G), TTA164CTT (671T>C 673A>T), TTG166CTA (677T>C 679G>A), TTG168ATC (683T>A 685G>C), TCC169CAG (686T>C 687C>A 688C>G), AAC170GCA (689A>G 690A>C 691C>A), CTA172CCA (696T>C), AAT174AAC (703T>C), GAT175GAA (706T>A), AGT179AGG (718T>G), GAA184AAG (731G>A 733A>G) | | | | | | | | |
| ToTV_sRNA2gp2 | 1 | 1199 | 69.3% | 5084 | 88.7% | 831 (100%) | 724 (87.1%) | 0/0/0/0 | 3 |
| Protein mutations: | M1T (703T>C), M2K (706T>A), V6G (718T>G), K11R (733A>G), S17A (750A>G 751G>C), Y48F (844A>T), E61D (884A>T), N70K (911T>G), L128M (1083C>A), V132M (1095G>A 1097T>G), Q211H (1334G>T), T228A (1383A>G 1385C>G), S230* (1390C>A), T232P (1395A>C 1397T>A), D233E (1400T>G), Q242L (1426A>T 1427G>T), D245N (1434G>A), S252Q (1455A>C 1456G>A 1457C>G), A254S (1461G>T), N481E (2142A>G 2144C>G), S487R (2162T>A), V488G (2164T>G), E489G (2167A>G), E490S (2169G>A 2170A>G 2171A>C), I491R (2173T>G 2174A>G), P492S (2175C>T 2177A>C), S493T (2178T>A), R524S (2271A>T 2272G>C 2273G>A), N526D (2277A>G 2279T>C), E527D (2282A>T), M529F (2286A>T 2288G>C), E531D (2294A>C), Q534E (2301C>G 2303A>G), N537M (2311A>T 2312T>G), N538E (2313A>G 2315T>A), E539D (2318A>T), G540A (2320G>C 2321T>C), A541P (2322G>C 2324G>C), T542I (2326C>T 2327A>T), E543Q (2328G>C 2330G>A), N555K (2366C>G), S556E (2367T>G 2368C>A 2369C>G), N559H (2376A>C), A566N (2397G>A 2398C>A 2399A>C), S571A (2412A>G 2413G>C 2414T>C), S572N (2415T>A 2416C>A 2417T>C), A575Q (2424G>C 2425C>A 2426T>A), V577L (2430G>C 2432G>A), S579A (2436T>G 2438A>T), G588S (2463G>A), S590K (2469T>A 2470C>A 2471C>G), M593T (2479T>C 2480G>T), R601K (2503G>A), L607S (2521T>C 2522A>C), I608L (2523A>C 2525T>G), M609I (2528G>T), Y610F (2530A>T), I622V (2565A>G 2567T>A), V635A (2605T>C), K639T (2617A>C 2618G>C), I643V (2628A>G 2630C>G), S647T (2640T>A), L648V (2643C>G 2645G>C), R654H (2661A>C 2662G>A 2663G>T), E665D (2696A>T), C667V (2700T>G 2701G>T 2702T>A), T671Q (2712A>C 2713C>A 2714G>A), K675T (2725A>C 2726A>C), V677I (2730G>A), S683A (2748T>G 2750G>T), A685V (2755C>T 2756T>G), L699F (2798G>T), S707C (2820A>T), I712V (2835A>G 2837C>T), Q715H (2846G>T), D740E (2921C>G), A761T (2982G>A), S763T (2989G>C), I771V (3012A>G 3014C>A), I809V (3126A>G 3128T>C), N827K (3182C>A), S829N (3187G>A), K864N (3293A>C), T868S (3303A>T 3305C>T), L891M (3372T>A), C912* (3437C>A), N936S (3508A>G), S948T (3543T>A), Q975A (3624C>G 3625A>C), V976L (3627G>C 3629G>T), P977G (3630C>G 3631C>G), D980E (3641C>A), L1001V (3702T>G), M1095V (3984A>G 3986G>T), S1096T (3988G>C), D1097T (3990G>A 3991A>C), H1105Q (4016T>A), I1120V (4059A>G 4061A>T), N1123R (4069A>G 4070T>G), A1124G (4072C>G), L1150I (4149C>A 4151G>T), L1153M (4158T>A), Q1163E (4188C>G), H1175Y (4224C>T), S1195G (4284A>G 4286T>C) | | | | | | | | |

| | Begin | End | Coverage | Score | Concordance | Matches | Identities | I/D/M/F* | Stop Codons |
|---|---|---|---|---|---|---|---|---|---|
| **NT** | **30** | **5329** | **72.7%** | **5298** | **67.5%** | **3927 (99.9%)** | **3293 (83.8%)** | **1/1** | |
| Codon mutations: | ATG1ACG (703T>C), ATG2AAG (706T>A), GTG6GGG (718T>G), AGG10AGA (731G>A), AAG11AGG (733A>G), AGT17GCT (750A>G 751G>C), GCA18GCC (755A>C), GCT19GCA (758T>A), GTA20GTC (761A>C), ACA29ACT (788A>T), GTT32GTG (797T>G), GGC35GGG (806C>G), CTG39CTC (818G>C), CTT40CTG (821T>G), AAA42AAG (827A>G), CTG43TTG (828C>T), GGC44GGA (833C>A), TAT48TTT (844A>T), GGG49GGC (848G>C), CTC51CTG (854C>G), GGC53GGA (860C>A), GCC54GCT (863C>T), CAA55CAG (866A>G), GAG56GAA (869G>A), GAA61GAT (884A>T), CTG62CTT (887G>T), AAT70AAG (911T>G), TCT76TCA (929T>A), TCA77TCG (932A>G), TTT78TTC (935T>C), TCG85TCC (956G>C), TTG95TTA (986G>A), GTT98GTC (995T>C), GTT103GTG (1010T>G), GAA110GAG (1031A>G), GGT111GGG (1034T>G), GTA113GTC (1040A>C), TTA117TTG (1052A>G), AGT120AGC (1061T>C), ACC124ACT (1073C>T), CTG128ATG (1083C>A), GTT132ATG (1095G>A 1097T>G), TTG133TTA (1100G>A), TCC136TCT (1109C>T), GGG138GGA (1115G>A), GCA139GCT (1118A>T), TTA143TTG (1130A>G), CCG146CCT (1139G>T), ACC147ACA (1142C>A), CTA150TTG (1149C>T 1151A>G), AAA154AAG (1163A>G), GGC156GGG (1169C>G), TTC159TTT (1178C>T), ACA162ACT (1187A>T), ACT168ACA (1205T>A), CCT173CCA (1220T>A), TCA175TCT (1226A>T), GCC178GCA (1235C>A), CAA182CAG (1247A>G), CGT187CGA (1262T>A), TCA192TCG (1277A>G), CTT204CTC (1313T>C), CAC205CAT (1316C>T), CGA206AGG (1317C>A 1319A>G), CCT207CCA (1322T>A), CAA208CAG (1325A>G), TTT209TTC (1328T>C), TTC210TTT (1331C>T), CAG211CAT (1334G>T), CAG214CAA (1343G>A), CAG218CAA (1355G>A), ATA220ATT (1361A>T), TCA221TCT (1364A>T), TCA222TCT (1367A>T), GTA223GTC (1370A>C), TCT225TCG (1376T>G), ACC228GCG (1383A>G 1385C>G), TCC229TCT (1388C>T), TCA230TAA (1390C>A), GCA231GCC (1394A>C), ACT232CCA (1395A>C 1397T>A), GAT233GAG (1400T>G), TCA234TCT (1403A>T), GTG235GTA (1406G>A), CTT236TTA (1407C>T 1409T>A), CCG237CCT (1412G>T), GTG239GTA (1418G>A), GGA240GGT (1421A>T), CAG242CTT (1426A>T 1427G>T), GGC244GGG (1433C>G), GAC245AAC (1434G>A), ACA246ACT (1439A>T), GTG250GTT (1451G>T), CCA251CCC (1454A>C), AGC252CAG (1455A>C 1456G>A 1457C>G), GAA253GAG (1460A>G), GCA254TCA (1461G>T), AAA480.GA (2140A>G), AAC481GAG (2142A>G 2144C>G), ACT486ACA (2159T>A), AGT487AGA (2162T>A), GTG488GGG (2164T>G), GAA489GGA (2167A>G), GAA490AGC (2169G>A 2170A>G 2171A>C), ATA491AGG (2173T>G 2174A>G), CCA492TCC (2175C>T 2177A>C), TCC493ACC (2178T>A), ACC494ACT (2183C>T), AGG524TCA (2271A>T 2272G>C 2273G>A), CCC525CCT (2276C>T), AAT526GAC (2277A>G 2279T>C), GAA527GAT (2282A>T), GTT528GTC (2285T>C), ATG529TTC (2286A>T 2288G>C), GAA531GAC (2294A>C), CAA534GAG (2301C>G 2303A>G), AAT537ATG (2311A>T 2312T>G), AAT538GAA (2313A>G 2315T>A), GAA539GAT (2318A>T), GGT540GCC (2320G>C 2321T>C), GCG541CCC (2322G>C 2324G>C), ACA542ATT (2326C>T 2327A>T), GAG543CAA (2328G>C 2330G>A), CTG544TTG (2331C>T), TTA546TTG (2339A>G), CAA548CAG (2345A>G), GTG552GTT (2357G>T), AGT554AGC (2363T>C), AAC555AAG (2366C>G), TCC556GAG (2367T>G 2368C>A 2369C>G), TTC558TTT (2375C>T), AAT559CAT (2376A>C), GTT560GTC (2381T>C), GGA561GGC (2384A>C), GTT562GTC (2387T>C), GAG564GAA (2393G>A), GCA566AAC (2397G>A 2398C>A 2399A>C), AGC570TCT (2409A>T 2410G>C 2411C>T), AGT571GCC (2412A>G 2413G>C 2414T>C), TCT572AAC (2415T>A 2416C>A 2417T>C), GTT573GTG (2420T>G), GCT575CAA (2424G>C 2425C>A 2426T>A), GTG577CTA (2430G>C 2432G>A), TCA579GCT (2436T>G 2438A>T), GCG581GCC (2444G>C), CTC582TTA (2445C>T 2447C>A), GCG584GCA (2453G>A), GCT585GCA (2456T>A), CTC586CTG (2459C>G), TTT587TTC (2462T>C), GGT588AGT (2463G>A), TCC590AAG (2469T>A 2470C>A 2471C>G), AAG591AAA (2474G>A), GAG592GAA (2477G>A), ATG593ACT (2479T>C 2480G>T), TCA594TCC (2483A>C), GGA596GGC (2489A>C), TCG597TCA (2492G>A), CAA598CAG (2495A>G), CTA600CTG (2501A>G), AGG601AAG (2503G>A), TAT602TAC (2507T>C), GCC605GCT (2516C>T), TTA607TCC (2521T>C 2522A>C), ATT608CTG (2523A>C 2525T>G), ATG609ATT (2528G>T), TAC610TTC (2530A>T), GTC612GTA (2537C>A), ATC613ATT (2540C>T), TAT615TAC (2546T>C), ATT616ATA (2549T>A), TCC617TCT (2552C>T), GGC618GGT (2555C>T), GGT620GGG (2561T>G), ATT622GTA (2565A>G 2567T>A), TCG623TCA (2570G>A), GGT624GGA (2573T>A), GCC627GCA (2582C>A), CTG628TTG (2583C>T), GTT629GTA (2588T>A), GAG632GAA (2597G>A), GTG635GCG (2605T>C), CTC636CTA (2609C>A), AGA638CGT (2613A>C 2615A>T), AAG639ACC (2617A>C 2618G>C), TTC642TTT (2627C>T), ATC643GTG (2628A>G 2630C>G), AAC644AAT (2633C>T), GCC646GCA (2639C>A), TCT647ACT (2640T>A), CTG648GTC (2643C>G 2645G>C), CAT653CAC (2660T>C), AGG654CAT (2661A>C 2662G>A 2663G>T), TCA657TCT (2672A>T), TCT659TCA (2678T>A), GAA660GAG (2681A>G), AGT662TCA (2685A>T 2686G>C 2687T>A), GGG664GGT (2693G>T), GAA665GAT (2696A>T), TGT667GTA (2700T>G 2701G>T 2702T>A), ACA669ACT (2708A>T), CCT670CCG (2711T>G), ACG671CAA (2712A>C 2713C>A 2714G>A), GGT672GGC (2717T>C), ATC673ATA (2720C>A), AAA675ACC (2725A>C 2726A>C), TTC676TTT (2729C>T), GTT677ATT (2730G>A), CCA678CCT (2735A>T), CTT679TTG (2736C>T 2738T>G), GCC682GCT (2747C>T), TCG683GCT (2748T>G 2750G>T), GGG684GGC (2753G>C), GCT685GTG (2755C>T 2756T>G), CAC697CAT (2792C>T), CCC698CCT (2795C>T), TTG699TTT (2798G>T), GCT700GCC (2801T>C), GAG706GAA (2819G>A), AGC707TGC (2820A>T), CCT709CCC (2828T>C), ATC712GTT (2835A>G 2837C>T), CTG714TTA (2841C>T 2843G>A), CAG715CAT (2846G>T), AAA717AAG (2852A>G), TCG720TCC (2861G>C), ACC721ACT (2864C>T), ATC723ATA (2870C>A), CCT726CCA (2879T>A), CCT727CCG (2882T>G), CGT728CGG (2885T>G), TTG729CTG (2886T>C), GCA731GCT (2894A>T), GCA733GCT (2900A>T), GGT736GGA (2909T>A), AAA738AAG (2915A>G), GAC740GAG (2921C>G), CCA745CCT (2936A>T), CGA746CGG (2939A>G), GTA752GTT (2957A>T), CTG753TTG (2958C>T), CAC755CAT (2966C>T), AAT757AAC (2972T>C), GGA759GGG (2978A>G), GTG760GTA (2981G>A), GCT761ACT (2982G>A), GCT762GCA (2987T>A), AGC763ACC (2989G>C), GGT765GGC (2996T>C), ACT766ACA (2999T>A), TTA768CTA (3003T>C), AGC770AGT (3011C>T), ATC771GTA (3012A>G 3014C>A), TCT773TCC (3020T>C), CCG774CCT (3023G>T), ATA777ATT (3032A>T), GAC781GAT (3044C>T), GGT782GGC (3047T>C), ACG783ACA (3050G>A), TCC787TCA (3062C>A), TTG788CTG (3063T>C), ATA792ATT (3077A>T), GCA793GCT (3080A>T), CGC794CGA (3083C>A), TGC796TGT (3089C>T), ACT800ACA (3101T>A), ACT802ACA (3107T>A), GTG804GTT (3113G>T), ATT807ATA (3122T>A), ATT809GTC (3126A>G 3128T>C), GAG810GAA (3131G>A), ACT812ACC (3137T>C), TTC814TTT (3143C>T), CAT815CAC (3146T>C), GGT817GGA (3152T>A), TTG819CTT (3156T>C 3158G>T), GGA822GGT (3167A>T), ACA825ACT (3176A>T), CTG826CTA (3179G>A), AAC827AAA (3182C>A), AGC829AAC (3187G>A), ACC832ACA (3197C>A), GAC835GAT (3206C>T), TTA837CTA (3210T>C), CCG840CCC (3221G>C), AAA864AAC (3293A>C), CTT866CTG (3299T>G), ACC868TCT (3303A>T 3305C>T), ACT869ACA (3308T>A), GGT870GGG (3311T>G), TCT874TCA (3323T>A), CTA875TTA (3324C>T), CCG876CCA (3329G>A), CCC878CCA (3335C>A), CAC883CAT (3350C>T), AGG885CGG (3354A>C), TTG886CTA (3357T>C 3359G>A), ACA889ACG (3368A>G), GTC890GTG (3371C>G), TTG891ATG (3372T>A), GTA895GTT (3386A>T), ATA899ATT (3398A>T), CTA901CTG (3404A>G), ACG903ACT (3410G>T), GTC904GTT (3413C>T), GTA906GTC (3419A>C), ACA907ACG (3422A>G), CAG909CAA (3428G>A), TGC912TGA (3437C>A), GAA914GAG (3443A>G), TTG916TTA (3449G>A), GTC917GTG (3452C>G), CTT918CTA (3455T>A), GGG919GGC (3458G>C), GGC920GGA (3461C>A), GTT924GTG (3473T>G), CCC926CCT (3479C>T), ATT927ATC (3482T>C), TAC928TAT (3485C>T), GGA929GGT (3488A>T), CAC930CAT (3491C>T), GGA934GGG (3503A>G), AAT936AGT (3508A>G), GCT937GCC (3512T>C), GTG938GTT (3515G>T), TTC940TTT (3521C>T), CTA941TTG (3522C>T 3524A>G), TTC942TTT (3527C>T), GAG944GAA (3533G>A), TCT948ACT (3543T>A), CGC950CGT (3551C>T), GAG954GAA (3563G>A), TTA956CTG (3567T>C 3569A>G), CGC957CGA (3572C>A), AAG958AAA (3575G>A), AAC960AAT (3581C>T), ACA963ACC (3590A>C), CCA972CCC (3617A>C), GTA974GTT (3623A>T), CAA975GCA (3624C>G 3625A>C), GTG976CTT (3627G>C 3629G>T), CCG977GGG (3630C>G 3631C>G), CTC978CTT (3635C>T), AGA979AGG (3638A>G), GAC980GAA (3641C>A), TCA983TCT (3650A>T), GGG985GGA (3656G>A), GTA987GTT (3662A>T), GTG991GTT (3674G>T), GAC996GAT (3689C>T), CGA999CGC (3698A>C), TTG1001GTG (3702T>G), TTA1003CTA (3708T>C), TGC1005TGT (3716C>T), CCC1007CCG (3722C>G), TCC1009TCT (3728C>T), GGT1094GGA (3983T>A), ATG1095GTT (3984A>G 3986G>T), AGC1096ACC (3988G>C), GAC1097ACC (3990G>A 3991A>C), AAC1098AAT (3995C>T), ATC1099ATT (3998C>T), CAT1105CAA (4016T>A), GAG1108GAA (4025G>A), TTT1110TTC (4031T>C), AGA1112CGG (4035A>C 4037A>G), CGC1113CGG (4040C>G), ACC1115ACT (4046C>T), AAA1116AAG (4049A>G), GCC1117GCA (4052C>A), ATA1120GTT (4059A>G 4061A>T), CCA1122CCT (4067A>T), AAT1123AGG (4069A>G 4070T>G), GCT1124GGT (4072C>G), TCA1125TCT (4076A>T), AGA1126CGT (4077A>C 4079A>T), ATA1127ATT (4082A>T), CAA1131CAG (4094A>G), CTA1134CTG (4103A>G), GGA1135GGT (4106A>T), GTG1146.TA (4139G>A), ATA1147ATT (4142A>T), TCT1149TCC (4148T>C), CTG1150ATT (4149C>A 4151G>T), TTG1153ATG (4158T>A), GTC1154GTT (4163C>T), GGA1157GGG (4172A>G), TTC1160TTT (4181C>T), TTG1162TTA (4187G>A), CAA1163GAA (4188C>G), GCT1165GCC (4196T>C), TCA1169TCG (4208A>G), GCA1170GCT (4211A>T), AAG1174AAA (4223G>A), CAT1175TAT (4224C>T), TTG1176CTT (4227T>C 4229G>T), ACC1183ACT (4250C>T), CTT1185CTA (4256T>A), ACT1187ACC (4262T>C), TTT1190TTC (4271T>C), CTA1193CTG (4280A>G), AGT1195GGC (4284A>G 4286T>C), TTT1196TTC (4289T>C) | | | | | | | | |

#### Proteins

| | Begin | End | Coverage | Score | Concordance | Matches | Identities | I/D/M/F* | Stop Codons |
|---|---|---|---|---|---|---|---|---|---|
| Orf1 (YP_001040017.1) | 52 | 188 | 58.0% | 561 | 81.1% | 109 (100%) | 86 (78.9%) | 0/0/0/0 | 1 |
| Protein mutations: | L57I (350C>A 352A>C), V59I (356G>A), N64T (372A>C), G84A (432G>C 433C>T), H93Q (460T>G), Q108H (505G>T), R109K (507G>A 508G>A), I110V (509A>G 511T>A), V112I (515G>A), R146P (618G>C 619C>T), L149V (626C>G 628C>A), K150N (631A>T), E151K (632G>A), L155A (644C>G 645T>C 646T>C), L158Q (654T>A), S162T (665T>A), L168I (683T>A 685G>C), S169Q (686T>C 687C>A 688C>G), N170A (689A>G 690A>C 691C>A), L172P (696T>C), D175E (706T>A), S179R (718T>G), E184K (731G>A 733A>G) | | | | | | | | |
| Codon mutations: | GCA53GCT (340A>T), CTA57ATC (350C>A 352A>C), TCT58TCC (355T>C), GTT59ATT (356G>A), AAC63AAT (370C>T), AAT64ACT (372A>C), AGC65AGT (376C>T), GTT69GTA (388T>A), TAT73TAC (400T>C), TTA74CTG (401T>C 403A>G), CTG75CTA (406G>A), ACT79ACG (418T>G), CCA81CCC (424A>C), ACT83ACA (430T>A), GGC84GCT (432G>C 433C>T), GTT85GTC (436T>C), AAG86AAA (439G>A), CTA87CTC (442A>C), TCT88TCA (445T>A), GGC89GGA (448C>A), TTA90TTG (451A>G), CAT93CAG (460T>G), AGA94CGA (461A>C), CGT97CGA (472T>A), ACC101ACG (484C>G), AGA102AGG (487A>G), CTA105CTT (496A>T), GCT107GCA (502T>A), CAG108CAT (505G>T), AGG109AAA (507G>A 508G>A), ATT110GTA (509A>G 511T>A), TCT111TCC (514T>C), GTA112ATA (515G>A), TCC113TCA (520C>A), GAC141..A (604C>A), GAG142GAA (607G>A), GCC143GCA (610C>A), AAA144AAG (613A>G), CGC146CCT (618G>C 619C>T), TTA147TTG (622A>G), CTC149GTA (626C>G 628C>A), AAA150AAT (631A>T), GAG151AAG (632G>A), AAA152AAG (637A>G), CTT155GCC (644C>G 645T>C 646T>C), GAA156GAG (649A>G), CTA158CAA (654T>A), TCC159TCT (658C>T), ATA161ATC (664A>C), TCC162ACC (665T>A), AAA163AAG (670A>G), TTA164CTT (671T>C 673A>T), TTG166CTA (677T>C 679G>A), TTG168ATC (683T>A 685G>C), TCC169CAG (686T>C 687C>A 688C>G), AAC170GCA (689A>G 690A>C 691C>A), CTA172CCA (696T>C), AAT174AAC (703T>C), GAT175GAA (706T>A), AGT179AGG (718T>G), GAA184AAG (731G>A 733A>G) | | | | | | | | |
| polyprotein (YP_001040018.1) | 1 | 1199 | 69.3% | 5084 | 88.7% | 831 (100%) | 724 (87.1%) | 0/0/0/0 | 3 |
| Protein mutations: | M1T (703T>C), M2K (706T>A), V6G (718T>G), K11R (733A>G), S17A (750A>G 751G>C), Y48F (844A>T), E61D (884A>T), N70K (911T>G), L128M (1083C>A), V132M (1095G>A 1097T>G), Q211H (1334G>T), T228A (1383A>G 1385C>G), S230* (1390C>A), T232P (1395A>C 1397T>A), D233E (1400T>G), Q242L (1426A>T 1427G>T), D245N (1434G>A), S252Q (1455A>C 1456G>A 1457C>G), A254S (1461G>T), N481E (2142A>G 2144C>G), S487R (2162T>A), V488G (2164T>G), E489G (2167A>G), E490S (2169G>A 2170A>G 2171A>C), I491R (2173T>G 2174A>G), P492S (2175C>T 2177A>C), S493T (2178T>A), R524S (2271A>T 2272G>C 2273G>A), N526D (2277A>G 2279T>C), E527D (2282A>T), M529F (2286A>T 2288G>C), E531D (2294A>C), Q534E (2301C>G 2303A>G), N537M (2311A>T 2312T>G), N538E (2313A>G 2315T>A), E539D (2318A>T), G540A (2320G>C 2321T>C), A541P (2322G>C 2324G>C), T542I (2326C>T 2327A>T), E543Q (2328G>C 2330G>A), N555K (2366C>G), S556E (2367T>G 2368C>A 2369C>G), N559H (2376A>C), A566N (2397G>A 2398C>A 2399A>C), S571A (2412A>G 2413G>C 2414T>C), S572N (2415T>A 2416C>A 2417T>C), A575Q (2424G>C 2425C>A 2426T>A), V577L (2430G>C 2432G>A), S579A (2436T>G 2438A>T), G588S (2463G>A), S590K (2469T>A 2470C>A 2471C>G), M593T (2479T>C 2480G>T), R601K (2503G>A), L607S (2521T>C 2522A>C), I608L (2523A>C 2525T>G), M609I (2528G>T), Y610F (2530A>T), I622V (2565A>G 2567T>A), V635A (2605T>C), K639T (2617A>C 2618G>C), I643V (2628A>G 2630C>G), S647T (2640T>A), L648V (2643C>G 2645G>C), R654H (2661A>C 2662G>A 2663G>T), E665D (2696A>T), C667V (2700T>G 2701G>T 2702T>A), T671Q (2712A>C 2713C>A 2714G>A), K675T (2725A>C 2726A>C), V677I (2730G>A), S683A (2748T>G 2750G>T), A685V (2755C>T 2756T>G), L699F (2798G>T), S707C (2820A>T), I712V (2835A>G 2837C>T), Q715H (2846G>T), D740E (2921C>G), A761T (2982G>A), S763T (2989G>C), I771V (3012A>G 3014C>A), I809V (3126A>G 3128T>C), N827K (3182C>A), S829N (3187G>A), K864N (3293A>C), T868S (3303A>T 3305C>T), L891M (3372T>A), C912* (3437C>A), N936S (3508A>G), S948T (3543T>A), Q975A (3624C>G 3625A>C), V976L (3627G>C 3629G>T), P977G (3630C>G 3631C>G), D980E (3641C>A), L1001V (3702T>G), M1095V (3984A>G 3986G>T), S1096T (3988G>C), D1097T (3990G>A 3991A>C), H1105Q (4016T>A), I1120V (4059A>G 4061A>T), N1123R (4069A>G 4070T>G), A1124G (4072C>G), L1150I (4149C>A 4151G>T), L1153M (4158T>A), Q1163E (4188C>G), H1175Y (4224C>T), S1195G (4284A>G 4286T>C) | | | | | | | | |

| | Begin | End | Coverage | Score | Concordance | Matches | Identities | I/D/M/F* | Stop Codons |
|---|---|---|---|---|---|---|---|---|---|
| **NT** | **30** | **5329** | **72.7%** | **5298** | **67.5%** | **3927 (99.9%)** | **3293 (83.8%)** | **1/1** | |
| Codon mutations: | ATG1ACG (703T>C), ATG2AAG (706T>A), GTG6GGG (718T>G), AGG10AGA (731G>A), AAG11AGG (733A>G), AGT17GCT (750A>G 751G>C), GCA18GCC (755A>C), GCT19GCA (758T>A), GTA20GTC (761A>C), ACA29ACT (788A>T), GTT32GTG (797T>G), GGC35GGG (806C>G), CTG39CTC (818G>C), CTT40CTG (821T>G), AAA42AAG (827A>G), CTG43TTG (828C>T), GGC44GGA (833C>A), TAT48TTT (844A>T), GGG49GGC (848G>C), CTC51CTG (854C>G), GGC53GGA (860C>A), GCC54GCT (863C>T), CAA55CAG (866A>G), GAG56GAA (869G>A), GAA61GAT (884A>T), CTG62CTT (887G>T), AAT70AAG (911T>G), TCT76TCA (929T>A), TCA77TCG (932A>G), TTT78TTC (935T>C), TCG85TCC (956G>C), TTG95TTA (986G>A), GTT98GTC (995T>C), GTT103GTG (1010T>G), GAA110GAG (1031A>G), GGT111GGG (1034T>G), GTA113GTC (1040A>C), TTA117TTG (1052A>G), AGT120AGC (1061T>C), ACC124ACT (1073C>T), CTG128ATG (1083C>A), GTT132ATG (1095G>A 1097T>G), TTG133TTA (1100G>A), TCC136TCT (1109C>T), GGG138GGA (1115G>A), GCA139GCT (1118A>T), TTA143TTG (1130A>G), CCG146CCT (1139G>T), ACC147ACA (1142C>A), CTA150TTG (1149C>T 1151A>G), AAA154AAG (1163A>G), GGC156GGG (1169C>G), TTC159TTT (1178C>T), ACA162ACT (1187A>T), ACT168ACA (1205T>A), CCT173CCA (1220T>A), TCA175TCT (1226A>T), GCC178GCA (1235C>A), CAA182CAG (1247A>G), CGT187CGA (1262T>A), TCA192TCG (1277A>G), CTT204CTC (1313T>C), CAC205CAT (1316C>T), CGA206AGG (1317C>A 1319A>G), CCT207CCA (1322T>A), CAA208CAG (1325A>G), TTT209TTC (1328T>C), TTC210TTT (1331C>T), CAG211CAT (1334G>T), CAG214CAA (1343G>A), CAG218CAA (1355G>A), ATA220ATT (1361A>T), TCA221TCT (1364A>T), TCA222TCT (1367A>T), GTA223GTC (1370A>C), TCT225TCG (1376T>G), ACC228GCG (1383A>G 1385C>G), TCC229TCT (1388C>T), TCA230TAA (1390C>A), GCA231GCC (1394A>C), ACT232CCA (1395A>C 1397T>A), GAT233GAG (1400T>G), TCA234TCT (1403A>T), GTG235GTA (1406G>A), CTT236TTA (1407C>T 1409T>A), CCG237CCT (1412G>T), GTG239GTA (1418G>A), GGA240GGT (1421A>T), CAG242CTT (1426A>T 1427G>T), GGC244GGG (1433C>G), GAC245AAC (1434G>A), ACA246ACT (1439A>T), GTG250GTT (1451G>T), CCA251CCC (1454A>C), AGC252CAG (1455A>C 1456G>A 1457C>G), GAA253GAG (1460A>G), GCA254TCA (1461G>T), AAA480.GA (2140A>G), AAC481GAG (2142A>G 2144C>G), ACT486ACA (2159T>A), AGT487AGA (2162T>A), GTG488GGG (2164T>G), GAA489GGA (2167A>G), GAA490AGC (2169G>A 2170A>G 2171A>C), ATA491AGG (2173T>G 2174A>G), CCA492TCC (2175C>T 2177A>C), TCC493ACC (2178T>A), ACC494ACT (2183C>T), AGG524TCA (2271A>T 2272G>C 2273G>A), CCC525CCT (2276C>T), AAT526GAC (2277A>G 2279T>C), GAA527GAT (2282A>T), GTT528GTC (2285T>C), ATG529TTC (2286A>T 2288G>C), GAA531GAC (2294A>C), CAA534GAG (2301C>G 2303A>G), AAT537ATG (2311A>T 2312T>G), AAT538GAA (2313A>G 2315T>A), GAA539GAT (2318A>T), GGT540GCC (2320G>C 2321T>C), GCG541CCC (2322G>C 2324G>C), ACA542ATT (2326C>T 2327A>T), GAG543CAA (2328G>C 2330G>A), CTG544TTG (2331C>T), TTA546TTG (2339A>G), CAA548CAG (2345A>G), GTG552GTT (2357G>T), AGT554AGC (2363T>C), AAC555AAG (2366C>G), TCC556GAG (2367T>G 2368C>A 2369C>G), TTC558TTT (2375C>T), AAT559CAT (2376A>C), GTT560GTC (2381T>C), GGA561GGC (2384A>C), GTT562GTC (2387T>C), GAG564GAA (2393G>A), GCA566AAC (2397G>A 2398C>A 2399A>C), AGC570TCT (2409A>T 2410G>C 2411C>T), AGT571GCC (2412A>G 2413G>C 2414T>C), TCT572AAC (2415T>A 2416C>A 2417T>C), GTT573GTG (2420T>G), GCT575CAA (2424G>C 2425C>A 2426T>A), GTG577CTA (2430G>C 2432G>A), TCA579GCT (2436T>G 2438A>T), GCG581GCC (2444G>C), CTC582TTA (2445C>T 2447C>A), GCG584GCA (2453G>A), GCT585GCA (2456T>A), CTC586CTG (2459C>G), TTT587TTC (2462T>C), GGT588AGT (2463G>A), TCC590AAG (2469T>A 2470C>A 2471C>G), AAG591AAA (2474G>A), GAG592GAA (2477G>A), ATG593ACT (2479T>C 2480G>T), TCA594TCC (2483A>C), GGA596GGC (2489A>C), TCG597TCA (2492G>A), CAA598CAG (2495A>G), CTA600CTG (2501A>G), AGG601AAG (2503G>A), TAT602TAC (2507T>C), GCC605GCT (2516C>T), TTA607TCC (2521T>C 2522A>C), ATT608CTG (2523A>C 2525T>G), ATG609ATT (2528G>T), TAC610TTC (2530A>T), GTC612GTA (2537C>A), ATC613ATT (2540C>T), TAT615TAC (2546T>C), ATT616ATA (2549T>A), TCC617TCT (2552C>T), GGC618GGT (2555C>T), GGT620GGG (2561T>G), ATT622GTA (2565A>G 2567T>A), TCG623TCA (2570G>A), GGT624GGA (2573T>A), GCC627GCA (2582C>A), CTG628TTG (2583C>T), GTT629GTA (2588T>A), GAG632GAA (2597G>A), GTG635GCG (2605T>C), CTC636CTA (2609C>A), AGA638CGT (2613A>C 2615A>T), AAG639ACC (2617A>C 2618G>C), TTC642TTT (2627C>T), ATC643GTG (2628A>G 2630C>G), AAC644AAT (2633C>T), GCC646GCA (2639C>A), TCT647ACT (2640T>A), CTG648GTC (2643C>G 2645G>C), CAT653CAC (2660T>C), AGG654CAT (2661A>C 2662G>A 2663G>T), TCA657TCT (2672A>T), TCT659TCA (2678T>A), GAA660GAG (2681A>G), AGT662TCA (2685A>T 2686G>C 2687T>A), GGG664GGT (2693G>T), GAA665GAT (2696A>T), TGT667GTA (2700T>G 2701G>T 2702T>A), ACA669ACT (2708A>T), CCT670CCG (2711T>G), ACG671CAA (2712A>C 2713C>A 2714G>A), GGT672GGC (2717T>C), ATC673ATA (2720C>A), AAA675ACC (2725A>C 2726A>C), TTC676TTT (2729C>T), GTT677ATT (2730G>A), CCA678CCT (2735A>T), CTT679TTG (2736C>T 2738T>G), GCC682GCT (2747C>T), TCG683GCT (2748T>G 2750G>T), GGG684GGC (2753G>C), GCT685GTG (2755C>T 2756T>G), CAC697CAT (2792C>T), CCC698CCT (2795C>T), TTG699TTT (2798G>T), GCT700GCC (2801T>C), GAG706GAA (2819G>A), AGC707TGC (2820A>T), CCT709CCC (2828T>C), ATC712GTT (2835A>G 2837C>T), CTG714TTA (2841C>T 2843G>A), CAG715CAT (2846G>T), AAA717AAG (2852A>G), TCG720TCC (2861G>C), ACC721ACT (2864C>T), ATC723ATA (2870C>A), CCT726CCA (2879T>A), CCT727CCG (2882T>G), CGT728CGG (2885T>G), TTG729CTG (2886T>C), GCA731GCT (2894A>T), GCA733GCT (2900A>T), GGT736GGA (2909T>A), AAA738AAG (2915A>G), GAC740GAG (2921C>G), CCA745CCT (2936A>T), CGA746CGG (2939A>G), GTA752GTT (2957A>T), CTG753TTG (2958C>T), CAC755CAT (2966C>T), AAT757AAC (2972T>C), GGA759GGG (2978A>G), GTG760GTA (2981G>A), GCT761ACT (2982G>A), GCT762GCA (2987T>A), AGC763ACC (2989G>C), GGT765GGC (2996T>C), ACT766ACA (2999T>A), TTA768CTA (3003T>C), AGC770AGT (3011C>T), ATC771GTA (3012A>G 3014C>A), TCT773TCC (3020T>C), CCG774CCT (3023G>T), ATA777ATT (3032A>T), GAC781GAT (3044C>T), GGT782GGC (3047T>C), ACG783ACA (3050G>A), TCC787TCA (3062C>A), TTG788CTG (3063T>C), ATA792ATT (3077A>T), GCA793GCT (3080A>T), CGC794CGA (3083C>A), TGC796TGT (3089C>T), ACT800ACA (3101T>A), ACT802ACA (3107T>A), GTG804GTT (3113G>T), ATT807ATA (3122T>A), ATT809GTC (3126A>G 3128T>C), GAG810GAA (3131G>A), ACT812ACC (3137T>C), TTC814TTT (3143C>T), CAT815CAC (3146T>C), GGT817GGA (3152T>A), TTG819CTT (3156T>C 3158G>T), GGA822GGT (3167A>T), ACA825ACT (3176A>T), CTG826CTA (3179G>A), AAC827AAA (3182C>A), AGC829AAC (3187G>A), ACC832ACA (3197C>A), GAC835GAT (3206C>T), TTA837CTA (3210T>C), CCG840CCC (3221G>C), AAA864AAC (3293A>C), CTT866CTG (3299T>G), ACC868TCT (3303A>T 3305C>T), ACT869ACA (3308T>A), GGT870GGG (3311T>G), TCT874TCA (3323T>A), CTA875TTA (3324C>T), CCG876CCA (3329G>A), CCC878CCA (3335C>A), CAC883CAT (3350C>T), AGG885CGG (3354A>C), TTG886CTA (3357T>C 3359G>A), ACA889ACG (3368A>G), GTC890GTG (3371C>G), TTG891ATG (3372T>A), GTA895GTT (3386A>T), ATA899ATT (3398A>T), CTA901CTG (3404A>G), ACG903ACT (3410G>T), GTC904GTT (3413C>T), GTA906GTC (3419A>C), ACA907ACG (3422A>G), CAG909CAA (3428G>A), TGC912TGA (3437C>A), GAA914GAG (3443A>G), TTG916TTA (3449G>A), GTC917GTG (3452C>G), CTT918CTA (3455T>A), GGG919GGC (3458G>C), GGC920GGA (3461C>A), GTT924GTG (3473T>G), CCC926CCT (3479C>T), ATT927ATC (3482T>C), TAC928TAT (3485C>T), GGA929GGT (3488A>T), CAC930CAT (3491C>T), GGA934GGG (3503A>G), AAT936AGT (3508A>G), GCT937GCC (3512T>C), GTG938GTT (3515G>T), TTC940TTT (3521C>T), CTA941TTG (3522C>T 3524A>G), TTC942TTT (3527C>T), GAG944GAA (3533G>A), TCT948ACT (3543T>A), CGC950CGT (3551C>T), GAG954GAA (3563G>A), TTA956CTG (3567T>C 3569A>G), CGC957CGA (3572C>A), AAG958AAA (3575G>A), AAC960AAT (3581C>T), ACA963ACC (3590A>C), CCA972CCC (3617A>C), GTA974GTT (3623A>T), CAA975GCA (3624C>G 3625A>C), GTG976CTT (3627G>C 3629G>T), CCG977GGG (3630C>G 3631C>G), CTC978CTT (3635C>T), AGA979AGG (3638A>G), GAC980GAA (3641C>A), TCA983TCT (3650A>T), GGG985GGA (3656G>A), GTA987GTT (3662A>T), GTG991GTT (3674G>T), GAC996GAT (3689C>T), CGA999CGC (3698A>C), TTG1001GTG (3702T>G), TTA1003CTA (3708T>C), TGC1005TGT (3716C>T), CCC1007CCG (3722C>G), TCC1009TCT (3728C>T), GGT1094GGA (3983T>A), ATG1095GTT (3984A>G 3986G>T), AGC1096ACC (3988G>C), GAC1097ACC (3990G>A 3991A>C), AAC1098AAT (3995C>T), ATC1099ATT (3998C>T), CAT1105CAA (4016T>A), GAG1108GAA (4025G>A), TTT1110TTC (4031T>C), AGA1112CGG (4035A>C 4037A>G), CGC1113CGG (4040C>G), ACC1115ACT (4046C>T), AAA1116AAG (4049A>G), GCC1117GCA (4052C>A), ATA1120GTT (4059A>G 4061A>T), CCA1122CCT (4067A>T), AAT1123AGG (4069A>G 4070T>G), GCT1124GGT (4072C>G), TCA1125TCT (4076A>T), AGA1126CGT (4077A>C 4079A>T), ATA1127ATT (4082A>T), CAA1131CAG (4094A>G), CTA1134CTG (4103A>G), GGA1135GGT (4106A>T), GTG1146.TA (4139G>A), ATA1147ATT (4142A>T), TCT1149TCC (4148T>C), CTG1150ATT (4149C>A 4151G>T), TTG1153ATG (4158T>A), GTC1154GTT (4163C>T), GGA1157GGG (4172A>G), TTC1160TTT (4181C>T), TTG1162TTA (4187G>A), CAA1163GAA (4188C>G), GCT1165GCC (4196T>C), TCA1169TCG (4208A>G), GCA1170GCT (4211A>T), AAG1174AAA (4223G>A), CAT1175TAT (4224C>T), TTG1176CTT (4227T>C 4229G>T), ACC1183ACT (4250C>T), CTT1185CTA (4256T>A), ACT1187ACC (4262T>C), TTT1190TTC (4271T>C), CTA1193CTG (4280A>G), AGT1195GGC (4284A>G 4286T>C), TTT1196TTC (4289T>C) | | | | | | | | |

*: Inserts / Deletes / Misaligned / Frameshifts

#### Analysis details

This analysis was performed with panviral2.64

### NGS Details (UN18_val): Torradovirus marchitezum (segment RNA 1)

#### Assembly

| | |
|---|---|
| **Coverage Length** | 5735 (1 contig(s)) |
| **Depth Of Coverage** | 116.5 |
| **Number Of Reads** | 5075 |
| **Reads Per Million** | 95.25 rpm (after QC) |
| **Ambiguities** | 0 |
| **Assembly Method** | de novo + reference guided assembly |
| **Consensus Caller** | Bcf Tools |

#### Coverage Map

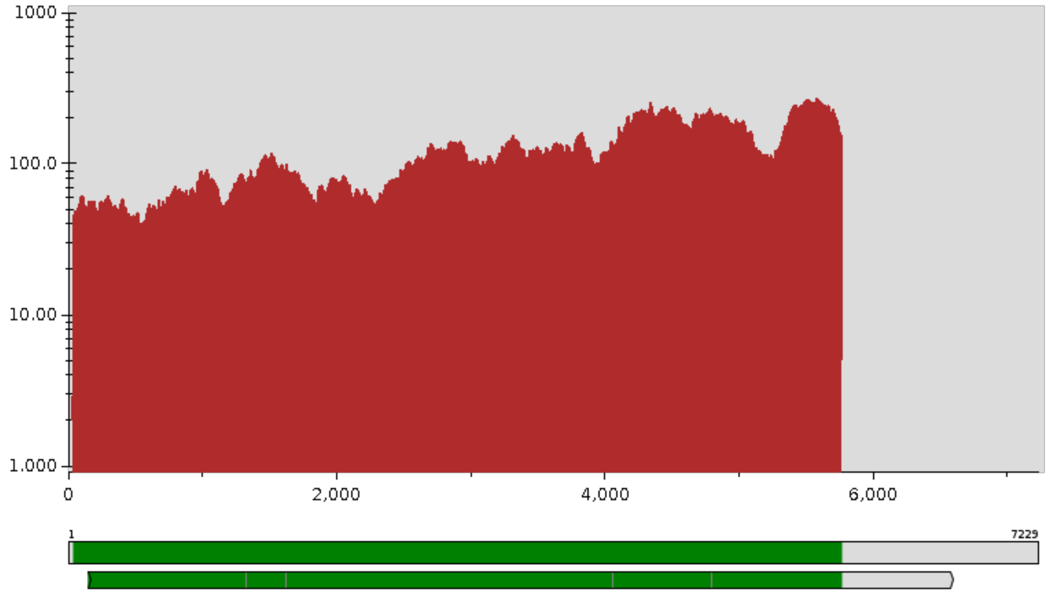

#### Assignment

| | |
|---|---|
| **Type** | Torradovirus marchitezum (Taxonomy ID: 3048376) |
| **Reference Genome** | NC_010987.1 |
| **NT Identity (%)** | 65.7968 |
| **AA Identity (%)** | 68.984 |
| **Number Of Stop Codons** | 0 |
| **Number Of CDS** | 1 |

#### Alignment

| | |
|---|---|
| **Alignment Score** | 3466.0 (NT) + 9403.0 (AA) = 12869.0 |
| **Concordance (%)** | 52.9392 |

| Alignment Method | Global, seeded, nucleotide + amino acids (AGA) |
|---|---|

#### Genome Region

Sequence starts at position 35 and ends at position 5769 relative to NC_010987.1 reference sequence.

#### Alignment Detailed Statistics

| | Begin | End | Coverage | Score | Concordance | Matches | Identities | I/D/M/F* | Stop Codons |
|---|---|---|---|---|---|---|---|---|---|
| NT | 35 | 5769 | 79.3% | 3466 | 30.8% | 5682 (98.6%) | 3757 (65.2%) | 28/53 | |

| | Begin | End | Coverage | Score | Concordance | Matches | Identities | I/D/M/F* | Stop Codons |
|---|---|---|---|---|---|---|---|---|---|
| NT | 35 | 5769 | 79.3% | 3466 | 30.8% | 5682 (98.6%) | 3757 (65.2%) | 28/53 | |

Mutations:

41A>C, 44G>T, 45A>T, 50G>C, 51C>T, 52C>T, 55T>A, 57T>A, 58T>A, 62T>C, 63G>C, 66A>T, 66_67insGTT, 70G>A, 74C>T, 75T>G, 79A>T, 80A>C, 81A>C, 82C>A, 88C>T, 88_89insG, 91C>T, 92A>T, 95T>C, 96T>C, 97C>A, 105C>T, 106T>C, 108A>G, 109C>G, 111C>T, 112G>T, 113A>T, 117T>C, 118T>A, 121T>A, 122T>C, 123A>C, 125A>C, 127T>A, 128_135delCCTCAATT, 139C>A, 146T>A, 155G>A, 158G>C, 159T>G, 161C>T, 162_191delTCCAAGGTTGCTGGTGTTGATGACGCCAGT, 198T>G, 200C>T, 203C>T, 204A>C, 209A>T, 212C>T, 213T>C, 216G>T, 217G>C, 221A>G, 222C>T, 224T>G, 227A>T, 228A>C, 230T>G, 231T>A, 233T>A, 234A>G, 236T>G, 239C>A, 240A>T, 241A>C, 243T>C, 245T>G, 248A>C, 251A>G, 252G>A, 254G>A, 255G>A, 257T>G, 259G>A, 260T>G, 262A>G, 266T>G, 268C>G, 269C>T, 271C>A, 272T>A, 275G>A, 278A>T, 279C>G, 284G>A, 285A>T, 287T>C, 292A>G, 293T>C, 297C>A, 302G>A, 305T>A, 306C>T, 308G>A, 314A>T, 317T>A, 318G>A, 320G>T, 321A>G, 322C>T, 324G>A, 326T>A, 327G>A, 328T>C, 329G>T, 330T>G, 332C>A, 333A>G, 334C>A, 335A>T, 338T>C, 342G>A, 343T>A, 344G>T, 345G>A, 346C>G, 347C>T, 348A>G, 349C>T, 359A>T, 360C>G, 362T>A, 363A>G, 365T>A, 369G>T, 371T>A, 374G>A, 376A>G, 377C>G, 385G>C, 386T>C, 387G>A, 389A>G, 390C>G, 395A>T, 398T>A, 401A>T, 405C>G, 412C>G, 413G>T, 414T>A, 415A>T, 417T>A, 418C>A, 425G>C, 426T>A, 427C>A, 428C>A, 437C>T, 439A>G, 441T>A, 443T>C, 446C>A, 448A>T, 449G>T, 459T>C, 461A>G, 465T>G, 467G>T, 468T>G, 471G>T, 472A>T, 474A>G, 476T>C, 477T>A, 479G>C, 480T>A, 481C>A, 482C>A, 488A>T, 489G>T, 493A>T, 494A>T, 495T>A, 496T>G, 502T>A, 504G>T, 506T>A, 510G>C, 515A>G, 516A>G, 518T>A, 520A>C, 523G>C, 524T>A, 526A>G, 527A>T, 530T>A, 531G>T, 532T>A, 533A>T, 537A>G, 539A>T, 542T>A, 545A>T, 546T>A, 547A>T, 548T>A, 550G>C, 557T>A, 560A>C, 564G>T, 566T>C, 567A>C, 569A>T, 570A>T, 573T>C, 575G>A, 578A>G, 579G>T, 581G>A, 582C>T, 584T>A, 586T>A, 588T>A, 590T>A, 591G>A, 593T>C, 600A>G, 601G>T, 602T>A, 611T>A, 614G>A, 615G>A, 616G>T, 617T>G, 621T>A, 622C>G, 623T>A, 625A>G, 626C>T, 627A>G, 628C>T, 629T>A, 632C>T, 633A>T, 634A>G, 635A>T, 641C>A, 645T>C, 647G>A, 648A>G, 650A>T, 653C>A, 656A>T, 660A>T, 662G>C, 665T>C, 668A>T, 669A>T, 670G>C, 674A>G, 677C>T, 681T>A, 683A>T, 684G>A, 685A>G, 695A>C, 696A>C, 697A>T, 698A>C, 702G>T, 704C>T, 707A>G, 708C>A, 709G>A, 713C>T, 714A>T, 716C>T, 717A>C, 718T>A, 720G>A, 725C>A, 728A>G, 732A>T, 733G>C, 735A>G, 740T>G, 746G>T, 749A>G, 750T>A, 751C>G, 752A>C, 753G>T, 755T>A, 756T>A, 757C>A, 758A>T, 770G>C, 771T>A, 772C>G, 776G>A, 780A>C, 785C>T, 786A>T, 788A>T, 789A>T, 791A>G, 792T>A, 794G>A, 797T>A, 803G>C, 806T>C, 809G>A, 818G>A, 821C>T, 822A>G, 824T>G, 830G>A, 839A>G, 849T>C, 851G>T, 853G>C, 854C>A, 860A>G, 861T>C, 863G>A, 873C>T, 875C>G, 881G>A, 882A>G, 884A>G, 887A>C, 893T>A, 899G>A, 914A>C, 918A>C, 919C>A, 920T>A, 921C>T, 923T>G, 926C>T, 927C>A, 928G>A, 929T>G, 932T>C, 938A>G, 947C>G, 962A>T, 968T>G, 969C>T, 971T>G, 977A>T, 978A>G, 980A>T, 982T>C, 983C>T, 985G>A, 986C>T, 988C>T, 989C>A, 992T>C, 995A>G, 998C>T, 1004T>C, 1010G>T, 1011T>C, 1016A>T, 1019T>A, 1022A>T, 1031G>A, 1032C>A, 1034G>T, 1043T>C, 1046G>C, 1049C>T, 1059C>A, 1060A>C, 1061G>T, 1064G>A, 1068G>T, 1069A>C, 1077T>G, 1078G>C, 1091G>A, 1092C>A, 1094C>G, 1100G>A, 1109C>T, 1113A>C, 1114A>C, 1115G>T, 1121C>T, 1122C>T, 1124T>A, 1126C>T, 1142G>T, 1144G>A, 1145G>A, 1148A>G, 1149G>C, 1151A>G, 1154T>A, 1163C>A, 1169T>C, 1172C>T, 1178A>T, 1187A>T, 1190T>G, 1199C>T, 1200A>T, 1205C>T, 1211G>A, 1215A>T, 1216G>C, 1221G>A, 1223C>A, 1228C>T, 1232G>A, 1235A>G, 1238A>T, 1242G>C, 1244T>C, 1245T>A, 1246A>T, 1247T>A, 1250T>A, 1255G>A, 1256T>C, 1260A>C, 1261G>A, 1262G>A, 1265C>T, 1267T>C, 1268G>A, 1271C>A, 1277C>T, 1283T>C, 1284C>A, 1286T>A, 1295T>C, 1304A>T, 1313C>T, 1316G>T, 1317T>G, 1318C>A, 1319C>G, 1325G>A, 1328T>C, 1331T>C, 1337T>C, 1340A>G, 1341T>A, 1342C>G, 1349C>A, 1350T>C, 1352A>G, 1355T>C, 1358T>C, 1367G>A, 1373C>T, 1374A>G, 1375G>C, 1376A>C, 1380G>C, 1381G>A, 1382T>A, 1388A>C, 1391A>G, 1394C>G, 1397T>C, 1398C>A, 1401C>T, 1403A>G, 1412A>T, 1419A>C, 1421G>C, 1427A>T, 1430T>C, 1436C>A, 1439C>T, 1463G>C, 1472C>T, 1478A>T, 1482T>C, 1487A>G, 1493G>A, 1499G>C, 1502G>A, 1508A>G, 1511T>C, 1517A>G, 1520T>C, 1523G>C, 1529T>C, 1538A>C, 1553C>A, 1556A>G, 1559A>C, 1565C>T, 1571T>A, 1574G>A, 1583C>G, 1584A>C, 1595C>A, 1598C>A, 1604T>C, 1616A>C, 1622T>C, 1623C>T, 1626A>G, 1628C>T, 1634T>A, 1637T>A, 1640C>A, 1646T>C, 1649T>G, 1652T>G, 1661A>T, 1673G>A, 1674C>A, 1679A>G, 1680A>C, 1688C>T, 1689C>T, 1697G>A, 1700T>C, 1701A>C, 1702T>G, 1706G>A, 1708A>T, 1712C>G, 1715G>A, 1718T>C, 1721C>A, 1724T>A, 1727A>G, 1728G>A, 1730T>C, 1733T>A, 1736G>A, 1738G>C, 1748A>C, 1751T>C, 1757A>C, 1758G>A, 1760G>T, 1762A>G, 1763G>A, 1769A>G, 1772A>G, 1781T>C, 1784C>A, 1787T>A, 1793G>A, 1794A>G, 1796T>C, 1797T>C, 1799G>C, 1802C>T, 1806A>T, 1807G>C, 1811T>C, 1812C>T, 1814T>G, 1820G>A, 1829A>G, 1835A>G, 1838T>C, 1839C>A, 1844A>C, 1847T>C, 1848G>C, 1849C>A, 1850A>C, 1854G>A, 1856A>T, 1857T>G, 1861A>C, 1862G>C, 1866T>C, 1868G>C, 1870T>A, 1871T>C, 1872G>A, 1874A>C, 1875G>C, 1876C>A, 1877T>G, 1878G>T, 1879A>C, 1881T>A, 1883T>A, 1886C>T, 1887A>T, 1888T>G, 1889A>T, 1890A>G, 1891G>A, 1892T>A, 1893C>G, 1894C>A, 1895T>G, 1898G>A, 1899A>G, 1901A>T, 1904T>C, 1910T>G, 1911C>G, 1921C>T, 1922A>T, 1925T>A, 1928T>A, 1934A>G, 1935G>A, 1937G>A, 1940T>C, 1941G>T, 1943G>T, 1944G>C, 1948G>A, 1949G>T, 1953A>C, 1954G>A, 1956T>A, 1958G>A, 1959C>A, 1962A>T, 1963C>T, 1964C>T, 1965A>G, 1967G>C, 1968C>G, 1972A>T, 1974G>A, 1975A>G, 1976A>C, 1979A>C, 1981T>G, 1982A>T, 1983T>G, 1985G>T, 1986_1991delGTGCCC, 1994T>A, 1998G>C, 2000G>C, 2001A>T, 2002G>C, 2006A>G, 2007A>G, 2019A>G, 2020T>G, 2021C>A, 2025G>T, 2028A>C, 2029G>A, 2030A>G, 2033G>A, 2036C>A, 2039G>T, 2042A>G, 2045G>A, 2049T>C, 2054A>G, 2060A>T, 2063C>T, 2066G>A, 2069C>A, 2070A>G, 2071G>C, 2072T>A, 2075G>A, 2076G>A, 2081T>A, 2082G>T, 2083A>T, 2084G>T, 2088C>T, 2089T>A, 2096T>C, 2102C>T, 2103C>G, 2104A>C, 2106C>T, 2107A>C, 2108T>A, 2117C>A, 2120G>T, 2122A>T, 2127T>G, 2128C>T, 2129T>C, 2132T>G, 2133G>T, 2134A>C, 2135T>A, 2136G>A, 2137A>G, 2142C>G, 2143A>C, 2144T>C, 2152T>G, 2153T>C, 2156C>T, 2158A>C, 2159C>T, 2160A>C, 2161A>G, 2162A>C, 2166A>G, 2167G>T, 2169G>A, 2170T>A, 2171C>T, 2172A>G, 2174C>G, 2178T>A, 2180G>T, 2183T>C, 2184A>G, 2185G>A, 2186A>G, 2189A>G, 2192A>T, 2193G>T, 2194A>C, 2195T>G, 2196G>A, 2200A>C, 2202C>G, 2203A>G, 2204G>A, 2208C>G, 2209G>C, 2210A>C, 2217G>A, 2219A>C, 2220C>A, 2222A>G, 2223A>G, 2224A>C, 2228T>C, 2231T>A, 2241A>G, 2243A>C, 2255G>A, 2258T>C, 2268G>A, 2269C>G, 2270A>T, 2271T>A, 2277A>G, 2279C>G, 2282T>A, 2285G>A, 2286A>G, 2287A>C, 2288G>T, 2289A>C, 2291A>C, 2292C>A, 2294A>G, 2296C>A, 2297A>T, 2300G>A, 2303G>A, 2305C>G, 2306A>G, 2307G>A, 2308A>T, 2310A>T, 2313A>T, 2314G>A, 2315G>C, 2317C>A, 2321A>C, 2322T>A, 2323T>G, 2324T>A, 2331T>A, 2332C>G, 2333T>A, 2334T>G, 2335G>C, 2336C>A, 2337A>C, 2339T>G, 2342T>A, 2345A>T, 2350T>C, 2357C>T, 2358C>T, 2360C>A, 2363A>G, 2369C>T, 2372G>A, 2375G>A, 2381A>T, 2385T>C, 2386T>A, 2390T>C, 2393G>A, 2396T>C, 2397T>C, 2402T>C, 2415A>C, 2417A>G, 2426_2427insAGAAGG, 2429C>G, 2431A>T, 2432A>G, 2435A>C, 2436T>A, 2438A>T, 2441A>G, 2444G>A, 2445C>A, 2448_2453delAAGGAA, 2456A>T, 2460A>G, 2462C>A, 2463A>C, 2466A>T, 2467T>C, 2468T>G, 2469T>C, 2470G>T, 2474C>T, 2475C>G, 2477T>C, 2478A>T, 2479A>C, 2480A>C, 2482T>C, 2483C>A, 2484G>A, 2488A>G, 2489A>G, 2490T>A, 2491A>G, 2492T>A, 2495C>G, 2496A>C, 2498G>A, 2501T>A, 2503C>T, 2504A>G, 2508T>A, 2509G>A, 2510G>T, 2511A>G, 2512G>A, 2513C>T, 2516T>G, 2522T>C, 2523G>A, 2524C>T, 2525A>T, 2526C>T, 2530G>A, 2537C>T, 2549T>G, 2556A>C, 2561G>A, 2567T>C, 2573A>G, 2574G>A, 2575T>C, 2577T>A, 2579T>G, 2582A>T, 2584G>C, 2586A>G, 2588T>A, 2591A>T, 2597T>C, 2598T>A, 2600G>T, 2601G>T, 2603G>T, 2610C>A, 2612G>A, 2615T>C, 2617A>G, 2620C>T, 2621T>G, 2622T>A, 2624C>T, 2629A>C, 2631G>T, 2633A>C, 2634T>C, 2635C>A, 2641G>A, 2651G>T, 2663C>G, 2669T>C, 2675C>A, 2678T>C, 2693T>A, 2695A>G, 2696T>A, 2696_2697insTCC, 2697A>C, 2699G>T, 2700C>G, 2702C>A, 2703A>T, 2704G>C, 2708T>G, 2709C>A, 2711T>G, 2714C>G, 2715G>T, 2718G>A, 2720T>C, 2729T>A, 2733T>C, 2736T>A, 2737T>A, 2738G>T, 2740C>A, 2741A>T, 2744A>G, 2744_2745insGGA, 2745G>A, 2747T>A, 2754G>T, 2756T>C, 2757G>A, 2758A>G, 2759A>G, 2765A>T, 2771A>G, 2774T>C, 2775C>G, 2777G>A, 2778A>T, 2779G>C, 2780C>T, 2783C>T, 2784A>G, 2785A>C, 2786G>C, 2798G>A, 2802T>C, 2807A>G, 2808C>G, 2810T>G, 2812G>C, 2813T>A, 2822G>T, 2829A>C, 2831A>T, 2834C>A, 2837C>T, 2846T>C, 2847T>A, 2848C>G, 2849T>C, 2851G>A, 2852T>A, 2864C>T, 2870T>A, 2873T>C, 2876A>T, 2880C>A, 2881A>C, 2882C>G, 2883A>C, 2885T>G, 2886G>C, 2887C>T, 2888T>C, 2891A>T, 2897T>A, 2900T>C, 2907A>T, 2915C>T, 2920C>A, 2921C>A, 2922A>C, 2928C>G, 2929G>T, 2930A>C, 2932C>G, 2933T>C, 2934A>C, 2936A>G, 2939T>G, 2948C>T, 2953G>A, 2955G>C, 2957A>T, 2958T>G, 2960A>C, 2962G>A, 2964A>T, 2967A>T, 2969T>G, 2970G>C, 2971A>C, 2972G>T, 2974G>A, 2975C>A, 2981C>T, 2982A>G, 2984C>T, 2985A>G, 2987T>A, 2988C>T, 2990G>A, 2991T>C, 2994C>A, 2996C>G, 3002T>C, 3003G>T, 3004A>G, 3008A>T, 3011T>C, 3014T>G, 3017A>T, 3020G>A, 3021C>G, 3022C>T, 3023T>C, 3030C>A, 3032T>A, 3035A>G, 3041C>T, 3042C>G, 3044A>T, 3045A>G, 3046G>A, 3055A>C, 3056G>A, 3071G>A, 3074C>A, 3077A>T, 3078C>G, 3083A>C, 3089C>T, 3090T>G, 3091G>T, 3092T>C, 3098T>A, 3101T>G, 3104G>A, 3107T>A, 3109C>A, 3116A>T, 3117C>G, 3118G>A, 3119T>G, 3120C>A, 3121T>A, 3122T>A, 3123G>C, 3124G>A, 3125C>G, 3128T>C, 3131A>G, 3134G>C, 3144A>T, 3145T>A, 3146G>T, 3147C>T, 3161A>C, 3164G>A, 3170C>T, 3175T>A, 3176T>G, 3182G>C, 3183G>A, 3184A>C, 3188T>C, 3189G>A, 3190A>G, 3191A>G, 3192C>A, 3196A>G, 3198T>A, 3200C>A, 3201T>A, 3202C>A, 3203C>T, 3204A>T, 3205G>C, 3206T>A, 3210A>C, 3212G>C, 3212_3213insTCA, 3215T>C, 3216T>A, 3218C>T, 3219T>G, 3225C>G, 3228T>G, 3230T>C, 3242C>T, 3243A>G, 3245A>C, 3249G>A, 3251G>C, 3263C>T, 3269C>A, 3272G>T, 3279C>T, 3281G>A, 3284A>G, 3288A>T, 3289G>C, 3291G>A, 3293T>C, 3296G>A, 3299T>C, 3308T>A, 3309G>T, 3310T>G, 3311G>C, 3314C>T, 3320C>T, 3323C>T, 3326C>T, 3332A>T, 3335A>C, 3343G>A, 3344C>A, 3346T>C, 3347A>T, 3351G>A, 3353T>A, 3356A>G, 3357C>T, 3358G>T, 3359T>G, 3366A>G, 3368C>A, 3369T>G, 3371T>A, 3380T>C, 3383A>T, 3387A>G, 3389C>A, 3398A>C, 3401A>G, 3404A>G, 3405G>C, 3407G>A, 3414C>G, 3416A>T, 3417A>C, 3418C>A, 3419T>A, 3425G>A, 3428C>T, 3432G>A, 3433C>T, 3434A>G, 3437A>C, 3443A>G, 3444C>A, 3446C>A, 3452T>C, 3453A>T, 3455T>G, 3458G>C, 3461G>A, 3462T>C, 3463C>A, 3464A>T, 3465A>G, 3468A>C, 3471A>T, 3472G>C, 3473C>A, 3482A>T, 3483C>G, 3484A>C, 3485G>T, 3488T>C, 3494G>C, 3500C>G, 3506G>T, 3507T>G, 3508A>C, 3509C>T, 3511A>T, 3512C>T, 3513C>T, 3515T>G, 3518A>C, 3521A>T, 3527A>T, 3530G>A, 3534T>A, 3539C>A, 3542C>A, 3545A>G, 3548T>G, 3551C>A, 3563T>C, 3566G>C, 3567C>A, 3577A>C, 3579T>C, 3587A>G, 3593C>T, 3596A>T, 3597C>G, 3598C>A, 3599T>A, 3603G>C, 3605G>A, 3608A>T, 3611A>T, 3614A>T, 3617C>A, 3618A>C, 3620T>G, 3623A>C, 3624A>T, 3626A>T, 3629T>G, 3630C>A, 3632T>A, 3635C>T, 3644T>G, 3650T>A, 3656C>T, 3659C>T, 3665G>T, 3671G>C, 3672G>A, 3674A>T, 3675A>G, 3676T>A, 3695T>C, 3697G>C, 3698T>A, 3699G>A, 3700G>A, 3701A>T, 3707T>G, 3710T>C, 3711G>A, 3712A>G, 3719C>A, 3720A>C, 3721A>G, 3722A>G, 3723C>G, 3724T>C, 3725A>T, 3731C>T, 3735A>G, 3736A>G, 3745G>C, 3746C>T, 3747T>G, 3749G>C, 3750C>A, 3752G>T, 3753C>T, 3758T>A, 3759A>G, 3761C>T, 3762A>C, 3764G>C, 3765C>A, 3766G>A, 3767T>G, 3770C>T, 3773G>A, 3775A>G, 3777A>G, 3779C>G, 3782T>A, 3785G>T, 3788A>T, 3789C>A, 3790G>T, 3791T>C, 3794A>C, 3795T>A, 3796G>C, 3797T>A, 3798G>A, 3799A>C, 3801A>C, 3803G>T, 3804G>T, 3805A>G, 3812T>C, 3813G>A, 3815A>C, 3818G>C, 3821C>G, 3827A>T, 3828A>G, 3830A>T, 3836A>T, 3838G>T, 3842T>C, 3844A>T, 3847A>T, 3852G>T, 3853G>C, 3860G>A, 3863G>C, 3872T>A, 3875C>T, 3881T>C, 3884C>T, 3890C>T, 3893G>A, 3894A>C, 3899G>T, 3900A>T, 3905G>T, 3907T>C, 3908G>C, 3911C>T, 3914G>A, 3917A>G, 3923T>C, 3926G>T, 3927C>G, 3930T>G, 3932C>A, 3933A>C, 3934C>A, 3935T>G, 3937G>A, 3938C>G, 3939G>A, 3940A>G, 3941A>T, 3942G>A, 3943G>C, 3944A>C, 3945G>C, 3947G>A, 3950A>G, 3953G>A, 3954G>C, 3955A>T, 3962A>G, 3966G>A, 3968C>T, 3969A>T, 3977A>T, 3981G>C, 3992A>G, 3995G>A, 3996C>T, 4005C>G, 4007G>A, 4010T>A, 4016A>G, 4019C>T, 4021T>A, 4023C>T, 4031T>A, 4032C>A, 4037C>T, 4043C>T, 4049C>T, 4052C>T, 4055A>G, 4058T>C, 4062C>A, 4064C>G, 4067C>A, 4070A>T, 4073T>G, 4076G>C, 4079A>G, 4082C>A, 4085T>C, 4088C>G, 4091G>A, 4093G>A, 4094A>G, 4097G>A, 4101A>C, 4103G>T, 4106C>T, 4115A>T, 4116C>T, 4118T>G, 4121T>A, 4130C>T, 4137T>C, 4142C>G, 4143A>C, 4145G>C, 4148G>A, 4154T>C, 4163C>T, 4166A>T, 4169C>A, 4170A>T, 4171C>G, 4172A>T, 4175G>A, 4182T>G, 4183T>C, 4184G>A, 4187C>T, 4192C>T, 4194A>C, 4196A>G, 4199A>C, 4208T>G, 4214T>C, 4215G>A, 4220T>C, 4222C>T, 4223A>T, 4224A>T, 4225G>C, 4226T>A, 4227T>C, 4228C>A, 4238A>C, 4239T>A, 4240T>C, 4241G>C, 4242T>C, 4250T>C, 4251G>C, 4252G>A, 4256T>C, 4257A>C, 4259A>G, 4260G>A, 4261C>G, 4268A>T, 4271C>T, 4274G>T, 4277A>T, 4283A>T, 4286T>A, 4289T>C, 4292T>C, 4310A>G, 4313A>G, 4316T>A, 4317G>A, 4319T>A, 4325T>A, 4331G>A, 4337T>A, 4354A>C, 4361C>T, 4364A>G, 4368A>G, 4369A>C, 4370G>T, 4373T>A, 4376C>A, 4379T>C, 4380C>G, 4381C>A, 4382T>G, 4388C>A, 4394T>A, 4395C>A, 4403A>T, 4406T>G, 4409A>G, 4418A>T, 4424T>C, 4433A>T, 4436G>A, 4449G>C, 4451G>C, 4454T>C, 4460G>A, 4466G>T, 4469C>A, 4475A>T, 4478A>T, 4487A>T, 4490A>C, 4493G>C, 4499C>A, 4502C>T, 4505T>C, 4508T>C, 4511C>T, 4512C>T, 4514C>A, 4517T>C, 4532A>G, 4538C>T, 4541T>C, 4547A>G, 4548T>C, 4550G>C, 4551T>C, 4553A>C, 4559A>G, 4565T>C, 4568T>C, 4569G>A, 4570C>A, 4571T>A, 4574C>A, 4576G>A, 4580T>A, 4581A>T, 4583A>G, 4586A>T, 4589T>A, 4592A>G, 4593C>T, 4604C>T, 4608C>G, 4610G>C, 4611C>T, 4616C>T, 4619A>T, 4622T>A, 4625T>G, 4631A>G, 4646T>G, 4652A>G, 4656T>A, 4658A>G, 4661T>C, 4664C>T, 4667G>A, 4673T>C, 4674T>G, 4675C>A, 4676T>G, 4677C>T, 4679G>A, 4682A>T, 4683A>G, 4684A>C, 4685T>A, 4694T>A, 4698T>C, 4705T>A, 4706G>A, 4709C>G, 4712T>C, 4715A>T, 4718C>T, 4721T>C, 4730T>C, 4736C>T, 4740T>A, 4741C>A, 4742G>C, 4746G>A, 4751T>C, 4757G>T, 4760A>C, 4761C>A, 4766T>C, 4769A>G, 4782C>T, 4784A>G, 4787T>G, 4791C>T, 4796T>C, 4802T>C, 4805T>C, 4808A>G, 4811A>C, 4818C>T, 4820T>G, 4823A>T, 4829C>A, 4830A>C, 4832G>A, 4835T>C, 4838T>C, 4853A>C, 4856C>T, 4857A>G, 4859T>G, 4862G>A, 4865C>A, 4877A>C, 4883T>A, 4886C>A, 4887C>G, 4888C>T, 4889A>T, 4892T>G, 4898T>A, 4901C>T, 4903C>T, 4904T>C, 4905C>A, 4906A>C, 4907G>T, 4914A>T, 4915T>A, 4916G>T, 4931A>G, 4934A>T, 4935A>G, 4938C>A, 4940T>A, 4943A>G, 4952T>G, 4955C>T, 4958T>A, 4960G>A, 4961T>G, 4964G>C, 4966C>T, 4975A>C, 4976G>A, 4977C>T, 4979T>G, 4980G>A, 4981T>C, 4982C>A, 4983T>G, 4984T>A, 4985A>C, 4986G>A, 4991T>C, 4994G>A, 4996C>A, 4997G>A, 5000G>A, 5001T>A, 5003G>A, 5004G>A, 5005A>T, 5006T>G, 5007G>C, 5009A>G, 5010A>C, 5012G>A, 5015T>C, 5018C>G, 5019A>C, 5020G>C, 5022G>A, 5023C>T, 5024T>C, 5028A>C, 5029T>A, 5030C>T, 5036T>A, 5039C>T, 5043T>G, 5044T>C, 5045G>T, 5049G>C, 5050T>A, 5051G>A, 5055C>A, 5056A>G, 5057G>T, 5060A>G, 5064A>C, 5066T>A, 5067G>C, 5068C>A, 5070A>G, 5071A>C, 5072G>C, 5075A>T, 5078A>T, 5081G>A, 5084T>C, 5085T>A, 5086C>G, 5087A>T, 5089G>A, 5090T>C, 5091G>A, 5092C>T, 5093C>A, 5094C>G, 5095T>G, 5097A>G, 5099A>T, 5100A>C, 5102T>G, 5103T>C, 5104G>T, 5105T>G, 5109A>C, 5112G>A, 5113C>G, 5114A>C, 5117A>G, 5119C>A, 5120A>T, 5127T>A, 5129C>T, 5132T>C, 5133A>G, 5134C>T, 5138A>T, 5139T>C, 5141G>T, 5142G>C, 5143A>C, 5144C>A, 5146C>A, 5150T>A, 5151G>A, 5153G>A, 5156G>A, 5159C>T, 5160A>G, 5161C>A, 5163A>G, 5169G>T, 5170A>C, 5171G>A, 5173G>A, 5174T>A, 5176A>G, 5178C>G, 5180G>C, 5181A>C, 5184T>G, 5185G>A, 5186G>A, 5190G>T, 5191C>G, 5192A>T, 5193C>T, 5195A>T, 5196A>G, 5197A>G, 5198C>A, 5199A>G, 5202G>T, 5203G>T, 5204T>C, 5207T>C, 5214G>T, 5216A>G, 5219T>A, 5222T>A, 5224C>A, 5227C>T, 5228C>T, 5229C>A, 5231C>T, 5232T>C, 5234C>T, 5236C>A, 5237G>A, 5246A>G, 5248G>A, 5249G>C, 5250T>A, 5251A>C, 5252C>A, 5256A>C, 5264C>T, 5267T>C, 5279G>A, 5282G>A, 5283T>A, 5285C>A, 5287A>C, 5288T>A, 5294T>G, 5339G>C, 5342C>T, 5345T>A, 5349T>G, 5354T>C, 5357A>C, 5359T>A, 5363T>C, 5364G>A, 5365C>A, 5366C>T, 5370A>C, 5372G>A, 5374A>G, 5376G>C, 5377T>G, 5378G>C, 5382G>T, 5385G>A, 5386G>T, 5387A>G, 5390A>G, 5391C>A, 5392A>T, 5395T>G, 5397G>A, 5399A>C, 5400A>C, 5401C>A, 5402C>A, 5405T>A, 5408C>T, 5411T>C, 5415G>A, 5417G>T, 5423A>G, 5429A>T, 5438C>G, 5439G>A, 5441C>T, 5444A>C, 5453A>T, 5456G>T, 5460A>T, 5462G>T, 5465C>A, 5469G>T, 5474A>C, 5476G>A, 5477G>T, 5480A>G, 5484T>A, 5486T>A, 5488T>C, 5489G>T, 5493G>C, 5495C>A, 5496A>C, 5498A>T, 5504T>C, 5511C>A, 5513C>A, 5517A>T, 5519T>G, 5521G>C, 5522A>G, 5523A>G, 5524A>G, 5525T>C, 5526G>T, 5531T>G, 5534C>T, 5535C>T, 5537C>G, 5541A>C, 5542C>A, 5543C>G, 5555C>T, 5559G>A, 5561G>T, 5564G>A, 5570T>C, 5573T>C, 5574C>A, 5575A>C, 5579_5580insGTC, 5584C>A, 5585T>G, 5586C>A, 5588T>A, 5589G>A, 5591A>T, 5592G>T, 5593A>G, 5597A>G, 5598G>A, 5599T>C, 5600G>A, 5607A>G, 5608G>C, 5610G>A, 5614C>G, 5615A>G, 5616_5618delTCG, 5620T>G, 5622C>G, 5624C>T, 5625A>G, 5627T>A, 5629G>T, 5630T>G, 5633T>G, 5635T>A, 5636G>C, 5642A>T, 5642_5643insCATGCT, 5644G>C, 5645G>T, 5647T>A, 5650A>C, 5651A>C, 5652T>C, 5654G>T, 5657C>A, 5663C>T, 5666C>A, 5669T>A, 5670A>G, 5672A>G, 5673C>G, 5674A>G, 5675T>C, 5676G>T, 5679G>T, 5680A>C, 5681T>A, 5682T>A, 5684A>T, 5685G>T, 5689A>G,

| | |
|---|---|
| | 5691C>A, 5695C>T, 5696A>G, 5700A>G, 5703C>A, 5705G>A, 5706A>T, 5711A>C, 5714T>C, 5715G>A, 5716T>C, 5717G>T, 5720A>T, 5723A>G, 5727C>A, 5728A>T, 5729T>G, 5732A>T, 5735A>T, 5740G>C, 5742A>G, 5746T>G, 5747T>C, 5750C>A, 5754C>A, 5755A>G, 5756G>A, 5757G>A |

#### CDS

| | | | | | | | | | |
|---|---|---|---|---|---|---|---|---|---|
| ToMarV_RNA1gp1 | 1 | 1877 | 87.2% | 9403 | 72.1% | 1862 (98.8%) | 1290 (68.4%) | 8/15/0/0 | 0 |
| Protein mutations: | M6I (158G>C), F7V (159T>G 161C>T), S8_S17del (162_191delTCCAAGGTTGCTGGTGTTGATGACGCCAGT), S20A (198T>G 200C>T), T22P (204A>C), F25L (213T>C), G26S (216G>T 217G>C), N30Q (228A>C 230T>G), S31T (231T>A 233T>A), I32V (234A>G 236T>G), K34S (240A>T 241A>C), F35L (243T>C 245T>G), A38T (252G>A 254G>A), V39M (255G>A 257T>G), S40K (259G>A 260T>G), N41S (262A>G), I42M (266T>G), T43S (268C>G 269C>T), T44K (271C>A 272T>A), M45I (275G>A), Q47E (279C>G), I49F (285A>T 287T>C), D51G (292A>G 293T>C), L53I (297C>A), D55E (305T>A), V60I (318G>A 320G>T), T61V (321A>G 322C>T), D62K (324G>A 326T>A), V63T (327G>A 328T>C 329G>T), S64A (330T>G 332C>A), T65D (333A>G 334C>A 335A>T), V68N (342G>A 343T>A 344G>T), A69S (345G>A 346C>G 347C>T), T70V (348A>G 349C>T), L74V (360C>G 362T>A), N75E (363A>G 365T>A), V77L (369G>T 371T>A), D79G (376A>G 377C>G), G82A (385G>C 386T>C), A83T (387G>A 389A>G), L84V (390C>G), D86E (398T>A), L89V (405C>G), A91G (412C>G 413G>T), Y92I (414T>A 415A>T), S93N (417T>A 418C>A), L95F (425G>C), S96K (426T>A 427C>A 428C>A), K100R (439A>G), S101T (441T>A 443T>C), K103I (448A>T 449G>T), L109V (465T>G 467G>T), F110V (468T>G), E111L (471G>T 472A>T), S112G (474A>G 476T>C), L113I (477T>A 479G>C), S114K (480T>A 481C>A 482C>A), K116N (488A>T), V117F (489G>T), K118I (493A>T 494A>T), L119R (495T>A 496T>G), F121Y (502T>A), A122S (504G>T 506T>A), V124L (510G>C), N126E (516A>G 518T>A), K127T (520A>C), S128T (523G>C 524T>A), E129G (526A>G 527A>T), D130E (530T>A), V131Y (531G>T 532T>A 533A>T), I133V (537A>G 539A>T), Y136I (546T>A 547A>T 548T>A), G137A (550G>C), A142S (564G>T 566T>C), I143L (567A>C 569A>T), T144S (570A>T), V147L (579G>T 581G>A), L149Q (586T>A), F150I (588T>A 590T>A), V151I (591G>A 593T>C), S154V (600A>G 601G>T 602T>A), G159M (615G>A 616G>T 617T>G), S161R (621T>A 622C>G 623T>A), N162S (625A>G 626C>T), T163V (627A>G 628C>T 629T>A), K165C (633A>T 634A>G 635A>T), I170V (648A>G 650A>T), M174F (660A>T 662G>C), L181I (681T>A 683A>T), D182S (684G>A 685A>G), K186L (696A>C 697A>T 698A>C), A188S (702G>T 704C>T), R190K (708C>A 709G>A), T192S (714A>T 716C>T), M193Q (717A>C 718T>A), V194M (720G>A), M199V (735A>G), V205L (753G>T 755T>A), S206N (756T>A 757C>A 758A>T), M214L (780A>C), T216S (786A>T 788A>T), I217L (789A>T 791A>G), L218I (792T>A 794G>A), I228V (822A>G 824T>G), S238T (853G>C 854C>A), T248A (882A>G 884A>G), T260Q (918A>C 919C>A 920T>A), R263K (927C>A 928G>A 929T>G), I280V (978A>G 980A>T), V281A (982T>C 983C>T), G282D (985G>A 986C>T), T283I (988C>T 989C>A), L298I (1032C>A 1034G>T), Q307T (1059C>A 1060A>C 1061G>T), D310S (1068G>T 1069A>C), C313A (1077T>G 1078G>C), K325P (1113A>C 1114A>C 1115G>T), P328S (1122C>T 1124T>A), T329I (1126C>T), R335K (1144G>A 1145G>A), E337Q (1149G>C 1151A>G), M354L (1200A>T), A361T (1221G>A 1223C>A), A363V (1228C>T), E366D (1238A>T), V368L (1242G>C 1244T>C), Y369I (1245T>A 1246A>T 1247T>A), S372N (1255G>A 1256T>C), R374Q (1260A>C 1261G>A 1262G>A), M376T (1267T>C 1268G>A), S393E (1317T>G 1318C>A 1319C>G), R412A (1374A>G 1375G>C 1376A>C), G414Q (1380G>C 1381G>A 1382T>A), I496V (1626A>G 1628C>T), M521R (1701A>C 1702T>G), E523V (1708A>T), D530N (1728G>A 1730T>C), G533A (1738G>C), R536S (1748A>C), V540I (1758G>A 1760G>T), E541G (1762A>G 1763G>A), N552D (1794A>G 1796T>C), Q567K (1839C>A), A570H (1848G>C 1849C>A 1850A>C), V572I (1854G>A 1856A>T), S573A (1857T>G), K574T (1861A>C 1862G>C), F577Y (1870T>A 1871T>C), E578N (1872G>A 1874A>C), A579Q (1875G>C 1876C>A 1877T>G), E580S (1878G>T 1879A>C), S581T (1881T>A 1883T>A), I583C (1887A>T 1888T>G 1889A>T), S584E (1890A>G 1891G>A 1892T>A), P585E (1893C>G 1894C>A 1895T>G), I587V (1899A>G 1901A>T), Q591E (1911C>G), A594V (1921C>T 1922A>T), D595E (1925T>A), D596E (1928T>A), I598M (1934A>G), A599T (1935G>A 1937G>A), V601F (1941G>T 1943G>T), D602H (1944G>C), G603D (1948G>A 1949G>T), R605Q (1953A>C 1954G>A), L606I (1956T>A 1958G>A), R607S (1959C>A), T608F (1962A>T 1963C>T 1964C>T), K609D (1965A>G 1967G>C), Q610E (1968C>G), D611V (1972A>T), E612S (1974G>A 1975A>G 1976A>C), I614S (1981T>G 1982A>T), L615V (1983T>G 1985G>T), V616_P617del (1986_1991delGTGCCC), V620L (1998G>C 2000G>C), N623D (2007A>G), I627G (2019A>G 2020T>G 2021C>A), A629S (2025G>T), R630Q (2028A>C 2029G>A 2030A>G), S644A (2070A>G 2071G>C 2072T>A), D646N (2076G>A), E648F (2082G>T 2083A>T 2084G>T), L650Y (2088C>T 2089T>A), H655A (2103C>G 2104A>C), H656S (2106C>T 2107A>C 2108T>A), M660I (2120G>T), Y661F (2122A>T), S663V (2127T>G 2128C>T 2129T>C), D665S (2133G>T 2134A>C 2135T>A), D666S (2136G>A 2137A>G), H668A (2142C>G 2143A>C 2144T>C), I671S (2152T>G 2153T>C), N673T (2158A>C 2159C>T), K674R (2160A>C 2161A>G 2162A>C), R676V (2166A>G 2167G>T), V677N (2169G>A 2170T>A 2171C>T), N678E (2172A>G 2174C>G), L680I (2178T>A 2180G>T), R682E (2184A>G 2185G>A 2186A>G), D685S (2193G>T 2194A>C 2195T>G), A686T (2196G>A), E687A (2200A>C), Q688G (2202C>G 2203A>G 2204G>A), R690A (2208C>G 2209G>C 2210A>C), V693I (2217G>A 2219A>C), Q694K (2220C>A 2222A>G), N695A (2223A>G 2224A>C), I701V (2241A>G 2243A>C), A710S (2268G>A 2269C>G 2270A>T), S711T (2271T>A), I713V (2277A>G 2279C>G), K716A (2286A>G 2287A>C 2288G>T), K717H (2289A>C 2291A>C), L718M (2292C>A 2294A>G), A719D (2296C>A 2297A>T), T722R (2305C>G 2306A>G), E723M (2307G>A 2308A>T), M724L (2310A>T), R725Y (2313A>T 2314G>A 2315G>C), T726N (2317C>A), F728R (2322T>A 2323T>G 2324T>A), S731R (2331T>A 2332C>G 2333T>A), C732A (2334T>G 2335G>C 2336C>A), I733L (2337A>C 2339T>G), D734E (2342T>A), V737A (2350T>C), I741M (2363A>G), F749H (2385T>C 2386T>A), C762_N763insRR (2426_2427insAGAAGG), N763K (2429C>G), K764M (2431A>T 2432A>G), S766T (2436T>A 2438A>T), P769T (2445C>A), K770_E771del (2448_2453delAAGGAA), I774V (2460A>G 2462C>A), K775Q (2463A>C), I776S (2466A>T 2467T>C 2468T>G), C777L (2469T>C 2470G>T), H779D (2475C>G 2477T>C), K780S (2478A>T 2479A>C 2480A>C), I781T (2482T>C 2483C>A), V782I (2484G>A), E783G (2488A>G 2489A>G), Y784R (2490T>A 2491A>G 2492T>A), I785M (2495C>G), K786Q (2496A>C 2498G>A), D787E (2501T>A), T788M (2503C>T 2504A>G), W790N (2508T>A 2509G>A 2510G>T), S791D (2511A>G 2512G>A 2513C>T), A795I (2523G>A 2524C>T 2525A>T), R797K (2530G>A), I806L (2556A>C), M807I (2561G>A), V812T (2574G>A 2575T>C), F813M (2577T>A 2579T>G), G815A (2584G>C), I816V (2586A>G 2588T>A), L820I (2598T>A 2600G>T), A821S (2601G>T 2603G>T), Q824K (2610C>A 2612G>A), N826S (2617A>G), S827L (2620C>T 2621T>G), F828I (2622T>A 2624C>T), N830T (2629A>C), A831S (2631G>T 2633A>C), S832Q (2634T>C 2635C>A), R834K (2641G>A), N852R (2695A>G 2696T>A), N852_K853insS (2696_2697insTCC), K853H (2697A>C 2699G>T), L854V (2700C>G 2702C>A), R855S (2703A>T 2704G>C), N856K (2708T>G), L857M (2709C>A 2711T>G), V859F (2715G>T), G860S (2718G>A 2720T>C), S865P (2733T>C), L866N (2736T>A 2737T>A 2738G>T), A867D (2740C>A 2741A>T), Q868_D869insG (2744_2745insGGA), D869K (2745G>A 2747T>A), A872S (2754G>T 2756T>C), E873R (2757G>A 2758A>G 2759A>G), Q879E (2775C>G 2777G>A), K882A (2784A>G 2785A>C 2786G>C), L890V (2808C>G 2810T>G), G891A (2812G>C 2813T>A), G904E (2851G>A 2852T>A), H914T (2880C>A 2881A>C 2882C>G), N915Q (2883A>C 2885T>G), A916L (2886G>C 2887C>T 2888T>C), T923S (2907A>T), A927E (2920C>A 2921C>A), N928H (2922A>C), R930V (2928C>G 2929G>T 2930A>C), S931C (2932C>G 2933T>C), K932Q (2934A>C 2936A>G), R938K (2953G>A), E939H (2955G>C 2957A>T), S940A (2958T>G 2960A>C), R941K (2962G>A), I942F (2964A>T), I943L (2967A>T 2969T>G), E944P (2970G>C 2971A>C 2972G>T), G945E (2974G>A 2975C>A), I948V (2982A>G 2984C>T), I949V (2985A>G 2987T>A), D955C (3003G>T 3004A>G), P961V (3021C>G 3022C>T 3023T>C), P968A (3042C>G 3044A>T), R969E (3045A>G 3046G>A), E972A (3055A>C 3056G>A), Q980E (3078C>G), C984V (3090T>G 3091G>T 3092T>C), T990K (3109C>A), R993E (3117C>G 3118G>A 3119T>G), L994K (3120C>A 3121T>A 3122T>A), G995Q (3123G>C 3124G>A 3125C>G), E998D (3134G>C), M1002Y (3144A>T 3145T>A 3146G>T), P1003S (3147C>T), V1012E (3175T>A 3176T>G), E1015T (3183G>A 3184A>C), E1017R (3189G>A 3190A>G 3191A>G), L1018M (3192C>A), N1019S (3196A>G), S1020T (3198T>A 3200C>A), S1021N (3201T>A 3202C>A 3203C>T), K1024H (3210A>C 3212G>C), K1024_I1025insS (3212_3213insTCA), C1026S (3216T>A 3218C>T), S1027A (3219T>G), Q1029E (3225C>G), S1030A (3228T>G 3230T>C), I1035V (3243A>G 3245A>C), G1037S (3249G>A 3251G>C), V1051I (3291G>A 3293T>C), V1057C (3309G>T 3310T>G 3311G>C), G1068E (3343G>A 3344C>A), V1069A (3346T>C 3347A>T), A1071T (3351G>A 3353T>A), R1073L (3357C>T 3358G>T 3359T>G), I1076V (3366A>G 3368C>A), S1077A (3369T>G 3371T>A), I1083V (3387A>G 3389C>A), E1086D (3398A>C), E1089Q (3405G>C 3407G>A), P1092A (3414C>G 3416A>T), T1093Q (3417A>C 3418C>A 3419T>A), A1098M (3432G>A 3433C>T 3434A>G), I1105L (3453A>T 3455T>G), S1108H (3462T>C 3463C>A 3464A>T), I1109V (3465A>G), Q1115A (3483C>G 3484A>C 3485G>T), E1118D (3494G>C), N1120K (3500C>G), Y1123A (3507T>G 3508A>C 3509C>T), Y1124F (3511A>T 3512C>T), L1132M (3534T>A), Q1146P (3577A>C), P1153E (3597C>G 3598C>A 3599T>A), E1155Q (3603G>C 3605G>A), I1160L (3618A>C 3620T>G), T1162S (3624A>T 3626A>T), H1164K (3630C>A 3632T>A), V1178I (3672G>A 3674A>T), M1179E (3675A>G 3676T>A), C1186S (3697G>C 3698T>A), G1187N (3699G>A 3700G>A 3701A>T), D1189E (3707T>G), D1191S (3711G>A 3712A>G), K1194R (3720A>C 3721A>G 3722A>G), L1195A (3723C>G 3724T>C 3725A>T), N1199G (3735A>G 3736A>G), C1202S (3745G>C 3746C>T), S1203A (3747T>G 3749G>C), L1204I (3750C>A 3752G>T), I1207V (3759A>G 3761C>T), M1208L (3762A>C 3764G>C), R1209K (3765C>A 3766G>A 3767T>G), K1212R (3775A>G), I1213V (3777A>G 3779C>G), R1217I (3789C>A 3790G>T 3791T>C), C1219T (3795T>A 3796G>C 3797T>A), E1220T (3798G>A 3799A>C), D1222C (3804G>T 3805A>G), V1225I (3813G>A 3815A>C), I1230V (3828A>G 3830A>T), C1233F (3838G>T), Y1235F (3844A>T), Y1236F (3847A>T), G1238S (3852G>T 3853G>C), M1252L (3894A>C), M1254L (3900A>T), M1256T (3907T>C 3908G>C), Q1263E (3927C>G), S1264A (3930T>G 3932C>A), T1265Q (3933A>C 3934C>A 3935T>G), G1266E (3937G>A 3938C>G), E1267S (3939G>A 3940A>G 3941A>T), G1268T (3942G>A 3943G>C 3944A>C), E1269Q (3945G>C 3947G>A), E1272L (3954G>C 3955A>T), D1276N (3966G>A 3968C>T), T1277S (3969A>T), E1281Q (3981G>C), Q1289E (4005C>G 4007G>A), I1294N (4021T>A), H1295Y (4023C>T), D1297E (4031T>A), L1298M (4032C>A), E1312D (4076G>C), R1318K (4093G>A 4094A>G), T1344C (4170A>T 4171C>G 4172A>T), L1348A (4182T>G 4183T>C 4184G>A), A1351V (4192C>T), I1352L (4194A>C 4196A>G), D1359N (4215G>A), T1361I (4222C>T 4223A>T), S1363H (4227T>C 4228C>A), L1367T (4239T>A 4240T>C 4241G>C), G1371Q (4251G>C 4252G>A), A1374S (4260G>A 4261C>G), V1393I (4317G>A 4319T>A), F1399L (4337T>A), E1405A (4354A>C), K1410A (4368A>G 4369A>C 4370G>T), P1414E (4380C>G 4381C>A 4382T>G), V1437L (4449G>C 4451G>C), A1477K (4569G>A 4570C>A 4571T>A), R1479K (4576G>A), I1481L (4581A>T 4583A>G), P1485S (4593C>T), Q1490D (4608C>G 4610G>C), L1506M (4656T>A 4658A>G), S1512E (4674T>G 4675C>A 4676T>G), N1515A (4683A>G 4684A>C 4685T>A), H1518Q (4694T>A), L1522Q (4705T>A 4706G>A), S1534N (4740T>A 4741C>A 4742G>C), D1536N (4746G>A), E1539D (4757G>T), Q1541K (4761C>A), H1563Q (4829C>A), M1564L (4830A>C 4832G>A), I1573V (4857A>G 4859T>G), P1583V (4887C>G 4888C>T 4889A>T), T1588I (4903C>T 4904T>C), Q1589T (4905C>A 4906A>C 4907G>T), M1592Y (4914A>T 4915T>A 4916G>T), I1599V (4935A>G), H1604Q (4952T>G), R1607Q (4960G>A 4961T>G), A1609V (4966C>T), K1612T (4975A>C 4976G>A), V1614T (4980G>A 4981T>C 4982C>A), L1615D (4983T>G 4984T>A 4985A>C), D1616N (4986G>A), T1619K (4996C>A 4997G>A), L1621I (5001T>A 5003G>A), D1622M (5004G>A 5005A>T 5006T>G), E1623Q (5007G>C 5009A>G), F1626L (5018C>G), R1627P (5019A>C 5020G>C), A1628I (5022G>A 5023C>T 5024T>C), I1630H (5028A>C 5029T>A 5030C>T), L1635A (5043T>G 5044T>C 5045G>T), V1637Q (5049G>C 5050T>A 5051G>A), Q1639S (5055C>A 5056A>G 5057G>T), I1642L (5064A>C 5066T>A), A1643Q (5067G>C 5068C>A), K1644A (5070A>G 5071A>C 5072G>C), S1650N (5089G>A 5090T>C), A1651I (5091G>A 5092C>T 5093C>A), L1652G (5094C>G 5095T>G), T1653A (5097A>G 5099A>T), I1654L (5100A>C 5102T>G), C1655L (5103T>C 5104G>T 5105T>G), M1657L (5109A>C), A1658S (5112G>A 5113C>G 5114A>C), A1660D (5119C>A 5120A>T), S1663T (5127T>A 5129C>T), T1665V (5133A>G 5134C>T), D1668P (5142G>C 5143A>C 5144C>A), A1669E (5146C>A), E1671K (5151G>A 5153G>A), T1674E (5160A>G 5161C>A), N1675D (5163A>G), E1677S (5169G>T 5170A>C 5171G>A), R1678Q (5173G>A 5174T>A), D1679G (5176A>G), L1680V (5178C>G 5180G>C), N1681H (5181A>C), W1682E (5184T>G 5185G>A 5186G>A), A1684C (5190G>T 5191C>G 5192A>T), P1685S (5193C>T 5195A>T), N1686G (5196A>G 5197A>G 5198C>A), I1687V (5199A>G), G1688F (5202G>T 5203G>T 5204T>C), A1692S (5214G>T 5216A>G), A1695E (5224C>A), A1696V (5227C>T 5228C>T), H1697N (5229C>A 5231C>T), Y1698H (5232T>C 5234C>T), T1699K (5236C>A 5237G>A), G1703D (5248G>A 5249G>C), Y1704T (5250T>A 5251A>C 5252C>A), I1706L (5256A>C), S1715T (5283T>A 5285C>A), N1716T (5287A>C 5288T>A), S1737A (5349T>G), M1740K (5359T>A), A1742N (5364G>A 5365C>A 5366C>T), K1744Q (5370A>C 5372G>A), N1745S (5374A>G), V1746R (5376G>C 5377T>G 5378G>C), A1748S (5382G>T), G1749M (5385G>A 5386G>T 5387A>G), H1751I (5391C>A 5392A>T), V1752G (5395T>G), A1753T (5397G>A 5399A>C), T1754Q (5400A>C 5401C>A 5402C>A), N1755K (5405T>A), V1759I (5415G>A 5417G>T), G1767S (5439G>A 5441C>T), M1774F (5460A>T 5462G>T), A1777S (5469G>T), R1779N (5476G>A 5477G>T), C1782R (5484T>A 5486T>A), V1783A (5488T>C 5489G>T), D1785Q (5493G>C 5495C>A), I1786L (5496A>C 5498A>T), I1793L (5517A>T 5519T>G), G1794A (5521G>C 5522A>G), N1795G (5523A>G 5524A>G 5525T>C), V1796L (5526G>T), T1801Q (5541A>C 5542C>A 5543C>G), V1807I (5559G>A 5561G>T), Q1812T (5574C>A 5575A>C), C1813_V1814insV (5579_5580insGTC), P1815Q (5584C>A 5585T>G), H1816K (5586C>A 5588T>A), G1817S (5589G>A 5591A>T), D1818C (5592G>T 5593A>G), V1820T (5598G>A 5599T>C 5600G>A), S1823A (5607A>G 5608G>C), G1824S (5610G>A), A1825G (5614C>G 5615A>G), S1826del (5616_5618delTCG), L1827R (5620T>G), H1828D (5622C>G 5624C>T), S1829G (5625A>G 5627T>A), S1830M (5629G>T 5630T>G), F1831L (5633T>G), M1832N (5635T>A 5636G>C), Q1834H (5642A>T), Q1834_G1835insHA (5642_5643insCATGCT), G1835A (5644G>C 5645G>T), F1836Y (5647T>A), E1837A (5650A>C 5651A>C), I1844V (5670A>G 5672A>G), H1845G (5673C>G 5674A>G 5675T>C), G1846C (5676G>T), D1847S (5679G>T 5680A>C 5681T>A), L1848I (5682T>A 5684A>T), A1849S (5685G>T), K1850R (5689A>G), Q1851K (5691C>A), T1852M (5695C>T 5696A>G), M1854V (5700A>G), M1856L (5706A>T), V1859T (5715G>A 5716T>C 5717G>T), H1863M (5727C>A 5728A>T 5729T>G), G1867A (5740G>C), N1868D (5742A>G), F1869C (5746T>G 5747T>C), Q1872R (5754C>A 5755A>G 5756G>A), D1873N (5757G>A) | | | | | | | | |

| | Begin | End | Coverage | Score | Concordance | Matches | Identities | I/D/M/F* | Stop Codons |
|---|---|---|---|---|---|---|---|---|---|
| **NT** | **35** | **5769** | **79.3%** | **3466** | **30.8%** | **5682 (98.6%)** | **3757 (65.2%)** | **28/53** | |

Codon mutations:

GCT2GCA (146T>A), AAG5AAA (155G>A), ATG6ATC (158G>C), TTC7GTT (159T>G 161C>T), TCC8_AGT17del (162_191delTCCAAGGTTGCTGGTGTTGATGACGCCAGT), TCC20GCT (198T>G 200C>T), ACC21ACT (203C>T), ACT22CCT (204A>C), TCA23TCT (209A>T), TTC24TTT (212C>T), TTT25CTT (213T>C), GGT26TCT (216G>T 217G>C), GAA27GAG (221A>G), CTT28TTG (222C>T 224T>G), TCA29TCT (227A>T), AAT30CAG (228A>C 230T>G), TCT31ACA (231T>A 233T>A), ATT32GTG (234A>G 236T>G), TCC33TCA (239C>A), AAA34TCA (240A>T 241A>C), TTT35CTG (243T>C 245T>G), TCA36TCC (248A>C), AAA37AAG (251A>G), GCG38ACA (252G>A 254G>A), GTT39ATG (255G>A 257T>G), AGT40AAG (259G>A 260T>G), AAT41AGT (262A>G), ATT42ATG (266T>G), ACC43AGT (268C>G 269C>T), ACT44AAA (271C>A 272T>A), ATG45ATA (275G>A), TCA46TCT (278A>T), CAG47GAG (279C>G), AAG48AAA (284G>A), ATT49TTC (285A>T 287T>C), GAT51GGC (292A>G 293T>C), CTT53ATT (297C>A), GAG54GAA (302G>A), GAT55GAA (305T>A), CTG56TTA (306C>T 308G>A), CCA58CCT (314A>T), TCT59TCA (317T>A), GTG60ATT (318G>A 320G>T), ACT61GTT (321A>G 322C>T), GAT62AAA (324G>A 326T>A), GTG63ACT (327G>A 328T>C 329G>T), TCC64GCA (330T>G 332C>A), ACA65GAT (333A>G 334C>A 335A>T), TCT66TCC (338T>C), GTG68AAT (342G>A 343T>A 344G>T), GCC69AGT (345G>A 346C>G 347C>T), ACT70GTT (348A>G 349C>T), TCA73TCT (359A>T), CTT74GTA (360C>G 362T>A), AAT75GAA (363A>G 365T>A), GTT77TTA (369G>T 371T>A), TTG78TTA (374G>A), GAC79GGG (376A>G 377C>G), GGT82GCC (385G>C 386T>C), GCA83ACG (387G>A 389A>G), CTT84GTT (390C>G), ATA85ATT (395A>T), GAT86GAA (398T>A), CCA87CCT (401A>T), CTT89GTT (405C>G), GCG91GGT (412C>G 413G>T), TAC92ATC (414T>A 415A>T), TCT93AAT (417T>A 418C>A), TTG95TTC (425G>C), TCC96AAA (426T>A 427C>A 428C>A), TAC99TAT (437C>T), AAA100AGA (439A>G), TCT101ACC (441T>A 443T>C), ATC102ATA (446C>A), AAG103ATT (448A>T 449G>T), TTA107CTG (459T>C 461A>G), TTG109GTT (465T>G 467G>T), TTT110GTT (468T>G), GAG111TTG (471G>T 472A>T), AGT112GGC (474A>G 476T>C), TTG113ATC (477T>A 479G>C), TCC114AAA (480T>A 481C>A 482C>A), AAA116AAT (488A>T), GTT117TTT (489G>T), AAA118ATT (493A>T 494A>T), TTA119AGA (495T>A 496T>G), TTT121TAT (502T>A), GCT122TCA (504G>T 506T>A), GTG124CTG (510G>C), TTA125TTG (515A>G), AAT126GAA (516A>G 518T>A), AAA127ACA (520A>C), AGT128ACA (523G>C 524T>A), GAA129GGT (526A>G 527A>T), GAT130GAA (530T>A), GTA131TAT (531G>T 532T>A 533A>T), ATA133GTT (537A>G 539A>T), GTT134GTA (542T>A), GTA135GTT (545A>T), TAT136ATA (546T>A 547A>T 548T>A), GGT137GCT (550G>C), CTT139CTA (557T>A), GTA140GTC (560A>C), GCT142TCC (564G>T 566T>C), ATA143CTT (567A>C 569A>T), ACA144TCA (570A>T), TTG145CTA (573T>C 575G>A), TTA146TTG (578A>G), GTG147TTA (579G>T 581G>A), CTT148TTA (582C>T 584T>A), CTA149CAA (586T>A), TTT150ATA (588T>A 590T>A), GTT151ATC (591G>A 593T>C), AGT154GTA (600A>G 601G>T 602T>A), GTT157GTA (611T>A), GAG158GAA (614G>A), GGT159ATG (615G>A 616G>T 617T>G), TCT161AGA (621T>A 622C>G 623T>A), AAC162AGT (625A>G 626C>T), ACT163GTA (627A>G 628C>T 629T>A), GTC164GTT (632C>T), AAA165TGT (633A>T 634A>G 635A>T), ATC167ATA (641C>A), TTG169CTA (645T>C 647G>A), ATA170GTT (648A>G 650A>T), GTC171GTA (653C>A), GGA172GGT (656A>T), ATG174TTC (660A>T 662G>C), TTT175TTC (665T>C), TCA176TCT (668A>T), AGT177TCT (669A>T 670G>C), CTA178CTG (674A>G), TAC179TAT (677C>T), TTA181ATT (681T>A 683A>T), GAT182AGT (684G>A 685A>G), CCA185CCC (695A>C), AAA186CTC (696A>C 697A>T 698A>C), GCC188TCT (702G>T 704C>T), GAA189GAG (707A>G), CGA190AAA (708C>A 709G>A), TTC191TTT (713C>T), ACC192TCT (714A>T 716C>T), ATG193CAG (717A>C 718T>A), GTG194ATG (720G>A), GCC195GCA (725C>A), CAA196CAG (728A>G), AGT198TCT (732A>T 733G>C), ATG199GTG (735A>G), CTT200CTG (740T>G), GGG202GGT (746G>T), GAA203GAG (749A>G), TCA204AGC (750T>A 751C>G 752A>C), GTT205TTA (753G>T 755T>A), TCA206AAT (756T>A 757C>A 758A>T), ACG210ACC (770G>C), TCT211AGT (771T>A 772C>G), CAG212CAA (776G>A), ATG214CTG (780A>C), TCC215TCT (785C>T), ACA216TCT (786A>T 788A>T), ATA217TTG (789A>T 791A>G), TTG218ATA (792T>A 794G>A), GCT219GCA (797T>A), GGG221GGC (803G>C), ATT222ATC (806T>C), TCG223TCA (809G>A), GTG226GTA (818G>A), TTC227TTT (821C>T), ATT228GTG (822A>G 824T>G), GTG230GTA (830G>A), AGA233AGG (839A>G), TTG237CTT (849T>C 851G>T), AGC238ACA (853G>C 854C>A), CCA240CCG (860A>G), TTG241CTA (861T>C 863G>A), CTC245TTG (873C>T 875C>G), TCG247TCA (881G>A), ACA248GCG (882A>G 884A>G), GGA249GGC (887A>C), GGT251GGA (893T>A), CAG253CAA (899G>A), CTA258CTC (914A>C), ACT260CAA (918A>C 919C>A 920T>A), CTT261TTG (921C>T 923T>G), TAC262TAT (926C>T), CGT263AAG (927C>A 928G>A 929T>G), AAT264AAC (932T>C), AAA266AAG (938A>G), ACC269ACG (947C>G), TCA274TCT (962A>T), GTT276GTG (968T>G), CTT277TTG (969C>T 971T>G), ATA279ATT (977A>T), ATA280GTT (978A>G 980A>T), GTC281GCT (982T>C 983C>T), GGC282GAT (985G>A 986C>T), ACC283ATA (988C>T 989C>A), TTT284TTC (992T>C), GGA285GGG (995A>G), TTC286TTT (998C>T), AAT288AAC (1004T>C), GTG290GTT (1010G>T), TTG291CTG (1011T>C), TCA292TCT (1016A>T), GCT293GCA (1019T>A), ATA294ATT (1022A>T), ACG297ACA (1031G>A), CTG298ATT (1032C>A 1034G>T), GAT301GAC (1043T>C), CTG302CTC (1046G>C), TTC303TTT (1049C>T), CAG307ACT (1059C>A 1060A>C 1061G>T), GAG308GAA (1064G>A), GAT310TCT (1068G>T 1069A>C), TGT313GCT (1077T>G 1078G>C), ACG317ACA (1091G>A), CGC318AGG (1092C>A 1094C>G), GAG320GAA (1100G>A), GCC323GCT (1109C>T), AAG325CCT (1113A>C 1114A>C 1115G>T), TTC327TTT (1121C>T), CCT328TCA (1122C>T 1124T>A), ACC329ATC (1126C>T), CTG334CTT (1142G>T), AGG335AAA (1144G>A 1145G>A), GAA336GAG (1148A>G), GAA337CAG (1149G>C 1151A>G), GCT338GCA (1154T>A), ATC341ATA (1163C>A), GCT343GCC (1169T>C), TAC344TAT (1172C>T), GCA346GCT (1178A>T), CCA349CCT (1187A>T), GTT350GTG (1190T>G), TTC353TTT (1199C>T), ATG354TTG (1200A>T), AGC355AGT (1205C>T), AGG357AGA (1211G>A), AGT359TCT (1215A>T 1216G>C), GCC361ACA (1221G>A 1223C>A), GCG363GTG (1228C>T), CAG364CAA (1232G>A), TTA365TTG (1235A>G), GAA366GAT (1238A>T), GTT368CTC (1242G>C 1244T>C), TAT369ATA (1245T>A 1246A>T 1247T>A), GCT370GCA (1250T>A), AGT372AAC (1255G>A 1256T>C), AGG374CAA (1260A>C 1261G>A 1262G>A), CAC375CAT (1265C>T), ATG376ACA (1267T>C 1268G>A), GGC377GGA (1271C>A), GGC379GGT (1277C>T), TAT381TAC (1283T>C), CGT382AGA (1284C>A 1286T>A), CCT385CCC (1295T>C), GTA388GTT (1304A>T), TTC391TTT (1313C>T), GGG392GGT (1316G>T), TCC393GAG (1317T>G 1318C>A 1319C>G), GGG395GGA (1325G>A), TGT396TGC (1328T>C), GGT397GGC (1331T>C), TCT399TCC (1337T>C), ACA400ACG (1340A>G), TCC401AGC (1341T>A 1342C>G), CGC403CGA (1349C>A), TTA404CTG (1350T>C 1352A>G), TTT405TTC (1355T>C), ATT406ATC (1358T>C), GTG409GTA (1367G>A), GAC411GAT (1373C>T), AGA412GCC (1374A>G 1375G>C 1376A>C), GGT414CAA (1380G>C 1381G>A 1382T>A), CCA416CCC (1388A>C), AAA417AAG (1391A>G), CTC418CTG (1394C>G), AAT419AAC (1397T>C), CGG420AGG (1398C>A), CTA421TTG (1401C>T 1403A>G), GTA424GTT (1412A>T), AGG427CGC (1419A>C 1421G>C), GCA429GCT (1427A>T), TAT430TAC (1430T>C), TCC432TCA (1436C>A), AAC433AAT (1439C>T), CTG441CTC (1463G>C), GAC444GAT (1472C>T), GGA446GGT (1478A>T), TTG448CTG (1482T>C), CGA449CGG (1487A>G), GGG451GGA (1493G>A), GGG453GGC (1499G>C), CAG454CAA (1502G>A), CAA456CAG (1508A>G), GAT457GAC (1511T>C), AAA459AAG (1517A>G), GAT460GAC (1520T>C), CTG461CTC (1523G>C), GAT463GAC (1529T>C), TCA466TCC (1538A>C), CCC471CCA (1553C>A), TTA472TTG (1556A>G), CCA473CCC (1559A>C), GCC475GCT (1565C>T), GTT477GTA (1571T>A), GAG478GAA (1574G>A), GGC481GGG (1583C>G), AGG482CGG (1584A>C), ACC485ACA (1595C>A), TCC486TCA (1598C>A), TAT488TAC (1604T>C), ACA492ACC (1616A>C), AAT494AAC (1622T>C), CTG495TTG (1623C>T), ATC496GTT (1626A>G 1628C>T), GCT498GCA (1634T>A), CCT499CCA (1637T>A), GCC500GCA (1640C>A), TGT502TGC (1646T>C), GGT503GGG (1649T>G), CTT504CTG (1652T>G), CCA507CCT (1661A>T), GAG511GAA (1673G>A), CGG512AGG (1674C>A), AGA513AGG (1679A>G), AGG514CGG (1680A>C), ATC516ATT (1688C>T), CTG517TTG (1689C>T), GAG519GAA (1697G>A), TGT520TGC (1700T>C), ATG521CGG (1701A>C 1702T>G), AAG522AAA (1706G>A), GAG523GTG (1708A>T), GGC524GGG (1712C>G), GAG525GAA (1715G>A), TTT526TTC (1718T>C), TCC527TCA (1721C>A), ACT528ACA (1724T>A), GAA529GAG (1727A>G), GAT530AAC (1728G>A 1730T>C), CCT531CCA (1733T>A), ACG532ACA (1736G>A), GGA533GCA (1738G>C), AGA536AGC (1748A>C), TTT537TTC (1751T>C), ATA539ATC (1757A>C), GTG540ATT (1758G>A 1760G>T), GAG541GGA (1762A>G 1763G>A), AGA543AGG (1769A>G), AGA544AGG (1772A>G), GCT547GCC (1781T>C), ATC548ATA (1784C>A), ACT549ACA (1787T>A), AGG551AGA (1793G>A), AAT552GAC (1794A>G 1796T>C), TTG553CTC (1797T>C 1799G>C), ACC554ACT (1802C>T), AGT556TCT (1806A>T 1807G>C), GAT557GAC (1811T>C), CTT558TTG (1812C>T 1814T>G), GAG560GAA (1820G>A), GTA563GTG (1829A>G), AAA565AAG (1835A>G), TGT566TGC (1838T>C), CAG567AAG (1839C>A), GTA568GTC (1844A>C), CAT569CAC (1847T>C), GCA570CAC (1848G>C 1849C>A 1850A>C), GTA572ATT (1854G>A 1856A>T), TCA573GCA (1857T>G), AAG574ACC (1861A>C 1862G>C), TTG576CTC (1866T>C 1868G>C), TTT577TAC (1870T>A 1871T>C), GAA578AAC (1872G>A 1874A>C), GCT579CAG (1875G>C 1876C>A 1877T>G), GAA580TCA (1878G>T 1879A>C), TCT581ACA (1881T>A 1883T>A), GGC582GGT (1886C>T), ATA583TGT (1887A>T 1888T>G 1889A>T), AGT584GAA (1890A>G 1891G>A 1892T>A), CCT585GAG (1893C>G 1894C>A 1895T>G), AAG586AAA (1898G>A), ATA587GTT (1899A>G 1901A>T), GCT588GCC (1904T>C), GTT590GTG (1910T>G), CAA591GAA (1911C>G), GCA594GTT (1921C>T 1922A>T), GAT595GAA (1925T>A), GAT596GAA (1928T>A), ATA598ATG (1934A>G), GCG599ACA (1935G>A 1937G>A), TCT600TCC (1940T>C), GTG601TTT (1941G>T 1943G>T), GAT602CAT (1944G>C), GGG603GAT (1948G>A 1949G>T), AGA605CAA (1953A>C 1954G>A), TTG606ATA (1956T>A 1958G>A), CGC607AGC (1959C>A), ACC608TTT (1962A>T 1963C>T 1964C>T), AAG609GAC (1965A>G 1967G>C), CAA610GAA (1968C>G), GAT611GTT (1972A>T), GAA612AGC (1974G>A 1975A>G 1976A>C), CCA613CCC (1979A>C), ATA614AGT (1981T>G 1982A>T), TTG615GTT (1983T>G 1985G>T), GTG616_CCC617del (1986_1991delGTGCCC), ACT618ACA (1994T>A), GTG620CTC (1998G>C 2000G>C), AGT621TCT (2001A>T 2002G>C), GAA622GAG (2006A>G), AAT623GAT (2007A>G), ATC627GGA (2019A>G 2020T>G 2021C>A), GCA629TCA (2025G>T), AGA630CAG (2028A>C 2029G>A 2030A>G), GAG631GAA (2033G>A), CTC632CTA (2036C>A), ACG633ACT (2039G>T), GTA634GTG (2042A>G), GAG635GAA (2045G>A), TTG637CTG (2049T>C), AAA638AAG (2054A>G), GCA640GCT (2060A>T), TAC641TAT (2063C>T), CAG642CAA (2066G>A), GGC643GGA (2069C>A), AGT644GCA (2070A>G 2071G>C 2072T>A), TTG645TTA (2075G>A), GAC646AAC (2076G>A), CCT647CCA (2081T>A), GAG648TTT (2082G>T 2083A>T 2084G>T), CTT650TAT (2088C>T 2089T>A), CCT652CCC (2096T>C), GAC654GAT (2102C>T), CAT655GCT (2103C>G 2104A>C), CAT656TCA (2106C>T 2107A>C 2108T>A), GCC659GCA (2117C>A), ATG660ATT (2120G>T), TAT661TTT (2122A>T), TCT663GTC (2127T>G 2128C>T 2129T>C), CTT664CTG (2132T>G), GAT665TCA (2133G>T 2134A>C 2135T>A), GAT666AGT (2136G>A 2137A>G), CAT668GCC (2142C>G 2143A>C 2144T>C), ATT671AGC (2152T>G 2153T>C), TTC672TTT (2156C>T), AAC673ACT (2158A>C 2159C>T), AAA674CGC (2160A>C 2161A>G 2162A>C), AGA676GTA (2166A>G 2167G>T), GTC677AAT (2169G>A 2170T>A 2171C>T), AAC678GAG (2172A>G 2174C>G), TTG680ATT (2178T>A 2180G>T), TAT681TAC (2183T>C), AGA682GAG (2184A>G 2185G>A 2186A>G), GGA683GGG (2189A>G), GCA684GCT (2192A>T), GAT685TCG (2193G>T 2194A>C 2195T>G), GCT686ACT (2196G>A), GAA687GCA (2200A>C), CAG688GGA (2202C>G 2203A>G 2204G>A), CGA690GCC (2208C>G 2209G>C 2210A>C), GTA693ATC (2217G>A 2219A>C), CAA694AAG (2220C>A 2222A>G), AAC695GCC (2223A>G 2224A>C), ATT696ATC (2228T>C), CCT697CCA (2231T>A), ATA701GTC (2241A>G 2243A>C), AAG705AAA (2255G>A), AGT706AGC (2258T>C), GCA710AGT (2268G>A 2269C>G 2270A>T), TCA711ACA (2271T>A), ATC713GTG (2277A>G 2279C>G), TCT714TCA (2282T>A), GAG715GAA (2285G>A), AAG716GCT (2286A>G 2287A>C 2288G>T), AAA717CAC (2289A>C 2291A>C), CTA718ATG (2292C>A 2294A>G), GCA719GAT (2296C>A 2297A>T), GTG720GTA (2300G>A), CAG721CAA (2303G>A), ACA722AGG (2305C>G 2306A>G), GAG723ATG (2307G>A 2308A>T), ATG724TTG (2310A>T), AGG725TAC (2313A>T 2314G>A 2315G>C), ACT726AAT (2317C>A), GGA727GGC (2321A>C), TTT728AGA (2322T>A 2323T>G 2324T>A), TCT731AGA (2331T>A 2332C>G 2333T>A), TGC732GCA (2334T>G 2335G>C 2336C>A), ATT733CTG (2337A>C 2339T>G), GAT734GAA (2342T>A), GCA735GCT (2345A>T), GTT737GCT (2350T>C), ACC739ACT (2357C>T), CTC740TTA (2358C>T 2360C>A), ATA741ATG (2363A>G), ATC743ATT (2369C>T), GAG744GAA (2372G>A), CAG745CAA (2375G>A), CCA747CCT (2381A>T), TTT749CAT (2385T>C 2386T>A), GTT750GTC (2390T>C), CAG751CAA (2393G>A), TTT752TTC (2396T>C), TTA753CTA (2397T>C), TAT754TAC (2402T>C), AGA759CGG (2415A>C 2417A>G), TGC762_AAC763insAGAAGG (2426_2427insAGAAGG), AAC763AAG (2429C>G), AAA764ATG (2431A>T 2432A>G), GTA765GTC (2435A>C), TCA766ACT (2436T>A 2438A>T), AAA767AAG (2441A>G), GAG768GAA (2444G>A), CCA769ACA (2445C>A), AAG770_GAA771del (2448_2453delAAGGAA), TCA772TCT (2456A>T), ATC774GTA (2460A>G 2462C>A), AAA775CAA (2463A>C), ATT776TCG (2466A>T 2467T>C 2468T>G), TGC777CTC (2469T>C 2470G>T), TAC778TAT (2474C>T), CAT779GAC (2475C>G 2477T>C), AAA780TCC (2478A>T 2479A>C 2480A>C), ATC781ACA (2482T>C 2483C>A), GTT782ATT (2484G>A), GAA783GGG (2488A>G 2489A>G), TAT784AGA (2490T>A 2491A>G 2492T>A), ATC785ATG (2495C>G), AAG786CAA (2496A>C 2498G>A), GAT787GAA (2501T>A), ACA788ATG (2503C>T 2504A>G), TGG790AAT (2508T>A 2509G>A 2510G>T), AGC791GAT (2511A>G 2512G>A 2513C>T), CTT792CTG (2516T>G), TAT794TAC (2522T>C), GCA795ATT (2523G>A 2524C>T 2525A>T), CTG796TTG (2526C>T), AGA797AAA (2530G>A), CTC799CTT (2537C>T), GGT803GGG (2549T>G), ATA806CTA (2556A>C), ATG807ATA (2561G>A), CTT809CTC (2567T>C), GGA811GGG (2573A>G), GTT812ACT (2574G>A 2575T>C), TTT813ATG (2577T>A 2579T>G), GGA814GGT (2582A>T), GGC815GCC (2584G>C), ATT816GTA (2586A>G 2588T>A), ACA817ACT (2591A>T), TTT819TTC (2597T>C), TTG820ATT (2598T>A 2600G>T), GCG821TCT (2601G>T 2603G>T), CAG824AAA (2610C>A 2612G>A), AGT825AGC (2615T>C), AAC826AGC (2617A>G), TCT827TTG (2620C>T 2621T>G), TTC828ATT (2622T>A 2624C>T), AAT830ACT (2629A>C), GCA831TCC (2631G>T 2633A>C), TCG832CAG (2634T>C 2635C>A), AGA834AAA (2641G>A), GTG837GTT (2651G>T), TCC841TCG (2663C>G), AGT843AGC (2669T>C), TCC845TCA (2675C>A), AGT846AGC (2678T>C), GGT851GGA (2693T>A), AAT852AGA (2695A>G 2696T>A), AAT852_AAG853insTCC (2696_2697insTCC), AAG853CAT (2697A>C 2699G>T), CTC854GTA (2700C>G 2702C>A), AGA855TCA (2703A>T 2704G>C), AAT856AAG (2708T>G), CTT857ATG (2709C>A 2711T>G), CTC858CTG (2714C>G), GTT859TTT (2715G>T), GGT860AGC (2718G>A 2720T>C), TCT863TCA (2729T>A), TCA865CCA (2733T>C), TTG866AAT (2736T>A 2737T>A 2738G>T), GCA867GAT (2740C>A 2741A>T), CAA868CAG (2744A>G), CAA868_GAT869insGGA (2744_2745insGGA), GAT869AAA (2745G>A 2747T>A), GCT872TCC (2754G>T 2756T>C), GAA873AGG (2757G>A 2758A>G 2759A>G), GGA875GGT (2765A>T), GTA877GTG (2771A>G), AAT878AAC (2774T>C), CAG879GAA (2775C>G 2777G>A), AGC880TCT (2778A>T 2779G>C 2780C>T), CTC881CTT (2783C>T), AAG882GCC (2784A>G 2785A>C 2786G>C), GTG886GTA (2798G>A), TTA888CTA (2802T>C), AGA889AGG (2807A>G), CTT890GTG (2808C>G 2810T>G), GGT891GCA (2812G>C 2813T>A), GTG894GTT (2822G>T), AGA897CGT (2829A>C 2831A>T), GGC898GGA (2834C>A), ACC899ACT (2837C>T), TGT902TGC (2846T>C), TCT903AGC (2847T>A 2848C>G 2849T>C), GGT904GAA (2851G>A 2852T>A), ACC908ACT (2864C>T), GCT910GCA (2870T>A), CAT911CAC (2873T>C), GCA912GCT (2876A>T), CAC914ACG (2880C>A 2881A>C 2882C>G), AAT915CAG (2883A>C 2885T>G), GCT916CTC (2886G>C 2887C>T 2888T>C), CGA917CGT (2891A>T), GGT919GGA (2897T>A), ACT920ACC (2900T>C), ACA923TCA (2907A>T), ATC925ATT (2915C>T), GCC927GAA (2920C>A 2921C>A), AAT928CAT (2922A>C), CGA930GTC (2928C>G 2929G>T 2930A>C), TCT931TGC (2932C>G 2933T>C), AAA932CAG (2934A>C 2936A>G), GTT933GTG (2939T>G), AAC936AAT (2948C>T), AGA938AAA (2953G>A), GAA939CAT (2955G>C 2957A>T), TCA940GCC (2958T>G 2960A>C), AGG941AAG (2962G>A), ATT942TTT (2964A>T), ATT943TTG (2967A>T 2969T>G), GAG944CCT (2970G>C 2971A>C 2972G>T), GGC945GAA (2974G>A 2975C>A), GAC947GAT (2981C>T), ATC948GTT (2982A>G 2984C>T), ATT949GTA (2985A>G 2987T>A), CTG950TTA (2988C>T 2990G>A), TTG951CTG (2991T>C), CGC952AGG (2994C>A 2996C>G), GGT954GGC (3002T>C), GAT955TGT (3003G>T 3004A>G), CCA956CCT (3008A>T), GAT957GAC (3011T>C), GGT958GGG (3014T>G), CCA959CCT (3017A>T), AAG960AAA (3020G>A), CCT961GTC (3021C>G 3022C>T 3023T>C), CGT964AGA (3030C>A 3032T>A), AAA965AAG (3035A>G), TTC967TTT (3041C>T), CCA968GCT (3042C>G 3044A>T), AGA969GAA (3045A>G 3046G>A), GAG972GCA (3055A>C 3056G>A), AAG977AAA

(3071G>A), GGC978GGA (3074C>A), TCA979TCT (3077A>T), CAA980GAA (3078C>G), GGA981GGC (3083A>C), TGC983TGT (3089C>T), TGT984GTC (3090T>G 3091G>T 3092T>C), GCT986GCA (3098T>A), GTT987GTG (3101T>G), GCG988GCA (3104G>A), TCT989TCA (3107T>A), ACA990AAA (3109C>A), CCA992CCT (3116A>T), CGT993GAG (3117C>G 3118G>A 3119T>G), CTT994AAA (3120C>A 3121T>A 3122T>A), GGC995CAG (3123G>C 3124G>A 3125C>G), AAT996AAC (3128T>C), TTA997TTG (3131A>G), GAG998GAC (3134G>C), ATG1002TAT (3144A>T 3145T>A 3146G>T), CCA1003TCA (3147C>T), TCA1007TCC (3161A>C), AAG1008AAA (3164G>A), TAC1010TAT (3170C>T), GTT1012GAG (3175T>A 3176T>G), GTG1014GTC (3182G>C), GAA1015ACA (3183G>A 3184A>C), TAT1016TAC (3188T>C), GAA1017AGG (3189G>A 3190A>G 3191A>G), CTG1018ATG (3192C>A), AAC1019AGC (3196A>G), TCC1020ACA (3198T>A 3200C>A), TCC1021AAT (3201T>A 3202C>A 3203C>T), AGT1022TCA (3204A>T 3205G>C 3206T>A), AAG1024CAC (3210A>C 3212G>C), AAG1024_ATT1025insTCA (3212_3213insTCA), ATT1025ATC (3215T>C), TGC1026AGT (3216T>A 3218C>T), TCT1027GCT (3219T>G), CAA1029GAA (3225C>G), TCT1030GCC (3228T>G 3230T>C), CAC1034CAT (3242C>T), ATA1035GTC (3243A>G 3245A>C), GGG1037AGC (3249G>A 3251G>C), GAC1041GAT (3263C>T), GGC1043GGA (3269C>A), ACG1044ACT (3272G>T), CTG1047TTA (3279C>T 3281G>A), TTA1048TTG (3284A>G), AGT1050TCT (3288A>T 3289G>C), GTT1051ATC (3291G>A 3293T>C), CAG1052CAA (3296G>A), AAT1053AAC (3299T>C), CCT1056CCA (3308T>A), GTG1057TGC (3309G>T 3310T>G 3311G>C), ATC1058ATT (3314C>T), GGC1060GGT (3320C>T), ATC1061ATT (3323C>T), CAC1062CAT (3326C>T), GCA1064GCT (3332A>T), TCA1065TCC (3335A>C), GGC1068GAA (3343G>A 3344C>A), GTA1069GCT (3346T>C 3347A>T), GCT1071ACA (3351G>A 3353T>A), GAA1072GAG (3356A>G), CGT1073TTG (3357C>T 3358G>T 3359T>G), ATC1076GTA (3366A>G 3368C>A), TCT1077GCA (3369T>G 3371T>A), GCT1080GCC (3380T>C), ACA1081ACT (3383A>T), ATC1083GTA (3387A>G 3389C>A), GAA1086GAC (3398A>C), CAA1087CAG (3401A>G), CTA1088CTG (3404A>G), GAG1089CAA (3405G>C 3407G>A), CCA1092GCT (3414C>G 3416A>T), ACT1093CAA (3417A>C 3418C>A 3419T>A), CCG1095CCA (3425G>A), GTC1096GTT (3428C>T), GCA1098ATG (3432G>A 3433C>T 3434A>G), GCA1099GCC (3437A>C), GTA1101GTG (3443A>G), CGC1102AGA (3444C>A 3446C>A), GAT1104GAC (3452T>C), ATT1105TTG (3453A>T 3455T>G), CTG1106CTC (3458G>C), AAG1107AAA (3461G>A), TCA1108CAT (3462T>C 3463C>A 3464A>T), ATT1109GTT (3465A>G), AGA1110CGA (3468A>C), AGC1111TCA (3471A>T 3472G>C 3473C>A), ACA1114ACT (3482A>T), CAG1115GCT (3483C>G 3484A>C 3485G>T), CTT1116CTC (3488T>C), GAG1118GAC (3494G>C), AAC1120AAG (3500C>G), GTG1122GTT (3506G>T), TAC1123GCT (3507T>G 3508A>C 3509C>T), TAC1124TTT (3511A>T 3512C>T), CTT1125TTG (3513C>T 3515T>G), GGA1126GGC (3518A>C), ACA1127ACT (3521A>T), CCA1129CCT (3527A>T), CAG1130CAA (3530G>A), TTG1132ATG (3534T>A), GCC1133GCA (3539C>A), GCC1134GCA (3542C>A), ACA1135ACG (3545A>G), GTT1136GTG (3548T>G), CCC1137CCA (3551C>A), ACT1141ACC (3563T>C), CTG1142CTC (3566G>C), CGG1143AGG (3567C>A), CAA1146CCA (3577A>C), TTG1147CTG (3579T>C), GAA1149GAG (3587A>G), TTC1151TTT (3593C>T), GGA1152GGT (3596A>T), CCT1153GAA (3597C>G 3598C>A 3599T>A), GAG1155CAA (3603G>C 3605G>A), ACA1156ACT (3608A>T), GCA1157GCT (3611A>T), CCA1158CCT (3614A>T), TCC1159TCA (3617C>A), ATT1160CTG (3618A>C 3620T>G), CTA1161CTC (3623A>C), ACA1162TCT (3624A>T 3626A>T), GTT1163GTG (3629T>G), CAT1164AAA (3630C>A 3632T>A), GAC1165GAT (3635C>T), GGT1168GGG (3644T>G), GGT1170GGA (3650T>A), GAC1172GAT (3656C>T), CCC1173CCT (3659C>T), GTG1175GTT (3665G>T), GGG1177GGC (3671G>C), GTA1178ATT (3672G>A 3674A>T), ATG1179GAG (3675A>G 3676T>A), GCT1185GCC (3695T>C), TGT1186TCA (3697G>C 3698T>A), GGA1187AAT (3699G>A 3700G>A 3701A>T), GAT1189GAG (3707T>G), GAT1190GAC (3710T>C), GAC1191AGC (3711G>A 3712A>G), GCC1193GCA (3719C>A), AAA1194CGG (3720A>C 3721A>G 3722A>G), CTA1195GCT (3723C>G 3724T>C 3725A>T), TTC1197TTT (3731C>T), AAT1199GGT (3735A>G 3736A>G), TGC1202TCT (3745G>C 3746C>T), TCG1203GCC (3747T>G 3749G>C), CTG1204ATT (3750C>A 3752G>T), CTA1205TTA (3753C>T), CCT1206CCA (3758T>A), ATC1207GTT (3759A>G 3761C>T), ATG1208CTC (3762A>C 3764G>C), CGT1209AAG (3765C>A 3766G>A 3767T>G), AGC1210AGT (3770C>T), CAG1211CAA (3773G>A), AAG1212AGG (3775A>G), ATC1213GTG (3777A>G 3779C>G), CCT1214CCA (3782T>A), GGG1215GGT (3785G>T), GGA1216GGT (3788A>T), CGT1217ATC (3789C>A 3790G>T 3791T>C), CCA1218CCC (3794A>C), TGT1219ACA (3795T>A 3796G>C 3797T>A), GAA1220ACA (3798G>A 3799A>C), AGG1221CGT (3801A>C 3803G>T), GAT1222TGT (3804G>T 3805A>G), GAT1224GAC (3812T>C), GTA1225ATC (3813G>A 3815A>C), GTG1226GTC (3818G>C), CTC1227CTG (3821C>G), GGA1229GGT (3827A>T), ATA1230GTT (3828A>G 3830A>T), GGA1232GGT (3836A>T), TGT1233TTT (3838G>T), GAT1234GAC (3842T>C), TAC1235TTC (3844A>T), TAT1236TTT (3847A>T), GGC1238TCC (3852G>T 3853G>C), GAG1240GAA (3860G>A), CTG1241CTC (3863G>C), TCT1244TCA (3872T>A), TGC1245TGT (3875C>T), TAT1247TAC (3881T>C), CCC1248CCT (3884C>T), AAC1250AAT (3890C>T), AAG1251AAA (3893G>A), ATG1252CTG (3894A>C), GGG1253GGT (3899G>T), ATG1254TTG (3900A>T), GGG1255GGT (3905G>T), ATG1256ACC (3907T>C 3908G>C), AAC1257AAT (3911C>T), AAG1258AAA (3914G>A), AGA1259AGG (3917A>G), TTT1261TTC (3923T>C), GTG1262GTT (3926G>T), CAA1263GAA (3927C>G), TCC1264GCA (3930T>G 3932C>A), ACT1265CAG (3933A>C 3934C>A 3935T>G), GGC1266GAG (3937G>A 3938C>G), GAA1267AGT (3939G>A 3940A>G 3941A>T), GGA1268ACC (3942G>A 3943G>C 3944A>C), GAG1269CAA (3945G>C 3947G>A), AGA1270AGG (3950A>G), GTG1271GTA (3953G>A), GAA1272CTA (3954G>C 3955A>T), AAA1274AAG (3962A>G), GAC1276AAT (3966G>A 3968C>T), ACT1277TCT (3969A>T), GTA1279GTT (3977A>T), GAA1281CAA (3981G>C), GAA1284GAG (3992A>G), GAG1285GAA (3995G>A), CTA1286TTA (3996C>T), CAG1289GAA (4005C>G 4007G>A), ATT1290ATA (4010T>A), AAA1292AAG (4016A>G), GGC1293GGT (4019C>T), ATC1294AAC (4021T>A), CAT1295TAT (4023C>T), GAT1297GAA (4031T>A), CTG1298ATG (4032C>A), GTC1299GTT (4037C>T), ACC1301ACT (4043C>T), TGC1303TGT (4049C>T), GCC1304GCT (4052C>T), AAA1305AAG (4055A>G), GAT1306GAC (4058T>C), CGC1308AGG (4062C>A 4064C>G), CTC1309CTA (4067C>A), CCA1310CCT (4070A>T), CTT1311CTG (4073T>G), GAG1312GAC (4076G>C), AAA1313AAG (4079A>G), ATC1314ATA (4082C>A), TAT1315TAC (4085T>C), GGC1316GGG (4088C>G), AAG1317AAA (4091G>A), AGA1318AAG (4093G>A 4094A>G), AAG1319AAA (4097G>A), AGG1321CGT (4101A>C 4103G>T), CTC1322CTT (4106C>T), ATA1325ATT (4115A>T), CTT1326TTG (4116C>T 4118T>G), CCT1327CCA (4121T>A), TAC1330TAT (4130C>T), TTG1333CTG (4137T>C), GTC1334GTG (4142C>G), AGG1335CGC (4143A>C 4145G>C), AAG1336AAA (4148G>A), TTT1338TTC (4154T>C), TTC1341TTT (4163C>T), TCA1342TCT (4166A>T), GCC1343GCA (4169C>A), ACA1344TGT (4170A>T 4171C>G 4172A>T), TTG1345TTA (4175G>A), TTG1348GCA (4182T>G 4183T>C 4184G>A), CAC1349CAT (4187C>T), GCT1351GTT (4192C>T), ATA1352CTG (4194A>C 4196A>G), CCA1353CCC (4199A>C), GTT1356GTG (4208T>G), ATT1358ATC (4214T>C), GAT1359AAT (4215G>A), CCT1360CCC (4220T>C), ACA1361ATT (4222C>T 4223A>T), AGT1362TCA (4224A>T 4225G>C 4226T>A), TCT1363CAT (4227T>C 4228C>A), ACA1366ACC (4238A>C), TTG1367ACC (4239T>A 4240T>C 4241G>C), TTG1368CTG (4242T>C), AAT1370AAC (4250T>C), GGG1371CAG (4251G>C 4252G>A), TTT1372TTC (4256T>C), AGA1373CGG (4257A>C 4259A>G), GCT1374AGT (4260G>A 4261C>G), TCA1376TCT (4268A>T), GAC1377GAT (4271C>T), GTG1378GTT (4274G>T), GGA1379GGT (4277A>T), TCA1381TCT (4283A>T), GCT1382GCA (4286T>A), GAT1383GAC (4289T>C), TAT1384TAC (4292T>C), AGA1390AGG (4310A>G), GCA1391GCG (4313A>G), CCT1392CCA (4316T>A), GTT1393ATA (4317G>A 4319T>A), GCT1395GCA (4325T>A), CAG1397CAA (4331G>A), TTT1399TTA (4337T>A), GAA1405GCA (4354A>C), TAC1407TAT (4361C>T), GGA1408GGG (4364A>G), AAG1410GCT (4368A>G 4369A>C 4370G>T), CCT1411CCA (4373T>A), GGC1412GGA (4376C>A), AGT1413AGC (4379T>C), CCT1414GAG (4380C>G 4381C>A 4382T>G), TCC1416TCA (4388C>A), GCT1418GCA (4394T>A), CGA1419AGA (4395C>A), GCA1421GCT (4403A>T), CTT1422CTG (4406T>G), TTA1423TTG (4409A>G), GCA1426GCT (4418A>T), TGT1428TGC (4424T>C), ACA1431ACT (4433A>T), CTG1432CTA (4436G>A), GTG1437CTC (4449G>C 4451G>C), TTT1438TTC (4454T>C), TTG1440TTA (4460G>A), GGG1442GGT (4466G>T), GGC1443GGA (4469C>A), CCA1445CCT (4475A>T), TCA1446TCT (4478A>T), GCA1449GCT (4487A>T), CTA1450CTC (4490A>C), ACG1451ACC (4493G>C), ATC1453ATA (4499C>A), TTC1454TTT (4502C>T), AAT1455AAC (4505T>C), TCT1456TCC (4508T>C), CTC1457CTT (4511C>T), CTC1458TTA (4512C>T 4514C>A), AAT1459AAC (4517T>C), CGA1464CGG (4532A>G), GCC1466GCT (4538C>T), TTT1467TTC (4541T>C), TCA1469TCG (4547A>G), TTG1470CTC (4548T>C 4550G>C), TTA1471CTC (4551T>C 4553A>C), AGA1473AGG (4559A>G), CAT1475CAC (4565T>C), ATT1476ATC (4568T>C), GCT1477AAA (4569G>A 4570C>A 4571T>A), GCC1478GCA (4574C>A), AGG1479AAG (4576G>A), GCT1480GCA (4580T>A), ATA1481TTG (4581A>T 4583A>G), GGA1482GGT (4586A>T), GTT1483GTA (4589T>A), AAA1484AAG (4592A>G), CCA1485TCA (4593C>T), TTC1488TTT (4604C>T), CAG1490GAC (4608C>G 4610G>C), CTA1491TTA (4611C>T), TTC1492TTT (4616C>T), ATA1493ATT (4619A>T), GCT1494GCA (4622T>A), GTT1495GTG (4625T>G), GGA1497GGG (4631A>G), GTT1502GTG (4646T>G), GTA1504GTG (4652A>G), TTA1506ATG (4656T>A 4658A>G), CAT1507CAC (4661T>C), CTC1508CTT (4664C>T), CAG1509CAA (4667G>A), TAT1511TAC (4673T>C), TCT1512GAG (4674T>G 4675C>A 4676T>G), CTG1513TTA (4677C>T 4679G>A), CCA1514CCT (4682A>T), AAT1515GCA (4683A>G 4684A>C 4685T>A), CAT1518CAA (4694T>A), TTA1520CTA (4698T>C), CTG1522CAA (4705T>A 4706G>A), GTC1523GTG (4709C>G), AAT1524AAC (4712T>C), GTA1525GTT (4715A>T), ATC1526ATT (4718C>T), ATT1527ATC (4721T>C), GGT1530GGC (4730T>C), GAC1532GAT (4736C>T), TCG1534AAC (4740T>A 4741C>A 4742G>C), GAT1536AAT (4746G>A), GTT1537GTC (4751T>C), GAG1539GAT (4757G>T), GTA1540GTC (4760A>C), CAA1541AAA (4761C>A), TTT1542TTC (4766T>C), CAA1543CAG (4769A>G), CTA1548TTG (4782C>T 4784A>G), ACT1549ACG (4787T>G), CTG1551TTG (4791C>T), AGT1552AGC (4796T>C), GGT1554GGC (4802T>C), TTT1555TTC (4805T>C), AAA1556AAG (4808A>G), CGA1557CGC (4811A>C), CTT1560TTG (4818C>T 4820T>G), GGA1561GGT (4823A>T), CAC1563CAA (4829C>A), ATG1564CTA (4830A>C 4832G>A), GCT1565GCC (4835T>C), CCT1566CCC (4838T>C), TCA1571TCC (4853A>C), ATC1572ATT (4856C>T), ATT1573GTG (4857A>G 4859T>G), GAG1574GAA (4862G>A), CCC1575CCA (4865C>A), ATA1579ATC (4877A>C), CCT1581CCA (4883T>A), GCC1582GCA (4886C>A), CCA1583GTT (4887C>G 4888C>T 4889A>T), GGT1584GGG (4892T>G), CCT1586CCA (4898T>A), GAC1587GAT (4901C>T), ACT1588ATC (4903C>T 4904T>C), CAG1589ACT (4905C>A 4906A>C 4907G>T), ATG1592TAT (4914A>T 4915T>A 4916G>T), ACA1597ACG (4931A>G), GGA1598GGT (4934A>T), ATA1599GTA (4935A>G), CGT1600AGA (4938C>A 4940T>A), GAA1601GAG (4943A>G), CAT1604CAG (4952T>G), CAC1605CAT (4955C>T), GGT1606GGA (4958T>A), CGT1607CAG (4960G>A 4961T>G), GTG1608GTC (4964G>C), GCT1609GTT (4966C>T), AAG1612ACA (4975A>C 4976G>A), CTT1613TTG (4977C>T 4979T>G), GTC1614ACA (4980G>A 4981T>C 4982C>A), TTA1615GAC (4983T>G 4984T>A 4985A>C), GAT1616AAT (4986G>A), GTT1617GTC (4991T>C), CAG1618CAA (4994G>A), ACG1619AAA (4996C>A 4997G>A), GCG1620GCA (5000G>A), TTG1621ATA (5001T>A 5003G>A), GAT1622ATG (5004G>A 5005A>T 5006T>G), GAA1623CAG (5007G>C 5009A>G), AGG1624CGA (5010A>C 5012G>A), GGT1625GGC (5015T>C), TTC1626TTG (5018C>G), AGA1627CCA (5019A>C 5020G>C), GCT1628ATC (5022G>A 5023C>T 5024T>C), ATC1630CAT (5028A>C 5029T>A 5030C>T), CCT1632CCA (5036T>A), TCC1633TCT (5039C>T), TTG1635GCT (5043T>G 5044T>C 5045G>T), GTG1637CAA (5049G>C 5050T>A 5051G>A), CAG1639AGT (5055C>A 5056A>G 5057G>T), GAA1640GAG (5060A>G), ATT1642CTA (5064A>C 5066T>A), GCA1643CAA (5067G>C 5068C>A), AAG1644GCC (5070A>G 5071A>C 5072G>C), GTA1645GTT (5075A>T), ACA1646ACT (5078A>T), GGG1647GGA (5081G>A), GAT1648GAC (5084T>C), TCA1649AGT (5085T>A 5086C>G 5087A>T), AGT1650AAC (5089G>A 5090T>C), GCC1651ATA (5091G>A 5092C>T 5093C>A), CTG1652GGG (5094C>G 5095T>G), ACA1653GCT (5097A>G 5099A>T), ATT1654CTG (5100A>C 5102T>G), TGT1655CTG (5103T>C 5104G>T 5105T>G), ATG1657CTG (5109A>C), GCA1658AGC (5112G>A 5113C>G 5114A>C), AAA1659AAG (5117A>G), GCA1660GAT (5119C>A 5120A>T), TCC1663ACT (5127T>A 5129C>T), TAT1664TAC (5132T>C), ACG1665GTG (5133A>G 5134C>T), CCA1666CCT (5138A>T), TTG1667CTT (5139T>C 5141G>T), GAC1668CCA (5142G>C 5143A>C 5144C>A), GCA1669GAA (5146C>A), GGT1670GGA (5150T>A), GAG1671AAA (5151G>A 5153G>A), AAG1672AAA (5156G>A), ATC1673ATT (5159C>T), ACA1674GAA (5160A>G 5161C>A), AAT1675GAT (5163A>G), GAG1677TCA (5169G>T 5170A>C 5171G>A), CGT1678CAA (5173G>A 5174T>A), GAT1679GGT (5176A>G), CTG1680GTC (5178C>G 5180G>C), AAT1681CAT (5181A>C), TGG1682GAA (5184T>G 5185G>A 5186G>A), GCA1684TGT (5190G>T 5191C>G 5192A>T), CCA1685TCT (5193C>T 5195A>T), AAC1686GGA (5196A>G 5197A>G 5198C>A), ATT1687GTT (5199A>G), GGT1688TTC (5202G>T 5203G>T 5204T>C), TTT1689TTC (5207T>C), GCA1692TCG (5214G>T 5216A>G), CGT1693CGA (5219T>A), ACT1694ACA (5222T>A), GCA1695GAA (5224C>A), GCC1696GTT (5227C>T 5228C>T), CAC1697AAT (5229C>A 5231C>T), TAC1698CAT (5232T>C 5234C>T), ACG1699AAA (5236C>A 5237G>A), GAA1702GAG (5246A>G), GGG1703GAC (5248G>A 5249G>C), TAC1704ACA (5250T>A 5251A>C 5252C>A), ATT1706CTT (5256A>C), AAC1708AAT (5264C>T), TGT1709TGC (5267T>C), AAG1713AAA (5279G>A), AAG1714AAA (5282G>A), TCC1715ACA (5283T>A 5285C>A), AAT1716ACA (5287A>C 5288T>A), GTT1718GTG (5294T>G), CCG1733CCC (5339G>C), TAC1734TAT (5342C>T), ACT1735ACA (5345T>A), TCA1737GCA (5349T>G), GCT1738GCC (5354T>C), ATA1739ATC (5357A>C), ATG1740AAG (5359T>A), GAT1741GAC (5363T>C), GCC1742AAT (5364G>A 5365C>A 5366C>T), AAG1744CAA (5370A>C 5372G>A), AAT1745AGT (5374A>G), GTG1746CGC (5376G>C 5377T>G 5378G>C), GCA1748TCA (5382G>T), GGA1749ATG (5385G>A 5386G>T 5387A>G), GGA1750GGG (5390A>G), CAT1751ATT (5391C>A 5392A>T), GTA1752GGA (5395T>G), GCA1753ACC (5397G>A 5399A>C), ACC1754CAA (5400A>C 5401C>A 5402C>A), AAT1755AAA (5405T>A), GTC1756GTT (5408C>T), GTT1757GTC (5411T>C), GTG1759ATT (5415G>A 5417G>T), GGA1761GGG (5423A>G), GGA1763GGT (5429A>T), GTC1766GTG (5438C>G), GGC1767AGT (5439G>A 5441C>T), CCA1768CCC (5444A>C), GCA1771GCT (5453A>T), GCG1772GCT (5456G>T), ATG1774TTT (5460A>T 5462G>T), GCC1775GCA (5465C>A), GCA1777TCA (5469G>T), ACA1778ACC (5474A>C), AGG1779AAT (5476G>A 5477G>T), CAA1780CAG (5480A>G), TGT1782AGA (5484T>A 5486T>A), GTG1783GCT (5488T>C 5489G>T), GAC1785CAA (5493G>C 5495C>A), ATA1786CTT (5496A>C 5498A>T), GTT1788GTC (5504T>C), CGC1791AGA (5511C>A 5513C>A), ATT1793TTG (5517A>T 5519T>G), GGA1794GCG (5521G>C 5522A>G), AAT1795GGC (5523A>G 5524A>G 5525T>C), GTG1796TTG (5526G>T), CTT1797CTG (5531T>G), GAC1798GAT (5534C>T), CTC1799TTG (5535C>T 5537C>G), ACC1801CAG (5541A>C 5542C>A 5543C>G), GGC1805GGT (5555C>T), GTG1807ATT (5559G>A 5561G>T), CAG1808CAA (5564G>A), TTT1810TTC (5570T>C), CTT1811CTC (5573T>C), CAA1812ACA (5574C>A 5575A>C), TGT1813_GTG1814insGTC (5579_5580insGTC), CCT1815CAG (5584C>A 5585T>G), CAT1816AAA (5586C>A 5588T>A), GGA1817AGT (5589G>A 5591A>T), GAC1818TGC (5592G>T 5593A>G), AAA1819AAG (5597A>G), GTG1820ACA (5598G>A 5599T>C 5600G>A), AGT1823GCT (5607A>G 5608G>C), GGT1824AGT (5610G>A), GCA1825GGG (5614C>G 5615A>G), TCG1826del (5616_5618delTCG), CTC1827CGC (5620T>G), CAC1828GAT (5622C>G 5624C>T), AGT1829GGA (5625A>G 5627T>A), AGT1830ATG (5629G>T 5630T>G), TTT1831TTG (5633T>G), ATG1832AAC (5635T>A 5636G>C), CAA1834CAT (5642A>T), CAA1834_GGG1835insCATGCT (5642_5643insCATGCT), GGG1835GCT (5644G>C 5645G>T), TTT1836TAT (5647T>A), GAA1837GCC (5650A>C 5651A>C), TTG1838CTT (5652T>C 5654G>T), GGC1839GGA (5657C>A), CTC1841CTT (5663C>T), CGC1842CGA (5666C>A), ATT1843ATA (5669T>A), ATA1844GTG (5670A>G 5672A>G), CAT1845GGC (5673C>G 5674A>G 5675T>C), GGT1846TGT (5676G>T), GAT1847TCA (5679G>T 5680A>C 5681T>A), TTA1848ATT (5682T>A 5684A>T), GCA1849TCA (5685G>T), AAA1850AGA (5689A>G), CAG1851AAG (5691C>A), ACA1852ATG (5695C>T 5696A>G), ATG1854GTG (5700A>G), CGG1855AGA (5703C>A 5705G>A), ATG1856TTG (5706A>T), CCA1857CCC (5711A>C), TAT1858TAC (5714T>C), GTG1859ACT (5715G>A 5716T>C 5717G>T), GTA1860GTT (5720A>T), GGA1861GGG (5723A>G), CAT1863ATG (5727C>A 5728A>T 5729T>G), GGA1864GGT (5732A>T), GGA1865GGT (5735A>T), GGA1867GCA (5740G>C), AAT1868GAT (5742A>G), TTT1869TGC (5746T>G 5747T>C), TCC1870TCA (5750C>A), CAG1872AGA (5754C>A 5755A>G 5756G>A), GAT1873AAT (5757G>A)

#### Proteins

| | | | | | | | | | |
|---|---|---|---|---|---|---|---|---|---|
| polyprotein (YP_001976147.1) | 1 | 1877 | 87.2% | 9403 | 72.1% | 1862 (98.8%) | 1290 (68.4%) | 8/15/0/0 | 0 |

| | Begin | End | Coverage | Score | Concordance | Matches | Identities | I/D/M/F* | Stop Codons |
|---|---|---|---|---|---|---|---|---|---|
| **NT** | **35** | **5769** | **79.3%** | **3466** | **30.8%** | **5682 (98.6%)** | **3757 (65.2%)** | **28/53** | |
| Protein mutations: | M6I (158G>C), F7V (159T>G 161C>T), S8_S17del (162_191delTCCAAGGTTGCTGGTGTTGATGACGCCAGT), S20A (198T>G 200C>T), T22P (204A>C), F25L (213T>C), G26S (216G>T 217G>C), N30Q (228A>C 230T>G), S31T (231T>A 233T>A), I32V (234A>G 236T>G), K34S (240A>T 241A>C), F35L (243T>C 245T>G), A38T (252G>A 254G>A), V39M (255G>A 257T>G), S40K (259G>A 260T>G), N41S (262A>G), I42M (266T>G), T43S (268C>G 269C>T), T44K (271C>A 272T>A), M45I (275G>A), Q47E (279C>G), I49F (285A>T 287T>C), D51G (292A>G 293T>C), L53I (297C>A), D55E (305T>A), V60I (318G>A 320G>T), T61V (321A>G 322C>T), D62K (324G>A 326T>A), V63T (327G>A 328T>C 329G>T), S64A (330T>G 332C>A), T65D (333A>G 334C>A 335A>T), V68N (342G>A 343T>A 344G>T), A69S (345G>A 346C>G 347C>T), T70V (348A>G 349C>T), L74V (360C>G 362T>A), N75E (363A>G 365T>A), V77L (369G>T 371T>A), D79G (376A>G 377C>G), G82A (385G>C 386T>C), A83T (387G>A 389A>G), L84V (390C>G), D86E (398T>A), L89V (405C>G), A91G (412C>G 413G>T), Y92I (414T>A 415A>T), S93N (417T>A 418C>A), L95F (425G>C), S96K (426T>A 427C>A 428C>A), K100R (439A>G), S101T (441T>A 443T>C), K103I (448A>T 449G>T), L109V (465T>G 467G>T), F110V (468T>G), E111L (471G>T 472A>T), S112G (474A>G 476T>C), L113I (477T>A 479G>C), S114K (480T>A 481C>A 482C>A), K116N (488A>T), V117F (489G>T), K118I (493A>T 494A>T), L119R (495T>A 496T>G), F121Y (502T>A), A122S (504G>T 506T>A), V124L (510G>C), N126E (516A>G 518T>A), K127T (520A>C), S128T (523G>C 524T>A), E129G (526A>G 527A>T), D130E (530T>A), V131Y (531G>T 532T>A 533A>T), I133V (537A>G 539A>T), Y136I (546T>A 547A>T 548T>A), G137A (550G>C), A142S (564G>T 566T>C), I143L (567A>C 569A>T), T144S (570A>T), V147L (579G>T 581G>A), L149Q (586T>A), F150I (588T>A 590T>A), V151I (591G>A 593T>C), S154V (600A>G 601G>T 602T>A), G159M (615G>A 616G>T 617T>G), S161R (621T>A 622C>G 623T>A), N162S (625A>G 626C>T), T163V (627A>G 628C>T 629T>A), K165C (633A>T 634A>G 635A>T), I170V (648A>G 650A>T), M174F (660A>T 662G>C), L181I (681T>A 683A>T), D182S (684G>A 685A>G), K186L (696A>C 697A>T 698A>C), A188S (702G>T 704C>T), R190K (708C>A 709G>A), T192S (714A>T 716C>T), M193Q (717A>C 718T>A), V194M (720G>A), M199V (735A>G), V205L (753G>T 755T>A), S206N (756T>A 757C>A 758A>T), M214L (780A>C), T216S (786A>T 788A>T), I217L (789A>T 791A>G), L218I (792T>A 794G>A), I228V (822A>G 824T>G), S238T (853G>C 854C>A), T248A (882A>G 884A>G), T260Q (918A>C 919C>A 920T>A), R263K (927C>A 928G>A 929T>G), I280V (978A>G 980A>T), V281A (982T>C 983C>T), G282D (985G>A 986C>T), T283I (988C>T 989C>A), L298I (1032C>A 1034G>T), Q307T (1059C>A 1060A>C 1061G>T), D310S (1068G>T 1069A>C), C313A (1077T>G 1078G>C), K325P (1113A>C 1114A>C 1115G>T), P328S (1122C>T 1124T>A), T329I (1126C>T), R335K (1144G>A 1145G>A), E337Q (1149G>C 1151A>G), M354L (1200A>T), A361T (1221G>A 1223C>A), A363V (1228C>T), E366D (1238A>T), V368L (1242G>C 1244T>C), Y369I (1245T>A 1246A>T 1247T>A), S372N (1255G>A 1256T>C), R374Q (1260A>C 1261G>A 1262G>A), M376T (1267T>C 1268G>A), S393E (1317T>G 1318C>A 1319C>G), R412A (1374A>G 1375G>C 1376A>C), G414Q (1380G>C 1381G>A 1382T>A), I496V (1626A>G 1628C>T), M521R (1701A>C 1702T>G), E523V (1708A>T), D530N (1728G>A 1730T>C), G533A (1738G>C), R536S (1748A>C), V540I (1758G>A 1760G>T), E541G (1762A>G 1763G>A), N552D (1794A>G 1796T>C), Q567K (1839C>A), A570H (1848G>C 1849C>A 1850A>C), V572I (1854G>A 1856A>T), S573A (1857T>G), K574T (1861A>C 1862G>C), F577Y (1870T>A 1871T>C), E578N (1872G>A 1874A>C), A579Q (1875G>C 1876C>A 1877T>G), E580S (1878G>T 1879A>C), S581T (1881T>A 1883T>A), I583C (1887A>T 1888T>G 1889A>T), S584E (1890A>G 1891G>A 1892T>A), P585E (1893C>G 1894C>A 1895T>G), I587V (1899A>G 1901A>T), Q591E (1911C>G), A594V (1921C>T 1922A>T), D595E (1925T>A), D596E (1928T>A), I598M (1934A>G), A599T (1935G>A 1937G>A), V601F (1941G>T 1943G>T), D602H (1944G>C), G603D (1948G>A 1949G>T), R605Q (1953A>C 1954G>A), L606I (1956T>A 1958G>A), R607S (1959C>A), T608F (1962A>T 1963C>T 1964C>T), K609D (1965A>G 1967G>C), Q610E (1968C>G), D611V (1972A>T), E612S (1974G>A 1975A>G 1976A>C), I614S (1981T>G 1982A>T), L615V (1983T>G 1985G>T), V616_P617del (1986_1991delGTGCCC), V620L (1998G>C 2000G>C), N623D (2007A>G), I627G (2019A>G 2020T>G 2021C>A), A629S (2025G>T), R630Q (2028A>C 2029G>A 2030A>G), S644A (2070A>G 2071G>C 2072T>A), D646N (2076G>A), E648F (2082G>T 2083A>T 2084G>T), L650Y (2088C>T 2089T>A), H655A (2103C>G 2104A>C), H656S (2106C>T 2107A>C 2108T>A), M660I (2120G>T), Y661F (2122A>T), S663V (2127T>G 2128C>T 2129T>C), D665S (2133G>T 2134A>C 2135T>A), D666S (2136G>A 2137A>G), H668A (2142C>G 2143A>C 2144T>C), I671S (2152T>G 2153T>C), N673T (2158A>C 2159C>T), K674R (2160A>C 2161A>G 2162A>C), R676V (2166A>G 2167G>T), V677N (2169G>A 2170T>A 2171C>T), N678E (2172A>G 2174C>G), L680I (2178T>A 2180G>T), R682E (2184A>G 2185G>A 2186A>G), D685S (2193G>T 2194A>C 2195T>G), A686T (2196G>A), E687A (2200A>C), Q688G (2202C>G 2203A>G 2204G>A), R690A (2208C>G 2209G>C 2210A>C), V693I (2217G>A 2219A>C), Q694K (2220C>A 2222A>G), N695A (2223A>G 2224A>C), I701V (2241A>G 2243A>C), A710S (2268G>A 2269C>G 2270A>T), S711T (2271T>A), I713V (2277A>G 2279C>G), K716A (2286A>G 2287A>C 2288G>T), K717H (2289A>C 2291A>C), L718M (2292C>A 2294A>G), A719D (2296C>A 2297A>T), T722R (2305C>G 2306A>G), E723M (2307G>A 2308A>T), M724L (2310A>T), R725Y (2313A>T 2314G>A 2315G>C), T726N (2317C>A), F728R (2322T>A 2323T>G 2324T>A), S731R (2331T>A 2332C>G 2333T>A), C732A (2334T>G 2335G>C 2336C>A), I733L (2337A>C 2339T>G), D734E (2342T>A), V737A (2350T>C), I741M (2363A>G), F749H (2385T>C 2386T>A), C762_N763insRR (2426_2427insAGAAGG), N763K (2429C>G), K764M (2431A>T 2432A>G), S766T (2436T>A 2438A>T), P769T (2445C>A), K770_E771del (2448_2453delAAGGAA), I774V (2460A>G 2462C>A), K775Q (2463A>C), I776S (2466A>T 2467T>C 2468T>G), C777L (2469T>C 2470G>T), H779D (2475C>G 2477T>C), K780S (2478A>T 2479A>C 2480A>C), I781T (2482T>C 2483C>A), V782I (2484G>A), E783G (2488A>G 2489A>G), Y784R (2490T>A 2491A>G 2492T>A), I785M (2495C>G), K786Q (2496A>C 2498G>A), D787E (2501T>A), T788M (2503C>T 2504A>G), W790N (2508T>A 2509G>A 2510G>T), S791D (2511A>G 2512G>A 2513C>T), A795I (2523G>A 2524C>T 2525A>T), R797K (2530G>A), I806L (2556A>C), M807I (2561G>A), V812T (2574G>A 2575T>C), F813M (2577T>A 2579T>G), G815A (2584G>C), I816V (2586A>G 2588T>A), L820I (2598T>A 2600G>T), A821S (2601G>T 2603G>T), Q824K (2610C>A 2612G>A), N826S (2617A>G), S827L (2620C>T 2621T>G), F828I (2622T>A 2624C>T), N830T (2629A>C), A831S (2631G>T 2633A>C), S832Q (2634T>C 2635C>A), R834K (2641G>A), N852R (2695A>G 2696T>A), N852_K853insS (2696_2697insTCC), K853H (2697A>C 2699G>T), L854V (2700C>G 2702C>A), R855S (2703A>T 2704G>C), N856K (2708T>G), L857M (2709C>A 2711T>G), V859F (2715G>T), G860S (2718G>A 2720T>C), S865P (2733T>C), L866N (2736T>A 2737T>A 2738G>T), A867D (2740C>A 2741A>T), Q868_D869insG (2744_2745insGGA), D869K (2745G>A 2747T>A), A872S (2754G>T 2756T>C), E873R (2757G>A 2758A>G 2759A>G), Q879E (2775C>G 2777G>A), K882A (2784A>G 2785A>C 2786G>C), L890V (2808C>G 2810T>G), G891A (2812G>C 2813T>A), G904E (2851G>A 2852T>A), H914T (2880C>A 2881A>C 2882C>G), N915Q (2883A>C 2885T>G), A916L (2886G>C 2887C>T 2888T>C), T923S (2907A>T), A927E (2920C>A 2921C>A), N928H (2922A>C), R930V (2928C>G 2929G>T 2930A>C), S931C (2932C>G 2933T>C), K932Q (2934A>C 2936A>G), R938K (2953G>A), E939H (2955G>C 2957A>T), S940A (2958T>G 2960A>C), R941K (2962G>A), I942F (2964A>T), I943L (2967A>T 2969T>G), E944P (2970G>C 2971A>C 2972G>T), G945E (2974G>A 2975C>A), I948V (2982A>G 2984C>T), I949V (2985A>G 2987T>A), D955C (3003G>T 3004A>G), P961V (3021C>G 3022C>T 3023T>C), P968A (3042C>G 3044A>T), R969E (3045A>G 3046G>A), E972A (3055A>C 3056G>A), Q980E (3078C>G), C984V (3090T>G 3091G>T 3092T>C), T990K (3109C>A), R993E (3117C>G 3118G>A 3119T>G), L994K (3120C>A 3121T>A 3122T>A), G995Q (3123G>C 3124G>A 3125C>G), E998D (3134G>C), M1002Y (3144A>T 3145T>A 3146G>T), P1003S (3147C>T), V1012E (3175T>A 3176T>G), E1015T (3183G>A 3184A>C), E1017R (3189G>A 3190A>G 3191A>G), L1018M (3192C>A), N1019S (3196A>G), S1020T (3198T>A 3200C>A), S1021N (3201T>A 3202C>A 3203C>T), K1024H (3210A>C 3212G>C), K1024_I1025insS (3212_3213insTCA), C1026S (3216T>A 3218C>T), S1027A (3219T>G), Q1029E (3225C>G), S1030A (3228T>G 3230T>C), I1035V (3243A>G 3245A>C), G1037S (3249G>A 3251G>C), V1051I (3291G>A 3293T>C), V1057C (3309G>T 3310T>G 3311G>C), G1068E (3343G>A 3344C>A), V1069A (3346T>C 3347A>T), A1071T (3351G>A 3353T>A), R1073L (3357C>T 3358G>T 3359T>G), I1076V (3366A>G 3368C>A), S1077A (3369T>G 3371T>A), I1083V (3387A>G 3389C>A), E1086D (3398A>C), E1089Q (3405G>C 3407G>A), P1092A (3414C>G 3416A>T), T1093Q (3417A>C 3418C>A 3419T>A), A1098M (3432G>A 3433C>T 3434A>G), I1105L (3453A>T 3455T>G), S1108H (3462T>C 3463C>A 3464A>T), I1109V (3465A>G), Q1115A (3483C>G 3484A>C 3485G>T), E1118D (3494G>C), N1120K (3500C>G), Y1123A (3507T>G 3508A>C 3509C>T), Y1124F (3511A>T 3512C>T), L1132M (3534T>A), Q1146P (3577A>C), P1153E (3597C>G 3598C>A 3599T>A), E1155Q (3603G>C 3605G>A), I1160L (3618A>C 3620T>G), T1162S (3624A>T 3626A>T), H1164K (3630C>A 3632T>A), V1178I (3672G>A 3674A>T), M1179E (3675A>G 3676T>A), C1186S (3697G>C 3698T>A), G1187N (3699G>A 3700G>A 3701A>T), D1189E (3707T>G), D1191S (3711G>A 3712A>G), K1194R (3720A>C 3721A>G 3722A>G), L1195A (3723C>G 3724T>C 3725A>T), N1199G (3735A>G 3736A>G), C1202S (3745G>C 3746C>T), S1203A (3747T>G 3749G>C), L1204I (3750C>A 3752G>T), I1207V (3759A>G 3761C>T), M1208L (3762A>C 3764G>C), R1209K (3765C>A 3766G>A 3767T>G), K1212R (3775A>G), I1213V (3777A>G 3779C>G), R1217I (3789C>A 3790G>T 3791T>C), C1219T (3795T>A 3796G>C 3797T>A), E1220T (3798G>A 3799A>C), D1222C (3804G>T 3805A>G), V1225I (3813G>A 3815A>C), I1230V (3828A>G 3830A>T), C1233F (3838G>T), Y1235F (3844A>T), Y1236F (3847A>T), G1238S (3852G>T 3853G>C), M1252L (3894A>C), M1254L (3900A>T), M1256T (3907T>C 3908G>C), Q1263E (3927C>G), S1264A (3930T>G 3932C>A), T1265Q (3933A>C 3934C>A 3935T>G), G1266E (3937G>A 3938C>G), E1267S (3939G>A 3940A>G 3941A>T), G1268T (3942G>A 3943G>C 3944A>C), E1269Q (3945G>C 3947G>A), E1272L (3954G>C 3955A>T), D1276N (3966G>A 3968C>T), T1277S (3969A>T), E1281Q (3981G>C), Q1289E (4005C>G 4007G>A), I1294N (4021T>A), H1295Y (4023C>T), D1297E (4031T>A), L1298M (4032C>A), E1312D (4076G>C), R1318K (4093G>A 4094A>G), T1344C (4170A>T 4171C>G 4172A>T), L1348A (4182T>G 4183T>C 4184G>A), A1351V (4192C>T), I1352L (4194A>C 4196A>G), D1359N (4215G>A), T1361I (4222C>T 4223A>T), S1363H (4227T>C 4228C>A), L1367T (4239T>A 4240T>C 4241G>C), G1371Q (4251G>C 4252G>A), A1374S (4260G>A 4261C>G), V1393I (4317G>A 4319T>A), F1399L (4337T>A), E1405A (4354A>C), K1410A (4368A>G 4369A>C 4370G>T), P1414E (4380C>G 4381C>A 4382T>G), V1437L (4449G>C 4451G>C), A1477K (4569G>A 4570C>A 4571T>A), R1479K (4576G>A), I1481L (4581A>T 4583A>G), P1485S (4593C>T), Q1490D (4608C>G 4610G>C), L1506M (4656T>A 4658A>G), S1512E (4674T>G 4675C>A 4676T>G), N1515A (4683A>G 4684A>C 4685T>A), H1518Q (4694T>A), L1522Q (4705T>A 4706G>A), S1534N (4740T>A 4741C>A 4742G>C), D1536N (4746G>A), E1539D (4757G>T), Q1541K (4761C>A), H1563Q (4829C>A), M1564L (4830A>C 4832G>A), I1573V (4857A>G 4859T>G), P1583V (4887C>G 4888C>T 4889A>T), T1588I (4903C>T 4904T>C), Q1589T (4905C>A 4906A>C 4907G>T), M1592Y (4914A>T 4915T>A 4916G>T), I1599V (4935A>G), H1604Q (4952T>G), R1607Q (4960G>A 4961T>G), A1609V (4966C>T), K1612T (4975A>C 4976G>A), V1614T (4980G>A 4981T>C 4982C>A), L1615D (4983T>G 4984T>A 4985A>C), D1616N (4986G>A), T1619K (4996C>A 4997G>A), L1621I (5001T>A 5003G>A), D1622M (5004G>A 5005A>T 5006T>G), E1623Q (5007G>C 5009A>G), F1626L (5018C>G), R1627P (5019A>C 5020G>C), A1628I (5022G>A 5023C>T 5024T>C), I1630H (5028A>C 5029T>A 5030C>T), L1635A (5043T>G 5044T>C 5045G>T), V1637Q (5049G>C 5050T>A 5051G>A), Q1639S (5055C>A 5056A>G 5057G>T), I1642L (5064A>C 5066T>A), A1643Q (5067G>C 5068C>A), K1644A (5070A>G 5071A>C 5072G>C), S1650N (5089G>A 5090T>C), A1651I (5091G>A 5092C>T 5093C>A), L1652G (5094C>G 5095T>G), T1653A (5097A>G 5099A>T), I1654L (5100A>C 5102T>G), C1655L (5103T>C 5104G>T 5105T>G), M1657L (5109A>C), A1658S (5112G>A 5113C>G 5114A>C), A1660D (5119C>A 5120A>T), S1663T (5127T>A 5129C>T), T1665V (5133A>G 5134C>T), D1668P (5142G>C 5143A>C 5144C>A), A1669E (5146C>A), E1671K (5151G>A 5153G>A), T1674E (5160A>G 5161C>A), N1675D (5163A>G), E1677S (5169G>T 5170A>C 5171G>A), R1678Q (5173G>A 5174T>A), D1679G (5176A>G), L1680V (5178C>G 5180G>C), N1681H (5181A>C), W1682E (5184T>G 5185G>A 5186G>A), A1684C (5190G>T 5191C>G 5192A>T), P1685S (5193C>T 5195A>T), N1686G (5196A>G 5197A>G 5198C>A), I1687V (5199A>G), G1688F (5202G>T 5203G>T 5204T>C), A1692S (5214G>T 5216A>G), A1695E (5224C>A), A1696V (5227C>T 5228C>T), H1697N (5229C>A 5231C>T), Y1698H (5232T>C 5234C>T), T1699K (5236C>A 5237G>A), G1703D (5248G>A 5249G>C), Y1704T (5250T>A 5251A>C 5252C>A), I1706L (5256A>C), S1715T (5283T>A 5285C>A), N1716T (5287A>C 5288T>A), S1737A (5349T>G), M1740K (5359T>A), A1742N (5364G>A 5365C>A 5366C>T), K1744Q (5370A>C 5372G>A), N1745S (5374A>G), V1746R (5376G>C 5377T>G 5378G>C), A1748S (5382G>T), G1749M (5385G>A 5386G>T 5387A>G), H1751I (5391C>A 5392A>T), V1752G (5395T>G), A1753T (5397G>A 5399A>C), T1754Q (5400A>C 5401C>A 5402C>A), N1755K (5405T>A), V1759I (5415G>A 5417G>T), G1767S (5439G>A 5441C>T), M1774F (5460A>T 5462G>T), A1777S (5469G>T), R1779N (5476G>A 5477G>T), C1782R (5484T>A 5486T>A), V1783A (5488T>C 5489G>T), D1785Q (5493G>C 5495C>A), I1786L (5496A>C 5498A>T), I1793L (5517A>T 5519T>G), G1794A (5521G>C 5522A>G), N1795G (5523A>G 5524A>G 5525T>C), V1796L (5526G>T), T1801Q (5541A>C 5542C>A 5543C>G), V1807I (5559G>A 5561G>T), Q1812T (5574C>A 5575A>C), C1813_V1814insV (5579_5580insGTC), P1815Q (5584C>A 5585T>G), H1816K (5586C>A 5588T>A), G1817S (5589G>A 5591A>T), D1818C (5592G>T 5593A>G), V1820T (5598G>A 5599T>C 5600G>A), S1823A (5607A>G 5608G>C), G1824S (5610G>A), A1825G (5614C>G 5615A>G), S1826del (5616_5618delTCG), L1827R (5620T>G), H1828D (5622C>G 5624C>T), S1829G (5625A>G 5627T>A), S1830M (5629G>T 5630T>G), F1831L (5633T>G), M1832N (5635T>A 5636G>C), Q1834H (5642A>T), Q1834_G1835insHA (5642_5643insCATGCT), G1835A (5644G>C 5645G>T), F1836Y (5647T>A), E1837A (5650A>C 5651A>C), I1844V (5670A>G 5672A>G), H1845G (5673C>G 5674A>G 5675T>C), G1846C (5676G>T), D1847S (5679G>T 5680A>C 5681T>A), L1848I (5682T>A 5684A>T), A1849S (5685G>T), K1850R (5689A>G), Q1851K (5691C>A), T1852M (5695C>T 5696A>G), M1854V (5700A>G), M1856L (5706A>T), V1859T (5715G>A 5716T>C 5717G>T), H1863M (5727C>A 5728A>T 5729T>G), G1867A (5740G>C), N1868D (5742A>G), F1869C (5746T>G 5747T>C), Q1872R (5754C>A 5755A>G 5756G>A), D1873N (5757G>A) | | | | | | | | |

| | Begin | End | Coverage | Score | Concordance | Matches | Identities | I/D/M/F* | Stop Codons |
|---|---|---|---|---|---|---|---|---|---|
| NT | 35 | 5769 | 79.3% | 3466 | 30.8% | 5682 (98.6%) | 3757 (65.2%) | 28/53 | |

Codon mutations:

GCT2GCA (146T>A), AAG5AAA (155G>A), ATG6ATC (158G>C), TTC7GTT (159T>G 161C>T), TCC8_AGT17del (162_191delTCCAAGGTTGCTGGTGTTGATGACGCCAGT), TCC20GCT (198T>G 200C>T), ACC21ACT (203C>T), ACT22CCT (204A>C), TCA23TCT (209A>T), TTC24TTT (212C>T), TTT25CTT (213T>C), GGT26TCT (216G>T 217G>C), GAA27GAG (221A>G), CTT28TTG (222C>T 224T>G), TCA29TCT (227A>T), AAT30CAG (228A>C 230T>G), TCT31ACA (231T>A 233T>A), ATT32GTG (234A>G 236T>G), TCC33TCA (239C>A), AAA34TCA (240A>T 241A>C), TTT35CTG (243T>C 245T>G), TCA36TCC (248A>C), AAA37AAG (251A>G), GCG38ACA (252G>A 254G>A), GTT39ATG (255G>A 257T>G), AGT40AAG (259G>A 260T>G), AAT41AGT (262A>G), ATT42ATG (266T>G), ACC43AGT (268C>G 269C>T), ACT44AAA (271C>A 272T>A), ATG45ATA (275G>A), TCA46TCT (278A>T), CAG47GAG (279C>G), AAG48AAA (284G>A), ATT49TTC (285A>T 287T>C), GAT51GGC (292A>G 293T>C), CTT53ATT (297C>A), GAG54GAA (302G>A), GAT55GAA (305T>A), CTG56TTA (306C>T 308G>A), CCA58CCT (314A>T), TCT59TCA (317T>A), GTG60ATT (318G>A 320G>T), ACT61GTT (321A>G 322C>T), GAT62AAA (324G>A 326T>A), GTG63ACT (327G>A 328T>C 329G>T), TCC64GCA (330T>G 332C>A), ACA65GAT (333A>G 334C>A 335A>T), TCT66TCC (338T>C), GTG68AAT (342G>A 343T>A 344G>T), GCC69AGT (345G>A 346C>G 347C>T), ACT70GTT (348A>G 349C>T), TCA73TCT (359A>T), CTT74GTA (360C>G 362T>A), AAT75GAA (363A>G 365T>A), GTT77TTA (369G>T 371T>A), TTG78TTA (374G>A), GAC79GGG (376A>G 377C>G), GGT82GCC (385G>C 386T>C), GCA83ACG (387G>A 389A>G), CTT84GTT (390C>G), ATA85ATT (395A>T), GAT86GAA (398T>A), CCA87CCT (401A>T), CTT89GTT (405C>G), GCG91GGT (412C>G 413G>T), TAC92ATC (414T>A 415A>T), TCT93AAT (417T>A 418C>A), TTG95TTC (425G>C), TCC96AAA (426T>A 427C>A 428C>A), TAC99TAT (437C>T), AAA100AGA (439A>G), TCT101ACC (441T>A 443T>C), ATC102ATA (446C>A), AAG103ATT (448A>T 449G>T), TTA107CTG (459T>C 461A>G), TTG109GTT (465T>G 467G>T), TTT110GTT (468T>G), GAG111TTG (471G>T 472A>T), AGT112GGC (474A>G 476T>C), TTG113ATC (477T>A 479G>C), TCC114AAA (480T>A 481C>A 482C>A), AAA116AAT (488A>T), GTT117TTT (489G>T), AAA118ATT (493A>T 494A>T), TTA119AGA (495T>A 496T>G), TTT121TAT (502T>A), GCT122TCA (504G>T 506T>A), GTG124CTG (510G>C), TTA125TTG (515A>G), AAT126GAA (516A>G 518T>A), AAA127ACA (520A>C), AGT128ACA (523G>C 524T>A), GAA129GGT (526A>G 527A>T), GAT130GAA (530T>A), GTA131TAT (531G>T 532T>A 533A>T), ATA133GTT (537A>G 539A>T), GTT134GTA (542T>A), GTA135GTT (545A>T), TAT136ATA (546T>A 547A>T 548T>A), GGT137GCT (550G>C), CTT139CTA (557T>A), GTA140GTC (560A>C), GCT142TCC (564G>T 566T>C), ATA143CTT (567A>C 569A>T), ACA144TCA (570A>T), TTG145CTA (573T>C 575G>A), TTA146TTG (578A>G), GTG147TTA (579G>T 581G>A), CTT148TTA (582C>T 584T>A), CTA149CAA (586T>A), TTT150ATA (588T>A 590T>A), GTT151ATC (591G>A 593T>C), AGT154GTA (600A>G 601G>T 602T>A), GTT157GTA (611T>A), GAG158GAA (614G>A), GGT159ATG (615G>A 616G>T 617T>G), TCT161AGA (621T>A 622C>G 623T>A), AAC162AGT (625A>G 626C>T), ACT163GTA (627A>G 628C>T 629T>A), GTC164GTT (632C>T), AAA165TGT (633A>T 634A>G 635A>T), ATC167ATA (641C>A), TTG169CTA (645T>C 647G>A), ATA170GTT (648A>G 650A>T), GTC171GTA (653C>A), GGA172GGT (656A>T), ATG174TTC (660A>T 662G>C), TTT175TTC (665T>C), TCA176TCT (668A>T), AGT177TCT (669A>T 670G>C), CTA178CTG (674A>G), TAC179TAT (677C>T), TTA181ATT (681T>A 683A>T), GAT182AGT (684G>A 685A>G), CCA185CCC (695A>C), AAA186CTC (696A>C 697A>T 698A>C), GCC188TCT (702G>T 704C>T), GAA189GAG (707A>G), CGA190AAA (708C>A 709G>A), TTC191TTT (713C>T), ACC192TCT (714A>T 716C>T), ATG193CAG (717A>C 718T>A), GTG194ATG (720G>A), GCC195GCA (725C>A), CAA196CAG (728A>G), AGT198TCT (732A>T 733G>C), ATG199GTG (735A>G), CTT200CTG (740T>G), GGG202GGT (746G>T), GAA203GAG (749A>G), TCA204AGC (750T>A 751C>G 752A>C), GTT205TTA (753G>T 755T>A), TCA206AAT (756T>A 757C>A 758A>T), ACG210ACC (770G>C), TCT211AGT (771T>A 772C>G), CAG212CAA (776G>A), ATG214CTG (780A>C), TCC215TCT (785C>T), ACA216TCT (786A>T 788A>T), ATA217TTG (789A>T 791A>G), TTG218ATA (792T>A 794G>A), GCT219GCA (797T>A), GGG221GGC (803G>C), ATT222ATC (806T>C), TCG223TCA (809G>A), GTG226GTA (818G>A), TTC227TTT (821C>T), ATT228GTG (822A>G 824T>G), GTG230GTA (830G>A), AGA233AGG (839A>G), TTG237CTT (849T>C 851G>T), AGC238ACA (853G>C 854C>A), CCA240CCG (860A>G), TTG241CTA (861T>C 863G>A), CTC245TTG (873C>T 875C>G), TCG247TCA (881G>A), ACA248GCG (882A>G 884A>G), GGA249GGC (887A>C), GGT251GGA (893T>A), CAG253CAA (899G>A), CTA258CTC (914A>C), ACT260CAA (918A>C 919C>A 920T>A), CTT261TTG (921C>T 923T>G), TAC262TAT (926C>T), CGT263AAG (927C>A 928G>A 929T>G), AAT264AAC (932T>C), AAA266AAG (938A>G), ACC269ACG (947C>G), TCA274TCT (962A>T), GTT276GTG (968T>G), CTT277TTG (969C>T 971T>G), ATA279ATT (977A>T), ATA280GTT (978A>G 980A>T), GTC281GCT (982T>C 983C>T), GGC282GAT (985G>A 986C>T), ACC283ATA (988C>T 989C>A), TTT284TTC (992T>C), GGA285GGG (995A>G), TTC286TTT (998C>T), AAT288AAC (1004T>C), GTG290GTT (1010G>T), TTG291CTG (1011T>C), TCA292TCT (1016A>T), GCT293GCA (1019T>A), ATA294ATT (1022A>T), ACG297ACA (1031G>A), CTG298ATT (1032C>A 1034G>T), GAT301GAC (1043T>C), CTG302CTC (1046G>C), TTC303TTT (1049C>T), CAG307ACT (1059C>A 1060A>C 1061G>T), GAG308GAA (1064G>A), GAT310TCT (1068G>T 1069A>C), TGT313GCT (1077T>G 1078G>C), ACG317ACA (1091G>A), CGC318AGG (1092C>A 1094C>G), GAG320GAA (1100G>A), GCC323GCT (1109C>T), AAG325CCT (1113A>C 1114A>C 1115G>T), TTC327TTT (1121C>T), CCT328TCA (1122C>T 1124T>A), ACC329ATC (1126C>T), CTG334CTT (1142G>T), AGG335AAA (1144G>A 1145G>A), GAA336GAG (1148A>G), GAA337CAG (1149G>C 1151A>G), GCT338GCA (1154T>A), ATC341ATA (1163C>A), GCT343GCC (1169T>C), TAC344TAT (1172C>T), GCA346GCT (1178A>T), CCA349CCT (1187A>T), GTT350GTG (1190T>G), TTC353TTT (1199C>T), ATG354TTG (1200A>T), AGC355AGT (1205C>T), AGG357AGA (1211G>A), AGT359TCT (1215A>T 1216G>C), GCC361ACA (1221G>A 1223C>A), GCG363GTG (1228C>T), CAG364CAA (1232G>A), TTA365TTG (1235A>G), GAA366GAT (1238A>T), GTT368CTC (1242G>C 1244T>C), TAT369ATA (1245T>A 1246A>T 1247T>A), GCT370GCA (1250T>A), AGT372AAC (1255G>A 1256T>C), AGG374CAA (1260A>C 1261G>A 1262G>A), CAC375CAT (1265C>T), ATG376ACA (1267T>C 1268G>A), GGC377GGA (1271C>A), GGC379GGT (1277C>T), TAT381TAC (1283T>C), CGT382AGA (1284C>A 1286T>A), CCT385CCC (1295T>C), GTA388GTT (1304A>T), TTC391TTT (1313C>T), GGG392GGT (1316G>T), TCC393GAG (1317T>G 1318C>A 1319C>G), GGG395GGA (1325G>A), TGT396TGC (1328T>C), GGT397GGC (1331T>C), TCT399TCC (1337T>C), ACA400ACG (1340A>G), TCC401AGC (1341T>A 1342C>G), CGC403CGA (1349C>A), TTA404CTG (1350T>C 1352A>G), TTT405TTC (1355T>C), ATT406ATC (1358T>C), GTG409GTA (1367G>A), GAC411GAT (1373C>T), AGA412GCC (1374A>G 1375G>C 1376A>C), GGT414CAA (1380G>C 1381G>A 1382T>A), CCA416CCC (1388A>C), AAA417AAG (1391A>G), CTC418CTG (1394C>G), AAT419AAC (1397T>C), CGG420AGG (1398C>A), CTA421TTG (1401C>T 1403A>G), GTA424GTT (1412A>T), AGG427CGC (1419A>C 1421G>C), GCA429GCT (1427A>T), TAT430TAC (1430T>C), TCC432TCA (1436C>A), AAC433AAT (1439C>T), CTG441CTC (1463G>C), GAC444GAT (1472C>T), GGA446GGT (1478A>T), TTG448CTG (1482T>C), CGA449CGG (1487A>G), GGG451GGA (1493G>A), GGG453GGC (1499G>C), CAG454CAA (1502G>A), CAA456CAG (1508A>G), GAT457GAC (1511T>C), AAA459AAG (1517A>G), GAT460GAC (1520T>C), CTG461CTC (1523G>C), GAT463GAC (1529T>C), TCA466TCC (1538A>C), CCC471CCA (1553C>A), TTA472TTG (1556A>G), CCA473CCC (1559A>C), GCC475GCT (1565C>T), GTT477GTA (1571T>A), GAG478GAA (1574G>A), GGC481GGG (1583C>G), AGG482CGG (1584A>C), ACC485ACA (1595C>A), TCC486TCA (1598C>A), TAT488TAC (1604T>C), ACA492ACC (1616A>C), AAT494AAC (1622T>C), CTG495TTG (1623C>T), ATC496GTT (1626A>G 1628C>T), GCT498GCA (1634T>A), CCT499CCA (1637T>A), GCC500GCA (1640C>A), TGT502TGC (1646T>C), GGT503GGG (1649T>G), CTT504CTG (1652T>G), CCA507CCT (1661A>T), GAG511GAA (1673G>A), CGG512AGG (1674C>A), AGA513AGG (1679A>G), AGG514CGG (1680A>C), ATC516ATT (1688C>T), CTG517TTG (1689C>T), GAG519GAA (1697G>A), TGT520TGC (1700T>C), ATG521CGG (1701A>C 1702T>G), AAG522AAA (1706G>A), GAG523GTG (1708A>T), GGC524GGG (1712C>G), GAG525GAA (1715G>A), TTT526TTC (1718T>C), TCC527TCA (1721C>A), ACT528ACA (1724T>A), GAA529GAG (1727A>G), GAT530AAC (1728G>A 1730T>C), CCT531CCA (1733T>A), ACG532ACA (1736G>A), GGA533GCA (1738G>C), AGA536AGC (1748A>C), TTT537TTC (1751T>C), ATA539ATC (1757A>C), GTG540ATT (1758G>A 1760G>T), GAG541GGA (1762A>G 1763G>A), AGA543AGG (1769A>G), AGA544AGG (1772A>G), GCT547GCC (1781T>C), ATC548ATA (1784C>A), ACT549ACA (1787T>A), AGG551AGA (1793G>A), AAT552GAC (1794A>G 1796T>C), TTG553CTC (1797T>C 1799G>C), ACC554ACT (1802C>T), AGT556TCT (1806A>T 1807G>C), GAT557GAC (1811T>C), CTT558TTG (1812C>T 1814T>G), GAG560GAA (1820G>A), GTA563GTG (1829A>G), AAA565AAG (1835A>G), TGT566TGC (1838T>C), CAG567AAG (1839C>A), GTA568GTC (1844A>C), CAT569CAC (1847T>C), GCA570CAC (1848G>C 1849C>A 1850A>C), GTA572ATT (1854G>A 1856A>T), TCA573GCA (1857T>G), AAG574ACC (1861A>C 1862G>C), TTG576CTC (1866T>C 1868G>C), TTT577TAC (1870T>A 1871T>C), GAA578AAC (1872G>A 1874A>C), GCT579CAG (1875G>C 1876C>A 1877T>G), GAA580TCA (1878G>T 1879A>C), TCT581ACA (1881T>A 1883T>A), GGC582GGT (1886C>T), ATA583TGT (1887A>T 1888T>G 1889A>T), AGT584GAA (1890A>G 1891G>A 1892T>A), CCT585GAG (1893C>G 1894C>A 1895T>G), AAG586AAA (1898G>A), ATA587GTT (1899A>G 1901A>T), GCT588GCC (1904T>C), GTT590GTG (1910T>G), CAA591GAA (1911C>G), GCA594GTT (1921C>T 1922A>T), GAT595GAA (1925T>A), GAT596GAA (1928T>A), ATA598ATG (1934A>G), GCG599ACA (1935G>A 1937G>A), TCT600TCC (1940T>C), GTG601TTT (1941G>T 1943G>T), GAT602CAT (1944G>C), GGG603GAT (1948G>A 1949G>T), AGA605CAA (1953A>C 1954G>A), TTG606ATA (1956T>A 1958G>A), CGC607AGC (1959C>A), ACC608TTT (1962A>T 1963C>T 1964C>T), AAG609GAC (1965A>G 1967G>C), CAA610GAA (1968C>G), GAT611GTT (1972A>T), GAA612AGC (1974G>A 1975A>G 1976A>C), CCA613CCC (1979A>C), ATA614AGT (1981T>G 1982A>T), TTG615GTT (1983T>G 1985G>T), GTG616_CCC617del (1986_1991delGTGCCC), ACT618ACA (1994T>A), GTG620CTC (1998G>C 2000G>C), AGT621TCT (2001A>T 2002G>C), GAA622GAG (2006A>G), AAT623GAT (2007A>G), ATC627GGA (2019A>G 2020T>G 2021C>A), GCA629TCA (2025G>T), AGA630CAG (2028A>C 2029G>A 2030A>G), GAG631GAA (2033G>A), CTC632CTA (2036C>A), ACG633ACT (2039G>T), GTA634GTG (2042A>G), GAG635GAA (2045G>A), TTG637CTG (2049T>C), AAA638AAG (2054A>G), GCA640GCT (2060A>T), TAC641TAT (2063C>T), CAG642CAA (2066G>A), GGC643GGA (2069C>A), AGT644GCA (2070A>G 2071G>C 2072T>A), TTG645TTA (2075G>A), GAC646AAC (2076G>A), CCT647CCA (2081T>A), GAG648TTT (2082G>T 2083A>T 2084G>T), CTT650TAT (2088C>T 2089T>A), CCT652CCC (2096T>C), GAC654GAT (2102C>T), CAT655GCT (2103C>G 2104A>C), CAT656TCA (2106C>T 2107A>C 2108T>A), GCC659GCA (2117C>A), ATG660ATT (2120G>T), TAT661TTT (2122A>T), TCT663GTC (2127T>G 2128C>T 2129T>C), CTT664CTG (2132T>G), GAT665TCA (2133G>T 2134A>C 2135T>A), GAT666AGT (2136G>A 2137A>G), CAT668GCC (2142C>G 2143A>C 2144T>C), ATT671AGC (2152T>G 2153T>C), TTC672TTT (2156C>T), AAC673ACT (2158A>C 2159C>T), AAA674CGC (2160A>C 2161A>G 2162A>C), AGA676GTA (2166A>G 2167G>T), GTC677AAT (2169G>A 2170T>A 2171C>T), AAC678GAG (2172A>G 2174C>G), TTG680ATT (2178T>A 2180G>T), TAT681TAC (2183T>C), AGA682GAG (2184A>G 2185G>A 2186A>G), GGA683GGG (2189A>G), GCA684GCT (2192A>T), GAT685TCG (2193G>T 2194A>C 2195T>G), GCT686ACT (2196G>A), GAA687GCA (2200A>C), CAG688GGA (2202C>G 2203A>G 2204G>A), CGA690GCC (2208C>G 2209G>C 2210A>C), GTA693ATC (2217G>A 2219A>C), CAA694AAG (2220C>A 2222A>G), AAC695GCC (2223A>G 2224A>C), ATT696ATC (2228T>C), CCT697CCA (2231T>A), ATA701GTC (2241A>G 2243A>C), AAG705AAA (2255G>A), AGT706AGC (2258T>C), GCA710AGT (2268G>A 2269C>G 2270A>T), TCA711ACA (2271T>A), ATC713GTG (2277A>G 2279C>G), TCT714TCA (2282T>A), GAG715GAA (2285G>A), AAG716GCT (2286A>G 2287A>C 2288G>T), AAA717CAC (2289A>C 2291A>C), CTA718ATG (2292C>A 2294A>G), GCA719GAT (2296C>A 2297A>T), GTG720GTA (2300G>A), CAG721CAA (2303G>A), ACA722AGG (2305C>G 2306A>G), GAG723ATG (2307G>A 2308A>T), ATG724TTG (2310A>T), AGG725TAC (2313A>T 2314G>A 2315G>C), ACT726AAT (2317C>A), GGA727GGC (2321A>C), TTT728AGA (2322T>A 2323T>G 2324T>A), TCT731AGA (2331T>A 2332C>G 2333T>A), TGC732GCA (2334T>G 2335G>C 2336C>A), ATT733CTG (2337A>C 2339T>G), GAT734GAA (2342T>A), GCA735GCT (2345A>T), GTT737GCT (2350T>C), ACC739ACT (2357C>T), CTC740TTA (2358C>T 2360C>A), ATA741ATG (2363A>G), ATC743ATT (2369C>T), GAG744GAA (2372G>A), CAG745CAA (2375G>A), CCA747CCT (2381A>T), TTT749CAT (2385T>C 2386T>A), GTT750GTC (2390T>C), CAG751CAA (2393G>A), TTT752TTC (2396T>C), TTA753CTA (2397T>C), TAT754TAC (2402T>C), AGA759CGG (2415A>C 2417A>G), TGC762_AAC763insAGAAGG (2426_2427insAGAAGG), AAC763AAG (2429C>G), AAA764ATG (2431A>T 2432A>G), GTA765GTC (2435A>C), TCA766ACT (2436T>A 2438A>T), AAA767AAG (2441A>G), GAG768GAA (2444G>A), CCA769ACA (2445C>A), AAG770_GAA771del (2448_2453delAAGGAA), TCA772TCT (2456A>T), ATC774GTA (2460A>G 2462C>A), AAA775CAA (2463A>C), ATT776TCG (2466A>T 2467T>C 2468T>G), TGC777CTC (2469T>C 2470G>T), TAC778TAT (2474C>T), CAT779GAC (2475C>G 2477T>C), AAA780TCC (2478A>T 2479A>C 2480A>C), ATC781ACA (2482T>C 2483C>A), GTT782ATT (2484G>A), GAA783GGG (2488A>G 2489A>G), TAT784AGA (2490T>A 2491A>G 2492T>A), ATC785ATG (2495C>G), AAG786CAA (2496A>C 2498G>A), GAT787GAA (2501T>A), ACA788ATG (2503C>T 2504A>G), TGG790AAT (2508T>A 2509G>A 2510G>T), AGC791GAT (2511A>G 2512G>A 2513C>T), CTT792CTG (2516T>G), TAT794TAC (2522T>C), GCA795ATT (2523G>A 2524C>T 2525A>T), CTG796TTG (2526C>T), AGA797AAA (2530G>A), CTC799CTT (2537C>T), GGT803GGG (2549T>G), ATA806CTA (2556A>C), ATG807ATA (2561G>A), CTT809CTC (2567T>C), GGA811GGG (2573A>G), GTT812ACT (2574G>A 2575T>C), TTT813ATG (2577T>A 2579T>G), GGA814GGT (2582A>T), GGC815GCC (2584G>C), ATT816GTA (2586A>G 2588T>A), ACA817ACT (2591A>T), TTT819TTC (2597T>C), TTG820ATT (2598T>A 2600G>T), GCG821TCT (2601G>T 2603G>T), CAG824AAA (2610C>A 2612G>A), AGT825AGC (2615T>C), AAC826AGC (2617A>G), TCT827TTG (2620C>T 2621T>G), TTC828ATT (2622T>A 2624C>T), AAT830ACT (2629A>C), GCA831TCC (2631G>T 2633A>C), TCG832CAG (2634T>C 2635C>A), AGA834AAA (2641G>A), GTG837GTT (2651G>T), TCC841TCG (2663C>G), AGT843AGC (2669T>C), TCC845TCA (2675C>A), AGT846AGC (2678T>C), GGT851GGA (2693T>A), AAT852AGA (2695A>G 2696T>A), AAT852_AAG853insTCC (2696_2697insTCC), AAG853CAT (2697A>C 2699G>T), CTC854GTA (2700C>G 2702C>A), AGA855TCA (2703A>T 2704G>C), AAT856AAG (2708T>G), CTT857ATG (2709C>A 2711T>G), CTC858CTG (2714C>G), GTT859TTT (2715G>T), GGT860AGC (2718G>A 2720T>C), TCT863TCA (2729T>A), TCA865CCA (2733T>C), TTG866AAT (2736T>A 2737T>A 2738G>T), GCA867GAT (2740C>A 2741A>T), CAA868CAG (2744A>G), CAA868_GAT869insGGA (2744_2745insGGA), GAT869AAA (2745G>A 2747T>A), GCT872TCC (2754G>T 2756T>C), GAA873AGG (2757G>A 2758A>G 2759A>G), GGA875GGT (2765A>T), GTA877GTG (2771A>G), AAT878AAC (2774T>C), CAG879GAA (2775C>G 2777G>A), AGC880TCT (2778A>T 2779G>C 2780C>T), CTC881CTT (2783C>T), AAG882GCC (2784A>G 2785A>C 2786G>C), GTG886GTA (2798G>A), TTA888CTA (2802T>C), AGA889AGG (2807A>G), CTT890GTG (2808C>G 2810T>G), GGT891GCA (2812G>C 2813T>A), GTG894GTT (2822G>T), AGA897CGT (2829A>C 2831A>T), GGC898GGA (2834C>A), ACC899ACT (2837C>T), TGT902TGC (2846T>C), TCT903AGC (2847T>A 2848C>G 2849T>C), GGT904GAA (2851G>A 2852T>A), ACC908ACT (2864C>T), GCT910GCA (2870T>A), CAT911CAC (2873T>C), GCA912GCT (2876A>T), CAC914ACG (2880C>A 2881A>C 2882C>G), AAT915CAG (2883A>C 2885T>G), GCT916CTC (2886G>C 2887C>T 2888T>C), CGA917CGT (2891A>T), GGT919GGA (2897T>A), ACT920ACC (2900T>C), ACA923TCA (2907A>T), ATC925ATT (2915C>T), GCC927GAA (2920C>A 2921C>A), AAT928CAT (2922A>C), CGA930GTC (2928C>G 2929G>T 2930A>C), TCT931TGC (2932C>G 2933T>C), AAA932CAG (2934A>C 2936A>G), GTT933GTG (2939T>G), AAC936AAT (2948C>T), AGA938AAA (2953G>A), GAA939CAT (2955G>C 2957A>T), TCA940GCC (2958T>G 2960A>C), AGG941AAG (2962G>A), ATT942TTT (2964A>T), ATT943TTG (2967A>T 2969T>G), GAG944CCT (2970G>C 2971A>C 2972G>T), GGC945GAA (2974G>A 2975C>A), GAC947GAT (2981C>T), ATC948GTT (2982A>G 2984C>T), ATT949GTA (2985A>G 2987T>A), CTG950TTA (2988C>T 2990G>A), TTG951CTG (2991T>C), CGC952AGG (2994C>A 2996C>G), GGT954GGC (3002T>C), GAT955TGT (3003G>T 3004A>G), CCA956CCT (3008A>T), GAT957GAC (3011T>C), GGT958GGG (3014T>G), CCA959CCT (3017A>T), AAG960AAA (3020G>A), CCT961GTC (3021C>G 3022C>T 3023T>C), CGT964AGA (3030C>A 3032T>A), AAA965AAG (3035A>G), TTC967TTT (3041C>T), CCA968CGT (3042C>G 3044A>T), AGA969GAA (3045A>G 3046G>A), GAG972GCA (3055A>C 3056G>A), AAG977AAA

| | | | | | | | | | |
|---|---|---|---|---|---|---|---|---|---|
| | (3071G>A), GGC978GGA (3074C>A), TCA979TCT (3077A>T), CAA980GAA (3078C>G), GGA981GGC (3083A>C), TGC983TGT (3089C>T), TGT984GTC (3090T>G 3091G>T 3092T>C), GCT986GCA (3098T>A), GTT987GTG (3101T>G), GCG988GCA (3104G>A), TCT989TCA (3107T>A), ACA990AAA (3109C>A), CCA992CCT (3116A>T), CGT993GAG (3117C>G 3118G>A 3119T>G), CTT994AAA (3120C>A 3121T>A 3122T>A), GGC995CAG (3123G>C 3124G>A 3125C>G), AAT996AAC (3128T>C), TTA997TTG (3131A>G), GAG998GAC (3134G>C), ATG1002TAT (3144A>T 3145T>A 3146G>T), CCA1003TCA (3147C>T), TCA1007TCC (3161A>C), AAG1008AAA (3164G>A), TAC1010TAT (3170C>T), GTT1012GAG (3175T>A 3176T>G), GTG1014GTC (3182G>C), GAA1015ACA (3183G>A 3184A>C), TAT1016TAC (3188T>C), GAA1017AGG (3189G>A 3190A>G 3191A>G), CTG1018ATG (3192C>A), AAC1019AGC (3196A>G), TCC1020ACA (3198T>A 3200C>A), TCC1021AAT (3201T>A 3202C>A 3203C>T), AGT1022TCA (3204A>T 3205G>C 3206T>A), AAG1024CAC (3210A>C 3212G>C), AAG1024_ATT1025insTCA (3212_3213insTCA), ATT1025ATC (3215T>C), TGC1026AGT (3216T>A 3218C>T), TCT1027GCT (3219T>G), CAA1029GAA (3225C>G), TCT1030GCC (3228T>G 3230T>C), CAC1034CAT (3242C>T), ATA1035GTC (3243A>G 3245A>C), GGG1037AGC (3249G>A 3251G>C), GAC1041GAT (3263C>T), GGC1043GGA (3269C>A), ACG1044ACT (3272G>T), CTG1047TTA (3279C>T 3281G>A), TTA1048TTG (3284A>G), AGT1050TCT (3288A>T 3289G>C), GTT1051ATC (3291G>A 3293T>C), CAG1052CAA (3296G>A), AAT1053AAC (3299T>C), CCT1056CCA (3308T>A), GTG1057TGC (3309G>T 3310T>G 3311G>C), ATC1058ATT (3314C>T), GGC1060GGT (3320C>T), ATC1061ATT (3323C>T), CAC1062CAT (3326C>T), GCA1064GCT (3332A>T), TCA1065TCC (3335A>C), GGC1068GAA (3343G>A 3344C>A), GTA1069GCT (3346T>C 3347A>T), GCT1071ACA (3351G>A 3353T>A), GAA1072GAG (3356A>G), CGT1073TTG (3357C>T 3358G>T 3359T>G), ATC1076GTA (3366A>G 3368C>A), TCT1077GCA (3369T>G 3371T>A), GCT1080GCC (3380T>C), ACA1081ACT (3383A>T), ATC1083GTA (3387A>G 3389C>A), GAA1086GAC (3398A>C), CAA1087CAG (3401A>G), CTA1088CTG (3404A>G), GAG1089CAA (3405G>C 3407G>A), CCA1092GCT (3414C>G 3416A>T), ACT1093CAA (3417A>C 3418C>A 3419T>A), CCG1095CCA (3425G>A), GTC1096GTT (3428C>T), GCA1098ATG (3432G>A 3433C>T 3434A>G), GCA1099GCC (3437A>C), GTA1101GTG (3443A>G), CGC1102AGA (3444C>A 3446C>A), GAT1104GAC (3452T>C), ATT1105TTG (3453A>T 3455T>G), CTG1106CTC (3458G>C), AAG1107AAA (3461G>A), TCA1108CAT (3462T>C 3463C>A 3464A>T), ATT1109GTT (3465A>G), AGA1110CGA (3468A>C), AGC1111TCA (3471A>T 3472G>C 3473C>A), ACA1114ACT (3482A>T), CAG1115GCT (3483C>G 3484A>C 3485G>T), CTT1116CTC (3488T>C), GAG1118GAC (3494G>C), AAC1120AAG (3500C>G), GTG1122GTT (3506G>T), TAC1123GCT (3507T>G 3508A>C 3509C>T), TAC1124TTT (3511A>T 3512C>T), CTT1125TTG (3513C>T 3515T>G), GGA1126GGC (3518A>C), ACA1127ACT (3521A>T), CCA1129CCT (3527A>T), CAG1130CAA (3530G>A), TTG1132ATG (3534T>A), GCC1133GCA (3539C>A), GCC1134GCA (3542C>A), ACA1135ACG (3545A>G), GTT1136GTG (3548T>G), CCC1137CCA (3551C>A), ACT1141ACC (3563T>C), CTG1142CTC (3566G>C), CGG1143AGG (3567C>A), CAA1146CCA (3577A>C), TTG1147CTG (3579T>C), GAA1149GAG (3587A>G), TTC1151TTT (3593C>T), GGA1152GGT (3596A>T), CCT1153GAA (3597C>G 3598C>A 3599T>A), GAG1155CAA (3603G>C 3605G>A), ACA1156ACT (3608A>T), GCA1157GCT (3611A>T), CCA1158CCT (3614A>T), TCC1159TCA (3617C>A), ATT1160CTG (3618A>C 3620T>G), CTA1161CTC (3623A>C), ACA1162TCT (3624A>T 3626A>T), GTT1163GTG (3629T>G), CAT1164AAA (3630C>A 3632T>A), GAC1165GAT (3635C>T), GGT1168GGG (3644T>G), GGT1170GGA (3650T>A), GAC1172GAT (3656C>T), CCC1173CCT (3659C>T), GTG1175GTT (3665G>T), GGG1177GGC (3671G>C), GTA1178ATT (3672G>A 3674A>T), ATG1179GAG (3675A>G 3676T>A), GCT1185GCC (3695T>C), TGT1186TCA (3697G>C 3698T>A), GGA1187AAT (3699G>A 3700G>A 3701A>T), GAT1189GAG (3707T>G), GAT1190GAC (3710T>C), GAC1191AGC (3711G>A 3712A>G), GCC1193GCA (3719C>A), AAA1194CGG (3720A>C 3721A>G 3722A>G), CTA1195GCT (3723C>G 3724T>C 3725A>T), TTC1197TTT (3731C>T), AAT1199GGT (3735A>G 3736A>G), TGC1202TCT (3745G>C 3746C>T), TCG1203GCC (3747T>G 3749G>C), CTG1204ATT (3750C>A 3752G>T), CTA1205TTA (3753C>T), CCT1206CCA (3758T>A), ATC1207GTT (3759A>G 3761C>T), ATG1208CTC (3762A>C 3764G>C), CGT1209AAG (3765C>A 3766G>A 3767T>G), AGC1210AGT (3770C>T), CAG1211CAA (3773G>A), AAG1212AGG (3775A>G), ATC1213GTG (3777A>G 3779C>G), CCT1214CCA (3782T>A), GGG1215GGT (3785G>T), GGA1216GGT (3788A>T), CGT1217ATC (3789C>A 3790G>T 3791T>C), CCA1218CCC (3794A>C), TGT1219ACA (3795T>A 3796G>C 3797T>A), GAA1220ACA (3798G>A 3799A>C), AGG1221CGT (3801A>C 3803G>T), GAT1222TGT (3804G>T 3805A>G), GAT1224GAC (3812T>C), GTA1225ATC (3813G>A 3815A>C), GTG1226GTC (3818G>C), CTC1227CTG (3821C>G), GGA1229GGT (3827A>T), ATA1230GTT (3828A>G 3830A>T), GGA1232GGT (3836A>T), TGT1233TTT (3838G>T), GAT1234GAC (3842T>C), TAC1235TTC (3844A>T), TAT1236TTT (3847A>T), GGC1238TCC (3852G>T 3853G>C), GAG1240GAA (3860G>A), CTG1241CTC (3863G>C), TCT1244TCA (3872T>A), TGC1245TGT (3875C>T), TAT1247TAC (3881T>C), CCC1248CCT (3884C>T), AAC1250AAT (3890C>T), AAG1251AAA (3893G>A), ATG1252CTG (3894A>C), GGG1253GGT (3899G>T), ATG1254TTG (3900A>T), GGG1255GGT (3905G>T), ATG1256ACC (3907T>C 3908G>C), AAC1257AAT (3911C>T), AAG1258AAA (3914G>A), AGA1259AGG (3917A>G), TTT1261TTC (3923T>C), GTG1262GTT (3926G>T), CAA1263GAA (3927C>G), TCC1264GCA (3930T>G 3932C>A), ACT1265CAG (3933A>C 3934C>A 3935T>G), GGC1266GAG (3937G>A 3938C>G), GAA1267AGT (3939G>A 3940A>G 3941A>T), GGA1268ACC (3942G>A 3943G>C 3944A>C), GAG1269CAA (3945G>C 3947G>A), AGA1270AGG (3950A>G), GTG1271GTA (3953G>A), GAA1272CTA (3954G>C 3955A>T), AAA1274AAG (3962A>G), GAC1276AAT (3966G>A 3968C>T), ACT1277TCT (3969A>T), GTA1279GTT (3977A>T), GAA1281CAA (3981G>C), GAA1284GAG (3992A>G), GAG1285GAA (3995G>A), CTA1286TTA (3996C>T), CAG1289GAA (4005C>G 4007G>A), ATT1290ATA (4010T>A), AAA1292AAG (4016A>G), GGC1293GGT (4019C>T), ATC1294AAC (4021T>A), CAT1295TAT (4023C>T), GAT1297GAA (4031T>A), CTG1298ATG (4032C>A), GTC1299GTT (4037C>T), ACC1301ACT (4043C>T), TGC1303TGT (4049C>T), GCC1304GCT (4052C>T), AAA1305AAG (4055A>G), GAT1306GAC (4058T>C), CGC1308AGG (4062C>A 4064C>G), CTC1309CTA (4067C>A), CCA1310CCT (4070A>T), CTT1311CTG (4073T>G), GAG1312GAC (4076G>C), AAA1313AAG (4079A>G), ATC1314ATA (4082C>A), TAT1315TAC (4085T>C), GGC1316GGG (4088C>G), AAG1317AAA (4091G>A), AGA1318AAG (4093G>A 4094A>G), AAG1319AAA (4097G>A), AGG1321CGT (4101A>C 4103G>T), CTC1322CTT (4106C>T), ATA1325ATT (4115A>T), CTT1326TTG (4116C>T 4118T>G), CCT1327CCA (4121T>A), TAC1330TAT (4130C>T), TTG1333CTG (4137T>C), GTC1334GTG (4142C>G), AGG1335CGC (4143A>C 4145G>C), AAG1336AAA (4148G>A), TTT1338TTC (4154T>C), TTC1341TTT (4163C>T), TCA1342TCT (4166A>T), GCC1343GCA (4169C>A), ACA1344TGT (4170A>T 4171C>G 4172A>T), TTG1345TTA (4175G>A), TTG1348GCA (4182T>G 4183T>C 4184G>A), CAC1349CAT (4187C>T), GCT1351GTT (4192C>T), ATA1352CTG (4194A>C 4196A>G), CCA1353CCC (4199A>C), GTT1356GTG (4208T>G), ATT1358ATC (4214T>C), GAT1359AAT (4215G>A), CCT1360CCC (4220T>C), ACA1361ATT (4222C>T 4223A>T), AGT1362TCA (4224A>T 4225G>C 4226T>A), TCT1363CAT (4227T>C 4228C>A), ACA1366ACC (4238A>C), TTG1367ACC (4239T>A 4240T>C 4241G>C), TTG1368CTG (4242T>C), AAT1370AAC (4250T>C), GGG1371CAG (4251G>C 4252G>A), TTT1372TTC (4256T>C), AGA1373CGG (4257A>C 4259A>G), GCT1374AGT (4260G>A 4261C>G), TCA1376TCT (4268A>T), GAC1377GAT (4271C>T), GTG1378GTT (4274G>T), GGA1379GGT (4277A>T), TCA1381TCT (4283A>T), GCT1382GCA (4286T>A), GAT1383GAC (4289T>C), TAT1384TAC (4292T>C), AGA1390AGG (4310A>G), GCA1391GCG (4313A>G), CCT1392CCA (4316T>A), GTT1393ATA (4317G>A 4319T>A), GCT1395GCA (4325T>A), CAG1397CAA (4331G>A), TTT1399TTA (4337T>A), GAA1405GCA (4354A>C), TAC1407TAT (4361C>T), GGA1408GGG (4364A>G), AAG1410GCT (4368A>G 4369A>C 4370G>T), CCT1411CCA (4373T>A), GGC1412GGA (4376C>A), AGT1413AGC (4379T>C), CCT1414GAG (4380C>G 4381C>A 4382T>G), TCC1416TCA (4388C>A), GCT1418GCA (4394T>A), CGA1419AGA (4395C>A), GCA1421GCT (4403A>T), CTT1422CTG (4406T>G), TTA1423TTG (4409A>G), GCA1426GCT (4418A>T), TGT1428TGC (4424T>C), ACA1431ACT (4433A>T), CTG1432CTA (4436G>A), GTG1437CTC (4449G>C 4451G>C), TTT1438TTC (4454T>C), TTG1440TTA (4460G>A), GGG1442GGT (4466G>T), GGC1443GGA (4469C>A), CCA1445CCT (4475A>T), TCA1446TCT (4478A>T), GCA1449GCT (4487A>T), CTA1450CTC (4490A>C), ACG1451ACC (4493G>C), ATC1453ATA (4499C>A), TTC1454TTT (4502C>T), AAT1455AAC (4505T>C), TCT1456TCC (4508T>C), CTC1457CTT (4511C>T), CTC1458TTA (4512C>T 4514C>A), AAT1459AAC (4517T>C), CGA1464CGG (4532A>G), GCC1466GCT (4538C>T), TTT1467TTC (4541T>C), TCA1469TCG (4547A>G), TTG1470CTC (4548T>C 4550G>C), TTA1471CTC (4551T>C 4553A>C), AGA1473AGG (4559A>G), CAT1475CAC (4565T>C), ATT1476ATC (4568T>C), GCT1477AAA (4569G>A 4570C>A 4571T>A), GCC1478GCA (4574C>A), AGG1479AAG (4576G>A), GCT1480GCA (4580T>A), ATA1481TTG (4581A>T 4583A>G), GGA1482GGT (4586A>T), GTT1483GTA (4589T>A), AAA1484AAG (4592A>G), CCA1485TCA (4593C>T), TTC1488TTT (4604C>T), CAG1490GAC (4608C>G 4610G>C), CTA1491TTA (4611C>T), TTC1492TTT (4616C>T), ATA1493ATT (4619A>T), GCT1494GCA (4622T>A), GTT1495GTG (4625T>G), GGA1497GGG (4631A>G), GTT1502GTG (4646T>G), GTA1504GTG (4652A>G), TTA1506ATG (4656T>A 4658A>G), CAT1507CAC (4661T>C), CTC1508CTT (4664C>T), CAG1509CAA (4667G>A), TAT1511TAC (4673T>C), TCT1512GAG (4674T>G 4675C>A 4676T>G), CTG1513TTA (4677C>T 4679G>A), CCA1514CCT (4682A>T), AAT1515GCA (4683A>G 4684A>C 4685T>A), CAT1518CAA (4694T>A), TTA1520CTA (4698T>C), CTG1522CAA (4705T>A 4706G>A), GTC1523GTG (4709C>G), AAT1524AAC (4712T>C), GTA1525GTT (4715A>T), ATC1526ATT (4718C>T), ATT1527ATC (4721T>C), GGT1530GGC (4730T>C), GAC1532GAT (4736C>T), TCG1534AAC (4740T>A 4741C>A 4742G>C), GAT1536AAT (4746G>A), GTT1537GTC (4751T>C), GAG1539GAT (4757G>T), GTA1540GTC (4760A>C), CAA1541AAA (4761C>A), TTT1542TTC (4766T>C), CAA1543CAG (4769A>G), CTA1548TTG (4782C>T 4784A>G), ACT1549ACG (4787T>G), CTG1551TTG (4791C>T), AGT1552AGC (4796T>C), GGT1554GGC (4802T>C), TTT1555TTC (4805T>C), AAA1556AAG (4808A>G), CGA1557CGC (4811A>C), CTT1560TTG (4818C>T 4820T>G), GGA1561GGT (4823A>T), CAC1563CAA (4829C>A), ATG1564CTA (4830A>C 4832G>A), GCT1565GCC (4835T>C), CCT1566CCC (4838T>C), TCA1571TCC (4853A>C), ATC1572ATT (4856C>T), ATT1573GTG (4857A>G 4859T>G), GAG1574GAA (4862G>A), CCC1575CCA (4865C>A), ATA1579ATC (4877A>C), CCT1581CCA (4883T>A), GCC1582GCA (4886C>A), CCA1583GTT (4887C>G 4888C>T 4889A>T), GGT1584GGG (4892T>G), CCT1586CCA (4898T>A), GAC1587GAT (4901C>T), ACT1588ATC (4903C>T 4904T>C), CAG1589ACT (4905C>A 4906A>C 4907G>T), ATG1592TAT (4914A>T 4915T>A 4916G>T), ACA1597ACG (4931A>G), GGA1598GGT (4934A>T), ATA1599GTA (4935A>G), CGT1600AGA (4938C>A 4940T>A), GAA1601GAG (4943A>G), CAT1604CAG (4952T>G), CAC1605CAT (4955C>T), GGT1606GGA (4958T>A), CGT1607CAG (4960G>A 4961T>G), GTG1608GTC (4964G>C), GCT1609GTT (4966C>T), AAG1612ACA (4975A>C 4976G>A), CTT1613TTG (4977C>T 4979T>G), GTC1614ACA (4980G>A 4981T>C 4982C>A), TTA1615GAC (4983T>G 4984T>A 4985A>C), GAT1616AAT (4986G>A), GTT1617GTC (4991T>C), CAG1618CAA (4994G>A), ACG1619AAA (4996C>A 4997G>A), GCG1620GCA (5000G>A), TTG1621ATA (5001T>A 5003G>A), GAT1622ATG (5004G>A 5005A>T 5006T>G), GAA1623CAG (5007G>C 5009A>G), AGG1624CGA (5010A>C 5012G>A), GGT1625GGC (5015T>C), TTC1626TTG (5018C>G), AGA1627CCA (5019A>C 5020G>C), GCT1628ATC (5022G>A 5023C>T 5024T>C), ATC1630CAT (5028A>C 5029T>A 5030C>T), CCT1632CCA (5036T>A), TCC1633TCT (5039C>T), TTG1635GCT (5043T>G 5044T>C 5045G>T), GTG1637CAA (5049G>C 5050T>A 5051G>A), CAG1639AGT (5055C>A 5056A>G 5057G>T), GAA1640GAG (5060A>G), ATT1642CTA (5064A>C 5066T>A), GCA1643CAA (5067G>C 5068C>A), AAG1644GCC (5070A>G 5071A>C 5072G>C), GTA1645GTT (5075A>T), ACA1646ACT (5078A>T), GGG1647GGA (5081G>A), GAT1648GAC (5084T>C), TCA1649AGT (5085T>A 5086C>G 5087A>T), AGT1650AAC (5089G>A 5090T>C), GCC1651ATA (5091G>A 5092C>T 5093C>A), CTG1652GGG (5094C>G 5095T>G), ACA1653GCT (5097A>G 5099A>T), ATT1654CTG (5100A>C 5102T>G), TGT1655CTG (5103T>C 5104G>T 5105T>G), ATG1657CTG (5109A>C), GCA1658AGC (5112G>A 5113C>G 5114A>C), AAA1659AAG (5117A>G), GCA1660GAT (5119C>A 5120A>T), TCC1663ACT (5127T>A 5129C>T), TAT1664TAC (5132T>C), ACG1665GTG (5133A>G 5134C>T), CCA1666CCT (5138A>T), TTG1667CTT (5139T>C 5141G>T), GAC1668CCA (5142G>C 5143A>C 5144C>A), GCA1669GAA (5146C>A), GGT1670GGA (5150T>A), GAG1671AAA (5151G>A 5153G>A), AAG1672AAA (5156G>A), ATC1673ATT (5159C>T), ACA1674GAA (5160A>G 5161C>A), AAT1675GAT (5163A>G), GAG1677TCA (5169G>T 5170A>C 5171G>A), CGT1678CAA (5173G>A 5174T>A), GAT1679GGT (5176A>G), CTG1680GTC (5178C>G 5180G>C), AAT1681CAT (5181A>C), TGG1682GAA (5184T>G 5185G>A 5186G>A), GCA1684TGT (5190G>T 5191C>G 5192A>T), CCA1685TCT (5193C>T 5195A>T), AAC1686GGA (5196A>G 5197A>G 5198C>A), ATT1687GTT (5199A>G), GGT1688TTC (5202G>T 5203G>T 5204T>C), TTT1689TTC (5207T>C), GCA1692TCG (5214G>T 5216A>G), CGT1693CGA (5219T>A), ACT1694ACA (5222T>A), GCA1695GAA (5224C>A), GCC1696GTT (5227C>T 5228C>T), CAC1697AAT (5229C>A 5231C>T), TAC1698CAT (5232T>C 5234C>T), ACG1699AAA (5236C>A 5237G>A), GAA1702GAG (5246A>G), GGG1703GAC (5248G>A 5249G>C), TAC1704ACA (5250T>A 5251A>C 5252C>A), ATT1706CTT (5256A>C), AAC1708AAT (5264C>T), TGT1709TGC (5267T>C), AAG1713AAA (5279G>A), AAG1714AAA (5282G>A), TCC1715ACA (5283T>A 5285C>A), AAT1716ACA (5287A>C 5288T>A), GTT1718GTG (5294T>G), CCG1733CCC (5339G>C), TAC1734TAT (5342C>T), ACT1735ACA (5345T>A), TCA1737GCA (5349T>G), GCT1738GCC (5354T>C), ATA1739ATC (5357A>C), ATG1740AAG (5359T>A), GAT1741GAC (5363T>C), GCC1742AAT (5364G>A 5365C>A 5366C>T), AAG1744CAA (5370A>C 5372G>A), AAT1745AGT (5374A>G), GTG1746CGC (5376G>C 5377T>G 5378G>C), GCA1748TCA (5382G>T), GGA1749ATG (5385G>A 5386G>T 5387A>G), GGA1750GGG (5390A>G), CAT1751ATT (5391C>A 5392A>T), GTA1752GGA (5395T>G), GCA1753ACC (5397G>A 5399A>C), ACC1754CAA (5400A>C 5401C>A 5402C>A), AAT1755AAA (5405T>A), GTC1756GTT (5408C>T), GTT1757GTC (5411T>C), GTG1759ATT (5415G>A 5417G>T), GGA1761GGG (5423A>G), GGA1763GGT (5429A>T), GTC1766GTG (5438C>G), GGC1767AGT (5439G>A 5441C>T), CCA1768CCC (5444A>C), GCA1771GCT (5453A>T), GCG1772GCT (5456G>T), ATG1774TTT (5460A>T 5462G>T), GCC1775GCA (5465C>A), GCA1777TCA (5469G>T), ACA1778ACC (5474A>C), AGG1779AAT (5476G>A 5477G>T), CAA1780CAG (5480A>G), TGT1782AGA (5484T>A 5486T>A), GTG1783GCT (5488T>C 5489G>T), GAC1785CAA (5493G>C 5495C>A), ATA1786CTT (5496A>C 5498A>T), GTT1788GTC (5504T>C), CGC1791AGA (5511C>A 5513C>A), ATT1793TTG (5517A>T 5519T>G), GGA1794GCG (5521G>C 5522A>G), AAT1795GGC (5523A>G 5524A>G 5525T>C), GTG1796TTG (5526G>T), CTT1797CTG (5531T>G), GAC1798GAT (5534C>T), CTC1799TTG (5535C>T 5537C>G), ACC1801CAG (5541A>C 5542C>A 5543C>G), GGC1805GGT (5555C>T), GTG1807ATT (5559G>A 5561G>T), CAG1808CAA (5564G>A), TTT1810TTC (5570T>C), CTT1811CTC (5573T>C), CAA1812ACA (5574C>A 5575A>C), TGT1813_GTG1814insGTC (5579_5580insGTC), CCT1815CAG (5584C>A 5585T>G), CAT1816AAA (5586C>A 5588T>A), GGA1817AGT (5589G>A 5591A>T), GAC1818TGC (5592G>T 5593A>G), AAA1819AAG (5597A>G), GTG1820ACA (5598G>A 5599T>C 5600G>A), AGT1823GCT (5607A>G 5608G>C), GGT1824AGT (5610G>A), GCA1825GGG (5614C>G 5615A>G), TCG1826del (5616_5618delTCG), CTC1827CGC (5620T>G), CAC1828GAT (5622C>G 5624C>T), AGT1829GGA (5625A>G 5627T>A), AGT1830ATG (5629G>T 5630T>G), TTT1831TTG (5633T>G), ATG1832AAC (5635T>A 5636G>C), CAA1834CAT (5642A>T), CAA1834_GGG1835insCATGCT (5642_5643insCATGCT), GGG1835GCT (5644G>C 5645G>T), TTT1836TAT (5647T>A), GAA1837GCC (5650A>C 5651A>C), TTG1838CTT (5652T>C 5654G>T), GGC1839GGA (5657C>A), CTC1841CTT (5663C>T), CGC1842CGA (5666C>A), ATT1843ATA (5669T>A), ATA1844GTG (5670A>G 5672A>G), CAT1845GGC (5673C>G 5674A>G 5675T>C), GGT1846TGT (5676G>T), GAT1847TCA (5679G>T 5680A>C 5681T>A), TTA1848ATT (5682T>A 5684A>T), GCA1849TCA (5685G>T), AAA1850AGA (5689A>G), CAG1851AAG (5691C>A), ACA1852ATG (5695C>T 5696A>G), ATG1854GTG (5700A>G), CGG1855AGA (5703C>A 5705G>A), ATG1856TTG (5706A>T), CCA1857CCC (5711A>C), TAT1858TAC (5714T>C), GTG1859ACT (5715G>A 5716T>C 5717G>T), GTA1860GTT (5720A>T), GGA1861GGG (5723A>G), CAT1863ATG (5727C>A 5728A>T 5729T>G), GGA1864GGT (5732A>T), GGA1865GGT (5735A>T), GGA1867GCA (5740G>C), AAT1868GAT (5742A>G), TTT1869TGC (5746T>G 5747T>C), TCC1870TCA (5750C>A), CAG1872AGA (5754C>A 5755A>G 5756G>A), GAT1873AAT (5757G>A) | | | | | | | | |
| helicase (YP_001976154.1) | 1 | 98 | 100% | 652 | 97.2% | 98 (100%) | 96 (98.0%) | 0/0/0/0 | 0 |
| Protein mutations: | R16A (1374A>G 1375G>C 1376A>C), G18Q (1380G>C 1381G>A 1382T>A) | | | | | | | | |

| | Begin | End | Coverage | Score | Concordance | Matches | Identities | I/D/M/F* | Stop Codons |
|---|---|---|---|---|---|---|---|---|---|
| **NT** | **35** | **5769** | **79.3%** | **3466** | **30.8%** | **5682 (98.6%)** | **3757 (65.2%)** | **28/53** | |
| Codon mutations: | GGT1GGC (1331T>C), TCT3TCC (1337T>C), ACA4ACG (1340A>G), TCC5AGC (1341T>A 1342C>G), CGC7CGA (1349C>A), TTA8CTG (1350T>C 1352A>G), TTT9TTC (1355T>C), ATT10ATC (1358T>C), GTG13GTA (1367G>A), GAC15GAT (1373C>T), AGA16GCC (1374A>G 1375G>C 1376A>C), GGT18CAA (1380G>C 1381G>A 1382T>A), CCA20CCC (1388A>C), AAA21AAG (1391A>G), CTC22CTG (1394C>G), AAT23AAC (1397T>C), CGG24AGG (1398C>A), CTA25TTG (1401C>T 1403A>G), GTA28GTT (1412A>T), AGG31CGC (1419A>C 1421G>C), GCA33GCT (1427A>T), TAT34TAC (1430T>C), TCC36TCA (1436C>A), AAC37AAT (1439C>T), CTG45CTC (1463G>C), GAC48GAT (1472C>T), GGA50GGT (1478A>T), TTG52CTG (1482T>C), CGA53CGG (1487A>G), GGG55GGA (1493G>A), GGG57GGC (1499G>C), CAG58CAA (1502G>A), CAA60CAG (1508A>G), GAT61GAC (1511T>C), AAA63AAG (1517A>G), GAT64GAC (1520T>C), CTG65CTC (1523G>C), GAT67GAC (1529T>C), TCA70TCC (1538A>C), CCC75CCA (1553C>A), TTA76TTG (1556A>G), CCA77CCC (1559A>C), GCC79GCT (1565C>T), GTT81GTA (1571T>A), GAG82GAA (1574G>A), GGC85GGG (1583C>G), AGG86CGG (1584A>C), ACC89ACA (1595C>A), TCC90TCA (1598C>A), TAT92TAC (1604T>C), ACA96ACC (1616A>C), AAT98AAC (1622T>C) | | | | | | | | |
| RNA-dependent RNA polymerase (YP_001976153.1) | 1 | 249 | 100% | 1561 | 88.9% | 249 (100%) | 217 (87.1%) | 0/0/0/0 | 0 |
| Protein mutations: | E8D (4076G>C), R14K (4093G>A 4094A>G), T40C (4170A>T 4171C>G 4172A>T), L44A (4182T>G 4183T>C 4184G>A), A47V (4192C>T), I48L (4194A>C 4196A>G), D55N (4215G>A), T57I (4222C>T 4223A>T), S59H (4227T>C 4228C>A), L63T (4239T>A 4240T>C 4241G>C), G67Q (4251G>C 4252G>A), A70S (4260G>A 4261C>G), V89I (4317G>A 4319T>A), F95L (4337T>A), E101A (4354A>C), K106A (4368A>G 4369A>C 4370G>T), P110E (4380C>G 4381C>A 4382T>G), V133L (4449G>C 4451G>C), A173K (4569G>A 4570C>A 4571T>A), R175K (4576G>A), I177L (4581A>T 4583A>G), P181S (4593C>T), Q186D (4608C>G 4610G>C), L202M (4656T>A 4658A>G), S208E (4674T>G 4675C>A 4676T>G), N211A (4683A>G 4684A>C 4685T>A), H214Q (4694T>A), L218Q (4705T>A 4706G>A), S230N (4740T>A 4741C>A 4742G>C), D232N (4746G>A), E235D (4757G>T), Q237K (4761C>A) | | | | | | | | |
| Codon mutations: | AAA1AAG (4055A>G), GAT2GAC (4058T>C), CGC4AGG (4062C>A 4064C>G), CTC5CTA (4067C>A), CCA6CCT (4070A>T), CTT7CTG (4073T>G), GAG8GAC (4076G>C), AAA9AAG (4079A>G), ATC10ATA (4082C>A), TAT11TAC (4085T>C), GGC12GGG (4088C>G), AAG13AAA (4091G>A), AGA14AAG (4093G>A 4094A>G), AAG15AAA (4097G>A), AGG17CGT (4101A>C 4103G>T), CTC18CTT (4106C>T), ATA21ATT (4115A>T), CTT22TTG (4116C>T 4118T>G), CCT23CCA (4121T>A), TAC26TAT (4130C>T), TTG29CTG (4137T>C), GTC30GTG (4142C>G), AGG31CGC (4143A>C 4145G>C), AAG32AAA (4148G>A), TTT34TTC (4154T>C), TTC37TTT (4163C>T), TCA38TCT (4166A>T), GCC39GCA (4169C>A), ACA40TGT (4170A>T 4171C>G 4172A>T), TTG41TTA (4175G>A), TTG44GCA (4182T>G 4183T>C 4184G>A), CAC45CAT (4187C>T), GCT47GTT (4192C>T), ATA48CTG (4194A>C 4196A>G), CCA49CCC (4199A>C), GTT52GTG (4208T>G), ATT54ATC (4214T>C), GAT55AAT (4215G>A), CCT56CCC (4220T>C), ACA57ATT (4222C>T 4223A>T), AGT58TCA (4224A>T 4225G>C 4226T>A), TCT59CAT (4227T>C 4228C>A), ACA62ACC (4238A>C), TTG63ACC (4239T>A 4240T>C 4241G>C), TTG64CTG (4242T>C), AAT66AAC (4250T>C), GGG67CAG (4251G>C 4252G>A), TTT68TTC (4256T>C), AGA69CGG (4257A>C 4259A>G), GCT70AGT (4260G>A 4261C>G), TCA72TCT (4268A>T), GAC73GAT (4271C>T), GTG74GTT (4274G>T), GGA75GGT (4277A>T), TCA77TCT (4283A>T), GCT78GCA (4286T>A), GAT79GAC (4289T>C), TAT80TAC (4292T>C), AGA86AGG (4310A>G), GCA87GCG (4313A>G), CCT88CCA (4316T>A), GTT89ATA (4317G>A 4319T>A), GCT91GCA (4325T>A), CAG93CAA (4331G>A), TTT95TTA (4337T>A), GAA101GCA (4354A>C), TAC103TAT (4361C>T), GGA104GGG (4364A>G), AAG106GCT (4368A>G 4369A>C 4370G>T), CCT107CCA (4373T>A), GGC108GGA (4376C>A), AGT109AGC (4379T>C), CCT110GAG (4380C>G 4381C>A 4382T>G), TCC112TCA (4388C>A), GCT114GCA (4394T>A), CGA115AGA (4395C>A), GCA117GCT (4403A>T), CTT118CTG (4406T>G), TTA119TTG (4409A>G), GCA122GCT (4418A>T), TGT124TGC (4424T>C), ACA127ACT (4433A>T), CTG128CTA (4436G>A), GTG133CTC (4449G>C 4451G>C), TTT134TTC (4454T>C), TTG136TTA (4460G>A), GGG138GGT (4466G>T), GGC139GGA (4469C>A), CCA141CCT (4475A>T), TCA142TCT (4478A>T), GCA145GCT (4487A>T), CTA146CTC (4490A>C), ACG147ACC (4493G>C), ATC149ATA (4499C>A), TTC150TTT (4502C>T), AAT151AAC (4505T>C), TCT152TCC (4508T>C), CTC153CTT (4511C>T), CTC154TTA (4512C>T 4514C>A), AAT155AAC (4517T>C), CGA160CGG (4532A>G), GCC162GCT (4538C>T), TTT163TTC (4541T>C), TCA165TCG (4547A>G), TTG166CTC (4548T>C 4550G>C), TTA167CTC (4551T>C 4553A>C), AGA169AGG (4559A>G), CAT171CAC (4565T>C), ATT172ATC (4568T>C), GCT173AAA (4569G>A 4570C>A 4571T>A), GCC174GCA (4574C>A), AGG175AAG (4576G>A), GCT176GCA (4580T>A), ATA177TTG (4581A>T 4583A>G), GGA178GGT (4586A>T), GTT179GTA (4589T>A), AAA180AAG (4592A>G), CCA181TCA (4593C>T), TTC184TTT (4604C>T), CAG186GAC (4608C>G 4610G>C), CTA187TTA (4611C>T), TTC188TTT (4616C>T), ATA189ATT (4619A>T), GCT190GCA (4622T>A), GTT191GTG (4625T>G), GGA193GGG (4631A>G), GTT198GTG (4646T>G), GTA200GTG (4652A>G), TTA202ATG (4656T>A 4658A>G), CAT203CAC (4661T>C), CTC204CTT (4664C>T), CAG205CAA (4667G>A), TAT207TAC (4673T>C), TCT208GAG (4674T>G 4675C>A 4676T>G), CTG209TTA (4677C>T 4679G>A), CCA210CCT (4682A>T), AAT211GCA (4683A>G 4684A>C 4685T>A), CAT214CAA (4694T>A), TTA216CTA (4698T>C), CTG218CAA (4705T>A 4706G>A), GTC219GTG (4709C>G), AAT220AAC (4712T>C), GTA221GTT (4715A>T), ATC222ATT (4718C>T), ATT223ATC (4721T>C), GGT226GGC (4730T>C), GAC228GAT (4736C>T), TCG230AAC (4740T>A 4741C>A 4742G>C), GAT232AAT (4746G>A), GTT233GTC (4751T>C), GAG235GAT (4757G>T), GTA236GTC (4760A>C), CAA237AAA (4761C>A), TTT238TTC (4766T>C), CAA239CAG (4769A>G), CTA244TTG (4782C>T 4784A>G), ACT245ACG (4787T>G), CTG247TTG (4791C>T), AGT248AGC (4796T>C) | | | | | | | | |

*: Inserts / Deletes / Misaligned / Frameshifts

#### Analysis details

This analysis was performed with panviral2.64

### NGS Details (UN18_val): Torradovirus marchitezum (segment RNA 2)

#### Assembly

| | |
|---|---|
| Coverage Length | 4645 (1 contig(s)) |
| Depth Of Coverage | 709.3 |
| Number Of Reads | 26259 |
| Reads Per Million | 492.84 rpm (after QC) |
| Ambiguities | 0 |
| Assembly Method | de novo + reference guided assembly |
| Consensus Caller | Bcf Tools |

#### Coverage Map

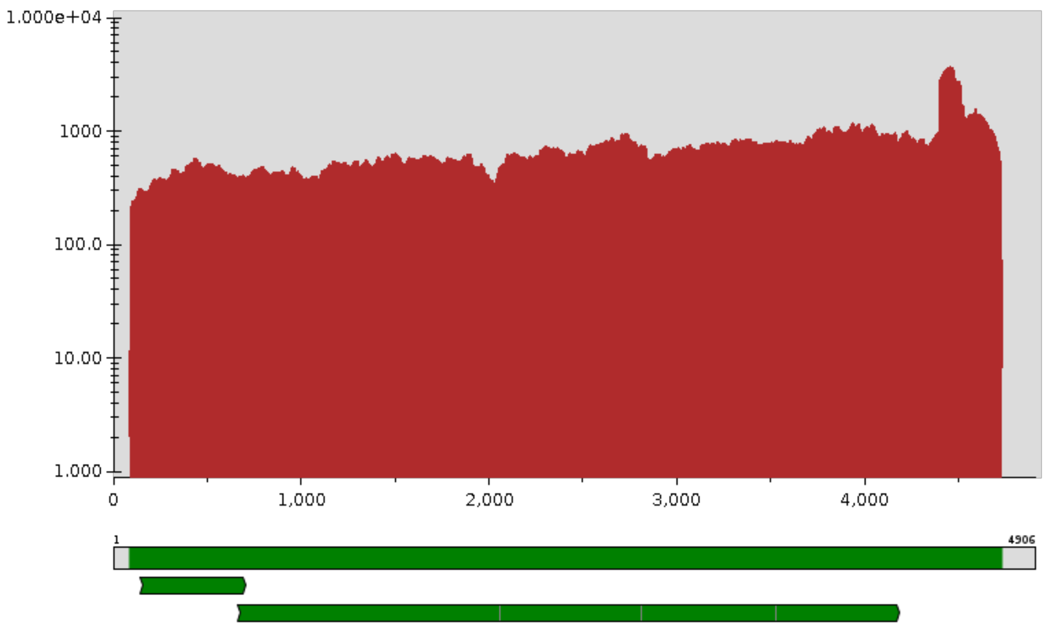

#### Assignment

| | |
|---|---|
| Type | Torradovirus marchitezum (Taxonomy ID: 3048376) |
| Reference Genome | NC_010988.1 |
| NT Identity (%) | 64.3495 |
| AA Identity (%) | 67.9739 |
| Number Of Stop Codons | 2 |
| Number Of CDS | 2 |

#### Alignment

| | |
|---|---|
| Alignment Score | 2570.0 (NT) + 6793.0 (AA) = 9363.0 |
| Concordance (%) | 50.7067 |

| **Alignment Method** | Global, seeded, nucleotide + amino acids (AGA) |
|---|---|

#### Genome Region

Sequence starts at position 89 and ends at position 4733 relative to NC_010988.1 reference sequence.

#### Alignment Detailed Statistics

| | Begin | End | Coverage | Score | Concordance | Matches | Identities | I/D/M/F* | Stop Codons |
|---|---|---|---|---|---|---|---|---|---|
| **NT** | **89** | **4733** | **94.7%** | **2570** | **28.4%** | **4597 (96.7%)** | **3027 (63.7%)** | **107/48** | |

Mutations:

101G>T, 106C>A, 109A>C, 110T>C, 111T>C, 118T>C, 120C>A, 121T>A, 122T>A, 124G>C, 128T>A, 129G>A, 130T>A, 132G>T, 133_134insCACTCT, 135C>T, 137T>G, 144A>T, 147C>T, 150T>C, 151G>A, 153G>T, 154C>A, 155G>A, 156T>G, 157T>C, 159G>C, 162T>C, 165A>C, 166T>C, 167C>A, 168T>A, 169G>A, 174C>G, 177A>T, 182C>A, 183T>A, 186T>C, 190C>A, 198T>A, 201C>A, 202A>G, 204C>A, 207G>T, 209A>C, 210T>C, 214A>G, 220T>A, 221C>G, 229G>A, 230T>C, 234C>T, 235A>G, 236C>T, 237C>T, 241T>A, 243G>T, 246A>T, 261C>T, 262C>T, 264T>A, 267C>T, 273A>G, 274G>A, 277G>T, 278T>G, 282T>C, 291T>A, 298G>A, 300C>T, 303A>G, 306T>G, 307T>C, 309A>T, 310A>T, 312C>T, 315T>G, 316T>A, 317C>G, 324G>A, 333G>T, 336A>C, 340T>G, 342A>T, 348T>A, 351A>C, 354C>T, 357T>C, 358T>C, 360G>A, 361T>C, 363G>A, 374A>C, 375T>G, 379C>G, 380C>A, 381C>T, 384T>C, 387G>A, 388A>G, 389A>C, 390C>A, 402T>A, 405T>C, 408A>G, 409C>A, 414C>G, 416A>G, 417T>A, 423A>C, 424G>A, 426G>T, 427C>A, 428G>A, 429T>A, 430C>T, 432T>G, 441C>A, 442A>C, 444A>G, 448G>T, 450C>G, 451C>T, 453A>G, 456G>A, 458A>C, 459A>C, 462C>T, 465G>A, 468T>G, 469T>C, 470C>A, 477A>T, 478G>T, 480A>T, 481T>A, 483A>C, 489A>T, 492T>A, 495T>C, 496C>A, 497C>A, 498T>C, 499T>C, 501T>C, 508_525delGACAGTAAAGGCAAGAGT, 527A>C, 529G>T, 532G>T, 533A>C, 534G>A, 535C>G, 538C>A, 540C>T, 541A>G, 544_549delCGAGAG, 550G>A, 551A>G, 553C>T, 555C>A, 557T>G, 558T>C, 559A>G, 560A>G, 564A>G, 565G>A, 566T>A, 567T>C, 568G>A, 572T>C, 573T>A, 576C>T, 579A>G, 580C>G, 581G>A, 582T>A, 584A>C, 585G>C, 586T>C, 588A>T, 591G>A, 592A>G, 593G>T, 594A>T, 595T>G, 596T>C, 597T>C, 598C>A, 599A>G, 600G>A, 606G>A, 607T>C, 610G>C, 611C>A, 613A>G, 614G>A, 615T>A, 616C>G, 617A>C, 619A>C, 620A>G, 621A>G, 629A>T, 631C>G, 632A>T, 633G>C, 639A>C, 640C>A, 641A>G, 642A>G, 643C>T, 645T>G, 649C>A, 651C>T, 652T>C, 653C>A, 654C>A, 655A>G, 656A>G, 661G>C, 662T>C, 663T>A, 666T>C, 669T>C, 670A>G, 672T>G, 675C>T, 676A>G, 681C>T, 682T>G, 687T>C, 693T>G, 694G>A, 695A>G, 696A>T, 697A>G, 698G>A, 702G>C, 706C>A, 710A>G, 712T>C, 714A>G, 716C>T, 717A>C, 718G>A, 719_721delCAG, 722G>A, 724G>A, 729C>T, 730C>A, 733C>T, 736G>C, 738G>A, 739G>A, 745A>T, 748A>T, 751A>G, 754A>C, 760T>G, 766G>T, 769G>A, 771C>A, 772C>A, 775A>T, 784T>C, 785T>C, 787G>T, 790C>T, 792G>A, 796G>A, 797C>T, 799T>G, 802C>A, 811C>A, 814C>T, 817G>C, 818G>A, 820T>A, 821C>T, 823T>G, 828G>C, 829T>A, 835A>G, 841G>A, 844A>G, 853G>T, 854T>C, 856G>T, 858T>A, 862C>T, 874A>T, 875T>A, 876C>G, 877A>T, 880G>A, 881A>G, 883C>G, 884A>G, 892T>A, 899A>G, 901A>T, 904C>T, 908C>A, 909A>G, 910T>G, 912A>C, 913G>C, 914T>A, 916T>A, 917G>A, 918T>C, 919T>A, 922G>A, 925T>A, 926A>C, 927A>G, 928G>T, 931C>T, 936C>G, 937A>G, 940C>T, 943G>A, 949T>C, 953C>T, 955G>A, 956T>C, 958T>C, 961T>G, 965C>T, 967T>G, 970C>T, 973C>T, 979C>T, 982A>C, 987T>C, 988T>A, 993G>A, 997G>A, 1003A>T, 1004A>G, 1005G>C, 1006G>C, 1009T>C, 1012G>A, 1015A>C, 1018T>C, 1019C>T, 1021T>G, 1024T>C, 1027A>T, 1030T>C, 1033T>C, 1040C>A, 1041T>C, 1042C>T, 1043T>G, 1044C>A, 1045A>T, 1048G>T, 1057A>G, 1063G>A, 1065C>T, 1066A>T, 1067C>T, 1071C>A, 1075G>T, 1076G>T, 1078C>T, 1080A>G, 1081T>A, 1087G>A, 1090G>A, 1093A>C, 1099C>A, 1105C>T, 1108C>T, 1111C>T, 1127C>T, 1129C>G, 1132A>G, 1135A>G, 1138G>C, 1156C>A, 1162G>A, 1165C>T, 1167A>T, 1168A>C, 1171T>C, 1174C>G, 1177G>A, 1180A>T, 1183G>A, 1189G>T, 1195C>T, 1198G>A, 1204C>A, 1207A>T, 1223G>A, 1224A>G, 1225C>T, 1226A>C, 1228A>T, 1229A>C, 1231G>T, 1237A>T, 1240T>A, 1244T>A, 1247C>G, 1249G>C, 1252A>T, 1255A>C, 1258A>T, 1262T>A, 1263C>G, 1270C>T, 1273A>T, 1276A>G, 1279A>T, 1282C>T, 1285T>C, 1286C>A, 1288T>A, 1291G>T, 1294A>G, 1306T>C, 1309A>G, 1315G>A, 1318T>A, 1321T>A, 1327T>C, 1330A>T, 1333T>A, 1339T>G, 1342T>C, 1345G>C, 1347G>A, 1351_1352insGTA, 1354A>G, 1357T>A, 1359C>G, 1363A>G, 1364A>C, 1367_1369delGCT, 1370C>T, 1372T>G, 1375A>G, 1380A>G, 1382G>C, 1384C>T, 1390T>G, 1393A>T, 1395A>C, 1396A>C, 1397C>G, 1398A>C, 1399G>A, 1405G>T, 1407G>A, 1408C>T, 1411A>C, 1418G>A, 1420C>T, 1423C>G, 1424A>C, 1425T>C, 1427A>C, 1429T>G, 1432A>G, 1433A>G, 1434G>C, 1435C>A, 1437C>A, 1442A>G, 1444C>G, 1447G>C, 1448A>T, 1449G>C, 1456T>A, 1462A>G, 1471T>C, 1474C>A, 1477A>G, 1480G>A, 1488A>G, 1492C>A, 1496C>G, 1498G>A, 1501C>A, 1508C>T, 1513G>T, 1513_1514insGACTTT, 1517G>T, 1518T>C, 1521G>C, 1523A>T, 1525A>T, 1526T>A, 1527C>G, 1529T>G, 1531C>G, 1533T>A, 1534A>T, 1536T>A, 1537C>G, 1538A>G, 1540C>A, 1541A>G, 1542C>T, 1543A>G, 1544C>A, 1546T>A, 1549A>G, 1555A>T, 1557T>G, 1559C>A, 1560A>T, 1565_1567delGAG, 1574T>A, 1575G>C, 1576T>G, 1577G>T, 1578G>C, 1579G>T, 1580G>C, 1581G>T, 1583T>C, 1591A>T, 1594A>C, 1595_1600delCCCAAG, 1603A>G, 1605A>T, 1606G>A, 1607G>T, 1608G>C, 1609C>A, 1615C>A, 1616T>G, 1617T>G, 1618C>A, 1620A>C, 1621C>A, 1622A>C, 1624G>C, 1625G>C, 1626G>A, 1627T>A, 1629C>A, 1630C>A, 1631A>G, 1633T>C, 1639T>C, 1644T>A, 1645G>T, 1651G>T, 1652G>C, 1654G>A, 1655T>A, 1656C>G, 1657G>T, 1660C>T, 1662C>A, 1664C>G, 1666C>T, 1667A>T, 1671A>C, 1672T>C, 1679G>A, 1680A>G, 1682G>A, 1684T>C, 1692A>C, 1693A>C, 1695G>T, 1696A>T, 1700A>G, 1702A>G, 1703G>C, 1705T>C, 1706C>G, 1711G>A, 1712G>A, 1714A>T, 1715C>G, 1717A>T, 1719T>A, 1721A>G, 1723C>T, 1726C>T, 1727T>G, 1729T>G, 1730A>C, 1731G>A, 1732T>G, 1733G>A, 1740A>G, 1742A>G, 1743G>C, 1744A>T, 1747A>T, 1750C>T, 1751G>C, 1753T>C, 1754A>T, 1755T>G, 1759G>C, 1764G>A, 1769G>A, 1773C>A, 1774A>G, 1776A>T, 1778C>A, 1781C>G, 1783T>C, 1784C>T, 1789T>C, 1790A>T, 1791A>G, 1795C>A, 1799A>T, 1800G>T, 1801G>T, 1804G>A, 1807C>T, 1810A>C, 1813C>T, 1818A>C, 1828A>T, 1831A>T, 1834A>T, 1835C>T, 1837T>G, 1839A>C, 1840A>G, 1843T>A, 1849T>A, 1858T>G, 1859A>G, 1861C>T, 1862C>G, 1863A>G, 1864A>G, 1865G>A, 1866G>T, 1867C>G, 1871A>G, 1872C>A, 1873T>A, 1874C>A, 1876C>A, 1879T>A, 1880T>C, 1881T>C, 1882C>A, 1891T>C, 1892G>T, 1895A>T, 1898T>A, 1899C>G, 1900T>C, 1903C>G, 1907G>C, 1915T>G, 1918G>T, 1924C>A, 1930T>C, 1931C>A, 1932A>T, 1933A>G, 1936A>G, 1939T>C, 1943T>C, 1944C>A, 1945C>A, 1948C>T, 1953T>G, 1954T>G, 1955C>T, 1957A>T, 1958G>T, 1961C>G, 1963G>T, 1964A>T, 1965C>T, 1966T>A, 1968A>C, 1969C>A, 1972T>C, 1975A>G, 1981G>T, 1982A>G, 1984T>C, 1991T>G, 1992G>T, 1993C>G, 2002A>T, 2003T>C, 2005A>T, 2008C>A, 2009T>C, 2012A>G, 2013A>C, 2017G>T, 2018C>A, 2020T>G, 2020_2021insGTGAAA, 2023G>T, 2024T>C, 2025T>C, 2026G>C, 2027A>T, 2029T>G, 2030G>A, 2033G>A, 2034A>G, 2035G>T, 2038G>T, 2040A>T, 2043T>C, 2044G>A, 2045G>T, 2046C>G, 2050_2051insTTGAAT, 2051A>C, 2053A>C, 2054A>T, 2055G>T, 2057T>G, 2058C>G, 2059T>A, 2060G>A, 2063A>C, 2064G>C, 2066C>A, 2067A>G, 2068A>C, 2069G>A, 2070C>T, 2071A>G, 2072C>A, 2073A>G, 2074G>T, 2078A>G, 2081G>C, 2082C>A, 2085G>C, 2086G>T, 2089C>A, 2091T>A, 2092A>T, 2095T>C, 2095_2096insAGC, 2096G>A, 2098A>G, 2099A>T, 2101T>G, 2104G>A, 2105C>A, 2112A>G, 2113C>A, 2115G>A, 2116A>G, 2120A>C, 2121C>T, 2122T>G, 2128C>A, 2134G>T, 2135T>A, 2140T>C, 2143C>A, 2144A>G, 2146G>A, 2147A>C, 2149G>T, 2152A>G, 2152_2153insAGT, 2155T>G, 2156C>G, 2157C>G, 2158C>T, 2160C>T, 2162A>C, 2164G>A, 2165C>A, 2166C>G, 2168G>A, 2170T>G, 2171A>G, 2174A>G, 2176A>G, 2177G>T, 2178A>C, 2184T>A, 2185T>G, 2188A>T, 2191G>A, 2193G>A, 2194A>T, 2195G>C, 2197A>G, 2202A>G, 2203A>G, 2204G>A, 2209T>A, 2210C>G, 2212A>G, 2216G>A, 2217G>A, 2221T>C, 2222G>A, 2225A>T, 2227T>C, 2233C>T, 2239A>G, 2240C>G, 2241C>A, 2242A>G, 2249G>A, 2250A>T, 2251C>G, 2252A>G, 2253C>G, 2254A>T, 2255T>G, 2256C>A, 2257C>T, 2258A>G, 2259T>C, 2260G>C, 2261G>C, 2263A>G, 2264G>T, 2266T>A, 2267G>A, 2269A>T, 2275T>G, 2276A>T, 2281G>A, 2284A>G, 2287G>A, 2289T>C, 2290G>A, 2296A>G, 2302C>T, 2308T>G, 2311C>T, 2323A>G, 2324G>A, 2335C>T, 2336G>C, 2338T>G, 2342G>A, 2344G>A, 2348T>A, 2349C>G, 2351G>A, 2352C>A, 2353T>C, 2365G>A, 2368A>G, 2371A>T, 2372A>C, 2374G>T, 2375A>G, 2376G>C, 2377C>A, 2378T>C, 2380G>A, 2381G>T, 2383T>C, 2384T>C, 2386G>C, 2389C>A, 2390G>A, 2391C>T, 2392T>G, 2395C>A, 2396C>T, 2398A>G, 2401C>T, 2402T>A, 2403C>G, 2404A>C, 2407A>G, 2409G>A, 2410C>G, 2413A>G, 2417A>G, 2422A>C, 2428T>G, 2429G>T, 2431G>C, 2440T>C, 2446T>C, 2449C>T, 2452T>C, 2455A>T, 2458T>A, 2459C>A, 2460T>A, 2461A>C, 2462A>C, 2464C>T, 2469A>T, 2470C>T, 2474A>G, 2476A>C, 2482T>G, 2485T>C, 2486G>A, 2495G>A, 2497T>G, 2503T>G, 2504A>G, 2506C>G, 2509T>G, 2512T>A, 2515A>G, 2516C>T, 2518G>A, 2521T>C, 2536A>G, 2539T>C, 2544T>C, 2545G>A, 2552C>A, 2556A>C, 2557G>C, 2572C>T, 2575C>A, 2581A>C, 2582T>A, 2584G>T, 2596G>A, 2600A>C, 2601C>A, 2602A>T, 2603T>C, 2608T>G, 2618C>G, 2624A>T, 2625A>C, 2631A>G, 2633G>A, 2639T>G, 2640G>T, 2641C>G, 2644T>C, 2645A>T, 2647C>A, 2650A>T, 2651A>C, 2652C>A, 2656G>T, 2662C>T, 2664A>C, 2665G>C, 2667A>T, 2668C>T, 2671G>A, 2674T>A, 2675C>T, 2681G>A, 2682A>C, 2683A>T, 2684G>T, 2685G>C, 2686T>A, 2687A>C, 2688C>T, 2689A>G, 2692A>T, 2694C>T, 2695C>T, 2696A>C, 2701T>A, 2702T>C, 2704A>T, 2707T>C, 2713G>T, 2728A>C, 2731C>T, 2737A>G, 2738T>G, 2740T>A, 2746A>C, 2752G>A, 2756A>G, 2758T>G, 2759A>T, 2760G>C, 2761T>A, 2762G>A, 2767A>T, 2770C>T, 2773T>C, 2774T>A, 2776G>A, 2780T>C, 2783C>T, 2784A>C, 2785G>C, 2791A>G, 2795T>C, 2797A>C, 2803C>A, 2810C>A, 2812T>G, 2822C>A, 2824A>G, 2825A>T, 2829T>G, 2832C>G, 2833A>G, 2836G>A, 2842A>G, 2844A>T, 2848C>A, 2852A>C, 2853A>G, 2855G>C, 2857G>A, 2860G>C, 2861C>A, 2862A>C, 2863G>A, 2866A>C, 2868A>T, 2869T>C, 2873C>A, 2874C>G, 2875A>T, 2876A>C, 2878G>C, 2881T>C, 2884A>T, 2887T>A, 2896T>G, 2897T>C, 2899G>T, 2900T>C, 2902A>G, 2905T>C, 2908T>C, 2915G>A, 2917G>T, 2918A>G, 2919C>T, 2921T>G, 2923A>T, 2924T>G, 2926C>A, 2927C>G, 2929A>C, 2941A>T, 2950C>T, 2951A>G, 2953A>C, 2956T>C, 2959A>C, 2971A>T, 2977G>A, 2986C>T, 2988C>T, 2992G>A, 2998G>A, 3001T>A, 3002T>C, 3010T>A, 3016A>T, 3019C>A, 3022G>A, 3031G>A, 3039C>G, 3040T>A, 3043A>C, 3046T>C, 3059A>G, 3061T>G, 3064C>T, 3065A>G, 3076C>T, 3077T>C, 3088A>T, 3091T>G, 3097G>T, 3100A>T, 3103T>A, 3106A>G, 3107C>T, 3109T>G, 3112A>C, 3115T>G, 3119A>G, 3120A>C, 3121C>A, 3124C>A, 3125A>G, 3127A>G, 3131A>T, 3135A>C, 3136T>A, 3139T>G, 3151T>G, 3160A>C, 3169A>G, 3172C>T, 3176C>T, 3178T>G, 3187T>G, 3192A>G, 3193A>G, 3198G>A, 3199G>A, 3203C>A, 3204G>A, 3205G>A, 3208T>C, 3211T>A, 3212A>C, 3214T>G, 3217A>C, 3229A>G, 3230A>T, 3231A>G, 3232A>C, 3235T>C, 3236T>C, 3238G>C, 3241T>G, 3244C>T, 3247A>C, 3250G>A, 3253A>C, 3256G>A, 3262C>A, 3265G>A, 3275C>A, 3280C>T, 3281T>C, 3282T>A, 3283T>A, 3286C>T, 3290T>A, 3295C>T, 3296T>C, 3301T>C, 3307G>T, 3310A>T, 3322C>G, 3331G>A, 3335A>T, 3337A>G, 3340T>C, 3347T>A, 3349C>T, 3355T>A, 3358C>A, 3367G>A, 3373G>A, 3374T>C, 3377C>A, 3383A>G, 3388T>C, 3391T>G, 3392T>C, 3394G>T, 3397T>G, 3400C>T, 3401A>T, 3403A>T, 3406A>G, 3409T>A, 3412G>T, 3415A>G, 3421A>T, 3427G>T, 3436A>C, 3439A>G, 3442C>G, 3444A>G, 3448C>T, 3449T>G, 3451A>T, 3454T>G, 3460C>T, 3463T>G, 3466C>T, 3476G>A, 3478T>C, 3479T>C, 3481A>G, 3482T>A, 3483C>A, 3484A>T, 3490T>C, 3497A>C, 3502A>G, 3505G>A, 3506T>C, 3514G>A, 3517C>A, 3520C>T, 3521A>G, 3526G>T, 3527G>A, 3528A>T, 3529G>C, 3531A>T, 3532C>T, 3533C>G, 3535G>C, 3536G>C, 3544C>G, 3548G>A, 3550T>C, 3551_3556delCCGCCC, 3561A>C, 3562G>A, 3563G>T, 3569A>G, 3570G>T, 3572A>T, 3573T>C, 3574T>C, 3575C>T, 3577G>C, 3578T>C, 3579C>A, 3580C>G, 3583A>C, 3586A>G, 3589G>A, 3592A>G, 3595T>C, 3596T>A, 3598C>A, 3604T>A, 3607T>A, 3610A>G, 3619C>T, 3622C>T, 3623A>G, 3624T>C, 3625G>T, 3627G>C, 3628T>A, 3638G>A, 3639A>C, 3640C>A, 3643A>C, 3650T>G, 3652G>A, 3655A>T, 3656A>T, 3658T>G, 3661T>A, 3667A>C, 3670A>T, 3676C>A, 3677A>G, 3679A>G, 3680A>C, 3682G>C, 3688A>T, 3690G>C, 3691G>A, 3692C>T, 3693A>C, 3700T>A, 3701G>A, 3706G>A, 3715C>A, 3721T>C, 3727T>C, 3733T>C, 3740A>T, 3742A>C, 3745A>T, 3760A>T, 3763T>A, 3770A>G, 3772T>A, 3776T>A, 3779A>T, 3784G>A, 3787A>G, 3791A>C, 3794G>A, 3795T>C, 3796A>T, 3799A>G, 3800T>A, 3801C>G, 3803A>T, 3804G>C, 3805T>C, 3806C>A, 3807C>A, 3808T>C, 3810A>G, 3812G>A, 3814T>A, 3817A>T, 3821G>A, 3823A>C, 3824C>T, 3826A>G, 3829T>C, 3830G>A, 3835T>A, 3836T>C, 3841T>C, 3842G>A, 3844C>A, 3850C>G, 3859T>C, 3861C>A, 3863G>T, 3864C>T, 3868G>T, 3869C>G, 3871T>A, 3872A>T, 3873A>C, 3875A>G, 3876A>C, 3877A>G, 3881A>G, 3882A>C, 3883T>A, 3884T>C, 3890G>T, 3901T>C, 3904A>T, 3907C>A, 3910A>T, 3925A>C, 3928C>T, 3931T>G, 3934G>T, 3935G>A, 3937A>C, 3940A>C, 3943T>C, 3945C>T, 3946G>C, 3952A>T, 3955C>T, 3956A>G, 3957C>T, 3958T>A, 3971G>C, 3976T>C, 3979C>T, 3982A>T, 3983A>C, 3985G>C, 3988G>T, 3989C>T, 3990A>T, 3991T>C, 3996G>A, 3998A>G, 3999T>C, 4000G>C, 4002G>A, 4003G>A, 4004G>T, 4005A>C, 4006A>C, 4008T>A, 4009C>T, 4010T>G, 4011C>A, 4012A>T, 4014G>A, 4015C>G, 4016A>G, 4018G>A, 4019C>A, 4020A>G, 4021A>T, 4024C>A, 4025C>A, 4027C>A, 4030C>T, 4031A>G, 4033G>T, 4036A>T, 4037C>T, 4039A>G, 4045T>C, 4047G>A, 4051T>A, 4054A>C, 4055A>T, 4056A>T, 4057T>G, 4058C>T, 4061A>C, 4063C>A, 4066A>C, 4070T>C, 4072G>T, 4075T>A, 4078T>A, 4079T>G, 4080C>T, 4081C>T, 4083C>A, 4084C>A, 4085A>C, 4087A>T, 4088G>A, 4090G>T, 4096C>A, 4102G>A, 4105A>C, 4106C>A, 4108T>A, 4109A>G, 4111A>C, 4112T>A, 4113C>A, 4114T>G, 4117T>A, 4120A>T, 4122T>C, 4129C>T, 4132G>A, 4144C>T, 4150T>C, 4153T>A, 4154T>C, 4155C>T, 4159C>A, 4163C>A, 4168A>G, 4171A>G, 4177T>C, 4180C>A, 4181A>G, 4189A>G, 4192T>G, 4195C>T, 4198C>T, 4210C>A, 4213A>T, 4228A>G, 4242T>A, 4247A>G, 4248A>C, 4259C>T, 4261A>G, 4267T>A, 4274T>C, 4276G>A, 4279A>T, 4280A>T, 4281A>T, 4283A>C, 4284C>T, 4285A>T, 4286C>T, 4287T>G, 4289A>T, 4291C>A, 4291_4292insCGTA, 4294C>T, 4304T>G, 4306G>A, 4309G>A, 4312G>T, 4312_4313insAGTAAATAAATAAAGT, 4317G>T, 4320G>T, 4324T>G, 4325T>G, 4330G>C, 4331C>A, 4333A>G, 4336A>T, 4339A>G, 4340G>T, 4342G>T, 4343A>T, 4345A>T, 4346G>C, 4349A>G, 4351G>A, 4363T>C, 4364C>T, 4365T>C, 4370A>T, 4372A>G, 4373G>A, 4374A>G, 4384C>T, 4385C>G, 4386T>A, 4391T>G, 4393C>T, 4394A>G, 4398A>T, 4399G>T, 4400A>G, 4404A>G, 4415T>C, 4420delA, 4422A>G, 4426G>A, 4437C>T, 4441T>C, 4449A>T, 4451T>G, 4472A>G, 4477_4478insTCCTCCACA, 4479A>C, 4480T>A, 4482T>A, 4486G>T, 4488T>A, 4489T>A, 4490A>C, 4491A>G, 4492C>G, 4493A>T, 4497C>G, 4499C>G, 4500G>T, 4501T>G, 4502T>A, 4506A>T, 4510T>A, 4510_4511insTCACTTA, 4518A>G, 4519G>A, 4520A>C, 4521C>A, 4523T>C, 4525A>G, 4526T>G, 4532C>G, 4534A>T, 4535A>T, 4539A>C, 4540_4541insAGTTAACGCGAACACAGGTTGTTTCCCTATAGAAACTA, 4542C>A, 4544T>C, 4548T>G, 4552C>G, 4553A>C, 4554T>A, 4556T>C, 4557A>T, 4559G>A, 4562A>G, 4563_4564delTA, 4566A>G, 4568G>A, 4569A>G, 4570G>C, 4581C>T, 4594T>C, 4595T>A, 4602C>G, 4606A>T, 4613T>C, 4614C>T, 4616C>T, 4618A>G, 4626T>C, 4628G>A, 4630G>A, 4632C>T, 4637T>A, 4639G>A, 4642A>T, 4643G>T, 4646C>A, 4654C>T, 4675G>C, 4680A>G, 4684T>A, 4685T>A, 4688C>T, 4689C>G, 4704G>C, 4706G>A, 4708T>A, 4722T>G

**CDS**

| | Begin | End | Coverage | Score | Concordance | Matches | Identities | I/D/M/F* | Stop Codons |
|---|---|---|---|---|---|---|---|---|---|
| **NT** | **89** | **4733** | **94.7%** | **2570** | **28.4%** | **4597 (96.7%)** | **3027 (63.7%)** | **107/48** | |
| ToMarV_RNA2gp1 | 1 | 191 | 100% | 784 | 66.7% | 183 (95.8%) | 116 (60.7%) | 0/8/0/0 | 1 |
| Protein mutations: | G5S (151G>A 153G>T), R6K (154C>A 155G>A 156T>G), S10Q (166T>C 167C>A 168T>A), V11I (169G>A), D12E (174C>G), E13D (177A>T), A15E (182C>A 183T>A), H18N (190C>A), T22A (202A>G 204C>A), N24T (209A>C 210T>C), I26V (214A>G), V31T (229G>A 230T>C), T33V (235A>G 236C>T 237C>T), L35I (241T>A 243G>T), V46I (274G>A), V47C (277G>T 278T>G), V54I (298G>A 300C>T), T58S (310A>T 312C>T), L68V (340T>G 342A>T), N79T (374A>C 375T>G), P81D (379C>G 380C>A 381C>T), N84A (388A>G 389A>C 390C>A), L91I (409C>A), H93R (416A>G 417T>A), V96I (424G>A 426G>T), R97K (427C>A 428G>A 429T>A), V104L (448G>T 450C>G), E107A (458A>C 459A>C), S111H (469T>C 470C>A), A114S (478G>T 480A>T), S115T (481T>A 483A>C), P120N (496C>A 497C>A 498T>C), S121P (499T>C 501T>C), D124_S129del (508_525delGACAGTAAAGGCAAGAGT), K130T (527A>C), V131L (529G>T), E132S (532G>T 533A>C 534G>A), Q133E (535C>G), P134T (538C>A 540C>T), T135A (541A>G), R136_E137del (544_549delCGAGAG), D138S (550G>A 551A>G), I140S (557T>G 558T>C), K141G (559A>G 560A>G), V143N (565G>A 566T>A 567T>C), E144K (568G>A), V145A (572T>C 573T>A), R148E (580C>G 581G>A 582T>A), E149A (584A>C 585G>C), R152V (592A>G 593G>T 594A>T), F153A (595T>G 596T>C 597T>C), Q154R (598C>A 599A>G 600G>A), A158Q (610G>C 611C>A), S159E (613A>G 614G>A 615T>A), Q160A (616C>G 617A>C), K161R (619A>C 620A>G 621A>G), N164I (629A>T), Q165V (631C>G 632A>T 633G>C), Q168R (640C>A 641A>G 642A>G), L171I (649C>A 651C>T), S172Q (652T>C 653C>A 654C>A), N173G (655A>G 656A>G), V175P (661G>C 662T>C 663T>A), N178E (670A>G 672T>G), I180V (676A>G), S182A (682T>G), E186S (694G>A 695A>G 696A>T), S187D (697A>G 698G>A), Q190K (706C>A) | | | | | | | | |
| Codon mutations: | TCA2TCT (144A>T), TTC3TTT (147C>T), ATT4ATC (150T>C), GGG5AGT (151G>A 153G>T), CGT6AAG (154C>A 155G>A 156T>G), TTG7CTC (157T>C 159G>C), AAT8AAC (162T>C), ACA9ACC (165A>C), TCT10CAA (166T>C 167C>A 168T>A), GTA11ATA (169G>A), GAC12GAG (174C>G), GAA13GAT (177A>T), GCT15GAA (182C>A 183T>A), TTT16TTC (186T>C), CAT18AAT (190C>A), GTT20GTA (198T>A), GCC21GCA (201C>A), ACC22GCA (202A>G 204C>A), TCG23TCT (207G>T), AAT24ACC (209A>C 210T>C), ATT26GTT (214A>G), TCT28AGT (220T>A 221C>G), GTT31ACT (229G>A 230T>C), GGC32GGT (234C>T), ACC33GTT (235A>G 236C>T 237C>T), TTG35ATT (241T>A 243G>T), ATA36ATT (246A>T), ACC41ACT (261C>T), CTT42TTA (262C>T 264T>A), GAC43GAT (267C>T), AAA45AAG (273A>G), GTT46ATT (274G>A), GTT47TGT (277G>T 278T>G), CCT48CCC (282T>C), GGT51GGA (291T>A), GTC54ATT (298G>A 300C>T), TCA55TCG (303A>G), GTT56GTG (306T>G), TTA57CTT (307T>C 309A>T), ACC58TCT (310A>T 312C>T), GTT59GTG (315T>G), TCT60AGT (316T>A 317C>G), GAG62GAA (324G>A), ACG65ACT (333G>T), CCA66CCC (336A>C), TTA68GTT (340T>G 342A>T), CCT70CCA (348T>A), GGA71GGC (351A>C), CAC72CAT (354C>T), TAT73TAC (357T>C), TTG74CTA (358T>C 360G>A), TTG75CTA (361T>C 363G>A), AAT79ACG (374A>C 375T>G), CCC81GAT (379C>G 380C>A 381C>T), ATT82ATC (384T>C), AAG83AAA (387G>A), AAC84GCA (388A>G 389A>C 390C>A), TCT88TCA (402T>A), GGT89GGC (405T>C), TTA90TTG (408A>G), CTT91ATT (409C>A), GTC92GTG (414C>G), CAT93CGA (416A>G 417T>A), TCA95TCC (423A>C), GTG96ATT (424G>A 426G>T), CGT97AAA (427C>A 428G>A 429T>A), CTT98TTG (430C>T 432T>G), ACC101ACA (441C>A), AGA102CGG (442A>C 444A>G), GTC104TTG (448G>T 450C>G), CTA105TTG (451C>T 453A>G), GAG106GAA (456G>A), GAA107GCC (458A>C 459A>C), AAC108AAT (462C>T), AAG109AAA (465G>A), GTT110GTG (468T>G), TCT111CAT (469T>C 470C>A), TCA113TCT (477A>T), GCA114TCT (478G>T 480A>T), TCA115ACC (481T>A 483A>C), TCA117TCT (489A>T), TCT118TCA (492T>A), TCT119TCC (495T>C), CCT120AAC (496C>A 497C>A 498T>C), TCT121CCC (499T>C 501T>C), GAC124_AGT129del (508_525delGACAGTAAAGGCAAGAGT), AAA130ACA (527A>C), GTA131TTA (529G>T), GAG132TCA (532G>T 533A>C 534G>A), CAA133GAA (535C>G), CCC134ACT (538C>A 540C>T), ACA135GCA (541A>G), CGA136_GAG137del (544_549delCGAGAG), GAT138AGT (550G>A 551A>G), CTC139TTA (553C>T 555C>A), ATT140AGC (557T>G 558T>C), AAA141GGA (559A>G 560A>G), GAA142GAG (564A>G), GTT143AAC (565G>A 566T>A 567T>C), GAG144AAG (568G>A), GTT145GCA (572T>C 573T>A), CTC146CTT (576C>T), AAA147AAG (579A>G), CGT148GAA (580C>G 581G>A 582T>A), GAG149GCC (584A>C 585G>C), TTA150CTT (586T>C 588A>T), GAG151GAA (591G>A), AGA152GTT (592A>G 593G>T 594A>T), TTT153GCC (595T>G 596T>C 597T>C), CAG154AGA (598C>A 599A>G 600G>A), GAG156GAA (606G>A), TTG157CTG (607T>C), GCA158CAA (610G>C 611C>A), AGT159GAA (613A>G 614G>A 615T>A), CAA160GCA (616C>G 617A>C), AAA161CGG (619A>C 620A>G 621A>G), AAT164ATT (629A>T), CAG165GTC (631C>G 632A>T 633G>C), CTA167CTC (639A>C), CAA168AGG (640C>A 641A>G 642A>G), CTT169TTG (643C>T 645T>G), CTC171ATT (649C>A 651C>T), TCC172CAA (652T>C 653C>A 654C>A), AAT173GGT (655A>G 656A>G), GTT175CCA (661G>C 662T>C 663T>A), AGT176AGC (666T>C), AAT177AAC (669T>C), AAT178GAG (670A>G 672T>G), GAC179GAT (675C>T), ATC180GTC (676A>G), TTC181TTT (681C>T), TCA182GCA (682T>G), GGT183GGC (687T>C), ACT185ACG (693T>G), GAA186AGT (694G>A 695A>G 696A>T), AGT187GAT (697A>G 698G>A), GGG188GGC (702G>C), CAG190AAG (706C>A), TAA191TGA (710A>G) | | | | | | | | |
| ToMarV_RNA2gp2 | 1 | 1192 | 100% | 6009 | 73.1% | 1185 (98.7%) | 820 (68.3%) | 9/7/0/0 | 1 |
| Protein mutations: | M1R (672T>G), T2M (675C>T 676A>G), S4L (681C>T 682T>G), V6A (687T>C), L8R (693T>G 694G>A), K9V (695A>G 696A>T 697A>G), V10M (698G>A), G11A (702G>C), N14D (710A>G 712T>C), H15R (714A>G), Q16S (716C>T 717A>C 718G>A), Q17del (719_721delCAG), E18K (722G>A 724G>A), A20V (729C>T 730C>A), E22D (736G>C), R23K (738G>A 739G>A), A34E (771C>A 772C>A), R41K (792G>A), V50I (818G>A 820T>A), G53A (828G>C 829T>A), E61D (853G>T), F63Y (858T>A), I71V (881A>G 883C>G), T72A (884A>G), T77A (899A>G 901A>T), H80R (908C>A 909A>G 910T>G), K81T (912A>C 913G>C), F82I (914T>A 916T>A), V83T (917G>A 918T>C 919T>A), K86R (926A>C 927A>G 928G>T), A89G (936C>G 937A>G), Y96H (956T>C 958T>C), V106A (987T>C 988T>A), S108N (993G>A), R112A (1004A>G 1005G>C 1006G>C), L124T (1040C>A 1041T>C 1042C>T), S125D (1043T>G 1044C>A 1045A>T), T132I (1065C>T 1066A>T), S134Y (1071C>A), L135F (1075G>T), A136S (1076G>T 1078C>T), D137G (1080A>G 1081T>A), Q166L (1167A>T 1168A>C), D185S (1223G>A 1224A>G 1225C>T), T186P (1226A>C 1228A>T), S192T (1244T>A), Q193D (1247C>G 1249G>C), Q194H (1252A>T), E195D (1255A>C), S226N (1347G>A), G227_Q228insV (1351_1352insGTA), S229R (1357T>A), T230S (1359C>G), M232L (1364A>C), A233del (1367_1369delGCT), K237R (1380A>G), A238P (1382G>C 1384C>T), E242A (1395A>C 1396A>C), Q243A (1397C>G 1398A>C 1399G>A), G246D (1407G>A 1408C>T), D250N (1418G>A 1420C>T), M252P (1424A>C 1425T>C), N253Q (1427A>C 1429T>G), S255A (1433A>G 1434G>C 1435C>A), T256N (1437C>A), I258V (1442A>G 1444C>G), K273R (1488A>G), P276A (1496C>G 1498G>A), P281_A282insDF (1513_1514insGACTTT), V283S (1517G>T 1518T>C), S284T (1521G>C), T285S (1523A>T 1525A>T), S286R (1526T>A 1527C>G), S287A (1529T>G 1531C>G), L288Y (1533T>A 1534A>T), L289Q (1536T>A 1537C>G), S290G (1538A>G 1540C>A), T291V (1541A>G 1542C>T 1543A>G), V296G (1557T>G), H297I (1559C>A 1560A>T), E299del (1565_1567delGAG), C302T (1574T>A 1575G>C 1576T>G), G303S (1577G>T 1578G>C 1579G>T), G304L (1580G>C 1581G>T), P309_K310del (1595_1600delCCCAAG), K312I (1605A>T 1606G>A), G313S (1607G>T 1608G>C 1609C>A), F316G (1616T>G 1617T>G 1618C>A), N317T (1620A>C 1621C>A), M318L (1622A>C 1624G>C), G319Q (1625G>C 1626G>A 1627T>A), A320E (1629C>A 1630C>A), I321V (1631A>G 1633T>C), L325Y (1644T>A 1645G>T), M327I (1651G>T), E328Q (1652G>C 1654G>A), A331E (1662C>A), H332D (1664C>G 1666C>T), I333F (1667A>T), D334A (1671A>C 1672T>C), D337S (1679G>A 1680A>G), D338N (1682G>A 1684T>C), E341A (1692A>C 1693A>C), R342L (1695G>T 1696A>T), I344V (1700A>G 1702A>G), A345P (1703G>C 1705T>C), Q346E (1706C>G), V348I (1712G>A 1714A>T), Q349D (1715C>G 1717A>T), F350Y (1719T>A), I351V (1721A>G 1723C>T), F353V (1727T>G 1729T>G), S354Q (1730A>C 1731G>A 1732T>G), E355K (1733G>A), K357R (1740A>G), R358A (1742A>G 1743G>C 1744A>T), V361L (1751G>C 1753T>C), I362C (1754A>T 1755T>G), R365K (1764G>A), V367I (1769G>A), A368E (1773C>A 1774A>G), K369M (1776A>T), H371D (1781C>G 1783T>C), N374C (1790A>T 1791A>G), R377F (1799A>T 1800G>T 1801G>T), D383A (1818A>C), E387D (1831A>T), E390A (1839A>C 1840A>G), N396K (1858T>G), I397V (1859A>G 1861C>T), Q398G (1862C>G 1863A>G 1864A>G), G399M (1865G>A 1866G>T 1867C>G), T401E (1871A>G 1872C>A 1873T>A), F404P (1880T>C 1881T>C 1882C>A), A408S (1892G>T), M409L (1895A>T), E413Q (1907G>C), Q421M (1931C>A 1932A>T 1933A>G), S425Q (1943T>C 1944C>A 1945C>A), I428R (1953T>G 1954T>G), Q429Y (1955C>T 1957A>T), G430W (1958G>T), Q431D (1961C>G 1963G>T), T432L (1964A>T 1965C>T 1966T>A), N433T (1968A>C 1969C>A), N438D (1982A>G 1984T>C), C441V (1991T>G 1992G>T 1993C>G), K448A (2012A>G 2013A>C), E449D (2017G>T), R450_Q451insVK (2020_2021insGTGAAA), Q451H (2023G>T), L452P (2024T>C 2025T>C 2026G>C), I453L (2027A>T 2029T>G), V454M (2030G>A), E455S (2033G>A 2034A>G 2035G>T), E456D (2038G>T), Q457L (2040A>T), V458A (2043T>C 2044G>A), A459C (2045G>T 2046C>G), Q460_R461insLN (2050_2051insTTGAAT), R462L (2054A>T 2055G>T), S463G (2057T>G 2058C>G 2059T>A), E464K (2060G>A), R465P (2063A>C 2064G>C), Q466S (2066C>A 2067A>G 2068A>C), A467M (2069G>A 2070C>T 2071A>G), Q468S (2072C>A 2073A>G 2074G>T), T470A (2078A>G), A471Q (2081G>C 2082C>A), R472T (2085G>C 2086G>T), I474N (2091T>A 2092A>T), A475_E476insS (2095_2096insAGC), E476K (2096G>A 2098A>G), S477W (2099A>T 2101T>G), P479T (2105C>A), D481G (2112A>G 2113C>A), R482K (2115G>A 2116A>G), T484L (2120A>C 2121C>T 2122T>G), S489T (2135T>A), T492A (2144A>G 2146G>A), M493L (2147A>C 2149G>T), E494_D495insS (2152_2153insAGT), D495E (2155T>G), P496G (2156C>G 2157C>G 2158C>T), T497I (2160C>T), K498Q (2162A>C 2164G>A), P499R (2165C>A 2166C>G), D500K (2168G>A 2170T>G), K501E (2171A>G), I502V (2174A>G 2176A>G), E503S (2177G>T 2178A>C), V505E (2184T>A 2185T>G), G508D (2193G>A 2194A>T), A509P (2195G>C 2197A>G), E511G (2202A>G 2203A>G), E512K (2204G>A), Q514E (2210C>G 2212A>G), G516N (2216G>A 2217G>A), V518I (2222G>A), I519F (2225A>T 2227T>C), P524E (2240C>G 2241C>A 2242A>G), D527M (2249G>A 2250A>T 2251C>G), T528G (2252A>G 2253C>G 2254A>T), S529D (2255T>G 2256C>A 2257C>T), M530A (2258A>G 2259T>C 2260G>C), A531P (2261G>C 2263A>G), V532L (2264G>T 2266T>A), E533N (2267G>A 2269A>T), D535E (2275T>G), M536L (2276A>T), V540A (2289T>C 2290G>A), D546E (2308T>G), V552I (2324G>A), V556L (2336G>C 2338T>G), E558K (2342G>A 2344G>A), A561N (2351G>A 2352C>A 2353T>C), M568L (2372A>C 2374G>T), S569A (2375A>G 2376G>C 2377C>A), A571S (2381G>T 2383T>C), A574M (2390G>A 2391C>T 2392T>G), S580K (2409G>A 2410C>G), T583A (2417A>G), A587S (2429G>T 2431G>C), L597N (2459C>A 2460T>A 2461A>C), I598L (2462A>C 2464C>T), Y600F (2469A>T 2470C>T), I602V (2474A>G 2476A>C), V606I (2486G>A), V609M (2495G>A 2497T>G), I612V (2504A>G 2506C>G), V625A (2544T>C 2545G>A), K629T (2556A>C 2557G>C), L638I (2582T>A 2584G>T), T644H (2600A>C 2601C>A 2602A>T), Q650E (2618C>G), N652S (2624A>T 2625A>C), E654G (2631A>G), E655K (2633G>A), C657V (2639T>G 2640G>T 2641C>G), T659S (2645A>T 2647C>A), T661Q (2651A>C 2652C>A), K665T (2664A>C 2665G>C), Y666F (2667A>T 2668C>T), E671T (2681G>A 2682A>C 2683A>T), G672S (2684G>T 2685G>C 2686T>A), T673L (2687A>C 2688C>T 2689A>G), A675V (2694C>T 2695C>T), T676P (2696A>C), D677E (2701T>A), S690A (2738T>G 2740T>A), N696E (2756A>G 2758T>G), V698I (2762G>A), L702I (2774T>A 2776G>A), Q705S (2783C>T 2784A>C 2785G>C), L714M (2810C>A 2812T>G), M719L (2825A>T), I720R (2829T>G), A721G (2832C>G 2833A>G), Y725F (2844A>T), K728R (2852A>C 2853A>G), A729P (2855G>C 2857G>A), Q731T (2861C>A 2862A>C 2863G>A), Y733F (2868A>T 2869T>C), P735S (2873C>A 2874C>G 2875A>T), G749S (2915G>A 2917G>T), T750V (2918A>G 2919C>T), S751A (2921T>G 2923A>T), S752A (2924T>G 2926C>A), P753A (2927C>G 2929A>C), I761V (2951A>G 2953A>C), T773I (2988C>T), T790R (3039C>G 3040T>A), I797V (3059A>G 3061T>G), I799V (3065A>G), N817A (3119A>G 3120A>C 3121C>A), K819E (3125A>G 3127A>G), T821S (3131A>T), N822T (3135A>C 3136T>A), F827L (3151T>G), K841R (3192A>G 3193A>G), R843Q (3198G>A 3199G>A), R845K (3203C>A 3204G>A 3205G>A), I848L (3212A>C 3214T>G), K854C (3230A>T 3231A>G 3232A>C), Q869K (3275C>A), F871Q (3281T>C 3282T>A 3283T>A), L874M (3290T>A), I889L (3335A>T 3337A>G), S893T (3347T>A 3349C>T), C902R (3374T>C), L903I (3377C>A), N905D (3383A>G), T911S (3401A>T 3403A>T), K925R (3444A>G), S927A (3449T>G 3451A>T), D936N (3476G>A 3478T>C), S938N (3482T>A 3483C>A 3484A>T), I943L (3497A>C), D949E (3517C>A), I951V (3521A>G), E952D (3526G>T), E953I (3527G>A 3528A>T 3529G>C), Y954F (3531A>T 3532C>T), Q955D (3533C>G 3535G>C), E956Q (3536G>C), D960N (3548G>A 3550T>C), P961_P962del (3551_3556delCCGCCC), K964T (3561A>C 3562G>A), A965S (3563G>T), S967V (3569A>G 3570G>T), I968S (3572A>T 3573T>C 3574T>C), L969F (3575C>T 3577G>C), S970Q (3578T>C 3579C>A 3580C>G), S976T (3596T>A 3598C>A), M985A (3623A>G 3624T>C 3625G>T), G986A (3627G>C 3628T>A), D990T (3638G>A 3639A>C 3640C>A), E991D (3643A>C), L994V (3650T>G 3652G>A), I996L (3656A>T 3658T>G), I1003V (3677A>G 3679A>G), E1006D (3688A>T), G1007A (3690G>C 3691G>A), H1008S (3692C>T 3693A>C), V1011I (3701G>A), T1024S (3740A>T 3742A>C), N1034E (3770A>G 3772T>A), S1036T (3776T>A), I1037F (3779A>T), I1039M (3787A>G), V1042T (3794G>A 3795T>C 3796A>T), P1046N (3806C>A 3807C>A 3808T>C), N1047S (3810A>G), V1048I (3812G>A 3814T>A), V1051I (3821G>A 3823A>C), V1054I (3830G>A), F1056L (3836T>C), A1058T (3842G>A 3844C>A), P1064Q (3861C>A), A1065F (3863G>T 3864C>T), L1067V (3869C>G 3871T>A), N1068S (3872A>T 3873A>C), K1069A (3875A>G 3876A>C 3877A>G), N1071A (3881A>G 3882A>C 3883T>A), Y1072H (3884T>C), V1074L (3890G>T), A1089T (3935G>A 3937A>C), T1092I (3945C>T 3946G>C), T1096V (3956A>G 3957C>T 3958T>A), E1101Q (3971G>C), H1107F (3989C>T 3990A>T 3991T>C), R1109K (3996G>A), M1110A (3998A>G 3999T>C 4000G>C), R1111K (4002G>A 4003G>A), E1112S (4004G>T 4005A>C 4006A>C), F1113Y (4008T>A 4009C>T), S1114D (4010T>G 4011C>A 4012A>T), S1115K (4014G>A 4015C>G), K1116E (4016A>G 4018G>A), Q1117S (4019C>A 4020A>G 4021A>T), M1121V (4031A>G 4033G>T), R1126K (4047G>A), N1129L (4055A>T 4056A>T 4057T>G), I1131L (4061A>C 4063C>A), S1137V (4079T>G 4080C>T 4081C>T), A1138E (4083C>A 4084C>A), I1139L (4085A>C 4087A>T), V1140I (4088G>A 4090G>T), L1146I (4106C>A 4108T>A), I1147V (4109A>G 4111A>C), S1148K (4112T>A 4113C>A 4114T>G), L1151P (4122T>C), S1162L (4154T>C 4155C>T), H1165N (4163C>A), N1171D (4181A>G), I1191N (4242T>A) | | | | | | | | |

| | Begin | End | Coverage | Score | Concordance | Matches | Identities | I/D/M/F* | Stop Codons |
|---|---|---|---|---|---|---|---|---|---|
| **NT** | **89** | **4733** | **94.7%** | **2570** | **28.4%** | **4597 (96.7%)** | **3027 (63.7%)** | **107/48** | |

Codon mutations:

ATG1AGG (672T>G), ACA2ATG (675C>T 676A>G), TCT4TTG (681C>T 682T>G), GTT6GCT (687T>C), CTG8CGA (693T>G 694G>A), AAA9GTG (695A>G 696A>T 697A>G), GTG10ATG (698G>A), GGC11GCC (702G>C), CCC12CCA (706C>A), AAT14GAC (710A>G 712T>C), CAC15CGC (714A>G), CAG16TCA (716C>T 717A>C 718G>A), CAG17del (719_721delCAG), GAG18AAA (722G>A 724G>A), GCC20GTA (729C>T 730C>A), TCC21TCT (733C>T), GAG22GAC (736G>C), AGG23AAA (738G>A 739G>A), ATA25ATT (745A>T), TCA26TCT (748A>T), CAA27CAG (751A>G), ATA28ATC (754A>C), GCT30GCG (760T>G), GTG32GTT (766G>T), GAG33GAA (769G>A), GCC34GAA (771C>A 772C>A), GGA35GGT (775A>T), AAT38AAC (784T>C), TTG39CTT (785T>C 787G>T), CTC40CTT (790C>T), AGA41AAA (792G>A), AAG42AAA (796G>A), CTT43TTG (797C>T 799T>G), GGC44GGA (802C>A), TCC47TCA (811C>A), TAC48TAT (814C>T), GGG49GGC (817G>C), GTT50ATA (818G>A 820T>A), CTT51TTG (821C>T 823T>G), GGT53GCA (828G>C 829T>A), CAA55CAG (835A>G), AAG57AAA (841G>A), AGA58AGG (844A>G), GAG61GAT (853G>T), TTG62CTT (854T>C 856G>T), TTT63TAT (858T>A), GAC64GAT (862C>T), GTA68GTT (874A>T), TCA69AGT (875T>A 876C>G 877A>T), AAG70AAA (880G>A), ATC71GTG (881A>G 883C>G), ACA72GCA (884A>G), CTT74CTA (892T>A), ACA77GCT (899A>G 901A>T), TTC78TTT (904C>T), CAT80AGG (908C>A 909A>G 910T>G), AAG81ACC (912A>C 913G>C), TTT82ATA (914T>A 916T>A), GTT83ACA (917G>A 918T>C 919T>A), GAG84GAA (922G>A), TCT85TCA (925T>A), AAG86CGT (926A>C 927A>G 928G>T), GAC87GAT (931C>T), GCA89GGG (936C>G 937A>G), AAC90AAT (940C>T), TTG91TTA (943G>A), TTT93TTC (949T>C), CTG95TTA (953C>T 955G>A), TAT96CAC (956T>C 958T>C), GGT97GGG (961T>G), CTT99TTG (965C>T 967T>G), TTC100TTT (970C>T), TTC101TTT (973C>T), GTC103GTT (979C>T), CCA104CCC (982A>C), GTT106GCA (987T>C 988T>A), AGT108AAT (993G>A), GGG109GGA (997G>A), GGA111GGT (1003A>T), AGG112GCC (1004A>G 1005G>C 1006G>C), GTT113GTC (1009T>C), AAG114AAA (1012G>A), ATA115ATC (1015A>C), AGT116AGC (1018T>C), CTT117TTG (1019C>T 1021T>G), TGT118TGC (1024T>C), TCA119TCT (1027A>T), AGT120AGC (1030T>C), AAT121AAC (1033T>C), CTC124ACT (1040C>A 1041T>C 1042C>T), TCA125GAT (1043T>G 1044C>A 1045A>T), CCG126CCT (1048G>T), CAA129CAG (1057A>G), AAG131AAA (1063G>A), ACA132ATT (1065C>T 1066A>T), CTG133TTG (1067C>T), TCC134TAC (1071C>A), TTG135TTT (1075G>T), GCC136TCT (1076G>T 1078C>T), GAT137GGA (1080A>G 1081T>A), GCG139GCA (1087G>A), CAG140CAA (1090G>A), GCA141GCC (1093A>C), CTC143CTA (1099C>A), AGC145AGT (1105C>T), CCC146CCT (1108C>T), AGC147AGT (1111C>T), CTC153TTG (1127C>T 1129C>G), AAA154AAG (1132A>G), AGA155AGG (1135A>G), GGG156GGC (1138G>C), ACC162ACA (1156C>A), GAG164GAA (1162G>A), TGC165TGT (1165C>T), CAA166CTC (1167A>T 1168A>C), AAT167AAC (1171T>C), ACC168ACG (1174C>G), AGG169AGA (1177G>A), GCA170GCT (1180A>T), CAG171CAA (1183G>A), CCG173CCT (1189G>T), TCC175TCT (1195C>T), GTG176GTA (1198G>A), GCC178GCA (1204C>A), ATA179ATT (1207A>T), GAC185AGT (1223G>A 1224A>G 1225C>T), ACA186CCT (1226A>C 1228A>T), AGG187CGT (1229A>C 1231G>T), GCA189GCT (1237A>T), GTT190GTA (1240T>A), TCA192ACA (1244T>A), CAG193GAC (1247C>G 1249G>C), CAA194CAT (1252A>T), GAA195GAC (1255A>C), ACA196ACT (1258A>T), TCT198AGT (1262T>A 1263C>G), GCC200GCT (1270C>T), ATA201ATT (1273A>T), GAA202GAG (1276A>G), GCA203GCT (1279A>T), CTC204CTT (1282C>T), AAT205AAC (1285T>C), CGT206AGA (1286C>A 1288T>A), CCG207CCT (1291G>T), CAA208CAG (1294A>G), GAT212GAC (1306T>C), AGA213AGG (1309A>G), GAG215GAA (1315G>A), GCT216GCA (1318T>A), GCT217GCA (1321T>A), TAT219TAC (1327T>C), ATA220ATT (1330A>T), GCT221GCA (1333T>A), GTT223GTG (1339T>G), TAT224TAC (1342T>C), TCG225TCC (1345G>C), AGT226AAT (1347G>A), GGG227_CAA228insGTA (1351_1352insGTA), CAA228CAG (1354A>G), AGT229AGA (1357T>A), ACT230AGT (1359C>G), CAA231CAG (1363A>G), ATG232CTG (1364A>C), GCT233del (1367_1369delGCT), CTT234TTG (1370C>T 1372T>G), GAA235GAG (1375A>G), AAA237AGA (1380A>G), GCC238CCT (1382G>C 1384C>T), GTT240GTG (1390T>G), GGA241GGT (1393A>T), GAA242GCC (1395A>C 1396A>C), CAG243GCA (1397C>G 1398A>C 1399G>A), GGG245GGT (1405G>T), GGC246GAT (1407G>A 1408C>T), ACA247ACC (1411A>C), GAC250AAT (1418G>A 1420C>T), GTC251GTG (1423C>G), ATG252CCG (1424A>C 1425T>C), AAT253CAG (1427A>C 1429T>G), GAA254GAG (1432A>G), AGC255GCA (1433A>G 1434G>C 1435C>A), ACT256AAT (1437C>A), ATC258GTG (1442A>G 1444C>G), CGG259CGC (1447G>C), AGT260TCT (1448A>T 1449G>C), TCT262TCA (1456T>A), AGA264AGG (1462A>G), ACT267ACC (1471T>C), CTC268CTA (1474C>A), AAA269AAG (1477A>G), GTG270GTA (1480G>A), AAG273AGG (1488A>G), CGC274CGA (1492C>A), CCG276GCA (1496C>G 1498G>A), TCC277TCA (1501C>A), CTG280TTG (1508C>T), CCG281CCT (1513G>T), CCG281_GCT282insGACTTT (1513_1514insGACTTT), GTA283TCA (1517G>T 1518T>C), AGT284ACT (1521G>C), ACA285TCT (1523A>T 1525A>T), TCA286AGA (1526T>A 1527C>G), TCC287GCG (1529T>G 1531C>G), TTA288TAT (1533T>A 1534A>T), CTC289CAG (1536T>A 1537C>G), AGC290GGA (1538A>G 1540C>A), ACA291GTG (1541A>G 1542C>T 1543A>G), CGT292AGA (1544C>A 1546T>A), GAA293GAG (1549A>G), ACA295ACT (1555A>T), GTT296GGT (1557T>G), CAT297ATT (1559C>A 1560A>T), GAG299del (1565_1567delGAG), TGT302ACG (1574T>A 1575G>C 1576T>G), GGG303TCT (1577G>T 1578G>C 1579G>T), GGA304CTA (1580G>C 1581G>T), TTG305CTG (1583T>C), CCA307CCT (1591A>T), GCA308GCC (1594A>C), CCC309_AAG310del (1595_1600delCCCAAG), AAA311AAG (1603A>G), AAG312ATA (1605A>T 1606G>A), GGC313TCA (1607G>T 1608G>C 1609C>A), GCC315GCA (1615C>A), TTC316GGA (1616T>G 1617T>G 1618C>A), AAC317ACA (1620A>C 1621C>A), ATG318CTC (1622A>C 1624G>C), GGT319CAA (1625G>C 1626G>A 1627T>A), GCC320GAA (1629C>A 1630C>A), ATT321GTC (1631A>G 1633T>C), GAT323GAC (1639T>C), TTG325TAT (1644T>A 1645G>T), ATG327ATT (1651G>T), GAG328CAA (1652G>C 1654G>A), TCG329AGT (1655T>A 1656C>G 1657G>T), TTC330TTT (1660C>T), GCA331GAA (1662C>A), CAC332GAT (1664C>G 1666C>T), ATT333TTT (1667A>T), GAT334GCC (1671A>C 1672T>C), GAT337AGT (1679G>A 1680A>G), GAT338AAC (1682G>A 1684T>C), GAA341GCC (1692A>C 1693A>C), CGA342CTT (1695G>T 1696A>T), ATA344GTG (1700A>G 1702A>G), GCT345CCC (1703G>C 1705T>C), CAG346GAG (1706C>G), CAG347CAA (1711G>A), GTA348ATT (1712G>A 1714A>T), CAA349GAT (1715C>G 1717A>T), TTT350TAT (1719T>A), ATC351GTT (1721A>G 1723C>T), CTC352CTT (1726C>T), TTT353GTG (1727T>G 1729T>G), AGT354CAG (1730A>C 1731G>A 1732T>G), GAG355AAG (1733G>A), AAA357AGA (1740A>G), AGA358GCT (1742A>G 1743G>C 1744A>T), GGA359GGT (1747A>T), AAC360AAT (1750C>T), GTT361CTC (1751G>C 1753T>C), ATT362TGT (1754A>T 1755T>G), GTG363GTC (1759G>C), AGA365AAA (1764G>A), GTT367ATT (1769G>A), GCA368GAG (1773C>A 1774A>G), AAG369ATG (1776A>T), CGG370AGG (1778C>A), CAT371GAC (1781C>G 1783T>C), CTG372TTG (1784C>T), CAT373CAC (1789T>C), AAT374TGT (1790A>T 1791A>G), ATC375ATA (1795C>A), AGG377TTT (1799A>T 1800G>T 1801G>T), GAG378GAA (1804G>A), CAC379CAT (1807C>T), ATA380ATC (1810A>C), ACC381ACT (1813C>T), GAC383GCC (1818A>C), GTA386GTT (1828A>T), GAA387GAT (1831A>T), ATA388ATT (1834A>T), CTT389TTG (1835C>T 1837T>G), GAA390GCG (1839A>C 1840A>G), GGT391GGA (1843T>A), GGT393GGA (1849T>A), AAT396AAG (1858T>G), ATC397GTT (1859A>G 1861C>T), CAA398GGG (1862C>G 1863A>G 1864A>G), GGC399ATG (1865G>A 1866G>T 1867C>G), ACT401GAA (1871A>G 1872C>A 1873T>A), CGC402AGA (1874C>A 1876C>A), ACT403ACA (1879T>A), TTC404CCA (1880T>C 1881T>C 1882C>A), TAT407TAC (1891T>C), GCT408TCT (1892G>T), ATG409TTG (1895A>T), TCT410AGC (1898T>A 1899C>G 1900T>C), CTC411CTG (1903C>G), GAG413CAG (1907G>C), GTT415GTG (1915T>G), GTG416GTT (1918G>T), CTC418CTA (1924C>A), CAT420CAC (1930T>C), CAA421ATG (1931C>A 1932A>T 1933A>G), AGA422AGG (1936A>G), GAT423GAC (1939T>C), TCC425CAA (1943T>C 1944C>A 1945C>A), TTC426TTT (1948C>T), ATT428AGG (1953T>G 1954T>G), CAA429TAT (1955C>T 1957A>T), GGG430TGG (1958G>T), CAG431GAT (1961C>G 1963G>T), ACT432TTA (1964A>T 1965C>T 1966T>A), AAC433ACA (1968A>C 1969C>A), AAT434AAC (1972T>C), CCA435CCG (1975A>G), CCG437CCT (1981G>T), AAT438GAC (1982A>G 1984T>C), TGC441GTG (1991T>G 1992G>T 1993C>G), GGA444GGT (2002A>T), TTA445CTT (2003T>C 2005A>T), ACC446ACA (2008C>A), TTG447CTG (2009T>C), AAA448GCA (2012A>G 2013A>C), GAG449GAT (2017G>T), CGT450AGG (2018C>A 2020T>G), CGT450_CAG451insGTGAAA (2020_2021insGTGAAA), CAG451CAT (2023G>T), TTG452CCC (2024T>C 2025T>C 2026G>C), ATT453TTG (2027A>T 2029T>G), GTG454ATG (2030G>A), GAG455AGT (2033G>A 2034A>G 2035G>T), GAG456GAT (2038G>T), CAA457CTA (2040A>T), GTG458GCA (2043T>C 2044G>A), GCC459TGC (2045G>T 2046C>G), CAA460_AGA461insTTGAAT (2050_2051insTTGAAT), AGA461CGC (2051A>C 2053A>C), AGG462TTG (2054A>T 2055G>T), TCT463GGA (2057T>G 2058C>G 2059T>A), GAA464AAA (2060G>A), AGA465CCA (2063A>C 2064G>C), CAA466AGC (2066C>A 2067A>G 2068A>C), GCA467ATG (2069G>A 2070C>T 2071A>G), CAG468AGT (2072C>A 2073A>G 2074G>T), ACA470GCA (2078A>G), GCA471CAA (2081G>C 2082C>A), AGG472ACT (2085G>C 2086G>T), GGC473GGA (2089C>A), ATA474AAT (2091T>A 2092A>T), GCT475GCC (2095T>C), GCT475_GAA476insAGC (2095_2096insAGC), GAA476AAG (2096G>A 2098A>G), AGT477TGG (2099A>T 2101T>G), CAG478CAA (2104G>A), CCT479ACT (2105C>A), GAC481GGA (2112A>G 2113C>A), AGA482AAG (2115G>A 2116A>G), ACT484CTG (2120A>C 2121C>T 2122T>G), TCC486TCA (2128C>A), GTG488GTT (2134G>T), TCA489ACA (2135T>A), ACT490ACC (2140T>C), ACC491ACA (2143C>A), ACG492GCA (2144A>G 2146G>A), ATG493CTT (2147A>C 2149G>T), GAA494GAG (2152A>G), GAA494_GAT495insAGT (2152_2153insAGT), GAT495GAG (2155T>G), CCC496GGT (2156C>G 2157C>G 2158C>T), ACC497ATC (2160C>T), AAG498CAA (2162A>C 2164G>A), CCA499AGA (2165C>A 2166C>G), GAT500AAG (2168G>A 2170T>G), AAG501GAG (2171A>G), ATA502GTG (2174A>G 2176A>G), GAA503TCA (2177G>T 2178A>C), GTT505GAG (2184T>A 2185T>G), GCA506GCT (2188A>T), GAG507GAA (2191G>A), GGA508GAT (2193G>A 2194A>T), GCA509CCG (2195G>C 2197A>G), GAA511GGG (2202A>G 2203A>G), GAA512AAA (2204G>A), ACT513ACA (2209T>A), CAA514GAG (2210C>G 2212A>G), GGT516AAT (2216G>A 2217G>A), GAT517GAC (2221T>C), GTT518ATT (2222G>A), ATT519TTC (2225A>T 2227T>C), GAC521GAT (2233C>T), GGA523GGG (2239A>G), CCA524GAG (2240C>G 2241C>A 2242A>G), GAC527ATG (2249G>A 2250A>T 2251C>G), ACA528GGT (2252A>G 2253C>G 2254A>T), TCC529GAT (2255T>G 2256C>A 2257C>T), ATG530GCC (2258A>G 2259T>C 2260G>C), GCA531CCG (2261G>C 2263A>G), GTT532TTA (2264G>T 2266T>A), GAA533AAT (2267G>A 2269A>T), GAT535GAG (2275T>G), ATG536TTG (2276A>T), CAG537CAA (2281G>A), CAA538CAG (2284A>G), CCG539CCA (2287G>A), GTG540GCA (2289T>C 2290G>A), GTA542GTG (2296A>G), AGC544AGT (2302C>T), GAT546GAG (2308T>G), TTC547TTT (2311C>T), GGA551GGG (2323A>G), GTT552ATT (2324G>A), TTC555TTT (2335C>T), GTT556CTG (2336G>C 2338T>G), GAG558AAA (2342G>A 2344G>A), TCC560AGC (2348T>A 2349C>G), GCT561AAC (2351G>A 2352C>A 2353T>C), GAG565GAA (2365G>A), CAA566CAG (2368A>G), GTA567GTT (2371A>T), ATG568CTT (2372A>C 2374G>T), AGC569GCA (2375A>G 2376G>C 2377C>A), TTG570CTA (2378T>C 2380G>A), GCT571TCC (2381G>T 2383T>C), TTG572CTC (2384T>C 2386G>C), CCC573CCA (2389C>A), GCT574ATG (2390G>A 2391C>T 2392T>G), GCC575GCA (2395C>A), CTA576TTG (2396C>T 2398A>G), TTC577TTT (2401C>T), TCA578AGC (2402T>A 2403C>G 2404A>C), AAA579AAG (2407A>G), AGC580AAG (2409G>A 2410C>G), AAA581AAG (2413A>G), ACT583GCT (2417A>G), TCA584TCC (2422A>C), GGT586GGG (2428T>G), GCG587TCC (2429G>T 2431G>C), CTT590CTC (2440T>C), TAT592TAC (2446T>C), TAC593TAT (2449C>T), GAT594GAC (2452T>C), GCA595GCT (2455A>T), GCT596GCA (2458T>A), CTA597AAC (2459C>A 2460T>A 2461A>C), ATC598CTT (2462A>C 2464C>T), TAC600TTT (2469A>T 2470C>T), ATA602GTC (2474A>G 2476A>C), CTT604CTG (2482T>G), TAT605TAC (2485T>C), GTT606ATT (2486G>A), GTT609ATG (2495G>A 2497T>G), GCT611GCG (2503T>G), ATC612GTG (2504A>G 2506C>G), TCT613TCG (2509T>G), GGT614GGA (2512T>A), CAA615CAG (2515A>G), CTG616TTA (2516C>T 2518G>A), GCT617GCC (2521T>C), GAA622GAG (2536A>G), TGT623TGC (2539T>C), GTG625GCA (2544T>C 2545G>A), CGA628AGA (2552C>A), AAG629ACC (2556A>C 2557G>C), AAC634AAT (2572C>T), ATC635ATA (2575C>A), ACA637ACC (2581A>C), TTG638ATT (2582T>A 2584G>T), AAG642AAA (2596G>A), ACA644CAT (2600A>C 2601C>A 2602A>T), TTG645CTG (2603T>C), GTT646GTG (2608T>G), CAA650GAA (2618C>G), AAC652TCC (2624A>T 2625A>C), GAG654GGG (2631A>G), GAG655AAG (2633G>A), TGC657GTG (2639T>G 2640G>T 2641C>G), TTT658TTC (2644T>C), ACC659TCA (2645A>T 2647C>A), CCA660CCT (2650A>T), ACA661CAA (2651A>C 2652C>A), GGG662GGT (2656G>T), GGC664GGT (2662C>T), AAG665ACC (2664A>C 2665G>C), TAC666TTT (2667A>T 2668C>T), GTG667GTA (2671G>A), CCT668CCA (2674T>A), CTG669TTG (2675C>T), GAA671ACT (2681G>A 2682A>C 2683A>T), GGT672TCA (2684G>T 2685G>C 2686T>A), ACA673CTG (2687A>C 2688C>T 2689A>G), GGA674GGT (2692A>T), GCC675GTT (2694C>T 2695C>T), ACT676CCT (2696A>C), GAT677GAA (2701T>A), TTA678CTT (2702T>C 2704A>T), GGT679GGC (2707T>C), GTG681GTT (2713G>T), ACA686ACC (2728A>C), CAC687CAT (2731C>T), TTA689TTG (2737A>G), TCT690GCA (2738T>G 2740T>A), GCA692GCC (2746A>C), GAG694GAA (2752G>A), AAT696GAG (2756A>G 2758T>G), AGT697TCA (2759A>T 2760G>C 2761T>A), GTA698ATA (2762G>A), CCA699CCT (2767A>T), TGC700TGT (2770C>T), CAT701CAC (2773T>C), TTG702ATA (2774T>A 2776G>A), TTA704CTA (2780T>C), CAG705TCC (2783C>T 2784A>C 2785G>C), AAA707AAG (2791A>G), TTA709CTC (2795T>C 2797A>C), ACC711ACA (2803C>A), CTT714ATG (2810C>A 2812T>G), CGA718AGG (2822C>A 2824A>G), ATG719TTG (2825A>T), ATA720AGA (2829T>G), GCA721GGG (2832C>G 2833A>G), CAG722CAA (2836G>A), CAA724CAG (2842A>G), TAT725TTT (2844A>T), GGC726GGA (2848C>A), AAG728CGG (2852A>C 2853A>G), GCG729CCA (2855G>C 2857G>A), GGG730GGC (2860G>C), CAG731ACA (2861C>A 2862A>C 2863G>A), ACA732ACC (2866A>C), TAT733TTC (2868A>T 2869T>C), CCA735AGT (2873C>A 2874C>G 2875A>T), AGG736CGC (2876A>C 2878G>C), TTT737TTC (2881T>C), CCA738CCT (2884A>T), ACT739ACA (2887T>A), GTT742GTG (2896T>G), TTG743CTT (2897T>C 2899G>T), TTA744CTG (2900T>C 2902A>G), CAT745CAC (2905T>C), TAT746TAC (2908T>C), GGG749AGT (2915G>A 2917G>T), ACA750GTA (2918A>G 2919C>T), TCA751GCT (2921T>G 2923A>T), TCC752GCA (2924T>G 2926C>A), CCA753GCC (2927C>G 2929A>C), ACA757ACT (2941A>T), AGC760AGT (2950C>T), ATA761GTC (2951A>G 2953A>C), TTT762TTC (2956T>C), TCA763TCC (2959A>C), ATA767ATT (2971A>T), GAG769GAA (2977G>A), GGC772GGT (2986C>T), ACC773ATC (2988C>T), TTG774TTA (2992G>A), CCG776CCA (2998G>A), TCT777TCA (3001T>A), TTG778CTG (3002T>C), GGT780GGA (3010T>A), ATA782ATT (3016A>T), GCC783GCA (3019C>A), AGG784AGA (3022G>A), AAG787AAA (3031G>A), ACT790AGA (3039C>G 3040T>A), GGA791GGC (3043A>C), ACT792ACC (3046T>C), ATT797GTG (3059A>G 3061T>G), TGC798TGT (3064C>T), ATT799GTT (3065A>G), ACC802ACT (3076C>T), TTG803CTG (3077T>C), TCA806TCT (3088A>T), GGT807GGG (3091T>G), CTG809CTT (3097G>T), GCA810GCT (3100A>T), ATT811ATA (3103T>A), GGA812GGG (3106A>G), CTT813TTG (3107C>T 3109T>G), GGA814GGC (3112A>C), ACT815ACG (3115T>G), AAC817GCA (3119A>G 3120A>C 3121C>A), ACC818ACA (3124C>A), AAA819GAG (3125A>G 3127A>G), ACC821TCC (3131A>T), AAT822ACA (3135A>C 3136T>A), GCT823GCG (3139T>G), TTT827TTG (3151T>G), CCA830CCC (3160A>C), GTA833GTG (3169A>G), TGC834TGT (3172C>T), CTT836TTG (3176C>T 3178T>G), GGT839GGG (3187T>G), AAA841AGG (3192A>G 3193A>G), CGG843CAA (3198G>A 3199G>A), CGG845AAA (3203C>A 3204G>A 3205G>A), TGT846TGC (3208T>C), TCT847TCA (3211T>A), ATT848CTG (3212A>C 3214T>G), ACA849ACC (3217A>C), GGA853GGG (3229A>G), AAA854TGC (3230A>T 3231A>G 3232A>C), AAT855AAC (3235T>C), TTG856CTC (3236T>C 3238G>C), CTT857CTG (3241T>G), TCC858TCT (3244C>T), ACA859ACC (3247A>C), GGG860GGA (3250G>A), CGA861CGC (3253A>C), AAG862AAA (3256G>A), TCC864TCA (3262C>A), TTG865TTA (3265G>A), CAG869AAG (3275C>A), CAC870CAT (3280C>T), TTT871CAA (3281T>C 3282T>A 3283T>A), TCC872TCT (3286C>T), TTG874ATG (3290T>A), CGC875CGT (3295C>T), TTG876CTG (3296T>C), TTT877TTC (3301T>C), ACG879ACT (3307G>T), GTA880GTT (3310A>T), CTC884CTG (3322C>G), ACG887ACA (3331G>A), ATA889TTG (3335A>T 3337A>G), CAT890CAC (3340T>C), TCC893ACT (3347T>A 3349C>T), GGT895GGA (3355T>A), GTC896GTA (3358C>A), CAG899CAA (3367G>A), AAG901AAA (3373G>A), TGC902CGC (3374T>C), CTT903ATT (3377C>A), AAT905GAT (3383A>G), CTT906CTC (3388T>C), ACT907ACG (3391T>G), TTG908CTT (3392T>C 3394G>T), GGT909GGG (3397T>G), GGC910GGT (3400C>T), ACA911TCT (3401A>T 3403A>T), GTA912GTG (3406A>G), TCT913TCA (3409T>A), GTG914GTT (3412G>T), AAA915AAG (3415A>G), ATA917ATT (3421A>T), GGG919GGT (3427G>T), ACA922ACC (3436A>C), AAA923AAG (3439A>G), GGC924GGG (3442C>G), AAA925AGA (3444A>G), AGC926AGT (3448C>T), TCA927GCT (3449T>G 3451A>T), GTT928GTG (3454T>G),

| | |
|---|---|
| | TTC930TTT (3460C>T), CTT931CTG (3463T>G), TTC932TTT (3466C>T), GAT936AAC (3476G>A 3478T>C), TTA937CTG (3479T>C 3481A>G), TCA938AAT (3482T>A 3483C>A 3484A>T), CGT940CGC (3490T>C), ATT943CTT (3497A>C), GAA944GAG (3502A>G), AAG945AAA (3505G>A), TTG946CTG (3506T>C), AAG948AAA (3514G>A), GAC949GAA (3517C>A), AAC950AAT (3520C>T), ATT951GTT (3521A>G), GAG952GAT (3526G>T), GAG953ATC (3527G>A 3528A>T 3529G>C), TAC954TTT (3531A>T 3532C>T), CAG955GAC (3533C>G 3535G>C), GAG956CAG (3536G>C), GGC958GGG (3544C>G), GAT960AAC (3548G>A 3550T>C), CCG961_CCC962del (3551_3556delCCGCCC), AAG964ACA (3561A>C 3562G>A), GCT965TCT (3563G>T), AGT967GTT (3569A>G 3570G>T), ATT968TCC (3572A>T 3573T>C 3574T>C), CTG969TTC (3575C>T 3577G>C), TCC970CAG (3578T>C 3579C>A 3580C>G), ATA971ATC (3583A>C), AGA972AGG (3586A>G), GAG973GAA (3589G>A), AAA974AAG (3592A>G), TTT975TTC (3595T>C), TCC976ACA (3596T>A 3598C>A), GGT978GGA (3604T>A), GCT979GCA (3607T>A), GTA980GTG (3610A>G), TTC983TTT (3619C>T), TGC984TGT (3622C>T), ATG985GCT (3623A>G 3624T>C 3625G>T), GGT986GCA (3627G>C 3628T>A), GAC990ACA (3638G>A 3639A>C 3640C>A), GAA991GAC (3643A>C), TTG994GTA (3650T>G 3652G>A), GTA995GTT (3655A>T), ATT996TTG (3656A>T 3658T>G), CCT997CCA (3661T>A), GCA999GCC (3667A>C), CCA1000CCT (3670A>T), TCC1002TCA (3676C>A), ATA1003GTG (3677A>G 3679A>G), AGG1004CGC (3680A>C 3682G>C), GAA1006GAT (3688A>T), GGG1007GCA (3690G>C 3691G>A), CAC1008TCC (3692C>T 3693A>C), CCT1010CCA (3700T>A), GTT1011ATT (3701G>A), AAG1012AAA (3706G>A), ATC1015ATA (3715C>A), TGT1017TGC (3721T>C), TTT1019TTC (3727T>C), GAT1021GAC (3733T>C), ACA1024TCC (3740A>T 3742A>C), TCA1025TCT (3745A>T), TCA1030TCT (3760A>T), GGT1031GGA (3763T>A), AAT1034GAA (3770A>G 3772T>A), TCA1036ACA (3776T>A), ATT1037TTT (3779A>T), GTG1038GTA (3784G>A), ATA1039ATG (3787A>G), AGA1041CGA (3791A>C), GTA1042ACT (3794G>A 3795T>C 3796A>T), CAA1043CAG (3799A>G), TCC1044AGC (3800T>A 3801C>G), AGT1045TCC (3803A>T 3804G>C 3805T>C), CCT1046AAC (3806C>A 3807C>A 3808T>C), AAT1047AGT (3810A>G), GTT1048ATA (3812G>A 3814T>A), GGA1049GGT (3817A>T), GTA1051ATC (3821G>A 3823A>C), CTA1052TTG (3824C>T 3826A>G), AAT1053AAC (3829T>C), GTT1054ATT (3830G>A), GCT1055GCA (3835T>A), TTT1056CTT (3836T>C), GAT1057GAC (3841T>C), GCC1058ACA (3842G>A 3844C>A), GGC1060GGG (3850C>G), TTT1063TTC (3859T>C), CCA1064CAA (3861C>A), GCT1065TTT (3863G>T 3864C>T), GGG1066GGT (3868G>T), CTT1067GTA (3869C>G 3871T>A), AAT1068TCT (3872A>T 3873A>C), AAA1069GCG (3875A>G 3876A>C 3877A>G), AAT1071GCA (3881A>G 3882A>C 3883T>A), TAT1072CAT (3884T>C), GTA1074TTA (3890G>T), GGT1077GGC (3901T>C), GGA1078GGT (3904A>T), GGC1079GGA (3907C>A), ACA1080ACT (3910A>T), TCA1085TCC (3925A>C), TAC1086TAT (3928C>T), GGT1087GGG (3931T>G), GTG1088GTT (3934G>T), GCA1089ACC (3935G>A 3937A>C), ACA1090ACC (3940A>C), AAT1091AAC (3943T>C), ACG1092ATC (3945C>T 3946G>C), TCA1094TCT (3952A>T), TTC1095TTT (3955C>T), ACT1096GTA (3956A>G 3957C>T 3958T>A), GAG1101CAG (3971G>C), TTT1102TTC (3976T>C), TTC1103TTT (3979C>T), CCA1104CCT (3982A>T), AGG1105CGC (3983A>C 3985G>C), CGG1106CGT (3988G>T), CAT1107TTC (3989C>T 3990A>T 3991T>C), AGG1109AAG (3996G>A), ATG1110GCC (3998A>G 3999T>C 4000G>C), AGG1111AAA (4002G>A 4003G>A), GAA1112TCC (4004G>T 4005A>C 4006A>C), TTC1113TAT (4008T>A 4009C>T), TCA1114GAT (4010T>G 4011C>A 4012A>T), AGC1115AAG (4014G>A 4015C>G), AAG1116GAA (4016A>G 4018G>A), CAA1117AGT (4019C>A 4020A>G 4021A>T), TCC1118TCA (4024C>A), CGC1119AGA (4025C>A 4027C>A), ATC1120ATT (4030C>T), ATG1121GTT (4031A>G 4033G>T), TCA1122TCT (4036A>T), CTA1123TTG (4037C>T 4039A>G), GAT1125GAC (4045T>C), AGG1126AAG (4047G>A), CTT1127CTA (4051T>A), GGA1128GGC (4054A>C), AAT1129TTG (4055A>T 4056A>T 4057T>G), CTG1130TTG (4058C>T), ATC1131CTA (4061A>C 4063C>A), ATA1132ATC (4066A>C), TTG1134CTT (4070T>C 4072G>T), CCT1135CCA (4075T>A), CCT1136CCA (4078T>A), TCC1137GTT (4079T>G 4080C>T 4081C>T), GCC1138GAA (4083C>A 4084C>A), ATA1139CTT (4085A>C 4087A>T), GTG1140ATT (4088G>A 4090G>T), TCC1142TCA (4096C>A), GAG1144GAA (4102G>A), ATA1145ATC (4105A>C), CTT1146ATA (4106C>A 4108T>A), ATA1147GTC (4109A>G 4111A>C), TCT1148AAG (4112T>A 4113C>A 4114T>G), CCT1149CCA (4117T>A), GGA1150GGT (4120A>T), CTT1151CCT (4122T>C), TTC1153TTT (4129C>T), AAG1154AAA (4132G>A), GCC1158GCT (4144C>T), CCT1160CCC (4150T>C), CCT1161CCA (4153T>A), TCT1162CTT (4154T>C 4155C>T), GCC1163GCA (4159C>A), CAT1165AAT (4163C>A), GAA1166GAG (4168A>G), AAA1167AAG (4171A>G), CTT1169CTC (4177T>C), GGC1170GGA (4180C>A), AAT1171GAT (4181A>G), CAA1173CAG (4189A>G), ACT1174ACG (4192T>G), CAC1175CAT (4195C>T), ACC1176ACT (4198C>T), ACC1180ACA (4210C>A), TCA1181TCT (4213A>T), CTA1186CTG (4228A>G), ATT1191AAT (4242T>A) |

#### Proteins

| | | | | | | | | | |
|---|---|---|---|---|---|---|---|---|---|
| hypothetical protein (YP_001976148.1) | 1 | 191 | 100% | 784 | 66.7% | 183 (95.8%) | 116 (60.7%) | 0/8/0/0 | 1 |
| Protein mutations: | G5S (151G>A 153G>T), R6K (154C>A 155G>A 156T>G), S10Q (166T>C 167C>A 168T>A), V11I (169G>A), D12E (174C>G), E13D (177A>T), A15E (182C>A 183T>A), H18N (190C>A), T22A (202A>G 204C>A), N24T (209A>C 210T>C), I26V (214A>G), V31T (229G>A 230T>C), T33V (235A>G 236C>T 237C>T), L35I (241T>A 243G>T), V46I (274G>A), V47C (277G>T 278T>G), V54I (298G>A 300C>T), T58S (310A>T 312C>T), L68V (340T>G 342A>T), N79T (374A>C 375T>G), P81D (379C>G 380C>A 381C>T), N84A (388A>G 389A>C 390C>A), L91I (409C>A), H93R (416A>G 417T>A), V96I (424G>A 426G>T), R97K (427C>A 428G>A 429T>A), V104L (448G>T 450C>G), E107A (458A>C 459A>C), S111H (469T>C 470C>A), A114S (478G>T 480A>T), S115T (481T>A 483A>C), P120N (496C>A 497C>A 498T>C), S121P (499T>C 501T>C), D124_S129del (508_525delGACAGTAAAGGCAAGAGT), K130T (527A>C), V131L (529G>T), E132S (532G>T 533A>C 534G>A), Q133E (535C>G), P134T (538C>A 540C>T), T135A (541A>G), R136_E137del (544_549delCGAGAG), D138S (550G>A 551A>G), I140S (557T>G 558T>C), K141G (559A>G 560A>G), V143N (565G>A 566T>A 567T>C), E144K (568G>A), V145A (572T>C 573T>A), R148E (580C>G 581G>A 582T>A), E149A (584A>C 585G>C), R152V (592A>G 593G>T 594A>T), F153A (595T>G 596T>C 597T>C), Q154R (598C>A 599A>G 600G>A), A158Q (610G>C 611C>A), S159E (613A>G 614G>A 615T>A), Q160A (616C>G 617A>C), K161R (619A>C 620A>G 621A>G), N164I (629A>T), Q165V (631C>G 632A>T 633G>C), Q168R (640C>A 641A>G 642A>G), L171I (649C>A 651C>T), S172Q (652T>C 653C>A 654C>A), N173G (655A>G 656A>G), V175P (661G>C 662T>C 663T>A), N178E (670A>G 672T>G), I180V (676A>G), S182A (682T>G), E186S (694G>A 695A>G 696A>T), S187D (697A>G 698G>A), Q190K (706C>A) | | | | | | | | |
| Codon mutations: | TCA2TCT (144A>T), TTC3TTT (147C>T), ATT4ATC (150T>C), GGG5AGT (151G>A 153G>T), CGT6AAG (154C>A 155G>A 156T>G), TTG7CTC (157T>C 159G>C), AAT8AAC (162T>C), ACA9ACC (165A>C), TCT10CAA (166T>C 167C>A 168T>A), GTA11ATA (169G>A), GAC12GAG (174C>G), GAA13GAT (177A>T), GCT15GAA (182C>A 183T>A), TTT16TTC (186T>C), CAT18AAT (190C>A), GTT20GTA (198T>A), GCC21GCA (201C>A), ACC22GCA (202A>G 204C>A), TCG23TCT (207G>T), AAT24ACC (209A>C 210T>C), ATT26GTT (214A>G), TCT28AGT (220T>A 221C>G), GTT31ACT (229G>A 230T>C), GGC32GGT (234C>T), ACC33GTT (235A>G 236C>T 237C>T), TTG35ATT (241T>A 243G>T), ATA36ATT (246A>T), ACC41ACT (261C>T), CTT42TTA (262C>T 264T>A), GAC43GAT (267C>T), AAA45AAG (273A>G), GTT46ATT (274G>A), GTT47TGT (277G>T 278T>G), CCT48CCC (282T>C), GGT51GGA (291T>A), GTC54ATT (298G>A 300C>T), TCA55TCG (303A>G), GTT56GTG (306T>G), TTA57CTT (307T>C 309A>T), ACC58TCT (310A>T 312C>T), GTT59GTG (315T>G), TCT60AGT (316T>A 317C>G), GAG62GAA (324G>A), ACG65ACT (333G>T), CCA66CCC (336A>C), TTA68GTT (340T>G 342A>T), CCT70CCA (348T>A), GGA71GGC (351A>C), CAC72CAT (354C>T), TAT73TAC (357T>C), TTG74CTA (358T>C 360G>A), TTG75CTA (361T>C 363G>A), AAT79ACG (374A>C 375T>G), CCC81GAT (379C>G 380C>A 381C>T), ATT82ATC (384T>C), AAG83AAA (387G>A), AAC84GCA (388A>G 389A>C 390C>A), TCT88TCA (402T>A), GGT89GGC (405T>C), TTA90TTG (408A>G), CTT91ATT (409C>A), GTC92GTG (414C>G), CAT93CGA (416A>G 417T>A), TCA95TCC (423A>C), GTG96ATT (424G>A 426G>T), CGT97AAA (427C>A 428G>A 429T>A), CTT98TTG (430C>T 432T>G), ACC101ACA (441C>A), AGA102CGG (442A>C 444A>G), GTC104TTG (448G>T 450C>G), CTA105TTG (451C>T 453A>G), GAG106GAA (456G>A), GAA107GCC (458A>C 459A>C), AAC108AAT (462C>T), AAG109AAA (465G>A), GTT110GTG (468T>G), TCT111CAT (469T>C 470C>A), TCA113TCT (477A>T), GCA114TCT (478G>T 480A>T), TCA115ACC (481T>A 483A>C), TCA117TCT (489A>T), TCT118TCA (492T>A), TCT119TCC (495T>C), CCT120AAC (496C>A 497C>A 498T>C), TCT121CCC (499T>C 501T>C), GAC124_AGT129del (508_525delGACAGTAAAGGCAAGAGT), AAA130ACA (527A>C), GTA131TTA (529G>T), GAG132TCA (532G>T 533A>C 534G>A), CAA133GAA (535C>G), CCC134ACT (538C>A 540C>T), ACA135GCA (541A>G), CGA136_GAG137del (544_549delCGAGAG), GAT138AGT (550G>A 551A>G), CTC139TTA (553C>T 555C>A), ATT140AGC (557T>G 558T>C), AAA141GGA (559A>G 560A>G), GAA142GAG (564A>G), GTT143AAC (565G>A 566T>A 567T>C), GAG144AAG (568G>A), GTT145GCA (572T>C 573T>A), CTC146CTT (576C>T), AAA147AAG (579A>G), CGT148GAA (580C>G 581G>A 582T>A), GAG149GCC (584A>C 585G>C), TTA150CTT (586T>C 588A>T), GAG151GAA (591G>A), AGA152GTT (592A>G 593G>T 594A>T), TTT153GCC (595T>G 596T>C 597T>C), CAG154AGA (598C>A 599A>G 600G>A), GAG156GAA (606G>A), TTG157CTG (607T>C), GCA158CAA (610G>C 611C>A), AGT159GAA (613A>G 614G>A 615T>A), CAA160GCA (616C>G 617A>C), AAA161CGG (619A>C 620A>G 621A>G), AAT164ATT (629A>T), CAG165GTC (631C>G 632A>T 633G>C), CTA167CTC (639A>C), CAA168AGG (640C>A 641A>G 642A>G), CTT169TTG (643C>T 645T>G), CTC171ATT (649C>A 651C>T), TCC172CAA (652T>C 653C>A 654C>A), AAT173GGT (655A>G 656A>G), GTT175CCA (661G>C 662T>C 663T>A), AGT176AGC (666T>C), AAT177AAC (669T>C), AAT178GAG (670A>G 672T>G), GAC179GAT (675C>T), ATC180GTC (676A>G), TTC181TTT (681C>T), TCA182GCA (682T>G), GGT183GGC (687T>C), ACT185ACG (693T>G), GAA186AGT (694G>A 695A>G 696A>T), AGT187GAT (697A>G 698G>A), GGG188GGC (702G>C), CAG190AAG (706C>A), TAA191TGA (710A>G) | | | | | | | | |
| polyprotein (YP_001976149.1) | 1 | 1192 | 100% | 6009 | 73.1% | 1185 (98.7%) | 820 (68.3%) | 9/7/0/0 | 1 |

| | Begin | End | Coverage | Score | Concordance | Matches | Identities | I/D/M/F* | Stop Codons |
|---|---|---|---|---|---|---|---|---|---|
| **NT** | **89** | **4733** | **94.7%** | **2570** | **28.4%** | **4597 (96.7%)** | **3027 (63.7%)** | **107/48** | |

Protein mutations:

M1R (672T>G), T2M (675C>T 676A>G), S4L (681C>T 682T>G), V6A (687T>C), L8R (693T>G 694G>A), K9V (695A>G 696A>T 697A>G), V10M (698G>A), G11A (702G>C), N14D (710A>G 712T>C), H15R (714A>G), Q16S (716C>T 717A>C 718G>A), Q17del (719_721delCAG), E18K (722G>A 724G>A), A20V (729C>T 730C>A), E22D (736G>C), R23K (738G>A 739G>A), A34E (771C>A 772C>A), R41K (792G>A), V50I (818G>A 820T>A), G53A (828G>C 829T>A), E61D (853G>T), F63Y (858T>A), I71V (881A>G 883C>G), T72A (884A>G), T77A (899A>G 901A>T), H80R (908C>A 909A>G 910T>G), K81T (912A>C 913G>C), F82I (914T>A 916T>A), V83T (917G>A 918T>C 919T>A), K86R (926A>C 927A>G 928G>T), A89G (936C>G 937A>G), Y96H (956T>C 958T>C), V106A (987T>C 988T>A), S108N (993G>A), R112A (1004A>G 1005G>C 1006G>C), L124T (1040C>A 1041T>C 1042C>T), S125D (1043T>G 1044C>A 1045A>T), T132I (1065C>T 1066A>T), S134Y (1071C>A), L135F (1075G>T), A136S (1076G>T 1078C>T), D137G (1080A>G 1081T>A), Q166L (1167A>T 1168A>C), D185S (1223G>A 1224A>G 1225C>T), T186P (1226A>C 1228A>T), S192T (1244T>A), Q193D (1247C>G 1249G>C), Q194H (1252A>T), E195D (1255A>C), S226N (1347G>A), G227_Q228insV (1351_1352insGTA), S229R (1357T>A), T230S (1359C>G), M232L (1364A>C), A233del (1367_1369delGCT), K237R (1380A>G), A238P (1382G>C 1384C>T), E242A (1395A>C 1396A>C), Q243A (1397C>G 1398A>C 1399G>A), G246D (1407G>A 1408C>T), D250N (1418G>A 1420C>T), M252P (1424A>C 1425T>C), N253Q (1427A>C 1429T>G), S255A (1433A>G 1434G>C 1435C>A), T256N (1437C>A), I258V (1442A>G 1444C>G), K273R (1488A>G), P276A (1496C>G 1498G>A), P281_A282insDF (1513_1514insGACTTT), V283S (1517G>T 1518T>C), S284T (1521G>C), T285S (1523A>T 1525A>T), S286R (1526T>A 1527C>G), S287A (1529T>G 1531C>G), L288Y (1533T>A 1534A>T), L289Q (1536T>A 1537C>G), S290G (1538A>G 1540C>A), T291V (1541A>G 1542C>T 1543A>G), V296G (1557T>G), H297I (1559C>A 1560A>T), E299del (1565_1567delGAG), C302T (1574T>A 1575G>C 1576T>G), G303S (1577G>T 1578G>C 1579G>T), G304L (1580G>C 1581G>T), P309_K310del (1595_1600delCCCAAG), K312I (1605A>T 1606G>A), G313S (1607G>T 1608G>C 1609C>A), F316G (1616T>G 1617T>G 1618C>A), N317T (1620A>C 1621C>A), M318L (1622A>C 1624G>C), G319Q (1625G>C 1626G>A 1627T>A), A320E (1629C>A 1630C>A), I321V (1631A>G 1633T>C), L325Y (1644T>A 1645G>T), M327I (1651G>T), E328Q (1652G>C 1654G>A), A331E (1662C>A), H332D (1664C>G 1666C>T), I333F (1667A>T), D334A (1671A>C 1672T>C), D337S (1679G>A 1680A>G), D338N (1682G>A 1684T>C), E341A (1692A>C 1693A>C), R342L (1695G>T 1696A>T), I344V (1700A>G 1702A>G), A345P (1703G>C 1705T>C), Q346E (1706C>G), V348I (1712G>A 1714A>T), Q349D (1715C>G 1717A>T), F350Y (1719T>A), I351V (1721A>G 1723C>T), F353V (1727T>G 1729T>G), S354Q (1730A>C 1731G>A 1732T>G), E355K (1733G>A), K357R (1740A>G), R358A (1742A>G 1743G>C 1744A>T), V361L (1751G>C 1753T>C), I362C (1754A>T 1755T>G), R365K (1764G>A), V367I (1769G>A), A368E (1773C>A 1774A>G), K369M (1776A>T), H371D (1781C>G 1783T>C), N374C (1790A>T 1791A>G), R377F (1799A>T 1800G>T 1801G>T), D383A (1818A>C), E387D (1831A>T), E390A (1839A>C 1840A>G), N396K (1858T>G), I397V (1859A>G 1861C>T), Q398G (1862C>G 1863A>G 1864A>G), G399M (1865G>A 1866G>T 1867C>G), T401E (1871A>G 1872C>A 1873T>A), F404P (1880T>C 1881T>C 1882C>A), A408S (1892G>T), M409L (1895A>T), E413Q (1907G>C), Q421M (1931C>A 1932A>T 1933A>G), S425Q (1943T>C 1944C>A 1945C>A), I428R (1953T>G 1954T>G), Q429Y (1955C>T 1957A>T), G430W (1958G>T), Q431D (1961C>G 1963G>T), T432L (1964A>T 1965C>T 1966T>A), N433T (1968A>C 1969C>A), N438D (1982A>G 1984T>C), C441V (1991T>G 1992G>T 1993C>G), K448A (2012A>G 2013A>C), E449D (2017G>T), R450_Q451insVK (2020_2021insGTGAAA), Q451H (2023G>T), L452P (2024T>C 2025T>C 2026G>C), I453L (2027A>T 2029T>G), V454M (2030G>A), E455S (2033G>A 2034A>G 2035G>T), E456D (2038G>T), Q457L (2040A>T), V458A (2043T>C 2044G>A), A459C (2045G>T 2046C>G), Q460_R461insLN (2050_2051insTTGAAT), R462L (2054A>T 2055G>T), S463G (2057T>G 2058C>G 2059T>A), E464K (2060G>A), R465P (2063A>C 2064G>C), Q466S (2066C>A 2067A>G 2068A>C), A467M (2069G>A 2070C>T 2071A>G), Q468S (2072C>A 2073A>G 2074G>T), T470A (2078A>G), A471Q (2081G>C 2082C>A), R472T (2085G>C 2086G>T), I474N (2091T>A 2092A>T), A475_E476insS (2095_2096insAGC), E476K (2096G>A 2098A>G), S477W (2099A>T 2101T>G), P479T (2105C>A), D481G (2112A>G 2113C>A), R482K (2115G>A 2116A>G), T484L (2120A>C 2121C>T 2122T>G), S489T (2135T>A), T492A (2144A>G 2146G>A), M493L (2147A>C 2149G>T), E494_D495insS (2152_2153insAGT), D495E (2155T>G), P496G (2156C>G 2157C>G 2158C>T), T497I (2160C>T), K498Q (2162A>C 2164G>A), P499R (2165C>A 2166C>G), D500K (2168G>A 2170T>G), K501E (2171A>G), I502V (2174A>G 2176A>G), E503S (2177G>T 2178A>C), V505E (2184T>A 2185T>G), G508D (2193G>A 2194A>T), A509P (2195G>C 2197A>G), E511G (2202A>G 2203A>G), E512K (2204G>A), Q514E (2210C>G 2212A>G), G516N (2216G>A 2217G>A), V518I (2222G>A), I519F (2225A>T 2227T>C), P524E (2240C>G 2241C>A 2242A>G), D527M (2249G>A 2250A>T 2251C>G), T528G (2252A>G 2253C>G 2254A>T), S529D (2255T>G 2256C>A 2257C>T), M530A (2258A>G 2259T>C 2260G>C), A531P (2261G>C 2263A>G), V532L (2264G>T 2266T>A), E533N (2267G>A 2269A>T), D535E (2275T>G), M536L (2276A>T), V540A (2289T>C 2290G>A), D546E (2308T>G), V552I (2324G>A), V556L (2336G>C 2338T>G), E558K (2342G>A 2344G>A), A561N (2351G>A 2352C>A 2353T>C), M568L (2372A>C 2374G>T), S569A (2375A>G 2376G>C 2377C>A), A571S (2381G>T 2383T>C), A574M (2390G>A 2391C>T 2392T>G), S580K (2409G>A 2410C>G), T583A (2417A>G), A587S (2429G>T 2431G>C), L597N (2459C>A 2460T>A 2461A>C), I598L (2462A>C 2464C>T), Y600F (2469A>T 2470C>T), I602V (2474A>G 2476A>C), V606I (2486G>A), V609M (2495G>A 2497T>G), I612V (2504A>G 2506C>G), V625A (2544T>C 2545G>A), K629T (2556A>C 2557G>C), L638I (2582T>A 2584G>T), T644H (2600A>C 2601C>A 2602A>T), Q650E (2618C>G), N652S (2624A>T 2625A>C), E654G (2631A>G), E655K (2633G>A), C657V (2639T>G 2640G>T 2641C>G), T659S (2645A>T 2647C>A), T661Q (2651A>C 2652C>A), K665T (2664A>C 2665G>C), Y666F (2667A>T 2668C>T), E671T (2681G>A 2682A>C 2683A>T), G672S (2684G>T 2685G>C 2686T>A), T673L (2687A>C 2688C>T 2689A>G), A675V (2694C>T 2695C>T), T676P (2696A>C), D677E (2701T>A), S690A (2738T>G 2740T>A), N696E (2756A>G 2758T>G), V698I (2762G>A), L702I (2774T>A 2776G>A), Q705S (2783C>T 2784A>C 2785G>C), L714M (2810C>A 2812T>G), M719L (2825A>T), I720R (2829T>G), A721G (2832C>G 2833A>G), Y725F (2844A>T), K728R (2852A>C 2853A>G), A729P (2855G>C 2857G>A), Q731T (2861C>A 2862A>C 2863G>A), Y733F (2868A>T 2869T>C), P735S (2873C>A 2874C>G 2875A>T), G749S (2915G>A 2917G>T), T750V (2918A>G 2919C>T), S751A (2921T>G 2923A>T), S752A (2924T>G 2926C>A), P753A (2927C>G 2929A>C), I761V (2951A>G 2953A>C), T773I (2988C>T), T790R (3039C>G 3040T>A), I797V (3059A>G 3061T>G), I799V (3065A>G), N817A (3119A>G 3120A>C 3121C>A), K819E (3125A>G 3127A>G), T821S (3131A>T), N822T (3135A>C 3136T>A), F827L (3151T>G), K841R (3192A>G 3193A>G), R843Q (3198G>A 3199G>A), R845K (3203C>A 3204G>A 3205G>A), I848L (3212A>C 3214T>G), K854C (3230A>T 3231A>G 3232A>C), Q869K (3275C>A), F871Q (3281T>C 3282T>A 3283T>A), L874M (3290T>A), I889L (3335A>T 3337A>G), S893T (3347T>A 3349C>T), C902R (3374T>C), L903I (3377C>A), N905D (3383A>G), T911S (3401A>T 3403A>T), K925R (3444A>G), S927A (3449T>G 3451A>T), D936N (3476G>A 3478T>C), S938N (3482T>A 3483C>A 3484A>T), I943L (3497A>C), D949E (3517C>A), I951V (3521A>G), E952D (3526G>T), E953I (3527G>A 3528A>T 3529G>C), Y954F (3531A>T 3532C>T), Q955D (3533C>G 3535G>C), E956Q (3536G>C), D960N (3548G>A 3550T>C), P961_P962del (3551_3556delCCGCCC), K964T (3561A>C 3562G>A), A965S (3563G>T), S967V (3569A>G 3570G>T), I968S (3572A>T 3573T>C 3574T>C), L969F (3575C>T 3577G>C), S970Q (3578T>C 3579C>A 3580C>G), S976T (3596T>A 3598C>A), M985A (3623A>G 3624T>C 3625G>T), G986A (3627G>C 3628T>A), D990T (3638G>A 3639A>C 3640C>A), E991D (3643A>C), L994V (3650T>G 3652G>A), I996L (3656A>T 3658T>G), I1003V (3677A>G 3679A>G), E1006D (3688A>T), G1007A (3690G>C 3691G>A), H1008S (3692C>T 3693A>C), V1011I (3701G>A), T1024S (3740A>T 3742A>C), N1034E (3770A>G 3772T>A), S1036T (3776T>A), I1037F (3779A>T), I1039M (3787A>G), V1042T (3794G>A 3795T>C 3796A>T), P1046N (3806C>A 3807C>A 3808T>C), N1047S (3810A>G), V1048I (3812G>A 3814T>A), V1051I (3821G>A 3823A>C), V1054I (3830G>A), F1056L (3836T>C), A1058T (3842G>A 3844C>A), P1064Q (3861C>A), A1065F (3863G>T 3864C>T), L1067V (3869C>G 3871T>A), N1068S (3872A>T 3873A>C), K1069A (3875A>G 3876A>C 3877A>G), N1071A (3881A>G 3882A>C 3883T>A), Y1072H (3884T>C), V1074L (3890G>T), A1089T (3935G>A 3937A>C), T1092I (3945C>T 3946G>C), T1096V (3956A>G 3957C>T 3958T>A), E1101Q (3971G>C), H1107F (3989C>T 3990A>T 3991T>C), R1109K (3996G>A), M1110A (3998A>G 3999T>C 4000G>C), R1111K (4002G>A 4003G>A), E1112S (4004G>T 4005A>C 4006A>C), F1113Y (4008T>A 4009C>T), S1114D (4010T>G 4011C>A 4012A>T), S1115K (4014G>A 4015C>G), K1116E (4016A>G 4018G>A), Q1117S (4019C>A 4020A>G 4021A>T), M1121V (4031A>G 4033G>T), R1126K (4047G>A), N1129L (4055A>T 4056A>T 4057T>G), I1131L (4061A>C 4063C>A), S1137V (4079T>G 4080C>T 4081C>T), A1138E (4083C>A 4084C>A), I1139L (4085A>C 4087A>T), V1140I (4088G>A 4090G>T), L1146I (4106C>A 4108T>A), I1147V (4109A>G 4111A>C), S1148K (4112T>A 4113C>A 4114T>G), L1151P (4122T>C), S1162L (4154T>C 4155C>T), H1165N (4163C>A), N1171D (4181A>G), I1191N (4242T>A)

| | Begin | End | Coverage | Score | Concordance | Matches | Identities | I/D/M/F* | Stop Codons |
|---|---|---|---|---|---|---|---|---|---|
| **NT** | **89** | **4733** | **94.7%** | **2570** | **28.4%** | **4597 (96.7%)** | **3027 (63.7%)** | **107/48** | |

Codon mutations:

ATG1AGG (672T>G), ACA2ATG (675C>T 676A>G), TCT4TTG (681C>T 682T>G), GTT6GCT (687T>C), CTG8CGA (693T>G 694G>A), AAA9GTG (695A>G 696A>T 697A>G), GTG10ATG (698G>A), GGC11GCC (702G>C), CCC12CCA (706C>A), AAT14GAC (710A>G 712T>C), CAC15CGC (714A>G), CAG16TCA (716C>T 717A>C 718G>A), CAG17del (719_721delCAG), GAG18AAA (722G>A 724G>A), GCC20GTA (729C>T 730C>A), TCC21TCT (733C>T), GAG22GAC (736G>C), AGG23AAA (738G>A 739G>A), ATA25ATT (745A>T), TCA26TCT (748A>T), CAA27CAG (751A>G), ATA28ATC (754A>C), GCT30GCG (760T>G), GTG32GTT (766G>T), GAG33GAA (769G>A), GCC34GAA (771C>A 772C>A), GGA35GGT (775A>T), AAT38AAC (784T>C), TTG39CTT (785T>C 787G>T), CTC40CTT (790C>T), AGA41AAA (792G>A), AAG42AAA (796G>A), CTT43TTG (797C>T 799T>G), GGC44GGA (802C>A), TCC47TCA (811C>A), TAC48TAT (814C>T), GGG49GGC (817G>C), GTT50ATA (818G>A 820T>A), CTT51TTG (821C>T 823T>G), GGT53GCA (828G>C 829T>A), CAA55CAG (835A>G), AAG57AAA (841G>A), AGA58AGG (844A>G), GAG61GAT (853G>T), TTG62CTT (854T>C 856G>T), TTT63TAT (858T>A), GAC64GAT (862C>T), GTA68GTT (874A>T), TCA69AGT (875T>A 876C>G 877A>T), AAG70AAA (880G>A), ATC71GTG (881A>G 883C>G), ACA72GCA (884A>G), CTT74CTA (892T>A), ACA77GCT (899A>G 901A>T), TTC78TTT (904C>T), CAT80AGG (908C>A 909A>G 910T>G), AAG81ACC (912A>C 913G>C), TTT82ATA (914T>A 916T>A), GTT83ACA (917G>A 918T>C 919T>A), GAG84GAA (922G>A), TCT85TCA (925T>A), AAG86CGT (926A>C 927A>G 928G>T), GAC87GAT (931C>T), GCA89GGG (936C>G 937A>G), AAC90AAT (940C>T), TTG91TTA (943G>A), TTT93TTC (949T>C), CTG95TTA (953C>T 955G>A), TAT96CAC (956T>C 958T>C), GGT97GGG (961T>G), CTT99TTG (965C>T 967T>G), TTC100TTT (970C>T), TTC101TTT (973C>T), GTC103GTT (979C>T), CCA104CCC (982A>C), GTT106GCA (987T>C 988T>A), AGT108AAT (993G>A), GGG109GGA (997G>A), GGA111GGT (1003A>T), AGG112GCC (1004A>G 1005G>C 1006G>C), GTT113GTC (1009T>C), AAG114AAA (1012G>A), ATA115ATC (1015A>C), AGT116AGC (1018T>C), CTT117TTG (1019C>T 1021T>G), TGT118TGC (1024T>C), TCA119TCT (1027A>T), AGT120AGC (1030T>C), AAT121AAC (1033T>C), CTC124ACT (1040C>A 1041T>C 1042C>T), TCA125GAT (1043T>G 1044C>A 1045A>T), CCG126CCT (1048G>T), CAA129CAG (1057A>G), AAG131AAA (1063G>A), ACA132ATT (1065C>T 1066A>T), CTG133TTG (1067C>T), TCC134TAC (1071C>A), TTG135TTT (1075G>T), GCC136TCT (1076G>T 1078C>T), GAT137GGA (1080A>G 1081T>A), GCG139GCA (1087G>A), CAG140CAA (1090G>A), GCA141GCC (1093A>C), CTC143CTA (1099C>A), AGC145AGT (1105C>T), CCC146CCT (1108C>T), AGC147AGT (1111C>T), CTC153TTG (1127C>T 1129C>G), AAA154AAG (1132A>G), AGA155AGG (1135A>G), GGG156GGC (1138G>C), ACC162ACA (1156C>A), GAG164GAA (1162G>A), TGC165TGT (1165C>T), CAA166CTC (1167A>T 1168A>C), AAT167AAC (1171T>C), ACC168ACG (1174C>G), AGG169AGA (1177G>A), GCA170GCT (1180A>T), CAG171CAA (1183G>A), CCG173CCT (1189G>T), TCC175TCT (1195C>T), GTG176GTA (1198G>A), GCC178GCA (1204C>A), ATA179ATT (1207A>T), GAC185AGT (1223G>A 1224A>G 1225C>T), ACA186CCT (1226A>C 1228A>T), AGG187CGT (1229A>C 1231G>T), GCA189GCT (1237A>T), GTT190GTA (1240T>A), TCA192ACA (1244T>A), CAG193GAC (1247C>G 1249G>C), CAA194CAT (1252A>T), GAA195GAC (1255A>C), ACA196ACT (1258A>T), TCT198AGT (1262T>A 1263C>G), GCC200GCT (1270C>T), ATA201ATT (1273A>T), GAA202GAG (1276A>G), GCA203GCT (1279A>T), CTC204CTT (1282C>T), AAT205AAC (1285T>C), CGT206AGA (1286C>A 1288T>A), CCG207CCT (1291G>T), CAA208CAG (1294A>G), GAT212GAC (1306T>C), AGA213AGG (1309A>G), GAG215GAA (1315G>A), GCT216GCA (1318T>A), GCT217GCA (1321T>A), TAT219TAC (1327T>C), ATA220ATT (1330A>T), GCT221GCA (1333T>A), GTT223GTG (1339T>G), TAT224TAC (1342T>C), TCG225TCC (1345G>C), AGT226AAT (1347G>A), GGG227_CAA228insGTA (1351_1352insGTA), CAA228CAG (1354A>G), AGT229AGA (1357T>A), ACT230AGT (1359C>G), CAA231CAG (1363A>G), ATG232CTG (1364A>C), GCT233del (1367_1369delGCT), CTT234TTG (1370C>T 1372T>G), GAA235GAG (1375A>G), AAA237AGA (1380A>G), GCC238CCT (1382G>C 1384C>T), GTT240GTG (1390T>G), GGA241GGT (1393A>T), GAA242GCC (1395A>C 1396A>C), CAG243GCA (1397C>G 1398A>C 1399G>A), GGG245GGT (1405G>T), GGC246GAT (1407G>A 1408C>T), ACA247ACC (1411A>C), GAC250AAT (1418G>A 1420C>T), GTC251GTG (1423C>G), ATG252CCG (1424A>C 1425T>C), AAT253CAG (1427A>C 1429T>G), GAA254GAG (1432A>G), AGC255GCA (1433A>G 1434G>C 1435C>A), ACT256AAT (1437C>A), ATC258GTG (1442A>G 1444C>G), CGG259CGC (1447G>C), AGT260TCT (1448A>T 1449G>C), TCT262TCA (1456T>A), AGA264AGG (1462A>G), ACT267ACC (1471T>C), CTC268CTA (1474C>A), AAA269AAG (1477A>G), GTG270GTA (1480G>A), AAG273AGG (1488A>G), CGC274CGA (1492C>A), CCG276GCA (1496C>G 1498G>A), TCC277TCA (1501C>A), CTG280TTG (1508C>T), CCG281CCT (1513G>T), CCG281_GCT282insGACTTT (1513_1514insGACTTT), GTA283TCA (1517G>T 1518T>C), AGT284ACT (1521G>C), ACA285TCT (1523A>T 1525A>T), TCA286AGA (1526T>A 1527C>G), TCC287GCG (1529T>G 1531C>G), TTA288TAT (1533T>A 1534A>T), CTC289CAG (1536T>A 1537C>G), AGC290GGA (1538A>G 1540C>A), ACA291GTG (1541A>G 1542C>T 1543A>G), CGT292AGA (1544C>A 1546T>A), GAA293GAG (1549A>G), ACA295ACT (1555A>T), GTT296GGT (1557T>G), CAT297ATT (1559C>A 1560A>T), GAG299del (1565_1567delGAG), TGT302ACG (1574T>A 1575G>C 1576T>G), GGG303TCT (1577G>T 1578G>C 1579G>T), GGA304CTA (1580G>C 1581G>T), TTG305CTG (1583T>C), CCA307CCT (1591A>T), GCA308GCC (1594A>C), CCC309_AAG310del (1595_1600delCCCAAG), AAA311AAG (1603A>G), AAG312ATA (1605A>T 1606G>A), GGC313TCA (1607G>T 1608G>C 1609C>A), GCC315GCA (1615C>A), TTC316GGA (1616T>G 1617T>G 1618C>A), AAC317ACA (1620A>C 1621C>A), ATG318CTC (1622A>C 1624G>C), GGT319CAA (1625G>C 1626G>A 1627T>A), GCC320GAA (1629C>A 1630C>A), ATT321GTC (1631A>G 1633T>C), GAT323GAC (1639T>C), TTG325TAT (1644T>A 1645G>T), ATG327ATT (1651G>T), GAG328CAA (1652G>C 1654G>A), TCG329AGT (1655T>A 1656C>G 1657G>T), TTC330TTT (1660C>T), GCA331GAA (1662C>A), CAC332GAT (1664C>G 1666C>T), ATT333TTT (1667A>T), GAT334GCC (1671A>C 1672T>C), GAT337AGT (1679G>A 1680A>G), GAT338AAC (1682G>A 1684T>C), GAA341GCC (1692A>C 1693A>C), CGA342CTT (1695G>T 1696A>T), ATA344GTG (1700A>G 1702A>G), GCT345CCC (1703G>C 1705T>C), CAG346GAG (1706C>G), CAG347CAA (1711G>A), GTA348ATT (1712G>A 1714A>T), CAA349GAT (1715C>G 1717A>T), TTT350TAT (1719T>A), ATC351GTT (1721A>G 1723C>T), CTC352CTT (1726C>T), TTT353GTG (1727T>G 1729T>G), AGT354CAG (1730A>C 1731G>A 1732T>G), GAG355AAG (1733G>A), AAA357AGA (1740A>G), AGA358GCT (1742A>G 1743G>C 1744A>T), GGA359GGT (1747A>T), AAC360AAT (1750C>T), GTT361CTC (1751G>C 1753T>C), ATT362TGT (1754A>T 1755T>G), GTG363GTC (1759G>C), AGA365AAA (1764G>A), GTT367ATT (1769G>A), GCA368GAG (1773C>A 1774A>G), AAG369ATG (1776A>T), CGG370AGG (1778C>A), CAT371GAC (1781C>G 1783T>C), CTG372TTG (1784C>T), CAT373CAC (1789T>C), AAT374TGT (1790A>T 1791A>G), ATC375ATA (1795C>A), AGG377TTT (1799A>T 1800G>T 1801G>T), GAG378GAA (1804G>A), CAC379CAT (1807C>T), ATA380ATC (1810A>C), ACC381ACT (1813C>T), GAC383GCC (1818A>C), GTA386GTT (1828A>T), GAA387GAT (1831A>T), ATA388ATT (1834A>T), CTT389TTG (1835C>T 1837T>G), GAA390GCG (1839A>C 1840A>G), GGT391GGA (1843T>A), GGT393GGA (1849T>A), AAT396AAG (1858T>G), ATC397GTT (1859A>G 1861C>T), CAA398GGG (1862C>G 1863A>G 1864A>G), GGC399ATG (1865G>A 1866G>T 1867C>G), ACT401GAA (1871A>G 1872C>A 1873T>A), CGC402AGA (1874C>A 1876C>A), ACT403ACA (1879T>A), TTC404CCA (1880T>C 1881T>C 1882C>A), TAT407TAC (1891T>C), GCT408TCT (1892G>T), ATG409TTG (1895A>T), TCT410AGC (1898T>A 1899C>G 1900T>C), CTC411CTG (1903C>G), GAG413CAG (1907G>C), GTT415GTG (1915T>G), GTG416GTT (1918G>T), CTC418CTA (1924C>A), CAT420CAC (1930T>C), CAA421ATG (1931C>A 1932A>T 1933A>G), AGA422AGG (1936A>G), GAT423GAC (1939T>C), TCC425CAA (1943T>C 1944C>A 1945C>A), TTC426TTT (1948C>T), ATT428AGG (1953T>G 1954T>G), CAA429TAT (1955C>T 1957A>T), GGG430TGG (1958G>T), CAG431GAT (1961C>G 1963G>T), ACT432TTA (1964A>T 1965C>T 1966T>A), AAC433ACA (1968A>C 1969C>A), AAT434AAC (1972T>C), CCA435CCG (1975A>G), CCG437CCT (1981G>T), AAT438GAC (1982A>G 1984T>C), TGC441GTG (1991T>G 1992G>T 1993C>G), GGA444GGT (2002A>T), TTA445CTT (2003T>C 2005A>T), ACC446ACA (2008C>A), TTG447CTG (2009T>C), AAA448GCA (2012A>G 2013A>C), GAG449GAT (2017G>T), CGT450AGG (2018C>A 2020T>G), CGT450_CAG451insGTGAAA (2020_2021insGTGAAA), CAG451CAT (2023G>T), TTG452CCC (2024T>C 2025T>C 2026G>C), ATT453TTG (2027A>T 2029T>G), GTG454ATG (2030G>A), GAG455AGT (2033G>A 2034A>G 2035G>T), GAG456GAT (2038G>T), CAA457CTA (2040A>T), GTG458GCA (2043T>C 2044G>A), GCC459TGC (2045G>T 2046C>G), CAA460_AGA461insTTGAAT (2050_2051insTTGAAT), AGA461CGC (2051A>C 2053A>C), AGG462TTG (2054A>T 2055G>T), TCT463GGA (2057T>G 2058C>G 2059T>A), GAA464AAA (2060G>A), AGA465CCA (2063A>C 2064G>C), CAA466AGC (2066C>A 2067A>G 2068A>C), GCA467ATG (2069G>A 2070C>T 2071A>G), CAG468AGT (2072C>A 2073A>G 2074G>T), ACA470GCA (2078A>G), GCA471CAA (2081G>C 2082C>A), AGG472ACT (2085G>C 2086G>T), GGC473GGA (2089C>A), ATA474AAT (2091T>A 2092A>T), GCT475GCC (2095T>C), GCT475_GAA476insAGC (2095_2096insAGC), GAA476AAG (2096G>A 2098A>G), AGT477TGG (2099A>T 2101T>G), CAG478CAA (2104G>A), CCT479ACT (2105C>A), GAC481GGA (2112A>G 2113C>A), AGA482AAG (2115G>A 2116A>G), ACT484CTG (2120A>C 2121C>T 2122T>G), TCC486TCA (2128C>A), GTG488GTT (2134G>T), TCA489ACA (2135T>A), ACT490ACC (2140T>C), ACC491ACA (2143C>A), ACG492GCA (2144A>G 2146G>A), ATG493CTT (2147A>C 2149G>T), GAA494GAG (2152A>G), GAA494_GAT495insAGT (2152_2153insAGT), GAT495GAG (2155T>G), CCC496GGT (2156C>G 2157C>G 2158C>T), ACC497ATC (2160C>T), AAG498CAA (2162A>C 2164G>A), CCA499AGA (2165C>A 2166C>G), GAT500AAG (2168G>A 2170T>G), AAG501GAG (2171A>G), ATA502GTG (2174A>G 2176A>G), GAA503TCA (2177G>T 2178A>C), GTT505GAG (2184T>A 2185T>G), GCA506GCT (2188A>T), GAG507GAA (2191G>A), GGA508GAT (2193G>A 2194A>T), GCA509CCG (2195G>C 2197A>G), GAA511GGG (2202A>G 2203A>G), GAA512AAA (2204G>A), ACT513ACA (2209T>A), CAA514GAG (2210C>G 2212A>G), GGT516AAT (2216G>A 2217G>A), GAT517GAC (2221T>C), GTT518ATT (2222G>A), ATT519TTC (2225A>T 2227T>C), GAC521GAT (2233C>T), GGA523GGG (2239A>G), CCA524GAG (2240C>G 2241C>A 2242A>G), GAC527ATG (2249G>A 2250A>T 2251C>G), ACA528GGT (2252A>G 2253C>G 2254A>T), TCC529GAT (2255T>G 2256C>A 2257C>T), ATG530GCC (2258A>G 2259T>C 2260G>C), GCA531CCG (2261G>C 2263A>G), GTT532TTA (2264G>T 2266T>A), GAA533AAT (2267G>A 2269A>T), GAT535GAG (2275T>G), ATG536TTG (2276A>T), CAG537CAA (2281G>A), CAA538CAG (2284A>G), CCG539CCA (2287G>A), GTG540GCA (2289T>C 2290G>A), GTA542GTG (2296A>G), AGC544AGT (2302C>T), GAT546GAG (2308T>G), TTC547TTT (2311C>T), GGA551GGG (2323A>G), GTT552ATT (2324G>A), TTC555TTT (2335C>T), GTT556CTG (2336G>C 2338T>G), GAG558AAA (2342G>A 2344G>A), TCC560AGC (2348T>A 2349C>G), GCT561AAC (2351G>A 2352C>A 2353T>C), GAG565GAA (2365G>A), CAA566CAG (2368A>G), GTA567GTT (2371A>T), ATG568CTT (2372A>C 2374G>T), AGC569GCA (2375A>G 2376G>C 2377C>A), TTG570CTA (2378T>C 2380G>A), GCT571TCC (2381G>T 2383T>C), TTG572CTC (2384T>C 2386G>C), CCC573CCA (2389C>A), GCT574ATG (2390G>A 2391C>T 2392T>G), GCC575GCA (2395C>A), CTA576TTG (2396C>T 2398A>G), TTC577TTT (2401C>T), TCA578AGC (2402T>A 2403C>G 2404A>C), AAA579AAG (2407A>G), AGC580AAG (2409G>A 2410C>G), AAA581AAG (2413A>G), ACT583GCT (2417A>G), TCA584TCC (2422A>C), GGT586GGG (2428T>G), GCG587TCC (2429G>T 2431G>C), CTT590CTC (2440T>C), TAT592TAC (2446T>C), TAC593TAT (2449C>T), GAT594GAC (2452T>C), GCA595GCT (2455A>T), GCT596GCA (2458T>A), CTA597AAC (2459C>A 2460T>A 2461A>C), ATC598CTT (2462A>C 2464C>T), TAC600TTT (2469A>T 2470C>T), ATA602GTC (2474A>G 2476A>C), CTT604CTG (2482T>G), TAT605TAC (2485T>C), GTT606ATT (2486G>A), GTT609ATG (2495G>A 2497T>G), GCT611GCG (2503T>G), ATC612GTG (2504A>G 2506C>G), TCT613TCG (2509T>G), GGT614GGA (2512T>A), CAA615CAG (2515A>G), CTG616TTA (2516C>T 2518G>A), GCT617GCC (2521T>C), GAA622GAG (2536A>G), TGT623TGC (2539T>C), GTG625GCA (2544T>C 2545G>A), CGA628AGA (2552C>A), AAG629ACC (2556A>C 2557G>C), AAC634AAT (2572C>T), ATC635ATA (2575C>A), ACA637ACC (2581A>C), TTG638ATT (2582T>A 2584G>T), AAG642AAA (2596G>A), ACA644CAT (2600A>C 2601C>A 2602A>T), TTG645CTG (2603T>C), GTT646GTG (2608T>G), CAA650GAA (2618C>G), AAC652TCC (2624A>T 2625A>C), GAG654GGG (2631A>G), GAG655AAG (2633G>A), TGC657GTG (2639T>G 2640G>T 2641C>G), TTT658TTC (2644T>C), ACC659TCA (2645A>T 2647C>A), CCA660CCT (2650A>T), ACA661CAA (2651A>C 2652C>A), GGG662GGT (2656G>T), GGC664GGT (2662C>T), AAG665ACC (2664A>C 2665G>C), TAC666TTT (2667A>T 2668C>T), GTG667GTA (2671G>A), CCT668CCA (2674T>A), CTG669TTG (2675C>T), GAA671ACT (2681G>A 2682A>C 2683A>T), GGT672TCA (2684G>T 2685G>C 2686T>A), ACA673CTG (2687A>C 2688C>T 2689A>G), GGA674GGT (2692A>T), GCC675GTT (2694C>T 2695C>T), ACT676CCT (2696A>C), GAT677GAA (2701T>A), TTA678CTT (2702T>C 2704A>T), GGT679GGC (2707T>C), GTG681GTT (2713G>T), ACA686ACC (2728A>C), CAC687CAT (2731C>T), TTA689TTG (2737A>G), TCT690GCA (2738T>G 2740T>A), GCA692GCC (2746A>C), GAG694GAA (2752G>A), AAT696GAG (2756A>G 2758T>G), AGT697TCA (2759A>T 2760G>C 2761T>A), GTA698ATA (2762G>A), CCA699CCT (2767A>T), TGC700TGT (2770C>T), CAT701CAC (2773T>C), TTG702ATA (2774T>A 2776G>A), TTA704CTA (2780T>C), CAG705TCC (2783C>T 2784A>C 2785G>C), AAA707AAG (2791A>G), TTA709CTC (2795T>C 2797A>C), ACC711ACA (2803C>A), CTT714ATG (2810C>A 2812T>G), CGA718AGG (2822C>A 2824A>G), ATG719TTG (2825A>T), ATA720AGA (2829T>G), GCA721GGG (2832C>G 2833A>G), CAG722CAA (2836G>A), CAA724CAG (2842A>G), TAT725TTT (2844A>T), GGC726GGA (2848C>A), AAG728CGG (2852A>C 2853A>G), GCG729CCA (2855G>C 2857G>A), GGG730GGC (2860G>C), CAG731ACA (2861C>A 2862A>C 2863G>A), ACA732ACC (2866A>C), TAT733TTC (2868A>T 2869T>C), CCA735AGT (2873C>A 2874C>G 2875A>T), AGG736CGC (2876A>C 2878G>C), TTT737TTC (2881T>C), CCA738CCT (2884A>T), ACT739ACA (2887T>A), GTT742GTG (2896T>G), TTG743CTT (2897T>C 2899G>T), TTA744CTG (2900T>C 2902A>G), CAT745CAC (2905T>C), TAT746TAC (2908T>C), GGG749AGT (2915G>A 2917G>T), ACA750GTA (2918A>G 2919C>T), TCA751GCT (2921T>G 2923A>T), TCC752GCA (2924T>G 2926C>A), CCA753GCC (2927C>G 2929A>C), ACA757ACT (2941A>T), AGC760AGT (2950C>T), ATA761GTC (2951A>G 2953A>C), TTT762TTC (2956T>C), TCA763TCC (2959A>C), ATA767ATT (2971A>T), GAG769GAA (2977G>A), GGC772GGT (2986C>T), ACC773ATC (2988C>T), TTG774TTA (2992G>A), CCG776CCA (2998G>A), TCT777TCA (3001T>A), TTG778CTG (3002T>C), GGT780GGA (3010T>A), ATA782ATT (3016A>T), GCC783GCA (3019C>A), AGG784AGA (3022G>A), AAG787AAA (3031G>A), ACT790AGA (3039C>G 3040T>A), GGA791GGC (3043A>C), ACT792ACC (3046T>C), ATT797GTG (3059A>G 3061T>G), TGC798TGT (3064C>T), ATT799GTT (3065A>G), ACC802ACT (3076C>T), TTG803CTG (3077T>C), TCA806TCT (3088A>T), GGT807GGG (3091T>G), CTG809CTT (3097G>T), GCA810GCT (3100A>T), ATT811ATA (3103T>A), GGA812GGG (3106A>G), CTT813TTG (3107C>T 3109T>G), GGA814GGC (3112A>C), ACT815ACG (3115T>G), AAC817GCA (3119A>G 3120A>C 3121C>A), ACC818ACA (3124C>A), AAA819GAG (3125A>G 3127A>G), ACC821TCC (3131A>T), AAT822ACA (3135A>C 3136T>A), GCT823GCG (3139T>G), TTT827TTG (3151T>G), CCA830CCC (3160A>C), GTA833GTG (3169A>G), TGC834TGT (3172C>T), CTT836TTG (3176C>T 3178T>G), GGT839GGG (3187T>G), AAA841AGG (3192A>G 3193A>G), CGG843CAA (3198G>A 3199G>A), CGG845AAA (3203C>A 3204G>A 3205G>A), TGT846TGC (3208T>C), TCT847TCA (3211T>A), ATT848CTG (3212A>C 3214T>G), ACA849ACC (3217A>C), GGA853GGG (3229A>G), AAA854TGC (3230A>T 3231A>G 3232A>C), AAT855AAC (3235T>C), TTG856CTC (3236T>C 3238G>C), CTT857CTG (3241T>G), TCC858TCT (3244C>T), ACA859ACC (3247A>C), GGG860GGA (3250G>A), CGA861CGC (3253A>C), AAG862AAA (3256G>A), TCC864TCA (3262C>A), TTG865TTA (3265G>A), CAG869AAG (3275C>A), CAC870CAT (3280C>T), TTT871CAA (3281T>C 3282T>A 3283T>A), TCC872TCT (3286C>T), TTG874ATG (3290T>A), CGC875CGT (3295C>T), TTG876CTG (3296T>C), TTT877TTC (3301T>C), ACG879ACT (3307G>T), GTA880GTT (3310A>T), CTC884CTG (3322C>G), ACG887ACA (3331G>A), ATA889TTG (3335A>T 3337A>G), CAT890CAC (3340T>C), TCC893ACT (3347T>A 3349C>T), GGT895GGA (3355T>A), GTC896GTA (3358C>A), CAG899CAA (3367G>A), AAG901AAA (3373G>A), TGC902CGC (3374T>C), CTT903ATT (3377C>A), AAT905GAT (3383A>G), CTT906CTC (3388T>C), ACT907ACG (3391T>G), TTG908CTT (3392T>C 3394G>T), GGT909GGG (3397T>G), GGC910GGT (3400C>T), ACA911TCT (3401A>T 3403A>T), GTA912GTG (3406A>G), TCT913TCA (3409T>A), GTG914GTT (3412G>T), AAA915AAG (3415A>G), ATA917ATT (3421A>T), GGG919GGT (3427G>T), ACA922ACC (3436A>C), AAA923AAG (3439A>G), GGC924GGG (3442C>G), AAA925AGA (3444A>G), AGC926AGT (3448C>T), TCA927GCT (3449T>G 3451A>T), GTT928GTG (3454T>G),

| | | | | | | | | | |
|---|---|---|---|---|---|---|---|---|---|
| | TTC930TTT (3460C>T), CTT931CTG (3463T>G), TTC932TTT (3466C>T), GAT936AAC (3476G>A 3478T>C), TTA937CTG (3479T>C 3481A>G), TCA938AAT (3482T>A 3483C>A 3484A>T), CGT940CGC (3490T>C), ATT943CTT (3497A>C), GAA944GAG (3502A>G), AAG945AAA (3505G>A), TTG946CTG (3506T>C), AAG948AAA (3514G>A), GAC949GAA (3517C>A), AAC950AAT (3520C>T), ATT951GTT (3521A>G), GAG952GAT (3526G>T), GAG953ATC (3527G>A 3528A>T 3529G>C), TAC954TTT (3531A>T 3532C>T), CAG955GAC (3533C>G 3535G>C), GAG956CAG (3536G>C), GGC958GGG (3544C>G), GAT960AAC (3548G>A 3550T>C), CCG961_CCC962del (3551_3556delCCGCCC), AAG964ACA (3561A>C 3562G>A), GCT965TCT (3563G>T), AGT967GTT (3569A>G 3570G>T), ATT968TCC (3572A>T 3573T>C 3574T>C), CTG969TTC (3575C>T 3577G>C), TCC970CAG (3578T>C 3579C>A 3580C>G), ATA971ATC (3583A>C), AGA972AGG (3586A>G), GAG973GAA (3589G>A), AAA974AAG (3592A>G), TTT975TTC (3595T>C), TCC976ACA (3596T>A 3598C>A), GGT978GGA (3604T>A), GCT979GCA (3607T>A), GTA980GTG (3610A>G), TTC983TTT (3619C>T), TGC984TGT (3622C>T), ATG985GCT (3623A>G 3624T>C 3625G>T), GGT986GCA (3627G>C 3628T>A), GAC990ACA (3638G>A 3639A>C 3640C>A), GAA991GAC (3643A>C), TTG994GTA (3650T>G 3652G>A), GTA995GTT (3655A>T), ATT996TTG (3656A>T 3658T>G), CCT997CCA (3661T>A), GCA999GCC (3667A>C), CCA1000CCT (3670A>T), TCC1002TCA (3676C>A), ATA1003GTG (3677A>G 3679A>G), AGG1004CGC (3680A>C 3682G>C), GAA1006GAT (3688A>T), GGG1007GCA (3690G>C 3691G>A), CAC1008TCC (3692C>T 3693A>C), CCT1010CCA (3700T>A), GTT1011ATT (3701G>A), AAG1012AAA (3706G>A), ATC1015ATA (3715C>A), TGT1017TGC (3721T>C), TTT1019TTC (3727T>C), GAT1021GAC (3733T>C), ACA1024TCC (3740A>T 3742A>C), TCA1025TCT (3745A>T), TCA1030TCT (3760A>T), GGT1031GGA (3763T>A), AAT1034GAA (3770A>G 3772T>A), TCA1036ACA (3776T>A), ATT1037TTT (3779A>T), GTG1038GTA (3784G>A), ATA1039ATG (3787A>G), AGA1041CGA (3791A>C), GTA1042ACT (3794G>A 3795T>C 3796A>T), CAA1043CAG (3799A>G), TCC1044AGC (3800T>A 3801C>G), AGT1045TCC (3803A>T 3804G>C 3805T>C), CCT1046AAC (3806C>A 3807C>A 3808T>C), AAT1047AGT (3810A>G), GTT1048ATA (3812G>A 3814T>A), GGA1049GGT (3817A>T), GTA1051ATC (3821G>A 3823A>C), CTA1052TTG (3824C>T 3826A>G), AAT1053AAC (3829T>C), GTT1054ATT (3830G>A), GCT1055GCA (3835T>A), TTT1056CTT (3836T>C), GAT1057GAC (3841T>C), GCC1058ACA (3842G>A 3844C>A), GGC1060GGG (3850C>G), TTT1063TTC (3859T>C), CCA1064CAA (3861C>A), GCT1065TTT (3863G>T 3864C>T), GGG1066GGT (3868G>T), CTT1067GTA (3869C>G 3871T>A), AAT1068TCT (3872A>T 3873A>C), AAA1069GCG (3875A>G 3876A>C 3877A>G), AAT1071GCA (3881A>G 3882A>C 3883T>A), TAT1072CAT (3884T>C), GTA1074TTA (3890G>T), GGT1077GGC (3901T>C), GGA1078GGT (3904A>T), GGC1079GGA (3907C>A), ACA1080ACT (3910A>T), TCA1085TCC (3925A>C), TAC1086TAT (3928C>T), GGT1087GGG (3931T>G), GTG1088GTT (3934G>T), GCA1089ACC (3935G>A 3937A>C), ACA1090ACC (3940A>C), AAT1091AAC (3943T>C), ACG1092ATC (3945C>T 3946G>C), TCA1094TCT (3952A>T), TTC1095TTT (3955C>T), ACT1096GTA (3956A>G 3957C>T 3958T>A), GAG1101CAG (3971G>C), TTT1102TTC (3976T>C), TTC1103TTT (3979C>T), CCA1104CCT (3982A>T), AGG1105CGC (3983A>C 3985G>C), CGG1106CGT (3988G>T), CAT1107TTC (3989C>T 3990A>T 3991T>C), AGG1109AAG (3996G>A), ATG1110GCC (3998A>G 3999T>C 4000G>C), AGG1111AAA (4002G>A 4003G>A), GAA1112TCC (4004G>T 4005A>C 4006A>C), TTC1113TAT (4008T>A 4009C>T), TCA1114GAT (4010T>G 4011C>A 4012A>T), AGC1115AAG (4014G>A 4015C>G), AAG1116GAA (4016A>G 4018G>A), CAA1117AGT (4019C>A 4020A>G 4021A>T), TCC1118TCA (4024C>A), CGC1119AGA (4025C>A 4027C>A), ATC1120ATT (4030C>T), ATG1121GTT (4031A>G 4033G>T), TCA1122TCT (4036A>T), CTA1123TTG (4037C>T 4039A>G), GAT1125GAC (4045T>C), AGG1126AAG (4047G>A), CTT1127CTA (4051T>A), GGA1128GGC (4054A>C), AAT1129TTG (4055A>T 4056A>T 4057T>G), CTG1130TTG (4058C>T), ATC1131CTA (4061A>C 4063C>A), ATA1132ATC (4066A>C), TTG1134CTT (4070T>C 4072G>T), CCT1135CCA (4075T>A), CCT1136CCA (4078T>A), TCC1137GTT (4079T>G 4080C>T 4081C>T), GCC1138GAA (4083C>A 4084C>A), ATA1139CTT (4085A>C 4087A>T), GTG1140ATT (4088G>A 4090G>T), TCC1142TCA (4096C>A), GAG1144GAA (4102G>A), ATA1145ATC (4105A>C), CTT1146ATA (4106C>A 4108T>A), ATA1147GTC (4109A>G 4111A>C), TCT1148AAG (4112T>A 4113C>A 4114T>G), CCT1149CCA (4117T>A), GGA1150GGT (4120A>T), CTT1151CCT (4122T>C), TTC1153TTT (4129C>T), AAG1154AAA (4132G>A), GCC1158GCT (4144C>T), CCT1160CCC (4150T>C), CCT1161CCA (4153T>A), TCT1162CTT (4154T>C 4155C>T), GCC1163GCA (4159C>A), CAT1165AAT (4163C>A), GAA1166GAG (4168A>G), AAA1167AAG (4171A>G), CTT1169CTC (4177T>C), GGC1170GGA (4180C>A), AAT1171GAT (4181A>G), CAA1173CAG (4189A>G), ACT1174ACG (4192T>G), CAC1175CAT (4195C>T), ACC1176ACT (4198C>T), ACC1180ACA (4210C>A), TCA1181TCT (4213A>T), CTA1186CTG (4228A>G), ATT1191AAT (4242T>A) | | | | | | | | |
| Coat protein A (YP_001976155.1) | 1 | 254 | 100% | 1189 | 70.1% | 254 (99.2%) | 165 (64.5%) | 2/0/0/0 | 0 |
| Protein mutations: | T2A (2078A>G), A3Q (2081G>C 2082C>A), R4T (2085G>C 2086G>T), I6N (2091T>A 2092A>T), A7_E8insS (2095_2096insAGC), E8K (2096G>A 2098A>G), S9W (2099A>T 2101T>G), P11T (2105C>A), D13G (2112A>G 2113C>A), R14K (2115G>A 2116A>G), T16L (2120A>C 2121C>T 2122T>G), S21T (2135T>A), T24A (2144A>G 2146G>A), M25L (2147A>C 2149G>T), E26_D27insS (2152_2153insAGT), D27E (2155T>G), P28G (2156C>G 2157C>G 2158C>T), T29I (2160C>T), K30Q (2162A>C 2164G>A), P31R (2165C>A 2166C>G), D32K (2168G>A 2170T>G), K33E (2171A>G), I34V (2174A>G 2176A>G), E35S (2177G>T 2178A>C), V37E (2184T>A 2185T>G), G40D (2193G>A 2194A>T), A41P (2195G>C 2197A>G), E43G (2202A>G 2203A>G), E44K (2204G>A), Q46E (2210C>G 2212A>G), G48N (2216G>A 2217G>A), V50I (2222G>A), I51F (2225A>T 2227T>C), P56E (2240C>G 2241C>A 2242A>G), D59M (2249G>A 2250A>T 2251C>G), T60G (2252A>G 2253C>G 2254A>T), S61D (2255T>G 2256C>A 2257C>T), M62A (2258A>G 2259T>C 2260G>C), A63P (2261G>C 2263A>G), V64L (2264G>T 2266T>A), E65N (2267G>A 2269A>T), D67E (2275T>G), M68L (2276A>T), V72A (2289T>C 2290G>A), D78E (2308T>G), V84I (2324G>A), V88L (2336G>C 2338T>G), E90K (2342G>A 2344G>A), A93N (2351G>A 2352C>A 2353T>C), M100L (2372A>C 2374G>T), S101A (2375A>G 2376G>C 2377C>A), A103S (2381G>T 2383T>C), A106M (2390G>A 2391C>T 2392T>G), S112K (2409G>A 2410C>G), T115A (2417A>G), A119S (2429G>T 2431G>C), L129N (2459C>A 2460T>A 2461A>C), I130L (2462A>C 2464C>T), Y132F (2469A>T 2470C>T), I134V (2474A>G 2476A>C), V138I (2486G>A), V141M (2495G>A 2497T>G), I144V (2504A>G 2506C>G), V157A (2544T>C 2545G>A), K161T (2556A>C 2557G>C), L170I (2582T>A 2584G>T), T176H (2600A>C 2601C>A 2602A>T), Q182E (2618C>G), N184S (2624A>T 2625A>C), E186G (2631A>G), E187K (2633G>A), C189V (2639T>G 2640G>T 2641C>G), T191S (2645A>T 2647C>A), T193Q (2651A>C 2652C>A), K197T (2664A>C 2665G>C), Y198F (2667A>T 2668C>T), E203T (2681G>A 2682A>C 2683A>T), G204S (2684G>T 2685G>C 2686T>A), T205L (2687A>C 2688C>T 2689A>G), A207V (2694C>T 2695C>T), T208P (2696A>C), D209E (2701T>A), S222A (2738T>G 2740T>A), N228E (2756A>G 2758T>G), V230I (2762G>A), L234I (2774T>A 2776G>A), Q237S (2783C>T 2784A>C 2785G>C), L246M (2810C>A 2812T>G), M251L (2825A>T), I252R (2829T>G), A253G (2832C>G 2833A>G) | | | | | | | | |
| Codon mutations: | ACA2GCA (2078A>G), GCA3CAA (2081G>C 2082C>A), AGG4ACT (2085G>C 2086G>T), GGC5GGA (2089C>A), ATA6AAT (2091T>A 2092A>T), GCT7GCC (2095T>C), GCT7_GAA8insAGC (2095_2096insAGC), GAA8AAG (2096G>A 2098A>G), AGT9TGG (2099A>T 2101T>G), CAG10CAA (2104G>A), CCT11ACT (2105C>A), GAC13GGA (2112A>G 2113C>A), AGA14AAG (2115G>A 2116A>G), ACT16CTG (2120A>C 2121C>T 2122T>G), TCC18TCA (2128C>A), GTG20GTT (2134G>T), TCA21ACA (2135T>A), ACT22ACC (2140T>C), ACC23ACA (2143C>A), ACG24GCA (2144A>G 2146G>A), ATG25CTT (2147A>C 2149G>T), GAA26GAG (2152A>G), GAA26_GAT27insAGT (2152_2153insAGT), GAT27GAG (2155T>G), CCC28GGT (2156C>G 2157C>G 2158C>T), ACC29ATC (2160C>T), AAG30CAA (2162A>C 2164G>A), CCA31AGA (2165C>A 2166C>G), GAT32AAG (2168G>A 2170T>G), AAG33GAG (2171A>G), ATA34GTG (2174A>G 2176A>G), GAA35TCA (2177G>T 2178A>C), GTT37GAG (2184T>A 2185T>G), GCA38GCT (2188A>T), GAG39GAA (2191G>A), GGA40GAT (2193G>A 2194A>T), GCA41CCG (2195G>C 2197A>G), GAA43GGG (2202A>G 2203A>G), GAA44AAA (2204G>A), ACT45ACA (2209T>A), CAA46GAG (2210C>G 2212A>G), GGT48AAT (2216G>A 2217G>A), GAT49GAC (2221T>C), GTT50ATT (2222G>A), ATT51TTC (2225A>T 2227T>C), GAC53GAT (2233C>T), GGA55GGG (2239A>G), CCA56GAG (2240C>G 2241C>A 2242A>G), GAC59ATG (2249G>A 2250A>T 2251C>G), ACA60GGT (2252A>G 2253C>G 2254A>T), TCC61GAT (2255T>G 2256C>A 2257C>T), ATG62GCC (2258A>G 2259T>C 2260G>C), GCA63CCG (2261G>C 2263A>G), GTT64TTA (2264G>T 2266T>A), GAA65AAT (2267G>A 2269A>T), GAT67GAG (2275T>G), ATG68TTG (2276A>T), CAG69CAA (2281G>A), CAA70CAG (2284A>G), CCG71CCA (2287G>A), GTG72GCA (2289T>C 2290G>A), GTA74GTG (2296A>G), AGC76AGT (2302C>T), GAT78GAG (2308T>G), TTC79TTT (2311C>T), GGA83GGG (2323A>G), GTT84ATT (2324G>A), TTC87TTT (2335C>T), GTT88CTG (2336G>C 2338T>G), GAG90AAA (2342G>A 2344G>A), TCC92AGC (2348T>A 2349C>G), GCT93AAC (2351G>A 2352C>A 2353T>C), GAG97GAA (2365G>A), CAA98CAG (2368A>G), GTA99GTT (2371A>T), ATG100CTT (2372A>C 2374G>T), AGC101GCA (2375A>G 2376G>C 2377C>A), TTG102CTA (2378T>C 2380G>A), GCT103TCC (2381G>T 2383T>C), TTG104CTC (2384T>C 2386G>C), CCC105CCA (2389C>A), GCT106ATG (2390G>A 2391C>T 2392T>G), GCC107GCA (2395C>A), CTA108TTG (2396C>T 2398A>G), TTC109TTT (2401C>T), TCA110AGC (2402T>A 2403C>G 2404A>C), AAA111AAG (2407A>G), AGC112AAG (2409G>A 2410C>G), AAA113AAG (2413A>G), ACT115GCT (2417A>G), TCA116TCC (2422A>C), GGT118GGG (2428T>G), GCG119TCC (2429G>T 2431G>C), CTT122CTC (2440T>C), TAT124TAC (2446T>C), TAC125TAT (2449C>T), GAT126GAC (2452T>C), GCA127GCT (2455A>T), GCT128GCA (2458T>A), CTA129AAC (2459C>A 2460T>A 2461A>C), ATC130CTT (2462A>C 2464C>T), TAC132TTT (2469A>T 2470C>T), ATA134GTC (2474A>G 2476A>C), CTT136CTG (2482T>G), TAT137TAC (2485T>C), GTT138ATT (2486G>A), GTT141ATG (2495G>A 2497T>G), GCT143GCG (2503T>G), ATC144GTG (2504A>G 2506C>G), TCT145TCG (2509T>G), GGT146GGA (2512T>A), CAA147CAG (2515A>G), CTG148TTA (2516C>T 2518G>A), GCT149GCC (2521T>C), GAA154GAG (2536A>G), TGT155TGC (2539T>C), GTG157GCA (2544T>C 2545G>A), CGA160AGA (2552C>A), AAG161ACC (2556A>C 2557G>C), AAC166AAT (2572C>T), ATC167ATA (2575C>A), ACA169ACC (2581A>C), TTG170ATT (2582T>A 2584G>T), AAG174AAA (2596G>A), ACA176CAT (2600A>C 2601C>A 2602A>T), TTG177CTG (2603T>C), GTT178GTG (2608T>G), CAA182GAA (2618C>G), AAC184TCC (2624A>T 2625A>C), GAG186GGG (2631A>G), GAG187AAG (2633G>A), TGC189GTG (2639T>G 2640G>T 2641C>G), TTT190TTC (2644T>C), ACC191TCA (2645A>T 2647C>A), CCA192CCT (2650A>T), ACA193CAA (2651A>C 2652C>A), GGG194GGT (2656G>T), GGC196GGT (2662C>T), AAG197ACC (2664A>C 2665G>C), TAC198TTT (2667A>T 2668C>T), GTG199GTA (2671G>A), CCT200CCA (2674T>A), CTG201TTG (2675C>T), GAA203ACT (2681G>A 2682A>C 2683A>T), GGT204TCA (2684G>T 2685G>C 2686T>A), ACA205CTG (2687A>C 2688C>T 2689A>G), GGA206GGT (2692A>T), GCC207GTT (2694C>T 2695C>T), ACT208CCT (2696A>C), GAT209GAA (2701T>A), TTA210CTT (2702T>C 2704A>T), GGT211GGC (2707T>C), GTG213GTT (2713G>T), ACA218ACC (2728A>C), CAC219CAT (2731C>T), TTA221TTG (2737A>G), TCT222GCA (2738T>G 2740T>A), GCA224GCC (2746A>C), GAG226GAA (2752G>A), AAT228GAG (2756A>G 2758T>G), AGT229TCA (2759A>T 2760G>C 2761T>A), GTA230ATA (2762G>A), CCA231CCT (2767A>T), TGC232TGT (2770C>T), CAT233CAC (2773T>C), TTG234ATA (2774T>A 2776G>A), TTA236CTA (2780T>C), CAG237TCC (2783C>T 2784A>C 2785G>C), AAA239AAG (2791A>G), TTA241CTC (2795T>C 2797A>C), ACC243ACA (2803C>A), CTT246ATG (2810C>A 2812T>G), CGA250AGG (2822C>A 2824A>G), ATG251TTG (2825A>T), ATA252AGA (2829T>G), GCA253GGG (2832C>G 2833A>G), CAG254CAA (2836G>A) | | | | | | | | |
| Coat protein B (YP_001976156.1) | 1 | 244 | 100% | 1422 | 83.4% | 242 (99.2%) | 192 (78.7%) | 0/2/0/0 | 0 |
| Protein mutations: | Y3F (2844A>T), K6R (2852A>C 2853A>G), A7P (2855G>C 2857G>A), Q9T (2861C>A 2862A>C 2863G>A), Y11F (2868A>T 2869T>C), P13S (2873C>A 2874C>G 2875A>T), G27S (2915G>A 2917G>T), T28V (2918A>G 2919C>T), S29A (2921T>G 2923A>T), S30A (2924T>G 2926C>A), P31A (2927C>G 2929A>C), I39V (2951A>G 2953A>C), T51I (2988C>T), T68R (3039C>G 3040T>A), I75V (3059A>G 3061T>G), I77V (3065A>G), N95A (3119A>G 3120A>C 3121C>A), K97E (3125A>G 3127A>G), T99S (3131A>T), N100T (3135A>C 3136T>A), F105L (3151T>G), K119R (3192A>G 3193A>G), R121Q (3198G>A 3199G>A), R123K (3203C>A 3204G>A 3205G>A), I126L (3212A>C 3214T>G), K132C (3230A>T 3231A>G 3232A>C), Q147K (3275C>A), F149Q (3281T>C 3282T>A 3283T>A), L152M (3290T>A), I167L (3335A>T 3337A>G), S171T (3347T>A 3349C>T), C180R (3374T>C), L181I (3377C>A), N183D (3383A>G), T189S (3401A>T 3403A>T), K203R (3444A>G), S205A (3449T>G 3451A>T), D214N (3476G>A 3478T>C), S216N (3482T>A 3483C>A 3484A>T), I221L (3497A>C), D227E (3517C>A), I229V (3521A>G), E230D (3526G>T), E231I (3527G>A 3528A>T 3529G>C), Y232F (3531A>T 3532C>T), Q233D (3533C>G 3535G>C), E234Q (3536G>C), D238N (3548G>A 3550T>C), P239_P240del (3551_3556delCCGCCC), K242T (3561A>C 3562G>A), A243S (3563G>T) | | | | | | | | |

| | Begin | End | Coverage | Score | Concordance | Matches | Identities | I/D/M/F* | Stop Codons |
|---|---|---|---|---|---|---|---|---|---|
| **NT** | **89** | **4733** | **94.7%** | **2570** | **28.4%** | **4597 (96.7%)** | **3027 (63.7%)** | **107/48** | |
| Codon mutations: | CAA2CAG (2842A>G), TAT3TTT (2844A>T), GGC4GGA (2848C>A), AAG6CGG (2852A>C 2853A>G), GCG7CCA (2855G>C 2857G>A), GGG8GGC (2860G>C), CAG9ACA (2861C>A 2862A>C 2863G>A), ACA10ACC (2866A>C), TAT11TTC (2868A>T 2869T>C), CCA13AGT (2873C>A 2874C>G 2875A>T), AGG14CGC (2876A>C 2878G>C), TTT15TTC (2881T>C), CCA16CCT (2884A>T), ACT17ACA (2887T>A), GTT20GTG (2896T>G), TTG21CTT (2897T>C 2899G>T), TTA22CTG (2900T>C 2902A>G), CAT23CAC (2905T>C), TAT24TAC (2908T>C), GGG27AGT (2915G>A 2917G>T), ACA28GTA (2918A>G 2919C>T), TCA29GCT (2921T>G 2923A>T), TCC30GCA (2924T>G 2926C>A), CCA31GCC (2927C>G 2929A>C), ACA35ACT (2941A>T), AGC38AGT (2950C>T), ATA39GTC (2951A>G 2953A>C), TTT40TTC (2956T>C), TCA41TCC (2959A>C), ATA45ATT (2971A>T), GAG47GAA (2977G>A), GGC50GGT (2986C>T), ACC51ATC (2988C>T), TTG52TTA (2992G>A), CCG54CCA (2998G>A), TCT55TCA (3001T>A), TTG56CTG (3002T>C), GGT58GGA (3010T>A), ATA60ATT (3016A>T), GCC61GCA (3019C>A), AGG62AGA (3022G>A), AAG65AAA (3031G>A), ACT68AGA (3039C>G 3040T>A), GGA69GGC (3043A>C), ACT70ACC (3046T>C), ATT75GTG (3059A>G 3061T>G), TGC76TGT (3064C>T), ATT77GTT (3065A>G), ACC80ACT (3076C>T), TTG81CTG (3077T>C), TCA84TCT (3088A>T), GGT85GGG (3091T>G), CTG87CTT (3097G>T), GCA88GCT (3100A>T), ATT89ATA (3103T>A), GGA90GGG (3106A>G), CTT91TTG (3107C>T 3109T>G), GGA92GGC (3112A>C), ACT93ACG (3115T>G), AAC95GCA (3119A>G 3120A>C 3121C>A), ACC96ACA (3124C>A), AAA97GAG (3125A>G 3127A>G), ACC99TCC (3131A>T), AAT100ACA (3135A>C 3136T>A), GCT101GCG (3139T>G), TTT105TTG (3151T>G), CCA108CCC (3160A>C), GTA111GTG (3169A>G), TGC112TGT (3172C>T), CTT114TTG (3176C>T 3178T>G), GGT117GGG (3187T>G), AAA119AGG (3192A>G 3193A>G), CGG121CAA (3198G>A 3199G>A), CGG123AAA (3203C>A 3204G>A 3205G>A), TGT124TGC (3208T>C), TCT125TCA (3211T>A), ATT126CTG (3212A>C 3214T>G), ACA127ACC (3217A>C), GGA131GGG (3229A>G), AAA132TGC (3230A>T 3231A>G 3232A>C), AAT133AAC (3235T>C), TTG134CTC (3236T>C 3238G>C), CTT135CTG (3241T>G), TCC136TCT (3244C>T), ACA137ACC (3247A>C), GGG138GGA (3250G>A), CGA139CGC (3253A>C), AAG140AAA (3256G>A), TCC142TCA (3262C>A), TTG143TTA (3265G>A), CAG147AAG (3275C>A), CAC148CAT (3280C>T), TTT149CAA (3281T>C 3282T>A 3283T>A), TCC150TCT (3286C>T), TTG152ATG (3290T>A), CGC153CGT (3295C>T), TTG154CTG (3296T>C), TTT155TTC (3301T>C), ACG157ACT (3307G>T), GTA158GTT (3310A>T), CTC162CTG (3322C>G), ACG165ACA (3331G>A), ATA167TTG (3335A>T 3337A>G), CAT168CAC (3340T>C), TCC171ACT (3347T>A 3349C>T), GGT173GGA (3355T>A), GTC174GTA (3358C>A), CAG177CAA (3367G>A), AAG179AAA (3373G>A), TGC180CGC (3374T>C), CTT181ATT (3377C>A), AAT183GAT (3383A>G), CTT184CTC (3388T>C), ACT185ACG (3391T>G), TTG186CTT (3392T>C 3394G>T), GGT187GGG (3397T>G), GGC188GGT (3400C>T), ACA189TCT (3401A>T 3403A>T), GTA190GTG (3406A>G), TCT191TCA (3409T>A), GTG192GTT (3412G>T), AAA193AAG (3415A>G), ATA195ATT (3421A>T), GGG197GGT (3427G>T), ACA200ACC (3436A>C), AAA201AAG (3439A>G), GGC202GGG (3442C>G), AAA203AGA (3444A>G), AGC204AGT (3448C>T), TCA205GCT (3449T>G 3451A>T), GTT206GTG (3454T>G), TTC208TTT (3460C>T), CTT209CTG (3463T>G), TTC210TTT (3466C>T), GAT214AAC (3476G>A 3478T>C), TTA215CTG (3479T>C 3481A>G), TCA216AAT (3482T>A 3483C>A 3484A>T), CGT218CGC (3490T>C), ATT221CTT (3497A>C), GAA222GAG (3502A>G), AAG223AAA (3505G>A), TTG224CTG (3506T>C), AAG226AAA (3514G>A), GAC227GAA (3517C>A), AAC228AAT (3520C>T), ATT229GTT (3521A>G), GAG230GAT (3526G>T), GAG231ATC (3527G>A 3528A>T 3529G>C), TAC232TTT (3531A>T 3532C>T), CAG233GAC (3533C>G 3535G>C), GAG234CAG (3536G>C), GGC236GGG (3544C>G), GAT238AAC (3548G>A 3550T>C), CCG239_CCC240del (3551_3556delCCGCCC), AAG242ACA (3561A>C 3562G>A), GCT243TCT (3563G>T) | | | | | | | | |
| Coat protein C (YP_001976157.1) | 1 | 225 | 100% | 1271 | 77.0% | 225 (100%) | 158 (70.2%) | 0/0/0/0 | 0 |
| Protein mutations: | S1V (3569A>G 3570G>T), I2S (3572A>T 3573T>C 3574T>C), L3F (3575C>T 3577G>C), S4Q (3578T>C 3579C>A 3580C>G), S10T (3596T>A 3598C>A), M19A (3623A>G 3624T>C 3625G>T), G20A (3627G>C 3628T>A), D24T (3638G>A 3639A>C 3640C>A), E25D (3643A>C), L28V (3650T>G 3652G>A), I30L (3656A>T 3658T>G), I37V (3677A>G 3679A>G), E40D (3688A>T), G41A (3690G>C 3691G>A), H42S (3692C>T 3693A>C), V45I (3701G>A), T58S (3740A>T 3742A>C), N68E (3770A>G 3772T>A), S70T (3776T>A), I71F (3779A>T), I73M (3787A>G), V76T (3794G>A 3795T>C 3796A>T), P80N (3806C>A 3807C>A 3808T>C), N81S (3810A>G), V82I (3812G>A 3814T>A), V85I (3821G>A 3823A>C), V88I (3830G>A), F90L (3836T>C), A92T (3842G>A 3844C>A), P98Q (3861C>A), A99F (3863G>T 3864C>T), L101V (3869C>G 3871T>A), N102S (3872A>T 3873A>C), K103A (3875A>G 3876A>C 3877A>G), N105A (3881A>G 3882A>C 3883T>A), Y106H (3884T>C), V108L (3890G>T), A123T (3935G>A 3937A>C), T126I (3945C>T 3946G>C), T130V (3956A>G 3957C>T 3958T>A), E135Q (3971G>C), H141F (3989C>T 3990A>T 3991T>C), R143K (3996G>A), M144A (3998A>G 3999T>C 4000G>C), R145K (4002G>A 4003G>A), E146S (4004G>T 4005A>C 4006A>C), F147Y (4008T>A 4009C>T), S148D (4010T>G 4011C>A 4012A>T), S149K (4014G>A 4015C>G), K150E (4016A>G 4018G>A), Q151S (4019C>A 4020A>G 4021A>T), M155V (4031A>G 4033G>T), R160K (4047G>A), N163L (4055A>T 4056A>T 4057T>G), I165L (4061A>C 4063C>A), S171V (4079T>G 4080C>T 4081C>T), A172E (4083C>A 4084C>A), I173L (4085A>C 4087A>T), V174I (4088G>A 4090G>T), L180I (4106C>A 4108T>A), I181V (4109A>G 4111A>C), S182K (4112T>A 4113C>A 4114T>G), L185P (4122T>C), S196L (4154T>C 4155C>T), H199N (4163C>A), N205D (4181A>G), I225N (4242T>A) | | | | | | | | |
| Codon mutations: | AGT1GTT (3569A>G 3570G>T), ATT2TCC (3572A>T 3573T>C 3574T>C), CTG3TTC (3575C>T 3577G>C), TCC4CAG (3578T>C 3579C>A 3580C>G), ATA5ATC (3583A>C), AGA6AGG (3586A>G), GAG7GAA (3589G>A), AAA8AAG (3592A>G), TTT9TTC (3595T>C), TCC10ACA (3596T>A 3598C>A), GGT12GGA (3604T>A), GCT13GCA (3607T>A), GTA14GTG (3610A>G), TTC17TTT (3619C>T), TGC18TGT (3622C>T), ATG19GCT (3623A>G 3624T>C 3625G>T), GGT20GCA (3627G>C 3628T>A), GAC24ACA (3638G>A 3639A>C 3640C>A), GAA25GAC (3643A>C), TTG28GTA (3650T>G 3652G>A), GTA29GTT (3655A>T), ATT30TTG (3656A>T 3658T>G), CCT31CCA (3661T>A), GCA33GCC (3667A>C), CCA34CCT (3670A>T), TCC36TCA (3676C>A), ATA37GTG (3677A>G 3679A>G), AGG38CGC (3680A>C 3682G>C), GAA40GAT (3688A>T), GGG41GCA (3690G>C 3691G>A), CAC42TCC (3692C>T 3693A>C), CCT44CCA (3700T>A), GTT45ATT (3701G>A), AAG46AAA (3706G>A), ATC49ATA (3715C>A), TGT51TGC (3721T>C), TTT53TTC (3727T>C), GAT55GAC (3733T>C), ACA58TCC (3740A>T 3742A>C), TCA59TCT (3745A>T), TCA64TCT (3760A>T), GGT65GGA (3763T>A), AAT68GAA (3770A>G 3772T>A), TCA70ACA (3776T>A), ATT71TTT (3779A>T), GTG72GTA (3784G>A), ATA73ATG (3787A>G), AGA75CGA (3791A>C), GTA76ACT (3794G>A 3795T>C 3796A>T), CAA77CAG (3799A>G), TCC78AGC (3800T>A 3801C>G), AGT79TCC (3803A>T 3804G>C 3805T>C), CCT80AAC (3806C>A 3807C>A 3808T>C), AAT81AGT (3810A>G), GTT82ATA (3812G>A 3814T>A), GGA83GGT (3817A>T), GTA85ATC (3821G>A 3823A>C), CTA86TTG (3824C>T 3826A>G), AAT87AAC (3829T>C), GTT88ATT (3830G>A), GCT89GCA (3835T>A), TTT90CTT (3836T>C), GAT91GAC (3841T>C), GCC92ACA (3842G>A 3844C>A), GGC94GGG (3850C>G), TTT97TTC (3859T>C), CCA98CAA (3861C>A), GCT99TTT (3863G>T 3864C>T), GGG100GGT (3868G>T), CTT101GTA (3869C>G 3871T>A), AAT102TCT (3872A>T 3873A>C), AAA103GCG (3875A>G 3876A>C 3877A>G), AAT105GCA (3881A>G 3882A>C 3883T>A), TAT106CAT (3884T>C), GTA108TTA (3890G>T), GGT111GGC (3901T>C), GGA112GGT (3904A>T), GGC113GGA (3907C>A), ACA114ACT (3910A>T), TCA119TCC (3925A>C), TAC120TAT (3928C>T), GGT121GGG (3931T>G), GTG122GTT (3934G>T), GCA123ACC (3935G>A 3937A>C), ACA124ACC (3940A>C), AAT125AAC (3943T>C), ACG126ATC (3945C>T 3946G>C), TCA128TCT (3952A>T), TTC129TTT (3955C>T), ACT130GTA (3956A>G 3957C>T 3958T>A), GAG135CAG (3971G>C), TTT136TTC (3976T>C), TTC137TTT (3979C>T), CCA138CCT (3982A>T), AGG139CGC (3983A>C 3985G>C), CGG140CGT (3988G>T), CAT141TTC (3989C>T 3990A>T 3991T>C), AGG143AAG (3996G>A), ATG144GCC (3998A>G 3999T>C 4000G>C), AGG145AAA (4002G>A 4003G>A), GAA146TCC (4004G>T 4005A>C 4006A>C), TTC147TAT (4008T>A 4009C>T), TCA148GAT (4010T>G 4011C>A 4012A>T), AGC149AAG (4014G>A 4015C>G), AAG150GAA (4016A>G 4018G>A), CAA151AGT (4019C>A 4020A>G 4021A>T), TCC152TCA (4024C>A), CGC153AGA (4025C>A 4027C>A), ATC154ATT (4030C>T), ATG155GTT (4031A>G 4033G>T), TCA156TCT (4036A>T), CTA157TTG (4037C>T 4039A>G), GAT159GAC (4045T>C), AGG160AAG (4047G>A), CTT161CTA (4051T>A), GGA162GGC (4054A>C), AAT163TTG (4055A>T 4056A>T 4057T>G), CTG164TTG (4058C>T), ATC165CTA (4061A>C 4063C>A), ATA166ATC (4066A>C), TTG168CTT (4070T>C 4072G>T), CCT169CCA (4075T>A), CCT170CCA (4078T>A), TCC171GTT (4079T>G 4080C>T 4081C>T), GCC172GAA (4083C>A 4084C>A), ATA173CTT (4085A>C 4087A>T), GTG174ATT (4088G>A 4090G>T), TCC176TCA (4096C>A), GAG178GAA (4102G>A), ATA179ATC (4105A>C), CTT180ATA (4106C>A 4108T>A), ATA181GTC (4109A>G 4111A>C), TCT182AAG (4112T>A 4113C>A 4114T>G), CCT183CCA (4117T>A), GGA184GGT (4120A>T), CTT185CCT (4122T>C), TTC187TTT (4129C>T), AAG188AAA (4132G>A), GCC192GCT (4144C>T), CCT194CCC (4150T>C), CCT195CCA (4153T>A), TCT196CTT (4154T>C 4155C>T), GCC197GCA (4159C>A), CAT199AAT (4163C>A), GAA200GAG (4168A>G), AAA201AAG (4171A>G), CTT203CTC (4177T>C), GGC204GGA (4180C>A), AAT205GAT (4181A>G), CAA207CAG (4189A>G), ACT208ACG (4192T>G), CAC209CAT (4195C>T), ACC210ACT (4198C>T), ACC214ACA (4210C>A), TCA215TCT (4213A>T), CTA220CTG (4228A>G), ATT225AAT (4242T>A) | | | | | | | | |

*: Inserts / Deletes / Misaligned / Frameshifts

#### Analysis details

This analysis was performed with panviral2.64

### NGS Details (UN18_val): Tomato necrotic dwarf virus (segment RNA1)

#### Assembly

| | |
|---|---|
| **Coverage Length** | 6096 (1 contig(s)) |
| **Depth Of Coverage** | 468.2 |
| **Number Of Reads** | 21581 |
| **Reads Per Million** | 405.04 rpm (after QC) |
| **Ambiguities** | 0 |
| **Assembly Method** | de novo + reference guided assembly |
| **Consensus Caller** | Bcf Tools |

#### Coverage Map

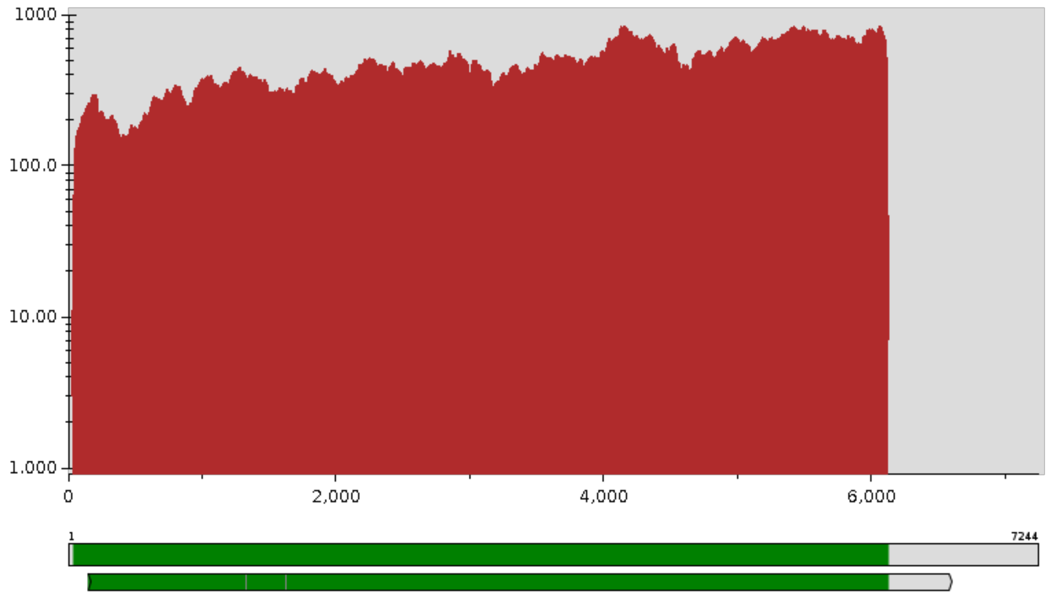

#### Assignment

| | |
|---|---|
| **Type** | Tomato necrotic dwarf virus (Taxonomy ID: 1481465) |
| **Reference Genome** | NC_027926.1 |
| **NT Identity (%)** | 64.5613 |
| **AA Identity (%)** | 66.0633 |
| **Number Of Stop Codons** | 0 |
| **Number Of CDS** | 1 |

#### Alignment

| | |
|---|---|
| **Alignment Score** | 3374.0 (NT) + 9636.0 (AA) = 13010.0 |
| **Concordance (%)** | 50.6383 |

| | |
|---|---|
| **Alignment Method** | Global, seeded, nucleotide + amino acids (AGA) |

#### Genome Region

Sequence starts at position 33 and ends at position 6128 relative to NC_027926.1 reference sequence.

#### Alignment Detailed Statistics

| | Begin | End | Coverage | Score | Concordance | Matches | Identities | I/D/M/F* | Stop Codons |
|---|---|---|---|---|---|---|---|---|---|
| **NT** | **33** | **6128** | **84.2%** | **3374** | **28.2%** | **6043 (98.8%)** | **3915 (64.0%)** | **21/53** | |

| | Begin | End | Coverage | Score | Concordance | Matches | Identities | I/D/M/F* | Stop Codons |
|---|---|---|---|---|---|---|---|---|---|
| **NT** | **33** | **6128** | **84.2%** | **3374** | **28.2%** | **6043 (98.8%)** | **3915 (64.0%)** | **21/53** | |

Mutations:

35T>C, 40G>A, 42C>T, 43C>T, 49T>C, 50C>T, 52A>C, 54T>C, 62A>C, 64T>C, 65T>A, 66G>T, 67G>A, 69A>C, 70T>A, 71T>A, 74A>C, 77G>T, 79T>C, 82A>T, 84G>T,
85_93delCTAAACTTC, 99C>A, 101T>C, 109_119delGAAACTTTTAA, 121A>C, 122C>T, 123A>C, 125T>A, 129T>C, 130T>C, 131C>T, 134C>G, 135T>A, 136T>A, 137A>T, 138T>C,
142G>A, 143T>C, 144G>A, 145A>T, 146T>A, 148A>C, 156T>A, 157C>T, 165A>G, 170T>G, 171T>G, 172G>A, 175T>C, 178T>C, 180A>T, 181T>G,
184_201delGAGAAGACTGGATCAGTT, 202C>T, 204C>T, 207T>A, 208A>C, 213T>A, 221G>C, 222T>A, 226C>G, 228G>T, 232G>T, 233G>C, 238A>T, 240C>T, 243A>T, 244A>T,
245A>C, 246A>T, 247T>C, 249C>T, 255A>G, 256G>A, 259A>C, 261T>C, 262A>T, 263G>C, 267C>T, 268A>G, 270C>T, 272C>A, 273A>T, 274T>A, 275C>A, 279A>T, 287A>C,
288A>C, 292T>G, 293C>T, 294T>A, 300T>C, 301T>A, 306T>G, 310T>G, 312G>C, 315A>G, 318A>G, 319T>C, 321T>A, 324G>T, 326C>T, 327T>G, 329A>C, 330T>A, 334T>G,
336T>A, 337T>A, 339T>A, 342T>A, 347T>C, 348T>C, 357T>C, 358T>A, 359C>A, 360C>T, 363A>T, 364T>G, 366G>A, 367A>G, 369T>A, 371A>G, 375G>T, 376C>T, 378T>A,
387A>T, 389G>C, 390T>A, 391T>A, 392C>A, 393C>A, 394T>A, 396G>A, 399T>G, 405T>C, 408T>C, 409C>A, 411G>C, 414A>G, 416C>G, 417A>T, 420C>T, 421T>G, 422C>A,
427T>G, 429G>A, 430G>A, 431C>A, 435C>T, 445T>A, 450C>A, 451A>G, 452A>T, 453G>A, 457A>C, 459G>T, 462T>A, 463T>C, 465A>T, 466C>A, 467G>A, 469T>A, 472T>G,
474C>T, 475G>T, 476A>T, 481T>G, 483G>C, 484T>A, 485C>A, 486C>G, 487A>G, 493G>T, 495T>C, 497G>T, 498A>C, 499A>C, 506T>A, 507T>C, 508C>G, 510G>A, 514G>T,
516T>A, 522G>A, 525G>A, 526A>G, 527G>T, 532G>T, 533A>C, 534T>A, 535G>T, 543T>A, 546A>G, 547G>T, 550T>A, 551A>T, 552T>A, 554G>C, 561G>A, 570A>T, 571A>T,
573A>G, 574A>G, 577C>A, 580C>T, 582T>G, 583G>C, 585T>A, 586T>C, 588G>A, 589T>C, 593G>A, 597T>A, 604T>G, 605C>T, 609C>T, 612A>T, 613A>G, 615T>A, 616G>A,
617A>C, 618A>G, 619G>C, 620G>C, 621A>T, 622G>T, 624A>T, 625T>A, 626C>A, 629G>A, 631A>G, 632C>T, 633C>A, 634A>G, 637A>G, 638A>T, 639A>T, 641A>C, 643A>T,
645T>G, 648C>T, 650T>C, 652A>G, 654A>T, 657A>T, 660G>A, 664A>T, 666G>T, 669T>C, 673A>T, 674G>C, 675T>A, 684C>T, 687G>A, 690C>T, 699C>A, 701A>C, 702G>A,
706G>T, 710A>G, 717C>T, 718A>T, 720A>C, 721A>C, 722T>A, 724G>C, 726A>G, 729A>G, 736A>G, 737G>C, 738T>A, 739A>G, 741G>A, 744A>T, 753G>A, 754T>A, 755C>G,
756T>C, 757G>T, 759C>G, 760T>A, 762A>G, 765C>T, 768T>A, 774A>T, 775T>A, 776C>G, 780G>A, 781T>C, 783G>T, 784A>T, 786G>A, 789A>C, 790A>T, 792T>A, 793A>T,
795T>G, 796A>T, 801A>T, 810T>C, 813C>A, 816G>T, 817C>T, 819C>A, 822G>A, 826A>G, 828A>T, 831A>T, 834G>T, 837A>T, 840A>G, 846C>A, 853T>C, 855G>T, 856A>T,
857G>C, 858C>A, 861T>C, 864C>A, 867C>G, 876T>C, 885A>T, 886A>G, 891G>A, 897G>T, 900T>G, 915A>G, 916T>C, 921T>C, 922A>C, 923C>A, 924A>G, 933A>G, 936T>C,
942A>G, 948C>T, 951T>A, 954A>T, 960A>C, 963T>C, 966T>A, 972G>T, 978A>G, 981T>A, 982A>G, 984A>G, 988G>A, 990T>C, 991G>A, 992C>T, 1002C>T, 1005G>A, 1014G>A,
1015C>T, 1017T>G, 1020T>A, 1023A>G, 1026T>A, 1028G>A, 1029C>T, 1032A>T, 1035A>C, 1038G>A, 1041T>C, 1044T>C, 1048T>C, 1050G>T, 1054G>C, 1056A>G, 1063C>A,
1064A>C, 1071G>T, 1072G>T, 1073A>C, 1074T>G, 1076C>A, 1077T>G, 1081T>G, 1082G>C, 1086T>A, 1090G>T, 1094C>T, 1095C>G, 1096A>C, 1098G>C, 1099T>C, 1101G>T,
1104G>A, 1107C>T, 1113C>A, 1117A>C, 1118A>C, 1119A>T, 1122C>A, 1125C>T, 1128T>A, 1129G>A, 1130A>T, 1131C>T, 1134A>G, 1138A>C, 1140T>A, 1143C>T, 1144C>T,
1146T>A, 1148G>A, 1149G>A, 1158A>C, 1159C>G, 1161G>C, 1164G>A, 1167T>A, 1173A>T, 1185A>C, 1188C>T, 1194T>C, 1197C>A, 1204A>T, 1212T>C, 1213C>A, 1215A>G,
1218G>T, 1219A>T, 1220G>C, 1224A>T, 1227T>C, 1230C>A, 1233T>G, 1239T>A, 1242A>T, 1245G>A, 1246G>C, 1248C>G, 1249T>G, 1250A>T, 1254G>T, 1258A>G, 1259G>C,
1260T>C, 1261T>A, 1264A>C, 1265G>A, 1270G>T, 1271T>C, 1278T>G, 1281T>C, 1284G>A, 1290A>G, 1296A>G, 1299C>T, 1308T>G, 1316A>T, 1320T>A, 1323C>T, 1329G>A,
1344A>G, 1345G>T, 1346C>G, 1347A>T, 1353A>C, 1354T>C, 1356G>T, 1372T>C, 1374G>C, 1377T>C, 1378C>G, 1379G>C, 1380C>A, 1386G>T, 1389G>A, 1392T>A, 1398T>G,
1402A>C, 1404A>C, 1405C>T, 1407T>G, 1413T>G, 1416T>A, 1419C>T, 1422G>A, 1423C>A, 1425C>A, 1431C>A, 1440T>C, 1443T>C, 1449C>A, 1452C>T, 1455G>A, 1461T>A,
1465T>C, 1467G>T, 1476C>T, 1482A>T, 1485A>T, 1486T>C, 1488G>T, 1489A>C, 1491A>G, 1503A>G, 1506A>G, 1512A>G, 1518T>A, 1536T>C, 1542C>T, 1545A>C, 1551A>T,
1557A>T, 1560G>T, 1563C>A, 1569A>T, 1572T>A, 1575G>C, 1578A>G, 1581C>T, 1584G>A, 1587G>A, 1593T>C, 1599T>A, 1608T>C, 1617A>C, 1626T>C, 1627C>T, 1629T>G,
1630G>A, 1635A>T, 1638A>T, 1641T>A, 1644C>T, 1656T>A, 1659C>G, 1671T>A, 1677G>A, 1678C>A, 1680T>G, 1683A>G, 1686G>A, 1692C>T, 1704T>C, 1706T>G, 1707C>A,
1712A>T, 1713A>G, 1716T>C, 1719A>T, 1723T>G, 1724C>A, 1726A>C, 1729G>A, 1731G>T, 1733A>C, 1734C>T, 1740C>A, 1742G>C, 1743A>C, 1749G>A, 1750A>G, 1752G>T,
1756A>G, 1758C>T, 1759A>G, 1761T>G, 1771C>A, 1773G>A, 1776G>A, 1788A>C, 1798C>G, 1799A>C, 1800C>T, 1804A>T, 1805G>C, 1806T>C, 1809C>T, 1810T>G, 1811C>A,
1812G>T, 1821A>G, 1822G>C, 1824C>A, 1836C>A, 1842T>C, 1844A>G, 1848T>G, 1851T>C, 1852G>T, 1853C>A, 1858A>G, 1860A>G, 1865A>G, 1866G>C, 1869G>A, 1872G>T,
1874T>A, 1875T>C, 1876G>A, 1878A>T, 1879G>A, 1881A>T, 1882G>A, 1883A>C, 1885A>T, 1887T>G, 1890C>A, 1891A>T, 1893T>G, 1894T>G, 1896A>T, 1897A>G, 1903A>G,
1904C>T, 1905A>T, 1908C>T, 1914T>C, 1915G>A, 1916C>A, 1920A>T, 1921T>G, 1924T>G, 1925C>T, 1929A>T, 1932G>A, 1935T>G, 1936G>C, 1940C>A, 1941A>T, 1945G>T,
1947T>C, 1948G>C, 1950T>A, 1952G>A, 1953G>T, 1956T>C, 1957_1959delCGC, 1962C>A, 1963A>C, 1966A>G, 1967C>T, 1968A>T, 1969A>T, 1970A>T, 1971A>G, 1973A>C,
1974A>T, 1975G>C, 1976A>C, 1979T>A, 1984A>G, 1985T>C, 1988T>C, 1989G>T, 1990A>T, 1991C>A, 1993C>G, 1994C>A, 1995C>G, 1996A>C, 1997C>A, 1998A>G, 1999G>A,
2000T>C, 2001T>A, 2004A>T, 2010G>A, 2012A>C, 2013A>T, 2016T>A, 2018G>A, 2020G>A, 2022G>T, 2023A>G, 2031T>C, 2032A>C, 2034A>G, 2038T>C, 2040A>C, 2046G>T,
2049A>G, 2055T>C, 2060T>A, 2067T>C, 2070G>A, 2073C>G, 2077T>A, 2080G>A, 2083C>A, 2084C>A, 2085G>A, 2086G>T, 2087A>T, 2088G>T, 2089A>G, 2090G>A, 2091T>G,
2094G>T, 2097T>C, 2100T>A, 2107T>C, 2108A>T, 2109T>C, 2110C>T, 2111A>C, 2112A>C, 2115A>G, 2118A>G, 2122C>A, 2126T>A, 2131T>G, 2132C>T, 2133C>G, 2137G>T,
2138A>C, 2140G>T, 2142A>T, 2145G>T, 2146C>G, 2147A>C, 2149C>A, 2151G>A, 2154A>G, 2155G>A, 2156T>A, 2157C>T, 2160C>T, 2161A>G, 2162T>A, 2163G>A, 2165G>T,
2166A>G, 2170A>G, 2172G>A, 2174C>A, 2178C>T, 2179A>T, 2182A>T, 2184C>G, 2185T>G, 2186A>T, 2193T>A, 2196C>A, 2198A>G, 2200G>C, 2203G>C, 2205A>G, 2208A>G,
2211C>T, 2212A>G, 2213G>C, 2214G>T, 2225A>G, 2226A>G, 2229T>C, 2245C>A, 2256T>C, 2259A>G, 2262C>T, 2268C>T, 2272G>A, 2273C>G, 2274T>C, 2275T>A, 2277C>A,
2283C>T, 2285G>C, 2290A>G, 2293A>C, 2294A>T, 2300C>A, 2304A>C, 2308A>C, 2309C>G, 2310A>C, 2311G>A, 2313G>T, 2314A>C, 2316G>T, 2317C>G, 2318G>A, 2319C>G,
2320A>C, 2321C>A, 2325A>G, 2326T>A, 2327T>A, 2328T>A, 2331A>G, 2332C>T, 2337T>G, 2338T>G, 2339G>C, 2340T>G, 2341A>C, 2344G>C, 2346T>G, 2354T>C, 2355T>A,
2362T>C, 2364G>T, 2366T>C, 2367T>G, 2370C>T, 2373C>T, 2374C>G, 2376G>A, 2389T>A, 2390T>G, 2391T>A, 2394G>A, 2400T>C, 2401T>C, 2409A>T, 2412C>T, 2421T>C,
2424T>C, 2432C>G, 2433C>G, 2440T>G, 2442C>A, 2442_2443insGGT, 2444A>C, 2445A>T, 2448G>C, 2449G>A, 2450C>A, 2451C>G, 2454A>G, 2455G>A, 2456A>G, 2457C>A,
2458T>A, 2459C>A, 2464G>A, 2466C>T, 2468A>T, 2469G>C, 2470A>G, 2471T>A, 2475T>C, 2476T>A, 2477A>T, 2478C>G, 2479C>A, 2480A>T, 2481C>T, 2482A>G, 2483A>G,
2484A>T, 2487T>C, 2488G>A, 2489T>A, 2493T>C, 2494T>A, 2496C>G, 2501A>T, 2502A>C, 2503G>C, 2505A>G, 2507C>T, 2508T>G, 2512T>A, 2513G>A, 2514G>T, 2515A>G,
2520A>G, 2523A>C, 2525A>T, 2526T>C, 2527G>A, 2528C>T, 2529A>C, 2530C>T, 2534G>A, 2553C>A, 2556A>G, 2560A>T, 2562C>G, 2565G>A, 2568G>A, 2577A>C, 2578T>A,
2579T>C, 2580G>C, 2581T>A, 2583T>G, 2584G>A, 2585G>A, 2586G>T, 2588G>C, 2589A>C, 2590A>G, 2592A>T, 2595A>T, 2601T>C, 2602C>A, 2605G>A, 2610T>C, 2611T>C,
2612G>T, 2613G>C, 2614C>A, 2616A>G, 2617A>G, 2618G>C, 2619C>A, 2620A>G, 2621A>C, 2622T>A, 2623T>A, 2627T>C, 2628T>C, 2631A>T, 2632A>T, 2633G>T, 2634T>A,
2638A>G, 2639G>A, 2643A>C, 2649T>G, 2652T>C, 2655C>A, 2658A>C, 2673C>T, 2679T>C, 2685A>G, 2688G>A, 2691C>T, 2694G>A, 2697C>G, 2698C>A, 2699C>A, 2700C>G,
2701T>A, 2705G>A, 2707C>T, 2709G>C, 2710A>G, 2711A>C, 2712G>A, 2715T>A, 2717T>C, 2718G>C, 2722G>C, 2724G>A, 2730G>A, 2731A>G, 2732G>C, 2733T>G, 2734A>G,
2735G>C, 2736C>A, 2738C>A, 2744T>A, 2745T>C, 2745_2746insCAT, 2746A>G, 2747C>T, 2748A>G, 2749C>G, 2751A>C, 2752G>C, 2753A>G, 2762C>A, 2763T>A, 2764G>C,
2766A>G, 2772G>A, 2778C>T, 2781C>T, 2782C>G, 2785G>A, 2786C>G, 2787A>T, 2790C>G, 2791A>C, 2793A>C, 2796G>A, 2799T>C, 2800A>T, 2803A>G, 2808A>T, 2809C>T,
2812C>A, 2814G>A, 2815C>A, 2819G>C, 2820A>T, 2823G>A, 2829G>C, 2832C>T, 2844A>G, 2850G>T, 2856A>T, 2859G>T, 2865A>T, 2866A>C, 2868G>C, 2883A>C, 2886C>T,
2887C>A, 2888A>G, 2889C>A, 2892T>C, 2893G>A, 2898A>G, 2904A>C, 2907A>T, 2914A>T, 2919T>C, 2922C>T, 2926C>G, 2927C>A, 2928T>G, 2929A>C, 2931C>T, 2934A>C,
2936G>T, 2937G>T, 2939C>G, 2940A>T, 2943G>T, 2951A>T, 2955C>T, 2959C>A, 2960G>C, 2961G>C, 2962G>C, 2967C>A, 2969A>G, 2971G>T, 2973C>G, 2976T>A, 2982A>T,
2989A>G, 2991C>G, 2995C>T, 2998C>T, 3000A>G, 3001C>A, 3003T>G, 3006G>A, 3009T>G, 3012C>T, 3015T>A, 3028C>T, 3030T>A, 3036T>A, 3037C>A, 3039C>A, 3042G>A,
3051A>T, 3063G>T, 3066C>G, 3069C>T, 3071T>A, 3073A>T, 3075C>T, 3078A>G, 3081T>C, 3084T>A, 3085C>A, 3086A>T, 3093A>G, 3096T>C, 3097T>G, 3098G>T, 3099T>G,
3102A>G, 3105C>T, 3108G>A, 3111G>T, 3116C>A, 3117C>G, 3123A>C, 3125G>A, 3126A>G, 3127C>A, 3128T>A, 3129G>A, 3130G>T, 3131G>C, 3135C>T, 3136C>T, 3138A>G,
3141A>G, 3142T>G, 3145T>C, 3147G>T, 3152T>A, 3168C>A, 3171A>G, 3177T>C, 3180A>C, 3182T>A, 3183C>A, 3186G>A, 3189G>C, 3190G>A, 3191A>C, 3192G>A, 3197A>G,
3198G>A, 3204C>T, 3207T>G, 3208G>A, 3209C>A, 3210C>T, 3211A>G, 3212G>C, 3213T>A, 3217A>G, 3218A>C, 3219A>T, 3220T>C, 3222G>T, 3225T>C, 3228G>A, 3231G>A,
3232C>G, 3233A>T, 3234A>T, 3235T>G, 3243A>G, 3246T>C, 3249T>C, 3252G>T, 3256G>A, 3258A>T, 3267T>A, 3276A>T, 3283T>C, 3285A>C, 3286C>T, 3288T>A, 3294C>A,
3296G>C, 3300T>G, 3309A>G, 3312A>G, 3315T>G, 3316G>T, 3317T>G, 3318T>C, 3324G>T, 3327C>A, 3330T>A, 3339T>A, 3342A>T, 3350G>A, 3353T>C, 3354T>C, 3357A>T,
3363A>G, 3369A>T, 3371T>A, 3373A>G, 3375T>G, 3376T>G, 3381C>A, 3384T>C, 3387C>A, 3390G>A, 3393C>A, 3394A>G, 3396C>G, 3399C>T, 3400C>A, 3402G>A, 3405C>T,
3408G>A, 3409C>T, 3411T>G, 3415A>G, 3417C>T, 3429G>A, 3432T>C, 3433G>A, 3440C>T, 3444A>T, 3453A>C, 3459C>A, 3460A>C, 3462T>G, 3463C>T, 3465C>G, 3466A>G,
3468A>G, 3469G>C, 3470C>A, 3472A>C, 3474T>G, 3477G>A, 3478A>T, 3479G>C, 3480T>C, 3482G>A, 3483A>G, 3486A>G, 3487A>G, 3488C>A, 3489C>A, 3491C>G, 3492T>A,
3493C>T, 3495T>C, 3501A>G, 3507C>A, 3510A>G, 3513C>G, 3514T>G, 3516C>T, 3519T>C, 3520C>T, 3522T>G, 3531T>A, 3534T>G, 3535C>T, 3536A>T, 3537A>G, 3541T>A,
3546C>T, 3549A>T, 3555A>G, 3558C>G, 3561T>C, 3567T>A, 3570A>C, 3573C>T, 3575G>A, 3577A>C, 3578A>G, 3579G>C, 3584A>C, 3585A>C, 3586T>C, 3597A>T, 3603T>G,
3604C>G, 3605C>A, 3606T>C, 3610G>C, 3612A>G, 3615C>T, 3618T>C, 3621C>A, 3624A>C, 3627C>T, 3630C>A, 3631A>T, 3633C>T, 3635T>C, 3639T>A, 3646C>A, 3654G>A,
3657A>G, 3660C>T, 3666A>C, 3675A>T, 3681G>C, 3682A>G, 3683T>A, 3687A>G, 3690C>T, 3699T>A, 3703C>T, 3704A>C, 3705T>C, 3706G>A, 3707G>A, 3708T>C, 3714T>G,
3717T>G, 3718G>C, 3719A>C, 3720T>A, 3726T>A, 3729A>G, 3730T>C, 3733T>G, 3741A>G, 3742A>G, 3743C>G, 3747C>T, 3751A>G, 3752G>C, 3753T>A, 3756A>C, 3757C>A,
3759G>T, 3762G>T, 3765A>T, 3766A>G, 3767C>T, 3768A>G, 3769A>C, 3771G>T, 3774A>G, 3776A>T, 3777C>G, 3780A>G, 3781A>C, 3782A>G, 3795C>G, 3801A>T, 3802C>G,
3803A>C, 3804G>A, 3805G>A, 3806A>T, 3807G>C, 3811G>A, 3822G>A, 3828G>C, 3837T>A, 3845G>T, 3854A>T, 3860G>C, 3861C>A, 3868C>T, 3870C>G, 3876A>G, 3879C>A,
3888C>T, 3894C>T, 3901A>T, 3906C>A, 3907A>C, 3909G>T, 3912A>T, 3913A>T, 3914T>C, 3915G>T, 3921G>A, 3922C>A, 3927G>A, 3933C>G, 3935T>A, 3937C>G, 3939A>C,
3940A>C, 3941C>A, 3943G>C, 3944G>C, 3946G>A, 3948A>C, 3949G>T, 3950G>C, 3951T>G, 3952G>C, 3954T>G, 3956A>G, 3957A>G, 3960A>T, 3961G>C, 3962A>T, 3963G>T,
3966T>C, 3968G>A, 3973G>A, 3981A>T, 3984G>C, 3990A>T, 3993T>A, 3999G>A, 4002A>G, 4003C>T, 4008C>T, 4009A>G, 4010T>C, 4011T>G, 4014A>G, 4015A>G, 4017T>A,
4023G>A, 4028T>C, 4030C>T, 4031A>T, 4035G>T, 4038T>C, 4041A>C, 4042A>G, 4047C>A, 4050A>C, 4053A>G, 4059C>G, 4062G>A, 4068G>A, 4071C>G, 4072C>T, 4074T>G,
4077T>A, 4083A>G, 4104A>G, 4107A>C, 4108A>C, 4110A>C, 4111T>C, 4113A>G, 4116T>C, 4119G>A, 4122A>C, 4137C>T, 4146G>A, 4149C>G, 4150A>C, 4161T>C, 4173C>T,
4176C>T, 4177A>T, 4178C>G, 4179C>T, 4188A>T, 4189T>G, 4190T>C, 4197T>C, 4200C>A, 4201A>C, 4206A>T, 4209T>C, 4215T>A, 4218A>T, 4221A>C, 4222G>A, 4230C>G,
4233T>C, 4234T>C, 4235C>A, 4243T>A, 4245T>A, 4246C>G, 4248T>G, 4251T>G, 4254C>T, 4256A>G, 4257C>T, 4259A>G, 4263C>T, 4267G>T, 4269T>A, 4279A>G, 4280C>A,
4281T>G, 4284G>A, 4287T>C, 4293T>C, 4302A>G, 4305T>C, 4311C>T, 4314G>A, 4315C>A, 4317C>A, 4323A>C, 4324G>A, 4326G>C, 4329C>T, 4332T>A, 4335T>C, 4338G>A,
4351T>C, 4356T>G, 4361A>C, 4362G>T, 4364A>T, 4370A>G, 4372A>G, 4373C>A, 4374C>T, 4375A>G, 4383A>G, 4384G>A, 4385A>G, 4386G>T, 4391C>A, 4392T>G, 4395C>A,
4401A>T, 4404A>G, 4407T>C, 4408G>A, 4410T>A, 4411C>T, 4413T>A, 4414C>T, 4416C>A, 4425A>C, 4428A>T, 4437C>T, 4441T>C, 4443A>C, 4450G>A, 4452T>C, 4455A>G,
4456G>T, 4458T>G, 4461C>T, 4464T>C, 4465T>C, 4467G>A, 4470T>G, 4476C>A, 4482G>C, 4494A>T, 4495C>T, 4497T>G, 4500C>T, 4509C>T, 4512C>T, 4516C>T, 4519T>C,
4521G>A, 4537C>A, 4542T>C, 4545T>G, 4548C>T, 4551C>A, 4557T>G, 4560A>G, 4563A>G, 4566G>A, 4569T>A, 4572C>T, 4573A>G, 4575T>C, 4576G>A, 4577C>A, 4578C>A,
4581T>C, 4583G>A, 4585G>T, 4588A>G, 4590A>G, 4593T>C, 4594G>A, 4596A>T, 4602T>A, 4603G>A, 4604G>A, 4608T>C, 4614C>T, 4618C>T, 4620T>G, 4638A>T, 4647T>C,
4648C>T, 4653G>T, 4656T>G, 4659G>T, 4662T>A, 4663C>A, 4669C>T, 4681T>G, 4682C>A, 4683A>T, 4686G>A, 4689T>A, 4690A>T, 4691A>C, 4692T>C, 4698C>A, 4699C>A,
4700A>G, 4701G>A, 4707T>G, 4710C>A, 4711A>G, 4712T>C, 4713G>T, 4716A>T, 4719T>C, 4728C>T, 4731A>G, 4737G>T, 4738C>T, 4743T>C, 4749T>A, 4759A>G, 4761C>G,
4762C>A, 4765G>A, 4766T>C, 4767G>A, 4768C>A, 4770G>A, 4773C>T, 4775A>T, 4779C>T, 4785A>T, 4788A>G, 4794T>C, 4803C>T, 4809A>T, 4812T>C, 4815A>G, 4818T>A,
4824C>A, 4825C>T, 4827T>G, 4830G>A, 4836C>A, 4857G>T, 4861A>G, 4863A>C, 4866T>C, 4872T>C, 4875G>T, 4878C>T, 4884C>T, 4887A>G, 4890T>A, 4894C>G, 4895C>T,
4899T>A, 4909A>G, 4910C>T, 4911T>A, 4912G>C, 4913T>A, 4914G>A, 4917A>T, 4921A>T, 4922T>A, 4923G>T, 4926G>A, 4930G>A, 4932G>T, 4938T>A, 4942C>A, 4944T>C,
4945C>A, 4947T>A, 4950G>A, 4956T>C, 4966A>C, 4967G>A, 4968A>G, 4970T>C, 4971T>C, 4972G>A, 4974A>T, 4980G>A, 4982A>G, 4983G>T, 4986A>T, 4987A>G, 4988C>T,
4989T>A, 4990T>A, 4999C>T, 5000A>T, 5001G>T, 5002T>G, 5004T>A, 5007C>T, 5012G>A, 5013A>G, 5014G>C, 5017C>A, 5019T>A, 5020G>A, 5021G>A, 5022A>T, 5026G>C,
5027G>C, 5029G>A, 5030C>T, 5031A>T, 5033C>T, 5034A>G, 5038T>A, 5040T>G, 5043T>C, 5044T>A, 5049T>C, 5050A>T, 5051T>A, 5052G>T, 5056G>C, 5058G>T, 5061G>A,
5062C>A, 5063A>G, 5064G>T, 5065G>C, 5071A>T, 5073C>G, 5074G>A, 5077A>G, 5093C>A, 5094A>G, 5095T>A, 5096C>G, 5097A>C, 5098G>A, 5099C>T, 5100G>T, 5101T>G,
5102T>G, 5104A>G, 5105C>T, 5106A>T, 5107A>T, 5109C>G, 5110T>A, 5111G>T, 5112T>G, 5115A>G, 5116A>C, 5118G>A, 5119G>T, 5121C>A, 5122A>G, 5124A>T, 5128T>A,
5129C>A, 5130T>A, 5132T>A, 5133C>T, 5136T>A, 5138T>A, 5142A>C, 5148G>A, 5151A>T, 5152C>G, 5158A>G, 5160T>A, 5164G>A, 5167_5169delGAA, 5178A>G, 5179A>G,
5180G>A, 5181A>G, 5183A>G, 5185C>G, 5188A>C, 5191T>G, 5192G>A, 5193G>A, 5198C>T, 5199C>G, 5202A>T, 5203A>G, 5205C>T, 5206A>G, 5208T>A, 5209G>T, 5210G>T,
5217C>T, 5218T>A, 5219C>G, 5220A>T, 5223C>T, 5226T>G, 5233G>T, 5235C>A, 5238G>C, 5239A>C, 5240C>A, 5241T>A, 5242G>C, 5248G>T, 5249A>C, 5250G>A, 5251C>T,
5253T>A, 5256G>T, 5257T>A, 5258G>A, 5262A>C, 5263G>C, 5265A>G, 5271C>T, 5274T>C, 5277A>T, 5280G>A, 5283G>C, 5284A>C, 5285A>G, 5286G>C, 5288A>G, 5289G>A,
5291G>C, 5292C>A, 5294A>C, 5301T>G, 5302C>A, 5304C>A, 5307A>G, 5313T>C, 5315G>A, 5325T>C, 5340G>A, 5346A>C, 5349T>C, 5352C>T, 5353A>T, 5356T>G, 5364T>C,
5366T>A, 5371G>C, 5372C>A, 5377A>C, 5379A>C, 5380G>A, 5381G>A, 5382C>G, 5384A>G, 5385A>T, 5386A>C, 5389T>G, 5390C>A, 5393A>T, 5394G>T, 5396A>G, 5397C>T,
5398A>G, 5399T>A, 5402C>G, 5403T>G, 5410G>T, 5411A>C, 5415T>C, 5416G>A, 5417T>C, 5418G>C, 5424G>T, 5430G>T, 5442T>A, 5443G>A, 5445G>A, 5446A>G, 5447G>C,
5448C>T, 5463T>A, 5466A>G, 5467A>T, 5469G>T, 5472T>A, 5473G>T, 5483G>A, 5487G>A, 5490T>C, 5491T>A, 5493C>G, 5494A>G, 5495T>C, 5496T>C, 5497G>A, 5498A>C,
5499G>A, 5502T>C, 5503A>C, 5505C>T, 5512A>C, 5514G>C, 5515T>C, 5517G>A, 5520C>A, 5523T>C, 5526T>G, 5528G>C, 5529C>T, 5530A>G, 5531A>G, 5532T>G, 5536T>C,
5550A>C, 5553T>C, 5556C>T, 5567T>C, 5568T>G, 5571G>A, 5577T>G, 5581C>A, 5589G>T, 5590C>G, 5591C>A, 5592T>A, 5593C>G, 5595G>C, 5595_5596insACC, 5596G>T,
5597A>C, 5599G>C, 5600G>A, 5601G>C, 5604A>G, 5605A>T, 5609C>G, 5614G>A, 5615T>G, 5620T>A, 5622G>A, 5623_5625delTCT, 5627T>A, 5628G>T, 5629C>G, 5630A>G,
5631A>C, 5633G>C, 5634T>C, 5636G>C, 5642T>A, 5643G>C, 5649A>T, 5649_5650insCACAGT, 5651G>C, 5652T>A, 5656G>T, 5657A>C, 5658G>A, 5659C>G, 5664C>A,

| | |
|---|---|
| | 5666A>T, 5667A>T, 5668A>C, 5670T>A, 5671C>A, 5673T>G, 5674G>A, 5676C>A, 5677A>G, 5679C>T, 5681A>C, 5682C>T, 5683G>T, 5684G>C, 5685T>C, 5688T>C, 5689T>C, 5691A>C, 5694A>T, 5695A>G, 5696T>A, 5697G>A, 5698G>A, 5700A>G, 5707T>G, 5709G>T, 5712C>T, 5713A>T, 5721T>C, 5722G>A, 5723T>C, 5724T>A, 5725G>A, 5727G>T, 5730C>T, 5733A>T, 5734C>G, 5736T>A, 5739T>A, 5742A>T, 5747G>C, 5749A>C, 5751C>G, 5753T>G, 5754T>C, 5755A>T, 5757C>A, 5760A>G, 5763A>G, 5766T>C, 5769T>G, 5770G>A, 5772T>G, 5775A>T, 5776T>G, 5777T>C, 5778G>T, 5781T>G, 5783C>A, 5784A>G, 5786G>C, 5787T>A, 5788C>T, 5790T>G, 5791G>T, 5792A>C, 5793A>T, 5794A>G, 5795A>T, 5796G>C, 5797T>A, 5806G>A, 5807A>C, 5808A>T, 5809T>C, 5811G>T, 5812C>G, 5813T>C, 5814T>C, 5819A>G, 5820G>A, 5821A>G, 5822A>G, 5823C>A, 5824A>G, 5825C>T, 5826G>C, 5830C>G, 5835C>A, 5845G>A, 5847G>A, 5848A>G, 5851A>T, 5853G>C, 5854A>T, 5856T>C, 5860A>C, 5862A>C, 5866G>T, 5867T>G, 5868G>T, 5870C>A, 5872C>A, 5873A>T, 5874A>G, 5878G>C, 5880C>T, 5881A>G, 5882G>A, 5883T>G, 5887G>C, 5889A>G, 5890A>G, 5893C>A, 5894A>G, 5895G>T, 5902T>G, 5904C>A, 5905T>G, 5906A>T, 5907T>A, 5908C>T, 5909C>T, 5912A>T, 5915C>G, 5916C>G, 5918C>G, 5919A>C, 5920A>G, 5921C>T, 5922C>T, 5923G>A, 5924T>A, 5926G>A, 5931G>A, 5932C>A, 5934C>G, 5935A>T, 5936A>T, 5938G>T, 5939T>C, 5940A>C, 5943T>C, 5944G>A, 5945C>T, 5946T>G, 5947G>A, 5950G>C, 5951C>A, 5952T>A, 5955T>G, 5955_5956insATG, 5956A>T, 5958C>A, 5960T>A, 5961T>A, 5962G>A, 5963T>C, 5965A>C, 5968A>G, 5969C>T, 5973T>C, 5975A>G, 5976A>C, 5979T>A, 5981T>G, 5982C>T, 5983C>T, 5985G>T, 5986_5988delGGA, 5991A>C, 5992G>A, 5993T>C, 5994T>A, 5997C>T, 5998C>A, 6000G>T, 6004A>G, 6005A>C, 6006T>A, 6008T>G, 6009T>G, 6012C>A, 6014G>C, 6016C>A, 6017C>A, 6018T>G, 6021G>C, 6022T>A, 6023G>T, 6024G>T, 6024_6025insGTC, 6026A>T, 6027A>G, 6029T>C, 6030T>A, 6031G>A, 6032G>A, 6037G>A, 6042C>G, 6043A>C, 6046_6048delATG, 6050C>G, 6052A>T, 6054A>T, 6055C>A, 6057C>T, 6058T>C, 6060A>T, 6066G>A, 6068G>C, 6069T>A, 6070C>G, 6072T>G, 6073C>T, 6077C>T, 6078A>T, 6086T>C, 6087G>T, 6090G>A, 6094T>C, 6095C>T, 6098C>T, 6099C>T, 6100A>G, 6104A>G, 6105G>A, 6109T>A, 6112C>T, 6114C>G, 6115A>C, 6119G>A |

**CDS**

| | | | | | | | | | |
|---|---|---|---|---|---|---|---|---|---|
| APL33_sRA11gp1 | 1 | 1993 | 92.7% | 9636 | 70.1% | 1982 (99.1%) | 1314 (65.7%) | 7/11/0/0 | 0 |

| | Begin | End | Coverage | Score | Concordance | Matches | Identities | I/D/M/F* | Stop Codons |
|---|---|---|---|---|---|---|---|---|---|
| **NT** | **33** | **6128** | **84.2%** | **3374** | **28.2%** | **6043 (98.8%)** | **3915 (64.0%)** | **21/53** | |
| Protein mutations: | L3F (157C>T), F7W (170T>G 171T>G), G8S (172G>A), S9P (175T>C), S10P (178T>C 180A>T), S11A (181T>G), E12_V17del (184_201delGAGAAGACTGGATCAGTT), P18S (202C>T 204C>T), T20P (208A>C), G24A (221G>C 222T>A), L26V (226C>G 228G>T), G28S (232G>T 233G>C), I30F (238A>T 240C>T), K32S (244A>T 245A>C 246A>T), F33L (247T>C 249C>T), A36T (256G>A), I37L (259A>C 261T>C), I40V (268A>G 270C>T), T41N (272C>A 273A>T), S42K (274T>A 275C>A), K46T (287A>C 288A>C), S48V (292T>G 293C>T 294T>A), L51I (301T>A), D52E (306T>G), L54V (310T>G 312G>C), S57P (319T>C 321T>A), T59M (326C>T 327T>G), D60A (329A>C 330T>A), S62A (334T>G 336T>A), S63T (337T>A 339T>A), V66A (347T>C 348T>C), S70N (358T>A 359C>A 360C>T), L72V (364T>G 366G>A), N73E (367A>G 369T>A), K74R (371A>G), G80A (389G>C 390T>A), S81K (391T>A 392C>A 393C>A), L82I (394T>A 396G>A), I83M (399T>G), L87I (409C>A 411G>C), A89G (416C>G 417A>T), S91D (421T>G 422C>A), L93V (427T>G 429G>A), A94K (430G>A 431C>A), S99T (445T>A), K101V (451A>G 452A>T 453G>A), M103L (457A>C 459G>T), R106K (466C>A 467G>A), L107M (469T>A), F108V (472T>G 474C>T), E109L (475G>T 476A>T), L111V (481T>G 483G>C), S112K (484T>A 485C>A 486C>G), N113D (487A>G), V115F (493G>T 495T>C), R116I (497G>T 498A>C), I117L (499A>C), F119Y (506T>A 507T>C), Q120E (508C>G 510G>A), V122L (514G>T 516T>A), S126V (526A>G 527G>T), D128S (532G>T 533A>C 534T>A), V129L (535G>T), V133F (547G>T), Y134I (550T>A 551A>T 552T>A), G135A (554G>C), I141L (571A>T 573A>G), T142A (574A>G), L143I (577C>A), V145L (583G>C 585T>A), C148Y (593G>A), S152V (604T>G 605C>T), I155V (613A>G 615T>A), E156T (616G>A 617A>C 618A>G), G157P (619G>C 620G>C 621A>T), V158F (622G>T 624A>T), S159N (625T>A 626C>A), S160N (629G>A), T161V (631A>G 632C>T 633C>A), I162V (634A>G), K163V (637A>G 638A>T 639A>T), N164T (641A>C), I165L (643A>T 645T>G), L167S (650T>C), I168V (652A>G 654A>T), M172F (664A>T 666G>T), K184T (701A>C 702G>A), A186S (706G>T), E187G (710A>G), T190S (718A>T 720A>C), M191Q (721A>C 722T>A), V192L (724G>C 726A>G), S196A (736A>G 737G>C 738T>A), M197V (739A>G 741G>A), V203L (757G>T 759C>G), S204T (760T>A 762A>G), M212L (784A>T 786G>A), T214S (790A>T 792T>A), I215L (793A>T 795T>G), M216L (796A>T), I226V (826A>G 828A>T), T246A (886A>G), T258Q (922A>C 923C>A 924A>G), I278V (982A>G 984A>G), G280S (988G>A 990T>C), A281I (991G>A 992C>T), S293N (1028G>A 1029C>T), E302Q (1054G>C 1056A>G), Q305T (1063C>A 1064A>C), D308S (1072G>T 1073A>C 1074T>G), A309E (1076C>A 1077T>G), C311A (1081T>G 1082G>C), D312E (1086T>A), A314S (1090G>T), T315M (1094C>T 1095C>G), K323P (1117A>C 1118A>C 1119A>T), D327I (1129G>A 1130A>T 1131C>T), N330Q (1138A>C 1140T>A), R333K (1148G>A 1149G>A), L337V (1159C>G 1161G>C), M352L (1204A>T), E364D (1242A>T), V366L (1246G>C 1248C>G), Y367V (1249T>G 1250A>T), S370A (1258A>G 1259G>C 1260T>C), C371S (1261T>A), R372Q (1264A>C 1265G>A), V374S (1270G>T 1271T>C), Y389F (1316A>T), A399C (1345G>T 1346C>G 1347A>T), R410A (1378C>G 1379G>C 1380C>A), V494I (1630G>A), I519R (1706T>G 1707C>A), E521V (1712A>T 1713A>G), E523D (1719A>T), S525D (1723T>G 1724C>A), T526P (1726A>C), E527N (1729G>A 1731G>T), D528A (1733A>C 1734C>T), G531A (1742G>C 1743A>C), M534V (1750A>G 1752G>T), N536D (1756A>G 1758C>T), I537V (1759A>G 1761T>G), H550A (1798C>G 1799A>C 1800C>T), S554D (1810T>G 1811C>A 1812G>T), D558Q (1822G>C 1824C>A), Q565R (1844A>G), A568Y (1852G>T 1853C>A), I570V (1858A>G 1860A>G), K572S (1865A>G 1866G>C), F575Y (1874T>A 1875T>C), E576N (1876G>A 1878A>T), A577T (1879G>A 1881A>T), E578T (1882G>A 1883A>C), T579S (1885A>T 1887T>G), I581L (1891A>T 1893T>G), S582A (1894T>G 1896A>T), N583D (1897A>G), T585V (1903A>G 1904C>T 1905A>T), A589N (1915G>A 1916C>A), S591A (1921T>G), S592V (1924T>G 1925C>T), E593D (1929A>T), V596L (1936G>C), A597D (1940C>A 1941A>T), V599F (1945G>T 1947T>C), D600Q (1948G>C 1950T>A), G601D (1952G>A 1953G>T), R603del (1957_1959delCGC), F604L (1962C>A), K605Q (1963A>C), T606V (1966A>G 1967C>T 1968A>T), K607L (1969A>T 1970A>T 1971A>G), Q608P (1973A>C 1974A>T), D609P (1975G>C 1976A>C), V610E (1979T>A), I612A (1984A>G 1985T>C), V613A (1988T>C 1989G>T), T614Y (1990A>T 1991C>A), P615E (1993C>G 1994C>A 1995C>G), T616Q (1996A>C 1997C>A 1998A>G), V617T (1999G>A 2000T>C 2001T>A), E621A (2012A>C 2013A>T), D622E (2016T>A), R623K (2018G>A), V624I (2020G>A 2022G>T), I625V (2023A>G), K628Q (2032A>C 2034A>G), F637Y (2060T>A), L643M (2077T>A), D644N (2080G>A), P645K (2083C>A 2084C>A 2085G>A), E646F (2086G>T 2087A>T 2088G>T), S647E (2089A>G 2090G>A 2091T>G), L648F (2094G>T), Y653L (2107T>C 2108A>T 2109T>C), Q654S (2110C>T 2111A>C 2112A>C), L658I (2122C>A), F659Y (2126T>A), S661V (2131T>G 2132C>T 2133C>G), D663S (2137G>T 2138A>C), A664S (2140G>T 2142A>T), E665D (2145G>T), H666A (2146C>G 2147A>C), Q667K (2149C>A 2151G>A), V669N (2155G>A 2156T>A 2157C>T), M671E (2161A>G 2162T>A 2163G>A), R672M (2165G>T 2166A>G), M674V (2170A>G 2172G>A), T675N (2174C>A), M677L (2179A>T), I678L (2182A>T 2184C>G), Y679V (2185T>G 2186A>T), N683S (2198A>G), A684P (2200G>C), E685Q (2203G>C 2205A>G), R688A (2212A>G 2213G>C 2214G>T), Q692R (2225A>G 2226A>G), L699I (2245C>A), A708S (2272G>A 2273C>G 2274T>C), S709T (2275T>A 2277C>A), C712S (2285G>C), K714E (2290A>G), K715L (2293A>C 2294A>T), A717D (2300C>A), T720R (2308A>C 2309C>G 2310A>C), E721N (2311G>A 2313G>T), M722L (2314A>C 2316G>T), R723E (2317C>G 2318G>A 2319C>G), T724Q (2320A>C 2321C>A), F726K (2326T>A 2327T>A 2328T>A), H728Y (2332C>T), N729K (2337T>G), C730A (2338T>G 2339G>C 2340T>G), I731L (2341A>C), D732Q (2344G>C 2346T>G), V735A (2354T>C 2355T>A), I739T (2366T>C 2367T>G), Q742E (2374C>G 2376G>A), F747R (2389T>A 2390T>G 2391T>A), T761R (2432C>G 2433C>G), S764A (2440T>G 2442C>A), S764_K765insG (2442_2443insGGT), K765T (2444A>C 2445A>T), E766D (2448G>C), A767K (2449G>A 2450C>A 2451C>G), D769R (2455G>A 2456A>G 2457C>A), S770N (2458T>A 2459C>A), V772I (2464G>A 2466C>T), K773I (2468A>T 2469G>C), I774E (2470A>G 2471T>A), Y776M (2476T>A 2477A>T 2478C>G), H777I (2479C>A 2480A>T 2481C>T), K778G (2482A>G 2483A>G 2484A>T), V780K (2488G>A 2489T>A), Y782K (2494T>A 2496C>G), K784I (2501A>T 2502A>C), E785Q (2503G>C 2505A>G), T786M (2507C>T 2508T>G), W788N (2512T>A 2513G>A 2514G>T), N789D (2515A>G), Y792F (2525A>T 2526T>C), A793I (2527G>A 2528C>T 2529A>C), R795K (2534G>A), I804L (2560A>T 2562C>G), M805I (2565G>A), L810T (2578T>A 2579T>C 2580G>C), F811M (2581T>A 2583T>G), G812N (2584G>A 2585G>A 2586G>T), G813A (2588G>C 2589A>C), I814V (2590A>G 2592A>T), L818I (2602C>A), A819T (2605G>A), W821L (2611T>C 2612G>T 2613G>C), Q822K (2614C>A 2616A>G), S823A (2617A>G 2618G>C 2619C>A), N824A (2620A>G 2621A>C 2622T>A), S825T (2623T>A), F826S (2627T>C 2628T>C), S828L (2632A>T 2633G>T 2634T>A), S830D (2638A>G 2639G>A), P850K (2698C>A 2699C>A 2700C>G), S851T (2701T>A), R852H (2705G>A), L853F (2707C>T 2709G>C), K854A (2710A>G 2711A>C 2712G>A), N855K (2715T>A), L856S (2717T>C 2718G>C), V858L (2722G>C 2724G>A), S861A (2731A>G 2732G>C 2733T>G), S862A (2734A>G 2735G>C 2736C>A), A863D (2738C>A), I865N (2744T>A 2745T>C), I865_T866insH (2745_2746insCAT), T866V (2746A>G 2747C>T 2748A>G), Q867D (2749C>G 2751A>C), E868R (2752G>C 2753A>G), A871E (2762C>A 2763T>A), E872Q (2764G>C 2766A>G), Q878E (2782C>G), A879S (2785G>A 2786C>G 2787A>T), K881H (2791A>C 2793A>C), M884L (2800A>T), I885V (2803A>G), L889I (2815C>A), G890A (2819G>C 2820A>T), M906L (2866A>C 2868G>C), H913R (2887C>A 2888A>G 2889C>A), V915M (2893G>A), T922S (2914A>T), P926E (2926C>G 2927C>A 2928T>G), N927H (2929A>C 2931C>T), R929I (2936G>T 2937G>T), S930C (2939C>G 2940A>T), K931N (2943G>T), Y934F (2951A>T), R937T (2959C>A 2960G>C 2961G>C), E938Q (2962G>C), K940R (2969A>G), V941L (2971G>T 2973C>G), I947V (2989A>G 2991C>G), P960S (3028C>T 3030T>A), E971D (3063G>T), F974Y (3071T>A), T975S (3073A>T 3075C>T), Q979M (3085C>A 3086A>T), C983V (3097T>G 3098G>T 3099T>G), T989K (3116C>A 3117C>G), R992K (3125G>A 3126A>G), L993K (3127C>A 3128T>A 3129G>A), G994S (3130G>T 3131G>C), F998V (3142T>G), F1001Y (3152T>A), V1011E (3182T>A 3183C>A), E1014T (3190G>A 3191A>C 3192G>A), E1016G (3197A>G 3198G>A), A1020N (3208G>A 3209C>A 3210C>T), S1021A (3211A>G 3212G>C 3213T>A), K1023A (3217A>G 3218A>C 3219A>T), Q1028V (3232C>G 3233A>T 3234A>T), S1029A (3235T>G), M1034I (3252G>T), G1036S (3256G>A 3258A>T), S1049T (3296G>C), V1056C (3316G>T 3317T>G 3318T>C), G1067D (3350G>A), I1068T (3353T>C 3354T>C), F1074Y (3371T>A), I1075V (3373A>G 3375T>G), S1076A (3376T>G), I1082V (3394A>G 3396C>G), N1089D (3415A>G 3417C>T), V1095I (3433G>A), A1097V (3440C>T), D1103E (3459C>A), I1104L (3460A>C 3462T>G), K1106E (3466A>G 3468A>G), A1107H (3469G>C 3470C>A), I1108L (3472A>C 3474T>G), R1111K (3482G>A 3483A>G), T1113E (3487A>G 3488C>A 3489C>A), T1114R (3491C>G 3492T>A), L1115F (3493C>T 3495T>C), N1119K (3507C>A), S1122A (3514T>G 3516C>T), Q1129L (3535C>T 3536A>T 3537A>G), L1131M (3541T>A), R1142K (3575G>A), K1143R (3577A>C 3578A>G 3579G>C), Q1145P (3584A>C 3585A>C), P1152D (3604C>G 3605C>A 3606T>C), E1154Q (3610G>C 3612A>G), T1161S (3631A>T 3633C>T), V1162A (3635T>C), N1163K (3639T>A), M1178E (3682A>G 3683T>A), H1185S (3703C>T 3704A>C 3705T>C), G1186N (3706G>A 3707G>A 3708T>C), D1188E (3714T>G), D1189E (3717T>G), D1190P (3718G>C 3719A>C 3720T>A), S1195A (3733T>G), T1198G (3742A>G 3743C>G), S1201A (3751A>G 3752G>C 3753T>A), L1203I (3757C>A 3759G>T), T1206V (3766A>G 3767C>T 3768A>G), M1207L (3769A>C 3771G>T), N1209M (3776A>T 3777C>G), K1211R (3781A>C 3782A>G), Q1218A (3802C>G 3803A>C 3804G>A), E1219I (3805G>A 3806A>T 3807G>C), D1221N (3811G>A), C1232F (3845G>T), Y1235F (3854A>T), G1237A (3860G>C 3861C>A), M1251L (3901A>T), M1253L (3907A>C 3909G>T), M1255S (3913A>T 3914T>C 3915G>T), V1262E (3935T>A), P1263A (3937C>G 3939A>C), T1264Q (3940A>C 3941C>A), G1265P (3943G>C 3944G>C), E1266N (3946G>A 3948A>C), G1267S (3949G>T 3950G>C 3951T>G), D1268Q (3952G>C 3954T>G), K1269R (3956A>G 3957A>G), E1271L (3961G>C 3962A>T 3963G>T), R1273K (3968G>A), D1275N (3973G>A), E1280D (3990A>T), I1287A (4009A>G 4010T>C 4011T>G), I1289V (4015A>G 4017T>A), I1293T (4028T>C), H1294F (4030C>T 4031A>T), I1298V (4042A>G), T1343C (4177A>T 4178C>G 4179C>T), L1347A (4189T>G 4190T>C), I1351L (4201A>C), D1358N (4222G>A), S1362H (4234T>C 4235C>A), S1365T (4243T>A 4245T>A), L1366V (4246C>G 4248T>G), N1369S (4256A>G 4257C>T), N1370S (4259A>G), A1373S (4267G>T 4269T>A), T1377E (4279A>G 4280C>A 4281T>G), V1392I (4324G>A 4326G>C), E1404A (4361A>C 4362G>T), Y1405F (4364A>T), D1407G (4370A>G), T1408D (4372A>G 4373C>A 4374C>T), K1409E (4375A>G), E1412S (4384G>A 4385A>G 4386G>T), A1414E (4391C>A 4392T>G), A1420T (4408G>A 4410T>A), D1434N (4450G>A 4452T>C), V1436L (4456G>T 4458T>G), I1475V (4573A>G 4575T>C), A1476K (4576G>A 4577C>A 4578C>A), R1478K (4583G>A), A1479S (4585G>T), I1480V (4588A>G 4590A>G), V1482I (4594G>A 4596A>T), G1485N (4603G>A 4604G>A), L1505M (4663C>A), S1511D (4681T>G 4682C>A 4683A>T), N1514S (4690A>T 4691A>C 4692T>C), Q1517R (4699C>A 4700A>G 4701G>A), D1520E (4710C>A), M1521A (4711A>G 4712T>C 4713G>T), N1537E (4759A>G 4761C>G), Q1538K (4762C>A), V1539T (4765G>A 4766T>C 4767G>A), Q1540K (4768C>A 4770G>A), Q1542L (4775A>T), H1562Q (4836C>A), I1571V (4861A>G 4863A>C), M1575I (4875G>T), P1582V (4894C>G 4895C>T), T1587V (4909A>G 4910C>T 4911T>A), V1588Q (4912G>C 4913T>A 4914G>A), M1591Y (4921A>T 4922T>A 4923G>T), V1594I (4930G>A 4932G>T), L1598I (4942C>A 4944T>C), R1606Q (4966A>C 4967G>A 4968A>G), V1607A (4970T>C 4971T>C), A1608T (4972G>A 4974A>T), K1611S (4982A>G 4983G>T), L1612F (4986A>T), T1613V (4987A>G 4988C>T 4989T>A), Y1614N (4990T>A), Q1617F (4999C>T 5000A>T 5001G>T), C1618G (5002T>G 5004T>A), R1621K (5012G>A 5013A>G), E1622Q (5014G>C), G1624N (5020G>A 5021G>A 5022A>T), G1626P (5026G>C 5027G>C), A1627I (5029G>A 5030C>T 5031A>T), T1628M (5033C>T 5034A>G), F1630M (5038T>A 5040T>G), S1632T (5044T>A), M1634Y (5050A>T 5051T>A 5052G>T), V1636L (5056G>C 5058G>T), Q1638S (5062C>A 5063A>G 5064G>T), E1639Q (5065G>C), I1641L (5071A>T 5073C>G), A1642T (5074G>A), K1643E (5077A>G), A1648E (5093C>A 5094A>G), A1650I (5098G>A 5099C>T 5100G>T), L1651G (5101T>G 5102T>G), T1652V (5104A>G 5105C>T 5106A>T), I1653L (5107A>T 5109C>G), C1654M (5110T>A 5111G>T 5112T>G), M1656L (5116A>C 5118G>A), A1657S (5119G>T 5121C>A), K1658D (5122A>G 5124A>T), S1660K (5128T>A 5129C>A 5130T>A), I1661N (5132T>A 5133C>T), F1663Y (5138T>A), P1668A (5152C>G), N1670E (5158A>G 5160T>A), V1672I (5164G>A), E1673del (5167_5169delGAA), R1677E (5179A>G 5180G>A 5181A>G), D1678G (5183A>G), L1679V (5185C>G), N1680H (5188A>C), W1681E (5191T>G 5192G>A 5193G>A), A1683V (5198C>T 5199C>G), N1685D (5203A>G 5205C>T), I1686V (5206A>G 5208T>A), G1687F (5209G>T 5210G>T), A1695S (5233G>T 5235C>A), K1696N (5238G>C), T1697Q (5239A>C 5240C>A 5241T>A), E1698Q (5242G>C), E1700S (5248G>T 5249A>C 5250G>A), P1701S (5251C>T 5253T>A), C1703N (5257T>A 5258G>A), V1705L (5263G>C 5265A>G), K1712R (5284A>C 5285A>G 5286G>C), K1713R (5288A>G 5289G>A), S1714T (5291G>C 5292C>A), K1715T (5294A>C), S1722N (5315G>A), M1730I (5340G>A), M1735L (5353A>T), S1736A (5356T>G), L1739Q (5366T>A), A1741H (5371G>C 5372C>A), G1744K (5380G>A 5381G>A 5382C>G), K1745S (5384A>G 5385A>T), M1746L (5386A>C), S1747E (5389T>G 5390C>A), E1748V (5393A>T 5394G>T), D1749G (5396A>G 5397C>T), I1750D (5398A>G 5399T>A), A1751G (5402C>G 5403T>G), D1754S (5410G>T 5411A>C), V1756T (5416G>A 5417T>C 5418G>C), V1765I (5443G>A 5445G>A), S1766A (5446A>G 5447G>C 5448C>T), M1773F (5467A>T 5469G>T), V1775L (5473G>T), R1778H (5483G>A), C1781R (5491T>A 5493C>G), I1782A (5494A>G 5495T>C 5496T>C), E1783T (5497G>A 5498A>C 5499G>A), I1785L (5503A>C 5505C>T), I1792M (5526T>G), G1793A (5528G>C 5529C>T), N1794G (5530A>G 5531A>G 5532T>G), V1806A (5567T>C 5568T>G), F1809L (5577T>G), Q1811K (5581C>A), P1814E (5590C>G 5591C>A 5592T>A), Q1815D (5593C>G 5595G>C), Q1815_E1816insT (5595_5596insACC), E1816S (5596G>T 5597A>C), G1817H (5599G>C 5600G>A 5601G>C), M1819L (5605A>T), A1820G (5609C>G), V1822S (5614G>A 5615T>G), S1824T (5620T>A 5622G>A), S1825del (5623_5625delTCT), L1826Y (5627T>A 5628G>T), Q1827G (5629C>G 5630A>G 5631A>C), S1828T (5633G>C 5634T>C), S1829T (5636G>C), M1831N (5642T>A 5643G>C), Q1833H (5649A>T), Q1833_G1834insHS (5649_5650insCACAGT), G1834A (5651G>C 5652T>A), E1836S (5656G>T 5657A>C 5658G>A), L1837V (5659C>G), Q1839L (5666A>T 5667A>T), I1840L (5668A>C 5670T>A), V1842I (5674G>A 5676C>A), I1843V (5677A>G 5679C>T), H1844P (5681A>C 5682C>T), G1845S (5683G>T 5684G>C 5685T>C), M1849E (5695A>G 5696T>A 5697G>A), E1850K (5698G>A 5700A>G), L1853V (5707T>G 5709G>T), M1855L (5713A>T), V1858T (5722G>A 5723T>C 5724T>A), V1859I (5725G>A 5727G>T), H1862E (5734C>G 5736T>A), G1866A (5747G>C), N1867Q (5749A>C 5751C>G), F1868C (5753T>G 5754T>C), T1869S (5755A>T 5757C>A), D1874K (5770G>A 5772T>G), Q1875H (5775A>T), L1876A (5776T>G 5777T>C 5778G>T), F1877L (5781T>G), A1878E (5783C>A 5784A>G), S1879T (5786G>C 5787T>A), E1881S (5791G>T 5792A>C 5793A>T), K1882V (5794A>G 5795A>T 5796G>C), C1883S (5797T>A), E1886T (5806G>A 5807A>C 5808A>T), L1888A (5812C>G 5813T>C 5814T>C), K1890R (5819A>G 5820G>A), N1891G (5821A>G 5822A>G 5823C>A), T1892V (5824A>G 5825C>T 5826G>C), L1894V (5830C>G), E1899K (5845G>A 5847G>A), K1900E (5848A>G), M1901F (5851A>T 5853G>C), T1902S (5854A>T 5856T>C), K1904H (5860A>C 5862A>C), V1906C (5866G>T 5867T>G 5868G>T), A1907E (5870C>A), Q1908M (5872C>A 5873A>T 5874A>G), V1910L (5878G>C 5880C>T), S1911E (5881A>G 5882G>A 5883T>G), V1913L (5887G>C 5889A>G), K1914E (5890A>G), Q1915S (5893C>A 5894A>G 5895G>T), F1918V (5902T>G 5904C>A), Y1919V (5905T>G 5906A>T 5907T>A), P1920F (5908C>T 5909C>T), K1921M (5912A>T), A1922G (5915C>G 5916C>G), T1923S (5918C>G 5919A>C), T1924V (5920A>G 5921C>T 5922C>T), V1925K (5923G>A 5924T>A), E1926K (5926G>A), L1928M (5932C>A 5934C>G), K1929L (5935A>T 5936A>T), V1930S (5938G>T 5939T>C 5940A>C), A1932M (5944G>A 5945C>T 5946T>G), D1933N (5947G>A), A1934Q (5950G>C 5951C>A 5952T>A), H1935Q (5955T>G), H1935_T1936insM (5955_5956insATG), T1936S (5956A>T 5958C>A), I1937K (5960T>A 5961T>A), V1938T (5962G>A 5963T>C), K1939Q (5965A>C), T1940V (5968A>G 5969C>T), K1942S (5975A>G 5976A>C), F1944C (5981T>G 5982C>T), R1945C (5983C>T 5985G>T), G1946del (5986_5988delGGA), V1948T (5992G>A 5993T>C 5994T>A), R1950S (5998C>A 6000G>T), N1952A (6004A>G 6005A>C 6006T>A), F1953W (6008T>G 6009T>G), S1955T (6014G>C), P1956K (6016C>A 6017C>A 6018T>G), E1957D (6021G>C), W1958I (6022T>A 6023G>T 6024G>T), W1958_K1959insV (6024_6025insGTC), K1959M (6026A>T 6027A>G), I1960T (6029T>C 6030T>A), G1961K (6031G>A 6032G>A), E1963K (6037G>A), N1964K (6042C>G), T1965P (6043A>C), M1966del (6046_6048delATG), A1967G (6050C>G), T1968S (6052A>T 6054A>T), L1969I (6055C>A 6057C>T), S1970P (6058T>C 6060A>T), S1973T (6068G>C 6069T>A), L1974V (6070C>G 6072T>G), L1975F (6073C>T), P1976L (6077C>T 6078A>T), L1979P (6086T>C 6087G>T), S1982L (6094T>C 6095C>T), A1983V (6098C>T 6099C>T), M1984V (6100A>G), K1985R (6104A>G 6105G>A), L1987M (6109T>A), S1989R (6115A>C), R1990K (6119G>A) | | | | | | | | |

| | Begin | End | Coverage | Score | Concordance | Matches | Identities | I/D/M/F* | Stop Codons |
|---|---|---|---|---|---|---|---|---|---|
| **NT** | **33** | **6128** | **84.2%** | **3374** | **28.2%** | **6043 (98.8%)** | **3915 (64.0%)** | **21/53** | |

GCT2GCA (156T>A), CTT3TTT (157C>T), AAA5AAG (165A>G), TTT7TGG (170T>G 171T>G), GGT8AGT (172G>A), TCT9CCT (175T>C), TCA10CCT (178T>C 180A>T), TCT11GCT (181T>G), GAG12_GTT17del (184_201delGAGAAGACTGGATCAGTT), CCC18TCT (202C>T 204C>T), TCT19TCA (207T>A), ACT20CCT (208A>C), TCT21TCA (213T>A), GGT24GCA (221G>C 222T>A), CTG26GTT (226C>G 228G>T), GGT28TCT (232G>T 233G>C), ATC30TTT (238A>T 240C>T), TCA31TCT (243A>T), AAA32TCT (244A>T 245A>C 246A>T), TTC33CTT (247T>C 249C>T), AAA35AAG (255A>G), GCT36ACT (256G>A), ATT37CTC (259A>C 261T>C), AGC38TCC (262A>T 263G>C), AAC39AAT (267C>T), ATC40GTT (268A>G 270C>T), ACA41AAT (272C>A 273A>T), TCA42AAA (274T>A 275C>A), GTA43GTT (279A>T), AAA46ACC (287A>C 288A>C), TCT48GTA (292T>G 293C>T 294T>A), CAT50CAC (300T>C), TTA51ATA (301T>A), GAT52GAG (306T>G), TTG54GTC (310T>G 312G>C), AAA55AAG (315A>G), CCA56CCG (318A>G), TCT57CCA (319T>C 321T>A), GTG58GTT (324G>T), ACT59ATG (326C>T 327T>G), GAT60GCA (329A>C 330T>A), TCT62GCA (334T>G 336T>A), TCT63ACA (337T>A 339T>A), TCT64TCA (342T>A), GTT66GCC (347T>C 348T>C), TGT69TGC (357T>C), TCC70AAT (358T>A 359C>A 360C>T), GCA71GCT (363A>T), TTG72GTA (364T>G 366G>A), AAT73GAA (367A>G 369T>A), AAA74AGA (371A>G), GTG75GTT (375G>T), CTT76TTA (376C>T 378T>A), ATA79ATT (387A>T), GGT80GCA (389G>C 390T>A), TCC81AAA (391T>A 392C>A 393C>A), TTG82ATA (394T>A 396G>A), ATT83ATG (399T>G), CCT85CCC (405T>C), TTT86TTC (408T>C), CTG87ATC (409C>A 411G>C), AAA88AAG (414A>G), GCA89GGT (416C>G 417A>T), TAC90TAT (420C>T), TCT91GAT (421T>G 422C>A), TTG93GTA (427T>G 429G>A), GCA94AAA (430G>A 431C>A), ACC95ACT (435C>T), TCT99ACT (445T>A), ATC100ATA (450C>A), AAG101GTA (451A>G 452A>T 453G>A), ATG103CTT (457A>C 459G>T), GTT104GTA (462T>A), TTA105CTT (463T>C 465A>T), CGG106AAG (466C>A 467G>A), TTG107ATG (469T>A), TTC108GTT (472T>G 474C>T), GAA109TTA (475G>T 476A>T), TTG111GTC (481T>G 483G>C), TCC112AAG (484T>A 485C>A 486C>G), AAT113GAT (487A>G), GTT115TTC (493G>T 495T>C), AGA116ATC (497G>T 498A>C), ATA117CTA (499A>C), TTT119TAC (506T>A 507T>C), CAG120GAA (508C>G 510G>A), GTT122TTA (514G>T 516T>A), GAG124GAA (522G>A), AAG125AAA (525G>A), AGT126GTT (526A>G 527G>T), GAT128TCA (532G>T 533A>C 534T>A), GTA129TTA (535G>T), ATT131ATA (543T>A), GTA132GTG (546A>G), GTT133TTT (547G>T), TAT134ATA (550T>A 551A>T 552T>A), GGT135GCT (554G>C), TTG137TTA (561G>A), GCA140GCT (570A>T), ATA141TTG (571A>T 573A>G), ACT142GCT (574A>G), CTT143ATT (577C>A), CTT144TTG (580C>T 582T>G), GTT145CTA (583G>C 585T>A), TTG146CTA (586T>C 588G>A), TTG147CTG (589T>C), TGT148TAT (593G>A), GTT149GTA (597T>A), TCA152GTA (604T>G 605C>T), GAC153GAT (609C>T), ATA154ATT (612A>T), ATT155GTA (613A>G 615T>A), GAA156ACG (616G>A 617A>C 618A>G), GGA157CCT (619G>C 620G>C 621A>T), GTA158TTT (622G>T 624A>T), TCT159AAT (625T>A 626C>A), AGT160AAT (629G>A), ACC161GTA (631A>G 632C>T 633C>A), ATT162GTT (634A>G), AAA163GTT (637A>G 638A>T 639A>T), AAT164ACT (641A>C), ATT165TTG (643A>T 645T>G), TTC166TTT (648C>T), TTA167TCA (650T>C), ATA168GTT (652A>G 654A>T), GTA169GTT (657A>T), GGG170GGA (660G>A), ATG172TTT (664A>T 666G>T), TTT173TTC (669T>C), AGT175TCA (673A>T 674G>C 675T>A), AAC178AAT (684C>T), TTG179TTA (687G>A), GAC180GAT (690C>T), CCC183CCA (699C>A), AAG184ACA (701A>C 702G>A), GCT186TCT (706G>T), GAG187GGG (710A>G), TTC189TTT (717C>T), ACA190TCC (718A>T 720A>C), ATG191CAG (721A>C 722T>A), GTA192CTG (724G>C 726A>G), GCA193GCG (729A>G), AGT196GCA (736A>G 737G>C 738T>A), ATG197GTA (739A>G 741G>A), CTA198CTT (744A>T), GAG201GAA (753G>A), TCT202AGC (754T>A 755C>G 756T>C), GTC203TTG (757G>T 759C>G), TCA204ACG (760T>A 762A>G), CAC205CAT (765C>T), ACT206ACA (768T>A), ACA208ACT (774A>T), TCC209AGC (775T>A 776C>G), CAG210CAA (780G>A), TTG211CTT (781T>C 783G>T), ATG212TTA (784A>T 786G>A), TCA213TCC (789A>C), ACT214TCA (790A>T 792T>A), ATT215TTG (793A>T 795T>G), ATG216TTG (796A>T), GCA217GCT (801A>T), ATT220ATC (810T>C), TCC221TCA (813C>A), ACG222ACT (816G>T), CTC223TTA (817C>T 819C>A), GTG224GTA (822G>A), ATA226GTT (826A>G 828A>T), GCA227GCT (831A>T), GTG228GTT (834G>T), CCA229CCT (837A>T), GGA230GGG (840A>G), CCC232CCA (846C>A), TTG235CTT (853T>C 855G>T), AGC236TCA (856A>T 857G>C 858C>A), AAT237AAC (861T>C), CCC238CCA (864C>A), CTC239CTG (867C>G), ATT242ATC (876T>C), TCA245TCT (885A>T), ACT246GCT (886A>G), GGG247GGA (891G>A), GGG249GGT (897G>T), GCT250GCG (900T>G), CAA255CAG (915A>G), TTG256CTG (916T>C), TTT257TTC (921T>C), ACA258CAG (922A>C 923C>A 924A>G), AGA261AGG (933A>G), AAT262AAC (936T>C), AAA264AAG (942A>G), TGC266TGT (948C>T), ACT267ACA (951T>A), TCA268TCT (954A>T), GCA270GCC (960A>C), TTT271TTC (963T>C), TCT272TCA (966T>A), GTG274GTT (972G>T), GAA276GAG (978A>G), ATT277ATA (981T>A), ATA278GTG (982A>G 984A>G), GGT280AGC (988G>A 990T>C), GCT281ATT (991G>A 992C>T), TTC284TTT (1002C>T), AAG285AAA (1005G>A), GTG288GTA (1014G>A), CTT289TTG (1015C>T 1017T>G), TCT290TCA (1020T>A), GCA291GCG (1023A>G), ATT292ATA (1026T>A), AGC293AAT (1028G>A 1029C>T), GCA294GCT (1032A>T), ACA295ACC (1035A>C), TTG296TTA (1038G>A), TCT297TCC (1041T>C), ACT298ACC (1044T>C), TTG300CTT (1048T>C 1050G>T), GAA302CAG (1054G>C 1056A>G), CAG305ACG (1063C>A 1064A>C), GTG307GTT (1071G>T), GAT308TCG (1072G>T 1073A>C 1074T>G), GCT309GAG (1076C>A 1077T>G), TGT311GCT (1081T>G 1082G>C), GAT312GAA (1086T>A), GCT314TCT (1090G>T), ACC315ATG (1094C>T 1095C>G), AGG316CGC (1096A>C 1098G>C), TTG317CTT (1099T>C 1101G>T), GAG318GAA (1104G>A), AAC319AAT (1107C>T), GCC321GCA (1113C>A), AAA323CCT (1117A>C 1118A>C 1119A>T), GCC324GCA (1122C>A), TTC325TTT (1125C>T), CCT326CCA (1128T>A), GAC327ATT (1129G>A 1130A>T 1131C>T), AAA328AAG (1134A>G), AAT330CAA (1138A>C 1140T>A), CAC331CAT (1143C>T), CTT332TTA (1144C>T 1146T>A), AGG333AAA (1148G>A 1149G>A), GCA336GCC (1158A>C), CTG337GTC (1159C>G 1161G>C), AAG338AAA (1164G>A), ATT339ATA (1167T>A), GCA341GCT (1173A>T), ACA345ACC (1185A>C), CAC346CAT (1188C>T), GTT348GTC (1194T>C), GCC349GCA (1197C>A), ATG352TTG (1204A>T), CAT354CAC (1212T>C), CGA355AGG (1213C>A 1215A>G), GTG356GTT (1218G>T), AGT357TCT (1219A>T 1220G>C), GCA358GCT (1224A>T), GCT359GCC (1227T>C), ATC360ATA (1230C>A), ACT361ACG (1233T>G), CTT363CTA (1239T>A), GAA364GAT (1242A>T), AAG365AAA (1245G>A), GTC366CTG (1246G>C 1248C>G), TAT367GTT (1249T>G 1250A>T), GCG368GCT (1254G>T), AGT370GCC (1258A>G 1259G>C 1260T>C), TGC371AGC (1261T>A), AGA372CAA (1264A>C 1265G>A), GTT374TCT (1270G>T 1271T>C), GTT376GTG (1278T>G), GGT377GGC (1281T>C), CAG378CAA (1284G>A), AGA380AGG (1290A>G), GAA382GAG (1296A>G), CCC383CCT (1299C>T), GTT386GTG (1308T>G), TAT389TTT (1316A>T), GGT390GGA (1320T>A), GCC391GCT (1323C>T), GGG393GGA (1329G>A), ACA398ACG (1344A>G), GCA399TGT (1345G>T 1346C>G 1347A>T), CGA401CGC (1353A>C), TTG402CTT (1354T>C 1356G>T), TTG408CTC (1372T>C 1374G>C), GAT409GAC (1377T>C), CGC410GCA (1378C>G 1379G>C 1380C>A), GGG412GGT (1386G>T), GAG413GAA (1389G>A), CCT414CCA (1392T>A), CTT416CTG (1398T>G), AGA418CGC (1402A>C 1404A>C), CTT419TTG (1405C>T 1407T>G), GCT421GCG (1413T>G), GTT422GTA (1416T>A), AGC423AGT (1419C>T), AAG424AAA (1422G>A), CGC425AGA (1423C>A 1425C>A), GCC427GCA (1431C>A), TCT430TCC (1440T>C), AAT431AAC (1443T>C), GCC433GCA (1449C>A), CAC434CAT (1452C>T), CAG435CAA (1455G>A), GCT437GCA (1461T>A), TTG439CTT (1465T>C 1467G>T), GAC442GAT (1476C>T), GGA444GGT (1482A>T), GCA445GCT (1485A>T), TTG446CTT (1486T>C 1488G>T), AGA447CGG (1489A>C 1491A>G), GGA451GGG (1503A>G), CAA452CAG (1506A>G), CAA454CAG (1512A>G), ATT456ATA (1518T>A), ATT462ATC (1536T>C), TCC464TCT (1542C>T), ACA465ACC (1545A>C), CCA467CCT (1551A>T), CCA469CCT (1557A>T), CTG470CTT (1560G>T), CCC471CCA (1563C>A), GCA473GCT (1569A>T), GCT474GCA (1572T>A), GTG475GTC (1575G>C), GAA476GAG (1578A>G), GAC477GAT (1581C>T), AAG478AAA (1584G>A), GGG479GGA (1587G>A), CAT481CAC (1593T>C), ACT483ACA (1599T>A), TAT486TAC (1608T>C), GCA489GCC (1617A>C), AAT492AAC (1626T>C), CTT493TTG (1627C>T 1629T>G), GTA494ATA (1630G>A), TCA495TCT (1635A>T), GCA496GCT (1638A>T), CCT497CCA (1641T>A), GCC498GCT (1644C>T), CTT502CTA (1656T>A), ACC503ACG (1659C>G), GCT507GCA (1671T>A), GAG509GAA (1677G>A), CGT510AGG (1678C>A 1680T>G), AGA511AGG (1683A>G), AGG512AGA (1686G>A), ATC514ATT (1692C>T), TGT518TGC (1704T>C), ATC519AGA (1706T>G 1707C>A), GAA521GTG (1712A>T 1713A>G), GGT522GGC (1716T>C), GAA523GAT (1719A>T), TCT525GAT (1723T>G 1724C>A), ACA526CCA (1726A>C), GAG527AAT (1729G>A 1731G>T), GAC528GCT (1733A>C 1734C>T), ACC530ACA (1740C>A), GGA531GCC (1742G>C 1743A>C), CTG533CTA (1749G>A), ATG534GTT (1750A>G 1752G>T), AAC536GAT (1756A>G 1758C>T), ATT537GTG (1759A>G 1761T>G), CGG541AGA (1771C>A 1773G>A), AGG542AGA (1776G>A), ATA546ATC (1788A>C), CAC550GCT (1798C>G 1799A>C 1800C>T), AGT552TCC (1804A>T 1805G>C 1806T>C), TAC553TAT (1809C>T), TCG554GAT (1810T>G 1811C>A 1812G>T), TTA557TTG (1821A>G), GAC558CAA (1822G>C 1824C>A), GCC562GCA (1836C>A), TGT564TGC (1842T>C), CAG565CGG (1844A>G), GTT566GTG (1848T>G), CAT567CAC (1851T>C), GCT568TAT (1852G>T 1853C>A), ATA570GTG (1858A>G 1860A>G), AAG572AGC (1865A>G 1866G>C), CAG573CAA (1869G>A), CTG574CTT (1872G>T), TTT575TAC (1874T>A 1875T>C), GAA576AAT (1876G>A 1878A>T), GCA577ACT (1879G>A 1881A>T), GAA578ACA (1882G>A 1883A>C), ACT579TCG (1885A>T 1887T>G), GGC580GGA (1890C>A), ATT581TTG (1891A>T 1893T>G), TCA582GCT (1894T>G 1896A>T), AAT583GAT (1897A>G), ACA585GTT (1903A>G 1904C>T 1905A>T), GCC586GCT (1908C>T), GTT588GTC (1914T>C), GCT589AAT (1915G>A 1916C>A), GTA590GTT (1920A>T), TCT591GCT (1921T>G), TCT592GTT (1924T>G 1925C>T), GAA593GAT (1929A>T), GAG594GAA (1932G>A), GTT595GTG (1935T>G), GTG596CTG (1936G>C), GCA597GAT (1940C>A 1941A>T), GTT599TTC (1945G>T 1947T>C), GAT600CAA (1948G>C 1950T>A), GGG601GAT (1952G>A 1953G>T), GCT602GCC (1956T>C), CGC603del (1957_1959delCGC), TTC604TTA (1962C>A), AAA605CAA (1963A>C), ACA606GTT (1966A>G 1967C>T 1968A>T), AAA607TTG (1969A>T 1970A>T 1971A>G), CAA608CCT (1973A>C 1974A>T), GAT609CCT (1975G>C 1976A>C), GTG610GAG (1979T>A), ATA612GCA (1984A>G 1985T>C), GTG613GCT (1988T>C 1989G>T), ACT614TAT (1990A>T 1991C>A), CCC615GAG (1993C>G 1994C>A 1995C>G), ACA616CAG (1996A>C 1997C>A 1998A>G), GTT617ACA (1999G>A 2000T>C 2001T>A), ATA618ATT (2004A>T), GAG620GAA (2010G>A), GAA621GCT (2012A>C 2013A>T), GAT622GAA (2016T>A), AGA623AAA (2018G>A), GTG624ATT (2020G>A 2022G>T), ATT625GTT (2023A>G), TCT627TCC (2031T>C), AAA628CAG (2032A>C 2034A>G), TTA630CTC (2038T>C 2040A>C), GTG632GTT (2046G>T), GAA633GAG (2049A>G), CTT635CTC (2055T>C), TTT637TAT (2060T>A), TAT639TAC (2067T>C), CAG640CAA (2070G>A), GGC641GGG (2073C>G), TTG643ATG (2077T>A), GAC644AAC (2080G>A), CCG645AAA (2083C>A 2084C>A 2085G>A), GAG646TTT (2086G>T 2087A>T 2088G>T), AGT647GAG (2089A>G 2090G>A 2091T>G), TTG648TTT (2094G>T), TTT649TTC (2097T>C), CCT650CCA (2100T>A), TAT653CTC (2107T>C 2108A>T 2109T>C), CAA654TCC (2110C>T 2111A>C 2112A>C), AAA655AAG (2115A>G), CAA656CAG (2118A>G), CTC658ATC (2122C>A), TTT659TAT (2126T>A), TCC661GTG (2131T>G 2132C>T 2133C>G), GAT663TCT (2137G>T 2138A>C), GCA664TCT (2140G>T 2142A>T), GAG665GAT (2145G>T), CAC666GCC (2146C>G 2147A>C), CAG667AAA (2149C>A 2151G>A), CAA668CAG (2154A>G), GTC669AAT (2155G>A 2156T>A 2157C>T), TTC670TTT (2160C>T), ATG671GAA (2161A>G 2162T>A 2163G>A), AGA672ATG (2165G>T 2166A>G), ATG674GTA (2170A>G 2172G>A), ACT675AAT (2174C>A), GAC676GAT (2178C>T), ATG677TTG (2179A>T), ATC678TTG (2182A>T 2184C>G), TAC679GTC (2185T>G 2186A>T), GGT681GGA (2193T>A), GCC682GCA (2196C>A), AAT683AGT (2198A>G), GCT684CCT (2200G>C), GAA685CAG (2203G>C 2205A>G), CAA686CAG (2208A>G), TAC687TAT (2211C>T), AGG688GCT (2212A>G 2213G>C 2214G>T), CAA692CGG (2225A>G 2226A>G), AAT693AAC (2229T>C), CTC699ATC (2245C>A), TTT702TTC (2256T>C), AAA703AAG (2259A>G), AGC704AGT (2262C>T), ATC706ATT (2268C>T), GCT708AGC (2272G>A 2273C>G 2274T>C), TCC709ACA (2275T>A 2277C>A), ATC711ATT (2283C>T), TGT712TCT (2285G>C), AAG714GAG (2290A>G), AAG715CTG (2293A>C 2294A>T), GCT717GAT (2300C>A), GTA718GTC (2304A>C), ACA720CGC (2308A>C 2309C>G 2310A>C), GAG721AAT (2311G>A 2313G>T), ATG722CTT (2314A>C 2316G>T), CGC723GAG (2317C>G 2318G>A 2319C>G), ACA724CAA (2320A>C 2321C>A), GGA725GGG (2325A>G), TTT726AAA (2326T>A 2327T>A 2328T>A), GCA727GCG (2331A>G), CAT728TAT (2332C>T), AAT729AAG (2337T>G), TGT730GCG (2338T>G 2339G>C 2340T>G), ATT731CTT (2341A>C), GAT732CAG (2344G>C 2346T>G), GTT735GCA (2354T>C 2355T>A), TTG738CTT (2362T>C 2364G>T), ATT739ACG (2366T>C 2367T>G), TGC740TGT (2370C>T), ATC741ATT (2373C>T), CAG742GAA (2374C>G 2376G>A), TTT747AGA (2389T>A 2390T>G 2391T>A), GTG748GTA (2394G>A), TTT750TTC (2400T>C), TTG751CTG (2401T>C), ACA753ACT (2409A>T), GCC754GCT (2412C>T), CGT757CGC (2421T>C), TAT758TAC (2424T>C), ACC761AGG (2432C>G 2433C>G), TCC764GCA (2440T>G 2442C>A), TCC764_AAA765insGGT (2442_2443insGGT), AAA765ACT (2444A>C 2445A>T), GAG766GAC (2448G>C), GCC767AAG (2449G>A 2450C>A 2451C>G), AAA768AAG (2454A>G), GAC769AGA (2455G>A 2456A>G 2457C>A), TCT770AAT (2458T>A 2459C>A), GTC772ATT (2464G>A 2466C>T), AAG773ATC (2468A>T 2469G>C), ATA774GAA (2470A>G 2471T>A), TGT775TGC (2475T>C), TAC776ATG (2476T>A 2477A>T 2478C>G), CAC777ATT (2479C>A 2480A>T 2481C>T), AAA778GGT (2482A>G 2483A>G 2484A>T), ATT779ATC (2487T>C), GTG780AAG (2488G>A 2489T>A), GAT781GAC (2493T>C), TAC782AAG (2494T>A 2496C>G), AAA784ATC (2501A>T 2502A>C), GAA785CAG (2503G>C 2505A>G), ACT786ATG (2507C>T 2508T>G), TGG788AAT (2512T>A 2513G>A 2514G>T), AAC789GAC (2515A>G), TTA790TTG (2520A>G), CCA791CCC (2523A>C), TAT792TTC (2525A>T 2526T>C), GCA793ATC (2527G>A 2528C>T 2529A>C), CTG794TTG (2530C>T), AGG795AAG (2534G>A), GGC801GGA (2553C>A), CTA802CTG (2556A>G), ATC804TTG (2560A>T 2562C>G), ATG805ATA (2565G>A), GCG806GCA (2568G>A), GGA809GGC (2577A>C), TTG810ACC (2578T>A 2579T>C 2580G>C), TTT811ATG (2581T>A 2583T>G), GGG812AAT (2584G>A 2585G>A 2586G>T), GGA813GCC (2588G>C 2589A>C), ATA814GTT (2590A>G 2592A>T), ACA815ACT (2595A>T), TTT817TTC (2601T>C), CTT818ATT (2602C>A), GCT819ACT (2605G>A), TGT820TGC (2610T>C), TGG821CTC (2611T>C 2612G>T 2613G>C), CAA822AAG (2614C>A 2616A>G), AGC823GCA (2617A>G 2618G>C 2619C>A), AAT824GCA (2620A>G 2621A>C 2622T>A), TCT825ACT (2623T>A), TTT826TCC (2627T>C 2628T>C), CCA827CCT (2631A>T), AGT828TTA (2632A>T 2633G>T 2634T>A), AGT830GAT (2638A>G 2639G>A), GGA831GGC (2643A>C), GGT833GGG (2649T>G), GGT834GGC (2652T>C), GTC835GTA (2655C>A), ACA836ACC (2658A>C), AGC841AGT (2673C>T), TCT843TCC (2679T>C), AAA845AAG (2685A>G), AAG846AAA (2688G>A), AAC847AAT (2691C>T), AAG848AAA (2694G>A), GGC849GGG (2697C>G), CCC850AAG (2698C>A 2699C>A 2700C>G), TCT851ACT (2701T>A), CGT852CAT (2705G>A), CTG853TTC (2707C>T 2709G>C), AAG854GCA (2710A>G 2711A>C 2712G>A), AAT855AAA (2715T>A), TTG856TCC (2717T>C 2718G>C), GTG858CTA (2722G>C 2724G>A), CAG860CAA (2730G>A), AGT861GCG (2731A>G 2732G>C 2733T>G), AGC862GCA (2734A>G 2735G>C 2736C>A), GCT863GAT (2738C>A), ATT865AAC (2744T>A 2745T>C), ATT865_ACA866insCAT (2745_2746insCAT), ACA866GTG (2746A>G 2747C>T 2748A>G), CAA867GAC (2749C>G 2751A>C), GAA868CGA (2752G>C 2753A>G), GCT871GAA (2762C>A 2763T>A), GAA872CAG (2764G>C 2766A>G), GGG874GGA (2772G>A), GTC876GTT (2778C>T), AAC877AAT (2781C>T), CAA878GAA (2782C>G), GCA879AGT (2785G>A 2786C>G 2787A>T), CTC880CTG (2790C>G), AAA881CAC (2791A>C 2793A>C), AAG882AAA (2796G>A), AAT883AAC (2799T>C), ATG884TTG (2800A>T), ATT885GTT (2803A>G), GTA886GTT (2808A>T), CTG887TTG (2809C>T), CGG888AGA (2812C>A 2814G>A), CTT889ATT (2815C>A), GGA890GCT (2819G>C 2820A>T), GAG891GAA (2823G>A), GTG893GTC (2829G>C), TAC894TAT (2832C>T), ACA898ACG (2844A>G), GTG900GTT (2850G>T), TCA902TCT (2856A>T), GGG903GGT (2859G>T), ATA905ATT (2865A>T), ATG906CTC (2866A>C 2868G>C), GCA911GCC (2883A>C), TTC912TTT (2886C>T), CAC913AGA (2887C>A 2888A>G 2889C>A), AAT914AAC (2892T>C), GTG915ATG (2893G>A), AGA916AGG (2898A>G), GGA918GGC (2904A>C), ACA919ACT (2907A>T), ACC922TCC (2914A>T), ATT923ATC (2919T>C), ATC924ATT (2922C>T), CCT926GAG (2926C>G 2927C>A 2928T>G), AAC927CAT (2929A>C 2931C>T), TCA928TCC (2934A>C), AGG929ATT (2936G>T 2937G>T), TCA930TGT (2939C>G 2940A>T), AAG931AAT (2943G>T), TAC934TTC (2951A>T), AAC935AAT (2955C>T), CGG937ACC (2959C>A 2960G>C 2961G>C), GAA938CAA (2962G>C), TCC939TCA (2967C>A), AAG940AGG (2969A>G), GTC941TTG (2971G>T 2973C>G), ATT942ATA (2976T>A), GGA944GGT (2982A>T), ATC947GTG (2989A>G 2991C>G), CTG949TTG (2995C>T), CTA950TTG

Codon mutations:

(2998C>T 3000A>G), CGT951AGG (3001C>A 3003T>G), GTG952GTA (3006G>A), GGT953GGG (3009T>G), GAC954GAT (3012C>T), CCT955CCA (3015T>A), CCT960TCA (3028C>T 3030T>A), ATT962ATA (3036T>A), CGC963AGA (3037C>A 3039C>A), AAG964AAA (3042G>A), CCA967CCT (3051A>T), GAG971GAT (3063G>T), GTC972GTG (3066C>G), TGC973TGT (3069C>T), TTC974TAC (3071T>A), ACC975TCT (3073A>T 3075C>T), AAA976AAG (3078A>G), GGT977GGC (3081T>C), TCT978TCA (3084T>A), CAG979ATG (3085C>A 3086A>T), TTA981TTG (3093A>G), TGT982TGC (3096T>C), TGT983GTG (3097T>G 3098G>T 3099T>G), AGA984AGG (3102A>G), GCC985GCT (3105C>T), GTG986GTA (3108G>A), GCG987GCT (3111G>T), ACC989AAG (3116C>A 3117C>G), CCA991CCC (3123A>C), AGA992AAG (3125G>A 3126A>G), CTG993AAA (3127C>A 3128T>A 3129G>A), GGC994TCC (3130G>T 3131G>C), AAC995AAT (3135C>T), CTA996TTG (3136C>T 3138A>G), GAA997GAG (3141A>G), TTT998GTT (3142T>G), TTG999CTT (3145T>C 3147G>T), TTT1001TAT (3152T>A), TCC1006TCA (3168C>A), AAA1007AAG (3171A>G), TAT1009TAC (3177T>C), ACA1010ACC (3180A>C), GTC1011GAA (3182T>A 3183C>A), AAG1012AAA (3186G>A), GTG1013GTC (3189G>C), GAG1014ACA (3190G>A 3191A>C 3192G>A), GAG1016GGA (3197A>G 3198G>A), GAC1018GAT (3204C>T), TCT1019TCG (3207T>G), GCC1020AAT (3208G>A 3209C>A 3210C>T), AGT1021GCA (3211A>G 3212G>C 3213T>A), AAA1023GCT (3217A>G 3218A>C 3219A>T), TTG1024CTT (3220T>C 3222G>T), AGT1025AGC (3225T>C), TCG1026TCA (3228G>A), CAG1027CAA (3231G>A), CAA1028GTT (3232C>G 3233A>T 3234A>T), TCA1029GCA (3235T>G), GAA1031GAG (3243A>G), TAT1032TAC (3246T>C), CAT1033CAC (3249T>C), ATG1034ATT (3252G>T), GGA1036AGT (3256G>A 3258A>T), GGT1039GGA (3267T>A), GGA1042GGT (3276A>T), TTA1045CTC (3283T>C 3285A>C), CTT1046TTA (3286C>T 3288T>A), CCC1048CCA (3294C>A), AGT1049ACT (3296G>C), GTT1050GTG (3300T>G), AAA1053AAG (3309A>G), CAA1054CAG (3312A>G), CCT1055CCG (3315T>G), GTT1056TGC (3316G>T 3317T>G 3318T>C), GTG1058GTT (3324G>T), GGC1059GGA (3327C>A), ATT1060ATA (3330T>A), GCT1063GCA (3339T>A), TCA1064TCT (3342A>T), GGT1067GAT (3350G>A), ATT1068ACC (3353T>C 3354T>C), GCA1069GCT (3357A>T), GAA1071GAG (3363A>G), GGA1073GGT (3369A>T), TTC1074TAC (3371T>A), ATT1075GTG (3373A>G 3375T>G), TCT1076GCT (3376T>G), TCC1077TCA (3381C>A), AAT1078AAC (3384T>C), GCC1079GCA (3387C>A), ACG1080ACA (3390G>A), GCC1081GCA (3393C>A), ATC1082GTG (3394A>G 3396C>G), TTC1083TTT (3399C>T), CGG1084AGA (3400C>A 3402G>A), GAC1085GAT (3405C>T), CAG1086CAA (3408G>A), CTT1087TTG (3409C>T 3411T>G), AAC1089GAT (3415A>G 3417C>T), GGG1093GGA (3429G>A), CCT1094CCC (3432T>C), GTT1095ATT (3433G>A), GCG1097GTG (3440C>T), GCA1098GCT (3444A>T), CGA1101CGC (3453A>C), GAC1103GAA (3459C>A), ATT1104CTG (3460A>C 3462T>G), CTC1105TTG (3463C>T 3465C>G), AAA1106GAG (3466A>G 3468A>G), GCC1107CAC (3469G>C 3470C>A), ATT1108CTG (3472A>C 3474T>G), AGG1109AGA (3477G>A), AGT1110TCC (3478A>T 3479G>C 3480T>C), AGA1111AAG (3482G>A 3483A>G), GAA1112GAG (3486A>G), ACC1113GAA (3487A>G 3488C>A 3489C>A), ACT1114AGA (3491C>G 3492T>A), CTT1115TTC (3493C>T 3495T>C), GAA1117GAG (3501A>G), AAC1119AAA (3507C>A), CAA1120CAG (3510A>G), GTC1121GTG (3513C>G), TCC1122GCT (3514T>G 3516C>T), TTT1123TTC (3519T>C), CTT1124TTG (3520C>T 3522T>G), GTT1127GTA (3531T>A), CCT1128CCG (3534T>G), CAA1129TTG (3535C>T 3536A>T 3537A>G), TTG1131ATG (3541T>A), GCC1132GCT (3546C>T), GCA1133GCT (3549A>T), GTA1135GTG (3555A>G), CCC1136CCG (3558C>G), CAT1137CAC (3561T>C), ACT1139ACA (3567T>A), ACA1140ACC (3570A>C), CTC1141CTT (3573C>T), AGA1142AAA (3575G>A), AAG1143CGC (3577A>C 3578A>G 3579G>C), CAA1145CCC (3584A>C 3585A>C), TTG1146CTG (3586T>C), GCA1149GCT (3597A>T), GGT1151GGG (3603T>G), CCT1152GAC (3604C>G 3605C>A 3606T>C), GAA1154CAG (3610G>C 3612A>G), ACC1155ACT (3615C>T), GCT1156GCC (3618T>C), CCC1157CCA (3621C>A), TCA1158TCC (3624A>C), ATC1159ATT (3627C>T), CTC1160CTA (3630C>A), ACC1161TCT (3631A>T 3633C>T), GTC1162GCC (3635T>C), AAT1163AAA (3639T>A), CGG1166AGG (3646C>A), GAG1168GAA (3654G>A), GGA1169GGG (3657A>G), TTC1170TTT (3660C>T), CCA1172CCC (3666A>C), GCA1175GCT (3675A>T), GTG1177GTC (3681G>C), ATG1178GAG (3682A>G 3683T>A), AAA1179AAG (3687A>G), TAC1180TAT (3690C>T), ACT1183ACA (3699T>A), CAT1185TCC (3703C>T 3704A>C 3705T>C), GGT1186AAC (3706G>A 3707G>A 3708T>C), GAT1188GAG (3714T>G), GAT1189GAG (3717T>G), GAT1190CCA (3718G>C 3719A>C 3720T>A), GCT1192GCA (3726T>A), AAA1193AAG (3729A>G), TTG1194CTG (3730T>C), TCC1195GCC (3733T>G), GAA1197GAG (3741A>G), ACT1198GGT (3742A>G 3743C>G), CTC1199CTT (3747C>T), AGT1201GCA (3751A>G 3752G>C 3753T>A), TCA1202TCC (3756A>C), CTG1203ATT (3757C>A 3759G>T), CTG1204CTT (3762G>T), CCA1205CCT (3765A>T), ACA1206GTG (3766A>G 3767C>T 3768A>G), ATG1207CTT (3769A>C 3771G>T), AAA1208AAG (3774A>G), AAC1209ATG (3776A>T 3777C>G), CAA1210CAG (3780A>G), AAA1211CGA (3781A>C 3782A>G), GGC1215GGG (3795C>G), CCA1217CCT (3801A>T), CAG1218GCA (3802C>G 3803A>C 3804G>A), GAG1219ATC (3805G>A 3806A>T 3807G>C), GAT1221AAT (3811G>A), GTG1224GTA (3822G>A), CTG1226CTC (3828G>C), ATT1229ATA (3837T>A), TGT1232TTT (3845G>T), TAT1235TTT (3854A>T), GGC1237GCA (3860G>C 3861C>A), CTC1240TTG (3868C>T 3870C>G), ACA1242ACG (3876A>G), TCC1243TCA (3879C>A), TAC1246TAT (3888C>T), TTC1248TTT (3894C>T), ATG1251TTG (3901A>T), GGC1252GGA (3906C>A), ATG1253CTT (3907A>C 3909G>T), GGA1254GGT (3912A>T), ATG1255TCT (3913A>T 3914T>C 3915G>T), AAG1257AAA (3921G>A), CGG1258AGG (3922C>A), GAG1259GAA (3927G>A), GTC1261GTG (3933C>G), GTA1262GAA (3935T>A), CCA1263GCC (3937C>G 3939A>C), ACA1264CAA (3940A>C 3941C>A), GGA1265CCA (3943G>C 3944G>C), GAA1266AAC (3946G>A 3948A>C), GGT1267TCG (3949G>T 3950G>C 3951T>G), GAT1268CAG (3952G>C 3954T>G), AAA1269AGG (3956A>G 3957A>G), GTA1270GTT (3960A>T), GAG1271CTT (3961G>C 3962A>T 3963G>T), CTT1272CTC (3966T>C), AGA1273AAA (3968G>A), GAC1275AAC (3973G>A), CCA1277CCT (3981A>T), GTG1278GTC (3984G>C), GAA1280GAT (3990A>T), GCT1281GCA (3993T>A), GAG1283GAA (3999G>A), GAA1284GAG (4002A>G), CTG1285TTG (4003C>T), GAC1286GAT (4008C>T), ATT1287GCG (4009A>G 4010T>C 4011T>G), CAA1288CAG (4014A>G), ATT1289GTA (4015A>G 4017T>A), CAG1291CAA (4023G>A), ATA1293ACA (4028T>C), CAT1294TTT (4030C>T 4031A>T), GTG1295GTT (4035G>T), GAT1296GAC (4038T>C), CTA1297CTC (4041A>C), ATC1298GTC (4042A>G), ACC1299ACA (4047C>A), ACA1300ACC (4050A>C), CAA1301CAG (4053A>G), GCC1303GCG (4059C>G), AAG1304AAA (4062G>A), GAG1306GAA (4068G>A), CGC1307CGG (4071C>G), CTT1308TTG (4072C>T 4074T>G), CCT1309CCA (4077T>A), GAA1311GAG (4083A>G), AAA1318AAG (4104A>G), ACA1319ACC (4107A>C), AGA1320CGC (4108A>C 4110A>C), TTA1321CTG (4111T>C 4113A>G), TTT1322TTC (4116T>C), GAG1323GAA (4119G>A), ATA1324ATC (4122A>C), TAC1329TAT (4137C>T), CTG1332CTA (4146G>A), GTC1333GTG (4149C>G), AGG1334CGG (4150A>C), TTT1337TTC (4161T>C), TCC1341TCT (4173C>T), GCC1342GCT (4176C>T), ACC1343TGT (4177A>T 4178C>G 4179C>T), GCA1346GCT (4188A>T), TTA1347GCA (4189T>G 4190T>C), AAT1349AAC (4197T>C), GTC1350GTA (4200C>A), ATC1351CTC (4201A>C), CCA1352CCT (4206A>T), TGT1353TGC (4209T>C), GTT1355GTA (4215T>A), GGA1356GGT (4218A>T), ATA1357ATC (4221A>C), GAT1358AAT (4222G>A), ACC1360ACG (4230C>G), AGT1361AGC (4233T>C), TCT1362CAT (4234T>C 4235C>A), TCT1365ACA (4243T>A 4245T>A), CTT1366GTG (4246C>G 4248T>G), CTT1367CTG (4251T>G), GCC1368GCT (4254C>T), AAC1369AGT (4256A>G 4257C>T), AAC1370AGC (4259A>G), TTC1371TTT (4263C>T), GCT1373TCA (4267G>T 4269T>A), ACT1377GAG (4279A>G 4280C>A 4281T>G), GGG1378GGA (4284G>A), TTT1379TTC (4287T>C), GCT1381GCC (4293T>C), TCA1384TCG (4302A>G), AGT1385AGC (4305T>C), GAC1387GAT (4311C>T), GGG1388GGA (4314G>A), CGC1389AGA (4315C>A 4317C>A), CCA1391CCC (4323A>C), GTG1392ATC (4324G>A 4326G>C), TTC1393TTT (4329C>T), GCT1394GCA (4332T>A), TTT1395TTC (4335T>C), CAG1396CAA (4338G>A), TTG1401CTG (4351T>C), GTT1402GTG (4356T>G), GAG1404GCT (4361A>C 4362G>T), TAT1405TTT (4364A>T), GAT1407GGT (4370A>G), ACC1408GAT (4372A>G 4373C>A 4374C>T), AAG1409GAG (4375A>G), GGA1411GGG (4383A>G), GAG1412AGT (4384G>A 4385A>G 4386G>T), GCT1414GAG (4391C>A 4392T>G), TCC1415TCA (4395C>A), GCA1417GCT (4401A>T), AGA1418AGG (4404A>G), CAT1419CAC (4407T>C), GCT1420ACA (4408G>A 4410T>A), CTT1421TTA (4411C>T 4413T>A), CTC1422TTA (4414C>T 4416C>A), GCA1425GCC (4425A>C), TCA1426TCT (4428A>T), TAC1429TAT (4437C>T), TTA1431CTC (4441T>C 4443A>C), GAT1434AAC (4450G>A 4452T>C), AAA1435AAG (4455A>G), GTT1436TTG (4456G>T 4458T>G), TTC1437TTT (4461C>T), CGT1438CGC (4464T>C), TTG1439CTA (4465T>C 4467G>A), GTT1440GTG (4470T>G), GGC1442GGA (4476C>A), CCG1444CCC (4482G>C), GCA1448GCT (4494A>T), CTT1449TTG (4495C>T 4497T>G), ACC1450ACT (4500C>T), TTC1453TTT (4509C>T), AAC1454AAT (4512C>T), CTG1456TTG (4516C>T), TTG1457CTA (4519T>C 4521G>A), CGG1463AGG (4537C>A), TAT1464TAC (4542T>C), GCT1465GCG (4545T>G), TTC1466TTT (4548C>T), ATC1467ATA (4551C>A), CTT1469CTG (4557T>G), TTA1470TTG (4560A>G), AGA1471AGG (4563A>G), AGG1472AGA (4566G>A), CCT1473CCA (4569T>A), CAC1474CAT (4572C>T), ATT1475GTC (4573A>G 4575T>C), GCC1476AAA (4576G>A 4577C>A 4578C>A), GCT1477GCC (4581T>C), AGA1478AAA (4583G>A), GCT1479TCT (4585G>T), ATA1480GTG (4588A>G 4590A>G), GGT1481GGC (4593T>C), GTA1482ATT (4594G>A 4596A>T), CCT1484CCA (4602T>A), GGT1485AAT (4603G>A 4604G>A), GAT1486GAC (4608T>C), AAC1488AAT (4614C>T), CTT1490TTG (4618C>T 4620T>G), GGA1496GGT (4638A>T), AAT1499AAC (4647T>C), CTG1500TTG (4648C>T), GTG1501GTT (4653G>T), GCT1502GCG (4656T>G), GTG1503GTT (4659G>T), CCT1504CCA (4662T>A), CTG1505ATG (4663C>A), CTG1507TTG (4669C>T), TCA1511GAT (4681T>G 4682C>A 4683A>T), CTG1512CTA (4686G>A), CCT1513CCA (4689T>A), AAT1514TCC (4690A>T 4691A>C 4692T>C), GCC1516GCA (4698C>A), CAG1517AGA (4699C>A 4700A>G 4701G>A), CTT1519CTG (4707T>G), GAC1520GAA (4710C>A), ATG1521GCT (4711A>G 4712T>C 4713G>T), GTA1522GTT (4716A>T), AAT1523AAC (4719T>C), ATC1526ATT (4728C>T), AAA1527AAG (4731A>G), GGG1529GGT (4737G>T), CTG1530TTG (4738C>T), GAT1531GAC (4743T>C), TCT1533TCA (4749T>A), AAC1537GAG (4759A>G 4761C>G), CAG1538AAG (4762C>A), GTG1539ACA (4765G>A 4766T>C 4767G>A), CAG1540AAA (4768C>A 4770G>A), TTC1541TTT (4773C>T), CAG1542CTG (4775A>T), GAC1543GAT (4779C>T), TCA1545TCT (4785A>T), GAA1546GAG (4788A>G), ACT1548ACC (4794T>C), AGC1551AGT (4803C>T), GGA1553GGT (4809A>T), TTT1554TTC (4812T>C), AAA1555AAG (4815A>G), CGT1556CGA (4818T>A), GCC1558GCA (4824C>A), CTT1559TTG (4825C>T 4827T>G), GGG1560GGA (4830G>A), CAC1562CAA (4836C>A), GTG1569GTT (4857G>T), ATA1571GTC (4861A>G 4863A>C), ATT1572ATC (4866T>C), CCT1574CCC (4872T>C), ATG1575ATT (4875G>T), TAC1576TAT (4878C>T), ATC1578ATT (4884C>T), AGA1579AGG (4887A>G), CCT1580CCA (4890T>A), CCG1582GTG (4894C>G 4895C>T), GGT1583GGA (4899T>A), ACT1587GTA (4909A>G 4910C>T 4911T>A), GTG1588CAA (4912G>C 4913T>A 4914G>A), GCA1589GCT (4917A>T), ATG1591TAT (4921A>T 4922T>A 4923G>T), GAG1592GAA (4926G>A), GTG1594ATT (4930G>A 4932G>T), ACT1596ACA (4938T>A), CTT1598ATC (4942C>A 4944T>C), CGT1599AGA (4945C>A 4947T>A), GAG1600GAA (4950G>A), TTT1602TTC (4956T>C), AGA1606CAG (4966A>C 4967G>A 4968A>G), GTT1607GCC (4970T>C 4971T>C), GCA1608ACT (4972G>A 4974A>T), GAG1610GAA (4980G>A), AAG1611AGT (4982A>G 4983G>T), TTA1612TTT (4986A>T), ACT1613GTA (4987A>G 4988C>T 4989T>A), TAT1614AAT (4990T>A), CAG1617TTT (4999C>T 5000A>T 5001G>T), TGT1618GGA (5002T>G 5004T>A), GCC1619GCT (5007C>T), AGA1621AAG (5012G>A 5013A>G), GAG1622CAG (5014G>C), CGT1623AGA (5017C>A 5019T>A), GGA1624AAT (5020G>A 5021G>A 5022A>T), GGC1626CCC (5026G>C 5027G>C), GCA1627ATT (5029G>A 5030C>T 5031A>T), ACA1628ATG (5033C>T 5034A>G), TTT1630ATG (5038T>A 5040T>G), CCT1631CCC (5043T>C), TCT1632ACT (5044T>A), TAT1633TAC (5049T>C), ATG1634TAT (5050A>T 5051T>A 5052G>T), GTG1636CTT (5056G>C 5058G>T), GAG1637GAA (5061G>A), CAG1638AGT (5062C>A 5063A>G 5064G>T), GAA1639CAA (5065G>C), ATC1641TTG (5071A>T 5073C>G), GCA1642ACA (5074G>A), AAG1643GAG (5077A>G), GCA1648GAG (5093C>A 5094A>G), TCA1649AGC (5095T>A 5096C>G 5097A>C), GCG1650ATT (5098G>A 5099C>T 5100G>T), TTG1651GGG (5101T>G 5102T>G), ACA1652GTT (5104A>G 5105C>T 5106A>T), ATC1653TTG (5107A>T 5109C>G), TGT1654ATG (5110T>A 5111G>T 5112T>G), GAA1655GAG (5115A>G), ATG1656CTA (5116A>C 5118G>A), GCC1657TCA (5119G>T 5121C>A), AAA1658GAT (5122A>G 5124A>T), TCT1660AAA (5128T>A 5129C>A 5130T>A), ATC1661AAT (5132T>A 5133C>T), ACT1662ACA (5136T>A), TTC1663TAC (5138T>A), ACA1664ACC (5142A>C), TTG1666TTA (5148G>A), CCA1667CCT (5151A>T), CCT1668GCT (5152C>G), AAT1670GAA (5158A>G 5160T>A), GTT1672ATT (5164G>A), GAA1673del (5167_5169delGAA), GAA1676GAG (5178A>G), AGA1677GAG (5179A>G 5180G>A 5181A>G), GAT1678GGT (5183A>G), CTC1679GTC (5185C>G), AAT1680CAT (5188A>C), TGG1681GAA (5191T>G 5192G>A 5193G>A), GCC1683GTG (5198C>T 5199C>G), CCA1684CCT (5202A>T), AAC1685GAT (5203A>G 5205C>T), ATT1686GTA (5206A>G 5208T>A), GGT1687TTT (5209G>T 5210G>T), TGC1689TGT (5217C>T), TCA1690AGT (5218T>A 5219C>G 5220A>T), GCC1691GCT (5223C>T), CGT1692CGG (5226T>G), GCC1695TCA (5233G>T 5235C>A), AAG1696AAC (5238G>C), ACT1697CAA (5239A>C 5240C>A 5241T>A), GAA1698CAA (5242G>C), GAG1700TCA (5248G>T 5249A>C 5250G>A), CCT1701TCA (5251C>T 5253T>A), GGG1702GGT (5256G>T), TGC1703AAC (5257T>A 5258G>A), ATA1704ATC (5262A>C), GTA1705CTG (5263G>C 5265A>G), AAC1707AAT (5271C>T), TGT1708TGC (5274T>C), ACA1709ACT (5277A>T), GGG1710GGA (5280G>A), GCG1711GCC (5283G>C), AAG1712CGC (5284A>C 5285A>G 5286G>C), AAG1713AGA (5288A>G 5289G>A), AGC1714ACA (5291G>C 5292C>A), AAA1715ACA (5294A>C), GTT1717GTG (5301T>G), CGC1718AGA (5302C>A 5304C>A), GGA1719GGG (5307A>G), GCT1721GCC (5313T>C), AGT1722AAT (5315G>A), GAT1725GAC (5325T>C), ATG1730ATA (5340G>A), CCA1732CCC (5346A>C), TAT1733TAC (5349T>C), ACC1734ACT (5352C>T), ATG1735TTG (5353A>T), TCA1736GCA (5356T>G), ATT1738ATC (5364T>C), CTG1739CAG (5366T>A), GCT1741CAT (5371G>C 5372C>A), AGA1743CGC (5377A>C 5379A>C), GGC1744AAG (5380G>A 5381G>A 5382C>G), AAA1745AGT (5384A>G 5385A>T), ATG1746CTG (5386A>C), TCA1747GAA (5389T>G 5390C>A), GAG1748GTT (5393A>T 5394G>T), GAC1749GGT (5396A>G 5397C>T), ATT1750GAT (5398A>G 5399T>A), GCT1751GGG (5402C>G 5403T>G), GAT1754TCT (5410G>T 5411A>C), GTT1755GTC (5415T>C), GTG1756ACC (5416G>A 5417T>C 5418G>C), GTG1758GTT (5424G>T), GGG1760GGT (5430G>T), GCT1764GCA (5442T>A), GTG1765ATA (5443G>A 5445G>A), AGC1766GCT (5446A>G 5447G>C 5448C>T), GCT1771GCA (5463T>A), TTA1772TTG (5466A>G), ATG1773TTT (5467A>T 5469G>T), GCT1774GCA (5472T>A), GTG1775TTG (5473G>T), CGT1778CAT (5483G>A), CAG1779CAA (5487G>A), TAT1780TAC (5490T>C), TGC1781AGG (5491T>A 5493C>G), ATT1782GCC (5494A>G 5495T>C 5496T>C), GAG1783ACA (5497G>A 5498A>C 5499G>A), GAT1784GAC (5502T>C), ATC1785CTT (5503A>C 5505C>T), AGG1788CGC (5512A>C 5514G>C), TTG1789CTA (5515T>C 5517G>A), CGC1790CGA (5520C>A), ACT1791ACC (5523T>C), ATT1792ATG (5526T>G), GGC1793GCT (5528G>C 5529C>T), AAT1794GGG (5530A>G 5531A>G 5532T>G), TTG1796CTG (5536T>C), ACA1800ACC (5550A>C), TAT1801TAC (5553T>C), CCC1802CCT (5556C>T), GTT1806GCG (5567T>C 5568T>G), CAG1807CAA (5571G>A), TTT1809TTG (5577T>G), CAA1811AAA (5581C>A), GTG1813GTT (5589G>T), CCT1814GAA (5590C>G 5591C>A 5592T>A), CAG1815GAC (5593C>G 5595G>C), CAG1815_GAA1816insACC (5595_5596insACC), GAA1816TCA (5596G>T 5597A>C), GGG1817CAC (5599G>C 5600G>A 5601G>C), AAA1818AAG (5604A>G), ATG1819TTG (5605A>T), GCA1820GGA (5609C>G), GTT1822AGT (5614G>A 5615T>G), TCG1824ACA (5620T>A 5622G>A), TCT1825del (5623_5625delTCT), TTG1826TAT (5627T>A 5628G>T), CAA1827GGC (5629C>G 5630A>G 5631A>C), AGT1828ACC (5633G>C 5634T>C), AGT1829ACT (5636G>C), ATG1831AAC (5642T>A 5643G>C), CAA1833CAT (5649A>T), CAA1833_GGT1834insCACAGT (5649_5650insCACAGT), GGT1834GCA (5651G>C 5652T>A), GAG1836TCA (5656G>T 5657A>C 5658G>A), CTT1837GTT (5659C>G), GGC1838GGA (5664C>A), CAA1839CTT (5666A>T 5667A>T), ATT1840CTA (5668A>C 5670T>A), CGT1841AGG (5671C>A 5673T>G), GTC1842ATA (5674G>A 5676C>A), ATC1843GTT (5677A>G 5679C>T), CAC1844CCT (5681A>C 5682C>T), GGT1845TCC (5683G>T 5684G>C 5685T>C), GAT1846GAC (5688T>C), TTA1847CTC (5689T>C 5691A>C), TCA1848TCT (5694A>T), ATG1849GAA (5695A>G 5696T>A 5697G>A), GAA1850AAG (5698G>A 5700A>G), TTG1853GTT (5707T>G 5709G>T), CGC1854CGT (5712C>T), ATG1855TTG (5713A>T), TAT1857TAC (5721T>C), GTT1858ACA (5722G>A 5723T>C 5724T>A), GTG1859ATT (5725G>A 5727G>T), GGC1860GGT (5730C>T), CCA1861CCT (5733A>T), CAT1862GAA (5734C>G 5736T>A), GGT1863GGA (5739T>A), GGA1864GGT (5742A>T), GGC1866GCC (5747G>C), AAC1867CAG (5749A>C 5751C>G), TTT1868TGC (5753T>G 5754T>C), ACC1869TCA (5755A>T 5757C>A), ACA1870ACG (5760A>G), AAA1871AAG (5763A>G), AAT1872AAC (5766T>C), CTT1873CTG (5769T>G), GAT1874AAG (5770G>A 5772T>G), CAA1875CAT (5775A>T), TTG1876GCT (5776T>G 5777T>C 5778G>T), TTT1877TTG (5781T>G), GCA1878GAG (5783C>A 5784A>G), AGT1879ACA (5786G>C 5787T>A), CTT1880TTG (5788C>T 5790T>G), GAA1881TCT (5791G>T 5792A>C 5793A>T), AAG1882GTC (5794A>G 5795A>T 5796G>C), TGT1883AGT (5797T>A), GAA1886ACT (5806G>A 5807A>C 5808A>T), TTG1887CTT (5809T>C 5811G>T), CTT1888GCC (5812C>G 5813T>C 5814T>C), AAG1890AGA (5819A>G 5820G>A), AAC1891GGA (5821A>G 5822A>G 5823C>A), ACG1892GTC (5824A>G 5825C>T 5826G>C), CTG1894GTG (5830C>G), ACC1895ACA (5835C>A), GAG1899AAA (5845G>A 5847G>A), AAA1900GAA (5848A>G), ATG1901TTC (5851A>T 5853G>C), ACT1902TCC (5854A>T 5856T>C), AAA1904CAC (5860A>C 5862A>C), GTG1906TGT (5866G>T 5867T>G 5868G>T), GCA1907GAA (5870C>A), CAA1908ATG (5872C>A 5873A>T 5874A>G), GTC1910CTT (5878G>C 5880C>T), AGT1911GAG (5881A>G 5882G>A 5883T>G), GTA1913CTG (5887G>C 5889A>G), AAA1914GAA (5890A>G), CAG1915AGT (5893C>A 5894A>G 5895G>T), TTC1918GTA

| | |
|---|---|
| | (5902T>G 5904C>A), TAT1919GTA (5905T>G 5906A>T 5907T>A), CCC1920TTC (5908C>T 5909C>T), AAG1921ATG (5912A>T), GCC1922GGG (5915C>G 5916C>G), ACA1923AGC (5918C>G 5919A>C), ACC1924GTT (5920A>G 5921C>T 5922C>T), GTG1925AAG (5923G>A 5924T>A), GAA1926AAA (5926G>A), CAG1927CAA (5931G>A), CTC1928ATG (5932C>A 5934C>G), AAG1929TTG (5935A>T 5936A>T), GTA1930TCC (5938G>T 5939T>C 5940A>C), TTT1931TTC (5943T>C), GCT1932ATG (5944G>A 5945C>T 5946T>G), GAT1933AAT (5947G>A), GCT1934CAA (5950G>C 5951C>A 5952T>A), CAT1935CAG (5955T>G), CAT1935_ACC1936insATG (5955_5956insATG), ACC1936TCA (5956A>T 5958C>A), ATT1937AAA (5960T>A 5961T>A), GTC1938ACC (5962G>A 5963T>C), AAA1939CAA (5965A>C), ACT1940GTT (5968A>G 5969C>T), GCT1941GCC (5973T>C), AAA1942AGC (5975A>G 5976A>C), CCT1943CCA (5979T>A), TTC1944TGT (5981T>G 5982C>T), CGG1945TGT (5983C>T 5985G>T), GGA1946del (5986_5988delGGA), GTA1947GTC (5991A>C), GTT1948ACA (5992G>A 5993T>C 5994T>A), TTC1949TTT (5997C>T), CGG1950AGT (5998C>A 6000G>T), AAT1952GCA (6004A>G 6005A>C 6006T>A), TTT1953TGG (6008T>G 6009T>G), CTC1954CTA (6012C>A), AGT1955ACT (6014G>C), CCT1956AAG (6016C>A 6017C>A 6018T>G), GAG1957GAC (6021G>C), TGG1958ATT (6022T>A 6023G>T 6024G>T), TGG1958_AAA1959insGTC (6024_6025insGTC), AAA1959ATG (6026A>T 6027A>G), ATT1960ACA (6029T>C 6030T>A), GGA1961AAA (6031G>A 6032G>A), GAG1963AAG (6037G>A), AAC1964AAG (6042C>G), ACA1965CCA (6043A>C), ATG1966del (6046_6048delATG), GCA1967GGA (6050C>G), ACA1968TCT (6052A>T 6054A>T), CTC1969ATT (6055C>A 6057C>T), TCA1970CCT (6058T>C 6060A>T), GAG1972GAA (6066G>A), AGT1973ACA (6068G>C 6069T>A), CTT1974GTG (6070C>G 6072T>G), CTT1975TTT (6073C>T), CCA1976CTT (6077C>T 6078A>T), CTG1979CCT (6086T>C 6087G>T), TCG1980TCA (6090G>A), TCA1982CTA (6094T>C 6095C>T), GCC1983GTT (6098C>T 6099C>T), ATG1984GTG (6100A>G), AAG1985AGA (6104A>G 6105G>A), TTG1987ATG (6109T>A), CTC1988TTG (6112C>T 6114C>G), AGC1989CGC (6115A>C), AGA1990AAA (6119G>A) |

**Proteins**

| | | | | | | | | | |
|---|---|---|---|---|---|---|---|---|---|
| polyprotein (YP_009165993.1) | 1 | 1993 | 92.7% | 9636 | 70.1% | 1982 (99.1%) | 1314 (65.7%) | 7/11/0/0 | 0 |

| | Begin | End | Coverage | Score | Concordance | Matches | Identities | I/D/M/F* | Stop Codons |
|---|---|---|---|---|---|---|---|---|---|
| **NT** | **33** | **6128** | **84.2%** | **3374** | **28.2%** | **6043 (98.8%)** | **3915 (64.0%)** | **21/53** | |

Protein mutations: L3F (157C>T), F7W (170T>G 171T>G), G8S (172G>A), S9P (175T>C), S10P (178T>C 180A>T), S11A (181T>G), E12_V17del (184_201delGAGAAGACTGGATCAGTT), P18S (202C>T 204C>T), T20P (208A>C), G24A (221G>C 222T>A), L26V (226C>G 228G>T), G28S (232G>T 233G>C), I30F (238A>T 240C>T), K32S (244A>T 245A>C 246A>T), F33L (247T>C 249C>T), A36T (256G>A), I37L (259A>C 261T>C), I40V (268A>G 270C>T), T41N (272C>A 273A>T), S42K (274T>A 275C>A), K46T (287A>C 288A>C), S48V (292T>G 293C>T 294T>A), L51I (301T>A), D52E (306T>G), L54V (310T>G 312G>C), S57P (319T>C 321T>A), T59M (326C>T 327T>G), D60A (329A>C 330T>A), S62A (334T>G 336T>A), S63T (337T>A 339T>A), V66A (347T>C 348T>C), S70N (358T>A 359C>A 360C>T), L72V (364T>G 366G>A), N73E (367A>G 369T>A), K74R (371A>G), G80A (389G>C 390T>A), S81K (391T>A 392C>A 393C>A), L82I (394T>A 396G>A), I83M (399T>G), L87I (409C>A 411G>C), A89G (416C>G 417A>T), S91D (421T>G 422C>A), L93V (427T>G 429G>A), A94K (430G>A 431C>A), S99T (445T>A), K101V (451A>G 452A>T 453G>A), M103L (457A>C 459G>T), R106K (466C>A 467G>A), L107M (469T>A), F108V (472T>G 474C>T), E109L (475G>T 476A>T), L111V (481T>G 483G>C), S112K (484T>A 485C>A 486C>G), N113D (487A>G), V115F (493G>T 495T>C), R116I (497G>T 498A>C), I117L (499A>C), F119Y (506T>A 507T>C), Q120E (508C>G 510G>A), V122L (514G>T 516T>A), S126V (526A>G 527G>T), D128S (532G>T 533A>C 534T>A), V129L (535G>T), V133F (547G>T), Y134I (550T>A 551A>T 552T>A), G135A (554G>C), I141L (571A>T 573A>G), T142A (574A>G), L143I (577C>A), V145L (583G>C 585T>A), C148Y (593G>A), S152V (604T>G 605C>T), I155V (613A>G 615T>A), E156T (616G>A 617A>C 618A>G), G157P (619G>C 620G>C 621A>T), V158F (622G>T 624A>T), S159N (625T>A 626C>A), S160N (629G>A), T161V (631A>G 632C>T 633C>A), I162V (634A>G), K163V (637A>G 638A>T 639A>T), N164T (641A>C), I165L (643A>T 645T>G), L167S (650T>C), I168V (652A>G 654A>T), M172F (664A>T 666G>T), K184T (701A>C 702G>A), A186S (706G>T), E187G (710A>G), T190S (718A>T 720A>C), M191Q (721A>C 722T>A), V192L (724G>C 726A>G), S196A (736A>G 737G>C 738T>A), M197V (739A>G 741G>A), V203L (757G>T 759C>G), S204T (760T>A 762A>G), M212L (784A>T 786G>A), T214S (790A>T 792T>A), I215L (793A>T 795T>G), M216L (796A>T), I226V (826A>G 828A>T), T246A (886A>G), T258Q (922A>C 923C>A 924A>G), I278V (982A>G 984A>G), G280S (988G>A 990T>C), A281I (991G>A 992C>T), S293N (1028G>A 1029C>T), E302Q (1054G>C 1056A>G), Q305T (1063C>A 1064A>C), D308S (1072G>T 1073A>C 1074T>G), A309E (1076C>A 1077T>G), C311A (1081T>G 1082G>C), D312E (1086T>A), A314S (1090G>T), T315M (1094C>T 1095C>G), K323P (1117A>C 1118A>C 1119A>T), D327I (1129G>A 1130A>T 1131C>T), N330Q (1138A>C 1140T>A), R333K (1148G>A 1149G>A), L337V (1159C>G 1161G>C), M352L (1204A>T), E364D (1242A>T), V366L (1246G>C 1248C>G), Y367V (1249T>G 1250A>T), S370A (1258A>G 1259G>C 1260T>C), C371S (1261T>A), R372Q (1264A>C 1265G>A), V374S (1270G>T 1271T>C), Y389F (1316A>T), A399C (1345G>T 1346C>G 1347A>T), R410A (1378C>G 1379G>C 1380C>A), V494I (1630G>A), I519R (1706T>G 1707C>A), E521V (1712A>T 1713A>G), E523D (1719A>T), S525D (1723T>G 1724C>A), T526P (1726A>C), E527N (1729G>A 1731G>T), D528A (1733A>C 1734C>T), G531A (1742G>C 1743A>C), M534V (1750A>G 1752G>T), N536D (1756A>G 1758C>T), I537V (1759A>G 1761T>G), H550A (1798C>G 1799A>C 1800C>T), S554D (1810T>G 1811C>A 1812G>T), D558Q (1822G>C 1824C>A), Q565R (1844A>G), A568Y (1852G>T 1853C>A), I570V (1858A>G 1860A>G), K572S (1865A>G 1866G>C), F575Y (1874T>A 1875T>C), E576N (1876G>A 1878A>T), A577T (1879G>A 1881A>T), E578T (1882G>A 1883A>C), T579S (1885A>T 1887T>G), I581L (1891A>T 1893T>G), S582A (1894T>G 1896A>T), N583D (1897A>G), T585V (1903A>G 1904C>T 1905A>T), A589N (1915G>A 1916C>A), S591A (1921T>G), S592V (1924T>G 1925C>T), E593D (1929A>T), V596L (1936G>C), A597D (1940C>A 1941A>T), V599F (1945G>T 1947T>C), D600Q (1948G>C 1950T>A), G601D (1952G>A 1953G>T), R603del (1957_1959delCGC), F604L (1962C>A), K605Q (1963A>C), T606V (1966A>G 1967C>T 1968A>T), K607L (1969A>T 1970A>T 1971A>G), Q608P (1973A>C 1974A>T), D609P (1975G>C 1976A>C), V610E (1979T>A), I612A (1984A>G 1985T>C), V613A (1988T>C 1989G>T), T614Y (1990A>T 1991C>A), P615E (1993C>G 1994C>A 1995C>G), T616Q (1996A>C 1997C>A 1998A>G), V617T (1999G>A 2000T>C 2001T>A), E621A (2012A>C 2013A>T), D622E (2016T>A), R623K (2018G>A), V624I (2020G>A 2022G>T), I625V (2023A>G), K628Q (2032A>C 2034A>G), F637Y (2060T>A), L643M (2077T>A), D644N (2080G>A), P645K (2083C>A 2084C>A 2085G>A), E646F (2086G>T 2087A>T 2088G>T), S647E (2089A>G 2090G>A 2091T>G), L648F (2094G>T), Y653L (2107T>C 2108A>T 2109T>C), Q654S (2110C>T 2111A>C 2112A>C), L658I (2122C>A), F659Y (2126T>A), S661V (2131T>G 2132C>T 2133C>G), D663S (2137G>T 2138A>C), A664S (2140G>T 2142A>T), E665D (2145G>T), H666A (2146C>G 2147A>C), Q667K (2149C>A 2151G>A), V669N (2155G>A 2156T>A 2157C>T), M671E (2161A>G 2162T>A 2163G>A), R672M (2165G>T 2166A>G), M674V (2170A>G 2172G>A), T675N (2174C>A), M677L (2179A>T), I678L (2182A>T 2184C>G), Y679V (2185T>G 2186A>T), N683S (2198A>G), A684P (2200G>C), E685Q (2203G>C 2205A>G), R688A (2212A>G 2213G>C 2214G>T), Q692R (2225A>G 2226A>G), L699I (2245C>A), A708S (2272G>A 2273C>G 2274T>C), S709T (2275T>A 2277C>A), C712S (2285G>C), K714E (2290A>G), K715L (2293A>C 2294A>T), A717D (2300C>A), T720R (2308A>C 2309C>G 2310A>C), E721N (2311G>A 2313G>T), M722L (2314A>C 2316G>T), R723E (2317C>G 2318G>A 2319C>G), T724Q (2320A>C 2321C>A), F726K (2326T>A 2327T>A 2328T>A), H728Y (2332C>T), N729K (2337T>G), C730A (2338T>G 2339G>C 2340T>G), I731L (2341A>C), D732Q (2344G>C 2346T>G), V735A (2354T>C 2355T>A), I739T (2366T>C 2367T>G), Q742E (2374C>G 2376G>A), F747R (2389T>A 2390T>G 2391T>A), T761R (2432C>G 2433C>G), S764A (2440T>G 2442C>A), S764_K765insG (2442_2443insGGT), K765T (2444A>C 2445A>T), E766D (2448G>C), A767K (2449G>A 2450C>A 2451C>G), D769R (2455G>A 2456A>G 2457C>A), S770N (2458T>A 2459C>A), V772I (2464G>A 2466C>T), K773I (2468A>T 2469G>C), I774E (2470A>G 2471T>A), Y776M (2476T>A 2477A>T 2478C>G), H777I (2479C>A 2480A>T 2481C>T), K778G (2482A>G 2483A>G 2484A>T), V780K (2488G>A 2489T>A), Y782K (2494T>A 2496C>G), K784I (2501A>T 2502A>C), E785Q (2503G>C 2505A>G), T786M (2507C>T 2508T>G), W788N (2512T>A 2513G>A 2514G>T), N789D (2515A>G), Y792F (2525A>T 2526T>C), A793I (2527G>A 2528C>T 2529A>C), R795K (2534G>A), I804L (2560A>T 2562C>G), M805I (2565G>A), L810T (2578T>A 2579T>C 2580G>C), F811M (2581T>A 2583T>G), G812N (2584G>A 2585G>A 2586G>T), G813A (2588G>C 2589A>C), I814V (2590A>G 2592A>T), L818I (2602C>A), A819T (2605G>A), W821L (2611T>C 2612G>T 2613G>C), Q822K (2614C>A 2616A>G), S823A (2617A>G 2618G>C 2619C>A), N824A (2620A>G 2621A>C 2622T>A), S825T (2623T>A), F826S (2627T>C 2628T>C), S828L (2632A>T 2633G>T 2634T>A), S830D (2638A>G 2639G>A), P850K (2698C>A 2699C>A 2700C>G), S851T (2701T>A), R852H (2705G>A), L853F (2707C>T 2709G>C), K854A (2710A>G 2711A>C 2712G>A), N855K (2715T>A), L856S (2717T>C 2718G>C), V858L (2722G>C 2724G>A), S861A (2731A>G 2732G>C 2733T>G), S862A (2734A>G 2735G>C 2736C>A), A863D (2738C>A), I865N (2744T>A 2745T>C), I865_T866insH (2745_2746insCAT), T866V (2746A>G 2747C>T 2748A>G), Q867D (2749C>G 2751A>C), E868R (2752G>C 2753A>G), A871E (2762C>A 2763T>A), E872Q (2764G>C 2766A>G), Q878E (2782C>G), A879S (2785G>A 2786C>G 2787A>T), K881H (2791A>C 2793A>C), M884L (2800A>T), I885V (2803A>G), L889I (2815C>A), G890A (2819G>C 2820A>T), M906L (2866A>C 2868G>C), H913R (2887C>A 2888A>G 2889C>A), V915M (2893G>A), T922S (2914A>T), P926E (2926C>G 2927C>A 2928T>G), N927H (2929A>C 2931C>T), R929I (2936G>T 2937G>T), S930C (2939C>G 2940A>T), K931N (2943G>T), Y934F (2951A>T), R937T (2959C>A 2960G>C 2961G>C), E938Q (2962G>C), K940R (2969A>G), V941L (2971G>T 2973C>G), I947V (2989A>G 2991C>G), P960S (3028C>T 3030T>A), E971D (3063G>T), F974Y (3071T>A), T975S (3073A>T 3075C>T), Q979M (3085C>A 3086A>T), C983V (3097T>G 3098G>T 3099T>G), T989K (3116C>A 3117C>G), R992K (3125G>A 3126A>G), L993K (3127C>A 3128T>A 3129G>A), G994S (3130G>T 3131G>C), F998V (3142T>G), F1001Y (3152T>A), V1011E (3182T>A 3183C>A), E1014T (3190G>A 3191A>C 3192G>A), E1016G (3197A>G 3198G>A), A1020N (3208G>A 3209C>A 3210C>T), S1021A (3211A>G 3212G>C 3213T>A), K1023A (3217A>G 3218A>C 3219A>T), Q1028V (3232C>G 3233A>T 3234A>T), S1029A (3235T>G), M1034I (3252G>T), G1036S (3256G>A 3258A>T), S1049T (3296G>C), V1056C (3316G>T 3317T>G 3318T>C), G1067D (3350G>A), I1068T (3353T>C 3354T>C), F1074Y (3371T>A), I1075V (3373A>G 3375T>G), S1076A (3376T>G), I1082V (3394A>G 3396C>G), N1089D (3415A>G 3417C>T), V1095I (3433G>A), A1097V (3440C>T), D1103E (3459C>A), I1104L (3460A>C 3462T>G), K1106E (3466A>G 3468A>G), A1107H (3469G>C 3470C>A), I1108L (3472A>C 3474T>G), R1111K (3482G>A 3483A>G), T1113E (3487A>G 3488C>A 3489C>A), T1114R (3491C>G 3492T>A), L1115F (3493C>T 3495T>C), N1119K (3507C>A), S1122A (3514T>G 3516C>T), Q1129L (3535C>T 3536A>T 3537A>G), L1131M (3541T>A), R1142K (3575G>A), K1143R (3577A>C 3578A>G 3579G>C), Q1145P (3584A>C 3585A>C), P1152D (3604C>G 3605C>A 3606T>C), E1154Q (3610G>C 3612A>G), T1161S (3631A>T 3633C>T), V1162A (3635T>C), N1163K (3639T>A), M1178E (3682A>G 3683T>A), H1185S (3703C>T 3704A>C 3705T>C), G1186N (3706G>A 3707G>A 3708T>C), D1188E (3714T>G), D1189E (3717T>G), D1190P (3718G>C 3719A>C 3720T>A), S1195A (3733T>G), T1198G (3742A>G 3743C>G), S1201A (3751A>G 3752G>C 3753T>A), L1203I (3757C>A 3759G>T), T1206V (3766A>G 3767C>T 3768A>G), M1207L (3769A>C 3771G>T), N1209M (3776A>T 3777C>G), K1211R (3781A>C 3782A>G), Q1218A (3802C>G 3803A>C 3804G>A), E1219I (3805G>A 3806A>T 3807G>C), D1221N (3811G>A), C1232F (3845G>T), Y1235F (3854A>T), G1237A (3860G>C 3861C>A), M1251L (3901A>T), M1253L (3907A>C 3909G>T), M1255S (3913A>T 3914T>C 3915G>T), V1262E (3935T>A), P1263A (3937C>G 3939A>C), T1264Q (3940A>C 3941C>A), G1265P (3943G>C 3944G>C), E1266N (3946G>A 3948A>C), G1267S (3949G>T 3950G>C 3951T>G), D1268Q (3952G>C 3954T>G), K1269R (3956A>G 3957A>G), E1271L (3961G>C 3962A>T 3963G>T), R1273K (3968G>A), D1275N (3973G>A), E1280D (3990A>T), I1287A (4009A>G 4010T>C 4011T>G), I1289V (4015A>G 4017T>A), I1293T (4028T>C), H1294F (4030C>T 4031A>T), I1298V (4042A>G), T1343C (4177A>T 4178C>G 4179C>T), L1347A (4189T>G 4190T>C), I1351L (4201A>C), D1358N (4222G>A), S1362H (4234T>C 4235C>A), S1365T (4243T>A 4245T>A), L1366V (4246C>G 4248T>G), N1369S (4256A>G 4257C>T), N1370S (4259A>G), A1373S (4267G>T 4269T>A), T1377E (4279A>G 4280C>A 4281T>G), V1392I (4324G>A 4326G>C), E1404A (4361A>C 4362G>T), Y1405F (4364A>T), D1407G (4370A>G), T1408D (4372A>G 4373C>A 4374C>T), K1409E (4375A>G), E1412S (4384G>A 4385A>G 4386G>T), A1414E (4391C>A 4392T>G), A1420T (4408G>A 4410T>A), D1434N (4450G>A 4452T>C), V1436L (4456G>T 4458T>G), I1475V (4573A>G 4575T>C), A1476K (4576G>A 4577C>A 4578C>A), R1478K (4583G>A), A1479S (4585G>T), I1480V (4588A>G 4590A>G), V1482I (4594G>A 4596A>T), G1485N (4603G>A 4604G>A), L1505M (4663C>A), S1511D (4681T>G 4682C>A 4683A>T), N1514S (4690A>T 4691A>C 4692T>C), Q1517R (4699C>A 4700A>G 4701G>A), D1520E (4710C>A), M1521A (4711A>G 4712T>C 4713G>T), N1537E (4759A>G 4761C>G), Q1538K (4762C>A), V1539T (4765G>A 4766T>C 4767G>A), Q1540K (4768C>A 4770G>A), Q1542L (4775A>T), H1562Q (4836C>A), I1571V (4861A>G 4863A>C), M1575I (4875G>T), P1582V (4894C>G 4895C>T), T1587V (4909A>G 4910C>T 4911T>A), V1588Q (4912G>C 4913T>A 4914G>A), M1591Y (4921A>T 4922T>A 4923G>T), V1594I (4930G>A 4932G>T), L1598I (4942C>A 4944T>C), R1606Q (4966A>C 4967G>A 4968A>G), V1607A (4970T>C 4971T>C), A1608T (4972G>A 4974A>T), K1611S (4982A>G 4983G>T), L1612F (4986A>T), T1613V (4987A>G 4988C>T 4989T>A), Y1614N (4990T>A), Q1617F (4999C>T 5000A>T 5001G>T), C1618G (5002T>G 5004T>A), R1621K (5012G>A 5013A>G), E1622Q (5014G>C), G1624N (5020G>A 5021G>A 5022A>T), G1626P (5026G>C 5027G>C), A1627I (5029G>A 5030C>T 5031A>T), T1628M (5033C>T 5034A>G), F1630M (5038T>A 5040T>G), S1632T (5044T>A), M1634Y (5050A>T 5051T>A 5052G>T), V1636L (5056G>C 5058G>T), Q1638S (5062C>A 5063A>G 5064G>T), E1639Q (5065G>C), I1641L (5071A>T 5073C>G), A1642T (5074G>A), K1643E (5077A>G), A1648E (5093C>A 5094A>G), A1650I (5098G>A 5099C>T 5100G>T), L1651G (5101T>G 5102T>G), T1652V (5104A>G 5105C>T 5106A>T), I1653L (5107A>T 5109C>G), C1654M (5110T>A 5111G>T 5112T>G), M1656L (5116A>C 5118G>A), A1657S (5119G>T 5121C>A), K1658D (5122A>G 5124A>T), S1660K (5128T>A 5129C>A 5130T>A), I1661N (5132T>A 5133C>T), F1663Y (5138T>A), P1668A (5152C>G), N1670E (5158A>G 5160T>A), V1672I (5164G>A), E1673del (5167_5169delGAA), R1677E (5179A>G 5180G>A 5181A>G), D1678G (5183A>G), L1679V (5185C>G), N1680H (5188A>C), W1681E (5191T>G 5192G>A 5193G>A), A1683V (5198C>T 5199C>G), N1685D (5203A>G 5205C>T), I1686V (5206A>G 5208T>A), G1687F (5209G>T 5210G>T), A1695S (5233G>T 5235C>A), K1696N (5238G>C), T1697Q (5239A>C 5240C>A 5241T>A), E1698Q (5242G>C), E1700S (5248G>T 5249A>C 5250G>A), P1701S (5251C>T 5253T>A), C1703N (5257T>A 5258G>A), V1705L (5263G>C 5265A>G), K1712R (5284A>C 5285A>G 5286G>C), K1713R (5288A>G 5289G>A), S1714T (5291G>C 5292C>A), K1715T (5294A>C), S1722N (5315G>A), M1730I (5340G>A), M1735L (5353A>T), S1736A (5356T>G), L1739Q (5366T>A), A1741H (5371G>C 5372C>A), G1744K (5380G>A 5381G>A 5382C>G), K1745S (5384A>G 5385A>T), M1746L (5386A>C), S1747E (5389T>G 5390C>A), E1748V (5393A>T 5394G>T), D1749G (5396A>G 5397C>T), I1750D (5398A>G 5399T>A), A1751G (5402C>G 5403T>G), D1754S (5410G>T 5411A>C), V1756T (5416G>A 5417T>C 5418G>C), V1765I (5443G>A 5445G>A), S1766A (5446A>G 5447G>C 5448C>T), M1773F (5467A>T 5469G>T), V1775L (5473G>T), R1778H (5483G>A), C1781R (5491T>A 5493C>G), I1782A (5494A>G 5495T>C 5496T>C), E1783T (5497G>A 5498A>C 5499G>A), I1785L (5503A>C 5505C>T), I1792M (5526T>G), G1793A (5528G>C 5529C>T), N1794G (5530A>G 5531A>G 5532T>G), V1806A (5567T>C 5568T>G), F1809L (5577T>G), Q1811K (5581C>A), P1814E (5590C>G 5591C>A 5592T>A), Q1815D (5593C>G 5595G>C), Q1815_E1816insT (5595_5596insACC), E1816S (5596G>T 5597A>C), G1817H (5599G>C 5600G>A 5601G>C), M1819L (5605A>T), A1820G (5609C>G), V1822S (5614G>A 5615T>G), S1824T (5620T>A 5622G>A), S1825del (5623_5625delTCT), L1826Y (5627T>A 5628G>T), Q1827G (5629C>G 5630A>G 5631A>C), S1828T (5633G>C 5634T>C), S1829T (5636G>C), M1831N (5642T>A 5643G>C), Q1833H (5649A>T), Q1833_G1834insHS (5649_5650insCACAGT), G1834A (5651G>C 5652T>A), E1836S (5656G>T 5657A>C 5658G>A), L1837V (5659C>G), Q1839L (5666A>T 5667A>T), I1840L (5668A>C 5670T>A), V1842I (5674G>A 5676C>A), I1843V (5677A>G 5679C>T), H1844P (5681A>C 5682C>T), G1845S (5683G>T 5684G>C 5685T>C), M1849E (5695A>G 5696T>A 5697G>A), E1850K (5698G>A 5700A>G), L1853V (5707T>G 5709G>T), M1855L (5713A>T), V1858T (5722G>A 5723T>C 5724T>A), V1859I (5725G>A 5727G>T), H1862E (5734C>G 5736T>A), G1866A (5747G>C), N1867Q (5749A>C 5751C>G), F1868C (5753T>G 5754T>C), T1869S (5755A>T 5757C>A), D1874K (5770G>A 5772T>G), Q1875H (5775A>T), L1876A (5776T>G 5777T>C 5778G>T), F1877L (5781T>G), A1878E (5783C>A 5784A>G), S1879T (5786G>C 5787T>A), E1881S (5791G>T 5792A>C 5793A>T), K1882V (5794A>G 5795A>T 5796G>C), C1883S (5797T>A), E1886T (5806G>A 5807A>C 5808A>T), L1888A (5812C>G 5813T>C 5814T>C), K1890R (5819A>G 5820G>A), N1891G (5821A>G 5822A>G 5823C>A), T1892V (5824A>G 5825C>T 5826G>C), L1894V (5830C>G), E1899K (5845G>A 5847G>A), K1900E (5848A>G), M1901F (5851A>T 5853G>C), T1902S (5854A>T 5856T>C), K1904H (5860A>C 5862A>C), V1906C (5866G>T 5867T>G 5868G>T), A1907E (5870C>A), Q1908M (5872C>A 5873A>T 5874A>G), V1910L (5878G>C 5880C>T), S1911E (5881A>G 5882G>A 5883T>G), V1913L (5887G>C 5889A>G), K1914E (5890A>G), Q1915S (5893C>A 5894A>G 5895G>T), F1918V (5902T>G 5904C>A), Y1919V (5905T>G 5906A>T 5907T>A), P1920F (5908C>T 5909C>T), K1921M (5912A>T), A1922G (5915C>G 5916C>G), T1923S (5918C>G 5919A>C), T1924V (5920A>G 5921C>T 5922C>T), V1925K (5923G>A 5924T>A), E1926K (5926G>A), L1928M (5932C>A 5934C>G), K1929L (5935A>T 5936A>T), V1930S (5938G>T 5939T>C 5940A>C), A1932M (5944G>A 5945C>T 5946T>G), D1933N (5947G>A), A1934Q (5950G>C 5951C>A 5952T>A), H1935Q (5955T>G), H1935_T1936insM (5955_5956insATG), T1936S (5956A>T 5958C>A), I1937K (5960T>A 5961T>A), V1938T (5962G>A 5963T>C), K1939Q (5965A>C), T1940V (5968A>G 5969C>T), K1942S (5975A>G 5976A>C), F1944C (5981T>G 5982C>T), R1945C (5983C>T 5985G>T), G1946del (5986_5988delGGA), V1948T (5992G>A 5993T>C 5994T>A), R1950S (5998C>A 6000G>T), N1952A (6004A>G 6005A>C 6006T>A), F1953W (6008T>G 6009T>G), S1955T (6014G>C), P1956K (6016C>A 6017C>A 6018T>G), E1957D (6021G>C), W1958I (6022T>A 6023G>T 6024G>T), W1958_K1959insV (6024_6025insGTC), K1959M (6026A>T 6027A>G), I1960T (6029T>C 6030T>A), G1961K (6031G>A 6032G>A), E1963K (6037G>A), N1964K (6042C>G), T1965P (6043A>C), M1966del (6046_6048delATG), A1967G (6050C>G), T1968S (6052A>T 6054A>T), L1969I (6055C>A 6057C>T), S1970P (6058T>C 6060A>T), S1973T (6068G>C 6069T>A), L1974V (6070C>G 6072T>G), L1975F (6073C>T), P1976L (6077C>T 6078A>T), L1979P (6086T>C 6087G>T), S1982L (6094T>C 6095C>T), A1983V (6098C>T 6099C>T), M1984V (6100A>G), K1985R (6104A>G 6105G>A), L1987M (6109T>A), S1989R (6115A>C), R1990K (6119G>A)

| | Begin | End | Coverage | Score | Concordance | Matches | Identities | I/D/M/F* | Stop Codons |
|---|---|---|---|---|---|---|---|---|---|
| **NT** | **33** | **6128** | **84.2%** | **3374** | **28.2%** | **6043 (98.8%)** | **3915 (64.0%)** | **21/53** | |

GCT2GCA (156T>A), CTT3TTT (157C>T), AAA5AAG (165A>G), TTT7TGG (170T>G 171T>G), GGT8AGT (172G>A), TCT9CCT (175T>C), TCA10CCT (178T>C 180A>T), TCT11GCT (181T>G), GAG12_GTT17del (184_201delGAGAAGACTGGATCAGTT), CCC18TCT (202C>T 204C>T), TCT19TCA (207T>A), ACT20CCT (208A>C), TCT21TCA (213T>A), GGT24GCA (221G>C 222T>A), CTG26GTT (226C>G 228G>T), GGT28TCT (232G>T 233G>C), ATC30TTT (238A>T 240C>T), TCA31TCT (243A>T), AAA32TCT (244A>T 245A>C 246A>T), TTC33CTT (247T>C 249C>T), AAA35AAG (255A>G), GCT36ACT (256G>A), ATT37CTC (259A>C 261T>C), AGC38TCC (262A>T 263G>C), AAC39AAT (267C>T), ATC40GTT (268A>G 270C>T), ACA41AAT (272C>A 273A>T), TCA42AAA (274T>A 275C>A), GTA43GTT (279A>T), AAA46ACC (287A>C 288A>C), TCT48GTA (292T>G 293C>T 294T>A), CAT50CAC (300T>C), TTA51ATA (301T>A), GAT52GAG (306T>G), TTG54GTC (310T>G 312G>C), AAA55AAG (315A>G), CCA56CCG (318A>G), TCT57CCA (319T>C 321T>A), GTG58GTT (324G>T), ACT59ATG (326C>T 327T>G), GAT60GCA (329A>C 330T>A), TCT62GCA (334T>G 336T>A), TCT63ACA (337T>A 339T>A), TCT64TCA (342T>A), GTT66GCC (347T>C 348T>C), TGT69TGC (357T>C), TCC70AAT (358T>A 359C>A 360C>T), GCA71GCT (363A>T), TTG72GTA (364T>G 366G>A), AAT73GAA (367A>G 369T>A), AAA74AGA (371A>G), GTG75GTT (375G>T), CTT76TTA (376C>T 378T>A), ATA79ATT (387A>T), GGT80GCA (389G>C 390T>A), TCC81AAA (391T>A 392C>A 393C>A), TTG82ATA (394T>A 396G>A), ATT83ATG (399T>G), CCT85CCC (405T>C), TTT86TTC (408T>C), CTG87ATC (409C>A 411G>C), AAA88AAG (414A>G), GCA89GGT (416C>G 417A>T), TAC90TAT (420C>T), TCT91GAT (421T>G 422C>A), TTG93GTA (427T>G 429G>A), GCA94AAA (430G>A 431C>A), ACC95ACT (435C>T), TCT99ACT (445T>A), ATC100ATA (450C>A), AAG101GTA (451A>G 452A>T 453G>A), ATG103CTT (457A>C 459G>T), GTT104GTA (462T>A), TTA105CTT (463T>C 465A>T), CGG106AAG (466C>A 467G>A), TTG107ATG (469T>A), TTC108GTT (472T>G 474C>T), GAA109TTA (475G>T 476A>T), TTG111GTC (481T>G 483G>C), TCC112AAG (484T>A 485C>A 486C>G), AAT113GAT (487A>G), GTT115TTC (493G>T 495T>C), AGA116ATC (497G>T 498A>C), ATA117CTA (499A>C), TTT119TAC (506T>A 507T>C), CAG120GAA (508C>G 510G>A), GTT122TTA (514G>T 516T>A), GAG124GAA (522G>A), AAG125AAA (525G>A), AGT126GTT (526A>G 527G>T), GAT128TCA (532G>T 533A>C 534T>A), GTA129TTA (535G>T), ATT131ATA (543T>A), GTA132GTG (546A>G), GTT133TTT (547G>T), TAT134ATA (550T>A 551A>T 552T>A), GGT135GCT (554G>C), TTG137TTA (561G>A), GCA140GCT (570A>T), ATA141TTG (571A>T 573A>G), ACT142GCT (574A>G), CTT143ATT (577C>A), CTT144TTG (580C>T 582T>G), GTT145CTA (583G>C 585T>A), TTG146CTA (586T>C 588G>A), TTG147CTG (589T>C), TGT148TAT (593G>A), GTT149GTA (597T>A), TCA152GTA (604T>G 605C>T), GAC153GAT (609C>T), ATA154ATT (612A>T), ATT155GTA (613A>G 615T>A), GAA156ACG (616G>A 617A>C 618A>G), GGA157CCT (619G>C 620G>C 621A>T), GTA158TTT (622G>T 624A>T), TCT159AAT (625T>A 626C>A), AGT160AAT (629G>A), ACC161GTA (631A>G 632C>T 633C>A), ATT162GTT (634A>G), AAA163GTT (637A>G 638A>T 639A>T), AAT164ACT (641A>C), ATT165TTG (643A>T 645T>G), TTC166TTT (648C>T), TTA167TCA (650T>C), ATA168GTT (652A>G 654A>T), GTA169GTT (657A>T), GGG170GGA (660G>A), ATG172TTT (664A>T 666G>T), TTT173TTC (669T>C), AGT175TCA (673A>T 674G>C 675T>A), AAC178AAT (684C>T), TTG179TTA (687G>A), GAC180GAT (690C>T), CCC183CCA (699C>A), AAG184ACA (701A>C 702G>A), GCT186TCT (706G>T), GAG187GGG (710A>G), TTC189TTT (717C>T), ACA190TCC (718A>T 720A>C), ATG191CAG (721A>C 722T>A), GTA192CTG (724G>C 726A>G), GCA193GCG (729A>G), AGT196GCA (736A>G 737G>C 738T>A), ATG197GTA (739A>G 741G>A), CTA198CTT (744A>T), GAG201GAA (753G>A), TCT202AGC (754T>A 755C>G 756T>C), GTC203TTG (757G>T 759C>G), TCA204ACG (760T>A 762A>G), CAC205CAT (765C>T), ACT206ACA (768T>A), ACA208ACT (774A>T), TCC209AGC (775T>A 776C>G), CAG210CAA (780G>A), TTG211CTT (781T>C 783G>T), ATG212TTA (784A>T 786G>A), TCA213TCC (789A>C), ACT214TCA (790A>T 792T>A), ATT215TTG (793A>T 795T>G), ATG216TTG (796A>T), GCA217GCT (801A>T), ATT220ATC (810T>C), TCC221TCA (813C>A), ACG222ACT (816G>T), CTC223TTA (817C>T 819C>A), GTG224GTA (822G>A), ATA226GTT (826A>G 828A>T), GCA227GCT (831A>T), GTG228GTT (834G>T), CCA229CCT (837A>T), GGA230GGG (840A>G), CCC232CCA (846C>A), TTG235CTT (853T>C 855G>T), AGC236TCA (856A>T 857G>C 858C>A), AAT237AAC (861T>C), CCC238CCA (864C>A), CTC239CTG (867C>G), ATT242ATC (876T>C), TCA245TCT (885A>T), ACT246GCT (886A>G), GGG247GGA (891G>A), GGG249GGT (897G>T), GCT250GCG (900T>G), CAA255CAG (915A>G), TTG256CTG (916T>C), TTT257TTC (921T>C), ACA258CAG (922A>C 923C>A 924A>G), AGA261AGG (933A>G), AAT262AAC (936T>C), AAA264AAG (942A>G), TGC266TGT (948C>T), ACT267ACA (951T>A), TCA268TCT (954A>T), GCA270GCC (960A>C), TTT271TTC (963T>C), TCT272TCA (966T>A), GTG274GTT (972G>T), GAA276GAG (978A>G), ATT277ATA (981T>A), ATA278GTG (982A>G 984A>G), GGT280AGC (988G>A 990T>C), GCT281ATT (991G>A 992C>T), TTC284TTT (1002C>T), AAG285AAA (1005G>A), GTG288GTA (1014G>A), CTT289TTG (1015C>T 1017T>G), TCT290TCA (1020T>A), GCA291GCG (1023A>G), ATT292ATA (1026T>A), AGC293AAT (1028G>A 1029C>T), GCA294GCT (1032A>T), ACA295ACC (1035A>C), TTG296TTA (1038G>A), TCT297TCC (1041T>C), ACT298ACC (1044T>C), TTG300CTT (1048T>C 1050G>T), GAA302CAG (1054G>C 1056A>G), CAG305ACG (1063C>A 1064A>C), GTG307GTT (1071G>T), GAT308TCG (1072G>T 1073A>C 1074T>G), GCT309GAG (1076C>A 1077T>G), TGT311GCT (1081T>G 1082G>C), GAT312GAA (1086T>A), GCT314TCT (1090G>T), ACC315ATG (1094C>T 1095C>G), AGG316CGC (1096A>C 1098G>C), TTG317CTT (1099T>C 1101G>T), GAG318GAA (1104G>A), AAC319AAT (1107C>T), GCC321GCA (1113C>A), AAA323CCT (1117A>C 1118A>C 1119A>T), GCC324GCA (1122C>A), TTC325TTT (1125C>T), CCT326CCA (1128T>A), GAC327ATT (1129G>A 1130A>T 1131C>T), AAA328AAG (1134A>G), AAT330CAA (1138A>C 1140T>A), CAC331CAT (1143C>T), CTT332TTA (1144C>T 1146T>A), AGG333AAA (1148G>A 1149G>A), GCA336GCC (1158A>C), CTG337GTC (1159C>G 1161G>C), AAG338AAA (1164G>A), ATT339ATA (1167T>A), GCA341GCT (1173A>T), ACA345ACC (1185A>C), CAC346CAT (1188C>T), GTT348GTC (1194T>C), GCC349GCA (1197C>A), ATG352TTG (1204A>T), CAT354CAC (1212T>C), CGA355AGG (1213C>A 1215A>G), GTG356GTT (1218G>T), AGT357TCT (1219A>T 1220G>C), GCA358GCT (1224A>T), GCT359GCC (1227T>C), ATC360ATA (1230C>A), ACT361ACG (1233T>G), CTT363CTA (1239T>A), GAA364GAT (1242A>T), AAG365AAA (1245G>A), GTC366CTG (1246G>C 1248C>G), TAT367GTT (1249T>G 1250A>T), GCG368GCT (1254G>T), AGT370GCC (1258A>G 1259G>C 1260T>C), TGC371AGC (1261T>A), AGA372CAA (1264A>C 1265G>A), GTT374TCT (1270G>T 1271T>C), GTT376GTG (1278T>G), GGT377GGC (1281T>C), CAG378CAA (1284G>A), AGA380AGG (1290A>G), GAA382GAG (1296A>G), CCC383CCT (1299C>T), GTT386GTG (1308T>G), TAT389TTT (1316A>T), GGT390GGA (1320T>A), GCC391GCT (1323C>T), GGG393GGA (1329G>A), ACA398ACG (1344A>G), GCA399TGT (1345G>T 1346C>G 1347A>T), CGA401CGC (1353A>C), TTG402CTT (1354T>C 1356G>T), TTG408CTC (1372T>C 1374G>C), GAT409GAC (1377T>C), CGC410GCA (1378C>G 1379G>C 1380C>A), GGG412GGT (1386G>T), GAG413GAA (1389G>A), CCT414CCA (1392T>A), CTT416CTG (1398T>G), AGA418CGC (1402A>C 1404A>C), CTT419TTG (1405C>T 1407T>G), GCT421GCG (1413T>G), GTT422GTA (1416T>A), AGC423AGT (1419C>T), AAG424AAA (1422G>A), CGC425AGA (1423C>A 1425C>A), GCC427GCA (1431C>A), TCT430TCC (1440T>C), AAT431AAC (1443T>C), GCC433GCA (1449C>A), CAC434CAT (1452C>T), CAG435CAA (1455G>A), GCT437GCA (1461T>A), TTG439CTT (1465T>C 1467G>T), GAC442GAT (1476C>T), GGA444GGT (1482A>T), GCA445GCT (1485A>T), TTG446CTT (1486T>C 1488G>T), AGA447CGG (1489A>C 1491A>G), GGA451GGG (1503A>G), CAA452CAG (1506A>G), CAA454CAG (1512A>G), ATT456ATA (1518T>A), ATT462ATC (1536T>C), TCC464TCT (1542C>T), ACA465ACC (1545A>C), CCA467CCT (1551A>T), CCA469CCT (1557A>T), CTG470CTT (1560G>T), CCC471CCA (1563C>A), GCA473GCT (1569A>T), GCT474GCA (1572T>A), GTG475GTC (1575G>C), GAA476GAG (1578A>G), GAC477GAT (1581C>T), AAG478AAA (1584G>A), GGG479GGA (1587G>A), CAT481CAC (1593T>C), ACT483ACA (1599T>A), TAT486TAC (1608T>C), GCA489GCC (1617A>C), AAT492AAC (1626T>C), CTT493TTG (1627C>T 1629T>G), GTA494ATA (1630G>A), TCA495TCT (1635A>T), GCA496GCT (1638A>T), CCT497CCA (1641T>A), GCC498GCT (1644C>T), CTT502CTA (1656T>A), ACC503ACG (1659C>G), GCT507GCA (1671T>A), GAG509GAA (1677G>A), CGT510AGG (1678C>A 1680T>G), AGA511AGG (1683A>G), AGG512AGA (1686G>A), ATC514ATT (1692C>T), TGT518TGC (1704T>C), ATC519AGA (1706T>G 1707C>A), GAA521GTG (1712A>T 1713A>G), GGT522GGC (1716T>C), GAA523GAT (1719A>T), TCT525GAT (1723T>G 1724C>A), ACA526CCA (1726A>C), GAG527AAT (1729G>A 1731G>T), GAC528GCT (1733A>C 1734C>T), ACC530ACA (1740C>A), GGA531GCC (1742G>C 1743A>C), CTG533CTA (1749G>A), ATG534GTT (1750A>G 1752G>T), AAC536GAT (1756A>G 1758C>T), ATT537GTG (1759A>G 1761T>G), CGG541AGA (1771C>A 1773G>A), AGG542AGA (1776G>A), ATA546ATC (1788A>C), CAC550GCT (1798C>G 1799A>C 1800C>T), AGT552TCC (1804A>T 1805G>C 1806T>C), TAC553TAT (1809C>T), TCG554GAT (1810T>G 1811C>A 1812G>T), TTA557TTG (1821A>G), GAC558CAA (1822G>C 1824C>A), GCC562GCA (1836C>A), TGT564TGC (1842T>C), CAG565CGG (1844A>G), GTT566GTG (1848T>G), CAT567CAC (1851T>C), GCT568TAT (1852G>T 1853C>A), ATA570GTG (1858A>G 1860A>G), AAG572AGC (1865A>G 1866G>C), CAG573CAA (1869G>A), CTG574CTT (1872G>T), TTT575TAC (1874T>A 1875T>C), GAA576AAT (1876G>A 1878A>T), GCA577ACT (1879G>A 1881A>T), GAA578ACA (1882G>A 1883A>C), ACT579TCG (1885A>T 1887T>G), GGC580GGA (1890C>A), ATT581TTG (1891A>T 1893T>G), TCA582GCT (1894T>G 1896A>T), AAT583GAT (1897A>G), ACA585GTT (1903A>G 1904C>T 1905A>T), GCC586GCT (1908C>T), GTT588GTC (1914T>C), GCT589AAT (1915G>A 1916C>A), GTA590GTT (1920A>T), TCT591GCT (1921T>G), TCT592GTT (1924T>G 1925C>T), GAA593GAT (1929A>T), GAG594GAA (1932G>A), GTT595GTG (1935T>G), GTG596CTG (1936G>C), GCA597GAT (1940C>A 1941A>T), GTT599TTC (1945G>T 1947T>C), GAT600CAA (1948G>C 1950T>A), GGG601GAT (1952G>A 1953G>T), GCT602GCC (1956T>C), CGC603del (1957_1959delCGC), TTC604TTA (1962C>A), AAA605CAA (1963A>C), ACA606GTT (1966A>G 1967C>T 1968A>T), AAA607TTG (1969A>T 1970A>T 1971A>G), CAA608CCT (1973A>C 1974A>T), GAT609CCT (1975G>C 1976A>C), GTG610GAG (1979T>A), ATA612GCA (1984A>G 1985T>C), GTG613GCT (1988T>C 1989G>T), ACT614TAT (1990A>T 1991C>A), CCC615GAG (1993C>G 1994C>A 1995C>G), ACA616CAG (1996A>C 1997C>A 1998A>G), GTT617ACA (1999G>A 2000T>C 2001T>A), ATA618ATT (2004A>T), GAG620GAA (2010G>A), GAA621GCT (2012A>C 2013A>T), GAT622GAA (2016T>A), AGA623AAA (2018G>A), GTG624ATT (2020G>A 2022G>T), ATT625GTT (2023A>G), TCT627TCC (2031T>C), AAA628CAG (2032A>C 2034A>G), TTA630CTC (2038T>C 2040A>C), GTG632GTT (2046G>T), GAA633GAG (2049A>G), CTT635CTC (2055T>C), TTT637TAT (2060T>A), TAT639TAC (2067T>C), CAG640CAA (2070G>A), GGC641GGG (2073C>G), TTG643ATG (2077T>A), GAC644AAC (2080G>A), CCG645AAA (2083C>A 2084C>A 2085G>A), GAG646TTT (2086G>T 2087A>T 2088G>T), AGT647GAG (2089A>G 2090G>A 2091T>G), TTG648TTT (2094G>T), TTT649TTC (2097T>C), CCT650CCA (2100T>A), TAT653CTC (2107T>C 2108A>T 2109T>C), CAA654TCC (2110C>T 2111A>C 2112A>C), AAA655AAG (2115A>G), CAA656CAG (2118A>G), CTC658ATC (2122C>A), TTT659TAT (2126T>A), TCC661GTG (2131T>G 2132C>T 2133C>G), GAT663TCT (2137G>T 2138A>C), GCA664TCT (2140G>T 2142A>T), GAG665GAT (2145G>T), CAC666GCC (2146C>G 2147A>C), CAG667AAA (2149C>A 2151G>A), CAA668CAG (2154A>G), GTC669AAT (2155G>A 2156T>A 2157C>T), TTC670TTT (2160C>T), ATG671GAA (2161A>G 2162T>A 2163G>A), AGA672ATG (2165G>T 2166A>G), ATG674GTA (2170A>G 2172G>A), ACT675AAT (2174C>A), GAC676GAT (2178C>T), ATG677TTG (2179A>T), ATC678TTG (2182A>T 2184C>G), TAC679GTC (2185T>G 2186A>T), GGT681GGA (2193T>A), GCC682GCA (2196C>A), AAT683AGT (2198A>G), GCT684CCT (2200G>C), GAA685CAG (2203G>C 2205A>G), CAA686CAG (2208A>G), TAC687TAT (2211C>T), AGG688GCT (2212A>G 2213G>C 2214G>T), CAA692CGG (2225A>G 2226A>G), AAT693AAC (2229T>C), CTC699ATC (2245C>A), TTT702TTC (2256T>C), AAA703AAG (2259A>G), AGC704AGT (2262C>T), ATC706ATT (2268C>T), GCT708AGC (2272G>A 2273C>G 2274T>C), TCC709ACA (2275T>A 2277C>A), ATC711ATT (2283C>T), TGT712TCT (2285G>C), AAG714GAG (2290A>G), AAG715CTG (2293A>C 2294A>T), GCT717GAT (2300C>A), GTA718GTC (2304A>C), ACA720CGC (2308A>C 2309C>G 2310A>C), GAG721AAT (2311G>A 2313G>T), ATG722CTT (2314A>C 2316G>T), CGC723GAG (2317C>G 2318G>A 2319C>G), ACA724CAA (2320A>C 2321C>A), GGA725GGG (2325A>G), TTT726AAA (2326T>A 2327T>A 2328T>A), GCA727GCG (2331A>G), CAT728TAT (2332C>T), AAT729AAG (2337T>G), TGT730GCG (2338T>G 2339G>C 2340T>G), ATT731CTT (2341A>C), GAT732CAG (2344G>C 2346T>G), GTT735GCA (2354T>C 2355T>A), TTG738CTT (2362T>C 2364G>T), ATT739ACG (2366T>C 2367T>G), TGC740TGT (2370C>T), ATC741ATT (2373C>T), CAG742GAA (2374C>G 2376G>A), TTT747AGA (2389T>A 2390T>G 2391T>A), GTG748GTA (2394G>A), TTT750TTC (2400T>C), TTG751CTG (2401T>C), ACA753ACT (2409A>T), GCC754GCT (2412C>T), CGT757CGC (2421T>C), TAT758TAC (2424T>C), ACC761AGG (2432C>G 2433C>G), TCC764GCA (2440T>G 2442C>A), TCC764_AAA765insGGT (2442_2443insGGT), AAA765ACT (2444A>C 2445A>T), GAG766GAC (2448G>C), GCC767AAG (2449G>A 2450C>A 2451C>G), AAA768AAG (2454A>G), GAC769AGA (2455G>A 2456A>G 2457C>A), TCT770AAT (2458T>A 2459C>A), GTC772ATT (2464G>A 2466C>T), AAG773ATC (2468A>T 2469G>C), ATA774GAA (2470A>G 2471T>A), TGT775TGC (2475T>C), TAC776ATG (2476T>A 2477A>T 2478C>G), CAC777ATT (2479C>A 2480A>T 2481C>T), AAA778GGT (2482A>G 2483A>G 2484A>T), ATT779ATC (2487T>C), GTG780AAG (2488G>A 2489T>A), GAT781GAC (2493T>C), TAC782AAG (2494T>A 2496C>G), AAA784ATC (2501A>T 2502A>C), GAA785CAG (2503G>C 2505A>G), ACT786ATG (2507C>T 2508T>G), TGG788AAT (2512T>A 2513G>A 2514G>T), AAC789GAC (2515A>G), TTA790TTG (2520A>G), CCA791CCC (2523A>C), TAT792TTC (2525A>T 2526T>C), GCA793ATC (2527G>A 2528C>T 2529A>C), CTG794TTG (2530C>T), AGG795AAG (2534G>A), GGC801GGA (2553C>A), CTA802CTG (2556A>G), ATC804TTG (2560A>T 2562C>G), ATG805ATA (2565G>A), GCG806GCA (2568G>A), GGA809GGC (2577A>C), TTG810ACC (2578T>A 2579T>C 2580G>C), TTT811ATG (2581T>A 2583T>G), GGG812AAT (2584G>A 2585G>A 2586G>T), GGA813GCC (2588G>C 2589A>C), ATA814GTT (2590A>G 2592A>T), ACA815ACT (2595A>T), TTT817TTC (2601T>C), CTT818ATT (2602C>A), GCT819ACT (2605G>A), TGT820TGC (2610T>C), TGG821CTC (2611T>C 2612G>T 2613G>C), CAA822AAG (2614C>A 2616A>G), AGC823GCA (2617A>G 2618G>C 2619C>A), AAT824GCA (2620A>G 2621A>C 2622T>A), TCT825ACT (2623T>A), TTT826TCC (2627T>C 2628T>C), CCA827CCT (2631A>T), AGT828TTA (2632A>T 2633G>T 2634T>A), AGT830GAT (2638A>G 2639G>A), GGA831GGC (2643A>C), GGT833GGG (2649T>G), GGT834GGC (2652T>C), GTC835GTA (2655C>A), ACA836ACC (2658A>C), AGC841AGT (2673C>T), TCT843TCC (2679T>C), AAA845AAG (2685A>G), AAG846AAA (2688G>A), AAC847AAT (2691C>T), AAG848AAA (2694G>A), GGC849GGG (2697C>G), CCC850AAG (2698C>A 2699C>A 2700C>G), TCT851ACT (2701T>A), CGT852CAT (2705G>A), CTG853TTC (2707C>T 2709G>C), AAG854GCA (2710A>G 2711A>C 2712G>A), AAT855AAA (2715T>A), TTG856TCC (2717T>C 2718G>C), GTG858CTA (2722G>C 2724G>A), CAG860CAA (2730G>A), AGT861GCG (2731A>G 2732G>C 2733T>G), AGC862GCA (2734A>G 2735G>C 2736C>A), GCT863GAT (2738C>A), ATT865AAC (2744T>A 2745T>C), ATT865_ACA866insCAT (2745_2746insCAT), ACA866GTG (2746A>G 2747C>T 2748A>G), CAA867GAC (2749C>G 2751A>C), GAA868CGA (2752G>C 2753A>G), GCT871GAA (2762C>A 2763T>A), GAA872CAG (2764G>C 2766A>G), GGG874GGA (2772G>A), GTC876GTT (2778C>T), AAC877AAT (2781C>T), CAA878GAA (2782C>G), GCA879AGT (2785G>A 2786C>G 2787A>T), CTC880CTG (2790C>G), AAA881CAC (2791A>C 2793A>C), AAG882AAA (2796G>A), AAT883AAC (2799T>C), ATG884TTG (2800A>T), ATT885GTT (2803A>G), GTA886GTT (2808A>T), CTG887TTG (2809C>T), CGG888AGA (2812C>A 2814G>A), CTT889ATT (2815C>A), GGA890GCT (2819G>C 2820A>T), GAG891GAA (2823G>A), GTG893GTC (2829G>C), TAC894TAT (2832C>T), ACA898ACG (2844A>G), GTG900GTT (2850G>T), TCA902TCT (2856A>T), GGG903GGT (2859G>T), ATA905ATT (2865A>T), ATG906CTC (2866A>C 2868G>C), GCA911GCC (2883A>C), TTC912TTT (2886C>T), CAC913AGA (2887C>A 2888A>G 2889C>A), AAT914AAC (2892T>C), GTG915ATG (2893G>A), AGA916AGG (2898A>G), GGA918GGC (2904A>C), ACA919ACT (2907A>T), ACC922TCC (2914A>T), ATT923ATC (2919T>C), ATC924ATT (2922C>T), CCT926GAG (2926C>G 2927C>A 2928T>G), AAC927CAT (2929A>C 2931C>T), TCA928TCC (2934A>C), AGG929ATT (2936G>T 2937G>T), TCA930TGT (2939C>G 2940A>T), AAG931AAT (2943G>T), TAC934TTC (2951A>T), AAC935AAT (2955C>T), CGG937ACC (2959C>A 2960G>C 2961G>C), GAA938CAA (2962G>C), TCC939TCA (2967C>A), AAG940AGG (2969A>G), GTC941TTG (2971G>T 2973C>G), ATT942ATA (2976T>A), GGA944GGT (2982A>T), ATC947GTG (2989A>G 2991C>G), CTG949TTG (2995C>T), CTA950TTG

Codon mutations:

(2998C>T 3000A>G), CGT951AGG (3001C>A 3003T>G), GTG952GTA (3006G>A), GGT953GGG (3009T>G), GAC954GAT (3012C>T), CCT955CCA (3015T>A), CCT960TCA (3028C>T 3030T>A), ATT962ATA (3036T>A), CGC963AGA (3037C>A 3039C>A), AAG964AAA (3042G>A), CCA967CCT (3051A>T), GAG971GAT (3063G>T), GTC972GTG (3066C>G), TGC973TGT (3069C>T), TTC974TAC (3071T>A), ACC975TCT (3073A>T 3075C>T), AAA976AAG (3078A>G), GGT977GGC (3081T>C), TCT978TCA (3084T>A), CAG979ATG (3085C>A 3086A>T), TTA981TTG (3093A>G), TGT982TGC (3096T>C), TGT983GTG (3097T>G 3098G>T 3099T>G), AGA984AGG (3102A>G), GCC985GCT (3105C>T), GTG986GTA (3108G>A), GCG987GCT (3111G>T), ACC989AAG (3116C>A 3117C>G), CCA991CCC (3123A>C), AGA992AAG (3125G>A 3126A>G), CTG993AAA (3127C>A 3128T>A 3129G>A), GGC994TCC (3130G>T 3131G>C), AAC995AAT (3135C>T), CTA996TTG (3136C>T 3138A>G), GAA997GAG (3141A>G), TTT998GTT (3142T>G), TTG999CTT (3145T>C 3147G>T), TTT1001TAT (3152T>A), TCC1006TCA (3168C>A), AAA1007AAG (3171A>G), TAT1009TAC (3177T>C), ACA1010ACC (3180A>C), GTC1011GAA (3182T>A 3183C>A), AAG1012AAA (3186G>A), GTG1013GTC (3189G>C), GAG1014ACA (3190G>A 3191A>C 3192G>A), GAG1016GGA (3197A>G 3198G>A), GAC1018GAT (3204C>T), TCT1019TCG (3207T>G), GCC1020AAT (3208G>A 3209C>A 3210C>T), AGT1021GCA (3211A>G 3212G>C 3213T>A), AAA1023GCT (3217A>G 3218A>C 3219A>T), TTG1024CTT (3220T>C 3222G>T), AGT1025AGC (3225T>C), TCG1026TCA (3228G>A), CAG1027CAA (3231G>A), CAA1028GTT (3232C>G 3233A>T 3234A>T), TCA1029GCA (3235T>G), GAA1031GAG (3243A>G), TAT1032TAC (3246T>C), CAT1033CAC (3249T>C), ATG1034ATT (3252G>T), GGA1036AGT (3256G>A 3258A>T), GGT1039GGA (3267T>A), GGA1042GGT (3276A>T), TTA1045CTC (3283T>C 3285A>C), CTT1046TTA (3286C>T 3288T>A), CCC1048CCA (3294C>A), AGT1049ACT (3296G>C), GTT1050GTG (3300T>G), AAA1053AAG (3309A>G), CAA1054CAG (3312A>G), CCT1055CCG (3315T>G), GTT1056TGC (3316G>T 3317T>G 3318T>C), GTG1058GTT (3324G>T), GGC1059GGA (3327C>A), ATT1060ATA (3330T>A), GCT1063GCA (3339T>A), TCA1064TCT (3342A>T), GGT1067GAT (3350G>A), ATT1068ACC (3353T>C 3354T>C), GCA1069GCT (3357A>T), GAA1071GAG (3363A>G), GGA1073GGT (3369A>T), TTC1074TAC (3371T>A), ATT1075GTG (3373A>G 3375T>G), TCT1076GCT (3376T>G), TCC1077TCA (3381C>A), AAT1078AAC (3384T>C), GCC1079GCA (3387C>A), ACG1080ACA (3390G>A), GCC1081GCA (3393C>A), ATC1082GTG (3394A>G 3396C>G), TTC1083TTT (3399C>T), CGG1084AGA (3400C>A 3402G>A), GAC1085GAT (3405C>T), CAG1086CAA (3408G>A), CTT1087TTG (3409C>T 3411T>G), AAC1089GAT (3415A>G 3417C>T), GGG1093GGA (3429G>A), CCT1094CCC (3432T>C), GTT1095ATT (3433G>A), GCG1097GTG (3440C>T), GCA1098GCT (3444A>T), CGA1101CGC (3453A>C), GAC1103GAA (3459C>A), ATT1104CTG (3460A>C 3462T>G), CTC1105TTG (3463C>T 3465C>G), AAA1106GAG (3466A>G 3468A>G), GCC1107CAC (3469G>C 3470C>A), ATT1108CTG (3472A>C 3474T>G), AGG1109AGA (3477G>A), AGT1110TCC (3478A>T 3479G>C 3480T>C), AGA1111AAG (3482G>A 3483A>G), GAA1112GAG (3486A>G), ACC1113GAA (3487A>G 3488C>A 3489C>A), ACT1114AGA (3491C>G 3492T>A), CTT1115TTC (3493C>T 3495T>C), GAA1117GAG (3501A>G), AAC1119AAA (3507C>A), CAA1120CAG (3510A>G), GTC1121GTG (3513C>G), TCC1122GCT (3514T>G 3516C>T), TTT1123TTC (3519T>C), CTT1124TTG (3520C>T 3522T>G), GTT1127GTA (3531T>A), CCT1128CCG (3534T>G), CAA1129TTG (3535C>T 3536A>T 3537A>G), TTG1131ATG (3541T>A), GCC1132GCT (3546C>T), GCA1133GCT (3549A>T), GTA1135GTG (3555A>G), CCC1136CCG (3558C>G), CAT1137CAC (3561T>C), ACT1139ACA (3567T>A), ACA1140ACC (3570A>C), CTC1141CTT (3573C>T), AGA1142AAA (3575G>A), AAG1143CGC (3577A>C 3578A>G 3579G>C), CAA1145CCC (3584A>C 3585A>C), TTG1146CTG (3586T>C), GCA1149GCT (3597A>T), GGT1151GGG (3603T>G), CCT1152GAC (3604C>G 3605C>A 3606T>C), GAA1154CAG (3610G>C 3612A>G), ACC1155ACT (3615C>T), GCT1156GCC (3618T>C), CCC1157CCA (3621C>A), TCA1158TCC (3624A>C), ATC1159ATT (3627C>T), CTC1160CTA (3630C>A), ACC1161TCT (3631A>T 3633C>T), GTC1162GCC (3635T>C), AAT1163AAA (3639T>A), CGG1166AGG (3646C>A), GAG1168GAA (3654G>A), GGA1169GGG (3657A>G), TTC1170TTT (3660C>T), CCA1172CCC (3666A>C), GCA1175GCT (3675A>T), GTG1177GTC (3681G>C), ATG1178GAG (3682A>G 3683T>A), AAA1179AAG (3687A>G), TAC1180TAT (3690C>T), ACT1183ACA (3699T>A), CAT1185TCC (3703C>T 3704A>C 3705T>C), GGT1186AAC (3706G>A 3707G>A 3708T>C), GAT1188GAG (3714T>G), GAT1189GAG (3717T>G), GAT1190CCA (3718G>C 3719A>C 3720T>A), GCT1192GCA (3726T>A), AAA1193AAG (3729A>G), TTG1194CTG (3730T>C), TCC1195GCC (3733T>G), GAA1197GAG (3741A>G), ACT1198GGT (3742A>G 3743C>G), CTC1199CTT (3747C>T), AGT1201GCA (3751A>G 3752G>C 3753T>A), TCA1202TCC (3756A>C), CTG1203ATT (3757C>A 3759G>T), CTG1204CTT (3762G>T), CCA1205CCT (3765A>T), ACA1206GTG (3766A>G 3767C>T 3768A>G), ATG1207CTT (3769A>C 3771G>T), AAA1208AAG (3774A>G), AAC1209ATG (3776A>T 3777C>G), CAA1210CAG (3780A>G), AAA1211CGA (3781A>C 3782A>G), GGC1215GGG (3795C>G), CCA1217CCT (3801A>T), CAG1218GCA (3802C>G 3803A>C 3804G>A), GAG1219ATC (3805G>A 3806A>T 3807G>C), GAT1221AAT (3811G>A), GTG1224GTA (3822G>A), CTG1226CTC (3828G>C), ATT1229ATA (3837T>A), TGT1232TTT (3845G>T), TAT1235TTT (3854A>T), GGC1237GCA (3860G>C 3861C>A), CTC1240TTG (3868C>T 3870C>G), ACA1242ACG (3876A>G), TCC1243TCA (3879C>A), TAC1246TAT (3888C>T), TTC1248TTT (3894C>T), ATG1251TTG (3901A>T), GGC1252GGA (3906C>A), ATG1253CTT (3907A>C 3909G>T), GGA1254GGT (3912A>T), ATG1255TCT (3913A>T 3914T>C 3915G>T), AAG1257AAA (3921G>A), CGG1258AGG (3922C>A), GAG1259GAA (3927G>A), GTC1261GTG (3933C>G), GTA1262GAA (3935T>A), CCA1263GCC (3937C>G 3939A>C), ACA1264CAA (3940A>C 3941C>A), GGA1265CCA (3943G>C 3944G>C), GAA1266AAC (3946G>A 3948A>C), GGT1267TCG (3949G>T 3950G>C 3951T>G), GAT1268CAG (3952G>C 3954T>G), AAA1269AGG (3956A>G 3957A>G), GTA1270GTT (3960A>T), GAG1271CTT (3961G>C 3962A>T 3963G>T), CTT1272CTC (3966T>C), AGA1273AAA (3968G>A), GAC1275AAC (3973G>A), CCA1277CCT (3981A>T), GTG1278GTC (3984G>C), GAA1280GAT (3990A>T), GCT1281GCA (3993T>A), GAG1283GAA (3999G>A), GAA1284GAG (4002A>G), CTG1285TTG (4003C>T), GAC1286GAT (4008C>T), ATT1287GCG (4009A>G 4010T>C 4011T>G), CAA1288CAG (4014A>G), ATT1289GTA (4015A>G 4017T>A), CAG1291CAA (4023G>A), ATA1293ACA (4028T>C), CAT1294TTT (4030C>T 4031A>T), GTG1295GTT (4035G>T), GAT1296GAC (4038T>C), CTA1297CTC (4041A>C), ATC1298GTC (4042A>G), ACC1299ACA (4047C>A), ACA1300ACC (4050A>C), CAA1301CAG (4053A>G), GCC1303GCG (4059C>G), AAG1304AAA (4062G>A), GAG1306GAA (4068G>A), CGC1307CGG (4071C>G), CTT1308TTG (4072C>T 4074T>G), CCT1309CCA (4077T>A), GAA1311GAG (4083A>G), AAA1318AAG (4104A>G), ACA1319ACC (4107A>C), AGA1320CGC (4108A>C 4110A>C), TTA1321CTG (4111T>C 4113A>G), TTT1322TTC (4116T>C), GAG1323GAA (4119G>A), ATA1324ATC (4122A>C), TAC1329TAT (4137C>T), CTG1332CTA (4146G>A), GTC1333GTG (4149C>G), AGG1334CGG (4150A>C), TTT1337TTC (4161T>C), TCC1341TCT (4173C>T), GCC1342GCT (4176C>T), ACC1343TGT (4177A>T 4178C>G 4179C>T), GCA1346GCT (4188A>T), TTA1347GCA (4189T>G 4190T>C), AAT1349AAC (4197T>C), GTC1350GTA (4200C>A), ATC1351CTC (4201A>C), CCA1352CCT (4206A>T), TGT1353TGC (4209T>C), GTT1355GTA (4215T>A), GGA1356GGT (4218A>T), ATA1357ATC (4221A>C), GAT1358AAT (4222G>A), ACC1360ACG (4230C>G), AGT1361AGC (4233T>C), TCT1362CAT (4234T>C 4235C>A), TCT1365ACA (4243T>A 4245T>A), CTT1366GTG (4246C>G 4248T>G), CTT1367CTG (4251T>G), GCC1368GCT (4254C>T), AAC1369AGT (4256A>G 4257C>T), AAC1370AGC (4259A>G), TTC1371TTT (4263C>T), GCT1373TCA (4267G>T 4269T>A), ACT1377GAG (4279A>G 4280C>A 4281T>G), GGG1378GGA (4284G>A), TTT1379TTC (4287T>C), GCT1381GCC (4293T>C), TCA1384TCG (4302A>G), AGT1385AGC (4305T>C), GAC1387GAT (4311C>T), GGG1388GGA (4314G>A), CGC1389AGA (4315C>A 4317C>A), CCA1391CCC (4323A>C), GTG1392ATC (4324G>A 4326G>C), TTC1393TTT (4329C>T), GCT1394GCA (4332T>A), TTT1395TTC (4335T>C), CAG1396CAA (4338G>A), TTG1401CTG (4351T>C), GTT1402GTG (4356T>G), GAG1404GCT (4361A>C 4362G>T), TAT1405TTT (4364A>T), GAT1407GGT (4370A>G), ACC1408GAT (4372A>G 4373C>A 4374C>T), AAG1409GAG (4375A>G), GGA1411GGG (4383A>G), GAG1412AGT (4384G>A 4385A>G 4386G>T), GCT1414GAG (4391C>A 4392T>G), TCC1415TCA (4395C>A), GCA1417GCT (4401A>T), AGA1418AGG (4404A>G), CAT1419CAC (4407T>C), GCT1420ACA (4408G>A 4410T>A), CTT1421TTA (4411C>T 4413T>A), CTC1422TTA (4414C>T 4416C>A), GCA1425GCC (4425A>C), TCA1426TCT (4428A>T), TAC1429TAT (4437C>T), TTA1431CTC (4441T>C 4443A>C), GAT1434AAC (4450G>A 4452T>C), AAA1435AAG (4455A>G), GTT1436TTG (4456G>T 4458T>G), TTC1437TTT (4461C>T), CGT1438CGC (4464T>C), TTG1439CTA (4465T>C 4467G>A), GTT1440GTG (4470T>G), GGC1442GGA (4476C>A), CCG1444CCC (4482G>C), GCA1448GCT (4494A>T), CTT1449TTG (4495C>T 4497T>G), ACC1450ACT (4500C>T), TTC1453TTT (4509C>T), AAC1454AAT (4512C>T), CTG1456TTG (4516C>T), TTG1457CTA (4519T>C 4521G>A), CGG1463AGG (4537C>A), TAT1464TAC (4542T>C), GCT1465GCG (4545T>G), TTC1466TTT (4548C>T), ATC1467ATA (4551C>A), CTT1469CTG (4557T>G), TTA1470TTG (4560A>G), AGA1471AGG (4563A>G), AGG1472AGA (4566G>A), CCT1473CCA (4569T>A), CAC1474CAT (4572C>T), ATT1475GTC (4573A>G 4575T>C), GCC1476AAA (4576G>A 4577C>A 4578C>A), GCT1477GCC (4581T>C), AGA1478AAA (4583G>A), GCT1479TCT (4585G>T), ATA1480GTG (4588A>G 4590A>G), GGT1481GGC (4593T>C), GTA1482ATT (4594G>A 4596A>T), CCT1484CCA (4602T>A), GGT1485AAT (4603G>A 4604G>A), GAT1486GAC (4608T>C), AAC1488AAT (4614C>T), CTT1490TTG (4618C>T 4620T>G), GGA1496GGT (4638A>T), AAT1499AAC (4647T>C), CTG1500TTG (4648C>T), GTG1501GTT (4653G>T), GCT1502GCG (4656T>G), GTG1503GTT (4659G>T), CCT1504CCA (4662T>A), CTG1505ATG (4663C>A), CTG1507TTG (4669C>T), TCA1511GAT (4681T>G 4682C>A 4683A>T), CTG1512CTA (4686G>A), CCT1513CCA (4689T>A), AAT1514TCC (4690A>T 4691A>C 4692T>C), GCC1516GCA (4698C>A), CAG1517AGA (4699C>A 4700A>G 4701G>A), CTT1519CTG (4707T>G), GAC1520GAA (4710C>A), ATG1521GCT (4711A>G 4712T>C 4713G>T), GTA1522GTT (4716A>T), AAT1523AAC (4719T>C), ATC1526ATT (4728C>T), AAA1527AAG (4731A>G), GGG1529GGT (4737G>T), CTG1530TTG (4738C>T), GAT1531GAC (4743T>C), TCT1533TCA (4749T>A), AAC1537GAG (4759A>G 4761C>G), CAG1538AAG (4762C>A), GTG1539ACA (4765G>A 4766T>C 4767G>A), CAG1540AAA (4768C>A 4770G>A), TTC1541TTT (4773C>T), CAG1542CTG (4775A>T), GAC1543GAT (4779C>T), TCA1545TCT (4785A>T), GAA1546GAG (4788A>G), ACT1548ACC (4794T>C), AGC1551AGT (4803C>T), GGA1553GGT (4809A>T), TTT1554TTC (4812T>C), AAA1555AAG (4815A>G), CGT1556CGA (4818T>A), GCC1558GCA (4824C>A), CTT1559TTG (4825C>T 4827T>G), GGG1560GGA (4830G>A), CAC1562CAA (4836C>A), GTG1569GTT (4857G>T), ATA1571GTC (4861A>G 4863A>C), ATT1572ATC (4866T>C), CCT1574CCC (4872T>C), ATG1575ATT (4875G>T), TAC1576TAT (4878C>T), ATC1578ATT (4884C>T), AGA1579AGG (4887A>G), CCT1580CCA (4890T>A), CCG1582GTG (4894C>G 4895C>T), GGT1583GGA (4899T>A), ACT1587GTA (4909A>G 4910C>T 4911T>A), GTG1588CAA (4912G>C 4913T>A 4914G>A), GCA1589GCT (4917A>T), ATG1591TAT (4921A>T 4922T>A 4923G>T), GAG1592GAA (4926G>A), GTG1594ATT (4930G>A 4932G>T), ACT1596ACA (4938T>A), CTT1598ATC (4942C>A 4944T>C), CGT1599AGA (4945C>A 4947T>A), GAG1600GAA (4950G>A), TTT1602TTC (4956T>C), AGA1606CAG (4966A>C 4967G>A 4968A>G), GTT1607GCC (4970T>C 4971T>C), GCA1608ACT (4972G>A 4974A>T), GAG1610GAA (4980G>A), AAG1611AGT (4982A>G 4983G>T), TTA1612TTT (4986A>T), ACT1613GTA (4987A>G 4988C>T 4989T>A), TAT1614AAT (4990T>A), CAG1617TTT (4999C>T 5000A>T 5001G>T), TGT1618GGA (5002T>G 5004T>A), GCC1619GCT (5007C>T), AGA1621AAG (5012G>A 5013A>G), GAG1622CAG (5014G>C), CGT1623AGA (5017C>A 5019T>A), GGA1624AAT (5020G>A 5021G>A 5022A>T), GGC1626CCC (5026G>C 5027G>C), GCA1627ATT (5029G>A 5030C>T 5031A>T), ACA1628ATG (5033C>T 5034A>G), TTT1630ATG (5038T>A 5040T>G), CCT1631CCC (5043T>C), TCT1632ACT (5044T>A), TAT1633TAC (5049T>C), ATG1634TAT (5050A>T 5051T>A 5052G>T), GTG1636CTT (5056G>C 5058G>T), GAG1637GAA (5061G>A), CAG1638AGT (5062C>A 5063A>G 5064G>T), GAA1639CAA (5065G>C), ATC1641TTG (5071A>T 5073C>G), GCA1642ACA (5074G>A), AAG1643GAG (5077A>G), GCA1648GAG (5093C>A 5094A>G), TCA1649AGC (5095T>A 5096C>G 5097A>C), GCG1650ATT (5098G>A 5099C>T 5100G>T), TTG1651GGG (5101T>G 5102T>G), ACA1652GTT (5104A>G 5105C>T 5106A>T), ATC1653TTG (5107A>T 5109C>G), TGT1654ATG (5110T>A 5111G>T 5112T>G), GAA1655GAG (5115A>G), ATG1656CTA (5116A>C 5118G>A), GCC1657TCA (5119G>T 5121C>A), AAA1658GAT (5122A>G 5124A>T), TCT1660AAA (5128T>A 5129C>A 5130T>A), ATC1661AAT (5132T>A 5133C>T), ACT1662ACA (5136T>A), TTC1663TAC (5138T>A), ACA1664ACC (5142A>C), TTG1666TTA (5148G>A), CCA1667CCT (5151A>T), CCT1668GCT (5152C>G), AAT1670GAA (5158A>G 5160T>A), GTT1672ATT (5164G>A), GAA1673del (5167_5169delGAA), GAA1676GAG (5178A>G), AGA1677GAG (5179A>G 5180G>A 5181A>G), GAT1678GGT (5183A>G), CTC1679GTC (5185C>G), AAT1680CAT (5188A>C), TGG1681GAA (5191T>G 5192G>A 5193G>A), GCC1683GTG (5198C>T 5199C>G), CCA1684CCT (5202A>T), AAC1685GAT (5203A>G 5205C>T), ATT1686GTA (5206A>G 5208T>A), GGT1687TTT (5209G>T 5210G>T), TGC1689TGT (5217C>T), TCA1690AGT (5218T>A 5219C>G 5220A>T), GCC1691GCT (5223C>T), CGT1692CGG (5226T>G), GCC1695TCA (5233G>T 5235C>A), AAG1696AAC (5238G>C), ACT1697CAA (5239A>C 5240C>A 5241T>A), GAA1698CAA (5242G>C), GAG1700TCA (5248G>T 5249A>C 5250G>A), CCT1701TCA (5251C>T 5253T>A), GGG1702GGT (5256G>T), TGC1703AAC (5257T>A 5258G>A), ATA1704ATC (5262A>C), GTA1705CTG (5263G>C 5265A>G), AAC1707AAT (5271C>T), TGT1708TGC (5274T>C), ACA1709ACT (5277A>T), GGG1710GGA (5280G>A), GCG1711GCC (5283G>C), AAG1712CGC (5284A>C 5285A>G 5286G>C), AAG1713AGA (5288A>G 5289G>A), AGC1714ACA (5291G>C 5292C>A), AAA1715ACA (5294A>C), GTT1717GTG (5301T>G), CGC1718AGA (5302C>A 5304C>A), GGA1719GGG (5307A>G), GCT1721GCC (5313T>C), AGT1722AAT (5315G>A), GAT1725GAC (5325T>C), ATG1730ATA (5340G>A), CCA1732CCC (5346A>C), TAT1733TAC (5349T>C), ACC1734ACT (5352C>T), ATG1735TTG (5353A>T), TCA1736GCA (5356T>G), ATT1738ATC (5364T>C), CTG1739CAG (5366T>A), GCT1741CAT (5371G>C 5372C>A), AGA1743CGC (5377A>C 5379A>C), GGC1744AAG (5380G>A 5381G>A 5382C>G), AAA1745AGT (5384A>G 5385A>T), ATG1746CTG (5386A>C), TCA1747GAA (5389T>G 5390C>A), GAG1748GTT (5393A>T 5394G>T), GAC1749GGT (5396A>G 5397C>T), ATT1750GAT (5398A>G 5399T>A), GCT1751GGG (5402C>G 5403T>G), GAT1754TCT (5410G>T 5411A>C), GTT1755GTC (5415T>C), GTG1756ACC (5416G>A 5417T>C 5418G>C), GTG1758GTT (5424G>T), GGG1760GGT (5430G>T), GCT1764GCA (5442T>A), GTG1765ATA (5443G>A 5445G>A), AGC1766GCT (5446A>G 5447G>C 5448C>T), GCT1771GCA (5463T>A), TTA1772TTG (5466A>G), ATG1773TTT (5467A>T 5469G>T), GCT1774GCA (5472T>A), GTG1775TTG (5473G>T), CGT1778CAT (5483G>A), CAG1779CAA (5487G>A), TAT1780TAC (5490T>C), TGC1781AGG (5491T>A 5493C>G), ATT1782GCC (5494A>G 5495T>C 5496T>C), GAG1783ACA (5497G>A 5498A>C 5499G>A), GAT1784GAC (5502T>C), ATC1785CTT (5503A>C 5505C>T), AGG1788CGC (5512A>C 5514G>C), TTG1789CTA (5515T>C 5517G>A), CGC1790CGA (5520C>A), ACT1791ACC (5523T>C), ATT1792ATG (5526T>G), GGC1793GCT (5528G>C 5529C>T), AAT1794GGG (5530A>G 5531A>G 5532T>G), TTG1796CTG (5536T>C), ACA1800ACC (5550A>C), TAT1801TAC (5553T>C), CCC1802CCT (5556C>T), GTT1806GCG (5567T>C 5568T>G), CAG1807CAA (5571G>A), TTT1809TTG (5577T>G), CAA1811AAA (5581C>A), GTG1813GTT (5589G>T), CCT1814GAA (5590C>G 5591C>A 5592T>A), CAG1815GAC (5593C>G 5595G>C), CAG1815_GAA1816insACC (5595_5596insACC), GAA1816TCA (5596G>T 5597A>C), GGG1817CAC (5599G>C 5600G>A 5601G>C), AAA1818AAG (5604A>G), ATG1819TTG (5605A>T), GCA1820GGA (5609C>G), GTT1822AGT (5614G>A 5615T>G), TCG1824ACA (5620T>A 5622G>A), TCT1825del (5623_5625delTCT), TTG1826TAT (5627T>A 5628G>T), CAA1827GGC (5629C>G 5630A>G 5631A>C), AGT1828ACC (5633G>C 5634T>C), AGT1829ACT (5636G>C), ATG1831AAC (5642T>A 5643G>C), CAA1833CAT (5649A>T), CAA1833_GGT1834insCACAGT (5649_5650insCACAGT), GGT1834GCA (5651G>C 5652T>A), GAG1836TCA (5656G>T 5657A>C 5658G>A), CTT1837GTT (5659C>G), GGC1838GGA (5664C>A), CAA1839CTT (5666A>T 5667A>T), ATT1840CTA (5668A>C 5670T>A), CGT1841AGG (5671C>A 5673T>G), GTC1842ATA (5674G>A 5676C>A), ATC1843GTT (5677A>G 5679C>T), CAC1844CCT (5681A>C 5682C>T), GGT1845TCC (5683G>T 5684G>C 5685T>C), GAT1846GAC (5688T>C), TTA1847CTC (5689T>C 5691A>C), TCA1848TCT (5694A>T), ATG1849GAA (5695A>G 5696T>A 5697G>A), GAA1850AAG (5698G>A 5700A>G), TTG1853GTT (5707T>G 5709G>T), CGC1854CGT (5712C>T), ATG1855TTG (5713A>T), TAT1857TAC (5721T>C), GTT1858ACA (5722G>A 5723T>C 5724T>A), GTG1859ATT (5725G>A 5727G>T), GGC1860GGT (5730C>T), CCA1861CCT (5733A>T), CAT1862GAA (5734C>G 5736T>A), GGT1863GGA (5739T>A), GGA1864GGT (5742A>T), GGC1866GCC (5747G>C), AAC1867CAG (5749A>C 5751C>G), TTT1868TGC (5753T>G 5754T>C), ACC1869TCA (5755A>T 5757C>A), ACA1870ACG (5760A>G), AAA1871AAG (5763A>G), AAT1872AAC (5766T>C), CTT1873CTG (5769T>G), GAT1874AAG (5770G>A 5772T>G), CAA1875CAT (5775A>T), TTG1876GCT (5776T>G 5777T>C 5778G>T), TTT1877TTG (5781T>G), GCA1878GAG (5783C>A 5784A>G), AGT1879ACA (5786G>C 5787T>A), CTT1880TTG (5788C>T 5790T>G), GAA1881TCT (5791G>T 5792A>C 5793A>T), AAG1882GTC (5794A>G 5795A>T 5796G>C), TGT1883AGT (5797T>A), GAA1886ACT (5806G>A 5807A>C 5808A>T), TTG1887CTT (5809T>C 5811G>T), CTT1888GCC (5812C>G 5813T>C 5814T>C), AAG1890AGA (5819A>G 5820G>A), AAC1891GGA (5821A>G 5822A>G 5823C>A), ACG1892GTC (5824A>G 5825C>T 5826G>C), CTG1894GTG (5830C>G), ACC1895ACA (5835C>A), GAG1899AAA (5845G>A 5847G>A), AAA1900GAA (5848A>G), ATG1901TTC (5851A>T 5853G>C), ACT1902TCC (5854A>T 5856T>C), AAA1904CAC (5860A>C 5862A>C), GTG1906TGT (5866G>T 5867T>G 5868G>T), GCA1907GAA (5870C>A), CAA1908ATG (5872C>A 5873A>T 5874A>G), GTC1910CTT (5878G>C 5880C>T), AGT1911GAG (5881A>G 5882G>A 5883T>G), GTA1913CTG (5887G>C 5889A>G), AAA1914GAA (5890A>G), CAG1915AGT (5893C>A 5894A>G 5895G>T), TTC1918GTA

| | | | | | | | | | |
|---|---|---|---|---|---|---|---|---|---|
| | (5902T>G 5904C>A), TAT1919GTA (5905T>G 5906A>T 5907T>A), CCC1920TTC (5908C>T 5909C>T), AAG1921ATG (5912A>T), GCC1922GGG (5915C>G 5916C>G), ACA1923AGC (5918C>G 5919A>C), ACC1924GTT (5920A>G 5921C>T 5922C>T), GTG1925AAG (5923G>A 5924T>A), GAA1926AAA (5926G>A), CAG1927CAA (5931G>A), CTC1928ATG (5932C>A 5934C>G), AAG1929TTG (5935A>T 5936A>T), GTA1930TCC (5938G>T 5939T>C 5940A>C), TTT1931TTC (5943T>C), GCT1932ATG (5944G>A 5945C>T 5946T>G), GAT1933AAT (5947G>A), GCT1934CAA (5950G>C 5951C>A 5952T>A), CAT1935CAG (5955T>G), CAT1935_ACC1936insATG (5955_5956insATG), ACC1936TCA (5956A>T 5958C>A), ATT1937AAA (5960T>A 5961T>A), GTC1938ACC (5962G>A 5963T>C), AAA1939CAA (5965A>C), ACT1940GTT (5968A>G 5969C>T), GCT1941GCC (5973T>C), AAA1942AGC (5975A>G 5976A>C), CCT1943CCA (5979T>A), TTC1944TGT (5981T>G 5982C>T), CGG1945TGT (5983C>T 5985G>T), GGA1946del (5986_5988delGGA), GTA1947GTC (5991A>C), GTT1948ACA (5992G>A 5993T>C 5994T>A), TTC1949TTT (5997C>T), CGG1950AGT (5998C>A 6000G>T), AAT1952GCA (6004A>G 6005A>C 6006T>A), TTT1953TGG (6008T>G 6009T>G), CTC1954CTA (6012C>A), AGT1955ACT (6014G>C), CCT1956AAG (6016C>A 6017C>A 6018T>G), GAG1957GAC (6021G>C), TGG1958ATT (6022T>A 6023G>T 6024G>T), TGG1958_AAA1959insGTC (6024_6025insGTC), AAA1959ATG (6026A>T 6027A>G), ATT1960ACA (6029T>C 6030T>A), GGA1961AAA (6031G>A 6032G>A), GAG1963AAG (6037G>A), AAC1964AAG (6042C>G), ACA1965CCA (6043A>C), ATG1966del (6046_6048delATG), GCA1967GGA (6050C>G), ACA1968TCT (6052A>T 6054A>T), CTC1969ATT (6055C>A 6057C>T), TCA1970CCT (6058T>C 6060A>T), GAG1972GAA (6066G>A), AGT1973ACA (6068G>C 6069T>A), CTT1974GTG (6070C>G 6072T>G), CTT1975TTT (6073C>T), CCA1976CTT (6077C>T 6078A>T), CTG1979CCT (6086T>C 6087G>T), TCG1980TCA (6090G>A), TCA1982CTA (6094T>C 6095C>T), GCC1983GTT (6098C>T 6099C>T), ATG1984GTG (6100A>G), AAG1985AGA (6104A>G 6105G>A), TTG1987ATG (6109T>A), CTC1988TTG (6112C>T 6114C>G), AGC1989CGC (6115A>C), AGA1990AAA (6119G>A) | | | | | | | | |
| helicase motif A-C (YP_009167369.1) | 1 | 98 | 100% | 654 | 97.6% | 98 (100%) | 96 (98.0%) | 0/0/0/0 | 0 |
| Protein mutations: | A5C (1345G>T 1346C>G 1347A>T), R16A (1378C>G 1379G>C 1380C>A) | | | | | | | | |
| Codon mutations: | ACA4ACG (1344A>G), GCA5TGT (1345G>T 1346C>G 1347A>T), CGA7CGC (1353A>C), TTG8CTT (1354T>C 1356G>T), TTG14CTC (1372T>C 1374G>C), GAT15GAC (1377T>C), CGC16GCA (1378C>G 1379G>C 1380C>A), GGG18GGT (1386G>T), GAG19GAA (1389G>A), CCT20CCA (1392T>A), CTT22CTG (1398T>G), AGA24CGC (1402A>C 1404A>C), CTT25TTG (1405C>T 1407T>G), GCT27GCG (1413T>G), GTT28GTA (1416T>A), AGC29AGT (1419C>T), AAG30AAA (1422G>A), CGC31AGA (1423C>A 1425C>A), GCC33GCA (1431C>A), TCT36TCC (1440T>C), AAT37AAC (1443T>C), GCC39GCA (1449C>A), CAC40CAT (1452C>T), CAG41CAA (1455G>A), GCT43GCA (1461T>A), TTG45CTT (1465T>C 1467G>T), GAC48GAT (1476C>T), GGA50GGT (1482A>T), GCA51GCT (1485A>T), TTG52CTT (1486T>C 1488G>T), AGA53CGG (1489A>C 1491A>G), GGA57GGG (1503A>G), CAA58CAG (1506A>G), CAA60CAG (1512A>G), ATT62ATA (1518T>A), ATT68ATC (1536T>C), TCC70TCT (1542C>T), ACA71ACC (1545A>C), CCA73CCT (1551A>T), CCA75CCT (1557A>T), CTG76CTT (1560G>T), CCC77CCA (1563C>A), GCA79GCT (1569A>T), GCT80GCA (1572T>A), GTG81GTC (1575G>C), GAA82GAG (1578A>G), GAC83GAT (1581C>T), AAG84AAA (1584G>A), GGG85GGA (1587G>A), CAT87CAC (1593T>C), ACT89ACA (1599T>A), TAT92TAC (1608T>C), GCA95GCC (1617A>C), AAT98AAC (1626T>C) | | | | | | | | |

*: Inserts / Deletes / Misaligned / Frameshifts

#### Analysis details

This analysis was performed with panviral2.64

#### NGS Details (UN18_val): Bracoviriform glomeratae (segment NC_043292.1)

##### Assembly

| | |
|---|---|
| **Coverage Length** | 323 (1 contig(s)) |
| **Depth Of Coverage** | 4199.5 |
| **Number Of Reads** | 11330 |
| **Reads Per Million** | 212.65 rpm (after QC) |
| **Ambiguities** | 0 |
| **Assembly Method** | de novo + reference guided assembly |
| **Consensus Caller** | Bcf Tools |

##### Coverage Map

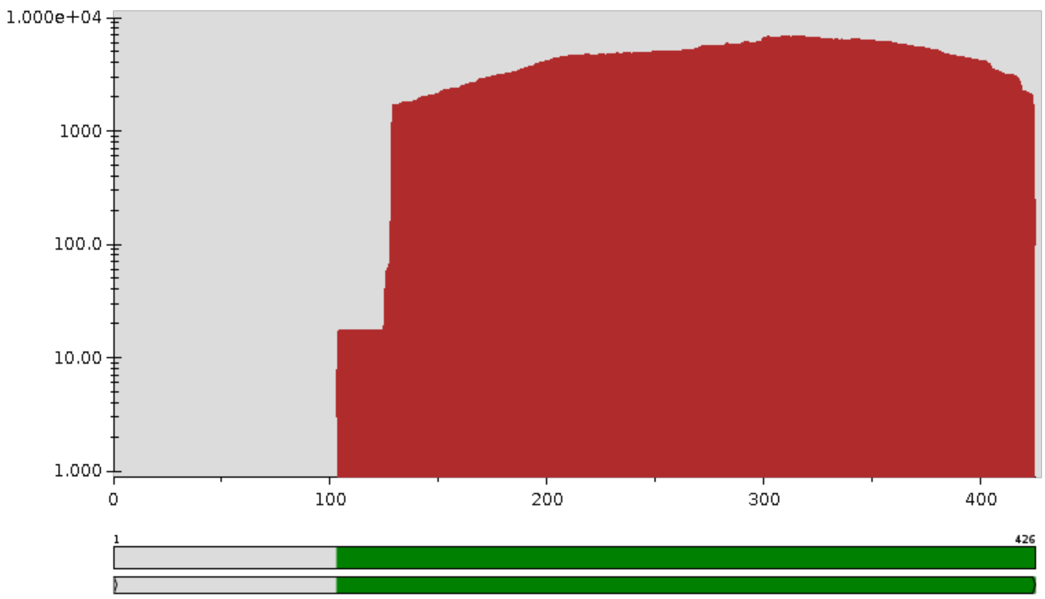

##### Assignment

| | |
|---|---|
| **Type** | Bracoviriform glomeratae (Taxonomy ID: 257816) |
| **Reference Genome** | NC_043292.1 |
| **NT Identity (%)** | 72.7554 |
| **AA Identity (%)** | 84.1121 |
| **Number Of Stop Codons** | 1 |
| **Number Of CDS** | 1 |

##### Alignment

| | |
|---|---|
| **Alignment Score** | 294.0 (NT) + 580.0 (AA) = 874.0 |
| **Concordance (%)** | 65.7637 |

| Alignment Method | Global, seeded, nucleotide + amino acids (AGA) |
|---|---|

#### Genome Region

Sequence starts at position 104 and ends at position 426 relative to NC_043292.1 reference sequence.

#### Alignment Detailed Statistics

| | Begin | End | Coverage | Score | Concordance | Matches | Identities | I/D/M/F* | Stop Codons |
|---|---|---|---|---|---|---|---|---|---|
| **NT** | **104** | **426** | **75.8%** | **294** | **45.5%** | **323 (100%)** | **235 (72.8%)** | **0/0** | |
| Mutations: | 105G>A, 108C>T, 111A>G, 113A>G, 117A>C, 119G>A, 129C>A, 132A>G, 136A>G, 137C>G, 138G>C, 150A>C, 153A>G, 155C>G, 156G>A, 159C>A, 166C>A, 168T>G, 169T>C, 172C>A, 174C>G, 177A>G, 183T>A, 184C>A, 189T>C, 198A>G, 201A>C, 207C>T, 214A>G, 219C>T, 222T>C, 223C>A, 225T>G, 228G>T, 235C>A, 237T>G, 243A>T, 246C>G, 255C>T, 261A>G, 264A>G, 265G>A, 273A>G, 278T>C, 279T>C, 284A>G, 285T>A, 288T>G, 289T>C, 300C>T, 301C>T, 303C>G, 309C>T, 312T>G, 315C>T, 322G>T, 329T>C, 333T>C, 336C>T, 339A>G, 345A>C, 347A>G, 348A>G, 349C>A, 351T>A, 360C>T, 363C>T, 366C>T, 375C>T, 378C>T, 384A>C, 385T>C, 387G>C, 390A>G, 391C>A, 392A>G, 393T>A, 394A>C, 396A>G, 399C>A, 400C>A, 402T>G, 404T>C, 405G>T, 406A>C, 414T>A, 416A>T, 420A>G | | | | | | | | |
| **CDS** | | | | | | | | | |
| FK954_p501 | 36 | 142 | 75.4% | 580 | 84.4% | 107 (100%) | 90 (84.1%) | 0/0/0/0 | 1 |
| Protein mutations: | E38G (113A>G), R40K (119G>A), T46G (136A>G 137C>G 138G>C), A52G (155C>G 156G>A), Y57H (169T>C), T72A (214A>G), V89I (265G>A), I93T (278T>C 279T>C), D95G (284A>G 285T>A), A108S (322G>T), I110T (329T>C), K116R (347A>G 348A>G), H131R (391C>A 392A>G 393T>A), K132Q (394A>C 396A>G), M135T (404T>C 405G>T), I136L (406A>C), Y139F (416A>T) | | | | | | | | |
| Codon mutations: | TTG35.TA (105G>A), GGC36GGT (108C>T), AAA37AAG (111A>G), GAA38GGA (113A>G), GGA39GGC (117A>C), AGA40AAA (119G>A), GGC43GGA (129C>A), AAA44AAG (132A>G), ACG46GGC (136A>G 137C>G 138G>C), GGA50GGC (150A>C), AAA51AAG (153A>G), GCG52GGA (155C>G 156G>A), GGC53GGA (159C>A), CGT56AGG (166C>A 168T>G), TAT57CAT (169T>C), CGC58AGG (172C>A 174C>G), AAA59AAG (177A>G), CTT61CTA (183T>A), CGA62AGA (184C>A), GAT63GAC (189T>C), CAA66CAG (198A>G), GGA67GGC (201A>C), ACC69ACT (207C>T), ACT72GCT (214A>G), ATC73ATT (219C>T), CGT74CGC (222T>C), CGT75AGG (223C>A 225T>G), CTG76CTT (228G>T), CGT79AGG (235C>A 237T>G), GGA81GGT (243A>T), GTC82GTG (246C>G), ATC85ATT (255C>T), GGA87GGG (261A>G), TTA88TTG (264A>G), GTC89ATC (265G>A), GAA91GAG (273A>G), ATT93ACC (278T>C 279T>C), GAT95GGA (284A>G 285T>A), GTT96GTG (288T>G), TTG97CTG (289T>C), TTC100TTT (300C>T), CTC101TTG (301C>T 303C>G), AAC103AAT (309C>T), GTT104GTG (312T>G), ATC105ATT (315C>T), GCT108TCT (322G>T), ATC110ACC (329T>C), TAT111TAC (333T>C), ACC112ACT (336C>T), GAA113GAG (339A>G), GCA115GCC (345A>C), AAA116AGG (347A>G 348A>G), CGT117AGA (349C>A 351T>A), GTC120GTT (360C>T), ACC121ACT (363C>T), GCC122GCT (366C>T), GTC125GTT (375C>T), GTC126GTT (378C>T), GCA128GCC (384A>C), TTG129CTC (385T>C 387G>C), AAA130AAG (390A>G), CAT131AGA (391C>A 392A>G 393T>A), AAA132CAG (394A>C 396A>G), GGC133GGA (399C>A), CGT134AGG (400C>A 402T>G), ATG135ACT (404T>C 405G>T), ATC136CTC (406A>C), GGT138GGA (414T>A), TAT139TTT (416A>T), GGA140GGG (420A>G) | | | | | | | | |
| **Proteins** | | | | | | | | | |
| putative histone 4 (YP_009665791.1) | 36 | 142 | 75.4% | 580 | 84.4% | 107 (100%) | 90 (84.1%) | 0/0/0/0 | 1 |
| Protein mutations: | E38G (113A>G), R40K (119G>A), T46G (136A>G 137C>G 138G>C), A52G (155C>G 156G>A), Y57H (169T>C), T72A (214A>G), V89I (265G>A), I93T (278T>C 279T>C), D95G (284A>G 285T>A), A108S (322G>T), I110T (329T>C), K116R (347A>G 348A>G), H131R (391C>A 392A>G 393T>A), K132Q (394A>C 396A>G), M135T (404T>C 405G>T), I136L (406A>C), Y139F (416A>T) | | | | | | | | |
| Codon mutations: | TTG35.TA (105G>A), GGC36GGT (108C>T), AAA37AAG (111A>G), GAA38GGA (113A>G), GGA39GGC (117A>C), AGA40AAA (119G>A), GGC43GGA (129C>A), AAA44AAG (132A>G), ACG46GGC (136A>G 137C>G 138G>C), GGA50GGC (150A>C), AAA51AAG (153A>G), GCG52GGA (155C>G 156G>A), GGC53GGA (159C>A), CGT56AGG (166C>A 168T>G), TAT57CAT (169T>C), CGC58AGG (172C>A 174C>G), AAA59AAG (177A>G), CTT61CTA (183T>A), CGA62AGA (184C>A), GAT63GAC (189T>C), CAA66CAG (198A>G), GGA67GGC (201A>C), ACC69ACT (207C>T), ACT72GCT (214A>G), ATC73ATT (219C>T), CGT74CGC (222T>C), CGT75AGG (223C>A 225T>G), CTG76CTT (228G>T), CGT79AGG (235C>A 237T>G), GGA81GGT (243A>T), GTC82GTG (246C>G), ATC85ATT (255C>T), GGA87GGG (261A>G), TTA88TTG (264A>G), GTC89ATC (265G>A), GAA91GAG (273A>G), ATT93ACC (278T>C 279T>C), GAT95GGA (284A>G 285T>A), GTT96GTG (288T>G), TTG97CTG (289T>C), TTC100TTT (300C>T), CTC101TTG (301C>T 303C>G), AAC103AAT (309C>T), GTT104GTG (312T>G), ATC105ATT (315C>T), GCT108TCT (322G>T), ATC110ACC (329T>C), TAT111TAC (333T>C), ACC112ACT (336C>T), GAA113GAG (339A>G), GCA115GCC (345A>C), AAA116AGG (347A>G 348A>G), CGT117AGA (349C>A 351T>A), GTC120GTT (360C>T), ACC121ACT (363C>T), GCC122GCT (366C>T), GTC125GTT (375C>T), GTC126GTT (378C>T), GCA128GCC (384A>C), TTG129CTC (385T>C 387G>C), AAA130AAG (390A>G), CAT131AGA (391C>A 392A>G 393T>A), AAA132CAG (394A>C 396A>G), GGC133GGA (399C>A), CGT134AGG (400C>A 402T>G), ATG135ACT (404T>C 405G>T), ATC136CTC (406A>C), GGT138GGA (414T>A), TAT139TTT (416A>T), GGA140GGG (420A>G) | | | | | | | | |

*: Inserts / Deletes / Misaligned / Frameshifts

#### Analysis details

This analysis was performed with panviral2.64

### NGS Details (UN18_val): Potato leafroll virus

#### Assembly

| | |
|---|---|
| **Coverage Length** | 5830 (1 contig(s)) |
| **Depth Of Coverage** | 98.0 |
| **Number Of Reads** | 4303 |
| **Reads Per Million** | 80.76 rpm (after QC) |
| **Ambiguities** | 0 |
| **Assembly Method** | read mapping against reference + variant calling |
| **Consensus Caller** | Bcf Tools |

#### Coverage Map

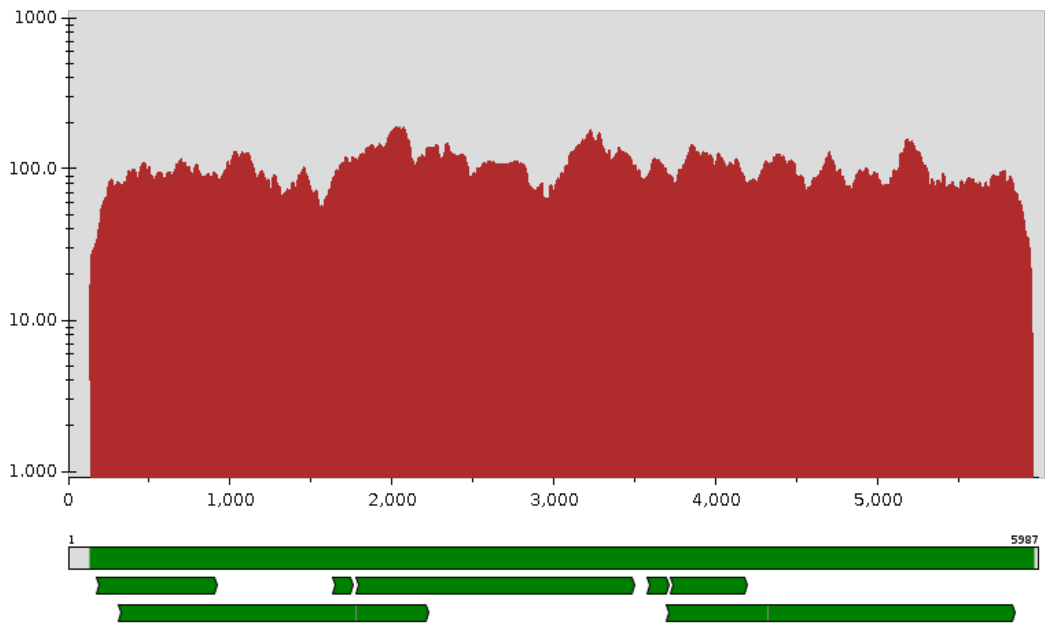

#### Assignment

| | |
|---|---|
| **Type** | Potato leafroll virus (Taxonomy ID: 12045) |
| **Reference Genome** | NC_001747.1 |
| **NT Identity (%)** | 98.0614 |
| **AA Identity (%)** | 97.3103 |
| **Number Of Stop Codons** | 9 |
| **Number Of CDS** | 8 |

#### Alignment

| | |
|---|---|
| **Alignment Score** | 11196.0 (NT) + 21113.0 (AA) = 32309.0 |
| **Concordance (%)** | 97.0444 |

| Alignment Method | Global, seeded, nucleotide + amino acids (AGA) |
|---|---|

#### Genome Region

Sequence starts at position 135 and ends at position 5964 relative to NC_001747.1 reference sequence.

#### Alignment Detailed Statistics

| | Begin | End | Coverage | Score | Concordance | Matches | Identities | I/D/M/F* | Stop Codons |
|---|---|---|---|---|---|---|---|---|---|
| **NT** | **135** | **5964** | **97.4%** | **11196** | **96.1%** | **5829 (99.9%)** | **5716 (98.0%)** | **0/1** | |
| Mutations: | 243C>T, 249C>T, 279G>T, 293T>C, 343C>T, 513C>T, 525C>T, 561C>T, 600T>C, 648T>C, 699A>C, 715C>T, 745G>A, 822C>T, 825T>G, 864C>T, 922G>A, 931A>T, 934A>G, 1079G>A, 1093A>G, 1117G>A, 1664G>A, 1674C>T, 1678T>A, 1755T>C, 1795C>A, 1815A>T, 1823G>A, 2007T>C, 2051G>A, 2079C>T, 2154C>T, 2265A>G, 2283G>A, 2313A>G, 2316T>G, 2325C>T, 2470G>A, 2580A>G, 2586A>G, 2613C>T, 2622T>C, 2637C>T, 2649C>T, 2697G>A, 2729A>G, 2811T>A, 2817C>T, 2856G>A, 2877C>T, 2898T>C, 2931T>C, 2982C>T, 3000T>C, 3033C>T, 3036C>T, 3069C>T, 3075T>A, 3102T>C, 3117C>T, 3321C>T, 3468T>C, 3510delA, 3525A>G, 3548C>T, 3585A>G, 3604T>C, 3605T>A, 3652T>A, 3721T>G, 3748T>G, 3750C>A, 3788T>G, 3876G>A, 3878T>A, 3911G>C, 3935C>T, 3939A>T, 4031A>G, 4034C>T, 4113G>A, 4166C>T, 4172T>C, 4214T>G, 4601A>G, 4608A>G, 4610C>T, 4616C>T, 4817T>C, 4883C>T, 4886T>C, 4895T>C, 4898C>T, 4979C>T, 5038C>T, 5042G>A, 5048C>T, 5062C>T, 5071A>G, 5072C>T, 5129G>A, 5199C>T, 5214C>A, 5215G>A, 5229G>A, 5385G>A, 5501C>T, 5508T>C, 5522C>T, 5781G>A, 5816A>G, 5824A>G, 5925T>C | | | | | | | | |
| **CDS** | | | | | | | | | |
| PLRVgp1 | 1 | 248 | 100% | 1695 | 97.9% | 248 (100%) | 243 (98.0%) | 0/0/0/0 | 1 |
| Protein mutations: | Q35H (279G>T), I40T (293T>C), H57Y (343C>T), Q175H (699A>C), V191I (745G>A) | | | | | | | | |
| Codon mutations: | TTC23TTT (243C>T), GTC25GTT (249C>T), CAG35CAT (279G>T), ATT40ACT (293T>C), CAT57TAT (343C>T), ACC113ACT (513C>T), TAC117TAT (525C>T), TAC129TAT (561C>T), CAT142CAC (600T>C), CGT158CGC (648T>C), CAA175CAC (699A>C), CTA181TTA (715C>T), GTT191ATT (745G>A), CGC216CGT (822C>T), GCT217GCG (825T>G), GTC230GTT (864C>T) | | | | | | | | |
| PLRVgp2 | 1 | 1063 | 100% | 7338 | 98.4% | 1063 (100%) | 1043 (98.1%) | 0/0/0/0 | 1 |
| Protein mutations: | P69L (513C>T), T73I (525C>T), T85I (561C>T), M98T (600T>C), V114A (648T>C), N131T (699A>C), A172V (822C>T), L173R (825T>G), S186F (864C>T), E208D (931A>T), E258K (1079G>A), E453K (1664G>A), T456I (1674C>T), N457K (1678T>A), L483P (1755T>C), Q497K (1795C>A), R506Q (1823G>A), G582E (2051G>A), A722T (2470G>A), N808S (2729A>G) | | | | | | | | |
| Codon mutations: | TTC12TTT (343C>T), CCG69CTG (513C>T), ACA73ATA (525C>T), ACA85ATA (561C>T), ATG98ACG (600T>C), GTG114GCG (648T>C), AAC131ACC (699A>C), CTC136CTT (715C>T), TTG146TTA (745G>A), GCG172GTG (822C>T), CTA173CGA (825T>G), TCC186TTC (864C>T), AGG205AGA (922G>A), GAA208GAT (931A>T), GGA209GGG (934A>G), GAA258AAA (1079G>A), GCA262GCG (1093A>G), AGG270AGA (1117G>A), GAG453AAG (1664G>A), ACA456ATA (1674C>T), AAT457AAA (1678T>A), CTA483CCA (1755T>C), CAA497AAA (1795C>A), CGA503CGT (1815A>T), CGA506CAA (1823G>A), TTT567TTC (2007T>C), GGA582GAA (2051G>A), TGC591TGT (2079C>T), CAC616CAT (2154C>T), TCA653TCG (2265A>G), GCG659GCA (2283G>A), ACA669ACG (2313A>G), GTT670GTG (2316T>G), TAC673TAT (2325C>T), GCA722ACA (2470G>A), GAA758GAG (2580A>G), CTA760CTG (2586A>G), ATC769ATT (2613C>T), TTT772TTC (2622T>C), CCC777CCT (2637C>T), AGC781AGT (2649C>T), GTG797GTA (2697G>A), AAT808AGT (2729A>G), GCT835GCA (2811T>A), TTC837TTT (2817C>T), GTG850GTA (2856G>A), CAC857CAT (2877C>T), ACT864ACC (2898T>C), TAT875TAC (2931T>C), AAC892AAT (2982C>T), CGT898CGC (3000T>C), AAC909AAT (3033C>T), TCC910TCT (3036C>T), GCC921GCT (3069C>T), ACT923ACA (3075T>A), AGT932AGC (3102T>C), TCC937TCT (3117C>T), AAC1005AAT (3321C>T), GTT1054GTC (3468T>C) | | | | | | | | |
| PLRVgp3 | 1 | 640 | 100% | 4167 | 97.1% | 640 (100%) | 619 (96.7%) | 0/0/0/0 | 1 |
| Protein mutations: | P69L (513C>T), T73I (525C>T), T85I (561C>T), M98T (600T>C), V114A (648T>C), N131T (699A>C), A172V (822C>T), L173R (825T>G), S186F (864C>T), E208D (931A>T), E258K (1079G>A), E453K (1664G>A), T456I (1674C>T), N457K (1678T>A), L483P (1755T>C), D503V (1815A>T), D506N (1823G>A), L567S (2007T>C), E582K (2051G>A), A591V (2079C>T), T616I (2154C>T) | | | | | | | | |
| Codon mutations: | TTC12TTT (343C>T), CCG69CTG (513C>T), ACA73ATA (525C>T), ACA85ATA (561C>T), ATG98ACG (600T>C), GTG114GCG (648T>C), AAC131ACC (699A>C), CTC136CTT (715C>T), TTG146TTA (745G>A), GCG172GTG (822C>T), CTA173CGA (825T>G), TCC186TTC (864C>T), AGG205AGA (922G>A), GAA208GAT (931A>T), GGA209GGG (934A>G), GAA258AAA (1079G>A), GCA262GCG (1093A>G), AGG270AGA (1117G>A), GAG453AAG (1664G>A), ACA456ATA (1674C>T), AAT457AAA (1678T>A), CTA483CCA (1755T>C), GCC496GCA (1795C>A), GAC503GTC (1815A>T), GAC506AAC (1823G>A), TTA567TCA (2007T>C), GAG582AAG (2051G>A), GCG591GTG (2079C>T), ACC616ATC (2154C>T) | | | | | | | | |
| Rap1 | 1 | 42 | 100% | 260 | 90.6% | 42 (100%) | 39 (92.9%) | 0/0/0/0 | 1 |
| Protein mutations: | Q15* (1674C>T), M16K (1678T>A), *42Q (1755T>C) | | | | | | | | |
| Codon mutations: | GAG11GAA (1664G>A), CAA15TAA (1674C>T), ATG16AAG (1678T>A), TAA42CAA (1755T>C) | | | | | | | | |
| ORF3a | 1 | 46 | 100% | 286 | 95.7% | 46 (100%) | 43 (93.5%) | 0/0/0/0 | 1 |
| Protein mutations: | K4R (3585A>G), L11I (3605T>A), F26L (3652T>A) | | | | | | | | |
| Codon mutations: | AAA4AGA (3585A>G), GCT10GCC (3604T>C), TTA11ATA (3605T>A), TTT26TTA (3652T>A) | | | | | | | | |
| PLRVgp4 | 1 | 718 | 100% | 4944 | 98.1% | 718 (100%) | 701 (97.6%) | 0/0/0/0 | 2 |
| Protein mutations: | V10G (3721T>G), M19R (3748T>G), V62I (3876G>A 3878T>A), T83S (3939A>T), V141I (4113G>A), I306V (4608A>G 4610C>T), P449L (5038C>T), A457V (5062C>T), H460R (5071A>G 5072C>T), P503S (5199C>T), R508K (5214C>A 5215G>A), V513I (5229G>A), E565K (5385G>A), S606P (5508T>C), E697K (5781G>A), I708M (5816A>G), K711R (5824A>G) | | | | | | | | |
| Codon mutations: | GTC10GGC (3721T>G), ATG19AGG (3748T>G), CGA20AGA (3750C>A), GTT32GTG (3788T>G), GTT62ATA (3876G>A 3878T>A), GTG73GTC (3911G>C), GGC81GGT (3935C>T), ACC83TCC (3939A>T), ACA113ACG (4031A>G), AGC114AGT (4034C>T), GTA141ATA (4113G>A), GCC158GCT (4166C>T), ACT160ACC (4172T>C), TCT174TCG (4214T>G), TCA303TCG (4601A>G), ATC306GTT (4608A>G 4610C>T), TGC308TGT (4616C>T), GGT375GGC (4817T>C), TTC397TTT (4883C>T), CTT398CTC (4886T>C), CCT401CCC (4895T>C), GCC402GCT (4898C>T), ACC429ACT (4979C>T), CCA449CTA (5038C>T), CGG450CGA (5042G>A), GGC452GGT (5048C>T), GCC457GTC (5062C>T), CAC460CGT (5071A>G 5072C>T), ACG479ACA (5129G>A), CCA503TCA (5199C>T), CGA508AAA (5214C>A 5215G>A), GTA513ATA (5229G>A), GAA565AAA (5385G>A), AAC603AAT (5501C>T), TCC606CCC (5508T>C), CCC610CCT (5522C>T), GAA697AAA (5781G>A), ATA708ATG (5816A>G), AAG711AGG (5824A>G) | | | | | | | | |
| PLRVgp5 | 1 | 209 | 100% | 1416 | 98.7% | 209 (100%) | 204 (97.6%) | 0/0/0/0 | 1 |
| Protein mutations: | V10G (3721T>G), M19R (3748T>G), V62I (3876G>A 3878T>A), T83S (3939A>T), V141I (4113G>A) | | | | | | | | |
| Codon mutations: | GTC10GGC (3721T>G), ATG19AGG (3748T>G), CGA20AGA (3750C>A), GTT32GTG (3788T>G), GTT62ATA (3876G>A 3878T>A), GTG73GTC (3911G>C), GGC81GGT (3935C>T), ACC83TCC (3939A>T), ACA113ACG (4031A>G), AGC114AGT (4034C>T), GTA141ATA (4113G>A), GCC158GCT (4166C>T), ACT160ACC (4172T>C), TCT174TCG (4214T>G) | | | | | | | | |
| PLRVgp6 | 1 | 157 | 100% | 1007 | 91.4% | 157 (100%) | 147 (93.6%) | 0/0/0/0 | 1 |
| Protein mutations: | S2A (3721T>G), C11G (3748T>G 3750C>A), F24C (3788T>G), F54Y (3878T>A), C65S (3911G>C), A73V (3935C>T), Q105R (4031A>G), A106V (4034C>T), P150L (4166C>T), L152P (4172T>C) | | | | | | | | |
| Codon mutations: | TCA2GCA (3721T>G), TGC11GGA (3748T>G 3750C>A), TTC24TGC (3788T>G), GAG53GAA (3876G>A), TTC54TAC (3878T>A), TGT65TCT (3911G>C), GCA73GTA (3935C>T), ACA74ACT (3939A>T), CAA105CGA (4031A>G), GCA106GTA (4034C>T), AAG132AAA (4113G>A), CCA150CTA (4166C>T), CTT152CCT (4172T>C) | | | | | | | | |
| **Proteins** | | | | | | | | | |
| P0 protein (NP_056746.1) | 1 | 248 | 100% | 1695 | 97.9% | 248 (100%) | 243 (98.0%) | 0/0/0/0 | 1 |
| Protein mutations: | Q35H (279G>T), I40T (293T>C), H57Y (343C>T), Q175H (699A>C), V191I (745G>A) | | | | | | | | |

| | Begin | End | Coverage | Score | Concordance | Matches | Identities | I/D/M/F* | Stop Codons |
|---|---|---|---|---|---|---|---|---|---|
| **NT** | **135** | **5964** | **97.4%** | **11196** | **96.1%** | **5829 (99.9%)** | **5716 (98.0%)** | **0/1** | |
| Codon mutations: | TTC23TTT (243C>T), GTC25GTT (249C>T), CAG35CAT (279G>T), ATT40ACT (293T>C), CAT57TAT (343C>T), ACC113ACT (513C>T), TAC117TAT (525C>T), TAC129TAT (561C>T), CAT142CAC (600T>C), CGT158CGC (648T>C), CAA175CAC (699A>C), CTA181TTA (715C>T), GTT191ATT (745G>A), CGC216CGT (822C>T), GCT217GCG (825T>G), GTC230GTT (864C>T) | | | | | | | | |
| RNA-dependent RNA polymerase (NP_056748.3) | 1 | 1063 | 100% | 7338 | 98.4% | 1063 (100%) | 1043 (98.1%) | 0/0/0/0 | 1 |
| Protein mutations: | P69L (513C>T), T73I (525C>T), T85I (561C>T), M98T (600T>C), V114A (648T>C), N131T (699A>C), A172V (822C>T), L173R (825T>G), S186F (864C>T), E208D (931A>T), E258K (1079G>A), E453K (1664G>A), T456I (1674C>T), N457K (1678T>A), L483P (1755T>C), Q497K (1795C>A), R506Q (1823G>A), G582E (2051G>A), A722T (2470G>A), N808S (2729A>G) | | | | | | | | |
| Codon mutations: | TTC12TTT (343C>T), CCG69CTG (513C>T), ACA73ATA (525C>T), ACA85ATA (561C>T), ATG98ACG (600T>C), GTG114GCG (648T>C), AAC131ACC (699A>C), CTC136CTT (715C>T), TTG146TTA (745G>A), GCG172GTG (822C>T), CTA173CGA (825T>G), TCC186TTC (864C>T), AGG205AGA (922G>A), GAA208GAT (931A>T), GGA209GGG (934A>G), GAA258AAA (1079G>A), GCA262GCG (1093A>G), AGG270AGA (1117G>A), GAG453AAG (1664G>A), ACA456ATA (1674C>T), AAT457AAA (1678T>A), CTA483CCA (1755T>C), CAA497AAA (1795C>A), CGA503CGT (1815A>T), CGA506CAA (1823G>A), TTT567TTC (2007T>C), GGA582GAA (2051G>A), TGC591TGT (2079C>T), CAC616CAT (2154C>T), TCA653TCG (2265A>G), GCG659GCA (2283G>A), ACA669ACG (2313A>G), GTT670GTG (2316T>G), TAC673TAT (2325C>T), GCA722ACA (2470G>A), GAA758GAG (2580A>G), CTA760CTG (2586A>G), ATC769ATT (2613C>T), TTT772TTC (2622T>C), CCC777CCT (2637C>T), AGC781AGT (2649C>T), GTG797GTA (2697G>A), AAT808AGT (2729A>G), GCT835GCA (2811T>A), TTC837TTT (2817C>T), GTG850GTA (2856G>A), CAC857CAT (2877C>T), ACT864ACC (2898T>C), TAT875TAC (2931T>C), AAC892AAT (2982C>T), CGT898CGC (3000T>C), AAC909AAT (3033C>T), TCC910TCT (3036C>T), GCC921GCT (3069C>T), ACT923ACA (3075T>A), AGT932AGC (3102T>C), TCC937TCT (3117C>T), AAC1005AAT (3321C>T), GTT1054GTC (3468T>C) | | | | | | | | |
| P1 protein (NP_056747.1) | 1 | 640 | 100% | 4167 | 97.1% | 640 (100%) | 619 (96.7%) | 0/0/0/0 | 1 |
| Protein mutations: | P69L (513C>T), T73I (525C>T), T85I (561C>T), M98T (600T>C), V114A (648T>C), N131T (699A>C), A172V (822C>T), L173R (825T>G), S186F (864C>T), E208D (931A>T), E258K (1079G>A), E453K (1664G>A), T456I (1674C>T), N457K (1678T>A), L483P (1755T>C), D503V (1815A>T), D506N (1823G>A), L567S (2007T>C), E582K (2051G>A), A591V (2079C>T), T616I (2154C>T) | | | | | | | | |
| Codon mutations: | TTC12TTT (343C>T), CCG69CTG (513C>T), ACA73ATA (525C>T), ACA85ATA (561C>T), ATG98ACG (600T>C), GTG114GCG (648T>C), AAC131ACC (699A>C), CTC136CTT (715C>T), TTG146TTA (745G>A), GCG172GTG (822C>T), CTA173CGA (825T>G), TCC186TTC (864C>T), AGG205AGA (922G>A), GAA208GAT (931A>T), GGA209GGG (934A>G), GAA258AAA (1079G>A), GCA262GCG (1093A>G), AGG270AGA (1117G>A), GAG453AAG (1664G>A), ACA456ATA (1674C>T), AAT457AAA (1678T>A), CTA483CCA (1755T>C), GCC496GCA (1795C>A), GAC503GTC (1815A>T), GAC506AAC (1823G>A), TTA567TCA (2007T>C), GAG582AAG (2051G>A), GCG591GTG (2079C>T), ACC616ATC (2154C>T) | | | | | | | | |
| Replication-associated protein (YP_006355442.1) | 1 | 42 | 100% | 260 | 90.6% | 42 (100%) | 39 (92.9%) | 0/0/0/0 | 1 |
| Protein mutations: | Q15* (1674C>T), M16K (1678T>A), *42Q (1755T>C) | | | | | | | | |
| Codon mutations: | GAG11GAA (1664G>A), CAA15TAA (1674C>T), ATG16AAG (1678T>A), TAA42CAA (1755T>C) | | | | | | | | |
| protein 3a (YP_009179365.2) | 1 | 46 | 100% | 286 | 95.7% | 46 (100%) | 43 (93.5%) | 0/0/0/0 | 1 |
| Protein mutations: | K4R (3585A>G), L11I (3605T>A), F26L (3652T>A) | | | | | | | | |
| Codon mutations: | AAA4AGA (3585A>G), GCT10GCC (3604T>C), TTA11ATA (3605T>A), TTT26TTA (3652T>A) | | | | | | | | |
| CP read-through protein (NP_056751.2) | 1 | 718 | 100% | 4944 | 98.1% | 718 (100%) | 701 (97.6%) | 0/0/0/0 | 2 |
| Protein mutations: | V10G (3721T>G), M19R (3748T>G), V62I (3876G>A 3878T>A), T83S (3939A>T), V141I (4113G>A), I306V (4608A>G 4610C>T), P449L (5038C>T), A457V (5062C>T), H460R (5071A>G 5072C>T), P503S (5199C>T), R508K (5214C>A 5215G>A), V513I (5229G>A), E565K (5385G>A), S606P (5508T>C), E697K (5781G>A), I708M (5816A>G), K711R (5824A>G) | | | | | | | | |
| Codon mutations: | GTC10GGC (3721T>G), ATG19AGG (3748T>G), CGA20AGA (3750C>A), GTT32GTG (3788T>G), GTT62ATA (3876G>A 3878T>A), GTG73GTC (3911G>C), GGC81GGT (3935C>T), ACC83TCC (3939A>T), ACA113ACG (4031A>G), AGC114AGT (4034C>T), GTA141ATA (4113G>A), GCC158GCT (4166C>T), ACT160ACC (4172T>C), TCT174TCG (4214T>G), TCA303TCG (4601A>G), ATC306GTT (4608A>G 4610C>T), TGC308TGT (4616C>T), GGT375GGC (4817T>C), TTC397TTT (4883C>T), CTT398CTC (4886T>C), CCT401CCC (4895T>C), GCC402GCT (4898C>T), ACC429ACT (4979C>T), CCA449CTA (5038C>T), CGG450CGA (5042G>A), GGC452GGT (5048C>T), GCC457GTC (5062C>T), CAC460CGT (5071A>G 5072C>T), ACG479ACA (5129G>A), CCA503TCA (5199C>T), CGA508AAA (5214C>A 5215G>A), GTA513ATA (5229G>A), GAA565AAA (5385G>A), AAC603AAT (5501C>T), TCC606CCC (5508T>C), CCC610CCT (5522C>T), GAA697AAA (5781G>A), ATA708ATG (5816A>G), AAG711AGG (5824A>G) | | | | | | | | |
| coat protein (NP_056749.1) | 1 | 209 | 100% | 1416 | 98.7% | 209 (100%) | 204 (97.6%) | 0/0/0/0 | 1 |
| Protein mutations: | V10G (3721T>G), M19R (3748T>G), V62I (3876G>A 3878T>A), T83S (3939A>T), V141I (4113G>A) | | | | | | | | |
| Codon mutations: | GTC10GGC (3721T>G), ATG19AGG (3748T>G), CGA20AGA (3750C>A), GTT32GTG (3788T>G), GTT62ATA (3876G>A 3878T>A), GTG73GTC (3911G>C), GGC81GGT (3935C>T), ACC83TCC (3939A>T), ACA113ACG (4031A>G), AGC114AGT (4034C>T), GTA141ATA (4113G>A), GCC158GCT (4166C>T), ACT160ACC (4172T>C), TCT174TCG (4214T>G) | | | | | | | | |
| movement protein (NP_056750.1) | 1 | 157 | 100% | 1007 | 91.4% | 157 (100%) | 147 (93.6%) | 0/0/0/0 | 1 |
| Protein mutations: | S2A (3721T>G), C11G (3748T>G 3750C>A), F24C (3788T>G), F54Y (3878T>A), C65S (3911G>C), A73V (3935C>T), Q105R (4031A>G), A106V (4034C>T), P150L (4166C>T), L152P (4172T>C) | | | | | | | | |
| Codon mutations: | TCA2GCA (3721T>G), TGC11GGA (3748T>G 3750C>A), TTC24TGC (3788T>G), GAG53GAA (3876G>A), TTC54TAC (3878T>A), TGT65TCT (3911G>C), GCA73GTA (3935C>T), ACA74ACT (3939A>T), CAA105CGA (4031A>G), GCA106GTA (4034C>T), AAG132AAA (4113G>A), CCA150CTA (4166C>T), CTT152CCT (4172T>C) | | | | | | | | |

*: Inserts / Deletes / Misaligned / Frameshifts

#### Analysis details

This analysis was performed with panviral2.64

### NGS Details (UN18_val): Duamitovirus soch1

#### Assembly

| | |
|---|---|
| **Coverage Length** | 2273 (1 contig(s)) |
| **Depth Of Coverage** | 45.1 |
| **Number Of Reads** | 853 |
| **Reads Per Million** | 16.01 rpm (after QC) |
| **Ambiguities** | 0 |
| **Assembly Method** | de novo + reference guided assembly |
| **Consensus Caller** | Bcf Tools |

#### Coverage Map

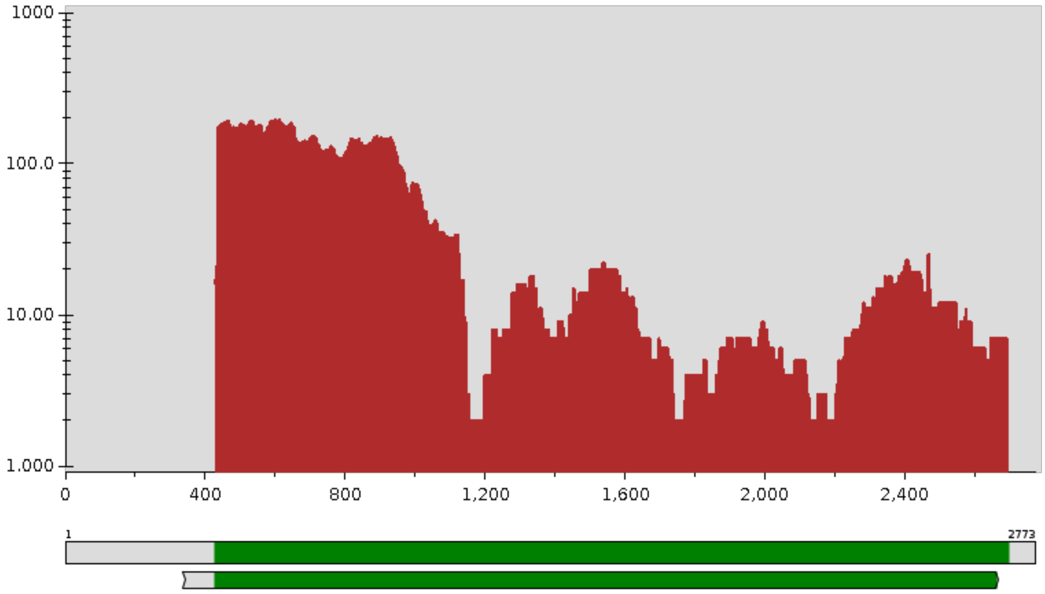

#### Assignment

| | |
|---|---|
| **Type** | Duamitovirus soch1 (Taxonomy ID: 2955838) |
| **Reference Genome** | NC_076524.1 |
| **NT Identity (%)** | 68.9124 |
| **AA Identity (%)** | 68.3646 |
| **Number Of Stop Codons** | 4 |
| **Number Of CDS** | 1 |

#### Alignment

| | |
|---|---|
| **Alignment Score** | 1680.0 (NT) + 3316.0 (AA) = 4996.0 |
| **Concordance (%)** | 52.8854 |

| | |
|---|---|
| **Alignment Method** | Global, seeded, nucleotide + amino acids (AGA) |

#### Genome Region

Sequence starts at position 428 and ends at position 2700 relative to NC_076524.1 reference sequence.

#### Alignment Detailed Statistics

| | Begin | End | Coverage | Score | Concordance | Matches | Identities | I/D/M/F* | Stop Codons |
|---|---|---|---|---|---|---|---|---|---|
| **NT** | **428** | **2700** | **82.0%** | **1680** | **37.3%** | **2266 (99.5%)** | **1565 (68.7%)** | **5/7** | |
| Mutations: | 437A>G, 441A>C, 443C>T, 447C>T, 459C>T, 466C>T, 470C>T, 471G>A, 472C>T, 473T>G, 477T>A, 480A>C, 481C>A, 482A>T, 485A>G, 489A>G, 494T>C, 497T>A, 500T>G, 503A>C, 506T>C, 509C>T, 512T>C, 513G>A, 516T>A, 517C>A, 518T>A, 520A>C, 524G>T, 525A>G, 526C>T, 527T>G, 529C>A, 530C>A, 531T>C, 533A>G, 534C>A, 535G>A, 536C>A, 539G>A, 540A>C, 541A>G, 542G>T, 545G>A, 548C>T, 550A>T, 556T>A, 557C>T, 560A>T, 569C>T, 575G>A, 581C>T, 582G>A, 587A>T, 589G>C, 591T>C, 593A>T, 599A>T, 602C>T, 612T>A, 613C>G, 614C>T, 616T>A, 618C>T, 619G>C, 620A>T, 624G>A, 626A>T, 629A>C, 630C>A, 632A>C, 633C>A, 635T>G, 638G>T, 641G>T, 644T>A, 647G>T, 651C>T, 653C>G, 656A>C, 657A>C, 661C>G, 662G>T, 665T>A, 668C>T, 672C>A, 673G>A, 674T>G, 680C>A, 683C>T, 684A>G, 685A>C, 686A>C, 692G>A, 695T>A, 699G>C, 700C>A, 701A>C, 702A>G, 705C>A, 707T>G, 708A>G, 709G>C, 710G>A, 713A>G, 714G>T, 715A>C, 719A>T, 722C>T, 737T>C, 746C>T, 752A>T, 761G>A, 765G>T, 767T>A, 770G>A, 771G>T, 772T>A, 773T>G, 774T>A, 776G>T, 777A>G, 778A>C, 779A>G, 782T>G, 796G>A, 797T>G, 803A>C, 809T>C, 810G>C, 815C>A, 818A>T, 821T>G, 824C>T, 827A>T, 831C>T, 833T>A, 841G>A, 842G>T, 850T>G, 851A>T, 857G>T, 860G>T, 862A>G, 869T>G, 875G>T, 880T>G, 881C>A, 885C>A, 887T>G, 892A>G, 905A>T, 908C>A, 909T>C, 910T>A, 916G>A, 918G>T, 919G>C, 923T>A, 926A>T, 927A>C, 929G>T, 934T>A, 935T>G, 938G>T, 942T>A, 943C>A, 948C>A, 971C>A, 972C>T, 975A>C, 976A>T, 977T>C, 983T>C, 990A>C, 992G>C, 993T>C, 994C>G, 995A>G, 997A>T, 998C>T, 999G>C, 1000G>A, 1001T>A, 1003T>A, 1004A>T, 1005G>T, 1007C>G, 1010A>G, 1016C>T, 1017A>G, 1018A>T, 1019A>T, 1022T>A, 1025T>C, 1026C>G, 1027A>G, 1030A>C, 1031G>A, 1034T>G, 1035A>G, 1036G>C, 1044T>G, 1046C>T, 1049G>A, 1050T>G, 1051T>A, 1052T>A, 1054C>A, 1055A>C, 1061T>C, 1067T>G, 1073G>A, 1076A>G, 1083C>A, 1088G>T, 1091C>G, 1099G>T, 1104A>G, 1105C>T, 1106A>G, 1115T>G, 1121A>C, 1122T>A, 1124C>T, 1127G>A, 1130C>T, 1133C>G, 1142A>C, 1145C>T, 1205A>T, 1211C>T, 1212T>G, 1213T>A, 1217T>C, 1220C>T, 1223A>T, 1224C>T, 1225A>T, 1226C>A, 1232G>A, 1235A>G, 1236A>T, 1237G>C, 1241C>A, 1244A>C, 1247A>C, 1249G>A, 1250G>C, 1254T>A, 1255C>G, 1256A>T, 1257A>T, 1259C>T, 1260T>C, 1262A>T, 1266A>G, 1268T>G, 1269C>G, 1274T>C, 1275C>A, 1276G>A, 1277A>G, 1280G>T, 1283G>T, 1286A>G, 1291T>C, 1292T>A, 1295T>G, 1298T>A, 1302C>T, 1304T>A, 1306G>C, 1308G>C, 1309C>A, 1322C>A, 1328A>T, 1335A>C, 1338G>A, 1340A>C, 1345T>C, 1349G>A, 1355C>T, 1359A>G, 1364_1365insAAAAA, 1369G>A, 1370A>G, 1371C>T, 1372T>G, 1373T>A, 1376C>G, 1377C>T, 1378A>T, 1379C>T, 1385T>C, 1389G>C, 1391T>A, 1397T>A, 1402C>A, 1409G>A, 1413A>C, 1414A>G, 1415A>C, 1416A>C, 1418A>T, 1419A>C, 1420A>C, 1422A>G, 1424A>G, 1427C>T, 1430G>A, 1431A>G, 1433T>A, 1445G>A, 1446A>C, 1447A>G, 1448A>T, 1451T>G, 1455T>G, 1457G>A, 1458G>C, 1459C>G, 1460T>A, 1464A>C, 1465A>G, 1466G>A, 1467C>T, 1468A>C, 1469A>T, 1470A>C, 1472G>A, 1475C>G, 1476T>A, 1477T>A, 1478T>A, 1479A>G, 1480A>G, 1481A>T, 1486G>C, 1489A>T, 1490T>C, 1502A>C, 1505G>A, 1508A>G, 1526G>A, 1538G>A, 1541T>C, 1545C>T, 1547T>C, 1553T>C, 1554A>T, 1555T>G, 1556G>C, 1559A>G, 1566C>A, 1567C>G, 1571C>A, 1572T>C, 1580C>A, 1589T>C, 1593_1597delAGCTC, 1598T>A, 1607T>C, 1613G>T, 1614A>T, 1617C>T, 1619T>G, 1620G>A, 1622T>C, 1623A>G, 1624A>T, 1626A>C, 1627A>C, 1628A>T, 1632T>A, 1634A>G, 1635A>G, 1636C>T, 1637A>C, 1643A>G, 1644G>A, 1647T>A, 1648A>G, 1664G>A, 1670G>A, 1676T>G, 1679G>A, 1682C>T, 1684T>A, 1686C>T, 1691T>A, 1694T>A, 1698G>T, 1700A>C, 1703T>A, 1707T>G, 1710T>C, 1712G>T, 1715T>C, 1736A>G, 1739A>T, 1744G>A, 1745C>A, 1746A>T, 1748A>T, 1754T>C, 1758G>A, 1759G>C, 1760A>G, 1761T>A, 1765A>C, 1766C>T, 1767T>A, 1769T>C, 1772A>T, 1777G>C, 1778G>T, 1787T>C, 1788T>C, 1790A>G, 1791C>T, 1819G>C, 1820T>A, 1821A>G, 1823A>G, 1829C>G, 1830A>C, 1831A>G, 1832T>G, 1833G>C, 1840A>G, 1847A>G, 1848T>C, 1856A>G, 1859C>T, 1860A>G, 1861A>G, 1865C>T, 1867A>C, 1871T>C, 1874G>T, 1875delA, 1878C>G, 1880A>T, 1886T>G, 1887T>A, 1888T>G, 1889G>A, 1895T>C, 1898T>G, 1901T>C, 1910T>C, 1919G>T, 1925G>A, 1934T>C, 1937A>G, 1938T>G, 1939C>A, 1940A>C, 1950G>T, 1952T>C, 1953T>C, 1955A>T, 1964T>C, 1967C>T, 1976T>A, 1980T>A, 1981T>C, 1982A>T, 1985T>C, 1989C>A, 1991T>G, 1994T>A, 2000T>G, 2003A>T, 2006T>G, 2009C>T, 2010G>C, 2011C>A, 2012C>A, 2020T>C, 2028G>T, 2029A>C, 2030T>G, 2034A>T, 2035A>C, 2036C>A, 2038T>C, 2042A>G, 2043G>A, 2050A>C, 2051A>C, 2053G>T, 2054A>T, 2057G>T, 2058G>T, 2063T>G, 2069G>A, 2071A>T, 2074A>G, 2075G>A, 2078A>T, 2082G>T, 2084G>T, 2087C>T, 2090A>G, 2091T>A, 2093C>G, 2094A>T, 2099C>T, 2103A>T, 2104G>C, 2105C>T, 2108G>A, 2117G>A, 2118C>A, 2120T>A, 2123A>G, 2129T>G, 2133C>G, 2134A>C, 2135T>A, 2136C>A, 2137T>G, 2138T>C, 2141G>A, 2145G>C, 2146T>A, 2147T>C, 2148C>A, 2156C>T, 2157T>C, 2160C>T, 2166T>C, 2167A>T, 2173G>A, 2174G>T, 2183A>G, 2186G>A, 2187G>T, 2188G>T, 2191T>A, 2192G>A, 2195C>T, 2204C>T, 2206A>T, 2213A>G, 2216G>A, 2223G>A, 2226A>G, 2227G>A, 2231C>T, 2236T>G, 2237T>G, 2240A>G, 2248A>C, 2250G>C, 2251T>C, 2259T>A, 2263A>G, 2264G>A, 2265A>C, 2276G>T, 2279T>G, 2280C>T, 2288C>T, 2297A>G, 2298G>A, 2299T>G, 2300T>A, 2306A>C, 2309A>T, 2315T>C, 2322C>T, 2326G>T, 2328C>A, 2329A>G, 2336A>G, 2337G>A, 2346T>C, 2348A>G, 2351G>A, 2355C>G, 2358A>T, 2359A>C, 2360C>T, 2372G>T, 2373A>G, 2375C>T, 2376T>G, 2377G>C, 2378T>A, 2379A>G, 2380T>C, 2381G>A, 2382A>T, 2383A>C, 2384C>A, 2385C>G, 2390T>A, 2394T>A, 2400A>G, 2402G>T, 2403A>C, 2404G>A, 2405T>G, 2406C>T, 2408C>G, 2411C>G, 2414C>A, 2415T>G, 2416C>T, 2417C>T, 2423A>G, 2426C>T, 2427G>A, 2428C>G, 2429A>T, 2431C>A, 2438G>A, 2444G>A, 2445C>A, 2446T>C, 2447C>A, 2448G>A, 2450A>C, 2451A>G, 2452C>A, 2453T>A, 2457T>C, 2465A>G, 2477C>T, 2478C>T, 2489A>G, 2492T>C, 2495C>T, 2504T>A, 2510T>C, 2514A>G, 2518C>T, 2519C>G, 2524C>T, 2525C>T, 2528T>A, 2532C>A, 2533A>G, 2536G>A, 2537G>T, 2543T>G, 2551A>G, 2552C>T, 2555T>C, 2556G>T, 2557T>C, 2561T>C, 2568A>G, 2571A>C, 2572A>G, 2582A>T, 2585G>A, 2597G>A, 2598A>G, 2599A>C, 2600C>T, 2601G>A, 2609T>A, 2613T>G, 2615T>A, 2618T>A, 2622G>T, 2624A>T, 2626A>T, 2627C>A, 2629A>C, 2631A>T, 2634A>C, 2635C>T, 2638G>T, 2639A>G, 2644A>G, 2646G>T, 2647A>G, 2648A>T, 2649G>A, 2650T>G, 2652G>T, 2653G>C, 2654C>A, 2656T>A, 2657A>T, 2658A>G, 2659G>A, 2660A>C, 2661C>T, 2664T>A, 2665A>G, 2667delG, 2674T>A, 2678C>T, 2681G>A, 2686T>A, 2689T>C, 2690G>C, 2694A>G, 2695T>A | | | | | | | | |

**CDS**

| | Begin | End | Coverage | Score | Concordance | Matches | Identities | I/D/M/F* | Stop Codons |
|---|---|---|---|---|---|---|---|---|---|
| RdRp | 32 | 777 | 95.9% | 3316 | 63.5% | 744 (99.6%) | 510 (68.3%) | 2/1/3/3 | 4 |
| Protein mutations: | I36L (441A>C 443C>T), L38F (447C>T), A44V (466C>T), A46M (471G>A 472C>T 473T>G), L48M (477T>A), T49H (480A>C 481C>A 482A>T), I52V (489A>G), V60I (513G>A), S61K (516T>A 517C>A 518T>A), K62T (520A>C), T64V (525A>G 526C>T 527T>G), T65K (529C>A 530C>A), R67K (534C>A 535G>A 536C>A), K69R (540A>C 541A>G 542G>T), Y72F (550A>T), F74Y (556T>A 557C>T), A83T (582G>A), C85S (589G>C), F94Y (616T>A), R95S (618C>T 619G>C 620A>T), E97N (624G>A 626A>T), E98D (629A>C), P99T (630C>A 632A>C), L100M (633C>A 635T>G), S109C (661C>G 662G>T), R113K (672C>A 673G>A 674T>G), K117A (684A>G 685A>C 686A>C), A122H (699G>C 700C>A 701A>C), I123V (702A>G), R125A (708A>G 709G>C 710G>A), D127S (714G>T 715A>C), E128D (719A>T), A144S (765G>T 767T>A), V146* (771G>T 772T>A 773T>G), L147I (774T>A 776G>T), K148A (777A>G 778A>C 779A>G), S154K (796G>A 797T>G), E159Q (810G>C), P166S (831C>T 833T>A), G169D (841G>A 842G>T), L172R (850T>G 851A>T), E176G (862A>G), I178M (869T>G), F182* (880T>G 881C>A), H184K (885C>A 887T>G), Q186R (892A>G), F192H (909T>C 910T>A), R194Q (916G>A), G195S (918G>T 919G>C), M198L (927A>C 929G>T), I200K (934T>A 935T>G), S203N (942T>A 943C>A), Q205K (948C>A), N214L (975A>C 976A>T 977T>C), S220R (993T>C 994C>G 995A>G), Y221F (997A>T 998C>T), G222Q (999G>C 1000G>A 1001T>A), L223Y (1003T>A 1004A>T), A224S (1005G>T 1007C>G), K228V (1017A>G 1018A>T 1019A>T), Q231G (1026C>G 1027A>G), E232A (1030A>C 1031G>A), S234A (1035A>G 1036G>C), F237V (1044T>G 1046C>T), F239E (1050T>G 1051T>A 1052T>A), P240H (1054C>A 1055A>C), L250I (1083C>A), W255L (1099G>T), T257V (1104A>G 1105C>T 1106A>G), I260M (1115T>G), F263I (1122T>A 1124C>T), L290F (1205A>T), L293E (1212T>G 1213T>A), E296D (1223A>T), H297L (1224C>T 1225A>T 1226C>A), W305Y (1249G>A 1250G>C), T308S (1257A>T 1259C>T), S311G (1266A>G 1268T>G), P312A (1269C>G), R314K (1275C>A 1276G>A 1277A>G), V319A (1291T>C 1292T>A), G324A (1306G>C), A325Q (1308G>C 1309C>A), V335I (1338G>A 1340A>C), V337A (1345T>C), M338I (1349G>A), I342V (1359A>G), K343_Q344insKX (1364_1365insAAAAA), R345K (1369G>A 1370A>G), L346* (1371C>T 1372T>G 1373T>A), H348F (1377C>T 1378A>T 1379C>T), D352Q (1389G>C 1391T>A), T356K (1402C>A), K360R (1413A>C 1414A>G 1415A>C), I361L (1416A>C 1418A>T), K362P (1419A>C 1420A>C), T363A (1422A>G 1424A>G), T366A (1431A>G 1433T>A), K371R (1446A>C 1447A>G 1448A>T), L374V (1455T>G 1457G>A), A375R (1458G>C 1459C>G 1460T>A), K377R (1464A>C 1465A>G 1466G>A), Q378S (1467C>T 1468A>C 1469A>T), K379Q (1470A>C 1472G>A), N380K (1475C>G), F381K (1476T>A 1477T>A 1478T>A), K382G (1479A>G 1480A>G 1481A>T), C384S (1486G>C), Y385F (1489A>T 1490T>C), L389F (1502A>C), M401I (1538G>A), L404F (1545C>T 1547T>C), M407C (1554A>T 1555T>G 1556G>C), I408M (1559A>G), P411S (1566C>A 1567C>G), S420del (1593_1597delAGCTC), I427F (1614A>T), V429I (1620G>A 1622T>C), N430V (1623A>G 1624A>T), K431P (1626A>C 1627A>C 1628A>T), L433M (1632T>A 1634A>G), T434V (1635A>G 1636C>T 1637A>C), V437M (1644G>A), Y438S (1647T>A 1648A>G), F450Y (1684T>A), H451Y (1686C>T), A455S (1698G>T 1700A>C), S458A (1707T>G), S470K (1744G>A 1745C>A), K471Y (1746A>T 1748A>T), G475T (1758G>A 1759G>C 1760A>G), C476S (1761T>A), N477T (1765A>C 1766C>T), S478T (1767T>A 1769T>C), G481A (1777G>C 1778G>T), S495T (1819G>C 1820T>A), K496E (1821A>G 1823A>G), N499R (1830A>C 1831A>G 1832T>G), E500Q (1833G>C), K502R (1840A>G), N509G (1860A>G 1861A>G), K511T (1867A>C), P515A (1878C>G 1880A>T), L518R (1887T>A 1888T>G 1889G>A), D521E (1898T>G), S535D (1938T>G 1939C>A 1940A>C), D539Y (1950G>T 1952T>C), L549T (1980T>A 1981T>C 1982A>T), A559Q (2010G>C 2011C>A 2012C>A), I562T (2020T>C), D565S (2028G>T 2029A>C 2030T>G), N567S (2034A>T 2035A>C 2036C>A), I568T (2038T>C), V570I (2043G>A), Q572P (2050A>C 2051A>C), R573I (2053G>T 2054A>T), L574F (2057G>T), G575W (2058G>T), Y579F (2071A>T), K580R (2074A>G 2075G>A), A583S (2082G>T 2084G>T), F586M (2091T>A 2093C>G), T587S (2094A>T), H600A (2133C>G 2134A>C 2135T>A), L601S (2136C>A 2137T>G 2138T>C), V604H (2145G>C 2146T>A 2147T>C), Y611L (2166T>C 2167A>T), W613Y (2173G>A 2174G>T), G618F (2187G>T 2188G>T), M619K (2191T>A 2192G>A), Y624F (2206A>T), V630I (2223G>A), S631D (2226A>G 2227G>A), L634R (2236T>G 2237T>G), K638T (2248A>C), V639P (2250G>C 2251T>C), L642I (2259T>A), Q643R (2263A>G 2264G>A), I644L (2265A>C), K647N (2276G>T), D648E (2279T>G), V655R (2298G>A 2299T>G 2300T>A), R664L (2326G>T), Q665R (2328C>A 2329A>G), E668K (2337G>A), L674V (2355C>G), N675S (2358A>T 2359A>C 2360C>T), I680V (2373A>G 2375C>T), C681A (2376T>G 2377G>C 2378T>A), M682A (2379A>G 2380T>C 2381G>A), N683S (2382A>T 2383A>C 2384C>A), P684A (2385C>G), D685E (2390T>A), S687T (2394T>A), K689D (2400A>G 2402G>T), S690Q (2403A>C 2404G>A 2405T>G), I692M (2411C>G), D693E (2414C>A), S694V (2415T>G 2416C>T 2417C>T), I696M (2423A>G), A698S (2427G>A 2428C>G 2429A>T), T699K (2431C>A), W701* (2438G>A), L704T (2445C>A 2446T>C 2447C>A), E705N (2448G>A 2450A>C), T706E (2451A>G 2452C>A 2453T>A), F708L (2457T>C), N727D (2514A>G), P728L (2518C>T 2519C>G), T730I (2524C>T 2525C>T), Q733R (2532C>A 2533A>G), W734Y (2536G>A 2537G>T), F736L (2543T>G), N739S (2551A>G 2552C>T), V741S (2556G>T 2557T>C), N745D (2568A>G), K746R (2571A>C 2572A>G), L749F (2582A>T), N755A (2598A>G 2599A>C 2600C>T), D756N (2601G>A), S760A (2613T>G 2615T>A), E763Y (2622G>T 2624A>T), Y764L (2626A>T 2627C>A), Q765P (2629A>C), T766S (2631A>T), T767L (2634A>C 2635C>T), G768V (2638G>T 2639A>G), E770G (2644A>G), E771C (2646G>T 2647A>G 2648A>T), V772R (2649G>A 2650T>G), G773S (2652G>T 2653G>C 2654C>A), L774Y (2656T>A 2657A>T), R775D (2658A>G 2659G>A 2660A>C), L776F (2661C>T), *777R (2664T>A 2665A>G) | | | | | | | | |

| | Begin | End | Coverage | Score | Concordance | Matches | Identities | I/D/M/F* | Stop Codons |
|---|---|---|---|---|---|---|---|---|---|
| **NT** | **428** | **2700** | **82.0%** | **1680** | **37.3%** | **2266 (99.5%)** | **1565 (68.7%)** | **5/7** | |

Codon mutations:

AAA34AAG (437A>G), ATC36CTT (441A>C 443C>T), CTT38TTT (447C>T), CTA42TTA (459C>T), GCC44GTC (466C>T), TCC45TCT (470C>T), GCT46ATG (471G>A 472C>T 473T>G), TTG48ATG (477T>A), ACA49CAT (480A>C 481C>A 482A>T), AAA50AAG (485A>G), ATC52GTC (489A>G), GCT53GCC (494T>C), CTT54CTA (497T>A), GCT55GCG (500T>G), GGA56GGC (503A>C), TGT57TGC (506T>C), GAC58GAT (509C>T), TTT59TTC (512T>C), GTA60ATA (513G>A), TCT61AAA (516T>A 517C>A 518T>A), AAA62ACA (520A>C), GTG63GTT (524G>T), ACT64GTG (525A>G 526C>T 527T>G), ACC65AAA (529C>A 530C>A), TTA66CTG (531T>C 533A>G), CGC67AAA (534C>A 535G>A 536C>A), CGG68CGA (539G>A), AAG69CGT (540A>C 541A>G 542G>T), TCG70TCA (545G>A), GGC71GGT (548C>T), TAT72TTT (550A>T), TTC74TAT (556T>A 557C>T), ACA75ACT (560A>T), TAC78TAT (569C>T), AAG80AAA (575G>A), TGC82TGT (581C>T), GCA83ACA (582G>A), GTA84GTT (587A>T), TGC85TCC (589G>C), TTA86CTT (591T>C 593A>T), CGA88CGT (599A>T), TAC89TAT (602C>T), TCC93AGT (612T>A 613C>G 614C>T), TTT94TAT (616T>A), CGA95TCT (618C>T 619G>C 620A>T), GAA97AAT (624G>A 626A>T), GAA98GAC (629A>C), CCA99ACC (630C>A 632A>C), CTT100ATG (633C>A 635T>G), TCG101TCT (638G>T), GTG102GTT (641G>T), CCT103CCA (644T>A), GTG104GTT (647G>T), CTC106TTG (651C>T 653C>G), ACA107ACC (656A>C), AGA108CGA (657A>C), TCG109TGT (661C>G 662G>T), GGT110GGA (665T>A), ATC111ATT (668C>T), CGT113AAG (672C>A 673G>A 674T>G), ATC115ATA (680C>A), CCC116CCT (683C>T), AAA117GCC (684A>G 685A>C 686A>C), TTG119TTA (692G>A), CGT120CGA (695T>A), GCA122CAC (699G>C 700C>A 701A>C), ATA123GTA (702A>G), CGT124AGG (705C>A 707T>G), AGG125GCA (708A>G 709G>C 710G>A), AAA126AAG (713A>G), GAT127TCT (714G>T 715A>C), GAA128GAT (719A>T), CAC129CAT (722C>T), GTT134GTC (737T>C), TAC137TAT (746C>T), TCA139TCT (752A>T), GGG142GGA (761G>A), GCT144TCA (765G>T 767T>A), AAG145AAA (770G>A), GTT146TAG (771G>T 772T>A 773T>G), TTG147ATT (774T>A 776G>T), AAA148GCG (777A>G 778A>C 779A>G), GTT149GTG (782T>G), AGT154AAG (796G>A 797T>G), GCA156GCC (803A>C), TTT158TTC (809T>C), GAA159CAA (810G>C), TCC160TCA (815C>A), ATA161ATT (818A>T), ACT162ACG (821T>G), ACC163ACT (824C>T), CCA164CCT (827A>T), CCT166TCA (831C>T 833T>A), GGG169GAT (841G>A 842G>T), CTA172CGT (850T>G 851A>T), GTG174GTT (857G>T), CTG175CTT (860G>T), GAA176GGA (862A>G), ATT178ATG (869T>G), ACG180ACT (875G>T), TTC182TGA (880T>G 881C>A), CAT184AAG (885C>A 887T>G), CAA186CGA (892A>G), CTA190CTT (905A>T), CCC191CCA (908C>A), TTT192CAT (909T>C 910T>A), CGA194CAA (916G>A), GGC195TCC (918G>T 919G>C), ATT196ATA (923T>A), CCA197CCT (926A>T), ATG198CTT (927A>C 929G>T), ATT200AAG (934T>A 935T>G), GGG201GGT (938G>T), TCT203AAT (942T>A 943C>A), CAA205AAA (948C>A), CCC212CCA (971C>A), CTA213TTA (972C>T), AAT214CTC (975A>C 976A>T 977T>C), AAT216AAC (983T>C), AGG219CGC (990A>C 992G>C), TCA220CGG (993T>C 994C>G 995A>G), TAC221TTT (997A>T 998C>T), GGT222CAA (999G>C 1000G>A 1001T>A), TTA223TAT (1003T>A 1004A>T), GCC224TCG (1005G>T 1007C>G), GTA225GTG (1010A>G), GAC227GAT (1016C>T), AAA228GTT (1017A>G 1018A>T 1019A>T), GTT229GTA (1022T>A), GAT230GAC (1025T>C), CAA231GGA (1026C>G 1027A>G), GAG232GCA (1030A>C 1031G>A), GCT233GCG (1034T>G), AGT234GCT (1035A>G 1036G>C), TTC237GTT (1044T>G 1046C>T), AAG238AAA (1049G>A), TTT239GAA (1050T>G 1051T>A 1052T>A), CCA240CAC (1054C>A 1055A>C), ATT242ATC (1061T>C), GTT244GTG (1067T>G), TTG246TTA (1073G>A), AAA247AAG (1076A>G), CTA250ATA (1083C>A), GCG251GCT (1088G>T), GCC252GCG (1091C>G), TGG255TTG (1099G>T), ACA257GTG (1104A>G 1105C>T 1106A>G), ATT260ATG (1115T>G), TCA262TCC (1121A>C), TTC263ATT (1122T>A 1124C>T), CAG264CAA (1127G>A), GAC265GAT (1130C>T), GGC266GGG (1133C>G), TCA269TCC (1142A>C), CCC270CCT (1145C>T), TTA290TTT (1205A>T), AAC292AAT (1211C>T), TTA293GAA (1212T>G 1213T>A), TCT294TCC (1217T>C), CTC295CTT (1220C>T), GAA296GAT (1223A>T), CAC297TTA (1224C>T 1225A>T 1226C>A), GAG299GAA (1232G>A), AAA300AAG (1235A>G), AGT301TCT (1236A>T 1237G>C), GTC302GTA (1241C>A), GGA303GGC (1244A>C), CCA304CCC (1247A>C), TGG305TAC (1249G>A 1250G>C), TCA307AGT (1254T>A 1255C>G 1256A>T), ACC308TCT (1257A>T 1259C>T), TTA309CTT (1260T>C 1262A>T), AGT311GGG (1266A>G 1268T>G), CCT312GCT (1269C>G), TAT313TAC (1274T>C), CGA314AAG (1275C>A 1276G>A 1277A>G), GGG315GGT (1280G>T), GTG316GTT (1283G>T), CCA317CCG (1286A>G), GTT319GCA (1291T>C 1292T>A), ACT320ACG (1295T>G), GGT321GGA (1298T>A), CTT323TTA (1302C>T 1304T>A), GGT324GCT (1306G>C), GCA325CAA (1308G>C 1309C>A), GGC329GGA (1322C>A), GGA331GGT (1328A>T), AGG334CGG (1335A>C), GTA335ATC (1338G>A 1340A>C), GTA337GCA (1345T>C), ATG338ATA (1349G>A), AAC340AAT (1355C>T), ATA342GTA (1359A>G), AAA343_CAA344insAAAAA- (1364_1365insAAAAA), AGA345AAG (1369G>A 1370A>G), CTT346TGA (1371C>T 1372T>G 1373T>A), CTC347CTG (1376C>G), CAC348TTT (1377C>T 1378A>T 1379C>T), GTT350GTC (1385T>C), GAT352CAA (1389G>C 1391T>A), GCT354GCA (1397T>A), ACA356AAA (1402C>A), CTG358CTA (1409G>A), AAA360CGC (1413A>C 1414A>G 1415A>C), ATA361CTT (1416A>C 1418A>T), AAG362CCG (1419A>C 1420A>C), ACA363GCG (1422A>G 1424A>G), GAC364GAT (1427C>T), GGG365GGA (1430G>A), ACT366GCA (1431A>G 1433T>A), GAG370GAA (1445G>A), AAA371CGT (1446A>C 1447A>G 1448A>T), CCT372CCG (1451T>G), TTG374GTA (1455T>G 1457G>A), GCT375CGA (1458G>C 1459C>G 1460T>A), AAG377CGA (1464A>C 1465A>G 1466G>A), CAA378TCT (1467C>T 1468A>C 1469A>T), AAG379CAA (1470A>C 1472G>A), AAC380AAG (1475C>G), TTT381AAA (1476T>A 1477T>A 1478T>A), AAA382GGT (1479A>G 1480A>G 1481A>T), TGT384TCT (1486G>C), TAT385TTC (1489A>T 1490T>C), TTA389TTC (1502A>C), AAG390AAA (1505G>A), TCA391TCG (1508A>G), CCG397CCA (1526G>A), ATG401ATA (1538G>A), TAT402TAC (1541T>C), CTT404TTC (1545C>T 1547T>C), TCT406TCC (1553T>C), ATG407TGC (1554A>T 1555T>G 1556G>C), ATA408ATG (1559A>G), CCT411AGT (1566C>A 1567C>G), ACC412ACA (1571C>A), TTG413CTG (1572T>C), TCC415TCA (1580C>A), GTT418GTC (1589T>C), AGC420del (1593_1597delAGCTC), TCT421--A (1593_1597delAGCTC 1598T>A), CTT424CTC (1607T>C), ACG426ACT (1613G>T), ATC427TTC (1614A>T), CTT428TTG (1617C>T 1619T>G), GTT429ATC (1620G>A 1622T>C), AAC430GTC (1623A>G 1624A>T), AAA431CCT (1626A>C 1627A>C 1628A>T), TTA433ATG (1632T>A 1634A>G), ACA434GTC (1635A>G 1636C>T 1637A>C), AGA436AGG (1643A>G), GTG437ATG (1644G>A), TAT438AGT (1647T>A 1648A>G), CTG443CTA (1664G>A), GGG445GGA (1670G>A), CCT447CCG (1676T>G), TTG448TTA (1679G>A), GGC449GGT (1682C>T), TTC450TAC (1684T>A), CAC451TAC (1686C>T), GGT452GGA (1691T>A), TCT453TCA (1694T>A), GCA455TCC (1698G>T 1700A>C), CTT456CTA (1703T>A), TCG458GCG (1707T>G), TTG459CTT (1710T>C 1712G>T), TCT460TCC (1715T>C), TTA467TTG (1736A>G), GCA468GCT (1739A>T), AGC470AAA (1744G>A 1745C>A), AAA471TAT (1746A>T 1748A>T), TAT473TAC (1754T>C), GGA475ACG (1758G>A 1759G>C 1760A>G), TGT476AGT (1761T>A), AAC477ACT (1765A>C 1766C>T), TCT478ACC (1767T>A 1769T>C), CCA479CCT (1772A>T), GGG481GCT (1777G>C 1778G>T), GCT484GCC (1787T>C), TTA485CTG (1788T>C 1790A>G), CTA486TTA (1791C>T), AGT495ACA (1819G>C 1820T>A), AAA496GAG (1821A>G 1823A>G), GCC498GCG (1829C>G), AAT499CGG (1830A>C 1831A>G 1832T>G), GAG500CAG (1833G>C), AAG502AGG (1840A>G), TTA504TTG (1847A>G), TTG505CTG (1848T>C), AGA507AGG (1856A>G), CTC508CTT (1859C>T), AAT509GGT (1860A>G 1861A>G), GTC510GTT (1865C>T), AAA511ACA (1867A>C), ATT512ATC (1871T>C), TCG513TCT (1874G>T), ATA514-TA (1875delA), CCA515GCT (1878C>G 1880A>T), TCT517TCG (1886T>G), TTG518AGA (1887T>A 1888T>G 1889G>A), TCT520TCC (1895T>C), GAT521GAG (1898T>G), AAT522AAC (1901T>C), ATT525ATC (1910T>C), GCG528GCT (1919G>T), AGG530AGA (1925G>A), ACT533ACC (1934T>C), AAA534AAG (1937A>G), TCA535GAC (1938T>G 1939C>A 1940A>C), GAT539TAC (1950G>T 1952T>C), TTA540CTT (1953T>C 1955A>T), ATT543ATC (1964T>C), TCC544TCT (1967C>T), GCT547GCA (1976T>A), TTA549ACT (1980T>A 1981T>C 1982A>T), TCT550TCC (1985T>C), CGT552AGG (1989C>A 1991T>G), ACT553ACA (1994T>A), GTT555GTG (2000T>G), GGA556GGT (2003A>T), CTT557CTG (2006T>G), TGC558TGT (2009C>T), GCC559CAA (2010G>C 2011C>A 2012C>A), ATC562ACC (2020T>C), GAT565TCG (2028G>T 2029A>C 2030T>G), AAC567TCA (2034A>T 2035A>C 2036C>A), ATA568ACA (2038T>C), TCA569TCG (2042A>G), GTT570ATT (2043G>A), CAA572CCC (2050A>C 2051A>C), AGA573ATT (2053G>T 2054A>T), TTG574TTT (2057G>T), GGG575TGG (2058G>T), GGT576GGG (2063T>G), GGG578GGA (2069G>A), TAC579TTC (2071A>T), AAG580AGA (2074A>G 2075G>A), GTA581GTT (2078A>T), GCG583TCT (2082G>T 2084G>T), CGC584CGT (2087C>T), TTA585TTG (2090A>G), TTC586ATG (2091T>A 2093C>G), ACT587TCT (2094A>T), ACC588ACT (2099C>T), AGC590TCT (2103A>T 2104G>C 2105C>T), AAG591AAA (2108G>A), GAG594GAA (2117G>A), CGT595AGA (2118C>A 2120T>A), CTA596CTG (2123A>G), GCT598GCG (2129T>G), CAT600GCA (2133C>G 2134A>C 2135T>A), CTT601AGC (2136C>A 2137T>G 2138T>C), AAG602AAA (2141G>A), GTT604CAC (2145G>C 2146T>A 2147T>C), CGG605AGG (2148C>A), CAC607CAT (2156C>T), TTA608CTA (2157T>C), CTA609TTA (2160C>T), TAC611CTC (2166T>C 2167A>T), TGG613TAT (2173G>A 2174G>T), GGA616GGG (2183A>G), AGG617AGA (2186G>A), GGT618TTT (2187G>T 2188G>T), ATG619AAA (2191T>A 2192G>A), CCC620CCT (2195C>T), CCC623CCT (2204C>T), TAT624TTT (2206A>T), AAA626AAG (2213A>G), GGG627GGA (2216G>A), GTA630ATA (2223G>A), AGT631GAT (2226A>G 2227G>A), TAC632TAT (2231C>T), CTT634CGG (2236T>G 2237T>G), AAA635AAG (2240A>G), AAA638ACA (2248A>C), GTT639CCT (2250G>C 2251T>C), TTA642ATA (2259T>A), CAG643CGA (2263A>G 2264G>A), ATC644CTC (2265A>C), AAG647AAT (2276G>T), GAT648GAG (2279T>G), CTG649TTG (2280C>T), TTC651TTT (2288C>T), GAA654GAG (2297A>G), GTT655AGA (2298G>A 2299T>G 2300T>A), ATA657ATC (2306A>C), CTA658CTT (2309A>T), CGT660CGC (2315T>C), CTA663TTA (2322C>T), CGC664CTC (2326G>T), CAA665AGA (2328C>A 2329A>G), GTA667GTG (2336A>G), GAA668AAA (2337G>A), TTA671CTG (2346T>C 2348A>G), AAG672AAA (2351G>A), CTC674GTC (2355C>G), AAC675TCT (2358A>T 2359A>C 2360C>T), ACG679ACT (2372G>T), ATC680GTT (2373A>G 2375C>T), TGT681GCA (2376T>G 2377G>C 2378T>A), ATG682GCA (2379A>G 2380T>C 2381G>A), AAC683TCA (2382A>T 2383A>C 2384C>A), CCA684GCA (2385C>G), GAT685GAA (2390T>A), TCC687ACC (2394T>A), AAG689GAT (2400A>G 2402G>T), AGT690CAG (2403A>C 2404G>A 2405T>G), CTC691TTG (2406C>T 2408C>G), ATC692ATG (2411C>G), GAC693GAA (2414C>A), TCC694GTT (2415T>G 2416C>T 2417C>T), ATA696ATG (2423A>G), TGC697TGT (2426C>T), GCA698AGT (2427G>A 2428C>G 2429A>T), ACA699AAA (2431C>A), TGG701TGA (2438G>A), AGG703AGA (2444G>A), CTC704ACA (2445C>A 2446T>C 2447C>A), GAA705AAC (2448G>A 2450A>C), ACT706GAA (2451A>G 2452C>A 2453T>A), TTT708CTT (2457T>C), TTA710TTG (2465A>G), GGC714GGT (2477C>T), CTA715TTA (2478C>T), AAA718AAG (2489A>G), TGT719TGC (2492T>C), TAC720TAT (2495C>T), GGT723GGA (2504T>A), GGT725GGC (2510T>C), AAT727GAT (2514A>G), CCC728CTG (2518C>T 2519C>G), ACC730ATT (2524C>T 2525C>T), ACT731ACA (2528T>A), CAG733AGG (2532C>A 2533A>G), TGG734TAT (2536G>A 2537G>T), TTT736TTG (2543T>G), AAC739AGT (2551A>G 2552C>T), ACT740ACC (2555T>C), GTT741TCT (2556G>T 2557T>C), ATT742ATC (2561T>C), AAT745GAT (2568A>G), AAA746CGA (2571A>C 2572A>G), TTA749TTT (2582A>T), GGG750GGA (2585G>A), GGG754GGA (2597G>A), AAC755GCT (2598A>G 2599A>C 2600C>T), GAT756AAT (2601G>A), ATT758ATA (2609T>A), TCT760GCA (2613T>G 2615T>A), CCT761CCA (2618T>A), GAA763TAT (2622G>T 2624A>T), TAC764TTA (2626A>T 2627C>A), CAA765CCA (2629A>C), ACT766TCT (2631A>T), ACA767CTA (2634A>C 2635C>T), GGA768GTG (2638G>T 2639A>G), GAG770GGG (2644A>G), GAA771TGT (2646G>T 2647A>G 2648A>T), GTG772AGG (2649G>A 2650T>G), GGC773TCA (2652G>T 2653G>C 2654C>A), TTA774TAT (2656T>A 2657A>T), AGA775GAC (2658A>G 2659G>A 2660A>C), CTT776TTT (2661C>T), TAG777AGG (2664T>A 2665A>G)

#### Proteins

| | | | | | | | | | |
|---|---|---|---|---|---|---|---|---|---|
| RNA-dependent RNA polymerase (YP_010798874.1) | 32 | 777 | 95.9% | 3316 | 63.5% | 744 (99.6%) | 510 (68.3%) | 2/1/3/3 | 4 |

| | Begin | End | Coverage | Score | Concordance | Matches | Identities | I/D/M/F* | Stop Codons |
|---|---|---|---|---|---|---|---|---|---|
| **NT** | **428** | **2700** | **82.0%** | **1680** | **37.3%** | **2266 (99.5%)** | **1565 (68.7%)** | **5/7** | |

Protein mutations:

I36L (441A>C 443C>T), L38F (447C>T), A44V (466C>T), A46M (471G>A 472C>T 473T>G), L48M (477T>A), T49H (480A>C 481C>A 482A>T), I52V (489A>G), V60I (513G>A), S61K (516T>A 517C>A 518T>A), K62T (520A>C), T64V (525A>G 526C>T 527T>G), T65K (529C>A 530C>A), R67K (534C>A 535G>A 536C>A), K69R (540A>C 541A>G 542G>T), Y72F (550A>T), F74Y (556T>A 557C>T), A83T (582G>A), C85S (589G>C), F94Y (616T>A), R95S (618C>T 619G>C 620A>T), E97N (624G>A 626A>T), E98D (629A>C), P99T (630C>A 632A>C), L100M (633C>A 635T>G), S109C (661C>G 662G>T), R113K (672C>A 673G>A 674T>G), K117A (684A>G 685A>C 686A>C), A122H (699G>C 700C>A 701A>C), I123V (702A>G), R125A (708A>G 709G>C 710G>A), D127S (714G>T 715A>C), E128D (719A>T), A144S (765G>T 767T>A), V146* (771G>T 772T>A 773T>G), L147I (774T>A 776G>T), K148A (777A>G 778A>C 779A>G), S154K (796G>A 797T>G), E159Q (810G>C), P166S (831C>T 833T>A), G169D (841G>A 842G>T), L172R (850T>G 851A>T), E176G (862A>G), I178M (869T>G), F182* (880T>G 881C>A), H184K (885C>A 887T>G), Q186R (892A>G), F192H (909T>C 910T>A), R194Q (916G>A), G195S (918G>T 919G>C), M198L (927A>C 929G>T), I200K (934T>A 935T>G), S203N (942T>A 943C>A), Q205K (948C>A), N214L (975A>C 976A>T 977T>C), S220R (993T>C 994C>G 995A>G), Y221F (997A>T 998C>T), G222Q (999G>C 1000G>A 1001T>A), L223Y (1003T>A 1004A>T), A224S (1005G>T 1007C>G), K228V (1017A>G 1018A>T 1019A>T), Q231G (1026C>G 1027A>G), E232A (1030A>C 1031G>A), S234A (1035A>G 1036G>C), F237V (1044T>G 1046C>T), F239E (1050T>G 1051T>A 1052T>A), P240H (1054C>A 1055A>C), L250I (1083C>A), W255L (1099G>T), T257V (1104A>G 1105C>T 1106A>G), I260M (1115T>G), F263I (1122T>A 1124C>T), L290F (1205A>T), L293E (1212T>G 1213T>A), E296D (1223A>T), H297L (1224C>T 1225A>T 1226C>A), W305Y (1249G>A 1250G>C), T308S (1257A>T 1259C>T), S311G (1266A>G 1268T>G), P312A (1269C>G), R314K (1275C>A 1276G>A 1277A>G), V319A (1291T>C 1292T>A), G324A (1306G>C), A325Q (1308G>C 1309C>A), V335I (1338G>A 1340A>C), V337A (1345T>C), M338I (1349G>A), I342V (1359A>G), K343_Q344insKX (1364_1365insAAAAA), R345K (1369G>A 1370A>G), L346* (1371C>T 1372T>G 1373T>A), H348F (1377C>T 1378A>T 1379C>T), D352Q (1389G>C 1391T>A), T356K (1402C>A), K360R (1413A>C 1414A>G 1415A>C), I361L (1416A>C 1418A>T), K362P (1419A>C 1420A>C), T363A (1422A>G 1424A>G), T366A (1431A>G 1433T>A), K371R (1446A>C 1447A>G 1448A>T), L374V (1455T>G 1457G>A), A375R (1458G>C 1459C>G 1460T>A), K377R (1464A>C 1465A>G 1466G>A), Q378S (1467C>T 1468A>C 1469A>T), K379Q (1470A>C 1472G>A), N380K (1475C>G), F381K (1476T>A 1477T>A 1478T>A), K382G (1479A>G 1480A>G 1481A>T), C384S (1486G>C), Y385F (1489A>T 1490T>C), L389F (1502A>C), M401I (1538G>A), L404F (1545C>T 1547T>C), M407C (1554A>T 1555T>G 1556G>C), I408M (1559A>G), P411S (1566C>A 1567C>G), S420del (1593_1597delAGCTC), I427F (1614A>T), V429I (1620G>A 1622T>C), N430V (1623A>G 1624A>T), K431P (1626A>C 1627A>C 1628A>T), L433M (1632T>A 1634A>G), T434V (1635A>G 1636C>T 1637A>C), V437M (1644G>A), Y438S (1647T>A 1648A>G), F450Y (1684T>A), H451Y (1686C>T), A455S (1698G>T 1700A>C), S458A (1707T>G), S470K (1744G>A 1745C>A), K471Y (1746A>T 1748A>T), G475T (1758G>A 1759G>C 1760A>G), C476S (1761T>A), N477T (1765A>C 1766C>T), S478T (1767T>A 1769T>C), G481A (1777G>C 1778G>T), S495T (1819G>C 1820T>A), K496E (1821A>G 1823A>G), N499R (1830A>C 1831A>G 1832T>G), E500Q (1833G>C), K502R (1840A>G), N509G (1860A>G 1861A>G), K511T (1867A>C), P515A (1878C>G 1880A>T), L518R (1887T>A 1888T>G 1889G>A), D521E (1898T>G), S535D (1938T>G 1939C>A 1940A>C), D539Y (1950G>T 1952T>C), L549T (1980T>A 1981T>C 1982A>T), A559Q (2010G>C 2011C>A 2012C>A), I562T (2020T>C), D565S (2028G>T 2029A>C 2030T>G), N567S (2034A>T 2035A>C 2036C>A), I568T (2038T>C), V570I (2043G>A), Q572P (2050A>C 2051A>C), R573I (2053G>T 2054A>T), L574F (2057G>T), G575W (2058G>T), Y579F (2071A>T), K580R (2074A>G 2075G>A), A583S (2082G>T 2084G>T), F586M (2091T>A 2093C>G), T587S (2094A>T), H600A (2133C>G 2134A>C 2135T>A), L601S (2136C>A 2137T>G 2138T>C), V604H (2145G>C 2146T>A 2147T>C), Y611L (2166T>C 2167A>T), W613Y (2173G>A 2174G>T), G618F (2187G>T 2188G>T), M619K (2191T>A 2192G>A), Y624F (2206A>T), V630I (2223G>A), S631D (2226A>G 2227G>A), L634R (2236T>G 2237T>G), K638T (2248A>C), V639P (2250G>C 2251T>C), L642I (2259T>A), Q643R (2263A>G 2264G>A), I644L (2265A>C), K647N (2276G>T), D648E (2279T>G), V655R (2298G>A 2299T>G 2300T>A), R664L (2326G>T), Q665R (2328C>A 2329A>G), E668K (2337G>A), L674V (2355C>G), N675S (2358A>T 2359A>C 2360C>T), I680V (2373A>G 2375C>T), C681A (2376T>G 2377G>C 2378T>A), M682A (2379A>G 2380T>C 2381G>A), N683S (2382A>T 2383A>C 2384C>A), P684A (2385C>G), D685E (2390T>A), S687T (2394T>A), K689D (2400A>G 2402G>T), S690Q (2403A>C 2404G>A 2405T>G), I692M (2411C>G), D693E (2414C>A), S694V (2415T>G 2416C>T 2417C>T), I696M (2423A>G), A698S (2427G>A 2428C>G 2429A>T), T699K (2431C>A), W701* (2438G>A), L704T (2445C>A 2446T>C 2447C>A), E705N (2448G>A 2450A>C), T706E (2451A>G 2452C>A 2453T>A), F708L (2457T>C), N727D (2514A>G), P728L (2518C>T 2519C>G), T730I (2524C>T 2525C>T), Q733R (2532C>A 2533A>G), W734Y (2536G>A 2537G>T), F736L (2543T>G), N739S (2551A>G 2552C>T), V741S (2556G>T 2557T>C), N745D (2568A>G), K746R (2571A>C 2572A>G), L749F (2582A>T), N755A (2598A>G 2599A>C 2600C>T), D756N (2601G>A), S760A (2613T>G 2615T>A), E763Y (2622G>T 2624A>T), Y764L (2626A>T 2627C>A), Q765P (2629A>C), T766S (2631A>T), T767L (2634A>C 2635C>T), G768V (2638G>T 2639A>G), E770G (2644A>G), E771C (2646G>T 2647A>G 2648A>T), V772R (2649G>A 2650T>G), G773S (2652G>T 2653G>C 2654C>A), L774Y (2656T>A 2657A>T), R775D (2658A>G 2659G>A 2660A>C), L776F (2661C>T), *777R (2664T>A 2665A>G)

| | Begin | End | Coverage | Score | Concordance | Matches | Identities | I/D/M/F* | Stop Codons |
|---|---|---|---|---|---|---|---|---|---|
| NT | 428 | 2700 | 82.0% | 1680 | 37.3% | 2266 (99.5%) | 1565 (68.7%) | 5/7 | |

Codon mutations: AAA34AAG (437A>G), ATC36CTT (441A>C 443C>T), CTT38TTT (447C>T), CTA42TTA (459C>T), GCC44GTC (466C>T), TCC45TCT (470C>T), GCT46ATG (471G>A 472C>T 473T>G), TTG48ATG (477T>A), ACA49CAT (480A>C 481C>A 482A>T), AAA50AAG (485A>G), ATC52GTC (489A>G), GCT53GCC (494T>C), CTT54CTA (497T>A), GCT55GCG (500T>G), GGA56GGC (503A>C), TGT57TGC (506T>C), GAC58GAT (509C>T), TTT59TTC (512T>C), GTA60ATA (513G>A), TCT61AAA (516T>A 517C>A 518T>A), AAA62ACA (520A>C), GTG63GTT (524G>T), ACT64GTG (525A>G 526C>T 527T>G), ACC65AAA (529C>A 530C>A), TTA66CTG (531T>C 533A>G), CGC67AAA (534C>A 535G>A 536C>A), CGG68CGA (539G>A), AAG69CGT (540A>C 541A>G 542G>T), TCG70TCA (545G>A), GGC71GGT (548C>T), TAT72TTT (550A>T), TTC74TAT (556T>A 557C>T), ACA75ACT (560A>T), TAC78TAT (569C>T), AAG80AAA (575G>A), TGC82TGT (581C>T), GCA83ACA (582G>A), GTA84GTT (587A>T), TGC85TCC (589G>C), TTA86CTT (591T>C 593A>T), CGA88CGT (599A>T), TAC89TAT (602C>T), TCC93AGT (612T>A 613C>G 614C>T), TTT94TAT (616T>A), CGA95TCT (618C>T 619G>C 620A>T), GAA97AAT (624G>A 626A>T), GAA98GAC (629A>C), CCA99ACC (630C>A 632A>C), CTT100ATG (633C>A 635T>G), TCG101TCT (638G>T), GTG102GTT (641G>T), CCT103CCA (644T>A), GTG104GTT (647G>T), CTC106TTG (651C>T 653C>G), ACA107ACC (656A>C), AGA108CGA (657A>C), TCG109TGT (661C>G 662G>T), GGT110GGA (665T>A), ATC111ATT (668C>T), CGT113AAG (672C>A 673G>A 674T>G), ATC115ATA (680C>A), CCC116CCT (683C>T), AAA117GCC (684A>G 685A>C 686A>C), TTG119TTA (692G>A), CGT120CGA (695T>A), GCA122CAC (699G>C 700C>A 701A>C), ATA123GTA (702A>G), CGT124AGG (705C>A 707T>G), AGG125GCA (708A>G 709G>C 710G>A), AAA126AAG (713A>G), GAT127TCT (714G>T 715A>C), GAA128GAT (719A>T), CAC129CAT (722C>T), GTT134GTC (737T>C), TAC137TAT (746C>T), TCA139TCT (752A>T), GGG142GGA (761G>A), GCT144TCA (765G>T 767T>A), AAG145AAA (770G>A), GTT146TAG (771G>T 772T>A 773T>G), TTG147ATT (774T>A 776G>T), AAA148GCG (777A>G 778A>C 779A>G), GTT149GTG (782T>G), AGT154AAG (796G>A 797T>G), GCA156GCC (803A>C), TTT158TTC (809T>C), GAA159CAA (810G>C), TCC160TCA (815C>A), ATA161ATT (818A>T), ACT162ACG (821T>G), ACC163ACT (824C>T), CCA164CCT (827A>T), CCT166TCA (831C>T 833T>A), GGG169GAT (841G>A 842G>T), CTA172CGT (850T>G 851A>T), GTG174GTT (857G>T), CTG175CTT (860G>T), GAA176GGA (862A>G), ATT178ATG (869T>G), ACG180ACT (875G>T), TTC182TGA (880T>G 881C>A), CAT184AAG (885C>A 887T>G), CAA186CGA (892A>G), CTA190CTT (905A>T), CCC191CCA (908C>A), TTT192CAT (909T>C 910T>A), CGA194CAA (916G>A), GGC195TCC (918G>T 919G>C), ATT196ATA (923T>A), CCA197CCT (926A>T), ATG198CTT (927A>C 929G>T), ATT200AAG (934T>A 935T>G), GGG201GGT (938G>T), TCT203AAT (942T>A 943C>A), CAA205AAA (948C>A), CCC212CCA (971C>A), CTA213TTA (972C>T), AAT214CTC (975A>C 976A>T 977T>C), AAT216AAC (983T>C), AGG219CGC (990A>C 992G>C), TCA220CGG (993T>C 994C>G 995A>G), TAC221TTT (997A>T 998C>T), GGT222CAA (999G>C 1000G>A 1001T>A), TTA223TAT (1003T>A 1004A>T), GCC224TCG (1005G>T 1007C>G), GTA225GTG (1010A>G), GAC227GAT (1016C>T), AAA228GTT (1017A>G 1018A>T 1019A>T), GTT229GTA (1022T>A), GAT230GAC (1025T>C), CAA231GGA (1026C>G 1027A>G), GAG232GCA (1030A>C 1031G>A), GCT233GCG (1034T>G), AGT234GCT (1035A>G 1036G>C), TTC237GTT (1044T>G 1046C>T), AAG238AAA (1049G>A), TTT239GAA (1050T>G 1051T>A 1052T>A), CCA240CAC (1054C>A 1055A>C), ATT242ATC (1061T>C), GTT244GTG (1067T>G), TTG246TTA (1073G>A), AAA247AAG (1076A>G), CTA250ATA (1083C>A), GCG251GCT (1088G>T), GCC252GCG (1091C>G), TGG255TTG (1099G>T), ACA257GTG (1104A>G 1105C>T 1106A>G), ATT260ATG (1115T>G), TCA262TCC (1121A>C), TTC263ATT (1122T>A 1124C>T), CAG264CAA (1127G>A), GAC265GAT (1130C>T), GGC266GGG (1133C>G), TCA269TCC (1142A>C), CCC270CCT (1145C>T), TTA290TTT (1205A>T), AAC292AAT (1211C>T), TTA293GAA (1212T>G 1213T>A), TCT294TCC (1217T>C), CTC295CTT (1220C>T), GAA296GAT (1223A>T), CAC297TTA (1224C>T 1225A>T 1226C>A), GAG299GAA (1232G>A), AAA300AAG (1235A>G), AGT301TCT (1236A>T 1237G>C), GTC302GTA (1241C>A), GGA303GGC (1244A>C), CCA304CCC (1247A>C), TGG305TAC (1249G>A 1250G>C), TCA307AGT (1254T>A 1255C>G 1256A>T), ACC308TCT (1257A>T 1259C>T), TTA309CTT (1260T>C 1262A>T), AGT311GGG (1266A>G 1268T>G), CCT312GCT (1269C>G), TAT313TAC (1274T>C), CGA314AAG (1275C>A 1276G>A 1277A>G), GGG315GGT (1280G>T), GTG316GTT (1283G>T), CCA317CCG (1286A>G), GTT319GCA (1291T>C 1292T>A), ACT320ACG (1295T>G), GGT321GGA (1298T>A), CTT323TTA (1302C>T 1304T>A), GGT324GCT (1306G>C), GCA325CAA (1308G>C 1309C>A), GGC329GGA (1322C>A), GGA331GGT (1328A>T), AGG334CGG (1335A>C), GTA335ATC (1338G>A 1340A>C), GTA337GCA (1345T>C), ATG338ATA (1349G>A), AAC340AAT (1355C>T), ATA342GTA (1359A>G), AAA343_CAA344insAAAAA- (1364_1365insAAAAA), AGA345AAG (1369G>A 1370A>G), CTT346TGA (1371C>T 1372T>G 1373T>A), CTC347CTG (1376C>G), CAC348TTT (1377C>T 1378A>T 1379C>T), GTT350GTC (1385T>C), GAT352CAA (1389G>C 1391T>A), GCT354GCA (1397T>A), ACA356AAA (1402C>A), CTG358CTA (1409G>A), AAA360CGC (1413A>C 1414A>G 1415A>C), ATA361CTT (1416A>C 1418A>T), AAG362CCG (1419A>C 1420A>C), ACA363GCG (1422A>G 1424A>G), GAC364GAT (1427C>T), GGG365GGA (1430G>A), ACT366GCA (1431A>G 1433T>A), GAG370GAA (1445G>A), AAA371CGT (1446A>C 1447A>G 1448A>T), CCT372CCG (1451T>G), TTG374GTA (1455T>G 1457G>A), GCT375CGA (1458G>C 1459C>G 1460T>A), AAG377CGA (1464A>C 1465A>G 1466G>A), CAA378TCT (1467C>T 1468A>C 1469A>T), AAG379CAA (1470A>C 1472G>A), AAC380AAG (1475C>G), TTT381AAA (1476T>A 1477T>A 1478T>A), AAA382GGT (1479A>G 1480A>G 1481A>T), TGT384TCT (1486G>C), TAT385TTC (1489A>T 1490T>C), TTA389TTC (1502A>C), AAG390AAA (1505G>A), TCA391TCG (1508A>G), CCG397CCA (1526G>A), ATG401ATA (1538G>A), TAT402TAC (1541T>C), CTT404TTC (1545C>T 1547T>C), TCT406TCC (1553T>C), ATG407TGC (1554A>T 1555T>G 1556G>C), ATA408ATG (1559A>G), CCT411AGT (1566C>A 1567C>G), ACC412ACA (1571C>A), TTG413CTG (1572T>C), TCC415TCA (1580C>A), GTT418GTC (1589T>C), AGC420del (1593_1597delAGCTC), TCT421--A (1593_1597delAGCTC 1598T>A), CTT424CTC (1607T>C), ACG426ACT (1613G>T), ATC427TTC (1614A>T), CTT428TTG (1617C>T 1619T>G), GTT429ATC (1620G>A 1622T>C), AAC430GTC (1623A>G 1624A>T), AAA431CCT (1626A>C 1627A>C 1628A>T), TTA433ATG (1632T>A 1634A>G), ACA434GTC (1635A>G 1636C>T 1637A>C), AGA436AGG (1643A>G), GTG437ATG (1644G>A), TAT438AGT (1647T>A 1648A>G), CTG443CTA (1664G>A), GGG445GGA (1670G>A), CCT447CCG (1676T>G), TTG448TTA (1679G>A), GGC449GGT (1682C>T), TTC450TAC (1684T>A), CAC451TAC (1686C>T), GGT452GGA (1691T>A), TCT453TCA (1694T>A), GCA455TCC (1698G>T 1700A>C), CTT456CTA (1703T>A), TCG458GCG (1707T>G), TTG459CTT (1710T>C 1712G>T), TCT460TCC (1715T>C), TTA467TTG (1736A>G), GCA468GCT (1739A>T), AGC470AAA (1744G>A 1745C>A), AAA471TAT (1746A>T 1748A>T), TAT473TAC (1754T>C), GGA475ACG (1758G>A 1759G>C 1760A>G), TGT476AGT (1761T>A), AAC477ACT (1765A>C 1766C>T), TCT478ACC (1767T>A 1769T>C), CCA479CCT (1772A>T), GGG481GCT (1777G>C 1778G>T), GCT484GCC (1787T>C), TTA485CTG (1788T>C 1790A>G), CTA486TTA (1791C>T), AGT495ACA (1819G>C 1820T>A), AAA496GAG (1821A>G 1823A>G), GCC498GCG (1829C>G), AAT499CGG (1830A>C 1831A>G 1832T>G), GAG500CAG (1833G>C), AAG502AGG (1840A>G), TTA504TTG (1847A>G), TTG505CTG (1848T>C), AGA507AGG (1856A>G), CTC508CTT (1859C>T), AAT509GGT (1860A>G 1861A>G), GTC510GTT (1865C>T), AAA511ACA (1867A>C), ATT512ATC (1871T>C), TCG513TCT (1874G>T), ATA514-TA (1875delA), CCA515GCT (1878C>G 1880A>T), TCT517TCG (1886T>G), TTG518AGA (1887T>A 1888T>G 1889G>A), TCT520TCC (1895T>C), GAT521GAG (1898T>G), AAT522AAC (1901T>C), ATT525ATC (1910T>C), GCG528GCT (1919G>T), AGG530AGA (1925G>A), ACT533ACC (1934T>C), AAA534AAG (1937A>G), TCA535GAC (1938T>G 1939C>A 1940A>C), GAT539TAC (1950G>T 1952T>C), TTA540CTT (1953T>C 1955A>T), ATT543ATC (1964T>C), TCC544TCT (1967C>T), GCT547GCA (1976T>A), TTA549ACT (1980T>A 1981T>C 1982A>T), TCT550TCC (1985T>C), CGT552AGG (1989C>A 1991T>G), ACT553ACA (1994T>A), GTT555GTG (2000T>G), GGA556GGT (2003A>T), CTT557CTG (2006T>G), TGC558TGT (2009C>T), GCC559CAA (2010G>C 2011C>A 2012C>A), ATC562ACC (2020T>C), GAT565TCG (2028G>T 2029A>C 2030T>G), AAC567TCA (2034A>T 2035A>C 2036C>A), ATA568ACA (2038T>C), TCA569TCG (2042A>G), GTT570ATT (2043G>A), CAA572CCC (2050A>C 2051A>C), AGA573ATT (2053G>T 2054A>T), TTG574TTT (2057G>T), GGG575TGG (2058G>T), GGT576GGG (2063T>G), GGG578GGA (2069G>A), TAC579TTC (2071A>T), AAG580AGA (2074A>G 2075G>A), GTA581GTT (2078A>T), GCG583TCT (2082G>T 2084G>T), CGC584CGT (2087C>T), TTA585TTG (2090A>G), TTC586ATG (2091T>A 2093C>G), ACT587TCT (2094A>T), ACC588ACT (2099C>T), AGC590TCT (2103A>T 2104G>C 2105C>T), AAG591AAA (2108G>A), GAG594GAA (2117G>A), CGT595AGA (2118C>A 2120T>A), CTA596CTG (2123A>G), GCT598GCG (2129T>G), CAT600GCA (2133C>G 2134A>C 2135T>A), CTT601AGC (2136C>A 2137T>G 2138T>C), AAG602AAA (2141G>A), GTT604CAC (2145G>C 2146T>A 2147T>C), CGG605AGG (2148C>A), CAC607CAT (2156C>T), TTA608CTA (2157T>C), CTA609TTA (2160C>T), TAC611CTC (2166T>C 2167A>T), TGG613TAT (2173G>A 2174G>T), GGA616GGG (2183A>G), AGG617AGA (2186G>A), GGT618TTT (2187G>T 2188G>T), ATG619AAA (2191T>A 2192G>A), CCC620CCT (2195C>T), CCC623CCT (2204C>T), TAT624TTT (2206A>T), AAA626AAG (2213A>G), GGG627GGA (2216G>A), GTA630ATA (2223G>A), AGT631GAT (2226A>G 2227G>A), TAC632TAT (2231C>T), CTT634CGG (2236T>G 2237T>G), AAA635AAG (2240A>G), AAA638ACA (2248A>C), GTT639CCT (2250G>C 2251T>C), TTA642ATA (2259T>A), CAG643CGA (2263A>G 2264G>A), ATC644CTC (2265A>C), AAG647AAT (2276G>T), GAT648GAG (2279T>G), CTG649TTG (2280C>T), TTC651TTT (2288C>T), GAA654GAG (2297A>G), GTT655AGA (2298G>A 2299T>G 2300T>A), ATA657ATC (2306A>C), CTA658CTT (2309A>T), CGT660CGC (2315T>C), CTA663TTA (2322C>T), CGC664CTC (2326G>T), CAA665AGA (2328C>A 2329A>G), GTA667GTG (2336A>G), GAA668AAA (2337G>A), TTA671CTG (2346T>C 2348A>G), AAG672AAA (2351G>A), CTC674GTC (2355C>G), AAC675TCT (2358A>T 2359A>C 2360C>T), ACG679ACT (2372G>T), ATC680GTT (2373A>G 2375C>T), TGT681GCA (2376T>G 2377G>C 2378T>A), ATG682GCA (2379A>G 2380T>C 2381G>A), AAC683TCA (2382A>T 2383A>C 2384C>A), CCA684GCA (2385C>G), GAT685GAA (2390T>A), TCC687ACC (2394T>A), AAG689GAT (2400A>G 2402G>T), AGT690CAG (2403A>C 2404G>A 2405T>G), CTC691TTG (2406C>T 2408C>G), ATC692ATG (2411C>G), GAC693GAA (2414C>A), TCC694GTT (2415T>G 2416C>T 2417C>T), ATA696ATG (2423A>G), TGC697TGT (2426C>T), GCA698AGT (2427G>A 2428C>G 2429A>T), ACA699AAA (2431C>A), TGG701TGA (2438G>A), AGG703AGA (2444G>A), CTC704ACA (2445C>A 2446T>C 2447C>A), GAA705AAC (2448G>A 2450A>C), ACT706GAA (2451A>G 2452C>A 2453T>A), TTT708CTT (2457T>C), TTA710TTG (2465A>G), GGC714GGT (2477C>T), CTA715TTA (2478C>T), AAA718AAG (2489A>G), TGT719TGC (2492T>C), TAC720TAT (2495C>T), GGT723GGA (2504T>A), GGT725GGC (2510T>C), AAT727GAT (2514A>G), CCC728CTG (2518C>T 2519C>G), ACC730ATT (2524C>T 2525C>T), ACT731ACA (2528T>A), CAG733AGG (2532C>A 2533A>G), TGG734TAT (2536G>A 2537G>T), TTT736TTG (2543T>G), AAC739AGT (2551A>G 2552C>T), ACT740ACC (2555T>C), GTT741TCT (2556G>T 2557T>C), ATT742ATC (2561T>C), AAT745GAT (2568A>G), AAA746CGA (2571A>C 2572A>G), TTA749TTT (2582A>T), GGG750GGA (2585G>A), GGG754GGA (2597G>A), AAC755GCT (2598A>G 2599A>C 2600C>T), GAT756AAT (2601G>A), ATT758ATA (2609T>A), TCT760GCA (2613T>G 2615T>A), CCT761CCA (2618T>A), GAA763TAT (2622G>T 2624A>T), TAC764TTA (2626A>T 2627C>A), CAA765CCA (2629A>C), ACT766TCT (2631A>T), ACA767CTA (2634A>C 2635C>T), GGA768GTG (2638G>T 2639A>G), GAG770GGG (2644A>G), GAA771TGT (2646G>T 2647A>G 2648A>T), GTG772AGG (2649G>A 2650T>G), GGC773TCA (2652G>T 2653G>C 2654C>A), TTA774TAT (2656T>A 2657A>T), AGA775GAC (2658A>G 2659G>A 2660A>C), CTT776TTT (2661C>T), TAG777AGG (2664T>A 2665A>G)

*: Inserts / Deletes / Misaligned / Frameshifts

#### Analysis details

This analysis was performed with panviral2.64

### NGS Details (UN18_val): Solendovirus venanicotianae

#### Assembly

| | |
|---|---|
| **Coverage Length** | 1662 (4 contig(s)) |
| **Depth Of Coverage** | 15.1 |
| **Number Of Reads** | 218 |
| **Reads Per Million** | 4.09 rpm (after QC) |
| **Ambiguities** | 0 |
| **Assembly Method** | de novo + reference guided assembly |
| **Consensus Caller** | Bcf Tools |

#### Coverage Map

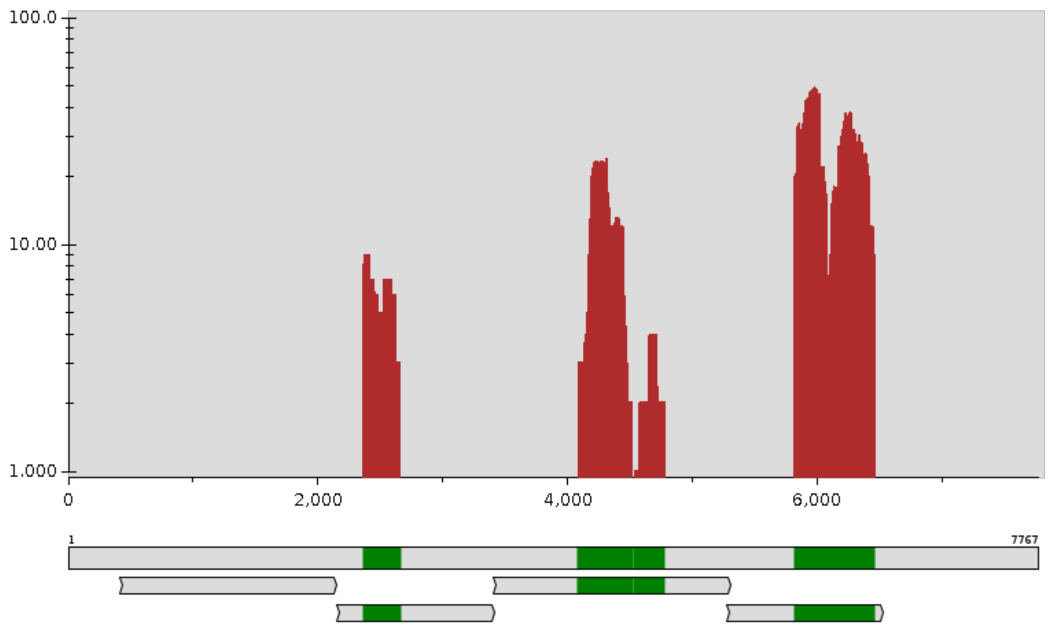

#### Assignment

| | |
|---|---|
| **Type** | Solendovirus venanicotianae (Taxonomy ID: 3048371) |
| **Reference Genome** | NC_003378.1 |
| **NT Identity (%)** | 77.9855 |
| **AA Identity (%)** | 73.9602 |
| **Number Of Stop Codons** | 2 |
| **Number Of CDS** | 4 |

#### Alignment

| | |
|---|---|
| **Alignment Score** | 1834.0 (NT) + 2746.0 (AA) = 4580.0 |
| **Concordance (%)** | 66.3971 |

| Alignment Method | Global, seeded, nucleotide + amino acids (AGA) |
|---|---|

#### Genome Region

Sequence starts at position 2358 and ends at position 6465 relative to NC_003378.1 reference sequence.

#### Alignment Detailed Statistics

| | Begin | End | Coverage | Score | Concordance | Matches | Identities | I/D/M/F* | Stop Codons |
|---|---|---|---|---|---|---|---|---|---|
| **NT** | **2358** | **6465** | **21.4%** | **1834** | **55.7%** | **1658 (99.8%)** | **1293 (77.8%)** | **0/4** | |
| Mutations: | 2358A>G, 2373T>A, 2376T>C, 2379C>T, 2388A>T, 2389G>A, 2390G>A, 2394T>A, 2397G>A, 2406A>T, 2426T>C, 2442A>T, 2454G>A, 2464C>T, 2469T>A, 2472G>A, 2482T>C, 2493T>A, 2503_2505delATA, 2535C>A, 2538T>A, 2545A>C, 2551C>T, 2555C>T, 2559T>C, 2562T>C, 2565A>G, 2568A>T, 2569A>T, 2571C>A, 2572C>G, 2589C>T, 2590A>G, 2595A>C, 2597G>C, 2598T>A, 2602A>G, 2606A>G, 2609G>A, 2613T>C, 2622T>C, 2628G>A, 2637T>C, 2638A>G, 2643A>C, 2646C>T, 2647G>C, 2649C>A, 2665C>T, 4086A>G, 4091T>A, 4097G>A, 4100T>G, 4103T>A, 4116G>A, 4118A>C, 4127C>T, 4139T>C, 4143G>A, 4161C>T, 4181A>T, 4187C>T, 4190G>A, 4196C>T, 4199A>T, 4241T>C, 4250C>A, 4256A>G, 4265G>A, 4292C>T, 4307C>T, 4313G>A, 4325T>C, 4328T>C, 4329T>C, 4331G>T, 4334A>G, 4340G>A, 4343T>A, 4357T>A, 4358A>T, 4362G>T, 4364T>A, 4367T>C, 4370A>T, 4380G>A, 4400T>A, 4411G>A, 4430A>T, 4436C>T, 4451C>T, 4454C>T, 4457T>C, 4467C>T, 4472A>T, 4482G>A, 4496T>C, 4503C>T, 4508T>C, 4513G>A, 4516C>T, 4550C>T, 4551G>A, 4552C>T, 4554C>G, 4568C>T, 4571T>A, 4574T>A, 4580C>G, 4587A>C, 4589A>G, 4591C>A, 4592C>A, 4595A>G, 4601A>C, 4604T>A, 4606T>A, 4607G>T, 4614C>G, 4616G>A, 4619A>T, 4626T>C, 4628A>C, 4634A>G, 4635C>A, 4636A>T, 4640A>T, 4641G>T, 4642A>C, 4643T>A, 4652A>C, 4655A>T, 4656A>G, 4659A>T, 4661G>A, 4670T>C, 4674G>T, 4675T>C, 4677C>A, 4682A>G, 4685A>C, 4686A>T, 4687T>G, 4688C>T, 4689A>G, 4691T>A, 4692C>T, 4695G>T, 4696A>C, 4697T>A, 4700A>T, 4703C>A, 4704A>C, 4706T>A, 4707G>A, 4709T>C, 4716A>C, 4718G>A, 4719A>G, 4722T>A, 4724A>C, 4728T>A, 4729C>G, 4730A>G, 4736A>T, 4740A>T, 4741T>G, 4742A>T, 4743G>T, 4748C>T, 4749C>T, 4751A>C, 4753T>C, 4757G>T, 4762A>T, 4769T>A, 4770A>G, 4772A>C, 4773T>C, 4779G>A, 5817T>C, 5820C>T, 5821C>A, 5824A>G, 5826T>A, 5831A>C, 5832C>A, 5833G>C, 5835A>G, 5838C>T, 5850C>A, 5851C>A, 5853T>A, 5857A>C, 5858C>A, 5859A>G, 5865C>T, 5868T>C, 5871A>G, 5872T>C, 5874G>A, 5877A>T, 5880T>A, 5881T>C, 5882T>A, 5889T>A, 5892A>T, 5894G>A, 5898A>T, 5901A>C, 5902T>C, 5907A>T, 5908G>A, 5909C>A, 5910C>A, 5913C>A, 5916C>T, 5921A>G, 5922C>T, 5925C>T, 5935G>A, 5936A>G, 5937T>C, 5940T>A, 5941G>A, 5950C>T, 5952G>T, 5953A>G, 5954C>A, 5955A>C, 5956G>T, 5959C>G, 5964G>A, 5968G>T, 5970C>A, 5972A>C, 5979T>A, 5991G>A, 5997C>T, 6001C>A, 6009G>A, 6012G>A, 6015A>T, 6018T>A, 6023G>A, 6024G>A, 6026C>A, 6027A>T, 6033C>T, 6036C>T, 6037G>A, 6040C>A, 6041G>A, 6045C>T, 6049T>A, 6051G>A, 6054T>A, 6055A>C, 6067C>T, 6075A>T, 6084G>A, 6087T>A, 6088A>G, 6099T>C, 6108T>A, 6112C>T, 6114C>G, 6117A>T, 6120G>A, 6123A>T, 6129C>T, 6132A>T, 6136G>T, 6139A>G, 6141G>T, 6144A>T, 6145G>A, 6146A>C, 6148G>C, 6150A>G, 6153A>G, 6154G>C, 6155A>C, 6156A>G, 6163G>C, 6164A>C, 6170T>A, 6172A>G, 6174T>G, 6177T>A, 6179C>T, 6181G>A, 6183A>T, 6186C>T, 6192T>C, 6201G>A, 6204A>T, 6207T>C, 6211A>C, 6214T>G, 6219T>A, 6223C>A, 6225A>T, 6232C>T, 6236C>G, 6240C>T, 6247C>A, 6253C>G, 6255G>A, 6258T>C, 6259G>C, 6264T>G, 6269G>T, 6270C>T, 6277T>A, 6282G>A, 6285A>T, 6288G>A, 6290C>T, 6308G>C, 6309C>A, 6315T>A, 6318A>G, 6319A>C, 6322G>A, 6325G>C, 6326C>A, 6334G>A, 6340C>G, 6341T>C, 6345A>G, 6352G>A, 6354C>A, 6355T>A, 6357A>G, 6359G>C, 6360T>A, 6363T>G, 6369A>T, 6371C>T, 6376C>A, 6377A>G, 6380C>A, 6381T>A, 6382A>C, 6384A>T, 6401G>A, 6402G>A, 6404G>A, 6406delA, 6407A>G, 6408T>A, 6411C>T, 6414C>T, 6417C>G, 6418C>G, 6419C>A, 6420A>T, 6423T>A, 6424C>T, 6426G>A, 6427T>C, 6446T>A, 6451A>G, 6453T>G, 6456C>A, 6462T>C | | | | | | | | |

##### CDS

| | Begin | End | Coverage | Score | Concordance | Matches | Identities | I/D/M/F* | Stop Codons |
|---|---|---|---|---|---|---|---|---|---|
| Tvvgp2 | 69 | 172 | 24.8% | 605 | 87.6% | 103 (99.0%) | 88 (84.6%) | 0/1/0/0 | 0 |
| Protein mutations: | G79K (2389G>A 2390G>A), I91T (2426T>C), P104S (2464C>T), I117del (2503_2505delATA), I131L (2545A>C), A134V (2555C>T), I139L (2569A>T 2571C>A), Q140E (2572C>G), I146V (2590A>G), S148T (2597G>C 2598T>A), I150V (2602A>G), K151R (2606A>G), G152E (2609G>A), I162V (2638A>G), A165P (2647G>C 2649C>A) | | | | | | | | |
| Codon mutations: | CAA68..G (2358A>G), ATT73ATA (2373T>A), GAT74GAC (2376T>C), TGC75TGT (2379C>T), ACA78ACT (2388A>T), GGA79AAA (2389G>A 2390G>A), GGT80GGA (2394T>A), AAG81AAA (2397G>A), ATA84ATT (2406A>T), ATA91ACA (2426T>C), ATA96ATT (2442A>T), AAG100AAA (2454G>A), CCA104TCA (2464C>T), ATT105ATA (2469T>A), AAG106AAA (2472G>A), TTA110CTA (2482T>C), ACT113ACA (2493T>A), ATA117del (2503_2505delATA), ACC127ACA (2535C>A), CCT128CCA (2538T>A), ATA131CTA (2545A>C), CTA133TTA (2551C>T), GCA134GTA (2555C>T), GAT135GAC (2559T>C), GAT136GAC (2562T>C), AGA137AGG (2565A>G), ATA138ATT (2568A>T), ATC139TTA (2569A>T 2571C>A), CAA140GAA (2572C>G), AGC145AGT (2589C>T), ATA146GTA (2590A>G), ATA147ATC (2595A>C), AGT148ACA (2597G>C 2598T>A), ATA150GTA (2602A>G), AAA151AGA (2606A>G), GGA152GAA (2609G>A), AAT153AAC (2613T>C), TAT156TAC (2622T>C), AAG158AAA (2628G>A), TTT161TTC (2637T>C), ATA162GTA (2638A>G), ATA163ATC (2643A>C), AGC164AGT (2646C>T), GCC165CCA (2647G>C 2649C>A), CTA171TTA (2665C>T) | | | | | | | | |
| Tvvgp3 | 225 | 458 | 36.5% | 1263 | 78.5% | 232 (100%) | 179 (77.2%) | 0/0/0/0 | 0 |
| Protein mutations: | K226E (4086A>G), D227E (4091T>A), D230E (4100T>G), N231K (4103T>A), E236N (4116G>A 4118A>C), A245T (4143G>A), H251Y (4161C>T), D311E (4343T>A), L316Y (4357T>A 4358A>T), A318S (4362G>T 4364T>A), G324R (4380G>A), G334E (4411G>A), E354D (4472A>T), V358I (4482G>A), R368K (4513G>A), T369I (4516C>T), A381I (4551G>A 4552C>T), H382D (4554C>G), S390R (4580C>G), K393Q (4587A>C 4589A>G), T394K (4591C>A 4592C>A), E397D (4601A>C), M399N (4606T>A 4607G>T), Q402E (4614C>G 4616G>A), I408M (4634A>G), Q409I (4635C>A 4636A>T), D411S (4641G>T 4642A>C 4643T>A), K416E (4656A>G), M417L (4659A>T 4661G>A), V422S (4674G>T 4675T>C), Q423K (4677C>A), I426C (4686A>T 4687T>G 4688C>T), N427E (4689A>G 4691T>A), L428F (4692C>T), D429S (4695G>T 4696A>C 4697T>A), E430D (4700A>T), N431K (4703C>A), I432L (4704A>C 4706T>A), D433N (4707G>A 4709T>C), K436Q (4716A>C 4718G>A), K437E (4719A>G), L438I (4722T>A 4724A>C), S440R (4728T>A 4729C>G 4730A>G), I444C (4740A>T 4741T>G 4742A>T), V445L (4743G>T), Q447Y (4749C>T 4751A>C), V448A (4753T>C), R449S (4757G>T), Y451F (4762A>T), K454D (4770A>G 4772A>C), E457K (4779G>A) | | | | | | | | |
| Codon mutations: | AAA226GAA (4086A>G), GAT227GAA (4091T>A), TTG229TTA (4097G>A), GAT230GAG (4100T>G), AAT231AAA (4103T>A), GAA236AAC (4116G>A 4118A>C), AGC239AGT (4127C>T), AGT243AGC (4139T>C), GCA245ACA (4143G>A), CAT251TAT (4161C>T), GGA257GGT (4181A>T), AGC259AGT (4187C>T), AGG260AGA (4190G>A), GTC262GTT (4196C>T), ATA263ATT (4199A>T), TAT277TAC (4241T>C), CCC280CCA (4250C>A), AAA282AAG (4256A>G), AAG285AAA (4265G>A), TAC294TAT (4292C>T), GAC299GAT (4307C>T), AAG301AAA (4313G>A), TAT305TAC (4325T>C), CAT306CAC (4328T>C), TTG307CTT (4329T>C 4331G>T), AAA308AAG (4334A>G), GAG310GAA (4340G>A), GAT311GAA (4343T>A), TTA316TAT (4357T>A 4358A>T), GCT318TCA (4362G>T 4364T>A), TTT319TTC (4367T>C), ACA320ACT (4370A>T), GGA324AGA (4380G>A), GTT330GTA (4400T>A), GGA334GAA (4411G>A), GGA340GGT (4430A>T), TAC342TAT (4436C>T), GAC347GAT (4451C>T), AAC348AAT (4454C>T), TAT349TAC (4457T>C), CTA353TTA (4467C>T), GAA354GAT (4472A>T), GTA358ATA (4482G>A), GAT362GAC (4496T>C), CTA365TTA (4503C>T), TAT366TAC (4508T>C), AGA368AAA (4513G>A), ACA369ATA (4516C>T), TTC380TTT (4550C>T), GCA381ATA (4551G>A 4552C>T), CAT382GAT (4554C>G), AAC386AAT (4568C>T), TCT387TCA (4571T>A), GGT388GGA (4574T>A), AGC390AGG (4580C>G), AAA393CAG (4587A>C 4589A>G), ACC394AAA (4591C>A 4592C>A), AAA395AAG (4595A>G), GAA397GAC (4601A>C), ATT398ATA (4604T>A), ATG399AAT (4606T>A 4607G>T), CAG402GAA (4614C>G 4616G>A), ATA403ATT (4619A>T), TTA406CTC (4626T>C 4628A>C), ATA408ATG (4634A>G), CAA409ATA (4635C>A 4636A>T), ATA410ATT (4640A>T), GAT411TCA (4641G>T 4642A>C 4643T>A), GGA414GGC (4652A>C), ATA415ATT (4655A>T), AAA416GAA (4656A>G), ATG417TTA (4659A>T 4661G>A), CAT420CAC (4670T>C), GTA422TCA (4674G>T 4675T>C), CAA423AAA (4677C>A), AAA424AAG (4682A>G), ATA425ATC (4685A>C), ATC426TGT (4686A>T 4687T>G 4688C>T), AAT427GAA (4689A>G 4691T>A), CTT428TTT (4692C>T), GAT429TCA (4695G>T 4696A>C 4697T>A), GAA430GAT (4700A>T), AAC431AAA (4703C>A), ATT432CTA (4704A>C 4706T>A), GAT433AAC (4707G>A 4709T>C), AAG436CAA (4716A>C 4718G>A), AAA437GAA (4719A>G), TTA438ATC (4722T>A 4724A>C), TCA440AGG (4728T>A 4729C>G 4730A>G), CTA442CTT (4736A>T), ATA444TGT (4740A>T 4741T>G 4742A>T), GTA445TTA (4743G>T), AAC446AAT (4748C>T), CAA447TAC (4749C>T 4751A>C), GTA448GCA (4753T>C), AGG449AGT (4757G>T), TAT451TTT (4762A>T), CCT453CCA (4769T>A), AAA454GAC (4770A>G 4772A>C), TTA455CTA (4773T>C), GAA457AAA (4779G>A) | | | | | | | | |
| Tvvgp4 | 180 | 397 | 52.2% | 878 | 59.0% | 218 (100%) | 142 (65.1%) | 0/0/1/1 | 2 |
| Protein mutations: | Q183K (5821C>A), N184E (5824A>G 5826T>A), N186T (5831A>C 5832C>A), E187Q (5833G>C 5835A>G), H192Q (5850C>A), H193K (5851C>A 5853T>A), T195Q (5857A>C 5858C>A 5859A>G), L203Q (5881T>C 5882T>A), N205K (5889T>A), S207N (5894G>A), K209N (5901A>C), A212K (5908G>A 5909C>A 5910C>A), N216S (5921A>G 5922C>T), D221S (5935G>A 5936A>G 5937T>C), V223I (5941G>A), Q226Y (5950C>T 5952G>T), T227D (5953A>G 5954C>A 5955A>C), G228* (5956G>T), Q229E (5959C>G), A232S (5968G>T 5970C>A), N233T (5972A>C), Q243K (6001C>A), G250E (6023G>A 6024G>A), T251N (6026C>A 6027A>T), V255I (6037G>A), R256K (6040C>A 6041G>A), S259T (6049T>A 6051G>A), T261P (6055A>C), E267D (6075A>T), I272V (6088A>G), E283D (6123A>T), E288* (6136G>T), K289D (6139A>G 6141G>T), E291T (6145G>A 6146A>C), E292Q (6148G>C 6150A>G), E294P (6154G>C 6155A>C 6156A>G), E297P (6163G>C 6164A>C), V299E (6170T>A), N300E (6172A>G 6174T>G), P302L (6179C>T), E303N (6181G>A 6183A>T), I313L (6211A>C), S314A (6214T>G), L317I (6223C>A 6225A>T), A321G (6236C>G), L325I (6247C>A), Q327E (6253C>G 6255G>A), A329P (6259G>C), I330M (6264T>G), S332I (6269G>T 6270C>T), S335T (6277T>A), E337D (6285A>T), T339I (6290C>T), C345S (6308G>C 6309C>A), D347E (6315T>A), I348M (6318A>G), E350K (6322G>A), A351Q (6325G>C 6326C>A), E354K (6334G>A), L356A (6340C>G 6341T>C), V360I (6352G>A 6354C>A), L361M (6355T>A 6357A>G), S362T (6359G>C 6360T>A), K365N (6369A>T), P366L (6371C>T), Q368R (6376C>A 6377A>G), P369Q (6380C>A 6381T>A), T370P (6382A>C 6384A>T), R376K (6401G>A 6402G>A), R377K (6404G>A), P382D (6418C>G 6419C>A 6420A>T), D383E (6423T>A), L391Q (6446T>A), S393G (6451A>G 6453T>G), H394Q (6456C>A) | | | | | | | | |

| | Begin | End | Coverage | Score | Concordance | Matches | Identities | I/D/M/F* | Stop Codons |
|---|---|---|---|---|---|---|---|---|---|
| **NT** | **2358** | **6465** | **21.4%** | **1834** | **55.7%** | **1658 (99.8%)** | **1293 (77.8%)** | **0/4** | |
| Codon mutations: | TCT181TCC (5817T>C), ACC182ACT (5820C>T), CAA183AAA (5821C>A), AAT184GAA (5824A>G 5826T>A), AAC186ACA (5831A>C 5832C>A), GAA187CAG (5833G>C 5835A>G), GAC188GAT (5838C>T), CAC192CAA (5850C>A), CAT193AAA (5851C>A 5853T>A), ACA195CAG (5857A>C 5858C>A 5859A>G), TAC197TAT (5865C>T), AAT198AAC (5868T>C), AAA199AAG (5871A>G), TTG200CTA (5872T>C 5874G>A), ATA201ATT (5877A>T), GCT202GCA (5880T>A), TTA203CAA (5881T>C 5882T>A), AAT205AAA (5889T>A), ACA206ACT (5892A>T), AGT207AAT (5894G>A), GCA208GCT (5898A>T), AAA209AAC (5901A>C), TTA210CTA (5902T>C), GTA211GTT (5907A>T), GCC212AAA (5908G>A 5909C>A 5910C>A), ACC213ACA (5913C>A), TGC214TGT (5916C>T), AAC216AGT (5921A>G 5922C>T), TAC217TAT (5925C>T), GAT221AGC (5935G>A 5936A>G 5937T>C), ACT222ACA (5940T>A), GTA223ATA (5941G>A), CAG226TAT (5950C>T 5952G>T), ACA227GAC (5953A>G 5954C>A 5955A>C), GGA228TGA (5956G>T), CAA229GAA (5959C>G), GAG230GAA (5964G>A), GCC232TCA (5968G>T 5970C>A), AAT233ACT (5972A>C), CCT235CCA (5979T>A), AAG239AAA (5991G>A), TTC241TTT (5997C>T), CAA243AAA (6001C>A), AAG245AAA (6009G>A), AGG246AGA (6012G>A), ATA247ATT (6015A>T), ACT248ACA (6018T>A), GGG250GAA (6023G>A 6024G>A), ACA251AAT (6026C>A 6027A>T), TTC253TTT (6033C>T), TAC254TAT (6036C>T), GTA255ATA (6037G>A), CGA256AAA (6040C>A 6041G>A), TTC257TTT (6045C>T), TCG259ACA (6049T>A 6051G>A), GCT260GCA (6054T>A), ACA261CCA (6055A>C), CTA265TTA (6067C>T), GAA267GAT (6075A>T), AAG270AAA (6084G>A), CCT271CCA (6087T>A), ATC272GTC (6088A>G), GTT275GTC (6099T>C), ATT278ATA (6108T>A), CTC280TTG (6112C>T 6114C>G), ACA281ACT (6117A>T), AGG282AGA (6120G>A), GAA283GAT (6123A>T), ATC285ATT (6129C>T), ATA286ATT (6132A>T), GAA288TAA (6136G>T), AAG289GAT (6139A>G 6141G>T), ATA290ATT (6144A>T), GAA291ACA (6145G>A 6146A>C), GAA292CAG (6148G>C 6150A>G), CAA293CAG (6153A>G), GAA294CCG (6154G>C 6155A>C 6156A>G), GAA297CCA (6163G>C 6164A>C), GTA299GAA (6170T>A), AAT300GAG (6172A>G 6174T>G), ATT301ATA (6177T>A), CCA302CTA (6179C>T), GAA303AAT (6181G>A 6183A>T), TTC304TTT (6186C>T), GCT306GCC (6192T>C), AGG309AGA (6201G>A), ATA310ATT (6204A>T), ATT311ATC (6207T>C), ATA313CTA (6211A>C), TCT314GCT (6214T>G), ACT315ACA (6219T>A), CTA317ATT (6223C>A 6225A>T), CTA320TTA (6232C>T), GCA321GGA (6236C>G), AAC322AAT (6240C>T), CTA325ATA (6247C>A), CAG327GAA (6253C>G 6255G>A), AAT328AAC (6258T>C), GCA329CCA (6259G>C), ATT330ATG (6264T>G), AGC332ATT (6269G>T 6270C>T), TCA335ACA (6277T>A), AGG336AGA (6282G>A), GAA337GAT (6285A>T), CAG338CAA (6288G>A), ACA339ATA (6290C>T), TGC345TCA (6308G>C 6309C>A), GAT347GAA (6315T>A), ATA348ATG (6318A>G), AGA349CGA (6319A>C), GAA350AAA (6322G>A), GCA351CAA (6325G>C 6326C>A), GAA354AAA (6334G>A), CTA356GCA (6340C>G 6341T>C), AGA357AGG (6345A>G), GTC360ATA (6352G>A 6354C>A), TTA361ATG (6355T>A 6357A>G), AGT362ACA (6359G>C 6360T>A), CTT363CTG (6363T>G), AAA365AAT (6369A>T), CCA366CTA (6371C>T), CAA368AGA (6376C>A 6377A>G), CCT369CAA (6380C>A 6381T>A), ACA370CCT (6382A>C 6384A>T), AGG376AAA (6401G>A 6402G>A), AGG377AAG (6404G>A), AAT378-GA (6406delA 6407A>G 6408T>A), TTC379TTT (6411C>T), ATC380ATT (6414C>T), TCC381TCG (6417C>G), CCA382GAT (6418C>G 6419C>A 6420A>T), GAT383GAA (6423T>A), CTG384TTA (6424C>T 6426G>A), TTA385CTA (6427T>C), CTA391CAA (6446T>A), AGT393GGG (6451A>G 6453T>G), CAC394CAA (6456C>A), TAT396TAC (6462T>C) | | | | | | | | |

**Proteins**

| | Begin | End | Coverage | Score | Concordance | Matches | Identities | I/D/M/F* | Stop Codons |
|---|---|---|---|---|---|---|---|---|---|
| putative cell-to-cell movement protein (NP_569140.1) | 69 | 172 | 24.8% | 605 | 87.6% | 103 (99.0%) | 88 (84.6%) | 0/1/0/0 | 0 |
| Protein mutations: | G79K (2389G>A 2390G>A), I91T (2426T>C), P104S (2464C>T), I117del (2503_2505delATA), I131L (2545A>C), A134V (2555C>T), I139L (2569A>T 2571C>A), Q140E (2572C>G), I146V (2590A>G), S148T (2597G>C 2598T>A), I150V (2602A>G), K151R (2606A>G), G152E (2609G>A), I162V (2638A>G), A165P (2647G>C 2649C>A) | | | | | | | | |
| Codon mutations: | CAA68..G (2358A>G), ATT73ATA (2373T>A), GAT74GAC (2376T>C), TGC75TGT (2379C>T), ACA78ACT (2388A>T), GGA79AAA (2389G>A 2390G>A), GGT80GGA (2394T>A), AAG81AAA (2397G>A), ATA84ATT (2406A>T), ATA91ACA (2426T>C), ATA96ATT (2442A>T), AAG100AAA (2454G>A), CCA104TCA (2464C>T), ATT105ATA (2469T>A), AAG106AAA (2472G>A), TTA110CTA (2482T>C), ACT113ACA (2493T>A), ATA117del (2503_2505delATA), ACC127ACA (2535C>A), CCT128CCA (2538T>A), ATA131CTA (2545A>C), CTA133TTA (2551C>T), GCA134GTA (2555C>T), GAT135GAC (2559T>C), GAT136GAC (2562T>C), AGA137AGG (2565A>G), ATA138ATT (2568A>T), ATC139TTA (2569A>T 2571C>A), CAA140GAA (2572C>G), AGC145AGT (2589C>T), ATA146GTA (2590A>G), ATA147ATC (2595A>C), AGT148ACA (2597G>C 2598T>A), ATA150GTA (2602A>G), AAA151AGA (2606A>G), GGA152GAA (2609G>A), AAT153AAC (2613T>C), TAT156TAC (2622T>C), AAG158AAA (2628G>A), TTT161TTC (2637T>C), ATA162GTA (2638A>G), ATA163ATC (2643A>C), AGC164AGT (2646C>T), GCC165CCA (2647G>C 2649C>A), CTA171TTA (2665C>T) | | | | | | | | |
| polyprotein (NP_569141.1) | 225 | 458 | 36.5% | 1263 | 78.5% | 232 (100%) | 179 (77.2%) | 0/0/0/0 | 0 |
| Protein mutations: | K226E (4086A>G), D227E (4091T>A), D230E (4100T>G), N231K (4103T>A), E236N (4116G>A 4118A>C), A245T (4143G>A), H251Y (4161C>T), D311E (4343T>A), L316Y (4357T>A 4358A>T), A318S (4362G>T 4364T>A), G324R (4380G>A), G334E (4411G>A), E354D (4472A>T), V358I (4482G>A), R368K (4513G>A), T369I (4516C>T), A381I (4551G>A 4552C>T), H382D (4554C>G), S390R (4580C>G), K393Q (4587A>C 4589A>G), T394K (4591C>A 4592C>A), E397D (4601A>C), M399N (4606T>A 4607G>T), Q402E (4614C>G 4616G>A), I408M (4634A>G), Q409I (4635C>A 4636A>T), D411S (4641G>T 4642A>C 4643T>A), K416E (4656A>G), M417L (4659A>T 4661G>A), V422S (4674G>T 4675T>C), Q423K (4677C>A), I426C (4686A>T 4687T>G 4688C>T), N427E (4689A>G 4691T>A), L428F (4692C>T), D429S (4695G>T 4696A>C 4697T>A), E430D (4700A>T), N431K (4703C>A), I432L (4704A>C 4706T>A), D433N (4707G>A 4709T>C), K436Q (4716A>C 4718G>A), K437E (4719A>G), L438I (4722T>A 4724A>C), S440R (4728T>A 4729C>G 4730A>G), I444C (4740A>T 4741T>G 4742A>T), V445L (4743G>T), Q447Y (4749C>T 4751A>C), V448A (4753T>C), R449S (4757G>T), Y451F (4762A>T), K454D (4770A>G 4772A>C), E457K (4779G>A) | | | | | | | | |
| Codon mutations: | AAA226GAA (4086A>G), GAT227GAA (4091T>A), TTG229TTA (4097G>A), GAT230GAG (4100T>G), AAT231AAA (4103T>A), GAA236AAC (4116G>A 4118A>C), AGC239AGT (4127C>T), AGT243AGC (4139T>C), GCA245ACA (4143G>A), CAT251TAT (4161C>T), GGA257GGT (4181A>T), AGC259AGT (4187C>T), AGG260AGA (4190G>A), GTC262GTT (4196C>T), ATA263ATT (4199A>T), TAT277TAC (4241T>C), CCC280CCA (4250C>A), AAA282AAG (4256A>G), AAG285AAA (4265G>A), TAC294TAT (4292C>T), GAC299GAT (4307C>T), AAG301AAA (4313G>A), TAT305TAC (4325T>C), CAT306CAC (4328T>C), TTG307CTT (4329T>C 4331G>T), AAA308AAG (4334A>G), GAG310GAA (4340G>A), GAT311GAA (4343T>A), TTA316TAT (4357T>A 4358A>T), GCT318TCA (4362G>T 4364T>A), TTT319TTC (4367T>C), ACA320ACT (4370A>T), GGA324AGA (4380G>A), GTT330GTA (4400T>A), GGA334GAA (4411G>A), GGA340GGT (4430A>T), TAC342TAT (4436C>T), GAC347GAT (4451C>T), AAC348AAT (4454C>T), TAT349TAC (4457T>C), CTA353TTA (4467C>T), GAA354GAT (4472A>T), GTA358ATA (4482G>A), GAT362GAC (4496T>C), CTA365TTA (4503C>T), TAT366TAC (4508T>C), AGA368AAA (4513G>A), ACA369ATA (4516C>T), TTC380TTT (4550C>T), GCA381ATA (4551G>A 4552C>T), CAT382GAT (4554C>G), AAC386AAT (4568C>T), TCT387TCA (4571T>A), GGT388GGA (4574T>A), AGC390AGG (4580C>G), AAA393CAG (4587A>C 4589A>G), ACC394AAA (4591C>A 4592C>A), AAA395AAG (4595A>G), GAA397GAC (4601A>C), ATT398ATA (4604T>A), ATG399AAT (4606T>A 4607G>T), CAG402GAA (4614C>G 4616G>A), ATA403ATT (4619A>T), TTA406CTC (4626T>C 4628A>C), ATA408ATG (4634A>G), CAA409ATA (4635C>A 4636A>T), ATA410ATT (4640A>T), GAT411TCA (4641G>T 4642A>C 4643T>A), GGA414GGC (4652A>C), ATA415ATT (4655A>T), AAA416GAA (4656A>G), ATG417TTA (4659A>T 4661G>A), CAT420CAC (4670T>C), GTA422TCA (4674G>T 4675T>C), CAA423AAA (4677C>A), AAA424AAG (4682A>G), ATA425ATC (4685A>C), ATC426TGT (4686A>T 4687T>G 4688C>T), AAT427GAA (4689A>G 4691T>A), CTT428TTT (4692C>T), GAT429TCA (4695G>T 4696A>C 4697T>A), GAA430GAT (4700A>T), AAC431AAA (4703C>A), ATT432CTA (4704A>C 4706T>A), GAT433AAC (4707G>A 4709T>C), AAG436CAA (4716A>C 4718G>A), AAA437GAA (4719A>G), TTA438ATC (4722T>A 4724A>C), TCA440AGG (4728T>A 4729C>G 4730A>G), CTA442CTT (4736A>T), ATA444TGT (4740A>T 4741T>G 4742A>T), GTA445TTA (4743G>T), AAC446AAT (4748C>T), CAA447TAC (4749C>T 4751A>C), GTA448GCA (4753T>C), AGG449AGT (4757G>T), TAT451TTT (4762A>T), CCT453CCA (4769T>A), AAA454GAC (4770A>G 4772A>C), TTA455CTA (4773T>C), GAA457AAA (4779G>A) | | | | | | | | |
| putative transactivator factor (NP_569142.1) | 180 | 397 | 52.2% | 878 | 59.0% | 218 (100%) | 142 (65.1%) | 0/0/1/1 | 2 |
| Protein mutations: | Q183K (5821C>A), N184E (5824A>G 5826T>A), N186T (5831A>C 5832C>A), E187Q (5833G>C 5835A>G), H192Q (5850C>A), H193K (5851C>A 5853T>A), T195Q (5857A>C 5858C>A 5859A>G), L203Q (5881T>C 5882T>A), N205K (5889T>A), S207N (5894G>A), K209N (5901A>C), A212K (5908G>A 5909C>A 5910C>A), N216S (5921A>G 5922C>T), D221S (5935G>A 5936A>G 5937T>C), V223I (5941G>A), Q226Y (5950C>T 5952G>T), T227D (5953A>G 5954C>A 5955A>C), G228* (5956G>T), Q229E (5959C>G), A232S (5968G>T 5970C>A), N233T (5972A>C), Q243K (6001C>A), G250E (6023G>A 6024G>A), T251N (6026C>A 6027A>T), V255I (6037G>A), R256K (6040C>A 6041G>A), S259T (6049T>A 6051G>A), T261P (6055A>C), E267D (6075A>T), I272V (6088A>G), E283D (6123A>T), E288* (6136G>T), K289D (6139A>G 6141G>T), E291T (6145G>A 6146A>C), E292Q (6148G>C 6150A>G), E294P (6154G>C 6155A>C 6156A>G), E297P (6163G>C 6164A>C), V299E (6170T>A), N300E (6172A>G 6174T>G), P302L (6179C>T), E303N (6181G>A 6183A>T), I313L (6211A>C), S314A (6214T>G), L317I (6223C>A 6225A>T), A321G (6236C>G), L325I (6247C>A), Q327E (6253C>G 6255G>A), A329P (6259G>C), I330M (6264T>G), S332I (6269G>T 6270C>T), S335T (6277T>A), E337D (6285A>T), T339I (6290C>T), C345S (6308G>C 6309C>A), D347E (6315T>A), I348M (6318A>G), E350K (6322G>A), A351Q (6325G>C 6326C>A), E354K (6334G>A), L356A (6340C>G 6341T>C), V360I (6352G>A 6354C>A), L361M (6355T>A 6357A>G), S362T (6359G>C 6360T>A), K365N (6369A>T), P366L (6371C>T), Q368R (6376C>A 6377A>G), P369Q (6380C>A 6381T>A), T370P (6382A>C 6384A>T), R376K (6401G>A 6402G>A), R377K (6404G>A), P382D (6418C>G 6419C>A 6420A>T), D383E (6423T>A), L391Q (6446T>A), S393G (6451A>G 6453T>G), H394Q (6456C>A) | | | | | | | | |

| | Begin | End | Coverage | Score | Concordance | Matches | Identities | I/D/M/F* | Stop Codons |
|---|---|---|---|---|---|---|---|---|---|
| **NT** | **2358** | **6465** | **21.4%** | **1834** | **55.7%** | **1658 (99.8%)** | **1293 (77.8%)** | **0/4** | |
| Codon mutations: | TCT181TCC (5817T>C), ACC182ACT (5820C>T), CAA183AAA (5821C>A), AAT184GAA (5824A>G 5826T>A), AAC186ACA (5831A>C 5832C>A), GAA187CAG (5833G>C 5835A>G), GAC188GAT (5838C>T), CAC192CAA (5850C>A), CAT193AAA (5851C>A 5853T>A), ACA195CAG (5857A>C 5858C>A 5859A>G), TAC197TAT (5865C>T), AAT198AAC (5868T>C), AAA199AAG (5871A>G), TTG200CTA (5872T>C 5874G>A), ATA201ATT (5877A>T), GCT202GCA (5880T>A), TTA203CAA (5881T>C 5882T>A), AAT205AAA (5889T>A), ACA206ACT (5892A>T), AGT207AAT (5894G>A), GCA208GCT (5898A>T), AAA209AAC (5901A>C), TTA210CTA (5902T>C), GTA211GTT (5907A>T), GCC212AAA (5908G>A 5909C>A 5910C>A), ACC213ACA (5913C>A), TGC214TGT (5916C>T), AAC216AGT (5921A>G 5922C>T), TAC217TAT (5925C>T), GAT221AGC (5935G>A 5936A>G 5937T>C), ACT222ACA (5940T>A), GTA223ATA (5941G>A), CAG226TAT (5950C>T 5952G>T), ACA227GAC (5953A>G 5954C>A 5955A>C), GGA228TGA (5956G>T), CAA229GAA (5959C>G), GAG230GAA (5964G>A), GCC232TCA (5968G>T 5970C>A), AAT233ACT (5972A>C), CCT235CCA (5979T>A), AAG239AAA (5991G>A), TTC241TTT (5997C>T), CAA243AAA (6001C>A), AAG245AAA (6009G>A), AGG246AGA (6012G>A), ATA247ATT (6015A>T), ACT248ACA (6018T>A), GGG250GAA (6023G>A 6024G>A), ACA251AAT (6026C>A 6027A>T), TTC253TTT (6033C>T), TAC254TAT (6036C>T), GTA255ATA (6037G>A), CGA256AAA (6040C>A 6041G>A), TTC257TTT (6045C>T), TCG259ACA (6049T>A 6051G>A), GCT260GCA (6054T>A), ACA261CCA (6055A>C), CTA265TTA (6067C>T), GAA267GAT (6075A>T), AAG270AAA (6084G>A), CCT271CCA (6087T>A), ATC272GTC (6088A>G), GTT275GTC (6099T>C), ATT278ATA (6108T>A), CTC280TTG (6112C>T 6114C>G), ACA281ACT (6117A>T), AGG282AGA (6120G>A), GAA283GAT (6123A>T), ATC285ATT (6129C>T), ATA286ATT (6132A>T), GAA288TAA (6136G>T), AAG289GAT (6139A>G 6141G>T), ATA290ATT (6144A>T), GAA291ACA (6145G>A 6146A>C), GAA292CAG (6148G>C 6150A>G), CAA293CAG (6153A>G), GAA294CCG (6154G>C 6155A>C 6156A>G), GAA297CCA (6163G>C 6164A>C), GTA299GAA (6170T>A), AAT300GAG (6172A>G 6174T>G), ATT301ATA (6177T>A), CCA302CTA (6179C>T), GAA303AAT (6181G>A 6183A>T), TTC304TTT (6186C>T), GCT306GCC (6192T>C), AGG309AGA (6201G>A), ATA310ATT (6204A>T), ATT311ATC (6207T>C), ATA313CTA (6211A>C), TCT314GCT (6214T>G), ACT315ACA (6219T>A), CTA317ATT (6223C>A 6225A>T), CTA320TTA (6232C>T), GCA321GGA (6236C>G), AAC322AAT (6240C>T), CTA325ATA (6247C>A), CAG327GAA (6253C>G 6255G>A), AAT328AAC (6258T>C), GCA329CCA (6259G>C), ATT330ATG (6264T>G), AGC332ATT (6269G>T 6270C>T), TCA335ACA (6277T>A), AGG336AGA (6282G>A), GAA337GAT (6285A>T), CAG338CAA (6288G>A), ACA339ATA (6290C>T), TGC345TCA (6308G>C 6309C>A), GAT347GAA (6315T>A), ATA348ATG (6318A>G), AGA349CGA (6319A>C), GAA350AAA (6322G>A), GCA351CAA (6325G>C 6326C>A), GAA354AAA (6334G>A), CTA356GCA (6340C>G 6341T>C), AGA357AGG (6345A>G), GTC360ATA (6352G>A 6354C>A), TTA361ATG (6355T>A 6357A>G), AGT362ACA (6359G>C 6360T>A), CTT363CTG (6363T>G), AAA365AAT (6369A>T), CCA366CTA (6371C>T), CAA368AGA (6376C>A 6377A>G), CCT369CAA (6380C>A 6381T>A), ACA370CCT (6382A>C 6384A>T), AGG376AAA (6401G>A 6402G>A), AGG377AAG (6404G>A), AAT378-GA (6406delA 6407A>G 6408T>A), TTC379TTT (6411C>T), ATC380ATT (6414C>T), TCC381TCG (6417C>G), CCA382GAT (6418C>G 6419C>A 6420A>T), GAT383GAA (6423T>A), CTG384TTA (6424C>T 6426G>A), TTA385CTA (6427T>C), CTA391CAA (6446T>A), AGT393GGG (6451A>G 6453T>G), CAC394CAA (6456C>A), TAT396TAC (6462T>C) | | | | | | | | |

*: Inserts / Deletes / Misaligned / Frameshifts

#### Analysis details

This analysis was performed with panviral2.64

### NGS Details (UN18_val): Potato virus Y

#### Assembly

| | |
|---|---|
| **Coverage Length** | 1875 (7 contig(s)) |
| **Depth Of Coverage** | 3.1 |
| **Number Of Reads** | 45 |
| **Reads Per Million** | 0.84 rpm (after QC) |
| **Ambiguities** | 0 |
| **Assembly Method** | de novo + reference guided assembly |
| **Consensus Caller** | Bcf Tools |

#### Coverage Map

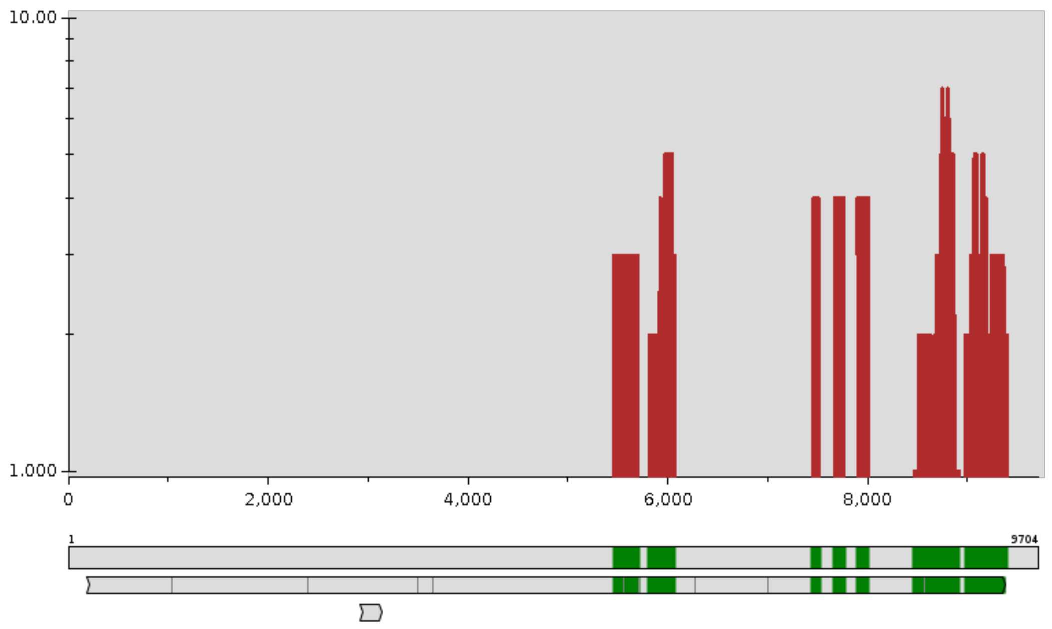

#### Assignment

| | |
|---|---|
| **Type** | Potato virus Y (Taxonomy ID: 12216) |
| **Reference Genome** | NC_001616.1 |
| **NT Identity (%)** | 85.1634 |
| **AA Identity (%)** | 91.1475 |
| **Number Of Stop Codons** | 1 |
| **Number Of CDS** | 2 |

#### Alignment

| | |
|---|---|
| **Alignment Score** | 2600.0 (NT) + 3847.0 (AA) = 6447.0 |
| **Concordance (%)** | 82.2321 |

| Alignment Method | Global, seeded, nucleotide + amino acids (AGA) |
|---|---|

#### Genome Region

Sequence starts at position 5448 and ends at position 9414 relative to NC_001616.1 reference sequence.

#### Alignment Detailed Statistics

| | Begin | End | Coverage | Score | Concordance | Matches | Identities | I/D/M/F* | Stop Codons |
|---|---|---|---|---|---|---|---|---|---|
| **NT** | **5448** | **9414** | **19.3%** | **2600** | **70.0%** | **1866 (99.5%)** | **1590 (84.8%)** | **1/9** | |
| Mutations: | 5452T>C, 5458G>A, 5467T>C, 5470A>G, 5471C>T, 5473A>G, 5482C>T, 5485A>C, 5488T>C, 5489T>C, 5491G>A, 5497T>C, 5506T>C, 5509G>A, 5512G>A, 5516T>C, 5530T>C, 5533G>A, 5536G>T, 5539G>A, 5548T>G, 5554C>T, 5560T>A, 5561G>A, 5563G>A, 5566G>C, 5569A>G, 5582_5590delCTCAAGTTG, 5591A>C, 5593G>A, 5596G>A, 5597A>G, 5605C>A, 5617G>T, 5619C>T, 5620T>G, 5629G>A, 5630A>C, 5641C>T, 5650A>T, 5662A>T, 5665A>G, 5689A>G, 5693G>A, 5698G>A, 5713A>G, 5716G>C, 5719A>G, 5803C>T, 5836G>A, 5839A>G, 5845A>G, 5848A>G, 5857T>C, 5860C>T, 5863A>T, 5869T>A, 5875T>C, 5884C>T, 5893C>T, 5894A>G, 5896C>G, 5899C>T, 5908G>A, 5911T>C, 5923G>A, 5938A>G, 5941C>T, 5944T>C, 5947T>C, 5959G>A, 5960C>G, 5961G>C, 5962G>C, 5974A>G, 5986T>A, 5989C>T, 5990A>G, 5998T>C, 6008A>C, 6010A>T, 6019A>T, 6022G>T, 6025A>T, 6030A>C, 6049C>A, 6058G>A, 6061A>G, 6064C>A, 6070T>C, 6072G>C, 6073T>C, 6079G>A, 6081C>A, 6085A>T, 7432T>C, 7441T>C, 7459G>A, 7462A>G, 7468A>T, 7471C>A, 7477A>G, 7480C>T, 7486G>T, 7495T>C, 7498A>G, 7501T>C, 7503T>C, 7504G>A, 7513T>A, 7519C>T, 7522A>G, 7531G>A, 7666A>G, 7672T>C, 7678A>T, 7684G>A, 7690T>C, 7696T>C, 7702T>G, 7709G>A, 7713A>G, 7714A>G, 7715C>T, 7726T>C, 7735G>A, 7744A>G, 7747C>T, 7762C>T, 7768G>A, 7771T>C, 7780A>G, 7783A>C, 7894A>T, 7897T>C, 7921C>T, 7927G>A, 7936T>C, 7939T>C, 7942T>C, 7966G>A, 7969T>C, 7975T>C, 7987C>T, 7990T>C, 8005T>C, 8458A>G, 8460G>A, 8464A>G, 8467G>A, 8480G>A, 8482T>A, 8497G>A, 8498C>T, 8500A>G, 8502G>A, 8503A>G, 8506C>T, 8515A>T, 8524C>T, 8527A>C, 8528T>C, 8530A>G, 8539G>C, 8546C>T, 8547T>G, 8552T>A, 8563A>G, 8574C>G, 8585A>G, 8598G>A, 8602C>T, 8603A>T, 8604A>C, 8605C>T, 8620A>G, 8622C>A, 8625A>G, 8626G>A, 8629G>A, 8632C>T, 8638C>T, 8641G>A, 8647C>T, 8648C>T, 8649C>T, 8650G>C, 8653C>T, 8656A>G, 8658G>A, 8664A>C, 8665T>G, 8674T>G, 8679C>T, 8680A>T, 8683C>A, 8692G>A, 8695A>T, 8698T>C, 8707G>A, 8708A>C, 8716G>A, 8722C>T, 8725G>A, 8731A>G, 8737A>G, 8743C>T, 8745C>A, 8746A>G, 8749C>T, 8752G>A, 8755A>T, 8770C>T, 8771T>C, 8776A>G, 8779T>C, 8782G>A, 8791T>C, 8797A>G, 8800A>G, 8803A>G, 8809T>C, 8812T>C, 8818T>C, 8824G>A, 8827A>G, 8848G>A, 8854T>C, 8856A>G, 8857G>A, 8863G>A, 8865G>A, 8867A>C, 8869G>T, 8872A>G, 8883G>A, 8888A>T, 8929C>T, 8983G>A, 8984A>G, 8986T>C, 8988A>T, 8998G>A, 9004G>A, 9007G>A, 9016C>T, 9037C>A, 9085A>T, 9097C>T, 9131A>G, 9136A>T, 9145G>T, 9149A>G, 9150T>G, 9151G>A, 9152G>A, 9157A>G, 9160G>T, 9163T>C, 9184G>A, 9196A>G, 9214G>A, 9217A>G, 9220G>A, 9244A>T, 9247G>A, 9256C>T, 9260C>T, 9262T>C, 9274G>A, 9280C>T, 9289C>T, 9304G>A, 9334C>T, 9364C>G, 9377_9378insT, 9387C>T, 9404G>A | | | | | | | | |

##### CDS

| | Begin | End | Coverage | Score | Concordance | Matches | Identities | I/D/M/F* | Stop Codons |
|---|---|---|---|---|---|---|---|---|---|
| PVYgp1 | 1756 | 3064 | 20.0% | 3847 | 91.5% | 610 (99.5%) | 556 (90.7%) | 0/3/0/0 | 1 |
| Protein mutations: | A1793T (5561G>A 5563G>A), L1800_L1802del (5582_5590delCTCAAGTTG), K1803Q (5591A>C 5593G>A), I1805V (5597A>G), N1807K (5605C>A), A1812V (5619C>T 5620T>G), I1816L (5630A>C), V1837I (5693G>A), I1904V (5894A>G 5896C>G), R1926A (5960C>G 5961G>C 5962G>C), I1936V (5990A>G), E1945D (6019A>T), K1949T (6030A>C), D1955E (6049C>A), M1958I (6058G>A), S1963T (6072G>C 6073T>C), T1966N (6081C>A), L2440S (7503T>C 7504G>A), D2509N (7709G>A), K2510R (7713A>G 7714A>G), R2759K (8460G>A), A2766T (8480G>A 8482T>A), R2773K (8502G>A 8503A>G), E2777D (8515A>T), E2785D (8539G>C), L2788C (8546C>T 8547T>G), S2790T (8552T>A), A2797G (8574C>G), I2801V (8585A>G), G2805E (8598G>A), N2807S (8603A>T 8604A>C 8605C>T), P2813Q (8622C>A), E2814G (8625A>G 8626G>A), P2822F (8648C>T 8649C>T 8650G>C), G2825E (8658G>A), D2827A (8664A>C 8665T>G), A2832V (8679C>T 8680A>T), T2854K (8745C>A 8746A>G), E2891G (8856A>G 8857G>A), R2894Q (8865G>A), M2895L (8867A>C 8869G>T), G2900E (8883G>A), T2902S (8888A>T), N2934D (8984A>G 8986T>C), E2935V (8988A>T), I2983V (9131A>G), M2989G (9149A>G 9150T>G 9151G>A), G2990S (9152G>A), P3026S (9260C>T 9262T>C) | | | | | | | | |
| Codon mutations: | CTT1756CTC (5452T>C), AAG1758AAA (5458G>A), AGT1761AGC (5467T>C), CAA1762CAG (5470A>G), CTA1763TTG (5471C>T 5473A>G), TTC1766TTT (5482C>T), TCA1767TCC (5485A>C), AAT1768AAC (5488T>C), TTG1769CTA (5489T>C 5491G>A), GGT1771GGC (5497T>C), TGT1774TGC (5506T>C), GAG1775GAA (5509G>A), GAG1776GAA (5512G>A), TTA1778CTA (5516T>C), TAT1782TAC (5530T>C), GAG1783GAA (5533G>A), TCG1784TCT (5536G>T), TTG1785TTA (5539G>A), GTT1788GTG (5548T>G), CAC1790CAT (5554C>T), GCT1792GCA (5560T>A), GCG1793ACA (5561G>A 5563G>A), ACG1794ACC (5566G>C), TCA1795TCG (5569A>G), CTC1800_TTG1802del (5582_5590delCTCAAGTTG), AAG1803CAA (5591A>C 5593G>A), GGG1804GGA (5596G>A), ATT1805GTT (5597A>G), AAC1807AAA (5605C>A), GTG1811GTT (5617G>T), GCT1812GTG (5619C>T 5620T>G), TTG1815TTA (5629G>A), ATC1816CTC (5630A>C), GGC1819GGT (5641C>T), GCA1822GCT (5650A>T), ATA1826ATT (5662A>T), GGA1827GGG (5665A>G), CAA1835CAG (5689A>G), GTT1837ATT (5693G>A), GAG1838GAA (5698G>A), CAA1843CAG (5713A>G), GGG1844GGC (5716G>C), AAA1845AAG (5719A>G), GAC1873GAT (5803C>T), AGG1884AGA (5836G>A), AAA1885AAG (5839A>G), GGA1887GGG (5845A>G), AAA1888AAG (5848A>G), GGT1891GGC (5857T>C), ACC1892ACT (5860C>T), ACA1893ACT (5863A>T), GGT1895GGA (5869T>A), GGT1897GGC (5875T>C), AGC1900AGT (5884C>T), TTC1903TTT (5893C>T), ATC1904GTG (5894A>G 5896C>G), AAC1905AAT (5899C>T), GGG1908GGA (5908G>A), TTT1909TTC (5911T>C), GAG1913GAA (5923G>A), CAA1918CAG (5938A>G), TTC1919TTT (5941C>T), GTT1920GTC (5944T>C), GAT1921GAC (5947T>C), GGG1925GGA (5959G>A), CGG1926GCC (5960C>G 5961G>C 5962G>C), GAA1930GAG (5974A>G), GCT1934GCA (5986T>A), GAC1935GAT (5989C>T), ATT1936GTT (5990A>G), GAT1938GAC (5998T>C), AGA1942CGT (6008A>C 6010A>T), GAA1945GAT (6019A>T), GTG1946GTT (6022G>T), CGA1947CGT (6025A>T), AAA1949ACA (6030A>C), GAC1955GAA (6049C>A), ATG1958ATA (6058G>A), CAA1959CAG (6061A>G), GCC1960GCA (6064C>A), GGT1962GGC (6070T>C), AGT1963ACC (6072G>C 6073T>C), ACG1965ACA (6079G>A), ACC1966AAC (6081C>A), ATA1967ATT (6085A>T), TAT2416..C (7432T>C), CAT2419CAC (7441T>C), AAG2425AAA (7459G>A), GAA2426GAG (7462A>G), ATA2428ATT (7468A>T), GTC2429GTA (7471C>A), CAA2431CAG (7477A>G), AGC2432AGT (7480C>T), CTG2434CTT (7486G>T), TAT2437TAC (7495T>C), AAA2438AAG (7498A>G), GGT2439GGC (7501T>C), TTG2440TCA (7503T>C 7504G>A), ATT2443ATA (7513T>A), AAC2445AAT (7519C>T), GGA2446GGG (7522A>G), AAG2449AAA (7531G>A), GAA2494GAG (7666A>G), TGT2496TGC (7672T>C), ACA2498ACT (7678A>T), GGG2500GGA (7684G>A), ACT2502ACC (7690T>C), TTT2504TTC (7696T>C), GGT2506GGG (7702T>G), GAT2509AAT (7709G>A), AAA2510AGG (7713A>G 7714A>G), CTG2511TTG (7715C>T), CGT2514CGC (7726T>C), GAG2517GAA (7735G>A), GTA2520GTG (7744A>G), TAC2521TAT (7747C>T), GGC2526GGT (7762C>T), CAG2528CAA (7768G>A), TTT2529TTC (7771T>C), TCA2532TCG (7780A>G), CTA2533CTC (7783A>C), CCA2570CCT (7894A>T), ATT2571ATC (7897T>C), GTC2579GTT (7921C>T), AAG2581AAA (7927G>A), GGT2584GGC (7936T>C), AAT2585AAC (7939T>C), AAT2586AAC (7942T>C), GTG2594GTA (7966G>A), GAT2595GAC (7969T>C), TCT2597TCC (7975T>C), GTC2601GTT (7987C>T), CTT2602CTC (7990T>C), GCT2607GCC (8005T>C), CTA2758CTG (8458A>G), AGG2759AAG (8460G>A), AAA2760AAG (8464A>G), CTG2761CTA (8467G>A), GCT2766ACA (8480G>A 8482T>A), GAG2771GAA (8497G>A), CTA2772TTG (8498C>T 8500A>G), AGA2773AAG (8502G>A 8503A>G), GCC2774GCT (8506C>T), GAA2777GAT (8515A>T), GTC2780GTT (8524C>T), GCA2781GCC (8527A>C), TTA2782CTG (8528T>C 8530A>G), GAG2785GAC (8539G>C), CTT2788TGT (8546C>T 8547T>G), TCT2790ACT (8552T>A), GTA2793GTG (8563A>G), GCA2797GGA (8574C>G), ATT2801GTT (8585A>G), GGA2805GAA (8598G>A), AGC2806AGT (8602C>T), AAC2807TCT (8603A>T 8604A>C 8605C>T), AAA2812AAG (8620A>G), CCA2813CAA (8622C>A), GAG2814GGA (8625A>G 8626G>A), CAG2815CAA (8629G>A), GGC2816GGT (8632C>T), ATC2818ATT (8638C>T), CAG2819CAA (8641G>A), AAC2821AAT (8647C>T), CCG2822TTC (8648C>T 8649C>T 8650G>C), AAC2823AAT (8653C>T), AAA2824AAG (8656A>G), GGA2825GAA (8658G>A), GAT2827GCG (8664A>C 8665T>G), GTT2830GTG (8674T>G), GCA2832GTT (8679C>T 8680A>T), GGC2833GGA (8683C>A), GGG2836GGA (8692G>A), ACA2837ACT (8695A>T), CAT2838CAC (8698T>C), CCG2841CCA (8707G>A), AGA2842CGA (8708A>C), AAG2844AAA (8716G>A), ATC2846ATT (8722C>T), ACG2847ACA (8725G>A), AAA2849AAG (8731A>G), AGA2851AGG (8737A>G), CCC2853CCT (8743C>T), ACA2854AAG (8745C>A 8746A>G), AGC2855AGT (8749C>T), AAG2856AAA (8752G>A), GGA2857GGT (8755A>T), AAC2862AAT (8770C>T), TTA2863CTA (8771T>C), GAA2864GAG (8776A>G), CAT2865CAC (8779T>C), TTG2866TTA (8782G>A), TAT2869TAC (8791T>C), CCA2871CCG (8797A>G), CAA2872CAG (8800A>G), CAA2873CAG (8803A>G), GAT2875GAC (8809T>C), ATT2876ATC (8812T>C), AAT2878AAC (8818T>C), CGG2880CGA (8824G>A), GCA2881GCG (8827A>G), ACG2888ACA (8848G>A), TAT2890TAC (8854T>C), GAG2891GGA (8856A>G 8857G>A), GTG2893GTA (8863G>A), CGG2894CAG (8865G>A), ATG2895CTT (8867A>C 8869G>T), GCA2896GCG (8872A>G), GGA2900GAA (8883G>A), ACT2902TCT (8888A>T), TGC2915TGT (8929C>T), GGG2933GGA (8983G>A), AAT2934GAC (8984A>G 8986T>C), GAA2935GTA (8988A>T), GAG2938GAA (8998G>A), CCG2940CCA (9004G>A), TTG2941TTA (9007G>A), ATC2944ATT (9016C>T), ACC2951ACA (9037C>A), ATA2967ATT (9085A>T), AAC2971AAT (9097C>T), ATT2983GTT (9131A>G), CGA2984CGT (9136A>T), CGG2987CGT (9145G>T), ATG2989GGA (9149A>G 9150T>G 9151G>A), GGT2990AGT (9152G>A), TTA2991TTG (9157A>G), GCG2992GCT (9160G>T), CGT2993CGC (9163T>C), GAG3000GAA (9184G>A), CGA3004CGG (9196A>G), AGG3010AGA (9214G>A), GAA3011GAG (9217A>G), GCG3012GCA (9220G>A), GCA3020GCT (9244A>T), TTG3021TTA (9247G>A), GCC3024GCT (9256C>T), CCT3026TCC (9260C>T 9262T>C), GGG3030GGA (9274G>A), GAC3032GAT (9280C>T), ATC3035ATT (9289C>T), GAG3040GAA (9304G>A), GTC3050GTT (9334C>T), GTC3060GTG (9364C>G) | | | | | | | | |

##### Proteins

| | Begin | End | Coverage | Score | Concordance | Matches | Identities | I/D/M/F* | Stop Codons |
|---|---|---|---|---|---|---|---|---|---|
| polyprotein (NP_056759.1) | 1756 | 3064 | 20.0% | 3847 | 91.5% | 610 (99.5%) | 556 (90.7%) | 0/3/0/0 | 1 |
| Protein mutations: | A1793T (5561G>A 5563G>A), L1800_L1802del (5582_5590delCTCAAGTTG), K1803Q (5591A>C 5593G>A), I1805V (5597A>G), N1807K (5605C>A), A1812V (5619C>T 5620T>G), I1816L (5630A>C), V1837I (5693G>A), I1904V (5894A>G 5896C>G), R1926A (5960C>G 5961G>C 5962G>C), I1936V (5990A>G), E1945D (6019A>T), K1949T (6030A>C), D1955E (6049C>A), M1958I (6058G>A), S1963T (6072G>C 6073T>C), T1966N (6081C>A), L2440S (7503T>C 7504G>A), D2509N (7709G>A), K2510R (7713A>G 7714A>G), R2759K (8460G>A), A2766T (8480G>A 8482T>A), R2773K (8502G>A 8503A>G), E2777D (8515A>T), E2785D (8539G>C), L2788C (8546C>T 8547T>G), S2790T (8552T>A), A2797G (8574C>G), I2801V (8585A>G), G2805E (8598G>A), N2807S (8603A>T 8604A>C 8605C>T), P2813Q (8622C>A), E2814G (8625A>G 8626G>A), P2822F (8648C>T 8649C>T 8650G>C), G2825E (8658G>A), D2827A (8664A>C 8665T>G), A2832V (8679C>T 8680A>T), T2854K (8745C>A 8746A>G), E2891G (8856A>G 8857G>A), R2894Q (8865G>A), M2895L (8867A>C 8869G>T), G2900E (8883G>A), T2902S (8888A>T), N2934D (8984A>G 8986T>C), E2935V (8988A>T), I2983V (9131A>G), M2989G (9149A>G 9150T>G 9151G>A), G2990S (9152G>A), P3026S (9260C>T 9262T>C) | | | | | | | | |

| | Begin | End | Coverage | Score | Concordance | Matches | Identities | I/D/M/F* | Stop Codons |
|---|---|---|---|---|---|---|---|---|---|
| **NT** | **5448** | **9414** | **19.3%** | **2600** | **70.0%** | **1866 (99.5%)** | **1590 (84.8%)** | **1/9** | |
| Codon mutations: | CTT1756CTC (5452T>C), AAG1758AAA (5458G>A), AGT1761AGC (5467T>C), CAA1762CAG (5470A>G), CTA1763TTG (5471C>T 5473A>G), TTC1766TTT (5482C>T), TCA1767TCC (5485A>C), AAT1768AAC (5488T>C), TTG1769CTA (5489T>C 5491G>A), GGT1771GGC (5497T>C), TGT1774TGC (5506T>C), GAG1775GAA (5509G>A), GAG1776GAA (5512G>A), TTA1778CTA (5516T>C), TAT1782TAC (5530T>C), GAG1783GAA (5533G>A), TCG1784TCT (5536G>T), TTG1785TTA (5539G>A), GTT1788GTG (5548T>G), CAC1790CAT (5554C>T), GCT1792GCA (5560T>A), GCG1793ACA (5561G>A 5563G>A), ACG1794ACC (5566G>C), TCA1795TCG (5569A>G), CTC1800_TTG1802del (5582_5590delCTCAAGTTG), AAG1803CAA (5591A>C 5593G>A), GGG1804GGA (5596G>A), ATT1805GTT (5597A>G), AAC1807AAA (5605C>A), GTG1811GTT (5617G>T), GCT1812GTG (5619C>T 5620T>G), TTG1815TTA (5629G>A), ATC1816CTC (5630A>C), GGC1819GGT (5641C>T), GCA1822GCT (5650A>T), ATA1826ATT (5662A>T), GGA1827GGG (5665A>G), CAA1835CAG (5689A>G), GTT1837ATT (5693G>A), GAG1838GAA (5698G>A), CAA1843CAG (5713A>G), GGG1844GGC (5716G>C), AAA1845AAG (5719A>G), GAC1873GAT (5803C>T), AGG1884AGA (5836G>A), AAA1885AAG (5839A>G), GGA1887GGG (5845A>G), AAA1888AAG (5848A>G), GGT1891GGC (5857T>C), ACC1892ACT (5860C>T), ACA1893ACT (5863A>T), GGT1895GGA (5869T>A), GGT1897GGC (5875T>C), AGC1900AGT (5884C>T), TTC1903TTT (5893C>T), ATC1904GTG (5894A>G 5896C>G), AAC1905AAT (5899C>T), GGG1908GGA (5908G>A), TTT1909TTC (5911T>C), GAG1913GAA (5923G>A), CAA1918CAG (5938A>G), TTC1919TTT (5941C>T), GTT1920GTC (5944T>C), GAT1921GAC (5947T>C), GGG1925GGA (5959G>A), CGG1926GCC (5960C>G 5961G>C 5962G>C), GAA1930GAG (5974A>G), GCT1934GCA (5986T>A), GAC1935GAT (5989C>T), ATT1936GTT (5990A>G), GAT1938GAC (5998T>C), AGA1942CGT (6008A>C 6010A>T), GAA1945GAT (6019A>T), GTG1946GTT (6022G>T), CGA1947CGT (6025A>T), AAA1949ACA (6030A>C), GAC1955GAA (6049C>A), ATG1958ATA (6058G>A), CAA1959CAG (6061A>G), GCC1960GCA (6064C>A), GGT1962GGC (6070T>C), AGT1963ACC (6072G>C 6073T>C), ACG1965ACA (6079G>A), ACC1966AAC (6081C>A), ATA1967ATT (6085A>T), TAT2416..C (7432T>C), CAT2419CAC (7441T>C), AAG2425AAA (7459G>A), GAA2426GAG (7462A>G), ATA2428ATT (7468A>T), GTC2429GTA (7471C>A), CAA2431CAG (7477A>G), AGC2432AGT (7480C>T), CTG2434CTT (7486G>T), TAT2437TAC (7495T>C), AAA2438AAG (7498A>G), GGT2439GGC (7501T>C), TTG2440TCA (7503T>C 7504G>A), ATT2443ATA (7513T>A), AAC2445AAT (7519C>T), GGA2446GGG (7522A>G), AAG2449AAA (7531G>A), GAA2494GAG (7666A>G), TGT2496TGC (7672T>C), ACA2498ACT (7678A>T), GGG2500GGA (7684G>A), ACT2502ACC (7690T>C), TTT2504TTC (7696T>C), GGT2506GGG (7702T>G), GAT2509AAT (7709G>A), AAA2510AGG (7713A>G 7714A>G), CTG2511TTG (7715C>T), CGT2514CGC (7726T>C), GAG2517GAA (7735G>A), GTA2520GTG (7744A>G), TAC2521TAT (7747C>T), GGC2526GGT (7762C>T), CAG2528CAA (7768G>A), TTT2529TTC (7771T>C), TCA2532TCG (7780A>G), CTA2533CTC (7783A>C), CCA2570CCT (7894A>T), ATT2571ATC (7897T>C), GTC2579GTT (7921C>T), AAG2581AAA (7927G>A), GGT2584GGC (7936T>C), AAT2585AAC (7939T>C), AAT2586AAC (7942T>C), GTG2594GTA (7966G>A), GAT2595GAC (7969T>C), TCT2597TCC (7975T>C), GTC2601GTT (7987C>T), CTT2602CTC (7990T>C), GCT2607GCC (8005T>C), CTA2758CTG (8458A>G), AGG2759AAG (8460G>A), AAA2760AAG (8464A>G), CTG2761CTA (8467G>A), GCT2766ACA (8480G>A 8482T>A), GAG2771GAA (8497G>A), CTA2772TTG (8498C>T 8500A>G), AGA2773AAG (8502G>A 8503A>G), GCC2774GCT (8506C>T), GAA2777GAT (8515A>T), GTC2780GTT (8524C>T), GCA2781GCC (8527A>C), TTA2782CTG (8528T>C 8530A>G), GAG2785GAC (8539G>C), CTT2788TGT (8546C>T 8547T>G), TCT2790ACT (8552T>A), GTA2793GTG (8563A>G), GCA2797GGA (8574C>G), ATT2801GTT (8585A>G), GGA2805GAA (8598G>A), AGC2806AGT (8602C>T), AAC2807TCT (8603A>T 8604A>C 8605C>T), AAA2812AAG (8620A>G), CCA2813CAA (8622C>A), GAG2814GGA (8625A>G 8626G>A), CAG2815CAA (8629G>A), GGC2816GGT (8632C>T), ATC2818ATT (8638C>T), CAG2819CAA (8641G>A), AAC2821AAT (8647C>T), CCG2822TTC (8648C>T 8649C>T 8650G>C), AAC2823AAT (8653C>T), AAA2824AAG (8656A>G), GGA2825GAA (8658G>A), GAT2827GCG (8664A>C 8665T>G), GTT2830GTG (8674T>G), GCA2832GTT (8679C>T 8680A>T), GGC2833GGA (8683C>A), GGG2836GGA (8692G>A), ACA2837ACT (8695A>T), CAT2838CAC (8698T>C), CCG2841CCA (8707G>A), AGA2842CGA (8708A>C), AAG2844AAA (8716G>A), ATC2846ATT (8722C>T), ACG2847ACA (8725G>A), AAA2849AAG (8731A>G), AGA2851AGG (8737A>G), CCC2853CCT (8743C>T), ACA2854AAG (8745C>A 8746A>G), AGC2855AGT (8749C>T), AAG2856AAA (8752G>A), GGA2857GGT (8755A>T), AAC2862AAT (8770C>T), TTA2863CTA (8771T>C), GAA2864GAG (8776A>G), CAT2865CAC (8779T>C), TTG2866TTA (8782G>A), TAT2869TAC (8791T>C), CCA2871CCG (8797A>G), CAA2872CAG (8800A>G), CAA2873CAG (8803A>G), GAT2875GAC (8809T>C), ATT2876ATC (8812T>C), AAT2878AAC (8818T>C), CGG2880CGA (8824G>A), GCA2881GCG (8827A>G), ACG2888ACA (8848G>A), TAT2890TAC (8854T>C), GAG2891GGA (8856A>G 8857G>A), GTG2893GTA (8863G>A), CGG2894CAG (8865G>A), ATG2895CTT (8867A>C 8869G>T), GCA2896GCG (8872A>G), GGA2900GAA (8883G>A), ACT2902TCT (8888A>T), TGC2915TGT (8929C>T), GGG2933GGA (8983G>A), AAT2934GAC (8984A>G 8986T>C), GAA2935GTA (8988A>T), GAG2938GAA (8998G>A), CCG2940CCA (9004G>A), TTG2941TTA (9007G>A), ATC2944ATT (9016C>T), ACC2951ACA (9037C>A), ATA2967ATT (9085A>T), AAC2971AAT (9097C>T), ATT2983GTT (9131A>G), CGA2984CGT (9136A>T), CGG2987CGT (9145G>T), ATG2989GGA (9149A>G 9150T>G 9151G>A), GGT2990AGT (9152G>A), TTA2991TTG (9157A>G), GCG2992GCT (9160G>T), CGT2993CGC (9163T>C), GAG3000GAA (9184G>A), CGA3004CGG (9196A>G), AGG3010AGA (9214G>A), GAA3011GAG (9217A>G), GCG3012GCA (9220G>A), GCA3020GCT (9244A>T), TTG3021TTA (9247G>A), GCC3024GCT (9256C>T), CCT3026TCC (9260C>T 9262T>C), GGG3030GGA (9274G>A), GAC3032GAT (9280C>T), ATC3035ATT (9289C>T), GAG3040GAA (9304G>A), GTC3050GTT (9334C>T), GTC3060GTG (9364C>G) | | | | | | | | |
| CI protein (NP_734246.1) | 599 | 634 | 5.7% | 245 | 98.4% | 36 (100%) | 36 (100%) | 0/0/0/0 | 0 |
| Protein mutations: | *none* | | | | | | | | |
| Codon mutations: | CTT599CTC (5452T>C), AAG601AAA (5458G>A), AGT604AGC (5467T>C), CAA605CAG (5470A>G), CTA606TTG (5471C>T 5473A>G), TTC609TTT (5482C>T), TCA610TCC (5485A>C), AAT611AAC (5488T>C), TTG612CTA (5489T>C 5491G>A), GGT614GGC (5497T>C), TGT617TGC (5506T>C), GAG618GAA (5509G>A), GAG619GAA (5512G>A), TTA621CTA (5516T>C), TAT625TAC (5530T>C), GAG626GAA (5533G>A), TCG627TCT (5536G>T), TTG628TTA (5539G>A), GTT631GTG (5548T>G), CAC633CAT (5554C>T) | | | | | | | | |
| 6K2 protein (NP_734247.1) | 1 | 52 | 100% | 277 | 91.7% | 49 (94.2%) | 42 (80.8%) | 0/3/0/0 | 0 |
| Protein mutations: | A2T (5561G>A 5563G>A), L9_L11del (5582_5590delCTCAAGTTG), K12Q (5591A>C 5593G>A), I14V (5597A>G), N16K (5605C>A), A21V (5619C>T 5620T>G), I25L (5630A>C), V46I (5693G>A) | | | | | | | | |
| Codon mutations: | GCT1GCA (5560T>A), GCG2ACA (5561G>A 5563G>A), ACG3ACC (5566G>C), TCA4TCG (5569A>G), CTC9_TTG11del (5582_5590delCTCAAGTTG), AAG12CAA (5591A>C 5593G>A), GGG13GGA (5596G>A), ATT14GTT (5597A>G), AAC16AAA (5605C>A), GTG20GTT (5617G>T), GCT21GTG (5619C>T 5620T>G), TTG24TTA (5629G>A), ATC25CTC (5630A>C), GGC28GGT (5641C>T), GCA31GCT (5650A>T), ATA35ATT (5662A>T), GGA36GGG (5665A>G), CAA44CAG (5689A>G), GTT46ATT (5693G>A), GAG47GAA (5698G>A), CAA52CAG (5713A>G) | | | | | | | | |
| NIa-VPg protein (NP_734252.1) | 1 | 125 | 54.3% | 631 | 91.2% | 102 (100%) | 91 (89.2%) | 0/0/0/0 | 0 |
| Protein mutations: | I61V (5894A>G 5896C>G), R83A (5960C>G 5961G>C 5962G>C), I93V (5990A>G), E102D (6019A>T), K106T (6030A>C), D112E (6049C>A), M115I (6058G>A), S120T (6072G>C 6073T>C), T123N (6081C>A) | | | | | | | | |
| Codon mutations: | GGG1GGC (5716G>C), AAA2AAG (5719A>G), GAC30GAT (5803C>T), AGG41AGA (5836G>A), AAA42AAG (5839A>G), GGA44GGG (5845A>G), AAA45AAG (5848A>G), GGT48GGC (5857T>C), ACC49ACT (5860C>T), ACA50ACT (5863A>T), GGT52GGA (5869T>A), GGT54GGC (5875T>C), AGC57AGT (5884C>T), TTC60TTT (5893C>T), ATC61GTG (5894A>G 5896C>G), AAC62AAT (5899C>T), GGG65GGA (5908G>A), TTT66TTC (5911T>C), GAG70GAA (5923G>A), CAA75CAG (5938A>G), TTC76TTT (5941C>T), GTT77GTC (5944T>C), GAT78GAC (5947T>C), GGG82GGA (5959G>A), CGG83GCC (5960C>G 5961G>C 5962G>C), GAA87GAG (5974A>G), GCT91GCA (5986T>A), GAC92GAT (5989C>T), ATT93GTT (5990A>G), GAT95GAC (5998T>C), AGA99CGT (6008A>C 6010A>T), GAA102GAT (6019A>T), GTG103GTT (6022G>T), CGA104CGT (6025A>T), AAA106ACA (6030A>C), GAC112GAA (6049C>A), ATG115ATA (6058G>A), CAA116CAG (6061A>G), GCC117GCA (6064C>A), GGT119GGC (6070T>C), AGT120ACC (6072G>C 6073T>C), ACG122ACA (6079G>A), ACC123AAC (6081C>A), ATA124ATT (6085A>T) | | | | | | | | |
| NIb protein (NP_734249.1) | 142 | 521 | 31.7% | 1109 | 92.3% | 165 (100%) | 152 (92.1%) | 0/0/0/0 | 0 |
| Protein mutations: | L165S (7503T>C 7504G>A), D234N (7709G>A), K235R (7713A>G 7714A>G), R484K (8460G>A), A491T (8480G>A 8482T>A), R498K (8502G>A 8503A>G), E502D (8515A>T), E510D (8539G>C), L513C (8546C>T 8547T>G), S515T (8552T>A) | | | | | | | | |
| Codon mutations: | TAT141..C (7432T>C), CAT144CAC (7441T>C), AAG150AAA (7459G>A), GAA151GAG (7462A>G), ATA153ATT (7468A>T), GTC154GTA (7471C>A), CAA156CAG (7477A>G), AGC157AGT (7480C>T), CTG159CTT (7486G>T), TAT162TAC (7495T>C), AAA163AAG (7498A>G), GGT164GGC (7501T>C), TTG165TCA (7503T>C 7504G>A), ATT168ATA (7513T>A), AAC170AAT (7519C>T), GGA171GGG (7522A>G), AAG174AAA (7531G>A), GAA219GAG (7666A>G), TGT221TGC (7672T>C), ACA223ACT (7678A>T), GGG225GGA (7684G>A), ACT227ACC (7690T>C), TTT229TTC (7696T>C), GGT231GGG (7702T>G), GAT234AAT (7709G>A), AAA235AGG (7713A>G 7714A>G), CTG236TTG (7715C>T), CGT239CGC (7726T>C), GAG242GAA (7735G>A), GTA245GTG (7744A>G), TAC246TAT (7747C>T), GGC251GGT (7762C>T), CAG253CAA (7768G>A), TTT254TTC (7771T>C), TCA257TCG (7780A>G), CTA258CTC (7783A>C), CCA295CCT (7894A>T), ATT296ATC (7897T>C), GTC304GTT (7921C>T), AAG306AAA (7927G>A), GGT309GGC (7936T>C), AAT310AAC (7939T>C), AAT311AAC (7942T>C), GTG319GTA (7966G>A), GAT320GAC (7969T>C), TCT322TCC (7975T>C), GTC326GTT (7987C>T), CTT327CTC (7990T>C), GCT332GCC (8005T>C), CTA483CTG (8458A>G), AGG484AAG (8460G>A), AAA485AAG (8464A>G), CTG486CTA (8467G>A), GCT491ACA (8480G>A 8482T>A), GAG496GAA (8497G>A), CTA497TTG (8498C>T 8500A>G), AGA498AAG (8502G>A 8503A>G), GCC499GCT (8506C>T), GAA502GAT (8515A>T), GTC505GTT (8524C>T), GCA506GCC (8527A>C), TTA507CTG (8528T>C 8530A>G), GAG510GAC (8539G>C), CTT513TGT (8546C>T 8547T>G), TCT515ACT (8552T>A), GTA518GTG (8563A>G) | | | | | | | | |
| coat protein (NP_734250.1) | 1 | 267 | 96.3% | 1584 | 90.1% | 257 (100%) | 234 (91.1%) | 0/0/0/0 | 0 |
| Protein mutations: | A1G (8574C>G), I5V (8585A>G), G9E (8598G>A), N11S (8603A>T 8604A>C 8605C>T), P17Q (8622C>A), E18G (8625A>G 8626G>A), P26F (8648C>T 8649C>T 8650G>C), G29E (8658G>A), D31A (8664A>C 8665T>G), A36V (8679C>T 8680A>T), T58K (8745C>A 8746A>G), E95G (8856A>G 8857G>A), R98Q (8865G>A), M99L (8867A>C 8869G>T), G104E (8883G>A), T106S (8888A>T), N138D (8984A>G 8986T>C), E139V (8988A>T), I187V (9131A>G), M193G (9149A>G 9150T>G 9151G>A), G194S (9152G>A), P230S (9260C>T 9262T>C) | | | | | | | | |

| | Begin | End | Coverage | Score | Concordance | Matches | Identities | I/D/M/F* | Stop Codons |
|---|---|---|---|---|---|---|---|---|---|
| **NT** | **5448** | **9414** | **19.3%** | **2600** | **70.0%** | **1866 (99.5%)** | **1590 (84.8%)** | **1/9** | |
| Codon mutations: | GCA1GGA (8574C>G), ATT5GTT (8585A>G), GGA9GAA (8598G>A), AGC10AGT (8602C>T), AAC11TCT (8603A>T 8604A>C 8605C>T), AAA16AAG (8620A>G), CCA17CAA (8622C>A), GAG18GGA (8625A>G 8626G>A), CAG19CAA (8629G>A), GGC20GGT (8632C>T), ATC22ATT (8638C>T), CAG23CAA (8641G>A), AAC25AAT (8647C>T), CCG26TTC (8648C>T 8649C>T 8650G>C), AAC27AAT (8653C>T), AAA28AAG (8656A>G), GGA29GAA (8658G>A), GAT31GCG (8664A>C 8665T>G), GTT34GTG (8674T>G), GCA36GTT (8679C>T 8680A>T), GGC37GGA (8683C>A), GGG40GGA (8692G>A), ACA41ACT (8695A>T), CAT42CAC (8698T>C), CCG45CCA (8707G>A), AGA46CGA (8708A>C), AAG48AAA (8716G>A), ATC50ATT (8722C>T), ACG51ACA (8725G>A), AAA53AAG (8731A>G), AGA55AGG (8737A>G), CCC57CCT (8743C>T), ACA58AAG (8745C>A 8746A>G), AGC59AGT (8749C>T), AAG60AAA (8752G>A), GGA61GGT (8755A>T), AAC66AAT (8770C>T), TTA67CTA (8771T>C), GAA68GAG (8776A>G), CAT69CAC (8779T>C), TTG70TTA (8782G>A), TAT73TAC (8791T>C), CCA75CCG (8797A>G), CAA76CAG (8800A>G), CAA77CAG (8803A>G), GAT79GAC (8809T>C), ATT80ATC (8812T>C), AAT82AAC (8818T>C), CGG84CGA (8824G>A), GCA85GCG (8827A>G), ACG92ACA (8848G>A), TAT94TAC (8854T>C), GAG95GGA (8856A>G 8857G>A), GTG97GTA (8863G>A), CGG98CAG (8865G>A), ATG99CTT (8867A>C 8869G>T), GCA100GCG (8872A>G), GGA104GAA (8883G>A), ACT106TCT (8888A>T), TGC119TGT (8929C>T), GGG137GGA (8983G>A), AAT138GAC (8984A>G 8986T>C), GAA139GTA (8988A>T), GAG142GAA (8998G>A), CCG144CCA (9004G>A), TTG145TTA (9007G>A), ATC148ATT (9016C>T), ACC155ACA (9037C>A), ATA171ATT (9085A>T), AAC175AAT (9097C>T), ATT187GTT (9131A>G), CGA188CGT (9136A>T), CGG191CGT (9145G>T), ATG193GGA (9149A>G 9150T>G 9151G>A), GGT194AGT (9152G>A), TTA195TTG (9157A>G), GCG196GCT (9160G>T), CGT197CGC (9163T>C), GAG204GAA (9184G>A), CGA208CGG (9196A>G), AGG214AGA (9214G>A), GAA215GAG (9217A>G), GCG216GCA (9220G>A), GCA224GCT (9244A>T), TTG225TTA (9247G>A), GCC228GCT (9256C>T), CCT230TCC (9260C>T 9262T>C), GGG234GGA (9274G>A), GAC236GAT (9280C>T), ATC239ATT (9289C>T), GAG244GAA (9304G>A), GTC254GTT (9334C>T), GTC264GTG (9364C>G) | | | | | | | | |

*: Inserts / Deletes / Misaligned / Frameshifts

#### Analysis details

This analysis was performed with panviral2.64

### NGS Details (UN18_val): Duamitovirus peex1

#### Assembly

| | |
|---|---|
| **Coverage Length** | 705 (3 contig(s)) |
| **Depth Of Coverage** | 5.3 |
| **Number Of Reads** | 32 |
| **Reads Per Million** | 0.60 rpm (after QC) |
| **Ambiguities** | 0 |
| **Assembly Method** | de novo + reference guided assembly |
| **Consensus Caller** | Bcf Tools |

#### Coverage Map

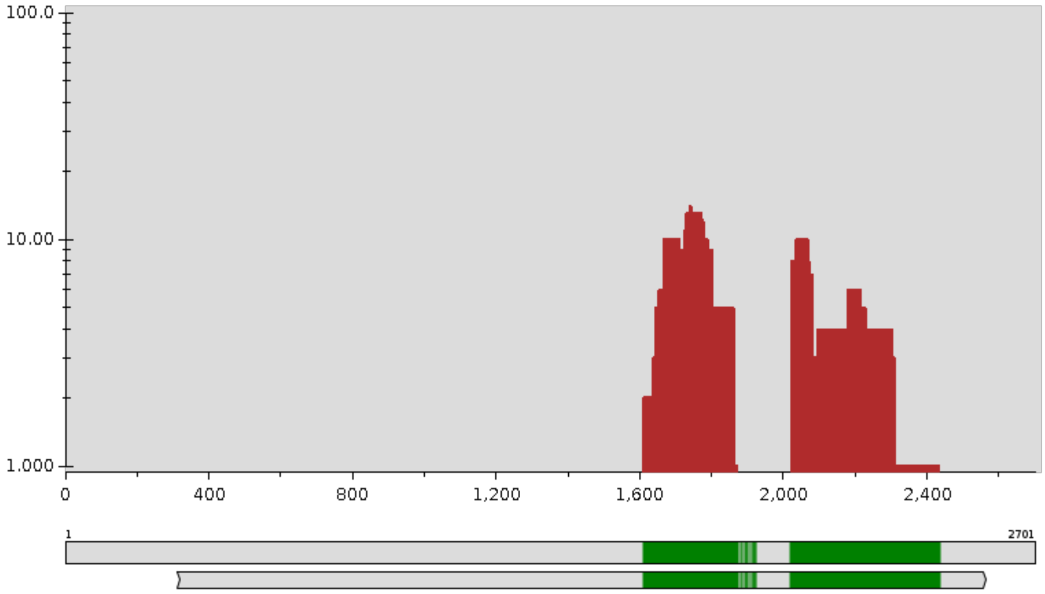

#### Assignment

| | |
|---|---|
| **Type** | Duamitovirus peex1 (Taxonomy ID: 2955799) |
| **Reference Genome** | NC_076525.1 |
| **NT Identity (%)** | 69.7443 |
| **AA Identity (%)** | 67.5325 |
| **Number Of Stop Codons** | 0 |
| **Number Of CDS** | 1 |

#### Alignment

| | |
|---|---|
| **Alignment Score** | 546.0 (NT) + 1021.0 (AA) = 1567.0 |
| **Concordance (%)** | 54.6484 |

| Alignment Method | Global, seeded, nucleotide + amino acids (AGA) |
|---|---|

#### Genome Region

Sequence starts at position 1609 and ends at position 2439 relative to NC_076525.1 reference sequence.

#### Alignment Detailed Statistics

| | Begin | End | Coverage | Score | Concordance | Matches | Identities | I/D/M/F* | Stop Codons |
|---|---|---|---|---|---|---|---|---|---|
| **NT** | **1609** | **2439** | **26.1%** | **546** | **39.1%** | **704 (99.9%)** | **491 (69.6%)** | **0/1** | |
| Mutations: | 1611G>A, 1613T>A, 1616G>A, 1622T>G, 1623C>T, 1628C>T, 1632C>T, 1633T>A, 1634G>C, 1655C>G, 1658G>T, 1661A>C, 1671T>A, 1676A>G, 1680T>C, 1682G>A, 1685G>T, 1689G>A, 1691G>A, 1692A>T, 1694G>T, 1696T>C, 1697C>A, 1703G>A, 1704G>A, 1705G>C, 1706C>G, 1707C>A, 1711G>C, 1713C>A, 1714G>C, 1715G>C, 1721C>T, 1722T>G, 1723G>C, 1724G>T, 1727A>T, 1730T>C, 1736T>G, 1737C>T, 1739C>A, 1751T>C, 1754C>A, 1757T>C, 1758G>A, 1764G>A, 1765A>C, 1766G>A, 1772C>G, 1774G>C, 1775C>G, 1776A>C, 1777A>G, 1781A>G, 1784T>C, 1785G>A, 1786C>G, 1788G>A, 1789A>T, 1790A>T, 1792A>T, 1793C>G, 1799G>T, 1800C>A, 1808A>T, 1814C>A, 1817T>C, 1820G>T, 1821delA, 1824C>G, 1825A>C, 1834T>G, 1841G>C, 1842A>G, 1844T>G, 1847T>C, 1850T>A, 1851T>A, 1853T>C, 1854C>A, 1856T>C, 1862C>T, 1865A>T, 1871G>A, 1880T>C, 1884T>G, 1885T>A, 1886A>C, 1893G>A, 1896G>T, 1898T>C, 1899T>C, 1901A>T, 1902A>T, 1903G>C, 1908G>A, 1914T>C, 1916A>T, 1918A>G, 1920T>G, 1922C>A, 2020G>A, 2024T>C, 2027A>T, 2033C>T, 2036T>G, 2040A>T, 2042G>T, 2051T>A, 2052A>C, 2053A>C, 2057C>A, 2066G>A, 2067A>C, 2069T>A, 2075T>G, 2077T>C, 2088T>C, 2090T>A, 2091T>A, 2095T>G, 2096A>G, 2098C>T, 2102G>T, 2104G>T, 2105G>A, 2109T>C, 2119G>T, 2120G>T, 2133G>T, 2134G>T, 2135A>T, 2136A>G, 2137A>G, 2138G>A, 2139C>A, 2145G>A, 2156A>G, 2158G>A, 2159A>G, 2161G>C, 2162T>G, 2165G>A, 2171C>A, 2173C>A, 2180T>G, 2184A>G, 2188A>C, 2189G>T, 2190T>G, 2195A>G, 2202G>C, 2210G>A, 2211T>A, 2213G>C, 2214A>T, 2216A>T, 2223C>G, 2225A>T, 2226C>T, 2231G>A, 2234A>T, 2240T>A, 2245A>G, 2246T>G, 2249G>A, 2255A>T, 2258A>G, 2261C>G, 2268G>A, 2270A>C, 2272G>T, 2274C>A, 2283G>A, 2292C>T, 2294T>G, 2295C>A, 2296G>C, 2297A>T, 2306A>T, 2315C>T, 2321G>A, 2325A>T, 2326A>T, 2327T>A, 2330C>A, 2333G>A, 2336A>T, 2345A>T, 2348C>T, 2354T>A, 2355A>T, 2357G>T, 2360A>T, 2363G>T, 2375G>C, 2390T>A, 2391A>T, 2393T>A, 2394C>A, 2396G>C, 2397A>G, 2398C>A, 2399T>A, 2403C>T, 2404T>A, 2406C>G, 2408T>A, 2412G>A, 2414G>C, 2425T>A, 2426A>C, 2427A>G, 2429C>T | | | | | | | | |

##### CDS

| | Begin | End | Coverage | Score | Concordance | Matches | Identities | I/D/M/F* | Stop Codons |
|---|---|---|---|---|---|---|---|---|---|
| RdRp | 434 | 710 | 30.8% | 1021 | 59.2% | 231 (100%) | 156 (67.5%) | 0/0/2/1 | 0 |
| Protein mutations: | A434T (1611G>A 1613T>A), L441Y (1632C>T 1633T>A 1634G>C), L454M (1671T>A), E460K (1689G>A 1691G>A), K461Y (1692A>T 1694G>T), V462A (1696T>C 1697C>A), G465T (1704G>A 1705G>C 1706C>G), R466S (1707C>A), S467T (1711G>C), R468T (1713C>A 1714G>C 1715G>C), W471A (1722T>G 1723G>C 1724G>T), E472D (1727A>T), A483T (1758G>A), E485T (1764G>A 1765A>C 1766G>A), G488A (1774G>C 1775C>G), K489R (1776A>C 1777A>G), A492R (1785G>A 1786C>G), E493I (1788G>A 1789A>T 1790A>T), Y494L (1792A>T 1793C>G), E496D (1799G>T), H505A (1824C>G 1825A>C), I508R (1834T>G), N511E (1842A>G 1844T>G), S514T (1851T>A 1853T>C), L515I (1854C>A 1856T>C), L525D (1884T>G 1885T>A 1886A>C), T577S (2040A>T 2042G>T), K581P (2052A>C 2053A>C), I586L (2067A>C 2069T>A), V589A (2077T>C), S593P (2088T>C 2090T>A), Y594N (2091T>A), I595R (2095T>G 2096A>G), S596L (2098C>T), Q597H (2102G>T), R598L (2104G>T 2105G>A), S600P (2109T>C), W603F (2119G>T 2120G>T), G608F (2133G>T 2134G>T 2135A>T), K609G (2136A>G 2137A>G 2138G>A), P610T (2139C>A), D612N (2145G>A), R616K (2158G>A 2159A>G), G617A (2161G>C 2162T>G), A621D (2173C>A), K625E (2184A>G), E626A (2188A>C 2189G>T), L627V (2190T>G), E631Q (2202G>C), L634I (2211T>A 2213G>C), I635F (2214A>T 2216A>T), Q638D (2223C>G 2225A>T), L641F (2234A>T), H645R (2245A>G 2246T>G), V653I (2268G>A 2270A>C), R654L (2272G>T), H655N (2274C>A), E658K (2283G>A), R662T (2295C>A 2296G>C 2297A>T), N672L (2325A>T 2326A>T 2327T>A), E675D (2336A>T), M682F (2355A>T 2357G>T), E683D (2360A>T), T694S (2391A>T 2393T>A), Q695N (2394C>A 2396G>C), T696E (2397A>G 2398C>A 2399T>A), L698Y (2403C>T 2404T>A), H699E (2406C>G 2408T>A), V701I (2412G>A 2414G>C), L705Y (2425T>A 2426A>C), I706V (2427A>G 2429C>T) | | | | | | | | |
| Codon mutations: | GCT434ACA (1611G>A 1613T>A), GGG435GGA (1616G>A), CCT437CCG (1622T>G), CTA438TTA (1623C>T), GGC439GGT (1628C>T), CTG441TAC (1632C>T 1633T>A 1634G>C), GCC448GCG (1655C>G), CTG449CTT (1658G>T), TCA450TCC (1661A>C), TTG454ATG (1671T>A), GTA455GTG (1676A>G), TTG457CTA (1680T>C 1682G>A), GCG458GCT (1685G>T), GAG460AAA (1689G>A 1691G>A), AAG461TAT (1692A>T 1694G>T), GTC462GCA (1696T>C 1697C>A), CCG464CCA (1703G>A), GGC465ACG (1704G>A 1705G>C 1706C>G), CGT466AGT (1707C>A), AGT467ACT (1711G>C), CGG468ACC (1713C>A 1714G>C 1715G>C), TTC470TTT (1721C>T), TGG471GCT (1722T>G 1723G>C 1724G>T), GAA472GAT (1727A>T), TAT473TAC (1730T>C), CTT475CTG (1736T>G), CTC476TTA (1737C>T 1739C>A), ATT480ATC (1751T>C), CTC481CTA (1754C>A), ATT482ATC (1757T>C), GCC483ACC (1758G>A), GAG485ACA (1764G>A 1765A>C 1766G>A), GTC487GTG (1772C>G), GGC488GCG (1774G>C 1775C>G), AAG489CGG (1776A>C 1777A>G), CAA490CAG (1781A>G), TAT491TAC (1784T>C), GCG492AGG (1785G>A 1786C>G), GAA493ATT (1788G>A 1789A>T 1790A>T), TAC494TTG (1792A>T 1793C>G), GAG496GAT (1799G>T), CGG497AGG (1800C>A), GGA499GGT (1808A>T), ACC501ACA (1814C>A), ATT502ATC (1817T>C), TCG503TCT (1820G>T), ATA504-TA (1821delA), CAT505GCT (1824C>G 1825A>C), ATA508AGA (1834T>G), TCG510TCC (1841G>C), AAT511GAG (1842A>G 1844T>G), AAT512AAC (1847T>C), GGT513GGA (1850T>A), TCT514ACC (1851T>A 1853T>C), CTT515ATC (1854C>A 1856T>C), TTC517TTT (1862C>T), GCA518GCT (1865A>T), AGG520AGA (1871G>A), ACT523..C (1880T>C), TTA525GAC (1884T>G 1885T>A 1886A>C), GTT528A.. (1893G>A), GAT529T.C (1896G>T 1898T>C), TTA530C.T (1899T>C 1901A>T), AGT531TC. (1902A>T 1903G>C), GTC533A.. (1908G>A), TTA535C.T (1914T>C 1916A>T), AAA536.G. (1918A>G), TCC537G.A (1920T>G 1922C>A), AGG570.AG (2020G>A), GTT571GTC (2024T>C), CGA572CGT (2027A>T), CGC574CGT (2033C>T), CTT575CTG (2036T>G), ACG577TCT (2040A>T 2042G>T), TCT580TCA (2051T>A), AAA581CCA (2052A>C 2053A>C), CGC582CGA (2057C>A), AGG585AGA (2066G>A), ATT586CTA (2067A>C 2069T>A), GCT588GCG (2075T>G), GTT589GCT (2077T>C), TCT593CCA (2088T>C 2090T>A), TAC594AAC (2091T>A), ATA595AGG (2095T>G 2096A>G), TCA596TTA (2098C>T), CAG597CAT (2102G>T), CGG598CTA (2104G>T 2105G>A), TCT600CCT (2109T>C), TGG603TTT (2119G>T 2120G>T), GGA608TTT (2133G>T 2134G>T 2135A>T), AAG609GGA (2136A>G 2137A>G 2138G>A), CCC610ACC (2139C>A), GAT612AAT (2145G>A), TTA615TTG (2156A>G), AGA616AAG (2158G>A 2159A>G), GGT617GCG (2161G>C 2162T>G), AAG618AAA (2165G>A), ATC620ATA (2171C>A), GCT621GAT (2173C>A), CTT623CTG (2180T>G), AAG625GAG (2184A>G), GAG626GCT (2188A>C 2189G>T), TTA627GTA (2190T>G), AAA628AAG (2195A>G), GAG631CAG (2202G>C), CAG633CAA (2210G>A), TTG634ATC (2211T>A 2213G>C), ATA635TTT (2214A>T 2216A>T), CAA638GAT (2223C>G 2225A>T), CTG639TTG (2226C>T), GTG640GTA (2231G>A), TTA641TTT (2234A>T), GGT643GGA (2240T>A), CAT645CGG (2245A>G 2246T>G), GAG646GAA (2249G>A), CTA648CTT (2255A>T), GAA649GAG (2258A>G), CGC650CGG (2261C>G), GTA653ATC (2268G>A 2270A>C), CGT654CTT (2272G>T), CAT655AAT (2274C>A), GAA658AAA (2283G>A), CTT661TTG (2292C>T 2294T>G), CGA662ACT (2295C>A 2296G>C 2297A>T), TCA665TCT (2306A>T), CAC668CAT (2315C>T), GTG670GTA (2321G>A), AAT672TTA (2325A>T 2326A>T 2327T>A), TCC673TCA (2330C>A), CCG674CCA (2333G>A), GAA675GAT (2336A>T), ATA678ATT (2345A>T), GAC679GAT (2348C>T), CTT681CTA (2354T>A), ATG682TTT (2355A>T 2357G>T), GAA683GAT (2360A>T), GTG684GTT (2363G>T), GCG688GCC (2375G>C), CGT693CGA (2390T>A), ACT694TCA (2391A>T 2393T>A), CAG695AAC (2394C>A 2396G>C), ACT696GAA (2397A>G 2398C>A 2399T>A), CTT698TAT (2403C>T 2404T>A), CAT699GAA (2406C>G 2408T>A), GTG701ATC (2412G>A 2414G>C), TTA705TAC (2425T>A 2426A>C), ATC706GTT (2427A>G 2429C>T) | | | | | | | | |

##### Proteins

| | Begin | End | Coverage | Score | Concordance | Matches | Identities | I/D/M/F* | Stop Codons |
|---|---|---|---|---|---|---|---|---|---|
| RNA-dependent RNA polymerase (YP_010798875.1) | 434 | 710 | 30.8% | 1021 | 59.2% | 231 (100%) | 156 (67.5%) | 0/0/2/1 | 0 |
| Protein mutations: | A434T (1611G>A 1613T>A), L441Y (1632C>T 1633T>A 1634G>C), L454M (1671T>A), E460K (1689G>A 1691G>A), K461Y (1692A>T 1694G>T), V462A (1696T>C 1697C>A), G465T (1704G>A 1705G>C 1706C>G), R466S (1707C>A), S467T (1711G>C), R468T (1713C>A 1714G>C 1715G>C), W471A (1722T>G 1723G>C 1724G>T), E472D (1727A>T), A483T (1758G>A), E485T (1764G>A 1765A>C 1766G>A), G488A (1774G>C 1775C>G), K489R (1776A>C 1777A>G), A492R (1785G>A 1786C>G), E493I (1788G>A 1789A>T 1790A>T), Y494L (1792A>T 1793C>G), E496D (1799G>T), H505A (1824C>G 1825A>C), I508R (1834T>G), N511E (1842A>G 1844T>G), S514T (1851T>A 1853T>C), L515I (1854C>A 1856T>C), L525D (1884T>G 1885T>A 1886A>C), T577S (2040A>T 2042G>T), K581P (2052A>C 2053A>C), I586L (2067A>C 2069T>A), V589A (2077T>C), S593P (2088T>C 2090T>A), Y594N (2091T>A), I595R (2095T>G 2096A>G), S596L (2098C>T), Q597H (2102G>T), R598L (2104G>T 2105G>A), S600P (2109T>C), W603F (2119G>T 2120G>T), G608F (2133G>T 2134G>T 2135A>T), K609G (2136A>G 2137A>G 2138G>A), P610T (2139C>A), D612N (2145G>A), R616K (2158G>A 2159A>G), G617A (2161G>C 2162T>G), A621D (2173C>A), K625E (2184A>G), E626A (2188A>C 2189G>T), L627V (2190T>G), E631Q (2202G>C), L634I (2211T>A 2213G>C), I635F (2214A>T 2216A>T), Q638D (2223C>G 2225A>T), L641F (2234A>T), H645R (2245A>G 2246T>G), V653I (2268G>A 2270A>C), R654L (2272G>T), H655N (2274C>A), E658K (2283G>A), R662T (2295C>A 2296G>C 2297A>T), N672L (2325A>T 2326A>T 2327T>A), E675D (2336A>T), M682F (2355A>T 2357G>T), E683D (2360A>T), T694S (2391A>T 2393T>A), Q695N (2394C>A 2396G>C), T696E (2397A>G 2398C>A 2399T>A), L698Y (2403C>T 2404T>A), H699E (2406C>G 2408T>A), V701I (2412G>A 2414G>C), L705Y (2425T>A 2426A>C), I706V (2427A>G 2429C>T) | | | | | | | | |

| | Begin | End | Coverage | Score | Concordance | Matches | Identities | I/D/M/F* | Stop Codons |
|---|---|---|---|---|---|---|---|---|---|
| **NT** | **1609** | **2439** | **26.1%** | **546** | **39.1%** | **704 (99.9%)** | **491 (69.6%)** | **0/1** | |
| Codon mutations: | GCT434ACA (1611G>A 1613T>A), GGG435GGA (1616G>A), CCT437CCG (1622T>G), CTA438TTA (1623C>T), GGC439GGT (1628C>T), CTG441TAC (1632C>T 1633T>A 1634G>C), GCC448GCG (1655C>G), CTG449CTT (1658G>T), TCA450TCC (1661A>C), TTG454ATG (1671T>A), GTA455GTG (1676A>G), TTG457CTA (1680T>C 1682G>A), GCG458GCT (1685G>T), GAG460AAA (1689G>A 1691G>A), AAG461TAT (1692A>T 1694G>T), GTC462GCA (1696T>C 1697C>A), CCG464CCA (1703G>A), GGC465ACG (1704G>A 1705G>C 1706C>G), CGT466AGT (1707C>A), AGT467ACT (1711G>C), CGG468ACC (1713C>A 1714G>C 1715G>C), TTC470TTT (1721C>T), TGG471GCT (1722T>G 1723G>C 1724G>T), GAA472GAT (1727A>T), TAT473TAC (1730T>C), CTT475CTG (1736T>G), CTC476TTA (1737C>T 1739C>A), ATT480ATC (1751T>C), CTC481CTA (1754C>A), ATT482ATC (1757T>C), GCC483ACC (1758G>A), GAG485ACA (1764G>A 1765A>C 1766G>A), GTC487GTG (1772C>G), GGC488GCG (1774G>C 1775C>G), AAG489CGG (1776A>C 1777A>G), CAA490CAG (1781A>G), TAT491TAC (1784T>C), GCG492AGG (1785G>A 1786C>G), GAA493ATT (1788G>A 1789A>T 1790A>T), TAC494TTG (1792A>T 1793C>G), GAG496GAT (1799G>T), CGG497AGG (1800C>A), GGA499GGT (1808A>T), ACC501ACA (1814C>A), ATT502ATC (1817T>C), TCG503TCT (1820G>T), ATA504-TA (1821delA), CAT505GCT (1824C>G 1825A>C), ATA508AGA (1834T>G), TCG510TCC (1841G>C), AAT511GAG (1842A>G 1844T>G), AAT512AAC (1847T>C), GGT513GGA (1850T>A), TCT514ACC (1851T>A 1853T>C), CTT515ATC (1854C>A 1856T>C), TTC517TTT (1862C>T), GCA518GCT (1865A>T), AGG520AGA (1871G>A), ACT523..C (1880T>C), TTA525GAC (1884T>G 1885T>A 1886A>C), GTT528A.. (1893G>A), GAT529T.C (1896G>T 1898T>C), TTA530C.T (1899T>C 1901A>T), AGT531TC. (1902A>T 1903G>C), GTC533A.. (1908G>A), TTA535C.T (1914T>C 1916A>T), AAA536.G. (1918A>G), TCC537G.A (1920T>G 1922C>A), AGG570.AG (2020G>A), GTT571GTC (2024T>C), CGA572CGT (2027A>T), CGC574CGT (2033C>T), CTT575CTG (2036T>G), ACG577TCT (2040A>T 2042G>T), TCT580TCA (2051T>A), AAA581CCA (2052A>C 2053A>C), CGC582CGA (2057C>A), AGG585AGA (2066G>A), ATT586CTA (2067A>C 2069T>A), GCT588GCG (2075T>G), GTT589GCT (2077T>C), TCT593CCA (2088T>C 2090T>A), TAC594AAC (2091T>A), ATA595AGG (2095T>G 2096A>G), TCA596TTA (2098C>T), CAG597CAT (2102G>T), CGG598CTA (2104G>T 2105G>A), TCT600CCT (2109T>C), TGG603TTT (2119G>T 2120G>T), GGA608TTT (2133G>T 2134G>T 2135A>T), AAG609GGA (2136A>G 2137A>G 2138G>A), CCC610ACC (2139C>A), GAT612AAT (2145G>A), TTA615TTG (2156A>G), AGA616AAG (2158G>A 2159A>G), GGT617GCG (2161G>C 2162T>G), AAG618AAA (2165G>A), ATC620ATA (2171C>A), GCT621GAT (2173C>A), CTT623CTG (2180T>G), AAG625GAG (2184A>G), GAG626GCT (2188A>C 2189G>T), TTA627GTA (2190T>G), AAA628AAG (2195A>G), GAG631CAG (2202G>C), CAG633CAA (2210G>A), TTG634ATC (2211T>A 2213G>C), ATA635TTT (2214A>T 2216A>T), CAA638GAT (2223C>G 2225A>T), CTG639TTG (2226C>T), GTG640GTA (2231G>A), TTA641TTT (2234A>T), GGT643GGA (2240T>A), CAT645CGG (2245A>G 2246T>G), GAG646GAA (2249G>A), CTA648CTT (2255A>T), GAA649GAG (2258A>G), CGC650CGG (2261C>G), GTA653ATC (2268G>A 2270A>C), CGT654CTT (2272G>T), CAT655AAT (2274C>A), GAA658AAA (2283G>A), CTT661TTG (2292C>T 2294T>G), CGA662ACT (2295C>A 2296G>C 2297A>T), TCA665TCT (2306A>T), CAC668CAT (2315C>T), GTG670GTA (2321G>A), AAT672TTA (2325A>T 2326A>T 2327T>A), TCC673TCA (2330C>A), CCG674CCA (2333G>A), GAA675GAT (2336A>T), ATA678ATT (2345A>T), GAC679GAT (2348C>T), CTT681CTA (2354T>A), ATG682TTT (2355A>T 2357G>T), GAA683GAT (2360A>T), GTG684GTT (2363G>T), GCG688GCC (2375G>C), CGT693CGA (2390T>A), ACT694TCA (2391A>T 2393T>A), CAG695AAC (2394C>A 2396G>C), ACT696GAA (2397A>G 2398C>A 2399T>A), CTT698TAT (2403C>T 2404T>A), CAT699GAA (2406C>G 2408T>A), GTG701ATC (2412G>A 2414G>C), TTA705TAC (2425T>A 2426A>C), ATC706GTT (2427A>G 2429C>T) | | | | | | | | |

*: Inserts / Deletes / Misaligned / Frameshifts

#### Analysis details

This analysis was performed with panviral2.64

### NGS Details (UN18_val): Harvey murine sarcoma virus

#### Assembly

| | |
|---|---|
| **Coverage Length** | 222 (1 contig(s)) |
| **Depth Of Coverage** | 2.2 |
| **Number Of Reads** | 6 |
| **Reads Per Million** | 0.11 rpm (after QC) |
| **Ambiguities** | 0 |
| **Assembly Method** | de novo + reference guided assembly |
| **Consensus Caller** | Bcf Tools |

#### Coverage Map

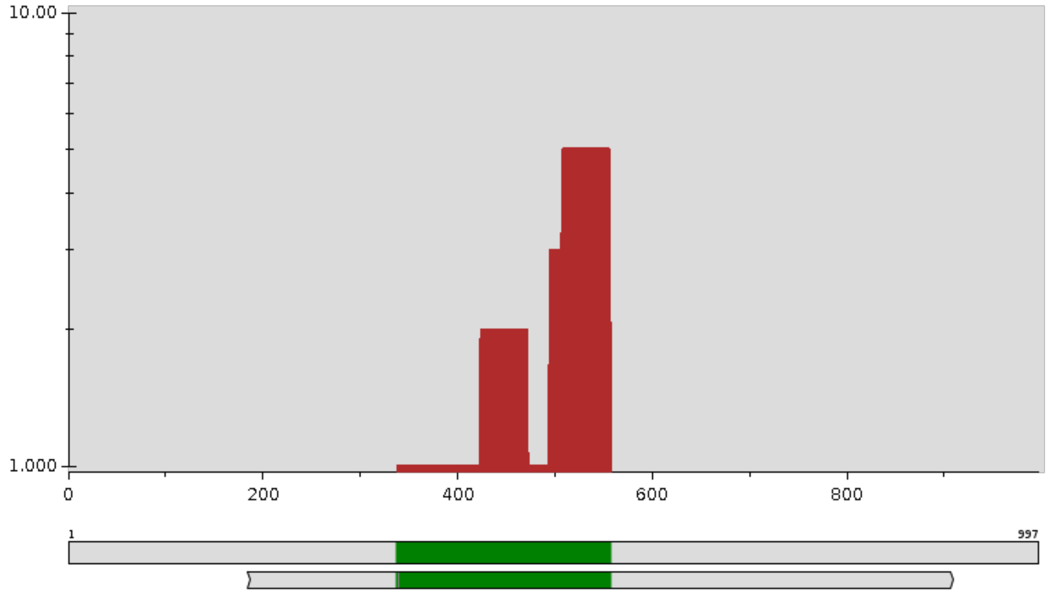

#### Assignment

| | |
|---|---|
| **Type** | Harvey murine sarcoma virus (Taxonomy ID: 11807) |
| **Reference Genome** | NC_038668.1 |
| **NT Identity (%)** | 73.4234 |
| **AA Identity (%)** | 85.034 |
| **Number Of Stop Codons** | 0 |
| **Number Of CDS** | 2 |

#### Alignment

| | |
|---|---|
| **Alignment Score** | 208.0 (NT) + 814.0 (AA) = 1022.0 |
| **Concordance (%)** | 71.669 |

| Alignment Method | Global, seeded, nucleotide + amino acids (AGA) |
|---|---|

#### Genome Region

Sequence starts at position 338 and ends at position 559 relative to NC_038668.1 reference sequence.

#### Alignment Detailed Statistics

| | Begin | End | Coverage | Score | Concordance | Matches | Identities | I/D/M/F* | Stop Codons |
|---|---|---|---|---|---|---|---|---|---|
| **NT** | **338** | **559** | **22.3%** | **208** | **46.8%** | **222 (100%)** | **163 (73.4%)** | **0/0** | |
| Mutations: | 346A>G, 358T>C, 364G>A, 367G>C, 370C>T, 372C>G, 374A>G, 376A>T, 379C>A, 382G>C, 392G>T, 393C>G, 394C>T, 395C>T, 409G>C, 417A>G, 424T>C, 427G>C, 433G>A, 439T>C, 442C>T, 445T>C, 448A>C, 451G>A, 457C>T, 461C>A, 466A>G, 469G>A, 470G>T, 471T>G, 472A>C, 475C>G, 478T>C, 483G>A, 488A>G, 489C>T, 490G>C, 491T>G, 492G>C, 493T>G, 494T>C, 496A>G, 499G>A, 503A>G, 505C>G, 506T>C, 508A>C, 514A>G, 515A>G, 517A>C, 520T>C, 523A>G, 526A>G, 532T>C, 533A>T, 534G>C, 544G>C, 547C>G, 550G>A | | | | | | | | |
| **CDS** | | | | | | | | | |
| D1R95_gp1 | 52 | 125 | 30.6% | 409 | 83.0% | 74 (100%) | 63 (85.1%) | 0/0/0/0 | 0 |
| Protein mutations: | A63G (372C>G), R64G (374A>G 376A>T), A70C (392G>T 393C>G 394C>T), N78S (417A>G), V96C (470G>T 471T>G 472A>C), G100E (483G>A), T102V (488A>G 489C>T 490G>C), C103A (491T>G 492G>C 493T>G), I107V (503A>G 505C>G), T111A (515A>G 517A>C), D121E (547C>G) | | | | | | | | |
| Codon mutations: | ACA54ACG (346A>G), CTT58CTC (358T>C), GTG60GTA (364G>A), GTG61GTC (367G>C), GGC62GGT (370C>T), GCT63GGT (372C>G), AGA64GGT (374A>G 376A>T), GGC65GGA (379C>A), GTG66GTC (382G>C), GCC70TGT (392G>T 393C>G 394C>T), CTG71TTG (395C>T), CTG75CTC (409G>C), AAC78AGC (417A>G), TTT80TTC (424T>C), GTG81GTC (427G>C), GAG83GAA (433G>A), GAT85GAC (439T>C), CCC86CCT (442C>T), ACT87ACC (445T>C), ATA88ATC (448A>C), GAG89GAA (451G>A), TCC91TCT (457C>T), CGG93AGG (461C>A), AAA94AAG (466A>G), CAG95CAA (469G>A), GTA96TGC (470G>T 471T>G 472A>C), GTC97GTG (475C>G), ATT98ATC (478T>C), GGG100GAG (483G>A), ACG102GTC (488A>G 489C>T 490G>C), TGT103GCG (491T>G 492G>C 493T>G), TTA104CTG (494T>C 496A>G), CTG105CTA (499G>A), ATC107GTG (503A>G 505C>G), TTA108CTC (506T>C 508A>C), ACA110ACG (514A>G), ACA111GCC (515A>G 517A>C), GGT112GGC (520T>C), CAA113CAG (523A>G), GAA114GAG (526A>G), TAT116TAC (532T>C), AGT117TCT (533A>T 534G>C), CGG120CGC (544G>C), GAC121GAG (547C>G), CAG122CAA (550G>A) | | | | | | | | |
| D1R95_gp2 | 1 | 73 | 38.4% | 405 | 82.8% | 73 (100%) | 62 (84.9%) | 0/0/0/0 | 0 |
| Protein mutations: | A11G (372C>G), R12G (374A>G 376A>T), A18C (392G>T 393C>G 394C>T), N26S (417A>G), V44C (470G>T 471T>G 472A>C), G48E (483G>A), T50V (488A>G 489C>T 490G>C), C51A (491T>G 492G>C 493T>G), I55V (503A>G 505C>G), T59A (515A>G 517A>C), D69E (547C>G) | | | | | | | | |
| Codon mutations: | ACA2ACG (346A>G), CTT6CTC (358T>C), GTG8GTA (364G>A), GTG9GTC (367G>C), GGC10GGT (370C>T), GCT11GGT (372C>G), AGA12GGT (374A>G 376A>T), GGC13GGA (379C>A), GTG14GTC (382G>C), GCC18TGT (392G>T 393C>G 394C>T), CTG19TTG (395C>T), CTG23CTC (409G>C), AAC26AGC (417A>G), TTT28TTC (424T>C), GTG29GTC (427G>C), GAG31GAA (433G>A), GAT33GAC (439T>C), CCC34CCT (442C>T), ACT35ACC (445T>C), ATA36ATC (448A>C), GAG37GAA (451G>A), TCC39TCT (457C>T), CGG41AGG (461C>A), AAA42AAG (466A>G), CAG43CAA (469G>A), GTA44TGC (470G>T 471T>G 472A>C), GTC45GTG (475C>G), ATT46ATC (478T>C), GGG48GAG (483G>A), ACG50GTC (488A>G 489C>T 490G>C), TGT51GCG (491T>G 492G>C 493T>G), TTA52CTG (494T>C 496A>G), CTG53CTA (499G>A), ATC55GTG (503A>G 505C>G), TTA56CTC (506T>C 508A>C), ACA58ACG (514A>G), ACA59GCC (515A>G 517A>C), GGT60GGC (520T>C), CAA61CAG (523A>G), GAA62GAG (526A>G), TAT64TAC (532T>C), AGT65TCT (533A>T 534G>C), CGG68CGC (544G>C), GAC69GAG (547C>G), CAG70CAA (550G>A) | | | | | | | | |
| **Proteins** | | | | | | | | | |
| hypothetical protein (YP_009507788.1) | 52 | 125 | 30.6% | 409 | 83.0% | 74 (100%) | 63 (85.1%) | 0/0/0/0 | 0 |
| Protein mutations: | A63G (372C>G), R64G (374A>G 376A>T), A70C (392G>T 393C>G 394C>T), N78S (417A>G), V96C (470G>T 471T>G 472A>C), G100E (483G>A), T102V (488A>G 489C>T 490G>C), C103A (491T>G 492G>C 493T>G), I107V (503A>G 505C>G), T111A (515A>G 517A>C), D121E (547C>G) | | | | | | | | |
| Codon mutations: | ACA54ACG (346A>G), CTT58CTC (358T>C), GTG60GTA (364G>A), GTG61GTC (367G>C), GGC62GGT (370C>T), GCT63GGT (372C>G), AGA64GGT (374A>G 376A>T), GGC65GGA (379C>A), GTG66GTC (382G>C), GCC70TGT (392G>T 393C>G 394C>T), CTG71TTG (395C>T), CTG75CTC (409G>C), AAC78AGC (417A>G), TTT80TTC (424T>C), GTG81GTC (427G>C), GAG83GAA (433G>A), GAT85GAC (439T>C), CCC86CCT (442C>T), ACT87ACC (445T>C), ATA88ATC (448A>C), GAG89GAA (451G>A), TCC91TCT (457C>T), CGG93AGG (461C>A), AAA94AAG (466A>G), CAG95CAA (469G>A), GTA96TGC (470G>T 471T>G 472A>C), GTC97GTG (475C>G), ATT98ATC (478T>C), GGG100GAG (483G>A), ACG102GTC (488A>G 489C>T 490G>C), TGT103GCG (491T>G 492G>C 493T>G), TTA104CTG (494T>C 496A>G), CTG105CTA (499G>A), ATC107GTG (503A>G 505C>G), TTA108CTC (506T>C 508A>C), ACA110ACG (514A>G), ACA111GCC (515A>G 517A>C), GGT112GGC (520T>C), CAA113CAG (523A>G), GAA114GAG (526A>G), TAT116TAC (532T>C), AGT117TCT (533A>T 534G>C), CGG120CGC (544G>C), GAC121GAG (547C>G), CAG122CAA (550G>A) | | | | | | | | |
| hypothetical protein (YP_009507789.1) | 1 | 73 | 38.4% | 405 | 82.8% | 73 (100%) | 62 (84.9%) | 0/0/0/0 | 0 |
| Protein mutations: | A11G (372C>G), R12G (374A>G 376A>T), A18C (392G>T 393C>G 394C>T), N26S (417A>G), V44C (470G>T 471T>G 472A>C), G48E (483G>A), T50V (488A>G 489C>T 490G>C), C51A (491T>G 492G>C 493T>G), I55V (503A>G 505C>G), T59A (515A>G 517A>C), D69E (547C>G) | | | | | | | | |
| Codon mutations: | ACA2ACG (346A>G), CTT6CTC (358T>C), GTG8GTA (364G>A), GTG9GTC (367G>C), GGC10GGT (370C>T), GCT11GGT (372C>G), AGA12GGT (374A>G 376A>T), GGC13GGA (379C>A), GTG14GTC (382G>C), GCC18TGT (392G>T 393C>G 394C>T), CTG19TTG (395C>T), CTG23CTC (409G>C), AAC26AGC (417A>G), TTT28TTC (424T>C), GTG29GTC (427G>C), GAG31GAA (433G>A), GAT33GAC (439T>C), CCC34CCT (442C>T), ACT35ACC (445T>C), ATA36ATC (448A>C), GAG37GAA (451G>A), TCC39TCT (457C>T), CGG41AGG (461C>A), AAA42AAG (466A>G), CAG43CAA (469G>A), GTA44TGC (470G>T 471T>G 472A>C), GTC45GTG (475C>G), ATT46ATC (478T>C), GGG48GAG (483G>A), ACG50GTC (488A>G 489C>T 490G>C), TGT51GCG (491T>G 492G>C 493T>G), TTA52CTG (494T>C 496A>G), CTG53CTA (499G>A), ATC55GTG (503A>G 505C>G), TTA56CTC (506T>C 508A>C), ACA58ACG (514A>G), ACA59GCC (515A>G 517A>C), GGT60GGC (520T>C), CAA61CAG (523A>G), GAA62GAG (526A>G), TAT64TAC (532T>C), AGT65TCT (533A>T 534G>C), CGG68CGC (544G>C), GAC69GAG (547C>G), CAG70CAA (550G>A) | | | | | | | | |

*: Inserts / Deletes / Misaligned / Frameshifts

#### Analysis details

This analysis was performed with panviral2.64

### NGS Details (UN18_val): Brazilian marseillevirus

#### Assembly

| | |
|---|---|
| **Coverage Length** | 240 (1 contig(s)) |
| **Depth Of Coverage** | 13067.1 |
| **Number Of Reads** | 31766 |
| **Reads Per Million** | 596.20 rpm (after QC) |
| **Ambiguities** | 0 |
| **Assembly Method** | de novo + reference guided assembly |
| **Consensus Caller** | Bcf Tools |

#### Coverage Map

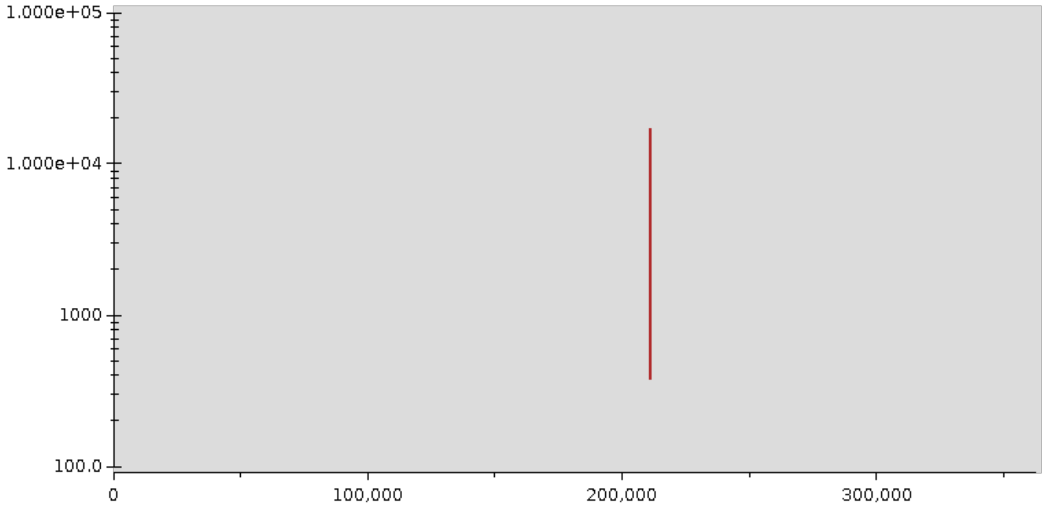

#### Assignment

| | |
|---|---|
| **Type** | Brazilian marseillevirus (Taxonomy ID: 1813599) |
| **Reference Genome** | NC_029692.1 |
| **NT Identity (%)** | 81.6667 |
| **AA Identity (%)** | 94.6667 |
| **Number Of Stop Codons** | 1 |
| **Number Of CDS** | 491 |

#### Alignment

| | |
|---|---|
| **Alignment Score** | 304.0 (NT) + 454.0 (AA) = 758.0 |
| **Concordance (%)** | 79.9578 |
| **Alignment Method** | Local, heuristic, nucleotide (BLASTN) |

#### Genome Region

Sequence starts at position 210884 and ends at position 211123 relative to NC_029692.1 reference sequence.

#### Alignment Detailed Statistics

| | Begin | End | Coverage | Score | Concordance | Matches | Identities | I/D/M/F* | Stop Codons |
|---|---|---|---|---|---|---|---|---|---|
| **NT** | **210884** | **211123** | **0.1%** | **304** | **63.3%** | **240 (100%)** | **196 (81.7%)** | **0/0** | |

Mutations: 210899T>A, 210901T>G, 210902T>G, 210905G>C, 210908A>C, 210910G>A, 210917A>G, 210926T>C, 210929T>C, 210932G>A, 210938A>G, 210953C>G, 210955T>G, 210956T>G, 210962T>C, 210965C>A, 210974G>A, 210977T>A, 210986A>C, 210988G>A, 210989G>C, 210995T>C, 211001A>T, 211004A>T, 211007A>G, 211009C>T, 211013T>C, 211031T>G, 211032T>G, 211033G>C, 211034T>C, 211037C>A, 211040G>A, 211043T>G, 211046G>A, 211061T>C, 211070G>C, 211079C>G, 211088C>G, 211091G>A, 211100T>A, 211103A>G, 211106C>G, 211108C>T

*: Inserts / Deletes / Misaligned / Frameshifts

#### Analysis details

This analysis was performed with panviral2.64

### NGS Details (UN18_val): Tomato chocolate spot virus (segment RNA 1)

#### Assembly

| | |
|---|---|
| **Coverage Length** | 775 (1 contig(s)) |
| **Depth Of Coverage** | 416.7 |
| **Number Of Reads** | 2065 |
| **Reads Per Million** | 38.76 rpm (after QC) |
| **Ambiguities** | 0 |
| **Assembly Method** | de novo + reference guided assembly |
| **Consensus Caller** | Bcf Tools |

#### Coverage Map

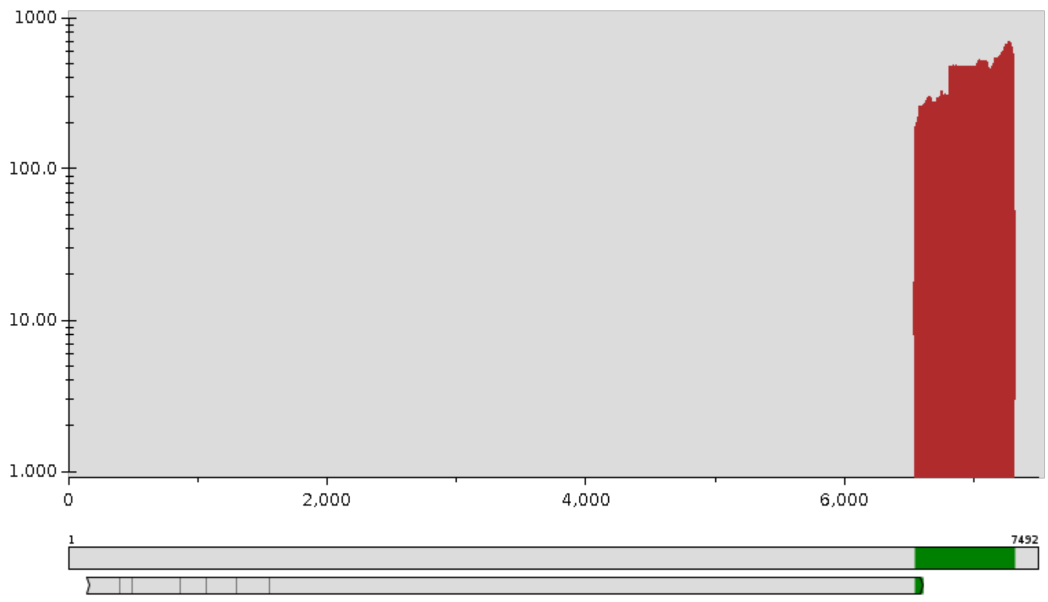

#### Assignment

| | |
|---|---|
| **Type** | Tomato chocolate spot virus (Taxonomy ID: 661101) |
| **Reference Genome** | NC_013075.1 |
| **NT Identity (%)** | 76.2393 |
| **AA Identity (%)** | 13.6364 |
| **Number Of Stop Codons** | 0 |
| **Number Of CDS** | 1 |

#### Alignment

| | |
|---|---|
| **Alignment Score** | 352.0 (NT) + 12.0 (AA) = 364.0 |
| **Concordance (%)** | 33.9869 |

| Alignment Method | Global, seeded, nucleotide + amino acids (AGA) |
|---|---|

#### Genome Region

Sequence starts at position 6539 and ends at position 7313 relative to NC_013075.1 reference sequence.

#### Alignment Detailed Statistics

| | Begin | End | Coverage | Score | Concordance | Matches | Identities | I/D/M/F* | Stop Codons |
|---|---|---|---|---|---|---|---|---|---|
| **NT** | **6539** | **7313** | **10.3%** | **352** | **38.5%** | **584 (75.3%)** | **446 (57.5%)** | **1/191** | |
| Mutations: | 6548T>A, 6550C>A, 6552G>A, 6554C>G, 6555C>T, 6556A>C, 6557C>T, 6558T>G, 6559T>G, 6560T>A, 6562G>A, 6564T>G, 6566G>C, 6569A>G, 6570C>A, 6571T>A, 6572G>T, 6574C>G, 6577T>A, 6579C>A, 6582C>G, 6583C>A, 6585T>G, 6586A>T, 6587C>T, 6589A>G, 6592T>G, 6593A>T, 6595C>A, 6598G>T, 6600G>C, 6604T>G, 6607G>T, 6608T>C, 6609C>G, 6610A>G, 6612G>C, 6615T>A, 6618delT, 6621C>T, 6622T>C, 6625G>T, 6626T>C, 6628T>A, 6629G>A, 6630T>C, 6633_6634insT, 6636T>A, 6637T>A, 6638A>G, 6639C>T, 6651A>G, 6653T>A, 6658_6660delGTG, 6662G>C, 6667G>A, 6668G>T, 6670A>G, 6671G>T, 6674T>A, 6675T>A, 6676G>T, 6677C>T, 6678A>C, 6683T>A, 6684_6688delATAAC, 6700T>C, 6711A>G, 6721G>T, 6723T>A, 6729T>A, 6735G>T, 6737A>G, 6740C>A, 6741G>A, 6753G>T, 6756T>C, 6758A>G, 6768A>G, 6777T>C, 6779T>A, 6783T>C, 6788T>C, 6795A>G, 6800G>A, 6806C>T, 6814C>T, 6815_6840delAATTGTTTGGTGTTGTCAACGCGCCG, 6846_6855delGGCCAAGAAA, 6861_6917delCCTTAGGCTTTATTGGTAAAGTCAAGATGTGGAGAAGGAATGCCAGCAAGCATAAGC, 6922_7010delTCTCTTAGGGTCGGGCTTAAGATGGCAGCCAGAATTAGAGCCTTTGAGTATGCTTGTGGGCATCTGGTTTTCACATCAACTCGTTTGGA, 7013A>T, 7014T>A, 7018C>G, 7019A>G, 7020A>T, 7023C>A, 7042G>T, 7044A>T, 7062G>A, 7070T>C, 7071C>T, 7084A>G, 7087T>C, 7098C>T, 7099T>C, 7107G>A, 7116C>T, 7118T>A, 7121G>A, 7122T>C, 7123C>T, 7124T>G, 7137A>T, 7138G>A, 7139A>G, 7140C>T, 7148A>G, 7151A>T, 7159A>T, 7162T>C, 7165G>A, 7171T>G, 7173C>T, 7177A>C, 7185G>A, 7199C>T, 7208A>T, 7209T>A, 7210C>T, 7211A>G, 7213C>T, 7214A>T, 7229T>C, 7245T>C, 7250G>A, 7253T>A, 7254G>A, 7256T>C, 7257G>A, 7258G>T, 7259C>G, 7274G>C, 7275T>C, 7292T>G, 7303T>A | | | | | | | | |
| **CDS** | | | | | | | | | |
| ToChSV_s1_gp1 | 2135 | 2156 | 1.0% | 12 | 7.5% | 22 (100%) | 3 (13.6%) | 0/0/0/0 | 0 |
| Protein mutations: | V2137D (6548T>A), Q2138K (6550C>A 6552G>A), S2139C (6554C>G 6555C>T), T2140L (6556A>C 6557C>T 6558T>G), F2141D (6559T>G 6560T>A), V2142M (6562G>A 6564T>G), R2143T (6566G>C), N2144R (6569A>G 6570C>A), C2145I (6571T>A 6572G>T), Q2146E (6574C>G), S2147T (6577T>A 6579C>A), I2148M (6582C>G), L2149M (6583C>A 6585T>G), T2150L (6586A>T 6587C>T), N2151D (6589A>G), Y2152V (6592T>G 6593A>T), Q2153K (6595C>A), E2154Y (6598G>T 6600G>C), *2156G (6604T>G) | | | | | | | | |
| Codon mutations: | GTC2137GAC (6548T>A), CAG2138AAA (6550C>A 6552G>A), TCC2139TGT (6554C>G 6555C>T), ACT2140CTG (6556A>C 6557C>T 6558T>G), TTT2141GAT (6559T>G 6560T>A), GTT2142ATG (6562G>A 6564T>G), AGA2143ACA (6566G>C), AAC2144AGA (6569A>G 6570C>A), TGT2145ATT (6571T>A 6572G>T), CAA2146GAA (6574C>G), TCC2147ACA (6577T>A 6579C>A), ATC2148ATG (6582C>G), CTT2149ATG (6583C>A 6585T>G), ACA2150TTA (6586A>T 6587C>T), AAT2151GAT (6589A>G), TAT2152GTT (6592T>G 6593A>T), CAG2153AAG (6595C>A), GAG2154TAC (6598G>T 6600G>C), TGA2156GGA (6604T>G) | | | | | | | | |
| **Proteins** | | | | | | | | | |
| polyprotein (YP_003097229.1) | 2135 | 2156 | 1.0% | 12 | 7.5% | 22 (100%) | 3 (13.6%) | 0/0/0/0 | 0 |
| Protein mutations: | V2137D (6548T>A), Q2138K (6550C>A 6552G>A), S2139C (6554C>G 6555C>T), T2140L (6556A>C 6557C>T 6558T>G), F2141D (6559T>G 6560T>A), V2142M (6562G>A 6564T>G), R2143T (6566G>C), N2144R (6569A>G 6570C>A), C2145I (6571T>A 6572G>T), Q2146E (6574C>G), S2147T (6577T>A 6579C>A), I2148M (6582C>G), L2149M (6583C>A 6585T>G), T2150L (6586A>T 6587C>T), N2151D (6589A>G), Y2152V (6592T>G 6593A>T), Q2153K (6595C>A), E2154Y (6598G>T 6600G>C), *2156G (6604T>G) | | | | | | | | |
| Codon mutations: | GTC2137GAC (6548T>A), CAG2138AAA (6550C>A 6552G>A), TCC2139TGT (6554C>G 6555C>T), ACT2140CTG (6556A>C 6557C>T 6558T>G), TTT2141GAT (6559T>G 6560T>A), GTT2142ATG (6562G>A 6564T>G), AGA2143ACA (6566G>C), AAC2144AGA (6569A>G 6570C>A), TGT2145ATT (6571T>A 6572G>T), CAA2146GAA (6574C>G), TCC2147ACA (6577T>A 6579C>A), ATC2148ATG (6582C>G), CTT2149ATG (6583C>A 6585T>G), ACA2150TTA (6586A>T 6587C>T), AAT2151GAT (6589A>G), TAT2152GTT (6592T>G 6593A>T), CAG2153AAG (6595C>A), GAG2154TAC (6598G>T 6600G>C), TGA2156GGA (6604T>G) | | | | | | | | |

*: Inserts / Deletes / Misaligned / Frameshifts

#### Analysis details

This analysis was performed with panviral2.64

### NGS Details (UN18_val): Tomato chocolate spot virus (segment RNA 1)

#### Assembly

| | |
|---|---|
| **Coverage Length** | 405 (2 contig(s)) |
| **Depth Of Coverage** | 1872.0 |
| **Number Of Reads** | 6736 |
| **Reads Per Million** | 126.42 rpm (after QC) |
| **Ambiguities** | 0 |
| **Assembly Method** | de novo + reference guided assembly |
| **Consensus Caller** | Bcf Tools |

#### Coverage Map

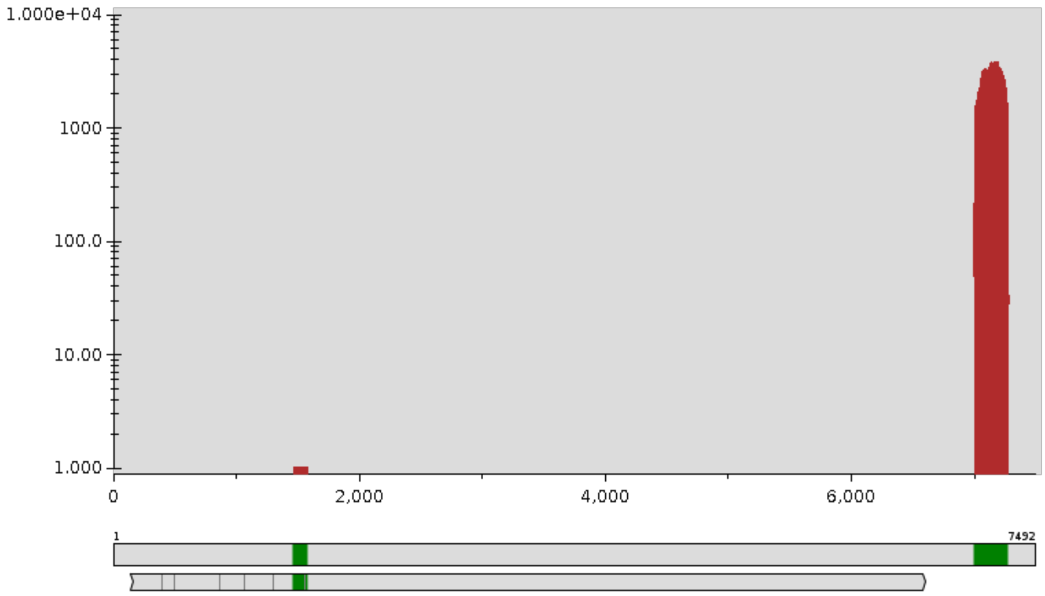

#### Assignment

| | |
|---|---|
| **Type** | Tomato chocolate spot virus (Taxonomy ID: 661101) |
| **Reference Genome** | NC_013075.1 |
| **NT Identity (%)** | 78.3251 |
| **AA Identity (%)** | 74.4186 |
| **Number Of Stop Codons** | 1 |
| **Number Of CDS** | 1 |

#### Alignment

| | |
|---|---|
| **Alignment Score** | 452.0 (NT) + 207.0 (AA) = 659.0 |
| **Concordance (%)** | 60.1277 |

| Alignment Method | Global, seeded, nucleotide + amino acids (AGA) |
|---|---|

#### Genome Region

Sequence starts at position 1458 and ends at position 7275 relative to NC_013075.1 reference sequence.

#### Alignment Detailed Statistics

| | Begin | End | Coverage | Score | Concordance | Matches | Identities | I/D/M/F* | Stop Codons |
|---|---|---|---|---|---|---|---|---|---|
| **NT** | **1458** | **7275** | **5.4%** | **452** | **55.8%** | **405 (99.8%)** | **318 (78.3%)** | **1/0** | |
| Mutations: | 1459A>G, 1461T>G, 1473C>T, 1479A>T, 1483T>C, 1485G>A, 1486C>A, 1488C>G, 1494T>A, 1497G>A, 1500T>G, 1503A>G, 1504T>A, 1506T>A, 1509A>G, 1512T>C, 1515C>A, 1518A>G, 1530C>A, 1536G>A, 1539A>T, 1542G>C, 1554G>T, 1555T>C, 1557A>T, 1559C>A, 1560T>A, 1562T>A, 1563G>C, 1564G>T, 1566T>A, 1567G>C, 1569T>A, 1570G>T, 1571T>A, 1572G>C, 1573G>T, 1576G>A, 1584G>A, 7012T>G, 7018C>T, 7019A>G, 7020A>G, 7020_7021insT, 7023C>A, 7042G>T, 7044A>T, 7047G>A, 7048C>G, 7066G>A, 7082G>A, 7083T>C, 7098C>T, 7099T>C, 7104T>A, 7107G>A, 7116C>G, 7121G>A, 7127A>C, 7140C>T, 7151A>T, 7152T>C, 7183C>T, 7184T>C, 7185G>T, 7186T>G, 7187T>C, 7188G>A, 7196C>T, 7197A>G, 7198A>C, 7199C>A, 7200A>G, 7201G>A, 7207A>T, 7208A>T, 7209T>G, 7210C>T, 7212T>A, 7219T>A, 7224T>C, 7242A>T, 7255A>T, 7257G>A, 7258G>C, 7260T>C, 7262C>G, 7264T>C | | | | | | | | |
| **CDS** | | | | | | | | | |
| ToChSV_s1_gp1 | 441 | 483 | 2.0% | 207 | 69.2% | 43 (100%) | 32 (74.4%) | 0/0/0/0 | 1 |
| Protein mutations: | I441V (1459A>G 1461T>G), C456R (1504T>A 1506T>A), D464E (1530C>A), P474Q (1559C>A 1560T>A), M475N (1562T>A 1563G>C), A476S (1564G>T 1566T>A), A477P (1567G>C 1569T>A), V478Y (1570G>T 1571T>A 1572G>C), E479* (1573G>T), D480N (1576G>A) | | | | | | | | |
| Codon mutations: | ATT441GTG (1459A>G 1461T>G), GAC445GAT (1473C>T), GGA447GGT (1479A>T), TTG449CTA (1483T>C 1485G>A), CGC450AGG (1486C>A 1488C>G), GGT452GGA (1494T>A), GCG453GCA (1497G>A), GGT454GGG (1500T>G), CAA455CAG (1503A>G), TGT456AGA (1504T>A 1506T>A), CAA457CAG (1509A>G), GAT458GAC (1512T>C), ATC459ATA (1515C>A), AAA460AAG (1518A>G), GAC464GAA (1530C>A), AAG466AAA (1536G>A), TCA467TCT (1539A>T), ACG468ACC (1542G>C), CCG472CCT (1554G>T), TTA473CTT (1555T>C 1557A>T), CCT474CAA (1559C>A 1560T>A), ATG475AAC (1562T>A 1563G>C), GCT476TCA (1564G>T 1566T>A), GCT477CCA (1567G>C 1569T>A), GTG478TAC (1570G>T 1571T>A 1572G>C), GAG479TAG (1573G>T), GAT480AAT (1576G>A), GGG482GGA (1584G>A) | | | | | | | | |
| **Proteins** | | | | | | | | | |
| polyprotein (YP_003097229.1) | 441 | 483 | 2.0% | 207 | 69.2% | 43 (100%) | 32 (74.4%) | 0/0/0/0 | 1 |
| Protein mutations: | I441V (1459A>G 1461T>G), C456R (1504T>A 1506T>A), D464E (1530C>A), P474Q (1559C>A 1560T>A), M475N (1562T>A 1563G>C), A476S (1564G>T 1566T>A), A477P (1567G>C 1569T>A), V478Y (1570G>T 1571T>A 1572G>C), E479* (1573G>T), D480N (1576G>A) | | | | | | | | |
| Codon mutations: | ATT441GTG (1459A>G 1461T>G), GAC445GAT (1473C>T), GGA447GGT (1479A>T), TTG449CTA (1483T>C 1485G>A), CGC450AGG (1486C>A 1488C>G), GGT452GGA (1494T>A), GCG453GCA (1497G>A), GGT454GGG (1500T>G), CAA455CAG (1503A>G), TGT456AGA (1504T>A 1506T>A), CAA457CAG (1509A>G), GAT458GAC (1512T>C), ATC459ATA (1515C>A), AAA460AAG (1518A>G), GAC464GAA (1530C>A), AAG466AAA (1536G>A), TCA467TCT (1539A>T), ACG468ACC (1542G>C), CCG472CCT (1554G>T), TTA473CTT (1555T>C 1557A>T), CCT474CAA (1559C>A 1560T>A), ATG475AAC (1562T>A 1563G>C), GCT476TCA (1564G>T 1566T>A), GCT477CCA (1567G>C 1569T>A), GTG478TAC (1570G>T 1571T>A 1572G>C), GAG479TAG (1573G>T), GAT480AAT (1576G>A), GGG482GGA (1584G>A) | | | | | | | | |
| RNA-dependent RNA polymerase (YP_003097234.1) | 52 | 84 | 39.3% | 201 | 85.9% | 33 (100%) | 30 (90.9%) | 0/0/0/0 | 0 |
| Protein mutations: | I52V (1459A>G 1461T>G), C67R (1504T>A 1506T>A), D75E (1530C>A) | | | | | | | | |
| Codon mutations: | ATT52GTG (1459A>G 1461T>G), GAC56GAT (1473C>T), GGA58GGT (1479A>T), TTG60CTA (1483T>C 1485G>A), CGC61AGG (1486C>A 1488C>G), GGT63GGA (1494T>A), GCG64GCA (1497G>A), GGT65GGG (1500T>G), CAA66CAG (1503A>G), TGT67AGA (1504T>A 1506T>A), CAA68CAG (1509A>G), GAT69GAC (1512T>C), ATC70ATA (1515C>A), AAA71AAG (1518A>G), GAC75GAA (1530C>A), AAG77AAA (1536G>A), TCA78TCT (1539A>T), ACG79ACC (1542G>C), CCG83CCT (1554G>T), TTA84CTT (1555T>C 1557A>T) | | | | | | | | |

*: Inserts / Deletes / Misaligned / Frameshifts

#### Analysis details

This analysis was performed with panviral2.64

### NGS Details (UN18_val): Tomato chocolate spot virus (segment RNA2)

#### Assembly

| | |
|---|---|
| **Coverage Length** | 1113 (2 contig(s)) |
| **Depth Of Coverage** | 1610.5 |
| **Number Of Reads** | 14899 |
| **Reads Per Million** | 279.63 rpm (after QC) |
| **Ambiguities** | 0 |
| **Assembly Method** | de novo + reference guided assembly |
| **Consensus Caller** | Bcf Tools |

#### Coverage Map

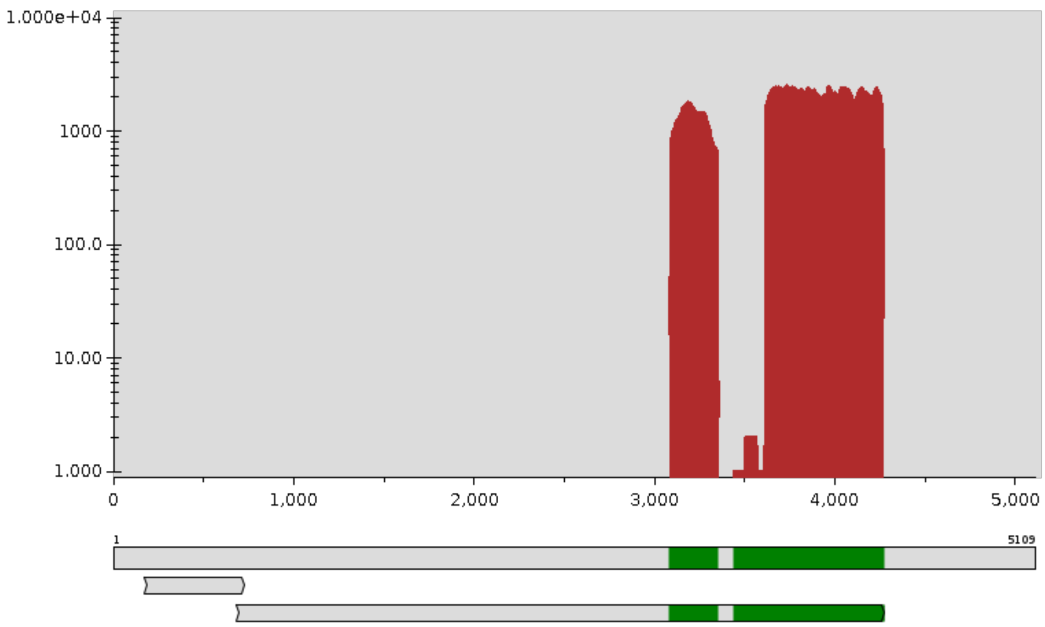

#### Assignment

| | |
|---|---|
| **Type** | Tomato chocolate spot virus (Taxonomy ID: 661101) |
| **Reference Genome** | NC_013076.1 |
| **NT Identity (%)** | 67.028 |
| **AA Identity (%)** | 74.7967 |
| **Number Of Stop Codons** | 1 |
| **Number Of CDS** | 2 |

#### Alignment

| | |
|---|---|
| **Alignment Score** | 739.0 (NT) + 2112.0 (AA) = 2851.0 |
| **Concordance (%)** | 58.9659 |

| Alignment Method | Global, seeded, nucleotide + amino acids (AGA) |
|---|---|

#### Genome Region

Sequence starts at position 3084 and ends at position 4276 relative to NC_013076.1 reference sequence.

#### Alignment Detailed Statistics

| | Begin | End | Coverage | Score | Concordance | Matches | Identities | I/D/M/F* | Stop Codons |
|---|---|---|---|---|---|---|---|---|---|
| **NT** | **3084** | **4276** | **21.8%** | **739** | **33.6%** | **1107 (99.5%)** | **742 (66.7%)** | **0/6** | |
| Mutations: | 3094T>A, 3098A>G, 3100T>C, 3103G>A, 3106A>G, 3109G>C, 3110C>T, 3112C>A, 3115C>T, 3121A>T, 3128T>C, 3130G>T, 3136A>T, 3139A>T, 3140C>T, 3142C>G, 3145G>T, 3148A>T, 3151G>A, 3152A>G, 3153A>C, 3154C>A, 3158A>G, 3164T>G, 3166A>C, 3168A>C, 3169T>A, 3172A>T, 3182T>C, 3184T>A, 3187C>T, 3193T>C, 3196T>C, 3199A>G, 3202G>A, 3211T>A, 3214G>A, 3215A>G, 3224A>C, 3225A>G, 3226G>T, 3231G>A, 3232C>G, 3238A>G, 3242A>T, 3244T>C, 3245G>C, 3247C>A, 3250G>A, 3253C>T, 3263A>T, 3264A>G, 3265G>C, 3268C>T, 3269T>C, 3272T>C, 3274G>C, 3277C>T, 3280T>A, 3283A>G, 3284A>C, 3289G>A, 3292C>T, 3296C>T, 3298G>A, 3301G>A, 3307G>A, 3308G>A, 3309G>A, 3310A>G, 3314T>C, 3315T>A, 3316T>A, 3317T>G, 3319C>T, 3323T>A, 3325A>G, 3328G>C, 3331T>A, 3337T>A, 3343T>G, 3442T>A, 3445G>T, 3454C>T, 3460A>T, 3463C>T, 3468C>A, 3469A>C, 3472A>T, 3475A>T, 3477A>G, 3478G>A, 3482G>T, 3484G>T, 3490C>T, 3499C>T, 3505G>A, 3509G>A, 3511T>C, 3512A>C, 3516C>A, 3517C>T, 3520G>T, 3522G>T, 3526G>T, 3530A>C, 3533G>T, 3535A>G, 3538G>A, 3541C>G, 3544G>A, 3547G>A, 3550T>A, 3553C>T, 3556G>T, 3559A>T, 3560C>A, 3561A>T, 3562A>C, 3564A>T, 3568T>C, 3569G>C, 3570C>A, 3571C>G, 3574G>A, 3577T>G, 3581C>A, 3583G>C, 3584_3589delCCAGTC, 3593C>A, 3594A>C, 3596A>T, 3597G>C, 3598C>T, 3601G>A, 3604G>T, 3605A>T, 3606A>C, 3607T>C, 3610G>C, 3611T>C, 3612C>A, 3613C>G, 3614A>C, 3617A>C, 3622G>A, 3628T>C, 3631A>T, 3637T>A, 3640A>T, 3643C>T, 3646G>A, 3652C>T, 3655C>T, 3656A>T, 3657T>C, 3658G>T, 3660G>C, 3671G>A, 3672A>C, 3673A>C, 3674G>A, 3675A>G, 3676A>C, 3677A>C, 3679G>C, 3680T>C, 3683C>G, 3685T>G, 3688A>T, 3689A>C, 3691A>T, 3703C>G, 3709A>T, 3712G>A, 3715C>A, 3718T>C, 3719G>T, 3720A>C, 3722G>C, 3724T>C, 3725C>A, 3726A>C, 3727G>C, 3731G>C, 3733T>C, 3745A>C, 3751C>G, 3757T>C, 3760C>T, 3772T>C, 3773A>T, 3775A>T, 3778C>A, 3784T>C, 3787C>T, 3793A>T, 3799C>T, 3802C>T, 3803A>G, 3805T>G, 3808T>C, 3809T>A, 3812A>C, 3814A>C, 3815G>A, 3817T>C, 3818G>A, 3820G>C, 3823C>T, 3824C>A, 3826A>G, 3829T>C, 3832G>A, 3833T>G, 3834C>G, 3836A>T, 3837G>C, 3838C>T, 3839C>A, 3840C>A, 3844T>C, 3845G>A, 3847C>T, 3850A>T, 3853C>T, 3854A>G, 3856T>A, 3859G>T, 3865A>G, 3869C>T, 3871T>G, 3875T>A, 3880T>A, 3883T>C, 3889A>T, 3893C>T, 3895A>T, 3896G>T, 3897C>A, 3898A>T, 3901C>T, 3902C>G, 3905A>G, 3906G>C, 3907T>A, 3908A>G, 3913A>G, 3914A>G, 3915A>C, 3916T>A, 3917T>C, 3922A>G, 3923G>C, 3934A>C, 3937A>T, 3940A>T, 3946G>A, 3952C>T, 3955T>C, 3961T>C, 3964T>A, 3970A>C, 3973A>T, 3979A>T, 3985T>C, 3989A>G, 3990C>T, 3994T>G, 4000T>C, 4004G>A, 4013T>C, 4014G>C, 4016A>C, 4021A>G, 4022C>T, 4024C>T, 4028C>A, 4029G>A, 4030C>G, 4031A>G, 4032T>C, 4033G>A, 4034A>C, 4039G>A, 4042C>T, 4043T>G, 4044C>A, 4045A>C, 4048A>T, 4050C>A, 4051T>G, 4052C>A, 4053A>G, 4054A>T, 4058A>C, 4060A>T, 4063A>T, 4064A>G, 4066G>T, 4070C>T, 4072T>G, 4078T>C, 4080G>A, 4081G>A, 4082C>T, 4084C>G, 4088A>C, 4089A>T, 4090C>T, 4091T>C, 4093A>C, 4094A>T, 4096A>G, 4099A>C, 4105T>A, 4108C>A, 4116G>A, 4117G>C, 4118C>T, 4120T>A, 4121G>A, 4123G>T, 4126T>C, 4129A>C, 4132A>T, 4135G>A, 4138T>C, 4139C>A, 4141C>G, 4142A>G, 4144C>T, 4145T>A, 4146C>A, 4147C>G, 4150G>A, 4153T>G, 4156T>A, 4159C>T, 4162C>T, 4164A>G, 4165A>G, 4166C>T, 4168G>A, 4172G>C, 4174G>T, 4177A>C, 4187T>C, 4188C>T, 4189T>G, 4192C>T, 4195C>T, 4198C>T, 4201G>A, 4204G>A, 4206T>A, 4213G>T, 4214A>G, 4216C>T, 4225A>G, 4228C>T, 4234T>C, 4255T>A, 4258G>A, 4261C>G, 4267T>C | | | | | | | | |

##### CDS

| | Begin | End | Coverage | Score | Concordance | Matches | Identities | I/D/M/F* | Stop Codons |
|---|---|---|---|---|---|---|---|---|---|
| ToChV_s2_gp2 | 801 | 1197 | 31.0% | 2112 | 79.8% | 369 (99.5%) | 276 (74.4%) | 0/2/0/0 | 1 |
| Protein mutations: | I805V (3098A>G 3100T>C), N823A (3152A>G 3153A>C 3154C>A), K825E (3158A>G), S827A (3164T>G 3166A>C), N828T (3168A>C 3169T>A), F833L (3182T>C 3184T>A), M844V (3215A>G), K847R (3224A>C 3225A>G 3226G>T), R849Q (3231G>A 3232C>G), T853S (3242A>T 3244T>C), V854L (3245G>C 3247C>A), K860C (3263A>T 3264A>G 3265G>C), G875K (3308G>A 3309G>A 3310A>G), F877Q (3314T>C 3315T>A 3316T>A), S878A (3317T>G 3319C>T), L880M (3323T>A 3325A>G), T928N (3468C>A 3469A>C), K929N (3472A>T), K931R (3477A>G 3478G>A), A933S (3482G>T 3484G>T), D942N (3509G>A 3511T>C), M943L (3512A>C), T944N (3516C>A 3517C>T), Q945H (3520G>T), R946L (3522G>T), K947N (3526G>T), I949L (3530A>C), E950* (3533G>T 3535A>G), D955E (3550T>A), E958D (3559A>T), Q959I (3560C>A 3561A>T 3562A>C), Y960F (3564A>T), A962Q (3569G>C 3570C>A 3571C>G), Q966N (3581C>A 3583G>C), P967_V968del (3584_3589delCCAGTC), Q970T (3593C>A 3594A>C), N974S (3605A>T 3606A>C 3607T>C), L975F (3610G>C), S976Q (3611T>C 3612C>A 3613C>G), I977L (3614A>C), M991S (3656A>T 3657T>C 3658G>T), G992A (3660G>C), E996T (3671G>A 3672A>C 3673A>C), E997S (3674G>A 3675A>G 3676A>C), L1000V (3683C>G 3685T>G), I1002L (3689A>C 3691A>T), E1012S (3719G>T 3720A>C), A1013P (3722G>C 3724T>C), Q1014T (3725C>A 3726A>C 3727G>C), A1016P (3731G>C 3733T>C), T1030S (3773A>T 3775A>T), N1040E (3803A>G 3805T>G), S1042T (3809T>A), I1043L (3812A>C 3814A>C), V1044I (3815G>A 3817T>C), V1045I (3818G>A 3820G>C), S1050G (3833T>G 3834C>G), P1052N (3839C>A 3840C>A), V1054I (3845G>A 3847C>T), I1057V (3854A>G 3856T>A), S1064T (3875T>A), P1070S (3893C>T 3895A>T), A1071Y (3896G>T 3897C>A 3898A>T), L1073V (3902C>G), S1074A (3905A>G 3906G>C 3907T>A), K1075E (3908A>G), N1077A (3914A>G 3915A>C 3916T>A), Y1078H (3917T>C), V1080L (3923G>C), T1102V (3989A>G 3990C>T), E1107K (4004G>A), C1110P (4013T>C 4014G>C), H1113Y (4022C>T 4024C>T), R1115K (4028C>A 4029G>A 4030C>G), M1116A (4031A>G 4032T>C 4033G>A), S1120D (4043T>G 4044C>A 4045A>C), K1121N (4048A>T), T1122K (4050C>A 4051T>G), Q1123S (4052C>A 4053A>G 4054A>T), M1127V (4064A>G 4066G>T), R1132K (4080G>A 4081G>A), N1135L (4088A>C 4089A>T 4090C>T), I1137L (4094A>T 4096A>G), G1144D (4116G>A 4117G>C), V1146I (4121G>A 4123G>T), L1152M (4139C>A 4141C>G), I1153V (4142A>G 4144C>T), S1154K (4145T>A 4146C>A 4147C>G), K1160R (4164A>G 4165A>G), V1163L (4172G>C 4174G>T), S1168L (4187T>C 4188C>T 4189T>G), F1174Y (4206T>A), N1177D (4214A>G 4216C>T) | | | | | | | | |
| Codon mutations: | ATT803ATA (3094T>A), ATT805GTC (3098A>G 3100T>C), GAG806GAA (3103G>A), AAA807AAG (3106A>G), ACG808ACC (3109G>C), CTC809TTA (3110C>T 3112C>A), TTC810TTT (3115C>T), TCA812TCT (3121A>T), TTG815CTT (3128T>C 3130G>T), ATA817ATT (3136A>T), GGA818GGT (3139A>T), CTC819TTG (3140C>T 3142C>G), GGG820GGT (3145G>T), ACA821ACT (3148A>T), CTG822CTA (3151G>A), AAC823GCA (3152A>G 3153A>C 3154C>A), AAA825GAA (3158A>G), TCA827GCC (3164T>G 3166A>C), AAT828ACA (3168A>C 3169T>A), GCA829GCT (3172A>T), TTT833CTA (3182T>C 3184T>A), AAC834AAT (3187C>T), CCT836CCC (3193T>C), CAT837CAC (3196T>C), GTA838GTG (3199A>G), GTG839GTA (3202G>A), CTT842CTA (3211T>A), GAG843GAA (3214G>A), ATG844GTG (3215A>G), AAG847CGT (3224A>C 3225A>G 3226G>T), CGC849CAG (3231G>A 3232C>G), AAA851AAG (3238A>G), ACT853TCC (3242A>T 3244T>C), GTC854CTA (3245G>C 3247C>A), ACG855ACA (3250G>A), AAC856AAT (3253C>T), AAG860TGC (3263A>T 3264A>G 3265G>C), AAC861AAT (3268C>T), TTG862CTG (3269T>C), TTG863CTC (3272T>C 3274G>C), TCC864TCT (3277C>T), ACT865ACA (3280T>A), GGA866GGG (3283A>G), AGG867CGG (3284A>C), AAG868AAA (3289G>A), AGC869AGT (3292C>T), CTG871TTA (3296C>T 3298G>A), CCG872CCA (3301G>A), CCG874CCA (3307G>A), GGA875AAG (3308G>A 3309G>A 3310A>G), TTT877CAA (3314T>C 3315T>A 3316T>A), TCC878GCT (3317T>G 3319C>T), TTA880ATG (3323T>A 3325A>G), CGG881CGC (3328G>C), CTT882CTA (3331T>A), GCT884GCA (3337T>A), GTT886GTG (3343T>G), TCT919TCA (3442T>A), GTG920GTT (3445G>T), ATC923ATT (3454C>T), GGA925GGT (3460A>T), CAC926CAT (3463C>T), ACA928AAC (3468C>A 3469A>C), AAA929AAT (3472A>T), GGA930GGT (3475A>T), AAG931AGA (3477A>G 3478G>A), GCG933TCT (3482G>T 3484G>T), GAC935GAT (3490C>T), TTC938TTT (3499C>T), GAG940GAA (3505G>A), GAT942AAC (3509G>A 3511T>C), ATG943CTG (3512A>C), ACC944AAT (3516C>A 3517C>T), CAG945CAT (3520G>T), CGC946CTC (3522G>T), AAG947AAT (3526G>T), ATT949CTT (3530A>C), GAA950TAG (3533G>T 3535A>G), AAG951AAA (3538G>A), CTC952CTG (3541C>G), AGG953AGA (3544G>A), AAG954AAA (3547G>A), GAT955GAA (3550T>A), AAC956AAT (3553C>T), GTG957GTT (3556G>T), GAA958GAT (3559A>T), CAA959ATC (3560C>A 3561A>T 3562A>C), TAT960TTT (3564A>T), GAT961GAC (3568T>C), GCC962CAG (3569G>C 3570C>A 3571C>G), AAG963AAA (3574G>A), GGT964GGG (3577T>G), CAG966AAC (3581C>A 3583G>C), CCA967_GTC968del (3584_3589delCCAGTC), CAA970ACA (3593C>A 3594A>C), AGC971TCT (3596A>T 3597G>C 3598C>T), CAG972CAA (3601G>A), GTG973GTT (3604G>T), AAT974TCC (3605A>T 3606A>C 3607T>C), TTG975TTC (3610G>C), TCC976CAG (3611T>C 3612C>A 3613C>G), ATT977CTT (3614A>C), AGG978CGG (3617A>C), GAG979GAA (3622G>A), TTT981TTC (3628T>C), TCA982TCT (3631A>T), GGT984GGA (3637T>A), GCA985GCT (3640A>T), GTC986GTT (3643C>T), CAG987CAA (3646G>A), TTC989TTT (3652C>T), TGC990TGT (3655C>T), ATG991TCT (3656A>T 3657T>C 3658G>T), GGC992GCC (3660G>C), GAA996ACC (3671G>A 3672A>C 3673A>C), GAA997AGC (3674G>A 3675A>G 3676A>C), AGG998CGC (3677A>C 3679G>C), TTG999CTG (3680T>C), CTT1000GTG (3683C>G 3685T>G), GTA1001GTT (3688A>T), ATA1002CTT (3689A>C 3691A>T), CCC1006CCG (3703C>G), TCA1008TCT (3709A>T), GTG1009GTA (3712G>A), CGC1010CGA (3715C>A), TTT1011TTC (3718T>C), GAA1012TCA (3719G>T 3720A>C), GCT1013CCC (3722G>C 3724T>C), CAG1014ACC (3725C>A 3726A>C 3727G>C), GCT1016CCC (3731G>C 3733T>C), GCA1020GCC (3745A>C), ACC1022ACG (3751C>G), CCT1024CCC (3757T>C), TTC1025TTT (3760C>T), TGT1029TGC (3772T>C), ACA1030TCT (3773A>T 3775A>T), TCC1031TCA (3778C>A), TGT1033TGC (3784T>C), TAC1034TAT (3787C>T), TCA1036TCT (3793A>T), AGC1038AGT (3799C>T), CTC1039CTT (3802C>T), AAT1040GAG (3803A>G 3805T>G), TAT1041TAC (3808T>C), TCC1042ACC (3809T>A), ATA1043CTC (3812A>C 3814A>C), GTT1044ATC (3815G>A 3817T>C), GTG1045ATC (3818G>A 3820G>C), CAC1046CAT (3823C>T), CGA1047AGG (3824C>A 3826A>G), GTT1048GTC (3829T>C), CAG1049CAA (3832G>A), TCC1050GGC (3833T>G 3834C>G), AGC1051TCT (3836A>T 3837G>C 3838C>T), CCT1052AAT (3839C>A 3840C>A), AAT1053AAC (3844T>C), GTC1054ATT (3845G>A 3847C>T), GGA1055GGT (3850A>T), GGC1056GGT (3853C>T), ATT1057GTA (3854A>G 3856T>A), CTG1058CTT (3859G>T), GTA1060GTG (3865A>G), CTT1062TTG (3869C>T 3871T>G), TCC1064ACC (3875T>A), TCT1065TCA (3880T>A), GGT1066GGC (3883T>C), CCA1068CCT (3889A>T), CCA1070TCT (3893C>T 3895A>T), GCA1071TAT (3896G>T 3897C>A 3898A>T), GGC1072GGT (3901C>T), CTG1073GTG (3902C>G), AGT1074GCA (3905A>G 3906G>C 3907T>A), AAG1075GAG (3908A>G), GGA1076GGG (3913A>G), AAT1077GCA (3914A>G 3915A>C 3916T>A), TAT1078CAT (3917T>C), GTA1079GTG (3922A>G), GTT1080CTT (3923G>C), GGA1083GGC (3934A>C), GGA1084GGT (3937A>T), GGA1085GGT (3940A>T), AAG1087AAA (3946G>A), AAC1089AAT (3952C>T), TTT1090TTC (3955T>C), TAT1092TAC (3961T>C), GGT1093GGA (3964T>A), ACA1095ACC (3970A>C), ACA1096ACT (3973A>T), ATA1098ATT (3979A>T), TCT1100TCC (3985T>C), ACG1102GTG (3989A>G 3990C>T), GTT1103GTG (3994T>G), GAT1105GAC (4000T>C), GAA1107AAA (4004G>A), TGT1110CCT (4013T>C 4014G>C), AGG1111CGG (4016A>C), AGA1112AGG (4021A>G), CAC1113TAT (4022C>T 4024C>T), CGC1115AAG (4028C>A 4029G>A 4030C>G), ATG1116GCA (4031A>G 4032T>C 4033G>A), AGA1117CGA (4034A>C), GAG1118GAA (4039G>A), TTC1119TTT (4042C>T), TCA1120GAC (4043T>G 4044C>A 4045A>C), AAA1121AAT (4048A>T), ACT1122AAG (4050C>A 4051T>G), CAA1123AGT (4052C>A 4053A>G 4054A>T), AGA1125CGT (4058A>C 4060A>T), ATA1126ATT (4063A>T), ATG1127GTT (4064A>G 4066G>T), CTT1129TTG (4070C>T 4072T>G), GAT1131GAC (4078T>C), AGG1132AAA (4080G>A 4081G>A), CTC1133TTG (4082C>T 4084C>G), AAC1135CTT (4088A>C 4089A>T 4090C>T), TTA1136CTC (4091T>C 4093A>C), ATA1137TTG (4094A>T 4096A>G), ATA1138ATC (4099A>C), CTT1140CTA (4105T>A), CCC1141CCA (4108C>A), GGG1144GAC (4116G>A 4117G>C), CTT1145TTA (4118C>T 4120T>A), GTG1146ATT (4121G>A 4123G>T), AGT1147AGC (4126T>C), TCA1148TCC (4129A>C), ATA1149ATT (4132A>T), GAG1150GAA (4135G>A), ATT1151ATC (4138T>C), CTC1152ATG (4139C>A 4141C>G), ATC1153GTT (4142A>G 4144C>T), TCC1154AAG (4145T>A 4146C>A 4147C>G), CCG1155CCA (4150G>A), GGT1156GGG (4153T>G), CCT1157CCA (4156T>A), GAC1158GAT (4159C>T), TTC1159TTT (4162C>T), AAA1160AGG (4164A>G 4165A>G), CTG1161TTA (4166C>T 4168G>A), GTG1163CTT (4172G>C 4174G>T), GCA1164GCC (4177A>C), TCT1168CTG (4187T>C 4188C>T 4189T>G), GCC1169GCT (4192C>T), AAC1170AAT (4195C>T), CAC1171CAT (4198C>T), GAG1172GAA (4201G>A), AAG1173AAA (4204G>A), TTT1174TAT (4206T>A), GGG1176GGT (4213G>T), AAC1177GAT (4214A>G 4216C>T), ACA1180ACG (4225A>G), CAC1181CAT (4228C>T), TAT1183TAC (4234T>C), TCT1190TCA (4255T>A), GAG1191GAA (4258G>A), CTC1192CTG (4261C>G), GAT1194GAC (4267T>C) | | | | | | | | |

##### Proteins

| | Begin | End | Coverage | Score | Concordance | Matches | Identities | I/D/M/F* | Stop Codons |
|---|---|---|---|---|---|---|---|---|---|
| polyprotein (YP_003097231.1) | 801 | 1197 | 31.0% | 2112 | 79.8% | 369 (99.5%) | 276 (74.4%) | 0/2/0/0 | 1 |

| | Begin | End | Coverage | Score | Concordance | Matches | Identities | I/D/M/F* | Stop Codons |
|---|---|---|---|---|---|---|---|---|---|
| **NT** | **3084** | **4276** | **21.8%** | **739** | **33.6%** | **1107 (99.5%)** | **742 (66.7%)** | **0/6** | |
| Protein mutations: | I805V (3098A>G 3100T>C), N823A (3152A>G 3153A>C 3154C>A), K825E (3158A>G), S827A (3164T>G 3166A>C), N828T (3168A>C 3169T>A), F833L (3182T>C 3184T>A), M844V (3215A>G), K847R (3224A>C 3225A>G 3226G>T), R849Q (3231G>A 3232C>G), T853S (3242A>T 3244T>C), V854L (3245G>C 3247C>A), K860C (3263A>T 3264A>G 3265G>C), G875K (3308G>A 3309G>A 3310A>G), F877Q (3314T>C 3315T>A 3316T>A), S878A (3317T>G 3319C>T), L880M (3323T>A 3325A>G), T928N (3468C>A 3469A>C), K929N (3472A>T), K931R (3477A>G 3478G>A), A933S (3482G>T 3484G>T), D942N (3509G>A 3511T>C), M943L (3512A>C), T944N (3516C>A 3517C>T), Q945H (3520G>T), R946L (3522G>T), K947N (3526G>T), I949L (3530A>C), E950* (3533G>T 3535A>G), D955E (3550T>A), E958D (3559A>T), Q959I (3560C>A 3561A>T 3562A>C), Y960F (3564A>T), A962Q (3569G>C 3570C>A 3571C>G), Q966N (3581C>A 3583G>C), P967_V968del (3584_3589delCCAGTC), Q970T (3593C>A 3594A>C), N974S (3605A>T 3606A>C 3607T>C), L975F (3610G>C), S976Q (3611T>C 3612C>A 3613C>G), I977L (3614A>C), M991S (3656A>T 3657T>C 3658G>T), G992A (3660G>C), E996T (3671G>A 3672A>C 3673A>C), E997S (3674G>A 3675A>G 3676A>C), L1000V (3683C>G 3685T>G), I1002L (3689A>C 3691A>T), E1012S (3719G>T 3720A>C), A1013P (3722G>C 3724T>C), Q1014T (3725C>A 3726A>C 3727G>C), A1016P (3731G>C 3733T>C), T1030S (3773A>T 3775A>T), N1040E (3803A>G 3805T>G), S1042T (3809T>A), I1043L (3812A>C 3814A>C), V1044I (3815G>A 3817T>C), V1045I (3818G>A 3820G>C), S1050G (3833T>G 3834C>G), P1052N (3839C>A 3840C>A), V1054I (3845G>A 3847C>T), I1057V (3854A>G 3856T>A), S1064T (3875T>A), P1070S (3893C>T 3895A>T), A1071Y (3896G>T 3897C>A 3898A>T), L1073V (3902C>G), S1074A (3905A>G 3906G>C 3907T>A), K1075E (3908A>G), N1077A (3914A>G 3915A>C 3916T>A), Y1078H (3917T>C), V1080L (3923G>C), T1102V (3989A>G 3990C>T), E1107K (4004G>A), C1110P (4013T>C 4014G>C), H1113Y (4022C>T 4024C>T), R1115K (4028C>A 4029G>A 4030C>G), M1116A (4031A>G 4032T>C 4033G>A), S1120D (4043T>G 4044C>A 4045A>C), K1121N (4048A>T), T1122K (4050C>A 4051T>G), Q1123S (4052C>A 4053A>G 4054A>T), M1127V (4064A>G 4066G>T), R1132K (4080G>A 4081G>A), N1135L (4088A>C 4089A>T 4090C>T), I1137L (4094A>T 4096A>G), G1144D (4116G>A 4117G>C), V1146I (4121G>A 4123G>T), L1152M (4139C>A 4141C>G), I1153V (4142A>G 4144C>T), S1154K (4145T>A 4146C>A 4147C>G), K1160R (4164A>G 4165A>G), V1163L (4172G>C 4174G>T), S1168L (4187T>C 4188C>T 4189T>G), F1174Y (4206T>A), N1177D (4214A>G 4216C>T) | | | | | | | | |
| Codon mutations: | ATT803ATA (3094T>A), ATT805GTC (3098A>G 3100T>C), GAG806GAA (3103G>A), AAA807AAG (3106A>G), ACG808ACC (3109G>C), CTC809TTA (3110C>T 3112C>A), TTC810TTT (3115C>T), TCA812TCT (3121A>T), TTG815CTT (3128T>C 3130G>T), ATA817ATT (3136A>T), GGA818GGT (3139A>T), CTC819TTG (3140C>T 3142C>G), GGG820GGT (3145G>T), ACA821ACT (3148A>T), CTG822CTA (3151G>A), AAC823GCA (3152A>G 3153A>C 3154C>A), AAA825GAA (3158A>G), TCA827GCC (3164T>G 3166A>C), AAT828ACA (3168A>C 3169T>A), GCA829GCT (3172A>T), TTT833CTA (3182T>C 3184T>A), AAC834AAT (3187C>T), CCT836CCC (3193T>C), CAT837CAC (3196T>C), GTA838GTG (3199A>G), GTG839GTA (3202G>A), CTT842CTA (3211T>A), GAG843GAA (3214G>A), ATG844GTG (3215A>G), AAG847CGT (3224A>C 3225A>G 3226G>T), CGC849CAG (3231G>A 3232C>G), AAA851AAG (3238A>G), ACT853TCC (3242A>T 3244T>C), GTC854CTA (3245G>C 3247C>A), ACG855ACA (3250G>A), AAC856AAT (3253C>T), AAG860TGC (3263A>T 3264A>G 3265G>C), AAC861AAT (3268C>T), TTG862CTG (3269T>C), TTG863CTC (3272T>C 3274G>C), TCC864TCT (3277C>T), ACT865ACA (3280T>A), GGA866GGG (3283A>G), AGG867CGG (3284A>C), AAG868AAA (3289G>A), AGC869AGT (3292C>T), CTG871TTA (3296C>T 3298G>A), CCG872CCA (3301G>A), CCG874CCA (3307G>A), GGA875AAG (3308G>A 3309G>A 3310A>G), TTT877CAA (3314T>C 3315T>A 3316T>A), TCC878GCT (3317T>G 3319C>T), TTA880ATG (3323T>A 3325A>G), CGG881CGC (3328G>C), CTT882CTA (3331T>A), GCT884GCA (3337T>A), GTT886GTG (3343T>G), TCT919TCA (3442T>A), GTG920GTT (3445G>T), ATC923ATT (3454C>T), GGA925GGT (3460A>T), CAC926CAT (3463C>T), ACA928AAC (3468C>A 3469A>C), AAA929AAT (3472A>T), GGA930GGT (3475A>T), AAG931AGA (3477A>G 3478G>A), GCG933TCT (3482G>T 3484G>T), GAC935GAT (3490C>T), TTC938TTT (3499C>T), GAG940GAA (3505G>A), GAT942AAC (3509G>A 3511T>C), ATG943CTG (3512A>C), ACC944AAT (3516C>A 3517C>T), CAG945CAT (3520G>T), CGC946CTC (3522G>T), AAG947AAT (3526G>T), ATT949CTT (3530A>C), GAA950TAG (3533G>T 3535A>G), AAG951AAA (3538G>A), CTC952CTG (3541C>G), AGG953AGA (3544G>A), AAG954AAA (3547G>A), GAT955GAA (3550T>A), AAC956AAT (3553C>T), GTG957GTT (3556G>T), GAA958GAT (3559A>T), CAA959ATC (3560C>A 3561A>T 3562A>C), TAT960TTT (3564A>T), GAT961GAC (3568T>C), GCC962CAG (3569G>C 3570C>A 3571C>G), AAG963AAA (3574G>A), GGT964GGG (3577T>G), CAG966AAC (3581C>A 3583G>C), CCA967_GTC968del (3584_3589delCCAGTC), CAA970ACA (3593C>A 3594A>C), AGC971TCT (3596A>T 3597G>C 3598C>T), CAG972CAA (3601G>A), GTG973GTT (3604G>T), AAT974TCC (3605A>T 3606A>C 3607T>C), TTG975TTC (3610G>C), TCC976CAG (3611T>C 3612C>A 3613C>G), ATT977CTT (3614A>C), AGG978CGG (3617A>C), GAG979GAA (3622G>A), TTT981TTC (3628T>C), TCA982TCT (3631A>T), GGT984GGA (3637T>A), GCA985GCT (3640A>T), GTC986GTT (3643C>T), CAG987CAA (3646G>A), TTC989TTT (3652C>T), TGC990TGT (3655C>T), ATG991TCT (3656A>T 3657T>C 3658G>T), GGC992GCC (3660G>C), GAA996ACC (3671G>A 3672A>C 3673A>C), GAA997AGC (3674G>A 3675A>G 3676A>C), AGG998CGC (3677A>C 3679G>C), TTG999CTG (3680T>C), CTT1000GTG (3683C>G 3685T>G), GTA1001GTT (3688A>T), ATA1002CTT (3689A>C 3691A>T), CCC1006CCG (3703C>G), TCA1008TCT (3709A>T), GTG1009GTA (3712G>A), CGC1010CGA (3715C>A), TTT1011TTC (3718T>C), GAA1012TCA (3719G>T 3720A>C), GCT1013CCC (3722G>C 3724T>C), CAG1014ACC (3725C>A 3726A>C 3727G>C), GCT1016CCC (3731G>C 3733T>C), GCA1020GCC (3745A>C), ACC1022ACG (3751C>G), CCT1024CCC (3757T>C), TTC1025TTT (3760C>T), TGT1029TGC (3772T>C), ACA1030TCT (3773A>T 3775A>T), TCC1031TCA (3778C>A), TGT1033TGC (3784T>C), TAC1034TAT (3787C>T), TCA1036TCT (3793A>T), AGC1038AGT (3799C>T), CTC1039CTT (3802C>T), AAT1040GAG (3803A>G 3805T>G), TAT1041TAC (3808T>C), TCC1042ACC (3809T>A), ATA1043CTC (3812A>C 3814A>C), GTT1044ATC (3815G>A 3817T>C), GTG1045ATC (3818G>A 3820G>C), CAC1046CAT (3823C>T), CGA1047AGG (3824C>A 3826A>G), GTT1048GTC (3829T>C), CAG1049CAA (3832G>A), TCC1050GGC (3833T>G 3834C>G), AGC1051TCT (3836A>T 3837G>C 3838C>T), CCT1052AAT (3839C>A 3840C>A), AAT1053AAC (3844T>C), GTC1054ATT (3845G>A 3847C>T), GGA1055GGT (3850A>T), GGC1056GGT (3853C>T), ATT1057GTA (3854A>G 3856T>A), CTG1058CTT (3859G>T), GTA1060GTG (3865A>G), CTT1062TTG (3869C>T 3871T>G), TCC1064ACC (3875T>A), TCT1065TCA (3880T>A), GGT1066GGC (3883T>C), CCA1068CCT (3889A>T), CCA1070TCT (3893C>T 3895A>T), GCA1071TAT (3896G>T 3897C>A 3898A>T), GGC1072GGT (3901C>T), CTG1073GTG (3902C>G), AGT1074GCA (3905A>G 3906G>C 3907T>A), AAG1075GAG (3908A>G), GGA1076GGG (3913A>G), AAT1077GCA (3914A>G 3915A>C 3916T>A), TAT1078CAT (3917T>C), GTA1079GTG (3922A>G), GTT1080CTT (3923G>C), GGA1083GGC (3934A>C), GGA1084GGT (3937A>T), GGA1085GGT (3940A>T), AAG1087AAA (3946G>A), AAC1089AAT (3952C>T), TTT1090TTC (3955T>C), TAT1092TAC (3961T>C), GGT1093GGA (3964T>A), ACA1095ACC (3970A>C), ACA1096ACT (3973A>T), ATA1098ATT (3979A>T), TCT1100TCC (3985T>C), ACG1102GTG (3989A>G 3990C>T), GTT1103GTG (3994T>G), GAT1105GAC (4000T>C), GAA1107AAA (4004G>A), TGT1110CCT (4013T>C 4014G>C), AGG1111CGG (4016A>C), AGA1112AGG (4021A>G), CAC1113TAT (4022C>T 4024C>T), CGC1115AAG (4028C>A 4029G>A 4030C>G), ATG1116GCA (4031A>G 4032T>C 4033G>A), AGA1117CGA (4034A>C), GAG1118GAA (4039G>A), TTC1119TTT (4042C>T), TCA1120GAC (4043T>G 4044C>A 4045A>C), AAA1121AAT (4048A>T), ACT1122AAG (4050C>A 4051T>G), CAA1123AGT (4052C>A 4053A>G 4054A>T), AGA1125CGT (4058A>C 4060A>T), ATA1126ATT (4063A>T), ATG1127GTT (4064A>G 4066G>T), CTT1129TTG (4070C>T 4072T>G), GAT1131GAC (4078T>C), AGG1132AAA (4080G>A 4081G>A), CTC1133TTG (4082C>T 4084C>G), AAC1135CTT (4088A>C 4089A>T 4090C>T), TTA1136CTC (4091T>C 4093A>C), ATA1137TTG (4094A>T 4096A>G), ATA1138ATC (4099A>C), CTT1140CTA (4105T>A), CCC1141CCA (4108C>A), GGG1144GAC (4116G>A 4117G>C), CTT1145TTA (4118C>T 4120T>A), GTG1146ATT (4121G>A 4123G>T), AGT1147AGC (4126T>C), TCA1148TCC (4129A>C), ATA1149ATT (4132A>T), GAG1150GAA (4135G>A), ATT1151ATC (4138T>C), CTC1152ATG (4139C>A 4141C>G), ATC1153GTT (4142A>G 4144C>T), TCC1154AAG (4145T>A 4146C>A 4147C>G), CCG1155CCA (4150G>A), GGT1156GGG (4153T>G), CCT1157CCA (4156T>A), GAC1158GAT (4159C>T), TTC1159TTT (4162C>T), AAA1160AGG (4164A>G 4165A>G), CTG1161TTA (4166C>T 4168G>A), GTG1163CTT (4172G>C 4174G>T), GCA1164GCC (4177A>C), TCT1168CTG (4187T>C 4188C>T 4189T>G), GCC1169GCT (4192C>T), AAC1170AAT (4195C>T), CAC1171CAT (4198C>T), GAG1172GAA (4201G>A), AAG1173AAA (4204G>A), TTT1174TAT (4206T>A), GGG1176GGT (4213G>T), AAC1177GAT (4214A>G 4216C>T), ACA1180ACG (4225A>G), CAC1181CAT (4228C>T), TAT1183TAC (4234T>C), TCT1190TCA (4255T>A), GAG1191GAA (4258G>A), CTC1192CTG (4261C>G), GAT1194GAC (4267T>C) | | | | | | | | |

*: Inserts / Deletes / Misaligned / Frameshifts

#### Analysis details

This analysis was performed with panviral2.64

### NGS Details (UN18_val): Tokyovirus A1

#### Assembly

| | |
|---|---|
| **Coverage Length** | 164 (1 contig(s)) |
| **Depth Of Coverage** | 4800.1 |
| **Number Of Reads** | 12764 |
| **Reads Per Million** | 239.56 rpm (after QC) |
| **Ambiguities** | 0 |
| **Assembly Method** | de novo + reference guided assembly |
| **Consensus Caller** | Bcf Tools |

#### Coverage Map

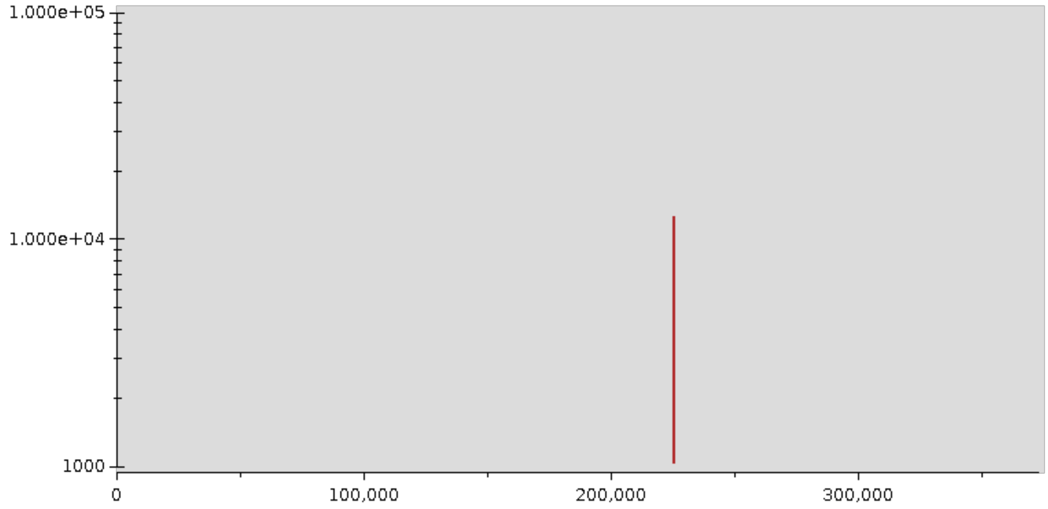

#### Assignment

| | |
|---|---|
| **Type** | Tokyovirus A1 (Taxonomy ID: 1826170) |
| **Reference Genome** | NC_030230.1 |
| **NT Identity (%)** | 76.8293 |
| **AA Identity (%)** | 90.7407 |
| **Number Of Stop Codons** | 0 |
| **Number Of CDS** | 470 |

#### Alignment

| | |
|---|---|
| **Alignment Score** | 176.0 (NT) + 332.0 (AA) = 508.0 |
| **Concordance (%)** | 74.4868 |
| **Alignment Method** | Local, heuristic, nucleotide (BLASTN) |

#### Genome Region

Sequence starts at position 225492 and ends at position 225655 relative to NC_030230.1 reference sequence.

#### Alignment Detailed Statistics

| | Begin | End | Coverage | Score | Concordance | Matches | Identities | I/D/M/F* | Stop Codons |
|---|---|---|---|---|---|---|---|---|---|
| **NT** | **225492** | **225655** | **0.1%** | **176** | **53.7%** | **164 (100%)** | **126 (76.8%)** | **0/0** | |

Mutations: 225498T>G, 225500T>G, 225507A>G, 225510G>A, 225513G>C, 225516A>T, 225522T>C, 225525T>C, 225528T>C, 225531G>A, 225533C>T, 225534A>G, 225540A>G, 225545C>A, 225552T>G, 225554T>G, 225555C>G, 225561T>C, 225564T>A, 225573C>G, 225585A>C, 225587G>A, 225588T>G, 225591T>C, 225594T>C, 225600G>T, 225603C>G, 225609A>G, 225618G>A, 225621T>C, 225630T>G, 225631T>C, 225632G>T, 225636C>G, 225640G>T, 225641C>T, 225642T>G, 225651G>A

*: Inserts / Deletes / Misaligned / Frameshifts

#### Analysis details

This analysis was performed with panviral2.64

### NGS Details (UN18_val): Leucotheavirus sp4

#### Assembly

| | |
|---|---|
| **Coverage Length** | 863 (1 contig(s)) |
| **Depth Of Coverage** | 880.3 |
| **Number Of Reads** | 6064 |
| **Reads Per Million** | 113.81 rpm (after QC) |
| **Ambiguities** | 0 |
| **Assembly Method** | de novo + reference guided assembly |
| **Consensus Caller** | Bcf Tools |

#### Coverage Map

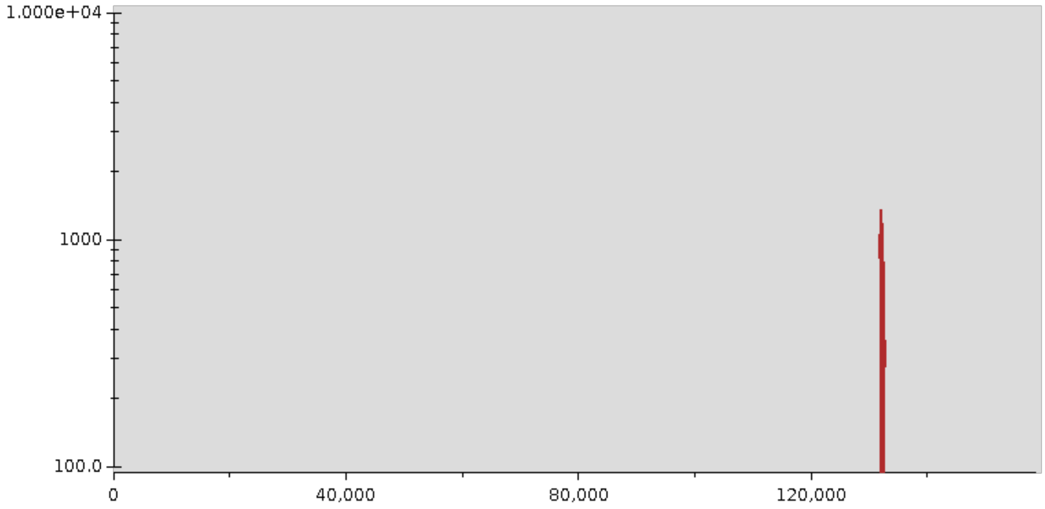

#### Assignment

| | |
|---|---|
| **Type** | Leucotheavirus sp4 (Taxonomy ID: 2734113) |
| **Reference Genome** | NC_048102.1 |
| **NT Identity (%)** | 75.898 |
| **AA Identity (%)** | 85.4167 |
| **Number Of Stop Codons** | 0 |
| **Number Of CDS** | 194 |

#### Alignment

| | |
|---|---|
| **Alignment Score** | 894.0 (NT) + 1851.0 (AA) = 2745.0 |
| **Concordance (%)** | 72.7731 |
| **Alignment Method** | Local, heuristic, nucleotide (BLASTN) |

#### Genome Region

Sequence starts at position 131800 and ends at position 132662 relative to NC_048102.1 reference sequence.

#### Alignment Detailed Statistics

| | Begin | End | Coverage | Score | Concordance | Matches | Identities | I/D/M/F* | Stop Codons |
|---|---|---|---|---|---|---|---|---|---|
| NT | 131800 | 132662 | 0.5% | 894 | 51.8% | 863 (100%) | 655 (75.9%) | 0/0 | |
| Mutations: | 131802T>A, 131816G>T, 131818T>A, 131821T>G, 131824A>T, 131827T>A, 131830G>T, 131833T>C, 131836T>A, 131842C>T, 131845C>T, 131851T>C, 131857T>C, 131858G>A, 131861G>A, 131868G>C, 131872A>T, 131873A>G, 131874A>C, 131875T>A, 131878A>T, 131881T>A, 131885C>T, 131887T>A, 131890C>T, 131896T>C, 131899C>A, 131902T>C, 131911A>G, 131915A>T, 131917C>T, 131918C>G, 131920C>T, 131926G>A, 131932G>A, 131938T>C, 131949T>A, 131950C>T, 131951C>G, 131956C>A, 131957G>A, 131959A>T, 131960A>G, 131962C>T, 131963T>C, 131965T>A, 131972C>T, 131974G>A, 131975A>C, 131980T>C, 131981A>G, 131983C>A, 131984T>G, 131985T>C, 131987G>T, 131988C>G, 131989A>T, 131998A>T, 132004A>G, 132010A>G, 132014T>A, 132015C>G, 132018A>T, 132019T>C, 132025T>G, 132034T>A, 132043C>T, 132044T>G, 132045G>C, 132055C>T, 132058T>A, 132061A>T, 132073A>T, 132076A>T, 132077T>A, 132085C>T, 132089C>T, 132092G>A, 132097T>C, 132100T>A, 132101T>A, 132112T>A, 132113T>A, 132114C>G, 132115G>T, 132118C>T, 132124C>T, 132126C>G, 132127A>T, 132136C>A, 132139C>A, 132142T>C, 132157C>T, 132159A>T, 132164C>A, 132166G>T, 132169G>A, 132175A>G, 132178A>T, 132193G>T, 132202C>A, 132205C>T, 132208T>C, 132212C>T, 132214C>A, 132220T>G, 132226C>T, 132235T>C, 132238A>C, 132241T>C, 132244G>A, 132253A>T, 132266C>T, 132271T>A, 132278T>A, 132279C>G, 132280A>T, 132281C>T, 132284G>A, 132286T>C, 132287C>A, 132289T>G, 132298T>A, 132303C>A, 132307G>A, 132310C>T, 132311C>G, 132312A>C, 132313G>T, 132316C>T, 132317T>G, 132319T>A, 132327A>G, 132328G>A, 132340A>G, 132346G>A, 132352C>T, 132355C>T, 132358T>C, 132361T>A, 132364T>C, 132367A>T, 132373A>T, 132376C>T, 132379C>T, 132382T>C, 132385T>A, 132386C>T, 132397G>A, 132403A>T, 132404T>A, 132405C>G, 132406G>T, 132418C>T, 132425C>T, 132427G>A, 132439T>A, 132445A>T, 132454C>A, 132457T>A, 132469C>T, 132472T>C, 132475A>T, 132476C>T, 132478T>A, 132482G>A, 132484T>C, 132490C>T, 132496A>T, 132505G>A, 132517C>T, 132526G>A, 132530A>G, 132532C>A, 132533C>G, 132538T>C, 132542T>C, 132543C>A, 132544T>G, 132548A>C, 132549A>G, 132550G>T, 132553T>A, 132554C>A, 132556G>T, 132557C>A, 132558C>A, 132559T>C, 132562C>T, 132568A>T, 132571C>T, 132572G>A, 132574T>C, 132575C>A, 132583C>T, 132587G>A, 132588G>A, 132589T>C, 132590T>C, 132592G>T, 132613G>A, 132619C>T, 132622A>T, 132625C>T, 132637C>A, 132643T>A, 132650G>A, 132652T>C | | | | | | | | |

*: Inserts / Deletes / Misaligned / Frameshifts

#### Analysis details

This analysis was performed with panviral2.64

### NGS Details (UN18_val): Noumeavirus

#### Assembly

| | |
|---|---|
| **Coverage Length** | 153 (1 contig(s)) |
| **Depth Of Coverage** | 2766.6 |
| **Number Of Reads** | 4184 |
| **Reads Per Million** | 78.53 rpm (after QC) |
| **Ambiguities** | 0 |
| **Assembly Method** | de novo + reference guided assembly |
| **Consensus Caller** | Bcf Tools |

#### Coverage Map

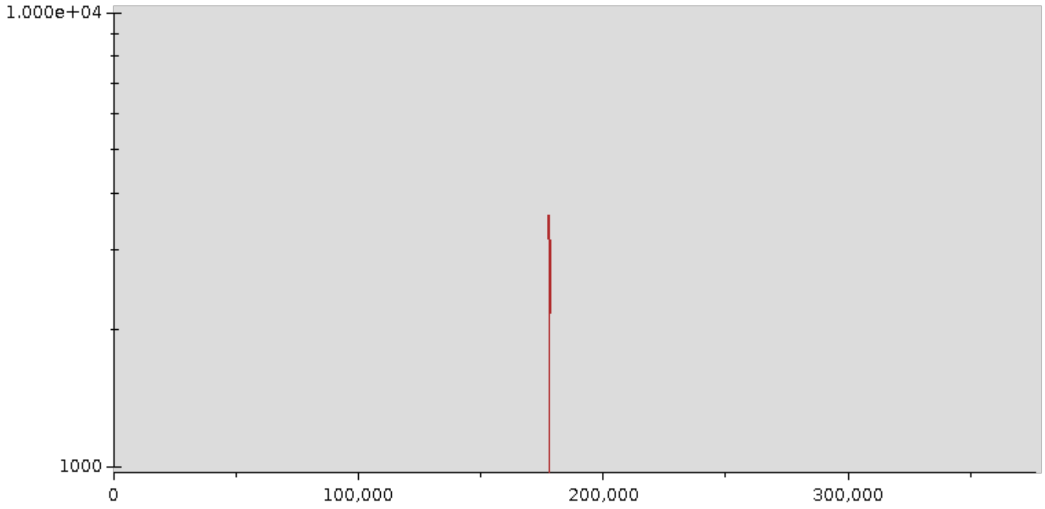

#### Assignment

| | |
|---|---|
| **Type** | Noumeavirus (Taxonomy ID: 1955558) |
| **Reference Genome** | NC_033775.1 |
| **NT Identity (%)** | 81.6993 |
| **AA Identity (%)** | 92.1569 |
| **Number Of Stop Codons** | 0 |
| **Number Of CDS** | 452 |

#### Alignment

| | |
|---|---|
| **Alignment Score** | 194.0 (NT) + 300.0 (AA) = 494.0 |
| **Concordance (%)** | 79.6774 |
| **Alignment Method** | Local, heuristic, nucleotide (BLASTN) |

#### Genome Region

Sequence starts at position 177941 and ends at position 178093 relative to NC_033775.1 reference sequence.

#### Alignment Detailed Statistics

| | Begin | End | Coverage | Score | Concordance | Matches | Identities | I/D/M/F* | Stop Codons |
|---|---|---|---|---|---|---|---|---|---|
| **NT** | **177941** | **178093** | **0.1%** | **194** | **63.4%** | **153 (100%)** | **125 (81.7%)** | **0/0** | |

Mutations: 177953A>G, 177956G>C, 177977C>T, 177980A>T, 177989C>T, 177992G>C, 177996G>A, 178001A>G, 178002C>A, 178003A>C, 178004A>C, 178010T>C, 178019A>G, 178022A>G, 178025A>G, 178028C>T, 178034G>A, 178037C>T, 178043A>G, 178044C>A, 178046A>G, 178049T>G, 178055C>T, 178058C>T, 178061A>G, 178064A>G, 178079A>T, 178080A>C

*: Inserts / Deletes / Misaligned / Frameshifts

#### Analysis details

This analysis was performed with panviral2.64

#### NGS Details (UN18_val): Betabaculovirus disaccharalis

##### Assembly

| | |
|---|---|
| **Coverage Length** | 125 (1 contig(s)) |
| **Depth Of Coverage** | 1580.3 |
| **Number Of Reads** | 2638 |
| **Reads Per Million** | 49.51 rpm (after QC) |
| **Ambiguities** | 0 |
| **Assembly Method** | de novo + reference guided assembly |
| **Consensus Caller** | Bcf Tools |

##### Coverage Map

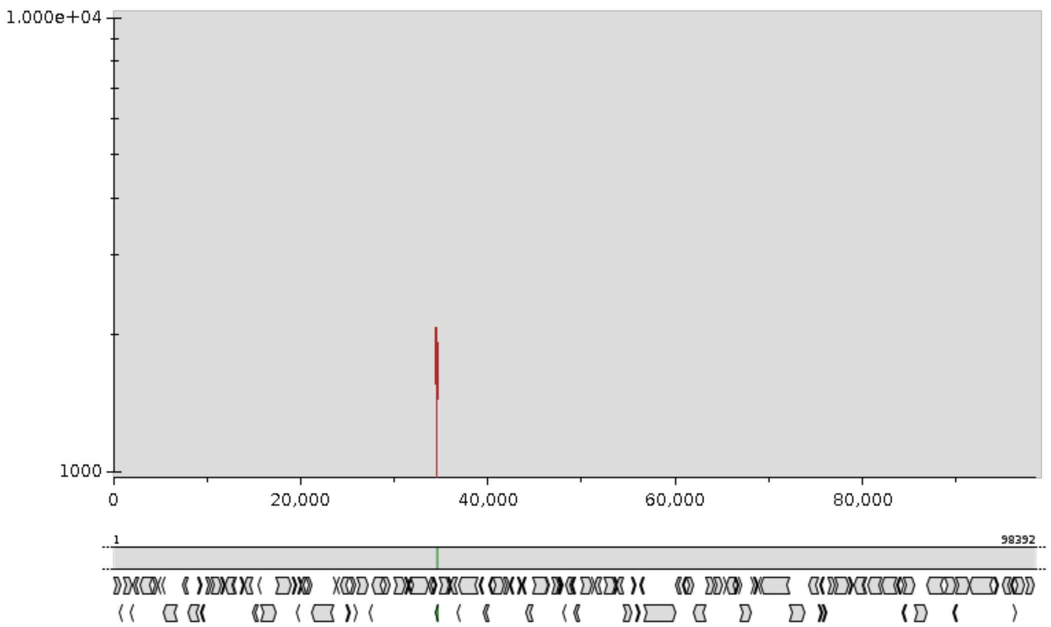

##### Assignment

| | |
|---|---|
| **Type** | Betabaculovirus disaccharalis (Taxonomy ID: 3047795) |
| **Reference Genome** | NC_028491.1 |
| **NT Identity (%)** | 82.4 |
| **AA Identity (%)** | 92.8571 |
| **Number Of Stop Codons** | 0 |
| **Number Of CDS** | 125 |

##### Alignment

| | |
|---|---|
| **Alignment Score** | 162.0 (NT) + 265.0 (AA) = 427.0 |
| **Concordance (%)** | 81.1787 |

| Alignment Method | Local, heuristic, nucleotide (BLASTN) |
|---|---|

#### Genome Region

Sequence starts at position 34471 and ends at position 34595 relative to NC_028491.1 reference sequence.

#### Alignment Detailed Statistics

| | Begin | End | Coverage | Score | Concordance | Matches | Identities | I/D/M/F* | Stop Codons |
|---|---|---|---|---|---|---|---|---|---|
| **NT** | **34471** | **34595** | **0.1%** | **162** | **64.8%** | **125 (100%)** | **103 (82.4%)** | **0/0** | |
| Mutations: | 34482G>T, 34485T>C, 34488A>C, 34490G>A, 34491A>C, 34512G>T, 34513C>G, 34514T>A, 34521T>C, 34524A>G, 34533A>G, 34538A>C, 34545T>C, 34547G>T, 34548C>G, 34549T>C, 34551G>A, 34554T>C, 34569G>A, 34575T>A, 34578T>C, 34580A>G | | | | | | | | |
| **CDS** | | | | | | | | | |
| v-ubq | 61 | 102 | 35.3% | 265 | 88.9% | 42 (100%) | 39 (92.9%) | 0/0/0/0 | 0 |
| Protein mutations: | E76G (34548C>G 34549T>C), S80A (34538A>C) | | | | | | | | |
| Codon mutations: | TTA66CTG (34578T>C 34580A>G), ATA67ATT (34575T>A), GCC69GCT (34569G>A), GAA74GAG (34554T>C), GAC75GAT (34551G>A), GAG76GGC (34548C>G 34549T>C), CGA77AGG (34545T>C 34547G>T), TCA80GCA (34538A>C), GAT81GAC (34533A>G), ATT84ATC (34524A>G), CAA85CAG (34521T>C), AGC88TCA (34512G>T 34513C>G 34514T>A), CGT95CGG (34491A>C), CTT96TTG (34488A>C 34490G>A), AGA97AGG (34485T>C), GGC98GGA (34482G>T) | | | | | | | | |
| **Proteins** | | | | | | | | | |
| ubiquitin-like protein (YP_009182246.1) | 61 | 102 | 35.3% | 265 | 88.9% | 42 (100%) | 39 (92.9%) | 0/0/0/0 | 0 |
| Protein mutations: | E76G (34548C>G 34549T>C), S80A (34538A>C) | | | | | | | | |
| Codon mutations: | TTA66CTG (34578T>C 34580A>G), ATA67ATT (34575T>A), GCC69GCT (34569G>A), GAA74GAG (34554T>C), GAC75GAT (34551G>A), GAG76GGC (34548C>G 34549T>C), CGA77AGG (34545T>C 34547G>T), TCA80GCA (34538A>C), GAT81GAC (34533A>G), ATT84ATC (34524A>G), CAA85CAG (34521T>C), AGC88TCA (34512G>T 34513C>G 34514T>A), CGT95CGG (34491A>C), CTT96TTG (34488A>C 34490G>A), AGA97AGG (34485T>C), GGC98GGA (34482G>T) | | | | | | | | |

*: Inserts / Deletes / Misaligned / Frameshifts

#### Analysis details

This analysis was performed with panviral2.64

### NGS Details (UN18_val): Lausannevirus

#### Assembly

| | |
|---|---|
| **Coverage Length** | 132 (1 contig(s)) |
| **Depth Of Coverage** | 1524.8 |
| **Number Of Reads** | 2565 |
| **Reads Per Million** | 48.14 rpm (after QC) |
| **Ambiguities** | 0 |
| **Assembly Method** | de novo + reference guided assembly |
| **Consensus Caller** | Bcf Tools |

#### Coverage Map

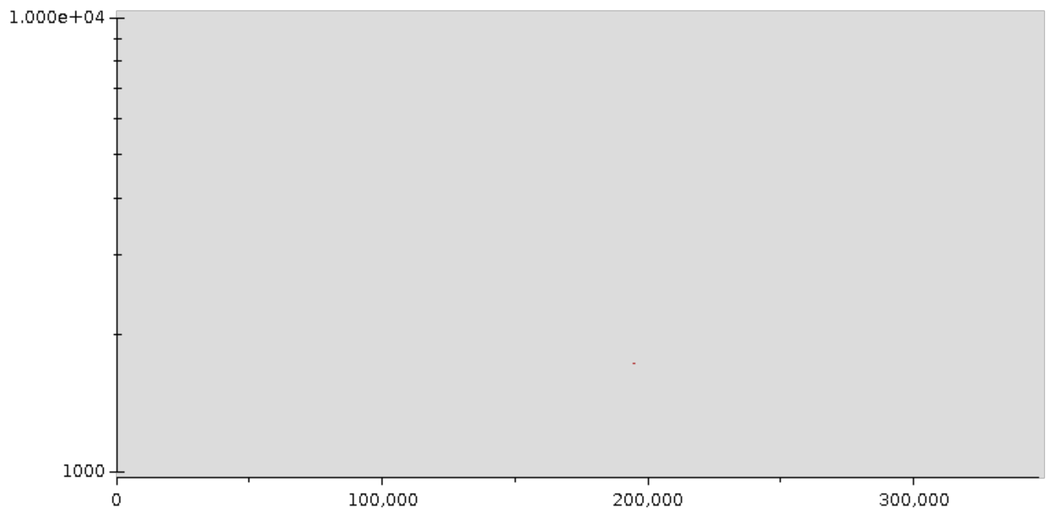

#### Assignment

| | |
|---|---|
| **Type** | Lausannevirus (Taxonomy ID: 999883) |
| **Reference Genome** | NC_015326.1 |
| **NT Identity (%)** | 83.3333 |
| **AA Identity (%)** | 100.0 |
| **Number Of Stop Codons** | 1 |
| **Number Of CDS** | 444 |

#### Alignment

| | |
|---|---|
| **Alignment Score** | 176.0 (NT) + 282.0 (AA) = 458.0 |
| **Concordance (%)** | 83.8828 |
| **Alignment Method** | Local, heuristic, nucleotide (BLASTN) |

#### Genome Region

Sequence starts at position 194913 and ends at position 195044 relative to NC_015326.1 reference sequence.

#### Alignment Detailed Statistics

| | Begin | End | Coverage | Score | Concordance | Matches | Identities | I/D/M/F* | Stop Codons |
|---|---|---|---|---|---|---|---|---|---|
| **NT** | **194913** | **195044** | **0.1%** | **176** | **66.7%** | **132 (100%)** | **110 (83.3%)** | **0/0** | |

Mutations: 194925T>C, 194930A>G, 194931A>G, 194937G>A, 194946G>A, 194955T>C, 194958A>G, 194967A>G, 194970G>T, 194976A>G, 194979T>G, 194981T>G, 194982C>G, 194988T>C, 194991G>A, 195003T>A, 195012A>G, 195015T>C, 195017G>T, 195018T>C, 195024G>A, 195030A>T

*: Inserts / Deletes / Misaligned / Frameshifts

#### Analysis details

This analysis was performed with panviral2.64

### NGS Details (UN18_val): Brazilian marseillevirus

#### Assembly

| | |
|---|---|
| **Coverage Length** | 228 (1 contig(s)) |
| **Depth Of Coverage** | 558.0 |
| **Number Of Reads** | 2433 |
| **Reads Per Million** | 45.66 rpm (after QC) |
| **Ambiguities** | 0 |
| **Assembly Method** | de novo + reference guided assembly |
| **Consensus Caller** | Bcf Tools |

#### Coverage Map

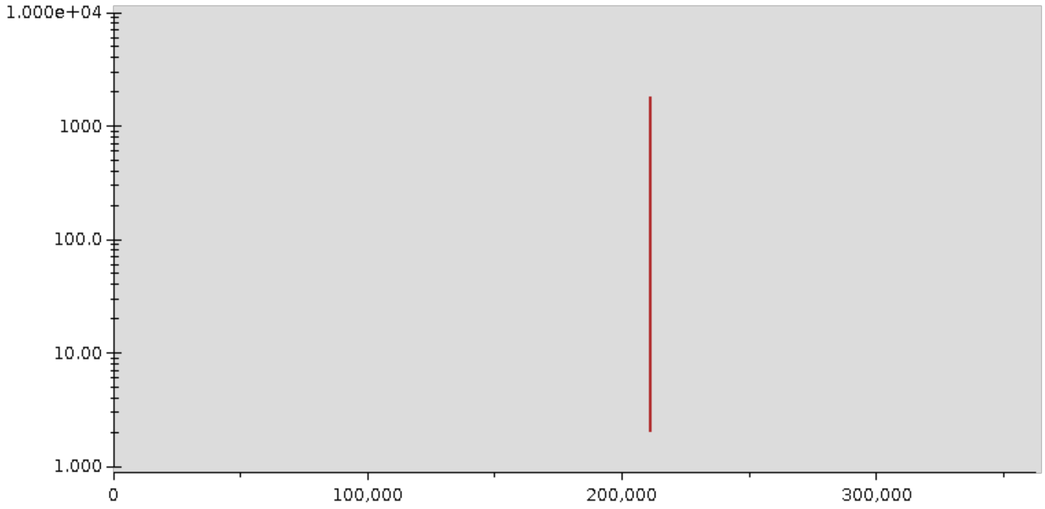

#### Assignment

| | |
|---|---|
| **Type** | Brazilian marseillevirus (Taxonomy ID: 1813599) |
| **Reference Genome** | NC_029692.1 |
| **NT Identity (%)** | 81.5789 |
| **AA Identity (%)** | 95.8904 |
| **Number Of Stop Codons** | 0 |
| **Number Of CDS** | 491 |

#### Alignment

| | |
|---|---|
| **Alignment Score** | 288.0 (NT) + 452.0 (AA) = 740.0 |
| **Concordance (%)** | 80.8743 |
| **Alignment Method** | Local, heuristic, nucleotide (BLASTN) |

#### Genome Region

Sequence starts at position 210896 and ends at position 211123 relative to NC_029692.1 reference sequence.

#### Alignment Detailed Statistics

| | Begin | End | Coverage | Score | Concordance | Matches | Identities | I/D/M/F* | Stop Codons |
|---|---|---|---|---|---|---|---|---|---|
| **NT** | **210896** | **211123** | **0.1%** | **288** | **63.2%** | **228 (100%)** | **186 (81.6%)** | **0/0** | |

Mutations: 210905G>C, 210908A>G, 210914A>G, 210917A>G, 210920G>C, 210926T>C, 210929T>C, 210938A>G, 210941A>G, 210944C>G, 210947A>G, 210953C>A, 210955T>G, 210956T>G, 210959A>G, 210962T>C, 210965C>G, 210977T>A, 210986A>G, 210989G>A, 210992T>C, 210995T>C, 211001A>G, 211004A>G, 211009C>T, 211010T>G, 211013T>C, 211031T>C, 211034T>C, 211043T>C, 211055A>G, 211056G>A, 211061T>C, 211064A>G, 211073A>G, 211079C>G, 211088C>G, 211094A>G, 211100T>G, 211103A>G, 211106C>G, 211108C>T

*: Inserts / Deletes / Misaligned / Frameshifts

#### Analysis details

This analysis was performed with panviral2.64

### NGS Details (UN18_val): Betabaculovirus disaccharalis

#### Assembly

| | |
|---|---|
| **Coverage Length** | 129 (1 contig(s)) |
| **Depth Of Coverage** | 1390.5 |
| **Number Of Reads** | 2060 |
| **Reads Per Million** | 38.66 rpm (after QC) |
| **Ambiguities** | 0 |
| **Assembly Method** | de novo + reference guided assembly |
| **Consensus Caller** | Bcf Tools |

#### Coverage Map

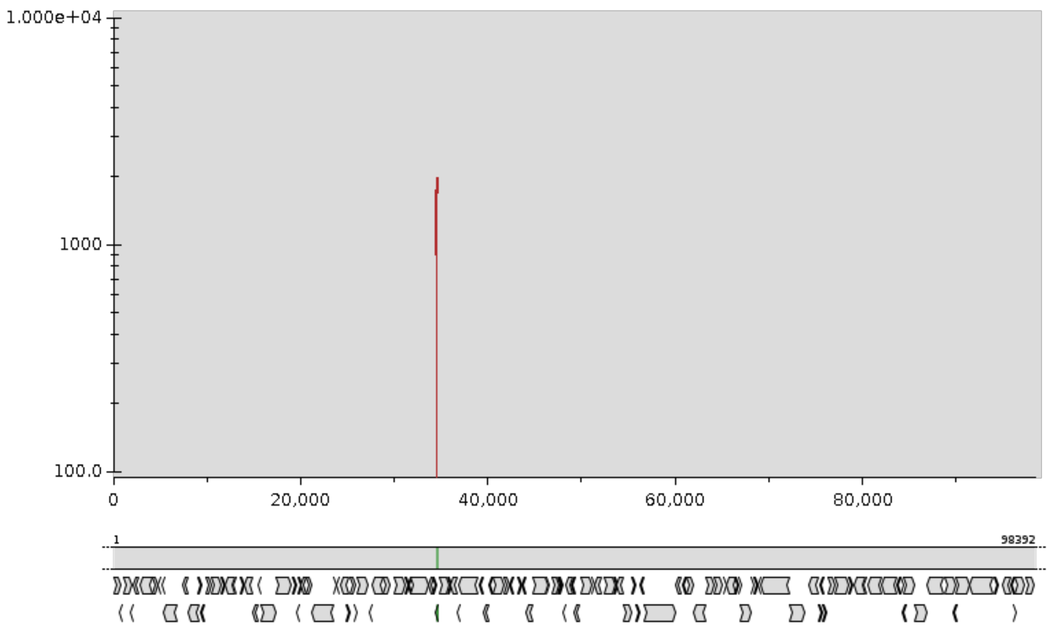

#### Assignment

| | |
|---|---|
| **Type** | Betabaculovirus disaccharalis (Taxonomy ID: 3047795) |
| **Reference Genome** | NC_028491.1 |
| **NT Identity (%)** | 83.7209 |
| **AA Identity (%)** | 88.3721 |
| **Number Of Stop Codons** | 0 |
| **Number Of CDS** | 125 |

#### Alignment

| | |
|---|---|
| **Alignment Score** | 174.0 (NT) + 257.0 (AA) = 431.0 |
| **Concordance (%)** | 79.0826 |

| Alignment Method | Local, heuristic, nucleotide (BLASTN) |
|---|---|

#### Genome Region

Sequence starts at position 34471 and ends at position 34599 relative to NC_028491.1 reference sequence.

#### Alignment Detailed Statistics

| | Begin | End | Coverage | Score | Concordance | Matches | Identities | I/D/M/F* | Stop Codons |
|---|---|---|---|---|---|---|---|---|---|
| **NT** | **34471** | **34599** | **0.1%** | **174** | **67.4%** | **129 (100%)** | **108 (83.7%)** | **0/0** | |
| Mutations: | 34482G>T, 34485T>C, 34488A>C, 34490G>A, 34503A>C, 34511T>G, 34513C>G, 34514T>A, 34521T>C, 34524A>G, 34533A>G, 34538A>C, 34542T>A, 34545T>C, 34547G>T, 34549T>C, 34554T>C, 34569G>A, 34575T>G, 34578T>C, 34587T>C | | | | | | | | |

##### CDS

| | Begin | End | Coverage | Score | Concordance | Matches | Identities | I/D/M/F* | Stop Codons |
|---|---|---|---|---|---|---|---|---|---|
| v-ubq | 60 | 102 | 36.1% | 257 | 81.8% | 43 (100%) | 38 (88.4%) | 0/0/0/0 | 0 |
| Protein mutations: | E76G (34549T>C), S80A (34538A>C), T89P (34511T>G), H91Q (34503A>C) | | | | | | | | |
| Codon mutations: | CAA63CAG (34587T>C), TTA66TTG (34578T>C), ATA67ATC (34575T>G), GCC69GCT (34569G>A), GAA74GAG (34554T>C), GAG76GGG (34549T>C), CGA77AGG (34545T>C 34547G>T), ACA78ACT (34542T>A), TCA80GCA (34538A>C), GAT81GAC (34533A>G), ATT84ATC (34524A>G), CAA85CAG (34521T>C), AGC88TCC (34513C>G 34514T>A), ACT89CCT (34511T>G), CAT91CAG (34503A>C), CTT96TTG (34488A>C 34490G>A), AGA97AGG (34485T>C), GGC98GGA (34482G>T) | | | | | | | | |

##### Proteins

| | Begin | End | Coverage | Score | Concordance | Matches | Identities | I/D/M/F* | Stop Codons |
|---|---|---|---|---|---|---|---|---|---|
| ubiquitin-like protein (YP_009182246.1) | 60 | 102 | 36.1% | 257 | 81.8% | 43 (100%) | 38 (88.4%) | 0/0/0/0 | 0 |
| Protein mutations: | E76G (34549T>C), S80A (34538A>C), T89P (34511T>G), H91Q (34503A>C) | | | | | | | | |
| Codon mutations: | CAA63CAG (34587T>C), TTA66TTG (34578T>C), ATA67ATC (34575T>G), GCC69GCT (34569G>A), GAA74GAG (34554T>C), GAG76GGG (34549T>C), CGA77AGG (34545T>C 34547G>T), ACA78ACT (34542T>A), TCA80GCA (34538A>C), GAT81GAC (34533A>G), ATT84ATC (34524A>G), CAA85CAG (34521T>C), AGC88TCC (34513C>G 34514T>A), ACT89CCT (34511T>G), CAT91CAG (34503A>C), CTT96TTG (34488A>C 34490G>A), AGA97AGG (34485T>C), GGC98GGA (34482G>T) | | | | | | | | |

*: Inserts / Deletes / Misaligned / Frameshifts

#### Analysis details

This analysis was performed with panviral2.64

### NGS Details (UN18_val): Bracoviriform congregatae (segment Circle 7)

#### Assembly

| | |
|---|---|
| **Coverage Length** | 172 (1 contig(s)) |
| **Depth Of Coverage** | 965.9 |
| **Number Of Reads** | 1498 |
| **Reads Per Million** | 28.12 rpm (after QC) |
| **Ambiguities** | 0 |
| **Assembly Method** | de novo + reference guided assembly |
| **Consensus Caller** | Bcf Tools |

#### Coverage Map

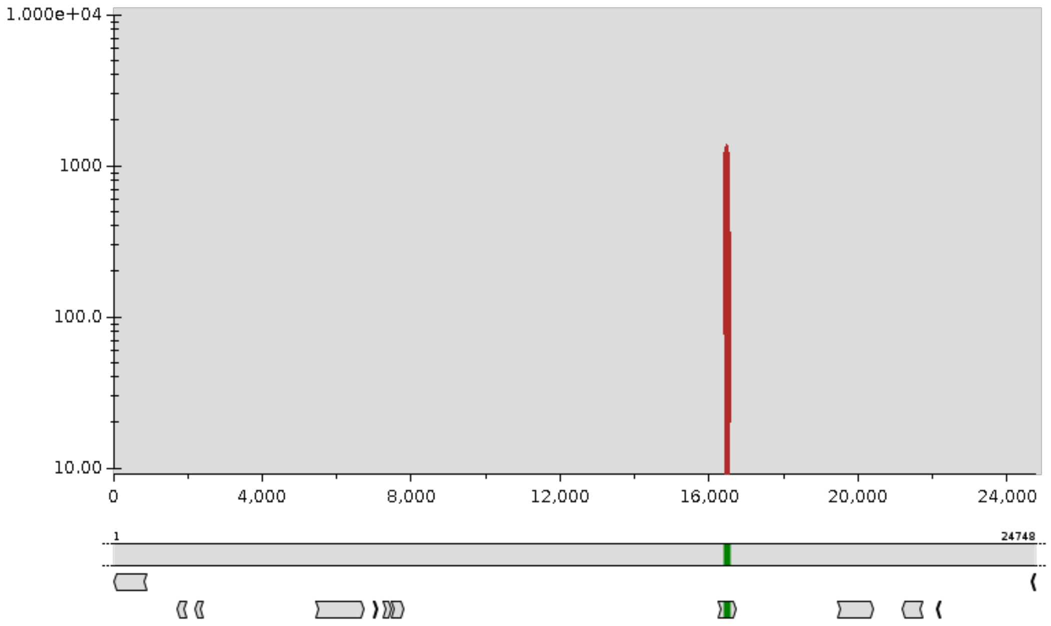

#### Assignment

| | |
|---|---|
| **Type** | Bracoviriform congregatae (Taxonomy ID: 39640) |
| **Reference Genome** | NC_006639.1 |
| **NT Identity (%)** | 77.3256 |
| **AA Identity (%)** | 91.2281 |
| **Number Of Stop Codons** | 0 |
| **Number Of CDS** | 5 |

#### Alignment

| | |
|---|---|
| **Alignment Score** | 188.0 (NT) + 350.0 (AA) = 538.0 |
| **Concordance (%)** | 75.3501 |

| Alignment Method | Global, seeded, nucleotide + amino acids (AGA) |
|---|---|

#### Genome Region

Sequence starts at position 16393 and ends at position 16564 relative to NC_006639.1 reference sequence.

#### Alignment Detailed Statistics

| | Begin | End | Coverage | Score | Concordance | Matches | Identities | I/D/M/F* | Stop Codons |
|---|---|---|---|---|---|---|---|---|---|
| **NT** | **16393** | **16564** | **0.7%** | **188** | **54.7%** | **172 (100%)** | **133 (77.3%)** | **0/0** | |
| Mutations: | 16402A>T, 16405A>G, 16411G>C, 16414A>G, 16423A>C, 16429G>A, 16432C>A, 16435T>C, 16439C>A, 16441T>G, 16444T>C, 16445C>A, 16447C>G, 16449G>A, 16450A>G, 16451T>G, 16452C>T, 16453T>G, 16456T>A, 16457C>A, 16459A>G, 16471A>G, 16474A>T, 16477C>T, 16480C>T, 16492C>T, 16496C>A, 16498T>G, 16501G>T, 16508C>A, 16510T>A, 16516A>T, 16519C>G, 16526G>A, 16528C>T, 16538G>A, 16540C>T, 16546A>G, 16551T>C | | | | | | | | |
| **CDS** | | | | | | | | | |
| Histone | 55 | 111 | 36.5% | 350 | 93.6% | 57 (100%) | 52 (91.2%) | 0/0/0/0 | 0 |
| Protein mutations: | R73K (16449G>A 16450A>G), S74V (16451T>G 16452C>T 16453T>G), V99I (16526G>A 16528C>T), V103I (16538G>A 16540C>T), I107T (16551T>C) | | | | | | | | |
| Codon mutations: | GTA57GTT (16402A>T), AAA58AAG (16405A>G), GGG60GGC (16411G>C), AAA61AAG (16414A>G), GGA64GGC (16423A>C), GGG66GGA (16429G>A), GGC67GGA (16432C>A), GCT68GCC (16435T>C), CGT70AGG (16439C>A 16441T>G), CAT71CAC (16444T>C), CGC72AGG (16445C>A 16447C>G), AGA73AAG (16449G>A 16450A>G), TCT74GTG (16451T>G 16452C>T 16453T>G), CTT75CTA (16456T>A), CGA76AGG (16457C>A 16459A>G), CAA80CAG (16471A>G), GGA81GGT (16474A>T), ATC82ATT (16477C>T), ACC83ACT (16480C>T), ATC87ATT (16492C>T), CGT89AGG (16496C>A 16498T>G), CTG90CTT (16501G>T), CGT93AGA (16508C>A 16510T>A), GGA95GGT (16516A>T), GTC96GTG (16519C>G), GTC99ATT (16526G>A 16528C>T), GTC103ATT (16538G>A 16540C>T), GAA105GAG (16546A>G), ATT107ACT (16551T>C) | | | | | | | | |
| **Proteins** | | | | | | | | | |
| hypothetical protein (YP_184795.1) | 55 | 111 | 36.5% | 350 | 93.6% | 57 (100%) | 52 (91.2%) | 0/0/0/0 | 0 |
| Protein mutations: | R73K (16449G>A 16450A>G), S74V (16451T>G 16452C>T 16453T>G), V99I (16526G>A 16528C>T), V103I (16538G>A 16540C>T), I107T (16551T>C) | | | | | | | | |
| Codon mutations: | GTA57GTT (16402A>T), AAA58AAG (16405A>G), GGG60GGC (16411G>C), AAA61AAG (16414A>G), GGA64GGC (16423A>C), GGG66GGA (16429G>A), GGC67GGA (16432C>A), GCT68GCC (16435T>C), CGT70AGG (16439C>A 16441T>G), CAT71CAC (16444T>C), CGC72AGG (16445C>A 16447C>G), AGA73AAG (16449G>A 16450A>G), TCT74GTG (16451T>G 16452C>T 16453T>G), CTT75CTA (16456T>A), CGA76AGG (16457C>A 16459A>G), CAA80CAG (16471A>G), GGA81GGT (16474A>T), ATC82ATT (16477C>T), ACC83ACT (16480C>T), ATC87ATT (16492C>T), CGT89AGG (16496C>A 16498T>G), CTG90CTT (16501G>T), CGT93AGA (16508C>A 16510T>A), GGA95GGT (16516A>T), GTC96GTG (16519C>G), GTC99ATT (16526G>A 16528C>T), GTC103ATT (16538G>A 16540C>T), GAA105GAG (16546A>G), ATT107ACT (16551T>C) | | | | | | | | |

*: Inserts / Deletes / Misaligned / Frameshifts

#### Analysis details

This analysis was performed with panviral2.64

### NGS Details (UN18_val): Tunisvirus fontaine2

#### Assembly

| | |
|---|---|
| **Coverage Length** | 175 (1 contig(s)) |
| **Depth Of Coverage** | 465.1 |
| **Number Of Reads** | 895 |
| **Reads Per Million** | 16.80 rpm (after QC) |
| **Ambiguities** | 0 |
| **Assembly Method** | de novo + reference guided assembly |
| **Consensus Caller** | Bcf Tools |

#### Coverage Map

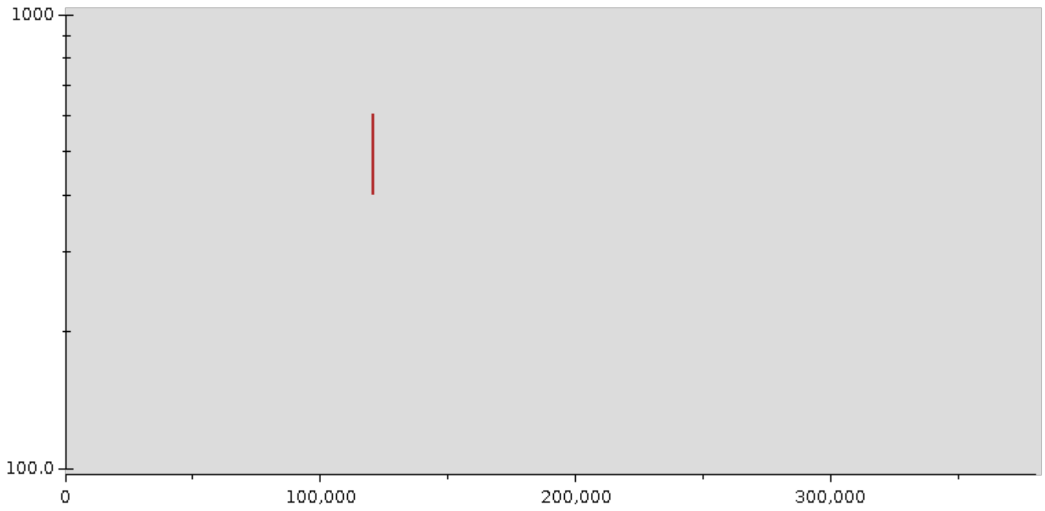

#### Assignment

| | |
|---|---|
| **Type** | Tunisvirus fontaine2 (Taxonomy ID: 1421067) |
| **Reference Genome** | NC_038511.1 |
| **NT Identity (%)** | 80.0 |
| **AA Identity (%)** | 91.3793 |
| **Number Of Stop Codons** | 1 |
| **Number Of CDS** | 484 |

#### Alignment

| | |
|---|---|
| **Alignment Score** | 210.0 (NT) + 342.0 (AA) = 552.0 |
| **Concordance (%)** | 77.2028 |
| **Alignment Method** | Local, heuristic, nucleotide (BLASTN) |

#### Genome Region

Sequence starts at position 120430 and ends at position 120604 relative to NC_038511.1 reference sequence.

#### Alignment Detailed Statistics

| | Begin | End | Coverage | Score | Concordance | Matches | Identities | I/D/M/F* | Stop Codons |
|---|---|---|---|---|---|---|---|---|---|
| **NT** | **120430** | **120604** | **0.1%** | **210** | **60.0%** | **175 (100%)** | **140 (80.0%)** | **0/0** | |

Mutations: 120440T>C, 120449A>C, 120458A>T, 120461G>T, 120470T>C, 120475C>T, 120476G>A, 120479A>G, 120485G>A, 120488A>C, 120494C>T, 120496T>G, 120497C>A, 120508G>A, 120509A>C, 120510C>T, 120511T>G, 120512C>T, 120515C>T, 120518G>A, 120521G>A, 120524G>A, 120527A>T, 120532G>T, 120536T>C, 120542C>A, 120548A>T, 120551G>C, 120560A>G, 120563T>C, 120572T>G, 120573T>G, 120574G>T, 120582C>T, 120584C>G

*: Inserts / Deletes / Misaligned / Frameshifts

#### Analysis details

This analysis was performed with panviral2.64

### NGS Details (UN18_val): Lowelvirus tuscon4d

#### Assembly

| | |
|---|---|
| **Coverage Length** | 619 (2 contig(s)) |
| **Depth Of Coverage** | 106.0 |
| **Number Of Reads** | 634 |
| **Reads Per Million** | 11.90 rpm (after QC) |
| **Ambiguities** | 0 |
| **Assembly Method** | de novo + reference guided assembly |
| **Consensus Caller** | Bcf Tools |

#### Coverage Map

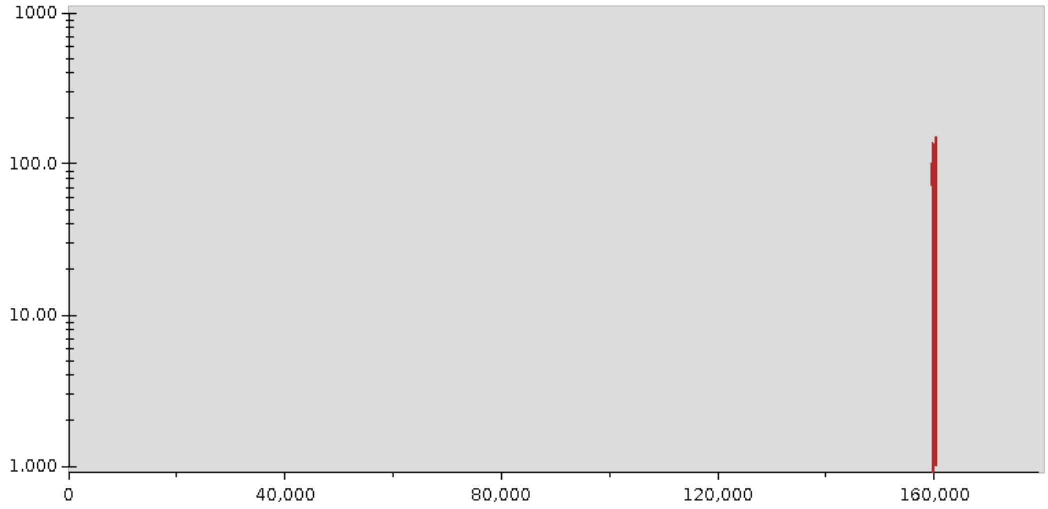

#### Assignment

| | |
|---|---|
| **Type** | Lowelvirus tuscon4d (Taxonomy ID: 2956131) |
| **Reference Genome** | NC_026923.1 |
| **NT Identity (%)** | 76.8233 |
| **AA Identity (%)** | 88.2353 |
| **Number Of Stop Codons** | 1 |
| **Number Of CDS** | 218 |

#### Alignment

| | |
|---|---|
| **Alignment Score** | 651.0 (NT) + 1264.0 (AA) = 1915.0 |
| **Concordance (%)** | 74.0275 |
| **Alignment Method** | Local, heuristic, nucleotide (BLASTN) |

#### Genome Region

Sequence starts at position 159602 and ends at position 160401 relative to NC_026923.1 reference sequence.

#### Alignment Detailed Statistics

| | Begin | End | Coverage | Score | Concordance | Matches | Identities | I/D/M/F* | Stop Codons |
|---|---|---|---|---|---|---|---|---|---|
| **NT** | **159602** | **160401** | **0.3%** | **651** | **53.2%** | **617 (99.7%)** | **474 (76.6%)** | **0/2** | |

Mutations: 159612T>A, 159624G>A, 159625T>G, 159627T>A, 159633C>A, 159636C>T, 159639C>T, 159640C>A, 159641A>C, 159642A>T, 159645C>T, 159651C>T, 159655C>T, 159657T>G, 159660A>G, 159663G>T, 159664A>C, 159666T>G, 159670T>A, 159672C>T, 159687C>T, 159690T>A, 159693A>T, 159696C>T, 159698C>G, 159699T>C, 159702C>A, 159711T>C, 159715C>T, 159720G>T, 159726T>C, 159729A>G, 159730A>C, 159738T>A, 159739C>T, 159740T>C, 159741A>T, 159745G>C, 159746G>A, 159747T>A, 159748A>T, 159750C>G, 159751C>A, 159753T>A, 159759C>T, 159765T>A, 159768T>C, 159771G>A, 159783T>A, 159798C>T, 159799A>T, 159800G>C, 159801C>T, 159807C>T, 159810T>G, 159813C>T, 159822G>A, 159832T>G, 159833C>G, 159834A>T, 159855C>T, 159858G>T, 159867A>T, 159870C>A, 159874A>C, 159880C>A, 159882G>C, 159883T>C, 159885G>A, 159888C>T, 159889C>T, 159894G>A, 159897T>G, 159900C>T, 159903C>T, 159906C>T, 159913C>T, 159915C>G, 159924C>T, 159942T>C, 159946A>G, 159949C>T, 159951A>G, 159954T>C, 159956G>C, 159957A>T, 159960A>T, 159961C>T, 159964T>C, 159966G>A, 159969T>C, 159972A>C, 159975C>T, 159987T>C, 159993G>A, 159996C>T, 159999A>T, 160002G>A, 160011T>C, 160014A>T, 160017A>T, 160018C>G, 160019A>G, 160020A>T, 160026C>T, 160032T>C, 160033A>C, 160034A>G, 160035G>T, 160038G>T, 160042G>A, 160044G>C, 160246A>C, 160248A>C, 160254A>G, 160270G>A, 160275C>T, 160287G>A, 160290C>T, 160291_160292delAT, 160293C>A, 160296A>C, 160299G>A, 160302T>C, 160308A>T, 160309C>A, 160326A>G, 160327C>G, 160330G>C, 160331C>A, 160332T>G, 160338A>G, 160341A>T, 160350C>T, 160356C>A, 160359C>T, 160368A>G, 160371A>T, 160377T>A, 160378A>C, 160380A>T, 160383T>A, 160389A>T, 160392G>T

*: Inserts / Deletes / Misaligned / Frameshifts

#### Analysis details

This analysis was performed with panviral2.64

#### NGS Details (UN18_val): Cassava brown streak virus

##### Assembly

| | |
|---|---|
| **Coverage Length** | 521 (1 contig(s)) |
| **Depth Of Coverage** | 103.8 |
| **Number Of Reads** | 450 |
| **Reads Per Million** | 8.45 rpm (after QC) |
| **Ambiguities** | 0 |
| **Assembly Method** | de novo + reference guided assembly |
| **Consensus Caller** | Bcf Tools |

##### Coverage Map

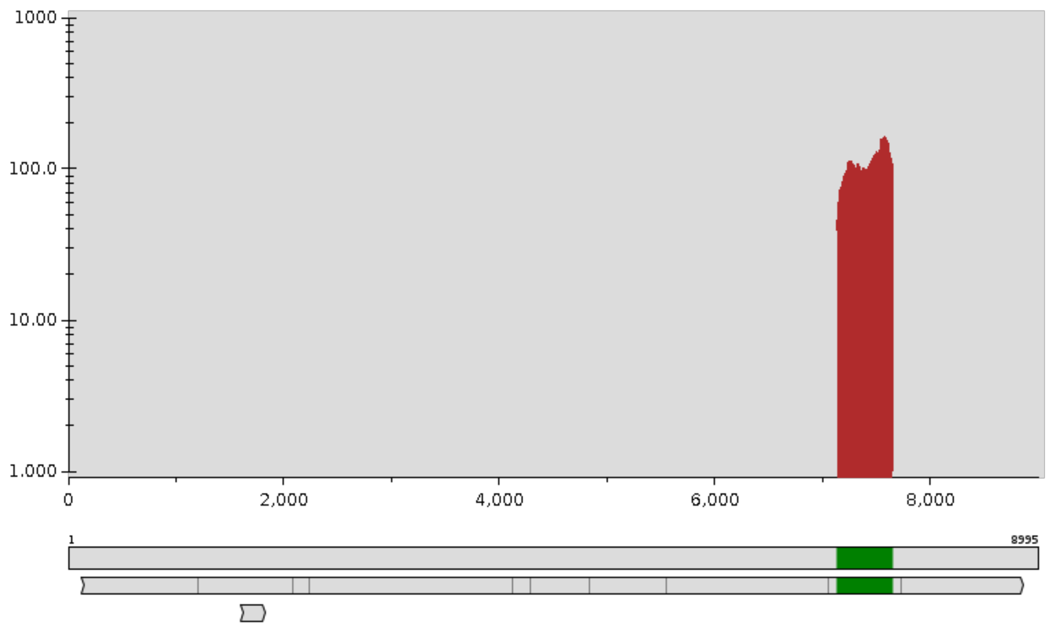

##### Assignment

| | |
|---|---|
| **Type** | Cassava brown streak virus (Taxonomy ID: 137758) |
| **Reference Genome** | NC_012698.2 |
| **NT Identity (%)** | 60.4607 |
| **AA Identity (%)** | 57.4713 |
| **Number Of Stop Codons** | 0 |
| **Number Of CDS** | 2 |

##### Alignment

| | |
|---|---|
| **Alignment Score** | 218.0 (NT) + 732.0 (AA) = 950.0 |
| **Concordance (%)** | 42.166 |

| **Alignment Method** | Global, seeded, nucleotide + amino acids (AGA) |
|---|---|

#### Genome Region

Sequence starts at position 7130 and ends at position 7650 relative to NC_012698.2 reference sequence.

#### Alignment Detailed Statistics

| | Begin | End | Coverage | Score | Concordance | Matches | Identities | I/D/M/F* | Stop Codons |
|---|---|---|---|---|---|---|---|---|---|
| **NT** | **7130** | **7650** | **5.8%** | **218** | **20.9%** | **521 (100%)** | **315 (60.5%)** | **0/0** | |
| Mutations: | 7130A>G, 7133A>G, 7134A>G, 7136T>G, 7142C>T, 7145A>T, 7148A>C, 7151G>A, 7157A>T, 7158C>A, 7163A>G, 7167A>G, 7175A>T, 7177A>G, 7178A>G, 7179C>G, 7180A>C, 7184A>T, 7185T>C, 7190T>C, 7192C>A, 7193C>G, 7194A>T, 7202A>C, 7203A>T, 7205C>T, 7206A>C, 7207T>A, 7208C>G, 7211T>C, 7213G>T, 7214G>T, 7217A>G, 7218G>C, 7229A>T, 7234C>T, 7235A>T, 7241A>G, 7242A>G, 7243C>A, 7244A>G, 7245G>C, 7246T>C, 7250G>A, 7253A>T, 7256T>A, 7257A>T, 7258T>C, 7270T>C, 7271A>T, 7272C>A, 7273G>A, 7274T>A, 7275G>A, 7277A>T, 7280G>T, 7281T>G, 7283T>C, 7284G>A, 7286G>A, 7287T>G, 7288T>A, 7290A>G, 7292T>G, 7293G>A, 7294G>A, 7295A>T, 7298G>A, 7304T>G, 7307C>A, 7310A>T, 7313A>G, 7319A>T, 7320A>T, 7323T>C, 7325G>C, 7331T>C, 7332G>A, 7334T>C, 7337T>C, 7340C>T, 7343T>G, 7355A>G, 7358A>T, 7361C>T, 7364T>C, 7374A>T, 7377G>C, 7380G>A, 7381G>A, 7385T>C, 7388A>T, 7390T>A, 7391A>T, 7397A>T, 7398T>C, 7400A>T, 7401T>A, 7403T>C, 7406G>C, 7409G>A, 7410G>T, 7413G>A, 7414A>T, 7416C>G, 7418G>T, 7422C>G, 7424A>G, 7425A>G, 7432T>C, 7433G>A, 7436T>A, 7437A>T, 7438G>A, 7442T>C, 7443C>A, 7445C>G, 7448C>T, 7449G>A, 7451G>C, 7455G>T, 7457T>A, 7458T>C, 7462T>C, 7463A>T, 7464A>C, 7465A>T, 7467A>G, 7468A>G, 7469A>G, 7470G>C, 7471T>C, 7472T>A, 7473G>A, 7474G>A, 7476G>A, 7477A>C, 7481T>G, 7484T>G, 7487A>G, 7489T>C, 7490C>T, 7494A>G, 7495A>T, 7496G>T, 7499T>A, 7500G>A, 7501T>A, 7502G>A, 7503C>A, 7504T>C, 7505A>G, 7506A>C, 7507G>T, 7508A>G, 7511T>A, 7512G>A, 7517T>A, 7520A>G, 7521A>C, 7522T>C, 7523G>A, 7524C>G, 7526A>T, 7527C>A, 7529C>G, 7535A>C, 7539T>A, 7540C>A, 7541A>T, 7544T>C, 7547G>A, 7553C>T, 7556A>G, 7559T>A, 7562C>T, 7568A>T, 7570T>A, 7572A>G, 7573A>G, 7576G>A, 7577G>T, 7578A>G, 7579G>A, 7580A>C, 7581A>C, 7586A>T, 7588T>A, 7589T>C, 7599A>C, 7600T>C, 7601G>C, 7602G>A, 7603C>A, 7604C>G, 7607G>A, 7610G>A, 7616T>C, 7618T>A, 7622A>T, 7628T>C, 7631A>C, 7632T>G, 7633T>G, 7635C>A, 7636G>A | | | | | | | | |

##### CDS

| | Begin | End | Coverage | Score | Concordance | Matches | Identities | I/D/M/F* | Stop Codons |
|---|---|---|---|---|---|---|---|---|---|
| CBSV_gp1 | 2336 | 2509 | 6.0% | 732 | 60.0% | 174 (100%) | 100 (57.5%) | 0/0/0/0 | 0 |
| Protein mutations: | I2337V (7134A>G 7136T>G), Q2345K (7158C>A), K2348E (7167A>G), K2351R (7177A>G 7178A>G), Q2352A (7179C>G 7180A>C), F2354L (7185T>C), P2356Q (7192C>A 7193C>G), T2357S (7194A>T), I2360F (7203A>T 7205C>T), I2361Q (7206A>C 7207T>A 7208C>G), R2363L (7213G>T 7214G>T), V2365L (7218G>C), P2370L (7234C>T 7235A>T), T2373E (7242A>G 7243C>A 7244A>G), V2374P (7245G>C 7246T>C), E2376D (7253A>T), I2378S (7257A>T 7258T>C), V2382A (7270T>C 7271A>T), R2383K (7272C>A 7273G>A 7274T>A), V2384I (7275G>A 7277A>T), S2386A (7281T>G 7283T>C), E2387K (7284G>A 7286G>A), L2388E (7287T>G 7288T>A), I2389V (7290A>G 7292T>G), G2390N (7293G>A 7294G>A 7295A>T), S2399C (7320A>T), D2403N (7332G>A 7334T>C), N2406K (7343T>G), M2417L (7374A>T), E2418Q (7377G>C), G2419K (7380G>A 7381G>A), L2422H (7390T>A 7391A>T), Y2426N (7401T>A 7403T>C), K2427N (7406G>C), V2429L (7410G>T), E2430M (7413G>A 7414A>T), P2431A (7416C>G 7418G>T), Q2433E (7422C>G 7424A>G), N2434D (7425A>G), M2436T (7432T>C 7433G>A), S2438Y (7437A>T 7438G>A), L2440M (7443C>A 7445C>G), V2442I (7449G>A 7451G>C), A2444S (7455G>T 7457T>A), F2445L (7458T>C), V2446A (7462T>C 7463A>T), N2447L (7464A>C 7465A>T), K2448G (7467A>G 7468A>G 7469A>G), V2449P (7470G>C 7471T>C 7472T>A), G2450N (7473G>A 7474G>A), D2451T (7476G>A 7477A>C), D2452E (7481T>G), I2454M (7487A>G), I2455T (7489T>C 7490C>T), K2457V (7494A>G 7495A>T 7496G>T), V2459K (7500G>A 7501T>A 7502G>A), L2460T (7503C>A 7504T>C 7505A>G), R2461L (7506A>C 7507G>T 7508A>G), E2463K (7512G>A), M2466P (7521A>C 7522T>C 7523G>A), P2467A (7524C>G 7526A>T), S2472N (7539T>A 7540C>A 7541A>T), L2482H (7570T>A), N2483G (7572A>G 7573A>G), W2484Y (7576G>A 7577G>T), R2485D (7578A>G 7579G>A 7580A>C), K2486Q (7581A>C), F2488Y (7588T>A 7589T>C), M2492P (7599A>C 7600T>C 7601G>C), A2493K (7602G>A 7603C>A 7604C>G), M2498K (7618T>A), F2503G (7632T>G 7633T>G), R2504K (7635C>A 7636G>A) | | | | | | | | |
| Codon mutations: | GCA2335..G (7130A>G), CCA2336CCG (7133A>G), ATT2337GTG (7134A>G 7136T>G), TTC2339TTT (7142C>T), GTA2340GTT (7145A>T), ACA2341ACC (7148A>C), GGG2342GGA (7151G>A), GCA2344GCT (7157A>T), CAA2345AAA (7158C>A), AAA2346AAG (7163A>G), AAG2348GAG (7167A>G), GTA2350GTT (7175A>T), AAA2351AGG (7177A>G 7178A>G), CAA2352GCA (7179C>G 7180A>C), ATA2353ATT (7184A>T), TTT2354CTT (7185T>C), GGT2355GGC (7190T>C), CCC2356CAG (7192C>A 7193C>G), ACT2357TCT (7194A>T), CCA2359CCC (7202A>C), ATC2360TTT (7203A>T 7205C>T), ATC2361CAG (7206A>C 7207T>A 7208C>G), TCT2362TCC (7211T>C), CGG2363CTT (7213G>T 7214G>T), AAA2364AAG (7217A>G), GTT2365CTT (7218G>C), CCA2368CCT (7229A>T), CCA2370CTT (7234C>T 7235A>T), GGA2372GGG (7241A>G), ACA2373GAG (7242A>G 7243C>A 7244A>G), GTT2374CCT (7245G>C 7246T>C), GAG2375GAA (7250G>A), GAA2376GAT (7253A>T), ATT2377ATA (7256T>A), ATC2378TCC (7257A>T 7258T>C), GTA2382GCT (7270T>C 7271A>T), CGT2383AAA (7272C>A 7273G>A 7274T>A), GTA2384ATT (7275G>A 7277A>T), GCG2385GCT (7280G>T), TCT2386GCC (7281T>G 7283T>C), GAG2387AAA (7284G>A 7286G>A), TTG2388GAG (7287T>G 7288T>A), ATT2389GTG (7290A>G 7292T>G), GGA2390AAT (7293G>A 7294G>A 7295A>T), GGG2391GGA (7298G>A), GTT2393GTG (7304T>G), CTC2394CTA (7307C>A), GTA2395GTT (7310A>T), GAA2396GAG (7313A>G), ACA2398ACT (7319A>T), AGT2399TGT (7320A>T), TTG2400CTC (7323T>C 7325G>C), TTT2402TTC (7331T>C), GAT2403AAC (7332G>A 7334T>C), GCT2404GCC (7337T>C), CTC2405CTT (7340C>T), AAT2406AAG (7343T>G), GGA2410GGG (7355A>G), CCA2411CCT (7358A>T), TAC2412TAT (7361C>T), ATT2413ATC (7364T>C), ATG2417TTG (7374A>T), GAA2418CAA (7377G>C), GGG2419AAG (7380G>A 7381G>A), ATT2420ATC (7385T>C), GGA2421GGT (7388A>T), CTA2422CAT (7390T>A 7391A>T), GGA2424GGT (7397A>T), TTA2425CTT (7398T>C 7400A>T), TAT2426AAC (7401T>A 7403T>C), AAG2427AAC (7406G>C), TTG2428TTA (7409G>A), GTG2429TTG (7410G>T), GAG2430ATG (7413G>A 7414A>T), CCG2431GCT (7416C>G 7418G>T), CAA2433GAG (7422C>G 7424A>G), AAT2434GAT (7425A>G), ATG2436ACA (7432T>C 7433G>A), GCT2437GCA (7436T>A), AGT2438TAT (7437A>T 7438G>A), GCT2439GCC (7442T>C), CTC2440ATG (7443C>A 7445C>G), TGC2441TGT (7448C>T), GTG2442ATC (7449G>A 7451G>C), GCT2444TCA (7455G>T 7457T>A), TTT2445CTT (7458T>C), GTA2446GCT (7462T>C 7463A>T), AAT2447CTT (7464A>C 7465A>T), AAA2448GGG (7467A>G 7468A>G 7469A>G), GTT2449CCA (7470G>C 7471T>C 7472T>A), GGT2450AAT (7473G>A 7474G>A), GAT2451ACT (7476G>A 7477A>C), GAT2452GAG (7481T>G), CCT2453CCG (7484T>G), ATA2454ATG (7487A>G), ATC2455ACT (7489T>C 7490C>T), AAG2457GTT (7494A>G 7495A>T 7496G>T), GGT2458GGA (7499T>A), GTG2459AAA (7500G>A 7501T>A 7502G>A), CTA2460ACG (7503C>A 7504T>C 7505A>G), AGA2461CTG (7506A>C 7507G>T 7508A>G), GGT2462GGA (7511T>A), GAG2463AAG (7512G>A), ATT2464ATA (7517T>A), GTA2465GTG (7520A>G), ATG2466CCA (7521A>C 7522T>C 7523G>A), CCA2467GCT (7524C>G 7526A>T), CGC2468AGG (7527C>A 7529C>G), CCA2470CCC (7535A>C), TCA2472AAT (7539T>A 7540C>A 7541A>T), TTT2473TTC (7544T>C), GGG2474GGA (7547G>A), GAC2476GAT (7553C>T), CCA2477CCG (7556A>G), ATT2478ATA (7559T>A), TTC2479TTT (7562C>T), CCA2481CCT (7568A>T), CTT2482CAT (7570T>A), AAC2483GGC (7572A>G 7573A>G), TGG2484TAT (7576G>A 7577G>T), AGA2485GAC (7578A>G 7579G>A 7580A>C), AAG2486CAG (7581A>C), ACA2487ACT (7586A>T), TTT2488TAC (7588T>A 7589T>C), ATG2492CCC (7599A>C 7600T>C 7601G>C), GCC2493AAG (7602G>A 7603C>A 7604C>G), GAG2494GAA (7607G>A), GAG2495GAA (7610G>A), AAT2497AAC (7616T>C), ATG2498AAG (7618T>A), ATA2499ATT (7622A>T), CAT2501CAC (7628T>C), CGA2502CGC (7631A>C), TTT2503GGT (7632T>G 7633T>G), CGA2504AAA (7635C>A 7636G>A) | | | | | | | | |

##### Proteins

| | Begin | End | Coverage | Score | Concordance | Matches | Identities | I/D/M/F* | Stop Codons |
|---|---|---|---|---|---|---|---|---|---|
| polyprotein (YP_007027011.1) | 2336 | 2509 | 6.0% | 732 | 60.0% | 174 (100%) | 100 (57.5%) | 0/0/0/0 | 0 |
| Protein mutations: | I2337V (7134A>G 7136T>G), Q2345K (7158C>A), K2348E (7167A>G), K2351R (7177A>G 7178A>G), Q2352A (7179C>G 7180A>C), F2354L (7185T>C), P2356Q (7192C>A 7193C>G), T2357S (7194A>T), I2360F (7203A>T 7205C>T), I2361Q (7206A>C 7207T>A 7208C>G), R2363L (7213G>T 7214G>T), V2365L (7218G>C), P2370L (7234C>T 7235A>T), T2373E (7242A>G 7243C>A 7244A>G), V2374P (7245G>C 7246T>C), E2376D (7253A>T), I2378S (7257A>T 7258T>C), V2382A (7270T>C 7271A>T), R2383K (7272C>A 7273G>A 7274T>A), V2384I (7275G>A 7277A>T), S2386A (7281T>G 7283T>C), E2387K (7284G>A 7286G>A), L2388E (7287T>G 7288T>A), I2389V (7290A>G 7292T>G), G2390N (7293G>A 7294G>A 7295A>T), S2399C (7320A>T), D2403N (7332G>A 7334T>C), N2406K (7343T>G), M2417L (7374A>T), E2418Q (7377G>C), G2419K (7380G>A 7381G>A), L2422H (7390T>A 7391A>T), Y2426N (7401T>A 7403T>C), K2427N (7406G>C), V2429L (7410G>T), E2430M (7413G>A 7414A>T), P2431A (7416C>G 7418G>T), Q2433E (7422C>G 7424A>G), N2434D (7425A>G), M2436T (7432T>C 7433G>A), S2438Y (7437A>T 7438G>A), L2440M (7443C>A 7445C>G), V2442I (7449G>A 7451G>C), A2444S (7455G>T 7457T>A), F2445L (7458T>C), V2446A (7462T>C 7463A>T), N2447L (7464A>C 7465A>T), K2448G (7467A>G 7468A>G 7469A>G), V2449P (7470G>C 7471T>C 7472T>A), G2450N (7473G>A 7474G>A), D2451T (7476G>A 7477A>C), D2452E (7481T>G), I2454M (7487A>G), I2455T (7489T>C 7490C>T), K2457V (7494A>G 7495A>T 7496G>T), V2459K (7500G>A 7501T>A 7502G>A), L2460T (7503C>A 7504T>C 7505A>G), R2461L (7506A>C 7507G>T 7508A>G), E2463K (7512G>A), M2466P (7521A>C 7522T>C 7523G>A), P2467A (7524C>G 7526A>T), S2472N (7539T>A 7540C>A 7541A>T), L2482H (7570T>A), N2483G (7572A>G 7573A>G), W2484Y (7576G>A 7577G>T), R2485D (7578A>G 7579G>A 7580A>C), K2486Q (7581A>C), F2488Y (7588T>A 7589T>C), M2492P (7599A>C 7600T>C 7601G>C), A2493K (7602G>A 7603C>A 7604C>G), M2498K (7618T>A), F2503G (7632T>G 7633T>G), R2504K (7635C>A 7636G>A) | | | | | | | | |

| | Begin | End | Coverage | Score | Concordance | Matches | Identities | I/D/M/F* | Stop Codons |
|---|---|---|---|---|---|---|---|---|---|
| **NT** | **7130** | **7650** | **5.8%** | **218** | **20.9%** | **521 (100%)** | **315 (60.5%)** | **0/0** | |
| Codon mutations: | GCA2335..G (7130A>G), CCA2336CCG (7133A>G), ATT2337GTG (7134A>G 7136T>G), TTC2339TTT (7142C>T), GTA2340GTT (7145A>T), ACA2341ACC (7148A>C), GGG2342GGA (7151G>A), GCA2344GCT (7157A>T), CAA2345AAA (7158C>A), AAA2346AAG (7163A>G), AAG2348GAG (7167A>G), GTA2350GTT (7175A>T), AAA2351AGG (7177A>G 7178A>G), CAA2352GCA (7179C>G 7180A>C), ATA2353ATT (7184A>T), TTT2354CTT (7185T>C), GGT2355GGC (7190T>C), CCC2356CAG (7192C>A 7193C>G), ACT2357TCT (7194A>T), CCA2359CCC (7202A>C), ATC2360TTT (7203A>T 7205C>T), ATC2361CAG (7206A>C 7207T>A 7208C>G), TCT2362TCC (7211T>C), CGG2363CTT (7213G>T 7214G>T), AAA2364AAG (7217A>G), GTT2365CTT (7218G>C), CCA2368CCT (7229A>T), CCA2370CTT (7234C>T 7235A>T), GGA2372GGG (7241A>G), ACA2373GAG (7242A>G 7243C>A 7244A>G), GTT2374CCT (7245G>C 7246T>C), GAG2375GAA (7250G>A), GAA2376GAT (7253A>T), ATT2377ATA (7256T>A), ATC2378TCC (7257A>T 7258T>C), GTA2382GCT (7270T>C 7271A>T), CGT2383AAA (7272C>A 7273G>A 7274T>A), GTA2384ATT (7275G>A 7277A>T), GCG2385GCT (7280G>T), TCT2386GCC (7281T>G 7283T>C), GAG2387AAA (7284G>A 7286G>A), TTG2388GAG (7287T>G 7288T>A), ATT2389GTG (7290A>G 7292T>G), GGA2390AAT (7293G>A 7294G>A 7295A>T), GGG2391GGA (7298G>A), GTT2393GTG (7304T>G), CTC2394CTA (7307C>A), GTA2395GTT (7310A>T), GAA2396GAG (7313A>G), ACA2398ACT (7319A>T), AGT2399TGT (7320A>T), TTG2400CTC (7323T>C 7325G>C), TTT2402TTC (7331T>C), GAT2403AAC (7332G>A 7334T>C), GCT2404GCC (7337T>C), CTC2405CTT (7340C>T), AAT2406AAG (7343T>G), GGA2410GGG (7355A>G), CCA2411CCT (7358A>T), TAC2412TAT (7361C>T), ATT2413ATC (7364T>C), ATG2417TTG (7374A>T), GAA2418CAA (7377G>C), GGG2419AAG (7380G>A 7381G>A), ATT2420ATC (7385T>C), GGA2421GGT (7388A>T), CTA2422CAT (7390T>A 7391A>T), GGA2424GGT (7397A>T), TTA2425CTT (7398T>C 7400A>T), TAT2426AAC (7401T>A 7403T>C), AAG2427AAC (7406G>C), TTG2428TTA (7409G>A), GTG2429TTG (7410G>T), GAG2430ATG (7413G>A 7414A>T), CCG2431GCT (7416C>G 7418G>T), CAA2433GAG (7422C>G 7424A>G), AAT2434GAT (7425A>G), ATG2436ACA (7432T>C 7433G>A), GCT2437GCA (7436T>A), AGT2438TAT (7437A>T 7438G>A), GCT2439GCC (7442T>C), CTC2440ATG (7443C>A 7445C>G), TGC2441TGT (7448C>T), GTG2442ATC (7449G>A 7451G>C), GCT2444TCA (7455G>T 7457T>A), TTT2445CTT (7458T>C), GTA2446GCT (7462T>C 7463A>T), AAT2447CTT (7464A>C 7465A>T), AAA2448GGG (7467A>G 7468A>G 7469A>G), GTT2449CCA (7470G>C 7471T>C 7472T>A), GGT2450AAT (7473G>A 7474G>A), GAT2451ACT (7476G>A 7477A>C), GAT2452GAG (7481T>G), CCT2453CCG (7484T>G), ATA2454ATG (7487A>G), ATC2455ACT (7489T>C 7490C>T), AAG2457GTT (7494A>G 7495A>T 7496G>T), GGT2458GGA (7499T>A), GTG2459AAA (7500G>A 7501T>A 7502G>A), CTA2460ACG (7503C>A 7504T>C 7505A>G), AGA2461CTG (7506A>C 7507G>T 7508A>G), GGT2462GGA (7511T>A), GAG2463AAG (7512G>A), ATT2464ATA (7517T>A), GTA2465GTG (7520A>G), ATG2466CCA (7521A>C 7522T>C 7523G>A), CCA2467GCT (7524C>G 7526A>T), CGC2468AGG (7527C>A 7529C>G), CCA2470CCC (7535A>C), TCA2472AAT (7539T>A 7540C>A 7541A>T), TTT2473TTC (7544T>C), GGG2474GGA (7547G>A), GAC2476GAT (7553C>T), CCA2477CCG (7556A>G), ATT2478ATA (7559T>A), TTC2479TTT (7562C>T), CCA2481CCT (7568A>T), CTT2482CAT (7570T>A), AAC2483GGC (7572A>G 7573A>G), TGG2484TAT (7576G>A 7577G>T), AGA2485GAC (7578A>G 7579G>A 7580A>C), AAG2486CAG (7581A>C), ACA2487ACT (7586A>T), TTT2488TAC (7588T>A 7589T>C), ATG2492CCC (7599A>C 7600T>C 7601G>C), GCC2493AAG (7602G>A 7603C>A 7604C>G), GAG2494GAA (7607G>A), GAG2495GAA (7610G>A), AAT2497AAC (7616T>C), ATG2498AAG (7618T>A), ATA2499ATT (7622A>T), CAT2501CAC (7628T>C), CGA2502CGC (7631A>C), TTT2503GGT (7632T>G 7633T>G), CGA2504AAA (7635C>A 7636G>A) | | | | | | | | |
| HAM1-like protein (YP_007032446.1) | 28 | 201 | 77.0% | 732 | 60.0% | 174 (100%) | 100 (57.5%) | 0/0/0/0 | 0 |
| Protein mutations: | I29V (7134A>G 7136T>G), Q37K (7158C>A), K40E (7167A>G), K43R (7177A>G 7178A>G), Q44A (7179C>G 7180A>C), F46L (7185T>C), P48Q (7192C>A 7193C>G), T49S (7194A>T), I52F (7203A>T 7205C>T), I53Q (7206A>C 7207T>A 7208C>G), R55L (7213G>T 7214G>T), V57L (7218G>C), P62L (7234C>T 7235A>T), T65E (7242A>G 7243C>A 7244A>G), V66P (7245G>C 7246T>C), E68D (7253A>T), I70S (7257A>T 7258T>C), V74A (7270T>C 7271A>T), R75K (7272C>A 7273G>A 7274T>A), V76I (7275G>A 7277A>T), S78A (7281T>G 7283T>C), E79K (7284G>A 7286G>A), L80E (7287T>G 7288T>A), I81V (7290A>G 7292T>G), G82N (7293G>A 7294G>A 7295A>T), S91C (7320A>T), D95N (7332G>A 7334T>C), N98K (7343T>G), M109L (7374A>T), E110Q (7377G>C), G111K (7380G>A 7381G>A), L114H (7390T>A 7391A>T), Y118N (7401T>A 7403T>C), K119N (7406G>C), V121L (7410G>T), E122M (7413G>A 7414A>T), P123A (7416C>G 7418G>T), Q125E (7422C>G 7424A>G), N126D (7425A>G), M128T (7432T>C 7433G>A), S130Y (7437A>T 7438G>A), L132M (7443C>A 7445C>G), V134I (7449G>A 7451G>C), A136S (7455G>T 7457T>A), F137L (7458T>C), V138A (7462T>C 7463A>T), N139L (7464A>C 7465A>T), K140G (7467A>G 7468A>G 7469A>G), V141P (7470G>C 7471T>C 7472T>A), G142N (7473G>A 7474G>A), D143T (7476G>A 7477A>C), D144E (7481T>G), I146M (7487A>G), I147T (7489T>C 7490C>T), K149V (7494A>G 7495A>T 7496G>T), V151K (7500G>A 7501T>A 7502G>A), L152T (7503C>A 7504T>C 7505A>G), R153L (7506A>C 7507G>T 7508A>G), E155K (7512G>A), M158P (7521A>C 7522T>C 7523G>A), P159A (7524C>G 7526A>T), S164N (7539T>A 7540C>A 7541A>T), L174H (7570T>A), N175G (7572A>G 7573A>G), W176Y (7576G>A 7577G>T), R177D (7578A>G 7579G>A 7580A>C), K178Q (7581A>C), F180Y (7588T>A 7589T>C), M184P (7599A>C 7600T>C 7601G>C), A185K (7602G>A 7603C>A 7604C>G), M190K (7618T>A), F195G (7632T>G 7633T>G), R196K (7635C>A 7636G>A) | | | | | | | | |
| Codon mutations: | GCA27..G (7130A>G), CCA28CCG (7133A>G), ATT29GTG (7134A>G 7136T>G), TTC31TTT (7142C>T), GTA32GTT (7145A>T), ACA33ACC (7148A>C), GGG34GGA (7151G>A), GCA36GCT (7157A>T), CAA37AAA (7158C>A), AAA38AAG (7163A>G), AAG40GAG (7167A>G), GTA42GTT (7175A>T), AAA43AGG (7177A>G 7178A>G), CAA44GCA (7179C>G 7180A>C), ATA45ATT (7184A>T), TTT46CTT (7185T>C), GGT47GGC (7190T>C), CCC48CAG (7192C>A 7193C>G), ACT49TCT (7194A>T), CCA51CCC (7202A>C), ATC52TTT (7203A>T 7205C>T), ATC53CAG (7206A>C 7207T>A 7208C>G), TCT54TCC (7211T>C), CGG55CTT (7213G>T 7214G>T), AAA56AAG (7217A>G), GTT57CTT (7218G>C), CCA60CCT (7229A>T), CCA62CTT (7234C>T 7235A>T), GGA64GGG (7241A>G), ACA65GAG (7242A>G 7243C>A 7244A>G), GTT66CCT (7245G>C 7246T>C), GAG67GAA (7250G>A), GAA68GAT (7253A>T), ATT69ATA (7256T>A), ATC70TCC (7257A>T 7258T>C), GTA74GCT (7270T>C 7271A>T), CGT75AAA (7272C>A 7273G>A 7274T>A), GTA76ATT (7275G>A 7277A>T), GCG77GCT (7280G>T), TCT78GCC (7281T>G 7283T>C), GAG79AAA (7284G>A 7286G>A), TTG80GAG (7287T>G 7288T>A), ATT81GTG (7290A>G 7292T>G), GGA82AAT (7293G>A 7294G>A 7295A>T), GGG83GGA (7298G>A), GTT85GTG (7304T>G), CTC86CTA (7307C>A), GTA87GTT (7310A>T), GAA88GAG (7313A>G), ACA90ACT (7319A>T), AGT91TGT (7320A>T), TTG92CTC (7323T>C 7325G>C), TTT94TTC (7331T>C), GAT95AAC (7332G>A 7334T>C), GCT96GCC (7337T>C), CTC97CTT (7340C>T), AAT98AAG (7343T>G), GGA102GGG (7355A>G), CCA103CCT (7358A>T), TAC104TAT (7361C>T), ATT105ATC (7364T>C), ATG109TTG (7374A>T), GAA110CAA (7377G>C), GGG111AAG (7380G>A 7381G>A), ATT112ATC (7385T>C), GGA113GGT (7388A>T), CTA114CAT (7390T>A 7391A>T), GGA116GGT (7397A>T), TTA117CTT (7398T>C 7400A>T), TAT118AAC (7401T>A 7403T>C), AAG119AAC (7406G>C), TTG120TTA (7409G>A), GTG121TTG (7410G>T), GAG122ATG (7413G>A 7414A>T), CCG123GCT (7416C>G 7418G>T), CAA125GAG (7422C>G 7424A>G), AAT126GAT (7425A>G), ATG128ACA (7432T>C 7433G>A), GCT129GCA (7436T>A), AGT130TAT (7437A>T 7438G>A), GCT131GCC (7442T>C), CTC132ATG (7443C>A 7445C>G), TGC133TGT (7448C>T), GTG134ATC (7449G>A 7451G>C), GCT136TCA (7455G>T 7457T>A), TTT137CTT (7458T>C), GTA138GCT (7462T>C 7463A>T), AAT139CTT (7464A>C 7465A>T), AAA140GGG (7467A>G 7468A>G 7469A>G), GTT141CCA (7470G>C 7471T>C 7472T>A), GGT142AAT (7473G>A 7474G>A), GAT143ACT (7476G>A 7477A>C), GAT144GAG (7481T>G), CCT145CCG (7484T>G), ATA146ATG (7487A>G), ATC147ACT (7489T>C 7490C>T), AAG149GTT (7494A>G 7495A>T 7496G>T), GGT150GGA (7499T>A), GTG151AAA (7500G>A 7501T>A 7502G>A), CTA152ACG (7503C>A 7504T>C 7505A>G), AGA153CTG (7506A>C 7507G>T 7508A>G), GGT154GGA (7511T>A), GAG155AAG (7512G>A), ATT156ATA (7517T>A), GTA157GTG (7520A>G), ATG158CCA (7521A>C 7522T>C 7523G>A), CCA159GCT (7524C>G 7526A>T), CGC160AGG (7527C>A 7529C>G), CCA162CCC (7535A>C), TCA164AAT (7539T>A 7540C>A 7541A>T), TTT165TTC (7544T>C), GGG166GGA (7547G>A), GAC168GAT (7553C>T), CCA169CCG (7556A>G), ATT170ATA (7559T>A), TTC171TTT (7562C>T), CCA173CCT (7568A>T), CTT174CAT (7570T>A), AAC175GGC (7572A>G 7573A>G), TGG176TAT (7576G>A 7577G>T), AGA177GAC (7578A>G 7579G>A 7580A>C), AAG178CAG (7581A>C), ACA179ACT (7586A>T), TTT180TAC (7588T>A 7589T>C), ATG184CCC (7599A>C 7600T>C 7601G>C), GCC185AAG (7602G>A 7603C>A 7604C>G), GAG186GAA (7607G>A), GAG187GAA (7610G>A), AAT189AAC (7616T>C), ATG190AAG (7618T>A), ATA191ATT (7622A>T), CAT193CAC (7628T>C), CGA194CGC (7631A>C), TTT195GGT (7632T>G 7633T>G), CGA196AAA (7635C>A 7636G>A) | | | | | | | | |

*: Inserts / Deletes / Misaligned / Frameshifts

#### Analysis details

This analysis was performed with panviral2.64

### NGS Details (UN18_val): Human gammaherpesvirus 8

#### Assembly

| | |
|---|---|
| **Coverage Length** | 119 (1 contig(s)) |
| **Depth Of Coverage** | 183.0 |
| **Number Of Reads** | 294 |
| **Reads Per Million** | 5.52 rpm (after QC) |
| **Ambiguities** | 0 |
| **Assembly Method** | de novo + reference guided assembly |
| **Consensus Caller** | Bcf Tools |

#### Coverage Map

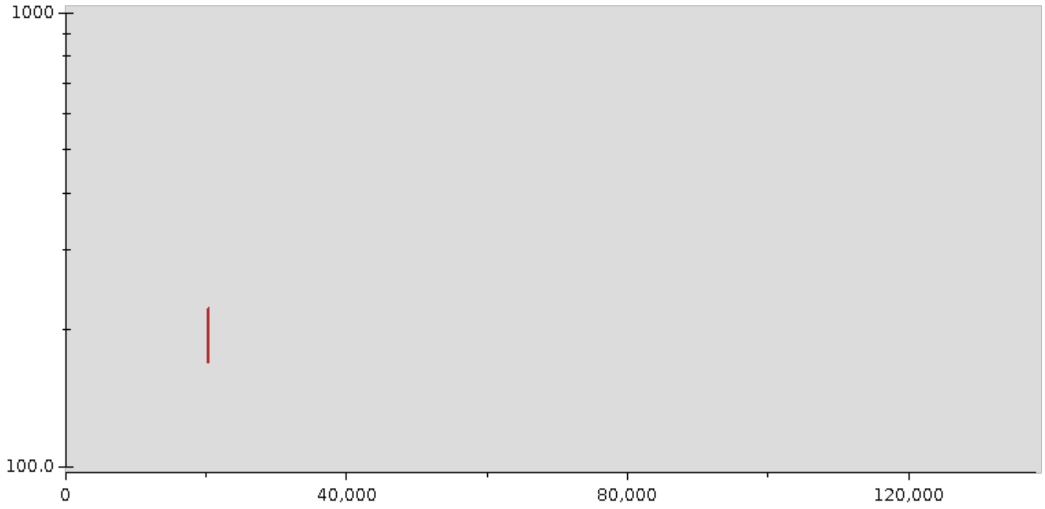

#### Assignment

| | |
|---|---|
| **Type** | Human gammaherpesvirus 8 (Taxonomy ID: 37296) |
| **Subtype** | Could not assign |
| **Reference Genome** | NC_009333.1 |
| **NT Identity (%)** | 83.1933 |
| **AA Identity (%)** | 84.6154 |
| **Number Of Stop Codons** | 0 |
| **Number Of CDS** | 86 |

#### Alignment

| | |
|---|---|
| **Alignment Score** | 158.0 (NT) + 260.0 (AA) = 418.0 |
| **Concordance (%)** | 77.4074 |
| **Alignment Method** | Local, heuristic, nucleotide (BLASTN) |

#### Genome Region

Sequence starts at position 20278 and ends at position 20396 relative to NC_009333.1 reference sequence.

#### Alignment Detailed Statistics

| | Begin | End | Coverage | Score | Concordance | Matches | Identities | I/D/M/F* | Stop Codons |
|---|---|---|---|---|---|---|---|---|---|
| **NT** | **20278** | **20396** | **0.1%** | **158** | **66.4%** | **119 (100%)** | **99 (83.2%)** | **0/0** | |

Mutations: 20293A>T, 20296A>C, 20311G>A, 20314T>G, 20315C>G, 20317C>A, 20320C>T, 20322T>G, 20331G>T, 20341A>C, 20347A>C, 20350G>A, 20352C>T, 20353A>G, 20369C>G, 20370A>C, 20371T>A, 20376A>T, 20383G>A, 20386C>T

*: Inserts / Deletes / Misaligned / Frameshifts

#### Analysis details

This analysis was performed with panviral2.64

### NGS Details (UN18_val): Cladosporium fulvum T-1 virus

#### Assembly

| | |
|---|---|
| Coverage Length | 1209 (4 contig(s)) |
| Depth Of Coverage | 15.0 |
| Number Of Reads | 179 |
| Reads Per Million | 3.36 rpm (after QC) |
| Ambiguities | 0 |
| Assembly Method | de novo + reference guided assembly |
| Consensus Caller | Bcf Tools |

#### Coverage Map

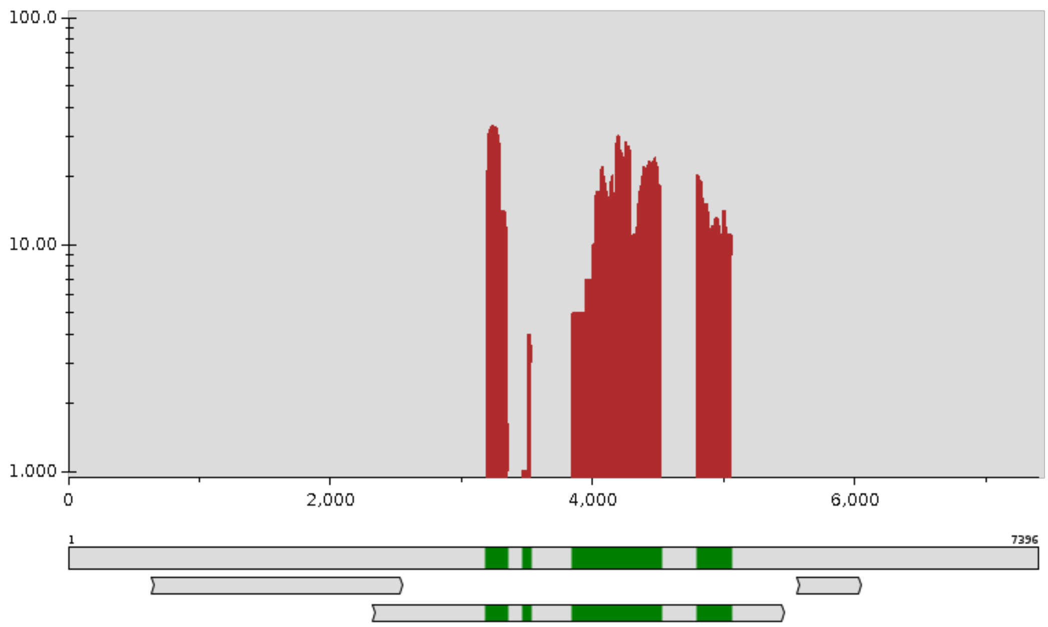

#### Assignment

| | |
|---|---|
| Type | Cladosporium fulvum T-1 virus (Taxonomy ID: 2052899) |
| Reference Genome | NC_043491.1 |
| NT Identity (%) | 53.1486 |
| AA Identity (%) | 42.5693 |
| Number Of Stop Codons | 0 |
| Number Of CDS | 3 |

#### Alignment

| | |
|---|---|
| Alignment Score | 96.0 (NT) + 1234.0 (AA) = 1330.0 |
| Concordance (%) | 26.2586 |

| **Alignment Method** | Global, seeded, nucleotide + amino acids (AGA) |
|---|---|

#### Genome Region

Sequence starts at position 3189 and ends at position 5061 relative to NC_043491.1 reference sequence.

#### Alignment Detailed Statistics

| | Begin | End | Coverage | Score | Concordance | Matches | Identities | I/D/M/F* | Stop Codons |
|---|---|---|---|---|---|---|---|---|---|
| **NT** | **3189** | **5061** | **16.3%** | **96** | **4.2%** | **1182 (97.0%)** | **633 (52.0%)** | **9/27** | |
| Mutations: | 3189G>C, 3190C>A, 3191A>G, 3194T>C, 3195G>T, 3197T>C, 3198C>T, 3200C>A, 3201G>A, 3204T>A, 3205C>G, 3207T>A, 3208T>C, 3210A>C, 3211A>T, 3214G>T, 3216A>T, 3217A>G, 3219G>T, 3220A>G, 3221G>C, 3222G>C, 3223C>A, 3224A>C, 3225A>C, 3226G>T, 3229C>G, 3230G>T, 3231C>T, 3232C>G, 3233T>C, 3234T>C, 3236C>A, 3237C>A, 3238T>G, 3243A>T, 3244C>G, 3246A>C, 3251T>A, 3253_3254insAGGGATATT, 3261A>G, 3262G>A, 3265A>T, 3266A>G, 3268C>G, 3269A>T, 3271T>G, 3272C>A, 3273A>T, 3280G>T, 3281A>T, 3282A>C, 3283A>T, 3284G>A, 3285A>C, 3289T>G, 3292A>C, 3294G>C, 3295G>T, 3296G>A, 3297G>T, 3298A>C, 3301C>T, 3303T>C, 3304A>T, 3307T>C, 3308C>A, 3309A>G, 3314T>G, 3317G>C, 3318A>C, 3319G>T, 3320A>G, 3321A>C, 3326C>T, 3328A>G, 3329C>G, 3330A>C, 3331G>T, 3332A>G, 3333C>A, 3334C>A, 3337A>G, 3338C>A, 3339G>A, 3340A>G, 3341G>A, 3343A>G, 3344T>C, 3345G>A, 3347C>T, 3457C>T, 3460A>G, 3461A>C, 3464T>C, 3469C>T, 3470G>A, 3473A>G, 3475C>A, 3478G>T, 3479A>G, 3481C>G, 3488C>A, 3489G>A, 3490A>G, 3496G>C, 3497C>A, 3499A>G, 3502C>T, 3504A>G, 3505C>A, 3508C>T, 3511A>T, 3514A>T, 3515G>T, 3516C>T, 3517A>G, 3518C>T, 3519A>T, 3520A>T, 3524A>C, 3525G>A, 3527T>C, 3529A>T, 3530A>C, 3531C>A, 3532C>G, 3839A>T, 3841G>T, 3842A>G, 3843C>T, 3844A>C, 3845G>A, 3846C>A, 3847A>G, 3848C>A, 3849C>T, 3850C>G, 3856A>G, 3859C>T, 3862A>G, 3865C>T, 3866C>G, 3867A>C, 3869A>C, 3871G>A, 3874A>G, 3880C>A, 3882A>C, 3883G>C, 3886T>C, 3887T>C, 3892C>G, 3893T>C, 3894T>A, 3898C>A, 3899A>G, 3901C>T, 3905A>G, 3906C>A, 3908A>G, 3909C>A, 3910A>C, 3912G>A, 3913G>A, 3914A>G, 3918C>G, 3922C>G, 3929G>A, 3930C>A, 3931A>G, 3934G>A, 3935A>G, 3936C>T, 3937A>G, 3940G>A, 3941T>G, 3943A>G, 3946C>T, 3948G>T, 3952A>T, 3957C>A, 3959G>A, 3960A>C, 3964G>C, 3970A>T, 3973C>G, 3975A>C, 3979T>A, 3980G>C, 3982A>T, 3983C>A, 3984A>G, 3985G>A, 3988A>T, 3992C>T, 3997A>G, 4000C>A, 4003C>T, 4009C>T, 4010A>T, 4025A>G, 4027G>T, 4029A>G, 4030C>A, 4033T>C, 4038A>C, 4041C>A, 4045A>C, 4048A>T, 4052A>C, 4057G>T, 4058A>G, 4059T>A, 4060G>T, 4061C>T, 4063T>G, 4064A>C, 4065C>T, 4066A>G, 4068G>A, 4069A>G, 4072A>G, 4075C>A, 4080A>C, 4081C>G, 4085A>G, 4091G>T, 4092G>C, 4093A>C, 4094A>G, 4097G>A, 4100C>T, 4101A>G, 4102G>C, 4103A>G, 4104C>A, 4105C>A, 4111G>T, 4115A>C, 4117A>G, 4119G>A, 4120A>T, 4129A>C, 4130C>G, 4131A>C, 4132G>A, 4133T>A, 4134G>T, 4135C>T, 4138T>A, 4143C>A, 4144C>A, 4149C>T, 4150G>A, 4153T>G, 4154C>A, 4155G>A, 4156A>G, 4159A>G, 4160T>C, 4161T>C, 4162C>A, 4165T>C, 4166G>T, 4167G>T, 4169A>G, 4170G>A, 4171C>G, 4172A>C, 4173A>T, 4174G>A, 4175G>C, 4176A>C, 4177A>T, 4178G>T, 4180C>T, 4181C>G, 4183C>G, 4184A>G, 4186C>A, 4187G>C, 4189G>C, 4192C>T, 4195C>T, 4201T>G, 4204T>C, 4206T>A, 4207G>A, 4210A>T, 4211A>G, 4213A>G, 4216C>A, 4218C>G, 4219A>C, 4220T>G, 4221G>T, 4222T>A, 4223C>T, 4225A>G, 4226A>G, 4227C>T, 4228A>G, 4231G>A, 4232_4237delACACAC, 4240T>A, 4243G>T, 4244A>G, 4248G>A, 4252C>T, 4255A>G, 4263A>T, 4265T>G, 4267T>A, 4268T>A, 4269C>G, 4271C>A, 4277A>T, 4279G>A, 4281C>A, 4283A>G, 4284C>A, 4285A>G, 4286G>A, 4288G>A, 4293A>G, 4294G>A, 4295A>C, 4296A>G, 4297C>G, 4301G>A, 4302A>C, 4303C>A, 4304A>G, 4309T>A, 4312C>G, 4319C>A, 4321T>G, 4322C>A, 4323T>C, 4330T>C, 4333T>C, 4334G>C, 4335C>A, 4337G>T, 4338C>G, 4340A>C, 4344A>G, 4345A>C, 4346C>A, 4347A>C, 4348T>C, 4355G>C, 4356T>A, 4357G>C, 4361G>T, 4363C>G, 4364G>C, 4365A>T, 4366G>T, 4369C>T, 4370C>T, 4372A>T, 4373_4375delCCG, 4381A>C, 4382A>T, 4383C>T, 4384G>A, 4385A>G, 4387T>C, 4388C>A, 4389T>A, 4390T>A, 4391T>A, 4393A>C, 4397C>A, 4400A>G, 4401A>T, 4403A>G, 4404A>C, 4405T>C, 4406C>A, 4407T>C, 4410C>G, 4411G>C, 4418A>C, 4419C>A, 4420G>A, 4421A>T, 4423G>T, 4424A>C, 4425C>A, 4426G>A, 4430G>A, 4438C>G, 4440G>C, 4441A>C, 4443G>A, 4444A>G, 4450C>A, 4451C>A, 4453C>G, 4457T>C, 4458C>A, 4459G>A, 4462G>T, 4463C>T, 4465G>C, 4466C>T, 4468T>G, 4470G>C, 4471G>A, 4472C>G, 4474G>A, 4476A>T, 4477C>T, 4478A>G, 4480G>C, 4482T>A, 4486A>G, 4487A>C, 4489C>G, 4490A>G, 4492A>G, 4497C>A, 4498T>G, 4504A>C, 4506C>A, 4507A>G, 4509A>G, 4510G>C, 4513C>T, 4515G>T, 4516C>T, 4517C>G, 4518C>T, 4519G>T, 4522A>C, 4789A>T, 4790C>G, 4795C>A, 4796A>G, 4797G>C, 4798C>T, 4800T>A, 4803C>A, 4804A>T, 4806T>G, 4807C>G, 4810A>C, 4813G>A, 4819A>G, 4820C>G, 4821T>A, 4823A>G, 4825G>T, 4829C>G, 4830T>A, 4831A>G, 4833G>T, 4838A>G, 4845A>C, 4846A>T, 4849C>T, 4850G>C, 4852A>T, 4853C>G, 4854A>T, 4855C>G, 4861A>G, 4865A>G, 4870A>G, 4872C>T, 4873T>G, 4875C>A, 4878G>A, 4879G>A, 4881A>G, 4882C>T, 4883G>C, 4885G>C, 4886A>G, 4887A>C, 4889T>G, 4890A>C, 4891C>T, 4896A>T, 4897C>A, 4901C>G, 4904T>C, 4905T>C, 4906C>A, 4907A>C, 4908G>T, 4910A>C, 4912A>C, 4913C>A, 4914C>T, 4916C>G, 4918A>C, 4919A>G, 4920C>A, 4923A>G, 4933C>A, 4934G>A, 4935A>G, 4936G>T, 4937G>A, 4940A>T, 4942G>C, 4954T>C, 4957G>A, 4958A>T, 4959A>G, 4960A>C, 4961C>T, 4963C>A, 4966G>A, 4968G>A, 4969G>A, 4973_4990delAAGGATCGAGTCACAGGA, 4991C>G, 4995C>G, 4996C>G, 4998A>G, 4999T>C, 5000G>A, 5001A>G, 5004T>A, 5005G>T, 5009C>A, 5011A>G, 5014C>T, 5015A>G, 5017G>C, 5020C>G, 5027C>T, 5029C>T, 5031C>G, 5032A>C, 5038T>C, 5040C>G, 5042C>G, 5043A>T, 5044C>A, 5050T>A, 5053T>A, 5055C>T, 5056A>C, 5057T>C, 5059A>G, 5060G>A, 5061A>C | | | | | | | | |
| **CDS** | | | | | | | | | |
| homologue_of_retroviral_POL_genes | 289 | 912 | 38.5% | 1234 | 44.2% | 394 (97.0%) | 169 (41.6%) | 3/9/0/0 | 0 |
| Protein mutations: | K289E (3191A>G), W290L (3194T>C 3195G>T), S291L (3197T>C 3198C>T), R292K (3200C>A 3201G>A), L293Q (3204T>A 3205C>G), F294Y (3207T>A 3208T>C), E295A (3210A>C 3211A>T), E296D (3214G>T), E297V (3216A>T 3217A>G), R298M (3219G>T 3220A>G), G299P (3221G>C 3222G>C 3223C>A), K300P (3224A>C 3225A>C 3226G>T), D301E (3229C>G), A302L (3230G>T 3231C>T 3232C>G), L303P (3233T>C 3234T>C), P304K (3236C>A 3237C>A 3238T>G), H306L (3243A>T 3244C>G), Q307P (3246A>C), W309R (3251T>A), W309_D310insRDI (3253_3254insAGGGATATT), K312R (3261A>G 3262G>A), N314E (3266A>G 3268C>G), I315L (3269A>T 3271T>G), Q316M (3272C>A 3273A>T), K319S (3281A>T 3282A>C 3283A>T), E320T (3284G>A 3285A>C), W323S (3294G>C 3295G>T), G324I (3296G>A 3297G>T 3298A>C), L326P (3303T>C 3304A>T), Q328R (3308C>A 3309A>G), S330A (3314T>G), E331P (3317G>C 3318A>C 3319G>T), K332A (3320A>G 3321A>C), Q335A (3329C>G 3330A>C 3331G>T), T336E (3332A>G 3333C>A 3334C>A), R338K (3338C>A 3339G>A 3340A>G), E339K (3341G>A 3343A>G), W340Q (3344T>C 3345G>A), K379Q (3461A>C), E382K (3470G>A), I383V (3473A>G 3475C>A), I385V (3479A>G 3481C>G), R388K (3488C>A 3489G>A 3490A>G), L391M (3497C>A 3499A>G), N393R (3504A>G 3505C>A), E395D (3511A>T), E396D (3514A>T), A397L (3515G>T 3516C>T 3517A>G), Q398F (3518C>T 3519A>T 3520A>T), R400Q (3524A>C 3525G>A), T402Q (3530A>C 3531C>A 3532C>G), K505Y (3839A>T 3841G>T), T506V (3842A>G 3843C>T 3844A>C), A507K (3845G>A 3846C>A 3847A>G), P508M (3848C>A 3849C>T 3850C>G), H514A (3866C>G 3867A>C), K515Q (3869A>C 3871G>A), K519T (3882A>C 3883G>C), F523H (3893T>C 3894T>A), I525V (3899A>G 3901C>T), T527E (3905A>G 3906C>A), T528D (3908A>G 3909C>A 3910A>C), G529E (3912G>A 3913G>A), I530V (3914A>G), T531R (3918C>G), I532M (3922C>G), A535K (3929G>A 3930C>A 3931A>G), T537V (3935A>G 3936C>T 3937A>G), S539A (3941T>G 3943A>G), R541I (3948G>T), E542D (3952A>T), P544Q (3957C>A), E545T (3959G>A 3960A>C), K550T (3975A>C), D551E (3979T>A), V552L (3980G>C 3982A>T), Q553R (3983C>A 3984A>G 3985G>A), L556F (3992C>T), N562Y (4010A>T), K567D (4025A>G 4027G>T), D568G (4029A>G 4030C>A), K571T (4038A>C), T572K (4041C>A), M576L (4052A>C), M578D (4058A>G 4059T>A 4060G>T), T580L (4064A>C 4065C>T 4066A>G), R581K (4068G>A 4069A>G), D583E (4075C>A), N585T (4080A>C 4081C>G), K587E (4085A>G), G589S (4091G>T 4092G>C 4093A>C), K590E (4094A>G), E591K (4097G>A), Q592C (4100C>T 4101A>G 4102G>C), T593E (4103A>G 4104C>A 4105C>A), K597Q (4115A>C 4117A>G), R598N (4119G>A 4120A>T), E601D (4129A>C), Q602A (4130C>G 4131A>C 4132G>A), C603I (4133T>A 4134G>T 4135C>T), A606E (4143C>A 4144C>A), T608I (4149C>T 4150G>A), R610K (4154C>A 4155G>A 4156A>G), F612P (4160T>C 4161T>C 4162C>A), G614F (4166G>T 4167G>T), S615E (4169A>G 4170G>A 4171C>G), K616L (4172A>C 4173A>T 4174G>A), E617P (4175G>C 4176A>C 4177A>T), V618F (4178G>T 4180C>T), H619E (4181C>G 4183C>G), I620V (4184A>G 4186C>A), E621H (4187G>C 4189G>C), M627K (4206T>A 4207G>A), I629V (4211A>G 4213A>G), A631G (4218C>G 4219A>C), C632V (4220T>G 4221G>T 4222T>A), T634V (4226A>G 4227C>T 4228A>G), T636_H637del (4232_4237delACACAC), D638E (4240T>A), K640E (4244A>G), R641K (4248G>A), Y646F (4263A>T), Y647E (4265T>G 4267T>A), M651L (4277A>T 4279G>A), T652N (4281C>A), T653E (4283A>G 4284C>A 4285A>G), A654T (4286G>A 4288G>A), Q656R (4293A>G 4294G>A), N657R (4295A>C 4296A>G 4297C>G), D659T (4301G>A 4302A>C 4303C>A), I660V (4304A>G), H661Q (4309T>A), D662E (4312C>G), L665M (4319C>A 4321T>G), L666T (4322C>A 4323T>C), A670H (4334G>C 4335C>A), A671C (4337G>T 4338C>G), M672L (4340A>C), Q673R (4344A>G 4345A>C), H674T (4346C>A 4347A>C 4348T>C), V677H (4355G>C 4356T>A 4357G>C), V679L (4361G>T 4363C>G), E680L (4364G>C 4365A>T 4366G>T), P682S (4370C>T 4372A>T), P683del (4373_4375delCCG), L685F (4381A>C), T686L (4382A>T 4383C>T 4384G>A), I687V (4385A>G 4387T>C), L688K (4388C>A 4389T>A 4390T>A), S689T (4391T>A 4393A>C), H691N (4397C>A), K692V (4400A>G 4401A>T), N693A (4403A>G 4404A>C 4405T>C), L694T (4406C>A 4407T>C), T695S (4410C>G 4411G>C), T698Q (4418A>C 4419C>A 4420G>A), T699S (4421A>T 4423G>T), T700Q (4424A>C 4425C>A 4426G>A), E702K (4430G>A), R705P (4440G>C 4441A>C), R706K (4443G>A 4444A>G), S711Q (4457T>C 4458C>A 4459G>A), E712D (4462G>T), L713F (4463C>T 4465G>C), G715A (4470G>C 4471G>A), Q716E (4472C>G 4474G>A), Y717F (4476A>T 4477C>T), K718D (4478A>G 4480G>C), F719Y (4482T>A), I721L (4487A>C 4489C>G), K722E (4490A>G 4492A>G), T724K (4497C>A 4498T>G), T727K (4506C>A 4507A>G), E728G (4509A>G 4510G>C), G730V (4515G>T 4516C>T), P731V (4517C>G 4518C>T 4519G>T), Q822E (4790C>G), S824A (4796A>G 4797G>C 4798C>T), F825Y (4800T>A), S826Y (4803C>A 4804A>T), F827W (4806T>G 4807C>G), L832D (4820C>G 4821T>A), K833D (4823A>G 4825G>T), L835E (4829C>G 4830T>A 4831A>G), R836L (4833G>T), I838V (4838A>G), K840T (4845A>C 4846A>T), V842L (4850G>C 4852A>T), H843V (4853C>G 4854A>T 4855C>G), N847D (4865A>G), A849V (4872C>T 4873T>G), A850E (4875C>A), R851Q (4878G>A 4879G>A), H852R (4881A>G 4882C>T), A853P (4883G>C 4885G>C), K854A (4886A>G 4887A>C), Y855A (4889T>G 4890A>C 4891C>T), H857L (4896A>T 4897C>A), Q859E (4901C>G), F860P (4904T>C 4905T>C 4906C>A), R861L (4907A>C 4908G>T), T862P (4910A>C 4912A>C), P863I (4913C>A 4914C>T), P864A (4916C>G 4918A>C), T865E (4919A>G 4920C>A), K866R (4923A>G), D869E (4933C>A), E870S (4934G>A 4935A>G 4936G>T), V871I (4937G>A), T872S (4940A>T 4942G>C), K878C (4958A>T 4959A>G 4960A>C), R881K (4968G>A 4969G>A), K883_G888del (4973_4990delAAGGATCGAGTCACAGGA), Q889E (4991C>G), A890G (4995C>G 4996C>G), Y891C (4998A>G 4999T>C), D892S (5000G>A 5001A>G), M893N (5004T>A 5005G>T), L895M (5009C>A 5011A>G), M897V (5015A>G 5017G>C), L901F (5027C>T 5029C>T), T902S (5031C>G 5032A>C), A905G (5040C>G), H906V (5042C>G 5043A>T 5044C>A), A910V (5055C>T 5056A>C), S911P (5057T>C 5059A>G) | | | | | | | | |

| | Begin | End | Coverage | Score | Concordance | Matches | Identities | I/D/M/F* | Stop Codons |
|---|---|---|---|---|---|---|---|---|---|
| **NT** | **3189** | **5061** | **16.3%** | **96** | **4.2%** | **1182 (97.0%)** | **633 (52.0%)** | **9/27** | |

Codon mutations:

CGC288.CA (3189G>C 3190C>A), AAG289GAG (3191A>G), TGG290CTG (3194T>C 3195G>T), TCG291CTG (3197T>C 3198C>T), CGA292AAA (3200C>A 3201G>A), CTC293CAG (3204T>A 3205C>G), TTT294TAC (3207T>A 3208T>C), GAA295GCT (3210A>C 3211A>T), GAG296GAT (3214G>T), GAA297GTG (3216A>T 3217A>G), AGA298ATG (3219G>T 3220A>G), GGC299CCA (3221G>C 3222G>C 3223C>A), AAG300CCT (3224A>C 3225A>C 3226G>T), GAC301GAG (3229C>G), GCC302TTG (3230G>T 3231C>T 3232C>G), TTA303CCA (3233T>C 3234T>C), CCT304AAG (3236C>A 3237C>A 3238T>G), CAC306CTG (3243A>T 3244C>G), CAA307CCA (3246A>C), TGG309AGG (3251T>A), TGG309_GAT310insAGGGATATT (3253_3254insAGGGATATT), AAG312AGA (3261A>G 3262G>A), ATA313ATT (3265A>T), AAC314GAG (3266A>G 3268C>G), ATT315TTG (3269A>T 3271T>G), CAG316ATG (3272C>A 3273A>T), GGG318GGT (3280G>T), AAA319TCT (3281A>T 3282A>C 3283A>T), GAG320ACG (3284G>A 3285A>C), CCT321CCG (3289T>G), CCA322CCC (3292A>C), TGG323TCT (3294G>C 3295G>T), GGA324ATC (3296G>A 3297G>T 3298A>C), CCC325CCT (3301C>T), CTA326CCT (3303T>C 3304A>T), TAT327TAC (3307T>C), CAA328AGA (3308C>A 3309A>G), TCT330GCT (3314T>G), GAG331CCT (3317G>C 3318A>C 3319G>T), AAA332GCA (3320A>G 3321A>C), CTA334TTG (3326C>T 3328A>G), CAG335GCT (3329C>G 3330A>C 3331G>T), ACC336GAA (3332A>G 3333C>A 3334C>A), CTA337CTG (3337A>G), CGA338AAG (3338C>A 3339G>A 3340A>G), GAA339AAG (3341G>A 3343A>G), TGG340CAG (3344T>C 3345G>A), CTG341TTG (3347C>T), TAC377TAT (3457C>T), CGA378CGG (3460A>G), AAG379CAG (3461A>C), TTG380CTG (3464T>C), AAC381AAT (3469C>T), GAG382AAG (3470G>A), ATC383GTA (3473A>G 3475C>A), ACG384ACT (3478G>T), ATC385GTG (3479A>G 3481C>G), CGA388AAG (3488C>A 3489G>A 3490A>G), CCG390CCC (3496G>C), CTA391ATG (3497C>A 3499A>G), CCC392CCT (3502C>T), AAC393AGA (3504A>G 3505C>A), ATC394ATT (3508C>T), GAA395GAT (3511A>T), GAA396GAT (3514A>T), GCA397TTG (3515G>T 3516C>T 3517A>G), CAA398TTT (3518C>T 3519A>T 3520A>T), AGA400CAA (3524A>C 3525G>A), TTA401CTT (3527T>C 3529A>T), ACC402CAG (3530A>C 3531C>A 3532C>G), AAG505TAT (3839A>T 3841G>T), ACA506GTC (3842A>G 3843C>T 3844A>C), GCA507AAG (3845G>A 3846C>A 3847A>G), CCC508ATG (3848C>A 3849C>T 3850C>G), AAA510AAG (3856A>G), TGC511TGT (3859C>T), GAA512GAG (3862A>G), TTC513TTT (3865C>T), CAC514GCC (3866C>G 3867A>C), AAG515CAA (3869A>C 3871G>A), AAA516AAG (3874A>G), GTC518GTA (3880C>A), AAG519ACC (3882A>C 3883G>C), TTT520TTC (3886T>C), TTA521CTA (3887T>C), GGC522GGG (3892C>G), TTT523CAT (3893T>C 3894T>A), ATC524ATA (3898C>A), ATC525GTT (3899A>G 3901C>T), ACA527GAA (3905A>G 3906C>A), ACA528GAC (3908A>G 3909C>A 3910A>C), GGG529GAA (3912G>A 3913G>A), ATA530GTA (3914A>G), ACG531AGG (3918C>G), ATC532ATG (3922C>G), GCA535AAG (3929G>A 3930C>A 3931A>G), AAG536AAA (3934G>A), ACA537GTG (3935A>G 3936C>T 3937A>G), CAG538CAA (3940G>A), TCA539GCG (3941T>G 3943A>G), ATC540ATT (3946C>T), AGA541ATA (3948G>T), GAA542GAT (3952A>T), CCA544CAA (3957C>A), GAA545ACA (3959G>A 3960A>C), CCG546CCC (3964G>C), ACA548ACT (3970A>T), GTC549GTG (3973C>G), AAG550ACG (3975A>C), GAT551GAA (3979T>A), GTA552CTT (3980G>C 3982A>T), CAG553AGA (3983C>A 3984A>G 3985G>A), TCA554TCT (3988A>T), CTT556TTT (3992C>T), GGA557GGG (3997A>G), CTC558CTA (4000C>A), GCC559GCT (4003C>T), TAC561TAT (4009C>T), AAC562TAC (4010A>T), AAG567GAT (4025A>G 4027G>T), GAC568GGA (4029A>G 4030C>A), TAT569TAC (4033T>C), AAG571ACG (4038A>C), ACA572AAA (4041C>A), GCA573GCC (4045A>C), GCA574GCT (4048A>T), ATG576CTG (4052A>C), ACG577ACT (4057G>T), ATG578GAT (4058A>G 4059T>A 4060G>T), CTT579TTG (4061C>T 4063T>G), ACA580CTG (4064A>C 4065C>T 4066A>G), AGA581AAG (4068G>A 4069A>G), AAA582AAG (4072A>G), GAC583GAA (4075C>A), AAC585ACG (4080A>C 4081C>G), AAA587GAA (4085A>G), GGA589TCC (4091G>T 4092G>C 4093A>C), AAA590GAA (4094A>G), GAA591AAA (4097G>A), CAG592TGC (4100C>T 4101A>G 4102G>C), ACC593GAA (4103A>G 4104C>A 4105C>A), GCG595GCT (4111G>T), AAA597CAG (4115A>C 4117A>G), AGA598AAT (4119G>A 4120A>T), GAA601GAC (4129A>C), CAG602GCA (4130C>G 4131A>C 4132G>A), TGC603ATT (4133T>A 4134G>T 4135C>T), GCT604GCA (4138T>A), GCC606GAA (4143C>A 4144C>A), ACG608ATA (4149C>T 4150G>A), CTT609CTG (4153T>G), CGA610AAG (4154C>A 4155G>A 4156A>G), CTA611CTG (4159A>G), TTC612CCA (4160T>C 4161T>C 4162C>A), GAT613GAC (4165T>C), GGT614TTT (4166G>T 4167G>T), AGC615GAG (4169A>G 4170G>A 4171C>G), AAG616CTA (4172A>C 4173A>T 4174G>A), GAA617CCT (4175G>C 4176A>C 4177A>T), GTC618TTT (4178G>T 4180C>T), CAC619GAG (4181C>G 4183C>G), ATC620GTA (4184A>G 4186C>A), GAG621CAC (4187G>C 4189G>C), ACC622ACT (4192C>T), GAC623GAT (4195C>T), TCT625TCG (4201T>G), GAT626GAC (4204T>C), ATG627AAA (4206T>A 4207G>A), GCA628GCT (4210A>T), ATA629GTG (4211A>G 4213A>G), GGC630GGA (4216C>A), GCA631GGC (4218C>G 4219A>C), TGT632GTA (4220T>G 4221G>T 4222T>A), CTA633TTG (4223C>T 4225A>G), ACA634GTG (4226A>G 4227C>T 4228A>G), CAG635CAA (4231G>A), ACA636_CAC637del (4232_4237delACACAC), GAT638GAA (4240T>A), GGG639GGT (4243G>T), AAA640GAA (4244A>G), AGA641AAA (4248G>A), CAC642CAT (4252C>T), CCA643CCG (4255A>G), TAT646TTT (4263A>T), TAT647GAA (4265T>G 4267T>A), TCC648AGC (4268T>A 4269C>G), CGG649AGG (4271C>A), ATG651TTA (4277A>T 4279G>A), ACC652AAC (4281C>A), ACA653GAG (4283A>G 4284C>A 4285A>G), GCG654ACA (4286G>A 4288G>A), CAG656CGA (4293A>G 4294G>A), AAC657CGG (4295A>C 4296A>G 4297C>G), GAC659ACA (4301G>A 4302A>C 4303C>A), ATC660GTC (4304A>G), CAT661CAA (4309T>A), GAC662GAG (4312C>G), CTT665ATG (4319C>A 4321T>G), CTA666ACA (4322C>A 4323T>C), ATT668ATC (4330T>C), GTT669GTC (4333T>C), GCC670CAC (4334G>C 4335C>A), GCC671TGC (4337G>T 4338C>G), ATG672CTG (4340A>C), CAA673CGC (4344A>G 4345A>C), CAT674ACC (4346C>A 4347A>C 4348T>C), GTG677CAC (4355G>C 4356T>A 4357G>C), GTC679TTG (4361G>T 4363C>G), GAG680CTT (4364G>C 4365A>T 4366G>T), GGC681GGT (4369C>T), CCA682TCT (4370C>T 4372A>T), CCG683del (4373_4375delCCG), TTA685TTC (4381A>C), ACG686TTA (4382A>T 4383C>T 4384G>A), ATT687GTC (4385A>G 4387T>C), CTT688AAA (4388C>A 4389T>A 4390T>A), TCA689ACC (4391T>A 4393A>C), CAC691AAC (4397C>A), AAG692GTG (4400A>G 4401A>T), AAT693GCC (4403A>G 4404A>C 4405T>C), CTC694ACC (4406C>A 4407T>C), ACG695AGC (4410C>G 4411G>C), ACG698CAA (4418A>C 4419C>A 4420G>A), ACG699TCT (4421A>T 4423G>T), ACG700CAA (4424A>C 4425C>A 4426G>A), GAA702AAA (4430G>A), ACC704ACG (4438C>G), CGA705CCC (4440G>C 4441A>C), AGA706AAG (4443G>A 4444A>G), GCC708GCA (4450C>A), CGC709AGG (4451C>A 4453C>G), TCG711CAA (4457T>C 4458C>A 4459G>A), GAG712GAT (4462G>T), CTG713TTC (4463C>T 4465G>C), CTT714TTG (4466C>T 4468T>G), GGG715GCA (4470G>C 4471G>A), CAG716GAA (4472C>G 4474G>A), TAC717TTT (4476A>T 4477C>T), AAG718GAC (4478A>G 4480G>C), TTC719TAC (4482T>A), GAA720GAG (4486A>G), ATC721CTG (4487A>C 4489C>G), AAA722GAG (4490A>G 4492A>G), ACT724AAG (4497C>A 4498T>G), GGA726GGC (4504A>C), ACA727AAG (4506C>A 4507A>G), GAG728GGC (4509A>G 4510G>C), AAC729AAT (4513C>T), GGC730GTT (4515G>T 4516C>T), CCG731GTT (4517C>G 4518C>T 4519G>T), GCA732GCC (4522A>C), ATA821ATT (4789A>T), CAG822GAG (4790C>G), CGC823CGA (4795C>A), AGC824GCT (4796A>G 4797G>C 4798C>T), TTC825TAC (4800T>A), TCA826TAT (4803C>A 4804A>T), TTC827TGG (4806T>G 4807C>G), CCA828CCC (4810A>C), CAG829CAA (4813G>A), AGA831AGG (4819A>G), CTT832GAT (4820C>G 4821T>A), AAG833GAT (4823A>G 4825G>T), CTA835GAG (4829C>G 4830T>A 4831A>G), CGC836CTC (4833G>T), ATC838GTC (4838A>G), AAA840ACT (4845A>C 4846A>T), TGC841TGT (4849C>T), GTA842CTT (4850G>C 4852A>T), CAC843GTG (4853C>G 4854A>T 4855C>G), CAA845CAG (4861A>G), AAC847GAC (4865A>G), AAA848AAG (4870A>G), GCT849GTG (4872C>T 4873T>G), GCA850GAA (4875C>A), CGG851CAA (4878G>A 4879G>A), CAC852CGT (4881A>G 4882C>T), GCG853CCC (4883G>C 4885G>C), AAA854GCA (4886A>G 4887A>C), TAC855GCT (4889T>G 4890A>C 4891C>T), CAC857CTA (4896A>T 4897C>A), CAG859GAG (4901C>G), TTC860CCA (4904T>C 4905T>C 4906C>A), AGG861CTG (4907A>C 4908G>T), ACA862CCC (4910A>C 4912A>C), CCA863ATA (4913C>A 4914C>T), CCA864GCC (4916C>G 4918A>C), ACG865GAG (4919A>G 4920C>A), AAA866AGA (4923A>G), GAC869GAA (4933C>A), GAG870AGT (4934G>A 4935A>G 4936G>T), GTT871ATT (4937G>A), ACG872TCC (4940A>T 4942G>C), ATT876ATC (4954T>C), ACG877ACA (4957G>A), AAA878TGC (4958A>T 4959A>G 4960A>C), CTC879TTA (4961C>T 4963C>A), CCG880CCA (4966G>A), AGG881AAA (4968G>A 4969G>A), AAG883_GGA888del (4973_4990delAAGGATCGAGTCACAGGA), CAA889GAA (4991C>G), GCC890GGG (4995C>G 4996C>G), TAT891TGC (4998A>G 4999T>C), GAC892AGC (5000G>A 5001A>G), ATG893AAT (5004T>A 5005G>T), CTA895ATG (5009C>A 5011A>G), GTC896GTT (5014C>T), ATG897GTC (5015A>G 5017G>C), GTC898GTG (5020C>G), CTC901TTT (5027C>T 5029C>T), ACA902AGC (5031C>G 5032A>C), TAT904TAC (5038T>C), GCA905GGA (5040C>G), CAC906GTA (5042C>G 5043A>T 5044C>A), ATT908ATA (5050T>A), CCT909CCA (5053T>A), GCA910GTC (5055C>T 5056A>C), TCA911CCG (5057T>C 5059A>G), GAA912AC. (5060G>A 5061A>C)

**Proteins**

| | Begin | End | Coverage | Score | Concordance | Matches | Identities | I/D/M/F* | Stop Codons |
|---|---|---|---|---|---|---|---|---|---|
| Reverse Transcriptase (YP_009666308.1) | 289 | 912 | 38.5% | 1234 | 44.2% | 394 (97.0%) | 169 (41.6%) | 3/9/0/0 | 0 |

| | Begin | End | Coverage | Score | Concordance | Matches | Identities | I/D/M/F* | Stop Codons |
|---|---|---|---|---|---|---|---|---|---|
| **NT** | **3189** | **5061** | **16.3%** | **96** | **4.2%** | **1182 (97.0%)** | **633 (52.0%)** | **9/27** | |
| Protein mutations: | K289E (3191A>G), W290L (3194T>C 3195G>T), S291L (3197T>C 3198C>T), R292K (3200C>A 3201G>A), L293Q (3204T>A 3205C>G), F294Y (3207T>A 3208T>C), E295A (3210A>C 3211A>T), E296D (3214G>T), E297V (3216A>T 3217A>G), R298M (3219G>T 3220A>G), G299P (3221G>C 3222G>C 3223C>A), K300P (3224A>C 3225A>C 3226G>T), D301E (3229C>G), A302L (3230G>T 3231C>T 3232C>G), L303P (3233T>C 3234T>C), P304K (3236C>A 3237C>A 3238T>G), H306L (3243A>T 3244C>G), Q307P (3246A>C), W309R (3251T>A), W309_D310insRDI (3253_3254insAGGGATATT), K312R (3261A>G 3262G>A), N314E (3266A>G 3268C>G), I315L (3269A>T 3271T>G), Q316M (3272C>A 3273A>T), K319S (3281A>T 3282A>C 3283A>T), E320T (3284G>A 3285A>C), W323S (3294G>C 3295G>T), G324I (3296G>A 3297G>T 3298A>C), L326P (3303T>C 3304A>T), Q328R (3308C>A 3309A>G), S330A (3314T>G), E331P (3317G>C 3318A>C 3319G>T), K332A (3320A>G 3321A>C), Q335A (3329C>G 3330A>C 3331G>T), T336E (3332A>G 3333C>A 3334C>A), R338K (3338C>A 3339G>A 3340A>G), E339K (3341G>A 3343A>G), W340Q (3344T>C 3345G>A), K379Q (3461A>C), E382K (3470G>A), I383V (3473A>G 3475C>A), I385V (3479A>G 3481C>G), R388K (3488C>A 3489G>A 3490A>G), L391M (3497C>A 3499A>G), N393R (3504A>G 3505C>A), E395D (3511A>T), E396D (3514A>T), A397L (3515G>T 3516C>T 3517A>G), Q398F (3518C>T 3519A>T 3520A>T), R400Q (3524A>C 3525G>A), T402Q (3530A>C 3531C>A 3532C>G), K505Y (3839A>T 3841G>T), T506V (3842A>G 3843C>T 3844A>C), A507K (3845G>A 3846C>A 3847A>G), P508M (3848C>A 3849C>T 3850C>G), H514A (3866C>G 3867A>C), K515Q (3869A>C 3871G>A), K519T (3882A>C 3883G>C), F523H (3893T>C 3894T>A), I525V (3899A>G 3901C>T), T527E (3905A>G 3906C>A), T528D (3908A>G 3909C>A 3910A>C), G529E (3912G>A 3913G>A), I530V (3914A>G), T531R (3918C>G), I532M (3922C>G), A535K (3929G>A 3930C>A 3931A>G), T537V (3935A>G 3936C>T 3937A>G), S539A (3941T>G 3943A>G), R541I (3948G>T), E542D (3952A>T), P544Q (3957C>A), E545T (3959G>A 3960A>C), K550T (3975A>C), D551E (3979T>A), V552L (3980G>C 3982A>T), Q553R (3983C>A 3984A>G 3985G>A), L556F (3992C>T), N562Y (4010A>T), K567D (4025A>G 4027G>T), D568G (4029A>G 4030C>A), K571T (4038A>C), T572K (4041C>A), M576L (4052A>C), M578D (4058A>G 4059T>A 4060G>T), T580L (4064A>C 4065C>T 4066A>G), R581K (4068G>A 4069A>G), D583E (4075C>A), N585T (4080A>C 4081C>G), K587E (4085A>G), G589S (4091G>T 4092G>C 4093A>C), K590E (4094A>G), E591K (4097G>A), Q592C (4100C>T 4101A>G 4102G>C), T593E (4103A>G 4104C>A 4105C>A), K597Q (4115A>C 4117A>G), R598N (4119G>A 4120A>T), E601D (4129A>C), Q602A (4130C>G 4131A>C 4132G>A), C603I (4133T>A 4134G>T 4135C>T), A606E (4143C>A 4144C>A), T608I (4149C>T 4150G>A), R610K (4154C>A 4155G>A 4156A>G), F612P (4160T>C 4161T>C 4162C>A), G614F (4166G>T 4167G>T), S615E (4169A>G 4170G>A 4171C>G), K616L (4172A>C 4173A>T 4174G>A), E617P (4175G>C 4176A>C 4177A>T), V618F (4178G>T 4180C>T), H619E (4181C>G 4183C>G), I620V (4184A>G 4186C>A), E621H (4187G>C 4189G>C), M627K (4206T>A 4207G>A), I629V (4211A>G 4213A>G), A631G (4218C>G 4219A>C), C632V (4220T>G 4221G>T 4222T>A), T634V (4226A>G 4227C>T 4228A>G), T636_H637del (4232_4237delACACAC), D638E (4240T>A), K640E (4244A>G), R641K (4248G>A), Y646F (4263A>T), Y647E (4265T>G 4267T>A), M651L (4277A>T 4279G>A), T652N (4281C>A), T653E (4283A>G 4284C>A 4285A>G), A654T (4286G>A 4288G>A), Q656R (4293A>G 4294G>A), N657R (4295A>C 4296A>G 4297C>G), D659T (4301G>A 4302A>C 4303C>A), I660V (4304A>G), H661Q (4309T>A), D662E (4312C>G), L665M (4319C>A 4321T>G), L666T (4322C>A 4323T>C), A670H (4334G>C 4335C>A), A671C (4337G>T 4338C>G), M672L (4340A>C), Q673R (4344A>G 4345A>C), H674T (4346C>A 4347A>C 4348T>C), V677H (4355G>C 4356T>A 4357G>C), V679L (4361G>T 4363C>G), E680L (4364G>C 4365A>T 4366G>T), P682S (4370C>T 4372A>T), P683del (4373_4375delCCG), L685F (4381A>C), T686L (4382A>T 4383C>T 4384G>A), I687V (4385A>G 4387T>C), L688K (4388C>A 4389T>A 4390T>A), S689T (4391T>A 4393A>C), H691N (4397C>A), K692V (4400A>G 4401A>T), N693A (4403A>G 4404A>C 4405T>C), L694T (4406C>A 4407T>C), T695S (4410C>G 4411G>C), T698Q (4418A>C 4419C>A 4420G>A), T699S (4421A>T 4423G>T), T700Q (4424A>C 4425C>A 4426G>A), E702K (4430G>A), R705P (4440G>C 4441A>C), R706K (4443G>A 4444A>G), S711Q (4457T>C 4458C>A 4459G>A), E712D (4462G>T), L713F (4463C>T 4465G>C), G715A (4470G>C 4471G>A), Q716E (4472C>G 4474G>A), Y717F (4476A>T 4477C>T), K718D (4478A>G 4480G>C), F719Y (4482T>A), I721L (4487A>C 4489C>G), K722E (4490A>G 4492A>G), T724K (4497C>A 4498T>G), T727K (4506C>A 4507A>G), E728G (4509A>G 4510G>C), G730V (4515G>T 4516C>T), P731V (4517C>G 4518C>T 4519G>T), Q822E (4790C>G), S824A (4796A>G 4797G>C 4798C>T), F825Y (4800T>A), S826Y (4803C>A 4804A>T), F827W (4806T>G 4807C>G), L832D (4820C>G 4821T>A), K833D (4823A>G 4825G>T), L835E (4829C>G 4830T>A 4831A>G), R836L (4833G>T), I838V (4838A>G), K840T (4845A>C 4846A>T), V842L (4850G>C 4852A>T), H843V (4853C>G 4854A>T 4855C>G), N847D (4865A>G), A849V (4872C>T 4873T>G), A850E (4875C>A), R851Q (4878G>A 4879G>A), H852R (4881A>G 4882C>T), A853P (4883G>C 4885G>C), K854A (4886A>G 4887A>C), Y855A (4889T>G 4890A>C 4891C>T), H857L (4896A>T 4897C>A), Q859E (4901C>G), F860P (4904T>C 4905T>C 4906C>A), R861L (4907A>C 4908G>T), T862P (4910A>C 4912A>C), P863I (4913C>A 4914C>T), P864A (4916C>G 4918A>C), T865E (4919A>G 4920C>A), K866R (4923A>G), D869E (4933C>A), E870S (4934G>A 4935A>G 4936G>T), V871I (4937G>A), T872S (4940A>T 4942G>C), K878C (4958A>T 4959A>G 4960A>C), R881K (4968G>A 4969G>A), K883_G888del (4973_4990delAAGGATCGAGTCACAGGA), Q889E (4991C>G), A890G (4995C>G 4996C>G), Y891C (4998A>G 4999T>C), D892S (5000G>A 5001A>G), M893N (5004T>A 5005G>T), L895M (5009C>A 5011A>G), M897V (5015A>G 5017G>C), L901F (5027C>T 5029C>T), T902S (5031C>G 5032A>C), A905G (5040C>G), H906V (5042C>G 5043A>T 5044C>A), A910V (5055C>T 5056A>C), S911P (5057T>C 5059A>G) | | | | | | | | |

| | Begin | End | Coverage | Score | Concordance | Matches | Identities | I/D/M/F* | Stop Codons |
|---|---|---|---|---|---|---|---|---|---|
| **NT** | **3189** | **5061** | **16.3%** | **96** | **4.2%** | **1182 (97.0%)** | **633 (52.0%)** | **9/27** | |

Codon mutations: CGC288.CA (3189G>C 3190C>A), AAG289GAG (3191A>G), TGG290CTG (3194T>C 3195G>T), TCG291CTG (3197T>C 3198C>T), CGA292AAA (3200C>A 3201G>A), CTC293CAG (3204T>A 3205C>G), TTT294TAC (3207T>A 3208T>C), GAA295GCT (3210A>C 3211A>T), GAG296GAT (3214G>T), GAA297GTG (3216A>T 3217A>G), AGA298ATG (3219G>T 3220A>G), GGC299CCA (3221G>C 3222G>C 3223C>A), AAG300CCT (3224A>C 3225A>C 3226G>T), GAC301GAG (3229C>G), GCC302TTG (3230G>T 3231C>T 3232C>G), TTA303CCA (3233T>C 3234T>C), CCT304AAG (3236C>A 3237C>A 3238T>G), CAC306CTG (3243A>T 3244C>G), CAA307CCA (3246A>C), TGG309AGG (3251T>A), TGG309_GAT310insAGGGATATT (3253_3254insAGGGATATT), AAG312AGA (3261A>G 3262G>A), ATA313ATT (3265A>T), AAC314GAG (3266A>G 3268C>G), ATT315TTG (3269A>T 3271T>G), CAG316ATG (3272C>A 3273A>T), GGG318GGT (3280G>T), AAA319TCT (3281A>T 3282A>C 3283A>T), GAG320ACG (3284G>A 3285A>C), CCT321CCG (3289T>G), CCA322CCC (3292A>C), TGG323TCT (3294G>C 3295G>T), GGA324ATC (3296G>A 3297G>T 3298A>C), CCC325CCT (3301C>T), CTA326CCT (3303T>C 3304A>T), TAT327TAC (3307T>C), CAA328AGA (3308C>A 3309A>G), TCT330GCT (3314T>G), GAG331CCT (3317G>C 3318A>C 3319G>T), AAA332GCA (3320A>G 3321A>C), CTA334TTG (3326C>T 3328A>G), CAG335GCT (3329C>G 3330A>C 3331G>T), ACC336GAA (3332A>G 3333C>A 3334C>A), CTA337CTG (3337A>G), CGA338AAG (3338C>A 3339G>A 3340A>G), GAA339AAG (3341G>A 3343A>G), TGG340CAG (3344T>C 3345G>A), CTG341TTG (3347C>T), TAC377TAT (3457C>T), CGA378CGG (3460A>G), AAG379CAG (3461A>C), TTG380CTG (3464T>C), AAC381AAT (3469C>T), GAG382AAG (3470G>A), ATC383GTA (3473A>G 3475C>A), ACG384ACT (3478G>T), ATC385GTG (3479A>G 3481C>G), CGA388AAG (3488C>A 3489G>A 3490A>G), CCG390CCC (3496G>C), CTA391ATG (3497C>A 3499A>G), CCC392CCT (3502C>T), AAC393AGA (3504A>G 3505C>A), ATC394ATT (3508C>T), GAA395GAT (3511A>T), GAA396GAT (3514A>T), GCA397TTG (3515G>T 3516C>T 3517A>G), CAA398TTT (3518C>T 3519A>T 3520A>T), AGA400CAA (3524A>C 3525G>A), TTA401CTT (3527T>C 3529A>T), ACC402CAG (3530A>C 3531C>A 3532C>G), AAG505TAT (3839A>T 3841G>T), ACA506GTC (3842A>G 3843C>T 3844A>C), GCA507AAG (3845G>A 3846C>A 3847A>G), CCC508ATG (3848C>A 3849C>T 3850C>G), AAA510AAG (3856A>G), TGC511TGT (3859C>T), GAA512GAG (3862A>G), TTC513TTT (3865C>T), CAC514GCC (3866C>G 3867A>C), AAG515CAA (3869A>C 3871G>A), AAA516AAG (3874A>G), GTC518GTA (3880C>A), AAG519ACC (3882A>C 3883G>C), TTT520TTC (3886T>C), TTA521CTA (3887T>C), GGC522GGG (3892C>G), TTT523CAT (3893T>C 3894T>A), ATC524ATA (3898C>A), ATC525GTT (3899A>G 3901C>T), ACA527GAA (3905A>G 3906C>A), ACA528GAC (3908A>G 3909C>A 3910A>C), GGG529GAA (3912G>A 3913G>A), ATA530GTA (3914A>G), ACG531AGG (3918C>G), ATC532ATG (3922C>G), GCA535AAG (3929G>A 3930C>A 3931A>G), AAG536AAA (3934G>A), ACA537GTG (3935A>G 3936C>T 3937A>G), CAG538CAA (3940G>A), TCA539GCG (3941T>G 3943A>G), ATC540ATT (3946C>T), AGA541ATA (3948G>T), GAA542GAT (3952A>T), CCA544CAA (3957C>A), GAA545ACA (3959G>A 3960A>C), CCG546CCC (3964G>C), ACA548ACT (3970A>T), GTC549GTG (3973C>G), AAG550ACG (3975A>C), GAT551GAA (3979T>A), GTA552CTT (3980G>C 3982A>T), CAG553AGA (3983C>A 3984A>G 3985G>A), TCA554TCT (3988A>T), CTT556TTT (3992C>T), GGA557GGG (3997A>G), CTC558CTA (4000C>A), GCC559GCT (4003C>T), TAC561TAT (4009C>T), AAC562TAC (4010A>T), AAG567GAT (4025A>G 4027G>T), GAC568GGA (4029A>G 4030C>A), TAT569TAC (4033T>C), AAG571ACG (4038A>C), ACA572AAA (4041C>A), GCA573GCC (4045A>C), GCA574GCT (4048A>T), ATG576CTG (4052A>C), ACG577ACT (4057G>T), ATG578GAT (4058A>G 4059T>A 4060G>T), CTT579TTG (4061C>T 4063T>G), ACA580CTG (4064A>C 4065C>T 4066A>G), AGA581AAG (4068G>A 4069A>G), AAA582AAG (4072A>G), GAC583GAA (4075C>A), AAC585ACG (4080A>C 4081C>G), AAA587GAA (4085A>G), GGA589TCC (4091G>T 4092G>C 4093A>C), AAA590GAA (4094A>G), GAA591AAA (4097G>A), CAG592TGC (4100C>T 4101A>G 4102G>C), ACC593GAA (4103A>G 4104C>A 4105C>A), GCG595GCT (4111G>T), AAA597CAG (4115A>C 4117A>G), AGA598AAT (4119G>A 4120A>T), GAA601GAC (4129A>C), CAG602GCA (4130C>G 4131A>C 4132G>A), TGC603ATT (4133T>A 4134G>T 4135C>T), GCT604GCA (4138T>A), GCC606GAA (4143C>A 4144C>A), ACG608ATA (4149C>T 4150G>A), CTT609CTG (4153T>G), CGA610AAG (4154C>A 4155G>A 4156A>G), CTA611CTG (4159A>G), TTC612CCA (4160T>C 4161T>C 4162C>A), GAT613GAC (4165T>C), GGT614TTT (4166G>T 4167G>T), AGC615GAG (4169A>G 4170G>A 4171C>G), AAG616CTA (4172A>C 4173A>T 4174G>A), GAA617CCT (4175G>C 4176A>C 4177A>T), GTC618TTT (4178G>T 4180C>T), CAC619GAG (4181C>G 4183C>G), ATC620GTA (4184A>G 4186C>A), GAG621CAC (4187G>C 4189G>C), ACC622ACT (4192C>T), GAC623GAT (4195C>T), TCT625TCG (4201T>G), GAT626GAC (4204T>C), ATG627AAA (4206T>A 4207G>A), GCA628GCT (4210A>T), ATA629GTG (4211A>G 4213A>G), GGC630GGA (4216C>A), GCA631GGC (4218C>G 4219A>C), TGT632GTA (4220T>G 4221G>T 4222T>A), CTA633TTG (4223C>T 4225A>G), ACA634GTG (4226A>G 4227C>T 4228A>G), CAG635CAA (4231G>A), ACA636_CAC637del (4232_4237delACACAC), GAT638GAA (4240T>A), GGG639GGT (4243G>T), AAA640GAA (4244A>G), AGA641AAA (4248G>A), CAC642CAT (4252C>T), CCA643CCG (4255A>G), TAT646TTT (4263A>T), TAT647GAA (4265T>G 4267T>A), TCC648AGC (4268T>A 4269C>G), CGG649AGG (4271C>A), ATG651TTA (4277A>T 4279G>A), ACC652AAC (4281C>A), ACA653GAG (4283A>G 4284C>A 4285A>G), GCG654ACA (4286G>A 4288G>A), CAG656CGA (4293A>G 4294G>A), AAC657CGG (4295A>C 4296A>G 4297C>G), GAC659ACA (4301G>A 4302A>C 4303C>A), ATC660GTC (4304A>G), CAT661CAA (4309T>A), GAC662GAG (4312C>G), CTT665ATG (4319C>A 4321T>G), CTA666ACA (4322C>A 4323T>C), ATT668ATC (4330T>C), GTT669GTC (4333T>C), GCC670CAC (4334G>C 4335C>A), GCC671TGC (4337G>T 4338C>G), ATG672CTG (4340A>C), CAA673CGC (4344A>G 4345A>C), CAT674ACC (4346C>A 4347A>C 4348T>C), GTG677CAC (4355G>C 4356T>A 4357G>C), GTC679TTG (4361G>T 4363C>G), GAG680CTT (4364G>C 4365A>T 4366G>T), GGC681GGT (4369C>T), CCA682TCT (4370C>T 4372A>T), CCG683del (4373_4375delCCG), TTA685TTC (4381A>C), ACG686TTA (4382A>T 4383C>T 4384G>A), ATT687GTC (4385A>G 4387T>C), CTT688AAA (4388C>A 4389T>A 4390T>A), TCA689ACC (4391T>A 4393A>C), CAC691AAC (4397C>A), AAG692GTG (4400A>G 4401A>T), AAT693GCC (4403A>G 4404A>C 4405T>C), CTC694ACC (4406C>A 4407T>C), ACG695AGC (4410C>G 4411G>C), ACG698CAA (4418A>C 4419C>A 4420G>A), ACG699TCT (4421A>T 4423G>T), ACG700CAA (4424A>C 4425C>A 4426G>A), GAA702AAA (4430G>A), ACC704ACG (4438C>G), CGA705CCC (4440G>C 4441A>C), AGA706AAG (4443G>A 4444A>G), GCC708GCA (4450C>A), CGC709AGG (4451C>A 4453C>G), TCG711CAA (4457T>C 4458C>A 4459G>A), GAG712GAT (4462G>T), CTG713TTC (4463C>T 4465G>C), CTT714TTG (4466C>T 4468T>G), GGG715GCA (4470G>C 4471G>A), CAG716GAA (4472C>G 4474G>A), TAC717TTT (4476A>T 4477C>T), AAG718GAC (4478A>G 4480G>C), TTC719TAC (4482T>A), GAA720GAG (4486A>G), ATC721CTG (4487A>C 4489C>G), AAA722GAG (4490A>G 4492A>G), ACT724AAG (4497C>A 4498T>G), GGA726GGC (4504A>C), ACA727AAG (4506C>A 4507A>G), GAG728GGC (4509A>G 4510G>C), AAC729AAT (4513C>T), GGC730GTT (4515G>T 4516C>T), CCG731GTT (4517C>G 4518C>T 4519G>T), GCA732GCC (4522A>C), ATA821ATT (4789A>T), CAG822GAG (4790C>G), CGC823CGA (4795C>A), AGC824GCT (4796A>G 4797G>C 4798C>T), TTC825TAC (4800T>A), TCA826TAT (4803C>A 4804A>T), TTC827TGG (4806T>G 4807C>G), CCA828CCC (4810A>C), CAG829CAA (4813G>A), AGA831AGG (4819A>G), CTT832GAT (4820C>G 4821T>A), AAG833GAT (4823A>G 4825G>T), CTA835GAG (4829C>G 4830T>A 4831A>G), CGC836CTC (4833G>T), ATC838GTC (4838A>G), AAA840ACT (4845A>C 4846A>T), TGC841TGT (4849C>T), GTA842CTT (4850G>C 4852A>T), CAC843GTG (4853C>G 4854A>T 4855C>G), CAA845CAG (4861A>G), AAC847GAC (4865A>G), AAA848AAG (4870A>G), GCT849GTG (4872C>T 4873T>G), GCA850GAA (4875C>A), CGG851CAA (4878G>A 4879G>A), CAC852CGT (4881A>G 4882C>T), GCG853CCC (4883G>C 4885G>C), AAA854GCA (4886A>G 4887A>C), TAC855GCT (4889T>G 4890A>C 4891C>T), CAC857CTA (4896A>T 4897C>A), CAG859GAG (4901C>G), TTC860CCA (4904T>C 4905T>C 4906C>A), AGG861CTG (4907A>C 4908G>T), ACA862CCC (4910A>C 4912A>C), CCA863ATA (4913C>A 4914C>T), CCA864GCC (4916C>G 4918A>C), ACG865GAG (4919A>G 4920C>A), AAA866AGA (4923A>G), GAC869GAA (4933C>A), GAG870AGT (4934G>A 4935A>G 4936G>T), GTT871ATT (4937G>A), ACG872TCC (4940A>T 4942G>C), ATT876ATC (4954T>C), ACG877ACA (4957G>A), AAA878TGC (4958A>T 4959A>G 4960A>C), CTC879TTA (4961C>T 4963C>A), CCG880CCA (4966G>A), AGG881AAA (4968G>A 4969G>A), AAG883_GGA888del (4973_4990delAAGGATCGAGTCACAGGA), CAA889GAA (4991C>G), GCC890GGG (4995C>G 4996C>G), TAT891TGC (4998A>G 4999T>C), GAC892AGC (5000G>A 5001A>G), ATG893AAT (5004T>A 5005G>T), CTA895ATG (5009C>A 5011A>G), GTC896GTT (5014C>T), ATG897GTC (5015A>G 5017G>C), GTC898GTG (5020C>G), CTC901TTT (5027C>T 5029C>T), ACA902AGC (5031C>G 5032A>C), TAT904TAC (5038T>C), GCA905GGA (5040C>G), CAC906GTA (5042C>G 5043A>T 5044C>A), ATT908ATA (5050T>A), CCT909CCA (5053T>A), GCA910GTC (5055C>T 5056A>C), TCA911CCG (5057T>C 5059A>G), GAA912AC. (5060G>A 5061A>C)

*: Inserts / Deletes / Misaligned / Frameshifts

#### Analysis details

This analysis was performed with panviral2.64

### NGS Details (UN18_val): Errantivirus

#### Assembly

| | |
|---|---|
| **Coverage Length** | 724 (3 contig(s)) |
| **Depth Of Coverage** | 16.0 |
| **Number Of Reads** | 149 |
| **Reads Per Million** | 2.80 rpm (after QC) |
| **Ambiguities** | 0 |
| **Assembly Method** | de novo + reference guided assembly |
| **Consensus Caller** | Bcf Tools |

#### Coverage Map

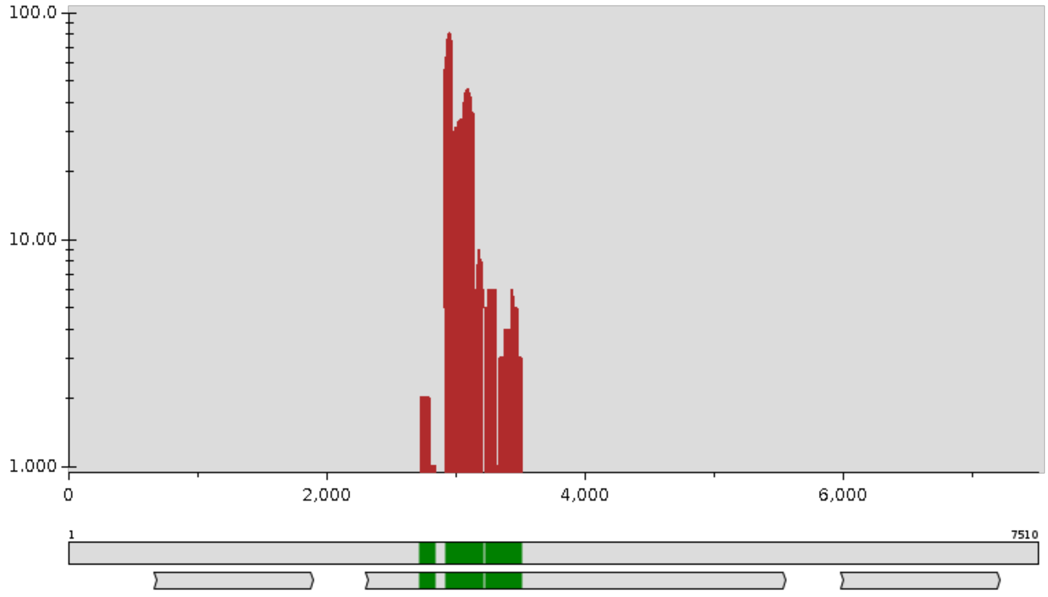

#### Assignment

| | |
|---|---|
| **Type** | Errantivirus (Taxonomy ID: 186666) |
| **Reference Genome** | NC_038512.1 |
| **NT Identity (%)** | 56.0773 |
| **AA Identity (%)** | 44.8133 |
| **Number Of Stop Codons** | 0 |
| **Number Of CDS** | 3 |

#### Alignment

| | |
|---|---|
| **Alignment Score** | 176.0 (NT) + 804.0 (AA) = 980.0 |
| **Concordance (%)** | 31.2799 |

| Alignment Method | Global, seeded, nucleotide + amino acids (AGA) |
|---|---|

#### Genome Region

Sequence starts at position 2720 and ends at position 3520 relative to NC_038512.1 reference sequence.

#### Alignment Detailed Statistics

| | Begin | End | Coverage | Score | Concordance | Matches | Identities | I/D/M/F* | Stop Codons |
|---|---|---|---|---|---|---|---|---|---|
| **NT** | **2720** | **3520** | **9.6%** | **176** | **12.2%** | **724 (100%)** | **406 (56.1%)** | **0/0** | |
| Mutations: | 2720G>A, 2722G>A, 2723T>A, 2727C>T, 2730A>G, 2732A>C, 2733G>C, 2734T>A, 2738C>A, 2746C>T, 2747T>A, 2748T>C, 2749C>T, 2752C>A, 2755T>G, 2756C>A, 2757G>A, 2758C>G, 2759C>G, 2761G>T, 2765G>A, 2768A>G, 2769G>A, 2770G>A, 2771G>A, 2773C>G, 2774C>A, 2775A>T, 2776A>G, 2777A>G, 2780A>G, 2781C>A, 2782G>T, 2783A>G, 2785A>G, 2786A>T, 2789T>C, 2795C>A, 2796A>C, 2801A>G, 2806A>T, 2809A>G, 2811C>A, 2812A>G, 2813T>A, 2814C>G, 2815A>T, 2816G>A, 2817A>T, 2818C>T, 2819T>A, 2820C>G, 2822G>C, 2827G>T, 2828A>T, 2829G>C, 2839A>T, 2917G>C, 2919A>T, 2920G>C, 2924A>G, 2926C>G, 2927G>A, 2928A>G, 2929T>G, 2932C>T, 2934A>G, 2935A>G, 2937A>T, 2941G>A, 2944A>C, 2949A>C, 2950C>T, 2953A>T, 2954A>G, 2955G>A, 2959C>G, 2960G>C, 2963C>T, 2968C>T, 2969A>G, 2972T>G, 2975G>A, 2976G>A, 2978A>G, 2979A>G, 2980G>C, 2981T>G, 2982G>C, 2983C>T, 2984C>G, 2985A>T, 2986A>T, 2989C>T, 2993A>T, 2995C>A, 2997C>A, 2998C>G, 2999T>C, 3001A>T, 3008G>T, 3009C>T, 3011A>T, 3012G>C, 3016G>A, 3018T>A, 3019T>C, 3020T>G, 3025G>A, 3026G>A, 3028G>T, 3029G>C, 3030A>G, 3032A>G, 3034G>T, 3035G>A, 3037C>G, 3038C>G, 3039C>T, 3040T>G, 3041C>G, 3042A>C, 3050T>C, 3052G>C, 3058C>T, 3061G>T, 3066A>G, 3067C>G, 3068G>A, 3069T>C, 3071G>C, 3073A>C, 3074C>G, 3076C>T, 3079G>T, 3084T>A, 3085T>C, 3091C>T, 3092C>T, 3094T>A, 3095C>G, 3096G>T, 3097A>C, 3104A>T, 3106G>T, 3109A>G, 3112A>G, 3113A>T, 3114A>C, 3115A>T, 3119T>G, 3121A>G, 3122C>T, 3124A>G, 3127T>C, 3130T>A, 3139A>C, 3140G>C, 3142T>C, 3146G>A, 3151T>G, 3152G>A, 3155C>T, 3157A>T, 3158A>C, 3159G>A, 3160A>C, 3161G>T, 3162G>C, 3164C>T, 3165T>A, 3168A>T, 3169A>C, 3171A>G, 3172T>A, 3175C>G, 3176A>T, 3179T>A, 3180G>T, 3182C>T, 3184C>G, 3187C>A, 3189A>T, 3190C>T, 3191C>T, 3193T>C, 3196C>T, 3202T>C, 3203A>C, 3205T>C, 3206G>A, 3215A>T, 3226G>A, 3229A>G, 3233C>A, 3235G>A, 3245G>A, 3246A>G, 3248C>A, 3249G>C, 3250A>T, 3253T>G, 3254T>C, 3256C>G, 3259A>G, 3260A>G, 3261G>T, 3262A>T, 3263C>T, 3265T>G, 3272A>G, 3273G>A, 3274T>A, 3275A>C, 3277C>G, 3280C>A, 3281A>T, 3283A>T, 3284A>G, 3285T>C, 3286T>A, 3287C>A, 3290A>T, 3291T>A, 3292G>T, 3295C>T, 3300C>G, 3301C>T, 3309T>G, 3311A>T, 3312A>T, 3313G>T, 3316C>G, 3320A>G, 3321C>T, 3322T>G, 3324C>A, 3325T>G, 3329C>T, 3331T>G, 3338A>T, 3340C>T, 3346C>T, 3347A>G, 3348G>C, 3349G>C, 3351A>C, 3352C>T, 3355T>A, 3356A>G, 3358C>T, 3359A>G, 3360A>C, 3361G>A, 3362C>A, 3365A>G, 3367C>T, 3371G>A, 3373T>G, 3379T>A, 3380T>G, 3381C>A, 3382C>T, 3384C>T, 3386A>G, 3389C>A, 3393A>C, 3396A>G, 3397T>G, 3398C>A, 3399T>C, 3400G>A, 3403T>A, 3408A>C, 3409G>A, 3413C>G, 3414C>T, 3415T>G, 3419G>C, 3422A>T, 3424A>G, 3426A>G, 3428C>G, 3429A>G, 3430A>C, 3433T>C, 3436A>T, 3439C>T, 3443C>G, 3444T>C, 3445C>A, 3451T>C, 3454C>T, 3455C>A, 3459A>G, 3467C>A, 3468C>G, 3471A>G, 3477C>G, 3478A>T, 3479C>G, 3480G>T, 3481A>T, 3482C>A, 3484C>T, 3485A>G, 3489A>G, 3498C>A, 3499A>T, 3500C>G, 3504G>T, 3505C>G, 3511A>G, 3515G>A, 3516G>A, 3517T>G, 3519G>T, 3520T>A | | | | | | | | |

**CDS**

| | Begin | End | Coverage | Score | Concordance | Matches | Identities | I/D/M/F* | Stop Codons |
|---|---|---|---|---|---|---|---|---|---|
| D1R33_gp2 | 139 | 405 | 22.2% | 804 | 47.2% | 241 (100%) | 108 (44.8%) | 0/0/0/0 | 0 |
| Protein mutations: | V139I (2720G>A 2722G>A), Y140N (2723T>A), T141I (2727C>T), K142R (2730A>G), S143P (2732A>C 2733G>C 2734T>A), F148T (2747T>A 2748T>C 2749C>T), H150Q (2755T>G), R151K (2756C>A 2757G>A 2758C>G), Q152D (2759C>G 2761G>T), V154I (2765G>A), R155E (2768A>G 2769G>A 2770G>A), D156K (2771G>A 2773C>G), Q157M (2774C>A 2775A>T 2776A>G), I158V (2777A>G), T159D (2780A>G 2781C>A 2782G>T), K160E (2783A>G 2785A>G), M161L (2786A>T), Q164T (2795C>A 2796A>C), I166V (2801A>G), P169Q (2811C>A 2812A>G), D171I (2816G>A 2817A>T 2818C>T), A173P (2822G>C), W174C (2827G>T), K205I (2919A>T 2920G>C), I207V (2924A>G 2926C>G), D208R (2927G>A 2928A>G 2929T>G), K210R (2934A>G 2935A>G), Y211F (2937A>T), N215T (2949A>C 2950C>T), S217D (2954A>G 2955G>A), D218E (2959C>G), V219L (2960G>C), L220F (2963C>T), K222E (2969A>G), L223V (2972T>G), G224N (2975G>A 2976G>A), K225G (2978A>G 2979A>G 2980G>C), C226A (2981T>G 2982G>C 2983C>T), Q227V (2984C>G 2985A>T 2986A>T), T230S (2993A>T 2995C>A), T231K (2997C>A 2998C>G), A235L (3008G>T 3009C>T), F238Y (3018T>A 3019T>C), Y239D (3020T>G), V241I (3026G>A 3028G>T), E242R (3029G>C 3030A>G), M243V (3032A>G 3034G>T), D244K (3035G>A 3037C>G), P245V (3038C>G 3039C>T 3040T>G), Q246A (3041C>G 3042A>C), S249P (3050T>C 3052G>C), N254R (3066A>G 3067C>G), V255T (3068G>A 3069T>C), E256H (3071G>C 3073A>C), H257D (3074C>G 3076C>T), F260Y (3084T>A 3085T>C), R264V (3095C>G 3096G>T 3097A>C), M267F (3104A>T 3106G>T), K270S (3113A>T 3114A>C 3115A>T), S272A (3119T>G 3121A>G), P273S (3122C>T 3124A>G), R278S (3139A>C), V279L (3140G>C 3142T>C), D281N (3146G>A), N282K (3151T>G), V283I (3152G>A), L284F (3155C>T 3157A>T), R285H (3158A>C 3159G>A 3160A>C), G286S (3161G>T 3162G>C), L287Y (3164C>T 3165T>A), Q288L (3168A>T 3169A>C), N289R (3171A>G 3172T>A), N290K (3175C>G), I291F (3176A>T), C292I (3179T>A 3180G>T), Y295F (3189A>T 3190C>T), L296F (3191C>T 3193T>C), I300L (3203A>C 3205T>C), V301I (3206G>A), L310I (3233C>A 3235G>A), E314R (3245G>A 3246A>G), R315T (3248C>A 3249G>C 3250A>T), F317L (3254T>C 3256C>G), R319V (3260A>G 3261G>T 3262A>T), S323E (3272A>G 3273G>A 3274T>A), N324Q (3275A>C 3277C>G), F325L (3280C>A), K326Y (3281A>T 3283A>T), I327A (3284A>G 3285T>C 3286T>A), Q328K (3287C>A), M329Y (3290A>T 3291T>A 3292G>T), S332C (3300C>G 3301C>T), L335W (3309T>G), K336F (3311A>T 3312A>T 3313G>T), T339V (3320A>G 3321C>T 3322T>G), A340E (3324C>A 3325T>G), I345F (3338A>T 3340C>T), R348A (3347A>G 3348G>C 3349G>C), D349A (3351A>C 3352C>T), I351V (3356A>G 3358C>T), K352A (3359A>G 3360A>C 3361G>A), P353T (3362C>A), N354D (3365A>G 3367C>T), D356K (3371G>A 3373T>G), S359D (3380T>G 3381C>A 3382C>T), A360V (3384C>T), I361V (3386A>G), Q362K (3389C>A), K363T (3393A>C), Y364W (3396A>G 3397T>G), L365T (3398C>A 3399T>C 3400G>A), K368T (3408A>C 3409G>A), P370V (3413C>G 3414C>T 3415T>G), E372Q (3419G>C), I373L (3422A>T 3424A>G), K374R (3426A>G), Q375G (3428C>G 3429A>G 3430A>C), L377F (3436A>T), L380A (3443C>G 3444T>C 3445C>A), K385R (3459A>G), P388R (3467C>A 3468C>G), D389G (3471A>G), A391G (3477C>G 3478A>T), R392V (3479C>G 3480G>T 3481A>T), L393I (3482C>A 3484C>T), T394A (3485A>G), K395R (3489A>G), T398N (3498C>A 3499A>T), Q399E (3500C>G), C400L (3504G>T 3505C>G), G404K (3515G>A 3516G>A 3517T>G), S405I (3519G>T 3520T>A) | | | | | | | | |
| Codon mutations: | GTG139ATA (2720G>A 2722G>A), TAC140AAC (2723T>A), ACC141ATC (2727C>T), AAA142AGA (2730A>G), AGT143CCA (2732A>C 2733G>C 2734T>A), CGG145AGG (2738C>A), CCC147CCT (2746C>T), TTC148ACT (2747T>A 2748T>C 2749C>T), ATC149ATA (2752C>A), CAT150CAG (2755T>G), CGC151AAG (2756C>A 2757G>A 2758C>G), CAG152GAT (2759C>G 2761G>T), GTT154ATT (2765G>A), AGG155GAA (2768A>G 2769G>A 2770G>A), GAC156AAG (2771G>A 2773C>G), CAA157ATG (2774C>A 2775A>T 2776A>G), ATC158GTC (2777A>G), ACG159GAT (2780A>G 2781C>A 2782G>T), AAA160GAG (2783A>G 2785A>G), ATG161TTG (2786A>T), TTG162CTG (2789T>C), CAA164ACA (2795C>A 2796A>C), ATT166GTT (2801A>G), ATA167ATT (2806A>T), AGA168AGG (2809A>G), CCA169CAG (2811C>A 2812A>G), TCA170AGT (2813T>A 2814C>G 2815A>T), GAC171ATT (2816G>A 2817A>T 2818C>T), TCT172AGT (2819T>A 2820C>G), GCA173CCA (2822G>C), TGG174TGT (2827G>T), AGC175TCC (2828A>T 2829G>C), ATA178ATT (2839A>T), GAG204..C (2917G>C), AAG205ATC (2919A>T 2920G>C), ATC207GTG (2924A>G 2926C>G), GAT208AGG (2927G>A 2928A>G 2929T>G), GAC209GAT (2932C>T), AAA210AGG (2934A>G 2935A>G), TAC211TTC (2937A>T), CCG212CCA (2941G>A), ATA213ATC (2944A>C), AAC215ACT (2949A>C 2950C>T), ATA216ATT (2953A>T), AGT217GAT (2954A>G 2955G>A), GAC218GAG (2959C>G), GTA219CTA (2960G>C), CTT220TTT (2963C>T), GAC221GAT (2968C>T), AAG222GAG (2969A>G), TTA223GTA (2972T>G), GGT224AAT (2975G>A 2976G>A), AAG225GGC (2978A>G 2979A>G 2980G>C), TGC226GCT (2981T>G 2982G>C 2983C>T), CAA227GTT (2984C>G 2985A>T 2986A>T), TAC228TAT (2989C>T), ACC230TCA (2993A>T 2995C>A), ACC231AAG (2997C>A 2998C>G), TTA232CTT (2999T>C 3001A>T), GCA235TTA (3008G>T 3009C>T), AGT236TCT (3011A>T 3012G>C), GGG237GGA (3016G>A), TTT238TAC (3018T>A 3019T>C), TAT239GAT (3020T>G), CAG240CAA (3025G>A), GTG241ATT (3026G>A 3028G>T), GAG242CGG (3029G>C 3030A>G), ATG243GTT (3032A>G 3034G>T), GAC244AAG (3035G>A 3037C>G), CCT245GTG (3038C>G 3039C>T 3040T>G), CAA246GCA (3041C>G 3042A>C), TCG249CCC (3050T>C 3052G>C), ACC251ACT (3058C>T), GCG252GCT (3061G>T), AAC254AGG (3066A>G 3067C>G), GTA255ACA (3068G>A 3069T>C), GAA256CAC (3071G>C 3073A>C), CAC257GAT (3074C>G 3076C>T), GGG258GGT (3079G>T), TTT260TAC (3084T>A 3085T>C), TTC262TTT (3091C>T), CTT263TTA (3092C>T 3094T>A), CGA264GTC (3095C>G 3096G>T 3097A>C), ATG267TTT (3104A>T 3106G>T), GGA268GGG (3109A>G), TTA269TTG (3112A>G), AAA270TCT (3113A>T 3114A>C 3115A>T), TCA272GCG (3119T>G 3121A>G), CCA273TCG (3122C>T 3124A>G), TCT274TCC (3127T>C), ACT275ACA (3130T>A), AGA278AGC (3139A>C), GTT279CTC (3140G>C 3142T>C), GAC281AAC (3146G>A), AAT282AAG (3151T>G), GTC283ATC (3152G>A), CTA284TTT (3155C>T 3157A>T), AGA285CAC (3158A>C 3159G>A 3160A>C), GGT286TCT (3161G>T 3162G>C), CTC287TAC (3164C>T 3165T>A), CAA288CTC (3168A>T 3169A>C), AAT289AGA (3171A>G 3172T>A), AAC290AAG (3175C>G), ATC291TTC (3176A>T), TGT292ATT (3179T>A 3180G>T), CTC293TTG (3182C>T 3184C>G), GTC294GTA (3187C>A), TAC295TTT (3189A>T 3190C>T), CTT296TTC (3191C>T 3193T>C), GAC297GAT (3196C>T), ATT299ATC (3202T>C), ATT300CTC (3203A>C 3205T>C), GTC301ATC (3206G>A), ACT304TC. (3215A>T), CAG307..A (3226G>A), GAA308GAG (3229A>G), CTG310ATA (3233C>A 3235G>A), GAA314AGA (3245G>A 3246A>G), CGA315ACT (3248C>A 3249G>C 3250A>T), GTT316GTG (3253T>G), TTC317CTG (3254T>C 3256C>G), CAA318CAG (3259A>G), AGA319GTT (3260A>G 3261G>T 3262A>T), CTT320TTG (3263C>T 3265T>G), AGT323GAA (3272A>G 3273G>A 3274T>A), AAC324CAG (3275A>C 3277C>G), TTC325TTA (3280C>A), AAA326TAT (3281A>T 3283A>T), ATT327GCA (3284A>G 3285T>C 3286T>A), CAA328AAA (3287C>A), ATG329TAT (3290A>T 3291T>A 3292G>T), GAC330GAT (3295C>T), TCC332TGT (3300C>G 3301C>T), TTG335TGG (3309T>G), AAG336TTT (3311A>T 3312A>T 3313G>T), CTC337CTG (3316C>G), ACT339GTG (3320A>G 3321C>T 3322T>G), GCT340GAG (3324C>A 3325T>G), CTT342TTG (3329C>T 3331T>G), ATC345TTT (3338A>T 3340C>T), AGC347AGT (3346C>T), AGG348GCC (3347A>G 3348G>C 3349G>C), GAC349GCT (3351A>C 3352C>T), GGT350GGA (3355T>A), ATC351GTT (3356A>G 3358C>T), AAG352GCA (3359A>G 3360A>C 3361G>A), CCT353ACT (3362C>A), AAC354GAT (3365A>G 3367C>T), GAT356AAG (3371G>A 3373T>G), ATT358ATA (3379T>A), TCC359GAT (3380T>G 3381C>A 3382C>T), GCT360GTT (3384C>T), ATT361GTT (3386A>G), CAA362AAA (3389C>A), AAA363ACA (3393A>C), TAT364TGG (3396A>G 3397T>G), CTG365ACA (3398C>A 3399T>C 3400G>A), ATT366ATA (3403T>A), AAG368ACA (3408A>C 3409G>A), CCT370GTG (3413C>G 3414C>T 3415T>G), GAA372CAA (3419G>C), ATA373TTG (3422A>T 3424A>G), AAA374AGA (3426A>G), CAA375GGC (3428C>G 3429A>G 3430A>C), TTT376TTC (3433T>C), TTA377TTT (3436A>T), GGC378GGT (3439C>T), CTC380GCA (3443C>G 3444T>C 3445C>A), TAT382TAC (3451T>C), TAC383TAT (3454C>T), CGA384AGA (3455C>A), AAA385AGA (3459A>G), CCA388AGA (3467C>A 3468C>G), GAT389GGT (3471A>G), GCA391GGT (3477C>G 3478A>T), CGA392GTT (3479C>G 3480G>T 3481A>T), CTC393ATT (3482C>A 3484C>T), ACA394GCA (3485A>G), AAA395AGA (3489A>G), ACA398AAT (3498C>A 3499A>T), CAG399GAG (3500C>G), TGC400TTG (3504G>T 3505C>G), AAA402AAG (3511A>G), GGT404AAG (3515G>A 3516G>A 3517T>G), AGT405ATA (3519G>T 3520T>A) | | | | | | | | |

**Proteins**

| | Begin | End | Coverage | Score | Concordance | Matches | Identities | I/D/M/F* | Stop Codons |
|---|---|---|---|---|---|---|---|---|---|
| ORF B (YP_009507248.1) | 139 | 405 | 22.2% | 804 | 47.2% | 241 (100%) | 108 (44.8%) | 0/0/0/0 | 0 |

| | Begin | End | Coverage | Score | Concordance | Matches | Identities | I/D/M/F* | Stop Codons |
|---|---|---|---|---|---|---|---|---|---|
| **NT** | **2720** | **3520** | **9.6%** | **176** | **12.2%** | **724 (100%)** | **406 (56.1%)** | **0/0** | |
| Protein mutations: | V139I (2720G>A 2722G>A), Y140N (2723T>A), T141I (2727C>T), K142R (2730A>G), S143P (2732A>C 2733G>C 2734T>A), F148T (2747T>A 2748T>C 2749C>T), H150Q (2755T>G), R151K (2756C>A 2757G>A 2758C>G), Q152D (2759C>G 2761G>T), V154I (2765G>A), R155E (2768A>G 2769G>A 2770G>A), D156K (2771G>A 2773C>G), Q157M (2774C>A 2775A>T 2776A>G), I158V (2777A>G), T159D (2780A>G 2781C>A 2782G>T), K160E (2783A>G 2785A>G), M161L (2786A>T), Q164T (2795C>A 2796A>C), I166V (2801A>G), P169Q (2811C>A 2812A>G), D171I (2816G>A 2817A>T 2818C>T), A173P (2822G>C), W174C (2827G>T), K205I (2919A>T 2920G>C), I207V (2924A>G 2926C>G), D208R (2927G>A 2928A>G 2929T>G), K210R (2934A>G 2935A>G), Y211F (2937A>T), N215T (2949A>C 2950C>T), S217D (2954A>G 2955G>A), D218E (2959C>G), V219L (2960G>C), L220F (2963C>T), K222E (2969A>G), L223V (2972T>G), G224N (2975G>A 2976G>A), K225G (2978A>G 2979A>G 2980G>C), C226A (2981T>G 2982G>C 2983C>T), Q227V (2984C>G 2985A>T 2986A>T), T230S (2993A>T 2995C>A), T231K (2997C>A 2998C>G), A235L (3008G>T 3009C>T), F238Y (3018T>A 3019T>C), Y239D (3020T>G), V241I (3026G>A 3028G>T), E242R (3029G>C 3030A>G), M243V (3032A>G 3034G>T), D244K (3035G>A 3037C>G), P245V (3038C>G 3039C>T 3040T>G), Q246A (3041C>G 3042A>C), S249P (3050T>C 3052G>C), N254R (3066A>G 3067C>G), V255T (3068G>A 3069T>C), E256H (3071G>C 3073A>C), H257D (3074C>G 3076C>T), F260Y (3084T>A 3085T>C), R264V (3095C>G 3096G>T 3097A>C), M267F (3104A>T 3106G>T), K270S (3113A>T 3114A>C 3115A>T), S272A (3119T>G 3121A>G), P273S (3122C>T 3124A>G), R278S (3139A>C), V279L (3140G>C 3142T>C), D281N (3146G>A), N282K (3151T>G), V283I (3152G>A), L284F (3155C>T 3157A>T), R285H (3158A>C 3159G>A 3160A>C), G286S (3161G>T 3162G>C), L287Y (3164C>T 3165T>A), Q288L (3168A>T 3169A>C), N289R (3171A>G 3172T>A), N290K (3175C>G), I291F (3176A>T), C292I (3179T>A 3180G>T), Y295F (3189A>T 3190C>T), L296F (3191C>T 3193T>C), I300L (3203A>C 3205T>C), V301I (3206G>A), L310I (3233C>A 3235G>A), E314R (3245G>A 3246A>G), R315T (3248C>A 3249G>C 3250A>T), F317L (3254T>C 3256C>G), R319V (3260A>G 3261G>T 3262A>T), S323E (3272A>G 3273G>A 3274T>A), N324Q (3275A>C 3277C>G), F325L (3280C>A), K326Y (3281A>T 3283A>T), I327A (3284A>G 3285T>C 3286T>A), Q328K (3287C>A), M329Y (3290A>T 3291T>A 3292G>T), S332C (3300C>G 3301C>T), L335W (3309T>G), K336F (3311A>T 3312A>T 3313G>T), T339V (3320A>G 3321C>T 3322T>G), A340E (3324C>A 3325T>G), I345F (3338A>T 3340C>T), R348A (3347A>G 3348G>C 3349G>C), D349A (3351A>C 3352C>T), I351V (3356A>G 3358C>T), K352A (3359A>G 3360A>C 3361G>A), P353T (3362C>A), N354D (3365A>G 3367C>T), D356K (3371G>A 3373T>G), S359D (3380T>G 3381C>A 3382C>T), A360V (3384C>T), I361V (3386A>G), Q362K (3389C>A), K363T (3393A>C), Y364W (3396A>G 3397T>G), L365T (3398C>A 3399T>C 3400G>A), K368T (3408A>C 3409G>A), P370V (3413C>G 3414C>T 3415T>G), E372Q (3419G>C), I373L (3422A>T 3424A>G), K374R (3426A>G), Q375G (3428C>G 3429A>G 3430A>C), L377F (3436A>T), L380A (3443C>G 3444T>C 3445C>A), K385R (3459A>G), P388R (3467C>A 3468C>G), D389G (3471A>G), A391G (3477C>G 3478A>T), R392V (3479C>G 3480G>T 3481A>T), L393I (3482C>A 3484C>T), T394A (3485A>G), K395R (3489A>G), T398N (3498C>A 3499A>T), Q399E (3500C>G), C400L (3504G>T 3505C>G), G404K (3515G>A 3516G>A 3517T>G), S405I (3519G>T 3520T>A) | | | | | | | | |
| Codon mutations: | GTG139ATA (2720G>A 2722G>A), TAC140AAC (2723T>A), ACC141ATC (2727C>T), AAA142AGA (2730A>G), AGT143CCA (2732A>C 2733G>C 2734T>A), CGG145AGG (2738C>A), CCC147CCT (2746C>T), TTC148ACT (2747T>A 2748T>C 2749C>T), ATC149ATA (2752C>A), CAT150CAG (2755T>G), CGC151AAG (2756C>A 2757G>A 2758C>G), CAG152GAT (2759C>G 2761G>T), GTT154ATT (2765G>A), AGG155GAA (2768A>G 2769G>A 2770G>A), GAC156AAG (2771G>A 2773C>G), CAA157ATG (2774C>A 2775A>T 2776A>G), ATC158GTC (2777A>G), ACG159GAT (2780A>G 2781C>A 2782G>T), AAA160GAG (2783A>G 2785A>G), ATG161TTG (2786A>T), TTG162CTG (2789T>C), CAA164ACA (2795C>A 2796A>C), ATT166GTT (2801A>G), ATA167ATT (2806A>T), AGA168AGG (2809A>G), CCA169CAG (2811C>A 2812A>G), TCA170AGT (2813T>A 2814C>G 2815A>T), GAC171ATT (2816G>A 2817A>T 2818C>T), TCT172AGT (2819T>A 2820C>G), GCA173CCA (2822G>C), TGG174TGT (2827G>T), AGC175TCC (2828A>T 2829G>C), ATA178ATT (2839A>T), GAG204..C (2917G>C), AAG205ATC (2919A>T 2920G>C), ATC207GTG (2924A>G 2926C>G), GAT208AGG (2927G>A 2928A>G 2929T>G), GAC209GAT (2932C>T), AAA210AGG (2934A>G 2935A>G), TAC211TTC (2937A>T), CCG212CCA (2941G>A), ATA213ATC (2944A>C), AAC215ACT (2949A>C 2950C>T), ATA216ATT (2953A>T), AGT217GAT (2954A>G 2955G>A), GAC218GAG (2959C>G), GTA219CTA (2960G>C), CTT220TTT (2963C>T), GAC221GAT (2968C>T), AAG222GAG (2969A>G), TTA223GTA (2972T>G), GGT224AAT (2975G>A 2976G>A), AAG225GGC (2978A>G 2979A>G 2980G>C), TGC226GCT (2981T>G 2982G>C 2983C>T), CAA227GTT (2984C>G 2985A>T 2986A>T), TAC228TAT (2989C>T), ACC230TCA (2993A>T 2995C>A), ACC231AAG (2997C>A 2998C>G), TTA232CTT (2999T>C 3001A>T), GCA235TTA (3008G>T 3009C>T), AGT236TCT (3011A>T 3012G>C), GGG237GGA (3016G>A), TTT238TAC (3018T>A 3019T>C), TAT239GAT (3020T>G), CAG240CAA (3025G>A), GTG241ATT (3026G>A 3028G>T), GAG242CGG (3029G>C 3030A>G), ATG243GTT (3032A>G 3034G>T), GAC244AAG (3035G>A 3037C>G), CCT245GTG (3038C>G 3039C>T 3040T>G), CAA246GCA (3041C>G 3042A>C), TCG249CCC (3050T>C 3052G>C), ACC251ACT (3058C>T), GCG252GCT (3061G>T), AAC254AGG (3066A>G 3067C>G), GTA255ACA (3068G>A 3069T>C), GAA256CAC (3071G>C 3073A>C), CAC257GAT (3074C>G 3076C>T), GGG258GGT (3079G>T), TTT260TAC (3084T>A 3085T>C), TTC262TTT (3091C>T), CTT263TTA (3092C>T 3094T>A), CGA264GTC (3095C>G 3096G>T 3097A>C), ATG267TTT (3104A>T 3106G>T), GGA268GGG (3109A>G), TTA269TTG (3112A>G), AAA270TCT (3113A>T 3114A>C 3115A>T), TCA272GCG (3119T>G 3121A>G), CCA273TCG (3122C>T 3124A>G), TCT274TCC (3127T>C), ACT275ACA (3130T>A), AGA278AGC (3139A>C), GTT279CTC (3140G>C 3142T>C), GAC281AAC (3146G>A), AAT282AAG (3151T>G), GTC283ATC (3152G>A), CTA284TTT (3155C>T 3157A>T), AGA285CAC (3158A>C 3159G>A 3160A>C), GGT286TCT (3161G>T 3162G>C), CTC287TAC (3164C>T 3165T>A), CAA288CTC (3168A>T 3169A>C), AAT289AGA (3171A>G 3172T>A), AAC290AAG (3175C>G), ATC291TTC (3176A>T), TGT292ATT (3179T>A 3180G>T), CTC293TTG (3182C>T 3184C>G), GTC294GTA (3187C>A), TAC295TTT (3189A>T 3190C>T), CTT296TTC (3191C>T 3193T>C), GAC297GAT (3196C>T), ATT299ATC (3202T>C), ATT300CTC (3203A>C 3205T>C), GTC301ATC (3206G>A), ACT304TC. (3215A>T), CAG307..A (3226G>A), GAA308GAG (3229A>G), CTG310ATA (3233C>A 3235G>A), GAA314AGA (3245G>A 3246A>G), CGA315ACT (3248C>A 3249G>C 3250A>T), GTT316GTG (3253T>G), TTC317CTG (3254T>C 3256C>G), CAA318CAG (3259A>G), AGA319GTT (3260A>G 3261G>T 3262A>T), CTT320TTG (3263C>T 3265T>G), AGT323GAA (3272A>G 3273G>A 3274T>A), AAC324CAG (3275A>C 3277C>G), TTC325TTA (3280C>A), AAA326TAT (3281A>T 3283A>T), ATT327GCA (3284A>G 3285T>C 3286T>A), CAA328AAA (3287C>A), ATG329TAT (3290A>T 3291T>A 3292G>T), GAC330GAT (3295C>T), TCC332TGT (3300C>G 3301C>T), TTG335TGG (3309T>G), AAG336TTT (3311A>T 3312A>T 3313G>T), CTC337CTG (3316C>G), ACT339GTG (3320A>G 3321C>T 3322T>G), GCT340GAG (3324C>A 3325T>G), CTT342TTG (3329C>T 3331T>G), ATC345TTT (3338A>T 3340C>T), AGC347AGT (3346C>T), AGG348GCC (3347A>G 3348G>C 3349G>C), GAC349GCT (3351A>C 3352C>T), GGT350GGA (3355T>A), ATC351GTT (3356A>G 3358C>T), AAG352GCA (3359A>G 3360A>C 3361G>A), CCT353ACT (3362C>A), AAC354GAT (3365A>G 3367C>T), GAT356AAG (3371G>A 3373T>G), ATT358ATA (3379T>A), TCC359GAT (3380T>G 3381C>A 3382C>T), GCT360GTT (3384C>T), ATT361GTT (3386A>G), CAA362AAA (3389C>A), AAA363ACA (3393A>C), TAT364TGG (3396A>G 3397T>G), CTG365ACA (3398C>A 3399T>C 3400G>A), ATT366ATA (3403T>A), AAG368ACA (3408A>C 3409G>A), CCT370GTG (3413C>G 3414C>T 3415T>G), GAA372CAA (3419G>C), ATA373TTG (3422A>T 3424A>G), AAA374AGA (3426A>G), CAA375GGC (3428C>G 3429A>G 3430A>C), TTT376TTC (3433T>C), TTA377TTT (3436A>T), GGC378GGT (3439C>T), CTC380GCA (3443C>G 3444T>C 3445C>A), TAT382TAC (3451T>C), TAC383TAT (3454C>T), CGA384AGA (3455C>A), AAA385AGA (3459A>G), CCA388AGA (3467C>A 3468C>G), GAT389GGT (3471A>G), GCA391GGT (3477C>G 3478A>T), CGA392GTT (3479C>G 3480G>T 3481A>T), CTC393ATT (3482C>A 3484C>T), ACA394GCA (3485A>G), AAA395AGA (3489A>G), ACA398AAT (3498C>A 3499A>T), CAG399GAG (3500C>G), TGC400TTG (3504G>T 3505C>G), AAA402AAG (3511A>G), GGT404AAG (3515G>A 3516G>A 3517T>G), AGT405ATA (3519G>T 3520T>A) | | | | | | | | |

*: Inserts / Deletes / Misaligned / Frameshifts

#### Analysis details

This analysis was performed with panviral2.64

### NGS Details (UN18_val): Marseillevirus marseillevirus

#### Assembly

| | |
|---|---|
| **Coverage Length** | 115 (1 contig(s)) |
| **Depth Of Coverage** | 52.2 |
| **Number Of Reads** | 75 |
| **Reads Per Million** | 1.41 rpm (after QC) |
| **Ambiguities** | 0 |
| **Assembly Method** | de novo + reference guided assembly |
| **Consensus Caller** | Bcf Tools |

#### Coverage Map

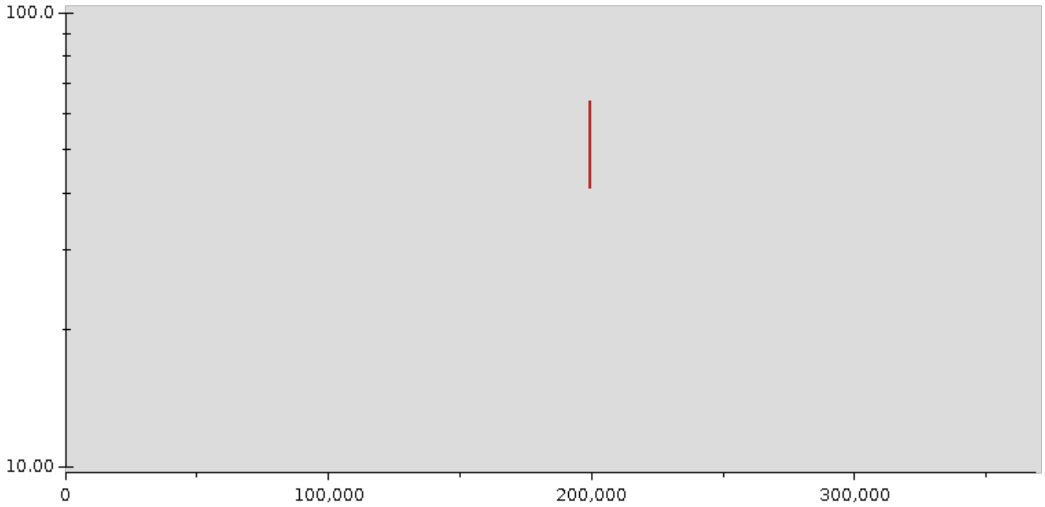

#### Assignment

| | |
|---|---|
| **Type** | Marseillevirus marseillevirus (Taxonomy ID: 694581) |
| **Reference Genome** | NC_013756.1 |
| **NT Identity (%)** | 81.7391 |
| **AA Identity (%)** | 97.3684 |
| **Number Of Stop Codons** | 1 |
| **Number Of CDS** | 428 |

#### Alignment

| | |
|---|---|
| **Alignment Score** | 146.0 (NT) + 245.0 (AA) = 391.0 |
| **Concordance (%)** | 82.1429 |
| **Alignment Method** | Local, heuristic, nucleotide (BLASTN) |

#### Genome Region

Sequence starts at position 199404 and ends at position 199518 relative to NC_013756.1 reference sequence.

#### Alignment Detailed Statistics

| | Begin | End | Coverage | Score | Concordance | Matches | Identities | I/D/M/F* | Stop Codons |
|---|---|---|---|---|---|---|---|---|---|
| NT | 199404 | 199518 | 0.1% | 146 | 63.5% | 115 (100%) | 94 (81.7%) | 0/0 | |

Mutations: 199411A>C, 199414T>A, 199416T>G, 199423A>G, 199432A>G, 199444T>C, 199449C>T, 199459A>C, 199462C>A, 199464A>G, 199465A>G, 199470T>G, 199471T>G, 199474A>G, 199477T>C, 199480C>G, 199489T>G, 199495G>A, 199501G>C, 199503G>A, 199516C>T

*: Inserts / Deletes / Misaligned / Frameshifts

#### Analysis details

This analysis was performed with panviral2.64

### NGS Details (UN18_val): Badnavirus occultipomeae

#### Assembly

| | |
|---|---|
| **Coverage Length** | 260 (1 contig(s)) |
| **Depth Of Coverage** | 25.5 |
| **Number Of Reads** | 66 |
| **Reads Per Million** | 1.24 rpm (after QC) |
| **Ambiguities** | 0 |
| **Assembly Method** | de novo + reference guided assembly |
| **Consensus Caller** | Bcf Tools |

#### Coverage Map

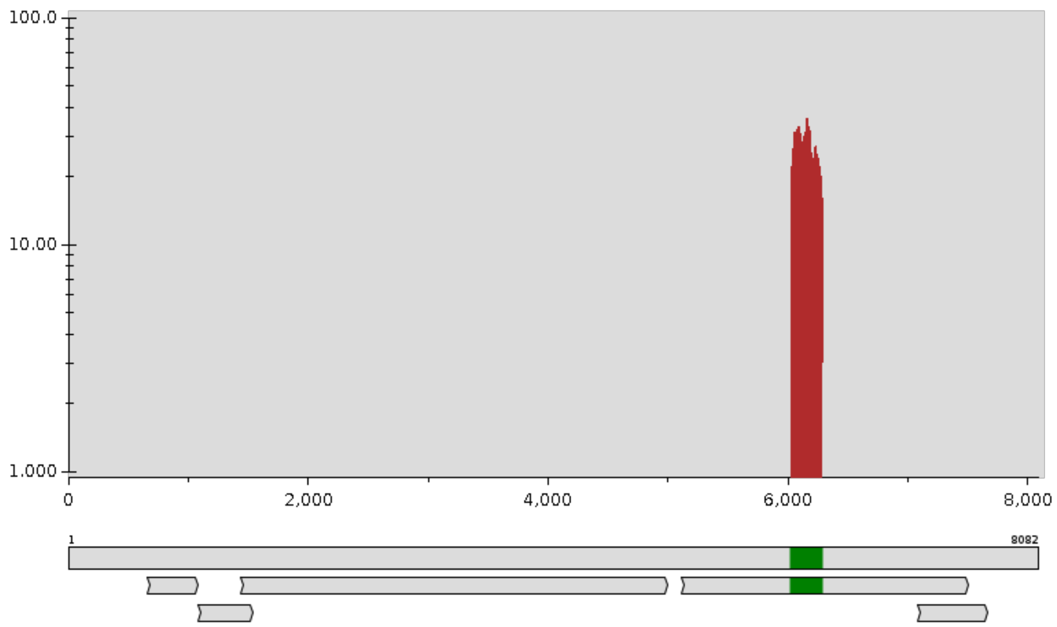

#### Assignment

| | |
|---|---|
| **Type** | Badnavirus occultipomeae (Taxonomy ID: 3048353) |
| **Reference Genome** | NC_015655.1 |
| **NT Identity (%)** | 58.7591 |
| **AA Identity (%)** | 49.4505 |
| **Number Of Stop Codons** | 1 |
| **Number Of CDS** | 5 |

#### Alignment

| | |
|---|---|
| **Alignment Score** | 101.0 (NT) + 190.0 (AA) = 291.0 |
| **Concordance (%)** | 25.2824 |

| Alignment Method | Global, seeded, nucleotide + amino acids (AGA) |
|---|---|

#### Genome Region

Sequence starts at position 6026 and ends at position 6285 relative to NC_015655.1 reference sequence.

#### Alignment Detailed Statistics

| | Begin | End | Coverage | Score | Concordance | Matches | Identities | I/D/M/F* | Stop Codons |
|---|---|---|---|---|---|---|---|---|---|
| **NT** | **6026** | **6285** | **3.2%** | **101** | **19.4%** | **260 (94.9%)** | **161 (58.8%)** | **14/0** | |
| Mutations: | 6033A>T, 6042A>G, 6043T>A, 6045T>A, 6048T>C, 6051C>A, 6053A>G, 6055G>T, 6062T>A, 6067C>T, 6070A>T, 6074G>A, 6075A>G, 6076A>G, 6078G>A, 6079G>A, 6081G>A, 6084A>G, 6085A>T, 6086A>G, 6087G>T, 6088T>G, 6089C>A, 6090C>T, 6092A>T, 6093A>C, 6097T>A, 6098G>A, 6099G>A, 6109T>C, 6111G>A, 6116C>A, 6117A>T, 6118G>T, 6120A>T, 6123G>C, 6124C>T, 6125T>A, 6126C>T, 6128A>T, 6132A>G, 6135T>C, 6136G>T, 6137A>T, 6138A>G, 6145C>T, 6153A>G, 6154C>T, 6158T>C, 6159G>T, 6168A>T, 6173A>C, 6174T>C, 6178C>A, 6179A>T, 6181A>G, 6182G>A, 6183A>T, 6184A>C, 6185A>T, 6186G>C, 6190G>A, 6194A>G, 6195T>A, 6197C>T, 6198A>G, 6201T>C, 6203G>A, 6205G>C, 6206G>C, 6207A>T, 6207_6208insTATTCAAGCCTTAT, 6208A>C, 6209C>T, 6210A>T, 6213T>C, 6214G>A, 6215C>T, 6216A>G, 6219T>A, 6220A>G, 6222C>T, 6223G>A, 6224C>T, 6225A>T, 6228A>C, 6230A>T, 6240C>T, 6243A>C, 6249A>C, 6251T>A, 6255T>C, 6256G>A, 6258A>G, 6264A>G, 6270G>A, 6271C>T, 6275A>C, 6276G>C, 6277G>T | | | | | | | | |
| **CDS** | | | | | | | | | |
| SPBVa_gp4 | 305 | 390 | 10.8% | 190 | 29.9% | 86 (94.5%) | 45 (49.5%) | 5/0/1/1 | 1 |
| Protein mutations: | F310I (6043T>A 6045T>A), K313R (6053A>G), A314S (6055G>T), F316Y (6062T>A), Q318* (6067C>T), I319L (6070A>T), R320K (6074G>A 6075A>G), M321V (6076A>G 6078G>A), E322K (6079G>A 6081G>A), K324C (6085A>T 6086A>G 6087G>T), S325D (6088T>G 6089C>A 6090C>T), K326I (6092A>T 6093A>C), W328K (6097T>A 6098G>A 6099G>A), W332R (6109T>C 6111G>A), P334H (6116C>A 6117A>T), E335Y (6118G>T 6120A>T), L337Y (6124C>T 6125T>A 6126C>T), Y338F (6128A>T), E341L (6136G>T 6137A>T 6138A>G), P344S (6145C>T), M348T (6158T>C 6159G>T), D353A (6173A>C 6174T>C), Q355M (6178C>A 6179A>T), R356D (6181A>G 6182G>A 6183A>T), K357L (6184A>C 6185A>T 6186G>C), D359N (6190G>A), N360R (6194A>G 6195T>A), A361V (6197C>T 6198A>G), R363K (6203G>A), G364P (6205G>C 6206G>C 6207A>T), G364_T365insYSSLX (6207_6208insTATTCAAGCCTTAT), T365L (6208A>C 6209C>T 6210A>T), A367M (6214G>A 6215C>T 6216A>G), F368L (6219T>A), I369V (6220A>G 6222C>T), A370I (6223G>A 6224C>T 6225A>T), Y372F (6230A>T), F379Y (6251T>A), E381K (6256G>A 6258A>G), H386Y (6271C>T), E387A (6275A>C 6276G>C), D388Y (6277G>T) | | | | | | | | |
| Codon mutations: | ATA306ATT (6033A>T), AAA309AAG (6042A>G), TTT310ATA (6043T>A 6045T>A), GAT311GAC (6048T>C), CTC312CTA (6051C>A), AAA313AGA (6053A>G), GCA314TCA (6055G>T), TTT316TAT (6062T>A), CAG318TAG (6067C>T), ATA319TTA (6070A>T), AGA320AAG (6074G>A 6075A>G), ATG321GTA (6076A>G 6078G>A), GAG322AAA (6079G>A 6081G>A), GAA323GAG (6084A>G), AAG324TGT (6085A>T 6086A>G 6087G>T), TCC325GAT (6088T>G 6089C>A 6090C>T), AAA326ATC (6092A>T 6093A>C), TGG328AAA (6097T>A 6098G>A 6099G>A), TGG332CGA (6109T>C 6111G>A), CCA334CAT (6116C>A 6117A>T), GAA335TAT (6118G>T 6120A>T), GGG336GGC (6123G>C), CTC337TAT (6124C>T 6125T>A 6126C>T), TAT338TTT (6128A>T), GAA339GAG (6132A>G), TTT340TTC (6135T>C), GAA341TTG (6136G>T 6137A>T 6138A>G), CCC344TCC (6145C>T), GGA346GGG (6153A>G), CTG347TTG (6154C>T), ATG348ACT (6158T>C 6159G>T), CCA351CCT (6168A>T), GAT353GCC (6173A>C 6174T>C), CAG355ATG (6178C>A 6179A>T), AGA356GAT (6181A>G 6182G>A 6183A>T), AAG357CTC (6184A>C 6185A>T 6186G>C), GAT359AAT (6190G>A), AAT360AGA (6194A>G 6195T>A), GCA361GTG (6197C>T 6198A>G), TTT362TTC (6201T>C), AGG363AAG (6203G>A), GGA364CCT (6205G>C 6206G>C 6207A>T), GGA364_ACA365insTATTCAAGCCTTAT-(6207_6208insTATTCAAGCCTTAT), ACA365CTT (6208A>C 6209C>T 6210A>T), GAT366GAC (6213T>C), GCA367ATG (6214G>A 6215C>T 6216A>G), TTT368TTA (6219T>A), ATC369GTT (6220A>G 6222C>T), GCA370ATT (6223G>A 6224C>T 6225A>T), GTA371GTC (6228A>C), TAT372TTT (6230A>T), GAC375GAT (6240C>T), ATA376ATC (6243A>C), GTA378GTC (6249A>C), TTC379TAC (6251T>A), TCT380TCC (6255T>C), GAA381AAG (6256G>A 6258A>G), GAA383GAG (6264A>G), GAG385GAA (6270G>A), CAT386TAT (6271C>T), GAG387GCC (6275A>C 6276G>C), GAT388TAT (6277G>T) | | | | | | | | |
| **Proteins** | | | | | | | | | |
| RNaseH/reverse transcriptase (YP_004581513.1) | 305 | 390 | 10.8% | 190 | 29.9% | 86 (94.5%) | 45 (49.5%) | 5/0/1/1 | 1 |
| Protein mutations: | F310I (6043T>A 6045T>A), K313R (6053A>G), A314S (6055G>T), F316Y (6062T>A), Q318* (6067C>T), I319L (6070A>T), R320K (6074G>A 6075A>G), M321V (6076A>G 6078G>A), E322K (6079G>A 6081G>A), K324C (6085A>T 6086A>G 6087G>T), S325D (6088T>G 6089C>A 6090C>T), K326I (6092A>T 6093A>C), W328K (6097T>A 6098G>A 6099G>A), W332R (6109T>C 6111G>A), P334H (6116C>A 6117A>T), E335Y (6118G>T 6120A>T), L337Y (6124C>T 6125T>A 6126C>T), Y338F (6128A>T), E341L (6136G>T 6137A>T 6138A>G), P344S (6145C>T), M348T (6158T>C 6159G>T), D353A (6173A>C 6174T>C), Q355M (6178C>A 6179A>T), R356D (6181A>G 6182G>A 6183A>T), K357L (6184A>C 6185A>T 6186G>C), D359N (6190G>A), N360R (6194A>G 6195T>A), A361V (6197C>T 6198A>G), R363K (6203G>A), G364P (6205G>C 6206G>C 6207A>T), G364_T365insYSSLX (6207_6208insTATTCAAGCCTTAT), T365L (6208A>C 6209C>T 6210A>T), A367M (6214G>A 6215C>T 6216A>G), F368L (6219T>A), I369V (6220A>G 6222C>T), A370I (6223G>A 6224C>T 6225A>T), Y372F (6230A>T), F379Y (6251T>A), E381K (6256G>A 6258A>G), H386Y (6271C>T), E387A (6275A>C 6276G>C), D388Y (6277G>T) | | | | | | | | |
| Codon mutations: | ATA306ATT (6033A>T), AAA309AAG (6042A>G), TTT310ATA (6043T>A 6045T>A), GAT311GAC (6048T>C), CTC312CTA (6051C>A), AAA313AGA (6053A>G), GCA314TCA (6055G>T), TTT316TAT (6062T>A), CAG318TAG (6067C>T), ATA319TTA (6070A>T), AGA320AAG (6074G>A 6075A>G), ATG321GTA (6076A>G 6078G>A), GAG322AAA (6079G>A 6081G>A), GAA323GAG (6084A>G), AAG324TGT (6085A>T 6086A>G 6087G>T), TCC325GAT (6088T>G 6089C>A 6090C>T), AAA326ATC (6092A>T 6093A>C), TGG328AAA (6097T>A 6098G>A 6099G>A), TGG332CGA (6109T>C 6111G>A), CCA334CAT (6116C>A 6117A>T), GAA335TAT (6118G>T 6120A>T), GGG336GGC (6123G>C), CTC337TAT (6124C>T 6125T>A 6126C>T), TAT338TTT (6128A>T), GAA339GAG (6132A>G), TTT340TTC (6135T>C), GAA341TTG (6136G>T 6137A>T 6138A>G), CCC344TCC (6145C>T), GGA346GGG (6153A>G), CTG347TTG (6154C>T), ATG348ACT (6158T>C 6159G>T), CCA351CCT (6168A>T), GAT353GCC (6173A>C 6174T>C), CAG355ATG (6178C>A 6179A>T), AGA356GAT (6181A>G 6182G>A 6183A>T), AAG357CTC (6184A>C 6185A>T 6186G>C), GAT359AAT (6190G>A), AAT360AGA (6194A>G 6195T>A), GCA361GTG (6197C>T 6198A>G), TTT362TTC (6201T>C), AGG363AAG (6203G>A), GGA364CCT (6205G>C 6206G>C 6207A>T), GGA364_ACA365insTATTCAAGCCTTAT-(6207_6208insTATTCAAGCCTTAT), ACA365CTT (6208A>C 6209C>T 6210A>T), GAT366GAC (6213T>C), GCA367ATG (6214G>A 6215C>T 6216A>G), TTT368TTA (6219T>A), ATC369GTT (6220A>G 6222C>T), GCA370ATT (6223G>A 6224C>T 6225A>T), GTA371GTC (6228A>C), TAT372TTT (6230A>T), GAC375GAT (6240C>T), ATA376ATC (6243A>C), GTA378GTC (6249A>C), TTC379TAC (6251T>A), TCT380TCC (6255T>C), GAA381AAG (6256G>A 6258A>G), GAA383GAG (6264A>G), GAG385GAA (6270G>A), CAT386TAT (6271C>T), GAG387GCC (6275A>C 6276G>C), GAT388TAT (6277G>T) | | | | | | | | |

*: Inserts / Deletes / Misaligned / Frameshifts

#### Analysis details

This analysis was performed with panviral2.64

### NGS Details (UN18_val): Caulimovirus venafragariae

#### Assembly

| | |
|---|---|
| **Coverage Length** | 570 (2 contig(s)) |
| **Depth Of Coverage** | 9.9 |
| **Number Of Reads** | 48 |
| **Reads Per Million** | 0.90 rpm (after QC) |
| **Ambiguities** | 0 |
| **Assembly Method** | de novo + reference guided assembly |
| **Consensus Caller** | Bcf Tools |

#### Coverage Map

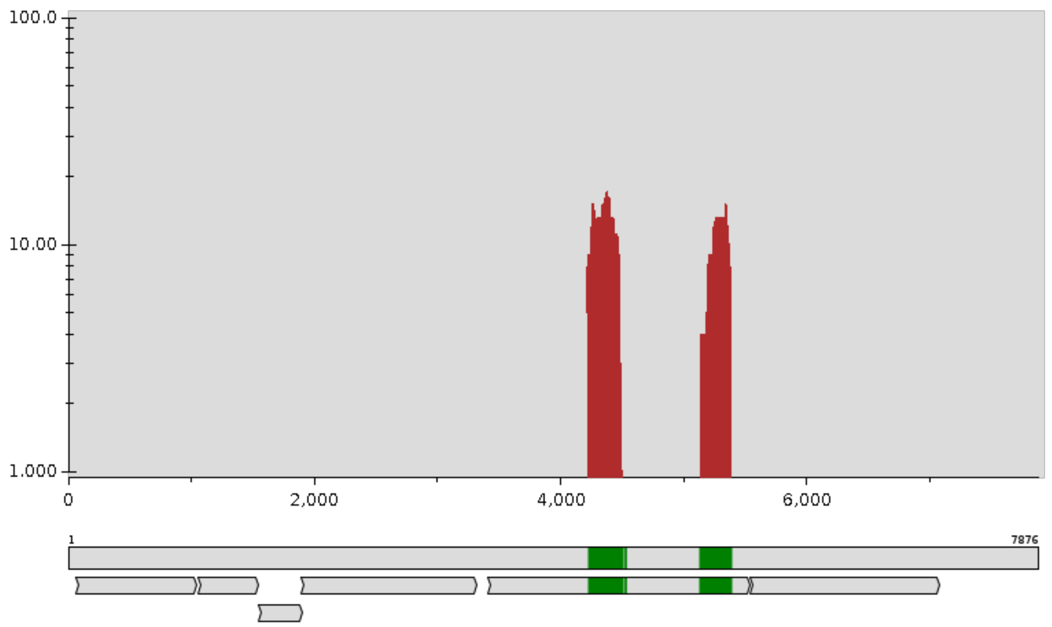

#### Assignment

| | |
|---|---|
| **Type** | Caulimovirus venafragariae (Taxonomy ID: 3048344) |
| **Reference Genome** | NC_001725.1 |
| **NT Identity (%)** | 58.2734 |
| **AA Identity (%)** | 54.8913 |
| **Number Of Stop Codons** | 0 |
| **Number Of CDS** | 6 |

#### Alignment

| | |
|---|---|
| **Alignment Score** | 152.0 (NT) + 606.0 (AA) = 758.0 |
| **Concordance (%)** | 35.4578 |

| Alignment Method | Global, seeded, nucleotide + amino acids (AGA) |
|---|---|

#### Genome Region

Sequence starts at position 4219 and ends at position 5394 relative to NC_001725.1 reference sequence.

#### Alignment Detailed Statistics

| | Begin | End | Coverage | Score | Concordance | Matches | Identities | I/D/M/F* | Stop Codons |
|---|---|---|---|---|---|---|---|---|---|
| **NT** | **4219** | **5394** | **7.2%** | **152** | **14.1%** | **556 (97.5%)** | **324 (56.8%)** | **0/14** | |
| Mutations: | 4219T>C, 4222T>A, 4223T>A, 4228A>C, 4230T>C, 4232A>G, 4235A>T, 4236A>C, 4242T>A, 4244A>G, 4245G>A, 4246T>G, 4247G>T, 4254A>T, 4259A>T, 4260G>A, 4263T>C, 4264T>A, 4265C>A, 4266A>T, 4267C>G, 4268C>A, 4269T>A, 4275C>T, 4276C>A, 4277G>A, 4279G>A, 4280A>C, 4281A>T, 4284G>A, 4292C>T, 4299C>T, 4300G>C, 4306C>T, 4309C>T, 4311C>A, 4314G>T, 4315C>A, 4317T>G, 4318G>A, 4319G>A, 4320T>A, 4321A>T, 4323C>A, 4326C>A, 4330_4331delCC, 4332T>C, 4335C>T, 4338G>A, 4341T>C, 4344A>G, 4348T>A, 4349C>G, 4350C>T, 4351T>A, 4352C>G, 4353A>T, 4359C>A, 4371G>A, 4374T>C, 4380C>G, 4381G>A, 4383G>A, 4384A>C, 4385T>A, 4386C>A, 4395C>A, 4401A>T, 4402A>C, 4413T>A, 4423A>G, 4426T>C, 4428A>T, 4432G>A, 4434C>A, 4435C>A, 4438A>T, 4439C>G, 4440A>T, 4441A>G, 4442A>T, 4443G>C, 4444G>T, 4445G>T, 4446A>T, 4452C>A, 4455C>T, 4456C>T, 4458A>T, 4459C>A, 4464T>A, 4470G>A, 4471G>A, 4473G>T, 4474C>A, 4476A>T, 4479G>T, 4480C>A, 4482T>A, 4483C>A, 4485A>T, 4486A>C, 4487G>T, 4489A>G, 4490T>C, 4492G>A, 4493G>A, 4497T>A, 4500G>A, 4505T>A, 4506T>C, 4508A>T, 4509C>T, 4512T>A, 4513T>A, 4514C>A, 4515C>A, 4521C>T, 4524T>C, 4528T>A, 4529C>G, 5126C>A, 5133G>A, 5139G>C, 5141C>T, 5142T>A, 5143A>G, 5145C>A, 5146A>T, 5148C>A, 5149G>C, 5152T>A, 5153G>C, 5154C>T, 5160A>T, 5163A>T, 5167G>T, 5169C>T, 5182A>T, 5184C>A, 5185C>T, 5187C>A, 5188A>C, 5191G>A, 5194A>G, 5196A>T, 5199T>A, 5200C>A, 5201C>A, 5202A>T, 5206_5217delGGTAAAGAGGTA, 5220A>T, 5223C>T, 5226G>A, 5230G>A, 5233T>A, 5234C>G, 5235A>T, 5247A>T, 5248C>G, 5249C>A, 5251G>A, 5253T>A, 5256G>A, 5258A>C, 5259G>T, 5261A>G, 5262T>G, 5266C>T, 5267A>T, 5268T>A, 5270G>C, 5271T>A, 5280G>A, 5284A>T, 5286C>A, 5289T>A, 5290T>G, 5292C>T, 5295T>A, 5296A>G, 5298T>A, 5300A>G, 5301G>A, 5303C>G, 5304A>C, 5307C>A, 5308A>C, 5309A>G, 5311G>A, 5312C>A, 5313T>G, 5317A>T, 5318G>C, 5319A>T, 5322T>A, 5324A>T, 5325T>C, 5326A>T, 5328C>A, 5331T>A, 5334T>A, 5335T>A, 5337T>A, 5338A>C, 5343T>C, 5344C>A, 5346T>C, 5347G>A, 5349A>C, 5352G>A, 5353A>T, 5355T>G, 5358T>C, 5364T>G, 5365A>C, 5367T>G, 5369C>T, 5370T>A, 5371G>T, 5373T>A, 5374T>G, 5375A>G, 5377T>C, 5379T>A, 5380G>A, 5382G>A, 5383A>T, 5384G>T, 5385A>C, 5387C>A, 5388A>T, 5391T>A, 5392A>C | | | | | | | | |

##### CDS

| | Begin | End | Coverage | Score | Concordance | Matches | Identities | I/D/M/F* | Stop Codons |
|---|---|---|---|---|---|---|---|---|---|
| ORF_V | 272 | 663 | 26.5% | 606 | 47.3% | 184 (97.9%) | 101 (53.7%) | 0/4/1/1 | 0 |
| Protein mutations: | L273K (4222T>A 4223T>A), T275P (4228A>C 4230T>C), N276S (4232A>G), K277I (4235A>T 4236A>C), K280R (4244A>G 4245G>A), C281V (4246T>G 4247G>T), Q285L (4259A>T 4260G>A), S287N (4264T>A 4265C>A 4266A>T), P288E (4267C>G 4268C>A 4269T>A), R291K (4276C>A 4277G>A), E292T (4279G>A 4280A>C 4281A>T), T296I (4292C>T), E299Q (4300G>C), K303N (4314G>T), L304M (4315C>A 4317T>G), G305K (4318G>A 4319G>A 4320T>A), I306L (4321A>T 4323C>A), E326K (4381G>A 4383G>A), I327Q (4384A>C 4385T>A 4386C>A), K340E (4423A>G), D343K (4432G>A 4434C>A), H344N (4435C>A), T345C (4438A>T 4439C>G 4440A>T), K346V (4441A>G 4442A>T 4443G>C), G347F (4444G>T 4445G>T 4446A>T), L351F (4456C>T 4458A>T), L352I (4459C>A), E356N (4471G>A 4473G>T), Q357N (4474C>A 4476A>T), L359I (4480C>A 4482T>A), Q360N (4483C>A 4485A>T), R361L (4486A>C 4487G>T), I362A (4489A>G 4490T>C), G363K (4492G>A 4493G>A), F367Y (4505T>A 4506T>C), Y368F (4508A>T 4509C>T), S370K (4513T>A 4514C>A 4515C>A), K578N (5139G>C), P579L (5141C>T 5142T>A), I580V (5143A>G 5145C>A), I581L (5146A>T 5148C>A), E582Q (5149G>C), C583T (5152T>A 5153G>C 5154C>T), D588Y (5167G>T 5169C>T), I593L (5182A>T 5184C>A), K595Q (5188A>C), A596T (5191G>A), K597D (5194A>G 5196A>T), P599N (5200C>A 5201C>A 5202A>T), G601_V604del (5206_5217delGGTAAAGAGGTA), A609T (5230G>A), K614N (5247A>T), P615E (5248C>G 5249C>A), A616T (5251G>A 5253T>A), K618T (5258A>C 5259G>T), N619R (5261A>G 5262T>G), H621L (5266C>T 5267A>T 5268T>A), S622T (5270G>C 5271T>A), I627L (5284A>T 5286C>A), S629A (5290T>G 5292C>T), I631V (5296A>G 5298T>A), K632R (5300A>G 5301G>A), A633G (5303C>G 5304A>C), K635R (5308A>C 5309A>G), A636K (5311G>A 5312C>A 5313T>G), R638S (5317A>T 5318G>C 5319A>T), Y640F (5324A>T 5325T>C), I641L (5326A>T 5328C>A), Y644K (5335T>A 5337T>A), K645Q (5338A>C), L647I (5344C>A 5346T>C), V648I (5347G>A 5349A>C), T650S (5353A>T 5355T>G), N654Q (5365A>C 5367T>G), A655V (5369C>T 5370T>A), A656S (5371G>T 5373T>A), Y657G (5374T>G 5375A>G), F658L (5377T>C 5379T>A), V659I (5380G>A 5382G>A), R660F (5383A>T 5384G>T 5385A>C), T661N (5387C>A 5388A>T), N662K (5391T>A), I663L (5392A>C) | | | | | | | | |
| Codon mutations: | TTA272CTA (4219T>C), TTA273AAA (4222T>A 4223T>A), ACT275CCC (4228A>C 4230T>C), AAC276AGC (4232A>G), AAA277ATC (4235A>T 4236A>C), ATT279ATA (4242T>A), AAG280AGA (4244A>G 4245G>A), TGT281GTT (4246T>G 4247G>T), CCA283CCT (4254A>T), CAG285CTA (4259A>T 4260G>A), TAT286TAC (4263T>C), TCA287AAT (4264T>A 4265C>A 4266A>T), CCT288GAA (4267C>G 4268C>A 4269T>A), GAC290GAT (4275C>T), CGA291AAA (4276C>A 4277G>A), GAA292ACT (4279G>A 4280A>C 4281A>T), GAG293GAA (4284G>A), ACT296ATT (4292C>T), ATC298ATT (4299C>T), GAA299CAA (4300G>C), CTA301TTA (4306C>T), CTC302TTA (4309C>T 4311C>A), AAG303AAT (4314G>T), CTT304ATG (4315C>A 4317T>G), GGT305AAA (4318G>A 4319G>A 4320T>A), ATC306TTA (4321A>T 4323C>A), ATC307ATA (4326C>A), CCT309--C (4330_4331delCC 4332T>C), AGC310AGT (4335C>T), AAG311AAA (4338G>A), AGT312AGC (4341T>C), CCA313CCG (4344A>G), TCC315AGT (4348T>A 4349C>G 4350C>T), TCA316AGT (4351T>A 4352C>G 4353A>T), GCC318GCA (4359C>A), AGG322AGA (4371G>A), AAT323AAC (4374T>C), GCC325GCG (4380C>G), GAG326AAA (4381G>A 4383G>A), ATC327CAA (4384A>C 4385T>A 4386C>A), GGC330GGA (4395C>A), GCA332GCT (4401A>T), AGA333CGA (4402A>C), ATT336ATA (4413T>A), AAG340GAG (4423A>G), TTA341CTT (4426T>C 4428A>T), GAC343AAA (4432G>A 4434C>A), CAT344AAT (4435C>A), ACA345TGT (4438A>T 4439C>G 4440A>T), AAG346GTC (4441A>G 4442A>T 4443G>C), GGA347TTT (4444G>T 4445G>T 4446A>T), GGC349GGA (4452C>A), TAC350TAT (4455C>T), CTA351TTT (4456C>T 4458A>T), CTT352ATT (4459C>A), CCT353CCA (4464T>A), AAG355AAA (4470G>A), GAG356AAT (4471G>A 4473G>T), CAA357AAT (4474C>A 4476A>T), CTG358CTT (4479G>T), CTT359ATA (4480C>A 4482T>A), CAA360AAT (4483C>A 4485A>T), AGA361CTA (4486A>C 4487G>T), ATC362GCC (4489A>G 4490T>C), GGA363AAA (4492G>A 4493G>A), GGT364GGA (4497T>A), AAG365AAA (4500G>A), TTT367TAC (4505T>A 4506T>C), TAC368TTT (4508A>T 4509C>T), TCT369TCA (4512T>A), TCC370AAA (4513T>A 4514C>A 4515C>A), GAC372..T (4521C>T), TGT373..C (4524T>C), TCT375AG. (4528T>A 4529C>G), TCA574.AA (5126C>A), GAG576GAA (5133G>A), AAG578AAC (5139G>C), CCT579CTA (5141C>T 5142T>A), ATC580GTA (5143A>G 5145C>A), ATC581TTA (5146A>T 5148C>A), GAA582CAA (5149G>C), TGC583ACT (5152T>A 5153G>C 5154C>T), GCA585GCT (5160A>T), TCA586TCT (5163A>T), GAC588TAT (5167G>T 5169C>T), ATC593TTA (5182A>T 5184C>A), CTC594TTA (5185C>T 5187C>A), AAA595CAA (5188A>C), GCA596ACA (5191G>A), AAA597GAT (5194A>G 5196A>T), CTT598CTA (5199T>A), CCA599AAT (5200C>A 5201C>A 5202A>T), GGT601_GTA604del (5206_5217delGGTAAAGAGGTA), ATA605ATT (5220A>T), TGC606TGT (5223C>T), AGG607AGA (5226G>A), GCT609ACT (5230G>A), TCA610AGT (5233T>A 5234C>G 5235A>T), AAA614AAT (5247A>T), CCA615GAA (5248C>G 5249C>A), GCT616ACA (5251G>A 5253T>A), GAG617GAA (5256G>A), AAG618ACT (5258A>C 5259G>T), AAT619AGG (5261A>G 5262T>G), CAT621TTA (5266C>T 5267A>T 5268T>A), AGT622ACA (5270G>C 5271T>A), AAG625AAA (5280G>A), ATC627TTA (5284A>T 5286C>A), CTT628CTA (5289T>A), TCC629GCT (5290T>G 5292C>T), ATT630ATA (5295T>A), ATT631GTA (5296A>G 5298T>A), AAG632AGA (5300A>G 5301G>A), GCA633GGC (5303C>G 5304A>C), ATC634ATA (5307C>A), AAA635CGA (5308A>C 5309A>G), GCT636AAG (5311G>A 5312C>A 5313T>G), AGA638TCT (5317A>T 5318G>C 5319A>T), GCT639GCA (5322T>A), TAT640TTC (5324A>T 5325T>C), ATC641TTA (5326A>T 5328C>A), CTT642CTA (5331T>A), CCT643CCA (5334T>A), TAT644AAA (5335T>A 5337T>A), AAA645CAA (5338A>C), TTT646TTC (5343T>C), CTT647ATC (5344C>A 5346T>C), GTA648ATC (5347G>A 5349A>C), AGG649AGA (5352G>A), ACT650TCG (5353A>T 5355T>G), GAT651GAC (5358T>C), ACT653ACG (5364T>G), AAT654CAG (5365A>C 5367T>G), GCT655GTA (5369C>T 5370T>A), GCT656TCA (5371G>T 5373T>A), TAT657GGT (5374T>G 5375A>G), TTT658CTA (5377T>C 5379T>A), GTG659ATA (5380G>A 5382G>A), AGA660TTC (5383A>T 5384G>T 5385A>C), ACA661AAT (5387C>A 5388A>T), AAT662AAA (5391T>A), ATT663CTT (5392A>C) | | | | | | | | |

##### Proteins

| | Begin | End | Coverage | Score | Concordance | Matches | Identities | I/D/M/F* | Stop Codons |
|---|---|---|---|---|---|---|---|---|---|
| hypothetical protein (NP_043933.1) | 272 | 663 | 26.5% | 606 | 47.3% | 184 (97.9%) | 101 (53.7%) | 0/4/1/1 | 0 |
| Protein mutations: | L273K (4222T>A 4223T>A), T275P (4228A>C 4230T>C), N276S (4232A>G), K277I (4235A>T 4236A>C), K280R (4244A>G 4245G>A), C281V (4246T>G 4247G>T), Q285L (4259A>T 4260G>A), S287N (4264T>A 4265C>A 4266A>T), P288E (4267C>G 4268C>A 4269T>A), R291K (4276C>A 4277G>A), E292T (4279G>A 4280A>C 4281A>T), T296I (4292C>T), E299Q (4300G>C), K303N (4314G>T), L304M (4315C>A 4317T>G), G305K (4318G>A 4319G>A 4320T>A), I306L (4321A>T 4323C>A), E326K (4381G>A 4383G>A), I327Q (4384A>C 4385T>A 4386C>A), K340E (4423A>G), D343K (4432G>A 4434C>A), H344N (4435C>A), T345C (4438A>T 4439C>G 4440A>T), K346V (4441A>G 4442A>T 4443G>C), G347F (4444G>T 4445G>T 4446A>T), L351F (4456C>T 4458A>T), L352I (4459C>A), E356N (4471G>A 4473G>T), Q357N (4474C>A 4476A>T), L359I (4480C>A 4482T>A), Q360N (4483C>A 4485A>T), R361L (4486A>C 4487G>T), I362A (4489A>G 4490T>C), G363K (4492G>A 4493G>A), F367Y (4505T>A 4506T>C), Y368F (4508A>T 4509C>T), S370K (4513T>A 4514C>A 4515C>A), K578N (5139G>C), P579L (5141C>T 5142T>A), I580V (5143A>G 5145C>A), I581L (5146A>T 5148C>A), E582Q (5149G>C), C583T (5152T>A 5153G>C 5154C>T), D588Y (5167G>T 5169C>T), I593L (5182A>T 5184C>A), K595Q (5188A>C), A596T (5191G>A), K597D (5194A>G 5196A>T), P599N (5200C>A 5201C>A 5202A>T), G601_V604del (5206_5217delGGTAAAGAGGTA), A609T (5230G>A), K614N (5247A>T), P615E (5248C>G 5249C>A), A616T (5251G>A 5253T>A), K618T (5258A>C 5259G>T), N619R (5261A>G 5262T>G), H621L (5266C>T 5267A>T 5268T>A), S622T (5270G>C 5271T>A), I627L (5284A>T 5286C>A), S629A (5290T>G 5292C>T), I631V (5296A>G 5298T>A), K632R (5300A>G 5301G>A), A633G (5303C>G 5304A>C), K635R (5308A>C 5309A>G), A636K (5311G>A 5312C>A 5313T>G), R638S (5317A>T 5318G>C 5319A>T), Y640F (5324A>T 5325T>C), I641L (5326A>T 5328C>A), Y644K (5335T>A 5337T>A), K645Q (5338A>C), L647I (5344C>A 5346T>C), V648I (5347G>A 5349A>C), T650S (5353A>T 5355T>G), N654Q (5365A>C 5367T>G), A655V (5369C>T 5370T>A), A656S (5371G>T 5373T>A), Y657G (5374T>G 5375A>G), F658L (5377T>C 5379T>A), V659I (5380G>A 5382G>A), R660F (5383A>T 5384G>T 5385A>C), T661N (5387C>A 5388A>T), N662K (5391T>A), I663L (5392A>C) | | | | | | | | |

| | Begin | End | Coverage | Score | Concordance | Matches | Identities | I/D/M/F* | Stop Codons |
|---|---|---|---|---|---|---|---|---|---|
| **NT** | **4219** | **5394** | **7.2%** | **152** | **14.1%** | **556 (97.5%)** | **324 (56.8%)** | **0/14** | |
| Codon mutations: | TTA272CTA (4219T>C), TTA273AAA (4222T>A 4223T>A), ACT275CCC (4228A>C 4230T>C), AAC276AGC (4232A>G), AAA277ATC (4235A>T 4236A>C), ATT279ATA (4242T>A), AAG280AGA (4244A>G 4245G>A), TGT281GTT (4246T>G 4247G>T), CCA283CCT (4254A>T), CAG285CTA (4259A>T 4260G>A), TAT286TAC (4263T>C), TCA287AAT (4264T>A 4265C>A 4266A>T), CCT288GAA (4267C>G 4268C>A 4269T>A), GAC290GAT (4275C>T), CGA291AAA (4276C>A 4277G>A), GAA292ACT (4279G>A 4280A>C 4281A>T), GAG293GAA (4284G>A), ACT296ATT (4292C>T), ATC298ATT (4299C>T), GAA299CAA (4300G>C), CTA301TTA (4306C>T), CTC302TTA (4309C>T 4311C>A), AAG303AAT (4314G>T), CTT304ATG (4315C>A 4317T>G), GGT305AAA (4318G>A 4319G>A 4320T>A), ATC306TTA (4321A>T 4323C>A), ATC307ATA (4326C>A), CCT309--C (4330_4331delCC 4332T>C), AGC310AGT (4335C>T), AAG311AAA (4338G>A), AGT312AGC (4341T>C), CCA313CCG (4344A>G), TCC315AGT (4348T>A 4349C>G 4350C>T), TCA316AGT (4351T>A 4352C>G 4353A>T), GCC318GCA (4359C>A), AGG322AGA (4371G>A), AAT323AAC (4374T>C), GCC325GCG (4380C>G), GAG326AAA (4381G>A 4383G>A), ATC327CAA (4384A>C 4385T>A 4386C>A), GGC330GGA (4395C>A), GCA332GCT (4401A>T), AGA333CGA (4402A>C), ATT336ATA (4413T>A), AAG340GAG (4423A>G), TTA341CTT (4426T>C 4428A>T), GAC343AAA (4432G>A 4434C>A), CAT344AAT (4435C>A), ACA345TGT (4438A>T 4439C>G 4440A>T), AAG346GTC (4441A>G 4442A>T 4443G>C), GGA347TTT (4444G>T 4445G>T 4446A>T), GGC349GGA (4452C>A), TAC350TAT (4455C>T), CTA351TTT (4456C>T 4458A>T), CTT352ATT (4459C>A), CCT353CCA (4464T>A), AAG355AAA (4470G>A), GAG356AAT (4471G>A 4473G>T), CAA357AAT (4474C>A 4476A>T), CTG358CTT (4479G>T), CTT359ATA (4480C>A 4482T>A), CAA360AAT (4483C>A 4485A>T), AGA361CTA (4486A>C 4487G>T), ATC362GCC (4489A>G 4490T>C), GGA363AAA (4492G>A 4493G>A), GGT364GGA (4497T>A), AAG365AAA (4500G>A), TTT367TAC (4505T>A 4506T>C), TAC368TTT (4508A>T 4509C>T), TCT369TCA (4512T>A), TCC370AAA (4513T>A 4514C>A 4515C>A), GAC372..T (4521C>T), TGT373..C (4524T>C), TCT375AG. (4528T>A 4529C>G), TCA574.AA (5126C>A), GAG576GAA (5133G>A), AAG578AAC (5139G>C), CCT579CTA (5141C>T 5142T>A), ATC580GTA (5143A>G 5145C>A), ATC581TTA (5146A>T 5148C>A), GAA582CAA (5149G>C), TGC583ACT (5152T>A 5153G>C 5154C>T), GCA585GCT (5160A>T), TCA586TCT (5163A>T), GAC588TAT (5167G>T 5169C>T), ATC593TTA (5182A>T 5184C>A), CTC594TTA (5185C>T 5187C>A), AAA595CAA (5188A>C), GCA596ACA (5191G>A), AAA597GAT (5194A>G 5196A>T), CTT598CTA (5199T>A), CCA599AAT (5200C>A 5201C>A 5202A>T), GGT601_GTA604del (5206_5217delGGTAAAGAGGTA), ATA605ATT (5220A>T), TGC606TGT (5223C>T), AGG607AGA (5226G>A), GCT609ACT (5230G>A), TCA610AGT (5233T>A 5234C>G 5235A>T), AAA614AAT (5247A>T), CCA615GAA (5248C>G 5249C>A), GCT616ACA (5251G>A 5253T>A), GAG617GAA (5256G>A), AAG618ACT (5258A>C 5259G>T), AAT619AGG (5261A>G 5262T>G), CAT621TTA (5266C>T 5267A>T 5268T>A), AGT622ACA (5270G>C 5271T>A), AAG625AAA (5280G>A), ATC627TTA (5284A>T 5286C>A), CTT628CTA (5289T>A), TCC629GCT (5290T>G 5292C>T), ATT630ATA (5295T>A), ATT631GTA (5296A>G 5298T>A), AAG632AGA (5300A>G 5301G>A), GCA633GGC (5303C>G 5304A>C), ATC634ATA (5307C>A), AAA635CGA (5308A>C 5309A>G), GCT636AAG (5311G>A 5312C>A 5313T>G), AGA638TCT (5317A>T 5318G>C 5319A>T), GCT639GCA (5322T>A), TAT640TTC (5324A>T 5325T>C), ATC641TTA (5326A>T 5328C>A), CTT642CTA (5331T>A), CCT643CCA (5334T>A), TAT644AAA (5335T>A 5337T>A), AAA645CAA (5338A>C), TTT646TTC (5343T>C), CTT647ATC (5344C>A 5346T>C), GTA648ATC (5347G>A 5349A>C), AGG649AGA (5352G>A), ACT650TCG (5353A>T 5355T>G), GAT651GAC (5358T>C), ACT653ACG (5364T>G), AAT654CAG (5365A>C 5367T>G), GCT655GTA (5369C>T 5370T>A), GCT656TCA (5371G>T 5373T>A), TAT657GGT (5374T>G 5375A>G), TTT658CTA (5377T>C 5379T>A), GTG659ATA (5380G>A 5382G>A), AGA660TTC (5383A>T 5384G>T 5385A>C), ACA661AAT (5387C>A 5388A>T), AAT662AAA (5391T>A), ATT663CTT (5392A>C) | | | | | | | | |

*: Inserts / Deletes / Misaligned / Frameshifts

#### Analysis details

This analysis was performed with panviral2.64

### NGS Details (UN18_val): Badnavirus occultipomeae

#### Assembly

| | |
|---|---|
| **Coverage Length** | 253 (1 contig(s)) |
| **Depth Of Coverage** | 14.4 |
| **Number Of Reads** | 38 |
| **Reads Per Million** | 0.71 rpm (after QC) |
| **Ambiguities** | 0 |
| **Assembly Method** | de novo + reference guided assembly |
| **Consensus Caller** | Bcf Tools |

#### Coverage Map

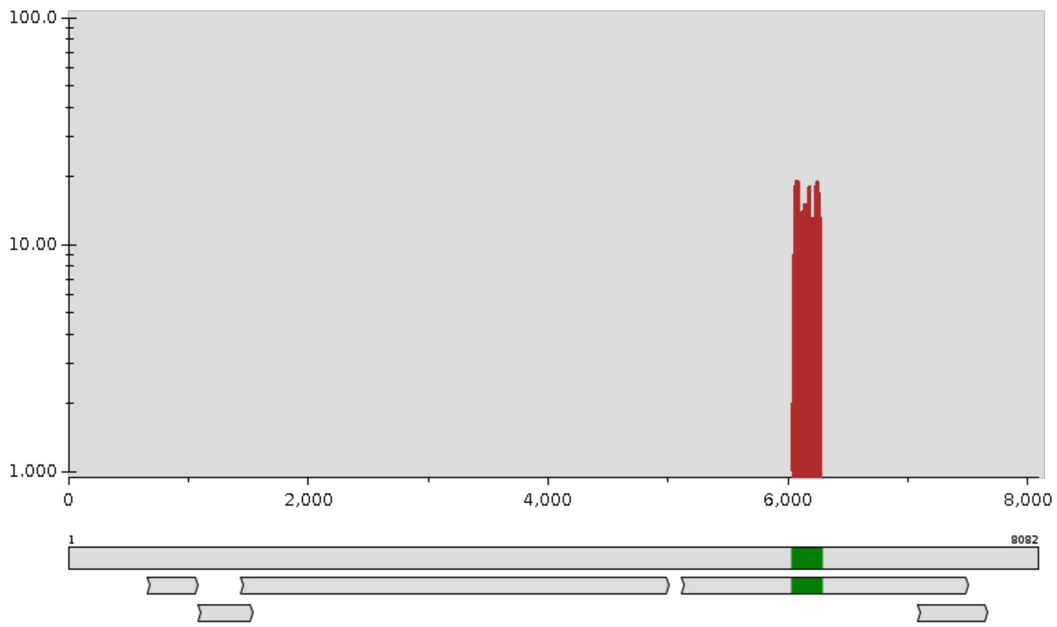

#### Assignment

| | |
|---|---|
| **Type** | Badnavirus occultipomeae (Taxonomy ID: 3048353) |
| **Reference Genome** | NC_015655.1 |
| **NT Identity (%)** | 60.5469 |
| **AA Identity (%)** | 51.7647 |
| **Number Of Stop Codons** | 0 |
| **Number Of CDS** | 5 |

#### Alignment

| | |
|---|---|
| **Alignment Score** | 102.0 (NT) + 312.0 (AA) = 414.0 |
| **Concordance (%)** | 36.7347 |

| Alignment Method | Global, seeded, nucleotide + amino acids (AGA) |
|---|---|

#### Genome Region

Sequence starts at position 6033 and ends at position 6285 relative to NC_015655.1 reference sequence.

#### Alignment Detailed Statistics

| | Begin | End | Coverage | Score | Concordance | Matches | Identities | I/D/M/F* | Stop Codons |
|---|---|---|---|---|---|---|---|---|---|
| **NT** | **6033** | **6285** | **3.1%** | **102** | **20.2%** | **253 (98.8%)** | **155 (60.5%)** | **3/0** | |
| Mutations: | 6033A>C, 6039A>T, 6042A>G, 6043T>A, 6045T>A, 6048T>C, 6053A>G, 6055G>T, 6062T>A, 6070A>T, 6072A>G, 6074G>A, 6076A>G, 6079G>A, 6080A>G, 6081G>A, 6085A>T, 6086A>G, 6087G>T, 6088T>G, 6089C>A, 6092A>T, 6093A>C, 6096A>G, 6097T>A, 6098G>A, 6102T>G, 6109T>C, 6114T>C, 6116C>G, 6117A>G, 6118G>T, 6120A>T, 6123G>T, 6125T>A, 6128A>T, 6134T>G, 6136G>T, 6137A>T, 6138A>G, 6141C>A, 6145C>T, 6150T>C, 6153A>T, 6154C>T, 6155T>C, 6158T>C, 6159G>A, 6168A>T, 6173A>C, 6174T>C, 6178C>A, 6179A>T, 6181A>G, 6182G>A, 6183A>T, 6184A>T, 6185A>T, 6190G>A, 6192T>C, 6194A>G, 6195T>A, 6197C>T, 6201T>C, 6204G>A, 6205G>C, 6206G>A, 6207A>G, 6207_6208insTAT, 6208A>C, 6209C>T, 6210A>T, 6213T>C, 6214G>A, 6215C>T, 6216A>G, 6219T>C, 6220A>G, 6222C>A, 6223G>A, 6224C>T, 6225A>C, 6228A>G, 6230A>T, 6234T>C, 6240C>T, 6243A>C, 6251T>A, 6252C>T, 6255T>C, 6256G>A, 6257A>G, 6259A>G, 6264A>G, 6267A>G, 6275A>T, 6276G>T, 6277G>A, 6282T>C | | | | | | | | |

**CDS**

| | Begin | End | Coverage | Score | Concordance | Matches | Identities | I/D/M/F* | Stop Codons |
|---|---|---|---|---|---|---|---|---|---|
| SPBVa_gp4 | 307 | 390 | 10.6% | 312 | 49.8% | 84 (98.8%) | 44 (51.8%) | 1/0/0/0 | 0 |
| Protein mutations: | F310I (6043T>A 6045T>A), K313R (6053A>G), A314S (6055G>T), F316Y (6062T>A), I319L (6070A>T 6072A>G), R320K (6074G>A), M321V (6076A>G), E322R (6079G>A 6080A>G 6081G>A), K324C (6085A>T 6086A>G 6087G>T), S325D (6088T>G 6089C>A), K326I (6092A>T 6093A>C), W328K (6097T>A 6098G>A), W332R (6109T>C), P334R (6116C>G 6117A>G), E335Y (6118G>T 6120A>T), L337H (6125T>A), Y338F (6128A>T), F340C (6134T>G), E341L (6136G>T 6137A>T 6138A>G), P344S (6145C>T), L347S (6154C>T 6155T>C), M348T (6158T>C 6159G>A), D353A (6173A>C 6174T>C), Q355M (6178C>A 6179A>T), R356D (6181A>G 6182G>A 6183A>T), K357L (6184A>T 6185A>T), D359N (6190G>A 6192T>C), N360R (6194A>G 6195T>A), A361V (6197C>T), G364Q (6205G>C 6206G>A 6207A>G), G364_T365insY (6207_6208insTAT), T365L (6208A>C 6209C>T 6210A>T), A367M (6214G>A 6215C>T 6216A>G), I369V (6220A>G 6222C>A), A370I (6223G>A 6224C>T 6225A>C), Y372F (6230A>T), F379Y (6251T>A 6252C>T), E381R (6256G>A 6257A>G), N382D (6259A>G), E387V (6275A>T 6276G>T), D388N (6277G>A) | | | | | | | | |
| Codon mutations: | ATA306..C (6033A>C), TCA308TCT (6039A>T), AAA309AAG (6042A>G), TTT310ATA (6043T>A 6045T>A), GAT311GAC (6048T>C), AAA313AGA (6053A>G), GCA314TCA (6055G>T), TTT316TAT (6062T>A), ATA319TTG (6070A>T 6072A>G), AGA320AAA (6074G>A), ATG321GTG (6076A>G), GAG322AGA (6079G>A 6080A>G 6081G>A), AAG324TGT (6085A>T 6086A>G 6087G>T), TCC325GAC (6088T>G 6089C>A), AAA326ATC (6092A>T 6093A>C), CCA327CCG (6096A>G), TGG328AAG (6097T>A 6098G>A), ACT329ACG (6102T>G), TGG332CGG (6109T>C), ACT333ACC (6114T>C), CCA334CGG (6116C>G 6117A>G), GAA335TAT (6118G>T 6120A>T), GGG336GGT (6123G>T), CTC337CAC (6125T>A), TAT338TTT (6128A>T), TTT340TGT (6134T>G), GAA341TTG (6136G>T 6137A>T 6138A>G), GTC342GTA (6141C>A), CCC344TCC (6145C>T), TTT345TTC (6150T>C), GGA346GGT (6153A>T), CTG347TCG (6154C>T 6155T>C), ATG348ACA (6158T>C 6159G>A), CCA351CCT (6168A>T), GAT353GCC (6173A>C 6174T>C), CAG355ATG (6178C>A 6179A>T), AGA356GAT (6181A>G 6182G>A 6183A>T), AAG357TTG (6184A>T 6185A>T), GAT359AAC (6190G>A 6192T>C), AAT360AGA (6194A>G 6195T>A), GCA361GTA (6197C>T), TTT362TTC (6201T>C), AGG363AGA (6204G>A), GGA364CAG (6205G>C 6206G>A 6207A>G), GGA364_ACA365insTAT (6207_6208insTAT), ACA365CTT (6208A>C 6209C>T 6210A>T), GAT366GAC (6213T>C), GCA367ATG (6214G>A 6215C>T 6216A>G), TTT368TTC (6219T>C), ATC369GTA (6220A>G 6222C>A), GCA370ATC (6223G>A 6224C>T 6225A>C), GTA371GTG (6228A>G), TAT372TTT (6230A>T), ATT373ATC (6234T>C), GAC375GAT (6240C>T), ATA376ATC (6243A>C), TTC379TAT (6251T>A 6252C>T), TCT380TCC (6255T>C), GAA381AGA (6256G>A 6257A>G), AAT382GAT (6259A>G), GAA383GAG (6264A>G), GAA384GAG (6267A>G), GAG387GTT (6275A>T 6276G>T), GAT388AAT (6277G>A), CAT389CAC (6282T>C) | | | | | | | | |

**Proteins**

| | Begin | End | Coverage | Score | Concordance | Matches | Identities | I/D/M/F* | Stop Codons |
|---|---|---|---|---|---|---|---|---|---|
| RNaseH/reverse transcriptase (YP_004581513.1) | 307 | 390 | 10.6% | 312 | 49.8% | 84 (98.8%) | 44 (51.8%) | 1/0/0/0 | 0 |
| Protein mutations: | F310I (6043T>A 6045T>A), K313R (6053A>G), A314S (6055G>T), F316Y (6062T>A), I319L (6070A>T 6072A>G), R320K (6074G>A), M321V (6076A>G), E322R (6079G>A 6080A>G 6081G>A), K324C (6085A>T 6086A>G 6087G>T), S325D (6088T>G 6089C>A), K326I (6092A>T 6093A>C), W328K (6097T>A 6098G>A), W332R (6109T>C), P334R (6116C>G 6117A>G), E335Y (6118G>T 6120A>T), L337H (6125T>A), Y338F (6128A>T), F340C (6134T>G), E341L (6136G>T 6137A>T 6138A>G), P344S (6145C>T), L347S (6154C>T 6155T>C), M348T (6158T>C 6159G>A), D353A (6173A>C 6174T>C), Q355M (6178C>A 6179A>T), R356D (6181A>G 6182G>A 6183A>T), K357L (6184A>T 6185A>T), D359N (6190G>A 6192T>C), N360R (6194A>G 6195T>A), A361V (6197C>T), G364Q (6205G>C 6206G>A 6207A>G), G364_T365insY (6207_6208insTAT), T365L (6208A>C 6209C>T 6210A>T), A367M (6214G>A 6215C>T 6216A>G), I369V (6220A>G 6222C>A), A370I (6223G>A 6224C>T 6225A>C), Y372F (6230A>T), F379Y (6251T>A 6252C>T), E381R (6256G>A 6257A>G), N382D (6259A>G), E387V (6275A>T 6276G>T), D388N (6277G>A) | | | | | | | | |
| Codon mutations: | ATA306..C (6033A>C), TCA308TCT (6039A>T), AAA309AAG (6042A>G), TTT310ATA (6043T>A 6045T>A), GAT311GAC (6048T>C), AAA313AGA (6053A>G), GCA314TCA (6055G>T), TTT316TAT (6062T>A), ATA319TTG (6070A>T 6072A>G), AGA320AAA (6074G>A), ATG321GTG (6076A>G), GAG322AGA (6079G>A 6080A>G 6081G>A), AAG324TGT (6085A>T 6086A>G 6087G>T), TCC325GAC (6088T>G 6089C>A), AAA326ATC (6092A>T 6093A>C), CCA327CCG (6096A>G), TGG328AAG (6097T>A 6098G>A), ACT329ACG (6102T>G), TGG332CGG (6109T>C), ACT333ACC (6114T>C), CCA334CGG (6116C>G 6117A>G), GAA335TAT (6118G>T 6120A>T), GGG336GGT (6123G>T), CTC337CAC (6125T>A), TAT338TTT (6128A>T), TTT340TGT (6134T>G), GAA341TTG (6136G>T 6137A>T 6138A>G), GTC342GTA (6141C>A), CCC344TCC (6145C>T), TTT345TTC (6150T>C), GGA346GGT (6153A>T), CTG347TCG (6154C>T 6155T>C), ATG348ACA (6158T>C 6159G>A), CCA351CCT (6168A>T), GAT353GCC (6173A>C 6174T>C), CAG355ATG (6178C>A 6179A>T), AGA356GAT (6181A>G 6182G>A 6183A>T), AAG357TTG (6184A>T 6185A>T), GAT359AAC (6190G>A 6192T>C), AAT360AGA (6194A>G 6195T>A), GCA361GTA (6197C>T), TTT362TTC (6201T>C), AGG363AGA (6204G>A), GGA364CAG (6205G>C 6206G>A 6207A>G), GGA364_ACA365insTAT (6207_6208insTAT), ACA365CTT (6208A>C 6209C>T 6210A>T), GAT366GAC (6213T>C), GCA367ATG (6214G>A 6215C>T 6216A>G), TTT368TTC (6219T>C), ATC369GTA (6220A>G 6222C>A), GCA370ATC (6223G>A 6224C>T 6225A>C), GTA371GTG (6228A>G), TAT372TTT (6230A>T), ATT373ATC (6234T>C), GAC375GAT (6240C>T), ATA376ATC (6243A>C), TTC379TAT (6251T>A 6252C>T), TCT380TCC (6255T>C), GAA381AGA (6256G>A 6257A>G), AAT382GAT (6259A>G), GAA383GAG (6264A>G), GAA384GAG (6267A>G), GAG387GTT (6275A>T 6276G>T), GAT388AAT (6277G>A), CAT389CAC (6282T>C) | | | | | | | | |

*: Inserts / Deletes / Misaligned / Frameshifts

#### Analysis details

This analysis was performed with panviral2.64

### NGS Details (UN18_val): Brazilian marseillevirus

#### Assembly

| | |
|---|---|
| **Coverage Length** | 172 (1 contig(s)) |
| **Depth Of Coverage** | 15.2 |
| **Number Of Reads** | 33 |
| **Reads Per Million** | 0.62 rpm (after QC) |
| **Ambiguities** | 0 |
| **Assembly Method** | de novo + reference guided assembly |
| **Consensus Caller** | Bcf Tools |

#### Coverage Map

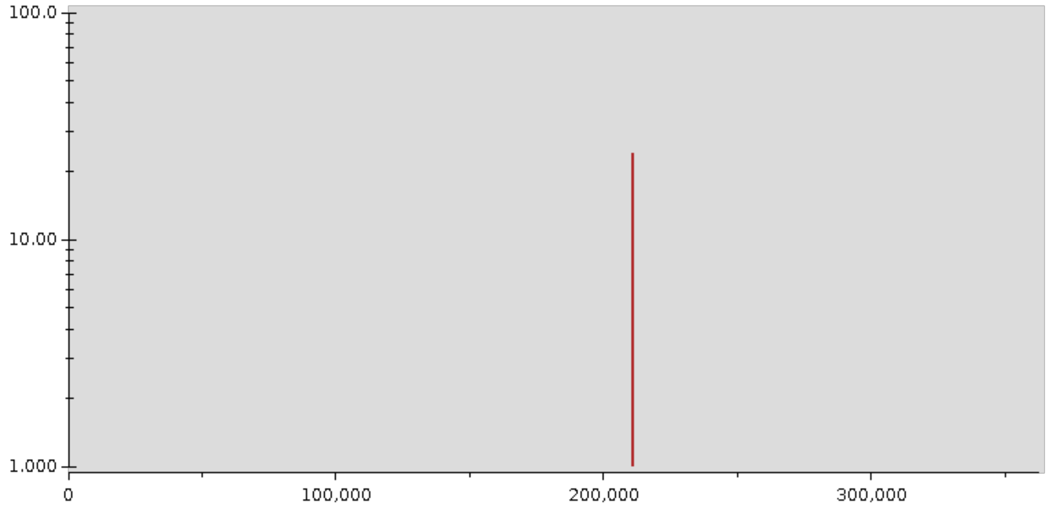

#### Assignment

| | |
|---|---|
| **Type** | Brazilian marseillevirus (Taxonomy ID: 1813599) |
| **Reference Genome** | NC_029692.1 |
| **NT Identity (%)** | 78.4884 |
| **AA Identity (%)** | 91.2281 |
| **Number Of Stop Codons** | 0 |
| **Number Of CDS** | 491 |

#### Alignment

| | |
|---|---|
| **Alignment Score** | 196.0 (NT) + 345.0 (AA) = 541.0 |
| **Concordance (%)** | 76.5205 |
| **Alignment Method** | Local, heuristic, nucleotide (BLASTN) |

#### Genome Region

Sequence starts at position 210894 and ends at position 211065 relative to NC_029692.1 reference sequence.

#### Alignment Detailed Statistics

| | Begin | End | Coverage | Score | Concordance | Matches | Identities | I/D/M/F* | Stop Codons |
|---|---|---|---|---|---|---|---|---|---|
| NT | 210894 | 211065 | 0.1% | 196 | 57.0% | 172 (100%) | 135 (78.5%) | 0/0 | |

Mutations: 210896A>G, 210901T>G, 210902T>C, 210904G>A, 210908A>G, 210914A>C, 210923C>T, 210926T>C, 210929T>C, 210932G>A, 210938A>G, 210941A>G, 210946C>A, 210953C>T, 210956T>A, 210959A>G, 210962T>C, 210965C>G, 210968C>T, 210971C>T, 210974G>A, 210977T>G, 210986A>G, 210989G>T, 210992T>C, 210995T>C, 211004A>T, 211007A>G, 211009C>T, 211013T>C, 211031T>G, 211032T>C, 211033G>T, 211034T>C, 211040G>A, 211043T>G, 211055A>C

*: Inserts / Deletes / Misaligned / Frameshifts

#### Analysis details

This analysis was performed with panviral2.64

### NGS Details (UN18_val): Rahariannevirus raharianne

#### Assembly

| | |
|---|---|
| **Coverage Length** | 1643 (6 contig(s)) |
| **Depth Of Coverage** | 2.4 |
| **Number Of Reads** | 31 |
| **Reads Per Million** | 0.58 rpm (after QC) |
| **Ambiguities** | 0 |
| **Assembly Method** | de novo + reference guided assembly |
| **Consensus Caller** | Bcf Tools |

#### Coverage Map

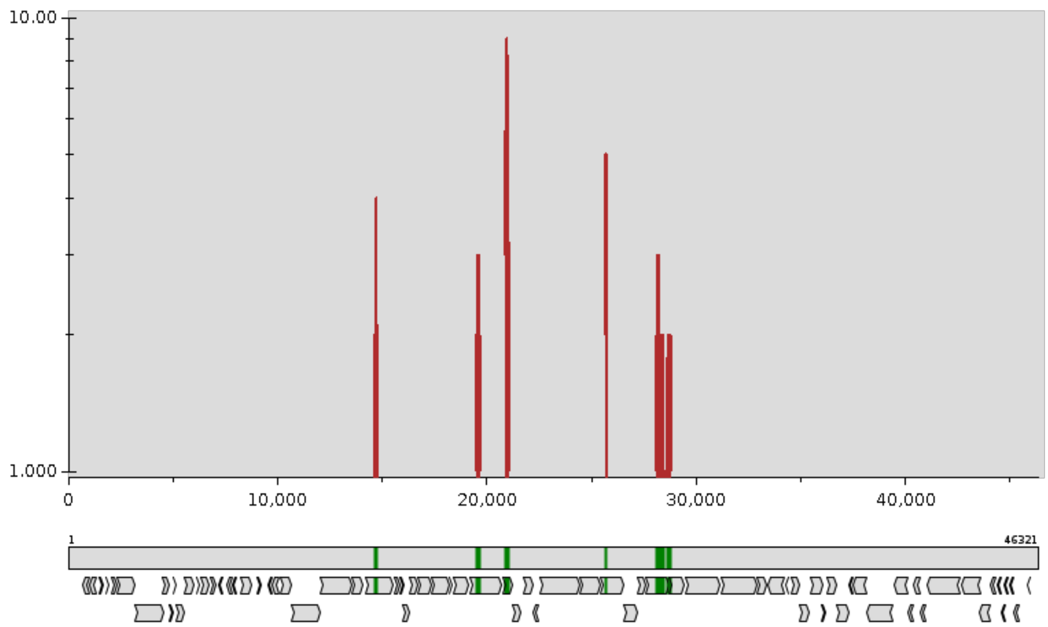

#### Assignment

| | |
|---|---|
| **Type** | Rahariannevirus raharianne (Taxonomy ID: 2846050) |
| **Reference Genome** | NC_054955.1 |
| **NT Identity (%)** | 69.7974 |
| **AA Identity (%)** | 65.3061 |
| **Number Of Stop Codons** | 1 |
| **Number Of CDS** | 75 |

#### Alignment

| | |
|---|---|
| **Alignment Score** | 1249.0 (NT) + 2380.0 (AA) = 3629.0 |
| **Concordance (%)** | 55.1939 |

| Alignment Method | Local, heuristic, nucleotide (BLASTN) |
|---|---|

#### Genome Region

Sequence starts at position 14591 and ends at position 28832 relative to NC_054955.1 reference sequence.

#### Alignment Detailed Statistics

| | Begin | End | Coverage | Score | Concordance | Matches | Identities | I/D/M/F* | Stop Codons |
|---|---|---|---|---|---|---|---|---|---|
| **NT** | **14591** | **28832** | **3.5%** | **1249** | **38.8%** | **1626 (98.8%)** | **1137 (69.1%)** | **3/17** | |
| Mutations: | 14595A>G, 14596A>C, 14597T>C, 14600C>G, 14603G>C, 14609T>C, 14640A>G, 14641C>T, 14642G>C, 14648G>C, 14652T>G, 14661G>C, 14662C>T, 14663C>G, 14666C>G, 14670T>A, 14671C>G, 14672G>C, 14678C>T, 14682A>G, 14684C>G, 14697A>C, 14699C>G, 14715C>G, 14716A>T, 14718T>A, 14723C>T, 14729G>C, 14734G>C, 14753G>C, 14759T>G, 14769A>G, 14780G>C, 14783T>C, 14788C>T, 14789A>G, 14794C>T, 14798C>T, 14802T>G, 14803C>G, 19482C>A, 19483G>A, 19484C>A, 19485T>G, 19486G>C, 19493C>G, 19496C>G, 19512A>G, 19513C>T, 19520G>C, 19521T>A, 19522C>G, 19523G>C, 19525C>T, 19526G>C, 19527A>G, 19528T>C, 19529C>G, 19534A>T, 19542G>T, 19545G>A, 19547G>C, 19550G>C, 19551C>T, 19553G>C, 19557G>T, 19559G>C, 19574G>C, 19575C>T, 19577G>C, 19584T>G, 19585C>G, 19586G>C, 19587G>A, 19589G>C, 19595C>G, 19596G>A, 19601C>G, 19602G>T, 19604C>G, 19607G>C, 19611A>T, 19612G>C, 19613C>G, 19614G>C, 19615C>A, 19616C>G, 19620A>T, 19621A>C, 19622T>G, 19626T>C, 19627G>A, 19633T>C, 19634A>C, 19635A>G, 19636C>T, 19640C>T, 19641G>T, 19650A>G, 19651C>G, 19652G>C, 19653C>T, 19655A>G, 19656A>G, 19657A>T, 19658C>G, 19666T>C, 19667T>C, 19683G>T, 19685T>G, 20839G>C, 20840G>A, 20841T>C, 20846C>G, 20847A>T, 20852T>G, 20854A>C, 20856C>G, 20857T>C, 20861A>C, 20866G>C, 20867C>A, 20869G>C, 20871C>T, 20874A>G, 20876A>C, 20877A>T, 20884C>G, 20885T>C, 20886C>A, 20887T>G, 20891A>C, 20894A>T, 20895G>C, 20896C>G, 20897A>G, 20898A>C, 20901A>C, 20902G>C, 20905T>C, 20906G>A, 20909A>G, 20914T>G, 20915G>A, 20917G>C, 20918C>A, 20919G>A, 20920C>G, 20921G>A, 20927C>G, 20929C>T, 20945G>C, 20946A>G, 20950C>T, 20952T>C, 20959T>C, 20972T>C, 20973C>G, 20974T>C, 20978T>A, 20979C>G, 20980G>C, 20984G>A, 20985C>A, 20986G>C, 20987A>G, 20989T>G, 20992A>G, 20998A>G, 21002G>A, 21003C>G, 21004G>C, 21005G>A, 21007G>A, 21014A>G, 21028A>G, 21034A>G, 21035T>G, 21037G>C, 21062T>C, 21063C>A, 21067C>G, 21071C>A, 21072A>C, 21076T>C, 21077T>A, 21078A>C, 21079T>C, 21085G>C, 25634T>C, 25635T>C, 25637C>G, 25650T>A, 25651C>G, 25653G>A, 25659A>G, 25662A>G, 25665T>C, 25666A>C, 25667C>G, 25672A>C, 25673G>C, 25675C>A, 25676G>C, 25679C>A, 25682C>G, 25683A>T, 25685C>G, 25691G>C, 25697T>C, 25700G>C, 25706G>C, 25709T>C, 25720A>T, 25724C>T, 25727C>G, 25733G>C, 25737C>A, 25744C>G, 25771C>T, 25772G>C, 28051T>G, 28053G>C, 28054C>G, 28062T>C, 28064G>A, 28065C>T, 28066T>C, 28068G>C, 28072C>G, 28073G>A, 28075A>C, 28076T>A, 28077C>A, 28078G>C, 28081C>G, 28082A>C, 28086T>G, 28088G>C, 28089G>A, 28090C>G, 28093A>C, 28096T>C, 28097T>G, 28099C>G, 28108T>C, 28114C>G, 28117G>C, 28123C>G, 28126A>G, 28137A>C, 28138C>G, 28141C>T, 28147T>C, 28154G>A, 28156A>G, 28159C>T, 28162T>C, 28165G>C, 28166C>T, 28168G>C, 28171T>C, 28172A>T, 28177C>G, 28181A>G, 28185C>G, 28190G>A, 28192A>C, 28193G>A, 28194C>A, 28195G>C, 28207G>A, 28213G>C, 28215A>G, 28216A>G, 28217T>G, 28219C>T, 28222T>C, 28223G>A, 28229T>A, 28238A>C, 28239C>T, 28240C>G, 28241T>G, 28243G>C, 28244C>A, 28246G>C, 28254G>A, 28255A>C, 28256C>A, 28258G>C, 28260G>A, 28264T>C, 28270G>C, 28271C>A, 28272T>C, 28273T>C, 28277A>G, 28278A>G, 28280C>T, 28282G>T, 28283A>G, 28285G>C, 28290C>G, 28296T>C, 28298A>G, 28304A>C, 28309T>C, 28310A>T, 28316G>T, 28317T>A, 28318G>C, 28321C>G, 28322A>G, 28324C>G, 28336A>C, 28339G>C, 28342C>T, 28343T>G, 28348A>G, 28351G>C, 28357G>C, 28358C>T, 28360T>C, 28366G>C, 28367T>A, 28369A>C, 28372T>C, 28373T>A, 28374C>G, 28375G>C, 28381T>C, 28390T>C, 28399A>G, 28400C>G, 28402A>C, 28403T>G, 28405C>T, 28406A>T, 28407G>C, 28408C>G, 28410G>C, 28426T>G, 28429G>T, 28433G>C, 28435G>T, 28438T>G, 28440A>C, 28441T>G, 28445T>A, 28446C>A, 28447T>C, 28457A>G, 28459G>C, 28460G>A, 28462G>C, 28465C>T, 28467T>C, 28468C>G, 28472G>A, 28473C>G, 28474G>C, 28480T>C, 28481G>A, 28484A>C, 28485C>A, 28486G>C, 28488T>C, 28489G>C, 28492G>C, 28494C>T, 28495G>C, 28496A>G, 28497A>T, 28498G>C, 28501G>T, 28502T>C, 28505A>G, 28507C>G, 28508G>C, 28509T>A, 28510G>A, 28511A>G, 28516A>C, 28517C>G, 28520A>C, 28523A>T, 28525G>C, 28528T>C, 28553G>A, 28555A>G, 28561G>C, 28564T>C, 28565T>A, 28567G>C, 28568C>A, 28569G>A, 28574G>T, 28576C>G, 28579A>G, 28580G>C, 28582G>C, 28585T>C, 28588T>C, 28590A>G, 28594T>G, 28600T>G, 28602G>C, 28603C>A, 28609G>C, 28613A>G, 28618G>C, 28622G>A, 28624G>C, 28634A>G, 28645G>T, 28648T>C, 28652G>A, 28654T>G, 28657G>C, 28660G>C, 28662G>A, 28665T>A, 28666T>G, 28669G>A, 28675A>C, 28678A>G, 28679T>G, 28680G>C, 28681G>C, 28682T>G, 28683C>G, 28684G>C, 28685G>A, 28687A>G, 28688G>A, 28691A>T, 28692G>C, 28693C>G, 28697G>A, 28699A>C, 28703T>G, 28706_28719delAAACAACTCTGGTG, 28724A>T, 28725A>C, 28729A>G, 28730G>C, 28731C>A, 28732G>A, 28734T>C, 28737C>T, 28740T>C, 28743A>G, 28744C>A, 28745A>G, 28746A>C, 28749T>C, 28750G>A, 28753T>A, 28754A>G, 28756C>A, 28757A>C, 28767C>G, 28768G>C, 28769G>A, 28770C>G, 28771T>C, 28772A>T, 28774C>A, 28776G>C, 28779T>G, 28780C>G, 28782_28783insGGG, 28789_28791delGGC, 28792G>C, 28794T>G, 28795G>C, 28796C>A, 28797G>A, 28799C>G, 28800G>C, 28802A>T, 28803G>C, 28804T>G, 28805C>T, 28809A>G, 28810G>A, 28814T>G, 28815G>C, 28816A>G, 28817C>A, 28830G>A | | | | | | | | |

##### CDS

| | Begin | End | Coverage | Score | Concordance | Matches | Identities | I/D/M/F* | Stop Codons |
|---|---|---|---|---|---|---|---|---|---|
| KMC43_gp27 | 133 | 207 | 16.9% | 411 | 85.8% | 75 (100%) | 62 (82.7%) | 0/0/0/0 | 0 |
| Protein mutations: | N134A (14595A>G 14596A>C 14597T>C), T149V (14640A>G 14641C>T 14642G>C), S153A (14652T>G), A156L (14661G>C 14662C>T 14663C>G), I163V (14682A>G 14684C>G), N168Q (14697A>C 14699C>G), Q174V (14715C>G 14716A>T), L175M (14718T>A), G180A (14734G>C), I192V (14769A>G), A198V (14788C>T 14789A>G), A200V (14794C>T), S203G (14802T>G 14803C>G) | | | | | | | | |
| Codon mutations: | AAT134GCC (14595A>G 14596A>C 14597T>C), GCC135GCG (14600C>G), CTG136CTC (14603G>C), AAT138AAC (14609T>C), ACG149GTC (14640A>G 14641C>T 14642G>C), CTG151CTC (14648G>C), TCG153GCG (14652T>G), GCC156CTG (14661G>C 14662C>T 14663C>G), ACC157ACG (14666C>G), TCG159AGC (14670T>A 14671C>G 14672G>C), GAC161GAT (14678C>T), ATC163GTG (14682A>G 14684C>G), AAC168CAG (14697A>C 14699C>G), CAG174GTG (14715C>G 14716A>T), TTG175ATG (14718T>A), GTC176GTT (14723C>T), GTG178GTC (14729G>C), GGC180GCC (14734G>C), GTG186GTC (14753G>C), GTT188GTG (14759T>G), ATC192GTC (14769A>G), CTG195CTC (14780G>C), ACT196ACC (14783T>C), GCA198GTG (14788C>T 14789A>G), GCC200GTC (14794C>T), GAC201GAT (14798C>T), TCC203GGC (14802T>G 14803C>G) | | | | | | | | |
| KMC43_gp36 | 89 | 162 | 14.5% | 317 | 67.3% | 74 (100%) | 49 (66.2%) | 0/0/0/0 | 0 |
| Protein mutations: | R92K (19482C>A 19483G>A 19484C>A), C93A (19485T>G 19486G>C), T102V (19512A>G 19513C>T), T106I (19525C>T 19526G>C), I107A (19527A>G 19528T>C 19529C>G), Q109L (19534A>T), G112C (19542G>T), V113I (19545G>A 19547G>C), L115F (19551C>T 19553G>C), A117S (19557G>T 19559G>C), L123F (19575C>T 19577G>C), S126G (19584T>G 19585C>G 19586G>C), V127I (19587G>A 19589G>C), A130T (19596G>A), A132S (19602G>T 19604C>G), A136Q (19614G>C 19615C>A 19616C>G), N138S (19620A>T 19621A>C 19622T>G), W140Q (19626T>C 19627G>A), V142A (19633T>C 19634A>C), T143V (19635A>G 19636C>T), A145S (19641G>T), T148G (19650A>G 19651C>G 19652G>C), N150V (19656A>G 19657A>T 19658C>G), V153A (19666T>C 19667T>C), A159S (19683G>T 19685T>G) | | | | | | | | |
| Codon mutations: | CGC92AAA (19482C>A 19483G>A 19484C>A), TGC93GCC (19485T>G 19486G>C), CGC95CGG (19493C>G), GTC96GTG (19496C>G), ACG102GTG (19512A>G 19513C>T), GCG104GCC (19520G>C), TCG105AGC (19521T>A 19522C>G 19523G>C), ACG106ATC (19525C>T 19526G>C), ATC107GCG (19527A>G 19528T>C 19529C>G), CAG109CTG (19534A>T), GGC112TGC (19542G>T), GTG113ATC (19545G>A 19547G>C), ACG114ACC (19550G>C), CTG115TTC (19551C>T 19553G>C), GCG117TCC (19557G>T 19559G>C), ACG122ACC (19574G>C), CTG123TTC (19575C>T 19577G>C), TCG126GGC (19584T>G 19585C>G 19586G>C), GTG127ATC (19587G>A 19589G>C), GTC129GTG (19595C>G), GCG130ACG (19596G>A), CTC131CTG (19601C>G), GCC132TCG (19602G>T 19604C>G), GCG133GCC (19607G>C), AGC135TCG (19611A>T 19612G>C 19613C>G), GCC136CAG (19614G>C 19615C>A 19616C>G), AAT138TCG (19620A>T 19621A>C 19622T>G), TGG140CAG (19626T>C 19627G>A), GTA142GCC (19633T>C 19634A>C), ACG143GTG (19635A>G 19636C>T), GTC144GTT (19640C>T), GCG145TCG (19641G>T), ACG148GGC (19650A>G 19651C>G 19652G>C), CTA149TTG (19653C>T 19655A>G), AAC150GTG (19656A>G 19657A>T 19658C>G), GTT153GCC (19666T>C 19667T>C), GCT159TCG (19683G>T 19685T>G) | | | | | | | | |
| KMC43_gp37 | 13 | 99 | 60.4% | 399 | 66.6% | 87 (100%) | 54 (62.1%) | 0/0/0/0 | 0 |
| Protein mutations: | V15T (20840G>A 20841T>C), Q17V (20846C>G 20847A>T), S19A (20852T>G 20854A>C), T20S (20856C>G 20857T>C), T22P (20861A>C), Q24N (20867C>A 20869G>C), P25L (20871C>T), N26S (20874A>G), K27L (20876A>C 20877A>T), S30Q (20885T>C 20886C>A 20887T>G), K32Q (20891A>C), K34A (20897A>G 20898A>C), Q35P (20901A>C 20902G>C), V37M (20906G>A), T38A (20909A>G), D39E (20914T>G), V40I (20915G>A 20917G>C), R41K (20918C>A 20919G>A 20920C>G), V42I (20921G>A), R44G (20927C>G 20929C>T), D50R (20945G>C 20946A>G), V52A (20952T>C), S59R (20972T>C 20973C>G 20974T>C), A63N (20984G>A 20985C>A 20986G>C), I64V (20987A>G 20989T>G), A69S (21002G>A 21003C>G 21004G>C), A70T (21005G>A 21007G>A), N73D (21014A>G), S80A (21035T>G 21037G>C), S89Q (21062T>C 21063C>A), Q92T (21071C>A 21072A>C), Y94T (21077T>A 21078A>C 21079T>C) | | | | | | | | |
| Codon mutations: | CTG14CTC (20839G>C), GTC15ACC (20840G>A 20841T>C), CAG17GTG (20846C>G 20847A>T), TCA19GCC (20852T>G 20854A>C), ACT20AGC (20856C>G 20857T>C), ACG22CCG (20861A>C), CTG23CTC (20866G>C), CAG24AAC (20867C>A 20869G>C), CCC25CTC (20871C>T), AAC26AGC (20874A>G), AAG27CTG (20876A>C 20877A>T), ACC29ACG (20884C>G), TCT30CAG (20885T>C 20886C>A 20887T>G), AAG32CAG (20891A>C), AGC33TCG (20894A>T 20895G>C 20896C>G), AAG34GCG (20897A>G 20898A>C), CAG35CCC (20901A>C 20902G>C), GAT36GAC (20905T>C), GTG37ATG (20906G>A), ACC38GCC (20909A>G), GAT39GAG (20914T>G), GTG40ATC (20915G>A 20917G>C), CGC41AAG (20918C>A 20919G>A 20920C>G), GTC42ATC (20921G>A), CGC44GGT (20927C>G 20929C>T), GAC50CGC (20945G>C 20946A>G), CAC51CAT (20950C>T), GTT52GCT (20952T>C), TTT54TTC (20959T>C), TCT59CGC (20972T>C 20973C>G 20974T>C), TCG61AGC (20978T>A 20979C>G 20980G>C), GCG63AAC (20984G>A 20985C>A 20986G>C), ATT64GTG (20987A>G 20989T>G), GAA65GAG (20992A>G), CAA67CAG (20998A>G), GCG69AGC (21002G>A 21003C>G 21004G>C), GCG70ACA (21005G>A 21007G>A), AAC73GAC (21014A>G), CAA77CAG (21028A>G), GAA79GAG (21034A>G), TCG80GCC (21035T>G 21037G>C), TCG89CAG (21062T>C 21063C>A), CCC90CCG (21067C>G), CAG92ACG (21071C>A 21072A>C), ATT93ATC (21076T>C), TAT94ACC (21077T>A 21078A>C 21079T>C), ACG96ACC (21085G>C) | | | | | | | | |
| KMC43_gp43 | 71 | 122 | 13.9% | 262 | 73.8% | 52 (100%) | 38 (73.1%) | 0/0/0/0 | 0 |
| Protein mutations: | F74L (25635T>C 25637C>G), D80N (25653G>A), T82A (25659A>G), N83D (25662A>G), Y84P (25665T>C 25666A>C 25667C>G), E86A (25672A>C 25673G>C), T87N (25675C>A 25676G>C), D88E (25679C>A), T90S (25683A>T 25685C>G), Y102F (25720A>T), L108M (25737C>A), T110S (25744C>G), A119V (25771C>T 25772G>C) | | | | | | | | |
| Codon mutations: | ATT73ATC (25634T>C), TTC74CTG (25635T>C 25637C>G), TCC79AGC (25650T>A 25651C>G), GAT80AAT (25653G>A), ACG82GCG (25659A>G), AAC83GAC (25662A>G), TAC84CCG (25665T>C 25666A>C 25667C>G), GAG86GCC (25672A>C 25673G>C), ACG87AAC (25675C>A 25676G>C), GAC88GAA (25679C>A), CTC89CTG (25682C>G), ACC90TCG (25683A>T 25685C>G), CTG92CTC (25691G>C), TAT94TAC (25697T>C), GGG95GGC (25700G>C), GTG97GTC (25706G>C), GAT98GAC (25709T>C), TAC102TTC (25720A>T), GAC103GAT (25724C>T), CCC104CCG (25727C>G), TCG106TCC (25733G>C), CTG108ATG (25737C>A), ACC110AGC (25744C>G), GCG119GTC (25771C>T 25772G>C) | | | | | | | | |
| KMC43_gp46 | 157 | 375 | 57.3% | 867 | 68.1% | 215 (100%) | 133 (61.9%) | 0/0/0/0 | 1 |

| | Begin | End | Coverage | Score | Concordance | Matches | Identities | I/D/M/F* | Stop Codons |
|---|---|---|---|---|---|---|---|---|---|
| **NT** | **14591** | **28832** | **3.5%** | **1249** | **38.8%** | **1626 (98.8%)** | **1137 (69.1%)** | **3/17** | |
| Protein mutations: | G159A (28053G>C 28054C>G), V162A (28062T>C), A163I (28064G>A 28065C>T 28066T>C), G164A (28068G>C), V166I (28073G>A 28075A>C), S167N (28076T>A 28077C>A 28078G>C), I169L (28082A>C), V170G (28086T>G), G171Q (28088G>C 28089G>A 28090C>G), S174A (28097T>G 28099C>G), N187T (28137A>C 28138C>G), A193T (28154G>A 28156A>G), L197F (28166C>T 28168G>C), T199S (28172A>T), I202V (28181A>G), A203G (28185C>G), V205I (28190G>A 28192A>C), A206N (28193G>A 28194C>A 28195G>C), K213R (28215A>G 28216A>G), S214A (28217T>G 28219C>T), V216I (28223G>A), Y218N (28229T>A), T221L (28238A>C 28239C>T 28240C>G), S222A (28241T>G 28243G>C), L223I (28244C>A 28246G>C), G226D (28254G>A 28255A>C), L227I (28256C>A 28258G>C), S228N (28260G>A), L232T (28271C>A 28272T>C 28273T>C), N234G (28277A>G 28278A>G), Q235Y (28280C>T 28282G>T), T236A (28283A>G 28285G>C), A238G (28290C>G), V240A (28296T>C), T241A (28298A>G), M243L (28304A>C), N245Y (28310A>T), V247Y (28316G>T 28317T>A 28318G>C), I249V (28322A>G 28324C>G), S256A (28343T>G), L261F (28358C>T 28360T>C), S264T (28367T>A 28369A>C), L275V (28400C>G 28402A>C), C276G (28403T>G 28405C>T), S278T (28410G>C), V286L (28433G>C 28435G>T), N288T (28440A>C 28441T>G), S290N (28445T>A 28446C>A 28447T>C), T294A (28457A>G 28459G>C), A295T (28460G>A 28462G>C), I297T (28467T>C 28468C>G), A299S (28472G>A 28473C>G 28474G>C), D302N (28481G>A), T303H (28484A>C 28485C>A 28486G>C), V304A (28488T>C 28489G>C), T306I (28494C>T 28495G>C), K307V (28496A>G 28497A>T 28498G>C), I310V (28505A>G 28507C>G), V311Q (28508G>C 28509T>A 28510G>A), T312A (28511A>G), L314V (28517C>G), K315Q (28520A>C), T316S (28523A>T 28525G>C), A326T (28553G>A 28555A>G), S330T (28565T>A 28567G>C), R331K (28568C>A 28569G>A), A333S (28574G>T 28576C>G), V335L (28580G>C 28582G>C), D338G (28590A>G), G342A (28602G>C 28603C>A), I346V (28613A>G), V349I (28622G>A 28624G>C), S353G (28634A>G), A359T (28652G>A 28654T>G), S362N (28662G>A), L363Q (28665T>A 28666T>G), W368A (28679T>G 28680G>C 28681G>C), S369G (28682T>G 28683C>G 28684G>C), A370T (28685G>A 28687A>G), V371I (28688G>A), A374T (28697G>A 28699A>C) | | | | | | | | |
| Codon mutations: | GCT158GCG (28051T>G), GGC159GCG (28053G>C 28054C>G), GTG162GCG (28062T>C), GCT163ATC (28064G>A 28065C>T 28066T>C), GGC164GCC (28068G>C), GCC165GCG (28072C>G), GTA166ATC (28073G>A 28075A>C), TCG167AAC (28076T>A 28077C>A 28078G>C), ACC168ACG (28081C>G), ATC169CTC (28082A>C), GTC170GGC (28086T>G), GGC171CAG (28088G>C 28089G>A 28090C>G), GCA172GCC (28093A>C), ATT173ATC (28096T>C), TCC174GCG (28097T>G 28099C>G), GAT177GAC (28108T>C), GTC179GTG (28114C>G), ACG180ACC (28117G>C), GCC182GCG (28123C>G), GCA183GCG (28126A>G), AAC187ACG (28137A>C 28138C>G), GGC188GGT (28141C>T), GAT190GAC (28147T>C), GCA193ACG (28154G>A 28156A>G), GAC194GAT (28159C>T), GCT195GCC (28162T>C), GCG196GCC (28165G>C), CTG197TTC (28166C>T 28168G>C), CGT198CGC (28171T>C), ACG199TCG (28172A>T), CGC200CGG (28177C>G), ATC202GTC (28181A>G), GCC203GGC (28185C>G), GTA205ATC (28190G>A 28192A>C), GCG206AAC (28193G>A 28194C>A 28195G>C), AAG210AAA (28207G>A), ACG212ACC (28213G>C), AAA213AGG (28215A>G 28216A>G), TCC214GCT (28217T>G 28219C>T), GCT215GCC (28222T>C), GTC216ATC (28223G>A), TAT218AAT (28229T>A), ACC221CTG (28238A>C 28239C>T 28240C>G), TCG222GCC (28241T>G 28243G>C), CTG223ATC (28244C>A 28246G>C), GGA226GAC (28254G>A 28255A>C), CTG227ATC (28256C>A 28258G>C), AGC228AAC (28260G>A), TAT229TAC (28264T>C), CTG231CTC (28270G>C), CTT232ACC (28271C>A 28272T>C 28273T>C), AAC234GGC (28277A>G 28278A>G), CAG235TAT (28280C>T 28282G>T), ACG236GCC (28283A>G 28285G>C), GCC238GGC (28290C>G), GTG240GCG (28296T>C), ACG241GCG (28298A>G), ATG243CTG (28304A>C), GGT244GGC (28309T>C), AAC245TAC (28310A>T), GTG247TAC (28316G>T 28317T>A 28318G>C), GTC248GTG (28321C>G), ATC249GTG (28322A>G 28324C>G), GGA253GGC (28336A>C), ACG254ACC (28339G>C), GGC255GGT (28342C>T), TCG256GCG (28343T>G), CCA257CCG (28348A>G), TCG258TCC (28351G>C), ACG260ACC (28357G>C), CTT261TTC (28358C>T 28360T>C), TCG263TCC (28366G>C), TCA264ACC (28367T>A 28369A>C), GTT265GTC (28372T>C), TCG266AGC (28373T>A 28374C>G 28375G>C), GCT268GCC (28381T>C), GCT271GCC (28390T>C), CCA274CCG (28399A>G), CTA275GTC (28400C>G 28402A>C), TGC276GGT (28403T>G 28405C>T), AGC277TCG (28406A>T 28407G>C 28408C>G), AGC278ACC (28410G>C), GCT283GCG (28426T>G), CCG284CCT (28429G>T), GTG286CTT (28433G>C 28435G>T), GTT287GTG (28438T>G), AAT288ACG (28440A>C 28441T>G), TCT290AAC (28445T>A 28446C>A 28447T>C), ACG294GCC (28457A>G 28459G>C), GCG295ACC (28460G>A 28462G>C), ACC296ACT (28465C>T), ATC297ACG (28467T>C 28468C>G), GCG299AGC (28472G>A 28473C>G 28474G>C), TAT301TAC (28480T>C), GAC302AAC (28481G>A), ACG303CAC (28484A>C 28485C>A 28486G>C), GTG304GCC (28488T>C 28489G>C), GCG305GCC (28492G>C), ACG306ATC (28494C>T 28495G>C), AAG307GTC (28496A>G 28497A>T 28498G>C), GCG308GCT (28501G>T), TTG309CTG (28502T>C), ATC310GTG (28505A>G 28507C>G), GTG311CAA (28508G>C 28509T>A 28510G>A), ACC312GCC (28511A>G), GCA313GCC (28516A>C), CTG314GTG (28517C>G), AAG315CAG (28520A>C), ACG316TCC (28523A>T 28525G>C), TAT317TAC (28528T>C), GCA326ACG (28553G>A 28555A>G), CCG328CCC (28561G>C), TAT329TAC (28564T>C), TCG330ACC (28565T>A 28567G>C), CGG331AAG (28568C>A 28569G>A), GCC333TCG (28574G>T 28576C>G), CAA334CAG (28579A>G), GTG335CTC (28580G>C 28582G>C), GCT336GCC (28585T>C), TAT337TAC (28588T>C), GAC338GGC (28590A>G), GCT339GCG (28594T>G), CCT341CCG (28600T>G), GGC342GCA (28602G>C 28603C>A), ACG344ACC (28609G>C), ATC346GTC (28613A>G), ACG347ACC (28618G>C), GTG349ATC (28622G>A 28624G>C), AGC353GGC (28634A>G), GCG356GCT (28645G>T), GAT357GAC (28648T>C), GCT359ACG (28652G>A 28654T>G), GCG360GCC (28657G>C), ACG361ACC (28660G>C), AGT362AAT (28662G>A), CTT363CAG (28665T>A 28666T>G), CAG364CAA (28669G>A), ATA366ATC (28675A>C), AAA367AAG (28678A>G), TGG368GCC (28679T>G 28680G>C 28681G>C), TCG369GGC (28682T>G 28683C>G 28684G>C), GCA370ACG (28685G>A 28687A>G), GTC371ATC (28688G>A), AGC372TCG (28691A>T 28692G>C 28693C>G), GCA374ACC (28697G>A 28699A>C) | | | | | | | | |
| KMC43_gp47 | 1 | 36 | 14.8% | 124 | 50.6% | 35 (94.6%) | 16 (43.2%) | 1/1/0/0 | 0 |
| Protein mutations: | S2A (28729A>G 28730G>C 28731C>A), V3I (28732G>A 28734T>C), Q7S (28744C>A 28745A>G 28746A>C), V9I (28750G>A), Y10S (28753T>A 28754A>G), Q11T (28756C>A 28757A>C), G15Q (28768G>C 28769G>A 28770C>G), Y16L (28771T>C 28772A>T), L17I (28774C>A 28776G>C), R19G (28780C>G), R19_W20insG (28782_28783insGGG), G22del (28789_28791delGGC), D23Q (28792G>C 28794T>G), A24Q (28795G>C 28796C>A 28797G>A), A25G (28799C>G 28800G>C), Q26L (28802A>T 28803G>C), S27V (28804T>G 28805C>T), V29I (28810G>A), L30R (28814T>G 28815G>C), T31D (28816A>G 28817C>A) | | | | | | | | |
| Codon mutations: | AGC2GCA (28729A>G 28730G>C 28731C>A), GTT3ATC (28732G>A 28734T>C), GGC4GGT (28737C>T), GAT5GAC (28740T>C), CAA6CAG (28743A>G), CAA7AGC (28744C>A 28745A>G 28746A>C), GAT8GAC (28749T>C), GTC9ATC (28750G>A), TAC10AGC (28753T>A 28754A>G), CAG11ACG (28756C>A 28757A>C), CGC14CGG (28767C>G), GGC15CAG (28768G>C 28769G>A 28770C>G), TAC16CTC (28771T>C 28772A>T), CTG17ATC (28774C>A 28776G>C), CCT18CCG (28779T>G), CGC19GGC (28780C>G), CGC19_TGG20insGGG (28782_28783insGGG), GGC22del (28789_28791delGGC), GAT23CAG (28792G>C 28794T>G), GCG24CAA (28795G>C 28796C>A 28797G>A), GCG25GGC (28799C>G 28800G>C), CAG26CTC (28802A>T 28803G>C), TCG27GTG (28804T>G 28805C>T), CCA28CCG (28809A>G), GTC29ATC (28810G>A), CTG30CGC (28814T>G 28815G>C), ACC31GAC (28816A>G 28817C>A), CAG35CAA (28830G>A) | | | | | | | | |

**Proteins**

| | Begin | End | Coverage | Score | Concordance | Matches | Identities | I/D/M/F* | Stop Codons |
|---|---|---|---|---|---|---|---|---|---|
| DUF4043 domain-containing protein (YP_010078151.1) | 133 | 207 | 16.9% | 411 | 85.8% | 75 (100%) | 62 (82.7%) | 0/0/0/0 | 0 |
| Protein mutations: | N134A (14595A>G 14596A>C 14597T>C), T149V (14640A>G 14641C>T 14642G>C), S153A (14652T>G), A156L (14661G>C 14662C>T 14663C>G), I163V (14682A>G 14684C>G), N168Q (14697A>C 14699C>G), Q174V (14715C>G 14716A>T), L175M (14718T>A), G180A (14734G>C), I192V (14769A>G), A198V (14788C>T 14789A>G), A200V (14794C>T), S203G (14802T>G 14803C>G) | | | | | | | | |
| Codon mutations: | AAT134GCC (14595A>G 14596A>C 14597T>C), GCC135GCG (14600C>G), CTG136CTC (14603G>C), AAT138AAC (14609T>C), ACG149GTC (14640A>G 14641C>T 14642G>C), CTG151CTC (14648G>C), TCG153GCG (14652T>G), GCC156CTG (14661G>C 14662C>T 14663C>G), ACC157ACG (14666C>G), TCG159AGC (14670T>A 14671C>G 14672G>C), GAC161GAT (14678C>T), ATC163GTG (14682A>G 14684C>G), AAC168CAG (14697A>C 14699C>G), CAG174GTG (14715C>G 14716A>T), TTG175ATG (14718T>A), GTC176GTT (14723C>T), GTG178GTC (14729G>C), GGC180GCC (14734G>C), GTG186GTC (14753G>C), GTT188GTG (14759T>G), ATC192GTC (14769A>G), CTG195CTC (14780G>C), ACT196ACC (14783T>C), GCA198GTG (14788C>T 14789A>G), GCC200GTC (14794C>T), GAC201GAT (14798C>T), TCC203GGC (14802T>G 14803C>G) | | | | | | | | |
| tail protein (YP_010078160.1) | 89 | 162 | 14.5% | 317 | 67.3% | 74 (100%) | 49 (66.2%) | 0/0/0/0 | 0 |
| Protein mutations: | R92K (19482C>A 19483G>A 19484C>A), C93A (19485T>G 19486G>C), T102V (19512A>G 19513C>T), T106I (19525C>T 19526G>C), I107A (19527A>G 19528T>C 19529C>G), Q109L (19534A>T), G112C (19542G>T), V113I (19545G>A 19547G>C), L115F (19551C>T 19553G>C), A117S (19557G>T 19559G>C), L123F (19575C>T 19577G>C), S126G (19584T>G 19585C>G 19586G>C), V127I (19587G>A 19589G>C), A130T (19596G>A), A132S (19602G>T 19604C>G), A136Q (19614G>C 19615C>A 19616C>G), N138S (19620A>T 19621A>C 19622T>G), W140Q (19626T>C 19627G>A), V142A (19633T>C 19634A>C), T143V (19635A>G 19636C>T), A145S (19641G>T), T148G (19650A>G 19651C>G 19652G>C), N150V (19656A>G 19657A>T 19658C>G), V153A (19666T>C 19667T>C), A159S (19683G>T 19685T>G) | | | | | | | | |
| Codon mutations: | CGC92AAA (19482C>A 19483G>A 19484C>A), TGC93GCC (19485T>G 19486G>C), CGC95CGG (19493C>G), GTC96GTG (19496C>G), ACG102GTG (19512A>G 19513C>T), GCG104GCC (19520G>C), TCG105AGC (19521T>A 19522C>G 19523G>C), ACG106ATC (19525C>T 19526G>C), ATC107GCG (19527A>G 19528T>C 19529C>G), CAG109CTG (19534A>T), GGC112TGC (19542G>T), GTG113ATC (19545G>A 19547G>C), ACG114ACC (19550G>C), CTG115TTC (19551C>T 19553G>C), GCG117TCC (19557G>T 19559G>C), ACG122ACC (19574G>C), CTG123TTC (19575C>T 19577G>C), TCG126GGC (19584T>G 19585C>G 19586G>C), GTG127ATC (19587G>A 19589G>C), GTC129GTG (19595C>G), GCG130ACG (19596G>A), CTC131CTG (19601C>G), GCC132TCG (19602G>T 19604C>G), GCG133GCC (19607G>C), AGC135TCG (19611A>T 19612G>C 19613C>G), GCC136CAG (19614G>C 19615C>A 19616C>G), AAT138TCG (19620A>T 19621A>C 19622T>G), TGG140CAG (19626T>C 19627G>A), GTA142GCC (19633T>C 19634A>C), ACG143GTG (19635A>G 19636C>T), GTC144GTT (19640C>T), GCG145TCG (19641G>T), ACG148GGC (19650A>G 19651C>G 19652G>C), CTA149TTG (19653C>T 19655A>G), AAC150GTG (19656A>G 19657A>T 19658C>G), GTT153GCC (19666T>C 19667T>C), GCT159TCG (19683G>T 19685T>G) | | | | | | | | |
| hypothetical protein (YP_010078161.1) | 13 | 99 | 60.4% | 399 | 66.6% | 87 (100%) | 54 (62.1%) | 0/0/0/0 | 0 |
| Protein mutations: | V15T (20840G>A 20841T>C), Q17V (20846C>G 20847A>T), S19A (20852T>G 20854A>C), T20S (20856C>G 20857T>C), T22P (20861A>C), Q24N (20867C>A 20869G>C), P25L (20871C>T), N26S (20874A>G), K27L (20876A>C 20877A>T), S30Q (20885T>C 20886C>A 20887T>G), K32Q (20891A>C), K34A (20897A>G 20898A>C), Q35P (20901A>C 20902G>C), V37M (20906G>A), T38A (20909A>G), D39E (20914T>G), V40I (20915G>A 20917G>C), R41K (20918C>A 20919G>A 20920C>G), V42I (20921G>A), R44G (20927C>G 20929C>T), D50R (20945G>C 20946A>G), V52A (20952T>C), S59R (20972T>C 20973C>G 20974T>C), A63N (20984G>A 20985C>A 20986G>C), I64V (20987A>G 20989T>G), A69S (21002G>A 21003C>G 21004G>C), A70T (21005G>A 21007G>A), N73D (21014A>G), S80A (21035T>G 21037G>C), S89Q (21062T>C 21063C>A), Q92T (21071C>A 21072A>C), Y94T (21077T>A 21078A>C 21079T>C) | | | | | | | | |

| | Begin | End | Coverage | Score | Concordance | Matches | Identities | I/D/M/F* | Stop Codons |
|---|---|---|---|---|---|---|---|---|---|
| **NT** | **14591** | **28832** | **3.5%** | **1249** | **38.8%** | **1626 (98.8%)** | **1137 (69.1%)** | **3/17** | |
| Codon mutations: | CTG14CTC (20839G>C), GTC15ACC (20840G>A 20841T>C), CAG17GTG (20846C>G 20847A>T), TCA19GCC (20852T>G 20854A>C), ACT20AGC (20856C>G 20857T>C), ACG22CCG (20861A>C), CTG23CTC (20866G>C), CAG24AAC (20867C>A 20869G>C), CCC25CTC (20871C>T), AAC26AGC (20874A>G), AAG27CTG (20876A>C 20877A>T), ACC29ACG (20884C>G), TCT30CAG (20885T>C 20886C>A 20887T>G), AAG32CAG (20891A>C), AGC33TCG (20894A>T 20895G>C 20896C>G), AAG34GCG (20897A>G 20898A>C), CAG35CCC (20901A>C 20902G>C), GAT36GAC (20905T>C), GTG37ATG (20906G>A), ACC38GCC (20909A>G), GAT39GAG (20914T>G), GTG40ATC (20915G>A 20917G>C), CGC41AAG (20918C>A 20919G>A 20920C>G), GTC42ATC (20921G>A), CGC44GGT (20927C>G 20929C>T), GAC50CGC (20945G>C 20946A>G), CAC51CAT (20950C>T), GTT52GCT (20952T>C), TTT54TTC (20959T>C), TCT59CGC (20972T>C 20973C>G 20974T>C), TCG61AGC (20978T>A 20979C>G 20980G>C), GCG63AAC (20984G>A 20985C>A 20986G>C), ATT64GTG (20987A>G 20989T>G), GAA65GAG (20992A>G), CAA67CAG (20998A>G), GCG69AGC (21002G>A 21003C>G 21004G>C), GCG70ACA (21005G>A 21007G>A), AAC73GAC (21014A>G), CAA77CAG (21028A>G), GAA79GAG (21034A>G), TCG80GCC (21035T>G 21037G>C), TCG89CAG (21062T>C 21063C>A), CCC90CCG (21067C>G), CAG92ACG (21071C>A 21072A>C), ATT93ATC (21076T>C), TAT94ACC (21077T>A 21078A>C 21079T>C), ACG96ACC (21085G>C) | | | | | | | | |
| hypothetical protein (YP_010078167.1) | 71 | 122 | 13.9% | 262 | 73.8% | 52 (100%) | 38 (73.1%) | 0/0/0/0 | 0 |
| Protein mutations: | F74L (25635T>C 25637C>G), D80N (25653G>A), T82A (25659A>G), N83D (25662A>G), Y84P (25665T>C 25666A>C 25667C>G), E86A (25672A>C 25673G>C), T87N (25675C>A 25676G>C), D88E (25679C>A), T90S (25683A>T 25685C>G), Y102F (25720A>T), L108M (25737C>A), T110S (25744C>G), A119V (25771C>T 25772G>C) | | | | | | | | |
| Codon mutations: | ATT73ATC (25634T>C), TTC74CTG (25635T>C 25637C>G), TCC79AGC (25650T>A 25651C>G), GAT80AAT (25653G>A), ACG82GCG (25659A>G), AAC83GAC (25662A>G), TAC84CCG (25665T>C 25666A>C 25667C>G), GAG86GCC (25672A>C 25673G>C), ACG87AAC (25675C>A 25676G>C), GAC88GAA (25679C>A), CTC89CTG (25682C>G), ACC90TCG (25683A>T 25685C>G), CTG92CTC (25691G>C), TAT94TAC (25697T>C), GGG95GGC (25700G>C), GTG97GTC (25706G>C), GAT98GAC (25709T>C), TAC102TTC (25720A>T), GAC103GAT (25724C>T), CCC104CCG (25727C>G), TCG106TCC (25733G>C), CTG108ATG (25737C>A), ACC110AGC (25744C>G), GCG119GTC (25771C>T 25772G>C) | | | | | | | | |
| hypothetical protein (YP_010078170.1) | 157 | 375 | 57.3% | 867 | 68.1% | 215 (100%) | 133 (61.9%) | 0/0/0/0 | 1 |
| Protein mutations: | G159A (28053G>C 28054C>G), V162A (28062T>C), A163I (28064G>A 28065C>T 28066T>C), G164A (28068G>C), V166I (28073G>A 28075A>C), S167N (28076T>A 28077C>A 28078G>C), I169L (28082A>C), V170G (28086T>G), G171Q (28088G>C 28089G>A 28090C>G), S174A (28097T>G 28099C>G), N187T (28137A>C 28138C>G), A193T (28154G>A 28156A>G), L197F (28166C>T 28168G>C), T199S (28172A>T), I202V (28181A>G), A203G (28185C>G), V205I (28190G>A 28192A>C), A206N (28193G>A 28194C>A 28195G>C), K213R (28215A>G 28216A>G), S214A (28217T>G 28219C>T), V216I (28223G>A), Y218N (28229T>A), T221L (28238A>C 28239C>T 28240C>G), S222A (28241T>G 28243G>C), L223I (28244C>A 28246G>C), G226D (28254G>A 28255A>C), L227I (28256C>A 28258G>C), S228N (28260G>A), L232T (28271C>A 28272T>C 28273T>C), N234G (28277A>G 28278A>G), Q235Y (28280C>T 28282G>T), T236A (28283A>G 28285G>C), A238G (28290C>G), V240A (28296T>C), T241A (28298A>G), M243L (28304A>C), N245Y (28310A>T), V247Y (28316G>T 28317T>A 28318G>C), I249V (28322A>G 28324C>G), S256A (28343T>G), L261F (28358C>T 28360T>C), S264T (28367T>A 28369A>C), L275V (28400C>G 28402A>C), C276G (28403T>G 28405C>T), S278T (28410G>C), V286L (28433G>C 28435G>T), N288T (28440A>C 28441T>G), S290N (28445T>A 28446C>A 28447T>C), T294A (28457A>G 28459G>C), A295T (28460G>A 28462G>C), I297T (28467T>C 28468C>G), A299S (28472G>A 28473C>G 28474G>C), D302N (28481G>A), T303H (28484A>C 28485C>A 28486G>C), V304A (28488T>C 28489G>C), T306I (28494C>T 28495G>C), K307V (28496A>G 28497A>T 28498G>C), I310V (28505A>G 28507C>G), V311Q (28508G>C 28509T>A 28510G>A), T312A (28511A>G), L314V (28517C>G), K315Q (28520A>C), T316S (28523A>T 28525G>C), A326T (28553G>A 28555A>G), S330T (28565T>A 28567G>C), R331K (28568C>A 28569G>A), A333S (28574G>T 28576C>G), V335L (28580G>C 28582G>C), D338G (28590A>G), G342A (28602G>C 28603C>A), I346V (28613A>G), V349I (28622G>A 28624G>C), S353G (28634A>G), A359T (28652G>A 28654T>G), S362N (28662G>A), L363Q (28665T>A 28666T>G), W368A (28679T>G 28680G>C 28681G>C), S369G (28682T>G 28683C>G 28684G>C), A370T (28685G>A 28687A>G), V371I (28688G>A), A374T (28697G>A 28699A>C) | | | | | | | | |
| Codon mutations: | GCT158GCG (28051T>G), GGC159GCG (28053G>C 28054C>G), GTG162GCG (28062T>C), GCT163ATC (28064G>A 28065C>T 28066T>C), GGC164GCC (28068G>C), GCC165GCG (28072C>G), GTA166ATC (28073G>A 28075A>C), TCG167AAC (28076T>A 28077C>A 28078G>C), ACC168ACG (28081C>G), ATC169CTC (28082A>C), GTC170GGC (28086T>G), GGC171CAG (28088G>C 28089G>A 28090C>G), GCA172GCC (28093A>C), ATT173ATC (28096T>C), TCC174GCG (28097T>G 28099C>G), GAT177GAC (28108T>C), GTC179GTG (28114C>G), ACG180ACC (28117G>C), GCC182GCG (28123C>G), GCA183GCG (28126A>G), AAC187ACG (28137A>C 28138C>G), GGC188GGT (28141C>T), GAT190GAC (28147T>C), GCA193ACG (28154G>A 28156A>G), GAC194GAT (28159C>T), GCT195GCC (28162T>C), GCG196GCC (28165G>C), CTG197TTC (28166C>T 28168G>C), CGT198CGC (28171T>C), ACG199TCG (28172A>T), CGC200CGG (28177C>G), ATC202GTC (28181A>G), GCC203GGC (28185C>G), GTA205ATC (28190G>A 28192A>C), GCG206AAC (28193G>A 28194C>A 28195G>C), AAG210AAA (28207G>A), ACG212ACC (28213G>C), AAA213AGG (28215A>G 28216A>G), TCC214GCT (28217T>G 28219C>T), GCT215GCC (28222T>C), GTC216ATC (28223G>A), TAT218AAT (28229T>A), ACC221CTG (28238A>C 28239C>T 28240C>G), TCG222GCC (28241T>G 28243G>C), CTG223ATC (28244C>A 28246G>C), GGA226GAC (28254G>A 28255A>C), CTG227ATC (28256C>A 28258G>C), AGC228AAC (28260G>A), TAT229TAC (28264T>C), CTG231CTC (28270G>C), CTT232ACC (28271C>A 28272T>C 28273T>C), AAC234GGC (28277A>G 28278A>G), CAG235TAT (28280C>T 28282G>T), ACG236GCC (28283A>G 28285G>C), GCC238GGC (28290C>G), GTG240GCG (28296T>C), ACG241GCG (28298A>G), ATG243CTG (28304A>C), GGT244GGC (28309T>C), AAC245TAC (28310A>T), GTG247TAC (28316G>T 28317T>A 28318G>C), GTC248GTG (28321C>G), ATC249GTG (28322A>G 28324C>G), GGA253GGC (28336A>C), ACG254ACC (28339G>C), GGC255GGT (28342C>T), TCG256GCG (28343T>G), CCA257CCG (28348A>G), TCG258TCC (28351G>C), ACG260ACC (28357G>C), CTT261TTC (28358C>T 28360T>C), TCG263TCC (28366G>C), TCA264ACC (28367T>A 28369A>C), GTT265GTC (28372T>C), TCG266AGC (28373T>A 28374C>G 28375G>C), GCT268GCC (28381T>C), GCT271GCC (28390T>C), CCA274CCG (28399A>G), CTA275GTC (28400C>G 28402A>C), TGC276GGT (28403T>G 28405C>T), AGC277TCG (28406A>T 28407G>C 28408C>G), AGC278ACC (28410G>C), GCT283GCG (28426T>G), CCG284CCT (28429G>T), GTG286CTT (28433G>C 28435G>T), GTT287GTG (28438T>G), AAT288ACG (28440A>C 28441T>G), TCT290AAC (28445T>A 28446C>A 28447T>C), ACG294GCC (28457A>G 28459G>C), GCG295ACC (28460G>A 28462G>C), ACC296ACT (28465C>T), ATC297ACG (28467T>C 28468C>G), GCG299AGC (28472G>A 28473C>G 28474G>C), TAT301TAC (28480T>C), GAC302AAC (28481G>A), ACG303CAC (28484A>C 28485C>A 28486G>C), GTG304GCC (28488T>C 28489G>C), GCG305GCC (28492G>C), ACG306ATC (28494C>T 28495G>C), AAG307GTC (28496A>G 28497A>T 28498G>C), GCG308GCT (28501G>T), TTG309CTG (28502T>C), ATC310GTG (28505A>G 28507C>G), GTG311CAA (28508G>C 28509T>A 28510G>A), ACC312GCC (28511A>G), GCA313GCC (28516A>C), CTG314GTG (28517C>G), AAG315CAG (28520A>C), ACG316TCC (28523A>T 28525G>C), TAT317TAC (28528T>C), GCA326ACG (28553G>A 28555A>G), CCG328CCC (28561G>C), TAT329TAC (28564T>C), TCG330ACC (28565T>A 28567G>C), CGG331AAG (28568C>A 28569G>A), GCC333TCG (28574G>T 28576C>G), CAA334CAG (28579A>G), GTG335CTC (28580G>C 28582G>C), GCT336GCC (28585T>C), TAT337TAC (28588T>C), GAC338GGC (28590A>G), GCT339GCG (28594T>G), CCT341CCG (28600T>G), GGC342GCA (28602G>C 28603C>A), ACG344ACC (28609G>C), ATC346GTC (28613A>G), ACG347ACC (28618G>C), GTG349ATC (28622G>A 28624G>C), AGC353GGC (28634A>G), GCG356GCT (28645G>T), GAT357GAC (28648T>C), GCT359ACG (28652G>A 28654T>G), GCG360GCC (28657G>C), ACG361ACC (28660G>C), AGT362AAT (28662G>A), CTT363CAG (28665T>A 28666T>G), CAG364CAA (28669G>A), ATA366ATC (28675A>C), AAA367AAG (28678A>G), TGG368GCC (28679T>G 28680G>C 28681G>C), TCG369GGC (28682T>G 28683C>G 28684G>C), GCA370ACG (28685G>A 28687A>G), GTC371ATC (28688G>A), AGC372TCG (28691A>T 28692G>C 28693C>G), GCA374ACC (28697G>A 28699A>C) | | | | | | | | |
| hypothetical protein (YP_010078171.1) | 1 | 36 | 14.8% | 124 | 50.6% | 35 (94.6%) | 16 (43.2%) | 1/1/0/0 | 0 |
| Protein mutations: | S2A (28729A>G 28730G>C 28731C>A), V3I (28732G>A 28734T>C), Q7S (28744C>A 28745A>G 28746A>C), V9I (28750G>A), Y10S (28753T>A 28754A>G), Q11T (28756C>A 28757A>C), G15Q (28768G>C 28769G>A 28770C>G), Y16L (28771T>C 28772A>T), L17I (28774C>A 28776G>C), R19G (28780C>G), R19_W20insG (28782_28783insGGG), G22del (28789_28791delGGC), D23Q (28792G>C 28794T>G), A24Q (28795G>C 28796C>A 28797G>A), A25G (28799C>G 28800G>C), Q26L (28802A>T 28803G>C), S27V (28804T>G 28805C>T), V29I (28810G>A), L30R (28814T>G 28815G>C), T31D (28816A>G 28817C>A) | | | | | | | | |
| Codon mutations: | AGC2GCA (28729A>G 28730G>C 28731C>A), GTT3ATC (28732G>A 28734T>C), GGC4GGT (28737C>T), GAT5GAC (28740T>C), CAA6CAG (28743A>G), CAA7AGC (28744C>A 28745A>G 28746A>C), GAT8GAC (28749T>C), GTC9ATC (28750G>A), TAC10AGC (28753T>A 28754A>G), CAG11ACG (28756C>A 28757A>C), CGC14CGG (28767C>G), GGC15CAG (28768G>C 28769G>A 28770C>G), TAC16CTC (28771T>C 28772A>T), CTG17ATC (28774C>A 28776G>C), CCT18CCG (28779T>G), CGC19GGC (28780C>G), CGC19_TGG20insGGG (28782_28783insGGG), GGC22del (28789_28791delGGC), GAT23CAG (28792G>C 28794T>G), GCG24CAA (28795G>C 28796C>A 28797G>A), GCG25GGC (28799C>G 28800G>C), CAG26CTC (28802A>T 28803G>C), TCG27GTG (28804T>G 28805C>T), CCA28CCG (28809A>G), GTC29ATC (28810G>A), CTG30CGC (28814T>G 28815G>C), ACC31GAC (28816A>G 28817C>A), CAG35CAA (28830G>A) | | | | | | | | |

*: Inserts / Deletes / Misaligned / Frameshifts

#### Analysis details

This analysis was performed with panviral2.64

### NGS Details (UN18_val): Caulimovirus venafragariae

#### Assembly

| | |
|---|---|
| **Coverage Length** | 838 (1 contig(s)) |
| **Depth Of Coverage** | 4.8 |
| **Number Of Reads** | 30 |
| **Reads Per Million** | 0.56 rpm (after QC) |
| **Ambiguities** | 0 |
| **Assembly Method** | de novo + reference guided assembly |
| **Consensus Caller** | Bcf Tools |

#### Coverage Map

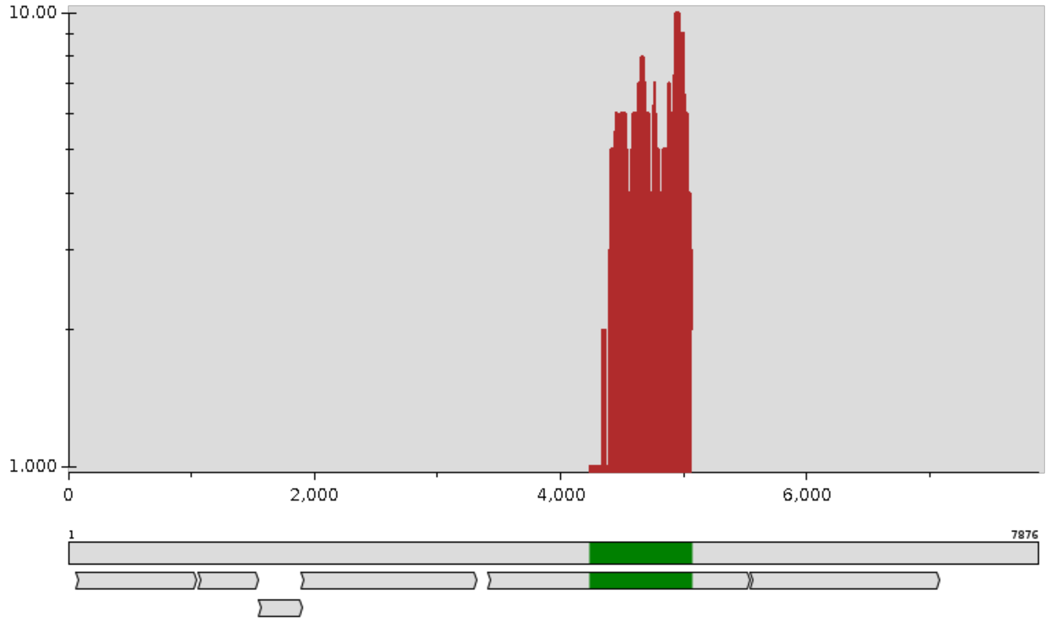

#### Assignment

| | |
|---|---|
| **Type** | Caulimovirus venafragariae (Taxonomy ID: 3048344) |
| **Reference Genome** | NC_001725.1 |
| **NT Identity (%)** | 58.0723 |
| **AA Identity (%)** | 52.8986 |
| **Number Of Stop Codons** | 1 |
| **Number Of CDS** | 6 |

#### Alignment

| | |
|---|---|
| **Alignment Score** | 242.0 (NT) + 916.0 (AA) = 1158.0 |
| **Concordance (%)** | 33.7413 |

| Alignment Method | Global, seeded, nucleotide + amino acids (AGA) |
|---|---|

#### Genome Region

Sequence starts at position 4232 and ends at position 5069 relative to NC_001725.1 reference sequence.

#### Alignment Detailed Statistics

| | Begin | End | Coverage | Score | Concordance | Matches | Identities | I/D/M/F* | Stop Codons |
|---|---|---|---|---|---|---|---|---|---|
| **NT** | **4232** | **5069** | **10.6%** | **242** | **14.8%** | **830 (99.0%)** | **482 (57.5%)** | **0/8** | |

Mutations: 4233C>T, 4235A>T, 4239T>C, 4242T>A, 4244A>G, 4245G>A, 4246T>G, 4247G>T, 4249A>C, 4254A>C, 4258C>T, 4259A>T, 4260G>T, 4264T>A, 4265C>A, 4266A>C, 4267C>G, 4268C>A, 4270C>G, 4275C>T, 4276C>A, 4277G>A, 4279G>A, 4281A>T, 4284G>A, 4287T>C, 4289A>G, 4292C>T, 4299C>T, 4300G>C, 4302A>T, 4308A>T, 4314G>A, 4315C>A, 4317T>G, 4318G>A, 4319G>A, 4320T>G, 4321A>T, 4323C>A, 4330C>G, 4332T>A, 4337A>G, 4338G>A, 4347T>C, 4348T>A, 4349C>G, 4351T>A, 4352C>G, 4353A>T, 4356A>C, 4359C>G, 4371G>A, 4374T>C, 4380C>G, 4383G>A, 4384A>C, 4385T>A, 4386C>A, 4395C>T, 4398A>G, 4401A>T, 4402A>C, 4410A>T, 4413T>A, 4423A>G, 4426T>C, 4428A>T, 4432G>A, 4434C>A, 4435C>A, 4437T>C, 4438A>T, 4439C>G, 4440A>T, 4442A>T, 4443G>T, 4444G>T, 4445G>T, 4446A>T, 4449T>C, 4452C>A, 4456C>T, 4458A>T, 4459C>A, 4461T>C, 4467C>T, 4470G>A, 4471G>A, 4472A>C, 4473G>T, 4474C>A, 4476A>T, 4479G>T, 4480C>A, 4482T>A, 4483C>A, 4485A>T, 4486A>C, 4487G>T, 4489A>G, 4492G>A, 4493G>A, 4496G>T, 4497T>A, 4500G>A, 4505T>A, 4506T>C, 4508A>T, 4509C>T, 4512T>C, 4513T>A, 4514C>A, 4515C>A, 4521C>T, 4528T>A, 4529C>G, 4543G>A, 4546C>A, 4548C>A, 4551T>C, 4552G>T, 4554T>A, 4555C>G, 4556C>A, 4557A>G, 4561A>T, 4568A>C, 4569G>T, 4575C>A, 4578T>A, 4585T>A, 4586G>C, 4590C>T, 4591C>A, 4593A>T, 4599C>T, 4602C>T, 4614C>A, 4618C>T, 4620T>A, 4633C>A, 4635A>T, 4638C>A, 4641T>A, 4642G>C, 4643C>A, 4644T>A, 4650C>T, 4656G>A, 4657C>A, 4659C>A, 4663G>T, 4665C>T, 4666G>A, 4670G>T, 4671T>A, 4672C>T, 4674C>T, 4675_4679delAGCAA, 4680C>T, 4681A>C, 4684T>A, 4686T>A, 4687C>A, 4688C>A, 4689A>G, 4690C>A, 4692G>A, 4697G>T, 4698T>G, 4699G>A, 4700C>T, 4701T>A, 4707T>C, 4708G>A, 4710T>A, 4719C>T, 4720A>T, 4722C>G, 4725C>A, 4727T>G, 4729A>T, 4730G>C, 4731C>A, 4733A>C, 4737T>A, 4738G>T, 4740A>T, 4741G>C, 4743A>G, 4750T>C, 4752A>T, 4753G>A, 4754G>C, 4759G>C, 4761C>T, 4765A>G, 4766T>A, 4767T>A, 4768G>T, 4771C>T, 4772T>C, 4776C>A, 4777A>C, 4778G>T, 4779A>C, 4782C>T, 4783A>G, 4784A>T, 4785A>C, 4786G>A, 4787C>A, 4788C>T, 4789C>A, 4790T>A, 4791A>T, 4793G>A, 4794A>C, 4800G>A, 4801C>T, 4803T>A, 4806C>T, 4807A>C, 4809A>G, 4812G>A, 4818C>A, 4819C>G, 4821G>A, 4822T>A, 4824G>A, 4825T>A, 4826G>A, 4827C>T, 4832C>A, 4833A>T, 4835C>A, 4836T>A, 4839C>A, 4840A>G, 4842T>A, 4851T>A, 4852C>A, 4854A>G, 4855G>A, 4857A>T, 4860A>T, 4861G>T, 4862A>C, 4865G>A, 4867G>A, 4868G>A, 4869A>T, 4870A>G, 4871A>G, 4872T>A, 4873C>A, 4875C>A, 4876A>G, 4879G>T, 4881C>A, 4884G>A, 4886G>C, 4890T>C, 4893C>A, 4894G>T, 4895G>C, 4896A>C, 4897C>A, 4898T>A, 4900C>A, 4902C>G, 4903T>A, 4906G>T, 4908T>A, 4910C>A, 4911T>G, 4917A>T, 4920T>C, 4921C>A, 4923A>G, 4924C>T, 4926T>A, 4928G>T, 4929T>A, 4930G>A, 4931A>C, 4932T>C, 4934G>A, 4935G>A, 4936A>C, 4938T>A, 4940C>A, 4941C>A, 4942T>A, 4954T>C, 4956A>T, 4959C>A, 4960C>T, 4961T>G, 4963C>T, 4968T>C, 4972A>G, 4973T>C, 4974C>A, 4975T>G, 4976C>G, 4977C>A, 4979C>A, 4980T>A, 4982A>T, 4983T>C, 4984T>A, 4986T>A, 4990A>G, 4993A>T, 5001C>G, 5002C>A, 5003T>A, 5004T>G, 5007G>A, 5008T>A, 5009C>A, 5010A>T, 5012C>T, 5014T>C, 5016A>T, 5019G>A, 5020_5022delGTT, 5028T>C, 5030A>T, 5031G>C, 5033A>G, 5035G>A, 5037G>A, 5038A>T, 5039T>C, 5042C>A, 5043T>C, 5044T>C, 5046G>A, 5048C>T, 5052G>A, 5054C>G, 5056G>T, 5057A>T, 5058G>T, 5059A>T, 5060A>T, 5061A>C

##### CDS

| | Begin | End | Coverage | Score | Concordance | Matches | Identities | I/D/M/F* | Stop Codons |
|---|---|---|---|---|---|---|---|---|---|
| ORF_V | 277 | 555 | 39.2% | 916 | 47.2% | 276 (99.3%) | 146 (52.5%) | 0/2/1/1 | 1 |

Protein mutations: K277I (4235A>T), K280R (4244A>G 4245G>A), C281V (4246T>G 4247G>T), K282Q (4249A>C), Q285F (4258C>T 4259A>T 4260G>T), S287N (4264T>A 4265C>A 4266A>C), P288D (4267C>G 4268C>A), Q289E (4270C>G), R291K (4276C>A 4277G>A), E292N (4279G>A 4281A>T), K295R (4289A>G), T296I (4292C>T), E299H (4300G>C 4302A>T), L304M (4315C>A 4317T>G), G305K (4318G>A 4319G>A 4320T>G), I306L (4321A>T 4323C>A), P309A (4330C>G 4332T>A), K311R (4337A>G 4338G>A), I327Q (4384A>C 4385T>A 4386C>A), K340E (4423A>G), D343K (4432G>A 4434C>A), H344N (4435C>A 4437T>C), T345C (4438A>T 4439C>G 4440A>T), K346I (4442A>T 4443G>T), G347F (4444G>T 4445G>T 4446A>T), L351F (4456C>T 4458A>T), L352I (4459C>A 4461T>C), E356T (4471G>A 4472A>C 4473G>T), Q357N (4474C>A 4476A>T), L359I (4480C>A 4482T>A), Q360N (4483C>A 4485A>T), R361L (4486A>C 4487G>T), I362V (4489A>G), G363K (4492G>A 4493G>A), G364V (4496G>T 4497T>A), F367Y (4505T>A 4506T>C), Y368F (4508A>T 4509C>T), S370K (4513T>A 4514C>A 4515C>A), V380I (4543G>A), A383S (4552G>T 4554T>A), P384E (4555C>G 4556C>A 4557A>G), T386S (4561A>T), Q388P (4568A>C 4569G>T), C394T (4585T>A 4586G>C), Q396N (4591C>A 4593A>T), P405S (4618C>T 4620T>A), Q410N (4633C>A 4635A>T), A413Q (4642G>C 4643C>A 4644T>A), H418K (4657C>A 4659C>A), D420Y (4663G>T 4665C>T), E421K (4666G>A), S422I (4670G>T 4671T>A), L423F (4672C>T 4674C>T), S424del (4675_4679delAGCAA), M426L (4681A>C), Y427K (4684T>A 4686T>A), P428K (4687C>A 4688C>A 4689A>G), Q429K (4690C>A 4692G>A), C431L (4697G>T 4698T>G), A432I (4699G>A 4700C>T 4701T>A), V435I (4708G>A 4710T>A), I439L (4720A>T 4722C>G), F441C (4727T>G), K443T (4733A>C), E445Y (4738G>T 4740A>T), E446Q (4741G>C 4743A>G), G450T (4753G>A 4754G>C), V452L (4759G>C 4761C>T), I454E (4765A>G 4766T>A 4767T>A), V455F (4768G>T), L456S (4771C>T 4772T>C), N457K (4776C>A), R458L (4777A>C 4778G>T 4779A>C), K460V (4783A>G 4784A>T 4785A>C), A461N (4786G>A 4787C>A 4788C>T), L462N (4789C>A 4790T>A 4791A>T), G463D (4793G>A 4794A>C), K468Q (4807A>C 4809A>G), Q472E (4819C>G 4821G>A), L473I (4822T>A 4824G>A), C474N (4825T>A 4826G>A 4827C>T), T476N (4832C>A 4833A>T), T477K (4835C>A 4836T>A), N479E (4840A>G 4842T>A), L483M (4852C>A 4854A>G), V484I (4855G>A 4857A>T), E486S (4861G>T 4862A>C), R487K (4865G>A), G488N (4867G>A 4868G>A 4869A>T), N489G (4870A>G 4871A>G 4872T>A), L490I (4873C>A 4875C>A), K491E (4876A>G), V492L (4879G>T 4881C>A), S494T (4886G>C), G497S (4894G>T 4895G>C 4896A>C), L498K (4897C>A 4898T>A), H499K (4900C>A 4902C>G), L500I (4903T>A), V501L (4906G>T 4908T>A), A502E (4910C>A 4911T>G), Q506K (4921C>A 4923A>G), S508I (4928G>T 4929T>A), D509T (4930G>A 4931A>C 4932T>C), R510K (4934G>A 4935G>A), N511Q (4936A>C 4938T>A), A512E (4940C>A 4941C>A), L513I (4942T>A), L519C (4960C>T 4961T>G), I523A (4972A>G 4973T>C 4974C>A), S524G (4975T>G 4976C>G 4977C>A), A525E (4979C>A 4980T>A), Y526F (4982A>T 4983T>C), F527I (4984T>A 4986T>A), K529E (4990A>G), I530L (4993A>T), N532K (5001C>G), L533K (5002C>A 5003T>A 5004T>G), S535N (5008T>A 5009C>A 5010A>T), P536L (5012C>T), V539del (5020_5022delGTT), K542I (5030A>T 5031G>C), K543R (5033A>G), E544K (5035G>A 5037G>A), I545S (5038A>T 5039T>C), T546N (5042C>A 5043T>C), W547R (5044T>C 5046G>A), S548F (5048C>T), W549* (5052G>A), T550S (5054C>G), E551F (5056G>T 5057A>T 5058G>T), K552F (5059A>T 5060A>T 5061A>C)

Codon mutations: AAC276.AT (4233C>T), AAA277ATA (4235A>T), ATT278ATC (4239T>C), ATT279ATA (4242T>A), AAG280AGA (4244A>G 4245G>A), TGT281GTT (4246T>G 4247G>T), AAA282CAA (4249A>C), CCA283CCC (4254A>C), CAG285TTT (4258C>T 4259A>T 4260G>T), TCA287AAC (4264T>A 4265C>A 4266A>C), CCT288GAT (4267C>G 4268C>A), CAA289GAA (4270C>G), GAC290GAT (4275C>T), CGA291AAA (4276C>A 4277G>A), GAA292AAT (4279G>A 4281A>T), GAG293GAA (4284G>A), TTT294TTC (4287T>C), AAA295AGA (4289A>G), ACT296ATT (4292C>T), ATC298ATT (4299C>T), GAA299CAT (4300G>C 4302A>T), CTA301CTT (4308A>T), AAG303AAA (4314G>A), CTT304ATG (4315C>A 4317T>G), GGT305AAG (4318G>A 4319G>A 4320T>G), ATC306TTA (4321A>T 4323C>A), CCT309GCA (4330C>G 4332T>A), AAG311AGA (4337A>G 4338G>A), CAT314CAC (4347T>C), TCC315AGC (4348T>A 4349C>G), TCA316AGT (4351T>A 4352C>G 4353A>T), CCA317CCC (4356A>C), GCC318GCG (4359C>G), AGG322AGA (4371G>A), AAT323AAC (4374T>C), GCC325GCG (4380C>G), GAG326GAA (4383G>A), ATC327CAA (4384A>C 4385T>A 4386C>A), GGC330GGT (4395C>T), AAA331AAG (4398A>G), GCA332GCT (4401A>T), AGA333CGA (4402A>C), GTA335GTT (4410A>T), ATT336ATA (4413T>A), AAG340GAG (4423A>G), TTA341CTT (4426T>C 4428A>T), GAC343AAA (4432G>A 4434C>A), CAT344AAC (4435C>A 4437T>C), ACA345TGT (4438A>T 4439C>G 4440A>T), AAG346ATT (4442A>T 4443G>T), GGA347TTT (4444G>T 4445G>T 4446A>T), GAT348GAC (4449T>C), GGC349GGA (4452C>A), CTA351TTT (4456C>T 4458A>T), CTT352ATC (4459C>A 4461T>C), AAC354AAT (4467C>T), AAG355AAA (4470G>A), GAG356ACT (4471G>A 4472A>C 4473G>T), CAA357AAT (4474C>A 4476A>T), CTG358CTT (4479G>T), CTT359ATA (4480C>A 4482T>A), CAA360AAT (4483C>A 4485A>T), AGA361CTA (4486A>C 4487G>T), ATC362GTC (4489A>G), GGA363AAA (4492G>A 4493G>A), GGT364GTA (4496G>T 4497T>A), AAG365AAA (4500G>A), TTT367TAC (4505T>A 4506T>C), TAC368TTT (4508A>T 4509C>T), TCT369TCC (4512T>C), TCC370AAA (4513T>A 4514C>A 4515C>A), GAC372GAT (4521C>T), TCT375AGT (4528T>A 4529C>G), GTA380ATA (4543G>A), CGC381AGA (4546C>A 4548C>A), CTT382CTC (4551T>C), GCT383TCA (4552G>T 4554T>A), CCA384GAG (4555C>G 4556C>A 4557A>G), ACA386TCA (4561A>T), CAG388CCT (4568A>C 4569G>T), ACC390ACA (4575C>A), GCT391GCA (4578T>A), TGT394ACT (4585T>A 4586G>C), CCC395CCT (4590C>T), CAA396AAT (4591C>A 4593A>T), CAC398CAT (4599C>T), TAC399TAT (4602C>T), GTC403GTA (4614C>A), CCT405TCA (4618C>T 4620T>A), CAA410AAT (4633C>A 4635A>T), GCC411GCA (4638C>A), CCT412CCA (4641T>A), GCT413CAA (4642G>C 4643C>A 4644T>A), TTC415TTT (4650C>T), AGG417AGA (4656G>A), CAC418AAA (4657C>A 4659C>A), GAC420TAT (4663G>T 4665C>T), GAA421AAA (4666G>A), AGT422ATA (4670G>T 4671T>A), CTC423TTT (4672C>T 4674C>T), AGC424del (4675_4679delAGCAA), AAC425--T (4675_4679delAGCAA 4680C>T), ATG426CTG (4681A>C), TAT427AAA (4684T>A 4686T>A), CCA428AAG (4687C>A 4688C>A 4689A>G), CAG429AAA (4690C>A 4692G>A), TGT431TTG (4697G>T 4698T>G), GCT432ATA (4699G>A 4700C>T 4701T>A), TAT434TAC (4707T>C), GTT435ATA (4708G>A 4710T>A), ATC438ATT (4719C>T), ATC439TTG (4720A>T 4722C>G), GTC440GTA (4725C>A), TTC441TGC (4727T>G), AGC442TCA (4729A>T 4730G>C 4731C>A), AAA443ACA (4733A>C), ACT444ACA (4737T>A), GAA445TAT (4738G>T 4740A>T), GAA446CAG (4741G>C 4743A>G), TTA449CTT (4750T>C 4752A>T), GGA450ACA (4753G>A 4754G>C), GTC452CTT (4759G>C 4761C>T), ATT454GAA (4765A>G 4766T>A 4767T>A), GTT455TTT (4768G>T), CTA456TCA (4771C>T 4772T>C), AAC457AAA (4776C>A), AGA458CTC (4777A>C 4778G>T 4779A>C), TGC459TGT (4782C>T), AAA460GTC (4783A>G 4784A>T 4785A>C), GCC461AAT (4786G>A 4787C>A 4788C>T), CTA462AAT (4789C>A 4790T>A 4791A>T), GGA463GAC (4793G>A 4794A>C), GTG465GTA (4800G>A), CTT466TTA (4801C>T 4803T>A), AGC467AGT (4806C>T), AAA468CAG (4807A>C 4809A>G), AAG469AAA (4812G>A), GCC471GCA (4818C>A), CAG472GAA (4819C>G 4821G>A), TTG473ATA (4822T>A 4824G>A), TGC474AAT (4825T>A 4826G>A 4827C>T), ACA476AAT (4832C>A 4833A>T), ACT477AAA (4835C>A 4836T>A), ATC478ATA (4839C>A), AAT479GAA (4840A>G 4842T>A), GGT482GGA (4851T>A), CTA483ATG (4852C>A 4854A>G), GTA484ATT (4855G>A 4857A>T), ATA485ATT (4860A>T), GAA486TCA (4861G>T 4862A>C), AGA487AAA (4865G>A), GGA488AAT (4867G>A 4868G>A 4869A>T), AAT489GGA (4870A>G 4871A>G 4872T>A), CTC490ATA (4873C>A 4875C>A), AAA491GAA (4876A>G), GTC492TTA (4879G>T 4881C>A), CAG493CAA (4884G>A), AGT494ACT (4886G>C), CAT495CAC (4890T>C), ATC496ATA (4893C>A), GGA497TCC (4894G>T 4895G>C 4896A>C), CTA498AAA (4897C>A 4898T>A), CAC499AAG (4900C>A 4902C>G), TTA500ATA (4903T>A), GTT501TTA (4906G>T 4908T>A), GCT502GAG (4910C>A 4911T>G), CCA504CCT (4917A>T), GAT505GAC (4920T>C), CAA506AAG (4921C>A 4923A>G), CTT507TTA (4924C>T 4926T>A), AGT508ATA (4928G>T 4929T>A), GAT509ACC (4930G>A 4931A>C 4932T>C), AGG510AAA (4934G>A 4935G>A), AAT511CAA (4936A>C 4938T>A), GCC512GAA (4940C>A 4941C>A), TTA513ATA (4942T>A), TTA517CTT (4954T>C 4956A>T), GGC518GGA (4959C>A), CTC519TGC (4960C>T 4961T>G), CTA520TTA (4963C>T), AAT521AAC (4968T>C), ATC523GCA (4972A>G 4973T>C 4974C>A), TCC524GGA (4975T>G 4976C>G 4977C>A), GCT525GAA (4979C>A 4980T>A), TAT526TTC (4982A>T 4983T>C), TTT527ATA (4984T>A 4986T>A), AAA529GAA (4990A>G), ATA530TTA (4993A>T), AAC532AAG (5001C>G), CTT533AAG (5002C>A 5003T>A 5004T>G), AGG534AGA (5007G>A), TCA535AAT (5008T>A 5009C>A 5010A>T), CCT536CTT (5012C>T), TTA537CTT (5014T>C 5016A>T), CAG538CAA (5019G>A), GTT539del (5020_5022delGTT), CTT541CTC (5028T>C), AAG542ATC (5030A>T 5031G>C), AAA543AGA (5033A>G), GAG544AAA (5035G>A 5037G>A), ATA545TCA (5038A>T 5039T>C), ACT546AAC (5042C>A 5043T>C), TGG547CGA (5044T>C 5046G>A), TCT548TTT (5048C>T), TGG549TGA (5052G>A), ACT550AGT (5054C>G), GAG551TTT (5056G>T 5057A>T 5058G>T), AAA552TTC (5059A>T 5060A>T 5061A>C)

##### Proteins

| | Begin | End | Coverage | Score | Concordance | Matches | Identities | I/D/M/F* | Stop Codons |
|---|---|---|---|---|---|---|---|---|---|
| hypothetical protein (NP_043933.1) | 277 | 555 | 39.2% | 916 | 47.2% | 276 (99.3%) | 146 (52.5%) | 0/2/1/1 | 1 |

| | Begin | End | Coverage | Score | Concordance | Matches | Identities | I/D/M/F* | Stop Codons |
|---|---|---|---|---|---|---|---|---|---|
| NT | 4232 | 5069 | 10.6% | 242 | 14.8% | 830 (99.0%) | 482 (57.5%) | 0/8 | |

| | |
|---|---|
| Protein mutations: | K277I (4235A>T), K280R (4244A>G 4245G>A), C281V (4246T>G 4247G>T), K282Q (4249A>C), Q285F (4258C>T 4259A>T 4260G>T), S287N (4264T>A 4265C>A 4266A>C), P288D (4267C>G 4268C>A), Q289E (4270C>G), R291K (4276C>A 4277G>A), E292N (4279G>A 4281A>T), K295R (4289A>G), T296I (4292C>T), E299H (4300G>C 4302A>T), L304M (4315C>A 4317T>G), G305K (4318G>A 4319G>A 4320T>G), I306L (4321A>T 4323C>A), P309A (4330C>G 4332T>A), K311R (4337A>G 4338G>A), I327Q (4384A>C 4385T>A 4386C>A), K340E (4423A>G), D343K (4432G>A 4434C>A), H344N (4435C>A 4437T>C), T345C (4438A>T 4439C>G 4440A>T), K346I (4442A>T 4443G>T), G347F (4444G>T 4445G>T 4446A>T), L351F (4456C>T 4458A>T), L352I (4459C>A 4461T>C), E356T (4471G>A 4472A>C 4473G>T), Q357N (4474C>A 4476A>T), L359I (4480C>A 4482T>A), Q360N (4483C>A 4485A>T), R361L (4486A>C 4487G>T), I362V (4489A>G), G363K (4492G>A 4493G>A), G364V (4496G>T 4497T>A), F367Y (4505T>A 4506T>C), Y368F (4508A>T 4509C>T), S370K (4513T>A 4514C>A 4515C>A), V380I (4543G>A), A383S (4552G>T 4554T>A), P384E (4555C>G 4556C>A 4557A>G), T386S (4561A>T), Q388P (4568A>C 4569G>T), C394T (4585T>A 4586G>C), Q396N (4591C>A 4593A>T), P405S (4618C>T 4620T>A), Q410N (4633C>A 4635A>T), A413Q (4642G>C 4643C>A 4644T>A), H418K (4657C>A 4659C>A), D420Y (4663G>T 4665C>T), E421K (4666G>A), S422I (4670G>T 4671T>A), L423F (4672C>T 4674C>T), S424del (4675_4679delAGCAA), M426L (4681A>C), Y427K (4684T>A 4686T>A), P428K (4687C>A 4688C>A 4689A>G), Q429K (4690C>A 4692G>A), C431L (4697G>T 4698T>G), A432I (4699G>A 4700C>T 4701T>A), V435I (4708G>A 4710T>A), I439L (4720A>T 4722C>G), F441C (4727T>G), K443T (4733A>C), E445Y (4738G>T 4740A>T), E446Q (4741G>C 4743A>G), G450T (4753G>A 4754G>C), V452L (4759G>C 4761C>T), I454E (4765A>G 4766T>A 4767T>A), V455F (4768G>T), L456S (4771C>T 4772T>C), N457K (4776C>A), R458L (4777A>C 4778G>T 4779A>C), K460V (4783A>G 4784A>T 4785A>C), A461N (4786G>A 4787C>A 4788C>T), L462N (4789C>A 4790T>A 4791A>T), G463D (4793G>A 4794A>C), K468Q (4807A>C 4809A>G), Q472E (4819C>G 4821G>A), L473I (4822T>A 4824G>A), C474N (4825T>A 4826G>A 4827C>T), T476N (4832C>A 4833A>T), T477K (4835C>A 4836T>A), N479E (4840A>G 4842T>A), L483M (4852C>A 4854A>G), V484I (4855G>A 4857A>T), E486S (4861G>T 4862A>C), R487K (4865G>A), G488N (4867G>A 4868G>A 4869A>T), N489G (4870A>G 4871A>G 4872T>A), L490I (4873C>A 4875C>A), K491E (4876A>G), V492L (4879G>T 4881C>A), S494T (4886G>C), G497S (4894G>T 4895G>C 4896A>C), L498K (4897C>A 4898T>A), H499K (4900C>A 4902C>G), L500I (4903T>A), V501L (4906G>T 4908T>A), A502E (4910C>A 4911T>G), Q506K (4921C>A 4923A>G), S508I (4928G>T 4929T>A), D509T (4930G>A 4931A>C 4932T>C), R510K (4934G>A 4935G>A), N511Q (4936A>C 4938T>A), A512E (4940C>A 4941C>A), L513I (4942T>A), L519C (4960C>T 4961T>G), I523A (4972A>G 4973T>C 4974C>A), S524G (4975T>G 4976C>G 4977C>A), A525E (4979C>A 4980T>A), Y526F (4982A>T 4983T>C), F527I (4984T>A 4986T>A), K529E (4990A>G), I530L (4993A>T), N532K (5001C>G), L533K (5002C>A 5003T>A 5004T>G), S535N (5008T>A 5009C>A 5010A>T), P536L (5012C>T), V539del (5020_5022delGTT), K542I (5030A>T 5031G>C), K543R (5033A>G), E544K (5035G>A 5037G>A), I545S (5038A>T 5039T>C), T546N (5042C>A 5043T>C), W547R (5044T>C 5046G>A), S548F (5048C>T), W549* (5052G>A), T550S (5054C>G), E551F (5056G>T 5057A>T 5058G>T), K552F (5059A>T 5060A>T 5061A>C) |
| Codon mutations: | AAC276.AT (4233C>T), AAA277ATA (4235A>T), ATT278ATC (4239T>C), ATT279ATA (4242T>A), AAG280AGA (4244A>G 4245G>A), TGT281GTT (4246T>G 4247G>T), AAA282CAA (4249A>C), CCA283CCC (4254A>C), CAG285TTT (4258C>T 4259A>T 4260G>T), TCA287AAC (4264T>A 4265C>A 4266A>C), CCT288GAT (4267C>G 4268C>A), CAA289GAA (4270C>G), GAC290GAT (4275C>T), CGA291AAA (4276C>A 4277G>A), GAA292AAT (4279G>A 4281A>T), GAG293GAA (4284G>A), TTT294TTC (4287T>C), AAA295AGA (4289A>G), ACT296ATT (4292C>T), ATC298ATT (4299C>T), GAA299CAT (4300G>C 4302A>T), CTA301CTT (4308A>T), AAG303AAA (4314G>A), CTT304ATG (4315C>A 4317T>G), GGT305AAG (4318G>A 4319G>A 4320T>G), ATC306TTA (4321A>T 4323C>A), CCT309GCA (4330C>G 4332T>A), AAG311AGA (4337A>G 4338G>A), CAT314CAC (4347T>C), TCC315AGC (4348T>A 4349C>G), TCA316AGT (4351T>A 4352C>G 4353A>T), CCA317CCC (4356A>C), GCC318GCG (4359C>G), AGG322AGA (4371G>A), AAT323AAC (4374T>C), GCC325GCG (4380C>G), GAG326GAA (4383G>A), ATC327CAA (4384A>C 4385T>A 4386C>A), GGC330GGT (4395C>T), AAA331AAG (4398A>G), GCA332GCT (4401A>T), AGA333CGA (4402A>C), GTA335GTT (4410A>T), ATT336ATA (4413T>A), AAG340GAG (4423A>G), TTA341CTT (4426T>C 4428A>T), GAC343AAA (4432G>A 4434C>A), CAT344AAC (4435C>A 4437T>C), ACA345TGT (4438A>T 4439C>G 4440A>T), AAG346ATT (4442A>T 4443G>T), GGA347TTT (4444G>T 4445G>T 4446A>T), GAT348GAC (4449T>C), GGC349GGA (4452C>A), CTA351TTT (4456C>T 4458A>T), CTT352ATC (4459C>A 4461T>C), AAC354AAT (4467C>T), AAG355AAA (4470G>A), GAG356ACT (4471G>A 4472A>C 4473G>T), CAA357AAT (4474C>A 4476A>T), CTG358CTT (4479G>T), CTT359ATA (4480C>A 4482T>A), CAA360AAT (4483C>A 4485A>T), AGA361CTA (4486A>C 4487G>T), ATC362GTC (4489A>G), GGA363AAA (4492G>A 4493G>A), GGT364GTA (4496G>T 4497T>A), AAG365AAA (4500G>A), TTT367TAC (4505T>A 4506T>C), TAC368TTT (4508A>T 4509C>T), TCT369TCC (4512T>C), TCC370AAA (4513T>A 4514C>A 4515C>A), GAC372GAT (4521C>T), TCT375AGT (4528T>A 4529C>G), GTA380ATA (4543G>A), CGC381AGA (4546C>A 4548C>A), CTT382CTC (4551T>C), GCT383TCA (4552G>T 4554T>A), CCA384GAG (4555C>G 4556C>A 4557A>G), ACA386TCA (4561A>T), CAG388CCT (4568A>C 4569G>T), ACC390ACA (4575C>A), GCT391GCA (4578T>A), TGT394ACT (4585T>A 4586G>C), CCC395CCT (4590C>T), CAA396AAT (4591C>A 4593A>T), CAC398CAT (4599C>T), TAC399TAT (4602C>T), GTC403GTA (4614C>A), CCT405TCA (4618C>T 4620T>A), CAA410AAT (4633C>A 4635A>T), GCC411GCA (4638C>A), CCT412CCA (4641T>A), GCT413CAA (4642G>C 4643C>A 4644T>A), TTC415TTT (4650C>T), AGG417AGA (4656G>A), CAC418AAA (4657C>A 4659C>A), GAC420TAT (4663G>T 4665C>T), GAA421AAA (4666G>A), AGT422ATA (4670G>T 4671T>A), CTC423TTT (4672C>T 4674C>T), AGC424del (4675_4679delAGCAA), AAC425--T (4675_4679delAGCAA 4680C>T), ATG426CTG (4681A>C), TAT427AAA (4684T>A 4686T>A), CCA428AAG (4687C>A 4688C>A 4689A>G), CAG429AAA (4690C>A 4692G>A), TGT431TTG (4697G>T 4698T>G), GCT432ATA (4699G>A 4700C>T 4701T>A), TAT434TAC (4707T>C), GTT435ATA (4708G>A 4710T>A), ATC438ATT (4719C>T), ATC439TTG (4720A>T 4722C>G), GTC440GTA (4725C>A), TTC441TGC (4727T>G), AGC442TCA (4729A>T 4730G>C 4731C>A), AAA443ACA (4733A>C), ACT444ACA (4737T>A), GAA445TAT (4738G>T 4740A>T), GAA446CAG (4741G>C 4743A>G), TTA449CTT (4750T>C 4752A>T), GGA450ACA (4753G>A 4754G>C), GTC452CTT (4759G>C 4761C>T), ATT454GAA (4765A>G 4766T>A 4767T>A), GTT455TTT (4768G>T), CTA456TCA (4771C>T 4772T>C), AAC457AAA (4776C>A), AGA458CTC (4777A>C 4778G>T 4779A>C), TGC459TGT (4782C>T), AAA460GTC (4783A>G 4784A>T 4785A>C), GCC461AAT (4786G>A 4787C>A 4788C>T), CTA462AAT (4789C>A 4790T>A 4791A>T), GGA463GAC (4793G>A 4794A>C), GTG465GTA (4800G>A), CTT466TTA (4801C>T 4803T>A), AGC467AGT (4806C>T), AAA468CAG (4807A>C 4809A>G), AAG469AAA (4812G>A), GCC471GCA (4818C>A), CAG472GAA (4819C>G 4821G>A), TTG473ATA (4822T>A 4824G>A), TGC474AAT (4825T>A 4826G>A 4827C>T), ACA476AAT (4832C>A 4833A>T), ACT477AAA (4835C>A 4836T>A), ATC478ATA (4839C>A), AAT479GAA (4840A>G 4842T>A), GGT482GGA (4851T>A), CTA483ATG (4852C>A 4854A>G), GTA484ATT (4855G>A 4857A>T), ATA485ATT (4860A>T), GAA486TCA (4861G>T 4862A>C), AGA487AAA (4865G>A), GGA488AAT (4867G>A 4868G>A 4869A>T), AAT489GGA (4870A>G 4871A>G 4872T>A), CTC490ATA (4873C>A 4875C>A), AAA491GAA (4876A>G), GTC492TTA (4879G>T 4881C>A), CAG493CAA (4884G>A), AGT494ACT (4886G>C), CAT495CAC (4890T>C), ATC496ATA (4893C>A), GGA497TCC (4894G>T 4895G>C 4896A>C), CTA498AAA (4897C>A 4898T>A), CAC499AAG (4900C>A 4902C>G), TTA500ATA (4903T>A), GTT501TTA (4906G>T 4908T>A), GCT502GAG (4910C>A 4911T>G), CCA504CCT (4917A>T), GAT505GAC (4920T>C), CAA506AAG (4921C>A 4923A>G), CTT507TTA (4924C>T 4926T>A), AGT508ATA (4928G>T 4929T>A), GAT509ACC (4930G>A 4931A>C 4932T>C), AGG510AAA (4934G>A 4935G>A), AAT511CAA (4936A>C 4938T>A), GCC512GAA (4940C>A 4941C>A), TTA513ATA (4942T>A), TTA517CTT (4954T>C 4956A>T), GGC518GGA (4959C>A), CTC519TGC (4960C>T 4961T>G), CTA520TTA (4963C>T), AAT521AAC (4968T>C), ATC523GCA (4972A>G 4973T>C 4974C>A), TCC524GGA (4975T>G 4976C>G 4977C>A), GCT525GAA (4979C>A 4980T>A), TAT526TTC (4982A>T 4983T>C), TTT527ATA (4984T>A 4986T>A), AAA529GAA (4990A>G), ATA530TTA (4993A>T), AAC532AAG (5001C>G), CTT533AAG (5002C>A 5003T>A 5004T>G), AGG534AGA (5007G>A), TCA535AAT (5008T>A 5009C>A 5010A>T), CCT536CTT (5012C>T), TTA537CTT (5014T>C 5016A>T), CAG538CAA (5019G>A), GTT539del (5020_5022delGTT), CTT541CTC (5028T>C), AAG542ATC (5030A>T 5031G>C), AAA543AGA (5033A>G), GAG544AAA (5035G>A 5037G>A), ATA545TCA (5038A>T 5039T>C), ACT546AAC (5042C>A 5043T>C), TGG547CGA (5044T>C 5046G>A), TCT548TTT (5048C>T), TGG549TGA (5052G>A), ACT550AGT (5054C>G), GAG551TTT (5056G>T 5057A>T 5058G>T), AAA552TTC (5059A>T 5060A>T 5061A>C) |

*: Inserts / Deletes / Misaligned / Frameshifts

#### Analysis details

This analysis was performed with panviral2.64

### NGS Details (UN18_val): Badnavirus maculapiperis

#### Assembly

| | |
|---|---|
| **Coverage Length** | 234 (1 contig(s)) |
| **Depth Of Coverage** | 13.7 |
| **Number Of Reads** | 30 |
| **Reads Per Million** | 0.56 rpm (after QC) |
| **Ambiguities** | 0 |
| **Assembly Method** | de novo + reference guided assembly |
| **Consensus Caller** | Bcf Tools |

#### Coverage Map

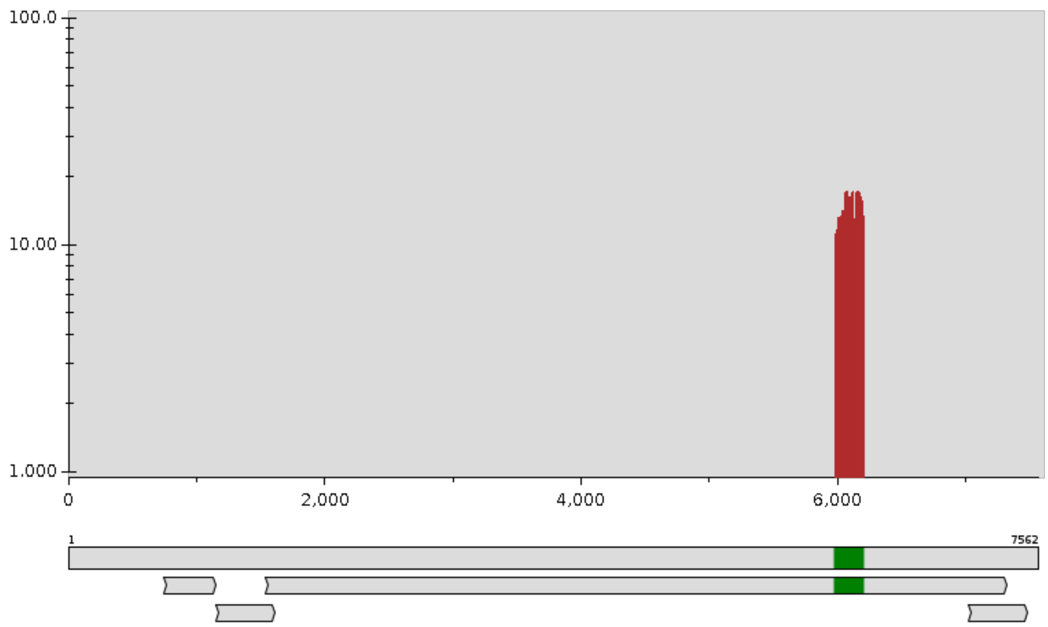

#### Assignment

| | |
|---|---|
| **Type** | Badnavirus maculapiperis (Taxonomy ID: 3048272) |
| **Reference Genome** | NC_022365.1 |
| **NT Identity (%)** | 57.8723 |
| **AA Identity (%)** | 53.1646 |
| **Number Of Stop Codons** | 0 |
| **Number Of CDS** | 4 |

#### Alignment

| | |
|---|---|
| **Alignment Score** | 57.0 (NT) + 188.0 (AA) = 245.0 |
| **Concordance (%)** | 29.5852 |

| Alignment Method | Global, seeded, nucleotide + amino acids (AGA) |
|---|---|

#### Genome Region

Sequence starts at position 5977 and ends at position 6210 relative to NC_022365.1 reference sequence.

#### Alignment Detailed Statistics

| | Begin | End | Coverage | Score | Concordance | Matches | Identities | I/D/M/F* | Stop Codons |
|---|---|---|---|---|---|---|---|---|---|
| **NT** | **5977** | **6210** | **3.1%** | **57** | **12.6%** | **232 (97.9%)** | **136 (57.4%)** | **3/2** | |
| Mutations: | 5989T>A, 5991A>C, 5994A>G, 5995T>C, 5997C>T, 6000C>T, 6003A>G, 6005A>G, 6009T>A, 6014T>A, 6018C>T, 6022G>A, 6024A>T, 6025G>A, 6026C>G, 6031G>A, 6032A>C, 6033C>A, 6034C>A, 6035C>A, 6036T>G, 6039A>G, 6040T>G, 6041C>A, 6043A>G, 6045A>C, 6046G>C, 6049T>A, 6050G>A, 6051G>A, 6055G>A, 6057C>A, 6060C>T, 6063C>G, 6066C>T, 6068C>A, 6070A>T, 6071A>C, 6075G>C, 6077T>A, 6078A>C, 6081C>T, 6084A>G, 6086G>A, 6087G>C, 6090T>A, 6093C>A, 6099A>C, 6105G>C, 6108T>C, 6110A>C, 6111A>T, 6121G>T, 6123T>A, 6124G>A, 6125T>G, 6126G>T, 6129C>T, 6133A>G, 6135A>G, 6136A>C, 6137A>T, 6138A>G, 6142G>A, 6144C>T, 6145A>C, 6147C>T, 6148T>G, 6149G>T, 6153C>T, 6156A>G, 6158G>A, 6159T>A, 6159_6160insCAC, 6160A>C, 6162G>T, 6163G>A, 6164A>G, 6165A>G, 6166G>A, 6168T>G, 6171T>C, 6174A>T, 6175G>T, 6176C>T, 6177T>G, 6180C>T, 6182A>T, 6183C>T, 6184_6185delAT, 6199G>A, 6201G>C, 6203T>A, 6205T>A, 6206C>G, 6207A>C, 6208G>A, 6210A>G | | | | | | | | |

##### CDS

| | Begin | End | Coverage | Score | Concordance | Matches | Identities | I/D/M/F* | Stop Codons |
|---|---|---|---|---|---|---|---|---|---|
| P878_gp3 | 1481 | 1558 | 4.0% | 188 | 33.4% | 78 (98.7%) | 42 (53.2%) | 1/0/1/1 | 0 |
| Protein mutations: | S1485T (5989T>A 5991A>C), F1487L (5995T>C 5997C>T), K1490R (6005A>G), F1493Y (6014T>A), V1496I (6022G>A 6024A>T), A1497R (6025G>A 6026C>G), D1499T (6031G>A 6032A>C 6033C>A), P1500K (6034C>A 6035C>A 6036T>G), S1502D (6040T>G 6041C>A), I1503V (6043A>G 6045A>C), E1504Q (6046G>C), W1505K (6049T>A 6050G>A 6051G>A), A1507T (6055G>A 6057C>A), S1509R (6063C>G), P1511H (6068C>A), N1512S (6070A>T 6071A>C), L1514H (6077T>A 6078A>C), W1517Y (6086G>A 6087G>C), K1525T (6110A>C 6111A>T), A1529S (6121G>T 6123T>A), V1530S (6124G>A 6125T>G 6126G>T), R1533G (6133A>G 6135A>G), K1534L (6136A>C 6137A>T 6138A>G), D1536N (6142G>A 6144C>T), N1537H (6145A>C 6147C>T), C1538V (6148T>G 6149G>T), G1541E (6158G>A 6159T>A), G1541_M1542insH (6159_6160insCAC), M1542L (6160A>C 6162G>T), E1543R (6163G>A 6164A>G 6165A>G), D1544K (6166G>A 6168T>G), A1547L (6175G>T 6176C>T 6177T>G), Y1549F (6182A>T 6183C>T), V1555I (6199G>A 6201G>C), F1556Y (6203T>A), E1558K (6208G>A 6210A>G) | | | | | | | | |
| Codon mutations: | TCA1485ACC (5989T>A 5991A>C), AAA1486AAG (5994A>G), TTC1487CTT (5995T>C 5997C>T), GAC1488GAT (6000C>T), CTA1489CTG (6003A>G), AAA1490AGA (6005A>G), TCT1491TCA (6009T>A), TTT1493TAT (6014T>A), CAC1494CAT (6018C>T), GTA1496ATT (6022G>A 6024A>T), GCG1497AGG (6025G>A 6026C>G), GAC1499ACA (6031G>A 6032A>C 6033C>A), CCT1500AAG (6034C>A 6035C>A 6036T>G), GAA1501GAG (6039A>G), TCT1502GAT (6040T>G 6041C>A), ATA1503GTC (6043A>G 6045A>C), GAG1504CAG (6046G>C), TGG1505AAA (6049T>A 6050G>A 6051G>A), GCC1507ACA (6055G>A 6057C>A), TTC1508TTT (6060C>T), AGC1509AGG (6063C>G), ACC1510ACT (6066C>T), CCT1511CAT (6068C>A), AAT1512TCT (6070A>T 6071A>C), GGG1513GGC (6075G>C), CTA1514CAC (6077T>A 6078A>C), TAC1515TAT (6081C>T), GAA1516GAG (6084A>G), TGG1517TAC (6086G>A 6087G>C), CTT1518CTA (6090T>A), GTC1519GTA (6093C>A), CCA1521CCC (6099A>C), GGG1523GGC (6105G>C), CTT1524CTC (6108T>C), AAA1525ACT (6110A>C 6111A>T), GCT1529TCA (6121G>T 6123T>A), GTG1530AGT (6124G>A 6125T>G 6126G>T), TTC1531TTT (6129C>T), AGA1533GGG (6133A>G 6135A>G), AAA1534CTG (6136A>C 6137A>T 6138A>G), GAC1536AAT (6142G>A 6144C>T), AAC1537CAT (6145A>C 6147C>T), TGT1538GTT (6148T>G 6149G>T), TTC1539TTT (6153C>T), AAA1540AAG (6156A>G), GGT1541GAA (6158G>A 6159T>A), GGT1541_ATG1542insCAC (6159_6160insCAC), ATG1542CTT (6160A>C 6162G>T), GAA1543AGG (6163G>A 6164A>G 6165A>G), GAT1544AAG (6166G>A 6168T>G), TTT1545TTC (6171T>C), ATA1546ATT (6174A>T), GCT1547TTG (6175G>T 6176C>T 6177T>G), GTC1548GTT (6180C>T), TAC1549TTT (6182A>T 6183C>T), ATT1550--T (6184_6185delAT), GTG1555ATC (6199G>A 6201G>C), TTC1556TAC (6203T>A), TCA1557AGC (6205T>A 6206C>G 6207A>C), GAA1558AAG (6208G>A 6210A>G) | | | | | | | | |

##### Proteins

| | Begin | End | Coverage | Score | Concordance | Matches | Identities | I/D/M/F* | Stop Codons |
|---|---|---|---|---|---|---|---|---|---|
| ORF3 (YP_008567619.1) | 1481 | 1558 | 4.0% | 188 | 33.4% | 78 (98.7%) | 42 (53.2%) | 1/0/1/1 | 0 |
| Protein mutations: | S1485T (5989T>A 5991A>C), F1487L (5995T>C 5997C>T), K1490R (6005A>G), F1493Y (6014T>A), V1496I (6022G>A 6024A>T), A1497R (6025G>A 6026C>G), D1499T (6031G>A 6032A>C 6033C>A), P1500K (6034C>A 6035C>A 6036T>G), S1502D (6040T>G 6041C>A), I1503V (6043A>G 6045A>C), E1504Q (6046G>C), W1505K (6049T>A 6050G>A 6051G>A), A1507T (6055G>A 6057C>A), S1509R (6063C>G), P1511H (6068C>A), N1512S (6070A>T 6071A>C), L1514H (6077T>A 6078A>C), W1517Y (6086G>A 6087G>C), K1525T (6110A>C 6111A>T), A1529S (6121G>T 6123T>A), V1530S (6124G>A 6125T>G 6126G>T), R1533G (6133A>G 6135A>G), K1534L (6136A>C 6137A>T 6138A>G), D1536N (6142G>A 6144C>T), N1537H (6145A>C 6147C>T), C1538V (6148T>G 6149G>T), G1541E (6158G>A 6159T>A), G1541_M1542insH (6159_6160insCAC), M1542L (6160A>C 6162G>T), E1543R (6163G>A 6164A>G 6165A>G), D1544K (6166G>A 6168T>G), A1547L (6175G>T 6176C>T 6177T>G), Y1549F (6182A>T 6183C>T), V1555I (6199G>A 6201G>C), F1556Y (6203T>A), E1558K (6208G>A 6210A>G) | | | | | | | | |
| Codon mutations: | TCA1485ACC (5989T>A 5991A>C), AAA1486AAG (5994A>G), TTC1487CTT (5995T>C 5997C>T), GAC1488GAT (6000C>T), CTA1489CTG (6003A>G), AAA1490AGA (6005A>G), TCT1491TCA (6009T>A), TTT1493TAT (6014T>A), CAC1494CAT (6018C>T), GTA1496ATT (6022G>A 6024A>T), GCG1497AGG (6025G>A 6026C>G), GAC1499ACA (6031G>A 6032A>C 6033C>A), CCT1500AAG (6034C>A 6035C>A 6036T>G), GAA1501GAG (6039A>G), TCT1502GAT (6040T>G 6041C>A), ATA1503GTC (6043A>G 6045A>C), GAG1504CAG (6046G>C), TGG1505AAA (6049T>A 6050G>A 6051G>A), GCC1507ACA (6055G>A 6057C>A), TTC1508TTT (6060C>T), AGC1509AGG (6063C>G), ACC1510ACT (6066C>T), CCT1511CAT (6068C>A), AAT1512TCT (6070A>T 6071A>C), GGG1513GGC (6075G>C), CTA1514CAC (6077T>A 6078A>C), TAC1515TAT (6081C>T), GAA1516GAG (6084A>G), TGG1517TAC (6086G>A 6087G>C), CTT1518CTA (6090T>A), GTC1519GTA (6093C>A), CCA1521CCC (6099A>C), GGG1523GGC (6105G>C), CTT1524CTC (6108T>C), AAA1525ACT (6110A>C 6111A>T), GCT1529TCA (6121G>T 6123T>A), GTG1530AGT (6124G>A 6125T>G 6126G>T), TTC1531TTT (6129C>T), AGA1533GGG (6133A>G 6135A>G), AAA1534CTG (6136A>C 6137A>T 6138A>G), GAC1536AAT (6142G>A 6144C>T), AAC1537CAT (6145A>C 6147C>T), TGT1538GTT (6148T>G 6149G>T), TTC1539TTT (6153C>T), AAA1540AAG (6156A>G), GGT1541GAA (6158G>A 6159T>A), GGT1541_ATG1542insCAC (6159_6160insCAC), ATG1542CTT (6160A>C 6162G>T), GAA1543AGG (6163G>A 6164A>G 6165A>G), GAT1544AAG (6166G>A 6168T>G), TTT1545TTC (6171T>C), ATA1546ATT (6174A>T), GCT1547TTG (6175G>T 6176C>T 6177T>G), GTC1548GTT (6180C>T), TAC1549TTT (6182A>T 6183C>T), ATT1550--T (6184_6185delAT), GTG1555ATC (6199G>A 6201G>C), TTC1556TAC (6203T>A), TCA1557AGC (6205T>A 6206C>G 6207A>C), GAA1558AAG (6208G>A 6210A>G) | | | | | | | | |

*: Inserts / Deletes / Misaligned / Frameshifts

#### Analysis details

This analysis was performed with panviral2.64

### NGS Details (UN18_val): Epiphyllum badnavirus 1

#### Assembly

| | |
|---|---|
| **Coverage Length** | 244 (1 contig(s)) |
| **Depth Of Coverage** | 12.9 |
| **Number Of Reads** | 30 |
| **Reads Per Million** | 0.56 rpm (after QC) |
| **Ambiguities** | 0 |
| **Assembly Method** | de novo + reference guided assembly |
| **Consensus Caller** | Bcf Tools |

#### Coverage Map

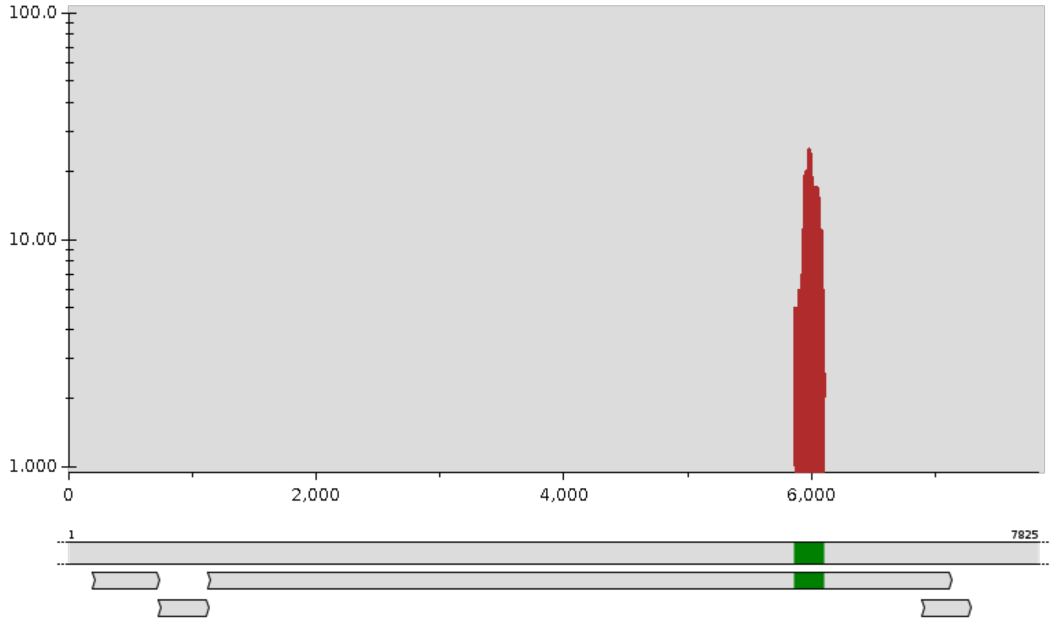

#### Assignment

| | |
|---|---|
| **Type** | Epiphyllum badnavirus 1 (Taxonomy ID: 2518008) |
| **Reference Genome** | NC_076247.1 |
| **NT Identity (%)** | 57.085 |
| **AA Identity (%)** | 49.3976 |
| **Number Of Stop Codons** | 0 |
| **Number Of CDS** | 4 |

#### Alignment

| | |
|---|---|
| **Alignment Score** | 64.0 (NT) + 291.0 (AA) = 355.0 |
| **Concordance (%)** | 32.5987 |

| **Alignment Method** | Global, seeded, nucleotide + amino acids (AGA) |
|---|---|

#### Genome Region

Sequence starts at position 5860 and ends at position 6103 relative to NC_076247.1 reference sequence.

#### Alignment Detailed Statistics

| | Begin | End | Coverage | Score | Concordance | Matches | Identities | I/D/M/F* | Stop Codons |
|---|---|---|---|---|---|---|---|---|---|
| **NT** | **5860** | **6103** | **3.1%** | **64** | **13.1%** | **244 (98.8%)** | **141 (57.1%)** | **3/0** | |
| Mutations: | 5865C>G, 5868A>C, 5870T>A, 5878A>T, 5880T>G, 5882T>A, 5883G>A, 5884A>G, 5886G>T, 5887G>C, 5888A>G, 5889C>G, 5893G>T, 5894A>C, 5896T>G, 5897C>A, 5898C>T, 5899A>G, 5904T>G, 5905T>A, 5906G>A, 5911G>A, 5916C>T, 5918T>G, 5922C>A, 5924C>G, 5925A>T, 5926G>T, 5931A>G, 5933T>A, 5934T>C, 5940A>G, 5942G>T, 5943G>C, 5944T>G, 5946A>G, 5949T>C, 5953C>T, 5955A>T, 5961A>G, 5962T>C, 5964G>T, 5966A>C, 5967G>T, 5981T>C, 5982G>T, 5985C>T, 5986C>A, 5987A>T, 5988A>G, 5989C>G, 5990G>A, 5991A>G, 5992A>T, 5993A>T, 5998G>A, 6000C>T, 6001A>C, 6002A>G, 6003C>A, 6004T>G, 6005G>T, 6006T>G, 6009C>T, 6011G>A, 6012A>G, 6012_6013insCCC, 6013G>T, 6016C>T, 6019T>G, 6020C>A, 6021A>C, 6022G>C, 6023G>A, 6024C>G, 6027T>C, 6030A>G, 6031G>A, 6032C>T, 6033A>T, 6036T>G, 6038A>T, 6039C>T, 6043G>A, 6048C>T, 6052C>T, 6057C>T, 6059T>A, 6060C>T, 6061A>T, 6062G>C, 6063T>C, 6066C>G, 6068C>G, 6072A>G, 6073G>A, 6074A>T, 6078C>T, 6084A>G, 6095A>G, 6097T>A, 6098C>T, 6100A>G, 6103C>T | | | | | | | | |
| **CDS** | | | | | | | | | |
| QKM20_gp3 | 1578 | 1659 | 4.1% | 291 | 48.1% | 82 (98.8%) | 41 (49.4%) | 1/0/0/0 | 0 |
| Protein mutations: | S1579R (5865C>G), F1581Y (5870T>A), I1584L (5878A>T 5880T>G), M1585K (5882T>A 5883G>A), M1586V (5884A>G 5886G>T), D1587R (5887G>C 5888A>G 5889C>G), E1589S (5893G>T 5894A>C), S1590D (5896T>G 5897C>A 5898C>T), I1591V (5899A>G), W1593K (5905T>A 5906G>A), A1595T (5911G>A), L1597R (5918T>G), P1599R (5924C>G 5925A>T), D1600Y (5926G>T), L1602H (5933T>A 5934T>C), W1605F (5942G>T 5943G>C), L1606V (5944T>G 5946A>G), P1609S (5953C>T 5955A>T), K1613T (5966A>C 5967G>T), V1618A (5981T>C 5982G>T), Q1620M (5986C>A 5987A>T 5988A>G), R1621E (5989C>G 5990G>A 5991A>G), K1622L (5992A>T 5993A>T), D1624N (5998G>A 6000C>T), N1625R (6001A>C 6002A>G 6003C>A), C1626V (6004T>G 6005G>T 6006T>G), R1628K (6011G>A 6012A>G), R1628_D1629insP (6012_6013insCCC), D1629Y (6013G>T), S1631D (6019T>G 6020C>A 6021A>C), G1632Q (6022G>C 6023G>A 6024C>G), A1635I (6031G>A 6032C>T 6033A>T), Y1637F (6038A>T 6039C>T), D1639N (6043G>A), F1644Y (6059T>A 6060C>T), N1646K (6066C>G), T1647S (6068C>G), D1649I (6073G>A 6074A>T), K1656R (6095A>G), S1657I (6097T>A 6098C>T), M1658V (6100A>G) | | | | | | | | |
| Codon mutations: | AGC1579AGG (5865C>G), GGA1580GGC (5868A>C), TTC1581TAC (5870T>A), ATT1584TTG (5878A>T 5880T>G), ATG1585AAA (5882T>A 5883G>A), ATG1586GTT (5884A>G 5886G>T), GAC1587CGG (5887G>C 5888A>G 5889C>G), GAA1589TCA (5893G>T 5894A>C), TCC1590GAT (5896T>G 5897C>A 5898C>T), ATC1591GTC (5899A>G), CCT1592CCG (5904T>G), TGG1593AAG (5905T>A 5906G>A), GCC1595ACC (5911G>A), TTC1596TTT (5916C>T), CTG1597CGG (5918T>G), ACC1598ACA (5922C>A), CCA1599CGT (5924C>G 5925A>T), GAT1600TAT (5926G>T), GGA1601GGG (5931A>G), CTT1602CAC (5933T>A 5934T>C), GAA1604GAG (5940A>G), TGG1605TTC (5942G>T 5943G>C), TTA1606GTG (5944T>G 5946A>G), GTT1607GTC (5949T>C), CCA1609TCT (5953C>T 5955A>T), GGA1611GGG (5961A>G), TTG1612CTT (5962T>C 5964G>T), AAG1613ACT (5966A>C 5967G>T), GTG1618GCT (5981T>C 5982G>T), TTC1619TTT (5985C>T), CAA1620ATG (5986C>A 5987A>T 5988A>G), CGA1621GAG (5989C>G 5990G>A 5991A>G), AAG1622TTG (5992A>T 5993A>T), GAC1624AAT (5998G>A 6000C>T), AAC1625CGA (6001A>C 6002A>G 6003C>A), TGT1626GTG (6004T>G 6005G>T 6006T>G), TTC1627TTT (6009C>T), AGA1628AAG (6011G>A 6012A>G), AGA1628_GAC1629insCCC (6012_6013insCCC), GAC1629TAC (6013G>T), CTG1630TTG (6016C>T), TCA1631GAC (6019T>G 6020C>A 6021A>C), GGC1632CAG (6022G>C 6023G>A 6024C>G), TTT1633TTC (6027T>C), GTA1634GTG (6030A>G), GCA1635ATT (6031G>A 6032C>T 6033A>T), GTT1636GTG (6036T>G), TAC1637TTT (6038A>T 6039C>T), GAT1639AAT (6043G>A), GAC1640GAT (6048C>T), CTG1642TTG (6052C>T), GTC1643GTT (6057C>T), TTC1644TAT (6059T>A 6060C>T), AGT1645TCC (6061A>T 6062G>C 6063T>C), AAC1646AAG (6066C>G), ACT1647AGT (6068C>G), GAA1648GAG (6072A>G), GAT1649ATT (6073G>A 6074A>T), GAC1650GAT (6078C>T), GCA1652GCG (6084A>G), AAG1656AGG (6095A>G), TCC1657ATC (6097T>A 6098C>T), ATG1658GTG (6100A>G), CTG1659T.. (6103C>T) | | | | | | | | |
| **Proteins** | | | | | | | | | |
| polyprotein (YP_010797894.1) | 1578 | 1659 | 4.1% | 291 | 48.1% | 82 (98.8%) | 41 (49.4%) | 1/0/0/0 | 0 |
| Protein mutations: | S1579R (5865C>G), F1581Y (5870T>A), I1584L (5878A>T 5880T>G), M1585K (5882T>A 5883G>A), M1586V (5884A>G 5886G>T), D1587R (5887G>C 5888A>G 5889C>G), E1589S (5893G>T 5894A>C), S1590D (5896T>G 5897C>A 5898C>T), I1591V (5899A>G), W1593K (5905T>A 5906G>A), A1595T (5911G>A), L1597R (5918T>G), P1599R (5924C>G 5925A>T), D1600Y (5926G>T), L1602H (5933T>A 5934T>C), W1605F (5942G>T 5943G>C), L1606V (5944T>G 5946A>G), P1609S (5953C>T 5955A>T), K1613T (5966A>C 5967G>T), V1618A (5981T>C 5982G>T), Q1620M (5986C>A 5987A>T 5988A>G), R1621E (5989C>G 5990G>A 5991A>G), K1622L (5992A>T 5993A>T), D1624N (5998G>A 6000C>T), N1625R (6001A>C 6002A>G 6003C>A), C1626V (6004T>G 6005G>T 6006T>G), R1628K (6011G>A 6012A>G), R1628_D1629insP (6012_6013insCCC), D1629Y (6013G>T), S1631D (6019T>G 6020C>A 6021A>C), G1632Q (6022G>C 6023G>A 6024C>G), A1635I (6031G>A 6032C>T 6033A>T), Y1637F (6038A>T 6039C>T), D1639N (6043G>A), F1644Y (6059T>A 6060C>T), N1646K (6066C>G), T1647S (6068C>G), D1649I (6073G>A 6074A>T), K1656R (6095A>G), S1657I (6097T>A 6098C>T), M1658V (6100A>G) | | | | | | | | |
| Codon mutations: | AGC1579AGG (5865C>G), GGA1580GGC (5868A>C), TTC1581TAC (5870T>A), ATT1584TTG (5878A>T 5880T>G), ATG1585AAA (5882T>A 5883G>A), ATG1586GTT (5884A>G 5886G>T), GAC1587CGG (5887G>C 5888A>G 5889C>G), GAA1589TCA (5893G>T 5894A>C), TCC1590GAT (5896T>G 5897C>A 5898C>T), ATC1591GTC (5899A>G), CCT1592CCG (5904T>G), TGG1593AAG (5905T>A 5906G>A), GCC1595ACC (5911G>A), TTC1596TTT (5916C>T), CTG1597CGG (5918T>G), ACC1598ACA (5922C>A), CCA1599CGT (5924C>G 5925A>T), GAT1600TAT (5926G>T), GGA1601GGG (5931A>G), CTT1602CAC (5933T>A 5934T>C), GAA1604GAG (5940A>G), TGG1605TTC (5942G>T 5943G>C), TTA1606GTG (5944T>G 5946A>G), GTT1607GTC (5949T>C), CCA1609TCT (5953C>T 5955A>T), GGA1611GGG (5961A>G), TTG1612CTT (5962T>C 5964G>T), AAG1613ACT (5966A>C 5967G>T), GTG1618GCT (5981T>C 5982G>T), TTC1619TTT (5985C>T), CAA1620ATG (5986C>A 5987A>T 5988A>G), CGA1621GAG (5989C>G 5990G>A 5991A>G), AAG1622TTG (5992A>T 5993A>T), GAC1624AAT (5998G>A 6000C>T), AAC1625CGA (6001A>C 6002A>G 6003C>A), TGT1626GTG (6004T>G 6005G>T 6006T>G), TTC1627TTT (6009C>T), AGA1628AAG (6011G>A 6012A>G), AGA1628_GAC1629insCCC (6012_6013insCCC), GAC1629TAC (6013G>T), CTG1630TTG (6016C>T), TCA1631GAC (6019T>G 6020C>A 6021A>C), GGC1632CAG (6022G>C 6023G>A 6024C>G), TTT1633TTC (6027T>C), GTA1634GTG (6030A>G), GCA1635ATT (6031G>A 6032C>T 6033A>T), GTT1636GTG (6036T>G), TAC1637TTT (6038A>T 6039C>T), GAT1639AAT (6043G>A), GAC1640GAT (6048C>T), CTG1642TTG (6052C>T), GTC1643GTT (6057C>T), TTC1644TAT (6059T>A 6060C>T), AGT1645TCC (6061A>T 6062G>C 6063T>C), AAC1646AAG (6066C>G), ACT1647AGT (6068C>G), GAA1648GAG (6072A>G), GAT1649ATT (6073G>A 6074A>T), GAC1650GAT (6078C>T), GCA1652GCG (6084A>G), AAG1656AGG (6095A>G), TCC1657ATC (6097T>A 6098C>T), ATG1658GTG (6100A>G), CTG1659T.. (6103C>T) | | | | | | | | |

*: Inserts / Deletes / Misaligned / Frameshifts

#### Analysis details

This analysis was performed with panviral2.64

### NGS Details (UN18_val): Badnavirus epsiloninflatheobromae

#### Assembly

| | |
|---|---|
| **Coverage Length** | 616 (2 contig(s)) |
| **Depth Of Coverage** | 3.9 |
| **Number Of Reads** | 23 |
| **Reads Per Million** | 0.43 rpm (after QC) |
| **Ambiguities** | 0 |
| **Assembly Method** | de novo + reference guided assembly |
| **Consensus Caller** | Bcf Tools |

#### Coverage Map

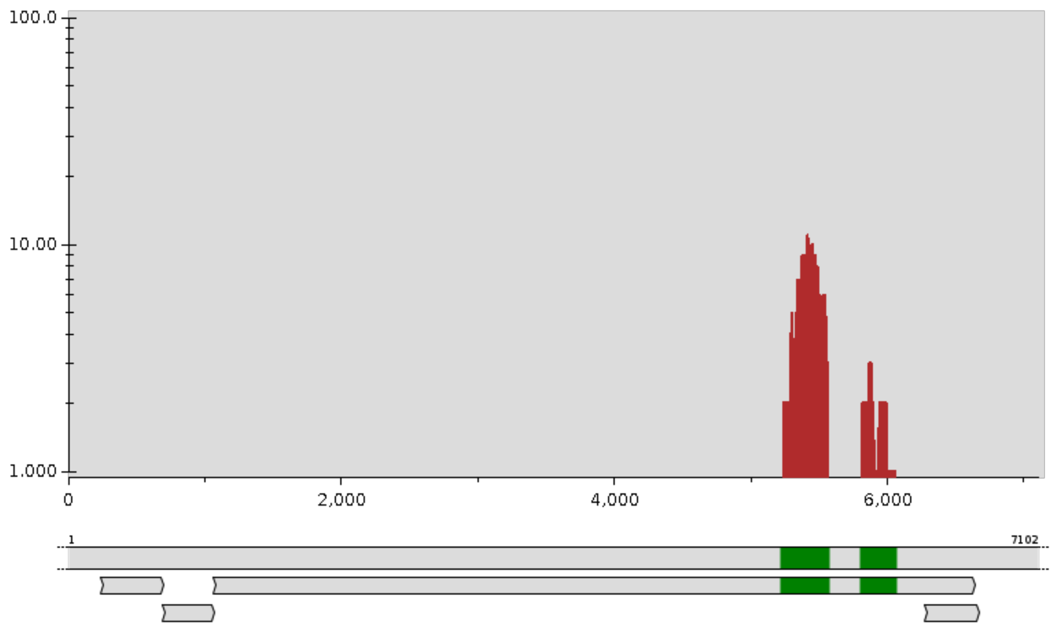

#### Assignment

| | |
|---|---|
| **Type** | Badnavirus epsiloninflatheobromae (Taxonomy ID: 3047711) |
| **Reference Genome** | NC_043535.1 |
| **NT Identity (%)** | 57.44 |
| **AA Identity (%)** | 48.5577 |
| **Number Of Stop Codons** | 0 |
| **Number Of CDS** | 4 |

#### Alignment

| | |
|---|---|
| **Alignment Score** | 177.0 (NT) + 779.0 (AA) = 956.0 |
| **Concordance (%)** | 34.8651 |

| Alignment Method | Global, seeded, nucleotide + amino acids (AGA) |
|---|---|

#### Genome Region

Sequence starts at position 5228 and ends at position 6068 relative to NC_043535.1 reference sequence.

#### Alignment Detailed Statistics

| | Begin | End | Coverage | Score | Concordance | Matches | Identities | I/D/M/F* | Stop Codons |
|---|---|---|---|---|---|---|---|---|---|
| **NT** | **5228** | **6068** | **8.7%** | **177** | **14.4%** | **616 (98.6%)** | **359 (57.4%)** | **9/0** | |
| Mutations: | 5228T>G, 5231A>G, 5233T>A, 5240G>A, 5243C>T, 5244G>A, 5245C>G, 5256G>C, 5257A>C, 5258G>C, 5259T>G, 5260C>A, 5261A>C, 5264C>T, 5267A>C, 5268T>A, 5269G>A, 5270G>A, 5273C>T, 5276G>C, 5280T>A, 5282G>A, 5283G>A, 5284T>C, 5285A>T, 5287C>A, 5288T>C, 5291A>T, 5294A>T, 5296T>C, 5303A>G, 5307C>T, 5312T>G, 5315G>T, 5318A>C, 5321T>C, 5325C>T, 5327C>A, 5329A>C, 5330A>C, 5333T>C, 5344T>C, 5345C>T, 5349C>T, 5350A>G, 5351A>C, 5353G>C, 5354A>G, 5355A>C, 5356A>T, 5361G>A, 5363T>A, 5364C>A, 5366T>G, 5367T>A, 5368G>T, 5369T>A, 5372_5373insCAT, 5373G>C, 5375A>T, 5376G>T, 5377G>A, 5378A>C, 5379A>C, 5380C>T, 5384A>T, 5385G>C, 5387G>A, 5390T>C, 5391A>G, 5393T>G, 5395C>T, 5396T>A, 5403A>T, 5405T>G, 5415A>G, 5420T>C, 5422T>A, 5429T>C, 5432A>C, 5433G>T, 5434A>T, 5439C>G, 5441G>A, 5444T>C, 5445A>G, 5446A>T, 5453T>C, 5454C>T, 5463T>G, 5465C>A, 5466C>T, 5468G>C, 5469G>C, 5471G>A, 5472A>G, 5473G>T, 5474A>C, 5476G>T, 5477C>A, 5478A>C, 5479A>G, 5481C>G, 5484C>A, 5488G>A, 5490T>C, 5492G>A, 5493G>T, 5494T>A, 5495G>C, 5496T>G, 5498A>C, 5500G>A, 5501C>G, 5503C>A, 5504A>G, 5505A>G, 5506C>A, 5507A>G, 5511A>T, 5512T>G, 5513G>C, 5514A>G, 5516A>G, 5517A>T, 5519A>T, 5521G>C, 5525G>A, 5526A>C, 5528A>T, 5529A>G, 5530G>A, 5531G>A, 5532A>G, 5534C>G, 5535G>C, 5537G>C, 5543A>G, 5547G>C, 5548C>A, 5549T>C, 5550G>T, 5552G>C, 5555C>T, 5557A>G, 5559G>C, 5564G>T, 5565A>G, 5568A>C, 5574C>A, 5576A>G, 5577C>G, 5804T>G, 5813G>A, 5814G>A, 5815A>G, 5816G>C, 5819C>A, 5824C>T, 5829G>T, 5830A>C, 5833C>A, 5834G>T, 5837C>T, 5838A>C, 5840C>A, 5846T>A, 5852T>A, 5859T>A, 5862A>C, 5864G>C, 5865G>C, 5878G>C, 5879A>T, 5880A>G, 5882C>A, 5884G>T, 5885T>A, 5888G>A, 5889T>G, 5890G>C, 5891G>A, 5893A>G, 5897T>A, 5898T>A, 5899T>A, 5900T>C, 5904A>T, 5905G>A, 5906G>C, 5907G>A, 5908A>G, 5909C>T, 5912A>T, 5913C>A, 5914G>A, 5921C>A, 5924G>A, 5927G>A, 5930A>T, 5936C>T, 5939T>C, 5940G>C, 5941C>A, 5948A>T, 5953T>A, 5956G>A, 5958C>G, 5959C>A, 5960_5961insAAGGGT, 5962T>A, 5965A>T, 5966A>G, 5967T>A, 5968C>G, 5969C>T, 5971C>G, 5972A>T, 5975C>A, 5980C>T, 5981T>C, 5984G>A, 5985A>G, 5988C>T, 5989A>T, 5990C>A, 5993G>T, 5994G>A, 5999G>A, 6000A>T, 6003A>G, 6005T>A, 6006T>C, 6011G>T, 6013A>G, 6014A>T, 6017T>C, 6020G>A, 6021C>T, 6027T>A, 6028A>T, 6034G>A, 6035T>C, 6039C>A, 6040C>G, 6041A>T, 6043C>A, 6044G>A, 6045G>A, 6047C>A, 6048A>G, 6049C>T, 6050T>A, 6051A>G, 6053C>A, 6056G>A, 6065T>C, 6066C>G, 6068G>A | | | | | | | | |

##### CDS

| | Begin | End | Coverage | Score | Concordance | Matches | Identities | I/D/M/F* | Stop Codons |
|---|---|---|---|---|---|---|---|---|---|
| FLA70_gp3 | 1390 | 1669 | 11.0% | 779 | 51.0% | 205 (98.6%) | 101 (48.6%) | 3/0/0/0 | 0 |
| Protein mutations: | F1391Y (5233T>A), A1395R (5244G>A 5245C>G), E1399P (5256G>C 5257A>C 5258G>C), S1400D (5259T>G 5260C>A 5261A>C), W1403K (5268T>A 5269G>A 5270G>A), W1407R (5280T>A 5282G>A), V1408T (5283G>A 5284T>C 5285A>T), P1409H (5287C>A 5288T>C), Q1410H (5291A>T), L1412P (5296T>C), M1418I (5315G>T), K1423T (5329A>C 5330A>C), I1428T (5344T>C 5345C>T), Q1430C (5349C>T 5350A>G 5351A>C), R1431T (5353G>C 5354A>G), K1432L (5355A>C 5356A>T), D1434K (5361G>A 5363T>A), H1435K (5364C>A 5366T>G), C1436I (5367T>A 5368G>T 5369T>A), F1437_A1438insH (5372_5373insCAT), A1438P (5373G>C 5375A>T), G1439Y (5376G>T 5377G>A 5378A>C), T1440L (5379A>C 5380C>T), E1441D (5384A>T), E1442Q (5385G>C 5387G>A), I1444V (5391A>G 5393T>G), A1445V (5395C>T 5396T>A), I1448L (5403A>T 5405T>G), I1452V (5415A>G), F1454Y (5422T>A), E1458L (5433G>T 5434A>T), Q1460E (5439C>G 5441G>A), K1462V (5445A>G 5446A>T), F1468V (5463T>G 5465C>A), L1469F (5466C>T 5468G>C), E1470Q (5469G>C 5471G>A), R1471V (5472A>G 5473G>T 5474A>C), C1472L (5476G>T 5477C>A), K1473R (5478A>C 5479A>G), Q1474E (5481C>G), H1475N (5484C>A), G1476E (5488G>A), V1478Y (5493G>T 5494T>A 5495G>C), L1479V (5496T>G 5498A>C), S1480K (5500G>A 5501C>G), P1481Q (5503C>A 5504A>G), T1482E (5505A>G 5506C>A 5507A>G), M1484C (5511A>T 5512T>G 5513G>C), K1485E (5514A>G 5516A>G), I1486F (5517A>T 5519A>T), G1487A (5521G>C), K1489H (5526A>C 5528A>T), R1490E (5529A>G 5530G>A 5531G>A), I1491V (5532A>G 5534C>G), E1492H (5535G>C 5537G>C), A1496H (5547G>C 5548C>A 5549T>C), V1497F (5550G>T 5552G>C), N1499S (5557A>G), E1500Q (5559G>C), K1502E (5565A>G), I1503L (5568A>C), L1505M (5574C>A 5576A>G), E1585S (5814G>A 5815A>G 5816G>C), P1588L (5824C>T), E1590S (5829G>T 5830A>C), A1591D (5833C>A 5834G>T), I1593L (5838A>C 5840C>A), C1600S (5859T>A), M1601L (5862A>C 5864G>C), E1602Q (5865G>C), G1606A (5878G>C 5879A>T), I1607V (5880A>G 5882C>A), C1608L (5884G>T 5885T>A), W1610A (5889T>G 5890G>C 5891G>A), K1611R (5893A>G), F1613N (5898T>A 5899T>A 5900T>C), R1615Y (5904A>T 5905G>A 5906G>C), D1616S (5907G>A 5908A>G 5909C>T), R1618K (5913C>A 5914G>A), D1620E (5921C>A), A1627Q (5940G>C 5941C>A), F1631Y (5953T>A), R1632K (5956G>A), P1633E (5958C>G 5959C>A), P1633_I1634insKG (5960_5961insAAGGGT), I1634N (5962T>A), K1635M (5965A>T 5966A>G), T1637S (5971C>G 5972A>T), A1640V (5980C>T 5981T>C), I1642V (5985A>G), H1643L (5988C>T 5989A>T 5990C>A), V1645I (5994G>A), M1646I (5999G>A), N1647Y (6000A>T), S1648G (6003A>G 6005T>A), K1650N (6011G>T), K1651S (6013A>G 6014A>T), Y1656I (6027T>A 6028A>T), S1658N (6034G>A 6035T>C), P1660S (6039C>A 6040C>G 6041A>T), A1661E (6043C>A 6044G>A), V1662I (6045G>A 6047C>A), T1663V (6048A>G 6049C>T 6050T>A), I1664V (6051A>G 6053C>A), Q1669E (6066C>G 6068G>A) | | | | | | | | |
| Codon mutations: | AGT1389..G (5228T>G), GGA1390GGG (5231A>G), TTT1391TAT (5233T>A), CAG1393CAA (5240G>A), ATC1394ATT (5243C>T), GCA1395AGA (5244G>A 5245C>G), GAG1399CCC (5256G>C 5257A>C 5258G>C), TCA1400GAC (5259T>G 5260C>A 5261A>C), ATC1401ATT (5264C>T), CCA1402CCC (5267A>C), TGG1403AAA (5268T>A 5269G>A 5270G>A), ACC1404ACT (5273C>T), GCG1405GCC (5276G>C), TGG1407AGA (5280T>A 5282G>A), GTA1408ACT (5283G>A 5284T>C 5285A>T), CCT1409CAC (5287C>A 5288T>C), CAA1410CAT (5291A>T), GGA1411GGT (5294A>T), CTT1412CCT (5296T>C), GAA1414GAG (5303A>G), CTG1416TTG (5307C>T), GTT1417GTG (5312T>G), ATG1418ATT (5315G>T), CCA1419CCC (5318A>C), TTT1420TTC (5321T>C), CTC1422TTA (5325C>T 5327C>A), AAA1423ACC (5329A>C 5330A>C), AAT1424AAC (5333T>C), ATC1428ACT (5344T>C 5345C>T), CAA1430TGC (5349C>T 5350A>G 5351A>C), AGA1431ACG (5353G>C 5354A>G), AAA1432CTA (5355A>C 5356A>T), GAT1434AAA (5361G>A 5363T>A), CAT1435AAG (5364C>A 5366T>G), TGT1436ATA (5367T>A 5368G>T 5369T>A), TTT1437_GCA1438insCAT (5372_5373insCAT), GCA1438CCT (5373G>C 5375A>T), GGA1439TAC (5376G>T 5377G>A 5378A>C), ACA1440CTA (5379A>C 5380C>T), GAA1441GAT (5384A>T), GAG1442CAA (5385G>C 5387G>A), TTT1443TTC (5390T>C), ATT1444GTG (5391A>G 5393T>G), GCT1445GTA (5395C>T 5396T>A), ATT1448TTG (5403A>T 5405T>G), ATC1452GTC (5415A>G), ATT1453ATC (5420T>C), TTC1454TAC (5422T>A), AAT1456AAC (5429T>C), ACA1457ACC (5432A>C), GAG1458TTG (5433G>T 5434A>T), CAG1460GAA (5439C>G 5441G>A), CAT1461CAC (5444T>C), AAG1462GTG (5445A>G 5446A>T), CAT1464CAC (5453T>C), CTG1465TTG (5454C>T), TTC1468GTA (5463T>G 5465C>A), CTG1469TTC (5466C>T 5468G>C), GAG1470CAA (5469G>C 5471G>A), AGA1471GTC (5472A>G 5473G>T 5474A>C), TGC1472TTA (5476G>T 5477C>A), AAG1473CGG (5478A>C 5479A>G), CAA1474GAA (5481C>G), CAT1475AAT (5484C>A), GGG1476GAG (5488G>A), TTG1477CTA (5490T>C 5492G>A), GTG1478TAC (5493G>T 5494T>A 5495G>C), TTA1479GTC (5496T>G 5498A>C), AGC1480AAG (5500G>A 5501C>G), CCA1481CAG (5503C>A 5504A>G), ACA1482GAG (5505A>G 5506C>A 5507A>G), ATG1484TGC (5511A>T 5512T>G 5513G>C), AAA1485GAG (5514A>G 5516A>G), ATA1486TTT (5517A>T 5519A>T), GGC1487GCC (5521G>C), CAG1488CAA (5525G>A), AAA1489CAT (5526A>C 5528A>T), AGG1490GAA (5529A>G 5530G>A 5531G>A), ATC1491GTG (5532A>G 5534C>G), GAG1492CAC (5535G>C 5537G>C), TTA1494TTG (5543A>G), GCT1496CAC (5547G>C 5548C>A 5549T>C), GTG1497TTC (5550G>T 5552G>C), ATC1498ATT (5555C>T), AAC1499AGC (5557A>G), GAA1500CAA (5559G>C), GGG1501GGT (5564G>T), AAG1502GAG (5565A>G), ATA1503CTA (5568A>C), CTA1505ATG (5574C>A 5576A>G), CAG1506GA. (5577C>G), CTT1581..G (5804T>G), TTG1584TTA (5813G>A), GAG1585AGC (5814G>A 5815A>G 5816G>C), ATC1586ATA (5819C>A), CCA1588CTA (5824C>T), GAA1590TCA (5829G>T 5830A>C), GCG1591GAT (5833C>A 5834G>T), TAC1592TAT (5837C>T), ATC1593CTA (5838A>C 5840C>A), ATT1595ATA (5846T>A), ACT1597ACA (5852T>A), TGC1600AGC (5859T>A), ATG1601CTC (5862A>C 5864G>C), GAA1602CAA (5865G>C), GGA1606GCT (5878G>C 5879A>T), ATC1607GTA (5880A>G 5882C>A), TGT1608TTA (5884G>T 5885T>A), AAG1609AAA (5888G>A), TGG1610GCA (5889T>G 5890G>C 5891G>A), AAA1611AGA (5893A>G), CCT1612CCA (5897T>A), TTT1613AAC (5898T>A 5899T>A 5900T>C), AGG1615TAC (5904A>T 5905G>A 5906G>C), GAC1616AGT (5907G>A 5908A>G 5909C>T), CCA1617CCT (5912A>T), CGA1618AAA (5913C>A 5914G>A), GAC1620GAA (5921C>A), GAG1621GAA (5924G>A), AAG1622AAA (5927G>A), ATA1623ATT (5930A>T), GCC1625GCT (5936C>T), TAT1626TAC (5939T>C), GCA1627CAA (5940G>C 5941C>A), GGA1629GGT (5948A>T), TTC1631TAC (5953T>A), AGA1632AAA (5956G>A), CCA1633GAA (5958C>G 5959C>A), CCA1633_ATC1634insAAGGGT (5960_5961insAAGGGT), ATC1634AAC (5962T>A), AAA1635ATG (5965A>T 5966A>G), TCC1636AGT (5967T>A 5968C>G 5969C>T), ACA1637AGT (5971C>G 5972A>T), ATC1638ATA (5975C>A), GCT1640GTC (5980C>T 5981T>C), GAG1641GAA (5984G>A), ATA1642GTA (5985A>G), CAC1643TTA (5988C>T 5989A>T 5990C>A), GCG1644GCT (5993G>T), GTA1645ATA (5994G>A), ATG1646ATA (5999G>A), AAT1647TAT (6000A>T), AGT1648GGA (6003A>G 6005T>A), TTA1649CTA (6006T>C), AAG1650AAT (6011G>T), AAA1651AGT (6013A>G 6014A>T), TTT1652TTC (6017T>C), AAG1653AAA (6020G>A), CTA1654TTA (6021C>T), TAT1656ATT (6027T>A 6028A>T), AGT1658AAC (6034G>A 6035T>C), CCA1660AGT (6039C>A 6040C>G 6041A>T), GCG1661GAA (6043C>A 6044G>A), GTC1662ATA (6045G>A 6047C>A), ACT1663GTA (6048A>G 6049C>T 6050T>A), ATC1664GTA (6051A>G 6053C>A), AGG1665AGA (6056G>A), TGT1668TGC (6065T>C), CAG1669GAA (6066C>G 6068G>A) | | | | | | | | |

##### Proteins

| | Begin | End | Coverage | Score | Concordance | Matches | Identities | I/D/M/F* | Stop Codons |
|---|---|---|---|---|---|---|---|---|---|
| ORF3 polyprotein (YP_009666830.1) | 1390 | 1669 | 11.0% | 779 | 51.0% | 205 (98.6%) | 101 (48.6%) | 3/0/0/0 | 0 |
| Protein mutations: | F1391Y (5233T>A), A1395R (5244G>A 5245C>G), E1399P (5256G>C 5257A>C 5258G>C), S1400D (5259T>G 5260C>A 5261A>C), W1403K (5268T>A 5269G>A 5270G>A), W1407R (5280T>A 5282G>A), V1408T (5283G>A 5284T>C 5285A>T), P1409H (5287C>A 5288T>C), Q1410H (5291A>T), L1412P (5296T>C), M1418I (5315G>T), K1423T (5329A>C 5330A>C), I1428T (5344T>C 5345C>T), Q1430C (5349C>T 5350A>G 5351A>C), R1431T (5353G>C 5354A>G), K1432L (5355A>C 5356A>T), D1434K (5361G>A 5363T>A), H1435K (5364C>A 5366T>G), C1436I (5367T>A 5368G>T 5369T>A), F1437_A1438insH (5372_5373insCAT), A1438P (5373G>C 5375A>T), G1439Y (5376G>T 5377G>A 5378A>C), T1440L (5379A>C 5380C>T), E1441D (5384A>T), E1442Q (5385G>C 5387G>A), I1444V (5391A>G 5393T>G), A1445V (5395C>T 5396T>A), I1448L (5403A>T 5405T>G), I1452V (5415A>G), F1454Y (5422T>A), E1458L (5433G>T 5434A>T), Q1460E (5439C>G 5441G>A), K1462V (5445A>G 5446A>T), F1468V (5463T>G 5465C>A), L1469F (5466C>T 5468G>C), E1470Q (5469G>C 5471G>A), R1471V (5472A>G 5473G>T 5474A>C), C1472L (5476G>T 5477C>A), K1473R (5478A>C 5479A>G), Q1474E (5481C>G), H1475N (5484C>A), G1476E (5488G>A), V1478Y (5493G>T 5494T>A 5495G>C), L1479V (5496T>G 5498A>C), S1480K (5500G>A 5501C>G), P1481Q (5503C>A 5504A>G), T1482E (5505A>G 5506C>A 5507A>G), M1484C (5511A>T 5512T>G 5513G>C), K1485E (5514A>G 5516A>G), I1486F (5517A>T 5519A>T), G1487A (5521G>C), K1489H (5526A>C 5528A>T), R1490E (5529A>G 5530G>A 5531G>A), I1491V (5532A>G 5534C>G), E1492H (5535G>C 5537G>C), A1496H (5547G>C 5548C>A 5549T>C), V1497F (5550G>T 5552G>C), N1499S (5557A>G), E1500Q (5559G>C), K1502E (5565A>G), I1503L (5568A>C), L1505M (5574C>A 5576A>G), E1585S (5814G>A 5815A>G 5816G>C), P1588L (5824C>T), E1590S (5829G>T 5830A>C), A1591D (5833C>A 5834G>T), I1593L (5838A>C 5840C>A), C1600S (5859T>A), M1601L (5862A>C 5864G>C), E1602Q (5865G>C), G1606A (5878G>C 5879A>T), I1607V (5880A>G 5882C>A), C1608L (5884G>T 5885T>A), W1610A (5889T>G 5890G>C 5891G>A), K1611R (5893A>G), F1613N (5898T>A 5899T>A 5900T>C), R1615Y (5904A>T 5905G>A 5906G>C), D1616S (5907G>A 5908A>G 5909C>T), R1618K (5913C>A 5914G>A), D1620E (5921C>A), A1627Q (5940G>C 5941C>A), F1631Y (5953T>A), R1632K (5956G>A), P1633E (5958C>G 5959C>A), P1633_I1634insKG (5960_5961insAAGGGT), I1634N (5962T>A), K1635M (5965A>T 5966A>G), T1637S (5971C>G 5972A>T), A1640V (5980C>T 5981T>C), I1642V (5985A>G), H1643L (5988C>T 5989A>T 5990C>A), V1645I (5994G>A), M1646I (5999G>A), N1647Y (6000A>T), S1648G (6003A>G 6005T>A), K1650N (6011G>T), K1651S (6013A>G 6014A>T), Y1656I (6027T>A 6028A>T), S1658N (6034G>A 6035T>C), P1660S (6039C>A 6040C>G 6041A>T), A1661E (6043C>A 6044G>A), V1662I (6045G>A 6047C>A), T1663V (6048A>G 6049C>T 6050T>A), I1664V (6051A>G 6053C>A), Q1669E (6066C>G 6068G>A) | | | | | | | | |

| | Begin | End | Coverage | Score | Concordance | Matches | Identities | I/D/M/F* | Stop Codons |
|---|---|---|---|---|---|---|---|---|---|
| **NT** | **5228** | **6068** | **8.7%** | **177** | **14.4%** | **616 (98.6%)** | **359 (57.4%)** | **9/0** | |
| Codon mutations: | AGT1389..G (5228T>G), GGA1390GGG (5231A>G), TTT1391TAT (5233T>A), CAG1393CAA (5240G>A), ATC1394ATT (5243C>T), GCA1395AGA (5244G>A 5245C>G), GAG1399CCC (5256G>C 5257A>C 5258G>C), TCA1400GAC (5259T>G 5260C>A 5261A>C), ATC1401ATT (5264C>T), CCA1402CCC (5267A>C), TGG1403AAA (5268T>A 5269G>A 5270G>A), ACC1404ACT (5273C>T), GCG1405GCC (5276G>C), TGG1407AGA (5280T>A 5282G>A), GTA1408ACT (5283G>A 5284T>C 5285A>T), CCT1409CAC (5287C>A 5288T>C), CAA1410CAT (5291A>T), GGA1411GGT (5294A>T), CTT1412CCT (5296T>C), GAA1414GAG (5303A>G), CTG1416TTG (5307C>T), GTT1417GTG (5312T>G), ATG1418ATT (5315G>T), CCA1419CCC (5318A>C), TTT1420TTC (5321T>C), CTC1422TTA (5325C>T 5327C>A), AAA1423ACC (5329A>C 5330A>C), AAT1424AAC (5333T>C), ATC1428ACT (5344T>C 5345C>T), CAA1430TGC (5349C>T 5350A>G 5351A>C), AGA1431ACG (5353G>C 5354A>G), AAA1432CTA (5355A>C 5356A>T), GAT1434AAA (5361G>A 5363T>A), CAT1435AAG (5364C>A 5366T>G), TGT1436ATA (5367T>A 5368G>T 5369T>A), TTT1437_GCA1438insCAT (5372_5373insCAT), GCA1438CCT (5373G>C 5375A>T), GGA1439TAC (5376G>T 5377G>A 5378A>C), ACA1440CTA (5379A>C 5380C>T), GAA1441GAT (5384A>T), GAG1442CAA (5385G>C 5387G>A), TTT1443TTC (5390T>C), ATT1444GTG (5391A>G 5393T>G), GCT1445GTA (5395C>T 5396T>A), ATT1448TTG (5403A>T 5405T>G), ATC1452GTC (5415A>G), ATT1453ATC (5420T>C), TTC1454TAC (5422T>A), AAT1456AAC (5429T>C), ACA1457ACC (5432A>C), GAG1458TTG (5433G>T 5434A>T), CAG1460GAA (5439C>G 5441G>A), CAT1461CAC (5444T>C), AAG1462GTG (5445A>G 5446A>T), CAT1464CAC (5453T>C), CTG1465TTG (5454C>T), TTC1468GTA (5463T>G 5465C>A), CTG1469TTC (5466C>T 5468G>C), GAG1470CAA (5469G>C 5471G>A), AGA1471GTC (5472A>G 5473G>T 5474A>C), TGC1472TTA (5476G>T 5477C>A), AAG1473CGG (5478A>C 5479A>G), CAA1474GAA (5481C>G), CAT1475AAT (5484C>A), GGG1476GAG (5488G>A), TTG1477CTA (5490T>C 5492G>A), GTG1478TAC (5493G>T 5494T>A 5495G>C), TTA1479GTC (5496T>G 5498A>C), AGC1480AAG (5500G>A 5501C>G), CCA1481CAG (5503C>A 5504A>G), ACA1482GAG (5505A>G 5506C>A 5507A>G), ATG1484TGC (5511A>T 5512T>G 5513G>C), AAA1485GAG (5514A>G 5516A>G), ATA1486TTT (5517A>T 5519A>T), GGC1487GCC (5521G>C), CAG1488CAA (5525G>A), AAA1489CAT (5526A>C 5528A>T), AGG1490GAA (5529A>G 5530G>A 5531G>A), ATC1491GTG (5532A>G 5534C>G), GAG1492CAC (5535G>C 5537G>C), TTA1494TTG (5543A>G), GCT1496CAC (5547G>C 5548C>A 5549T>C), GTG1497TTC (5550G>T 5552G>C), ATC1498ATT (5555C>T), AAC1499AGC (5557A>G), GAA1500CAA (5559G>C), GGG1501GGT (5564G>T), AAG1502GAG (5565A>G), ATA1503CTA (5568A>C), CTA1505ATG (5574C>A 5576A>G), CAG1506GA. (5577C>G), CTT1581..G (5804T>G), TTG1584TTA (5813G>A), GAG1585AGC (5814G>A 5815A>G 5816G>C), ATC1586ATA (5819C>A), CCA1588CTA (5824C>T), GAA1590TCA (5829G>T 5830A>C), GCG1591GAT (5833C>A 5834G>T), TAC1592TAT (5837C>T), ATC1593CTA (5838A>C 5840C>A), ATT1595ATA (5846T>A), ACT1597ACA (5852T>A), TGC1600AGC (5859T>A), ATG1601CTC (5862A>C 5864G>C), GAA1602CAA (5865G>C), GGA1606GCT (5878G>C 5879A>T), ATC1607GTA (5880A>G 5882C>A), TGT1608TTA (5884G>T 5885T>A), AAG1609AAA (5888G>A), TGG1610GCA (5889T>G 5890G>C 5891G>A), AAA1611AGA (5893A>G), CCT1612CCA (5897T>A), TTT1613AAC (5898T>A 5899T>A 5900T>C), AGG1615TAC (5904A>T 5905G>A 5906G>C), GAC1616AGT (5907G>A 5908A>G 5909C>T), CCA1617CCT (5912A>T), CGA1618AAA (5913C>A 5914G>A), GAC1620GAA (5921C>A), GAG1621GAA (5924G>A), AAG1622AAA (5927G>A), ATA1623ATT (5930A>T), GCC1625GCT (5936C>T), TAT1626TAC (5939T>C), GCA1627CAA (5940G>C 5941C>A), GGA1629GGT (5948A>T), TTC1631TAC (5953T>A), AGA1632AAA (5956G>A), CCA1633GAA (5958C>G 5959C>A), CCA1633_ATC1634insAAGGGT (5960_5961insAAGGGT), ATC1634AAC (5962T>A), AAA1635ATG (5965A>T 5966A>G), TCC1636AGT (5967T>A 5968C>G 5969C>T), ACA1637AGT (5971C>G 5972A>T), ATC1638ATA (5975C>A), GCT1640GTC (5980C>T 5981T>C), GAG1641GAA (5984G>A), ATA1642GTA (5985A>G), CAC1643TTA (5988C>T 5989A>T 5990C>A), GCG1644GCT (5993G>T), GTA1645ATA (5994G>A), ATG1646ATA (5999G>A), AAT1647TAT (6000A>T), AGT1648GGA (6003A>G 6005T>A), TTA1649CTA (6006T>C), AAG1650AAT (6011G>T), AAA1651AGT (6013A>G 6014A>T), TTT1652TTC (6017T>C), AAG1653AAA (6020G>A), CTA1654TTA (6021C>T), TAT1656ATT (6027T>A 6028A>T), AGT1658AAC (6034G>A 6035T>C), CCA1660AGT (6039C>A 6040C>G 6041A>T), GCG1661GAA (6043C>A 6044G>A), GTC1662ATA (6045G>A 6047C>A), ACT1663GTA (6048A>G 6049C>T 6050T>A), ATC1664GTA (6051A>G 6053C>A), AGG1665AGA (6056G>A), TGT1668TGC (6065T>C), CAG1669GAA (6066C>G 6068G>A) | | | | | | | | |

*: Inserts / Deletes / Misaligned / Frameshifts

#### Analysis details

This analysis was performed with panviral2.64

#### NGS Details (UN18_val): Dioscovirus dioscoreae

##### Assembly

| | |
|---|---|
| **Coverage Length** | 915 (1 contig(s)) |
| **Depth Of Coverage** | 3.2 |
| **Number Of Reads** | 23 |
| **Reads Per Million** | 0.43 rpm (after QC) |
| **Ambiguities** | 0 |
| **Assembly Method** | de novo + reference guided assembly |
| **Consensus Caller** | Bcf Tools |

##### Coverage Map

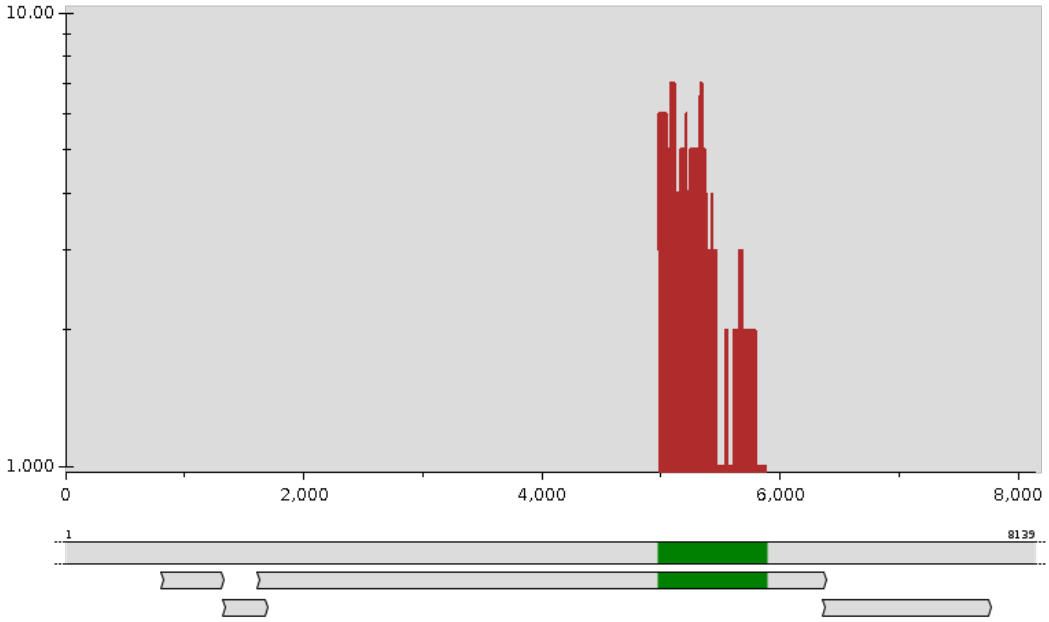

##### Assignment

| | |
|---|---|
| **Type** | Dioscovirus dioscoreae (Taxonomy ID: 3052184) |
| **Reference Genome** | NC_040712.1 |
| **NT Identity (%)** | 61.8421 |
| **AA Identity (%)** | 52.6316 |
| **Number Of Stop Codons** | 0 |
| **Number Of CDS** | 4 |

##### Alignment

| | |
|---|---|
| **Alignment Score** | 420.0 (NT) + 1186.0 (AA) = 1606.0 |
| **Concordance (%)** | 40.8444 |

| Alignment Method | Global, seeded, nucleotide + amino acids (AGA) |
|---|---|

#### Genome Region

Sequence starts at position 4980 and ends at position 5894 relative to NC_040712.1 reference sequence.

#### Alignment Detailed Statistics

| | Begin | End | Coverage | Score | Concordance | Matches | Identities | I/D/M/F* | Stop Codons |
|---|---|---|---|---|---|---|---|---|---|
| **NT** | **4980** | **5894** | **11.2%** | **420** | **23.2%** | **912 (99.7%)** | **564 (61.6%)** | **0/3** | |
| Mutations: | 4983C>A, 4986C>A, 4988T>A, 4989A>T, 4990A>T, 4992T>C, 4994G>A, 4997A>T, 5001A>T, 5003C>A, 5004C>G, 5009T>A, 5012T>A, 5015G>A, 5016A>G, 5018T>A, 5019G>A, 5021C>A, 5024A>C, 5025T>C, 5029A>T, 5030G>A, 5031T>C, 5038A>G, 5040A>G, 5041A>C, 5042G>C, 5046C>A, 5047A>G, 5048G>A, 5051T>A, 5053G>T, 5054A>G, 5057C>T, 5063G>A, 5066C>T, 5069C>A, 5078T>A, 5081T>A, 5082C>T, 5083T>A, 5091C>A, 5093G>A, 5094G>A, 5095A>T, 5096C>A, 5097A>G, 5098A>C, 5099G>A, 5100C>G, 5102T>A, 5105T>A, 5106A>G, 5107T>A, 5109C>G, 5112T>A, 5115C>A, 5116C>A, 5117A>T, 5118G>C, 5121A>G, 5122T>A, 5123G>C, 5124G>A, 5125C>T, 5126G>A, 5127A>G, 5132A>T, 5134A>T, 5136G>C, 5138A>G, 5140A>T, 5141G>A, 5144T>A, 5147T>A, 5149C>A, 5157C>T, 5159G>A, 5161C>A, 5162A>T, 5163C>T, 5165C>A, 5166A>G, 5167A>G, 5168G>A, 5171T>A, 5174C>A, 5178C>A, 5179C>G, 5180T>A, 5183C>A, 5184C>A, 5185A>C, 5187A>T, 5188G>C, 5190C>A, 5192C>A, 5195T>C, 5198G>A, 5199A>T, 5207T>A, 5213T>G, 5216C>A, 5219C>A, 5222A>C, 5226T>A, 5227C>A, 5228T>C, 5232C>A, 5233A>T, 5237A>C, 5243T>A, 5247A>G, 5248G>C, 5249T>C, 5250C>A, 5252T>A, 5259T>A, 5260A>T, 5261C>A, 5271C>A, 5276A>C, 5288A>T, 5289T>A, 5290A>G, 5291C>A, 5293A>C, 5294A>G, 5295G>T, 5297C>T, 5298C>G, 5299A>C, 5300A>T, 5303C>T, 5305C>A, 5306A>G, 5309C>A, 5313A>G, 5314G>A, 5317T>A, 5318T>A, 5319G>A, 5320A>G, 5321C>T, 5322T>G, 5324T>A, 5325C>T, 5327C>A, 5328C>A, 5331C>A, 5332T>A, 5335A>G, 5339T>A, 5340A>C, 5344A>G, 5345C>T, 5350T>C, 5351T>A, 5354C>A, 5356A>T, 5359G>A, 5360C>T, 5370C>T, 5371T>G, 5372A>T, 5375G>A, 5383T>A, 5384C>T, 5385C>T, 5386A>G, 5387C>G, 5390G>A, 5391A>G, 5395T>G, 5396G>A, 5400G>C, 5403C>G, 5404C>A, 5406C>G, 5408G>T, 5410G>A, 5424G>A, 5429T>C, 5430G>A, 5431T>C, 5432C>T, 5438A>T, 5439C>G, 5441G>A, 5444T>A, 5457A>T, 5471C>T, 5475C>T, 5477C>G, 5480A>G, 5482A>C, 5489T>A, 5490T>C, 5492A>T, 5493G>A, 5502C>A, 5507A>G, 5513T>C, 5519A>T, 5522C>T, 5523A>G, 5524A>G, 5526A>G, 5528A>T, 5531T>C, 5532T>A, 5533C>A, 5535G>A, 5536A>G, 5544T>C, 5545G>T, 5546T>A, 5549T>A, 5552C>T, 5553A>G, 5556G>T, 5558C>T, 5561T>C, 5564T>C, 5565T>C, 5567G>A, 5568A>G, 5573T>C, 5574T>A, 5575C>G, 5577G>A, 5578A>G, 5580A>G, 5581G>A, 5585A>G, 5589C>G, 5595G>C, 5598C>G, 5607T>C, 5608T>A, 5610C>A, 5611A>C, 5612G>A, 5613T>A, 5615C>T, 5618C>T, 5619C>A, 5620A>G, 5621A>G, 5622G>T, 5626G>T, 5627T>C, 5628A>G, 5629A>T, 5630G>A, 5631G>A, 5632A>G, 5633A>T, 5634G>A, 5639G>A, 5640C>A, 5642T>A, 5645T>A, 5646T>A, 5648G>A, 5649T>A, 5651A>C, 5656C>A, 5657A>G, 5661T>A, 5664A>G, 5667A>T, 5670G>T, 5672A>T, 5673G>A, 5674T>A, 5676G>A, 5679A>C, 5685G>A, 5687A>C, 5691C>T, 5693T>A, 5696C>A, 5697C>G, 5700G>A, 5701A>C, 5702A>T, 5703A>T, 5711A>T, 5714A>G, 5717A>G, 5718G>A, 5720T>A, 5721C>A, 5726C>A, 5739T>G, 5740T>C, 5741A>T, 5748A>G, 5756A>T, 5757T>A, 5759T>G, 5762T>A, 5763_5765delGAA, 5768C>T, 5769C>A, 5777A>T, 5779C>A, 5781T>A, 5782T>C, 5786G>A, 5788G>A, 5790C>T, 5795G>A, 5798G>A, 5807G>A, 5808A>T, 5811T>G, 5813A>G, 5816C>T, 5822C>T, 5828T>C, 5830A>T, 5834C>A, 5836C>A, 5843G>A, 5844A>G, 5846T>A, 5850T>A, 5851A>T, 5853A>G, 5855T>A, 5856A>G, 5859A>C, 5860T>C, 5864T>A, 5868A>G, 5869A>C, 5870T>A, 5873G>A, 5874T>A, 5875G>T, 5876T>A, 5877T>G, 5878C>G, 5879C>A, 5882C>T, 5889G>C, 5893G>A | | | | | | | | |

##### CDS

| | Begin | End | Coverage | Score | Concordance | Matches | Identities | I/D/M/F* | Stop Codons |
|---|---|---|---|---|---|---|---|---|---|
| EXK67_gp3 | 1124 | 1428 | 19.1% | 1186 | 55.9% | 304 (99.7%) | 160 (52.5%) | 0/1/0/0 | 0 |
| Protein mutations: | L1125I (4983C>A), H1126K (4986C>A 4988T>A), K1127L (4989A>T 4990A>T), E1129D (4997A>T), I1131L (5001A>T 5003C>A), Q1132E (5004C>G), N1136E (5016A>G 5018T>A), D1137K (5019G>A 5021C>A), K1140I (5029A>T 5030G>A), Y1141H (5031T>C), E1143G (5038A>G), K1144A (5040A>G 5041A>C 5042G>C), Q1146R (5046C>A 5047A>G 5048G>A), R1148M (5053G>T 5054A>G), D1157E (5081T>A), L1158Y (5082C>T 5083T>A), Q1161K (5091C>A 5093G>A), D1162I (5094G>A 5095A>T 5096C>A), K1163A (5097A>G 5098A>C 5099G>A), P1164A (5100C>G 5102T>A), I1166E (5106A>G 5107T>A), P1167A (5109C>G), S1168T (5112T>A), P1169N (5115C>A 5116C>A 5117A>T), E1170Q (5118G>C), M1171D (5121A>G 5122T>A 5123G>C), A1172I (5124G>A 5125C>T 5126G>A), K1173E (5127A>G), E1174D (5132A>T), Y1175F (5134A>T), E1176Q (5136G>C 5138A>G), K1177I (5140A>T 5141G>A), H1178Q (5144T>A), T1180K (5149C>A), A1184D (5161C>A 5162A>T), K1186G (5166A>G 5167A>G 5168G>A), P1190R (5178C>A 5179C>G 5180T>A), Q1192T (5184C>A 5185A>C), T1197S (5199A>T), I1201M (5213T>G), N1203K (5219C>A), K1204N (5222A>C), S1206N (5226T>A 5227C>A 5228T>C), Q1208I (5232C>A 5233A>T), S1213A (5247A>G 5248G>C 5249T>C), Y1217I (5259T>A 5260A>T 5261C>A), Y1227R (5289T>A 5290A>G 5291C>A), K1228T (5293A>C 5294A>G), D1229Y (5295G>T 5297C>T), Q1230A (5298C>G 5299A>C 5300A>T), T1232K (5305C>A 5306A>G), S1235D (5313A>G 5314G>A), I1236K (5317T>A 5318T>A), D1237S (5319G>A 5320A>G 5321C>T), Y1238E (5322T>G 5324T>A), L1240I (5328C>A), L1241N (5331C>A 5332T>A), K1242R (5335A>G), K1244Q (5340A>C), D1245G (5344A>G 5345C>T), I1247T (5350T>C 5351T>A), Y1249F (5356A>T), S1250N (5359G>A 5360C>T), L1254C (5370C>T 5371T>G 5372A>T), F1258Y (5383T>A 5384C>T), H1259W (5385C>T 5386A>G 5387C>G), I1261V (5391A>G), M1262R (5395T>G 5396G>A), D1264H (5400G>C), P1265E (5403C>G 5404C>A), Q1266D (5406C>G 5408G>T), S1267N (5410G>A), A1272T (5424G>A), V1274T (5430G>A 5431T>C 5432C>T), Q1277E (5439C>G 5441G>A), I1283L (5457A>T), N1291T (5482A>C), S1294P (5490T>C 5492A>T), V1295I (5493G>A), K1305G (5523A>G 5524A>G), K1306D (5526A>G 5528A>T), S1308K (5532T>A 5533C>A), E1309R (5535G>A 5536A>G), C1312L (5544T>C 5545G>T 5546T>A), I1315V (5553A>G), D1316Y (5556G>T 5558C>T), I1320V (5568A>G), E1323R (5577G>A 5578A>G), S1324D (5580A>G 5581G>A), I1325M (5585A>G), Q1327E (5589C>G), V1329L (5595G>C), Q1330E (5598C>G), L1333Q (5607T>C 5608T>A), Q1334T (5610C>A 5611A>C 5612G>A), F1335I (5613T>A 5615C>T), Q1337R (5619C>A 5620A>G 5621A>G), V1338L (5622G>T), C1339F (5626G>T 5627T>C), K1340V (5628A>G 5629A>T 5630G>A), E1341S (5631G>A 5632A>G 5633A>T), E1342K (5634G>A), L1344I (5640C>A 5642T>A), L1346I (5646T>A 5648G>A), S1347T (5649T>A 5651A>C), T1349K (5656C>A 5657A>G), L1351M (5661T>A), K1352E (5664A>G), I1353L (5667A>T), G1354C (5670G>T 5672A>T), V1355K (5673G>A 5674T>A), A1356T (5676G>A), N1357H (5679A>C), E1359N (5685G>A 5687A>C), L1363V (5697C>G), E1364T (5700G>A 5701A>C 5702A>T), I1365L (5703A>T), E1367D (5711A>T), V1370I (5718G>A 5720T>A), Q1371K (5721C>A), L1377A (5739T>G 5740T>C 5741A>T), I1380V (5748A>G), E1382D (5756A>T), F1383M (5757T>A 5759T>G), E1385del (5763_5765delGAA), Q1387K (5769C>A), E1389D (5777A>T), T1390K (5779C>A), L1391T (5781T>A 5782T>C), G1393D (5788G>A), I1400L (5808A>T), L1401V (5811T>G 5813A>G), Y1407F (5830A>T), P1409Q (5836C>A), S1412G (5844A>G 5846T>A), Y1414I (5850T>A 5851A>T), T1415A (5853A>G 5855T>A), R1416G (5856A>G), I1417P (5859A>C 5860T>C), F1418L (5864T>A), N1420A (5868A>G 5869A>C 5870T>A), C1422I (5874T>A 5875G>T 5876T>A), S1423G (5877T>G 5878C>G 5879C>A), E1427Q (5889G>C), R1428K (5893G>A) | | | | | | | | |
| Codon mutations: | CTA1125ATA (4983C>A), CAT1126AAA (4986C>A 4988T>A), AAA1127TTA (4989A>T 4990A>T), TTG1128CTA (4992T>C 4994G>A), GAA1129GAT (4997A>T), ATC1131TTA (5001A>T 5003C>A), CAA1132GAA (5004C>G), ATT1133ATA (5009T>A), ATT1134ATA (5012T>A), GGG1135GGA (5015G>A), AAT1136GAA (5016A>G 5018T>A), GAC1137AAA (5019G>A 5021C>A), CCA1138CCC (5024A>C), TTA1139CTA (5025T>C), AAG1140ATA (5029A>T 5030G>A), TAT1141CAT (5031T>C), GAA1143GGA (5038A>G), AAG1144GCC (5040A>G 5041A>C 5042G>C), CAG1146AGA (5046C>A 5047A>G 5048G>A), ATT1147ATA (5051T>A), AGA1148ATG (5053G>T 5054A>G), TGC1149TGT (5057C>T), CTG1151CTA (5063G>A), GAC1152GAT (5066C>T), ATC1153ATA (5069C>A), CCT1156CCA (5078T>A), GAT1157GAA (5081T>A), CTT1158TAT (5082C>T 5083T>A), CAG1161AAA (5091C>A 5093G>A), GAC1162ATA (5094G>A 5095A>T 5096C>A), AAG1163GCA (5097A>G 5098A>C 5099G>A), CCT1164GCA (5100C>G 5102T>A), ATT1165ATA (5105T>A), ATA1166GAA (5106A>G 5107T>A), CCT1167GCT (5109C>G), TCA1168ACA (5112T>A), CCA1169AAT (5115C>A 5116C>A 5117A>T), GAA1170CAA (5118G>C), ATG1171GAC (5121A>G 5122T>A 5123G>C), GCG1172ATA (5124G>A 5125C>T 5126G>A), AAA1173GAA (5127A>G), GAA1174GAT (5132A>T), TAT1175TTT (5134A>T), GAA1176CAG (5136G>C 5138A>G), AAG1177ATA (5140A>T 5141G>A), CAT1178CAA (5144T>A), ATT1179ATA (5147T>A), ACA1180AAA (5149C>A), CTG1183TTA (5157C>T 5159G>A), GCA1184GAT (5161C>A 5162A>T), CTC1185TTA (5163C>T 5165C>A), AAG1186GGA (5166A>G 5167A>G 5168G>A), GTT1187GTA (5171T>A), ATC1188ATA (5174C>A), CCT1190AGA (5178C>A 5179C>G 5180T>A), TCC1191TCA (5183C>A), CAA1192ACA (5184C>A 5185A>C), AGT1193TCT (5187A>T 5188G>C), CGC1194AGA (5190C>A 5192C>A), CAT1195CAC (5195T>C), AGG1196AGA (5198G>A), ACA1197TCA (5199A>T), GCT1199GCA (5207T>A), ATT1201ATG (5213T>G), GTC1202GTA (5216C>A), AAC1203AAA (5219C>A), AAA1204AAC (5222A>C), TCT1206AAC (5226T>A 5227C>A 5228T>C), CAA1208ATA (5232C>A 5233A>T), GTA1209GTC (5237A>C), GGT1211GGA (5243T>A), AGT1213GCC (5247A>G 5248G>C 5249T>C), CGT1214AGA (5250C>A 5252T>A), TAC1217ATA (5259T>A 5260A>T 5261C>A), CGA1221AGA (5271C>A), CTA1222CTC (5276A>C), ACA1226ACT (5288A>T), TAC1227AGA (5289T>A 5290A>G 5291C>A), AAA1228ACG (5293A>C 5294A>G), GAC1229TAT (5295G>T 5297C>T), CAA1230GCT (5298C>G 5299A>C 5300A>T), TAC1231TAT (5303C>T), ACA1232AAG (5305C>A 5306A>G), CTC1233CTA (5309C>A), AGT1235GAT (5313A>G 5314G>A), ATT1236AAA (5317T>A 5318T>A), GAC1237AGT (5319G>A 5320A>G 5321C>T), TAT1238GAA (5322T>G 5324T>A), CTC1239TTA (5325C>T 5327C>A), CTA1240ATA (5328C>A), CTT1241AAT (5331C>A 5332T>A), AAA1242AGA (5335A>G), ATT1243ATA (5339T>A), AAA1244CAA (5340A>C), GAC1245GGT (5344A>G 5345C>T), ATT1247ACA (5350T>C 5351T>A), GTC1248GTA (5354C>A), TAC1249TTC (5356A>T), AGC1250AAT (5359G>A 5360C>T), CTA1254TGT (5370C>T 5371T>G 5372A>T), AAG1255AAA (5375G>A), TTC1258TAT (5383T>A 5384C>T), CAC1259TGG (5385C>T 5386A>G 5387C>G), CAG1260CAA (5390G>A), ATT1261GTT (5391A>G), ATG1262AGA (5395T>G 5396G>A), GAT1264CAT (5400G>C), CCA1265GAA (5403C>G 5404C>A), CAG1266GAT (5406C>G 5408G>T), AGT1267AAT (5410G>A), GCA1272ACA (5424G>A), TTT1273TTC (5429T>C), GTC1274ACT (5430G>A 5431T>C 5432C>T), CCA1276CCT (5438A>T), CAG1277GAA (5439C>G 5441G>A), GGT1278GGA (5444T>A), ATA1283TTA (5457A>T), TTC1287TTT (5471C>T), CTC1289TTG (5475C>T 5477C>G), AAA1290AAG (5480A>G), AAT1291ACT (5482A>C), CCT1293CCA (5489T>A), TCA1294CCT (5490T>C 5492A>T), GTA1295ATA (5493G>A), CGA1298AGA (5502C>A), AAA1299AAG (5507A>G), GAT1301GAC (5513T>C), ATA1303ATT (5519A>T), TTC1304TTT (5522C>T), AAA1305GGA (5523A>G 5524A>G), AAA1306GAT (5526A>G 5528A>T), TAT1307TAC (5531T>C), TCA1308AAA (5532T>A 5533C>A), GAA1309AGA (5535G>A 5536A>G), TGT1312CTA (5544T>C 5545G>T 5546T>A), GTT1313GTA (5549T>A), TAC1314TAT (5552C>T), ATA1315GTA (5553A>G), GAC1316TAT (5556G>T 5558C>T), GAT1317GAC (5561T>C), ATT1318ATC (5564T>C), TTG1319CTA (5565T>C 5567G>A), ATA1320GTA (5568A>G), TTT1321TTC (5573T>C), TCT1322AGT (5574T>A 5575C>G), GAA1323AGA (5577G>A 5578A>G), AGT1324GAT (5580A>G 5581G>A), ATA1325ATG (5585A>G), CAA1327GAA (5589C>G), GTA1329CTA (5595G>C), CAA1330GAA (5598C>G), TTA1333CAA (5607T>C 5608T>A), CAG1334ACA (5610C>A 5611A>C 5612G>A), TTC1335ATT (5613T>A 5615C>T), TTC1336TTT (5618C>T), CAA1337AGG (5619C>A 5620A>G 5621A>G), GTA1338TTA (5622G>T), TGT1339TTC (5626G>T 5627T>C), AAG1340GTA (5628A>G 5629A>T 5630G>A), GAA1341AGT (5631G>A 5632A>G 5633A>T), GAA1342AAA (5634G>A), GGG1343GGA (5639G>A), CTT1344ATA (5640C>A 5642T>A), ATT1345ATA (5645T>A), TTG1346ATA (5646T>A 5648G>A), TCA1347ACC (5649T>A 5651A>C), ACA1349AAG (5656C>A 5657A>G), TTG1351ATG (5661T>A), AAA1352GAA (5664A>G), ATA1353TTA (5667A>T), GGA1354TGT (5670G>T 5672A>T), GTA1355AAA (5673G>A 5674T>A), GCA1356ACA (5676G>A), AAT1357CAT (5679A>C), GAA1359AAC (5685G>A 5687A>C), CTT1361TTA (5691C>T 5693T>A), GGC1362GGA (5696C>A), CTA1363GTA (5697C>G), GAA1364ACT (5700G>A 5701A>C 5702A>T), ATA1365TTA (5703A>T), GAA1367GAT (5711A>T), GGA1368GGG (5714A>G), AAA1369AAG (5717A>G), GTT1370ATA (5718G>A 5720T>A), CAA1371AAA (5721C>A), CTC1372CTA (5726C>A), TTA1377GCT (5739T>G 5740T>C 5741A>T), ATA1380GTA (5748A>G), GAA1382GAT (5756A>T), TTT1383ATG (5757T>A 5759T>G), CCT1384CCA (5762T>A), GAA1385del (5763_5765delGAA), GAC1386GAT (5768C>T), CAA1387AAA (5769C>A), GAA1389GAT (5777A>T), ACA1390AAA (5779C>A), TTA1391ACA (5781T>A 5782T>C), AAG1392AAA (5786G>A), GGT1393GAT (5788G>A), CTA1394TTA (5790C>T), CAG1395CAA (5795G>A), AAG1396AAA (5798G>A), GGG1399GGA (5807G>A), ATA1400TTA (5808A>T), TTA1401GTG (5811T>G 5813A>G), AAC1402AAT (5816C>T), GCC1404GCT (5822C>T), AAT1406AAC (5828T>C), TAT1407TTT (5830A>T), ATC1408ATA (5834C>A), CCA1409CAA (5836C>A), TTG1411TTA (5843G>A), AGT1412GGA (5844A>G 5846T>A), TAT1414ATT (5850T>A 5851A>T), ACT1415GCA (5853A>G 5855T>A), AGA1416GGA (5856A>G), ATA1417CCA (5859A>C 5860T>C), TTT1418TTA (5864T>A), AAT1420GCA (5868A>G 5869A>C 5870T>A), AAG1421AAA (5873G>A), TGT1422ATA (5874T>A 5875G>T 5876T>A), TCC1423GGA (5877T>G 5878C>G 5879C>A), AGC1424AGT (5882C>T), GAA1427CAA (5889G>C), AGA1428AAA (5893G>A) | | | | | | | | |

##### Proteins

| | Begin | End | Coverage | Score | Concordance | Matches | Identities | I/D/M/F* | Stop Codons |
|---|---|---|---|---|---|---|---|---|---|
| **NT** | **4980** | **5894** | **11.2%** | **420** | **23.2%** | **912 (99.7%)** | **564 (61.6%)** | **0/3** | |
| ORF3 (YP_009553219.1) | 1124 | 1428 | 19.1% | 1186 | 55.9% | 304 (99.7%) | 160 (52.5%) | 0/1/0/0 | 0 |
| Protein mutations: | L1125I (4983C>A), H1126K (4986C>A 4988T>A), K1127L (4989A>T 4990A>T), E1129D (4997A>T), I1131L (5001A>T 5003C>A), Q1132E (5004C>G), N1136E (5016A>G 5018T>A), D1137K (5019G>A 5021C>A), K1140I (5029A>T 5030G>A), Y1141H (5031T>C), E1143G (5038A>G), K1144A (5040A>G 5041A>C 5042G>C), Q1146R (5046C>A 5047A>G 5048G>A), R1148M (5053G>T 5054A>G), D1157E (5081T>A), L1158Y (5082C>T 5083T>A), Q1161K (5091C>A 5093G>A), D1162I (5094G>A 5095A>T 5096C>A), K1163A (5097A>G 5098A>C 5099G>A), P1164A (5100C>G 5102T>A), I1166E (5106A>G 5107T>A), P1167A (5109C>G), S1168T (5112T>A), P1169N (5115C>A 5116C>A 5117A>T), E1170Q (5118G>C), M1171D (5121A>G 5122T>A 5123G>C), A1172I (5124G>A 5125C>T 5126G>A), K1173E (5127A>G), E1174D (5132A>T), Y1175F (5134A>T), E1176Q (5136G>C 5138A>G), K1177I (5140A>T 5141G>A), H1178Q (5144T>A), T1180K (5149C>A), A1184D (5161C>A 5162A>T), K1186G (5166A>G 5167A>G 5168G>A), P1190R (5178C>A 5179C>G 5180T>A), Q1192T (5184C>A 5185A>C), T1197S (5199A>T), I1201M (5213T>G), N1203K (5219C>A), K1204N (5222A>C), S1206N (5226T>A 5227C>A 5228T>C), Q1208I (5232C>A 5233A>T), S1213A (5247A>G 5248G>C 5249T>C), Y1217I (5259T>A 5260A>T 5261C>A), Y1227R (5289T>A 5290A>G 5291C>A), K1228T (5293A>C 5294A>G), D1229Y (5295G>T 5297C>T), Q1230A (5298C>G 5299A>C 5300A>T), T1232K (5305C>A 5306A>G), S1235D (5313A>G 5314G>A), I1236K (5317T>A 5318T>A), D1237S (5319G>A 5320A>G 5321C>T), Y1238E (5322T>G 5324T>A), L1240I (5328C>A), L1241N (5331C>A 5332T>A), K1242R (5335A>G), K1244Q (5340A>C), D1245G (5344A>G 5345C>T), I1247T (5350T>C 5351T>A), Y1249F (5356A>T), S1250N (5359G>A 5360C>T), L1254C (5370C>T 5371T>G 5372A>T), F1258Y (5383T>A 5384C>T), H1259W (5385C>T 5386A>G 5387C>G), I1261V (5391A>G), M1262R (5395T>G 5396G>A), D1264H (5400G>C), P1265E (5403C>G 5404C>A), Q1266D (5406C>G 5408G>T), S1267N (5410G>A), A1272T (5424G>A), V1274T (5430G>A 5431T>C 5432C>T), Q1277E (5439C>G 5441G>A), I1283L (5457A>T), N1291T (5482A>C), S1294P (5490T>C 5492A>T), V1295I (5493G>A), K1305G (5523A>G 5524A>G), K1306D (5526A>G 5528A>T), S1308K (5532T>A 5533C>A), E1309R (5535G>A 5536A>G), C1312L (5544T>C 5545G>T 5546T>A), I1315V (5553A>G), D1316Y (5556G>T 5558C>T), I1320V (5568A>G), E1323R (5577G>A 5578A>G), S1324D (5580A>G 5581G>A), I1325M (5585A>G), Q1327E (5589C>G), V1329L (5595G>C), Q1330E (5598C>G), L1333Q (5607T>C 5608T>A), Q1334T (5610C>A 5611A>C 5612G>A), F1335I (5613T>A 5615C>T), Q1337R (5619C>A 5620A>G 5621A>G), V1338L (5622G>T), C1339F (5626G>T 5627T>C), K1340V (5628A>G 5629A>T 5630G>A), E1341S (5631G>A 5632A>G 5633A>T), E1342K (5634G>A), L1344I (5640C>A 5642T>A), L1346I (5646T>A 5648G>A), S1347T (5649T>A 5651A>C), T1349K (5656C>A 5657A>G), L1351M (5661T>A), K1352E (5664A>G), I1353L (5667A>T), G1354C (5670G>T 5672A>T), V1355K (5673G>A 5674T>A), A1356T (5676G>A), N1357H (5679A>C), E1359N (5685G>A 5687A>C), L1363V (5697C>G), E1364T (5700G>A 5701A>C 5702A>T), I1365L (5703A>T), E1367D (5711A>T), V1370I (5718G>A 5720T>A), Q1371K (5721C>A), L1377A (5739T>G 5740T>C 5741A>T), I1380V (5748A>G), E1382D (5756A>T), F1383M (5757T>A 5759T>G), E1385del (5763_5765delGAA), Q1387K (5769C>A), E1389D (5777A>T), T1390K (5779C>A), L1391T (5781T>A 5782T>C), G1393D (5788G>A), I1400L (5808A>T), L1401V (5811T>G 5813A>G), Y1407F (5830A>T), P1409Q (5836C>A), S1412G (5844A>G 5846T>A), Y1414I (5850T>A 5851A>T), T1415A (5853A>G 5855T>A), R1416G (5856A>G), I1417P (5859A>C 5860T>C), F1418L (5864T>A), N1420A (5868A>G 5869A>C 5870T>A), C1422I (5874T>A 5875G>T 5876T>A), S1423G (5877T>G 5878C>G 5879C>A), E1427Q (5889G>C), R1428K (5893G>A) | | | | | | | | |
| Codon mutations: | CTA1125ATA (4983C>A), CAT1126AAA (4986C>A 4988T>A), AAA1127TTA (4989A>T 4990A>T), TTG1128CTA (4992T>C 4994G>A), GAA1129GAT (4997A>T), ATC1131TTA (5001A>T 5003C>A), CAA1132GAA (5004C>G), ATT1133ATA (5009T>A), ATT1134ATA (5012T>A), GGG1135GGA (5015G>A), AAT1136GAA (5016A>G 5018T>A), GAC1137AAA (5019G>A 5021C>A), CCA1138CCC (5024A>C), TTA1139CTA (5025T>C), AAG1140ATA (5029A>T 5030G>A), TAT1141CAT (5031T>C), GAA1143GGA (5038A>G), AAG1144GCC (5040A>G 5041A>C 5042G>C), CAG1146AGA (5046C>A 5047A>G 5048G>A), ATT1147ATA (5051T>A), AGA1148ATG (5053G>T 5054A>G), TGC1149TGT (5057C>T), CTG1151CTA (5063G>A), GAC1152GAT (5066C>T), ATC1153ATA (5069C>A), CCT1156CCA (5078T>A), GAT1157GAA (5081T>A), CTT1158TAT (5082C>T 5083T>A), CAG1161AAA (5091C>A 5093G>A), GAC1162ATA (5094G>A 5095A>T 5096C>A), AAG1163GCA (5097A>G 5098A>C 5099G>A), CCT1164GCA (5100C>G 5102T>A), ATT1165ATA (5105T>A), ATA1166GAA (5106A>G 5107T>A), CCT1167GCT (5109C>G), TCA1168ACA (5112T>A), CCA1169AAT (5115C>A 5116C>A 5117A>T), GAA1170CAA (5118G>C), ATG1171GAC (5121A>G 5122T>A 5123G>C), GCG1172ATA (5124G>A 5125C>T 5126G>A), AAA1173GAA (5127A>G), GAA1174GAT (5132A>T), TAT1175TTT (5134A>T), GAA1176CAG (5136G>C 5138A>G), AAG1177ATA (5140A>T 5141G>A), CAT1178CAA (5144T>A), ATT1179ATA (5147T>A), ACA1180AAA (5149C>A), CTG1183TTA (5157C>T 5159G>A), GCA1184GAT (5161C>A 5162A>T), CTC1185TTA (5163C>T 5165C>A), AAG1186GGA (5166A>G 5167A>G 5168G>A), GTT1187GTA (5171T>A), ATC1188ATA (5174C>A), CCT1190AGA (5178C>A 5179C>G 5180T>A), TCC1191TCA (5183C>A), CAA1192ACA (5184C>A 5185A>C), AGT1193TCT (5187A>T 5188G>C), CGC1194AGA (5190C>A 5192C>A), CAT1195CAC (5195T>C), AGG1196AGA (5198G>A), ACA1197TCA (5199A>T), GCT1199GCA (5207T>A), ATT1201ATG (5213T>G), GTC1202GTA (5216C>A), AAC1203AAA (5219C>A), AAA1204AAC (5222A>C), TCT1206AAC (5226T>A 5227C>A 5228T>C), CAA1208ATA (5232C>A 5233A>T), GTA1209GTC (5237A>C), GGT1211GGA (5243T>A), AGT1213GCC (5247A>G 5248G>C 5249T>C), CGT1214AGA (5250C>A 5252T>A), TAC1217ATA (5259T>A 5260A>T 5261C>A), CGA1221AGA (5271C>A), CTA1222CTC (5276A>C), ACA1226ACT (5288A>T), TAC1227AGA (5289T>A 5290A>G 5291C>A), AAA1228ACG (5293A>C 5294A>G), GAC1229TAT (5295G>T 5297C>T), CAA1230GCT (5298C>G 5299A>C 5300A>T), TAC1231TAT (5303C>T), ACA1232AAG (5305C>A 5306A>G), CTC1233CTA (5309C>A), AGT1235GAT (5313A>G 5314G>A), ATT1236AAA (5317T>A 5318T>A), GAC1237AGT (5319G>A 5320A>G 5321C>T), TAT1238GAA (5322T>G 5324T>A), CTC1239TTA (5325C>T 5327C>A), CTA1240ATA (5328C>A), CTT1241AAT (5331C>A 5332T>A), AAA1242AGA (5335A>G), ATT1243ATA (5339T>A), AAA1244CAA (5340A>C), GAC1245GGT (5344A>G 5345C>T), ATT1247ACA (5350T>C 5351T>A), GTC1248GTA (5354C>A), TAC1249TTC (5356A>T), AGC1250AAT (5359G>A 5360C>T), CTA1254TGT (5370C>T 5371T>G 5372A>T), AAG1255AAA (5375G>A), TTC1258TAT (5383T>A 5384C>T), CAC1259TGG (5385C>T 5386A>G 5387C>G), CAG1260CAA (5390G>A), ATT1261GTT (5391A>G), ATG1262AGA (5395T>G 5396G>A), GAT1264CAT (5400G>C), CCA1265GAA (5403C>G 5404C>A), CAG1266GAT (5406C>G 5408G>T), AGT1267AAT (5410G>A), GCA1272ACA (5424G>A), TTT1273TTC (5429T>C), GTC1274ACT (5430G>A 5431T>C 5432C>T), CCA1276CCT (5438A>T), CAG1277GAA (5439C>G 5441G>A), GGT1278GGA (5444T>A), ATA1283TTA (5457A>T), TTC1287TTT (5471C>T), CTC1289TTG (5475C>T 5477C>G), AAA1290AAG (5480A>G), AAT1291ACT (5482A>C), CCT1293CCA (5489T>A), TCA1294CCT (5490T>C 5492A>T), GTA1295ATA (5493G>A), CGA1298AGA (5502C>A), AAA1299AAG (5507A>G), GAT1301GAC (5513T>C), ATA1303ATT (5519A>T), TTC1304TTT (5522C>T), AAA1305GGA (5523A>G 5524A>G), AAA1306GAT (5526A>G 5528A>T), TAT1307TAC (5531T>C), TCA1308AAA (5532T>A 5533C>A), GAA1309AGA (5535G>A 5536A>G), TGT1312CTA (5544T>C 5545G>T 5546T>A), GTT1313GTA (5549T>A), TAC1314TAT (5552C>T), ATA1315GTA (5553A>G), GAC1316TAT (5556G>T 5558C>T), GAT1317GAC (5561T>C), ATT1318ATC (5564T>C), TTG1319CTA (5565T>C 5567G>A), ATA1320GTA (5568A>G), TTT1321TTC (5573T>C), TCT1322AGT (5574T>A 5575C>G), GAA1323AGA (5577G>A 5578A>G), AGT1324GAT (5580A>G 5581G>A), ATA1325ATG (5585A>G), CAA1327GAA (5589C>G), GTA1329CTA (5595G>C), CAA1330GAA (5598C>G), TTA1333CAA (5607T>C 5608T>A), CAG1334ACA (5610C>A 5611A>C 5612G>A), TTC1335ATT (5613T>A 5615C>T), TTC1336TTT (5618C>T), CAA1337AGG (5619C>A 5620A>G 5621A>G), GTA1338TTA (5622G>T), TGT1339TTC (5626G>T 5627T>C), AAG1340GTA (5628A>G 5629A>T 5630G>A), GAA1341AGT (5631G>A 5632A>G 5633A>T), GAA1342AAA (5634G>A), GGG1343GGA (5639G>A), CTT1344ATA (5640C>A 5642T>A), ATT1345ATA (5645T>A), TTG1346ATA (5646T>A 5648G>A), TCA1347ACC (5649T>A 5651A>C), ACA1349AAG (5656C>A 5657A>G), TTG1351ATG (5661T>A), AAA1352GAA (5664A>G), ATA1353TTA (5667A>T), GGA1354TGT (5670G>T 5672A>T), GTA1355AAA (5673G>A 5674T>A), GCA1356ACA (5676G>A), AAT1357CAT (5679A>C), GAA1359AAC (5685G>A 5687A>C), CTT1361TTA (5691C>T 5693T>A), GGC1362GGA (5696C>A), CTA1363GTA (5697C>G), GAA1364ACT (5700G>A 5701A>C 5702A>T), ATA1365TTA (5703A>T), GAA1367GAT (5711A>T), GGA1368GGG (5714A>G), AAA1369AAG (5717A>G), GTT1370ATA (5718G>A 5720T>A), CAA1371AAA (5721C>A), CTC1372CTA (5726C>A), TTA1377GCT (5739T>G 5740T>C 5741A>T), ATA1380GTA (5748A>G), GAA1382GAT (5756A>T), TTT1383ATG (5757T>A 5759T>G), CCT1384CCA (5762T>A), GAA1385del (5763_5765delGAA), GAC1386GAT (5768C>T), CAA1387AAA (5769C>A), GAA1389GAT (5777A>T), ACA1390AAA (5779C>A), TTA1391ACA (5781T>A 5782T>C), AAG1392AAA (5786G>A), GGT1393GAT (5788G>A), CTA1394TTA (5790C>T), CAG1395CAA (5795G>A), AAG1396AAA (5798G>A), GGG1399GGA (5807G>A), ATA1400TTA (5808A>T), TTA1401GTG (5811T>G 5813A>G), AAC1402AAT (5816C>T), GCC1404GCT (5822C>T), AAT1406AAC (5828T>C), TAT1407TTT (5830A>T), ATC1408ATA (5834C>A), CCA1409CAA (5836C>A), TTG1411TTA (5843G>A), AGT1412GGA (5844A>G 5846T>A), TAT1414ATT (5850T>A 5851A>T), ACT1415GCA (5853A>G 5855T>A), AGA1416GGA (5856A>G), ATA1417CCA (5859A>C 5860T>C), TTT1418TTA (5864T>A), AAT1420GCA (5868A>G 5869A>C 5870T>A), AAG1421AAA (5873G>A), TGT1422ATA (5874T>A 5875G>T 5876T>A), TCC1423GGA (5877T>G 5878C>G 5879C>A), AGC1424AGT (5882C>T), GAA1427CAA (5889G>C), AGA1428AAA (5893G>A) | | | | | | | | |

*: Inserts / Deletes / Misaligned / Frameshifts

#### Analysis details

This analysis was performed with panviral2.64

### NGS Details (UN18_val): Caulimovirus minutangelicae

#### Assembly

| | |
|---|---|
| **Coverage Length** | 391 (1 contig(s)) |
| **Depth Of Coverage** | 6.0 |
| **Number Of Reads** | 17 |
| **Reads Per Million** | 0.32 rpm (after QC) |
| **Ambiguities** | 0 |
| **Assembly Method** | de novo + reference guided assembly |
| **Consensus Caller** | Bcf Tools |

#### Coverage Map

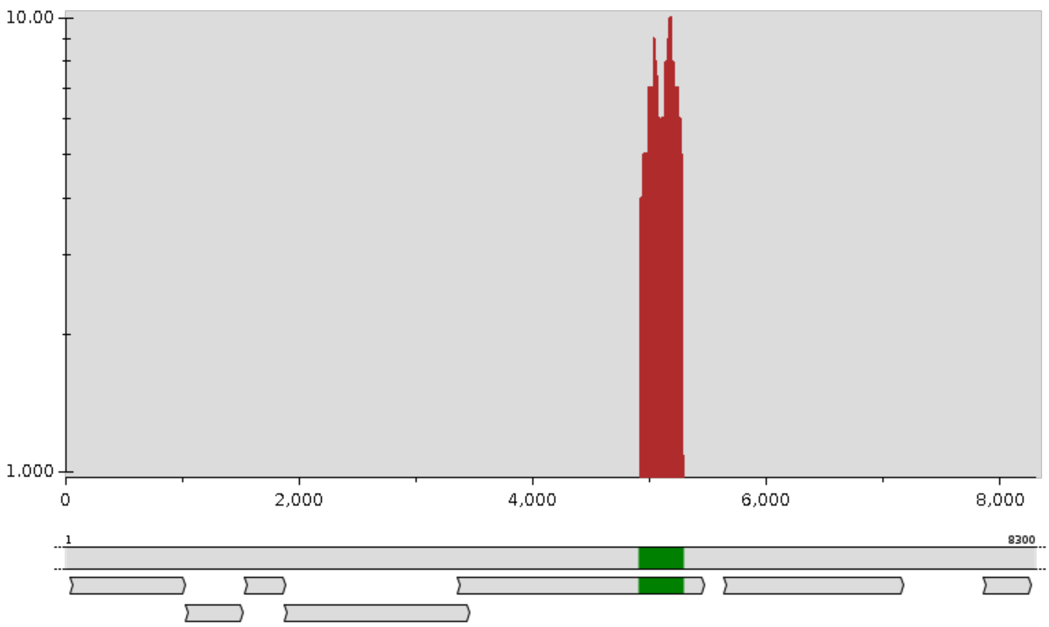

#### Assignment

| | |
|---|---|
| **Type** | Caulimovirus minutangelicae (Taxonomy ID: 3047363) |
| **Reference Genome** | NC_043523.1 |
| **NT Identity (%)** | 54.8813 |
| **AA Identity (%)** | 46.4567 |
| **Number Of Stop Codons** | 0 |
| **Number Of CDS** | 7 |

#### Alignment

| | |
|---|---|
| **Alignment Score** | 44.0 (NT) + 416.0 (AA) = 460.0 |
| **Concordance (%)** | 29.2435 |

| Alignment Method | Global, seeded, nucleotide + amino acids (AGA) |
|---|---|

#### Genome Region

Sequence starts at position 4910 and ends at position 5300 relative to NC_043523.1 reference sequence.

#### Alignment Detailed Statistics

| | Begin | End | Coverage | Score | Concordance | Matches | Identities | I/D/M/F* | Stop Codons |
|---|---|---|---|---|---|---|---|---|---|
| **NT** | **4910** | **5300** | **4.7%** | **44** | **6.0%** | **379 (96.9%)** | **208 (53.2%)** | **0/12** | |
| Mutations: | 4912T>C, 4915T>A, 4916T>A, 4917C>G, 4921T>A, 4923A>T, 4924C>T, 4927T>A, 4932G>A, 4933T>C, 4940T>A, 4941C>A, 4942C>A, 4945A>G, 4949G>A, 4950C>A, 4951A>T, 4953C>T, 4955C>T, 4960G>A, 4961G>A, 4962T>G, 4963A>G, 4964A>C, 4965A>T, 4966A>T, 4967C>A, 4969C>T, 4970A>C, 4971A>G, 4975G>A, 4976G>T, 4977A>C, 4978T>C, 4980T>A, 4981A>C, 4982T>C, 4983C>G, 4984T>A, 4985T>A, 4986G>T, 4988A>G, 4989A>G, 4990T>A, 4998C>A, 5000T>G, 5001C>A, 5003G>C, 5005C>T, 5008T>C, 5009T>A, 5010C>A, 5013A>G, 5014C>T, 5017G>T, 5018G>C, 5019C>A, 5020T>A, 5022A>G, 5023G>C, 5024T>C, 5029G>A, 5030A>C, 5034G>A, 5035T>A, 5037T>G, 5038G>T, 5039A>T, 5040A>C, 5041T>C, 5044T>A, 5047C>A, 5050G>A, 5053A>G, 5054T>C, 5057T>A, 5058A>G, 5059T>A, 5061A>T, 5066C>G, 5067T>A, 5068T>A, 5071T>G, 5074T>A, 5077T>C, 5080G>C, 5081C>T, 5084A>G, 5086C>T, 5087A>C, 5089C>A, 5090G>C, 5101A>T, 5104T>A, 5105C>A, 5107A>T, 5108G>T, 5110T>C, 5111T>C, 5121G>C, 5122C>A, 5123A>T, 5125T>A, 5128G>A, 5129A>C, 5131G>A, 5132G>A, 5134A>T, 5136C>A, 5137A>C, 5138A>C, 5139A>T, 5141_5143delACA, 5148G>A, 5150_5158delGAAGAGCTC, 5167A>G, 5170C>T, 5171G>A, 5173T>A, 5174T>A, 5175C>G, 5176A>T, 5179A>T, 5181G>C, 5182C>A, 5188A>T, 5189T>G, 5190C>A, 5192G>A, 5194A>C, 5201C>A, 5202G>A, 5206T>C, 5207C>T, 5208A>C, 5209T>A, 5210T>A, 5212T>A, 5221A>G, 5224G>A, 5226A>T, 5227T>A, 5230A>G, 5233T>C, 5234G>A, 5236T>A, 5237A>G, 5239T>A, 5244T>G, 5245T>C, 5248T>A, 5249A>C, 5250A>G, 5257C>T, 5260G>T, 5261A>G, 5262T>C, 5265A>T, 5270A>T, 5271C>T, 5272T>A, 5276G>A, 5277T>A, 5278A>G, 5281T>A, 5284T>C, 5286C>T, 5287T>C, 5288G>A, 5290T>C, 5294A>T, 5296A>G, 5299T>C | | | | | | | | |

##### CDS

| | Begin | End | Coverage | Score | Concordance | Matches | Identities | I/D/M/F* | Stop Codons |
|---|---|---|---|---|---|---|---|---|---|
| FLA58_gp5 | 517 | 647 | 18.6% | 416 | 48.8% | 127 (96.9%) | 59 (45.0%) | 0/4/0/0 | 0 |
| Protein mutations: | D520E (4921T>A), Y521F (4923A>T 4924C>T), G524D (4932G>A 4933T>C), S527K (4940T>A 4941C>A 4942C>A), A530N (4949G>A 4950C>A 4951A>T), P531L (4953C>T), V534R (4961G>A 4962T>G 4963A>G), K535L (4964A>C 4965A>T 4966A>T), L536I (4967C>A 4969C>T), K537R (4970A>C 4971A>G), D539S (4976G>T 4977A>C 4978T>C), I540N (4980T>A 4981A>C), S541R (4982T>C 4983C>G 4984T>A), W542M (4985T>A 4986G>T), N543G (4988A>G 4989A>G 4990T>A), A546D (4998C>A), S547E (5000T>G 5001C>A), D548H (5003G>C 5005C>T), S550K (5009T>A 5010C>A), Y551C (5013A>G 5014C>T), A553Q (5018G>C 5019C>A 5020T>A), K554S (5022A>G 5023G>C), K557Q (5030A>C), G558E (5034G>A 5035T>A), L559C (5037T>G 5038G>T), N560S (5039A>T 5040A>C 5041T>C), N561K (5044T>A), F562L (5047C>A), Y566R (5057T>A 5058A>G 5059T>A), H567L (5061A>T), L569E (5066C>G 5067T>A 5068T>A), D571E (5074T>A), K573N (5080G>C), I575V (5084A>G 5086C>T), I576L (5087A>C 5089C>A), E577Q (5090G>C), Q582N (5105C>A 5107A>T), D583Y (5108G>T 5110T>C), Y584H (5111T>C), G587A (5121G>C 5122C>A), I588L (5123A>T 5125T>A), K590Q (5129A>C 5131G>A), A591T (5132G>A 5134A>T), A592D (5136C>A 5137A>C), N593L (5138A>C 5139A>T), T594del (5141_5143delACA), G596E (5148G>A), E597_L599del (5150_5158delGAAGAGCTC), A604T (5171G>A 5173T>A), S607T (5181G>C 5182C>A), K609N (5188A>T), S610E (5189T>G 5190C>A), A611T (5192G>A 5194A>C), R614K (5201C>A 5202G>A), H616S (5207C>T 5208A>C 5209T>A), S617T (5210T>A 5212T>A), Y622L (5226A>T 5227T>A), V625I (5234G>A 5236T>A), I626V (5237A>G 5239T>A), V628G (5244T>G 5245T>C), K630R (5249A>C 5250A>G), I634A (5261A>G 5262T>C), Y635F (5265A>T), T637L (5270A>T 5271C>T 5272T>A), V639K (5276G>A 5277T>A 5278A>G), H640Q (5281T>A), T642I (5286C>T 5287T>C), V643I (5288G>A 5290T>C), T645S (5294A>T 5296A>G) | | | | | | | | |
| Codon mutations: | TAT517TAC (4912T>C), GCT518GCA (4915T>A), TCT519AGT (4916T>A 4917C>G), GAT520GAA (4921T>A), TAC521TTT (4923A>T 4924C>T), ATT522ATA (4927T>A), GGT524GAC (4932G>A 4933T>C), TCC527AAA (4940T>A 4941C>A 4942C>A), AAA528AAG (4945A>G), GCA530AAT (4949G>A 4950C>A 4951A>T), CCT531CTT (4953C>T), CTA532TTA (4955C>T), CAG533CAA (4960G>A), GTA534AGG (4961G>A 4962T>G 4963A>G), AAA535CTT (4964A>C 4965A>T 4966A>T), CTC536ATT (4967C>A 4969C>T), AAA537CGA (4970A>C 4971A>G), AAG538AAA (4975G>A), GAT539TCC (4976G>T 4977A>C 4978T>C), ATA540AAC (4980T>A 4981A>C), TCT541CGA (4982T>C 4983C>G 4984T>A), TGG542ATG (4985T>A 4986G>T), AAT543GGA (4988A>G 4989A>G 4990T>A), GCT546GAT (4998C>A), TCA547GAA (5000T>G 5001C>A), GAC548CAT (5003G>C 5005C>T), ACT549ACC (5008T>C), TCA550AAA (5009T>A 5010C>A), TAC551TGT (5013A>G 5014C>T), GTG552GTT (5017G>T), GCT553CAA (5018G>C 5019C>A 5020T>A), AAG554AGC (5022A>G 5023G>C), TTA555CTA (5024T>C), AAG556AAA (5029G>A), AAA557CAA (5030A>C), GGT558GAA (5034G>A 5035T>A), TTG559TGT (5037T>G 5038G>T), AAT560TCC (5039A>T 5040A>C 5041T>C), AAT561AAA (5044T>A), TTC562TTA (5047C>A), CCG563CCA (5050G>A), AAA564AAG (5053A>G), TTA565CTA (5054T>C), TAT566AGA (5057T>A 5058A>G 5059T>A), CAT567CTT (5061A>T), CTT569GAA (5066C>G 5067T>A 5068T>A), CCT570CCG (5071T>G), GAT571GAA (5074T>A), GAT572GAC (5077T>C), AAG573AAC (5080G>C), CTA574TTA (5081C>T), ATC575GTT (5084A>G 5086C>T), ATC576CTA (5087A>C 5089C>A), GAA577CAA (5090G>C), GCA580GCT (5101A>T), TCT581TCA (5104T>A), CAA582AAT (5105C>A 5107A>T), GAT583TAC (5108G>T 5110T>C), TAT584CAT (5111T>C), GGC587GCA (5121G>C 5122C>A), ATT588TTA (5123A>T 5125T>A), TTG589TTA (5128G>A), AAG590CAA (5129A>C 5131G>A), GCA591ACT (5132G>A 5134A>T), GCA592GAC (5136C>A 5137A>C), AAT593CTT (5138A>C 5139A>T), ACA594del (5141_5143delACA), GGA596GAA (5148G>A), GAA597_CTC599del (5150_5158delGAAGAGCTC), AGA602AGG (5167A>G), TAC603TAT (5170C>T), GCT604ACA (5171G>A 5173T>A), TCA605AGT (5174T>A 5175C>G 5176A>T), GGA606GGT (5179A>T), AGC607ACA (5181G>C 5182C>A), AAA609AAT (5188A>T), TCA610GAA (5189T>G 5190C>A), GCA611ACC (5192G>A 5194A>C), CGA614AAA (5201C>A 5202G>A), TAT615TAC (5206T>C), CAT616TCA (5207C>T 5208A>C 5209T>A), TCT617ACA (5210T>A 5212T>A), AAA620AAG (5221A>G), GAG621GAA (5224G>A), TAT622TTA (5226A>T 5227T>A), TTA623TTG (5230A>G), GCT624GCC (5233T>C), GTT625ATA (5234G>A 5236T>A), ATT626GTA (5237A>G 5239T>A), GTT628GGC (5244T>G 5245T>C), ATT629ATA (5248T>A), AAA630CGA (5249A>C 5250A>G), TTC632TTT (5257C>T), TCG633TCT (5260G>T), ATT634GCT (5261A>G 5262T>C), TAT635TTT (5265A>T), ACT637TTA (5270A>T 5271C>T 5272T>A), GTA639AAG (5276G>A 5277T>A 5278A>G), CAT640CAA (5281T>A), TTT641TTC (5284T>C), ACT642ATC (5286C>T 5287T>C), GTT643ATC (5288G>A 5290T>C), ACA645TCG (5294A>T 5296A>G), GAT646GAC (5299T>C) | | | | | | | | |

##### Proteins

| | Begin | End | Coverage | Score | Concordance | Matches | Identities | I/D/M/F* | Stop Codons |
|---|---|---|---|---|---|---|---|---|---|
| reverse transcriptase (YP_009666503.1) | 517 | 647 | 18.6% | 416 | 48.8% | 127 (96.9%) | 59 (45.0%) | 0/4/0/0 | 0 |
| Protein mutations: | D520E (4921T>A), Y521F (4923A>T 4924C>T), G524D (4932G>A 4933T>C), S527K (4940T>A 4941C>A 4942C>A), A530N (4949G>A 4950C>A 4951A>T), P531L (4953C>T), V534R (4961G>A 4962T>G 4963A>G), K535L (4964A>C 4965A>T 4966A>T), L536I (4967C>A 4969C>T), K537R (4970A>C 4971A>G), D539S (4976G>T 4977A>C 4978T>C), I540N (4980T>A 4981A>C), S541R (4982T>C 4983C>G 4984T>A), W542M (4985T>A 4986G>T), N543G (4988A>G 4989A>G 4990T>A), A546D (4998C>A), S547E (5000T>G 5001C>A), D548H (5003G>C 5005C>T), S550K (5009T>A 5010C>A), Y551C (5013A>G 5014C>T), A553Q (5018G>C 5019C>A 5020T>A), K554S (5022A>G 5023G>C), K557Q (5030A>C), G558E (5034G>A 5035T>A), L559C (5037T>G 5038G>T), N560S (5039A>T 5040A>C 5041T>C), N561K (5044T>A), F562L (5047C>A), Y566R (5057T>A 5058A>G 5059T>A), H567L (5061A>T), L569E (5066C>G 5067T>A 5068T>A), D571E (5074T>A), K573N (5080G>C), I575V (5084A>G 5086C>T), I576L (5087A>C 5089C>A), E577Q (5090G>C), Q582N (5105C>A 5107A>T), D583Y (5108G>T 5110T>C), Y584H (5111T>C), G587A (5121G>C 5122C>A), I588L (5123A>T 5125T>A), K590Q (5129A>C 5131G>A), A591T (5132G>A 5134A>T), A592D (5136C>A 5137A>C), N593L (5138A>C 5139A>T), T594del (5141_5143delACA), G596E (5148G>A), E597_L599del (5150_5158delGAAGAGCTC), A604T (5171G>A 5173T>A), S607T (5181G>C 5182C>A), K609N (5188A>T), S610E (5189T>G 5190C>A), A611T (5192G>A 5194A>C), R614K (5201C>A 5202G>A), H616S (5207C>T 5208A>C 5209T>A), S617T (5210T>A 5212T>A), Y622L (5226A>T 5227T>A), V625I (5234G>A 5236T>A), I626V (5237A>G 5239T>A), V628G (5244T>G 5245T>C), K630R (5249A>C 5250A>G), I634A (5261A>G 5262T>C), Y635F (5265A>T), T637L (5270A>T 5271C>T 5272T>A), V639K (5276G>A 5277T>A 5278A>G), H640Q (5281T>A), T642I (5286C>T 5287T>C), V643I (5288G>A 5290T>C), T645S (5294A>T 5296A>G) | | | | | | | | |
| Codon mutations: | TAT517TAC (4912T>C), GCT518GCA (4915T>A), TCT519AGT (4916T>A 4917C>G), GAT520GAA (4921T>A), TAC521TTT (4923A>T 4924C>T), ATT522ATA (4927T>A), GGT524GAC (4932G>A 4933T>C), TCC527AAA (4940T>A 4941C>A 4942C>A), AAA528AAG (4945A>G), GCA530AAT (4949G>A 4950C>A 4951A>T), CCT531CTT (4953C>T), CTA532TTA (4955C>T), CAG533CAA (4960G>A), GTA534AGG (4961G>A 4962T>G 4963A>G), AAA535CTT (4964A>C 4965A>T 4966A>T), CTC536ATT (4967C>A 4969C>T), AAA537CGA (4970A>C 4971A>G), AAG538AAA (4975G>A), GAT539TCC (4976G>T 4977A>C 4978T>C), ATA540AAC (4980T>A 4981A>C), TCT541CGA (4982T>C 4983C>G 4984T>A), TGG542ATG (4985T>A 4986G>T), AAT543GGA (4988A>G 4989A>G 4990T>A), GCT546GAT (4998C>A), TCA547GAA (5000T>G 5001C>A), GAC548CAT (5003G>C 5005C>T), ACT549ACC (5008T>C), TCA550AAA (5009T>A 5010C>A), TAC551TGT (5013A>G 5014C>T), GTG552GTT (5017G>T), GCT553CAA (5018G>C 5019C>A 5020T>A), AAG554AGC (5022A>G 5023G>C), TTA555CTA (5024T>C), AAG556AAA (5029G>A), AAA557CAA (5030A>C), GGT558GAA (5034G>A 5035T>A), TTG559TGT (5037T>G 5038G>T), AAT560TCC (5039A>T 5040A>C 5041T>C), AAT561AAA (5044T>A), TTC562TTA (5047C>A), CCG563CCA (5050G>A), AAA564AAG (5053A>G), TTA565CTA (5054T>C), TAT566AGA (5057T>A 5058A>G 5059T>A), CAT567CTT (5061A>T), CTT569GAA (5066C>G 5067T>A 5068T>A), CCT570CCG (5071T>G), GAT571GAA (5074T>A), GAT572GAC (5077T>C), AAG573AAC (5080G>C), CTA574TTA (5081C>T), ATC575GTT (5084A>G 5086C>T), ATC576CTA (5087A>C 5089C>A), GAA577CAA (5090G>C), GCA580GCT (5101A>T), TCT581TCA (5104T>A), CAA582AAT (5105C>A 5107A>T), GAT583TAC (5108G>T 5110T>C), TAT584CAT (5111T>C), GGC587GCA (5121G>C 5122C>A), ATT588TTA (5123A>T 5125T>A), TTG589TTA (5128G>A), AAG590CAA (5129A>C 5131G>A), GCA591ACT (5132G>A 5134A>T), GCA592GAC (5136C>A 5137A>C), AAT593CTT (5138A>C 5139A>T), ACA594del (5141_5143delACA), GGA596GAA (5148G>A), GAA597_CTC599del (5150_5158delGAAGAGCTC), AGA602AGG (5167A>G), TAC603TAT (5170C>T), GCT604ACA (5171G>A 5173T>A), TCA605AGT (5174T>A 5175C>G 5176A>T), GGA606GGT (5179A>T), AGC607ACA (5181G>C 5182C>A), AAA609AAT (5188A>T), TCA610GAA (5189T>G 5190C>A), GCA611ACC (5192G>A 5194A>C), CGA614AAA (5201C>A 5202G>A), TAT615TAC (5206T>C), CAT616TCA (5207C>T 5208A>C 5209T>A), TCT617ACA (5210T>A 5212T>A), AAA620AAG (5221A>G), GAG621GAA (5224G>A), TAT622TTA (5226A>T 5227T>A), TTA623TTG (5230A>G), GCT624GCC (5233T>C), GTT625ATA (5234G>A 5236T>A), ATT626GTA (5237A>G 5239T>A), GTT628GGC (5244T>G 5245T>C), ATT629ATA (5248T>A), AAA630CGA (5249A>C 5250A>G), TTC632TTT (5257C>T), TCG633TCT (5260G>T), ATT634GCT (5261A>G 5262T>C), TAT635TTT (5265A>T), ACT637TTA (5270A>T 5271C>T 5272T>A), GTA639AAG (5276G>A 5277T>A 5278A>G), CAT640CAA (5281T>A), TTT641TTC (5284T>C), ACT642ATC (5286C>T 5287T>C), GTT643ATC (5288G>A 5290T>C), ACA645TCG (5294A>T 5296A>G), GAT646GAC (5299T>C) | | | | | | | | |

*: Inserts / Deletes / Misaligned / Frameshifts

#### Analysis details

This analysis was performed with panviral2.64

### NGS Details (UN18_val): Caulimovirus venafragariae

#### Assembly

| | |
|---|---|
| **Coverage Length** | 607 (1 contig(s)) |
| **Depth Of Coverage** | 3.8 |
| **Number Of Reads** | 17 |
| **Reads Per Million** | 0.32 rpm (after QC) |
| **Ambiguities** | 0 |
| **Assembly Method** | de novo + reference guided assembly |
| **Consensus Caller** | Bcf Tools |

#### Coverage Map

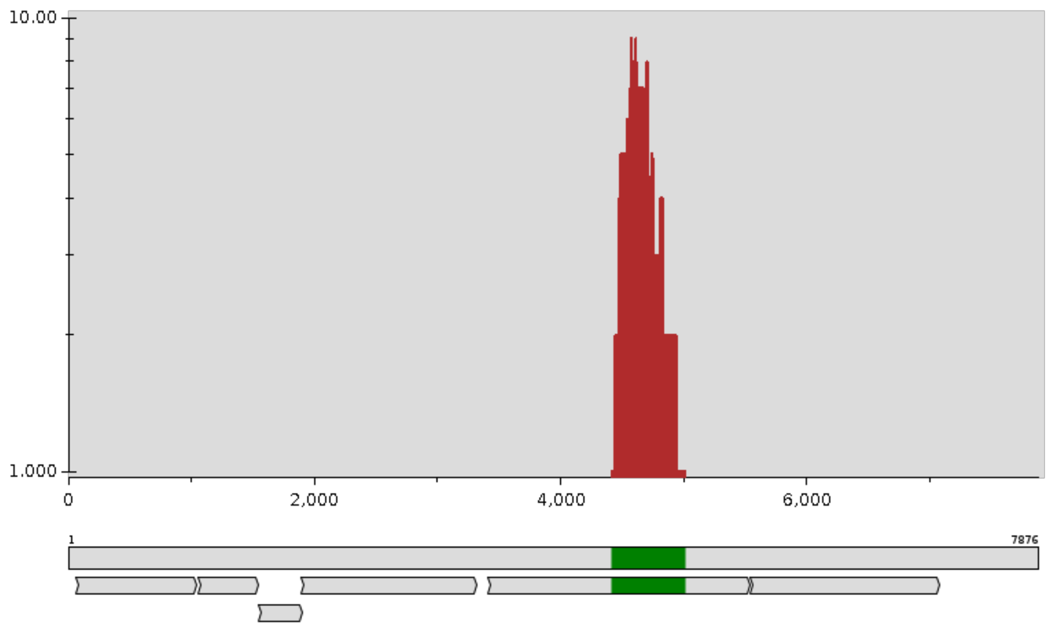

#### Assignment

| | |
|---|---|
| **Type** | Caulimovirus venafragariae (Taxonomy ID: 3048344) |
| **Reference Genome** | NC_001725.1 |
| **NT Identity (%)** | 56.0732 |
| **AA Identity (%)** | 51.2438 |
| **Number Of Stop Codons** | 0 |
| **Number Of CDS** | 6 |

#### Alignment

| | |
|---|---|
| **Alignment Score** | 131.0 (NT) + 766.0 (AA) = 897.0 |
| **Concordance (%)** | 34.66 |

| Alignment Method | Global, seeded, nucleotide + amino acids (AGA) |
|---|---|

#### Genome Region

Sequence starts at position 4411 and ends at position 5017 relative to NC_001725.1 reference sequence.

#### Alignment Detailed Statistics

| | Begin | End | Coverage | Score | Concordance | Matches | Identities | I/D/M/F* | Stop Codons |
|---|---|---|---|---|---|---|---|---|---|
| **NT** | **4411** | **5017** | **7.7%** | **131** | **11.0%** | **601 (99.0%)** | **337 (55.5%)** | **0/6** | |
| Mutations: | 4413T>A, 4423A>G, 4426T>C, 4428A>T, 4432G>A, 4434C>A, 4435C>A, 4438A>T, 4439C>G, 4440A>T, 4441A>G, 4442A>T, 4443G>C, 4444G>T, 4445G>T, 4446A>C, 4449T>C, 4452C>A, 4455C>T, 4456C>T, 4458A>C, 4459C>A, 4461T>C, 4464T>A, 4467C>T, 4470G>A, 4471G>A, 4473G>T, 4474C>A, 4476A>C, 4479G>T, 4480C>A, 4482T>A, 4483C>A, 4485A>T, 4486A>C, 4487G>T, 4488A>T, 4489A>G, 4490T>C, 4492G>A, 4493G>A, 4497T>A, 4500G>A, 4505T>A, 4506T>C, 4508A>T, 4509C>T, 4512T>A, 4513T>A, 4514C>A, 4515C>A, 4521C>T, 4524T>C, 4528T>A, 4529C>G, 4542G>A, 4543G>A, 4546C>A, 4547G>A, 4548C>A, 4551T>A, 4553C>A, 4554T>A, 4555C>G, 4556C>A, 4558G>A, 4560A>T, 4561A>T, 4563A>T, 4566T>A, 4567C>G, 4568A>C, 4569G>T, 4570C>T, 4575C>A, 4578T>A, 4584C>T, 4585T>A, 4586G>C, 4587T>A, 4590C>A, 4591C>A, 4593A>T, 4614C>G, 4633C>A, 4635A>T, 4642G>C, 4643C>A, 4644T>A, 4647T>C, 4650C>T, 4653A>G, 4657C>A, 4659C>A, 4665C>T, 4666G>A, 4670G>T, 4671T>A, 4672C>T, 4674C>T, 4675A>T, 4676G>C, 4678A>C, 4680C>A, 4682T>A, 4684T>A, 4687_4692delCCACAG, 4696T>C, 4697G>T, 4698T>A, 4699G>A, 4700C>T, 4701T>C, 4704A>C, 4707T>C, 4708G>A, 4710T>A, 4716C>T, 4719C>A, 4720A>T, 4722C>A, 4723G>A, 4725C>A, 4727T>G, 4728C>T, 4729A>T, 4730G>C, 4734A>C, 4735A>T, 4737T>A, 4738G>T, 4740A>T, 4741G>C, 4753G>A, 4754G>C, 4755A>T, 4759G>C, 4761C>T, 4765A>C, 4766T>A, 4767T>A, 4768G>T, 4771C>T, 4772T>C, 4773A>T, 4774A>C, 4776C>A, 4778G>T, 4782C>T, 4783A>T, 4784A>G, 4785A>C, 4787C>A, 4788C>A, 4789C>A, 4790T>A, 4791A>T, 4797A>T, 4801C>T, 4803T>A, 4807A>C, 4809A>G, 4812G>A, 4815A>G, 4818C>A, 4819C>G, 4821G>C, 4822T>A, 4824G>A, 4825T>A, 4826G>A, 4827C>T, 4832C>A, 4833A>T, 4834A>G, 4835C>A, 4836T>A, 4839C>T, 4840A>G, 4842T>A, 4845C>T, 4848T>C, 4851T>A, 4852C>A, 4854A>G, 4855G>A, 4860A>T, 4861G>T, 4862A>C, 4865G>A, 4866A>G, 4867G>A, 4868G>A, 4869A>T, 4870A>G, 4871A>G, 4872T>C, 4873C>A, 4875C>T, 4876A>G, 4879G>T, 4881C>A, 4884G>A, 4886G>C, 4893C>A, 4894G>T, 4895G>C, 4897C>A, 4898T>A, 4900C>A, 4902C>G, 4903T>A, 4905A>C, 4906G>T, 4907T>G, 4910C>A, 4911T>A, 4921C>A, 4926T>A, 4928G>A, 4930G>A, 4931A>C, 4932T>A, 4934G>A, 4935G>A, 4936A>C, 4938T>A, 4940C>A, 4941C>A, 4942T>A, 4944A>C, 4950A>G, 4954T>C, 4956A>T, 4959C>A, 4960C>T, 4961T>G, 4962C>T, 4963C>T, 4971T>C, 4972A>G, 4973T>C, 4974C>A, 4975T>A, 4976C>G, 4977C>T, 4979C>A, 4980T>A, 4982A>T, 4983T>C, 4984T>A, 4986T>A, 4990A>G, 4992A>C, 4993A>T, 5001C>A, 5002C>A, 5003T>A, 5004T>G, 5007G>A, 5008T>A, 5009C>A, 5010A>T, 5012C>T | | | | | | | | |

##### CDS

| | Begin | End | Coverage | Score | Concordance | Matches | Identities | I/D/M/F* | Stop Codons |
|---|---|---|---|---|---|---|---|---|---|
| ORF_V | 336 | 538 | 28.6% | 766 | 54.4% | 201 (99.0%) | 103 (50.7%) | 0/2/0/0 | 0 |
| Protein mutations: | K340E (4423A>G), D343K (4432G>A 4434C>A), H344N (4435C>A), T345C (4438A>T 4439C>G 4440A>T), K346V (4441A>G 4442A>T 4443G>C), G347F (4444G>T 4445G>T 4446A>C), L351F (4456C>T 4458A>C), L352I (4459C>A 4461T>C), E356N (4471G>A 4473G>T), Q357N (4474C>A 4476A>C), L359I (4480C>A 4482T>A), Q360N (4483C>A 4485A>T), R361L (4486A>C 4487G>T 4488A>T), I362A (4489A>G 4490T>C), G363K (4492G>A 4493G>A), F367Y (4505T>A 4506T>C), Y368F (4508A>T 4509C>T), S370K (4513T>A 4514C>A 4515C>A), V380I (4543G>A), R381K (4546C>A 4547G>A 4548C>A), A383E (4553C>A 4554T>A), P384E (4555C>G 4556C>A), E385N (4558G>A 4560A>T), T386S (4561A>T 4563A>T), Q388A (4567C>G 4568A>C 4569G>T), C394T (4585T>A 4586G>C 4587T>A), Q396N (4591C>A 4593A>T), Q410N (4633C>A 4635A>T), A413Q (4642G>C 4643C>A 4644T>A), H418K (4657C>A 4659C>A), E421K (4666G>A), S422I (4670G>T 4671T>A), L423F (4672C>T 4674C>T), N425Q (4678A>C 4680C>A), M426K (4682T>A), Y427N (4684T>A), P428_Q429del (4687_4692delCCACAG), C431L (4696T>C 4697G>T 4698T>A), A432I (4699G>A 4700C>T 4701T>C), V435I (4708G>A 4710T>A), I439L (4720A>T 4722C>A), V440I (4723G>A 4725C>A), F441C (4727T>G 4728C>T), K443N (4734A>C), T444S (4735A>T 4737T>A), E445Y (4738G>T 4740A>T), E446Q (4741G>C), G450T (4753G>A 4754G>C 4755A>T), V452L (4759G>C 4761C>T), I454Q (4765A>C 4766T>A 4767T>A), V455F (4768G>T), L456S (4771C>T 4772T>C 4773A>T), N457Q (4774A>C 4776C>A), R458I (4778G>T), K460C (4783A>T 4784A>G 4785A>C), A461E (4787C>A 4788C>A), L462N (4789C>A 4790T>A 4791A>T), K468Q (4807A>C 4809A>G), Q472D (4819C>G 4821G>C), L473I (4822T>A 4824G>A), C474N (4825T>A 4826G>A 4827C>T), T476N (4832C>A 4833A>T), T477E (4834A>G 4835C>A 4836T>A), N479E (4840A>G 4842T>A), L483M (4852C>A 4854A>G), V484I (4855G>A), E486S (4861G>T 4862A>C), R487K (4865G>A 4866A>G), G488N (4867G>A 4868G>A 4869A>T), N489G (4870A>G 4871A>G 4872T>C), L490I (4873C>A 4875C>T), K491E (4876A>G), V492L (4879G>T 4881C>A), S494T (4886G>C), G497S (4894G>T 4895G>C), L498K (4897C>A 4898T>A), H499K (4900C>A 4902C>G), L500I (4903T>A 4905A>C), V501C (4906G>T 4907T>G), A502E (4910C>A 4911T>A), Q506K (4921C>A), S508N (4928G>A), D509T (4930G>A 4931A>C 4932T>A), R510K (4934G>A 4935G>A), N511Q (4936A>C 4938T>A), A512E (4940C>A 4941C>A), L513I (4942T>A 4944A>C), L519C (4960C>T 4961T>G 4962C>T), I523A (4972A>G 4973T>C 4974C>A), A525E (4979C>A 4980T>A), Y526F (4982A>T 4983T>C), F527I (4984T>A 4986T>A), K529D (4990A>G 4992A>C), I530L (4993A>T), N532K (5001C>A), L533K (5002C>A 5003T>A 5004T>G), S535N (5008T>A 5009C>A 5010A>T), P536L (5012C>T) | | | | | | | | |
| Codon mutations: | ATT336ATA (4413T>A), AAG340GAG (4423A>G), TTA341CTT (4426T>C 4428A>T), GAC343AAA (4432G>A 4434C>A), CAT344AAT (4435C>A), ACA345TGT (4438A>T 4439C>G 4440A>T), AAG346GTC (4441A>G 4442A>T 4443G>C), GGA347TTC (4444G>T 4445G>T 4446A>C), GAT348GAC (4449T>C), GGC349GGA (4452C>A), TAC350TAT (4455C>T), CTA351TTC (4456C>T 4458A>C), CTT352ATC (4459C>A 4461T>C), CCT353CCA (4464T>A), AAC354AAT (4467C>T), AAG355AAA (4470G>A), GAG356AAT (4471G>A 4473G>T), CAA357AAC (4474C>A 4476A>C), CTG358CTT (4479G>T), CTT359ATA (4480C>A 4482T>A), CAA360AAT (4483C>A 4485A>T), AGA361CTT (4486A>C 4487G>T 4488A>T), ATC362GCC (4489A>G 4490T>C), GGA363AAA (4492G>A 4493G>A), GGT364GGA (4497T>A), AAG365AAA (4500G>A), TTT367TAC (4505T>A 4506T>C), TAC368TTT (4508A>T 4509C>T), TCT369TCA (4512T>A), TCC370AAA (4513T>A 4514C>A 4515C>A), GAC372GAT (4521C>T), TGT373TGC (4524T>C), TCT375AGT (4528T>A 4529C>G), CAG379CAA (4542G>A), GTA380ATA (4543G>A), CGC381AAA (4546C>A 4547G>A 4548C>A), CTT382CTA (4551T>A), GCT383GAA (4553C>A 4554T>A), CCA384GAA (4555C>G 4556C>A), GAA385AAT (4558G>A 4560A>T), ACA386TCT (4561A>T 4563A>T), ATT387ATA (4566T>A), CAG388GCT (4567C>G 4568A>C 4569G>T), CTA389TTA (4570C>T), ACC390ACA (4575C>A), GCT391GCA (4578T>A), AGC393AGT (4584C>T), TGT394ACA (4585T>A 4586G>C 4587T>A), CCC395CCA (4590C>A), CAA396AAT (4591C>A 4593A>T), GTC403GTG (4614C>G), CAA410AAT (4633C>A 4635A>T), GCT413CAA (4642G>C 4643C>A 4644T>A), ATT414ATC (4647T>C), TTC415TTT (4650C>T), CAA416CAG (4653A>G), CAC418AAA (4657C>A 4659C>A), GAC420GAT (4665C>T), GAA421AAA (4666G>A), AGT422ATA (4670G>T 4671T>A), CTC423TTT (4672C>T 4674C>T), AGC424TCC (4675A>T 4676G>C), AAC425CAA (4678A>C 4680C>A), ATG426AAG (4682T>A), TAT427AAT (4684T>A), CCA428_CAG429del (4687_4692delCCACAG), TGT431CTA (4696T>C 4697G>T 4698T>A), GCT432ATC (4699G>A 4700C>T 4701T>C), GTA433GTC (4704A>C), TAT434TAC (4707T>C), GTT435ATA (4708G>A 4710T>A), GAC437GAT (4716C>T), ATC438ATA (4719C>A), ATC439TTA (4720A>T 4722C>A), GTC440ATA (4723G>A 4725C>A), TTC441TGT (4727T>G 4728C>T), AGC442TCC (4729A>T 4730G>C), AAA443AAC (4734A>C), ACT444TCA (4735A>T 4737T>A), GAA445TAT (4738G>T 4740A>T), GAA446CAA (4741G>C), GGA450ACT (4753G>A 4754G>C 4755A>T), GTC452CTT (4759G>C 4761C>T), ATT454CAA (4765A>C 4766T>A 4767T>A), GTT455TTT (4768G>T), CTA456TCT (4771C>T 4772T>C 4773A>T), AAC457CAA (4774A>C 4776C>A), AGA458ATA (4778G>T), TGC459TGT (4782C>T), AAA460TGC (4783A>T 4784A>G 4785A>C), GCC461GAA (4787C>A 4788C>A), CTA462AAT (4789C>A 4790T>A 4791A>T), ATA464ATT (4797A>T), CTT466TTA (4801C>T 4803T>A), AAA468CAG (4807A>C 4809A>G), AAG469AAA (4812G>A), AAA470AAG (4815A>G), GCC471GCA (4818C>A), CAG472GAC (4819C>G 4821G>C), TTG473ATA (4822T>A 4824G>A), TGC474AAT (4825T>A 4826G>A 4827C>T), ACA476AAT (4832C>A 4833A>T), ACT477GAA (4834A>G 4835C>A 4836T>A), ATC478ATT (4839C>T), AAT479GAA (4840A>G 4842T>A), TTC480TTT (4845C>T), CTT481CTC (4848T>C), GGT482GGA (4851T>A), CTA483ATG (4852C>A 4854A>G), GTA484ATA (4855G>A), ATA485ATT (4860A>T), GAA486TCA (4861G>T 4862A>C), AGA487AAG (4865G>A 4866A>G), GGA488AAT (4867G>A 4868G>A 4869A>T), AAT489GGC (4870A>G 4871A>G 4872T>C), CTC490ATT (4873C>A 4875C>T), AAA491GAA (4876A>G), GTC492TTA (4879G>T 4881C>A), CAG493CAA (4884G>A), AGT494ACT (4886G>C), ATC496ATA (4893C>A), GGA497TCA (4894G>T 4895G>C), CTA498AAA (4897C>A 4898T>A), CAC499AAG (4900C>A 4902C>G), TTA500ATC (4903T>A 4905A>C), GTT501TGT (4906G>T 4907T>G), GCT502GAA (4910C>A 4911T>A), CAA506AAA (4921C>A), CTT507CTA (4926T>A), AGT508AAT (4928G>A), GAT509ACA (4930G>A 4931A>C 4932T>A), AGG510AAA (4934G>A 4935G>A), AAT511CAA (4936A>C 4938T>A), GCC512GAA (4940C>A 4941C>A), TTA513ATC (4942T>A 4944A>C), AGA515AGG (4950A>G), TTA517CTT (4954T>C 4956A>T), GGC518GGA (4959C>A), CTC519TGT (4960C>T 4961T>G 4962C>T), CTA520TTA (4963C>T), TAT522TAC (4971T>C), ATC523GCA (4972A>G 4973T>C 4974C>A), TCC524AGT (4975T>A 4976C>G 4977C>T), GCT525GAA (4979C>A 4980T>A), TAT526TTC (4982A>T 4983T>C), TTT527ATA (4984T>A 4986T>A), AAA529GAC (4990A>G 4992A>C), ATA530TTA (4993A>T), AAC532AAA (5001C>A), CTT533AAG (5002C>A 5003T>A 5004T>G), AGG534AGA (5007G>A), TCA535AAT (5008T>A 5009C>A 5010A>T), CCT536CTT (5012C>T) | | | | | | | | |

##### Proteins

| | Begin | End | Coverage | Score | Concordance | Matches | Identities | I/D/M/F* | Stop Codons |
|---|---|---|---|---|---|---|---|---|---|
| hypothetical protein (NP_043933.1) | 336 | 538 | 28.6% | 766 | 54.4% | 201 (99.0%) | 103 (50.7%) | 0/2/0/0 | 0 |
| Protein mutations: | K340E (4423A>G), D343K (4432G>A 4434C>A), H344N (4435C>A), T345C (4438A>T 4439C>G 4440A>T), K346V (4441A>G 4442A>T 4443G>C), G347F (4444G>T 4445G>T 4446A>C), L351F (4456C>T 4458A>C), L352I (4459C>A 4461T>C), E356N (4471G>A 4473G>T), Q357N (4474C>A 4476A>C), L359I (4480C>A 4482T>A), Q360N (4483C>A 4485A>T), R361L (4486A>C 4487G>T 4488A>T), I362A (4489A>G 4490T>C), G363K (4492G>A 4493G>A), F367Y (4505T>A 4506T>C), Y368F (4508A>T 4509C>T), S370K (4513T>A 4514C>A 4515C>A), V380I (4543G>A), R381K (4546C>A 4547G>A 4548C>A), A383E (4553C>A 4554T>A), P384E (4555C>G 4556C>A), E385N (4558G>A 4560A>T), T386S (4561A>T 4563A>T), Q388A (4567C>G 4568A>C 4569G>T), C394T (4585T>A 4586G>C 4587T>A), Q396N (4591C>A 4593A>T), Q410N (4633C>A 4635A>T), A413Q (4642G>C 4643C>A 4644T>A), H418K (4657C>A 4659C>A), E421K (4666G>A), S422I (4670G>T 4671T>A), L423F (4672C>T 4674C>T), N425Q (4678A>C 4680C>A), M426K (4682T>A), Y427N (4684T>A), P428_Q429del (4687_4692delCCACAG), C431L (4696T>C 4697G>T 4698T>A), A432I (4699G>A 4700C>T 4701T>C), V435I (4708G>A 4710T>A), I439L (4720A>T 4722C>A), V440I (4723G>A 4725C>A), F441C (4727T>G 4728C>T), K443N (4734A>C), T444S (4735A>T 4737T>A), E445Y (4738G>T 4740A>T), E446Q (4741G>C), G450T (4753G>A 4754G>C 4755A>T), V452L (4759G>C 4761C>T), I454Q (4765A>C 4766T>A 4767T>A), V455F (4768G>T), L456S (4771C>T 4772T>C 4773A>T), N457Q (4774A>C 4776C>A), R458I (4778G>T), K460C (4783A>T 4784A>G 4785A>C), A461E (4787C>A 4788C>A), L462N (4789C>A 4790T>A 4791A>T), K468Q (4807A>C 4809A>G), Q472D (4819C>G 4821G>C), L473I (4822T>A 4824G>A), C474N (4825T>A 4826G>A 4827C>T), T476N (4832C>A 4833A>T), T477E (4834A>G 4835C>A 4836T>A), N479E (4840A>G 4842T>A), L483M (4852C>A 4854A>G), V484I (4855G>A), E486S (4861G>T 4862A>C), R487K (4865G>A 4866A>G), G488N (4867G>A 4868G>A 4869A>T), N489G (4870A>G 4871A>G 4872T>C), L490I (4873C>A 4875C>T), K491E (4876A>G), V492L (4879G>T 4881C>A), S494T (4886G>C), G497S (4894G>T 4895G>C), L498K (4897C>A 4898T>A), H499K (4900C>A 4902C>G), L500I (4903T>A 4905A>C), V501C (4906G>T 4907T>G), A502E (4910C>A 4911T>A), Q506K (4921C>A), S508N (4928G>A), D509T (4930G>A 4931A>C 4932T>A), R510K (4934G>A 4935G>A), N511Q (4936A>C 4938T>A), A512E (4940C>A 4941C>A), L513I (4942T>A 4944A>C), L519C (4960C>T 4961T>G 4962C>T), I523A (4972A>G 4973T>C 4974C>A), A525E (4979C>A 4980T>A), Y526F (4982A>T 4983T>C), F527I (4984T>A 4986T>A), K529D (4990A>G 4992A>C), I530L (4993A>T), N532K (5001C>A), L533K (5002C>A 5003T>A 5004T>G), S535N (5008T>A 5009C>A 5010A>T), P536L (5012C>T) | | | | | | | | |

| | Begin | End | Coverage | Score | Concordance | Matches | Identities | I/D/M/F* | Stop Codons |
|---|---|---|---|---|---|---|---|---|---|
| **NT** | **4411** | **5017** | **7.7%** | **131** | **11.0%** | **601 (99.0%)** | **337 (55.5%)** | **0/6** | |
| Codon mutations: | ATT336ATA (4413T>A), AAG340GAG (4423A>G), TTA341CTT (4426T>C 4428A>T), GAC343AAA (4432G>A 4434C>A), CAT344AAT (4435C>A), ACA345TGT (4438A>T 4439C>G 4440A>T), AAG346GTC (4441A>G 4442A>T 4443G>C), GGA347TTC (4444G>T 4445G>T 4446A>C), GAT348GAC (4449T>C), GGC349GGA (4452C>A), TAC350TAT (4455C>T), CTA351TTC (4456C>T 4458A>C), CTT352ATC (4459C>A 4461T>C), CCT353CCA (4464T>A), AAC354AAT (4467C>T), AAG355AAA (4470G>A), GAG356AAT (4471G>A 4473G>T), CAA357AAC (4474C>A 4476A>C), CTG358CTT (4479G>T), CTT359ATA (4480C>A 4482T>A), CAA360AAT (4483C>A 4485A>T), AGA361CTT (4486A>C 4487G>T 4488A>T), ATC362GCC (4489A>G 4490T>C), GGA363AAA (4492G>A 4493G>A), GGT364GGA (4497T>A), AAG365AAA (4500G>A), TTT367TAC (4505T>A 4506T>C), TAC368TTT (4508A>T 4509C>T), TCT369TCA (4512T>A), TCC370AAA (4513T>A 4514C>A 4515C>A), GAC372GAT (4521C>T), TGT373TGC (4524T>C), TCT375AGT (4528T>A 4529C>G), CAG379CAA (4542G>A), GTA380ATA (4543G>A), CGC381AAA (4546C>A 4547G>A 4548C>A), CTT382CTA (4551T>A), GCT383GAA (4553C>A 4554T>A), CCA384GAA (4555C>G 4556C>A), GAA385AAT (4558G>A 4560A>T), ACA386TCT (4561A>T 4563A>T), ATT387ATA (4566T>A), CAG388GCT (4567C>G 4568A>C 4569G>T), CTA389TTA (4570C>T), ACC390ACA (4575C>A), GCT391GCA (4578T>A), AGC393AGT (4584C>T), TGT394ACA (4585T>A 4586G>C 4587T>A), CCC395CCA (4590C>A), CAA396AAT (4591C>A 4593A>T), GTC403GTG (4614C>G), CAA410AAT (4633C>A 4635A>T), GCT413CAA (4642G>C 4643C>A 4644T>A), ATT414ATC (4647T>C), TTC415TTT (4650C>T), CAA416CAG (4653A>G), CAC418AAA (4657C>A 4659C>A), GAC420GAT (4665C>T), GAA421AAA (4666G>A), AGT422ATA (4670G>T 4671T>A), CTC423TTT (4672C>T 4674C>T), AGC424TCC (4675A>T 4676G>C), AAC425CAA (4678A>C 4680C>A), ATG426AAG (4682T>A), TAT427AAT (4684T>A), CCA428_CAG429del (4687_4692delCCACAG), TGT431CTA (4696T>C 4697G>T 4698T>A), GCT432ATC (4699G>A 4700C>T 4701T>C), GTA433GTC (4704A>C), TAT434TAC (4707T>C), GTT435ATA (4708G>A 4710T>A), GAC437GAT (4716C>T), ATC438ATA (4719C>A), ATC439TTA (4720A>T 4722C>A), GTC440ATA (4723G>A 4725C>A), TTC441TGT (4727T>G 4728C>T), AGC442TCC (4729A>T 4730G>C), AAA443AAC (4734A>C), ACT444TCA (4735A>T 4737T>A), GAA445TAT (4738G>T 4740A>T), GAA446CAA (4741G>C), GGA450ACT (4753G>A 4754G>C 4755A>T), GTC452CTT (4759G>C 4761C>T), ATT454CAA (4765A>C 4766T>A 4767T>A), GTT455TTT (4768G>T), CTA456TCT (4771C>T 4772T>C 4773A>T), AAC457CAA (4774A>C 4776C>A), AGA458ATA (4778G>T), TGC459TGT (4782C>T), AAA460TGC (4783A>T 4784A>G 4785A>C), GCC461GAA (4787C>A 4788C>A), CTA462AAT (4789C>A 4790T>A 4791A>T), ATA464ATT (4797A>T), CTT466TTA (4801C>T 4803T>A), AAA468CAG (4807A>C 4809A>G), AAG469AAA (4812G>A), AAA470AAG (4815A>G), GCC471GCA (4818C>A), CAG472GAC (4819C>G 4821G>C), TTG473ATA (4822T>A 4824G>A), TGC474AAT (4825T>A 4826G>A 4827C>T), ACA476AAT (4832C>A 4833A>T), ACT477GAA (4834A>G 4835C>A 4836T>A), ATC478ATT (4839C>T), AAT479GAA (4840A>G 4842T>A), TTC480TTT (4845C>T), CTT481CTC (4848T>C), GGT482GGA (4851T>A), CTA483ATG (4852C>A 4854A>G), GTA484ATA (4855G>A), ATA485ATT (4860A>T), GAA486TCA (4861G>T 4862A>C), AGA487AAG (4865G>A 4866A>G), GGA488AAT (4867G>A 4868G>A 4869A>T), AAT489GGC (4870A>G 4871A>G 4872T>C), CTC490ATT (4873C>A 4875C>T), AAA491GAA (4876A>G), GTC492TTA (4879G>T 4881C>A), CAG493CAA (4884G>A), AGT494ACT (4886G>C), ATC496ATA (4893C>A), GGA497TCA (4894G>T 4895G>C), CTA498AAA (4897C>A 4898T>A), CAC499AAG (4900C>A 4902C>G), TTA500ATC (4903T>A 4905A>C), GTT501TGT (4906G>T 4907T>G), GCT502GAA (4910C>A 4911T>A), CAA506AAA (4921C>A), CTT507CTA (4926T>A), AGT508AAT (4928G>A), GAT509ACA (4930G>A 4931A>C 4932T>A), AGG510AAA (4934G>A 4935G>A), AAT511CAA (4936A>C 4938T>A), GCC512GAA (4940C>A 4941C>A), TTA513ATC (4942T>A 4944A>C), AGA515AGG (4950A>G), TTA517CTT (4954T>C 4956A>T), GGC518GGA (4959C>A), CTC519TGT (4960C>T 4961T>G 4962C>T), CTA520TTA (4963C>T), TAT522TAC (4971T>C), ATC523GCA (4972A>G 4973T>C 4974C>A), TCC524AGT (4975T>A 4976C>G 4977C>T), GCT525GAA (4979C>A 4980T>A), TAT526TTC (4982A>T 4983T>C), TTT527ATA (4984T>A 4986T>A), AAA529GAC (4990A>G 4992A>C), ATA530TTA (4993A>T), AAC532AAA (5001C>A), CTT533AAG (5002C>A 5003T>A 5004T>G), AGG534AGA (5007G>A), TCA535AAT (5008T>A 5009C>A 5010A>T), CCT536CTT (5012C>T) | | | | | | | | |

*: Inserts / Deletes / Misaligned / Frameshifts

#### Analysis details

This analysis was performed with panviral2.64

### NGS Details (UN18_val): Badnavirus maculasmallanthi

#### Assembly

| | |
|---|---|
| **Coverage Length** | 281 (1 contig(s)) |
| **Depth Of Coverage** | 5.8 |
| **Number Of Reads** | 13 |
| **Reads Per Million** | 0.24 rpm (after QC) |
| **Ambiguities** | 0 |
| **Assembly Method** | de novo + reference guided assembly |
| **Consensus Caller** | Bcf Tools |

#### Coverage Map

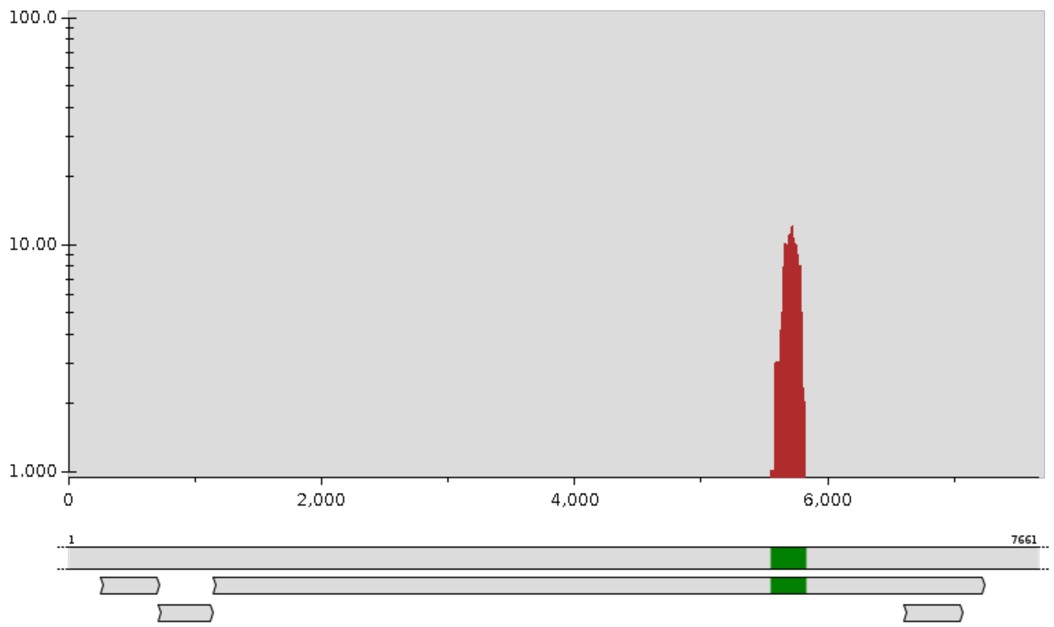

#### Assignment

| | |
|---|---|
| **Type** | Badnavirus maculasmallanthi (Taxonomy ID: 3048453) |
| **Reference Genome** | NC_026472.1 |
| **NT Identity (%)** | 56.338 |
| **AA Identity (%)** | 48.9362 |
| **Number Of Stop Codons** | 1 |
| **Number Of CDS** | 4 |

#### Alignment

| | |
|---|---|
| **Alignment Score** | 66.0 (NT) + 305.0 (AA) = 371.0 |
| **Concordance (%)** | 30.261 |

| Alignment Method | Global, seeded, nucleotide + amino acids (AGA) |
|---|---|

#### Genome Region

Sequence starts at position 5549 and ends at position 5829 relative to NC_026472.1 reference sequence.

#### Alignment Detailed Statistics

| | Begin | End | Coverage | Score | Concordance | Matches | Identities | I/D/M/F* | Stop Codons |
|---|---|---|---|---|---|---|---|---|---|
| **NT** | **5549** | **5829** | **3.7%** | **66** | **11.7%** | **281 (98.9%)** | **160 (56.3%)** | **3/0** | |
| Mutations: | 5549G>A, 5551A>T, 5554G>C, 5557A>G, 5558A>G, 5562A>T, 5563A>T, 5564A>T, 5566A>G, 5568A>G, 5574A>C, 5578T>A, 5587A>C, 5589G>T, 5595A>G, 5597T>A, 5601T>C, 5604A>G, 5605G>T, 5607T>G, 5608G>A, 5609C>G, 5610C>A, 5611A>G, 5613G>T, 5614G>T, 5615A>G, 5619A>G, 5620G>T, 5621A>C, 5623T>G, 5624C>A, 5625C>T, 5628A>C, 5630C>T, 5631C>A, 5632T>A, 5633G>A, 5634G>A, 5637G>T, 5638G>A, 5640C>T, 5644T>A, 5645T>G, 5646A>G, 5648T>C, 5649C>A, 5651C>G, 5652A>G, 5653G>T, 5654G>A, 5655A>C, 5658A>G, 5659T>C, 5660T>A, 5661A>T, 5664C>T, 5667A>G, 5669G>T, 5670G>T, 5671C>G, 5673T>G, 5676C>T, 5680C>T, 5682A>C, 5685C>T, 5686G>A, 5691C>A, 5693A>C, 5694A>T, 5703C>T, 5706T>C, 5707A>G, 5708T>C, 5709T>A, 5713C>A, 5714A>T, 5716A>G, 5717G>A, 5718A>G, 5719A>T, 5720A>T, 5721A>G, 5725G>A, 5727T>C, 5728A>C, 5729A>G, 5731T>G, 5732G>T, 5733C>G, 5736C>T, 5739_5740insCCC, 5740G>T, 5743A>T, 5744C>T, 5745A>G, 5748A>C, 5749G>T, 5750A>C, 5755A>G, 5757C>G, 5758G>A, 5759C>T, 5760C>T, 5763A>G, 5765A>T, 5775C>T, 5779C>T, 5787C>T, 5793C>G, 5799A>G, 5801C>A, 5805T>C, 5810A>T, 5811A>C, 5812A>C, 5814G>T, 5817T>C, 5818T>C, 5822A>G, 5825T>C, 5827A>G | | | | | | | | |

##### CDS

| | Begin | End | Coverage | Score | Concordance | Matches | Identities | I/D/M/F* | Stop Codons |
|---|---|---|---|---|---|---|---|---|---|
| UF61_gp3 | 1470 | 1562 | 4.6% | 305 | 45.4% | 93 (98.9%) | 46 (48.9%) | 1/0/0/0 | 1 |
| Protein mutations: | I1470L (5551A>T), G1471R (5554G>C), N1472G (5557A>G 5558A>G), K1474L (5563A>T 5564A>T), I1475V (5566A>G 5568A>G), F1479I (5578T>A), K1482H (5587A>C 5589G>T), F1485Y (5597T>A), V1488L (5605G>T 5607T>G), A1489R (5608G>A 5609C>G 5610C>A), M1490V (5611A>G 5613G>T), E1491* (5614G>T 5615A>G), E1493S (5620G>T 5621A>C), S1494D (5623T>G 5624C>A 5625C>T), P1496L (5630C>T 5631C>A), W1497K (5632T>A 5633G>A 5634G>A), A1499T (5638G>A 5640C>T), L1501R (5644T>A 5645T>G 5646A>G), I1502T (5648T>C 5649C>A), P1503R (5651C>G 5652A>G), G1504Y (5653G>T 5654G>A 5655A>C), L1506H (5659T>C 5660T>A 5661A>T), W1509F (5669G>T 5670G>T), L1510V (5671C>G 5673T>G), P1513S (5680C>T 5682A>C), G1515R (5686G>A), K1517T (5693A>C 5694A>T), I1522A (5707A>G 5708T>C 5709T>A), Q1524M (5713C>A 5714A>T), R1525E (5716A>G 5717G>A 5718A>G), K1526L (5719A>T 5720A>T 5721A>G), D1528N (5725G>A 5727T>C), K1529R (5728A>C 5729A>G), C1530V (5731T>G 5732G>T 5733C>G), K1532_D1533insP (5739_5740insCCC), D1533Y (5740G>T), T1534L (5743A>T 5744C>T 5745A>G), E1535D (5748A>C), E1536S (5749G>T 5750A>C), I1538V (5755A>G 5757C>G), A1539I (5758G>A 5759C>T 5760C>T), Y1541F (5765A>T), N1550K (5793C>G), A1553E (5801C>A), E1556V (5810A>T 5811A>C), K1560R (5822A>G), I1561T (5825T>C), M1562V (5827A>G) | | | | | | | | |
| Codon mutations: | AGA1469.AA (5549G>A), ATA1470TTA (5551A>T), GGG1471CGG (5554G>C), AAT1472GGT (5557A>G 5558A>G), GCA1473GCT (5562A>T), AAG1474TTG (5563A>T 5564A>T), ATA1475GTG (5566A>G 5568A>G), TCA1477TCC (5574A>C), TTT1479ATT (5578T>A), AAG1482CAT (5587A>C 5589G>T), GGA1484GGG (5595A>G), TTT1485TAT (5597T>A), CAT1486CAC (5601T>C), CAA1487CAG (5604A>G), GTT1488TTG (5605G>T 5607T>G), GCC1489AGA (5608G>A 5609C>G 5610C>A), ATG1490GTT (5611A>G 5613G>T), GAA1491TGA (5614G>T 5615A>G), GAA1492GAG (5619A>G), GAA1493TCA (5620G>T 5621A>C), TCC1494GAT (5623T>G 5624C>A 5625C>T), ATA1495ATC (5628A>C), CCC1496CTA (5630C>T 5631C>A), TGG1497AAA (5632T>A 5633G>A 5634G>A), ACG1498ACT (5637G>T), GCC1499ACT (5638G>A 5640C>T), TTA1501AGG (5644T>A 5645T>G 5646A>G), ATC1502ACA (5648T>C 5649C>A), CCA1503CGG (5651C>G 5652A>G), GGA1504TAC (5653G>T 5654G>A 5655A>C), GGA1505GGG (5658A>G), TTA1506CAT (5659T>C 5660T>A 5661A>T), TAC1507TAT (5664C>T), GAA1508GAG (5667A>G), TGG1509TTT (5669G>T 5670G>T), CTT1510GTG (5671C>G 5673T>G), GTC1511GTT (5676C>T), CCA1513TCC (5680C>T 5682A>C), TTC1514TTT (5685C>T), GGA1515AGA (5686G>A), CTC1516CTA (5691C>A), AAA1517ACT (5693A>C 5694A>T), CCC1520CCT (5703C>T), GCT1521GCC (5706T>C), ATT1522GCA (5707A>G 5708T>C 5709T>A), CAG1524ATG (5713C>A 5714A>T), AGA1525GAG (5716A>G 5717G>A 5718A>G), AAA1526TTG (5719A>T 5720A>T 5721A>G), GAT1528AAC (5725G>A 5727T>C), AAA1529CGA (5728A>C 5729A>G), TGC1530GTG (5731T>G 5732G>T 5733C>G), TTC1531TTT (5736C>T), AAA1532_GAC1533insCCC (5739_5740insCCC), GAC1533TAC (5740G>T), ACA1534TTG (5743A>T 5744C>T 5745A>G), GAA1535GAC (5748A>C), GAG1536TCG (5749G>T 5750A>C), ATC1538GTG (5755A>G 5757C>G), GCC1539ATT (5758G>A 5759C>T 5760C>T), GTA1540GTG (5763A>G), TAC1541TTC (5765A>T), GAC1544GAT (5775C>T), CTG1546TTG (5779C>T), TAC1548TAT (5787C>T), AAC1550AAG (5793C>G), GAA1552GAG (5799A>G), GCG1553GAG (5801C>A), GAT1554GAC (5805T>C), GAA1556GTC (5810A>T 5811A>C), AGG1557CGT (5812A>C 5814G>T), CAT1558CAC (5817T>C), TTA1559CTA (5818T>C), AAG1560AGG (5822A>G), ATT1561ACT (5825T>C), ATG1562GTG (5827A>G) | | | | | | | | |

##### Proteins

| | Begin | End | Coverage | Score | Concordance | Matches | Identities | I/D/M/F* | Stop Codons |
|---|---|---|---|---|---|---|---|---|---|
| ORF3 (YP_009121747.1) | 1470 | 1562 | 4.6% | 305 | 45.4% | 93 (98.9%) | 46 (48.9%) | 1/0/0/0 | 1 |
| Protein mutations: | I1470L (5551A>T), G1471R (5554G>C), N1472G (5557A>G 5558A>G), K1474L (5563A>T 5564A>T), I1475V (5566A>G 5568A>G), F1479I (5578T>A), K1482H (5587A>C 5589G>T), F1485Y (5597T>A), V1488L (5605G>T 5607T>G), A1489R (5608G>A 5609C>G 5610C>A), M1490V (5611A>G 5613G>T), E1491* (5614G>T 5615A>G), E1493S (5620G>T 5621A>C), S1494D (5623T>G 5624C>A 5625C>T), P1496L (5630C>T 5631C>A), W1497K (5632T>A 5633G>A 5634G>A), A1499T (5638G>A 5640C>T), L1501R (5644T>A 5645T>G 5646A>G), I1502T (5648T>C 5649C>A), P1503R (5651C>G 5652A>G), G1504Y (5653G>T 5654G>A 5655A>C), L1506H (5659T>C 5660T>A 5661A>T), W1509F (5669G>T 5670G>T), L1510V (5671C>G 5673T>G), P1513S (5680C>T 5682A>C), G1515R (5686G>A), K1517T (5693A>C 5694A>T), I1522A (5707A>G 5708T>C 5709T>A), Q1524M (5713C>A 5714A>T), R1525E (5716A>G 5717G>A 5718A>G), K1526L (5719A>T 5720A>T 5721A>G), D1528N (5725G>A 5727T>C), K1529R (5728A>C 5729A>G), C1530V (5731T>G 5732G>T 5733C>G), K1532_D1533insP (5739_5740insCCC), D1533Y (5740G>T), T1534L (5743A>T 5744C>T 5745A>G), E1535D (5748A>C), E1536S (5749G>T 5750A>C), I1538V (5755A>G 5757C>G), A1539I (5758G>A 5759C>T 5760C>T), Y1541F (5765A>T), N1550K (5793C>G), A1553E (5801C>A), E1556V (5810A>T 5811A>C), K1560R (5822A>G), I1561T (5825T>C), M1562V (5827A>G) | | | | | | | | |
| Codon mutations: | AGA1469.AA (5549G>A), ATA1470TTA (5551A>T), GGG1471CGG (5554G>C), AAT1472GGT (5557A>G 5558A>G), GCA1473GCT (5562A>T), AAG1474TTG (5563A>T 5564A>T), ATA1475GTG (5566A>G 5568A>G), TCA1477TCC (5574A>C), TTT1479ATT (5578T>A), AAG1482CAT (5587A>C 5589G>T), GGA1484GGG (5595A>G), TTT1485TAT (5597T>A), CAT1486CAC (5601T>C), CAA1487CAG (5604A>G), GTT1488TTG (5605G>T 5607T>G), GCC1489AGA (5608G>A 5609C>G 5610C>A), ATG1490GTT (5611A>G 5613G>T), GAA1491TGA (5614G>T 5615A>G), GAA1492GAG (5619A>G), GAA1493TCA (5620G>T 5621A>C), TCC1494GAT (5623T>G 5624C>A 5625C>T), ATA1495ATC (5628A>C), CCC1496CTA (5630C>T 5631C>A), TGG1497AAA (5632T>A 5633G>A 5634G>A), ACG1498ACT (5637G>T), GCC1499ACT (5638G>A 5640C>T), TTA1501AGG (5644T>A 5645T>G 5646A>G), ATC1502ACA (5648T>C 5649C>A), CCA1503CGG (5651C>G 5652A>G), GGA1504TAC (5653G>T 5654G>A 5655A>C), GGA1505GGG (5658A>G), TTA1506CAT (5659T>C 5660T>A 5661A>T), TAC1507TAT (5664C>T), GAA1508GAG (5667A>G), TGG1509TTT (5669G>T 5670G>T), CTT1510GTG (5671C>G 5673T>G), GTC1511GTT (5676C>T), CCA1513TCC (5680C>T 5682A>C), TTC1514TTT (5685C>T), GGA1515AGA (5686G>A), CTC1516CTA (5691C>A), AAA1517ACT (5693A>C 5694A>T), CCC1520CCT (5703C>T), GCT1521GCC (5706T>C), ATT1522GCA (5707A>G 5708T>C 5709T>A), CAG1524ATG (5713C>A 5714A>T), AGA1525GAG (5716A>G 5717G>A 5718A>G), AAA1526TTG (5719A>T 5720A>T 5721A>G), GAT1528AAC (5725G>A 5727T>C), AAA1529CGA (5728A>C 5729A>G), TGC1530GTG (5731T>G 5732G>T 5733C>G), TTC1531TTT (5736C>T), AAA1532_GAC1533insCCC (5739_5740insCCC), GAC1533TAC (5740G>T), ACA1534TTG (5743A>T 5744C>T 5745A>G), GAA1535GAC (5748A>C), GAG1536TCG (5749G>T 5750A>C), ATC1538GTG (5755A>G 5757C>G), GCC1539ATT (5758G>A 5759C>T 5760C>T), GTA1540GTG (5763A>G), TAC1541TTC (5765A>T), GAC1544GAT (5775C>T), CTG1546TTG (5779C>T), TAC1548TAT (5787C>T), AAC1550AAG (5793C>G), GAA1552GAG (5799A>G), GCG1553GAG (5801C>A), GAT1554GAC (5805T>C), GAA1556GTC (5810A>T 5811A>C), AGG1557CGT (5812A>C 5814G>T), CAT1558CAC (5817T>C), TTA1559CTA (5818T>C), AAG1560AGG (5822A>G), ATT1561ACT (5825T>C), ATG1562GTG (5827A>G) | | | | | | | | |

*: Inserts / Deletes / Misaligned / Frameshifts

#### Analysis details

This analysis was performed with panviral2.64

### NGS Details (UN18_val): Pinus nigra virus 1

#### Assembly

| | |
|---|---|
| **Coverage Length** | 437 (1 contig(s)) |
| **Depth Of Coverage** | 3.8 |
| **Number Of Reads** | 13 |
| **Reads Per Million** | 0.24 rpm (after QC) |
| **Ambiguities** | 0 |
| **Assembly Method** | de novo + reference guided assembly |
| **Consensus Caller** | Bcf Tools |

#### Coverage Map

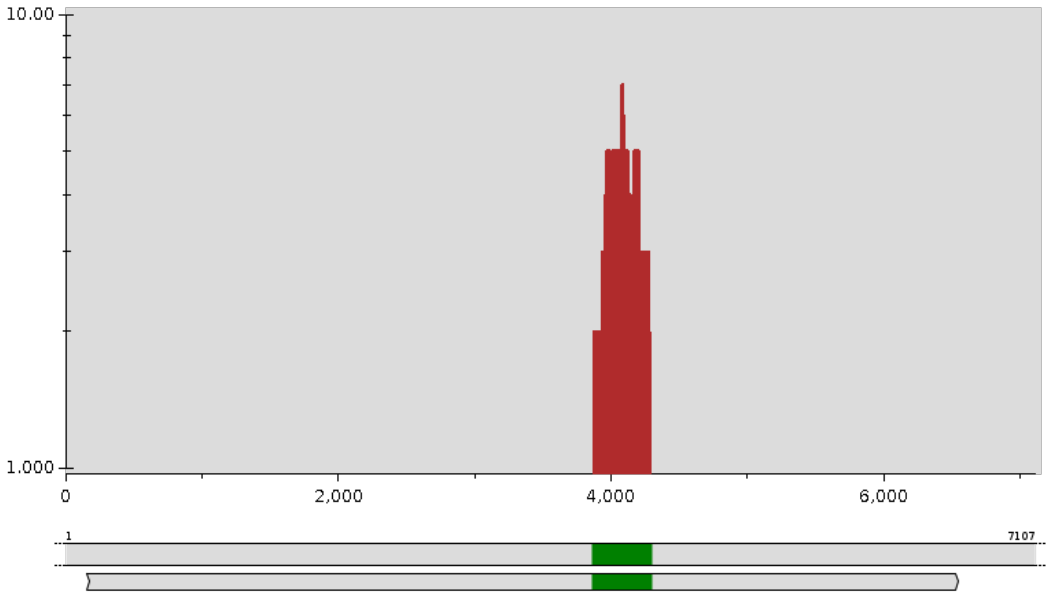

#### Assignment

| | |
|---|---|
| **Type** | Pinus nigra virus 1 (Taxonomy ID: 2267679) |
| **Reference Genome** | NC_040841.1 |
| **NT Identity (%)** | 57.9545 |
| **AA Identity (%)** | 48.2993 |
| **Number Of Stop Codons** | 0 |
| **Number Of CDS** | 1 |

#### Alignment

| | |
|---|---|
| **Alignment Score** | 134.0 (NT) + 536.0 (AA) = 670.0 |
| **Concordance (%)** | 34.4473 |

| Alignment Method | Global, seeded, nucleotide + amino acids (AGA) |
|---|---|

#### Genome Region

Sequence starts at position 3866 and ends at position 4302 relative to NC_040841.1 reference sequence.

#### Alignment Detailed Statistics

| | Begin | End | Coverage | Score | Concordance | Matches | Identities | I/D/M/F* | Stop Codons |
|---|---|---|---|---|---|---|---|---|---|
| **NT** | **3866** | **4302** | **6.1%** | **134** | **15.3%** | **437 (99.3%)** | **255 (58.0%)** | **3/0** | |
| Mutations: | 3868A>C, 3871T>C, 3873T>A, 3874A>T, 3877A>T, 3879G>A, 3880G>T, 3883T>A, 3884C>A, 3885A>C, 3887T>G, 3892G>A, 3894T>A, 3897T>A, 3904T>C, 3905A>T, 3906A>C, 3907A>C, 3910A>T, 3913A>G, 3914A>T, 3916G>A, 3919A>T, 3922A>T, 3931A>G, 3935C>T, 3937T>A, 3941G>T, 3943C>T, 3944A>G, 3946C>T, 3947A>T, 3948C>T, 3949T>A, 3950C>A, 3952G>A, 3953A>T, 3954A>G, 3955T>G, 3956T>A, 3960A>G, 3961T>A, 3967A>G, 3968T>A, 3970G>T, 3973A>T, 3975G>A, 3976A>T, 3977C>A, 3978C>A, 3980G>A, 3981A>T, 3982A>G, 3984T>A, 3985G>T, 3986A>C, 3989A>T, 3992C>A, 3996A>G, 3999T>C, 4001C>T, 4002A>T, 4003G>T, 4004C>A, 4006T>A, 4008G>C, 4009T>C, 4012G>T, 4013G>A, 4015G>T, 4017T>A, 4021C>T, 4024G>A, 4030C>T, 4036G>A, 4042G>T, 4044A>T, 4045T>C, 4047A>G, 4048T>G, 4051G>A, 4054A>T, 4061C>T, 4062A>C, 4064C>A, 4065C>A, 4067G>A, 4074G>A, 4076C>T, 4078C>T, 4084A>T, 4092T>A, 4093A>C, 4094T>G, 4095G>T, 4096T>C, 4099A>C, 4100G>T, 4101C>T, 4102A>C, 4106T>C, 4107T>A, 4108C>A, 4112C>G, 4114G>A, 4120A>T, 4124G>A, 4126T>G, 4129A>C, 4135T>A, 4138G>A, 4141G>A, 4147G>C, 4150A>T, 4151G>T, 4153C>T, 4154T>G, 4155T>A, 4156C>A, 4164G>A, 4165A>C, 4166A>G, 4167G>T, 4168G>T, 4172G>A, 4176A>C, 4180C>T, 4184G>A, 4185C>A, 4187A>C, 4189A>C, 4190T>A, 4191A>T, 4192T>C, 4193G>A, 4194A>G, 4195T>C, 4196T>A, 4197T>A, 4198T>C, 4201A>C, 4201_4202insTCT, 4202G>A, 4205A>G, 4206C>T, 4207A>G, 4210C>T, 4213A>T, 4220A>G, 4225A>G, 4228T>A, 4229C>T, 4230A>T, 4232A>T, 4233G>C, 4235C>G, 4236C>A, 4237A>G, 4241G>A, 4243A>T, 4244C>G, 4246G>T, 4247A>T, 4248A>C, 4255G>T, 4258G>A, 4261C>T, 4265G>A, 4267A>T, 4269T>C, 4270T>C, 4273T>C, 4274C>T, 4276T>C, 4277G>A, 4280G>A, 4281A>T, 4282A>T, 4288G>A, 4290A>G, 4292C>A, 4297A>T, 4298A>C | | | | | | | | |

##### CDS

| | Begin | End | Coverage | Score | Concordance | Matches | Identities | I/D/M/F* | Stop Codons |
|---|---|---|---|---|---|---|---|---|---|
| EXL67_gp1 | 1238 | 1383 | 6.9% | 536 | 49.8% | 146 (99.3%) | 71 (48.3%) | 1/0/0/0 | 0 |
| Protein mutations: | L1240Y (3873T>A 3874A>T), R1242N (3879G>A 3880G>T), N1243K (3883T>A), H1244T (3884C>A 3885A>C), S1245A (3887T>G), I1247K (3894T>A), V1248E (3897T>A), K1251S (3905A>T 3906A>C 3907A>C), M1254L (3914A>T 3916G>A), A1263S (3941G>T 3943C>T), I1264V (3944A>G 3946C>T), T1265L (3947A>T 3948C>T 3949T>A), Q1266K (3950C>A 3952G>A), N1267W (3953A>T 3954A>G 3955T>G), F1268I (3956T>A), N1269R (3960A>G 3961T>A), L1272I (3968T>A 3970G>T), R1274N (3975G>A 3976A>T), P1275K (3977C>A 3978C>A), E1276M (3980G>A 3981A>T 3982A>G), V1277D (3984T>A 3985G>T), I1278L (3986A>C), M1279L (3989A>T), Q1280K (3992C>A), K1281R (3996A>G), I1282T (3999T>C), Q1283F (4001C>T 4002A>T 4003G>T), H1284K (4004C>A 4006T>A), S1285T (4008G>C 4009T>C), K1286N (4012G>T), V1287I (4013G>A 4015G>T), F1288Y (4017T>A), Y1297F (4044A>T 4045T>C), Y1298W (4047A>G 4048T>G), Q1303S (4061C>T 4062A>C), P1304K (4064C>A 4065C>A), E1305K (4067G>A), R1307K (4074G>A), H1308Y (4076C>T 4078C>T), I1313N (4092T>A 4093A>C), C1314V (4094T>G 4095G>T 4096T>C), A1316F (4100G>T 4101C>T 4102A>C), F1318Q (4106T>C 4107T>A 4108C>A), Q1320E (4112C>G 4114G>A), K1322N (4120A>T), V1324M (4124G>A 4126T>G), A1333S (4151G>T 4153C>T), F1334E (4154T>G 4155T>A 4156C>A), R1337N (4164G>A 4165A>C), R1338V (4166A>G 4167G>T 4168G>T), D1340N (4172G>A), Y1341S (4176A>C), A1344N (4184G>A 4185C>A), K1345H (4187A>C 4189A>C), Y1346I (4190T>A 4191A>T 4192T>C), D1347S (4193G>A 4194A>G 4195T>C), F1348N (4196T>A 4197T>A 4198T>C), I1349_V1350insS (4201_4202insTCT), V1350I (4202G>A), T1351V (4205A>G 4206C>T 4207A>G), I1356V (4220A>G), H1359F (4229C>T 4230A>T), P1361E (4235C>G 4236C>A 4237A>G), V1363I (4241G>A 4243A>T), Q1364D (4244C>G 4246G>T), N1365S (4247A>T 4248A>C), L1367F (4255G>T), E1371N (4265G>A 4267A>T), I1372T (4269T>C 4270T>C), L1374F (4274C>T 4276T>C), E1375K (4277G>A), E1376I (4280G>A 4281A>T 4282A>T), K1379R (4290A>G), H1380N (4292C>A), I1382L (4298A>C) | | | | | | | | |
| Codon mutations: | GCA1238GCC (3868A>C), TTT1239TTC (3871T>C), TTA1240TAT (3873T>A 3874A>T), GTA1241GTT (3877A>T), AGG1242AAT (3879G>A 3880G>T), AAT1243AAA (3883T>A), CAT1244ACT (3884C>A 3885A>C), TCT1245GCT (3887T>G), GAG1246GAA (3892G>A), ATA1247AAA (3894T>A), GTA1248GAA (3897T>A), GGT1250GGC (3904T>C), AAA1251TCC (3905A>T 3906A>C 3907A>C), CCA1252CCT (3910A>T), AGA1253AGG (3913A>G), ATG1254TTA (3914A>T 3916G>A), GTA1255GTT (3919A>T), ATA1256ATT (3922A>T), AAA1259AAG (3931A>G), CTT1261TTA (3935C>T 3937T>A), GCC1263TCT (3941G>T 3943C>T), ATC1264GTT (3944A>G 3946C>T), ACT1265TTA (3947A>T 3948C>T 3949T>A), CAG1266AAA (3950C>A 3952G>A), AAT1267TGG (3953A>T 3954A>G 3955T>G), TTT1268ATT (3956T>A), AAT1269AGA (3960A>G 3961T>A), CCA1271CCG (3967A>G), TTG1272ATT (3968T>A 3970G>T), CCA1273CCT (3973A>T), AGA1274AAT (3975G>A 3976A>T), CCA1275AAA (3977C>A 3978C>A), GAA1276ATG (3980G>A 3981A>T 3982A>G), GTG1277GAT (3984T>A 3985G>T), ATT1278CTT (3986A>C), ATG1279TTG (3989A>T), CAA1280AAA (3992C>A), AAA1281AGA (3996A>G), ATT1282ACT (3999T>C), CAG1283TTT (4001C>T 4002A>T 4003G>T), CAT1284AAA (4004C>A 4006T>A), AGT1285ACC (4008G>C 4009T>C), AAG1286AAT (4012G>T), GTG1287ATT (4013G>A 4015G>T), TTC1288TAC (4017T>A), AGC1289AGT (4021C>T), AAG1290AAA (4024G>A), GAC1292GAT (4030C>T), AAG1294AAA (4036G>A), GGG1296GGT (4042G>T), TAT1297TTC (4044A>T 4045T>C), TAT1298TGG (4047A>G 4048T>G), CAG1299CAA (4051G>A), ATA1300ATT (4054A>T), CAG1303TCG (4061C>T 4062A>C), CCA1304AAA (4064C>A 4065C>A), GAA1305AAA (4067G>A), AGA1307AAA (4074G>A), CAC1308TAT (4076C>T 4078C>T), ACA1310ACT (4084A>T), ATA1313AAC (4092T>A 4093A>C), TGT1314GTC (4094T>G 4095G>T 4096T>C), CCA1315CCC (4099A>C), GCA1316TTC (4100G>T 4101C>T 4102A>C), TTC1318CAA (4106T>C 4107T>A 4108C>A), CAG1320GAA (4112C>G 4114G>A), AAA1322AAT (4120A>T), GTT1324ATG (4124G>A 4126T>G), CCA1325CCC (4129A>C), GGT1327GGA (4135T>A), TTG1328TTA (4138G>A), AAG1329AAA (4141G>A), GCG1331GCC (4147G>C), CCA1332CCT (4150A>T), GCC1333TCT (4151G>T 4153C>T), TTC1334GAA (4154T>G 4155T>A 4156C>A), AGA1337AAC (4164G>A 4165A>C), AGG1338GTT (4166A>G 4167G>T 4168G>T), GAT1340AAT (4172G>A), TAT1341TCT (4176A>C), ATC1342ATT (4180C>T), GCT1344AAT (4184G>A 4185C>A), AAA1345CAC (4187A>C 4189A>C), TAT1346ATC (4190T>A 4191A>T 4192T>C), GAT1347AGC (4193G>A 4194A>G 4195T>C), TTT1348AAC (4196T>A 4197T>A 4198T>C), ATA1349ATC (4201A>C), ATA1349_GTT1350insTCT (4201_4202insTCT), GTT1350ATT (4202G>A), ACA1351GTG (4205A>G 4206C>T 4207A>G), TAC1352TAT (4210C>T), ATA1353ATT (4213A>T), ATT1356GTT (4220A>G), TTA1357TTG (4225A>G), ATT1358ATA (4228T>A), CAT1359TTT (4229C>T 4230A>T), AGT1360TCT (4232A>T 4233G>C), CCA1361GAG (4235C>G 4236C>A 4237A>G), GTA1363ATT (4241G>A 4243A>T), CAG1364GAT (4244C>G 4246G>T), AAT1365TCT (4247A>T 4248A>C), TTG1367TTT (4255G>T), AAG1368AAA (4258G>A), CAC1369CAT (4261C>T), GAA1371AAT (4265G>A 4267A>T), ATT1372ACC (4269T>C 4270T>C), TTT1373TTC (4273T>C), CTT1374TTC (4274C>T 4276T>C), GAA1375AAA (4277G>A), GAA1376ATT (4280G>A 4281A>T 4282A>T), AAG1378AAA (4288G>A), AAA1379AGA (4290A>G), CAT1380AAT (4292C>A), GGA1381GGT (4297A>T), ATT1382CTT (4298A>C) | | | | | | | | |

##### Proteins

| | Begin | End | Coverage | Score | Concordance | Matches | Identities | I/D/M/F* | Stop Codons |
|---|---|---|---|---|---|---|---|---|---|
| polyprotein (YP_009553669.1) | 1238 | 1383 | 6.9% | 536 | 49.8% | 146 (99.3%) | 71 (48.3%) | 1/0/0/0 | 0 |
| Protein mutations: | L1240Y (3873T>A 3874A>T), R1242N (3879G>A 3880G>T), N1243K (3883T>A), H1244T (3884C>A 3885A>C), S1245A (3887T>G), I1247K (3894T>A), V1248E (3897T>A), K1251S (3905A>T 3906A>C 3907A>C), M1254L (3914A>T 3916G>A), A1263S (3941G>T 3943C>T), I1264V (3944A>G 3946C>T), T1265L (3947A>T 3948C>T 3949T>A), Q1266K (3950C>A 3952G>A), N1267W (3953A>T 3954A>G 3955T>G), F1268I (3956T>A), N1269R (3960A>G 3961T>A), L1272I (3968T>A 3970G>T), R1274N (3975G>A 3976A>T), P1275K (3977C>A 3978C>A), E1276M (3980G>A 3981A>T 3982A>G), V1277D (3984T>A 3985G>T), I1278L (3986A>C), M1279L (3989A>T), Q1280K (3992C>A), K1281R (3996A>G), I1282T (3999T>C), Q1283F (4001C>T 4002A>T 4003G>T), H1284K (4004C>A 4006T>A), S1285T (4008G>C 4009T>C), K1286N (4012G>T), V1287I (4013G>A 4015G>T), F1288Y (4017T>A), Y1297F (4044A>T 4045T>C), Y1298W (4047A>G 4048T>G), Q1303S (4061C>T 4062A>C), P1304K (4064C>A 4065C>A), E1305K (4067G>A), R1307K (4074G>A), H1308Y (4076C>T 4078C>T), I1313N (4092T>A 4093A>C), C1314V (4094T>G 4095G>T 4096T>C), A1316F (4100G>T 4101C>T 4102A>C), F1318Q (4106T>C 4107T>A 4108C>A), Q1320E (4112C>G 4114G>A), K1322N (4120A>T), V1324M (4124G>A 4126T>G), A1333S (4151G>T 4153C>T), F1334E (4154T>G 4155T>A 4156C>A), R1337N (4164G>A 4165A>C), R1338V (4166A>G 4167G>T 4168G>T), D1340N (4172G>A), Y1341S (4176A>C), A1344N (4184G>A 4185C>A), K1345H (4187A>C 4189A>C), Y1346I (4190T>A 4191A>T 4192T>C), D1347S (4193G>A 4194A>G 4195T>C), F1348N (4196T>A 4197T>A 4198T>C), I1349_V1350insS (4201_4202insTCT), V1350I (4202G>A), T1351V (4205A>G 4206C>T 4207A>G), I1356V (4220A>G), H1359F (4229C>T 4230A>T), P1361E (4235C>G 4236C>A 4237A>G), V1363I (4241G>A 4243A>T), Q1364D (4244C>G 4246G>T), N1365S (4247A>T 4248A>C), L1367F (4255G>T), E1371N (4265G>A 4267A>T), I1372T (4269T>C 4270T>C), L1374F (4274C>T 4276T>C), E1375K (4277G>A), E1376I (4280G>A 4281A>T 4282A>T), K1379R (4290A>G), H1380N (4292C>A), I1382L (4298A>C) | | | | | | | | |
| Codon mutations: | GCA1238GCC (3868A>C), TTT1239TTC (3871T>C), TTA1240TAT (3873T>A 3874A>T), GTA1241GTT (3877A>T), AGG1242AAT (3879G>A 3880G>T), AAT1243AAA (3883T>A), CAT1244ACT (3884C>A 3885A>C), TCT1245GCT (3887T>G), GAG1246GAA (3892G>A), ATA1247AAA (3894T>A), GTA1248GAA (3897T>A), GGT1250GGC (3904T>C), AAA1251TCC (3905A>T 3906A>C 3907A>C), CCA1252CCT (3910A>T), AGA1253AGG (3913A>G), ATG1254TTA (3914A>T 3916G>A), GTA1255GTT (3919A>T), ATA1256ATT (3922A>T), AAA1259AAG (3931A>G), CTT1261TTA (3935C>T 3937T>A), GCC1263TCT (3941G>T 3943C>T), ATC1264GTT (3944A>G 3946C>T), ACT1265TTA (3947A>T 3948C>T 3949T>A), CAG1266AAA (3950C>A 3952G>A), AAT1267TGG (3953A>T 3954A>G 3955T>G), TTT1268ATT (3956T>A), AAT1269AGA (3960A>G 3961T>A), CCA1271CCG (3967A>G), TTG1272ATT (3968T>A 3970G>T), CCA1273CCT (3973A>T), AGA1274AAT (3975G>A 3976A>T), CCA1275AAA (3977C>A 3978C>A), GAA1276ATG (3980G>A 3981A>T 3982A>G), GTG1277GAT (3984T>A 3985G>T), ATT1278CTT (3986A>C), ATG1279TTG (3989A>T), CAA1280AAA (3992C>A), AAA1281AGA (3996A>G), ATT1282ACT (3999T>C), CAG1283TTT (4001C>T 4002A>T 4003G>T), CAT1284AAA (4004C>A 4006T>A), AGT1285ACC (4008G>C 4009T>C), AAG1286AAT (4012G>T), GTG1287ATT (4013G>A 4015G>T), TTC1288TAC (4017T>A), AGC1289AGT (4021C>T), AAG1290AAA (4024G>A), GAC1292GAT (4030C>T), AAG1294AAA (4036G>A), GGG1296GGT (4042G>T), TAT1297TTC (4044A>T 4045T>C), TAT1298TGG (4047A>G 4048T>G), CAG1299CAA (4051G>A), ATA1300ATT (4054A>T), CAG1303TCG (4061C>T 4062A>C), CCA1304AAA (4064C>A 4065C>A), GAA1305AAA (4067G>A), AGA1307AAA (4074G>A), CAC1308TAT (4076C>T 4078C>T), ACA1310ACT (4084A>T), ATA1313AAC (4092T>A 4093A>C), TGT1314GTC (4094T>G 4095G>T 4096T>C), CCA1315CCC (4099A>C), GCA1316TTC (4100G>T 4101C>T 4102A>C), TTC1318CAA (4106T>C 4107T>A 4108C>A), CAG1320GAA (4112C>G 4114G>A), AAA1322AAT (4120A>T), GTT1324ATG (4124G>A 4126T>G), CCA1325CCC (4129A>C), GGT1327GGA (4135T>A), TTG1328TTA (4138G>A), AAG1329AAA (4141G>A), GCG1331GCC (4147G>C), CCA1332CCT (4150A>T), GCC1333TCT (4151G>T 4153C>T), TTC1334GAA (4154T>G 4155T>A 4156C>A), AGA1337AAC (4164G>A 4165A>C), AGG1338GTT (4166A>G 4167G>T 4168G>T), GAT1340AAT (4172G>A), TAT1341TCT (4176A>C), ATC1342ATT (4180C>T), GCT1344AAT (4184G>A 4185C>A), AAA1345CAC (4187A>C 4189A>C), TAT1346ATC (4190T>A 4191A>T 4192T>C), GAT1347AGC (4193G>A 4194A>G 4195T>C), TTT1348AAC (4196T>A 4197T>A 4198T>C), ATA1349ATC (4201A>C), ATA1349_GTT1350insTCT (4201_4202insTCT), GTT1350ATT (4202G>A), ACA1351GTG (4205A>G 4206C>T 4207A>G), TAC1352TAT (4210C>T), ATA1353ATT (4213A>T), ATT1356GTT (4220A>G), TTA1357TTG (4225A>G), ATT1358ATA (4228T>A), CAT1359TTT (4229C>T 4230A>T), AGT1360TCT (4232A>T 4233G>C), CCA1361GAG (4235C>G 4236C>A 4237A>G), GTA1363ATT (4241G>A 4243A>T), CAG1364GAT (4244C>G 4246G>T), AAT1365TCT (4247A>T 4248A>C), TTG1367TTT (4255G>T), AAG1368AAA (4258G>A), CAC1369CAT (4261C>T), GAA1371AAT (4265G>A 4267A>T), ATT1372ACC (4269T>C 4270T>C), TTT1373TTC (4273T>C), CTT1374TTC (4274C>T 4276T>C), GAA1375AAA (4277G>A), GAA1376ATT (4280G>A 4281A>T 4282A>T), AAG1378AAA (4288G>A), AAA1379AGA (4290A>G), CAT1380AAT (4292C>A), GGA1381GGT (4297A>T), ATT1382CTT (4298A>C) | | | | | | | | |

*: Inserts / Deletes / Misaligned / Frameshifts

#### Analysis details

This analysis was performed with panviral2.64

### NGS Details (UN18_val): Hibiscus bacilliform virus GD1

#### Assembly

| | |
|---|---|
| **Coverage Length** | 235 (1 contig(s)) |
| **Depth Of Coverage** | 6.1 |
| **Number Of Reads** | 12 |
| **Reads Per Million** | 0.23 rpm (after QC) |
| **Ambiguities** | 0 |
| **Assembly Method** | de novo + reference guided assembly |
| **Consensus Caller** | Bcf Tools |

#### Coverage Map

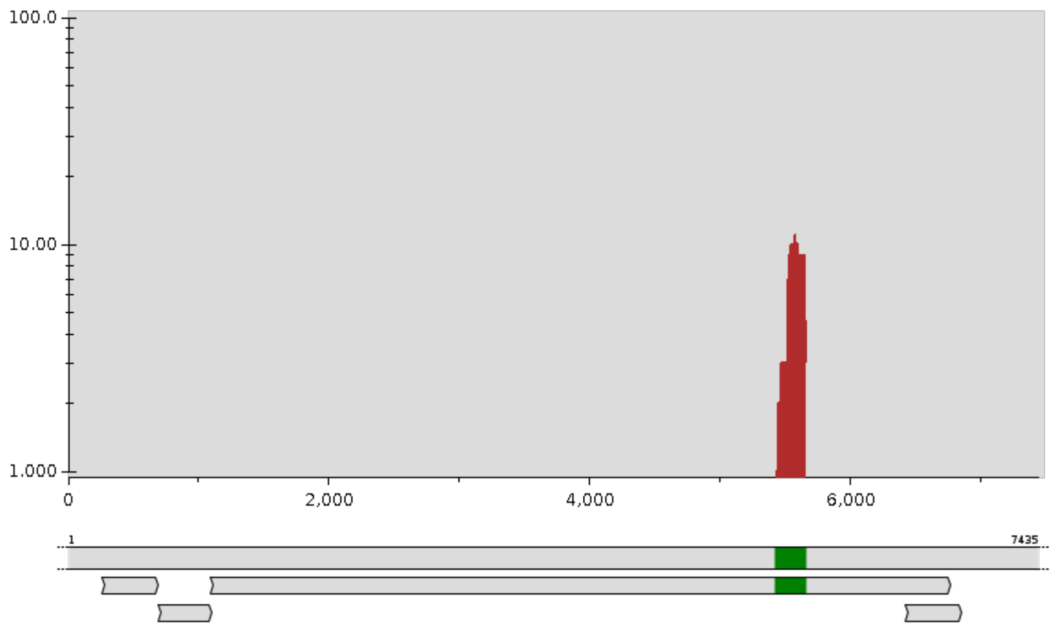

#### Assignment

| | |
|---|---|
| **Type** | Hibiscus bacilliform virus GD1 (Taxonomy ID: 1459800) |
| **Reference Genome** | NC_023485.1 |
| **NT Identity (%)** | 56.3025 |
| **AA Identity (%)** | 47.5 |
| **Number Of Stop Codons** | 0 |
| **Number Of CDS** | 4 |

#### Alignment

| | |
|---|---|
| **Alignment Score** | 54.0 (NT) + 274.0 (AA) = 328.0 |
| **Concordance (%)** | 31.5688 |

| Alignment Method | Global, seeded, nucleotide + amino acids (AGA) |
|---|---|

#### Genome Region

Sequence starts at position 5421 and ends at position 5655 relative to NC_023485.1 reference sequence.

#### Alignment Detailed Statistics

| | Begin | End | Coverage | Score | Concordance | Matches | Identities | I/D/M/F* | Stop Codons |
|---|---|---|---|---|---|---|---|---|---|
| **NT** | **5421** | **5655** | **3.2%** | **54** | **11.5%** | **235 (98.7%)** | **134 (56.3%)** | **3/0** | |
| Mutations: | 5422C>A, 5426C>T, 5427A>C, 5430T>A, 5431G>A, 5435T>A, 5436G>A, 5441C>T, 5442T>C, 5445G>A, 5446T>C, 5447T>C, 5449C>A, 5450A>T, 5451G>T, 5453T>C, 5456A>C, 5458T>A, 5459G>C, 5465A>G, 5467G>T, 5468G>T, 5470T>C, 5480A>T, 5486A>G, 5489T>A, 5491A>C, 5492A>T, 5495C>T, 5502G>A, 5503C>A, 5505G>A, 5506T>C, 5511C>A, 5512A>T, 5514C>G, 5515G>A, 5516A>T, 5517A>C, 5518A>T, 5523G>A, 5524A>C, 5525C>T, 5526A>C, 5527A>G, 5528C>G, 5531C>G, 5534C>T, 5536A>G, 5537A>G, 5537_5538insCCA, 5538G>T, 5539G>A, 5541A>C, 5542C>T, 5546A>T, 5547G>A, 5552T>C, 5555C>T, 5556G>A, 5557C>T, 5561A>G, 5563A>T, 5564C>T, 5573C>T, 5576A>T, 5579C>G, 5584T>A, 5588C>T, 5590A>G, 5591G>C, 5592T>A, 5593C>G, 5595G>C, 5601G>C, 5603A>G, 5607A>G, 5608T>C, 5609C>T, 5610A>C, 5615C>T, 5616A>C, 5618A>T, 5619C>A, 5622G>A, 5624C>T, 5625A>G, 5626T>C, 5627G>A, 5630A>G, 5631G>C, 5633A>G, 5635A>C, 5637T>C, 5638G>T, 5640C>A, 5642G>A, 5646A>C, 5648T>C, 5649G>A, 5654T>A, 5655G>T | | | | | | | | |

##### CDS

| | Begin | End | Coverage | Score | Concordance | Matches | Identities | I/D/M/F* | Stop Codons |
|---|---|---|---|---|---|---|---|---|---|
| BU35_gp3 | 1444 | 1522 | 4.2% | 274 | 47.7% | 79 (98.8%) | 38 (47.5%) | 1/0/0/0 | 0 |
| Protein mutations: | S1444Y (5422C>A), K1446Q (5427A>C), W1447K (5430T>A 5431G>A), A1449T (5436G>A), W1451R (5442T>C), V1452T (5445G>A 5446T>C 5447T>C), P1453H (5449C>A 5450A>T), D1454Y (5451G>T 5453T>C), L1456H (5458T>A 5459G>C), W1459F (5467G>T 5468G>T), L1460P (5470T>C), K1467T (5491A>C 5492A>T), A1471K (5502G>A 5503C>A), V1472T (5505G>A 5506T>C), Q1474M (5511C>A 5512A>T), R1475D (5514C>G 5515G>A 5516A>T), K1476L (5517A>C 5518A>T), D1478T (5523G>A 5524A>C 5525C>T), N1479R (5526A>C 5527A>G 5528C>G), K1482R (5536A>G 5537A>G), K1482_G1483insP (5537_5538insCCA), G1483Y (5538G>T 5539G>A), T1484L (5541A>C 5542C>T), E1485D (5546A>T), A1486T (5547G>A), A1489I (5556G>A 5557C>T), Y1491F (5563A>T 5564C>T), F1498Y (5584T>A), Q1500R (5590A>G 5591G>C), E1502Q (5595G>C), E1504Q (5601G>C 5603A>G), I1506A (5607A>G 5608T>C 5609C>T), K1507Q (5610A>C), I1509L (5616A>C 5618A>T), V1511I (5622G>A 5624C>T), M1512A (5625A>G 5626T>C 5627G>A), E1514Q (5631G>C 5633A>G), K1515T (5635A>C), C1516L (5637T>C 5638G>T), N1519H (5646A>C 5648T>C), G1520R (5649G>A) | | | | | | | | |
| Codon mutations: | TCT1444TAT (5422C>A), ATC1445ATT (5426C>T), AAG1446CAG (5427A>C), TGG1447AAG (5430T>A 5431G>A), ACT1448ACA (5435T>A), GCT1449ACT (5436G>A), TTC1450TTT (5441C>T), TGG1451CGG (5442T>C), GTT1452ACC (5445G>A 5446T>C 5447T>C), CCA1453CAT (5449C>A 5450A>T), GAT1454TAC (5451G>T 5453T>C), GGA1455GGC (5456A>C), CTG1456CAC (5458T>A 5459G>C), GAA1458GAG (5465A>G), TGG1459TTT (5467G>T 5468G>T), CTA1460CCA (5470T>C), CCA1463CCT (5480A>T), GGA1465GGG (5486A>G), CTT1466CTA (5489T>A), AAA1467ACT (5491A>C 5492A>T), AAC1468AAT (5495C>T), GCA1471AAA (5502G>A 5503C>A), GTA1472ACA (5505G>A 5506T>C), CAG1474ATG (5511C>A 5512A>T), CGA1475GAT (5514C>G 5515G>A 5516A>T), AAG1476CTG (5517A>C 5518A>T), GAC1478ACT (5523G>A 5524A>C 5525C>T), AAC1479CGG (5526A>C 5527A>G 5528C>G), GTC1480GTG (5531C>G), TTC1481TTT (5534C>T), AAA1482AGG (5536A>G 5537A>G), AAA1482_GGT1483insCCA (5537_5538insCCA), GGT1483TAT (5538G>T 5539G>A), ACT1484CTT (5541A>C 5542C>T), GAA1485GAT (5546A>T), GCC1486ACC (5547G>A), TTT1487TTC (5552T>C), ATC1488ATT (5555C>T), GCT1489ATT (5556G>A 5557C>T), GTA1490GTG (5561A>G), TAC1491TTT (5563A>T 5564C>T), GAC1494GAT (5573C>T), ATA1495ATT (5576A>T), CTC1496CTG (5579C>G), TTT1498TAT (5584T>A), TCC1499TCT (5588C>T), CAG1500CGC (5590A>G 5591G>C), TCT1501AGT (5592T>A 5593C>G), GAG1502CAG (5595G>C), GAA1504CAG (5601G>C 5603A>G), ATC1506GCT (5607A>G 5608T>C 5609C>T), AAG1507CAG (5610A>C), CAC1508CAT (5615C>T), ATA1509CTT (5616A>C 5618A>T), CGG1510AGG (5619C>A), GTC1511ATT (5622G>A 5624C>T), ATG1512GCA (5625A>G 5626T>C 5627G>A), TTA1513TTG (5630A>G), GAA1514CAG (5631G>C 5633A>G), AAA1515ACA (5635A>C), TGC1516CTC (5637T>C 5638G>T), CGG1517AGA (5640C>A 5642G>A), AAT1519CAC (5646A>C 5648T>C), GGG1520AGG (5649G>A), CTT1521CTA (5654T>A), GTT1522T.. (5655G>T) | | | | | | | | |

##### Proteins

| | Begin | End | Coverage | Score | Concordance | Matches | Identities | I/D/M/F* | Stop Codons |
|---|---|---|---|---|---|---|---|---|---|
| polyprotein (YP_009002585.1) | 1444 | 1522 | 4.2% | 274 | 47.7% | 79 (98.8%) | 38 (47.5%) | 1/0/0/0 | 0 |
| Protein mutations: | S1444Y (5422C>A), K1446Q (5427A>C), W1447K (5430T>A 5431G>A), A1449T (5436G>A), W1451R (5442T>C), V1452T (5445G>A 5446T>C 5447T>C), P1453H (5449C>A 5450A>T), D1454Y (5451G>T 5453T>C), L1456H (5458T>A 5459G>C), W1459F (5467G>T 5468G>T), L1460P (5470T>C), K1467T (5491A>C 5492A>T), A1471K (5502G>A 5503C>A), V1472T (5505G>A 5506T>C), Q1474M (5511C>A 5512A>T), R1475D (5514C>G 5515G>A 5516A>T), K1476L (5517A>C 5518A>T), D1478T (5523G>A 5524A>C 5525C>T), N1479R (5526A>C 5527A>G 5528C>G), K1482R (5536A>G 5537A>G), K1482_G1483insP (5537_5538insCCA), G1483Y (5538G>T 5539G>A), T1484L (5541A>C 5542C>T), E1485D (5546A>T), A1486T (5547G>A), A1489I (5556G>A 5557C>T), Y1491F (5563A>T 5564C>T), F1498Y (5584T>A), Q1500R (5590A>G 5591G>C), E1502Q (5595G>C), E1504Q (5601G>C 5603A>G), I1506A (5607A>G 5608T>C 5609C>T), K1507Q (5610A>C), I1509L (5616A>C 5618A>T), V1511I (5622G>A 5624C>T), M1512A (5625A>G 5626T>C 5627G>A), E1514Q (5631G>C 5633A>G), K1515T (5635A>C), C1516L (5637T>C 5638G>T), N1519H (5646A>C 5648T>C), G1520R (5649G>A) | | | | | | | | |
| Codon mutations: | TCT1444TAT (5422C>A), ATC1445ATT (5426C>T), AAG1446CAG (5427A>C), TGG1447AAG (5430T>A 5431G>A), ACT1448ACA (5435T>A), GCT1449ACT (5436G>A), TTC1450TTT (5441C>T), TGG1451CGG (5442T>C), GTT1452ACC (5445G>A 5446T>C 5447T>C), CCA1453CAT (5449C>A 5450A>T), GAT1454TAC (5451G>T 5453T>C), GGA1455GGC (5456A>C), CTG1456CAC (5458T>A 5459G>C), GAA1458GAG (5465A>G), TGG1459TTT (5467G>T 5468G>T), CTA1460CCA (5470T>C), CCA1463CCT (5480A>T), GGA1465GGG (5486A>G), CTT1466CTA (5489T>A), AAA1467ACT (5491A>C 5492A>T), AAC1468AAT (5495C>T), GCA1471AAA (5502G>A 5503C>A), GTA1472ACA (5505G>A 5506T>C), CAG1474ATG (5511C>A 5512A>T), CGA1475GAT (5514C>G 5515G>A 5516A>T), AAG1476CTG (5517A>C 5518A>T), GAC1478ACT (5523G>A 5524A>C 5525C>T), AAC1479CGG (5526A>C 5527A>G 5528C>G), GTC1480GTG (5531C>G), TTC1481TTT (5534C>T), AAA1482AGG (5536A>G 5537A>G), AAA1482_GGT1483insCCA (5537_5538insCCA), GGT1483TAT (5538G>T 5539G>A), ACT1484CTT (5541A>C 5542C>T), GAA1485GAT (5546A>T), GCC1486ACC (5547G>A), TTT1487TTC (5552T>C), ATC1488ATT (5555C>T), GCT1489ATT (5556G>A 5557C>T), GTA1490GTG (5561A>G), TAC1491TTT (5563A>T 5564C>T), GAC1494GAT (5573C>T), ATA1495ATT (5576A>T), CTC1496CTG (5579C>G), TTT1498TAT (5584T>A), TCC1499TCT (5588C>T), CAG1500CGC (5590A>G 5591G>C), TCT1501AGT (5592T>A 5593C>G), GAG1502CAG (5595G>C), GAA1504CAG (5601G>C 5603A>G), ATC1506GCT (5607A>G 5608T>C 5609C>T), AAG1507CAG (5610A>C), CAC1508CAT (5615C>T), ATA1509CTT (5616A>C 5618A>T), CGG1510AGG (5619C>A), GTC1511ATT (5622G>A 5624C>T), ATG1512GCA (5625A>G 5626T>C 5627G>A), TTA1513TTG (5630A>G), GAA1514CAG (5631G>C 5633A>G), AAA1515ACA (5635A>C), TGC1516CTC (5637T>C 5638G>T), CGG1517AGA (5640C>A 5642G>A), AAT1519CAC (5646A>C 5648T>C), GGG1520AGG (5649G>A), CTT1521CTA (5654T>A), GTT1522T.. (5655G>T) | | | | | | | | |

*: Inserts / Deletes / Misaligned / Frameshifts

#### Analysis details

This analysis was performed with panviral2.64

#### NGS Details (UN18_val): Caulimovirus tessellomirabilis

##### Assembly

| | |
|---|---|
| **Coverage Length** | 467 (1 contig(s)) |
| **Depth Of Coverage** | 3.1 |
| **Number Of Reads** | 11 |
| **Reads Per Million** | 0.21 rpm (after QC) |
| **Ambiguities** | 0 |
| **Assembly Method** | de novo + reference guided assembly |
| **Consensus Caller** | Bcf Tools |

##### Coverage Map

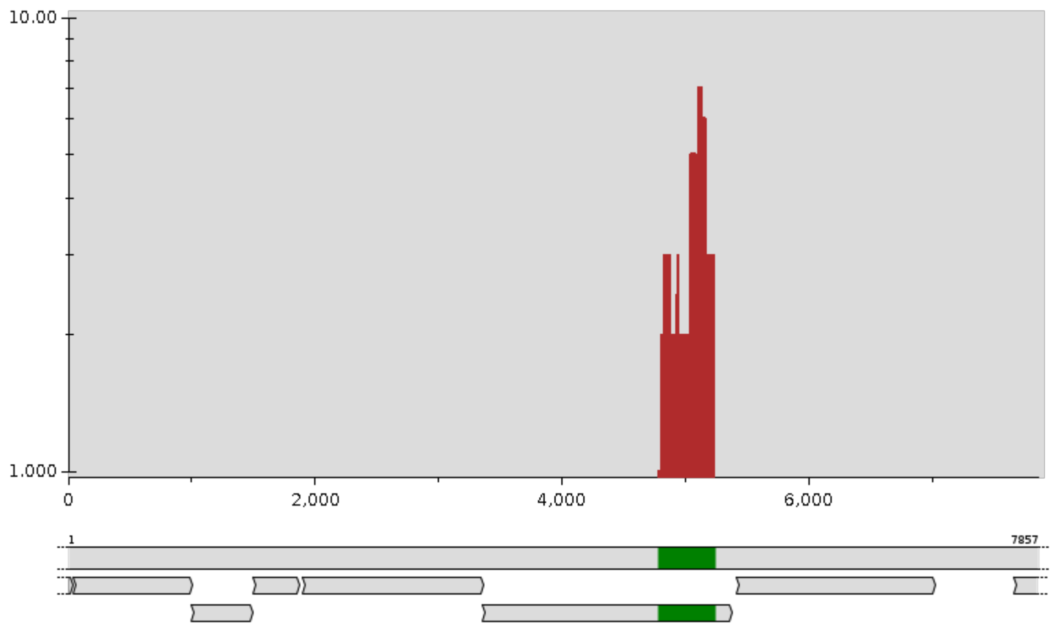

##### Assignment

| | |
|---|---|
| **Type** | Caulimovirus tessellomirabilis (Taxonomy ID: 3048203) |
| **Reference Genome** | NC_004036.1 |
| **NT Identity (%)** | 58.1858 |
| **AA Identity (%)** | 47.6821 |
| **Number Of Stop Codons** | 0 |
| **Number Of CDS** | 7 |

##### Alignment

| | |
|---|---|
| **Alignment Score** | 106.0 (NT) + 464.0 (AA) = 570.0 |
| **Concordance (%)** | 30.8108 |

| Alignment Method | Global, seeded, nucleotide + amino acids (AGA) |
|---|---|

#### Genome Region

Sequence starts at position 4777 and ends at position 5243 relative to NC_004036.1 reference sequence.

#### Alignment Detailed Statistics

| | Begin | End | Coverage | Score | Concordance | Matches | Identities | I/D/M/F* | Stop Codons |
|---|---|---|---|---|---|---|---|---|---|
| **NT** | **4777** | **5243** | **5.9%** | **106** | **12.3%** | **452 (96.8%)** | **263 (56.3%)** | **0/15** | |
| Mutations: | 4780G>A, 4790G>T, 4791T>G, 4792T>C, 4793C>T, 4797C>A, 4798A>C, 4806A>G, 4807T>A, 4808A>G, 4809G>A, 4810C>A, 4812A>T, 4813T>C, 4816C>A, 4820A>G, 4822G>A, 4823C>T, 4825T>A, 4829G>A, 4834A>G, 4835C>A, 4837G>A, 4840A>T, 4842C>T, 4843G>T, 4844T>C, 4849G>A, 4850G>A, 4851T>A, 4853A>C, 4854A>T, 4855G>T, 4856T>A, 4858G>C, 4860A>G, 4861G>A, 4865G>T, 4866A>C, 4867T>A, 4868C>A, 4870G>T, 4871G>C, 4872T>G, 4875G>T, 4877A>G, 4878T>G, 4879C>A, 4885G>A, 4886C>A, 4889T>A, 4890C>A, 4892G>C, 4898G>C, 4899A>T, 4902A>G, 4903C>T, 4907A>C, 4912G>A, 4913A>C, 4915A>C, 4918G>A, 4919A>G, 4923G>A, 4926T>G, 4927A>T, 4928A>G, 4929T>C, 4933T>A, 4934T>C, 4946T>C, 4947A>G, 4948T>A, 4949C>T, 4955A>G, 4957G>A, 4958A>C, 4959A>C, 4966C>T, 4968G>A, 4972A>G, 4976A>T, 4978T>A, 4979G>C, 4990T>C, 4993T>C, 4997C>T, 4999C>T, 5000T>C, 5001T>A, 5002C>T, 5008T>A, 5010G>C, 5012G>T, 5014C>A, 5015C>T, 5017T>A, 5018_5023delAAGGCC, 5029C>A, 5030A>G, 5031C>A, 5033G>C, 5034A>T, 5036G>A, 5037G>A, 5042_5047delGAATTA, 5050C>T, 5053C>T, 5056G>A, 5059T>C, 5060T>A, 5062T>A, 5063T>A, 5064C>G, 5065A>T, 5071A>C, 5074C>T, 5077A>C, 5078C>G, 5079C>A, 5087T>G, 5089G>T, 5095C>T, 5096C>T, 5097A>C, 5098T>C, 5100G>C, 5101T>A, 5107G>A, 5117C>T, 5122G>A, 5126A>G, 5127A>T, 5129C>A, 5130A>G, 5131G>A, 5133T>G, 5134G>A, 5137T>A, 5138A>C, 5139C>G, 5140T>A, 5147A>T, 5148G>C, 5149T>A, 5151T>C, 5152T>A, 5154A>C, 5155T>C, 5156C>T, 5159A>T, 5160C>T, 5161C>A, 5163C>A, 5166T>A, 5167T>A, 5168T>C, 5169G>C, 5170T>A, 5174A>G, 5175C>T, 5176A>T, 5177G>A, 5179C>T, 5181G>A, 5182G>A, 5184C>T, 5192G>A, 5193T>C, 5195A>C, 5197T>A, 5198_5200delCTT, 5201T>A, 5204A>T, 5205A>C, 5212T>G, 5215G>A, 5216A>T, 5217A>T, 5221A>T, 5224G>A, 5225A>C, 5227T>C, 5228A>C, 5231G>T, 5232G>C, 5236C>A, 5237A>C, 5238G>C, 5240A>C, 5241A>C, 5242A>T | | | | | | | | |

##### CDS

| | Begin | End | Coverage | Score | Concordance | Matches | Identities | I/D/M/F* | Stop Codons |
|---|---|---|---|---|---|---|---|---|---|
| MiMVgp6 | 475 | 630 | 23.1% | 464 | 46.2% | 151 (96.8%) | 72 (46.2%) | 0/5/0/0 | 0 |
| Protein mutations: | V479C (4790G>T 4791T>G 4792T>C), T481N (4797C>A 4798A>C), D484G (4806A>G 4807T>A), S485E (4808A>G 4809G>A 4810C>A), Y486F (4812A>T 4813T>C), K489E (4820A>G 4822G>A), E492K (4829G>A), K495N (4840A>T), P496L (4842C>T 4843G>T), V499K (4850G>A 4851T>A), K500L (4853A>C 4854A>T 4855G>T), L501I (4856T>A 4858G>C), K502R (4860A>G 4861G>A), D504S (4865G>T 4866A>C 4867T>A), Q505N (4868C>A 4870G>T), V506R (4871G>C 4872T>G), W507L (4875G>T), I508G (4877A>G 4878T>G 4879C>A), Q511K (4886C>A), S512K (4889T>A 4890C>A), D513H (4892G>C), D515L (4898G>C 4899A>T), Y516C (4902A>G 4903C>T), K518Q (4907A>C), I520L (4913A>C 4915A>C), K522E (4919A>G), G523E (4923G>A), L524C (4926T>G 4927A>T), I525A (4928A>G 4929T>C), N526K (4933T>A), F527L (4934T>C), Y531R (4946T>C 4947A>G 4948T>A), K534E (4955A>G 4957G>A), K535P (4958A>C 4959A>C), S538N (4968G>A), I541L (4976A>T 4978T>A), E542Q (4979G>C), H548Y (4997C>T 4999C>T), F549H (5000T>C 5001T>A 5002C>T), G552A (5010G>C), V553L (5012G>T 5014C>A), K555_A556del (5018_5023delAAGGCC), T559D (5030A>G 5031C>A), E560L (5033G>C 5034A>T), G561N (5036G>A 5037G>A), E563_L564del (5042_5047delGAATTA), S569T (5060T>A 5062T>A), K574N (5077A>C), P575E (5078C>G 5079C>A), L578V (5087T>G 5089G>T), H581S (5096C>T 5097A>C 5098T>C), S582T (5100G>C 5101T>A), K591V (5126A>G 5127A>T), Q592R (5129C>A 5130A>G 5131G>A), V593G (5133T>G 5134G>A), T595R (5138A>C 5139C>G 5140T>A), I599T (5151T>C 5152T>A), Y600S (5154A>C 5155T>C), T602L (5159A>T 5160C>T 5161C>A), P603H (5163C>A), V604E (5166T>A 5167T>A), C605P (5168T>C 5169G>C 5170T>A), T607V (5174A>G 5175C>T 5176A>T), V608I (5177G>A 5179C>T), R609K (5181G>A 5182G>A), T610I (5184C>T), V613T (5192G>A 5193T>C), N614Q (5195A>C 5197T>A), L615del (5198_5200delCTT), L616I (5201T>A), K617S (5204A>T 5205A>C), F619L (5212T>G), M620I (5215G>A), N621F (5216A>T 5217A>T), K622N (5221A>T), I624L (5225A>C 5227T>C), T625P (5228A>C), G626S (5231G>T 5232G>C), D627E (5236C>A), S628P (5237A>C 5238G>C), K629P (5240A>C 5241A>C 5242A>T) | | | | | | | | |
| Codon mutations: | AGG475AGA (4780G>A), GTT479TGC (4790G>T 4791T>G 4792T>C), CTA480TTA (4793C>T), ACA481AAC (4797C>A 4798A>C), GAT484GGA (4806A>G 4807T>A), AGC485GAA (4808A>G 4809G>A 4810C>A), TAT486TTC (4812A>T 4813T>C), ATC487ATA (4816C>A), AAG489GAA (4820A>G 4822G>A), CTT490TTA (4823C>T 4825T>A), GAA492AAA (4829G>A), AAA493AAG (4834A>G), CGG494AGA (4835C>A 4837G>A), AAA495AAT (4840A>T), CCG496CTT (4842C>T 4843G>T), TTA497CTA (4844T>C), CAG498CAA (4849G>A), GTA499AAA (4850G>A 4851T>A), AAG500CTT (4853A>C 4854A>T 4855G>T), TTG501ATC (4856T>A 4858G>C), AAG502AGA (4860A>G 4861G>A), GAT504TCA (4865G>T 4866A>C 4867T>A), CAG505AAT (4868C>A 4870G>T), GTA506CGA (4871G>C 4872T>G), TGG507TTG (4875G>T), ATC508GGA (4877A>G 4878T>G 4879C>A), ACG510ACA (4885G>A), CAA511AAA (4886C>A), TCA512AAA (4889T>A 4890C>A), GAT513CAT (4892G>C), GAT515CTT (4898G>C 4899A>T), TAC516TGT (4902A>G 4903C>T), AAA518CAA (4907A>C), AAG519AAA (4912G>A), ATA520CTC (4913A>C 4915A>C), AAG521AAA (4918G>A), AAA522GAA (4919A>G), GGA523GAA (4923G>A), TTA524TGT (4926T>G 4927A>T), ATT525GCT (4928A>G 4929T>C), AAT526AAA (4933T>A), TTT527CTT (4934T>C), TAT531CGA (4946T>C 4947A>G 4948T>A), CTA532TTA (4949C>T), AAG534GAA (4955A>G 4957G>A), AAA535CCA (4958A>C 4959A>C), GAC537GAT (4966C>T), AGT538AAT (4968G>A), TTA539TTG (4972A>G), ATT541TTA (4976A>T 4978T>A), GAA542CAA (4979G>C), GCT545GCC (4990T>C), TCT546TCC (4993T>C), CAC548TAT (4997C>T 4999C>T), TTC549CAT (5000T>C 5001T>A 5002C>T), GGT551GGA (5008T>A), GGA552GCA (5010G>C), GTC553TTA (5012G>T 5014C>A), CTT554TTA (5015C>T 5017T>A), AAG555_GCC556del (5018_5023delAAGGCC), ACC558ACA (5029C>A), ACT559GAT (5030A>G 5031C>A), GAA560CTA (5033G>C 5034A>T), GGT561AAT (5036G>A 5037G>A), GAA563_TTA564del (5042_5047delGAATTA), ATC565ATT (5050C>T), TGC566TGT (5053C>T), AGG567AGA (5056G>A), TAT568TAC (5059T>C), TCT569ACA (5060T>A 5062T>A), TCA570AGT (5063T>A 5064C>G 5065A>T), ACA572ACC (5071A>C), TTC573TTT (5074C>T), AAA574AAC (5077A>C), CCA575GAA (5078C>G 5079C>A), TTG578GTT (5087T>G 5089G>T), TAC580TAT (5095C>T), CAT581TCC (5096C>T 5097A>C 5098T>C), AGT582ACA (5100G>C 5101T>A), GAG584GAA (5107G>A), CTA588TTA (5117C>T), GCG589GCA (5122G>A), AAA591GTA (5126A>G 5127A>T), CAG592AGA (5129C>A 5130A>G 5131G>A), GTG593GGA (5133T>G 5134G>A), ATT594ATA (5137T>A), ACT595CGA (5138A>C 5139C>G 5140T>A), AGT598TCA (5147A>T 5148G>C 5149T>A), ATT599ACA (5151T>C 5152T>A), TAT600TCC (5154A>C 5155T>C), CTA601TTA (5156C>T), ACC602TTA (5159A>T 5160C>T 5161C>A), CCT603CAT (5163C>A), GTT604GAA (5166T>A 5167T>A), TGT605CCA (5168T>C 5169G>C 5170T>A), ACA607GTT (5174A>G 5175C>T 5176A>T), GTC608ATT (5177G>A 5179C>T), AGG609AAA (5181G>A 5182G>A), ACA610ATA (5184C>T), GTA613ACA (5192G>A 5193T>C), AAT614CAA (5195A>C 5197T>A), CTT615del (5198_5200delCTT), TTA616ATA (5201T>A), AAA617TCA (5204A>T 5205A>C), TTT619TTG (5212T>G), ATG620ATA (5215G>A), AAT621TTT (5216A>T 5217A>T), AAA622AAT (5221A>T), AAG623AAA (5224G>A), ATT624CTC (5225A>C 5227T>C), ACT625CCT (5228A>C), GGT626TCT (5231G>T 5232G>C), GAC627GAA (5236C>A), AGT628CCT (5237A>C 5238G>C), AAA629CCT (5240A>C 5241A>C 5242A>T) | | | | | | | | |

##### Proteins

| | Begin | End | Coverage | Score | Concordance | Matches | Identities | I/D/M/F* | Stop Codons |
|---|---|---|---|---|---|---|---|---|---|
| hypothetical protein (NP_659397.1) | 475 | 630 | 23.1% | 464 | 46.2% | 151 (96.8%) | 72 (46.2%) | 0/5/0/0 | 0 |
| Protein mutations: | V479C (4790G>T 4791T>G 4792T>C), T481N (4797C>A 4798A>C), D484G (4806A>G 4807T>A), S485E (4808A>G 4809G>A 4810C>A), Y486F (4812A>T 4813T>C), K489E (4820A>G 4822G>A), E492K (4829G>A), K495N (4840A>T), P496L (4842C>T 4843G>T), V499K (4850G>A 4851T>A), K500L (4853A>C 4854A>T 4855G>T), L501I (4856T>A 4858G>C), K502R (4860A>G 4861G>A), D504S (4865G>T 4866A>C 4867T>A), Q505N (4868C>A 4870G>T), V506R (4871G>C 4872T>G), W507L (4875G>T), I508G (4877A>G 4878T>G 4879C>A), Q511K (4886C>A), S512K (4889T>A 4890C>A), D513H (4892G>C), D515L (4898G>C 4899A>T), Y516C (4902A>G 4903C>T), K518Q (4907A>C), I520L (4913A>C 4915A>C), K522E (4919A>G), G523E (4923G>A), L524C (4926T>G 4927A>T), I525A (4928A>G 4929T>C), N526K (4933T>A), F527L (4934T>C), Y531R (4946T>C 4947A>G 4948T>A), K534E (4955A>G 4957G>A), K535P (4958A>C 4959A>C), S538N (4968G>A), I541L (4976A>T 4978T>A), E542Q (4979G>C), H548Y (4997C>T 4999C>T), F549H (5000T>C 5001T>A 5002C>T), G552A (5010G>C), V553L (5012G>T 5014C>A), K555_A556del (5018_5023delAAGGCC), T559D (5030A>G 5031C>A), E560L (5033G>C 5034A>T), G561N (5036G>A 5037G>A), E563_L564del (5042_5047delGAATTA), S569T (5060T>A 5062T>A), K574N (5077A>C), P575E (5078C>G 5079C>A), L578V (5087T>G 5089G>T), H581S (5096C>T 5097A>C 5098T>C), S582T (5100G>C 5101T>A), K591V (5126A>G 5127A>T), Q592R (5129C>A 5130A>G 5131G>A), V593G (5133T>G 5134G>A), T595R (5138A>C 5139C>G 5140T>A), I599T (5151T>C 5152T>A), Y600S (5154A>C 5155T>C), T602L (5159A>T 5160C>T 5161C>A), P603H (5163C>A), V604E (5166T>A 5167T>A), C605P (5168T>C 5169G>C 5170T>A), T607V (5174A>G 5175C>T 5176A>T), V608I (5177G>A 5179C>T), R609K (5181G>A 5182G>A), T610I (5184C>T), V613T (5192G>A 5193T>C), N614Q (5195A>C 5197T>A), L615del (5198_5200delCTT), L616I (5201T>A), K617S (5204A>T 5205A>C), F619L (5212T>G), M620I (5215G>A), N621F (5216A>T 5217A>T), K622N (5221A>T), I624L (5225A>C 5227T>C), T625P (5228A>C), G626S (5231G>T 5232G>C), D627E (5236C>A), S628P (5237A>C 5238G>C), K629P (5240A>C 5241A>C 5242A>T) | | | | | | | | |
| Codon mutations: | AGG475AGA (4780G>A), GTT479TGC (4790G>T 4791T>G 4792T>C), CTA480TTA (4793C>T), ACA481AAC (4797C>A 4798A>C), GAT484GGA (4806A>G 4807T>A), AGC485GAA (4808A>G 4809G>A 4810C>A), TAT486TTC (4812A>T 4813T>C), ATC487ATA (4816C>A), AAG489GAA (4820A>G 4822G>A), CTT490TTA (4823C>T 4825T>A), GAA492AAA (4829G>A), AAA493AAG (4834A>G), CGG494AGA (4835C>A 4837G>A), AAA495AAT (4840A>T), CCG496CTT (4842C>T 4843G>T), TTA497CTA (4844T>C), CAG498CAA (4849G>A), GTA499AAA (4850G>A 4851T>A), AAG500CTT (4853A>C 4854A>T 4855G>T), TTG501ATC (4856T>A 4858G>C), AAG502AGA (4860A>G 4861G>A), GAT504TCA (4865G>T 4866A>C 4867T>A), CAG505AAT (4868C>A 4870G>T), GTA506CGA (4871G>C 4872T>G), TGG507TTG (4875G>T), ATC508GGA (4877A>G 4878T>G 4879C>A), ACG510ACA (4885G>A), CAA511AAA (4886C>A), TCA512AAA (4889T>A 4890C>A), GAT513CAT (4892G>C), GAT515CTT (4898G>C 4899A>T), TAC516TGT (4902A>G 4903C>T), AAA518CAA (4907A>C), AAG519AAA (4912G>A), ATA520CTC (4913A>C 4915A>C), AAG521AAA (4918G>A), AAA522GAA (4919A>G), GGA523GAA (4923G>A), TTA524TGT (4926T>G 4927A>T), ATT525GCT (4928A>G 4929T>C), AAT526AAA (4933T>A), TTT527CTT (4934T>C), TAT531CGA (4946T>C 4947A>G 4948T>A), CTA532TTA (4949C>T), AAG534GAA (4955A>G 4957G>A), AAA535CCA (4958A>C 4959A>C), GAC537GAT (4966C>T), AGT538AAT (4968G>A), TTA539TTG (4972A>G), ATT541TTA (4976A>T 4978T>A), GAA542CAA (4979G>C), GCT545GCC (4990T>C), TCT546TCC (4993T>C), CAC548TAT (4997C>T 4999C>T), TTC549CAT (5000T>C 5001T>A 5002C>T), GGT551GGA (5008T>A), GGA552GCA (5010G>C), GTC553TTA (5012G>T 5014C>A), CTT554TTA (5015C>T 5017T>A), AAG555_GCC556del (5018_5023delAAGGCC), ACC558ACA (5029C>A), ACT559GAT (5030A>G 5031C>A), GAA560CTA (5033G>C 5034A>T), GGT561AAT (5036G>A 5037G>A), GAA563_TTA564del (5042_5047delGAATTA), ATC565ATT (5050C>T), TGC566TGT (5053C>T), AGG567AGA (5056G>A), TAT568TAC (5059T>C), TCT569ACA (5060T>A 5062T>A), TCA570AGT (5063T>A 5064C>G 5065A>T), ACA572ACC (5071A>C), TTC573TTT (5074C>T), AAA574AAC (5077A>C), CCA575GAA (5078C>G 5079C>A), TTG578GTT (5087T>G 5089G>T), TAC580TAT (5095C>T), CAT581TCC (5096C>T 5097A>C 5098T>C), AGT582ACA (5100G>C 5101T>A), GAG584GAA (5107G>A), CTA588TTA (5117C>T), GCG589GCA (5122G>A), AAA591GTA (5126A>G 5127A>T), CAG592AGA (5129C>A 5130A>G 5131G>A), GTG593GGA (5133T>G 5134G>A), ATT594ATA (5137T>A), ACT595CGA (5138A>C 5139C>G 5140T>A), AGT598TCA (5147A>T 5148G>C 5149T>A), ATT599ACA (5151T>C 5152T>A), TAT600TCC (5154A>C 5155T>C), CTA601TTA (5156C>T), ACC602TTA (5159A>T 5160C>T 5161C>A), CCT603CAT (5163C>A), GTT604GAA (5166T>A 5167T>A), TGT605CCA (5168T>C 5169G>C 5170T>A), ACA607GTT (5174A>G 5175C>T 5176A>T), GTC608ATT (5177G>A 5179C>T), AGG609AAA (5181G>A 5182G>A), ACA610ATA (5184C>T), GTA613ACA (5192G>A 5193T>C), AAT614CAA (5195A>C 5197T>A), CTT615del (5198_5200delCTT), TTA616ATA (5201T>A), AAA617TCA (5204A>T 5205A>C), TTT619TTG (5212T>G), ATG620ATA (5215G>A), AAT621TTT (5216A>T 5217A>T), AAA622AAT (5221A>T), AAG623AAA (5224G>A), ATT624CTC (5225A>C 5227T>C), ACT625CCT (5228A>C), GGT626TCT (5231G>T 5232G>C), GAC627GAA (5236C>A), AGT628CCT (5237A>C 5238G>C), AAA629CCT (5240A>C 5241A>C 5242A>T) | | | | | | | | |

*: Inserts / Deletes / Misaligned / Frameshifts

#### Analysis details

This analysis was performed with panviral2.64

### NGS Details (UN18_val): Badnavirus tessellocastaneae

#### Assembly

| | |
|---|---|
| **Coverage Length** | 263 (1 contig(s)) |
| **Depth Of Coverage** | 2.8 |
| **Number Of Reads** | 6 |
| **Reads Per Million** | 0.11 rpm (after QC) |
| **Ambiguities** | 0 |
| **Assembly Method** | de novo + reference guided assembly |
| **Consensus Caller** | Bcf Tools |

#### Coverage Map

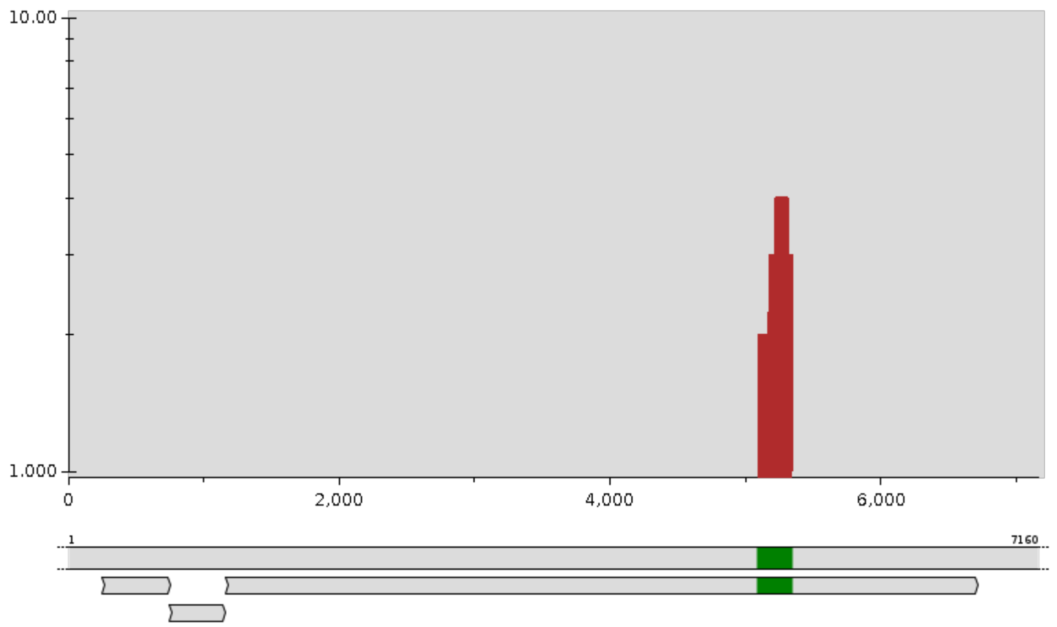

#### Assignment

| | |
|---|---|
| **Type** | Badnavirus tessellocastaneae (Taxonomy ID: 3051987) |
| **Reference Genome** | NC_076885.1 |
| **NT Identity (%)** | 55.8935 |
| **AA Identity (%)** | 54.5455 |
| **Number Of Stop Codons** | 0 |
| **Number Of CDS** | 3 |

#### Alignment

| | |
|---|---|
| **Alignment Score** | 62.0 (NT) + 353.0 (AA) = 415.0 |
| **Concordance (%)** | 36.1813 |

| Alignment Method | Global, seeded, nucleotide + amino acids (AGA) |
|---|---|

#### Genome Region

Sequence starts at position 5087 and ends at position 5349 relative to NC_076885.1 reference sequence.

#### Alignment Detailed Statistics

| | Begin | End | Coverage | Score | Concordance | Matches | Identities | I/D/M/F* | Stop Codons |
|---|---|---|---|---|---|---|---|---|---|
| **NT** | **5087** | **5349** | **3.7%** | **62** | **11.8%** | **263 (100%)** | **147 (55.9%)** | **0/0** | |
| Mutations: | 5087C>G, 5092T>C, 5098A>G, 5100C>A, 5102C>T, 5104C>A, 5109T>C, 5110G>C, 5113T>C, 5116C>T, 5117C>G, 5118A>T, 5122A>G, 5125C>T, 5126C>A, 5130A>T, 5132T>C, 5134C>A, 5135C>A, 5137C>T, 5142G>T, 5143A>T, 5146A>T, 5148C>A, 5149A>G, 5150T>G, 5151C>A, 5152C>A, 5153A>T, 5155C>A, 5156A>T, 5158C>A, 5159A>G, 5161G>T, 5162A>G, 5163G>A, 5164A>G, 5165G>T, 5167T>G, 5168G>C, 5169C>A, 5170A>T, 5174T>G, 5176A>C, 5180G>T, 5181T>A, 5182C>T, 5188C>T, 5192T>C, 5194T>A, 5197T>C, 5200A>G, 5202A>G, 5203G>A, 5204A>T, 5205G>C, 5209G>T, 5211T>A, 5215T>C, 5219G>A, 5221G>A, 5222G>A, 5223T>G, 5228G>T, 5229A>T, 5230A>T, 5234G>T, 5235A>C, 5236G>T, 5237T>A, 5238C>A, 5239A>T, 5245A>T, 5246T>A, 5247G>A, 5248G>A, 5251A>G, 5252G>A, 5254G>A, 5258G>A, 5259T>G, 5260G>A, 5261G>A, 5262T>C, 5263A>C, 5265C>A, 5266A>T, 5267G>C, 5268G>A, 5272A>G, 5273T>C, 5274T>A, 5275A>C, 5278C>T, 5283G>T, 5284G>C, 5285C>T, 5287G>A, 5290T>G, 5296A>C, 5302G>C, 5305A>T, 5307G>C, 5308A>C, 5314A>C, 5317A>C, 5320A>C, 5323C>A, 5326C>T, 5329A>G, 5332G>C, 5333A>C, 5334A>T, 5335G>C, 5339G>A, 5341C>T | | | | | | | | |

##### CDS

| | Begin | End | Coverage | Score | Concordance | Matches | Identities | I/D/M/F* | Stop Codons |
|---|---|---|---|---|---|---|---|---|---|
| ORF3 | 1309 | 1396 | 4.7% | 353 | 55.9% | 88 (100%) | 48 (54.5%) | 0/0/0/0 | 0 |
| Protein mutations: | L1309V (5087C>G), A1313E (5100C>A), M1316T (5109T>C 5110G>C), H1319V (5117C>G 5118A>T), Q1322K (5126C>A), Y1323F (5130A>T), S1324P (5132T>C 5134C>A), L1325I (5135C>A 5137C>T), G1327V (5142G>T 5143A>T), T1329K (5148C>A 5149A>G), S1330E (5150T>G 5151C>A 5152C>A), I1331L (5153A>T 5155C>A), I1332L (5156A>T 5158C>A), K1333D (5159A>G 5161G>T), R1334E (5162A>G 5163G>A 5164A>G), V1335L (5165G>T 5167T>G), A1336H (5168G>C 5169C>A 5170A>T), S1338A (5174T>G 5176A>C), V1340Y (5180G>T 5181T>A 5182C>T), F1344L (5192T>C 5194T>A), K1347R (5202A>G 5203G>A), F1350Y (5211T>A), V1353I (5219G>A 5221G>A), V1354R (5222G>A 5223T>G), E1356F (5228G>T 5229A>T 5230A>T), E1358S (5234G>T 5235A>C 5236G>T), S1359N (5237T>A 5238C>A 5239A>T), W1362K (5246T>A 5247G>A 5248G>A), A1364T (5252G>A 5254G>A), V1366R (5258G>A 5259T>G 5260G>A), V1367T (5261G>A 5262T>C 5263A>C), P1368H (5265C>A 5266A>T), G1369Q (5267G>C 5268G>A), L1371H (5273T>C 5274T>A 5275A>C), W1374F (5283G>T 5284G>C), R1382T (5307G>C 5308A>C), R1390S (5332G>C), K1391L (5333A>C 5334A>T 5335G>C), D1393N (5339G>A 5341C>T) | | | | | | | | |
| Codon mutations: | CTA1309GTA (5087C>G), GAT1310GAC (5092T>C), AGA1312AGG (5098A>G), GCA1313GAA (5100C>A), CTC1314TTA (5102C>T 5104C>A), ATG1316ACC (5109T>C 5110G>C), AAT1317AAC (5113T>C), ACC1318ACT (5116C>T), CAC1319GTC (5117C>G 5118A>T), AAA1320AAG (5122A>G), GAC1321GAT (5125C>T), CAG1322AAG (5126C>A), TAT1323TTT (5130A>T), TCC1324CCA (5132T>C 5134C>A), CTC1325ATT (5135C>A 5137C>T), GGA1327GTT (5142G>T 5143A>T), ATA1328ATT (5146A>T), ACA1329AAG (5148C>A 5149A>G), TCC1330GAA (5150T>G 5151C>A 5152C>A), ATC1331TTA (5153A>T 5155C>A), ATC1332TTA (5156A>T 5158C>A), AAG1333GAT (5159A>G 5161G>T), AGA1334GAG (5162A>G 5163G>A 5164A>G), GTT1335TTG (5165G>T 5167T>G), GCA1336CAT (5168G>C 5169C>A 5170A>T), TCA1338GCC (5174T>G 5176A>C), GTC1340TAT (5180G>T 5181T>A 5182C>T), TCC1342TCT (5188C>T), TTT1344CTA (5192T>C 5194T>A), GAT1345GAC (5197T>C), TTA1346TTG (5200A>G), AAG1347AGA (5202A>G 5203G>A), AGT1348TCT (5204A>T 5205G>C), GGG1349GGT (5209G>T), TTC1350TAC (5211T>A), CAT1351CAC (5215T>C), GTG1353ATA (5219G>A 5221G>A), GTA1354AGA (5222G>A 5223T>G), GAA1356TTT (5228G>T 5229A>T 5230A>T), GAG1358TCT (5234G>T 5235A>C 5236G>T), TCA1359AAT (5237T>A 5238C>A 5239A>T), CCA1361CCT (5245A>T), TGG1362AAA (5246T>A 5247G>A 5248G>A), ACA1363ACG (5251A>G), GCG1364ACA (5252G>A 5254G>A), GTG1366AGA (5258G>A 5259T>G 5260G>A), GTA1367ACC (5261G>A 5262T>C 5263A>C), CCA1368CAT (5265C>A 5266A>T), GGA1369CAA (5267G>C 5268G>A), GGA1370GGG (5272A>G), TTA1371CAC (5273T>C 5274T>A 5275A>C), TAC1372TAT (5278C>T), TGG1374TTC (5283G>T 5284G>C), CTG1375TTA (5285C>T 5287G>A), GTT1376GTG (5290T>G), CCA1378CCC (5296A>C), GGG1380GGC (5302G>C), CTA1381CTT (5305A>T), AGA1382ACC (5307G>C 5308A>C), GCA1384GCC (5314A>C), CCA1385CCC (5317A>C), TCA1386TCC (5320A>C), ACC1387ACA (5323C>A), TTC1388TTT (5326C>T), CAA1389CAG (5329A>G), AGG1390AGC (5332G>C), AAG1391CTC (5333A>C 5334A>T 5335G>C), GAC1393AAT (5339G>A 5341C>T) | | | | | | | | |

##### Proteins

| | Begin | End | Coverage | Score | Concordance | Matches | Identities | I/D/M/F* | Stop Codons |
|---|---|---|---|---|---|---|---|---|---|
| polyprotein (YP_010800602.1) | 1309 | 1396 | 4.7% | 353 | 55.9% | 88 (100%) | 48 (54.5%) | 0/0/0/0 | 0 |
| Protein mutations: | L1309V (5087C>G), A1313E (5100C>A), M1316T (5109T>C 5110G>C), H1319V (5117C>G 5118A>T), Q1322K (5126C>A), Y1323F (5130A>T), S1324P (5132T>C 5134C>A), L1325I (5135C>A 5137C>T), G1327V (5142G>T 5143A>T), T1329K (5148C>A 5149A>G), S1330E (5150T>G 5151C>A 5152C>A), I1331L (5153A>T 5155C>A), I1332L (5156A>T 5158C>A), K1333D (5159A>G 5161G>T), R1334E (5162A>G 5163G>A 5164A>G), V1335L (5165G>T 5167T>G), A1336H (5168G>C 5169C>A 5170A>T), S1338A (5174T>G 5176A>C), V1340Y (5180G>T 5181T>A 5182C>T), F1344L (5192T>C 5194T>A), K1347R (5202A>G 5203G>A), F1350Y (5211T>A), V1353I (5219G>A 5221G>A), V1354R (5222G>A 5223T>G), E1356F (5228G>T 5229A>T 5230A>T), E1358S (5234G>T 5235A>C 5236G>T), S1359N (5237T>A 5238C>A 5239A>T), W1362K (5246T>A 5247G>A 5248G>A), A1364T (5252G>A 5254G>A), V1366R (5258G>A 5259T>G 5260G>A), V1367T (5261G>A 5262T>C 5263A>C), P1368H (5265C>A 5266A>T), G1369Q (5267G>C 5268G>A), L1371H (5273T>C 5274T>A 5275A>C), W1374F (5283G>T 5284G>C), R1382T (5307G>C 5308A>C), R1390S (5332G>C), K1391L (5333A>C 5334A>T 5335G>C), D1393N (5339G>A 5341C>T) | | | | | | | | |
| Codon mutations: | CTA1309GTA (5087C>G), GAT1310GAC (5092T>C), AGA1312AGG (5098A>G), GCA1313GAA (5100C>A), CTC1314TTA (5102C>T 5104C>A), ATG1316ACC (5109T>C 5110G>C), AAT1317AAC (5113T>C), ACC1318ACT (5116C>T), CAC1319GTC (5117C>G 5118A>T), AAA1320AAG (5122A>G), GAC1321GAT (5125C>T), CAG1322AAG (5126C>A), TAT1323TTT (5130A>T), TCC1324CCA (5132T>C 5134C>A), CTC1325ATT (5135C>A 5137C>T), GGA1327GTT (5142G>T 5143A>T), ATA1328ATT (5146A>T), ACA1329AAG (5148C>A 5149A>G), TCC1330GAA (5150T>G 5151C>A 5152C>A), ATC1331TTA (5153A>T 5155C>A), ATC1332TTA (5156A>T 5158C>A), AAG1333GAT (5159A>G 5161G>T), AGA1334GAG (5162A>G 5163G>A 5164A>G), GTT1335TTG (5165G>T 5167T>G), GCA1336CAT (5168G>C 5169C>A 5170A>T), TCA1338GCC (5174T>G 5176A>C), GTC1340TAT (5180G>T 5181T>A 5182C>T), TCC1342TCT (5188C>T), TTT1344CTA (5192T>C 5194T>A), GAT1345GAC (5197T>C), TTA1346TTG (5200A>G), AAG1347AGA (5202A>G 5203G>A), AGT1348TCT (5204A>T 5205G>C), GGG1349GGT (5209G>T), TTC1350TAC (5211T>A), CAT1351CAC (5215T>C), GTG1353ATA (5219G>A 5221G>A), GTA1354AGA (5222G>A 5223T>G), GAA1356TTT (5228G>T 5229A>T 5230A>T), GAG1358TCT (5234G>T 5235A>C 5236G>T), TCA1359AAT (5237T>A 5238C>A 5239A>T), CCA1361CCT (5245A>T), TGG1362AAA (5246T>A 5247G>A 5248G>A), ACA1363ACG (5251A>G), GCG1364ACA (5252G>A 5254G>A), GTG1366AGA (5258G>A 5259T>G 5260G>A), GTA1367ACC (5261G>A 5262T>C 5263A>C), CCA1368CAT (5265C>A 5266A>T), GGA1369CAA (5267G>C 5268G>A), GGA1370GGG (5272A>G), TTA1371CAC (5273T>C 5274T>A 5275A>C), TAC1372TAT (5278C>T), TGG1374TTC (5283G>T 5284G>C), CTG1375TTA (5285C>T 5287G>A), GTT1376GTG (5290T>G), CCA1378CCC (5296A>C), GGG1380GGC (5302G>C), CTA1381CTT (5305A>T), AGA1382ACC (5307G>C 5308A>C), GCA1384GCC (5314A>C), CCA1385CCC (5317A>C), TCA1386TCC (5320A>C), ACC1387ACA (5323C>A), TTC1388TTT (5326C>T), CAA1389CAG (5329A>G), AGG1390AGC (5332G>C), AAG1391CTC (5333A>C 5334A>T 5335G>C), GAC1393AAT (5339G>A 5341C>T) | | | | | | | | |

*: Inserts / Deletes / Misaligned / Frameshifts

#### Analysis details

This analysis was performed with panviral2.64

### NGS Details (UN18_val): Betabaculovirus erellonis

#### Assembly

| | |
|---|---|
| **Coverage Length** | 91 (1 contig(s)) |
| **Depth Of Coverage** | 5.9 |
| **Number Of Reads** | 6 |
| **Reads Per Million** | 0.11 rpm (after QC) |
| **Ambiguities** | 0 |
| **Assembly Method** | de novo + reference guided assembly |
| **Consensus Caller** | Bcf Tools |

#### Coverage Map

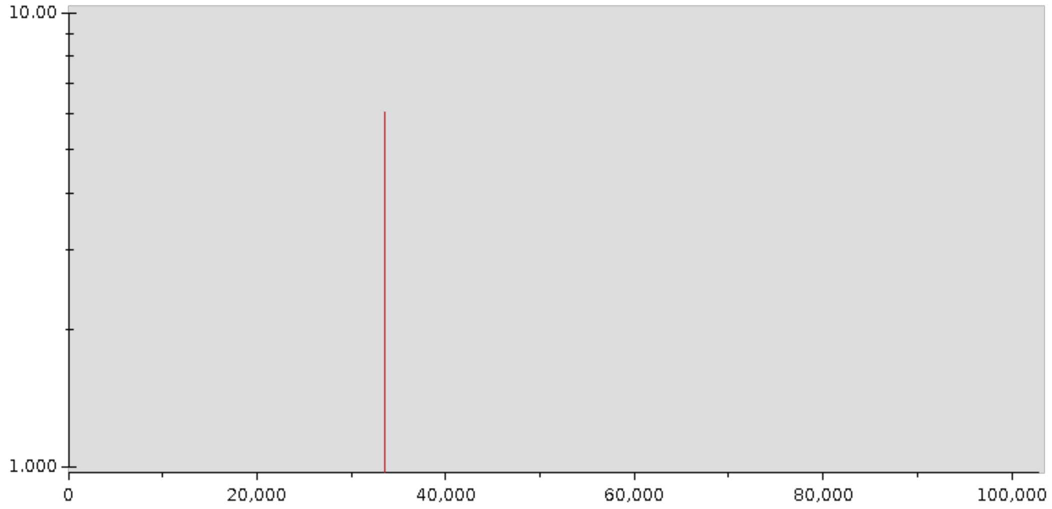

#### Assignment

| | |
|---|---|
| **Type** | Betabaculovirus erellonis (Taxonomy ID: 3047828) |
| **Reference Genome** | NC_025257.1 |
| **NT Identity (%)** | 82.4176 |
| **AA Identity (%)** | 93.3333 |
| **Number Of Stop Codons** | 0 |
| **Number Of CDS** | 130 |

#### Alignment

| | |
|---|---|
| **Alignment Score** | 118.0 (NT) + 178.0 (AA) = 296.0 |
| **Concordance (%)** | 80.4348 |
| **Alignment Method** | Local, heuristic, nucleotide (BLASTN) |

#### Genome Region

Sequence starts at position 33523 and ends at position 33613 relative to NC_025257.1 reference sequence.

#### Alignment Detailed Statistics

| | Begin | End | Coverage | Score | Concordance | Matches | Identities | I/D/M/F* | Stop Codons |
|---|---|---|---|---|---|---|---|---|---|
| **NT** | **33523** | **33613** | **0.1%** | **118** | **64.8%** | **91 (100%)** | **75 (82.4%)** | **0/0** | |

Mutations: 33525A>T, 33537T>A, 33539A>G, 33540C>G, 33549T>C, 33556C>G, 33557T>A, 33561T>C, 33567T>G, 33576A>G, 33580G>C, 33581A>T, 33582G>C, 33585T>A, 33592T>C, 33606T>C

*: Inserts / Deletes / Misaligned / Frameshifts

#### Analysis details

This analysis was performed with panviral2.64

### NGS Details (UN18_val): Gihfavirus pelohabitans

#### Assembly

| | |
|---|---|
| **Coverage Length** | 155 (1 contig(s)) |
| **Depth Of Coverage** | 2.3 |
| **Number Of Reads** | 4 |
| **Reads Per Million** | 0.08 rpm (after QC) |
| **Ambiguities** | 0 |
| **Assembly Method** | de novo + reference guided assembly |
| **Consensus Caller** | Bcf Tools |

#### Coverage Map

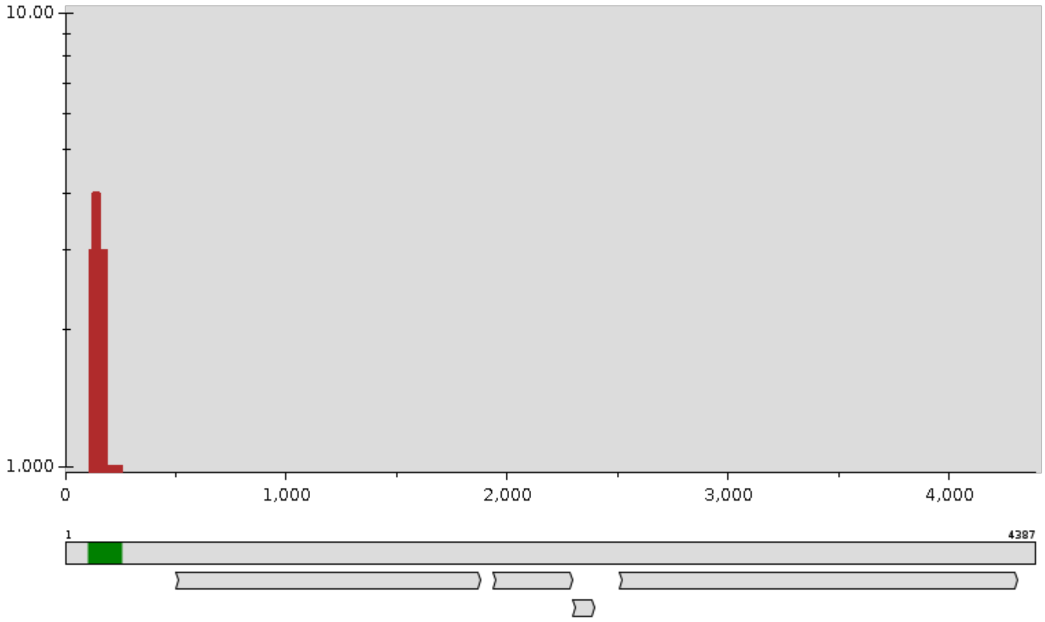

#### Assignment

| | |
|---|---|
| **Type** | Gihfavirus pelohabitans (Taxonomy ID: 2844652) |
| **Reference Genome** | NC_074583.1 |
| **NT Identity (%)** | 97.4194 |
| **AA Identity (%)** | 0.0 |
| **Number Of Stop Codons** | 0 |
| **Number Of CDS** | 4 |

#### Alignment

| | |
|---|---|
| **Alignment Score** | 294.0 (NT) + 0.0 (AA) = 294.0 |
| **Concordance (%)** | 94.8387 |

| Alignment Method | Global, seeded, nucleotide + amino acids (AGA) |
|---|---|

#### Genome Region

Sequence starts at position 106 and ends at position 260 relative to NC_074583.1 reference sequence.

#### Alignment Detailed Statistics

| | Begin | End | Coverage | Score | Concordance | Matches | Identities | I/D/M/F* | Stop Codons |
|---|---|---|---|---|---|---|---|---|---|
| **NT** | **106** | **260** | **3.5%** | **294** | **94.8%** | **155 (100%)** | **151 (97.4%)** | **0/0** | |
| Mutations: | 106G>T, 169A>C, 178T>G, 182T>C | | | | | | | | |

**CDS**

**Proteins**

*: Inserts / Deletes / Misaligned / Frameshifts

#### Analysis details

This analysis was performed with panviral2.64

### NGS Details (UN18_val): Duamitovirus dapi1

#### Assembly

| | |
|---|---|
| **Coverage Length** | 134 (1 contig(s)) |
| **Depth Of Coverage** | 3.8 |
| **Number Of Reads** | 4 |
| **Reads Per Million** | 0.08 rpm (after QC) |
| **Ambiguities** | 0 |
| **Assembly Method** | de novo + reference guided assembly |
| **Consensus Caller** | Bcf Tools |

#### Coverage Map

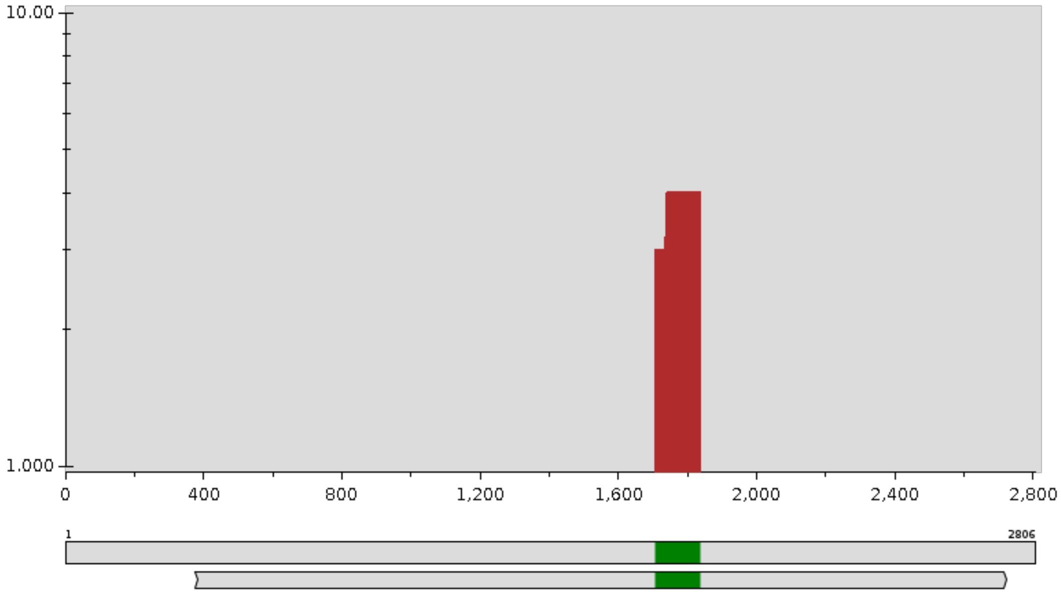

#### Assignment

| | |
|---|---|
| **Type** | Duamitovirus dapi1 (Taxonomy ID: 2955778) |
| **Reference Genome** | NC_076528.1 |
| **NT Identity (%)** | 69.403 |
| **AA Identity (%)** | 66.6667 |
| **Number Of Stop Codons** | 0 |
| **Number Of CDS** | 1 |

#### Alignment

| | |
|---|---|
| **Alignment Score** | 104.0 (NT) + 249.0 (AA) = 353.0 |
| **Concordance (%)** | 57.6797 |

| Alignment Method | Global, seeded, nucleotide + amino acids (AGA) |
|---|---|

#### Genome Region

Sequence starts at position 1707 and ends at position 1840 relative to NC_076528.1 reference sequence.

#### Alignment Detailed Statistics

| | Begin | End | Coverage | Score | Concordance | Matches | Identities | I/D/M/F* | Stop Codons |
|---|---|---|---|---|---|---|---|---|---|
| **NT** | **1707** | **1840** | **4.8%** | **104** | **38.8%** | **134 (100%)** | **93 (69.4%)** | **0/0** | |
| Mutations: | 1710C>T, 1713A>G, 1715G>A, 1725T>C, 1728G>C, 1734C>T, 1736A>T, 1737C>G, 1738T>A, 1739C>G, 1743C>T, 1749G>T, 1750T>C, 1751C>T, 1756G>A, 1763C>A, 1774G>C, 1776A>T, 1783C>A, 1785G>A, 1788G>T, 1789G>A, 1791C>T, 1792T>G, 1793G>A, 1794G>A, 1800A>G, 1804C>G, 1805C>G, 1806T>C, 1809T>G, 1810C>A, 1811A>C, 1816T>C, 1819A>T, 1820A>G, 1821G>C, 1827C>T, 1830T>C, 1831C>A, 1839T>C | | | | | | | | |
| **CDS** | | | | | | | | | |
| RdRp | 445 | 489 | 5.7% | 249 | 68.8% | 45 (100%) | 30 (66.7%) | 0/0/0/0 | 0 |
| Protein mutations: | R447K (1715G>A), Y454L (1736A>T 1737C>G), S459L (1750T>C 1751C>T), A461T (1756G>A), T463K (1763C>A), V467L (1774G>C 1776A>T), L470I (1783C>A 1785G>A), A472T (1789G>A 1791C>T), W473E (1792T>G 1793G>A 1794G>A), P477G (1804C>G 1805C>G 1806T>C), Q479T (1810C>A 1811A>C), F481L (1816T>C), K482C (1819A>T 1820A>G 1821G>C), L486I (1831C>A) | | | | | | | | |
| Codon mutations: | TTC445TTT (1710C>T), ACA446ACG (1713A>G), AGG447AAG (1715G>A), CCT450CCC (1725T>C), CTG451CTC (1728G>C), TTC453TTT (1734C>T), TAC454TTG (1736A>T 1737C>G), TCT455AGT (1738T>A 1739C>G), TCC456TCT (1743C>T), CCG458CCT (1749G>T), TCA459CTA (1750T>C 1751C>T), GCA461ACA (1756G>A), ACA463AAA (1763C>A), GTA467CTT (1774G>C 1776A>T), CTG470ATA (1783C>A 1785G>A), GCG471GCT (1788G>T), GCC472ACT (1789G>A 1791C>T), TGG473GAA (1792T>G 1793G>A 1794G>A), GTA475GTG (1800A>G), CCT477GGC (1804C>G 1805C>G 1806T>C), ACT478ACG (1809T>G), CAG479ACG (1810C>A 1811A>C), TTT481CTT (1816T>C), AAG482TGC (1819A>T 1820A>G 1821G>C), TAC484TAT (1827C>T), GCT485GCC (1830T>C), CTT486ATT (1831C>A), GGT488GGC (1839T>C) | | | | | | | | |
| **Proteins** | | | | | | | | | |
| RNA-dependent RNA polymerase (YP_010798878.1) | 445 | 489 | 5.7% | 249 | 68.8% | 45 (100%) | 30 (66.7%) | 0/0/0/0 | 0 |
| Protein mutations: | R447K (1715G>A), Y454L (1736A>T 1737C>G), S459L (1750T>C 1751C>T), A461T (1756G>A), T463K (1763C>A), V467L (1774G>C 1776A>T), L470I (1783C>A 1785G>A), A472T (1789G>A 1791C>T), W473E (1792T>G 1793G>A 1794G>A), P477G (1804C>G 1805C>G 1806T>C), Q479T (1810C>A 1811A>C), F481L (1816T>C), K482C (1819A>T 1820A>G 1821G>C), L486I (1831C>A) | | | | | | | | |
| Codon mutations: | TTC445TTT (1710C>T), ACA446ACG (1713A>G), AGG447AAG (1715G>A), CCT450CCC (1725T>C), CTG451CTC (1728G>C), TTC453TTT (1734C>T), TAC454TTG (1736A>T 1737C>G), TCT455AGT (1738T>A 1739C>G), TCC456TCT (1743C>T), CCG458CCT (1749G>T), TCA459CTA (1750T>C 1751C>T), GCA461ACA (1756G>A), ACA463AAA (1763C>A), GTA467CTT (1774G>C 1776A>T), CTG470ATA (1783C>A 1785G>A), GCG471GCT (1788G>T), GCC472ACT (1789G>A 1791C>T), TGG473GAA (1792T>G 1793G>A 1794G>A), GTA475GTG (1800A>G), CCT477GGC (1804C>G 1805C>G 1806T>C), ACT478ACG (1809T>G), CAG479ACG (1810C>A 1811A>C), TTT481CTT (1816T>C), AAG482TGC (1819A>T 1820A>G 1821G>C), TAC484TAT (1827C>T), GCT485GCC (1830T>C), CTT486ATT (1831C>A), GGT488GGC (1839T>C) | | | | | | | | |

*: Inserts / Deletes / Misaligned / Frameshifts

#### Analysis details

This analysis was performed with panviral2.64

### NGS Details (UN18_val): Fusarium graminearum mycotymovirus 1

#### Assembly

| | |
|---|---|
| **Coverage Length** | 269 (2 contig(s)) |
| **Depth Of Coverage** | 2.0 |
| **Number Of Reads** | 4 |
| **Reads Per Million** | 0.08 rpm (after QC) |
| **Ambiguities** | 0 |
| **Assembly Method** | de novo + reference guided assembly |
| **Consensus Caller** | Bcf Tools |

#### Coverage Map

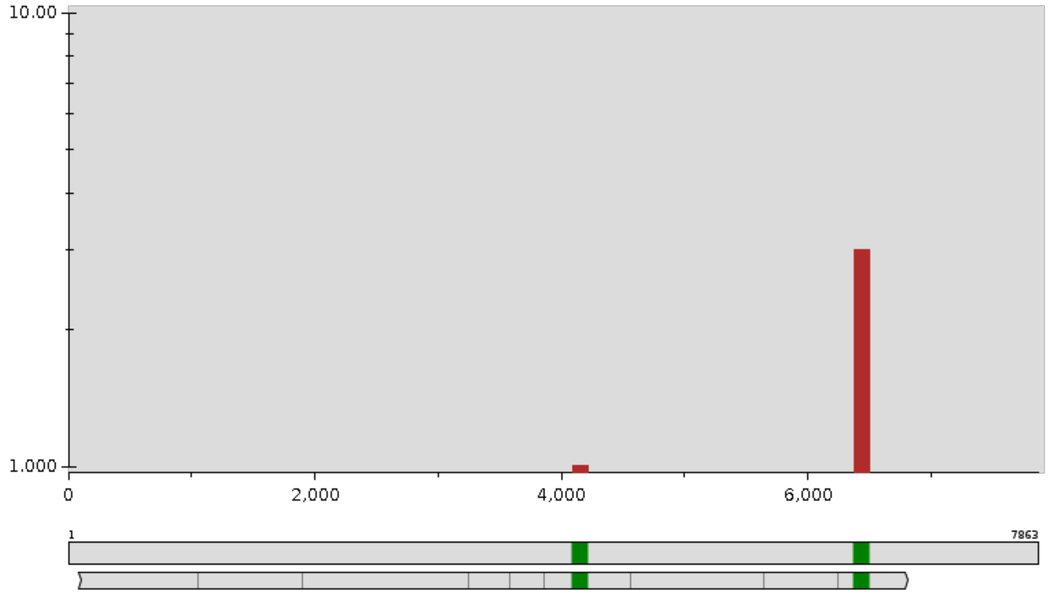

#### Assignment

| | |
|---|---|
| **Type** | Fusarium graminearum mycotymovirus 1 (Taxonomy ID: 1809243) |
| **Reference Genome** | NC_040784.1 |
| **NT Identity (%)** | 69.5167 |
| **AA Identity (%)** | 76.6667 |
| **Number Of Stop Codons** | 0 |
| **Number Of CDS** | 1 |

#### Alignment

| | |
|---|---|
| **Alignment Score** | 210.0 (NT) + 483.0 (AA) = 693.0 |
| **Concordance (%)** | 59.7929 |

| Alignment Method | Global, seeded, nucleotide + amino acids (AGA) |
|---|---|

#### Genome Region

Sequence starts at position 4085 and ends at position 6503 relative to NC_040784.1 reference sequence.

#### Alignment Detailed Statistics

| | Begin | End | Coverage | Score | Concordance | Matches | Identities | I/D/M/F* | Stop Codons |
|---|---|---|---|---|---|---|---|---|---|
| **NT** | **4085** | **6503** | **3.4%** | **210** | **39.0%** | **269 (100%)** | **187 (69.5%)** | **0/0** | |
| Mutations: | 4085T>A, 4087T>G, 4093T>C, 4100A>G, 4105C>T, 4106G>T, 4107C>T, 4108A>C, 4111T>C, 4114G>C, 4123C>T, 4124A>C, 4125G>A, 4126C>G, 4129G>C, 4138A>C, 4141G>T, 4142C>G, 4144G>T, 4145A>G, 4150T>A, 4153C>T, 4156C>T, 4159T>C, 4162A>G, 4165G>A, 4168A>T, 4170G>A, 4171T>G, 4177T>C, 4183A>C, 4184G>A, 4186C>T, 4187T>G, 4188G>C, 4189C>T, 4192T>G, 4193T>A, 4194C>G, 4201A>T, 4205T>A, 4207C>T, 4213C>T, 4215G>C, 6370A>C, 6376C>T, 6379A>G, 6385G>T, 6388A>T, 6397T>C, 6406C>T, 6407A>C, 6409G>C, 6415C>T, 6421C>T, 6424T>C, 6425G>A, 6426G>A, 6427C>T, 6434G>A, 6435C>G, 6436G>C, 6438A>C, 6439A>C, 6440T>A, 6450A>G, 6454G>T, 6455C>T, 6457C>A, 6458A>G, 6459A>C, 6460C>T, 6463A>G, 6466T>C, 6472G>C, 6478C>T, 6481C>T, 6482C>G, 6484C>G, 6493C>T, 6496C>T, 6499C>T | | | | | | | | |
| **CDS** | | | | | | | | | |
| EXL10_gp1 | 1335 | 2141 | 4.0% | 483 | 75.0% | 90 (100%) | 69 (76.7%) | 0/0/0/0 | 0 |
| Protein mutations: | F1335M (4085T>A 4087T>G), I1340V (4100A>G), A1342F (4106G>T 4107C>T 4108A>C), S1348Q (4124A>C 4125G>A 4126C>G), L1354V (4142C>G 4144G>T), I1355V (4145A>G), G1363E (4170G>A 4171T>G), D1368N (4184G>A 4186C>T), C1369A (4187T>G 4188G>C 4189C>T), D1370E (4192T>G), S1375T (4205T>A 4207C>T), C1378S (4215G>C), M2109L (6407A>C 6409G>C), G2115N (6425G>A 6426G>A 6427C>T), A2118S (6434G>A 6435C>G 6436G>C), K2119T (6438A>C 6439A>C), S2120T (6440T>A), N2123S (6450A>G), N2126A (6458A>G 6459A>C 6460C>T), L2134V (6482C>G 6484C>G) | | | | | | | | |
| Codon mutations: | TTT1335ATG (4085T>A 4087T>G), CCT1337CCC (4093T>C), ATC1340GTC (4100A>G), GAC1341GAT (4105C>T), GCA1342TTC (4106G>T 4107C>T 4108A>C), TGT1343TGC (4111T>C), CTG1344CTC (4114G>C), GAC1347GAT (4123C>T), AGC1348CAG (4124A>C 4125G>A 4126C>G), ACG1349ACC (4129G>C), TCA1352TCC (4138A>C), GTG1353GTT (4141G>T), CTG1354GTT (4142C>G 4144G>T), ATC1355GTC (4145A>G), ATT1356ATA (4150T>A), GGC1357GGT (4153C>T), GAC1358GAT (4156C>T), GTT1359GTC (4159T>C), ACA1360ACG (4162A>G), CAG1361CAA (4165G>A), GGA1362GGT (4168A>T), GGT1363GAG (4170G>A 4171T>G), CAT1365CAC (4177T>C), TCA1367TCC (4183A>C), GAC1368AAT (4184G>A 4186C>T), TGC1369GCT (4187T>G 4188G>C 4189C>T), GAT1370GAG (4192T>G), TCC1371AGC (4193T>A 4194C>G), CTA1373CTT (4201A>T), TCC1375ACT (4205T>A 4207C>T), GGC1377GGT (4213C>T), TGT1378TCT (4215G>C), GGA2096..C (6370A>C), ATC2098ATT (6376C>T), AAA2099AAG (6379A>G), CCG2101CCT (6385G>T), CGA2102CGT (6388A>T), TTT2105TTC (6397T>C), CTC2108CTT (6406C>T), ATG2109CTC (6407A>C 6409G>C), GCC2111GCT (6415C>T), GAC2113GAT (6421C>T), CGT2114CGC (6424T>C), GGC2115AAT (6425G>A 6426G>A 6427C>T), GCG2118AGC (6434G>A 6435C>G 6436G>C), AAA2119ACC (6438A>C 6439A>C), TCT2120ACT (6440T>A), AAC2123AGC (6450A>G), CTG2124CTT (6454G>T), CTC2125TTA (6455C>T 6457C>A), AAC2126GCT (6458A>G 6459A>C 6460C>T), GAA2127GAG (6463A>G), GTT2128GTC (6466T>C), GTG2130GTC (6472G>C), TAC2132TAT (6478C>T), CAC2133CAT (6481C>T), CTC2134GTG (6482C>G 6484C>G), CAC2137CAT (6493C>T), GTC2138GTT (6496C>T), TAC2139TAT (6499C>T) | | | | | | | | |
| **Proteins** | | | | | | | | | |
| replication-associated polyprotein (YP_009553357.1) | 1335 | 2141 | 4.0% | 483 | 75.0% | 90 (100%) | 69 (76.7%) | 0/0/0/0 | 0 |
| Protein mutations: | F1335M (4085T>A 4087T>G), I1340V (4100A>G), A1342F (4106G>T 4107C>T 4108A>C), S1348Q (4124A>C 4125G>A 4126C>G), L1354V (4142C>G 4144G>T), I1355V (4145A>G), G1363E (4170G>A 4171T>G), D1368N (4184G>A 4186C>T), C1369A (4187T>G 4188G>C 4189C>T), D1370E (4192T>G), S1375T (4205T>A 4207C>T), C1378S (4215G>C), M2109L (6407A>C 6409G>C), G2115N (6425G>A 6426G>A 6427C>T), A2118S (6434G>A 6435C>G 6436G>C), K2119T (6438A>C 6439A>C), S2120T (6440T>A), N2123S (6450A>G), N2126A (6458A>G 6459A>C 6460C>T), L2134V (6482C>G 6484C>G) | | | | | | | | |
| Codon mutations: | TTT1335ATG (4085T>A 4087T>G), CCT1337CCC (4093T>C), ATC1340GTC (4100A>G), GAC1341GAT (4105C>T), GCA1342TTC (4106G>T 4107C>T 4108A>C), TGT1343TGC (4111T>C), CTG1344CTC (4114G>C), GAC1347GAT (4123C>T), AGC1348CAG (4124A>C 4125G>A 4126C>G), ACG1349ACC (4129G>C), TCA1352TCC (4138A>C), GTG1353GTT (4141G>T), CTG1354GTT (4142C>G 4144G>T), ATC1355GTC (4145A>G), ATT1356ATA (4150T>A), GGC1357GGT (4153C>T), GAC1358GAT (4156C>T), GTT1359GTC (4159T>C), ACA1360ACG (4162A>G), CAG1361CAA (4165G>A), GGA1362GGT (4168A>T), GGT1363GAG (4170G>A 4171T>G), CAT1365CAC (4177T>C), TCA1367TCC (4183A>C), GAC1368AAT (4184G>A 4186C>T), TGC1369GCT (4187T>G 4188G>C 4189C>T), GAT1370GAG (4192T>G), TCC1371AGC (4193T>A 4194C>G), CTA1373CTT (4201A>T), TCC1375ACT (4205T>A 4207C>T), GGC1377GGT (4213C>T), TGT1378TCT (4215G>C), GGA2096..C (6370A>C), ATC2098ATT (6376C>T), AAA2099AAG (6379A>G), CCG2101CCT (6385G>T), CGA2102CGT (6388A>T), TTT2105TTC (6397T>C), CTC2108CTT (6406C>T), ATG2109CTC (6407A>C 6409G>C), GCC2111GCT (6415C>T), GAC2113GAT (6421C>T), CGT2114CGC (6424T>C), GGC2115AAT (6425G>A 6426G>A 6427C>T), GCG2118AGC (6434G>A 6435C>G 6436G>C), AAA2119ACC (6438A>C 6439A>C), TCT2120ACT (6440T>A), AAC2123AGC (6450A>G), CTG2124CTT (6454G>T), CTC2125TTA (6455C>T 6457C>A), AAC2126GCT (6458A>G 6459A>C 6460C>T), GAA2127GAG (6463A>G), GTT2128GTC (6466T>C), GTG2130GTC (6472G>C), TAC2132TAT (6478C>T), CAC2133CAT (6481C>T), CTC2134GTG (6482C>G 6484C>G), CAC2137CAT (6493C>T), GTC2138GTT (6496C>T), TAC2139TAT (6499C>T) | | | | | | | | |
| viral RNA helicase (YP_009553722.1) | 76 | 120 | 19.1% | 236 | 70.9% | 45 (100%) | 33 (73.3%) | 0/0/0/0 | 0 |
| Protein mutations: | F76M (4085T>A 4087T>G), I81V (4100A>G), A83F (4106G>T 4107C>T 4108A>C), S89Q (4124A>C 4125G>A 4126C>G), L95V (4142C>G 4144G>T), I96V (4145A>G), G104E (4170G>A 4171T>G), D109N (4184G>A 4186C>T), C110A (4187T>G 4188G>C 4189C>T), D111E (4192T>G), S116T (4205T>A 4207C>T), C119S (4215G>C) | | | | | | | | |
| Codon mutations: | TTT76ATG (4085T>A 4087T>G), CCT78CCC (4093T>C), ATC81GTC (4100A>G), GAC82GAT (4105C>T), GCA83TTC (4106G>T 4107C>T 4108A>C), TGT84TGC (4111T>C), CTG85CTC (4114G>C), GAC88GAT (4123C>T), AGC89CAG (4124A>C 4125G>A 4126C>G), ACG90ACC (4129G>C), TCA93TCC (4138A>C), GTG94GTT (4141G>T), CTG95GTT (4142C>G 4144G>T), ATC96GTC (4145A>G), ATT97ATA (4150T>A), GGC98GGT (4153C>T), GAC99GAT (4156C>T), GTT100GTC (4159T>C), ACA101ACG (4162A>G), CAG102CAA (4165G>A), GGA103GGT (4168A>T), GGT104GAG (4170G>A 4171T>G), CAT106CAC (4177T>C), TCA108TCC (4183A>C), GAC109AAT (4184G>A 4186C>T), TGC110GCT (4187T>G 4188G>C 4189C>T), GAT111GAG (4192T>G), TCC112AGC (4193T>A 4194C>G), CTA114CTT (4201A>T), TCC116ACT (4205T>A 4207C>T), GGC118GGT (4213C>T), TGT119TCT (4215G>C) | | | | | | | | |

*: Inserts / Deletes / Misaligned / Frameshifts

#### Analysis details

This analysis was performed with panviral2.64

### NGS Details (UN18_val): Tokyovirus A1

#### Assembly

| | |
|---|---|
| **Coverage Length** | 135 (1 contig(s)) |
| **Depth Of Coverage** | 2.9 |
| **Number Of Reads** | 4 |
| **Reads Per Million** | 0.08 rpm (after QC) |
| **Ambiguities** | 0 |
| **Assembly Method** | de novo + reference guided assembly |
| **Consensus Caller** | Bcf Tools |

#### Coverage Map

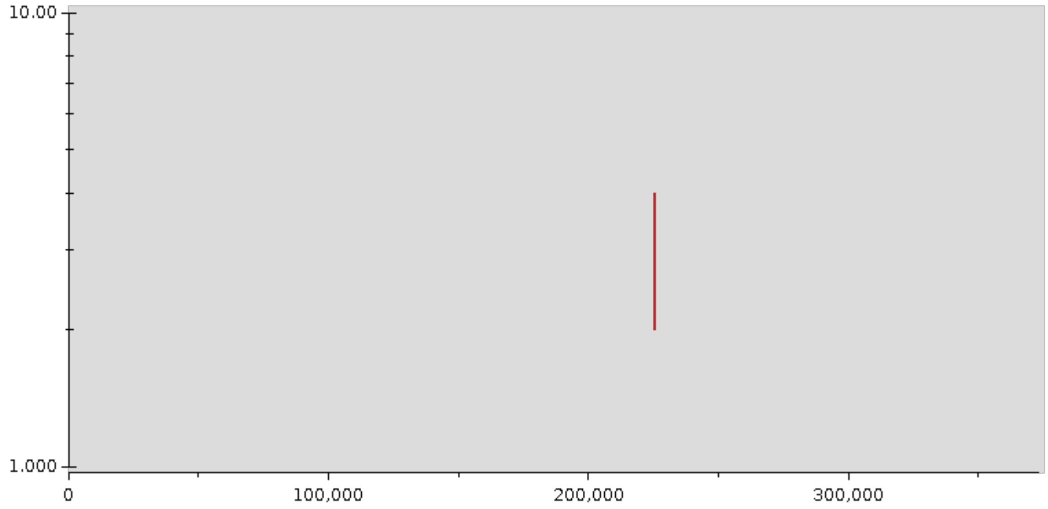

#### Assignment

| | |
|---|---|
| **Type** | Tokyovirus A1 (Taxonomy ID: 1826170) |
| **Reference Genome** | NC_030230.1 |
| **NT Identity (%)** | 77.037 |
| **AA Identity (%)** | 93.3333 |
| **Number Of Stop Codons** | 0 |
| **Number Of CDS** | 470 |

#### Alignment

| | |
|---|---|
| **Alignment Score** | 146.0 (NT) + 291.0 (AA) = 437.0 |
| **Concordance (%)** | 77.3451 |
| **Alignment Method** | Local, heuristic, nucleotide (BLASTN) |

#### Genome Region

Sequence starts at position 225496 and ends at position 225630 relative to NC_030230.1 reference sequence.

#### Alignment Detailed Statistics

| | Begin | End | Coverage | Score | Concordance | Matches | Identities | I/D/M/F* | Stop Codons |
|---|---|---|---|---|---|---|---|---|---|
| NT | 225496 | 225630 | 0.1% | 146 | 54.1% | 135 (100%) | 104 (77.0%) | 0/0 | |

Mutations: 225503G>A, 225504G>C, 225507A>C, 225509G>A, 225513G>C, 225515G>A, 225516A>G, 225519G>A, 225520G>C, 225521A>T, 225522T>C, 225525T>C, 225528T>C, 225531G>A, 225533C>T, 225534A>G, 225540A>G, 225544G>C, 225545C>T, 225552T>C, 225555C>T, 225558A>G, 225564T>G, 225567C>T, 225570C>T, 225573C>T, 225585A>G, 225597G>A, 225603C>A, 225606A>G, 225621T>C

*: Inserts / Deletes / Misaligned / Frameshifts

#### Analysis details

This analysis was performed with panviral2.64

### NGS Details (UN18_val): Botryotinia fuckeliana partitivirus 1 (segment RNA 1)

#### Assembly

| | |
|---|---|
| **Coverage Length** | 135 (1 contig(s)) |
| **Depth Of Coverage** | 3.6 |
| **Number Of Reads** | 4 |
| **Reads Per Million** | 0.08 rpm (after QC) |
| **Ambiguities** | 0 |
| **Assembly Method** | de novo + reference guided assembly |
| **Consensus Caller** | Bcf Tools |

#### Coverage Map

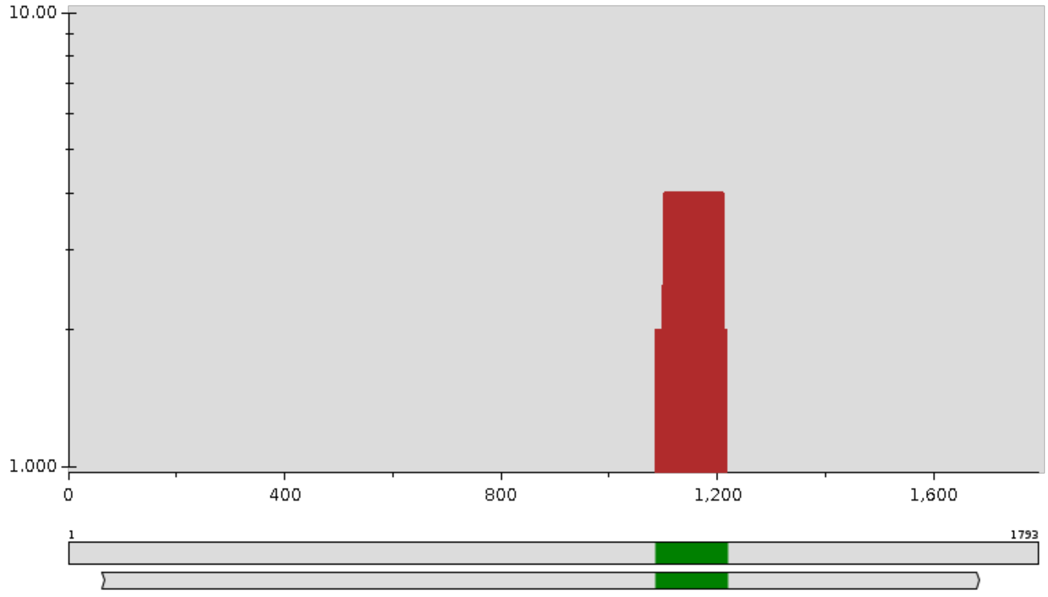

#### Assignment

| | |
|---|---|
| **Type** | Botryotinia fuckeliana partitivirus 1 (Taxonomy ID: 425010) |
| **Reference Genome** | NC_010349.1 |
| **NT Identity (%)** | 68.1481 |
| **AA Identity (%)** | 71.1111 |
| **Number Of Stop Codons** | 0 |
| **Number Of CDS** | 1 |

#### Alignment

| | |
|---|---|
| **Alignment Score** | 98.0 (NT) + 243.0 (AA) = 341.0 |
| **Concordance (%)** | 55.6281 |

| Alignment Method | Global, seeded, nucleotide + amino acids (AGA) |
|---|---|

#### Genome Region

Sequence starts at position 1086 and ends at position 1220 relative to NC_010349.1 reference sequence.

#### Alignment Detailed Statistics

| | Begin | End | Coverage | Score | Concordance | Matches | Identities | I/D/M/F* | Stop Codons |
|---|---|---|---|---|---|---|---|---|---|
| **NT** | **1086** | **1220** | **7.5%** | **98** | **36.3%** | **135 (100%)** | **92 (68.1%)** | **0/0** | |
| Mutations: | 1096A>T, 1102T>C, 1103C>A, 1105T>A, 1110T>A, 1111C>T, 1112A>G, 1114G>A, 1117T>A, 1123T>A, 1129A>T, 1130T>A, 1132G>C, 1134G>A, 1135G>C, 1137G>T, 1138G>C, 1141G>C, 1144G>A, 1145A>C, 1147G>C, 1150T>C, 1156T>C, 1165T>C, 1166C>A, 1167A>C, 1171A>T, 1172C>G, 1174A>T, 1177T>C, 1178G>A, 1183T>C, 1184T>C, 1186G>C, 1187G>A, 1192C>T, 1195C>T, 1201T>C, 1202G>A, 1204A>G, 1205G>A, 1206A>C, 1210C>T | | | | | | | | |
| **CDS** | | | | | | | | | |
| RdRp | 343 | 387 | 8.3% | 243 | 67.9% | 45 (100%) | 32 (71.1%) | 0/0/0/0 | 0 |
| Protein mutations: | F350Y (1110T>A 1111C>T), R351G (1112A>G 1114G>A), S357T (1130T>A 1132G>C), W358Y (1134G>A 1135G>C), W359F (1137G>T 1138G>C), M362L (1145A>C 1147G>C), H369T (1166C>A 1167A>C), L371V (1172C>G 1174A>T), D373N (1178G>A), A376T (1187G>A), V381M (1202G>A 1204A>G), E382T (1205G>A 1206A>C) | | | | | | | | |
| Codon mutations: | CGA345CGT (1096A>T), TTT347TTC (1102T>C), CGT348AGA (1103C>A 1105T>A), TTC350TAT (1110T>A 1111C>T), AGG351GGA (1112A>G 1114G>A), GGT352GGA (1117T>A), CCT354CCA (1123T>A), GGA356GGT (1129A>T), TCG357ACC (1130T>A 1132G>C), TGG358TAC (1134G>A 1135G>C), TGG359TTC (1137G>T 1138G>C), ACG360ACC (1141G>C), CAG361CAA (1144G>A), ATG362CTC (1145A>C 1147G>C), ATT363ATC (1150T>C), AGT365AGC (1156T>C), AAT368AAC (1165T>C), CAC369ACC (1166C>A 1167A>C), ATA370ATT (1171A>T), CTA371GTT (1172C>G 1174A>T), ATT372ATC (1177T>C), GAT373AAT (1178G>A), TAT374TAC (1183T>C), TTG375CTC (1184T>C 1186G>C), GCA376ACA (1187G>A), GAC377GAT (1192C>T), TGC378TGT (1195C>T), GAT380GAC (1201T>C), GTA381ATG (1202G>A 1204A>G), GAG382ACG (1205G>A 1206A>C), ATC383ATT (1210C>T) | | | | | | | | |
| **Proteins** | | | | | | | | | |
| RNA-dependent RNA polymerase (YP_001686789.1) | 343 | 387 | 8.3% | 243 | 67.9% | 45 (100%) | 32 (71.1%) | 0/0/0/0 | 0 |
| Protein mutations: | F350Y (1110T>A 1111C>T), R351G (1112A>G 1114G>A), S357T (1130T>A 1132G>C), W358Y (1134G>A 1135G>C), W359F (1137G>T 1138G>C), M362L (1145A>C 1147G>C), H369T (1166C>A 1167A>C), L371V (1172C>G 1174A>T), D373N (1178G>A), A376T (1187G>A), V381M (1202G>A 1204A>G), E382T (1205G>A 1206A>C) | | | | | | | | |
| Codon mutations: | CGA345CGT (1096A>T), TTT347TTC (1102T>C), CGT348AGA (1103C>A 1105T>A), TTC350TAT (1110T>A 1111C>T), AGG351GGA (1112A>G 1114G>A), GGT352GGA (1117T>A), CCT354CCA (1123T>A), GGA356GGT (1129A>T), TCG357ACC (1130T>A 1132G>C), TGG358TAC (1134G>A 1135G>C), TGG359TTC (1137G>T 1138G>C), ACG360ACC (1141G>C), CAG361CAA (1144G>A), ATG362CTC (1145A>C 1147G>C), ATT363ATC (1150T>C), AGT365AGC (1156T>C), AAT368AAC (1165T>C), CAC369ACC (1166C>A 1167A>C), ATA370ATT (1171A>T), CTA371GTT (1172C>G 1174A>T), ATT372ATC (1177T>C), GAT373AAT (1178G>A), TAT374TAC (1183T>C), TTG375CTC (1184T>C 1186G>C), GCA376ACA (1187G>A), GAC377GAT (1192C>T), TGC378TGT (1195C>T), GAT380GAC (1201T>C), GTA381ATG (1202G>A 1204A>G), GAG382ACG (1205G>A 1206A>C), ATC383ATT (1210C>T) | | | | | | | | |

*: Inserts / Deletes / Misaligned / Frameshifts

#### Analysis details

This analysis was performed with panviral2.64

### NGS Details (UN18_val): Gammaflexivirus PaGFV-1

#### Assembly

| | |
|---|---|
| **Coverage Length** | 135 (1 contig(s)) |
| **Depth Of Coverage** | 2.0 |
| **Number Of Reads** | 2 |
| **Reads Per Million** | 0.04 rpm (after QC) |
| **Ambiguities** | 0 |
| **Assembly Method** | de novo + reference guided assembly |
| **Consensus Caller** | Bcf Tools |

#### Coverage Map

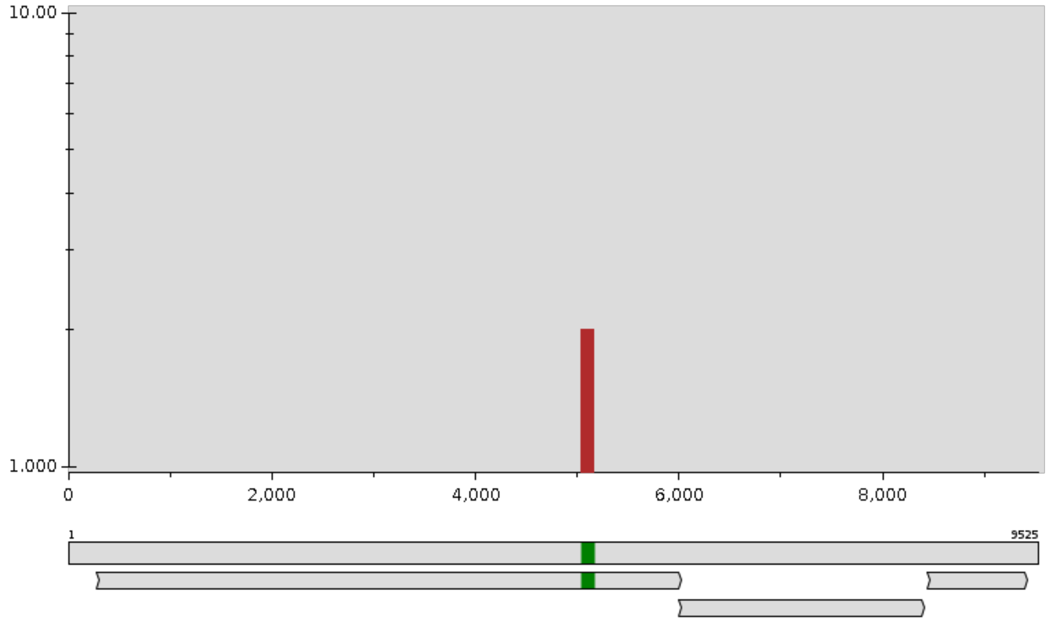

#### Assignment

| | |
|---|---|
| **Type** | Gammaflexivirus PaGFV-1 (Taxonomy ID: 2955928) |
| **Reference Genome** | NC_076696.1 |
| **NT Identity (%)** | 80.0 |
| **AA Identity (%)** | 86.6667 |
| **Number Of Stop Codons** | 0 |
| **Number Of CDS** | 3 |

#### Alignment

| | |
|---|---|
| **Alignment Score** | 162.0 (NT) + 246.0 (AA) = 408.0 |
| **Concordance (%)** | 73.7794 |

| Alignment Method | Global, seeded, nucleotide + amino acids (AGA) |
|---|---|

#### Genome Region

Sequence starts at position 5038 and ends at position 5172 relative to NC_076696.1 reference sequence.

#### Alignment Detailed Statistics

| | Begin | End | Coverage | Score | Concordance | Matches | Identities | I/D/M/F* | Stop Codons |
|---|---|---|---|---|---|---|---|---|---|
| **NT** | **5038** | **5172** | **1.4%** | **162** | **60.0%** | **135 (100%)** | **108 (80.0%)** | **0/0** | |
| Mutations: | 5040T>A, 5042T>A, 5045G>C, 5048C>T, 5052C>G, 5053A>G, 5054C>T, 5057G>A, 5063G>A, 5072T>C, 5078A>T, 5084T>C, 5093G>A, 5096T>C, 5102C>G, 5103C>G, 5111T>G, 5112G>A, 5114T>A, 5126C>T, 5138T>C, 5147G>C, 5150T>G, 5159C>T, 5160T>G, 5163A>C, 5165A>C | | | | | | | | |

##### CDS

| | Begin | End | Coverage | Score | Concordance | Matches | Identities | I/D/M/F* | Stop Codons |
|---|---|---|---|---|---|---|---|---|---|
| QKQ70_gp1 | 1588 | 1632 | 2.3% | 246 | 80.4% | 45 (100%) | 39 (86.7%) | 0/0/0/0 | 0 |
| Protein mutations: | S1588T (5040T>A 5042T>A), H1592G (5052C>G 5053A>G 5054C>T), Q1609E (5103C>G), D1612K (5112G>A 5114T>A), S1628A (5160T>G) | | | | | | | | |
| Codon mutations: | TCT1588ACA (5040T>A 5042T>A), CTG1589CTC (5045G>C), GAC1590GAT (5048C>T), CAC1592GGT (5052C>G 5053A>G 5054C>T), AAG1593AAA (5057G>A), GAG1595GAA (5063G>A), CTT1598CTC (5072T>C), TCA1600TCT (5078A>T), GAT1602GAC (5084T>C), AAG1605AAA (5093G>A), CTT1606CTC (5096T>C), ACC1608ACG (5102C>G), CAG1609GAG (5103C>G), CGT1611CGG (5111T>G), GAT1612AAA (5112G>A 5114T>A), GGC1616GGT (5126C>T), ACT1620ACC (5138T>C), GCG1623GCC (5147G>C), GCT1624GCG (5150T>G), AAC1627AAT (5159C>T), TCT1628GCT (5160T>G), AGA1629CGC (5163A>C 5165A>C) | | | | | | | | |

##### Proteins

| | Begin | End | Coverage | Score | Concordance | Matches | Identities | I/D/M/F* | Stop Codons |
|---|---|---|---|---|---|---|---|---|---|
| replicase (YP_010799909.1) | 1588 | 1632 | 2.3% | 246 | 80.4% | 45 (100%) | 39 (86.7%) | 0/0/0/0 | 0 |
| Protein mutations: | S1588T (5040T>A 5042T>A), H1592G (5052C>G 5053A>G 5054C>T), Q1609E (5103C>G), D1612K (5112G>A 5114T>A), S1628A (5160T>G) | | | | | | | | |
| Codon mutations: | TCT1588ACA (5040T>A 5042T>A), CTG1589CTC (5045G>C), GAC1590GAT (5048C>T), CAC1592GGT (5052C>G 5053A>G 5054C>T), AAG1593AAA (5057G>A), GAG1595GAA (5063G>A), CTT1598CTC (5072T>C), TCA1600TCT (5078A>T), GAT1602GAC (5084T>C), AAG1605AAA (5093G>A), CTT1606CTC (5096T>C), ACC1608ACG (5102C>G), CAG1609GAG (5103C>G), CGT1611CGG (5111T>G), GAT1612AAA (5112G>A 5114T>A), GGC1616GGT (5126C>T), ACT1620ACC (5138T>C), GCG1623GCC (5147G>C), GCT1624GCG (5150T>G), AAC1627AAT (5159C>T), TCT1628GCT (5160T>G), AGA1629CGC (5163A>C 5165A>C) | | | | | | | | |

*: Inserts / Deletes / Misaligned / Frameshifts

#### Analysis details

This analysis was performed with panviral2.64

#### NGS Details (UN18_val): Sugarcane chlorotic streak virus

##### Assembly

| | |
|---|---|
| **Coverage Length** | 121 (1 contig(s)) |
| **Depth Of Coverage** | 1.9 |
| **Number Of Reads** | 2 |
| **Reads Per Million** | 0.04 rpm (after QC) |
| **Ambiguities** | 0 |
| **Assembly Method** | de novo + reference guided assembly |
| **Consensus Caller** | Bcf Tools |

##### Coverage Map

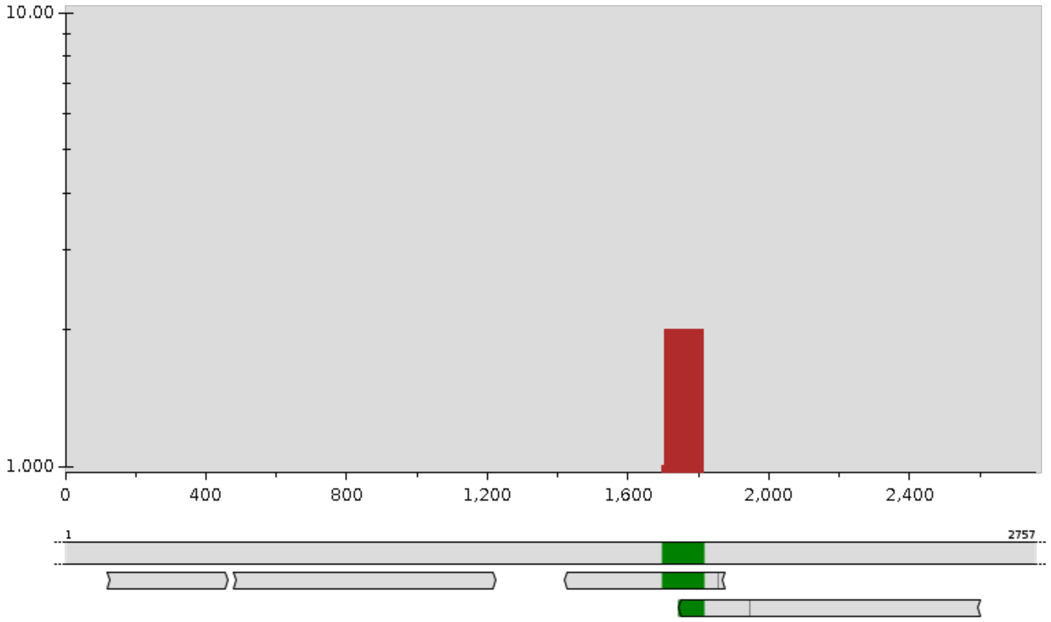

##### Assignment

| | |
|---|---|
| **Type** | Sugarcane chlorotic streak virus (Taxonomy ID: 1919062) |
| **Reference Genome** | NC_032004.1 |
| **NT Identity (%)** | 62.8099 |
| **AA Identity (%)** | 55.7692 |
| **Number Of Stop Codons** | 1 |
| **Number Of CDS** | 5 |

##### Alignment

| | |
|---|---|
| **Alignment Score** | 62.0 (NT) + 441.0 (AA) = 503.0 |
| **Concordance (%)** | 50.1996 |

| Alignment Method | Global, seeded, nucleotide + amino acids (AGA) |
|---|---|

#### Genome Region

Sequence starts at position 1696 and ends at position 1816 relative to NC_032004.1 reference sequence.

#### Alignment Detailed Statistics

| | Begin | End | Coverage | Score | Concordance | Matches | Identities | I/D/M/F* | Stop Codons |
|---|---|---|---|---|---|---|---|---|---|
| **NT** | **1696** | **1816** | **4.4%** | **62** | **25.6%** | **121 (100%)** | **76 (62.8%)** | **0/0** | |
| Mutations: | 1701A>G, 1704A>T, 1710G>A, 1713G>A, 1714A>T, 1715C>A, 1716G>C, 1718C>A, 1720G>C, 1721C>T, 1722C>G, 1723T>A, 1724C>T, 1725T>A, 1734A>G, 1735G>T, 1736A>T, 1737G>A, 1738G>C, 1740C>G, 1741C>A, 1744G>T, 1755T>G, 1756T>C, 1757G>A, 1758C>A, 1759C>A, 1761G>A, 1767A>G, 1776C>A, 1779G>A, 1784T>G, 1785A>T, 1791A>T, 1793A>T, 1794T>C, 1796A>T, 1797C>T, 1800T>A, 1803T>A, 1806C>T, 1809T>A, 1811T>A, 1812T>A, 1815G>C | | | | | | | | |
| **CDS** | | | | | | | | | |
| BS327_gp3 | 234 | 273 | 11.0% | 206 | 65.8% | 40 (100%) | 25 (62.5%) | 0/0/0/0 | 0 |
| Protein mutations: | T235S (1809T>A 1811T>A), S240T (1794T>C 1796A>T), S241T (1791A>T 1793A>T), W252F (1758C>A 1759C>A), Q253C (1755T>G 1756T>C 1757G>A), A257D (1744G>T), W258F (1740C>G 1741C>A), T259S (1737G>A 1738G>C), S260N (1734A>G 1735G>T 1736A>T), E263D (1725T>A), E264I (1722C>G 1723T>A 1724C>T), A265R (1720G>C 1721C>T), V266L (1716G>C 1718C>A), V267Y (1713G>A 1714A>T 1715C>A) | | | | | | | | |
| Codon mutations: | GGC233.GG (1815G>C), CCA234CCT (1812T>A), ACA235TCT (1809T>A 1811T>A), AGG236AGA (1806C>T), ACA237ACT (1803T>A), GGA238GGT (1800T>A), AAG239AAA (1797C>T), TCA240ACG (1794T>C 1796A>T), TCT241ACA (1791A>T 1793A>T), GCT243GCA (1785A>T), AGA244CGA (1784T>G), AGC245AGT (1779G>A), CTG246CTT (1776C>A), CAT249CAC (1767A>G), TAC251TAT (1761G>A), TGG252TTT (1758C>A 1759C>A), CAA253TGC (1755T>G 1756T>C 1757G>A), GCC257GAC (1744G>T), TGG258TTC (1740C>G 1741C>A), ACC259AGT (1737G>A 1738G>C), TCT260AAC (1734A>G 1735G>T 1736A>T), GAA263GAT (1725T>A), GAG264ATC (1722C>G 1723T>A 1724C>T), GCA265AGA (1720G>C 1721C>T), GTC266TTG (1716G>C 1718C>A), GTC267TAT (1713G>A 1714A>T 1715C>A), AAC268AAT (1710G>A), ATT270ATA (1704A>T), GAT271GAC (1701A>G) | | | | | | | | |
| BS327_gp4 | 22 | 61 | 26.3% | 206 | 65.8% | 40 (100%) | 25 (62.5%) | 0/0/0/0 | 0 |
| Protein mutations: | T23S (1809T>A 1811T>A), S28T (1794T>C 1796A>T), S29T (1791A>T 1793A>T), W40F (1758C>A 1759C>A), Q41C (1755T>G 1756T>C 1757G>A), A45D (1744G>T), W46F (1740C>G 1741C>A), T47S (1737G>A 1738G>C), S48N (1734A>G 1735G>T 1736A>T), E51D (1725T>A), E52I (1722C>G 1723T>A 1724C>T), A53R (1720G>C 1721C>T), V54L (1716G>C 1718C>A), V55Y (1713G>A 1714A>T 1715C>A) | | | | | | | | |
| Codon mutations: | GGC21.GG (1815G>C), CCA22CCT (1812T>A), ACA23TCT (1809T>A 1811T>A), AGG24AGA (1806C>T), ACA25ACT (1803T>A), GGA26GGT (1800T>A), AAG27AAA (1797C>T), TCA28ACG (1794T>C 1796A>T), TCT29ACA (1791A>T 1793A>T), GCT31GCA (1785A>T), AGA32CGA (1784T>G), AGC33AGT (1779G>A), CTG34CTT (1776C>A), CAT37CAC (1767A>G), TAC39TAT (1761G>A), TGG40TTT (1758C>A 1759C>A), CAA41TGC (1755T>G 1756T>C 1757G>A), GCC45GAC (1744G>T), TGG46TTC (1740C>G 1741C>A), ACC47AGT (1737G>A 1738G>C), TCT48AAC (1734A>G 1735G>T 1736A>T), GAA51GAT (1725T>A), GAG52ATC (1722C>G 1723T>A 1724C>T), GCA53AGA (1720G>C 1721C>T), GTC54TTG (1716G>C 1718C>A), GTC55TAT (1713G>A 1714A>T 1715C>A), AAC56AAT (1710G>A), ATT58ATA (1704A>T), GAT59GAC (1701A>G) | | | | | | | | |
| BS327_gp5 | 264 | 287 | 8.4% | 29 | 17.9% | 24 (100%) | 8 (33.3%) | 0/0/0/0 | 1 |
| Protein mutations: | A264G (1815G>C), Q265L (1811T>A 1812T>A), Q266L (1809T>A), G267E (1806C>T), Q268L (1803T>A), E269V (1800T>A), S270K (1796A>T 1797C>T), H271R (1793A>T 1794T>C), L272H (1791A>T), L274H (1784T>G 1785A>T), A276V (1779G>A), W277L (1776C>A), I280T (1767A>G), T282I (1761G>A), G283F (1757G>A 1758C>A 1759C>A), K284A (1755T>G 1756T>C) | | | | | | | | |
| Codon mutations: | GCC264GGC (1815G>C), CAA265CTT (1811T>A 1812T>A), CAA266CTA (1809T>A), GGA267GAA (1806C>T), CAG268CTG (1803T>A), GAA269GTA (1800T>A), AGT270AAA (1796A>T 1797C>T), CAT271CGA (1793A>T 1794T>C), CTT272CAT (1791A>T), CTA274CAC (1784T>G 1785A>T), GCC276GTC (1779G>A), TGG277TTG (1776C>A), ATA280ACA (1767A>G), ACT282ATT (1761G>A), GGC283TTT (1757G>A 1758C>A 1759C>A), AAA284GCA (1755T>G 1756T>C) | | | | | | | | |
| **Proteins** | | | | | | | | | |
| Rep-associated protein (YP_009325925.1) | 234 | 273 | 11.0% | 206 | 65.8% | 40 (100%) | 25 (62.5%) | 0/0/0/0 | 0 |
| Protein mutations: | T235S (1809T>A 1811T>A), S240T (1794T>C 1796A>T), S241T (1791A>T 1793A>T), W252F (1758C>A 1759C>A), Q253C (1755T>G 1756T>C 1757G>A), A257D (1744G>T), W258F (1740C>G 1741C>A), T259S (1737G>A 1738G>C), S260N (1734A>G 1735G>T 1736A>T), E263D (1725T>A), E264I (1722C>G 1723T>A 1724C>T), A265R (1720G>C 1721C>T), V266L (1716G>C 1718C>A), V267Y (1713G>A 1714A>T 1715C>A) | | | | | | | | |
| Codon mutations: | GGC233.GG (1815G>C), CCA234CCT (1812T>A), ACA235TCT (1809T>A 1811T>A), AGG236AGA (1806C>T), ACA237ACT (1803T>A), GGA238GGT (1800T>A), AAG239AAA (1797C>T), TCA240ACG (1794T>C 1796A>T), TCT241ACA (1791A>T 1793A>T), GCT243GCA (1785A>T), AGA244CGA (1784T>G), AGC245AGT (1779G>A), CTG246CTT (1776C>A), CAT249CAC (1767A>G), TAC251TAT (1761G>A), TGG252TTT (1758C>A 1759C>A), CAA253TGC (1755T>G 1756T>C 1757G>A), GCC257GAC (1744G>T), TGG258TTC (1740C>G 1741C>A), ACC259AGT (1737G>A 1738G>C), TCT260AAC (1734A>G 1735G>T 1736A>T), GAA263GAT (1725T>A), GAG264ATC (1722C>G 1723T>A 1724C>T), GCA265AGA (1720G>C 1721C>T), GTC266TTG (1716G>C 1718C>A), GTC267TAT (1713G>A 1714A>T 1715C>A), AAC268AAT (1710G>A), ATT270ATA (1704A>T), GAT271GAC (1701A>G) | | | | | | | | |
| RepB (YP_009325926.1) | 22 | 61 | 26.3% | 206 | 65.8% | 40 (100%) | 25 (62.5%) | 0/0/0/0 | 0 |
| Protein mutations: | T23S (1809T>A 1811T>A), S28T (1794T>C 1796A>T), S29T (1791A>T 1793A>T), W40F (1758C>A 1759C>A), Q41C (1755T>G 1756T>C 1757G>A), A45D (1744G>T), W46F (1740C>G 1741C>A), T47S (1737G>A 1738G>C), S48N (1734A>G 1735G>T 1736A>T), E51D (1725T>A), E52I (1722C>G 1723T>A 1724C>T), A53R (1720G>C 1721C>T), V54L (1716G>C 1718C>A), V55Y (1713G>A 1714A>T 1715C>A) | | | | | | | | |
| Codon mutations: | GGC21.GG (1815G>C), CCA22CCT (1812T>A), ACA23TCT (1809T>A 1811T>A), AGG24AGA (1806C>T), ACA25ACT (1803T>A), GGA26GGT (1800T>A), AAG27AAA (1797C>T), TCA28ACG (1794T>C 1796A>T), TCT29ACA (1791A>T 1793A>T), GCT31GCA (1785A>T), AGA32CGA (1784T>G), AGC33AGT (1779G>A), CTG34CTT (1776C>A), CAT37CAC (1767A>G), TAC39TAT (1761G>A), TGG40TTT (1758C>A 1759C>A), CAA41TGC (1755T>G 1756T>C 1757G>A), GCC45GAC (1744G>T), TGG46TTC (1740C>G 1741C>A), ACC47AGT (1737G>A 1738G>C), TCT48AAC (1734A>G 1735G>T 1736A>T), GAA51GAT (1725T>A), GAG52ATC (1722C>G 1723T>A 1724C>T), GCA53AGA (1720G>C 1721C>T), GTC54TTG (1716G>C 1718C>A), GTC55TAT (1713G>A 1714A>T 1715C>A), AAC56AAT (1710G>A), ATT58ATA (1704A>T), GAT59GAC (1701A>G) | | | | | | | | |
| RepA (YP_009325927.1) | 264 | 287 | 8.4% | 29 | 17.9% | 24 (100%) | 8 (33.3%) | 0/0/0/0 | 1 |
| Protein mutations: | A264G (1815G>C), Q265L (1811T>A 1812T>A), Q266L (1809T>A), G267E (1806C>T), Q268L (1803T>A), E269V (1800T>A), S270K (1796A>T 1797C>T), H271R (1793A>T 1794T>C), L272H (1791A>T), L274H (1784T>G 1785A>T), A276V (1779G>A), W277L (1776C>A), I280T (1767A>G), T282I (1761G>A), G283F (1757G>A 1758C>A 1759C>A), K284A (1755T>G 1756T>C) | | | | | | | | |
| Codon mutations: | GCC264GGC (1815G>C), CAA265CTT (1811T>A 1812T>A), CAA266CTA (1809T>A), GGA267GAA (1806C>T), CAG268CTG (1803T>A), GAA269GTA (1800T>A), AGT270AAA (1796A>T 1797C>T), CAT271CGA (1793A>T 1794T>C), CTT272CAT (1791A>T), CTA274CAC (1784T>G 1785A>T), GCC276GTC (1779G>A), TGG277TTG (1776C>A), ATA280ACA (1767A>G), ACT282ATT (1761G>A), GGC283TTT (1757G>A 1758C>A 1759C>A), AAA284GCA (1755T>G 1756T>C) | | | | | | | | |

*: Inserts / Deletes / Misaligned / Frameshifts

#### Analysis details

This analysis was performed with panviral2.64

### NGS Details (UN18_val): Seussvirus seuss

#### Assembly

| | |
|---|---|
| **Coverage Length** | 232 (1 contig(s)) |
| **Depth Of Coverage** | 1.1 |
| **Number Of Reads** | 2 |
| **Reads Per Million** | 0.04 rpm (after QC) |
| **Ambiguities** | 0 |
| **Assembly Method** | de novo + reference guided assembly |
| **Consensus Caller** | Bcf Tools |

#### Coverage Map

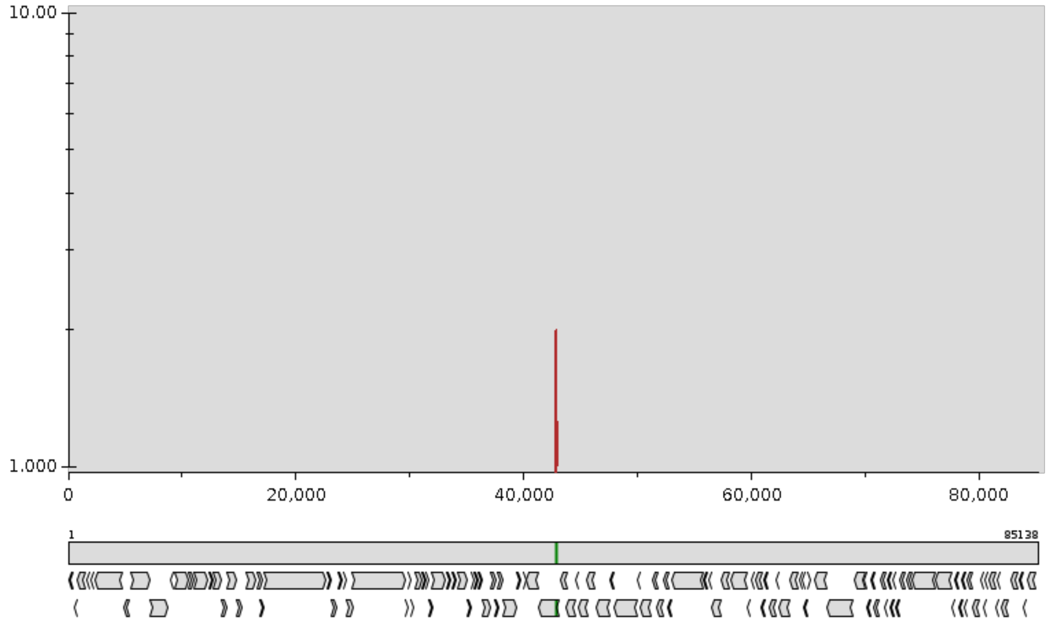

#### Assignment

| | |
|---|---|
| **Type** | Seussvirus seuss (Taxonomy ID: 2734278) |
| **Reference Genome** | NC_047757.1 |
| **NT Identity (%)** | 76.2931 |
| **AA Identity (%)** | 79.4872 |
| **Number Of Stop Codons** | 0 |
| **Number Of CDS** | 120 |

#### Alignment

| | |
|---|---|
| **Alignment Score** | 244.0 (NT) + 499.0 (AA) = 743.0 |
| **Concordance (%)** | 70.4265 |

| Alignment Method | Local, heuristic, nucleotide (BLASTN) |
|---|---|

#### Genome Region

Sequence starts at position 42730 and ends at position 42961 relative to NC_047757.1 reference sequence.

#### Alignment Detailed Statistics

| | Begin | End | Coverage | Score | Concordance | Matches | Identities | I/D/M/F* | Stop Codons |
|---|---|---|---|---|---|---|---|---|---|
| **NT** | **42730** | **42961** | **0.3%** | **244** | **52.6%** | **232 (100%)** | **177 (76.3%)** | **0/0** | |
| Mutations: | 42734T>G, 42737C>T, 42747G>C, 42748A>C, 42752G>C, 42756T>C, 42757C>T, 42764G>A, 42782A>G, 42786T>C, 42787T>G, 42788G>C, 42789C>G, 42790T>A, 42791G>T, 42794C>G, 42801G>A, 42802T>C, 42820T>C, 42821C>G, 42822T>A, 42823G>C, 42830C>G, 42831G>T, 42832C>T, 42837T>C, 42842C>G, 42843T>G, 42844C>G, 42845G>C, 42848G>C, 42849C>G, 42850C>A, 42851G>A, 42854C>G, 42858A>T, 42860A>G, 42872A>C, 42886C>T, 42893A>G, 42899G>C, 42902G>C, 42904T>C, 42908G>C, 42928G>A, 42929C>G, 42930T>C, 42933G>C, 42934A>T, 42935C>G, 42938G>A, 42941G>A, 42944A>G, 42956G>C, 42959G>A | | | | | | | | |

##### CDS

| | Begin | End | Coverage | Score | Concordance | Matches | Identities | I/D/M/F* | Stop Codons |
|---|---|---|---|---|---|---|---|---|---|
| HOR08_gp052 | 48 | 125 | 13.1% | 499 | 83.2% | 78 (100%) | 62 (79.5%) | 0/0/0/0 | 0 |
| Protein mutations: | M56I (42935C>G), Q58R (42929C>G 42930T>C), I67V (42902G>C 42904T>C), A73T (42886C>T), F82Y (42858A>T), G85S (42848G>C 42849C>G 42850C>A), E87P (42842C>G 42843T>G 42844C>G), N89S (42837T>C), A91N (42830C>G 42831G>T 42832C>T), Q94V (42821C>G 42822T>A 42823G>C), N95D (42820T>C), T101V (42801G>A 42802T>C), N106R (42786T>C 42787T>G), D116S (42756T>C 42757C>T), S119G (42747G>C 42748A>C) | | | | | | | | |
| Codon mutations: | CAC48CAT (42959G>A), GCC49GCG (42956G>C), TAT53TAC (42944A>G), GAC54GAT (42941G>A), TAC55TAT (42938G>A), ATG56ATC (42935C>G), TCC57AGC (42933G>C 42934A>T), CAG58CGC (42929C>G 42930T>C), CTG59TTG (42928G>A), ACC65ACG (42908G>C), ATC67GTG (42902G>C 42904T>C), CTC68CTG (42899G>C), AAT70AAC (42893A>G), GCG73ACG (42886C>T), CTT77CTG (42872A>C), TGT81TGC (42860A>G), TTC82TAC (42858A>T), CTG83CTC (42854C>G), AAC84AAT (42851G>A), GGC85TCG (42848G>C 42849C>G 42850C>A), GTC86GTG (42845G>C), GAG87CCC (42842C>G 42843T>G 42844C>G), AAC89AGC (42837T>C), GCG91AAC (42830C>G 42831G>T 42832C>T), CAG94GTC (42821C>G 42822T>A 42823G>C), AAC95GAC (42820T>C), ACC101GTC (42801G>A 42802T>C), CTG103CTC (42794C>G), GCC104GCA (42791G>T), AGC105TCG (42788G>C 42789C>G 42790T>A), AAC106CGC (42786T>C 42787T>G), GGT107GGC (42782A>G), TAC113TAT (42764G>A), GAC116AGC (42756T>C 42757C>T), GTC117GTG (42752G>C), TCC119GGC (42747G>C 42748A>C), GAG122GAA (42737C>T), CCA123CCC (42734T>G) | | | | | | | | |

##### Proteins

| | Begin | End | Coverage | Score | Concordance | Matches | Identities | I/D/M/F* | Stop Codons |
|---|---|---|---|---|---|---|---|---|---|
| ribonucleotide reductase large subunit (YP_009785562.1) | 48 | 125 | 13.1% | 499 | 83.2% | 78 (100%) | 62 (79.5%) | 0/0/0/0 | 0 |
| Protein mutations: | M56I (42935C>G), Q58R (42929C>G 42930T>C), I67V (42902G>C 42904T>C), A73T (42886C>T), F82Y (42858A>T), G85S (42848G>C 42849C>G 42850C>A), E87P (42842C>G 42843T>G 42844C>G), N89S (42837T>C), A91N (42830C>G 42831G>T 42832C>T), Q94V (42821C>G 42822T>A 42823G>C), N95D (42820T>C), T101V (42801G>A 42802T>C), N106R (42786T>C 42787T>G), D116S (42756T>C 42757C>T), S119G (42747G>C 42748A>C) | | | | | | | | |
| Codon mutations: | CAC48CAT (42959G>A), GCC49GCG (42956G>C), TAT53TAC (42944A>G), GAC54GAT (42941G>A), TAC55TAT (42938G>A), ATG56ATC (42935C>G), TCC57AGC (42933G>C 42934A>T), CAG58CGC (42929C>G 42930T>C), CTG59TTG (42928G>A), ACC65ACG (42908G>C), ATC67GTG (42902G>C 42904T>C), CTC68CTG (42899G>C), AAT70AAC (42893A>G), GCG73ACG (42886C>T), CTT77CTG (42872A>C), TGT81TGC (42860A>G), TTC82TAC (42858A>T), CTG83CTC (42854C>G), AAC84AAT (42851G>A), GGC85TCG (42848G>C 42849C>G 42850C>A), GTC86GTG (42845G>C), GAG87CCC (42842C>G 42843T>G 42844C>G), AAC89AGC (42837T>C), GCG91AAC (42830C>G 42831G>T 42832C>T), CAG94GTC (42821C>G 42822T>A 42823G>C), AAC95GAC (42820T>C), ACC101GTC (42801G>A 42802T>C), CTG103CTC (42794C>G), GCC104GCA (42791G>T), AGC105TCG (42788G>C 42789C>G 42790T>A), AAC106CGC (42786T>C 42787T>G), GGT107GGC (42782A>G), TAC113TAT (42764G>A), GAC116AGC (42756T>C 42757C>T), GTC117GTG (42752G>C), TCC119GGC (42747G>C 42748A>C), GAG122GAA (42737C>T), CCA123CCC (42734T>G) | | | | | | | | |

*: Inserts / Deletes / Misaligned / Frameshifts

#### Analysis details

This analysis was performed with panviral2.64

### NGS Details (UN18_val): Fusarium poae narnavirus 2

#### Assembly

| | |
|---|---|
| **Coverage Length** | 246 (2 contig(s)) |
| **Depth Of Coverage** | 1.0 |
| **Number Of Reads** | 2 |
| **Reads Per Million** | 0.04 rpm (after QC) |
| **Ambiguities** | 0 |
| **Assembly Method** | de novo + reference guided assembly |
| **Consensus Caller** | Bcf Tools |

#### Coverage Map

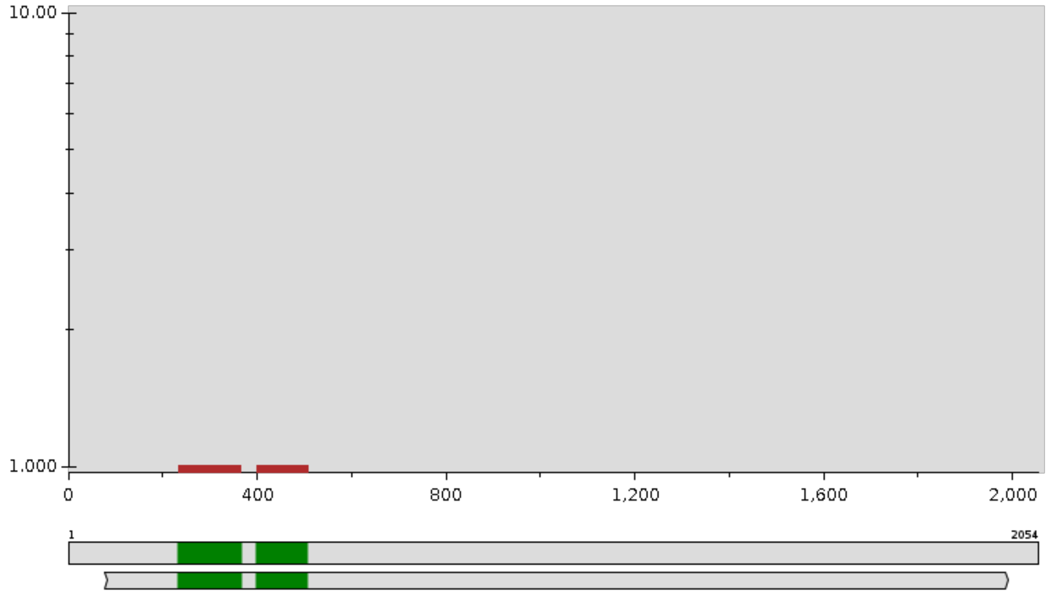

#### Assignment

| | |
|---|---|
| **Type** | Fusarium poae narnavirus 2 (Taxonomy ID: 1849532) |
| **Reference Genome** | NC_030866.1 |
| **NT Identity (%)** | 74.7967 |
| **AA Identity (%)** | 74.3902 |
| **Number Of Stop Codons** | 0 |
| **Number Of CDS** | 1 |

#### Alignment

| | |
|---|---|
| **Alignment Score** | 244.0 (NT) + 507.0 (AA) = 751.0 |
| **Concordance (%)** | 66.7556 |

| | |
|---|---|
| **Alignment Method** | Global, seeded, nucleotide + amino acids (AGA) |

#### Genome Region

Sequence starts at position 233 and ends at position 509 relative to NC_030866.1 reference sequence.

#### Alignment Detailed Statistics

| | Begin | End | Coverage | Score | Concordance | Matches | Identities | I/D/M/F* | Stop Codons |
|---|---|---|---|---|---|---|---|---|---|
| **NT** | **233** | **509** | **12.0%** | **244** | **49.6%** | **246 (100%)** | **184 (74.8%)** | **0/0** | |
| Mutations: | 234T>C, 238C>A, 240G>A, 242G>A, 243A>G, 244A>G, 245A>G, 249C>T, 250A>G, 252T>C, 255G>A, 256C>G, 257C>A, 258T>A, 263C>A, 264A>C, 268T>C, 269T>A, 270G>C, 272A>G, 273C>G, 279T>C, 282G>T, 283T>C, 285T>G, 288C>T, 289C>T, 290G>C, 294G>C, 301C>A, 304C>G, 309C>T, 312C>A, 315T>C, 321C>T, 332A>G, 334A>T, 335G>T, 336C>G, 337A>G, 338T>A, 339A>T, 355A>G, 399G>A, 403C>T, 405T>A, 406A>T, 408T>A, 411A>G, 420T>C, 423A>G, 429T>G, 447G>C, 450C>A, 468C>G, 471G>A, 475A>T, 476A>C, 477C>T, 480G>C, 484C>T, 486T>C | | | | | | | | |

##### CDS

| | Begin | End | Coverage | Score | Concordance | Matches | Identities | I/D/M/F* | Stop Codons |
|---|---|---|---|---|---|---|---|---|---|
| BHR96_gp1 | 53 | 144 | 12.9% | 507 | 77.2% | 82 (100%) | 61 (74.4%) | 0/0/0/0 | 0 |
| Protein mutations: | R55K (242G>A 243A>G), K56G (244A>G 245A>G), I58V (250A>G 252T>C), P60E (256C>G 257C>A 258T>A), A62D (263C>A 264A>C), L64H (268T>C 269T>A 270G>C), D65G (272A>G 273C>G), S69P (283T>C 285T>G), R71S (289C>T 290G>C), E72D (294G>C), Q75K (301C>A), L76V (304C>G), D85G (332A>G), S86L (334A>T 335G>T 336C>G), I87D (337A>G 338T>A 339A>T), I93V (355A>G), T110S (406A>T 408T>A), N133S (475A>T 476A>C 477C>T), H136Y (484C>T 486T>C) | | | | | | | | |
| Codon mutations: | TCT52.CC (234T>C), CGG54AGA (238C>A 240G>A), AGA55AAG (242G>A 243A>G), AAA56GGA (244A>G 245A>G), TTC57TTT (249C>T), ATT58GTC (250A>G 252T>C), GCG59GCA (255G>A), CCT60GAA (256C>G 257C>A 258T>A), GCA62GAC (263C>A 264A>C), TTG64CAC (268T>C 269T>A 270G>C), GAC65GGG (272A>G 273C>G), CAT67CAC (279T>C), CGG68CGT (282G>T), TCT69CCG (283T>C 285T>G), TTC70TTT (288C>T), CGC71TCC (289C>T 290G>C), GAG72GAC (294G>C), CAA75AAA (301C>A), CTC76GTC (304C>G), GTC77GTT (309C>T), ATC78ATA (312C>A), GAT79GAC (315T>C), TCC81TCT (321C>T), GAT85GGT (332A>G), AGC86TTG (334A>T 335G>T 336C>G), ATA87GAT (337A>G 338T>A 339A>T), ATC93GTC (355A>G), CGG107..A (399G>A), CTT109TTA (403C>T 405T>A), ACT110TCA (406A>T 408T>A), GGA111GGG (411A>G), AAT114AAC (420T>C), CGA115CGG (423A>G), CTT117CTG (429T>G), TCG123TCC (447G>C), ATC124ATA (450C>A), GTC130GTG (468C>G), CTG131CTA (471G>A), AAC133TCT (475A>T 476A>C 477C>T), GGG134GGC (480G>C), CAT136TAC (484C>T 486T>C) | | | | | | | | |

##### Proteins

| | Begin | End | Coverage | Score | Concordance | Matches | Identities | I/D/M/F* | Stop Codons |
|---|---|---|---|---|---|---|---|---|---|
| RNA-dependent RNA polymerase (YP_009272903.1) | 53 | 144 | 12.9% | 507 | 77.2% | 82 (100%) | 61 (74.4%) | 0/0/0/0 | 0 |
| Protein mutations: | R55K (242G>A 243A>G), K56G (244A>G 245A>G), I58V (250A>G 252T>C), P60E (256C>G 257C>A 258T>A), A62D (263C>A 264A>C), L64H (268T>C 269T>A 270G>C), D65G (272A>G 273C>G), S69P (283T>C 285T>G), R71S (289C>T 290G>C), E72D (294G>C), Q75K (301C>A), L76V (304C>G), D85G (332A>G), S86L (334A>T 335G>T 336C>G), I87D (337A>G 338T>A 339A>T), I93V (355A>G), T110S (406A>T 408T>A), N133S (475A>T 476A>C 477C>T), H136Y (484C>T 486T>C) | | | | | | | | |
| Codon mutations: | TCT52.CC (234T>C), CGG54AGA (238C>A 240G>A), AGA55AAG (242G>A 243A>G), AAA56GGA (244A>G 245A>G), TTC57TTT (249C>T), ATT58GTC (250A>G 252T>C), GCG59GCA (255G>A), CCT60GAA (256C>G 257C>A 258T>A), GCA62GAC (263C>A 264A>C), TTG64CAC (268T>C 269T>A 270G>C), GAC65GGG (272A>G 273C>G), CAT67CAC (279T>C), CGG68CGT (282G>T), TCT69CCG (283T>C 285T>G), TTC70TTT (288C>T), CGC71TCC (289C>T 290G>C), GAG72GAC (294G>C), CAA75AAA (301C>A), CTC76GTC (304C>G), GTC77GTT (309C>T), ATC78ATA (312C>A), GAT79GAC (315T>C), TCC81TCT (321C>T), GAT85GGT (332A>G), AGC86TTG (334A>T 335G>T 336C>G), ATA87GAT (337A>G 338T>A 339A>T), ATC93GTC (355A>G), CGG107..A (399G>A), CTT109TTA (403C>T 405T>A), ACT110TCA (406A>T 408T>A), GGA111GGG (411A>G), AAT114AAC (420T>C), CGA115CGG (423A>G), CTT117CTG (429T>G), TCG123TCC (447G>C), ATC124ATA (450C>A), GTC130GTG (468C>G), CTG131CTA (471G>A), AAC133TCT (475A>T 476A>C 477C>T), GGG134GGC (480G>C), CAT136TAC (484C>T 486T>C) | | | | | | | | |

*: Inserts / Deletes / Misaligned / Frameshifts

#### Analysis details

This analysis was performed with panviral2.64

### Analysis of UN24

| | |
|---|---|
| **Technology** | Paired-end short reads |
| **Input Files** | UN24_R1.fq.gz (1.61 GB), UN24_R2.fq.gz (1.67 GB) |
| **Submitted On** | 2023-09-26 13:12:56 UTC |
| **Duration** | 3h 59m 33s |
| **Tool Version** | panviral2.64 |
| **Location** | https://www.genomedetective.com/db/ui/analysis/d87ef240-8dcb-4afa-b65b-122051e282e6 |

#### Statistics

| | |
|---|---|
| **Original Read Length** | 20 - 150 |
| **Trimmed Read Length** | 50 - 135 |

| | # Reads | % of Reads |
|---|---|---|
| **Input file** | 50312498 | 100.0% |
| **After QC** | 49956410 | 99.3% |
| **After filtering** | 1799410 | 3.6% |
| **Mapped back to contigs** | 79354 | 0.2% |

#### Assignments

| Assignment | No. of Reads | Depth of Coverage | Identity | | Genome Coverage | |
|---|---|---|---|---|---|---|
| | | | NT | AA | | |
| Diachasmimorpha longicaudata entomopoxvirus (segment NC_043455.1) | 10408 | 1124.4 | 60.2% | 55.7% | 84.0% | 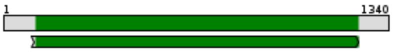 |
| Bracoviriform glomeratae (segment NC_043292.1) | 5765 | 2222.6 | 72.9% | 84.5% | 72.8% | 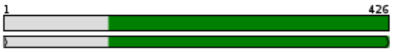 |
| Duamitovirus soch1 | 434 | 27.9 | 68.6% | 70.1% | 67.7% | 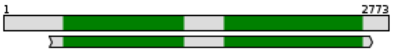 |
| Colombian datura virus | 252 | 5.7 | 76.3% | 85.0% | 60.8% | 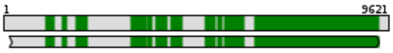 |
| Solendovirus venanicotianae | 201 | 11.2 | 78.2% | 75.3% | 26.1% | 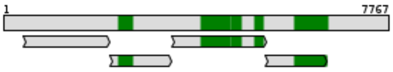 |
| Potato virus Y | 77 | 3.3 | 84.5% | 90.8% | 31.6% | 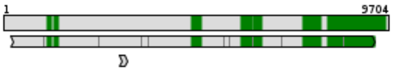 |
| Torradovirus lycopersici (2 segments out of 2) | 62 | 2.5 | 86.3% | 92.5% | 24.8% | |
| Torradovirus lycopersici (segment RNA 1) | 16 | 2.6 | 87.5% | 97.8% | 10.6% | 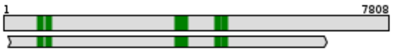 |

| Assignment | No. of Reads | Depth of Coverage | Identity | | Genome Coverage | |
|---|---|---|---|---|---|---|
| | | | NT | AA | | |
| Torradovirus lycopersici (segment RNA 2) | 46 | 2.4 | 85.9% | 89.9% | 45.3% | 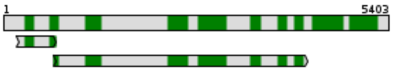  |

#### Discoveries

| Similar to | No. of Reads | Depth of Coverage | Identity | | Genome Coverage | |
|---|---|---|---|---|---|---|
| | | | NT | AA | | |
| Lausannevirus | 20011 | 11097.3 | 80.5% | 96.6% | 0.1% | 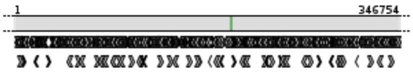  |
| Noumeavirus | 14282 | 5688.3 | 82.0% | 95.8% | 0.1% | 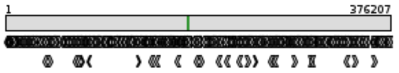  |
| Makelovirus prm1 | 7006 | 894.1 | 76.7% | 87.2% | 0.7% | 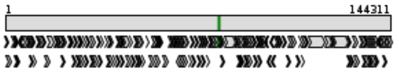  |
| Brazilian marseillevirus | 4824 | 2168.2 | 82.2% | 93.3% | 0.1% | 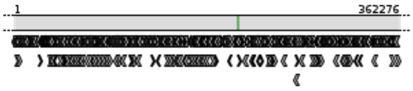  |
| Tokyovirus A1 | 3678 | 1769.2 | 80.6% | 93.1% | 0.0% | 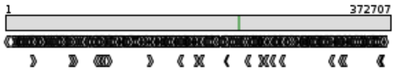  |
| Marseillevirus marseillevirus | 2867 | 2178.8 | 83.6% | 94.3% | 0.0% | 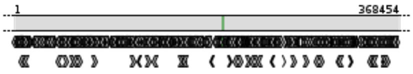  |
| Tunisvirus fontaine2 | 1907 | 946.0 | 78.2% | 90.5% | 0.1% | 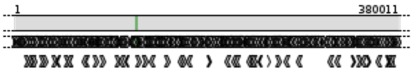  |
| Yellowstone lake phycodnavirus 1 | 1645 | 856.7 | 81.6% | 90.3% | 0.1% | 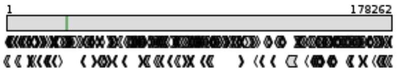  |
| Amsacta moorei entomopoxvirus | 1423 | 860.5 | 80.2% | 81.8% | 0.1% | 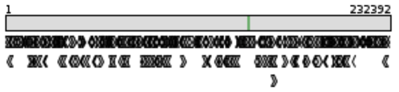  |
| Yellowstone lake phycodnavirus 1 | 1029 | 648.7 | 81.9% | 83.3% | 0.1% | 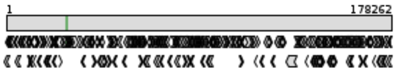  |
| Cladosporium fulvum T-1 virus | 749 | 56.9 | 53.1% | 47.6% | 18.9% | 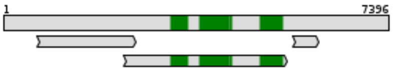  |
| Errantivirus | 503 | 36.2 | 52.1% | 41.3% | 14.8% | 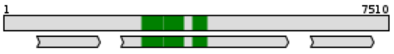  |
| Cassava brown streak virus | 421 | 94.4 | 60.8% | 57.7% | 5.8% | 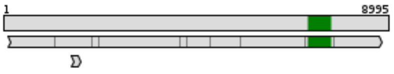  |
| Nodensvirus spm2 | 374 | 194.4 | 81.4% | 86.8% | 0.1% | 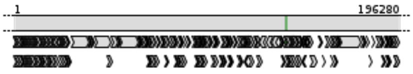  |

| Similar to | No. of Reads | Depth of Coverage | Identity | | Genome Coverage | |
|---|---|---|---|---|---|---|
| | | | NT | AA | | |
| Rahariannevirus raharianne | 176 | 5.0 | 70.8% | 68.1% | 9.4% | 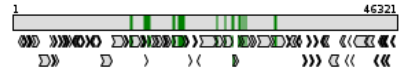 |
| Caulimovirus venafragariae | 168 | 21.5 | 58.5% | 54.4% | 10.9% | 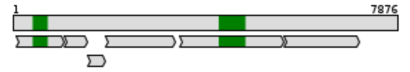 |
| Dioscovirus dioscoreae | 161 | 12.2 | 59.7% | 49.6% | 16.9% | 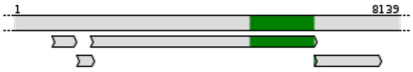 |
| Badnavirus occultipomeae | 157 | 53.6 | 58.1% | 52.3% | 3.9% | 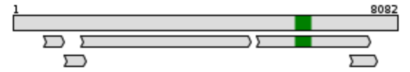 |
| Pseudomonas phage phi297 | 122 | 7.3 | 72.7% | 67.2% | 4.3% | 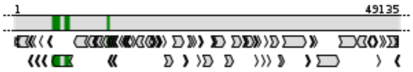 |
| Epiphyllum badnavirus 1 | 90 | 35.8 | 53.9% | 51.1% | 3.6% | 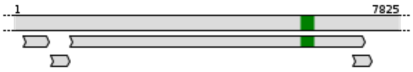 |
| Caulimovirus venafragariae | 74 | 31.5 | 57.6% | 47.1% | 3.3% | 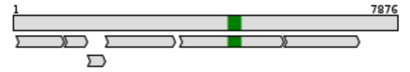 |
| Badnavirus volubetulae | 63 | 27.0 | 54.1% | 49.4% | 3.1% | 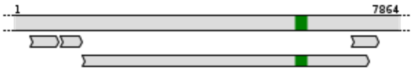 |
| Yellowstone lake phycodnavirus 1 | 55 | 35.6 | 83.7% | 93.3% | 0.1% | 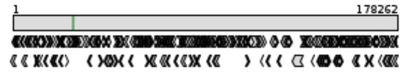 |
| Caulimovirus tessellobrassicae | 32 | 4.8 | 55.4% | 44.9% | 10.1% | 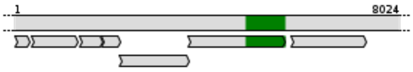 |
| Bracoviriform congregatae (2 segments out of 30) | 25 | 8.2 | 78.9% | 86.2% | N/A | |
| Bracoviriform congregatae (segment Circle 7) | 20 | 12.2 | 77.8% | 84.4% | 0.5% | 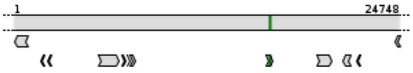 |
| Bracoviriform congregatae (segment Circle 7) | 5 | 4.4 | 80.0% | 87.8% | 0.6% | 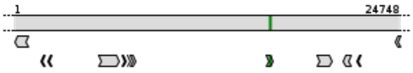 |
| Pinus nigra virus 1 | 24 | 10.8 | 54.4% | 40.5% | 3.5% | 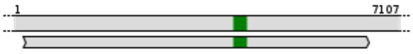 |
| Badnavirus venatheobromae | 21 | 8.9 | 58.0% | 48.9% | 3.7% | 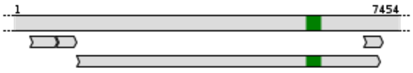 |
| Unknown | 21 | 7.4 | 58.0% | 55.8% | 3.2% | 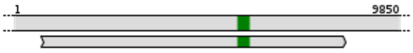 |
| Petuvirus venapetuniae | 20 | 6.3 | 61.1% | 54.3% | 4.4% | 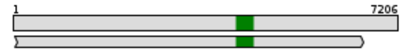 |

| Similar to | No. of Reads | Depth of Coverage | Identity | | Genome Coverage |
|---|---|---|---|---|---|
| | | | NT | AA | |
| Marseillevirus marseillevirus | 18 | 11.4 | 82.1% | 92.3% | 0.0% |
| Metaplexis yellow mottle-associated virus | 17 | 6.8 | 58.7% | 45.9% | 3.8% |
| Cavemovirus venamanihotis | 15 | 3.3 | 57.2% | 47.4% | 7.1% |
| Caulimovirus tessellobrassicae | 13 | 4.5 | 52.4% | 48.4% | 4.8% |
| Duamitovirus peex1 | 12 | 8.0 | 70.4% | 66.7% | 5.0% |
| Caulimovirus tessellobrassicae | 12 | 4.9 | 58.3% | 45.6% | 4.1% |
| Badnavirus venaribis | 12 | 6.4 | 53.9% | 48.1% | 3.1% |
| Badnavirus rutilanscamelliae | 11 | 3.5 | 56.0% | 53.7% | 4.4% |
| Soymovirus virgarachidis | 10 | 3.9 | 59.4% | 48.1% | 4.1% |
| Dioscovirus dioscoreae | 10 | 2.6 | 63.9% | 57.3% | 6.3% |
| Caulimovirus minutangelicae | 9 | 3.2 | 56.7% | 50.0% | 3.3% |
| Badnavirus maculaucubae | 9 | 3.6 | 56.7% | 45.7% | 3.1% |
| Caulimovirus venafragariae | 8 | 3.8 | 61.5% | 58.0% | 3.4% |
| Caulimovirus venafragariae | 8 | 3.5 | 62.3% | 64.7% | 3.3% |
| Petuvirus venapetuniae | 8 | 1.9 | 59.6% | 51.5% | 5.6% |
| Epiphyllum badnavirus 1 | 8 | 3.9 | 53.8% | 46.6% | 3.3% |
| Chinaberry tree badnavirus 1 | 7 | 2.7 | 59.8% | 47.7% | 3.7% |

| Similar to | No. of Reads | Depth of Coverage | Identity | | Genome Coverage | |
|---|---|---|---|---|---|---|
| | | | NT | AA | | |
| Gihfavirus pelohabitans | 6 | 2.4 | 98.2% | 0.0% | 6.3% | 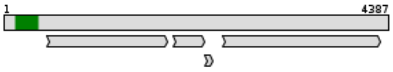 |
| Duamitovirus peex1 | 6 | 3.1 | 72.3% | 64.8% | 9.8% | 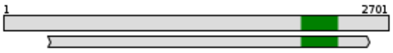 |
| Badnavirus venatheobromae | 6 | 2.3 | 54.1% | 42.9% | 4.7% | 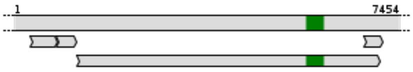 |
| Petuvirus venapetuniae | 6 | 2.5 | 62.5% | 53.2% | 3.9% | 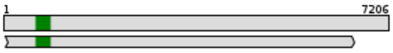 |
| Marseillevirus marseillevirus | 6 | 4.7 | 80.9% | 94.6% | 0.0% | 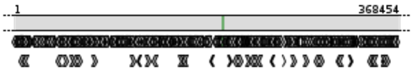 |
| Punavirus P1 | 4 | 2.2 | 100.0% | 98.0% | 0.2% | 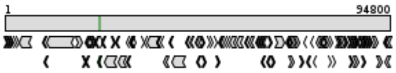 |
| Sugarcane chlorotic streak virus | 2 | 2.0 | 64.9% | 58.4% | 4.9% | 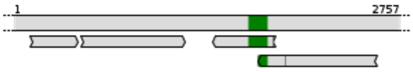 |
| Caulimovirus venafragariae | 2 | 1.1 | 59.4% | 51.9% | 3.1% | 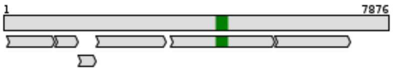 |
| Tomato associated geminivirus 1 | 2 | 1.0 | 76.0% | 79.1% | 9.9% | 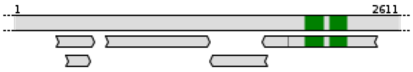 |

### NGS Details (UN24): Torradovirus lycopersici (segment RNA 1)

#### Assembly

| | |
|---|---|
| **Coverage Length** | 826 (5 contig(s)) |
| **Depth Of Coverage** | 2.6 |
| **Number Of Reads** | 16 |
| **Reads Per Million** | 0.32 rpm (after QC) |
| **Ambiguities** | 0 |
| **Assembly Method** | de novo + reference guided assembly |
| **Consensus Caller** | Bcf Tools |

#### Coverage Map

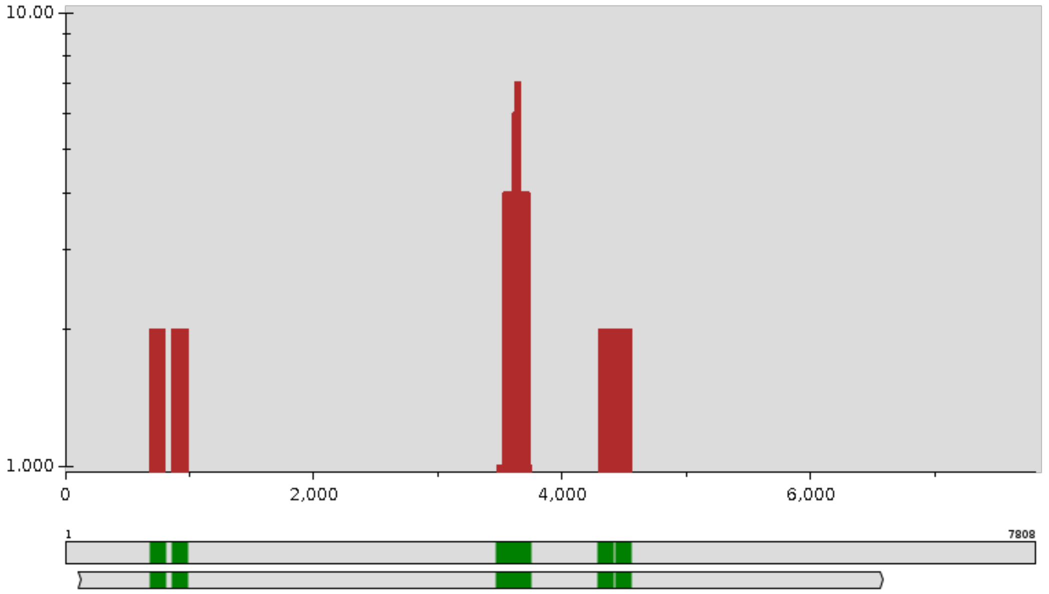

#### Assignment

| | |
|---|---|
| **Type** | Torradovirus lycopersici (Taxonomy ID: 3048378) |
| **Reference Genome** | NC_009013.1 |
| **NT Identity (%)** | 87.5303 |
| **AA Identity (%)** | 97.8182 |
| **Number Of Stop Codons** | 0 |
| **Number Of CDS** | 1 |

#### Alignment

| | |
|---|---|
| **Alignment Score** | 1240.0 (NT) + 1909.0 (AA) = 3149.0 |
| **Concordance (%)** | 87.9363 |

| | |
|---|---|
| **Alignment Method** | Global, seeded, nucleotide + amino acids (AGA) |

#### Genome Region

Sequence starts at position 677 and ends at position 4563 relative to NC_009013.1 reference sequence.

#### Alignment Detailed Statistics

| | Begin | End | Coverage | Score | Concordance | Matches | Identities | I/D/M/F* | Stop Codons |
|---|---|---|---|---|---|---|---|---|---|
| **NT** | **677** | **4563** | **10.6%** | **1240** | **75.1%** | **826 (100%)** | **723 (87.5%)** | **0/0** | |
| Mutations: | 694A>G, 707C>T, 715G>A, 718A>G, 719T>G, 721C>T, 722A>G, 727A>T, 730C>T, 739T>G, 748G>T, 751A>T, 754A>G, 758C>T, 760C>A, 763A>T, 766C>T, 772G>C, 775T>A, 778T>A, 781C>A, 790C>T, 793G>A, 796T>A, 799A>T, 802T>G, 805T>C, 806A>C, 808A>T, 862T>A, 865T>C, 877T>C, 881T>C, 883G>T, 886T>C, 889C>A, 895C>T, 898G>A, 901T>C, 913T>C, 949A>T, 958T>A, 964T>A, 967T>C, 976A>T, 977G>A, 979T>A, 988T>A, 3481C>T, 3484A>T, 3508G>A, 3534G>A, 3538T>C, 3580T>C, 3583C>T, 3610A>G, 3625C>T, 3628G>A, 3643A>G, 3676C>T, 3685G>A, 3701C>T, 3710G>A, 3739A>T, 3751A>G, 3754G>A, 4288C>T, 4291T>C, 4294A>G, 4309T>G, 4312G>T, 4318C>T, 4324T>C, 4327A>T, 4345A>T, 4348C>T, 4357C>G, 4358A>C, 4369T>A, 4381T>A, 4384A>C, 4399T>C, 4414T>G, 4429A>G, 4441A>T, 4444A>G, 4453C>G, 4459T>C, 4471T>C, 4486T>C, 4489T>C, 4493C>A, 4495T>G, 4501A>T, 4507T>C, 4510T>A, 4514T>C, 4525A>C, 4528T>C, 4531A>T, 4537T>A, 4549G>A, 4561T>G | | | | | | | | |

##### CDS

| | Begin | End | Coverage | Score | Concordance | Matches | Identities | I/D/M/F* | Stop Codons |
|---|---|---|---|---|---|---|---|---|---|
| ToTV_sRNA1gp1 | 191 | 1486 | 12.7% | 1909 | 97.8% | 275 (100%) | 269 (97.8%) | 0/0/0/0 | 0 |
| Protein mutations: | S205A (719T>G 721C>T), M206V (722A>G), V291I (977G>A 979T>A), R1143K (3534G>A), A1202T (3710G>A) | | | | | | | | |
| Codon mutations: | GCA196GCG (694A>G), CTA201TTA (707C>T), GGG203GGA (715G>A), GAA204GAG (718A>G), TCC205GCT (719T>G 721C>T), ATG206GTG (722A>G), TCA207TCT (727A>T), CAC208CAT (730C>T), CTT211CTG (739T>G), GTG214GTT (748G>T), GTA215GTT (751A>T), GCA216GCG (754A>G), CTC218TTA (758C>T 760C>A), ATA219ATT (763A>T), GCC220GCT (766C>T), GGG222GGC (772G>C), ATT223ATA (775T>A), TCT224TCA (778T>A), ACC225ACA (781C>A), TTC228TTT (790C>T), GTG229GTA (793G>A), GCT230GCA (796T>A), GTA231GTT (799A>T), CCT232CCG (802T>G), GGT233GGC (805T>C), AGA234CGT (806A>C 808A>T), GGT252GGA (862T>A), GCT253GCC (865T>C), AAT257AAC (877T>C), TTG259CTT (881T>C 883G>T), TTT260TTC (886T>C), ACC261ACA (889C>A), TTC263TTT (895C>T), AGG264AGA (898G>A), AAT265AAC (901T>C), TGT269TGC (913T>C), ATA281ATT (949A>T), ATT284ATA (958T>A), GGT286GGA (964T>A), TTT287TTC (967T>C), CCA290CCT (976A>T), GTT291ATA (977G>A 979T>A), GCT294GCA (988T>A), GGC1125GGT (3481C>T), ACA1126ACT (3484A>T), ACG1134ACA (3508G>A), AGG1143AAG (3534G>A), AGT1144AGC (3538T>C), TCT1158TCC (3580T>C), ATC1159ATT (3583C>T), GAA1168GAG (3610A>G), TAC1173TAT (3625C>T), GTG1174GTA (3628G>A), AAA1179AAG (3643A>G), GAC1190GAT (3676C>T), AGG1193AGA (3685G>A), CTA1199TTA (3701C>T), GCA1202ACA (3710G>A), CGA1211CGT (3739A>T), GGA1215GGG (3751A>G), AAG1216AAA (3754G>A), GCC1394GCT (4288C>T), TTT1395TTC (4291T>C), CAA1396CAG (4294A>G), CTT1401CTG (4309T>G), GTG1402GTT (4312G>T), GAC1404GAT (4318C>T), TAT1406TAC (4324T>C), GGA1407GGT (4327A>T), CCA1413CCT (4345A>T), GAC1414GAT (4348C>T), GCC1417GCG (4357C>G), AGA1418CGA (4358A>C), CTT1421CTA (4369T>A), GCT1425GCA (4381T>A), TCA1426TCC (4384A>C), ATT1431ATC (4399T>C), GTT1436GTG (4414T>G), GGA1441..G (4429A>G), TCA1445TCT (4441A>T), GGA1446GGG (4444A>G), CTC1449CTG (4453C>G), GTT1451GTC (4459T>C), TCT1455TCC (4471T>C), TTT1460TTC (4486T>C), TAT1461TAC (4489T>C), CGT1463AGG (4493C>A 4495T>G), GCA1465GCT (4501A>T), ATT1467ATC (4507T>C), TCT1468TCA (4510T>A), TTG1470CTG (4514T>C), CCA1473CCC (4525A>C), CAT1474CAC (4528T>C), ATA1475ATT (4531A>T), GCT1477GCA (4537T>A), GGG1481GGA (4549G>A), TCT1485TCG (4561T>G) | | | | | | | | |

##### Proteins

| | Begin | End | Coverage | Score | Concordance | Matches | Identities | I/D/M/F* | Stop Codons |
|---|---|---|---|---|---|---|---|---|---|
| polyprotein (YP_001039627.1) | 191 | 1486 | 12.7% | 1909 | 97.8% | 275 (100%) | 269 (97.8%) | 0/0/0/0 | 0 |
| Protein mutations: | S205A (719T>G 721C>T), M206V (722A>G), V291I (977G>A 979T>A), R1143K (3534G>A), A1202T (3710G>A) | | | | | | | | |
| Codon mutations: | GCA196GCG (694A>G), CTA201TTA (707C>T), GGG203GGA (715G>A), GAA204GAG (718A>G), TCC205GCT (719T>G 721C>T), ATG206GTG (722A>G), TCA207TCT (727A>T), CAC208CAT (730C>T), CTT211CTG (739T>G), GTG214GTT (748G>T), GTA215GTT (751A>T), GCA216GCG (754A>G), CTC218TTA (758C>T 760C>A), ATA219ATT (763A>T), GCC220GCT (766C>T), GGG222GGC (772G>C), ATT223ATA (775T>A), TCT224TCA (778T>A), ACC225ACA (781C>A), TTC228TTT (790C>T), GTG229GTA (793G>A), GCT230GCA (796T>A), GTA231GTT (799A>T), CCT232CCG (802T>G), GGT233GGC (805T>C), AGA234CGT (806A>C 808A>T), GGT252GGA (862T>A), GCT253GCC (865T>C), AAT257AAC (877T>C), TTG259CTT (881T>C 883G>T), TTT260TTC (886T>C), ACC261ACA (889C>A), TTC263TTT (895C>T), AGG264AGA (898G>A), AAT265AAC (901T>C), TGT269TGC (913T>C), ATA281ATT (949A>T), ATT284ATA (958T>A), GGT286GGA (964T>A), TTT287TTC (967T>C), CCA290CCT (976A>T), GTT291ATA (977G>A 979T>A), GCT294GCA (988T>A), GGC1125GGT (3481C>T), ACA1126ACT (3484A>T), ACG1134ACA (3508G>A), AGG1143AAG (3534G>A), AGT1144AGC (3538T>C), TCT1158TCC (3580T>C), ATC1159ATT (3583C>T), GAA1168GAG (3610A>G), TAC1173TAT (3625C>T), GTG1174GTA (3628G>A), AAA1179AAG (3643A>G), GAC1190GAT (3676C>T), AGG1193AGA (3685G>A), CTA1199TTA (3701C>T), GCA1202ACA (3710G>A), CGA1211CGT (3739A>T), GGA1215GGG (3751A>G), AAG1216AAA (3754G>A), GCC1394GCT (4288C>T), TTT1395TTC (4291T>C), CAA1396CAG (4294A>G), CTT1401CTG (4309T>G), GTG1402GTT (4312G>T), GAC1404GAT (4318C>T), TAT1406TAC (4324T>C), GGA1407GGT (4327A>T), CCA1413CCT (4345A>T), GAC1414GAT (4348C>T), GCC1417GCG (4357C>G), AGA1418CGA (4358A>C), CTT1421CTA (4369T>A), GCT1425GCA (4381T>A), TCA1426TCC (4384A>C), ATT1431ATC (4399T>C), GTT1436GTG (4414T>G), GGA1441..G (4429A>G), TCA1445TCT (4441A>T), GGA1446GGG (4444A>G), CTC1449CTG (4453C>G), GTT1451GTC (4459T>C), TCT1455TCC (4471T>C), TTT1460TTC (4486T>C), TAT1461TAC (4489T>C), CGT1463AGG (4493C>A 4495T>G), GCA1465GCT (4501A>T), ATT1467ATC (4507T>C), TCT1468TCA (4510T>A), TTG1470CTG (4514T>C), CCA1473CCC (4525A>C), CAT1474CAC (4528T>C), ATA1475ATT (4531A>T), GCT1477GCA (4537T>A), GGG1481GGA (4549G>A), TCT1485TCG (4561T>G) | | | | | | | | |

*: Inserts / Deletes / Misaligned / Frameshifts

#### Analysis details

This analysis was performed with panviral2.64

### NGS Details (UN24): Torradovirus lycopersici (segment RNA 2)

#### Assembly

| | |
|---|---|
| **Coverage Length** | 2450 (10 contig(s)) |
| **Depth Of Coverage** | 2.4 |
| **Number Of Reads** | 46 |
| **Reads Per Million** | 0.92 rpm (after QC) |
| **Ambiguities** | 0 |
| **Assembly Method** | de novo + reference guided assembly |
| **Consensus Caller** | Bcf Tools |

#### Coverage Map

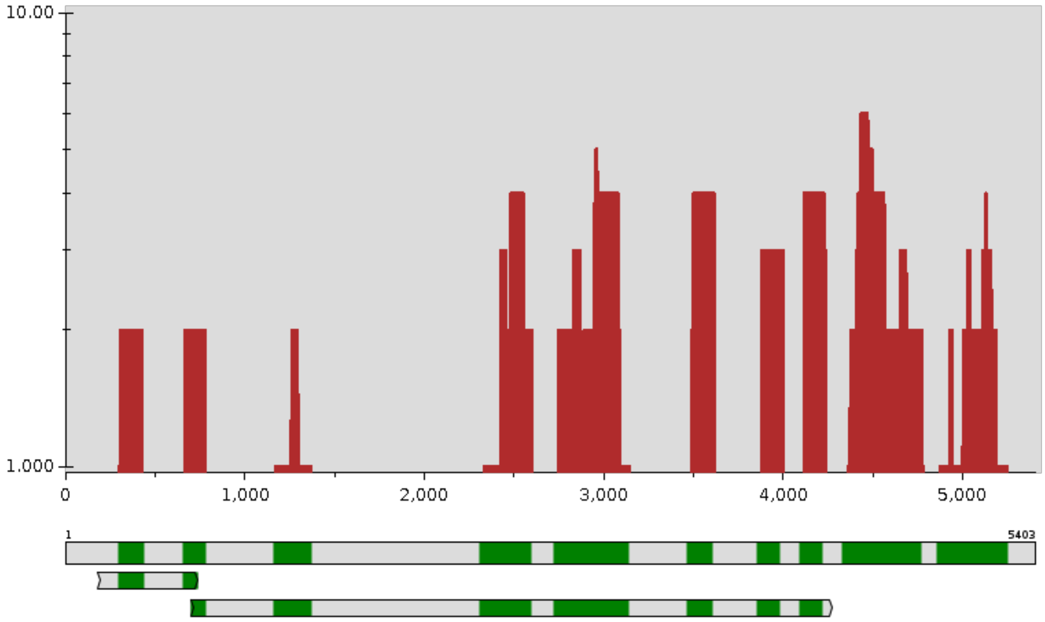

#### Assignment

| | |
|---|---|
| **Type** | Torradovirus lycopersici (Taxonomy ID: 3048378) |
| **Reference Genome** | NC_009032.1 |
| **NT Identity (%)** | 85.9457 |
| **AA Identity (%)** | 89.9281 |
| **Number Of Stop Codons** | 1 |
| **Number Of CDS** | 2 |

#### Alignment

| | |
|---|---|
| **Alignment Score** | 3470.0 (NT) + 3541.0 (AA) = 7011.0 |
| **Concordance (%)** | 80.7998 |

| Alignment Method | Global, seeded, nucleotide + amino acids (AGA) |
|---|---|

#### Genome Region

Sequence starts at position 294 and ends at position 5255 relative to NC_009032.1 reference sequence.

#### Alignment Detailed Statistics

| | Begin | End | Coverage | Score | Concordance | Matches | Identities | I/D/M/F* | Stop Codons |
|---|---|---|---|---|---|---|---|---|---|
| **NT** | **294** | **5255** | **45.3%** | **3470** | **71.6%** | **2439 (98.3%)** | **2122 (85.6%)** | **30/11** | |
| Mutations: | 301A>T, 304T>C, 305C>T, 307A>G, 313C>T, 316G>A, 325C>T, 329A>G, 334T>A, 340A>T, 352A>T, 356G>A, 370C>T, 372A>C, 376C>T, 382A>G, 383C>T, 385G>A, 388T>A, 400T>C, 401T>C, 403A>C, 404C>T, 409A>C, 415A>T, 418T>G, 428A>T, 433C>A, 658C>A, 661A>G, 665T>G, 670A>G, 673A>G, 675G>A, 678T>C, 679G>A, 683T>A, 685G>A, 686T>G, 688C>T, 689A>T, 690A>C, 694A>G, 696T>C, 697A>T, 699G>C, 700T>A, 704G>A, 715C>T, 721C>T, 733A>G, 742G>A, 748A>G, 749A>G, 755A>T, 758T>A, 760T>C, 773A>G, 788A>T, 1166A>G, 1178C>T, 1188A>C, 1196C>T, 1199T>C, 1200G>A, 1201G>A, 1205T>G, 1206C>A, 1208G>A, 1223C>T, 1226A>T, 1232G>T, 1235C>A, 1254G>A, 1255A>G, 1257A>C, 1258T>C, 1259C>T, 1268C>T, 1269A>G, 1271T>A, 1274C>T, 1275T>A, 1278A>G, 1280G>C, 1283G>T, 1286A>C, 1289A>T, 1293T>A, 1294C>G, 1314C>A, 1317C>A, 1325A>G, 1337T>C, 1340A>G, 1346G>A, 1355G>A, 1361A>T, 1362T>G, 1367A>T, 1370A>G, 1373T>C, 1376T>C, 2326C>T, 2336A>T, 2339A>G, 2351C>A, 2354C>T, 2357G>T, 2387T>G, 2393G>A, 2405G>A, 2408A>G, 2414T>C, 2433C>T, 2465T>C, 2501A>C, 2503G>A, 2519A>G, 2573T>C, 2612C>T, 2792C>T, 2819G>A, 2840T>C, 2841C>T, 2885T>C, 2919G>A, 2939A>G, 2968A>C, 2969C>T, 3003T>C, 3050G>A, 3104C>T, 3122T>C, 3128T>C, 3131G>A, 3485C>T, 3488A>G, 3491C>T, 3497T>C, 3503A>G, 3508A>G, 3515G>T, 3522C>T, 3524A>G, 3527C>T, 3530T>A, 3533G>A, 3539C>T, 3540T>C, 3542G>C, 3545T>A, 3555G>A, 3556A>C, 3557A>C, 3560T>C, 3564A>C, 3567T>A, 3569A>G, 3575G>A, 3578A>G, 3581C>T, 3590A>C, 3605A>G, 3611G>A, 3617A>T, 3620G>A, 3623A>T, 3629G>C, 3631C>A, 3878G>C, 3881C>T, 3882A>G, 3884C>A, 3885A>G, 3890A>G, 3893T>C, 3896A>T, 3911T>C, 3914T>A, 3920A>T, 3923C>T, 3926A>C, 3927A>C, 3928A>G, 3929G>A, 3932C>A, 3933A>G, 3935C>T, 3941T>G, 3944C>A, 3947T>C, 3950T>A, 3953T>A, 3960G>A, 3971T>A, 3983T>A, 3984A>G, 3986G>T, 3988G>A, 3989C>T, 3990G>A, 3991A>G, 3995C>T, 3998C>T, 4001T>C, 4112C>G, 4124A>G, 4133A>G, 4142A>C, 4145C>T, 4149C>A, 4151G>A, 4154A>G, 4157C>A, 4166G>A, 4181C>T, 4187G>A, 4188C>G, 4196T>A, 4202T>C, 4205T>A, 4208A>T, 4211A>T, 4214T>C, 4223G>A, 4224C>T, 4225A>T, 4227T>C, 4229G>T, 4232T>C, 4241A>G, 4244G>A, 4354G>A, 4355T>G, 4381A>G, 4382A>C, 4383C>T, 4391A>T, 4392T>C, 4397A>G, 4399T>A, 4410_4411insTAACGGAGA, 4424C>T, 4429T>C, 4447T>A, 4448_4449insT, 4457A>G, 4468T>C, 4496T>A, 4506A>T, 4525T>C, 4567_4568insT, 4605T>C, 4671C>T, 4749T>A, 4888G>T, 4889A>T, 4894A>T, 4896G>A, 4908T>A, 4915G>T, 4916C>T, 4917G>T, 4928A>T, 4930G>A, 4940C>A, 4941C>G, 4943A>T, 4946_4947insC, 4951_4952insGAGCACTGCCATTA, 4955A>C, 4957G>A, 4958A>C, 4959A>C, 4961C>T, 4969A>T, 4972T>G, 4973T>G, 4974T>A, 4976A>T, 4977A>T, 4978G>A, 4993G>A, 4994C>A, 4996A>C, 5000C>T, 5003A>T, 5004G>T, 5019A>T, 5020C>T, 5031T>C, 5032G>A, 5040C>T, 5041A>G, 5053A>G, 5054T>C, 5056G>A, 5099C>T, 5101A>G, 5102T>C, 5107_5108insT, 5109A>C, 5110A>T, 5111A>C, 5121_5122insG, 5122T>A, 5122_5123insG, 5141T>A, 5150A>G, 5185T>A, 5196T>G, 5197T>A, 5198T>G, 5199_5200insC, 5206C>T, 5208G>A, 5211A>G, 5212A>C, 5213A>T, 5214C>T, 5221_5227delACTGCTT, 5233_5236delGTTG, 5241T>A, 5242T>A, 5244G>T, 5246T>A, 5249A>T | | | | | | | | |

##### CDS

| | Begin | End | Coverage | Score | Concordance | Matches | Identities | I/D/M/F* | Stop Codons |
|---|---|---|---|---|---|---|---|---|---|
| ToTV_sRNA2gp1 | 39 | 188 | 41.5% | 490 | 88.3% | 78 (100%) | 65 (83.3%) | 0/0/0/0 | 1 |
| Protein mutations: | T50A (329A>G), V59I (356G>A), N64T (372A>C), T83S (428A>T), S162A (665T>G), R165K (675G>A), L166S (678T>C 679G>A), L168I (683T>A 685G>A), S169A (686T>G 688C>T), N170S (689A>T 690A>C), L172P (696T>C 697A>T), S173T (699G>C 700T>A), D175N (704G>A) | | | | | | | | |
| Codon mutations: | CCA40CCT (301A>T), ACT41ACC (304T>C), CTA42TTG (305C>T 307A>G), TTC44TTT (313C>T), AAG45AAA (316G>A), CCC48CCT (325C>T), ACA50GCA (329A>G), GGT51GGA (334T>A), GCA53GCT (340A>T), CTA57CTT (352A>T), GTT59ATT (356G>A), AAC63AAT (370C>T), AAT64ACT (372A>C), AGC65AGT (376C>T), CAA67CAG (382A>G), CTG68TTA (383C>T 385G>A), GTT69GTA (388T>A), TAT73TAC (400T>C), TTA74CTC (401T>C 403A>C), CTG75TTG (404C>T), CGA76CGC (409A>C), GGA78GGT (415A>T), ACT79ACG (418T>G), ACT83TCT (428A>T), GGC84GGA (433C>A), TCC159TCA (658C>A), GAA160GAG (661A>G), TCC162GCC (665T>G), AAA163AAG (670A>G), TTA164TTG (673A>G), AGA165AAA (675G>A), TTG166TCA (678T>C 679G>A), TTG168ATA (683T>A 685G>A), TCC169GCT (686T>G 688C>T), AAC170TCC (689A>T 690A>C), CAA171CAG (694A>G), CTA172CCT (696T>C 697A>T), AGT173ACA (699G>C 700T>A), GAT175AAT (704G>A), TTC178TTT (715C>T), GGC180GGT (721C>T), GAA184GAG (733A>G), AAG187AAA (742G>A) | | | | | | | | |
| ToTV_sRNA2gp2 | 1 | 1182 | 39.9% | 3051 | 91.6% | 478 (100%) | 435 (91.0%) | 0/0/0/0 | 0 |
| Protein mutations: | M1I (704G>A), S5L (715C>T), A7V (721C>T), K11R (733A>G), S14N (742G>A), K16R (748A>G 749A>G), V20A (760T>C), M163L (1188A>C), G167N (1200G>A 1201G>A), D185S (1254G>A 1255A>G), I186P (1257A>C 1258T>C 1259C>T), I190V (1269A>G 1271T>A), S192T (1275T>A), K193D (1278A>G 1280G>C), Q194H (1283G>T), E195D (1286A>C), H205N (1314C>A), S221A (1362T>G), T542I (2326C>T), E545D (2336A>T), R601K (2503G>A), D740N (2919G>A), Y756S (2968A>C 2969C>T), N936S (3508A>G), E952T (3555G>A 3556A>C 3557A>C), K955Q (3564A>C), L956M (3567T>A 3569A>G), P977Q (3631C>A), I1061V (3882A>G 3884C>A), T1062A (3885A>G), K1076R (3927A>C 3928A>G 3929G>A), T1078A (3933A>G 3935C>T), A1087T (3960G>A), M1095V (3984A>G 3986G>T), S1096N (3988G>A 3989C>T), D1097S (3990G>A 3991A>G), L1150I (4149C>A 4151G>A), Q1163E (4188C>G), H1175F (4224C>T 4225A>T) | | | | | | | | |
| Codon mutations: | ATG1ATA (704G>A), TCA5TTA (715C>T), GCT7GTT (721C>T), AAG11AGG (733A>G), AGT14AAT (742G>A), AAA16AGG (748A>G 749A>G), GCA18GCT (755A>T), GCT19GCA (758T>A), GTA20GCA (760T>C), TTA24TTG (773A>G), ACA29ACT (788A>T), AGA155AGG (1166A>G), TTC159TTT (1178C>T), ATG163CTG (1188A>C), TGC165TGT (1196C>T), CTT166CTC (1199T>C), GGC167AAC (1200G>A 1201G>A), ACT168ACG (1205T>G), CGG169AGA (1206C>A 1208G>A), TGC174TGT (1223C>T), TCA175TCT (1226A>T), GTG177GTT (1232G>T), GCC178GCA (1235C>A), GAT185AGT (1254G>A 1255A>G), ATC186CCT (1257A>C 1258T>C 1259C>T), GCC189GCT (1268C>T), ATT190GTA (1269A>G 1271T>A), TAC191TAT (1274C>T), TCA192ACA (1275T>A), AAG193GAC (1278A>G 1280G>C), CAG194CAT (1283G>T), GAA195GAC (1286A>C), ACA196ACT (1289A>T), TCT198AGT (1293T>A 1294C>G), CAC205AAC (1314C>A), CGA206AGA (1317C>A), CAA208CAG (1325A>G), GAT212GAC (1337T>C), AGA213AGG (1340A>G), GAG215GAA (1346G>A), CAG218CAA (1355G>A), ATA220ATT (1361A>T), TCA221GCA (1362T>G), TCA222TCT (1367A>T), GTA223GTG (1370A>G), TAT224TAC (1373T>C), TCT225TCC (1376T>C), ACA542ATA (2326C>T), GAA545GAT (2336A>T), TTA546TTG (2339A>G), GCC550GCA (2351C>A), TGC551TGT (2354C>T), GTG552GTT (2357G>T), GTT562GTG (2387T>G), GAG564GAA (2393G>A), AAG568AAA (2405G>A), AAA569AAG (2408A>G), AGT571AGC (2414T>C), CTA578TTA (2433C>T), GGT588GGC (2465T>C), CTA600CTC (2501A>C), AGG601AAG (2503G>A), GCA606GCG (2519A>G), GGT624GGC (2573T>C), AAC637AAT (2612C>T), CAC697CAT (2792C>T), GAG706GAA (2819G>A), CAT713CAC (2840T>C), CTG714TTG (2841C>T), CGT728CGC (2885T>C), GAC740AAC (2919G>A), CGA746CGG (2939A>G), TAC756TCT (2968A>C 2969C>T), TTA768CTA (3003T>C), ACG783ACA (3050G>A), GGC801GGT (3104C>T), ATT807ATC (3122T>C), ATT809ATC (3128T>C), GAG810GAA (3131G>A), TAC928..T (3485C>T), GGA929GGG (3488A>G), CAC930CAT (3491C>T), ACT932ACC (3497T>C), GGA934GGG (3503A>G), AAT936AGT (3508A>G), GTG938GTT (3515G>T), CTA941TTG (3522C>T 3524A>G), TTC942TTT (3527C>T), TCT943TCA (3530T>A), GAG944GAA (3533G>A), GAC946GAT (3539C>T), TTG947CTC (3540T>C 3542G>C), TCT948TCA (3545T>A), GAA952ACC (3555G>A 3556A>C 3557A>C), ATT953ATC (3560T>C), AAA955CAA (3564A>C), TTA956ATG (3567T>A 3569A>G), AAG958AAA (3575G>A), GAA959GAG (3578A>G), AAC960AAT (3581C>T), ACA963ACC (3590A>C), GGA968GGG (3605A>G), AAG970AAA (3611G>A), CCA972CCT (3617A>T), CAG973CAA (3620G>A), GTA974GTT (3623A>T), GTG976GTC (3629G>C), CCG977CAG (3631C>A), CTG1059CTC (3878G>C), AAC1060AAT (3881C>T), ATC1061GTA (3882A>G 3884C>A), ACC1062GCC (3885A>G), TTA1063TTG (3890A>G), GAT1064GAC (3893T>C), TCA1065TCT (3896A>T), TTT1070TTC (3911T>C), CCT1071CCA (3914T>A), GGA1073GGT (3920A>T), ATC1074ATT (3923C>T), TCA1075TCC (3926A>C), AAG1076CGA (3927A>C 3928A>G 3929G>A), GGC1077GGA (3932C>A), ACC1078GCT (3933A>G 3935C>T), GTT1080GTG (3941T>G), GTC1081GTA (3944C>A), TCT1082TCC (3947T>C), GCT1083GCA (3950T>A), GGT1084GGA (3953T>A), GCA1087ACA (3960G>A), GCT1090GCA (3971T>A), GGT1094GGA (3983T>A), ATG1095GTT (3984A>G 3986G>T), AGC1096AAT (3988G>A 3989C>T), GAC1097AGC (3990G>A 3991A>G), AAC1098AAT (3995C>T), ATC1099ATT (3998C>T), TTT1100TTC (4001T>C), CTC1137..G (4112C>G), TTA1141TTG (4124A>G), AAA1144AAG (4133A>G), ATA1147ATC (4142A>C), AGC1148AGT (4145C>T), CTG1150ATA (4149C>A 4151G>A), GAA1151GAG (4154A>G), ATC1152ATA (4157C>A), AAG1155AAA (4166G>A), TTC1160TTT (4181C>T), TTG1162TTA (4187G>A), CAA1163GAA (4188C>G), GCT1165GCA (4196T>A), GCT1167GCC (4202T>C), CCT1168CCA (4205T>A), TCA1169TCT (4208A>T), GCA1170GCT (4211A>T), AAT1171AAC (4214T>C), AAG1174AAA (4223G>A), CAT1175TTT (4224C>T 4225A>T), TTG1176CTT (4227T>C 4229G>T), GGT1177GGC (4232T>C), CAA1180CAG (4241A>G), ACG1181ACA (4244G>A) | | | | | | | | |

##### Proteins

| | Begin | End | Coverage | Score | Concordance | Matches | Identities | I/D/M/F* | Stop Codons |
|---|---|---|---|---|---|---|---|---|---|
| Orf1 (YP_001040017.1) | 39 | 188 | 41.5% | 490 | 88.3% | 78 (100%) | 65 (83.3%) | 0/0/0/0 | 1 |
| Protein mutations: | T50A (329A>G), V59I (356G>A), N64T (372A>C), T83S (428A>T), S162A (665T>G), R165K (675G>A), L166S (678T>C 679G>A), L168I (683T>A 685G>A), S169A (686T>G 688C>T), N170S (689A>T 690A>C), L172P (696T>C 697A>T), S173T (699G>C 700T>A), D175N (704G>A) | | | | | | | | |
| Codon mutations: | CCA40CCT (301A>T), ACT41ACC (304T>C), CTA42TTG (305C>T 307A>G), TTC44TTT (313C>T), AAG45AAA (316G>A), CCC48CCT (325C>T), ACA50GCA (329A>G), GGT51GGA (334T>A), GCA53GCT (340A>T), CTA57CTT (352A>T), GTT59ATT (356G>A), AAC63AAT (370C>T), AAT64ACT (372A>C), AGC65AGT (376C>T), CAA67CAG (382A>G), CTG68TTA (383C>T 385G>A), GTT69GTA (388T>A), TAT73TAC (400T>C), TTA74CTC (401T>C 403A>C), CTG75TTG (404C>T), CGA76CGC (409A>C), GGA78GGT (415A>T), ACT79ACG (418T>G), ACT83TCT (428A>T), GGC84GGA (433C>A), TCC159TCA (658C>A), GAA160GAG (661A>G), TCC162GCC (665T>G), AAA163AAG (670A>G), TTA164TTG (673A>G), AGA165AAA (675G>A), TTG166TCA (678T>C 679G>A), TTG168ATA (683T>A 685G>A), TCC169GCT (686T>G 688C>T), AAC170TCC (689A>T 690A>C), CAA171CAG (694A>G), CTA172CCT (696T>C 697A>T), AGT173ACA (699G>C 700T>A), GAT175AAT (704G>A), TTC178TTT (715C>T), GGC180GGT (721C>T), GAA184GAG (733A>G), AAG187AAA (742G>A) | | | | | | | | |
| polyprotein (YP_001040018.1) | 1 | 1182 | 39.9% | 3051 | 91.6% | 478 (100%) | 435 (91.0%) | 0/0/0/0 | 0 |

| | Begin | End | Coverage | Score | Concordance | Matches | Identities | I/D/M/F* | Stop Codons |
|---|---|---|---|---|---|---|---|---|---|
| **NT** | **294** | **5255** | **45.3%** | **3470** | **71.6%** | **2439 (98.3%)** | **2122 (85.6%)** | **30/11** | |

| | |
|---|---|
| Protein mutations: | M1I (704G>A), S5L (715C>T), A7V (721C>T), K11R (733A>G), S14N (742G>A), K16R (748A>G 749A>G), V20A (760T>C), M163L (1188A>C), G167N (1200G>A 1201G>A), D185S (1254G>A 1255A>G), I186P (1257A>C 1258T>C 1259C>T), I190V (1269A>G 1271T>A), S192T (1275T>A), K193D (1278A>G 1280G>C), Q194H (1283G>T), E195D (1286A>C), H205N (1314C>A), S221A (1362T>G), T542I (2326C>T), E545D (2336A>T), R601K (2503G>A), D740N (2919G>A), Y756S (2968A>C 2969C>T), N936S (3508A>G), E952T (3555G>A 3556A>C 3557A>C), K955Q (3564A>C), L956M (3567T>A 3569A>G), P977Q (3631C>A), I1061V (3882A>G 3884C>A), T1062A (3885A>G), K1076R (3927A>C 3928A>G 3929G>A), T1078A (3933A>G 3935C>T), A1087T (3960G>A), M1095V (3984A>G 3986G>T), S1096N (3988G>A 3989C>T), D1097S (3990G>A 3991A>G), L1150I (4149C>A 4151G>A), Q1163E (4188C>G), H1175F (4224C>T 4225A>T) |
| Codon mutations: | ATG1ATA (704G>A), TCA5TTA (715C>T), GCT7GTT (721C>T), AAG11AGG (733A>G), AGT14AAT (742G>A), AAA16AGG (748A>G 749A>G), GCA18GCT (755A>T), GCT19GCA (758T>A), GTA20GCA (760T>C), TTA24TTG (773A>G), ACA29ACT (788A>T), AGA155AGG (1166A>G), TTC159TTT (1178C>T), ATG163CTG (1188A>C), TGC165TGT (1196C>T), CTT166CTC (1199T>C), GGC167AAC (1200G>A 1201G>A), ACT168ACG (1205T>G), CGG169AGA (1206C>A 1208G>A), TGC174TGT (1223C>T), TCA175TCT (1226A>T), GTG177GTT (1232G>T), GCC178GCA (1235C>A), GAT185AGT (1254G>A 1255A>G), ATC186CCT (1257A>C 1258T>C 1259C>T), GCC189GCT (1268C>T), ATT190GTA (1269A>G 1271T>A), TAC191TAT (1274C>T), TCA192ACA (1275T>A), AAG193GAC (1278A>G 1280G>C), CAG194CAT (1283G>T), GAA195GAC (1286A>C), ACA196ACT (1289A>T), TCT198AGT (1293T>A 1294C>G), CAC205AAC (1314C>A), CGA206AGA (1317C>A), CAA208CAG (1325A>G), GAT212GAC (1337T>C), AGA213AGG (1340A>G), GAG215GAA (1346G>A), CAG218CAA (1355G>A), ATA220ATT (1361A>T), TCA221GCA (1362T>G), TCA222TCT (1367A>T), GTA223GTG (1370A>G), TAT224TAC (1373T>C), TCT225TCC (1376T>C), ACA542ATA (2326C>T), GAA545GAT (2336A>T), TTA546TTG (2339A>G), GCC550GCA (2351C>A), TGC551TGT (2354C>T), GTG552GTT (2357G>T), GTT562GTG (2387T>G), GAG564GAA (2393G>A), AAG568AAA (2405G>A), AAA569AAG (2408A>G), AGT571AGC (2414T>C), CTA578TTA (2433C>T), GGT588GGC (2465T>C), CTA600CTC (2501A>C), AGG601AAG (2503G>A), GCA606GCG (2519A>G), GGT624GGC (2573T>C), AAC637AAT (2612C>T), CAC697CAT (2792C>T), GAG706GAA (2819G>A), CAT713CAC (2840T>C), CTG714TTG (2841C>T), CGT728CGC (2885T>C), GAC740AAC (2919G>A), CGA746CGG (2939A>G), TAC756TCT (2968A>C 2969C>T), TTA768CTA (3003T>C), ACG783ACA (3050G>A), GGC801GGT (3104C>T), ATT807ATC (3122T>C), ATT809ATC (3128T>C), GAG810GAA (3131G>A), TAC928..T (3485C>T), GGA929GGG (3488A>G), CAC930CAT (3491C>T), ACT932ACC (3497T>C), GGA934GGG (3503A>G), AAT936AGT (3508A>G), GTG938GTT (3515G>T), CTA941TTG (3522C>T 3524A>G), TTC942TTT (3527C>T), TCT943TCA (3530T>A), GAG944GAA (3533G>A), GAC946GAT (3539C>T), TTG947CTC (3540T>C 3542G>C), TCT948TCA (3545T>A), GAA952ACC (3555G>A 3556A>C 3557A>C), ATT953ATC (3560T>C), AAA955CAA (3564A>C), TTA956ATG (3567T>A 3569A>G), AAG958AAA (3575G>A), GAA959GAG (3578A>G), AAC960AAT (3581C>T), ACA963ACC (3590A>C), GGA968GGG (3605A>G), AAG970AAA (3611G>A), CCA972CCT (3617A>T), CAG973CAA (3620G>A), GTA974GTT (3623A>T), GTG976GTC (3629G>C), CCG977CAG (3631C>A), CTG1059CTC (3878G>C), AAC1060AAT (3881C>T), ATC1061GTA (3882A>G 3884C>A), ACC1062GCC (3885A>G), TTA1063TTG (3890A>G), GAT1064GAC (3893T>C), TCA1065TCT (3896A>T), TTT1070TTC (3911T>C), CCT1071CCA (3914T>A), GGA1073GGT (3920A>T), ATC1074ATT (3923C>T), TCA1075TCC (3926A>C), AAG1076CGA (3927A>C 3928A>G 3929G>A), GGC1077GGA (3932C>A), ACC1078GCT (3933A>G 3935C>T), GTT1080GTG (3941T>G), GTC1081GTA (3944C>A), TCT1082TCC (3947T>C), GCT1083GCA (3950T>A), GGT1084GGA (3953T>A), GCA1087ACA (3960G>A), GCT1090GCA (3971T>A), GGT1094GGA (3983T>A), ATG1095GTT (3984A>G 3986G>T), AGC1096AAT (3988G>A 3989C>T), GAC1097AGC (3990G>A 3991A>G), AAC1098AAT (3995C>T), ATC1099ATT (3998C>T), TTT1100TTC (4001T>C), CTC1137..G (4112C>G), TTA1141TTG (4124A>G), AAA1144AAG (4133A>G), ATA1147ATC (4142A>C), AGC1148AGT (4145C>T), CTG1150ATA (4149C>A 4151G>A), GAA1151GAG (4154A>G), ATC1152ATA (4157C>A), AAG1155AAA (4166G>A), TTC1160TTT (4181C>T), TTG1162TTA (4187G>A), CAA1163GAA (4188C>G), GCT1165GCA (4196T>A), GCT1167GCC (4202T>C), CCT1168CCA (4205T>A), TCA1169TCT (4208A>T), GCA1170GCT (4211A>T), AAT1171AAC (4214T>C), AAG1174AAA (4223G>A), CAT1175TTT (4224C>T 4225A>T), TTG1176CTT (4227T>C 4229G>T), GGT1177GGC (4232T>C), CAA1180CAG (4241A>G), ACG1181ACA (4244G>A) |

*: Inserts / Deletes / Misaligned / Frameshifts

#### Analysis details

This analysis was performed with panviral2.64

### NGS Details (UN24): Diachasmimorpha longicaudata entomopoxvirus (segment NC_043455.1)

#### Assembly

| | |
|---|---|
| **Coverage Length** | 1126 (1 contig(s)) |
| **Depth Of Coverage** | 1124.4 |
| **Number Of Reads** | 10408 |
| **Reads Per Million** | 208.34 rpm (after QC) |
| **Ambiguities** | 0 |
| **Assembly Method** | de novo + reference guided assembly |
| **Consensus Caller** | Bcf Tools |

#### Coverage Map

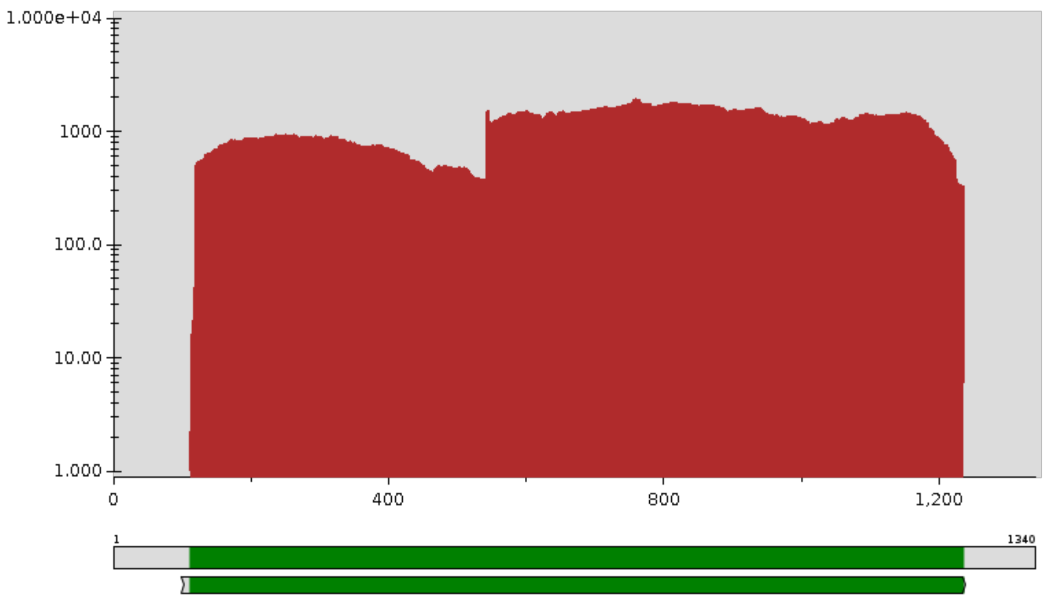

#### Assignment

| | |
|---|---|
| **Type** | Diachasmimorpha longicaudata entomopoxvirus (Taxonomy ID: 109981) |
| **Reference Genome** | NC_043455.1 |
| **NT Identity (%)** | 60.2131 |
| **AA Identity (%)** | 55.7333 |
| **Number Of Stop Codons** | 0 |
| **Number Of CDS** | 1 |

#### Alignment

| | |
|---|---|
| **Alignment Score** | 442.0 (NT) + 1481.0 (AA) = 1923.0 |
| **Concordance (%)** | 40.7934 |

| Alignment Method | Global, seeded, nucleotide + amino acids (AGA) |
|---|---|

#### Genome Region

Sequence starts at position 112 and ends at position 1237 relative to NC_043455.1 reference sequence.

#### Alignment Detailed Statistics

| | Begin | End | Coverage | Score | Concordance | Matches | Identities | I/D/M/F* | Stop Codons |
|---|---|---|---|---|---|---|---|---|---|
| **NT** | **112** | **1237** | **84.0%** | **442** | **19.8%** | **1123 (99.5%)** | **678 (60.1%)** | **3/3** | |
| Mutations: | 121A>T, 126A>C, 127C>T, 128C>G, 130T>G, 136C>G, 137C>A, 139C>T, 142G>A, 143A>G, 145A>T, 146A>G, 148T>C, 149A>C, 151C>T, 154A>T, 156A>G, 157G>A, 160T>G, 163A>T, 166T>C, 167T>C, 168C>A, 169T>G, 171G>A, 172T>C, 175A>G, 178T>C, 187T>A, 190A>T, 191A>G, 193T>C, 203A>C, 204A>G, 208C>T, 209A>G, 211T>A, 214T>G, 217A>T, 218T>A, 219G>T, 223T>A, 226T>G, 230A>C, 231A>G, 232A>T, 238G>C, 241A>T, 243T>C, 244C>T, 247A>G, 256G>C, 259A>T, 262A>C, 271A>C, 272G>T, 274A>C, 276C>T, 277A>G, 278T>A, 279A>T, 283A>T, 284A>C, 286C>T, 288G>C, 289T>C, 294T>G, 295A>C, 298G>A, 299C>G, 300A>T, 301A>T, 302A>G, 304A>G, 310T>C, 311T>A, 312C>A, 313T>A, 320A>G, 322T>G, 323A>G, 325C>A, 331T>A, 337C>A, 341A>T, 343C>T, 346A>T, 352T>G, 355A>G, 356T>C, 358G>T, 361T>A, 362C>G, 363T>C, 364C>T, 368G>A, 370T>G, 371C>G, 374C>A, 375G>A, 376T>G, 379A>G, 380T>A, 382G>T, 383C>T, 384A>T, 385A>G, 386A>G, 394G>A, 395A>G, 400T>C, 401C>A, 403A>T, 404_406delTAT, 409T>C, 410T>A, 412T>A, 413A>C, 415A>G, 416T>G, 417G>C, 418T>A, 421A>T, 423T>C, 424T>G, 438C>A, 442T>C, 443A>G, 445C>G, 446A>G, 447A>G, 448G>T, 452A>G, 453G>A, 454C>T, 455C>A, 456A>T, 457G>C, 458G>A, 459A>G, 462C>A, 463T>G, 467A>G, 469_470insCAT, 470A>G, 471A>G, 474C>T, 475C>A, 478G>A, 482T>G, 484G>C, 485A>T, 486T>C, 500C>A, 503A>G, 505G>C, 506A>T, 507T>G, 511T>C, 512C>A, 514A>G, 515C>A, 517C>T, 519C>A, 520C>A, 521C>A, 523A>G, 525A>G, 528G>C, 530A>T, 532C>A, 533G>C, 534A>G, 535T>A, 540A>G, 541A>G, 544A>C, 547T>C, 550A>G, 553A>G, 554G>T, 559A>T, 560A>T, 562A>G, 574T>C, 577G>A, 581T>C, 583G>C, 584A>T, 585T>C, 586A>T, 587G>A, 588A>G, 589T>A, 590A>G, 591A>G, 595T>C, 596T>A, 597T>A, 602A>C, 604A>G, 607A>T, 608C>T, 610A>T, 616T>A, 620G>C, 622A>G, 623T>C, 628T>G, 629G>C, 630A>C, 632A>C, 633G>C, 634T>A, 635C>A, 637T>A, 638G>A, 646T>G, 647A>G, 648T>G, 650T>G, 652A>T, 655A>C, 658A>T, 661A>C, 664C>T, 665G>A, 667T>G, 671T>C, 674A>G, 675G>A, 678T>C, 680A>C, 682C>T, 683A>G, 685T>G, 687C>T, 688A>T, 689T>A, 692C>A, 693A>G, 694G>A, 695G>A, 696T>A, 697C>G, 700T>C, 705G>A, 706A>C, 707G>A, 709T>G, 715A>G, 717A>G, 718A>G, 724G>T, 727A>G, 730A>G, 731A>C, 732A>G, 735C>A, 739A>G, 740T>C, 742G>C, 745A>T, 748C>T, 754C>T, 759G>A, 765A>T, 770A>G, 772A>C, 778A>C, 779A>G, 781G>T, 784A>G, 785A>G, 787T>A, 790T>G, 792T>G, 793T>G, 796A>G, 797G>C, 798C>T, 799A>T, 810T>G, 820T>C, 823C>G, 824C>A, 825A>C, 826C>T, 829A>G, 830A>G, 831G>C, 833C>A, 838T>C, 841A>G, 843C>G, 844A>T, 845C>G, 847A>C, 850A>C, 854T>G, 855G>T, 859T>C, 862A>T, 863C>A, 865T>G, 866A>C, 869C>A, 878G>T, 879T>G, 880A>G, 885T>C, 886G>T, 889A>T, 890T>A, 891G>A, 892T>G, 893T>A, 895A>G, 896A>C, 897C>G, 898A>C, 900A>G, 901T>C, 902A>C, 904A>T, 905A>G, 908T>C, 909T>A, 915C>T, 919A>T, 920A>G, 921G>C, 922C>A, 924T>C, 925C>T, 934T>C, 938T>G, 939C>A, 943A>G, 944C>A, 946A>C, 947G>A, 948A>C, 949A>T, 950C>A, 952T>A, 956T>A, 958T>C, 965A>C, 966A>G, 967A>G, 970A>G, 973T>C, 974A>C, 977G>T, 978A>C, 982T>C, 983A>T, 984A>C, 986A>T, 989A>C, 991A>T, 992A>G, 994A>G, 998C>A, 1000T>C, 1001T>A, 1007A>G, 1012A>C, 1013T>C, 1015A>T, 1018A>T, 1019A>C, 1021A>T, 1024C>T, 1033T>A, 1036A>G, 1040A>G, 1042A>C, 1045T>C, 1046T>C, 1048A>T, 1051T>C, 1054C>T, 1057T>C, 1063T>C, 1064T>C, 1070C>A, 1071A>C, 1073A>C, 1075T>A, 1076A>C, 1077G>C, 1087T>C, 1088A>C, 1090A>G, 1103A>C, 1105A>T, 1111T>A, 1112A>C, 1120C>A, 1121C>A, 1123A>G, 1126A>G, 1129G>T, 1130A>G, 1141T>C, 1145A>G, 1147A>C, 1150T>C, 1151G>A, 1153A>G, 1156C>T, 1159A>T, 1160T>G, 1161T>A, 1163C>A, 1164C>G, 1165A>G, 1167A>T, 1172G>T, 1173C>T, 1177A>C, 1178T>A, 1181G>C, 1183A>G, 1185C>A, 1186G>A, 1192T>C, 1195T>C, 1196A>G, 1197C>T, 1199A>G, 1200A>T, 1204A>T, 1207T>G, 1210A>G, 1211A>C, 1213G>C, 1218A>C, 1219G>C, 1223A>G, 1227T>C, 1232T>C, 1234G>C, 1235A>C, 1237A>G | | | | | | | | |

##### CDS

| | Begin | End | Coverage | Score | Concordance | Matches | Identities | I/D/M/F* | Stop Codons |
|---|---|---|---|---|---|---|---|---|---|
| FLA14_p101 | 5 | 379 | 98.7% | 1481 | 59.6% | 374 (99.5%) | 209 (55.6%) | 1/1/0/0 | 0 |
| Protein mutations: | E7D (121A>T), D9A (126A>C 127C>T), H10E (128C>G 130T>G), L13I (137C>A 139C>T), K15D (143A>G 145A>T), N16D (146A>G 148T>C), I17L (149A>C 151C>T), K19R (156A>G 157G>A), S23Q (167T>C 168C>A 169T>G), C24Y (171G>A 172T>C), T31A (191A>G 193T>C), K35R (203A>C 204A>G), I37V (209A>G 211T>A), F38L (214T>G), C40I (218T>A 219G>T), K44R (230A>C 231A>G 232A>T), V48A (243T>C 244C>T), A58S (272G>T 274A>C), T59M (276C>T 277A>G), Y60I (278T>A 279A>T), I62L (284A>C 286C>T), S63T (288G>C 289T>C), L65C (294T>G 295A>C), Q67V (299C>G 300A>T 301A>T), I68V (302A>G 304A>G), S71K (311T>A 312C>A 313T>A), N74E (320A>G 322T>G), I75V (323A>G 325C>A), T81S (341A>T 343C>T), L88A (362C>G 363T>C 364C>T), A90T (368G>A 370T>G), Q91E (371C>G), R92K (374C>A 375G>A 376T>G), L94I (380T>A 382G>T), Q95L (383C>T 384A>T 385A>G), T96A (386A>G), N99D (395A>G), L101I (401C>A 403A>T), Y102del (404_406delTAT), F104I (410T>A 412T>A), K105Q (413A>C 415A>G), C106A (416T>G 417G>C 418T>A), Q107H (421A>T), V108A (423T>C 424T>G), T113K (438C>A), I115V (443A>G 445C>G), K116G (446A>G 447A>G 448G>T), S118D (452A>G 453G>A 454C>T), Q119I (455C>A 456A>T 457G>C), E120R (458G>A 459A>G), T121K (462C>A 463T>G), K123E (467A>G), K123_K124insH (469_470insCAT), K124G (470A>G 471A>G), A125V (474C>T 475C>A), L128V (482T>G 484G>C), I129S (485A>T 486T>C), M135V (503A>G 505G>C), I136C (506A>T 507T>G), L138M (512C>A 514A>G), L139I (515C>A 517C>T), T140K (519C>A 520C>A), K142R (525A>G), S143T (528G>C), I144L (530A>T 532C>A), D145R (533G>C 534A>G 535T>A), K147R (540A>G 541A>G), I151M (553A>G), V152F (554G>T), I154L (560A>T 562A>G), I162S (584A>T 585T>C 586A>T), D163R (587G>A 588A>G 589T>A), N164G (590A>G 591A>G), L166K (596T>A 597T>A), K168Q (602A>C 604A>G), Q170Y (608C>T 610A>T), E174Q (620G>C 622A>G), F175L (623T>C), F176L (628T>G), E177P (629G>C 630A>C), S178P (632A>C 633G>C 634T>A), H179K (635C>A 637T>A), V180I (638G>A), I183G (647A>G 648T>G), L184V (650T>G 652A>T), L185F (655A>C), V189M (665G>A 667T>G), S191P (671T>C), R192E (674A>G 675G>A), V193A (678T>C), I194L (680A>C 682C>T), N195E (683A>G 685T>G), T196I (687C>T 688A>T), S197T (689T>A), Q198R (692C>A 693A>G 694G>A), V199K (695G>A 696T>A 697C>G), R202N (705G>A 706A>C), D203K (707G>A 709T>G), K206R (717A>G 718A>G), N211R (731A>C 732A>G), A212D (735C>A), R220K (759G>A), Y222F (765A>T), I224V (770A>G 772A>C), K227D (779A>G 781G>T), N229E (785A>G 787T>A), D230E (790T>G), F231W (792T>G 793T>G), A233L (797G>C 798C>T 799A>T), F237C (810T>G), D241E (823C>G), H242T (824C>A 825A>C 826C>T), S244A (830A>G 831G>C), L245I (833C>A), T248S (843C>G 844A>T), L249V (845C>G 847A>C), C252V (854T>G 855G>T), Q257K (869C>A), V260W (878G>T 879T>G 880A>G), M262T (885T>C 886G>T), E263D (889A>T), C264K (890T>A 891G>A 892T>G), L265M (893T>A 895A>G), T266R (896A>C 897C>G 898A>C), N267S (900A>G 901T>C), N269D (905A>G), F270H (908T>C 909T>A), A272V (915C>T), S274A (920A>G 921G>C 922C>A), I275T (924T>C 925C>T), S280D (938T>G 939C>A), Q282N (944C>A 946A>C), E283T (947G>A 948A>C 949A>T), F286I (956T>A 958T>C), K289R (965A>C 966A>G 967A>G), D293S (977G>T 978A>C), K295S (983A>T 984A>C), T296S (986A>T), I298V (992A>G 994A>G), L300I (998C>A 1000T>C), S301T (1001T>A), N303D (1007A>G), I314V (1040A>G 1042A>C), H324T (1070C>A 1071A>C), N325Q (1073A>C 1075T>A), R326P (1076A>C 1077G>C), I330L (1088A>C 1090A>G), I344V (1130A>G), I349V (1145A>G 1147A>C), E351K (1151G>A 1153A>G), E353D (1159A>T), L354E (1160T>G 1161T>A), P355R (1163C>A 1164C>G 1165A>G), K356M (1167A>T), A358F (1172G>T 1173C>T), E359D (1177A>C), L360I (1178T>A), E361Q (1181G>C 1183A>G), T362K (1185C>A 1186G>A), T366V (1196A>G 1197C>T), K367V (1199A>G 1200A>T), D369E (1207T>G), M371L (1211A>C 1213G>C), E373A (1218A>C 1219G>C), I375V (1223A>G), V376A (1227T>C), I379L (1235A>C 1237A>G) | | | | | | | | |

| | Begin | End | Coverage | Score | Concordance | Matches | Identities | I/D/M/F* | Stop Codons |
|---|---|---|---|---|---|---|---|---|---|
| **NT** | **112** | **1237** | **84.0%** | **442** | **19.8%** | **1123 (99.5%)** | **678 (60.1%)** | **3/3** | |
| Codon mutations: | GAA7GAT (121A>T), GAC9GCT (126A>C 127C>T), CAT10GAG (128C>G 130T>G), GGC12GGG (136C>G), CTC13ATT (137C>A 139C>T), AAG14AAA (142G>A), AAA15GAT (143A>G 145A>T), AAT16GAC (146A>G 148T>C), ATC17CTT (149A>C 151C>T), CTA18CTT (154A>T), AAG19AGA (156A>G 157G>A), GGT20GGG (160T>G), ATA21ATT (163A>T), TAT22TAC (166T>C), TCT23CAG (167T>C 168C>A 169T>G), TGT24TAC (171G>A 172T>C), GGA25GGG (175A>G), TTT26TTC (178T>C), CCT29CCA (187T>A), TCA30TCT (190A>T), ACT31GCC (191A>G 193T>C), AAA35CGA (203A>C 204A>G), GCC36GCT (208C>T), ATT37GTA (209A>G 211T>A), TTT38TTG (214T>G), CCA39CCT (217A>T), TGT40ATT (218T>A 219G>T), ATT41ATA (223T>A), TCT42TCG (226T>G), AAA44CGT (230A>C 231A>G 232A>T), GTG46GTC (238G>C), ATA47ATT (241A>T), GTC48GCT (243T>C 244C>T), CAA49CAG (247A>G), TCG52TCC (256G>C), GGA53GGT (259A>T), ACA54ACC (262A>C), ACA57ACC (271A>C), GCA58TCC (272G>T 274A>C), ACA59ATG (276C>T 277A>G), TAT60ATT (278T>A 279A>T), GCA61GCT (283A>T), ATC62CTT (284A>C 286C>T), AGT63ACC (288G>C 289T>C), TTA65TGC (294T>G 295A>C), CAG66CAA (298G>A), CAA67GTT (299C>G 300A>T 301A>T), ATA68GTG (302A>G 304A>G), ACT70ACC (310T>C), TCT71AAA (311T>A 312C>A 313T>A), AAT74GAG (320A>G 322T>G), ATC75GTA (323A>G 325C>A), GCT77GCA (331T>A), ATC79ATA (337C>A), ACC81TCT (341A>T 343C>T), CCA82CCT (346A>T), CGT84CGG (352T>G), GAA85GAG (355A>G), TTG86CTT (356T>C 358G>T), GCT87GCA (361T>A), CTC88GCT (362C>G 363T>C 364C>T), GCT90ACG (368G>A 370T>G), CAA91GAA (371C>G), CGT92AAG (374C>A 375G>A 376T>G), GTA93GTG (379A>G), TTG94ATT (380T>A 382G>T), CAA95TTG (383C>T 384A>T 385A>G), ACA96GCA (386A>G), GGG98GGA (394G>A), AAT99GAT (395A>G), TAT100TAC (400T>C), CTA101ATT (401C>A 403A>T), TAT102del (404_406delTAT), AAT103AAC (409T>C), TTT104ATA (410T>A 412T>A), AAA105CAG (413A>C 415A>G), TGT106GCA (416T>G 417G>C 418T>A), CAA107CAT (421A>T), GTT108GCG (423T>C 424T>G), ACA113AAA (438C>A), AGT114AGC (442T>C), ATC115GTG (443A>G 445C>G), AAG116GGT (446A>G 447A>G 448G>T), AGC118GAT (452A>G 453G>A 454C>T), CAG119ATC (455C>A 456A>T 457G>C), GAA120AGA (458G>A 459A>G), ACT121AAG (462C>A 463T>G), AAG123GAG (467A>G), AAG123_AAA124insCAT (469_470insCAT), AAA124GGA (470A>G 471A>G), GCC125GTA (474C>T 475C>A), CAG126CAA (478G>A), TTG128GTC (482T>G 484G>C), ATT129TCT (485A>T 486T>C), CGA134AGA (500C>A), ATG135GTC (503A>G 505G>C), ATT136TGT (506A>T 507T>G), GAT137GAC (511T>C), CTA138ATG (512C>A 514A>G), CTC139ATT (515C>A 517C>T), ACC140AAA (519C>A 520C>A), CGA141AGG (521C>A 523A>G), AAA142AGA (525A>G), AGT143ACT (528G>C), ATC144TTA (530A>T 532C>A), GAT145CGA (533G>C 534A>G 535T>A), AAA147AGG (540A>G 541A>G), GCA148GCC (544A>C), ATT149ATC (547T>C), AAA150AAG (550A>G), ATA151ATG (553A>G), GTT152TTT (554G>T), GTA153GTT (559A>T), ATA154TTG (560A>T 562A>G), GAT158GAC (574T>C), GAG159GAA (577G>A), TTG161CTC (581T>C 583G>C), ATA162TCT (584A>T 585T>C 586A>T), GAT163AGA (587G>A 588A>G 589T>A), AAT164GGT (590A>G 591A>G), TTT165TTC (595T>C), TTG166AAG (596T>A 597T>A), AAA168CAG (602A>C 604A>G), ATA169ATT (607A>T), CAA170TAT (608C>T 610A>T), ATT172ATA (616T>A), GAA174CAG (620G>C 622A>G), TTT175CTT (623T>C), TTT176TTG (628T>G), GAA177CCA (629G>C 630A>C), AGT178CCA (632A>C 633G>C 634T>A), CAT179AAA (635C>A 637T>A), GTC180ATC (638G>A), GTT182GTG (646T>G), ATT183GGT (647A>G 648T>G), TTA184GTT (650T>G 652A>T), TTA185TTC (655A>C), TCA186TCT (658A>T), GCA187GCC (661A>C), ACC188ACT (664C>T), GTT189ATG (665G>A 667T>G), TCA191CCA (671T>C), AGG192GAG (674A>G 675G>A), GTT193GCT (678T>C), ATC194CTT (680A>C 682C>T), AAT195GAG (683A>G 685T>G), ACA196ATT (687C>T 688A>T), TCT197ACT (689T>A), CAG198AGA (692C>A 693A>G 694G>A), GTC199AAG (695G>A 696T>A 697C>G), TTT200TTC (700T>C), AGA202AAC (705G>A 706A>C), GAT203AAG (707G>A 709T>G), GTA205GTG (715A>G), AAA206AGG (717A>G 718A>G), CTG208CTT (724G>T), GTA209GTG (727A>G), AAA210AAG (730A>G), AAT211CGT (731A>C 732A>G), GCT212GAT (735C>A), GAA213GAG (739A>G), TTG214CTC (740T>C 742G>C), ACA215ACT (745A>T), CTC216CTT (748C>T), GGC218GGT (754C>T), AGG220AAG (759G>A), TAT222TTT (765A>T), ATA224GTC (770A>G 772A>C), GTA226GTC (778A>C), AAG227GAT (779A>G 781G>T), AAA228AAG (784A>G), AAT229GAA (785A>G 787T>A), GAT230GAG (790T>G), TTT231TGG (792T>G 793T>G), AAA232AAG (796A>G), GCA233CTT (797G>C 798C>T 799A>T), TTT237TGT (810T>G), TAT240TAC (820T>C), GAC241GAG (823C>G), CAC242ACT (824C>A 825A>C 826C>T), TTA243TTG (829A>G), AGC244GCC (830A>G 831G>C), CTC245ATC (833C>A), ACT246ACC (838T>C), CAA247CAG (841A>G), ACA248AGT (843C>G 844A>T), CTA249GTC (845C>G 847A>C), ATA250ATC (850A>C), TGT252GTT (854T>G 855G>T), AAT253AAC (859T>C), ACA254ACT (862A>T), CGT255AGG (863C>A 865T>G), AGA256CGA (866A>C), CAA257AAA (869C>A), GTA260TGG (878G>T 879T>G 880A>G), ATG262ACT (885T>C 886G>T), GAA263GAT (889A>T), TGT264AAG (890T>A 891G>A 892T>G), TTA265ATG (893T>A 895A>G), ACA266CGC (896A>C 897C>G 898A>C), AAT267AGC (900A>G 901T>C), AGA268CGT (902A>C 904A>T), AAT269GAT (905A>G), TTC270CAC (908T>C 909T>A), GCA272GTA (915C>T), TCA273TCT (919A>T), AGC274GCA (920A>G 921G>C 922C>A), ATC275ACT (924T>C 925C>T), GAT278GAC (934T>C), TCC280GAC (938T>G 939C>A), CAA281CAG (943A>G), CAA282AAC (944C>A 946A>C), GAA283ACT (947G>A 948A>C 949A>T), CGT284AGA (950C>A 952T>A), TTT286ATC (956T>A 958T>C), AAA289CGG (965A>C 966A>G 967A>G), GAA290GAG (970A>G), TTT291TTC (973T>C), AGA292CGA (974A>C), GAT293TCT (977G>T 978A>C), GGT294GGC (982T>C), AAA295TCA (983A>T 984A>C), ACT296TCT (986A>T), AGA297CGT (989A>C 991A>T), ATA298GTG (992A>G 994A>G), CTT300ATC (998C>A 1000T>C), TCA301ACA (1001T>A), AAT303GAT (1007A>G), CTA304CTC (1012A>C), TTA305CTT (1013T>C 1015A>T), GCA306GCT (1018A>T), AGA307CGT (1019A>C 1021A>T), GGC308GGT (1024C>T), GTT311GTA (1033T>A), CAA312CAG (1036A>G), ATA314GTC (1040A>G 1042A>C), TCT315TCC (1045T>C), TTA316CTT (1046T>C 1048A>T), GTT317GTC (1051T>C), ATC318ATT (1054C>T), AAT319AAC (1057T>C), GAT321GAC (1063T>C), TTG322CTG (1064T>C), CAT324ACT (1070C>A 1071A>C), AAT325CAA (1073A>C 1075T>A), AGA326CCA (1076A>C 1077G>C), TAT329TAC (1087T>C), ATA330CTG (1088A>C 1090A>G), AGA335CGT (1103A>C 1105A>T), GGT337GGA (1111T>A), AGA338CGA (1112A>C), GGC340GGA (1120C>A), CGA341AGG (1121C>A 1123A>G), AAA342AAG (1126A>G), GGG343GGT (1129G>T), ATT344GTT (1130A>G), AAT347AAC (1141T>C), ATA349GTC (1145A>G 1147A>C), ACT350ACC (1150T>C), GAA351AAG (1151G>A 1153A>G), GAC352GAT (1156C>T), GAA353GAT (1159A>T), TTA354GAA (1160T>G 1161T>A), CCA355AGG (1163C>A 1164C>G 1165A>G), AAG356ATG (1167A>T), GCT358TTT (1172G>T 1173C>T), GAA359GAC (1177A>C), TTA360ATA (1178T>A), GAA361CAG (1181G>C 1183A>G), ACG362AAA (1185C>A 1186G>A), TAT364TAC (1192T>C), AAT365AAC (1195T>C), ACC366GTC (1196A>G 1197C>T), AAA367GTA (1199A>G 1200A>T), ATA368ATT (1204A>T), GAT369GAG (1207T>G), GAA370GAG (1210A>G), ATG371CTC (1211A>C 1213G>C), GAG373GCC (1218A>C 1219G>C), ATT375GTT (1223A>G), GTT376GCT (1227T>C), TTG378CTC (1232T>C 1234G>C), ATA379CTG (1235A>C 1237A>G) | | | | | | | | |

#### Proteins

| | Begin | End | Coverage | Score | Concordance | Matches | Identities | I/D/M/F* | Stop Codons |
|---|---|---|---|---|---|---|---|---|---|
| hypothetical protein (YP_009666148.1) | 5 | 379 | 98.7% | 1481 | 59.6% | 374 (99.5%) | 209 (55.6%) | 1/1/0/0 | 0 |
| Protein mutations: | E7D (121A>T), D9A (126A>C 127C>T), H10E (128C>G 130T>G), L13I (137C>A 139C>T), K15D (143A>G 145A>T), N16D (146A>G 148T>C), I17L (149A>C 151C>T), K19R (156A>G 157G>A), S23Q (167T>C 168C>A 169T>G), C24Y (171G>A 172T>C), T31A (191A>G 193T>C), K35R (203A>C 204A>G), I37V (209A>G 211T>A), F38L (214T>G), C40I (218T>A 219G>T), K44R (230A>C 231A>G 232A>T), V48A (243T>C 244C>T), A58S (272G>T 274A>C), T59M (276C>T 277A>G), Y60I (278T>A 279A>T), I62L (284A>C 286C>T), S63T (288G>C 289T>C), L65C (294T>G 295A>C), Q67V (299C>G 300A>T 301A>T), I68V (302A>G 304A>G), S71K (311T>A 312C>A 313T>A), N74E (320A>G 322T>G), I75V (323A>G 325C>A), T81S (341A>T 343C>T), L88A (362C>G 363T>C 364C>T), A90T (368G>A 370T>G), Q91E (371C>G), R92K (374C>A 375G>A 376T>G), L94I (380T>A 382G>T), Q95L (383C>T 384A>T 385A>G), T96A (386A>G), N99D (395A>G), L101I (401C>A 403A>T), Y102del (404_406delTAT), F104I (410T>A 412T>A), K105Q (413A>C 415A>G), C106A (416T>G 417G>C 418T>A), Q107H (421A>T), V108A (423T>C 424T>G), T113K (438C>A), I115V (443A>G 445C>G), K116G (446A>G 447A>G 448G>T), S118D (452A>G 453G>A 454C>T), Q119I (455C>A 456A>T 457G>C), E120R (458G>A 459A>G), T121K (462C>A 463T>G), K123E (467A>G), K123_K124insH (469_470insCAT), K124G (470A>G 471A>G), A125V (474C>T 475C>A), L128V (482T>G 484G>C), I129S (485A>T 486T>C), M135V (503A>G 505G>C), I136C (506A>T 507T>G), L138M (512C>A 514A>G), L139I (515C>A 517C>T), T140K (519C>A 520C>A), K142R (525A>G), S143T (528G>C), I144L (530A>T 532C>A), D145R (533G>C 534A>G 535T>A), K147R (540A>G 541A>G), I151M (553A>G), V152F (554G>T), I154L (560A>T 562A>G), I162S (584A>T 585T>C 586A>T), D163R (587G>A 588A>G 589T>A), N164G (590A>G 591A>G), L166K (596T>A 597T>A), K168Q (602A>C 604A>G), Q170Y (608C>T 610A>T), E174Q (620G>C 622A>G), F175L (623T>C), F176L (628T>G), E177P (629G>C 630A>C), S178P (632A>C 633G>C 634T>A), H179K (635C>A 637T>A), V180I (638G>A), I183G (647A>G 648T>G), L184V (650T>G 652A>T), L185F (655A>C), V189M (665G>A 667T>G), S191P (671T>C), R192E (674A>G 675G>A), V193A (678T>C), I194L (680A>C 682C>T), N195E (683A>G 685T>G), T196I (687C>T 688A>T), S197T (689T>A), Q198R (692C>A 693A>G 694G>A), V199K (695G>A 696T>A 697C>G), R202N (705G>A 706A>C), D203K (707G>A 709T>G), K206R (717A>G 718A>G), N211R (731A>C 732A>G), A212D (735C>A), R220K (759G>A), Y222F (765A>T), I224V (770A>G 772A>C), K227D (779A>G 781G>T), N229E (785A>G 787T>A), D230E (790T>G), F231W (792T>G 793T>G), A233L (797G>C 798C>T 799A>T), F237C (810T>G), D241E (823C>G), H242T (824C>A 825A>C 826C>T), S244A (830A>G 831G>C), L245I (833C>A), T248S (843C>G 844A>T), L249V (845C>G 847A>C), C252V (854T>G 855G>T), Q257K (869C>A), V260W (878G>T 879T>G 880A>G), M262T (885T>C 886G>T), E263D (889A>T), C264K (890T>A 891G>A 892T>G), L265M (893T>A 895A>G), T266R (896A>C 897C>G 898A>C), N267S (900A>G 901T>C), N269D (905A>G), F270H (908T>C 909T>A), A272V (915C>T), S274A (920A>G 921G>C 922C>A), I275T (924T>C 925C>T), S280D (938T>G 939C>A), Q282N (944C>A 946A>C), E283T (947G>A 948A>C 949A>T), F286I (956T>A 958T>C), K289R (965A>C 966A>G 967A>G), D293S (977G>T 978A>C), K295S (983A>T 984A>C), T296S (986A>T), I298V (992A>G 994A>G), L300I (998C>A 1000T>C), S301T (1001T>A), N303D (1007A>G), I314V (1040A>G 1042A>C), H324T (1070C>A 1071A>C), N325Q (1073A>C 1075T>A), R326P (1076A>C 1077G>C), I330L (1088A>C 1090A>G), I344V (1130A>G), I349V (1145A>G 1147A>C), E351K (1151G>A 1153A>G), E353D (1159A>T), L354E (1160T>G 1161T>A), P355R (1163C>A 1164C>G 1165A>G), K356M (1167A>T), A358F (1172G>T 1173C>T), E359D (1177A>C), L360I (1178T>A), E361Q (1181G>C 1183A>G), T362K (1185C>A 1186G>A), T366V (1196A>G 1197C>T), K367V (1199A>G 1200A>T), D369E (1207T>G), M371L (1211A>C 1213G>C), E373A (1218A>C 1219G>C), I375V (1223A>G), V376A (1227T>C), I379L (1235A>C 1237A>G) | | | | | | | | |

| | Begin | End | Coverage | Score | Concordance | Matches | Identities | I/D/M/F* | Stop Codons |
|---|---|---|---|---|---|---|---|---|---|
| **NT** | **112** | **1237** | **84.0%** | **442** | **19.8%** | **1123 (99.5%)** | **678 (60.1%)** | **3/3** | |
| Codon mutations: | GAA7GAT (121A>T), GAC9GCT (126A>C 127C>T), CAT10GAG (128C>G 130T>G), GGC12GGG (136C>G), CTC13ATT (137C>A 139C>T), AAG14AAA (142G>A), AAA15GAT (143A>G 145A>T), AAT16GAC (146A>G 148T>C), ATC17CTT (149A>C 151C>T), CTA18CTT (154A>T), AAG19AGA (156A>G 157G>A), GGT20GGG (160T>G), ATA21ATT (163A>T), TAT22TAC (166T>C), TCT23CAG (167T>C 168C>A 169T>G), TGT24TAC (171G>A 172T>C), GGA25GGG (175A>G), TTT26TTC (178T>C), CCT29CCA (187T>A), TCA30TCT (190A>T), ACT31GCC (191A>G 193T>C), AAA35CGA (203A>C 204A>G), GCC36GCT (208C>T), ATT37GTA (209A>G 211T>A), TTT38TTG (214T>G), CCA39CCT (217A>T), TGT40ATT (218T>A 219G>T), ATT41ATA (223T>A), TCT42TCG (226T>G), AAA44CGT (230A>C 231A>G 232A>T), GTG46GTC (238G>C), ATA47ATT (241A>T), GTC48GCT (243T>C 244C>T), CAA49CAG (247A>G), TCG52TCC (256G>C), GGA53GGT (259A>T), ACA54ACC (262A>C), ACA57ACC (271A>C), GCA58TCC (272G>T 274A>C), ACA59ATG (276C>T 277A>G), TAT60ATT (278T>A 279A>T), GCA61GCT (283A>T), ATC62CTT (284A>C 286C>T), AGT63ACC (288G>C 289T>C), TTA65TGC (294T>G 295A>C), CAG66CAA (298G>A), CAA67GTT (299C>G 300A>T 301A>T), ATA68GTG (302A>G 304A>G), ACT70ACC (310T>C), TCT71AAA (311T>A 312C>A 313T>A), AAT74GAG (320A>G 322T>G), ATC75GTA (323A>G 325C>A), GCT77GCA (331T>A), ATC79ATA (337C>A), ACC81TCT (341A>T 343C>T), CCA82CCT (346A>T), CGT84CGG (352T>G), GAA85GAG (355A>G), TTG86CTT (356T>C 358G>T), GCT87GCA (361T>A), CTC88GCT (362C>G 363T>C 364C>T), GCT90ACG (368G>A 370T>G), CAA91GAA (371C>G), CGT92AAG (374C>A 375G>A 376T>G), GTA93GTG (379A>G), TTG94ATT (380T>A 382G>T), CAA95TTG (383C>T 384A>T 385A>G), ACA96GCA (386A>G), GGG98GGA (394G>A), AAT99GAT (395A>G), TAT100TAC (400T>C), CTA101ATT (401C>A 403A>T), TAT102del (404_406delTAT), AAT103AAC (409T>C), TTT104ATA (410T>A 412T>A), AAA105CAG (413A>C 415A>G), TGT106GCA (416T>G 417G>C 418T>A), CAA107CAT (421A>T), GTT108GCG (423T>C 424T>G), ACA113AAA (438C>A), AGT114AGC (442T>C), ATC115GTG (443A>G 445C>G), AAG116GGT (446A>G 447A>G 448G>T), AGC118GAT (452A>G 453G>A 454C>T), CAG119ATC (455C>A 456A>T 457G>C), GAA120AGA (458G>A 459A>G), ACT121AAG (462C>A 463T>G), AAG123GAG (467A>G), AAG123_AAA124insCAT (469_470insCAT), AAA124GGA (470A>G 471A>G), GCC125GTA (474C>T 475C>A), CAG126CAA (478G>A), TTG128GTC (482T>G 484G>C), ATT129TCT (485A>T 486T>C), CGA134AGA (500C>A), ATG135GTC (503A>G 505G>C), ATT136TGT (506A>T 507T>G), GAT137GAC (511T>C), CTA138ATG (512C>A 514A>G), CTC139ATT (515C>A 517C>T), ACC140AAA (519C>A 520C>A), CGA141AGG (521C>A 523A>G), AAA142AGA (525A>G), AGT143ACT (528G>C), ATC144TTA (530A>T 532C>A), GAT145CGA (533G>C 534A>G 535T>A), AAA147AGG (540A>G 541A>G), GCA148GCC (544A>C), ATT149ATC (547T>C), AAA150AAG (550A>G), ATA151ATG (553A>G), GTT152TTT (554G>T), GTA153GTT (559A>T), ATA154TTG (560A>T 562A>G), GAT158GAC (574T>C), GAG159GAA (577G>A), TTG161CTC (581T>C 583G>C), ATA162TCT (584A>T 585T>C 586A>T), GAT163AGA (587G>A 588A>G 589T>A), AAT164GGT (590A>G 591A>G), TTT165TTC (595T>C), TTG166AAG (596T>A 597T>A), AAA168CAG (602A>C 604A>G), ATA169ATT (607A>T), CAA170TAT (608C>T 610A>T), ATT172ATA (616T>A), GAA174CAG (620G>C 622A>G), TTT175CTT (623T>C), TTT176TTG (628T>G), GAA177CCA (629G>C 630A>C), AGT178CCA (632A>C 633G>C 634T>A), CAT179AAA (635C>A 637T>A), GTC180ATC (638G>A), GTT182GTG (646T>G), ATT183GGT (647A>G 648T>G), TTA184GTT (650T>G 652A>T), TTA185TTC (655A>C), TCA186TCT (658A>T), GCA187GCC (661A>C), ACC188ACT (664C>T), GTT189ATG (665G>A 667T>G), TCA191CCA (671T>C), AGG192GAG (674A>G 675G>A), GTT193GCT (678T>C), ATC194CTT (680A>C 682C>T), AAT195GAG (683A>G 685T>G), ACA196ATT (687C>T 688A>T), TCT197ACT (689T>A), CAG198AGA (692C>A 693A>G 694G>A), GTC199AAG (695G>A 696T>A 697C>G), TTT200TTC (700T>C), AGA202AAC (705G>A 706A>C), GAT203AAG (707G>A 709T>G), GTA205GTG (715A>G), AAA206AGG (717A>G 718A>G), CTG208CTT (724G>T), GTA209GTG (727A>G), AAA210AAG (730A>G), AAT211CGT (731A>C 732A>G), GCT212GAT (735C>A), GAA213GAG (739A>G), TTG214CTC (740T>C 742G>C), ACA215ACT (745A>T), CTC216CTT (748C>T), GGC218GGT (754C>T), AGG220AAG (759G>A), TAT222TTT (765A>T), ATA224GTC (770A>G 772A>C), GTA226GTC (778A>C), AAG227GAT (779A>G 781G>T), AAA228AAG (784A>G), AAT229GAA (785A>G 787T>A), GAT230GAG (790T>G), TTT231TGG (792T>G 793T>G), AAA232AAG (796A>G), GCA233CTT (797G>C 798C>T 799A>T), TTT237TGT (810T>G), TAT240TAC (820T>C), GAC241GAG (823C>G), CAC242ACT (824C>A 825A>C 826C>T), TTA243TTG (829A>G), AGC244GCC (830A>G 831G>C), CTC245ATC (833C>A), ACT246ACC (838T>C), CAA247CAG (841A>G), ACA248AGT (843C>G 844A>T), CTA249GTC (845C>G 847A>C), ATA250ATC (850A>C), TGT252GTT (854T>G 855G>T), AAT253AAC (859T>C), ACA254ACT (862A>T), CGT255AGG (863C>A 865T>G), AGA256CGA (866A>C), CAA257AAA (869C>A), GTA260TGG (878G>T 879T>G 880A>G), ATG262ACT (885T>C 886G>T), GAA263GAT (889A>T), TGT264AAG (890T>A 891G>A 892T>G), TTA265ATG (893T>A 895A>G), ACA266CGC (896A>C 897C>G 898A>C), AAT267AGC (900A>G 901T>C), AGA268CGT (902A>C 904A>T), AAT269GAT (905A>G), TTC270CAC (908T>C 909T>A), GCA272GTA (915C>T), TCA273TCT (919A>T), AGC274GCA (920A>G 921G>C 922C>A), ATC275ACT (924T>C 925C>T), GAT278GAC (934T>C), TCC280GAC (938T>G 939C>A), CAA281CAG (943A>G), CAA282AAC (944C>A 946A>C), GAA283ACT (947G>A 948A>C 949A>T), CGT284AGA (950C>A 952T>A), TTT286ATC (956T>A 958T>C), AAA289CGG (965A>C 966A>G 967A>G), GAA290GAG (970A>G), TTT291TTC (973T>C), AGA292CGA (974A>C), GAT293TCT (977G>T 978A>C), GGT294GGC (982T>C), AAA295TCA (983A>T 984A>C), ACT296TCT (986A>T), AGA297CGT (989A>C 991A>T), ATA298GTG (992A>G 994A>G), CTT300ATC (998C>A 1000T>C), TCA301ACA (1001T>A), AAT303GAT (1007A>G), CTA304CTC (1012A>C), TTA305CTT (1013T>C 1015A>T), GCA306GCT (1018A>T), AGA307CGT (1019A>C 1021A>T), GGC308GGT (1024C>T), GTT311GTA (1033T>A), CAA312CAG (1036A>G), ATA314GTC (1040A>G 1042A>C), TCT315TCC (1045T>C), TTA316CTT (1046T>C 1048A>T), GTT317GTC (1051T>C), ATC318ATT (1054C>T), AAT319AAC (1057T>C), GAT321GAC (1063T>C), TTG322CTG (1064T>C), CAT324ACT (1070C>A 1071A>C), AAT325CAA (1073A>C 1075T>A), AGA326CCA (1076A>C 1077G>C), TAT329TAC (1087T>C), ATA330CTG (1088A>C 1090A>G), AGA335CGT (1103A>C 1105A>T), GGT337GGA (1111T>A), AGA338CGA (1112A>C), GGC340GGA (1120C>A), CGA341AGG (1121C>A 1123A>G), AAA342AAG (1126A>G), GGG343GGT (1129G>T), ATT344GTT (1130A>G), AAT347AAC (1141T>C), ATA349GTC (1145A>G 1147A>C), ACT350ACC (1150T>C), GAA351AAG (1151G>A 1153A>G), GAC352GAT (1156C>T), GAA353GAT (1159A>T), TTA354GAA (1160T>G 1161T>A), CCA355AGG (1163C>A 1164C>G 1165A>G), AAG356ATG (1167A>T), GCT358TTT (1172G>T 1173C>T), GAA359GAC (1177A>C), TTA360ATA (1178T>A), GAA361CAG (1181G>C 1183A>G), ACG362AAA (1185C>A 1186G>A), TAT364TAC (1192T>C), AAT365AAC (1195T>C), ACC366GTC (1196A>G 1197C>T), AAA367GTA (1199A>G 1200A>T), ATA368ATT (1204A>T), GAT369GAG (1207T>G), GAA370GAG (1210A>G), ATG371CTC (1211A>C 1213G>C), GAG373GCC (1218A>C 1219G>C), ATT375GTT (1223A>G), GTT376GCT (1227T>C), TTG378CTC (1232T>C 1234G>C), ATA379CTG (1235A>C 1237A>G) | | | | | | | | |

*: Inserts / Deletes / Misaligned / Frameshifts

#### Analysis details

This analysis was performed with panviral2.64

### NGS Details (UN24): Bracoviriform glomeratae (segment NC_043292.1)

#### Assembly

| | |
|---|---|
| **Coverage Length** | 310 (1 contig(s)) |
| **Depth Of Coverage** | 2222.6 |
| **Number Of Reads** | 5765 |
| **Reads Per Million** | 115.40 rpm (after QC) |
| **Ambiguities** | 0 |
| **Assembly Method** | de novo + reference guided assembly |
| **Consensus Caller** | Bcf Tools |

#### Coverage Map

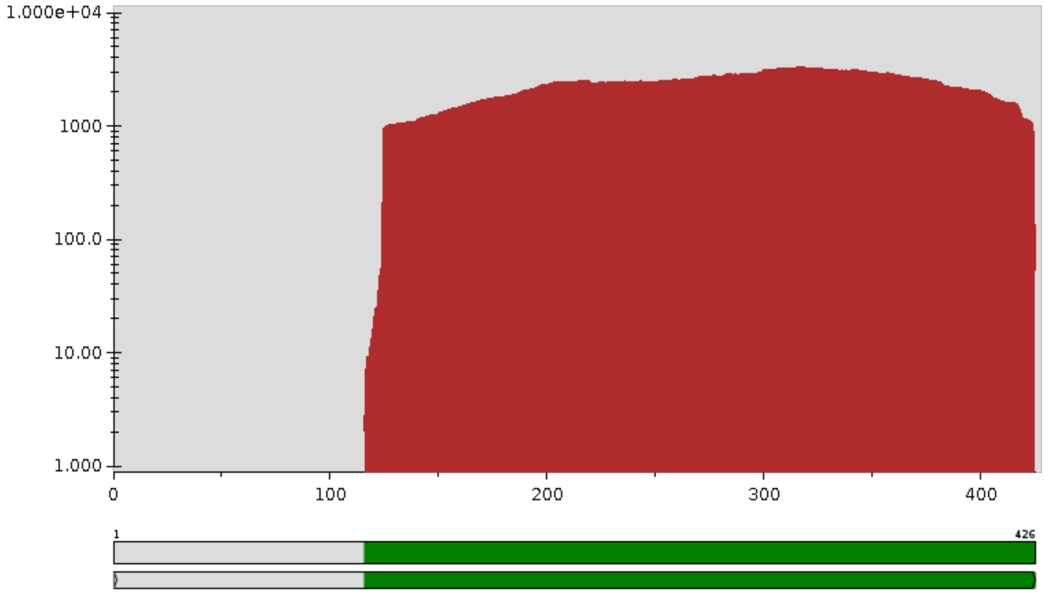

#### Assignment

| | |
|---|---|
| **Type** | Bracoviriform glomeratae (Taxonomy ID: 257816) |
| **Reference Genome** | NC_043292.1 |
| **NT Identity (%)** | 72.9032 |
| **AA Identity (%)** | 84.466 |
| **Number Of Stop Codons** | 1 |
| **Number Of CDS** | 1 |

#### Alignment

| | |
|---|---|
| **Alignment Score** | 284.0 (NT) + 567.0 (AA) = 851.0 |
| **Concordance (%)** | 66.6406 |

| **Alignment Method** | Global, seeded, nucleotide + amino acids (AGA) |
|---|---|

#### Genome Region

Sequence starts at position 117 and ends at position 426 relative to NC_043292.1 reference sequence.

#### Alignment Detailed Statistics

| | Begin | End | Coverage | Score | Concordance | Matches | Identities | I/D/M/F* | Stop Codons |
|---|---|---|---|---|---|---|---|---|---|
| **NT** | **117** | **426** | **72.8%** | **284** | **45.8%** | **310 (100%)** | **226 (72.9%)** | **0/0** | |
| Mutations: | 126G>T, 129C>A, 132A>G, 136A>G, 137C>G, 138G>C, 150A>C, 153A>G, 155C>G, 156G>A, 159C>A, 166C>A, 168T>G, 169T>C, 171T>C, 172C>A, 174C>G, 177A>G, 183T>A, 184C>A, 189T>C, 198A>G, 201A>C, 207C>T, 214A>G, 219C>T, 222T>C, 223C>A, 225T>G, 228G>T, 235C>A, 237T>G, 243A>T, 246C>G, 255C>T, 261A>G, 262T>C, 264A>G, 265G>A, 273A>G, 278T>C, 279T>C, 284A>G, 285T>A, 288T>G, 289T>C, 300C>T, 301C>T, 303C>G, 309C>T, 312T>G, 315C>T, 322G>T, 329T>C, 333T>C, 336C>T, 339A>G, 345A>C, 347A>G, 348A>G, 349C>A, 351T>A, 360C>T, 363C>T, 366C>T, 375C>T, 378C>T, 384A>C, 385T>C, 387G>C, 390A>G, 391C>A, 392A>G, 393T>A, 394A>C, 399C>A, 400C>A, 402T>G, 404T>C, 405G>T, 406A>C, 414T>A, 416A>T, 420A>T | | | | | | | | |
| **CDS** | | | | | | | | | |
| FK954_p501 | 40 | 142 | 72.5% | 567 | 85.3% | 103 (100%) | 87 (84.5%) | 0/0/0/0 | 1 |
| Protein mutations: | L42F (126G>T), T46G (136A>G 137C>G 138G>C), A52G (155C>G 156G>A), Y57H (169T>C 171T>C), T72A (214A>G), V89I (265G>A), I93T (278T>C 279T>C), D95G (284A>G 285T>A), A108S (322G>T), I110T (329T>C), K116R (347A>G 348A>G), H131R (391C>A 392A>G 393T>A), K132Q (394A>C), M135T (404T>C 405G>T), I136L (406A>C), Y139F (416A>T) | | | | | | | | |
| Codon mutations: | TTG42TTT (126G>T), GGC43GGA (129C>A), AAA44AAG (132A>G), ACG46GGC (136A>G 137C>G 138G>C), GGA50GGC (150A>C), AAA51AAG (153A>G), GCG52GGA (155C>G 156G>A), GGC53GGA (159C>A), CGT56AGG (166C>A 168T>G), TAT57CAC (169T>C 171T>C), CGC58AGG (172C>A 174C>G), AAA59AAG (177A>G), CTT61CTA (183T>A), CGA62AGA (184C>A), GAT63GAC (189T>C), CAA66CAG (198A>G), GGA67GGC (201A>C), ACC69ACT (207C>T), ACT72GCT (214A>G), ATC73ATT (219C>T), CGT74CGC (222T>C), CGT75AGG (223C>A 225T>G), CTG76CTT (228G>T), CGT79AGG (235C>A 237T>G), GGA81GGT (243A>T), GTC82GTG (246C>G), ATC85ATT (255C>T), GGA87GGG (261A>G), TTA88CTG (262T>C 264A>G), GTC89ATC (265G>A), GAA91GAG (273A>G), ATT93ACC (278T>C 279T>C), GAT95GGA (284A>G 285T>A), GTT96GTG (288T>G), TTG97CTG (289T>C), TTC100TTT (300C>T), CTC101TTG (301C>T 303C>G), AAC103AAT (309C>T), GTT104GTG (312T>G), ATC105ATT (315C>T), GCT108TCT (322G>T), ATC110ACC (329T>C), TAT111TAC (333T>C), ACC112ACT (336C>T), GAA113GAG (339A>G), GCA115GCC (345A>C), AAA116AGG (347A>G 348A>G), CGT117AGA (349C>A 351T>A), GTC120GTT (360C>T), ACC121ACT (363C>T), GCC122GCT (366C>T), GTC125GTT (375C>T), GTC126GTT (378C>T), GCA128GCC (384A>C), TTG129CTC (385T>C 387G>C), AAA130AAG (390A>G), CAT131AGA (391C>A 392A>G 393T>A), AAA132CAA (394A>C), GGC133GGA (399C>A), CGT134AGG (400C>A 402T>G), ATG135ACT (404T>C 405G>T), ATC136CTC (406A>C), GGT138GGA (414T>A), TAT139TTT (416A>T), GGA140GGT (420A>T) | | | | | | | | |
| **Proteins** | | | | | | | | | |
| putative histone 4 (YP_009665791.1) | 40 | 142 | 72.5% | 567 | 85.3% | 103 (100%) | 87 (84.5%) | 0/0/0/0 | 1 |
| Protein mutations: | L42F (126G>T), T46G (136A>G 137C>G 138G>C), A52G (155C>G 156G>A), Y57H (169T>C 171T>C), T72A (214A>G), V89I (265G>A), I93T (278T>C 279T>C), D95G (284A>G 285T>A), A108S (322G>T), I110T (329T>C), K116R (347A>G 348A>G), H131R (391C>A 392A>G 393T>A), K132Q (394A>C), M135T (404T>C 405G>T), I136L (406A>C), Y139F (416A>T) | | | | | | | | |
| Codon mutations: | TTG42TTT (126G>T), GGC43GGA (129C>A), AAA44AAG (132A>G), ACG46GGC (136A>G 137C>G 138G>C), GGA50GGC (150A>C), AAA51AAG (153A>G), GCG52GGA (155C>G 156G>A), GGC53GGA (159C>A), CGT56AGG (166C>A 168T>G), TAT57CAC (169T>C 171T>C), CGC58AGG (172C>A 174C>G), AAA59AAG (177A>G), CTT61CTA (183T>A), CGA62AGA (184C>A), GAT63GAC (189T>C), CAA66CAG (198A>G), GGA67GGC (201A>C), ACC69ACT (207C>T), ACT72GCT (214A>G), ATC73ATT (219C>T), CGT74CGC (222T>C), CGT75AGG (223C>A 225T>G), CTG76CTT (228G>T), CGT79AGG (235C>A 237T>G), GGA81GGT (243A>T), GTC82GTG (246C>G), ATC85ATT (255C>T), GGA87GGG (261A>G), TTA88CTG (262T>C 264A>G), GTC89ATC (265G>A), GAA91GAG (273A>G), ATT93ACC (278T>C 279T>C), GAT95GGA (284A>G 285T>A), GTT96GTG (288T>G), TTG97CTG (289T>C), TTC100TTT (300C>T), CTC101TTG (301C>T 303C>G), AAC103AAT (309C>T), GTT104GTG (312T>G), ATC105ATT (315C>T), GCT108TCT (322G>T), ATC110ACC (329T>C), TAT111TAC (333T>C), ACC112ACT (336C>T), GAA113GAG (339A>G), GCA115GCC (345A>C), AAA116AGG (347A>G 348A>G), CGT117AGA (349C>A 351T>A), GTC120GTT (360C>T), ACC121ACT (363C>T), GCC122GCT (366C>T), GTC125GTT (375C>T), GTC126GTT (378C>T), GCA128GCC (384A>C), TTG129CTC (385T>C 387G>C), AAA130AAG (390A>G), CAT131AGA (391C>A 392A>G 393T>A), AAA132CAA (394A>C), GGC133GGA (399C>A), CGT134AGG (400C>A 402T>G), ATG135ACT (404T>C 405G>T), ATC136CTC (406A>C), GGT138GGA (414T>A), TAT139TTT (416A>T), GGA140GGT (420A>T) | | | | | | | | |

*: Inserts / Deletes / Misaligned / Frameshifts

#### Analysis details

This analysis was performed with panviral2.64

#### NGS Details (UN24): Duamitovirus soch1

##### Assembly

| | |
|---|---|
| **Coverage Length** | 1876 (2 contig(s)) |
| **Depth Of Coverage** | 27.9 |
| **Number Of Reads** | 434 |
| **Reads Per Million** | 8.69 rpm (after QC) |
| **Ambiguities** | 0 |
| **Assembly Method** | de novo + reference guided assembly |
| **Consensus Caller** | Bcf Tools |

##### Coverage Map

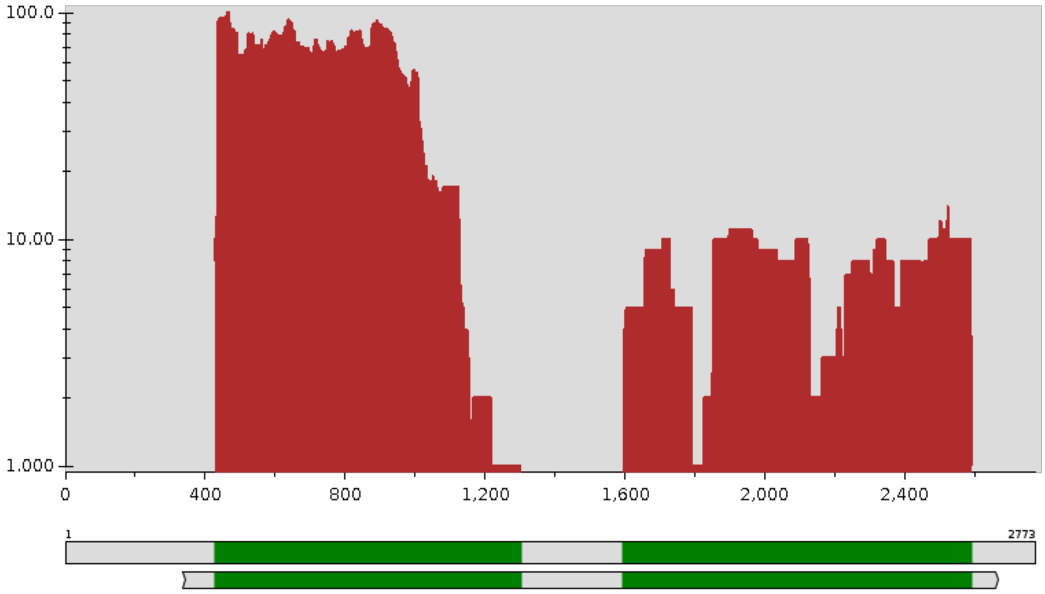

##### Assignment

| | |
|---|---|
| **Type** | Duamitovirus soch1 (Taxonomy ID: 2955838) |
| **Reference Genome** | NC_076524.1 |
| **NT Identity (%)** | 68.6065 |
| **AA Identity (%)** | 70.1278 |
| **Number Of Stop Codons** | 3 |
| **Number Of CDS** | 1 |

##### Alignment

| | |
|---|---|
| **Alignment Score** | 1364.0 (NT) + 2912.0 (AA) = 4276.0 |
| **Concordance (%)** | 53.93 |

| Alignment Method | Global, seeded, nucleotide + amino acids (AGA) |
|---|---|

#### Genome Region

Sequence starts at position 428 and ends at position 2593 relative to NC_076524.1 reference sequence.

#### Alignment Detailed Statistics

| | Begin | End | Coverage | Score | Concordance | Matches | Identities | I/D/M/F* | Stop Codons |
|---|---|---|---|---|---|---|---|---|---|
| **NT** | **428** | **2593** | **67.7%** | **1364** | **36.6%** | **1872 (99.7%)** | **1285 (68.5%)** | **1/4** | |
| Mutations: | 437A>G, 441A>C, 443C>T, 447C>T, 459C>T, 466C>T, 470C>T, 471G>A, 472C>T, 473T>G, 477T>A, 480A>C, 481C>A, 482A>T, 485A>G, 489A>G, 494T>C, 497T>A, 500T>G, 503A>C, 506T>C, 509C>T, 512T>C, 513G>A, 516T>A, 517C>A, 518T>A, 520A>C, 524G>T, 525A>G, 526C>T, 527T>G, 529C>A, 530C>A, 531T>C, 533A>G, 534C>A, 535G>A, 536C>A, 539G>A, 540A>C, 541A>G, 542G>T, 545G>A, 548C>T, 550A>T, 556T>A, 557C>T, 560A>T, 569C>T, 575G>A, 581C>T, 582G>A, 587A>T, 589G>C, 591T>C, 593A>T, 599A>T, 602C>T, 612T>A, 613C>G, 614C>T, 616T>A, 618C>T, 619G>C, 620A>T, 624G>A, 626A>T, 629A>C, 630C>A, 632A>C, 633C>A, 635T>G, 638G>T, 641G>T, 644T>A, 647G>T, 651C>T, 653C>G, 656A>C, 657A>C, 661C>G, 662G>T, 665T>A, 668C>T, 672C>A, 673G>A, 674T>G, 680C>A, 683C>T, 684A>G, 685A>C, 686A>C, 692G>A, 695T>A, 699G>C, 700C>A, 701A>C, 702A>G, 705C>A, 707T>G, 708A>G, 709G>C, 710G>A, 713A>G, 714G>T, 715A>C, 719A>T, 722C>T, 737T>C, 746C>T, 752A>T, 761G>A, 765G>T, 767T>A, 770G>A, 771G>T, 772T>A, 773T>G, 774T>A, 776G>T, 777A>G, 778A>C, 779A>G, 782T>G, 796G>A, 797T>G, 803A>C, 809T>C, 810G>C, 815C>A, 818A>T, 821T>G, 824C>T, 827A>T, 831C>T, 833T>A, 841G>A, 842G>T, 850T>G, 851A>T, 857G>T, 860G>T, 862A>G, 869T>G, 875G>T, 880T>G, 881C>A, 885C>A, 887T>G, 892A>G, 905A>T, 908C>A, 909T>C, 910T>A, 916G>A, 918G>T, 919G>C, 923T>A, 926A>T, 927A>C, 929G>T, 934T>A, 935T>G, 938G>T, 942T>A, 943C>A, 948C>A, 971C>A, 972C>T, 975A>C, 976A>T, 977T>C, 983T>C, 990A>C, 992G>C, 993T>C, 994C>G, 995A>G, 997A>T, 998C>T, 999G>C, 1000G>A, 1001T>A, 1003T>A, 1004A>T, 1005G>T, 1007C>G, 1010A>G, 1016C>T, 1017A>G, 1018A>T, 1019A>T, 1022T>A, 1025T>C, 1026C>G, 1027A>G, 1030A>C, 1031G>A, 1034T>G, 1035A>G, 1036G>C, 1044T>G, 1046C>T, 1049G>A, 1050T>G, 1051T>A, 1052T>A, 1054C>A, 1055A>C, 1061T>C, 1067T>G, 1073G>A, 1076A>G, 1083C>A, 1088G>T, 1091C>G, 1099G>T, 1104A>G, 1105C>T, 1106A>G, 1115T>G, 1121A>C, 1122T>A, 1124C>T, 1127G>A, 1130C>T, 1133C>G, 1142A>C, 1145C>T, 1148G>T, 1151G>T, 1152T>C, 1154G>T, 1160C>T, 1161A>C, 1164T>A, 1165C>T, 1166T>G, 1169C>T, 1172C>A, 1174T>A, 1175T>C, 1178T>A, 1181A>T, 1184C>T, 1185A>T, 1186G>C, 1187G>T, 1188G>A, 1189C>A, 1191G>A, 1192C>A, 1193A>T, 1196T>C, 1197_1199delGAA, 1205A>T, 1211C>T, 1212T>G, 1213T>A, 1217T>C, 1220C>T, 1223A>T, 1224C>T, 1225A>T, 1226C>A, 1232G>A, 1236A>C, 1237G>A, 1238T>A, 1241C>A, 1247A>C, 1248T>C, 1249G>T, 1250G>T, 1253T>C, 1256A>T, 1258C>G, 1265G>A, 1266A>G, 1268T>G, 1269C>G, 1271T>C, 1274T>C, 1276G>A, 1280G>A, 1283G>T, 1292T>G, 1596T>A, 1598T>A, 1607T>C, 1613G>T, 1614A>T, 1617C>T, 1619T>G, 1620G>A, 1622T>C, 1623A>G, 1624A>T, 1626A>C, 1627A>C, 1628A>T, 1632T>A, 1634A>G, 1635A>C, 1636C>T, 1637A>C, 1643A>G, 1647T>A, 1648A>G, 1655A>T, 1661T>C, 1662C>T, 1664G>A, 1667T>A, 1676T>G, 1679G>A, 1682C>T, 1684T>A, 1686C>T, 1688C>T, 1691T>A, 1694T>A, 1699C>T, 1700A>T, 1700_1701insC, 1703T>A, 1706C>T, 1707T>G, 1712G>T, 1715T>C, 1721C>T, 1727A>G, 1734T>C, 1739A>T, 1744G>A, 1745C>A, 1746A>T, 1748A>T, 1754T>C, 1758G>A, 1759G>C, 1760A>G, 1761T>A, 1765A>C, 1766C>T, 1767T>A, 1769T>C, 1772A>T, 1777G>C, 1778G>T, 1787T>C, 1788T>C, 1790A>G, 1791C>T, 1819G>C, 1820T>A, 1821A>G, 1823A>G, 1829C>G, 1830A>C, 1831A>G, 1832T>G, 1833G>C, 1840A>G, 1847A>G, 1848T>C, 1856A>G, 1859C>T, 1860A>G, 1861A>G, 1865C>T, 1867A>C, 1871T>C, 1874G>T, 1875delA, 1878C>G, 1880A>T, 1886T>G, 1887T>A, 1888T>G, 1889G>A, 1895T>C, 1898T>G, 1901T>C, 1910T>C, 1919G>T, 1925G>A, 1934T>C, 1937A>G, 1938T>G, 1939C>A, 1940A>C, 1950G>T, 1952T>C, 1953T>C, 1955A>T, 1964T>C, 1967C>T, 1976T>A, 1980T>A, 1981T>C, 1982A>T, 1985T>C, 1989C>A, 1991T>G, 1994T>A, 2000T>G, 2003A>T, 2006T>G, 2009C>T, 2010G>C, 2011C>A, 2012C>A, 2020T>C, 2028G>T, 2029A>C, 2030T>G, 2034A>T, 2035A>C, 2036C>A, 2038T>C, 2042A>G, 2043G>A, 2050A>C, 2051A>C, 2053G>T, 2054A>T, 2057G>T, 2058G>T, 2063T>G, 2069G>A, 2071A>T, 2074A>G, 2075G>A, 2078A>T, 2082G>T, 2084G>T, 2087C>T, 2090A>G, 2091T>A, 2093C>G, 2094A>T, 2099C>T, 2103A>T, 2104G>C, 2105C>T, 2108G>A, 2117G>A, 2118C>A, 2120T>A, 2123A>G, 2129T>G, 2133C>A, 2134A>G, 2135T>G, 2138T>C, 2141G>A, 2146T>G, 2148C>A, 2152C>T, 2158T>C, 2167A>T, 2168C>A, 2173G>T, 2174G>T, 2186G>A, 2191T>A, 2192G>A, 2195C>T, 2204C>T, 2208T>C, 2210G>A, 2213A>G, 2225A>G, 2230A>T, 2231C>T, 2234C>T, 2237T>G, 2239A>G, 2240A>G, 2243A>G, 2250G>C, 2251T>C, 2252T>C, 2264G>T, 2273A>T, 2274A>G, 2276G>A, 2280C>A, 2285G>A, 2288C>T, 2298G>C, 2299T>G, 2300T>G, 2303A>G, 2306A>C, 2309A>T, 2315T>C, 2322C>T, 2326G>T, 2328C>A, 2330A>T, 2336A>G, 2337G>A, 2346T>C, 2348A>T, 2349A>C, 2350A>G, 2355C>G, 2357C>T, 2358A>T, 2359A>C, 2360C>T, 2369T>C, 2372G>T, 2377G>A, 2378T>G, 2382A>T, 2383A>C, 2384C>T, 2385C>T, 2386C>T, 2387A>T, 2390T>A, 2396C>T, 2399T>C, 2400A>G, 2402G>A, 2403A>C, 2404G>A, 2405T>A, 2406C>T, 2408C>G, 2411C>G, 2414C>T, 2415T>G, 2417C>T, 2420T>C, 2423A>T, 2431C>A, 2432A>G, 2435C>A, 2441A>G, 2442A>C, 2444G>A, 2445C>T, 2446T>C, 2447C>G, 2448G>A, 2450A>T, 2451A>G, 2452C>A, 2453T>A, 2456T>C, 2457T>C, 2459T>G, 2460G>A, 2462T>A, 2465A>G, 2466A>G, 2477C>T, 2478C>T, 2489A>G, 2492T>C, 2495C>T, 2504T>A, 2510T>C, 2514A>G, 2518C>T, 2519C>G, 2525C>T, 2528T>A, 2532C>A, 2533A>G, 2536G>A, 2537G>T, 2541T>C, 2543T>C, 2551A>G, 2552C>T, 2555T>C, 2556G>T, 2557T>C, 2561T>C, 2568A>G, 2571A>C, 2572A>G, 2582A>G, 2585G>A | | | | | | | | |

##### CDS

| | Begin | End | Coverage | Score | Concordance | Matches | Identities | I/D/M/F* | Stop Codons |
|---|---|---|---|---|---|---|---|---|---|
| RdRp | 32 | 753 | 80.6% | 2912 | 66.2% | 625 (99.7%) | 439 (70.0%) | 1/1/2/2 | 3 |
| Protein mutations: | I36L (441A>C 443C>T), L38F (447C>T), A44V (466C>T), A46M (471G>A 472C>T 473T>G), L48M (477T>A), T49H (480A>C 481C>A 482A>T), I52V (489A>G), V60I (513G>A), S61K (516T>A 517C>A 518T>A), K62T (520A>C), T64V (525A>G 526C>T 527T>G), T65K (529C>A 530C>A), R67K (534C>A 535G>A 536C>A), K69R (540A>C 541A>G 542G>T), Y72F (550A>T), F74Y (556T>A 557C>T), A83T (582G>A), C85S (589G>C), F94Y (616T>A), R95S (618C>T 619G>C 620A>T), E97N (624G>A 626A>T), E98D (629A>C), P99T (630C>A 632A>C), L100M (633C>A 635T>G), S109C (661C>G 662G>T), R113K (672C>A 673G>A 674T>G), K117A (684A>G 685A>C 686A>C), A122H (699G>C 700C>A 701A>C), I123V (702A>G), R125A (708A>G 709G>C 710G>A), D127S (714G>T 715A>C), E128D (719A>T), A144S (765G>T 767T>A), V146* (771G>T 772T>A 773T>G), L147I (774T>A 776G>T), K148A (777A>G 778A>C 779A>G), S154K (796G>A 797T>G), E159Q (810G>C), P166S (831C>T 833T>A), G169D (841G>A 842G>T), L172R (850T>G 851A>T), E176G (862A>G), I178M (869T>G), F182* (880T>G 881C>A), H184K (885C>A 887T>G), Q186R (892A>G), F192H (909T>C 910T>A), R194Q (916G>A), G195S (918G>T 919G>C), M198L (927A>C 929G>T), I200K (934T>A 935T>G), S203N (942T>A 943C>A), Q205K (948C>A), N214L (975A>C 976A>T 977T>C), S220R (993T>C 994C>G 995A>G), Y221F (997A>T 998C>T), G222Q (999G>C 1000G>A 1001T>A), L223Y (1003T>A 1004A>T), A224S (1005G>T 1007C>G), K228V (1017A>G 1018A>T 1019A>T), Q231G (1026C>G 1027A>G), E232A (1030A>C 1031G>A), S234A (1035A>G 1036G>C), F237V (1044T>G 1046C>T), F239E (1050T>G 1051T>A 1052T>A), P240H (1054C>A 1055A>C), L250I (1083C>A), W255L (1099G>T), T257V (1104A>G 1105C>T 1106A>G), I260M (1115T>G), F263I (1122T>A 1124C>T), M272I (1151G>T), K276Q (1161A>C), S277M (1164T>A 1165C>T 1166T>G), F279L (1172C>A), F280Y (1174T>A 1175T>C), L282F (1181A>T), R284S (1185A>T 1186G>C 1187G>T), A285N (1188G>A 1189C>A), A286N (1191G>A 1192C>A 1193A>T), E288del (1197_1199delGAA), L290F (1205A>T), L293E (1212T>G 1213T>A), E296D (1223A>T), H297L (1224C>T 1225A>T 1226C>A), S301Q (1236A>C 1237G>A 1238T>A), W305L (1248T>C 1249G>T 1250G>T), T308S (1258C>G), S311G (1266A>G 1268T>G), P312A (1269C>G 1271T>C), R314Q (1276G>A), S421T (1596T>A 1598T>A), I427F (1614A>T), V429I (1620G>A 1622T>C), N430V (1623A>G 1624A>T), K431P (1626A>C 1627A>C 1628A>T), L433M (1632T>A 1634A>G), T434L (1635A>C 1636C>T 1637A>C), Y438S (1647T>A 1648A>G), F450Y (1684T>A), H451Y (1686C>T 1688C>T), A455V (1699C>T 1700A>T), A455_L456insX (1700_1701insC), S458A (1707T>G), L459F (1712G>T), I464M (1727A>G), S470K (1744G>A 1745C>A), K471Y (1746A>T 1748A>T), G475T (1758G>A 1759G>C 1760A>G), C476S (1761T>A), N477T (1765A>C 1766C>T), S478T (1767T>A 1769T>C), G481A (1777G>C 1778G>T), S495T (1819G>C 1820T>A), K496E (1821A>G 1823A>G), N499R (1830A>C 1831A>G 1832T>G), E500Q (1833G>C), K502R (1840A>G), N509G (1860A>G 1861A>G), K511T (1867A>C), P515A (1878C>G 1880A>T), L518R (1887T>A 1888T>G 1889G>A), D521E (1898T>G), S535D (1938T>G 1939C>A 1940A>C), D539Y (1950G>T 1952T>C), L549T (1980T>A 1981T>C 1982A>T), A559Q (2010G>C 2011C>A 2012C>A), I562T (2020T>C), D565S (2028G>T 2029A>C 2030T>G), N567S (2034A>T 2035A>C 2036C>A), I568T (2038T>C), V570I (2043G>A), Q572P (2050A>C 2051A>C), R573I (2053G>T 2054A>T), L574F (2057G>T), G575W (2058G>T), Y579F (2071A>T), K580R (2074A>G 2075G>A), A583S (2082G>T 2084G>T), F586M (2091T>A 2093C>G), T587S (2094A>T), H600R (2133C>A 2134A>G 2135T>G), V604G (2146T>G), S606L (2152C>T), L608S (2158T>C), Y611L (2167A>T 2168C>A), W613F (2173G>T 2174G>T), M619K (2191T>A 2192G>A), Y632F (2230A>T 2231C>T), K635R (2239A>G 2240A>G), V639P (2250G>C 2251T>C 2252T>C), Q643H (2264G>T), K647E (2274A>G 2276G>A), L649M (2280C>A), V655R (2298G>C 2299T>G 2300T>G), R664L (2326G>T), Q665N (2328C>A 2330A>T), E668K (2337G>A), K672R (2349A>C 2350A>G), L674V (2355C>G 2357C>T), N675S (2358A>T 2359A>C 2360C>T), C681* (2377G>A 2378T>G), N683S (2382A>T 2383A>C 2384C>T), P684F (2385C>T 2386C>T 2387A>T), D685E (2390T>A), K689E (2400A>G 2402G>A), S690Q (2403A>C 2404G>A 2405T>A), I692M (2411C>G), S694A (2415T>G 2417C>T), T699K (2431C>A 2432A>G), L704S (2445C>T 2446T>C 2447C>G), E705N (2448G>A 2450A>T), T706E (2451A>G 2452C>A 2453T>A), F708L (2457T>C 2459T>G), D709K (2460G>A 2462T>A), I711V (2466A>G), N727D (2514A>G), P728L (2518C>T 2519C>G), Q733R (2532C>A 2533A>G), W734Y (2536G>A 2537G>T), F736L (2541T>C 2543T>C), N739S (2551A>G 2552C>T), V741S (2556G>T 2557T>C), N745D (2568A>G), K746R (2571A>C 2572A>G) | | | | | | | | |

| | Begin | End | Coverage | Score | Concordance | Matches | Identities | I/D/M/F* | Stop Codons |
|---|---|---|---|---|---|---|---|---|---|
| **NT** | **428** | **2593** | **67.7%** | **1364** | **36.6%** | **1872 (99.7%)** | **1285 (68.5%)** | **1/4** | |

Codon mutations:

AAA34AAG (437A>G), ATC36CTT (441A>C 443C>T), CTT38TTT (447C>T), CTA42TTA (459C>T), GCC44GTC (466C>T), TCC45TCT (470C>T), GCT46ATG (471G>A 472C>T 473T>G), TTG48ATG (477T>A), ACA49CAT (480A>C 481C>A 482A>T), AAA50AAG (485A>G), ATC52GTC (489A>G), GCT53GCC (494T>C), CTT54CTA (497T>A), GCT55GCG (500T>G), GGA56GGC (503A>C), TGT57TGC (506T>C), GAC58GAT (509C>T), TTT59TTC (512T>C), GTA60ATA (513G>A), TCT61AAA (516T>A 517C>A 518T>A), AAA62ACA (520A>C), GTG63GTT (524G>T), ACT64GTG (525A>G 526C>T 527T>G), ACC65AAA (529C>A 530C>A), TTA66CTG (531T>C 533A>G), CGC67AAA (534C>A 535G>A 536C>A), CGG68CGA (539G>A), AAG69CGT (540A>C 541A>G 542G>T), TCG70TCA (545G>A), GGC71GGT (548C>T), TAT72TTT (550A>T), TTC74TAT (556T>A 557C>T), ACA75ACT (560A>T), TAC78TAT (569C>T), AAG80AAA (575G>A), TGC82TGT (581C>T), GCA83ACA (582G>A), GTA84GTT (587A>T), TGC85TCC (589G>C), TTA86CTT (591T>C 593A>T), CGA88CGT (599A>T), TAC89TAT (602C>T), TCC93AGT (612T>A 613C>G 614C>T), TTT94TAT (616T>A), CGA95TCT (618C>T 619G>C 620A>T), GAA97AAT (624G>A 626A>T), GAA98GAC (629A>C), CCA99ACC (630C>A 632A>C), CTT100ATG (633C>A 635T>G), TCG101TCT (638G>T), GTG102GTT (641G>T), CCT103CCA (644T>A), GTG104GTT (647G>T), CTC106TTG (651C>T 653C>G), ACA107ACC (656A>C), AGA108CGA (657A>C), TCG109TGT (661C>G 662G>T), GGT110GGA (665T>A), ATC111ATT (668C>T), CGT113AAG (672C>A 673G>A 674T>G), ATC115ATA (680C>A), CCC116CCT (683C>T), AAA117GCC (684A>G 685A>C 686A>C), TTG119TTA (692G>A), CGT120CGA (695T>A), GCA122CAC (699G>C 700C>A 701A>C), ATA123GTA (702A>G), CGT124AGG (705C>A 707T>G), AGG125GCA (708A>G 709G>C 710G>A), AAA126AAG (713A>G), GAT127TCT (714G>T 715A>C), GAA128GAT (719A>T), CAC129CAT (722C>T), GTT134GTC (737T>C), TAC137TAT (746C>T), TCA139TCT (752A>T), GGG142GGA (761G>A), GCT144TCA (765G>T 767T>A), AAG145AAA (770G>A), GTT146TAG (771G>T 772T>A 773T>G), TTG147ATT (774T>A 776G>T), AAA148GCG (777A>G 778A>C 779A>G), GTT149GTG (782T>G), AGT154AAG (796G>A 797T>G), GCA156GCC (803A>C), TTT158TTC (809T>C), GAA159CAA (810G>C), TCC160TCA (815C>A), ATA161ATT (818A>T), ACT162ACG (821T>G), ACC163ACT (824C>T), CCA164CCT (827A>T), CCT166TCA (831C>T 833T>A), GGG169GAT (841G>A 842G>T), CTA172CGT (850T>G 851A>T), GTG174GTT (857G>T), CTG175CTT (860G>T), GAA176GGA (862A>G), ATT178ATG (869T>G), ACG180ACT (875G>T), TTC182TGA (880T>G 881C>A), CAT184AAG (885C>A 887T>G), CAA186CGA (892A>G), CTA190CTT (905A>T), CCC191CCA (908C>A), TTT192CAT (909T>C 910T>A), CGA194CAA (916G>A), GGC195TCC (918G>T 919G>C), ATT196ATA (923T>A), CCA197CCT (926A>T), ATG198CTT (927A>C 929G>T), ATT200AAG (934T>A 935T>G), GGG201GGT (938G>T), TCT203AAT (942T>A 943C>A), CAA205AAA (948C>A), CCC212CCA (971C>A), CTA213TTA (972C>T), AAT214CTC (975A>C 976A>T 977T>C), AAT216AAC (983T>C), AGG219CGC (990A>C 992G>C), TCA220CGG (993T>C 994C>G 995A>G), TAC221TTT (997A>T 998C>T), GGT222CAA (999G>C 1000G>A 1001T>A), TTA223TAT (1003T>A 1004A>T), GCC224TCG (1005G>T 1007C>G), GTA225GTG (1010A>G), GAC227GAT (1016C>T), AAA228GTT (1017A>G 1018A>T 1019A>T), GTT229GTA (1022T>A), GAT230GAC (1025T>C), CAA231GGA (1026C>G 1027A>G), GAG232GCA (1030A>C 1031G>A), GCT233GCG (1034T>G), AGT234GCT (1035A>G 1036G>C), TTC237GTT (1044T>G 1046C>T), AAG238AAA (1049G>A), TTT239GAA (1050T>G 1051T>A 1052T>A), CCA240CAC (1054C>A 1055A>C), ATT242ATC (1061T>C), GTT244GTG (1067T>G), TTG246TTA (1073G>A), AAA247AAG (1076A>G), CTA250ATA (1083C>A), GCG251GCT (1088G>T), GCC252GCG (1091C>G), TGG255TTG (1099G>T), ACA257GTG (1104A>G 1105C>T 1106A>G), ATT260ATG (1115T>G), TCA262TCC (1121A>C), TTC263ATT (1122T>A 1124C>T), CAG264CAA (1127G>A), GAC265GAT (1130C>T), GGC266GGG (1133C>G), TCA269TCC (1142A>C), CCC270CCT (1145C>T), GGG271GGT (1148G>T), ATG272ATT (1151G>T), TTG273CTT (1152T>C 1154G>T), TCC275TCT (1160C>T), AAA276CAA (1161A>C), TCT277ATG (1164T>A 1165C>T 1166T>G), GTC278GTT (1169C>T), TTC279TTA (1172C>A), TTT280TAC (1174T>A 1175T>C), CCT281CCA (1178T>A), TTA282TTT (1181A>T), GAC283GAT (1184C>T), AGG284TCT (1185A>T 1186G>C 1187G>T), GCT285AAT (1188G>A 1189C>A), GCA286AAT (1191G>A 1192C>A 1193A>T), ACT287ACC (1196T>C), GAA288del (1197_1199delGAA), TTA290TTT (1205A>T), AAC292AAT (1211C>T), TTA293GAA (1212T>G 1213T>A), TCT294TCC (1217T>C), CTC295CTT (1220C>T), GAA296GAT (1223A>T), CAC297TTA (1224C>T 1225A>T 1226C>A), GAG299GAA (1232G>A), AGT301CAA (1236A>C 1237G>A 1238T>A), GTC302GTA (1241C>A), CCA304CCC (1247A>C), TGG305CTT (1248T>C 1249G>T 1250G>T), TTT306TTC (1253T>C), TCA307TCT (1256A>T), ACC308AGC (1258C>G), TTG310TTA (1265G>A), AGT311GGG (1266A>G 1268T>G), CCT312GCC (1269C>G 1271T>C), TAT313TAC (1274T>C), CGA314CAA (1276G>A), GGG315GGA (1280G>A), GTG316GTT (1283G>T), GTT319GTG (1292T>G), TCT421ACA (1596T>A 1598T>A), CTT424CTC (1607T>C), ACG426ACT (1613G>T), ATC427TTC (1614A>T), CTT428TTG (1617C>T 1619T>G), GTT429ATC (1620G>A 1622T>C), AAC430GTC (1623A>G 1624A>T), AAA431CCT (1626A>C 1627A>C 1628A>T), TTA433ATG (1632T>A 1634A>G), ACA434CTC (1635A>C 1636C>T 1637A>C), AGA436AGG (1643A>G), TAT438AGT (1647T>A 1648A>G), ATA440ATT (1655A>T), TTT442TTC (1661T>C), CTG443TTA (1662C>T 1664G>A), ACT444ACA (1667T>A), CCT447CCG (1676T>G), TTG448TTA (1679G>A), GGC449GGT (1682C>T), TTC450TAC (1684T>A), CAC451TAT (1686C>T 1688C>T), GGT452GGA (1691T>A), TCT453TCA (1694T>A), GCA455GTT (1699C>T 1700A>T), GCA455_CTT456insC-- (1700_1701insC), CTT456CTA (1703T>A), TTC457TTT (1706C>T), TCG458GCG (1707T>G), TTG459TTT (1712G>T), TCT460TCC (1715T>C), CAC462CAT (1721C>T), ATA464ATG (1727A>G), TTA467CTA (1734T>C), GCA468GCT (1739A>T), AGC470AAA (1744G>A 1745C>A), AAA471TAT (1746A>T 1748A>T), TAT473TAC (1754T>C), GGA475ACG (1758G>A 1759G>C 1760A>G), TGT476AGT (1761T>A), AAC477ACT (1765A>C 1766C>T), TCT478ACC (1767T>A 1769T>C), CCA479CCT (1772A>T), GGG481GCT (1777G>C 1778G>T), GCT484GCC (1787T>C), TTA485CTG (1788T>C 1790A>G), CTA486TTA (1791C>T), AGT495ACA (1819G>C 1820T>A), AAA496GAG (1821A>G 1823A>G), GCC498GCG (1829C>G), AAT499CGG (1830A>C 1831A>G 1832T>G), GAG500CAG (1833G>C), AAG502AGG (1840A>G), TTA504TTG (1847A>G), TTG505CTG (1848T>C), AGA507AGG (1856A>G), CTC508CTT (1859C>T), AAT509GGT (1860A>G 1861A>G), GTC510GTT (1865C>T), AAA511ACA (1867A>C), ATT512ATC (1871T>C), TCG513TCT (1874G>T), ATA514-TA (1875delA), CCA515GCT (1878C>G 1880A>T), TCT517TCG (1886T>G), TTG518AGA (1887T>A 1888T>G 1889G>A), TCT520TCC (1895T>C), GAT521GAG (1898T>G), AAT522AAC (1901T>C), ATT525ATC (1910T>C), GCG528GCT (1919G>T), AGG530AGA (1925G>A), ACT533ACC (1934T>C), AAA534AAG (1937A>G), TCA535GAC (1938T>G 1939C>A 1940A>C), GAT539TAC (1950G>T 1952T>C), TTA540CTT (1953T>C 1955A>T), ATT543ATC (1964T>C), TCC544TCT (1967C>T), GCT547GCA (1976T>A), TTA549ACT (1980T>A 1981T>C 1982A>T), TCT550TCC (1985T>C), CGT552AGG (1989C>A 1991T>G), ACT553ACA (1994T>A), GTT555GTG (2000T>G), GGA556GGT (2003A>T), CTT557CTG (2006T>G), TGC558TGT (2009C>T), GCC559CAA (2010G>C 2011C>A 2012C>A), ATC562ACC (2020T>C), GAT565TCG (2028G>T 2029A>C 2030T>G), AAC567TCA (2034A>T 2035A>C 2036C>A), ATA568ACA (2038T>C), TCA569TCG (2042A>G), GTT570ATT (2043G>A), CAA572CCC (2050A>C 2051A>C), AGA573ATT (2053G>T 2054A>T), TTG574TTT (2057G>T), GGG575TGG (2058G>T), GGT576GGG (2063T>G), GGG578GGA (2069G>A), TAC579TTC (2071A>T), AAG580AGA (2074A>G 2075G>A), GTA581GTT (2078A>T), GCG583TCT (2082G>T 2084G>T), CGC584CGT (2087C>T), TTA585TTG (2090A>G), TTC586ATG (2091T>A 2093C>G), ACT587TCT (2094A>T), ACC588ACT (2099C>T), AGC590TCT (2103A>T 2104G>C 2105C>T), AAG591AAA (2108G>A), GAG594GAA (2117G>A), CGT595AGA (2118C>A 2120T>A), CTA596CTG (2123A>G), GCT598GCG (2129T>G), CAT600AGG (2133C>A 2134A>G 2135T>G), CTT601CTC (2138T>C), AAG602AAA (2141G>A), GTT604GGT (2146T>G), CGG605AGG (2148C>A), TCA606TTA (2152C>T), TTA608TCA (2158T>C), TAC611TTA (2167A>T 2168C>A), TGG613TTT (2173G>T 2174G>T), AGG617AGA (2186G>A), ATG619AAA (2191T>A 2192G>A), CCC620CCT (2195C>T), CCC623CCT (2204C>T), TTG625CTA (2208T>C 2210G>A), AAA626AAG (2213A>G), GTA630GTG (2225A>G), TAC632TTT (2230A>T 2231C>T), CTC633CTT (2234C>T), CTT634CTG (2237T>G), AAA635AGG (2239A>G 2240A>G), GAA636GAG (2243A>G), GTT639CCC (2250G>C 2251T>C 2252T>C), CAG643CAT (2264G>T), CCA646CCT (2273A>T), AAG647GAA (2274A>G 2276G>A), CTG649ATG (2280C>A), GTG650GTA (2285G>A), TTC651TTT (2288C>T), GTT655CGG (2298G>C 2299T>G 2300T>G), GAA656GAG (2303A>G), ATA657ATC (2306A>C), CTA658CTT (2309A>T), CGT660CGC (2315T>C), CTA663TTA (2322C>T), CGC664CTC (2326G>T), CAA665AAT (2328C>A 2330A>T), GTA667GTG (2336A>G), GAA668AAA (2337G>A), TTA671CTT (2346T>C 2348A>T), AAG672CGG (2349A>C 2350A>G), CTC674GTT (2355C>G 2357C>T), AAC675TCT (2358A>T 2359A>C 2360C>T), CAT678CAC (2369T>C), ACG679ACT (2372G>T), TGT681TAG (2377G>A 2378T>G), AAC683TCT (2382A>T 2383A>C 2384C>T), CCA684TTT (2385C>T 2386C>T 2387A>T), GAT685GAA (2390T>A), TCC687TCT (2396C>T), ATT688ATC (2399T>C), AAG689GAA (2400A>G 2402G>A), AGT690CAA (2403A>C 2404G>A 2405T>A), CTC691TTG (2406C>T 2408C>G), ATC692ATG (2411C>G), GAC693GAT (2414C>T), TCC694GCT (2415T>G 2417C>T), CCT695CCC (2420T>C), ATA696ATT (2423A>T), ACA699AAG (2431C>A 2432A>G), TCC700TCA (2435C>A), AAA702AAG (2441A>G), AGG703CGA (2442A>C 2444G>A), CTC704TCG (2445C>T 2446T>C 2447C>G), GAA705AAT (2448G>A 2450A>T), ACT706GAA (2451A>G 2452C>A 2453T>A), GAT707GAC (2456T>C), TTT708CTG (2457T>C 2459T>G), GAT709AAA (2460G>A 2462T>A), TTA710TTG (2465A>G), ATC711GTC (2466A>G), GGC714GGT (2477C>T), CTA715TTA (2478C>T), AAA718AAG (2489A>G), TGT719TGC (2492T>C), TAC720TAT (2495C>T), GGT723GGA (2504T>A), GGT725GGC (2510T>C), AAT727GAT (2514A>G), CCC728CTG (2518C>T 2519C>G), ACC730ACT (2525C>T), ACT731ACA (2528T>A), CAG733AGG (2532C>A 2533A>G), TGG734TAT (2536G>A 2537G>T), TTT736CTC (2541T>C 2543T>C), AAC739AGT (2551A>G 2552C>T), ACT740ACC (2555T>C), GTT741TCT (2556G>T 2557T>C), ATT742ATC (2561T>C), AAT745GAT (2568A>G), AAA746CGA (2571A>C 2572A>G), TTA749TTG (2582A>G), GGG750GGA (2585G>A)

#### Proteins

| | Begin | End | Coverage | Score | Concordance | Matches | Identities | I/D/M/F* | Stop Codons |
|---|---|---|---|---|---|---|---|---|---|
| RNA-dependent RNA polymerase (YP_010798874.1) | 32 | 753 | 80.6% | 2912 | 66.2% | 625 (99.7%) | 439 (70.0%) | 1/1/2/2 | 3 |

Protein mutations:

I36L (441A>C 443C>T), L38F (447C>T), A44V (466C>T), A46M (471G>A 472C>T 473T>G), L48M (477T>A), T49H (480A>C 481C>A 482A>T), I52V (489A>G), V60I (513G>A), S61K (516T>A 517C>A 518T>A), K62T (520A>C), T64V (525A>G 526C>T 527T>G), T65K (529C>A 530C>A), R67K (534C>A 535G>A 536C>A), K69R (540A>C 541A>G 542G>T), Y72F (550A>T), F74Y (556T>A 557C>T), A83T (582G>A), C85S (589G>C), F94Y (616T>A), R95S (618C>T 619G>C 620A>T), E97N (624G>A 626A>T), E98D (629A>C), P99T (630C>A 632A>C), L100M (633C>A 635T>G), S109C (661C>G 662G>T), R113K (672C>A 673G>A 674T>G), K117A (684A>G 685A>C 686A>C), A122H (699G>C 700C>A 701A>C), I123V (702A>G), R125A (708A>G 709G>C 710G>A), D127S (714G>T 715A>C), E128D (719A>T), A144S (765G>T 767T>A), V146* (771G>T 772T>A 773T>G), L147I (774T>A 776G>T), K148A (777A>G 778A>C 779A>G), S154K (796G>A 797T>G), E159Q (810G>C), P166S (831C>T 833T>A), G169D (841G>A 842G>T), L172R (850T>G 851A>T), E176G (862A>G), I178M (869T>G), F182* (880T>G 881C>A), H184K (885C>A 887T>G), Q186R (892A>G), F192H (909T>C 910T>A), R194Q (916G>A), G195S (918G>T 919G>C), M198L (927A>C 929G>T), I200K (934T>A 935T>G), S203N (942T>A 943C>A), Q205K (948C>A), N214L (975A>C 976A>T 977T>C), S220R (993T>C 994C>G 995A>G), Y221F (997A>T 998C>T), G222Q (999G>C 1000G>A 1001T>A), L223Y (1003T>A 1004A>T), A224S (1005G>T 1007C>G), K228V (1017A>G 1018A>T 1019A>T), Q231G (1026C>G 1027A>G), E232A (1030A>C 1031G>A), S234A (1035A>G 1036G>C), F237V (1044T>G 1046C>T), F239E (1050T>G 1051T>A 1052T>A), P240H (1054C>A 1055A>C), L250I (1083C>A), W255L (1099G>T), T257V (1104A>G 1105C>T 1106A>G), I260M (1115T>G), F263I (1122T>A 1124C>T), M272I (1151G>T), K276Q (1161A>C), S277M (1164T>A 1165C>T 1166T>G), F279L (1172C>A), F280Y (1174T>A 1175T>C), L282F (1181A>T), R284S (1185A>T 1186G>C 1187G>T), A285N (1188G>A 1189C>A), A286N (1191G>A 1192C>A 1193A>T), E288del (1197_1199delGAA), L290F (1205A>T), L293E (1212T>G 1213T>A), E296D (1223A>T), H297L (1224C>T 1225A>T 1226C>A), S301Q (1236A>C 1237G>A 1238T>A), W305L (1248T>C 1249G>T 1250G>T), T308S (1258C>G), S311G (1266A>G 1268T>G), P312A (1269C>G 1271T>C), R314Q (1276G>A), S421T (1596T>A 1598T>A), I427F (1614A>T), V429I (1620G>A 1622T>C), N430V (1623A>G 1624A>T), K431P (1626A>C 1627A>C 1628A>T), L433M (1632T>A 1634A>G), T434L (1635A>C 1636C>T 1637A>C), Y438S (1647T>A 1648A>G), F450Y (1684T>A), H451Y (1686C>T 1688C>T), A455V (1699C>T 1700A>T), A455_L456insX (1700_1701insC), S458A (1707T>G), L459F (1712G>T), I464M (1727A>G), S470K (1744G>A 1745C>A), K471Y (1746A>T 1748A>T), G475T (1758G>A 1759G>C 1760A>G), C476S (1761T>A), N477T (1765A>C 1766C>T), S478T (1767T>A 1769T>C), G481A (1777G>C 1778G>T), S495T (1819G>C 1820T>A), K496E (1821A>G 1823A>G), N499R (1830A>C 1831A>G 1832T>G), E500Q (1833G>C), K502R (1840A>G), N509G (1860A>G 1861A>G), K511T (1867A>C), P515A (1878C>G 1880A>T), L518R (1887T>A 1888T>G 1889G>A), D521E (1898T>G), S535D (1938T>G 1939C>A 1940A>C), D539Y (1950G>T 1952T>C), L549T (1980T>A 1981T>C 1982A>T), A559Q (2010G>C 2011C>A 2012C>A), I562T (2020T>C), D565S (2028G>T 2029A>C 2030T>G), N567S (2034A>T 2035A>C 2036C>A), I568T (2038T>C), V570I (2043G>A), Q572P (2050A>C 2051A>C), R573I (2053G>T 2054A>T), L574F (2057G>T), G575W (2058G>T), Y579F (2071A>T), K580R (2074A>G 2075G>A), A583S (2082G>T 2084G>T), F586M (2091T>A 2093C>G), T587S (2094A>T), H600R (2133C>A 2134A>G 2135T>G), V604G (2146T>G), S606L (2152C>T), L608S (2158T>C), Y611L (2167A>T 2168C>A), W613F (2173G>T 2174G>T), M619K (2191T>A 2192G>A), Y632F (2230A>T 2231C>T), K635R (2239A>G 2240A>G), V639P (2250G>C 2251T>C 2252T>C), Q643H (2264G>T), K647E (2274A>G 2276G>A), L649M (2280C>A), V655R (2298G>C 2299T>G 2300T>G), R664L (2326G>T), Q665N (2328C>A 2330A>T), E668K (2337G>A), K672R (2349A>C 2350A>G), L674V (2355C>G 2357C>T), N675S (2358A>T 2359A>C 2360C>T), C681* (2377G>A 2378T>G), N683S (2382A>T 2383A>C 2384C>T), P684F (2385C>T 2386C>T 2387A>T), D685E (2390T>A), K689E (2400A>G 2402G>A), S690Q (2403A>C 2404G>A 2405T>A), I692M (2411C>G), S694A (2415T>G 2417C>T), T699K (2431C>A 2432A>G), L704S (2445C>T 2446T>C 2447C>G), E705N (2448G>A 2450A>T), T706E (2451A>G 2452C>A 2453T>A), F708L (2457T>C 2459T>G), D709K (2460G>A 2462T>A), I711V (2466A>G), N727D (2514A>G), P728L (2518C>T 2519C>G), Q733R (2532C>A 2533A>G), W734Y (2536G>A 2537G>T), F736L (2541T>C 2543T>C), N739S (2551A>G 2552C>T), V741S (2556G>T 2557T>C), N745D (2568A>G), K746R (2571A>C 2572A>G)

| | Begin | End | Coverage | Score | Concordance | Matches | Identities | I/D/M/F* | Stop Codons |
|---|---|---|---|---|---|---|---|---|---|
| **NT** | **428** | **2593** | **67.7%** | **1364** | **36.6%** | **1872 (99.7%)** | **1285 (68.5%)** | **1/4** | |
| Codon mutations: | AAA34AAG (437A>G), ATC36CTT (441A>C 443C>T), CTT38TTT (447C>T), CTA42TTA (459C>T), GCC44GTC (466C>T), TCC45TCT (470C>T), GCT46ATG (471G>A 472C>T 473T>G), TTG48ATG (477T>A), ACA49CAT (480A>C 481C>A 482A>T), AAA50AAG (485A>G), ATC52GTC (489A>G), GCT53GCC (494T>C), CTT54CTA (497T>A), GCT55GCG (500T>G), GGA56GGC (503A>C), TGT57TGC (506T>C), GAC58GAT (509C>T), TTT59TTC (512T>C), GTA60ATA (513G>A), TCT61AAA (516T>A 517C>A 518T>A), AAA62ACA (520A>C), GTG63GTT (524G>T), ACT64GTG (525A>G 526C>T 527T>G), ACC65AAA (529C>A 530C>A), TTA66CTG (531T>C 533A>G), CGC67AAA (534C>A 535G>A 536C>A), CGG68CGA (539G>A), AAG69CGT (540A>C 541A>G 542G>T), TCG70TCA (545G>A), GGC71GGT (548C>T), TAT72TTT (550A>T), TTC74TAT (556T>A 557C>T), ACA75ACT (560A>T), TAC78TAT (569C>T), AAG80AAA (575G>A), TGC82TGT (581C>T), GCA83ACA (582G>A), GTA84GTT (587A>T), TGC85TCC (589G>C), TTA86CTT (591T>C 593A>T), CGA88CGT (599A>T), TAC89TAT (602C>T), TCC93AGT (612T>A 613C>G 614C>T), TTT94TAT (616T>A), CGA95TCT (618C>T 619G>C 620A>T), GAA97AAT (624G>A 626A>T), GAA98GAC (629A>C), CCA99ACC (630C>A 632A>C), CTT100ATG (633C>A 635T>G), TCG101TCT (638G>T), GTG102GTT (641G>T), CCT103CCA (644T>A), GTG104GTT (647G>T), CTC106TTG (651C>T 653C>G), ACA107ACC (656A>C), AGA108CGA (657A>C), TCG109TGT (661C>G 662G>T), GGT110GGA (665T>A), ATC111ATT (668C>T), CGT113AAG (672C>A 673G>A 674T>G), ATC115ATA (680C>A), CCC116CCT (683C>T), AAA117GCC (684A>G 685A>C 686A>C), TTG119TTA (692G>A), CGT120CGA (695T>A), GCA122CAC (699G>C 700C>A 701A>C), ATA123GTA (702A>G), CGT124AGG (705C>A 707T>G), AGG125GCA (708A>G 709G>C 710G>A), AAA126AAG (713A>G), GAT127TCT (714G>T 715A>C), GAA128GAT (719A>T), CAC129CAT (722C>T), GTT134GTC (737T>C), TAC137TAT (746C>T), TCA139TCT (752A>T), GGG142GGA (761G>A), GCT144TCA (765G>T 767T>A), AAG145AAA (770G>A), GTT146TAG (771G>T 772T>A 773T>G), TTG147ATT (774T>A 776G>T), AAA148GCG (777A>G 778A>C 779A>G), GTT149GTG (782T>G), AGT154AAG (796G>A 797T>G), GCA156GCC (803A>C), TTT158TTC (809T>C), GAA159CAA (810G>C), TCC160TCA (815C>A), ATA161ATT (818A>T), ACT162ACG (821T>G), ACC163ACT (824C>T), CCA164CCT (827A>T), CCT166TCA (831C>T 833T>A), GGG169GAT (841G>A 842G>T), CTA172CGT (850T>G 851A>T), GTG174GTT (857G>T), CTG175CTT (860G>T), GAA176GGA (862A>G), ATT178ATG (869T>G), ACG180ACT (875G>T), TTC182TGA (880T>G 881C>A), CAT184AAG (885C>A 887T>G), CAA186CGA (892A>G), CTA190CTT (905A>T), CCC191CCA (908C>A), TTT192CAT (909T>C 910T>A), CGA194CAA (916G>A), GGC195TCC (918G>T 919G>C), ATT196ATA (923T>A), CCA197CCT (926A>T), ATG198CTT (927A>C 929G>T), ATT200AAG (934T>A 935T>G), GGG201GGT (938G>T), TCT203AAT (942T>A 943C>A), CAA205AAA (948C>A), CCC212CCA (971C>A), CTA213TTA (972C>T), AAT214CTC (975A>C 976A>T 977T>C), AAT216AAC (983T>C), AGG219CGC (990A>C 992G>C), TCA220CGG (993T>C 994C>G 995A>G), TAC221TTT (997A>T 998C>T), GGT222CAA (999G>C 1000G>A 1001T>A), TTA223TAT (1003T>A 1004A>T), GCC224TCG (1005G>T 1007C>G), GTA225GTG (1010A>G), GAC227GAT (1016C>T), AAA228GTT (1017A>G 1018A>T 1019A>T), GTT229GTA (1022T>A), GAT230GAC (1025T>C), CAA231GGA (1026C>G 1027A>G), GAG232GCA (1030A>C 1031G>A), GCT233GCG (1034T>G), AGT234GCT (1035A>G 1036G>C), TTC237GTT (1044T>G 1046C>T), AAG238AAA (1049G>A), TTT239GAA (1050T>G 1051T>A 1052T>A), CCA240CAC (1054C>A 1055A>C), ATT242ATC (1061T>C), GTT244GTG (1067T>G), TTG246TTA (1073G>A), AAA247AAG (1076A>G), CTA250ATA (1083C>A), GCG251GCT (1088G>T), GCC252GCG (1091C>G), TGG255TTG (1099G>T), ACA257GTG (1104A>G 1105C>T 1106A>G), ATT260ATG (1115T>G), TCA262TCC (1121A>C), TTC263ATT (1122T>A 1124C>T), CAG264CAA (1127G>A), GAC265GAT (1130C>T), GGC266GGG (1133C>G), TCA269TCC (1142A>C), CCC270CCT (1145C>T), GGG271GGT (1148G>T), ATG272ATT (1151G>T), TTG273CTT (1152T>C 1154G>T), TCC275TCT (1160C>T), AAA276CAA (1161A>C), TCT277ATG (1164T>A 1165C>T 1166T>G), GTC278GTT (1169C>T), TTC279TTA (1172C>A), TTT280TAC (1174T>A 1175T>C), CCT281CCA (1178T>A), TTA282TTT (1181A>T), GAC283GAT (1184C>T), AGG284TCT (1185A>T 1186G>C 1187G>T), GCT285AAT (1188G>A 1189C>A), GCA286AAT (1191G>A 1192C>A 1193A>T), ACT287ACC (1196T>C), GAA288del (1197_1199delGAA), TTA290TTT (1205A>T), AAC292AAT (1211C>T), TTA293GAA (1212T>G 1213T>A), TCT294TCC (1217T>C), CTC295CTT (1220C>T), GAA296GAT (1223A>T), CAC297TTA (1224C>T 1225A>T 1226C>A), GAG299GAA (1232G>A), AGT301CAA (1236A>C 1237G>A 1238T>A), GTC302GTA (1241C>A), CCA304CCC (1247A>C), TGG305CTT (1248T>C 1249G>T 1250G>T), TTT306TTC (1253T>C), TCA307TCT (1256A>T), ACC308AGC (1258C>G), TTG310TTA (1265G>A), AGT311GGG (1266A>G 1268T>G), CCT312GCC (1269C>G 1271T>C), TAT313TAC (1274T>C), CGA314CAA (1276G>A), GGG315GGA (1280G>A), GTG316GTT (1283G>T), GTT319GTG (1292T>G), TCT421ACA (1596T>A 1598T>A), CTT424CTC (1607T>C), ACG426ACT (1613G>T), ATC427TTC (1614A>T), CTT428TTG (1617C>T 1619T>G), GTT429ATC (1620G>A 1622T>C), AAC430GTC (1623A>G 1624A>T), AAA431CCT (1626A>C 1627A>C 1628A>T), TTA433ATG (1632T>A 1634A>G), ACA434CTC (1635A>C 1636C>T 1637A>C), AGA436AGG (1643A>G), TAT438AGT (1647T>A 1648A>G), ATA440ATT (1655A>T), TTT442TTC (1661T>C), CTG443TTA (1662C>T 1664G>A), ACT444ACA (1667T>A), CCT447CCG (1676T>G), TTG448TTA (1679G>A), GGC449GGT (1682C>T), TTC450TAC (1684T>A), CAC451TAT (1686C>T 1688C>T), GGT452GGA (1691T>A), TCT453TCA (1694T>A), GCA455GTT (1699C>T 1700A>T), GCA455_CTT456insC-- (1700_1701insC), CTT456CTA (1703T>A), TTC457TTT (1706C>T), TCG458GCG (1707T>G), TTG459TTT (1712G>T), TCT460TCC (1715T>C), CAC462CAT (1721C>T), ATA464ATG (1727A>G), TTA467CTA (1734T>C), GCA468GCT (1739A>T), AGC470AAA (1744G>A 1745C>A), AAA471TAT (1746A>T 1748A>T), TAT473TAC (1754T>C), GGA475ACG (1758G>A 1759G>C 1760A>G), TGT476AGT (1761T>A), AAC477ACT (1765A>C 1766C>T), TCT478ACC (1767T>A 1769T>C), CCA479CCT (1772A>T), GGG481GCT (1777G>C 1778G>T), GCT484GCC (1787T>C), TTA485CTG (1788T>C 1790A>G), CTA486TTA (1791C>T), AGT495ACA (1819G>C 1820T>A), AAA496GAG (1821A>G 1823A>G), GCC498GCG (1829C>G), AAT499CGG (1830A>C 1831A>G 1832T>G), GAG500CAG (1833G>C), AAG502AGG (1840A>G), TTA504TTG (1847A>G), TTG505CTG (1848T>C), AGA507AGG (1856A>G), CTC508CTT (1859C>T), AAT509GGT (1860A>G 1861A>G), GTC510GTT (1865C>T), AAA511ACA (1867A>C), ATT512ATC (1871T>C), TCG513TCT (1874G>T), ATA514-TA (1875delA), CCA515GCT (1878C>G 1880A>T), TCT517TCG (1886T>G), TTG518AGA (1887T>A 1888T>G 1889G>A), TCT520TCC (1895T>C), GAT521GAG (1898T>G), AAT522AAC (1901T>C), ATT525ATC (1910T>C), GCG528GCT (1919G>T), AGG530AGA (1925G>A), ACT533ACC (1934T>C), AAA534AAG (1937A>G), TCA535GAC (1938T>G 1939C>A 1940A>C), GAT539TAC (1950G>T 1952T>C), TTA540CTT (1953T>C 1955A>T), ATT543ATC (1964T>C), TCC544TCT (1967C>T), GCT547GCA (1976T>A), TTA549ACT (1980T>A 1981T>C 1982A>T), TCT550TCC (1985T>C), CGT552AGG (1989C>A 1991T>G), ACT553ACA (1994T>A), GTT555GTG (2000T>G), GGA556GGT (2003A>T), CTT557CTG (2006T>G), TGC558TGT (2009C>T), GCC559CAA (2010G>C 2011C>A 2012C>A), ATC562ACC (2020T>C), GAT565TCG (2028G>T 2029A>C 2030T>G), AAC567TCA (2034A>T 2035A>C 2036C>A), ATA568ACA (2038T>C), TCA569TCG (2042A>G), GTT570ATT (2043G>A), CAA572CCC (2050A>C 2051A>C), AGA573ATT (2053G>T 2054A>T), TTG574TTT (2057G>T), GGG575TGG (2058G>T), GGT576GGG (2063T>G), GGG578GGA (2069G>A), TAC579TTC (2071A>T), AAG580AGA (2074A>G 2075G>A), GTA581GTT (2078A>T), GCG583TCT (2082G>T 2084G>T), CGC584CGT (2087C>T), TTA585TTG (2090A>G), TTC586ATG (2091T>A 2093C>G), ACT587TCT (2094A>T), ACC588ACT (2099C>T), AGC590TCT (2103A>T 2104G>C 2105C>T), AAG591AAA (2108G>A), GAG594GAA (2117G>A), CGT595AGA (2118C>A 2120T>A), CTA596CTG (2123A>G), GCT598GCG (2129T>G), CAT600AGG (2133C>A 2134A>G 2135T>G), CTT601CTC (2138T>C), AAG602AAA (2141G>A), GTT604GGT (2146T>G), CGG605AGG (2148C>A), TCA606TTA (2152C>T), TTA608TCA (2158T>C), TAC611TTA (2167A>T 2168C>A), TGG613TTT (2173G>T 2174G>T), AGG617AGA (2186G>A), ATG619AAA (2191T>A 2192G>A), CCC620CCT (2195C>T), CCC623CCT (2204C>T), TTG625CTA (2208T>C 2210G>A), AAA626AAG (2213A>G), GTA630GTG (2225A>G), TAC632TTT (2230A>T 2231C>T), CTC633CTT (2234C>T), CTT634CTG (2237T>G), AAA635AGG (2239A>G 2240A>G), GAA636GAG (2243A>G), GTT639CCC (2250G>C 2251T>C 2252T>C), CAG643CAT (2264G>T), CCA646CCT (2273A>T), AAG647GAA (2274A>G 2276G>A), CTG649ATG (2280C>A), GTG650GTA (2285G>A), TTC651TTT (2288C>T), GTT655CGG (2298G>C 2299T>G 2300T>G), GAA656GAG (2303A>G), ATA657ATC (2306A>C), CTA658CTT (2309A>T), CGT660CGC (2315T>C), CTA663TTA (2322C>T), CGC664CTC (2326G>T), CAA665AAT (2328C>A 2330A>T), GTA667GTG (2336A>G), GAA668AAA (2337G>A), TTA671CTT (2346T>C 2348A>T), AAG672CGG (2349A>C 2350A>G), CTC674GTT (2355C>G 2357C>T), AAC675TCT (2358A>T 2359A>C 2360C>T), CAT678CAC (2369T>C), ACG679ACT (2372G>T), TGT681TAG (2377G>A 2378T>G), AAC683TCT (2382A>T 2383A>C 2384C>T), CCA684TTT (2385C>T 2386C>T 2387A>T), GAT685GAA (2390T>A), TCC687TCT (2396C>T), ATT688ATC (2399T>C), AAG689GAA (2400A>G 2402G>A), AGT690CAA (2403A>C 2404G>A 2405T>A), CTC691TTG (2406C>T 2408C>G), ATC692ATG (2411C>G), GAC693GAT (2414C>T), TCC694GCT (2415T>G 2417C>T), CCT695CCC (2420T>C), ATA696ATT (2423A>T), ACA699AAG (2431C>A 2432A>G), TCC700TCA (2435C>A), AAA702AAG (2441A>G), AGG703CGA (2442A>C 2444G>A), CTC704TCG (2445C>T 2446T>C 2447C>G), GAA705AAT (2448G>A 2450A>T), ACT706GAA (2451A>G 2452C>A 2453T>A), GAT707GAC (2456T>C), TTT708CTG (2457T>C 2459T>G), GAT709AAA (2460G>A 2462T>A), TTA710TTG (2465A>G), ATC711GTC (2466A>G), GGC714GGT (2477C>T), CTA715TTA (2478C>T), AAA718AAG (2489A>G), TGT719TGC (2492T>C), TAC720TAT (2495C>T), GGT723GGA (2504T>A), GGT725GGC (2510T>C), AAT727GAT (2514A>G), CCC728CTG (2518C>T 2519C>G), ACC730ACT (2525C>T), ACT731ACA (2528T>A), CAG733AGG (2532C>A 2533A>G), TGG734TAT (2536G>A 2537G>T), TTT736CTC (2541T>C 2543T>C), AAC739AGT (2551A>G 2552C>T), ACT740ACC (2555T>C), GTT741TCT (2556G>T 2557T>C), ATT742ATC (2561T>C), AAT745GAT (2568A>G), AAA746CGA (2571A>C 2572A>G), TTA749TTG (2582A>G), GGG750GGA (2585G>A) | | | | | | | | |

*: Inserts / Deletes / Misaligned / Frameshifts

#### Analysis details

This analysis was performed with panviral2.64

#### NGS Details (UN24): Colombian datura virus

##### Assembly

| | |
|---|---|
| **Coverage Length** | 5854 (11 contig(s)) |
| **Depth Of Coverage** | 5.7 |
| **Number Of Reads** | 252 |
| **Reads Per Million** | 5.04 rpm (after QC) |
| **Ambiguities** | 0 |
| **Assembly Method** | de novo + reference guided assembly |
| **Consensus Caller** | Bcf Tools |

##### Coverage Map

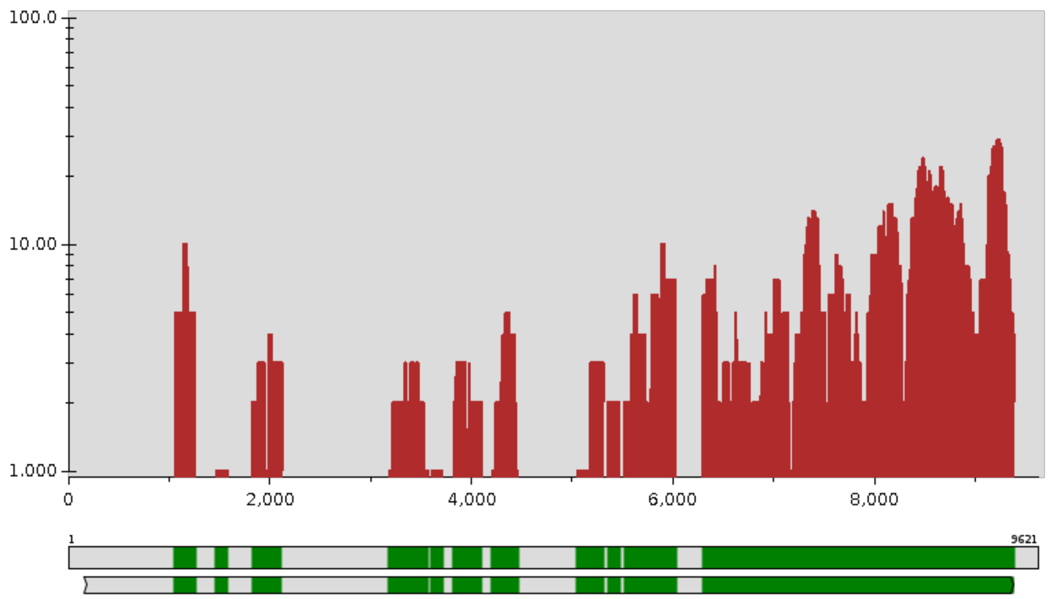

##### Assignment

| | |
|---|---|
| **Type** | Colombian datura virus (Taxonomy ID: 91613) |
| **Reference Genome** | NC_020072.1 |
| **NT Identity (%)** | 76.3496 |
| **AA Identity (%)** | 85.0 |
| **Number Of Stop Codons** | 1 |
| **Number Of CDS** | 1 |

##### Alignment

| | |
|---|---|
| **Alignment Score** | 6095.0 (NT) + 11399.0 (AA) = 17494.0 |
| **Concordance (%)** | 71.3983 |

| Alignment Method | Global, seeded, nucleotide + amino acids (AGA) |
|---|---|

#### Genome Region

Sequence starts at position 1045 and ends at position 9396 relative to NC_020072.1 reference sequence.

#### Alignment Detailed Statistics

| | Begin | End | Coverage | Score | Concordance | Matches | Identities | I/D/M/F* | Stop Codons |
|---|---|---|---|---|---|---|---|---|---|
| **NT** | **1045** | **9396** | **60.8%** | **6095** | **52.5%** | **5835 (99.7%)** | **4455 (76.1%)** | **0/19** | |

Mutations:

1045T>A, 1048T>C, 1052T>A, 1057T>C, 1063T>C, 1064C>T, 1066T>A, 1072C>T, 1079A>G, 1080A>C, 1081G>A, 1084T>C, 1087T>C, 1090A>T, 1099A>G, 1102T>A, 1105A>G, 1118C>T, 1120G>A, 1129T>C, 1132A>G, 1138C>T, 1142G>A, 1144G>A, 1147A>G, 1149G>A, 1156A>T, 1157A>T, 1159T>A, 1160G>A, 1186G>C, 1189T>A, 1192T>C, 1193T>C, 1194T>A, 1198T>C, 1201T>A, 1207C>T, 1210G>T, 1216A>G, 1219G>A, 1225G>A, 1226T>C, 1237A>G, 1243C>G, 1255C>T, 1257C>T, 1258A>T, 1463C>A, 1465G>A, 1468C>T, 1471G>A, 1477C>A, 1489T>C, 1490C>T, 1492T>G, 1504T>A, 1505C>T, 1506T>G, 1507A>T, 1508T>C, 1510A>T, 1511T>C, 1516G>A, 1520T>A, 1522T>A, 1523T>A, 1524T>C, 1526A>T, 1529A>G, 1530G>A, 1540G>A, 1546T>C, 1547G>A, 1556A>G, 1562T>C, 1564G>A, 1567T>G, 1571G>A, 1576A>T, 1577C>A, 1583T>C, 1585G>C, 1588C>T, 1823A>C, 1825A>C, 1828A>G, 1829T>C, 1831A>T, 1834A>G, 1840C>A, 1844T>C, 1846G>T, 1847A>G, 1849T>G, 1868T>A, 1870T>A, 1871C>T, 1873T>A, 1876G>A, 1888C>T, 1891G>T, 1894G>A, 1896A>G, 1900A>T, 1901A>G, 1902C>A, 1903A>G, 1904G>T, 1910C>G, 1912C>A, 1933C>T, 1937C>A, 1939T>A, 1945T>C, 1951C>T, 1952T>C, 1954A>T, 1957C>T, 1966C>T, 1975T>A, 1987C>G, 1994T>C, 1996G>T, 1999A>T, 2006T>C, 2008G>T, 2020C>A, 2026G>A, 2029T>C, 2030C>A, 2032T>A, 2041T>C, 2045T>A, 2047T>C, 2050T>A, 2053T>C, 2054G>T, 2056A>T, 2059A>G, 2062T>C, 2065A>T, 2068T>C, 2069C>T, 2071T>A, 2077G>C, 2078G>T, 2079A>C, 2081A>T, 2082T>C, 2084A>G, 2085G>A, 2086T>A, 2087G>C, 2089A>G, 2092G>A, 2097A>T, 2098C>T, 2110A>G, 2113A>T, 3172T>A, 3174G>A, 3177T>A, 3181C>A, 3184C>T, 3186A>G, 3189A>C, 3190A>G, 3199A>T, 3200G>A, 3201C>T, 3203G>C, 3208T>A, 3211T>C, 3214T>C, 3215A>G, 3232A>G, 3241G>A, 3244T>G, 3245T>C, 3249G>C, 3257C>T, 3259G>A, 3263C>T, 3265A>G, 3268C>A, 3274C>A, 3280A>T, 3286T>G, 3289T>C, 3295C>T, 3299G>A, 3301G>C, 3309G>A, 3310G>A, 3313T>C, 3320G>A, 3328G>A, 3336A>C, 3337C>A, 3338A>C, 3340G>T, 3343A>T, 3347G>A, 3348C>T, 3352C>T, 3361A>G, 3362A>G, 3367T>G, 3373C>T, 3376C>T, 3385C>T, 3391C>T, 3395A>G, 3397T>A, 3400T>G, 3401C>A, 3403T>G, 3404T>C, 3405T>C, 3406A>C, 3409G>C, 3412T>A, 3416T>G, 3418T>A, 3421A>G, 3426G>A, 3428A>G, 3429G>A, 3430C>G, 3431A>C, 3432A>G, 3433G>A, 3434G>A, 3436G>C, 3439C>T, 3440T>C, 3445T>A, 3446G>A, 3448T>C, 3450C>A, 3452C>G, 3453A>C, 3454G>T, 3457A>C, 3460C>G, 3461A>G, 3463G>T, 3467A>G, 3469T>G, 3470G>A, 3475T>C, 3481T>C, 3482C>T, 3483A>G, 3484G>C, 3492A>G, 3493A>T, 3496T>G, 3497A>C, 3498C>A, 3499T>A, 3505G>A, 3508G>A, 3511T>A, 3515C>T, 3516A>C, 3517G>T, 3523C>T, 3526C>A, 3528A>T, 3538C>A, 3544A>T, 3548G>A, 3550C>T, 3551A>G, 3556T>A, 3559G>T, 3562T>C, 3563G>A, 3565A>C, 3571T>C, 3598T>C, 3600A>T, 3602A>C, 3603A>G, 3608C>T, 3610C>G, 3612G>A, 3616A>G, 3619T>C, 3622A>G, 3625A>C, 3626T>C, 3628A>G, 3631G>T, 3634T>C, 3637T>A, 3646G>A, 3649A>G, 3650A>G, 3652T>G, 3655T>A, 3656C>A, 3658T>A, 3659T>C, 3661G>T, 3664G>A, 3665A>T, 3666G>C, 3667T>A, 3670T>A, 3671C>T, 3673T>G, 3679C>T, 3691C>A, 3694A>T, 3700T>C, 3706A>G, 3712A>C, 3715G>T, 3812C>A, 3814T>G, 3817A>T, 3820C>T, 3823C>T, 3824C>A, 3826C>G, 3832A>G, 3835A>C, 3836A>C, 3838G>A, 3844T>A, 3850T>C, 3853T>C, 3854C>A, 3862T>A, 3868C>T, 3871A>T, 3874G>A, 3883G>A, 3886T>C, 3895A>T, 3896G>A, 3897A>G, 3898G>T, 3899A>G, 3900C>T, 3901C>G, 3904G>A, 3911T>C, 3922T>A, 3931T>A, 3934T>A, 3937T>G, 3946T>A, 3961T>C, 3962T>C, 3965T>A, 3970G>A, 3973A>G, 3982C>T, 3991G>A, 4001A>C, 4006T>C, 4009G>A, 4015T>C, 4021G>T, 4023T>A, 4024T>C, 4027G>A, 4030A>G, 4031T>C, 4033G>A, 4035A>G, 4036G>A, 4039A>T, 4042G>A, 4053C>T, 4054T>A, 4057T>C, 4060A>C, 4064T>C, 4066G>A, 4072A>C, 4073A>C, 4081A>G, 4084G>C, 4093T>A, 4108C>A, 4111T>C, 4195C>T, 4201G>A, 4213T>A, 4216A>G, 4222T>A, 4231T>C, 4232T>C, 4234G>C, 4235C>T, 4237T>G, 4240T>C, 4252T>C, 4260G>A, 4265T>C, 4267G>T, 4273G>T, 4285G>A, 4288A>T, 4291C>G, 4303A>G, 4307T>A, 4309A>C, 4312A>T, 4318T>C, 4328A>G, 4330C>G, 4336T>G, 4339A>G, 4344A>G, 4348A>C, 4351T>A, 4354C>T, 4357G>A, 4366G>T, 4367T>G, 4378T>G, 4384G>A, 4387A>T, 4392C>T, 4393T>G, 4400G>A, 4405A>T, 4411A>T, 4414C>T, 4420A>T, 4421T>C, 4423A>C, 4438T>C, 4450T>G, 4453T>C, 4454T>A, 4455C>A, 4456A>T, 4457C>T, 4459T>A, 4465A>G, 4466T>C, 4468A>G, 4469C>T, 4471T>G, 5038G>C, 5045C>T, 5050A>T, 5056A>C, 5057C>A, 5059C>T, 5062G>A, 5063A>G, 5065C>G, 5068T>A, 5069G>A, 5070G>A, 5071A>T, 5072G>T, 5073T>C, 5074T>A, 5080T>C, 5083A>G, 5092T>A, 5095T>C, 5096A>C, 5098A>T, 5099A>C, 5101G>C, 5102A>C, 5103C>T, 5104A>T, 5107C>A, 5113T>A, 5116T>A, 5117G>C, 5125A>G, 5128T>C, 5137T>A, 5138G>C, 5140T>A, 5146C>T, 5147A>T, 5155A>T, 5158C>A, 5161G>A, 5166A>G, 5170T>G, 5171_5178delCACAGGGA, 5179G>C, 5182A>C, 5191A>T, 5194T>G, 5195T>A, 5196G>A, 5224T>A, 5225A>C, 5227A>C, 5228C>T, 5233T>A, 5236T>C, 5239C>T, 5245T>C, 5248A>C, 5254G>C, 5260T>C, 5266G>A, 5269G>A, 5272T>A, 5284A>T, 5287T>C, 5288G>T, 5291A>C, 5293G>A, 5302T>A, 5308A>C, 5311C>T, 5314G>A, 5356T>A, 5365T>C, 5371A>T, 5372A>T, 5373C>G, 5374A>T, 5386C>T, 5392G>T, 5393C>T, 5395A>G, 5404C>T, 5416T>C, 5419A>T, 5425A>G, 5431T>A, 5432A>C, 5434G>T, 5446G>A, 5453T>C, 5455G>T, 5515G>A, 5530G>T, 5531C>A, 5533C>A, 5534G>T, 5536C>T, 5539T>C, 5545C>A, 5555G>T, 5557A>C, 5566C>T, 5569A>G, 5573T>C, 5574C>A, 5580G>A, 5585T>G, 5588A>C, 5589A>G, 5590G>C, 5593T>C, 5597G>A, 5598G>A, 5599T>A, 5600T>C, 5602A>T, 5608A>T, 5611A>G, 5619A>G, 5620G>T, 5623T>C, 5624T>C, 5626G>C, 5629C>T, 5632T>A, 5635A>G, 5639G>A, 5640C>T, 5641T>C, 5645A>G, 5647A>T, 5648A>G, 5650A>T, 5668A>T, 5674T>A, 5681C>T, 5683T>A, 5686C>T, 5687A>T, 5688A>C, 5689G>T, 5696C>A, 5698A>G, 5699A>G, 5700G>A, 5701T>G, 5703A>G, 5704T>G, 5705T>G, 5706C>T, 5707T>A, 5709A>G, 5710A>T, 5713A>G, 5714G>A, 5715A>C, 5716A>G, 5724T>C, 5725G>C, 5728A>G, 5734G>A, 5744G>A, 5746A>C, 5750_5754delAAGTT, 5755G>A, 5758A>C, 5764A>T, 5770C>A, 5776G>A, 5779G>A, 5782A>G, 5788T>C, 5794C>T, 5797C>T, 5800A>G, 5803A>C, 5812C>A, 5815C>A, 5818C>A, 5821A>T, 5824C>T, 5827T>C, 5839A>T, 5845C>A, 5846T>A, 5847C>A, 5848A>G, 5851A>G, 5863A>G, 5869A>T, 5878A>C, 5887T>C, 5890G>A, 5896A>G, 5899G>A, 5902C>T, 5917A>T, 5926T>A, 5929C>T, 5932C>T, 5935T>C, 5938A>G, 5944T>A, 5947T>A, 5953G>A, 5959A>T, 5962A>G, 5968T>A, 5969C>A, 5970G>A, 5971C>G, 5976C>T, 5977A>G, 5983A>G, 5986T>A, 5989G>A, 5995T>A, 5998C>T, 6016G>A, 6025A>C, 6028A>G, 6031A>C, 6036C>A, 6037T>A, 6289G>A, 6295A>G, 6296G>T, 6298G>A, 6301A>T, 6302C>T, 6304T>A, 6307A>G, 6313A>G, 6316T>G, 6319A>T, 6334C>T, 6337T>A, 6340T>C, 6341G>T, 6346T>C, 6349T>C, 6352A>G, 6355G>A, 6358A>C, 6364T>C, 6373T>A, 6376G>T, 6377A>C, 6379G>A, 6382C>G, 6391T>A, 6394C>G, 6400T>A, 6406A>G, 6411T>A, 6412C>T, 6415A>T, 6418T>C, 6421G>A, 6427G>A, 6433A>G, 6436C>T, 6437A>C, 6439A>T, 6451T>G, 6455A>C, 6460A>G, 6461A>G, 6463C>T, 6472A>T, 6478T>A, 6479T>C, 6481G>T, 6490T>C, 6491C>A, 6493C>A, 6497C>T, 6499T>G, 6502G>A, 6505G>A, 6506T>C, 6508G>T, 6520G>A, 6526G>T, 6527A>G, 6528A>G, 6529C>T, 6535C>T, 6541A>T, 6542A>T, 6547T>C, 6550A>G, 6562T>C, 6565T>C, 6577T>C, 6586G>A, 6589G>A, 6592C>T, 6593C>A, 6595A>G, 6596C>A, 6598T>G, 6610A>G, 6613G>A, 6616A>G, 6619C>A, 6626G>A, 6628G>T, 6643G>A, 6646A>C, 6652C>T, 6653A>C, 6663A>T, 6667T>G, 6681C>T, 6683A>G, 6685A>T, 6688T>A, 6691A>T, 6700A>G, 6703A>T, 6718T>C, 6724C>T, 6725A>T, 6739C>G, 6740C>A, 6742G>T, 6748T>G, 6752_6756delCCATT, 6757A>C, 6763T>A, 6766G>A, 6769A>G, 6778G>C, 6784T>A, 6787A>G, 6790T>A, 6799T>A, 6802G>A, 6809T>C, 6814A>G, 6820T>C, 6823C>T, 6829C>A, 6832C>T, 6835T>C, 6841A>G, 6850A>T, 6851A>G, 6853T>C, 6854T>A, 6856G>C, 6857A>T, 6859A>G, 6861T>A, 6862G>A, 6866G>T, 6869C>A, 6871C>G, 6874A>T, 6875A>C, 6877G>T, 6880T>G, 6886G>T, 6898A>G, 6901C>T, 6904T>C, 6905T>C, 6907G>A, 6910T>C, 6919C>T, 6925C>T, 6929C>T, 6931C>G, 6934A>G, 6935A>C, 6937T>C, 6940A>G, 6955G>A, 6956C>G, 6958G>A, 6970T>A, 6972A>C, 6973G>A, 6979T>A, 6980G>A, 6982T>A, 6988T>C, 6989C>T, 6991T>G, 6992T>C, 6994T>C, 6997T>A, 7000G>A, 7005T>C, 7006T>A, 7010A>G, 7011T>C, 7016G>T, 7018C>T, 7019A>C, 7021T>A, 7024G>T, 7027A>G, 7030T>G, 7036G>T, 7038A>T, 7039C>T, 7048T>A, 7049T>C, 7051T>G, 7054C>T, 7058T>C, 7060G>T, 7063C>A, 7066A>T, 7067A>G, 7069T>C, 7070G>T, 7072T>C, 7073A>T, 7075A>C, 7078C>T, 7088T>C, 7090A>T, 7093C>T, 7099A>G, 7105T>G, 7108T>G, 7111G>A, 7117G>A, 7121A>C, 7122A>C, 7123T>A, 7124C>T, 7129C>T, 7132C>T, 7133T>G, 7135G>C, 7138T>C, 7139T>C, 7141A>G, 7144C>T, 7151G>A, 7152A>G, 7153A>T, 7156G>T, 7158A>C, 7159G>A, 7162A>G, 7163A>C, 7168T>C, 7172C>A, 7177A>T, 7178C>T, 7180A>G, 7186A>C, 7192T>C, 7194T>A, 7201C>T, 7202A>G, 7203A>C, 7205T>C, 7207G>A, 7212G>A, 7213A>G, 7216G>A, 7219A>T, 7225G>T, 7234A>T, 7240A>G, 7243C>T, 7246A>G, 7248G>A, 7253A>G, 7254C>T, 7258G>A, 7264A>T, 7265T>G, 7270G>C, 7276A>T, 7282C>T, 7285T>C, 7287A>G, 7289A>C, 7290G>A, 7291A>G, 7294A>C, 7297G>C, 7299A>C, 7304T>G, 7306G>C, 7309T>A, 7312T>C, 7318C>T, 7320A>T, 7321A>T, 7330T>C, 7351G>T, 7361A>C, 7366G>A, 7369A>C, 7372C>T, 7378T>A, 7381A>G, 7384T>C, 7396A>T, 7405T>C, 7408A>T, 7414A>G, 7423A>G, 7426A>G, 7429T>C, 7432C>T, 7438C>A, 7439A>T, 7440A>C, 7443G>C, 7444C>A, 7445T>A, 7453T>C, 7456T>C, 7457T>C, 7459G>T, 7462A>G, 7464A>G, 7465T>G, 7467T>A, 7471G>A, 7473G>T, 7474A>G, 7477C>A, 7483T>C, 7484G>A, 7486G>A, 7487A>C, 7489A>C, 7492C>T, 7494T>A, 7498T>C, 7501C>T, 7507T>C, 7510G>A, 7513G>A, 7522C>A, 7525A>C, 7528T>A, 7537A>G, 7538T>C, 7541C>A, 7543G>A, 7547A>T, 7549T>G, 7555G>A, 7558T>G, 7559G>C, 7561G>A, 7567C>T, 7570G>A, 7571A>T, 7576A>C, 7582T>C, 7591A>T, 7595G>A, 7597T>A, 7600T>C, 7604T>C, 7606A>T, 7609G>A, 7612T>A, 7615A>G, 7627G>A, 7636T>C, 7645A>G, 7648C>T, 7651T>C, 7654T>A, 7655C>T, 7657C>A, 7661T>C, 7663G>C, 7664C>A, 7672C>A, 7675A>T, 7681C>T, 7684A>T, 7693A>G, 7696T>C, 7700A>G, 7705G>T, 7711T>C, 7714A>G, 7715T>C, 7717G>T, 7723A>G, 7724G>A, 7726G>C, 7727C>T, 7729T>A, 7732T>G, 7742A>G, 7743C>T, 7750T>C, 7756T>A, 7759C>T, 7768G>A, 7774T>C, 7781C>T, 7783A>G, 7789A>T, 7795T>C, 7796T>A, 7798G>C, 7805A>C, 7807T>C, 7810T>C, 7814G>A, 7820C>T, 7822C>A, 7824A>T, 7825C>T, 7834A>G, 7843T>C, 7846T>A, 7853G>A, 7854T>C, 7855G>C, 7863A>G, 7864G>A, 7873C>T, 7876C>T, 7896T>A, 7897A>C, 7900C>A, 7915T>G, 7918A>T, 7921A>T, 7930T>C, 7936C>T, 7939T>C, 7942T>C, 7944G>C, 7948A>G, 7951A>G, 7957T>A, 7960A>T, 7963C>T, 7966T>G, 7969C>T, 7978G>T, 7984T>A, 7987C>T, 7988A>C, 7999T>C, 8002C>T, 8008G>T, 8011C>T, 8014A>T, 8015C>A, 8017T>A, 8018G>A, 8023T>C, 8026G>T, 8028A>T, 8029A>T, 8032G>T, 8033T>A, 8034C>T, 8035C>G, 8041A>G, 8044T>C, 8046A>T, 8050T>A, 8056A>G, 8063C>T, 8074T>A, 8077A>G, 8080G>A, 8084C>T, 8085A>C, 8086G>T, 8095C>T, 8099T>C, 8101G>T, 8104G>A, 8110T>C, 8113A>G, 8114G>T, 8116T>A, 8119A>G, 8124T>G, 8125A>T, 8134T>A, 8135C>A, 8137T>G, 8138A>T, 8139T>C, 8140T>A, 8146T>A, 8161A>T, 8163A>G, 8164A>G, 8167C>T, 8168C>A, 8170C>A, 8171A>C, 8172G>T, 8173T>A, 8179G>T, 8191A>C, 8194C>T, 8200T>A, 8201G>A, 8203T>G, 8206C>G, 8215T>A, 8216A>G, 8218T>C, 8221C>T, 8224A>C, 8227T>A, 8242G>A, 8243C>A, 8245G>A, 8251T>A, 8254C>A, 8260T>C, 8269C>T, 8270A>C, 8275A>T, 8278A>T, 8281A>G, 8284A>C, 8285T>A, 8291C>A, 8294C>T, 8299A>G, 8323G>A, 8341A>G, 8342C>T, 8347T>A, 8349A>T, 8353G>A, 8357A>C, 8369C>T, 8371T>G, 8383G>A, 8395C>T, 8411G>C, 8416G>A, 8418G>A, 8422T>A, 8428C>T, 8431A>G, 8434T>C, 8437A>G, 8440T>G, 8443T>A, 8444C>T, 8455A>G, 8461T>A, 8464T>C, 8465G>A, 8467T>C, 8473C>A, 8476T>G, 8479A>T, 8480T>A, 8481T>A, 8482G>A, 8485A>G, 8488G>A, 8489C>T, 8491T>G, 8499C>T, 8500A>T, 8503G>T, 8506C>T, 8507A>G, 8515T>A, 8518T>G, 8520A>C, 8524T>C, 8525G>T, 8526A>T, 8527T>G, 8528A>G, 8529G>A, 8530C>T, 8531T>G, 8533G>A, 8535C>A, 8537T>C, 8538C>A, 8539T>A, 8542A>G, 8543G>C, 8545T>C, 8548A>G, 8549T>A, 8550A>T, 8551T>A, 8552C>T, 8560T>A, 8563T>C, 8565C>A, 8578T>A, 8581T>C, 8587A>C, 8591G>A, 8596A>T, 8599G>A, 8601A>G, 8602C>T, 8606T>G, 8607C>G, 8624A>C, 8625G>T, 8626T>C, 8628T>C, 8634C>A, 8635G>A, 8636A>C, 8640G>A, 8641A>C, 8642G>C, 8644G>A, 8647A>G, 8650G>C, 8654C>T, 8655T>C, 8658T>C, 8668A>G, 8680G>T, 8686T>A, 8701C>G, 8710A>T, 8722A>G, 8728T>A, 8731T>C, 8734T>A, 8746A>C, 8761T>G, 8763A>G, 8770T>C, 8773T>C, 8775G>A, 8779G>T, 8782C>T, 8786C>T, 8791G>C, 8792A>C, 8793A>T, 8794C>G, 8803A>T, 8815T>C, 8821A>T, 8839C>A, 8851A>G, 8861A>G, 8864C>A, 8865G>A, 8866T>A, 8869A>G, 8870C>A, 8887T>C, 8890T>A, 8896A>G, 8899G>A, 8905G>A, 8911C>T, 8921T>C, 8923A>T, 8938C>T, 8959C>T, 8962A>T, 8968T>C, 8977T>C, 8992G>C, 9013A>G, 9025T>A, 9028A>G, 9040C>A, 9046T>C, 9052A>G, 9061G>A, 9064T>C, 9073T>C, 9074C>T, 9109T>C, 9124A>T, 9125C>A, 9127T>A, 9142C>T, 9156T>A, 9157G>T, 9160T>C, 9163G>T, 9181C>T, 9191G>A, 9193T>C, 9196T>C, 9197C>T, 9200C>A, 9211C>T, 9244T>A, 9256A>T, 9261A>C, 9262C>A, 9265A>G, 9268G>C, 9271G>A, 9272T>C, 9274A>C, 9283G>C, 9307A>G, 9316C>A, 9325C>T, 9332C>A, 9333A>C, 9334A>T, 9340T>G, 9346A>G, 9347G>A, 9348G>A, 9349T>C, 9364T>A, 9382G>A, 9383A>T, 9384G>T, 9387delT

**CDS**

| | Begin | End | Coverage | Score | Concordance | Matches | Identities | I/D/M/F* | Stop Codons |
|---|---|---|---|---|---|---|---|---|---|
| G357_gp1 | 298 | 3076 | 63.2% | 11399 | 85.2% | 1940 (99.8%) | 1649 (84.8%) | 0/4/3/3 | 1 |

| | Begin | End | Coverage | Score | Concordance | Matches | Identities | I/D/M/F* | Stop Codons |
|---|---|---|---|---|---|---|---|---|---|
| **NT** | **1045** | **9396** | **60.8%** | **6095** | **52.5%** | **5835 (99.7%)** | **4455 (76.1%)** | **0/19** | |
| Protein mutations: | L300M (1052T>A), K309A (1079A>G 1080A>C 1081G>A), A330T (1142G>A 1144G>A), R332K (1149G>A), E334D (1156A>T), I335L (1157A>T 1159T>A), D336N (1160G>A), L347Q (1193T>C 1194T>A), T368I (1257C>T 1258A>T), D450E (1504T>A), L451C (1505C>T 1506T>G 1507A>T), S456T (1520T>A 1522T>A), L457T (1523T>A 1524T>C), M458L (1526A>T), S459D (1529A>G 1530G>A), V465I (1547G>A), S468G (1556A>G), V473I (1571G>A), I565V (1847A>G 1849T>G), S572T (1868T>A 1870T>A), K581R (1896A>G), T583E (1901A>G 1902C>A 1903A>G), A584S (1904G>T), P586A (1910C>G 1912C>A), L626I (2030C>A 2032T>A), S631T (2045T>A 2047T>C), A634S (2054G>T 2056A>T), K641N (2077G>C), E642S (2078G>T 2079A>C), I643S (2081A>T 2082T>C), S644E (2084A>G 2085G>A 2086T>A), E645Q (2087G>C 2089A>G), Y648F (2097A>T 2098C>T), R1007K (3174G>A), F1008Y (3177T>A), K1011R (3186A>G), K1012T (3189A>C 3190A>G), A1016I (3200G>A 3201C>T), V1017L (3203G>C), S1018R (3208T>A), I1021V (3215A>G), S1032T (3249G>C), V1049I (3299G>A 3301G>C), R1052K (3309G>A 3310G>A), A1056T (3320G>A), N1061T (3336A>C 3337C>A), K1062H (3338A>C 3340G>T), A1065M (3347G>A 3348C>T), I1070V (3362A>G), N1081E (3395A>G 3397T>A), L1084P (3404T>C 3405T>C 3406A>C), S1088A (3416T>G 3418T>A), R1091K (3426G>A), S1092E (3428A>G 3429G>A 3430C>G), K1093R (3431A>C 3432A>G 3433G>A), V1094I (3434G>A 3436G>C), S1096P (3440T>C), D1098N (3446G>A 3448T>C), T1099K (3450C>A), Q1100A (3452C>G 3453A>C 3454G>T), I1102M (3460C>G), K1103D (3461A>G 3463G>T), I1105V (3467A>G 3469T>G), D1106N (3470G>A), Q1110C (3482C>T 3483A>G 3484G>C), E1113G (3492A>G 3493A>T), T1115Q (3497A>C 3498C>A 3499T>A), Q1121S (3515C>T 3516A>C 3517G>T), Y1125F (3528A>T), V1132I (3548G>A 3550C>T), I1133V (3551A>G), V1137I (3563G>A 3565A>C), Y1149F (3600A>T), K1150R (3602A>C 3603A>G), S1153N (3612G>A), T1166A (3650A>G 3652T>G), K1228Q (3836A>C 3838G>A), E1248S (3896G>A 3897A>G 3898G>T), T1249V (3899A>G 3900C>T 3901C>G), C1271S (3965T>A), F1290Y (4023T>A 4024T>C), K1294R (4035A>G 4036G>A), A1300V (4053C>T 4054T>A), R1369K (4260G>A), S1385T (4307T>A 4309A>C), I1392V (4328A>G 4330C>G), I1394M (4336T>G), N1397S (4344A>G), S1405A (4367T>G), A1413V (4392C>T 4393T>G), V1416I (4400G>A), S1434N (4454T>A 4455C>A 4456A>T), H1635N (5057C>A 5059C>T), S1637G (5063A>G 5065C>G), G1639N (5069G>A 5070G>A 5071A>T), V1640S (5072G>T 5073T>C 5074T>A), T1650L (5102A>C 5103C>T 5104A>T), H1653Q (5113T>A), E1655Q (5117G>C), A1662P (5138G>C 5140T>A), T1665S (5147A>T), E1667D (5155A>T), K1671R (5166A>G), H1673_R1674del (5171_5178delCACAGGGA), C1681N (5195T>A 5196G>A), A1712S (5288G>T), T1740C (5372A>T 5373C>G 5374A>T), L1793I (5531C>A 5533C>A), A1794S (5534G>T 5536C>T), A1801S (5555G>T 5557A>C), S1807Q (5573T>C 5574C>A), G1809E (5580G>A), S1811A (5585T>G), K1812R (5588A>C 5589A>G 5590G>C), G1815K (5597G>A 5598G>A 5599T>A), K1822S (5619A>G 5620G>T), A1829I (5639G>A 5640C>T 5641T>C), I1831V (5645A>G 5647A>T), I1832V (5648A>G 5650A>T), K1845S (5687A>T 5688A>C 5689G>T), Q1848K (5696C>A 5698A>G), S1849E (5699A>G 5700G>A 5701T>G), N1850R (5703A>G 5704T>G), S1851V (5705T>G 5706C>T 5707T>A), K1852S (5709A>G 5710A>T), E1854T (5714G>A 5715A>C 5716A>G), L1857P (5724T>C 5725G>C), A1864T (5744G>A 5746A>C), K1866del (5750_5754delAAGTT), S1898K (5846T>A 5847C>A 5848A>G), R1939K (5969C>A 5970G>A 5971C>G), A1941V (5976C>T 5977A>G), A1961E (6036C>A 6037T>A), A2048S (6296G>T 6298G>A), A2063S (6341G>T), I2076M (6382C>G), F2086Y (6411T>A 6412C>T), M2101L (6455A>C), I2103V (6461A>G 6463C>T), Q2106H (6472A>T), E2124D (6526G>T), N2125G (6527A>G 6528A>G 6529C>T), I2130L (6542A>T), L2148M (6596C>A 6598T>G), V2158I (6626G>A 6628G>T), M2167L (6653A>C), Q2170L (6663A>T), T2176I (6681C>T), I2177V (6683A>G 6685A>T), T2191S (6725A>T), Q2196N (6740C>A 6742G>T), P2200del (6752_6756delCCATT), K2208N (6778G>C), Q2232H (6850A>T), N2233D (6851A>G 6853T>C), L2234I (6854T>A 6856G>C), I2235L (6857A>T 6859A>G), M2236K (6861T>A 6862G>A), A2238S (6866G>T), L2239M (6869C>A 6871C>G), E2240D (6874A>T), M2241L (6875A>C 6877G>T), D2242E (6880T>G), I2261L (6935A>C 6937T>C), Q2268E (6956C>G 6958G>A), E2273A (6972A>C 6973G>A), V2276I (6980G>A 6982T>A), F2280L (6992T>C 6994T>C), D2281E (6997T>A), I2284T (7005T>C 7006T>A), I2286A (7010A>G 7011T>C), A2288S (7016G>T 7018C>T), N2289Q (7019A>C 7021T>A), M2290I (7024G>T), N2292K (7030T>G), Y2295F (7038A>T 7039C>T), C2299R (7049T>C 7051T>G), I2305V (7067A>G 7069T>C), A2306S (7070G>T 7072T>C), T2307S (7073A>T 7075A>C), N2323P (7121A>C 7122A>C 7123T>A), L2327V (7133T>G 7135G>C), E2333S (7151G>A 7152A>G 7153A>T), E2334D (7156G>T), E2335A (7158A>C 7159G>A), K2337Q (7163A>C), L2347Q (7194T>A), K2350A (7202A>G 7203A>C), R2353K (7212G>A 7213A>G), S2365N (7248G>A), T2367V (7253A>G 7254C>T), S2371A (7265T>G), D2378G (7287A>G), R2379Q (7289A>C 7290G>A 7291A>G), N2382T (7299A>C), L2384V (7304T>G 7306G>C), E2389V (7320A>T 7321A>T), M2403L (7361A>C), N2429S (7439A>T 7440A>C), C2430S (7443G>C 7444C>A), S2431T (7445T>A), N2437R (7464A>G 7465T>G), F2438Y (7467T>A), R2440M (7473G>T 7474A>G), E2444K (7484G>A 7486G>A), F2447Y (7494T>A), I2465L (7547A>T 7549T>G), E2469Q (7559G>C 7561G>A), T2473S (7571A>T), V2481I (7595G>A 7597T>A), Q2504K (7664C>A), R2516G (7700A>G), G2524S (7724G>A 7726G>C), T2530V (7742A>G 7743C>T), L2548I (7796T>A 7798G>C), I2551L (7805A>C 7807T>C), V2554I (7814G>A), Y2557F (7824A>T 7825C>T), V2567T (7853G>A 7854T>C 7855G>C), K2570R (7863A>G 7864G>A), I2581N (7896T>A 7897A>C), S2597T (7944G>C), I2612L (7988A>C), M2618I (8008G>T), D2622N (8018G>A), K2625I (8028A>T 8029A>T), E2626D (8032G>T), S2627M (8033T>A 8034C>T 8035C>G), Y2631F (8046A>T), I2641M (8077A>G), Q2644S (8084C>T 8085A>C 8086G>T), A2654S (8114G>T 8116T>A), L2657C (8124T>G 8125A>T), L2661M (8135C>A 8137T>G), I2662S (8138A>T 8139T>C 8140T>A), K2670R (8163A>G 8164A>G), S2673L (8171A>C 8172G>T 8173T>A), V2683M (8201G>A 8203T>G), I2688V (8216A>G 8218T>C), S2711T (8285T>A), K2732M (8349A>T), E2753Q (8411G>C), R2755K (8418G>A), A2771T (8465G>A 8467T>C), E2775D (8479A>T), L2776K (8480T>A 8481T>A 8482G>A), A2782V (8499C>T 8500A>T), I2785V (8507A>G), D2788E (8518T>G), Y2789S (8520A>C), D2791L (8525G>T 8526A>T 8527T>G), S2792D (8528A>G 8529G>A 8530C>T), L2793V (8531T>G 8533G>A), A2794E (8535C>A), S2795Q (8537T>C 8538C>A 8539T>A), D2797H (8543G>C 8545T>C), I2798M (8548A>G), Y2799I (8549T>A 8550A>T 8551T>A), L2800F (8552C>T), T2804K (8565C>A), A2813T (8591G>A), N2816S (8601A>G 8602C>T), S2818G (8606T>G 8607C>G), S2824L (8624A>C 8625G>T 8626T>C), V2825A (8628T>C), T2827K (8634C>A 8635G>A), T2828P (8636A>C), G2829D (8640G>A 8641A>C), E2830Q (8642G>C 8644G>A), K2832N (8650G>C), L2834S (8654C>T 8655T>C), V2835A (8658T>C), N2869K (8761T>G), K2870R (8763A>G), S2874N (8775G>A), N2880L (8792A>C 8793A>T 8794C>G), H2895Q (8839C>A), N2903D (8861A>G), R2904K (8864C>A 8865G>A 8866T>A), E2946D (8992G>C), M3001N (9156T>A 9157G>T), V3013I (9191G>A 9193T>C), P3015S (9197C>T), N3036T (9261A>C 9262C>A), Q3060T (9332C>A 9333A>C 9334A>T), G3065N (9347G>A 9348G>A 9349T>C) | | | | | | | | |

| | Begin | End | Coverage | Score | Concordance | Matches | Identities | I/D/M/F* | Stop Codons |
|---|---|---|---|---|---|---|---|---|---|
| NT | 1045 | 9396 | 60.8% | 6095 | 52.5% | 5835 (99.7%) | 4455 (76.1%) | 0/19 | |

Codon mutations:

CGT297..A (1045T>A), GAT298GAC (1048T>C), TTG300ATG (1052T>A), TGT301TGC (1057T>C), GAT303GAC (1063T>C), CTT304TTA (1064C>T 1066T>A), TAC306TAT (1072C>T), AAG309GCA (1079A>G 1080A>C 1081G>A), CAT310CAC (1084T>C), TTT311TTC (1087T>C), TCA312TCT (1090A>T), CAA315CAG (1099A>G), ATT316ATA (1102T>A), TCA317TCG (1105A>G), CTG322TTA (1118C>T 1120G>A), TGT325TGC (1129T>C), AAA326AAG (1132A>G), TAC328TAT (1138C>T), GCG330ACA (1142G>A 1144G>A), GGA331GGG (1147A>G), AGA332AAA (1149G>A), GAA334GAT (1156A>T), ATT335TTA (1157A>T 1159T>A), GAT336AAT (1160G>A), TCG344TCC (1186G>C), GTT345GTA (1189T>A), TCT346TCC (1192T>C), TTA347CAA (1193T>C 1194T>A), TGT348TGC (1198T>C), GGT349GGA (1201T>A), GTC351GTT (1207C>T), GCG352GCT (1210G>T), TTA354TTG (1216A>G), GTG355GTA (1219G>A), CAG357CAA (1225G>A), TTA358CTA (1226T>C), CCA361CCG (1237A>G), GGC363GGG (1243C>G), TGC367TGT (1255C>T), ACA368ATT (1257C>T 1258A>T), CGG437AGA (1463C>A 1465G>A), AAC438AAT (1468C>T), AAG439AAA (1471G>A), CCC441CCA (1477C>A), GTT445GTC (1489T>C), CTT446TTG (1490C>T 1492T>G), GAT450GAA (1504T>A), CTA451TGT (1505C>T 1506T>G 1507A>T), TTA452CTT (1508T>C 1510A>T), TTG453CTG (1511T>C), AAG454AAA (1516G>A), TCT456ACA (1520T>A 1522T>A), TTA457ACA (1523T>A 1524T>C), ATG458TTG (1526A>T), AGT459GAT (1529A>G 1530G>A), GAG462GAA (1540G>A), GGT464GGC (1546T>C), GTA465ATA (1547G>A), AGT468GGT (1556A>G), TTG470CTA (1562T>C 1564G>A), CTT471CTG (1567T>G), GTT473ATT (1571G>A), GCA474GCT (1576A>T), CGA475AGA (1577C>A), TTG477CTC (1583T>C 1585G>C), AAC478AAT (1588C>T), AGA557CGC (1823A>C 1825A>C), AAA558AAG (1828A>G), TTA559CTT (1829T>C 1831A>T), GCA560GCG (1834A>G), GGC562GGA (1840C>A), TTG564CTT (1844T>C 1846G>T), ATT565GTG (1847A>G 1849T>G), TCT572ACA (1868T>A 1870T>A), CTT573TTA (1871C>T 1873T>A), CGG574CGA (1876G>A), GTC578GTT (1888C>T), GGG579GGT (1891G>T), GAG580GAA (1894G>A), AAG581AGG (1896A>G), ATA582ATT (1900A>T), ACA583GAG (1901A>G 1902C>A 1903A>G), GCA584TCA (1904G>T), CCC586GCA (1910C>G 1912C>A), AGC593AGT (1933C>T), CGT595AGA (1937C>A 1939T>A), GAT597GAC (1945T>C), TTC599TTT (1951C>T), TTA600CTT (1952T>C 1954A>T), TAC601TAT (1957C>T), TGC604TGT (1966C>T), ACT607ACA (1975T>A), GGC611GGG (1987C>G), TTG614CTT (1994T>C 1996G>T), CTA615CTT (1999A>T), TTG618CTT (2006T>C 2008G>T), ACC622ACA (2020C>A), AAG624AAA (2026G>A), CAT625CAC (2029T>C), CTT626ATA (2030C>A 2032T>A), GGT629GGC (2041T>C), TCT631ACC (2045T>A 2047T>C), GGT632GGA (2050T>A), GAT633GAC (2053T>C), GCA634TCT (2054G>T 2056A>T), AAA635AAG (2059A>G), TTT636TTC (2062T>C), ATA637ATT (2065A>T), GAT638GAC (2068T>C), CTT639TTA (2069C>T 2071T>A), AAG641AAC (2077G>C), GAA642TCA (2078G>T 2079A>C), ATT643TCT (2081A>T 2082T>C), AGT644GAA (2084A>G 2085G>A 2086T>A), GAA645CAG (2087G>C 2089A>G), GGG646GGA (2092G>A), TAC648TTT (2097A>T 2098C>T), GAA652GAG (2110A>G), GGA653GGT (2113A>T), GCT1006..A (3172T>A), AGG1007AAG (3174G>A), TTC1008TAC (3177T>A), ACC1009ACA (3181C>A), AAC1010AAT (3184C>T), AAG1011AGG (3186A>G), AAA1012ACG (3189A>C 3190A>G), TCA1015TCT (3199A>T), GCT1016ATT (3200G>A 3201C>T), GTT1017CTT (3203G>C), AGT1018AGA (3208T>A), TAT1019TAC (3211T>C), TGT1020TGC (3214T>C), ATA1021GTA (3215A>G), AAA1026AAG (3232A>G), ACG1029ACA (3241G>A), GTT1030GTG (3244T>G), TTG1031CTG (3245T>C), AGT1032ACT (3249G>C), CTG1035TTA (3257C>T 3259G>A), CTA1037TTG (3263C>T 3265A>G), ACC1038ACA (3268C>A), GTC1040GTA (3274C>A), CGA1042CGT (3280A>T), GTT1044GTG (3286T>G), CAT1045CAC (3289T>C), TAC1047TAT (3295C>T), GTG1049ATC (3299G>A 3301G>C), AGG1052AAA (3309G>A 3310G>A), AAT1053AAC (3313T>C), GCA1056ACA (3320G>A), GAG1058GAA (3328G>A), AAC1061ACA (3336A>C 3337C>A), AAG1062CAT (3338A>C 3340G>T), GCA1063GCT (3343A>T), GCG1065ATG (3347G>A 3348C>T), TGC1066TGT (3352C>T), GAA1069GAG (3361A>G), ATT1070GTT (3362A>G), GGT1071GGG (3367T>G), ATC1073ATT (3373C>T), ATC1074ATT (3376C>T), TTC1077TTT (3385C>T), TTC1079TTT (3391C>T), AAT1081GAA (3395A>G 3397T>A), GGT1082GGG (3400T>G), CGT1083AGG (3401C>A 3403T>G), TTA1084CCC (3404T>C 3405T>C 3406A>C), CCG1085CCC (3409G>C), ACT1086ACA (3412T>A), TCT1088GCA (3416T>G 3418T>A), GAA1089GAG (3421A>G), AGA1091AAA (3426G>A), AGC1092GAG (3428A>G 3429G>A 3430C>G), AAG1093CGA (3431A>C 3432A>G 3433G>A), GTG1094ATC (3434G>A 3436G>C), AAC1095AAT (3439C>T), TCA1096CCA (3440T>C), TCT1097TCA (3445T>A), GAT1098AAC (3446G>A 3448T>C), ACA1099AAA (3450C>A), CAG1100GCT (3452C>G 3453A>C 3454G>T), CTA1101CTC (3457A>C), ATC1102ATG (3460C>G), AAG1103GAT (3461A>G 3463G>T), ATT1105GTG (3467A>G 3469T>G), GAC1106AAC (3470G>A), CAT1107CAC (3475T>C), TAT1109TAC (3481T>C), CAG1110TGC (3482C>T 3483A>G 3484G>C), GAA1113GGT (3492A>G 3493A>T), ACT1114ACG (3496T>G), ACT1115CAA (3497A>C 3498C>A 3499T>A), CAG1117CAA (3505G>A), AAG1118AAA (3508G>A), GGT1119GGA (3511T>A), CAG1121TCT (3515C>T 3516A>C 3517G>T), TTC1123TTT (3523C>T), GCC1124GCA (3526C>A), TAC1125TTC (3528A>T), CGC1128CGA (3538C>A), ATA1130ATT (3544A>T), GTC1132ATT (3548G>A 3550C>T), ATA1133GTA (3551A>G), ACT1134ACA (3556T>A), CTG1135CTT (3559G>T), CTT1136CTC (3562T>C), GTA1137ATC (3563G>A 3565A>C), GCT1139GCC (3571T>C), GTT1148GTC (3598T>C), TAC1149TTC (3600A>T), AAA1150CGA (3602A>C 3603A>G), CTC1152TTG (3608C>T 3610C>G), AGC1153AAC (3612G>A), AAA1154AAG (3616A>G), TTT1155TTC (3619T>C), AAA1156AAG (3622A>G), GGA1157GGC (3625A>C), TTA1158CTG (3626T>C 3628A>G), GTG1159GTT (3631G>T), GTT1160GTC (3634T>C), TCT1161TCA (3637T>A), AAG1164AAA (3646G>A), GAA1165GAG (3649A>G), ACT1166GCG (3650A>G 3652T>G), ACT1167ACA (3655T>A), CGT1168AGA (3656C>A 3658T>A), TTG1169CTT (3659T>C 3661G>T), GAG1170GAA (3664G>A), AGT1171TCA (3665A>T 3666G>C 3667T>A), TCT1172TCA (3670T>A), CTT1173TTG (3671C>T 3673T>G), GAC1175GAT (3679C>T), ATC1179ATA (3691C>A), GCA1180GCT (3694A>T), AAT1182AAC (3700T>C), CAA1184CAG (3706A>G), ACA1186ACC (3712A>C), GTG1187GTT (3715G>T), CGT1220AGG (3812C>A 3814T>G), CCA1221CCT (3817A>T), CAC1222CAT (3820C>T), TAC1223TAT (3823C>T), CGC1224AGG (3824C>A 3826C>G), GAA1226GAG (3832A>G), GGA1227GGC (3835A>C), AAG1228CAA (3836A>C 3838G>A), ATT1230ATA (3844T>A), TTT1232TTC (3850T>C), ACT1233ACC (3853T>C), CGA1234AGA (3854C>A), ACT1236ACA (3862T>A), AGC1238AGT (3868C>T), TCA1239TCT (3871A>T), GTG1240GTA (3874G>A), GAG1243GAA (3883G>A), ATT1244ATC (3886T>C), TCA1247TCT (3895A>T), GAG1248AGT (3896G>A 3897A>G 3898G>T), ACC1249GTG (3899A>G 3900C>T 3901C>G), AAG1250AAA (3904G>A), TTA1253CTA (3911T>C), GGT1256GGA (3922T>A), GGT1259GGA (3931T>A), TCT1260TCA (3934T>A), GGT1261GGG (3937T>G), ACT1264ACA (3946T>A), AAT1269AAC (3961T>C), TTG1270CTG (3962T>C), TGC1271AGC (3965T>A), AAG1272AAA (3970G>A), AGA1273AGG (3973A>G), GTC1276GTT (3982C>T), CTG1279CTA (3991G>A), AGA1283CGA (4001A>C), CCT1284CCC (4006T>C), TTG1285TTA (4009G>A), AAT1287AAC (4015T>C), GTG1289GTT (4021G>T), TTT1290TAC (4023T>A 4024T>C), CAG1291CAA (4027G>A), CAA1292CAG (4030A>G), TTG1293CTA (4031T>C 4033G>A), AAG1294AGA (4035A>G 4036G>A), GCA1295GCT (4039A>T), CAG1296CAA (4042G>A), GCT1300GTA (4053C>T 4054T>A), AGT1301AGC (4057T>C), CCA1302CCC (4060A>C), TTG1304CTA (4064T>C 4066G>A), ATA1306ATC (4072A>C), AGA1307CGA (4073A>C), AAA1309AAG (4081A>G), TCG1310TCC (4084G>C), GGT1313GGA (4093T>A), ACC1318ACA (4108C>A), ATT1319ATC (4111T>C), TGC1347TGT (4195C>T), GTG1349GTA (4201G>A), TCT1353TCA (4213T>A), GGA1354GGG (4216A>G), GCT1356GCA (4222T>A), AAT1359AAC (4231T>C), TTG1360CTC (4232T>C 4234G>C), CTT1361TTG (4235C>T 4237T>G), TTT1362TTC (4240T>C), TTT1366TTC (4252T>C), AGA1369AAA (4260G>A), TTG1371CTT (4265T>C 4267G>T), GTG1373GTT (4273G>T), CCG1377CCA (4285G>A), CCA1378CCT (4288A>T), GGC1379GGG (4291C>G), GAA1383GAG (4303A>G), TCA1385ACC (4307T>A 4309A>C), ACA1386ACT (4312A>T), TAT1388TAC (4318T>C), ATC1392GTG (4328A>G 4330C>G), ATT1394ATG (4336T>G), GAA1395GAG (4339A>G), AAC1397AGC (4344A>G), CTA1398CTC (4348A>C), TCT1399TCA (4351T>A), TTC1400TTT (4354C>T), CAG1401CAA (4357G>A), GTG1404GTT (4366G>T), TCA1405GCA (4367T>G), GGT1408GGG (4378T>G), CAG1410CAA (4384G>A), TCA1411TCT (4387A>T), GCT1413GTG (4392C>T 4393T>G), GTC1416ATC (4400G>A), CGA1417CGT (4405A>T), GGA1419GGT (4411A>T), AAC1420AAT (4414C>T), ATA1422ATT (4420A>T), TTA1423CTC (4421T>C 4423A>C), AGT1428AGC (4438T>C), GTT1432GTG (4450T>G), GAT1433GAC (4453T>C), TCA1434AAT (4454T>A 4455C>A 4456A>T), CTT1435TTA (4457C>T 4459T>A), AAA1437AAG (4465A>G), TTA1438CTG (4466T>C 4468A>G), CTT1439TTG (4469C>T 4471T>G), TTG1628..C (5038G>C), CTG1631TTG (5045C>T), GCA1632GCT (5050A>T), CCA1634CCC (5056A>C), CAC1635AAT (5057C>A 5059C>T), AAG1636AAA (5062G>A), AGC1637GGG (5063A>G 5065C>G), ATT1638ATA (5068T>A), GGA1639AAT (5069G>A 5070G>A 5071A>T), GTT1640TCA (5072G>T 5073T>C 5074T>A), ATT1642ATC (5080T>C), CCA1643CCG (5083A>G), ACT1646ACA (5092T>A), TAT1647TAC (5095T>C), AGA1648CGT (5096A>C 5098A>T), AGG1649CGC (5099A>C 5101G>C), ACA1650CTT (5102A>C 5103C>T 5104A>T), GGC1651GGA (5107C>A), CAT1653CAA (5113T>A), ACT1654ACA (5116T>A), GAA1655CAA (5117G>C), GAA1657GAG (5125A>G), AAT1658AAC (5128T>C), ATT1661ATA (5137T>A), GCT1662CCA (5138G>C 5140T>A), CTC1664CTT (5146C>T), ACA1665TCA (5147A>T), GAA1667GAT (5155A>T), GTC1668GTA (5158C>A), CCG1669CCA (5161G>A), AAA1671AGA (5166A>G), GTT1672GTG (5170T>G), CAC1673_AGG1674del (5171_5178delCACAGGGA), GAG1675--C (5171_5178delCACAGGGA 5179G>C), GTA1676GTC (5182A>C), GCA1679GCT (5191A>T), GTT1680GTG (5194T>G), TGT1681AAT (5195T>A 5196G>A), GGT1690GGA (5224T>A), AGA1691CGC (5225A>C 5227A>C), CTA1692TTA (5228C>T), ACT1693ACA (5233T>A), AGT1694AGC (5236T>C), CAC1695CAT (5239C>T), GTT1697GTC (5245T>C), TCA1698TCC (5248A>C), GTG1700GTC (5254G>C), TAT1702TAC (5260T>C), TTG1704TTA (5266G>A), CAG1705CAA (5269G>A), ACT1706ACA (5272T>A), TCA1710TCT (5284A>T), ATT1711ATC (5287T>C), GCA1712TCA (5288G>T), AGG1713CGA (5291A>C 5293G>A), ACT1716ACA (5302T>A), ATA1718ATC (5308A>C), AAC1719AAT (5311C>T), CAG1720CAA (5314G>A), GCT1734GCA (5356T>A), AGT1737AGC (5365T>C), ACA1739ACT (5371A>T), ACA1740TGT (5372A>T 5373C>G 5374A>T), AAC1744AAT (5386C>T), TCG1746TCT (5392G>T), CTA1747TTG (5393C>T 5395A>G), GTC1750GTT (5404C>T), ATT1754ATC (5416T>C), CGA1755CGT (5419A>T), CGA1757CGG (5425A>G), GCT1759GCA (5431T>A), AGG1760CGT (5432A>C 5434G>T), CAG1764CAA (5446G>A), TTG1767CTT (5453T>C 5455G>T), CGG1787CGA (5515G>A), GTG1792GTT (5530G>T), CTC1793ATA (5531C>A 5533C>A), GCC1794TCT (5534G>T 5536C>T), AAT1795AAC (5539T>C), GGC1797GGA (5545C>A), GCA1801TCC (5555G>T 5557A>C), TTC1804TTT (5566C>T), CAA1805CAG (5569A>G), TCA1807CAA (5573T>C 5574C>A), GGA1809GAA (5580G>A), TCA1811GCA (5585T>G), AAG1812CGC (5588A>C 5589A>G 5590G>C), TTT1813TTC (5593T>C), GGT1815AAA (5597G>A 5598G>A 5599T>A), TTA1816CTT (5600T>C 5602A>T), GGA1818GGT (5608A>T), CAA1819CAG (5611A>G), AAG1822AGT (5619A>G 5620G>T), TCT1823TCC (5623T>C), TTG1824CTC (5624T>C 5626G>C), ATC1825ATT (5629C>T), ACT1826ACA (5632T>A), AGA1827AGG (5635A>G), GCT1829ATC (5639G>A 5640C>T 5641T>C), ATA1831GTT (5645A>G 5647A>T), ATA1832GTT (5648A>G 5650A>T), GGA1838GGT (5668A>T), GCT1840GCA (5674T>A), CTT1843TTA (5681C>T 5683T>A), TAC1844TAT (5686C>T), AAG1845TCT (5687A>T 5688A>C 5689G>T), CAA1848AAG (5696C>A 5698A>G), AGT1849GAG (5699A>G 5700G>A 5701T>G), AAT1850AGG (5703A>G 5704T>G), TCT1851GTA (5705T>G 5706C>T 5707T>A), AAA1852AGT (5709A>G 5710A>T), GAA1853GAG (5713A>G), GAA1854ACG (5714G>A 5715A>C 5716A>G), CTG1857CCC (5724T>C 5725G>C), CAA1858CAG (5728A>G), AGG1860AGA (5734G>A), GCA1864ACC (5744G>A 5746A>C), AAG1866del (5750_5754delAAGTT), TTG1867--A (5750_5754delAAGTT 5755G>A), CGA1868CGC (5758A>C), GCA1870GCT (5764A>T), TCC1872TCA (5770C>A), GCG1874GCA (5776G>A), GAG1875GAA (5779G>A), AAA1876AAG (5782A>G), AAT1878AAC (5788T>C), GTC1880GTT (5794C>T), GTC1881GTT (5797C>T), GAA1882GAG (5800A>G), GGA1883GGC (5803A>C), GCC1886GCA (5812C>A), GCC1887GCA (5815C>A), CTC1888CTA (5818C>A), ACA1889ACT (5821A>T), CAC1890CAT (5824C>T), TAT1891TAC (5827T>C), GCA1895GCT (5839A>T), GCC1897GCA (5845C>A), TCA1898AAG (5846T>A 5847C>A 5848A>G), AAA1899AAG (5851A>G), AAA1903AAG (5863A>G), ACA1905ACT (5869A>T), GGA1908GGC (5878A>C), CAT1911CAC (5887T>C), AAG1912AAA (5890G>A), AGA1914AGG (5896A>G), AAG1915AAA (5899G>A), TTC1916TTT (5902C>T), GGA1921GGT (5917A>T), CCT1924CCA (5926T>A), ACC1925ACT (5929C>T), GAC1926GAT (5932C>T), TTT1927TTC (5935T>C), CAA1928CAG (5938A>G), GTT1930GTA (5944T>A), CGT1931CGA (5947T>A), GTG1933GTA (5953G>A), CCA1935CCT (5959A>T), TTA1936TTG (5962A>G), GGT1938GGA (5968T>A), CGC1939AAG (5969C>A 5970G>A 5971C>G), GCA1941GTG (5976C>T 5977A>G), GAA1943GAG (5983A>G), TCT1944TCA (5986T>A), CCG1945CCA (5989G>A), ACT1947ACA (5995T>A), GAC1948GAT (5998C>T), GAG1954GAA (6016G>A), GGA1957GGC (6025A>C), AAA1958AAG (6028A>G), ATA1959ATC (6031A>C), GCT1961GAA (6036C>A 6037T>A), TTG2045TTA (6289G>A), GGA2047GGG (6295A>G), GCG2048TCA (6296G>T 6298G>A), TCA2049TCT (6301A>T), CTT2050TTA (6302C>T 6304T>A), GCA2051GCG (6307A>G), GGA2053GGG (6313A>G), GTT2054GTG (6316T>G), CGA2055CGT (6319A>T), ATC2060ATT (6334C>T), TCT2061TCA (6337T>A), GCT2062GCC (6340T>C), GCA2063TCA (6341G>T), CTT2064CTC (6346T>C), TGT2065TGC (6349T>C), AAA2066AAG (6352A>G), CTG2067CTA (6355G>A), ACA2068ACC (6358A>C), GAT2070GAC (6364T>C), GGT2073GGA (6373T>A), GTG2074GTT (6376G>T), AGG2075CGA (6377A>C 6379G>A), ATC2076ATG (6382C>G), TCT2079TCA (6391T>A), GGC2080GGG (6394C>G), GGT2082GGA (6400T>A), GGA2084GGG (6406A>G), TTC2086TAT (6411T>A 6412C>T), ATA2087ATT (6415A>T), ATT2088ATC (6418T>C), ACG2089ACA (6421G>A), AAG2091AAA (6427G>A), TTA2093TTG (6433A>G), TTC2094TTT (6436C>T), AGA2095CGT (6437A>C 6439A>T), GGT2099GGG (6451T>G), ATG2101CTG (6455A>C), GAA2102GAG (6460A>G), ATC2103GTT (6461A>G 6463C>T), CAA2106CAT (6472A>T), GGT2108GGA (6478T>A), TTG2109CTT (6479T>C 6481G>T), ATT2112ATC (6490T>C), CGC2113AGA (6491C>A 6493C>A), CTT2115TTG (6497C>T 6499T>G), AAG2116AAA (6502G>A), ACG2117ACA (6505G>A), TTG2118CTT (6506T>C 6508G>T), TTG2122TTA (6520G>A), GAG2124GAT (6526G>T), AAC2125GGT (6527A>G 6528A>G 6529C>T), GAC2127GAT (6535C>T), CTA2129CTT (6541A>T), ATA2130TTA (6542A>T), ATT2131ATC (6547T>C), AAA2132AAG (6550A>G), GAT2136GAC (6562T>C), TTT2137TTC (6565T>C), GCT2141GCC (6577T>C), TTG2144TTA (6586G>A), AAG2145AAA (6589G>A), TTC2146TTT (6592C>T), CGA2147AGG (6593C>A 6595A>G), CTT2148ATG (6596C>A 6598T>G), GAA2152GAG (6610A>G), GAG2153GAA (6613G>A), AAA2154AAG (6616A>G), GCC2155GCA (6619C>A), GTG2158ATT (6626G>A 6628G>T), CAG2163CAA (6643G>A), ACA2164ACC (6646A>C), AGC2166AGT (6652C>T), ATG2167CTG (6653A>C), CAA2170CTA (6663A>T), GTT2171GTG (6667T>G), ACA2176ATA (6681C>T), ATA2177GTT (6683A>G 6685A>T), GTT2178GTA (6688T>A), CCA2179CCT (6691A>T), AAA2182AAG (6700A>G), TCA2183TCT (6703A>T), CAT2188CAC (6718T>C), ATC2190ATT (6724C>T), ACA2191TCA (6725A>T), GGC2195GGG (6739C>G), CAG2196AAT (6740C>A 6742G>T), GGT2198GGG (6748T>G), CCA2200del (6752_6756delCCATT), TTA2201--C (6752_6756delCCATT 6757A>C), GCT2203GCA (6763T>A), TTG2204TTA (6766G>A), AAA2205AAG (6769A>G), AAG2208AAC (6778G>C), CTT2210CTA (6784T>A), GGA2211GGG (6787A>G), ATT2212ATA (6790T>A), GGT2215GGA (6799T>A), TCG2216TCA (6802G>A), TTA2219CTA (6809T>C), ACA2220ACG (6814A>G), AAT2222AAC (6820T>C), CAC2223CAT (6823C>T), ACC2225ACA (6829C>A), TAC2226TAT (6832C>T), ATT2227ATC (6835T>C), GAA2229GAG (6841A>G), CAA2232CAT (6850A>T), AAT2233GAC (6851A>G 6853T>C), TTG2234ATC (6854T>A 6856G>C), ATA2235TTG (6857A>T 6859A>G), ATG2236AAA (6861T>A 6862G>A), GCA2238TCA (6866G>T), CTC2239ATG (6869C>A 6871C>G), GAA2240GAT (6874A>T), ATG2241CTT (6875A>C 6877G>T), GAT2242GAG (6880T>G), GTG2244GTT (6886G>T), AAA2248AAG (6898A>G), TTC2249TTT (6901C>T), AAT2250AAC (6904T>C), TTG2251CTA (6905T>C 6907G>A), AAT2252AAC (6910T>C), AGC2255AGT (6919C>T), GGC2257GGT (6925C>T), CTC2259TTG (6929C>T 6931C>G), AAA2260AAG (6934A>G), ATT2261CTC (6935A>C 6937T>C), CAA2262CAG (6940A>G), GAG2267GAA (6955G>A), CAG2268GAA (6956C>G 6958G>A), ACT2272ACA (6970T>A), GAG2273GCA (6972A>C 6973G>A), ATT2275ATA (6979T>A), GTT2276ATA (6980G>A 6982T>A), GAT2278GAC (6988T>C), CTT2279TTG (6989C>T 6991T>G), TTT2280CTC (6992T>C 6994T>C), GAT2281GAA (6997T>A), GAG2282GAA (7000G>A), ATT2284ACA (7005T>C 7006T>A), ATA2286GCA (7010A>G 7011T>C), GCC2288TCT (7016G>T 7018C>T), AAT2289CAA (7019A>C 7021T>A), ATG2290ATT (7024G>T), CAA2291CAG (7027A>G), AAT2292AAG (7030T>G), GTG2294GTT (7036G>T), TAC2295TTT (7038A>T 7039C>T), ATT2298ATA (7048T>A), TGT2299CGG (7049T>C

7051T>G), GAC2300GAT (7054C>T), TTG2302CTT (7058T>C 7060G>T), GTC2303GTA (7063C>A), CCA2304CCT (7066A>T), ATT2305GTC (7067A>G 7069T>C), GCT2306TCC (7070G>T 7072T>C), ACA2307TCC (7073A>T 7075A>C), TGC2308TGT (7078C>T), TTA2312CTT (7088T>C 7090A>T), GTC2313GTT (7093C>T), AAA2315AAG (7099A>G), GTT2317GTG (7105T>G), GTT2318GTG (7108T>G), AAG2319AAA (7111G>A), AAG2321AAA (7117G>A), AAT2323CCA (7121A>C 7122A>C 7123T>A), CTG2324TTG (7124C>T), TTC2325TTT (7129C>T), AGC2326AGT (7132C>T), TTG2327GTC (7133T>G 7135G>C), TAT2328TAC (7138T>C), TTA2329CTG (7139T>C 7141A>G), GCC2330GCT (7144C>T), GAA2333AGT (7151G>A 7152A>G 7153A>T), GAG2334GAT (7156G>T), GAG2335GCA (7158A>C 7159G>A), GAA2336GAG (7162A>G), AAA2337CAA (7163A>C), TAT2338TAC (7168T>C), CGA2340AGA (7172C>A), CCA2341CCT (7177A>T), CTA2342TTG (7178C>T 7180A>G), GGA2344GGC (7186A>C), TAT2346TAC (7192T>C), CTA2347CAA (7194T>A), AGC2349AGT (7201C>T), AAA2350GCA (7202A>G 7203A>C), TTG2351CTA (7205T>C 7207G>A), AGA2353AAG (7212G>A 7213A>G), GAG2354GAA (7216G>A), GCA2355GCT (7219A>T), CTG2357CTT (7225G>T), ATA2360ATT (7234A>T), AAA2362AAG (7240A>G), TAC2363TAT (7243C>T), TCA2364TCG (7246A>G), AGT2365AAT (7248G>A), ACA2367GTA (7253A>G 7254C>T), GAG2368GAA (7258G>A), GGA2370GGT (7264A>T), TCT2371GCT (7265T>G), GTG2372GTC (7270G>C), ACA2374ACT (7276A>T), ATC2376ATT (7282C>T), TTT2377TTC (7285T>C), GAT2378GGT (7287A>G), AGA2379CAG (7289A>C 7290G>A 7291A>G), GCA2380GCC (7294A>C), GTG2381GTC (7297G>C), AAT2382ACT (7299A>C), TTG2384GTC (7304T>G 7306G>C), ATT2385ATA (7309T>A), AAT2386AAC (7312T>C), CTC2388CTT (7318C>T), GAA2389GTT (7320A>T 7321A>T), GGT2392GGC (7330T>C), GTG2399GTT (7351G>T), ATG2403CTG (7361A>C), CAG2404CAA (7366G>A), ATA2405ATC (7369A>C), TTC2406TTT (7372C>T), GCT2408GCA (7378T>A), TTA2409TTG (7381A>G), AAT2410AAC (7384T>C), GCA2414GCT (7396A>T), GCT2417GCC (7405T>C), CTA2418CTT (7408A>T), AAA2420AAG (7414A>G), AAA2423AAG (7423A>G), AGA2424AGG (7426A>G), GAT2425GAC (7429T>C), TAC2426TAT (7432C>T), GCC2428GCA (7438C>A), AAT2429TCT (7439A>T 7440A>C), TGC2430TCA (7443G>C 7444C>A), TCA2431ACA (7445T>A), GCT2433GCC (7453T>C), GAT2434GAC (7456T>C), TTG2435CTT (7457T>C 7459G>T), GAA2436GAG (7462A>G), AAT2437AGG (7464A>G 7465T>G), TTC2438TAC (7467T>A), TTG2439TTA (7471G>A), AGA2440ATG (7473G>T 7474A>G), GCC2441GCA (7477C>A), TGT2443TGC (7483T>C), GAG2444AAA (7484G>A 7486G>A), AGA2445CGC (7487A>C 7489A>C), CTC2446CTT (7492C>T), TTT2447TAT (7494T>A), CTT2448CTC (7498T>C), GGC2449GGT (7501C>T), CTT2451CTC (7507T>C), GGG2452GGA (7510G>A), TTG2453TTA (7513G>A), GGC2456GGA (7522C>A), TCA2457TCC (7525A>C), CTT2458CTA (7528T>A), GAA2461GAG (7537A>G), TTG2462CTG (7538T>C), CGG2463AGA (7541C>A 7543G>A), ATT2465TTG (7547A>T 7549T>G), AAG2467AAA (7555G>A), GTT2468GTG (7558T>G), GAG2469CAA (7559G>C 7561G>A), AAC2471AAT (7567C>T), AAG2472AAA (7570G>A), ACA2473TCA (7571A>T), CGA2474CGC (7576A>C), TTT2476TTC (7582T>C), GCA2479GCT (7591A>T), GTT2481ATA (7595G>A 7597T>A), GAT2482GAC (7600T>C), TTA2484CTT (7604T>C 7606A>T), CTG2485CTA (7609G>A), GCT2486GCA (7612T>A), GGA2487GGG (7615A>G), GTG2491GTA (7627G>A), TTT2494TTC (7636T>C), CAA2497CAG (7645A>G), TTC2498TTT (7648C>T), TAT2499TAC (7651T>C), TCT2500TCA (7654T>A), CTC2501TTA (7655C>T 7657C>A), TTG2503CTC (7661T>C 7663G>C), CAG2504AAG (7664C>A), CCC2506CCA (7672C>A), TCA2507TCT (7675A>T), GTC2509GTT (7681C>T), GGA2510GGT (7684A>T), AAA2513AAG (7693A>G), TTT2514TTC (7696T>C), AGA2516GGA (7700A>G), GGG2517GGT (7705G>T), GAT2519GAC (7711T>C), AAA2520AAG (7714A>G), TTG2521CTT (7715T>C 7717G>T), AGA2523AGG (7723A>G), GGG2524AGC (7724G>A 7726G>C), CTT2525TTA (7727C>T 7729T>A), CCT2526CCG (7732T>G), ACT2530GTT (7742A>G 7743C>T), TGT2532TGC (7750T>C), GCT2534GCA (7756T>A), GAC2535GAT (7759C>T), AGG2538AGA (7768G>A), GAT2540GAC (7774T>C), CTA2543TTG (7781C>T 7783A>G), CCA2545CCT (7789A>T), CTT2547CTC (7795T>C), TTG2548ATC (7796T>A 7798G>C), ATT2551CTC (7805A>C 7807T>C), ATT2552ATC (7810T>C), GTC2554ATC (7814G>A), CTC2556TTA (7820C>T 7822C>A), TAC2557TTT (7824A>T 7825C>T), GAA2560GAG (7834A>G), GAT2563GAC (7843T>C), ATT2564ATA (7846T>A), GTG2567ACC (7853G>A 7854T>C 7855G>C), AAG2570AGA (7863A>G 7864G>A), TAC2573TAT (7873C>T), ACC2574ACT (7876C>T), ATA2581AAC (7896T>A 7897A>C), TCC2582TCA (7900C>A), ACT2587ACG (7915T>G), GTA2588GTT (7918A>T), ATA2589ATT (7921A>T), CAT2592CAC (7930T>C), GGC2594GGT (7936C>T), AAT2595AAC (7939T>C), AAT2596AAC (7942T>C), AGT2597ACT (7944G>C), GGA2598GGG (7948A>G), CAA2599CAG (7951A>G), TCT2601TCA (7957T>A), ACA2602ACT (7960A>T), GTC2603GTT (7963C>T), GTT2604GTG (7966T>G), GAC2605GAT (7969C>T), CTG2608CTT (7978G>T), GTT2610GTA (7984T>A), AAC2611AAT (7987C>T), ATA2612CTA (7988A>C), TTT2615TTC (7999T>C), TAC2616TAT (8002C>T), ATG2618ATT (8008G>T), CAC2619CAT (8011C>T), TCA2620TCT (8014A>T), CGT2621AGA (8015C>A 8017T>A), GAC2622AAC (8018G>A), ATT2623ATC (8023T>C), CCG2624CCT (8026G>T), AAA2625ATT (8028A>T 8029A>T), GAG2626GAT (8032G>T), TCC2627ATG (8033T>A 8034C>T 8035C>G), AGA2629AGG (8041A>G), TTT2630TTC (8044T>C), TAT2631TTT (8046A>T), GTT2632GTA (8050T>A), GGA2634GGG (8056A>G), CTG2637TTG (8063C>T), GCT2640GCA (8074T>A), ATA2641ATG (8077A>G), GAG2642GAA (8080G>A), CAG2644TCT (8084C>T 8085A>C 8086G>T), GAC2647GAT (8095C>T), TTG2649CTT (8099T>C 8101G>T), AAG2650AAA (8104G>A), TTT2652TTC (8110T>C), CAA2653CAG (8113A>G), GCT2654TCA (8114G>T 8116T>A), TTA2655TTG (8119A>G), TTA2657TGT (8124T>G 8125A>T), GGT2660GGA (8134T>A), CTT2661ATG (8135C>A 8137T>G), ATT2662TCA (8138A>T 8139T>C 8140T>A), ACT2664ACA (8146T>A), ACA2669ACT (8161A>T), AAA2670AGG (8163A>G 8164A>G), GAC2671GAT (8167C>T), CGC2672AGA (8168C>A 8170C>A), AGT2673CTA (8171A>C 8172G>T 8173T>A), GTG2675GTT (8179G>T), TCA2679TCC (8191A>C), CAC2680CAT (8194C>T), GCT2682GCA (8200T>A), GTT2683ATG (8201G>A 8203T>G), CTC2684CTG (8206C>G), GGT2687GGA (8215T>A), ATT2688GTC (8216A>G 8218T>C), TAC2689TAT (8221C>T), ATA2690ATC (8224A>C), CCT2691CCA (8227T>A), GAG2696GAA (8242G>A), CGG2697AGA (8243C>A 8245G>A), GTT2699GTA (8251T>A), TCC2700TCA (8254C>A), CTT2702CTC (8260T>C), GAC2705GAT (8269C>T), AGA2706CGA (8270A>C), TCA2707TCT (8275A>T), GGA2708GGT (8278A>T), GAA2709GAG (8281A>G), CCA2710CCC (8284A>C), TCT2711ACT (8285T>A), CGG2713AGG (8291C>A), CTG2714TTG (8294C>T), GAA2715GAG (8299A>G), GAG2723GAA (8323G>A), GAA2729GAG (8341A>G), CTG2730TTG (8342C>T), ACT2731ACA (8347T>A), AAG2732ATG (8349A>T), AGG2733AGA (8353G>A), AGA2735CGA (8357A>C), CTT2739TTG (8369C>T 8371T>G), GAG2743GAA (8383G>A), TAC2747TAT (8395C>T), GAA2753CAA (8411G>C), GGG2754GGA (8416G>A), AGA2755AAA (8418G>A), GCT2756GCA (8422T>A), TAC2758TAT (8428C>T), CTA2759CTG (8431A>G), GCT2760GCC (8434T>C), GAA2761GAG (8437A>G), ACT2762ACG (8440T>G), GCT2763GCA (8443T>A), CTG2764TTG (8444C>T), TTA2767TTG (8455A>G), CTT2769CTA (8461T>A), GAT2770GAC (8464T>C), GCT2771ACC (8465G>A 8467T>C), CCC2773CCA (8473C>A), ACT2774ACG (8476T>G), GAA2775GAT (8479A>T), TTG2776AAA (8480T>A 8481T>A 8482G>A), GAA2777GAG (8485A>G), TTG2778TTA (8488G>A), CTT2779TTG (8489C>T 8491T>G), GCA2782GTT (8499C>T 8500A>T), GGG2783GGT (8503G>T), TGC2784TGT (8506C>T), ATC2785GTC (8507A>G), GGT2787GGA (8515T>A), GAT2788GAG (8518T>G), TAT2789TCT (8520A>C), GAT2790GAC (8524T>C), GAT2791TTG (8525G>T 8526A>T 8527T>G), AGC2792GAT (8528A>G 8529G>A 8530C>T), TTG2793GTA (8531T>G 8533G>A), GCA2794GAA (8535C>A), TCT2795CAA (8537T>C 8538C>A 8539T>A), GAA2796GAG (8542A>G), GAT2797CAC (8543G>C 8545T>C), ATA2798ATG (8548A>G), TAT2799ATA (8549T>A 8550A>T 8551T>A), CTC2800TTC (8552C>T), GCT2802GCA (8560T>A), GAT2803GAC (8563T>C), ACA2804AAA (8565C>A), GGT2808GGA (8578T>A), AGT2809AGC (8581T>C), ACA2811ACC (8587A>C), GCC2813ACC (8591G>A), GGA2814GGT (8596A>T), AAG2815AAA (8599G>A), AAC2816AGT (8601A>G 8602C>T), TCA2818GGA (8606T>G 8607C>G), AGT2824CTC (8624A>C 8625G>T 8626T>C), GTT2825GCT (8628T>C), ACG2827AAA (8634C>A 8635G>A), ACT2828CCT (8636A>C), GGA2829GAC (8640G>A 8641A>C), GAG2830CAA (8642G>C 8644G>A), AAA2831AAG (8647A>G), AAG2832AAC (8650G>C), CTT2834TCT (8654C>T 8655T>C), GTC2835GCC (8658T>C), CAA2838CAG (8668A>G), GTG2842GTT (8680G>T), GCT2844GCA (8686T>A), ACC2849ACG (8701C>G), GTA2852GTT (8710A>T), AAA2856AAG (8722A>G), ATT2858ATA (8728T>A), CCT2859CCC (8731T>C), ACT2860ACA (8734T>A), CTA2864CTC (8746A>C), AAT2869AAG (8761T>G), AAA2870AGA (8763A>G), GTT2872GTC (8770T>C), ATT2873ATC (8773T>C), AGT2874AAT (8775G>A), CTG2875CTT (8779G>T), GAC2876GAT (8782C>T), CTG2878TTG (8786C>T), CTG2879CTC (8791G>C), AAC2880CTG (8792A>C 8793A>T 8794C>G), CCA2883CCT (8803A>T), GAT2887GAC (8815T>C), TCA2889TCT (8821A>T), CAC2895CAA (8839C>A), GAA2899GAG (8851A>G), AAT2903GAT (8861A>G), CGT2904AAA (8864C>A 8865G>A 8866T>A), GTA2905GTG (8869A>G), CGA2906AGA (8870C>A), GTT2911GTC (8887T>C), ACT2912ACA (8890T>A), GAA2914GAG (8896A>G), CAG2915CAA (8899G>A), CAG2917CAA (8905G>A), ATC2919ATT (8911C>T), TTA2923CTT (8921T>C 8923A>T), ATC2928ATT (8938C>T), AAC2935AAT (8959C>T), ATA2936ATT (8962A>T), GGT2938GGC (8968T>C), ACT2941ACC (8977T>C), GAG2946GAC (8992G>C), TTA2953TTG (9013A>G), ATT2957ATA (9025T>A), GAA2958GAG (9028A>G), CCC2962CCA (9040C>A), TTT2964TTC (9046T>C), CAA2966CAG (9052A>G), GCG2969GCA (9061G>A), CAT2970CAC (9064T>C), GAT2973GAC (9073T>C), CTA2974TTA (9074C>T), GAT2985GAC (9109T>C), CCA2990CCT (9124A>T), CGT2991AGA (9125C>A 9127T>A), CGC2996CGT (9142C>T), ATG3001AAT (9156T>A 9157G>T), AGT3002AGC (9160T>C), CTG3003CTT (9163G>T), GAC3009GAT (9181C>T), GTT3013ATC (9191G>A 9193T>C), ACT3014ACC (9196T>C), CCG3015TCG (9197C>T), CGG3016AGG (9200C>A), ACC3019ACT (9211C>T), GCT3030GCA (9244T>A), CGA3034CGT (9256A>T), AAC3036ACA (9261A>C 9262C>A), ACA3037ACG (9265A>G), ACG3038ACC (9268G>C), AAG3039AAA (9271G>A), TTA3040CTC (9272T>C 9274A>C), CTG3043CTC (9283G>C), GAA3051GAG (9307A>G), ACC3054ACA (9316C>A), CAC3057CAT (9325C>T), CAA3060ACT (9332C>A 9333A>C 9334A>T), GTT3062GTG (9340T>G), AGA3064AGG (9346A>G), GGT3065AAC (9347G>A 9348G>A 9349T>C), CTT3070CTA (9364T>A), TAG3076TAA (9382G>A)

#### Proteins

| | | | | | | | | | |
|---|---|---|---|---|---|---|---|---|---|
| polyprotein (YP_007346986.1) | 298 | 3076 | 63.2% | 11399 | 85.2% | 1940 (99.8%) | 1649 (84.8%) | 0/4/3/3 | 1 |

| | Begin | End | Coverage | Score | Concordance | Matches | Identities | I/D/M/F* | Stop Codons |
|---|---|---|---|---|---|---|---|---|---|
| **NT** | **1045** | **9396** | **60.8%** | **6095** | **52.5%** | **5835 (99.7%)** | **4455 (76.1%)** | **0/19** | |
| Protein mutations: | L300M (1052T>A), K309A (1079A>G 1080A>C 1081G>A), A330T (1142G>A 1144G>A), R332K (1149G>A), E334D (1156A>T), I335L (1157A>T 1159T>A), D336N (1160G>A), L347Q (1193T>C 1194T>A), T368I (1257C>T 1258A>T), D450E (1504T>A), L451C (1505C>T 1506T>G 1507A>T), S456T (1520T>A 1522T>A), L457T (1523T>A 1524T>C), M458L (1526A>T), S459D (1529A>G 1530G>A), V465I (1547G>A), S468G (1556A>G), V473I (1571G>A), I565V (1847A>G 1849T>G), S572T (1868T>A 1870T>A), K581R (1896A>G), T583E (1901A>G 1902C>A 1903A>G), A584S (1904G>T), P586A (1910C>G 1912C>A), L626I (2030C>A 2032T>A), S631T (2045T>A 2047T>C), A634S (2054G>T 2056A>T), K641N (2077G>C), E642S (2078G>T 2079A>C), I643S (2081A>T 2082T>C), S644E (2084A>G 2085G>A 2086T>A), E645Q (2087G>C 2089A>G), Y648F (2097A>T 2098C>T), R1007K (3174G>A), F1008Y (3177T>A), K1011R (3186A>G), K1012T (3189A>C 3190A>G), A1016I (3200G>A 3201C>T), V1017L (3203G>C), S1018R (3208T>A), I1021V (3215A>G), S1032T (3249G>C), V1049I (3299G>A 3301G>C), R1052K (3309G>A 3310G>A), A1056T (3320G>A), N1061T (3336A>C 3337C>A), K1062H (3338A>C 3340G>T), A1065M (3347G>A 3348C>T), I1070V (3362A>G), N1081E (3395A>G 3397T>A), L1084P (3404T>C 3405T>C 3406A>C), S1088A (3416T>G 3418T>A), R1091K (3426G>A), S1092E (3428A>G 3429G>A 3430C>G), K1093R (3431A>C 3432A>G 3433G>A), V1094I (3434G>A 3436G>C), S1096P (3440T>C), D1098N (3446G>A 3448T>C), T1099K (3450C>A), Q1100A (3452C>G 3453A>C 3454G>T), I1102M (3460C>G), K1103D (3461A>G 3463G>T), I1105V (3467A>G 3469T>G), D1106N (3470G>A), Q1110C (3482C>T 3483A>G 3484G>C), E1113G (3492A>G 3493A>T), T1115Q (3497A>C 3498C>A 3499T>A), Q1121S (3515C>T 3516A>C 3517G>T), Y1125F (3528A>T), V1132I (3548G>A 3550C>T), I1133V (3551A>G), V1137I (3563G>A 3565A>C), Y1149F (3600A>T), K1150R (3602A>C 3603A>G), S1153N (3612G>A), T1166A (3650A>G 3652T>G), K1228Q (3836A>C 3838G>A), E1248S (3896G>A 3897A>G 3898G>T), T1249V (3899A>G 3900C>T 3901C>G), C1271S (3965T>A), F1290Y (4023T>A 4024T>C), K1294R (4035A>G 4036G>A), A1300V (4053C>T 4054T>A), R1369K (4260G>A), S1385T (4307T>A 4309A>C), I1392V (4328A>G 4330C>G), I1394M (4336T>G), N1397S (4344A>G), S1405A (4367T>G), A1413V (4392C>T 4393T>G), V1416I (4400G>A), S1434N (4454T>A 4455C>A 4456A>T), H1635N (5057C>A 5059C>T), S1637G (5063A>G 5065C>G), G1639N (5069G>A 5070G>A 5071A>T), V1640S (5072G>T 5073T>C 5074T>A), T1650L (5102A>C 5103C>T 5104A>T), H1653Q (5113T>A), E1655Q (5117G>C), A1662P (5138G>C 5140T>A), T1665S (5147A>T), E1667D (5155A>T), K1671R (5166A>G), H1673_R1674del (5171_5178delCACAGGGA), C1681N (5195T>A 5196G>A), A1712S (5288G>T), T1740C (5372A>T 5373C>G 5374A>T), L1793I (5531C>A 5533C>A), A1794S (5534G>T 5536C>T), A1801S (5555G>T 5557A>C), S1807Q (5573T>C 5574C>A), G1809E (5580G>A), S1811A (5585T>G), K1812R (5588A>C 5589A>G 5590G>C), G1815K (5597G>A 5598G>A 5599T>A), K1822S (5619A>G 5620G>T), A1829I (5639G>A 5640C>T 5641T>C), I1831V (5645A>G 5647A>T), I1832V (5648A>G 5650A>T), K1845S (5687A>T 5688A>C 5689G>T), Q1848K (5696C>A 5698A>G), S1849E (5699A>G 5700G>A 5701T>G), N1850R (5703A>G 5704T>G), S1851V (5705T>G 5706C>T 5707T>A), K1852S (5709A>G 5710A>T), E1854T (5714G>A 5715A>C 5716A>G), L1857P (5724T>C 5725G>C), A1864T (5744G>A 5746A>C), K1866del (5750_5754delAAGTT), S1898K (5846T>A 5847C>A 5848A>G), R1939K (5969C>A 5970G>A 5971C>G), A1941V (5976C>T 5977A>G), A1961E (6036C>A 6037T>A), A2048S (6296G>T 6298G>A), A2063S (6341G>T), I2076M (6382C>G), F2086Y (6411T>A 6412C>T), M2101L (6455A>C), I2103V (6461A>G 6463C>T), Q2106H (6472A>T), E2124D (6526G>T), N2125G (6527A>G 6528A>G 6529C>T), I2130L (6542A>T), L2148M (6596C>A 6598T>G), V2158I (6626G>A 6628G>T), M2167L (6653A>C), Q2170L (6663A>T), T2176I (6681C>T), I2177V (6683A>G 6685A>T), T2191S (6725A>T), Q2196N (6740C>A 6742G>T), P2200del (6752_6756delCCATT), K2208N (6778G>C), Q2232H (6850A>T), N2233D (6851A>G 6853T>C), L2234I (6854T>A 6856G>C), I2235L (6857A>T 6859A>G), M2236K (6861T>A 6862G>A), A2238S (6866G>T), L2239M (6869C>A 6871C>G), E2240D (6874A>T), M2241L (6875A>C 6877G>T), D2242E (6880T>G), I2261L (6935A>C 6937T>C), Q2268E (6956C>G 6958G>A), E2273A (6972A>C 6973G>A), V2276I (6980G>A 6982T>A), F2280L (6992T>C 6994T>C), D2281E (6997T>A), I2284T (7005T>C 7006T>A), I2286A (7010A>G 7011T>C), A2288S (7016G>T 7018C>T), N2289Q (7019A>C 7021T>A), M2290I (7024G>T), N2292K (7030T>G), Y2295F (7038A>T 7039C>T), C2299R (7049T>C 7051T>G), I2305V (7067A>G 7069T>C), A2306S (7070G>T 7072T>C), T2307S (7073A>T 7075A>C), N2323P (7121A>C 7122A>C 7123T>A), L2327V (7133T>G 7135G>C), E2333S (7151G>A 7152A>G 7153A>T), E2334D (7156G>T), E2335A (7158A>C 7159G>A), K2337Q (7163A>C), L2347Q (7194T>A), K2350A (7202A>G 7203A>C), R2353K (7212G>A 7213A>G), S2365N (7248G>A), T2367V (7253A>G 7254C>T), S2371A (7265T>G), D2378G (7287A>G), R2379Q (7289A>C 7290G>A 7291A>G), N2382T (7299A>C), L2384V (7304T>G 7306G>C), E2389V (7320A>T 7321A>T), M2403L (7361A>C), N2429S (7439A>T 7440A>C), C2430S (7443G>C 7444C>A), S2431T (7445T>A), N2437R (7464A>G 7465T>G), F2438Y (7467T>A), R2440M (7473G>T 7474A>G), E2444K (7484G>A 7486G>A), F2447Y (7494T>A), I2465L (7547A>T 7549T>G), E2469Q (7559G>C 7561G>A), T2473S (7571A>T), V2481I (7595G>A 7597T>A), Q2504K (7664C>A), R2516G (7700A>G), G2524S (7724G>A 7726G>C), T2530V (7742A>G 7743C>T), L2548I (7796T>A 7798G>C), I2551L (7805A>C 7807T>C), V2554I (7814G>A), Y2557F (7824A>T 7825C>T), V2567T (7853G>A 7854T>C 7855G>C), K2570R (7863A>G 7864G>A), I2581N (7896T>A 7897A>C), S2597T (7944G>C), I2612L (7988A>C), M2618I (8008G>T), D2622N (8018G>A), K2625I (8028A>T 8029A>T), E2626D (8032G>T), S2627M (8033T>A 8034C>T 8035C>G), Y2631F (8046A>T), I2641M (8077A>G), Q2644S (8084C>T 8085A>C 8086G>T), A2654S (8114G>T 8116T>A), L2657C (8124T>G 8125A>T), L2661M (8135C>A 8137T>G), I2662S (8138A>T 8139T>C 8140T>A), K2670R (8163A>G 8164A>G), S2673L (8171A>C 8172G>T 8173T>A), V2683M (8201G>A 8203T>G), I2688V (8216A>G 8218T>C), S2711T (8285T>A), K2732M (8349A>T), E2753Q (8411G>C), R2755K (8418G>A), A2771T (8465G>A 8467T>C), E2775D (8479A>T), L2776K (8480T>A 8481T>A 8482G>A), A2782V (8499C>T 8500A>T), I2785V (8507A>G), D2788E (8518T>G), Y2789S (8520A>C), D2791L (8525G>T 8526A>T 8527T>G), S2792D (8528A>G 8529G>A 8530C>T), L2793V (8531T>G 8533G>A), A2794E (8535C>A), S2795Q (8537T>C 8538C>A 8539T>A), D2797H (8543G>C 8545T>C), I2798M (8548A>G), Y2799I (8549T>A 8550A>T 8551T>A), L2800F (8552C>T), T2804K (8565C>A), A2813T (8591G>A), N2816S (8601A>G 8602C>T), S2818G (8606T>G 8607C>G), S2824L (8624A>C 8625G>T 8626T>C), V2825A (8628T>C), T2827K (8634C>A 8635G>A), T2828P (8636A>C), G2829D (8640G>A 8641A>C), E2830Q (8642G>C 8644G>A), K2832N (8650G>C), L2834S (8654C>T 8655T>C), V2835A (8658T>C), N2869K (8761T>G), K2870R (8763A>G), S2874N (8775G>A), N2880L (8792A>C 8793A>T 8794C>G), H2895Q (8839C>A), N2903D (8861A>G), R2904K (8864C>A 8865G>A 8866T>A), E2946D (8992G>C), M3001N (9156T>A 9157G>T), V3013I (9191G>A 9193T>C), P3015S (9197C>T), N3036T (9261A>C 9262C>A), Q3060T (9332C>A 9333A>C 9334A>T), G3065N (9347G>A 9348G>A 9349T>C) | | | | | | | | |

| | Begin | End | Coverage | Score | Concordance | Matches | Identities | I/D/M/F* | Stop Codons |
|---|---|---|---|---|---|---|---|---|---|
| NT | 1045 | 9396 | 60.8% | 6095 | 52.5% | 5835 (99.7%) | 4455 (76.1%) | 0/19 | |

Codon mutations:

CGT297..A (1045T>A), GAT298GAC (1048T>C), TTG300ATG (1052T>A), TGT301TGC (1057T>C), GAT303GAC (1063T>C), CTT304TTA (1064C>T 1066T>A), TAC306TAT (1072C>T), AAG309GCA (1079A>G 1080A>C 1081G>A), CAT310CAC (1084T>C), TTT311TTC (1087T>C), TCA312TCT (1090A>T), CAA315CAG (1099A>G), ATT316ATA (1102T>A), TCA317TCG (1105A>G), CTG322TTA (1118C>T 1120G>A), TGT325TGC (1129T>C), AAA326AAG (1132A>G), TAC328TAT (1138C>T), GCG330ACA (1142G>A 1144G>A), GGA331GGG (1147A>G), AGA332AAA (1149G>A), GAA334GAT (1156A>T), ATT335TTA (1157A>T 1159T>A), GAT336AAT (1160G>A), TCG344TCC (1186G>C), GTT345GTA (1189T>A), TCT346TCC (1192T>C), TTA347CAA (1193T>C 1194T>A), TGT348TGC (1198T>C), GGT349GGA (1201T>A), GTC351GTT (1207C>T), GCG352GCT (1210G>T), TTA354TTG (1216A>G), GTG355GTA (1219G>A), CAG357CAA (1225G>A), TTA358CTA (1226T>C), CCA361CCG (1237A>G), GGC363GGG (1243C>G), TGC367TGT (1255C>T), ACA368ATT (1257C>T 1258A>T), CGG437AGA (1463C>A 1465G>A), AAC438AAT (1468C>T), AAG439AAA (1471G>A), CCC441CCA (1477C>A), GTT445GTC (1489T>C), CTT446TTG (1490C>T 1492T>G), GAT450GAA (1504T>A), CTA451TGT (1505C>T 1506T>G 1507A>T), TTA452CTT (1508T>C 1510A>T), TTG453CTG (1511T>C), AAG454AAA (1516G>A), TCT456ACA (1520T>A 1522T>A), TTA457ACA (1523T>A 1524T>C), ATG458TTG (1526A>T), AGT459GAT (1529A>G 1530G>A), GAG462GAA (1540G>A), GGT464GGC (1546T>C), GTA465ATA (1547G>A), AGT468GGT (1556A>G), TTG470CTA (1562T>C 1564G>A), CTT471CTG (1567T>G), GTT473ATT (1571G>A), GCA474GCT (1576A>T), CGA475AGA (1577C>A), TTG477CTC (1583T>C 1585G>C), AAC478AAT (1588C>T), AGA557CGC (1823A>C 1825A>C), AAA558AAG (1828A>G), TTA559CTT (1829T>C 1831A>T), GCA560GCG (1834A>G), GGC562GGA (1840C>A), TTG564CTT (1844T>C 1846G>T), ATT565GTG (1847A>G 1849T>G), TCT572ACA (1868T>A 1870T>A), CTT573TTA (1871C>T 1873T>A), CGG574CGA (1876G>A), GTC578GTT (1888C>T), GGG579GGT (1891G>T), GAG580GAA (1894G>A), AAG581AGG (1896A>G), ATA582ATT (1900A>T), ACA583GAG (1901A>G 1902C>A 1903A>G), GCA584TCA (1904G>T), CCC586GCA (1910C>G 1912C>A), AGC593AGT (1933C>T), CGT595AGA (1937C>A 1939T>A), GAT597GAC (1945T>C), TTC599TTT (1951C>T), TTA600CTT (1952T>C 1954A>T), TAC601TAT (1957C>T), TGC604TGT (1966C>T), ACT607ACA (1975T>A), GGC611GGG (1987C>G), TTG614CTT (1994T>C 1996G>T), CTA615CTT (1999A>T), TTG618CTT (2006T>C 2008G>T), ACC622ACA (2020C>A), AAG624AAA (2026G>A), CAT625CAC (2029T>C), CTT626ATA (2030C>A 2032T>A), GGT629GGC (2041T>C), TCT631ACC (2045T>A 2047T>C), GGT632GGA (2050T>A), GAT633GAC (2053T>C), GCA634TCT (2054G>T 2056A>T), AAA635AAG (2059A>G), TTT636TTC (2062T>C), ATA637ATT (2065A>T), GAT638GAC (2068T>C), CTT639TTA (2069C>T 2071T>A), AAG641AAC (2077G>C), GAA642TCA (2078G>T 2079A>C), ATT643TCT (2081A>T 2082T>C), AGT644GAA (2084A>G 2085G>A 2086T>A), GAA645CAG (2087G>C 2089A>G), GGG646GGA (2092G>A), TAC648TTT (2097A>T 2098C>T), GAA652GAG (2110A>G), GGA653GGT (2113A>T), GCT1006..A (3172T>A), AGG1007AAG (3174G>A), TTC1008TAC (3177T>A), ACC1009ACA (3181C>A), AAC1010AAT (3184C>T), AAG1011AGG (3186A>G), AAA1012ACG (3189A>C 3190A>G), TCA1015TCT (3199A>T), GCT1016ATT (3200G>A 3201C>T), GTT1017CTT (3203G>C), AGT1018AGA (3208T>A), TAT1019TAC (3211T>C), TGT1020TGC (3214T>C), ATA1021GTA (3215A>G), AAA1026AAG (3232A>G), ACG1029ACA (3241G>A), GTT1030GTG (3244T>G), TTG1031CTG (3245T>C), AGT1032ACT (3249G>C), CTG1035TTA (3257C>T 3259G>A), CTA1037TTG (3263C>T 3265A>G), ACC1038ACA (3268C>A), GTC1040GTA (3274C>A), CGA1042CGT (3280A>T), GTT1044GTG (3286T>G), CAT1045CAC (3289T>C), TAC1047TAT (3295C>T), GTG1049ATC (3299G>A 3301G>C), AGG1052AAA (3309G>A 3310G>A), AAT1053AAC (3313T>C), GCA1056ACA (3320G>A), GAG1058GAA (3328G>A), AAC1061ACA (3336A>C 3337C>A), AAG1062CAT (3338A>C 3340G>T), GCA1063GCT (3343A>T), GCG1065ATG (3347G>A 3348C>T), TGC1066TGT (3352C>T), GAA1069GAG (3361A>G), ATT1070GTT (3362A>G), GGT1071GGG (3367T>G), ATC1073ATT (3373C>T), ATC1074ATT (3376C>T), TTC1077TTT (3385C>T), TTC1079TTT (3391C>T), AAT1081GAA (3395A>G 3397T>A), GGT1082GGG (3400T>G), CGT1083AGG (3401C>A 3403T>G), TTA1084CCC (3404T>C 3405T>C 3406A>C), CCG1085CCC (3409G>C), ACT1086ACA (3412T>A), TCT1088GCA (3416T>G 3418T>A), GAA1089GAG (3421A>G), AGA1091AAA (3426G>A), AGC1092GAG (3428A>G 3429G>A 3430C>G), AAG1093CGA (3431A>C 3432A>G 3433G>A), GTG1094ATC (3434G>A 3436G>C), AAC1095AAT (3439C>T), TCA1096CCA (3440T>C), TCT1097TCA (3445T>A), GAT1098AAC (3446G>A 3448T>C), ACA1099AAA (3450C>A), CAG1100GCT (3452C>G 3453A>C 3454G>T), CTA1101CTC (3457A>C), ATC1102ATG (3460C>G), AAG1103GAT (3461A>G 3463G>T), ATT1105GTG (3467A>G 3469T>G), GAC1106AAC (3470G>A), CAT1107CAC (3475T>C), TAT1109TAC (3481T>C), CAG1110TGC (3482C>T 3483A>G 3484G>C), GAA1113GGT (3492A>G 3493A>T), ACT1114ACG (3496T>G), ACT1115CAA (3497A>C 3498C>A 3499T>A), CAG1117CAA (3505G>A), AAG1118AAA (3508G>A), GGT1119GGA (3511T>A), CAG1121TCT (3515C>T 3516A>C 3517G>T), TTC1123TTT (3523C>T), GCC1124GCA (3526C>A), TAC1125TTC (3528A>T), CGC1128CGA (3538C>A), ATA1130ATT (3544A>T), GTC1132ATT (3548G>A 3550C>T), ATA1133GTA (3551A>G), ACT1134ACA (3556T>A), CTG1135CTT (3559G>T), CTT1136CTC (3562T>C), GTA1137ATC (3563G>A 3565A>C), GCT1139GCC (3571T>C), GTT1148GTC (3598T>C), TAC1149TTC (3600A>T), AAA1150CGA (3602A>C 3603A>G), CTC1152TTG (3608C>T 3610C>G), AGC1153AAC (3612G>A), AAA1154AAG (3616A>G), TTT1155TTC (3619T>C), AAA1156AAG (3622A>G), GGA1157GGC (3625A>C), TTA1158CTG (3626T>C 3628A>G), GTG1159GTT (3631G>T), GTT1160GTC (3634T>C), TCT1161TCA (3637T>A), AAG1164AAA (3646G>A), GAA1165GAG (3649A>G), ACT1166GCG (3650A>G 3652T>G), ACT1167ACA (3655T>A), CGT1168AGA (3656C>A 3658T>A), TTG1169CTT (3659T>C 3661G>T), GAG1170GAA (3664G>A), AGT1171TCA (3665A>T 3666G>C 3667T>A), TCT1172TCA (3670T>A), CTT1173TTG (3671C>T 3673T>G), GAC1175GAT (3679C>T), ATC1179ATA (3691C>A), GCA1180GCT (3694A>T), AAT1182AAC (3700T>C), CAA1184CAG (3706A>G), ACA1186ACC (3712A>C), GTG1187GTT (3715G>T), CGT1220AGG (3812C>A 3814T>G), CCA1221CCT (3817A>T), CAC1222CAT (3820C>T), TAC1223TAT (3823C>T), CGC1224AGG (3824C>A 3826C>G), GAA1226GAG (3832A>G), GGA1227GGC (3835A>C), AAG1228CAA (3836A>C 3838G>A), ATT1230ATA (3844T>A), TTT1232TTC (3850T>C), ACT1233ACC (3853T>C), CGA1234AGA (3854C>A), ACT1236ACA (3862T>A), AGC1238AGT (3868C>T), TCA1239TCT (3871A>T), GTG1240GTA (3874G>A), GAG1243GAA (3883G>A), ATT1244ATC (3886T>C), TCA1247TCT (3895A>T), GAG1248AGT (3896G>A 3897A>G 3898G>T), ACC1249GTG (3899A>G 3900C>T 3901C>G), AAG1250AAA (3904G>A), TTA1253CTA (3911T>C), GGT1256GGA (3922T>A), GGT1259GGA (3931T>A), TCT1260TCA (3934T>A), GGT1261GGG (3937T>G), ACT1264ACA (3946T>A), AAT1269AAC (3961T>C), TTG1270CTG (3962T>C), TGC1271AGC (3965T>A), AAG1272AAA (3970G>A), AGA1273AGG (3973A>G), GTC1276GTT (3982C>T), CTG1279CTA (3991G>A), AGA1283CGA (4001A>C), CCT1284CCC (4006T>C), TTG1285TTA (4009G>A), AAT1287AAC (4015T>C), GTG1289GTT (4021G>T), TTT1290TAC (4023T>A 4024T>C), CAG1291CAA (4027G>A), CAA1292CAG (4030A>G), TTG1293CTA (4031T>C 4033G>A), AAG1294AGA (4035A>G 4036G>A), GCA1295GCT (4039A>T), CAG1296CAA (4042G>A), GCT1300GTA (4053C>T 4054T>A), AGT1301AGC (4057T>C), CCA1302CCC (4060A>C), TTG1304CTA (4064T>C 4066G>A), ATA1306ATC (4072A>C), AGA1307CGA (4073A>C), AAA1309AAG (4081A>G), TCG1310TCC (4084G>C), GGT1313GGA (4093T>A), ACC1318ACA (4108C>A), ATT1319ATC (4111T>C), TGC1347TGT (4195C>T), GTG1349GTA (4201G>A), TCT1353TCA (4213T>A), GGA1354GGG (4216A>G), GCT1356GCA (4222T>A), AAT1359AAC (4231T>C), TTG1360CTC (4232T>C 4234G>C), CTT1361TTG (4235C>T 4237T>G), TTT1362TTC (4240T>C), TTT1366TTC (4252T>C), AGA1369AAA (4260G>A), TTG1371CTT (4265T>C 4267G>T), GTG1373GTT (4273G>T), CCG1377CCA (4285G>A), CCA1378CCT (4288A>T), GGC1379GGG (4291C>G), GAA1383GAG (4303A>G), TCA1385ACC (4307T>A 4309A>C), ACA1386ACT (4312A>T), TAT1388TAC (4318T>C), ATC1392GTG (4328A>G 4330C>G), ATT1394ATG (4336T>G), GAA1395GAG (4339A>G), AAC1397AGC (4344A>G), CTA1398CTC (4348A>C), TCT1399TCA (4351T>A), TTC1400TTT (4354C>T), CAG1401CAA (4357G>A), GTG1404GTT (4366G>T), TCA1405GCA (4367T>G), GGT1408GGG (4378T>G), CAG1410CAA (4384G>A), TCA1411TCT (4387A>T), GCT1413GTG (4392C>T 4393T>G), GTC1416ATC (4400G>A), CGA1417CGT (4405A>T), GGA1419GGT (4411A>T), AAC1420AAT (4414C>T), ATA1422ATT (4420A>T), TTA1423CTC (4421T>C 4423A>C), AGT1428AGC (4438T>C), GTT1432GTG (4450T>G), GAT1433GAC (4453T>C), TCA1434AAT (4454T>A 4455C>A 4456A>T), CTT1435TTA (4457C>T 4459T>A), AAA1437AAG (4465A>G), TTA1438CTG (4466T>C 4468A>G), CTT1439TTG (4469C>T 4471T>G), TTG1628..C (5038G>C), CTG1631TTG (5045C>T), GCA1632GCT (5050A>T), CCA1634CCC (5056A>C), CAC1635AAT (5057C>A 5059C>T), AAG1636AAA (5062G>A), AGC1637GGG (5063A>G 5065C>G), ATT1638ATA (5068T>A), GGA1639AAT (5069G>A 5070G>A 5071A>T), GTT1640TCA (5072G>T 5073T>C 5074T>A), ATT1642ATC (5080T>C), CCA1643CCG (5083A>G), ACT1646ACA (5092T>A), TAT1647TAC (5095T>C), AGA1648CGT (5096A>C 5098A>T), AGG1649CGC (5099A>C 5101G>C), ACA1650CTT (5102A>C 5103C>T 5104A>T), GGC1651GGA (5107C>A), CAT1653CAA (5113T>A), ACT1654ACA (5116T>A), GAA1655CAA (5117G>C), GAA1657GAG (5125A>G), AAT1658AAC (5128T>C), ATT1661ATA (5137T>A), GCT1662CCA (5138G>C 5140T>A), CTC1664CTT (5146C>T), ACA1665TCA (5147A>T), GAA1667GAT (5155A>T), GTC1668GTA (5158C>A), CCG1669CCA (5161G>A), AAA1671AGA (5166A>G), GTT1672GTG (5170T>G), CAC1673_AGG1674del (5171_5178delCACAGGGA), GAG1675--C (5171_5178delCACAGGGA 5179G>C), GTA1676GTC (5182A>C), GCA1679GCT (5191A>T), GTT1680GTG (5194T>G), TGT1681AAT (5195T>A 5196G>A), GGT1690GGA (5224T>A), AGA1691CGC (5225A>C 5227A>C), CTA1692TTA (5228C>T), ACT1693ACA (5233T>A), AGT1694AGC (5236T>C), CAC1695CAT (5239C>T), GTT1697GTC (5245T>C), TCA1698TCC (5248A>C), GTG1700GTC (5254G>C), TAT1702TAC (5260T>C), TTG1704TTA (5266G>A), CAG1705CAA (5269G>A), ACT1706ACA (5272T>A), TCA1710TCT (5284A>T), ATT1711ATC (5287T>C), GCA1712TCA (5288G>T), AGG1713CGA (5291A>C 5293G>A), ACT1716ACA (5302T>A), ATA1718ATC (5308A>C), AAC1719AAT (5311C>T), CAG1720CAA (5314G>A), GCT1734GCA (5356T>A), AGT1737AGC (5365T>C), ACA1739ACT (5371A>T), ACA1740TGT (5372A>T 5373C>G 5374A>T), AAC1744AAT (5386C>T), TCG1746TCT (5392G>T), CTA1747TTG (5393C>T 5395A>G), GTC1750GTT (5404C>T), ATT1754ATC (5416T>C), CGA1755CGT (5419A>T), CGA1757CGG (5425A>G), GCT1759GCA (5431T>A), AGG1760CGT (5432A>C 5434G>T), CAG1764CAA (5446G>A), TTG1767CTT (5453T>C 5455G>T), CGG1787CGA (5515G>A), GTG1792GTT (5530G>T), CTC1793ATA (5531C>A 5533C>A), GCC1794TCT (5534G>T 5536C>T), AAT1795AAC (5539T>C), GGC1797GGA (5545C>A), GCA1801TCC (5555G>T 5557A>C), TTC1804TTT (5566C>T), CAA1805CAG (5569A>G), TCA1807CAA (5573T>C 5574C>A), GGA1809GAA (5580G>A), TCA1811GCA (5585T>G), AAG1812CGC (5588A>C 5589A>G 5590G>C), TTT1813TTC (5593T>C), GGT1815AAA (5597G>A 5598G>A 5599T>A), TTA1816CTT (5600T>C 5602A>T), GGA1818GGT (5608A>T), CAA1819CAG (5611A>G), AAG1822AGT (5619A>G 5620G>T), TCT1823TCC (5623T>C), TTG1824CTC (5624T>C 5626G>C), ATC1825ATT (5629C>T), ACT1826ACA (5632T>A), AGA1827AGG (5635A>G), GCT1829ATC (5639G>A 5640C>T 5641T>C), ATA1831GTT (5645A>G 5647A>T), ATA1832GTT (5648A>G 5650A>T), GGA1838GGT (5668A>T), GCT1840GCA (5674T>A), CTT1843TTA (5681C>T 5683T>A), TAC1844TAT (5686C>T), AAG1845TCT (5687A>T 5688A>C 5689G>T), CAA1848AAG (5696C>A 5698A>G), AGT1849GAG (5699A>G 5700G>A 5701T>G), AAT1850AGG (5703A>G 5704T>G), TCT1851GTA (5705T>G 5706C>T 5707T>A), AAA1852AGT (5709A>G 5710A>T), GAA1853GAG (5713A>G), GAA1854ACG (5714G>A 5715A>C 5716A>G), CTG1857CCC (5724T>C 5725G>C), CAA1858CAG (5728A>G), AGG1860AGA (5734G>A), GCA1864ACC (5744G>A 5746A>C), AAG1866del (5750_5754delAAGTT), TTG1867--A (5750_5754delAAGTT 5755G>A), CGA1868CGC (5758A>C), GCA1870GCT (5764A>T), TCC1872TCA (5770C>A), GCG1874GCA (5776G>A), GAG1875GAA (5779G>A), AAA1876AAG (5782A>G), AAT1878AAC (5788T>C), GTC1880GTT (5794C>T), GTC1881GTT (5797C>T), GAA1882GAG (5800A>G), GGA1883GGC (5803A>C), GCC1886GCA (5812C>A), GCC1887GCA (5815C>A), CTC1888CTA (5818C>A), ACA1889ACT (5821A>T), CAC1890CAT (5824C>T), TAT1891TAC (5827T>C), GCA1895GCT (5839A>T), GCC1897GCA (5845C>A), TCA1898AAG (5846T>A 5847C>A 5848A>G), AAA1899AAG (5851A>G), AAA1903AAG (5863A>G), ACA1905ACT (5869A>T), GGA1908GGC (5878A>C), CAT1911CAC (5887T>C), AAG1912AAA (5890G>A), AGA1914AGG (5896A>G), AAG1915AAA (5899G>A), TTC1916TTT (5902C>T), GGA1921GGT (5917A>T), CCT1924CCA (5926T>A), ACC1925ACT (5929C>T), GAC1926GAT (5932C>T), TTT1927TTC (5935T>C), CAA1928CAG (5938A>G), GTT1930GTA (5944T>A), CGT1931CGA (5947T>A), GTG1933GTA (5953G>A), CCA1935CCT (5959A>T), TTA1936TTG (5962A>G), GGT1938GGA (5968T>A), CGC1939AAG (5969C>A 5970G>A 5971C>G), GCA1941GTG (5976C>T 5977A>G), GAA1943GAG (5983A>G), TCT1944TCA (5986T>A), CCG1945CCA (5989G>A), ACT1947ACA (5995T>A), GAC1948GAT (5998C>T), GAG1954GAA (6016G>A), GGA1957GGC (6025A>C), AAA1958AAG (6028A>G), ATA1959ATC (6031A>C), GCT1961GAA (6036C>A 6037T>A), TTG2045TTA (6289G>A), GGA2047GGG (6295A>G), GCG2048TCA (6296G>T 6298G>A), TCA2049TCT (6301A>T), CTT2050TTA (6302C>T 6304T>A), GCA2051GCG (6307A>G), GGA2053GGG (6313A>G), GTT2054GTG (6316T>G), CGA2055CGT (6319A>T), ATC2060ATT (6334C>T), TCT2061TCA (6337T>A), GCT2062GCC (6340T>C), GCA2063TCA (6341G>T), CTT2064CTC (6346T>C), TGT2065TGC (6349T>C), AAA2066AAG (6352A>G), CTG2067CTA (6355G>A), ACA2068ACC (6358A>C), GAT2070GAC (6364T>C), GGT2073GGA (6373T>A), GTG2074GTT (6376G>T), AGG2075CGA (6377A>C 6379G>A), ATC2076ATG (6382C>G), TCT2079TCA (6391T>A), GGC2080GGG (6394C>G), GGT2082GGA (6400T>A), GGA2084GGG (6406A>G), TTC2086TAT (6411T>A 6412C>T), ATA2087ATT (6415A>T), ATT2088ATC (6418T>C), ACG2089ACA (6421G>A), AAG2091AAA (6427G>A), TTA2093TTG (6433A>G), TTC2094TTT (6436C>T), AGA2095CGT (6437A>C 6439A>T), GGT2099GGG (6451T>G), ATG2101CTG (6455A>C), GAA2102GAG (6460A>G), ATC2103GTT (6461A>G 6463C>T), CAA2106CAT (6472A>T), GGT2108GGA (6478T>A), TTG2109CTT (6479T>C 6481G>T), ATT2112ATC (6490T>C), CGC2113AGA (6491C>A 6493C>A), CTT2115TTG (6497C>T 6499T>G), AAG2116AAA (6502G>A), ACG2117ACA (6505G>A), TTG2118CTT (6506T>C 6508G>T), TTG2122TTA (6520G>A), GAG2124GAT (6526G>T), AAC2125GGT (6527A>G 6528A>G 6529C>T), GAC2127GAT (6535C>T), CTA2129CTT (6541A>T), ATA2130TTA (6542A>T), ATT2131ATC (6547T>C), AAA2132AAG (6550A>G), GAT2136GAC (6562T>C), TTT2137TTC (6565T>C), GCT2141GCC (6577T>C), TTG2144TTA (6586G>A), AAG2145AAA (6589G>A), TTC2146TTT (6592C>T), CGA2147AGG (6593C>A 6595A>G), CTT2148ATG (6596C>A 6598T>G), GAA2152GAG (6610A>G), GAG2153GAA (6613G>A), AAA2154AAG (6616A>G), GCC2155GCA (6619C>A), GTG2158ATT (6626G>A 6628G>T), CAG2163CAA (6643G>A), ACA2164ACC (6646A>C), AGC2166AGT (6652C>T), ATG2167CTG (6653A>C), CAA2170CTA (6663A>T), GTT2171GTG (6667T>G), ACA2176ATA (6681C>T), ATA2177GTT (6683A>G 6685A>T), GTT2178GTA (6688T>A), CCA2179CCT (6691A>T), AAA2182AAG (6700A>G), TCA2183TCT (6703A>T), CAT2188CAC (6718T>C), ATC2190ATT (6724C>T), ACA2191TCA (6725A>T), GGC2195GGG (6739C>G), CAG2196AAT (6740C>A 6742G>T), GGT2198GGG (6748T>G), CCA2200del (6752_6756delCCATT), TTA2201--C (6752_6756delCCATT 6757A>C), GCT2203GCA (6763T>A), TTG2204TTA (6766G>A), AAA2205AAG (6769A>G), AAG2208AAC (6778G>C), CTT2210CTA (6784T>A), GGA2211GGG (6787A>G), ATT2212ATA (6790T>A), GGT2215GGA (6799T>A), TCG2216TCA (6802G>A), TTA2219CTA (6809T>C), ACA2220ACG (6814A>G), AAT2222AAC (6820T>C), CAC2223CAT (6823C>T), ACC2225ACA (6829C>A), TAC2226TAT (6832C>T), ATT2227ATC (6835T>C), GAA2229GAG (6841A>G), CAA2232CAT (6850A>T), AAT2233GAC (6851A>G 6853T>C), TTG2234ATC (6854T>A 6856G>C), ATA2235TTG (6857A>T 6859A>G), ATG2236AAA (6861T>A 6862G>A), GCA2238TCA (6866G>T), CTC2239ATG (6869C>A 6871C>G), GAA2240GAT (6874A>T), ATG2241CTT (6875A>C 6877G>T), GAT2242GAG (6880T>G), GTG2244GTT (6886G>T), AAA2248AAG (6898A>G), TTC2249TTT (6901C>T), AAT2250AAC (6904T>C), TTG2251CTA (6905T>C 6907G>A), AAT2252AAC (6910T>C), AGC2255AGT (6919C>T), GGC2257GGT (6925C>T), CTC2259TTG (6929C>T 6931C>G), AAA2260AAG (6934A>G), ATT2261CTC (6935A>C 6937T>C), CAA2262CAG (6940A>G), GAG2267GAA (6955G>A), CAG2268GAA (6956C>G 6958G>A), ACT2272ACA (6970T>A), GAG2273GCA (6972A>C 6973G>A), ATT2275ATA (6979T>A), GTT2276ATA (6980G>A 6982T>A), GAT2278GAC (6988T>C), CTT2279TTG (6989C>T 6991T>G), TTT2280CTC (6992T>C 6994T>C), GAT2281GAA (6997T>A), GAG2282GAA (7000G>A), ATT2284ACA (7005T>C 7006T>A), ATA2286GCA (7010A>G 7011T>C), GCC2288TCT (7016G>T 7018C>T), AAT2289CAA (7019A>C 7021T>A), ATG2290ATT (7024G>T), CAA2291CAG (7027A>G), AAT2292AAG (7030T>G), GTG2294GTT (7036G>T), TAC2295TTT (7038A>T 7039C>T), ATT2298ATA (7048T>A), TGT2299CGG (7049T>C

7051T>G), GAC2300GAT (7054C>T), TTG2302CTT (7058T>C 7060G>T), GTC2303GTA (7063C>A), CCA2304CCT (7066A>T), ATT2305GTC (7067A>G 7069T>C), GCT2306TCC (7070G>T 7072T>C), ACA2307TCC (7073A>T 7075A>C), TGC2308TGT (7078C>T), TTA2312CTT (7088T>C 7090A>T), GTC2313GTT (7093C>T), AAA2315AAG (7099A>G), GTT2317GTG (7105T>G), GTT2318GTG (7108T>G), AAG2319AAA (7111G>A), AAG2321AAA (7117G>A), AAT2323CCA (7121A>C 7122A>C 7123T>A), CTG2324TTG (7124C>T), TTC2325TTT (7129C>T), AGC2326AGT (7132C>T), TTG2327GTC (7133T>G 7135G>C), TAT2328TAC (7138T>C), TTA2329CTG (7139T>C 7141A>G), GCC2330GCT (7144C>T), GAA2333AGT (7151G>A 7152A>G 7153A>T), GAG2334GAT (7156G>T), GAG2335GCA (7158A>C 7159G>A), GAA2336GAG (7162A>G), AAA2337CAA (7163A>C), TAT2338TAC (7168T>C), CGA2340AGA (7172C>A), CCA2341CCT (7177A>T), CTA2342TTG (7178C>T 7180A>G), GGA2344GGC (7186A>C), TAT2346TAC (7192T>C), CTA2347CAA (7194T>A), AGC2349AGT (7201C>T), AAA2350GCA (7202A>G 7203A>C), TTG2351CTA (7205T>C 7207G>A), AGA2353AAG (7212G>A 7213A>G), GAG2354GAA (7216G>A), GCA2355GCT (7219A>T), CTG2357CTT (7225G>T), ATA2360ATT (7234A>T), AAA2362AAG (7240A>G), TAC2363TAT (7243C>T), TCA2364TCG (7246A>G), AGT2365AAT (7248G>A), ACA2367GTA (7253A>G 7254C>T), GAG2368GAA (7258G>A), GGA2370GGT (7264A>T), TCT2371GCT (7265T>G), GTG2372GTC (7270G>C), ACA2374ACT (7276A>T), ATC2376ATT (7282C>T), TTT2377TTC (7285T>C), GAT2378GGT (7287A>G), AGA2379CAG (7289A>C 7290G>A 7291A>G), GCA2380GCC (7294A>C), GTG2381GTC (7297G>C), AAT2382ACT (7299A>C), TTG2384GTC (7304T>G 7306G>C), ATT2385ATA (7309T>A), AAT2386AAC (7312T>C), CTC2388CTT (7318C>T), GAA2389GTT (7320A>T 7321A>T), GGT2392GGC (7330T>C), GTG2399GTT (7351G>T), ATG2403CTG (7361A>C), CAG2404CAA (7366G>A), ATA2405ATC (7369A>C), TTC2406TTT (7372C>T), GCT2408GCA (7378T>A), TTA2409TTG (7381A>G), AAT2410AAC (7384T>C), GCA2414GCT (7396A>T), GCT2417GCC (7405T>C), CTA2418CTT (7408A>T), AAA2420AAG (7414A>G), AAA2423AAG (7423A>G), AGA2424AGG (7426A>G), GAT2425GAC (7429T>C), TAC2426TAT (7432C>T), GCC2428GCA (7438C>A), AAT2429TCT (7439A>T 7440A>C), TGC2430TCA (7443G>C 7444C>A), TCA2431ACA (7445T>A), GCT2433GCC (7453T>C), GAT2434GAC (7456T>C), TTG2435CTT (7457T>C 7459G>T), GAA2436GAG (7462A>G), AAT2437AGG (7464A>G 7465T>G), TTC2438TAC (7467T>A), TTG2439TTA (7471G>A), AGA2440ATG (7473G>T 7474A>G), GCC2441GCA (7477C>A), TGT2443TGC (7483T>C), GAG2444AAA (7484G>A 7486G>A), AGA2445CGC (7487A>C 7489A>C), CTC2446CTT (7492C>T), TTT2447TAT (7494T>A), CTT2448CTC (7498T>C), GGC2449GGT (7501C>T), CTT2451CTC (7507T>C), GGG2452GGA (7510G>A), TTG2453TTA (7513G>A), GGC2456GGA (7522C>A), TCA2457TCC (7525A>C), CTT2458CTA (7528T>A), GAA2461GAG (7537A>G), TTG2462CTG (7538T>C), CGG2463AGA (7541C>A 7543G>A), ATT2465TTG (7547A>T 7549T>G), AAG2467AAA (7555G>A), GTT2468GTG (7558T>G), GAG2469CAA (7559G>C 7561G>A), AAC2471AAT (7567C>T), AAG2472AAA (7570G>A), ACA2473TCA (7571A>T), CGA2474CGC (7576A>C), TTT2476TTC (7582T>C), GCA2479GCT (7591A>T), GTT2481ATA (7595G>A 7597T>A), GAT2482GAC (7600T>C), TTA2484CTT (7604T>C 7606A>T), CTG2485CTA (7609G>A), GCT2486GCA (7612T>A), GGA2487GGG (7615A>G), GTG2491GTA (7627G>A), TTT2494TTC (7636T>C), CAA2497CAG (7645A>G), TTC2498TTT (7648C>T), TAT2499TAC (7651T>C), TCT2500TCA (7654T>A), CTC2501TTA (7655C>T 7657C>A), TTG2503CTC (7661T>C 7663G>C), CAG2504AAG (7664C>A), CCC2506CCA (7672C>A), TCA2507TCT (7675A>T), GTC2509GTT (7681C>T), GGA2510GGT (7684A>T), AAA2513AAG (7693A>G), TTT2514TTC (7696T>C), AGA2516GGA (7700A>G), GGG2517GGT (7705G>T), GAT2519GAC (7711T>C), AAA2520AAG (7714A>G), TTG2521CTT (7715T>C 7717G>T), AGA2523AGG (7723A>G), GGG2524AGC (7724G>A 7726G>C), CTT2525TTA (7727C>T 7729T>A), CCT2526CCG (7732T>G), ACT2530GTT (7742A>G 7743C>T), TGT2532TGC (7750T>C), GCT2534GCA (7756T>A), GAC2535GAT (7759C>T), AGG2538AGA (7768G>A), GAT2540GAC (7774T>C), CTA2543TTG (7781C>T 7783A>G), CCA2545CCT (7789A>T), CTT2547CTC (7795T>C), TTG2548ATC (7796T>A 7798G>C), ATT2551CTC (7805A>C 7807T>C), ATT2552ATC (7810T>C), GTC2554ATC (7814G>A), CTC2556TTA (7820C>T 7822C>A), TAC2557TTT (7824A>T 7825C>T), GAA2560GAG (7834A>G), GAT2563GAC (7843T>C), ATT2564ATA (7846T>A), GTG2567ACC (7853G>A 7854T>C 7855G>C), AAG2570AGA (7863A>G 7864G>A), TAC2573TAT (7873C>T), ACC2574ACT (7876C>T), ATA2581AAC (7896T>A 7897A>C), TCC2582TCA (7900C>A), ACT2587ACG (7915T>G), GTA2588GTT (7918A>T), ATA2589ATT (7921A>T), CAT2592CAC (7930T>C), GGC2594GGT (7936C>T), AAT2595AAC (7939T>C), AAT2596AAC (7942T>C), AGT2597ACT (7944G>C), GGA2598GGG (7948A>G), CAA2599CAG (7951A>G), TCT2601TCA (7957T>A), ACA2602ACT (7960A>T), GTC2603GTT (7963C>T), GTT2604GTG (7966T>G), GAC2605GAT (7969C>T), CTG2608CTT (7978G>T), GTT2610GTA (7984T>A), AAC2611AAT (7987C>T), ATA2612CTA (7988A>C), TTT2615TTC (7999T>C), TAC2616TAT (8002C>T), ATG2618ATT (8008G>T), CAC2619CAT (8011C>T), TCA2620TCT (8014A>T), CGT2621AGA (8015C>A 8017T>A), GAC2622AAC (8018G>A), ATT2623ATC (8023T>C), CCG2624CCT (8026G>T), AAA2625ATT (8028A>T 8029A>T), GAG2626GAT (8032G>T), TCC2627ATG (8033T>A 8034C>T 8035C>G), AGA2629AGG (8041A>G), TTT2630TTC (8044T>C), TAT2631TTT (8046A>T), GTT2632GTA (8050T>A), GGA2634GGG (8056A>G), CTG2637TTG (8063C>T), GCT2640GCA (8074T>A), ATA2641ATG (8077A>G), GAG2642GAA (8080G>A), CAG2644TCT (8084C>T 8085A>C 8086G>T), GAC2647GAT (8095C>T), TTG2649CTT (8099T>C 8101G>T), AAG2650AAA (8104G>A), TTT2652TTC (8110T>C), CAA2653CAG (8113A>G), GCT2654TCA (8114G>T 8116T>A), TTA2655TTG (8119A>G), TTA2657TGT (8124T>G 8125A>T), GGT2660GGA (8134T>A), CTT2661ATG (8135C>A 8137T>G), ATT2662TCA (8138A>T 8139T>C 8140T>A), ACT2664ACA (8146T>A), ACA2669ACT (8161A>T), AAA2670AGG (8163A>G 8164A>G), GAC2671GAT (8167C>T), CGC2672AGA (8168C>A 8170C>A), AGT2673CTA (8171A>C 8172G>T 8173T>A), GTG2675GTT (8179G>T), TCA2679TCC (8191A>C), CAC2680CAT (8194C>T), GCT2682GCA (8200T>A), GTT2683ATG (8201G>A 8203T>G), CTC2684CTG (8206C>G), GGT2687GGA (8215T>A), ATT2688GTC (8216A>G 8218T>C), TAC2689TAT (8221C>T), ATA2690ATC (8224A>C), CCT2691CCA (8227T>A), GAG2696GAA (8242G>A), CGG2697AGA (8243C>A 8245G>A), GTT2699GTA (8251T>A), TCC2700TCA (8254C>A), CTT2702CTC (8260T>C), GAC2705GAT (8269C>T), AGA2706CGA (8270A>C), TCA2707TCT (8275A>T), GGA2708GGT (8278A>T), GAA2709GAG (8281A>G), CCA2710CCC (8284A>C), TCT2711ACT (8285T>A), CGG2713AGG (8291C>A), CTG2714TTG (8294C>T), GAA2715GAG (8299A>G), GAG2723GAA (8323G>A), GAA2729GAG (8341A>G), CTG2730TTG (8342C>T), ACT2731ACA (8347T>A), AAG2732ATG (8349A>T), AGG2733AGA (8353G>A), AGA2735CGA (8357A>C), CTT2739TTG (8369C>T 8371T>G), GAG2743GAA (8383G>A), TAC2747TAT (8395C>T), GAA2753CAA (8411G>C), GGG2754GGA (8416G>A), AGA2755AAA (8418G>A), GCT2756GCA (8422T>A), TAC2758TAT (8428C>T), CTA2759CTG (8431A>G), GCT2760GCC (8434T>C), GAA2761GAG (8437A>G), ACT2762ACG (8440T>G), GCT2763GCA (8443T>A), CTG2764TTG (8444C>T), TTA2767TTG (8455A>G), CTT2769CTA (8461T>A), GAT2770GAC (8464T>C), GCT2771ACC (8465G>A 8467T>C), CCC2773CCA (8473C>A), ACT2774ACG (8476T>G), GAA2775GAT (8479A>T), TTG2776AAA (8480T>A 8481T>A 8482G>A), GAA2777GAG (8485A>G), TTG2778TTA (8488G>A), CTT2779TTG (8489C>T 8491T>G), GCA2782GTT (8499C>T 8500A>T), GGG2783GGT (8503G>T), TGC2784TGT (8506C>T), ATC2785GTC (8507A>G), GGT2787GGA (8515T>A), GAT2788GAG (8518T>G), TAT2789TCT (8520A>C), GAT2790GAC (8524T>C), GAT2791TTG (8525G>T 8526A>T 8527T>G), AGC2792GAT (8528A>G 8529G>A 8530C>T), TTG2793GTA (8531T>G 8533G>A), GCA2794GAA (8535C>A), TCT2795CAA (8537T>C 8538C>A 8539T>A), GAA2796GAG (8542A>G), GAT2797CAC (8543G>C 8545T>C), ATA2798ATG (8548A>G), TAT2799ATA (8549T>A 8550A>T 8551T>A), CTC2800TTC (8552C>T), GCT2802GCA (8560T>A), GAT2803GAC (8563T>C), ACA2804AAA (8565C>A), GGT2808GGA (8578T>A), AGT2809AGC (8581T>C), ACA2811ACC (8587A>C), GCC2813ACC (8591G>A), GGA2814GGT (8596A>T), AAG2815AAA (8599G>A), AAC2816AGT (8601A>G 8602C>T), TCA2818GGA (8606T>G 8607C>G), AGT2824CTC (8624A>C 8625G>T 8626T>C), GTT2825GCT (8628T>C), ACG2827AAA (8634C>A 8635G>A), ACT2828CCT (8636A>C), GGA2829GAC (8640G>A 8641A>C), GAG2830CAA (8642G>C 8644G>A), AAA2831AAG (8647A>G), AAG2832AAC (8650G>C), CTT2834TCT (8654C>T 8655T>C), GTC2835GCC (8658T>C), CAA2838CAG (8668A>G), GTG2842GTT (8680G>T), GCT2844GCA (8686T>A), ACC2849ACG (8701C>G), GTA2852GTT (8710A>T), AAA2856AAG (8722A>G), ATT2858ATA (8728T>A), CCT2859CCC (8731T>C), ACT2860ACA (8734T>A), CTA2864CTC (8746A>C), AAT2869AAG (8761T>G), AAA2870AGA (8763A>G), GTT2872GTC (8770T>C), ATT2873ATC (8773T>C), AGT2874AAT (8775G>A), CTG2875CTT (8779G>T), GAC2876GAT (8782C>T), CTG2878TTG (8786C>T), CTG2879CTC (8791G>C), AAC2880CTG (8792A>C 8793A>T 8794C>G), CCA2883CCT (8803A>T), GAT2887GAC (8815T>C), TCA2889TCT (8821A>T), CAC2895CAA (8839C>A), GAA2899GAG (8851A>G), AAT2903GAT (8861A>G), CGT2904AAA (8864C>A 8865G>A 8866T>A), GTA2905GTG (8869A>G), CGA2906AGA (8870C>A), GTT2911GTC (8887T>C), ACT2912ACA (8890T>A), GAA2914GAG (8896A>G), CAG2915CAA (8899G>A), CAG2917CAA (8905G>A), ATC2919ATT (8911C>T), TTA2923CTT (8921T>C 8923A>T), ATC2928ATT (8938C>T), AAC2935AAT (8959C>T), ATA2936ATT (8962A>T), GGT2938GGC (8968T>C), ACT2941ACC (8977T>C), GAG2946GAC (8992G>C), TTA2953TTG (9013A>G), ATT2957ATA (9025T>A), GAA2958GAG (9028A>G), CCC2962CCA (9040C>A), TTT2964TTC (9046T>C), CAA2966CAG (9052A>G), GCG2969GCA (9061G>A), CAT2970CAC (9064T>C), GAT2973GAC (9073T>C), CTA2974TTA (9074C>T), GAT2985GAC (9109T>C), CCA2990CCT (9124A>T), CGT2991AGA (9125C>A 9127T>A), CGC2996CGT (9142C>T), ATG3001AAT (9156T>A 9157G>T), AGT3002AGC (9160T>C), CTG3003CTT (9163G>T), GAC3009GAT (9181C>T), GTT3013ATC (9191G>A 9193T>C), ACT3014ACC (9196T>C), CCG3015TCG (9197C>T), CGG3016AGG (9200C>A), ACC3019ACT (9211C>T), GCT3030GCA (9244T>A), CGA3034CGT (9256A>T), AAC3036ACA (9261A>C 9262C>A), ACA3037ACG (9265A>G), ACG3038ACC (9268G>C), AAG3039AAA (9271G>A), TTA3040CTC (9272T>C 9274A>C), CTG3043CTC (9283G>C), GAA3051GAG (9307A>G), ACC3054ACA (9316C>A), CAC3057CAT (9325C>T), CAA3060ACT (9332C>A 9333A>C 9334A>T), GTT3062GTG (9340T>G), AGA3064AGG (9346A>G), GGT3065AAC (9347G>A 9348G>A 9349T>C), CTT3070CTA (9364T>A), TAG3076TAA (9382G>A)

*: Inserts / Deletes / Misaligned / Frameshifts

#### Analysis details

This analysis was performed with panviral2.64

### NGS Details (UN24): Solendovirus venanicotianae

#### Assembly

| | |
|---|---|
| **Coverage Length** | 2027 (5 contig(s)) |
| **Depth Of Coverage** | 11.2 |
| **Number Of Reads** | 201 |
| **Reads Per Million** | 4.02 rpm (after QC) |
| **Ambiguities** | 26 |
| **Assembly Method** | de novo + reference guided assembly |
| **Consensus Caller** | Bcf Tools |

#### Coverage Map

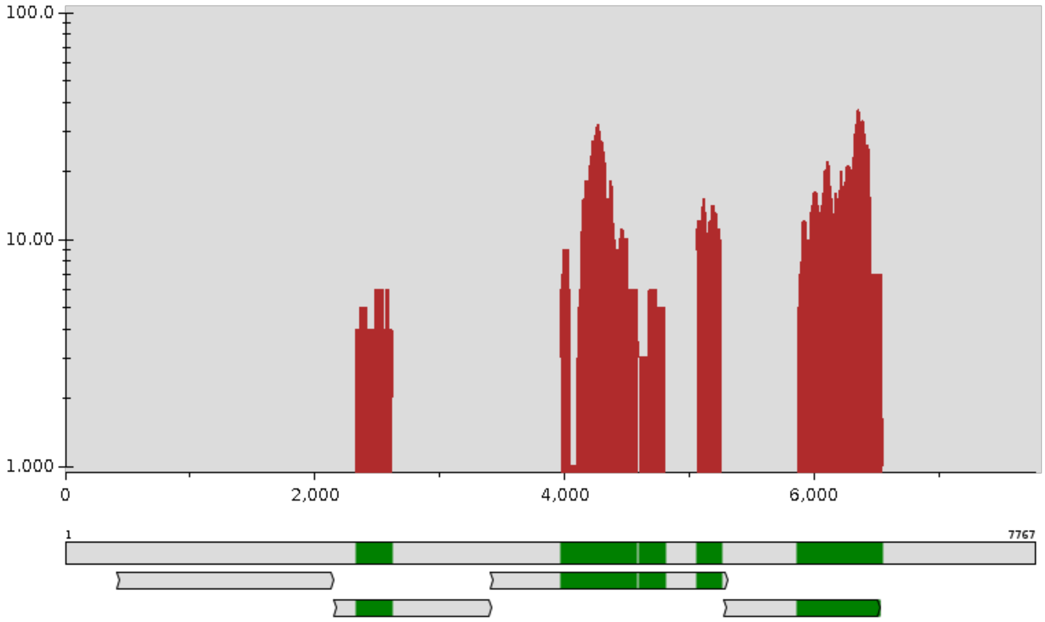

#### Assignment

| | |
|---|---|
| **Type** | Solendovirus venanicotianae (Taxonomy ID: 3048371) |
| **Reference Genome** | NC_003378.1 |
| **NT Identity (%)** | 78.1728 |
| **AA Identity (%)** | 75.2994 |
| **Number Of Stop Codons** | 3 |
| **Number Of CDS** | 4 |

#### Alignment

| | |
|---|---|
| **Alignment Score** | 2323.0 (NT) + 3342.0 (AA) = 5665.0 |
| **Concordance (%)** | 66.0564 |

| **Alignment Method** | Global, seeded, nucleotide + amino acids (AGA) |
|---|---|

#### Genome Region

Sequence starts at position 2326 and ends at position 6548 relative to NC_003378.1 reference sequence.

#### Alignment Detailed Statistics

| | Begin | End | Coverage | Score | Concordance | Matches | Identities | I/D/M/F* | Stop Codons |
|---|---|---|---|---|---|---|---|---|---|
| **NT** | **2326** | **6548** | **26.1%** | **2323** | **57.5%** | **2025 (99.9%)** | **1583 (78.1%)** | **0/2** | |
| Mutations: | 2326A>T, 2329C>A, 2333A>T, 2347G>A, 2350G>C, 2373T>A, 2379C>T, 2381A>G, 2385T>C, 2388A>T, 2389G>A, 2394T>A, 2397G>A, 2400C>A, 2409A>W, 2410A>C, 2412A>W, 2415A>R, 2417A>S, 2418C>Y, 2420A>R, 2421A>R, 2423G>A, 2424A>R, 2425A>R, 2426T>Y, 2428A>R, 2432A>M, 2433T>M, 2434A>M, 2442A>W, 2443C>R, 2446G>R, 2448T>C, 2454G>A, 2457A>T, 2461A>G, 2463C>A, 2469T>A, 2472G>A, 2475T>C, 2481C>T, 2482T>C, 2493T>C, 2500T>C, 2501T>A, 2505A>T, 2538T>A, 2541A>M, 2545A>M, 2550T>Y, 2562T>Y, 2565A>R, 2569A>R, 2571C>A, 2572C>K, 2574A>W, 2577T>W, 2589C>T, 2602A>G, 2610A>T, 2616T>A, 3974G>T, 3977G>T, 3989A>T, 3991A>C, 4010T>A, 4011G>A, 4013A>T, 4014G>A, 4016G>A, 4017A>G, 4025A>T, 4037A>G, 4040T>A, 4045A>G, 4056A>G, 4059G>A, 4061T>C, 4065A>G, 4066C>A, 4076A>G, 4088A>T, 4091T>A, 4094A>G, 4098G>A, 4100T>A, 4103T>G, 4104A>G, 4105A>G, 4115A>G, 4116G>A, 4121T>C, 4122A>G, 4124T>A, 4127C>T, 4133T>C, 4181A>T, 4187C>T, 4190G>A, 4196C>T, 4199A>T, 4212T>C, 4241T>C, 4247A>T, 4250C>A, 4253T>C, 4265G>A, 4274A>G, 4286T>C, 4292C>T, 4298T>C, 4307C>T, 4313G>A, 4325T>C, 4328T>C, 4331G>A, 4338G>A, 4340G>A, 4362G>A, 4364T>A, 4388T>C, 4400T>A, 4406A>T, 4409T>C, 4415T>C, 4421T>C, 4436C>T, 4451C>T, 4456A>G, 4460T>C, 4481A>T, 4484A>T, 4500C>T, 4503C>T, 4508T>C, 4510C>A, 4513G>A, 4516C>T, 4517A>G, 4518C>A, 4519A>T, 4523T>A, 4524G>C, 4532A>C, 4533A>T, 4534A>T, 4542G>C, 4551G>A, 4552C>T, 4559T>A, 4567A>T, 4568C>A, 4569T>G, 4571T>G, 4574T>A, 4596G>T, 4599G>A, 4604T>A, 4612A>C, 4613T>A, 4616G>A, 4619A>G, 4626T>C, 4637A>G, 4652A>G, 4666C>A, 4667T>A, 4670T>C, 4688C>T, 4692C>T, 4693T>C, 4694T>A, 4695G>A, 4697T>C, 4701A>G, 4703C>T, 4706T>A, 4709T>C, 4722T>C, 4730A>C, 4733T>C, 4734C>T, 4740A>C, 4742A>G, 4745A>C, 4752G>A, 4755A>C, 4757G>T, 4763T>C, 4769T>A, 4784C>T, 4785T>C, 4791C>T, 4793T>A, 4794T>A, 4797C>T, 4799G>T, 4806C>T, 5076T>C, 5077T>A, 5079G>T, 5094C>T, 5117A>T, 5120T>G, 5123C>T, 5125C>A, 5128G>A, 5135T>A, 5136G>A, 5147C>T, 5150T>C, 5156A>G, 5174T>A, 5180A>G, 5189T>C, 5195C>T, 5211C>A, 5217C>T, 5219G>A, 5230T>A, 5234A>G, 5237C>T, 5242C>A, 5256A>T, 5859A>G, 5865C>T, 5868T>C, 5872T>C, 5874G>T, 5877A>T, 5880T>A, 5881T>C, 5882T>A, 5886A>T, 5889T>A, 5892A>C, 5894G>A, 5901A>C, 5902T>C, 5908G>A, 5909C>A, 5910C>A, 5913C>A, 5916C>T, 5921A>G, 5925C>T, 5929C>T, 5931A>G, 5932C>A, 5935G>A, 5937T>C, 5950C>T, 5952G>T, 5953A>G, 5954C>A, 5955A>C, 5956G>A, 5959C>G, 5964G>A, 5968G>T, 5970C>G, 5972A>C, 5973T>C, 5978C>T, 5979T>A, 5986T>C, 5991G>A, 5997C>T, 6001C>A, 6002A>T, 6009G>A, 6012G>A, 6015A>T, 6018T>A, 6022G>A, 6023G>A, 6024G>T, 6026C>A, 6027A>T, 6028T>C, 6033C>T, 6036C>T, 6037G>A, 6040C>A, 6041G>A, 6045C>T, 6049T>A, 6051G>A, 6054T>A, 6055A>C, 6067C>T, 6075A>T, 6078G>A, 6081A>C, 6084G>A, 6087T>A, 6090C>A, 6105A>G, 6108T>A, 6112C>T, 6114C>G, 6120G>A, 6123A>T, 6127A>T, 6129C>A, 6139A>G, 6141G>C, 6145G>A, 6146A>G, 6148G>C, 6150A>G, 6154G>C, 6155A>C, 6157G>A, 6163G>C, 6164A>C, 6169G>A, 6171A>T, 6172A>G, 6174T>G, 6177T>A, 6181G>A, 6182A>G, 6183A>T, 6186C>T, 6189T>C, 6201G>A, 6204A>T, 6211A>C, 6219T>G, 6223C>A, 6225A>C, 6232C>T, 6246T>C, 6247C>A, 6248T>C, 6253C>G, 6254A>G, 6258T>C, 6259G>C, 6264T>G, 6269G>A, 6270C>T, 6271T>C, 6277T>A, 6279A>C, 6281G>A, 6282G>A, 6285A>C, 6290C>T, 6299A>T, 6300T>C, 6301T>G, 6302C>A, 6305A>C, 6306T>G, 6307T>G, 6308G>C, 6309C>A, 6316A>T, 6319A>C, 6321A>G, 6322G>C, 6325G>A, 6340C>G, 6341T>C, 6352G>A, 6354C>A, 6355T>A, 6357A>G, 6359G>C, 6360T>A, 6361C>T, 6363T>G, 6369A>T, 6375A>G, 6376C>A, 6380C>A, 6381T>A, 6382A>C, 6384A>T, 6386C>T, 6387C>T, 6389C>T, 6401G>A, 6402G>A, 6404G>A, 6405G>T, 6406A>G, 6407A>G, 6408T>A, 6411C>T, 6414C>T, 6417C>G, 6418C>G, 6419C>A, 6420A>C, 6423T>A, 6424C>T, 6426G>A, 6427T>C, 6442A>T, 6443A>C, 6444G>A, 6445C>A, 6447A>T, 6450T>C, 6451A>G, 6453T>A, 6454C>A, 6455A>G, 6456C>A, 6459A>T, 6463C>G, 6464C>A, 6465A>T, 6468C>A, 6471C>T, 6472C>A, 6478T>A, 6479C>A, 6480A>C, 6482A>G, 6488A>G, 6489A>T, 6493G>A, 6494A>C, 6495A>C, 6496G>A, 6497A>G, 6501C>T, 6502A>T, 6503T>A, 6505G>A, 6513C>A, 6514G>A, 6516A>C, 6517C>A, 6519A>C, 6520C>T, 6529T>A, 6530_6531delTC | | | | | | | | |
| **CDS** | | | | | | | | | |
| Tvvgp2 | 58 | 156 | 23.6% | 310 | 46.3% | 99 (100%) | 74 (74.7%) | 0/0/10/0 | 1 |
| Protein mutations: | R58* (2326A>T), Q59K (2329C>A), K60I (2333A>T), V65I (2347G>A), V66L (2350G>C), K76R (2381A>G), G79R (2389G>A), I86L (2410A>C 2412A>W), R90K (2423G>A 2424A>R), D98B (2446G>R 2448T>C), N103E (2461A>G 2463C>A), L116Q (2500T>C 2501T>A), I131J (2545A>M), I150V (2602A>G) | | | | | | | | |
| Codon mutations: | AGA58TGA (2326A>T), CAA59AAA (2329C>A), AAA60ATA (2333A>T), GTA65ATA (2347G>A), GTT66CTT (2350G>C), ATT73ATA (2373T>A), TGC75TGT (2379C>T), AAA76AGA (2381A>G), TAT77TAC (2385T>C), ACA78ACT (2388A>T), GGA79AGA (2389G>A), GGT80GGA (2394T>A), AAG81AAA (2397G>A), GCC82GCA (2400C>A), CCA85CCW (2409A>W), ATA86CTW (2410A>C 2412A>W), ATA87ATR (2415A>R), AAC88ASY (2417A>S 2418C>Y), AAA89ARR (2420A>R 2421A>R), AGA90AAR (2423G>A 2424A>R), ATA91RYA (2425A>R 2426T>Y), ATA92RTA (2428A>R), AAT93AMM (2432A>M 2433T>M), AAA94MAA (2434A>M), ATA96ATW (2442A>W), CAA97RAA (2443C>R), GAT98RAC (2446G>R 2448T>C), AAG100AAA (2454G>A), GCA101GCT (2457A>T), AAC103GAA (2461A>G 2463C>A), ATT105ATA (2469T>A), AAG106AAA (2472G>A), TAT107TAC (2475T>C), CAC109CAT (2481C>T), TTA110CTA (2482T>C), ACT113ACC (2493T>C), TTA116CAA (2500T>C 2501T>A), ATA117ATT (2505A>T), CCT128CCA (2538T>A), ATA129ATM (2541A>M), ATA131MTA (2545A>M), TAT132TAY (2550T>Y), GAT136GAY (2562T>Y), AGA137AGR (2565A>R), ATC139RTA (2569A>R 2571C>A), CAA140KAW (2572C>K 2574A>W), CCT141CCW (2577T>W), AGC145AGT (2589C>T), ATA150GTA (2602A>G), GGA152GGT (2610A>T), CTT154CTA (2616T>A) | | | | | | | | |
| Tvvgp3 | 188 | 617 | 54.4% | 2026 | 85.1% | 346 (100%) | 293 (84.7%) | 0/0/0/0 | 1 |
| Protein mutations: | W188C (3974G>T), E189D (3977G>T), K194T (3991A>C), E201N (4011G>A 4013A>T), E202K (4014G>A 4016G>A), N203D (4017A>G), K212R (4045A>G), N216D (4056A>G), D217N (4059G>A 4061T>C), T219E (4065A>G 4066C>A), K226N (4088A>T), D227E (4091T>A), D230K (4098G>A 4100T>A), N231K (4103T>G), K232G (4104A>G 4105A>G), E236K (4116G>A), N238E (4122A>G 4124T>A), E310K (4338G>A 4340G>A), A318T (4362G>A 4364T>A), Y349C (4456A>G), S367* (4510C>A), R368K (4513G>A), T369M (4516C>T 4517A>G), Q370I (4518C>A 4519A>T), D371E (4523T>A), E372Q (4524G>C), K375L (4533A>T 4534A>T), E378Q (4542G>C), A381I (4551G>A 4552C>T), N386I (4567A>T 4568C>A), S387A (4569T>G 4571T>G), A396S (4596G>T), E397K (4599G>A), N401T (4612A>C 4613T>A), I403M (4619A>G), T419K (4666C>A 4667T>A), L428S (4692C>T 4693T>C 4694T>A), D429N (4695G>A 4697T>C), N431D (4701A>G 4703C>T), I444L (4740A>C 4742A>G), V448I (4752G>A), P461S (4791C>T 4793T>A), L462I (4794T>A), Q463Y (4797C>T 4799G>T), F556H (5076T>C 5077T>A), A557S (5079G>T), T572N (5125C>A), R573K (5128G>A), V576I (5136G>A), I607K (5230T>A), T611K (5242C>A), I616L (5256A>T) | | | | | | | | |
| Codon mutations: | TGG188TGT (3974G>T), GAG189GAT (3977G>T), ACA193ACT (3989A>T), AAA194ACA (3991A>C), ATT200ATA (4010T>A), GAA201AAT (4011G>A 4013A>T), GAG202AAA (4014G>A 4016G>A), AAT203GAT (4017A>G), ATA205ATT (4025A>T), AAA209AAG (4037A>G), CCT210CCA (4040T>A), AAA212AGA (4045A>G), AAT216GAT (4056A>G), GAT217AAC (4059G>A 4061T>C), ACA219GAA (4065A>G 4066C>A), AAA222AAG (4076A>G), AAA226AAT (4088A>T), GAT227GAA (4091T>A), TTA228TTG (4094A>G), GAT230AAA (4098G>A 4100T>A), AAT231AAG (4103T>G), AAA232GGA (4104A>G 4105A>G), CAA235CAG (4115A>G), GAA236AAA (4116G>A), AGT237AGC (4121T>C), AAT238GAA (4122A>G 4124T>A), AGC239AGT (4127C>T), CAT241CAC (4133T>C), GGA257GGT (4181A>T), AGC259AGT (4187C>T), AGG260AGA (4190G>A), GTC262GTT (4196C>T), ATA263ATT (4199A>T), TTA268CTA (4212T>C), TAT277TAC (4241T>C), ATA279ATT (4247A>T), CCC280CCA (4250C>A), AAT281AAC (4253T>C), AAG285AAA (4265G>A), CAA288CAG (4274A>G), TAT292TAC (4286T>C), TAC294TAT (4292C>T), AGT296AGC (4298T>C), GAC299GAT (4307C>T), AAG301AAA (4313G>A), TAT305TAC (4325T>C), CAT306CAC (4328T>C), TTG307TTA (4331G>A), GAG310AAA (4338G>A 4340G>A), GCT318ACA (4362G>A 4364T>A), TAT326TAC (4388T>C), GTT330GTA (4400T>A), CCA332CCT (4406A>T), TTT333TTC (4409T>C), TAT335TAC (4415T>C), AAT337AAC (4421T>C), TAC342TAT (4436C>T), GAC347GAT (4451C>T), TAT349TGT (4456A>G), TTT350TTC (4460T>C), ATA357ATT (4481A>T), GTA358GTT (4484A>T), CTA364TTA (4500C>T), CTA365TTA (4503C>T), TAT366TAC (4508T>C), TCA367TAA (4510C>A), AGA368AAA (4513G>A), ACA369ATG (4516C>T 4517A>G), CAA370ATA (4518C>A 4519A>T), GAT371GAA (4523T>A), GAA372CAA (4524G>C), ATA374ATC (4532A>C), AAA375TTA (4533A>T 4534A>T), GAA378CAA (4542G>C), GCA381ATA (4551G>A 4552C>T), ATT383ATA (4559T>A), AAC386ATA (4567A>T 4568C>A), TCT387GCG (4569T>G 4571T>G), GGT388GGA (4574T>A), GCA396TCA (4596G>T), GAA397AAA (4599G>A), ATT398ATA (4604T>A), AAT401ACA (4612A>C 4613T>A), CAG402CAA (4616G>A), ATA403ATG (4619A>G), TTA406CTA (4626T>C), CAA409CAG (4637A>G), GGA414GGG (4652A>G), ACT419AAA (4666C>A 4667T>A), CAT420CAC (4670T>C), ATC426ATT (4688C>T), CTT428TCA (4692C>T 4693T>C 4694T>A), GAT429AAC (4695G>A 4697T>C), AAC431GAT (4701A>G 4703C>T), ATT432ATA (4706T>A), GAT433GAC (4709T>C), TTA438CTA (4722T>C), TCA440TCC (4730A>C), TTT441TTC (4733T>C), CTA442TTA (4734C>T), ATA444CTG (4740A>C 4742A>G), GTA445GTC (4745A>C), GTA448ATA (4752G>A), AGG449CGT (4755A>C 4757G>T), TAT451TAC (4763T>C), CCT453CCA (4769T>A), AAC458AAT (4784C>T), TTA459CTA (4785T>C), CCT461TCA (4791C>T 4793T>A), TTA462ATA (4794T>A), CAG463TAT (4797C>T 4799G>T), CTA466TTA (4806C>T), TTT556CAT (5076T>C 5077T>A), GCA557TCA (5079G>T), CTA562TTA (5094C>T), ATA569ATT (5117A>T), GTT570GTG (5120T>G), TAC571TAT (5123C>T), ACC572AAC (5125C>A), AGA573AAA (5128G>A), ATT575ATA (5135T>A), GTA576ATA (5136G>A), GAC579GAT (5147C>T), AAT580AAC (5150T>C), CAA582CAG (5156A>G), ACT588ACA (5174T>A), AAA590AAG (5180A>G), GAT593GAC (5189T>C), GTC595GTT (5195C>T), CGG601AGG (5211C>A), CTG603TTA (5217C>T 5219G>A), ATA607AAA (5230T>A), TTA608TTG (5234A>G), AAC609AAT (5237C>T), ACA611AAA (5242C>A), ATA616TTA (5256A>T) | | | | | | | | |
| Tvvgp4 | 196 | 418 | 53.3% | 1006 | 66.1% | 223 (100%) | 136 (61.0%) | 0/0/0/0 | 1 |
| Protein mutations: | L203Q (5881T>C 5882T>A), N205K (5889T>A), S207N (5894G>A), K209N (5901A>C), A212K (5908G>A 5909C>A 5910C>A), N216S (5921A>G), L220I (5932C>A), D221N (5935G>A 5937T>C), Q226Y (5950C>T 5952G>T), T227D (5953A>G 5954C>A 5955A>C), G228R (5956G>A), Q229E (5959C>G), A232S (5968G>T 5970C>G), N233T (5972A>C 5973T>C), P235L (5978C>T 5979T>A), Y238H (5986T>C), Q243I (6001C>A 6002A>T), G250N (6022G>A 6023G>A 6024G>T), T251N (6026C>A 6027A>T), V255I (6037G>A), R256K (6040C>A 6041G>A), S259T (6049T>A 6051G>A), T261P (6055A>C), E267D (6075A>T), E283D (6123A>T), I285L (6127A>T 6129C>A), K289D (6139A>G 6141G>C), E291R (6145G>A 6146A>G), E292Q (6148G>C 6150A>G), E294P (6154G>C 6155A>C), E295K (6157G>A), E297P (6163G>C 6164A>C), V299I (6169G>A 6171A>T), N300E (6172A>G 6174T>G), E303S (6181G>A 6182A>G 6183A>T), I313L (6211A>C), L317I (6223C>A 6225A>C), L325T (6247C>A 6248T>C), Q327G (6253C>G 6254A>G), A329P (6259G>C), I330M (6264T>G), S332N (6269G>A 6270C>T), Y333H (6271T>C), S335T (6277T>A 6279A>C), R336K (6281G>A 6282G>A), E337D (6285A>C), T339I (6290C>T), Y342F (6299A>T 6300T>C), S343E (6301T>G 6302C>A), N344T (6305A>C 6306T>G), C345A (6307T>G 6308G>C 6309C>A), I348L (6316A>T), E350Q (6322G>C), A351T (6325G>A), L356A (6340C>G 6341T>C), V360I (6352G>A 6354C>A), L361M (6355T>A 6357A>G), S362T (6359G>C 6360T>A), K365N (6369A>T), Q368K (6376C>A), P369Q (6380C>A 6381T>A), T370P (6382A>C 6384A>T), T371I (6386C>T 6387C>T), T372I (6389C>T), R376K (6401G>A 6402G>A), R377N (6404G>A 6405G>T), N378G (6406A>G 6407A>G 6408T>A), P382D (6418C>G 6419C>A 6420A>C), D383E (6423T>A), K390S (6442A>T 6443A>C 6444G>A), L391I (6445C>A 6447A>T), S393G (6451A>G 6453T>A), H394R (6454C>A 6455A>G 6456C>A), K395N (6459A>T), P397D (6463C>G 6464C>A 6465A>T), D398E (6468C>A), L400I (6472C>A), S402N (6478T>A 6479C>A 6480A>C), K403R (6482A>G), K405S (6488A>G 6489A>T), E407T (6493G>A 6494A>C 6495A>C), D408S (6496G>A 6497A>G), I410Y (6502A>T 6503T>A), V411I (6505G>A), D413E (6513C>A), V414I (6514G>A 6516A>C), Q415N (6517C>A 6519A>C) | | | | | | | | |

| | Begin | End | Coverage | Score | Concordance | Matches | Identities | I/D/M/F* | Stop Codons |
|---|---|---|---|---|---|---|---|---|---|
| **NT** | **2326** | **6548** | **26.1%** | **2323** | **57.5%** | **2025 (99.9%)** | **1583 (78.1%)** | **0/2** | |
| Codon mutations: | ACA195..G (5859A>G), TAC197TAT (5865C>T), AAT198AAC (5868T>C), TTG200CTT (5872T>C 5874G>T), ATA201ATT (5877A>T), GCT202GCA (5880T>A), TTA203CAA (5881T>C 5882T>A), CCA204CCT (5886A>T), AAT205AAA (5889T>A), ACA206ACC (5892A>C), AGT207AAT (5894G>A), AAA209AAC (5901A>C), TTA210CTA (5902T>C), GCC212AAA (5908G>A 5909C>A 5910C>A), ACC213ACA (5913C>A), TGC214TGT (5916C>T), AAC216AGC (5921A>G), TAC217TAT (5925C>T), CTA219TTG (5929C>T 5931A>G), CTA220ATA (5932C>A), GAT221AAC (5935G>A 5937T>C), CAG226TAT (5950C>T 5952G>T), ACA227GAC (5953A>G 5954C>A 5955A>C), GGA228AGA (5956G>A), CAA229GAA (5959C>G), GAG230GAA (5964G>A), GCC232TCG (5968G>T 5970C>G), AAT233ACC (5972A>C 5973T>C), CCT235CTA (5978C>T 5979T>A), TAC238CAC (5986T>C), AAG239AAA (5991G>A), TTC241TTT (5997C>T), CAA243ATA (6001C>A 6002A>T), AAG245AAA (6009G>A), AGG246AGA (6012G>A), ATA247ATT (6015A>T), ACT248ACA (6018T>A), GGG250AAT (6022G>A 6023G>A 6024G>T), ACA251AAT (6026C>A 6027A>T), TTA252CTA (6028T>C), TTC253TTT (6033C>T), TAC254TAT (6036C>T), GTA255ATA (6037G>A), CGA256AAA (6040C>A 6041G>A), TTC257TTT (6045C>T), TCG259ACA (6049T>A 6051G>A), GCT260GCA (6054T>A), ACA261CCA (6055A>C), CTA265TTA (6067C>T), GAA267GAT (6075A>T), GAG268GAA (6078G>A), ATA269ATC (6081A>C), AAG270AAA (6084G>A), CCT271CCA (6087T>A), ATC272ATA (6090C>A), AAA277AAG (6105A>G), ATT278ATA (6108T>A), CTC280TTG (6112C>T 6114C>G), AGG282AGA (6120G>A), GAA283GAT (6123A>T), ATC285TTA (6127A>T 6129C>A), AAG289GAC (6139A>G 6141G>C), GAA291AGA (6145G>A 6146A>G), GAA292CAG (6148G>C 6150A>G), GAA294CCA (6154G>C 6155A>C), GAG295AAG (6157G>A), GAA297CCA (6163G>C 6164A>C), GTA299ATT (6169G>A 6171A>T), AAT300GAG (6172A>G 6174T>G), ATT301ATA (6177T>A), GAA303AGT (6181G>A 6182A>G 6183A>T), TTC304TTT (6186C>T), TAT305TAC (6189T>C), AGG309AGA (6201G>A), ATA310ATT (6204A>T), ATA313CTA (6211A>C), ACT315ACG (6219T>G), CTA317ATC (6223C>A 6225A>C), CTA320TTA (6232C>T), TAT324TAC (6246T>C), CTA325ACA (6247C>A 6248T>C), CAG327GGG (6253C>G 6254A>G), AAT328AAC (6258T>C), GCA329CCA (6259G>C), ATT330ATG (6264T>G), AGC332AAT (6269G>A 6270C>T), TAT333CAT (6271T>C), TCA335ACC (6277T>A 6279A>C), AGG336AAA (6281G>A 6282G>A), GAA337GAC (6285A>C), ACA339ATA (6290C>T), TAT342TTC (6299A>T 6300T>C), TCA343GAA (6301T>G 6302C>A), AAT344ACG (6305A>C 6306T>G), TGC345GCA (6307T>G 6308G>C 6309C>A), ATA348TTA (6316A>T), AGA349CGG (6319A>C 6321A>G), GAA350CAA (6322G>C), GCA351ACA (6325G>A), CTA356GCA (6340C>G 6341T>C), GTC360ATA (6352G>A 6354C>A), TTA361ATG (6355T>A 6357A>G), AGT362ACA (6359G>C 6360T>A), CTT363TTG (6361C>T 6363T>G), AAA365AAT (6369A>T), GAA367GAG (6375A>G), CAA368AAA (6376C>A), CCT369CAA (6380C>A 6381T>A), ACA370CCT (6382A>C 6384A>T), ACC371ATT (6386C>T 6387C>T), ACA372ATA (6389C>T), AGG376AAA (6401G>A 6402G>A), AGG377AAT (6404G>A 6405G>T), AAT378GGA (6406A>G 6407A>G 6408T>A), TTC379TTT (6411C>T), ATC380ATT (6414C>T), TCC381TCG (6417C>G), CCA382GAC (6418C>G 6419C>A 6420A>C), GAT383GAA (6423T>A), CTG384TTA (6424C>T 6426G>A), TTA385CTA (6427T>C), AAG390TCA (6442A>T 6443A>C 6444G>A), CTA391ATT (6445C>A 6447A>T), ATT392ATC (6450T>C), AGT393GGA (6451A>G 6453T>A), CAC394AGA (6454C>A 6455A>G 6456C>A), AAA395AAT (6459A>T), CCA397GAT (6463C>G 6464C>A 6465A>T), GAC398GAA (6468C>A), CAC399CAT (6471C>T), CTA400ATA (6472C>A), TCA402AAC (6478T>A 6479C>A 6480A>C), AAA403AGA (6482A>G), AAA405AGT (6488A>G 6489A>T), GAA407ACC (6493G>A 6494A>C 6495A>C), GAT408AGT (6496G>A 6497A>G), AAC409AAT (6501C>T), ATT410TAT (6502A>T 6503T>A), GTT411ATT (6505G>A), GAC413GAA (6513C>A), GTA414ATC (6514G>A 6516A>C), CAA415AAC (6517C>A 6519A>C), CTG416TTG (6520C>T) | | | | | | | | |

#### Proteins

| | Begin | End | Coverage | Score | Concordance | Matches | Identities | I/D/M/F* | Stop Codons |
|---|---|---|---|---|---|---|---|---|---|
| putative cell-to-cell movement protein (NP_569140.1) | 58 | 156 | 23.6% | 310 | 46.3% | 99 (100%) | 74 (74.7%) | 0/0/10/0 | 1 |
| Protein mutations: | R58* (2326A>T), Q59K (2329C>A), K60I (2333A>T), V65I (2347G>A), V66L (2350G>C), K76R (2381A>G), G79R (2389G>A), I86L (2410A>C 2412A>W), R90K (2423G>A 2424A>R), D98B (2446G>R 2448T>C), N103E (2461A>G 2463C>A), L116Q (2500T>C 2501T>A), I131J (2545A>M), I150V (2602A>G) | | | | | | | | |
| Codon mutations: | AGA58TGA (2326A>T), CAA59AAA (2329C>A), AAA60ATA (2333A>T), GTA65ATA (2347G>A), GTT66CTT (2350G>C), ATT73ATA (2373T>A), TGC75TGT (2379C>T), AAA76AGA (2381A>G), TAT77TAC (2385T>C), ACA78ACT (2388A>T), GGA79AGA (2389G>A), GGT80GGA (2394T>A), AAG81AAA (2397G>A), GCC82GCA (2400C>A), CCA85CCW (2409A>W), ATA86CTW (2410A>C 2412A>W), ATA87ATR (2415A>R), AAC88ASY (2417A>S 2418C>Y), AAA89ARR (2420A>R 2421A>R), AGA90AAR (2423G>A 2424A>R), ATA91RYA (2425A>R 2426T>Y), ATA92RTA (2428A>R), AAT93AMM (2432A>M 2433T>M), AAA94MAA (2434A>M), ATA96ATW (2442A>W), CAA97RAA (2443C>R), GAT98RAC (2446G>R 2448T>C), AAG100AAA (2454G>A), GCA101GCT (2457A>T), AAC103GAA (2461A>G 2463C>A), ATT105ATA (2469T>A), AAG106AAA (2472G>A), TAT107TAC (2475T>C), CAC109CAT (2481C>T), TTA110CTA (2482T>C), ACT113ACC (2493T>C), TTA116CAA (2500T>C 2501T>A), ATA117ATT (2505A>T), CCT128CCA (2538T>A), ATA129ATM (2541A>M), ATA131MTA (2545A>M), TAT132TAY (2550T>Y), GAT136GAY (2562T>Y), AGA137AGR (2565A>R), ATC139RTA (2569A>R 2571C>A), CAA140KAW (2572C>K 2574A>W), CCT141CCW (2577T>W), AGC145AGT (2589C>T), ATA150GTA (2602A>G), GGA152GGT (2610A>T), CTT154CTA (2616T>A) | | | | | | | | |
| polyprotein (NP_569141.1) | 188 | 617 | 54.4% | 2026 | 85.1% | 346 (100%) | 293 (84.7%) | 0/0/0/0 | 1 |
| Protein mutations: | W188C (3974G>T), E189D (3977G>T), K194T (3991A>C), E201N (4011G>A 4013A>T), E202K (4014G>A 4016G>A), N203D (4017A>G), K212R (4045A>G), N216D (4056A>G), D217N (4059G>A 4061T>C), T219E (4065A>G 4066C>A), K226N (4088A>T), D227E (4091T>A), D230K (4098G>A 4100T>A), N231K (4103T>G), K232G (4104A>G 4105A>G), E236K (4116G>A), N238E (4122A>G 4124T>A), E310K (4338G>A 4340G>A), A318T (4362G>A 4364T>A), Y349C (4456A>G), S367* (4510C>A), R368K (4513G>A), T369M (4516C>T 4517A>G), Q370I (4518C>A 4519A>T), D371E (4523T>A), E372Q (4524G>C), K375L (4533A>T 4534A>T), E378Q (4542G>C), A381I (4551G>A 4552C>T), N386I (4567A>T 4568C>A), S387A (4569T>G 4571T>G), A396S (4596G>T), E397K (4599G>A), N401T (4612A>C 4613T>A), I403M (4619A>G), T419K (4666C>A 4667T>A), L428S (4692C>T 4693T>C 4694T>A), D429N (4695G>A 4697T>C), N431D (4701A>G 4703C>T), I444L (4740A>C 4742A>G), V448I (4752G>A), P461S (4791C>T 4793T>A), L462I (4794T>A), Q463Y (4797C>T 4799G>T), F556H (5076T>C 5077T>A), A557S (5079G>T), T572N (5125C>A), R573K (5128G>A), V576I (5136G>A), I607K (5230T>A), T611K (5242C>A), I616L (5256A>T) | | | | | | | | |
| Codon mutations: | TGG188TGT (3974G>T), GAG189GAT (3977G>T), ACA193ACT (3989A>T), AAA194ACA (3991A>C), ATT200ATA (4010T>A), GAA201AAT (4011G>A 4013A>T), GAG202AAA (4014G>A 4016G>A), AAT203GAT (4017A>G), ATA205ATT (4025A>T), AAA209AAG (4037A>G), CCT210CCA (4040T>A), AAA212AGA (4045A>G), AAT216GAT (4056A>G), GAT217AAC (4059G>A 4061T>C), ACA219GAA (4065A>G 4066C>A), AAA222AAG (4076A>G), AAA226AAT (4088A>T), GAT227GAA (4091T>A), TTA228TTG (4094A>G), GAT230AAA (4098G>A 4100T>A), AAT231AAG (4103T>G), AAA232GGA (4104A>G 4105A>G), CAA235CAG (4115A>G), GAA236AAA (4116G>A), AGT237AGC (4121T>C), AAT238GAA (4122A>G 4124T>A), AGC239AGT (4127C>T), CAT241CAC (4133T>C), GGA257GGT (4181A>T), AGC259AGT (4187C>T), AGG260AGA (4190G>A), GTC262GTT (4196C>T), ATA263ATT (4199A>T), TTA268CTA (4212T>C), TAT277TAC (4241T>C), ATA279ATT (4247A>T), CCC280CCA (4250C>A), AAT281AAC (4253T>C), AAG285AAA (4265G>A), CAA288CAG (4274A>G), TAT292TAC (4286T>C), TAC294TAT (4292C>T), AGT296AGC (4298T>C), GAC299GAT (4307C>T), AAG301AAA (4313G>A), TAT305TAC (4325T>C), CAT306CAC (4328T>C), TTG307TTA (4331G>A), GAG310AAA (4338G>A 4340G>A), GCT318ACA (4362G>A 4364T>A), TAT326TAC (4388T>C), GTT330GTA (4400T>A), CCA332CCT (4406A>T), TTT333TTC (4409T>C), TAT335TAC (4415T>C), AAT337AAC (4421T>C), TAC342TAT (4436C>T), GAC347GAT (4451C>T), TAT349TGT (4456A>G), TTT350TTC (4460T>C), ATA357ATT (4481A>T), GTA358GTT (4484A>T), CTA364TTA (4500C>T), CTA365TTA (4503C>T), TAT366TAC (4508T>C), TCA367TAA (4510C>A), AGA368AAA (4513G>A), ACA369ATG (4516C>T 4517A>G), CAA370ATA (4518C>A 4519A>T), GAT371GAA (4523T>A), GAA372CAA (4524G>C), ATA374ATC (4532A>C), AAA375TTA (4533A>T 4534A>T), GAA378CAA (4542G>C), GCA381ATA (4551G>A 4552C>T), ATT383ATA (4559T>A), AAC386ATA (4567A>T 4568C>A), TCT387GCG (4569T>G 4571T>G), GGT388GGA (4574T>A), GCA396TCA (4596G>T), GAA397AAA (4599G>A), ATT398ATA (4604T>A), AAT401ACA (4612A>C 4613T>A), CAG402CAA (4616G>A), ATA403ATG (4619A>G), TTA406CTA (4626T>C), CAA409CAG (4637A>G), GGA414GGG (4652A>G), ACT419AAA (4666C>A 4667T>A), CAT420CAC (4670T>C), ATC426ATT (4688C>T), CTT428TCA (4692C>T 4693T>C 4694T>A), GAT429AAC (4695G>A 4697T>C), AAC431GAT (4701A>G 4703C>T), ATT432ATA (4706T>A), GAT433GAC (4709T>C), TTA438CTA (4722T>C), TCA440TCC (4730A>C), TTT441TTC (4733T>C), CTA442TTA (4734C>T), ATA444CTG (4740A>C 4742A>G), GTA445GTC (4745A>C), GTA448ATA (4752G>A), AGG449CGT (4755A>C 4757G>T), TAT451TAC (4763T>C), CCT453CCA (4769T>A), AAC458AAT (4784C>T), TTA459CTA (4785T>C), CCT461TCA (4791C>T 4793T>A), TTA462ATA (4794T>A), CAG463TAT (4797C>T 4799G>T), CTA466TTA (4806C>T), TTT556CAT (5076T>C 5077T>A), GCA557TCA (5079G>T), CTA562TTA (5094C>T), ATA569ATT (5117A>T), GTT570GTG (5120T>G), TAC571TAT (5123C>T), ACC572AAC (5125C>A), AGA573AAA (5128G>A), ATT575ATA (5135T>A), GTA576ATA (5136G>A), GAC579GAT (5147C>T), AAT580AAC (5150T>C), CAA582CAG (5156A>G), ACT588ACA (5174T>A), AAA590AAG (5180A>G), GAT593GAC (5189T>C), GTC595GTT (5195C>T), CGG601AGG (5211C>A), CTG603TTA (5217C>T 5219G>A), ATA607AAA (5230T>A), TTA608TTG (5234A>G), AAC609AAT (5237C>T), ACA611AAA (5242C>A), ATA616TTA (5256A>T) | | | | | | | | |
| putative transactivator factor (NP_569142.1) | 196 | 418 | 53.3% | 1006 | 66.1% | 223 (100%) | 136 (61.0%) | 0/0/0/0 | 1 |
| Protein mutations: | L203Q (5881T>C 5882T>A), N205K (5889T>A), S207N (5894G>A), K209N (5901A>C), A212K (5908G>A 5909C>A 5910C>A), N216S (5921A>G), L220I (5932C>A), D221N (5935G>A 5937T>C), Q226Y (5950C>T 5952G>T), T227D (5953A>G 5954C>A 5955A>C), G228R (5956G>A), Q229E (5959C>G), A232S (5968G>T 5970C>G), N233T (5972A>C 5973T>C), P235L (5978C>T 5979T>A), Y238H (5986T>C), Q243I (6001C>A 6002A>T), G250N (6022G>A 6023G>A 6024G>T), T251N (6026C>A 6027A>T), V255I (6037G>A), R256K (6040C>A 6041G>A), S259T (6049T>A 6051G>A), T261P (6055A>C), E267D (6075A>T), E283D (6123A>T), I285L (6127A>T 6129C>A), K289D (6139A>G 6141G>C), E291R (6145G>A 6146A>G), E292Q (6148G>C 6150A>G), E294P (6154G>C 6155A>C), E295K (6157G>A), E297P (6163G>C 6164A>C), V299I (6169G>A 6171A>T), N300E (6172A>G 6174T>G), E303S (6181G>A 6182A>G 6183A>T), I313L (6211A>C), L317I (6223C>A 6225A>C), L325T (6247C>A 6248T>C), Q327G (6253C>G 6254A>G), A329P (6259G>C), I330M (6264T>G), S332N (6269G>A 6270C>T), Y333H (6271T>C), S335T (6277T>A 6279A>C), R336K (6281G>A 6282G>A), E337D (6285A>C), T339I (6290C>T), Y342F (6299A>T 6300T>C), S343E (6301T>G 6302C>A), N344T (6305A>C 6306T>G), C345A (6307T>G 6308G>C 6309C>A), I348L (6316A>T), E350Q (6322G>C), A351T (6325G>A), L356A (6340C>G 6341T>C), V360I (6352G>A 6354C>A), L361M (6355T>A 6357A>G), S362T (6359G>C 6360T>A), K365N (6369A>T), Q368K (6376C>A), P369Q (6380C>A 6381T>A), T370P (6382A>C 6384A>T), T371I (6386C>T 6387C>T), T372I (6389C>T), R376K (6401G>A 6402G>A), R377N (6404G>A 6405G>T), N378G (6406A>G 6407A>G 6408T>A), P382D (6418C>G 6419C>A 6420A>C), D383E (6423T>A), K390S (6442A>T 6443A>C 6444G>A), L391I (6445C>A 6447A>T), S393G (6451A>G 6453T>A), H394R (6454C>A 6455A>G 6456C>A), K395N (6459A>T), P397D (6463C>G 6464C>A 6465A>T), D398E (6468C>A), L400I (6472C>A), S402N (6478T>A 6479C>A 6480A>C), K403R (6482A>G), K405S (6488A>G 6489A>T), E407T (6493G>A 6494A>C 6495A>C), D408S (6496G>A 6497A>G), I410Y (6502A>T 6503T>A), V411I (6505G>A), D413E (6513C>A), V414I (6514G>A 6516A>C), Q415N (6517C>A 6519A>C) | | | | | | | | |

| | Begin | End | Coverage | Score | Concordance | Matches | Identities | I/D/M/F* | Stop Codons |
|---|---|---|---|---|---|---|---|---|---|
| **NT** | **2326** | **6548** | **26.1%** | **2323** | **57.5%** | **2025 (99.9%)** | **1583 (78.1%)** | **0/2** | |

| | |
|---|---|
| Codon mutations: | ACA195..G (5859A>G), TAC197TAT (5865C>T), AAT198AAC (5868T>C), TTG200CTT (5872T>C 5874G>T), ATA201ATT (5877A>T), GCT202GCA (5880T>A), TTA203CAA (5881T>C 5882T>A), CCA204CCT (5886A>T), AAT205AAA (5889T>A), ACA206ACC (5892A>C), AGT207AAT (5894G>A), AAA209AAC (5901A>C), TTA210CTA (5902T>C), GCC212AAA (5908G>A 5909C>A 5910C>A), ACC213ACA (5913C>A), TGC214TGT (5916C>T), AAC216AGC (5921A>G), TAC217TAT (5925C>T), CTA219TTG (5929C>T 5931A>G), CTA220ATA (5932C>A), GAT221AAC (5935G>A 5937T>C), CAG226TAT (5950C>T 5952G>T), ACA227GAC (5953A>G 5954C>A 5955A>C), GGA228AGA (5956G>A), CAA229GAA (5959C>G), GAG230GAA (5964G>A), GCC232TCG (5968G>T 5970C>G), AAT233ACC (5972A>C 5973T>C), CCT235CTA (5978C>T 5979T>A), TAC238CAC (5986T>C), AAG239AAA (5991G>A), TTC241TTT (5997C>T), CAA243ATA (6001C>A 6002A>T), AAG245AAA (6009G>A), AGG246AGA (6012G>A), ATA247ATT (6015A>T), ACT248ACA (6018T>A), GGG250AAT (6022G>A 6023G>A 6024G>T), ACA251AAT (6026C>A 6027A>T), TTA252CTA (6028T>C), TTC253TTT (6033C>T), TAC254TAT (6036C>T), GTA255ATA (6037G>A), CGA256AAA (6040C>A 6041G>A), TTC257TTT (6045C>T), TCG259ACA (6049T>A 6051G>A), GCT260GCA (6054T>A), ACA261CCA (6055A>C), CTA265TTA (6067C>T), GAA267GAT (6075A>T), GAG268GAA (6078G>A), ATA269ATC (6081A>C), AAG270AAA (6084G>A), CCT271CCA (6087T>A), ATC272ATA (6090C>A), AAA277AAG (6105A>G), ATT278ATA (6108T>A), CTC280TTG (6112C>T 6114C>G), AGG282AGA (6120G>A), GAA283GAT (6123A>T), ATC285TTA (6127A>T 6129C>A), AAG289GAC (6139A>G 6141G>C), GAA291AGA (6145G>A 6146A>G), GAA292CAG (6148G>C 6150A>G), GAA294CCA (6154G>C 6155A>C), GAG295AAG (6157G>A), GAA297CCA (6163G>C 6164A>C), GTA299ATT (6169G>A 6171A>T), AAT300GAG (6172A>G 6174T>G), ATT301ATA (6177T>A), GAA303AGT (6181G>A 6182A>G 6183A>T), TTC304TTT (6186C>T), TAT305TAC (6189T>C), AGG309AGA (6201G>A), ATA310ATT (6204A>T), ATA313CTA (6211A>C), ACT315ACG (6219T>G), CTA317ATC (6223C>A 6225A>C), CTA320TTA (6232C>T), TAT324TAC (6246T>C), CTA325ACA (6247C>A 6248T>C), CAG327GGG (6253C>G 6254A>G), AAT328AAC (6258T>C), GCA329CCA (6259G>C), ATT330ATG (6264T>G), AGC332AAT (6269G>A 6270C>T), TAT333CAT (6271T>C), TCA335ACC (6277T>A 6279A>C), AGG336AAA (6281G>A 6282G>A), GAA337GAC (6285A>C), ACA339ATA (6290C>T), TAT342TTC (6299A>T 6300T>C), TCA343GAA (6301T>G 6302C>A), AAT344ACG (6305A>C 6306T>G), TGC345GCA (6307T>G 6308G>C 6309C>A), ATA348TTA (6316A>T), AGA349CGG (6319A>C 6321A>G), GAA350CAA (6322G>C), GCA351ACA (6325G>A), CTA356GCA (6340C>G 6341T>C), GTC360ATA (6352G>A 6354C>A), TTA361ATG (6355T>A 6357A>G), AGT362ACA (6359G>C 6360T>A), CTT363TTG (6361C>T 6363T>G), AAA365AAT (6369A>T), GAA367GAG (6375A>G), CAA368AAA (6376C>A), CCT369CAA (6380C>A 6381T>A), ACA370CCT (6382A>C 6384A>T), ACC371ATT (6386C>T 6387C>T), ACA372ATA (6389C>T), AGG376AAA (6401G>A 6402G>A), AGG377AAT (6404G>A 6405G>T), AAT378GGA (6406A>G 6407A>G 6408T>A), TTC379TTT (6411C>T), ATC380ATT (6414C>T), TCC381TCG (6417C>G), CCA382GAC (6418C>G 6419C>A 6420A>C), GAT383GAA (6423T>A), CTG384TTA (6424C>T 6426G>A), TTA385CTA (6427T>C), AAG390TCA (6442A>T 6443A>C 6444G>A), CTA391ATT (6445C>A 6447A>T), ATT392ATC (6450T>C), AGT393GGA (6451A>G 6453T>A), CAC394AGA (6454C>A 6455A>G 6456C>A), AAA395AAT (6459A>T), CCA397GAT (6463C>G 6464C>A 6465A>T), GAC398GAA (6468C>A), CAC399CAT (6471C>T), CTA400ATA (6472C>A), TCA402AAC (6478T>A 6479C>A 6480A>C), AAA403AGA (6482A>G), AAA405AGT (6488A>G 6489A>T), GAA407ACC (6493G>A 6494A>C 6495A>C), GAT408AGT (6496G>A 6497A>G), AAC409AAT (6501C>T), ATT410TAT (6502A>T 6503T>A), GTT411ATT (6505G>A), GAC413GAA (6513C>A), GTA414ATC (6514G>A 6516A>C), CAA415AAC (6517C>A 6519A>C), CTG416TTG (6520C>T) |

*: Inserts / Deletes / Misaligned / Frameshifts

#### Analysis details

This analysis was performed with panviral2.64

### NGS Details (UN24): Potato virus Y

#### Assembly

| | |
|---|---|
| **Coverage Length** | 3067 (7 contig(s)) |
| **Depth Of Coverage** | 3.3 |
| **Number Of Reads** | 77 |
| **Reads Per Million** | 1.54 rpm (after QC) |
| **Ambiguities** | 0 |
| **Assembly Method** | de novo + reference guided assembly |
| **Consensus Caller** | Bcf Tools |

#### Coverage Map

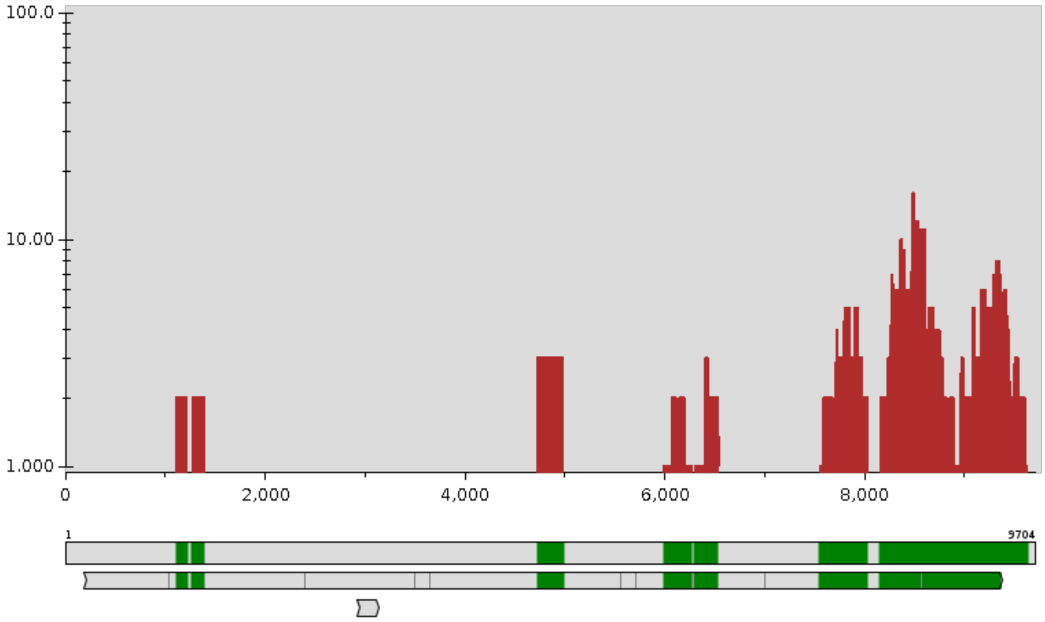

#### Assignment

| | |
|---|---|
| **Type** | Potato virus Y (Taxonomy ID: 12216) |
| **Reference Genome** | NC_001616.1 |
| **NT Identity (%)** | 84.5098 |
| **AA Identity (%)** | 90.7824 |
| **Number Of Stop Codons** | 1 |
| **Number Of CDS** | 2 |

#### Alignment

| | |
|---|---|
| **Alignment Score** | 4154.0 (NT) + 6023.0 (AA) = 10177.0 |
| **Concordance (%)** | 81.0916 |

| **Alignment Method** | Global, seeded, nucleotide + amino acids (AGA) |
|---|---|

#### Genome Region

Sequence starts at position 1103 and ends at position 9641 relative to NC_001616.1 reference sequence.

#### Alignment Detailed Statistics

| | Begin | End | Coverage | Score | Concordance | Matches | Identities | I/D/M/F* | Stop Codons |
|---|---|---|---|---|---|---|---|---|---|
| **NT** | **1103** | **9641** | **31.6%** | **4154** | **68.5%** | **3057 (99.6%)** | **2586 (84.2%)** | **3/10** | |
| Mutations: | 1105C>T, 1108A>G, 1114A>G, 1117T>G, 1123A>G, 1126T>A, 1129C>T, 1143G>A, 1147A>T, 1150T>A, 1154T>A, 1156G>A, 1165C>T, 1168T>C, 1172C>T, 1174T>A, 1201C>T, 1219C>T, 1221G>A, 1265A>G, 1267A>G, 1273T>C, 1276G>A, 1278A>G, 1283T>C, 1290C>T, 1291G>A, 1298G>A, 1299A>C, 1300C>T, 1306T>C, 1307A>G, 1309A>G, 1312T>C, 1316A>G, 1318T>A, 1327G>A, 1329T>C, 1333G>C, 1336A>G, 1337G>A, 1343C>T, 1345A>G, 1354G>A, 1357G>T, 1359A>T, 1360C>T, 1361C>T, 1363G>A, 1367C>T, 1369C>G, 1373C>A, 1378C>T, 1384G>A, 1385A>G, 1393A>G, 1396C>T, 4738A>G, 4744A>T, 4747A>G, 4748T>C, 4765T>C, 4796G>A, 4798C>T, 4803C>G, 4804T>C, 4807G>A, 4810G>T, 4813T>G, 4816T>C, 4831T>C, 4834C>T, 4840A>T, 4843G>A, 4852A>G, 4853_4858delGGCGTT, 4859T>G, 4864T>C, 4867T>C, 4870G>C, 4873T>A, 4882T>C, 4903A>T, 4909G>A, 4919T>C, 4921G>A, 4924T>C, 4927C>G, 4933T>C, 4936C>A, 4939G>A, 4942T>C, 4945C>T, 4963A>G, 4972T>C, 4990T>C, 5986T>A, 5989C>T, 5990A>G, 6008A>C, 6010A>T, 6019A>T, 6022G>T, 6025A>T, 6030A>C, 6049C>A, 6058G>A, 6061A>G, 6064C>A, 6070T>C, 6072G>A, 6073T>C, 6074A>C, 6079G>A, 6081C>A, 6085A>T, 6091A>T, 6094C>T, 6111G>C, 6112T>C, 6118A>G, 6124G>A, 6127G>A, 6128A>G, 6130T>C, 6133T>C, 6136A>G, 6148C>T, 6157A>G, 6158G>A, 6163T>C, 6169A>G, 6170A>T, 6172A>G, 6178C>T, 6187A>G, 6190T>C, 6196G>A, 6203C>T, 6205C>T, 6208A>G, 6209C>T, 6211A>G, 6217G>A, 6229T>A, 6230G>A, 6238C>T, 6247G>A, 6253A>T, 6257G>A, 6258C>A, 6274T>C, 6298C>T, 6301G>A, 6304A>G, 6307C>T, 6313C>T, 6322C>T, 6328A>T, 6331T>G, 6334T>C, 6337G>A, 6338C>T, 6346A>G, 6358T>C, 6361G>A, 6362G>A, 6367A>C, 6370G>A, 6376C>T, 6379T>G, 6385A>C, 6391A>T, 6394A>G, 6397C>T, 6401G>A, 6405C>T, 6406G>A, 6412C>T, 6418A>G, 6421T>C, 6429A>T, 6436T>A, 6445G>A, 6450A>G, 6454C>A, 6460C>T, 6463T>A, 6466A>G, 6469C>T, 6472G>A, 6475G>A, 6481T>C, 6485C>A, 6487C>T, 6490T>C, 6496C>T, 6502G>A, 6514T>C, 6517G>A, 6527C>A, 6532C>A, 7546T>C, 7570G>A, 7573G>A, 7579G>A, 7588T>C, 7595C>T, 7604T>C, 7606G>A, 7615T>A, 7624T>C, 7633C>T, 7636C>T, 7639T>C, 7642T>C, 7648T>C, 7666A>G, 7672T>C, 7678A>T, 7684G>A, 7690T>C, 7696T>C, 7702T>G, 7709G>A, 7713A>G, 7714A>G, 7715C>T, 7726T>C, 7735G>A, 7744A>G, 7747C>T, 7762C>T, 7768G>A, 7771T>C, 7783A>C, 7786T>C, 7793C>T, 7795A>G, 7798C>T, 7804T>C, 7812C>T, 7834A>G, 7837C>T, 7844G>T, 7846G>A, 7855G>A, 7861G>A, 7870A>G, 7873C>T, 7876T>A, 7879G>A, 7882T>A, 7883G>A, 7894A>T, 7897T>C, 7921C>T, 7927G>A, 7936T>C, 7939T>C, 7942T>C, 7954T>C, 7966G>A, 7969T>C, 7975T>C, 7987C>T, 7990T>C, 8005T>C, 8032A>G, 8038C>T, 8151A>G, 8152C>T, 8164G>A, 8178G>A, 8182G>A, 8191G>A, 8197C>T, 8208G>A, 8213C>T, 8218A>G, 8221C>T, 8224G>A, 8239A>G, 8260T>C, 8263A>G, 8281T>C, 8284A>G, 8294C>G, 8296A>G, 8299G>A, 8302C>T, 8311A>G, 8317T>C, 8320C>T, 8326T>A, 8335G>A, 8338C>T, 8349C>T, 8353A>G, 8362C>T, 8365A>G, 8368C>T, 8380C>T, 8383A>G, 8389A>G, 8404T>C, 8408G>T, 8413A>G, 8419G>T, 8434T>C, 8440T>C, 8452G>A, 8458A>G, 8460G>A, 8464A>G, 8467G>A, 8480G>A, 8482T>A, 8497G>A, 8498C>T, 8500A>G, 8502G>A, 8503A>G, 8506C>T, 8515A>T, 8524C>T, 8527A>C, 8528T>C, 8530A>G, 8539G>C, 8546C>T, 8547T>G, 8552T>A, 8563A>G, 8574C>G, 8585A>G, 8598G>A, 8602C>T, 8603A>T, 8604A>C, 8605C>T, 8620A>G, 8622C>A, 8625A>G, 8626G>A, 8629G>A, 8632C>T, 8638C>T, 8641G>A, 8647C>T, 8648C>T, 8649C>T, 8650G>C, 8653C>T, 8656A>G, 8658G>A, 8664A>C, 8665T>G, 8674T>G, 8679C>T, 8680A>T, 8683C>A, 8692G>A, 8695A>T, 8707G>A, 8708A>C, 8716G>A, 8722C>T, 8725G>A, 8731A>G, 8737A>G, 8743C>T, 8745C>A, 8746A>G, 8749C>T, 8752G>A, 8755A>T, 8770C>T, 8771T>C, 8776A>G, 8779T>C, 8782G>A, 8791T>C, 8797A>G, 8800A>G, 8803A>G, 8809T>C, 8812T>C, 8818T>C, 8827A>G, 8838A>C, 8848G>A, 8854T>C, 8857G>A, 8863G>A, 8865G>A, 8867A>C, 8869G>T, 8872A>G, 8882G>A, 8883G>A, 8888A>T, 8929C>T, 8954G>A, 8959C>T, 8962A>T, 8983G>A, 8984A>G, 8988A>T, 8998G>A, 9004G>A, 9007G>A, 9016C>T, 9037C>A, 9085A>T, 9097C>T, 9131A>G, 9136A>T, 9145G>T, 9149A>G, 9150T>G, 9151G>A, 9152G>A, 9157A>G, 9160G>T, 9163T>C, 9184G>A, 9196A>G, 9214G>A, 9217A>G, 9220G>A, 9244A>T, 9247G>A, 9256C>T, 9260C>T, 9262T>C, 9274G>A, 9280C>T, 9289C>T, 9304G>A, 9334C>T, 9364C>G, 9377_9378insT, 9387C>T, 9404G>A, 9411C>T, 9412A>G, 9414A>T, 9423G>A, 9427T>G, 9454A>G, 9458T>C, 9467G>A, 9470_9471insT, 9482G>A, 9483C>T, 9496T>C, 9516T>G, 9517delA, 9533G>A, 9534G>T, 9548A>T, 9551C>T, 9553_9554delTA, 9556A>T, 9557T>A, 9558A>T, 9574T>A, 9581G>T, 9582T>G, 9583G>C, 9584_9585insG, 9590C>T, 9591T>C, 9607C>T, 9610T>C, 9615delC, 9620A>G | | | | | | | | |

##### CDS

| | | | | | | | | | |
|---|---|---|---|---|---|---|---|---|---|
| PVYgp1 | 307 | 3064 | 30.5% | 6023 | 91.8% | 933 (99.8%) | 847 (90.6%) | 0/2/0/0 | 1 |
| Protein mutations: | R320K (1143G>A), L324I (1154T>A 1156G>A), S346N (1221G>A), R361G (1265A>G 1267A>G), N365S (1278A>G), A369V (1290C>T 1291G>A), D372T (1298G>A 1299A>C 1300C>T), I375V (1307A>G 1309A>G), N378E (1316A>G 1318T>A), I382T (1329T>C), E385K (1337G>A), D392V (1359A>T 1360C>T), L397I (1373C>A), I401V (1385A>G), V1538I (4796G>A 4798C>T), T1540S (4803C>G 4804T>C), G1557_V1558del (4853_4858delGGCGTT), S1559A (4859T>G), I1936V (5990A>G), E1945D (6019A>T), K1949T (6030A>C), D1955E (6049C>A), M1958I (6058G>A), S1963N (6072G>A 6073T>C), N1964H (6074A>C), T1966N (6081C>A), C1976S (6111G>C 6112T>C), I1982V (6128A>G 6130T>C), V1992I (6158G>A), T1996S (6170A>T 6172A>G), L2007F (6203C>T 6205C>T), V2016I (6230G>A), A2025K (6257G>A 6258C>A), A2060T (6362G>A), V2073I (6401G>A), A2074V (6405C>T 6406G>A), Y2082F (6429A>T), Q2089R (6450A>G), H2101N (6485C>A 6487C>T), L2115I (6527C>A), D2509N (7709G>A), K2510R (7713A>G 7714A>G), T2543I (7812C>T), V2554L (7844G>T 7846G>A), V2567I (7883G>A), N2656S (8151A>G 8152C>T), R2665K (8178G>A), R2675K (8208G>A), P2704A (8294C>G 8296A>G), S2722F (8349C>T), A2742S (8408G>T), M2756I (8452G>A), R2759K (8460G>A), A2766T (8480G>A 8482T>A), R2773K (8502G>A 8503A>G), E2777D (8515A>T), E2785D (8539G>C), L2788C (8546C>T 8547T>G), S2790T (8552T>A), A2797G (8574C>G), I2801V (8585A>G), G2805E (8598G>A), N2807S (8603A>T 8604A>C 8605C>T), P2813Q (8622C>A), E2814G (8625A>G 8626G>A), P2822F (8648C>T 8649C>T 8650G>C), G2825E (8658G>A), D2827A (8664A>C 8665T>G), A2832V (8679C>T 8680A>T), T2854K (8745C>A 8746A>G), Q2885P (8838A>C), R2894Q (8865G>A), M2895L (8867A>C 8869G>T), G2900K (8882G>A 8883G>A), T2902S (8888A>T), V2924I (8954G>A), N2934D (8984A>G), E2935V (8988A>T), I2983V (9131A>G), M2989G (9149A>G 9150T>G 9151G>A), G2990S (9152G>A), P3026S (9260C>T 9262T>C) | | | | | | | | |

| | Begin | End | Coverage | Score | Concordance | Matches | Identities | I/D/M/F* | Stop Codons |
|---|---|---|---|---|---|---|---|---|---|
| **NT** | **1103** | **9641** | **31.6%** | **4154** | **68.5%** | **3057 (99.6%)** | **2586 (84.2%)** | **3/10** | |
| Codon mutations: | CAC307CAT (1105C>T), ACA308ACG (1108A>G), GTA310GTG (1114A>G), GCT311GCG (1117T>G), TTA313TTG (1123A>G), CCT314CCA (1126T>A), GTC315GTT (1129C>T), AGG320AAG (1143G>A), GTA321GTT (1147A>T), GCT322GCA (1150T>A), TTG324ATA (1154T>A 1156G>A), CAC327CAT (1165C>T), AGT328AGC (1168T>C), CTT330TTA (1172C>T 1174T>A), ACC339ACT (1201C>T), GCC345GCT (1219C>T), AGC346AAC (1221G>A), AGA361GGG (1265A>G 1267A>G), GGT363GGC (1273T>C), TTG364TTA (1276G>A), AAT365AGT (1278A>G), TTG367CTG (1283T>C), GCG369GTA (1290C>T 1291G>A), GAC372ACT (1298G>A 1299A>C 1300C>T), TTT374TTC (1306T>C), ATA375GTG (1307A>G 1309A>G), CAT376CAC (1312T>C), AAT378GAA (1316A>G 1318T>A), TTG381TTA (1327G>A), ATA382ACA (1329T>C), GCG383GCC (1333G>C), TTA384TTG (1336A>G), GAG385AAG (1337G>A), CTA387TTG (1343C>T 1345A>G), CCG390CCA (1354G>A), GTG391GTT (1357G>T), GAC392GTT (1359A>T 1360C>T), CTG393TTA (1361C>T 1363G>A), CTC395TTG (1367C>T 1369C>G), CTT397ATT (1373C>A), TTC398TTT (1378C>T), GAG400GAA (1384G>A), ATA401GTA (1385A>G), AAA403AAG (1393A>G), TCC404TCT (1396C>T), AAA1518AAG (4738A>G), GTA1520GTT (4744A>T), GCA1521GCG (4747A>G), TTG1522CTG (4748T>C), ACT1527ACC (4765T>C), GTC1538ATT (4796G>A 4798C>T), ACT1540AGC (4803C>G 4804T>C), GAG1541GAA (4807G>A), GCG1542GCT (4810G>T), GCT1543GCG (4813T>G), CTT1544CTC (4816T>C), TAT1549TAC (4831T>C), AAC1550AAT (4834C>T), CCA1552CCT (4840A>T), GTG1553GTA (4843G>A), GGA1556GGG (4852A>G), GGC1557_GTT1558del (4853_4858delGGCGTT), TCA1559GCA (4859T>G), ACT1560ACC (4864T>C), AGT1561AGC (4867T>C), CTG1562CTC (4870G>C), ATT1563ATA (4873T>A), TGT1566TGC (4882T>C), ACA1573ACT (4903A>T), CAG1575CAA (4909G>A), TTG1579CTA (4919T>C 4921G>A), AGT1580AGC (4924T>C), CCC1581CCG (4927C>G), TTT1583TTC (4933T>C), ATC1584ATA (4936C>A), CAG1585CAA (4939G>A), AAT1586AAC (4942T>C), TTC1587TTT (4945C>T), TCA1593TCG (4963A>G), CCT1596CCC (4972T>C), CTT1602CTC (4990T>C), GCT1934GCA (5986T>A), GAC1935GAT (5989C>T), ATT1936GTT (5990A>G), AGA1942CGT (6008A>C 6010A>T), GAA1945GAT (6019A>T), GTG1946GTT (6022G>T), CGA1947CGT (6025A>T), AAA1949ACA (6030A>C), GAC1955GAA (6049C>A), ATG1958ATA (6058G>A), CAA1959CAG (6061A>G), GCC1960GCA (6064C>A), GGT1962GGC (6070T>C), AGT1963AAC (6072G>A 6073T>C), AAC1964CAC (6074A>C), ACG1965ACA (6079G>A), ACC1966AAC (6081C>A), ATA1967ATT (6085A>T), GCA1969GCT (6091A>T), TAC1970TAT (6094C>T), TGT1976TCC (6111G>C 6112T>C), AAA1978AAG (6118A>G), TTG1980TTA (6124G>A), AAG1981AAA (6127G>A), ATT1982GTC (6128A>G 6130T>C), GAT1983GAC (6133T>C), TTA1984TTG (6136A>G), AAC1988AAT (6148C>T), AAA1991AAG (6157A>G), GTT1992ATT (6158G>A), TGT1993TGC (6163T>C), AAA1995AAG (6169A>G), ACA1996TCG (6170A>T 6172A>G), GGC1998GGT (6178C>T), AAA2001AAG (6187A>G), TTT2002TTC (6190T>C), GAG2004GAA (6196G>A), CTC2007TTT (6203C>T 6205C>T), GAA2008GAG (6208A>G), CTA2009TTG (6209C>T 6211A>G), CAG2011CAA (6217G>A), GCT2015GCA (6229T>A), GTA2016ATA (6230G>A), GTC2018GTT (6238C>T), AAG2021AAA (6247G>A), ATA2023ATT (6253A>T), GCA2025AAA (6257G>A 6258C>A), CAT2030CAC (6274T>C), GGC2038GGT (6298C>T), TTG2039TTA (6301G>A), AGA2040AGG (6304A>G), GAC2041GAT (6307C>T), AAC2043AAT (6313C>T), GCC2046GCT (6322C>T), ACA2048ACT (6328A>T), GTT2049GTG (6331T>G), TGT2050TGC (6334T>C), AGG2051AGA (6337G>A), CTG2052TTG (6338C>T), GTA2054GTG (6346A>G), TAT2058TAC (6358T>C), GGG2059GGA (6361G>A), GCA2060ACA (6362G>A), TCA2061TCC (6367A>C), GAG2062GAA (6370G>A), TAC2064TAT (6376C>T), GGT2065GGG (6379T>G), GGA2067GGC (6385A>C), GGA2069GGT (6391A>T), GCA2070GCG (6394A>G), TAC2071TAT (6397C>T), GTA2073ATA (6401G>A), GCG2074GTA (6405C>T 6406G>A), CAC2076CAT (6412C>T), TTA2078TTG (6418A>G), TTT2079TTC (6421T>C), TAC2082TTC (6429A>T), GGT2084GGA (6436T>A), GAG2087GAA (6445G>A), CAA2089CGA (6450A>G), TCC2090TCA (6454C>A), CAC2092CAT (6460C>T), GGT2093GGA (6463T>A), ACA2094ACG (6466A>G), TTC2095TTT (6469C>T), AGG2096AGA (6472G>A), GTG2097GTA (6475G>A), AAT2099AAC (6481T>C), CAC2101AAT (6485C>A 6487C>T), AGT2102AGC (6490T>C), AGC2104AGT (6496C>T), CTG2106CTA (6502G>A), GGT2110GGC (6514T>C), AGG2111AGA (6517G>A), CTC2115ATC (6527C>A), ATC2116ATA (6532C>A), TGT2454..C (7546T>C), AAG2462AAA (7570G>A), ACG2463ACA (7573G>A), ACG2465ACA (7579G>A), GCT2468GCC (7588T>C), CTA2471TTA (7595C>T), TTG2474CTA (7604T>C 7606G>A), GGT2477GGA (7615T>A), TGT2480TGC (7624T>C), GAC2483GAT (7633C>T), TTC2484TTT (7636C>T), AAT2485AAC (7639T>C), AAT2486AAC (7642T>C), TTT2488TTC (7648T>C), GAA2494GAG (7666A>G), TGT2496TGC (7672T>C), ACA2498ACT (7678A>T), GGG2500GGA (7684G>A), ACT2502ACC (7690T>C), TTT2504TTC (7696T>C), GGT2506GGG (7702T>G), GAT2509AAT (7709G>A), AAA2510AGG (7713A>G 7714A>G), CTG2511TTG (7715C>T), CGT2514CGC (7726T>C), GAG2517GAA (7735G>A), GTA2520GTG (7744A>G), TAC2521TAT (7747C>T), GGC2526GGT (7762C>T), CAG2528CAA (7768G>A), TTT2529TTC (7771T>C), CTA2533CTC (7783A>C), ACT2534ACC (7786T>C), CTA2537TTG (7793C>T 7795A>G), ATC2538ATT (7798C>T), GCT2540GCC (7804T>C), ACC2543ATC (7812C>T), GAA2550GAG (7834A>G), GAC2551GAT (7837C>T), GTG2554TTA (7844G>T 7846G>A), CAG2557CAA (7855G>A), CTG2559CTA (7861G>A), TTA2562TTG (7870A>G), TAC2563TAT (7873C>T), ACT2564ACA (7876T>A), GAG2565GAA (7879G>A), ATT2566ATA (7882T>A), GTT2567ATT (7883G>A), CCA2570CCT (7894A>T), ATT2571ATC (7897T>C), GTC2579GTT (7921C>T), AAG2581AAA (7927G>A), GGT2584GGC (7936T>C), AAT2585AAC (7939T>C), AAT2586AAC (7942T>C), CCT2590CCC (7954T>C), GTG2594GTA (7966G>A), GAT2595GAC (7969T>C), TCT2597TCC (7975T>C), GTC2601GTT (7987C>T), CTT2602CTC (7990T>C), GCT2607GCC (8005T>C), GAA2616GAG (8032A>G), ATC2618ATT (8038C>T), AAC2656AGT (8151A>G 8152C>T), TCG2660TCA (8164G>A), AGG2665AAG (8178G>A), AAG2666AAA (8182G>A), TTG2669TTA (8191G>A), TTC2671TTT (8197C>T), AGA2675AAA (8208G>A), CTG2677TTG (8213C>T), CTA2678CTG (8218A>G), ATC2679ATT (8221C>T), GAG2680GAA (8224G>A), CCA2685CCG (8239A>G), ATT2692ATC (8260T>C), GTA2693GTG (8263A>G), GAT2699GAC (8281T>C), AGA2700AGG (8284A>G), CCA2704GCG (8294C>G 8296A>G), GAG2705GAA (8299G>A), CAC2706CAT (8302C>T), GAA2709GAG (8311A>G), ATT2711ATC (8317T>C), TGC2712TGT (8320C>T), GCT2714GCA (8326T>A), GAG2717GAA (8335G>A), TCC2718TCT (8338C>T), TCT2722TTT (8349C>T), GAA2723GAG (8353A>G), CAC2726CAT (8362C>T), CAA2727CAG (8365A>G), ATC2728ATT (8368C>T), TAC2732TAT (8380C>T), TCA2733TCG (8383A>G), TTA2735TTG (8389A>G), CCT2740CCC (8404T>C), GCA2742TCA (8408G>T), ACA2743ACG (8413A>G), GCG2745GCT (8419G>T), GCT2750GCC (8434T>C), TAT2752TAC (8440T>C), ATG2756ATA (8452G>A), CTA2758CTG (8458A>G), AGG2759AAG (8460G>A), AAA2760AAG (8464A>G), CTG2761CTA (8467G>A), GCT2766ACA (8480G>A 8482T>A), GAG2771GAA (8497G>A), CTA2772TTG (8498C>T 8500A>G), AGA2773AAG (8502G>A 8503A>G), GCC2774GCT (8506C>T), GAA2777GAT (8515A>T), GTC2780GTT (8524C>T), GCA2781GCC (8527A>C), TTA2782CTG (8528T>C 8530A>G), GAG2785GAC (8539G>C), CTT2788TGT (8546C>T 8547T>G), TCT2790ACT (8552T>A), GTA2793GTG (8563A>G), GCA2797GGA (8574C>G), ATT2801GTT (8585A>G), GGA2805GAA (8598G>A), AGC2806AGT (8602C>T), AAC2807TCT (8603A>T 8604A>C 8605C>T), AAA2812AAG (8620A>G), CCA2813CAA (8622C>A), GAG2814GGA (8625A>G 8626G>A), CAG2815CAA (8629G>A), GGC2816GGT (8632C>T), ATC2818ATT (8638C>T), CAG2819CAA (8641G>A), AAC2821AAT (8647C>T), CCG2822TTC (8648C>T 8649C>T 8650G>C), AAC2823AAT (8653C>T), AAA2824AAG (8656A>G), GGA2825GAA (8658G>A), GAT2827GCG (8664A>C 8665T>G), GTT2830GTG (8674T>G), GCA2832GTT (8679C>T 8680A>T), GGC2833GGA (8683C>A), GGG2836GGA (8692G>A), ACA2837ACT (8695A>T), CCG2841CCA (8707G>A), AGA2842CGA (8708A>C), AAG2844AAA (8716G>A), ATC2846ATT (8722C>T), ACG2847ACA (8725G>A), AAA2849AAG (8731A>G), AGA2851AGG (8737A>G), CCC2853CCT (8743C>T), ACA2854AAG (8745C>A 8746A>G), AGC2855AGT (8749C>T), AAG2856AAA (8752G>A), GGA2857GGT (8755A>T), AAC2862AAT (8770C>T), TTA2863CTA (8771T>C), GAA2864GAG (8776A>G), CAT2865CAC (8779T>C), TTG2866TTA (8782G>A), TAT2869TAC (8791T>C), CCA2871CCG (8797A>G), CAA2872CAG (8800A>G), CAA2873CAG (8803A>G), GAT2875GAC (8809T>C), ATT2876ATC (8812T>C), AAT2878AAC (8818T>C), GCA2881GCG (8827A>G), CAG2885CCG (8838A>C), ACG2888ACA (8848G>A), TAT2890TAC (8854T>C), GAG2891GAA (8857G>A), GTG2893GTA (8863G>A), CGG2894CAG (8865G>A), ATG2895CTT (8867A>C 8869G>T), GCA2896GCG (8872A>G), GGA2900AAA (8882G>A 8883G>A), ACT2902TCT (8888A>T), TGC2915TGT (8929C>T), GTC2924ATC (8954G>A), AAC2925AAT (8959C>T), GGA2926GGT (8962A>T), GGG2933GGA (8983G>A), AAT2934GAT (8984A>G), GAA2935GTA (8988A>T), GAG2938GAA (8998G>A), CCG2940CCA (9004G>A), TTG2941TTA (9007G>A), ATC2944ATT (9016C>T), ACC2951ACA (9037C>A), ATA2967ATT (9085A>T), AAC2971AAT (9097C>T), ATT2983GTT (9131A>G), CGA2984CGT (9136A>T), CGG2987CGT (9145G>T), ATG2989GGA (9149A>G 9150T>G 9151G>A), GGT2990AGT (9152G>A), TTA2991TTG (9157A>G), GCG2992GCT (9160G>T), CGT2993CGC (9163T>C), GAG3000GAA (9184G>A), CGA3004CGG (9196A>G), AGG3010AGA (9214G>A), GAA3011GAG (9217A>G), GCG3012GCA (9220G>A), GCA3020GCT (9244A>T), TTG3021TTA (9247G>A), GCC3024GCT (9256C>T), CCT3026TCC (9260C>T 9262T>C), GGG3030GGA (9274G>A), GAC3032GAT (9280C>T), ATC3035ATT (9289C>T), GAG3040GAA (9304G>A), GTC3050GTT (9334C>T), GTC3060GTG (9364C>G) | | | | | | | | |

#### Proteins

| | Begin | End | Coverage | Score | Concordance | Matches | Identities | I/D/M/F* | Stop Codons |
|---|---|---|---|---|---|---|---|---|---|
| polyprotein (NP_056759.1) | 307 | 3064 | 30.5% | 6023 | 91.8% | 933 (99.8%) | 847 (90.6%) | 0/2/0/0 | 1 |
| Protein mutations: | R320K (1143G>A), L324I (1154T>A 1156G>A), S346N (1221G>A), R361G (1265A>G 1267A>G), N365S (1278A>G), A369V (1290C>T 1291G>A), D372T (1298G>A 1299A>C 1300C>T), I375V (1307A>G 1309A>G), N378E (1316A>G 1318T>A), I382T (1329T>C), E385K (1337G>A), D392V (1359A>T 1360C>T), L397I (1373C>A), I401V (1385A>G), V1538I (4796G>A 4798C>T), T1540S (4803C>G 4804T>C), G1557_V1558del (4853_4858delGGCGTT), S1559A (4859T>G), I1936V (5990A>G), E1945D (6019A>T), K1949T (6030A>C), D1955E (6049C>A), M1958I (6058G>A), S1963N (6072G>A 6073T>C), N1964H (6074A>C), T1966N (6081C>A), C1976S (6111G>C 6112T>C), I1982V (6128A>G 6130T>C), V1992I (6158G>A), T1996S (6170A>T 6172A>G), L2007F (6203C>T 6205C>T), V2016I (6230G>A), A2025K (6257G>A 6258C>A), A2060T (6362G>A), V2073I (6401G>A), A2074V (6405C>T 6406G>A), Y2082F (6429A>T), Q2089R (6450A>G), H2101N (6485C>A 6487C>T), L2115I (6527C>A), D2509N (7709G>A), K2510R (7713A>G 7714A>G), T2543I (7812C>T), V2554L (7844G>T 7846G>A), V2567I (7883G>A), N2656S (8151A>G 8152C>T), R2665K (8178G>A), R2675K (8208G>A), P2704A (8294C>G 8296A>G), S2722F (8349C>T), A2742S (8408G>T), M2756I (8452G>A), R2759K (8460G>A), A2766T (8480G>A 8482T>A), R2773K (8502G>A 8503A>G), E2777D (8515A>T), E2785D (8539G>C), L2788C (8546C>T 8547T>G), S2790T (8552T>A), A2797G (8574C>G), I2801V (8585A>G), G2805E (8598G>A), N2807S (8603A>T 8604A>C 8605C>T), P2813Q (8622C>A), E2814G (8625A>G 8626G>A), P2822F (8648C>T 8649C>T 8650G>C), G2825E (8658G>A), D2827A (8664A>C 8665T>G), A2832V (8679C>T 8680A>T), T2854K (8745C>A 8746A>G), Q2885P (8838A>C), R2894Q (8865G>A), M2895L (8867A>C 8869G>T), G2900K (8882G>A 8883G>A), T2902S (8888A>T), V2924I (8954G>A), N2934D (8984A>G), E2935V (8988A>T), I2983V (9131A>G), M2989G (9149A>G 9150T>G 9151G>A), G2990S (9152G>A), P3026S (9260C>T 9262T>C) | | | | | | | | |

| | Begin | End | Coverage | Score | Concordance | Matches | Identities | I/D/M/F* | Stop Codons |
|---|---|---|---|---|---|---|---|---|---|
| **NT** | **1103** | **9641** | **31.6%** | **4154** | **68.5%** | **3057 (99.6%)** | **2586 (84.2%)** | **3/10** | |
| Codon mutations: | CAC307CAT (1105C>T), ACA308ACG (1108A>G), GTA310GTG (1114A>G), GCT311GCG (1117T>G), TTA313TTG (1123A>G), CCT314CCA (1126T>A), GTC315GTT (1129C>T), AGG320AAG (1143G>A), GTA321GTT (1147A>T), GCT322GCA (1150T>A), TTG324ATA (1154T>A 1156G>A), CAC327CAT (1165C>T), AGT328AGC (1168T>C), CTT330TTA (1172C>T 1174T>A), ACC339ACT (1201C>T), GCC345GCT (1219C>T), AGC346AAC (1221G>A), AGA361GGG (1265A>G 1267A>G), GGT363GGC (1273T>C), TTG364TTA (1276G>A), AAT365AGT (1278A>G), TTG367CTG (1283T>C), GCG369GTA (1290C>T 1291G>A), GAC372ACT (1298G>A 1299A>C 1300C>T), TTT374TTC (1306T>C), ATA375GTG (1307A>G 1309A>G), CAT376CAC (1312T>C), AAT378GAA (1316A>G 1318T>A), TTG381TTA (1327G>A), ATA382ACA (1329T>C), GCG383GCC (1333G>C), TTA384TTG (1336A>G), GAG385AAG (1337G>A), CTA387TTG (1343C>T 1345A>G), CCG390CCA (1354G>A), GTG391GTT (1357G>T), GAC392GTT (1359A>T 1360C>T), CTG393TTA (1361C>T 1363G>A), CTC395TTG (1367C>T 1369C>G), CTT397ATT (1373C>A), TTC398TTT (1378C>T), GAG400GAA (1384G>A), ATA401GTA (1385A>G), AAA403AAG (1393A>G), TCC404TCT (1396C>T), AAA1518AAG (4738A>G), GTA1520GTT (4744A>T), GCA1521GCG (4747A>G), TTG1522CTG (4748T>C), ACT1527ACC (4765T>C), GTC1538ATT (4796G>A 4798C>T), ACT1540AGC (4803C>G 4804T>C), GAG1541GAA (4807G>A), GCG1542GCT (4810G>T), GCT1543GCG (4813T>G), CTT1544CTC (4816T>C), TAT1549TAC (4831T>C), AAC1550AAT (4834C>T), CCA1552CCT (4840A>T), GTG1553GTA (4843G>A), GGA1556GGG (4852A>G), GGC1557_GTT1558del (4853_4858delGGCGTT), TCA1559GCA (4859T>G), ACT1560ACC (4864T>C), AGT1561AGC (4867T>C), CTG1562CTC (4870G>C), ATT1563ATA (4873T>A), TGT1566TGC (4882T>C), ACA1573ACT (4903A>T), CAG1575CAA (4909G>A), TTG1579CTA (4919T>C 4921G>A), AGT1580AGC (4924T>C), CCC1581CCG (4927C>G), TTT1583TTC (4933T>C), ATC1584ATA (4936C>A), CAG1585CAA (4939G>A), AAT1586AAC (4942T>C), TTC1587TTT (4945C>T), TCA1593TCG (4963A>G), CCT1596CCC (4972T>C), CTT1602CTC (4990T>C), GCT1934GCA (5986T>A), GAC1935GAT (5989C>T), ATT1936GTT (5990A>G), AGA1942CGT (6008A>C 6010A>T), GAA1945GAT (6019A>T), GTG1946GTT (6022G>T), CGA1947CGT (6025A>T), AAA1949ACA (6030A>C), GAC1955GAA (6049C>A), ATG1958ATA (6058G>A), CAA1959CAG (6061A>G), GCC1960GCA (6064C>A), GGT1962GGC (6070T>C), AGT1963AAC (6072G>A 6073T>C), AAC1964CAC (6074A>C), ACG1965ACA (6079G>A), ACC1966AAC (6081C>A), ATA1967ATT (6085A>T), GCA1969GCT (6091A>T), TAC1970TAT (6094C>T), TGT1976TCC (6111G>C 6112T>C), AAA1978AAG (6118A>G), TTG1980TTA (6124G>A), AAG1981AAA (6127G>A), ATT1982GTC (6128A>G 6130T>C), GAT1983GAC (6133T>C), TTA1984TTG (6136A>G), AAC1988AAT (6148C>T), AAA1991AAG (6157A>G), GTT1992ATT (6158G>A), TGT1993TGC (6163T>C), AAA1995AAG (6169A>G), ACA1996TCG (6170A>T 6172A>G), GGC1998GGT (6178C>T), AAA2001AAG (6187A>G), TTT2002TTC (6190T>C), GAG2004GAA (6196G>A), CTC2007TTT (6203C>T 6205C>T), GAA2008GAG (6208A>G), CTA2009TTG (6209C>T 6211A>G), CAG2011CAA (6217G>A), GCT2015GCA (6229T>A), GTA2016ATA (6230G>A), GTC2018GTT (6238C>T), AAG2021AAA (6247G>A), ATA2023ATT (6253A>T), GCA2025AAA (6257G>A 6258C>A), CAT2030CAC (6274T>C), GGC2038GGT (6298C>T), TTG2039TTA (6301G>A), AGA2040AGG (6304A>G), GAC2041GAT (6307C>T), AAC2043AAT (6313C>T), GCC2046GCT (6322C>T), ACA2048ACT (6328A>T), GTT2049GTG (6331T>G), TGT2050TGC (6334T>C), AGG2051AGA (6337G>A), CTG2052TTG (6338C>T), GTA2054GTG (6346A>G), TAT2058TAC (6358T>C), GGG2059GGA (6361G>A), GCA2060ACA (6362G>A), TCA2061TCC (6367A>C), GAG2062GAA (6370G>A), TAC2064TAT (6376C>T), GGT2065GGG (6379T>G), GGA2067GGC (6385A>C), GGA2069GGT (6391A>T), GCA2070GCG (6394A>G), TAC2071TAT (6397C>T), GTA2073ATA (6401G>A), GCG2074GTA (6405C>T 6406G>A), CAC2076CAT (6412C>T), TTA2078TTG (6418A>G), TTT2079TTC (6421T>C), TAC2082TTC (6429A>T), GGT2084GGA (6436T>A), GAG2087GAA (6445G>A), CAA2089CGA (6450A>G), TCC2090TCA (6454C>A), CAC2092CAT (6460C>T), GGT2093GGA (6463T>A), ACA2094ACG (6466A>G), TTC2095TTT (6469C>T), AGG2096AGA (6472G>A), GTG2097GTA (6475G>A), AAT2099AAC (6481T>C), CAC2101AAT (6485C>A 6487C>T), AGT2102AGC (6490T>C), AGC2104AGT (6496C>T), CTG2106CTA (6502G>A), GGT2110GGC (6514T>C), AGG2111AGA (6517G>A), CTC2115ATC (6527C>A), ATC2116ATA (6532C>A), TGT2454..C (7546T>C), AAG2462AAA (7570G>A), ACG2463ACA (7573G>A), ACG2465ACA (7579G>A), GCT2468GCC (7588T>C), CTA2471TTA (7595C>T), TTG2474CTA (7604T>C 7606G>A), GGT2477GGA (7615T>A), TGT2480TGC (7624T>C), GAC2483GAT (7633C>T), TTC2484TTT (7636C>T), AAT2485AAC (7639T>C), AAT2486AAC (7642T>C), TTT2488TTC (7648T>C), GAA2494GAG (7666A>G), TGT2496TGC (7672T>C), ACA2498ACT (7678A>T), GGG2500GGA (7684G>A), ACT2502ACC (7690T>C), TTT2504TTC (7696T>C), GGT2506GGG (7702T>G), GAT2509AAT (7709G>A), AAA2510AGG (7713A>G 7714A>G), CTG2511TTG (7715C>T), CGT2514CGC (7726T>C), GAG2517GAA (7735G>A), GTA2520GTG (7744A>G), TAC2521TAT (7747C>T), GGC2526GGT (7762C>T), CAG2528CAA (7768G>A), TTT2529TTC (7771T>C), CTA2533CTC (7783A>C), ACT2534ACC (7786T>C), CTA2537TTG (7793C>T 7795A>G), ATC2538ATT (7798C>T), GCT2540GCC (7804T>C), ACC2543ATC (7812C>T), GAA2550GAG (7834A>G), GAC2551GAT (7837C>T), GTG2554TTA (7844G>T 7846G>A), CAG2557CAA (7855G>A), CTG2559CTA (7861G>A), TTA2562TTG (7870A>G), TAC2563TAT (7873C>T), ACT2564ACA (7876T>A), GAG2565GAA (7879G>A), ATT2566ATA (7882T>A), GTT2567ATT (7883G>A), CCA2570CCT (7894A>T), ATT2571ATC (7897T>C), GTC2579GTT (7921C>T), AAG2581AAA (7927G>A), GGT2584GGC (7936T>C), AAT2585AAC (7939T>C), AAT2586AAC (7942T>C), CCT2590CCC (7954T>C), GTG2594GTA (7966G>A), GAT2595GAC (7969T>C), TCT2597TCC (7975T>C), GTC2601GTT (7987C>T), CTT2602CTC (7990T>C), GCT2607GCC (8005T>C), GAA2616GAG (8032A>G), ATC2618ATT (8038C>T), AAC2656AGT (8151A>G 8152C>T), TCG2660TCA (8164G>A), AGG2665AAG (8178G>A), AAG2666AAA (8182G>A), TTG2669TTA (8191G>A), TTC2671TTT (8197C>T), AGA2675AAA (8208G>A), CTG2677TTG (8213C>T), CTA2678CTG (8218A>G), ATC2679ATT (8221C>T), GAG2680GAA (8224G>A), CCA2685CCG (8239A>G), ATT2692ATC (8260T>C), GTA2693GTG (8263A>G), GAT2699GAC (8281T>C), AGA2700AGG (8284A>G), CCA2704GCG (8294C>G 8296A>G), GAG2705GAA (8299G>A), CAC2706CAT (8302C>T), GAA2709GAG (8311A>G), ATT2711ATC (8317T>C), TGC2712TGT (8320C>T), GCT2714GCA (8326T>A), GAG2717GAA (8335G>A), TCC2718TCT (8338C>T), TCT2722TTT (8349C>T), GAA2723GAG (8353A>G), CAC2726CAT (8362C>T), CAA2727CAG (8365A>G), ATC2728ATT (8368C>T), TAC2732TAT (8380C>T), TCA2733TCG (8383A>G), TTA2735TTG (8389A>G), CCT2740CCC (8404T>C), GCA2742TCA (8408G>T), ACA2743ACG (8413A>G), GCG2745GCT (8419G>T), GCT2750GCC (8434T>C), TAT2752TAC (8440T>C), ATG2756ATA (8452G>A), CTA2758CTG (8458A>G), AGG2759AAG (8460G>A), AAA2760AAG (8464A>G), CTG2761CTA (8467G>A), GCT2766ACA (8480G>A 8482T>A), GAG2771GAA (8497G>A), CTA2772TTG (8498C>T 8500A>G), AGA2773AAG (8502G>A 8503A>G), GCC2774GCT (8506C>T), GAA2777GAT (8515A>T), GTC2780GTT (8524C>T), GCA2781GCC (8527A>C), TTA2782CTG (8528T>C 8530A>G), GAG2785GAC (8539G>C), CTT2788TGT (8546C>T 8547T>G), TCT2790ACT (8552T>A), GTA2793GTG (8563A>G), GCA2797GGA (8574C>G), ATT2801GTT (8585A>G), GGA2805GAA (8598G>A), AGC2806AGT (8602C>T), AAC2807TCT (8603A>T 8604A>C 8605C>T), AAA2812AAG (8620A>G), CCA2813CAA (8622C>A), GAG2814GGA (8625A>G 8626G>A), CAG2815CAA (8629G>A), GGC2816GGT (8632C>T), ATC2818ATT (8638C>T), CAG2819CAA (8641G>A), AAC2821AAT (8647C>T), CCG2822TTC (8648C>T 8649C>T 8650G>C), AAC2823AAT (8653C>T), AAA2824AAG (8656A>G), GGA2825GAA (8658G>A), GAT2827GCG (8664A>C 8665T>G), GTT2830GTG (8674T>G), GCA2832GTT (8679C>T 8680A>T), GGC2833GGA (8683C>A), GGG2836GGA (8692G>A), ACA2837ACT (8695A>T), CCG2841CCA (8707G>A), AGA2842CGA (8708A>C), AAG2844AAA (8716G>A), ATC2846ATT (8722C>T), ACG2847ACA (8725G>A), AAA2849AAG (8731A>G), AGA2851AGG (8737A>G), CCC2853CCT (8743C>T), ACA2854AAG (8745C>A 8746A>G), AGC2855AGT (8749C>T), AAG2856AAA (8752G>A), GGA2857GGT (8755A>T), AAC2862AAT (8770C>T), TTA2863CTA (8771T>C), GAA2864GAG (8776A>G), CAT2865CAC (8779T>C), TTG2866TTA (8782G>A), TAT2869TAC (8791T>C), CCA2871CCG (8797A>G), CAA2872CAG (8800A>G), CAA2873CAG (8803A>G), GAT2875GAC (8809T>C), ATT2876ATC (8812T>C), AAT2878AAC (8818T>C), GCA2881GCG (8827A>G), CAG2885CCG (8838A>C), ACG2888ACA (8848G>A), TAT2890TAC (8854T>C), GAG2891GAA (8857G>A), GTG2893GTA (8863G>A), CGG2894CAG (8865G>A), ATG2895CTT (8867A>C 8869G>T), GCA2896GCG (8872A>G), GGA2900AAA (8882G>A 8883G>A), ACT2902TCT (8888A>T), TGC2915TGT (8929C>T), GTC2924ATC (8954G>A), AAC2925AAT (8959C>T), GGA2926GGT (8962A>T), GGG2933GGA (8983G>A), AAT2934GAT (8984A>G), GAA2935GTA (8988A>T), GAG2938GAA (8998G>A), CCG2940CCA (9004G>A), TTG2941TTA (9007G>A), ATC2944ATT (9016C>T), ACC2951ACA (9037C>A), ATA2967ATT (9085A>T), AAC2971AAT (9097C>T), ATT2983GTT (9131A>G), CGA2984CGT (9136A>T), CGG2987CGT (9145G>T), ATG2989GGA (9149A>G 9150T>G 9151G>A), GGT2990AGT (9152G>A), TTA2991TTG (9157A>G), GCG2992GCT (9160G>T), CGT2993CGC (9163T>C), GAG3000GAA (9184G>A), CGA3004CGG (9196A>G), AGG3010AGA (9214G>A), GAA3011GAG (9217A>G), GCG3012GCA (9220G>A), GCA3020GCT (9244A>T), TTG3021TTA (9247G>A), GCC3024GCT (9256C>T), CCT3026TCC (9260C>T 9262T>C), GGG3030GGA (9274G>A), GAC3032GAT (9280C>T), ATC3035ATT (9289C>T), GAG3040GAA (9304G>A), GTC3050GTT (9334C>T), GTC3060GTG (9364C>G) | | | | | | | | |
| HC-Pro protein (NP_734244.1) | 23 | 121 | 19.1% | 538 | 84.1% | 87 (100%) | 71 (81.6%) | 0/0/0/0 | 0 |
| Protein mutations: | R36K (1143G>A), L40I (1154T>A 1156G>A), S62N (1221G>A), R77G (1265A>G 1267A>G), N81S (1278A>G), A85V (1290C>T 1291G>A), D88T (1298G>A 1299A>C 1300C>T), I91V (1307A>G 1309A>G), N94E (1316A>G 1318T>A), I98T (1329T>C), E101K (1337G>A), D108V (1359A>T 1360C>T), L113I (1373C>A), I117V (1385A>G) | | | | | | | | |
| Codon mutations: | CAC23CAT (1105C>T), ACA24ACG (1108A>G), GTA26GTG (1114A>G), GCT27GCG (1117T>G), TTA29TTG (1123A>G), CCT30CCA (1126T>A), GTC31GTT (1129C>T), AGG36AAG (1143G>A), GTA37GTT (1147A>T), GCT38GCA (1150T>A), TTG40ATA (1154T>A 1156G>A), CAC43CAT (1165C>T), AGT44AGC (1168T>C), CTT46TTA (1172C>T 1174T>A), ACC55ACT (1201C>T), GCC61GCT (1219C>T), AGC62AAC (1221G>A), AGA77GGG (1265A>G 1267A>G), GGT79GGC (1273T>C), TTG80TTA (1276G>A), AAT81AGT (1278A>G), TTG83CTG (1283T>C), GCG85GTA (1290C>T 1291G>A), GAC88ACT (1298G>A 1299A>C 1300C>T), TTT90TTC (1306T>C), ATA91GTG (1307A>G 1309A>G), CAT92CAC (1312T>C), AAT94GAA (1316A>G 1318T>A), TTG97TTA (1327G>A), ATA98ACA (1329T>C), GCG99GCC (1333G>C), TTA100TTG (1336A>G), GAG101AAG (1337G>A), CTA103TTG (1343C>T 1345A>G), CCG106CCA (1354G>A), GTG107GTT (1357G>T), GAC108GTT (1359A>T 1360C>T), CTG109TTA (1361C>T 1363G>A), CTC111TTG (1367C>T 1369C>G), CTT113ATT (1373C>A), TTC114TTT (1378C>T), GAG116GAA (1384G>A), ATA117GTA (1385A>G), AAA119AAG (1393A>G), TCC120TCT (1396C>T) | | | | | | | | |
| CI protein (NP_734246.1) | 356 | 447 | 14.5% | 600 | 98.2% | 90 (97.8%) | 86 (93.5%) | 0/2/0/0 | 0 |
| Protein mutations: | V381I (4796G>A 4798C>T), T383S (4803C>G 4804T>C), G400_V401del (4853_4858delGGCGTT), S402A (4859T>G) | | | | | | | | |
| Codon mutations: | AAA361AAG (4738A>G), GTA363GTT (4744A>T), GCA364GCG (4747A>G), TTG365CTG (4748T>C), ACT370ACC (4765T>C), GTC381ATT (4796G>A 4798C>T), ACT383AGC (4803C>G 4804T>C), GAG384GAA (4807G>A), GCG385GCT (4810G>T), GCT386GCG (4813T>G), CTT387CTC (4816T>C), TAT392TAC (4831T>C), AAC393AAT (4834C>T), CCA395CCT (4840A>T), GTG396GTA (4843G>A), GGA399GGG (4852A>G), GGC400_GTT401del (4853_4858delGGCGTT), TCA402GCA (4859T>G), ACT403ACC (4864T>C), AGT404AGC (4867T>C), CTG405CTC (4870G>C), ATT406ATA (4873T>A), TGT409TGC (4882T>C), ACA416ACT (4903A>T), CAG418CAA (4909G>A), TTG422CTA (4919T>C 4921G>A), AGT423AGC (4924T>C), CCC424CCG (4927C>G), TTT426TTC (4933T>C), ATC427ATA (4936C>A), CAG428CAA (4939G>A), AAT429AAC (4942T>C), TTC430TTT (4945C>T), TCA436TCG (4963A>G), CCT439CCC (4972T>C), CTT445CTC (4990T>C) | | | | | | | | |
| NIa-VPg protein (NP_734252.1) | 91 | 188 | 52.1% | 619 | 88.2% | 98 (100%) | 83 (84.7%) | 0/0/0/0 | 0 |
| Protein mutations: | I93V (5990A>G), E102D (6019A>T), K106T (6030A>C), D112E (6049C>A), M115I (6058G>A), S120N (6072G>A 6073T>C), N121H (6074A>C), T123N (6081C>A), C133S (6111G>C 6112T>C), I139V (6128A>G 6130T>C), V149I (6158G>A), T153S (6170A>T 6172A>G), L164F (6203C>T 6205C>T), V173I (6230G>A), A182K (6257G>A 6258C>A) | | | | | | | | |
| Codon mutations: | GCT91GCA (5986T>A), GAC92GAT (5989C>T), ATT93GTT (5990A>G), AGA99CGT (6008A>C 6010A>T), GAA102GAT (6019A>T), GTG103GTT (6022G>T), CGA104CGT (6025A>T), AAA106ACA (6030A>C), GAC112GAA (6049C>A), ATG115ATA (6058G>A), CAA116CAG (6061A>G), GCC117GCA (6064C>A), GGT119GGC (6070T>C), AGT120AAC (6072G>A 6073T>C), AAC121CAC (6074A>C), ACG122ACA (6079G>A), ACC123AAC (6081C>A), ATA124ATT (6085A>T), GCA126GCT (6091A>T), TAC127TAT (6094C>T), TGT133TCC (6111G>C 6112T>C), AAA135AAG (6118A>G), TTG137TTA (6124G>A), AAG138AAA (6127G>A), ATT139GTC (6128A>G 6130T>C), GAT140GAC (6133T>C), TTA141TTG (6136A>G), AAC145AAT (6148C>T), AAA148AAG (6157A>G), GTT149ATT (6158G>A), TGT150TGC (6163T>C), AAA152AAG (6169A>G), ACA153TCG (6170A>T 6172A>G), GGC155GGT (6178C>T), AAA158AAG (6187A>G), TTT159TTC (6190T>C), GAG161GAA (6196G>A), CTC164TTT (6203C>T 6205C>T), GAA165GAG (6208A>G), CTA166TTG (6209C>T 6211A>G), CAG168CAA (6217G>A), GCT172GCA (6229T>A), GTA173ATA (6230G>A), GTC175GTT (6238C>T), AAG178AAA (6247G>A), ATA180ATT (6253A>T), GCA182AAA (6257G>A 6258C>A), CAT187CAC (6274T>C) | | | | | | | | |
| NIa-Pro protein (NP_734248.1) | 1 | 88 | 34.4% | 538 | 92.0% | 84 (100%) | 76 (90.5%) | 0/0/0/0 | 0 |
| Protein mutations: | A29T (6362G>A), V42I (6401G>A), A43V (6405C>T 6406G>A), Y51F (6429A>T), Q58R (6450A>G), H70N (6485C>A 6487C>T), L84I (6527C>A) | | | | | | | | |

| | Begin | End | Coverage | Score | Concordance | Matches | Identities | I/D/M/F* | Stop Codons |
|---|---|---|---|---|---|---|---|---|---|
| **NT** | **1103** | **9641** | **31.6%** | **4154** | **68.5%** | **3057 (99.6%)** | **2586 (84.2%)** | **3/10** | |
| Codon mutations: | GGC7GGT (6298C>T), TTG8TTA (6301G>A), AGA9AGG (6304A>G), GAC10GAT (6307C>T), AAC12AAT (6313C>T), GCC15GCT (6322C>T), ACA17ACT (6328A>T), GTT18GTG (6331T>G), TGT19TGC (6334T>C), AGG20AGA (6337G>A), CTG21TTG (6338C>T), GTA23GTG (6346A>G), TAT27TAC (6358T>C), GGG28GGA (6361G>A), GCA29ACA (6362G>A), TCA30TCC (6367A>C), GAG31GAA (6370G>A), TAC33TAT (6376C>T), GGT34GGG (6379T>G), GGA36GGC (6385A>C), GGA38GGT (6391A>T), GCA39GCG (6394A>G), TAC40TAT (6397C>T), GTA42ATA (6401G>A), GCG43GTA (6405C>T 6406G>A), CAC45CAT (6412C>T), TTA47TTG (6418A>G), TTT48TTC (6421T>C), TAC51TTC (6429A>T), GGT53GGA (6436T>A), GAG56GAA (6445G>A), CAA58CGA (6450A>G), TCC59TCA (6454C>A), CAC61CAT (6460C>T), GGT62GGA (6463T>A), ACA63ACG (6466A>G), TTC64TTT (6469C>T), AGG65AGA (6472G>A), GTG66GTA (6475G>A), AAT68AAC (6481T>C), CAC70AAT (6485C>A 6487C>T), AGT71AGC (6490T>C), AGC73AGT (6496C>T), CTG75CTA (6502G>A), GGT79GGC (6514T>C), AGG80AGA (6517G>A), CTC84ATC (6527C>A), ATC85ATA (6532C>A) | | | | | | | | |
| NIb protein (NP_734249.1) | 180 | 521 | 58.7% | 2066 | 94.0% | 306 (100%) | 286 (93.5%) | 0/0/0/0 | 0 |
| Protein mutations: | D234N (7709G>A), K235R (7713A>G 7714A>G), T268I (7812C>T), V279L (7844G>T 7846G>A), V292I (7883G>A), N381S (8151A>G 8152C>T), R390K (8178G>A), R400K (8208G>A), P429A (8294C>G 8296A>G), S447F (8349C>T), A467S (8408G>T), M481I (8452G>A), R484K (8460G>A), A491T (8480G>A 8482T>A), R498K (8502G>A 8503A>G), E502D (8515A>T), E510D (8539G>C), L513C (8546C>T 8547T>G), S515T (8552T>A) | | | | | | | | |
| Codon mutations: | TGT179..C (7546T>C), AAG187AAA (7570G>A), ACG188ACA (7573G>A), ACG190ACA (7579G>A), GCT193GCC (7588T>C), CTA196TTA (7595C>T), TTG199CTA (7604T>C 7606G>A), GGT202GGA (7615T>A), TGT205TGC (7624T>C), GAC208GAT (7633C>T), TTC209TTT (7636C>T), AAT210AAC (7639T>C), AAT211AAC (7642T>C), TTT213TTC (7648T>C), GAA219GAG (7666A>G), TGT221TGC (7672T>C), ACA223ACT (7678A>T), GGG225GGA (7684G>A), ACT227ACC (7690T>C), TTT229TTC (7696T>C), GGT231GGG (7702T>G), GAT234AAT (7709G>A), AAA235AGG (7713A>G 7714A>G), CTG236TTG (7715C>T), CGT239CGC (7726T>C), GAG242GAA (7735G>A), GTA245GTG (7744A>G), TAC246TAT (7747C>T), GGC251GGT (7762C>T), CAG253CAA (7768G>A), TTT254TTC (7771T>C), CTA258CTC (7783A>C), ACT259ACC (7786T>C), CTA262TTG (7793C>T 7795A>G), ATC263ATT (7798C>T), GCT265GCC (7804T>C), ACC268ATC (7812C>T), GAA275GAG (7834A>G), GAC276GAT (7837C>T), GTG279TTA (7844G>T 7846G>A), CAG282CAA (7855G>A), CTG284CTA (7861G>A), TTA287TTG (7870A>G), TAC288TAT (7873C>T), ACT289ACA (7876T>A), GAG290GAA (7879G>A), ATT291ATA (7882T>A), GTT292ATT (7883G>A), CCA295CCT (7894A>T), ATT296ATC (7897T>C), GTC304GTT (7921C>T), AAG306AAA (7927G>A), GGT309GGC (7936T>C), AAT310AAC (7939T>C), AAT311AAC (7942T>C), CCT315CCC (7954T>C), GTG319GTA (7966G>A), GAT320GAC (7969T>C), TCT322TCC (7975T>C), GTC326GTT (7987C>T), CTT327CTC (7990T>C), GCT332GCC (8005T>C), GAA341GAG (8032A>G), ATC343ATT (8038C>T), AAC381AGT (8151A>G 8152C>T), TCG385TCA (8164G>A), AGG390AAG (8178G>A), AAG391AAA (8182G>A), TTG394TTA (8191G>A), TTC396TTT (8197C>T), AGA400AAA (8208G>A), CTG402TTG (8213C>T), CTA403CTG (8218A>G), ATC404ATT (8221C>T), GAG405GAA (8224G>A), CCA410CCG (8239A>G), ATT417ATC (8260T>C), GTA418GTG (8263A>G), GAT424GAC (8281T>C), AGA425AGG (8284A>G), CCA429GCG (8294C>G 8296A>G), GAG430GAA (8299G>A), CAC431CAT (8302C>T), GAA434GAG (8311A>G), ATT436ATC (8317T>C), TGC437TGT (8320C>T), GCT439GCA (8326T>A), GAG442GAA (8335G>A), TCC443TCT (8338C>T), TCT447TTT (8349C>T), GAA448GAG (8353A>G), CAC451CAT (8362C>T), CAA452CAG (8365A>G), ATC453ATT (8368C>T), TAC457TAT (8380C>T), TCA458TCG (8383A>G), TTA460TTG (8389A>G), CCT465CCC (8404T>C), GCA467TCA (8408G>T), ACA468ACG (8413A>G), GCG470GCT (8419G>T), GCT475GCC (8434T>C), TAT477TAC (8440T>C), ATG481ATA (8452G>A), CTA483CTG (8458A>G), AGG484AAG (8460G>A), AAA485AAG (8464A>G), CTG486CTA (8467G>A), GCT491ACA (8480G>A 8482T>A), GAG496GAA (8497G>A), CTA497TTG (8498C>T 8500A>G), AGA498AAG (8502G>A 8503A>G), GCC499GCT (8506C>T), GAA502GAT (8515A>T), GTC505GTT (8524C>T), GCA506GCC (8527A>C), TTA507CTG (8528T>C 8530A>G), GAG510GAC (8539G>C), CTT513TGT (8546C>T 8547T>G), TCT515ACT (8552T>A), GTA518GTG (8563A>G) | | | | | | | | |
| coat protein (NP_734250.1) | 1 | 267 | 100% | 1661 | 91.1% | 267 (100%) | 244 (91.4%) | 0/0/0/0 | 0 |
| Protein mutations: | A1G (8574C>G), I5V (8585A>G), G9E (8598G>A), N11S (8603A>T 8604A>C 8605C>T), P17Q (8622C>A), E18G (8625A>G 8626G>A), P26F (8648C>T 8649C>T 8650G>C), G29E (8658G>A), D31A (8664A>C 8665T>G), A36V (8679C>T 8680A>T), T58K (8745C>A 8746A>G), Q89P (8838A>C), R98Q (8865G>A), M99L (8867A>C 8869G>T), G104K (8882G>A 8883G>A), T106S (8888A>T), V128I (8954G>A), N138D (8984A>G), E139V (8988A>T), I187V (9131A>G), M193G (9149A>G 9150T>G 9151G>A), G194S (9152G>A), P230S (9260C>T 9262T>C) | | | | | | | | |
| Codon mutations: | GCA1GGA (8574C>G), ATT5GTT (8585A>G), GGA9GAA (8598G>A), AGC10AGT (8602C>T), AAC11TCT (8603A>T 8604A>C 8605C>T), AAA16AAG (8620A>G), CCA17CAA (8622C>A), GAG18GGA (8625A>G 8626G>A), CAG19CAA (8629G>A), GGC20GGT (8632C>T), ATC22ATT (8638C>T), CAG23CAA (8641G>A), AAC25AAT (8647C>T), CCG26TTC (8648C>T 8649C>T 8650G>C), AAC27AAT (8653C>T), AAA28AAG (8656A>G), GGA29GAA (8658G>A), GAT31GCG (8664A>C 8665T>G), GTT34GTG (8674T>G), GCA36GTT (8679C>T 8680A>T), GGC37GGA (8683C>A), GGG40GGA (8692G>A), ACA41ACT (8695A>T), CCG45CCA (8707G>A), AGA46CGA (8708A>C), AAG48AAA (8716G>A), ATC50ATT (8722C>T), ACG51ACA (8725G>A), AAA53AAG (8731A>G), AGA55AGG (8737A>G), CCC57CCT (8743C>T), ACA58AAG (8745C>A 8746A>G), AGC59AGT (8749C>T), AAG60AAA (8752G>A), GGA61GGT (8755A>T), AAC66AAT (8770C>T), TTA67CTA (8771T>C), GAA68GAG (8776A>G), CAT69CAC (8779T>C), TTG70TTA (8782G>A), TAT73TAC (8791T>C), CCA75CCG (8797A>G), CAA76CAG (8800A>G), CAA77CAG (8803A>G), GAT79GAC (8809T>C), ATT80ATC (8812T>C), AAT82AAC (8818T>C), GCA85GCG (8827A>G), CAG89CCG (8838A>C), ACG92ACA (8848G>A), TAT94TAC (8854T>C), GAG95GAA (8857G>A), GTG97GTA (8863G>A), CGG98CAG (8865G>A), ATG99CTT (8867A>C 8869G>T), GCA100GCG (8872A>G), GGA104AAA (8882G>A 8883G>A), ACT106TCT (8888A>T), TGC119TGT (8929C>T), GTC128ATC (8954G>A), AAC129AAT (8959C>T), GGA130GGT (8962A>T), GGG137GGA (8983G>A), AAT138GAT (8984A>G), GAA139GTA (8988A>T), GAG142GAA (8998G>A), CCG144CCA (9004G>A), TTG145TTA (9007G>A), ATC148ATT (9016C>T), ACC155ACA (9037C>A), ATA171ATT (9085A>T), AAC175AAT (9097C>T), ATT187GTT (9131A>G), CGA188CGT (9136A>T), CGG191CGT (9145G>T), ATG193GGA (9149A>G 9150T>G 9151G>A), GGT194AGT (9152G>A), TTA195TTG (9157A>G), GCG196GCT (9160G>T), CGT197CGC (9163T>C), GAG204GAA (9184G>A), CGA208CGG (9196A>G), AGG214AGA (9214G>A), GAA215GAG (9217A>G), GCG216GCA (9220G>A), GCA224GCT (9244A>T), TTG225TTA (9247G>A), GCC228GCT (9256C>T), CCT230TCC (9260C>T 9262T>C), GGG234GGA (9274G>A), GAC236GAT (9280C>T), ATC239ATT (9289C>T), GAG244GAA (9304G>A), GTC254GTT (9334C>T), GTC264GTG (9364C>G) | | | | | | | | |

*: Inserts / Deletes / Misaligned / Frameshifts

#### Analysis details

This analysis was performed with panviral2.64

### NGS Details (UN24): Lausannevirus

#### Assembly

| | |
|---|---|
| **Coverage Length** | 174 (1 contig(s)) |
| **Depth Of Coverage** | 11097.3 |
| **Number Of Reads** | 20011 |
| **Reads Per Million** | 400.57 rpm (after QC) |
| **Ambiguities** | 0 |
| **Assembly Method** | de novo + reference guided assembly |
| **Consensus Caller** | Bcf Tools |

#### Coverage Map

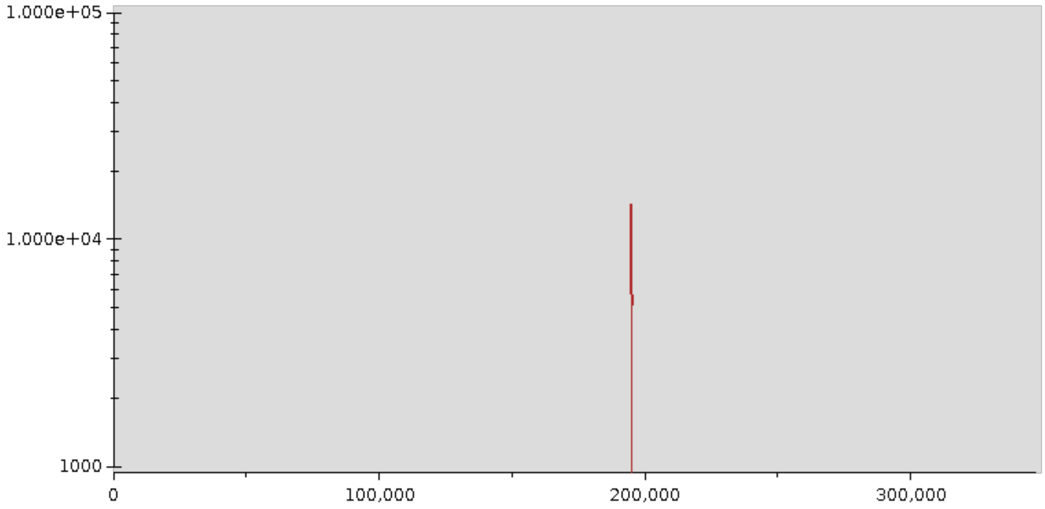

#### Assignment

| | |
|---|---|
| **Type** | Lausannevirus (Taxonomy ID: 999883) |
| **Reference Genome** | NC_015326.1 |
| **NT Identity (%)** | 80.4598 |
| **AA Identity (%)** | 96.5517 |
| **Number Of Stop Codons** | 0 |
| **Number Of CDS** | 444 |

#### Alignment

| | |
|---|---|
| **Alignment Score** | 212.0 (NT) + 362.0 (AA) = 574.0 |
| **Concordance (%)** | 79.5014 |
| **Alignment Method** | Local, heuristic, nucleotide (BLASTN) |

#### Genome Region

Sequence starts at position 194931 and ends at position 195104 relative to NC_015326.1 reference sequence.

#### Alignment Detailed Statistics

| | Begin | End | Coverage | Score | Concordance | Matches | Identities | I/D/M/F* | Stop Codons |
|---|---|---|---|---|---|---|---|---|---|
| **NT** | **194931** | **195104** | **0.1%** | **212** | **60.9%** | **174 (100%)** | **140 (80.5%)** | **0/0** | |
| Mutations: | 194931A>G, 194940A>C, 194946G>T, 194952T>C, 194955T>C, 194958A>G, 194967A>G, 194970G>C, 194979T>G, 194981T>G, 194982C>G, 194988T>C, 195003T>A, 195012A>C, 195015T>C, 195017G>T, 195018T>C, 195021T>C, 195033A>G, 195036G>C, 195042T>C, 195048T>C, 195054T>C, 195057T>A, 195058T>G, 195059G>C, 195060C>T, 195063G>C, 195066G>A, 195069C>G, 195072A>G, 195081C>A, 195087T>C, 195093T>C | | | | | | | | |

*: Inserts / Deletes / Misaligned / Frameshifts

#### Analysis details

This analysis was performed with panviral2.64

### NGS Details (UN24): Noumeavirus

#### Assembly

| | |
|---|---|
| **Coverage Length** | 217 (1 contig(s)) |
| **Depth Of Coverage** | 5688.3 |
| **Number Of Reads** | 14282 |
| **Reads Per Million** | 285.89 rpm (after QC) |
| **Ambiguities** | 0 |
| **Assembly Method** | de novo + reference guided assembly |
| **Consensus Caller** | Bcf Tools |

#### Coverage Map

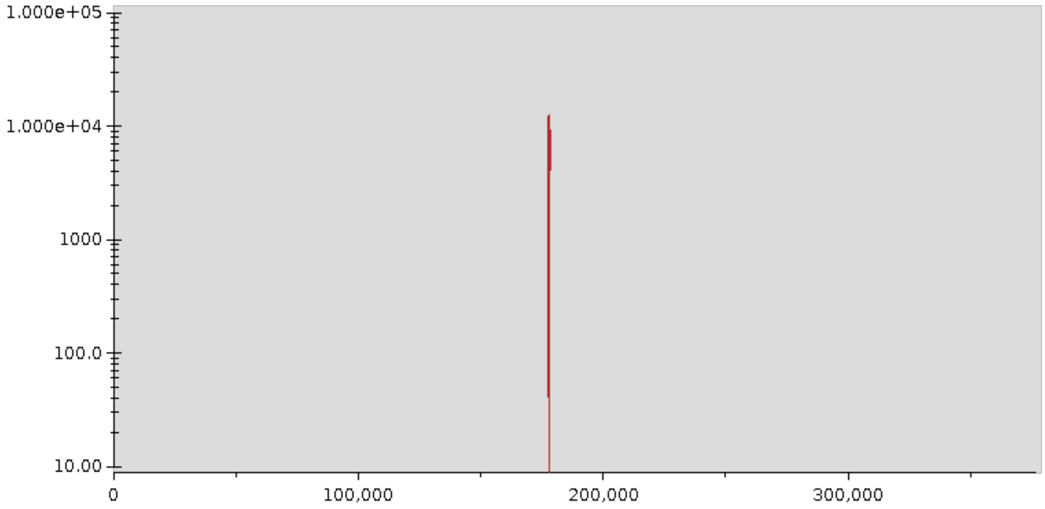

#### Assignment

| | |
|---|---|
| **Type** | Noumeavirus (Taxonomy ID: 1955558) |
| **Reference Genome** | NC_033775.1 |
| **NT Identity (%)** | 82.0276 |
| **AA Identity (%)** | 95.7746 |
| **Number Of Stop Codons** | 0 |
| **Number Of CDS** | 452 |

#### Alignment

| | |
|---|---|
| **Alignment Score** | 278.0 (NT) + 435.0 (AA) = 713.0 |
| **Concordance (%)** | 80.839 |
| **Alignment Method** | Local, heuristic, nucleotide (BLASTN) |

#### Genome Region

Sequence starts at position 177915 and ends at position 178136 relative to NC_033775.1 reference sequence.

#### Alignment Detailed Statistics

| | Begin | End | Coverage | Score | Concordance | Matches | Identities | I/D/M/F* | Stop Codons |
|---|---|---|---|---|---|---|---|---|---|
| NT | 177915 | 178136 | 0.1% | 278 | 64.1% | 217 (100%) | 178 (82.0%) | 0/0 | |

Mutations: 177926A>G, 177929T>C, 177941T>A, 177947A>C, 177950A>G, 177953A>G, 177980A>C, 177989C>T, 177992G>C, 177995C>T, 178001A>G, 178002C>G, 178003A>C, 178004A>T, 178019A>G, 178025A>G, 178028C>T, 178034G>A, 178043A>G, 178044C>A, 178047C>T, 178049T>G, 178055C>T, 178064A>G, 178070C>T, 178079A>C, 178080A>C, 178082G>T, 178088T>C, 178091C>T, 178097T>C, 178101G>A, 178106A>G, 178112A>G, 178118A>T, 178121G>C, 178124T>C, 178127T>G, 178136A>G

*: Inserts / Deletes / Misaligned / Frameshifts

#### Analysis details

This analysis was performed with panviral2.64

### NGS Details (UN24): Makelovirus prm1

#### Assembly

| | |
|---|---|
| **Coverage Length** | 1010 (1 contig(s)) |
| **Depth Of Coverage** | 894.1 |
| **Number Of Reads** | 7006 |
| **Reads Per Million** | 140.24 rpm (after QC) |
| **Ambiguities** | 0 |
| **Assembly Method** | de novo + reference guided assembly |
| **Consensus Caller** | Bcf Tools |

#### Coverage Map

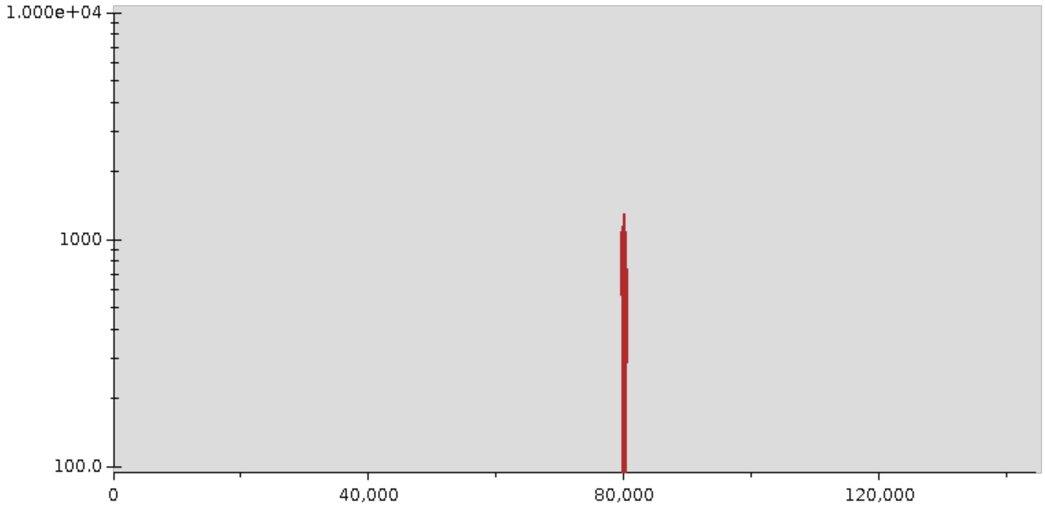

#### Assignment

| | |
|---|---|
| **Type** | Makelovirus prm1 (Taxonomy ID: 2956181) |
| **Reference Genome** | NC_055761.1 |
| **NT Identity (%)** | 76.7327 |
| **AA Identity (%)** | 87.2404 |
| **Number Of Stop Codons** | 0 |
| **Number Of CDS** | 190 |

#### Alignment

| | |
|---|---|
| **Alignment Score** | 1080.0 (NT) + 2246.0 (AA) = 3326.0 |
| **Concordance (%)** | 74.3406 |
| **Alignment Method** | Local, heuristic, nucleotide (BLASTN) |

#### Genome Region

Sequence starts at position 79508 and ends at position 80517 relative to NC_055761.1 reference sequence.

#### Alignment Detailed Statistics

| | Begin | End | Coverage | Score | Concordance | Matches | Identities | I/D/M/F* | Stop Codons |
|---|---|---|---|---|---|---|---|---|---|
| **NT** | **79508** | **80517** | **0.7%** | **1080** | **53.5%** | **1010 (100%)** | **775 (76.7%)** | **0/0** | |

Mutations: 79519A>G, 79520A>T, 79522A>G, 79523A>C, 79529C>T, 79530G>A, 79532G>C, 79536G>A, 79541G>T, 79542T>A, 79543C>G, 79544T>C, 79547A>T, 79548A>G, 79550C>A, 79563G>A, 79565C>T, 79568T>A, 79574C>T, 79580G>T, 79581C>T, 79592A>T, 79595T>C, 79596C>T, 79598G>A, 79602G>A, 79604A>G, 79610C>T, 79611G>T, 79612T>C, 79613C>T, 79614T>G, 79615G>T, 79616C>A, 79619C>T, 79624C>T, 79628G>C, 79632G>A, 79643C>T, 79646T>A, 79649G>A, 79655C>T, 79658C>T, 79664C>T, 79673A>T, 79676C>T, 79677G>T, 79679T>A, 79682T>G, 79688G>A, 79689A>C, 79691G>T, 79694T>C, 79703C>T, 79712T>C, 79718T>C, 79719G>A, 79722G>A, 79724A>T, 79728T>A, 79733C>T, 79734A>G, 79735A>C, 79736C>A, 79739A>T, 79742T>A, 79746C>T, 79748T>A, 79751C>T, 79754C>T, 79760C>A, 79763T>C, 79772T>G, 79779C>G, 79791C>T, 79793C>A, 79799T>C, 79811C>T, 79812C>G, 79817G>A, 79818G>A, 79820C>T, 79823A>T, 79824T>C, 79826T>A, 79833C>T, 79835G>A, 79836A>C, 79838C>T, 79842A>G, 79844C>A, 79845T>G, 79846T>C, 79847C>T, 79850C>T, 79871A>G, 79874C>T, 79875T>A, 79876C>G, 79879A>T, 79880T>C, 79895T>A, 79898C>T, 79904C>T, 79905T>G, 79906G>C, 79907C>T, 79916C>T, 79919T>A, 79922A>T, 79931T>A, 79937A>T, 79938T>A, 79940T>C, 79946C>T, 79950C>T, 79953G>A, 79958T>C, 79961T>A, 79962T>A, 79973T>A, 79974T>A, 79975C>G, 79976G>T, 79979C>T, 79985C>T, 79994C>T, 79997C>A, 80000C>A, 80018C>T, 80025C>A, 80030C>A, 80039A>T, 80042A>G, 80054G>T, 80063C>A, 80066C>T, 80069T>C, 80073C>T, 80075T>A, 80078T>C, 80090C>A, 80096T>C, 80099T>C, 80105G>A, 80114A>T, 80120C>T, 80126T>C, 80127C>T, 80132T>A, 80139T>A, 80140C>G, 80141G>T, 80142C>T, 80145G>A, 80147T>C, 80148C>A, 80150T>G, 80159C>A, 80162G>A, 80165C>T, 80168G>A, 80171C>T, 80172C>G, 80173A>C, 80174G>T, 80177C>T, 80178T>G, 80180T>A, 80188A>G, 80189G>A, 80201A>G, 80207G>A, 80210C>T, 80213C>T, 80216C>T, 80222G>A, 80225T>C, 80237C>T, 80240C>T, 80243T>C, 80246C>A, 80247C>T, 80261C>T, 80265T>A, 80266C>G, 80267C>T, 80279C>T, 80286C>T, 80288G>A, 80291T>C, 80300T>A, 80306A>T, 80315T>A, 80318C>A, 80333T>C, 80336A>T, 80337C>T, 80339T>A, 80343G>A, 80346T>A, 80347C>G, 80351C>T, 80357A>T, 80366G>A, 80369C>T, 80378C>T, 80387G>A, 80390C>T, 80391A>G, 80393C>A, 80394A>G, 80396G>T, 80399T>C, 80400G>A, 80409A>C, 80410A>G, 80411A>T, 80414C>A, 80415C>A, 80417C>T, 80433G>A, 80435T>C, 80436C>A, 80438A>C, 80447C>T, 80453G>T, 80456G>T, 80462G>A, 80465A>T, 80471C>T, 80474G>A, 80486C>T, 80498G>A, 80504G>A

*: Inserts / Deletes / Misaligned / Frameshifts

#### Analysis details

This analysis was performed with panviral2.64

#### NGS Details (UN24): Brazilian marseillevirus

##### Assembly

| | |
|---|---|
| **Coverage Length** | 242 (1 contig(s)) |
| **Depth Of Coverage** | 2168.2 |
| **Number Of Reads** | 4824 |
| **Reads Per Million** | 96.56 rpm (after QC) |
| **Ambiguities** | 0 |
| **Assembly Method** | de novo + reference guided assembly |
| **Consensus Caller** | Bcf Tools |

##### Coverage Map

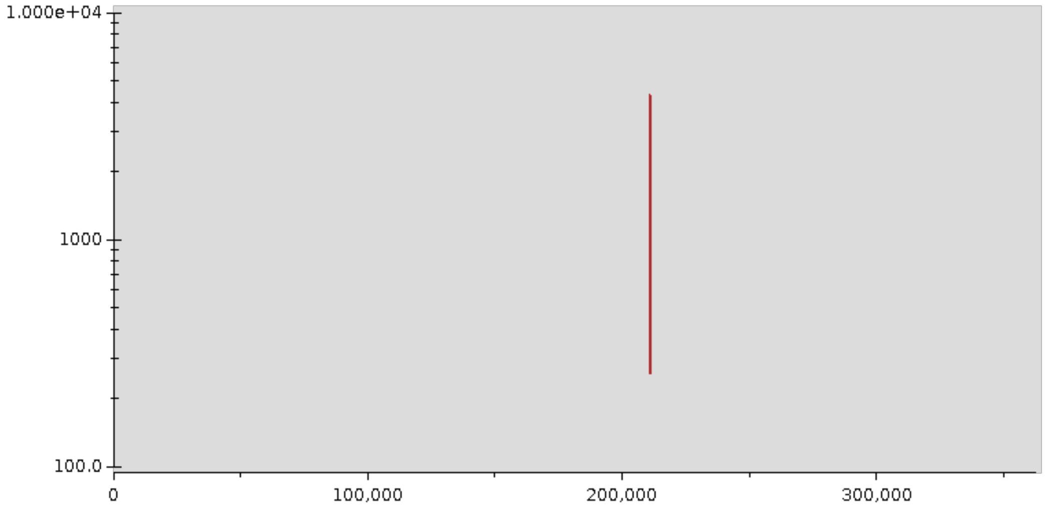

##### Assignment

| | |
|---|---|
| **Type** | Brazilian marseillevirus (Taxonomy ID: 1813599) |
| **Reference Genome** | NC_029692.1 |
| **NT Identity (%)** | 82.2314 |
| **AA Identity (%)** | 93.3333 |
| **Number Of Stop Codons** | 1 |
| **Number Of CDS** | 491 |

##### Alignment

| | |
|---|---|
| **Alignment Score** | 312.0 (NT) + 452.0 (AA) = 764.0 |
| **Concordance (%)** | 80.2521 |
| **Alignment Method** | Local, heuristic, nucleotide (BLASTN) |

##### Genome Region

Sequence starts at position 210884 and ends at position 211125 relative to NC_029692.1 reference sequence.

#### Alignment Detailed Statistics

| | Begin | End | Coverage | Score | Concordance | Matches | Identities | I/D/M/F* | Stop Codons |
|---|---|---|---|---|---|---|---|---|---|
| NT | 210884 | 211125 | 0.1% | 312 | 64.5% | 242 (100%) | 199 (82.2%) | 0/0 | |

Mutations: 210899T>G, 210901T>G, 210902T>G, 210917A>G, 210926T>C, 210938A>G, 210941A>G, 210944C>G, 210950G>T, 210953C>G, 210955T>G, 210956T>G, 210965C>G, 210974G>A, 210977T>G, 210986A>C, 210989G>C, 210991G>T, 210992T>C, 210995T>C, 211001A>T, 211004A>G, 211007A>G, 211009C>T, 211010T>C, 211025G>A, 211031T>A, 211032T>G, 211033G>C, 211040G>A, 211043T>G, 211058G>A, 211061T>C, 211064A>C, 211070G>A, 211072G>A, 211079C>G, 211088C>G, 211094A>C, 211100T>A, 211106C>G, 211108C>T, 211115T>C

*: Inserts / Deletes / Misaligned / Frameshifts

#### Analysis details

This analysis was performed with panviral2.64

### NGS Details (UN24): Tokyovirus A1

#### Assembly

| | |
|---|---|
| **Coverage Length** | 180 (1 contig(s)) |
| **Depth Of Coverage** | 1769.2 |
| **Number Of Reads** | 3678 |
| **Reads Per Million** | 73.62 rpm (after QC) |
| **Ambiguities** | 0 |
| **Assembly Method** | de novo + reference guided assembly |
| **Consensus Caller** | Bcf Tools |

#### Coverage Map

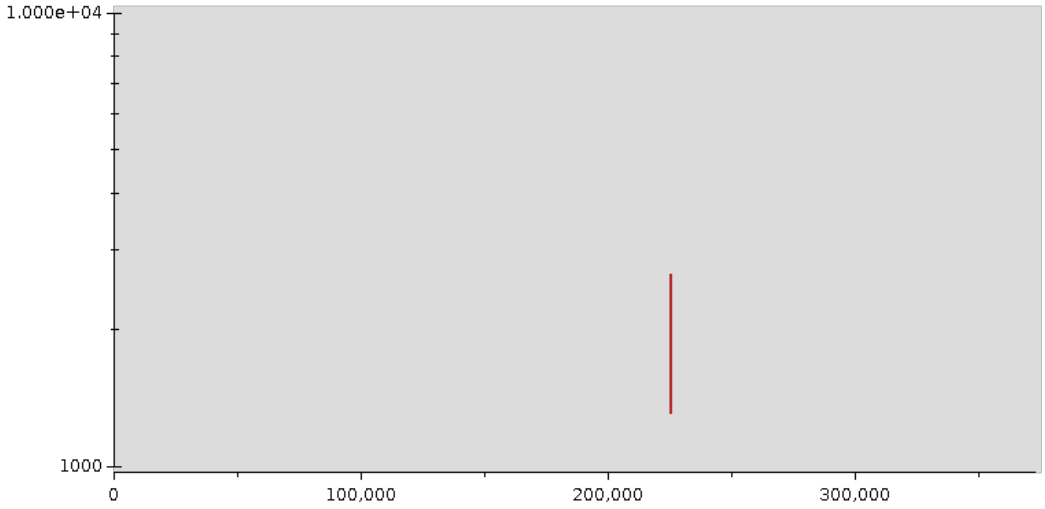

#### Assignment

| | |
|---|---|
| **Type** | Tokyovirus A1 (Taxonomy ID: 1826170) |
| **Reference Genome** | NC_030230.1 |
| **NT Identity (%)** | 80.5556 |
| **AA Identity (%)** | 93.1034 |
| **Number Of Stop Codons** | 1 |
| **Number Of CDS** | 470 |

#### Alignment

| | |
|---|---|
| **Alignment Score** | 220.0 (NT) + 353.0 (AA) = 573.0 |
| **Concordance (%)** | 78.4932 |
| **Alignment Method** | Local, heuristic, nucleotide (BLASTN) |

#### Genome Region

Sequence starts at position 225483 and ends at position 225662 relative to NC_030230.1 reference sequence.

#### Alignment Detailed Statistics

| | Begin | End | Coverage | Score | Concordance | Matches | Identities | I/D/M/F* | Stop Codons |
|---|---|---|---|---|---|---|---|---|---|
| **NT** | **225483** | **225662** | **0.1%** | **220** | **61.1%** | **180 (100%)** | **145 (80.6%)** | **0/0** | |

Mutations: 225498T>C, 225504G>C, 225507A>C, 225510G>A, 225513G>T, 225519G>T, 225528T>C, 225533C>T, 225534A>G, 225543G>C, 225549A>G, 225552T>G, 225554T>G, 225555C>G, 225564T>G, 225573C>A, 225579G>A, 225582G>A, 225585A>C, 225590G>T, 225594T>C, 225600G>C, 225603C>T, 225609A>T, 225621T>C, 225624G>T, 225630T>G, 225631T>G, 225632G>C, 225636C>G, 225640G>T, 225641C>T, 225642T>G, 225648G>T, 225651G>A

*: Inserts / Deletes / Misaligned / Frameshifts

#### Analysis details

This analysis was performed with panviral2.64

#### NGS Details (UN24): Marseillevirus marseillevirus

##### Assembly

| | |
|---|---|
| **Coverage Length** | 110 (1 contig(s)) |
| **Depth Of Coverage** | 2178.8 |
| **Number Of Reads** | 2867 |
| **Reads Per Million** | 57.39 rpm (after QC) |
| **Ambiguities** | 0 |
| **Assembly Method** | de novo + reference guided assembly |
| **Consensus Caller** | Bcf Tools |

##### Coverage Map

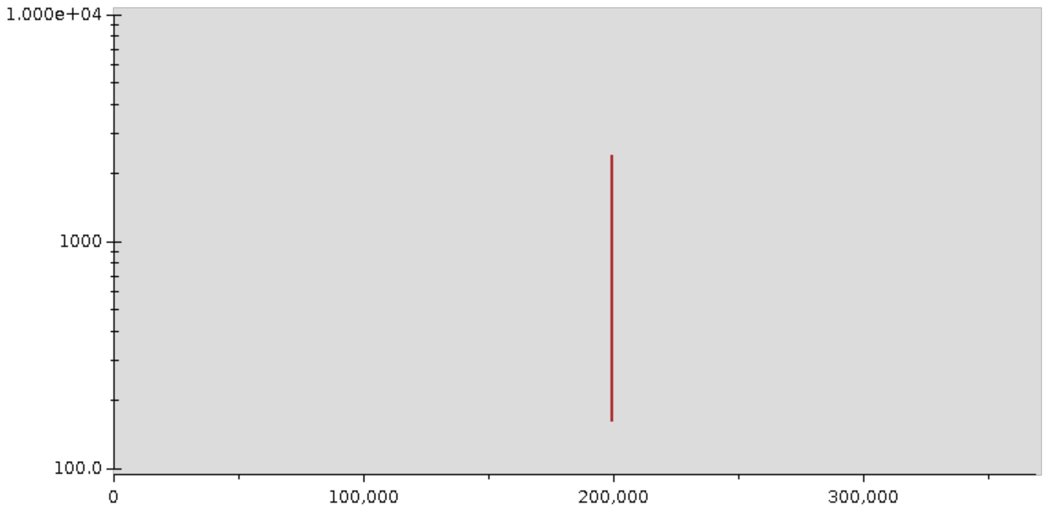

##### Assignment

| | |
|---|---|
| **Type** | Marseillevirus marseillevirus (Taxonomy ID: 694581) |
| **Reference Genome** | NC_013756.1 |
| **NT Identity (%)** | 83.6364 |
| **AA Identity (%)** | 94.2857 |
| **Number Of Stop Codons** | 1 |
| **Number Of CDS** | 428 |

##### Alignment

| | |
|---|---|
| **Alignment Score** | 148.0 (NT) + 204.0 (AA) = 352.0 |
| **Concordance (%)** | 80.3653 |
| **Alignment Method** | Local, heuristic, nucleotide (BLASTN) |

##### Genome Region

Sequence starts at position 199400 and ends at position 199509 relative to NC_013756.1 reference sequence.

#### Alignment Detailed Statistics

| | Begin | End | Coverage | Score | Concordance | Matches | Identities | I/D/M/F* | Stop Codons |
|---|---|---|---|---|---|---|---|---|---|
| NT | 199400 | 199509 | 0.1% | 148 | 67.3% | 110 (100%) | 92 (83.6%) | 0/0 | |

Mutations: 199411A>G, 199414T>A, 199416T>G, 199417C>G, 199421A>T, 199429G>C, 199431G>A, 199432A>T, 199435C>G, 199448A>T, 199449C>T, 199459A>C, 199464A>G, 199471T>A, 199489T>C, 199492G>T, 199495G>A, 199498G>A

*: Inserts / Deletes / Misaligned / Frameshifts

#### Analysis details

This analysis was performed with panviral2.64

#### NGS Details (UN24): Tunisvirus fontaine2

##### Assembly

| | |
|---|---|
| **Coverage Length** | 197 (1 contig(s)) |
| **Depth Of Coverage** | 946.0 |
| **Number Of Reads** | 1907 |
| **Reads Per Million** | 38.17 rpm (after QC) |
| **Ambiguities** | 0 |
| **Assembly Method** | de novo + reference guided assembly |
| **Consensus Caller** | Bcf Tools |

##### Coverage Map

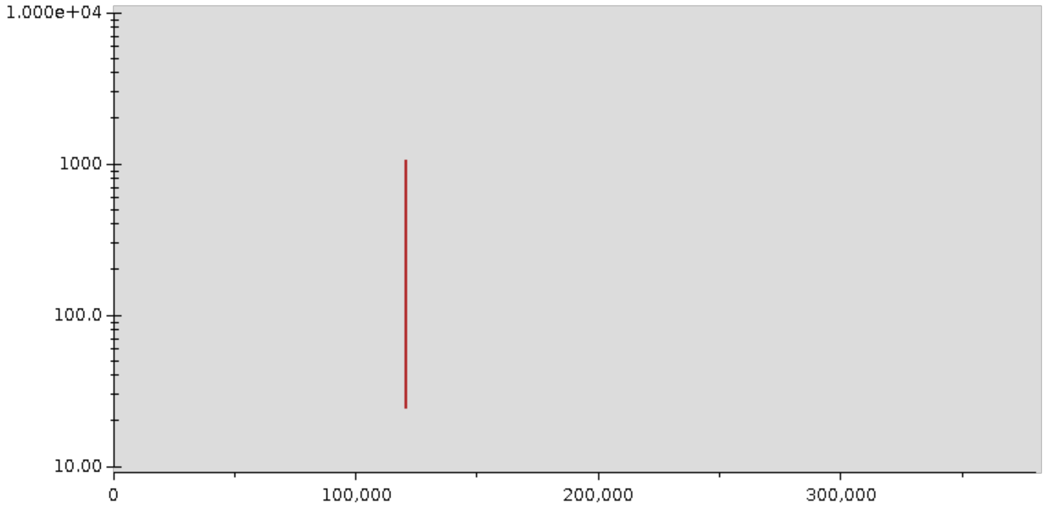

##### Assignment

| | |
|---|---|
| **Type** | Tunisvirus fontaine2 (Taxonomy ID: 1421067) |
| **Reference Genome** | NC_038511.1 |
| **NT Identity (%)** | 78.1726 |
| **AA Identity (%)** | 90.4762 |
| **Number Of Stop Codons** | 1 |
| **Number Of CDS** | 484 |

##### Alignment

| | |
|---|---|
| **Alignment Score** | 222.0 (NT) + 378.0 (AA) = 600.0 |
| **Concordance (%)** | 74.813 |
| **Alignment Method** | Local, heuristic, nucleotide (BLASTN) |

##### Genome Region

Sequence starts at position 120425 and ends at position 120621 relative to NC_038511.1 reference sequence.

#### Alignment Detailed Statistics

| | Begin | End | Coverage | Score | Concordance | Matches | Identities | I/D/M/F* | Stop Codons |
|---|---|---|---|---|---|---|---|---|---|
| **NT** | **120425** | **120621** | **0.1%** | **222** | **56.3%** | **197 (100%)** | **154 (78.2%)** | **0/0** | |

Mutations: 120440T>C, 120449A>C, 120451G>A, 120452G>A, 120455A>C, 120457G>A, 120461G>T, 120464T>C, 120470T>C, 120475C>T, 120482A>G, 120485G>A, 120488A>T, 120494C>A, 120496T>G, 120497C>T, 120506C>G, 120509A>C, 120510C>T, 120511T>G, 120515C>T, 120527A>C, 120529G>A, 120536T>C, 120545A>T, 120551G>A, 120557T>C, 120569T>C, 120572T>A, 120573T>G, 120574G>C, 120575C>T, 120582C>T, 120584C>G, 120587G>T, 120590G>A, 120596C>A, 120597G>C, 120598A>T, 120600C>G, 120601T>A, 120605C>A, 120608A>C

*: Inserts / Deletes / Misaligned / Frameshifts

#### Analysis details

This analysis was performed with panviral2.64

### NGS Details (UN24): Yellowstone lake phycodnavirus 1

#### Assembly

| | |
|---|---|
| **Coverage Length** | 185 (1 contig(s)) |
| **Depth Of Coverage** | 856.7 |
| **Number Of Reads** | 1645 |
| **Reads Per Million** | 32.93 rpm (after QC) |
| **Ambiguities** | 0 |
| **Assembly Method** | de novo + reference guided assembly |
| **Consensus Caller** | Bcf Tools |

#### Coverage Map

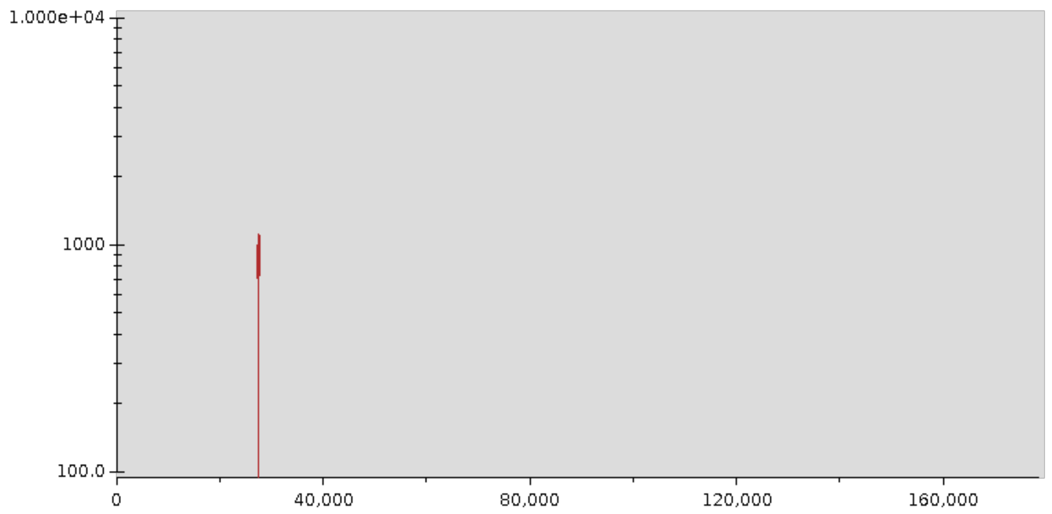

#### Assignment

| | |
|---|---|
| **Type** | Yellowstone lake phycodnavirus 1 (Taxonomy ID: 1586713) |
| **Reference Genome** | NC_028112.1 |
| **NT Identity (%)** | 81.6216 |
| **AA Identity (%)** | 90.3226 |
| **Number Of Stop Codons** | 0 |
| **Number Of CDS** | 248 |

#### Alignment

| | |
|---|---|
| **Alignment Score** | 234.0 (NT) + 371.0 (AA) = 605.0 |
| **Concordance (%)** | 79.5007 |
| **Alignment Method** | Local, heuristic, nucleotide (BLASTN) |

#### Genome Region

Sequence starts at position 27457 and ends at position 27641 relative to NC_028112.1 reference sequence.

#### Alignment Detailed Statistics

| | Begin | End | Coverage | Score | Concordance | Matches | Identities | I/D/M/F* | Stop Codons |
|---|---|---|---|---|---|---|---|---|---|
| **NT** | **27457** | **27641** | **0.1%** | **234** | **63.2%** | **185 (100%)** | **151 (81.6%)** | **0/0** | |

Mutations: 27470G>C, 27476C>A, 27477A>T, 27481T>G, 27489T>A, 27492A>G, 27501C>T, 27510A>T, 27513T>C, 27516T>A, 27529C>A, 27531T>G, 27534A>G, 27552A>G, 27555C>G, 27563A>G, 27564T>G, 27567T>C, 27573A>T, 27576A>T, 27579C>T, 27585T>C, 27586G>A, 27588C>T, 27591A>G, 27594G>A, 27600C>A, 27609T>C, 27612A>T, 27615C>T, 27616T>C, 27618G>T, 27624T>C, 27627C>T

*: Inserts / Deletes / Misaligned / Frameshifts

#### Analysis details

This analysis was performed with panviral2.64

### NGS Details (UN24): Amsacta moorei entomopoxvirus

#### Assembly

| | |
|---|---|
| **Coverage Length** | 131 (1 contig(s)) |
| **Depth Of Coverage** | 860.5 |
| **Number Of Reads** | 1423 |
| **Reads Per Million** | 28.48 rpm (after QC) |
| **Ambiguities** | 0 |
| **Assembly Method** | de novo + reference guided assembly |
| **Consensus Caller** | Bcf Tools |

#### Coverage Map

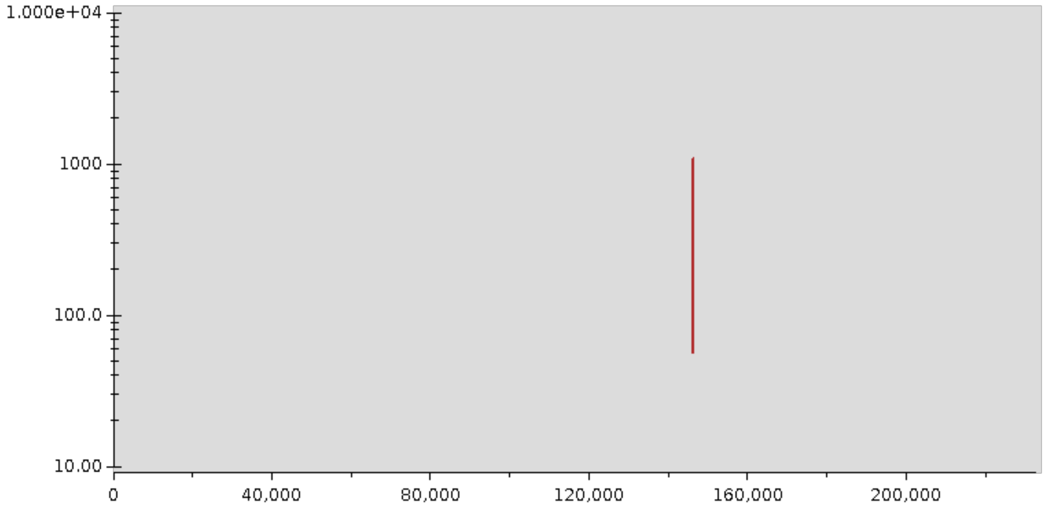

#### Assignment

| | |
|---|---|
| **Type** | Amsacta moorei entomopoxvirus (Taxonomy ID: 28321) |
| **Reference Genome** | NC_002520.1 |
| **NT Identity (%)** | 80.1527 |
| **AA Identity (%)** | 81.8182 |
| **Number Of Stop Codons** | 0 |
| **Number Of CDS** | 294 |

#### Alignment

| | |
|---|---|
| **Alignment Score** | 158.0 (NT) + 251.0 (AA) = 409.0 |
| **Concordance (%)** | 74.4991 |
| **Alignment Method** | Local, heuristic, nucleotide (BLASTN) |

#### Genome Region

Sequence starts at position 146093 and ends at position 146223 relative to NC_002520.1 reference sequence.

#### Alignment Detailed Statistics

| | Begin | End | Coverage | Score | Concordance | Matches | Identities | I/D/M/F* | Stop Codons |
|---|---|---|---|---|---|---|---|---|---|
| **NT** | **146093** | **146223** | **0.1%** | **158** | **60.3%** | **131 (100%)** | **105 (80.2%)** | **0/0** | |

Mutations: 146104A>G, 146107T>A, 146109A>C, 146113A>T, 146119G>T, 146121T>C, 146125A>C, 146128T>C, 146131T>C, 146140A>C, 146143A>G, 146144A>T, 146146A>G, 146155T>C, 146179T>C, 146188T>G, 146194A>T, 146195T>G, 146196T>C, 146197T>C, 146200A>G, 146202T>C, 146206T>G, 146207G>T, 146208A>C, 146209T>A

*: Inserts / Deletes / Misaligned / Frameshifts

#### Analysis details

This analysis was performed with panviral2.64

### NGS Details (UN24): Yellowstone lake phycodnavirus 1

#### Assembly

| | |
|---|---|
| **Coverage Length** | 144 (1 contig(s)) |
| **Depth Of Coverage** | 648.7 |
| **Number Of Reads** | 1029 |
| **Reads Per Million** | 20.60 rpm (after QC) |
| **Ambiguities** | 0 |
| **Assembly Method** | de novo + reference guided assembly |
| **Consensus Caller** | Bcf Tools |

#### Coverage Map

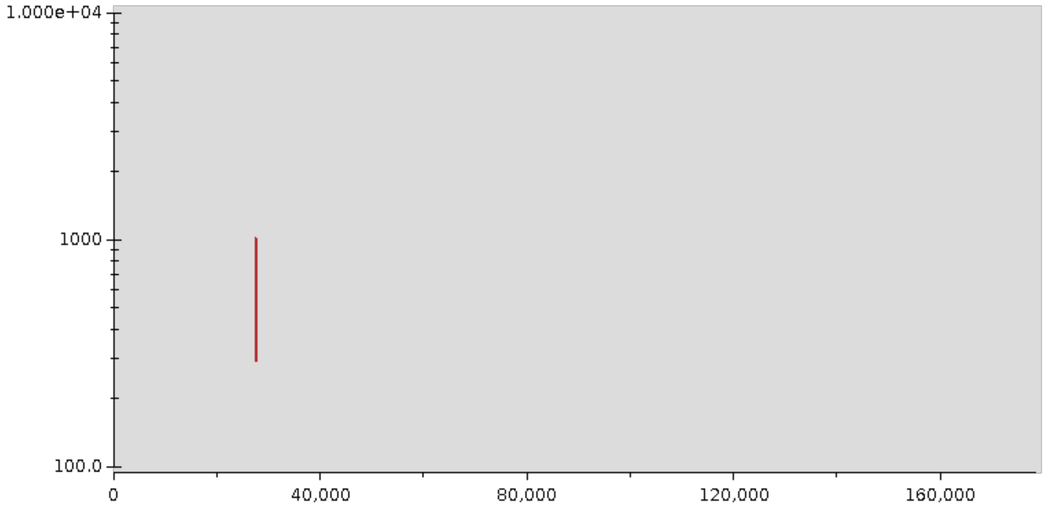

#### Assignment

| | |
|---|---|
| **Type** | Yellowstone lake phycodnavirus 1 (Taxonomy ID: 1586713) |
| **Reference Genome** | NC_028112.1 |
| **NT Identity (%)** | 81.9444 |
| **AA Identity (%)** | 83.3333 |
| **Number Of Stop Codons** | 0 |
| **Number Of CDS** | 248 |

#### Alignment

| | |
|---|---|
| **Alignment Score** | 184.0 (NT) + 267.0 (AA) = 451.0 |
| **Concordance (%)** | 75.2922 |
| **Alignment Method** | Local, heuristic, nucleotide (BLASTN) |

#### Genome Region

Sequence starts at position 27486 and ends at position 27629 relative to NC_028112.1 reference sequence.

#### Alignment Detailed Statistics

| | Begin | End | Coverage | Score | Concordance | Matches | Identities | I/D/M/F* | Stop Codons |
|---|---|---|---|---|---|---|---|---|---|
| **NT** | **27486** | **27629** | **0.1%** | **184** | **63.9%** | **144 (100%)** | **118 (81.9%)** | **0/0** | |

Mutations: 27498G>A, 27508G>T, 27510A>T, 27516T>C, 27534A>G, 27549G>A, 27550C>A, 27555C>T, 27558G>T, 27563A>G, 27564T>C, 27565C>A, 27567T>G, 27570C>A, 27573A>T, 27574G>T, 27585T>C, 27586G>A, 27588C>A, 27591A>G, 27597A>T, 27600C>A, 27606T>C, 27610C>T, 27612A>G, 27615C>G

*: Inserts / Deletes / Misaligned / Frameshifts

#### Analysis details

This analysis was performed with panviral2.64

### NGS Details (UN24): Cladosporium fulvum T-1 virus

#### Assembly

| | |
|---|---|
| **Coverage Length** | 1401 (3 contig(s)) |
| **Depth Of Coverage** | 56.9 |
| **Number Of Reads** | 749 |
| **Reads Per Million** | 14.99 rpm (after QC) |
| **Ambiguities** | 0 |
| **Assembly Method** | de novo + reference guided assembly |
| **Consensus Caller** | Bcf Tools |

#### Coverage Map

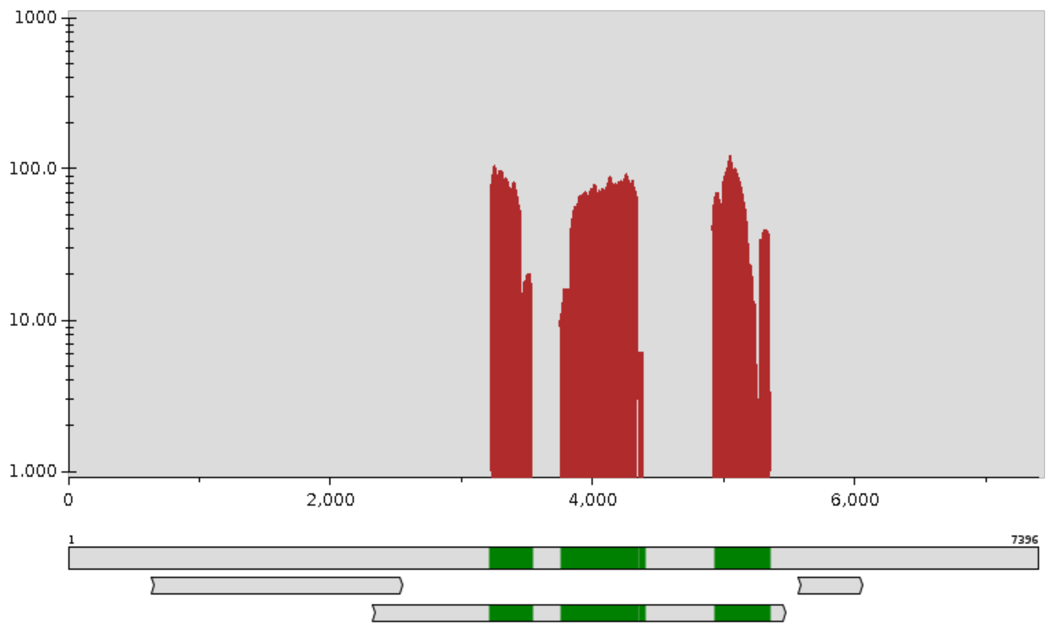

#### Assignment

| | |
|---|---|
| **Type** | Cladosporium fulvum T-1 virus (Taxonomy ID: 2052899) |
| **Reference Genome** | NC_043491.1 |
| **NT Identity (%)** | 53.0786 |
| **AA Identity (%)** | 47.5584 |
| **Number Of Stop Codons** | 1 |
| **Number Of CDS** | 3 |

#### Alignment

| | |
|---|---|
| **Alignment Score** | 123.0 (NT) + 1606.0 (AA) = 1729.0 |
| **Concordance (%)** | 29.2505 |

| Alignment Method | Global, seeded, nucleotide + amino acids (AGA) |
|---|---|

#### Genome Region

Sequence starts at position 3223 and ends at position 5354 relative to NC_043491.1 reference sequence.

#### Alignment Detailed Statistics

| | Begin | End | Coverage | Score | Concordance | Matches | Identities | I/D/M/F* | Stop Codons |
|---|---|---|---|---|---|---|---|---|---|
| **NT** | **3223** | **5354** | **18.9%** | **123** | **4.5%** | **1383 (96.6%)** | **750 (52.4%)** | **30/18** | |
| Mutations: | 3223C>G, 3224A>C, 3225A>T, 3229C>T, 3231C>T, 3232C>G, 3233T>A, 3235A>G, 3238T>A, 3238_3239insCCTGAGTTGCCAAAG, 3243A>T, 3244C>G, 3246A>C, 3251T>A, 3253_3254insAGGGATATT, 3261A>G, 3262G>A, 3265A>T, 3266A>G, 3268C>G, 3269A>T, 3271T>G, 3272C>A, 3273A>T, 3280G>T, 3281A>T, 3282A>C, 3283A>T, 3284G>A, 3285A>C, 3289T>G, 3292A>T, 3293T>G, 3294G>C, 3295G>A, 3296G>C, 3297G>A, 3299C>G, 3301C>T, 3303T>C, 3304A>T, 3307T>C, 3308C>A, 3309A>G, 3314T>G, 3317G>C, 3318A>C, 3319G>T, 3326C>T, 3328A>G, 3329C>G, 3330A>C, 3331G>T, 3332A>G, 3333C>A, 3334C>A, 3337A>G, 3338C>A, 3340A>G, 3341G>A, 3343A>G, 3344T>C, 3345G>A, 3352G>C, 3355G>A, 3356A>T, 3357A>T, 3361A>T, 3363C>A, 3364C>T, 3365A>T, 3366A>C, 3371T>C, 3372G>T, 3376A>T, 3378G>A, 3379A>G, 3381G>C, 3382A>T, 3385C>T, 3387C>A, 3388C>G, 3389T>G, 3391A>T, 3392A>C, 3393G>C, 3394T>C, 3395G>T, 3396C>A, 3397A>T, 3400A>T, 3401A>G, 3403T>A, 3406A>T, 3407T>G, 3408G>T, 3409C>T, 3410A>T, 3412G>A, 3415C>T, 3416G>C, 3417T>A, 3418T>G, 3419C>A, 3420C>A, 3425G>C, 3426C>A, 3427A>G, 3428A>G, 3430C>T, 3434A>T, 3435A>C, 3437C>T, 3440C>A, 3442A>G, 3443C>A, 3445C>G, 3446G>T, 3447T>G, 3448A>C, 3449C>A, 3450A>T, 3451A>T, 3460A>G, 3461A>C, 3464T>C, 3470G>A, 3473A>G, 3475C>G, 3478G>T, 3481C>A, 3487C>T, 3488C>A, 3489G>A, 3490A>G, 3493T>C, 3497C>T, 3499A>G, 3502C>T, 3504A>G, 3505C>G, 3508C>T, 3511A>T, 3514A>T, 3515G>T, 3516C>T, 3517A>G, 3518C>T, 3519A>T, 3520A>T, 3523C>T, 3524A>C, 3525G>A, 3529A>T, 3530A>C, 3531C>A, 3532C>G, 3536T>G, 3538A>T, 3751C>T, 3757A>T, 3758C>G, 3760G>C, 3761G>A, 3763C>A, 3766C>T, 3768C>G, 3769A>T, 3774G>A, 3775A>T, 3776T>A, 3778C>G, 3779C>T, 3781C>G, 3782C>G, 3784G>T, 3791A>G, 3792C>T, 3793C>G, 3796G>C, 3799A>T, 3800G>T, 3802T>G, 3803C>A, 3804A>G, 3805A>G, 3806G>T, 3807A>T, 3808T>G, 3811G>A, 3814C>A, 3815G>C, 3819G>T, 3820A>T, 3821C>T, 3823C>A, 3825C>G, 3827A>G, 3829G>A, 3831C>A, 3833G>A, 3834G>A, 3835A>G, 3838C>G, 3839A>T, 3841G>T, 3842A>G, 3843C>T, 3844A>T, 3845G>A, 3846C>A, 3848C>A, 3849C>T, 3850C>G, 3856A>G, 3859C>T, 3862A>G, 3865C>T, 3866C>G, 3867A>C, 3869A>C, 3871G>A, 3880C>A, 3886T>C, 3887T>C, 3893T>C, 3894T>A, 3898C>A, 3899A>G, 3901C>T, 3905A>G, 3906C>A, 3908A>G, 3909C>A, 3910A>C, 3912G>A, 3913G>A, 3914A>G, 3918C>G, 3922C>G, 3925C>T, 3929G>A, 3930C>A, 3931A>G, 3934G>A, 3935A>G, 3936C>T, 3937A>G, 3940G>A, 3941T>G, 3943A>C, 3946C>T, 3947A>G, 3948G>T, 3952A>T, 3957C>A, 3960A>C, 3964G>C, 3967G>A, 3970A>T, 3973C>G, 3975A>C, 3980G>C, 3982A>G, 3983C>A, 3984A>G, 3988A>T, 3992C>T, 3994T>G, 3997A>G, 4000C>A, 4003C>T, 4010A>T, 4012C>T, 4025A>G, 4027G>T, 4029A>G, 4030C>A, 4033T>C, 4038A>C, 4041C>T, 4048A>T, 4052A>C, 4057G>T, 4058A>G, 4059T>A, 4060G>T, 4063T>G, 4064A>C, 4065C>T, 4066A>G, 4068G>A, 4069A>G, 4072A>G, 4073G>A, 4075C>A, 4081C>G, 4085A>G, 4086A>T, 4091G>T, 4092G>C, 4093A>G, 4094A>G, 4095A>T, 4097G>A, 4098A>G, 4100C>T, 4101A>G, 4102G>C, 4103A>G, 4104C>A, 4105C>A, 4111G>T, 4114C>T, 4115A>C, 4117A>G, 4120A>T, 4121C>T, 4123C>G, 4130C>G, 4131A>C, 4132G>T, 4133T>A, 4134G>T, 4135C>T, 4138T>A, 4143C>A, 4144C>A, 4149C>T, 4150G>A, 4153T>G, 4154C>A, 4155G>A, 4156A>G, 4159A>G, 4159_4160insCCAAAT, 4162C>T, 4165T>G, 4166G>C, 4167G>T, 4168T>A, 4169A>C, 4170G>C, 4171C>T, 4172A>T, 4173A>T, 4174G>T, 4177A>G, 4180C>A, 4184A>G, 4186C>G, 4187G>C, 4192C>T, 4195C>T, 4201T>G, 4204T>C, 4206T>A, 4207G>A, 4210A>T, 4211A>G, 4213A>G, 4216C>A, 4218C>G, 4219A>C, 4220T>G, 4221G>T, 4222T>A, 4223C>T, 4225A>G, 4226A>G, 4227C>T, 4228A>G, 4231G>A, 4232_4237delACACAC, 4240T>A, 4243G>T, 4244_4249delAAAAGA, 4252C>T, 4255A>G, 4263A>T, 4265T>G, 4267T>A, 4268T>A, 4269C>G, 4271C>A, 4277A>T, 4279G>A, 4281C>A, 4282C>T, 4283A>G, 4284C>A, 4285A>T, 4288G>T, 4291A>G, 4296A>G, 4297C>G, 4301G>T, 4302A>C, 4303C>T, 4305T>C, 4306C>T, 4312C>A, 4319C>A, 4321T>G, 4322C>G, 4324A>T, 4327C>A, 4328A>G, 4330T>G, 4333T>G, 4334G>C, 4335C>A, 4337G>T, 4338C>G, 4340A>C, 4346C>A, 4347A>C, 4348T>C, 4355G>C, 4356T>A, 4357G>C, 4361G>T, 4363C>G, 4364G>C, 4365A>T, 4366G>T, 4369C>T, 4370C>T, 4372A>T, 4373_4375delCCG, 4381A>C, 4382A>T, 4383C>T, 4384G>A, 4385A>G, 4387T>C, 4388C>A, 4389T>A, 4390T>A, 4391T>A, 4393A>C, 4915A>T, 4918A>T, 4921G>T, 4922A>T, 4924A>G, 4925C>A, 4926C>A, 4927A>G, 4933C>A, 4939T>G, 4941C>A, 4942G>T, 4951C>T, 4952A>G, 4955A>G, 4956C>T, 4957G>A, 4958A>G, 4959A>G, 4961C>T, 4963C>G, 4966G>A, 4968G>A, 4972A>T, 4974A>G, 4976G>A, 4977A>G, 4978T>G, 4980G>A, 4982G>A, 4983T>C, 4984C>T, 4986C>G, 4988G>A, 4990A>G, 4994_4996delGCC, 4997T>C, 5002C>T, 5003A>T, 5004T>C, 5005G>T, 5009C>T, 5010T>G, 5011A>G, 5014C>G, 5017G>A, 5020C>T, 5023C>T, 5024A>C, 5026A>G, 5027C>A, 5029C>G, 5032A>G, 5037A>C, 5038T>A, 5041A>C, 5044C>T, 5047C>T, 5048A>C, 5050T>A, 5053T>A, 5055C>T, 5056A>C, 5057T>A, 5058C>A, 5059A>G, 5060G>A, 5061A>C, 5062A>T, 5064T>C, 5065A>T, 5067A>T, 5070C>G, 5074A>T, 5077G>A, 5078C>G, 5080G>T, 5081C>T, 5082T>A, 5083C>T, 5085G>C, 5086A>C, 5087T>A, 5089C>A, 5092C>G, 5093G>T, 5094T>A, 5095A>C, 5096C>A, 5098G>T, 5099G>C, 5100A>G, 5101C>T, 5102A>G, 5103G>A, 5104A>G, 5108A>G, 5110C>T, 5114T>C, 5115A>T, 5119C>T, 5122A>T, 5123T>A, 5125C>T, 5128G>T, 5129G>T, 5130A>T, 5131A>G, 5132G>T, 5133T>C, 5134A>C, 5135T>A, 5137C>T, 5141A>T, 5143A>T, 5146C>T, 5151A>G, 5152C>T, 5153A>G, 5154A>C, 5155G>T, 5157T>A, 5158C>G, 5161C>T, 5164A>T, 5168A>C, 5172A>T, 5173C>T, 5178A>G, 5179G>A, 5180A>T, 5182G>A, 5183C>T, 5185C>T, 5186A>C, 5187T>A, 5188G>T, 5189G>A, 5190G>A, 5191A>G, 5192A>G, 5193C>G, 5194G>A, 5195A>C, 5200A>T, 5202T>C, 5207C>G, 5208A>T, 5209C>A, 5216T>A, 5217C>G, 5218A>C, 5221A>C, 5224A>T, 5226A>T, 5227C>T, 5233A>T, 5234G>C, 5236G>A, 5239G>T, 5245G>A, 5251G>A, 5257A>G, 5260G>C, 5262A>T, 5263C>T, 5272C>T, 5276C>G, 5278A>C, 5284A>T, 5285C>A, 5292A>T, 5293C>T, 5294A>G, 5296C>G, 5298A>G, 5301A>C, 5302C>T, 5303G>C, 5304C>A, 5305A>T, 5309G>A, 5311C>A, 5312A>G, 5314C>T, 5319T>C, 5321T>A, 5322C>A, 5323A>G, 5326A>G, 5329G>A, 5330C>G, 5331C>A, 5332A>T, 5333A>G, 5338G>C, 5341G>A, 5342A>T, 5345G>T | | | | | | | | |

**CDS**

| | | | | | | | | | |
|---|---|---|---|---|---|---|---|---|---|
| homologue_of_retroviral_POL_genes | 300 | 1010 | 44.6% | 1606 | 49.8% | 461 (96.6%) | 224 (47.0%) | 10/6/0/0 | 1 |

| | Begin | End | Coverage | Score | Concordance | Matches | Identities | I/D/M/F* | Stop Codons |
|---|---|---|---|---|---|---|---|---|---|
| **NT** | **3223** | **5354** | **18.9%** | **123** | **4.5%** | **1383 (96.6%)** | **750 (52.4%)** | **30/18** | |
| Protein mutations: | K300L (3224A>C 3225A>T), A302V (3231C>T 3232C>G), L303M (3233T>A 3235A>G), P304_K305insPELPK (3238_3239insCCTGAGTTGCCAAAG), H306L (3243A>T 3244C>G), Q307P (3246A>C), W309R (3251T>A), W309_D310insRDI (3253_3254insAGGGATATT), K312R (3261A>G 3262G>A), N314E (3266A>G 3268C>G), I315L (3269A>T 3271T>G), Q316M (3272C>A 3273A>T), K319S (3281A>T 3282A>C 3283A>T), E320T (3284G>A 3285A>C), W323A (3293T>G 3294G>C 3295G>A), G324Q (3296G>C 3297G>A), P325A (3299C>G 3301C>T), L326P (3303T>C 3304A>T), Q328R (3308C>A 3309A>G), S330A (3314T>G), E331P (3317G>C 3318A>C 3319G>T), Q335A (3329C>G 3330A>C 3331G>T), T336E (3332A>G 3333C>A 3334C>A), E339K (3341G>A 3343A>G), W340Q (3344T>C 3345G>A), K342N (3352G>C), K344L (3356A>T 3357A>T), A346D (3363C>A 3364C>T), K347S (3365A>T 3366A>C), W349L (3371T>C 3372G>T), R351Q (3378G>A 3379A>G), R352P (3381G>C 3382A>T), T354K (3387C>A 3388C>G), S355A (3389T>G 3391A>T), S356P (3392A>C 3393G>C 3394T>C), A357Y (3395G>T 3396C>A 3397A>T), T359A (3401A>G 3403T>A), C361V (3407T>G 3408G>T 3409C>T), M362L (3410A>T 3412G>A), V364Q (3416G>C 3417T>A 3418T>G), P365K (3419C>A 3420C>A), A367Q (3425G>C 3426C>A 3427A>G), N368D (3428A>G 3430C>T), K370S (3434A>T 3435A>C), L373M (3443C>A 3445C>G), V374C (3446G>T 3447T>G 3448A>C), Q375I (3449C>A 3450A>T 3451A>T), K379Q (3461A>C), E382K (3470G>A), I383V (3473A>G 3475C>G), R388K (3488C>A 3489G>A 3490A>G), N393R (3504A>G 3505C>G), E395D (3511A>T), E396D (3514A>T), A397L (3515G>T 3516C>T 3517A>G), Q398F (3518C>T 3519A>T 3520A>T), R400Q (3524A>C 3525G>A), L401F (3529A>T), T402Q (3530A>C 3531C>A 3532C>G), S404A (3536T>G 3538A>T), L478V (3758C>G 3760G>C), V479I (3761G>A 3763C>A), T481S (3768C>G 3769A>T), G483D (3774G>A 3775A>T), S484T (3776T>A 3778C>G), Q486D (3782C>G 3784G>T), T489V (3791A>G 3792C>T 3793C>G), K490N (3796G>C), Q491H (3799A>T), V492L (3800G>T 3802T>G), Q493R (3803C>A 3804A>G 3805A>G), D494L (3806G>T 3807A>T 3808T>G), F496L (3814C>A), E497Q (3815G>C), R498L (3819G>T 3820A>T), T500R (3825C>G), K501E (3827A>G 3829G>A), S502Y (3831C>A), G503K (3833G>A 3834G>A 3835A>G), F504L (3838C>G), K505Y (3839A>T 3841G>T), T506V (3842A>G 3843C>T 3844A>T), A507K (3845G>A 3846C>A), P508M (3848C>A 3849C>T 3850C>G), H514A (3866C>G 3867A>C), K515Q (3869A>C 3871G>A), F523H (3893T>C 3894T>A), I525V (3899A>G 3901C>T), T527E (3905A>G 3906C>A), T528D (3908A>G 3909C>A 3910A>C), G529E (3912G>A 3913G>A), I530V (3914A>G), T531R (3918C>G), I532M (3922C>G), A535K (3929G>A 3930C>A 3931A>G), T537V (3935A>G 3936C>T 3937A>G), S539A (3941T>G 3943A>C), R541V (3947A>G 3948G>T), E542D (3952A>T), P544Q (3957C>A), E545A (3960A>C), K550T (3975A>C), V552L (3980G>C 3982A>G), Q553R (3983C>A 3984A>G), N562Y (4010A>T 4012C>T), K567D (4025A>G 4027G>T), D568G (4029A>G 4030C>A), K571T (4038A>C), T572I (4041C>T), M576L (4052A>C), M578D (4058A>G 4059T>A 4060G>T), T580L (4064A>C 4065C>T 4066A>G), R581K (4068G>A 4069A>G), D583K (4073G>A 4075C>A), N585K (4081C>G), K587V (4085A>G 4086A>T), G589S (4091G>T 4092G>C 4093A>G), K590V (4094A>G 4095A>T), E591R (4097G>A 4098A>G), Q592C (4100C>T 4101A>G 4102G>C), T593E (4103A>G 4104C>A 4105C>A), K597Q (4115A>C 4117A>G), R598S (4120A>T), Q602A (4130C>G 4131A>C 4132G>T), C603I (4133T>A 4134G>T 4135C>T), A606E (4143C>A 4144C>A), T608I (4149C>T 4150G>A), R610K (4154C>A 4155G>A 4156A>G), L611_F612insPN (4159_4160insCCAAAT), D613E (4165T>G), G614L (4166G>C 4167G>T 4168T>A), S615P (4169A>C 4170G>C 4171C>T), K616F (4172A>T 4173A>T 4174G>T), I620V (4184A>G 4186C>G), E621Q (4187G>C), M627K (4206T>A 4207G>A), I629V (4211A>G 4213A>G), A631G (4218C>G 4219A>C), C632V (4220T>G 4221G>T 4222T>A), T634V (4226A>G 4227C>T 4228A>G), T636_H637del (4232_4237delACACAC), D638E (4240T>A), K640_R641del (4244_4249delAAAAGA), Y646F (4263A>T), Y647E (4265T>G 4267T>A), M651L (4277A>T 4279G>A), T652N (4281C>A 4282C>T), T653D (4283A>G 4284C>A 4285A>T), N657R (4296A>G 4297C>G), D659S (4301G>T 4302A>C 4303C>T), I660T (4305T>C 4306C>T), D662E (4312C>A), L665M (4319C>A 4321T>G), L666V (4322C>G 4324A>T), I668V (4328A>G 4330T>G), A670H (4334G>C 4335C>A), A671C (4337G>T 4338C>G), M672L (4340A>C), H674T (4346C>A 4347A>C 4348T>C), V677H (4355G>C 4356T>A 4357G>C), V679L (4361G>T 4363C>G), E680L (4364G>C 4365A>T 4366G>T), P682S (4370C>T 4372A>T), P683del (4373_4375delCCG), L685F (4381A>C), T686L (4382A>T 4383C>T 4384G>A), I687V (4385A>G 4387T>C), L688K (4388C>A 4389T>A 4390T>A), S689T (4391T>A 4393A>C), K866* (4922A>T 4924A>G), P867K (4925C>A 4926C>A 4927A>G), D869E (4933C>A), T872N (4941C>A 4942G>T), I876V (4952A>G), T877V (4955A>G 4956C>T 4957G>A), K878G (4958A>G 4959A>G), R881K (4968G>A), K883R (4974A>G), D884R (4976G>A 4977A>G 4978T>G), R885Q (4980G>A), V886T (4982G>A 4983T>C 4984C>T), T887R (4986C>G), G888R (4988G>A 4990A>G), A890del (4994_4996delGCC), Y891H (4997T>C), M893S (5003A>T 5004T>C 5005G>T), L895W (5009C>T 5010T>G 5011A>G), M897I (5017G>A), L901M (5027C>A 5029C>G), Y904S (5037A>C 5038T>A), I908L (5048A>C 5050T>A), A910V (5055C>T 5056A>C), S911K (5057T>A 5058C>A 5059A>G), E912T (5060G>A 5061A>C 5062A>T), I913T (5064T>C 5065A>T), Y914F (5067A>T), T915S (5070C>G), Q918D (5078C>G 5080G>T), L919Y (5081C>T 5082T>A 5083C>T), G920A (5085G>C 5086A>C), Y921K (5087T>A 5089C>A), V923Y (5093G>T 5094T>A 5095A>C), L924I (5096C>A 5098G>T), D925R (5099G>C 5100A>G 5101C>T), R926E (5102A>G 5103G>A 5104A>G), I928V (5108A>G 5110C>T), Y930L (5114T>C 5115A>T), F933I (5123T>A 5125C>T), E935L (5129G>T 5130A>T 5131A>G), V936S (5132G>T 5133T>C 5134A>C), F937I (5135T>A 5137C>T), T939S (5141A>T 5143A>T), D942G (5151A>G 5152C>T), K943A (5153A>G 5154A>C 5155G>T), L944Q (5157T>A 5158C>G), N948H (5168A>C), Y949F (5172A>T 5173C>T), K951R (5178A>G 5179G>A), T952S (5180A>T 5182G>A), L953F (5183C>T 5185C>T), M954H (5186A>C 5187T>A 5188G>T), G955K (5189G>A 5190G>A 5191A>G), T956G (5192A>G 5193C>G 5194G>A), I957L (5195A>C), I959T (5202T>C), H961V (5207C>G 5208A>T 5209C>A), Y967F (5226A>T 5227C>T), E970Q (5234G>C 5236G>A), N979I (5262A>T 5263C>T), Q984D (5276C>G 5278A>C), Y989F (5292A>T 5293C>T), I990V (5294A>G 5296C>G), N991S (5298A>G), Y992S (5301A>C 5302C>T), A993H (5303G>C 5304C>A 5305A>T), D995K (5309G>A 5311C>A), N996D (5312A>G 5314C>T), V998A (5319T>C), S999K (5321T>A 5322C>A 5323A>G), P1002D (5330C>G 5331C>A 5332A>T), M1003V (5333A>G), I1006F (5342A>T), A1007S (5345G>T) | | | | | | | | |

| | Begin | End | Coverage | Score | Concordance | Matches | Identities | I/D/M/F* | Stop Codons |
|---|---|---|---|---|---|---|---|---|---|
| **NT** | **3223** | **5354** | **18.9%** | **123** | **4.5%** | **1383 (96.6%)** | **750 (52.4%)** | **30/18** | |
| Codon mutations: | GGC299..G (3223C>G), AAG300CTG (3224A>C 3225A>T), GAC301GAT (3229C>T), GCC302GTG (3231C>T 3232C>G), TTA303ATG (3233T>A 3235A>G), CCT304CCA (3238T>A), CCT304_AAG305insCCTGAGTTGCCAAAG (3238_3239insCCTGAGTTGCCAAAG), CAC306CTG (3243A>T 3244C>G), CAA307CCA (3246A>C), TGG309AGG (3251T>A), TGG309_GAT310insAGGGATATT (3253_3254insAGGGATATT), AAG312AGA (3261A>G 3262G>A), ATA313ATT (3265A>T), AAC314GAG (3266A>G 3268C>G), ATT315TTG (3269A>T 3271T>G), CAG316ATG (3272C>A 3273A>T), GGG318GGT (3280G>T), AAA319TCT (3281A>T 3282A>C 3283A>T), GAG320ACG (3284G>A 3285A>C), CCT321CCG (3289T>G), CCA322CCT (3292A>T), TGG323GCA (3293T>G 3294G>C 3295G>A), GGA324CAA (3296G>C 3297G>A), CCC325GCT (3299C>G 3301C>T), CTA326CCT (3303T>C 3304A>T), TAT327TAC (3307T>C), CAA328AGA (3308C>A 3309A>G), TCT330GCT (3314T>G), GAG331CCT (3317G>C 3318A>C 3319G>T), CTA334TTG (3326C>T 3328A>G), CAG335GCT (3329C>G 3330A>C 3331G>T), ACC336GAA (3332A>G 3333C>A 3334C>A), CTA337CTG (3337A>G), CGA338AGG (3338C>A 3340A>G), GAA339AAG (3341G>A 3343A>G), TGG340CAG (3344T>C 3345G>A), AAG342AAC (3352G>C), GAG343GAA (3355G>A), AAG344TTG (3356A>T 3357A>T), CTA345CTT (3361A>T), GCC346GAT (3363C>A 3364C>T), AAA347TCA (3365A>T 3366A>C), TGG349CTG (3371T>C 3372G>T), ATA350ATT (3376A>T), CGA351CAG (3378G>A 3379A>G), CGA352CCT (3381G>C 3382A>T), TCC353TCT (3385C>T), ACC354AAG (3387C>A 3388C>G), TCA355GCT (3389T>G 3391A>T), AGT356CCC (3392A>C 3393G>C 3394T>C), GCA357TAT (3395G>T 3396C>A 3397A>T), GGA358GGT (3400A>T), ACT359GCA (3401A>G 3403T>A), CCA360CCT (3406A>T), TGC361GTT (3407T>G 3408G>T 3409C>T), ATG362TTA (3410A>T 3412G>A), TTC363TTT (3415C>T), GTT364CAG (3416G>C 3417T>A 3418T>G), CCA365AAA (3419C>A 3420C>A), GCA367CAG (3425G>C 3426C>A 3427A>G), AAC368GAT (3428A>G 3430C>T), AAA370TCA (3434A>T 3435A>C), CTA371TTA (3437C>T), CGA372AGG (3440C>A 3442A>G), CTC373ATG (3443C>A 3445C>G), GTA374TGC (3446G>T 3447T>G 3448A>C), CAA375ATT (3449C>A 3450A>T 3451A>T), CGA378CGG (3460A>G), AAG379CAG (3461A>C), TTG380CTG (3464T>C), GAG382AAG (3470G>A), ATC383GTG (3473A>G 3475C>G), ACG384ACT (3478G>T), ATC385ATA (3481C>A), AAC387AAT (3487C>T), CGA388AAG (3488C>A 3489G>A 3490A>G), TAT389TAC (3493T>C), CTA391TTG (3497C>T 3499A>G), CCC392CCT (3502C>T), AAC393AGG (3504A>G 3505C>G), ATC394ATT (3508C>T), GAA395GAT (3511A>T), GAA396GAT (3514A>T), GCA397TTG (3515G>T 3516C>T 3517A>G), CAA398TTT (3518C>T 3519A>T 3520A>T), GAC399GAT (3523C>T), AGA400CAA (3524A>C 3525G>A), TTA401TTT (3529A>T), ACC402CAG (3530A>C 3531C>A 3532C>G), TCA404GCT (3536T>G 3538A>T), GAC475GAT (3751C>T), ATA477ATT (3757A>T), CTG478GTC (3758C>G 3760G>C), GTC479ATA (3761G>A 3763C>A), TAC480TAT (3766C>T), ACA481AGT (3768C>G 3769A>T), GGA483GAT (3774G>A 3775A>T), TCC484ACG (3776T>A 3778C>G), CTC485TTG (3779C>T 3781C>G), CAG486GAT (3782C>G 3784G>T), ACC489GTG (3791A>G 3792C>T 3793C>G), AAG490AAC (3796G>C), CAA491CAT (3799A>T), GTT492TTG (3800G>T 3802T>G), CAA493AGG (3803C>A 3804A>G 3805A>G), GAT494TTG (3806G>T 3807A>T 3808T>G), GTG495GTA (3811G>A), TTC496TTA (3814C>A), GAA497CAA (3815G>C), CGA498CTT (3819G>T 3820A>T), CTC499TTA (3821C>T 3823C>A), ACG500AGG (3825C>G), AAG501GAA (3827A>G 3829G>A), TCC502TAC (3831C>A), GGA503AAG (3833G>A 3834G>A 3835A>G), TTC504TTG (3838C>G), AAG505TAT (3839A>T 3841G>T), ACA506GTT (3842A>G 3843C>T 3844A>T), GCA507AAA (3845G>A 3846C>A), CCC508ATG (3848C>A 3849C>T 3850C>G), AAA510AAG (3856A>G), TGC511TGT (3859C>T), GAA512GAG (3862A>G), TTC513TTT (3865C>T), CAC514GCC (3866C>G 3867A>C), AAG515CAA (3869A>C 3871G>A), GTC518GTA (3880C>A), TTT520TTC (3886T>C), TTA521CTA (3887T>C), TTT523CAT (3893T>C 3894T>A), ATC524ATA (3898C>A), ATC525GTT (3899A>G 3901C>T), ACA527GAA (3905A>G 3906C>A), ACA528GAC (3908A>G 3909C>A 3910A>C), GGG529GAA (3912G>A 3913G>A), ATA530GTA (3914A>G), ACG531AGG (3918C>G), ATC532ATG (3922C>G), GAC533GAT (3925C>T), GCA535AAG (3929G>A 3930C>A 3931A>G), AAG536AAA (3934G>A), ACA537GTG (3935A>G 3936C>T 3937A>G), CAG538CAA (3940G>A), TCA539GCC (3941T>G 3943A>C), ATC540ATT (3946C>T), AGA541GTA (3947A>G 3948G>T), GAA542GAT (3952A>T), CCA544CAA (3957C>A), GAA545GCA (3960A>C), CCG546CCC (3964G>C), AAG547AAA (3967G>A), ACA548ACT (3970A>T), GTC549GTG (3973C>G), AAG550ACG (3975A>C), GTA552CTG (3980G>C 3982A>G), CAG553AGG (3983C>A 3984A>G), TCA554TCT (3988A>T), CTT556TTG (3992C>T 3994T>G), GGA557GGG (3997A>G), CTC558CTA (4000C>A), GCC559GCT (4003C>T), AAC562TAT (4010A>T 4012C>T), AAG567GAT (4025A>G 4027G>T), GAC568GGA (4029A>G 4030C>A), TAT569TAC (4033T>C), AAG571ACG (4038A>C), ACA572ATA (4041C>T), GCA574GCT (4048A>T), ATG576CTG (4052A>C), ACG577ACT (4057G>T), ATG578GAT (4058A>G 4059T>A 4060G>T), CTT579CTG (4063T>G), ACA580CTG (4064A>C 4065C>T 4066A>G), AGA581AAG (4068G>A 4069A>G), AAA582AAG (4072A>G), GAC583AAA (4073G>A 4075C>A), AAC585AAG (4081C>G), AAA587GTA (4085A>G 4086A>T), GGA589TCG (4091G>T 4092G>C 4093A>G), AAA590GTA (4094A>G 4095A>T), GAA591AGA (4097G>A 4098A>G), CAG592TGC (4100C>T 4101A>G 4102G>C), ACC593GAA (4103A>G 4104C>A 4105C>A), GCG595GCT (4111G>T), TTC596TTT (4114C>T), AAA597CAG (4115A>C 4117A>G), AGA598AGT (4120A>T), CTC599TTG (4121C>T 4123C>G), CAG602GCT (4130C>G 4131A>C 4132G>T), TGC603ATT (4133T>A 4134G>T 4135C>T), GCT604GCA (4138T>A), GCC606GAA (4143C>A 4144C>A), ACG608ATA (4149C>T 4150G>A), CTT609CTG (4153T>G), CGA610AAG (4154C>A 4155G>A 4156A>G), CTA611CTG (4159A>G), CTA611_TTC612insCCAAAT (4159_4160insCCAAAT), TTC612TTT (4162C>T), GAT613GAG (4165T>G), GGT614CTA (4166G>C 4167G>T 4168T>A), AGC615CCT (4169A>C 4170G>C 4171C>T), AAG616TTT (4172A>T 4173A>T 4174G>T), GAA617GAG (4177A>G), GTC618GTA (4180C>A), ATC620GTG (4184A>G 4186C>G), GAG621CAG (4187G>C), ACC622ACT (4192C>T), GAC623GAT (4195C>T), TCT625TCG (4201T>G), GAT626GAC (4204T>C), ATG627AAA (4206T>A 4207G>A), GCA628GCT (4210A>T), ATA629GTG (4211A>G 4213A>G), GGC630GGA (4216C>A), GCA631GGC (4218C>G 4219A>C), TGT632GTA (4220T>G 4221G>T 4222T>A), CTA633TTG (4223C>T 4225A>G), ACA634GTG (4226A>G 4227C>T 4228A>G), CAG635CAA (4231G>A), ACA636_CAC637del (4232_4237delACACAC), GAT638GAA (4240T>A), GGG639GGT (4243G>T), AAA640_AGA641del (4244_4249delAAAAGA), CAC642CAT (4252C>T), CCA643CCG (4255A>G), TAT646TTT (4263A>T), TAT647GAA (4265T>G 4267T>A), TCC648AGC (4268T>A 4269C>G), CGG649AGG (4271C>A), ATG651TTA (4277A>T 4279G>A), ACC652AAT (4281C>A 4282C>T), ACA653GAT (4283A>G 4284C>A 4285A>T), GCG654GCT (4288G>T), GAA655GAG (4291A>G), AAC657AGG (4296A>G 4297C>G), GAC659TCT (4301G>T 4302A>C 4303C>T), ATC660ACT (4305T>C 4306C>T), GAC662GAA (4312C>A), CTT665ATG (4319C>A 4321T>G), CTA666GTT (4322C>G 4324A>T), GCC667GCA (4327C>A), ATT668GTG (4328A>G 4330T>G), GTT669GTG (4333T>G), GCC670CAC (4334G>C 4335C>A), GCC671TGC (4337G>T 4338C>G), ATG672CTG (4340A>C), CAT674ACC (4346C>A 4347A>C 4348T>C), GTG677CAC (4355G>C 4356T>A 4357G>C), GTC679TTG (4361G>T 4363C>G), GAG680CTT (4364G>C 4365A>T 4366G>T), GGC681GGT (4369C>T), CCA682TCT (4370C>T 4372A>T), CCG683del (4373_4375delCCG), TTA685TTC (4381A>C), ACG686TTA (4382A>T 4383C>T 4384G>A), ATT687GTC (4385A>G 4387T>C), CTT688AAA (4388C>A 4389T>A 4390T>A), TCA689ACC (4391T>A 4393A>C), CCA863..T (4915A>T), CCA864CCT (4918A>T), ACG865ACT (4921G>T), AAA866TAG (4922A>T 4924A>G), CCA867AAG (4925C>A 4926C>A 4927A>G), GAC869GAA (4933C>A), GTT871GTG (4939T>G), ACG872AAT (4941C>A 4942G>T), TTC875TTT (4951C>T), ATT876GTT (4952A>G), ACG877GTA (4955A>G 4956C>T 4957G>A), AAA878GGA (4958A>G 4959A>G), CTC879TTG (4961C>T 4963C>G), CCG880CCA (4966G>A), AGG881AAG (4968G>A), TCA882TCT (4972A>T), AAG883AGG (4974A>G), GAT884AGG (4976G>A 4977A>G 4978T>G), CGA885CAA (4980G>A), GTC886ACT (4982G>A 4983T>C 4984C>T), ACA887AGA (4986C>G), GGA888AGG (4988G>A 4990A>G), GCC890del (4994_4996delGCC), TAT891CAT (4997T>C), GAC892GAT (5002C>T), ATG893TCT (5003A>T 5004T>C 5005G>T), CTA895TGG (5009C>T 5010T>G 5011A>G), GTC896GTG (5014C>G), ATG897ATA (5017G>A), GTC898GTT (5020C>T), GAC899GAT (5023C>T), AGA900CGG (5024A>C 5026A>G), CTC901ATG (5027C>A 5029C>G), ACA902ACG (5032A>G), TAT904TCA (5037A>C 5038T>A), GCA905GCC (5041A>C), CAC906CAT (5044C>T), TTC907TTT (5047C>T), ATT908CTA (5048A>C 5050T>A), CCT909CCA (5053T>A), GCA910GTC (5055C>T 5056A>C), TCA911AAG (5057T>A 5058C>A 5059A>G), GAA912ACT (5060G>A 5061A>C 5062A>T), ATA913ACT (5064T>C 5065A>T), TAC914TTC (5067A>T), ACT915AGT (5070C>G), GCA916GCT (5074A>T), GAG917GAA (5077G>A), CAG918GAT (5078C>G 5080G>T), CTC919TAT (5081C>T 5082T>A 5083C>T), GGA920GCC (5085G>C 5086A>C), TAC921AAA (5087T>A 5089C>A), CTC922CTG (5092C>G), GTA923TAC (5093G>T 5094T>A 5095A>C), CTG924ATT (5096C>A 5098G>T), GAC925CGT (5099G>C 5100A>G 5101C>T), AGA926GAG (5102A>G 5103G>A 5104A>G), ATC928GTT (5108A>G 5110C>T), TAT930CTT (5114T>C 5115A>T), CAC931CAT (5119C>T), GGA932GGT (5122A>T), TTC933ATT (5123T>A 5125C>T), CCG934CCT (5128G>T), GAA935TTG (5129G>T 5130A>T 5131A>G), GTA936TCC (5132G>T 5133T>C 5134A>C), TTC937ATT (5135T>A 5137C>T), ACA939TCT (5141A>T 5143A>T), GAC940GAT (5146C>T), GAC942GGT (5151A>G 5152C>T), AAG943GCT (5153A>G 5154A>C 5155G>T), CTC944CAG (5157T>A 5158C>G), TTC945TTT (5161C>T), ACA946ACT (5164A>T), AAC948CAC (5168A>C), TAC949TTT (5172A>T 5173C>T), AAG951AGA (5178A>G 5179G>A), ACG952TCA (5180A>T 5182G>A), CTC953TTT (5183C>T 5185C>T), ATG954CAT (5186A>C 5187T>A 5188G>T), GGA955AAG (5189G>A 5190G>A 5191A>G), ACG956GGA (5192A>G 5193C>G 5194G>A), ATT957CTT (5195A>C), GGA958GGT (5200A>T), ATC959ACC (5202T>C), CAC961GTA (5207C>G 5208A>T 5209C>A), TCA964AGC (5216T>A 5217C>G 5218A>C), ACA965ACC (5221A>C), GCA966GCT (5224A>T), TAC967TTT (5226A>T 5227C>T), CCA969CCT (5233A>T), GAG970CAA (5234G>C 5236G>A), ACG971ACT (5239G>T), GGG973GGA (5245G>A), ACG975ACA (5251G>A), AGA977AGG (5257A>G), ACG978ACC (5260G>C), AAC979ATT (5262A>T 5263C>T), CTC982CTT (5272C>T), CAA984GAC (5276C>G 5278A>C), CTA986CTT (5284A>T), CGG987AGG (5285C>A), TAC989TTT (5292A>T 5293C>T), ATC990GTG (5294A>G 5296C>G), AAC991AGC (5298A>G), TAC992TCT (5301A>C 5302C>T), GCA993CAT (5303G>C 5304C>A 5305A>T), GAC995AAA (5309G>A 5311C>A), AAC996GAT (5312A>G 5314C>T), GTT998GCT (5319T>C), TCA999AAG (5321T>A 5322C>A 5323A>G), TTA1000TTG (5326A>G), CTG1001CTA (5329G>A), CCA1002GAT (5330C>G 5331C>A 5332A>T), ATG1003GTG (5333A>G), GCG1004GCC (5338G>C), CAG1005CAA (5341G>A), ATC1006TTC (5342A>T), GCA1007TCA (5345G>T) | | | | | | | | |

#### Proteins

| | Begin | End | Coverage | Score | Concordance | Matches | Identities | I/D/M/F* | Stop Codons |
|---|---|---|---|---|---|---|---|---|---|
| Reverse Transcriptase (YP_009666308.1) | 300 | 1010 | 44.6% | 1606 | 49.8% | 461 (96.6%) | 224 (47.0%) | 10/6/0/0 | 1 |

| | Begin | End | Coverage | Score | Concordance | Matches | Identities | I/D/M/F* | Stop Codons |
|---|---|---|---|---|---|---|---|---|---|
| **NT** | **3223** | **5354** | **18.9%** | **123** | **4.5%** | **1383 (96.6%)** | **750 (52.4%)** | **30/18** | |
| Protein mutations: | K300L (3224A>C 3225A>T), A302V (3231C>T 3232C>G), L303M (3233T>A 3235A>G), P304_K305insPELPK (3238_3239insCCTGAGTTGCCAAAG), H306L (3243A>T 3244C>G), Q307P (3246A>C), W309R (3251T>A), W309_D310insRDI (3253_3254insAGGGATATT), K312R (3261A>G 3262G>A), N314E (3266A>G 3268C>G), I315L (3269A>T 3271T>G), Q316M (3272C>A 3273A>T), K319S (3281A>T 3282A>C 3283A>T), E320T (3284G>A 3285A>C), W323A (3293T>G 3294G>C 3295G>A), G324Q (3296G>C 3297G>A), P325A (3299C>G 3301C>T), L326P (3303T>C 3304A>T), Q328R (3308C>A 3309A>G), S330A (3314T>G), E331P (3317G>C 3318A>C 3319G>T), Q335A (3329C>G 3330A>C 3331G>T), T336E (3332A>G 3333C>A 3334C>A), E339K (3341G>A 3343A>G), W340Q (3344T>C 3345G>A), K342N (3352G>C), K344L (3356A>T 3357A>T), A346D (3363C>A 3364C>T), K347S (3365A>T 3366A>C), W349L (3371T>C 3372G>T), R351Q (3378G>A 3379A>G), R352P (3381G>C 3382A>T), T354K (3387C>A 3388C>G), S355A (3389T>G 3391A>T), S356P (3392A>C 3393G>C 3394T>C), A357Y (3395G>T 3396C>A 3397A>T), T359A (3401A>G 3403T>A), C361V (3407T>G 3408G>T 3409C>T), M362L (3410A>T 3412G>A), V364Q (3416G>C 3417T>A 3418T>G), P365K (3419C>A 3420C>A), A367Q (3425G>C 3426C>A 3427A>G), N368D (3428A>G 3430C>T), K370S (3434A>T 3435A>C), L373M (3443C>A 3445C>G), V374C (3446G>T 3447T>G 3448A>C), Q375I (3449C>A 3450A>T 3451A>T), K379Q (3461A>C), E382K (3470G>A), I383V (3473A>G 3475C>G), R388K (3488C>A 3489G>A 3490A>G), N393R (3504A>G 3505C>G), E395D (3511A>T), E396D (3514A>T), A397L (3515G>T 3516C>T 3517A>G), Q398F (3518C>T 3519A>T 3520A>T), R400Q (3524A>C 3525G>A), L401F (3529A>T), T402Q (3530A>C 3531C>A 3532C>G), S404A (3536T>G 3538A>T), L478V (3758C>G 3760G>C), V479I (3761G>A 3763C>A), T481S (3768C>G 3769A>T), G483D (3774G>A 3775A>T), S484T (3776T>A 3778C>G), Q486D (3782C>G 3784G>T), T489V (3791A>G 3792C>T 3793C>G), K490N (3796G>C), Q491H (3799A>T), V492L (3800G>T 3802T>G), Q493R (3803C>A 3804A>G 3805A>G), D494L (3806G>T 3807A>T 3808T>G), F496L (3814C>A), E497Q (3815G>C), R498L (3819G>T 3820A>T), T500R (3825C>G), K501E (3827A>G 3829G>A), S502Y (3831C>A), G503K (3833G>A 3834G>A 3835A>G), F504L (3838C>G), K505Y (3839A>T 3841G>T), T506V (3842A>G 3843C>T 3844A>T), A507K (3845G>A 3846C>A), P508M (3848C>A 3849C>T 3850C>G), H514A (3866C>G 3867A>C), K515Q (3869A>C 3871G>A), F523H (3893T>C 3894T>A), I525V (3899A>G 3901C>T), T527E (3905A>G 3906C>A), T528D (3908A>G 3909C>A 3910A>C), G529E (3912G>A 3913G>A), I530V (3914A>G), T531R (3918C>G), I532M (3922C>G), A535K (3929G>A 3930C>A 3931A>G), T537V (3935A>G 3936C>T 3937A>G), S539A (3941T>G 3943A>C), R541V (3947A>G 3948G>T), E542D (3952A>T), P544Q (3957C>A), E545A (3960A>C), K550T (3975A>C), V552L (3980G>C 3982A>G), Q553R (3983C>A 3984A>G), N562Y (4010A>T 4012C>T), K567D (4025A>G 4027G>T), D568G (4029A>G 4030C>A), K571T (4038A>C), T572I (4041C>T), M576L (4052A>C), M578D (4058A>G 4059T>A 4060G>T), T580L (4064A>C 4065C>T 4066A>G), R581K (4068G>A 4069A>G), D583K (4073G>A 4075C>A), N585K (4081C>G), K587V (4085A>G 4086A>T), G589S (4091G>T 4092G>C 4093A>G), K590V (4094A>G 4095A>T), E591R (4097G>A 4098A>G), Q592C (4100C>T 4101A>G 4102G>C), T593E (4103A>G 4104C>A 4105C>A), K597Q (4115A>C 4117A>G), R598S (4120A>T), Q602A (4130C>G 4131A>C 4132G>T), C603I (4133T>A 4134G>T 4135C>T), A606E (4143C>A 4144C>A), T608I (4149C>T 4150G>A), R610K (4154C>A 4155G>A 4156A>G), L611_F612insPN (4159_4160insCCAAAT), D613E (4165T>G), G614L (4166G>C 4167G>T 4168T>A), S615P (4169A>C 4170G>C 4171C>T), K616F (4172A>T 4173A>T 4174G>T), I620V (4184A>G 4186C>G), E621Q (4187G>C), M627K (4206T>A 4207G>A), I629V (4211A>G 4213A>G), A631G (4218C>G 4219A>C), C632V (4220T>G 4221G>T 4222T>A), T634V (4226A>G 4227C>T 4228A>G), T636_H637del (4232_4237delACACAC), D638E (4240T>A), K640_R641del (4244_4249delAAAAGA), Y646F (4263A>T), Y647E (4265T>G 4267T>A), M651L (4277A>T 4279G>A), T652N (4281C>A 4282C>T), T653D (4283A>G 4284C>A 4285A>T), N657R (4296A>G 4297C>G), D659S (4301G>T 4302A>C 4303C>T), I660T (4305T>C 4306C>T), D662E (4312C>A), L665M (4319C>A 4321T>G), L666V (4322C>G 4324A>T), I668V (4328A>G 4330T>G), A670H (4334G>C 4335C>A), A671C (4337G>T 4338C>G), M672L (4340A>C), H674T (4346C>A 4347A>C 4348T>C), V677H (4355G>C 4356T>A 4357G>C), V679L (4361G>T 4363C>G), E680L (4364G>C 4365A>T 4366G>T), P682S (4370C>T 4372A>T), P683del (4373_4375delCCG), L685F (4381A>C), T686L (4382A>T 4383C>T 4384G>A), I687V (4385A>G 4387T>C), L688K (4388C>A 4389T>A 4390T>A), S689T (4391T>A 4393A>C), K866* (4922A>T 4924A>G), P867K (4925C>A 4926C>A 4927A>G), D869E (4933C>A), T872N (4941C>A 4942G>T), I876V (4952A>G), T877V (4955A>G 4956C>T 4957G>A), K878G (4958A>G 4959A>G), R881K (4968G>A), K883R (4974A>G), D884R (4976G>A 4977A>G 4978T>G), R885Q (4980G>A), V886T (4982G>A 4983T>C 4984C>T), T887R (4986C>G), G888R (4988G>A 4990A>G), A890del (4994_4996delGCC), Y891H (4997T>C), M893S (5003A>T 5004T>C 5005G>T), L895W (5009C>T 5010T>G 5011A>G), M897I (5017G>A), L901M (5027C>A 5029C>G), Y904S (5037A>C 5038T>A), I908L (5048A>C 5050T>A), A910V (5055C>T 5056A>C), S911K (5057T>A 5058C>A 5059A>G), E912T (5060G>A 5061A>C 5062A>T), I913T (5064T>C 5065A>T), Y914F (5067A>T), T915S (5070C>G), Q918D (5078C>G 5080G>T), L919Y (5081C>T 5082T>A 5083C>T), G920A (5085G>C 5086A>C), Y921K (5087T>A 5089C>A), V923Y (5093G>T 5094T>A 5095A>C), L924I (5096C>A 5098G>T), D925R (5099G>C 5100A>G 5101C>T), R926E (5102A>G 5103G>A 5104A>G), I928V (5108A>G 5110C>T), Y930L (5114T>C 5115A>T), F933I (5123T>A 5125C>T), E935L (5129G>T 5130A>T 5131A>G), V936S (5132G>T 5133T>C 5134A>C), F937I (5135T>A 5137C>T), T939S (5141A>T 5143A>T), D942G (5151A>G 5152C>T), K943A (5153A>G 5154A>C 5155G>T), L944Q (5157T>A 5158C>G), N948H (5168A>C), Y949F (5172A>T 5173C>T), K951R (5178A>G 5179G>A), T952S (5180A>T 5182G>A), L953F (5183C>T 5185C>T), M954H (5186A>C 5187T>A 5188G>T), G955K (5189G>A 5190G>A 5191A>G), T956G (5192A>G 5193C>G 5194G>A), I957L (5195A>C), I959T (5202T>C), H961V (5207C>G 5208A>T 5209C>A), Y967F (5226A>T 5227C>T), E970Q (5234G>C 5236G>A), N979I (5262A>T 5263C>T), Q984D (5276C>G 5278A>C), Y989F (5292A>T 5293C>T), I990V (5294A>G 5296C>G), N991S (5298A>G), Y992S (5301A>C 5302C>T), A993H (5303G>C 5304C>A 5305A>T), D995K (5309G>A 5311C>A), N996D (5312A>G 5314C>T), V998A (5319T>C), S999K (5321T>A 5322C>A 5323A>G), P1002D (5330C>G 5331C>A 5332A>T), M1003V (5333A>G), I1006F (5342A>T), A1007S (5345G>T) | | | | | | | | |

| | Begin | End | Coverage | Score | Concordance | Matches | Identities | I/D/M/F* | Stop Codons |
|---|---|---|---|---|---|---|---|---|---|
| **NT** | **3223** | **5354** | **18.9%** | **123** | **4.5%** | **1383 (96.6%)** | **750 (52.4%)** | **30/18** | |
| Codon mutations: | GGC299..G (3223C>G), AAG300CTG (3224A>C 3225A>T), GAC301GAT (3229C>T), GCC302GTG (3231C>T 3232C>G), TTA303ATG (3233T>A 3235A>G), CCT304CCA (3238T>A), CCT304_AAG305insCCTGAGTTGCCAAAG (3238_3239insCCTGAGTTGCCAAAG), CAC306CTG (3243A>T 3244C>G), CAA307CCA (3246A>C), TGG309AGG (3251T>A), TGG309_GAT310insAGGGATATT (3253_3254insAGGGATATT), AAG312AGA (3261A>G 3262G>A), ATA313ATT (3265A>T), AAC314GAG (3266A>G 3268C>G), ATT315TTG (3269A>T 3271T>G), CAG316ATG (3272C>A 3273A>T), GGG318GGT (3280G>T), AAA319TCT (3281A>T 3282A>C 3283A>T), GAG320ACG (3284G>A 3285A>C), CCT321CCG (3289T>G), CCA322CCT (3292A>T), TGG323GCA (3293T>G 3294G>C 3295G>A), GGA324CAA (3296G>C 3297G>A), CCC325GCT (3299C>G 3301C>T), CTA326CCT (3303T>C 3304A>T), TAT327TAC (3307T>C), CAA328AGA (3308C>A 3309A>G), TCT330GCT (3314T>G), GAG331CCT (3317G>C 3318A>C 3319G>T), CTA334TTG (3326C>T 3328A>G), CAG335GCT (3329C>G 3330A>C 3331G>T), ACC336GAA (3332A>G 3333C>A 3334C>A), CTA337CTG (3337A>G), CGA338AGG (3338C>A 3340A>G), GAA339AAG (3341G>A 3343A>G), TGG340CAG (3344T>C 3345G>A), AAG342AAC (3352G>C), GAG343GAA (3355G>A), AAG344TTG (3356A>T 3357A>T), CTA345CTT (3361A>T), GCC346GAT (3363C>A 3364C>T), AAA347TCA (3365A>T 3366A>C), TGG349CTG (3371T>C 3372G>T), ATA350ATT (3376A>T), CGA351CAG (3378G>A 3379A>G), CGA352CCT (3381G>C 3382A>T), TCC353TCT (3385C>T), ACC354AAG (3387C>A 3388C>G), TCA355GCT (3389T>G 3391A>T), AGT356CCC (3392A>C 3393G>C 3394T>C), GCA357TAT (3395G>T 3396C>A 3397A>T), GGA358GGT (3400A>T), ACT359GCA (3401A>G 3403T>A), CCA360CCT (3406A>T), TGC361GTT (3407T>G 3408G>T 3409C>T), ATG362TTA (3410A>T 3412G>A), TTC363TTT (3415C>T), GTT364CAG (3416G>C 3417T>A 3418T>G), CCA365AAA (3419C>A 3420C>A), GCA367CAG (3425G>C 3426C>A 3427A>G), AAC368GAT (3428A>G 3430C>T), AAA370TCA (3434A>T 3435A>C), CTA371TTA (3437C>T), CGA372AGG (3440C>A 3442A>G), CTC373ATG (3443C>A 3445C>G), GTA374TGC (3446G>T 3447T>G 3448A>C), CAA375ATT (3449C>A 3450A>T 3451A>T), CGA378CGG (3460A>G), AAG379CAG (3461A>C), TTG380CTG (3464T>C), GAG382AAG (3470G>A), ATC383GTG (3473A>G 3475C>G), ACG384ACT (3478G>T), ATC385ATA (3481C>A), AAC387AAT (3487C>T), CGA388AAG (3488C>A 3489G>A 3490A>G), TAT389TAC (3493T>C), CTA391TTG (3497C>T 3499A>G), CCC392CCT (3502C>T), AAC393AGG (3504A>G 3505C>G), ATC394ATT (3508C>T), GAA395GAT (3511A>T), GAA396GAT (3514A>T), GCA397TTG (3515G>T 3516C>T 3517A>G), CAA398TTT (3518C>T 3519A>T 3520A>T), GAC399GAT (3523C>T), AGA400CAA (3524A>C 3525G>A), TTA401TTT (3529A>T), ACC402CAG (3530A>C 3531C>A 3532C>G), TCA404GCT (3536T>G 3538A>T), GAC475GAT (3751C>T), ATA477ATT (3757A>T), CTG478GTC (3758C>G 3760G>C), GTC479ATA (3761G>A 3763C>A), TAC480TAT (3766C>T), ACA481AGT (3768C>G 3769A>T), GGA483GAT (3774G>A 3775A>T), TCC484ACG (3776T>A 3778C>G), CTC485TTG (3779C>T 3781C>G), CAG486GAT (3782C>G 3784G>T), ACC489GTG (3791A>G 3792C>T 3793C>G), AAG490AAC (3796G>C), CAA491CAT (3799A>T), GTT492TTG (3800G>T 3802T>G), CAA493AGG (3803C>A 3804A>G 3805A>G), GAT494TTG (3806G>T 3807A>T 3808T>G), GTG495GTA (3811G>A), TTC496TTA (3814C>A), GAA497CAA (3815G>C), CGA498CTT (3819G>T 3820A>T), CTC499TTA (3821C>T 3823C>A), ACG500AGG (3825C>G), AAG501GAA (3827A>G 3829G>A), TCC502TAC (3831C>A), GGA503AAG (3833G>A 3834G>A 3835A>G), TTC504TTG (3838C>G), AAG505TAT (3839A>T 3841G>T), ACA506GTT (3842A>G 3843C>T 3844A>T), GCA507AAA (3845G>A 3846C>A), CCC508ATG (3848C>A 3849C>T 3850C>G), AAA510AAG (3856A>G), TGC511TGT (3859C>T), GAA512GAG (3862A>G), TTC513TTT (3865C>T), CAC514GCC (3866C>G 3867A>C), AAG515CAA (3869A>C 3871G>A), GTC518GTA (3880C>A), TTT520TTC (3886T>C), TTA521CTA (3887T>C), TTT523CAT (3893T>C 3894T>A), ATC524ATA (3898C>A), ATC525GTT (3899A>G 3901C>T), ACA527GAA (3905A>G 3906C>A), ACA528GAC (3908A>G 3909C>A 3910A>C), GGG529GAA (3912G>A 3913G>A), ATA530GTA (3914A>G), ACG531AGG (3918C>G), ATC532ATG (3922C>G), GAC533GAT (3925C>T), GCA535AAG (3929G>A 3930C>A 3931A>G), AAG536AAA (3934G>A), ACA537GTG (3935A>G 3936C>T 3937A>G), CAG538CAA (3940G>A), TCA539GCC (3941T>G 3943A>C), ATC540ATT (3946C>T), AGA541GTA (3947A>G 3948G>T), GAA542GAT (3952A>T), CCA544CAA (3957C>A), GAA545GCA (3960A>C), CCG546CCC (3964G>C), AAG547AAA (3967G>A), ACA548ACT (3970A>T), GTC549GTG (3973C>G), AAG550ACG (3975A>C), GTA552CTG (3980G>C 3982A>G), CAG553AGG (3983C>A 3984A>G), TCA554TCT (3988A>T), CTT556TTG (3992C>T 3994T>G), GGA557GGG (3997A>G), CTC558CTA (4000C>A), GCC559GCT (4003C>T), AAC562TAT (4010A>T 4012C>T), AAG567GAT (4025A>G 4027G>T), GAC568GGA (4029A>G 4030C>A), TAT569TAC (4033T>C), AAG571ACG (4038A>C), ACA572ATA (4041C>T), GCA574GCT (4048A>T), ATG576CTG (4052A>C), ACG577ACT (4057G>T), ATG578GAT (4058A>G 4059T>A 4060G>T), CTT579CTG (4063T>G), ACA580CTG (4064A>C 4065C>T 4066A>G), AGA581AAG (4068G>A 4069A>G), AAA582AAG (4072A>G), GAC583AAA (4073G>A 4075C>A), AAC585AAG (4081C>G), AAA587GTA (4085A>G 4086A>T), GGA589TCG (4091G>T 4092G>C 4093A>G), AAA590GTA (4094A>G 4095A>T), GAA591AGA (4097G>A 4098A>G), CAG592TGC (4100C>T 4101A>G 4102G>C), ACC593GAA (4103A>G 4104C>A 4105C>A), GCG595GCT (4111G>T), TTC596TTT (4114C>T), AAA597CAG (4115A>C 4117A>G), AGA598AGT (4120A>T), CTC599TTG (4121C>T 4123C>G), CAG602GCT (4130C>G 4131A>C 4132G>T), TGC603ATT (4133T>A 4134G>T 4135C>T), GCT604GCA (4138T>A), GCC606GAA (4143C>A 4144C>A), ACG608ATA (4149C>T 4150G>A), CTT609CTG (4153T>G), CGA610AAG (4154C>A 4155G>A 4156A>G), CTA611CTG (4159A>G), CTA611_TTC612insCCAAAT (4159_4160insCCAAAT), TTC612TTT (4162C>T), GAT613GAG (4165T>G), GGT614CTA (4166G>C 4167G>T 4168T>A), AGC615CCT (4169A>C 4170G>C 4171C>T), AAG616TTT (4172A>T 4173A>T 4174G>T), GAA617GAG (4177A>G), GTC618GTA (4180C>A), ATC620GTG (4184A>G 4186C>G), GAG621CAG (4187G>C), ACC622ACT (4192C>T), GAC623GAT (4195C>T), TCT625TCG (4201T>G), GAT626GAC (4204T>C), ATG627AAA (4206T>A 4207G>A), GCA628GCT (4210A>T), ATA629GTG (4211A>G 4213A>G), GGC630GGA (4216C>A), GCA631GGC (4218C>G 4219A>C), TGT632GTA (4220T>G 4221G>T 4222T>A), CTA633TTG (4223C>T 4225A>G), ACA634GTG (4226A>G 4227C>T 4228A>G), CAG635CAA (4231G>A), ACA636_CAC637del (4232_4237delACACAC), GAT638GAA (4240T>A), GGG639GGT (4243G>T), AAA640_AGA641del (4244_4249delAAAAGA), CAC642CAT (4252C>T), CCA643CCG (4255A>G), TAT646TTT (4263A>T), TAT647GAA (4265T>G 4267T>A), TCC648AGC (4268T>A 4269C>G), CGG649AGG (4271C>A), ATG651TTA (4277A>T 4279G>A), ACC652AAT (4281C>A 4282C>T), ACA653GAT (4283A>G 4284C>A 4285A>T), GCG654GCT (4288G>T), GAA655GAG (4291A>G), AAC657AGG (4296A>G 4297C>G), GAC659TCT (4301G>T 4302A>C 4303C>T), ATC660ACT (4305T>C 4306C>T), GAC662GAA (4312C>A), CTT665ATG (4319C>A 4321T>G), CTA666GTT (4322C>G 4324A>T), GCC667GCA (4327C>A), ATT668GTG (4328A>G 4330T>G), GTT669GTG (4333T>G), GCC670CAC (4334G>C 4335C>A), GCC671TGC (4337G>T 4338C>G), ATG672CTG (4340A>C), CAT674ACC (4346C>A 4347A>C 4348T>C), GTG677CAC (4355G>C 4356T>A 4357G>C), GTC679TTG (4361G>T 4363C>G), GAG680CTT (4364G>C 4365A>T 4366G>T), GGC681GGT (4369C>T), CCA682TCT (4370C>T 4372A>T), CCG683del (4373_4375delCCG), TTA685TTC (4381A>C), ACG686TTA (4382A>T 4383C>T 4384G>A), ATT687GTC (4385A>G 4387T>C), CTT688AAA (4388C>A 4389T>A 4390T>A), TCA689ACC (4391T>A 4393A>C), CCA863..T (4915A>T), CCA864CCT (4918A>T), ACG865ACT (4921G>T), AAA866TAG (4922A>T 4924A>G), CCA867AAG (4925C>A 4926C>A 4927A>G), GAC869GAA (4933C>A), GTT871GTG (4939T>G), ACG872AAT (4941C>A 4942G>T), TTC875TTT (4951C>T), ATT876GTT (4952A>G), ACG877GTA (4955A>G 4956C>T 4957G>A), AAA878GGA (4958A>G 4959A>G), CTC879TTG (4961C>T 4963C>G), CCG880CCA (4966G>A), AGG881AAG (4968G>A), TCA882TCT (4972A>T), AAG883AGG (4974A>G), GAT884AGG (4976G>A 4977A>G 4978T>G), CGA885CAA (4980G>A), GTC886ACT (4982G>A 4983T>C 4984C>T), ACA887AGA (4986C>G), GGA888AGG (4988G>A 4990A>G), GCC890del (4994_4996delGCC), TAT891CAT (4997T>C), GAC892GAT (5002C>T), ATG893TCT (5003A>T 5004T>C 5005G>T), CTA895TGG (5009C>T 5010T>G 5011A>G), GTC896GTG (5014C>G), ATG897ATA (5017G>A), GTC898GTT (5020C>T), GAC899GAT (5023C>T), AGA900CGG (5024A>C 5026A>G), CTC901ATG (5027C>A 5029C>G), ACA902ACG (5032A>G), TAT904TCA (5037A>C 5038T>A), GCA905GCC (5041A>C), CAC906CAT (5044C>T), TTC907TTT (5047C>T), ATT908CTA (5048A>C 5050T>A), CCT909CCA (5053T>A), GCA910GTC (5055C>T 5056A>C), TCA911AAG (5057T>A 5058C>A 5059A>G), GAA912ACT (5060G>A 5061A>C 5062A>T), ATA913ACT (5064T>C 5065A>T), TAC914TTC (5067A>T), ACT915AGT (5070C>G), GCA916GCT (5074A>T), GAG917GAA (5077G>A), CAG918GAT (5078C>G 5080G>T), CTC919TAT (5081C>T 5082T>A 5083C>T), GGA920GCC (5085G>C 5086A>C), TAC921AAA (5087T>A 5089C>A), CTC922CTG (5092C>G), GTA923TAC (5093G>T 5094T>A 5095A>C), CTG924ATT (5096C>A 5098G>T), GAC925CGT (5099G>C 5100A>G 5101C>T), AGA926GAG (5102A>G 5103G>A 5104A>G), ATC928GTT (5108A>G 5110C>T), TAT930CTT (5114T>C 5115A>T), CAC931CAT (5119C>T), GGA932GGT (5122A>T), TTC933ATT (5123T>A 5125C>T), CCG934CCT (5128G>T), GAA935TTG (5129G>T 5130A>T 5131A>G), GTA936TCC (5132G>T 5133T>C 5134A>C), TTC937ATT (5135T>A 5137C>T), ACA939TCT (5141A>T 5143A>T), GAC940GAT (5146C>T), GAC942GGT (5151A>G 5152C>T), AAG943GCT (5153A>G 5154A>C 5155G>T), CTC944CAG (5157T>A 5158C>G), TTC945TTT (5161C>T), ACA946ACT (5164A>T), AAC948CAC (5168A>C), TAC949TTT (5172A>T 5173C>T), AAG951AGA (5178A>G 5179G>A), ACG952TCA (5180A>T 5182G>A), CTC953TTT (5183C>T 5185C>T), ATG954CAT (5186A>C 5187T>A 5188G>T), GGA955AAG (5189G>A 5190G>A 5191A>G), ACG956GGA (5192A>G 5193C>G 5194G>A), ATT957CTT (5195A>C), GGA958GGT (5200A>T), ATC959ACC (5202T>C), CAC961GTA (5207C>G 5208A>T 5209C>A), TCA964AGC (5216T>A 5217C>G 5218A>C), ACA965ACC (5221A>C), GCA966GCT (5224A>T), TAC967TTT (5226A>T 5227C>T), CCA969CCT (5233A>T), GAG970CAA (5234G>C 5236G>A), ACG971ACT (5239G>T), GGG973GGA (5245G>A), ACG975ACA (5251G>A), AGA977AGG (5257A>G), ACG978ACC (5260G>C), AAC979ATT (5262A>T 5263C>T), CTC982CTT (5272C>T), CAA984GAC (5276C>G 5278A>C), CTA986CTT (5284A>T), CGG987AGG (5285C>A), TAC989TTT (5292A>T 5293C>T), ATC990GTG (5294A>G 5296C>G), AAC991AGC (5298A>G), TAC992TCT (5301A>C 5302C>T), GCA993CAT (5303G>C 5304C>A 5305A>T), GAC995AAA (5309G>A 5311C>A), AAC996GAT (5312A>G 5314C>T), GTT998GCT (5319T>C), TCA999AAG (5321T>A 5322C>A 5323A>G), TTA1000TTG (5326A>G), CTG1001CTA (5329G>A), CCA1002GAT (5330C>G 5331C>A 5332A>T), ATG1003GTG (5333A>G), GCG1004GCC (5338G>C), CAG1005CAA (5341G>A), ATC1006TTC (5342A>T), GCA1007TCA (5345G>T) | | | | | | | | |

*: Inserts / Deletes / Misaligned / Frameshifts

#### Analysis details

This analysis was performed with panviral2.64

#### NGS Details (UN24): Errantivirus

##### Assembly

| | |
|---|---|
| **Coverage Length** | 1115 (2 contig(s)) |
| **Depth Of Coverage** | 36.2 |
| **Number Of Reads** | 503 |
| **Reads Per Million** | 10.07 rpm (after QC) |
| **Ambiguities** | 0 |
| **Assembly Method** | de novo + reference guided assembly |
| **Consensus Caller** | Bcf Tools |

##### Coverage Map

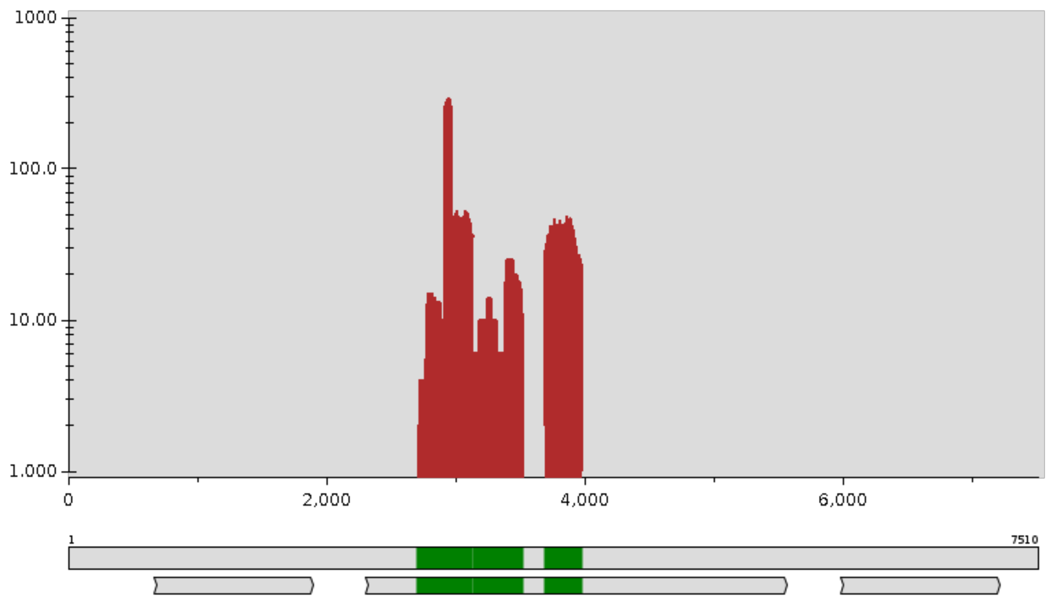

##### Assignment

| | |
|---|---|
| **Type** | Errantivirus (Taxonomy ID: 186666) |
| **Reference Genome** | NC_038512.1 |
| **NT Identity (%)** | 52.1429 |
| **AA Identity (%)** | 41.2869 |
| **Number Of Stop Codons** | 3 |
| **Number Of CDS** | 3 |

##### Alignment

| | |
|---|---|
| **Alignment Score** | 74.0 (NT) + 958.0 (AA) = 1032.0 |
| **Concordance (%)** | 22.3726 |

| **Alignment Method** | Global, seeded, nucleotide + amino acids (AGA) |
|---|---|

#### Genome Region

Sequence starts at position 2706 and ends at position 3983 relative to NC_038512.1 reference sequence.

#### Alignment Detailed Statistics

| | Begin | End | Coverage | Score | Concordance | Matches | Identities | I/D/M/F* | Stop Codons |
|---|---|---|---|---|---|---|---|---|---|
| **NT** | **2706** | **3983** | **14.8%** | **74** | **3.4%** | **1109 (98.5%)** | **584 (51.9%)** | **11/6** | |
| Mutations: | 2708G>T, 2710C>G, 2712A>T, 2715T>G, 2719C>A, 2722G>T, 2724A>C, 2725C>A, 2726A>G, 2727C>T, 2730A>G, 2731A>G, 2732A>C, 2733G>T, 2737C>T, 2738C>A, 2740G>A, 2744_2749delCCCTTC, 2751T>A, 2756C>T, 2757G>T, 2761G>A, 2762G>A, 2764A>G, 2765G>C, 2767T>A, 2768A>G, 2769G>A, 2771G>C, 2772A>T, 2772_2773insGA, 2773C>G, 2774C>A, 2777A>T, 2779C>G, 2780A>G, 2781C>A, 2782G>A, 2783A>G, 2785A>T, 2786A>T, 2788G>A, 2792G>A, 2794C>G, 2795C>A, 2797A>G, 2799G>A, 2800A>G, 2801A>T, 2806A>T, 2807A>T, 2812A>T, 2813T>A, 2814C>G, 2815A>T, 2817A>G, 2818C>A, 2821T>A, 2822G>C, 2824A>T, 2828A>G, 2830C>T, 2831T>G, 2836C>A, 2837A>G, 2841G>T, 2843G>T, 2846G>A, 2849C>A, 2850C>G, 2851C>A, 2857A>G, 2859T>A, 2860C>G, 2863C>T, 2865C>G, 2866T>G, 2869T>G, 2872G>T, 2874A>T, 2876C>A, 2877A>T, 2880A>G, 2882T>A, 2883G>T, 2885C>T, 2888C>A, 2890C>G, 2891G>T, 2892T>A, 2893A>T, 2894G>A, 2899C>T, 2901T>A, 2902C>T, 2903C>T, 2905T>A, 2907A>G, 2915G>A, 2917G>A, 2919A>T, 2920G>C, 2924A>G, 2926C>G, 2927G>A, 2928A>G, 2929T>G, 2932C>T, 2934A>G, 2935A>G, 2937A>T, 2941G>A, 2944A>C, 2949A>C, 2950C>T, 2953A>T, 2954A>G, 2955G>A, 2959C>G, 2960G>C, 2963C>T, 2968C>T, 2969A>G, 2972T>G, 2975G>A, 2976G>A, 2978A>G, 2979A>G, 2980G>C, 2981T>G, 2982G>C, 2983C>T, 2984C>G, 2985A>T, 2986A>T, 2989C>T, 2993A>T, 2995C>A, 2997C>A, 2998C>G, 2999T>C, 3001A>T, 3008G>T, 3009C>T, 3011A>T, 3012G>C, 3016G>A, 3018T>A, 3019T>C, 3020T>G, 3025G>A, 3026G>A, 3028G>T, 3029G>C, 3030A>G, 3032A>G, 3034G>T, 3035G>A, 3037C>G, 3038C>G, 3039C>T, 3040T>G, 3041C>G, 3042A>C, 3050T>C, 3052G>C, 3058C>T, 3061G>T, 3066A>G, 3067C>G, 3068G>A, 3069T>C, 3071G>C, 3073A>C, 3074C>G, 3076C>T, 3079G>T, 3084T>A, 3085T>C, 3091C>T, 3092C>T, 3094T>A, 3095C>G, 3096G>T, 3097A>C, 3104A>T, 3106G>T, 3109A>G, 3112A>G, 3113A>T, 3114A>C, 3115A>T, 3119T>G, 3121A>G, 3122C>T, 3124A>G, 3127T>C, 3130T>A, 3142T>G, 3146G>A, 3154C>T, 3155C>T, 3157A>T, 3159G>A, 3161G>C, 3162G>C, 3164C>T, 3166C>T, 3168A>T, 3169A>C, 3171A>G, 3172T>G, 3175C>A, 3176A>T, 3178C>T, 3179T>A, 3180G>T, 3181T>A, 3182C>T, 3184C>G, 3187C>A, 3189A>T, 3190C>T, 3191C>T, 3196C>T, 3202T>A, 3203A>C, 3206G>A, 3215A>G, 3216C>T, 3217T>A, 3220C>A, 3221C>T, 3224C>A, 3226G>A, 3229A>G, 3232C>T, 3233C>G, 3235G>T, 3238G>A, 3239A>C, 3242C>T, 3244G>A, 3246A>T, 3247A>T, 3248C>G, 3249G>C, 3250A>T, 3256C>T, 3257C>G, 3259A>T, 3262A>C, 3263C>T, 3265T>G, 3268A>G, 3269G>A, 3271A>C, 3272A>C, 3273G>A, 3274T>C, 3275A>T, 3276A>C, 3277C>T, 3280C>G, 3281A>T, 3283A>T, 3284A>G, 3285T>C, 3287C>A, 3291T>G, 3292G>A, 3293G>T, 3294A>C, 3295C>T, 3298G>A, 3300C>G, 3301C>T, 3302G>T, 3303A>C, 3304A>T, 3307C>T, 3308T>G, 3309T>G, 3310G>A, 3311A>C, 3313G>A, 3315T>C, 3316C>T, 3317G>A, 3319A>G, 3320A>G, 3321C>T, 3324C>A, 3325T>G, 3328T>C, 3329C>T, 3331T>A, 3337C>T, 3338A>G, 3340C>T, 3343A>T, 3345G>C, 3347A>G, 3349G>A, 3352C>A, 3355T>A, 3356A>G, 3358C>G, 3360A>G, 3361G>T, 3362C>A, 3365A>G, 3367C>T, 3372A>C, 3376G>A, 3379T>C, 3380T>G, 3381C>A, 3382C>A, 3388T>G, 3389C>A, 3394A>C, 3396A>G, 3397T>G, 3399T>C, 3400G>A, 3402T>G, 3403T>A, 3406A>T, 3409G>A, 3412C>A, 3417A>T, 3421A>G, 3422A>G, 3424A>T, 3425A>C, 3426A>G, 3427A>T, 3428C>A, 3429A>G, 3430A>T, 3434T>C, 3436A>T, 3439C>A, 3440C>T, 3442T>G, 3443C>G, 3445C>A, 3454C>T, 3455C>A, 3459A>T, 3463C>T, 3464A>G, 3466T>G, 3467C>G, 3468C>A, 3471A>G, 3472T>G, 3476G>T, 3478A>T, 3479C>T, 3480G>C, 3482C>A, 3484C>T, 3485A>T, 3487A>T, 3488A>G, 3489A>C, 3493C>T, 3494C>T, 3496T>A, 3500C>A, 3503T>C, 3504G>T, 3505C>A, 3506T>A, 3507T>C, 3509A>C, 3511A>C, 3514A>G, 3517T>A, 3518A>G, 3519G>C, 3520T>C, 3524G>T, 3526A>T, 3527A>C, 3528C>A, 3694A>G, 3694_3695insATCTTTCAT, 3695C>T, 3698G>A, 3702G>A, 3706C>T, 3710T>A, 3711C>G, 3712T>C, 3713C>A, 3714G>A, 3715A>G, 3718A>C, 3719C>T, 3721C>G, 3722A>G, 3726A>C, 3727G>A, 3728A>G, 3729G>C, 3731G>C, 3733A>G, 3734C>G, 3735T>C, 3742T>C, 3744G>C, 3745C>T, 3746A>G, 3747C>T, 3748A>T, 3750T>C, 3751A>T, 3754G>A, 3757A>G, 3760A>G, 3761T>A, 3763A>G, 3766G>A, 3769T>C, 3770A>G, 3775T>G, 3777G>T, 3778G>T, 3781T>C, 3782A>T, 3783C>T, 3784A>T, 3785A>G, 3787A>C, 3788T>A, 3790T>A, 3793T>C, 3799C>A, 3802C>T, 3803T>C, 3805A>G, 3806T>G, 3808C>G, 3813G>C, 3814A>C, 3818T>G, 3820T>C, 3822A>T, 3823G>T, 3824A>G, 3826A>T, 3828T>A, 3829G>T, 3835C>T, 3838C>T, 3839A>G, 3840A>C, 3841A>T, 3842C>G, 3844A>T, 3847A>T, 3848C>A, 3849A>G, 3852G>A, 3853G>C, 3854A>C, 3856G>A, 3857A>T, 3859G>T, 3861A>G, 3863T>A, 3864T>A, 3865A>G, 3868A>T, 3871C>T, 3872C>G, 3877C>A, 3878T>C, 3881C>A, 3883A>G, 3884A>C, 3888C>T, 3896C>A, 3897G>T, 3898A>T, 3899C>T, 3903G>T, 3907A>C, 3908A>C, 3909G>A, 3910T>A, 3915A>T, 3916T>C, 3920T>A, 3922C>T, 3923T>A, 3924C>A, 3925T>G, 3928A>T, 3929G>A, 3930T>A, 3931G>A, 3932T>G, 3936A>G, 3937G>A, 3940A>G, 3943A>G, 3944A>T, 3945A>G, 3946G>T, 3947T>G, 3948C>A, 3949T>G, 3952T>C, 3953A>C, 3954C>A, 3955C>A, 3957A>T, 3958C>T, 3964T>C, 3965G>C, 3966C>A, 3970T>A, 3973T>C, 3974C>A | | | | | | | | |

##### CDS

| | | | | | | | | | |
|---|---|---|---|---|---|---|---|---|---|
| D1R33_gp2 | 135 | 560 | 34.2% | 958 | 37.4% | 369 (98.4%) | 154 (41.1%) | 4/2/1/1 | 3 |
| Protein mutations: | D135* (2708G>T 2710C>G), E136V (2712A>T), V137G (2715T>G), Y140S (2724A>C 2725C>A), T141V (2726A>G 2727C>T), K142R (2730A>G 2731A>G), S143L (2732A>C 2733G>T), P147_F148del (2744_2749delCCCTTC), I149N (2751T>A), R151F (2756C>T 2757G>T), E153K (2762G>A 2764A>G), V154L (2765G>C 2767T>A), R155E (2768A>G 2769G>A), D156L (2771G>C 2772A>T 2772_2773insGA), D156_Q157insX (2772_2773insGA 2773C>G), Q157K (2774C>A), I158L (2777A>T 2779C>G), T159E (2780A>G 2781C>A 2782G>A), K160D (2783A>G 2785A>T), M161L (2786A>T 2788G>A), D163K (2792G>A 2794C>G), Q164K (2795C>A 2797A>G), G165E (2799G>A 2800A>G), I166F (2801A>T), R168* (2807A>T), D171G (2817A>G 2818C>A), A173P (2822G>C 2824A>T), S175G (2828A>G 2830C>T), S176A (2831T>G), I178V (2837A>G), W179L (2841G>T), V180F (2843G>T), V181M (2846G>A), P182R (2849C>A 2850C>G 2851C>A), I185K (2859T>A 2860C>G), A187G (2865C>G 2866T>G), K190I (2874A>T), Q191I (2876C>A 2877A>T), K192R (2880A>G), W193M (2882T>A 2883G>T), R194C (2885C>T), L195M (2888C>A 2890C>G), V196Y (2891G>T 2892T>A 2893A>T), V197I (2894G>A), F199Y (2901T>A 2902C>T), R200* (2903C>T 2905T>A), K201R (2907A>G), E204K (2915G>A 2917G>A), K205I (2919A>T 2920G>C), I207V (2924A>G 2926C>G), D208R (2927G>A 2928A>G 2929T>G), K210R (2934A>G 2935A>G), Y211F (2937A>T), N215T (2949A>C 2950C>T), S217D (2954A>G 2955G>A), D218E (2959C>G), V219L (2960G>C), L220F (2963C>T), K222E (2969A>G), L223V (2972T>G), G224N (2975G>A 2976G>A), K225G (2978A>G 2979A>G 2980G>C), C226A (2981T>G 2982G>C 2983C>T), Q227V (2984C>G 2985A>T 2986A>T), T230S (2993A>T 2995C>A), T231K (2997C>A 2998C>G), A235L (3008G>T 3009C>T), F238Y (3018T>A 3019T>C), Y239D (3020T>G), V241I (3026G>A 3028G>T), E242R (3029G>C 3030A>G), M243V (3032A>G 3034G>T), D244K (3035G>A 3037C>G), P245V (3038C>G 3039C>T 3040T>G), Q246A (3041C>G 3042A>C), S249P (3050T>C 3052G>C), N254R (3066A>G 3067C>G), V255T (3068G>A 3069T>C), E256H (3071G>C 3073A>C), H257D (3074C>G 3076C>T), F260Y (3084T>A 3085T>C), R264V (3095C>G 3096G>T 3097A>C), M267F (3104A>T 3106G>T), K270S (3113A>T 3114A>C 3115A>T), S272A (3119T>G 3121A>G), P273S (3122C>T 3124A>G), D281N (3146G>A), L284F (3155C>T 3157A>T), R285K (3159G>A), G286P (3161G>C 3162G>C), L287F (3164C>T 3166C>T), Q288L (3168A>T 3169A>C), N289R (3171A>G 3172T>G), N290K (3175C>A), I291F (3176A>T 3178C>T), C292I (3179T>A 3180G>T 3181T>A), Y295F (3189A>T 3190C>T), L296F (3191C>T), I300L (3203A>C), V301I (3206G>A), T304V (3215A>G 3216C>T 3217T>A), Q307K (3224C>A 3226G>A), L310V (3233C>G 3235G>T), N312H (3239A>C), E314V (3246A>T 3247A>T), R315A (3248C>G 3249G>C 3250A>T), Q318D (3257C>G 3259A>T), R319S (3262A>C), E322N (3269G>A 3271A>C), S323H (3272A>C 3273G>A 3274T>C), N324S (3275A>T 3276A>C 3277C>T), F325L (3280C>G), K326Y (3281A>T 3283A>T), I327A (3284A>G 3285T>C), Q328K (3287C>A), M329R (3291T>G 3292G>A), D330S (3293G>T 3294A>C 3295C>T), S332C (3300C>G 3301C>T), E333S (3302G>T 3303A>C 3304A>T), L335G (3308T>G 3309T>G 3310G>A), K336Q (3311A>C 3313G>A), L337P (3315T>C 3316C>T), E338K (3317G>A 3319A>G), T339V (3320A>G 3321C>T), A340E (3324C>A 3325T>G), I345V (3338A>G 3340C>T), S347T (3345G>C), R348G (3347A>G 3349G>A), D349E (3352C>A), I351V (3356A>G 3358C>G), K352S (3360A>G 3361G>T), P353T (3362C>A), N354D (3365A>G 3367C>T), D356A (3372A>C), S359E (3380T>G 3381C>A 3382C>A), I361M (3388T>G), Q362K (3389C>A), K363N (3394A>C), Y364W (3396A>G 3397T>G), L365P (3399T>C 3400G>A), I366R (3402T>G 3403T>A), K371M (3417A>T), I373V (3422A>G 3424A>T), K374R (3425A>C 3426A>G 3427A>T), Q375S (3428C>A 3429A>G 3430A>T), L380V (3443C>G 3445C>A), K385I (3459A>T), I387V (3464A>G 3466T>G), P388E (3467C>G 3468C>A), D389G (3471A>G 3472T>G), A391S (3476G>T 3478A>T), R392S (3479C>T 3480G>C), L393I (3482C>A 3484C>T), T394S (3485A>T 3487A>T), K395A (3488A>G 3489A>C), Q399K (3500C>A), C400L (3503T>C 3504G>T 3505C>A), L401T (3506T>A 3507T>C), K402H (3509A>C 3511A>C), S405A (3518A>G 3519G>C 3520T>C), V407F (3524G>T 3526A>T), K463_P464insIFH (3694_3695insATCTTTCAT), P464S (3695C>T), V465I (3698G>A), C466Y (3702G>A), R470K (3713C>A 3714G>A 3715A>G), N473D (3722A>G), E474A (3726A>C 3727G>A), S475A (3728A>G 3729G>C), E476Q (3731G>C 3733A>G), L477A (3734C>G 3735T>C), S480T (3744G>C 3745C>T), T481V (3746A>G 3747C>T 3748A>T), I482T (3750T>C 3751A>T), L486M (3761T>A 3763A>G), I489V (3770A>G), W491F (3777G>T 3778G>T), T493F (3782A>T 3783C>T 3784A>T), K494D (3785A>G 3787A>C), Y495K (3788T>A 3790T>A), F501V (3806T>G 3808C>G), R503T (3813G>C 3814A>C), F505V (3818T>G 3820T>C), K506I (3822A>T 3823G>T), I507V (3824A>G 3826A>T), L508Y (3828T>A 3829G>T), K512A (3839A>G 3840A>C 3841A>T), P513A (3842C>G 3844A>T), Q515R (3848C>A 3849A>G), W516Y (3852G>A 3853G>C), M517L (3854A>C 3856G>A), M518F (3857A>T 3859G>T), N519S (3861A>G), L520K (3863T>A 3864T>A 3865A>G), K521N (3868A>T), P523A (3872C>G), N524K (3877C>A), S525P (3878T>C), M527L (3884A>C), T528I (3888C>T), R531I (3896C>A 3897G>T 3898A>T), R533L (3903G>T), S535Q (3908A>C 3909G>A 3910T>A), Y537F (3915A>T 3916T>C), F539I (3920T>A 3922C>T), S540K (3923T>A 3924C>A 3925T>G), V542K (3929G>A 3930T>A 3931G>A), Y543D (3932T>G), K544R (3936A>G 3937G>A), K547C (3944A>T 3945A>G 3946G>T), S548E (3947T>G 3948C>A 3949T>G), T550Q (3953A>C 3954C>A 3955C>A), N551I (3957A>T 3958C>T), A554H (3965G>C 3966C>A), R557S (3974C>A) | | | | | | | | |

| | Begin | End | Coverage | Score | Concordance | Matches | Identities | I/D/M/F* | Stop Codons |
|---|---|---|---|---|---|---|---|---|---|
| **NT** | **2706** | **3983** | **14.8%** | **74** | **3.4%** | **1109 (98.5%)** | **584 (51.9%)** | **11/6** | |

Codon mutations:

GAC135TAG (2708G>T 2710C>G), GAA136GTA (2712A>T), GTA137GGA (2715T>G), CCC138CCA (2719C>A), GTG139GTT (2722G>T), TAC140TCA (2724A>C 2725C>A), ACC141GTC (2726A>G 2727C>T), AAA142AGG (2730A>G 2731A>G), AGT143CTT (2732A>C 2733G>T), TAC144TAT (2737C>T), CGG145AGA (2738C>A 2740G>A), CCC147_TTC148del (2744_2749delCCCTTC), ATC149AAC (2751T>A), CGC151TTC (2756C>T 2757G>T), CAG152CAA (2761G>A), GAA153AAG (2762G>A 2764A>G), GTT154CTA (2765G>C 2767T>A), AGG155GAG (2768A>G 2769G>A), GAC156CTG (2771G>C 2772A>T 2772_2773insGA), GAC156_CAA157insA-G (2772_2773insGA 2773C>G), CAA157AAA (2774C>A), ATC158TTG (2777A>T 2779C>G), ACG159GAA (2780A>G 2781C>A 2782G>A), AAA160GAT (2783A>G 2785A>T), ATG161TTA (2786A>T 2788G>A), GAC163AAG (2792G>A 2794C>G), CAA164AAG (2795C>A 2797A>G), GGA165GAG (2799G>A 2800A>G), ATT166TTT (2801A>T), ATA167ATT (2806A>T), AGA168TGA (2807A>T), CCA169CCT (2812A>T), TCA170AGT (2813T>A 2814C>G 2815A>T), GAC171GGA (2817A>G 2818C>A), TCT172TCA (2821T>A), GCA173CCT (2822G>C 2824A>T), AGC175GGT (2828A>G 2830C>T), TCA176GCA (2831T>G), CCC177CCA (2836C>A), ATA178GTA (2837A>G), TGG179TTG (2841G>T), GTT180TTT (2843G>T), GTG181ATG (2846G>A), CCC182AGA (2849C>A 2850C>G 2851C>A), AAA184AAG (2857A>G), ATC185AAG (2859T>A 2860C>G), GAC186GAT (2863C>T), GCT187GGG (2865C>G 2866T>G), TCT188TCG (2869T>G), GGG189GGT (2872G>T), AAA190ATA (2874A>T), CAA191ATA (2876C>A 2877A>T), AAG192AGG (2880A>G), TGG193ATG (2882T>A 2883G>T), CGT194TGT (2885C>T), CTC195ATG (2888C>A 2890C>G), GTA196TAT (2891G>T 2892T>A 2893A>T), GTT197ATT (2894G>A), GAC198GAT (2899C>T), TTC199TAT (2901T>A 2902C>T), CGT200TGA (2903C>T 2905T>A), AAG201AGG (2907A>G), GAG204AAA (2915G>A 2917G>A), AAG205ATC (2919A>T 2920G>C), ATC207GTG (2924A>G 2926C>G), GAT208AGG (2927G>A 2928A>G 2929T>G), GAC209GAT (2932C>T), AAA210AGG (2934A>G 2935A>G), TAC211TTC (2937A>T), CCG212CCA (2941G>A), ATA213ATC (2944A>C), AAC215ACT (2949A>C 2950C>T), ATA216ATT (2953A>T), AGT217GAT (2954A>G 2955G>A), GAC218GAG (2959C>G), GTA219CTA (2960G>C), CTT220TTT (2963C>T), GAC221GAT (2968C>T), AAG222GAG (2969A>G), TTA223GTA (2972T>G), GGT224AAT (2975G>A 2976G>A), AAG225GGC (2978A>G 2979A>G 2980G>C), TGC226GCT (2981T>G 2982G>C 2983C>T), CAA227GTT (2984C>G 2985A>T 2986A>T), TAC228TAT (2989C>T), ACC230TCA (2993A>T 2995C>A), ACC231AAG (2997C>A 2998C>G), TTA232CTT (2999T>C 3001A>T), GCA235TTA (3008G>T 3009C>T), AGT236TCT (3011A>T 3012G>C), GGG237GGA (3016G>A), TTT238TAC (3018T>A 3019T>C), TAT239GAT (3020T>G), CAG240CAA (3025G>A), GTG241ATT (3026G>A 3028G>T), GAG242CGG (3029G>C 3030A>G), ATG243GTT (3032A>G 3034G>T), GAC244AAG (3035G>A 3037C>G), CCT245GTG (3038C>G 3039C>T 3040T>G), CAA246GCA (3041C>G 3042A>C), TCG249CCC (3050T>C 3052G>C), ACC251ACT (3058C>T), GCG252GCT (3061G>T), AAC254AGG (3066A>G 3067C>G), GTA255ACA (3068G>A 3069T>C), GAA256CAC (3071G>C 3073A>C), CAC257GAT (3074C>G 3076C>T), GGG258GGT (3079G>T), TTT260TAC (3084T>A 3085T>C), TTC262TTT (3091C>T), CTT263TTA (3092C>T 3094T>A), CGA264GTC (3095C>G 3096G>T 3097A>C), ATG267TTT (3104A>T 3106G>T), GGA268GGG (3109A>G), TTA269TTG (3112A>G), AAA270TCT (3113A>T 3114A>C 3115A>T), TCA272GCG (3119T>G 3121A>G), CCA273TCG (3122C>T 3124A>G), TCT274TCC (3127T>C), ACT275ACA (3130T>A), GTT279.TG (3142T>G), GAC281AAC (3146G>A), GTC283GTT (3154C>T), CTA284TTT (3155C>T 3157A>T), AGA285AAA (3159G>A), GGT286CCT (3161G>C 3162G>C), CTC287TTT (3164C>T 3166C>T), CAA288CTC (3168A>T 3169A>C), AAT289AGG (3171A>G 3172T>G), AAC290AAA (3175C>A), ATC291TTT (3176A>T 3178C>T), TGT292ATA (3179T>A 3180G>T 3181T>A), CTC293TTG (3182C>T 3184C>G), GTC294GTA (3187C>A), TAC295TTT (3189A>T 3190C>T), CTT296TTT (3191C>T), GAC297GAT (3196C>T), ATT299ATA (3202T>A), ATT300CTT (3203A>C), GTC301ATC (3206G>A), ACT304GTA (3215A>G 3216C>T 3217T>A), TCC305TCA (3220C>A), CTA306TTA (3221C>T), CAG307AAA (3224C>A 3226G>A), GAA308GAG (3229A>G), CAC309CAT (3232C>T), CTG310GTT (3233C>G 3235G>T), GAG311GAA (3238G>A), AAC312CAC (3239A>C), CTG313TTA (3242C>T 3244G>A), GAA314GTT (3246A>T 3247A>T), CGA315GCT (3248C>G 3249G>C 3250A>T), TTC317TTT (3256C>T), CAA318GAT (3257C>G 3259A>T), AGA319AGC (3262A>C), CTT320TTG (3263C>T 3265T>G), AGA321AGG (3268A>G), GAA322AAC (3269G>A 3271A>C), AGT323CAC (3272A>C 3273G>A 3274T>C), AAC324TCT (3275A>T 3276A>C 3277C>T), TTC325TTG (3280C>G), AAA326TAT (3281A>T 3283A>T), ATT327GCT (3284A>G 3285T>C), CAA328AAA (3287C>A), ATG329AGA (3291T>G 3292G>A), GAC330TCT (3293G>T 3294A>C 3295C>T), AAG331AAA (3298G>A), TCC332TGT (3300C>G 3301C>T), GAA333TCT (3302G>T 3303A>C 3304A>T), TTC334TTT (3307C>T), TTG335GGA (3308T>G 3309T>G 3310G>A), AAG336CAA (3311A>C 3313G>A), CTC337CCT (3315T>C 3316C>T), GAA338AAG (3317G>A 3319A>G), ACT339GTT (3320A>G 3321C>T), GCT340GAG (3324C>A 3325T>G), TAT341TAC (3328T>C), CTT342TTA (3329C>T 3331T>A), CAC344CAT (3337C>T), ATC345GTT (3338A>G 3340C>T), ATA346ATT (3343A>T), AGC347ACC (3345G>C), AGG348GGA (3347A>G 3349G>A), GAC349GAA (3352C>A), GGT350GGA (3355T>A), ATC351GTG (3356A>G 3358C>G), AAG352AGT (3360A>G 3361G>T), CCT353ACT (3362C>A), AAC354GAT (3365A>G 3367C>T), GAT356GCT (3372A>C), AAG357AAA (3376G>A), ATT358ATC (3379T>C), TCC359GAA (3380T>G 3381C>A 3382C>A), ATT361ATG (3388T>G), CAA362AAA (3389C>A), AAA363AAC (3394A>C), TAT364TGG (3396A>G 3397T>G), CTG365CCA (3399T>C 3400G>A), ATT366AGA (3402T>G 3403T>A), CCA367CCT (3406A>T), AAG368AAA (3409G>A), ACC369ACA (3412C>A), AAG371ATG (3417A>T), GAA372GAG (3421A>G), ATA373GTT (3422A>G 3424A>T), AAA374CGT (3425A>C 3426A>G 3427A>T), CAA375AGT (3428C>A 3429A>G 3430A>T), TTA377CTT (3434T>C 3436A>T), GGC378GGA (3439C>A), CTT379TTG (3440C>T 3442T>G), CTC380GTA (3443C>G 3445C>A), TAC383TAT (3454C>T), CGA384AGA (3455C>A), AAA385ATA (3459A>T), TTC386TTT (3463C>T), ATT387GTG (3464A>G 3466T>G), CCA388GAA (3467C>G 3468C>A), GAT389GGG (3471A>G 3472T>G), GCA391TCT (3476G>T 3478A>T), CGA392TCA (3479C>T 3480G>C), CTC393ATT (3482C>A 3484C>T), ACA394TCT (3485A>T 3487A>T), AAA395GCA (3488A>G 3489A>C), CCC396CCT (3493C>T), CTT397TTA (3494C>T 3496T>A), CAG399AAG (3500C>A), TGC400CTA (3503T>C 3504G>T 3505C>A), TTA401ACA (3506T>A 3507T>C), AAA402CAC (3509A>C 3511A>C), AAA403AAG (3514A>G), GGT404GGA (3517T>A), AGT405GCC (3518A>G 3519G>C 3520T>C), GTA407TTT (3524G>T 3526A>T), ACT408CA. (3527A>C 3528C>A), AAA463AAG (3694A>G), AAA463_CCC464insATCTTTCAT (3694_3695insATCTTTCAT), CCC464TCC (3695C>T), GTC465ATC (3698G>A), TGT466TAT (3702G>A), TAC467TAT (3706C>T), TCT469AGC (3710T>A 3711C>G 3712T>C), CGA470AAG (3713C>A 3714G>A 3715A>G), ACA471ACC (3718A>C), CTC472TTG (3719C>T 3721C>G), AAT473GAT (3722A>G), GAG474GCA (3726A>C 3727G>A), AGC475GCC (3728A>G 3729G>C), GAA476CAG (3731G>C 3733A>G), CTA477GCA (3734C>G 3735T>C), TAT479TAC (3742T>C), AGC480ACT (3744G>C 3745C>T), ACA481GTT (3746A>G 3747C>T 3748A>T), ATA482ACT (3750T>C 3751A>T), GAG483GAA (3754G>A), AAA484AAG (3757A>G), GAA485GAG (3760A>G), TTA486ATG (3761T>A 3763A>G), CTG487CTA (3766G>A), GCT488GCC (3769T>C), ATA489GTA (3770A>G), GTT490GTG (3775T>G), TGG491TTT (3777G>T 3778G>T), GCT492GCC (3781T>C), ACA493TTT (3782A>T 3783C>T 3784A>T), AAA494GAC (3785A>G 3787A>C), TAT495AAA (3788T>A 3790T>A), TTT496TTC (3793T>C), CCC498CCA (3799C>A), TAC499TAT (3802C>T), TTA500CTG (3803T>C 3805A>G), TTC501GTG (3806T>G 3808C>G), AGA503ACC (3813G>C 3814A>C), TTT505GTC (3818T>G 3820T>C), AAG506ATT (3822A>T 3823G>T), ATA507GTT (3824A>G 3826A>T), TTG508TAT (3828T>A 3829G>T), GAC510GAT (3835C>T), CAC511CAT (3838C>T), AAA512GCT (3839A>G 3840A>C 3841A>T), CCA513GCT (3842C>G 3844A>T), CTA514CTT (3847A>T), CAG515AGG (3848C>A 3849A>G), TGG516TAC (3852G>A 3853G>C), ATG517CTA (3854A>C 3856G>A), ATG518TTT (3857A>T 3859G>T), AAC519AGC (3861A>G), TTA520AAG (3863T>A 3864T>A 3865A>G), AAA521AAT (3868A>T), GAC522GAT (3871C>T), CCA523GCA (3872C>G), AAC524AAA (3877C>A), TCA525CCA (3878T>C), CGA526AGG (3881C>A 3883A>G), ATG527CTG (3884A>C), ACT528ATT (3888C>T), CGA531ATT (3896C>A 3897G>T 3898A>T), CTA532TTA (3899C>T), CGA533CTA (3903G>T), CTA534CTC (3907A>C), AGT535CAA (3908A>C 3909G>A 3910T>A), TAT537TTC (3915A>T 3916T>C), TTC539ATT (3920T>A 3922C>T), TCT540AAG (3923T>A 3924C>A 3925T>G), GTA541GTT (3928A>T), GTG542AAA (3929G>A 3930T>A 3931G>A), TAC543GAC (3932T>G), AAG544AGA (3936A>G 3937G>A), AAA545AAG (3940A>G), GGA546GGG (3943A>G), AAG547TGT (3944A>T 3945A>G 3946G>T), TCT548GAG (3947T>G 3948C>A 3949T>G), AAT549AAC (3952T>C), ACC550CAA (3953A>C 3954C>A 3955C>A), AAC551ATT (3957A>T 3958C>T), GAT553GAC (3964T>C), GCC554CAC (3965G>C 3966C>A), CTT555CTA (3970T>A), TCT556TCC (3973T>C), CGT557AGT (3974C>A)

#### Proteins

| | Begin | End | Coverage | Score | Concordance | Matches | Identities | I/D/M/F* | Stop Codons |
|---|---|---|---|---|---|---|---|---|---|
| ORF B (YP_009507248.1) | 135 | 560 | 34.2% | 958 | 37.4% | 369 (98.4%) | 154 (41.1%) | 4/2/1/1 | 3 |

Protein mutations:

D135* (2708G>T 2710C>G), E136V (2712A>T), V137G (2715T>G), Y140S (2724A>C 2725C>A), T141V (2726A>G 2727C>T), K142R (2730A>G 2731A>G), S143L (2732A>C 2733G>T), P147_F148del (2744_2749delCCCTTC), I149N (2751T>A), R151F (2756C>T 2757G>T), E153K (2762G>A 2764A>G), V154L (2765G>C 2767T>A), R155E (2768A>G 2769G>A), D156L (2771G>C 2772A>T 2772_2773insGA), D156_Q157insX (2772_2773insGA 2773C>G), Q157K (2774C>A), I158L (2777A>T 2779C>G), T159E (2780A>G 2781C>A 2782G>A), K160D (2783A>G 2785A>T), M161L (2786A>T 2788G>A), D163K (2792G>A 2794C>G), Q164K (2795C>A 2797A>G), G165E (2799G>A 2800A>G), I166F (2801A>T), R168* (2807A>T), D171G (2817A>G 2818C>A), A173P (2822G>C 2824A>T), S175G (2828A>G 2830C>T), S176A (2831T>G), I178V (2837A>G), W179L (2841G>T), V180F (2843G>T), V181M (2846G>A), P182R (2849C>A 2850C>G 2851C>A), I185K (2859T>A 2860C>G), A187G (2865C>G 2866T>G), K190I (2874A>T), Q191I (2876C>A 2877A>T), K192R (2880A>G), W193M (2882T>A 2883G>T), R194C (2885C>T), L195M (2888C>A 2890C>G), V196Y (2891G>T 2892T>A 2893A>T), V197I (2894G>A), F199Y (2901T>A 2902C>T), R200* (2903C>T 2905T>A), K201R (2907A>G), E204K (2915G>A 2917G>A), K205I (2919A>T 2920G>C), I207V (2924A>G 2926C>G), D208R (2927G>A 2928A>G 2929T>G), K210R (2934A>G 2935A>G), Y211F (2937A>T), N215T (2949A>C 2950C>T), S217D (2954A>G 2955G>A), D218E (2959C>G), V219L (2960G>C), L220F (2963C>T), K222E (2969A>G), L223V (2972T>G), G224N (2975G>A 2976G>A), K225G (2978A>G 2979A>G 2980G>C), C226A (2981T>G 2982G>C 2983C>T), Q227V (2984C>G 2985A>T 2986A>T), T230S (2993A>T 2995C>A), T231K (2997C>A 2998C>G), A235L (3008G>T 3009C>T), F238Y (3018T>A 3019T>C), Y239D (3020T>G), V241I (3026G>A 3028G>T), E242R (3029G>C 3030A>G), M243V (3032A>G 3034G>T), D244K (3035G>A 3037C>G), P245V (3038C>G 3039C>T 3040T>G), Q246A (3041C>G 3042A>C), S249P (3050T>C 3052G>C), N254R (3066A>G 3067C>G), V255T (3068G>A 3069T>C), E256H (3071G>C 3073A>C), H257D (3074C>G 3076C>T), F260Y (3084T>A 3085T>C), R264V (3095C>G 3096G>T 3097A>C), M267F (3104A>T 3106G>T), K270S (3113A>T 3114A>C 3115A>T), S272A (3119T>G 3121A>G), P273S (3122C>T 3124A>G), D281N (3146G>A), L284F (3155C>T 3157A>T), R285K (3159G>A), G286P (3161G>C 3162G>C), L287F (3164C>T 3166C>T), Q288L (3168A>T 3169A>C), N289R (3171A>G 3172T>G), N290K (3175C>A), I291F (3176A>T 3178C>T), C292I (3179T>A 3180G>T 3181T>A), Y295F (3189A>T 3190C>T), L296F (3191C>T), I300L (3203A>C), V301I (3206G>A), T304V (3215A>G 3216C>T 3217T>A), Q307K (3224C>A 3226G>A), L310V (3233C>G 3235G>T), N312H (3239A>C), E314V (3246A>T 3247A>T), R315A (3248C>G 3249G>C 3250A>T), Q318D (3257C>G 3259A>T), R319S (3262A>C), E322N (3269G>A 3271A>C), S323H (3272A>C 3273G>A 3274T>C), N324S (3275A>T 3276A>C 3277C>T), F325L (3280C>G), K326Y (3281A>T 3283A>T), I327A (3284A>G 3285T>C), Q328K (3287C>A), M329R (3291T>G 3292G>A), D330S (3293G>T 3294A>C 3295C>T), S332C (3300C>G 3301C>T), E333S (3302G>T 3303A>C 3304A>T), L335G (3308T>G 3309T>G 3310G>A), K336Q (3311A>C 3313G>A), L337P (3315T>C 3316C>T), E338K (3317G>A 3319A>G), T339V (3320A>G 3321C>T), A340E (3324C>A 3325T>G), I345V (3338A>G 3340C>T), S347T (3345G>C), R348G (3347A>G 3349G>A), D349E (3352C>A), I351V (3356A>G 3358C>G), K352S (3360A>G 3361G>T), P353T (3362C>A), N354D (3365A>G 3367C>T), D356A (3372A>C), S359E (3380T>G 3381C>A 3382C>A), I361M (3388T>G), Q362K (3389C>A), K363N (3394A>C), Y364W (3396A>G 3397T>G), L365P (3399T>C 3400G>A), I366R (3402T>G 3403T>A), K371M (3417A>T), I373V (3422A>G 3424A>T), K374R (3425A>C 3426A>G 3427A>T), Q375S (3428C>A 3429A>G 3430A>T), L380V (3443C>G 3445C>A), K385I (3459A>T), I387V (3464A>G 3466T>G), P388E (3467C>G 3468C>A), D389G (3471A>G 3472T>G), A391S (3476G>T 3478A>T), R392S (3479C>T 3480G>C), L393I (3482C>A 3484C>T), T394S (3485A>T 3487A>T), K395A (3488A>G 3489A>C), Q399K (3500C>A), C400L (3503T>C 3504G>T 3505C>A), L401T (3506T>A 3507T>C), K402H (3509A>C 3511A>C), S405A (3518A>G 3519G>C 3520T>C), V407F (3524G>T 3526A>T), K463_P464insIFH (3694_3695insATCTTTCAT), P464S (3695C>T), V465I (3698G>A), C466Y (3702G>A), R470K (3713C>A 3714G>A 3715A>G), N473D (3722A>G), E474A (3726A>C 3727G>A), S475A (3728A>G 3729G>C), E476Q (3731G>C 3733A>G), L477A (3734C>G 3735T>C), S480T (3744G>C 3745C>T), T481V (3746A>G 3747C>T 3748A>T), I482T (3750T>C 3751A>T), L486M (3761T>A 3763A>G), I489V (3770A>G), W491F (3777G>T 3778G>T), T493F (3782A>T 3783C>T 3784A>T), K494D (3785A>G 3787A>C), Y495K (3788T>A 3790T>A), F501V (3806T>G 3808C>G), R503T (3813G>C 3814A>C), F505V (3818T>G 3820T>C), K506I (3822A>T 3823G>T), I507V (3824A>G 3826A>T), L508Y (3828T>A 3829G>T), K512A (3839A>G 3840A>C 3841A>T), P513A (3842C>G 3844A>T), Q515R (3848C>A 3849A>G), W516Y (3852G>A 3853G>C), M517L (3854A>C 3856G>A), M518F (3857A>T 3859G>T), N519S (3861A>G), L520K (3863T>A 3864T>A 3865A>G), K521N (3868A>T), P523A (3872C>G), N524K (3877C>A), S525P (3878T>C), M527L (3884A>C), T528I (3888C>T), R531I (3896C>A 3897G>T 3898A>T), R533L (3903G>T), S535Q (3908A>C 3909G>A 3910T>A), Y537F (3915A>T 3916T>C), F539I (3920T>A 3922C>T), S540K (3923T>A 3924C>A 3925T>G), V542K (3929G>A 3930T>A 3931G>A), Y543D (3932T>G), K544R (3936A>G 3937G>A), K547C (3944A>T 3945A>G 3946G>T), S548E (3947T>G 3948C>A 3949T>G), T550Q (3953A>C 3954C>A 3955C>A), N551I (3957A>T 3958C>T), A554H (3965G>C 3966C>A), R557S (3974C>A)

| | Begin | End | Coverage | Score | Concordance | Matches | Identities | I/D/M/F* | Stop Codons |
|---|---|---|---|---|---|---|---|---|---|
| **NT** | **2706** | **3983** | **14.8%** | **74** | **3.4%** | **1109 (98.5%)** | **584 (51.9%)** | **11/6** | |

Codon mutations: GAC135TAG (2708G>T 2710C>G), GAA136GTA (2712A>T), GTA137GGA (2715T>G), CCC138CCA (2719C>A), GTG139GTT (2722G>T), TAC140TCA (2724A>C 2725C>A), ACC141GTC (2726A>G 2727C>T), AAA142AGG (2730A>G 2731A>G), AGT143CTT (2732A>C 2733G>T), TAC144TAT (2737C>T), CGG145AGA (2738C>A 2740G>A), CCC147_TTC148del (2744_2749delCCCTTC), ATC149AAC (2751T>A), CGC151TTC (2756C>T 2757G>T), CAG152CAA (2761G>A), GAA153AAG (2762G>A 2764A>G), GTT154CTA (2765G>C 2767T>A), AGG155GAG (2768A>G 2769G>A), GAC156CTG (2771G>C 2772A>T 2772_2773insGA), GAC156_CAA157insA-G (2772_2773insGA 2773C>G), CAA157AAA (2774C>A), ATC158TTG (2777A>T 2779C>G), ACG159GAA (2780A>G 2781C>A 2782G>A), AAA160GAT (2783A>G 2785A>T), ATG161TTA (2786A>T 2788G>A), GAC163AAG (2792G>A 2794C>G), CAA164AAG (2795C>A 2797A>G), GGA165GAG (2799G>A 2800A>G), ATT166TTT (2801A>T), ATA167ATT (2806A>T), AGA168TGA (2807A>T), CCA169CCT (2812A>T), TCA170AGT (2813T>A 2814C>G 2815A>T), GAC171GGA (2817A>G 2818C>A), TCT172TCA (2821T>A), GCA173CCT (2822G>C 2824A>T), AGC175GGT (2828A>G 2830C>T), TCA176GCA (2831T>G), CCC177CCA (2836C>A), ATA178GTA (2837A>G), TGG179TTG (2841G>T), GTT180TTT (2843G>T), GTG181ATG (2846G>A), CCC182AGA (2849C>A 2850C>G 2851C>A), AAA184AAG (2857A>G), ATC185AAG (2859T>A 2860C>G), GAC186GAT (2863C>T), GCT187GGG (2865C>G 2866T>G), TCT188TCG (2869T>G), GGG189GGT (2872G>T), AAA190ATA (2874A>T), CAA191ATA (2876C>A 2877A>T), AAG192AGG (2880A>G), TGG193ATG (2882T>A 2883G>T), CGT194TGT (2885C>T), CTC195ATG (2888C>A 2890C>G), GTA196TAT (2891G>T 2892T>A 2893A>T), GTT197ATT (2894G>A), GAC198GAT (2899C>T), TTC199TAT (2901T>A 2902C>T), CGT200TGA (2903C>T 2905T>A), AAG201AGG (2907A>G), GAG204AAA (2915G>A 2917G>A), AAG205ATC (2919A>T 2920G>C), ATC207GTG (2924A>G 2926C>G), GAT208AGG (2927G>A 2928A>G 2929T>G), GAC209GAT (2932C>T), AAA210AGG (2934A>G 2935A>G), TAC211TTC (2937A>T), CCG212CCA (2941G>A), ATA213ATC (2944A>C), AAC215ACT (2949A>C 2950C>T), ATA216ATT (2953A>T), AGT217GAT (2954A>G 2955G>A), GAC218GAG (2959C>G), GTA219CTA (2960G>C), CTT220TTT (2963C>T), GAC221GAT (2968C>T), AAG222GAG (2969A>G), TTA223GTA (2972T>G), GGT224AAT (2975G>A 2976G>A), AAG225GGC (2978A>G 2979A>G 2980G>C), TGC226GCT (2981T>G 2982G>C 2983C>T), CAA227GTT (2984C>G 2985A>T 2986A>T), TAC228TAT (2989C>T), ACC230TCA (2993A>T 2995C>A), ACC231AAG (2997C>A 2998C>G), TTA232CTT (2999T>C 3001A>T), GCA235TTA (3008G>T 3009C>T), AGT236TCT (3011A>T 3012G>C), GGG237GGA (3016G>A), TTT238TAC (3018T>A 3019T>C), TAT239GAT (3020T>G), CAG240CAA (3025G>A), GTG241ATT (3026G>A 3028G>T), GAG242CGG (3029G>C 3030A>G), ATG243GTT (3032A>G 3034G>T), GAC244AAG (3035G>A 3037C>G), CCT245GTG (3038C>G 3039C>T 3040T>G), CAA246GCA (3041C>G 3042A>C), TCG249CCC (3050T>C 3052G>C), ACC251ACT (3058C>T), GCG252GCT (3061G>T), AAC254AGG (3066A>G 3067C>G), GTA255ACA (3068G>A 3069T>C), GAA256CAC (3071G>C 3073A>C), CAC257GAT (3074C>G 3076C>T), GGG258GGT (3079G>T), TTT260TAC (3084T>A 3085T>C), TTC262TTT (3091C>T), CTT263TTA (3092C>T 3094T>A), CGA264GTC (3095C>G 3096G>T 3097A>C), ATG267TTT (3104A>T 3106G>T), GGA268GGG (3109A>G), TTA269TTG (3112A>G), AAA270TCT (3113A>T 3114A>C 3115A>T), TCA272GCG (3119T>G 3121A>G), CCA273TCG (3122C>T 3124A>G), TCT274TCC (3127T>C), ACT275ACA (3130T>A), GTT279.TG (3142T>G), GAC281AAC (3146G>A), GTC283GTT (3154C>T), CTA284TTT (3155C>T 3157A>T), AGA285AAA (3159G>A), GGT286CCT (3161G>C 3162G>C), CTC287TTT (3164C>T 3166C>T), CAA288CTC (3168A>T 3169A>C), AAT289AGG (3171A>G 3172T>G), AAC290AAA (3175C>A), ATC291TTT (3176A>T 3178C>T), TGT292ATA (3179T>A 3180G>T 3181T>A), CTC293TTG (3182C>T 3184C>G), GTC294GTA (3187C>A), TAC295TTT (3189A>T 3190C>T), CTT296TTT (3191C>T), GAC297GAT (3196C>T), ATT299ATA (3202T>A), ATT300CTT (3203A>C), GTC301ATC (3206G>A), ACT304GTA (3215A>G 3216C>T 3217T>A), TCC305TCA (3220C>A), CTA306TTA (3221C>T), CAG307AAA (3224C>A 3226G>A), GAA308GAG (3229A>G), CAC309CAT (3232C>T), CTG310GTT (3233C>G 3235G>T), GAG311GAA (3238G>A), AAC312CAC (3239A>C), CTG313TTA (3242C>T 3244G>A), GAA314GTT (3246A>T 3247A>T), CGA315GCT (3248C>G 3249G>C 3250A>T), TTC317TTT (3256C>T), CAA318GAT (3257C>G 3259A>T), AGA319AGC (3262A>C), CTT320TTG (3263C>T 3265T>G), AGA321AGG (3268A>G), GAA322AAC (3269G>A 3271A>C), AGT323CAC (3272A>C 3273G>A 3274T>C), AAC324TCT (3275A>T 3276A>C 3277C>T), TTC325TTG (3280C>G), AAA326TAT (3281A>T 3283A>T), ATT327GCT (3284A>G 3285T>C), CAA328AAA (3287C>A), ATG329AGA (3291T>G 3292G>A), GAC330TCT (3293G>T 3294A>C 3295C>T), AAG331AAA (3298G>A), TCC332TGT (3300C>G 3301C>T), GAA333TCT (3302G>T 3303A>C 3304A>T), TTC334TTT (3307C>T), TTG335GGA (3308T>G 3309T>G 3310G>A), AAG336CAA (3311A>C 3313G>A), CTC337CCT (3315T>C 3316C>T), GAA338AAG (3317G>A 3319A>G), ACT339GTT (3320A>G 3321C>T), GCT340GAG (3324C>A 3325T>G), TAT341TAC (3328T>C), CTT342TTA (3329C>T 3331T>A), CAC344CAT (3337C>T), ATC345GTT (3338A>G 3340C>T), ATA346ATT (3343A>T), AGC347ACC (3345G>C), AGG348GGA (3347A>G 3349G>A), GAC349GAA (3352C>A), GGT350GGA (3355T>A), ATC351GTG (3356A>G 3358C>G), AAG352AGT (3360A>G 3361G>T), CCT353ACT (3362C>A), AAC354GAT (3365A>G 3367C>T), GAT356GCT (3372A>C), AAG357AAA (3376G>A), ATT358ATC (3379T>C), TCC359GAA (3380T>G 3381C>A 3382C>A), ATT361ATG (3388T>G), CAA362AAA (3389C>A), AAA363AAC (3394A>C), TAT364TGG (3396A>G 3397T>G), CTG365CCA (3399T>C 3400G>A), ATT366AGA (3402T>G 3403T>A), CCA367CCT (3406A>T), AAG368AAA (3409G>A), ACC369ACA (3412C>A), AAG371ATG (3417A>T), GAA372GAG (3421A>G), ATA373GTT (3422A>G 3424A>T), AAA374CGT (3425A>C 3426A>G 3427A>T), CAA375AGT (3428C>A 3429A>G 3430A>T), TTA377CTT (3434T>C 3436A>T), GGC378GGA (3439C>A), CTT379TTG (3440C>T 3442T>G), CTC380GTA (3443C>G 3445C>A), TAC383TAT (3454C>T), CGA384AGA (3455C>A), AAA385ATA (3459A>T), TTC386TTT (3463C>T), ATT387GTG (3464A>G 3466T>G), CCA388GAA (3467C>G 3468C>A), GAT389GGG (3471A>G 3472T>G), GCA391TCT (3476G>T 3478A>T), CGA392TCA (3479C>T 3480G>C), CTC393ATT (3482C>A 3484C>T), ACA394TCT (3485A>T 3487A>T), AAA395GCA (3488A>G 3489A>C), CCC396CCT (3493C>T), CTT397TTA (3494C>T 3496T>A), CAG399AAG (3500C>A), TGC400CTA (3503T>C 3504G>T 3505C>A), TTA401ACA (3506T>A 3507T>C), AAA402CAC (3509A>C 3511A>C), AAA403AAG (3514A>G), GGT404GGA (3517T>A), AGT405GCC (3518A>G 3519G>C 3520T>C), GTA407TTT (3524G>T 3526A>T), ACT408CA. (3527A>C 3528C>A), AAA463AAG (3694A>G), AAA463_CCC464insATCTTTCAT (3694_3695insATCTTTCAT), CCC464TCC (3695C>T), GTC465ATC (3698G>A), TGT466TAT (3702G>A), TAC467TAT (3706C>T), TCT469AGC (3710T>A 3711C>G 3712T>C), CGA470AAG (3713C>A 3714G>A 3715A>G), ACA471ACC (3718A>C), CTC472TTG (3719C>T 3721C>G), AAT473GAT (3722A>G), GAG474GCA (3726A>C 3727G>A), AGC475GCC (3728A>G 3729G>C), GAA476CAG (3731G>C 3733A>G), CTA477GCA (3734C>G 3735T>C), TAT479TAC (3742T>C), AGC480ACT (3744G>C 3745C>T), ACA481GTT (3746A>G 3747C>T 3748A>T), ATA482ACT (3750T>C 3751A>T), GAG483GAA (3754G>A), AAA484AAG (3757A>G), GAA485GAG (3760A>G), TTA486ATG (3761T>A 3763A>G), CTG487CTA (3766G>A), GCT488GCC (3769T>C), ATA489GTA (3770A>G), GTT490GTG (3775T>G), TGG491TTT (3777G>T 3778G>T), GCT492GCC (3781T>C), ACA493TTT (3782A>T 3783C>T 3784A>T), AAA494GAC (3785A>G 3787A>C), TAT495AAA (3788T>A 3790T>A), TTT496TTC (3793T>C), CCC498CCA (3799C>A), TAC499TAT (3802C>T), TTA500CTG (3803T>C 3805A>G), TTC501GTG (3806T>G 3808C>G), AGA503ACC (3813G>C 3814A>C), TTT505GTC (3818T>G 3820T>C), AAG506ATT (3822A>T 3823G>T), ATA507GTT (3824A>G 3826A>T), TTG508TAT (3828T>A 3829G>T), GAC510GAT (3835C>T), CAC511CAT (3838C>T), AAA512GCT (3839A>G 3840A>C 3841A>T), CCA513GCT (3842C>G 3844A>T), CTA514CTT (3847A>T), CAG515AGG (3848C>A 3849A>G), TGG516TAC (3852G>A 3853G>C), ATG517CTA (3854A>C 3856G>A), ATG518TTT (3857A>T 3859G>T), AAC519AGC (3861A>G), TTA520AAG (3863T>A 3864T>A 3865A>G), AAA521AAT (3868A>T), GAC522GAT (3871C>T), CCA523GCA (3872C>G), AAC524AAA (3877C>A), TCA525CCA (3878T>C), CGA526AGG (3881C>A 3883A>G), ATG527CTG (3884A>C), ACT528ATT (3888C>T), CGA531ATT (3896C>A 3897G>T 3898A>T), CTA532TTA (3899C>T), CGA533CTA (3903G>T), CTA534CTC (3907A>C), AGT535CAA (3908A>C 3909G>A 3910T>A), TAT537TTC (3915A>T 3916T>C), TTC539ATT (3920T>A 3922C>T), TCT540AAG (3923T>A 3924C>A 3925T>G), GTA541GTT (3928A>T), GTG542AAA (3929G>A 3930T>A 3931G>A), TAC543GAC (3932T>G), AAG544AGA (3936A>G 3937G>A), AAA545AAG (3940A>G), GGA546GGG (3943A>G), AAG547TGT (3944A>T 3945A>G 3946G>T), TCT548GAG (3947T>G 3948C>A 3949T>G), AAT549AAC (3952T>C), ACC550CAA (3953A>C 3954C>A 3955C>A), AAC551ATT (3957A>T 3958C>T), GAT553GAC (3964T>C), GCC554CAC (3965G>C 3966C>A), CTT555CTA (3970T>A), TCT556TCC (3973T>C), CGT557AGT (3974C>A)

*: Inserts / Deletes / Misaligned / Frameshifts

#### Analysis details

This analysis was performed with panviral2.64

### NGS Details (UN24): Cassava brown streak virus

#### Assembly

| | |
|---|---|
| **Coverage Length** | 525 (1 contig(s)) |
| **Depth Of Coverage** | 94.4 |
| **Number Of Reads** | 421 |
| **Reads Per Million** | 8.43 rpm (after QC) |
| **Ambiguities** | 0 |
| **Assembly Method** | de novo + reference guided assembly |
| **Consensus Caller** | Bcf Tools |

#### Coverage Map

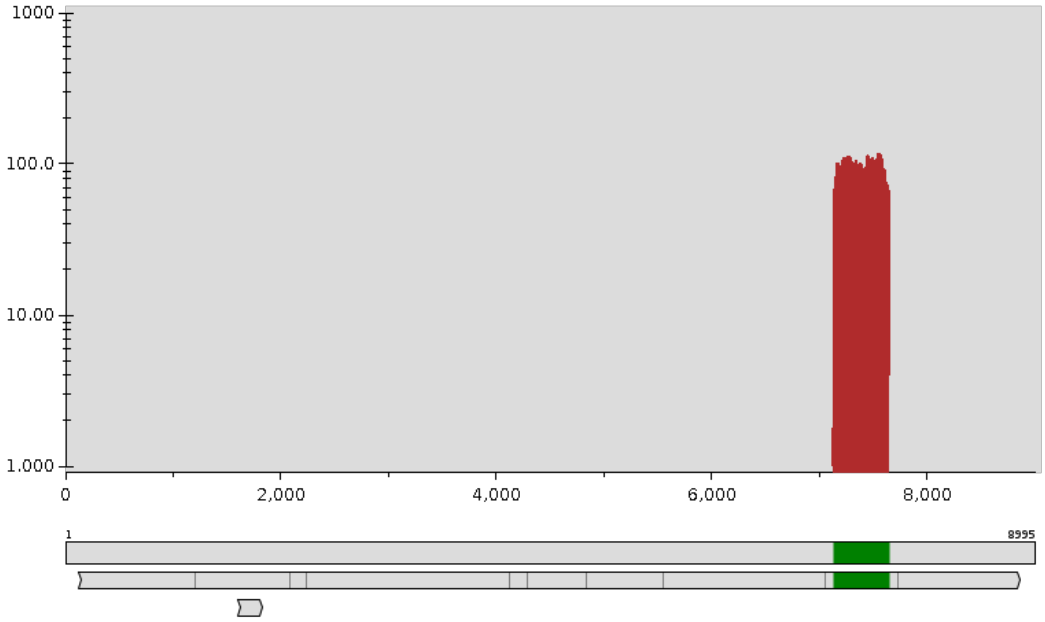

#### Assignment

| | |
|---|---|
| **Type** | Cassava brown streak virus (Taxonomy ID: 137758) |
| **Reference Genome** | NC_012698.2 |
| **NT Identity (%)** | 60.7619 |
| **AA Identity (%)** | 57.7143 |
| **Number Of Stop Codons** | 0 |
| **Number Of CDS** | 2 |

#### Alignment

| | |
|---|---|
| **Alignment Score** | 226.0 (NT) + 736.0 (AA) = 962.0 |
| **Concordance (%)** | 42.4724 |

| Alignment Method | Global, seeded, nucleotide + amino acids (AGA) |
|---|---|

#### Genome Region

Sequence starts at position 7126 and ends at position 7650 relative to NC_012698.2 reference sequence.

#### Alignment Detailed Statistics

| | Begin | End | Coverage | Score | Concordance | Matches | Identities | I/D/M/F* | Stop Codons |
|---|---|---|---|---|---|---|---|---|---|
| **NT** | **7126** | **7650** | **5.8%** | **226** | **21.5%** | **525 (100%)** | **319 (60.8%)** | **0/0** | |
| Mutations: | 7130A>G, 7133A>G, 7134A>G, 7136T>G, 7142C>T, 7145A>T, 7148A>C, 7151G>A, 7157A>T, 7158C>A, 7163A>G, 7167A>G, 7175A>T, 7177A>G, 7178A>G, 7179C>G, 7180A>C, 7184A>T, 7185T>C, 7190T>C, 7192C>A, 7193C>G, 7194A>T, 7202A>C, 7203A>T, 7205C>T, 7206A>C, 7207T>A, 7208C>G, 7211T>C, 7213G>T, 7214G>T, 7217A>G, 7218G>C, 7229A>T, 7234C>T, 7235A>T, 7241A>G, 7242A>G, 7243C>A, 7244A>G, 7245G>C, 7246T>C, 7250G>A, 7253A>T, 7256T>A, 7257A>T, 7258T>C, 7270T>C, 7271A>T, 7272C>A, 7273G>A, 7274T>A, 7275G>A, 7277A>T, 7280G>T, 7281T>G, 7283T>C, 7284G>A, 7286G>A, 7287T>G, 7288T>A, 7290A>G, 7292T>G, 7293G>A, 7294G>A, 7295A>T, 7298G>A, 7304T>G, 7307C>A, 7310A>T, 7313A>G, 7319A>T, 7320A>T, 7323T>C, 7325G>C, 7331T>C, 7332G>A, 7334T>C, 7337T>C, 7340C>T, 7343T>G, 7355A>G, 7358A>T, 7361C>T, 7364T>C, 7374A>T, 7377G>C, 7380G>A, 7381G>A, 7385T>C, 7388A>T, 7390T>A, 7391A>T, 7397A>T, 7398T>C, 7400A>T, 7401T>A, 7403T>C, 7406G>C, 7409G>A, 7410G>T, 7413G>A, 7414A>T, 7416C>G, 7418G>T, 7422C>G, 7424A>G, 7425A>G, 7432T>C, 7433G>A, 7436T>A, 7437A>T, 7438G>A, 7442T>C, 7443C>A, 7445C>G, 7448C>T, 7449G>A, 7451G>C, 7455G>T, 7457T>A, 7458T>C, 7462T>C, 7463A>T, 7464A>C, 7465A>T, 7467A>G, 7468A>G, 7469A>G, 7470G>C, 7471T>C, 7472T>A, 7473G>A, 7474G>A, 7476G>A, 7477A>C, 7481T>G, 7484T>G, 7487A>G, 7489T>C, 7490C>T, 7494A>G, 7495A>T, 7496G>T, 7499T>A, 7500G>A, 7501T>A, 7502G>A, 7503C>A, 7504T>C, 7505A>G, 7506A>C, 7507G>T, 7508A>G, 7511T>A, 7512G>A, 7517T>A, 7520A>G, 7521A>C, 7522T>C, 7523G>A, 7524C>G, 7526A>T, 7527C>A, 7529C>G, 7535A>C, 7539T>A, 7540C>A, 7541A>T, 7544T>C, 7547G>A, 7553C>T, 7556A>G, 7559T>A, 7562C>T, 7568A>T, 7570T>A, 7572A>G, 7573A>G, 7576G>A, 7577G>T, 7578A>G, 7579G>A, 7580A>C, 7581A>C, 7586A>T, 7588T>A, 7589T>C, 7599A>C, 7600T>C, 7601G>C, 7602G>A, 7603C>A, 7604C>G, 7607G>A, 7610G>A, 7616T>C, 7618T>A, 7622A>T, 7628T>C, 7631A>C, 7632T>G, 7633T>G, 7635C>A, 7636G>A | | | | | | | | |

##### CDS

| | Begin | End | Coverage | Score | Concordance | Matches | Identities | I/D/M/F* | Stop Codons |
|---|---|---|---|---|---|---|---|---|---|
| CBSV_gp1 | 2335 | 2509 | 6.0% | 736 | 60.0% | 175 (100%) | 101 (57.7%) | 0/0/0/0 | 0 |
| Protein mutations: | I2337V (7134A>G 7136T>G), Q2345K (7158C>A), K2348E (7167A>G), K2351R (7177A>G 7178A>G), Q2352A (7179C>G 7180A>C), F2354L (7185T>C), P2356Q (7192C>A 7193C>G), T2357S (7194A>T), I2360F (7203A>T 7205C>T), I2361Q (7206A>C 7207T>A 7208C>G), R2363L (7213G>T 7214G>T), V2365L (7218G>C), P2370L (7234C>T 7235A>T), T2373E (7242A>G 7243C>A 7244A>G), V2374P (7245G>C 7246T>C), E2376D (7253A>T), I2378S (7257A>T 7258T>C), V2382A (7270T>C 7271A>T), R2383K (7272C>A 7273G>A 7274T>A), V2384I (7275G>A 7277A>T), S2386A (7281T>G 7283T>C), E2387K (7284G>A 7286G>A), L2388E (7287T>G 7288T>A), I2389V (7290A>G 7292T>G), G2390N (7293G>A 7294G>A 7295A>T), S2399C (7320A>T), D2403N (7332G>A 7334T>C), N2406K (7343T>G), M2417L (7374A>T), E2418Q (7377G>C), G2419K (7380G>A 7381G>A), L2422H (7390T>A 7391A>T), Y2426N (7401T>A 7403T>C), K2427N (7406G>C), V2429L (7410G>T), E2430M (7413G>A 7414A>T), P2431A (7416C>G 7418G>T), Q2433E (7422C>G 7424A>G), N2434D (7425A>G), M2436T (7432T>C 7433G>A), S2438Y (7437A>T 7438G>A), L2440M (7443C>A 7445C>G), V2442I (7449G>A 7451G>C), A2444S (7455G>T 7457T>A), F2445L (7458T>C), V2446A (7462T>C 7463A>T), N2447L (7464A>C 7465A>T), K2448G (7467A>G 7468A>G 7469A>G), V2449P (7470G>C 7471T>C 7472T>A), G2450N (7473G>A 7474G>A), D2451T (7476G>A 7477A>C), D2452E (7481T>G), I2454M (7487A>G), I2455T (7489T>C 7490C>T), K2457V (7494A>G 7495A>T 7496G>T), V2459K (7500G>A 7501T>A 7502G>A), L2460T (7503C>A 7504T>C 7505A>G), R2461L (7506A>C 7507G>T 7508A>G), E2463K (7512G>A), M2466P (7521A>C 7522T>C 7523G>A), P2467A (7524C>G 7526A>T), S2472N (7539T>A 7540C>A 7541A>T), L2482H (7570T>A), N2483G (7572A>G 7573A>G), W2484Y (7576G>A 7577G>T), R2485D (7578A>G 7579G>A 7580A>C), K2486Q (7581A>C), F2488Y (7588T>A 7589T>C), M2492P (7599A>C 7600T>C 7601G>C), A2493K (7602G>A 7603C>A 7604C>G), M2498K (7618T>A), F2503G (7632T>G 7633T>G), R2504K (7635C>A 7636G>A) | | | | | | | | |
| Codon mutations: | GCA2335GCG (7130A>G), CCA2336CCG (7133A>G), ATT2337GTG (7134A>G 7136T>G), TTC2339TTT (7142C>T), GTA2340GTT (7145A>T), ACA2341ACC (7148A>C), GGG2342GGA (7151G>A), GCA2344GCT (7157A>T), CAA2345AAA (7158C>A), AAA2346AAG (7163A>G), AAG2348GAG (7167A>G), GTA2350GTT (7175A>T), AAA2351AGG (7177A>G 7178A>G), CAA2352GCA (7179C>G 7180A>C), ATA2353ATT (7184A>T), TTT2354CTT (7185T>C), GGT2355GGC (7190T>C), CCC2356CAG (7192C>A 7193C>G), ACT2357TCT (7194A>T), CCA2359CCC (7202A>C), ATC2360TTT (7203A>T 7205C>T), ATC2361CAG (7206A>C 7207T>A 7208C>G), TCT2362TCC (7211T>C), CGG2363CTT (7213G>T 7214G>T), AAA2364AAG (7217A>G), GTT2365CTT (7218G>C), CCA2368CCT (7229A>T), CCA2370CTT (7234C>T 7235A>T), GGA2372GGG (7241A>G), ACA2373GAG (7242A>G 7243C>A 7244A>G), GTT2374CCT (7245G>C 7246T>C), GAG2375GAA (7250G>A), GAA2376GAT (7253A>T), ATT2377ATA (7256T>A), ATC2378TCC (7257A>T 7258T>C), GTA2382GCT (7270T>C 7271A>T), CGT2383AAA (7272C>A 7273G>A 7274T>A), GTA2384ATT (7275G>A 7277A>T), GCG2385GCT (7280G>T), TCT2386GCC (7281T>G 7283T>C), GAG2387AAA (7284G>A 7286G>A), TTG2388GAG (7287T>G 7288T>A), ATT2389GTG (7290A>G 7292T>G), GGA2390AAT (7293G>A 7294G>A 7295A>T), GGG2391GGA (7298G>A), GTT2393GTG (7304T>G), CTC2394CTA (7307C>A), GTA2395GTT (7310A>T), GAA2396GAG (7313A>G), ACA2398ACT (7319A>T), AGT2399TGT (7320A>T), TTG2400CTC (7323T>C 7325G>C), TTT2402TTC (7331T>C), GAT2403AAC (7332G>A 7334T>C), GCT2404GCC (7337T>C), CTC2405CTT (7340C>T), AAT2406AAG (7343T>G), GGA2410GGG (7355A>G), CCA2411CCT (7358A>T), TAC2412TAT (7361C>T), ATT2413ATC (7364T>C), ATG2417TTG (7374A>T), GAA2418CAA (7377G>C), GGG2419AAG (7380G>A 7381G>A), ATT2420ATC (7385T>C), GGA2421GGT (7388A>T), CTA2422CAT (7390T>A 7391A>T), GGA2424GGT (7397A>T), TTA2425CTT (7398T>C 7400A>T), TAT2426AAC (7401T>A 7403T>C), AAG2427AAC (7406G>C), TTG2428TTA (7409G>A), GTG2429TTG (7410G>T), GAG2430ATG (7413G>A 7414A>T), CCG2431GCT (7416C>G 7418G>T), CAA2433GAG (7422C>G 7424A>G), AAT2434GAT (7425A>G), ATG2436ACA (7432T>C 7433G>A), GCT2437GCA (7436T>A), AGT2438TAT (7437A>T 7438G>A), GCT2439GCC (7442T>C), CTC2440ATG (7443C>A 7445C>G), TGC2441TGT (7448C>T), GTG2442ATC (7449G>A 7451G>C), GCT2444TCA (7455G>T 7457T>A), TTT2445CTT (7458T>C), GTA2446GCT (7462T>C 7463A>T), AAT2447CTT (7464A>C 7465A>T), AAA2448GGG (7467A>G 7468A>G 7469A>G), GTT2449CCA (7470G>C 7471T>C 7472T>A), GGT2450AAT (7473G>A 7474G>A), GAT2451ACT (7476G>A 7477A>C), GAT2452GAG (7481T>G), CCT2453CCG (7484T>G), ATA2454ATG (7487A>G), ATC2455ACT (7489T>C 7490C>T), AAG2457GTT (7494A>G 7495A>T 7496G>T), GGT2458GGA (7499T>A), GTG2459AAA (7500G>A 7501T>A 7502G>A), CTA2460ACG (7503C>A 7504T>C 7505A>G), AGA2461CTG (7506A>C 7507G>T 7508A>G), GGT2462GGA (7511T>A), GAG2463AAG (7512G>A), ATT2464ATA (7517T>A), GTA2465GTG (7520A>G), ATG2466CCA (7521A>C 7522T>C 7523G>A), CCA2467GCT (7524C>G 7526A>T), CGC2468AGG (7527C>A 7529C>G), CCA2470CCC (7535A>C), TCA2472AAT (7539T>A 7540C>A 7541A>T), TTT2473TTC (7544T>C), GGG2474GGA (7547G>A), GAC2476GAT (7553C>T), CCA2477CCG (7556A>G), ATT2478ATA (7559T>A), TTC2479TTT (7562C>T), CCA2481CCT (7568A>T), CTT2482CAT (7570T>A), AAC2483GGC (7572A>G 7573A>G), TGG2484TAT (7576G>A 7577G>T), AGA2485GAC (7578A>G 7579G>A 7580A>C), AAG2486CAG (7581A>C), ACA2487ACT (7586A>T), TTT2488TAC (7588T>A 7589T>C), ATG2492CCC (7599A>C 7600T>C 7601G>C), GCC2493AAG (7602G>A 7603C>A 7604C>G), GAG2494GAA (7607G>A), GAG2495GAA (7610G>A), AAT2497AAC (7616T>C), ATG2498AAG (7618T>A), ATA2499ATT (7622A>T), CAT2501CAC (7628T>C), CGA2502CGC (7631A>C), TTT2503GGT (7632T>G 7633T>G), CGA2504AAA (7635C>A 7636G>A) | | | | | | | | |

##### Proteins

| | Begin | End | Coverage | Score | Concordance | Matches | Identities | I/D/M/F* | Stop Codons |
|---|---|---|---|---|---|---|---|---|---|
| polyprotein (YP_007027011.1) | 2335 | 2509 | 6.0% | 736 | 60.0% | 175 (100%) | 101 (57.7%) | 0/0/0/0 | 0 |
| Protein mutations: | I2337V (7134A>G 7136T>G), Q2345K (7158C>A), K2348E (7167A>G), K2351R (7177A>G 7178A>G), Q2352A (7179C>G 7180A>C), F2354L (7185T>C), P2356Q (7192C>A 7193C>G), T2357S (7194A>T), I2360F (7203A>T 7205C>T), I2361Q (7206A>C 7207T>A 7208C>G), R2363L (7213G>T 7214G>T), V2365L (7218G>C), P2370L (7234C>T 7235A>T), T2373E (7242A>G 7243C>A 7244A>G), V2374P (7245G>C 7246T>C), E2376D (7253A>T), I2378S (7257A>T 7258T>C), V2382A (7270T>C 7271A>T), R2383K (7272C>A 7273G>A 7274T>A), V2384I (7275G>A 7277A>T), S2386A (7281T>G 7283T>C), E2387K (7284G>A 7286G>A), L2388E (7287T>G 7288T>A), I2389V (7290A>G 7292T>G), G2390N (7293G>A 7294G>A 7295A>T), S2399C (7320A>T), D2403N (7332G>A 7334T>C), N2406K (7343T>G), M2417L (7374A>T), E2418Q (7377G>C), G2419K (7380G>A 7381G>A), L2422H (7390T>A 7391A>T), Y2426N (7401T>A 7403T>C), K2427N (7406G>C), V2429L (7410G>T), E2430M (7413G>A 7414A>T), P2431A (7416C>G 7418G>T), Q2433E (7422C>G 7424A>G), N2434D (7425A>G), M2436T (7432T>C 7433G>A), S2438Y (7437A>T 7438G>A), L2440M (7443C>A 7445C>G), V2442I (7449G>A 7451G>C), A2444S (7455G>T 7457T>A), F2445L (7458T>C), V2446A (7462T>C 7463A>T), N2447L (7464A>C 7465A>T), K2448G (7467A>G 7468A>G 7469A>G), V2449P (7470G>C 7471T>C 7472T>A), G2450N (7473G>A 7474G>A), D2451T (7476G>A 7477A>C), D2452E (7481T>G), I2454M (7487A>G), I2455T (7489T>C 7490C>T), K2457V (7494A>G 7495A>T 7496G>T), V2459K (7500G>A 7501T>A 7502G>A), L2460T (7503C>A 7504T>C 7505A>G), R2461L (7506A>C 7507G>T 7508A>G), E2463K (7512G>A), M2466P (7521A>C 7522T>C 7523G>A), P2467A (7524C>G 7526A>T), S2472N (7539T>A 7540C>A 7541A>T), L2482H (7570T>A), N2483G (7572A>G 7573A>G), W2484Y (7576G>A 7577G>T), R2485D (7578A>G 7579G>A 7580A>C), K2486Q (7581A>C), F2488Y (7588T>A 7589T>C), M2492P (7599A>C 7600T>C 7601G>C), A2493K (7602G>A 7603C>A 7604C>G), M2498K (7618T>A), F2503G (7632T>G 7633T>G), R2504K (7635C>A 7636G>A) | | | | | | | | |

| | Begin | End | Coverage | Score | Concordance | Matches | Identities | I/D/M/F* | Stop Codons |
|---|---|---|---|---|---|---|---|---|---|
| **NT** | **7126** | **7650** | **5.8%** | **226** | **21.5%** | **525 (100%)** | **319 (60.8%)** | **0/0** | |
| Codon mutations: | GCA2335GCG (7130A>G), CCA2336CCG (7133A>G), ATT2337GTG (7134A>G 7136T>G), TTC2339TTT (7142C>T), GTA2340GTT (7145A>T), ACA2341ACC (7148A>C), GGG2342GGA (7151G>A), GCA2344GCT (7157A>T), CAA2345AAA (7158C>A), AAA2346AAG (7163A>G), AAG2348GAG (7167A>G), GTA2350GTT (7175A>T), AAA2351AGG (7177A>G 7178A>G), CAA2352GCA (7179C>G 7180A>C), ATA2353ATT (7184A>T), TTT2354CTT (7185T>C), GGT2355GGC (7190T>C), CCC2356CAG (7192C>A 7193C>G), ACT2357TCT (7194A>T), CCA2359CCC (7202A>C), ATC2360TTT (7203A>T 7205C>T), ATC2361CAG (7206A>C 7207T>A 7208C>G), TCT2362TCC (7211T>C), CGG2363CTT (7213G>T 7214G>T), AAA2364AAG (7217A>G), GTT2365CTT (7218G>C), CCA2368CCT (7229A>T), CCA2370CTT (7234C>T 7235A>T), GGA2372GGG (7241A>G), ACA2373GAG (7242A>G 7243C>A 7244A>G), GTT2374CCT (7245G>C 7246T>C), GAG2375GAA (7250G>A), GAA2376GAT (7253A>T), ATT2377ATA (7256T>A), ATC2378TCC (7257A>T 7258T>C), GTA2382GCT (7270T>C 7271A>T), CGT2383AAA (7272C>A 7273G>A 7274T>A), GTA2384ATT (7275G>A 7277A>T), GCG2385GCT (7280G>T), TCT2386GCC (7281T>G 7283T>C), GAG2387AAA (7284G>A 7286G>A), TTG2388GAG (7287T>G 7288T>A), ATT2389GTG (7290A>G 7292T>G), GGA2390AAT (7293G>A 7294G>A 7295A>T), GGG2391GGA (7298G>A), GTT2393GTG (7304T>G), CTC2394CTA (7307C>A), GTA2395GTT (7310A>T), GAA2396GAG (7313A>G), ACA2398ACT (7319A>T), AGT2399TGT (7320A>T), TTG2400CTC (7323T>C 7325G>C), TTT2402TTC (7331T>C), GAT2403AAC (7332G>A 7334T>C), GCT2404GCC (7337T>C), CTC2405CTT (7340C>T), AAT2406AAG (7343T>G), GGA2410GGG (7355A>G), CCA2411CCT (7358A>T), TAC2412TAT (7361C>T), ATT2413ATC (7364T>C), ATG2417TTG (7374A>T), GAA2418CAA (7377G>C), GGG2419AAG (7380G>A 7381G>A), ATT2420ATC (7385T>C), GGA2421GGT (7388A>T), CTA2422CAT (7390T>A 7391A>T), GGA2424GGT (7397A>T), TTA2425CTT (7398T>C 7400A>T), TAT2426AAC (7401T>A 7403T>C), AAG2427AAC (7406G>C), TTG2428TTA (7409G>A), GTG2429TTG (7410G>T), GAG2430ATG (7413G>A 7414A>T), CCG2431GCT (7416C>G 7418G>T), CAA2433GAG (7422C>G 7424A>G), AAT2434GAT (7425A>G), ATG2436ACA (7432T>C 7433G>A), GCT2437GCA (7436T>A), AGT2438TAT (7437A>T 7438G>A), GCT2439GCC (7442T>C), CTC2440ATG (7443C>A 7445C>G), TGC2441TGT (7448C>T), GTG2442ATC (7449G>A 7451G>C), GCT2444TCA (7455G>T 7457T>A), TTT2445CTT (7458T>C), GTA2446GCT (7462T>C 7463A>T), AAT2447CTT (7464A>C 7465A>T), AAA2448GGG (7467A>G 7468A>G 7469A>G), GTT2449CCA (7470G>C 7471T>C 7472T>A), GGT2450AAT (7473G>A 7474G>A), GAT2451ACT (7476G>A 7477A>C), GAT2452GAG (7481T>G), CCT2453CCG (7484T>G), ATA2454ATG (7487A>G), ATC2455ACT (7489T>C 7490C>T), AAG2457GTT (7494A>G 7495A>T 7496G>T), GGT2458GGA (7499T>A), GTG2459AAA (7500G>A 7501T>A 7502G>A), CTA2460ACG (7503C>A 7504T>C 7505A>G), AGA2461CTG (7506A>C 7507G>T 7508A>G), GGT2462GGA (7511T>A), GAG2463AAG (7512G>A), ATT2464ATA (7517T>A), GTA2465GTG (7520A>G), ATG2466CCA (7521A>C 7522T>C 7523G>A), CCA2467GCT (7524C>G 7526A>T), CGC2468AGG (7527C>A 7529C>G), CCA2470CCC (7535A>C), TCA2472AAT (7539T>A 7540C>A 7541A>T), TTT2473TTC (7544T>C), GGG2474GGA (7547G>A), GAC2476GAT (7553C>T), CCA2477CCG (7556A>G), ATT2478ATA (7559T>A), TTC2479TTT (7562C>T), CCA2481CCT (7568A>T), CTT2482CAT (7570T>A), AAC2483GGC (7572A>G 7573A>G), TGG2484TAT (7576G>A 7577G>T), AGA2485GAC (7578A>G 7579G>A 7580A>C), AAG2486CAG (7581A>C), ACA2487ACT (7586A>T), TTT2488TAC (7588T>A 7589T>C), ATG2492CCC (7599A>C 7600T>C 7601G>C), GCC2493AAG (7602G>A 7603C>A 7604C>G), GAG2494GAA (7607G>A), GAG2495GAA (7610G>A), AAT2497AAC (7616T>C), ATG2498AAG (7618T>A), ATA2499ATT (7622A>T), CAT2501CAC (7628T>C), CGA2502CGC (7631A>C), TTT2503GGT (7632T>G 7633T>G), CGA2504AAA (7635C>A 7636G>A) | | | | | | | | |
| HAM1-like protein (YP_007032446.1) | 27 | 201 | 77.4% | 736 | 60.0% | 175 (100%) | 101 (57.7%) | 0/0/0/0 | 0 |
| Protein mutations: | I29V (7134A>G 7136T>G), Q37K (7158C>A), K40E (7167A>G), K43R (7177A>G 7178A>G), Q44A (7179C>G 7180A>C), F46L (7185T>C), P48Q (7192C>A 7193C>G), T49S (7194A>T), I52F (7203A>T 7205C>T), I53Q (7206A>C 7207T>A 7208C>G), R55L (7213G>T 7214G>T), V57L (7218G>C), P62L (7234C>T 7235A>T), T65E (7242A>G 7243C>A 7244A>G), V66P (7245G>C 7246T>C), E68D (7253A>T), I70S (7257A>T 7258T>C), V74A (7270T>C 7271A>T), R75K (7272C>A 7273G>A 7274T>A), V76I (7275G>A 7277A>T), S78A (7281T>G 7283T>C), E79K (7284G>A 7286G>A), L80E (7287T>G 7288T>A), I81V (7290A>G 7292T>G), G82N (7293G>A 7294G>A 7295A>T), S91C (7320A>T), D95N (7332G>A 7334T>C), N98K (7343T>G), M109L (7374A>T), E110Q (7377G>C), G111K (7380G>A 7381G>A), L114H (7390T>A 7391A>T), Y118N (7401T>A 7403T>C), K119N (7406G>C), V121L (7410G>T), E122M (7413G>A 7414A>T), P123A (7416C>G 7418G>T), Q125E (7422C>G 7424A>G), N126D (7425A>G), M128T (7432T>C 7433G>A), S130Y (7437A>T 7438G>A), L132M (7443C>A 7445C>G), V134I (7449G>A 7451G>C), A136S (7455G>T 7457T>A), F137L (7458T>C), V138A (7462T>C 7463A>T), N139L (7464A>C 7465A>T), K140G (7467A>G 7468A>G 7469A>G), V141P (7470G>C 7471T>C 7472T>A), G142N (7473G>A 7474G>A), D143T (7476G>A 7477A>C), D144E (7481T>G), I146M (7487A>G), I147T (7489T>C 7490C>T), K149V (7494A>G 7495A>T 7496G>T), V151K (7500G>A 7501T>A 7502G>A), L152T (7503C>A 7504T>C 7505A>G), R153L (7506A>C 7507G>T 7508A>G), E155K (7512G>A), M158P (7521A>C 7522T>C 7523G>A), P159A (7524C>G 7526A>T), S164N (7539T>A 7540C>A 7541A>T), L174H (7570T>A), N175G (7572A>G 7573A>G), W176Y (7576G>A 7577G>T), R177D (7578A>G 7579G>A 7580A>C), K178Q (7581A>C), F180Y (7588T>A 7589T>C), M184P (7599A>C 7600T>C 7601G>C), A185K (7602G>A 7603C>A 7604C>G), M190K (7618T>A), F195G (7632T>G 7633T>G), R196K (7635C>A 7636G>A) | | | | | | | | |
| Codon mutations: | GCA27GCG (7130A>G), CCA28CCG (7133A>G), ATT29GTG (7134A>G 7136T>G), TTC31TTT (7142C>T), GTA32GTT (7145A>T), ACA33ACC (7148A>C), GGG34GGA (7151G>A), GCA36GCT (7157A>T), CAA37AAA (7158C>A), AAA38AAG (7163A>G), AAG40GAG (7167A>G), GTA42GTT (7175A>T), AAA43AGG (7177A>G 7178A>G), CAA44GCA (7179C>G 7180A>C), ATA45ATT (7184A>T), TTT46CTT (7185T>C), GGT47GGC (7190T>C), CCC48CAG (7192C>A 7193C>G), ACT49TCT (7194A>T), CCA51CCC (7202A>C), ATC52TTT (7203A>T 7205C>T), ATC53CAG (7206A>C 7207T>A 7208C>G), TCT54TCC (7211T>C), CGG55CTT (7213G>T 7214G>T), AAA56AAG (7217A>G), GTT57CTT (7218G>C), CCA60CCT (7229A>T), CCA62CTT (7234C>T 7235A>T), GGA64GGG (7241A>G), ACA65GAG (7242A>G 7243C>A 7244A>G), GTT66CCT (7245G>C 7246T>C), GAG67GAA (7250G>A), GAA68GAT (7253A>T), ATT69ATA (7256T>A), ATC70TCC (7257A>T 7258T>C), GTA74GCT (7270T>C 7271A>T), CGT75AAA (7272C>A 7273G>A 7274T>A), GTA76ATT (7275G>A 7277A>T), GCG77GCT (7280G>T), TCT78GCC (7281T>G 7283T>C), GAG79AAA (7284G>A 7286G>A), TTG80GAG (7287T>G 7288T>A), ATT81GTG (7290A>G 7292T>G), GGA82AAT (7293G>A 7294G>A 7295A>T), GGG83GGA (7298G>A), GTT85GTG (7304T>G), CTC86CTA (7307C>A), GTA87GTT (7310A>T), GAA88GAG (7313A>G), ACA90ACT (7319A>T), AGT91TGT (7320A>T), TTG92CTC (7323T>C 7325G>C), TTT94TTC (7331T>C), GAT95AAC (7332G>A 7334T>C), GCT96GCC (7337T>C), CTC97CTT (7340C>T), AAT98AAG (7343T>G), GGA102GGG (7355A>G), CCA103CCT (7358A>T), TAC104TAT (7361C>T), ATT105ATC (7364T>C), ATG109TTG (7374A>T), GAA110CAA (7377G>C), GGG111AAG (7380G>A 7381G>A), ATT112ATC (7385T>C), GGA113GGT (7388A>T), CTA114CAT (7390T>A 7391A>T), GGA116GGT (7397A>T), TTA117CTT (7398T>C 7400A>T), TAT118AAC (7401T>A 7403T>C), AAG119AAC (7406G>C), TTG120TTA (7409G>A), GTG121TTG (7410G>T), GAG122ATG (7413G>A 7414A>T), CCG123GCT (7416C>G 7418G>T), CAA125GAG (7422C>G 7424A>G), AAT126GAT (7425A>G), ATG128ACA (7432T>C 7433G>A), GCT129GCA (7436T>A), AGT130TAT (7437A>T 7438G>A), GCT131GCC (7442T>C), CTC132ATG (7443C>A 7445C>G), TGC133TGT (7448C>T), GTG134ATC (7449G>A 7451G>C), GCT136TCA (7455G>T 7457T>A), TTT137CTT (7458T>C), GTA138GCT (7462T>C 7463A>T), AAT139CTT (7464A>C 7465A>T), AAA140GGG (7467A>G 7468A>G 7469A>G), GTT141CCA (7470G>C 7471T>C 7472T>A), GGT142AAT (7473G>A 7474G>A), GAT143ACT (7476G>A 7477A>C), GAT144GAG (7481T>G), CCT145CCG (7484T>G), ATA146ATG (7487A>G), ATC147ACT (7489T>C 7490C>T), AAG149GTT (7494A>G 7495A>T 7496G>T), GGT150GGA (7499T>A), GTG151AAA (7500G>A 7501T>A 7502G>A), CTA152ACG (7503C>A 7504T>C 7505A>G), AGA153CTG (7506A>C 7507G>T 7508A>G), GGT154GGA (7511T>A), GAG155AAG (7512G>A), ATT156ATA (7517T>A), GTA157GTG (7520A>G), ATG158CCA (7521A>C 7522T>C 7523G>A), CCA159GCT (7524C>G 7526A>T), CGC160AGG (7527C>A 7529C>G), CCA162CCC (7535A>C), TCA164AAT (7539T>A 7540C>A 7541A>T), TTT165TTC (7544T>C), GGG166GGA (7547G>A), GAC168GAT (7553C>T), CCA169CCG (7556A>G), ATT170ATA (7559T>A), TTC171TTT (7562C>T), CCA173CCT (7568A>T), CTT174CAT (7570T>A), AAC175GGC (7572A>G 7573A>G), TGG176TAT (7576G>A 7577G>T), AGA177GAC (7578A>G 7579G>A 7580A>C), AAG178CAG (7581A>C), ACA179ACT (7586A>T), TTT180TAC (7588T>A 7589T>C), ATG184CCC (7599A>C 7600T>C 7601G>C), GCC185AAG (7602G>A 7603C>A 7604C>G), GAG186GAA (7607G>A), GAG187GAA (7610G>A), AAT189AAC (7616T>C), ATG190AAG (7618T>A), ATA191ATT (7622A>T), CAT193CAC (7628T>C), CGA194CGC (7631A>C), TTT195GGT (7632T>G 7633T>G), CGA196AAA (7635C>A 7636G>A) | | | | | | | | |

*: Inserts / Deletes / Misaligned / Frameshifts

#### Analysis details

This analysis was performed with panviral2.64

### NGS Details (UN24): Nodensvirus spm2

#### Assembly

| | |
|---|---|
| **Coverage Length** | 169 (1 contig(s)) |
| **Depth Of Coverage** | 194.4 |
| **Number Of Reads** | 374 |
| **Reads Per Million** | 7.49 rpm (after QC) |
| **Ambiguities** | 0 |
| **Assembly Method** | de novo + reference guided assembly |
| **Consensus Caller** | Bcf Tools |

#### Coverage Map

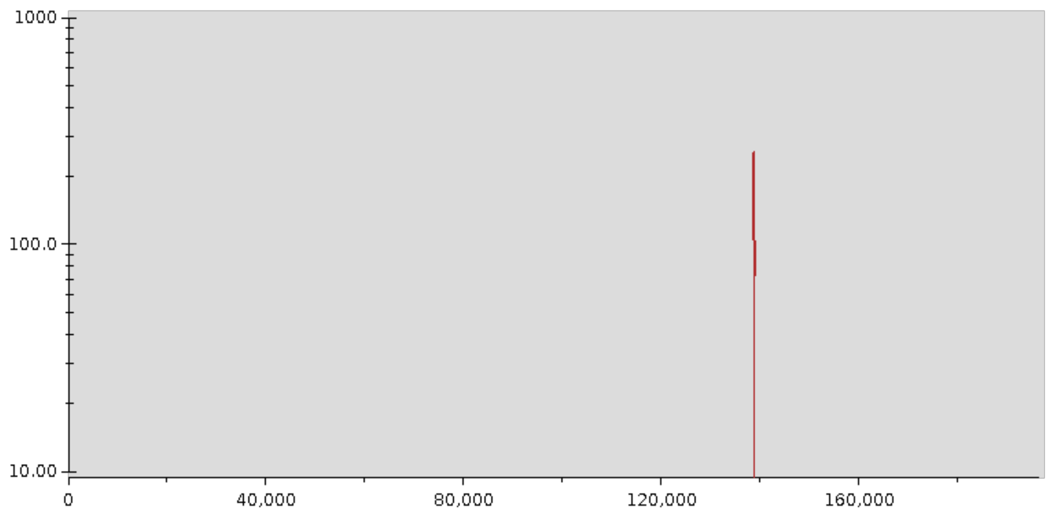

#### Assignment

| | |
|---|---|
| **Type** | Nodensvirus spm2 (Taxonomy ID: 2734126) |
| **Reference Genome** | NC_006820.1 |
| **NT Identity (%)** | 81.4371 |
| **AA Identity (%)** | 86.7925 |
| **Number Of Stop Codons** | 1 |
| **Number Of CDS** | 243 |

#### Alignment

| | |
|---|---|
| **Alignment Score** | 199.0 (NT) + 223.0 (AA) = 422.0 |
| **Concordance (%)** | 73.9635 |
| **Alignment Method** | Local, heuristic, nucleotide (BLASTN) |

#### Genome Region

Sequence starts at position 138722 and ends at position 138890 relative to NC_006820.1 reference sequence.

#### Alignment Detailed Statistics

| | Begin | End | Coverage | Score | Concordance | Matches | Identities | I/D/M/F* | Stop Codons |
|---|---|---|---|---|---|---|---|---|---|
| **NT** | **138722** | **138890** | **0.1%** | **199** | **61.6%** | **167 (98.8%)** | **136 (80.5%)** | **0/2** | |

Mutations: 138734T>C, 138740A>G, 138755C>T, 138756G>A, 138761C>T, 138773G>A, 138776C>T, 138778T>C, 138783_138784delCT, 138785G>A, 138788T>C, 138795C>A, 138797G>T, 138803C>T, 138812T>G, 138813C>G, 138816G>C, 138817T>A, 138824A>G, 138830C>T, 138836C>T, 138840G>A, 138842T>A, 138851A>G, 138854A>G, 138857A>T, 138858T>C, 138860G>A, 138864A>C, 138866A>T, 138869C>A, 138878G>T

*: Inserts / Deletes / Misaligned / Frameshifts

#### Analysis details

This analysis was performed with panviral2.64

### NGS Details (UN24): Rahariannevirus raharianne

#### Assembly

| | |
|---|---|
| **Coverage Length** | 4366 (11 contig(s)) |
| **Depth Of Coverage** | 5.0 |
| **Number Of Reads** | 176 |
| **Reads Per Million** | 3.52 rpm (after QC) |
| **Ambiguities** | 0 |
| **Assembly Method** | de novo + reference guided assembly |
| **Consensus Caller** | Bcf Tools |

#### Coverage Map

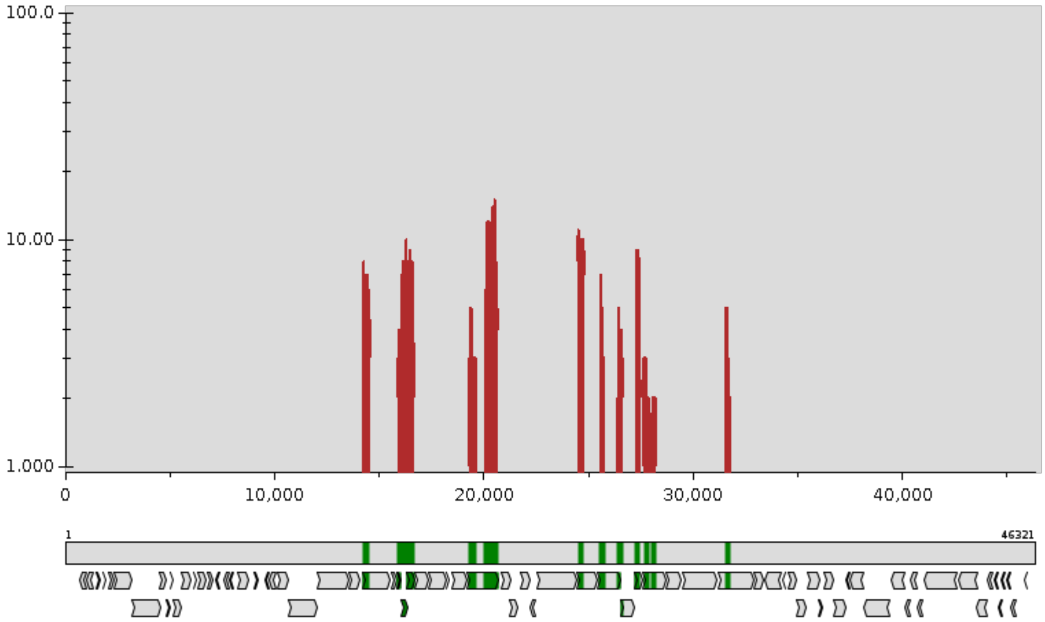

#### Assignment

| | |
|---|---|
| **Type** | Rahariannevirus raharianne (Taxonomy ID: 2846050) |
| **Reference Genome** | NC_054955.1 |
| **NT Identity (%)** | 70.758 |
| **AA Identity (%)** | 68.0603 |
| **Number Of Stop Codons** | 3 |
| **Number Of CDS** | 75 |

#### Alignment

| | |
|---|---|
| **Alignment Score** | 3582.0 (NT) + 6867.0 (AA) = 10449.0 |
| **Concordance (%)** | 57.8415 |

| Alignment Method | Local, heuristic, nucleotide (BLASTN) |
|---|---|

#### Genome Region

Sequence starts at position 14196 and ends at position 31794 relative to NC_054955.1 reference sequence.

#### Alignment Detailed Statistics

| | Begin | End | Coverage | Score | Concordance | Matches | Identities | I/D/M/F* | Stop Codons |
|---|---|---|---|---|---|---|---|---|---|
| NT | 14196 | 31794 | 9.4% | 3582 | 41.3% | 4358 (99.6%) | 3090 (70.6%) | 9/8 | |

Mutations:

14204C>T, 14205A>C, 14206G>A, 14207C>A, 14210T>C, 14219C>T, 14222T>C, 14231G>C, 14264T>C, 14270T>A, 14272C>G, 14285C>G, 14288C>G, 14289G>A, 14293T>A, 14294C>T, 14298C>T, 14299A>C, 14306A>C, 14309T>C, 14317G>A, 14318T>G, 14336T>C, 14339A>G, 14342C>A, 14357T>C, 14363T>C, 14366G>C, 14372G>T, 14375G>C, 14378C>G, 14381C>G, 14394C>G, 14397T>G, 14399G>C, 14400G>A, 14401C>G, 14402A>C, 14427T>A, 14432G>C, 14433T>A, 14434C>G, 14435G>C, 14447G>C, 14457T>A, 14462C>G, 14486G>C, 14500C>T, 14504G>C, 14516C>G, 14525G>C, 15865C>G, 15868T>C, 15870C>T, 15871C>G, 15872A>G, 15875C>G, 15878T>C, 15883C>G, 15884G>C, 15885A>G, 15886G>A, 15887T>G, 15888C>A, 15890A>T, 15895G>C, 15897G>T, 15898G>C, 15900T>G, 15901G>C, 15903A>T, 15904A>C, 15911A>C, 15913C>G, 15919A>C, 15920C>G, 15921A>T, 15924T>G, 15925C>G, 15926C>G, 15927A>C, 15930A>C, 15931A>T, 15934T>C, 15936C>A, 15939A>C, 15941G>A, 15947A>G, 15949C>A, 15950C>G, 15951C>T, 15952T>G, 15953G>C, 15954T>G, 15959G>A, 15961G>C, 15962G>C, 15963C>G, 15966C>A, 15974C>G, 15975A>C, 15976G>C, 15977C>A, 15981G>A, 15987G>C, 15988T>G, 15988_15989insATCAAGGCC, 15992T>C, 15993T>G, 15998G>A, 16001delA, 16002G>T, 16003C>G, 16014C>T, 16018G>A, 16025T>C, 16026T>C, 16035C>G, 16036A>C, 16037A>G, 16038A>C, 16040G>A, 16045A>G, 16047C>G, 16053T>C, 16057A>G, 16059G>T, 16061C>G, 16062C>G, 16065G>C, 16066C>A, 16067A>C, 16071G>C, 16074T>C, 16078T>A, 16086C>A, 16087A>G, 16088A>C, 16089G>C, 16091A>G, 16095T>C, 16099A>C, 16101C>G, 16113A>C, 16126A>C, 16128T>G, 16131T>G, 16134C>G, 16137T>G, 16140G>C, 16143G>C, 16146T>C, 16147G>A, 16149G>T, 16150G>C, 16151G>C, 16158C>G, 16161G>C, 16164C>T, 16170G>A, 16176G>C, 16177A>G, 16178C>T, 16179G>C, 16182C>T, 16183A>G, 16184C>G, 16190C>T, 16191T>C, 16198A>G, 16199A>T, 16204A>G, 16208G>C, 16209C>G, 16212G>C, 16218G>C, 16227G>C, 16231T>A, 16233A>C, 16239C>G, 16245T>C, 16251G>C, 16255C>T, 16258A>G, 16259A>C, 16260C>A, 16263C>G, 16270A>G, 16278T>C, 16282G>A, 16284T>C, 16293C>G, 16296T>C, 16297T>G, 16302T>C, 16305T>C, 16317C>T, 16319G>C, 16321G>C, 16323delG, 16335A>C, 16338C>G, 16339G>C, 16340C>A, 16341C>G, 16345A>C, 16346A>G, 16365T>C, 16368C>T, 16369G>T, 16371A>G, 16378A>G, 16380G>C, 16382T>A, 16383C>T, 16389G>C, 16395A>C, 16396T>A, 16397C>G, 16398G>T, 16399T>G, 16413T>C, 16414C>A, 16428G>C, 16431C>T, 16440T>G, 16443A>G, 16450A>C, 16453A>T, 16454A>C, 16455C>G, 16459A>C, 16464G>C, 16465G>C, 16467C>G, 16473A>G, 16478C>G, 16479T>C, 16483G>A, 16485G>C, 16494G>C, 16497T>C, 16500G>C, 16503T>C, 16512C>G, 16515T>G, 16518C>G, 16524G>C, 16527A>C, 16534T>G, 16535C>G, 16536T>C, 16537G>A, 16540A>T, 16541G>C, 16542C>G, 16545G>A, 16551C>T, 16557A>G, 16561C>A, 16566G>C, 16569C>G, 16576G>A, 16577T>C, 16578G>C, 16593A>G, 16594G>C, 16599A>C, 16602C>T, 16605C>T, 16608C>T, 16617T>C, 16618G>A, 16619C>G, 16629G>C, 16631G>T, 16632C>G, 16642G>C, 16648A>C, 16650G>C, 16654A>G, 16656T>G, 16662C>G, 16665G>C, 16668C>G, 16670G>C, 16671G>C, 16672C>T, 19263T>C, 19271A>G, 19274A>G, 19280G>C, 19283T>G, 19286A>G, 19290G>A, 19291T>C, 19292T>C, 19293T>C, 19294C>A, 19295C>G, 19296C>T, 19297T>A, 19298G>C, 19301C>G, 19319T>C, 19320G>A, 19322G>C, 19323C>T, 19325T>G, 19328T>C, 19329G>A, 19331T>C, 19334G>C, 19340C>G, 19343A>G, 19344A>T, 19352C>G, 19358C>G, 19362T>A, 19363C>G, 19364G>C, 19367G>C, 19370G>C, 19371C>A, 19372T>C, 19374A>G, 19381A>G, 19383A>C, 19384T>C, 19388G>T, 19389A>C, 19390T>A, 19392T>G, 19393A>C, 19397C>G, 19398C>G, 19399A>C, 19400G>C, 19401G>A, 19402C>T, 19410G>C, 19411C>A, 19413A>G, 19414T>C, 19424T>C, 19426C>A, 19427C>G, 19430T>C, 19434A>C, 19437G>A, 19438G>T, 19439C>G, 19445T>C, 19446G>A, 19448G>C, 19449G>T, 19452A>G, 19457A>G, 19458G>A, 19459T>C, 19460G>C, 19461C>A, 19462A>T, 19472G>C, 19482C>A, 19483G>A, 19484C>A, 19485T>G, 19486G>C, 19493C>G, 19496C>G, 19512A>G, 19513C>T, 19520G>C, 19521T>A, 19522C>G, 19523G>C, 19525C>T, 19526G>C, 19527A>G, 19528T>C, 19529C>G, 19534A>T, 19542G>T, 19545G>A, 19547G>C, 19550G>C, 19551C>T, 19553G>C, 19557G>T, 19559G>C, 19574G>C, 19575C>T, 19577G>C, 19584T>G, 19585C>G, 19586G>C, 19587G>A, 19589G>C, 19595C>G, 19596G>A, 19601C>G, 19602G>T, 19604C>G, 19607G>C, 19611A>T, 19612G>C, 19613C>G, 19614G>C, 19615C>A, 19616C>G, 19620A>T, 19621A>C, 19622T>G, 19626T>C, 19627G>A, 19633T>C, 19634A>C, 19635A>G, 19636C>T, 19640C>T, 19641G>T, 20005T>C, 20006C>G, 20013A>G, 20016G>A, 20017G>T, 20018C>G, 20019G>A, 20020T>C, 20028G>A, 20029C>G, 20039G>C, 20043A>G, 20044G>C, 20045C>G, 20051C>G, 20053C>T, 20056C>G, 20057G>C, 20058A>G, 20064G>A, 20065C>G, 20076A>G, 20078C>G, 20091T>G, 20092T>C, 20102G>T, 20108T>C, 20115G>T, 20116T>G, 20117G>C, 20124T>C, 20132G>C, 20133G>T, 20134T>A, 20139A>C, 20140G>A, 20141C>G, 20142G>A, 20144T>C, 20145A>C, 20146C>T, 20147A>T, 20151C>T, 20157T>A, 20158C>G, 20159G>C, 20165A>G, 20166G>T, 20168G>C, 20174C>G, 20177T>C, 20180T>C, 20187T>G, 20189G>C, 20211A>T, 20229T>C, 20231T>G, 20235G>A, 20237G>C, 20240G>C, 20253T>A, 20254C>G, 20255G>C, 20268A>C, 20269C>A, 20279C>G, 20282C>T, 20286T>C, 20292T>C, 20293C>A, 20298A>G, 20299T>G, 20301C>A, 20302A>T, 20303G>C, 20306G>C, 20309T>C, 20336C>G, 20343T>A, 20344C>G, 20345G>C, 20355T>G, 20356G>C, 20357C>G, 20360G>C, 20362C>T, 20363G>C, 20369T>C, 20385G>C, 20388C>G, 20394C>A, 20396G>C, 20411C>G, 20433A>T, 20434G>A, 20439G>C, 20444T>C, 20461A>T, 20463G>A, 20468G>T, 20477A>G, 20478T>A, 20479C>G, 20480G>C, 20481G>T, 20483C>G, 20489G>C, 20493G>A, 20495G>T, 20501G>C, 20502G>A, 20503C>T, 20504C>G, 20511C>A, 20513G>C, 20514T>A, 20515C>G, 20516G>C, 20517A>G, 20518A>C, 20526T>G, 20527C>A, 20530G>A, 20532A>C, 20535G>C, 20546C>A, 20547A>C, 20557C>A, 20559G>C, 20560T>C, 20561C>G, 20564T>C, 20571C>G, 20573A>G, 20574T>G, 20576G>C, 20578T>A, 20580T>A, 20581C>G, 20582G>C, 20588A>G, 20589A>G, 20591C>G, 20597T>G, 20598A>G, 20599A>C, 20610A>G, 20611T>C, 20613A>T, 20614A>C, 20615C>G, 20618G>C, 20624G>C, 20627T>G, 20634C>A, 20636T>G, 20642T>C, 20648C>G, 20656T>A, 20657T>G, 20664T>G, 20666G>C, 20669T>C, 20682A>C, 20685G>A, 20687G>C, 20690T>C, 20691C>A, 20693T>C, 24489A>G, 24491C>T, 24492G>C, 24495A>T, 24496G>C, 24500A>G, 24502T>G, 24505C>G, 24507A>C, 24508G>C, 24509T>G, 24510C>A, 24515C>G, 24516C>A, 24526G>C, 24527A>G, 24529C>G, 24532A>G, 24541G>C, 24542C>G, 24543G>T, 24544C>G, 24550C>G, 24553T>G, 24554G>A, 24562A>C, 24568G>C, 24572G>A, 24580G>C, 24587G>C, 24588C>G, 24591T>C, 24592C>G, 24593A>G, 24594C>A, 24595G>T, 24596G>C, 24597A>G, 24601C>G, 24602A>C, 24605A>G, 24606A>G, 24613T>C, 24616C>G, 24617T>A, 24618G>T, 24620C>A, 24625T>C, 24628C>T, 24629C>G, 24630A>T, 24631A>C, 24633A>G, 24634T>C, 24637T>C, 24638T>A, 24642C>A, 24643G>C, 24646A>C, 24651G>T, 24653C>T, 24655G>C, 24658T>G, 24661T>C, 24663C>G, 24664G>C, 24665A>C, 24670G>C, 24671G>A, 24673T>C, 24675C>G, 24676G>C, 24680A>C, 24681C>A, 24682C>G, 24685C>G, 24688G>A, 24690T>A, 24691G>A, 24695C>A, 24696A>G, 24697G>C, 24701T>G, 24702C>G, 24703A>C, 24704G>C, 24707T>C, 24710T>C, 24711C>G, 24714T>A, 24715T>C, 24716T>A, 24717C>A, 24718C>G, 24721A>G, 24722G>A, 24724G>C, 24725A>G, 24730G>C, 24733C>G, 24737C>T, 24738G>C, 24739C>G, 24743G>A, 24744A>C, 24745T>C, 24747T>A, 24748C>T, 24754T>C, 24755T>G, 24756T>C, 24757C>T, 24760T>C, 24775A>C, 24779A>G, 24780C>T, 24781G>C, 24787G>A, 25498C>G, 25506A>T, 25507G>C, 25508T>C, 25510T>G, 25511T>G, 25517C>G, 25520A>C, 25521A>G, 25522A>C, 25523C>G, 25531T>A, 25535T>C, 25536C>G, 25541T>C, 25553T>C, 25565A>C, 25566G>A, 25568G>T, 25571T>G, 25575C>A, 25578C>G, 25580G>C, 25582C>A, 25583G>C, 25584G>C, 25589A>C, 25590_25592delGAC, 25595G>C, 25596A>G, 25604C>T, 25607C>T, 25608A>C, 25610C>G, 25614A>G, 25615A>C, 25616A>C, 25620G>A, 25621C>T, 25622G>C, 25624T>A, 25634T>C, 25635T>C, 25637C>G, 25650T>A, 25651C>G, 25653G>A, 25659A>G, 25662A>G, 25665T>C, 25666A>C, 25667C>G, 25672A>C, 25673G>C, 25675C>A, 25676G>C, 25679C>A, 25682C>G, 25683A>T, 25685C>G, 25691G>C, 25697T>C, 25700G>C, 25706G>C, 25709T>C, 25720A>T, 25724C>T, 25727C>G, 25733G>C, 25737C>A, 25744C>G, 25771C>T, 25772G>C, 26324G>A, 26329C>T, 26330G>C, 26331C>G, 26332A>C, 26336C>T, 26339A>G, 26343A>C, 26344A>G, 26345G>C, 26346C>A, 26354G>C, 26366A>C, 26373A>C, 26375C>G, 26379A>C, 26380C>A, 26382C>G, 26383A>C, 26384G>C, 26386C>A, 26388G>T, 26390C>G, 26391A>G, 26393G>C, 26394C>A, 26396T>C, 26397C>A, 26398G>A, 26400C>G, 26403T>C, 26404C>A, 26405T>G, 26408G>C, 26418A>G, 26419A>C, 26420G>T, 26426T>C, 26430G>A, 26431A>C, 26437T>A, 26439G>C, 26440T>C, 26441A>C, 26442G>A, 26443A>T, 26444C>G, 26450T>C, 26453G>C, 26456G>A, 26457T>A, 26458C>T, 26466C>A, 26468C>G, 26471T>C, 26480T>C, 26481G>A, 26482T>C, 26483G>C, 26487G>T, 26488A>T, 26489A>G, 26493T>G, 26494C>A, 26505A>C, 26506T>G, 26510T>C, 26513T>C, 26516A>C, 26520_26522delGTG, 26525C>G, 26526T>C, 26527C>T, 26528C>G, 26529A>C, 26530T>A, 26537C>G, 26538A>G, 26539G>A, 26541A>C, 26542T>A, 26543T>C, 26547G>A, 26548A>G, 26551T>C, 26552C>G, 26553A>C, 26558G>C, 26559C>G, 26560T>C, 26563C>G, 26564G>C, 26565C>A, 26566G>A, 26569C>G, 26571A>T, 26572A>G, 26574T>G, 26575A>G, 26578G>C, 26579C>A, 26580A>C, 26582G>T, 26584T>C, 26586G>A, 26587C>T, 26589G>A, 26590C>G, 26593T>C, 26595A>C, 26597A>G, 26598G>C, 26599C>A, 26602G>C, 26605T>G, 26608C>G, 26611T>C, 26612G>A, 26616C>G, 26626C>G, 26629A>C, 26631G>C, 26632C>G, 26635G>C, 26636G>T, 26637C>T, 26639T>A, 26640C>G, 26641T>C, 26643C>T, 26644T>A, 26650C>G, 26652G>A, 26654A>C, 26656T>G, 26659G>A, 27222C>G, 27225T>C, 27228G>T, 27229T>C, 27230G>T, 27231G>T, 27232G>T, 27233T>G, 27234T>G, 27237T>C, 27244A>C, 27246C>G, 27249C>A, 27251C>G, 27252G>C, 27253T>A, 27254C>G, 27256T>C, 27258A>C, 27260C>G, 27261G>C, 27267T>C, 27268C>A, 27270G>C, 27272G>C, 27273C>G, 27274A>C, 27275C>T, 27277G>T, 27281A>G, 27284C>G, 27286G>A, 27287A>C, 27288C>G, 27289A>C, 27290C>A, 27295A>G, 27296C>G, 27297A>C, 27307A>G, 27309A>G, 27310G>C, 27318C>G, 27321T>G, 27322G>C, 27324G>C, 27327G>C, 27330T>C, 27333C>G, 27339G>C, 27343A>G, 27345C>A, 27351C>A, 27354T>G, 27355G>A, 27356A>G, 27357G>C, 27360T>C, 27379A>T, 27381G>C, 27382A>G, 27387T>C, 27391A>C, 27392C>T, 27399C>T, 27402G>C, 27406G>A, 27407C>A, 27409A>G, 27411T>G, 27412C>G, 27413G>T, 27416G>C, 27417A>C, 27425T>G, 27426G>C, 27432G>A, 27433G>A, 27436A>C, 27437G>T, 27439A>C, 27447G>C, 27448G>A, 27453T>G, 27456G>C, 27457C>A, 27459A>G, 27466G>C, 27467A>C, 27474A>G, 27475G>A, 27477G>C, 27478G>A, 27641T>G, 27642A>C, 27643C>A, 27646T>G, 27650G>C, 27651C>A, 27652C>G, 27653T>C, 27655A>C, 27658T>C, 27662C>A, 27668G>A, 27676G>C, 27677A>G, 27679C>G, 27685C>G, 27689G>T, 27691G>T, 27701A>G, 27702A>T, 27703T>G, 27706T>G, 27709C>A, 27710G>A, 27712C>T, 27714T>C, 27716A>C, 27731C>A, 27734A>G, 27735T>C, 27736C>G, 27737A>C, 27742A>G, 27743G>A, 27745G>T, 27746C>G, 27747T>C, 27748G>A, 27749G>T, 27752A>C, 27754C>G, 27761G>T, 27762C>T, 27763G>T, 27766T>G, 27769G>C, 27770G>A, 27771C>G, 27772G>C, 27774C>A, 27775C>T, 27779A>G, 27781G>C, 27785C>G, 27786T>C, 27787T>G, 27792C>G, 27793G>T, 27798T>C, 27802G>C, 27812G>T, 27814G>C, 27815A>G, 27820T>C, 27823T>G, 27826T>G, 27829G>C, 27830G>C, 27831T>A, 27833G>C, 27834C>A, 27841T>C, 27844A>C, 27845A>C, 27846C>A, 27850C>G, 27851G>A, 27857T>G, 27868G>T, 27869A>T, 27874C>G, 27875C>A, 27876A>C, 27877G>C, 27883C>T, 27886T>G, 27895T>C, 27896A>G, 27898A>G, 27899G>T, 27901C>T, 27904T>C, 27913A>G, 27922G>C, 27923A>T, 27929C>A, 27931A>G, 27933T>A, 27964T>C, 27967T>C, 27971T>G, 27976A>G, 27985C>G, 27988C>T, 27994T>G, 27997T>G, 28000G>C, 28006G>C, 28007G>A, 28009G>C, 28012C>T, 28013G>T, 28014T>G, 28015G>C, 28016A>G, 28017A>T, 28018T>G, 28020T>C, 28021A>G, 28022C>A, 28024G>C, 28028G>A, 28029T>A, 28031G>T, 28036G>C, 28038C>A, 28039G>C, 28042C>A, 28044G>C, 28045C>G, 28051T>G, 28053G>C, 28054C>G, 28062T>C, 28064G>A, 28065C>T, 28066T>C, 28068G>C, 28072C>G, 28073G>A, 28075A>C, 28076T>A, 28077C>A, 28078G>C, 28081C>G, 28082A>C, 28086T>G, 28088G>C, 28089G>A, 28090C>G, 28093A>C, 28096T>C, 28097T>G, 28099C>G, 28108T>C, 28114C>G, 28117G>C, 28123C>G, 28126A>G, 28137A>C, 28138C>G, 28141C>T, 28147T>C, 28154G>A, 28156A>G, 28159C>T, 28162T>C, 28165G>C, 28166C>T, 28168G>C, 28171T>C, 28172A>T, 28177C>G, 28181A>G, 28185C>G, 28190G>A, 28192A>C, 28193G>A, 28194C>A, 28195G>C, 28207G>A, 28213G>C, 28215A>G, 28216A>G, 28217T>G, 28219C>T, 28222T>C, 28223G>A, 31501T>C, 31504G>C, 31508C>T, 31510G>C, 31528T>C, 31531G>A, 31535A>G, 31537A>C, 31540C>T, 31543A>G, 31546T>C, 31547C>A, 31548A>G, 31549G>C, 31556G>A, 31559C>T, 31562A>G, 31563C>T, 31564T>G, 31565G>A, 31605G>A, 31606G>T, 31607A>C, 31608G>A, 31609C>G, 31616A>G, 31617A>C, 31618C>G, 31619A>G, 31620A>G, 31621C>A, 31623G>A, 31631G>A, 31632C>G, 31633A>T, 31639G>C, 31643A>G, 31660C>G, 31663T>G, 31672C>G, 31678T>A, 31682G>A, 31684A>C, 31687G>C, 31690T>A, 31693C>G, 31699C>A, 31701G>C, 31702C>G, 31708C>G, 31711C>G, 31714A>G, 31717C>A, 31724T>G, 31725C>T, 31729C>T, 31733A>G, 31734C>T, 31735C>G, 31738G>T, 31741G>C, 31743G>A, 31744G>T, 31745G>T, 31746T>C, 31753C>G, 31756C>T, 31760A>T, 31765T>C, 31771G>C, 31774C>G, 31783G>C, 31786G>C, 31789C>T

##### CDS

| | | | | | | | | | |
|---|---|---|---|---|---|---|---|---|---|
| KMC43_gp27 | 1 | 117 | 26.4% | 715 | 90.2% | 117 (100%) | 103 (88.0%) | 0/0/0/0 | 0 |

Protein mutations: S4Q (14205A>C 14206G>A 14207C>A), D25E (14270T>A), A26G (14272C>G), I30M (14285C>G), G32S (14289G>A), F33Y (14293T>A 14294C>T), Q35S (14298C>T 14299A>C), R41Q (14317G>A 14318T>G), P67A (14394C>G), S68A (14397T>G 14399G>C), A69S (14400G>A 14401C>G 14402A>C), S78T (14427T>A), S88T (14457T>A), T102M (14500C>T)

| | Begin | End | Coverage | Score | Concordance | Matches | Identities | I/D/M/F* | Stop Codons |
|---|---|---|---|---|---|---|---|---|---|
| **NT** | **14196** | **31794** | **9.4%** | **3582** | **41.3%** | **4358 (99.6%)** | **3090 (70.6%)** | **9/8** | |
| Codon mutations: | ATC3ATT (14204C>T), AGC4CAA (14205A>C 14206G>A 14207C>A), AAT5AAC (14210T>C), GCC8GCT (14219C>T), GCT9GCC (14222T>C), CCG12CCC (14231G>C), TTT23TTC (14264T>C), GAT25GAA (14270T>A), GCC26GGC (14272C>G), ATC30ATG (14285C>G), CTC31CTG (14288C>G), GGC32AGC (14289G>A), TTC33TAT (14293T>A 14294C>T), CAG35TCG (14298C>T 14299A>C), GCA37GCC (14306A>C), CGT38CGC (14309T>C), CGT41CAG (14317G>A 14318T>G), GGT47GGC (14336T>C), GAA48GAG (14339A>G), ACC49ACA (14342C>A), CGT54CGC (14357T>C), GGT56GGC (14363T>C), CTG57CTC (14366G>C), GCG59GCT (14372G>T), CCG60CCC (14375G>C), GTC61GTG (14378C>G), ACC62ACG (14381C>G), CCG67GCG (14394C>G), TCG68GCC (14397T>G 14399G>C), GCA69AGC (14400G>A 14401C>G 14402A>C), TCG78ACG (14427T>A), CCG79CCC (14432G>C), TCG80AGC (14433T>A 14434C>G 14435G>C), GTG84GTC (14447G>C), TCG88ACG (14457T>A), CTC89CTG (14462C>G), ACG97ACC (14486G>C), ACG102ATG (14500C>T), GTG103GTC (14504G>C), GTC107GTG (14516C>G), GTG110GTC (14525G>C) | | | | | | | | |
| KMC43_gp29 | 15 | 64 | 78.1% | -12 | -3.5% | 50 (94.3%) | 19 (35.8%) | 3/0/1/1 | 1 |
| Protein mutations: | S18L (15870C>T 15871C>G), I19V (15872A>G), L20V (15875C>G), D22E (15883C>G), E23R (15884G>C 15885A>G 15886G>A), S24D (15887T>G 15888C>A), I25F (15890A>T), W27F (15897G>T 15898G>C), M28S (15900T>G 15901G>C), E29V (15903A>T 15904A>C), I32L (15911A>C 15913C>G), Q35V (15920C>G 15921A>T), F36W (15924T>G 15925C>G), H37A (15926C>G 15927A>C), E38A (15930A>C 15931A>T), T40K (15936C>A), H41P (15939A>C), V42I (15941G>A), N44E (15947A>G 15949C>A), P45V (15950C>G 15951C>T 15952T>G), V46R (15953G>C 15954T>G), V48I (15959G>A 15961G>C), A49R (15962G>C 15963C>G), A50E (15966C>A), Q53A (15974C>G 15975A>C 15976G>C), H54N (15977C>A), G55D (15981G>A), R57P (15987G>C 15988T>G), R57_Y58insIKA (15988_15989insATCAAGGCC), L59R (15992T>C 15993T>G), V61I (15998G>A) | | | | | | | | |
| Codon mutations: | CCC16CCG (15865C>G), GAT17GAC (15868T>C), TCC18TTG (15870C>T 15871C>G), ATC19GTC (15872A>G), CTG20GTG (15875C>G), TTG21CTG (15878T>C), GAC22GAG (15883C>G), GAG23CGA (15884G>C 15885A>G 15886G>A), TCC24GAC (15887T>G 15888C>A), ATC25TTC (15890A>T), GGG26GGC (15895G>C), TGG27TTC (15897G>T 15898G>C), ATG28AGC (15900T>G 15901G>C), GAA29GTC (15903A>T 15904A>C), ATC32CTG (15911A>C 15913C>G), CGA34CGC (15919A>C), CAG35GTG (15920C>G 15921A>T), TTC36TGG (15924T>G 15925C>G), CAC37GCC (15926C>G 15927A>C), GAA38GCT (15930A>C 15931A>T), GGT39GGC (15934T>C), ACG40AAG (15936C>A), CAC41CCC (15939A>C), GTC42ATC (15941G>A), AAC44GAA (15947A>G 15949C>A), CCT45GTG (15950C>G 15951C>T 15952T>G), GTC46CGC (15953G>C 15954T>G), GTG48ATC (15959G>A 15961G>C), GCC49CGC (15962G>C 15963C>G), GCG50GAG (15966C>A), CAG53GCC (15974C>G 15975A>C 15976G>C), CAC54AAC (15977C>A), GGC55GAC (15981G>A), CGT57CCG (15987G>C 15988T>G), CGT57_TAC58insATCAAGGCC (15988_15989insATCAAGGCC), TTG59CGG (15992T>C 15993T>G), GTC61ATC (15998G>A), AGC62-TG (16001delA 16002G>T 16003C>G) | | | | | | | | |
| KMC43_gp30 | 1 | 103 | 100% | 538 | 80.9% | 103 (100%) | 77 (74.8%) | 0/0/0/0 | 1 |
| Protein mutations: | V5I (16018G>A), I7T (16025T>C 16026T>C), K11R (16036A>C 16037A>G 16038A>C), S12N (16040G>A), T14A (16045A>G 16047C>G), T18A (16057A>G 16059G>T), A19G (16061C>G 16062C>G), Q21T (16066C>A 16067A>C), S25T (16078T>A), K28A (16087A>G 16088A>C 16089G>C), D29G (16091A>G), I32L (16099A>C 16101C>G), N41Q (16126A>C 16128T>G), A48T (16147G>A 16149G>T), G49P (16150G>C 16151G>C), T58V (16177A>G 16178C>T 16179G>C), T60G (16183A>G 16184C>G), A62V (16190C>T 16191T>C), N65V (16198A>G 16199A>T), I67V (16204A>G), G68A (16208G>C 16209C>G), S76T (16231T>A 16233A>C), N85A (16258A>G 16259A>C 16260C>A), I89V (16270A>G), A93T (16282G>A 16284T>C), S98A (16297T>G) | | | | | | | | |
| Codon mutations: | GAC3GAT (16014C>T), GTC5ATC (16018G>A), ATT7ACC (16025T>C 16026T>C), GGC10GGG (16035C>G), AAA11CGC (16036A>C 16037A>G 16038A>C), AGC12AAC (16040G>A), ACC14GCG (16045A>G 16047C>G), AAT16AAC (16053T>C), ACG18GCT (16057A>G 16059G>T), GCC19GGG (16061C>G 16062C>G), GCG20GCC (16065G>C), CAG21ACG (16066C>A 16067A>C), GTG22GTC (16071G>C), GTT23GTC (16074T>C), TCG25ACG (16078T>A), CCC27CCA (16086C>A), AAG28GCC (16087A>G 16088A>C 16089G>C), GAC29GGC (16091A>G), TTT30TTC (16095T>C), ATC32CTG (16099A>C 16101C>G), CGA36CGC (16113A>C), AAT41CAG (16126A>C 16128T>G), GTT42GTG (16131T>G), CTC43CTG (16134C>G), GTT44GTG (16137T>G), GCG45GCC (16140G>C), GGG46GGC (16143G>C), ACT47ACC (16146T>C), GCG48ACT (16147G>A 16149G>T), GGG49CCG (16150G>C 16151G>C), GCC51GCG (16158C>G), GCG52GCC (16161G>C), TAC53TAT (16164C>T), GCG55GCA (16170G>A), ACG57ACC (16176G>C), ACG58GTC (16177A>G 16178C>T 16179G>C), GCC59GCT (16182C>T), ACC60GGC (16183A>G 16184C>G), GCT62GTC (16190C>T 16191T>C), AAC65GTC (16198A>G 16199A>T), ATC67GTC (16204A>G), GGC68GCG (16208G>C 16209C>G), GCG69GCC (16212G>C), CCG71CCC (16218G>C), GTG74GTC (16227G>C), TCA76ACC (16231T>A 16233A>C), CTC78CTG (16239C>G), GAT80GAC (16245T>C), CCG82CCC (16251G>C), CTG84TTG (16255C>T), AAC85GCA (16258A>G 16259A>C 16260C>A), GGC86GGG (16263C>G), ATC89GTC (16270A>G), CCT91CCC (16278T>C), GCT93ACC (16282G>A 16284T>C), ACC96ACG (16293C>G), GTT97GTC (16296T>C), TCT98GCT (16297T>G), GTT99GTC (16302T>C), TCT100TCC (16305T>C) | | | | | | | | |
| KMC43_gp31 | 1 | 117 | 91.4% | 710 | 86.3% | 117 (100%) | 93 (79.5%) | 0/0/0/0 | 0 |
| Protein mutations: | A6Q (16339G>C 16340C>A 16341C>G), K8R (16345A>C 16346A>G), G16W (16369G>T 16371A>G), M19V (16378A>G 16380G>C), F20Y (16382T>A 16383C>T), S26A (16399T>G), M43L (16450A>C), N44S (16453A>T 16454A>C 16455C>G), M46L (16459A>C), A48P (16465G>C 16467C>G), A52G (16478C>G 16479T>C), V54I (16483G>A 16485G>C), S71G (16534T>G 16535C>G 16536T>C), A72T (16537G>A), Q80K (16561C>A), V85T (16576G>A 16577T>C 16578G>C), V91L (16594G>C), A99S (16618G>A 16619C>G), R103L (16631G>T 16632C>G), E107Q (16642G>C), M109L (16648A>C 16650G>C), I111V (16654A>G 16656T>G), G116A (16670G>C 16671G>C) | | | | | | | | |
| Codon mutations: | ACA4ACC (16335A>C), ACC5ACG (16338C>G), GCC6CAG (16339G>C 16340C>A 16341C>G), AAG8CGG (16345A>C 16346A>G), TAT14TAC (16365T>C), TGC15TGT (16368C>T), GGA16TGG (16369G>T 16371A>G), ATG19GTC (16378A>G 16380G>C), TTC20TAT (16382T>A 16383C>T), GGG22GGC (16389G>C), CCA24CCC (16395A>C), TCG25AGT (16396T>A 16397C>G 16398G>T), TCG26GCG (16399T>G), TAT30TAC (16413T>C), CGG31AGG (16414C>A), GCG35GCC (16428G>C), TAC36TAT (16431C>T), CTT39CTG (16440T>G), GAA40GAG (16443A>G), ATG43CTG (16450A>C), AAC44TCG (16453A>T 16454A>C 16455C>G), ATG46CTG (16459A>C), CTG47CTC (16464G>C), GCC48CCG (16465G>C 16467C>G), GAA50GAG (16473A>G), GCT52GGC (16478C>G 16479T>C), GTG54ATC (16483G>A 16485G>C), ACG57ACC (16494G>C), TAT58TAC (16497T>C), CTG59CTC (16500G>C), GCT60GCC (16503T>C), ACC63ACG (16512C>G), ACT64ACG (16515T>G), CTC65CTG (16518C>G), ACG67ACC (16524G>C), GCA68GCC (16527A>C), TCT71GGC (16534T>G 16535C>G 16536T>C), GCG72ACG (16537G>A), AGC73TCG (16540A>T 16541G>C 16542C>G), GCG74GCA (16545G>A), CTC76CTT (16551C>T), ACA78ACG (16557A>G), CAG80AAG (16561C>A), GCG81GCC (16566G>C), GCC82GCG (16569C>G), GTG85ACC (16576G>A 16577T>C 16578G>C), GAA90GAG (16593A>G), GTG91CTG (16594G>C), GCA92GCC (16599A>C), GAC93GAT (16602C>T), CGC94CGT (16605C>T), GAC95GAT (16608C>T), TTT98TTC (16617T>C), GCC99AGC (16618G>A 16619C>G), CGG102CGC (16629G>C), CGC103CTG (16631G>T 16632C>G), GAG107CAG (16642G>C), ATG109CTC (16648A>C 16650G>C), ATT111GTG (16654A>G 16656T>G), CCC113CCG (16662C>G), GGG114GGC (16665G>C), CCC115CCG (16668C>G), GGG116GCC (16670G>C 16671G>C), CTC117T.. (16672C>T) | | | | | | | | |
| KMC43_gp36 | 17 | 497 | 72.0% | 1825 | 76.6% | 368 (100%) | 262 (71.2%) | 0/0/0/0 | 0 |
| Protein mutations: | V28T (19290G>A 19291T>C 19292T>C), S29Q (19293T>C 19294C>A 19295C>G), L30Y (19296C>T 19297T>A 19298G>C), V38I (19320G>A 19322G>C), V41I (19329G>A 19331T>C), T46S (19344A>T), L55T (19371C>A 19372T>C), I56V (19374A>G), N58S (19381A>G), M59P (19383A>C 19384T>C), M61Q (19389A>C 19390T>A), Y62A (19392T>G 19393A>C), Q64A (19398C>G 19399A>C 19400G>C), A65I (19401G>A 19402C>T), A68Q (19410G>C 19411C>A), I69A (19413A>G 19414T>C), T73K (19426C>A 19427C>G), M76L (19434A>C), G77M (19437G>A 19438G>T 19439C>G), V80I (19446G>A 19448G>C), A81S (19449G>T), I82V (19452A>G), V84T (19458G>A 19459T>C 19460G>C), Q85M (19461C>A 19462A>T), R92K (19482C>A 19483G>A 19484C>A), C93A (19485T>G 19486G>C), T102V (19512A>G 19513C>T), T106I (19525C>T 19526G>C), I107A (19527A>G 19528T>C 19529C>G), Q109L (19534A>T), G112C (19542G>T), V113I (19545G>A 19547G>C), L115F (19551C>T 19553G>C), A117S (19557G>T 19559G>C), L123F (19575C>T 19577G>C), S126G (19584T>G 19585C>G 19586G>C), V127I (19587G>A 19589G>C), A130T (19596G>A), A132S (19602G>T 19604C>G), A136Q (19614G>C 19615C>A 19616C>G), N138S (19620A>T 19621A>C 19622T>G), W140Q (19626T>C 19627G>A), V142A (19633T>C 19634A>C), T143V (19635A>G 19636C>T), A145S (19641G>T), I266T (20005T>C 20006C>G), I269V (20013A>G), G270M (20016G>A 20017G>T 20018C>G), V271T (20019G>A 20020T>C), A274S (20028G>A 20029C>G), S279A (20043A>G 20044G>C 20045C>G), T282I (20053C>T), T283S (20056C>G 20057G>C), T284A (20058A>G), A286S (20064G>A 20065C>G), I290V (20076A>G 20078C>G), F295A (20091T>G 20092T>C), V303C (20115G>T 20116T>G 20117G>C), V309Y (20133G>T 20134T>A), S311Q (20139A>C 20140G>A 20141C>G), V312I (20142G>A 20144T>C), T313L (20145A>C 20146C>T 20147A>T), A320S (20166G>T 20168G>C), S327A (20187T>G 20189G>C), T335S (20211A>T), Y341Q (20229T>C 20231T>G), V343I (20235G>A 20237G>C), T354Q (20268A>C 20269C>A), S362Q (20292T>C 20293C>A), I364G (20298A>G 20299T>G), Q365I (20301C>A 20302A>T 20303G>C), C383A (20355T>G 20356G>C 20357C>G), S385F (20362C>T 20363G>C), A393P (20385G>C), L394V (20388C>G), Q396N (20394C>A 20396G>C), S409Y (20433A>T 20434G>A), V411L (20439G>C), Y418F (20461A>T), V419I (20463G>A), A425S (20481G>T 20483C>G), A429T (20493G>A 20495G>T), A432M (20502G>A 20503C>T 20504C>G), L435I (20511C>A 20513G>C), N437A (20517A>G 20518A>C), S440D (20526T>G 20527C>A), S441N (20530G>A), M442L (20532A>C), E443Q (20535G>C), M447L (20547A>C), A450D (20557C>A), V451P (20559G>C 20560T>C 20561C>G), P455A (20571C>G 20573A>G), S456A (20574T>G 20576G>C), F457Y (20578T>A), I461V (20589A>G 20591C>G), N464A (20598A>G 20599A>C), M468A (20610A>G 20611T>C), N469S (20613A>T 20614A>C 20615C>G), L476M (20634C>A 20636T>G), I483K (20656T>A 20657T>G), S486A (20664T>G 20666G>C), I492L (20682A>C), V493I (20685G>A 20687G>C), L495I (20691C>A 20693T>C) | | | | | | | | |

| | Begin | End | Coverage | Score | Concordance | Matches | Identities | I/D/M/F* | Stop Codons |
|---|---|---|---|---|---|---|---|---|---|
| **NT** | **14196** | **31794** | **9.4%** | **3582** | **41.3%** | **4358 (99.6%)** | **3090 (70.6%)** | **9/8** | |
| Codon mutations: | TTG19CTG (19263T>C), GTA21GTG (19271A>G), CAA22CAG (19274A>G), GTG24GTC (19280G>C), CCT25CCG (19283T>G), CCA26CCG (19286A>G), GTT28ACC (19290G>A 19291T>C 19292T>C), TCC29CAG (19293T>C 19294C>A 19295C>G), CTG30TAC (19296C>T 19297T>A 19298G>C), CTC31CTG (19301C>G), AAT37AAC (19319T>C), GTG38ATC (19320G>A 19322G>C), CTT39TTG (19323C>T 19325T>G), GGT40GGC (19328T>C), GTT41ATC (19329G>A 19331T>C), GTG42GTC (19334G>C), ACC44ACG (19340C>G), GCA45GCG (19343A>G), ACC46TCC (19344A>T), GGC48GGG (19352C>G), ACC50ACG (19358C>G), TCG52AGC (19362T>A 19363C>G 19364G>C), CCG53CCC (19367G>C), ACG54ACC (19370G>C), CTG55ACG (19371C>A 19372T>C), ATC56GTC (19374A>G), AAC58AGC (19381A>G), ATG59CCG (19383A>C 19384T>C), GCG60GCT (19388G>T), ATG61CAG (19389A>C 19390T>A), TAC62GCC (19392T>G 19393A>C), GCC63GCG (19397C>G), CAG64GCC (19398C>G 19399A>C 19400G>C), GCC65ATC (19401G>A 19402C>T), GCG68CAG (19410G>C 19411C>A), ATC69GCC (19413A>G 19414T>C), CGT72CGC (19424T>C), ACC73AAG (19426C>A 19427C>G), TAT74TAC (19430T>C), ATG76CTG (19434A>C), GGC77ATG (19437G>A 19438G>T 19439C>G), GCT79GCC (19445T>C), GTG80ATC (19446G>A 19448G>C), GCG81TCG (19449G>T), ATC82GTC (19452A>G), GCA83GCG (19457A>G), GTG84ACC (19458G>A 19459T>C 19460G>C), CAG85ATG (19461C>A 19462A>T), GCG88GCC (19472G>C), CGC92AAA (19482C>A 19483G>A 19484C>A), TGC93GCC (19485T>G 19486G>C), CGC95CGG (19493C>G), GTC96GTG (19496C>G), ACG102GTG (19512A>G 19513C>T), GCG104GCC (19520G>C), TCG105AGC (19521T>A 19522C>G 19523G>C), ACG106ATC (19525C>T 19526G>C), ATC107GCG (19527A>G 19528T>C 19529C>G), CAG109CTG (19534A>T), GGC112TGC (19542G>T), GTG113ATC (19545G>A 19547G>C), ACG114ACC (19550G>C), CTG115TTC (19551C>T 19553G>C), GCG117TCC (19557G>T 19559G>C), ACG122ACC (19574G>C), CTG123TTC (19575C>T 19577G>C), TCG126GGC (19584T>G 19585C>G 19586G>C), GTG127ATC (19587G>A 19589G>C), GTC129GTG (19595C>G), GCG130ACG (19596G>A), CTC131CTG (19601C>G), GCC132TCG (19602G>T 19604C>G), GCG133GCC (19607G>C), AGC135TCG (19611A>T 19612G>C 19613C>G), GCC136CAG (19614G>C 19615C>A 19616C>G), AAT138TCG (19620A>T 19621A>C 19622T>G), TGG140CAG (19626T>C 19627G>A), GTA142GCC (19633T>C 19634A>C), ACG143GTG (19635A>G 19636C>T), GTC144GTT (19640C>T), GCG145TCG (19641G>T), ATC266ACG (20005T>C 20006C>G), ATC269GTC (20013A>G), GGC270ATG (20016G>A 20017G>T 20018C>G), GTC271ACC (20019G>A 20020T>C), GCT274AGT (20028G>A 20029C>G), ACG277ACC (20039G>C), AGC279GCG (20043A>G 20044G>C 20045C>G), GCC281GCG (20051C>G), ACC282ATC (20053C>T), ACG283AGC (20056C>G 20057G>C), ACC284GCC (20058A>G), GCC286AGC (20064G>A 20065C>G), ATC290GTG (20076A>G 20078C>G), TTC295GCC (20091T>G 20092T>C), CTG298CTT (20102G>T), GGT300GGC (20108T>C), GTG303TGC (20115G>T 20116T>G 20117G>C), TTG306CTG (20124T>C), ACG308ACC (20132G>C), GTC309TAC (20133G>T 20134T>A), AGC311CAG (20139A>C 20140G>A 20141C>G), GTT312ATC (20142G>A 20144T>C), ACA313CTT (20145A>C 20146C>T 20147A>T), CTG315TTG (20151C>T), TCG317AGC (20157T>A 20158C>G 20159G>C), CAA319CAG (20165A>G), GCG320TCC (20166G>T 20168G>C), GTC322GTG (20174C>G), GCT323GCC (20177T>C), GGT324GGC (20180T>C), TCG327GCC (20187T>G 20189G>C), ACG335TCG (20211A>T), TAT341CAG (20229T>C 20231T>G), GTG343ATC (20235G>A 20237G>C), GTG344GTC (20240G>C), TCG349AGC (20253T>A 20254C>G 20255G>C), ACG354CAG (20268A>C 20269C>A), TCC357TCG (20279C>G), GCC358GCT (20282C>T), TTG360CTG (20286T>C), TCG362CAG (20292T>C 20293C>A), ATC364GGC (20298A>G 20299T>G), CAG365ATC (20301C>A 20302A>T 20303G>C), GCG366GCC (20306G>C), GGT367GGC (20309T>C), CCC376CCG (20336C>G), TCG379AGC (20343T>A 20344C>G 20345G>C), TGC383GCG (20355T>G 20356G>C 20357C>G), CGG384CGC (20360G>C), TCG385TTC (20362C>T 20363G>C), CAT387CAC (20369T>C), GCG393CCG (20385G>C), CTG394GTG (20388C>G), CAG396AAC (20394C>A 20396G>C), ACC401ACG (20411C>G), AGC409TAC (20433A>T 20434G>A), GTC411CTC (20439G>C), AAT412AAC (20444T>C), TAC418TTC (20461A>T), GTC419ATC (20463G>A), GGG420GGT (20468G>T), CAA423CAG (20477A>G), TCG424AGC (20478T>A 20479C>G 20480G>C), GCC425TCG (20481G>T 20483C>G), GTG427GTC (20489G>C), GCG429ACT (20493G>A 20495G>T), GCG431GCC (20501G>C), GCC432ATG (20502G>A 20503C>T 20504C>G), CTG435ATC (20511C>A 20513G>C), TCG436AGC (20514T>A 20515C>G 20516G>C), AAC437GCC (20517A>G 20518A>C), TCC440GAC (20526T>G 20527C>A), AGC441AAC (20530G>A), ATG442CTG (20532A>C), GAG443CAG (20535G>C), GGC446GGA (20546C>A), ATG447CTG (20547A>C), GCC450GAC (20557C>A), GTC451CCG (20559G>C 20560T>C 20561C>G), AAT452AAC (20564T>C), CCA455GCG (20571C>G 20573A>G), TCG456GCC (20574T>G 20576G>C), TTC457TAC (20578T>A), TCG458AGC (20580T>A 20581C>G 20582G>C), CAA460CAG (20588A>G), ATC461GTG (20589A>G 20591C>G), GCT463GCG (20597T>G), AAC464GCC (20598A>G 20599A>C), ATG468GCG (20610A>G 20611T>C), AAC469TCG (20613A>T 20614A>C 20615C>G), CGG470CGC (20618G>C), GCG472GCC (20624G>C), CTT473CTG (20627T>G), CTT476ATG (20634C>A 20636T>G), GCT478GCC (20642T>C), GTC480GTG (20648C>G), ATT483AAG (20656T>A 20657T>G), TCG486GCC (20664T>G 20666G>C), GTT487GTC (20669T>C), ATC492CTC (20682A>C), GTG493ATC (20685G>A 20687G>C), AAT494AAC (20690T>C), CTT495ATC (20691C>A 20693T>C) | | | | | | | | |
| KMC43_gp42 | 12 | 115 | 32.4% | 453 | 59.9% | 104 (100%) | 57 (54.8%) | 0/0/0/0 | 0 |
| Protein mutations: | D14G (24489A>G), R15S (24491C>T 24492G>C), Q16L (24495A>T 24496G>C), I18V (24500A>G 24502T>G), E20A (24507A>C 24508G>C), S21E (24509T>G 24510C>A), P23E (24515C>G 24516C>A), T27A (24527A>G 24529C>G), R32V (24542C>G 24543G>T 24544C>G), D34E (24550C>G), V36I (24554G>A), V42I (24572G>A), A47R (24587G>C 24588C>G), F48S (24591T>C 24592C>G), T49D (24593A>G 24594C>A 24595G>T), E50R (24596G>C 24597A>G), I52L (24602A>C), N53G (24605A>G 24606A>G), W57M (24617T>A 24618G>T), L58M (24620C>A), Q61V (24629C>G 24630A>T 24631A>C), D62G (24633A>G 24634T>C), L64M (24638T>A), A65D (24642C>A 24643G>C), R68L (24651G>T), Q69Y (24653C>T 24655G>C), A72G (24663C>G 24664G>C), M73L (24665A>C), V75I (24671G>A 24673T>C), A76G (24675C>G 24676G>C), T78Q (24680A>C 24681C>A 24682C>G), V81E (24690T>A 24691G>A), Q83S (24695C>A 24696A>G 24697G>C), S85G (24701T>G 24702C>G 24703A>C), E86Q (24704G>C), S88R (24710T>C 24711C>G), F89Y (24714T>A 24715T>C), S90K (24716T>A 24717C>A 24718C>G), V92I (24722G>A 24724G>C), I93V (24725A>G), D95E (24733C>G), R97S (24737C>T 24738G>C 24739C>G), D99T (24743G>A 24744A>C 24745T>C), F100Y (24747T>A 24748C>T), F103A (24755T>G 24756T>C 24757C>T), T111V (24779A>G 24780C>T 24781G>C) | | | | | | | | |
| Codon mutations: | GAC14GGC (24489A>G), CGG15TCG (24491C>T 24492G>C), CAG16CTC (24495A>T 24496G>C), ATT18GTG (24500A>G 24502T>G), CCC19CCG (24505C>G), GAG20GCC (24507A>C 24508G>C), TCG21GAG (24509T>G 24510C>A), CCG23GAG (24515C>G 24516C>A), GGG26GGC (24526G>C), ACC27GCG (24527A>G 24529C>G), CAA28CAG (24532A>G), GCG31GCC (24541G>C), CGC32GTG (24542C>G 24543G>T 24544C>G), GAC34GAG (24550C>G), CTT35CTG (24553T>G), GTC36ATC (24554G>A), GGA38GGC (24562A>C), CGG40CGC (24568G>C), GTC42ATC (24572G>A), GCG44GCC (24580G>C), GCC47CGC (24587G>C 24588C>G), TTC48TCG (24591T>C 24592C>G), ACG49GAT (24593A>G 24594C>A 24595G>T), GAG50CGG (24596G>C 24597A>G), CCC51CCG (24601C>G), ATC52CTC (24602A>C), AAC53GGC (24605A>G 24606A>G), TCT55TCC (24613T>C), GGC56GGG (24616C>G), TGG57ATG (24617T>A 24618G>T), CTG58ATG (24620C>A), CTT59CTC (24625T>C), GGC60GGT (24628C>T), CAA61GTC (24629C>G 24630A>T 24631A>C), GAT62GGC (24633A>G 24634T>C), GCT63GCC (24637T>C), TTG64ATG (24638T>A), GCG65GAC (24642C>A 24643G>C), CGA66CGC (24646A>C), CGG68CTG (24651G>T), CAG69TAC (24653C>T 24655G>C), CTT70CTG (24658T>G), GAT71GAC (24661T>C), GCG72GGC (24663C>G 24664G>C), ATG73CTG (24665A>C), CGG74CGC (24670G>C), GTT75ATC (24671G>A 24673T>C), GCG76GGC (24675C>G 24676G>C), ACC78CAG (24680A>C 24681C>A 24682C>G), GTC79GTG (24685C>G), CAG80CAA (24688G>A), GTG81GAA (24690T>A 24691G>A), CAG83AGC (24695C>A 24696A>G 24697G>C), TCA85GGC (24701T>G 24702C>G 24703A>C), GAG86CAG (24704G>C), TTG87CTG (24707T>C), TCC88CGC (24710T>C 24711C>G), TTT89TAC (24714T>A 24715T>C), TCC90AAG (24716T>A 24717C>A 24718C>G), GTA91GTG (24721A>G), GTG92ATC (24722G>A 24724G>C), ATT93GTT (24725A>G), CGG94CGC (24730G>C), GAC95GAG (24733C>G), CGC97TCG (24737C>T 24738G>C 24739C>G), GAT99ACC (24743G>A 24744A>C 24745T>C), TTC100TAT (24747T>A 24748C>T), CGT102CGC (24754T>C), TTC103GCT (24755T>G 24756T>C 24757C>T), TAT104TAC (24760T>C), CGA109CGC (24775A>C), ACG111GTC (24779A>G 24780C>T 24781G>C), GAG113GAA (24787G>A) | | | | | | | | |
| KMC43_gp43 | 25 | 373 | 46.4% | 790 | 70.5% | 171 (98.8%) | 114 (65.9%) | 0/2/0/0 | 1 |
| Protein mutations: | A28G (25498C>G), F32W (25510T>G 25511T>G), N36A (25521A>G 25522A>C 25523C>G), F39Y (25531T>A), Q41E (25536C>G), A51T (25566G>A 25568G>T), L54M (25575C>A), P55A (25578C>G 25580G>C), A56D (25582C>A 25583G>C), E57Q (25584G>C), D59del (25590_25592delGAC), N61D (25596A>G), N65Q (25608A>C 25610C>G), K67A (25614A>G 25615A>C 25616A>C), A69I (25620G>A 25621C>T 25622G>C), F70Y (25624T>A), F74L (25635T>C 25637C>G), D80N (25653G>A), T82A (25659A>G), N83D (25662A>G), Y84P (25665T>C 25666A>C 25667C>G), E86A (25672A>C 25673G>C), T87N (25675C>A 25676G>C), D88E (25679C>A), T90S (25683A>T 25685C>G), Y102F (25720A>T), L108M (25737C>A), T110S (25744C>G), A119V (25771C>T 25772G>C), T305I (26329C>T 26330G>C), Q306A (26331C>G 26332A>C), K310R (26343A>C 26344A>G 26345G>C), L311M (26346C>A), I320L (26373A>C 26375C>G), T322Q (26379A>C 26380C>A), Q323A (26382C>G 26383A>C 26384G>C), T324K (26386C>A), A325S (26388G>T 26390C>G), M326V (26391A>G 26393G>C), L327I (26394C>A 26396T>C), R328K (26397C>A 26398G>A), L329V (26400C>G), S330Q (26403T>C 26404C>A 26405T>G), K335A (26418A>G 26419A>C 26420G>T), D339T (26430G>A 26431A>C), F341Y (26437T>A), V342P (26439G>C 26440T>C 26441A>C), D343M (26442G>A 26443A>T 26444C>G), S348M (26457T>A 26458C>T), L351M (26466C>A 26468C>G), V356T (26481G>A 26482T>C 26483G>C), E358L (26487G>T 26488A>T 26489A>G), S360D (26493T>G 26494C>A), I364R (26505A>C 26506T>G), E367D (26516A>C), V369del (26520_26522delGTG), S371L (26526T>C 26527C>T 26528C>G), I372Q (26529A>C 26530T>A) | | | | | | | | |
| Codon mutations: | GCC28GGC (25498C>G), AGT31TCC (25506A>T 25507G>C 25508T>C), TTT32TGG (25510T>G 25511T>G), GTC34GTG (25517C>G), ACA35ACC (25520A>C), AAC36GCG (25521A>G 25522A>C 25523C>G), TTC39TAC (25531T>A), TAT40TAC (25535T>C), CAG41GAG (25536C>G), GCT42GCC (25541T>C), CGT46CGC (25553T>C), GCA50GCC (25565A>C), GCG51ACT (25566G>A 25568G>T), TCT52TCG (25571T>G), CTG54ATG (25575C>A), CCG55GCC (25578C>G 25580G>C), GCG56GAC (25582C>A 25583G>C), GAG57CAG (25584G>C), ACA58ACC (25589A>C), GAC59del (25590_25592delGAC), GCG60GCC (25595G>C), AAC61GAC (25596A>G), TTC63TTT (25604C>T), TCC64TCT (25607C>T), AAC65CAG (25608A>C 25610C>G), AAA67GCC (25614A>G 25615A>C 25616A>C), GCG69ATC (25620G>A 25621C>T 25622G>C), TTC70TAC (25624T>A), ATT73ATC (25634T>C), TTC74CTG (25635T>C 25637C>G), TCC79AGC (25650T>A 25651C>G), GAT80AAT (25653G>A), ACG82GCG (25659A>G), AAC83GAC (25662A>G), TAC84CCG (25665T>C 25666A>C 25667C>G), GAG86GCC (25672A>C 25673G>C), ACG87AAC (25675C>A 25676G>C), GAC88GAA (25679C>A), CTC89CTG (25682C>G), ACC90TCG (25683A>T 25685C>G), CTG92CTC (25691G>C), TAT94TAC (25697T>C), GGG95GGC (25700G>C), GTG97GTC (25706G>C), GAT98GAC (25709T>C), TAC102TTC (25720A>T), GAC103GAT (25724C>T), CCC104CCG (25727C>G), TCG106TCC (25733G>C), CTG108ATG (25737C>A), ACC110AGC (25744C>G), GCG119GTC (25771C>T 25772G>C), GAG303GAA (26324G>A), ACG305ATC (26329C>T 26330G>C), CAG306GCG (26331C>G 26332A>C), CAC307CAT (26336C>T), GAA308GAG (26339A>G), AAG310CGC (26343A>C 26344A>G 26345G>C), CTG311ATG (26346C>A), GCG313GCC (26354G>C), GCA317GCC (26366A>C), ATC320CTG (26373A>C 26375C>G), ACG322CAG (26379A>C 26380C>A), CAG323GCC (26382C>G 26383A>C 26384G>C), ACG324AAG (26386C>A), GCC325TCG (26388G>T 26390C>G), ATG326GTC (26391A>G 26393G>C), CTT327ATC (26394C>A 26396T>C), CGG328AAG (26397C>A 26398G>A), CTG329GTG (26400C>G), TCT330CAG (26403T>C 26404C>A 26405T>G), GGG331GGC (26408G>C), AAG335GCT (26418A>G 26419A>C 26420G>T), GAT337GAC (26426T>C), GAC339ACC (26430G>A 26431A>C), TTC341TAC (26437T>A), GTA342CCC (26439G>C 26440T>C 26441A>C), GAC343ATG (26442G>A 26443A>T 26444C>G), ATT345ATC (26450T>C), ACG346ACC (26453G>C), CGG347CGA (26456G>A), TCG348ATG (26457T>A 26458C>T), CTC351ATG (26466C>A 26468C>G), GAT352GAC (26471T>C), TAT355TAC (26480T>C), GTG356ACC (26481G>A 26482T>C 26483G>C), GAA358TTG (26487G>T 26488A>T 26489A>G), TCC360GAC (26493T>G 26494C>A), ATC364CGC (26505A>C 26506T>G), AAT365AAC (26510T>C), CCT366CCC (26513T>C), GAA367GAC (26516A>C), GTG369del (26520_26522delGTG), CCC370CCG (26525C>G), TCC371CTG (26526T>C 26527C>T 26528C>G), ATA372CAA (26529A>C 26530T>A) | | | | | | | | |
| KMC43_gp44 | 1 | 44 | 19.7% | 136 | 47.1% | 44 (100%) | 20 (45.5%) | 0/0/0/0 | 0 |
| Protein mutations: | Q3G (26537C>G 26538A>G 26539G>A), H4P (26541A>C 26542T>A), F5L (26543T>C), R6Q (26547G>A 26548A>G), Q8A (26552C>G 26553A>C), A10R (26558G>C 26559C>G 26560T>C), A12Q (26564G>C 26565C>A 26566G>A), Q14L (26571A>T 26572A>G), L15R (26574T>G 26575A>G), Q17T (26579C>A 26580A>C), A18S (26582G>T 26584T>C), G19D (26586G>A 26587C>T), R20Q (26589G>A 26590C>G), E22A (26595A>C), S23A (26597A>G 26598G>C 26599C>A), V28I (26612G>A), T29S (26616C>G), D32E (26626C>G), G34A (26631G>C 26632C>G), A36F (26636G>T 26637C>T), A38V (26643C>T 26644T>A), R41K (26652G>A), I42L (26654A>C 26656T>G) | | | | | | | | |
| Codon mutations: | CAG3GGA (26537C>G 26538A>G 26539G>A), CAT4CCA (26541A>C 26542T>A), TTC5CTC (26543T>C), CGA6CAG (26547G>A 26548A>G), AAT7AAC (26551T>C), CAG8GCG (26552C>G 26553A>C), GCT10CGC (26558G>C 26559C>G 26560T>C), CTC11CTG (26563C>G), GCG12CAA (26564G>C 26565C>A 26566G>A), GCC13GCG (26569C>G), CAA14CTG (26571A>T 26572A>G), CTA15CGG (26574T>G 26575A>G), GCG16GCC (26578G>C), CAG17ACG (26579C>A 26580A>C), GCT18TCC (26582G>T 26584T>C), GGC19GAT (26586G>A 26587C>T), CGC20CAG (26589G>A 26590C>G), GCT21GCC (26593T>C), GAG22GCG (26595A>C), AGC23GCA (26597A>G 26598G>C 26599C>A), CGG24CGC (26602G>C), GTT25GTG (26605T>G), GGC26GGG (26608C>G), ATT27ATC (26611T>C), GTC28ATC (26612G>A), ACC29AGC (26616C>G), GAC32GAG (26626C>G), CCA33CCC (26629A>C), GGC34GCG (26631G>C 26632C>G), ACG35ACC (26635G>C), GCC36TTC (26636G>T 26637C>T), TCT37AGC (26639T>A 26640C>G 26641T>C), GCT38GTA (26643C>T 26644T>A), GTC40GTG (26650C>G), AGG41AAG (26652G>A), ATT42CTG (26654A>C 26656T>G), CAG43CAA (26659G>A) | | | | | | | | |
| KMC43_gp45 | 8 | 96 | 70.1% | 381 | 60.8% | 89 (100%) | 48 (53.9%) | 0/0/0/0 | 0 |

| | Begin | End | Coverage | Score | Concordance | Matches | Identities | I/D/M/F* | Stop Codons |
|---|---|---|---|---|---|---|---|---|---|
| **NT** | **14196** | **31794** | **9.4%** | **3582** | **41.3%** | **4358 (99.6%)** | **3090 (70.6%)** | **9/8** | |
| Protein mutations: | Q11H (27228G>T), W12L (27229T>C 27230G>T 27231G>T), V13W (27232G>T 27233T>G 27234T>G), I17L (27244A>C 27246C>G), T19S (27251C>G 27252G>C), S21P (27256T>C 27258A>C), T22S (27260C>G 27261G>C), L25I (27268C>A 27270G>C), G26A (27272G>C 27273C>G), T27L (27274A>C 27275C>T), A28S (27277G>T), N29S (27281A>G), A30G (27284C>G), D31T (27286G>A 27287A>C 27288C>G), T32Q (27289A>C 27290C>A), T34G (27295A>G 27296C>G 27297A>C), I38V (27307A>G 27309A>G), V39L (27310G>C), V43L (27322G>C 27324G>C), E48D (27339G>C), I50V (27343A>G 27345C>A), H52Q (27351C>A), E54S (27355G>A 27356A>G 27357G>C), K62Y (27379A>T 27381G>C), I63V (27382A>G), T66L (27391A>C 27392C>T), A71K (27406G>A 27407C>A), I72V (27409A>G 27411T>G), R73V (27412C>G 27413G>T), G74A (27416G>C 27417A>C), L77R (27425T>G 27426G>C), V80I (27433G>A), R81L (27436A>C 27437G>T), M82L (27439A>C), E84D (27447G>C), G85S (27448G>A), Q88K (27457C>A 27459A>G), E91P (27466G>C 27467A>C), V94I (27475G>A 27477G>C), D95N (27478G>A) | | | | | | | | |
| Codon mutations: | CTC9CTG (27222C>G), AAT10AAC (27225T>C), CAG11CAT (27228G>T), TGG12CTT (27229T>C 27230G>T 27231G>T), GTT13TGG (27232G>T 27233T>G 27234T>G), GGT14GGC (27237T>C), ATC17CTG (27244A>C 27246C>G), GGC18GGA (27249C>A), ACG19AGC (27251C>G 27252G>C), TCC20AGC (27253T>A 27254C>G), TCA21CCC (27256T>C 27258A>C), ACG22AGC (27260C>G 27261G>C), GAT24GAC (27267T>C), CTG25ATC (27268C>A 27270G>C), GGC26GCG (27272G>C 27273C>G), ACC27CTC (27274A>C 27275C>T), GCC28TCC (27277G>T), AAC29AGC (27281A>G), GCC30GGC (27284C>G), GAC31ACG (27286G>A 27287A>C 27288C>G), ACG32CAG (27289A>C 27290C>A), ACA34GGC (27295A>G 27296C>G 27297A>C), ATA38GTG (27307A>G 27309A>G), GTG39CTG (27310G>C), CGC41CGG (27318C>G), CTT42CTG (27321T>G), GTG43CTC (27322G>C 27324G>C), ACG44ACC (27327G>C), AAT45AAC (27330T>C), CCC46CCG (27333C>G), GAG48GAC (27339G>C), ATC50GTA (27343A>G 27345C>A), CAC52CAA (27351C>A), CCT53CCG (27354T>G), GAG54AGC (27355G>A 27356A>G 27357G>C), TAT55TAC (27360T>C), AAG62TAC (27379A>T 27381G>C), ATC63GTC (27382A>G), GGT64GGC (27387T>C), ACG66CTG (27391A>C 27392C>T), GAC68GAT (27399C>T), GTG69GTC (27402G>C), GCG71AAG (27406G>A 27407C>A), ATT72GTG (27409A>G 27411T>G), CGC73GTC (27412C>G 27413G>T), GGA74GCC (27416G>C 27417A>C), CTG77CGC (27425T>G 27426G>C), CAG79CAA (27432G>A), GTC80ATC (27433G>A), AGG81CTG (27436A>C 27437G>T), ATG82CTG (27439A>C), GAG84GAC (27447G>C), GGC85AGC (27448G>A), GTT86GTG (27453T>G), GCG87GCC (27456G>C), CAA88AAG (27457C>A 27459A>G), GAG91CCG (27466G>C 27467A>C), CAA93CAG (27474A>G), GTG94ATC (27475G>A 27477G>C), GAC95AAC (27478G>A) | | | | | | | | |
| KMC43_gp46 | 18 | 218 | 51.5% | 814 | 71.2% | 193 (100%) | 121 (62.7%) | 0/0/0/0 | 0 |
| Protein mutations: | Y22A (27641T>G 27642A>C 27643C>A), A25Q (27650G>C 27651C>A 27652C>G), L29M (27662C>A), V31I (27668G>A), I34V (27677A>G 27679C>G), V38F (27689G>T 27691G>T), N42V (27701A>G 27702A>T 27703T>G), V45I (27710G>A 27712C>T), V46A (27714T>C), I47L (27716A>C), L52I (27731C>A), I53A (27734A>G 27735T>C 27736C>G), M54L (27737A>C), V56I (27743G>A 27745G>T), L57A (27746C>G 27747T>C 27748G>A), A58S (27749G>T), I59L (27752A>C 27754C>G), A62F (27761G>T 27762C>T 27763G>T), A65S (27770G>A 27771C>G 27772G>C), T66N (27774C>A 27775C>T), T68A (27779A>G 27781G>C), L70A (27785C>G 27786T>C 27787T>G), T72S (27792C>G 27793G>T), V74A (27798T>C), V79F (27812G>T 27814G>C), T80A (27815A>G), V85Q (27830G>C 27831T>A), A86Q (27833G>C 27834C>A), T90Q (27845A>C 27846C>A), A92T (27851G>A), S94A (27857T>G), T98S (27869A>T), Q100T (27875C>A 27876A>C 27877G>C), T107A (27896A>G 27898A>G), A108S (27899G>T 27901C>T), T116S (27923A>T), Q118K (27929C>A 27931A>G), F119Y (27933T>A), S132A (27971T>G), I139M (27994T>G), A144T (28007G>A 28009G>C), V146C (28013G>T 28014T>G 28015G>C), N147V (28016A>G 28017A>T 28018T>G), V148A (28020T>C 28021A>G), P149T (28022C>A 28024G>C), V151K (28028G>A 28029T>A), A152S (28031G>T), T154N (28038C>A 28039G>C), G156A (28044G>C 28045C>G), G159A (28053G>C 28054C>G), V162A (28062T>C), A163I (28064G>A 28065C>T 28066T>C), G164A (28068G>C), V166I (28073G>A 28075A>C), S167N (28076T>A 28077C>A 28078G>C), I169L (28082A>C), V170G (28086T>G), G171Q (28088G>C 28089G>A 28090C>G), S174A (28097T>G 28099C>G), N187T (28137A>C 28138C>G), A193T (28154G>A 28156A>G), L197F (28166C>T 28168G>C), T199S (28172A>T), I202V (28181A>G), A203G (28185C>G), V205I (28190G>A 28192A>C), A206N (28193G>A 28194C>A 28195G>C), K213R (28215A>G 28216A>G), S214A (28217T>G 28219C>T), V216I (28223G>A) | | | | | | | | |
| Codon mutations: | TAC22GCA (27641T>G 27642A>C 27643C>A), TCT23TCG (27646T>G), GCC25CAG (27650G>C 27651C>A 27652C>G), TTA26CTC (27653T>C 27655A>C), GTT27GTC (27658T>C), CTG29ATG (27662C>A), GTC31ATC (27668G>A), TCG33TCC (27676G>C), ATC34GTG (27677A>G 27679C>G), CGC36CGG (27685C>G), GTG38TTT (27689G>T 27691G>T), AAT42GTG (27701A>G 27702A>T 27703T>G), GCT43GCG (27706T>G), GCC44GCA (27709C>A), GTC45ATT (27710G>A 27712C>T), GTG46GCG (27714T>C), ATC47CTC (27716A>C), CTC52ATC (27731C>A), ATC53GCG (27734A>G 27735T>C 27736C>G), ATG54CTG (27737A>C), CAA55CAG (27742A>G), GTG56ATT (27743G>A 27745G>T), CTG57GCA (27746C>G 27747T>C 27748G>A), GCG58TCG (27749G>T), ATC59CTG (27752A>C 27754C>G), GCG62TTT (27761G>T 27762C>T 27763G>T), GCT63GCG (27766T>G), ACG64ACC (27769G>C), GCG65AGC (27770G>A 27771C>G 27772G>C), ACC66AAT (27774C>A 27775C>T), ACG68GCC (27779A>G 27781G>C), CTT70GCG (27785C>G 27786T>C 27787T>G), ACG72AGT (27792C>G 27793G>T), GTG74GCG (27798T>C), GCG75GCC (27802G>C), GTG79TTC (27812G>T 27814G>C), ACG80GCG (27815A>G), CGT81CGC (27820T>C), CTT82CTG (27823T>G), GCT83GCG (27826T>G), GCG84GCC (27829G>C), GTG85CAG (27830G>C 27831T>A), GCG86CAG (27833G>C 27834C>A), ACT88ACC (27841T>C), GGA89GGC (27844A>C), ACG90CAG (27845A>C 27846C>A), GTC91GTG (27850C>G), GCC92ACC (27851G>A), TCG94GCG (27857T>G), ACG97ACT (27868G>T), ACG98TCG (27869A>T), ACC99ACG (27874C>G), CAG100ACC (27875C>A 27876A>C 27877G>C), GTC102GTT (27883C>T), GTT103GTG (27886T>G), CTT106CTC (27895T>C), ACA107GCG (27896A>G 27898A>G), GCC108TCT (27899G>T 27901C>T), GTT109GTC (27904T>C), ACA112ACG (27913A>G), GGG115GGC (27922G>C), ACG116TCG (27923A>T), CAA118AAG (27929C>A 27931A>G), TTC119TAC (27933T>A), GCT129GCC (27964T>C), TAT130TAC (27967T>C), TCG132GCG (27971T>G), ACA133ACG (27976A>G), GGC136GGG (27985C>G), TAC137TAT (27988C>T), ATT139ATG (27994T>G), GCT140GCG (27997T>G), GCG141GCC (28000G>C), GTG143GTC (28006G>C), GCG144ACC (28007G>A 28009G>C), AGC145AGT (28012C>T), GTG146TGC (28013G>T 28014T>G 28015G>C), AAT147GTG (28016A>G 28017A>T 28018T>G), GTA148GCG (28020T>C 28021A>G), CCG149ACC (28022C>A 28024G>C), GTG151AAG (28028G>A 28029T>A), GCA152TCA (28031G>T), GTG153GTC (28036G>C), ACG154AAC (28038C>A 28039G>C), GCC155GCA (28042C>A), GGC156GCG (28044G>C 28045C>G), GCT158GCG (28051T>G), GGC159GCG (28053G>C 28054C>G), GTG162GCG (28062T>C), GCT163ATC (28064G>A 28065C>T 28066T>C), GGC164GCC (28068G>C), GCC165GCG (28072C>G), GTA166ATC (28073G>A 28075A>C), TCG167AAC (28076T>A 28077C>A 28078G>C), ACC168ACG (28081C>G), ATC169CTC (28082A>C), GTC170GGC (28086T>G), GGC171CAG (28088G>C 28089G>A 28090C>G), GCA172GCC (28093A>C), ATT173ATC (28096T>C), TCC174GCG (28097T>G 28099C>G), GAT177GAC (28108T>C), GTC179GTG (28114C>G), ACG180ACC (28117G>C), GCC182GCG (28123C>G), GCA183GCG (28126A>G), AAC187ACG (28137A>C 28138C>G), GGC188GGT (28141C>T), GAT190GAC (28147T>C), GCA193ACG (28154G>A 28156A>G), GAC194GAT (28159C>T), GCT195GCC (28162T>C), GCG196GCC (28165G>C), CTG197TTC (28166C>T 28168G>C), CGT198CGC (28171T>C), ACG199TCG (28172A>T), CGC200CGG (28177C>G), ATC202GTC (28181A>G), GCC203GGC (28185C>G), GTA205ATC (28190G>A 28192A>C), GCG206AAC (28193G>A 28194C>A 28195G>C), AAG210AAA (28207G>A), ACG212ACC (28213G>C), AAA213AGG (28215A>G 28216A>G), TCC214GCT (28217T>G 28219C>T), GCT215GCC (28222T>C), GTC216ATC (28223G>A) | | | | | | | | |
| KMC43_gp49 | 105 | 204 | 17.3% | 517 | 79.4% | 100 (100%) | 79 (79.0%) | 0/0/0/0 | 0 |
| Protein mutations: | Q109Y (31508C>T 31510G>C), T118A (31535A>G 31537A>C), Q122S (31547C>A 31548A>G 31549G>C), D125N (31556G>A), T127V (31562A>G 31563C>T 31564T>G), A128T (31565G>A), W141Y (31605G>A 31606G>T), S142Q (31607A>C 31608G>A 31609C>G), N145A (31616A>G 31617A>C 31618C>G), N146G (31619A>G 31620A>G 31621C>A), S147N (31623G>A), A150S (31631G>A 31632C>G 31633A>T), T154A (31643A>G), V167I (31682G>A 31684A>C), S173T (31701G>C 31702C>G), S181V (31724T>G 31725C>T), T184V (31733A>G 31734C>T 31735C>G), W187Y (31743G>A 31744G>T), V188S (31745G>T 31746T>C), N193Y (31760A>T) | | | | | | | | |
| Codon mutations: | ACT106ACC (31501T>C), GTG107GTC (31504G>C), CAG109TAC (31508C>T 31510G>C), ATT115ATC (31528T>C), CAG116CAA (31531G>A), ACA118GCC (31535A>G 31537A>C), TAC119TAT (31540C>T), CAA120CAG (31543A>G), GAT121GAC (31546T>C), CAG122AGC (31547C>A 31548A>G 31549G>C), GAC125AAC (31556G>A), CTG126TTG (31559C>T), ACT127GTG (31562A>G 31563C>T 31564T>G), GCG128ACG (31565G>A), TGG141TAT (31605G>A 31606G>T), AGC142CAG (31607A>C 31608G>A 31609C>G), AAC145GCG (31616A>G 31617A>C 31618C>G), AAC146GGA (31619A>G 31620A>G 31621C>A), AGC147AAC (31623G>A), GCA150AGT (31631G>A 31632C>G 31633A>T), GCG152GCC (31639G>C), ACT154GCT (31643A>G), GTC159GTG (31660C>G), GTT160GTG (31663T>G), GCC163GCG (31672C>G), GCT165GCA (31678T>A), GTA167ATC (31682G>A 31684A>C), GCG168GCC (31687G>C), GCT169GCA (31690T>A), ACC170ACG (31693C>G), GGC172GGA (31699C>A), AGC173ACG (31701G>C 31702C>G), ACC175ACG (31708C>G), ACC176ACG (31711C>G), CCA177CCG (31714A>G), GCC178GCA (31717C>A), TCC181GTC (31724T>G 31725C>T), GGC182GGT (31729C>T), ACC184GTG (31733A>G 31734C>T 31735C>G), GGG185GGT (31738G>T), CTG186CTC (31741G>C), TGG187TAT (31743G>A 31744G>T), GTG188TCG (31745G>T 31746T>C), ACC190ACG (31753C>G), GTC191GTT (31756C>T), AAT193TAT (31760A>T), GGT194GGC (31765T>C), ACG196ACC (31771G>C), ACC197ACG (31774C>G), GCG200GCC (31783G>C), CCG201CCC (31786G>C), AAC202AAT (31789C>T) | | | | | | | | |

**Proteins**

| | Begin | End | Coverage | Score | Concordance | Matches | Identities | I/D/M/F* | Stop Codons |
|---|---|---|---|---|---|---|---|---|---|
| DUF4043 domain-containing protein (YP_010078151.1) | 1 | 117 | 26.4% | 715 | 90.2% | 117 (100%) | 103 (88.0%) | 0/0/0/0 | 0 |
| Protein mutations: | S4Q (14205A>C 14206G>A 14207C>A), D25E (14270T>A), A26G (14272C>G), I30M (14285C>G), G32S (14289G>A), F33Y (14293T>A 14294C>T), Q35S (14298C>T 14299A>C), R41Q (14317G>A 14318T>G), P67A (14394C>G), S68A (14397T>G 14399G>C), A69S (14400G>A 14401C>G 14402A>C), S78T (14427T>A), S88T (14457T>A), T102M (14500C>T) | | | | | | | | |
| Codon mutations: | ATC3ATT (14204C>T), AGC4CAA (14205A>C 14206G>A 14207C>A), AAT5AAC (14210T>C), GCC8GCT (14219C>T), GCT9GCC (14222T>C), CCG12CCC (14231G>C), TTT23TTC (14264T>C), GAT25GAA (14270T>A), GCC26GGC (14272C>G), ATC30ATG (14285C>G), CTC31CTG (14288C>G), GGC32AGC (14289G>A), TTC33TAT (14293T>A 14294C>T), CAG35TCG (14298C>T 14299A>C), GCA37GCC (14306A>C), CGT38CGC (14309T>C), CGT41CAG (14317G>A 14318T>G), GGT47GGC (14336T>C), GAA48GAG (14339A>G), ACC49ACA (14342C>A), CGT54CGC (14357T>C), GGT56GGC (14363T>C), CTG57CTC (14366G>C), GCG59GCT (14372G>T), CCG60CCC (14375G>C), GTC61GTG (14378C>G), ACC62ACG (14381C>G), CCG67GCG (14394C>G), TCG68GCC (14397T>G 14399G>C), GCA69AGC (14400G>A 14401C>G 14402A>C), TCG78ACG (14427T>A), CCG79CCC (14432G>C), TCG80AGC (14433T>A 14434C>G 14435G>C), GTG84GTC (14447G>C), TCG88ACG (14457T>A), CTC89CTG (14462C>G), ACG97ACC (14486G>C), ACG102ATG (14500C>T), GTG103GTC (14504G>C), GTC107GTG (14516C>G), GTG110GTC (14525G>C) | | | | | | | | |
| hypothetical protein (YP_010078153.1) | 15 | 64 | 78.1% | -12 | -3.5% | 50 (94.3%) | 19 (35.8%) | 3/0/1/1 | 1 |
| Protein mutations: | S18L (15870C>T 15871C>G), I19V (15872A>G), L20V (15875C>G), D22E (15883C>G), E23R (15884G>C 15885A>G 15886G>A), S24D (15887T>G 15888C>A), I25F (15890A>T), W27F (15897G>T 15898G>C), M28S (15900T>G 15901G>C), E29V (15903A>T 15904A>C), I32L (15911A>C 15913C>G), Q35V (15920C>G 15921A>T), F36W (15924T>G 15925C>G), H37A (15926C>G 15927A>C), E38A (15930A>C 15931A>T), T40K (15936C>A), H41P (15939A>C), V42I (15941G>A), N44E (15947A>G 15949C>A), P45V (15950C>G 15951C>T 15952T>G), V46R (15953G>C 15954T>G), V48I (15959G>A 15961G>C), A49R (15962G>C 15963C>G), A50E (15966C>A), Q53A (15974C>G 15975A>C 15976G>C), H54N (15977C>A), G55D (15981G>A), R57P (15987G>C 15988T>G), R57_Y58insIKA (15988_15989insATCAAGGCC), L59R (15992T>C 15993T>G), V61I (15998G>A) | | | | | | | | |

| | Begin | End | Coverage | Score | Concordance | Matches | Identities | I/D/M/F* | Stop Codons |
|---|---|---|---|---|---|---|---|---|---|
| **NT** | **14196** | **31794** | **9.4%** | **3582** | **41.3%** | **4358 (99.6%)** | **3090 (70.6%)** | **9/8** | |
| Codon mutations: | CCC16CCG (15865C>G), GAT17GAC (15868T>C), TCC18TTG (15870C>T 15871C>G), ATC19GTC (15872A>G), CTG20GTG (15875C>G), TTG21CTG (15878T>C), GAC22GAG (15883C>G), GAG23CGA (15884G>C 15885A>G 15886G>A), TCC24GAC (15887T>G 15888C>A), ATC25TTC (15890A>T), GGG26GGC (15895G>C), TGG27TTC (15897G>T 15898G>C), ATG28AGC (15900T>G 15901G>C), GAA29GTC (15903A>T 15904A>C), ATC32CTG (15911A>C 15913C>G), CGA34CGC (15919A>C), CAG35GTG (15920C>G 15921A>T), TTC36TGG (15924T>G 15925C>G), CAC37GCC (15926C>G 15927A>C), GAA38GCT (15930A>C 15931A>T), GGT39GGC (15934T>C), ACG40AAG (15936C>A), CAC41CCC (15939A>C), GTC42ATC (15941G>A), AAC44GAA (15947A>G 15949C>A), CCT45GTG (15950C>G 15951C>T 15952T>G), GTC46CGC (15953G>C 15954T>G), GTG48ATC (15959G>A 15961G>C), GCC49CGC (15962G>C 15963C>G), GCG50GAG (15966C>A), CAG53GCC (15974C>G 15975A>C 15976G>C), CAC54AAC (15977C>A), GGC55GAC (15981G>A), CGT57CCG (15987G>C 15988T>G), CGT57_TAC58insATCAAGGCC (15988_15989insATCAAGGCC), TTG59CGG (15992T>C 15993T>G), GTC61ATC (15998G>A), AGC62-TG (16001delA 16002G>T 16003C>G) | | | | | | | | |
| hypothetical protein (YP_010078154.1) | 1 | 103 | 100% | 538 | 80.9% | 103 (100%) | 77 (74.8%) | 0/0/0/0 | 1 |
| Protein mutations: | V5I (16018G>A), I7T (16025T>C 16026T>C), K11R (16036A>C 16037A>G 16038A>C), S12N (16040G>A), T14A (16045A>G 16047C>G), T18A (16057A>G 16059G>T), A19G (16061C>G 16062C>G), Q21T (16066C>A 16067A>C), S25T (16078T>A), K28A (16087A>G 16088A>C 16089G>C), D29G (16091A>G), I32L (16099A>C 16101C>G), N41Q (16126A>C 16128T>G), A48T (16147G>A 16149G>T), G49P (16150G>C 16151G>C), T58V (16177A>G 16178C>T 16179G>C), T60G (16183A>G 16184C>G), A62V (16190C>T 16191T>C), N65V (16198A>G 16199A>T), I67V (16204A>G), G68A (16208G>C 16209C>G), S76T (16231T>A 16233A>C), N85A (16258A>G 16259A>C 16260C>A), I89V (16270A>G), A93T (16282G>A 16284T>C), S98A (16297T>G) | | | | | | | | |
| Codon mutations: | GAC3GAT (16014C>T), GTC5ATC (16018G>A), ATT7ACC (16025T>C 16026T>C), GGC10GGG (16035C>G), AAA11CGC (16036A>C 16037A>G 16038A>C), AGC12AAC (16040G>A), ACC14GCG (16045A>G 16047C>G), AAT16AAC (16053T>C), ACG18GCT (16057A>G 16059G>T), GCC19GGG (16061C>G 16062C>G), GCG20GCC (16065G>C), CAG21ACG (16066C>A 16067A>C), GTG22GTC (16071G>C), GTT23GTC (16074T>C), TCG25ACG (16078T>A), CCC27CCA (16086C>A), AAG28GCC (16087A>G 16088A>C 16089G>C), GAC29GGC (16091A>G), TTT30TTC (16095T>C), ATC32CTG (16099A>C 16101C>G), CGA36CGC (16113A>C), AAT41CAG (16126A>C 16128T>G), GTT42GTG (16131T>G), CTC43CTG (16134C>G), GTT44GTG (16137T>G), GCG45GCC (16140G>C), GGG46GGC (16143G>C), ACT47ACC (16146T>C), GCG48ACT (16147G>A 16149G>T), GGG49CCG (16150G>C 16151G>C), GCC51GCG (16158C>G), GCG52GCC (16161G>C), TAC53TAT (16164C>T), GCG55GCA (16170G>A), ACG57ACC (16176G>C), ACG58GTC (16177A>G 16178C>T 16179G>C), GCC59GCT (16182C>T), ACC60GGC (16183A>G 16184C>G), GCT62GTC (16190C>T 16191T>C), AAC65GTC (16198A>G 16199A>T), ATC67GTC (16204A>G), GGC68GCG (16208G>C 16209C>G), GCG69GCC (16212G>C), CCG71CCC (16218G>C), GTG74GTC (16227G>C), TCA76ACC (16231T>A 16233A>C), CTC78CTG (16239C>G), GAT80GAC (16245T>C), CCG82CCC (16251G>C), CTG84TTG (16255C>T), AAC85GCA (16258A>G 16259A>C 16260C>A), GGC86GGG (16263C>G), ATC89GTC (16270A>G), CCT91CCC (16278T>C), GCT93ACC (16282G>A 16284T>C), ACC96ACG (16293C>G), GTT97GTC (16296T>C), TCT98GCT (16297T>G), GTT99GTC (16302T>C), TCT100TCC (16305T>C) | | | | | | | | |
| hypothetical protein (YP_010078155.1) | 1 | 117 | 91.4% | 710 | 86.3% | 117 (100%) | 93 (79.5%) | 0/0/0/0 | 0 |
| Protein mutations: | A6Q (16339G>C 16340C>A 16341C>G), K8R (16345A>C 16346A>G), G16W (16369G>T 16371A>G), M19V (16378A>G 16380G>C), F20Y (16382T>A 16383C>T), S26A (16399T>G), M43L (16450A>C), N44S (16453A>T 16454A>C 16455C>G), M46L (16459A>C), A48P (16465G>C 16467C>G), A52G (16478C>G 16479T>C), V54I (16483G>A 16485G>C), S71G (16534T>G 16535C>G 16536T>C), A72T (16537G>A), Q80K (16561C>A), V85T (16576G>A 16577T>C 16578G>C), V91L (16594G>C), A99S (16618G>A 16619C>G), R103L (16631G>T 16632C>G), E107Q (16642G>C), M109L (16648A>C 16650G>C), I111V (16654A>G 16656T>G), G116A (16670G>C 16671G>C) | | | | | | | | |
| Codon mutations: | ACA4ACC (16335A>C), ACC5ACG (16338C>G), GCC6CAG (16339G>C 16340C>A 16341C>G), AAG8CGG (16345A>C 16346A>G), TAT14TAC (16365T>C), TGC15TGT (16368C>T), GGA16TGG (16369G>T 16371A>G), ATG19GTC (16378A>G 16380G>C), TTC20TAT (16382T>A 16383C>T), GGG22GGC (16389G>C), CCA24CCC (16395A>C), TCG25AGT (16396T>A 16397C>G 16398G>T), TCG26GCG (16399T>G), TAT30TAC (16413T>C), CGG31AGG (16414C>A), GCG35GCC (16428G>C), TAC36TAT (16431C>T), CTT39CTG (16440T>G), GAA40GAG (16443A>G), ATG43CTG (16450A>C), AAC44TCG (16453A>T 16454A>C 16455C>G), ATG46CTG (16459A>C), CTG47CTC (16464G>C), GCC48CCG (16465G>C 16467C>G), GAA50GAG (16473A>G), GCT52GGC (16478C>G 16479T>C), GTG54ATC (16483G>A 16485G>C), ACG57ACC (16494G>C), TAT58TAC (16497T>C), CTG59CTC (16500G>C), GCT60GCC (16503T>C), ACC63ACG (16512C>G), ACT64ACG (16515T>G), CTC65CTG (16518C>G), ACG67ACC (16524G>C), GCA68GCC (16527A>C), TCT71GGC (16534T>G 16535C>G 16536T>C), GCG72ACG (16537G>A), AGC73TCG (16540A>T 16541G>C 16542C>G), GCG74GCA (16545G>A), CTC76CTT (16551C>T), ACA78ACG (16557A>G), CAG80AAG (16561C>A), GCG81GCC (16566G>C), GCC82GCG (16569C>G), GTG85ACC (16576G>A 16577T>C 16578G>C), GAA90GAG (16593A>G), GTG91CTG (16594G>C), GCA92GCC (16599A>C), GAC93GAT (16602C>T), CGC94CGT (16605C>T), GAC95GAT (16608C>T), TTT98TTC (16617T>C), GCC99AGC (16618G>A 16619C>G), CGG102CGC (16629G>C), CGC103CTG (16631G>T 16632C>G), GAG107CAG (16642G>C), ATG109CTC (16648A>C 16650G>C), ATT111GTG (16654A>G 16656T>G), CCC113CCG (16662C>G), GGG114GGC (16665G>C), CCC115CCG (16668C>G), GGG116GCC (16670G>C 16671G>C), CTC117T.. (16672C>T) | | | | | | | | |
| tail protein (YP_010078160.1) | 17 | 497 | 72.0% | 1825 | 76.6% | 368 (100%) | 262 (71.2%) | 0/0/0/0 | 0 |
| Protein mutations: | V28T (19290G>A 19291T>C 19292T>C), S29Q (19293T>C 19294C>A 19295C>G), L30Y (19296C>T 19297T>A 19298G>C), V38I (19320G>A 19322G>C), V41I (19329G>A 19331T>C), T46S (19344A>T), L55T (19371C>A 19372T>C), I56V (19374A>G), N58S (19381A>G), M59P (19383A>C 19384T>C), M61Q (19389A>C 19390T>A), Y62A (19392T>G 19393A>C), Q64A (19398C>G 19399A>C 19400G>C), A65I (19401G>A 19402C>T), A68Q (19410G>C 19411C>A), I69A (19413A>G 19414T>C), T73K (19426C>A 19427C>G), M76L (19434A>C), G77M (19437G>A 19438G>T 19439C>G), V80I (19446G>A 19448G>C), A81S (19449G>T), I82V (19452A>G), V84T (19458G>A 19459T>C 19460G>C), Q85M (19461C>A 19462A>T), R92K (19482C>A 19483G>A 19484C>A), C93A (19485T>G 19486G>C), T102V (19512A>G 19513C>T), T106I (19525C>T 19526G>C), I107A (19527A>G 19528T>C 19529C>G), Q109L (19534A>T), G112C (19542G>T), V113I (19545G>A 19547G>C), L115F (19551C>T 19553G>C), A117S (19557G>T 19559G>C), L123F (19575C>T 19577G>C), S126G (19584T>G 19585C>G 19586G>C), V127I (19587G>A 19589G>C), A130T (19596G>A), A132S (19602G>T 19604C>G), A136Q (19614G>C 19615C>A 19616C>G), N138S (19620A>T 19621A>C 19622T>G), W140Q (19626T>C 19627G>A), V142A (19633T>C 19634A>C), T143V (19635A>G 19636C>T), A145S (19641G>T), I266T (20005T>C 20006C>G), I269V (20013A>G), G270M (20016G>A 20017G>T 20018C>G), V271T (20019G>A 20020T>C), A274S (20028G>A 20029C>G), S279A (20043A>G 20044G>C 20045C>G), T282I (20053C>T), T283S (20056C>G 20057G>C), T284A (20058A>G), A286S (20064G>A 20065C>G), I290V (20076A>G 20078C>G), F295A (20091T>G 20092T>C), V303C (20115G>T 20116T>G 20117G>C), V309Y (20133G>T 20134T>A), S311Q (20139A>C 20140G>A 20141C>G), V312I (20142G>A 20144T>C), T313L (20145A>C 20146C>T 20147A>T), A320S (20166G>T 20168G>C), S327A (20187T>G 20189G>C), T335S (20211A>T), Y341Q (20229T>C 20231T>G), V343I (20235G>A 20237G>C), T354Q (20268A>C 20269C>A), S362Q (20292T>C 20293C>A), I364G (20298A>G 20299T>G), Q365I (20301C>A 20302A>T 20303G>C), C383A (20355T>G 20356G>C 20357C>G), S385F (20362C>T 20363G>C), A393P (20385G>C), L394V (20388C>G), Q396N (20394C>A 20396G>C), S409Y (20433A>T 20434G>A), V411L (20439G>C), Y418F (20461A>T), V419I (20463G>A), A425S (20481G>T 20483C>G), A429T (20493G>A 20495G>T), A432M (20502G>A 20503C>T 20504C>G), L435I (20511C>A 20513G>C), N437A (20517A>G 20518A>C), S440D (20526T>G 20527C>A), S441N (20530G>A), M442L (20532A>C), E443Q (20535G>C), M447L (20547A>C), A450D (20557C>A), V451P (20559G>C 20560T>C 20561C>G), P455A (20571C>G 20573A>G), S456A (20574T>G 20576G>C), F457Y (20578T>A), I461V (20589A>G 20591C>G), N464A (20598A>G 20599A>C), M468A (20610A>G 20611T>C), N469S (20613A>T 20614A>C 20615C>G), L476M (20634C>A 20636T>G), I483K (20656T>A 20657T>G), S486A (20664T>G 20666G>C), I492L (20682A>C), V493I (20685G>A 20687G>C), L495I (20691C>A 20693T>C) | | | | | | | | |

| | Begin | End | Coverage | Score | Concordance | Matches | Identities | I/D/M/F* | Stop Codons |
|---|---|---|---|---|---|---|---|---|---|
| **NT** | **14196** | **31794** | **9.4%** | **3582** | **41.3%** | **4358 (99.6%)** | **3090 (70.6%)** | **9/8** | |
| Codon mutations: | TTG19CTG (19263T>C), GTA21GTG (19271A>G), CAA22CAG (19274A>G), GTG24GTC (19280G>C), CCT25CCG (19283T>G), CCA26CCG (19286A>G), GTT28ACC (19290G>A 19291T>C 19292T>C), TCC29CAG (19293T>C 19294C>A 19295C>G), CTG30TAC (19296C>T 19297T>A 19298G>C), CTC31CTG (19301C>G), AAT37AAC (19319T>C), GTG38ATC (19320G>A 19322G>C), CTT39TTG (19323C>T 19325T>G), GGT40GGC (19328T>C), GTT41ATC (19329G>A 19331T>C), GTG42GTC (19334G>C), ACC44ACG (19340C>G), GCA45GCG (19343A>G), ACC46TCC (19344A>T), GGC48GGG (19352C>G), ACC50ACG (19358C>G), TCG52AGC (19362T>A 19363C>G 19364G>C), CCG53CCC (19367G>C), ACG54ACC (19370G>C), CTG55ACG (19371C>A 19372T>C), ATC56GTC (19374A>G), AAC58AGC (19381A>G), ATG59CCG (19383A>C 19384T>C), GCG60GCT (19388G>T), ATG61CAG (19389A>C 19390T>A), TAC62GCC (19392T>G 19393A>C), GCC63GCG (19397C>G), CAG64GCC (19398C>G 19399A>C 19400G>C), GCC65ATC (19401G>A 19402C>T), GCG68CAG (19410G>C 19411C>A), ATC69GCC (19413A>G 19414T>C), CGT72CGC (19424T>C), ACC73AAG (19426C>A 19427C>G), TAT74TAC (19430T>C), ATG76CTG (19434A>C), GGC77ATG (19437G>A 19438G>T 19439C>G), GCT79GCC (19445T>C), GTG80ATC (19446G>A 19448G>C), GCG81TCG (19449G>T), ATC82GTC (19452A>G), GCA83GCG (19457A>G), GTG84ACC (19458G>A 19459T>C 19460G>C), CAG85ATG (19461C>A 19462A>T), GCG88GCC (19472G>C), CGC92AAA (19482C>A 19483G>A 19484C>A), TGC93GCC (19485T>G 19486G>C), CGC95CGG (19493C>G), GTC96GTG (19496C>G), ACG102GTG (19512A>G 19513C>T), GCG104GCC (19520G>C), TCG105AGC (19521T>A 19522C>G 19523G>C), ACG106ATC (19525C>T 19526G>C), ATC107GCG (19527A>G 19528T>C 19529C>G), CAG109CTG (19534A>T), GGC112TGC (19542G>T), GTG113ATC (19545G>A 19547G>C), ACG114ACC (19550G>C), CTG115TTC (19551C>T 19553G>C), GCG117TCC (19557G>T 19559G>C), ACG122ACC (19574G>C), CTG123TTC (19575C>T 19577G>C), TCG126GGC (19584T>G 19585C>G 19586G>C), GTG127ATC (19587G>A 19589G>C), GTC129GTG (19595C>G), GCG130ACG (19596G>A), CTC131CTG (19601C>G), GCC132TCG (19602G>T 19604C>G), GCG133GCC (19607G>C), AGC135TCG (19611A>T 19612G>C 19613C>G), GCC136CAG (19614G>C 19615C>A 19616C>G), AAT138TCG (19620A>T 19621A>C 19622T>G), TGG140CAG (19626T>C 19627G>A), GTA142GCC (19633T>C 19634A>C), ACG143GTG (19635A>G 19636C>T), GTC144GTT (19640C>T), GCG145TCG (19641G>T), ATC266ACG (20005T>C 20006C>G), ATC269GTC (20013A>G), GGC270ATG (20016G>A 20017G>T 20018C>G), GTC271ACC (20019G>A 20020T>C), GCT274AGT (20028G>A 20029C>G), ACG277ACC (20039G>C), AGC279GCG (20043A>G 20044G>C 20045C>G), GCC281GCG (20051C>G), ACC282ATC (20053C>T), ACG283AGC (20056C>G 20057G>C), ACC284GCC (20058A>G), GCC286AGC (20064G>A 20065C>G), ATC290GTG (20076A>G 20078C>G), TTC295GCC (20091T>G 20092T>C), CTG298CTT (20102G>T), GGT300GGC (20108T>C), GTG303TGC (20115G>T 20116T>G 20117G>C), TTG306CTG (20124T>C), ACG308ACC (20132G>C), GTC309TAC (20133G>T 20134T>A), AGC311CAG (20139A>C 20140G>A 20141C>G), GTT312ATC (20142G>A 20144T>C), ACA313CTT (20145A>C 20146C>T 20147A>T), CTG315TTG (20151C>T), TCG317AGC (20157T>A 20158C>G 20159G>C), CAA319CAG (20165A>G), GCG320TCC (20166G>T 20168G>C), GTC322GTG (20174C>G), GCT323GCC (20177T>C), GGT324GGC (20180T>C), TCG327GCC (20187T>G 20189G>C), ACG335TCG (20211A>T), TAT341CAG (20229T>C 20231T>G), GTG343ATC (20235G>A 20237G>C), GTG344GTC (20240G>C), TCG349AGC (20253T>A 20254C>G 20255G>C), ACG354CAG (20268A>C 20269C>A), TCC357TCG (20279C>G), GCC358GCT (20282C>T), TTG360CTG (20286T>C), TCG362CAG (20292T>C 20293C>A), ATC364GGC (20298A>G 20299T>G), CAG365ATC (20301C>A 20302A>T 20303G>C), GCG366GCC (20306G>C), GGT367GGC (20309T>C), CCC376CCG (20336C>G), TCG379AGC (20343T>A 20344C>G 20345G>C), TGC383GCG (20355T>G 20356G>C 20357C>G), CGG384CGC (20360G>C), TCG385TTC (20362C>T 20363G>C), CAT387CAC (20369T>C), GCG393CCG (20385G>C), CTG394GTG (20388C>G), CAG396AAC (20394C>A 20396G>C), ACC401ACG (20411C>G), AGC409TAC (20433A>T 20434G>A), GTC411CTC (20439G>C), AAT412AAC (20444T>C), TAC418TTC (20461A>T), GTC419ATC (20463G>A), GGG420GGT (20468G>T), CAA423CAG (20477A>G), TCG424AGC (20478T>A 20479C>G 20480G>C), GCC425TCG (20481G>T 20483C>G), GTG427GTC (20489G>C), GCG429ACT (20493G>A 20495G>T), GCG431GCC (20501G>C), GCC432ATG (20502G>A 20503C>T 20504C>G), CTG435ATC (20511C>A 20513G>C), TCG436AGC (20514T>A 20515C>G 20516G>C), AAC437GCC (20517A>G 20518A>C), TCC440GAC (20526T>G 20527C>A), AGC441AAC (20530G>A), ATG442CTG (20532A>C), GAG443CAG (20535G>C), GGC446GGA (20546C>A), ATG447CTG (20547A>C), GCC450GAC (20557C>A), GTC451CCG (20559G>C 20560T>C 20561C>G), AAT452AAC (20564T>C), CCA455GCG (20571C>G 20573A>G), TCG456GCC (20574T>G 20576G>C), TTC457TAC (20578T>A), TCG458AGC (20580T>A 20581C>G 20582G>C), CAA460CAG (20588A>G), ATC461GTG (20589A>G 20591C>G), GCT463GCG (20597T>G), AAC464GCC (20598A>G 20599A>C), ATG468GCG (20610A>G 20611T>C), AAC469TCG (20613A>T 20614A>C 20615C>G), CGG470CGC (20618G>C), GCG472GCC (20624G>C), CTT473CTG (20627T>G), CTT476ATG (20634C>A 20636T>G), GCT478GCC (20642T>C), GTC480GTG (20648C>G), ATT483AAG (20656T>A 20657T>G), TCG486GCC (20664T>G 20666G>C), GTT487GTC (20669T>C), ATC492CTC (20682A>C), GTG493ATC (20685G>A 20687G>C), AAT494AAC (20690T>C), CTT495ATC (20691C>A 20693T>C) | | | | | | | | |
| hypothetical protein (YP_010078166.1) | 12 | 115 | 32.4% | 453 | 59.9% | 104 (100%) | 57 (54.8%) | 0/0/0/0 | 0 |
| Protein mutations: | D14G (24489A>G), R15S (24491C>T 24492G>C), Q16L (24495A>T 24496G>C), I18V (24500A>G 24502T>G), E20A (24507A>C 24508G>C), S21E (24509T>G 24510C>A), P23E (24515C>G 24516C>A), T27A (24527A>G 24529C>G), R32V (24542C>G 24543G>T 24544C>G), D34E (24550C>G), V36I (24554G>A), V42I (24572G>A), A47R (24587G>C 24588C>G), F48S (24591T>C 24592C>G), T49D (24593A>G 24594C>A 24595G>T), E50R (24596G>C 24597A>G), I52L (24602A>C), N53G (24605A>G 24606A>G), W57M (24617T>A 24618G>T), L58M (24620C>A), Q61V (24629C>G 24630A>T 24631A>C), D62G (24633A>G 24634T>C), L64M (24638T>A), A65D (24642C>A 24643G>C), R68L (24651G>T), Q69Y (24653C>T 24655G>C), A72G (24663C>G 24664G>C), M73L (24665A>C), V75I (24671G>A 24673T>C), A76G (24675C>G 24676G>C), T78Q (24680A>C 24681C>A 24682C>G), V81E (24690T>A 24691G>A), Q83S (24695C>A 24696A>G 24697G>C), S85G (24701T>G 24702C>G 24703A>C), E86Q (24704G>C), S88R (24710T>C 24711C>G), F89Y (24714T>A 24715T>C), S90K (24716T>A 24717C>A 24718C>G), V92I (24722G>A 24724G>C), I93V (24725A>G), D95E (24733C>G), R97S (24737C>T 24738G>C 24739C>G), D99T (24743G>A 24744A>C 24745T>C), F100Y (24747T>A 24748C>T), F103A (24755T>G 24756T>C 24757C>T), T111V (24779A>G 24780C>T 24781G>C) | | | | | | | | |
| Codon mutations: | GAC14GGC (24489A>G), CGG15TCG (24491C>T 24492G>C), CAG16CTC (24495A>T 24496G>C), ATT18GTG (24500A>G 24502T>G), CCC19CCG (24505C>G), GAG20GCC (24507A>C 24508G>C), TCG21GAG (24509T>G 24510C>A), CCG23GAG (24515C>G 24516C>A), GGG26GGC (24526G>C), ACC27GCG (24527A>G 24529C>G), CAA28CAG (24532A>G), GCG31GCC (24541G>C), CGC32GTG (24542C>G 24543G>T 24544C>G), GAC34GAG (24550C>G), CTT35CTG (24553T>G), GTC36ATC (24554G>A), GGA38GGC (24562A>C), CGG40CGC (24568G>C), GTC42ATC (24572G>A), GCG44GCC (24580G>C), GCC47CGC (24587G>C 24588C>G), TTC48TCG (24591T>C 24592C>G), ACG49GAT (24593A>G 24594C>A 24595G>T), GAG50CGG (24596G>C 24597A>G), CCC51CCG (24601C>G), ATC52CTC (24602A>C), AAC53GGC (24605A>G 24606A>G), TCT55TCC (24613T>C), GGC56GGG (24616C>G), TGG57ATG (24617T>A 24618G>T), CTG58ATG (24620C>A), CTT59CTC (24625T>C), GGC60GGT (24628C>T), CAA61GTC (24629C>G 24630A>T 24631A>C), GAT62GGC (24633A>G 24634T>C), GCT63GCC (24637T>C), TTG64ATG (24638T>A), GCG65GAC (24642C>A 24643G>C), CGA66CGC (24646A>C), CGG68CTG (24651G>T), CAG69TAC (24653C>T 24655G>C), CTT70CTG (24658T>G), GAT71GAC (24661T>C), GCG72GGC (24663C>G 24664G>C), ATG73CTG (24665A>C), CGG74CGC (24670G>C), GTT75ATC (24671G>A 24673T>C), GCG76GGC (24675C>G 24676G>C), ACC78CAG (24680A>C 24681C>A 24682C>G), GTC79GTG (24685C>G), CAG80CAA (24688G>A), GTG81GAA (24690T>A 24691G>A), CAG83AGC (24695C>A 24696A>G 24697G>C), TCA85GGC (24701T>G 24702C>G 24703A>C), GAG86CAG (24704G>C), TTG87CTG (24707T>C), TCC88CGC (24710T>C 24711C>G), TTT89TAC (24714T>A 24715T>C), TCC90AAG (24716T>A 24717C>A 24718C>G), GTA91GTG (24721A>G), GTG92ATC (24722G>A 24724G>C), ATT93GTT (24725A>G), CGG94CGC (24730G>C), GAC95GAG (24733C>G), CGC97TCG (24737C>T 24738G>C 24739C>G), GAT99ACC (24743G>A 24744A>C 24745T>C), TTC100TAT (24747T>A 24748C>T), CGT102CGC (24754T>C), TTC103GCT (24755T>G 24756T>C 24757C>T), TAT104TAC (24760T>C), CGA109CGC (24775A>C), ACG111GTC (24779A>G 24780C>T 24781G>C), GAG113GAA (24787G>A) | | | | | | | | |
| hypothetical protein (YP_010078167.1) | 25 | 373 | 46.4% | 790 | 70.5% | 171 (98.8%) | 114 (65.9%) | 0/2/0/0 | 1 |
| Protein mutations: | A28G (25498C>G), F32W (25510T>G 25511T>G), N36A (25521A>G 25522A>C 25523C>G), F39Y (25531T>A), Q41E (25536C>G), A51T (25566G>A 25568G>T), L54M (25575C>A), P55A (25578C>G 25580G>C), A56D (25582C>A 25583G>C), E57Q (25584G>C), D59del (25590_25592delGAC), N61D (25596A>G), N65Q (25608A>C 25610C>G), K67A (25614A>G 25615A>C 25616A>C), A69I (25620G>A 25621C>T 25622G>C), F70Y (25624T>A), F74L (25635T>C 25637C>G), D80N (25653G>A), T82A (25659A>G), N83D (25662A>G), Y84P (25665T>C 25666A>C 25667C>G), E86A (25672A>C 25673G>C), T87N (25675C>A 25676G>C), D88E (25679C>A), T90S (25683A>T 25685C>G), Y102F (25720A>T), L108M (25737C>A), T110S (25744C>G), A119V (25771C>T 25772G>C), T305I (26329C>T 26330G>C), Q306A (26331C>G 26332A>C), K310R (26343A>C 26344A>G 26345G>C), L311M (26346C>A), I320L (26373A>C 26375C>G), T322Q (26379A>C 26380C>A), Q323A (26382C>G 26383A>C 26384G>C), T324K (26386C>A), A325S (26388G>T 26390C>G), M326V (26391A>G 26393G>C), L327I (26394C>A 26396T>C), R328K (26397C>A 26398G>A), L329V (26400C>G), S330Q (26403T>C 26404C>A 26405T>G), K335A (26418A>G 26419A>C 26420G>T), D339T (26430G>A 26431A>C), F341Y (26437T>A), V342P (26439G>C 26440T>C 26441A>C), D343M (26442G>A 26443A>T 26444C>G), S348M (26457T>A 26458C>T), L351M (26466C>A 26468C>G), V356T (26481G>A 26482T>C 26483G>C), E358L (26487G>T 26488A>T 26489A>G), S360D (26493T>G 26494C>A), I364R (26505A>C 26506T>G), E367D (26516A>C), V369del (26520_26522delGTG), S371L (26526T>C 26527C>T 26528C>G), I372Q (26529A>C 26530T>A) | | | | | | | | |
| Codon mutations: | GCC28GGC (25498C>G), AGT31TCC (25506A>T 25507G>C 25508T>C), TTT32TGG (25510T>G 25511T>G), GTC34GTG (25517C>G), ACA35ACC (25520A>C), AAC36GCG (25521A>G 25522A>C 25523C>G), TTC39TAC (25531T>A), TAT40TAC (25535T>C), CAG41GAG (25536C>G), GCT42GCC (25541T>C), CGT46CGC (25553T>C), GCA50GCC (25565A>C), GCG51ACT (25566G>A 25568G>T), TCT52TCG (25571T>G), CTG54ATG (25575C>A), CCG55GCC (25578C>G 25580G>C), GCG56GAC (25582C>A 25583G>C), GAG57CAG (25584G>C), ACA58ACC (25589A>C), GAC59del (25590_25592delGAC), GCG60GCC (25595G>C), AAC61GAC (25596A>G), TTC63TTT (25604C>T), TCC64TCT (25607C>T), AAC65CAG (25608A>C 25610C>G), AAA67GCC (25614A>G 25615A>C 25616A>C), GCG69ATC (25620G>A 25621C>T 25622G>C), TTC70TAC (25624T>A), ATT73ATC (25634T>C), TTC74CTG (25635T>C 25637C>G), TCC79AGC (25650T>A 25651C>G), GAT80AAT (25653G>A), ACG82GCG (25659A>G), AAC83GAC (25662A>G), TAC84CCG (25665T>C 25666A>C 25667C>G), GAG86GCC (25672A>C 25673G>C), ACG87AAC (25675C>A 25676G>C), GAC88GAA (25679C>A), CTC89CTG (25682C>G), ACC90TCG (25683A>T 25685C>G), CTG92CTC (25691G>C), TAT94TAC (25697T>C), GGG95GGC (25700G>C), GTG97GTC (25706G>C), GAT98GAC (25709T>C), TAC102TTC (25720A>T), GAC103GAT (25724C>T), CCC104CCG (25727C>G), TCG106TCC (25733G>C), CTG108ATG (25737C>A), ACC110AGC (25744C>G), GCG119GTC (25771C>T 25772G>C), GAG303GAA (26324G>A), ACG305ATC (26329C>T 26330G>C), CAG306GCG (26331C>G 26332A>C), CAC307CAT (26336C>T), GAA308GAG (26339A>G), AAG310CGC (26343A>C 26344A>G 26345G>C), CTG311ATG (26346C>A), GCG313GCC (26354G>C), GCA317GCC (26366A>C), ATC320CTG (26373A>C 26375C>G), ACG322CAG (26379A>C 26380C>A), CAG323GCC (26382C>G 26383A>C 26384G>C), ACG324AAG (26386C>A), GCC325TCG (26388G>T 26390C>G), ATG326GTC (26391A>G 26393G>C), CTT327ATC (26394C>A 26396T>C), CGG328AAG (26397C>A 26398G>A), CTG329GTG (26400C>G), TCT330CAG (26403T>C 26404C>A 26405T>G), GGG331GGC (26408G>C), AAG335GCT (26418A>G 26419A>C 26420G>T), GAT337GAC (26426T>C), GAC339ACC (26430G>A 26431A>C), TTC341TAC (26437T>A), GTA342CCC (26439G>C 26440T>C 26441A>C), GAC343ATG (26442G>A 26443A>T 26444C>G), ATT345ATC (26450T>C), ACG346ACC (26453G>C), CGG347CGA (26456G>A), TCG348ATG (26457T>A 26458C>T), CTC351ATG (26466C>A 26468C>G), GAT352GAC (26471T>C), TAT355TAC (26480T>C), GTG356ACC (26481G>A 26482T>C 26483G>C), GAA358TTG (26487G>T 26488A>T 26489A>G), TCC360GAC (26493T>G 26494C>A), ATC364CGC (26505A>C 26506T>G), AAT365AAC (26510T>C), CCT366CCC (26513T>C), GAA367GAC (26516A>C), GTG369del (26520_26522delGTG), CCC370CCG (26525C>G), TCC371CTG (26526T>C 26527C>T 26528C>G), ATA372CAA (26529A>C 26530T>A) | | | | | | | | |
| baseplate assembly protein V (YP_010078168.1) | 1 | 44 | 19.7% | 136 | 47.1% | 44 (100%) | 20 (45.5%) | 0/0/0/0 | 0 |
| Protein mutations: | Q3G (26537C>G 26538A>G 26539G>A), H4P (26541A>C 26542T>A), F5L (26543T>C), R6Q (26547G>A 26548A>G), Q8A (26552C>G 26553A>C), A10R (26558G>C 26559C>G 26560T>C), A12Q (26564G>C 26565C>A 26566G>A), Q14L (26571A>T 26572A>G), L15R (26574T>G 26575A>G), Q17T (26579C>A 26580A>C), A18S (26582G>T 26584T>C), G19D (26586G>A 26587C>T), R20Q (26589G>A 26590C>G), E22A (26595A>C), S23A (26597A>G 26598G>C 26599C>A), V28I (26612G>A), T29S (26616C>G), D32E (26626C>G), G34A (26631G>C 26632C>G), A36F (26636G>T 26637C>T), A38V (26643C>T 26644T>A), R41K (26652G>A), I42L (26654A>C 26656T>G) | | | | | | | | |

| | Begin | End | Coverage | Score | Concordance | Matches | Identities | I/D/M/F* | Stop Codons |
|---|---|---|---|---|---|---|---|---|---|
| **NT** | **14196** | **31794** | **9.4%** | **3582** | **41.3%** | **4358 (99.6%)** | **3090 (70.6%)** | **9/8** | |
| Codon mutations: | CAG3GGA (26537C>G 26538A>G 26539G>A), CAT4CCA (26541A>C 26542T>A), TTC5CTC (26543T>C), CGA6CAG (26547G>A 26548A>G), AAT7AAC (26551T>C), CAG8GCG (26552C>G 26553A>C), GCT10CGC (26558G>C 26559C>G 26560T>C), CTC11CTG (26563C>G), GCG12CAA (26564G>C 26565C>A 26566G>A), GCC13GCG (26569C>G), CAA14CTG (26571A>T 26572A>G), CTA15CGG (26574T>G 26575A>G), GCG16GCC (26578G>C), CAG17ACG (26579C>A 26580A>C), GCT18TCC (26582G>T 26584T>C), GGC19GAT (26586G>A 26587C>T), CGC20CAG (26589G>A 26590C>G), GCT21GCC (26593T>C), GAG22GCG (26595A>C), AGC23GCA (26597A>G 26598G>C 26599C>A), CGG24CGC (26602G>C), GTT25GTG (26605T>G), GGC26GGG (26608C>G), ATT27ATC (26611T>C), GTC28ATC (26612G>A), ACC29AGC (26616C>G), GAC32GAG (26626C>G), CCA33CCC (26629A>C), GGC34GCG (26631G>C 26632C>G), ACG35ACC (26635G>C), GCC36TTC (26636G>T 26637C>T), TCT37AGC (26639T>A 26640C>G 26641T>C), GCT38GTA (26643C>T 26644T>A), GTC40GTG (26650C>G), AGG41AAG (26652G>A), ATT42CTG (26654A>C 26656T>G), CAG43CAA (26659G>A) | | | | | | | | |
| tail protein (YP_010078169.1) | 8 | 96 | 70.1% | 381 | 60.8% | 89 (100%) | 48 (53.9%) | 0/0/0/0 | 0 |
| Protein mutations: | Q11H (27228G>T), W12L (27229T>C 27230G>T 27231G>T), V13W (27232G>T 27233T>G 27234T>G), I17L (27244A>C 27246C>G), T19S (27251C>G 27252G>C), S21P (27256T>C 27258A>C), T22S (27260C>G 27261G>C), L25I (27268C>A 27270G>C), G26A (27272G>C 27273C>G), T27L (27274A>C 27275C>T), A28S (27277G>T), N29S (27281A>G), A30G (27284C>G), D31T (27286G>A 27287A>C 27288C>G), T32Q (27289A>C 27290C>A), T34G (27295A>G 27296C>G 27297A>C), I38V (27307A>G 27309A>G), V39L (27310G>C), V43L (27322G>C 27324G>C), E48D (27339G>C), I50V (27343A>G 27345C>A), H52Q (27351C>A), E54S (27355G>A 27356A>G 27357G>C), K62Y (27379A>T 27381G>C), I63V (27382A>G), T66L (27391A>C 27392C>T), A71K (27406G>A 27407C>A), I72V (27409A>G 27411T>G), R73V (27412C>G 27413G>T), G74A (27416G>C 27417A>C), L77R (27425T>G 27426G>C), V80I (27433G>A), R81L (27436A>C 27437G>T), M82L (27439A>C), E84D (27447G>C), G85S (27448G>A), Q88K (27457C>A 27459A>G), E91P (27466G>C 27467A>C), V94I (27475G>A 27477G>C), D95N (27478G>A) | | | | | | | | |
| Codon mutations: | CTC9CTG (27222C>G), AAT10AAC (27225T>C), CAG11CAT (27228G>T), TGG12CTT (27229T>C 27230G>T 27231G>T), GTT13TGG (27232G>T 27233T>G 27234T>G), GGT14GGC (27237T>C), ATC17CTG (27244A>C 27246C>G), GGC18GGA (27249C>A), ACG19AGC (27251C>G 27252G>C), TCC20AGC (27253T>A 27254C>G), TCA21CCC (27256T>C 27258A>C), ACG22AGC (27260C>G 27261G>C), GAT24GAC (27267T>C), CTG25ATC (27268C>A 27270G>C), GGC26GCG (27272G>C 27273C>G), ACC27CTC (27274A>C 27275C>T), GCC28TCC (27277G>T), AAC29AGC (27281A>G), GCC30GGC (27284C>G), GAC31ACG (27286G>A 27287A>C 27288C>G), ACG32CAG (27289A>C 27290C>A), ACA34GGC (27295A>G 27296C>G 27297A>C), ATA38GTG (27307A>G 27309A>G), GTG39CTG (27310G>C), CGC41CGG (27318C>G), CTT42CTG (27321T>G), GTG43CTC (27322G>C 27324G>C), ACG44ACC (27327G>C), AAT45AAC (27330T>C), CCC46CCG (27333C>G), GAG48GAC (27339G>C), ATC50GTA (27343A>G 27345C>A), CAC52CAA (27351C>A), CCT53CCG (27354T>G), GAG54AGC (27355G>A 27356A>G 27357G>C), TAT55TAC (27360T>C), AAG62TAC (27379A>T 27381G>C), ATC63GTC (27382A>G), GGT64GGC (27387T>C), ACG66CTG (27391A>C 27392C>T), GAC68GAT (27399C>T), GTG69GTC (27402G>C), GCG71AAG (27406G>A 27407C>A), ATT72GTG (27409A>G 27411T>G), CGC73GTC (27412C>G 27413G>T), GGA74GCC (27416G>C 27417A>C), CTG77CGC (27425T>G 27426G>C), CAG79CAA (27432G>A), GTC80ATC (27433G>A), AGG81CTG (27436A>C 27437G>T), ATG82CTG (27439A>C), GAG84GAC (27447G>C), GGC85AGC (27448G>A), GTT86GTG (27453T>G), GCG87GCC (27456G>C), CAA88AAG (27457C>A 27459A>G), GAG91CCG (27466G>C 27467A>C), CAA93CAG (27474A>G), GTG94ATC (27475G>A 27477G>C), GAC95AAC (27478G>A) | | | | | | | | |
| hypothetical protein (YP_010078170.1) | 18 | 218 | 51.5% | 814 | 71.2% | 193 (100%) | 121 (62.7%) | 0/0/0/0 | 0 |
| Protein mutations: | Y22A (27641T>G 27642A>C 27643C>A), A25Q (27650G>C 27651C>A 27652C>G), L29M (27662C>A), V31I (27668G>A), I34V (27677A>G 27679C>G), V38F (27689G>T 27691G>T), N42V (27701A>G 27702A>T 27703T>G), V45I (27710G>A 27712C>T), V46A (27714T>C), I47L (27716A>C), L52I (27731C>A), I53A (27734A>G 27735T>C 27736C>G), M54L (27737A>C), V56I (27743G>A 27745G>T), L57A (27746C>G 27747T>C 27748G>A), A58S (27749G>T), I59L (27752A>C 27754C>G), A62F (27761G>T 27762C>T 27763G>T), A65S (27770G>A 27771C>G 27772G>C), T66N (27774C>A 27775C>T), T68A (27779A>G 27781G>C), L70A (27785C>G 27786T>C 27787T>G), T72S (27792C>G 27793G>T), V74A (27798T>C), V79F (27812G>T 27814G>C), T80A (27815A>G), V85Q (27830G>C 27831T>A), A86Q (27833G>C 27834C>A), T90Q (27845A>C 27846C>A), A92T (27851G>A), S94A (27857T>G), T98S (27869A>T), Q100T (27875C>A 27876A>C 27877G>C), T107A (27896A>G 27898A>G), A108S (27899G>T 27901C>T), T116S (27923A>T), Q118K (27929C>A 27931A>G), F119Y (27933T>A), S132A (27971T>G), I139M (27994T>G), A144T (28007G>A 28009G>C), V146C (28013G>T 28014T>G 28015G>C), N147V (28016A>G 28017A>T 28018T>G), V148A (28020T>C 28021A>G), P149T (28022C>A 28024G>C), V151K (28028G>A 28029T>A), A152S (28031G>T), T154N (28038C>A 28039G>C), G156A (28044G>C 28045C>G), G159A (28053G>C 28054C>G), V162A (28062T>C), A163I (28064G>A 28065C>T 28066T>C), G164A (28068G>C), V166I (28073G>A 28075A>C), S167N (28076T>A 28077C>A 28078G>C), I169L (28082A>C), V170G (28086T>G), G171Q (28088G>C 28089G>A 28090C>G), S174A (28097T>G 28099C>G), N187T (28137A>C 28138C>G), A193T (28154G>A 28156A>G), L197F (28166C>T 28168G>C), T199S (28172A>T), I202V (28181A>G), A203G (28185C>G), V205I (28190G>A 28192A>C), A206N (28193G>A 28194C>A 28195G>C), K213R (28215A>G 28216A>G), S214A (28217T>G 28219C>T), V216I (28223G>A) | | | | | | | | |
| Codon mutations: | TAC22GCA (27641T>G 27642A>C 27643C>A), TCT23TCG (27646T>G), GCC25CAG (27650G>C 27651C>A 27652C>G), TTA26CTC (27653T>C 27655A>C), GTT27GTC (27658T>C), CTG29ATG (27662C>A), GTC31ATC (27668G>A), TCG33TCC (27676G>C), ATC34GTG (27677A>G 27679C>G), CGC36CGG (27685C>G), GTG38TTT (27689G>T 27691G>T), AAT42GTG (27701A>G 27702A>T 27703T>G), GCT43GCG (27706T>G), GCC44GCA (27709C>A), GTC45ATT (27710G>A 27712C>T), GTG46GCG (27714T>C), ATC47CTC (27716A>C), CTC52ATC (27731C>A), ATC53GCG (27734A>G 27735T>C 27736C>G), ATG54CTG (27737A>C), CAA55CAG (27742A>G), GTG56ATT (27743G>A 27745G>T), CTG57GCA (27746C>G 27747T>C 27748G>A), GCG58TCG (27749G>T), ATC59CTG (27752A>C 27754C>G), GCG62TTT (27761G>T 27762C>T 27763G>T), GCT63GCG (27766T>G), ACG64ACC (27769G>C), GCG65AGC (27770G>A 27771C>G 27772G>C), ACC66AAT (27774C>A 27775C>T), ACG68GCC (27779A>G 27781G>C), CTT70GCG (27785C>G 27786T>C 27787T>G), ACG72AGT (27792C>G 27793G>T), GTG74GCG (27798T>C), GCG75GCC (27802G>C), GTG79TTC (27812G>T 27814G>C), ACG80GCG (27815A>G), CGT81CGC (27820T>C), CTT82CTG (27823T>G), GCT83GCG (27826T>G), GCG84GCC (27829G>C), GTG85CAG (27830G>C 27831T>A), GCG86CAG (27833G>C 27834C>A), ACT88ACC (27841T>C), GGA89GGC (27844A>C), ACG90CAG (27845A>C 27846C>A), GTC91GTG (27850C>G), GCC92ACC (27851G>A), TCG94GCG (27857T>G), ACG97ACT (27868G>T), ACG98TCG (27869A>T), ACC99ACG (27874C>G), CAG100ACC (27875C>A 27876A>C 27877G>C), GTC102GTT (27883C>T), GTT103GTG (27886T>G), CTT106CTC (27895T>C), ACA107GCG (27896A>G 27898A>G), GCC108TCT (27899G>T 27901C>T), GTT109GTC (27904T>C), ACA112ACG (27913A>G), GGG115GGC (27922G>C), ACG116TCG (27923A>T), CAA118AAG (27929C>A 27931A>G), TTC119TAC (27933T>A), GCT129GCC (27964T>C), TAT130TAC (27967T>C), TCG132GCG (27971T>G), ACA133ACG (27976A>G), GGC136GGG (27985C>G), TAC137TAT (27988C>T), ATT139ATG (27994T>G), GCT140GCG (27997T>G), GCG141GCC (28000G>C), GTG143GTC (28006G>C), GCG144ACC (28007G>A 28009G>C), AGC145AGT (28012C>T), GTG146TGC (28013G>T 28014T>G 28015G>C), AAT147GTG (28016A>G 28017A>T 28018T>G), GTA148GCG (28020T>C 28021A>G), CCG149ACC (28022C>A 28024G>C), GTG151AAG (28028G>A 28029T>A), GCA152TCA (28031G>T), GTG153GTC (28036G>C), ACG154AAC (28038C>A 28039G>C), GCC155GCA (28042C>A), GGC156GCG (28044G>C 28045C>G), GCT158GCG (28051T>G), GGC159GCG (28053G>C 28054C>G), GTG162GCG (28062T>C), GCT163ATC (28064G>A 28065C>T 28066T>C), GGC164GCC (28068G>C), GCC165GCG (28072C>G), GTA166ATC (28073G>A 28075A>C), TCG167AAC (28076T>A 28077C>A 28078G>C), ACC168ACG (28081C>G), ATC169CTC (28082A>C), GTC170GGC (28086T>G), GGC171CAG (28088G>C 28089G>A 28090C>G), GCA172GCC (28093A>C), ATT173ATC (28096T>C), TCC174GCG (28097T>G 28099C>G), GAT177GAC (28108T>C), GTC179GTG (28114C>G), ACG180ACC (28117G>C), GCC182GCG (28123C>G), GCA183GCG (28126A>G), AAC187ACG (28137A>C 28138C>G), GGC188GGT (28141C>T), GAT190GAC (28147T>C), GCA193ACG (28154G>A 28156A>G), GAC194GAT (28159C>T), GCT195GCC (28162T>C), GCG196GCC (28165G>C), CTG197TTC (28166C>T 28168G>C), CGT198CGC (28171T>C), ACG199TCG (28172A>T), CGC200CGG (28177C>G), ATC202GTC (28181A>G), GCC203GGC (28185C>G), GTA205ATC (28190G>A 28192A>C), GCG206AAC (28193G>A 28194C>A 28195G>C), AAG210AAA (28207G>A), ACG212ACC (28213G>C), AAA213AGG (28215A>G 28216A>G), TCC214GCT (28217T>G 28219C>T), GCT215GCC (28222T>C), GTC216ATC (28223G>A) | | | | | | | | |
| hypothetical protein (YP_010078173.1) | 105 | 204 | 17.3% | 517 | 79.4% | 100 (100%) | 79 (79.0%) | 0/0/0/0 | 0 |
| Protein mutations: | Q109Y (31508C>T 31510G>C), T118A (31535A>G 31537A>C), Q122S (31547C>A 31548A>G 31549G>C), D125N (31556G>A), T127V (31562A>G 31563C>T 31564T>G), A128T (31565G>A), W141Y (31605G>A 31606G>T), S142Q (31607A>C 31608G>A 31609C>G), N145A (31616A>G 31617A>C 31618C>G), N146G (31619A>G 31620A>G 31621C>A), S147N (31623G>A), A150S (31631G>A 31632C>G 31633A>T), T154A (31643A>G), V167I (31682G>A 31684A>C), S173T (31701G>C 31702C>G), S181V (31724T>G 31725C>T), T184V (31733A>G 31734C>T 31735C>G), W187Y (31743G>A 31744G>T), V188S (31745G>T 31746T>C), N193Y (31760A>T) | | | | | | | | |
| Codon mutations: | ACT106ACC (31501T>C), GTG107GTC (31504G>C), CAG109TAC (31508C>T 31510G>C), ATT115ATC (31528T>C), CAG116CAA (31531G>A), ACA118GCC (31535A>G 31537A>C), TAC119TAT (31540C>T), CAA120CAG (31543A>G), GAT121GAC (31546T>C), CAG122AGC (31547C>A 31548A>G 31549G>C), GAC125AAC (31556G>A), CTG126TTG (31559C>T), ACT127GTG (31562A>G 31563C>T 31564T>G), GCG128ACG (31565G>A), TGG141TAT (31605G>A 31606G>T), AGC142CAG (31607A>C 31608G>A 31609C>G), AAC145GCG (31616A>G 31617A>C 31618C>G), AAC146GGA (31619A>G 31620A>G 31621C>A), AGC147AAC (31623G>A), GCA150AGT (31631G>A 31632C>G 31633A>T), GCG152GCC (31639G>C), ACT154GCT (31643A>G), GTC159GTG (31660C>G), GTT160GTG (31663T>G), GCC163GCG (31672C>G), GCT165GCA (31678T>A), GTA167ATC (31682G>A 31684A>C), GCG168GCC (31687G>C), GCT169GCA (31690T>A), ACC170ACG (31693C>G), GGC172GGA (31699C>A), AGC173ACG (31701G>C 31702C>G), ACC175ACG (31708C>G), ACC176ACG (31711C>G), CCA177CCG (31714A>G), GCC178GCA (31717C>A), TCC181GTC (31724T>G 31725C>T), GGC182GGT (31729C>T), ACC184GTG (31733A>G 31734C>T 31735C>G), GGG185GGT (31738G>T), CTG186CTC (31741G>C), TGG187TAT (31743G>A 31744G>T), GTG188TCG (31745G>T 31746T>C), ACC190ACG (31753C>G), GTC191GTT (31756C>T), AAT193TAT (31760A>T), GGT194GGC (31765T>C), ACG196ACC (31771G>C), ACC197ACG (31774C>G), GCG200GCC (31783G>C), CCG201CCC (31786G>C), AAC202AAT (31789C>T) | | | | | | | | |

*: Inserts / Deletes / Misaligned / Frameshifts

#### Analysis details

This analysis was performed with panviral2.64

### NGS Details (UN24): Caulimovirus venafragariae

#### Assembly

| | |
|---|---|
| **Coverage Length** | 857 (2 contig(s)) |
| **Depth Of Coverage** | 21.5 |
| **Number Of Reads** | 168 |
| **Reads Per Million** | 3.36 rpm (after QC) |
| **Ambiguities** | 0 |
| **Assembly Method** | de novo + reference guided assembly |
| **Consensus Caller** | Bcf Tools |

#### Coverage Map

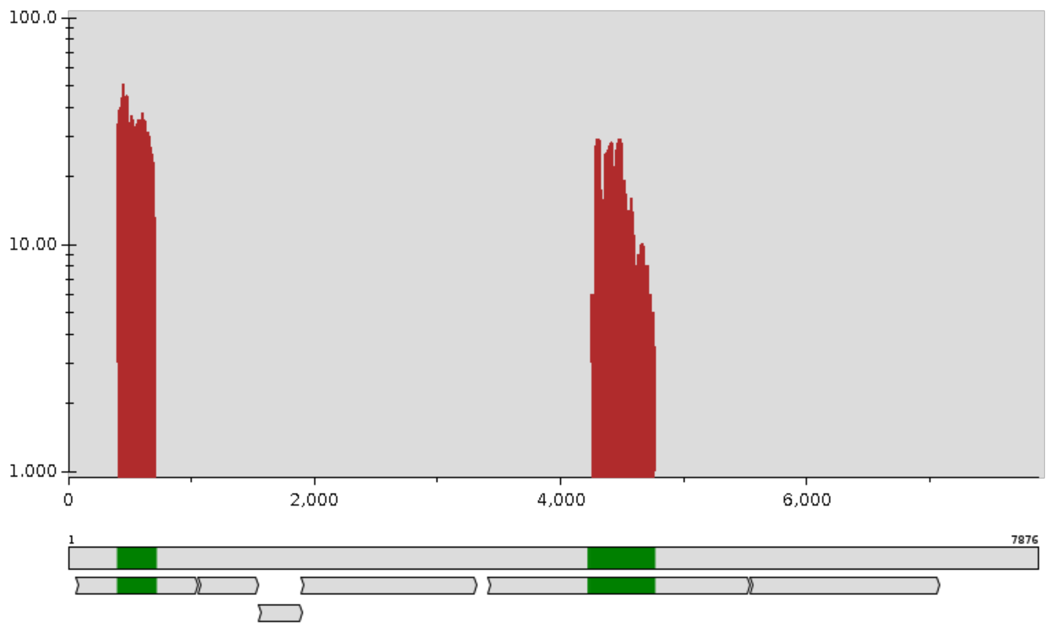

#### Assignment

| | |
|---|---|
| **Type** | Caulimovirus venafragariae (Taxonomy ID: 3048344) |
| **Reference Genome** | NC_001725.1 |
| **NT Identity (%)** | 58.4597 |
| **AA Identity (%)** | 54.4484 |
| **Number Of Stop Codons** | 1 |
| **Number Of CDS** | 6 |

#### Alignment

| | |
|---|---|
| **Alignment Score** | 272.0 (NT) + 1125.0 (AA) = 1397.0 |
| **Concordance (%)** | 38.159 |

| Alignment Method | Global, seeded, nucleotide + amino acids (AGA) |
|---|---|

#### Genome Region

Sequence starts at position 396 and ends at position 4770 relative to NC_001725.1 reference sequence.

#### Alignment Detailed Statistics

| | Begin | End | Coverage | Score | Concordance | Matches | Identities | I/D/M/F* | Stop Codons |
|---|---|---|---|---|---|---|---|---|---|
| **NT** | **396** | **4770** | **10.9%** | **272** | **16.0%** | **854 (99.3%)** | **501 (58.3%)** | **3/3** | |
| Mutations: | 403A>C, 405A>T, 411A>G, 412G>A, 416T>A, 418A>G, 424A>T, 426A>G, 430T>G, 431C>G, 432T>A, 433G>A, 435T>A, 436A>C, 441T>C, 445A>G, 447C>G, 451A>T, 452G>T, 456C>T, 459T>C, 461G>A, 462G>A, 463A>G, 464C>A, 468A>T, 472G>A, 474C>T, 476C>G, 483T>A, 485G>T, 486T>A, 487G>A, 490G>A, 491C>A, 492A>C, 495C>T, 496C>T, 497T>C, 501C>T, 502C>G, 503G>A, 504G>A, 508A>T, 510G>T, 512A>T, 513A>T, 516C>T, 517G>A, 521A>G, 525C>A, 527C>G, 529G>A, 530T>A, 531C>T, 532T>C, 534C>T, 537C>A, 538G>A, 539G>T, 544A>G, 552C>T, 553C>T, 555T>A, 556T>G, 558C>A, 562G>A, 563G>C, 564G>T, 567A>G, 568C>T, 570T>A, 571A>C, 573C>T, 579C>A, 582C>T, 583A>T, 585C>T, 588A>C, 590A>G, 592A>T, 593T>A, 594C>T, 596C>G, 598G>A, 600T>A, 601A>T, 602G>C, 603T>A, 604C>T, 606T>A, 608G>A, 612C>T, 613C>G, 617C>A, 618G>C, 619A>T, 624C>T, 625A>G, 627G>A, 628A>G, 630T>A, 633A>G, 635C>G, 636A>T, 637C>T, 639A>G, 640_642delGCT, 648C>T, 648_649insCAG, 649T>G, 651T>C, 652G>A, 656A>G, 658G>A, 663A>T, 666G>A, 667A>T, 669G>C, 670C>A, 672T>G, 673G>C, 674A>C, 675G>A, 678A>T, 685C>A, 686C>T, 687C>T, 688T>A, 689A>T, 690T>G, 692C>G, 693T>C, 696A>T, 698C>A, 699C>T, 702C>T, 703A>C, 705A>G, 706A>G, 707T>C, 4219T>C, 4222T>A, 4223T>A, 4227C>T, 4228A>C, 4230T>C, 4232A>G, 4235A>T, 4236A>C, 4239T>A, 4242T>A, 4244A>G, 4245G>A, 4246T>G, 4247G>T, 4249A>C, 4254A>C, 4258C>T, 4259A>T, 4260G>T, 4264T>A, 4265C>A, 4266A>C, 4267C>G, 4268C>A, 4270C>G, 4275C>T, 4276C>A, 4277G>A, 4279G>A, 4281A>T, 4282G>A, 4284G>A, 4287T>C, 4289A>G, 4292C>T, 4299C>T, 4300G>C, 4308A>T, 4314G>A, 4315C>T, 4317T>A, 4318G>A, 4319G>A, 4320T>A, 4321A>T, 4322T>A, 4323C>A, 4326C>A, 4330C>G, 4331C>A, 4332T>A, 4335C>T, 4348T>A, 4349C>G, 4351T>A, 4352C>G, 4353A>T, 4356A>T, 4359C>A, 4362C>T, 4366G>A, 4371G>A, 4380C>G, 4383G>A, 4384A>C, 4385T>A, 4386C>A, 4390C>A, 4395C>A, 4398A>G, 4401A>T, 4402A>C, 4410A>T, 4413T>A, 4423A>G, 4426T>C, 4428A>T, 4432G>A, 4434C>A, 4435C>A, 4437T>C, 4438A>T, 4439C>G, 4440A>T, 4442A>T, 4443G>T, 4444G>T, 4445G>T, 4446A>T, 4449T>C, 4452C>A, 4456C>T, 4458A>T, 4459C>A, 4461T>C, 4467C>T, 4470G>A, 4471G>A, 4472A>C, 4473G>T, 4474C>A, 4476A>T, 4479G>T, 4480C>A, 4482T>A, 4483C>A, 4485A>T, 4486A>C, 4487G>T, 4489A>G, 4492G>A, 4493G>A, 4496G>T, 4497T>A, 4500G>A, 4505T>A, 4506T>C, 4508A>T, 4509C>T, 4512T>C, 4513T>A, 4514C>A, 4515C>A, 4521C>T, 4528T>A, 4529C>G, 4543G>A, 4546C>A, 4548C>A, 4551T>C, 4552G>T, 4554T>A, 4555C>G, 4556C>A, 4557A>G, 4561A>T, 4568A>C, 4569G>T, 4575C>A, 4578T>A, 4585T>A, 4586G>C, 4590C>T, 4591C>A, 4593A>T, 4599C>T, 4602C>T, 4614C>A, 4618C>T, 4620T>A, 4633C>A, 4635A>T, 4638C>A, 4641T>A, 4642G>C, 4643C>A, 4644T>A, 4650C>T, 4656G>A, 4657C>A, 4659C>A, 4663G>T, 4665C>T, 4666G>A, 4670G>T, 4671T>A, 4672C>T, 4674C>T, 4675A>T, 4676G>C, 4677C>T, 4678A>G, 4680C>T, 4681A>C, 4684T>A, 4686T>A, 4687C>A, 4688C>A, 4689A>G, 4690C>A, 4692G>A, 4697G>T, 4698T>G, 4699G>A, 4700C>T, 4701T>A, 4707T>C, 4708G>A, 4710T>A, 4719C>T, 4720A>T, 4722C>G, 4725C>A, 4727T>G, 4729A>T, 4730G>C, 4731C>A, 4733A>C, 4737T>A, 4738G>T, 4740A>T, 4741G>C, 4743A>G, 4750T>C, 4752A>T, 4753G>A, 4754G>C, 4759G>C, 4761C>T, 4765A>G, 4766T>A, 4767T>A, 4768G>T | | | | | | | | |
| **CDS** | | | | | | | | | |
| ORF_I | 111 | 216 | 32.2% | 349 | 49.8% | 105 (98.1%) | 46 (43.0%) | 1/1/0/0 | 0 |
| Protein mutations: | K113H (403A>C 405A>T), I115M (411A>G), D116N (412G>A), F117Y (416T>A), I118V (418A>G), I120L (424A>T 426A>G), S122G (430T>G 431C>G 432T>A), V123I (433G>A 435T>A), K124Q (436A>C), I127V (445A>G 447C>G), S129F (451A>T 452G>T), R132K (461G>A 462G>A), T133E (463A>G 464C>A), D136N (472G>A 474C>T), S137C (476C>G), S140I (485G>T 486T>A), V141I (487G>A), A142N (490G>A 491C>A 492A>C), L144S (496C>T 497T>C), R146E (502C>G 503G>A 504G>A), M148F (508A>T 510G>T), K149I (512A>T 513A>T), A151T (517G>A), K152R (521A>G), D153E (525C>A), A154G (527C>G), V155N (529G>A 530T>A 531C>T), F156L (532T>C 534C>T), G158I (538G>A 539G>T), K160E (544A>G), S164A (556T>G 558C>A), G166T (562G>A 563G>C 564G>T), I169L (571A>C 573C>T), N173Y (583A>T 585C>T), K175R (590A>G), I176Y (592A>T 593T>A 594C>T), S177C (596C>G), V178I (598G>A 600T>A), R181K (608G>A), P183A (613C>G), T184N (617C>A 618G>C), I185F (619A>T), K187E (625A>G 627G>A), T188A (628A>G 630T>A), T190S (635C>G 636A>T), A192del (640_642delGCT), F194_F195insQ (648_649insCAG), F195V (649T>G 651T>C), E196K (652G>A), K197R (656A>G), E198K (658G>A), E199D (663A>T), M201F (667A>T 669G>C), H202K (670C>A 672T>G), E203P (673G>C 674A>C 675G>A), P207I (685C>A 686C>T 687C>T), Y208M (688T>A 689A>T 690T>G), T209S (692C>G 693T>C), S211Y (698C>A 699C>T), K213Q (703A>C 705A>G), I214A (706A>G 707T>C) | | | | | | | | |
| Codon mutations: | AAA113CAT (403A>C 405A>T), ATA115ATG (411A>G), GAT116AAT (412G>A), TTT117TAT (416T>A), ATT118GTT (418A>G), ATA120TTG (424A>T 426A>G), TCT122GGA (430T>G 431C>G 432T>A), GTT123ATA (433G>A 435T>A), AAA124CAA (436A>C), ATT125ATC (441T>C), ATC127GTG (445A>G 447C>G), AGC129TTC (451A>T 452G>T), ACC130ACT (456C>T), TTT131TTC (459T>C), AGG132AAA (461G>A 462G>A), ACA133GAA (463A>G 464C>A), GGA134GGT (468A>T), GAC136AAT (472G>A 474C>T), TCT137TGT (476C>G), ATT139ATA (483T>A), AGT140ATA (485G>T 486T>A), GTA141ATA (487G>A), GCA142AAC (490G>A 491C>A 492A>C), CTC143CTT (495C>T), CTT144TCT (496C>T 497T>C), GAC145GAT (501C>T), CGG146GAA (502C>G 503G>A 504G>A), ATG148TTT (508A>T 510G>T), AAA149ATT (512A>T 513A>T), AAC150AAT (516C>T), GCT151ACT (517G>A), AAA152AGA (521A>G), GAC153GAA (525C>A), GCA154GGA (527C>G), GTC155AAT (529G>A 530T>A 531C>T), TTC156CTT (532T>C 534C>T), GGC157GGA (537C>A), GGA158ATA (538G>A 539G>T), AAA160GAA (544A>G), AAC162AAT (552C>T), CTT163TTA (553C>T 555T>A), TCC164GCA (556T>G 558C>A), GGG166ACT (562G>A 563G>C 564G>T), AAA167AAG (567A>G), CTT168TTA (568C>T 570T>A), ATC169CTT (571A>C 573C>T), ACC171ACA (579C>A), TAC172TAT (582C>T), AAC173TAT (583A>T 585C>T), CCA174CCC (588A>C), AAA175AGA (590A>G), ATC176TAT (592A>T 593T>A 594C>T), TCT177TGT (596C>G), GTT178ATA (598G>A 600T>A), AGT179TCA (601A>T 602G>C 603T>A), CTT180TTA (604C>T 606T>A), AGA181AAA (608G>A), GAC182GAT (612C>T), CCT183GCT (613C>G), ACG184AAC (617C>A 618G>C), ATT185TTT (619A>T), AAC186AAT (624C>T), AAG187GAA (625A>G 627G>A), ACT188GCA (628A>G 630T>A), TTA189TTG (633A>G), ACA190AGT (635C>G 636A>T), CTA191TTG (637C>T 639A>G), GCT192del (640_642delGCT), TTC194TTT (648C>T), TTC194_TTT195insCAG (648_649insCAG), TTT195GTC (649T>G 651T>C), GAA196AAA (652G>A), AAA197AGA (656A>G), GAA198AAA (658G>A), GAA199GAT (663A>T), TTG200TTA (666G>A), ATG201TTC (667A>T 669G>C), CAT202AAG (670C>A 672T>G), GAG203CCA (673G>C 674A>C 675G>A), GGA204GGT (678A>T), CCC207ATT (685C>A 686C>T 687C>T), TAT208ATG (688T>A 689A>T 690T>G), ACT209AGC (692C>G 693T>C), ATA210ATT (696A>T), TCC211TAT (698C>A 699C>T), TAC212TAT (702C>T), AAA213CAG (703A>C 705A>G), ATA214GCA (706A>G 707T>C) | | | | | | | | |
| ORF_V | 281 | 455 | 24.7% | 776 | 58.8% | 175 (100%) | 107 (61.1%) | 0/0/0/0 | 1 |
| Protein mutations: | C281V (4246T>G 4247G>T), K282Q (4249A>C), Q285F (4258C>T 4259A>T 4260G>T), S287N (4264T>A 4265C>A 4266A>C), P288D (4267C>G 4268C>A), Q289E (4270C>G), R291K (4276C>A 4277G>A), E292N (4279G>A 4281A>T), E293K (4282G>A 4284G>A), K295R (4289A>G), T296I (4292C>T), E299Q (4300G>C), G305K (4318G>A 4319G>A 4320T>A), I306* (4321A>T 4322T>A 4323C>A), P309E (4330C>G 4331C>A 4332T>A), V321I (4366G>A), I327Q (4384A>C 4385T>A 4386C>A), K340E (4423A>G), D343K (4432G>A 4434C>A), H344N (4435C>A 4437T>C), T345C (4438A>T 4439C>G 4440A>T), K346I (4442A>T 4443G>T), G347F (4444G>T 4445G>T 4446A>T), L351F (4456C>T 4458A>T), L352I (4459C>A 4461T>C), E356T (4471G>A 4472A>C 4473G>T), Q357N (4474C>A 4476A>T), L359I (4480C>A 4482T>A), Q360N (4483C>A 4485A>T), R361L (4486A>C 4487G>T), I362V (4489A>G), G363K (4492G>A 4493G>A), G364V (4496G>T 4497T>A), F367Y (4505T>A 4506T>C), Y368F (4508A>T 4509C>T), S370K (4513T>A 4514C>A 4515C>A), V380I (4543G>A), A383S (4552G>T 4554T>A), P384E (4555C>G 4556C>A 4557A>G), T386S (4561A>T), Q388P (4568A>C 4569G>T), C394T (4585T>A 4586G>C), Q396N (4591C>A 4593A>T), P405S (4618C>T 4620T>A), Q410N (4633C>A 4635A>T), A413Q (4642G>C 4643C>A 4644T>A), H418K (4657C>A 4659C>A), D420Y (4663G>T 4665C>T), E421K (4666G>A), S422I (4670G>T 4671T>A), L423F (4672C>T 4674C>T), N425D (4678A>G 4680C>T), M426L (4681A>C), Y427K (4684T>A 4686T>A), P428K (4687C>A 4688C>A 4689A>G), Q429K (4690C>A 4692G>A), C431L (4697G>T 4698T>G), A432I (4699G>A 4700C>T 4701T>A), V435I (4708G>A 4710T>A), I439L (4720A>T 4722C>G), F441C (4727T>G), K443T (4733A>C), E445Y (4738G>T 4740A>T), E446Q (4741G>C 4743A>G), G450T (4753G>A 4754G>C), V452L (4759G>C 4761C>T), I454E (4765A>G 4766T>A 4767T>A), V455F (4768G>T) | | | | | | | | |
| Codon mutations: | TTA272C.. (4219T>C), TTA273AA. (4222T>A 4223T>A), GAC274..T (4227C>T), ACT275C.C (4228A>C 4230T>C), AAC276.G. (4232A>G), AAA277.TC (4235A>T 4236A>C), ATT278..A (4239T>A), ATT279..A (4242T>A), AAG280.GA (4244A>G 4245G>A), TGT281GTT (4246T>G 4247G>T), AAA282CAA (4249A>C), CCA283CCC (4254A>C), CAG285TTT (4258C>T 4259A>T 4260G>T), TCA287AAC (4264T>A 4265C>A 4266A>C), CCT288GAT (4267C>G 4268C>A), CAA289GAA (4270C>G), GAC290GAT (4275C>T), CGA291AAA (4276C>A 4277G>A), GAA292AAT (4279G>A 4281A>T), GAG293AAA (4282G>A 4284G>A), TTT294TTC (4287T>C), AAA295AGA (4289A>G), ACT296ATT (4292C>T), ATC298ATT (4299C>T), GAA299CAA (4300G>C), CTA301CTT (4308A>T), AAG303AAA (4314G>A), CTT304TTA (4315C>T 4317T>A), GGT305AAA (4318G>A 4319G>A 4320T>A), ATC306TAA (4321A>T 4322T>A 4323C>A), ATC307ATA (4326C>A), CCT309GAA (4330C>G 4331C>A 4332T>A), AGC310AGT (4335C>T), TCC315AGC (4348T>A 4349C>G), TCA316AGT (4351T>A 4352C>G 4353A>T), CCA317CCT (4356A>T), GCC318GCA (4359C>A), TTC319TTT (4362C>T), GTA321ATA (4366G>A), AGG322AGA (4371G>A), GCC325GCG (4380C>G), GAG326GAA (4383G>A), ATC327CAA (4384A>C 4385T>A 4386C>A), CGA329AGA (4390C>A), GGC330GGA (4395C>A), AAA331AAG (4398A>G), GCA332GCT (4401A>T), AGA333CGA (4402A>C), GTA335GTT (4410A>T), ATT336ATA (4413T>A), AAG340GAG (4423A>G), TTA341CTT (4426T>C 4428A>T), GAC343AAA (4432G>A 4434C>A), CAT344AAC (4435C>A 4437T>C), ACA345TGT (4438A>T 4439C>G 4440A>T), AAG346ATT (4442A>T 4443G>T), GGA347TTT (4444G>T 4445G>T 4446A>T), GAT348GAC (4449T>C), GGC349GGA (4452C>A), CTA351TTT (4456C>T 4458A>T), CTT352ATC (4459C>A 4461T>C), AAC354AAT (4467C>T), AAG355AAA (4470G>A), GAG356ACT (4471G>A 4472A>C 4473G>T), CAA357AAT (4474C>A 4476A>T), CTG358CTT (4479G>T), CTT359ATA (4480C>A 4482T>A), CAA360AAT (4483C>A 4485A>T), AGA361CTA (4486A>C 4487G>T), ATC362GTC (4489A>G), GGA363AAA (4492G>A 4493G>A), GGT364GTA (4496G>T 4497T>A), AAG365AAA (4500G>A), TTT367TAC (4505T>A 4506T>C), TAC368TTT (4508A>T 4509C>T), TCT369TCC (4512T>C), TCC370AAA (4513T>A 4514C>A 4515C>A), GAC372GAT (4521C>T), TCT375AGT (4528T>A 4529C>G), GTA380ATA (4543G>A), CGC381AGA (4546C>A 4548C>A), CTT382CTC (4551T>C), GCT383TCA (4552G>T 4554T>A), CCA384GAG (4555C>G 4556C>A 4557A>G), ACA386TCA (4561A>T), CAG388CCT (4568A>C 4569G>T), ACC390ACA (4575C>A), GCT391GCA (4578T>A), TGT394ACT (4585T>A 4586G>C), CCC395CCT (4590C>T), CAA396AAT (4591C>A 4593A>T), CAC398CAT (4599C>T), TAC399TAT (4602C>T), GTC403GTA (4614C>A), CCT405TCA (4618C>T 4620T>A), CAA410AAT (4633C>A 4635A>T), GCC411GCA (4638C>A), CCT412CCA (4641T>A), GCT413CAA (4642G>C 4643C>A 4644T>A), TTC415TTT (4650C>T), AGG417AGA (4656G>A), CAC418AAA (4657C>A 4659C>A), GAC420TAT (4663G>T 4665C>T), GAA421AAA (4666G>A), AGT422ATA (4670G>T 4671T>A), CTC423TTT (4672C>T 4674C>T), AGC424TCT (4675A>T 4676G>C 4677C>T), AAC425GAT (4678A>G 4680C>T), ATG426CTG (4681A>C), TAT427AAA (4684T>A 4686T>A), CCA428AAG (4687C>A 4688C>A 4689A>G), CAG429AAA (4690C>A 4692G>A), TGT431TTG (4697G>T 4698T>G), GCT432ATA (4699G>A 4700C>T 4701T>A), TAT434TAC (4707T>C), GTT435ATA (4708G>A 4710T>A), ATC438ATT (4719C>T), ATC439TTG (4720A>T 4722C>G), GTC440GTA (4725C>A), TTC441TGC (4727T>G), AGC442TCA (4729A>T 4730G>C 4731C>A), AAA443ACA (4733A>C), ACT444ACA (4737T>A), GAA445TAT (4738G>T 4740A>T), GAA446CAG (4741G>C 4743A>G), TTA449CTT (4750T>C 4752A>T), GGA450ACA (4753G>A 4754G>C), GTC452CTT (4759G>C 4761C>T), ATT454GAA (4765A>G 4766T>A 4767T>A), GTT455TTT (4768G>T) | | | | | | | | |
| **Proteins** | | | | | | | | | |
| hypothetical protein (NP_043929.1) | 111 | 216 | 32.2% | 349 | 49.8% | 105 (98.1%) | 46 (43.0%) | 1/1/0/0 | 0 |

| | Begin | End | Coverage | Score | Concordance | Matches | Identities | I/D/M/F* | Stop Codons |
|---|---|---|---|---|---|---|---|---|---|
| **NT** | **396** | **4770** | **10.9%** | **272** | **16.0%** | **854 (99.3%)** | **501 (58.3%)** | **3/3** | |
| Protein mutations: | K113H (403A>C 405A>T), I115M (411A>G), D116N (412G>A), F117Y (416T>A), I118V (418A>G), I120L (424A>T 426A>G), S122G (430T>G 431C>G 432T>A), V123I (433G>A 435T>A), K124Q (436A>C), I127V (445A>G 447C>G), S129F (451A>T 452G>T), R132K (461G>A 462G>A), T133E (463A>G 464C>A), D136N (472G>A 474C>T), S137C (476C>G), S140I (485G>T 486T>A), V141I (487G>A), A142N (490G>A 491C>A 492A>C), L144S (496C>T 497T>C), R146E (502C>G 503G>A 504G>A), M148F (508A>T 510G>T), K149I (512A>T 513A>T), A151T (517G>A), K152R (521A>G), D153E (525C>A), A154G (527C>G), V155N (529G>A 530T>A 531C>T), F156L (532T>C 534C>T), G158I (538G>A 539G>T), K160E (544A>G), S164A (556T>G 558C>A), G166T (562G>A 563G>C 564G>T), I169L (571A>C 573C>T), N173Y (583A>T 585C>T), K175R (590A>G), I176Y (592A>T 593T>A 594C>T), S177C (596C>G), V178I (598G>A 600T>A), R181K (608G>A), P183A (613C>G), T184N (617C>A 618G>C), I185F (619A>T), K187E (625A>G 627G>A), T188A (628A>G 630T>A), T190S (635C>G 636A>T), A192del (640_642delGCT), F194_F195insQ (648_649insCAG), F195V (649T>G 651T>C), E196K (652G>A), K197R (656A>G), E198K (658G>A), E199D (663A>T), M201F (667A>T 669G>C), H202K (670C>A 672T>G), E203P (673G>C 674A>C 675G>A), P207I (685C>A 686C>T 687C>T), Y208M (688T>A 689A>T 690T>G), T209S (692C>G 693T>C), S211Y (698C>A 699C>T), K213Q (703A>C 705A>G), I214A (706A>G 707T>C) | | | | | | | | |
| Codon mutations: | AAA113CAT (403A>C 405A>T), ATA115ATG (411A>G), GAT116AAT (412G>A), TTT117TAT (416T>A), ATT118GTT (418A>G), ATA120TTG (424A>T 426A>G), TCT122GGA (430T>G 431C>G 432T>A), GTT123ATA (433G>A 435T>A), AAA124CAA (436A>C), ATT125ATC (441T>C), ATC127GTG (445A>G 447C>G), AGC129TTC (451A>T 452G>T), ACC130ACT (456C>T), TTT131TTC (459T>C), AGG132AAA (461G>A 462G>A), ACA133GAA (463A>G 464C>A), GGA134GGT (468A>T), GAC136AAT (472G>A 474C>T), TCT137TGT (476C>G), ATT139ATA (483T>A), AGT140ATA (485G>T 486T>A), GTA141ATA (487G>A), GCA142AAC (490G>A 491C>A 492A>C), CTC143CTT (495C>T), CTT144TCT (496C>T 497T>C), GAC145GAT (501C>T), CGG146GAA (502C>G 503G>A 504G>A), ATG148TTT (508A>T 510G>T), AAA149ATT (512A>T 513A>T), AAC150AAT (516C>T), GCT151ACT (517G>A), AAA152AGA (521A>G), GAC153GAA (525C>A), GCA154GGA (527C>G), GTC155AAT (529G>A 530T>A 531C>T), TTC156CTT (532T>C 534C>T), GGC157GGA (537C>A), GGA158ATA (538G>A 539G>T), AAA160GAA (544A>G), AAC162AAT (552C>T), CTT163TTA (553C>T 555T>A), TCC164GCA (556T>G 558C>A), GGG166ACT (562G>A 563G>C 564G>T), AAA167AAG (567A>G), CTT168TTA (568C>T 570T>A), ATC169CTT (571A>C 573C>T), ACC171ACA (579C>A), TAC172TAT (582C>T), AAC173TAT (583A>T 585C>T), CCA174CCC (588A>C), AAA175AGA (590A>G), ATC176TAT (592A>T 593T>A 594C>T), TCT177TGT (596C>G), GTT178ATA (598G>A 600T>A), AGT179TCA (601A>T 602G>C 603T>A), CTT180TTA (604C>T 606T>A), AGA181AAA (608G>A), GAC182GAT (612C>T), CCT183GCT (613C>G), ACG184AAC (617C>A 618G>C), ATT185TTT (619A>T), AAC186AAT (624C>T), AAG187GAA (625A>G 627G>A), ACT188GCA (628A>G 630T>A), TTA189TTG (633A>G), ACA190AGT (635C>G 636A>T), CTA191TTG (637C>T 639A>G), GCT192del (640_642delGCT), TTC194TTT (648C>T), TTC194_TTT195insCAG (648_649insCAG), TTT195GTC (649T>G 651T>C), GAA196AAA (652G>A), AAA197AGA (656A>G), GAA198AAA (658G>A), GAA199GAT (663A>T), TTG200TTA (666G>A), ATG201TTC (667A>T 669G>C), CAT202AAG (670C>A 672T>G), GAG203CCA (673G>C 674A>C 675G>A), GGA204GGT (678A>T), CCC207ATT (685C>A 686C>T 687C>T), TAT208ATG (688T>A 689A>T 690T>G), ACT209AGC (692C>G 693T>C), ATA210ATT (696A>T), TCC211TAT (698C>A 699C>T), TAC212TAT (702C>T), AAA213CAG (703A>C 705A>G), ATA214GCA (706A>G 707T>C) | | | | | | | | |
| hypothetical protein (NP_043933.1) | 281 | 455 | 24.7% | 776 | 58.8% | 175 (100%) | 107 (61.1%) | 0/0/0/0 | 1 |
| Protein mutations: | C281V (4246T>G 4247G>T), K282Q (4249A>C), Q285F (4258C>T 4259A>T 4260G>T), S287N (4264T>A 4265C>A 4266A>C), P288D (4267C>G 4268C>A), Q289E (4270C>G), R291K (4276C>A 4277G>A), E292N (4279G>A 4281A>T), E293K (4282G>A 4284G>A), K295R (4289A>G), T296I (4292C>T), E299Q (4300G>C), G305K (4318G>A 4319G>A 4320T>A), I306* (4321A>T 4322T>A 4323C>A), P309E (4330C>G 4331C>A 4332T>A), V321I (4366G>A), I327Q (4384A>C 4385T>A 4386C>A), K340E (4423A>G), D343K (4432G>A 4434C>A), H344N (4435C>A 4437T>C), T345C (4438A>T 4439C>G 4440A>T), K346I (4442A>T 4443G>T), G347F (4444G>T 4445G>T 4446A>T), L351F (4456C>T 4458A>T), L352I (4459C>A 4461T>C), E356T (4471G>A 4472A>C 4473G>T), Q357N (4474C>A 4476A>T), L359I (4480C>A 4482T>A), Q360N (4483C>A 4485A>T), R361L (4486A>C 4487G>T), I362V (4489A>G), G363K (4492G>A 4493G>A), G364V (4496G>T 4497T>A), F367Y (4505T>A 4506T>C), Y368F (4508A>T 4509C>T), S370K (4513T>A 4514C>A 4515C>A), V380I (4543G>A), A383S (4552G>T 4554T>A), P384E (4555C>G 4556C>A 4557A>G), T386S (4561A>T), Q388P (4568A>C 4569G>T), C394T (4585T>A 4586G>C), Q396N (4591C>A 4593A>T), P405S (4618C>T 4620T>A), Q410N (4633C>A 4635A>T), A413Q (4642G>C 4643C>A 4644T>A), H418K (4657C>A 4659C>A), D420Y (4663G>T 4665C>T), E421K (4666G>A), S422I (4670G>T 4671T>A), L423F (4672C>T 4674C>T), N425D (4678A>G 4680C>T), M426L (4681A>C), Y427K (4684T>A 4686T>A), P428K (4687C>A 4688C>A 4689A>G), Q429K (4690C>A 4692G>A), C431L (4697G>T 4698T>G), A432I (4699G>A 4700C>T 4701T>A), V435I (4708G>A 4710T>A), I439L (4720A>T 4722C>G), F441C (4727T>G), K443T (4733A>C), E445Y (4738G>T 4740A>T), E446Q (4741G>C 4743A>G), G450T (4753G>A 4754G>C), V452L (4759G>C 4761C>T), I454E (4765A>G 4766T>A 4767T>A), V455F (4768G>T) | | | | | | | | |
| Codon mutations: | TTA272C.. (4219T>C), TTA273AA. (4222T>A 4223T>A), GAC274..T (4227C>T), ACT275C.C (4228A>C 4230T>C), AAC276.G. (4232A>G), AAA277.TC (4235A>T 4236A>C), ATT278..A (4239T>A), ATT279..A (4242T>A), AAG280.GA (4244A>G 4245G>A), TGT281GTT (4246T>G 4247G>T), AAA282CAA (4249A>C), CCA283CCC (4254A>C), CAG285TTT (4258C>T 4259A>T 4260G>T), TCA287AAC (4264T>A 4265C>A 4266A>C), CCT288GAT (4267C>G 4268C>A), CAA289GAA (4270C>G), GAC290GAT (4275C>T), CGA291AAA (4276C>A 4277G>A), GAA292AAT (4279G>A 4281A>T), GAG293AAA (4282G>A 4284G>A), TTT294TTC (4287T>C), AAA295AGA (4289A>G), ACT296ATT (4292C>T), ATC298ATT (4299C>T), GAA299CAA (4300G>C), CTA301CTT (4308A>T), AAG303AAA (4314G>A), CTT304TTA (4315C>T 4317T>A), GGT305AAA (4318G>A 4319G>A 4320T>A), ATC306TAA (4321A>T 4322T>A 4323C>A), ATC307ATA (4326C>A), CCT309GAA (4330C>G 4331C>A 4332T>A), AGC310AGT (4335C>T), TCC315AGC (4348T>A 4349C>G), TCA316AGT (4351T>A 4352C>G 4353A>T), CCA317CCT (4356A>T), GCC318GCA (4359C>A), TTC319TTT (4362C>T), GTA321ATA (4366G>A), AGG322AGA (4371G>A), GCC325GCG (4380C>G), GAG326GAA (4383G>A), ATC327CAA (4384A>C 4385T>A 4386C>A), CGA329AGA (4390C>A), GGC330GGA (4395C>A), AAA331AAG (4398A>G), GCA332GCT (4401A>T), AGA333CGA (4402A>C), GTA335GTT (4410A>T), ATT336ATA (4413T>A), AAG340GAG (4423A>G), TTA341CTT (4426T>C 4428A>T), GAC343AAA (4432G>A 4434C>A), CAT344AAC (4435C>A 4437T>C), ACA345TGT (4438A>T 4439C>G 4440A>T), AAG346ATT (4442A>T 4443G>T), GGA347TTT (4444G>T 4445G>T 4446A>T), GAT348GAC (4449T>C), GGC349GGA (4452C>A), CTA351TTT (4456C>T 4458A>T), CTT352ATC (4459C>A 4461T>C), AAC354AAT (4467C>T), AAG355AAA (4470G>A), GAG356ACT (4471G>A 4472A>C 4473G>T), CAA357AAT (4474C>A 4476A>T), CTG358CTT (4479G>T), CTT359ATA (4480C>A 4482T>A), CAA360AAT (4483C>A 4485A>T), AGA361CTA (4486A>C 4487G>T), ATC362GTC (4489A>G), GGA363AAA (4492G>A 4493G>A), GGT364GTA (4496G>T 4497T>A), AAG365AAA (4500G>A), TTT367TAC (4505T>A 4506T>C), TAC368TTT (4508A>T 4509C>T), TCT369TCC (4512T>C), TCC370AAA (4513T>A 4514C>A 4515C>A), GAC372GAT (4521C>T), TCT375AGT (4528T>A 4529C>G), GTA380ATA (4543G>A), CGC381AGA (4546C>A 4548C>A), CTT382CTC (4551T>C), GCT383TCA (4552G>T 4554T>A), CCA384GAG (4555C>G 4556C>A 4557A>G), ACA386TCA (4561A>T), CAG388CCT (4568A>C 4569G>T), ACC390ACA (4575C>A), GCT391GCA (4578T>A), TGT394ACT (4585T>A 4586G>C), CCC395CCT (4590C>T), CAA396AAT (4591C>A 4593A>T), CAC398CAT (4599C>T), TAC399TAT (4602C>T), GTC403GTA (4614C>A), CCT405TCA (4618C>T 4620T>A), CAA410AAT (4633C>A 4635A>T), GCC411GCA (4638C>A), CCT412CCA (4641T>A), GCT413CAA (4642G>C 4643C>A 4644T>A), TTC415TTT (4650C>T), AGG417AGA (4656G>A), CAC418AAA (4657C>A 4659C>A), GAC420TAT (4663G>T 4665C>T), GAA421AAA (4666G>A), AGT422ATA (4670G>T 4671T>A), CTC423TTT (4672C>T 4674C>T), AGC424TCT (4675A>T 4676G>C 4677C>T), AAC425GAT (4678A>G 4680C>T), ATG426CTG (4681A>C), TAT427AAA (4684T>A 4686T>A), CCA428AAG (4687C>A 4688C>A 4689A>G), CAG429AAA (4690C>A 4692G>A), TGT431TTG (4697G>T 4698T>G), GCT432ATA (4699G>A 4700C>T 4701T>A), TAT434TAC (4707T>C), GTT435ATA (4708G>A 4710T>A), ATC438ATT (4719C>T), ATC439TTG (4720A>T 4722C>G), GTC440GTA (4725C>A), TTC441TGC (4727T>G), AGC442TCA (4729A>T 4730G>C 4731C>A), AAA443ACA (4733A>C), ACT444ACA (4737T>A), GAA445TAT (4738G>T 4740A>T), GAA446CAG (4741G>C 4743A>G), TTA449CTT (4750T>C 4752A>T), GGA450ACA (4753G>A 4754G>C), GTC452CTT (4759G>C 4761C>T), ATT454GAA (4765A>G 4766T>A 4767T>A), GTT455TTT (4768G>T) | | | | | | | | |

*: Inserts / Deletes / Misaligned / Frameshifts

#### Analysis details

This analysis was performed with panviral2.64

### NGS Details (UN24): Dioscovirus dioscoreae

#### Assembly

| | |
|---|---|
| **Coverage Length** | 1376 (1 contig(s)) |
| **Depth Of Coverage** | 12.2 |
| **Number Of Reads** | 161 |
| **Reads Per Million** | 3.22 rpm (after QC) |
| **Ambiguities** | 0 |
| **Assembly Method** | de novo + reference guided assembly |
| **Consensus Caller** | Bcf Tools |

#### Coverage Map

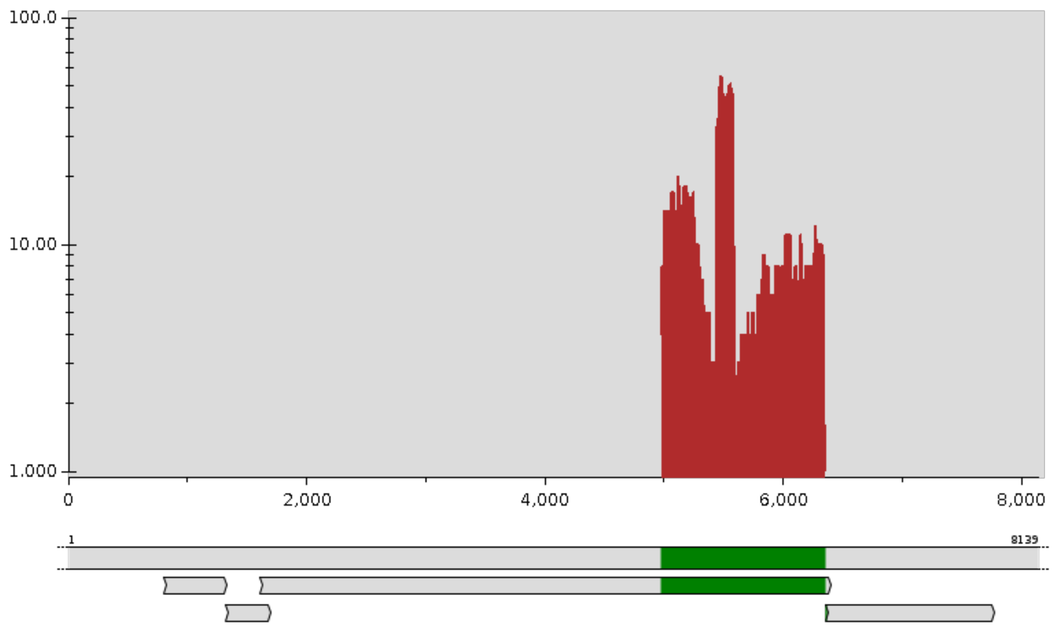

#### Assignment

| | |
|---|---|
| **Type** | Dioscovirus dioscoreae (Taxonomy ID: 3052184) |
| **Reference Genome** | NC_040712.1 |
| **NT Identity (%)** | 59.7242 |
| **AA Identity (%)** | 49.5652 |
| **Number Of Stop Codons** | 0 |
| **Number Of CDS** | 4 |

#### Alignment

| | |
|---|---|
| **Alignment Score** | 493.0 (NT) + 1519.0 (AA) = 2012.0 |
| **Concordance (%)** | 35.1314 |

| Alignment Method | Global, seeded, nucleotide + amino acids (AGA) |
|---|---|

#### Genome Region

Sequence starts at position 4980 and ends at position 6355 relative to NC_040712.1 reference sequence.

#### Alignment Detailed Statistics

| | Begin | End | Coverage | Score | Concordance | Matches | Identities | I/D/M/F* | Stop Codons |
|---|---|---|---|---|---|---|---|---|---|
| **NT** | **4980** | **6355** | **16.9%** | **493** | **18.2%** | **1369 (98.8%)** | **823 (59.4%)** | **9/7** | |
| Mutations: | 4983C>A, 4986C>A, 4988T>A, 4989A>T, 4990A>T, 4992T>C, 4994G>A, 4997A>T, 5001A>T, 5003C>A, 5004C>G, 5009T>A, 5012T>A, 5015G>A, 5016A>G, 5018T>A, 5019G>A, 5021C>A, 5024A>C, 5025T>C, 5029A>T, 5030G>A, 5031T>C, 5038A>G, 5040A>G, 5041A>C, 5042G>C, 5046C>A, 5047A>G, 5048G>A, 5051T>A, 5053G>T, 5054A>G, 5057C>T, 5063G>A, 5066C>T, 5069C>A, 5078T>A, 5081T>A, 5082C>T, 5083T>A, 5091C>A, 5093G>A, 5094G>A, 5095A>T, 5096C>A, 5097A>G, 5098A>C, 5099G>A, 5100C>G, 5102T>A, 5105T>A, 5106A>G, 5107T>A, 5109C>G, 5112T>A, 5115C>A, 5116C>A, 5117A>T, 5118G>C, 5121A>G, 5122T>A, 5123G>C, 5124G>A, 5125C>T, 5126G>A, 5127A>G, 5132A>T, 5134A>T, 5136G>C, 5138A>G, 5140A>T, 5141G>A, 5144T>A, 5147T>A, 5149C>A, 5157C>T, 5159G>A, 5161C>A, 5162A>T, 5163C>T, 5165C>A, 5166A>G, 5167A>G, 5168G>A, 5171T>A, 5174C>A, 5178C>A, 5179C>G, 5180T>A, 5183C>A, 5184C>A, 5185A>C, 5187A>T, 5188G>C, 5190C>A, 5192C>A, 5195T>C, 5198G>A, 5199A>T, 5207T>A, 5213T>G, 5216C>A, 5219C>A, 5220delA, 5222A>C, 5226T>A, 5227C>A, 5228T>C, 5232C>A, 5233A>T, 5237A>C, 5243T>A, 5247A>G, 5248G>C, 5249T>C, 5250C>A, 5252T>A, 5259T>A, 5260A>T, 5261C>A, 5271C>A, 5276A>C, 5288A>T, 5289T>A, 5290A>G, 5291C>A, 5293A>T, 5294A>G, 5297C>T, 5298C>G, 5299A>C, 5300A>T, 5303C>T, 5305C>A, 5307C>T, 5309C>A, 5313A>G, 5314G>A, 5315T>C, 5317T>A, 5318T>A, 5319G>A, 5321C>T, 5322T>G, 5324T>A, 5327C>A, 5331C>A, 5332T>A, 5335A>G, 5339T>A, 5340A>C, 5344A>G, 5345C>A, 5348G>A, 5350T>G, 5351T>A, 5352G>A, 5354C>T, 5356A>T, 5360C>T, 5369T>C, 5370C>T, 5371T>G, 5372A>T, 5375G>A, 5381A>G, 5383T>A, 5384C>T, 5385C>T, 5386A>G, 5387C>G, 5390G>A, 5391A>G, 5393T>A, 5395T>A, 5396G>A, 5400G>C, 5402T>C, 5405A>T, 5406C>G, 5408G>T, 5424G>A, 5430G>A, 5431T>C, 5432C>T, 5435C>T, 5439C>G, 5441G>A, 5444T>A, 5447T>C, 5449T>A, 5455G>T, 5456G>T, 5457A>C, 5459A>G, 5462A>T, 5471C>T, 5474A>C, 5477C>T, 5479A>C, 5492A>C, 5493G>T, 5494T>C, 5498C>T, 5501A>G, 5502C>A, 5503G>A, 5504A>T, 5505A>C, 5506A>T, 5507A>T, 5511G>A, 5514A>G, 5515A>C, 5517A>G, 5519A>C, 5522C>T, 5526A>C, 5527A>C, 5528_5529insCTC, 5529T>C, 5530A>T, 5532T>A, 5533C>G, 5535G>A, 5536A>G, 5539T>C, 5540T>A, 5543C>A, 5545G>T, 5546T>G, 5549T>C, 5551A>T, 5553A>T, 5555A>T, 5558C>T, 5564T>A, 5568A>G, 5570A>T, 5572T>A, 5573T>C, 5574T>A, 5575C>G, 5576T>C, 5577G>A, 5579A>G, 5580A>G, 5581G>A, 5585A>G, 5589C>G, 5595G>C, 5598C>G, 5607T>C, 5608T>A, 5610C>A, 5611A>C, 5612G>A, 5613T>A, 5615C>T, 5618C>T, 5619C>A, 5620A>G, 5621A>G, 5622G>T, 5626G>T, 5627T>C, 5628A>G, 5629A>T, 5630G>A, 5631G>A, 5632A>G, 5633A>T, 5634G>A, 5639G>A, 5640C>A, 5642T>A, 5645T>A, 5646T>A, 5648G>A, 5649T>A, 5651A>C, 5656C>A, 5657A>G, 5661T>A, 5664A>G, 5667A>T, 5670G>T, 5672A>T, 5673G>A, 5674T>A, 5676G>A, 5679A>C, 5685G>A, 5687A>C, 5691C>T, 5693T>A, 5696C>A, 5697C>G, 5700G>A, 5701A>C, 5702A>T, 5703A>T, 5711A>T, 5714A>G, 5717A>G, 5718G>A, 5720T>A, 5721C>A, 5726C>A, 5739T>G, 5740T>C, 5741A>T, 5748A>G, 5756A>T, 5757T>A, 5759T>G, 5762T>A, 5763_5765delGAA, 5768C>T, 5769C>A, 5777A>T, 5779C>A, 5781T>A, 5782T>C, 5786G>A, 5788G>A, 5790C>T, 5795G>A, 5798G>A, 5807G>A, 5808A>T, 5811T>G, 5813A>G, 5816C>T, 5822C>T, 5828T>C, 5830A>T, 5834C>A, 5836C>A, 5843G>A, 5844A>G, 5846T>A, 5850T>A, 5851A>T, 5853A>G, 5855T>A, 5856A>G, 5859A>C, 5860T>C, 5864T>A, 5868A>G, 5869A>C, 5870T>A, 5873G>A, 5874T>A, 5875G>T, 5876T>A, 5877T>G, 5878C>G, 5879C>A, 5882C>T, 5889G>C, 5893G>A, 5895A>C, 5897A>C, 5903T>C, 5904T>A, 5906T>A, 5907C>G, 5910G>A, 5912C>T, 5913T>A, 5914G>T, 5915G>T, 5918G>A, 5919A>C, 5924T>A, 5925A>C, 5926G>A, 5928A>C, 5933C>T, 5940G>A, 5941T>A, 5945T>A, 5947C>A, 5948C>A, 5951A>C, 5952T>A, 5957C>A, 5958C>G, 5959C>A, 5960T>C, 5963G>A, 5965T>G, 5966T>C, 5967T>A, 5973A>C, 5974A>T, 5976C>G, 5977C>A, 5979C>T, 5980A>C, 5982A>G, 5983G>A, 5988A>C, 5990T>A, 5999G>A, 6005T>C, 6009T>A, 6010C>G, 6011T>C, 6014A>C, 6015G>C, 6026G>A, 6028G>C, 6029A>T, 6030A>G, 6032T>A, 6039T>G, 6040G>C, 6041G>A, 6047C>A, 6048T>A, 6049C>A, 6050T>C, 6051C>A, 6054C>T, 6056A>C, 6057G>A, 6058A>G, 6060T>C, 6062A>T, 6063C>A, 6064C>A, 6065T>A, 6066T>A, 6067C>A, 6069A>G, 6070C>A, 6082C>G, 6083A>T, 6084C>G, 6085G>C, 6086G>T, 6089T>C, 6090T>C, 6091G>A, 6092T>A, 6100G>A, 6101C>A, 6104T>C, 6107_6108insGAAAAG, 6108T>G, 6109C>G, 6111G>A, 6112C>A, 6116C>G, 6117T>A, 6118C>G, 6120G>A, 6121C>G, 6122A>T, 6130C>T, 6131T>C, 6135A>G, 6138A>T, 6140G>A, 6144T>A, 6145G>T, 6146T>A, 6154T>G, 6156T>C, 6159G>A, 6163A>G, 6164G>T, 6167T>C, 6171A>T, 6175T>A, 6177T>A, 6179A>T, 6181A>T, 6182T>A, 6183G>A, 6185G>C, 6190A>G, 6191A>T, 6195T>A, 6197T>A, 6198A>G, 6199C>T, 6200C>A, 6201C>G, 6203T>A, 6206G>A, 6212C>T, 6215T>C, 6216A>G, 6217T>A, 6218T>A, 6221A>T, 6224T>A, 6230T>A, 6234T>C, 6237A>C, 6239C>A, 6243C>A, 6245T>A, 6246A>G, 6248T>C, 6249A>C, 6251T>G, 6257A>T, 6258T>A, 6259T>G, 6260G>T, 6262A>G, 6265T>A, 6268A>G, 6269T>A, 6270C>A, 6272T>A, 6276G>C, 6284C>T, 6285T>A, 6286G>T, 6287T>A, 6290T>C, 6291T>A, 6292T>C, 6293A>C, 6297A>T, 6299A>T, 6300G>A, 6301G>T, 6304T>A, 6305A>C, 6306G>C, 6307G>C, 6309_6311delTTA, 6313C>A, 6318A>G, 6319A>T, 6320A>T, 6321A>T, 6323A>C, 6327C>T, 6332C>A, 6333C>A, 6335G>A, 6338T>A | | | | | | | | |

##### CDS

| | Begin | End | Coverage | Score | Concordance | Matches | Identities | I/D/M/F* | Stop Codons |
|---|---|---|---|---|---|---|---|---|---|
| EXK67_gp3 | 1124 | 1582 | 28.8% | 1519 | 47.7% | 457 (98.9%) | 228 (49.4%) | 3/2/1/1 | 0 |
| Protein mutations: | L1125I (4983C>A), H1126K (4986C>A 4988T>A), K1127L (4989A>T 4990A>T), E1129D (4997A>T), I1131L (5001A>T 5003C>A), Q1132E (5004C>G), N1136E (5016A>G 5018T>A), D1137K (5019G>A 5021C>A), K1140I (5029A>T 5030G>A), Y1141H (5031T>C), E1143G (5038A>G), K1144A (5040A>G 5041A>C 5042G>C), Q1146R (5046C>A 5047A>G 5048G>A), R1148M (5053G>T 5054A>G), D1157E (5081T>A), L1158Y (5082C>T 5083T>A), Q1161K (5091C>A 5093G>A), D1162I (5094G>A 5095A>T 5096C>A), K1163A (5097A>G 5098A>C 5099G>A), P1164A (5100C>G 5102T>A), I1166E (5106A>G 5107T>A), P1167A (5109C>G), S1168T (5112T>A), P1169N (5115C>A 5116C>A 5117A>T), E1170Q (5118G>C), M1171D (5121A>G 5122T>A 5123G>C), A1172I (5124G>A 5125C>T 5126G>A), K1173E (5127A>G), E1174D (5132A>T), Y1175F (5134A>T), E1176Q (5136G>C 5138A>G), K1177I (5140A>T 5141G>A), H1178Q (5144T>A), T1180K (5149C>A), A1184D (5161C>A 5162A>T), K1186G (5166A>G 5167A>G 5168G>A), P1190R (5178C>A 5179C>G 5180T>A), Q1192T (5184C>A 5185A>C), T1197S (5199A>T), I1201M (5213T>G), N1203K (5219C>A), S1206N (5226T>A 5227C>A 5228T>C), Q1208I (5232C>A 5233A>T), S1213A (5247A>G 5248G>C 5249T>C), Y1217I (5259T>A 5260A>T 5261C>A), Y1227R (5289T>A 5290A>G 5291C>A), K1228M (5293A>T 5294A>G), Q1230A (5298C>G 5299A>C 5300A>T), T1232K (5305C>A), S1235D (5313A>G 5314G>A 5315T>C), I1236K (5317T>A 5318T>A), D1237N (5319G>A 5321C>T), Y1238E (5322T>G 5324T>A), L1241N (5331C>A 5332T>A), K1242R (5335A>G), K1244Q (5340A>C), D1245G (5344A>G 5345C>A), I1247R (5350T>G 5351T>A), V1248I (5352G>A 5354C>T), Y1249F (5356A>T), L1254C (5370C>T 5371T>G 5372A>T), F1258Y (5383T>A 5384C>T), H1259W (5385C>T 5386A>G 5387C>G), I1261V (5391A>G 5393T>A), M1262K (5395T>A 5396G>A), D1264H (5400G>C 5402T>C), Q1266D (5406C>G 5408G>T), A1272T (5424G>A), V1274T (5430G>A 5431T>C 5432C>T), Q1277E (5439C>G 5441G>A), F1280Y (5449T>A), W1282F (5455G>T 5456G>T), I1283L (5457A>C 5459A>G), K1290T (5479A>C), V1295S (5493G>T 5494T>C), R1298N (5502C>A 5503G>A 5504A>T), K1299L (5505A>C 5506A>T 5507A>T), D1301N (5511G>A), N1302A (5514A>G 5515A>C), I1303V (5517A>G 5519A>C), K1306P (5526A>C 5527A>C), K1306_Y1307insL (5528_5529insCTC), Y1307L (5529T>C 5530A>T), S1308R (5532T>A 5533C>G), E1309R (5535G>A 5536A>G), F1310S (5539T>C 5540T>A), C1312L (5545G>T 5546T>G), Y1314F (5551A>T), I1315F (5553A>T 5555A>T), I1320V (5568A>G 5570A>T), F1321Y (5572T>A 5573T>C), E1323K (5577G>A 5579A>G), S1324D (5580A>G 5581G>A), I1325M (5585A>G), Q1327E (5589C>G), V1329L (5595G>C), Q1330E (5598C>G), L1333Q (5607T>C 5608T>A), Q1334T (5610C>A 5611A>C 5612G>A), F1335I (5613T>A 5615C>T), Q1337R (5619C>A 5620A>G 5621A>G), V1338L (5622G>T), C1339F (5626G>T 5627T>C), K1340V (5628A>G 5629A>T 5630G>A), E1341S (5631G>A 5632A>G 5633A>T), E1342K (5634G>A), L1344I (5640C>A 5642T>A), L1346I (5646T>A 5648G>A), S1347T (5649T>A 5651A>C), T1349K (5656C>A 5657A>G), L1351M (5661T>A), K1352E (5664A>G), I1353L (5667A>T), G1354C (5670G>T 5672A>T), V1355K (5673G>A 5674T>A), A1356T (5676G>A), N1357H (5679A>C), E1359N (5685G>A 5687A>C), L1363V (5697C>G), E1364T (5700G>A 5701A>C 5702A>T), I1365L (5703A>T), E1367D (5711A>T), V1370I (5718G>A 5720T>A), Q1371K (5721C>A), L1377A (5739T>G 5740T>C 5741A>T), I1380V (5748A>G), E1382D (5756A>T), F1383M (5757T>A 5759T>G), E1385del (5763_5765delGAA), Q1387K (5769C>A), E1389D (5777A>T), T1390K (5779C>A), L1391T (5781T>A 5782T>C), G1393D (5788G>A), I1400L (5808A>T), L1401V (5811T>G 5813A>G), Y1407F (5830A>T), P1409Q (5836C>A), S1412G (5844A>G 5846T>A), Y1414I (5850T>A 5851A>T), T1415A (5853A>G 5855T>A), R1416G (5856A>G), I1417P (5859A>C 5860T>C), F1418L (5864T>A), N1420A (5868A>G 5869A>C 5870T>A), C1422I (5874T>A 5875G>T 5876T>A), S1423G (5877T>G 5878C>G 5879C>A), E1427Q (5889G>C), R1428K (5893G>A), K1429H (5895A>C 5897A>C), S1432T (5904T>A 5906T>A), Q1433E (5907C>G), D1434N (5910G>A 5912C>T), W1435I (5913T>A 5914G>T 5915G>T), M1437L (5919A>C), R1439Q (5925A>C 5926G>A), K1440Q (5928A>C), V1444K (5940G>A 5941T>A), T1446K (5947C>A 5948C>A), K1447N (5951A>C), L1448I (5952T>A), P1450D (5958C>G 5959C>A 5960T>C), I1452S (5965T>G 5966T>C), L1453I (5967T>A), K1455L (5973A>C 5974A>T), P1456E (5976C>G 5977C>A), Q1457S (5979C>T 5980A>C), S1458D (5982A>G 5983G>A), I1460L (5988A>C 5990T>A), E1469Q (6015G>C), G1473A (6028G>C 6029A>T), I1474V (6030A>G 6032T>A), W1477A (6039T>G 6040G>C 6041G>A), S1480N (6048T>A 6049C>A 6050T>C), Q1481K (6051C>A), Q1482Y (6054C>T 6056A>C), D1483S (6057G>A 6058A>G), S1484P (6060T>C 6062A>T), P1485K (6063C>A 6064C>A 6065T>A), S1486K (6066T>A 6067C>A), T1487E (6069A>G 6070C>A), S1491C (6082C>G 6083A>T), R1492A (6084C>G 6085G>C 6086G>T), C1494Q (6090T>C 6091G>A 6092T>A), S1497K (6100G>A 6101C>A), K1499_S1500insEK (6107_6108insGAAAAG), S1500G (6108T>G 6109C>G), A1501N (6111G>A 6112C>A), I1502M (6116C>G), A1504S (6120G>A 6121C>G 6122A>T), A1507V (6130C>T 6131T>C), I1509V (6135A>G), M1510L (6138A>T 6140G>A), C1512I (6144T>A 6145G>T 6146T>A), V1515G (6154T>G), D1517N (6159G>A), K1518S (6163A>G 6164G>T), I1521L (6171A>T), F1522Y (6175T>A), L1523I (6177T>A 6179A>T), Y1524L (6181A>T 6182T>A), E1525N (6183G>A 6185G>C), K1527S (6190A>G 6191A>T), F1529I (6195T>A 6197T>A), T1530V (6198A>G 6199C>T 6200C>A), L1531V (6201C>G 6203T>A), I1536E (6216A>G 6217T>A 6218T>A), N1540K (6230T>A), Y1542H (6234T>C), N1543Q (6237A>C 6239C>A), L1545I (6243C>A 6245T>A), N1546D (6246A>G 6248T>C), N1547Q (6249A>C 6251T>G), K1549N (6257A>T), L1550S (6258T>A 6259T>G 6260G>T), N1551S (6262A>G), I1552K (6265T>A), N1553R (6268A>G 6269T>A), V1556L (6276G>C), C1559I (6285T>A 6286G>T 6287T>A), L1561T (6291T>A 6292T>C 6293A>C), T1563S (6297A>T 6299A>T), G1564I (6300G>A 6301G>T), L1565Y (6304T>A 6305A>C), G1566P (6306G>C 6307G>C), L1567del (6309_6311delTTA), T1568N (6313C>A), K1570V (6318A>G 6319A>T 6320A>T), I1571F (6321A>T 6323A>C), H1573Y (6327C>T), Q1575K (6333C>A 6335G>A) | | | | | | | | |

| | Begin | End | Coverage | Score | Concordance | Matches | Identities | I/D/M/F* | Stop Codons |
|---|---|---|---|---|---|---|---|---|---|
| **NT** | **4980** | **6355** | **16.9%** | **493** | **18.2%** | **1369 (98.8%)** | **823 (59.4%)** | **9/7** | |

Codon mutations:

CTA1125ATA (4983C>A), CAT1126AAA (4986C>A 4988T>A), AAA1127TTA (4989A>T 4990A>T), TTG1128CTA (4992T>C 4994G>A), GAA1129GAT (4997A>T), ATC1131TTA (5001A>T 5003C>A), CAA1132GAA (5004C>G), ATT1133ATA (5009T>A), ATT1134ATA (5012T>A), GGG1135GGA (5015G>A), AAT1136GAA (5016A>G 5018T>A), GAC1137AAA (5019G>A 5021C>A), CCA1138CCC (5024A>C), TTA1139CTA (5025T>C), AAG1140ATA (5029A>T 5030G>A), TAT1141CAT (5031T>C), GAA1143GGA (5038A>G), AAG1144GCC (5040A>G 5041A>C 5042G>C), CAG1146AGA (5046C>A 5047A>G 5048G>A), ATT1147ATA (5051T>A), AGA1148ATG (5053G>T 5054A>G), TGC1149TGT (5057C>T), CTG1151CTA (5063G>A), GAC1152GAT (5066C>T), ATC1153ATA (5069C>A), CCT1156CCA (5078T>A), GAT1157GAA (5081T>A), CTT1158TAT (5082C>T 5083T>A), CAG1161AAA (5091C>A 5093G>A), GAC1162ATA (5094G>A 5095A>T 5096C>A), AAG1163GCA (5097A>G 5098A>C 5099G>A), CCT1164GCA (5100C>G 5102T>A), ATT1165ATA (5105T>A), ATA1166GAA (5106A>G 5107T>A), CCT1167GCT (5109C>G), TCA1168ACA (5112T>A), CCA1169AAT (5115C>A 5116C>A 5117A>T), GAA1170CAA (5118G>C), ATG1171GAC (5121A>G 5122T>A 5123G>C), GCG1172ATA (5124G>A 5125C>T 5126G>A), AAA1173GAA (5127A>G), GAA1174GAT (5132A>T), TAT1175TTT (5134A>T), GAA1176CAG (5136G>C 5138A>G), AAG1177ATA (5140A>T 5141G>A), CAT1178CAA (5144T>A), ATT1179ATA (5147T>A), ACA1180AAA (5149C>A), CTG1183TTA (5157C>T 5159G>A), GCA1184GAT (5161C>A 5162A>T), CTC1185TTA (5163C>T 5165C>A), AAG1186GGA (5166A>G 5167A>G 5168G>A), GTT1187GTA (5171T>A), ATC1188ATA (5174C>A), CCT1190AGA (5178C>A 5179C>G 5180T>A), TCC1191TCA (5183C>A), CAA1192ACA (5184C>A 5185A>C), AGT1193TCT (5187A>T 5188G>C), CGC1194AGA (5190C>A 5192C>A), CAT1195CAC (5195T>C), AGG1196AGA (5198G>A), ACA1197TCA (5199A>T), GCT1199GCA (5207T>A), ATT1201ATG (5213T>G), GTC1202GTA (5216C>A), AAC1203AAA (5219C>A), AAA1204-AC (5220delA 5222A>C), TCT1206AAC (5226T>A 5227C>A 5228T>C), CAA1208ATA (5232C>A 5233A>T), GTA1209GTC (5237A>C), GGT1211GGA (5243T>A), AGT1213GCC (5247A>G 5248G>C 5249T>C), CGT1214AGA (5250C>A 5252T>A), TAC1217ATA (5259T>A 5260A>T 5261C>A), CGA1221AGA (5271C>A), CTA1222CTC (5276A>C), ACA1226ACT (5288A>T), TAC1227AGA (5289T>A 5290A>G 5291C>A), AAA1228ATG (5293A>T 5294A>G), GAC1229GAT (5297C>T), CAA1230GCT (5298C>G 5299A>C 5300A>T), TAC1231TAT (5303C>T), ACA1232AAA (5305C>A), CTC1233TTA (5307C>T 5309C>A), AGT1235GAC (5313A>G 5314G>A 5315T>C), ATT1236AAA (5317T>A 5318T>A), GAC1237AAT (5319G>A 5321C>T), TAT1238GAA (5322T>G 5324T>A), CTC1239CTA (5327C>A), CTT1241AAT (5331C>A 5332T>A), AAA1242AGA (5335A>G), ATT1243ATA (5339T>A), AAA1244CAA (5340A>C), GAC1245GGA (5344A>G 5345C>A), AAG1246AAA (5348G>A), ATT1247AGA (5350T>G 5351T>A), GTC1248ATT (5352G>A 5354C>T), TAC1249TTC (5356A>T), AGC1250AGT (5360C>T), GAT1253GAC (5369T>C), CTA1254TGT (5370C>T 5371T>G 5372A>T), AAG1255AAA (5375G>A), GGA1257GGG (5381A>G), TTC1258TAT (5383T>A 5384C>T), CAC1259TGG (5385C>T 5386A>G 5387C>G), CAG1260CAA (5390G>A), ATT1261GTA (5391A>G 5393T>A), ATG1262AAA (5395T>A 5396G>A), GAT1264CAC (5400G>C 5402T>C), CCA1265CCT (5405A>T), CAG1266GAT (5406C>G 5408G>T), GCA1272ACA (5424G>A), GTC1274ACT (5430G>A 5431T>C 5432C>T), TGC1275TGT (5435C>T), CAG1277GAA (5439C>G 5441G>A), GGT1278GGA (5444T>A), CAT1279CAC (5447T>C), TTT1280TAT (5449T>A), TGG1282TTT (5455G>T 5456G>T), ATA1283CTG (5457A>C 5459A>G), GTA1284GTT (5462A>T), TTC1287TTT (5471C>T), GGA1288GGC (5474A>C), CTC1289CTT (5477C>T), AAA1290ACA (5479A>C), TCA1294TCC (5492A>C), GTA1295TCA (5493G>T 5494T>C), TTC1296TTT (5498C>T), CAA1297CAG (5501A>G), CGA1298AAT (5502C>A 5503G>A 5504A>T), AAA1299CTT (5505A>C 5506A>T 5507A>T), GAT1301AAT (5511G>A), AAT1302GCT (5514A>G 5515A>C), ATA1303GTC (5517A>G 5519A>C), TTC1304TTT (5522C>T), AAA1306CCA (5526A>C 5527A>C), AAA1306_TAT1307insCTC (5528_5529insCTC), TAT1307CTT (5529T>C 5530A>T), TCA1308AGA (5532T>A 5533C>G), GAA1309AGA (5535G>A 5536A>G), TTT1310TCA (5539T>C 5540T>A), GTC1311GTA (5543C>A), TGT1312TTG (5545G>T 5546T>G), GTT1313GTC (5549T>C), TAC1314TTC (5551A>T), ATA1315TTT (5553A>T 5555A>T), GAC1316GAT (5558C>T), ATT1318ATA (5564T>A), ATA1320GTT (5568A>G 5570A>T), TTT1321TAC (5572T>A 5573T>C), TCT1322AGC (5574T>A 5575C>G 5576T>C), GAA1323AAG (5577G>A 5579A>G), AGT1324GAT (5580A>G 5581G>A), ATA1325ATG (5585A>G), CAA1327GAA (5589C>G), GTA1329CTA (5595G>C), CAA1330GAA (5598C>G), TTA1333CAA (5607T>C 5608T>A), CAG1334ACA (5610C>A 5611A>C 5612G>A), TTC1335ATT (5613T>A 5615C>T), TTC1336TTT (5618C>T), CAA1337AGG (5619C>A 5620A>G 5621A>G), GTA1338TTA (5622G>T), TGT1339TTC (5626G>T 5627T>C), AAG1340GTA (5628A>G 5629A>T 5630G>A), GAA1341AGT (5631G>A 5632A>G 5633A>T), GAA1342AAA (5634G>A), GGG1343GGA (5639G>A), CTT1344ATA (5640C>A 5642T>A), ATT1345ATA (5645T>A), TTG1346ATA (5646T>A 5648G>A), TCA1347ACC (5649T>A 5651A>C), ACA1349AAG (5656C>A 5657A>G), TTG1351ATG (5661T>A), AAA1352GAA (5664A>G), ATA1353TTA (5667A>T), GGA1354TGT (5670G>T 5672A>T), GTA1355AAA (5673G>A 5674T>A), GCA1356ACA (5676G>A), AAT1357CAT (5679A>C), GAA1359AAC (5685G>A 5687A>C), CTT1361TTA (5691C>T 5693T>A), GGC1362GGA (5696C>A), CTA1363GTA (5697C>G), GAA1364ACT (5700G>A 5701A>C 5702A>T), ATA1365TTA (5703A>T), GAA1367GAT (5711A>T), GGA1368GGG (5714A>G), AAA1369AAG (5717A>G), GTT1370ATA (5718G>A 5720T>A), CAA1371AAA (5721C>A), CTC1372CTA (5726C>A), TTA1377GCT (5739T>G 5740T>C 5741A>T), ATA1380GTA (5748A>G), GAA1382GAT (5756A>T), TTT1383ATG (5757T>A 5759T>G), CCT1384CCA (5762T>A), GAA1385del (5763_5765delGAA), GAC1386GAT (5768C>T), CAA1387AAA (5769C>A), GAA1389GAT (5777A>T), ACA1390AAA (5779C>A), TTA1391ACA (5781T>A 5782T>C), AAG1392AAA (5786G>A), GGT1393GAT (5788G>A), CTA1394TTA (5790C>T), CAG1395CAA (5795G>A), AAG1396AAA (5798G>A), GGG1399GGA (5807G>A), ATA1400TTA (5808A>T), TTA1401GTG (5811T>G 5813A>G), AAC1402AAT (5816C>T), GCC1404GCT (5822C>T), AAT1406AAC (5828T>C), TAT1407TTT (5830A>T), ATC1408ATA (5834C>A), CCA1409CAA (5836C>A), TTG1411TTA (5843G>A), AGT1412GGA (5844A>G 5846T>A), TAT1414ATT (5850T>A 5851A>T), ACT1415GCA (5853A>G 5855T>A), AGA1416GGA (5856A>G), ATA1417CCA (5859A>C 5860T>C), TTT1418TTA (5864T>A), AAT1420GCA (5868A>G 5869A>C 5870T>A), AAG1421AAA (5873G>A), TGT1422ATA (5874T>A 5875G>T 5876T>A), TCC1423GGA (5877T>G 5878C>G 5879C>A), AGC1424AGT (5882C>T), GAA1427CAA (5889G>C), AGA1428AAA (5893G>A), AAA1429CAC (5895A>C 5897A>C), AAT1431AAC (5903T>C), TCT1432ACA (5904T>A 5906T>A), CAA1433GAA (5907C>G), GAC1434AAT (5910G>A 5912C>T), TGG1435ATT (5913T>A 5914G>T 5915G>T), AAG1436AAA (5918G>A), ATG1437CTG (5919A>C), GTT1438GTA (5924T>A), AGA1439CAA (5925A>C 5926G>A), AAA1440CAA (5928A>C), ATC1441ATT (5933C>T), GTA1444AAA (5940G>A 5941T>A), ATT1445ATA (5945T>A), ACC1446AAA (5947C>A 5948C>A), AAA1447AAC (5951A>C), TTA1448ATA (5952T>A), CCC1449CCA (5957C>A), CCT1450GAC (5958C>G 5959C>A 5960T>C), TTG1451TTA (5963G>A), ATT1452AGC (5965T>G 5966T>C), TTA1453ATA (5967T>A), AAA1455CTA (5973A>C 5974A>T), CCA1456GAA (5976C>G 5977C>A), CAA1457TCA (5979C>T 5980A>C), AGT1458GAT (5982A>G 5983G>A), ATT1460CTA (5988A>C 5990T>A), GAG1463GAA (5999G>A), GAT1465GAC (6005T>C), TCT1467AGC (6009T>A 6010C>G 6011T>C), CTA1468CTC (6014A>C), GAA1469CAA (6015G>C), GGG1472GGA (6026G>A), GGA1473GCT (6028G>C 6029A>T), ATT1474GTA (6030A>G 6032T>A), TGG1477GCA (6039T>G 6040G>C 6041G>A), CCC1479CCA (6047C>A), TCT1480AAC (6048T>A 6049C>A 6050T>C), CAA1481AAA (6051C>A), CAA1482TAC (6054C>T 6056A>C), GAT1483AGT (6057G>A 6058A>G), TCA1484CCT (6060T>C 6062A>T), CCT1485AAA (6063C>A 6064C>A 6065T>A), TCA1486AAA (6066T>A 6067C>A), ACA1487GAA (6069A>G 6070C>A), TCA1491TGT (6082C>G 6083A>T), CGG1492GCT (6084C>G 6085G>C 6086G>T), TAT1493TAC (6089T>C), TGT1494CAA (6090T>C 6091G>A 6092T>A), AGC1497AAA (6100G>A 6101C>A), TAT1498TAC (6104T>C), AAA1499_TCT1500insGAAAAG (6107_6108insGAAAAG), TCT1500GGT (6108T>G 6109C>G), GCC1501AAC (6111G>A 6112C>A), ATC1502ATG (6116C>G), TCT1503AGT (6117T>A 6118C>G), GCA1504AGT (6120G>A 6121C>G 6122A>T), GCT1507GTC (6130C>T 6131T>C), ATA1509GTA (6135A>G), ATG1510TTA (6138A>T 6140G>A), TGT1512ATA (6144T>A 6145G>T 6146T>A), GTA1515GGA (6154T>G), TTA1516CTA (6156T>C), GAT1517AAT (6159G>A), AAG1518AGT (6163A>G 6164G>T), TTT1519TTC (6167T>C), ATA1521TTA (6171A>T), TTT1522TAT (6175T>A), TTA1523ATT (6177T>A 6179A>T), TAT1524TTA (6181A>T 6182T>A), GAG1525AAC (6183G>A 6185G>C), AAA1527AGT (6190A>G 6191A>T), TTT1529ATA (6195T>A 6197T>A), ACC1530GTA (6198A>G 6199C>T 6200C>A), CTT1531GTA (6201C>G 6203T>A), AGG1532AGA (6206G>A), GAC1534GAT (6212C>T), TGT1535TGC (6215T>C), ATT1536GAA (6216A>G 6217T>A 6218T>A), GCA1537GCT (6221A>T), ATT1538ATA (6224T>A), AAT1540AAA (6230T>A), TAT1542CAT (6234T>C), AAC1543CAA (6237A>C 6239C>A), CTT1545ATA (6243C>A 6245T>A), AAT1546GAC (6246A>G 6248T>C), AAT1547CAG (6249A>C 6251T>G), AAA1549AAT (6257A>T), TTG1550AGT (6258T>A 6259T>G 6260G>T), AAT1551AGT (6262A>G), ATA1552AAA (6265T>A), AAT1553AGA (6268A>G 6269T>A), CGT1554AGA (6270C>A 6272T>A), GTA1556CTA (6276G>C), TTC1558TTT (6284C>T), TGT1559ATA (6285T>A 6286G>T 6287T>A), GAT1560GAC (6290T>C), TTA1561ACC (6291T>A 6292T>C 6293A>C), ACA1563TCT (6297A>T 6299A>T), GGA1564ATA (6300G>A 6301G>T), TTA1565TAC (6304T>A 6305A>C), GGT1566CCT (6306G>C 6307G>C), TTA1567del (6309_6311delTTA), ACT1568AAT (6313C>A), AAA1570GTT (6318A>G 6319A>T 6320A>T), ATA1571TTC (6321A>T 6323A>C), CAT1573TAT (6327C>T), ATC1574ATA (6332C>A), CAG1575AAA (6333C>A 6335G>A), GGT1576GGA (6338T>A)

#### Proteins

| | Begin | End | Coverage | Score | Concordance | Matches | Identities | I/D/M/F* | Stop Codons |
|---|---|---|---|---|---|---|---|---|---|
| ORF3 (YP_009553219.1) | 1124 | 1582 | 28.8% | 1519 | 47.7% | 457 (98.9%) | 228 (49.4%) | 3/2/1/1 | 0 |

Protein mutations:

L1125I (4983C>A), H1126K (4986C>A 4988T>A), K1127L (4989A>T 4990A>T), E1129D (4997A>T), I1131L (5001A>T 5003C>A), Q1132E (5004C>G), N1136E (5016A>G 5018T>A), D1137K (5019G>A 5021C>A), K1140I (5029A>T 5030G>A), Y1141H (5031T>C), E1143G (5038A>G), K1144A (5040A>G 5041A>C 5042G>C), Q1146R (5046C>A 5047A>G 5048G>A), R1148M (5053G>T 5054A>G), D1157E (5081T>A), L1158Y (5082C>T 5083T>A), Q1161K (5091C>A 5093G>A), D1162I (5094G>A 5095A>T 5096C>A), K1163A (5097A>G 5098A>C 5099G>A), P1164A (5100C>G 5102T>A), I1166E (5106A>G 5107T>A), P1167A (5109C>G), S1168T (5112T>A), P1169N (5115C>A 5116C>A 5117A>T), E1170Q (5118G>C), M1171D (5121A>G 5122T>A 5123G>C), A1172I (5124G>A 5125C>T 5126G>A), K1173E (5127A>G), E1174D (5132A>T), Y1175F (5134A>T), E1176Q (5136G>C 5138A>G), K1177I (5140A>T 5141G>A), H1178Q (5144T>A), T1180K (5149C>A), A1184D (5161C>A 5162A>T), K1186G (5166A>G 5167A>G 5168G>A), P1190R (5178C>A 5179C>G 5180T>A), Q1192T (5184C>A 5185A>C), T1197S (5199A>T), I1201M (5213T>G), N1203K (5219C>A), S1206N (5226T>A 5227C>A 5228T>C), Q1208I (5232C>A 5233A>T), S1213A (5247A>G 5248G>C 5249T>C), Y1217I (5259T>A 5260A>T 5261C>A), Y1227R (5289T>A 5290A>G 5291C>A), K1228M (5293A>T 5294A>G), Q1230A (5298C>G 5299A>C 5300A>T), T1232K (5305C>A), S1235D (5313A>G 5314G>A 5315T>C), I1236K (5317T>A 5318T>A), D1237N (5319G>A 5321C>T), Y1238E (5322T>G 5324T>A), L1241N (5331C>A 5332T>A), K1242R (5335A>G), K1244Q (5340A>C), D1245G (5344A>G 5345C>A), I1247R (5350T>G 5351T>A), V1248I (5352G>A 5354C>T), Y1249F (5356A>T), L1254C (5370C>T 5371T>G 5372A>T), F1258Y (5383T>A 5384C>T), H1259W (5385C>T 5386A>G 5387C>G), I1261V (5391A>G 5393T>A), M1262K (5395T>A 5396G>A), D1264H (5400G>C 5402T>C), Q1266D (5406C>G 5408G>T), A1272T (5424G>A), V1274T (5430G>A 5431T>C 5432C>T), Q1277E (5439C>G 5441G>A), F1280Y (5449T>A), W1282F (5455G>T 5456G>T), I1283L (5457A>C 5459A>G), K1290T (5479A>C), V1295S (5493G>T 5494T>C), R1298N (5502C>A 5503G>A 5504A>T), K1299L (5505A>C 5506A>T 5507A>T), D1301N (5511G>A), N1302A (5514A>G 5515A>C), I1303V (5517A>G 5519A>C), K1306P (5526A>C 5527A>C), K1306_Y1307insL (5528_5529insCTC), Y1307L (5529T>C 5530A>T), S1308R (5532T>A 5533C>G), E1309R (5535G>A 5536A>G), F1310S (5539T>C 5540T>A), C1312L (5545G>T 5546T>G), Y1314F (5551A>T), I1315F (5553A>T 5555A>T), I1320V (5568A>G 5570A>T), F1321Y (5572T>A 5573T>C), E1323K (5577G>A 5579A>G), S1324D (5580A>G 5581G>A), I1325M (5585A>G), Q1327E (5589C>G), V1329L (5595G>C), Q1330E (5598C>G), L1333Q (5607T>C 5608T>A), Q1334T (5610C>A 5611A>C 5612G>A), F1335I (5613T>A 5615C>T), Q1337R (5619C>A 5620A>G 5621A>G), V1338L (5622G>T), C1339F (5626G>T 5627T>C), K1340V (5628A>G 5629A>T 5630G>A), E1341S (5631G>A 5632A>G 5633A>T), E1342K (5634G>A), L1344I (5640C>A 5642T>A), L1346I (5646T>A 5648G>A), S1347T (5649T>A 5651A>C), T1349K (5656C>A 5657A>G), L1351M (5661T>A), K1352E (5664A>G), I1353L (5667A>T), G1354C (5670G>T 5672A>T), V1355K (5673G>A 5674T>A), A1356T (5676G>A), N1357H (5679A>C), E1359N (5685G>A 5687A>C), L1363V (5697C>G), E1364T (5700G>A 5701A>C 5702A>T), I1365L (5703A>T), E1367D (5711A>T), V1370I (5718G>A 5720T>A), Q1371K (5721C>A), L1377A (5739T>G 5740T>C 5741A>T), I1380V (5748A>G), E1382D (5756A>T), F1383M (5757T>A 5759T>G), E1385del (5763_5765delGAA), Q1387K (5769C>A), E1389D (5777A>T), T1390K (5779C>A), L1391T (5781T>A 5782T>C), G1393D (5788G>A), I1400L (5808A>T), L1401V (5811T>G 5813A>G), Y1407F (5830A>T), P1409Q (5836C>A), S1412G (5844A>G 5846T>A), Y1414I (5850T>A 5851A>T), T1415A (5853A>G 5855T>A), R1416G (5856A>G), I1417P (5859A>C 5860T>C), F1418L (5864T>A), N1420A (5868A>G 5869A>C 5870T>A), C1422I (5874T>A 5875G>T 5876T>A), S1423G (5877T>G 5878C>G 5879C>A), E1427Q (5889G>C), R1428K (5893G>A), K1429H (5895A>C 5897A>C), S1432T (5904T>A 5906T>A), Q1433E (5907C>G), D1434N (5910G>A 5912C>T), W1435I (5913T>A 5914G>T 5915G>T), M1437L (5919A>C), R1439Q (5925A>C 5926G>A), K1440Q (5928A>C), V1444K (5940G>A 5941T>A), T1446K (5947C>A 5948C>A), K1447N (5951A>C), L1448I (5952T>A), P1450D (5958C>G 5959C>A 5960T>C), I1452S (5965T>G 5966T>C), L1453I (5967T>A), K1455L (5973A>C 5974A>T), P1456E (5976C>G 5977C>A), Q1457S (5979C>T 5980A>C), S1458D (5982A>G 5983G>A), I1460L (5988A>C 5990T>A), E1469Q (6015G>C), G1473A (6028G>C 6029A>T), I1474V (6030A>G 6032T>A), W1477A (6039T>G 6040G>C 6041G>A), S1480N (6048T>A 6049C>A 6050T>C), Q1481K (6051C>A), Q1482Y (6054C>T 6056A>C), D1483S (6057G>A 6058A>G), S1484P (6060T>C 6062A>T), P1485K (6063C>A 6064C>A 6065T>A), S1486K (6066T>A 6067C>A), T1487E (6069A>G 6070C>A), S1491C (6082C>G 6083A>T), R1492A (6084C>G 6085G>C 6086G>T), C1494Q (6090T>C 6091G>A 6092T>A), S1497K (6100G>A 6101C>A), K1499_S1500insEK (6107_6108insGAAAAG), S1500G (6108T>G 6109C>G), A1501N (6111G>A 6112C>A), I1502M (6116C>G), A1504S (6120G>A 6121C>G 6122A>T), A1507V (6130C>T 6131T>C), I1509V (6135A>G), M1510L (6138A>T 6140G>A), C1512I (6144T>A 6145G>T 6146T>A), V1515G (6154T>G), D1517N (6159G>A), K1518S (6163A>G 6164G>T), I1521L (6171A>T), F1522Y (6175T>A), L1523I (6177T>A 6179A>T), Y1524L (6181A>T 6182T>A), E1525N (6183G>A 6185G>C), K1527S (6190A>G 6191A>T), F1529I (6195T>A 6197T>A), T1530V (6198A>G 6199C>T 6200C>A), L1531V (6201C>G 6203T>A), I1536E (6216A>G 6217T>A 6218T>A), N1540K (6230T>A), Y1542H (6234T>C), N1543Q (6237A>C 6239C>A), L1545I (6243C>A 6245T>A), N1546D (6246A>G 6248T>C), N1547Q (6249A>C 6251T>G), K1549N (6257A>T), L1550S (6258T>A 6259T>G 6260G>T), N1551S (6262A>G), I1552K (6265T>A), N1553R (6268A>G 6269T>A), V1556L (6276G>C), C1559I (6285T>A 6286G>T 6287T>A), L1561T (6291T>A 6292T>C 6293A>C), T1563S (6297A>T 6299A>T), G1564I (6300G>A 6301G>T), L1565Y (6304T>A 6305A>C), G1566P (6306G>C 6307G>C), L1567del (6309_6311delTTA), T1568N (6313C>A), K1570V (6318A>G 6319A>T 6320A>T), I1571F (6321A>T 6323A>C), H1573Y (6327C>T), Q1575K (6333C>A 6335G>A)

| | Begin | End | Coverage | Score | Concordance | Matches | Identities | I/D/M/F* | Stop Codons |
|---|---|---|---|---|---|---|---|---|---|
| **NT** | **4980** | **6355** | **16.9%** | **493** | **18.2%** | **1369 (98.8%)** | **823 (59.4%)** | **9/7** | |
| Codon mutations: | CTA1125ATA (4983C>A), CAT1126AAA (4986C>A 4988T>A), AAA1127TTA (4989A>T 4990A>T), TTG1128CTA (4992T>C 4994G>A), GAA1129GAT (4997A>T), ATC1131TTA (5001A>T 5003C>A), CAA1132GAA (5004C>G), ATT1133ATA (5009T>A), ATT1134ATA (5012T>A), GGG1135GGA (5015G>A), AAT1136GAA (5016A>G 5018T>A), GAC1137AAA (5019G>A 5021C>A), CCA1138CCC (5024A>C), TTA1139CTA (5025T>C), AAG1140ATA (5029A>T 5030G>A), TAT1141CAT (5031T>C), GAA1143GGA (5038A>G), AAG1144GCC (5040A>G 5041A>C 5042G>C), CAG1146AGA (5046C>A 5047A>G 5048G>A), ATT1147ATA (5051T>A), AGA1148ATG (5053G>T 5054A>G), TGC1149TGT (5057C>T), CTG1151CTA (5063G>A), GAC1152GAT (5066C>T), ATC1153ATA (5069C>A), CCT1156CCA (5078T>A), GAT1157GAA (5081T>A), CTT1158TAT (5082C>T 5083T>A), CAG1161AAA (5091C>A 5093G>A), GAC1162ATA (5094G>A 5095A>T 5096C>A), AAG1163GCA (5097A>G 5098A>C 5099G>A), CCT1164GCA (5100C>G 5102T>A), ATT1165ATA (5105T>A), ATA1166GAA (5106A>G 5107T>A), CCT1167GCT (5109C>G), TCA1168ACA (5112T>A), CCA1169AAT (5115C>A 5116C>A 5117A>T), GAA1170CAA (5118G>C), ATG1171GAC (5121A>G 5122T>A 5123G>C), GCG1172ATA (5124G>A 5125C>T 5126G>A), AAA1173GAA (5127A>G), GAA1174GAT (5132A>T), TAT1175TTT (5134A>T), GAA1176CAG (5136G>C 5138A>G), AAG1177ATA (5140A>T 5141G>A), CAT1178CAA (5144T>A), ATT1179ATA (5147T>A), ACA1180AAA (5149C>A), CTG1183TTA (5157C>T 5159G>A), GCA1184GAT (5161C>A 5162A>T), CTC1185TTA (5163C>T 5165C>A), AAG1186GGA (5166A>G 5167A>G 5168G>A), GTT1187GTA (5171T>A), ATC1188ATA (5174C>A), CCT1190AGA (5178C>A 5179C>G 5180T>A), TCC1191TCA (5183C>A), CAA1192ACA (5184C>A 5185A>C), AGT1193TCT (5187A>T 5188G>C), CGC1194AGA (5190C>A 5192C>A), CAT1195CAC (5195T>C), AGG1196AGA (5198G>A), ACA1197TCA (5199A>T), GCT1199GCA (5207T>A), ATT1201ATG (5213T>G), GTC1202GTA (5216C>A), AAC1203AAA (5219C>A), AAA1204-AC (5220delA 5222A>C), TCT1206AAC (5226T>A 5227C>A 5228T>C), CAA1208ATA (5232C>A 5233A>T), GTA1209GTC (5237A>C), GGT1211GGA (5243T>A), AGT1213GCC (5247A>G 5248G>C 5249T>C), CGT1214AGA (5250C>A 5252T>A), TAC1217ATA (5259T>A 5260A>T 5261C>A), CGA1221AGA (5271C>A), CTA1222CTC (5276A>C), ACA1226ACT (5288A>T), TAC1227AGA (5289T>A 5290A>G 5291C>A), AAA1228ATG (5293A>T 5294A>G), GAC1229GAT (5297C>T), CAA1230GCT (5298C>G 5299A>C 5300A>T), TAC1231TAT (5303C>T), ACA1232AAA (5305C>A), CTC1233TTA (5307C>T 5309C>A), AGT1235GAC (5313A>G 5314G>A 5315T>C), ATT1236AAA (5317T>A 5318T>A), GAC1237AAT (5319G>A 5321C>T), TAT1238GAA (5322T>G 5324T>A), CTC1239CTA (5327C>A), CTT1241AAT (5331C>A 5332T>A), AAA1242AGA (5335A>G), ATT1243ATA (5339T>A), AAA1244CAA (5340A>C), GAC1245GGA (5344A>G 5345C>A), AAG1246AAA (5348G>A), ATT1247AGA (5350T>G 5351T>A), GTC1248ATT (5352G>A 5354C>T), TAC1249TTC (5356A>T), AGC1250AGT (5360C>T), GAT1253GAC (5369T>C), CTA1254TGT (5370C>T 5371T>G 5372A>T), AAG1255AAA (5375G>A), GGA1257GGG (5381A>G), TTC1258TAT (5383T>A 5384C>T), CAC1259TGG (5385C>T 5386A>G 5387C>G), CAG1260CAA (5390G>A), ATT1261GTA (5391A>G 5393T>A), ATG1262AAA (5395T>A 5396G>A), GAT1264CAC (5400G>C 5402T>C), CCA1265CCT (5405A>T), CAG1266GAT (5406C>G 5408G>T), GCA1272ACA (5424G>A), GTC1274ACT (5430G>A 5431T>C 5432C>T), TGC1275TGT (5435C>T), CAG1277GAA (5439C>G 5441G>A), GGT1278GGA (5444T>A), CAT1279CAC (5447T>C), TTT1280TAT (5449T>A), TGG1282TTT (5455G>T 5456G>T), ATA1283CTG (5457A>C 5459A>G), GTA1284GTT (5462A>T), TTC1287TTT (5471C>T), GGA1288GGC (5474A>C), CTC1289CTT (5477C>T), AAA1290ACA (5479A>C), TCA1294TCC (5492A>C), GTA1295TCA (5493G>T 5494T>C), TTC1296TTT (5498C>T), CAA1297CAG (5501A>G), CGA1298AAT (5502C>A 5503G>A 5504A>T), AAA1299CTT (5505A>C 5506A>T 5507A>T), GAT1301AAT (5511G>A), AAT1302GCT (5514A>G 5515A>C), ATA1303GTC (5517A>G 5519A>C), TTC1304TTT (5522C>T), AAA1306CCA (5526A>C 5527A>C), AAA1306_TAT1307insCTC (5528_5529insCTC), TAT1307CTT (5529T>C 5530A>T), TCA1308AGA (5532T>A 5533C>G), GAA1309AGA (5535G>A 5536A>G), TTT1310TCA (5539T>C 5540T>A), GTC1311GTA (5543C>A), TGT1312TTG (5545G>T 5546T>G), GTT1313GTC (5549T>C), TAC1314TTC (5551A>T), ATA1315TTT (5553A>T 5555A>T), GAC1316GAT (5558C>T), ATT1318ATA (5564T>A), ATA1320GTT (5568A>G 5570A>T), TTT1321TAC (5572T>A 5573T>C), TCT1322AGC (5574T>A 5575C>G 5576T>C), GAA1323AAG (5577G>A 5579A>G), AGT1324GAT (5580A>G 5581G>A), ATA1325ATG (5585A>G), CAA1327GAA (5589C>G), GTA1329CTA (5595G>C), CAA1330GAA (5598C>G), TTA1333CAA (5607T>C 5608T>A), CAG1334ACA (5610C>A 5611A>C 5612G>A), TTC1335ATT (5613T>A 5615C>T), TTC1336TTT (5618C>T), CAA1337AGG (5619C>A 5620A>G 5621A>G), GTA1338TTA (5622G>T), TGT1339TTC (5626G>T 5627T>C), AAG1340GTA (5628A>G 5629A>T 5630G>A), GAA1341AGT (5631G>A 5632A>G 5633A>T), GAA1342AAA (5634G>A), GGG1343GGA (5639G>A), CTT1344ATA (5640C>A 5642T>A), ATT1345ATA (5645T>A), TTG1346ATA (5646T>A 5648G>A), TCA1347ACC (5649T>A 5651A>C), ACA1349AAG (5656C>A 5657A>G), TTG1351ATG (5661T>A), AAA1352GAA (5664A>G), ATA1353TTA (5667A>T), GGA1354TGT (5670G>T 5672A>T), GTA1355AAA (5673G>A 5674T>A), GCA1356ACA (5676G>A), AAT1357CAT (5679A>C), GAA1359AAC (5685G>A 5687A>C), CTT1361TTA (5691C>T 5693T>A), GGC1362GGA (5696C>A), CTA1363GTA (5697C>G), GAA1364ACT (5700G>A 5701A>C 5702A>T), ATA1365TTA (5703A>T), GAA1367GAT (5711A>T), GGA1368GGG (5714A>G), AAA1369AAG (5717A>G), GTT1370ATA (5718G>A 5720T>A), CAA1371AAA (5721C>A), CTC1372CTA (5726C>A), TTA1377GCT (5739T>G 5740T>C 5741A>T), ATA1380GTA (5748A>G), GAA1382GAT (5756A>T), TTT1383ATG (5757T>A 5759T>G), CCT1384CCA (5762T>A), GAA1385del (5763_5765delGAA), GAC1386GAT (5768C>T), CAA1387AAA (5769C>A), GAA1389GAT (5777A>T), ACA1390AAA (5779C>A), TTA1391ACA (5781T>A 5782T>C), AAG1392AAA (5786G>A), GGT1393GAT (5788G>A), CTA1394TTA (5790C>T), CAG1395CAA (5795G>A), AAG1396AAA (5798G>A), GGG1399GGA (5807G>A), ATA1400TTA (5808A>T), TTA1401GTG (5811T>G 5813A>G), AAC1402AAT (5816C>T), GCC1404GCT (5822C>T), AAT1406AAC (5828T>C), TAT1407TTT (5830A>T), ATC1408ATA (5834C>A), CCA1409CAA (5836C>A), TTG1411TTA (5843G>A), AGT1412GGA (5844A>G 5846T>A), TAT1414ATT (5850T>A 5851A>T), ACT1415GCA (5853A>G 5855T>A), AGA1416GGA (5856A>G), ATA1417CCA (5859A>C 5860T>C), TTT1418TTA (5864T>A), AAT1420GCA (5868A>G 5869A>C 5870T>A), AAG1421AAA (5873G>A), TGT1422ATA (5874T>A 5875G>T 5876T>A), TCC1423GGA (5877T>G 5878C>G 5879C>A), AGC1424AGT (5882C>T), GAA1427CAA (5889G>C), AGA1428AAA (5893G>A), AAA1429CAC (5895A>C 5897A>C), AAT1431AAC (5903T>C), TCT1432ACA (5904T>A 5906T>A), CAA1433GAA (5907C>G), GAC1434AAT (5910G>A 5912C>T), TGG1435ATT (5913T>A 5914G>T 5915G>T), AAG1436AAA (5918G>A), ATG1437CTG (5919A>C), GTT1438GTA (5924T>A), AGA1439CAA (5925A>C 5926G>A), AAA1440CAA (5928A>C), ATC1441ATT (5933C>T), GTA1444AAA (5940G>A 5941T>A), ATT1445ATA (5945T>A), ACC1446AAA (5947C>A 5948C>A), AAA1447AAC (5951A>C), TTA1448ATA (5952T>A), CCC1449CCA (5957C>A), CCT1450GAC (5958C>G 5959C>A 5960T>C), TTG1451TTA (5963G>A), ATT1452AGC (5965T>G 5966T>C), TTA1453ATA (5967T>A), AAA1455CTA (5973A>C 5974A>T), CCA1456GAA (5976C>G 5977C>A), CAA1457TCA (5979C>T 5980A>C), AGT1458GAT (5982A>G 5983G>A), ATT1460CTA (5988A>C 5990T>A), GAG1463GAA (5999G>A), GAT1465GAC (6005T>C), TCT1467AGC (6009T>A 6010C>G 6011T>C), CTA1468CTC (6014A>C), GAA1469CAA (6015G>C), GGG1472GGA (6026G>A), GGA1473GCT (6028G>C 6029A>T), ATT1474GTA (6030A>G 6032T>A), TGG1477GCA (6039T>G 6040G>C 6041G>A), CCC1479CCA (6047C>A), TCT1480AAC (6048T>A 6049C>A 6050T>C), CAA1481AAA (6051C>A), CAA1482TAC (6054C>T 6056A>C), GAT1483AGT (6057G>A 6058A>G), TCA1484CCT (6060T>C 6062A>T), CCT1485AAA (6063C>A 6064C>A 6065T>A), TCA1486AAA (6066T>A 6067C>A), ACA1487GAA (6069A>G 6070C>A), TCA1491TGT (6082C>G 6083A>T), CGG1492GCT (6084C>G 6085G>C 6086G>T), TAT1493TAC (6089T>C), TGT1494CAA (6090T>C 6091G>A 6092T>A), AGC1497AAA (6100G>A 6101C>A), TAT1498TAC (6104T>C), AAA1499_TCT1500insGAAAAG (6107_6108insGAAAAG), TCT1500GGT (6108T>G 6109C>G), GCC1501AAC (6111G>A 6112C>A), ATC1502ATG (6116C>G), TCT1503AGT (6117T>A 6118C>G), GCA1504AGT (6120G>A 6121C>G 6122A>T), GCT1507GTC (6130C>T 6131T>C), ATA1509GTA (6135A>G), ATG1510TTA (6138A>T 6140G>A), TGT1512ATA (6144T>A 6145G>T 6146T>A), GTA1515GGA (6154T>G), TTA1516CTA (6156T>C), GAT1517AAT (6159G>A), AAG1518AGT (6163A>G 6164G>T), TTT1519TTC (6167T>C), ATA1521TTA (6171A>T), TTT1522TAT (6175T>A), TTA1523ATT (6177T>A 6179A>T), TAT1524TTA (6181A>T 6182T>A), GAG1525AAC (6183G>A 6185G>C), AAA1527AGT (6190A>G 6191A>T), TTT1529ATA (6195T>A 6197T>A), ACC1530GTA (6198A>G 6199C>T 6200C>A), CTT1531GTA (6201C>G 6203T>A), AGG1532AGA (6206G>A), GAC1534GAT (6212C>T), TGT1535TGC (6215T>C), ATT1536GAA (6216A>G 6217T>A 6218T>A), GCA1537GCT (6221A>T), ATT1538ATA (6224T>A), AAT1540AAA (6230T>A), TAT1542CAT (6234T>C), AAC1543CAA (6237A>C 6239C>A), CTT1545ATA (6243C>A 6245T>A), AAT1546GAC (6246A>G 6248T>C), AAT1547CAG (6249A>C 6251T>G), AAA1549AAT (6257A>T), TTG1550AGT (6258T>A 6259T>G 6260G>T), AAT1551AGT (6262A>G), ATA1552AAA (6265T>A), AAT1553AGA (6268A>G 6269T>A), CGT1554AGA (6270C>A 6272T>A), GTA1556CTA (6276G>C), TTC1558TTT (6284C>T), TGT1559ATA (6285T>A 6286G>T 6287T>A), GAT1560GAC (6290T>C), TTA1561ACC (6291T>A 6292T>C 6293A>C), ACA1563TCT (6297A>T 6299A>T), GGA1564ATA (6300G>A 6301G>T), TTA1565TAC (6304T>A 6305A>C), GGT1566CCT (6306G>C 6307G>C), TTA1567del (6309_6311delTTA), ACT1568AAT (6313C>A), AAA1570GTT (6318A>G 6319A>T 6320A>T), ATA1571TTC (6321A>T 6323A>C), CAT1573TAT (6327C>T), ATC1574ATA (6332C>A), CAG1575AAA (6333C>A 6335G>A), GGT1576GGA (6338T>A) | | | | | | | | |

*: Inserts / Deletes / Misaligned / Frameshifts

#### Analysis details

This analysis was performed with panviral2.64

### NGS Details (UN24): Badnavirus occultipomeae

#### Assembly

| | |
|---|---|
| **Coverage Length** | 319 (1 contig(s)) |
| **Depth Of Coverage** | 53.6 |
| **Number Of Reads** | 157 |
| **Reads Per Million** | 3.14 rpm (after QC) |
| **Ambiguities** | 0 |
| **Assembly Method** | de novo + reference guided assembly |
| **Consensus Caller** | Bcf Tools |

#### Coverage Map

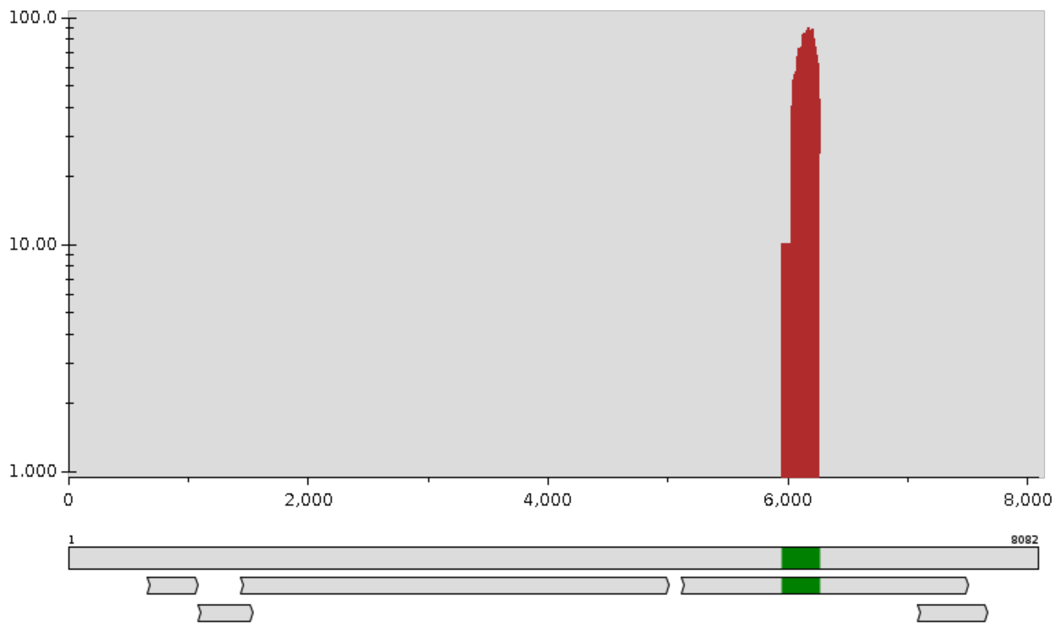

#### Assignment

| | |
|---|---|
| **Type** | Badnavirus occultipomeae (Taxonomy ID: 3048353) |
| **Reference Genome** | NC_015655.1 |
| **NT Identity (%)** | 58.0745 |
| **AA Identity (%)** | 52.3364 |
| **Number Of Stop Codons** | 0 |
| **Number Of CDS** | 5 |

#### Alignment

| | |
|---|---|
| **Alignment Score** | 98.0 (NT) + 408.0 (AA) = 506.0 |
| **Concordance (%)** | 36.5871 |

| **Alignment Method** | Global, seeded, nucleotide + amino acids (AGA) |
|---|---|

#### Genome Region

Sequence starts at position 5946 and ends at position 6264 relative to NC_015655.1 reference sequence.

#### Alignment Detailed Statistics

| | Begin | End | Coverage | Score | Concordance | Matches | Identities | I/D/M/F* | Stop Codons |
|---|---|---|---|---|---|---|---|---|---|
| **NT** | **5946** | **6264** | **3.9%** | **98** | **15.4%** | **319 (99.1%)** | **187 (58.1%)** | **3/0** | |
| Mutations: | 5949G>A, 5950C>A, 5951G>A, 5952C>A, 5955C>T, 5961C>T, 5962A>G, 5963A>C, 5967A>C, 5968G>C, 5970A>G, 5973A>G, 5976T>C, 5979G>C, 5982T>C, 5983T>C, 5985C>A, 5986C>G, 5988A>G, 5991G>C, 5992G>T, 5993G>T, 5994A>C, 5997C>T, 6002C>A, 6003T>A, 6006T>G, 6007G>C, 6009C>T, 6010A>G, 6011G>A, 6012T>C, 6016A>T, 6018A>G, 6019T>G, 6024A>G, 6025A>C, 6028A>G, 6029A>C, 6031A>G, 6033A>G, 6039A>T, 6042A>G, 6043T>A, 6048T>C, 6049C>T, 6051C>G, 6053A>G, 6054A>G, 6055G>T, 6057A>G, 6060C>T, 6062T>A, 6070A>T, 6072A>G, 6076A>G, 6078G>T, 6079G>C, 6080A>G, 6084A>G, 6085A>T, 6086A>C, 6087G>T, 6088T>G, 6089C>A, 6090C>T, 6092A>T, 6093A>C, 6096A>G, 6097T>A, 6098G>A, 6109T>C, 6114T>A, 6116C>G, 6117A>C, 6118G>T, 6120A>T, 6123G>T, 6125T>A, 6126C>T, 6137A>T, 6138A>G, 6141C>T, 6145C>T, 6153A>G, 6156G>T, 6158T>C, 6159G>C, 6165A>C, 6168A>T, 6171A>G, 6172G>A, 6173A>C, 6174T>A, 6177T>C, 6178C>A, 6179A>T, 6181A>G, 6182G>A, 6183A>G, 6184A>T, 6185A>T, 6190G>A, 6193A>C, 6194A>G, 6195T>G, 6197C>T, 6198A>G, 6203G>A, 6205G>T, 6206G>C, 6207A>C, 6207_6208insTAT, 6208A>T, 6209C>T, 6210A>G, 6214G>C, 6215C>A, 6216A>G, 6220A>G, 6222C>G, 6223G>A, 6224C>T, 6225A>T, 6228A>G, 6230A>T, 6234T>A, 6243A>C, 6249A>G, 6251T>A, 6255T>C, 6256G>A, 6264A>G | | | | | | | | |

##### CDS

| | Begin | End | Coverage | Score | Concordance | Matches | Identities | I/D/M/F* | Stop Codons |
|---|---|---|---|---|---|---|---|---|---|
| SPBVa_gp4 | 278 | 383 | 13.3% | 408 | 54.1% | 106 (99.1%) | 56 (52.3%) | 1/0/0/0 | 0 |
| Protein mutations: | R279K (5950C>A 5951G>A 5952C>A), N283A (5962A>G 5963A>C), E285Q (5968G>C 5970A>G), Q288H (5979G>C), S290P (5983T>C 5985C>A), L291V (5986C>G 5988A>G), G293F (5992G>T 5993G>T 5994A>C), T296K (6002C>A 6003T>A), I297M (6006T>G), V298L (6007G>C 6009C>T), S299D (6010A>G 6011G>A 6012T>C), I301L (6016A>T 6018A>G), S302A (6019T>G), K304Q (6025A>C), K305A (6028A>G 6029A>C), I306V (6031A>G 6033A>G), F310I (6043T>A), K313R (6053A>G 6054A>G), A314S (6055G>T 6057A>G), F316Y (6062T>A), I319L (6070A>T 6072A>G), M321V (6076A>G 6078G>T), E322R (6079G>C 6080A>G), K324S (6085A>T 6086A>C 6087G>T), S325D (6088T>G 6089C>A 6090C>T), K326I (6092A>T 6093A>C), W328K (6097T>A 6098G>A), W332R (6109T>C), P334R (6116C>G 6117A>C), E335Y (6118G>T 6120A>T), L337H (6125T>A 6126C>T), E341V (6137A>T 6138A>G), P344S (6145C>T), M348T (6158T>C 6159G>C), D353T (6172G>A 6173A>C 6174T>A), Q355M (6178C>A 6179A>T), R356E (6181A>G 6182G>A 6183A>G), K357L (6184A>T 6185A>T), D359N (6190G>A), N360R (6193A>C 6194A>G 6195T>G), A361V (6197C>T 6198A>G), R363K (6203G>A), G364S (6205G>T 6206G>C 6207A>C), G364_T365insY (6207_6208insTAT), T365L (6208A>T 6209C>T 6210A>G), A367Q (6214G>C 6215C>A 6216A>G), I369V (6220A>G 6222C>G), A370I (6223G>A 6224C>T 6225A>T), Y372F (6230A>T), F379Y (6251T>A), E381K (6256G>A) | | | | | | | | |
| Codon mutations: | AAG278AAA (5949G>A), CGC279AAA (5950C>A 5951G>A 5952C>A), CTC280CTT (5955C>T), GAC282GAT (5961C>T), AAT283GCT (5962A>G 5963A>C), ACA284ACC (5967A>C), GAA285CAG (5968G>C 5970A>G), AAA286AAG (5973A>G), GAT287GAC (5976T>C), CAG288CAC (5979G>C), TAT289TAC (5982T>C), TCC290CCA (5983T>C 5985C>A), CTA291GTG (5986C>G 5988A>G), CCG292CCC (5991G>C), GGA293TTC (5992G>T 5993G>T 5994A>C), ATC294ATT (5997C>T), ACT296AAA (6002C>A 6003T>A), ATT297ATG (6006T>G), GTC298CTT (6007G>C 6009C>T), AGT299GAC (6010A>G 6011G>A 6012T>C), ATA301TTG (6016A>T 6018A>G), TCT302GCT (6019T>G), GGA303GGG (6024A>G), AAA304CAA (6025A>C), AAG305GCG (6028A>G 6029A>C), ATA306GTG (6031A>G 6033A>G), TCA308TCT (6039A>T), AAA309AAG (6042A>G), TTT310ATT (6043T>A), GAT311GAC (6048T>C), CTC312TTG (6049C>T 6051C>G), AAA313AGG (6053A>G 6054A>G), GCA314TCG (6055G>T 6057A>G), GGC315GGT (6060C>T), TTT316TAT (6062T>A), ATA319TTG (6070A>T 6072A>G), ATG321GTT (6076A>G 6078G>T), GAG322CGG (6079G>C 6080A>G), GAA323GAG (6084A>G), AAG324TCT (6085A>T 6086A>C 6087G>T), TCC325GAT (6088T>G 6089C>A 6090C>T), AAA326ATC (6092A>T 6093A>C), CCA327CCG (6096A>G), TGG328AAG (6097T>A 6098G>A), TGG332CGG (6109T>C), ACT333ACA (6114T>A), CCA334CGC (6116C>G 6117A>C), GAA335TAT (6118G>T 6120A>T), GGG336GGT (6123G>T), CTC337CAT (6125T>A 6126C>T), GAA341GTG (6137A>T 6138A>G), GTC342GTT (6141C>T), CCC344TCC (6145C>T), GGA346GGG (6153A>G), CTG347CTT (6156G>T), ATG348ACC (6158T>C 6159G>C), GCA350GCC (6165A>C), CCA351CCT (6168A>T), GCA352GCG (6171A>G), GAT353ACA (6172G>A 6173A>C 6174T>A), TTT354TTC (6177T>C), CAG355ATG (6178C>A 6179A>T), AGA356GAG (6181A>G 6182G>A 6183A>G), AAG357TTG (6184A>T 6185A>T), GAT359AAT (6190G>A), AAT360CGG (6193A>C 6194A>G 6195T>G), GCA361GTG (6197C>T 6198A>G), AGG363AAG (6203G>A), GGA364TCC (6205G>T 6206G>C 6207A>C), GGA364_ACA365insTAT (6207_6208insTAT), ACA365TTG (6208A>T 6209C>T 6210A>G), GCA367CAG (6214G>C 6215C>A 6216A>G), ATC369GTG (6220A>G 6222C>G), GCA370ATT (6223G>A 6224C>T 6225A>T), GTA371GTG (6228A>G), TAT372TTT (6230A>T), ATT373ATA (6234T>A), ATA376ATC (6243A>C), GTA378GTG (6249A>G), TTC379TAC (6251T>A), TCT380TCC (6255T>C), GAA381AAA (6256G>A), GAA383GAG (6264A>G) | | | | | | | | |

##### Proteins

| | Begin | End | Coverage | Score | Concordance | Matches | Identities | I/D/M/F* | Stop Codons |
|---|---|---|---|---|---|---|---|---|---|
| RNaseH/reverse transcriptase (YP_004581513.1) | 278 | 383 | 13.3% | 408 | 54.1% | 106 (99.1%) | 56 (52.3%) | 1/0/0/0 | 0 |
| Protein mutations: | R279K (5950C>A 5951G>A 5952C>A), N283A (5962A>G 5963A>C), E285Q (5968G>C 5970A>G), Q288H (5979G>C), S290P (5983T>C 5985C>A), L291V (5986C>G 5988A>G), G293F (5992G>T 5993G>T 5994A>C), T296K (6002C>A 6003T>A), I297M (6006T>G), V298L (6007G>C 6009C>T), S299D (6010A>G 6011G>A 6012T>C), I301L (6016A>T 6018A>G), S302A (6019T>G), K304Q (6025A>C), K305A (6028A>G 6029A>C), I306V (6031A>G 6033A>G), F310I (6043T>A), K313R (6053A>G 6054A>G), A314S (6055G>T 6057A>G), F316Y (6062T>A), I319L (6070A>T 6072A>G), M321V (6076A>G 6078G>T), E322R (6079G>C 6080A>G), K324S (6085A>T 6086A>C 6087G>T), S325D (6088T>G 6089C>A 6090C>T), K326I (6092A>T 6093A>C), W328K (6097T>A 6098G>A), W332R (6109T>C), P334R (6116C>G 6117A>C), E335Y (6118G>T 6120A>T), L337H (6125T>A 6126C>T), E341V (6137A>T 6138A>G), P344S (6145C>T), M348T (6158T>C 6159G>C), D353T (6172G>A 6173A>C 6174T>A), Q355M (6178C>A 6179A>T), R356E (6181A>G 6182G>A 6183A>G), K357L (6184A>T 6185A>T), D359N (6190G>A), N360R (6193A>C 6194A>G 6195T>G), A361V (6197C>T 6198A>G), R363K (6203G>A), G364S (6205G>T 6206G>C 6207A>C), G364_T365insY (6207_6208insTAT), T365L (6208A>T 6209C>T 6210A>G), A367Q (6214G>C 6215C>A 6216A>G), I369V (6220A>G 6222C>G), A370I (6223G>A 6224C>T 6225A>T), Y372F (6230A>T), F379Y (6251T>A), E381K (6256G>A) | | | | | | | | |
| Codon mutations: | AAG278AAA (5949G>A), CGC279AAA (5950C>A 5951G>A 5952C>A), CTC280CTT (5955C>T), GAC282GAT (5961C>T), AAT283GCT (5962A>G 5963A>C), ACA284ACC (5967A>C), GAA285CAG (5968G>C 5970A>G), AAA286AAG (5973A>G), GAT287GAC (5976T>C), CAG288CAC (5979G>C), TAT289TAC (5982T>C), TCC290CCA (5983T>C 5985C>A), CTA291GTG (5986C>G 5988A>G), CCG292CCC (5991G>C), GGA293TTC (5992G>T 5993G>T 5994A>C), ATC294ATT (5997C>T), ACT296AAA (6002C>A 6003T>A), ATT297ATG (6006T>G), GTC298CTT (6007G>C 6009C>T), AGT299GAC (6010A>G 6011G>A 6012T>C), ATA301TTG (6016A>T 6018A>G), TCT302GCT (6019T>G), GGA303GGG (6024A>G), AAA304CAA (6025A>C), AAG305GCG (6028A>G 6029A>C), ATA306GTG (6031A>G 6033A>G), TCA308TCT (6039A>T), AAA309AAG (6042A>G), TTT310ATT (6043T>A), GAT311GAC (6048T>C), CTC312TTG (6049C>T 6051C>G), AAA313AGG (6053A>G 6054A>G), GCA314TCG (6055G>T 6057A>G), GGC315GGT (6060C>T), TTT316TAT (6062T>A), ATA319TTG (6070A>T 6072A>G), ATG321GTT (6076A>G 6078G>T), GAG322CGG (6079G>C 6080A>G), GAA323GAG (6084A>G), AAG324TCT (6085A>T 6086A>C 6087G>T), TCC325GAT (6088T>G 6089C>A 6090C>T), AAA326ATC (6092A>T 6093A>C), CCA327CCG (6096A>G), TGG328AAG (6097T>A 6098G>A), TGG332CGG (6109T>C), ACT333ACA (6114T>A), CCA334CGC (6116C>G 6117A>C), GAA335TAT (6118G>T 6120A>T), GGG336GGT (6123G>T), CTC337CAT (6125T>A 6126C>T), GAA341GTG (6137A>T 6138A>G), GTC342GTT (6141C>T), CCC344TCC (6145C>T), GGA346GGG (6153A>G), CTG347CTT (6156G>T), ATG348ACC (6158T>C 6159G>C), GCA350GCC (6165A>C), CCA351CCT (6168A>T), GCA352GCG (6171A>G), GAT353ACA (6172G>A 6173A>C 6174T>A), TTT354TTC (6177T>C), CAG355ATG (6178C>A 6179A>T), AGA356GAG (6181A>G 6182G>A 6183A>G), AAG357TTG (6184A>T 6185A>T), GAT359AAT (6190G>A), AAT360CGG (6193A>C 6194A>G 6195T>G), GCA361GTG (6197C>T 6198A>G), AGG363AAG (6203G>A), GGA364TCC (6205G>T 6206G>C 6207A>C), GGA364_ACA365insTAT (6207_6208insTAT), ACA365TTG (6208A>T 6209C>T 6210A>G), GCA367CAG (6214G>C 6215C>A 6216A>G), ATC369GTG (6220A>G 6222C>G), GCA370ATT (6223G>A 6224C>T 6225A>T), GTA371GTG (6228A>G), TAT372TTT (6230A>T), ATT373ATA (6234T>A), ATA376ATC (6243A>C), GTA378GTG (6249A>G), TTC379TAC (6251T>A), TCT380TCC (6255T>C), GAA381AAA (6256G>A), GAA383GAG (6264A>G) | | | | | | | | |

*: Inserts / Deletes / Misaligned / Frameshifts

#### Analysis details

This analysis was performed with panviral2.64

### NGS Details (UN24): Pseudomonas phage phi297

#### Assembly

| | |
|---|---|
| **Coverage Length** | 2114 (3 contig(s)) |
| **Depth Of Coverage** | 7.3 |
| **Number Of Reads** | 122 |
| **Reads Per Million** | 2.44 rpm (after QC) |
| **Ambiguities** | 0 |
| **Assembly Method** | de novo + reference guided assembly |
| **Consensus Caller** | Bcf Tools |

#### Coverage Map

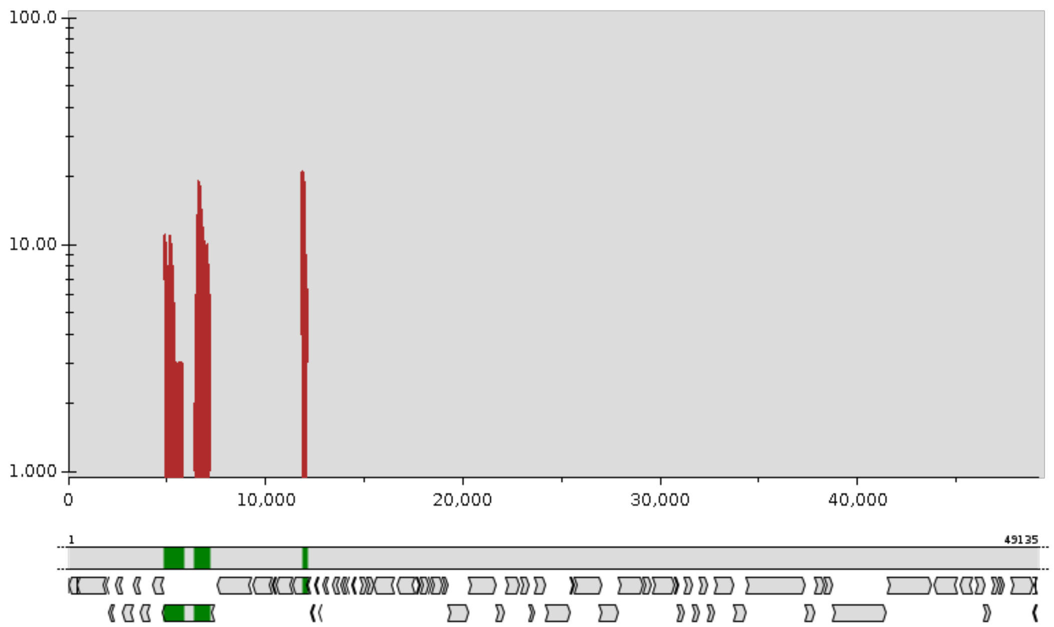

#### Assignment

| | |
|---|---|
| **Type** | Pseudomonas phage phi297 (Taxonomy ID: 1129145) |
| **Reference Genome** | NC_016762.1 |
| **NT Identity (%)** | 72.6802 |
| **AA Identity (%)** | 67.1853 |
| **Number Of Stop Codons** | 0 |
| **Number Of CDS** | 69 |

#### Alignment

| | |
|---|---|
| **Alignment Score** | 1899.0 (NT) + 3530.0 (AA) = 5429.0 |
| **Concordance (%)** | 58.7872 |

| Alignment Method | Local, heuristic, nucleotide (BLASTN) |
|---|---|

#### Genome Region

Sequence starts at position 4836 and ends at position 12136 relative to NC_016762.1 reference sequence.

#### Alignment Detailed Statistics

| | Begin | End | Coverage | Score | Concordance | Matches | Identities | I/D/M/F* | Stop Codons |
|---|---|---|---|---|---|---|---|---|---|
| **NT** | **4836** | **12136** | **4.3%** | **1899** | **45.2%** | **2108 (99.0%)** | **1543 (72.5%)** | **15/6** | |
| Mutations: | 4841A>T, 4842G>C, 4843C>A, 4847G>T, 4849G>A, 4855G>C, 4856T>G, 4857C>A, 4858G>C, 4862A>G, 4864G>C, 4866C>A, 4871T>G, 4872C>A, 4873T>G, 4874T>C, 4875C>G, 4877T>A, 4879C>G, 4882G>C, 4883G>T, 4884A>G, 4887T>C, 4891A>G, 4898G>A, 4900A>G, 4903G>A, 4916C>G, 4918G>C, 4919G>T, 4920A>T, 4922T>G, 4923T>C, 4927T>C, 4931T>C, 4932G>A, 4934G>C, 4940G>C, 4941T>C, 4942T>G, 4944T>G, 4945G>C, 4948C>G, 4949T>C, 4950T>C, 4951G>C, 4955T>A, 4956T>G, 4959G>C, 4960C>A, 4963G>C, 4965A>G, 4969C>G, 4972G>C, 4975G>C, 4981G>C, 4983T>G, 4984C>G, 4996A>G, 5000A>T, 5006A>T, 5007G>C, 5008G>C, 5011T>G, 5014T>C, 5016C>T, 5029C>G, 5030A>G, 5031C>T, 5035G>C, 5044G>C, 5045A>G, 5047G>A, 5061G>T, 5077T>C, 5080C>G, 5081T>C, 5083G>C, 5086C>G, 5087T>G, 5088T>C, 5089G>T, 5090C>G, 5091T>C, 5092G>C, 5093T>G, 5099T>G, 5100G>C, 5102G>C, 5103C>T, 5113C>G, 5122G>C, 5124C>T, 5125G>C, 5137T>G, 5146G>A, 5153G>C, 5154A>T, 5161T>C, 5162G>T, 5177C>G, 5178C>T, 5189T>A, 5191C>G, 5195G>C, 5196T>G, 5197G>C, 5198A>T, 5200C>G, 5201T>G, 5208A>C, 5209T>C, 5221G>C, 5226G>C, 5246G>C, 5247T>G, 5248G>C, 5249G>T, 5250A>C, 5254G>C, 5255C>G, 5256T>A, 5259A>G, 5260G>C, 5262C>T, 5267T>A, 5278C>G, 5281C>G, 5282A>T, 5285T>G, 5287G>A, 5290C>G, 5292G>T, 5293A>G, 5295T>G, 5302G>C, 5304A>G, 5305G>C, 5309G>T, 5312T>G, 5313C>G, 5314A>C, 5321G>A, 5323A>C, 5328T>C, 5332G>C, 5341G>C, 5343T>C, 5345A>T, 5359C>G, 5371T>C, 5373C>G, 5465G>T, 5470G>C, 5476C>G, 5479A>C, 5481T>C, 5486C>G, 5488G>A, 5492G>C, 5495T>C, 5496T>G, 5498G>C, 5499A>T, 5501T>G, 5502T>C, 5509G>C, 5512G>T, 5514G>C, 5540G>C, 5541G>A, 5550C>A, 5551A>G, 5557C>G, 5558T>C, 5559T>G, 5560G>C, 5562T>A, 5566G>A, 5575C>T, 5578T>C, 5581C>G, 5582G>C, 5584C>G, 5585C>T, 5587C>G, 5588G>C, 5589A>C, 5590G>C, 5606G>T, 5607C>T, 5617A>C, 5641C>G, 5642A>C, 5643T>G, 5649T>C, 5650A>G, 5651T>A, 5654G>T, 5655C>G, 5656C>G, 5657C>A, 5662A>G, 5663T>G, 5664G>C, 5666T>C, 5667T>G, 5669A>C, 5670C>G, 5675G>C, 5676C>A, 5677T>G, 5678C>G, 5679G>C, 5680A>G, 5681G>A, 5683C>G, 5684A>G, 5685C>T, 5686T>G, 5691T>C, 5694G>C, 5704G>C, 5705A>T, 5719A>G, 5721G>C, 5724_5725insCTC, 5725G>C, 5726G>A, 5727T>C, 5731T>G, 5736C>T, 5740A>C, 5742T>C, 5749A>G, 5751C>T, 5755C>G, 5765C>G, 5766A>C, 5772G>C, 5773A>C, 5775G>T, 5776C>G, 5779G>C, 5782G>C, 5785G>C, 5786G>A, 5787T>G, 5800T>C, 5806G>C, 5808T>C, 5812A>G, 5813T>A, 5826C>G, 5843T>C, 5844C>T, 6398A>G, 6399T>C, 6403G>C, 6406G>C, 6412A>G, 6416G>C, 6417C>G, 6424C>G, 6428T>C, 6433T>C, 6436C>A, 6437G>T, 6442G>C, 6448A>G, 6457G>C, 6459T>C, 6460C>A, 6466A>G, 6469A>G, 6470T>G, 6472T>C, 6475A>G, 6476G>T, 6484C>G, 6487G>C, 6489T>G, 6498G>T, 6499T>C, 6505G>C, 6507T>C, 6508C>G, 6511G>C, 6515G>T, 6516G>C, 6517A>C, 6523C>G, 6535T>G, 6536G>C, 6537C>G, 6538G>C, 6541G>C, 6548T>A, 6549G>C, 6556A>G, 6562G>C, 6563G>T, 6564T>G, 6566T>C, 6567T>G, 6568G>C, 6569C>T, 6570G>T, 6571A>G, 6573G>A, 6576G>T, 6577G>C, 6578C>T, 6580G>C, 6581C>G, 6591T>C, 6592A>C, 6593C>A, 6594G>T, 6595C>A, 6597C>T, 6598A>G, 6599T>G, 6600G>T, 6601G>A, 6609G>C, 6610C>G, 6612A>C, 6619G>C, 6621A>C, 6625T>G, 6627C>T, 6628T>C, 6646C>G, 6648G>T, 6649T>G, 6650T>G, 6651C>G, 6655T>C, 6659G>T, 6667T>G, 6677A>T, 6679C>G, 6688G>C, 6689C>G, 6690T>A, 6694G>C, 6695C>T, 6696G>T, 6700A>C, 6706G>C, 6708T>G, 6709C>G, 6716G>C, 6717A>T, 6718C>T, 6729T>G, 6736C>G, 6741C>G, 6745T>G, 6746T>C, 6747T>G, 6748G>C, 6749T>G, 6750G>C, 6751G>T, 6753C>G, 6754A>G, 6755T>C, 6756G>T, 6757G>C, 6759T>G, 6760G>C, 6763G>C, 6766T>G, 6768G>T, 6781A>G, 6782G>C, 6783A>T, 6790T>C, 6799C>A, 6801T>G, 6802G>C, 6817G>A, 6820A>G, 6832G>C, 6841C>G, 6842G>C, 6843A>T, 6847A>G, 6853T>C, 6861T>C, 6862G>A, 6863T>A, 6865T>C, 6871C>T, 6874G>A, 6877T>G, 6886A>C, 6896T>A, 6898C>G, 6901G>A, 6905C>G, 6907G>C, 6914T>G, 6918C>T, 6919G>C, 6920C>T, 6921G>T, 6922G>C, 6927C>G, 6929G>T, 6930C>T, 6933_6934insCATGTCGCG, 6934G>C, 6937A>G, 6943A>G, 6946A>G, 6955A>C, 6973C>G, 6997A>C, 7003A>T, 7004C>G, 7005T>C, 7006G>C, 7009T>C, 7021T>G, 7034C>A, 7036G>C, 7037C>A, 7039A>C, 7045G>C, 7053C>G, 7062C>T, 7063G>C, 7066C>T, 7072A>G, 7078C>G, 7088G>C, 7089T>C, 7091G>T, 7093G>C, 7102T>G, 7103A>G, 7105C>T, 7108G>C, 7113A>T, 7114A>G, 7116T>G, 7127C>T, 7128G>T, 7132G>C, 7141G>A, 7144C>G, 7145A>C, 7146T>G, 7147C>T, 7150G>C, 7160T>A, 7161A>T, 11839C>G, 11840T>C, 11841T>G, 11842C>G, 11853T>C, 11857T>C, 11858C>G, 11861G>A, 11864G>C, 11865A>G, 11872T>G, 11881G>C, 11882C>G, 11883T>C, 11885A>G, 11887G>T, 11890G>T, 11891G>T, 11892A>C, 11893_11898delGCCGCG, 11902G>A, 11911G>T, 11912A>T, 11913A>C, 11914C>T, 11917C>G, 11922G>T, 11923C>G, 11926C>G, 11928G>T, 11931T>C, 11934G>C, 11938C>G, 11940G>C, 11941C>A, 11942A>G, 11943G>C, 11944T>G, 11945T>C, 11947C>G, 11949A>C, 11956G>A, 11958A>C, 11965G>C, 11967T>A, 11968C>G, 11970_11971insAGT, 11971C>T, 11972T>C, 11977G>C, 11978T>G, 11979T>G, 11980G>C, 11982C>G, 11983C>G, 11984C>T, 11986T>G, 11988C>A, 11990G>T, 11991C>G, 11998C>G, 11999G>A, 12000C>G, 12009T>C, 12016G>C, 12018T>C, 12022G>C, 12025A>G, 12028G>C, 12032A>T, 12037C>G, 12038C>G, 12039G>C, 12042G>C, 12044G>C, 12045A>T, 12046T>G, 12047G>C, 12048A>G, 12049A>C, 12054T>C, 12061C>G, 12062T>C, 12063T>C, 12069G>T, 12073T>C, 12074A>T, 12075C>T, 12076G>C, 12077C>G, 12079C>T, 12086T>C, 12088G>T, 12089A>G, 12090A>C, 12091C>A, 12093C>G, 12107G>T, 12110C>G, 12113C>A, 12114A>G, 12115G>C, 12117A>T, 12118C>G, 12119G>C, 12120A>T, 12127C>G, 12129C>T, 12132T>G | | | | | | | | |
| **CDS** | | | | | | | | | |
| phi297_00010 | 89 | 865 | 68.5% | 3140 | 72.2% | 601 (99.3%) | 423 (69.9%) | 4/0/0/0 | 0 |
| Protein mutations: | Y90I (7160T>A 7161A>T), M95R (7144C>G 7145A>C 7146T>G), R101K (7127C>T 7128G>T), S105R (7114A>G 7116T>G), F106I (7113A>T), D107E (7108G>C), V109A (7102T>G 7103A>G), T113N (7091G>T), T114G (7088G>C 7089T>C), V123I (7062C>T), G126R (7053C>G), R131L (7036G>C 7037C>A), R132L (7034C>A), S142A (7003A>T 7004C>G 7005T>C), R165_A166insRDM (6933_6934insCATGTCGCG), A167N (6929G>T 6930C>T), G168R (6927C>G), R170K (6919G>C 6920C>T 6921G>T), G171S (6918C>T), D172A (6914T>G), R175P (6905C>G), Y178F (6896T>A), Y189F (6862G>A 6863T>A), I190V (6861T>C), Q221N (6766T>G 6768G>T), I224L (6757G>C 6759T>G), H225S (6754A>G 6755T>C 6756G>T), A226P (6751G>T 6753C>G), H227A (6748G>C 6749T>G 6750G>C), K228R (6745T>G 6746T>C 6747T>G), G230R (6741C>G), E231D (6736C>G), M234L (6729T>G), I241L (6706G>C 6708T>G), R245K (6694G>C 6695C>T 6696G>T), F251Y (6677A>T), E254D (6667T>G), A257E (6659G>T), E260P (6649T>G 6650T>G 6651C>G), L261I (6646C>G 6648G>T), V268I (6625T>G 6627C>T), S270A (6619G>C 6621A>C), S273A (6610C>G 6612A>C), H274D (6609G>C), H277T (6598A>G 6599T>G 6600G>T), E278N (6595C>A 6597C>T), R279M (6592A>C 6593C>A 6594G>T), N280D (6591T>C), G283A (6580G>C 6581C>G), R284Q (6577G>C 6578C>T), L285M (6576G>T), L286F (6571A>G 6573G>A), R287K (6568G>C 6569C>T 6570G>T), N288R (6566T>C 6567T>G), T289Q (6562G>C 6563G>T 6564T>G), Q294V (6548T>A 6549G>C), A298R (6535T>G 6536G>C 6537C>G), E302D (6523C>G), P305D (6515G>T 6516G>C), I308V (6505G>C 6507T>C), L311M (6498G>T), I314L (6487G>C 6489T>G), A318D (6475A>G 6476G>T), D320A (6469A>G 6470T>G), I324V (6457G>C 6459T>C), A331D (6436C>A 6437G>T), H334R (6428T>C), A338R (6416G>C 6417C>G), I344A (6398A>G 6399T>C), D529S (5843T>C 5844C>T), E535Q (5826C>G), Y539F (5812A>G 5813T>A), I541V (5806G>C 5808T>C), T548L (5785G>C 5786G>A 5787T>G), I549M (5782G>C), L553V (5772G>C), C555A (5765C>G 5766A>C), D560N (5749A>G 5751C>T), I563V (5740A>C 5742T>C), G565S (5736C>T), T568V (5725G>C 5726G>A 5727T>C), T568_G569insE (5724_5725insCTC), L570V (5719A>G 5721G>C), V575E (5704G>C 5705A>T), H579D (5694G>C), M580V (5691T>C), E581D (5686T>G), V582T (5683C>G 5684A>G 5685C>T), T583I (5680A>G 5681G>A), R584A (5677T>G 5678C>G 5679G>C), A585C (5675G>C 5676C>A), V587R (5669A>C 5670C>G), K588R (5666T>C 5667T>G), H589A (5662A>G 5663T>G 5664G>C), W591F (5656C>G 5657C>A), A592H (5654G>T 5655C>G), Y593F (5650A>G 5651T>A), M594V (5649T>C), M596R (5641C>G 5642A>C 5643T>G), A608N (5606G>T 5607C>T), S614G (5587C>G 5588G>C 5589A>C), W615Y (5584C>G 5585C>T), T616S (5581C>G 5582G>C), T623S (5560G>C 5562T>A), K624R (5557C>G 5558T>C 5559T>G), V627F (5550C>A), P630C (5540G>C 5541G>A), P639A (5512G>T 5514G>C), N643A (5501T>G 5502T>C), N645R (5495T>C 5496T>G), A646G (5492G>C), G648A (5486C>G), I650V (5479A>C 5481T>C), A655D (5465G>T), A686P (5371T>C 5373C>G), L690F (5359C>G), F695Y (5345A>T), T696A (5341G>C 5343T>C), N699K (5332G>C), N701D (5328T>C), P703L (5321G>A), E706P (5312T>G 5313C>G), A707E (5309G>T), D708E (5305G>C), F709L (5302G>C 5304A>G), I712L (5293A>G 5295T>G), P713T (5290C>G 5292G>T), D715A (5285T>G), V716D (5281C>G 5282A>T), M717I (5278C>G), Y721F (5267T>A), D723K (5260G>C 5262C>T), Y724H (5259A>G), S727E (5248G>C 5249G>T 5250A>C), T728R (5246G>C 5247T>G), H735D (5226G>C), S741A (5208A>C), E743A (5200C>G 5201T>G), V744E (5197G>C 5198A>T), T745R (5195G>C 5196T>G), K747M (5189T>A), G751T (5177C>G 5178C>T), P756Q (5161T>C 5162G>T), A769T (5122G>C 5124C>T), A776S (5102G>C 5103C>T), Q777A (5099T>G 5100G>C), Y779S (5092G>C 5093T>G), S780A (5089G>T 5090C>G 5091T>C), K781A (5086C>G 5087T>G 5088T>C), Q783R (5080C>G 5081T>C), L790M (5061G>T), V795A (5044G>C 5045A>G), V800T (5029C>G 5030A>G 5031C>T), E805K (5014T>C 5016C>T), D807E (5008G>C), L808E (5006A>T 5007G>C), F810Y (5000A>T), I816L (4981G>C 4983T>G), C822R (4963G>C 4965A>G), L824V (4959G>C), K825L (4955T>A 4956T>G), K827G (4948C>G 4949T>C 4950T>C), T830G (4940G>C 4941T>C), A832G (4934G>C), H833C (4931T>C 4932G>A), N836A (4922T>G 4923T>C), S837K (4918G>C 4919G>T 4920A>T), R838P (4916C>G), A844V (4898G>A), K848E (4887T>C), S849Q (4882G>C 4883G>T 4884A>G), E851V (4877T>A), E852R (4873T>G 4874T>C 4875C>G), E853S (4871T>G 4872C>A), A855S (4864G>C 4866C>A), M856T (4862A>G), D858S (4855G>C 4856T>G 4857C>A), A861D (4847G>T), L863D (4841A>T 4842G>C) | | | | | | | | |

| | Begin | End | Coverage | Score | Concordance | Matches | Identities | I/D/M/F* | Stop Codons |
|---|---|---|---|---|---|---|---|---|---|
| **NT** | **4836** | **12136** | **4.3%** | **1899** | **45.2%** | **2108 (99.0%)** | **1543 (72.5%)** | **15/6** | |
| Codon mutations: | TAC90ATC (7160T>A 7161A>T), GCC93GCG (7150G>C), GCG94GCA (7147C>T), ATG95CGC (7144C>G 7145A>C 7146T>G), CTC96CTT (7141G>A), ACC99ACG (7132G>C), CGG101AAG (7127C>T 7128G>T), AGT105CGC (7114A>G 7116T>G), TTC106ATC (7113A>T), GAC107GAG (7108G>C), GAG108GAA (7105C>T), GTA109GCC (7102T>G 7103A>G), CCC112CCG (7093G>C), ACC113AAC (7091G>T), ACC114GGC (7088G>C 7089T>C), CTG117CTC (7078C>G), AAT119AAC (7072A>G), GAG121GAA (7066C>T), ACC122ACG (7063G>C), GTC123ATC (7062C>T), GGC126CGC (7053C>G), CTC128CTG (7045G>C), CCT130CCG (7039A>C), CGC131CTG (7036G>C 7037C>A), CGG132CTG (7034C>A), GCA136GCC (7021T>G), GAA140GAG (7009T>C), GCC141GCG (7006G>C), AGT142GCA (7003A>T 7004C>G 7005T>C), CTT144CTG (6997A>C), ACG152ACC (6973C>G), GCT158GCG (6955A>C), GCT161GCC (6946A>G), GGT162GGC (6943A>G), GAT164GAC (6937A>G), CGC165CGG (6934G>C), CGC165_GCC166insCGCGACATG (6933_6934insCATGTCGCG), GCC167AAC (6929G>T 6930C>T), GGC168CGC (6927C>G), GTC169GTG (6922G>C), CGC170AAG (6919G>C 6920C>T 6921G>T), GGC171AGC (6918C>T), GAT172GCT (6914T>G), GTC174GTG (6907G>C), CGG175CCG (6905C>G), TAC176TAT (6901G>A), CGG177CGC (6898C>G), TAC178TTC (6896T>A), ACT181ACG (6886A>C), CCA184CCC (6877T>G), AGC185AGT (6874G>A), CCG186CCA (6871C>T), GAA188GAG (6865T>C), TAC189TTT (6862G>A 6863T>A), ATC190GTC (6861T>C), CTA192CTG (6853T>C), GCT194GCC (6847A>G), TCG196AGC (6841C>G 6842G>C 6843A>T), CTC199CTG (6832G>C), GAT203GAC (6820A>G), GTC204GTT (6817G>A), ACC209ACG (6802G>C), AGG210CGT (6799C>A 6801T>G), AAA213AAG (6790T>C), TCT216AGC (6781A>G 6782G>C 6783A>T), CAA221AAC (6766T>G 6768G>T), CTC222CTG (6763G>C), ACC223ACG (6760G>C), ATC224CTG (6757G>C 6759T>G), CAT225AGC (6754A>G 6755T>C 6756G>T), GCC226CCA (6751G>T 6753C>G), CAC227GCG (6748G>C 6749T>G 6750G>C), AAA228CGC (6745T>G 6746T>C 6747T>G), GGC230CGC (6741C>G), GAG231GAC (6736C>G), ATG234CTG (6729T>G), GCG237GCA (6718C>T), TCC238AGC (6716G>C 6717A>T), GCG240GCC (6709C>G), ATC241CTG (6706G>C 6708T>G), GTT243GTG (6700A>C), CGC245AAG (6694G>C 6695C>T 6696G>T), AGC247TCG (6688G>C 6689C>G 6690T>A), GGG250GGC (6679C>G), TTC251TAC (6677A>T), GAA254GAC (6667T>G), GCG257GAG (6659G>T), CTA258CTG (6655T>C), GAA260CCC (6649T>G 6650T>G 6651C>G), CTG261ATC (6646C>G 6648G>T), GAA267GAG (6628T>C), GTA268ATC (6625T>G 6627C>T), TCC270GCG (6619G>C 6621A>C), TCG273GCC (6610C>G 6612A>C), CAC274GAC (6609G>C), GGC276GGT (6601G>A), CAT277ACC (6598A>G 6599T>G 6600G>T), GAG278AAT (6595C>A 6597C>T), CGT279ATG (6592A>C 6593C>A 6594G>T), AAC280GAC (6591T>C), GGC283GCG (6580G>C 6581C>G), CGC284CAG (6577G>C 6578C>T), CTG285ATG (6576G>T), CTT286TTC (6571A>G 6573G>A), CGC287AAG (6568G>C 6569C>T 6570G>T), AAC288CGC (6566T>C 6567T>G), ACC289CAG (6562G>C 6563G>T 6564T>G), ATT291ATC (6556A>G), CAG294GTG (6548T>A 6549G>C), GCC296GCG (6541G>C), GCC297GCG (6538G>C), GCA298CGC (6535T>G 6536G>C 6537C>G), GAG302GAC (6523C>G), CTT304CTG (6517A>C), CCT305GAT (6515G>T 6516G>C), GCC306GCG (6511G>C), CGG307CGC (6508C>G), ATC308GTG (6505G>C 6507T>C), AAA310AAG (6499T>C), CTG311ATG (6498G>T), ATC314CTG (6487G>C 6489T>G), CGG315CGC (6484C>G), GCT318GAC (6475A>G 6476G>T), CCA319CCG (6472T>C), GAT320GCC (6469A>G 6470T>G), GCT321GCC (6466A>G), CGG323CGT (6460C>A), ATC324GTG (6457G>C 6459T>C), CAT327CAC (6448A>G), CTC329CTG (6442G>C), GCG331GAT (6436C>A 6437G>T), GAA332GAG (6433T>C), CAC334CGC (6428T>C), GCG335GCC (6424C>G), GCC338CGC (6416G>C 6417C>G), GCT339GCC (6412A>G), CCC341CCG (6406G>C), ACC342ACG (6403G>C), ATC344GCC (6398A>G 6399T>C), GAC529AGC (5843T>C 5844C>T), GAG535CAG (5826C>G), TAT539TTC (5812A>G 5813T>A), ATC541GTG (5806G>C 5808T>C), GAA543GAG (5800T>C), ACC548CTG (5785G>C 5786G>A 5787T>G), ATC549ATG (5782G>C), CCC550CCG (5779G>C), GGG551GGC (5776C>G), CGT552AGG (5773A>C 5775G>T), CTG553GTG (5772G>C), TGC555GCC (5765C>G 5766A>C), GTG558GTC (5755C>G), GAT560AAC (5749A>G 5751C>T), ATT563GTG (5740A>C 5742T>C), GGC565AGC (5736C>T), GGA566GGC (5731T>G), ACC568GTG (5725G>C 5726G>A 5727T>C), ACC568_GGC569insGAG (5724_5725insCTC), CTT570GTC (5719A>G 5721G>C), GTC575GAG (5704G>C 5705A>T), CAC579GAC (5694G>C), ATG580GTG (5691T>C), GAA581GAC (5686T>G), GTG582ACC (5683C>G 5684A>G 5685C>T), ACT583ATC (5680A>G 5681G>A), CGA584GCC (5677T>G 5678C>G 5679G>C), GCC585TGC (5675G>C 5676C>A), GTC587CGC (5669A>C 5670C>G), AAG588CGG (5666T>C 5667T>G), CAT589GCC (5662A>G 5663T>G 5664G>C), TGG591TTC (5656C>G 5657C>A), GCC592CAC (5654G>T 5655C>G), TAT593TTC (5650A>G 5651T>A), ATG594GTG (5649T>C), ATG596CGC (5641C>G 5642A>C 5643T>G), GTT604GTG (5617A>C), GCC608AAC (5606G>T 5607C>T), CTC613CTG (5590G>C), TCG614GGC (5587C>G 5588G>C 5589A>C), TGG615TAC (5584C>G 5585C>T), ACG616AGC (5581C>G 5582G>C), GAA617GAG (5578T>C), CAG618CAA (5575C>T), GAC621GAT (5566G>A), ACC623TCG (5560G>C 5562T>A), AAG624CGC (5557C>G 5558T>C 5559T>G), GGT626GGC (5551A>G), GTC627TTC (5550C>A), CCC630TGC (5540G>C 5541G>A), CCC639GCA (5512G>T 5514G>C), CCC640CCG (5509G>C), AAC643GCC (5501T>G 5502T>C), TCC644AGC (5498G>C 5499A>T), AAC645CGC (5495T>C 5496T>G), GCC646GGC (5492G>C), TAC647TAT (5488G>A), GGC648GCC (5486C>G), ATT650GTG (5479A>C 5481T>C), CCG651CCC (5476C>G), GTC653GTG (5470G>C), GCC655GAC (5465G>T), GCA686CCG (5371T>C 5373C>G), TTG690TTC (5359C>G), TTC695TAC (5345A>T), ACC696GCG (5341G>C 5343T>C), AAC699AAG (5332G>C), AAC701GAC (5328T>C), CGT702CGG (5323A>C), CCG703CTG (5321G>A), ACT705ACG (5314A>C), GAA706CCA (5312T>G 5313C>G), GCG707GAG (5309G>T), GAC708GAG (5305G>C), TTC709CTG (5302G>C 5304A>G), ATT712CTC (5293A>G 5295T>G), CCG713ACC (5290C>G 5292G>T), CAC714CAT (5287G>A), GAC715GCC (5285T>G), GTG716GAC (5281C>G 5282A>T), ATG717ATC (5278C>G), TAC721TTC (5267T>A), GAC723AAG (5260G>C 5262C>T), TAC724CAC (5259A>G), AGC725TCG (5254G>C 5255C>G 5256T>A), TCC727GAG (5248G>C 5249G>T 5250A>C), ACC728CGC (5246G>C 5247T>G), CAC735GAC (5226G>C), GTC736GTG (5221G>C), GAA740GAG (5209T>C), TCG741GCG (5208A>C), GAG743GCC (5200C>G 5201T>G), GTC744GAG (5197G>C 5198A>T), ACC745CGC (5195G>C 5196T>G), GGG746GGC (5191C>G), AAG747ATG (5189T>A), GGC751ACC (5177C>G 5178C>T), CCA756CAG (5161T>C 5162G>T), TCC759AGC (5153G>C 5154A>T), CAC761CAT (5146G>A), GTA764GTC (5137T>G), GTC768GTG (5125G>C), GCC769ACG (5122G>C 5124C>T), CGG772CGC (5113C>G), GCC776AGC (5102G>C 5103C>T), CAG777GCG (5099T>G 5100G>C), TAC779TCG (5092G>C 5093T>G), AGC780GCA (5089G>T 5090C>G 5091T>C), AAG781GCC (5086C>G 5087T>G 5088T>C), GGC782GGG (5083G>C), CAG783CGC (5080C>G 5081T>C), GAA784GAG (5077T>C), CTG790ATG (5061G>T), ATC794ATT (5047G>A), GTC795GCG (5044G>C 5045A>G), GCC798GCG (5035G>C), GTG800ACC (5029C>G 5030A>G 5031C>T), GAA805AAG (5014T>C 5016C>T), GGA806GGC (5011T>G), GAC807GAG (5008G>C), CTG808GAG (5006A>T 5007G>C), TTC810TAC (5000A>T), GAT811GAC (4996A>G), GGG815GGC (4984C>G), ATC816CTG (4981G>C 4983T>G), ACC818ACG (4975G>C), GTC819GTG (4972G>C), CCG820CCC (4969C>G), TGC822CGG (4963G>C 4965A>G), GCG823GCT (4960C>A), CTG824GTG (4959G>C), AAG825CTG (4955T>A 4956T>G), CTC826CTG (4951G>C), AAG827GGC (4948C>G 4949T>C 4950T>C), CGC828CGG (4945G>C), AGA829CGC (4942T>G 4944T>G), ACC830GGC (4940G>C 4941T>C), GCC832GGC (4934G>C), CAC833TGC (4931T>C 4932G>A), GAA834GAG (4927T>C), AAC836GCC (4922T>G 4923T>C), TCC837AAG (4918G>C 4919G>T 4920A>T), CGC838CCC (4916C>G), GAC842GAT (4903G>A), GGT843GGC (4900A>G), GCC844GTC (4898G>A), TAT846TAC (4891A>G), AAG848GAG (4887T>C), TCC849CAG (4882G>C 4883G>T 4884A>G), GCG850GCC (4879C>G), GAG851GTG (4877T>A), GAA852CGC (4873T>G 4874T>C 4875C>G), GAG853TCG (4871T>G 4872C>A), GCC855TCG (4864G>C 4866C>A), ATG856ACG (4862A>G), CCC857CCG (4858G>C), GAC858TCG (4855G>C 4856T>G 4857C>A), TTC860TTT (4849G>A), GCC861GAC (4847G>T), CTG862CTT (4843C>A), CTC863GAC (4841A>T 4842G>C) | | | | | | | | |
| phi297_00015 | 4 | 106 | 38.4% | 390 | 55.2% | 101 (97.1%) | 52 (50.0%) | 1/2/0/0 | 0 |
| Protein mutations: | T5P (12132T>G), V6I (12127C>G 12129C>T), S10T (12115G>C 12117A>T), W11L (12113C>A 12114A>G), G12A (12110C>G), A13E (12107G>T), A18P (12091C>A 12093C>G), F19A (12088G>T 12089A>G 12090A>C), K20R (12086T>C), G23A (12076G>C 12077C>G), V24K (12073T>C 12074A>T 12075C>T), L26I (12069G>T), K28G (12061C>G 12062T>C 12063T>C), S31G (12054T>C), S33R (12046T>G 12047G>C 12048A>G), P35A (12042G>C), R36A (12037C>G 12038C>G 12039G>C), L38Q (12032A>T), I43V (12016G>C 12018T>C), S46G (12009T>C), A49L (11998C>G 11999G>A 12000C>G), A52Q (11990G>T 11991C>G), A53S (11986T>G 11988C>A), R54H (11983C>G 11984C>T), V55L (11980G>C 11982C>G), N56P (11977G>C 11978T>G 11979T>G), E58G (11971C>T 11972T>C), E58_P59insT (11970_11971insAGT), I60L (11965G>C 11967T>A), Y63D (11956G>A 11958A>C), S66A (11947C>G 11949A>C), E67G (11944T>G 11945T>C), L68A (11941C>A 11942A>G 11943G>C), L69V (11938C>G 11940G>C), H71D (11934G>C), T72A (11931T>C), L73I (11926C>G 11928G>T), Q74H (11923C>G), Q75K (11922G>T), F78E (11911G>T 11912A>T 11913A>C), R83_G84del (11893_11898delGCCGCG), S85E (11890G>T 11891G>T 11892A>C), D86E (11887G>T), I87T (11885A>G), S88A (11881G>C 11882C>G 11883T>C), E91D (11872T>G), S94R (11864G>C 11865A>G), T95I (11861G>A), G96A (11857T>C 11858C>G), T98A (11853T>C), K102R (11839C>G 11840T>C 11841T>G) | | | | | | | | |
| Codon mutations: | ACC5CCC (12132T>G), GTG6ATC (12127C>G 12129C>T), TCG9AGC (12118C>G 12119G>C 12120A>T), TCC10ACG (12115G>C 12117A>T), TGG11CTG (12113C>A 12114A>G), GGC12GCC (12110C>G), GCG13GAG (12107G>T), GCG18CCT (12091C>A 12093C>G), TTC19GCA (12088G>T 12089A>G 12090A>C), AAG20AGG (12086T>C), GAG22GAA (12079C>T), GGC23GCG (12076G>C 12077C>G), GTA24AAG (12073T>C 12074A>T 12075C>T), CTC26ATC (12069G>T), AAG28GGC (12061C>G 12062T>C 12063T>C), AGC31GGC (12054T>C), CCT32CCG (12049A>C), TCA33CGC (12046T>G 12047G>C 12048A>G), TCC34AGC (12044G>C 12045A>T), CCC35GCC (12042G>C), CGG36GCC (12037C>G 12038C>G 12039G>C), CTG38CAG (12032A>T), CTC39CTG (12028G>C), GGT40GGC (12025A>G), ACC41ACG (12022G>C), ATC43GTG (12016G>C 12018T>C), AGC46GGC (12009T>C), GCG49CTC (11998C>G 11999G>A 12000C>G), GCG52CAG (11990G>T 11991C>G), GCA53TCC (11986T>G 11988C>A), CGG54CAC (11983C>G 11984C>T), GTC55CTG (11980G>C 11982C>G), AAC56CCG (11977G>C 11978T>G 11979T>G), GAG58GGA (11971C>T 11972T>C), GAG58_CCG59insACT (11970_11971insAGT), CCG59CCC (11968C>G), ATC60TTG (11965G>C 11967T>A), TAC63GAT (11956G>A 11958A>C), TCG66GCC (11947C>G 11949A>C), GAA67GGC (11944T>G 11945T>C), CTG68GCT (11941C>A 11942A>G 11943G>C), CTG69GTC (11938C>G 11940G>C), CAC71GAC (11934G>C), ACG72GCG (11931T>C), CTG73ATC (11926C>G 11928G>T), CAG74CAC (11923C>G), CAG75AAG (11922G>T), CCG76CCC (11917C>G), GAG77GAA (11914C>T), TTC78GAA (11911G>T 11912A>T 11913A>C), GAC81GAT (11902G>A), CGC83_GGC84del (11893_11898delGCCGCG), TCC85GAA (11890G>T 11891G>T 11892A>C), GAC86GAA (11887G>T), ATC87ACC (11885A>G), AGC88GCG (11881G>C 11882C>G 11883T>C), GAA91GAC (11872T>G), TCC94CGC (11864G>C 11865A>G), ACC95ATC (11861G>A), GGA96GCG (11857T>C 11858C>G), ACG98GCG (11853T>C), ACG101ACC (11842C>G), AAG102CGC (11839C>G 11840T>C 11841T>G) | | | | | | | | |

**Proteins**

| | | | | | | | | | |
|---|---|---|---|---|---|---|---|---|---|
| putative DNA methylase N-4/N-6 domain protein (YP_005098037.1) | 89 | 865 | 68.5% | 3140 | 72.2% | 601 (99.3%) | 423 (69.9%) | 4/0/0/0 | 0 |

| | Begin | End | Coverage | Score | Concordance | Matches | Identities | I/D/M/F* | Stop Codons |
|---|---|---|---|---|---|---|---|---|---|
| **NT** | **4836** | **12136** | **4.3%** | **1899** | **45.2%** | **2108 (99.0%)** | **1543 (72.5%)** | **15/6** | |
| Protein mutations: | Y90I (7160T>A 7161A>T), M95R (7144C>G 7145A>C 7146T>G), R101K (7127C>T 7128G>T), S105R (7114A>G 7116T>G), F106I (7113A>T), D107E (7108G>C), V109A (7102T>G 7103A>G), T113N (7091G>T), T114G (7088G>C 7089T>C), V123I (7062C>T), G126R (7053C>G), R131L (7036G>C 7037C>A), R132L (7034C>A), S142A (7003A>T 7004C>G 7005T>C), R165_A166insRDM (6933_6934insCATGTCGCG), A167N (6929G>T 6930C>T), G168R (6927C>G), R170K (6919G>C 6920C>T 6921G>T), G171S (6918C>T), D172A (6914T>G), R175P (6905C>G), Y178F (6896T>A), Y189F (6862G>A 6863T>A), I190V (6861T>C), Q221N (6766T>G 6768G>T), I224L (6757G>C 6759T>G), H225S (6754A>G 6755T>C 6756G>T), A226P (6751G>T 6753C>G), H227A (6748G>C 6749T>G 6750G>C), K228R (6745T>G 6746T>C 6747T>G), G230R (6741C>G), E231D (6736C>G), M234L (6729T>G), I241L (6706G>C 6708T>G), R245K (6694G>C 6695C>T 6696G>T), F251Y (6677A>T), E254D (6667T>G), A257E (6659G>T), E260P (6649T>G 6650T>G 6651C>G), L261I (6646C>G 6648G>T), V268I (6625T>G 6627C>T), S270A (6619G>C 6621A>C), S273A (6610C>G 6612A>C), H274D (6609G>C), H277T (6598A>G 6599T>G 6600G>T), E278N (6595C>A 6597C>T), R279M (6592A>C 6593C>A 6594G>T), N280D (6591T>C), G283A (6580G>C 6581C>G), R284Q (6577G>C 6578C>T), L285M (6576G>T), L286F (6571A>G 6573G>A), R287K (6568G>C 6569C>T 6570G>T), N288R (6566T>C 6567T>G), T289Q (6562G>C 6563G>T 6564T>G), Q294V (6548T>A 6549G>C), A298R (6535T>G 6536G>C 6537C>G), E302D (6523C>G), P305D (6515G>T 6516G>C), I308V (6505G>C 6507T>C), L311M (6498G>T), I314L (6487G>C 6489T>G), A318D (6475A>G 6476G>T), D320A (6469A>G 6470T>G), I324V (6457G>C 6459T>C), A331D (6436C>A 6437G>T), H334R (6428T>C), A338R (6416G>C 6417C>G), I344A (6398A>G 6399T>C), D529S (5843T>C 5844C>T), E535Q (5826C>G), Y539F (5812A>G 5813T>A), I541V (5806G>C 5808T>C), T548L (5785G>C 5786G>A 5787T>G), I549M (5782G>C), L553V (5772G>C), C555A (5765C>G 5766A>C), D560N (5749A>G 5751C>T), I563V (5740A>C 5742T>C), G565S (5736C>T), T568V (5725G>C 5726G>A 5727T>C), T568_G569insE (5724_5725insCTC), L570V (5719A>G 5721G>C), V575E (5704G>C 5705A>T), H579D (5694G>C), M580V (5691T>C), E581D (5686T>G), V582T (5683C>G 5684A>G 5685C>T), T583I (5680A>G 5681G>A), R584A (5677T>G 5678C>G 5679G>C), A585C (5675G>C 5676C>A), V587R (5669A>C 5670C>G), K588R (5666T>C 5667T>G), H589A (5662A>G 5663T>G 5664G>C), W591F (5656C>G 5657C>A), A592H (5654G>T 5655C>G), Y593F (5650A>G 5651T>A), M594V (5649T>C), M596R (5641C>G 5642A>C 5643T>G), A608N (5606G>T 5607C>T), S614G (5587C>G 5588G>C 5589A>C), W615Y (5584C>G 5585C>T), T616S (5581C>G 5582G>C), T623S (5560G>C 5562T>A), K624R (5557C>G 5558T>C 5559T>G), V627F (5550C>A), P630C (5540G>C 5541G>A), P639A (5512G>T 5514G>C), N643A (5501T>G 5502T>C), N645R (5495T>C 5496T>G), A646G (5492G>C), G648A (5486C>G), I650V (5479A>C 5481T>C), A655D (5465G>T), A686P (5371T>C 5373C>G), L690F (5359C>G), F695Y (5345A>T), T696A (5341G>C 5343T>C), N699K (5332G>C), N701D (5328T>C), P703L (5321G>A), E706P (5312T>G 5313C>G), A707E (5309G>T), D708E (5305G>C), F709L (5302G>C 5304A>G), I712L (5293A>G 5295T>G), P713T (5290C>G 5292G>T), D715A (5285T>G), V716D (5281C>G 5282A>T), M717I (5278C>G), Y721F (5267T>A), D723K (5260G>C 5262C>T), Y724H (5259A>G), S727E (5248G>C 5249G>T 5250A>C), T728R (5246G>C 5247T>G), H735D (5226G>C), S741A (5208A>C), E743A (5200C>G 5201T>G), V744E (5197G>C 5198A>T), T745R (5195G>C 5196T>G), K747M (5189T>A), G751T (5177C>G 5178C>T), P756Q (5161T>C 5162G>T), A769T (5122G>C 5124C>T), A776S (5102G>C 5103C>T), Q777A (5099T>G 5100G>C), Y779S (5092G>C 5093T>G), S780A (5089G>T 5090C>G 5091T>C), K781A (5086C>G 5087T>G 5088T>C), Q783R (5080C>G 5081T>C), L790M (5061G>T), V795A (5044G>C 5045A>G), V800T (5029C>G 5030A>G 5031C>T), E805K (5014T>C 5016C>T), D807E (5008G>C), L808E (5006A>T 5007G>C), F810Y (5000A>T), I816L (4981G>C 4983T>G), C822R (4963G>C 4965A>G), L824V (4959G>C), K825L (4955T>A 4956T>G), K827G (4948C>G 4949T>C 4950T>C), T830G (4940G>C 4941T>C), A832G (4934G>C), H833C (4931T>C 4932G>A), N836A (4922T>G 4923T>C), S837K (4918G>C 4919G>T 4920A>T), R838P (4916C>G), A844V (4898G>A), K848E (4887T>C), S849Q (4882G>C 4883G>T 4884A>G), E851V (4877T>A), E852R (4873T>G 4874T>C 4875C>G), E853S (4871T>G 4872C>A), A855S (4864G>C 4866C>A), M856T (4862A>G), D858S (4855G>C 4856T>G 4857C>A), A861D (4847G>T), L863D (4841A>T 4842G>C) | | | | | | | | |
| Codon mutations: | TAC90ATC (7160T>A 7161A>T), GCC93GCG (7150G>C), GCG94GCA (7147C>T), ATG95CGC (7144C>G 7145A>C 7146T>G), CTC96CTT (7141G>A), ACC99ACG (7132G>C), CGG101AAG (7127C>T 7128G>T), AGT105CGC (7114A>G 7116T>G), TTC106ATC (7113A>T), GAC107GAG (7108G>C), GAG108GAA (7105C>T), GTA109GCC (7102T>G 7103A>G), CCC112CCG (7093G>C), ACC113AAC (7091G>T), ACC114GGC (7088G>C 7089T>C), CTG117CTC (7078C>G), AAT119AAC (7072A>G), GAG121GAA (7066C>T), ACC122ACG (7063G>C), GTC123ATC (7062C>T), GGC126CGC (7053C>G), CTC128CTG (7045G>C), CCT130CCG (7039A>C), CGC131CTG (7036G>C 7037C>A), CGG132CTG (7034C>A), GCA136GCC (7021T>G), GAA140GAG (7009T>C), GCC141GCG (7006G>C), AGT142GCA (7003A>T 7004C>G 7005T>C), CTT144CTG (6997A>C), ACG152ACC (6973C>G), GCT158GCG (6955A>C), GCT161GCC (6946A>G), GGT162GGC (6943A>G), GAT164GAC (6937A>G), CGC165CGG (6934G>C), CGC165_GCC166insCGCGACATG (6933_6934insCATGTCGCG), GCC167AAC (6929G>T 6930C>T), GGC168CGC (6927C>G), GTC169GTG (6922G>C), CGC170AAG (6919G>C 6920C>T 6921G>T), GGC171AGC (6918C>T), GAT172GCT (6914T>G), GTC174GTG (6907G>C), CGG175CCG (6905C>G), TAC176TAT (6901G>A), CGG177CGC (6898C>G), TAC178TTC (6896T>A), ACT181ACG (6886A>C), CCA184CCC (6877T>G), AGC185AGT (6874G>A), CCG186CCA (6871C>T), GAA188GAG (6865T>C), TAC189TTT (6862G>A 6863T>A), ATC190GTC (6861T>C), CTA192CTG (6853T>C), GCT194GCC (6847A>G), TCG196AGC (6841C>G 6842G>C 6843A>T), CTC199CTG (6832G>C), GAT203GAC (6820A>G), GTC204GTT (6817G>A), ACC209ACG (6802G>C), AGG210CGT (6799C>A 6801T>G), AAA213AAG (6790T>C), TCT216AGC (6781A>G 6782G>C 6783A>T), CAA221AAC (6766T>G 6768G>T), CTC222CTG (6763G>C), ACC223ACG (6760G>C), ATC224CTG (6757G>C 6759T>G), CAT225AGC (6754A>G 6755T>C 6756G>T), GCC226CCA (6751G>T 6753C>G), CAC227GCG (6748G>C 6749T>G 6750G>C), AAA228CGC (6745T>G 6746T>C 6747T>G), GGC230CGC (6741C>G), GAG231GAC (6736C>G), ATG234CTG (6729T>G), GCG237GCA (6718C>T), TCC238AGC (6716G>C 6717A>T), GCG240GCC (6709C>G), ATC241CTG (6706G>C 6708T>G), GTT243GTG (6700A>C), CGC245AAG (6694G>C 6695C>T 6696G>T), AGC247TCG (6688G>C 6689C>G 6690T>A), GGG250GGC (6679C>G), TTC251TAC (6677A>T), GAA254GAC (6667T>G), GCG257GAG (6659G>T), CTA258CTG (6655T>C), GAA260CCC (6649T>G 6650T>G 6651C>G), CTG261ATC (6646C>G 6648G>T), GAA267GAG (6628T>C), GTA268ATC (6625T>G 6627C>T), TCC270GCG (6619G>C 6621A>C), TCG273GCC (6610C>G 6612A>C), CAC274GAC (6609G>C), GGC276GGT (6601G>A), CAT277ACC (6598A>G 6599T>G 6600G>T), GAG278AAT (6595C>A 6597C>T), CGT279ATG (6592A>C 6593C>A 6594G>T), AAC280GAC (6591T>C), GGC283GCG (6580G>C 6581C>G), CGC284CAG (6577G>C 6578C>T), CTG285ATG (6576G>T), CTT286TTC (6571A>G 6573G>A), CGC287AAG (6568G>C 6569C>T 6570G>T), AAC288CGC (6566T>C 6567T>G), ACC289CAG (6562G>C 6563G>T 6564T>G), ATT291ATC (6556A>G), CAG294GTG (6548T>A 6549G>C), GCC296GCG (6541G>C), GCC297GCG (6538G>C), GCA298CGC (6535T>G 6536G>C 6537C>G), GAG302GAC (6523C>G), CTT304CTG (6517A>C), CCT305GAT (6515G>T 6516G>C), GCC306GCG (6511G>C), CGG307CGC (6508C>G), ATC308GTG (6505G>C 6507T>C), AAA310AAG (6499T>C), CTG311ATG (6498G>T), ATC314CTG (6487G>C 6489T>G), CGG315CGC (6484C>G), GCT318GAC (6475A>G 6476G>T), CCA319CCG (6472T>C), GAT320GCC (6469A>G 6470T>G), GCT321GCC (6466A>G), CGG323CGT (6460C>A), ATC324GTG (6457G>C 6459T>C), CAT327CAC (6448A>G), CTC329CTG (6442G>C), GCG331GAT (6436C>A 6437G>T), GAA332GAG (6433T>C), CAC334CGC (6428T>C), GCG335GCC (6424C>G), GCC338CGC (6416G>C 6417C>G), GCT339GCC (6412A>G), CCC341CCG (6406G>C), ACC342ACG (6403G>C), ATC344GCC (6398A>G 6399T>C), GAC529AGC (5843T>C 5844C>T), GAG535CAG (5826C>G), TAT539TTC (5812A>G 5813T>A), ATC541GTG (5806G>C 5808T>C), GAA543GAG (5800T>C), ACC548CTG (5785G>C 5786G>A 5787T>G), ATC549ATG (5782G>C), CCC550CCG (5779G>C), GGG551GGC (5776C>G), CGT552AGG (5773A>C 5775G>T), CTG553GTG (5772G>C), TGC555GCC (5765C>G 5766A>C), GTG558GTC (5755C>G), GAT560AAC (5749A>G 5751C>T), ATT563GTG (5740A>C 5742T>C), GGC565AGC (5736C>T), GGA566GGC (5731T>G), ACC568GTG (5725G>C 5726G>A 5727T>C), ACC568_GGC569insGAG (5724_5725insCTC), CTT570GTC (5719A>G 5721G>C), GTC575GAG (5704G>C 5705A>T), CAC579GAC (5694G>C), ATG580GTG (5691T>C), GAA581GAC (5686T>G), GTG582ACC (5683C>G 5684A>G 5685C>T), ACT583ATC (5680A>G 5681G>A), CGA584GCC (5677T>G 5678C>G 5679G>C), GCC585TGC (5675G>C 5676C>A), GTC587CGC (5669A>C 5670C>G), AAG588CGG (5666T>C 5667T>G), CAT589GCC (5662A>G 5663T>G 5664G>C), TGG591TTC (5656C>G 5657C>A), GCC592CAC (5654G>T 5655C>G), TAT593TTC (5650A>G 5651T>A), ATG594GTG (5649T>C), ATG596CGC (5641C>G 5642A>C 5643T>G), GTT604GTG (5617A>C), GCC608AAC (5606G>T 5607C>T), CTC613CTG (5590G>C), TCG614GGC (5587C>G 5588G>C 5589A>C), TGG615TAC (5584C>G 5585C>T), ACG616AGC (5581C>G 5582G>C), GAA617GAG (5578T>C), CAG618CAA (5575C>T), GAC621GAT (5566G>A), ACC623TCG (5560G>C 5562T>A), AAG624CGC (5557C>G 5558T>C 5559T>G), GGT626GGC (5551A>G), GTC627TTC (5550C>A), CCC630TGC (5540G>C 5541G>A), CCC639GCA (5512G>T 5514G>C), CCC640CCG (5509G>C), AAC643GCC (5501T>G 5502T>C), TCC644AGC (5498G>C 5499A>T), AAC645CGC (5495T>C 5496T>G), GCC646GGC (5492G>C), TAC647TAT (5488G>A), GGC648GCC (5486C>G), ATT650GTG (5479A>C 5481T>C), CCG651CCC (5476C>G), GTC653GTG (5470G>C), GCC655GAC (5465G>T), GCA686CCG (5371T>C 5373C>G), TTG690TTC (5359C>G), TTC695TAC (5345A>T), ACC696GCG (5341G>C 5343T>C), AAC699AAG (5332G>C), AAC701GAC (5328T>C), CGT702CGG (5323A>C), CCG703CTG (5321G>A), ACT705ACG (5314A>C), GAA706CCA (5312T>G 5313C>G), GCG707GAG (5309G>T), GAC708GAG (5305G>C), TTC709CTG (5302G>C 5304A>G), ATT712CTC (5293A>G 5295T>G), CCG713ACC (5290C>G 5292G>T), CAC714CAT (5287G>A), GAC715GCC (5285T>G), GTG716GAC (5281C>G 5282A>T), ATG717ATC (5278C>G), TAC721TTC (5267T>A), GAC723AAG (5260G>C 5262C>T), TAC724CAC (5259A>G), AGC725TCG (5254G>C 5255C>G 5256T>A), TCC727GAG (5248G>C 5249G>T 5250A>C), ACC728CGC (5246G>C 5247T>G), CAC735GAC (5226G>C), GTC736GTG (5221G>C), GAA740GAG (5209T>C), TCG741GCG (5208A>C), GAG743GCC (5200C>G 5201T>G), GTC744GAG (5197G>C 5198A>T), ACC745CGC (5195G>C 5196T>G), GGG746GGC (5191C>G), AAG747ATG (5189T>A), GGC751ACC (5177C>G 5178C>T), CCA756CAG (5161T>C 5162G>T), TCC759AGC (5153G>C 5154A>T), CAC761CAT (5146G>A), GTA764GTC (5137T>G), GTC768GTG (5125G>C), GCC769ACG (5122G>C 5124C>T), CGG772CGC (5113C>G), GCC776AGC (5102G>C 5103C>T), CAG777GCG (5099T>G 5100G>C), TAC779TCG (5092G>C 5093T>G), AGC780GCA (5089G>T 5090C>G 5091T>C), AAG781GCC (5086C>G 5087T>G 5088T>C), GGC782GGG (5083G>C), CAG783CGC (5080C>G 5081T>C), GAA784GAG (5077T>C), CTG790ATG (5061G>T), ATC794ATT (5047G>A), GTC795GCG (5044G>C 5045A>G), GCC798GCG (5035G>C), GTG800ACC (5029C>G 5030A>G 5031C>T), GAA805AAG (5014T>C 5016C>T), GGA806GGC (5011T>G), GAC807GAG (5008G>C), CTG808GAG (5006A>T 5007G>C), TTC810TAC (5000A>T), GAT811GAC (4996A>G), GGG815GGC (4984C>G), ATC816CTG (4981G>C 4983T>G), ACC818ACG (4975G>C), GTC819GTG (4972G>C), CCG820CCC (4969C>G), TGC822CGG (4963G>C 4965A>G), GCG823GCT (4960C>A), CTG824GTG (4959G>C), AAG825CTG (4955T>A 4956T>G), CTC826CTG (4951G>C), AAG827GGC (4948C>G 4949T>C 4950T>C), CGC828CGG (4945G>C), AGA829CGC (4942T>G 4944T>G), ACC830GGC (4940G>C 4941T>C), GCC832GGC (4934G>C), CAC833TGC (4931T>C 4932G>A), GAA834GAG (4927T>C), AAC836GCC (4922T>G 4923T>C), TCC837AAG (4918G>C 4919G>T 4920A>T), CGC838CCC (4916C>G), GAC842GAT (4903G>A), GGT843GGC (4900A>G), GCC844GTC (4898G>A), TAT846TAC (4891A>G), AAG848GAG (4887T>C), TCC849CAG (4882G>C 4883G>T 4884A>G), GCG850GCC (4879C>G), GAG851GTG (4877T>A), GAA852CGC (4873T>G 4874T>C 4875C>G), GAG853TCG (4871T>G 4872C>A), GCC855TCG (4864G>C 4866C>A), ATG856ACG (4862A>G), CCC857CCG (4858G>C), GAC858TCG (4855G>C 4856T>G 4857C>A), TTC860TTT (4849G>A), GCC861GAC (4847G>T), CTG862CTT (4843C>A), CTC863GAC (4841A>T 4842G>C) | | | | | | | | |
| PD-(D/E)XK nuclease family protein (YP_005098042.1) | 4 | 106 | 38.4% | 390 | 55.2% | 101 (97.1%) | 52 (50.0%) | 1/2/0/0 | 0 |
| Protein mutations: | T5P (12132T>G), V6I (12127C>G 12129C>T), S10T (12115G>C 12117A>T), W11L (12113C>A 12114A>G), G12A (12110C>G), A13E (12107G>T), A18P (12091C>A 12093C>G), F19A (12088G>T 12089A>G 12090A>C), K20R (12086T>C), G23A (12076G>C 12077C>G), V24K (12073T>C 12074A>T 12075C>T), L26I (12069G>T), K28G (12061C>G 12062T>C 12063T>C), S31G (12054T>C), S33R (12046T>G 12047G>C 12048A>G), P35A (12042G>C), R36A (12037C>G 12038C>G 12039G>C), L38Q (12032A>T), I43V (12016G>C 12018T>C), S46G (12009T>C), A49L (11998C>G 11999G>A 12000C>G), A52Q (11990G>T 11991C>G), A53S (11986T>G 11988C>A), R54H (11983C>G 11984C>T), V55L (11980G>C 11982C>G), N56P (11977G>C 11978T>G 11979T>G), E58G (11971C>T 11972T>C), E58_P59insT (11970_11971insAGT), I60L (11965G>C 11967T>A), Y63D (11956G>A 11958A>C), S66A (11947C>G 11949A>C), E67G (11944T>G 11945T>C), L68A (11941C>A 11942A>G 11943G>C), L69V (11938C>G 11940G>C), H71D (11934G>C), T72A (11931T>C), L73I (11926C>G 11928G>T), Q74H (11923C>G), Q75K (11922G>T), F78E (11911G>T 11912A>T 11913A>C), R83_G84del (11893_11898delGCCGCG), S85E (11890G>T 11891G>T 11892A>C), D86E (11887G>T), I87T (11885A>G), S88A (11881G>C 11882C>G 11883T>C), E91D (11872T>G), S94R (11864G>C 11865A>G), T95I (11861G>A), G96A (11857T>C 11858C>G), T98A (11853T>C), K102R (11839C>G 11840T>C 11841T>G) | | | | | | | | |

| | Begin | End | Coverage | Score | Concordance | Matches | Identities | I/D/M/F* | Stop Codons |
|---|---|---|---|---|---|---|---|---|---|
| **NT** | **4836** | **12136** | **4.3%** | **1899** | **45.2%** | **2108 (99.0%)** | **1543 (72.5%)** | **15/6** | |
| Codon mutations: | ACC5CCC (12132T>G), GTG6ATC (12127C>G 12129C>T), TCG9AGC (12118C>G 12119G>C 12120A>T), TCC10ACG (12115G>C 12117A>T), TGG11CTG (12113C>A 12114A>G), GGC12GCC (12110C>G), GCG13GAG (12107G>T), GCG18CCT (12091C>A 12093C>G), TTC19GCA (12088G>T 12089A>G 12090A>C), AAG20AGG (12086T>C), GAG22GAA (12079C>T), GGC23GCG (12076G>C 12077C>G), GTA24AAG (12073T>C 12074A>T 12075C>T), CTC26ATC (12069G>T), AAG28GGC (12061C>G 12062T>C 12063T>C), AGC31GGC (12054T>C), CCT32CCG (12049A>C), TCA33CGC (12046T>G 12047G>C 12048A>G), TCC34AGC (12044G>C 12045A>T), CCC35GCC (12042G>C), CGG36GCC (12037C>G 12038C>G 12039G>C), CTG38CAG (12032A>T), CTC39CTG (12028G>C), GGT40GGC (12025A>G), ACC41ACG (12022G>C), ATC43GTG (12016G>C 12018T>C), AGC46GGC (12009T>C), GCG49CTC (11998C>G 11999G>A 12000C>G), GCG52CAG (11990G>T 11991C>G), GCA53TCC (11986T>G 11988C>A), CGG54CAC (11983C>G 11984C>T), GTC55CTG (11980G>C 11982C>G), AAC56CCG (11977G>C 11978T>G 11979T>G), GAG58GGA (11971C>T 11972T>C), GAG58_CCG59insACT (11970_11971insAGT), CCG59CCC (11968C>G), ATC60TTG (11965G>C 11967T>A), TAC63GAT (11956G>A 11958A>C), TCG66GCC (11947C>G 11949A>C), GAA67GGC (11944T>G 11945T>C), CTG68GCT (11941C>A 11942A>G 11943G>C), CTG69GTC (11938C>G 11940G>C), CAC71GAC (11934G>C), ACG72GCG (11931T>C), CTG73ATC (11926C>G 11928G>T), CAG74CAC (11923C>G), CAG75AAG (11922G>T), CCG76CCC (11917C>G), GAG77GAA (11914C>T), TTC78GAA (11911G>T 11912A>T 11913A>C), GAC81GAT (11902G>A), CGC83_GGC84del (11893_11898delGCCGCG), TCC85GAA (11890G>T 11891G>T 11892A>C), GAC86GAA (11887G>T), ATC87ACC (11885A>G), AGC88GCG (11881G>C 11882C>G 11883T>C), GAA91GAC (11872T>G), TCC94CGC (11864G>C 11865A>G), ACC95ATC (11861G>A), GGA96GCG (11857T>C 11858C>G), ACG98GCG (11853T>C), ACG101ACC (11842C>G), AAG102CGC (11839C>G 11840T>C 11841T>G) | | | | | | | | |

*: Inserts / Deletes / Misaligned / Frameshifts

#### Analysis details

This analysis was performed with panviral2.64

### NGS Details (UN24): Epiphyllum badnavirus 1

#### Assembly

| | |
|---|---|
| Coverage Length | 279 (1 contig(s)) |
| Depth Of Coverage | 35.8 |
| Number Of Reads | 90 |
| Reads Per Million | 1.80 rpm (after QC) |
| Ambiguities | 0 |
| Assembly Method | de novo + reference guided assembly |
| Consensus Caller | Bcf Tools |

#### Coverage Map

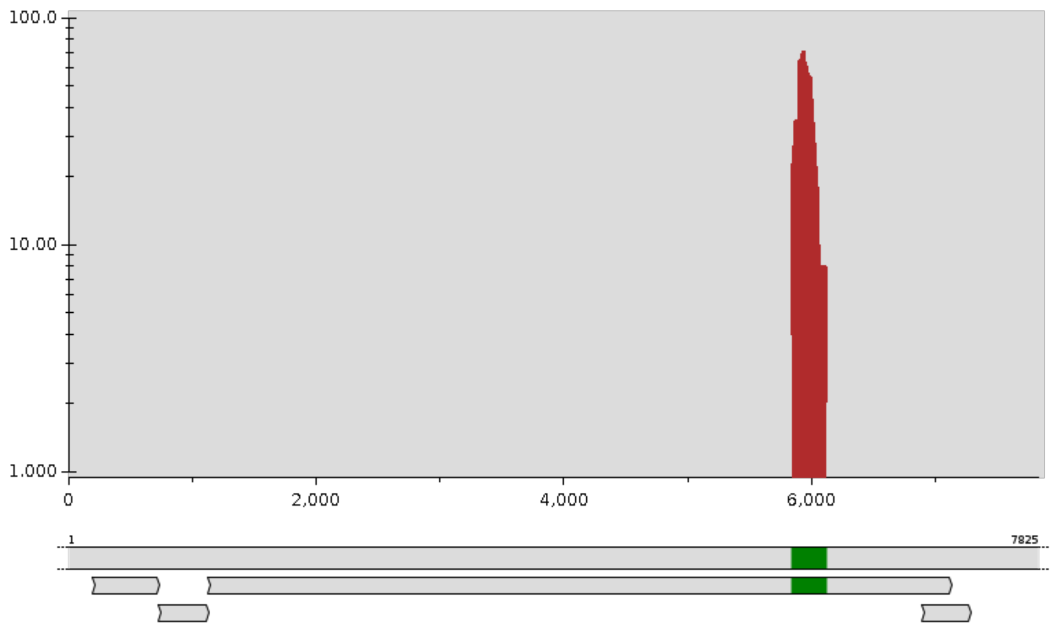

#### Assignment

| | |
|---|---|
| Type | Epiphyllum badnavirus 1 (Taxonomy ID: 2518008) |
| Reference Genome | NC_076247.1 |
| NT Identity (%) | 53.9007 |
| AA Identity (%) | 51.0638 |
| Number Of Stop Codons | 0 |
| Number Of CDS | 4 |

#### Alignment

| | |
|---|---|
| Alignment Score | 38.0 (NT) + 339.0 (AA) = 377.0 |
| Concordance (%) | 30.4278 |

| Alignment Method | Global, seeded, nucleotide + amino acids (AGA) |
|---|---|

#### Genome Region

Sequence starts at position 5840 and ends at position 6118 relative to NC_076247.1 reference sequence.

#### Alignment Detailed Statistics

| | Begin | End | Coverage | Score | Concordance | Matches | Identities | I/D/M/F* | Stop Codons |
|---|---|---|---|---|---|---|---|---|---|
| **NT** | **5840** | **6118** | **3.6%** | **38** | **6.8%** | **279 (98.9%)** | **152 (53.9%)** | **3/0** | |
| Mutations: | 5841C>T, 5847C>A, 5851T>A, 5853C>A, 5859G>T, 5860A>C, 5861A>G, 5862G>T, 5863A>T, 5864G>C, 5865C>G, 5868A>T, 5870T>A, 5871C>T, 5878A>T, 5880T>G, 5882T>A, 5884A>G, 5886G>T, 5887G>A, 5889C>G, 5892A>G, 5893G>T, 5894A>G, 5895A>T, 5896T>G, 5897C>A, 5898C>T, 5901C>T, 5904T>C, 5905T>A, 5906G>A, 5910C>A, 5913C>A, 5918T>G, 5922C>G, 5923C>A, 5924C>G, 5925A>G, 5926G>T, 5933T>A, 5936A>T, 5937C>T, 5938G>A, 5942G>T, 5943G>C, 5944T>C, 5946A>T, 5953C>T, 5955A>C, 5958C>T, 5961A>T, 5964G>A, 5966A>C, 5967G>T, 5970T>C, 5973C>T, 5976T>G, 5977G>A, 5982G>A, 5985C>T, 5986C>A, 5987A>T, 5988A>G, 5989C>G, 5990G>A, 5991A>C, 5992A>C, 5993A>T, 5994G>T, 5998G>A, 6000C>T, 6002A>G, 6003C>G, 6004T>A, 6005G>T, 6006T>A, 6011G>A, 6012A>G, 6013G>C, 6014A>C, 6015_6016insTAT, 6018G>C, 6019T>G, 6020C>A, 6021A>C, 6022G>A, 6023G>T, 6024C>G, 6026T>A, 6030A>G, 6032C>T, 6033A>T, 6036T>G, 6038A>T, 6059T>A, 6060C>T, 6061A>T, 6062G>C, 6064A>C, 6065A>G, 6066C>A, 6068C>G, 6069T>C, 6072A>G, 6073G>C, 6075T>A, 6078C>T, 6081T>C, 6082G>A, 6083C>T, 6084A>G, 6085C>G, 6087G>A, 6090C>T, 6091C>T, 6095A>G, 6097T>A, 6099C>G, 6100A>T, 6102G>A, 6103C>T, 6108G>A, 6113G>T, 6114T>G, 6115G>A, 6116A>G, 6118A>G | | | | | | | | |

##### CDS

| | Begin | End | Coverage | Score | Concordance | Matches | Identities | I/D/M/F* | Stop Codons |
|---|---|---|---|---|---|---|---|---|---|
| QKM20_gp3 | 1572 | 1664 | 4.6% | 339 | 48.8% | 93 (98.9%) | 48 (51.1%) | 1/0/0/0 | 0 |
| Protein mutations: | F1575I (5851T>A 5853C>A), K1578R (5860A>C 5861A>G 5862G>T), F1581Y (5870T>A 5871C>T), I1584L (5878A>T 5880T>G), M1585K (5882T>A), M1586V (5884A>G 5886G>T), D1587K (5887G>A 5889C>G), E1589C (5893G>T 5894A>G 5895A>T), S1590D (5896T>G 5897C>A 5898C>T), W1593K (5905T>A 5906G>A), L1597R (5918T>G), P1599R (5923C>A 5924C>G 5925A>G), D1600Y (5926G>T), L1602H (5933T>A), Y1603F (5936A>T 5937C>T), E1604K (5938G>A), W1605F (5942G>T 5943G>C), P1609S (5953C>T 5955A>C), K1613T (5966A>C 5967G>T), A1617T (5977G>A), Q1620M (5986C>A 5987A>T 5988A>G), R1621D (5989C>G 5990G>A 5991A>C), K1622L (5992A>C 5993A>T 5994G>T), D1624N (5998G>A 6000C>T), N1625R (6002A>G 6003C>G), C1626I (6004T>A 6005G>T 6006T>A), R1628K (6011G>A 6012A>G), D1629P (6013G>C 6014A>C), D1629_L1630insY (6015_6016insTAT), S1631D (6019T>G 6020C>A 6021A>C), G1632M (6022G>A 6023G>T 6024C>G), F1633Y (6026T>A), A1635V (6032C>T 6033A>T), Y1637F (6038A>T), F1644Y (6059T>A 6060C>T), N1646R (6064A>C 6065A>G 6066C>A), T1647S (6068C>G 6069T>C), D1649Q (6073G>C 6075T>A), A1652M (6082G>A 6083C>T 6084A>G), Q1653E (6085C>G 6087G>A), K1656R (6095A>G), S1657T (6097T>A 6099C>G), M1658L (6100A>T 6102G>A), C1662L (6113G>T 6114T>G), E1663R (6115G>A 6116A>G) | | | | | | | | |
| Codon mutations: | GTC1571.TT (5841C>T), TCC1573TCA (5847C>A), TTC1575ATA (5851T>A 5853C>A), CTG1577CTT (5859G>T), AAG1578CGT (5860A>C 5861A>G 5862G>T), AGC1579TCG (5863A>T 5864G>C 5865C>G), GGA1580GGT (5868A>T), TTC1581TAT (5870T>A 5871C>T), ATT1584TTG (5878A>T 5880T>G), ATG1585AAG (5882T>A), ATG1586GTT (5884A>G 5886G>T), GAC1587AAG (5887G>A 5889C>G), GAA1588GAG (5892A>G), GAA1589TGT (5893G>T 5894A>G 5895A>T), TCC1590GAT (5896T>G 5897C>A 5898C>T), ATC1591ATT (5901C>T), CCT1592CCC (5904T>C), TGG1593AAG (5905T>A 5906G>A), ACC1594ACA (5910C>A), GCC1595GCA (5913C>A), CTG1597CGG (5918T>G), ACC1598ACG (5922C>G), CCA1599AGG (5923C>A 5924C>G 5925A>G), GAT1600TAT (5926G>T), CTT1602CAT (5933T>A), TAC1603TTT (5936A>T 5937C>T), GAA1604AAA (5938G>A), TGG1605TTC (5942G>T 5943G>C), TTA1606CTT (5944T>C 5946A>T), CCA1609TCC (5953C>T 5955A>C), TTC1610TTT (5958C>T), GGA1611GGT (5961A>T), TTG1612TTA (5964G>A), AAG1613ACT (5966A>C 5967G>T), AAT1614AAC (5970T>C), GCC1615GCT (5973C>T), CCT1616CCG (5976T>G), GCT1617ACT (5977G>A), GTG1618GTA (5982G>A), TTC1619TTT (5985C>T), CAA1620ATG (5986C>A 5987A>T 5988A>G), CGA1621GAC (5989C>G 5990G>A 5991A>C), AAG1622CTT (5992A>C 5993A>T 5994G>T), GAC1624AAT (5998G>A 6000C>T), AAC1625AGG (6002A>G 6003C>G), TGT1626ATA (6004T>A 6005G>T 6006T>A), AGA1628AAG (6011G>A 6012A>G), GAC1629CCC (6013G>C 6014A>C), GAC1629_CTG1630insTAT (6015_6016insTAT), CTG1630CTC (6018G>C), TCA1631GAC (6019T>G 6020C>A 6021A>C), GGC1632ATG (6022G>A 6023G>T 6024C>G), TTT1633TAT (6026T>A), GTA1634GTG (6030A>G), GCA1635GTT (6032C>T 6033A>T), GTT1636GTG (6036T>G), TAC1637TTC (6038A>T), TTC1644TAT (6059T>A 6060C>T), AGT1645TCT (6061A>T 6062G>C), AAC1646CGA (6064A>C 6065A>G 6066C>A), ACT1647AGC (6068C>G 6069T>C), GAA1648GAG (6072A>G), GAT1649CAA (6073G>C 6075T>A), GAC1650GAT (6078C>T), CAT1651CAC (6081T>C), GCA1652ATG (6082G>A 6083C>T 6084A>G), CAG1653GAA (6085C>G 6087G>A), CAC1654CAT (6090C>T), CTG1655TTG (6091C>T), AAG1656AGG (6095A>G), TCC1657ACG (6097T>A 6099C>G), ATG1658TTA (6100A>T 6102G>A), CTG1659TTG (6103C>T), CAG1660CAA (6108G>A), TGT1662TTG (6113G>T 6114T>G), GAG1663AGG (6115G>A 6116A>G), AGA1664G.. (6118A>G) | | | | | | | | |

##### Proteins

| | Begin | End | Coverage | Score | Concordance | Matches | Identities | I/D/M/F* | Stop Codons |
|---|---|---|---|---|---|---|---|---|---|
| polyprotein (YP_010797894.1) | 1572 | 1664 | 4.6% | 339 | 48.8% | 93 (98.9%) | 48 (51.1%) | 1/0/0/0 | 0 |
| Protein mutations: | F1575I (5851T>A 5853C>A), K1578R (5860A>C 5861A>G 5862G>T), F1581Y (5870T>A 5871C>T), I1584L (5878A>T 5880T>G), M1585K (5882T>A), M1586V (5884A>G 5886G>T), D1587K (5887G>A 5889C>G), E1589C (5893G>T 5894A>G 5895A>T), S1590D (5896T>G 5897C>A 5898C>T), W1593K (5905T>A 5906G>A), L1597R (5918T>G), P1599R (5923C>A 5924C>G 5925A>G), D1600Y (5926G>T), L1602H (5933T>A), Y1603F (5936A>T 5937C>T), E1604K (5938G>A), W1605F (5942G>T 5943G>C), P1609S (5953C>T 5955A>C), K1613T (5966A>C 5967G>T), A1617T (5977G>A), Q1620M (5986C>A 5987A>T 5988A>G), R1621D (5989C>G 5990G>A 5991A>C), K1622L (5992A>C 5993A>T 5994G>T), D1624N (5998G>A 6000C>T), N1625R (6002A>G 6003C>G), C1626I (6004T>A 6005G>T 6006T>A), R1628K (6011G>A 6012A>G), D1629P (6013G>C 6014A>C), D1629_L1630insY (6015_6016insTAT), S1631D (6019T>G 6020C>A 6021A>C), G1632M (6022G>A 6023G>T 6024C>G), F1633Y (6026T>A), A1635V (6032C>T 6033A>T), Y1637F (6038A>T), F1644Y (6059T>A 6060C>T), N1646R (6064A>C 6065A>G 6066C>A), T1647S (6068C>G 6069T>C), D1649Q (6073G>C 6075T>A), A1652M (6082G>A 6083C>T 6084A>G), Q1653E (6085C>G 6087G>A), K1656R (6095A>G), S1657T (6097T>A 6099C>G), M1658L (6100A>T 6102G>A), C1662L (6113G>T 6114T>G), E1663R (6115G>A 6116A>G) | | | | | | | | |
| Codon mutations: | GTC1571.TT (5841C>T), TCC1573TCA (5847C>A), TTC1575ATA (5851T>A 5853C>A), CTG1577CTT (5859G>T), AAG1578CGT (5860A>C 5861A>G 5862G>T), AGC1579TCG (5863A>T 5864G>C 5865C>G), GGA1580GGT (5868A>T), TTC1581TAT (5870T>A 5871C>T), ATT1584TTG (5878A>T 5880T>G), ATG1585AAG (5882T>A), ATG1586GTT (5884A>G 5886G>T), GAC1587AAG (5887G>A 5889C>G), GAA1588GAG (5892A>G), GAA1589TGT (5893G>T 5894A>G 5895A>T), TCC1590GAT (5896T>G 5897C>A 5898C>T), ATC1591ATT (5901C>T), CCT1592CCC (5904T>C), TGG1593AAG (5905T>A 5906G>A), ACC1594ACA (5910C>A), GCC1595GCA (5913C>A), CTG1597CGG (5918T>G), ACC1598ACG (5922C>G), CCA1599AGG (5923C>A 5924C>G 5925A>G), GAT1600TAT (5926G>T), CTT1602CAT (5933T>A), TAC1603TTT (5936A>T 5937C>T), GAA1604AAA (5938G>A), TGG1605TTC (5942G>T 5943G>C), TTA1606CTT (5944T>C 5946A>T), CCA1609TCC (5953C>T 5955A>C), TTC1610TTT (5958C>T), GGA1611GGT (5961A>T), TTG1612TTA (5964G>A), AAG1613ACT (5966A>C 5967G>T), AAT1614AAC (5970T>C), GCC1615GCT (5973C>T), CCT1616CCG (5976T>G), GCT1617ACT (5977G>A), GTG1618GTA (5982G>A), TTC1619TTT (5985C>T), CAA1620ATG (5986C>A 5987A>T 5988A>G), CGA1621GAC (5989C>G 5990G>A 5991A>C), AAG1622CTT (5992A>C 5993A>T 5994G>T), GAC1624AAT (5998G>A 6000C>T), AAC1625AGG (6002A>G 6003C>G), TGT1626ATA (6004T>A 6005G>T 6006T>A), AGA1628AAG (6011G>A 6012A>G), GAC1629CCC (6013G>C 6014A>C), GAC1629_CTG1630insTAT (6015_6016insTAT), CTG1630CTC (6018G>C), TCA1631GAC (6019T>G 6020C>A 6021A>C), GGC1632ATG (6022G>A 6023G>T 6024C>G), TTT1633TAT (6026T>A), GTA1634GTG (6030A>G), GCA1635GTT (6032C>T 6033A>T), GTT1636GTG (6036T>G), TAC1637TTC (6038A>T), TTC1644TAT (6059T>A 6060C>T), AGT1645TCT (6061A>T 6062G>C), AAC1646CGA (6064A>C 6065A>G 6066C>A), ACT1647AGC (6068C>G 6069T>C), GAA1648GAG (6072A>G), GAT1649CAA (6073G>C 6075T>A), GAC1650GAT (6078C>T), CAT1651CAC (6081T>C), GCA1652ATG (6082G>A 6083C>T 6084A>G), CAG1653GAA (6085C>G 6087G>A), CAC1654CAT (6090C>T), CTG1655TTG (6091C>T), AAG1656AGG (6095A>G), TCC1657ACG (6097T>A 6099C>G), ATG1658TTA (6100A>T 6102G>A), CTG1659TTG (6103C>T), CAG1660CAA (6108G>A), TGT1662TTG (6113G>T 6114T>G), GAG1663AGG (6115G>A 6116A>G), AGA1664G.. (6118A>G) | | | | | | | | |

*: Inserts / Deletes / Misaligned / Frameshifts

#### Analysis details

This analysis was performed with panviral2.64

### NGS Details (UN24): Caulimovirus venafragariae

#### Assembly

| | |
|---|---|
| **Coverage Length** | 262 (1 contig(s)) |
| **Depth Of Coverage** | 31.5 |
| **Number Of Reads** | 74 |
| **Reads Per Million** | 1.48 rpm (after QC) |
| **Ambiguities** | 0 |
| **Assembly Method** | de novo + reference guided assembly |
| **Consensus Caller** | Bcf Tools |

#### Coverage Map

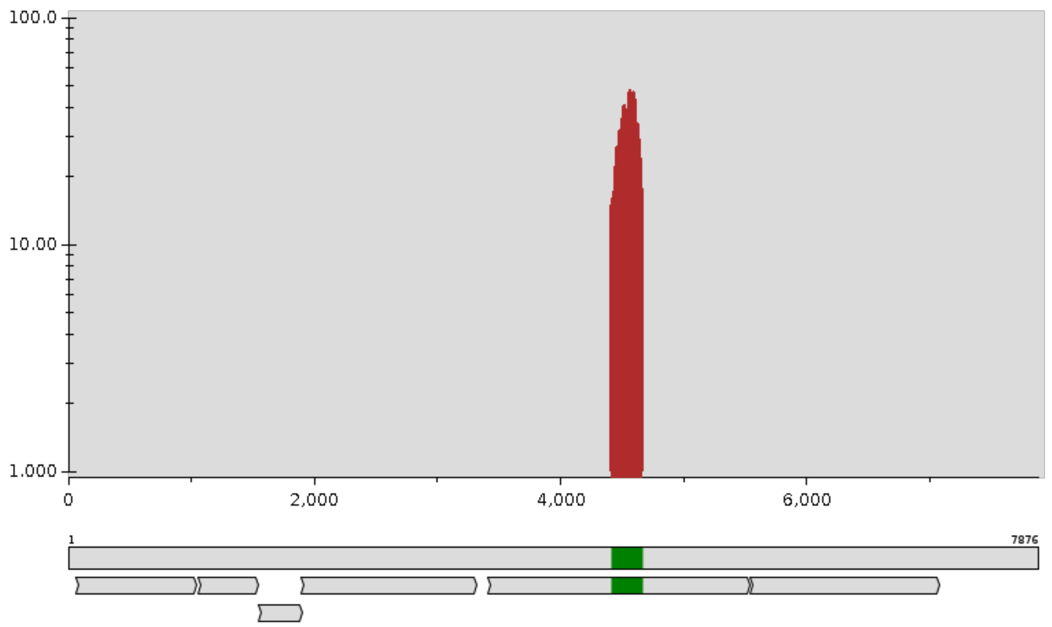

#### Assignment

| | |
|---|---|
| **Type** | Caulimovirus venafragariae (Taxonomy ID: 3048344) |
| **Reference Genome** | NC_001725.1 |
| **NT Identity (%)** | 57.6336 |
| **AA Identity (%)** | 47.1264 |
| **Number Of Stop Codons** | 0 |
| **Number Of CDS** | 6 |

#### Alignment

| | |
|---|---|
| **Alignment Score** | 80.0 (NT) + 295.0 (AA) = 375.0 |
| **Concordance (%)** | 31.9693 |

| Alignment Method | Global, seeded, nucleotide + amino acids (AGA) |
|---|---|

#### Genome Region

Sequence starts at position 4409 and ends at position 4670 relative to NC_001725.1 reference sequence.

#### Alignment Detailed Statistics

| | Begin | End | Coverage | Score | Concordance | Matches | Identities | I/D/M/F* | Stop Codons |
|---|---|---|---|---|---|---|---|---|---|
| **NT** | **4409** | **4670** | **3.3%** | **80** | **15.3%** | **262 (100%)** | **151 (57.6%)** | **0/0** | |
| Mutations: | 4410A>C, 4413T>G, 4414A>G, 4419T>C, 4421A>G, 4426T>C, 4428A>T, 4434C>T, 4435C>G, 4436A>C, 4442A>G, 4443G>A, 4444G>A, 4445G>A, 4450G>C, 4451G>A, 4457T>C, 4459C>G, 4461T>A, 4464T>C, 4465A>T, 4466A>T, 4469A>T, 4470G>T, 4472A>G, 4473G>T, 4477C>A, 4482T>A, 4483C>G, 4485A>C, 4489A>T, 4491C>G, 4493G>C, 4494A>T, 4497T>G, 4498A>C, 4500G>A, 4501A>G, 4502C>A, 4503C>T, 4505T>A, 4509C>T, 4511C>G, 4512T>C, 4514C>T, 4516T>C, 4518T>A, 4521C>T, 4522T>G, 4524T>G, 4525A>T, 4527A>C, 4530T>A, 4535T>A, 4537T>A, 4538G>A, 4539G>T, 4543G>A, 4545A>C, 4546C>A, 4547G>C, 4548C>A, 4549C>A, 4554T>G, 4560A>G, 4561A>G, 4562C>A, 4563A>C, 4564A>C, 4565T>A, 4566T>G, 4567C>G, 4569G>A, 4570C>A, 4571T>A, 4572A>G, 4576G>A, 4583G>C, 4584C>T, 4587T>C, 4590C>T, 4591C>T, 4593A>T, 4597C>A, 4598A>C, 4599C>A, 4602C>T, 4604A>C, 4607G>T, 4608G>C, 4609T>A, 4610T>A, 4611A>G, 4612G>C, 4613T>G, 4614C>G, 4626A>G, 4630A>T, 4631A>G, 4632A>C, 4633C>A, 4635A>T, 4638C>A, 4641T>G, 4644T>G, 4646T>C, 4647T>A, 4650C>T, 4656G>A, 4657C>T, 4658A>G | | | | | | | | |
| **CDS** | | | | | | | | | |
| ORF_V | 336 | 422 | 12.3% | 295 | 44.8% | 87 (100%) | 41 (47.1%) | 0/0/0/0 | 0 |
| Protein mutations: | I336M (4413T>G), N337D (4414A>G), K339R (4421A>G), H344A (4435C>G 4436A>C), K346R (4442A>G 4443G>A), G347K (4444G>A 4445G>A), G349H (4450G>C 4451G>A), L351P (4457T>C), L352V (4459C>G 4461T>A), N354F (4465A>T 4466A>T), K355I (4469A>T 4470G>T), E356G (4472A>G 4473G>T), L358M (4477C>A), Q360D (4483C>G 4485A>C), I362L (4489A>T 4491C>G), G363A (4493G>C 4494A>T), K365Q (4498A>C 4500G>A), T366D (4501A>G 4502C>A 4503C>T), F367Y (4505T>A), S369C (4511C>G 4512T>C), S370F (4514C>T), F371L (4516T>C 4518T>A), C373G (4522T>G 4524T>G), K374Y (4525A>T 4527A>C), F377Y (4535T>A), W378N (4537T>A 4538G>A 4539G>T), V380I (4543G>A 4545A>C), R381T (4546C>A 4547G>C 4548C>A), L382I (4549C>A), T386D (4561A>G 4562C>A 4563A>C), I387Q (4564A>C 4565T>A 4566T>G), Q388E (4567C>G 4569G>A), L389K (4570C>A 4571T>A 4572A>G), A391T (4576G>A), S393T (4583G>C 4584C>T), Q396Y (4591C>T 4593A>T), H398T (4597C>A 4598A>C 4599C>A), E400A (4604A>C), W401F (4607G>T 4608G>C), L402K (4609T>A 4610T>A 4611A>G), V403R (4612G>C 4613T>G 4614C>G), K409C (4630A>T 4631A>G 4632A>C), Q410N (4633C>A 4635A>T), I414T (4646T>C 4647T>A), H418C (4657C>T 4658A>G) | | | | | | | | |
| Codon mutations: | GTA335.TC (4410A>C), ATT336ATG (4413T>G), AAC337GAC (4414A>G), TAT338TAC (4419T>C), AAA339AGA (4421A>G), TTA341CTT (4426T>C 4428A>T), GAC343GAT (4434C>T), CAT344GCT (4435C>G 4436A>C), AAG346AGA (4442A>G 4443G>A), GGA347AAA (4444G>A 4445G>A), GGC349CAC (4450G>C 4451G>A), CTA351CCA (4457T>C), CTT352GTA (4459C>G 4461T>A), CCT353CCC (4464T>C), AAC354TTC (4465A>T 4466A>T), AAG355ATT (4469A>T 4470G>T), GAG356GGT (4472A>G 4473G>T), CTG358ATG (4477C>A), CTT359CTA (4482T>A), CAA360GAC (4483C>G 4485A>C), ATC362TTG (4489A>T 4491C>G), GGA363GCT (4493G>C 4494A>T), GGT364GGG (4497T>G), AAG365CAA (4498A>C 4500G>A), ACC366GAT (4501A>G 4502C>A 4503C>T), TTT367TAT (4505T>A), TAC368TAT (4509C>T), TCT369TGC (4511C>G 4512T>C), TCC370TTC (4514C>T), TTT371CTA (4516T>C 4518T>A), GAC372GAT (4521C>T), TGT373GGG (4522T>G 4524T>G), AAA374TAC (4525A>T 4527A>C), TCT375TCA (4530T>A), TTT377TAT (4535T>A), TGG378AAT (4537T>A 4538G>A 4539G>T), GTA380ATC (4543G>A 4545A>C), CGC381ACA (4546C>A 4547G>C 4548C>A), CTT382ATT (4549C>A), GCT383GCG (4554T>G), GAA385GAG (4560A>G), ACA386GAC (4561A>G 4562C>A 4563A>C), ATT387CAG (4564A>C 4565T>A 4566T>G), CAG388GAA (4567C>G 4569G>A), CTA389AAG (4570C>A 4571T>A 4572A>G), GCT391ACT (4576G>A), AGC393ACT (4583G>C 4584C>T), TGT394TGC (4587T>C), CCC395CCT (4590C>T), CAA396TAT (4591C>T 4593A>T), CAC398ACA (4597C>A 4598A>C 4599C>A), TAC399TAT (4602C>T), GAA400GCA (4604A>C), TGG401TTC (4607G>T 4608G>C), TTA402AAG (4609T>A 4610T>A 4611A>G), GTC403CGG (4612G>C 4613T>G 4614C>G), GGA407GGG (4626A>G), AAA409TGC (4630A>T 4631A>G 4632A>C), CAA410AAT (4633C>A 4635A>T), GCC411GCA (4638C>A), CCT412CCG (4641T>G), GCT413GCG (4644T>G), ATT414ACA (4646T>C 4647T>A), TTC415TTT (4650C>T), AGG417AGA (4656G>A), CAC418TGC (4657C>T 4658A>G) | | | | | | | | |
| **Proteins** | | | | | | | | | |
| hypothetical protein (NP_043933.1) | 336 | 422 | 12.3% | 295 | 44.8% | 87 (100%) | 41 (47.1%) | 0/0/0/0 | 0 |
| Protein mutations: | I336M (4413T>G), N337D (4414A>G), K339R (4421A>G), H344A (4435C>G 4436A>C), K346R (4442A>G 4443G>A), G347K (4444G>A 4445G>A), G349H (4450G>C 4451G>A), L351P (4457T>C), L352V (4459C>G 4461T>A), N354F (4465A>T 4466A>T), K355I (4469A>T 4470G>T), E356G (4472A>G 4473G>T), L358M (4477C>A), Q360D (4483C>G 4485A>C), I362L (4489A>T 4491C>G), G363A (4493G>C 4494A>T), K365Q (4498A>C 4500G>A), T366D (4501A>G 4502C>A 4503C>T), F367Y (4505T>A), S369C (4511C>G 4512T>C), S370F (4514C>T), F371L (4516T>C 4518T>A), C373G (4522T>G 4524T>G), K374Y (4525A>T 4527A>C), F377Y (4535T>A), W378N (4537T>A 4538G>A 4539G>T), V380I (4543G>A 4545A>C), R381T (4546C>A 4547G>C 4548C>A), L382I (4549C>A), T386D (4561A>G 4562C>A 4563A>C), I387Q (4564A>C 4565T>A 4566T>G), Q388E (4567C>G 4569G>A), L389K (4570C>A 4571T>A 4572A>G), A391T (4576G>A), S393T (4583G>C 4584C>T), Q396Y (4591C>T 4593A>T), H398T (4597C>A 4598A>C 4599C>A), E400A (4604A>C), W401F (4607G>T 4608G>C), L402K (4609T>A 4610T>A 4611A>G), V403R (4612G>C 4613T>G 4614C>G), K409C (4630A>T 4631A>G 4632A>C), Q410N (4633C>A 4635A>T), I414T (4646T>C 4647T>A), H418C (4657C>T 4658A>G) | | | | | | | | |
| Codon mutations: | GTA335.TC (4410A>C), ATT336ATG (4413T>G), AAC337GAC (4414A>G), TAT338TAC (4419T>C), AAA339AGA (4421A>G), TTA341CTT (4426T>C 4428A>T), GAC343GAT (4434C>T), CAT344GCT (4435C>G 4436A>C), AAG346AGA (4442A>G 4443G>A), GGA347AAA (4444G>A 4445G>A), GGC349CAC (4450G>C 4451G>A), CTA351CCA (4457T>C), CTT352GTA (4459C>G 4461T>A), CCT353CCC (4464T>C), AAC354TTC (4465A>T 4466A>T), AAG355ATT (4469A>T 4470G>T), GAG356GGT (4472A>G 4473G>T), CTG358ATG (4477C>A), CTT359CTA (4482T>A), CAA360GAC (4483C>G 4485A>C), ATC362TTG (4489A>T 4491C>G), GGA363GCT (4493G>C 4494A>T), GGT364GGG (4497T>G), AAG365CAA (4498A>C 4500G>A), ACC366GAT (4501A>G 4502C>A 4503C>T), TTT367TAT (4505T>A), TAC368TAT (4509C>T), TCT369TGC (4511C>G 4512T>C), TCC370TTC (4514C>T), TTT371CTA (4516T>C 4518T>A), GAC372GAT (4521C>T), TGT373GGG (4522T>G 4524T>G), AAA374TAC (4525A>T 4527A>C), TCT375TCA (4530T>A), TTT377TAT (4535T>A), TGG378AAT (4537T>A 4538G>A 4539G>T), GTA380ATC (4543G>A 4545A>C), CGC381ACA (4546C>A 4547G>C 4548C>A), CTT382ATT (4549C>A), GCT383GCG (4554T>G), GAA385GAG (4560A>G), ACA386GAC (4561A>G 4562C>A 4563A>C), ATT387CAG (4564A>C 4565T>A 4566T>G), CAG388GAA (4567C>G 4569G>A), CTA389AAG (4570C>A 4571T>A 4572A>G), GCT391ACT (4576G>A), AGC393ACT (4583G>C 4584C>T), TGT394TGC (4587T>C), CCC395CCT (4590C>T), CAA396TAT (4591C>T 4593A>T), CAC398ACA (4597C>A 4598A>C 4599C>A), TAC399TAT (4602C>T), GAA400GCA (4604A>C), TGG401TTC (4607G>T 4608G>C), TTA402AAG (4609T>A 4610T>A 4611A>G), GTC403CGG (4612G>C 4613T>G 4614C>G), GGA407GGG (4626A>G), AAA409TGC (4630A>T 4631A>G 4632A>C), CAA410AAT (4633C>A 4635A>T), GCC411GCA (4638C>A), CCT412CCG (4641T>G), GCT413GCG (4644T>G), ATT414ACA (4646T>C 4647T>A), TTC415TTT (4650C>T), AGG417AGA (4656G>A), CAC418TGC (4657C>T 4658A>G) | | | | | | | | |

*: Inserts / Deletes / Misaligned / Frameshifts

#### Analysis details

This analysis was performed with panviral2.64

### NGS Details (UN24): Badnavirus volubetulae

#### Assembly

| | |
|---|---|
| **Coverage Length** | 244 (1 contig(s)) |
| **Depth Of Coverage** | 27.0 |
| **Number Of Reads** | 63 |
| **Reads Per Million** | 1.26 rpm (after QC) |
| **Ambiguities** | 0 |
| **Assembly Method** | de novo + reference guided assembly |
| **Consensus Caller** | Bcf Tools |

#### Coverage Map

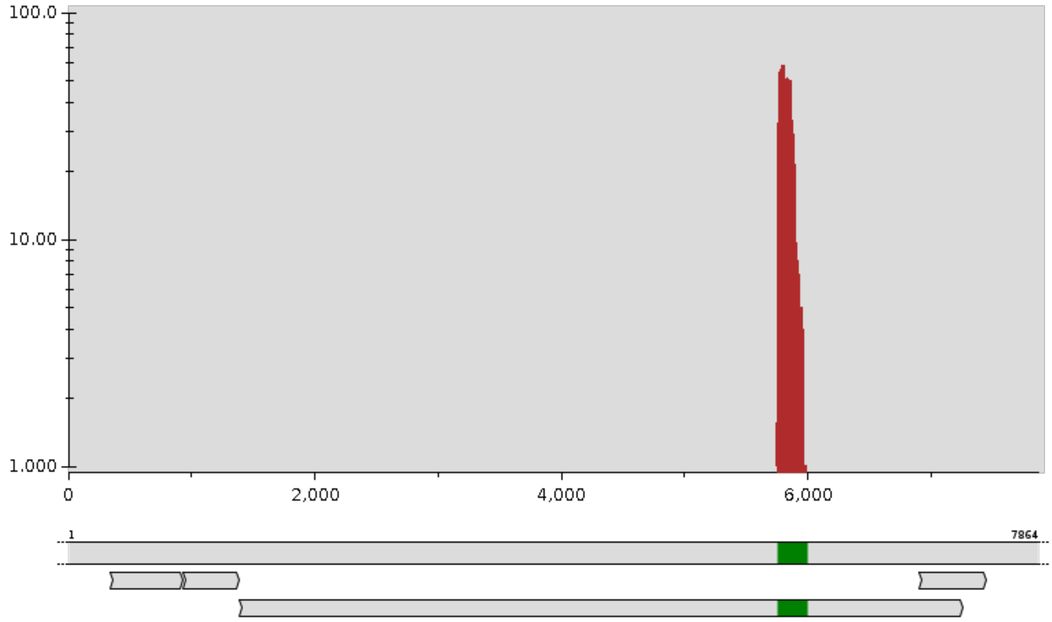

#### Assignment

| | |
|---|---|
| **Type** | Badnavirus volubetulae (Taxonomy ID: 3047683) |
| **Reference Genome** | NC_040635.1 |
| **NT Identity (%)** | 54.0984 |
| **AA Identity (%)** | 49.3827 |
| **Number Of Stop Codons** | 1 |
| **Number Of CDS** | 4 |

#### Alignment

| | |
|---|---|
| **Alignment Score** | 40.0 (NT) + 267.0 (AA) = 307.0 |
| **Concordance (%)** | 29.0445 |

| Alignment Method | Global, seeded, nucleotide + amino acids (AGA) |
|---|---|

#### Genome Region

Sequence starts at position 5753 and ends at position 5996 relative to NC_040635.1 reference sequence.

#### Alignment Detailed Statistics

| | Begin | End | Coverage | Score | Concordance | Matches | Identities | I/D/M/F* | Stop Codons |
|---|---|---|---|---|---|---|---|---|---|
| **NT** | **5753** | **5996** | **3.1%** | **40** | **8.2%** | **244 (100%)** | **132 (54.1%)** | **0/0** | |
| Mutations: | 5755G>T, 5756T>A, 5757C>T, 5758C>A, 5760C>T, 5770A>C, 5772T>G, 5773C>T, 5779C>A, 5781G>A, 5782A>G, 5783A>T, 5784C>G, 5788C>A, 5789A>T, 5790C>A, 5794G>A, 5797C>A, 5799A>G, 5802C>T, 5803A>C, 5804G>C, 5806C>T, 5808A>G, 5811A>G, 5812G>C, 5814A>T, 5817C>T, 5820C>T, 5821A>G, 5822C>A, 5823T>C, 5824A>T, 5826T>G, 5827G>T, 5829C>T, 5830C>G, 5832G>T, 5834G>A, 5835C>G, 5836A>C, 5839G>C, 5840G>A, 5841T>G, 5842A>G, 5843A>G, 5844T>A, 5847A>T, 5848A>T, 5849A>C, 5850G>C, 5853A>G, 5855T>C, 5859A>T, 5862A>G, 5863T>A, 5868T>C, 5869C>T, 5871T>G, 5873A>G, 5875A>T, 5876G>C, 5877T>G, 5880C>T, 5882T>A, 5889A>G, 5890G>T, 5891T>A, 5892C>G, 5894C>G, 5895C>T, 5896A>G, 5898G>T, 5899G>C, 5900A>G, 5901T>G, 5904A>G, 5905G>T, 5906A>C, 5908T>G, 5909C>A, 5910A>T, 5913A>C, 5916A>G, 5917T>A, 5918G>A, 5922A>T, 5923G>A, 5925A>T, 5928C>T, 5930T>G, 5931C>G, 5935C>T, 5936C>G, 5937A>C, 5938G>T, 5943G>T, 5944T>C, 5945T>A, 5946G>T, 5949C>T, 5954G>T, 5955G>T, 5956C>A, 5958A>G, 5965C>T, 5967A>C, 5973T>G, 5974C>T, 5978A>C, 5979A>C, 5988A>T | | | | | | | | |

##### CDS

| | Begin | End | Coverage | Score | Concordance | Matches | Identities | I/D/M/F* | Stop Codons |
|---|---|---|---|---|---|---|---|---|---|
| BLRaV_gp3 | 1456 | 1536 | 4.1% | 267 | 45.6% | 81 (100%) | 40 (49.4%) | 0/0/0/0 | 1 |
| Protein mutations: | V1456Y (5755G>T 5756T>A 5757C>T), L1457I (5758C>A 5760C>T), S1461R (5770A>C 5772T>G), Q1464K (5779C>A 5781G>A), N1465V (5782A>G 5783A>T 5784C>G), H1467I (5788C>A 5789A>T 5790C>A), D1469N (5794G>A), Q1470K (5797C>A 5799A>G), S1472P (5803A>C 5804G>C), G1475R (5812G>C 5814A>T), T1478D (5821A>G 5822C>A 5823T>C), I1479L (5824A>T 5826T>G), V1480F (5827G>T 5829C>T), Q1481D (5830C>G 5832G>T), R1482Q (5834G>A 5835C>G), I1483L (5836A>C), G1484Q (5839G>C 5840G>A 5841T>G), N1485G (5842A>G 5843A>G 5844T>A), K1487S (5848A>T 5849A>C 5850G>C), F1489S (5855T>C), F1492I (5863T>A), K1495R (5873A>G), F1498Y (5882T>A), V1501* (5890G>T 5891T>A 5892C>G), T1502S (5894C>G 5895C>T), M1503V (5896A>G 5898G>T), D1504R (5899G>C 5900A>G 5901T>G), E1506S (5905G>T 5906A>C), S1507D (5908T>G 5909C>A 5910A>T), W1510K (5917T>A 5918G>A), A1512T (5923G>A 5925A>T), L1514R (5930T>G 5931C>G), P1516C (5935C>T 5936C>G 5937A>C), D1517Y (5938G>T), L1519H (5944T>C 5945T>A 5946G>T), W1522F (5954G>T 5955G>T), L1523M (5956C>A 5958A>G), P1526S (5965C>T 5967A>C), L1529F (5974C>T), K1530T (5978A>C 5979A>C) | | | | | | | | |
| Codon mutations: | GTC1456TAT (5755G>T 5756T>A 5757C>T), CTC1457ATT (5758C>A 5760C>T), AGT1461CGG (5770A>C 5772T>G), CTG1462TTG (5773C>T), CAG1464AAA (5779C>A 5781G>A), AAC1465GTG (5782A>G 5783A>T 5784C>G), CAC1467ATA (5788C>A 5789A>T 5790C>A), GAT1469AAT (5794G>A), CAA1470AAG (5797C>A 5799A>G), TAC1471TAT (5802C>T), AGC1472CCC (5803A>C 5804G>C), CTA1473TTG (5806C>T 5808A>G), CCA1474CCG (5811A>G), GGA1475CGT (5812G>C 5814A>T), ATC1476ATT (5817C>T), AAC1477AAT (5820C>T), ACT1478GAC (5821A>G 5822C>A 5823T>C), ATT1479TTG (5824A>T 5826T>G), GTC1480TTT (5827G>T 5829C>T), CAG1481GAT (5830C>G 5832G>T), CGC1482CAG (5834G>A 5835C>G), ATA1483CTA (5836A>C), GGT1484CAG (5839G>C 5840G>A 5841T>G), AAT1485GGA (5842A>G 5843A>G 5844T>A), GCA1486GCT (5847A>T), AAG1487TCC (5848A>T 5849A>C 5850G>C), GTA1488GTG (5853A>G), TTC1489TCC (5855T>C), TCA1490TCT (5859A>T), AAA1491AAG (5862A>G), TTC1492ATC (5863T>A), GAT1493GAC (5868T>C), CTT1494TTG (5869C>T 5871T>G), AAG1495AGG (5873A>G), AGT1496TCG (5875A>T 5876G>C 5877T>G), GGC1497GGT (5880C>T), TTT1498TAT (5882T>A), CAA1500CAG (5889A>G), GTC1501TAG (5890G>T 5891T>A 5892C>G), ACC1502AGT (5894C>G 5895C>T), ATG1503GTT (5896A>G 5898G>T), GAT1504CGG (5899G>C 5900A>G 5901T>G), GAA1505GAG (5904A>G), GAA1506TCA (5905G>T 5906A>C), TCA1507GAT (5908T>G 5909C>A 5910A>T), ATA1508ATC (5913A>C), CCA1509CCG (5916A>G), TGG1510AAG (5917T>A 5918G>A), ACA1511ACT (5922A>T), GCA1512ACT (5923G>A 5925A>T), TTC1513TTT (5928C>T), CTC1514CGG (5930T>G 5931C>G), CCA1516TGC (5935C>T 5936C>G 5937A>C), GAT1517TAT (5938G>T), GGG1518GGT (5943G>T), TTG1519CAT (5944T>C 5945T>A 5946G>T), TAC1520TAT (5949C>T), TGG1522TTT (5954G>T 5955G>T), CTA1523ATG (5956C>A 5958A>G), CCA1526TCC (5965C>T 5967A>C), GGT1528GGG (5973T>G), CTT1529TTT (5974C>T), AAA1530ACC (5978A>C 5979A>C), CCA1533CCT (5988A>T) | | | | | | | | |

##### Proteins

| | Begin | End | Coverage | Score | Concordance | Matches | Identities | I/D/M/F* | Stop Codons |
|---|---|---|---|---|---|---|---|---|---|
| polyprotein (YP_009552737.1) | 1456 | 1536 | 4.1% | 267 | 45.6% | 81 (100%) | 40 (49.4%) | 0/0/0/0 | 1 |
| Protein mutations: | V1456Y (5755G>T 5756T>A 5757C>T), L1457I (5758C>A 5760C>T), S1461R (5770A>C 5772T>G), Q1464K (5779C>A 5781G>A), N1465V (5782A>G 5783A>T 5784C>G), H1467I (5788C>A 5789A>T 5790C>A), D1469N (5794G>A), Q1470K (5797C>A 5799A>G), S1472P (5803A>C 5804G>C), G1475R (5812G>C 5814A>T), T1478D (5821A>G 5822C>A 5823T>C), I1479L (5824A>T 5826T>G), V1480F (5827G>T 5829C>T), Q1481D (5830C>G 5832G>T), R1482Q (5834G>A 5835C>G), I1483L (5836A>C), G1484Q (5839G>C 5840G>A 5841T>G), N1485G (5842A>G 5843A>G 5844T>A), K1487S (5848A>T 5849A>C 5850G>C), F1489S (5855T>C), F1492I (5863T>A), K1495R (5873A>G), F1498Y (5882T>A), V1501* (5890G>T 5891T>A 5892C>G), T1502S (5894C>G 5895C>T), M1503V (5896A>G 5898G>T), D1504R (5899G>C 5900A>G 5901T>G), E1506S (5905G>T 5906A>C), S1507D (5908T>G 5909C>A 5910A>T), W1510K (5917T>A 5918G>A), A1512T (5923G>A 5925A>T), L1514R (5930T>G 5931C>G), P1516C (5935C>T 5936C>G 5937A>C), D1517Y (5938G>T), L1519H (5944T>C 5945T>A 5946G>T), W1522F (5954G>T 5955G>T), L1523M (5956C>A 5958A>G), P1526S (5965C>T 5967A>C), L1529F (5974C>T), K1530T (5978A>C 5979A>C) | | | | | | | | |
| Codon mutations: | GTC1456TAT (5755G>T 5756T>A 5757C>T), CTC1457ATT (5758C>A 5760C>T), AGT1461CGG (5770A>C 5772T>G), CTG1462TTG (5773C>T), CAG1464AAA (5779C>A 5781G>A), AAC1465GTG (5782A>G 5783A>T 5784C>G), CAC1467ATA (5788C>A 5789A>T 5790C>A), GAT1469AAT (5794G>A), CAA1470AAG (5797C>A 5799A>G), TAC1471TAT (5802C>T), AGC1472CCC (5803A>C 5804G>C), CTA1473TTG (5806C>T 5808A>G), CCA1474CCG (5811A>G), GGA1475CGT (5812G>C 5814A>T), ATC1476ATT (5817C>T), AAC1477AAT (5820C>T), ACT1478GAC (5821A>G 5822C>A 5823T>C), ATT1479TTG (5824A>T 5826T>G), GTC1480TTT (5827G>T 5829C>T), CAG1481GAT (5830C>G 5832G>T), CGC1482CAG (5834G>A 5835C>G), ATA1483CTA (5836A>C), GGT1484CAG (5839G>C 5840G>A 5841T>G), AAT1485GGA (5842A>G 5843A>G 5844T>A), GCA1486GCT (5847A>T), AAG1487TCC (5848A>T 5849A>C 5850G>C), GTA1488GTG (5853A>G), TTC1489TCC (5855T>C), TCA1490TCT (5859A>T), AAA1491AAG (5862A>G), TTC1492ATC (5863T>A), GAT1493GAC (5868T>C), CTT1494TTG (5869C>T 5871T>G), AAG1495AGG (5873A>G), AGT1496TCG (5875A>T 5876G>C 5877T>G), GGC1497GGT (5880C>T), TTT1498TAT (5882T>A), CAA1500CAG (5889A>G), GTC1501TAG (5890G>T 5891T>A 5892C>G), ACC1502AGT (5894C>G 5895C>T), ATG1503GTT (5896A>G 5898G>T), GAT1504CGG (5899G>C 5900A>G 5901T>G), GAA1505GAG (5904A>G), GAA1506TCA (5905G>T 5906A>C), TCA1507GAT (5908T>G 5909C>A 5910A>T), ATA1508ATC (5913A>C), CCA1509CCG (5916A>G), TGG1510AAG (5917T>A 5918G>A), ACA1511ACT (5922A>T), GCA1512ACT (5923G>A 5925A>T), TTC1513TTT (5928C>T), CTC1514CGG (5930T>G 5931C>G), CCA1516TGC (5935C>T 5936C>G 5937A>C), GAT1517TAT (5938G>T), GGG1518GGT (5943G>T), TTG1519CAT (5944T>C 5945T>A 5946G>T), TAC1520TAT (5949C>T), TGG1522TTT (5954G>T 5955G>T), CTA1523ATG (5956C>A 5958A>G), CCA1526TCC (5965C>T 5967A>C), GGT1528GGG (5973T>G), CTT1529TTT (5974C>T), AAA1530ACC (5978A>C 5979A>C), CCA1533CCT (5988A>T) | | | | | | | | |

*: Inserts / Deletes / Misaligned / Frameshifts

#### Analysis details

This analysis was performed with panviral2.64

### NGS Details (UN24): Yellowstone lake phycodnavirus 1

#### Assembly

| | |
|---|---|
| **Coverage Length** | 135 (1 contig(s)) |
| **Depth Of Coverage** | 35.6 |
| **Number Of Reads** | 55 |
| **Reads Per Million** | 1.10 rpm (after QC) |
| **Ambiguities** | 0 |
| **Assembly Method** | de novo + reference guided assembly |
| **Consensus Caller** | Bcf Tools |

#### Coverage Map

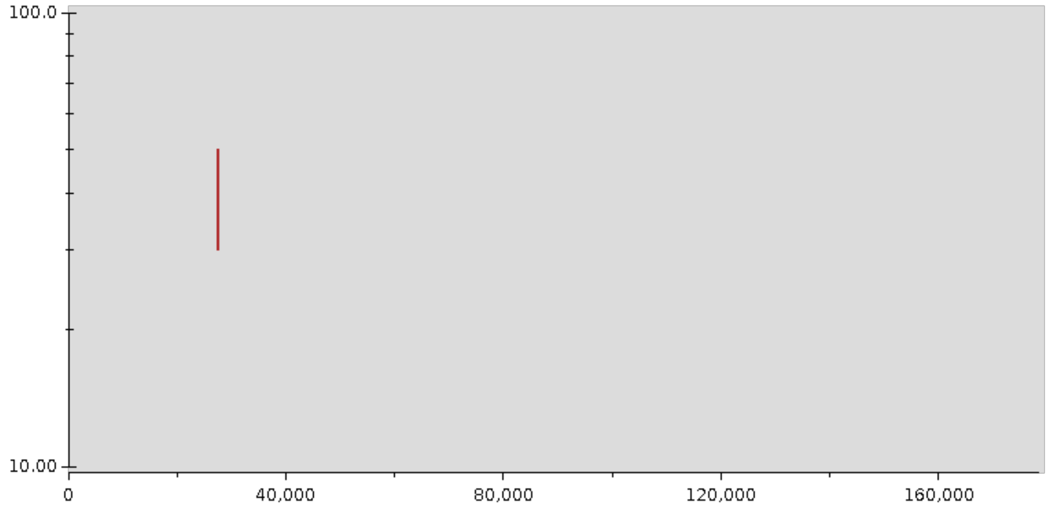

#### Assignment

| | |
|---|---|
| **Type** | Yellowstone lake phycodnavirus 1 (Taxonomy ID: 1586713) |
| **Reference Genome** | NC_028112.1 |
| **NT Identity (%)** | 83.7037 |
| **AA Identity (%)** | 93.3333 |
| **Number Of Stop Codons** | 0 |
| **Number Of CDS** | 248 |

#### Alignment

| | |
|---|---|
| **Alignment Score** | 182.0 (NT) + 273.0 (AA) = 455.0 |
| **Concordance (%)** | 82.13 |
| **Alignment Method** | Local, heuristic, nucleotide (BLASTN) |

#### Genome Region

Sequence starts at position 27473 and ends at position 27607 relative to NC_028112.1 reference sequence.

#### Alignment Detailed Statistics

| | Begin | End | Coverage | Score | Concordance | Matches | Identities | I/D/M/F* | Stop Codons |
|---|---|---|---|---|---|---|---|---|---|
| **NT** | **27473** | **27607** | **0.1%** | **182** | **67.4%** | **135 (100%)** | **113 (83.7%)** | **0/0** | |

Mutations: 27486G>A, 27492A>G, 27507G>A, 27510A>G, 27513T>C, 27529C>A, 27531T>G, 27534A>G, 27537T>C, 27540T>C, 27552A>G, 27563A>G, 27564T>C, 27567T>C, 27570C>T, 27573A>T, 27576A>G, 27582T>C, 27586G>A, 27591A>G, 27597A>G, 27600C>A

*: Inserts / Deletes / Misaligned / Frameshifts

#### Analysis details

This analysis was performed with panviral2.64

### NGS Details (UN24): Caulimovirus tessellobrassicae

#### Assembly

| | |
|---|---|
| **Coverage Length** | 812 (1 contig(s)) |
| **Depth Of Coverage** | 4.8 |
| **Number Of Reads** | 32 |
| **Reads Per Million** | 0.64 rpm (after QC) |
| **Ambiguities** | 0 |
| **Assembly Method** | de novo + reference guided assembly |
| **Consensus Caller** | Bcf Tools |

#### Coverage Map

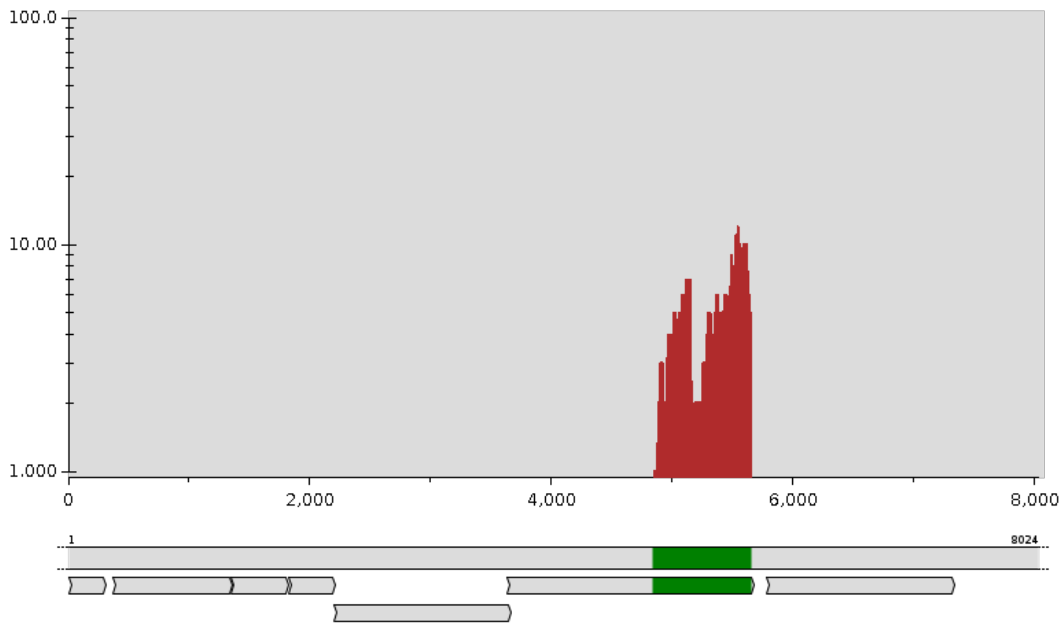

#### Assignment

| | |
|---|---|
| **Type** | Caulimovirus tessellobrassicae (Taxonomy ID: 3047738) |
| **Reference Genome** | NC_001497.2 |
| **NT Identity (%)** | 55.4293 |
| **AA Identity (%)** | 44.9057 |
| **Number Of Stop Codons** | 1 |
| **Number Of CDS** | 7 |

#### Alignment

| | |
|---|---|
| **Alignment Score** | 116.0 (NT) + 668.0 (AA) = 784.0 |
| **Concordance (%)** | 23.3403 |

| **Alignment Method** | Global, seeded, nucleotide + amino acids (AGA) |
|---|---|

#### Genome Region

Sequence starts at position 4845 and ends at position 5656 relative to NC_001497.2 reference sequence.

#### Alignment Detailed Statistics

| | Begin | End | Coverage | Score | Concordance | Matches | Identities | I/D/M/F* | Stop Codons |
|---|---|---|---|---|---|---|---|---|---|
| **NT** | **4845** | **5656** | **10.1%** | **116** | **7.6%** | **788 (96.6%)** | **439 (53.8%)** | **4/24** | |

Mutations: 4849C>T, 4851C>G, 4856T>G, 4858A>T, 4859G>C, 4860T>A, 4862A>C, 4863C>A, 4865A>C, 4866C>A, 4867G>T, 4869A>T, 4870G>C, 4872A>G, 4875T>A, 4881A>T, 4882C>A, 4883T>C, 4884T>A, 4887C>T, 4888G>C, 4890A>T, 4891G>A, 4892C>A, 4894A>G, 4895T>A, 4896G>A, 4897A>T, 4899C>T, 4901T>C, 4903C>A, 4906A>C, 4907A>T, 4908G>C, 4912A>G, 4913A>T, 4914T>C, 4915C>A, 4917A>T, 4918C>A, 4922G>A, 4923A>C, 4926T>A, 4927A>G, 4929C>A, 4930C>T, 4932T>A, 4933T>A, 4934C>G, 4935C>T, 4936A>C, 4941G>A, 4948C>G, 4951C>A, 4953C>A, 4954T>A, 4955T>A, 4956C>T, 4959G>A, 4962G>T, 4965G>A, 4969A>G, 4971C>A, 4980T>A, 4981C>A, 4983A>G, 4984G>A, 4985A>T, 4986A>T, 4989A>T, 4990G>T, 4991A>C, 4992T>A, 4993G>A, 4996G>A, 4997G>A, 4998A>T, 4999A>G, 5000C>G, 5002C>A, 5003A>T, 5004T>A, 5005A>G, 5007G>A, 5008C>T, 5009C>T, 5010T>A, 5014G>A, 5015G>C, 5016A>T, 5019T>C, 5022C>A, 5024T>C, 5025G>C, 5026G>A, 5029C>A, 5031C>G, 5034C>A, 5035A>T, 5036A>T, 5037C>A, 5038A>G, 5043C>T, 5046C>T, 5049T>C, 5051C>A, 5052C>G, 5053C>T, 5055T>A, 5056G>A, 5057A>T, 5059G>A, 5060A>C, 5064G>A, 5065A>C, 5067G>A, 5068C>G, 5071C>A, 5073T>A, 5076G>A, 5082C>T, 5083T>C, 5085A>T, 5088C>A, 5089A>T, 5090T>G, 5091A>C, 5092C>T, 5096C>A, 5097A>C, 5103C>A, 5104T>G, 5105C>G, 5106G>A, 5109T>A, 5111A>T, 5115C>A, 5118G>A, 5119A>G, 5121G>A, 5122C>T, 5127T>A, 5128C>A, 5130A>G, 5132T>A, 5133C>G, 5139G>T, 5141C>T, 5145G>T, 5149_5151delGCC, 5154G>A, 5155C>A, 5156T>C, 5158A>C, 5161G>C, 5166C>A, 5167G>A, 5169T>C, 5170C>A, 5171C>A, 5174G>T, 5175_5176insATTA, 5176A>G, 5187A>T, 5190G>A, 5191G>C, 5196C>T, 5199C>T, 5201A>G, 5202C>T, 5203A>G, 5205G>A, 5206C>A, 5211G>A, 5212G>C, 5214G>C, 5217G>A, 5218A>G, 5221A>G, 5223T>A, 5224C>T, 5225T>G, 5226G>T, 5227C>A, 5228A>T, 5229A>G, 5230G>A, 5231G>A, 5232A>G, 5233T>C, 5238T>A, 5239C>A, 5240C>A, 5241A>G, 5242C>A, 5244A>T, 5245C>A, 5246A>G, 5247T>A, 5248C>T, 5249A>C, 5250T>A, 5253C>A, 5254T>G, 5255T>A, 5262G>A, 5265G>C, 5268G>T, 5272A>G, 5274C>A, 5275A>T, 5277C>A, 5278G>C, 5280G>A, 5283C>A, 5289A>C, 5292A>C, 5295C>T, 5296G>T, 5298C>T, 5299T>C, 5301C>T, 5309G>C, 5310T>A, 5311A>T, 5314T>C, 5317A>C, 5320G>A, 5322T>A, 5323_5325delATC, 5326A>G, 5328A>C, 5329A>C, 5331T>A, 5338_5355delGGTACTAATACTGAGTTA, 5361C>T, 5363G>A, 5364A>G, 5367C>T, 5368G>A, 5371T>A, 5372C>G, 5378G>C, 5379C>T, 5385A>C, 5387C>A, 5388T>A, 5392G>A, 5395A>G, 5396A>T, 5397G>T, 5403C>T, 5404C>T, 5405A>C, 5408G>C, 5409C>A, 5415C>A, 5421G>A, 5422A>T, 5423C>T, 5427G>A, 5430G>A, 5434A>G, 5438A>G, 5439T>A, 5440A>G, 5441C>G, 5442T>A, 5446A>C, 5448G>A, 5455A>T, 5456G>C, 5457T>G, 5458A>G, 5459T>C, 5462A>G, 5463T>C, 5464C>T, 5467A>T, 5468C>T, 5469T>A, 5472T>A, 5473G>A, 5474T>A, 5475T>A, 5476C>T, 5477A>C, 5478T>A, 5482C>G, 5484G>T, 5489G>A, 5490G>A, 5505T>A, 5506T>A, 5508C>A, 5509A>T, 5510A>C, 5511G>A, 5512A>G, 5514T>G, 5517C>G, 5518G>A, 5520T>A, 5521A>T, 5522A>T, 5524C>A, 5525T>A, 5529T>A, 5530T>C, 5531A>T, 5533A>C, 5534A>C, 5535A>T, 5536G>T, 5537G>C, 5538A>C, 5541T>A, 5542T>C, 5544G>C, 5545A>C, 5548C>T, 5549T>A, 5551G>A, 5557A>C, 5558A>T, 5559C>G, 5560A>C, 5561T>A, 5562C>T, 5568G>A, 5573C>T, 5574A>T, 5576G>T, 5581A>T, 5582G>C, 5583C>G, 5584C>T, 5585A>T, 5592A>T, 5596G>A, 5598T>A, 5599G>A, 5605C>A, 5607C>T, 5610T>A, 5619C>T, 5622C>T, 5625C>T, 5626C>T, 5627A>T, 5628C>T, 5634G>A, 5637C>T

##### CDS

| | Begin | End | Coverage | Score | Concordance | Matches | Identities | I/D/M/F* | Stop Codons |
|---|---|---|---|---|---|---|---|---|---|
| CaMVgp6 | 405 | 675 | 39.9% | 668 | 36.1% | 263 (96.3%) | 119 (43.6%) | 2/8/1/1 | 1 |

Protein mutations: F408C (4856T>G), N410T (4862A>C 4863C>A), N411T (4865A>C 4866C>A), E412Y (4867G>T 4869A>T), E413Q (4870G>C 4872A>G), D414E (4875T>A), L417T (4882C>A 4883T>C 4884T>A), V419L (4888G>C 4890A>T), A420K (4891G>A 4892C>A), M421E (4894A>G 4895T>A 4896G>A), I422F (4897A>T 4899C>T), L423S (4901T>C), Q424K (4903C>A), K425L (4906A>C 4907A>T 4908G>C), N427V (4912A>G 4913A>T 4914T>C), Q428N (4915C>A 4917A>T), H429N (4918C>A), G430D (4922G>A 4923A>C), I432V (4927A>G 4929C>A), K435Q (4936A>C), Q439E (4948C>G), L440I (4951C>A 4953C>A), F441N (4954T>A 4955T>A 4956C>T), K443N (4962G>T), N446E (4969A>G 4971C>A), L450M (4981C>A 4983A>G), E451I (4984G>A 4985A>T 4986A>T), D453S (4990G>T 4991A>C 4992T>A), E454K (4993G>A), G455N (4996G>A 4997G>A 4998A>T), T456G (4999A>G 5000C>G), H457I (5002C>A 5003A>T 5004T>A), K458E (5005A>G 5007G>A), P459L (5008C>T 5009C>T 5010T>A), G461T (5014G>A 5015G>C 5016A>T), L464S (5024T>C 5025G>C), E465K (5026G>A), H466K (5029C>A 5031C>G), N468L (5035A>T 5036A>T 5037C>A), K469E (5038A>G), T473K (5051C>A 5052C>G), E475I (5056G>A 5057A>T), D476T (5059G>A 5060A>C), K478Q (5065A>C 5067G>A), Q479E (5068C>G), L480I (5071C>A 5073T>A), I486C (5089A>T 5090T>G 5091A>C), T488N (5096C>A 5097A>C), S491G (5104T>G 5105C>G 5106G>A), D492E (5109T>A), Y493F (5111A>T), K496E (5119A>G 5121G>A), Q499K (5128C>A 5130A>G), I500K (5132T>A 5133C>G), K502N (5139G>T), P503L (5141C>T), A506del (5149_5151delGCC), L508T (5155C>A 5156T>C), K509Q (5158A>C), E510Q (5161G>C), N511K (5166C>A), V512I (5167G>A 5169T>C), P513K (5170C>A 5171C>A), W514L (5174G>T), W514_R515insIX (5175_5176insATTA), R515G (5176A>G), K518N (5187A>T), D520H (5191G>C), Y523C (5201A>G 5202C>T), M524V (5203A>G 5205G>A), Q525K (5206C>A), V527L (5212G>C 5214G>C), K529E (5218A>G), N530E (5221A>G 5223T>A), L531C (5224C>T 5225T>G 5226G>T), Q532M (5227C>A 5228A>T 5229A>G), G533K (5230G>A 5231G>A 5232A>G), F534L (5233T>C), P536K (5239C>A 5240C>A 5241A>G), L537I (5242C>A 5244A>T), H538R (5245C>A 5246A>G 5247T>A), H539S (5248C>T 5249A>C 5250T>A), L541E (5254T>G 5255T>A), E544D (5265G>C), K545N (5268G>T), I547V (5272A>G 5274C>A), I548L (5275A>T 5277C>A), E549Q (5278G>C 5280G>A), D555Y (5296G>T 5298C>T), Y556H (5299T>C 5301C>T), G559A (5309G>C 5310T>A), M560L (5311A>T), K562Q (5317A>C), A563T (5320G>A 5322T>A), I564del (5323_5325delATC), K565D (5326A>G 5328A>C), I566L (5329A>C 5331T>A), G569_L574del (5338_5355delGGTACTAATACTGAGTTA), R577K (5363G>A 5364A>G), A579T (5368G>A), S582T (5378G>C 5379C>T), K584N (5385A>C), A585E (5387C>A 5388T>A), E587K (5392G>A), K588V (5395A>G 5396A>T 5397G>T), H591S (5404C>T 5405A>C), S592T (5408G>C 5409C>A), D594E (5415C>A), T597L (5422A>T 5423C>T), I601V (5434A>G), N602R (5438A>G 5439T>A), T603G (5440A>G 5441C>G 5442T>A), K605Q (5446A>C 5448G>A), I609A (5458A>G 5459T>C), Y610C (5462A>G 5463T>C), T612L (5467A>T 5468C>T 5469T>A), V614K (5473G>A 5474T>A 5475T>A), H615S (5476C>T 5477A>C 5478T>A), L617V (5482C>G 5484G>T), R619K (5489G>A 5490G>A), H624Q (5505T>A), F625I (5506T>A 5508C>A), K626S (5509A>T 5510A>C 5511G>A), S627G (5512A>G 5514T>G), F628L (5517C>G), V629I (5518G>A 5520T>A), N630F (5521A>T 5522A>T), L631N (5524C>A 5525T>A), N632K (5529T>A), Y633L (5530T>C 5531A>T), K634P (5533A>C 5534A>C 5535A>T), G635S (5536G>T 5537G>C 5538A>C), D636E (5541T>A), S637P (5542T>C 5544G>C), K638Q (5545A>C), L639Y (5548C>T 5549T>A), G640R (5551G>A), N642L (5557A>C 5558A>T 5559C>G), I643H (5560A>C 5561T>A 5562C>T), W645* (5568G>A), A647V (5573C>T 5574A>T), W648L (5576G>T), H651F (5584C>T 5585A>T), D655K (5596G>A 5598T>A), V656I (5599G>A), H658N (5605C>A 5607C>T), H665F (5626C>T 5627A>T 5628C>T)

Codon mutations: CTC406TTG (4849C>T 4851C>G), TTC408TGC (4856T>G), AGT409TCA (4858A>T 4859G>C 4860T>A), AAC410ACA (4862A>C 4863C>A), AAC411ACA (4865A>C 4866C>A), GAA412TAT (4867G>T 4869A>T), GAA413CAG (4870G>C 4872A>G), GAT414GAA (4875T>A), CTA416CTT (4881A>T), CTT417ACA (4882C>A 4883T>C 4884T>A), CAC418CAT (4887C>T), GTA419CTT (4888G>C 4890A>T), GCA420AAA (4891G>A 4892C>A), ATG421GAA (4894A>G 4895T>A 4896G>A), ATC422TTT (4897A>T 4899C>T), TTA423TCA (4901T>C), CAA424AAA (4903C>A), AAG425CTC (4906A>C 4907A>T 4908G>C), AAT427GTC (4912A>G 4913A>T 4914T>C), CAA428AAT (4915C>A 4917A>T), CAT429AAT (4918C>A), GGA430GAC (4922G>A 4923A>C), ATT431ATA (4926T>A), ATC432GTA (4927A>G 4929C>A), CTT433TTA (4930C>T 4932T>A), TCC434AGT (4933T>A 4934C>G 4935C>T), AAG435CAG (4936A>C), AAG436AAA (4941G>A), CAA439GAA (4948C>G), CTC440ATA (4951C>A 4953C>A), TTC441AAT (4954T>A 4955T>A 4956C>T), AAG442AAA (4959G>A), AAG443AAT (4962G>T), AAG444AAA (4965G>A), AAC446GAA (4969A>G 4971C>A), GGT449GGA (4980T>A), CTA450ATG (4981C>A 4983A>G), GAA451ATT (4984G>A 4985A>T 4986A>T), ATA452ATT (4989A>T), GAT453TCA (4990G>T 4991A>C 4992T>A), GAA454AAA (4993G>A), GGA455AAT (4996G>A 4997G>A 4998A>T), ACA456GGA (4999A>G 5000C>G), CAT457ATA (5002C>A 5003A>T 5004T>A), AAG458GAA (5005A>G 5007G>A), CCT459TTA (5008C>T 5009C>T 5010T>A), GGA461ACT (5014G>A 5015G>C 5016A>T), CAT462CAC (5019T>C), ATC463ATA (5022C>A), TTG464TCC (5024T>C 5025G>C), GAA465AAA (5026G>A), CAC466AAG (5029C>A 5031C>G), ATC467ATA (5034C>A), AAC468TTA (5035A>T 5036A>T 5037C>A), AAG469GAG (5038A>G), TTC470TTT (5043C>T), CCC471CCT (5046C>T), GAT472GAC (5049T>C), ACC473AAG (5051C>A 5052C>G), CTT474TTA (5053C>T 5055T>A), GAA475ATA (5056G>A 5057A>T), GAC476ACC (5059G>A 5060A>C), AAG477AAA (5064G>A), AAG478CAA (5065A>C 5067G>A), CAA479GAA (5068C>G), CTT480ATA (5071C>A 5073T>A), CAG481CAA (5076G>A), TTC483TTT (5082C>T), TTA484CTT (5083T>C 5085A>T), GGC485GGA (5088C>A), ATA486TGC (5089A>T 5090T>G 5091A>C), CTA487TTA (5092C>T), ACA488AAC (5096C>A 5097A>C), GCC490GCA (5103C>A), TCG491GGA (5104T>G 5105C>G 5106G>A), GAT492GAA (5109T>A), TAC493TTC (5111A>T), ATC494ATA (5115C>A), CCG495CCA (5118G>A), AAG496GAA (5119A>G 5121G>A), CTA497TTA (5122C>T), GCT498GCA (5127T>A), CAA499AAG (5128C>A 5130A>G), ATC500AAG (5132T>A 5133C>G), AAG502AAT (5139G>T), CCT503CTT (5141C>T), CTG504CTT (5145G>T), GCC506del (5149_5151delGCC), AAG507AAA (5154G>A), CTT508ACT (5155C>A 5156T>C), AAA509CAA (5158A>C), GAA510CAA (5161G>C), AAC511AAA (5166C>A), GTT512ATC (5167G>A 5169T>C), CCA513AAA (5170C>A 5171C>A), TGG514TTG (5174G>T), TGG514_AGA515insATTA-- (5175_5176insATTA), AGA515GGA (5176A>G), AAA518AAT (5187A>T), GAG519GAA (5190G>A), GAT520CAT (5191G>C), ACC521ACT (5196C>T), CTC522CTT (5199C>T), TAC523TGT (5201A>G 5202C>T), ATG524GTA (5203A>G 5205G>A), CAA525AAA (5206C>A), AAG526AAA (5211G>A), GTG527CTC (5212G>C 5214G>C), AAG528AAA (5217G>A), AAA529GAA (5218A>G), AAT530GAA (5221A>G 5223T>A), CTG531TGT (5224C>T 5225T>G 5226G>T), CAA532ATG (5227C>A 5228A>T 5229A>G), GGA533AAG (5230G>A 5231G>A 5232A>G), TTT534CTT (5233T>C), CCT535CCA (5238T>A), CCA536AAG (5239C>A 5240C>A 5241A>G), CTA537ATT (5242C>A 5244A>T), CAT538AGA (5245C>A 5246A>G 5247T>A), CAT539TCA (5248C>T 5249A>C 5250T>A), CCC540CCA (5253C>A), TTA541GAA (5254T>G 5255T>A), GAG543GAA (5262G>A), GAG544GAC (5265G>C), AAG545AAT (5268G>T), ATC547GTA (5272A>G 5274C>A), ATC548TTA (5275A>T 5277C>A), GAG549CAA (5278G>C 5280G>A), ACC550ACA (5283C>A), GCA552GCC (5289A>C), TCA553TCC (5292A>C), GAC554GAT (5295C>T), GAC555TAT (5296G>T 5298C>T), TAC556CAT (5299T>C 5301C>T), GGT559GCA (5309G>C 5310T>A), ATG560TTG (5311A>T), TTA561CTA (5314T>C), AAA562CAA (5317A>C), GCT563ACA (5320G>A 5322T>A), ATC564del (5323_5325delATC), AAA565GAC (5326A>G 5328A>C), ATT566CTA (5329A>C 5331T>A), GGT569_TTA574del (5338_5355delGGTACTAATACTGAGTTA), TGC576TGT (5361C>T), AGA577AAG (5363G>A 5364A>G), TAC578TAT (5367C>T), GCA579ACA (5368G>A), TCT580AGT (5371T>A 5372C>G), AGC582ACT (5378G>C 5379C>T), AAA584AAC (5385A>C), GCT585GAA (5387C>A 5388T>A), GAA587AAA (5392G>A), AAG588GTT (5395A>G 5396A>T 5397G>T), TAC590TAT (5403C>T), CAC591TCC (5404C>T 5405A>C), AGC592ACA (5408G>C 5409C>A), GAC594GAA (5415C>A), GAG596GAA (5421G>A), ACA597TTA (5422A>T 5423C>T), TTG598TTA (5427G>A), GCG599GCA (5430G>A), ATA601GTA (5434A>G), AAT602AGA (5438A>G 5439T>A), ACT603GGA (5440A>G 5441C>G 5442T>A), AAG605CAA (5446A>C 5448G>A), AGT608TCG (5455A>T 5456G>C 5457T>G), ATT609GCT (5458A>G 5459T>C), TAT610TGC (5462A>G 5463T>C), CTA611TTA (5464C>T), ACT612TTA (5467A>T 5468C>T 5469T>A), CCT613CCA (5472T>A), GTT614AAA (5473G>A 5474T>A 5475T>A), CAT615TCA (5476C>T 5477A>C 5478T>A), CTG617GTT (5482C>G 5484G>T), AGG619AAA (5489G>A 5490G>A), CAT624CAA (5505T>A), TTC625ATA (5506T>A 5508C>A), AAG626TCA (5509A>T 5510A>C 5511G>A), AGT627GGG (5512A>G 5514T>G), TTC628TTG (5517C>G), GTT629ATA (5518G>A 5520T>A), AAT630TTT (5521A>T 5522A>T), CTC631AAC (5524C>A 5525T>A), AAT632AAA (5529T>A), TAC633CTC (5530T>C 5531A>T), AAA634CCT (5533A>C 5534A>C 5535A>T), GGA635TCC (5536G>T 5537G>C 5538A>C), GAT636GAA (5541T>A), TCG637CCC (5542T>C 5544G>C), AAA638CAA (5545A>C), CTT639TAT (5548C>T 5549T>A), GGA640AGA (5551G>A), AAC642CTG (5557A>C 5558A>T 5559C>G), ATC643CAT (5560A>C 5561T>A 5562C>T), TGG645TGA (5568G>A), GCA647GTT (5573C>T 5574A>T), TGG648TTG (5576G>T), AGC650TCG (5581A>T 5582G>C 5583C>G), CAC651TTC (5584C>T 5585A>T), TCA653TCT (5592A>T), GAT655AAA (5596G>A 5598T>A), GTT656ATT (5599G>A), CAC658AAT (5605C>A 5607C>T), ATT659ATA (5610T>A), ACC662ACT (5619C>T), GAC663GAT (5622C>T), AAC664AAT (5625C>T), CAC665TTT (5626C>T 5627A>T 5628C>T), GCG667GCA (5634G>A), GAC668GAT (5637C>T)

##### Proteins

| | Begin | End | Coverage | Score | Concordance | Matches | Identities | I/D/M/F* | Stop Codons |
|---|---|---|---|---|---|---|---|---|---|
| reverse transcriptase (NP_056728.1) | 405 | 675 | 39.9% | 668 | 36.1% | 263 (96.3%) | 119 (43.6%) | 2/8/1/1 | 1 |

| | Begin | End | Coverage | Score | Concordance | Matches | Identities | I/D/M/F* | Stop Codons |
|---|---|---|---|---|---|---|---|---|---|
| **NT** | **4845** | **5656** | **10.1%** | **116** | **7.6%** | **788 (96.6%)** | **439 (53.8%)** | **4/24** | |
| Protein mutations: | F408C (4856T>G), N410T (4862A>C 4863C>A), N411T (4865A>C 4866C>A), E412Y (4867G>T 4869A>T), E413Q (4870G>C 4872A>G), D414E (4875T>A), L417T (4882C>A 4883T>C 4884T>A), V419L (4888G>C 4890A>T), A420K (4891G>A 4892C>A), M421E (4894A>G 4895T>A 4896G>A), I422F (4897A>T 4899C>T), L423S (4901T>C), Q424K (4903C>A), K425L (4906A>C 4907A>T 4908G>C), N427V (4912A>G 4913A>T 4914T>C), Q428N (4915C>A 4917A>T), H429N (4918C>A), G430D (4922G>A 4923A>C), I432V (4927A>G 4929C>A), K435Q (4936A>C), Q439E (4948C>G), L440I (4951C>A 4953C>A), F441N (4954T>A 4955T>A 4956C>T), K443N (4962G>T), N446E (4969A>G 4971C>A), L450M (4981C>A 4983A>G), E451I (4984G>A 4985A>T 4986A>T), D453S (4990G>T 4991A>C 4992T>A), E454K (4993G>A), G455N (4996G>A 4997G>A 4998A>T), T456G (4999A>G 5000C>G), H457I (5002C>A 5003A>T 5004T>A), K458E (5005A>G 5007G>A), P459L (5008C>T 5009C>T 5010T>A), G461T (5014G>A 5015G>C 5016A>T), L464S (5024T>C 5025G>C), E465K (5026G>A), H466K (5029C>A 5031C>G), N468L (5035A>T 5036A>T 5037C>A), K469E (5038A>G), T473K (5051C>A 5052C>G), E475I (5056G>A 5057A>T), D476T (5059G>A 5060A>C), K478Q (5065A>C 5067G>A), Q479E (5068C>G), L480I (5071C>A 5073T>A), I486C (5089A>T 5090T>G 5091A>C), T488N (5096C>A 5097A>C), S491G (5104T>G 5105C>G 5106G>A), D492E (5109T>A), Y493F (5111A>T), K496E (5119A>G 5121G>A), Q499K (5128C>A 5130A>G), I500K (5132T>A 5133C>G), K502N (5139G>T), P503L (5141C>T), A506del (5149_5151delGCC), L508T (5155C>A 5156T>C), K509Q (5158A>C), E510Q (5161G>C), N511K (5166C>A), V512I (5167G>A 5169T>C), P513K (5170C>A 5171C>A), W514L (5174G>T), W514_R515insIX (5175_5176insATTA), R515G (5176A>G), K518N (5187A>T), D520H (5191G>C), Y523C (5201A>G 5202C>T), M524V (5203A>G 5205G>A), Q525K (5206C>A), V527L (5212G>C 5214G>C), K529E (5218A>G), N530E (5221A>G 5223T>A), L531C (5224C>T 5225T>G 5226G>T), Q532M (5227C>A 5228A>T 5229A>G), G533K (5230G>A 5231G>A 5232A>G), F534L (5233T>C), P536K (5239C>A 5240C>A 5241A>G), L537I (5242C>A 5244A>T), H538R (5245C>A 5246A>G 5247T>A), H539S (5248C>T 5249A>C 5250T>A), L541E (5254T>G 5255T>A), E544D (5265G>C), K545N (5268G>T), I547V (5272A>G 5274C>A), I548L (5275A>T 5277C>A), E549Q (5278G>C 5280G>A), D555Y (5296G>T 5298C>T), Y556H (5299T>C 5301C>T), G559A (5309G>C 5310T>A), M560L (5311A>T), K562Q (5317A>C), A563T (5320G>A 5322T>A), I564del (5323_5325delATC), K565D (5326A>G 5328A>C), I566L (5329A>C 5331T>A), G569_L574del (5338_5355delGGTACTAATACTGAGTTA), R577K (5363G>A 5364A>G), A579T (5368G>A), S582T (5378G>C 5379C>T), K584N (5385A>C), A585E (5387C>A 5388T>A), E587K (5392G>A), K588V (5395A>G 5396A>T 5397G>T), H591S (5404C>T 5405A>C), S592T (5408G>C 5409C>A), D594E (5415C>A), T597L (5422A>T 5423C>T), I601V (5434A>G), N602R (5438A>G 5439T>A), T603G (5440A>G 5441C>G 5442T>A), K605Q (5446A>C 5448G>A), I609A (5458A>G 5459T>C), Y610C (5462A>G 5463T>C), T612L (5467A>T 5468C>T 5469T>A), V614K (5473G>A 5474T>A 5475T>A), H615S (5476C>T 5477A>C 5478T>A), L617V (5482C>G 5484G>T), R619K (5489G>A 5490G>A), H624Q (5505T>A), F625I (5506T>A 5508C>A), K626S (5509A>T 5510A>C 5511G>A), S627G (5512A>G 5514T>G), F628L (5517C>G), V629I (5518G>A 5520T>A), N630F (5521A>T 5522A>T), L631N (5524C>A 5525T>A), N632K (5529T>A), Y633L (5530T>C 5531A>T), K634P (5533A>C 5534A>C 5535A>T), G635S (5536G>T 5537G>C 5538A>C), D636E (5541T>A), S637P (5542T>C 5544G>C), K638Q (5545A>C), L639Y (5548C>T 5549T>A), G640R (5551G>A), N642L (5557A>C 5558A>T 5559C>G), I643H (5560A>C 5561T>A 5562C>T), W645* (5568G>A), A647V (5573C>T 5574A>T), W648L (5576G>T), H651F (5584C>T 5585A>T), D655K (5596G>A 5598T>A), V656I (5599G>A), H658N (5605C>A 5607C>T), H665F (5626C>T 5627A>T 5628C>T) | | | | | | | | |
| Codon mutations: | CTC406TTG (4849C>T 4851C>G), TTC408TGC (4856T>G), AGT409TCA (4858A>T 4859G>C 4860T>A), AAC410ACA (4862A>C 4863C>A), AAC411ACA (4865A>C 4866C>A), GAA412TAT (4867G>T 4869A>T), GAA413CAG (4870G>C 4872A>G), GAT414GAA (4875T>A), CTA416CTT (4881A>T), CTT417ACA (4882C>A 4883T>C 4884T>A), CAC418CAT (4887C>T), GTA419CTT (4888G>C 4890A>T), GCA420AAA (4891G>A 4892C>A), ATG421GAA (4894A>G 4895T>A 4896G>A), ATC422TTT (4897A>T 4899C>T), TTA423TCA (4901T>C), CAA424AAA (4903C>A), AAG425CTC (4906A>C 4907A>T 4908G>C), AAT427GTC (4912A>G 4913A>T 4914T>C), CAA428AAT (4915C>A 4917A>T), CAT429AAT (4918C>A), GGA430GAC (4922G>A 4923A>C), ATT431ATA (4926T>A), ATC432GTA (4927A>G 4929C>A), CTT433TTA (4930C>T 4932T>A), TCC434AGT (4933T>A 4934C>G 4935C>T), AAG435CAG (4936A>C), AAG436AAA (4941G>A), CAA439GAA (4948C>G), CTC440ATA (4951C>A 4953C>A), TTC441AAT (4954T>A 4955T>A 4956C>T), AAG442AAA (4959G>A), AAG443AAT (4962G>T), AAG444AAA (4965G>A), AAC446GAA (4969A>G 4971C>A), GGT449GGA (4980T>A), CTA450ATG (4981C>A 4983A>G), GAA451ATT (4984G>A 4985A>T 4986A>T), ATA452ATT (4989A>T), GAT453TCA (4990G>T 4991A>C 4992T>A), GAA454AAA (4993G>A), GGA455AAT (4996G>A 4997G>A 4998A>T), ACA456GGA (4999A>G 5000C>G), CAT457ATA (5002C>A 5003A>T 5004T>A), AAG458GAA (5005A>G 5007G>A), CCT459TTA (5008C>T 5009C>T 5010T>A), GGA461ACT (5014G>A 5015G>C 5016A>T), CAT462CAC (5019T>C), ATC463ATA (5022C>A), TTG464TCC (5024T>C 5025G>C), GAA465AAA (5026G>A), CAC466AAG (5029C>A 5031C>G), ATC467ATA (5034C>A), AAC468TTA (5035A>T 5036A>T 5037C>A), AAG469GAG (5038A>G), TTC470TTT (5043C>T), CCC471CCT (5046C>T), GAT472GAC (5049T>C), ACC473AAG (5051C>A 5052C>G), CTT474TTA (5053C>T 5055T>A), GAA475ATA (5056G>A 5057A>T), GAC476ACC (5059G>A 5060A>C), AAG477AAA (5064G>A), AAG478CAA (5065A>C 5067G>A), CAA479GAA (5068C>G), CTT480ATA (5071C>A 5073T>A), CAG481CAA (5076G>A), TTC483TTT (5082C>T), TTA484CTT (5083T>C 5085A>T), GGC485GGA (5088C>A), ATA486TGC (5089A>T 5090T>G 5091A>C), CTA487TTA (5092C>T), ACA488AAC (5096C>A 5097A>C), GCC490GCA (5103C>A), TCG491GGA (5104T>G 5105C>G 5106G>A), GAT492GAA (5109T>A), TAC493TTC (5111A>T), ATC494ATA (5115C>A), CCG495CCA (5118G>A), AAG496GAA (5119A>G 5121G>A), CTA497TTA (5122C>T), GCT498GCA (5127T>A), CAA499AAG (5128C>A 5130A>G), ATC500AAG (5132T>A 5133C>G), AAG502AAT (5139G>T), CCT503CTT (5141C>T), CTG504CTT (5145G>T), GCC506del (5149_5151delGCC), AAG507AAA (5154G>A), CTT508ACT (5155C>A 5156T>C), AAA509CAA (5158A>C), GAA510CAA (5161G>C), AAC511AAA (5166C>A), GTT512ATC (5167G>A 5169T>C), CCA513AAA (5170C>A 5171C>A), TGG514TTG (5174G>T), TGG514_AGA515insATTA-- (5175_5176insATTA), AGA515GGA (5176A>G), AAA518AAT (5187A>T), GAG519GAA (5190G>A), GAT520CAT (5191G>C), ACC521ACT (5196C>T), CTC522CTT (5199C>T), TAC523TGT (5201A>G 5202C>T), ATG524GTA (5203A>G 5205G>A), CAA525AAA (5206C>A), AAG526AAA (5211G>A), GTG527CTC (5212G>C 5214G>C), AAG528AAA (5217G>A), AAA529GAA (5218A>G), AAT530GAA (5221A>G 5223T>A), CTG531TGT (5224C>T 5225T>G 5226G>T), CAA532ATG (5227C>A 5228A>T 5229A>G), GGA533AAG (5230G>A 5231G>A 5232A>G), TTT534CTT (5233T>C), CCT535CCA (5238T>A), CCA536AAG (5239C>A 5240C>A 5241A>G), CTA537ATT (5242C>A 5244A>T), CAT538AGA (5245C>A 5246A>G 5247T>A), CAT539TCA (5248C>T 5249A>C 5250T>A), CCC540CCA (5253C>A), TTA541GAA (5254T>G 5255T>A), GAG543GAA (5262G>A), GAG544GAC (5265G>C), AAG545AAT (5268G>T), ATC547GTA (5272A>G 5274C>A), ATC548TTA (5275A>T 5277C>A), GAG549CAA (5278G>C 5280G>A), ACC550ACA (5283C>A), GCA552GCC (5289A>C), TCA553TCC (5292A>C), GAC554GAT (5295C>T), GAC555TAT (5296G>T 5298C>T), TAC556CAT (5299T>C 5301C>T), GGT559GCA (5309G>C 5310T>A), ATG560TTG (5311A>T), TTA561CTA (5314T>C), AAA562CAA (5317A>C), GCT563ACA (5320G>A 5322T>A), ATC564del (5323_5325delATC), AAA565GAC (5326A>G 5328A>C), ATT566CTA (5329A>C 5331T>A), GGT569_TTA574del (5338_5355delGGTACTAATACTGAGTTA), TGC576TGT (5361C>T), AGA577AAG (5363G>A 5364A>G), TAC578TAT (5367C>T), GCA579ACA (5368G>A), TCT580AGT (5371T>A 5372C>G), AGC582ACT (5378G>C 5379C>T), AAA584AAC (5385A>C), GCT585GAA (5387C>A 5388T>A), GAA587AAA (5392G>A), AAG588GTT (5395A>G 5396A>T 5397G>T), TAC590TAT (5403C>T), CAC591TCC (5404C>T 5405A>C), AGC592ACA (5408G>C 5409C>A), GAC594GAA (5415C>A), GAG596GAA (5421G>A), ACA597TTA (5422A>T 5423C>T), TTG598TTA (5427G>A), GCG599GCA (5430G>A), ATA601GTA (5434A>G), AAT602AGA (5438A>G 5439T>A), ACT603GGA (5440A>G 5441C>G 5442T>A), AAG605CAA (5446A>C 5448G>A), AGT608TCG (5455A>T 5456G>C 5457T>G), ATT609GCT (5458A>G 5459T>C), TAT610TGC (5462A>G 5463T>C), CTA611TTA (5464C>T), ACT612TTA (5467A>T 5468C>T 5469T>A), CCT613CCA (5472T>A), GTT614AAA (5473G>A 5474T>A 5475T>A), CAT615TCA (5476C>T 5477A>C 5478T>A), CTG617GTT (5482C>G 5484G>T), AGG619AAA (5489G>A 5490G>A), CAT624CAA (5505T>A), TTC625ATA (5506T>A 5508C>A), AAG626TCA (5509A>T 5510A>C 5511G>A), AGT627GGG (5512A>G 5514T>G), TTC628TTG (5517C>G), GTT629ATA (5518G>A 5520T>A), AAT630TTT (5521A>T 5522A>T), CTC631AAC (5524C>A 5525T>A), AAT632AAA (5529T>A), TAC633CTC (5530T>C 5531A>T), AAA634CCT (5533A>C 5534A>C 5535A>T), GGA635TCC (5536G>T 5537G>C 5538A>C), GAT636GAA (5541T>A), TCG637CCC (5542T>C 5544G>C), AAA638CAA (5545A>C), CTT639TAT (5548C>T 5549T>A), GGA640AGA (5551G>A), AAC642CTG (5557A>C 5558A>T 5559C>G), ATC643CAT (5560A>C 5561T>A 5562C>T), TGG645TGA (5568G>A), GCA647GTT (5573C>T 5574A>T), TGG648TTG (5576G>T), AGC650TCG (5581A>T 5582G>C 5583C>G), CAC651TTC (5584C>T 5585A>T), TCA653TCT (5592A>T), GAT655AAA (5596G>A 5598T>A), GTT656ATT (5599G>A), CAC658AAT (5605C>A 5607C>T), ATT659ATA (5610T>A), ACC662ACT (5619C>T), GAC663GAT (5622C>T), AAC664AAT (5625C>T), CAC665TTT (5626C>T 5627A>T 5628C>T), GCG667GCA (5634G>A), GAC668GAT (5637C>T) | | | | | | | | |

*: Inserts / Deletes / Misaligned / Frameshifts

#### Analysis details

This analysis was performed with panviral2.64

### NGS Details (UN24): Pinus nigra virus 1

#### Assembly

| | |
|---|---|
| **Coverage Length** | 246 (1 contig(s)) |
| **Depth Of Coverage** | 10.8 |
| **Number Of Reads** | 24 |
| **Reads Per Million** | 0.48 rpm (after QC) |
| **Ambiguities** | 0 |
| **Assembly Method** | de novo + reference guided assembly |
| **Consensus Caller** | Bcf Tools |

#### Coverage Map

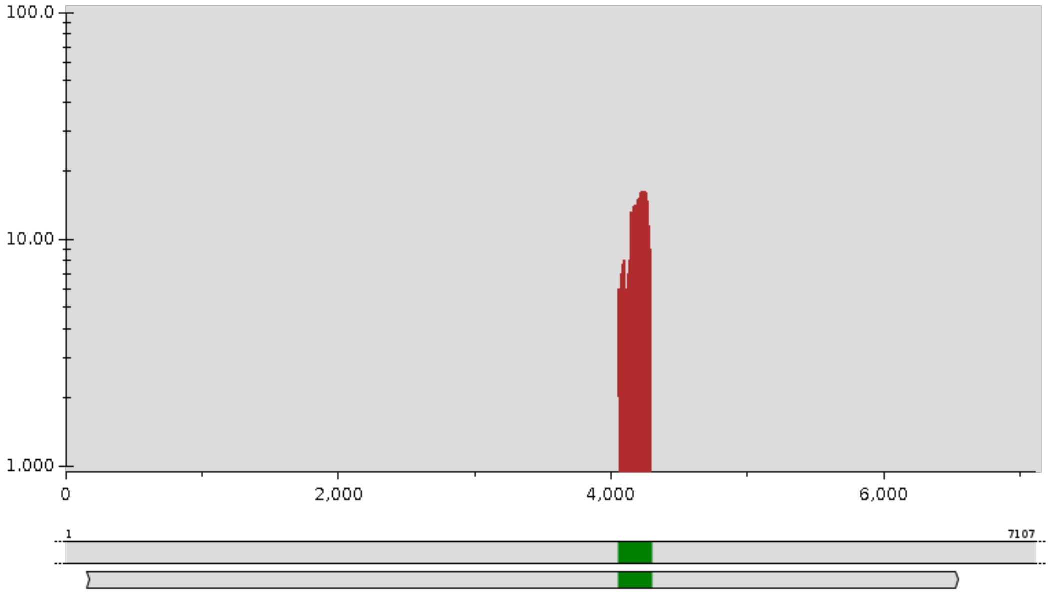

#### Assignment

| | |
|---|---|
| **Type** | Pinus nigra virus 1 (Taxonomy ID: 2267679) |
| **Reference Genome** | NC_040841.1 |
| **NT Identity (%)** | 54.3651 |
| **AA Identity (%)** | 40.4762 |
| **Number Of Stop Codons** | 0 |
| **Number Of CDS** | 1 |

#### Alignment

| | |
|---|---|
| **Alignment Score** | 41.0 (NT) + 281.0 (AA) = 322.0 |
| **Concordance (%)** | 28.9308 |

| Alignment Method | Global, seeded, nucleotide + amino acids (AGA) |
|---|---|

#### Genome Region

Sequence starts at position 4054 and ends at position 4299 relative to NC_040841.1 reference sequence.

#### Alignment Detailed Statistics

| | Begin | End | Coverage | Score | Concordance | Matches | Identities | I/D/M/F* | Stop Codons |
|---|---|---|---|---|---|---|---|---|---|
| **NT** | **4054** | **4299** | **3.5%** | **41** | **8.3%** | **246 (97.6%)** | **137 (54.4%)** | **6/0** | |
| Mutations: | 4056A>G, 4060T>G, 4061C>A, 4064C>T, 4065C>T, 4066A>G, 4073A>G, 4074G>T, 4075A>G, 4084A>T, 4090C>T, 4092T>G, 4094T>A, 4095G>C, 4098C>A, 4099A>T, 4100G>A, 4101C>T, 4102A>G, 4106T>C, 4107T>A, 4111T>C, 4112C>G, 4116G>T, 4117G>C, 4120A>G, 4123T>C, 4124G>C, 4126T>G, 4132T>C, 4135T>A, 4138G>A, 4140A>C, 4141G>A, 4147G>C, 4153C>A, 4154T>A, 4155T>C, 4156C>A, 4159T>C, 4163A>G, 4164G>C, 4165A>C, 4166A>T, 4167G>T, 4172G>A, 4175T>C, 4177T>G, 4178A>G, 4180C>A, 4184G>C, 4185C>A, 4186T>A, 4187A>C, 4188A>C, 4189A>C, 4192_4193insCTCAGA, 4193G>A, 4195T>G, 4199A>G, 4201A>G, 4202G>C, 4204T>G, 4205A>G, 4206C>T, 4207A>T, 4209A>T, 4211A>T, 4213A>T, 4219T>C, 4226A>G, 4228T>G, 4229C>T, 4231T>C, 4234T>C, 4235C>T, 4236C>T, 4237A>G, 4238G>T, 4239A>C, 4240T>C, 4241G>C, 4244C>G, 4246G>A, 4247A>G, 4253T>A, 4255G>T, 4256A>C, 4262T>C, 4264A>C, 4265G>A, 4266A>T, 4267A>T, 4268A>G, 4271T>G, 4273T>G, 4274C>T, 4277G>T, 4278A>C, 4279A>T, 4280G>A, 4281A>C, 4282A>C, 4283G>T, 4285C>G, 4287A>G, 4289A>G, 4295G>T, 4296G>C, 4298A>T | | | | | | | | |

##### CDS

| | Begin | End | Coverage | Score | Concordance | Matches | Identities | I/D/M/F* | Stop Codons |
|---|---|---|---|---|---|---|---|---|---|
| EXL67_gp1 | 1301 | 1382 | 3.8% | 281 | 44.4% | 82 (97.6%) | 34 (40.5%) | 2/0/0/0 | 0 |
| Protein mutations: | Q1301R (4056A>G), I1302M (4060T>G), Q1303K (4061C>A), P1304L (4064C>T 4065C>T 4066A>G), R1307V (4073A>G 4074G>T 4075A>G), I1313R (4092T>G), C1314T (4094T>A 4095G>C), P1315H (4098C>A 4099A>T), A1316M (4100G>A 4101C>T 4102A>G), F1318H (4106T>C 4107T>A), Q1320E (4112C>G), W1321F (4116G>T 4117G>C), V1324L (4124G>C 4126T>G), K1329T (4140A>C 4141G>A), F1334T (4154T>A 4155T>C 4156C>A), R1337A (4163A>G 4164G>C 4165A>C), R1338L (4166A>T 4167G>T), D1340N (4172G>A), Y1341Q (4175T>C 4177T>G), I1342V (4178A>G 4180C>A), A1344Q (4184G>C 4185C>A 4186T>A), K1345P (4187A>C 4188A>C 4189A>C), Y1346_D1347insLR (4192_4193insCTCAGA), D1347K (4193G>A 4195T>G), I1349V (4199A>G 4201A>G), V1350L (4202G>C 4204T>G), T1351V (4205A>G 4206C>T 4207A>T), Y1352F (4209A>T), I1353F (4211A>T 4213A>T), I1358V (4226A>G 4228T>G), H1359Y (4229C>T 4231T>C), P1361L (4235C>T 4236C>T 4237A>G), D1362S (4238G>T 4239A>C 4240T>C), V1363L (4241G>C), Q1364E (4244C>G 4246G>A), N1365D (4247A>G), L1367I (4253T>A 4255G>T), K1368Q (4256A>C), E1371I (4265G>A 4266A>T 4267A>T), I1372V (4268A>G), F1373V (4271T>G 4273T>G), L1374F (4274C>T), E1375S (4277G>T 4278A>C 4279A>T), E1376T (4280G>A 4281A>C 4282A>C), V1377L (4283G>T 4285C>G), K1378R (4287A>G), K1379E (4289A>G), G1381S (4295G>T 4296G>C) | | | | | | | | |
| Codon mutations: | CAA1301CGA (4056A>G), ATT1302ATG (4060T>G), CAG1303AAG (4061C>A), CCA1304TTG (4064C>T 4065C>T 4066A>G), AGA1307GTG (4073A>G 4074G>T 4075A>G), ACA1310ACT (4084A>T), TTC1312TTT (4090C>T), ATA1313AGA (4092T>G), TGT1314ACT (4094T>A 4095G>C), CCA1315CAT (4098C>A 4099A>T), GCA1316ATG (4100G>A 4101C>T 4102A>G), TTC1318CAC (4106T>C 4107T>A), TAT1319TAC (4111T>C), CAG1320GAG (4112C>G), TGG1321TTC (4116G>T 4117G>C), AAA1322AAG (4120A>G), GTT1323GTC (4123T>C), GTT1324CTG (4124G>C 4126T>G), TTT1326TTC (4132T>C), GGT1327GGA (4135T>A), TTG1328TTA (4138G>A), AAG1329ACA (4140A>C 4141G>A), GCG1331GCC (4147G>C), GCC1333GCA (4153C>A), TTC1334ACA (4154T>A 4155T>C 4156C>A), TTT1335TTC (4159T>C), AGA1337GCC (4163A>G 4164G>C 4165A>C), AGG1338TTG (4166A>T 4167G>T), GAT1340AAT (4172G>A), TAT1341CAG (4175T>C 4177T>G), ATC1342GTA (4178A>G 4180C>A), GCT1344CAA (4184G>C 4185C>A 4186T>A), AAA1345CCC (4187A>C 4188A>C 4189A>C), TAT1346_GAT1347insCTCAGA (4192_4193insCTCAGA), GAT1347AAG (4193G>A 4195T>G), ATA1349GTG (4199A>G 4201A>G), GTT1350CTG (4202G>C 4204T>G), ACA1351GTT (4205A>G 4206C>T 4207A>T), TAC1352TTC (4209A>T), ATA1353TTT (4211A>T 4213A>T), GAT1355GAC (4219T>C), ATT1358GTG (4226A>G 4228T>G), CAT1359TAC (4229C>T 4231T>C), AGT1360AGC (4234T>C), CCA1361TTG (4235C>T 4236C>T 4237A>G), GAT1362TCC (4238G>T 4239A>C 4240T>C), GTA1363CTA (4241G>C), CAG1364GAA (4244C>G 4246G>A), AAT1365GAT (4247A>G), TTG1367ATT (4253T>A 4255G>T), AAG1368CAG (4256A>C), TTA1370CTC (4262T>C 4264A>C), GAA1371ATT (4265G>A 4266A>T 4267A>T), ATT1372GTT (4268A>G), TTT1373GTG (4271T>G 4273T>G), CTT1374TTT (4274C>T), GAA1375TCT (4277G>T 4278A>C 4279A>T), GAA1376ACC (4280G>A 4281A>C 4282A>C), GTC1377TTG (4283G>T 4285C>G), AAG1378AGG (4287A>G), AAA1379GAA (4289A>G), GGA1381TCA (4295G>T 4296G>C), ATT1382TT. (4298A>T) | | | | | | | | |

##### Proteins

| | Begin | End | Coverage | Score | Concordance | Matches | Identities | I/D/M/F* | Stop Codons |
|---|---|---|---|---|---|---|---|---|---|
| polyprotein (YP_009553669.1) | 1301 | 1382 | 3.8% | 281 | 44.4% | 82 (97.6%) | 34 (40.5%) | 2/0/0/0 | 0 |
| Protein mutations: | Q1301R (4056A>G), I1302M (4060T>G), Q1303K (4061C>A), P1304L (4064C>T 4065C>T 4066A>G), R1307V (4073A>G 4074G>T 4075A>G), I1313R (4092T>G), C1314T (4094T>A 4095G>C), P1315H (4098C>A 4099A>T), A1316M (4100G>A 4101C>T 4102A>G), F1318H (4106T>C 4107T>A), Q1320E (4112C>G), W1321F (4116G>T 4117G>C), V1324L (4124G>C 4126T>G), K1329T (4140A>C 4141G>A), F1334T (4154T>A 4155T>C 4156C>A), R1337A (4163A>G 4164G>C 4165A>C), R1338L (4166A>T 4167G>T), D1340N (4172G>A), Y1341Q (4175T>C 4177T>G), I1342V (4178A>G 4180C>A), A1344Q (4184G>C 4185C>A 4186T>A), K1345P (4187A>C 4188A>C 4189A>C), Y1346_D1347insLR (4192_4193insCTCAGA), D1347K (4193G>A 4195T>G), I1349V (4199A>G 4201A>G), V1350L (4202G>C 4204T>G), T1351V (4205A>G 4206C>T 4207A>T), Y1352F (4209A>T), I1353F (4211A>T 4213A>T), I1358V (4226A>G 4228T>G), H1359Y (4229C>T 4231T>C), P1361L (4235C>T 4236C>T 4237A>G), D1362S (4238G>T 4239A>C 4240T>C), V1363L (4241G>C), Q1364E (4244C>G 4246G>A), N1365D (4247A>G), L1367I (4253T>A 4255G>T), K1368Q (4256A>C), E1371I (4265G>A 4266A>T 4267A>T), I1372V (4268A>G), F1373V (4271T>G 4273T>G), L1374F (4274C>T), E1375S (4277G>T 4278A>C 4279A>T), E1376T (4280G>A 4281A>C 4282A>C), V1377L (4283G>T 4285C>G), K1378R (4287A>G), K1379E (4289A>G), G1381S (4295G>T 4296G>C) | | | | | | | | |
| Codon mutations: | CAA1301CGA (4056A>G), ATT1302ATG (4060T>G), CAG1303AAG (4061C>A), CCA1304TTG (4064C>T 4065C>T 4066A>G), AGA1307GTG (4073A>G 4074G>T 4075A>G), ACA1310ACT (4084A>T), TTC1312TTT (4090C>T), ATA1313AGA (4092T>G), TGT1314ACT (4094T>A 4095G>C), CCA1315CAT (4098C>A 4099A>T), GCA1316ATG (4100G>A 4101C>T 4102A>G), TTC1318CAC (4106T>C 4107T>A), TAT1319TAC (4111T>C), CAG1320GAG (4112C>G), TGG1321TTC (4116G>T 4117G>C), AAA1322AAG (4120A>G), GTT1323GTC (4123T>C), GTT1324CTG (4124G>C 4126T>G), TTT1326TTC (4132T>C), GGT1327GGA (4135T>A), TTG1328TTA (4138G>A), AAG1329ACA (4140A>C 4141G>A), GCG1331GCC (4147G>C), GCC1333GCA (4153C>A), TTC1334ACA (4154T>A 4155T>C 4156C>A), TTT1335TTC (4159T>C), AGA1337GCC (4163A>G 4164G>C 4165A>C), AGG1338TTG (4166A>T 4167G>T), GAT1340AAT (4172G>A), TAT1341CAG (4175T>C 4177T>G), ATC1342GTA (4178A>G 4180C>A), GCT1344CAA (4184G>C 4185C>A 4186T>A), AAA1345CCC (4187A>C 4188A>C 4189A>C), TAT1346_GAT1347insCTCAGA (4192_4193insCTCAGA), GAT1347AAG (4193G>A 4195T>G), ATA1349GTG (4199A>G 4201A>G), GTT1350CTG (4202G>C 4204T>G), ACA1351GTT (4205A>G 4206C>T 4207A>T), TAC1352TTC (4209A>T), ATA1353TTT (4211A>T 4213A>T), GAT1355GAC (4219T>C), ATT1358GTG (4226A>G 4228T>G), CAT1359TAC (4229C>T 4231T>C), AGT1360AGC (4234T>C), CCA1361TTG (4235C>T 4236C>T 4237A>G), GAT1362TCC (4238G>T 4239A>C 4240T>C), GTA1363CTA (4241G>C), CAG1364GAA (4244C>G 4246G>A), AAT1365GAT (4247A>G), TTG1367ATT (4253T>A 4255G>T), AAG1368CAG (4256A>C), TTA1370CTC (4262T>C 4264A>C), GAA1371ATT (4265G>A 4266A>T 4267A>T), ATT1372GTT (4268A>G), TTT1373GTG (4271T>G 4273T>G), CTT1374TTT (4274C>T), GAA1375TCT (4277G>T 4278A>C 4279A>T), GAA1376ACC (4280G>A 4281A>C 4282A>C), GTC1377TTG (4283G>T 4285C>G), AAG1378AGG (4287A>G), AAA1379GAA (4289A>G), GGA1381TCA (4295G>T 4296G>C), ATT1382TT. (4298A>T) | | | | | | | | |

*: Inserts / Deletes / Misaligned / Frameshifts

#### Analysis details

This analysis was performed with panviral2.64

### NGS Details (UN24): Badnavirus venatheobromae

#### Assembly

| | |
|---|---|
| **Coverage Length** | 273 (1 contig(s)) |
| **Depth Of Coverage** | 8.9 |
| **Number Of Reads** | 21 |
| **Reads Per Million** | 0.42 rpm (after QC) |
| **Ambiguities** | 0 |
| **Assembly Method** | de novo + reference guided assembly |
| **Consensus Caller** | Bcf Tools |

#### Coverage Map

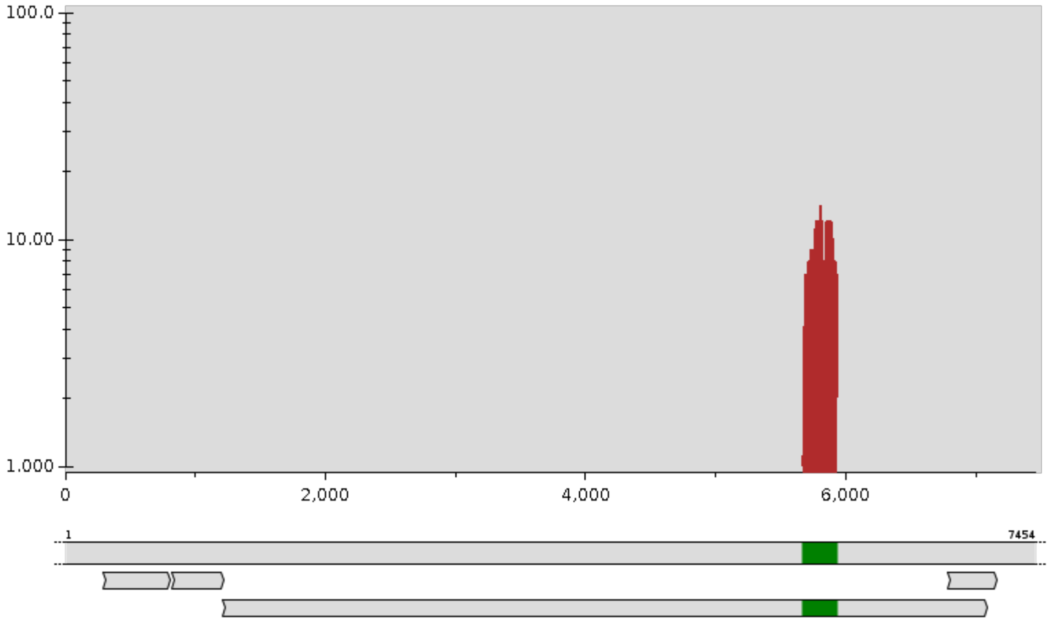

#### Assignment

| | |
|---|---|
| **Type** | Badnavirus venatheobromae (Taxonomy ID: 3047715) |
| **Reference Genome** | NC_033739.1 |
| **NT Identity (%)** | 57.971 |
| **AA Identity (%)** | 48.913 |
| **Number Of Stop Codons** | 1 |
| **Number Of CDS** | 4 |

#### Alignment

| | |
|---|---|
| **Alignment Score** | 82.0 (NT) + 324.0 (AA) = 406.0 |
| **Concordance (%)** | 33.1699 |

| Alignment Method | Global, seeded, nucleotide + amino acids (AGA) |
|---|---|

#### Genome Region

Sequence starts at position 5666 and ends at position 5938 relative to NC_033739.1 reference sequence.

#### Alignment Detailed Statistics

| | Begin | End | Coverage | Score | Concordance | Matches | Identities | I/D/M/F* | Stop Codons |
|---|---|---|---|---|---|---|---|---|---|
| **NT** | **5666** | **5938** | **3.7%** | **82** | **15.0%** | **273 (98.9%)** | **160 (58.0%)** | **3/0** | |
| Mutations: | 5666G>C, 5669T>C, 5670T>A, 5674C>G, 5675C>A, 5677A>G, 5679T>A, 5685T>C, 5687A>T, 5688A>C, 5689A>G, 5692C>T, 5698T>A, 5706G>T, 5708A>G, 5709G>A, 5710C>A, 5711A>G, 5712A>G, 5715G>A, 5720A>G, 5722C>G, 5723A>T, 5724T>G, 5725C>A, 5726A>T, 5732A>C, 5733T>A, 5734G>A, 5738A>G, 5741G>C, 5745T>C, 5747G>A, 5748G>A, 5749T>C, 5750C>T, 5751C>A, 5752C>G, 5753T>A, 5754G>T, 5759A>T, 5761T>A, 5762A>T, 5764A>T, 5770G>T, 5771G>C, 5772C>T, 5774A>G, 5777C>T, 5781C>T, 5783G>A, 5786C>T, 5789A>T, 5792A>G, 5794A>C, 5795G>C, 5801T>A, 5804A>G, 5807C>G, 5810T>A, 5814C>A, 5815A>T, 5816A>G, 5817A>G, 5818G>A, 5819A>C, 5820A>C, 5821A>T, 5822G>C, 5826G>A, 5828C>T, 5830A>G, 5831C>G, 5832T>G, 5833G>T, 5837T>C, 5841G>C, 5842G>C, 5843A>C, 5843_5844insTTC, 5844A>C, 5845C>T, 5846A>T, 5849A>T, 5850G>A, 5851A>T, 5852A>G, 5856A>G, 5858A>T, 5860C>T, 5861A>G, 5864T>A, 5866A>T, 5880C>T, 5882A>G, 5883G>A, 5887T>A, 5888C>T, 5891A>C, 5893A>G, 5896C>A, 5906A>G, 5909T>C, 5910G>A, 5912C>G, 5913C>A, 5915T>G, 5919C>T, 5921A>G, 5922C>A, 5926G>T, 5928T>G, 5930C>G, 5933C>G | | | | | | | | |

##### CDS

| | Begin | End | Coverage | Score | Concordance | Matches | Identities | I/D/M/F* | Stop Codons |
|---|---|---|---|---|---|---|---|---|---|
| B1U04_gp3 | 1486 | 1576 | 4.6% | 324 | 47.1% | 91 (98.9%) | 45 (48.9%) | 1/0/0/0 | 1 |
| Protein mutations: | Y1487N (5670T>A), S1488* (5674C>G 5675C>A), K1489R (5677A>G), F1490I (5679T>A), K1493R (5688A>C 5689A>G), S1494F (5692C>T), F1496Y (5698T>A), V1499L (5706G>T 5708A>G), A1500K (5709G>A 5710C>A 5711A>G), M1501V (5712A>G), E1502K (5715G>A), S1504C (5722C>G 5723A>T), S1505D (5724T>G 5725C>A 5726A>T), W1508K (5733T>A 5734G>A), W1512R (5745T>C 5747G>A), V1513T (5748G>A 5749T>C 5750C>T), P1514R (5751C>A 5752C>G 5753T>A), D1515Y (5754G>T), L1517H (5761T>A 5762A>T), Y1518F (5764A>T), W1520F (5770G>T 5771G>C), P1524S (5781C>T 5783G>A), K1528T (5794A>C 5795G>C), Q1535M (5814C>A 5815A>T 5816A>G), R1536D (5817A>G 5818G>A 5819A>C), K1537L (5820A>C 5821A>T 5822G>C), D1539N (5826G>A 5828C>T), N1540R (5830A>G 5831C>G), C1541V (5832T>G 5833G>T), G1544P (5841G>C 5842G>C 5843A>C), G1544_T1545insF (5843_5844insTTC), T1545L (5844A>C 5845C>T 5846A>T), E1546D (5849A>T), E1547M (5850G>A 5851A>T 5852A>G), I1549V (5856A>G 5858A>T), A1550V (5860C>T 5861A>G), Y1552F (5866A>T), V1558I (5883G>A), F1559Y (5887T>A 5888C>T), Q1561R (5893A>G), T1562N (5896C>A), V1567M (5910G>A 5912C>G), Q1571K (5922C>A), R1572M (5926G>T), F1573V (5928T>G 5930C>G), F1574L (5933C>G) | | | | | | | | |
| Codon mutations: | AAG1485..C (5666G>C), ATT1486ATC (5669T>C), TAT1487AAT (5670T>A), TCC1488TGA (5674C>G 5675C>A), AAG1489AGG (5677A>G), TTT1490ATT (5679T>A), TTA1492CTT (5685T>C 5687A>T), AAG1493CGG (5688A>C 5689A>G), TCC1494TTC (5692C>T), TTT1496TAT (5698T>A), GTA1499TTG (5706G>T 5708A>G), GCA1500AAG (5709G>A 5710C>A 5711A>G), ATG1501GTG (5712A>G), GAG1502AAG (5715G>A), GAA1503GAG (5720A>G), TCA1504TGT (5722C>G 5723A>T), TCA1505GAT (5724T>G 5725C>A 5726A>T), CCA1507CCC (5732A>C), TGG1508AAG (5733T>A 5734G>A), ACA1509ACG (5738A>G), GCG1510GCC (5741G>C), TGG1512CGA (5745T>C 5747G>A), GTC1513ACT (5748G>A 5749T>C 5750C>T), CCT1514AGA (5751C>A 5752C>G 5753T>A), GAT1515TAT (5754G>T), GGA1516GGT (5759A>T), CTA1517CAT (5761T>A 5762A>T), TAT1518TTT (5764A>T), TGG1520TTC (5770G>T 5771G>C), CTA1521TTG (5772C>T 5774A>G), GTC1522GTT (5777C>T), CCG1524TCA (5781C>T 5783G>A), TTC1525TTT (5786C>T), GGA1526GGT (5789A>T), TTA1527TTG (5792A>G), AAG1528ACC (5794A>C 5795G>C), GCT1530GCA (5801T>A), CCA1531CCG (5804A>G), GCC1532GCG (5807C>G), ATT1533ATA (5810T>A), CAA1535ATG (5814C>A 5815A>T 5816A>G), AGA1536GAC (5817A>G 5818G>A 5819A>C), AAG1537CTC (5820A>C 5821A>T 5822G>C), GAC1539AAT (5826G>A 5828C>T), AAC1540AGG (5830A>G 5831C>G), TGC1541GTC (5832T>G 5833G>T), TTT1542TTC (5837T>C), GGA1544CCC (5841G>C 5842G>C 5843A>C), GGA1544_ACA1545insTTC (5843_5844insTTC), ACA1545CTT (5844A>C 5845C>T 5846A>T), GAA1546GAT (5849A>T), GAA1547ATG (5850G>A 5851A>T 5852A>G), ATA1549GTT (5856A>G 5858A>T), GCA1550GTG (5860C>T 5861A>G), GTT1551GTA (5864T>A), TAT1552TTT (5866A>T), CTA1557TTG (5880C>T 5882A>G), GTC1558ATC (5883G>A), TTC1559TAT (5887T>A 5888C>T), TCA1560TCC (5891A>C), CAG1561CGG (5893A>G), ACT1562AAT (5896C>A), GAA1565GAG (5906A>G), CAT1566CAC (5909T>C), GTC1567ATG (5910G>A 5912C>G), CGT1568AGG (5913C>A 5915T>G), CTA1570TTG (5919C>T 5921A>G), CAA1571AAA (5922C>A), AGG1572ATG (5926G>T), TTC1573GTG (5928T>G 5930C>G), TTC1574TTG (5933C>G) | | | | | | | | |

##### Proteins

| | Begin | End | Coverage | Score | Concordance | Matches | Identities | I/D/M/F* | Stop Codons |
|---|---|---|---|---|---|---|---|---|---|
| polyprotein (YP_009345075.1) | 1486 | 1576 | 4.6% | 324 | 47.1% | 91 (98.9%) | 45 (48.9%) | 1/0/0/0 | 1 |
| Protein mutations: | Y1487N (5670T>A), S1488* (5674C>G 5675C>A), K1489R (5677A>G), F1490I (5679T>A), K1493R (5688A>C 5689A>G), S1494F (5692C>T), F1496Y (5698T>A), V1499L (5706G>T 5708A>G), A1500K (5709G>A 5710C>A 5711A>G), M1501V (5712A>G), E1502K (5715G>A), S1504C (5722C>G 5723A>T), S1505D (5724T>G 5725C>A 5726A>T), W1508K (5733T>A 5734G>A), W1512R (5745T>C 5747G>A), V1513T (5748G>A 5749T>C 5750C>T), P1514R (5751C>A 5752C>G 5753T>A), D1515Y (5754G>T), L1517H (5761T>A 5762A>T), Y1518F (5764A>T), W1520F (5770G>T 5771G>C), P1524S (5781C>T 5783G>A), K1528T (5794A>C 5795G>C), Q1535M (5814C>A 5815A>T 5816A>G), R1536D (5817A>G 5818G>A 5819A>C), K1537L (5820A>C 5821A>T 5822G>C), D1539N (5826G>A 5828C>T), N1540R (5830A>G 5831C>G), C1541V (5832T>G 5833G>T), G1544P (5841G>C 5842G>C 5843A>C), G1544_T1545insF (5843_5844insTTC), T1545L (5844A>C 5845C>T 5846A>T), E1546D (5849A>T), E1547M (5850G>A 5851A>T 5852A>G), I1549V (5856A>G 5858A>T), A1550V (5860C>T 5861A>G), Y1552F (5866A>T), V1558I (5883G>A), F1559Y (5887T>A 5888C>T), Q1561R (5893A>G), T1562N (5896C>A), V1567M (5910G>A 5912C>G), Q1571K (5922C>A), R1572M (5926G>T), F1573V (5928T>G 5930C>G), F1574L (5933C>G) | | | | | | | | |
| Codon mutations: | AAG1485..C (5666G>C), ATT1486ATC (5669T>C), TAT1487AAT (5670T>A), TCC1488TGA (5674C>G 5675C>A), AAG1489AGG (5677A>G), TTT1490ATT (5679T>A), TTA1492CTT (5685T>C 5687A>T), AAG1493CGG (5688A>C 5689A>G), TCC1494TTC (5692C>T), TTT1496TAT (5698T>A), GTA1499TTG (5706G>T 5708A>G), GCA1500AAG (5709G>A 5710C>A 5711A>G), ATG1501GTG (5712A>G), GAG1502AAG (5715G>A), GAA1503GAG (5720A>G), TCA1504TGT (5722C>G 5723A>T), TCA1505GAT (5724T>G 5725C>A 5726A>T), CCA1507CCC (5732A>C), TGG1508AAG (5733T>A 5734G>A), ACA1509ACG (5738A>G), GCG1510GCC (5741G>C), TGG1512CGA (5745T>C 5747G>A), GTC1513ACT (5748G>A 5749T>C 5750C>T), CCT1514AGA (5751C>A 5752C>G 5753T>A), GAT1515TAT (5754G>T), GGA1516GGT (5759A>T), CTA1517CAT (5761T>A 5762A>T), TAT1518TTT (5764A>T), TGG1520TTC (5770G>T 5771G>C), CTA1521TTG (5772C>T 5774A>G), GTC1522GTT (5777C>T), CCG1524TCA (5781C>T 5783G>A), TTC1525TTT (5786C>T), GGA1526GGT (5789A>T), TTA1527TTG (5792A>G), AAG1528ACC (5794A>C 5795G>C), GCT1530GCA (5801T>A), CCA1531CCG (5804A>G), GCC1532GCG (5807C>G), ATT1533ATA (5810T>A), CAA1535ATG (5814C>A 5815A>T 5816A>G), AGA1536GAC (5817A>G 5818G>A 5819A>C), AAG1537CTC (5820A>C 5821A>T 5822G>C), GAC1539AAT (5826G>A 5828C>T), AAC1540AGG (5830A>G 5831C>G), TGC1541GTC (5832T>G 5833G>T), TTT1542TTC (5837T>C), GGA1544CCC (5841G>C 5842G>C 5843A>C), GGA1544_ACA1545insTTC (5843_5844insTTC), ACA1545CTT (5844A>C 5845C>T 5846A>T), GAA1546GAT (5849A>T), GAA1547ATG (5850G>A 5851A>T 5852A>G), ATA1549GTT (5856A>G 5858A>T), GCA1550GTG (5860C>T 5861A>G), GTT1551GTA (5864T>A), TAT1552TTT (5866A>T), CTA1557TTG (5880C>T 5882A>G), GTC1558ATC (5883G>A), TTC1559TAT (5887T>A 5888C>T), TCA1560TCC (5891A>C), CAG1561CGG (5893A>G), ACT1562AAT (5896C>A), GAA1565GAG (5906A>G), CAT1566CAC (5909T>C), GTC1567ATG (5910G>A 5912C>G), CGT1568AGG (5913C>A 5915T>G), CTA1570TTG (5919C>T 5921A>G), CAA1571AAA (5922C>A), AGG1572ATG (5926G>T), TTC1573GTG (5928T>G 5930C>G), TTC1574TTG (5933C>G) | | | | | | | | |

*: Inserts / Deletes / Misaligned / Frameshifts

#### Analysis details

This analysis was performed with panviral2.64

### NGS Details (UN24): Unknown

#### Assembly

| | |
|---|---|
| **Coverage Length** | 314 (1 contig(s)) |
| **Depth Of Coverage** | 7.4 |
| **Number Of Reads** | 21 |
| **Reads Per Million** | 0.42 rpm (after QC) |
| **Ambiguities** | 0 |
| **Assembly Method** | de novo + reference guided assembly |
| **Consensus Caller** | Bcf Tools |

#### Coverage Map

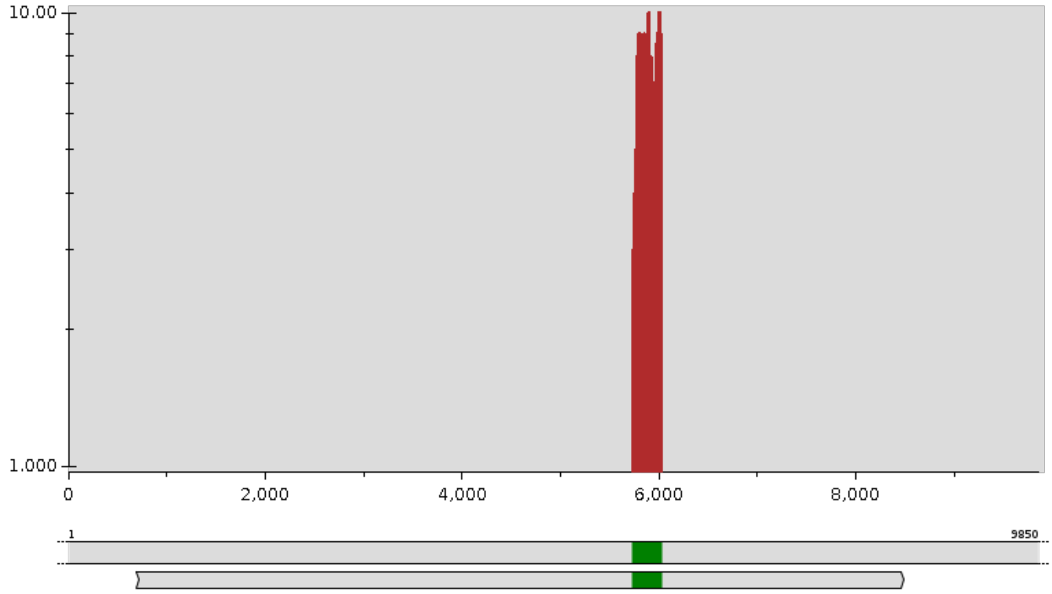

#### Assignment

| | |
|---|---|
| **Type** | Unknown (Taxonomy ID: -1) |
| **Reference Genome** | NC_028462.1 |
| **NT Identity (%)** | 57.9618 |
| **AA Identity (%)** | 55.7692 |
| **Number Of Stop Codons** | 0 |
| **Number Of CDS** | 1 |

#### Alignment

| | |
|---|---|
| **Alignment Score** | 100.0 (NT) + 411.0 (AA) = 511.0 |
| **Concordance (%)** | 37.8238 |

| **Alignment Method** | Global, seeded, nucleotide + amino acids (AGA) |
|---|---|

#### Genome Region

Sequence starts at position 5722 and ends at position 6035 relative to NC_028462.1 reference sequence.

#### Alignment Detailed Statistics

| | Begin | End | Coverage | Score | Concordance | Matches | Identities | I/D/M/F* | Stop Codons |
|---|---|---|---|---|---|---|---|---|---|
| **NT** | **5722** | **6035** | **3.2%** | **100** | **15.9%** | **314 (100%)** | **182 (58.0%)** | **0/0** | |
| Mutations: | 5727A>T, 5729C>A, 5730C>G, 5731A>C, 5735G>A, 5741T>A, 5743T>A, 5744C>A, 5745A>T, 5747C>T, 5750A>T, 5751A>G, 5752C>A, 5753C>A, 5754T>A, 5755A>G, 5756T>A, 5759T>A, 5760G>A, 5761A>G, 5762C>G, 5763A>T, 5766C>A, 5768A>T, 5771C>A, 5774G>A, 5780G>C, 5781T>A, 5782A>G, 5783T>C, 5784G>A, 5785C>T, 5786A>C, 5787C>A, 5789C>A, 5793C>A, 5795T>A, 5798A>T, 5801G>A, 5804C>T, 5809C>T, 5815G>A, 5818C>A, 5819T>A, 5820G>C, 5826G>A, 5829G>A, 5834G>A, 5838G>A, 5842G>T, 5843A>T, 5846A>G, 5847C>A, 5849C>T, 5850G>C, 5855G>A, 5856C>T, 5857A>T, 5859C>T, 5861C>A, 5864A>T, 5865C>A, 5868G>A, 5871C>T, 5873C>A, 5876T>A, 5877C>A, 5878A>G, 5879G>A, 5881C>A, 5882C>T, 5888G>A, 5898T>A, 5899C>G, 5900C>T, 5906A>C, 5912C>T, 5913T>A, 5914G>T, 5915T>G, 5918C>A, 5924T>C, 5929C>G, 5930T>C, 5933G>A, 5934A>C, 5935T>A, 5936C>A, 5939G>A, 5940A>C, 5942A>T, 5943C>G, 5944A>G, 5945G>A, 5951C>T, 5954C>A, 5957T>G, 5960C>A, 5963C>A, 5969C>T, 5975A>G, 5978C>T, 5982C>A, 5983G>A, 5985G>A, 5986T>A, 5987C>T, 5988A>T, 5989C>G, 5990C>T, 5991A>G, 5992A>T, 5993A>C, 5994G>T, 5995A>T, 5999T>C, 6002C>A, 6005C>T, 6008T>C, 6009C>A, 6011A>C, 6014C>A, 6018T>A, 6019T>A, 6020G>A, 6021G>A, 6023G>T, 6025C>A, 6027T>C, 6029A>T, 6030G>A, 6033T>A | | | | | | | | |
| **CDS** | | | | | | | | | |
| ATL80_gp1 | 1678 | 1781 | 4.0% | 411 | 56.2% | 104 (100%) | 58 (55.8%) | 0/0/0/0 | 0 |
| Protein mutations: | I1679L (5727A>T 5729C>A), H1680A (5730C>G 5731A>C), D1683E (5741T>A), V1684E (5743T>A 5744C>A), T1685S (5745A>T 5747C>T), Q1686H (5750A>T), T1687E (5751A>G 5752C>A 5753C>A), Y1688R (5754T>A 5755A>G 5756T>A), D1690R (5760G>A 5761A>G 5762C>G), I1691L (5763A>T), Q1692N (5766C>A 5768A>T), Y1697S (5781T>A 5782A>G 5783T>C), A1698I (5784G>A 5785C>T 5786A>C), L1699I (5787C>A 5789C>A), P1706L (5809C>T), S1708N (5815G>A), A1709E (5818C>A 5819T>A), E1710Q (5820G>C), E1712K (5826G>A), A1713T (5829G>A), E1716K (5838G>A), R1717I (5842G>T 5843A>T), L1719I (5847C>A 5849C>T), E1720Q (5850G>C), Q1722L (5856C>T 5857A>T), K1724N (5864A>T), L1725M (5865C>A), E1726K (5868G>A), Q1729R (5877C>A 5878A>G 5879G>A), P1730H (5881C>A 5882C>T), C1741M (5913T>A 5914G>T 5915T>G), A1746G (5929C>G 5930T>C), I1748Q (5934A>C 5935T>A 5936C>A), Q1751G (5943C>G 5944A>G 5945G>A), I1755M (5957T>G), R1764K (5982C>A 5983G>A), V1765N (5985G>A 5986T>A 5987C>T), T1766C (5988A>T 5989C>G 5990C>T), K1767V (5991A>G 5992A>T 5993A>C), D1768F (5994G>T 5995A>T), L1773I (6009C>A 6011A>C), L1776K (6018T>A 6019T>A 6020G>A), E1777N (6021G>A 6023G>T), T1778N (6025C>A), V1780I (6030G>A), Y1781N (6033T>A) | | | | | | | | |
| Codon mutations: | ATC1679TTA (5727A>T 5729C>A), CAC1680GCC (5730C>G 5731A>C), AAG1681AAA (5735G>A), GAT1683GAA (5741T>A), GTC1684GAA (5743T>A 5744C>A), ACC1685TCT (5745A>T 5747C>T), CAA1686CAT (5750A>T), ACC1687GAA (5751A>G 5752C>A 5753C>A), TAT1688AGA (5754T>A 5755A>G 5756T>A), GCT1689GCA (5759T>A), GAC1690AGG (5760G>A 5761A>G 5762C>G), ATA1691TTA (5763A>T), CAA1692AAT (5766C>A 5768A>T), CTC1693CTA (5771C>A), AAG1694AAA (5774G>A), CCG1696CCC (5780G>C), TAT1697AGC (5781T>A 5782A>G 5783T>C), GCA1698ATC (5784G>A 5785C>T 5786A>C), CTC1699ATA (5787C>A 5789C>A), CGT1701AGA (5793C>A 5795T>A), GTA1702GTT (5798A>T), AAG1703AAA (5801G>A), CCC1704CCT (5804C>T), CCA1706CTA (5809C>T), AGT1708AAT (5815G>A), GCT1709GAA (5818C>A 5819T>A), GAA1710CAA (5820G>C), GAA1712AAA (5826G>A), GCT1713ACT (5829G>A), GAG1714GAA (5834G>A), GAA1716AAA (5838G>A), AGA1717ATT (5842G>T 5843A>T), CAA1718CAG (5846A>G), CTC1719ATT (5847C>A 5849C>T), GAA1720CAA (5850G>C), GAG1721GAA (5855G>A), CAA1722TTA (5856C>T 5857A>T), CTC1723TTA (5859C>T 5861C>A), AAA1724AAT (5864A>T), CTG1725ATG (5865C>A), GAA1726AAA (5868G>A), CTC1727TTA (5871C>T 5873C>A), ATT1728ATA (5876T>A), CAG1729AGA (5877C>A 5878A>G 5879G>A), CCC1730CAT (5881C>A 5882C>T), AAG1732AAA (5888G>A), TCC1736AGT (5898T>A 5899C>G 5900C>T), CCA1738CCC (5906A>C), TTC1740TTT (5912C>T), TGT1741ATG (5913T>A 5914G>T 5915T>G), GTC1742GTA (5918C>A), AAT1744AAC (5924T>C), GCT1746GGC (5929C>G 5930T>C), GAG1747GAA (5933G>A), ATC1748CAA (5934A>C 5935T>A 5936C>A), AAG1749AAA (5939G>A), AGA1750CGT (5940A>C 5942A>T), CAG1751GGA (5943C>G 5944A>G 5945G>A), GCC1753GCT (5951C>T), CGC1754CGA (5954C>A), ATT1755ATG (5957T>G), GTC1756GTA (5960C>A), ATC1757ATA (5963C>A), TAC1759TAT (5969C>T), GAA1761GAG (5975A>G), CTC1762CTT (5978C>T), CGA1764AAA (5982C>A 5983G>A), GTC1765AAT (5985G>A 5986T>A 5987C>T), ACC1766TGT (5988A>T 5989C>G 5990C>T), AAA1767GTC (5991A>G 5992A>T 5993A>C), GAC1768TTC (5994G>T 5995A>T), GAT1769GAC (5999T>C), GGC1770GGA (6002C>A), TAC1771TAT (6005C>T), TTT1772TTC (6008T>C), CTA1773ATC (6009C>A 6011A>C), CCC1774CCA (6014C>A), TTG1776AAA (6018T>A 6019T>A 6020G>A), GAG1777AAT (6021G>A 6023G>T), ACC1778AAC (6025C>A), TTA1779CTT (6027T>C 6029A>T), GTA1780ATA (6030G>A), TAT1781AAT (6033T>A) | | | | | | | | |
| **Proteins** | | | | | | | | | |
| polyprotein (YP_009182100.1) | 1678 | 1781 | 4.0% | 411 | 56.2% | 104 (100%) | 58 (55.8%) | 0/0/0/0 | 0 |
| Protein mutations: | I1679L (5727A>T 5729C>A), H1680A (5730C>G 5731A>C), D1683E (5741T>A), V1684E (5743T>A 5744C>A), T1685S (5745A>T 5747C>T), Q1686H (5750A>T), T1687E (5751A>G 5752C>A 5753C>A), Y1688R (5754T>A 5755A>G 5756T>A), D1690R (5760G>A 5761A>G 5762C>G), I1691L (5763A>T), Q1692N (5766C>A 5768A>T), Y1697S (5781T>A 5782A>G 5783T>C), A1698I (5784G>A 5785C>T 5786A>C), L1699I (5787C>A 5789C>A), P1706L (5809C>T), S1708N (5815G>A), A1709E (5818C>A 5819T>A), E1710Q (5820G>C), E1712K (5826G>A), A1713T (5829G>A), E1716K (5838G>A), R1717I (5842G>T 5843A>T), L1719I (5847C>A 5849C>T), E1720Q (5850G>C), Q1722L (5856C>T 5857A>T), K1724N (5864A>T), L1725M (5865C>A), E1726K (5868G>A), Q1729R (5877C>A 5878A>G 5879G>A), P1730H (5881C>A 5882C>T), C1741M (5913T>A 5914G>T 5915T>G), A1746G (5929C>G 5930T>C), I1748Q (5934A>C 5935T>A 5936C>A), Q1751G (5943C>G 5944A>G 5945G>A), I1755M (5957T>G), R1764K (5982C>A 5983G>A), V1765N (5985G>A 5986T>A 5987C>T), T1766C (5988A>T 5989C>G 5990C>T), K1767V (5991A>G 5992A>T 5993A>C), D1768F (5994G>T 5995A>T), L1773I (6009C>A 6011A>C), L1776K (6018T>A 6019T>A 6020G>A), E1777N (6021G>A 6023G>T), T1778N (6025C>A), V1780I (6030G>A), Y1781N (6033T>A) | | | | | | | | |
| Codon mutations: | ATC1679TTA (5727A>T 5729C>A), CAC1680GCC (5730C>G 5731A>C), AAG1681AAA (5735G>A), GAT1683GAA (5741T>A), GTC1684GAA (5743T>A 5744C>A), ACC1685TCT (5745A>T 5747C>T), CAA1686CAT (5750A>T), ACC1687GAA (5751A>G 5752C>A 5753C>A), TAT1688AGA (5754T>A 5755A>G 5756T>A), GCT1689GCA (5759T>A), GAC1690AGG (5760G>A 5761A>G 5762C>G), ATA1691TTA (5763A>T), CAA1692AAT (5766C>A 5768A>T), CTC1693CTA (5771C>A), AAG1694AAA (5774G>A), CCG1696CCC (5780G>C), TAT1697AGC (5781T>A 5782A>G 5783T>C), GCA1698ATC (5784G>A 5785C>T 5786A>C), CTC1699ATA (5787C>A 5789C>A), CGT1701AGA (5793C>A 5795T>A), GTA1702GTT (5798A>T), AAG1703AAA (5801G>A), CCC1704CCT (5804C>T), CCA1706CTA (5809C>T), AGT1708AAT (5815G>A), GCT1709GAA (5818C>A 5819T>A), GAA1710CAA (5820G>C), GAA1712AAA (5826G>A), GCT1713ACT (5829G>A), GAG1714GAA (5834G>A), GAA1716AAA (5838G>A), AGA1717ATT (5842G>T 5843A>T), CAA1718CAG (5846A>G), CTC1719ATT (5847C>A 5849C>T), GAA1720CAA (5850G>C), GAG1721GAA (5855G>A), CAA1722TTA (5856C>T 5857A>T), CTC1723TTA (5859C>T 5861C>A), AAA1724AAT (5864A>T), CTG1725ATG (5865C>A), GAA1726AAA (5868G>A), CTC1727TTA (5871C>T 5873C>A), ATT1728ATA (5876T>A), CAG1729AGA (5877C>A 5878A>G 5879G>A), CCC1730CAT (5881C>A 5882C>T), AAG1732AAA (5888G>A), TCC1736AGT (5898T>A 5899C>G 5900C>T), CCA1738CCC (5906A>C), TTC1740TTT (5912C>T), TGT1741ATG (5913T>A 5914G>T 5915T>G), GTC1742GTA (5918C>A), AAT1744AAC (5924T>C), GCT1746GGC (5929C>G 5930T>C), GAG1747GAA (5933G>A), ATC1748CAA (5934A>C 5935T>A 5936C>A), AAG1749AAA (5939G>A), AGA1750CGT (5940A>C 5942A>T), CAG1751GGA (5943C>G 5944A>G 5945G>A), GCC1753GCT (5951C>T), CGC1754CGA (5954C>A), ATT1755ATG (5957T>G), GTC1756GTA (5960C>A), ATC1757ATA (5963C>A), TAC1759TAT (5969C>T), GAA1761GAG (5975A>G), CTC1762CTT (5978C>T), CGA1764AAA (5982C>A 5983G>A), GTC1765AAT (5985G>A 5986T>A 5987C>T), ACC1766TGT (5988A>T 5989C>G 5990C>T), AAA1767GTC (5991A>G 5992A>T 5993A>C), GAC1768TTC (5994G>T 5995A>T), GAT1769GAC (5999T>C), GGC1770GGA (6002C>A), TAC1771TAT (6005C>T), TTT1772TTC (6008T>C), CTA1773ATC (6009C>A 6011A>C), CCC1774CCA (6014C>A), TTG1776AAA (6018T>A 6019T>A 6020G>A), GAG1777AAT (6021G>A 6023G>T), ACC1778AAC (6025C>A), TTA1779CTT (6027T>C 6029A>T), GTA1780ATA (6030G>A), TAT1781AAT (6033T>A) | | | | | | | | |

*: Inserts / Deletes / Misaligned / Frameshifts

#### Analysis details

This analysis was performed with panviral2.64

### NGS Details (UN24): Bracoviriform congregatae (segment Circle 7)

#### Assembly

| | |
|---|---|
| **Coverage Length** | 145 (1 contig(s)) |
| **Depth Of Coverage** | 4.4 |
| **Number Of Reads** | 5 |
| **Reads Per Million** | 0.10 rpm (after QC) |
| **Ambiguities** | 0 |
| **Assembly Method** | de novo + reference guided assembly |
| **Consensus Caller** | Bcf Tools |

#### Coverage Map

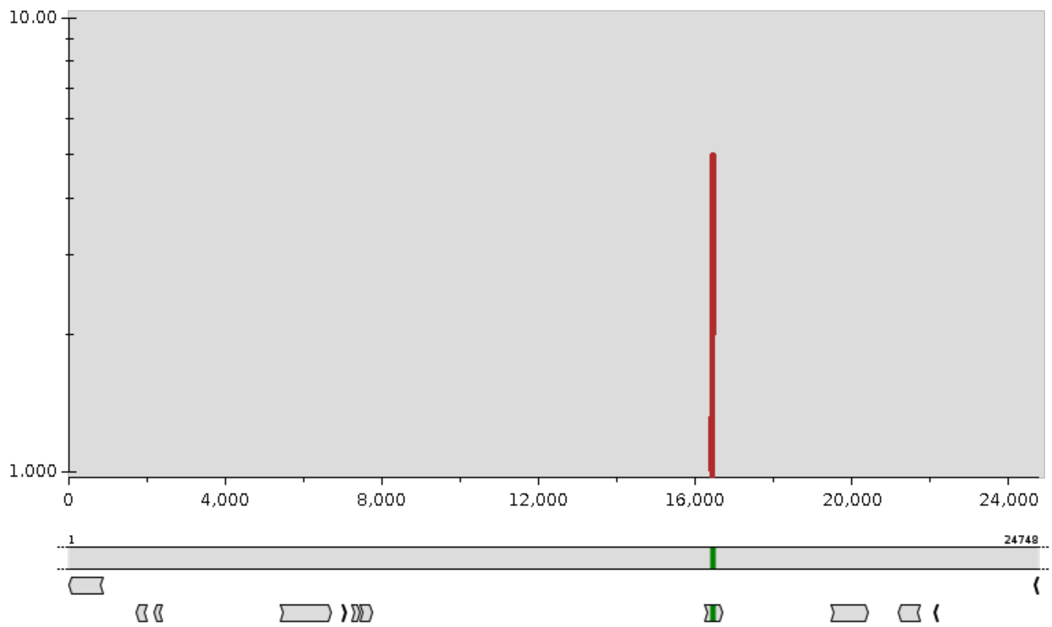

#### Assignment

| | |
|---|---|
| **Type** | Bracoviriform congregatae (Taxonomy ID: 39640) |
| **Reference Genome** | NC_006639.1 |
| **NT Identity (%)** | 80.0 |
| **AA Identity (%)** | 87.7551 |
| **Number Of Stop Codons** | 0 |
| **Number Of CDS** | 5 |

#### Alignment

| | |
|---|---|
| **Alignment Score** | 174.0 (NT) + 285.0 (AA) = 459.0 |
| **Concordance (%)** | 75.9934 |

| Alignment Method | Global, seeded, nucleotide + amino acids (AGA) |
|---|---|

#### Genome Region

Sequence starts at position 16385 and ends at position 16529 relative to NC_006639.1 reference sequence.

#### Alignment Detailed Statistics

| | Begin | End | Coverage | Score | Concordance | Matches | Identities | I/D/M/F* | Stop Codons |
|---|---|---|---|---|---|---|---|---|---|
| **NT** | **16385** | **16529** | **0.6%** | **174** | **60.0%** | **145 (100%)** | **116 (80.0%)** | **0/0** | |
| Mutations: | 16393A>T, 16397T>C, 16398C>G, 16399G>T, 16401T>G, 16408A>T, 16411G>A, 16414A>G, 16418T>C, 16420G>C, 16426A>G, 16429G>A, 16432C>A, 16435T>C, 16444T>C, 16449G>A, 16450A>G, 16451T>A, 16452C>T, 16453T>C, 16459A>T, 16462T>C, 16471A>G, 16474A>T, 16477C>T, 16486A>C, 16492C>T, 16501G>C, 16516A>T | | | | | | | | |
| **CDS** | | | | | | | | | |
| Histone | 52 | 100 | 31.4% | 285 | 89.6% | 49 (100%) | 43 (87.8%) | 0/0/0/0 | 0 |
| Protein mutations: | K54N (16393A>T), S56R (16397T>C 16398C>G 16399G>T), V57G (16401T>G), R73K (16449G>A 16450A>G), S74I (16451T>A 16452C>T 16453T>C) | | | | | | | | |
| Codon mutations: | AAA54AAT (16393A>T), TCG56CGT (16397T>C 16398C>G 16399G>T), GTA57GGA (16401T>G), GGA59GGT (16408A>T), GGG60GGA (16411G>A), AAA61AAG (16414A>G), TTG63CTC (16418T>C 16420G>C), AAA65AAG (16426A>G), GGG66GGA (16429G>A), GGC67GGA (16432C>A), GCT68GCC (16435T>C), CAT71CAC (16444T>C), AGA73AAG (16449G>A 16450A>G), TCT74ATC (16451T>A 16452C>T 16453T>C), CGA76CGT (16459A>T), GAT77GAC (16462T>C), CAA80CAG (16471A>G), GGA81GGT (16474A>T), ATC82ATT (16477C>T), CCA85CCC (16486A>C), ATC87ATT (16492C>T), CTG90CTC (16501G>C), GGA95GGT (16516A>T) | | | | | | | | |
| **Proteins** | | | | | | | | | |
| hypothetical protein (YP_184795.1) | 52 | 100 | 31.4% | 285 | 89.6% | 49 (100%) | 43 (87.8%) | 0/0/0/0 | 0 |
| Protein mutations: | K54N (16393A>T), S56R (16397T>C 16398C>G 16399G>T), V57G (16401T>G), R73K (16449G>A 16450A>G), S74I (16451T>A 16452C>T 16453T>C) | | | | | | | | |
| Codon mutations: | AAA54AAT (16393A>T), TCG56CGT (16397T>C 16398C>G 16399G>T), GTA57GGA (16401T>G), GGA59GGT (16408A>T), GGG60GGA (16411G>A), AAA61AAG (16414A>G), TTG63CTC (16418T>C 16420G>C), AAA65AAG (16426A>G), GGG66GGA (16429G>A), GGC67GGA (16432C>A), GCT68GCC (16435T>C), CAT71CAC (16444T>C), AGA73AAG (16449G>A 16450A>G), TCT74ATC (16451T>A 16452C>T 16453T>C), CGA76CGT (16459A>T), GAT77GAC (16462T>C), CAA80CAG (16471A>G), GGA81GGT (16474A>T), ATC82ATT (16477C>T), CCA85CCC (16486A>C), ATC87ATT (16492C>T), CTG90CTC (16501G>C), GGA95GGT (16516A>T) | | | | | | | | |

*: Inserts / Deletes / Misaligned / Frameshifts

#### Analysis details

This analysis was performed with panviral2.64

### NGS Details (UN24): Bracoviriform congregatae (segment Circle 7)

#### Assembly

| | |
|---|---|
| **Coverage Length** | 135 (1 contig(s)) |
| **Depth Of Coverage** | 12.2 |
| **Number Of Reads** | 20 |
| **Reads Per Million** | 0.40 rpm (after QC) |
| **Ambiguities** | 0 |
| **Assembly Method** | de novo + reference guided assembly |
| **Consensus Caller** | Bcf Tools |

#### Coverage Map

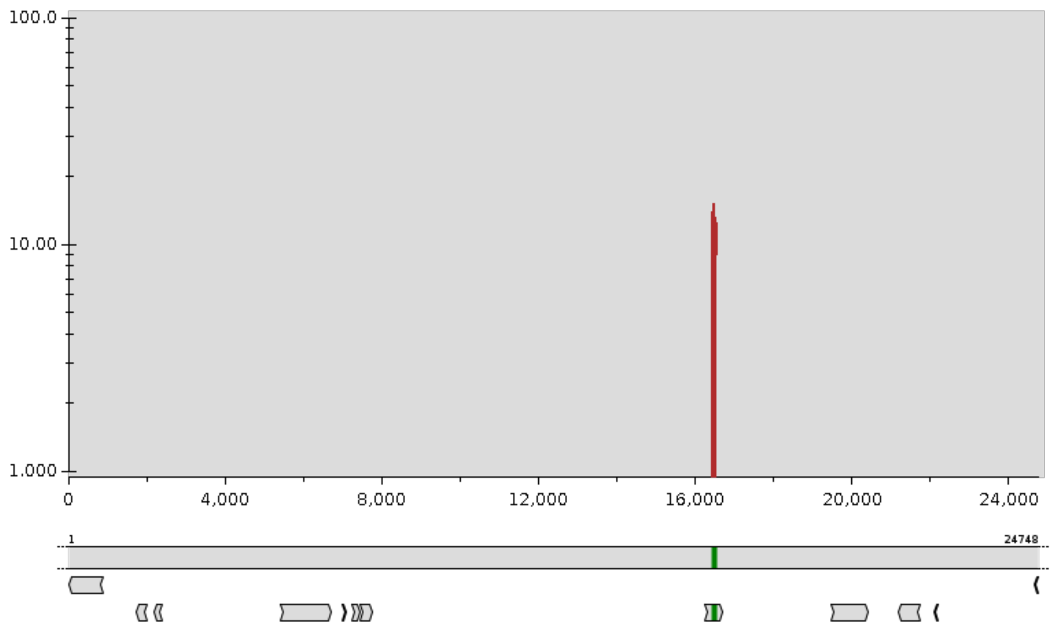

#### Assignment

| | |
|---|---|
| **Type** | Bracoviriform congregatae (Taxonomy ID: 39640) |
| **Reference Genome** | NC_006639.1 |
| **NT Identity (%)** | 77.7778 |
| **AA Identity (%)** | 84.4444 |
| **Number Of Stop Codons** | 0 |
| **Number Of CDS** | 5 |

#### Alignment

| | |
|---|---|
| **Alignment Score** | 150.0 (NT) + 266.0 (AA) = 416.0 |
| **Concordance (%)** | 74.2857 |

| Alignment Method | Global, seeded, nucleotide + amino acids (AGA) |
|---|---|

#### Genome Region

Sequence starts at position 16414 and ends at position 16548 relative to NC_006639.1 reference sequence.

#### Alignment Detailed Statistics

| | Begin | End | Coverage | Score | Concordance | Matches | Identities | I/D/M/F* | Stop Codons |
|---|---|---|---|---|---|---|---|---|---|
| **NT** | **16414** | **16548** | **0.5%** | **150** | **55.6%** | **135 (100%)** | **105 (77.8%)** | **0/0** | |
| Mutations: | 16414A>G, 16417A>T, 16418T>C, 16420G>C, 16423A>C, 16426A>G, 16429G>C, 16435T>C, 16444T>C, 16449G>A, 16450A>G, 16451T>A, 16452C>T, 16459A>T, 16462T>C, 16471A>G, 16474A>T, 16486A>C, 16495T>C, 16501G>C, 16516A>T, 16526G>A, 16528C>T, 16531T>A, 16533G>C, 16534A>C, 16535T>A, 16538G>A, 16543T>C, 16546A>G | | | | | | | | |
| **CDS** | | | | | | | | | |
| Histone | 62 | 106 | 28.8% | 266 | 88.4% | 45 (100%) | 38 (84.4%) | 0/0/0/0 | 0 |
| Protein mutations: | R73K (16449G>A 16450A>G), S74I (16451T>A 16452C>T), V99I (16526G>A 16528C>T), G101A (16533G>C 16534A>C), L102M (16535T>A), V103I (16538G>A) | | | | | | | | |
| Codon mutations: | AAA61..G (16414A>G), GGA62GGT (16417A>T), TTG63CTC (16418T>C 16420G>C), GGA64GGC (16423A>C), AAA65AAG (16426A>G), GGG66GGC (16429G>C), GCT68GCC (16435T>C), CAT71CAC (16444T>C), AGA73AAG (16449G>A 16450A>G), TCT74ATT (16451T>A 16452C>T), CGA76CGT (16459A>T), GAT77GAC (16462T>C), CAA80CAG (16471A>G), GGA81GGT (16474A>T), CCA85CCC (16486A>C), CGT88CGC (16495T>C), CTG90CTC (16501G>C), GGA95GGT (16516A>T), GTC99ATT (16526G>A 16528C>T), TCT100TCA (16531T>A), GGA101GCC (16533G>C 16534A>C), TTG102ATG (16535T>A), GTC103ATC (16538G>A), TAT104TAC (16543T>C), GAA105GAG (16546A>G) | | | | | | | | |
| **Proteins** | | | | | | | | | |
| hypothetical protein (YP_184795.1) | 62 | 106 | 28.8% | 266 | 88.4% | 45 (100%) | 38 (84.4%) | 0/0/0/0 | 0 |
| Protein mutations: | R73K (16449G>A 16450A>G), S74I (16451T>A 16452C>T), V99I (16526G>A 16528C>T), G101A (16533G>C 16534A>C), L102M (16535T>A), V103I (16538G>A) | | | | | | | | |
| Codon mutations: | AAA61..G (16414A>G), GGA62GGT (16417A>T), TTG63CTC (16418T>C 16420G>C), GGA64GGC (16423A>C), AAA65AAG (16426A>G), GGG66GGC (16429G>C), GCT68GCC (16435T>C), CAT71CAC (16444T>C), AGA73AAG (16449G>A 16450A>G), TCT74ATT (16451T>A 16452C>T), CGA76CGT (16459A>T), GAT77GAC (16462T>C), CAA80CAG (16471A>G), GGA81GGT (16474A>T), CCA85CCC (16486A>C), CGT88CGC (16495T>C), CTG90CTC (16501G>C), GGA95GGT (16516A>T), GTC99ATT (16526G>A 16528C>T), TCT100TCA (16531T>A), GGA101GCC (16533G>C 16534A>C), TTG102ATG (16535T>A), GTC103ATC (16538G>A), TAT104TAC (16543T>C), GAA105GAG (16546A>G) | | | | | | | | |

*: Inserts / Deletes / Misaligned / Frameshifts

#### Analysis details

This analysis was performed with panviral2.64

### NGS Details (UN24): Petuvirus venapetuniae

#### Assembly

| | |
|---|---|
| **Coverage Length** | 316 (1 contig(s)) |
| **Depth Of Coverage** | 6.3 |
| **Number Of Reads** | 20 |
| **Reads Per Million** | 0.40 rpm (after QC) |
| **Ambiguities** | 0 |
| **Assembly Method** | de novo + reference guided assembly |
| **Consensus Caller** | Bcf Tools |

#### Coverage Map

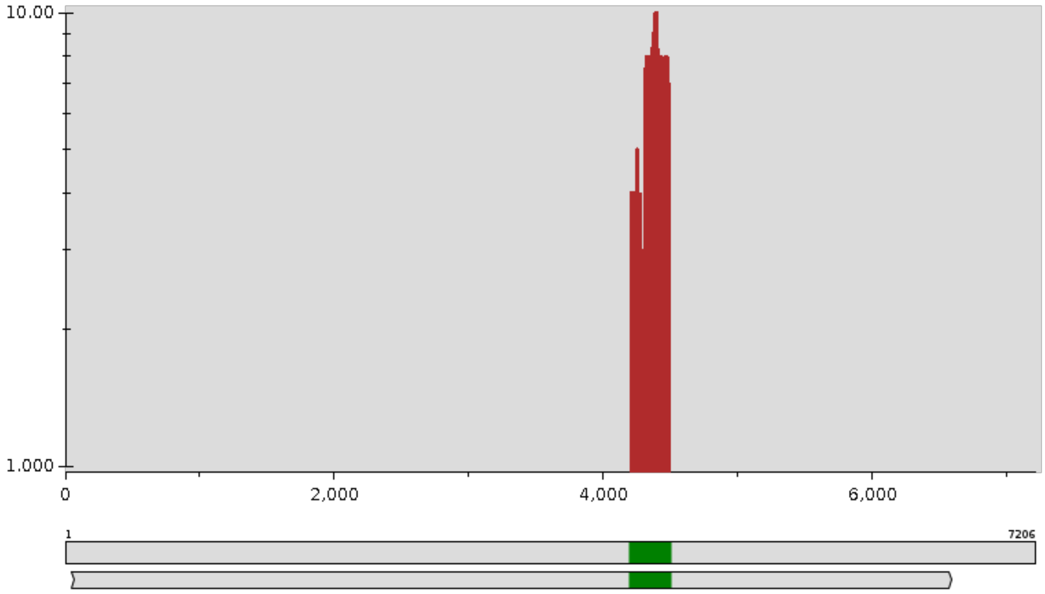

#### Assignment

| | |
|---|---|
| **Type** | Petuvirus venapetuniae (Taxonomy ID: 3048265) |
| **Reference Genome** | NC_001839.2 |
| **NT Identity (%)** | 61.0759 |
| **AA Identity (%)** | 54.2857 |
| **Number Of Stop Codons** | 0 |
| **Number Of CDS** | 1 |

#### Alignment

| | |
|---|---|
| **Alignment Score** | 140.0 (NT) + 387.0 (AA) = 527.0 |
| **Concordance (%)** | 37.8864 |

| | |
|---|---|
| **Alignment Method** | Global, seeded, nucleotide + amino acids (AGA) |

#### Genome Region

Sequence starts at position 4190 and ends at position 4505 relative to NC_001839.2 reference sequence.

#### Alignment Detailed Statistics

| | Begin | End | Coverage | Score | Concordance | Matches | Identities | I/D/M/F* | Stop Codons |
|---|---|---|---|---|---|---|---|---|---|
| **NT** | **4190** | **4505** | **4.4%** | **140** | **22.2%** | **316 (100%)** | **193 (61.1%)** | **0/0** | |
| Mutations: | 4196C>T, 4197A>G, 4198T>A, 4199C>T, 4201A>T, 4208A>T, 4211G>A, 4212G>T, 4215A>C, 4217C>T, 4219A>C, 4220T>C, 4222C>G, 4224G>C, 4225G>A, 4233C>G, 4234C>T, 4235A>T, 4239C>T, 4240A>T, 4241T>A, 4242C>G, 4244C>G, 4245C>G, 4247G>A, 4250A>C, 4251G>T, 4252C>G, 4253A>C, 4255T>A, 4259G>A, 4260G>A, 4262A>G, 4263T>A, 4264G>T, 4269G>A, 4271G>T, 4275C>T, 4276A>T, 4281T>A, 4282T>A, 4283T>G, 4285A>G, 4287C>T, 4292C>A, 4293G>A, 4298C>A, 4301T>A, 4302G>A, 4304T>G, 4307A>T, 4309A>C, 4310A>T, 4314G>T, 4319T>C, 4320G>A, 4321A>C, 4324C>T, 4334C>T, 4337C>T, 4340G>T, 4341A>G, 4342G>C, 4343G>T, 4344T>A, 4345C>G, 4346A>T, 4349G>A, 4350C>A, 4352A>G, 4354T>A, 4355G>A, 4356A>C, 4361A>T, 4363A>T, 4364A>T, 4365T>C, 4367A>C, 4368A>C, 4370G>A, 4376C>T, 4386C>A, 4397C>T, 4398C>A, 4400T>A, 4403C>T, 4406C>T, 4407C>A, 4410G>T, 4411A>G, 4412T>G, 4413G>A, 4414A>T, 4415C>T, 4417A>G, 4420T>A, 4430A>G, 4436G>A, 4437C>A, 4438T>G, 4439C>G, 4440A>G, 4441C>A, 4442A>T, 4443C>T, 4445C>A, 4448C>A, 4452C>A, 4453A>G, 4454C>A, 4459C>A, 4460C>T, 4461A>G, 4463A>T, 4466A>C, 4469G>T, 4470C>A, 4472C>A, 4481G>T, 4488C>T, 4490T>A, 4493G>A, 4501T>A | | | | | | | | |

##### CDS

| | Begin | End | Coverage | Score | Concordance | Matches | Identities | I/D/M/F* | Stop Codons |
|---|---|---|---|---|---|---|---|---|---|
| PVCVgp1 | 1381 | 1485 | 4.8% | 387 | 50.5% | 105 (100%) | 57 (54.3%) | 0/0/0/0 | 0 |
| Protein mutations: | I1383D (4197A>G 4198T>A 4199C>T), N1384I (4201A>T), A1388S (4212G>T), S1389R (4215A>C 4217C>T), H1390P (4219A>C 4220T>C), S1391C (4222C>G), G1392Q (4224G>C 4225G>A), P1395V (4233C>G 4234C>T 4235A>T), H1397L (4239C>T 4240A>T 4241T>A), L1398V (4242C>G 4244C>G), Q1399E (4245C>G 4247G>A), L1400F (4250A>C), A1401C (4251G>T 4252C>G 4253A>C), I1402K (4255T>A), E1404K (4260G>A 4262A>G), C1405I (4263T>A 4264G>T), E1407N (4269G>A 4271G>T), Q1409L (4275C>T 4276A>T), F1411K (4281T>A 4282T>A 4283T>G), D1412G (4285A>G), E1415K (4293G>A), D1418K (4302G>A 4304T>G), Q1420P (4309A>C 4310A>T), A1422S (4314G>T), E1424T (4320G>A 4321A>C), A1425V (4324C>T), K1430N (4340G>T), R1431A (4341A>G 4342G>C 4343G>T), Q1434K (4350C>A 4352A>G), V1435E (4354T>A 4355G>A), K1438I (4363A>T 4364A>T), Q1446K (4386C>A), H1450K (4398C>A 4400T>A), Q1453K (4407C>A), D1454W (4410G>T 4411A>G 4412T>G), D1455I (4413G>A 4414A>T 4415C>T), K1456R (4417A>G), F1457Y (4420T>A), L1463R (4437C>A 4438T>G 4439C>G), T1464D (4440A>G 4441C>A 4442A>T), F1466L (4448C>A), H1468R (4452C>A 4453A>G 4454C>A), S1470Y (4459C>A 4460C>T), K1471D (4461A>G 4463A>T), K1473N (4469G>T), L1474I (4470C>A 4472C>A), K1477N (4481G>T), F1484Y (4501T>A) | | | | | | | | |
| Codon mutations: | AAC1382AAT (4196C>T), ATC1383GAT (4197A>G 4198T>A 4199C>T), AAT1384ATT (4201A>T), ACA1386ACT (4208A>T), AAG1387AAA (4211G>A), GCA1388TCA (4212G>T), AGC1389CGT (4215A>C 4217C>T), CAT1390CCC (4219A>C 4220T>C), TCT1391TGT (4222C>G), GGA1392CAA (4224G>C 4225G>A), CCA1395GTT (4233C>G 4234C>T 4235A>T), CAT1397TTA (4239C>T 4240A>T 4241T>A), CTC1398GTG (4242C>G 4244C>G), CAG1399GAA (4245C>G 4247G>A), TTA1400TTC (4250A>C), GCA1401TGC (4251G>T 4252C>G 4253A>C), ATA1402AAA (4255T>A), AAG1403AAA (4259G>A), GAA1404AAG (4260G>A 4262A>G), TGT1405ATT (4263T>A 4264G>T), GAG1407AAT (4269G>A 4271G>T), CAA1409TTA (4275C>T 4276A>T), TTT1411AAG (4281T>A 4282T>A 4283T>G), GAT1412GGT (4285A>G), CTA1413TTA (4287C>T), ATC1414ATA (4292C>A), GAA1415AAA (4293G>A), CCC1416CCA (4298C>A), TCT1417TCA (4301T>A), GAT1418AAG (4302G>A 4304T>G), TCA1419TCT (4307A>T), CAA1420CCT (4309A>C 4310A>T), GCA1422TCA (4314G>T), TGT1423TGC (4319T>C), GAA1424ACA (4320G>A 4321A>C), GCT1425GTT (4324C>T), GTC1428GTT (4334C>T), AAC1429AAT (4337C>T), AAG1430AAT (4340G>T), AGG1431GCT (4341A>G 4342G>C 4343G>T), TCA1432AGT (4344T>A 4345C>G 4346A>T), GAG1433GAA (4349G>A), CAA1434AAG (4350C>A 4352A>G), GTG1435GAA (4354T>A 4355G>A), AGA1436CGA (4356A>C), GGA1437GGT (4361A>T), AAA1438ATT (4363A>T 4364A>T), TTA1439CTC (4365T>C 4367A>C), AGG1440CGA (4368A>C 4370G>A), GTC1442GTT (4376C>T), CAA1446AAA (4386C>A), AAC1449AAT (4397C>T), CAT1450AAA (4398C>A 4400T>A), TTC1451TTT (4403C>T), CTC1452CTT (4406C>T), CAA1453AAA (4407C>A), GAT1454TGG (4410G>T 4411A>G 4412T>G), GAC1455ATT (4413G>A 4414A>T 4415C>T), AAG1456AGG (4417A>G), TTT1457TAT (4420T>A), CCA1460CCG (4430A>G), AAG1462AAA (4436G>A), CTC1463AGG (4437C>A 4438T>G 4439C>G), ACA1464GAT (4440A>G 4441C>A 4442A>T), CTC1465TTA (4443C>T 4445C>A), TTC1466TTA (4448C>A), CAC1468AGA (4452C>A 4453A>G 4454C>A), TCC1470TAT (4459C>A 4460C>T), AAA1471GAT (4461A>G 4463A>T), GCA1472GCC (4466A>C), AAG1473AAT (4469G>T), CTC1474ATA (4470C>A 4472C>A), AAG1477AAT (4481G>T), CTT1480TTA (4488C>T 4490T>A), AAG1481AAA (4493G>A), TTT1484TAT (4501T>A) | | | | | | | | |

##### Proteins

| | Begin | End | Coverage | Score | Concordance | Matches | Identities | I/D/M/F* | Stop Codons |
|---|---|---|---|---|---|---|---|---|---|
| ORF I polyprotein (NP_127504.1) | 1381 | 1485 | 4.8% | 387 | 50.5% | 105 (100%) | 57 (54.3%) | 0/0/0/0 | 0 |
| Protein mutations: | I1383D (4197A>G 4198T>A 4199C>T), N1384I (4201A>T), A1388S (4212G>T), S1389R (4215A>C 4217C>T), H1390P (4219A>C 4220T>C), S1391C (4222C>G), G1392Q (4224G>C 4225G>A), P1395V (4233C>G 4234C>T 4235A>T), H1397L (4239C>T 4240A>T 4241T>A), L1398V (4242C>G 4244C>G), Q1399E (4245C>G 4247G>A), L1400F (4250A>C), A1401C (4251G>T 4252C>G 4253A>C), I1402K (4255T>A), E1404K (4260G>A 4262A>G), C1405I (4263T>A 4264G>T), E1407N (4269G>A 4271G>T), Q1409L (4275C>T 4276A>T), F1411K (4281T>A 4282T>A 4283T>G), D1412G (4285A>G), E1415K (4293G>A), D1418K (4302G>A 4304T>G), Q1420P (4309A>C 4310A>T), A1422S (4314G>T), E1424T (4320G>A 4321A>C), A1425V (4324C>T), K1430N (4340G>T), R1431A (4341A>G 4342G>C 4343G>T), Q1434K (4350C>A 4352A>G), V1435E (4354T>A 4355G>A), K1438I (4363A>T 4364A>T), Q1446K (4386C>A), H1450K (4398C>A 4400T>A), Q1453K (4407C>A), D1454W (4410G>T 4411A>G 4412T>G), D1455I (4413G>A 4414A>T 4415C>T), K1456R (4417A>G), F1457Y (4420T>A), L1463R (4437C>A 4438T>G 4439C>G), T1464D (4440A>G 4441C>A 4442A>T), F1466L (4448C>A), H1468R (4452C>A 4453A>G 4454C>A), S1470Y (4459C>A 4460C>T), K1471D (4461A>G 4463A>T), K1473N (4469G>T), L1474I (4470C>A 4472C>A), K1477N (4481G>T), F1484Y (4501T>A) | | | | | | | | |
| Codon mutations: | AAC1382AAT (4196C>T), ATC1383GAT (4197A>G 4198T>A 4199C>T), AAT1384ATT (4201A>T), ACA1386ACT (4208A>T), AAG1387AAA (4211G>A), GCA1388TCA (4212G>T), AGC1389CGT (4215A>C 4217C>T), CAT1390CCC (4219A>C 4220T>C), TCT1391TGT (4222C>G), GGA1392CAA (4224G>C 4225G>A), CCA1395GTT (4233C>G 4234C>T 4235A>T), CAT1397TTA (4239C>T 4240A>T 4241T>A), CTC1398GTG (4242C>G 4244C>G), CAG1399GAA (4245C>G 4247G>A), TTA1400TTC (4250A>C), GCA1401TGC (4251G>T 4252C>G 4253A>C), ATA1402AAA (4255T>A), AAG1403AAA (4259G>A), GAA1404AAG (4260G>A 4262A>G), TGT1405ATT (4263T>A 4264G>T), GAG1407AAT (4269G>A 4271G>T), CAA1409TTA (4275C>T 4276A>T), TTT1411AAG (4281T>A 4282T>A 4283T>G), GAT1412GGT (4285A>G), CTA1413TTA (4287C>T), ATC1414ATA (4292C>A), GAA1415AAA (4293G>A), CCC1416CCA (4298C>A), TCT1417TCA (4301T>A), GAT1418AAG (4302G>A 4304T>G), TCA1419TCT (4307A>T), CAA1420CCT (4309A>C 4310A>T), GCA1422TCA (4314G>T), TGT1423TGC (4319T>C), GAA1424ACA (4320G>A 4321A>C), GCT1425GTT (4324C>T), GTC1428GTT (4334C>T), AAC1429AAT (4337C>T), AAG1430AAT (4340G>T), AGG1431GCT (4341A>G 4342G>C 4343G>T), TCA1432AGT (4344T>A 4345C>G 4346A>T), GAG1433GAA (4349G>A), CAA1434AAG (4350C>A 4352A>G), GTG1435GAA (4354T>A 4355G>A), AGA1436CGA (4356A>C), GGA1437GGT (4361A>T), AAA1438ATT (4363A>T 4364A>T), TTA1439CTC (4365T>C 4367A>C), AGG1440CGA (4368A>C 4370G>A), GTC1442GTT (4376C>T), CAA1446AAA (4386C>A), AAC1449AAT (4397C>T), CAT1450AAA (4398C>A 4400T>A), TTC1451TTT (4403C>T), CTC1452CTT (4406C>T), CAA1453AAA (4407C>A), GAT1454TGG (4410G>T 4411A>G 4412T>G), GAC1455ATT (4413G>A 4414A>T 4415C>T), AAG1456AGG (4417A>G), TTT1457TAT (4420T>A), CCA1460CCG (4430A>G), AAG1462AAA (4436G>A), CTC1463AGG (4437C>A 4438T>G 4439C>G), ACA1464GAT (4440A>G 4441C>A 4442A>T), CTC1465TTA (4443C>T 4445C>A), TTC1466TTA (4448C>A), CAC1468AGA (4452C>A 4453A>G 4454C>A), TCC1470TAT (4459C>A 4460C>T), AAA1471GAT (4461A>G 4463A>T), GCA1472GCC (4466A>C), AAG1473AAT (4469G>T), CTC1474ATA (4470C>A 4472C>A), AAG1477AAT (4481G>T), CTT1480TTA (4488C>T 4490T>A), AAG1481AAA (4493G>A), TTT1484TAT (4501T>A) | | | | | | | | |

*: Inserts / Deletes / Misaligned / Frameshifts

#### Analysis details

This analysis was performed with panviral2.64

### NGS Details (UN24): Marseillevirus marseillevirus

#### Assembly

| | |
|---|---|
| **Coverage Length** | 117 (1 contig(s)) |
| **Depth Of Coverage** | 11.4 |
| **Number Of Reads** | 18 |
| **Reads Per Million** | 0.36 rpm (after QC) |
| **Ambiguities** | 0 |
| **Assembly Method** | de novo + reference guided assembly |
| **Consensus Caller** | Bcf Tools |

#### Coverage Map

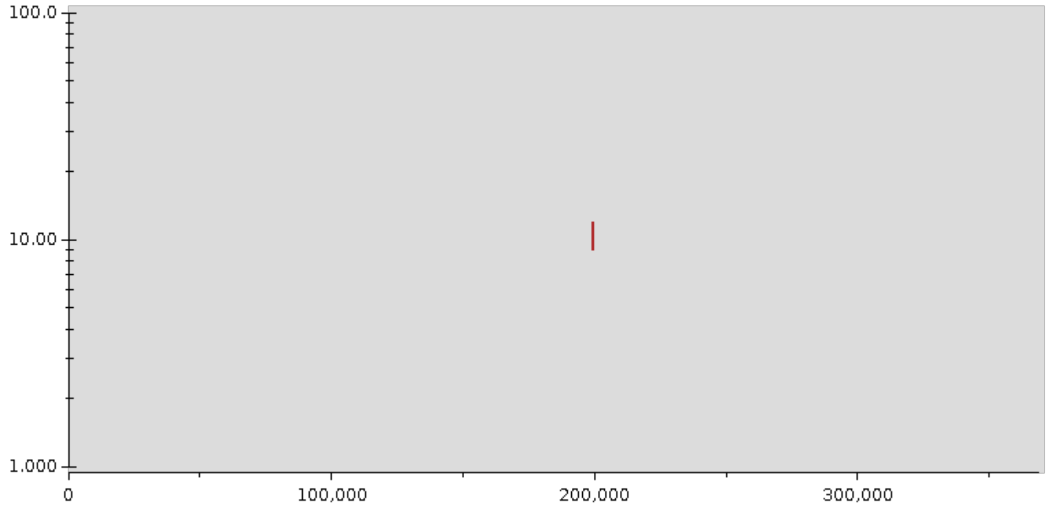

#### Assignment

| | |
|---|---|
| **Type** | Marseillevirus marseillevirus (Taxonomy ID: 694581) |
| **Reference Genome** | NC_013756.1 |
| **NT Identity (%)** | 82.0513 |
| **AA Identity (%)** | 92.3077 |
| **Number Of Stop Codons** | 0 |
| **Number Of CDS** | 428 |

#### Alignment

| | |
|---|---|
| **Alignment Score** | 150.0 (NT) + 248.0 (AA) = 398.0 |
| **Concordance (%)** | 81.893 |
| **Alignment Method** | Local, heuristic, nucleotide (BLASTN) |

#### Genome Region

Sequence starts at position 199409 and ends at position 199525 relative to NC_013756.1 reference sequence.

#### Alignment Detailed Statistics

| | Begin | End | Coverage | Score | Concordance | Matches | Identities | I/D/M/F* | Stop Codons |
|---|---|---|---|---|---|---|---|---|---|
| **NT** | **199409** | **199525** | **0.1%** | **150** | **64.1%** | **117 (100%)** | **96 (82.1%)** | **0/0** | |

Mutations: 199414T>A, 199416T>G, 199417C>G, 199420G>A, 199423A>C, 199425G>A, 199429G>A, 199432A>C, 199444T>C, 199449C>T, 199461C>A, 199462C>G, 199464A>G, 199468C>G, 199470T>G, 199471T>G, 199477T>C, 199480C>G, 199489T>G, 199513G>A, 199516C>G

*: Inserts / Deletes / Misaligned / Frameshifts

#### Analysis details

This analysis was performed with panviral2.64

#### NGS Details (UN24): Metaplexis yellow mottle-associated virus

##### Assembly

| | |
|---|---|
| **Coverage Length** | 293 (1 contig(s)) |
| **Depth Of Coverage** | 6.8 |
| **Number Of Reads** | 17 |
| **Reads Per Million** | 0.34 rpm (after QC) |
| **Ambiguities** | 0 |
| **Assembly Method** | de novo + reference guided assembly |
| **Consensus Caller** | Bcf Tools |

##### Coverage Map

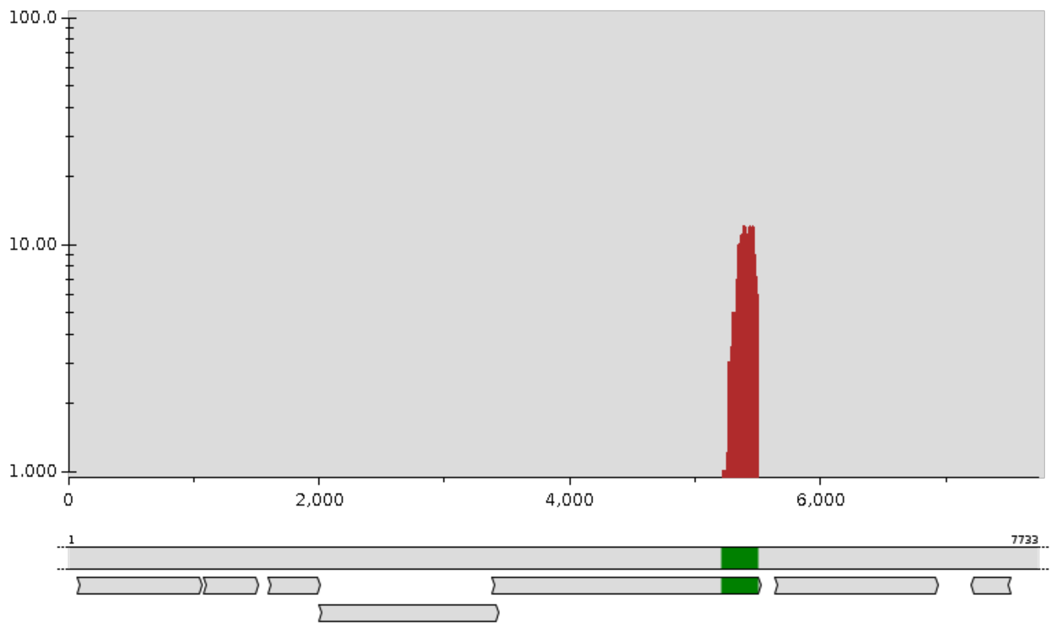

##### Assignment

| | |
|---|---|
| **Type** | Metaplexis yellow mottle-associated virus (Taxonomy ID: 2878269) |
| **Reference Genome** | NC_077108.1 |
| **NT Identity (%)** | 58.7031 |
| **AA Identity (%)** | 45.9184 |
| **Number Of Stop Codons** | 0 |
| **Number Of CDS** | 7 |

##### Alignment

| | |
|---|---|
| **Alignment Score** | 102.0 (NT) + 323.0 (AA) = 425.0 |
| **Concordance (%)** | 34.219 |

| **Alignment Method** | Global, seeded, nucleotide + amino acids (AGA) |
|---|---|

#### Genome Region

Sequence starts at position 5214 and ends at position 5506 relative to NC_077108.1 reference sequence.

#### Alignment Detailed Statistics

| | Begin | End | Coverage | Score | Concordance | Matches | Identities | I/D/M/F* | Stop Codons |
|---|---|---|---|---|---|---|---|---|---|
| **NT** | **5214** | **5506** | **3.8%** | **102** | **17.4%** | **293 (100%)** | **172 (58.7%)** | **0/0** | |
| Mutations: | 5217A>T, 5219C>T, 5226T>A, 5227G>C, 5229T>A, 5230C>G, 5231C>T, 5233G>A, 5234T>C, 5235T>A, 5243A>T, 5244C>G, 5246G>A, 5247G>A, 5252G>A, 5254A>C, 5255A>T, 5258T>A, 5262C>T, 5263A>C, 5264T>A, 5266G>C, 5267T>A, 5273G>A, 5276A>G, 5280A>T, 5285A>G, 5288T>C, 5289T>A, 5291G>A, 5292A>G, 5294T>A, 5298A>G, 5299C>G, 5300A>C, 5303T>A, 5304A>C, 5305A>G, 5306G>A, 5307G>A, 5308C>A, 5309T>A, 5313A>T, 5314A>C, 5315A>T, 5318A>T, 5320A>T, 5324G>A, 5331G>A, 5332T>A, 5333A>G, 5334A>C, 5335G>A, 5339T>C, 5342T>C, 5343G>A, 5344C>T, 5345T>C, 5349A>T, 5351A>G, 5354T>C, 5360T>A, 5361A>C, 5363T>G, 5366T>A, 5367A>T, 5368A>C, 5370T>G, 5371A>G, 5375C>G, 5378T>A, 5379A>T, 5380G>T, 5383C>A, 5387C>A, 5388A>C, 5390C>T, 5391T>C, 5394G>T, 5395G>C, 5399T>A, 5400T>C, 5401A>C, 5402C>T, 5403A>C, 5406C>T, 5408A>T, 5409G>A, 5410C>G, 5414A>G, 5415C>T, 5417T>A, 5418G>C, 5419T>A, 5420G>T, 5430C>G, 5431A>T, 5434A>T, 5435C>G, 5436C>T, 5438T>A, 5439A>T, 5440A>C, 5443A>T, 5444C>T, 5447C>T, 5449A>C, 5454G>A, 5461T>A, 5463C>A, 5465C>T, 5475A>T, 5476A>C, 5478A>G, 5479C>A, 5487C>T, 5495C>T, 5496A>T, 5499A>C, 5501G>A, 5504C>A | | | | | | | | |

##### CDS

| | Begin | End | Coverage | Score | Concordance | Matches | Identities | I/D/M/F* | Stop Codons |
|---|---|---|---|---|---|---|---|---|---|
| QKU89_gp5 | 611 | 708 | 13.7% | 323 | 48.6% | 98 (100%) | 45 (45.9%) | 0/0/0/0 | 0 |
| Protein mutations: | S612C (5217A>T 5219C>T), C615T (5226T>A 5227G>C), G617D (5233G>A 5234T>C), S618T (5235T>A), K620N (5243A>T), Q621E (5244C>G 5246G>A), A622T (5247G>A), K624T (5254A>C 5255A>T), N625K (5258T>A), H627S (5262C>T 5263A>C 5264T>A), S628T (5266G>C 5267T>A), I633L (5280A>T), L636I (5289T>A 5291G>A), I637V (5292A>G 5294T>A), T639G (5298A>G 5299C>G 5300A>C), K641R (5304A>C 5305A>G 5306G>A), A642K (5307G>A 5308C>A 5309T>A), K644S (5313A>T 5314A>C 5315A>T), Y646F (5320A>T), V650K (5331G>A 5332T>A 5333A>G), R651Q (5334A>C 5335G>A), A654I (5343G>A 5344C>T 5345T>C), T656S (5349A>T 5351A>G), N660Q (5361A>C 5363T>G), K662S (5367A>T 5368A>C), Y663G (5370T>G 5371A>G), F664L (5375C>G), S666F (5379A>T 5380G>T), T667N (5383C>A), N668K (5387C>A), I669L (5388A>C 5390C>T), S670P (5391T>C), G671S (5394G>T 5395G>C), D672E (5399T>A), Y673P (5400T>C 5401A>C 5402C>T), K674Q (5403A>C), Q675Y (5406C>T 5408A>T), A676R (5409G>A 5410C>G), V679H (5418G>C 5419T>A 5420G>T), Q683V (5430C>G 5431A>T), Y684L (5434A>T 5435C>G), N686S (5439A>T 5440A>C), Y687F (5443A>T 5444C>T), N689T (5449A>C), E691K (5454G>A), V693E (5461T>A), H694N (5463C>A 5465C>T), N698S (5475A>T 5476A>C), T699D (5478A>G 5479C>A), L702F (5487C>T), I705F (5496A>T), M706L (5499A>C 5501G>A) | | | | | | | | |
| Codon mutations: | AGC612TGT (5217A>T 5219C>T), TGT615ACT (5226T>A 5227G>C), TCC616AGT (5229T>A 5230C>G 5231C>T), GGT617GAC (5233G>A 5234T>C), TCA618ACA (5235T>A), AAA620AAT (5243A>T), CAG621GAA (5244C>G 5246G>A), GCC622ACC (5247G>A), GAG623GAA (5252G>A), AAA624ACT (5254A>C 5255A>T), AAT625AAA (5258T>A), CAT627TCA (5262C>T 5263A>C 5264T>A), AGT628ACA (5266G>C 5267T>A), GAG630GAA (5273G>A), AAA631AAG (5276A>G), ATA633TTA (5280A>T), TTA634TTG (5285A>G), GCT635GCC (5288T>C), TTG636ATA (5289T>A 5291G>A), ATT637GTA (5292A>G 5294T>A), ACA639GGC (5298A>G 5299C>G 5300A>C), ATT640ATA (5303T>A), AAG641CGA (5304A>C 5305A>G 5306G>A), GCT642AAA (5307G>A 5308C>A 5309T>A), AAA644TCT (5313A>T 5314A>C 5315A>T), GCA645GCT (5318A>T), TAT646TTT (5320A>T), CTG647CTA (5324G>A), GTA650AAG (5331G>A 5332T>A 5333A>G), AGA651CAA (5334A>C 5335G>A), TTT652TTC (5339T>C), ATT653ATC (5342T>C), GCT654ATC (5343G>A 5344C>T 5345T>C), ACA656TCG (5349A>T 5351A>G), GAT657GAC (5354T>C), ACT659ACA (5360T>A), AAT660CAG (5361A>C 5363T>G), GTT661GTA (5366T>A), AAA662TCA (5367A>T 5368A>C), TAT663GGT (5370T>G 5371A>G), TTC664TTG (5375C>G), ATT665ATA (5378T>A), AGT666TTT (5379A>T 5380G>T), ACT667AAT (5383C>A), AAC668AAA (5387C>A), ATC669CTT (5388A>C 5390C>T), TCA670CCA (5391T>C), GGT671TCT (5394G>T 5395G>C), GAT672GAA (5399T>A), TAC673CCT (5400T>C 5401A>C 5402C>T), AAA674CAA (5403A>C), CAA675TAT (5406C>T 5408A>T), GCA676AGA (5409G>A 5410C>G), AGA677AGG (5414A>G), CTT678TTA (5415C>T 5417T>A), GTG679CAT (5418G>C 5419T>A 5420G>T), CAA683GTA (5430C>G 5431A>T), TAC684TTG (5434A>T 5435C>G), CTT685TTA (5436C>T 5438T>A), AAT686TCT (5439A>T 5440A>C), TAC687TTT (5443A>T 5444C>T), TAC688TAT (5447C>T), AAT689ACT (5449A>C), GAA691AAA (5454G>A), GTA693GAA (5461T>A), CAC694AAT (5463C>A 5465C>T), AAT698TCT (5475A>T 5476A>C), ACT699GAT (5478A>G 5479C>A), CTT702TTT (5487C>T), GAC704GAT (5495C>T), ATC705TTC (5496A>T), ATG706CTA (5499A>C 5501G>A), TCC707TCA (5504C>A) | | | | | | | | |

##### Proteins

| | Begin | End | Coverage | Score | Concordance | Matches | Identities | I/D/M/F* | Stop Codons |
|---|---|---|---|---|---|---|---|---|---|
| polymerase polyprotein (YP_010805169.1) | 611 | 708 | 13.7% | 323 | 48.6% | 98 (100%) | 45 (45.9%) | 0/0/0/0 | 0 |
| Protein mutations: | S612C (5217A>T 5219C>T), C615T (5226T>A 5227G>C), G617D (5233G>A 5234T>C), S618T (5235T>A), K620N (5243A>T), Q621E (5244C>G 5246G>A), A622T (5247G>A), K624T (5254A>C 5255A>T), N625K (5258T>A), H627S (5262C>T 5263A>C 5264T>A), S628T (5266G>C 5267T>A), I633L (5280A>T), L636I (5289T>A 5291G>A), I637V (5292A>G 5294T>A), T639G (5298A>G 5299C>G 5300A>C), K641R (5304A>C 5305A>G 5306G>A), A642K (5307G>A 5308C>A 5309T>A), K644S (5313A>T 5314A>C 5315A>T), Y646F (5320A>T), V650K (5331G>A 5332T>A 5333A>G), R651Q (5334A>C 5335G>A), A654I (5343G>A 5344C>T 5345T>C), T656S (5349A>T 5351A>G), N660Q (5361A>C 5363T>G), K662S (5367A>T 5368A>C), Y663G (5370T>G 5371A>G), F664L (5375C>G), S666F (5379A>T 5380G>T), T667N (5383C>A), N668K (5387C>A), I669L (5388A>C 5390C>T), S670P (5391T>C), G671S (5394G>T 5395G>C), D672E (5399T>A), Y673P (5400T>C 5401A>C 5402C>T), K674Q (5403A>C), Q675Y (5406C>T 5408A>T), A676R (5409G>A 5410C>G), V679H (5418G>C 5419T>A 5420G>T), Q683V (5430C>G 5431A>T), Y684L (5434A>T 5435C>G), N686S (5439A>T 5440A>C), Y687F (5443A>T 5444C>T), N689T (5449A>C), E691K (5454G>A), V693E (5461T>A), H694N (5463C>A 5465C>T), N698S (5475A>T 5476A>C), T699D (5478A>G 5479C>A), L702F (5487C>T), I705F (5496A>T), M706L (5499A>C 5501G>A) | | | | | | | | |
| Codon mutations: | AGC612TGT (5217A>T 5219C>T), TGT615ACT (5226T>A 5227G>C), TCC616AGT (5229T>A 5230C>G 5231C>T), GGT617GAC (5233G>A 5234T>C), TCA618ACA (5235T>A), AAA620AAT (5243A>T), CAG621GAA (5244C>G 5246G>A), GCC622ACC (5247G>A), GAG623GAA (5252G>A), AAA624ACT (5254A>C 5255A>T), AAT625AAA (5258T>A), CAT627TCA (5262C>T 5263A>C 5264T>A), AGT628ACA (5266G>C 5267T>A), GAG630GAA (5273G>A), AAA631AAG (5276A>G), ATA633TTA (5280A>T), TTA634TTG (5285A>G), GCT635GCC (5288T>C), TTG636ATA (5289T>A 5291G>A), ATT637GTA (5292A>G 5294T>A), ACA639GGC (5298A>G 5299C>G 5300A>C), ATT640ATA (5303T>A), AAG641CGA (5304A>C 5305A>G 5306G>A), GCT642AAA (5307G>A 5308C>A 5309T>A), AAA644TCT (5313A>T 5314A>C 5315A>T), GCA645GCT (5318A>T), TAT646TTT (5320A>T), CTG647CTA (5324G>A), GTA650AAG (5331G>A 5332T>A 5333A>G), AGA651CAA (5334A>C 5335G>A), TTT652TTC (5339T>C), ATT653ATC (5342T>C), GCT654ATC (5343G>A 5344C>T 5345T>C), ACA656TCG (5349A>T 5351A>G), GAT657GAC (5354T>C), ACT659ACA (5360T>A), AAT660CAG (5361A>C 5363T>G), GTT661GTA (5366T>A), AAA662TCA (5367A>T 5368A>C), TAT663GGT (5370T>G 5371A>G), TTC664TTG (5375C>G), ATT665ATA (5378T>A), AGT666TTT (5379A>T 5380G>T), ACT667AAT (5383C>A), AAC668AAA (5387C>A), ATC669CTT (5388A>C 5390C>T), TCA670CCA (5391T>C), GGT671TCT (5394G>T 5395G>C), GAT672GAA (5399T>A), TAC673CCT (5400T>C 5401A>C 5402C>T), AAA674CAA (5403A>C), CAA675TAT (5406C>T 5408A>T), GCA676AGA (5409G>A 5410C>G), AGA677AGG (5414A>G), CTT678TTA (5415C>T 5417T>A), GTG679CAT (5418G>C 5419T>A 5420G>T), CAA683GTA (5430C>G 5431A>T), TAC684TTG (5434A>T 5435C>G), CTT685TTA (5436C>T 5438T>A), AAT686TCT (5439A>T 5440A>C), TAC687TTT (5443A>T 5444C>T), TAC688TAT (5447C>T), AAT689ACT (5449A>C), GAA691AAA (5454G>A), GTA693GAA (5461T>A), CAC694AAT (5463C>A 5465C>T), AAT698TCT (5475A>T 5476A>C), ACT699GAT (5478A>G 5479C>A), CTT702TTT (5487C>T), GAC704GAT (5495C>T), ATC705TTC (5496A>T), ATG706CTA (5499A>C 5501G>A), TCC707TCA (5504C>A) | | | | | | | | |

*: Inserts / Deletes / Misaligned / Frameshifts

#### Analysis details

This analysis was performed with panviral2.64

### NGS Details (UN24): Cavemovirus venamanihotis

#### Assembly

| | |
|---|---|
| **Coverage Length** | 577 (1 contig(s)) |
| **Depth Of Coverage** | 3.3 |
| **Number Of Reads** | 15 |
| **Reads Per Million** | 0.30 rpm (after QC) |
| **Ambiguities** | 0 |
| **Assembly Method** | de novo + reference guided assembly |
| **Consensus Caller** | Bcf Tools |

#### Coverage Map

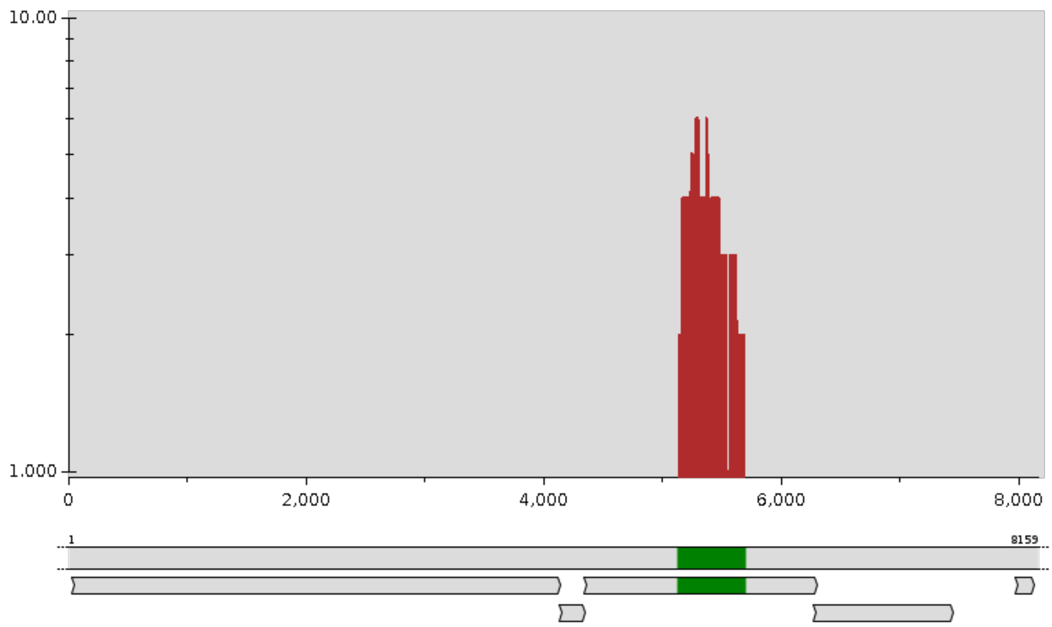

#### Assignment

| | |
|---|---|
| **Type** | Cavemovirus venamanihotis (Taxonomy ID: 3047736) |
| **Reference Genome** | NC_001648.1 |
| **NT Identity (%)** | 57.1924 |
| **AA Identity (%)** | 47.3958 |
| **Number Of Stop Codons** | 0 |
| **Number Of CDS** | 5 |

#### Alignment

| | |
|---|---|
| **Alignment Score** | 148.0 (NT) + 625.0 (AA) = 773.0 |
| **Concordance (%)** | 31.929 |

| **Alignment Method** | Global, seeded, nucleotide + amino acids (AGA) |
|---|---|

#### Genome Region

Sequence starts at position 5128 and ends at position 5704 relative to NC_001648.1 reference sequence.

#### Alignment Detailed Statistics

| | Begin | End | Coverage | Score | Concordance | Matches | Identities | I/D/M/F* | Stop Codons |
|---|---|---|---|---|---|---|---|---|---|
| **NT** | **5128** | **5704** | **7.1%** | **148** | **13.2%** | **568 (96.9%)** | **330 (56.3%)** | **9/9** | |
| Mutations: | 5130A>C, 5132G>T, 5135A>T, 5139G>A, 5148G>C, 5149A>C, 5151T>C, 5153A>G, 5156T>C, 5158A>C, 5159A>T, 5160A>G, 5161A>C, 5162A>T, 5163G>C, 5164C>T, 5165T>A, 5169G>T, 5170T>G, 5171A>G, 5172G>A, 5176A>G, 5180T>C, 5183T>G, 5186C>T, 5189A>T, 5196G>A, 5197A>G, 5198T>A, 5199A>G, 5200C>A, 5201A>T, 5202T>C, 5204A>T, 5205A>T, 5208C>A, 5210T>A, 5211A>C, 5215G>C, 5216T>C, 5217A>T, 5219A>T, 5220C>A, 5222G>A, 5224C>A, 5225A>C, 5227G>A, 5228A>T, 5229T>A, 5230A>T, 5231C>T, 5246T>C, 5247T>A, 5248G>T, 5249T>G, 5258A>C, 5263A>G, 5264T>G, 5267T>A, 5270A>C, 5271A>C, 5273G>A, 5274T>A, 5276A>T, 5277G>T, 5278A>T, 5279A>C, 5283G>A, 5285T>G, 5286A>G, 5287G>A, 5292A>T, 5294A>T, 5295T>A, 5297T>A, 5300A>T, 5306T>C, 5308C>A, 5309A>T, 5312T>C, 5316C>T, 5317A>T, 5318G>T, 5322T>C, 5324T>A, 5328C>G, 5330G>A, 5336A>T, 5338T>C, 5339A>T, 5340T>A, 5342A>G, 5352T>C, 5353A>T, 5354T>A, 5355C>A, 5357T>A, 5361T>G, 5363A>C, 5366A>T, 5367A>T, 5368G>T, 5369T>C, 5370A>G, 5371T>A, 5372T>A, 5375C>T, 5379C>A, 5381A>T, 5382T>A, 5384T>C, 5388G>A, 5392G>A, 5393A>T, 5396A>T, 5401G>A, 5402A>T, 5410A>T, 5411T>A, 5420T>C, 5426G>T, 5438T>C, 5441A>T, 5444A>G, 5445G>A, 5447C>T, 5449T>A, 5456G>A, 5459T>G, 5460A>T, 5466G>A, 5467A>T, 5473A>T, 5474A>C, 5476T>A, 5481A>C, 5485C>A, 5486A>T, 5488A>T, 5489A>C, 5492T>C, 5494G>A, 5496G>A, 5498T>A, 5501A>T, 5502A>G, 5503C>T, 5504A>T, 5505C>A, 5507T>A, 5508G>C, 5509C>A, 5515G>A, 5516A>T, 5519A>G, 5520A>G, 5522T>A, 5526A>T, 5527G>C, 5531G>A, 5532A>C, 5533A>C, 5539C>T, 5540A>G, 5541G>A, 5542A>G, 5543A>T, 5548G>T, 5550A>C, 5553G>A, 5554A>C, 5556A>G, 5558A>T, 5559A>G, 5561A>C, 5562G>C, 5563A>G, 5564T>A, 5567C>T, 5570T>A, 5574G>C, 5575T>A, 5576C>T, 5577C>A, 5579A>T, 5582A>T, 5583G>T, 5584A>G, 5585A>C, 5590G>A, 5592G>A, 5593G>A, 5594T>A, 5598_5606delGAATTACAG, 5607C>A, 5609T>C, 5615_5616insCAGAGATCT, 5619A>G, 5622A>T, 5623A>T, 5624A>T, 5625A>G, 5626T>C, 5629T>C, 5631G>A, 5634A>T, 5635A>T, 5636A>T, 5638A>T, 5640A>G, 5641C>A, 5642G>T, 5644A>T, 5645A>T, 5650A>C, 5651A>T, 5652A>G, 5654T>C, 5656A>G, 5657A>G, 5661G>A, 5662A>T, 5663A>T, 5664T>C, 5666A>C, 5671G>C, 5672T>A, 5673A>T, 5675A>T, 5678A>G, 5682T>A, 5683T>A, 5684A>T, 5690C>T, 5691C>T, 5693G>T, 5697A>T, 5698G>T, 5699A>T, 5701A>T | | | | | | | | |

##### CDS

| | | | | | | | | | |
|---|---|---|---|---|---|---|---|---|---|
| CsVMVgp3 | 263 | 454 | 29.4% | 625 | 47.3% | 189 (96.9%) | 91 (46.7%) | 3/3/0/0 | 0 |
| Protein mutations: | M263L (5130A>C 5132G>T), D266N (5139G>A), D269P (5148G>C 5149A>C), K272T (5158A>C 5159A>T), K273A (5160A>G 5161A>C 5162A>T), A274L (5163G>C 5164C>T 5165T>A), V276W (5169G>T 5170T>G 5171A>G), V277I (5172G>A), K278R (5176A>G), D285R (5196G>A 5197A>G 5198T>A), T286D (5199A>G 5200C>A 5201A>T), I288L (5205A>T), H289K (5208C>A 5210T>A), S291T (5215G>C 5216T>C), I292F (5217A>T 5219A>T), Q293K (5220C>A 5222G>A), A294D (5224C>A 5225A>C), R295N (5227G>A 5228A>T), Y296I (5229T>A 5230A>T 5231C>T), C302M (5247T>A 5248G>T 5249T>G), Y307W (5263A>G 5264T>G), H308Q (5267T>A), K310Q (5271A>C 5273G>A), L311I (5274T>A 5276A>T), E312F (5277G>T 5278A>T 5279A>C), D314K (5283G>A 5285T>G), S315D (5286A>G 5287G>A), K317Y (5292A>T 5294A>T), Y318K (5295T>A 5297T>A), T322N (5308C>A 5309A>T), Q325F (5316C>T 5317A>T 5318G>T), Y327Q (5322T>C 5324T>A), Q329E (5328C>G 5330G>A), K331N (5336A>T), V332A (5338T>C 5339A>T), L333M (5340T>A 5342A>G), Y337L (5352T>C 5353A>T 5354T>A), H338K (5355C>A 5357T>A), S340A (5361T>G 5363A>C), S342F (5367A>T 5368G>T 5369T>C), I343E (5370A>G 5371T>A 5372T>A), Q346N (5379C>A 5381A>T), F347I (5382T>A 5384T>C), D349N (5388G>A), R350N (5392G>A 5393A>T), R353N (5401G>A 5402A>T), Y356L (5410A>T 5411T>A), V368I (5445G>A 5447C>T), F369Y (5449T>A), I373F (5460A>T), E375M (5466G>A 5467A>T), K377I (5473A>T 5474A>C), I378K (5476T>A), I380L (5481A>C), A381D (5485C>A 5486A>T), K382I (5488A>T 5489A>C), R384K (5494G>A), D385K (5496G>A 5498T>A), T387V (5502A>G 5503C>T 5504A>T), L388I (5505C>A 5507T>A), A389Q (5508G>C 5509C>A), G391D (5515G>A 5516A>T), I393V (5520A>G 5522T>A), K397P (5532A>C 5533A>C), T399M (5539C>T 5540A>G), E400S (5541G>A 5542A>G 5543A>T), C402F (5548G>T), K403Q (5550A>C), E404T (5553G>A 5554A>C), K405D (5556A>G 5558A>T), I406V (5559A>G 5561A>C), D407R (5562G>C 5563A>G 5564T>A), V411H (5574G>C 5575T>A 5576C>T), Q412N (5577C>A 5579A>T), E414C (5583G>T 5584A>G 5585A>C), G416E (5590G>A), G417K (5592G>A 5593G>A 5594T>A), E419_Q421del (5598_5606delGAATTACAG), P422T (5607C>A 5609T>C), I424_I425insQRS (5615_5616insCAGAGATCT), N426D (5619A>G), K427F (5622A>T 5623A>T 5624A>T), I428A (5625A>G 5626T>C), L429S (5629T>C), E430K (5631G>A), K431F (5634A>T 5635A>T 5636A>T), H432L (5638A>T), T433D (5640A>G 5641C>A 5642G>T), K434I (5644A>T 5645A>T), K436T (5650A>C 5651A>T), N437D (5652A>G 5654T>C), K438R (5656A>G 5657A>G), E440I (5661G>A 5662A>T 5663A>T), S443T (5671G>C 5672T>A), I444F (5673A>T 5675A>T), L447N (5682T>A 5683T>A 5684A>T), Q450Y (5691C>T 5693G>T), R452F (5697A>T 5698G>T 5699A>T), H453L (5701A>T) | | | | | | | | |
| Codon mutations: | ATG263CTT (5130A>C 5132G>T), GTA264GTT (5135A>T), GAT266AAT (5139G>A), GAT269CCT (5148G>C 5149A>C), TTA270CTG (5151T>C 5153A>G), AAT271AAC (5156T>C), AAA272ACT (5158A>C 5159A>T), AAA273GCT (5160A>G 5161A>C 5162A>T), GCT274CTA (5163G>C 5164C>T 5165T>A), GTA276TGG (5169G>T 5170T>G 5171A>G), GTA277ATA (5172G>A), AAA278AGA (5176A>G), TAT279TAC (5180T>C), CCT280CCG (5183T>G), ATC281ATT (5186C>T), CCA282CCT (5189A>T), GAT285AGA (5196G>A 5197A>G 5198T>A), ACA286GAT (5199A>G 5200C>A 5201A>T), TTA287CTT (5202T>C 5204A>T), ATA288TTA (5205A>T), CAT289AAA (5208C>A 5210T>A), AGA290CGA (5211A>C), AGT291ACC (5215G>C 5216T>C), ATA292TTT (5217A>T 5219A>T), CAG293AAA (5220C>A 5222G>A), GCA294GAC (5224C>A 5225A>C), AGA295AAT (5227G>A 5228A>T), TAC296ATT (5229T>A 5230A>T 5231C>T), GAT301GAC (5246T>C), TGT302ATG (5247T>A 5248G>T 5249T>G), GGA305GGC (5258A>C), TAT307TGG (5263A>G 5264T>G), CAT308CAA (5267T>A), ATA309ATC (5270A>C), AAG310CAA (5271A>C 5273G>A), TTA311ATT (5274T>A 5276A>T), GAA312TTC (5277G>T 5278A>T 5279A>C), GAT314AAG (5283G>A 5285T>G), AGT315GAT (5286A>G 5287G>A), AAA317TAT (5292A>T 5294A>T), TAT318AAA (5295T>A 5297T>A), ACA319ACT (5300A>T), TTT321TTC (5306T>C), ACA322AAT (5308C>A 5309A>T), GTT323GTC (5312T>C), CAG325TTT (5316C>T 5317A>T 5318G>T), TAT327CAA (5322T>C 5324T>A), CAG329GAA (5328C>G 5330G>A), AAA331AAT (5336A>T), GTA332GCT (5338T>C 5339A>T), TTA333ATG (5340T>A 5342A>G), TAT337CTA (5352T>C 5353A>T 5354T>A), CAT338AAA (5355C>A 5357T>A), TCA340GCC (5361T>G 5363A>C), CCA341CCT (5366A>T), AGT342TTC (5367A>T 5368G>T 5369T>C), ATT343GAA (5370A>G 5371T>A 5372T>A), TTC344TTT (5375C>T), CAA346AAT (5379C>A 5381A>T), TTT347ATC (5382T>A 5384T>C), GAT349AAT (5388G>A), AGA350AAT (5392G>A 5393A>T), ATA351ATT (5396A>T), AGA353AAT (5401G>A 5402A>T), TAT356TTA (5410A>T 5411T>A), ATT359ATC (5420T>C), GTG361GTT (5426G>T), GAT365GAC (5438T>C), ATA366ATT (5441A>T), TTA367TTG (5444A>G), GTC368ATT (5445G>A 5447C>T), TTT369TAT (5449T>A), AAG371AAA (5456G>A), ACT372ACG (5459T>G), ATT373TTT (5460A>T), GAG375ATG (5466G>A 5467A>T), AAA377ATC (5473A>T 5474A>C), ATA378AAA (5476T>A), ATT380CTT (5481A>C), GCA381GAT (5485C>A 5486A>T), AAA382ATC (5488A>T 5489A>C), TTT383TTC (5492T>C), AGA384AAA (5494G>A), GAT385AAA (5496G>A 5498T>A), ATA386ATT (5501A>T), ACA387GTT (5502A>G 5503C>T 5504A>T), CTT388ATA (5505C>A 5507T>A), GCA389CAA (5508G>C 5509C>A), GGA391GAT (5515G>A 5516A>T), TTA392TTG (5519A>G), ATT393GTA (5520A>G 5522T>A), AGT395TCT (5526A>T 5527G>C), AAG396AAA (5531G>A), AAA397CCA (5532A>C 5533A>C), ACA399ATG (5539C>T 5540A>G), GAA400AGT (5541G>A 5542A>G 5543A>T), TGT402TTT (5548G>T), AAA403CAA (5550A>C), GAA404ACA (5553G>A 5554A>C), AAA405GAT (5556A>G 5558A>T), ATA406GTC (5559A>G 5561A>C), GAT407CGA (5562G>C 5563A>G 5564T>A), TTC408TTT (5567C>T), CTT409CTA (5570T>A), GTC411CAT (5574G>C 5575T>A 5576C>T), CAA412AAT (5577C>A 5579A>T), ATA413ATT (5582A>T), GAA414TGC (5583G>T 5584A>G 5585A>C), GGA416GAA (5590G>A), GGT417AAA (5592G>A 5593G>A 5594T>A), GAA419_CAG421del (5598_5606delGAATTACAG), CCT422ACC (5607C>A 5609T>C), ATT424_ATT425insCAGAGATCT (5615_5616insCAGAGATCT), AAT426GAT (5619A>G), AAA427TTT (5622A>T 5623A>T 5624A>T), ATA428GCA (5625A>G 5626T>C), TTA429TCA (5629T>C), GAA430AAA (5631G>A), AAA431TTT (5634A>T 5635A>T 5636A>T), CAT432CTT (5638A>T), ACG433GAT (5640A>G 5641C>A 5642G>T), AAA434ATT (5644A>T 5645A>T), AAA436ACT (5650A>C 5651A>T), AAT437GAC (5652A>G 5654T>C), AAA438AGG (5656A>G 5657A>G), GAA440ATT (5661G>A 5662A>T 5663A>T), TTA441CTC (5664T>C 5666A>C), AGT443ACA (5671G>C 5672T>A), ATA444TTT (5673A>T 5675A>T), TTA445TTG (5678A>G), TTA447AAT (5682T>A 5683T>A 5684A>T), AAC449AAT (5690C>T), CAG450TAT (5691C>T 5693G>T), AGA452TTT (5697A>T 5698G>T 5699A>T), CAT453CTT (5701A>T) | | | | | | | | |

##### Proteins

| | | | | | | | | | |
|---|---|---|---|---|---|---|---|---|---|
| aspartic protease/reverse transcriptase (NP_056848.1) | 263 | 454 | 29.4% | 625 | 47.3% | 189 (96.9%) | 91 (46.7%) | 3/3/0/0 | 0 |
| Protein mutations: | M263L (5130A>C 5132G>T), D266N (5139G>A), D269P (5148G>C 5149A>C), K272T (5158A>C 5159A>T), K273A (5160A>G 5161A>C 5162A>T), A274L (5163G>C 5164C>T 5165T>A), V276W (5169G>T 5170T>G 5171A>G), V277I (5172G>A), K278R (5176A>G), D285R (5196G>A 5197A>G 5198T>A), T286D (5199A>G 5200C>A 5201A>T), I288L (5205A>T), H289K (5208C>A 5210T>A), S291T (5215G>C 5216T>C), I292F (5217A>T 5219A>T), Q293K (5220C>A 5222G>A), A294D (5224C>A 5225A>C), R295N (5227G>A 5228A>T), Y296I (5229T>A 5230A>T 5231C>T), C302M (5247T>A 5248G>T 5249T>G), Y307W (5263A>G 5264T>G), H308Q (5267T>A), K310Q (5271A>C 5273G>A), L311I (5274T>A 5276A>T), E312F (5277G>T 5278A>T 5279A>C), D314K (5283G>A 5285T>G), S315D (5286A>G 5287G>A), K317Y (5292A>T 5294A>T), Y318K (5295T>A 5297T>A), T322N (5308C>A 5309A>T), Q325F (5316C>T 5317A>T 5318G>T), Y327Q (5322T>C 5324T>A), Q329E (5328C>G 5330G>A), K331N (5336A>T), V332A (5338T>C 5339A>T), L333M (5340T>A 5342A>G), Y337L (5352T>C 5353A>T 5354T>A), H338K (5355C>A 5357T>A), S340A (5361T>G 5363A>C), S342F (5367A>T 5368G>T 5369T>C), I343E (5370A>G 5371T>A 5372T>A), Q346N (5379C>A 5381A>T), F347I (5382T>A 5384T>C), D349N (5388G>A), R350N (5392G>A 5393A>T), R353N (5401G>A 5402A>T), Y356L (5410A>T 5411T>A), V368I (5445G>A 5447C>T), F369Y (5449T>A), I373F (5460A>T), E375M (5466G>A 5467A>T), K377I (5473A>T 5474A>C), I378K (5476T>A), I380L (5481A>C), A381D (5485C>A 5486A>T), K382I (5488A>T 5489A>C), R384K (5494G>A), D385K (5496G>A 5498T>A), T387V (5502A>G 5503C>T 5504A>T), L388I (5505C>A 5507T>A), A389Q (5508G>C 5509C>A), G391D (5515G>A 5516A>T), I393V (5520A>G 5522T>A), K397P (5532A>C 5533A>C), T399M (5539C>T 5540A>G), E400S (5541G>A 5542A>G 5543A>T), C402F (5548G>T), K403Q (5550A>C), E404T (5553G>A 5554A>C), K405D (5556A>G 5558A>T), I406V (5559A>G 5561A>C), D407R (5562G>C 5563A>G 5564T>A), V411H (5574G>C 5575T>A 5576C>T), Q412N (5577C>A 5579A>T), E414C (5583G>T 5584A>G 5585A>C), G416E (5590G>A), G417K (5592G>A 5593G>A 5594T>A), E419_Q421del (5598_5606delGAATTACAG), P422T (5607C>A 5609T>C), I424_I425insQRS (5615_5616insCAGAGATCT), N426D (5619A>G), K427F (5622A>T 5623A>T 5624A>T), I428A (5625A>G 5626T>C), L429S (5629T>C), E430K (5631G>A), K431F (5634A>T 5635A>T 5636A>T), H432L (5638A>T), T433D (5640A>G 5641C>A 5642G>T), K434I (5644A>T 5645A>T), K436T (5650A>C 5651A>T), N437D (5652A>G 5654T>C), K438R (5656A>G 5657A>G), E440I (5661G>A 5662A>T 5663A>T), S443T (5671G>C 5672T>A), I444F (5673A>T 5675A>T), L447N (5682T>A 5683T>A 5684A>T), Q450Y (5691C>T 5693G>T), R452F (5697A>T 5698G>T 5699A>T), H453L (5701A>T) | | | | | | | | |

| | Begin | End | Coverage | Score | Concordance | Matches | Identities | I/D/M/F* | Stop Codons |
|---|---|---|---|---|---|---|---|---|---|
| **NT** | **5128** | **5704** | **7.1%** | **148** | **13.2%** | **568 (96.9%)** | **330 (56.3%)** | **9/9** | |
| Codon mutations: | ATG263CTT (5130A>C 5132G>T), GTA264GTT (5135A>T), GAT266AAT (5139G>A), GAT269CCT (5148G>C 5149A>C), TTA270CTG (5151T>C 5153A>G), AAT271AAC (5156T>C), AAA272ACT (5158A>C 5159A>T), AAA273GCT (5160A>G 5161A>C 5162A>T), GCT274CTA (5163G>C 5164C>T 5165T>A), GTA276TGG (5169G>T 5170T>G 5171A>G), GTA277ATA (5172G>A), AAA278AGA (5176A>G), TAT279TAC (5180T>C), CCT280CCG (5183T>G), ATC281ATT (5186C>T), CCA282CCT (5189A>T), GAT285AGA (5196G>A 5197A>G 5198T>A), ACA286GAT (5199A>G 5200C>A 5201A>T), TTA287CTT (5202T>C 5204A>T), ATA288TTA (5205A>T), CAT289AAA (5208C>A 5210T>A), AGA290CGA (5211A>C), AGT291ACC (5215G>C 5216T>C), ATA292TTT (5217A>T 5219A>T), CAG293AAA (5220C>A 5222G>A), GCA294GAC (5224C>A 5225A>C), AGA295AAT (5227G>A 5228A>T), TAC296ATT (5229T>A 5230A>T 5231C>T), GAT301GAC (5246T>C), TGT302ATG (5247T>A 5248G>T 5249T>G), GGA305GGC (5258A>C), TAT307TGG (5263A>G 5264T>G), CAT308CAA (5267T>A), ATA309ATC (5270A>C), AAG310CAA (5271A>C 5273G>A), TTA311ATT (5274T>A 5276A>T), GAA312TTC (5277G>T 5278A>T 5279A>C), GAT314AAG (5283G>A 5285T>G), AGT315GAT (5286A>G 5287G>A), AAA317TAT (5292A>T 5294A>T), TAT318AAA (5295T>A 5297T>A), ACA319ACT (5300A>T), TTT321TTC (5306T>C), ACA322AAT (5308C>A 5309A>T), GTT323GTC (5312T>C), CAG325TTT (5316C>T 5317A>T 5318G>T), TAT327CAA (5322T>C 5324T>A), CAG329GAA (5328C>G 5330G>A), AAA331AAT (5336A>T), GTA332GCT (5338T>C 5339A>T), TTA333ATG (5340T>A 5342A>G), TAT337CTA (5352T>C 5353A>T 5354T>A), CAT338AAA (5355C>A 5357T>A), TCA340GCC (5361T>G 5363A>C), CCA341CCT (5366A>T), AGT342TTC (5367A>T 5368G>T 5369T>C), ATT343GAA (5370A>G 5371T>A 5372T>A), TTC344TTT (5375C>T), CAA346AAT (5379C>A 5381A>T), TTT347ATC (5382T>A 5384T>C), GAT349AAT (5388G>A), AGA350AAT (5392G>A 5393A>T), ATA351ATT (5396A>T), AGA353AAT (5401G>A 5402A>T), TAT356TTA (5410A>T 5411T>A), ATT359ATC (5420T>C), GTG361GTT (5426G>T), GAT365GAC (5438T>C), ATA366ATT (5441A>T), TTA367TTG (5444A>G), GTC368ATT (5445G>A 5447C>T), TTT369TAT (5449T>A), AAG371AAA (5456G>A), ACT372ACG (5459T>G), ATT373TTT (5460A>T), GAG375ATG (5466G>A 5467A>T), AAA377ATC (5473A>T 5474A>C), ATA378AAA (5476T>A), ATT380CTT (5481A>C), GCA381GAT (5485C>A 5486A>T), AAA382ATC (5488A>T 5489A>C), TTT383TTC (5492T>C), AGA384AAA (5494G>A), GAT385AAA (5496G>A 5498T>A), ATA386ATT (5501A>T), ACA387GTT (5502A>G 5503C>T 5504A>T), CTT388ATA (5505C>A 5507T>A), GCA389CAA (5508G>C 5509C>A), GGA391GAT (5515G>A 5516A>T), TTA392TTG (5519A>G), ATT393GTA (5520A>G 5522T>A), AGT395TCT (5526A>T 5527G>C), AAG396AAA (5531G>A), AAA397CCA (5532A>C 5533A>C), ACA399ATG (5539C>T 5540A>G), GAA400AGT (5541G>A 5542A>G 5543A>T), TGT402TTT (5548G>T), AAA403CAA (5550A>C), GAA404ACA (5553G>A 5554A>C), AAA405GAT (5556A>G 5558A>T), ATA406GTC (5559A>G 5561A>C), GAT407CGA (5562G>C 5563A>G 5564T>A), TTC408TTT (5567C>T), CTT409CTA (5570T>A), GTC411CAT (5574G>C 5575T>A 5576C>T), CAA412AAT (5577C>A 5579A>T), ATA413ATT (5582A>T), GAA414TGC (5583G>T 5584A>G 5585A>C), GGA416GAA (5590G>A), GGT417AAA (5592G>A 5593G>A 5594T>A), GAA419_CAG421del (5598_5606delGAATTACAG), CCT422ACC (5607C>A 5609T>C), ATT424_ATT425insCAGAGATCT (5615_5616insCAGAGATCT), AAT426GAT (5619A>G), AAA427TTT (5622A>T 5623A>T 5624A>T), ATA428GCA (5625A>G 5626T>C), TTA429TCA (5629T>C), GAA430AAA (5631G>A), AAA431TTT (5634A>T 5635A>T 5636A>T), CAT432CTT (5638A>T), ACG433GAT (5640A>G 5641C>A 5642G>T), AAA434ATT (5644A>T 5645A>T), AAA436ACT (5650A>C 5651A>T), AAT437GAC (5652A>G 5654T>C), AAA438AGG (5656A>G 5657A>G), GAA440ATT (5661G>A 5662A>T 5663A>T), TTA441CTC (5664T>C 5666A>C), AGT443ACA (5671G>C 5672T>A), ATA444TTT (5673A>T 5675A>T), TTA445TTG (5678A>G), TTA447AAT (5682T>A 5683T>A 5684A>T), AAC449AAT (5690C>T), CAG450TAT (5691C>T 5693G>T), AGA452TTT (5697A>T 5698G>T 5699A>T), CAT453CTT (5701A>T) | | | | | | | | |

*: Inserts / Deletes / Misaligned / Frameshifts

#### Analysis details

This analysis was performed with panviral2.64

#### NGS Details (UN24): Caulimovirus tessellobrassicae

##### Assembly

| | |
|---|---|
| Coverage Length | 385 (1 contig(s)) |
| Depth Of Coverage | 4.5 |
| Number Of Reads | 13 |
| Reads Per Million | 0.26 rpm (after QC) |
| Ambiguities | 0 |
| Assembly Method | de novo + reference guided assembly |
| Consensus Caller | Bcf Tools |

##### Coverage Map

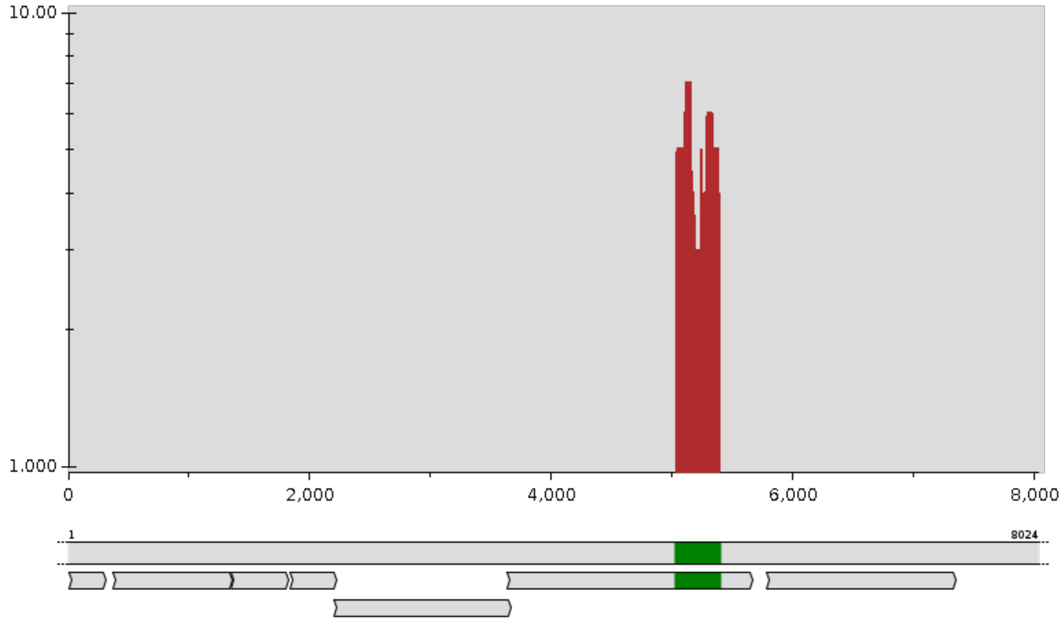

##### Assignment

| | |
|---|---|
| Type | Caulimovirus tessellobrassicae (Taxonomy ID: 3047738) |
| Reference Genome | NC_001497.2 |
| NT Identity (%) | 52.3936 |
| AA Identity (%) | 48.4127 |
| Number Of Stop Codons | 1 |
| Number Of CDS | 7 |

##### Alignment

| | |
|---|---|
| Alignment Score | 0.0 (NT) + 371.0 (AA) = 371.0 |
| Concordance (%) | 23.7973 |

| Alignment Method | Global, seeded, nucleotide + amino acids (AGA) |
|---|---|

#### Genome Region

Sequence starts at position 5020 and ends at position 5404 relative to NC_001497.2 reference sequence.

#### Alignment Detailed Statistics

| | Begin | End | Coverage | Score | Concordance | Matches | Identities | I/D/M/F* | Stop Codons |
|---|---|---|---|---|---|---|---|---|---|
| **NT** | **5020** | **5404** | **4.8%** | **0** | **0%** | **376 (97.7%)** | **197 (51.2%)** | **0/9** | |
| Mutations: | 5020A>G, 5021T>C, 5022C>T, 5023T>A, 5025G>T, 5029C>T, 5030A>T, 5032A>T, 5033T>C, 5034C>T, 5035A>G, 5036A>G, 5037C>T, 5043C>T, 5046C>T, 5047G>A, 5050A>G, 5051C>A, 5052C>A, 5053C>A, 5055T>A, 5056G>A, 5057A>C, 5058A>T, 5061C>T, 5064G>A, 5066A>C, 5067G>T, 5070A>G, 5071C>T, 5073T>G, 5079A>G, 5088C>A, 5090T>G, 5091A>C, 5094A>T, 5096C>A, 5097A>C, 5100T>C, 5102C>T, 5103C>T, 5104T>G, 5106G>A, 5109T>C, 5111A>T, 5112C>T, 5115C>T, 5118G>A, 5122C>G, 5125G>A, 5126C>G, 5127T>G, 5134A>T, 5136A>C, 5139G>A, 5142T>C, 5143C>T, 5145G>A, 5146C>T, 5148A>T, 5150C>A, 5153A>G, 5154G>A, 5155C>T, 5157T>G, 5160A>G, 5161G>A, 5163A>G, 5166C>T, 5167_5169delGTT, 5172A>G, 5173T>G, 5174G>T, 5175G>T, 5176A>C, 5177G>C, 5178A>C, 5179T>C, 5183C>A, 5187A>G, 5190G>A, 5193T>G, 5196C>T, 5199C>G, 5200T>G, 5201A>C, 5202C>T, 5203A>G, 5205G>C, 5214G>T, 5217G>A, 5218A>G, 5220A>T, 5222A>T, 5223T>G, 5224C>G, 5226G>C, 5227C>A, 5230G>A, 5231G>A, 5232A>C, 5233T>C, 5239C>T, 5240C>G, 5241A>T, 5242C>T, 5244A>G, 5245C>G, 5246A>G, 5247T>A, 5248C>A, 5249A>T, 5253C>T, 5254T>A, 5255T>A, 5256A>C, 5259A>C, 5262G>T, 5263G>T, 5265G>T, 5266A>T, 5267A>T, 5268G>T, 5269C>A, 5274C>T, 5275A>G, 5277C>T, 5280G>A, 5283C>T, 5289A>T, 5292A>C, 5296G>A, 5297A>T, 5298C>T, 5299T>G, 5301C>T, 5303G>A, 5304G>C, 5310T>A, 5313G>T, 5314T>C, 5320G>C, 5321C>A, 5322T>A, 5324T>A, 5325C>G, 5330T>C, 5332A>C, 5333A>C, 5334C>A, 5335G>A, 5338G>T, 5339G>C, 5340T>A, 5341A>G, 5343T>C, 5344A>G, 5346T>A, 5347_5349delACT, 5350G>C, 5352G>A, 5356A>G, 5358T>A, 5359_5361delTGC, 5362A>C, 5364A>G, 5367C>T, 5368G>A, 5370A>T, 5373T>G, 5377A>G, 5378G>T, 5379C>T, 5381T>G, 5382T>G, 5385A>T, 5386G>T, 5387C>T, 5392G>T, 5400T>C, 5403C>T, 5404C>A | | | | | | | | |

##### CDS

| | Begin | End | Coverage | Score | Concordance | Matches | Identities | I/D/M/F* | Stop Codons |
|---|---|---|---|---|---|---|---|---|---|
| CaMVgp6 | 463 | 591 | 19.0% | 371 | 43.2% | 126 (97.7%) | 61 (47.3%) | 0/3/0/0 | 1 |
| Protein mutations: | I463A (5020A>G 5021T>C 5022C>T), L464I (5023T>A 5025G>T), H466F (5029C>T 5030A>T), I467S (5032A>T 5033T>C 5034C>T), N468G (5035A>G 5036A>G 5037C>T), D472N (5047G>A), T473E (5050A>G 5051C>A 5052C>A), L474I (5053C>A 5055T>A), E475T (5056G>A 5057A>C 5058A>T), K478T (5066A>C 5067G>T), I486S (5090T>G 5091A>C), T488N (5096C>A 5097A>C), A490V (5102C>T 5103C>T), S491A (5104T>G 5106G>A), Y493F (5111A>T 5112C>T), L497V (5122C>G), A498R (5125G>A 5126C>G 5127T>G), R501C (5134A>T 5136A>C), Q505Y (5146C>T 5148A>T), A506D (5150C>A), K507R (5153A>G 5154G>A), E510K (5161G>A 5163A>G), V512del (5167_5169delGTT), W514V (5173T>G 5174G>T 5175G>T), R515P (5176A>C 5177G>C 5178A>C), W516R (5179T>C), T517K (5183C>A), D520E (5193T>G), Y523A (5200T>G 5201A>C 5202C>T), M524V (5203A>G 5205G>C), K529D (5218A>G 5220A>T), N530M (5222A>T 5223T>G), L531V (5224C>G 5226G>C), Q532K (5227C>A), G533N (5230G>A 5231G>A 5232A>C), F534L (5233T>C), P536C (5239C>T 5240C>G 5241A>T), H538G (5245C>G 5246A>G 5247T>A), H539I (5248C>A 5249A>T), L541N (5254T>A 5255T>A 5256A>C), E543D (5262G>T), E544Y (5263G>T 5265G>T), K545F (5266A>T 5267A>T 5268G>T), L546M (5269C>A), I548V (5275A>G 5277C>T), D555I (5296G>A 5297A>T 5298C>T), Y556D (5299T>G 5301C>T), W557Y (5303G>A 5304G>C), M560I (5313G>T), A563Q (5320G>C 5321C>A 5322T>A), I564K (5324T>A 5325C>G), I566T (5330T>C), N567P (5332A>C 5333A>C 5334C>A), E568K (5335G>A), G569S (5338G>T 5339G>C 5340T>A), T570A (5341A>G 5343T>C), N571E (5344A>G 5346T>A), T572del (5347_5349delACT), E573Q (5350G>C 5352G>A), I575V (5356A>G 5358T>A), C576del (5359_5361delTGC), A579T (5368G>A 5370A>T), S582V (5377A>G 5378G>T 5379C>T), F583W (5381T>G 5382T>G), K584N (5385A>T), A585F (5386G>T 5387C>T), E587* (5392G>T) | | | | | | | | |
| Codon mutations: | ATC463GCT (5020A>G 5021T>C 5022C>T), TTG464ATT (5023T>A 5025G>T), CAC466TTC (5029C>T 5030A>T), ATC467TCT (5032A>T 5033T>C 5034C>T), AAC468GGT (5035A>G 5036A>G 5037C>T), TTC470TTT (5043C>T), CCC471CCT (5046C>T), GAT472AAT (5047G>A), ACC473GAA (5050A>G 5051C>A 5052C>A), CTT474ATA (5053C>A 5055T>A), GAA475ACT (5056G>A 5057A>C 5058A>T), GAC476GAT (5061C>T), AAG477AAA (5064G>A), AAG478ACT (5066A>C 5067G>T), CAA479CAG (5070A>G), CTT480TTG (5071C>T 5073T>G), AGA482AGG (5079A>G), GGC485GGA (5088C>A), ATA486AGC (5090T>G 5091A>C), CTA487CTT (5094A>T), ACA488AAC (5096C>A 5097A>C), TAT489TAC (5100T>C), GCC490GTT (5102C>T 5103C>T), TCG491GCA (5104T>G 5106G>A), GAT492GAC (5109T>C), TAC493TTT (5111A>T 5112C>T), ATC494ATT (5115C>T), CCG495CCA (5118G>A), CTA497GTA (5122C>G), GCT498AGG (5125G>A 5126C>G 5127T>G), AGA501TGC (5134A>T 5136A>C), AAG502AAA (5139G>A), CCT503CCC (5142T>C), CTG504TTA (5143C>T 5145G>A), CAA505TAT (5146C>T 5148A>T), GCC506GAC (5150C>A), AAG507AGA (5153A>G 5154G>A), CTT508TTG (5155C>T 5157T>G), AAA509AAG (5160A>G), GAA510AAG (5161G>A 5163A>G), AAC511AAT (5166C>T), GTT512del (5167_5169delGTT), CCA513CCG (5172A>G), TGG514GTT (5173T>G 5174G>T 5175G>T), AGA515CCC (5176A>C 5177G>C 5178A>C), TGG516CGG (5179T>C), ACA517AAA (5183C>A), AAA518AAG (5187A>G), GAG519GAA (5190G>A), GAT520GAG (5193T>G), ACC521ACT (5196C>T), CTC522CTG (5199C>G), TAC523GCT (5200T>G 5201A>C 5202C>T), ATG524GTC (5203A>G 5205G>C), GTG527GTT (5214G>T), AAG528AAA (5217G>A), AAA529GAT (5218A>G 5220A>T), AAT530ATG (5222A>T 5223T>G), CTG531GTC (5224C>G 5226G>C), CAA532AAA (5227C>A), GGA533AAC (5230G>A 5231G>A 5232A>C), TTT534CTT (5233T>C), CCA536TGT (5239C>T 5240C>G 5241A>T), CTA537TTG (5242C>T 5244A>G), CAT538GGA (5245C>G 5246A>G 5247T>A), CAT539ATT (5248C>A 5249A>T), CCC540CCT (5253C>T), TTA541AAC (5254T>A 5255T>A 5256A>C), CCA542CCC (5259A>C), GAG543GAT (5262G>T), GAG544TAT (5263G>T 5265G>T), AAG545TTT (5266A>T 5267A>T 5268G>T), CTG546ATG (5269C>A), ATC547ATT (5274C>T), ATC548GTT (5275A>G 5277C>T), GAG549GAA (5280G>A), ACC550ACT (5283C>T), GCA552GCT (5289A>T), TCA553TCC (5292A>C), GAC555ATT (5296G>A 5297A>T 5298C>T), TAC556GAT (5299T>G 5301C>T), TGG557TAC (5303G>A 5304G>C), GGT559GGA (5310T>A), ATG560ATT (5313G>T), TTA561CTA (5314T>C), GCT563CAA (5320G>C 5321C>A 5322T>A), ATC564AAG (5324T>A 5325C>G), ATT566ACT (5330T>C), AAC567CCA (5332A>C 5333A>C 5334C>A), GAA568AAA (5335G>A), GGT569TCA (5338G>T 5339G>C 5340T>A), ACT570GCC (5341A>G 5343T>C), AAT571GAA (5344A>G 5346T>A), ACT572del (5347_5349delACT), GAG573CAA (5350G>C 5352G>A), ATT575GTA (5356A>G 5358T>A), TGC576del (5359_5361delTGC), AGA577CGG (5362A>C 5364A>G), TAC578TAT (5367C>T), GCA579ACT (5368G>A 5370A>T), TCT580TCG (5373T>G), AGC582GTT (5377A>G 5378G>T 5379C>T), TTT583TGG (5381T>G 5382T>G), AAA584AAT (5385A>T), GCT585TTT (5386G>T 5387C>T), GAA587TAA (5392G>T), AAT589AAC (5400T>C), TAC590TAT (5403C>T), CAC591A.. (5404C>A) | | | | | | | | |

##### Proteins

| | Begin | End | Coverage | Score | Concordance | Matches | Identities | I/D/M/F* | Stop Codons |
|---|---|---|---|---|---|---|---|---|---|
| reverse transcriptase (NP_056728.1) | 463 | 591 | 19.0% | 371 | 43.2% | 126 (97.7%) | 61 (47.3%) | 0/3/0/0 | 1 |
| Protein mutations: | I463A (5020A>G 5021T>C 5022C>T), L464I (5023T>A 5025G>T), H466F (5029C>T 5030A>T), I467S (5032A>T 5033T>C 5034C>T), N468G (5035A>G 5036A>G 5037C>T), D472N (5047G>A), T473E (5050A>G 5051C>A 5052C>A), L474I (5053C>A 5055T>A), E475T (5056G>A 5057A>C 5058A>T), K478T (5066A>C 5067G>T), I486S (5090T>G 5091A>C), T488N (5096C>A 5097A>C), A490V (5102C>T 5103C>T), S491A (5104T>G 5106G>A), Y493F (5111A>T 5112C>T), L497V (5122C>G), A498R (5125G>A 5126C>G 5127T>G), R501C (5134A>T 5136A>C), Q505Y (5146C>T 5148A>T), A506D (5150C>A), K507R (5153A>G 5154G>A), E510K (5161G>A 5163A>G), V512del (5167_5169delGTT), W514V (5173T>G 5174G>T 5175G>T), R515P (5176A>C 5177G>C 5178A>C), W516R (5179T>C), T517K (5183C>A), D520E (5193T>G), Y523A (5200T>G 5201A>C 5202C>T), M524V (5203A>G 5205G>C), K529D (5218A>G 5220A>T), N530M (5222A>T 5223T>G), L531V (5224C>G 5226G>C), Q532K (5227C>A), G533N (5230G>A 5231G>A 5232A>C), F534L (5233T>C), P536C (5239C>T 5240C>G 5241A>T), H538G (5245C>G 5246A>G 5247T>A), H539I (5248C>A 5249A>T), L541N (5254T>A 5255T>A 5256A>C), E543D (5262G>T), E544Y (5263G>T 5265G>T), K545F (5266A>T 5267A>T 5268G>T), L546M (5269C>A), I548V (5275A>G 5277C>T), D555I (5296G>A 5297A>T 5298C>T), Y556D (5299T>G 5301C>T), W557Y (5303G>A 5304G>C), M560I (5313G>T), A563Q (5320G>C 5321C>A 5322T>A), I564K (5324T>A 5325C>G), I566T (5330T>C), N567P (5332A>C 5333A>C 5334C>A), E568K (5335G>A), G569S (5338G>T 5339G>C 5340T>A), T570A (5341A>G 5343T>C), N571E (5344A>G 5346T>A), T572del (5347_5349delACT), E573Q (5350G>C 5352G>A), I575V (5356A>G 5358T>A), C576del (5359_5361delTGC), A579T (5368G>A 5370A>T), S582V (5377A>G 5378G>T 5379C>T), F583W (5381T>G 5382T>G), K584N (5385A>T), A585F (5386G>T 5387C>T), E587* (5392G>T) | | | | | | | | |
| Codon mutations: | ATC463GCT (5020A>G 5021T>C 5022C>T), TTG464ATT (5023T>A 5025G>T), CAC466TTC (5029C>T 5030A>T), ATC467TCT (5032A>T 5033T>C 5034C>T), AAC468GGT (5035A>G 5036A>G 5037C>T), TTC470TTT (5043C>T), CCC471CCT (5046C>T), GAT472AAT (5047G>A), ACC473GAA (5050A>G 5051C>A 5052C>A), CTT474ATA (5053C>A 5055T>A), GAA475ACT (5056G>A 5057A>C 5058A>T), GAC476GAT (5061C>T), AAG477AAA (5064G>A), AAG478ACT (5066A>C 5067G>T), CAA479CAG (5070A>G), CTT480TTG (5071C>T 5073T>G), AGA482AGG (5079A>G), GGC485GGA (5088C>A), ATA486AGC (5090T>G 5091A>C), CTA487CTT (5094A>T), ACA488AAC (5096C>A 5097A>C), TAT489TAC (5100T>C), GCC490GTT (5102C>T 5103C>T), TCG491GCA (5104T>G 5106G>A), GAT492GAC (5109T>C), TAC493TTT (5111A>T 5112C>T), ATC494ATT (5115C>T), CCG495CCA (5118G>A), CTA497GTA (5122C>G), GCT498AGG (5125G>A 5126C>G 5127T>G), AGA501TGC (5134A>T 5136A>C), AAG502AAA (5139G>A), CCT503CCC (5142T>C), CTG504TTA (5143C>T 5145G>A), CAA505TAT (5146C>T 5148A>T), GCC506GAC (5150C>A), AAG507AGA (5153A>G 5154G>A), CTT508TTG (5155C>T 5157T>G), AAA509AAG (5160A>G), GAA510AAG (5161G>A 5163A>G), AAC511AAT (5166C>T), GTT512del (5167_5169delGTT), CCA513CCG (5172A>G), TGG514GTT (5173T>G 5174G>T 5175G>T), AGA515CCC (5176A>C 5177G>C 5178A>C), TGG516CGG (5179T>C), ACA517AAA (5183C>A), AAA518AAG (5187A>G), GAG519GAA (5190G>A), GAT520GAG (5193T>G), ACC521ACT (5196C>T), CTC522CTG (5199C>G), TAC523GCT (5200T>G 5201A>C 5202C>T), ATG524GTC (5203A>G 5205G>C), GTG527GTT (5214G>T), AAG528AAA (5217G>A), AAA529GAT (5218A>G 5220A>T), AAT530ATG (5222A>T 5223T>G), CTG531GTC (5224C>G 5226G>C), CAA532AAA (5227C>A), GGA533AAC (5230G>A 5231G>A 5232A>C), TTT534CTT (5233T>C), CCA536TGT (5239C>T 5240C>G 5241A>T), CTA537TTG (5242C>T 5244A>G), CAT538GGA (5245C>G 5246A>G 5247T>A), CAT539ATT (5248C>A 5249A>T), CCC540CCT (5253C>T), TTA541AAC (5254T>A 5255T>A 5256A>C), CCA542CCC (5259A>C), GAG543GAT (5262G>T), GAG544TAT (5263G>T 5265G>T), AAG545TTT (5266A>T 5267A>T 5268G>T), CTG546ATG (5269C>A), ATC547ATT (5274C>T), ATC548GTT (5275A>G 5277C>T), GAG549GAA (5280G>A), ACC550ACT (5283C>T), GCA552GCT (5289A>T), TCA553TCC (5292A>C), GAC555ATT (5296G>A 5297A>T 5298C>T), TAC556GAT (5299T>G 5301C>T), TGG557TAC (5303G>A 5304G>C), GGT559GGA (5310T>A), ATG560ATT (5313G>T), TTA561CTA (5314T>C), GCT563CAA (5320G>C 5321C>A 5322T>A), ATC564AAG (5324T>A 5325C>G), ATT566ACT (5330T>C), AAC567CCA (5332A>C 5333A>C 5334C>A), GAA568AAA (5335G>A), GGT569TCA (5338G>T 5339G>C 5340T>A), ACT570GCC (5341A>G 5343T>C), AAT571GAA (5344A>G 5346T>A), ACT572del (5347_5349delACT), GAG573CAA (5350G>C 5352G>A), ATT575GTA (5356A>G 5358T>A), TGC576del (5359_5361delTGC), AGA577CGG (5362A>C 5364A>G), TAC578TAT (5367C>T), GCA579ACT (5368G>A 5370A>T), TCT580TCG (5373T>G), AGC582GTT (5377A>G 5378G>T 5379C>T), TTT583TGG (5381T>G 5382T>G), AAA584AAT (5385A>T), GCT585TTT (5386G>T 5387C>T), GAA587TAA (5392G>T), AAT589AAC (5400T>C), TAC590TAT (5403C>T), CAC591A.. (5404C>A) | | | | | | | | |

*: Inserts / Deletes / Misaligned / Frameshifts

#### Analysis details

This analysis was performed with panviral2.64

#### NGS Details (UN24): Duamitovirus peex1

##### Assembly

| | |
|---|---|
| **Coverage Length** | 135 (1 contig(s)) |
| **Depth Of Coverage** | 8.0 |
| **Number Of Reads** | 12 |
| **Reads Per Million** | 0.24 rpm (after QC) |
| **Ambiguities** | 0 |
| **Assembly Method** | de novo + reference guided assembly |
| **Consensus Caller** | Bcf Tools |

##### Coverage Map

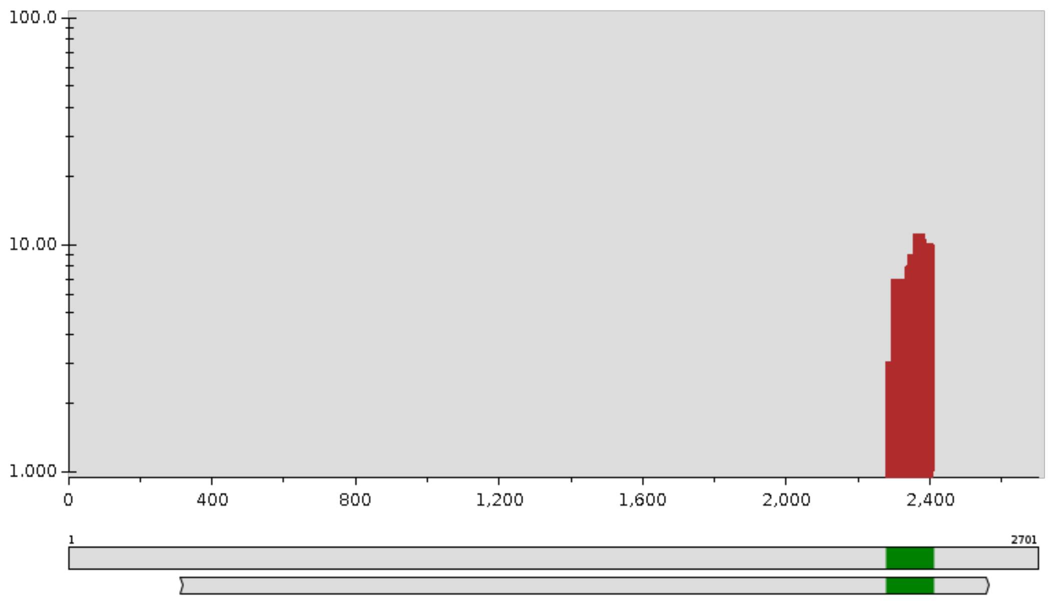

##### Assignment

| | |
|---|---|
| **Type** | Duamitovirus peex1 (Taxonomy ID: 2955799) |
| **Reference Genome** | NC_076525.1 |
| **NT Identity (%)** | 70.3704 |
| **AA Identity (%)** | 66.6667 |
| **Number Of Stop Codons** | 0 |
| **Number Of CDS** | 1 |

##### Alignment

| | |
|---|---|
| **Alignment Score** | 110.0 (NT) + 251.0 (AA) = 361.0 |
| **Concordance (%)** | 58.4142 |

| Alignment Method | Global, seeded, nucleotide + amino acids (AGA) |
|---|---|

#### Genome Region

Sequence starts at position 2278 and ends at position 2412 relative to NC_076525.1 reference sequence.

#### Alignment Detailed Statistics

| | Begin | End | Coverage | Score | Concordance | Matches | Identities | I/D/M/F* | Stop Codons |
|---|---|---|---|---|---|---|---|---|---|
| **NT** | **2278** | **2412** | **5.0%** | **110** | **40.7%** | **135 (100%)** | **95 (70.4%)** | **0/0** | |
| Mutations: | 2278G>A, 2281T>A, 2283G>A, 2294T>G, 2295C>A, 2296G>A, 2301C>G, 2303T>C, 2306A>T, 2311A>G, 2312T>C, 2315C>T, 2321G>T, 2325A>G, 2326A>C, 2327T>A, 2330C>A, 2331C>G, 2333G>T, 2339G>A, 2345A>T, 2348C>T, 2351A>G, 2352C>T, 2354T>C, 2363G>T, 2373G>A, 2374C>G, 2375G>T, 2377C>A, 2378T>A, 2379T>A, 2388C>A, 2390T>A, 2393T>A, 2394C>A, 2396G>C, 2398C>A, 2399T>A, 2406C>G | | | | | | | | |

**CDS**

| | Begin | End | Coverage | Score | Concordance | Matches | Identities | I/D/M/F* | Stop Codons |
|---|---|---|---|---|---|---|---|---|---|
| RdRp | 657 | 701 | 6.0% | 251 | 67.3% | 45 (100%) | 30 (66.7%) | 0/0/0/0 | 0 |
| Protein mutations: | M657K (2281T>A), E658K (2283G>A), R662K (2295C>A 2296G>A), L664V (2301C>G 2303T>C), Y667C (2311A>G 2312T>C), N672A (2325A>G 2326A>C 2327T>A), P674A (2331C>G 2333G>T), L681F (2352C>T 2354T>C), A688S (2373G>A 2374C>G 2375G>T), T689K (2377C>A 2378T>A), S690T (2379T>A), Q695N (2394C>A 2396G>C), T696K (2398C>A 2399T>A), H699D (2406C>G) | | | | | | | | |
| Codon mutations: | TGG656.AG (2278G>A), ATG657AAG (2281T>A), GAA658AAA (2283G>A), CTT661CTG (2294T>G), CGA662AAA (2295C>A 2296G>A), CTT664GTC (2301C>G 2303T>C), TCA665TCT (2306A>T), TAT667TGC (2311A>G 2312T>C), CAC668CAT (2315C>T), GTG670GTT (2321G>T), AAT672GCA (2325A>G 2326A>C 2327T>A), TCC673TCA (2330C>A), CCG674GCT (2331C>G 2333G>T), GTG676GTA (2339G>A), ATA678ATT (2345A>T), GAC679GAT (2348C>T), CAA680CAG (2351A>G), CTT681TTC (2352C>T 2354T>C), GTG684GTT (2363G>T), GCG688AGT (2373G>A 2374C>G 2375G>T), ACT689AAA (2377C>A 2378T>A), TCT690ACT (2379T>A), CGT693AGA (2388C>A 2390T>A), ACT694ACA (2393T>A), CAG695AAC (2394C>A 2396G>C), ACT696AAA (2398C>A 2399T>A), CAT699GAT (2406C>G) | | | | | | | | |

**Proteins**

| | Begin | End | Coverage | Score | Concordance | Matches | Identities | I/D/M/F* | Stop Codons |
|---|---|---|---|---|---|---|---|---|---|
| RNA-dependent RNA polymerase (YP_010798875.1) | 657 | 701 | 6.0% | 251 | 67.3% | 45 (100%) | 30 (66.7%) | 0/0/0/0 | 0 |
| Protein mutations: | M657K (2281T>A), E658K (2283G>A), R662K (2295C>A 2296G>A), L664V (2301C>G 2303T>C), Y667C (2311A>G 2312T>C), N672A (2325A>G 2326A>C 2327T>A), P674A (2331C>G 2333G>T), L681F (2352C>T 2354T>C), A688S (2373G>A 2374C>G 2375G>T), T689K (2377C>A 2378T>A), S690T (2379T>A), Q695N (2394C>A 2396G>C), T696K (2398C>A 2399T>A), H699D (2406C>G) | | | | | | | | |
| Codon mutations: | TGG656.AG (2278G>A), ATG657AAG (2281T>A), GAA658AAA (2283G>A), CTT661CTG (2294T>G), CGA662AAA (2295C>A 2296G>A), CTT664GTC (2301C>G 2303T>C), TCA665TCT (2306A>T), TAT667TGC (2311A>G 2312T>C), CAC668CAT (2315C>T), GTG670GTT (2321G>T), AAT672GCA (2325A>G 2326A>C 2327T>A), TCC673TCA (2330C>A), CCG674GCT (2331C>G 2333G>T), GTG676GTA (2339G>A), ATA678ATT (2345A>T), GAC679GAT (2348C>T), CAA680CAG (2351A>G), CTT681TTC (2352C>T 2354T>C), GTG684GTT (2363G>T), GCG688AGT (2373G>A 2374C>G 2375G>T), ACT689AAA (2377C>A 2378T>A), TCT690ACT (2379T>A), CGT693AGA (2388C>A 2390T>A), ACT694ACA (2393T>A), CAG695AAC (2394C>A 2396G>C), ACT696AAA (2398C>A 2399T>A), CAT699GAT (2406C>G) | | | | | | | | |

*: Inserts / Deletes / Misaligned / Frameshifts

#### Analysis details

This analysis was performed with panviral2.64

### NGS Details (UN24): Caulimovirus tessellobrassicae

#### Assembly

| | |
|---|---|
| **Coverage Length** | 330 (1 contig(s)) |
| **Depth Of Coverage** | 4.9 |
| **Number Of Reads** | 12 |
| **Reads Per Million** | 0.24 rpm (after QC) |
| **Ambiguities** | 0 |
| **Assembly Method** | de novo + reference guided assembly |
| **Consensus Caller** | Bcf Tools |

#### Coverage Map

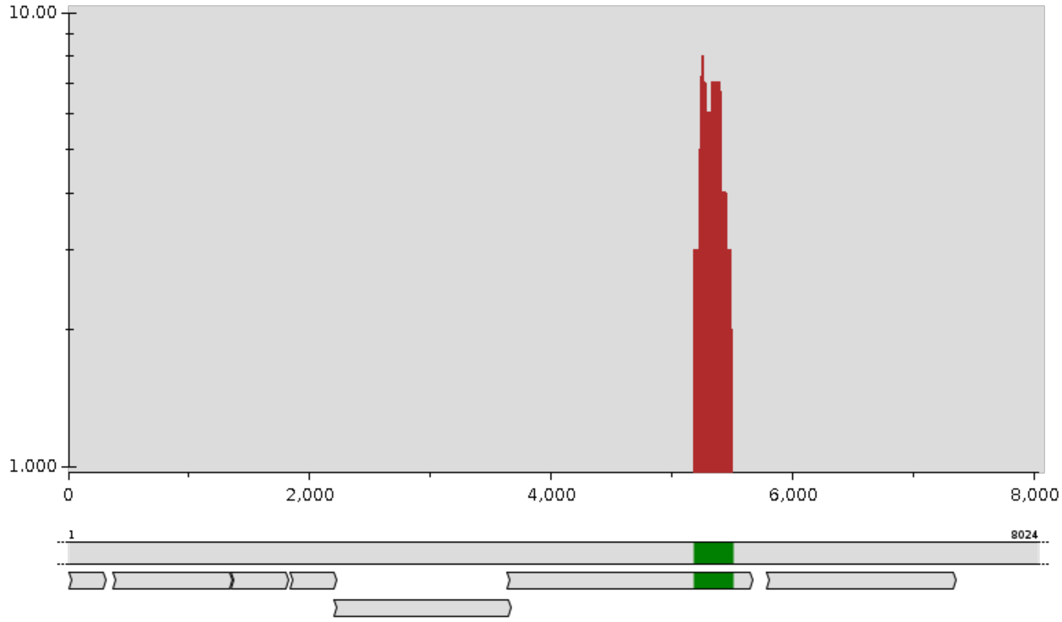

#### Assignment

| | |
|---|---|
| **Type** | Caulimovirus tessellobrassicae (Taxonomy ID: 3047738) |
| **Reference Genome** | NC_001497.2 |
| **NT Identity (%)** | 58.2524 |
| **AA Identity (%)** | 45.6311 |
| **Number Of Stop Codons** | 0 |
| **Number Of CDS** | 7 |

#### Alignment

| | |
|---|---|
| **Alignment Score** | 63.0 (NT) + 343.0 (AA) = 406.0 |
| **Concordance (%)** | 32.0189 |

| **Alignment Method** | Global, seeded, nucleotide + amino acids (AGA) |
|---|---|

#### Genome Region

Sequence starts at position 5174 and ends at position 5503 relative to NC_001497.2 reference sequence.

#### Alignment Detailed Statistics

| | Begin | End | Coverage | Score | Concordance | Matches | Identities | I/D/M/F* | Stop Codons |
|---|---|---|---|---|---|---|---|---|---|
| **NT** | **5174** | **5503** | **4.1%** | **63** | **10.9%** | **309 (93.6%)** | **180 (54.5%)** | **0/21** | |
| Mutations: | 5174G>T, 5176A>G, 5185A>G, 5187A>T, 5190G>A, 5191G>C, 5197C>A, 5198T>A, 5199C>A, 5201A>G, 5202C>T, 5203A>G, 5205G>T, 5210A>G, 5211G>C, 5212G>C, 5214G>A, 5217G>A, 5218A>C, 5221A>G, 5223T>A, 5224C>T, 5225T>G, 5226G>T, 5227C>T, 5228A>C, 5229A>C, 5230G>A, 5231G>A, 5235T>A, 5238T>A, 5239C>A, 5240C>A, 5241A>G, 5245C>A, 5246A>G, 5247T>A, 5249A>T, 5253C>A, 5254T>G, 5255T>A, 5259A>G, 5262G>A, 5265G>C, 5268G>C, 5269C>T, 5271G>A, 5272A>G, 5274C>T, 5275A>C, 5277C>A, 5278G>C, 5280G>A, 5283C>A, 5289A>T, 5293G>A, 5295C>T, 5296G>T, 5299T>C, 5301C>T, 5309G>C, 5310T>A, 5311A>T, 5313G>A, 5317A>C, 5320G>A, 5323_5325delATC, 5326A>G, 5328A>C, 5329A>C, 5334C>T, 5338_5355delGGTACTAATACTGAGTTA, 5361C>T, 5364A>G, 5367C>T, 5368G>A, 5371T>A, 5372C>G, 5376A>T, 5378G>C, 5379C>A, 5385A>T, 5387C>A, 5388T>A, 5389G>A, 5391A>C, 5396A>C, 5397G>T, 5400T>A, 5404C>T, 5405A>C, 5406C>A, 5408G>C, 5409C>A, 5415C>A, 5421G>A, 5422A>T, 5423C>T, 5430G>C, 5431G>A, 5434A>G, 5438A>G, 5439T>A, 5440A>G, 5441C>G, 5442T>C, 5446A>C, 5447A>G, 5448G>A, 5455A>T, 5456G>C, 5458A>G, 5459T>C, 5462A>T, 5463T>C, 5467A>C, 5468C>T, 5469T>A, 5472T>A, 5473G>A, 5474T>A, 5475T>G, 5478T>A, 5481T>C, 5482C>A, 5484G>C, 5487T>C, 5490G>A, 5491A>T, 5496T>C, 5502T>A | | | | | | | | |
| **CDS** | | | | | | | | | |
| CaMVgp6 | 515 | 624 | 16.2% | 343 | 47.4% | 103 (93.6%) | 47 (42.7%) | 0/7/0/0 | 0 |
| Protein mutations: | R515G (5176A>G), K518D (5185A>G 5187A>T), D520H (5191G>C), L522K (5197C>A 5198T>A 5199C>A), Y523C (5201A>G 5202C>T), M524V (5203A>G 5205G>T), K526S (5210A>G 5211G>C), V527L (5212G>C 5214G>A), K529Q (5218A>C), N530E (5221A>G 5223T>A), L531C (5224C>T 5225T>G 5226G>T), Q532S (5227C>T 5228A>C 5229A>C), G533K (5230G>A 5231G>A), F534L (5235T>A), P536K (5239C>A 5240C>A 5241A>G), H538R (5245C>A 5246A>G 5247T>A), H539L (5249A>T), L541E (5254T>G 5255T>A), E544D (5265G>C), K545N (5268G>C), I547V (5272A>G 5274C>T), I548L (5275A>C 5277C>A), E549Q (5278G>C 5280G>A), D554N (5293G>A 5295C>T), D555Y (5296G>T), Y556H (5299T>C 5301C>T), G559A (5309G>C 5310T>A), M560L (5311A>T 5313G>A), K562Q (5317A>C), A563T (5320G>A), I564del (5323_5325delATC), K565D (5326A>G 5328A>C), I566L (5329A>C), G569_L574del (5338_5355delGGTACTAATACTGAGTTA), A579T (5368G>A), S582T (5378G>C 5379C>A), K584N (5385A>T), A585E (5387C>A 5388T>A), A586T (5389G>A 5391A>C), K588T (5396A>C 5397G>T), N589K (5400T>A), H591S (5404C>T 5405A>C 5406C>A), S592T (5408G>C 5409C>A), D594E (5415C>A), T597L (5422A>T 5423C>T), V600I (5431G>A), I601V (5434A>G), N602R (5438A>G 5439T>A), T603G (5440A>G 5441C>G 5442T>C), K605R (5446A>C 5447A>G 5448G>A), I609A (5458A>G 5459T>C), Y610F (5462A>T 5463T>C), T612L (5467A>C 5468C>T 5469T>A), V614K (5473G>A 5474T>A 5475T>G), H615Q (5478T>A), L617I (5482C>A 5484G>C), T620S (5491A>T) | | | | | | | | |
| Codon mutations: | TGG514.TG (5174G>T), AGA515GGA (5176A>G), AAA518GAT (5185A>G 5187A>T), GAG519GAA (5190G>A), GAT520CAT (5191G>C), CTC522AAA (5197C>A 5198T>A 5199C>A), TAC523TGT (5201A>G 5202C>T), ATG524GTT (5203A>G 5205G>T), AAG526AGC (5210A>G 5211G>C), GTG527CTA (5212G>C 5214G>A), AAG528AAA (5217G>A), AAA529CAA (5218A>C), AAT530GAA (5221A>G 5223T>A), CTG531TGT (5224C>T 5225T>G 5226G>T), CAA532TCC (5227C>T 5228A>C 5229A>C), GGA533AAA (5230G>A 5231G>A), TTT534TTA (5235T>A), CCT535CCA (5238T>A), CCA536AAG (5239C>A 5240C>A 5241A>G), CAT538AGA (5245C>A 5246A>G 5247T>A), CAT539CTT (5249A>T), CCC540CCA (5253C>A), TTA541GAA (5254T>G 5255T>A), CCA542CCG (5259A>G), GAG543GAA (5262G>A), GAG544GAC (5265G>C), AAG545AAC (5268G>C), CTG546TTA (5269C>T 5271G>A), ATC547GTT (5272A>G 5274C>T), ATC548CTA (5275A>C 5277C>A), GAG549CAA (5278G>C 5280G>A), ACC550ACA (5283C>A), GCA552GCT (5289A>T), GAC554AAT (5293G>A 5295C>T), GAC555TAC (5296G>T), TAC556CAT (5299T>C 5301C>T), GGT559GCA (5309G>C 5310T>A), ATG560TTA (5311A>T 5313G>A), AAA562CAA (5317A>C), GCT563ACT (5320G>A), ATC564del (5323_5325delATC), AAA565GAC (5326A>G 5328A>C), ATT566CTT (5329A>C), AAC567AAT (5334C>T), GGT569_TTA574del (5338_5355delGGTACTAATACTGAGTTA), TGC576TGT (5361C>T), AGA577AGG (5364A>G), TAC578TAT (5367C>T), GCA579ACA (5368G>A), TCT580AGT (5371T>A 5372C>G), GGA581GGT (5376A>T), AGC582ACA (5378G>C 5379C>A), AAA584AAT (5385A>T), GCT585GAA (5387C>A 5388T>A), GCA586ACC (5389G>A 5391A>C), AAG588ACT (5396A>C 5397G>T), AAT589AAA (5400T>A), CAC591TCA (5404C>T 5405A>C 5406C>A), AGC592ACA (5408G>C 5409C>A), GAC594GAA (5415C>A), GAG596GAA (5421G>A), ACA597TTA (5422A>T 5423C>T), GCG599GCC (5430G>C), GTA600ATA (5431G>A), ATA601GTA (5434A>G), AAT602AGA (5438A>G 5439T>A), ACT603GGC (5440A>G 5441C>G 5442T>C), AAG605CGA (5446A>C 5447A>G 5448G>A), AGT608TCT (5455A>T 5456G>C), ATT609GCT (5458A>G 5459T>C), TAT610TTC (5462A>T 5463T>C), ACT612CTA (5467A>C 5468C>T 5469T>A), CCT613CCA (5472T>A), GTT614AAG (5473G>A 5474T>A 5475T>G), CAT615CAA (5478T>A), TTT616TTC (5481T>C), CTG617ATC (5482C>A 5484G>C), ATT618ATC (5487T>C), AGG619AGA (5490G>A), ACA620TCA (5491A>T), GAT621GAC (5496T>C), ACT623ACA (5502T>A) | | | | | | | | |
| **Proteins** | | | | | | | | | |
| reverse transcriptase (NP_056728.1) | 515 | 624 | 16.2% | 343 | 47.4% | 103 (93.6%) | 47 (42.7%) | 0/7/0/0 | 0 |
| Protein mutations: | R515G (5176A>G), K518D (5185A>G 5187A>T), D520H (5191G>C), L522K (5197C>A 5198T>A 5199C>A), Y523C (5201A>G 5202C>T), M524V (5203A>G 5205G>T), K526S (5210A>G 5211G>C), V527L (5212G>C 5214G>A), K529Q (5218A>C), N530E (5221A>G 5223T>A), L531C (5224C>T 5225T>G 5226G>T), Q532S (5227C>T 5228A>C 5229A>C), G533K (5230G>A 5231G>A), F534L (5235T>A), P536K (5239C>A 5240C>A 5241A>G), H538R (5245C>A 5246A>G 5247T>A), H539L (5249A>T), L541E (5254T>G 5255T>A), E544D (5265G>C), K545N (5268G>C), I547V (5272A>G 5274C>T), I548L (5275A>C 5277C>A), E549Q (5278G>C 5280G>A), D554N (5293G>A 5295C>T), D555Y (5296G>T), Y556H (5299T>C 5301C>T), G559A (5309G>C 5310T>A), M560L (5311A>T 5313G>A), K562Q (5317A>C), A563T (5320G>A), I564del (5323_5325delATC), K565D (5326A>G 5328A>C), I566L (5329A>C), G569_L574del (5338_5355delGGTACTAATACTGAGTTA), A579T (5368G>A), S582T (5378G>C 5379C>A), K584N (5385A>T), A585E (5387C>A 5388T>A), A586T (5389G>A 5391A>C), K588T (5396A>C 5397G>T), N589K (5400T>A), H591S (5404C>T 5405A>C 5406C>A), S592T (5408G>C 5409C>A), D594E (5415C>A), T597L (5422A>T 5423C>T), V600I (5431G>A), I601V (5434A>G), N602R (5438A>G 5439T>A), T603G (5440A>G 5441C>G 5442T>C), K605R (5446A>C 5447A>G 5448G>A), I609A (5458A>G 5459T>C), Y610F (5462A>T 5463T>C), T612L (5467A>C 5468C>T 5469T>A), V614K (5473G>A 5474T>A 5475T>G), H615Q (5478T>A), L617I (5482C>A 5484G>C), T620S (5491A>T) | | | | | | | | |
| Codon mutations: | TGG514.TG (5174G>T), AGA515GGA (5176A>G), AAA518GAT (5185A>G 5187A>T), GAG519GAA (5190G>A), GAT520CAT (5191G>C), CTC522AAA (5197C>A 5198T>A 5199C>A), TAC523TGT (5201A>G 5202C>T), ATG524GTT (5203A>G 5205G>T), AAG526AGC (5210A>G 5211G>C), GTG527CTA (5212G>C 5214G>A), AAG528AAA (5217G>A), AAA529CAA (5218A>C), AAT530GAA (5221A>G 5223T>A), CTG531TGT (5224C>T 5225T>G 5226G>T), CAA532TCC (5227C>T 5228A>C 5229A>C), GGA533AAA (5230G>A 5231G>A), TTT534TTA (5235T>A), CCT535CCA (5238T>A), CCA536AAG (5239C>A 5240C>A 5241A>G), CAT538AGA (5245C>A 5246A>G 5247T>A), CAT539CTT (5249A>T), CCC540CCA (5253C>A), TTA541GAA (5254T>G 5255T>A), CCA542CCG (5259A>G), GAG543GAA (5262G>A), GAG544GAC (5265G>C), AAG545AAC (5268G>C), CTG546TTA (5269C>T 5271G>A), ATC547GTT (5272A>G 5274C>T), ATC548CTA (5275A>C 5277C>A), GAG549CAA (5278G>C 5280G>A), ACC550ACA (5283C>A), GCA552GCT (5289A>T), GAC554AAT (5293G>A 5295C>T), GAC555TAC (5296G>T), TAC556CAT (5299T>C 5301C>T), GGT559GCA (5309G>C 5310T>A), ATG560TTA (5311A>T 5313G>A), AAA562CAA (5317A>C), GCT563ACT (5320G>A), ATC564del (5323_5325delATC), AAA565GAC (5326A>G 5328A>C), ATT566CTT (5329A>C), AAC567AAT (5334C>T), GGT569_TTA574del (5338_5355delGGTACTAATACTGAGTTA), TGC576TGT (5361C>T), AGA577AGG (5364A>G), TAC578TAT (5367C>T), GCA579ACA (5368G>A), TCT580AGT (5371T>A 5372C>G), GGA581GGT (5376A>T), AGC582ACA (5378G>C 5379C>A), AAA584AAT (5385A>T), GCT585GAA (5387C>A 5388T>A), GCA586ACC (5389G>A 5391A>C), AAG588ACT (5396A>C 5397G>T), AAT589AAA (5400T>A), CAC591TCA (5404C>T 5405A>C 5406C>A), AGC592ACA (5408G>C 5409C>A), GAC594GAA (5415C>A), GAG596GAA (5421G>A), ACA597TTA (5422A>T 5423C>T), GCG599GCC (5430G>C), GTA600ATA (5431G>A), ATA601GTA (5434A>G), AAT602AGA (5438A>G 5439T>A), ACT603GGC (5440A>G 5441C>G 5442T>C), AAG605CGA (5446A>C 5447A>G 5448G>A), AGT608TCT (5455A>T 5456G>C), ATT609GCT (5458A>G 5459T>C), TAT610TTC (5462A>T 5463T>C), ACT612CTA (5467A>C 5468C>T 5469T>A), CCT613CCA (5472T>A), GTT614AAG (5473G>A 5474T>A 5475T>G), CAT615CAA (5478T>A), TTT616TTC (5481T>C), CTG617ATC (5482C>A 5484G>C), ATT618ATC (5487T>C), AGG619AGA (5490G>A), ACA620TCA (5491A>T), GAT621GAC (5496T>C), ACT623ACA (5502T>A) | | | | | | | | |

*: Inserts / Deletes / Misaligned / Frameshifts

#### Analysis details

This analysis was performed with panviral2.64

### NGS Details (UN24): Badnavirus venaribis

#### Assembly

| | |
|---|---|
| **Coverage Length** | 238 (1 contig(s)) |
| **Depth Of Coverage** | 6.4 |
| **Number Of Reads** | 12 |
| **Reads Per Million** | 0.24 rpm (after QC) |
| **Ambiguities** | 0 |
| **Assembly Method** | de novo + reference guided assembly |
| **Consensus Caller** | Bcf Tools |

#### Coverage Map

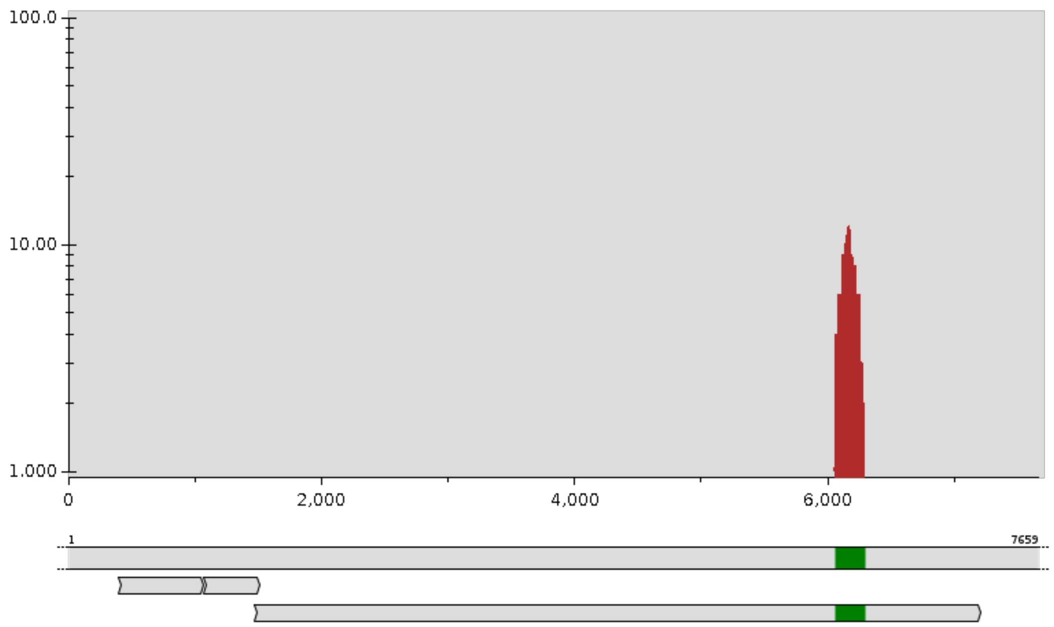

#### Assignment

| | |
|---|---|
| **Type** | Badnavirus venaribis (Taxonomy ID: 3047843) |
| **Reference Genome** | NC_018105.1 |
| **NT Identity (%)** | 53.9419 |
| **AA Identity (%)** | 48.1481 |
| **Number Of Stop Codons** | 0 |
| **Number Of CDS** | 3 |

#### Alignment

| | |
|---|---|
| **Alignment Score** | 32.0 (NT) + 266.0 (AA) = 298.0 |
| **Concordance (%)** | 27.9288 |

| Alignment Method | Global, seeded, nucleotide + amino acids (AGA) |
|---|---|

#### Genome Region

Sequence starts at position 6058 and ends at position 6295 relative to NC_018105.1 reference sequence.

#### Alignment Detailed Statistics

| | Begin | End | Coverage | Score | Concordance | Matches | Identities | I/D/M/F* | Stop Codons |
|---|---|---|---|---|---|---|---|---|---|
| **NT** | **6058** | **6295** | **3.1%** | **32** | **6.7%** | **238 (98.8%)** | **130 (53.9%)** | **3/0** | |
| Mutations: | 6061C>G, 6064T>A, 6065G>A, 6072G>C, 6076A>C, 6077T>G, 6078C>T, 6081G>C, 6083C>A, 6086T>A, 6087A>T, 6090A>C, 6092T>A, 6093C>T, 6101G>T, 6102G>T, 6113C>T, 6114T>G, 6117C>T, 6120T>A, 6121A>T, 6123T>G, 6124G>A, 6126T>A, 6129C>T, 6132T>C, 6135A>T, 6136G>T, 6139G>A, 6141T>A, 6144C>T, 6147G>A, 6148A>G, 6149G>C, 6150G>T, 6152A>C, 6153A>T, 6157G>A, 6158A>C, 6160A>G, 6162C>A, 6163T>A, 6164G>T, 6165C>T, 6168C>T, 6171C>T, 6171_6172insCCA, 6172A>C, 6174G>T, 6175C>A, 6178A>C, 6180A>T, 6181G>C, 6182A>G, 6183G>C, 6186C>T, 6189C>T, 6190G>T, 6191C>T, 6195T>A, 6197A>T, 6199A>T, 6201C>T, 6204C>T, 6207C>T, 6211C>T, 6213C>G, 6218T>A, 6219C>T, 6222C>T, 6223A>G, 6224A>C, 6225T>A, 6228C>T, 6229C>T, 6230T>G, 6234A>T, 6237A>T, 6241G>T, 6242A>T, 6244A>G, 6245G>A, 6246C>A, 6253C>G, 6254G>C, 6255G>A, 6256C>G, 6257A>T, 6258A>G, 6259A>G, 6261G>A, 6262C>T, 6265G>C, 6267A>G, 6268G>T, 6270T>G, 6272G>T, 6273C>G, 6277A>C, 6280A>C, 6283G>C, 6284G>A, 6285G>A, 6286T>C, 6288G>T, 6292C>G, 6293T>C, 6294G>C, 6295T>A | | | | | | | | |

##### CDS

| | Begin | End | Coverage | Score | Concordance | Matches | Identities | I/D/M/F* | Stop Codons |
|---|---|---|---|---|---|---|---|---|---|
| B227_gp3 | 1531 | 1610 | 4.2% | 266 | 44.7% | 80 (98.8%) | 39 (48.1%) | 1/0/0/0 | 0 |
| Protein mutations: | P1532A (6061C>G), W1533K (6064T>A 6065G>A), I1537R (6076A>C 6077T>G 6078C>T), P1539H (6083C>A), V1540D (6086T>A 6087A>T), F1542Y (6092T>A 6093C>T), W1545F (6101G>T 6102G>T), P1549L (6113C>T 6114T>G), I1552L (6121A>T 6123T>G), A1553T (6124G>A 6126T>A), A1557S (6136G>T), V1558I (6139G>A 6141T>A), R1561A (6148A>G 6149G>C 6150G>T), K1562T (6152A>C 6153A>T), D1564T (6157G>A 6158A>C), N1565E (6160A>G 6162C>A), C1566I (6163T>A 6164G>T 6165C>T), H1568_K1569insP (6171_6172insCCA), K1569H (6172A>C 6174G>T), L1570M (6175C>A), E1572R (6181G>C 6182A>G 6183G>C), A1575L (6190G>T 6191C>T), Y1577F (6197A>T), I1578F (6199A>T 6201C>T), F1584Y (6218T>A 6219C>T), N1586A (6223A>G 6224A>C 6225T>A), L1588W (6229C>T 6230T>G), Q1589H (6234A>T), E1590D (6237A>T), E1592L (6241G>T 6242A>T), S1593E (6244A>G 6245G>A 6246C>A), R1596A (6253C>G 6254G>C 6255G>A), Q1597V (6256C>G 6257A>T 6258A>G), M1598V (6259A>G 6261G>A), E1600Q (6265G>C 6267A>G), V1601L (6268G>T 6270T>G), C1602L (6272G>T 6273C>G), K1604Q (6277A>C), N1605H (6280A>C), G1606Q (6283G>C 6284G>A 6285G>A), L1609A (6292C>G 6293T>C 6294G>C) | | | | | | | | |
| Codon mutations: | CCT1532GCT (6061C>G), TGG1533AAG (6064T>A 6065G>A), GCG1535GCC (6072G>C), ATC1537CGT (6076A>C 6077T>G 6078C>T), ACG1538ACC (6081G>C), CCC1539CAC (6083C>A), GTA1540GAT (6086T>A 6087A>T), GGA1541GGC (6090A>C), TTC1542TAT (6092T>A 6093C>T), TGG1545TTT (6101G>T 6102G>T), CCT1549CTG (6113C>T 6114T>G), TTC1550TTT (6117C>T), GGT1551GGA (6120T>A), ATT1552TTG (6121A>T 6123T>G), GCT1553ACA (6124G>A 6126T>A), AAC1554AAT (6129C>T), GCT1555GCC (6132T>C), CCA1556CCT (6135A>T), GCG1557TCG (6136G>T), GTT1558ATA (6139G>A 6141T>A), TTC1559TTT (6144C>T), CAG1560CAA (6147G>A), AGG1561GCT (6148A>G 6149G>C 6150G>T), AAA1562ACT (6152A>C 6153A>T), GAT1564ACT (6157G>A 6158A>C), AAC1565GAA (6160A>G 6162C>A), TGC1566ATT (6163T>A 6164G>T 6165C>T), TTC1567TTT (6168C>T), CAC1568CAT (6171C>T), CAC1568_AAG1569insCCA (6171_6172insCCA), AAG1569CAT (6172A>C 6174G>T), CTG1570ATG (6175C>A), AGA1571CGT (6178A>C 6180A>T), GAG1572CGC (6181G>C 6182A>G 6183G>C), TTC1573TTT (6186C>T), GTC1574GTT (6189C>T), GCG1575TTG (6190G>T 6191C>T), GTT1576GTA (6195T>A), TAT1577TTT (6197A>T), ATC1578TTT (6199A>T 6201C>T), GAC1579GAT (6204C>T), GAC1580GAT (6207C>T), CTC1582TTG (6211C>T 6213C>G), TTC1584TAT (6218T>A 6219C>T), AGC1585AGT (6222C>T), AAT1586GCA (6223A>G 6224A>C 6225T>A), TCC1587TCT (6228C>T), CTG1588TGG (6229C>T 6230T>G), CAA1589CAT (6234A>T), GAA1590GAT (6237A>T), GAG1592TTG (6241G>T 6242A>T), AGC1593GAA (6244A>G 6245G>A 6246C>A), CGG1596GCA (6253C>G 6254G>C 6255G>A), CAA1597GTG (6256C>G 6257A>T 6258A>G), ATG1598GTA (6259A>G 6261G>A), CTG1599TTG (6262C>T), GAA1600CAG (6265G>C 6267A>G), GTT1601TTG (6268G>T 6270T>G), TGC1602TTG (6272G>T 6273C>G), AAG1604CAG (6277A>C), AAT1605CAT (6280A>C), GGG1606CAA (6283G>C 6284G>A 6285G>A), TTG1607CTT (6286T>C 6288G>T), CTG1609GCC (6292C>G 6293T>C 6294G>C), TCA1610A.. (6295T>A) | | | | | | | | |

##### Proteins

| | Begin | End | Coverage | Score | Concordance | Matches | Identities | I/D/M/F* | Stop Codons |
|---|---|---|---|---|---|---|---|---|---|
| polyprotein (YP_006495799.1) | 1531 | 1610 | 4.2% | 266 | 44.7% | 80 (98.8%) | 39 (48.1%) | 1/0/0/0 | 0 |
| Protein mutations: | P1532A (6061C>G), W1533K (6064T>A 6065G>A), I1537R (6076A>C 6077T>G 6078C>T), P1539H (6083C>A), V1540D (6086T>A 6087A>T), F1542Y (6092T>A 6093C>T), W1545F (6101G>T 6102G>T), P1549L (6113C>T 6114T>G), I1552L (6121A>T 6123T>G), A1553T (6124G>A 6126T>A), A1557S (6136G>T), V1558I (6139G>A 6141T>A), R1561A (6148A>G 6149G>C 6150G>T), K1562T (6152A>C 6153A>T), D1564T (6157G>A 6158A>C), N1565E (6160A>G 6162C>A), C1566I (6163T>A 6164G>T 6165C>T), H1568_K1569insP (6171_6172insCCA), K1569H (6172A>C 6174G>T), L1570M (6175C>A), E1572R (6181G>C 6182A>G 6183G>C), A1575L (6190G>T 6191C>T), Y1577F (6197A>T), I1578F (6199A>T 6201C>T), F1584Y (6218T>A 6219C>T), N1586A (6223A>G 6224A>C 6225T>A), L1588W (6229C>T 6230T>G), Q1589H (6234A>T), E1590D (6237A>T), E1592L (6241G>T 6242A>T), S1593E (6244A>G 6245G>A 6246C>A), R1596A (6253C>G 6254G>C 6255G>A), Q1597V (6256C>G 6257A>T 6258A>G), M1598V (6259A>G 6261G>A), E1600Q (6265G>C 6267A>G), V1601L (6268G>T 6270T>G), C1602L (6272G>T 6273C>G), K1604Q (6277A>C), N1605H (6280A>C), G1606Q (6283G>C 6284G>A 6285G>A), L1609A (6292C>G 6293T>C 6294G>C) | | | | | | | | |
| Codon mutations: | CCT1532GCT (6061C>G), TGG1533AAG (6064T>A 6065G>A), GCG1535GCC (6072G>C), ATC1537CGT (6076A>C 6077T>G 6078C>T), ACG1538ACC (6081G>C), CCC1539CAC (6083C>A), GTA1540GAT (6086T>A 6087A>T), GGA1541GGC (6090A>C), TTC1542TAT (6092T>A 6093C>T), TGG1545TTT (6101G>T 6102G>T), CCT1549CTG (6113C>T 6114T>G), TTC1550TTT (6117C>T), GGT1551GGA (6120T>A), ATT1552TTG (6121A>T 6123T>G), GCT1553ACA (6124G>A 6126T>A), AAC1554AAT (6129C>T), GCT1555GCC (6132T>C), CCA1556CCT (6135A>T), GCG1557TCG (6136G>T), GTT1558ATA (6139G>A 6141T>A), TTC1559TTT (6144C>T), CAG1560CAA (6147G>A), AGG1561GCT (6148A>G 6149G>C 6150G>T), AAA1562ACT (6152A>C 6153A>T), GAT1564ACT (6157G>A 6158A>C), AAC1565GAA (6160A>G 6162C>A), TGC1566ATT (6163T>A 6164G>T 6165C>T), TTC1567TTT (6168C>T), CAC1568CAT (6171C>T), CAC1568_AAG1569insCCA (6171_6172insCCA), AAG1569CAT (6172A>C 6174G>T), CTG1570ATG (6175C>A), AGA1571CGT (6178A>C 6180A>T), GAG1572CGC (6181G>C 6182A>G 6183G>C), TTC1573TTT (6186C>T), GTC1574GTT (6189C>T), GCG1575TTG (6190G>T 6191C>T), GTT1576GTA (6195T>A), TAT1577TTT (6197A>T), ATC1578TTT (6199A>T 6201C>T), GAC1579GAT (6204C>T), GAC1580GAT (6207C>T), CTC1582TTG (6211C>T 6213C>G), TTC1584TAT (6218T>A 6219C>T), AGC1585AGT (6222C>T), AAT1586GCA (6223A>G 6224A>C 6225T>A), TCC1587TCT (6228C>T), CTG1588TGG (6229C>T 6230T>G), CAA1589CAT (6234A>T), GAA1590GAT (6237A>T), GAG1592TTG (6241G>T 6242A>T), AGC1593GAA (6244A>G 6245G>A 6246C>A), CGG1596GCA (6253C>G 6254G>C 6255G>A), CAA1597GTG (6256C>G 6257A>T 6258A>G), ATG1598GTA (6259A>G 6261G>A), CTG1599TTG (6262C>T), GAA1600CAG (6265G>C 6267A>G), GTT1601TTG (6268G>T 6270T>G), TGC1602TTG (6272G>T 6273C>G), AAG1604CAG (6277A>C), AAT1605CAT (6280A>C), GGG1606CAA (6283G>C 6284G>A 6285G>A), TTG1607CTT (6286T>C 6288G>T), CTG1609GCC (6292C>G 6293T>C 6294G>C), TCA1610A.. (6295T>A) | | | | | | | | |

*: Inserts / Deletes / Misaligned / Frameshifts

#### Analysis details

This analysis was performed with panviral2.64

### NGS Details (UN24): Badnavirus rutilanscamelliae

#### Assembly

| | |
|---|---|
| **Coverage Length** | 364 (1 contig(s)) |
| **Depth Of Coverage** | 3.5 |
| **Number Of Reads** | 11 |
| **Reads Per Million** | 0.22 rpm (after QC) |
| **Ambiguities** | 0 |
| **Assembly Method** | de novo + reference guided assembly |
| **Consensus Caller** | Bcf Tools |

#### Coverage Map

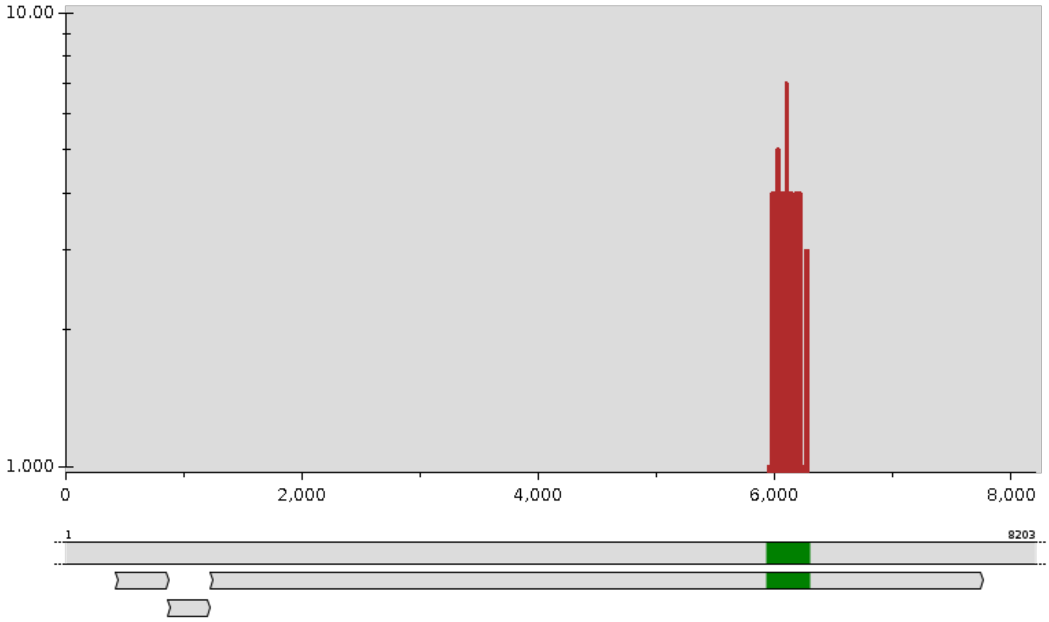

#### Assignment

| | |
|---|---|
| **Type** | Badnavirus rutilanscamelliae (Taxonomy ID: 3047719) |
| **Reference Genome** | NC_055598.1 |
| **NT Identity (%)** | 56.044 |
| **AA Identity (%)** | 53.719 |
| **Number Of Stop Codons** | 1 |
| **Number Of CDS** | 3 |

#### Alignment

| | |
|---|---|
| **Alignment Score** | 88.0 (NT) + 446.0 (AA) = 534.0 |
| **Concordance (%)** | 33.1883 |

| Alignment Method | Global, seeded, nucleotide + amino acids (AGA) |
|---|---|

#### Genome Region

Sequence starts at position 5938 and ends at position 6301 relative to NC_055598.1 reference sequence.

#### Alignment Detailed Statistics

| | Begin | End | Coverage | Score | Concordance | Matches | Identities | I/D/M/F* | Stop Codons |
|---|---|---|---|---|---|---|---|---|---|
| **NT** | **5938** | **6301** | **4.4%** | **88** | **12.1%** | **364 (100%)** | **204 (56.0%)** | **0/0** | |
| Mutations: | 5938C>A, 5941A>C, 5944C>T, 5946A>T, 5950G>A, 5953G>A, 5956C>T, 5959C>T, 5960C>T, 5962G>A, 5965G>A, 5971G>A, 5973T>A, 5975C>T, 5976A>G, 5977T>G, 5981G>A, 5984G>C, 5985C>A, 5986T>G, 5989G>C, 5990G>T, 5991A>T, 5993C>A, 5994C>A, 5995T>A, 5996C>G, 5998A>G, 6001C>T, 6003T>C, 6005G>T, 6007A>T, 6008T>A, 6010G>A, 6013C>T, 6020T>A, 6021G>A, 6022G>T, 6023A>G, 6024C>T, 6025T>C, 6029G>T, 6031G>T, 6034C>A, 6035T>C, 6036T>A, 6046G>A, 6047C>A, 6048T>A, 6049G>T, 6064G>T, 6067T>A, 6070G>A, 6072A>T, 6079A>T, 6080G>T, 6082A>T, 6084T>A, 6091A>G, 6093G>A, 6094G>A, 6096A>T, 6097G>C, 6101G>A, 6103C>T, 6104C>G, 6105G>A, 6106A>T, 6109C>T, 6121C>T, 6122G>A, 6123C>T, 6124T>A, 6125G>A, 6127A>T, 6130T>C, 6133T>A, 6134G>A, 6135C>T, 6139G>C, 6151T>C, 6154C>A, 6155C>T, 6157C>G, 6160A>T, 6162T>A, 6163C>T, 6164A>T, 6165G>C, 6166C>T, 6167C>A, 6168C>A, 6172A>T, 6173C>T, 6174C>T, 6179G>A, 6180C>T, 6181C>G, 6186A>T, 6187A>C, 6193C>T, 6196G>A, 6198C>A, 6199C>A, 6200C>A, 6201A>T, 6202A>C, 6205C>T, 6206G>A, 6207T>A, 6209G>A, 6211G>A, 6214A>T, 6215T>G, 6216G>T, 6217C>T, 6219A>T, 6220G>A, 6221G>C, 6224C>A, 6229G>T, 6230C>T, 6232T>G, 6235C>A, 6236T>A, 6238G>A, 6239A>T, 6240G>C, 6242C>A, 6243C>A, 6245A>C, 6250G>A, 6255A>G, 6256G>T, 6257G>T, 6259G>A, 6260G>T, 6261C>T, 6265G>A, 6266G>A, 6268C>A, 6269C>G, 6272A>G, 6275G>C, 6276A>G, 6277C>A, 6280T>C, 6283G>A, 6286C>A, 6287T>C, 6288C>A, 6289A>T, 6291C>A, 6292A>T, 6295T>A, 6296A>T, 6297C>G, 6298A>T, 6299A>C, 6301T>A | | | | | | | | |

##### CDS

| | Begin | End | Coverage | Score | Concordance | Matches | Identities | I/D/M/F* | Stop Codons |
|---|---|---|---|---|---|---|---|---|---|
| KM754_gp3 | 1573 | 1693 | 5.6% | 446 | 50.4% | 121 (100%) | 65 (53.7%) | 0/0/0/0 | 1 |
| Protein mutations: | K1573N (5941A>C), Y1575F (5946A>T), F1584Y (5973T>A), H1585W (5975C>T 5976A>G 5977T>G), V1587I (5981G>A), A1588Q (5984G>C 5985C>A 5986T>G), M1589I (5989G>C), D1590F (5990G>T 5991A>T), P1591K (5993C>A 5994C>A 5995T>A), Q1592E (5996C>G 5998A>G), I1594T (6003T>C), E1595Y (6005G>T 6007A>T), W1596R (6008T>A 6010G>A), W1600N (6020T>A 6021G>A 6022G>T), T1601V (6023A>G 6024C>T 6025T>C), V1603F (6029G>T 6031G>T), L1605Q (6035T>C 6036T>A), W1608* (6046G>A), L1609N (6047C>A 6048T>A 6049G>T), N1617I (6072A>T), A1620S (6080G>T 6082A>T), V1621E (6084T>A), R1624K (6093G>A 6094G>A), K1625I (6096A>T 6097G>C), D1627N (6101G>A 6103C>T), R1628D (6104C>G 6105G>A 6106A>T), A1634I (6122G>A 6123C>T 6124T>A), G1635S (6125G>A 6127A>T), A1638I (6134G>A 6135C>T), F1647Y (6162T>A 6163C>T), P1649K (6167C>A 6168C>A), P1651F (6173C>T 6174C>T), A1653M (6179G>A 6180C>T 6181C>G), K1655I (6186A>T 6187A>C), A1659E (6198C>A 6199C>A), Q1660I (6200C>A 6201A>T 6202A>C), V1662K (6206G>A 6207T>A), E1663K (6209G>A 6211G>A), C1665V (6215T>G 6216G>T 6217C>T), K1666I (6219A>T 6220G>A), E1667Q (6221G>C), H1668N (6224C>A), L1672I (6236T>A 6238G>A), P1674K (6242C>A 6243C>A), T1675P (6245A>C), K1678S (6255A>G 6256G>T), V1679L (6257G>T 6259G>A), A1680F (6260G>T 6261C>T), A1682T (6266G>A 6268C>A), Q1683E (6269C>G), I1684V (6272A>G), D1685R (6275G>C 6276A>G 6277C>A), S1689H (6287T>C 6288C>A 6289A>T), T1690N (6291C>A 6292A>T), T1692C (6296A>T 6297C>G 6298A>T), N1693Q (6299A>C 6301T>A) | | | | | | | | |
| Codon mutations: | AGC1572..A (5938C>A), AAA1573AAC (5941A>C), ATC1574ATT (5944C>T), TAT1575TTT (5946A>T), TCG1576TCA (5950G>A), AAG1577AAA (5953G>A), TTC1578TTT (5956C>T), GAC1579GAT (5959C>T), CTG1580TTA (5960C>T 5962G>A), AAG1581AAA (5965G>A), GGG1583GGA (5971G>A), TTT1584TAT (5973T>A), CAT1585TGG (5975C>T 5976A>G 5977T>G), GTT1587ATT (5981G>A), GCT1588CAG (5984G>C 5985C>A 5986T>G), ATG1589ATC (5989G>C), GAT1590TTT (5990G>T 5991A>T), CCT1591AAA (5993C>A 5994C>A 5995T>A), CAA1592GAG (5996C>G 5998A>G), CAC1593CAT (6001C>T), ATT1594ACT (6003T>C), GAA1595TAT (6005G>T 6007A>T), TGG1596AGA (6008T>A 6010G>A), ACC1597ACT (6013C>T), TGG1600AAT (6020T>A 6021G>A 6022G>T), ACT1601GTC (6023A>G 6024C>T 6025T>C), GTG1603TTT (6029G>T 6031G>T), GGC1604GGA (6034C>A), TTA1605CAA (6035T>C 6036T>A), TGG1608TGA (6046G>A), CTG1609AAT (6047C>A 6048T>A 6049G>T), GGG1614GGT (6064G>T), CTT1615CTA (6067T>A), AAG1616AAA (6070G>A), AAT1617ATT (6072A>T), CCA1619CCT (6079A>T), GCA1620TCT (6080G>T 6082A>T), GTA1621GAA (6084T>A), CAA1623CAG (6091A>G), AGG1624AAA (6093G>A 6094G>A), AAG1625ATC (6096A>T 6097G>C), GAC1627AAT (6101G>A 6103C>T), CGA1628GAT (6104C>G 6105G>A 6106A>T), ATC1629ATT (6109C>T), TAC1633TAT (6121C>T), GCT1634ATA (6122G>A 6123C>T 6124T>A), GGA1635AGT (6125G>A 6127A>T), TTT1636TTC (6130T>C), ATT1637ATA (6133T>A), GCT1638ATT (6134G>A 6135C>T), GTG1639GTC (6139G>C), GAT1643GAC (6151T>C), ATC1644ATA (6154C>A), CTC1645TTG (6155C>T 6157C>G), ATA1646ATT (6160A>T), TTC1647TAT (6162T>A 6163C>T), AGC1648TCT (6164A>T 6165G>C 6166C>T), CCA1649AAA (6167C>A 6168C>A), ACA1650ACT (6172A>T), CCT1651TTT (6173C>T 6174C>T), GCC1653ATG (6179G>A 6180C>T 6181C>G), AAA1655ATC (6186A>T 6187A>C), CAC1657CAT (6193C>T), TTG1658TTA (6196G>A), GCC1659GAA (6198C>A 6199C>A), CAA1660ATC (6200C>A 6201A>T 6202A>C), TTC1661TTT (6205C>T), GTG1662AAG (6206G>A 6207T>A), GAG1663AAA (6209G>A 6211G>A), ATA1664ATT (6214A>T), TGC1665GTT (6215T>G 6216G>T 6217C>T), AAG1666ATA (6219A>T 6220G>A), GAA1667CAA (6221G>C), CAT1668AAT (6224C>A), GGG1669GGT (6229G>T), CTT1670TTG (6230C>T 6232T>G), GTC1671GTA (6235C>A), TTG1672ATA (6236T>A 6238G>A), AGT1673TCT (6239A>T 6240G>C), CCA1674AAA (6242C>A 6243C>A), ACA1675CCA (6245A>C), AAG1676AAA (6250G>A), AAG1678AGT (6255A>G 6256G>T), GTG1679TTA (6257G>T 6259G>A), GCC1680TTC (6260G>T 6261C>T), CAG1681CAA (6265G>A), GCC1682ACA (6266G>A 6268C>A), CAA1683GAA (6269C>G), ATT1684GTT (6272A>G), GAC1685CGA (6275G>C 6276A>G 6277C>A), TTT1686TTC (6280T>C), TTG1687TTA (6283G>A), GGC1688GGA (6286C>A), TCA1689CAT (6287T>C 6288C>A 6289A>T), ACA1690AAT (6291C>A 6292A>T), ATT1691ATA (6295T>A), ACA1692TGT (6296A>T 6297C>G 6298A>T), AAT1693CAA (6299A>C 6301T>A) | | | | | | | | |

##### Proteins

| | Begin | End | Coverage | Score | Concordance | Matches | Identities | I/D/M/F* | Stop Codons |
|---|---|---|---|---|---|---|---|---|---|
| polyprotein (YP_010087856.1) | 1573 | 1693 | 5.6% | 446 | 50.4% | 121 (100%) | 65 (53.7%) | 0/0/0/0 | 1 |
| Protein mutations: | K1573N (5941A>C), Y1575F (5946A>T), F1584Y (5973T>A), H1585W (5975C>T 5976A>G 5977T>G), V1587I (5981G>A), A1588Q (5984G>C 5985C>A 5986T>G), M1589I (5989G>C), D1590F (5990G>T 5991A>T), P1591K (5993C>A 5994C>A 5995T>A), Q1592E (5996C>G 5998A>G), I1594T (6003T>C), E1595Y (6005G>T 6007A>T), W1596R (6008T>A 6010G>A), W1600N (6020T>A 6021G>A 6022G>T), T1601V (6023A>G 6024C>T 6025T>C), V1603F (6029G>T 6031G>T), L1605Q (6035T>C 6036T>A), W1608* (6046G>A), L1609N (6047C>A 6048T>A 6049G>T), N1617I (6072A>T), A1620S (6080G>T 6082A>T), V1621E (6084T>A), R1624K (6093G>A 6094G>A), K1625I (6096A>T 6097G>C), D1627N (6101G>A 6103C>T), R1628D (6104C>G 6105G>A 6106A>T), A1634I (6122G>A 6123C>T 6124T>A), G1635S (6125G>A 6127A>T), A1638I (6134G>A 6135C>T), F1647Y (6162T>A 6163C>T), P1649K (6167C>A 6168C>A), P1651F (6173C>T 6174C>T), A1653M (6179G>A 6180C>T 6181C>G), K1655I (6186A>T 6187A>C), A1659E (6198C>A 6199C>A), Q1660I (6200C>A 6201A>T 6202A>C), V1662K (6206G>A 6207T>A), E1663K (6209G>A 6211G>A), C1665V (6215T>G 6216G>T 6217C>T), K1666I (6219A>T 6220G>A), E1667Q (6221G>C), H1668N (6224C>A), L1672I (6236T>A 6238G>A), P1674K (6242C>A 6243C>A), T1675P (6245A>C), K1678S (6255A>G 6256G>T), V1679L (6257G>T 6259G>A), A1680F (6260G>T 6261C>T), A1682T (6266G>A 6268C>A), Q1683E (6269C>G), I1684V (6272A>G), D1685R (6275G>C 6276A>G 6277C>A), S1689H (6287T>C 6288C>A 6289A>T), T1690N (6291C>A 6292A>T), T1692C (6296A>T 6297C>G 6298A>T), N1693Q (6299A>C 6301T>A) | | | | | | | | |
| Codon mutations: | AGC1572..A (5938C>A), AAA1573AAC (5941A>C), ATC1574ATT (5944C>T), TAT1575TTT (5946A>T), TCG1576TCA (5950G>A), AAG1577AAA (5953G>A), TTC1578TTT (5956C>T), GAC1579GAT (5959C>T), CTG1580TTA (5960C>T 5962G>A), AAG1581AAA (5965G>A), GGG1583GGA (5971G>A), TTT1584TAT (5973T>A), CAT1585TGG (5975C>T 5976A>G 5977T>G), GTT1587ATT (5981G>A), GCT1588CAG (5984G>C 5985C>A 5986T>G), ATG1589ATC (5989G>C), GAT1590TTT (5990G>T 5991A>T), CCT1591AAA (5993C>A 5994C>A 5995T>A), CAA1592GAG (5996C>G 5998A>G), CAC1593CAT (6001C>T), ATT1594ACT (6003T>C), GAA1595TAT (6005G>T 6007A>T), TGG1596AGA (6008T>A 6010G>A), ACC1597ACT (6013C>T), TGG1600AAT (6020T>A 6021G>A 6022G>T), ACT1601GTC (6023A>G 6024C>T 6025T>C), GTG1603TTT (6029G>T 6031G>T), GGC1604GGA (6034C>A), TTA1605CAA (6035T>C 6036T>A), TGG1608TGA (6046G>A), CTG1609AAT (6047C>A 6048T>A 6049G>T), GGG1614GGT (6064G>T), CTT1615CTA (6067T>A), AAG1616AAA (6070G>A), AAT1617ATT (6072A>T), CCA1619CCT (6079A>T), GCA1620TCT (6080G>T 6082A>T), GTA1621GAA (6084T>A), CAA1623CAG (6091A>G), AGG1624AAA (6093G>A 6094G>A), AAG1625ATC (6096A>T 6097G>C), GAC1627AAT (6101G>A 6103C>T), CGA1628GAT (6104C>G 6105G>A 6106A>T), ATC1629ATT (6109C>T), TAC1633TAT (6121C>T), GCT1634ATA (6122G>A 6123C>T 6124T>A), GGA1635AGT (6125G>A 6127A>T), TTT1636TTC (6130T>C), ATT1637ATA (6133T>A), GCT1638ATT (6134G>A 6135C>T), GTG1639GTC (6139G>C), GAT1643GAC (6151T>C), ATC1644ATA (6154C>A), CTC1645TTG (6155C>T 6157C>G), ATA1646ATT (6160A>T), TTC1647TAT (6162T>A 6163C>T), AGC1648TCT (6164A>T 6165G>C 6166C>T), CCA1649AAA (6167C>A 6168C>A), ACA1650ACT (6172A>T), CCT1651TTT (6173C>T 6174C>T), GCC1653ATG (6179G>A 6180C>T 6181C>G), AAA1655ATC (6186A>T 6187A>C), CAC1657CAT (6193C>T), TTG1658TTA (6196G>A), GCC1659GAA (6198C>A 6199C>A), CAA1660ATC (6200C>A 6201A>T 6202A>C), TTC1661TTT (6205C>T), GTG1662AAG (6206G>A 6207T>A), GAG1663AAA (6209G>A 6211G>A), ATA1664ATT (6214A>T), TGC1665GTT (6215T>G 6216G>T 6217C>T), AAG1666ATA (6219A>T 6220G>A), GAA1667CAA (6221G>C), CAT1668AAT (6224C>A), GGG1669GGT (6229G>T), CTT1670TTG (6230C>T 6232T>G), GTC1671GTA (6235C>A), TTG1672ATA (6236T>A 6238G>A), AGT1673TCT (6239A>T 6240G>C), CCA1674AAA (6242C>A 6243C>A), ACA1675CCA (6245A>C), AAG1676AAA (6250G>A), AAG1678AGT (6255A>G 6256G>T), GTG1679TTA (6257G>T 6259G>A), GCC1680TTC (6260G>T 6261C>T), CAG1681CAA (6265G>A), GCC1682ACA (6266G>A 6268C>A), CAA1683GAA (6269C>G), ATT1684GTT (6272A>G), GAC1685CGA (6275G>C 6276A>G 6277C>A), TTT1686TTC (6280T>C), TTG1687TTA (6283G>A), GGC1688GGA (6286C>A), TCA1689CAT (6287T>C 6288C>A 6289A>T), ACA1690AAT (6291C>A 6292A>T), ATT1691ATA (6295T>A), ACA1692TGT (6296A>T 6297C>G 6298A>T), AAT1693CAA (6299A>C 6301T>A) | | | | | | | | |

*: Inserts / Deletes / Misaligned / Frameshifts

#### Analysis details

This analysis was performed with panviral2.64

#### NGS Details (UN24): Soymovirus virgarachidis

##### Assembly

| | |
|---|---|
| **Coverage Length** | 334 (1 contig(s)) |
| **Depth Of Coverage** | 3.9 |
| **Number Of Reads** | 10 |
| **Reads Per Million** | 0.20 rpm (after QC) |
| **Ambiguities** | 0 |
| **Assembly Method** | de novo + reference guided assembly |
| **Consensus Caller** | Bcf Tools |

##### Coverage Map

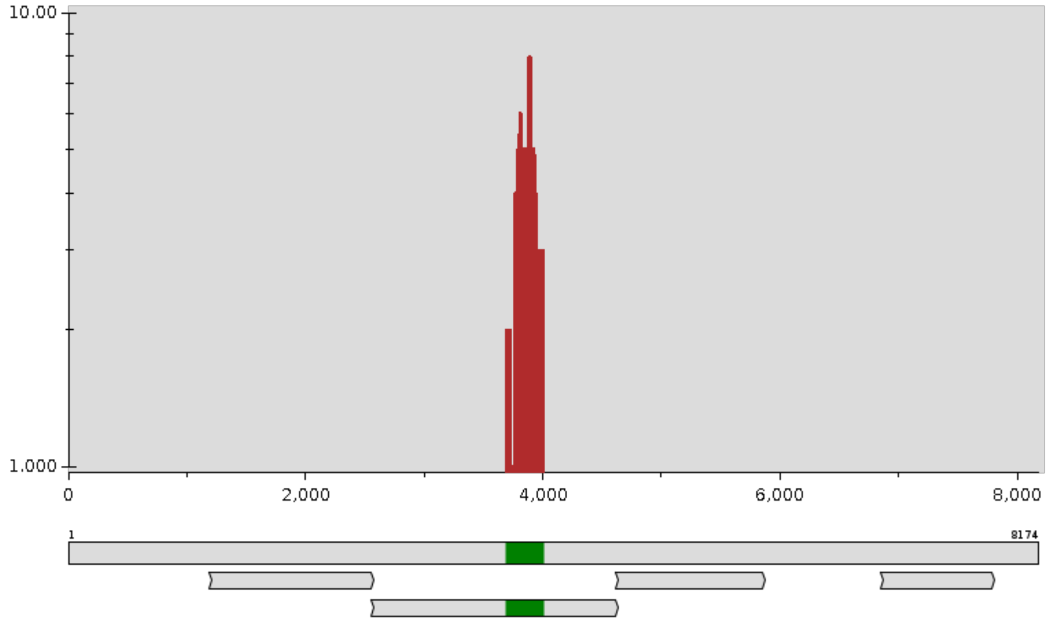

##### Assignment

| | |
|---|---|
| **Type** | Soymovirus virgarachidis (Taxonomy ID: 3048252) |
| **Reference Genome** | NC_001634.1 |
| **NT Identity (%)** | 59.4427 |
| **AA Identity (%)** | 48.1481 |
| **Number Of Stop Codons** | 2 |
| **Number Of CDS** | 4 |

##### Alignment

| | |
|---|---|
| **Alignment Score** | 84.0 (NT) + 215.0 (AA) = 299.0 |
| **Concordance (%)** | 25.4902 |

| **Alignment Method** | Global, seeded, nucleotide + amino acids (AGA) |
|---|---|

#### Genome Region

Sequence starts at position 3684 and ends at position 4017 relative to NC_001634.1 reference sequence.

#### Alignment Detailed Statistics

| | Begin | End | Coverage | Score | Concordance | Matches | Identities | I/D/M/F* | Stop Codons |
|---|---|---|---|---|---|---|---|---|---|
| **NT** | **3684** | **4017** | **4.1%** | **84** | **13.8%** | **323 (96.7%)** | **192 (57.5%)** | **0/11** | |
| Mutations: | 3684G>C, 3689G>A, 3693A>G, 3695G>C, 3696C>T, 3697A>G, 3701A>T, 3702T>A, 3703C>A, 3704G>T, 3706T>A, 3707T>A, 3708T>A, 3709T>A, 3710G>A, 3712A>T, 3718T>A, 3721G>A, 3724T>C, 3727C>T, 3730T>A, 3731T>A, 3733A>G, 3735A>T, 3736G>T, 3741T>C, 3742G>A, 3751A>G, 3752_3754delTTT, 3757T>A, 3760A>G, 3761C>T, 3762C>T, 3766G>A, 3768A>C, 3769A>T, 3775T>C, 3776_3777delTT, 3778A>C, 3779G>A, 3784G>A, 3788G>T, 3789C>T, 3790T>A, 3799A>T, 3803C>A, 3805G>A, 3806C>T, 3809C>A, 3810A>C, 3812G>A, 3813A>C, 3816G>A, 3818A>C, 3820G>A, 3821C>G, 3826T>A, 3829G>A, 3831A>G, 3838C>T, 3852A>T, 3853C>T, 3854A>G, 3855T>C, 3856T>A, 3863_3868delAAAGGA, 3871T>C, 3872T>A, 3873T>C, 3874T>A, 3875A>C, 3876A>C, 3877G>A, 3890G>A, 3892A>G, 3898A>T, 3899G>C, 3902T>C, 3907G>A, 3911A>C, 3913G>C, 3914C>A, 3916T>C, 3917T>A, 3918C>G, 3920G>A, 3923A>T, 3924A>C, 3926C>A, 3927T>A, 3930C>G, 3933G>T, 3935A>T, 3936A>G, 3942A>C, 3943T>C, 3944G>A, 3947T>G, 3948T>A, 3950G>C, 3951C>A, 3952C>T, 3957T>G, 3958G>T, 3959G>T, 3960C>G, 3961A>T, 3964T>A, 3965A>C, 3969G>A, 3970G>A, 3971T>C, 3973A>C, 3977C>G, 3979G>A, 3981T>A, 3982T>A, 3985C>T, 3986A>G, 3987A>T, 3988A>T, 3991C>G, 3992T>C, 3994A>T, 3999G>A, 4003G>T, 4004T>A, 4005A>G, 4006T>G, 4007G>C, 4009T>A, 4010G>C, 4013A>G, 4015G>T | | | | | | | | |
| **CDS** | | | | | | | | | |
| PCSVgp2 | 378 | 488 | 16.0% | 215 | 30.5% | 108 (97.3%) | 52 (46.8%) | 0/3/1/1 | 2 |
| Protein mutations: | E379K (3689G>A), K380R (3693A>G), A381L (3695G>C 3696C>T 3697A>G), I383* (3701A>T 3702T>A 3703C>A), A384S (3704G>T 3706T>A), F385K (3707T>A 3708T>A 3709T>A), E386N (3710G>A 3712A>T), L393M (3731T>A 3733A>G), K394I (3735A>T 3736G>T), L396S (3741T>C 3742G>A), F400del (3752_3754delTTT), P403L (3761C>T 3762C>T), K405T (3768A>C 3769A>T), E409K (3779G>A), A412L (3788G>T 3789C>T 3790T>A), Q417K (3803C>A 3805G>A), Q419T (3809C>A 3810A>C), D420T (3812G>A 3813A>C), R421K (3816G>A), K422Q (3818A>C 3820G>A), Q423E (3821C>G), K426R (3831A>G), Y433F (3852A>T 3853C>T), I434A (3854A>G 3855T>C 3856T>A), K437_G438del (3863_3868delAAAGGA), F440T (3872T>A 3873T>C 3874T>A), K441P (3875A>C 3876A>C 3877G>A), E446K (3890G>A 3892A>G), K448N (3898A>T), V449L (3899G>C), M453L (3911A>C 3913G>C), L454I (3914C>A 3916T>C), S455R (3917T>A 3918C>G), E456K (3920G>A), K457S (3923A>T 3924A>C), L458N (3926C>A 3927T>A), P459R (3930C>G), W460L (3933G>T), K461* (3935A>T 3936A>G), N463T (3942A>C 3943T>C), D464N (3944G>A), L465E (3947T>G 3948T>A), A466H (3950G>C 3951C>A 3952C>T), L468R (3957T>G 3958G>T), A469C (3959G>T 3960C>G 3961A>T), K471Q (3965A>C), R472K (3969G>A 3970G>A), Q475E (3977C>G 3979G>A), V476E (3981T>A 3982T>A), K478V (3986A>G 3987A>T 3988A>T), N479K (3991C>G), R482K (3999G>A), Y484R (4004T>A 4005A>G 4006T>G), V485L (4007G>C 4009T>A), A486P (4010G>C), K487D (4013A>G 4015G>T) | | | | | | | | |
| Codon mutations: | AGC377.CC (3684G>C), GAA379AAA (3689G>A), AAG380AGG (3693A>G), GCA381CTG (3695G>C 3696C>T 3697A>G), ATC383TAA (3701A>T 3702T>A 3703C>A), GCT384TCA (3704G>T 3706T>A), TTT385AAA (3707T>A 3708T>A 3709T>A), GAA386AAT (3710G>A 3712A>T), ATT388ATA (3718T>A), GAG389GAA (3721G>A), TTT390TTC (3724T>C), CTC391CTT (3727C>T), GGT392GGA (3730T>A), TTA393ATG (3731T>A 3733A>G), AAG394ATT (3735A>T 3736G>T), TTG396TCA (3741T>C 3742G>A), GGA399GGG (3751A>G), TTT400del (3752_3754delTTT), ATT401ATA (3757T>A), GAA402GAG (3760A>G), CCA403TTA (3761C>T 3762C>T), CAG404CAA (3766G>A), AAA405ACT (3768A>C 3769A>T), CTT407CTC (3775T>C), TTA408--C (3776_3777delTT 3778A>C), GAA409AAA (3779G>A), AAG410AAA (3784G>A), GCT412TTA (3788G>T 3789C>T 3790T>A), CCA415CCT (3799A>T), CAG417AAA (3803C>A 3805G>A), CTA418TTA (3806C>T), CAA419ACA (3809C>A 3810A>C), GAC420ACC (3812G>A 3813A>C), AGA421AAA (3816G>A), AAG422CAA (3818A>C 3820G>A), CAA423GAA (3821C>G), ATT424ATA (3826T>A), CAG425CAA (3829G>A), AAA426AGA (3831A>G), CTC428CTT (3838C>T), TAC433TTT (3852A>T 3853C>T), ATT434GCA (3854A>G 3855T>C 3856T>A), AAA437_GGA438del (3863_3868delAAAGGA), TTT439TTC (3871T>C), TTT440ACA (3872T>A 3873T>C 3874T>A), AAG441CCA (3875A>C 3876A>C 3877G>A), GAA446AAG (3890G>A 3892A>G), AAA448AAT (3898A>T), GTT449CTT (3899G>C), TTA450CTA (3902T>C), CAG451CAA (3907G>A), ATG453CTC (3911A>C 3913G>C), CTT454ATC (3914C>A 3916T>C), TCA455AGA (3917T>A 3918C>G), GAA456AAA (3920G>A), AAA457TCA (3923A>T 3924A>C), CTT458AAT (3926C>A 3927T>A), CCA459CGA (3930C>G), TGG460TTG (3933G>T), AAA461TGA (3935A>T 3936A>G), AAT463ACC (3942A>C 3943T>C), GAT464AAT (3944G>A), TTA465GAA (3947T>G 3948T>A), GCC466CAT (3950G>C 3951C>A 3952C>T), CTG468CGT (3957T>G 3958G>T), GCA469TGT (3959G>T 3960C>G 3961A>T), GTT470GTA (3964T>A), AAA471CAA (3965A>C), AGG472AAA (3969G>A 3970G>A), TTA473CTC (3971T>C 3973A>C), CAG475GAA (3977C>G 3979G>A), GTT476GAA (3981T>A 3982T>A), TGC477TGT (3985C>T), AAA478GTT (3986A>G 3987A>T 3988A>T), AAC479AAG (3991C>G), TTA480CTT (3992T>C 3994A>T), AGG482AAG (3999G>A), CTG483CTT (4003G>T), TAT484AGG (4004T>A 4005A>G 4006T>G), GTT485CTA (4007G>C 4009T>A), GCA486CCA (4010G>C), AAG487GAT (4013A>G 4015G>T) | | | | | | | | |
| **Proteins** | | | | | | | | | |
| reverse transcriptase (NP_042513.1) | 378 | 488 | 16.0% | 215 | 30.5% | 108 (97.3%) | 52 (46.8%) | 0/3/1/1 | 2 |
| Protein mutations: | E379K (3689G>A), K380R (3693A>G), A381L (3695G>C 3696C>T 3697A>G), I383* (3701A>T 3702T>A 3703C>A), A384S (3704G>T 3706T>A), F385K (3707T>A 3708T>A 3709T>A), E386N (3710G>A 3712A>T), L393M (3731T>A 3733A>G), K394I (3735A>T 3736G>T), L396S (3741T>C 3742G>A), F400del (3752_3754delTTT), P403L (3761C>T 3762C>T), K405T (3768A>C 3769A>T), E409K (3779G>A), A412L (3788G>T 3789C>T 3790T>A), Q417K (3803C>A 3805G>A), Q419T (3809C>A 3810A>C), D420T (3812G>A 3813A>C), R421K (3816G>A), K422Q (3818A>C 3820G>A), Q423E (3821C>G), K426R (3831A>G), Y433F (3852A>T 3853C>T), I434A (3854A>G 3855T>C 3856T>A), K437_G438del (3863_3868delAAAGGA), F440T (3872T>A 3873T>C 3874T>A), K441P (3875A>C 3876A>C 3877G>A), E446K (3890G>A 3892A>G), K448N (3898A>T), V449L (3899G>C), M453L (3911A>C 3913G>C), L454I (3914C>A 3916T>C), S455R (3917T>A 3918C>G), E456K (3920G>A), K457S (3923A>T 3924A>C), L458N (3926C>A 3927T>A), P459R (3930C>G), W460L (3933G>T), K461* (3935A>T 3936A>G), N463T (3942A>C 3943T>C), D464N (3944G>A), L465E (3947T>G 3948T>A), A466H (3950G>C 3951C>A 3952C>T), L468R (3957T>G 3958G>T), A469C (3959G>T 3960C>G 3961A>T), K471Q (3965A>C), R472K (3969G>A 3970G>A), Q475E (3977C>G 3979G>A), V476E (3981T>A 3982T>A), K478V (3986A>G 3987A>T 3988A>T), N479K (3991C>G), R482K (3999G>A), Y484R (4004T>A 4005A>G 4006T>G), V485L (4007G>C 4009T>A), A486P (4010G>C), K487D (4013A>G 4015G>T) | | | | | | | | |
| Codon mutations: | AGC377.CC (3684G>C), GAA379AAA (3689G>A), AAG380AGG (3693A>G), GCA381CTG (3695G>C 3696C>T 3697A>G), ATC383TAA (3701A>T 3702T>A 3703C>A), GCT384TCA (3704G>T 3706T>A), TTT385AAA (3707T>A 3708T>A 3709T>A), GAA386AAT (3710G>A 3712A>T), ATT388ATA (3718T>A), GAG389GAA (3721G>A), TTT390TTC (3724T>C), CTC391CTT (3727C>T), GGT392GGA (3730T>A), TTA393ATG (3731T>A 3733A>G), AAG394ATT (3735A>T 3736G>T), TTG396TCA (3741T>C 3742G>A), GGA399GGG (3751A>G), TTT400del (3752_3754delTTT), ATT401ATA (3757T>A), GAA402GAG (3760A>G), CCA403TTA (3761C>T 3762C>T), CAG404CAA (3766G>A), AAA405ACT (3768A>C 3769A>T), CTT407CTC (3775T>C), TTA408--C (3776_3777delTT 3778A>C), GAA409AAA (3779G>A), AAG410AAA (3784G>A), GCT412TTA (3788G>T 3789C>T 3790T>A), CCA415CCT (3799A>T), CAG417AAA (3803C>A 3805G>A), CTA418TTA (3806C>T), CAA419ACA (3809C>A 3810A>C), GAC420ACC (3812G>A 3813A>C), AGA421AAA (3816G>A), AAG422CAA (3818A>C 3820G>A), CAA423GAA (3821C>G), ATT424ATA (3826T>A), CAG425CAA (3829G>A), AAA426AGA (3831A>G), CTC428CTT (3838C>T), TAC433TTT (3852A>T 3853C>T), ATT434GCA (3854A>G 3855T>C 3856T>A), AAA437_GGA438del (3863_3868delAAAGGA), TTT439TTC (3871T>C), TTT440ACA (3872T>A 3873T>C 3874T>A), AAG441CCA (3875A>C 3876A>C 3877G>A), GAA446AAG (3890G>A 3892A>G), AAA448AAT (3898A>T), GTT449CTT (3899G>C), TTA450CTA (3902T>C), CAG451CAA (3907G>A), ATG453CTC (3911A>C 3913G>C), CTT454ATC (3914C>A 3916T>C), TCA455AGA (3917T>A 3918C>G), GAA456AAA (3920G>A), AAA457TCA (3923A>T 3924A>C), CTT458AAT (3926C>A 3927T>A), CCA459CGA (3930C>G), TGG460TTG (3933G>T), AAA461TGA (3935A>T 3936A>G), AAT463ACC (3942A>C 3943T>C), GAT464AAT (3944G>A), TTA465GAA (3947T>G 3948T>A), GCC466CAT (3950G>C 3951C>A 3952C>T), CTG468CGT (3957T>G 3958G>T), GCA469TGT (3959G>T 3960C>G 3961A>T), GTT470GTA (3964T>A), AAA471CAA (3965A>C), AGG472AAA (3969G>A 3970G>A), TTA473CTC (3971T>C 3973A>C), CAG475GAA (3977C>G 3979G>A), GTT476GAA (3981T>A 3982T>A), TGC477TGT (3985C>T), AAA478GTT (3986A>G 3987A>T 3988A>T), AAC479AAG (3991C>G), TTA480CTT (3992T>C 3994A>T), AGG482AAG (3999G>A), CTG483CTT (4003G>T), TAT484AGG (4004T>A 4005A>G 4006T>G), GTT485CTA (4007G>C 4009T>A), GCA486CCA (4010G>C), AAG487GAT (4013A>G 4015G>T) | | | | | | | | |

*: Inserts / Deletes / Misaligned / Frameshifts

#### Analysis details

This analysis was performed with panviral2.64

### NGS Details (UN24): Dioscovirus dioscoreae

#### Assembly

| | |
|---|---|
| **Coverage Length** | 513 (1 contig(s)) |
| **Depth Of Coverage** | 2.6 |
| **Number Of Reads** | 10 |
| **Reads Per Million** | 0.20 rpm (after QC) |
| **Ambiguities** | 0 |
| **Assembly Method** | de novo + reference guided assembly |
| **Consensus Caller** | Bcf Tools |

#### Coverage Map

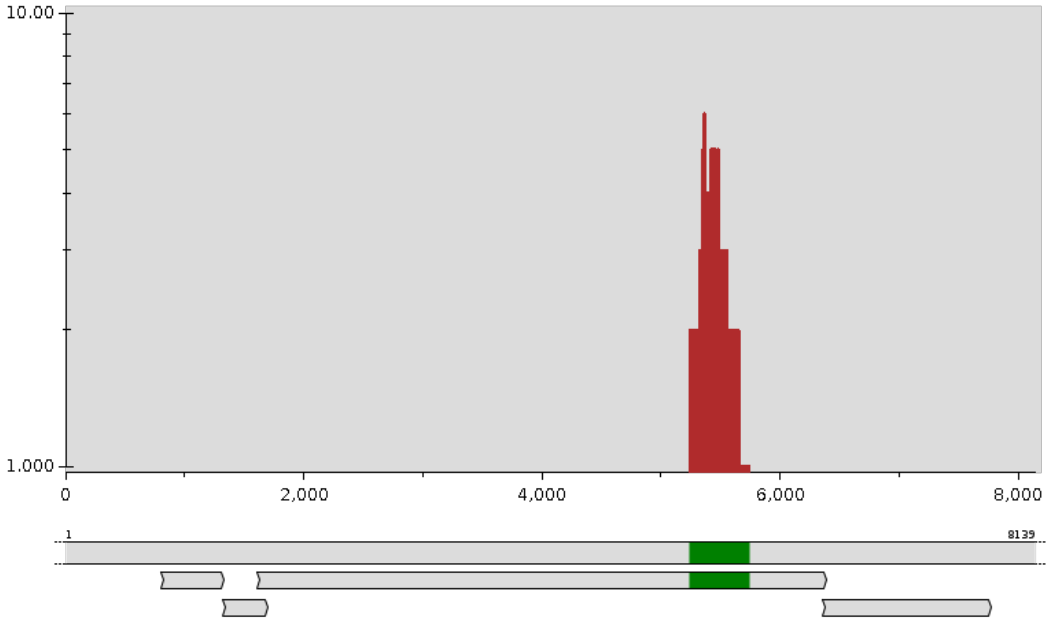

#### Assignment

| | |
|---|---|
| **Type** | Dioscovirus dioscoreae (Taxonomy ID: 3052184) |
| **Reference Genome** | NC_040712.1 |
| **NT Identity (%)** | 63.9376 |
| **AA Identity (%)** | 57.3099 |
| **Number Of Stop Codons** | 0 |
| **Number Of CDS** | 4 |

#### Alignment

| | |
|---|---|
| **Alignment Score** | 286.0 (NT) + 755.0 (AA) = 1041.0 |
| **Concordance (%)** | 46.7026 |

| Alignment Method | Global, seeded, nucleotide + amino acids (AGA) |
|---|---|

#### Genome Region

Sequence starts at position 5237 and ends at position 5749 relative to NC_040712.1 reference sequence.

#### Alignment Detailed Statistics

| | Begin | End | Coverage | Score | Concordance | Matches | Identities | I/D/M/F* | Stop Codons |
|---|---|---|---|---|---|---|---|---|---|
| **NT** | **5237** | **5749** | **6.3%** | **286** | **27.9%** | **513 (100%)** | **328 (63.9%)** | **0/0** | |
| Mutations: | 5237A>T, 5243T>A, 5247A>G, 5248G>C, 5249T>C, 5250C>A, 5252T>A, 5258A>C, 5259T>A, 5260A>T, 5261C>T, 5264T>A, 5270G>A, 5271C>A, 5274C>T, 5289T>G, 5291C>A, 5293A>T, 5294A>G, 5298C>G, 5299A>G, 5303C>T, 5305C>A, 5309C>A, 5313A>G, 5314G>A, 5315T>C, 5317T>A, 5318T>A, 5319G>A, 5320A>C, 5321C>A, 5322T>A, 5324T>A, 5325C>T, 5327C>A, 5328C>A, 5331C>A, 5332T>A, 5335A>G, 5339T>A, 5342A>T, 5344A>G, 5345C>A, 5348G>A, 5350T>G, 5351T>A, 5352G>A, 5354C>T, 5356A>T, 5357C>T, 5363A>G, 5370C>T, 5371T>G, 5372A>T, 5381A>C, 5383T>A, 5384C>T, 5385C>T, 5386A>G, 5387C>G, 5390G>A, 5391A>G, 5393T>A, 5395T>A, 5396G>A, 5400G>C, 5406C>G, 5408G>C, 5414A>T, 5417A>G, 5426A>C, 5430G>T, 5431T>A, 5432C>T, 5439C>G, 5441G>A, 5444T>A, 5450T>C, 5457A>T, 5461T>C, 5471C>T, 5475C>T, 5477C>A, 5482A>C, 5486A>T, 5489T>A, 5490T>C, 5493G>A, 5495A>T, 5498C>T, 5501A>G, 5502C>A, 5519A>T, 5522C>T, 5523A>G, 5524A>G, 5527A>G, 5532T>A, 5533C>A, 5535G>C, 5537A>T, 5541G>A, 5543C>T, 5545G>T, 5546T>A, 5552C>T, 5558C>T, 5561T>C, 5565T>C, 5567G>A, 5568A>G, 5570A>T, 5574T>A, 5575C>G, 5577G>A, 5579A>T, 5581G>C, 5582T>C, 5585A>G, 5589C>G, 5591A>G, 5595G>T, 5598C>G, 5599A>G, 5604T>C, 5606A>C, 5607T>C, 5608T>A, 5610C>A, 5613T>G, 5615C>T, 5618C>T, 5619C>A, 5621A>T, 5622G>C, 5626G>T, 5629A>T, 5630G>A, 5633A>C, 5634G>A, 5636A>T, 5639G>T, 5640C>T, 5642T>A, 5646T>A, 5648G>A, 5649T>A, 5650C>G, 5651A>T, 5656C>A, 5657A>G, 5660A>G, 5661T>A, 5663G>A, 5664A>G, 5666A>G, 5667A>T, 5670G>T, 5672A>C, 5673G>A, 5674T>A, 5676G>A, 5679A>T, 5684A>T, 5685G>A, 5687A>T, 5691C>T, 5693T>A, 5696C>A, 5697C>A, 5700G>A, 5701A>C, 5702A>C, 5718G>A, 5720T>A, 5721C>A, 5724C>T, 5726C>A, 5732C>T, 5739T>G, 5740T>C, 5741A>T, 5743A>G, 5747T>G | | | | | | | | |

##### CDS

| | Begin | End | Coverage | Score | Concordance | Matches | Identities | I/D/M/F* | Stop Codons |
|---|---|---|---|---|---|---|---|---|---|
| EXK67_gp3 | 1210 | 1380 | 10.7% | 755 | 62.2% | 171 (100%) | 98 (57.3%) | 0/0/0/0 | 0 |
| Protein mutations: | S1213A (5247A>G 5248G>C 5249T>C), Y1217I (5259T>A 5260A>T 5261C>T), N1218K (5264T>A), Y1227E (5289T>G 5291C>A), K1228M (5293A>T 5294A>G), Q1230G (5298C>G 5299A>G), T1232K (5305C>A), S1235D (5313A>G 5314G>A 5315T>C), I1236K (5317T>A 5318T>A), D1237T (5319G>A 5320A>C 5321C>A), Y1238K (5322T>A 5324T>A), L1240I (5328C>A), L1241N (5331C>A 5332T>A), K1242R (5335A>G), K1244N (5342A>T), D1245G (5344A>G 5345C>A), I1247R (5350T>G 5351T>A), V1248I (5352G>A 5354C>T), Y1249F (5356A>T 5357C>T), L1254C (5370C>T 5371T>G 5372A>T), F1258Y (5383T>A 5384C>T), H1259W (5385C>T 5386A>G 5387C>G), I1261V (5391A>G 5393T>A), M1262K (5395T>A 5396G>A), D1264H (5400G>C), Q1266D (5406C>G 5408G>C), V1274Y (5430G>T 5431T>A 5432C>T), Q1277E (5439C>G 5441G>A), I1283L (5457A>T), V1284A (5461T>C), N1291T (5482A>C), S1294P (5490T>C), V1295I (5493G>A 5495A>T), K1305G (5523A>G 5524A>G), K1306R (5527A>G), S1308K (5532T>A 5533C>A), E1309H (5535G>C 5537A>T), V1311I (5541G>A 5543C>T), C1312L (5545G>T 5546T>A), I1320V (5568A>G 5570A>T), E1323N (5577G>A 5579A>T), S1324T (5581G>C 5582T>C), I1325M (5585A>G), Q1327E (5589C>G 5591A>G), V1329L (5595G>T), Q1330G (5598C>G 5599A>G), L1333Q (5607T>C 5608T>A), Q1334K (5610C>A), F1335V (5613T>G 5615C>T), Q1337N (5619C>A 5621A>T), V1338L (5622G>C), C1339F (5626G>T), K1340I (5629A>T 5630G>A), E1341D (5633A>C), E1342N (5634G>A 5636A>T), L1346I (5646T>A 5648G>A), T1349K (5656C>A 5657A>G), L1351I (5661T>A 5663G>A), K1352E (5664A>G 5666A>G), I1353L (5667A>T), G1354C (5670G>T 5672A>C), V1355K (5673G>A 5674T>A), A1356T (5676G>A), N1357Y (5679A>T), E1359N (5685G>A 5687A>T), L1363I (5697C>A), E1364T (5700G>A 5701A>C 5702A>C), V1370I (5718G>A 5720T>A), Q1371K (5721C>A), L1377A (5739T>G 5740T>C 5741A>T), K1378R (5743A>G), N1379K (5747T>G) | | | | | | | | |
| Codon mutations: | GTA1209..T (5237A>T), GGT1211GGA (5243T>A), AGT1213GCC (5247A>G 5248G>C 5249T>C), CGT1214AGA (5250C>A 5252T>A), GTA1216GTC (5258A>C), TAC1217ATT (5259T>A 5260A>T 5261C>T), AAT1218AAA (5264T>A), AAG1220AAA (5270G>A), CGA1221AGA (5271C>A), CTA1222TTA (5274C>T), TAC1227GAA (5289T>G 5291C>A), AAA1228ATG (5293A>T 5294A>G), CAA1230GGA (5298C>G 5299A>G), TAC1231TAT (5303C>T), ACA1232AAA (5305C>A), CTC1233CTA (5309C>A), AGT1235GAC (5313A>G 5314G>A 5315T>C), ATT1236AAA (5317T>A 5318T>A), GAC1237ACA (5319G>A 5320A>C 5321C>A), TAT1238AAA (5322T>A 5324T>A), CTC1239TTA (5325C>T 5327C>A), CTA1240ATA (5328C>A), CTT1241AAT (5331C>A 5332T>A), AAA1242AGA (5335A>G), ATT1243ATA (5339T>A), AAA1244AAT (5342A>T), GAC1245GGA (5344A>G 5345C>A), AAG1246AAA (5348G>A), ATT1247AGA (5350T>G 5351T>A), GTC1248ATT (5352G>A 5354C>T), TAC1249TTT (5356A>T 5357C>T), AAA1251AAG (5363A>G), CTA1254TGT (5370C>T 5371T>G 5372A>T), GGA1257GGC (5381A>C), TTC1258TAT (5383T>A 5384C>T), CAC1259TGG (5385C>T 5386A>G 5387C>G), CAG1260CAA (5390G>A), ATT1261GTA (5391A>G 5393T>A), ATG1262AAA (5395T>A 5396G>A), GAT1264CAT (5400G>C), CAG1266GAC (5406C>G 5408G>C), ATA1268ATT (5414A>T), GAA1269GAG (5417A>G), GCA1272GCC (5426A>C), GTC1274TAT (5430G>T 5431T>A 5432C>T), CAG1277GAA (5439C>G 5441G>A), GGT1278GGA (5444T>A), TTT1280TTC (5450T>C), ATA1283TTA (5457A>T), GTA1284GCA (5461T>C), TTC1287TTT (5471C>T), CTC1289TTA (5475C>T 5477C>A), AAT1291ACT (5482A>C), GCA1292GCT (5486A>T), CCT1293CCA (5489T>A), TCA1294CCA (5490T>C), GTA1295ATT (5493G>A 5495A>T), TTC1296TTT (5498C>T), CAA1297CAG (5501A>G), CGA1298AGA (5502C>A), ATA1303ATT (5519A>T), TTC1304TTT (5522C>T), AAA1305GGA (5523A>G 5524A>G), AAA1306AGA (5527A>G), TCA1308AAA (5532T>A 5533C>A), GAA1309CAT (5535G>C 5537A>T), GTC1311ATT (5541G>A 5543C>T), TGT1312TTA (5545G>T 5546T>A), TAC1314TAT (5552C>T), GAC1316GAT (5558C>T), GAT1317GAC (5561T>C), TTG1319CTA (5565T>C 5567G>A), ATA1320GTT (5568A>G 5570A>T), TCT1322AGT (5574T>A 5575C>G), GAA1323AAT (5577G>A 5579A>T), AGT1324ACC (5581G>C 5582T>C), ATA1325ATG (5585A>G), CAA1327GAG (5589C>G 5591A>G), GTA1329TTA (5595G>T), CAA1330GGA (5598C>G 5599A>G), TTA1332CTC (5604T>C 5606A>C), TTA1333CAA (5607T>C 5608T>A), CAG1334AAG (5610C>A), TTC1335GTT (5613T>G 5615C>T), TTC1336TTT (5618C>T), CAA1337AAT (5619C>A 5621A>T), GTA1338CTA (5622G>C), TGT1339TTT (5626G>T), AAG1340ATA (5629A>T 5630G>A), GAA1341GAC (5633A>C), GAA1342AAT (5634G>A 5636A>T), GGG1343GGT (5639G>T), CTT1344TTA (5640C>T 5642T>A), TTG1346ATA (5646T>A 5648G>A), TCA1347AGT (5649T>A 5650C>G 5651A>T), ACA1349AAG (5656C>A 5657A>G), AAA1350AAG (5660A>G), TTG1351ATA (5661T>A 5663G>A), AAA1352GAG (5664A>G 5666A>G), ATA1353TTA (5667A>T), GGA1354TGC (5670G>T 5672A>C), GTA1355AAA (5673G>A 5674T>A), GCA1356ACA (5676G>A), AAT1357TAT (5679A>T), ATA1358ATT (5684A>T), GAA1359AAT (5685G>A 5687A>T), CTT1361TTA (5691C>T 5693T>A), GGC1362GGA (5696C>A), CTA1363ATA (5697C>A), GAA1364ACC (5700G>A 5701A>C 5702A>C), GTT1370ATA (5718G>A 5720T>A), CAA1371AAA (5721C>A), CTC1372TTA (5724C>T 5726C>A), CCC1374CCT (5732C>T), TTA1377GCT (5739T>G 5740T>C 5741A>T), AAA1378AGA (5743A>G), AAT1379AAG (5747T>G) | | | | | | | | |

##### Proteins

| | Begin | End | Coverage | Score | Concordance | Matches | Identities | I/D/M/F* | Stop Codons |
|---|---|---|---|---|---|---|---|---|---|
| ORF3 (YP_009553219.1) | 1210 | 1380 | 10.7% | 755 | 62.2% | 171 (100%) | 98 (57.3%) | 0/0/0/0 | 0 |
| Protein mutations: | S1213A (5247A>G 5248G>C 5249T>C), Y1217I (5259T>A 5260A>T 5261C>T), N1218K (5264T>A), Y1227E (5289T>G 5291C>A), K1228M (5293A>T 5294A>G), Q1230G (5298C>G 5299A>G), T1232K (5305C>A), S1235D (5313A>G 5314G>A 5315T>C), I1236K (5317T>A 5318T>A), D1237T (5319G>A 5320A>C 5321C>A), Y1238K (5322T>A 5324T>A), L1240I (5328C>A), L1241N (5331C>A 5332T>A), K1242R (5335A>G), K1244N (5342A>T), D1245G (5344A>G 5345C>A), I1247R (5350T>G 5351T>A), V1248I (5352G>A 5354C>T), Y1249F (5356A>T 5357C>T), L1254C (5370C>T 5371T>G 5372A>T), F1258Y (5383T>A 5384C>T), H1259W (5385C>T 5386A>G 5387C>G), I1261V (5391A>G 5393T>A), M1262K (5395T>A 5396G>A), D1264H (5400G>C), Q1266D (5406C>G 5408G>C), V1274Y (5430G>T 5431T>A 5432C>T), Q1277E (5439C>G 5441G>A), I1283L (5457A>T), V1284A (5461T>C), N1291T (5482A>C), S1294P (5490T>C), V1295I (5493G>A 5495A>T), K1305G (5523A>G 5524A>G), K1306R (5527A>G), S1308K (5532T>A 5533C>A), E1309H (5535G>C 5537A>T), V1311I (5541G>A 5543C>T), C1312L (5545G>T 5546T>A), I1320V (5568A>G 5570A>T), E1323N (5577G>A 5579A>T), S1324T (5581G>C 5582T>C), I1325M (5585A>G), Q1327E (5589C>G 5591A>G), V1329L (5595G>T), Q1330G (5598C>G 5599A>G), L1333Q (5607T>C 5608T>A), Q1334K (5610C>A), F1335V (5613T>G 5615C>T), Q1337N (5619C>A 5621A>T), V1338L (5622G>C), C1339F (5626G>T), K1340I (5629A>T 5630G>A), E1341D (5633A>C), E1342N (5634G>A 5636A>T), L1346I (5646T>A 5648G>A), T1349K (5656C>A 5657A>G), L1351I (5661T>A 5663G>A), K1352E (5664A>G 5666A>G), I1353L (5667A>T), G1354C (5670G>T 5672A>C), V1355K (5673G>A 5674T>A), A1356T (5676G>A), N1357Y (5679A>T), E1359N (5685G>A 5687A>T), L1363I (5697C>A), E1364T (5700G>A 5701A>C 5702A>C), V1370I (5718G>A 5720T>A), Q1371K (5721C>A), L1377A (5739T>G 5740T>C 5741A>T), K1378R (5743A>G), N1379K (5747T>G) | | | | | | | | |
| Codon mutations: | GTA1209..T (5237A>T), GGT1211GGA (5243T>A), AGT1213GCC (5247A>G 5248G>C 5249T>C), CGT1214AGA (5250C>A 5252T>A), GTA1216GTC (5258A>C), TAC1217ATT (5259T>A 5260A>T 5261C>T), AAT1218AAA (5264T>A), AAG1220AAA (5270G>A), CGA1221AGA (5271C>A), CTA1222TTA (5274C>T), TAC1227GAA (5289T>G 5291C>A), AAA1228ATG (5293A>T 5294A>G), CAA1230GGA (5298C>G 5299A>G), TAC1231TAT (5303C>T), ACA1232AAA (5305C>A), CTC1233CTA (5309C>A), AGT1235GAC (5313A>G 5314G>A 5315T>C), ATT1236AAA (5317T>A 5318T>A), GAC1237ACA (5319G>A 5320A>C 5321C>A), TAT1238AAA (5322T>A 5324T>A), CTC1239TTA (5325C>T 5327C>A), CTA1240ATA (5328C>A), CTT1241AAT (5331C>A 5332T>A), AAA1242AGA (5335A>G), ATT1243ATA (5339T>A), AAA1244AAT (5342A>T), GAC1245GGA (5344A>G 5345C>A), AAG1246AAA (5348G>A), ATT1247AGA (5350T>G 5351T>A), GTC1248ATT (5352G>A 5354C>T), TAC1249TTT (5356A>T 5357C>T), AAA1251AAG (5363A>G), CTA1254TGT (5370C>T 5371T>G 5372A>T), GGA1257GGC (5381A>C), TTC1258TAT (5383T>A 5384C>T), CAC1259TGG (5385C>T 5386A>G 5387C>G), CAG1260CAA (5390G>A), ATT1261GTA (5391A>G 5393T>A), ATG1262AAA (5395T>A 5396G>A), GAT1264CAT (5400G>C), CAG1266GAC (5406C>G 5408G>C), ATA1268ATT (5414A>T), GAA1269GAG (5417A>G), GCA1272GCC (5426A>C), GTC1274TAT (5430G>T 5431T>A 5432C>T), CAG1277GAA (5439C>G 5441G>A), GGT1278GGA (5444T>A), TTT1280TTC (5450T>C), ATA1283TTA (5457A>T), GTA1284GCA (5461T>C), TTC1287TTT (5471C>T), CTC1289TTA (5475C>T 5477C>A), AAT1291ACT (5482A>C), GCA1292GCT (5486A>T), CCT1293CCA (5489T>A), TCA1294CCA (5490T>C), GTA1295ATT (5493G>A 5495A>T), TTC1296TTT (5498C>T), CAA1297CAG (5501A>G), CGA1298AGA (5502C>A), ATA1303ATT (5519A>T), TTC1304TTT (5522C>T), AAA1305GGA (5523A>G 5524A>G), AAA1306AGA (5527A>G), TCA1308AAA (5532T>A 5533C>A), GAA1309CAT (5535G>C 5537A>T), GTC1311ATT (5541G>A 5543C>T), TGT1312TTA (5545G>T 5546T>A), TAC1314TAT (5552C>T), GAC1316GAT (5558C>T), GAT1317GAC (5561T>C), TTG1319CTA (5565T>C 5567G>A), ATA1320GTT (5568A>G 5570A>T), TCT1322AGT (5574T>A 5575C>G), GAA1323AAT (5577G>A 5579A>T), AGT1324ACC (5581G>C 5582T>C), ATA1325ATG (5585A>G), CAA1327GAG (5589C>G 5591A>G), GTA1329TTA (5595G>T), CAA1330GGA (5598C>G 5599A>G), TTA1332CTC (5604T>C 5606A>C), TTA1333CAA (5607T>C 5608T>A), CAG1334AAG (5610C>A), TTC1335GTT (5613T>G 5615C>T), TTC1336TTT (5618C>T), CAA1337AAT (5619C>A 5621A>T), GTA1338CTA (5622G>C), TGT1339TTT (5626G>T), AAG1340ATA (5629A>T 5630G>A), GAA1341GAC (5633A>C), GAA1342AAT (5634G>A 5636A>T), GGG1343GGT (5639G>T), CTT1344TTA (5640C>T 5642T>A), TTG1346ATA (5646T>A 5648G>A), TCA1347AGT (5649T>A 5650C>G 5651A>T), ACA1349AAG (5656C>A 5657A>G), AAA1350AAG (5660A>G), TTG1351ATA (5661T>A 5663G>A), AAA1352GAG (5664A>G 5666A>G), ATA1353TTA (5667A>T), GGA1354TGC (5670G>T 5672A>C), GTA1355AAA (5673G>A 5674T>A), GCA1356ACA (5676G>A), AAT1357TAT (5679A>T), ATA1358ATT (5684A>T), GAA1359AAT (5685G>A 5687A>T), CTT1361TTA (5691C>T 5693T>A), GGC1362GGA (5696C>A), CTA1363ATA (5697C>A), GAA1364ACC (5700G>A 5701A>C 5702A>C), GTT1370ATA (5718G>A 5720T>A), CAA1371AAA (5721C>A), CTC1372TTA (5724C>T 5726C>A), CCC1374CCT (5732C>T), TTA1377GCT (5739T>G 5740T>C 5741A>T), AAA1378AGA (5743A>G), AAT1379AAG (5747T>G) | | | | | | | | |

*: Inserts / Deletes / Misaligned / Frameshifts

#### Analysis details

This analysis was performed with panviral2.64

### NGS Details (UN24): Caulimovirus minutangelicae

#### Assembly

| | |
|---|---|
| **Coverage Length** | 270 (1 contig(s)) |
| **Depth Of Coverage** | 3.2 |
| **Number Of Reads** | 9 |
| **Reads Per Million** | 0.18 rpm (after QC) |
| **Ambiguities** | 0 |
| **Assembly Method** | de novo + reference guided assembly |
| **Consensus Caller** | Bcf Tools |

#### Coverage Map

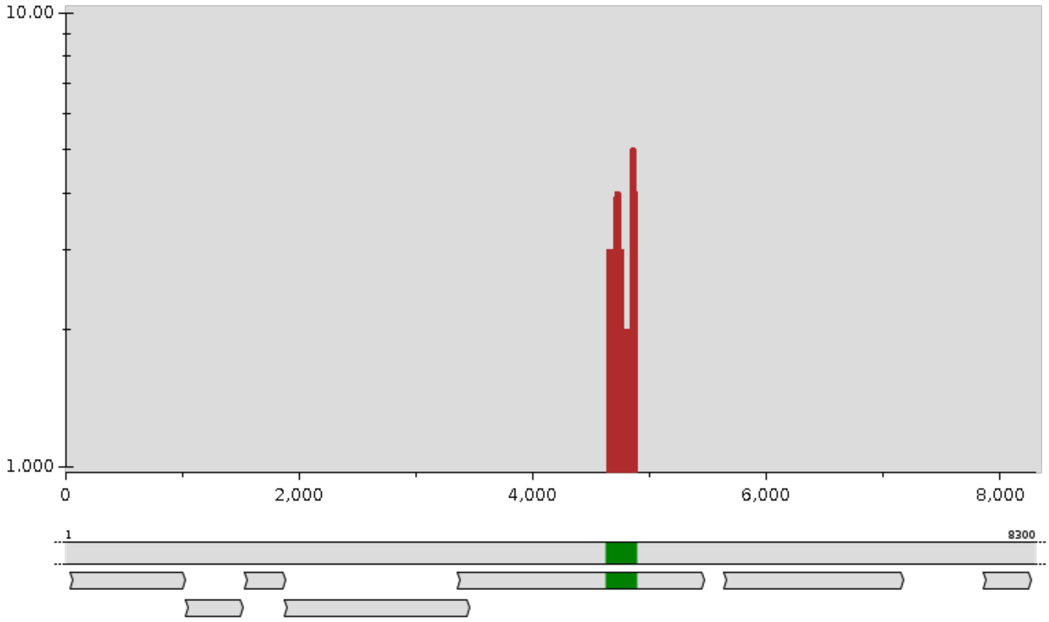

#### Assignment

| | |
|---|---|
| **Type** | Caulimovirus minutangelicae (Taxonomy ID: 3047363) |
| **Reference Genome** | NC_043523.1 |
| **NT Identity (%)** | 56.6667 |
| **AA Identity (%)** | 50.0 |
| **Number Of Stop Codons** | 0 |
| **Number Of CDS** | 7 |

#### Alignment

| | |
|---|---|
| **Alignment Score** | 72.0 (NT) + 314.0 (AA) = 386.0 |
| **Concordance (%)** | 32.7674 |

| Alignment Method | Global, seeded, nucleotide + amino acids (AGA) |
|---|---|

#### Genome Region

Sequence starts at position 4631 and ends at position 4900 relative to NC_043523.1 reference sequence.

#### Alignment Detailed Statistics

| | Begin | End | Coverage | Score | Concordance | Matches | Identities | I/D/M/F* | Stop Codons |
|---|---|---|---|---|---|---|---|---|---|
| **NT** | **4631** | **4900** | **3.3%** | **72** | **13.3%** | **270 (100%)** | **153 (56.7%)** | **0/0** | |
| Mutations: | 4631A>C, 4633A>T, 4638G>C, 4640T>A, 4641G>T, 4645T>C, 4648T>C, 4649G>A, 4651A>T, 4657T>C, 4658A>G, 4661C>T, 4666C>A, 4675T>A, 4676A>G, 4679G>A, 4680A>T, 4681A>T, 4684A>T, 4685G>T, 4686A>C, 4690C>T, 4691C>T, 4692A>T, 4693G>C, 4696A>G, 4702A>C, 4703G>A, 4704C>A, 4705A>T, 4708A>T, 4709G>T, 4711T>C, 4714A>C, 4717G>A, 4719A>C, 4720A>T, 4721T>A, 4722G>T, 4723C>T, 4724G>A, 4727C>A, 4728A>T, 4729A>C, 4730C>A, 4731T>A, 4732G>C, 4735A>C, 4736A>C, 4739A>G, 4741T>C, 4742C>A, 4745T>A, 4746C>G, 4747A>T, 4748A>G, 4749A>C, 4757G>A, 4758C>T, 4759T>A, 4760C>A, 4763T>C, 4765A>T, 4768C>T, 4769A>C, 4774A>T, 4775A>C, 4777A>G, 4780T>A, 4782A>G, 4783T>A, 4787T>C, 4789A>T, 4792T>A, 4794T>A, 4795A>T, 4798A>T, 4799A>T, 4801T>A, 4802G>T, 4810A>G, 4814C>T, 4817C>A, 4818G>A, 4819T>G, 4823C>A, 4824A>T, 4825A>T, 4826A>T, 4827A>G, 4829C>A, 4830A>G, 4831T>A, 4833T>C, 4834C>T, 4835C>A, 4837A>T, 4841C>T, 4842A>T, 4844A>G, 4845T>C, 4846T>A, 4848C>A, 4849A>T, 4852G>A, 4855C>T, 4858G>T, 4863A>C, 4867T>C, 4868G>A, 4869A>C, 4870A>T, 4873C>T, 4878A>C, 4894C>T, 4895T>C, 4897A>T | | | | | | | | |

##### CDS

| | Begin | End | Coverage | Score | Concordance | Matches | Identities | I/D/M/F* | Stop Codons |
|---|---|---|---|---|---|---|---|---|---|
| FLA58_gp5 | 424 | 513 | 12.8% | 314 | 49.2% | 90 (100%) | 45 (50.0%) | 0/0/0/0 | 0 |
| Protein mutations: | K424H (4631A>C 4633A>T), C426S (4638G>C), C427I (4640T>A 4641G>T), V430I (4649G>A 4651A>T), I433V (4658A>G), D438E (4675T>A), N439D (4676A>G), E440I (4679G>A 4680A>T 4681A>T), E441D (4684A>T), D442S (4685G>T 4686A>C), Q444F (4691C>T 4692A>T 4693G>C), A448N (4703G>A 4704C>A 4705A>T), V450F (4709G>T 4711T>C), L451F (4714A>C), K453T (4719A>C 4720A>T), C454I (4721T>A 4722G>T 4723C>T), E455K (4724G>A), Q456I (4727C>A 4728A>T 4729A>C), L457N (4730C>A 4731T>A 4732G>C), I459L (4736A>C), I460V (4739A>G 4741T>C), L461I (4742C>A), K463A (4748A>G 4749A>C), A466I (4757G>A 4758C>T 4759T>A), Q467K (4760C>A), K470Q (4769A>C), K472Q (4775A>C 4777A>G), N474R (4782A>G 4783T>A), L478H (4794T>A 4795A>T), E479D (4798A>T), I480L (4799A>T 4801T>A), D481Y (4802G>T), H485Y (4814C>T), R486K (4817C>A 4818G>A 4819T>G), Q488I (4823C>A 4824A>T 4825A>T), N489C (4826A>T 4827A>G), H490R (4829C>A 4830A>G 4831T>A), I491T (4833T>C 4834C>T), L492I (4835C>A 4837A>T), H494F (4841C>T 4842A>T), I495A (4844A>G 4845T>C 4846T>A), A496D (4848C>A 4849A>T), K501T (4863A>C), E503T (4868G>A 4869A>C 4870A>T), K506T (4878A>C) | | | | | | | | |
| Codon mutations: | AAA424CAT (4631A>C 4633A>T), TGT426TCT (4638G>C), TGT427ATT (4640T>A 4641G>T), GTT428GTC (4645T>C), TAT429TAC (4648T>C), GTA430ATT (4649G>A 4651A>T), GAT432GAC (4657T>C), ATT433GTT (4658A>G), CTA434TTA (4661C>T), ATC435ATA (4666C>A), GAT438GAA (4675T>A), AAT439GAT (4676A>G), GAA440ATT (4679G>A 4680A>T 4681A>T), GAA441GAT (4684A>T), GAT442TCT (4685G>T 4686A>C), CAC443CAT (4690C>T), CAG444TTC (4691C>T 4692A>T 4693G>C), AAA445AAG (4696A>G), CTA447CTC (4702A>C), GCA448AAT (4703G>A 4704C>A 4705A>T), GCA449GCT (4708A>T), GTT450TTC (4709G>T 4711T>C), TTA451TTC (4714A>C), AAG452AAA (4717G>A), AAA453ACT (4719A>C 4720A>T), TGC454ATT (4721T>A 4722G>T 4723C>T), GAA455AAA (4724G>A), CAA456ATC (4727C>A 4728A>T 4729A>C), CTG457AAC (4730C>A 4731T>A 4732G>C), GGA458GGC (4735A>C), ATT459CTT (4736A>C), ATT460GTC (4739A>G 4741T>C), CTT461ATT (4742C>A), TCA462AGT (4745T>A 4746C>G 4747A>T), AAA463GCA (4748A>G 4749A>C), GCT466ATA (4757G>A 4758C>T 4759T>A), CAG467AAG (4760C>A), TTA468CTT (4763T>C 4765A>T), TTC469TTT (4768C>T), AAA470CAA (4769A>C), ACA471ACT (4774A>T), AAA472CAG (4775A>C 4777A>G), ATT473ATA (4780T>A), AAT474AGA (4782A>G 4783T>A), TTA476CTT (4787T>C 4789A>T), GGT477GGA (4792T>A), CTA478CAT (4794T>A 4795A>T), GAA479GAT (4798A>T), ATT480TTA (4799A>T 4801T>A), GAT481TAT (4802G>T), GGA483GGG (4810A>G), CAT485TAT (4814C>T), CGT486AAG (4817C>A 4818G>A 4819T>G), CAA488ATT (4823C>A 4824A>T 4825A>T), AAC489TGC (4826A>T 4827A>G), CAT490AGA (4829C>A 4830A>G 4831T>A), ATC491ACT (4833T>C 4834C>T), CTA492ATT (4835C>A 4837A>T), CAT494TTT (4841C>T 4842A>T), ATT495GCA (4844A>G 4845T>C 4846T>A), GCA496GAT (4848C>A 4849A>T), AAG497AAA (4852G>A), TTC498TTT (4855C>T), CCG499CCT (4858G>T), AAA501ACA (4863A>C), ATT502ATC (4867T>C), GAA503ACT (4868G>A 4869A>C 4870A>T), GAC504GAT (4873C>T), AAA506ACA (4878A>C), TTC511TTT (4894C>T), TTA512CTT (4895T>C 4897A>T) | | | | | | | | |

##### Proteins

| | Begin | End | Coverage | Score | Concordance | Matches | Identities | I/D/M/F* | Stop Codons |
|---|---|---|---|---|---|---|---|---|---|
| reverse transcriptase (YP_009666503.1) | 424 | 513 | 12.8% | 314 | 49.2% | 90 (100%) | 45 (50.0%) | 0/0/0/0 | 0 |
| Protein mutations: | K424H (4631A>C 4633A>T), C426S (4638G>C), C427I (4640T>A 4641G>T), V430I (4649G>A 4651A>T), I433V (4658A>G), D438E (4675T>A), N439D (4676A>G), E440I (4679G>A 4680A>T 4681A>T), E441D (4684A>T), D442S (4685G>T 4686A>C), Q444F (4691C>T 4692A>T 4693G>C), A448N (4703G>A 4704C>A 4705A>T), V450F (4709G>T 4711T>C), L451F (4714A>C), K453T (4719A>C 4720A>T), C454I (4721T>A 4722G>T 4723C>T), E455K (4724G>A), Q456I (4727C>A 4728A>T 4729A>C), L457N (4730C>A 4731T>A 4732G>C), I459L (4736A>C), I460V (4739A>G 4741T>C), L461I (4742C>A), K463A (4748A>G 4749A>C), A466I (4757G>A 4758C>T 4759T>A), Q467K (4760C>A), K470Q (4769A>C), K472Q (4775A>C 4777A>G), N474R (4782A>G 4783T>A), L478H (4794T>A 4795A>T), E479D (4798A>T), I480L (4799A>T 4801T>A), D481Y (4802G>T), H485Y (4814C>T), R486K (4817C>A 4818G>A 4819T>G), Q488I (4823C>A 4824A>T 4825A>T), N489C (4826A>T 4827A>G), H490R (4829C>A 4830A>G 4831T>A), I491T (4833T>C 4834C>T), L492I (4835C>A 4837A>T), H494F (4841C>T 4842A>T), I495A (4844A>G 4845T>C 4846T>A), A496D (4848C>A 4849A>T), K501T (4863A>C), E503T (4868G>A 4869A>C 4870A>T), K506T (4878A>C) | | | | | | | | |
| Codon mutations: | AAA424CAT (4631A>C 4633A>T), TGT426TCT (4638G>C), TGT427ATT (4640T>A 4641G>T), GTT428GTC (4645T>C), TAT429TAC (4648T>C), GTA430ATT (4649G>A 4651A>T), GAT432GAC (4657T>C), ATT433GTT (4658A>G), CTA434TTA (4661C>T), ATC435ATA (4666C>A), GAT438GAA (4675T>A), AAT439GAT (4676A>G), GAA440ATT (4679G>A 4680A>T 4681A>T), GAA441GAT (4684A>T), GAT442TCT (4685G>T 4686A>C), CAC443CAT (4690C>T), CAG444TTC (4691C>T 4692A>T 4693G>C), AAA445AAG (4696A>G), CTA447CTC (4702A>C), GCA448AAT (4703G>A 4704C>A 4705A>T), GCA449GCT (4708A>T), GTT450TTC (4709G>T 4711T>C), TTA451TTC (4714A>C), AAG452AAA (4717G>A), AAA453ACT (4719A>C 4720A>T), TGC454ATT (4721T>A 4722G>T 4723C>T), GAA455AAA (4724G>A), CAA456ATC (4727C>A 4728A>T 4729A>C), CTG457AAC (4730C>A 4731T>A 4732G>C), GGA458GGC (4735A>C), ATT459CTT (4736A>C), ATT460GTC (4739A>G 4741T>C), CTT461ATT (4742C>A), TCA462AGT (4745T>A 4746C>G 4747A>T), AAA463GCA (4748A>G 4749A>C), GCT466ATA (4757G>A 4758C>T 4759T>A), CAG467AAG (4760C>A), TTA468CTT (4763T>C 4765A>T), TTC469TTT (4768C>T), AAA470CAA (4769A>C), ACA471ACT (4774A>T), AAA472CAG (4775A>C 4777A>G), ATT473ATA (4780T>A), AAT474AGA (4782A>G 4783T>A), TTA476CTT (4787T>C 4789A>T), GGT477GGA (4792T>A), CTA478CAT (4794T>A 4795A>T), GAA479GAT (4798A>T), ATT480TTA (4799A>T 4801T>A), GAT481TAT (4802G>T), GGA483GGG (4810A>G), CAT485TAT (4814C>T), CGT486AAG (4817C>A 4818G>A 4819T>G), CAA488ATT (4823C>A 4824A>T 4825A>T), AAC489TGC (4826A>T 4827A>G), CAT490AGA (4829C>A 4830A>G 4831T>A), ATC491ACT (4833T>C 4834C>T), CTA492ATT (4835C>A 4837A>T), CAT494TTT (4841C>T 4842A>T), ATT495GCA (4844A>G 4845T>C 4846T>A), GCA496GAT (4848C>A 4849A>T), AAG497AAA (4852G>A), TTC498TTT (4855C>T), CCG499CCT (4858G>T), AAA501ACA (4863A>C), ATT502ATC (4867T>C), GAA503ACT (4868G>A 4869A>C 4870A>T), GAC504GAT (4873C>T), AAA506ACA (4878A>C), TTC511TTT (4894C>T), TTA512CTT (4895T>C 4897A>T) | | | | | | | | |

*: Inserts / Deletes / Misaligned / Frameshifts

#### Analysis details

This analysis was performed with panviral2.64

### NGS Details (UN24): Badnavirus maculaucubae

#### Assembly

| | |
|---|---|
| **Coverage Length** | 279 (1 contig(s)) |
| **Depth Of Coverage** | 3.6 |
| **Number Of Reads** | 9 |
| **Reads Per Million** | 0.18 rpm (after QC) |
| **Ambiguities** | 0 |
| **Assembly Method** | de novo + reference guided assembly |
| **Consensus Caller** | Bcf Tools |

#### Coverage Map

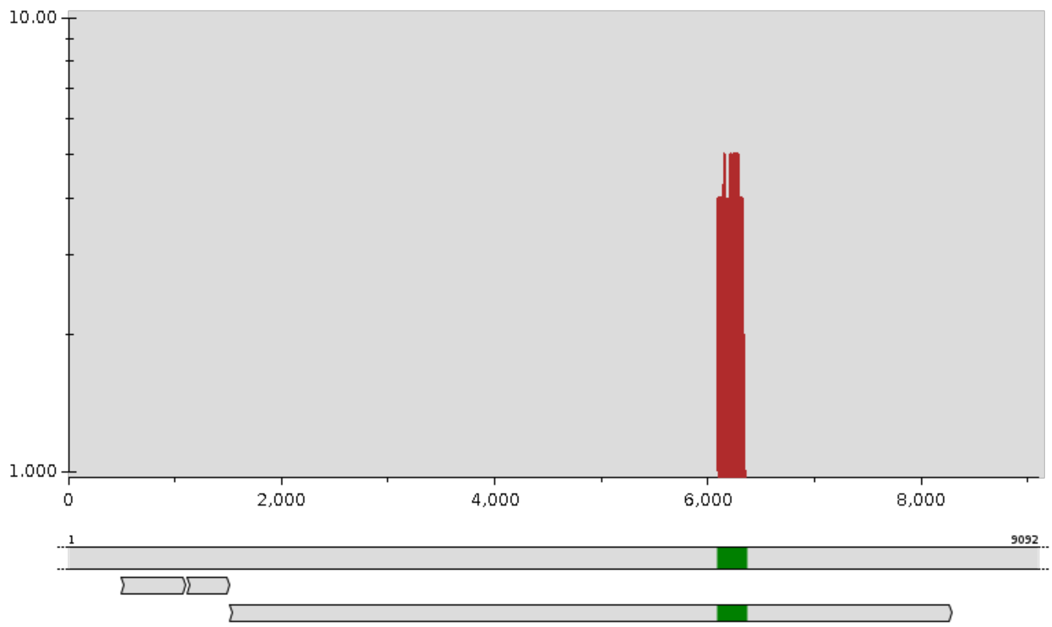

#### Assignment

| | |
|---|---|
| **Type** | Badnavirus maculaucubae (Taxonomy ID: 3051986) |
| **Reference Genome** | NC_076606.1 |
| **NT Identity (%)** | 56.7376 |
| **AA Identity (%)** | 45.7447 |
| **Number Of Stop Codons** | 1 |
| **Number Of CDS** | 3 |

#### Alignment

| | |
|---|---|
| **Alignment Score** | 70.0 (NT) + 300.0 (AA) = 370.0 |
| **Concordance (%)** | 30.3777 |

| Alignment Method | Global, seeded, nucleotide + amino acids (AGA) |
|---|---|

#### Genome Region

Sequence starts at position 6090 and ends at position 6368 relative to NC_076606.1 reference sequence.

#### Alignment Detailed Statistics

| | Begin | End | Coverage | Score | Concordance | Matches | Identities | I/D/M/F* | Stop Codons |
|---|---|---|---|---|---|---|---|---|---|
| **NT** | **6090** | **6368** | **3.1%** | **70** | **12.5%** | **279 (98.9%)** | **160 (56.7%)** | **3/0** | |
| Mutations: | 6093A>T, 6100A>T, 6101G>C, 6106T>A, 6116A>G, 6119C>T, 6120T>C, 6125T>A, 6131A>G, 6133G>C, 6135G>A, 6136G>A, 6137C>A, 6141G>A, 6142G>C, 6143A>G, 6144A>G, 6146A>C, 6148T>G, 6151A>G, 6152G>A, 6153C>T, 6156T>G, 6159A>T, 6160T>A, 6161G>A, 6162G>A, 6164C>T, 6165G>T, 6171C>T, 6172A>T, 6173T>G, 6177T>A, 6178C>A, 6179C>G, 6180A>G, 6181G>A, 6185G>A, 6186A>G, 6188T>A, 6189A>C, 6192C>T, 6195A>G, 6197G>T, 6198G>C, 6199C>T, 6201A>G, 6208C>T, 6210A>T, 6216T>A, 6219A>T, 6221A>C, 6222G>A, 6228A>C, 6234A>T, 6235G>A, 6236T>C, 6237G>T, 6241C>A, 6242A>T, 6243A>G, 6246G>C, 6247A>C, 6248A>T, 6249A>T, 6253G>A, 6255C>A, 6256A>G, 6257A>G, 6258C>A, 6259T>A, 6260G>T, 6261T>C, 6264C>T, 6266G>A, 6267_6268insCCA, 6269G>A, 6271A>T, 6272C>T, 6273A>G, 6274G>A, 6276A>T, 6277G>C, 6278A>T, 6279T>C, 6282C>T, 6285C>T, 6286G>A, 6287C>T, 6291T>G, 6293A>T, 6294C>T, 6306T>A, 6307C>G, 6310G>A, 6312A>C, 6314T>A, 6318T>A, 6319G>A, 6321A>G, 6323A>G, 6324T>C, 6325C>A, 6326T>G, 6327T>A, 6328C>A, 6333G>A, 6339G>A, 6345T>C, 6346C>T, 6348C>G, 6350A>G, 6354A>T, 6355A>G, 6357G>T, 6360G>A, 6362A>T, 6363A>T, 6364A>C, 6367T>G | | | | | | | | |
| **CDS** | | | | | | | | | |
| QKP79_gp3 | 1528 | 1620 | 4.1% | 300 | 43.9% | 93 (98.9%) | 43 (45.7%) | 1/0/0/0 | 1 |
| Protein mutations: | K1528N (6093A>T), F1533I (6106T>A), K1536R (6116A>G), S1537F (6119C>T 6120T>C), F1539Y (6125T>A), Q1541R (6131A>G), V1542L (6133G>C 6135G>A), A1543K (6136G>A 6137C>A), M1544I (6141G>A), E1545R (6142G>C 6143A>G 6144A>G), E1546A (6146A>C), S1547A (6148T>G), S1548D (6151A>G 6152G>A 6153C>T), I1549M (6156T>G), W1551K (6160T>A 6161G>A 6162G>A), T1552I (6164C>T 6165G>T), I1555* (6172A>T 6173T>G), P1557R (6178C>A 6179C>G 6180A>G), D1558N (6181G>A), G1559E (6185G>A 6186A>G), L1560H (6188T>A 6189A>C), W1563F (6197G>T 6198G>C), P1567S (6208C>T 6210A>T), K1571T (6221A>C 6222G>A), V1576T (6235G>A 6236T>C 6237G>T), Q1578M (6241C>A 6242A>T 6243A>G), R1579S (6246G>C), K1580L (6247A>C 6248A>T 6249A>T), D1582K (6253G>A 6255C>A), N1583G (6256A>G 6257A>G 6258C>A), C1584I (6259T>A 6260G>T 6261T>C), R1586K (6266G>A), R1586_G1587insP (6267_6268insCCA), G1587D (6269G>A), T1588L (6271A>T 6272C>T 6273A>G), E1589N (6274G>A 6276A>T), D1590L (6277G>C 6278A>T 6279T>C), A1593I (6286G>A 6287C>T), Y1595F (6293A>T 6294C>T), L1600V (6307C>G), V1601I (6310G>A 6312A>C), F1602Y (6314T>A), E1604K (6319G>A 6321A>G), N1605S (6323A>G 6324T>C), L1606R (6325C>A 6326T>G 6327T>A), Q1607K (6328C>A), K1614R (6350A>G), M1616V (6355A>G 6357G>T), E1618V (6362A>T 6363A>T), I1619L (6364A>C) | | | | | | | | |
| Codon mutations: | AAA1528AAT (6093A>T), AGC1531TCC (6100A>T 6101G>C), TTT1533ATT (6106T>A), AAA1536AGA (6116A>G), TCT1537TTC (6119C>T 6120T>C), TTT1539TAT (6125T>A), CAG1541CGG (6131A>G), GTG1542CTA (6133G>C 6135G>A), GCA1543AAA (6136G>A 6137C>A), ATG1544ATA (6141G>A), GAA1545CGG (6142G>C 6143A>G 6144A>G), GAA1546GCA (6146A>C), TCA1547GCA (6148T>G), AGC1548GAT (6151A>G 6152G>A 6153C>T), ATT1549ATG (6156T>G), CCA1550CCT (6159A>T), TGG1551AAA (6160T>A 6161G>A 6162G>A), ACG1552ATT (6164C>T 6165G>T), TTC1554TTT (6171C>T), ATA1555TGA (6172A>T 6173T>G), ACT1556ACA (6177T>A), CCA1557AGG (6178C>A 6179C>G 6180A>G), GAT1558AAT (6181G>A), GGA1559GAG (6185G>A 6186A>G), CTA1560CAC (6188T>A 6189A>C), TAC1561TAT (6192C>T), GAA1562GAG (6195A>G), TGG1563TTC (6197G>T 6198G>C), CTA1564TTG (6199C>T 6201A>G), CCA1567TCT (6208C>T 6210A>T), GGT1569GGA (6216T>A), CTA1570CTT (6219A>T), AAG1571ACA (6221A>C 6222G>A), GCA1573GCC (6228A>C), GCA1575GCT (6234A>T), GTG1576ACT (6235G>A 6236T>C 6237G>T), CAA1578ATG (6241C>A 6242A>T 6243A>G), AGG1579AGC (6246G>C), AAA1580CTT (6247A>C 6248A>T 6249A>T), GAC1582AAA (6253G>A 6255C>A), AAC1583GGA (6256A>G 6257A>G 6258C>A), TGT1584ATC (6259T>A 6260G>T 6261T>C), TTC1585TTT (6264C>T), AGG1586AAG (6266G>A), AGG1586_GGT1587insCCA (6267_6268insCCA), GGT1587GAT (6269G>A), ACA1588TTG (6271A>T 6272C>T 6273A>G), GAA1589AAT (6274G>A 6276A>T), GAT1590CTC (6277G>C 6278A>T 6279T>C), TTC1591TTT (6282C>T), ATC1592ATT (6285C>T), GCT1593ATT (6286G>A 6287C>T), GTT1594GTG (6291T>G), TAC1595TTT (6293A>T 6294C>T), ATT1599ATA (6306T>A), CTG1600GTG (6307C>G), GTA1601ATC (6310G>A 6312A>C), TTC1602TAC (6314T>A), TCT1603TCA (6318T>A), GAA1604AAG (6319G>A 6321A>G), AAT1605AGC (6323A>G 6324T>C), CTT1606AGA (6325C>A 6326T>G 6327T>A), CAA1607AAA (6328C>A), GAG1608GAA (6333G>A), GAG1610GAA (6339G>A), CAT1612CAC (6345T>C), CTC1613TTG (6346C>T 6348C>G), AAA1614AGA (6350A>G), ATA1615ATT (6354A>T), ATG1616GTT (6355A>G 6357G>T), CTG1617CTA (6360G>A), GAA1618GTT (6362A>T 6363A>T), ATA1619CTA (6364A>C), TGC1620GG. (6367T>G) | | | | | | | | |
| **Proteins** | | | | | | | | | |
| hypothetical protein (YP_010799265.1) | 1528 | 1620 | 4.1% | 300 | 43.9% | 93 (98.9%) | 43 (45.7%) | 1/0/0/0 | 1 |
| Protein mutations: | K1528N (6093A>T), F1533I (6106T>A), K1536R (6116A>G), S1537F (6119C>T 6120T>C), F1539Y (6125T>A), Q1541R (6131A>G), V1542L (6133G>C 6135G>A), A1543K (6136G>A 6137C>A), M1544I (6141G>A), E1545R (6142G>C 6143A>G 6144A>G), E1546A (6146A>C), S1547A (6148T>G), S1548D (6151A>G 6152G>A 6153C>T), I1549M (6156T>G), W1551K (6160T>A 6161G>A 6162G>A), T1552I (6164C>T 6165G>T), I1555* (6172A>T 6173T>G), P1557R (6178C>A 6179C>G 6180A>G), D1558N (6181G>A), G1559E (6185G>A 6186A>G), L1560H (6188T>A 6189A>C), W1563F (6197G>T 6198G>C), P1567S (6208C>T 6210A>T), K1571T (6221A>C 6222G>A), V1576T (6235G>A 6236T>C 6237G>T), Q1578M (6241C>A 6242A>T 6243A>G), R1579S (6246G>C), K1580L (6247A>C 6248A>T 6249A>T), D1582K (6253G>A 6255C>A), N1583G (6256A>G 6257A>G 6258C>A), C1584I (6259T>A 6260G>T 6261T>C), R1586K (6266G>A), R1586_G1587insP (6267_6268insCCA), G1587D (6269G>A), T1588L (6271A>T 6272C>T 6273A>G), E1589N (6274G>A 6276A>T), D1590L (6277G>C 6278A>T 6279T>C), A1593I (6286G>A 6287C>T), Y1595F (6293A>T 6294C>T), L1600V (6307C>G), V1601I (6310G>A 6312A>C), F1602Y (6314T>A), E1604K (6319G>A 6321A>G), N1605S (6323A>G 6324T>C), L1606R (6325C>A 6326T>G 6327T>A), Q1607K (6328C>A), K1614R (6350A>G), M1616V (6355A>G 6357G>T), E1618V (6362A>T 6363A>T), I1619L (6364A>C) | | | | | | | | |
| Codon mutations: | AAA1528AAT (6093A>T), AGC1531TCC (6100A>T 6101G>C), TTT1533ATT (6106T>A), AAA1536AGA (6116A>G), TCT1537TTC (6119C>T 6120T>C), TTT1539TAT (6125T>A), CAG1541CGG (6131A>G), GTG1542CTA (6133G>C 6135G>A), GCA1543AAA (6136G>A 6137C>A), ATG1544ATA (6141G>A), GAA1545CGG (6142G>C 6143A>G 6144A>G), GAA1546GCA (6146A>C), TCA1547GCA (6148T>G), AGC1548GAT (6151A>G 6152G>A 6153C>T), ATT1549ATG (6156T>G), CCA1550CCT (6159A>T), TGG1551AAA (6160T>A 6161G>A 6162G>A), ACG1552ATT (6164C>T 6165G>T), TTC1554TTT (6171C>T), ATA1555TGA (6172A>T 6173T>G), ACT1556ACA (6177T>A), CCA1557AGG (6178C>A 6179C>G 6180A>G), GAT1558AAT (6181G>A), GGA1559GAG (6185G>A 6186A>G), CTA1560CAC (6188T>A 6189A>C), TAC1561TAT (6192C>T), GAA1562GAG (6195A>G), TGG1563TTC (6197G>T 6198G>C), CTA1564TTG (6199C>T 6201A>G), CCA1567TCT (6208C>T 6210A>T), GGT1569GGA (6216T>A), CTA1570CTT (6219A>T), AAG1571ACA (6221A>C 6222G>A), GCA1573GCC (6228A>C), GCA1575GCT (6234A>T), GTG1576ACT (6235G>A 6236T>C 6237G>T), CAA1578ATG (6241C>A 6242A>T 6243A>G), AGG1579AGC (6246G>C), AAA1580CTT (6247A>C 6248A>T 6249A>T), GAC1582AAA (6253G>A 6255C>A), AAC1583GGA (6256A>G 6257A>G 6258C>A), TGT1584ATC (6259T>A 6260G>T 6261T>C), TTC1585TTT (6264C>T), AGG1586AAG (6266G>A), AGG1586_GGT1587insCCA (6267_6268insCCA), GGT1587GAT (6269G>A), ACA1588TTG (6271A>T 6272C>T 6273A>G), GAA1589AAT (6274G>A 6276A>T), GAT1590CTC (6277G>C 6278A>T 6279T>C), TTC1591TTT (6282C>T), ATC1592ATT (6285C>T), GCT1593ATT (6286G>A 6287C>T), GTT1594GTG (6291T>G), TAC1595TTT (6293A>T 6294C>T), ATT1599ATA (6306T>A), CTG1600GTG (6307C>G), GTA1601ATC (6310G>A 6312A>C), TTC1602TAC (6314T>A), TCT1603TCA (6318T>A), GAA1604AAG (6319G>A 6321A>G), AAT1605AGC (6323A>G 6324T>C), CTT1606AGA (6325C>A 6326T>G 6327T>A), CAA1607AAA (6328C>A), GAG1608GAA (6333G>A), GAG1610GAA (6339G>A), CAT1612CAC (6345T>C), CTC1613TTG (6346C>T 6348C>G), AAA1614AGA (6350A>G), ATA1615ATT (6354A>T), ATG1616GTT (6355A>G 6357G>T), CTG1617CTA (6360G>A), GAA1618GTT (6362A>T 6363A>T), ATA1619CTA (6364A>C), TGC1620GG. (6367T>G) | | | | | | | | |

*: Inserts / Deletes / Misaligned / Frameshifts

#### Analysis details

This analysis was performed with panviral2.64

### NGS Details (UN24): Caulimovirus venafragariae

#### Assembly

| | |
|---|---|
| **Coverage Length** | 265 (1 contig(s)) |
| **Depth Of Coverage** | 3.8 |
| **Number Of Reads** | 8 |
| **Reads Per Million** | 0.16 rpm (after QC) |
| **Ambiguities** | 0 |
| **Assembly Method** | de novo + reference guided assembly |
| **Consensus Caller** | Bcf Tools |

#### Coverage Map

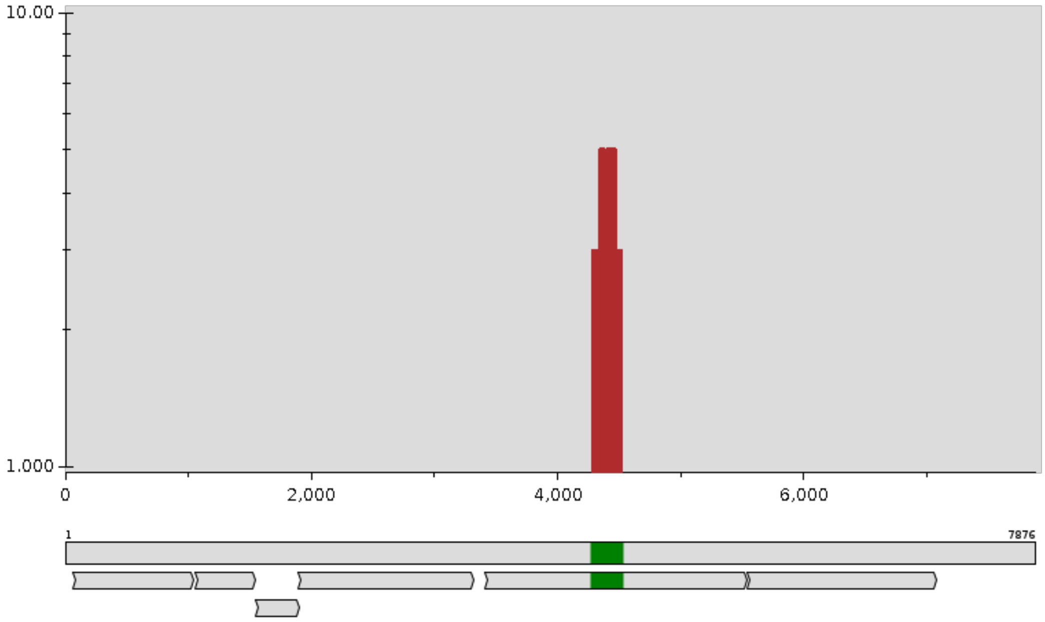

#### Assignment

| | |
|---|---|
| **Type** | Caulimovirus venafragariae (Taxonomy ID: 3048344) |
| **Reference Genome** | NC_001725.1 |
| **NT Identity (%)** | 61.5094 |
| **AA Identity (%)** | 57.9545 |
| **Number Of Stop Codons** | 1 |
| **Number Of CDS** | 6 |

#### Alignment

| | |
|---|---|
| **Alignment Score** | 122.0 (NT) + 343.0 (AA) = 465.0 |
| **Concordance (%)** | 41.5179 |

| Alignment Method | Global, seeded, nucleotide + amino acids (AGA) |
|---|---|

#### Genome Region

Sequence starts at position 4268 and ends at position 4532 relative to NC_001725.1 reference sequence.

#### Alignment Detailed Statistics

| | Begin | End | Coverage | Score | Concordance | Matches | Identities | I/D/M/F* | Stop Codons |
|---|---|---|---|---|---|---|---|---|---|
| **NT** | **4268** | **4532** | **3.4%** | **122** | **23.0%** | **265 (100%)** | **163 (61.5%)** | **0/0** | |
| Mutations: | 4268C>A, 4275C>T, 4276C>A, 4277G>T, 4284G>C, 4287T>C, 4290A>C, 4293T>A, 4296G>A, 4299C>T, 4300G>A, 4306C>T, 4309C>T, 4311C>A, 4314G>A, 4315C>T, 4317T>A, 4320T>A, 4321A>G, 4323C>T, 4326C>A, 4330C>A, 4331C>G, 4332T>A, 4333A>T, 4334G>C, 4337A>T, 4338G>T, 4339A>T, 4340G>C, 4342C>A, 4343C>A, 4348T>A, 4349C>G, 4350C>A, 4353A>T, 4354C>G, 4357G>A, 4359C>A, 4362C>T, 4370G>A, 4371G>A, 4378G>A, 4379C>G, 4388A>T, 4390C>A, 4394G>A, 4395C>A, 4397A>G, 4401A>T, 4407G>A, 4413T>A, 4416C>T, 4424A>G, 4425G>A, 4426T>C, 4428A>T, 4432G>A, 4434C>T, 4435C>A, 4440A>T, 4442A>G, 4443G>A, 4444G>A, 4445G>C, 4446A>G, 4451G>C, 4452C>A, 4455C>T, 4456C>A, 4457T>A, 4459C>T, 4461T>A, 4464T>A, 4465A>G, 4467C>T, 4470G>A, 4471G>A, 4472A>G, 4473G>T, 4474C>G, 4476A>T, 4477C>T, 4479G>A, 4480C>A, 4483C>A, 4485A>T, 4491C>G, 4492G>T, 4493G>A, 4496G>A, 4500G>A, 4502C>A, 4503C>A, 4506T>A, 4508A>T, 4510T>A, 4511C>G, 4513T>A, 4514C>A, 4515C>A, 4521C>T | | | | | | | | |
| **CDS** | | | | | | | | | |
| ORF_V | 289 | 376 | 12.4% | 343 | 56.2% | 88 (100%) | 51 (58.0%) | 0/0/0/0 | 1 |
| Protein mutations: | R291I (4276C>A 4277G>T), E293D (4284G>C), K295N (4290A>C), E299K (4300G>A), I306V (4321A>G 4323C>T), P309R (4330C>A 4331C>G 4332T>A), K311I (4337A>T 4338G>T), P313K (4342C>A 4343C>A), S315R (4348T>A 4349C>G 4350C>A), P317A (4354C>G), A318T (4357G>A 4359C>A), R322K (4370G>A 4371G>A), A325S (4378G>A 4379C>G), K328I (4388A>T), G330E (4394G>A 4395C>A), K331R (4397A>G), M334I (4407G>A), K340R (4424A>G 4425G>A), D343N (4432G>A 4434C>T), H344N (4435C>A), K346R (4442A>G 4443G>A), G347T (4444G>A 4445G>C 4446A>G), G349A (4451G>C 4452C>A), L351K (4456C>A 4457T>A), N354D (4465A>G 4467C>T), E356S (4471G>A 4472A>G 4473G>T), Q357D (4474C>G 4476A>T), L359I (4480C>A), Q360N (4483C>A 4485A>T), I362M (4491C>G), G363* (4492G>T 4493G>A), G364D (4496G>A), T366K (4502C>A 4503C>A), F367L (4506T>A), Y368F (4508A>T), S370K (4513T>A 4514C>A 4515C>A) | | | | | | | | |
| Codon mutations: | CCT288.AT (4268C>A), GAC290GAT (4275C>T), CGA291ATA (4276C>A 4277G>T), GAG293GAC (4284G>C), TTT294TTC (4287T>C), AAA295AAC (4290A>C), ACT296ACA (4293T>A), CAG297CAA (4296G>A), ATC298ATT (4299C>T), GAA299AAA (4300G>A), CTA301TTA (4306C>T), CTC302TTA (4309C>T 4311C>A), AAG303AAA (4314G>A), CTT304TTA (4315C>T 4317T>A), GGT305GGA (4320T>A), ATC306GTT (4321A>G 4323C>T), ATC307ATA (4326C>A), CCT309AGA (4330C>A 4331C>G 4332T>A), AGC310TCC (4333A>T 4334G>C), AAG311ATT (4337A>T 4338G>T), AGT312TCT (4339A>T 4340G>C), CCA313AAA (4342C>A 4343C>A), TCC315AGA (4348T>A 4349C>G 4350C>A), TCA316TCT (4353A>T), CCA317GCA (4354C>G), GCC318ACA (4357G>A 4359C>A), TTC319TTT (4362C>T), AGG322AAA (4370G>A 4371G>A), GCC325AGC (4378G>A 4379C>G), AAA328ATA (4388A>T), CGA329AGA (4390C>A), GGC330GAA (4394G>A 4395C>A), AAA331AGA (4397A>G), GCA332GCT (4401A>T), ATG334ATA (4407G>A), ATT336ATA (4413T>A), AAC337AAT (4416C>T), AAG340AGA (4424A>G 4425G>A), TTA341CTT (4426T>C 4428A>T), GAC343AAT (4432G>A 4434C>T), CAT344AAT (4435C>A), ACA345ACT (4440A>T), AAG346AGA (4442A>G 4443G>A), GGA347ACG (4444G>A 4445G>C 4446A>G), GGC349GCA (4451G>C 4452C>A), TAC350TAT (4455C>T), CTA351AAA (4456C>A 4457T>A), CTT352TTA (4459C>T 4461T>A), CCT353CCA (4464T>A), AAC354GAT (4465A>G 4467C>T), AAG355AAA (4470G>A), GAG356AGT (4471G>A 4472A>G 4473G>T), CAA357GAT (4474C>G 4476A>T), CTG358TTA (4477C>T 4479G>A), CTT359ATT (4480C>A), CAA360AAT (4483C>A 4485A>T), ATC362ATG (4491C>G), GGA363TAA (4492G>T 4493G>A), GGT364GAT (4496G>A), AAG365AAA (4500G>A), ACC366AAA (4502C>A 4503C>A), TTT367TTA (4506T>A), TAC368TTC (4508A>T), TCT369AGT (4510T>A 4511C>G), TCC370AAA (4513T>A 4514C>A 4515C>A), GAC372GAT (4521C>T) | | | | | | | | |
| **Proteins** | | | | | | | | | |
| hypothetical protein (NP_043933.1) | 289 | 376 | 12.4% | 343 | 56.2% | 88 (100%) | 51 (58.0%) | 0/0/0/0 | 1 |
| Protein mutations: | R291I (4276C>A 4277G>T), E293D (4284G>C), K295N (4290A>C), E299K (4300G>A), I306V (4321A>G 4323C>T), P309R (4330C>A 4331C>G 4332T>A), K311I (4337A>T 4338G>T), P313K (4342C>A 4343C>A), S315R (4348T>A 4349C>G 4350C>A), P317A (4354C>G), A318T (4357G>A 4359C>A), R322K (4370G>A 4371G>A), A325S (4378G>A 4379C>G), K328I (4388A>T), G330E (4394G>A 4395C>A), K331R (4397A>G), M334I (4407G>A), K340R (4424A>G 4425G>A), D343N (4432G>A 4434C>T), H344N (4435C>A), K346R (4442A>G 4443G>A), G347T (4444G>A 4445G>C 4446A>G), G349A (4451G>C 4452C>A), L351K (4456C>A 4457T>A), N354D (4465A>G 4467C>T), E356S (4471G>A 4472A>G 4473G>T), Q357D (4474C>G 4476A>T), L359I (4480C>A), Q360N (4483C>A 4485A>T), I362M (4491C>G), G363* (4492G>T 4493G>A), G364D (4496G>A), T366K (4502C>A 4503C>A), F367L (4506T>A), Y368F (4508A>T), S370K (4513T>A 4514C>A 4515C>A) | | | | | | | | |
| Codon mutations: | CCT288.AT (4268C>A), GAC290GAT (4275C>T), CGA291ATA (4276C>A 4277G>T), GAG293GAC (4284G>C), TTT294TTC (4287T>C), AAA295AAC (4290A>C), ACT296ACA (4293T>A), CAG297CAA (4296G>A), ATC298ATT (4299C>T), GAA299AAA (4300G>A), CTA301TTA (4306C>T), CTC302TTA (4309C>T 4311C>A), AAG303AAA (4314G>A), CTT304TTA (4315C>T 4317T>A), GGT305GGA (4320T>A), ATC306GTT (4321A>G 4323C>T), ATC307ATA (4326C>A), CCT309AGA (4330C>A 4331C>G 4332T>A), AGC310TCC (4333A>T 4334G>C), AAG311ATT (4337A>T 4338G>T), AGT312TCT (4339A>T 4340G>C), CCA313AAA (4342C>A 4343C>A), TCC315AGA (4348T>A 4349C>G 4350C>A), TCA316TCT (4353A>T), CCA317GCA (4354C>G), GCC318ACA (4357G>A 4359C>A), TTC319TTT (4362C>T), AGG322AAA (4370G>A 4371G>A), GCC325AGC (4378G>A 4379C>G), AAA328ATA (4388A>T), CGA329AGA (4390C>A), GGC330GAA (4394G>A 4395C>A), AAA331AGA (4397A>G), GCA332GCT (4401A>T), ATG334ATA (4407G>A), ATT336ATA (4413T>A), AAC337AAT (4416C>T), AAG340AGA (4424A>G 4425G>A), TTA341CTT (4426T>C 4428A>T), GAC343AAT (4432G>A 4434C>T), CAT344AAT (4435C>A), ACA345ACT (4440A>T), AAG346AGA (4442A>G 4443G>A), GGA347ACG (4444G>A 4445G>C 4446A>G), GGC349GCA (4451G>C 4452C>A), TAC350TAT (4455C>T), CTA351AAA (4456C>A 4457T>A), CTT352TTA (4459C>T 4461T>A), CCT353CCA (4464T>A), AAC354GAT (4465A>G 4467C>T), AAG355AAA (4470G>A), GAG356AGT (4471G>A 4472A>G 4473G>T), CAA357GAT (4474C>G 4476A>T), CTG358TTA (4477C>T 4479G>A), CTT359ATT (4480C>A), CAA360AAT (4483C>A 4485A>T), ATC362ATG (4491C>G), GGA363TAA (4492G>T 4493G>A), GGT364GAT (4496G>A), AAG365AAA (4500G>A), ACC366AAA (4502C>A 4503C>A), TTT367TTA (4506T>A), TAC368TTC (4508A>T), TCT369AGT (4510T>A 4511C>G), TCC370AAA (4513T>A 4514C>A 4515C>A), GAC372GAT (4521C>T) | | | | | | | | |

*: Inserts / Deletes / Misaligned / Frameshifts

#### Analysis details

This analysis was performed with panviral2.64

### NGS Details (UN24): Caulimovirus venafragariae

#### Assembly

| | |
|---|---|
| **Coverage Length** | 257 (1 contig(s)) |
| **Depth Of Coverage** | 3.5 |
| **Number Of Reads** | 8 |
| **Reads Per Million** | 0.16 rpm (after QC) |
| **Ambiguities** | 0 |
| **Assembly Method** | de novo + reference guided assembly |
| **Consensus Caller** | Bcf Tools |

#### Coverage Map

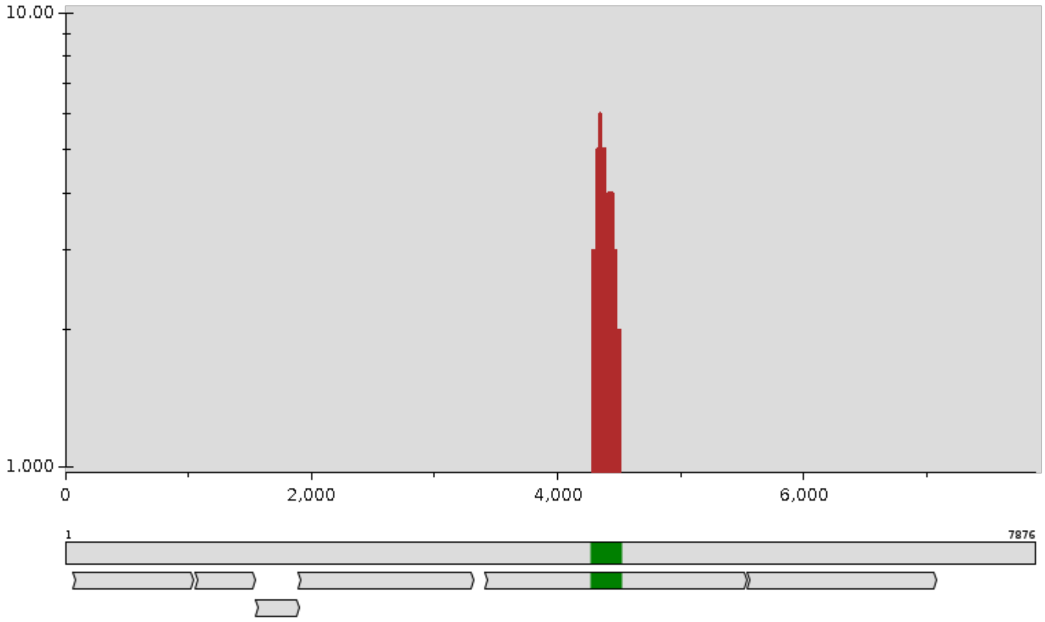

#### Assignment

| | |
|---|---|
| **Type** | Caulimovirus venafragariae (Taxonomy ID: 3048344) |
| **Reference Genome** | NC_001725.1 |
| **NT Identity (%)** | 62.2568 |
| **AA Identity (%)** | 64.7059 |
| **Number Of Stop Codons** | 0 |
| **Number Of CDS** | 6 |

#### Alignment

| | |
|---|---|
| **Alignment Score** | 126.0 (NT) + 389.0 (AA) = 515.0 |
| **Concordance (%)** | 46.9462 |

| | |
|---|---|
| **Alignment Method** | Global, seeded, nucleotide + amino acids (AGA) |

#### Genome Region

Sequence starts at position 4268 and ends at position 4524 relative to NC_001725.1 reference sequence.

#### Alignment Detailed Statistics

| | Begin | End | Coverage | Score | Concordance | Matches | Identities | I/D/M/F* | Stop Codons |
|---|---|---|---|---|---|---|---|---|---|
| **NT** | **4268** | **4524** | **3.3%** | **126** | **24.5%** | **257 (100%)** | **160 (62.3%)** | **0/0** | |
| Mutations: | 4268C>A, 4276C>G, 4277G>T, 4284G>T, 4289A>G, 4290A>T, 4292C>A, 4296G>A, 4299C>T, 4300G>A, 4311C>A, 4313A>C, 4314G>T, 4315C>T, 4317T>A, 4320T>A, 4321A>G, 4323C>T, 4326C>A, 4330C>A, 4331C>G, 4332T>A, 4333A>T, 4334G>C, 4335C>T, 4337A>C, 4338G>A, 4339A>T, 4340G>C, 4341T>A, 4342C>A, 4343C>A, 4348T>A, 4349C>G, 4350C>A, 4354C>G, 4362C>T, 4368A>T, 4371G>A, 4378G>A, 4379C>G, 4383G>A, 4386C>A, 4387A>G, 4388A>T, 4390C>A, 4395C>T, 4398A>G, 4402A>C, 4413T>A, 4419T>C, 4424A>G, 4425G>A, 4426T>C, 4428A>T, 4434C>T, 4435C>A, 4437T>C, 4440A>T, 4442A>G, 4443G>A, 4444G>A, 4445G>C, 4446A>G, 4451G>C, 4455C>T, 4456C>A, 4457T>A, 4458A>G, 4461T>A, 4464T>A, 4465A>G, 4467C>T, 4470G>A, 4471G>A, 4472A>G, 4473G>T, 4474C>G, 4477C>T, 4479G>A, 4480C>A, 4482T>A, 4483C>A, 4485A>T, 4491C>A, 4492G>C, 4493G>A, 4503C>A, 4504T>G, 4506T>A, 4508A>T, 4510T>A, 4511C>A, 4513T>A, 4514C>A, 4515C>A, 4521C>T | | | | | | | | |
| **CDS** | | | | | | | | | |
| ORF_V | 289 | 373 | 12.0% | 389 | 65.5% | 85 (100%) | 55 (64.7%) | 0/0/0/0 | 0 |
| Protein mutations: | R291V (4276C>G 4277G>T), E293D (4284G>T), K295S (4289A>G 4290A>T), T296N (4292C>A), E299K (4300G>A), K303T (4313A>C 4314G>T), I306V (4321A>G 4323C>T), P309R (4330C>A 4331C>G 4332T>A), K311T (4337A>C 4338G>A), P313K (4342C>A 4343C>A), S315R (4348T>A 4349C>G 4350C>A), P317A (4354C>G), A325S (4378G>A 4379C>G), K328V (4387A>G 4388A>T), K340R (4424A>G 4425G>A), H344N (4435C>A 4437T>C), K346R (4442A>G 4443G>A), G347T (4444G>A 4445G>C 4446A>G), G349A (4451G>C), L351K (4456C>A 4457T>A 4458A>G), N354D (4465A>G 4467C>T), E356S (4471G>A 4472A>G 4473G>T), Q357E (4474C>G), L359I (4480C>A 4482T>A), Q360N (4483C>A 4485A>T), G363Q (4492G>C 4493G>A), F367V (4504T>G 4506T>A), Y368F (4508A>T), S369N (4510T>A 4511C>A), S370K (4513T>A 4514C>A 4515C>A) | | | | | | | | |
| Codon mutations: | CCT288.AT (4268C>A), CGA291GTA (4276C>G 4277G>T), GAG293GAT (4284G>T), AAA295AGT (4289A>G 4290A>T), ACT296AAT (4292C>A), CAG297CAA (4296G>A), ATC298ATT (4299C>T), GAA299AAA (4300G>A), CTC302CTA (4311C>A), AAG303ACT (4313A>C 4314G>T), CTT304TTA (4315C>T 4317T>A), GGT305GGA (4320T>A), ATC306GTT (4321A>G 4323C>T), ATC307ATA (4326C>A), CCT309AGA (4330C>A 4331C>G 4332T>A), AGC310TCT (4333A>T 4334G>C 4335C>T), AAG311ACA (4337A>C 4338G>A), AGT312TCA (4339A>T 4340G>C 4341T>A), CCA313AAA (4342C>A 4343C>A), TCC315AGA (4348T>A 4349C>G 4350C>A), CCA317GCA (4354C>G), TTC319TTT (4362C>T), GTA321GTT (4368A>T), AGG322AGA (4371G>A), GCC325AGC (4378G>A 4379C>G), GAG326GAA (4383G>A), ATC327ATA (4386C>A), AAA328GTA (4387A>G 4388A>T), CGA329AGA (4390C>A), GGC330GGT (4395C>T), AAA331AAG (4398A>G), AGA333CGA (4402A>C), ATT336ATA (4413T>A), TAT338TAC (4419T>C), AAG340AGA (4424A>G 4425G>A), TTA341CTT (4426T>C 4428A>T), GAC343GAT (4434C>T), CAT344AAC (4435C>A 4437T>C), ACA345ACT (4440A>T), AAG346AGA (4442A>G 4443G>A), GGA347ACG (4444G>A 4445G>C 4446A>G), GGC349GCC (4451G>C), TAC350TAT (4455C>T), CTA351AAG (4456C>A 4457T>A 4458A>G), CTT352CTA (4461T>A), CCT353CCA (4464T>A), AAC354GAT (4465A>G 4467C>T), AAG355AAA (4470G>A), GAG356AGT (4471G>A 4472A>G 4473G>T), CAA357GAA (4474C>G), CTG358TTA (4477C>T 4479G>A), CTT359ATA (4480C>A 4482T>A), CAA360AAT (4483C>A 4485A>T), ATC362ATA (4491C>A), GGA363CAA (4492G>C 4493G>A), ACC366ACA (4503C>A), TTT367GTA (4504T>G 4506T>A), TAC368TTC (4508A>T), TCT369AAT (4510T>A 4511C>A), TCC370AAA (4513T>A 4514C>A 4515C>A), GAC372GAT (4521C>T) | | | | | | | | |
| **Proteins** | | | | | | | | | |
| hypothetical protein (NP_043933.1) | 289 | 373 | 12.0% | 389 | 65.5% | 85 (100%) | 55 (64.7%) | 0/0/0/0 | 0 |
| Protein mutations: | R291V (4276C>G 4277G>T), E293D (4284G>T), K295S (4289A>G 4290A>T), T296N (4292C>A), E299K (4300G>A), K303T (4313A>C 4314G>T), I306V (4321A>G 4323C>T), P309R (4330C>A 4331C>G 4332T>A), K311T (4337A>C 4338G>A), P313K (4342C>A 4343C>A), S315R (4348T>A 4349C>G 4350C>A), P317A (4354C>G), A325S (4378G>A 4379C>G), K328V (4387A>G 4388A>T), K340R (4424A>G 4425G>A), H344N (4435C>A 4437T>C), K346R (4442A>G 4443G>A), G347T (4444G>A 4445G>C 4446A>G), G349A (4451G>C), L351K (4456C>A 4457T>A 4458A>G), N354D (4465A>G 4467C>T), E356S (4471G>A 4472A>G 4473G>T), Q357E (4474C>G), L359I (4480C>A 4482T>A), Q360N (4483C>A 4485A>T), G363Q (4492G>C 4493G>A), F367V (4504T>G 4506T>A), Y368F (4508A>T), S369N (4510T>A 4511C>A), S370K (4513T>A 4514C>A 4515C>A) | | | | | | | | |
| Codon mutations: | CCT288.AT (4268C>A), CGA291GTA (4276C>G 4277G>T), GAG293GAT (4284G>T), AAA295AGT (4289A>G 4290A>T), ACT296AAT (4292C>A), CAG297CAA (4296G>A), ATC298ATT (4299C>T), GAA299AAA (4300G>A), CTC302CTA (4311C>A), AAG303ACT (4313A>C 4314G>T), CTT304TTA (4315C>T 4317T>A), GGT305GGA (4320T>A), ATC306GTT (4321A>G 4323C>T), ATC307ATA (4326C>A), CCT309AGA (4330C>A 4331C>G 4332T>A), AGC310TCT (4333A>T 4334G>C 4335C>T), AAG311ACA (4337A>C 4338G>A), AGT312TCA (4339A>T 4340G>C 4341T>A), CCA313AAA (4342C>A 4343C>A), TCC315AGA (4348T>A 4349C>G 4350C>A), CCA317GCA (4354C>G), TTC319TTT (4362C>T), GTA321GTT (4368A>T), AGG322AGA (4371G>A), GCC325AGC (4378G>A 4379C>G), GAG326GAA (4383G>A), ATC327ATA (4386C>A), AAA328GTA (4387A>G 4388A>T), CGA329AGA (4390C>A), GGC330GGT (4395C>T), AAA331AAG (4398A>G), AGA333CGA (4402A>C), ATT336ATA (4413T>A), TAT338TAC (4419T>C), AAG340AGA (4424A>G 4425G>A), TTA341CTT (4426T>C 4428A>T), GAC343GAT (4434C>T), CAT344AAC (4435C>A 4437T>C), ACA345ACT (4440A>T), AAG346AGA (4442A>G 4443G>A), GGA347ACG (4444G>A 4445G>C 4446A>G), GGC349GCC (4451G>C), TAC350TAT (4455C>T), CTA351AAG (4456C>A 4457T>A 4458A>G), CTT352CTA (4461T>A), CCT353CCA (4464T>A), AAC354GAT (4465A>G 4467C>T), AAG355AAA (4470G>A), GAG356AGT (4471G>A 4472A>G 4473G>T), CAA357GAA (4474C>G), CTG358TTA (4477C>T 4479G>A), CTT359ATA (4480C>A 4482T>A), CAA360AAT (4483C>A 4485A>T), ATC362ATA (4491C>A), GGA363CAA (4492G>C 4493G>A), ACC366ACA (4503C>A), TTT367GTA (4504T>G 4506T>A), TAC368TTC (4508A>T), TCT369AAT (4510T>A 4511C>A), TCC370AAA (4513T>A 4514C>A 4515C>A), GAC372GAT (4521C>T) | | | | | | | | |

*: Inserts / Deletes / Misaligned / Frameshifts

#### Analysis details

This analysis was performed with panviral2.64

### NGS Details (UN24): Petuvirus venapetuniae

#### Assembly

| | |
|---|---|
| **Coverage Length** | 403 (2 contig(s)) |
| **Depth Of Coverage** | 1.9 |
| **Number Of Reads** | 8 |
| **Reads Per Million** | 0.16 rpm (after QC) |
| **Ambiguities** | 0 |
| **Assembly Method** | de novo + reference guided assembly |
| **Consensus Caller** | Bcf Tools |

#### Coverage Map

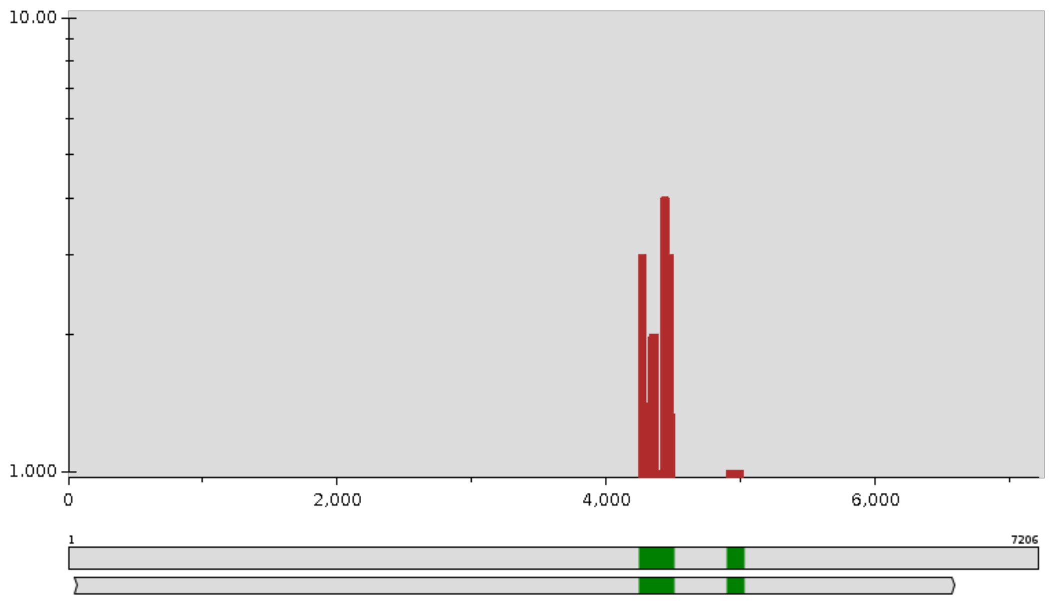

#### Assignment

| | |
|---|---|
| **Type** | Petuvirus venapetuniae (Taxonomy ID: 3048265) |
| **Reference Genome** | NC_001839.2 |
| **NT Identity (%)** | 59.5533 |
| **AA Identity (%)** | 51.4925 |
| **Number Of Stop Codons** | 1 |
| **Number Of CDS** | 1 |

#### Alignment

| | |
|---|---|
| **Alignment Score** | 154.0 (NT) + 522.0 (AA) = 676.0 |
| **Concordance (%)** | 38.106 |

| Alignment Method | Global, seeded, nucleotide + amino acids (AGA) |
|---|---|

#### Genome Region

Sequence starts at position 4238 and ends at position 5024 relative to NC_001839.2 reference sequence.

#### Alignment Detailed Statistics

| | Begin | End | Coverage | Score | Concordance | Matches | Identities | I/D/M/F* | Stop Codons |
|---|---|---|---|---|---|---|---|---|---|
| **NT** | **4238** | **5024** | **5.6%** | **154** | **19.1%** | **403 (100%)** | **240 (59.6%)** | **0/0** | |
| Mutations: | 4240A>T, 4242C>A, 4244C>G, 4245C>G, 4247G>A, 4250A>T, 4251G>T, 4252C>G, 4253A>T, 4255T>A, 4263T>A, 4264G>T, 4265T>C, 4266G>A, 4269G>T, 4270A>C, 4271G>T, 4275C>T, 4276A>T, 4278C>G, 4281T>A, 4282T>A, 4283T>C, 4284G>A, 4286T>A, 4287C>A, 4289A>T, 4293G>A, 4294A>G, 4298C>T, 4302G>A, 4304T>A, 4307A>T, 4309A>C, 4310A>T, 4314G>A, 4315C>G, 4316A>C, 4320G>T, 4321A>C, 4325T>C, 4328T>C, 4334C>T, 4337C>T, 4340G>A, 4342G>C, 4343G>T, 4344T>G, 4346A>T, 4349G>A, 4350C>A, 4354T>A, 4355G>A, 4361A>C, 4362A>T, 4363A>C, 4364A>C, 4365T>C, 4366T>C, 4367A>T, 4376C>T, 4379A>T, 4386C>A, 4388A>G, 4391A>T, 4397C>T, 4398C>T, 4399A>C, 4401T>G, 4403C>T, 4404C>T, 4406C>A, 4407C>A, 4410G>T, 4411A>G, 4412T>G, 4413G>A, 4414A>T, 4415C>T, 4417A>G, 4418G>A, 4420T>A, 4421T>C, 4424C>G, 4430A>T, 4436G>A, 4437C>A, 4439C>G, 4440A>G, 4441C>A, 4442A>T, 4445C>T, 4448C>G, 4449T>A, 4450C>A, 4451T>A, 4452C>A, 4453A>G, 4454C>A, 4455T>A, 4456T>C, 4457A>T, 4459C>T, 4460C>T, 4466A>C, 4469G>T, 4470C>A, 4472C>T, 4474T>A, 4475T>C, 4476T>A, 4477C>G, 4478A>T, 4481G>A, 4488C>A, 4490T>G, 4493G>A, 4499A>T, 4502T>C, 4894C>T, 4896G>A, 4898A>C, 4904G>A, 4905A>C, 4907A>C, 4908C>T, 4910A>G, 4916T>C, 4922G>T, 4923A>G, 4925T>G, 4926G>A, 4935A>G, 4938A>C, 4939G>A, 4952A>T, 4953G>T, 4955G>T, 4956G>A, 4958T>C, 4959A>T, 4961T>G, 4962G>A, 4965C>T, 4973C>T, 4976T>A, 4981G>T, 4982T>G, 4983G>A, 4985T>G, 4988G>T, 4992A>C, 4994G>A, 5001C>T, 5002C>A, 5003A>T, 5006G>T, 5008C>T, 5009A>T, 5015A>G, 5018A>G, 5019T>G, 5020G>A | | | | | | | | |

##### CDS

| | Begin | End | Coverage | Score | Concordance | Matches | Identities | I/D/M/F* | Stop Codons |
|---|---|---|---|---|---|---|---|---|---|
| PVCVgp1 | 1397 | 1658 | 6.1% | 522 | 52.9% | 134 (100%) | 69 (51.5%) | 0/0/0/0 | 1 |
| Protein mutations: | H1397L (4240A>T), L1398M (4242C>A 4244C>G), Q1399E (4245C>G 4247G>A), L1400F (4250A>T), A1401C (4251G>T 4252C>G 4253A>T), I1402K (4255T>A), C1405I (4263T>A 4264G>T 4265T>C), D1406N (4266G>A), E1407S (4269G>T 4270A>C 4271G>T), Q1409L (4275C>T 4276A>T), Q1410E (4278C>G), F1411N (4281T>A 4282T>A 4283T>C), D1412K (4284G>A 4286T>A), L1413I (4287C>A 4289A>T), E1415R (4293G>A 4294A>G), D1418K (4302G>A 4304T>A), Q1420P (4309A>C 4310A>T), A1422S (4314G>A 4315C>G 4316A>C), E1424S (4320G>T 4321A>C), R1431T (4342G>C 4343G>T), S1432A (4344T>G 4346A>T), Q1434K (4350C>A), V1435E (4354T>A 4355G>A), K1438S (4362A>T 4363A>C 4364A>C), L1439P (4365T>C 4366T>C 4367A>T), Q1446K (4386C>A 4388A>G), H1450S (4398C>T 4399A>C), F1451V (4401T>G 4403C>T), Q1453K (4407C>A), D1454W (4410G>T 4411A>G 4412T>G), D1455I (4413G>A 4414A>T 4415C>T), K1456R (4417A>G 4418G>A), F1457Y (4420T>A 4421T>C), L1463M (4437C>A 4439C>G), T1464D (4440A>G 4441C>A 4442A>T), F1466L (4448C>G), S1467K (4449T>A 4450C>A 4451T>A), H1468R (4452C>A 4453A>G 4454C>A), L1469T (4455T>A 4456T>C 4457A>T), S1470F (4459C>T 4460C>T), K1473N (4469G>T), L1474I (4470C>A 4472C>T), F1475Y (4474T>A 4475T>C), L1480M (4488C>A 4490T>G), S1615L (4894C>T), V1616I (4896G>A 4898A>C), I1619L (4905A>C 4907A>C), Q1620* (4908C>T 4910A>G), I1625V (4923A>G 4925T>G), V1626I (4926G>A), I1629V (4935A>G), R1630Q (4938A>C 4939G>A), E1635Y (4953G>T 4955G>T), V1636I (4956G>A 4958T>C), T1637S (4959A>T 4961T>G), E1638K (4962G>A), H1639Y (4965C>T), S1644M (4981G>T 4982T>G), D1645K (4983G>A 4985T>G), M1646I (4988G>T), K1648Q (4992A>C 4994G>A), P1651Y (5001C>T 5002C>A 5003A>T), A1653V (5008C>T 5009A>T), C1657D (5019T>G 5020G>A) | | | | | | | | |
| Codon mutations: | CAT1397CTT (4240A>T), CTC1398ATG (4242C>A 4244C>G), CAG1399GAA (4245C>G 4247G>A), TTA1400TTT (4250A>T), GCA1401TGT (4251G>T 4252C>G 4253A>T), ATA1402AAA (4255T>A), TGT1405ATC (4263T>A 4264G>T 4265T>C), GAT1406AAT (4266G>A), GAG1407TCT (4269G>T 4270A>C 4271G>T), CAA1409TTA (4275C>T 4276A>T), CAA1410GAA (4278C>G), TTT1411AAC (4281T>A 4282T>A 4283T>C), GAT1412AAA (4284G>A 4286T>A), CTA1413ATT (4287C>A 4289A>T), GAA1415AGA (4293G>A 4294A>G), CCC1416CCT (4298C>T), GAT1418AAA (4302G>A 4304T>A), TCA1419TCT (4307A>T), CAA1420CCT (4309A>C 4310A>T), GCA1422AGC (4314G>A 4315C>G 4316A>C), GAA1424TCA (4320G>T 4321A>C), GCT1425GCC (4325T>C), TTT1426TTC (4328T>C), GTC1428GTT (4334C>T), AAC1429AAT (4337C>T), AAG1430AAA (4340G>A), AGG1431ACT (4342G>C 4343G>T), TCA1432GCT (4344T>G 4346A>T), GAG1433GAA (4349G>A), CAA1434AAA (4350C>A), GTG1435GAA (4354T>A 4355G>A), GGA1437GGC (4361A>C), AAA1438TCC (4362A>T 4363A>C 4364A>C), TTA1439CCT (4365T>C 4366T>C 4367A>T), GTC1442GTT (4376C>T), ATA1443ATT (4379A>T), CAA1446AAG (4386C>A 4388A>G), CCA1447CCT (4391A>T), AAC1449AAT (4397C>T), CAT1450TCT (4398C>T 4399A>C), TTC1451GTT (4401T>G 4403C>T), CTC1452TTA (4404C>T 4406C>A), CAA1453AAA (4407C>A), GAT1454TGG (4410G>T 4411A>G 4412T>G), GAC1455ATT (4413G>A 4414A>T 4415C>T), AAG1456AGA (4417A>G 4418G>A), TTT1457TAC (4420T>A 4421T>C), CCC1458CCG (4424C>G), CCA1460CCT (4430A>T), AAG1462AAA (4436G>A), CTC1463ATG (4437C>A 4439C>G), ACA1464GAT (4440A>G 4441C>A 4442A>T), CTC1465CTT (4445C>T), TTC1466TTG (4448C>G), TCT1467AAA (4449T>A 4450C>A 4451T>A), CAC1468AGA (4452C>A 4453A>G 4454C>A), TTA1469ACT (4455T>A 4456T>C 4457A>T), TCC1470TTT (4459C>T 4460C>T), GCA1472GCC (4466A>C), AAG1473AAT (4469G>T), CTC1474ATT (4470C>A 4472C>T), TTT1475TAC (4474T>A 4475T>C), TCA1476AGT (4476T>A 4477C>G 4478A>T), AAG1477AAA (4481G>A), CTT1480ATG (4488C>A 4490T>G), AAG1481AAA (4493G>A), GGA1483GGT (4499A>T), TTT1484TTC (4502T>C), TCG1615TTG (4894C>T), GTA1616ATC (4896G>A 4898A>C), CAG1618CAA (4904G>A), ATA1619CTC (4905A>C 4907A>C), CAA1620TAG (4908C>T 4910A>G), TTT1622TTC (4916T>C), GGG1624GGT (4922G>T), ATT1625GTG (4923A>G 4925T>G), GTT1626ATT (4926G>A), ATC1629GTC (4935A>G), AGA1630CAA (4938A>C 4939G>A), CCA1634CCT (4952A>T), GAG1635TAT (4953G>T 4955G>T), GTT1636ATC (4956G>A 4958T>C), ACT1637TCG (4959A>T 4961T>G), GAG1638AAG (4962G>A), CAT1639TAT (4965C>T), TCC1641TCT (4973C>T), CCT1642CCA (4976T>A), AGT1644ATG (4981G>T 4982T>G), GAT1645AAG (4983G>A 4985T>G), ATG1646ATT (4988G>T), AAG1648CAA (4992A>C 4994G>A), CCA1651TAT (5001C>T 5002C>A 5003A>T), CCG1652CCT (5006G>T), GCA1653GTT (5008C>T 5009A>T), GGA1655GGG (5015A>G), AAA1656AAG (5018A>G), TGT1657GAT (5019T>G 5020G>A) | | | | | | | | |

##### Proteins

| | Begin | End | Coverage | Score | Concordance | Matches | Identities | I/D/M/F* | Stop Codons |
|---|---|---|---|---|---|---|---|---|---|
| ORF I polyprotein (NP_127504.1) | 1397 | 1658 | 6.1% | 522 | 52.9% | 134 (100%) | 69 (51.5%) | 0/0/0/0 | 1 |
| Protein mutations: | H1397L (4240A>T), L1398M (4242C>A 4244C>G), Q1399E (4245C>G 4247G>A), L1400F (4250A>T), A1401C (4251G>T 4252C>G 4253A>T), I1402K (4255T>A), C1405I (4263T>A 4264G>T 4265T>C), D1406N (4266G>A), E1407S (4269G>T 4270A>C 4271G>T), Q1409L (4275C>T 4276A>T), Q1410E (4278C>G), F1411N (4281T>A 4282T>A 4283T>C), D1412K (4284G>A 4286T>A), L1413I (4287C>A 4289A>T), E1415R (4293G>A 4294A>G), D1418K (4302G>A 4304T>A), Q1420P (4309A>C 4310A>T), A1422S (4314G>A 4315C>G 4316A>C), E1424S (4320G>T 4321A>C), R1431T (4342G>C 4343G>T), S1432A (4344T>G 4346A>T), Q1434K (4350C>A), V1435E (4354T>A 4355G>A), K1438S (4362A>T 4363A>C 4364A>C), L1439P (4365T>C 4366T>C 4367A>T), Q1446K (4386C>A 4388A>G), H1450S (4398C>T 4399A>C), F1451V (4401T>G 4403C>T), Q1453K (4407C>A), D1454W (4410G>T 4411A>G 4412T>G), D1455I (4413G>A 4414A>T 4415C>T), K1456R (4417A>G 4418G>A), F1457Y (4420T>A 4421T>C), L1463M (4437C>A 4439C>G), T1464D (4440A>G 4441C>A 4442A>T), F1466L (4448C>G), S1467K (4449T>A 4450C>A 4451T>A), H1468R (4452C>A 4453A>G 4454C>A), L1469T (4455T>A 4456T>C 4457A>T), S1470F (4459C>T 4460C>T), K1473N (4469G>T), L1474I (4470C>A 4472C>T), F1475Y (4474T>A 4475T>C), L1480M (4488C>A 4490T>G), S1615L (4894C>T), V1616I (4896G>A 4898A>C), I1619L (4905A>C 4907A>C), Q1620* (4908C>T 4910A>G), I1625V (4923A>G 4925T>G), V1626I (4926G>A), I1629V (4935A>G), R1630Q (4938A>C 4939G>A), E1635Y (4953G>T 4955G>T), V1636I (4956G>A 4958T>C), T1637S (4959A>T 4961T>G), E1638K (4962G>A), H1639Y (4965C>T), S1644M (4981G>T 4982T>G), D1645K (4983G>A 4985T>G), M1646I (4988G>T), K1648Q (4992A>C 4994G>A), P1651Y (5001C>T 5002C>A 5003A>T), A1653V (5008C>T 5009A>T), C1657D (5019T>G 5020G>A) | | | | | | | | |
| Codon mutations: | CAT1397CTT (4240A>T), CTC1398ATG (4242C>A 4244C>G), CAG1399GAA (4245C>G 4247G>A), TTA1400TTT (4250A>T), GCA1401TGT (4251G>T 4252C>G 4253A>T), ATA1402AAA (4255T>A), TGT1405ATC (4263T>A 4264G>T 4265T>C), GAT1406AAT (4266G>A), GAG1407TCT (4269G>T 4270A>C 4271G>T), CAA1409TTA (4275C>T 4276A>T), CAA1410GAA (4278C>G), TTT1411AAC (4281T>A 4282T>A 4283T>C), GAT1412AAA (4284G>A 4286T>A), CTA1413ATT (4287C>A 4289A>T), GAA1415AGA (4293G>A 4294A>G), CCC1416CCT (4298C>T), GAT1418AAA (4302G>A 4304T>A), TCA1419TCT (4307A>T), CAA1420CCT (4309A>C 4310A>T), GCA1422AGC (4314G>A 4315C>G 4316A>C), GAA1424TCA (4320G>T 4321A>C), GCT1425GCC (4325T>C), TTT1426TTC (4328T>C), GTC1428GTT (4334C>T), AAC1429AAT (4337C>T), AAG1430AAA (4340G>A), AGG1431ACT (4342G>C 4343G>T), TCA1432GCT (4344T>G 4346A>T), GAG1433GAA (4349G>A), CAA1434AAA (4350C>A), GTG1435GAA (4354T>A 4355G>A), GGA1437GGC (4361A>C), AAA1438TCC (4362A>T 4363A>C 4364A>C), TTA1439CCT (4365T>C 4366T>C 4367A>T), GTC1442GTT (4376C>T), ATA1443ATT (4379A>T), CAA1446AAG (4386C>A 4388A>G), CCA1447CCT (4391A>T), AAC1449AAT (4397C>T), CAT1450TCT (4398C>T 4399A>C), TTC1451GTT (4401T>G 4403C>T), CTC1452TTA (4404C>T 4406C>A), CAA1453AAA (4407C>A), GAT1454TGG (4410G>T 4411A>G 4412T>G), GAC1455ATT (4413G>A 4414A>T 4415C>T), AAG1456AGA (4417A>G 4418G>A), TTT1457TAC (4420T>A 4421T>C), CCC1458CCG (4424C>G), CCA1460CCT (4430A>T), AAG1462AAA (4436G>A), CTC1463ATG (4437C>A 4439C>G), ACA1464GAT (4440A>G 4441C>A 4442A>T), CTC1465CTT (4445C>T), TTC1466TTG (4448C>G), TCT1467AAA (4449T>A 4450C>A 4451T>A), CAC1468AGA (4452C>A 4453A>G 4454C>A), TTA1469ACT (4455T>A 4456T>C 4457A>T), TCC1470TTT (4459C>T 4460C>T), GCA1472GCC (4466A>C), AAG1473AAT (4469G>T), CTC1474ATT (4470C>A 4472C>T), TTT1475TAC (4474T>A 4475T>C), TCA1476AGT (4476T>A 4477C>G 4478A>T), AAG1477AAA (4481G>A), CTT1480ATG (4488C>A 4490T>G), AAG1481AAA (4493G>A), GGA1483GGT (4499A>T), TTT1484TTC (4502T>C), TCG1615TTG (4894C>T), GTA1616ATC (4896G>A 4898A>C), CAG1618CAA (4904G>A), ATA1619CTC (4905A>C 4907A>C), CAA1620TAG (4908C>T 4910A>G), TTT1622TTC (4916T>C), GGG1624GGT (4922G>T), ATT1625GTG (4923A>G 4925T>G), GTT1626ATT (4926G>A), ATC1629GTC (4935A>G), AGA1630CAA (4938A>C 4939G>A), CCA1634CCT (4952A>T), GAG1635TAT (4953G>T 4955G>T), GTT1636ATC (4956G>A 4958T>C), ACT1637TCG (4959A>T 4961T>G), GAG1638AAG (4962G>A), CAT1639TAT (4965C>T), TCC1641TCT (4973C>T), CCT1642CCA (4976T>A), AGT1644ATG (4981G>T 4982T>G), GAT1645AAG (4983G>A 4985T>G), ATG1646ATT (4988G>T), AAG1648CAA (4992A>C 4994G>A), CCA1651TAT (5001C>T 5002C>A 5003A>T), CCG1652CCT (5006G>T), GCA1653GTT (5008C>T 5009A>T), GGA1655GGG (5015A>G), AAA1656AAG (5018A>G), TGT1657GAT (5019T>G 5020G>A) | | | | | | | | |

*: Inserts / Deletes / Misaligned / Frameshifts

#### Analysis details

This analysis was performed with panviral2.64

### NGS Details (UN24): Epiphyllum badnavirus 1

#### Assembly

| | |
|---|---|
| Coverage Length | 261 (1 contig(s)) |
| Depth Of Coverage | 3.9 |
| Number Of Reads | 8 |
| Reads Per Million | 0.16 rpm (after QC) |
| Ambiguities | 0 |
| Assembly Method | de novo + reference guided assembly |
| Consensus Caller | Bcf Tools |

#### Coverage Map

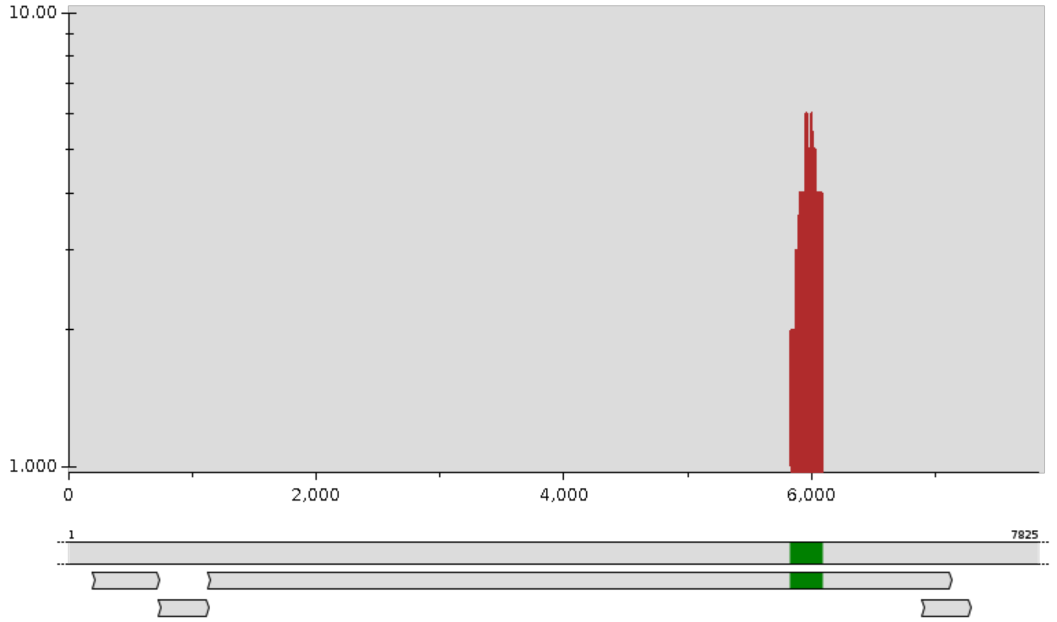

#### Assignment

| | |
|---|---|
| Type | Epiphyllum badnavirus 1 (Taxonomy ID: 2518008) |
| Reference Genome | NC_076247.1 |
| NT Identity (%) | 53.7879 |
| AA Identity (%) | 46.5909 |
| Number Of Stop Codons | 1 |
| Number Of CDS | 4 |

#### Alignment

| | |
|---|---|
| Alignment Score | 34.0 (NT) + 271.0 (AA) = 305.0 |
| Concordance (%) | 26.3158 |

| **Alignment Method** | Global, seeded, nucleotide + amino acids (AGA) |
|---|---|

#### Genome Region

Sequence starts at position 5831 and ends at position 6091 relative to NC_076247.1 reference sequence.

#### Alignment Detailed Statistics

| | Begin | End | Coverage | Score | Concordance | Matches | Identities | I/D/M/F* | Stop Codons |
|---|---|---|---|---|---|---|---|---|---|
| **NT** | **5831** | **6091** | **3.3%** | **34** | **6.5%** | **261 (98.9%)** | **142 (53.8%)** | **3/0** | |
| Mutations: | 5832T>G, 5835C>T, 5839G>C, 5840T>A, 5850G>A, 5851T>A, 5853C>T, 5859G>T, 5860A>T, 5863A>T, 5864G>C, 5865C>T, 5868A>C, 5870T>A, 5871C>T, 5874C>T, 5876A>T, 5878A>T, 5880T>G, 5882T>G, 5884A>G, 5887G>A, 5888A>G, 5889C>G, 5892A>G, 5893G>A, 5894A>G, 5895A>T, 5896T>G, 5897C>A, 5898C>T, 5901C>T, 5904T>A, 5905T>A, 5906G>A, 5910C>A, 5912C>T, 5913C>A, 5916C>T, 5918T>A, 5922C>G, 5923C>A, 5924C>G, 5926G>T, 5931A>C, 5933T>A, 5936A>T, 5937C>T, 5940A>G, 5942G>T, 5943G>C, 5944T>C, 5947G>A, 5949T>C, 5953C>T, 5955A>C, 5958C>T, 5960G>A, 5961A>G, 5964G>A, 5966A>C, 5967G>C, 5973C>T, 5976T>G, 5979T>A, 5982G>T, 5986C>A, 5987A>T, 5989C>G, 5990G>A, 5991A>C, 5992A>C, 5993A>T, 5994G>T, 5998G>A, 6000C>T, 6002A>G, 6003C>G, 6004T>G, 6005G>T, 6006T>G, 6011G>A, 6012A>T, 6012_6013insTTC, 6013G>T, 6015C>T, 6018G>C, 6019T>A, 6020C>A, 6021A>T, 6023G>C, 6024C>A, 6028G>A, 6030A>G, 6032C>T, 6033A>G, 6038A>T, 6042T>A, 6046G>T, 6051C>T, 6052C>T, 6055G>A, 6059T>A, 6061A>T, 6062G>C, 6063T>C, 6064A>C, 6065A>G, 6066C>T, 6068C>G, 6072A>G, 6075T>A, 6078C>A, 6082G>A, 6083C>T, 6084A>G, 6085C>A, 6086A>G, 6087G>T, 6091C>T | | | | | | | | |

##### CDS

| | Begin | End | Coverage | Score | Concordance | Matches | Identities | I/D/M/F* | Stop Codons |
|---|---|---|---|---|---|---|---|---|---|
| QKM20_gp3 | 1569 | 1655 | 4.3% | 271 | 41.8% | 87 (98.9%) | 41 (46.6%) | 1/0/0/0 | 1 |
| Protein mutations: | V1571H (5839G>C 5840T>A), F1575I (5851T>A 5853C>T), K1578* (5860A>T), F1581Y (5870T>A 5871C>T), Q1583L (5876A>T), I1584L (5878A>T 5880T>G), M1585R (5882T>G), M1586V (5884A>G), D1587R (5887G>A 5888A>G 5889C>G), E1589S (5893G>A 5894A>G 5895A>T), S1590D (5896T>G 5897C>A 5898C>T), W1593K (5905T>A 5906G>A), A1595V (5912C>T 5913C>A), L1597Q (5918T>A), P1599R (5923C>A 5924C>G), D1600Y (5926G>T), L1602H (5933T>A), Y1603F (5936A>T 5937C>T), W1605F (5942G>T 5943G>C), V1607I (5947G>A 5949T>C), P1609S (5953C>T 5955A>C), G1611E (5960G>A 5961A>G), K1613T (5966A>C 5967G>C), Q1620I (5986C>A 5987A>T), R1621D (5989C>G 5990G>A 5991A>C), K1622L (5992A>C 5993A>T 5994G>T), D1624N (5998G>A 6000C>T), N1625R (6002A>G 6003C>G), C1626V (6004T>G 6005G>T 6006T>G), R1628N (6011G>A 6012A>T), R1628_D1629insF (6012_6013insTTC), D1629Y (6013G>T 6015C>T), S1631N (6019T>A 6020C>A 6021A>T), G1632A (6023G>C 6024C>A), V1634M (6028G>A 6030A>G), A1635V (6032C>T 6033A>G), Y1637F (6038A>T), D1640Y (6046G>T), V1643I (6055G>A), F1644Y (6059T>A), N1646R (6064A>C 6065A>G 6066C>T), T1647S (6068C>G), D1649E (6075T>A), D1650E (6078C>A), A1652M (6082G>A 6083C>T 6084A>G), Q1653S (6085C>A 6086A>G 6087G>T) | | | | | | | | |
| Codon mutations: | AAT1568.AG (5832T>G), GCC1569GCT (5835C>T), GTC1571CAC (5839G>C 5840T>A), AAG1574AAA (5850G>A), TTC1575ATT (5851T>A 5853C>T), CTG1577CTT (5859G>T), AAG1578TAG (5860A>T), AGC1579TCT (5863A>T 5864G>C 5865C>T), GGA1580GGC (5868A>C), TTC1581TAT (5870T>A 5871C>T), CAC1582CAT (5874C>T), CAG1583CTG (5876A>T), ATT1584TTG (5878A>T 5880T>G), ATG1585AGG (5882T>G), ATG1586GTG (5884A>G), GAC1587AGG (5887G>A 5888A>G 5889C>G), GAA1588GAG (5892A>G), GAA1589AGT (5893G>A 5894A>G 5895A>T), TCC1590GAT (5896T>G 5897C>A 5898C>T), ATC1591ATT (5901C>T), CCT1592CCA (5904T>A), TGG1593AAG (5905T>A 5906G>A), ACC1594ACA (5910C>A), GCC1595GTA (5912C>T 5913C>A), TTC1596TTT (5916C>T), CTG1597CAG (5918T>A), ACC1598ACG (5922C>G), CCA1599AGA (5923C>A 5924C>G), GAT1600TAT (5926G>T), GGA1601GGC (5931A>C), CTT1602CAT (5933T>A), TAC1603TTT (5936A>T 5937C>T), GAA1604GAG (5940A>G), TGG1605TTC (5942G>T 5943G>C), TTA1606CTA (5944T>C), GTT1607ATC (5947G>A 5949T>C), CCA1609TCC (5953C>T 5955A>C), TTC1610TTT (5958C>T), GGA1611GAG (5960G>A 5961A>G), TTG1612TTA (5964G>A), AAG1613ACC (5966A>C 5967G>C), GCC1615GCT (5973C>T), CCT1616CCG (5976T>G), GCT1617GCA (5979T>A), GTG1618GTT (5982G>T), CAA1620ATA (5986C>A 5987A>T), CGA1621GAC (5989C>G 5990G>A 5991A>C), AAG1622CTT (5992A>C 5993A>T 5994G>T), GAC1624AAT (5998G>A 6000C>T), AAC1625AGG (6002A>G 6003C>G), TGT1626GTG (6004T>G 6005G>T 6006T>G), AGA1628AAT (6011G>A 6012A>T), AGA1628_GAC1629insTTC (6012_6013insTTC), GAC1629TAT (6013G>T 6015C>T), CTG1630CTC (6018G>C), TCA1631AAT (6019T>A 6020C>A 6021A>T), GGC1632GCA (6023G>C 6024C>A), GTA1634ATG (6028G>A 6030A>G), GCA1635GTG (6032C>T 6033A>G), TAC1637TTC (6038A>T), ATT1638ATA (6042T>A), GAC1640TAC (6046G>T), ATC1641ATT (6051C>T), CTG1642TTG (6052C>T), GTC1643ATC (6055G>A), TTC1644TAC (6059T>A), AGT1645TCC (6061A>T 6062G>C 6063T>C), AAC1646CGT (6064A>C 6065A>G 6066C>T), ACT1647AGT (6068C>G), GAA1648GAG (6072A>G), GAT1649GAA (6075T>A), GAC1650GAA (6078C>A), GCA1652ATG (6082G>A 6083C>T 6084A>G), CAG1653AGT (6085C>A 6086A>G 6087G>T), CTG1655T.. (6091C>T) | | | | | | | | |

##### Proteins

| | Begin | End | Coverage | Score | Concordance | Matches | Identities | I/D/M/F* | Stop Codons |
|---|---|---|---|---|---|---|---|---|---|
| polyprotein (YP_010797894.1) | 1569 | 1655 | 4.3% | 271 | 41.8% | 87 (98.9%) | 41 (46.6%) | 1/0/0/0 | 1 |
| Protein mutations: | V1571H (5839G>C 5840T>A), F1575I (5851T>A 5853C>T), K1578* (5860A>T), F1581Y (5870T>A 5871C>T), Q1583L (5876A>T), I1584L (5878A>T 5880T>G), M1585R (5882T>G), M1586V (5884A>G), D1587R (5887G>A 5888A>G 5889C>G), E1589S (5893G>A 5894A>G 5895A>T), S1590D (5896T>G 5897C>A 5898C>T), W1593K (5905T>A 5906G>A), A1595V (5912C>T 5913C>A), L1597Q (5918T>A), P1599R (5923C>A 5924C>G), D1600Y (5926G>T), L1602H (5933T>A), Y1603F (5936A>T 5937C>T), W1605F (5942G>T 5943G>C), V1607I (5947G>A 5949T>C), P1609S (5953C>T 5955A>C), G1611E (5960G>A 5961A>G), K1613T (5966A>C 5967G>C), Q1620I (5986C>A 5987A>T), R1621D (5989C>G 5990G>A 5991A>C), K1622L (5992A>C 5993A>T 5994G>T), D1624N (5998G>A 6000C>T), N1625R (6002A>G 6003C>G), C1626V (6004T>G 6005G>T 6006T>G), R1628N (6011G>A 6012A>T), R1628_D1629insF (6012_6013insTTC), D1629Y (6013G>T 6015C>T), S1631N (6019T>A 6020C>A 6021A>T), G1632A (6023G>C 6024C>A), V1634M (6028G>A 6030A>G), A1635V (6032C>T 6033A>G), Y1637F (6038A>T), D1640Y (6046G>T), V1643I (6055G>A), F1644Y (6059T>A), N1646R (6064A>C 6065A>G 6066C>T), T1647S (6068C>G), D1649E (6075T>A), D1650E (6078C>A), A1652M (6082G>A 6083C>T 6084A>G), Q1653S (6085C>A 6086A>G 6087G>T) | | | | | | | | |
| Codon mutations: | AAT1568.AG (5832T>G), GCC1569GCT (5835C>T), GTC1571CAC (5839G>C 5840T>A), AAG1574AAA (5850G>A), TTC1575ATT (5851T>A 5853C>T), CTG1577CTT (5859G>T), AAG1578TAG (5860A>T), AGC1579TCT (5863A>T 5864G>C 5865C>T), GGA1580GGC (5868A>C), TTC1581TAT (5870T>A 5871C>T), CAC1582CAT (5874C>T), CAG1583CTG (5876A>T), ATT1584TTG (5878A>T 5880T>G), ATG1585AGG (5882T>G), ATG1586GTG (5884A>G), GAC1587AGG (5887G>A 5888A>G 5889C>G), GAA1588GAG (5892A>G), GAA1589AGT (5893G>A 5894A>G 5895A>T), TCC1590GAT (5896T>G 5897C>A 5898C>T), ATC1591ATT (5901C>T), CCT1592CCA (5904T>A), TGG1593AAG (5905T>A 5906G>A), ACC1594ACA (5910C>A), GCC1595GTA (5912C>T 5913C>A), TTC1596TTT (5916C>T), CTG1597CAG (5918T>A), ACC1598ACG (5922C>G), CCA1599AGA (5923C>A 5924C>G), GAT1600TAT (5926G>T), GGA1601GGC (5931A>C), CTT1602CAT (5933T>A), TAC1603TTT (5936A>T 5937C>T), GAA1604GAG (5940A>G), TGG1605TTC (5942G>T 5943G>C), TTA1606CTA (5944T>C), GTT1607ATC (5947G>A 5949T>C), CCA1609TCC (5953C>T 5955A>C), TTC1610TTT (5958C>T), GGA1611GAG (5960G>A 5961A>G), TTG1612TTA (5964G>A), AAG1613ACC (5966A>C 5967G>C), GCC1615GCT (5973C>T), CCT1616CCG (5976T>G), GCT1617GCA (5979T>A), GTG1618GTT (5982G>T), CAA1620ATA (5986C>A 5987A>T), CGA1621GAC (5989C>G 5990G>A 5991A>C), AAG1622CTT (5992A>C 5993A>T 5994G>T), GAC1624AAT (5998G>A 6000C>T), AAC1625AGG (6002A>G 6003C>G), TGT1626GTG (6004T>G 6005G>T 6006T>G), AGA1628AAT (6011G>A 6012A>T), AGA1628_GAC1629insTTC (6012_6013insTTC), GAC1629TAT (6013G>T 6015C>T), CTG1630CTC (6018G>C), TCA1631AAT (6019T>A 6020C>A 6021A>T), GGC1632GCA (6023G>C 6024C>A), GTA1634ATG (6028G>A 6030A>G), GCA1635GTG (6032C>T 6033A>G), TAC1637TTC (6038A>T), ATT1638ATA (6042T>A), GAC1640TAC (6046G>T), ATC1641ATT (6051C>T), CTG1642TTG (6052C>T), GTC1643ATC (6055G>A), TTC1644TAC (6059T>A), AGT1645TCC (6061A>T 6062G>C 6063T>C), AAC1646CGT (6064A>C 6065A>G 6066C>T), ACT1647AGT (6068C>G), GAA1648GAG (6072A>G), GAT1649GAA (6075T>A), GAC1650GAA (6078C>A), GCA1652ATG (6082G>A 6083C>T 6084A>G), CAG1653AGT (6085C>A 6086A>G 6087G>T), CTG1655T.. (6091C>T) | | | | | | | | |

*: Inserts / Deletes / Misaligned / Frameshifts

#### Analysis details

This analysis was performed with panviral2.64

### NGS Details (UN24): Chinaberry tree badnavirus 1

#### Assembly

| | |
|---|---|
| **Coverage Length** | 261 (1 contig(s)) |
| **Depth Of Coverage** | 2.7 |
| **Number Of Reads** | 7 |
| **Reads Per Million** | 0.14 rpm (after QC) |
| **Ambiguities** | 0 |
| **Assembly Method** | de novo + reference guided assembly |
| **Consensus Caller** | Bcf Tools |

#### Coverage Map

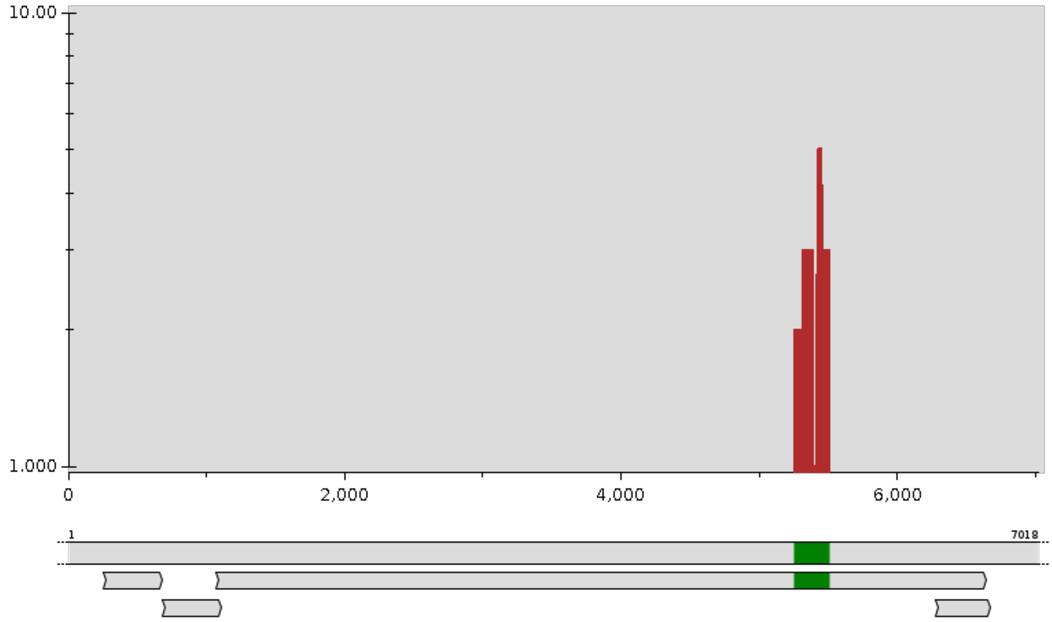

#### Assignment

| | |
|---|---|
| **Type** | Chinaberry tree badnavirus 1 (Taxonomy ID: 2908099) |
| **Reference Genome** | NC_077165.1 |
| **NT Identity (%)** | 59.8485 |
| **AA Identity (%)** | 47.7273 |
| **Number Of Stop Codons** | 1 |
| **Number Of CDS** | 4 |

#### Alignment

| | |
|---|---|
| **Alignment Score** | 98.0 (NT) + 310.0 (AA) = 408.0 |
| **Concordance (%)** | 34.9914 |

| **Alignment Method** | Global, seeded, nucleotide + amino acids (AGA) |
|---|---|

#### Genome Region

Sequence starts at position 5253 and ends at position 5513 relative to NC_077165.1 reference sequence.

#### Alignment Detailed Statistics

| | Begin | End | Coverage | Score | Concordance | Matches | Identities | I/D/M/F* | Stop Codons |
|---|---|---|---|---|---|---|---|---|---|
| **NT** | **5253** | **5513** | **3.7%** | **98** | **18.8%** | **261 (98.9%)** | **158 (59.8%)** | **3/0** | |
| Mutations: | 5255G>A, 5256T>A, 5264G>T, 5265A>T, 5266A>G, 5270G>C, 5273C>T, 5275T>A, 5282A>G, 5283G>T, 5285A>C, 5286G>A, 5287C>G, 5288C>G, 5289A>G, 5292G>A, 5293A>G, 5294C>G, 5295C>G, 5296C>A, 5298G>T, 5299A>G, 5300G>C, 5301T>A, 5302C>A, 5304G>A, 5306A>T, 5307G>C, 5308A>T, 5309A>C, 5310T>A, 5311G>A, 5316G>A, 5325G>A, 5326T>C, 5327T>A, 5328C>A, 5329C>G, 5331G>T, 5336G>C, 5338T>A, 5341A>T, 5347G>T, 5348G>T, 5349C>T, 5357T>G, 5358C>T, 5360A>G, 5363C>T, 5369A>G, 5371A>C, 5374A>C, 5380C>T, 5381A>C, 5387A>C, 5390C>T, 5391C>A, 5392A>T, 5393A>G, 5395G>A, 5396A>C, 5397A>T, 5398A>T, 5403G>A, 5405C>T, 5407A>G, 5409T>A, 5410G>T, 5411C>T, 5417G>A, 5418G>T, 5419G>C, 5420_5421insTAT, 5421A>C, 5422C>T, 5424G>T, 5426G>T, 5428A>T, 5429G>A, 5432C>T, 5433A>G, 5435T>G, 5437C>T, 5443A>T, 5444C>T, 5447A>T, 5453C>T, 5456C>A, 5460G>A, 5462C>T, 5464T>A, 5469A>C, 5471G>A, 5474C>T, 5480G>A, 5487G>A, 5488C>T, 5489A>G, 5492C>T, 5496G>T, 5498A>G, 5503A>T, 5504A>T, 5511G>C | | | | | | | | |

##### CDS

| | Begin | End | Coverage | Score | Concordance | Matches | Identities | I/D/M/F* | Stop Codons |
|---|---|---|---|---|---|---|---|---|---|
| QKV46_gp3 | 1395 | 1481 | 4.7% | 310 | 48.1% | 87 (98.9%) | 42 (47.7%) | 1/0/0/0 | 1 |
| Protein mutations: | F1396I (5256T>A), K1399* (5265A>T 5266A>G), F1402Y (5275T>A), V1405F (5283G>T 5285A>C), A1406R (5286G>A 5287C>G 5288C>G), M1407V (5289A>G), D1408R (5292G>A 5293A>G 5294C>G), P1409E (5295C>G 5296C>A), E1410C (5298G>T 5299A>G 5300G>C), S1411N (5301T>A 5302C>A), V1412I (5304G>A 5306A>T), E1413L (5307G>C 5308A>T 5309A>C), W1414K (5310T>A 5311G>A), A1416T (5316G>A), V1419T (5325G>A 5326T>C 5327T>A), P1420R (5328C>A 5329C>G), D1421Y (5331G>T), L1423H (5338T>A), Y1424F (5341A>T), W1426F (5347G>T 5348G>T), I1429M (5357T>G), P1430S (5358C>T 5360A>G), K1434T (5371A>C), N1435T (5374A>C), P1437L (5380C>T 5381A>C), Q1441M (5391C>A 5392A>T 5393A>G), R1442N (5395G>A 5396A>C), K1443L (5397A>T 5398A>T), D1445N (5403G>A 5405C>T), Q1446R (5407A>G), C1447I (5409T>A 5410G>T 5411C>T), G1450S (5418G>T 5419G>C), G1450_T1451insY (5420_5421insTAT), T1451L (5421A>C 5422C>T), E1452Y (5424G>T 5426G>T), K1453I (5428A>T 5429G>A), I1455V (5433A>G 5435T>G), A1456V (5437C>T), Y1458F (5443A>T 5444C>T), V1464I (5460G>A 5462C>T), F1465Y (5464T>A), K1467Q (5469A>C 5471G>A), A1473M (5487G>A 5488C>T 5489A>G), V1476L (5496G>T 5498A>G), K1478I (5503A>T 5504A>T), E1481Q (5511G>C) | | | | | | | | |
| Codon mutations: | AAG1395AAA (5255G>A), TTT1396ATT (5256T>A), CTG1398CTT (5264G>T), AAA1399TGA (5265A>T 5266A>G), TCG1400TCC (5270G>C), GGC1401GGT (5273C>T), TTT1402TAT (5275T>A), CAA1404CAG (5282A>G), GTA1405TTC (5283G>T 5285A>C), GCC1406AGG (5286G>A 5287C>G 5288C>G), ATG1407GTG (5289A>G), GAC1408AGG (5292G>A 5293A>G 5294C>G), CCG1409GAG (5295C>G 5296C>A), GAG1410TGC (5298G>T 5299A>G 5300G>C), TCT1411AAT (5301T>A 5302C>A), GTA1412ATT (5304G>A 5306A>T), GAA1413CTC (5307G>C 5308A>T 5309A>C), TGG1414AAG (5310T>A 5311G>A), GCA1416ACA (5316G>A), GTT1419ACA (5325G>A 5326T>C 5327T>A), CCA1420AGA (5328C>A 5329C>G), GAT1421TAT (5331G>T), GGG1422GGC (5336G>C), CTT1423CAT (5338T>A), TAT1424TTT (5341A>T), TGG1426TTT (5347G>T 5348G>T), CTG1427TTG (5349C>T), ATT1429ATG (5357T>G), CCA1430TCG (5358C>T 5360A>G), TTC1431TTT (5363C>T), TTA1433TTG (5369A>G), AAA1434ACA (5371A>C), AAT1435ACT (5374A>C), CCA1437CTC (5380C>T 5381A>C), ATA1439ATC (5387A>C), TTC1440TTT (5390C>T), CAA1441ATG (5391C>A 5392A>T 5393A>G), AGA1442AAC (5395G>A 5396A>C), AAG1443TTG (5397A>T 5398A>T), GAC1445AAT (5403G>A 5405C>T), CAA1446CGA (5407A>G), TGC1447ATT (5409T>A 5410G>T 5411C>T), AAG1449AAA (5417G>A), GGA1450TCA (5418G>T 5419G>C), GGA1450_ACA1451insTAT (5420_5421insTAT), ACA1451CTA (5421A>C 5422C>T), GAG1452TAT (5424G>T 5426G>T), AAG1453ATA (5428A>T 5429G>A), TTC1454TTT (5432C>T), ATT1455GTG (5433A>G 5435T>G), GCG1456GTG (5437C>T), TAC1458TTT (5443A>T 5444C>T), ATA1459ATT (5447A>T), GAC1461GAT (5453C>T), ATC1462ATA (5456C>A), GTC1464ATT (5460G>A 5462C>T), TTT1465TAT (5464T>A), AAG1467CAA (5469A>C 5471G>A), AAC1468AAT (5474C>T), GAG1470GAA (5480G>A), GCA1473ATG (5487G>A 5488C>T 5489A>G), GGC1474GGT (5492C>T), GTA1476TTG (5496G>T 5498A>G), AAA1478ATT (5503A>T 5504A>T), GAA1481CAA (5511G>C) | | | | | | | | |

##### Proteins

| | Begin | End | Coverage | Score | Concordance | Matches | Identities | I/D/M/F* | Stop Codons |
|---|---|---|---|---|---|---|---|---|---|
| ORF3 (YP_010805662.1) | 1395 | 1481 | 4.7% | 310 | 48.1% | 87 (98.9%) | 42 (47.7%) | 1/0/0/0 | 1 |
| Protein mutations: | F1396I (5256T>A), K1399* (5265A>T 5266A>G), F1402Y (5275T>A), V1405F (5283G>T 5285A>C), A1406R (5286G>A 5287C>G 5288C>G), M1407V (5289A>G), D1408R (5292G>A 5293A>G 5294C>G), P1409E (5295C>G 5296C>A), E1410C (5298G>T 5299A>G 5300G>C), S1411N (5301T>A 5302C>A), V1412I (5304G>A 5306A>T), E1413L (5307G>C 5308A>T 5309A>C), W1414K (5310T>A 5311G>A), A1416T (5316G>A), V1419T (5325G>A 5326T>C 5327T>A), P1420R (5328C>A 5329C>G), D1421Y (5331G>T), L1423H (5338T>A), Y1424F (5341A>T), W1426F (5347G>T 5348G>T), I1429M (5357T>G), P1430S (5358C>T 5360A>G), K1434T (5371A>C), N1435T (5374A>C), P1437L (5380C>T 5381A>C), Q1441M (5391C>A 5392A>T 5393A>G), R1442N (5395G>A 5396A>C), K1443L (5397A>T 5398A>T), D1445N (5403G>A 5405C>T), Q1446R (5407A>G), C1447I (5409T>A 5410G>T 5411C>T), G1450S (5418G>T 5419G>C), G1450_T1451insY (5420_5421insTAT), T1451L (5421A>C 5422C>T), E1452Y (5424G>T 5426G>T), K1453I (5428A>T 5429G>A), I1455V (5433A>G 5435T>G), A1456V (5437C>T), Y1458F (5443A>T 5444C>T), V1464I (5460G>A 5462C>T), F1465Y (5464T>A), K1467Q (5469A>C 5471G>A), A1473M (5487G>A 5488C>T 5489A>G), V1476L (5496G>T 5498A>G), K1478I (5503A>T 5504A>T), E1481Q (5511G>C) | | | | | | | | |
| Codon mutations: | AAG1395AAA (5255G>A), TTT1396ATT (5256T>A), CTG1398CTT (5264G>T), AAA1399TGA (5265A>T 5266A>G), TCG1400TCC (5270G>C), GGC1401GGT (5273C>T), TTT1402TAT (5275T>A), CAA1404CAG (5282A>G), GTA1405TTC (5283G>T 5285A>C), GCC1406AGG (5286G>A 5287C>G 5288C>G), ATG1407GTG (5289A>G), GAC1408AGG (5292G>A 5293A>G 5294C>G), CCG1409GAG (5295C>G 5296C>A), GAG1410TGC (5298G>T 5299A>G 5300G>C), TCT1411AAT (5301T>A 5302C>A), GTA1412ATT (5304G>A 5306A>T), GAA1413CTC (5307G>C 5308A>T 5309A>C), TGG1414AAG (5310T>A 5311G>A), GCA1416ACA (5316G>A), GTT1419ACA (5325G>A 5326T>C 5327T>A), CCA1420AGA (5328C>A 5329C>G), GAT1421TAT (5331G>T), GGG1422GGC (5336G>C), CTT1423CAT (5338T>A), TAT1424TTT (5341A>T), TGG1426TTT (5347G>T 5348G>T), CTG1427TTG (5349C>T), ATT1429ATG (5357T>G), CCA1430TCG (5358C>T 5360A>G), TTC1431TTT (5363C>T), TTA1433TTG (5369A>G), AAA1434ACA (5371A>C), AAT1435ACT (5374A>C), CCA1437CTC (5380C>T 5381A>C), ATA1439ATC (5387A>C), TTC1440TTT (5390C>T), CAA1441ATG (5391C>A 5392A>T 5393A>G), AGA1442AAC (5395G>A 5396A>C), AAG1443TTG (5397A>T 5398A>T), GAC1445AAT (5403G>A 5405C>T), CAA1446CGA (5407A>G), TGC1447ATT (5409T>A 5410G>T 5411C>T), AAG1449AAA (5417G>A), GGA1450TCA (5418G>T 5419G>C), GGA1450_ACA1451insTAT (5420_5421insTAT), ACA1451CTA (5421A>C 5422C>T), GAG1452TAT (5424G>T 5426G>T), AAG1453ATA (5428A>T 5429G>A), TTC1454TTT (5432C>T), ATT1455GTG (5433A>G 5435T>G), GCG1456GTG (5437C>T), TAC1458TTT (5443A>T 5444C>T), ATA1459ATT (5447A>T), GAC1461GAT (5453C>T), ATC1462ATA (5456C>A), GTC1464ATT (5460G>A 5462C>T), TTT1465TAT (5464T>A), AAG1467CAA (5469A>C 5471G>A), AAC1468AAT (5474C>T), GAG1470GAA (5480G>A), GCA1473ATG (5487G>A 5488C>T 5489A>G), GGC1474GGT (5492C>T), GTA1476TTG (5496G>T 5498A>G), AAA1478ATT (5503A>T 5504A>T), GAA1481CAA (5511G>C) | | | | | | | | |

*: Inserts / Deletes / Misaligned / Frameshifts

#### Analysis details

This analysis was performed with panviral2.64

### NGS Details (UN24): Gihfavirus pelohabitans

#### Assembly

| | |
|---|---|
| **Coverage Length** | 278 (1 contig(s)) |
| **Depth Of Coverage** | 2.4 |
| **Number Of Reads** | 6 |
| **Reads Per Million** | 0.12 rpm (after QC) |
| **Ambiguities** | 0 |
| **Assembly Method** | de novo + reference guided assembly |
| **Consensus Caller** | Bcf Tools |

#### Coverage Map

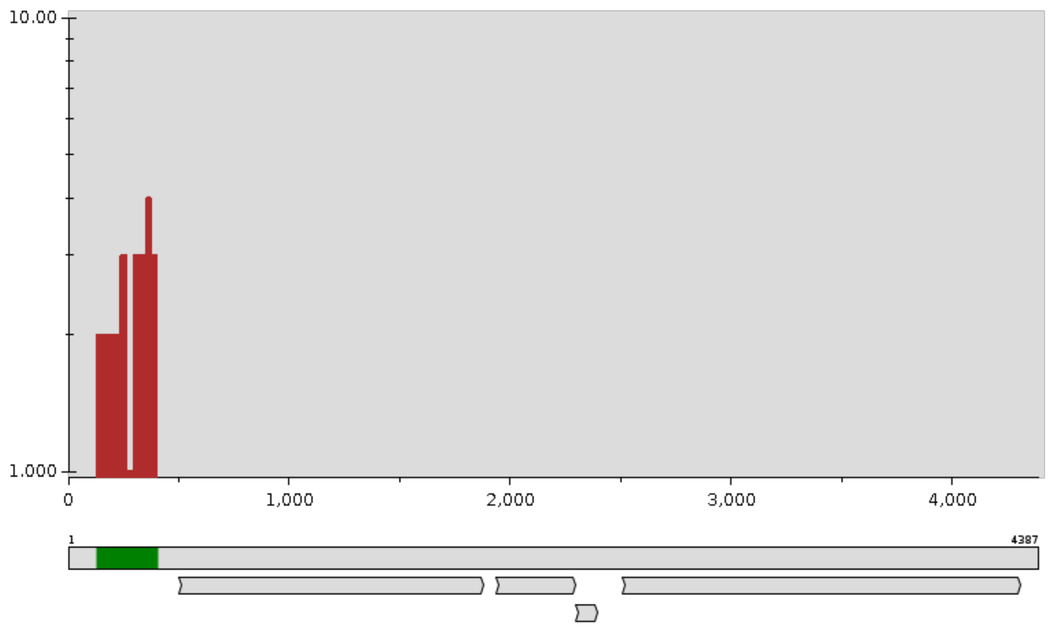

#### Assignment

| | |
|---|---|
| **Type** | Gihfavirus pelohabitans (Taxonomy ID: 2844652) |
| **Reference Genome** | NC_074583.1 |
| **NT Identity (%)** | 98.2014 |
| **AA Identity (%)** | 0.0 |
| **Number Of Stop Codons** | 0 |
| **Number Of CDS** | 4 |

#### Alignment

| | |
|---|---|
| **Alignment Score** | 536.0 (NT) + 0.0 (AA) = 536.0 |
| **Concordance (%)** | 96.4029 |

| Alignment Method | Global, seeded, nucleotide + amino acids (AGA) |
|---|---|

#### Genome Region

Sequence starts at position 127 and ends at position 404 relative to NC_074583.1 reference sequence.

#### Alignment Detailed Statistics

| | Begin | End | Coverage | Score | Concordance | Matches | Identities | I/D/M/F* | Stop Codons |
|---|---|---|---|---|---|---|---|---|---|
| NT | 127 | 404 | 6.3% | 536 | 96.4% | 278 (100%) | 273 (98.2%) | 0/0 | |
| Mutations: | 168G>A, 169A>C, 178T>G, 179C>T, 182T>C | | | | | | | | |

**CDS**

**Proteins**

*: Inserts / Deletes / Misaligned / Frameshifts

#### Analysis details

This analysis was performed with panviral2.64

### NGS Details (UN24): Duamitovirus peex1

#### Assembly

| | |
|---|---|
| Coverage Length | 264 (1 contig(s)) |
| Depth Of Coverage | 3.1 |
| Number Of Reads | 6 |
| Reads Per Million | 0.12 rpm (after QC) |
| Ambiguities | 0 |
| Assembly Method | de novo + reference guided assembly |
| Consensus Caller | Bcf Tools |

#### Coverage Map

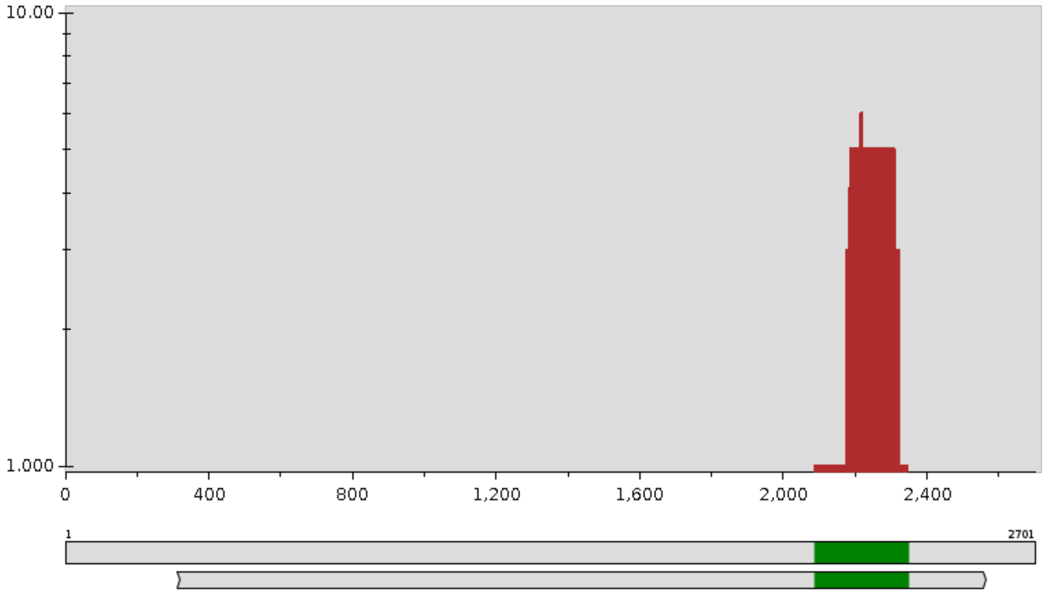

#### Assignment

| | |
|---|---|
| Type | Duamitovirus peex1 (Taxonomy ID: 2955799) |
| Reference Genome | NC_076525.1 |
| NT Identity (%) | 72.3485 |
| AA Identity (%) | 64.7727 |
| Number Of Stop Codons | 0 |
| Number Of CDS | 1 |

#### Alignment

| | |
|---|---|
| Alignment Score | 236.0 (NT) + 421.0 (AA) = 657.0 |
| Concordance (%) | 55.7725 |

| Alignment Method | Global, seeded, nucleotide + amino acids (AGA) |
|---|---|

#### Genome Region

Sequence starts at position 2087 and ends at position 2350 relative to NC_076525.1 reference sequence.

#### Alignment Detailed Statistics

| | Begin | End | Coverage | Score | Concordance | Matches | Identities | I/D/M/F* | Stop Codons |
|---|---|---|---|---|---|---|---|---|---|
| **NT** | **2087** | **2350** | **9.8%** | **236** | **44.7%** | **264 (100%)** | **191 (72.3%)** | **0/0** | |
| Mutations: | 2091T>A, 2095T>G, 2096A>G, 2098C>T, 2102G>T, 2104G>T, 2105G>A, 2109T>C, 2111T>A, 2119G>T, 2120G>T, 2133G>T, 2134G>T, 2135A>T, 2136A>G, 2137A>G, 2138G>A, 2139C>A, 2145G>A, 2156A>G, 2158G>A, 2159A>G, 2161G>C, 2162T>G, 2165G>A, 2171C>A, 2173C>A, 2180T>G, 2184A>G, 2188A>C, 2189G>T, 2190T>G, 2195A>G, 2202G>C, 2210G>A, 2211T>A, 2213G>C, 2214A>T, 2216A>T, 2223C>G, 2225A>T, 2226C>T, 2231G>A, 2232T>A, 2234A>T, 2240T>A, 2245A>G, 2246T>G, 2249G>A, 2255A>T, 2258A>G, 2261C>G, 2268G>A, 2270A>C, 2272G>T, 2274C>A, 2283G>A, 2292C>T, 2294T>G, 2295C>A, 2296G>C, 2297A>T, 2306A>T, 2315C>T, 2321G>A, 2325A>T, 2326A>T, 2327T>A, 2330C>A, 2333G>A, 2336A>T, 2345A>T, 2348C>T | | | | | | | | |
| **CDS** | | | | | | | | | |
| RdRp | 593 | 680 | 11.7% | 421 | 63.6% | 88 (100%) | 57 (64.8%) | 0/0/0/0 | 0 |
| Protein mutations: | Y594N (2091T>A), I595R (2095T>G 2096A>G), S596L (2098C>T), Q597H (2102G>T), R598L (2104G>T 2105G>A), S600P (2109T>C 2111T>A), W603F (2119G>T 2120G>T), G608F (2133G>T 2134G>T 2135A>T), K609G (2136A>G 2137A>G 2138G>A), P610T (2139C>A), D612N (2145G>A), R616K (2158G>A 2159A>G), G617A (2161G>C 2162T>G), A621D (2173C>A), K625E (2184A>G), E626A (2188A>C 2189G>T), L627V (2190T>G), E631Q (2202G>C), L634I (2211T>A 2213G>C), I635F (2214A>T 2216A>T), Q638D (2223C>G 2225A>T), L641I (2232T>A 2234A>T), H645R (2245A>G 2246T>G), V653I (2268G>A 2270A>C), R654L (2272G>T), H655N (2274C>A), E658K (2283G>A), R662T (2295C>A 2296G>C 2297A>T), N672L (2325A>T 2326A>T 2327T>A), E675D (2336A>T) | | | | | | | | |
| Codon mutations: | TAC594AAC (2091T>A), ATA595AGG (2095T>G 2096A>G), TCA596TTA (2098C>T), CAG597CAT (2102G>T), CGG598CTA (2104G>T 2105G>A), TCT600CCA (2109T>C 2111T>A), TGG603TTT (2119G>T 2120G>T), GGA608TTT (2133G>T 2134G>T 2135A>T), AAG609GGA (2136A>G 2137A>G 2138G>A), CCC610ACC (2139C>A), GAT612AAT (2145G>A), TTA615TTG (2156A>G), AGA616AAG (2158G>A 2159A>G), GGT617GCG (2161G>C 2162T>G), AAG618AAA (2165G>A), ATC620ATA (2171C>A), GCT621GAT (2173C>A), CTT623CTG (2180T>G), AAG625GAG (2184A>G), GAG626GCT (2188A>C 2189G>T), TTA627GTA (2190T>G), AAA628AAG (2195A>G), GAG631CAG (2202G>C), CAG633CAA (2210G>A), TTG634ATC (2211T>A 2213G>C), ATA635TTT (2214A>T 2216A>T), CAA638GAT (2223C>G 2225A>T), CTG639TTG (2226C>T), GTG640GTA (2231G>A), TTA641ATT (2232T>A 2234A>T), GGT643GGA (2240T>A), CAT645CGG (2245A>G 2246T>G), GAG646GAA (2249G>A), CTA648CTT (2255A>T), GAA649GAG (2258A>G), CGC650CGG (2261C>G), GTA653ATC (2268G>A 2270A>C), CGT654CTT (2272G>T), CAT655AAT (2274C>A), GAA658AAA (2283G>A), CTT661TTG (2292C>T 2294T>G), CGA662ACT (2295C>A 2296G>C 2297A>T), TCA665TCT (2306A>T), CAC668CAT (2315C>T), GTG670GTA (2321G>A), AAT672TTA (2325A>T 2326A>T 2327T>A), TCC673TCA (2330C>A), CCG674CCA (2333G>A), GAA675GAT (2336A>T), ATA678ATT (2345A>T), GAC679GAT (2348C>T) | | | | | | | | |
| **Proteins** | | | | | | | | | |
| RNA-dependent RNA polymerase (YP_010798875.1) | 593 | 680 | 11.7% | 421 | 63.6% | 88 (100%) | 57 (64.8%) | 0/0/0/0 | 0 |
| Protein mutations: | Y594N (2091T>A), I595R (2095T>G 2096A>G), S596L (2098C>T), Q597H (2102G>T), R598L (2104G>T 2105G>A), S600P (2109T>C 2111T>A), W603F (2119G>T 2120G>T), G608F (2133G>T 2134G>T 2135A>T), K609G (2136A>G 2137A>G 2138G>A), P610T (2139C>A), D612N (2145G>A), R616K (2158G>A 2159A>G), G617A (2161G>C 2162T>G), A621D (2173C>A), K625E (2184A>G), E626A (2188A>C 2189G>T), L627V (2190T>G), E631Q (2202G>C), L634I (2211T>A 2213G>C), I635F (2214A>T 2216A>T), Q638D (2223C>G 2225A>T), L641I (2232T>A 2234A>T), H645R (2245A>G 2246T>G), V653I (2268G>A 2270A>C), R654L (2272G>T), H655N (2274C>A), E658K (2283G>A), R662T (2295C>A 2296G>C 2297A>T), N672L (2325A>T 2326A>T 2327T>A), E675D (2336A>T) | | | | | | | | |
| Codon mutations: | TAC594AAC (2091T>A), ATA595AGG (2095T>G 2096A>G), TCA596TTA (2098C>T), CAG597CAT (2102G>T), CGG598CTA (2104G>T 2105G>A), TCT600CCA (2109T>C 2111T>A), TGG603TTT (2119G>T 2120G>T), GGA608TTT (2133G>T 2134G>T 2135A>T), AAG609GGA (2136A>G 2137A>G 2138G>A), CCC610ACC (2139C>A), GAT612AAT (2145G>A), TTA615TTG (2156A>G), AGA616AAG (2158G>A 2159A>G), GGT617GCG (2161G>C 2162T>G), AAG618AAA (2165G>A), ATC620ATA (2171C>A), GCT621GAT (2173C>A), CTT623CTG (2180T>G), AAG625GAG (2184A>G), GAG626GCT (2188A>C 2189G>T), TTA627GTA (2190T>G), AAA628AAG (2195A>G), GAG631CAG (2202G>C), CAG633CAA (2210G>A), TTG634ATC (2211T>A 2213G>C), ATA635TTT (2214A>T 2216A>T), CAA638GAT (2223C>G 2225A>T), CTG639TTG (2226C>T), GTG640GTA (2231G>A), TTA641ATT (2232T>A 2234A>T), GGT643GGA (2240T>A), CAT645CGG (2245A>G 2246T>G), GAG646GAA (2249G>A), CTA648CTT (2255A>T), GAA649GAG (2258A>G), CGC650CGG (2261C>G), GTA653ATC (2268G>A 2270A>C), CGT654CTT (2272G>T), CAT655AAT (2274C>A), GAA658AAA (2283G>A), CTT661TTG (2292C>T 2294T>G), CGA662ACT (2295C>A 2296G>C 2297A>T), TCA665TCT (2306A>T), CAC668CAT (2315C>T), GTG670GTA (2321G>A), AAT672TTA (2325A>T 2326A>T 2327T>A), TCC673TCA (2330C>A), CCG674CCA (2333G>A), GAA675GAT (2336A>T), ATA678ATT (2345A>T), GAC679GAT (2348C>T) | | | | | | | | |

*: Inserts / Deletes / Misaligned / Frameshifts

#### Analysis details

This analysis was performed with panviral2.64

### NGS Details (UN24): Badnavirus venatheobromae

#### Assembly

| | |
|---|---|
| **Coverage Length** | 354 (1 contig(s)) |
| **Depth Of Coverage** | 2.3 |
| **Number Of Reads** | 6 |
| **Reads Per Million** | 0.12 rpm (after QC) |
| **Ambiguities** | 0 |
| **Assembly Method** | de novo + reference guided assembly |
| **Consensus Caller** | Bcf Tools |

#### Coverage Map

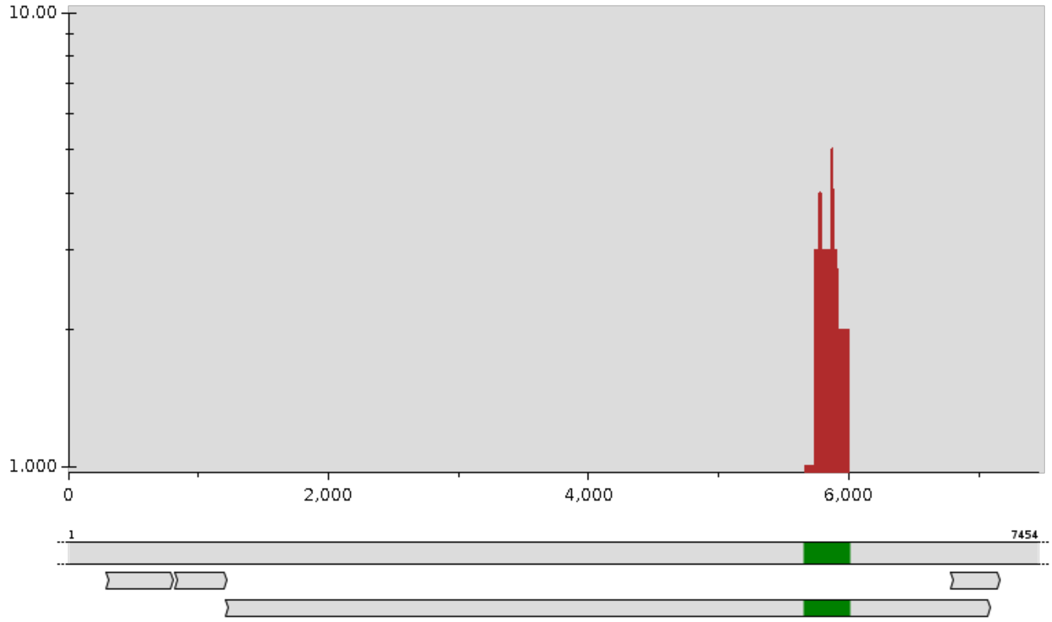

#### Assignment

| | |
|---|---|
| **Type** | Badnavirus venatheobromae (Taxonomy ID: 3047715) |
| **Reference Genome** | NC_033739.1 |
| **NT Identity (%)** | 54.0616 |
| **AA Identity (%)** | 42.8571 |
| **Number Of Stop Codons** | 0 |
| **Number Of CDS** | 4 |

#### Alignment

| | |
|---|---|
| **Alignment Score** | 52.0 (NT) + 369.0 (AA) = 421.0 |
| **Concordance (%)** | 26.9526 |

| Alignment Method | Global, seeded, nucleotide + amino acids (AGA) |
|---|---|

#### Genome Region

Sequence starts at position 5656 and ends at position 6009 relative to NC_033739.1 reference sequence.

#### Alignment Detailed Statistics

| | Begin | End | Coverage | Score | Concordance | Matches | Identities | I/D/M/F* | Stop Codons |
|---|---|---|---|---|---|---|---|---|---|
| **NT** | **5656** | **6009** | **4.7%** | **52** | **7.3%** | **354 (99.2%)** | **193 (54.1%)** | **3/0** | |
| Mutations: | 5659A>G, 5663A>C, 5666G>A, 5667A>G, 5673T>A, 5679T>C, 5681T>G, 5689A>G, 5690G>A, 5693C>A, 5696T>C, 5698T>A, 5702T>C, 5705A>G, 5706G>A, 5709G>A, 5710C>G, 5716A>C, 5717G>T, 5718G>C, 5721T>G, 5724T>G, 5725C>A, 5726A>T, 5727A>G, 5729T>G, 5731C>A, 5733T>A, 5734G>A, 5738A>T, 5741G>C, 5742T>A, 5744C>G, 5745T>A, 5748G>A, 5749T>C, 5750C>A, 5752C>A, 5753T>C, 5754G>T, 5755A>C, 5756T>A, 5759A>T, 5761T>A, 5762A>C, 5763T>A, 5765T>C, 5766G>A, 5768A>G, 5770G>A, 5771G>C, 5777C>G, 5783G>T, 5786C>T, 5789A>T, 5790T>C, 5792A>C, 5794A>C, 5795G>A, 5801T>C, 5805G>T, 5807C>T, 5809T>G, 5810T>C, 5813T>C, 5816A>G, 5817A>G, 5820A>C, 5821A>T, 5822G>C, 5826G>A, 5829A>C, 5831C>T, 5832T>G, 5833G>T, 5837T>C, 5840A>G, 5842G>A, 5843A>C, 5843_5844insCAT, 5844A>C, 5845C>T, 5846A>T, 5847G>A, 5848A>G, 5849A>G, 5850G>A, 5855T>C, 5858A>C, 5859G>C, 5860C>T, 5864T>A, 5866A>T, 5867T>C, 5868A>G, 5869T>C, 5876T>C, 5882A>C, 5885C>T, 5887T>A, 5888C>T, 5889T>A, 5890C>G, 5891A>T, 5896C>A, 5898G>A, 5899A>T, 5903A>G, 5907C>T, 5910G>C, 5912C>A, 5914G>A, 5915T>A, 5918T>C, 5919C>T, 5921A>G, 5922C>A, 5923A>G, 5926G>T, 5927G>T, 5928T>G, 5934C>A, 5936G>C, 5937A>C, 5939A>G, 5941G>T, 5942C>G, 5943C>A, 5944A>T, 5945G>C, 5948G>A, 5951T>C, 5952G>C, 5953G>A, 5955T>C, 5957A>C, 5958G>T, 5960C>A, 5961C>G, 5963G>C, 5966T>G, 5969G>A, 5970G>A, 5972T>C, 5976A>T, 5977T>G, 5978G>T, 5979G>T, 5981A>T, 5982A>T, 5984A>T, 5986C>G, 5989T>C, 5990C>T, 5991C>T, 5993A>T, 5994C>A, 5996A>G, 5997A>G, 5999T>G, 6002G>A, 6004T>A | | | | | | | | |

**CDS**

| | Begin | End | Coverage | Score | Concordance | Matches | Identities | I/D/M/F* | Stop Codons |
|---|---|---|---|---|---|---|---|---|---|
| B1U04_gp3 | 1483 | 1600 | 6.0% | 369 | 42.4% | 118 (99.2%) | 51 (42.9%) | 1/0/0/0 | 0 |
| Protein mutations: | N1483S (5659A>G), I1486V (5667A>G), S1488T (5673T>A), F1490L (5679T>C 5681T>G), K1493R (5689A>G 5690G>A), F1496Y (5698T>A), V1499I (5706G>A), A1500R (5709G>A 5710C>G), E1502A (5716A>C 5717G>T), E1503Q (5718G>C), S1504A (5721T>G), S1505D (5724T>G 5725C>A 5726A>T), I1506V (5727A>G 5729T>G), P1507Q (5731C>A), W1508K (5733T>A 5734G>A), F1511M (5742T>A 5744C>G), W1512R (5745T>A), V1513T (5748G>A 5749T>C 5750C>A), P1514H (5752C>A 5753T>C), D1515S (5754G>T 5755A>C 5756T>A), L1517H (5761T>A 5762A>C), Y1518N (5763T>A 5765T>C), E1519K (5766G>A 5768A>G), W1520Y (5770G>A 5771G>C), K1528T (5794A>C 5795G>A), A1532S (5805G>T 5807C>T), I1533S (5809T>G 5810T>C), R1536G (5817A>G), K1537L (5820A>C 5821A>T 5822G>C), D1539N (5826G>A), N1540H (5829A>C 5831C>T), C1541V (5832T>G 5833G>T), G1544D (5842G>A 5843A>C), G1544_T1545insH (5843_5844insCAT), T1545L (5844A>C 5845C>T 5846A>T), E1546R (5847G>A 5848A>G 5849A>G), E1547K (5850G>A), A1550L (5859G>C 5860C>T), Y1552F (5866A>T 5867T>C), I1553A (5868A>G 5869T>C), F1559Y (5887T>A 5888C>T), T1562N (5896C>A), E1563M (5898G>A 5899A>T), H1566Y (5907C>T), V1567L (5910G>C 5912C>A), R1568Q (5914G>A 5915T>A), Q1571R (5922C>A 5923A>G), R1572I (5926G>T 5927G>T), F1573V (5928T>G), Q1575N (5934C>A 5936G>C), I1576L (5937A>C 5939A>G), C1577L (5941G>T 5942C>G), Q1578I (5943C>A 5944A>T 5945G>C), G1581Q (5952G>C 5953G>A), V1583L (5958G>T 5960C>A), L1584V (5961C>G 5963G>C), S1585R (5966T>G), D1587N (5970G>A 5972T>C), M1589C (5976A>T 5977T>G 5978G>T), A1590S (5979G>T 5981A>T), I1591F (5982A>T 5984A>T), A1592G (5986C>G), V1593A (5989T>C 5990C>T), P1594S (5991C>T 5993A>T), Q1595K (5994C>A 5996A>G), I1596V (5997A>G 5999T>G), F1598Y (6004T>A) | | | | | | | | |
| Codon mutations: | AAT1483AGT (5659A>G), TCA1484TCC (5663A>C), AAG1485AAA (5666G>A), ATT1486GTT (5667A>G), TCC1488ACC (5673T>A), TTT1490CTG (5679T>C 5681T>G), AAG1493AGA (5689A>G 5690G>A), TCC1494TCA (5693C>A), GGT1495GGC (5696T>C), TTT1496TAT (5698T>A), CAT1497CAC (5702T>C), CAA1498CAG (5705A>G), GTA1499ATA (5706G>A), GCA1500AGA (5709G>A 5710C>G), GAG1502GCT (5716A>C 5717G>T), GAA1503CAA (5718G>C), TCA1504GCA (5721T>G), TCA1505GAT (5724T>G 5725C>A 5726A>T), ATT1506GTG (5727A>G 5729T>G), CCA1507CAA (5731C>A), TGG1508AAG (5733T>A 5734G>A), ACA1509ACT (5738A>T), GCG1510GCC (5741G>C), TTC1511ATG (5742T>A 5744C>G), TGG1512AGG (5745T>A), GTC1513ACA (5748G>A 5749T>C 5750C>A), CCT1514CAC (5752C>A 5753T>C), GAT1515TCA (5754G>T 5755A>C 5756T>A), GGA1516GGT (5759A>T), CTA1517CAC (5761T>A 5762A>C), TAT1518AAC (5763T>A 5765T>C), GAA1519AAG (5766G>A 5768A>G), TGG1520TAC (5770G>A 5771G>C), GTC1522GTG (5777C>G), CCG1524CCT (5783G>T), TTC1525TTT (5786C>T), GGA1526GGT (5789A>T), TTA1527CTC (5790T>C 5792A>C), AAG1528ACA (5794A>C 5795G>A), GCT1530GCC (5801T>C), GCC1532TCT (5805G>T 5807C>T), ATT1533AGC (5809T>G 5810T>C), TTT1534TTC (5813T>C), CAA1535CAG (5816A>G), AGA1536GGA (5817A>G), AAG1537CTC (5820A>C 5821A>T 5822G>C), GAC1539AAC (5826G>A), AAC1540CAT (5829A>C 5831C>T), TGC1541GTC (5832T>G 5833G>T), TTT1542TTC (5837T>C), AAA1543AAG (5840A>G), GGA1544GAC (5842G>A 5843A>C), GGA1544_ACA1545insCAT (5843_5844insCAT), ACA1545CTT (5844A>C 5845C>T 5846A>T), GAA1546AGG (5847G>A 5848A>G 5849A>G), GAA1547AAA (5850G>A), TTT1548TTC (5855T>C), ATA1549ATC (5858A>C), GCA1550CTA (5859G>C 5860C>T), GTT1551GTA (5864T>A), TAT1552TTC (5866A>T 5867T>C), ATT1553GCT (5868A>G 5869T>C), GAT1555GAC (5876T>C), CTA1557CTC (5882A>C), GTC1558GTT (5885C>T), TTC1559TAT (5887T>A 5888C>T), TCA1560AGT (5889T>A 5890C>G 5891A>T), ACT1562AAT (5896C>A), GAG1563ATG (5898G>A 5899A>T), GAA1564GAG (5903A>G), CAT1566TAT (5907C>T), GTC1567CTA (5910G>C 5912C>A), CGT1568CAA (5914G>A 5915T>A), CAT1569CAC (5918T>C), CTA1570TTG (5919C>T 5921A>G), CAA1571AGA (5922C>A 5923A>G), AGG1572ATT (5926G>T 5927G>T), TTC1573GTC (5928T>G), CAG1575AAC (5934C>A 5936G>C), ATA1576CTG (5937A>C 5939A>G), TGC1577TTG (5941G>T 5942C>G), CAG1578ATC (5943C>A 5944A>T 5945G>C), AAG1579AAA (5948G>A), CAT1580CAC (5951T>C), GGG1581CAG (5952G>C 5953G>A), TTA1582CTC (5955T>C 5957A>C), GTC1583TTA (5958G>T 5960C>A), CTG1584GTC (5961C>G 5963G>C), AGT1585AGG (5966T>G), AAG1586AAA (5969G>A), GAT1587AAC (5970G>A 5972T>C), ATG1589TGT (5976A>T 5977T>G 5978G>T), GCA1590TCT (5979G>T 5981A>T), ATA1591TTT (5982A>T 5984A>T), GCA1592GGA (5986C>G), GTC1593GCT (5989T>C 5990C>T), CCA1594TCT (5991C>T 5993A>T), CAA1595AAG (5994C>A 5996A>G), ATT1596GTG (5997A>G 5999T>G), GAG1597GAA (6002G>A), TTC1598TAC (6004T>A) | | | | | | | | |

**Proteins**

| | Begin | End | Coverage | Score | Concordance | Matches | Identities | I/D/M/F* | Stop Codons |
|---|---|---|---|---|---|---|---|---|---|
| polyprotein (YP_009345075.1) | 1483 | 1600 | 6.0% | 369 | 42.4% | 118 (99.2%) | 51 (42.9%) | 1/0/0/0 | 0 |
| Protein mutations: | N1483S (5659A>G), I1486V (5667A>G), S1488T (5673T>A), F1490L (5679T>C 5681T>G), K1493R (5689A>G 5690G>A), F1496Y (5698T>A), V1499I (5706G>A), A1500R (5709G>A 5710C>G), E1502A (5716A>C 5717G>T), E1503Q (5718G>C), S1504A (5721T>G), S1505D (5724T>G 5725C>A 5726A>T), I1506V (5727A>G 5729T>G), P1507Q (5731C>A), W1508K (5733T>A 5734G>A), F1511M (5742T>A 5744C>G), W1512R (5745T>A), V1513T (5748G>A 5749T>C 5750C>A), P1514H (5752C>A 5753T>C), D1515S (5754G>T 5755A>C 5756T>A), L1517H (5761T>A 5762A>C), Y1518N (5763T>A 5765T>C), E1519K (5766G>A 5768A>G), W1520Y (5770G>A 5771G>C), K1528T (5794A>C 5795G>A), A1532S (5805G>T 5807C>T), I1533S (5809T>G 5810T>C), R1536G (5817A>G), K1537L (5820A>C 5821A>T 5822G>C), D1539N (5826G>A), N1540H (5829A>C 5831C>T), C1541V (5832T>G 5833G>T), G1544D (5842G>A 5843A>C), G1544_T1545insH (5843_5844insCAT), T1545L (5844A>C 5845C>T 5846A>T), E1546R (5847G>A 5848A>G 5849A>G), E1547K (5850G>A), A1550L (5859G>C 5860C>T), Y1552F (5866A>T 5867T>C), I1553A (5868A>G 5869T>C), F1559Y (5887T>A 5888C>T), T1562N (5896C>A), E1563M (5898G>A 5899A>T), H1566Y (5907C>T), V1567L (5910G>C 5912C>A), R1568Q (5914G>A 5915T>A), Q1571R (5922C>A 5923A>G), R1572I (5926G>T 5927G>T), F1573V (5928T>G), Q1575N (5934C>A 5936G>C), I1576L (5937A>C 5939A>G), C1577L (5941G>T 5942C>G), Q1578I (5943C>A 5944A>T 5945G>C), G1581Q (5952G>C 5953G>A), V1583L (5958G>T 5960C>A), L1584V (5961C>G 5963G>C), S1585R (5966T>G), D1587N (5970G>A 5972T>C), M1589C (5976A>T 5977T>G 5978G>T), A1590S (5979G>T 5981A>T), I1591F (5982A>T 5984A>T), A1592G (5986C>G), V1593A (5989T>C 5990C>T), P1594S (5991C>T 5993A>T), Q1595K (5994C>A 5996A>G), I1596V (5997A>G 5999T>G), F1598Y (6004T>A) | | | | | | | | |
| Codon mutations: | AAT1483AGT (5659A>G), TCA1484TCC (5663A>C), AAG1485AAA (5666G>A), ATT1486GTT (5667A>G), TCC1488ACC (5673T>A), TTT1490CTG (5679T>C 5681T>G), AAG1493AGA (5689A>G 5690G>A), TCC1494TCA (5693C>A), GGT1495GGC (5696T>C), TTT1496TAT (5698T>A), CAT1497CAC (5702T>C), CAA1498CAG (5705A>G), GTA1499ATA (5706G>A), GCA1500AGA (5709G>A 5710C>G), GAG1502GCT (5716A>C 5717G>T), GAA1503CAA (5718G>C), TCA1504GCA (5721T>G), TCA1505GAT (5724T>G 5725C>A 5726A>T), ATT1506GTG (5727A>G 5729T>G), CCA1507CAA (5731C>A), TGG1508AAG (5733T>A 5734G>A), ACA1509ACT (5738A>T), GCG1510GCC (5741G>C), TTC1511ATG (5742T>A 5744C>G), TGG1512AGG (5745T>A), GTC1513ACA (5748G>A 5749T>C 5750C>A), CCT1514CAC (5752C>A 5753T>C), GAT1515TCA (5754G>T 5755A>C 5756T>A), GGA1516GGT (5759A>T), CTA1517CAC (5761T>A 5762A>C), TAT1518AAC (5763T>A 5765T>C), GAA1519AAG (5766G>A 5768A>G), TGG1520TAC (5770G>A 5771G>C), GTC1522GTG (5777C>G), CCG1524CCT (5783G>T), TTC1525TTT (5786C>T), GGA1526GGT (5789A>T), TTA1527CTC (5790T>C 5792A>C), AAG1528ACA (5794A>C 5795G>A), GCT1530GCC (5801T>C), GCC1532TCT (5805G>T 5807C>T), ATT1533AGC (5809T>G 5810T>C), TTT1534TTC (5813T>C), CAA1535CAG (5816A>G), AGA1536GGA (5817A>G), AAG1537CTC (5820A>C 5821A>T 5822G>C), GAC1539AAC (5826G>A), AAC1540CAT (5829A>C 5831C>T), TGC1541GTC (5832T>G 5833G>T), TTT1542TTC (5837T>C), AAA1543AAG (5840A>G), GGA1544GAC (5842G>A 5843A>C), GGA1544_ACA1545insCAT (5843_5844insCAT), ACA1545CTT (5844A>C 5845C>T 5846A>T), GAA1546AGG (5847G>A 5848A>G 5849A>G), GAA1547AAA (5850G>A), TTT1548TTC (5855T>C), ATA1549ATC (5858A>C), GCA1550CTA (5859G>C 5860C>T), GTT1551GTA (5864T>A), TAT1552TTC (5866A>T 5867T>C), ATT1553GCT (5868A>G 5869T>C), GAT1555GAC (5876T>C), CTA1557CTC (5882A>C), GTC1558GTT (5885C>T), TTC1559TAT (5887T>A 5888C>T), TCA1560AGT (5889T>A 5890C>G 5891A>T), ACT1562AAT (5896C>A), GAG1563ATG (5898G>A 5899A>T), GAA1564GAG (5903A>G), CAT1566TAT (5907C>T), GTC1567CTA (5910G>C 5912C>A), CGT1568CAA (5914G>A 5915T>A), CAT1569CAC (5918T>C), CTA1570TTG (5919C>T 5921A>G), CAA1571AGA (5922C>A 5923A>G), AGG1572ATT (5926G>T 5927G>T), TTC1573GTC (5928T>G), CAG1575AAC (5934C>A 5936G>C), ATA1576CTG (5937A>C 5939A>G), TGC1577TTG (5941G>T 5942C>G), CAG1578ATC (5943C>A 5944A>T 5945G>C), AAG1579AAA (5948G>A), CAT1580CAC (5951T>C), GGG1581CAG (5952G>C 5953G>A), TTA1582CTC (5955T>C 5957A>C), GTC1583TTA (5958G>T 5960C>A), CTG1584GTC (5961C>G 5963G>C), AGT1585AGG (5966T>G), AAG1586AAA (5969G>A), GAT1587AAC (5970G>A 5972T>C), ATG1589TGT (5976A>T 5977T>G 5978G>T), GCA1590TCT (5979G>T 5981A>T), ATA1591TTT (5982A>T 5984A>T), GCA1592GGA (5986C>G), GTC1593GCT (5989T>C 5990C>T), CCA1594TCT (5991C>T 5993A>T), CAA1595AAG (5994C>A 5996A>G), ATT1596GTG (5997A>G 5999T>G), GAG1597GAA (6002G>A), TTC1598TAC (6004T>A) | | | | | | | | |

*: Inserts / Deletes / Misaligned / Frameshifts

#### Analysis details

This analysis was performed with panviral2.64

### NGS Details (UN24): Petuvirus venapetuniae

#### Assembly

| | |
|---|---|
| **Coverage Length** | 281 (1 contig(s)) |
| **Depth Of Coverage** | 2.5 |
| **Number Of Reads** | 6 |
| **Reads Per Million** | 0.12 rpm (after QC) |
| **Ambiguities** | 0 |
| **Assembly Method** | de novo + reference guided assembly |
| **Consensus Caller** | Bcf Tools |

#### Coverage Map

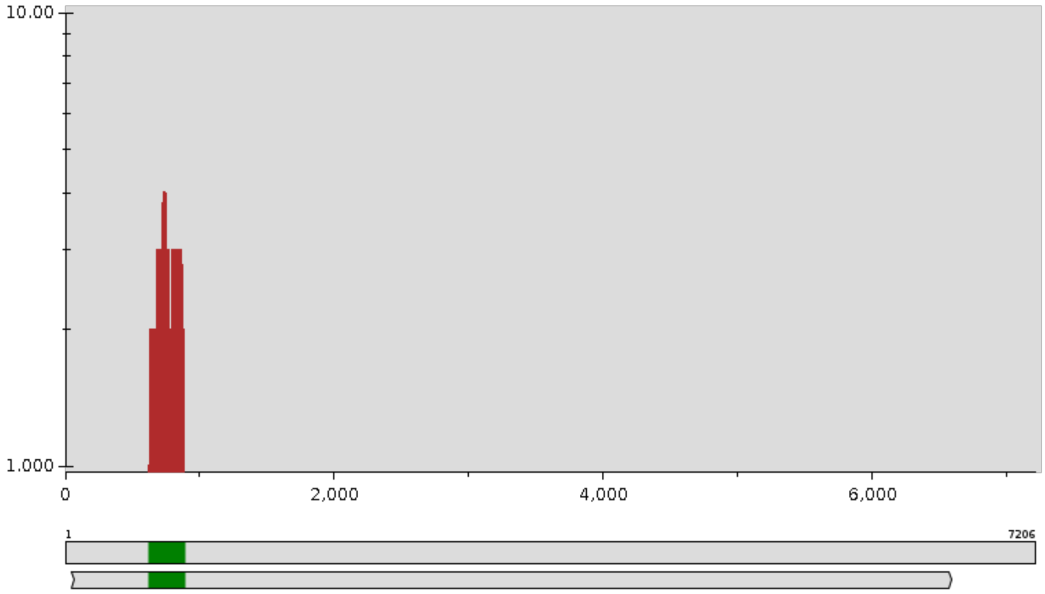

#### Assignment

| | |
|---|---|
| **Type** | Petuvirus venapetuniae (Taxonomy ID: 3048265) |
| **Reference Genome** | NC_001839.2 |
| **NT Identity (%)** | 62.5 |
| **AA Identity (%)** | 53.1915 |
| **Number Of Stop Codons** | 5 |
| **Number Of CDS** | 1 |

#### Alignment

| | |
|---|---|
| **Alignment Score** | 130.0 (NT) + 219.0 (AA) = 349.0 |
| **Concordance (%)** | 34.4186 |

| Alignment Method | Global, seeded, nucleotide + amino acids (AGA) |
|---|---|

#### Genome Region

Sequence starts at position 614 and ends at position 894 relative to NC_001839.2 reference sequence.

#### Alignment Detailed Statistics

| | Begin | End | Coverage | Score | Concordance | Matches | Identities | I/D/M/F* | Stop Codons |
|---|---|---|---|---|---|---|---|---|---|
| **NT** | **614** | **894** | **3.9%** | **130** | **23.6%** | **280 (99.6%)** | **175 (62.3%)** | **0/1** | |
| Mutations: | 615G>A, 618G>C, 620C>A, 621A>C, 627T>A, 629C>A, 630A>G, 631C>A, 632A>G, 635C>A, 636T>A, 638A>G, 641A>G, 644C>A, 645C>T, 647A>G, 656C>A, 658A>T, 659A>T, 668C>G, 670T>A, 672A>G, 674A>T, 676A>G, 680C>T, 681T>G, 682C>T, 686C>A, 693delA, 694C>A, 696T>C, 698A>T, 701T>C, 704C>T, 705C>T, 707A>G, 708A>C, 710T>A, 712C>T, 717C>A, 719T>A, 722C>T, 723C>T, 725A>G, 726A>G, 728T>C, 729C>A, 733C>T, 734T>C, 740T>C, 741T>C, 743G>A, 744A>T, 749C>T, 752T>A, 755A>T, 758T>C, 759G>A, 761G>T, 764A>T, 765G>T, 767C>T, 774C>T, 776C>A, 777A>C, 779A>C, 782T>A, 786G>T, 789G>A, 790G>T, 793G>A, 798G>A, 800C>T, 804C>T, 806A>G, 809T>C, 815A>G, 816G>A, 818C>G, 819C>T, 821T>A, 823G>A, 827G>C, 828T>A, 830G>A, 835A>G, 836A>G, 839A>C, 840G>A, 842T>G, 845A>T, 846A>T, 849A>T, 851A>G, 854A>T, 855T>A, 857G>C, 861G>A, 863T>C, 866T>A, 872A>C, 875A>T, 878T>A, 884A>G, 890A>G, 893G>A | | | | | | | | |
| **CDS** | | | | | | | | | |
| PVCVgp1 | 189 | 282 | 4.3% | 219 | 34.4% | 94 (100%) | 50 (53.2%) | 0/0/1/1 | 5 |
| Protein mutations: | D189N (615G>A), A190P (618G>C 620C>A), N191H (621A>C), S193T (627T>A 629C>A), T194E (630A>G 631C>A 632A>G), L196M (636T>A 638A>G), Q199* (645C>T 647A>G), Q203L (658A>T 659A>T), L207H (670T>A), T208A (672A>G 674A>T), K209R (676A>G), S211V (681T>G 682C>T), Q219* (705C>T 707A>G), I220L (708A>C 710T>A), A221V (712C>T), Q225* (723C>T 725A>G), N226D (726A>G 728T>C), H227N (729C>A), A228V (733C>T 734T>C), T232S (744A>T), E237N (759G>A 761G>T), E238D (764A>T), A239S (765G>T 767C>T), K243H (777A>C 779A>C), A246S (786G>T), G247I (789G>A 790G>T), S248N (793G>A), A250T (798G>A 800C>T), Q252* (804C>T 806A>G), V256M (816G>A 818C>G), P257S (819C>T 821T>A), R258K (823G>A), Q259H (827G>C), L260I (828T>A 830G>A), K262R (835A>G 836A>G), E263D (839A>C), D264K (840G>A 842T>G), I266F (846A>T), K267* (849A>T 851A>G), L269I (855T>A 857G>C), D271N (861G>A 863T>C), N276K (878T>A) | | | | | | | | |
| Codon mutations: | GAC189AAC (615G>A), GCC190CCA (618G>C 620C>A), AAT191CAT (621A>C), TCC193ACA (627T>A 629C>A), ACA194GAG (630A>G 631C>A 632A>G), GCC195GCA (635C>A), TTA196ATG (636T>A 638A>G), AAA197AAG (641A>G), ATC198ATA (644C>A), CAA199TAG (645C>T 647A>G), ATC202ATA (656C>A), CAA203CTT (658A>T 659A>T), CCC206CCG (668C>G), CTT207CAT (670T>A), ACA208GCT (672A>G 674A>T), AAA209AGA (676A>G), GAC210GAT (680C>T), TCC211GTC (681T>G 682C>T), ATC212ATA (686C>A), ACT215-AT (693delA 694C>A), TTA216CTT (696T>C 698A>T), CAT217CAC (701T>C), TAC218TAT (704C>T), CAA219TAG (705C>T 707A>G), ATT220CTA (708A>C 710T>A), GCT221GTT (712C>T), CGT223AGA (717C>A 719T>A), GTC224GTT (722C>T), CAA225TAG (723C>T 725A>G), AAT226GAC (726A>G 728T>C), CAT227AAT (729C>A), GCT228GTC (733C>T 734T>C), GAT230GAC (740T>C), TTG231CTA (741T>C 743G>A), ACT232TCT (744A>T), CTC233CTT (749C>T), CCT234CCA (752T>A), GGA235GGT (755A>T), GGT236GGC (758T>C), GAG237AAT (759G>A 761G>T), GAA238GAT (764A>T), GCC239TCT (765G>T 767C>T), CTC242TTA (774C>T 776C>A), AAA243CAC (777A>C 779A>C), ATT244ATA (782T>A), GCC246TCC (786G>T), GGA247ATA (789G>A 790G>T), AGT248AAT (793G>A), GCC250ACT (798G>A 800C>T), CAA252TAG (804C>T 806A>G), TGT253TGC (809T>C), CAA255CAG (815A>G), GTC256ATG (816G>A 818C>G), CCT257TCA (819C>T 821T>A), AGA258AAA (823G>A), CAG259CAC (827G>C), TTG260ATA (828T>A 830G>A), AAA262AGG (835A>G 836A>G), GAA263GAC (839A>C), GAT264AAG (840G>A 842T>G), CTA265CTT (845A>T), ATC266TTC (846A>T), AAA267TAG (849A>T 851A>G), ATA268ATT (854A>T), TTG269ATC (855T>A 857G>C), GAT271AAC (861G>A 863T>C), TCT272TCA (866T>A), GTA274GTC (872A>C), ACA275ACT (875A>T), AAT276AAA (878T>A), GAA278GAG (884A>G), TTA280TTG (890A>G), AGG281AGA (893G>A) | | | | | | | | |
| **Proteins** | | | | | | | | | |
| ORF I polyprotein (NP_127504.1) | 189 | 282 | 4.3% | 219 | 34.4% | 94 (100%) | 50 (53.2%) | 0/0/1/1 | 5 |
| Protein mutations: | D189N (615G>A), A190P (618G>C 620C>A), N191H (621A>C), S193T (627T>A 629C>A), T194E (630A>G 631C>A 632A>G), L196M (636T>A 638A>G), Q199* (645C>T 647A>G), Q203L (658A>T 659A>T), L207H (670T>A), T208A (672A>G 674A>T), K209R (676A>G), S211V (681T>G 682C>T), Q219* (705C>T 707A>G), I220L (708A>C 710T>A), A221V (712C>T), Q225* (723C>T 725A>G), N226D (726A>G 728T>C), H227N (729C>A), A228V (733C>T 734T>C), T232S (744A>T), E237N (759G>A 761G>T), E238D (764A>T), A239S (765G>T 767C>T), K243H (777A>C 779A>C), A246S (786G>T), G247I (789G>A 790G>T), S248N (793G>A), A250T (798G>A 800C>T), Q252* (804C>T 806A>G), V256M (816G>A 818C>G), P257S (819C>T 821T>A), R258K (823G>A), Q259H (827G>C), L260I (828T>A 830G>A), K262R (835A>G 836A>G), E263D (839A>C), D264K (840G>A 842T>G), I266F (846A>T), K267* (849A>T 851A>G), L269I (855T>A 857G>C), D271N (861G>A 863T>C), N276K (878T>A) | | | | | | | | |
| Codon mutations: | GAC189AAC (615G>A), GCC190CCA (618G>C 620C>A), AAT191CAT (621A>C), TCC193ACA (627T>A 629C>A), ACA194GAG (630A>G 631C>A 632A>G), GCC195GCA (635C>A), TTA196ATG (636T>A 638A>G), AAA197AAG (641A>G), ATC198ATA (644C>A), CAA199TAG (645C>T 647A>G), ATC202ATA (656C>A), CAA203CTT (658A>T 659A>T), CCC206CCG (668C>G), CTT207CAT (670T>A), ACA208GCT (672A>G 674A>T), AAA209AGA (676A>G), GAC210GAT (680C>T), TCC211GTC (681T>G 682C>T), ATC212ATA (686C>A), ACT215-AT (693delA 694C>A), TTA216CTT (696T>C 698A>T), CAT217CAC (701T>C), TAC218TAT (704C>T), CAA219TAG (705C>T 707A>G), ATT220CTA (708A>C 710T>A), GCT221GTT (712C>T), CGT223AGA (717C>A 719T>A), GTC224GTT (722C>T), CAA225TAG (723C>T 725A>G), AAT226GAC (726A>G 728T>C), CAT227AAT (729C>A), GCT228GTC (733C>T 734T>C), GAT230GAC (740T>C), TTG231CTA (741T>C 743G>A), ACT232TCT (744A>T), CTC233CTT (749C>T), CCT234CCA (752T>A), GGA235GGT (755A>T), GGT236GGC (758T>C), GAG237AAT (759G>A 761G>T), GAA238GAT (764A>T), GCC239TCT (765G>T 767C>T), CTC242TTA (774C>T 776C>A), AAA243CAC (777A>C 779A>C), ATT244ATA (782T>A), GCC246TCC (786G>T), GGA247ATA (789G>A 790G>T), AGT248AAT (793G>A), GCC250ACT (798G>A 800C>T), CAA252TAG (804C>T 806A>G), TGT253TGC (809T>C), CAA255CAG (815A>G), GTC256ATG (816G>A 818C>G), CCT257TCA (819C>T 821T>A), AGA258AAA (823G>A), CAG259CAC (827G>C), TTG260ATA (828T>A 830G>A), AAA262AGG (835A>G 836A>G), GAA263GAC (839A>C), GAT264AAG (840G>A 842T>G), CTA265CTT (845A>T), ATC266TTC (846A>T), AAA267TAG (849A>T 851A>G), ATA268ATT (854A>T), TTG269ATC (855T>A 857G>C), GAT271AAC (861G>A 863T>C), TCT272TCA (866T>A), GTA274GTC (872A>C), ACA275ACT (875A>T), AAT276AAA (878T>A), GAA278GAG (884A>G), TTA280TTG (890A>G), AGG281AGA (893G>A) | | | | | | | | |

*: Inserts / Deletes / Misaligned / Frameshifts

#### Analysis details

This analysis was performed with panviral2.64

### NGS Details (UN24): Marseillevirus marseillevirus

#### Assembly

| | |
|---|---|
| **Coverage Length** | 110 (1 contig(s)) |
| **Depth Of Coverage** | 4.7 |
| **Number Of Reads** | 6 |
| **Reads Per Million** | 0.12 rpm (after QC) |
| **Ambiguities** | 0 |
| **Assembly Method** | de novo + reference guided assembly |
| **Consensus Caller** | Bcf Tools |

#### Coverage Map

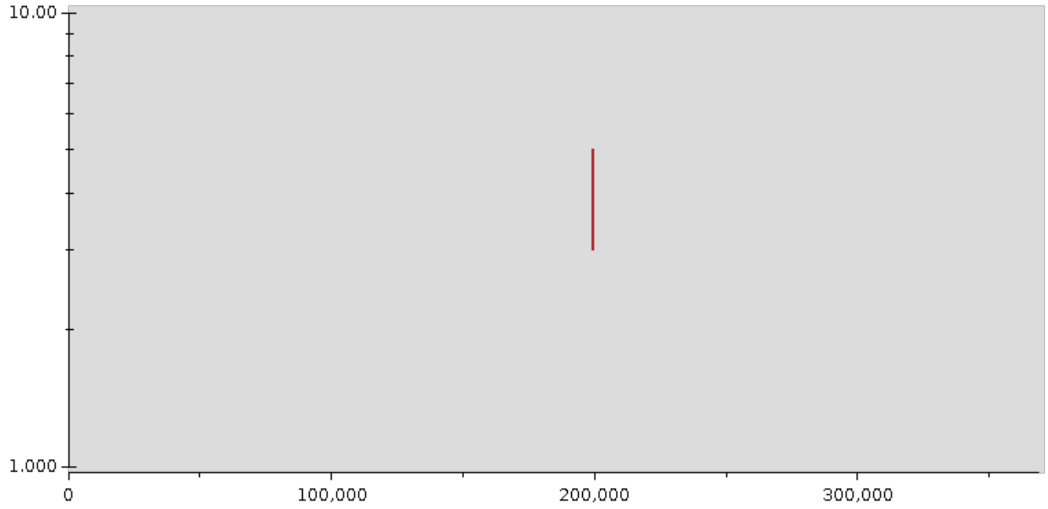

#### Assignment

| | |
|---|---|
| **Type** | Marseillevirus marseillevirus (Taxonomy ID: 694581) |
| **Reference Genome** | NC_013756.1 |
| **NT Identity (%)** | 80.9091 |
| **AA Identity (%)** | 94.5946 |
| **Number Of Stop Codons** | 0 |
| **Number Of CDS** | 428 |

#### Alignment

| | |
|---|---|
| **Alignment Score** | 136.0 (NT) + 235.0 (AA) = 371.0 |
| **Concordance (%)** | 81.3596 |
| **Alignment Method** | Local, heuristic, nucleotide (BLASTN) |

#### Genome Region

Sequence starts at position 199409 and ends at position 199518 relative to NC_013756.1 reference sequence.

#### Alignment Detailed Statistics

| | Begin | End | Coverage | Score | Concordance | Matches | Identities | I/D/M/F* | Stop Codons |
|---|---|---|---|---|---|---|---|---|---|
| **NT** | **199409** | **199518** | **0.1%** | **136** | **61.8%** | **110 (100%)** | **89 (80.9%)** | **0/0** | |

Mutations: 199414T>C, 199419G>A, 199420G>A, 199423A>C, 199425G>A, 199426G>A, 199429G>C, 199431G>A, 199435C>T, 199441C>T, 199444T>C, 199449C>T, 199453G>A, 199462C>T, 199464A>G, 199468C>A, 199470T>G, 199480C>G, 199501G>C, 199503G>A, 199504A>T

*: Inserts / Deletes / Misaligned / Frameshifts

#### Analysis details

This analysis was performed with panviral2.64

#### NGS Details (UN24): Punavirus P1

##### Assembly

| | |
|---|---|
| **Coverage Length** | 211 (1 contig(s)) |
| **Depth Of Coverage** | 2.2 |
| **Number Of Reads** | 4 |
| **Reads Per Million** | 0.08 rpm (after QC) |
| **Ambiguities** | 0 |
| **Assembly Method** | de novo + reference guided assembly |
| **Consensus Caller** | Bcf Tools |

##### Coverage Map

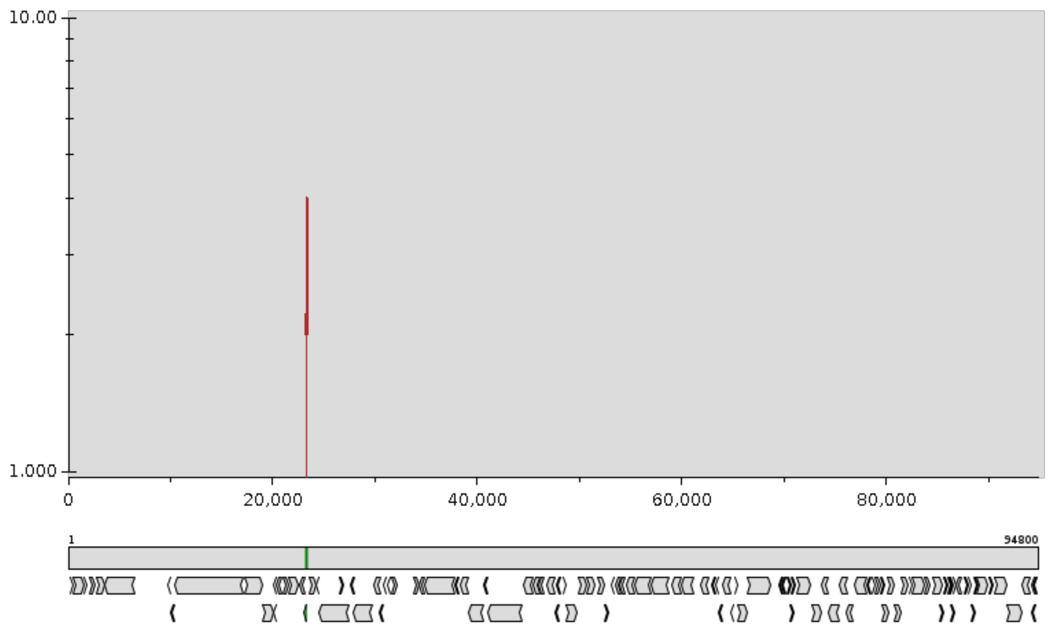

##### Assignment

| | |
|---|---|
| **Type** | Punavirus P1 (Taxonomy ID: 10678) |
| **Reference Genome** | NC_005856.1 |
| **NT Identity (%)** | 100.0 |
| **AA Identity (%)** | 98.0392 |
| **Number Of Stop Codons** | 0 |
| **Number Of CDS** | 110 |

##### Alignment

| | |
|---|---|
| **Alignment Score** | 422.0 (NT) + 377.0 (AA) = 799.0 |
| **Concordance (%)** | 100.0 |

| **Alignment Method** | Local, heuristic, nucleotide (BLASTN) |
|---|---|

#### Genome Region

Sequence starts at position 23187 and ends at position 23397 relative to NC_005856.1 reference sequence.

#### Alignment Detailed Statistics

| | Begin | End | Coverage | Score | Concordance | Matches | Identities | I/D/M/F* | Stop Codons |
|---|---|---|---|---|---|---|---|---|---|
| **NT** | **23187** | **23397** | **0.2%** | **422** | **100%** | **211 (100%)** | **211 (100%)** | **0/0** | |
| Mutations: | *none* | | | | | | | | |
| **CDS** | | | | | | | | | |
| insA | 1 | 51 | 55.4% | 377 | 97.7% | 51 (100%) | 50 (98.0%) | 0/0/0/0 | 0 |
| **Proteins** | | | | | | | | | |
| IS1 family transposase (YP_006488.1) | 1 | 51 | 55.4% | 377 | 97.7% | 51 (100%) | 50 (98.0%) | 0/0/0/0 | 0 |
| Protein mutations: | *none* | | | | | | | | |
| Codon mutations: | *none* | | | | | | | | |

*: Inserts / Deletes / Misaligned / Frameshifts

#### Analysis details

This analysis was performed with panviral2.64

#### NGS Details (UN24): Sugarcane chlorotic streak virus

##### Assembly

| | |
|---|---|
| **Coverage Length** | 134 (1 contig(s)) |
| **Depth Of Coverage** | 2.0 |
| **Number Of Reads** | 2 |
| **Reads Per Million** | 0.04 rpm (after QC) |
| **Ambiguities** | 0 |
| **Assembly Method** | de novo + reference guided assembly |
| **Consensus Caller** | Bcf Tools |

##### Coverage Map

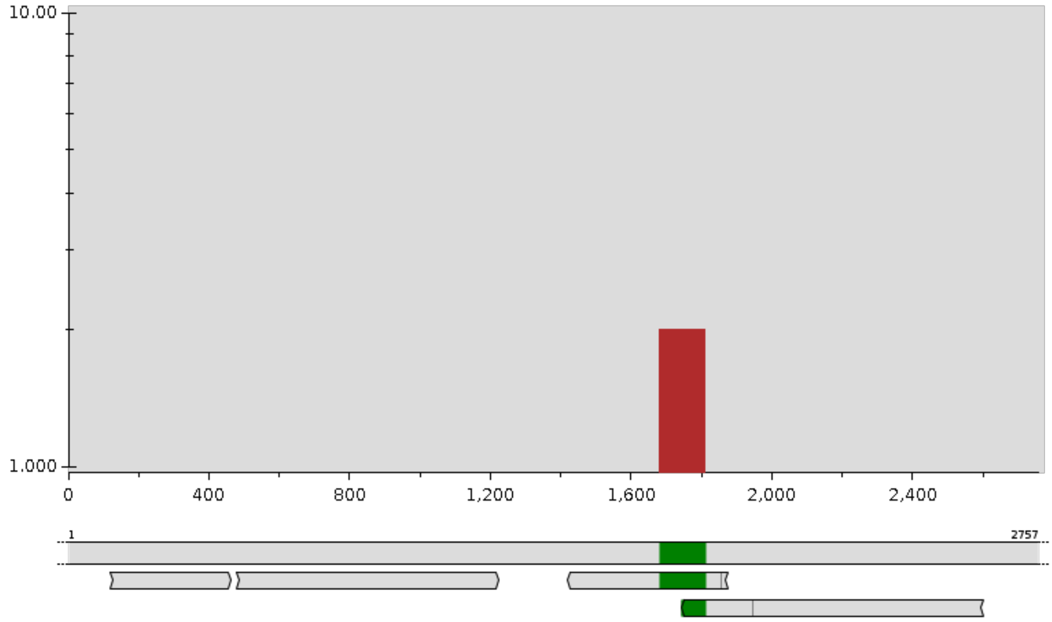

##### Assignment

| | |
|---|---|
| **Type** | Sugarcane chlorotic streak virus (Taxonomy ID: 1919062) |
| **Reference Genome** | NC_032004.1 |
| **NT Identity (%)** | 64.9254 |
| **AA Identity (%)** | 58.4071 |
| **Number Of Stop Codons** | 1 |
| **Number Of CDS** | 5 |

##### Alignment

| | |
|---|---|
| **Alignment Score** | 80.0 (NT) + 509.0 (AA) = 589.0 |
| **Concordance (%)** | 53.5455 |

| Alignment Method | Global, seeded, nucleotide + amino acids (AGA) |
|---|---|

#### Genome Region

Sequence starts at position 1681 and ends at position 1814 relative to NC_032004.1 reference sequence.

#### Alignment Detailed Statistics

| | Begin | End | Coverage | Score | Concordance | Matches | Identities | I/D/M/F* | Stop Codons |
|---|---|---|---|---|---|---|---|---|---|
| **NT** | **1681** | **1814** | **4.9%** | **80** | **29.9%** | **134 (100%)** | **87 (64.9%)** | **0/0** | |
| Mutations: | 1684T>A, 1689G>A, 1692A>C, 1695T>G, 1701A>G, 1704A>T, 1710G>A, 1713G>A, 1714A>T, 1715C>A, 1716G>C, 1718C>A, 1721C>T, 1722C>G, 1723T>A, 1724C>T, 1725T>A, 1734A>G, 1735G>T, 1736A>T, 1737G>A, 1738G>C, 1740C>G, 1741C>A, 1744G>T, 1755T>G, 1756T>C, 1757G>A, 1758C>A, 1759C>A, 1761G>A, 1767A>G, 1776C>A, 1779G>A, 1784T>G, 1785A>T, 1791A>T, 1793A>T, 1794T>C, 1796A>T, 1797C>T, 1800T>A, 1803T>A, 1806C>T, 1809T>A, 1811T>A, 1812T>A | | | | | | | | |
| **CDS** | | | | | | | | | |
| BS327_gp3 | 234 | 278 | 12.4% | 240 | 67.4% | 45 (100%) | 29 (64.4%) | 0/0/0/0 | 0 |
| Protein mutations: | T235S (1809T>A 1811T>A), S240T (1794T>C 1796A>T), S241T (1791A>T 1793A>T), W252F (1758C>A 1759C>A), Q253C (1755T>G 1756T>C 1757G>A), A257D (1744G>T), W258F (1740C>G 1741C>A), T259S (1737G>A 1738G>C), S260N (1734A>G 1735G>T 1736A>T), E263D (1725T>A), E264I (1722C>G 1723T>A 1724C>T), A265T (1721C>T), V266L (1716G>C 1718C>A), V267Y (1713G>A 1714A>T 1715C>A), Y277F (1684T>A) | | | | | | | | |
| Codon mutations: | CCA234CCT (1812T>A), ACA235TCT (1809T>A 1811T>A), AGG236AGA (1806C>T), ACA237ACT (1803T>A), GGA238GGT (1800T>A), AAG239AAA (1797C>T), TCA240ACG (1794T>C 1796A>T), TCT241ACA (1791A>T 1793A>T), GCT243GCA (1785A>T), AGA244CGA (1784T>G), AGC245AGT (1779G>A), CTG246CTT (1776C>A), CAT249CAC (1767A>G), TAC251TAT (1761G>A), TGG252TTT (1758C>A 1759C>A), CAA253TGC (1755T>G 1756T>C 1757G>A), GCC257GAC (1744G>T), TGG258TTC (1740C>G 1741C>A), ACC259AGT (1737G>A 1738G>C), TCT260AAC (1734A>G 1735G>T 1736A>T), GAA263GAT (1725T>A), GAG264ATC (1722C>G 1723T>A 1724C>T), GCA265ACA (1721C>T), GTC266TTG (1716G>C 1718C>A), GTC267TAT (1713G>A 1714A>T 1715C>A), AAC268AAT (1710G>A), ATT270ATA (1704A>T), GAT271GAC (1701A>G), ATA273ATC (1695T>G), CCT274CCG (1692A>C), TTC275TTT (1689G>A), TAC277TTC (1684T>A) | | | | | | | | |
| BS327_gp4 | 22 | 66 | 29.6% | 240 | 67.4% | 45 (100%) | 29 (64.4%) | 0/0/0/0 | 0 |
| Protein mutations: | T23S (1809T>A 1811T>A), S28T (1794T>C 1796A>T), S29T (1791A>T 1793A>T), W40F (1758C>A 1759C>A), Q41C (1755T>G 1756T>C 1757G>A), A45D (1744G>T), W46F (1740C>G 1741C>A), T47S (1737G>A 1738G>C), S48N (1734A>G 1735G>T 1736A>T), E51D (1725T>A), E52I (1722C>G 1723T>A 1724C>T), A53T (1721C>T), V54L (1716G>C 1718C>A), V55Y (1713G>A 1714A>T 1715C>A), Y65F (1684T>A) | | | | | | | | |
| Codon mutations: | CCA22CCT (1812T>A), ACA23TCT (1809T>A 1811T>A), AGG24AGA (1806C>T), ACA25ACT (1803T>A), GGA26GGT (1800T>A), AAG27AAA (1797C>T), TCA28ACG (1794T>C 1796A>T), TCT29ACA (1791A>T 1793A>T), GCT31GCA (1785A>T), AGA32CGA (1784T>G), AGC33AGT (1779G>A), CTG34CTT (1776C>A), CAT37CAC (1767A>G), TAC39TAT (1761G>A), TGG40TTT (1758C>A 1759C>A), CAA41TGC (1755T>G 1756T>C 1757G>A), GCC45GAC (1744G>T), TGG46TTC (1740C>G 1741C>A), ACC47AGT (1737G>A 1738G>C), TCT48AAC (1734A>G 1735G>T 1736A>T), GAA51GAT (1725T>A), GAG52ATC (1722C>G 1723T>A 1724C>T), GCA53ACA (1721C>T), GTC54TTG (1716G>C 1718C>A), GTC55TAT (1713G>A 1714A>T 1715C>A), AAC56AAT (1710G>A), ATT58ATA (1704A>T), GAT59GAC (1701A>G), ATA61ATC (1695T>G), CCT62CCG (1692A>C), TTC63TTT (1689G>A), TAC65TTC (1684T>A) | | | | | | | | |
| BS327_gp5 | 265 | 287 | 8.0% | 29 | 17.9% | 23 (100%) | 8 (34.8%) | 0/0/0/0 | 1 |
| Protein mutations: | Q265L (1811T>A 1812T>A), Q266L (1809T>A), G267E (1806C>T), Q268L (1803T>A), E269V (1800T>A), S270K (1796A>T 1797C>T), H271R (1793A>T 1794T>C), L272H (1791A>T), L274H (1784T>G 1785A>T), A276V (1779G>A), W277L (1776C>A), I280T (1767A>G), T282I (1761G>A), G283F (1757G>A 1758C>A 1759C>A), K284A (1755T>G 1756T>C) | | | | | | | | |
| Codon mutations: | CAA265CTT (1811T>A 1812T>A), CAA266CTA (1809T>A), GGA267GAA (1806C>T), CAG268CTG (1803T>A), GAA269GTA (1800T>A), AGT270AAA (1796A>T 1797C>T), CAT271CGA (1793A>T 1794T>C), CTT272CAT (1791A>T), CTA274CAC (1784T>G 1785A>T), GCC276GTC (1779G>A), TGG277TTG (1776C>A), ATA280ACA (1767A>G), ACT282ATT (1761G>A), GGC283TTT (1757G>A 1758C>A 1759C>A), AAA284GCA (1755T>G 1756T>C) | | | | | | | | |
| **Proteins** | | | | | | | | | |
| Rep-associated protein (YP_009325925.1) | 234 | 278 | 12.4% | 240 | 67.4% | 45 (100%) | 29 (64.4%) | 0/0/0/0 | 0 |
| Protein mutations: | T235S (1809T>A 1811T>A), S240T (1794T>C 1796A>T), S241T (1791A>T 1793A>T), W252F (1758C>A 1759C>A), Q253C (1755T>G 1756T>C 1757G>A), A257D (1744G>T), W258F (1740C>G 1741C>A), T259S (1737G>A 1738G>C), S260N (1734A>G 1735G>T 1736A>T), E263D (1725T>A), E264I (1722C>G 1723T>A 1724C>T), A265T (1721C>T), V266L (1716G>C 1718C>A), V267Y (1713G>A 1714A>T 1715C>A), Y277F (1684T>A) | | | | | | | | |
| Codon mutations: | CCA234CCT (1812T>A), ACA235TCT (1809T>A 1811T>A), AGG236AGA (1806C>T), ACA237ACT (1803T>A), GGA238GGT (1800T>A), AAG239AAA (1797C>T), TCA240ACG (1794T>C 1796A>T), TCT241ACA (1791A>T 1793A>T), GCT243GCA (1785A>T), AGA244CGA (1784T>G), AGC245AGT (1779G>A), CTG246CTT (1776C>A), CAT249CAC (1767A>G), TAC251TAT (1761G>A), TGG252TTT (1758C>A 1759C>A), CAA253TGC (1755T>G 1756T>C 1757G>A), GCC257GAC (1744G>T), TGG258TTC (1740C>G 1741C>A), ACC259AGT (1737G>A 1738G>C), TCT260AAC (1734A>G 1735G>T 1736A>T), GAA263GAT (1725T>A), GAG264ATC (1722C>G 1723T>A 1724C>T), GCA265ACA (1721C>T), GTC266TTG (1716G>C 1718C>A), GTC267TAT (1713G>A 1714A>T 1715C>A), AAC268AAT (1710G>A), ATT270ATA (1704A>T), GAT271GAC (1701A>G), ATA273ATC (1695T>G), CCT274CCG (1692A>C), TTC275TTT (1689G>A), TAC277TTC (1684T>A) | | | | | | | | |
| RepB (YP_009325926.1) | 22 | 66 | 29.6% | 240 | 67.4% | 45 (100%) | 29 (64.4%) | 0/0/0/0 | 0 |
| Protein mutations: | T23S (1809T>A 1811T>A), S28T (1794T>C 1796A>T), S29T (1791A>T 1793A>T), W40F (1758C>A 1759C>A), Q41C (1755T>G 1756T>C 1757G>A), A45D (1744G>T), W46F (1740C>G 1741C>A), T47S (1737G>A 1738G>C), S48N (1734A>G 1735G>T 1736A>T), E51D (1725T>A), E52I (1722C>G 1723T>A 1724C>T), A53T (1721C>T), V54L (1716G>C 1718C>A), V55Y (1713G>A 1714A>T 1715C>A), Y65F (1684T>A) | | | | | | | | |
| Codon mutations: | CCA22CCT (1812T>A), ACA23TCT (1809T>A 1811T>A), AGG24AGA (1806C>T), ACA25ACT (1803T>A), GGA26GGT (1800T>A), AAG27AAA (1797C>T), TCA28ACG (1794T>C 1796A>T), TCT29ACA (1791A>T 1793A>T), GCT31GCA (1785A>T), AGA32CGA (1784T>G), AGC33AGT (1779G>A), CTG34CTT (1776C>A), CAT37CAC (1767A>G), TAC39TAT (1761G>A), TGG40TTT (1758C>A 1759C>A), CAA41TGC (1755T>G 1756T>C 1757G>A), GCC45GAC (1744G>T), TGG46TTC (1740C>G 1741C>A), ACC47AGT (1737G>A 1738G>C), TCT48AAC (1734A>G 1735G>T 1736A>T), GAA51GAT (1725T>A), GAG52ATC (1722C>G 1723T>A 1724C>T), GCA53ACA (1721C>T), GTC54TTG (1716G>C 1718C>A), GTC55TAT (1713G>A 1714A>T 1715C>A), AAC56AAT (1710G>A), ATT58ATA (1704A>T), GAT59GAC (1701A>G), ATA61ATC (1695T>G), CCT62CCG (1692A>C), TTC63TTT (1689G>A), TAC65TTC (1684T>A) | | | | | | | | |
| RepA (YP_009325927.1) | 265 | 287 | 8.0% | 29 | 17.9% | 23 (100%) | 8 (34.8%) | 0/0/0/0 | 1 |
| Protein mutations: | Q265L (1811T>A 1812T>A), Q266L (1809T>A), G267E (1806C>T), Q268L (1803T>A), E269V (1800T>A), S270K (1796A>T 1797C>T), H271R (1793A>T 1794T>C), L272H (1791A>T), L274H (1784T>G 1785A>T), A276V (1779G>A), W277L (1776C>A), I280T (1767A>G), T282I (1761G>A), G283F (1757G>A 1758C>A 1759C>A), K284A (1755T>G 1756T>C) | | | | | | | | |
| Codon mutations: | CAA265CTT (1811T>A 1812T>A), CAA266CTA (1809T>A), GGA267GAA (1806C>T), CAG268CTG (1803T>A), GAA269GTA (1800T>A), AGT270AAA (1796A>T 1797C>T), CAT271CGA (1793A>T 1794T>C), CTT272CAT (1791A>T), CTA274CAC (1784T>G 1785A>T), GCC276GTC (1779G>A), TGG277TTG (1776C>A), ATA280ACA (1767A>G), ACT282ATT (1761G>A), GGC283TTT (1757G>A 1758C>A 1759C>A), AAA284GCA (1755T>G 1756T>C) | | | | | | | | |

*: Inserts / Deletes / Misaligned / Frameshifts

#### Analysis details

This analysis was performed with panviral2.64

### NGS Details (UN24): Caulimovirus venafragariae

#### Assembly

| | |
|---|---|
| **Coverage Length** | 244 (1 contig(s)) |
| **Depth Of Coverage** | 1.1 |
| **Number Of Reads** | 2 |
| **Reads Per Million** | 0.04 rpm (after QC) |
| **Ambiguities** | 0 |
| **Assembly Method** | de novo + reference guided assembly |
| **Consensus Caller** | Bcf Tools |

#### Coverage Map

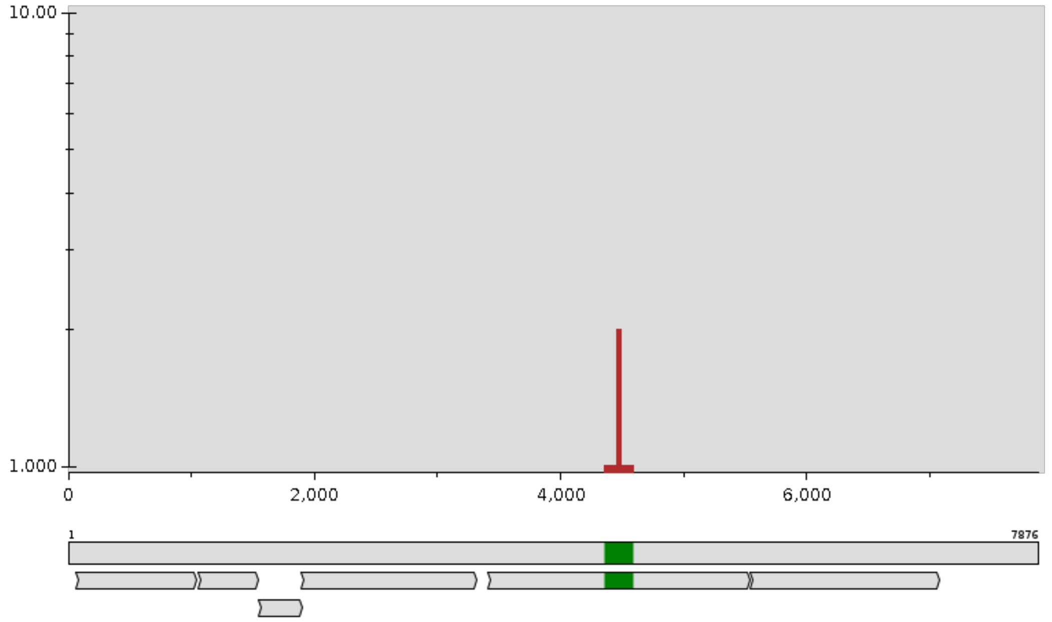

#### Assignment

| | |
|---|---|
| **Type** | Caulimovirus venafragariae (Taxonomy ID: 3048344) |
| **Reference Genome** | NC_001725.1 |
| **NT Identity (%)** | 59.4262 |
| **AA Identity (%)** | 51.8519 |
| **Number Of Stop Codons** | 3 |
| **Number Of CDS** | 6 |

#### Alignment

| | |
|---|---|
| **Alignment Score** | 92.0 (NT) + 295.0 (AA) = 387.0 |
| **Concordance (%)** | 36.4407 |

| Alignment Method | Global, seeded, nucleotide + amino acids (AGA) |
|---|---|

#### Genome Region

Sequence starts at position 4352 and ends at position 4595 relative to NC_001725.1 reference sequence.

#### Alignment Detailed Statistics

| | Begin | End | Coverage | Score | Concordance | Matches | Identities | I/D/M/F* | Stop Codons |
|---|---|---|---|---|---|---|---|---|---|
| **NT** | **4352** | **4595** | **3.1%** | **92** | **18.9%** | **244 (100%)** | **145 (59.4%)** | **0/0** | |
| Mutations: | 4354C>A, 4357G>A, 4359C>A, 4362C>T, 4371G>A, 4378G>A, 4379C>G, 4380C>T, 4383G>A, 4386C>A, 4387A>G, 4388A>T, 4390C>A, 4395C>A, 4398A>G, 4399G>A, 4407G>A, 4413T>A, 4416C>T, 4424A>G, 4425G>A, 4426T>C, 4428A>T, 4432G>A, 4434C>T, 4435C>A, 4440A>T, 4442A>G, 4443G>A, 4444G>A, 4445G>C, 4447G>T, 4451G>C, 4452C>A, 4455C>T, 4456C>A, 4457T>A, 4458A>T, 4459C>T, 4461T>A, 4462C>T, 4464T>A, 4465A>G, 4467C>T, 4470G>A, 4471G>A, 4472A>G, 4473G>T, 4474C>G, 4477C>T, 4479G>A, 4480C>A, 4483C>A, 4485A>T, 4491C>A, 4492G>C, 4493G>A, 4497T>A, 4502C>A, 4503C>G, 4504T>C, 4506T>G, 4508A>T, 4510T>A, 4511C>G, 4513T>A, 4514C>A, 4515C>A, 4530T>A, 4533A>G, 4535T>A, 4538G>A, 4539G>A, 4540C>T, 4542G>A, 4545A>C, 4548C>A, 4549C>A, 4551T>G, 4552G>C, 4553C>A, 4555C>A, 4556C>A, 4560A>T, 4562C>G, 4563A>C, 4566T>C, 4567C>G, 4569G>A, 4570C>T, 4571T>G, 4575C>G, 4576G>A, 4578T>A, 4583G>T, 4584C>G, 4587T>C, 4590C>T, 4591C>A | | | | | | | | |
| **CDS** | | | | | | | | | |
| ORF_V | 317 | 397 | 11.4% | 295 | 50.3% | 81 (100%) | 42 (51.9%) | 0/0/0/0 | 3 |
| Protein mutations: | P317T (4354C>A), A318T (4357G>A 4359C>A), A325S (4378G>A 4379C>G 4380C>T), K328V (4387A>G 4388A>T), A332T (4399G>A), M334I (4407G>A), K340R (4424A>G 4425G>A), D343N (4432G>A 4434C>T), H344N (4435C>A), K346R (4442A>G 4443G>A), G347T (4444G>A 4445G>C), D348Y (4447G>T), G349A (4451G>C 4452C>A), L351N (4456C>A 4457T>A 4458A>T), P353S (4462C>T 4464T>A), N354D (4465A>G 4467C>T), E356S (4471G>A 4472A>G 4473G>T), Q357E (4474C>G), L359I (4480C>A), Q360N (4483C>A 4485A>T), G363Q (4492G>C 4493G>A), T366K (4502C>A 4503C>G), F367L (4504T>C 4506T>G), Y368F (4508A>T), S370K (4513T>A 4514C>A 4515C>A), F377Y (4535T>A), W378* (4538G>A 4539G>A), Q379* (4540C>T 4542G>A), L382M (4549C>A 4551T>G), A383H (4552G>C 4553C>A), P384K (4555C>A 4556C>A), E385D (4560A>T), T386S (4562C>G 4563A>C), Q388E (4567C>G 4569G>A), L389* (4570C>T 4571T>G), A391T (4576G>A 4578T>A), S393M (4583G>T 4584C>G), Q396K (4591C>A) | | | | | | | | |
| Codon mutations: | CCA317ACA (4354C>A), GCC318ACA (4357G>A 4359C>A), TTC319TTT (4362C>T), AGG322AGA (4371G>A), GCC325AGT (4378G>A 4379C>G 4380C>T), GAG326GAA (4383G>A), ATC327ATA (4386C>A), AAA328GTA (4387A>G 4388A>T), CGA329AGA (4390C>A), GGC330GGA (4395C>A), AAA331AAG (4398A>G), GCA332ACA (4399G>A), ATG334ATA (4407G>A), ATT336ATA (4413T>A), AAC337AAT (4416C>T), AAG340AGA (4424A>G 4425G>A), TTA341CTT (4426T>C 4428A>T), GAC343AAT (4432G>A 4434C>T), CAT344AAT (4435C>A), ACA345ACT (4440A>T), AAG346AGA (4442A>G 4443G>A), GGA347ACA (4444G>A 4445G>C), GAT348TAT (4447G>T), GGC349GCA (4451G>C 4452C>A), TAC350TAT (4455C>T), CTA351AAT (4456C>A 4457T>A 4458A>T), CTT352TTA (4459C>T 4461T>A), CCT353TCA (4462C>T 4464T>A), AAC354GAT (4465A>G 4467C>T), AAG355AAA (4470G>A), GAG356AGT (4471G>A 4472A>G 4473G>T), CAA357GAA (4474C>G), CTG358TTA (4477C>T 4479G>A), CTT359ATT (4480C>A), CAA360AAT (4483C>A 4485A>T), ATC362ATA (4491C>A), GGA363CAA (4492G>C 4493G>A), GGT364GGA (4497T>A), ACC366AAG (4502C>A 4503C>G), TTT367CTG (4504T>C 4506T>G), TAC368TTC (4508A>T), TCT369AGT (4510T>A 4511C>G), TCC370AAA (4513T>A 4514C>A 4515C>A), TCT375TCA (4530T>A), GGA376GGG (4533A>G), TTT377TAT (4535T>A), TGG378TAA (4538G>A 4539G>A), CAG379TAA (4540C>T 4542G>A), GTA380GTC (4545A>C), CGC381CGA (4548C>A), CTT382ATG (4549C>A 4551T>G), GCT383CAT (4552G>C 4553C>A), CCA384AAA (4555C>A 4556C>A), GAA385GAT (4560A>T), ACA386AGC (4562C>G 4563A>C), ATT387ATC (4566T>C), CAG388GAA (4567C>G 4569G>A), CTA389TGA (4570C>T 4571T>G), ACC390ACG (4575C>G), GCT391ACA (4576G>A 4578T>A), AGC393ATG (4583G>T 4584C>G), TGT394TGC (4587T>C), CCC395CCT (4590C>T), CAA396AAA (4591C>A) | | | | | | | | |
| **Proteins** | | | | | | | | | |
| hypothetical protein (NP_043933.1) | 317 | 397 | 11.4% | 295 | 50.3% | 81 (100%) | 42 (51.9%) | 0/0/0/0 | 3 |
| Protein mutations: | P317T (4354C>A), A318T (4357G>A 4359C>A), A325S (4378G>A 4379C>G 4380C>T), K328V (4387A>G 4388A>T), A332T (4399G>A), M334I (4407G>A), K340R (4424A>G 4425G>A), D343N (4432G>A 4434C>T), H344N (4435C>A), K346R (4442A>G 4443G>A), G347T (4444G>A 4445G>C), D348Y (4447G>T), G349A (4451G>C 4452C>A), L351N (4456C>A 4457T>A 4458A>T), P353S (4462C>T 4464T>A), N354D (4465A>G 4467C>T), E356S (4471G>A 4472A>G 4473G>T), Q357E (4474C>G), L359I (4480C>A), Q360N (4483C>A 4485A>T), G363Q (4492G>C 4493G>A), T366K (4502C>A 4503C>G), F367L (4504T>C 4506T>G), Y368F (4508A>T), S370K (4513T>A 4514C>A 4515C>A), F377Y (4535T>A), W378* (4538G>A 4539G>A), Q379* (4540C>T 4542G>A), L382M (4549C>A 4551T>G), A383H (4552G>C 4553C>A), P384K (4555C>A 4556C>A), E385D (4560A>T), T386S (4562C>G 4563A>C), Q388E (4567C>G 4569G>A), L389* (4570C>T 4571T>G), A391T (4576G>A 4578T>A), S393M (4583G>T 4584C>G), Q396K (4591C>A) | | | | | | | | |
| Codon mutations: | CCA317ACA (4354C>A), GCC318ACA (4357G>A 4359C>A), TTC319TTT (4362C>T), AGG322AGA (4371G>A), GCC325AGT (4378G>A 4379C>G 4380C>T), GAG326GAA (4383G>A), ATC327ATA (4386C>A), AAA328GTA (4387A>G 4388A>T), CGA329AGA (4390C>A), GGC330GGA (4395C>A), AAA331AAG (4398A>G), GCA332ACA (4399G>A), ATG334ATA (4407G>A), ATT336ATA (4413T>A), AAC337AAT (4416C>T), AAG340AGA (4424A>G 4425G>A), TTA341CTT (4426T>C 4428A>T), GAC343AAT (4432G>A 4434C>T), CAT344AAT (4435C>A), ACA345ACT (4440A>T), AAG346AGA (4442A>G 4443G>A), GGA347ACA (4444G>A 4445G>C), GAT348TAT (4447G>T), GGC349GCA (4451G>C 4452C>A), TAC350TAT (4455C>T), CTA351AAT (4456C>A 4457T>A 4458A>T), CTT352TTA (4459C>T 4461T>A), CCT353TCA (4462C>T 4464T>A), AAC354GAT (4465A>G 4467C>T), AAG355AAA (4470G>A), GAG356AGT (4471G>A 4472A>G 4473G>T), CAA357GAA (4474C>G), CTG358TTA (4477C>T 4479G>A), CTT359ATT (4480C>A), CAA360AAT (4483C>A 4485A>T), ATC362ATA (4491C>A), GGA363CAA (4492G>C 4493G>A), GGT364GGA (4497T>A), ACC366AAG (4502C>A 4503C>G), TTT367CTG (4504T>C 4506T>G), TAC368TTC (4508A>T), TCT369AGT (4510T>A 4511C>G), TCC370AAA (4513T>A 4514C>A 4515C>A), TCT375TCA (4530T>A), GGA376GGG (4533A>G), TTT377TAT (4535T>A), TGG378TAA (4538G>A 4539G>A), CAG379TAA (4540C>T 4542G>A), GTA380GTC (4545A>C), CGC381CGA (4548C>A), CTT382ATG (4549C>A 4551T>G), GCT383CAT (4552G>C 4553C>A), CCA384AAA (4555C>A 4556C>A), GAA385GAT (4560A>T), ACA386AGC (4562C>G 4563A>C), ATT387ATC (4566T>C), CAG388GAA (4567C>G 4569G>A), CTA389TGA (4570C>T 4571T>G), ACC390ACG (4575C>G), GCT391ACA (4576G>A 4578T>A), AGC393ATG (4583G>T 4584C>G), TGT394TGC (4587T>C), CCC395CCT (4590C>T), CAA396AAA (4591C>A) | | | | | | | | |

*: Inserts / Deletes / Misaligned / Frameshifts

#### Analysis details

This analysis was performed with panviral2.64

### NGS Details (UN24): Tomato associated geminivirus 1

#### Assembly

| | |
|---|---|
| **Coverage Length** | 258 (2 contig(s)) |
| **Depth Of Coverage** | 1.0 |
| **Number Of Reads** | 2 |
| **Reads Per Million** | 0.04 rpm (after QC) |
| **Ambiguities** | 0 |
| **Assembly Method** | de novo + reference guided assembly |
| **Consensus Caller** | Bcf Tools |

#### Coverage Map

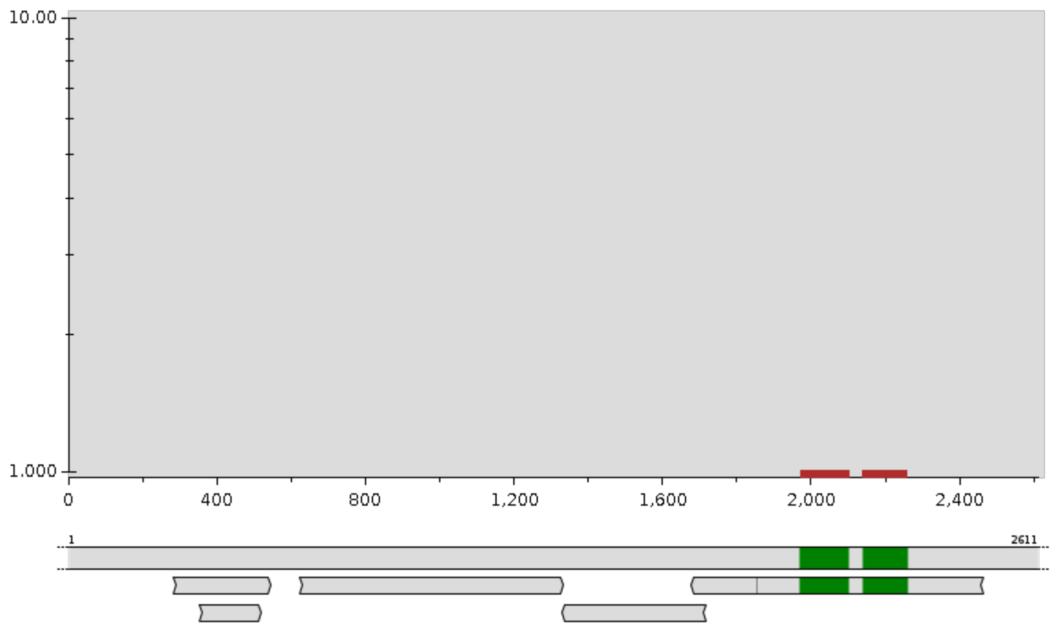

#### Assignment

| | |
|---|---|
| **Type** | Tomato associated geminivirus 1 (Taxonomy ID: 2027645) |
| **Reference Genome** | NC_035676.1 |
| **NT Identity (%)** | 75.969 |
| **AA Identity (%)** | 79.0698 |
| **Number Of Stop Codons** | 4 |
| **Number Of CDS** | 5 |

#### Alignment

| | |
|---|---|
| **Alignment Score** | 268.0 (NT) + 976.0 (AA) = 1244.0 |
| **Concordance (%)** | 71.8245 |

| Alignment Method | Global, seeded, nucleotide + amino acids (AGA) |
|---|---|

#### Genome Region

Sequence starts at position 1970 and ends at position 2261 relative to NC_035676.1 reference sequence.

#### Alignment Detailed Statistics

| | Begin | End | Coverage | Score | Concordance | Matches | Identities | I/D/M/F* | Stop Codons |
|---|---|---|---|---|---|---|---|---|---|
| **NT** | **1970** | **2261** | **9.9%** | **268** | **51.9%** | **258 (100%)** | **196 (76.0%)** | **0/0** | |
| Mutations: | 1981C>T, 1989G>A, 1992G>A, 1998C>G, 1999G>C, 2001G>A, 2007C>T, 2018G>T, 2019T>G, 2024T>G, 2027T>A, 2034G>A, 2037A>G, 2043G>T, 2049G>T, 2055T>A, 2056G>T, 2057A>T, 2061A>T, 2065C>G, 2067T>A, 2073G>T, 2076A>T, 2078C>A, 2079G>A, 2082T>C, 2084T>G, 2086T>G, 2087T>A, 2088C>T, 2094C>A, 2102T>A, 2139A>C, 2150A>C, 2151A>G, 2157G>A, 2160C>G, 2163A>G, 2175C>T, 2177T>A, 2190G>A, 2191A>C, 2192T>A, 2196T>G, 2205A>G, 2209C>G, 2210C>T, 2216C>T, 2220A>T, 2223G>A, 2227G>A, 2228T>C, 2229G>A, 2241C>G, 2243T>G, 2244C>T, 2247G>A, 2251T>C, 2252G>T, 2253C>G, 2256C>G, 2258G>T | | | | | | | | |

##### CDS

| | Begin | End | Coverage | Score | Concordance | Matches | Identities | I/D/M/F* | Stop Codons |
|---|---|---|---|---|---|---|---|---|---|
| CK833_gp4 | 69 | 166 | 25.8% | 488 | 78.0% | 86 (100%) | 68 (79.1%) | 0/0/0/0 | 2 |
| Protein mutations: | Q70N (2256C>G 2258G>T), Q72R (2251T>C 2252G>T), T80V (2227G>A 2228T>C), G84R (2216C>T), G86T (2209C>G 2210C>T), I92C (2190G>A 2191A>C 2192T>A), R97* (2175C>T 2177T>A), S106A (2150A>C), R122* (2102T>A), N127S (2086T>G 2087T>A), A130S (2076A>T 2078C>A), R134T (2065C>G), S137N (2055T>A 2056G>T 2057A>T), T147S (2027T>A), L150M (2018G>T), T156S (1998C>G 1999G>C), R162K (1981C>T) | | | | | | | | |
| Codon mutations: | CAG70AAC (2256C>G 2258G>T), ACG71ACC (2253C>G), CAG72AGG (2251T>C 2252G>T), AAC73AAT (2247G>A), CAG74CAA (2244C>T), AGG75CGC (2241C>G 2243T>G), ATC79ATT (2229G>A), ACC80GTC (2227G>A 2228T>C), GAC81GAT (2223G>A), CCT82CCA (2220A>T), GGA84AGA (2216C>T), GGA86ACA (2209C>G 2210C>T), ATT87ATC (2205A>G), CCA90CCC (2196T>G), ATC92TGT (2190G>A 2191A>C 2192T>A), AGG97TGA (2175C>T 2177T>A), GTT101GTC (2163A>G), GCG102GCC (2160C>G), GAC103GAT (2157G>A), ATT105ATC (2151A>G), TCC106GCC (2150A>C), GGT109GGG (2139A>C), AGA122TGA (2102T>A), CCG124CCT (2094C>A), AAG126AAA (2088C>T), AAT127TCT (2086T>G 2087T>A), AGA128CGG (2082T>C 2084T>G), GAC129GAT (2079G>A), GCT130TCA (2076A>T 2078C>A), ATC131ATA (2073G>T), ACA133ACT (2067T>A), AGA134ACA (2065C>G), ATT135ATA (2061A>T), TCA137AAT (2055T>A 2056G>T 2057A>T), TCC139TCA (2049G>T), TCC141TCA (2043G>T), TCT143TCC (2037A>G), GAC144GAT (2034G>A), ACC147TCC (2027T>A), AGA148CGA (2024T>G), GTA149GTC (2019T>G), CTG150ATG (2018G>T), CAG153CAA (2007C>T), TAC155TAT (2001G>A), ACG156AGC (1998C>G 1999G>C), GCC158GCT (1992G>A), AAC159AAT (1989G>A), AGA162AAA (1981C>T) | | | | | | | | |
| CK833_gp5 | 69 | 166 | 32.8% | 488 | 78.0% | 86 (100%) | 68 (79.1%) | 0/0/0/0 | 2 |
| Protein mutations: | Q70N (2256C>G 2258G>T), Q72R (2251T>C 2252G>T), T80V (2227G>A 2228T>C), G84R (2216C>T), G86T (2209C>G 2210C>T), I92C (2190G>A 2191A>C 2192T>A), R97* (2175C>T 2177T>A), S106A (2150A>C), R122* (2102T>A), N127S (2086T>G 2087T>A), A130S (2076A>T 2078C>A), R134T (2065C>G), S137N (2055T>A 2056G>T 2057A>T), T147S (2027T>A), L150M (2018G>T), T156S (1998C>G 1999G>C), R162K (1981C>T) | | | | | | | | |
| Codon mutations: | CAG70AAC (2256C>G 2258G>T), ACG71ACC (2253C>G), CAG72AGG (2251T>C 2252G>T), AAC73AAT (2247G>A), CAG74CAA (2244C>T), AGG75CGC (2241C>G 2243T>G), ATC79ATT (2229G>A), ACC80GTC (2227G>A 2228T>C), GAC81GAT (2223G>A), CCT82CCA (2220A>T), GGA84AGA (2216C>T), GGA86ACA (2209C>G 2210C>T), ATT87ATC (2205A>G), CCA90CCC (2196T>G), ATC92TGT (2190G>A 2191A>C 2192T>A), AGG97TGA (2175C>T 2177T>A), GTT101GTC (2163A>G), GCG102GCC (2160C>G), GAC103GAT (2157G>A), ATT105ATC (2151A>G), TCC106GCC (2150A>C), GGT109GGG (2139A>C), AGA122TGA (2102T>A), CCG124CCT (2094C>A), AAG126AAA (2088C>T), AAT127TCT (2086T>G 2087T>A), AGA128CGG (2082T>C 2084T>G), GAC129GAT (2079G>A), GCT130TCA (2076A>T 2078C>A), ATC131ATA (2073G>T), ACA133ACT (2067T>A), AGA134ACA (2065C>G), ATT135ATA (2061A>T), TCA137AAT (2055T>A 2056G>T 2057A>T), TCC139TCA (2049G>T), TCC141TCA (2043G>T), TCT143TCC (2037A>G), GAC144GAT (2034G>A), ACC147TCC (2027T>A), AGA148CGA (2024T>G), GTA149GTC (2019T>G), CTG150ATG (2018G>T), CAG153CAA (2007C>T), TAC155TAT (2001G>A), ACG156AGC (1998C>G 1999G>C), GCC158GCT (1992G>A), AAC159AAT (1989G>A), AGA162AAA (1981C>T) | | | | | | | | |

##### Proteins

| | Begin | End | Coverage | Score | Concordance | Matches | Identities | I/D/M/F* | Stop Codons |
|---|---|---|---|---|---|---|---|---|---|
| replication-associated protein (YP_009417307.1) | 69 | 166 | 25.8% | 488 | 78.0% | 86 (100%) | 68 (79.1%) | 0/0/0/0 | 2 |
| Protein mutations: | Q70N (2256C>G 2258G>T), Q72R (2251T>C 2252G>T), T80V (2227G>A 2228T>C), G84R (2216C>T), G86T (2209C>G 2210C>T), I92C (2190G>A 2191A>C 2192T>A), R97* (2175C>T 2177T>A), S106A (2150A>C), R122* (2102T>A), N127S (2086T>G 2087T>A), A130S (2076A>T 2078C>A), R134T (2065C>G), S137N (2055T>A 2056G>T 2057A>T), T147S (2027T>A), L150M (2018G>T), T156S (1998C>G 1999G>C), R162K (1981C>T) | | | | | | | | |
| Codon mutations: | CAG70AAC (2256C>G 2258G>T), ACG71ACC (2253C>G), CAG72AGG (2251T>C 2252G>T), AAC73AAT (2247G>A), CAG74CAA (2244C>T), AGG75CGC (2241C>G 2243T>G), ATC79ATT (2229G>A), ACC80GTC (2227G>A 2228T>C), GAC81GAT (2223G>A), CCT82CCA (2220A>T), GGA84AGA (2216C>T), GGA86ACA (2209C>G 2210C>T), ATT87ATC (2205A>G), CCA90CCC (2196T>G), ATC92TGT (2190G>A 2191A>C 2192T>A), AGG97TGA (2175C>T 2177T>A), GTT101GTC (2163A>G), GCG102GCC (2160C>G), GAC103GAT (2157G>A), ATT105ATC (2151A>G), TCC106GCC (2150A>C), GGT109GGG (2139A>C), AGA122TGA (2102T>A), CCG124CCT (2094C>A), AAG126AAA (2088C>T), AAT127TCT (2086T>G 2087T>A), AGA128CGG (2082T>C 2084T>G), GAC129GAT (2079G>A), GCT130TCA (2076A>T 2078C>A), ATC131ATA (2073G>T), ACA133ACT (2067T>A), AGA134ACA (2065C>G), ATT135ATA (2061A>T), TCA137AAT (2055T>A 2056G>T 2057A>T), TCC139TCA (2049G>T), TCC141TCA (2043G>T), TCT143TCC (2037A>G), GAC144GAT (2034G>A), ACC147TCC (2027T>A), AGA148CGA (2024T>G), GTA149GTC (2019T>G), CTG150ATG (2018G>T), CAG153CAA (2007C>T), TAC155TAT (2001G>A), ACG156AGC (1998C>G 1999G>C), GCC158GCT (1992G>A), AAC159AAT (1989G>A), AGA162AAA (1981C>T) | | | | | | | | |
| RepA (YP_009417308.1) | 69 | 166 | 32.8% | 488 | 78.0% | 86 (100%) | 68 (79.1%) | 0/0/0/0 | 2 |
| Protein mutations: | Q70N (2256C>G 2258G>T), Q72R (2251T>C 2252G>T), T80V (2227G>A 2228T>C), G84R (2216C>T), G86T (2209C>G 2210C>T), I92C (2190G>A 2191A>C 2192T>A), R97* (2175C>T 2177T>A), S106A (2150A>C), R122* (2102T>A), N127S (2086T>G 2087T>A), A130S (2076A>T 2078C>A), R134T (2065C>G), S137N (2055T>A 2056G>T 2057A>T), T147S (2027T>A), L150M (2018G>T), T156S (1998C>G 1999G>C), R162K (1981C>T) | | | | | | | | |
| Codon mutations: | CAG70AAC (2256C>G 2258G>T), ACG71ACC (2253C>G), CAG72AGG (2251T>C 2252G>T), AAC73AAT (2247G>A), CAG74CAA (2244C>T), AGG75CGC (2241C>G 2243T>G), ATC79ATT (2229G>A), ACC80GTC (2227G>A 2228T>C), GAC81GAT (2223G>A), CCT82CCA (2220A>T), GGA84AGA (2216C>T), GGA86ACA (2209C>G 2210C>T), ATT87ATC (2205A>G), CCA90CCC (2196T>G), ATC92TGT (2190G>A 2191A>C 2192T>A), AGG97TGA (2175C>T 2177T>A), GTT101GTC (2163A>G), GCG102GCC (2160C>G), GAC103GAT (2157G>A), ATT105ATC (2151A>G), TCC106GCC (2150A>C), GGT109GGG (2139A>C), AGA122TGA (2102T>A), CCG124CCT (2094C>A), AAG126AAA (2088C>T), AAT127TCT (2086T>G 2087T>A), AGA128CGG (2082T>C 2084T>G), GAC129GAT (2079G>A), GCT130TCA (2076A>T 2078C>A), ATC131ATA (2073G>T), ACA133ACT (2067T>A), AGA134ACA (2065C>G), ATT135ATA (2061A>T), TCA137AAT (2055T>A 2056G>T 2057A>T), TCC139TCA (2049G>T), TCC141TCA (2043G>T), TCT143TCC (2037A>G), GAC144GAT (2034G>A), ACC147TCC (2027T>A), AGA148CGA (2024T>G), GTA149GTC (2019T>G), CTG150ATG (2018G>T), CAG153CAA (2007C>T), TAC155TAT (2001G>A), ACG156AGC (1998C>G 1999G>C), GCC158GCT (1992G>A), AAC159AAT (1989G>A), AGA162AAA (1981C>T) | | | | | | | | |

*: Inserts / Deletes / Misaligned / Frameshifts

#### Analysis details

This analysis was performed with panviral2.64

### Analysis of UN59

| | |
|---|---|
| **Technology** | Paired-end short reads |
| **Input Files** | UN59_R1.fq.gz (1.47 GB), UN59_R2.fq.gz (1.52 GB) |
| **Submitted On** | 2023-09-26 13:15:02 UTC |
| **Duration** | 2h 52m 57s |
| **Tool Version** | panviral2.64 |
| **Location** | https://www.genomedetective.com/db/ui/analysis/2e79870b-faad-478b-bd65-8d012eb0827f |

#### Statistics

| | |
|---|---|
| **Original Read Length** | 20 - 150 |
| **Trimmed Read Length** | 50 - 135 |

| | # Reads | % of Reads |
|---|---|---|
| **Input file** | 46314422 | 100.0% |
| **After QC** | 46000056 | 99.3% |
| **After filtering** | 2552086 | 5.5% |
| **Mapped back to contigs** | 725798 | 1.6% |

#### Assignments

| Assignment | No. of Reads | Depth of Coverage | Identity | | Genome Coverage | |
|---|---|---|---|---|---|---|
| | | | NT | AA | | |
| Torradovirus lycopersici (2 segments out of 2) | 415162 | 4267.6 | 89.2% | 95.0% | 99.8% | |
| Torradovirus lycopersici (segment RNA 2) | 251860 | 6024.7 | 85.2% | 89.6% | 99.6% | 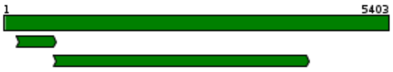 |
| Torradovirus lycopersici (segment RNA 1) | 163302 | 3055.8 | 92.0% | 98.4% | 99.9% | 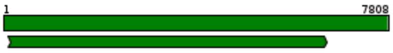 |
| Torradovirus marchitezum (2 segments out of 2) | 188453 | 2391.1 | 65.6% | 69.1% | 84.2% | |
| Torradovirus marchitezum (segment RNA 2) | 128651 | 3539.8 | 64.5% | 68.0% | 94.9% | 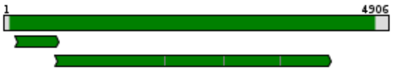 |
| Torradovirus marchitezum (segment RNA 1) | 59802 | 1430.4 | 66.5% | 69.9% | 77.0% | 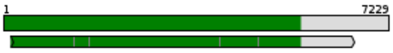 |
| Bracoviriform glomeratae (segment NC_043292.1) | 12058 | 4640.4 | 72.7% | 84.3% | 72.3% | 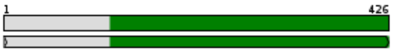 |
| Diachasmimorpha longicaudata entomopoxvirus (segment NC_043455.1) | 7928 | 876.9 | 59.1% | 55.3% | 83.7% | 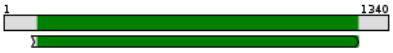 |

| Assignment | No. of Reads | Depth of Coverage | Identity | | Genome Coverage | |
|---|---|---|---|---|---|---|
| | | | NT | AA | | |
| Potato virus Y | 757 | 11.7 | 83.3% | 90.2% | 89.3% | 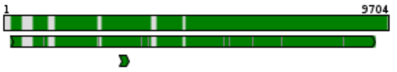 |
| Duamitovirus soch1 | 734 | 39.0 | 67.5% | 68.0% | 82.0% | 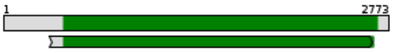 |
| Solendovirus venanicotianae | 285 | 14.6 | 79.8% | 76.8% | 29.4% | 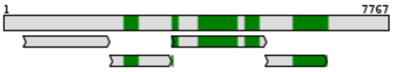 |

#### Discoveries

| Similar to | No. of Reads | Depth of Coverage | Identity | | Genome Coverage | |
|---|---|---|---|---|---|---|
| | | | NT | AA | | |
| Tomato chocolate spot virus (3 segments out of 2) | 71567 | 4220.5 | 70.5% | 73.2% | 19.7% | |
| Tomato chocolate spot virus (segment RNA 1) | 56205 | 7250.8 | 67.5% | 66.2% | 16.2% | 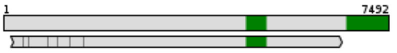 |
| Tomato chocolate spot virus (segment RNA2) | 9765 | 1360.0 | 69.7% | 75.4% | 15.9% | 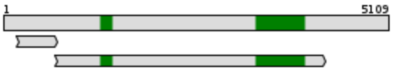 |
| Tomato chocolate spot virus (segment RNA 1) | 5597 | 1311.6 | 79.5% | 80.4% | 6.2% | 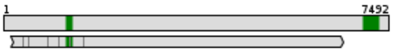 |
| Lausannevirus | 18309 | 9728.0 | 81.8% | 94.9% | 0.1% | 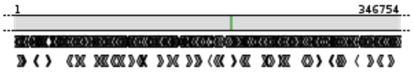 |
| Makelovirus prm1 | 4406 | 557.0 | 76.8% | 87.6% | 0.7% | 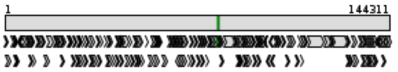 |
| Lausannevirus | 1051 | 709.4 | 79.8% | 95.3% | 0.0% | 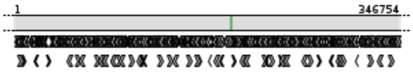 |
| Betabaculovirus disaccharalis | 905 | 510.6 | 83.9% | 91.3% | 0.1% | 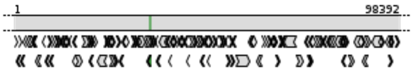 |
| Betabaculovirus disaccharalis | 820 | 486.0 | 82.8% | 88.9% | 0.1% | 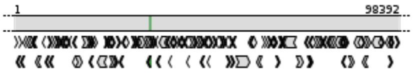 |
| Cladosporium fulvum T-1 virus | 529 | 42.1 | 54.1% | 47.2% | 19.8% | 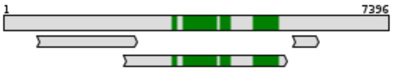 |
| Epiphyllum badnavirus 1 | 512 | 143.5 | 52.5% | 46.8% | 5.3% | 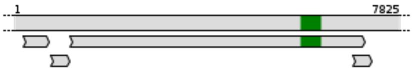 |
| Human gammaherpesvirus 8 (subtype: Could not assign) | 421 | 265.0 | 83.7% | 84.4% | 0.1% | 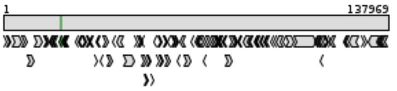 |
| Tomato necrotic dwarf virus (segment RNA1) | 407 | 176.1 | 75.2% | 86.1% | 4.2% | 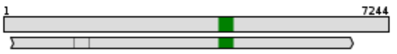 |

| Similar to | No. of Reads | Depth of Coverage | Identity | | Genome Coverage | |
|---|---|---|---|---|---|---|
| | | | NT | AA | | |
| Badnavirus maculasmallanthi | 339 | 121.8 | 56.5% | 53.5% | 3.9% | 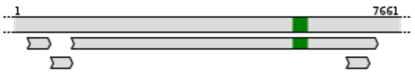 |
| Cassava brown streak virus | 273 | 61.7 | 61.1% | 58.2% | 5.9% | 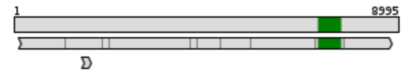 |
| Caulimovirus venafragariae | 134 | 12.6 | 56.6% | 48.5% | 15.5% | 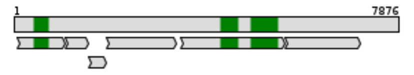 |
| Dioscovirus dioscoreae | 81 | 8.0 | 62.0% | 52.2% | 14.7% | 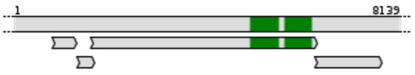 |
| Badnavirus occultipomeae | 80 | 30.9 | 58.1% | 50.0% | 3.2% | 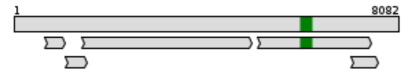 |
| Badnavirus venabougainvilleae | 73 | 25.3 | 55.2% | 46.9% | 3.3% | 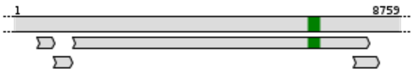 |
| Badnavirus occultipomeae | 50 | 18.2 | 57.8% | 49.0% | 3.7% | 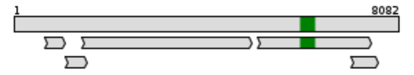 |
| Dioscorea bacilliform virus | 35 | 14.2 | 55.2% | 45.2% | 3.8% | 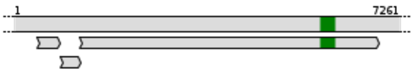 |
| Dioscorea bacilliform virus | 11 | 3.5 | 53.1% | 39.7% | 5.8% | 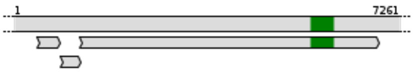 |
| Badnavirus venaribis | 36 | 9.7 | 52.9% | 45.9% | 5.8% | 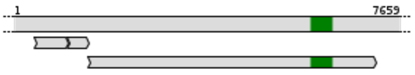 |
| Errantivirus | 32 | 8.5 | 31.4% | 30.3% | 6.6% | 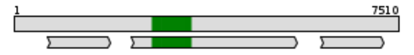 |
| Pinus nigra virus 1 | 27 | 6.5 | 56.6% | 43.5% | 7.5% | 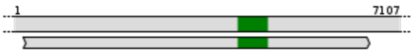 |
| Duamitovirus peex1 | 26 | 6.5 | 71.9% | 65.9% | 14.4% | 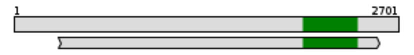 |
| Caulimovirus latensarmoraciae | 26 | 9.8 | 59.9% | 56.1% | 4.3% | 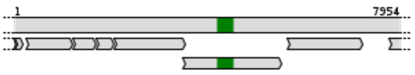 |
| Badnavirus volubetulae | 25 | 7.7 | 50.9% | 47.1% | 5.0% | 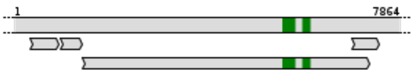 |
| Petuvirus venapetuniae | 25 | 5.3 | 51.9% | 42.9% | 8.5% | 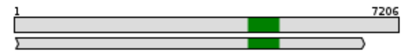 |
| Badnavirus maculasmallanthi | 23 | 5.8 | 58.7% | 60.4% | 6.3% | 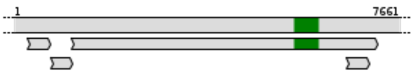 |

| Similar to | No. of Reads | Depth of Coverage | Identity | | Genome Coverage | |
|---|---|---|---|---|---|---|
| | | | NT | AA | | |
| Epiphyllum badnavirus 1 | 22 | 7.4 | 51.0% | 42.0% | 4.5% | 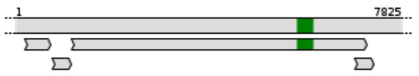  |
| Badnavirus rutilanscamelliae | 21 | 9.1 | 52.4% | 38.5% | 3.4% | 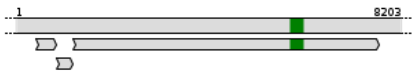  |
| Unknown | 20 | 5.6 | 56.7% | 53.6% | 4.3% | 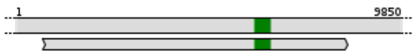  |
| Badnavirus venabougainvilleae | 18 | 6.9 | 53.3% | 46.1% | 3.5% | 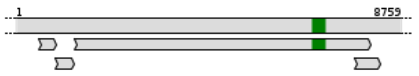  |
| Epiphyllum badnavirus 1 | 16 | 6.7 | 58.0% | 50.0% | 3.5% | 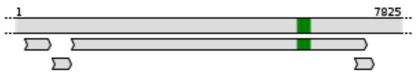  |
| Cavemovirus deltaepiphylli | 15 | 4.4 | 60.0% | 52.9% | 5.7% | 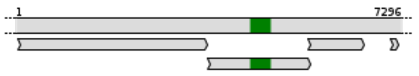  |
| Alphabaculovirus lydisparis | 8 | 4.3 | 78.6% | 89.1% | 0.1% | 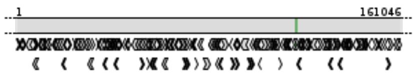  |
| Trialeurodes vaporariorum mononega-like virus 2 | 8 | 4.0 | 99.6% | 97.8% | 2.4% | 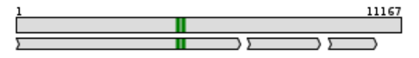  |
| Caulimovirus venafragariae | 7 | 2.2 | 65.1% | 60.9% | 5.0% | 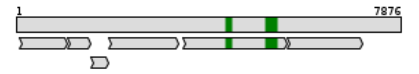  |
| Burkholderia phage Magia | 4 | 2.6 | 86.8% | 84.8% | 0.4% | 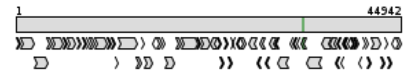  |
| Sugarcane chlorotic streak virus | 3 | 1.6 | 59.9% | 56.2% | 7.3% | 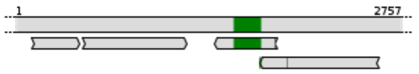  |
| Alphafusarivirus pleosporae | 2 | 1.1 | 75.2% | 85.7% | 3.8% | 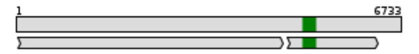  |
| Potato virus X | 2 | 1.2 | 79.1% | 97.3% | 3.5% | 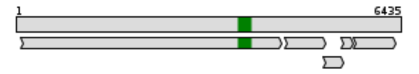  |
| Colombian datura virus | 2 | 1.0 | 98.1% | 97.8% | 2.8% | 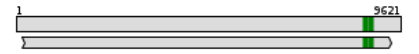  |
| Rahariannevirus raharianne | 2 | 1.0 | 82.1% | 82.1% | 0.5% | 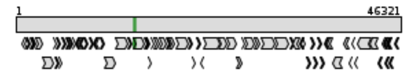  |
| Euphorbia caput-medusae latent virus | 1 | 1.0 | 64.2% | 66.3% | 4.6% | 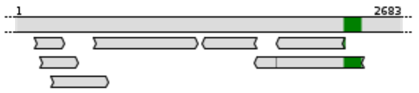  |
| Escherichia virus DE3 | 1 | 1.0 | 100.0% | 96.3% | 0.3% | 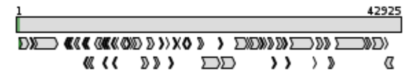  |

### NGS Details (UN59): Torradovirus lycopersici (segment RNA 1)

#### Assembly

| | |
|---|---|
| **Coverage Length** | 7799 (1 contig(s)) |
| **Depth Of Coverage** | 3055.8 |
| **Number Of Reads** | 163302 |
| **Reads Per Million** | 3550.04 rpm (after QC) |
| **Ambiguities** | 0 |
| **Assembly Method** | de novo + reference guided assembly |
| **Consensus Caller** | Bcf Tools |

#### Coverage Map

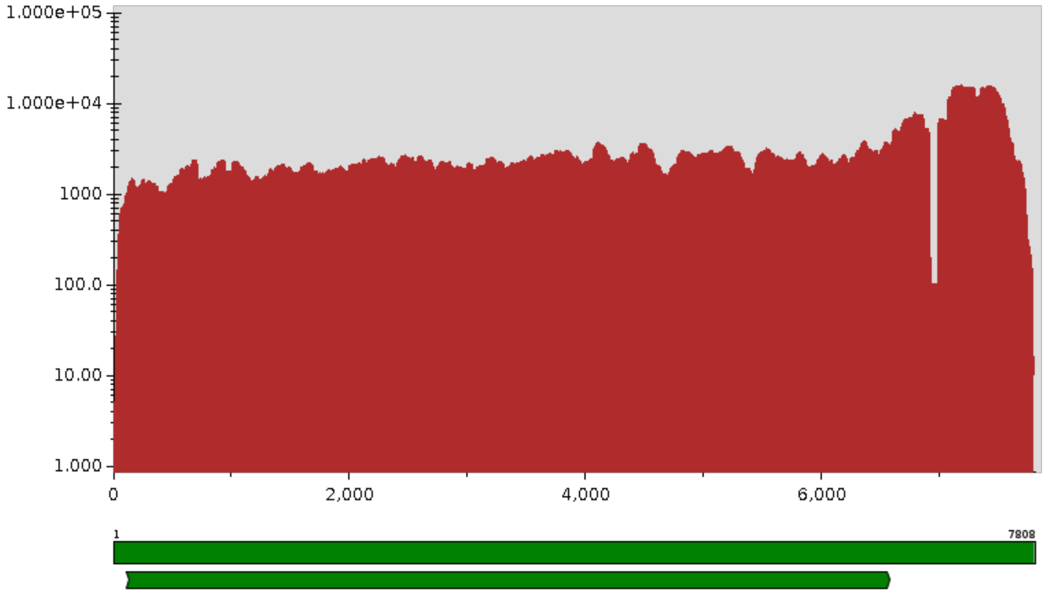

#### Assignment

| | |
|---|---|
| **Type** | Torradovirus lycopersici (Taxonomy ID: 3048378) |
| **Reference Genome** | NC_009013.1 |
| **NT Identity (%)** | 92.0064 |
| **AA Identity (%)** | 98.4252 |
| **Number Of Stop Codons** | 1 |
| **Number Of CDS** | 1 |

#### Alignment

| | |
|---|---|
| **Alignment Score** | 12080.0 (NT) + 14905.0 (AA) = 26985.0 |
| **Concordance (%)** | 91.0702 |

| Alignment Method | Global, seeded, nucleotide + amino acids (AGA) |
|---|---|

#### Genome Region

Sequence starts at position 1 and ends at position 7799 relative to NC_009013.1 reference sequence.

#### Alignment Detailed Statistics

| | Begin | End | Coverage | Score | Concordance | Matches | Identities | I/D/M/F* | Stop Codons |
|---|---|---|---|---|---|---|---|---|---|
| **NT** | **1** | **7799** | **99.9%** | **12080** | **83.0%** | **7519 (96.3%)** | **6929 (88.7%)** | **12/280** | |
| Mutations: | 2T>C, 12A>C, 15T>A, 76_77insT, 83C>T, 83_84insT, 87C>T, 115T>C, 127C>T, 128C>T, 130C>T, 135T>C, 148T>C, 158G>A, 160T>C, 161A>G, 187A>T, 199T>A, 208T>C, 220C>T, 224C>T, 253A>G, 259T>C, 262T>A, 265C>T, 340G>A, 349G>A, 373C>T, 376C>T, 445C>T, 487C>T, 502G>A, 541T>C, 559C>T, 578C>T, 628C>T, 655C>T, 676A>G, 694A>G, 736A>G, 842C>T, 880A>G, 898G>A, 925C>T, 952A>G, 964T>G, 970G>A, 997C>T, 1006T>C, 1033G>A, 1060T>C, 1078A>T, 1081C>T, 1093T>C, 1105A>G, 1108T>C, 1109C>T, 1165C>T, 1180G>A, 1196A>G, 1231G>A, 1240C>T, 1255G>A, 1267T>C, 1276A>G, 1300G>A, 1351A>G, 1372C>T, 1411T>C, 1417T>C, 1429C>T, 1450T>C, 1451C>T, 1480C>T, 1481A>G, 1489T>C, 1522C>T, 1552C>T, 1555C>T, 1570A>G, 1573T>C, 1582C>T, 1585A>T, 1591C>T, 1594T>C, 1607G>T, 1612G>A, 1618T>G, 1624C>A, 1645A>G, 1669C>T, 1681T>A, 1684G>A, 1699T>C, 1705T>C, 1737A>G, 1738A>G, 1771C>T, 1781T>C, 1789A>G, 1897A>G, 1910T>C, 1954T>C, 1960A>G, 1984C>T, 2014A>G, 2023T>C, 2056T>C, 2077C>A, 2082T>A, 2092A>G, 2107A>G, 2128T>C, 2138C>T, 2143T>C, 2149T>C, 2155T>C, 2174C>T, 2179A>C, 2182G>A, 2194T>A, 2200T>C, 2215G>A, 2218C>T, 2224C>T, 2236A>G, 2241A>G, 2250G>A, 2251C>T, 2257C>T, 2263G>A, 2275T>C, 2287G>A, 2290C>T, 2314C>T, 2323A>T, 2347C>T, 2350A>G, 2356C>T, 2359G>A, 2362C>T, 2377G>A, 2380T>C, 2392A>G, 2407T>C, 2409G>A, 2443T>C, 2446G>A, 2479G>A, 2485C>T, 2497G>A, 2500T>C, 2503G>A, 2509C>T, 2512T>C, 2542G>A, 2548C>T, 2557C>T, 2620G>A, 2626T>C, 2650G>A, 2665T>C, 2675T>C, 2683A>G, 2692G>A, 2702C>T, 2710A>G, 2719T>C, 2725T>C, 2731C>T, 2734C>T, 2743A>G, 2744T>C, 2765T>C, 2773A>G, 2809C>T, 2812G>A, 2827A>G, 2869C>T, 2881T>C, 2929C>T, 2930T>C, 2935G>A, 2941A>G, 2950C>T, 2951C>T, 3037C>T, 3040T>C, 3049A>G, 3064A>G, 3109T>C, 3163T>C, 3169C>T, 3202T>C, 3241T>C, 3253C>T, 3256G>A, 3265A>G, 3313T>C, 3406A>C, 3418C>A, 3421T>C, 3463A>G, 3466G>A, 3481C>T, 3484A>T, 3534G>A, 3541C>T, 3580T>C, 3583C>T, 3610A>G, 3613T>C, 3625C>T, 3628G>A, 3643A>G, 3646C>T, 3661A>G, 3664T>C, 3673G>A, 3676C>T, 3686C>T, 3706T>C, 3716T>C, 3721T>A, 3739A>T, 3751A>G, 3754G>A, 3758G>A, 3766G>A, 3808T>C, 3814C>T, 3817G>C, 3824T>C, 3835T>C, 3844T>C, 3857T>C, 3859G>A, 3868C>T, 3871T>C, 3883G>A, 3898T>C, 3907A>T, 3916C>T, 3925A>G, 3928G>A, 3949C>T, 3955G>A, 3961G>A, 3970A>G, 3973T>C, 4034T>C, 4066T>C, 4075A>G, 4084C>T, 4100T>C, 4118C>T, 4129C>T, 4147C>A, 4159G>A, 4162A>T, 4168A>G, 4186A>T, 4195A>G, 4207G>A, 4210T>C, 4228C>T, 4294A>G, 4339T>C, 4343C>T, 4348C>T, 4354T>C, 4396T>C, 4408C>T, 4414T>G, 4417T>C, 4420G>A, 4459T>C, 4462C>T, 4465C>T, 4468C>T, 4471T>C, 4474T>C, 4591T>C, 4636T>C, 4640C>T, 4660A>G, 4684A>C, 4696C>T, 4702G>A, 4705C>T, 4714T>C, 4735C>T, 4747A>G, 4759C>T, 4768T>C, 4774G>A, 4777C>T, 4783A>G, 4786A>G, 4789C>T, 4813A>G, 4816T>C, 4843G>A, 4898G>A, 4906G>A, 4969G>A, 4988A>G, 4993A>T, 5005T>C, 5012T>C, 5029G>A, 5065C>T, 5074T>C, 5077T>C, 5079T>C, 5102T>C, 5118C>T, 5119T>C, 5120G>A, 5125G>C, 5146T>C, 5148C>A, 5200G>A, 5218T>G, 5244G>A, 5266C>T, 5281C>T, 5329C>T, 5341T>A, 5344A>G, 5347C>T, 5369G>A, 5371G>A, 5386C>T, 5392G>A, 5407A>G, 5413C>T, 5419A>T, 5437T>C, 5440C>T, 5461T>C, 5471T>C, 5482A>G, 5533T>A, 5572T>C, 5578T>C, 5584T>C, 5617A>T, 5641T>C, 5647T>C, 5686G>A, 5690T>C, 5710T>A, 5716G>T, 5734T>C, 5737T>C, 5739G>A, 5746T>C, 5750T>C, 5758C>T, 5773G>A, 5784T>C, 5786T>C, 5797A>G, 5806C>T, 5836A>T, 5849T>C, 5850T>A, 5851G>A, 5878G>T, 5884T>C, 5887C>T, 5890C>T, 5914T>C, 5923T>C, 5927C>T, 5932C>T, 5955A>G, 5965G>A, 5998A>G, 6004C>T, 6016A>G, 6028C>T, 6031A>T, 6033C>T, 6035G>T, 6055T>A, 6067C>A, 6070T>C, 6094T>G, 6103G>A, 6106G>A, 6139T>C, 6145T>C, 6148T>C, 6154C>T, 6184A>G, 6187T>C, 6193C>T, 6196A>G, 6208A>G, 6226G>A, 6230A>G, 6232A>G, 6235G>A, 6256C>A, 6262A>G, 6265G>A, 6292T>A, 6298A>G, 6304A>G, 6308T>C, 6313T>C, 6319A>C, 6322G>A, 6323A>G, 6358T>C, 6372C>T, 6487T>C, 6493T>C, 6517C>T, 6545C>T, 6571C>T, 6587G>A, 6660C>T, 6661T>C, 6670C>T, 6685A>G, 6742A>T, 6766delC, 6786T>C, 6798_6799insA, 6806_6852delTAACGGAGAGTACTGACTTTAACTAGTTGGGAGTCCGGCTCCATTTG, 6854A>T, 6857A>G, 6858C>A, 6859C>T, 6860A>G, 6861A>G, 6864A>T, 6867T>A, 6871_6894delATTGGTTAAAGAGATTTGCACGCC, 6897T>A, 6905T>G, 6907T>A, 6908G>T, 6911T>C, 6912C>G, 6915G>C, 6918C>T, 6921T>G, 6923C>G, 6924T>A, 6927G>T, 6928T>G, 6931G>T, 6933_6935delAAG, 6938C>A, 6947A>C, 6948_6951delGGTT, 6957A>T, 6959_6968delTACGGGGAGC, 6984T>C, 6984_6985insG, 6988G>A, 6994G>A, 6997A>T, 6998T>C, 7002A>T, 7003T>A, 7005_7012delATAGTCCT, 7014T>A, 7020_7026delAAGGGTT, 7031T>A, 7033G>A, 7035T>G, 7037T>A, 7038G>C, 7043T>C, 7045T>G, 7047_7060delGCCGTGATGAAAGA, 7063A>T, 7069C>A, 7071T>A, 7074C>A, 7077T>C, 7083G>T, 7085A>G, 7086G>T, 7087_7099delGCTGAAACTACAC, 7101T>G, 7108T>A, 7109G>T, 7113T>G, 7117C>T, 7118T>G, 7120A>T, 7123_7125delCCT, 7129_7133delATCAG, 7136C>A, 7137G>C, 7141G>A, 7144_7169delATTCGATAATTTTCCGTAGCTTGCGT, 7173_7177delGCTGC, 7179C>G, 7182G>A, 7183T>C, 7184T>C, 7185A>C, 7189G>A, 7194_7216delAGTGAAGATGCGCCGTACCACGT, 7221C>T, 7224_7270delGGCAATGCCAGTGGTTCAGAGCGGGCCCTCAGAATAGAGGTTAAAAC, 7275_7285delGTGATGGTGTA, 7288T>A, 7294T>G, 7295A>C, 7297A>C, 7300T>G, 7303C>A, 7304_7315delACCCGGGTTGTG, 7320delG, 7325_7326delGT, 7332C>G, 7333G>T, 7336C>T, 7337A>G, 7339_7348delAGGTCCCACC, 7350T>A, 7351A>T, 7353A>T, 7354G>A, 7358A>C, 7360G>A, 7361A>C, 7362A>C, 7364C>T, 7368G>A, 7372A>T, 7375T>G, 7376T>G, 7377T>A, 7379A>T, 7388C>T, 7397G>A, 7398C>A, 7399G>A, 7400A>T, 7404C>T, 7407A>T, 7408G>T, 7423A>T, 7425G>A, 7435T>C, 7436G>A, 7444C>T, 7445A>G, 7457A>G, 7458T>C, 7460G>A, 7498C>T, 7503C>T, 7505A>G, 7511_7512insC, 7513A>C, 7514A>T, 7515A>C, 7522T>C, 7525_7526insG, 7526T>A, 7526_7527insG, 7545T>A, 7554A>G, 7569G>A, 7585T>C, 7589T>A, 7600T>G, 7601T>A, 7602T>G, 7603_7604insC, 7610C>T, 7612G>A, 7615A>G, 7616A>C, 7617A>T, 7618C>T, 7625A>T, 7628_7631delGCTT, 7637G>T, 7641T>A, 7644G>T, 7647T>G, 7648G>T, 7649T>C, 7657G>C, 7660T>A, 7665G>T, 7666T>C, 7667G>T, 7678C>T, 7686T>G, 7687T>A, 7689G>T, 7698G>A, 7717A>C, 7719T>A, 7723T>C, 7739T>A, 7742_7743insC, 7749T>C, 7751T>C, 7752G>A, 7754_7755insG, 7756_7757insT, 7762C>T, 7763A>C, 7764A>T, 7767C>T, 7773G>T, 7775G>A, 7778T>A, 7782A>T, 7783A>C, 7784_7785insT | | | | | | | | |

##### CDS

| | Begin | End | Coverage | Score | Concordance | Matches | Identities | I/D/M/F* | Stop Codons |
|---|---|---|---|---|---|---|---|---|---|
| ToTV_sRNA1gp1 | 1 | 2159 | 100% | 14905 | 98.9% | 2159 (100%) | 2125 (98.4%) | 0/0/0/0 | 1 |
| Protein mutations: | P8S (128C>T 130C>T), F10S (135T>C), A18T (158G>A 160T>C), T19A (161A>G), T364A (1196A>G), I459V (1481A>G), A501S (1607G>T), K544R (1737A>G 1738A>G), F659Y (2082T>A), N712S (2241A>G), S715N (2250G>A 2251C>T), R768K (2409G>A), R1143K (3534G>A), V1218I (3758G>A), P1413S (4343C>T), V1598I (4898G>A), T1628A (4988A>G), I1658T (5079T>C), T1671I (5118C>T 5119T>C), V1672I (5120G>A), T1681N (5148C>A), R1713K (5244G>A), V1755I (5369G>A 5371G>A), I1792M (5482A>G), R1878K (5739G>A), I1893T (5784T>C), L1915Q (5849T>C 5850T>A 5851G>A), N1950S (5955A>G), T1976I (6033C>T), A1977S (6035G>T), I2042V (6230A>G 6232A>G), I2073V (6323A>G), A2089V (6372C>T), L2147F (6545C>T) | | | | | | | | |

| | Begin | End | Coverage | Score | Concordance | Matches | Identities | I/D/M/F* | Stop Codons |
|---|---|---|---|---|---|---|---|---|---|
| **NT** | **1** | **7799** | **99.9%** | **12080** | **83.0%** | **7519 (96.3%)** | **6929 (88.7%)** | **12/280** | |
| Codon mutations: | TTT3TTC (115T>C), TTC7TTT (127C>T), CCC8TCT (128C>T 130C>T), TTC10TCC (135T>C), ACT14ACC (148T>C), GCT18ACC (158G>A 160T>C), ACC19GCC (161A>G), TCA27TCT (187A>T), GCT31GCA (199T>A), AGT34AGC (208T>C), CGC38CGT (220C>T), CTG40TTG (224C>T), CAA49CAG (253A>G), TCT51TCC (259T>C), TCT52TCA (262T>A), CAC53CAT (265C>T), TTG78TTA (340G>A), AAG81AAA (349G>A), TTC89TTT (373C>T), ATC90ATT (376C>T), AGC113AGT (445C>T), GAC127GAT (487C>T), GTG132GTA (502G>A), TCT145TCC (541T>C), TTC151TTT (559C>T), CTA158TTA (578C>T), AAC174AAT (628C>T), GAC183GAT (655C>T), CAA190CAG (676A>G), GCA196GCG (694A>G), CCA210CCG (736A>G), CTG246TTG (842C>T), CAA258CAG (880A>G), AGG264AGA (898G>A), GCC273GCT (925C>T), GTA282GTG (952A>G), GGT286GGG (964T>G), AAG288AAA (970G>A), GCC297GCT (997C>T), TCT300TCC (1006T>C), GAG309GAA (1033G>A), CAT318CAC (1060T>C), GCA324GCT (1078A>T), AAC325AAT (1081C>T), ACT329ACC (1093T>C), CAA333CAG (1105A>G), CAT334CAC (1108T>C), CTA335TTA (1109C>T), GCC353GCT (1165C>T), AGG358AGA (1180G>A), ACC364GCC (1196A>G), CAG375CAA (1231G>A), GGC378GGT (1240C>T), AGG383AGA (1255G>A), TTT387TTC (1267T>C), CAA390CAG (1276A>G), GGG398GGA (1300G>A), GAA415GAG (1351A>G), CTC422CTT (1372C>T), TAT435TAC (1411T>C), CAT437CAC (1417T>C), ATC441ATT (1429C>T), GCT448GCC (1450T>C), CTG449TTG (1451C>T), GAC458GAT (1480C>T), ATT459GTT (1481A>G), GAT461GAC (1489T>C), CCC472CCT (1522C>T), GGC482GGT (1552C>T), CGC483CGT (1555C>T), CGA488CGG (1570A>G), TAT489TAC (1573T>C), GCC492GCT (1582C>T), ACA493ACT (1585A>T), AAC495AAT (1591C>T), CTT496CTC (1594T>C), GCT501TCT (1607G>T), CAG502CAA (1612G>A), GGT504GGG (1618T>G), ACC506ACA (1624C>A), AGA513AGG (1645A>G), TGC521TGT (1669C>T), GGT525GGA (1681T>A), GAG526GAA (1684G>A), GCT531GCC (1699T>C), ACT533ACC (1705T>C), AAA544AGG (1737A>G 1738A>G), AGC555AGT (1771C>T), TTG559CTG (1781T>C), GAA561GAG (1789A>G), GCA597GCG (1897A>G), TTG602CTG (1910T>C), GTT616GTC (1954T>C), GTA618GTG (1960A>G), TAC626TAT (1984C>T), AAA636AAG (2014A>G), TAT639TAC (2023T>C), CCT650CCC (2056T>C), GCC657GCA (2077C>A), TTT659TAT (2082T>A), CTA662CTG (2092A>G), AAA667AAG (2107A>G), GTT674GTC (2128T>C), CTG678TTG (2138C>T), TAT679TAC (2143T>C), GGT681GGC (2149T>C), TGT683TGC (2155T>C), CTG690TTG (2174C>T), ATA691ATC (2179A>C), AAG692AAA (2182G>A), GCT696GCA (2194T>A), TAT698TAC (2200T>C), AAG703AAA (2215G>A), AGC704AGT (2218C>T), ATC706ATT (2224C>T), ACA710ACG (2236A>G), AAT712AGT (2241A>G), AGC715AAT (2250G>A 2251C>T), GAC717GAT (2257C>T), CAG719CAA (2263G>A), CCT723CCC (2275T>C), GCG727GCA (2287G>A), CAC728CAT (2290C>T), GAC736GAT (2314C>T), ATA739ATT (2323A>T), CAC747CAT (2347C>T), GTA748GTG (2350A>G), TTC750TTT (2356C>T), TTG751TTA (2359G>A), TAC752TAT (2362C>T), AGG757AGA (2377G>A), TAT758TAC (2380T>C), AGA762AGG (2392A>G), CCT767CCC (2407T>C), AGG768AAG (2409G>A), ATT779ATC (2443T>C), GTG780GTA (2446G>A), CCG791CCA (2479G>A), ATC793ATT (2485C>T), CTG797CTA (2497G>A), ATT798ATC (2500T>C), AAG799AAA (2503G>A), GGC801GGT (2509C>T), CTT802CTC (2512T>C), GGG812GGA (2542G>A), GTC814GTT (2548C>T), TTC817TTT (2557C>T), GAG838GAA (2620G>A), AAT840AAC (2626T>C), AAG848AAA (2650G>A), TTT853TTC (2665T>C), TTG857CTG (2675T>C), GCA859GCG (2683A>G), AAG862AAA (2692G>A), CTG866TTG (2702C>T), AAA868AAG (2710A>G), AGT871AGC (2719T>C), GAT873GAC (2725T>C), TTC875TTT (2731C>T), ATC876ATT (2734C>T), GGA879GGG (2743A>G), TTG880CTG (2744T>C), TTG887CTG (2765T>C), CTA889CTG (2773A>G), TGC901TGT (2809C>T), TCG902TCA (2812G>A), ACA907ACG (2827A>G), TTC921TTT (2869C>T), CAT925CAC (2881T>C), TTC941TTT (2929C>T), TTG942CTG (2930T>C), AAG943AAA (2935G>A), CAA945CAG (2941A>G), GTC948GTT (2950C>T), CTG949TTG (2951C>T), GGC977GGT (3037C>T), ACT978ACC (3040T>C), GTA981GTG (3049A>G), GTA986GTG (3064A>G), TTT1001TTC (3109T>C), TCT1019TCC (3163T>C), TCC1021TCT (3169C>T), TAT1032TAC (3202T>C), CTT1045CTC (3241T>C), AAC1049AAT (3253C>T), GTG1050GTA (3256G>A), AAA1053AAG (3265A>G), GCT1069GCC (3313T>C), GTA1100GTC (3406A>C), CTC1104CTA (3418C>A), CTT1105CTC (3421T>C), AAA1119AAG (3463A>G), CAG1120CAA (3466G>A), GGC1125GGT (3481C>T), ACA1126ACT (3484A>T), AGG1143AAG (3534G>A), GGC1145GGT (3541C>T), TCT1158TCC (3580T>C), ATC1159ATT (3583C>T), GAA1168GAG (3610A>G), GGT1169GGC (3613T>C), TAC1173TAT (3625C>T), GTG1174GTA (3628G>A), AAA1179AAG (3643A>G), TAC1180TAT (3646C>T), CAA1185CAG (3661A>G), AAT1186AAC (3664T>C), GAG1189GAA (3673G>A), GAC1190GAT (3676C>T), CTA1194TTA (3686C>T), CGT1200CGC (3706T>C), TTG1204CTG (3716T>C), CCT1205CCA (3721T>A), CGA1211CGT (3739A>T), GGA1215GGG (3751A>G), AAG1216AAA (3754G>A), GTC1218ATC (3758G>A), CAG1220CAA (3766G>A), TAT1234TAC (3808T>C), GAC1236GAT (3814C>T), GGG1237GGC (3817G>C), TTG1240CTG (3824T>C), TCT1243TCC (3835T>C), TAT1246TAC (3844T>C), TTG1251CTA (3857T>C 3859G>A), GGC1254GGT (3868C>T), ACT1255ACC (3871T>C), GAG1259GAA (3883G>A), AGT1264AGC (3898T>C), GGA1267GGT (3907A>T), GTC1270GTT (3916C>T), AAA1273AAG (3925A>G), AGG1274AGA (3928G>A), GAC1281GAT (3949C>T), GAG1283GAA (3955G>A), TTG1285TTA (3961G>A), GAA1288GAG (3970A>G), ATT1289ATC (3973T>C), TTG1310CTG (4034T>C), CGT1320CGC (4066T>C), GAA1323GAG (4075A>G), CCC1326CCT (4084C>T), TTG1332CTG (4100T>C), CTG1338TTG (4118C>T), TCC1341TCT (4129C>T), TCC1347TCA (4147C>A), CTG1351CTA (4159G>A), CCA1352CCT (4162A>T), AAA1354AAG (4168A>G), GGA1360GGT (4186A>T), GAA1363GAG (4195A>G), TTG1367TTA (4207G>A), GCT1368GCC (4210T>C), GTC1374GTT (4228C>T), CAA1396CAG (4294A>G), GGT1411GGC (4339T>C), CCA1413TCA (4343C>T), GAC1414GAT (4348C>T), AAT1416AAC (4354T>C), ACT1430ACC (4396T>C), AAC1434AAT (4408C>T), GTT1436GTG (4414T>G), TTT1437TTC (4417T>C), AGG1438AGA (4420G>A), GTT1451GTC (4459T>C), ATC1452ATT (4462C>T), TTC1453TTT (4465C>T), AAC1454AAT (4468C>T), TCT1455TCC (4471T>C), CTT1456CTC (4474T>C), TAT1495TAC (4591T>C), TAT1510TAC (4636T>C), CTG1512TTG (4640C>T), GAA1518GAG (4660A>G), ATA1526ATC (4684A>C), ATC1530ATT (4696C>T), AAG1532AAA (4702G>A), AAC1533AAT (4705C>T), GTT1536GTC (4714T>C), GAC1543GAT (4735C>T), CTA1547CTG (4747A>G), AGC1551AGT (4759C>T), TTT1554TTC (4768T>C), AGG1556AGA (4774G>A), CAC1557CAT (4777C>T), CTA1559CTG (4783A>G), GGA1560GGG (4786A>G), TAC1561TAT (4789C>T), GTA1569GTG (4813A>G), TCT1570TCC (4816T>C), AGG1579AGA (4843G>A), GTT1598ATT (4898G>A), GAG1600GAA (4906G>A), AAG1621AAA (4969G>A), ACC1628GCC (4988A>G), ACA1629ACT (4993A>T), TAT1633TAC (5005T>C), TTG1636CTG (5012T>C), CTG1641CTA (5029G>A), ATC1653ATT (5065C>T), CTT1656CTC (5074T>C), CAT1657CAC (5077T>C), ATT1658ACT (5079T>C), TTG1666CTG (5102T>C), ACT1671ATC (5118C>T 5119T>C), GTT1672ATT (5120G>A), CTG1673CTC (5125G>C), CAT1680CAC (5146T>C), ACT1681AAT (5148C>A), CAG1698CAA (5200G>A), GTT1704GTG (5218T>G), AGA1713AAA (5244G>A), CCC1720CCT (5266C>T), GAC1725GAT (5281C>T), CAC1741CAT (5329C>T), ATT1745ATA (5341T>A), GTA1746GTG (5344A>G), ACC1747ACT (5347C>T), GTG1755ATA (5369G>A 5371G>A), GGC1760GGT (5386C>T), GGG1762GGA (5392G>A), CCA1767CCG (5407A>G), TGC1769TGT (5413C>T), GCA1771GCT (5419A>T), ACT1777ACC (5437T>C), GGC1778GGT (5440C>T), GTT1785GTC (5461T>C), TTG1789CTG (5471T>C), ATA1792ATG (5482A>G), CTT1809CTA (5533T>A), GAT1822GAC (5572T>C), TGT1824TGC (5578T>C), ATT1826ATC (5584T>C), ATA1837ATT (5617A>T), GGT1845GGC (5641T>C), TTT1847TTC (5647T>C), CAG1860CAA (5686G>A), TTG1862CTG (5690T>C), CCT1868CCA (5710T>A), GGG1870GGT (5716G>T), ACT1876ACC (5734T>C), ACT1877ACC (5737T>C), AGA1878AAA (5739G>A), CTT1880CTC (5746T>C), TTG1882CTG (5750T>C), CTC1884CTT (5758C>T), AAG1889AAA (5773G>A), ATT1893ACT (5784T>C), TTA1894CTA (5786T>C), AGA1897AGG (5797A>G), TTC1900TTT (5806C>T), TCA1910TCT (5836A>T), TTG1915CAA (5849T>C 5850T>A 5851G>A), GGG1924GGT (5878G>T), TTT1926TTC (5884T>C), CCC1927CCT (5887C>T), CGC1928CGT (5890C>T), CTT1936CTC (5914T>C), TTT1939TTC (5923T>C), CTG1941TTG (5927C>T), AGC1942AGT (5932C>T), AAC1950AGC (5955A>G), ACG1953ACA (5965G>A), AAA1964AAG (5998A>G), ACC1966ACT (6004C>T), AAA1970AAG (6016A>G), TCC1974TCT (6028C>T), GCA1975GCT (6031A>T), ACA1976ATA (6033C>T), GCT1977TCT (6035G>T), GGT1983GGA (6055T>A), TCC1987TCA (6067C>A), GCT1988GCC (6070T>C), CTT1996CTG (6094T>G), CAG1999CAA (6103G>A), AAG2000AAA (6106G>A), TTT2011TTC (6139T>C), ATT2013ATC (6145T>C), TAT2014TAC (6148T>C), GTC2016GTT (6154C>T), GTA2026GTG (6184A>G), ACT2027ACC (6187T>C), TTC2029TTT (6193C>T), CAA2030CAG (6196A>G), CAA2034CAG (6208A>G), GGG2040GGA (6226G>A), ATA2042GTG (6230A>G 6232A>G), GAG2043GAA (6235G>A), ACC2050ACA (6256C>A), ACA2052ACG (6262A>G), CTG2053CTA (6265G>A), GTT2062GTA (6292T>A), GTA2064GTG (6298A>G), TTA2066TTG (6304A>G), TTG2068CTG (6308T>C), GAT2069GAC (6313T>C), ATA2071ATC (6319A>C), GAG2072GAA (6322G>A), ATC2073GTC (6323A>G), AAT2084AAC (6358T>C), GCA2089GTA (6372C>T), CAT2127CAC (6487T>C), CAT2129CAC (6493T>C), TCC2137TCT (6517C>T), CTT2147TTT (6545C>T), TAC2155TAT (6571C>T) | | | | | | | | |

#### Proteins

| | | | | | | | | | |
|---|---|---|---|---|---|---|---|---|---|
| polyprotein (YP_001039627.1) | 1 | 2159 | 100% | 14905 | 98.9% | 2159 (100%) | 2125 (98.4%) | 0/0/0/0 | 1 |
| Protein mutations: | P8S (128C>T 130C>T), F10S (135T>C), A18T (158G>A 160T>C), T19A (161A>G), T364A (1196A>G), I459V (1481A>G), A501S (1607G>T), K544R (1737A>G 1738A>G), F659Y (2082T>A), N712S (2241A>G), S715N (2250G>A 2251C>T), R768K (2409G>A), R1143K (3534G>A), V1218I (3758G>A), P1413S (4343C>T), V1598I (4898G>A), T1628A (4988A>G), I1658T (5079T>C), T1671I (5118C>T 5119T>C), V1672I (5120G>A), T1681N (5148C>A), R1713K (5244G>A), V1755I (5369G>A 5371G>A), I1792M (5482A>G), R1878K (5739G>A), I1893T (5784T>C), L1915Q (5849T>C 5850T>A 5851G>A), N1950S (5955A>G), T1976I (6033C>T), A1977S (6035G>T), I2042V (6230A>G 6232A>G), I2073V (6323A>G), A2089V (6372C>T), L2147F (6545C>T) | | | | | | | | |

| | Begin | End | Coverage | Score | Concordance | Matches | Identities | I/D/M/F* | Stop Codons |
|---|---|---|---|---|---|---|---|---|---|
| **NT** | **1** | **7799** | **99.9%** | **12080** | **83.0%** | **7519 (96.3%)** | **6929 (88.7%)** | **12/280** | |
| Codon mutations: | TTT3TTC (115T>C), TTC7TTT (127C>T), CCC8TCT (128C>T 130C>T), TTC10TCC (135T>C), ACT14ACC (148T>C), GCT18ACC (158G>A 160T>C), ACC19GCC (161A>G), TCA27TCT (187A>T), GCT31GCA (199T>A), AGT34AGC (208T>C), CGC38CGT (220C>T), CTG40TTG (224C>T), CAA49CAG (253A>G), TCT51TCC (259T>C), TCT52TCA (262T>A), CAC53CAT (265C>T), TTG78TTA (340G>A), AAG81AAA (349G>A), TTC89TTT (373C>T), ATC90ATT (376C>T), AGC113AGT (445C>T), GAC127GAT (487C>T), GTG132GTA (502G>A), TCT145TCC (541T>C), TTC151TTT (559C>T), CTA158TTA (578C>T), AAC174AAT (628C>T), GAC183GAT (655C>T), CAA190CAG (676A>G), GCA196GCG (694A>G), CCA210CCG (736A>G), CTG246TTG (842C>T), CAA258CAG (880A>G), AGG264AGA (898G>A), GCC273GCT (925C>T), GTA282GTG (952A>G), GGT286GGG (964T>G), AAG288AAA (970G>A), GCC297GCT (997C>T), TCT300TCC (1006T>C), GAG309GAA (1033G>A), CAT318CAC (1060T>C), GCA324GCT (1078A>T), AAC325AAT (1081C>T), ACT329ACC (1093T>C), CAA333CAG (1105A>G), CAT334CAC (1108T>C), CTA335TTA (1109C>T), GCC353GCT (1165C>T), AGG358AGA (1180G>A), ACC364GCC (1196A>G), CAG375CAA (1231G>A), GGC378GGT (1240C>T), AGG383AGA (1255G>A), TTT387TTC (1267T>C), CAA390CAG (1276A>G), GGG398GGA (1300G>A), GAA415GAG (1351A>G), CTC422CTT (1372C>T), TAT435TAC (1411T>C), CAT437CAC (1417T>C), ATC441ATT (1429C>T), GCT448GCC (1450T>C), CTG449TTG (1451C>T), GAC458GAT (1480C>T), ATT459GTT (1481A>G), GAT461GAC (1489T>C), CCC472CCT (1522C>T), GGC482GGT (1552C>T), CGC483CGT (1555C>T), CGA488CGG (1570A>G), TAT489TAC (1573T>C), GCC492GCT (1582C>T), ACA493ACT (1585A>T), AAC495AAT (1591C>T), CTT496CTC (1594T>C), GCT501TCT (1607G>T), CAG502CAA (1612G>A), GGT504GGG (1618T>G), ACC506ACA (1624C>A), AGA513AGG (1645A>G), TGC521TGT (1669C>T), GGT525GGA (1681T>A), GAG526GAA (1684G>A), GCT531GCC (1699T>C), ACT533ACC (1705T>C), AAA544AGG (1737A>G 1738A>G), AGC555AGT (1771C>T), TTG559CTG (1781T>C), GAA561GAG (1789A>G), GCA597GCG (1897A>G), TTG602CTG (1910T>C), GTT616GTC (1954T>C), GTA618GTG (1960A>G), TAC626TAT (1984C>T), AAA636AAG (2014A>G), TAT639TAC (2023T>C), CCT650CCC (2056T>C), GCC657GCA (2077C>A), TTT659TAT (2082T>A), CTA662CTG (2092A>G), AAA667AAG (2107A>G), GTT674GTC (2128T>C), CTG678TTG (2138C>T), TAT679TAC (2143T>C), GGT681GGC (2149T>C), TGT683TGC (2155T>C), CTG690TTG (2174C>T), ATA691ATC (2179A>C), AAG692AAA (2182G>A), GCT696GCA (2194T>A), TAT698TAC (2200T>C), AAG703AAA (2215G>A), AGC704AGT (2218C>T), ATC706ATT (2224C>T), ACA710ACG (2236A>G), AAT712AGT (2241A>G), AGC715AAT (2250G>A 2251C>T), GAC717GAT (2257C>T), CAG719CAA (2263G>A), CCT723CCC (2275T>C), GCG727GCA (2287G>A), CAC728CAT (2290C>T), GAC736GAT (2314C>T), ATA739ATT (2323A>T), CAC747CAT (2347C>T), GTA748GTG (2350A>G), TTC750TTT (2356C>T), TTG751TTA (2359G>A), TAC752TAT (2362C>T), AGG757AGA (2377G>A), TAT758TAC (2380T>C), AGA762AGG (2392A>G), CCT767CCC (2407T>C), AGG768AAG (2409G>A), ATT779ATC (2443T>C), GTG780GTA (2446G>A), CCG791CCA (2479G>A), ATC793ATT (2485C>T), CTG797CTA (2497G>A), ATT798ATC (2500T>C), AAG799AAA (2503G>A), GGC801GGT (2509C>T), CTT802CTC (2512T>C), GGG812GGA (2542G>A), GTC814GTT (2548C>T), TTC817TTT (2557C>T), GAG838GAA (2620G>A), AAT840AAC (2626T>C), AAG848AAA (2650G>A), TTT853TTC (2665T>C), TTG857CTG (2675T>C), GCA859GCG (2683A>G), AAG862AAA (2692G>A), CTG866TTG (2702C>T), AAA868AAG (2710A>G), AGT871AGC (2719T>C), GAT873GAC (2725T>C), TTC875TTT (2731C>T), ATC876ATT (2734C>T), GGA879GGG (2743A>G), TTG880CTG (2744T>C), TTG887CTG (2765T>C), CTA889CTG (2773A>G), TGC901TGT (2809C>T), TCG902TCA (2812G>A), ACA907ACG (2827A>G), TTC921TTT (2869C>T), CAT925CAC (2881T>C), TTC941TTT (2929C>T), TTG942CTG (2930T>C), AAG943AAA (2935G>A), CAA945CAG (2941A>G), GTC948GTT (2950C>T), CTG949TTG (2951C>T), GGC977GGT (3037C>T), ACT978ACC (3040T>C), GTA981GTG (3049A>G), GTA986GTG (3064A>G), TTT1001TTC (3109T>C), TCT1019TCC (3163T>C), TCC1021TCT (3169C>T), TAT1032TAC (3202T>C), CTT1045CTC (3241T>C), AAC1049AAT (3253C>T), GTG1050GTA (3256G>A), AAA1053AAG (3265A>G), GCT1069GCC (3313T>C), GTA1100GTC (3406A>C), CTC1104CTA (3418C>A), CTT1105CTC (3421T>C), AAA1119AAG (3463A>G), CAG1120CAA (3466G>A), GGC1125GGT (3481C>T), ACA1126ACT (3484A>T), AGG1143AAG (3534G>A), GGC1145GGT (3541C>T), TCT1158TCC (3580T>C), ATC1159ATT (3583C>T), GAA1168GAG (3610A>G), GGT1169GGC (3613T>C), TAC1173TAT (3625C>T), GTG1174GTA (3628G>A), AAA1179AAG (3643A>G), TAC1180TAT (3646C>T), CAA1185CAG (3661A>G), AAT1186AAC (3664T>C), GAG1189GAA (3673G>A), GAC1190GAT (3676C>T), CTA1194TTA (3686C>T), CGT1200CGC (3706T>C), TTG1204CTG (3716T>C), CCT1205CCA (3721T>A), CGA1211CGT (3739A>T), GGA1215GGG (3751A>G), AAG1216AAA (3754G>A), GTC1218ATC (3758G>A), CAG1220CAA (3766G>A), TAT1234TAC (3808T>C), GAC1236GAT (3814C>T), GGG1237GGC (3817G>C), TTG1240CTG (3824T>C), TCT1243TCC (3835T>C), TAT1246TAC (3844T>C), TTG1251CTA (3857T>C 3859G>A), GGC1254GGT (3868C>T), ACT1255ACC (3871T>C), GAG1259GAA (3883G>A), AGT1264AGC (3898T>C), GGA1267GGT (3907A>T), GTC1270GTT (3916C>T), AAA1273AAG (3925A>G), AGG1274AGA (3928G>A), GAC1281GAT (3949C>T), GAG1283GAA (3955G>A), TTG1285TTA (3961G>A), GAA1288GAG (3970A>G), ATT1289ATC (3973T>C), TTG1310CTG (4034T>C), CGT1320CGC (4066T>C), GAA1323GAG (4075A>G), CCC1326CCT (4084C>T), TTG1332CTG (4100T>C), CTG1338TTG (4118C>T), TCC1341TCT (4129C>T), TCC1347TCA (4147C>A), CTG1351CTA (4159G>A), CCA1352CCT (4162A>T), AAA1354AAG (4168A>G), GGA1360GGT (4186A>T), GAA1363GAG (4195A>G), TTG1367TTA (4207G>A), GCT1368GCC (4210T>C), GTC1374GTT (4228C>T), CAA1396CAG (4294A>G), GGT1411GGC (4339T>C), CCA1413TCA (4343C>T), GAC1414GAT (4348C>T), AAT1416AAC (4354T>C), ACT1430ACC (4396T>C), AAC1434AAT (4408C>T), GTT1436GTG (4414T>G), TTT1437TTC (4417T>C), AGG1438AGA (4420G>A), GTT1451GTC (4459T>C), ATC1452ATT (4462C>T), TTC1453TTT (4465C>T), AAC1454AAT (4468C>T), TCT1455TCC (4471T>C), CTT1456CTC (4474T>C), TAT1495TAC (4591T>C), TAT1510TAC (4636T>C), CTG1512TTG (4640C>T), GAA1518GAG (4660A>G), ATA1526ATC (4684A>C), ATC1530ATT (4696C>T), AAG1532AAA (4702G>A), AAC1533AAT (4705C>T), GTT1536GTC (4714T>C), GAC1543GAT (4735C>T), CTA1547CTG (4747A>G), AGC1551AGT (4759C>T), TTT1554TTC (4768T>C), AGG1556AGA (4774G>A), CAC1557CAT (4777C>T), CTA1559CTG (4783A>G), GGA1560GGG (4786A>G), TAC1561TAT (4789C>T), GTA1569GTG (4813A>G), TCT1570TCC (4816T>C), AGG1579AGA (4843G>A), GTT1598ATT (4898G>A), GAG1600GAA (4906G>A), AAG1621AAA (4969G>A), ACC1628GCC (4988A>G), ACA1629ACT (4993A>T), TAT1633TAC (5005T>C), TTG1636CTG (5012T>C), CTG1641CTA (5029G>A), ATC1653ATT (5065C>T), CTT1656CTC (5074T>C), CAT1657CAC (5077T>C), ATT1658ACT (5079T>C), TTG1666CTG (5102T>C), ACT1671ATC (5118C>T 5119T>C), GTT1672ATT (5120G>A), CTG1673CTC (5125G>C), CAT1680CAC (5146T>C), ACT1681AAT (5148C>A), CAG1698CAA (5200G>A), GTT1704GTG (5218T>G), AGA1713AAA (5244G>A), CCC1720CCT (5266C>T), GAC1725GAT (5281C>T), CAC1741CAT (5329C>T), ATT1745ATA (5341T>A), GTA1746GTG (5344A>G), ACC1747ACT (5347C>T), GTG1755ATA (5369G>A 5371G>A), GGC1760GGT (5386C>T), GGG1762GGA (5392G>A), CCA1767CCG (5407A>G), TGC1769TGT (5413C>T), GCA1771GCT (5419A>T), ACT1777ACC (5437T>C), GGC1778GGT (5440C>T), GTT1785GTC (5461T>C), TTG1789CTG (5471T>C), ATA1792ATG (5482A>G), CTT1809CTA (5533T>A), GAT1822GAC (5572T>C), TGT1824TGC (5578T>C), ATT1826ATC (5584T>C), ATA1837ATT (5617A>T), GGT1845GGC (5641T>C), TTT1847TTC (5647T>C), CAG1860CAA (5686G>A), TTG1862CTG (5690T>C), CCT1868CCA (5710T>A), GGG1870GGT (5716G>T), ACT1876ACC (5734T>C), ACT1877ACC (5737T>C), AGA1878AAA (5739G>A), CTT1880CTC (5746T>C), TTG1882CTG (5750T>C), CTC1884CTT (5758C>T), AAG1889AAA (5773G>A), ATT1893ACT (5784T>C), TTA1894CTA (5786T>C), AGA1897AGG (5797A>G), TTC1900TTT (5806C>T), TCA1910TCT (5836A>T), TTG1915CAA (5849T>C 5850T>A 5851G>A), GGG1924GGT (5878G>T), TTT1926TTC (5884T>C), CCC1927CCT (5887C>T), CGC1928CGT (5890C>T), CTT1936CTC (5914T>C), TTT1939TTC (5923T>C), CTG1941TTG (5927C>T), AGC1942AGT (5932C>T), AAC1950AGC (5955A>G), ACG1953ACA (5965G>A), AAA1964AAG (5998A>G), ACC1966ACT (6004C>T), AAA1970AAG (6016A>G), TCC1974TCT (6028C>T), GCA1975GCT (6031A>T), ACA1976ATA (6033C>T), GCT1977TCT (6035G>T), GGT1983GGA (6055T>A), TCC1987TCA (6067C>A), GCT1988GCC (6070T>C), CTT1996CTG (6094T>G), CAG1999CAA (6103G>A), AAG2000AAA (6106G>A), TTT2011TTC (6139T>C), ATT2013ATC (6145T>C), TAT2014TAC (6148T>C), GTC2016GTT (6154C>T), GTA2026GTG (6184A>G), ACT2027ACC (6187T>C), TTC2029TTT (6193C>T), CAA2030CAG (6196A>G), CAA2034CAG (6208A>G), GGG2040GGA (6226G>A), ATA2042GTG (6230A>G 6232A>G), GAG2043GAA (6235G>A), ACC2050ACA (6256C>A), ACA2052ACG (6262A>G), CTG2053CTA (6265G>A), GTT2062GTA (6292T>A), GTA2064GTG (6298A>G), TTA2066TTG (6304A>G), TTG2068CTG (6308T>C), GAT2069GAC (6313T>C), ATA2071ATC (6319A>C), GAG2072GAA (6322G>A), ATC2073GTC (6323A>G), AAT2084AAC (6358T>C), GCA2089GTA (6372C>T), CAT2127CAC (6487T>C), CAT2129CAC (6493T>C), TCC2137TCT (6517C>T), CTT2147TTT (6545C>T), TAC2155TAT (6571C>T) | | | | | | | | |

*: Inserts / Deletes / Misaligned / Frameshifts

#### Analysis details

This analysis was performed with panviral2.64

### NGS Details (UN59): Torradovirus lycopersici (segment RNA 2)

#### Assembly

| | |
|---|---|
| **Coverage Length** | 5379 (1 contig(s)) |
| **Depth Of Coverage** | 6024.7 |
| **Number Of Reads** | 251860 |
| **Reads Per Million** | 5475.21 rpm (after QC) |
| **Ambiguities** | 0 |
| **Assembly Method** | de novo + reference guided assembly |
| **Consensus Caller** | Bcf Tools |

#### Coverage Map

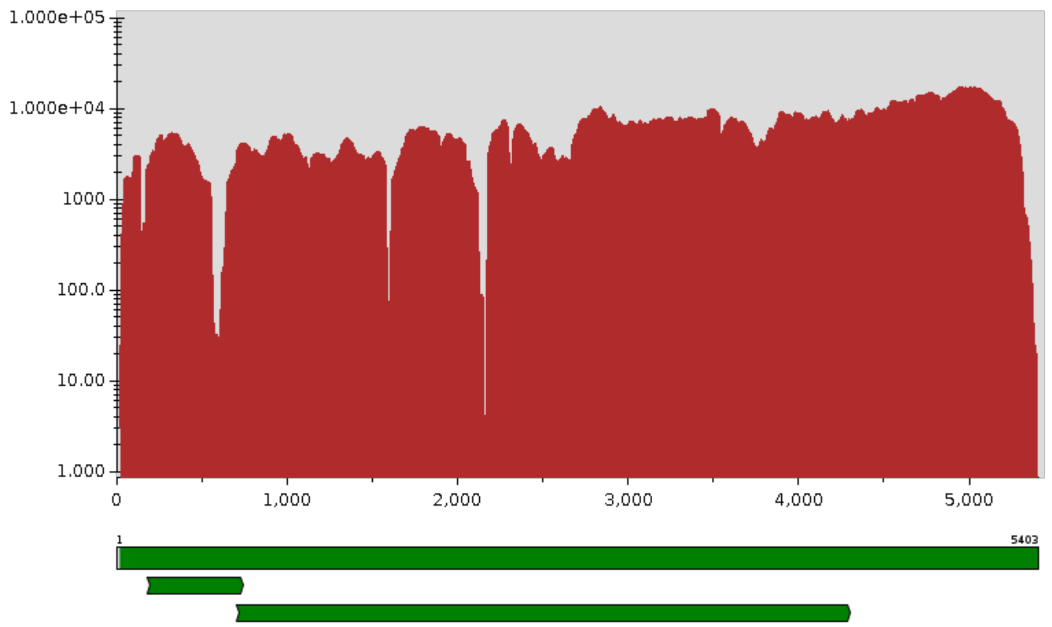

#### Assignment

| | |
|---|---|
| **Type** | Torradovirus lycopersici (Taxonomy ID: 3048378) |
| **Reference Genome** | NC_009032.1 |
| **NT Identity (%)** | 85.17 |
| **AA Identity (%)** | 89.6179 |
| **Number Of Stop Codons** | 5 |
| **Number Of CDS** | 2 |

#### Alignment

| | |
|---|---|
| **Alignment Score** | 7554.0 (NT) + 8679.0 (AA) = 16233.0 |
| **Concordance (%)** | 80.1828 |

| **Alignment Method** | Global, seeded, nucleotide + amino acids (AGA) |
|---|---|

#### Genome Region

Sequence starts at position 25 and ends at position 5403 relative to NC_009032.1 reference sequence.

#### Alignment Detailed Statistics

| | Begin | End | Coverage | Score | Concordance | Matches | Identities | I/D/M/F* | Stop Codons |
|---|---|---|---|---|---|---|---|---|---|
| **NT** | **25** | **5403** | **99.6%** | **7554** | **70.2%** | **5379 (99.9%)** | **4583 (85.2%)** | **2/0** | |

Mutations: 27T>G, 52C>T, 58T>C, 60G>A, 69A>G, 79A>G, 91T>A, 96C>T, 99T>G, 103G>C, 104A>G, 105G>T, 107G>A, 108C>G, 111C>A, 112G>A, 117C>T, 120A>G, 121A>C, 132C>T, 137T>C, 142G>A, 143G>T, 149G>T, 161A>T, 164T>A, 170G>T, 181T>C, 187A>C, 194T>G, 202G>T, 212C>G, 223G>A, 232C>T, 235G>A, 238A>G, 244A>T, 247C>T, 265G>A, 268T>A, 277C>T, 280T>C, 286A>C, 289A>C, 292C>T, 301A>T, 304T>C, 305C>T, 307A>G, 313C>T, 316G>A, 325C>T, 329A>G, 334T>A, 338G>A, 340A>T, 352A>T, 356G>A, 370C>T, 372A>C, 376C>T, 388T>A, 400T>C, 401T>C, 403A>C, 418T>G, 433C>T, 439G>A, 445T>A, 448C>A, 451A>T, 461A>C, 481C>T, 484C>A, 487A>G, 496A>G, 502T>A, 505G>C, 507G>A, 508G>A, 509A>G, 511T>A, 515G>A, 520C>T, 541C>T, 547T>G, 552C>A, 553T>G, 554G>A, 555C>A, 564A>G, 565A>T, 568A>T, 570A>G, 576C>A, 578C>G, 586A>G, 587T>C, 589A>T, 595G>T, 599C>T, 601T>A, 603A>C, 604C>T, 607G>A, 610C>T, 613A>G, 618G>A, 619C>A, 622A>G, 631A>G, 632G>T, 637A>G, 644C>G, 645T>C, 654T>A, 658C>A, 665T>G, 670A>G, 673A>G, 679G>A, 683T>A, 685G>A, 689A>T, 690A>C, 696T>C, 706T>C, 731G>A, 733A>G, 750A>G, 755A>T, 758T>A, 761A>G, 771T>C, 782G>A, 788A>T, 791T>A, 818G>A, 821T>C, 827A>G, 833C>T, 844A>T, 854C>A, 860C>G, 863C>T, 866A>G, 878C>T, 884A>G, 887G>T, 893C>T, 911T>C, 929T>A, 932A>T, 968A>G, 1010T>G, 1061T>C, 1073C>A, 1076T>C, 1083C>T, 1085G>A, 1095G>A, 1100G>A, 1109C>T, 1115G>A, 1118A>C, 1130A>G, 1139G>A, 1142C>T, 1151A>G, 1163A>G, 1169C>G, 1178C>T, 1187A>T, 1205T>A, 1220T>C, 1235C>A, 1247A>G, 1262T>C, 1277A>G, 1313T>C, 1316C>T, 1317C>A, 1319A>G, 1325A>G, 1328T>C, 1331C>T, 1343G>A, 1355G>A, 1361A>T, 1367A>T, 1370A>T, 1376T>A, 1383A>G, 1385C>T, 1388C>T, 1390C>G, 1394A>T, 1395A>T, 1397T>A, 1400T>C, 1403A>T, 1406G>A, 1407C>T, 1409T>A, 1412G>T, 1418G>A, 1426A>C, 1427G>T, 1434G>A, 1439A>C, 1451G>T, 1454A>T, 1456G>C, 1457C>T, 1460A>G, 1461G>T, 1465G>A, 1472C>G, 1475G>C, 1478C>G, 1481C>T, 1487C>G, 1490G>A, 1493G>A, 1496C>T, 1502C>T, 1514T>C, 1524T>C, 1526C>T, 1529A>T, 1538A>T, 1543C>T, 1545A>G, 1547C>T, 1553T>C, 1559C>A, 1560C>T, 1562C>A, 1568C>G, 1574G>A, 1575C>A, 1577T>G, 1589A>G, 1593C>G, 1595G>T, 1603G>A, 1605A>G, 1608T>C, 1610C>G, 1613A>G, 1616G>A, 1621G>C, 1622T>A, 1626G>T, 1627A>C, 1628T>C, 1631G>A, 1634T>A, 1646T>C, 1651A>G, 1655T>A, 1661T>C, 1662C>T, 1664G>T, 1667C>T, 1676T>C, 1677G>A, 1679G>T, 1685T>C, 1694T>C, 1700G>A, 1721G>C, 1724C>T, 1727A>G, 1730C>T, 1740C>T, 1745G>A, 1748G>A, 1749A>C, 1754G>A, 1757A>G, 1760T>G, 1763T>A, 1766A>G, 1769A>T, 1778A>C, 1784A>T, 1790G>A, 1793T>A, 1794G>A, 1797T>G, 1799C>G, 1805C>T, 1811A>C, 1814T>C, 1817A>T, 1819T>A, 1820C>G, 1823C>T, 1832G>C, 1841C>T, 1843A>G, 1845T>C, 1847G>T, 1850G>A, 1856C>T, 1863C>T, 1865C>T, 1875G>A, 1878A>G, 1892C>T, 1898T>G, 1899G>A, 1900T>C, 1901G>A, 1906C>T, 1907C>G, 1913T>A, 1928A>C, 1931C>T, 1938A>G, 1941C>T, 1943T>G, 1946C>A, 1947T>C, 1952A>C, 1955G>A, 1958A>G, 1967G>A, 1970A>G, 1979A>G, 1986G>A, 1994T>C, 2002T>A, 2012C>T, 2024T>C, 2027T>A, 2049T>A, 2051A>T, 2054G>T, 2059A>G, 2060A>C, 2066C>G, 2074T>A, 2083T>C, 2084G>A, 2087A>C, 2094C>G, 2096G>C, 2098C>A, 2099G>A, 2100C>A, 2101G>T, 2102C>G, 2106A>G, 2107G>A, 2111T>G, 2114C>T, 2115A>C, 2116A>G, 2119A>C, 2120G>A, 2123G>T, 2125C>T, 2126C>T, 2127A>G, 2128G>T, 2130G>A, 2132T>C, 2135G>A, 2136G>C, 2138G>T, 2144C>G, 2159T>A, 2162T>C, 2164T>G, 2170A>T, 2171A>T, 2174A>T, 2177A>T, 2178T>G, 2183C>T, 2186A>C, 2189C>T, 2195A>C, 2198G>A, 2199G>A, 2200T>C, 2204C>T, 2205A>G, 2210G>A, 2216C>G, 2219T>C, 2225T>A, 2228C>T, 2229A>G, 2230T>A, 2231G>T, 2240T>A, 2243T>C, 2252A>C, 2256G>A, 2259A>C, 2261T>A, 2262G>C, 2270A>G, 2285T>C, 2294A>C, 2303A>G, 2312T>C, 2315T>A, 2320G>C, 2321T>C, 2324G>C, 2326C>T, 2339A>G, 2345A>G, 2375C>T, 2393G>A, 2414T>C, 2420T>G, 2429G>A, 2432G>A, 2436T>G, 2444G>C, 2447C>A, 2453G>A, 2459C>G, 2469T>A, 2471C>G, 2474G>A, 2489A>C, 2492G>A, 2495A>G, 2501A>G, 2503G>A, 2507T>C, 2516C>T, 2523A>C, 2546T>C, 2552C>T, 2561T>G, 2567T>A, 2570G>A, 2573T>A, 2588T>A, 2609C>A, 2627C>T, 2630C>G, 2645G>C, 2660T>C, 2672A>T, 2693G>T, 2702T>A, 2708A>G, 2717T>C, 2720C>A, 2729C>T, 2747C>T, 2753G>T, 2774A>T, 2792C>T, 2795C>G, 2798G>A, 2799G>A, 2804T>C, 2835A>G, 2837C>T, 2840T>C, 2841C>T, 2846G>A, 2852A>G, 2861G>A, 2864C>A, 2867C>T, 2870C>A, 2879T>A, 2882T>A, 2886T>C, 2888G>A, 2891A>G, 2894A>G, 2900A>T, 2909T>C, 2915A>G, 2921C>T, 2927A>G, 2936A>C, 2939A>G, 2948A>G, 2954G>A, 2957A>T, 2960G>A, 2966C>T, 2972T>C, 2981G>T, 2982G>A, 2996T>G, 2999T>A, 3003T>C, 3011C>T, 3012A>G, 3014C>T, 3020T>C, 3023G>C, 3029C>T, 3032A>T, 3044C>T, 3047T>A, 3050G>C, 3051C>T, 3059A>T, 3062C>T, 3063T>C, 3065G>A, 3071A>G, 3077A>T, 3083C>A, 3101T>G, 3104C>T, 3107T>G, 3113G>C, 3122T>G, 3128T>C, 3131G>A, 3143C>T, 3146T>C, 3152T>A, 3158G>A, 3161C>A, 3167A>T, 3176A>C, 3179G>T, 3182C>T, 3187G>A, 3194C>T, 3195A>G, 3197C>T, 3200T>C, 3206C>T, 3212A>G, 3227T>G, 3230T>A, 3236C>T, 3239T>G, 3242G>A, 3248A>C, 3251A>T, 3253A>G, 3265G>A, 3269T>C, 3270A>T, 3272G>T, 3278C>T, 3293A>G, 3299T>A, 3303A>T, 3305C>T, 3311T>A, 3324C>T, 3329G>T, 3350C>T, 3354A>C, 3359G>A, 3368A>G, 3371C>G, 3372T>C, 3386A>T, 3404A>G, 3410G>A, 3413C>T, 3422A>G, 3428G>A, 3431T>G, 3437C>T, 3446T>C, 3449G>A, 3452C>A, 3455T>A, 3458G>A, 3461C>T, 3470T>C, 3473T>C, 3479C>T, 3482T>C, 3485C>T, 3488A>G, 3491C>T, 3503A>G, 3508A>G, 3512T>C, 3515G>T, 3522C>T, 3524A>G, 3527C>T, 3533G>A, 3543T>A, 3551C>T, 3560T>C, 3564A>C, 3567T>A, 3569A>G, 3575G>A, 3578A>G, 3581C>T, 3590A>C, 3605A>G, 3611G>A, 3617A>T, 3620G>A, 3623A>T, 3629G>C, 3631C>A, 3634T>C, 3635C>T, 3638A>G, 3641C>T, 3650A>T, 3656G>T, 3659T>C, 3662A>G, 3674G>C, 3689C>T, 3698A>C, 3702T>C, 3708T>C, 3716C>T, 3719A>G, 3722C>A, 3728C>G, 3742A>C, 3743G>C, 3746G>A, 3749A>T, 3752G>A, 3755A>G, 3770A>C, 3773C>T, 3776A>T, 3779C>T, 3800C>T, 3815A>T, 3827C>T, 3833T>A, 3839A>G, 3845A>G, 3848G>A, 3851G>A, 3852A>G, 3854T>G, 3857C>A, 3866A>C, 3869A>T, 3872A>T, 3878G>C, 3881C>T, 3882A>G, 3884C>A, 3885A>G, 3890A>G, 3893T>C, 3896A>T, 3920A>T, 3923C>T, 3927A>C, 3932C>A, 3933A>G, 3935C>T, 3941T>G, 3944C>A, 3947T>C, 3950T>A, 3953T>A, 3960G>A, 3971T>A, 3983T>A, 3984A>G, 3986G>T, 3988G>A, 3989C>T, 3990G>A, 3991A>G, 3995C>T, 3998C>T, 4001T>C, 4016T>G, 4025G>A, 4031T>C, 4034T>G, 4035A>C, 4037A>T, 4040C>T, 4046C>T, 4052C>T, 4059A>G, 4061A>T, 4067A>T, 4069A>G, 4070T>C, 4072C>T, 4077A>C, 4100A>C, 4103A>T, 4106A>C, 4112C>T, 4142A>C, 4149C>A, 4151G>A, 4181C>T, 4187G>A, 4188C>G, 4196T>A, 4208A>T, 4211A>T, 4223G>A, 4224C>T, 4227T>C, 4229G>T, 4250C>T, 4253C>T, 4256T>G, 4262T>C, 4265T>A, 4274A>T, 4281A>C, 4284A>G, 4286T>A, 4289T>C, 4300T>C, 4302T>G, 4316T>C, 4354G>A, 4355T>G, 4382A>T, 4383C>T, 4388A>T, 4391A>C, 4393A>G, 4424C>T, 4425T>C, 4429T>C, 4447T>A, 4457A>G, 4468T>C, 4496T>A, 4506A>T, 4525T>C, 4567_4568insT, 4605T>C, 4633T>A, 4634G>A, 4643G>A, 4648G>A, 4650T>C, 4651G>A, 4659A>G, 4672C>T, 4675A>T, 4812C>T, 4813C>T, 4860T>C, 4915G>A, 5020C>T, 5068T>C, 5118T>C, 5139T>C, 5141T>C, 5206C>T, 5213A>G, 5214C>T, 5218C>T, 5222C>T, 5224G>A, 5225C>A, 5226T>G, 5231A>T, 5233G>T, 5235T>A, 5236G>A, 5239_5240insA, 5242T>G, 5244G>A, 5249A>G, 5258A>G, 5259T>A, 5262A>T, 5282T>C, 5311C>T, 5313A>T, 5315T>C, 5325G>A, 5343C>T, 5350A>G, 5358C>T, 5370T>A, 5371G>A, 5377G>C, 5379A>G

##### CDS

| | Begin | End | Coverage | Score | Concordance | Matches | Identities | I/D/M/F* | Stop Codons |
|---|---|---|---|---|---|---|---|---|---|
| ToTV_sRNA2gp1 | 1 | 188 | 100% | 1072 | 88.7% | 188 (100%) | 159 (84.6%) | 0/0/0/0 | 2 |

Protein mutations: S5A (194T>G), L7F (202G>T), L11V (212C>G), T50A (329A>G), A53T (338G>A 340A>T), V59I (356G>A), N64T (372A>C), L90F (451A>T), Q108H (505G>C), R109K (507G>A 508G>A), I110V (509A>G 511T>A), V112I (515G>A), A124E (552C>A 553T>G), A125K (554G>A 555C>A), Q128R (564A>G 565A>T), Q130R (570A>G), T132K (576C>A), L133V (578C>G), E138D (595G>T), D141A (603A>C 604C>T), R146Q (618G>A 619C>A), E151* (632G>T), L155A (644C>G 645T>C), L158Q (654T>A), S162A (665T>G), L168I (683T>A 685G>A), N170S (689A>T 690A>C), L172P (696T>C), E184K (731G>A 733A>G)

Codon mutations: TCA2TCC (187A>C), TCT5GCT (194T>G), TTG7TTT (202G>T), CTT11GTT (212C>G), GAG14GAA (223G>A), CAC17CAT (232C>T), AAG18AAA (235G>A), CAA19CAG (238A>G), GCA21GCT (244A>T), GAC22GAT (247C>T), TCG28TCA (265G>A), GTT29GTA (268T>A), GGC32GGT (277C>T), TCT33TCC (280T>C), ATA35ATC (286A>C), ATA36ATC (289A>C), AAC37AAT (292C>T), CCA40CCT (301A>T), ACT41ACC (304T>C), CTA42TTG (305C>T 307A>G), TTC44TTT (313C>T), AAG45AAA (316G>A), CCC48CCT (325C>T), ACA50GCA (329A>G), GGT51GGA (334T>A), GCA53ACT (338G>A 340A>T), CTA57CTT (352A>T), GTT59ATT (356G>A), AAC63AAT (370C>T), AAT64ACT (372A>C), AGC65AGT (376C>T), GTT69GTA (388T>A), TAT73TAC (400T>C), TTA74CTC (401T>C 403A>C), ACT79ACG (418T>G), GGC84GGT (433C>T), AAG86AAA (439G>A), TCT88TCA (445T>A), GGC89GGA (448C>A), TTA90TTT (451A>T), AGA94CGA (461A>C), ACC100ACT (481C>T), ACC101ACA (484C>A), AGA102AGG (487A>G), CTA105CTG (496A>G), GCT107GCA (502T>A), CAG108CAC (505G>C), AGG109AAA (507G>A 508G>A), ATT110GTA (509A>G 511T>A), GTA112ATA (515G>A), TCC113TCT (520C>T), TCC120TCT (541C>T), GCT122GCG (547T>G), GCT124GAG (552C>A 553T>G), GCA125AAA (554G>A 555C>A), CAA128CGT (564A>G 565A>T), CCA129CCT (568A>T), CAA130CGA (570A>G), ACG132AAG (576C>A), CTT133GTT (578C>G), CAA135CAG (586A>G), TTA136CTT (587T>C 589A>T), GAG138GAT (595G>T), CTT140TTA (599C>T 601T>A), GAC141GCT (603A>C 604C>T), GAG142GAA (607G>A), GCC143GCT (610C>T), AAA144AAG (613A>G), CGC146CAA (618G>A 619C>A), TTA147TTG (622A>G), AAA150AAG (631A>G), GAG151TAG (632G>T), AAA152AAG (637A>G), CTT155GCT (644C>G 645T>C), CTA158CAA (654T>A), TCC159TCA (658C>A), TCC162GCC (665T>G), AAA163AAG (670A>G), TTA164TTG (673A>G), TTG166TTA (679G>A), TTG168ATA (683T>A 685G>A), AAC170TCC (689A>T 690A>C), CTA172CCA (696T>C), GAT175GAC (706T>C), GAA184AAG (731G>A 733A>G)

| | Begin | End | Coverage | Score | Concordance | Matches | Identities | I/D/M/F* | Stop Codons |
|---|---|---|---|---|---|---|---|---|---|
| ToTV_sRNA2gp2 | 1 | 1199 | 100% | 7607 | 91.9% | 1199 (100%) | 1084 (90.4%) | 0/0/0/0 | 3 |

Protein mutations: M2T (706T>C), K11R (733A>G), S17G (750A>G), Y48F (844A>T), V132I (1095G>A), T228A (1383A>G 1385C>T), S230* (1390C>G), T232S (1395A>T 1397T>A), Q242P (1426A>C 1427G>T), D245N (1434G>A), S252T (1456G>C 1457C>T), A254S (1461G>T), R255K (1465G>A), I257M (1472C>G), S275P (1524T>C 1526C>T), S281F (1543C>T), T282A (1545A>G 1547C>T), H286Q (1559C>A), Q298D (1593C>G 1595G>T), S301N (1603G>A), R302G (1605A>G), Y303Q (1608T>C 1610C>G), G307A (1621G>C 1622T>A), D309S (1626G>T 1627A>C 1628T>C), K317R (1651A>G), L321F (1662C>T 1664G>T), V326I (1677G>A 1679G>T), V365I (1794G>A), C366G (1797T>G 1799C>G), K370N (1811A>C), I373K (1819T>A 1820C>G), K377N (1832G>C), H381R (1843A>G), H388Y (1863C>T 1865C>T), V392M (1875G>A), I393V (1878A>G), I399M (1898T>G), V400T (1899G>A 1900T>C 1901G>A), A402V (1906C>T 1907C>G), E409D (1928A>C), N413D (1938A>G), D429N (1986G>A), F434Y (2002T>A), L450I (2049T>A 2051A>T), K453S (2059A>G 2060A>C), L458Q (2074T>A), V461A (2083T>C 2084G>A), Q465D (2094C>G 2096G>C), A466E (2098C>A 2099G>A), R467M (2100C>A 2101G>T 2102C>G), R469E (2106A>G 2107G>A), F470L (2111T>G), K472R (2115A>C 2116A>G), E473A (2119A>C 2120G>A), A475V (2125C>T 2126C>T), R476V (2127A>G 2128G>T), A477T (2130G>A 2132T>C), V479L (2136G>C 2138G>T), N481K (2144C>G), V488G (2164T>G), E490V (2170A>T 2171A>T), S493A (2178T>G), V500T (2199G>A 2200T>C), M502V (2205A>G), D508E (2225T>A), M510D (2229A>G 2230T>A 2231G>T), E517D (2252A>C), A519T (2256G>A), N520Q (2259A>C 2261T>A), E521Q (2262G>C), E531D (2294A>C), N538K (2315T>A), G540A (2320G>C 2321T>C), T542I (2326C>T), S579A (2436T>G), S590T (2469T>A 2471C>G), R601K (2503G>A), I608L (2523A>C), I643M (2630C>G), C667* (2702T>A), A700T (2799G>A), I712V (2835A>G 2837C>T), A761T (2982G>A), I771V (3012A>G 3014C>T), I807M (3122T>G), S829N (3187G>A), T832A (3195A>G 3197C>T), K851R (3253A>G), R855K (3265G>A), T857S (3270A>T 3272G>T), T868S (3303A>T 3305C>T), N936S (3508A>G), S948T (3543T>A), K955Q (3564A>C), L956M (3567T>A 3569A>G), P977Q (3631C>A), L978P (3634T>C 3635C>T), Q1014P (3742A>C 3743G>C), T1051A (3852A>G 3854T>G), I1061V (3882A>G 3884C>A), T1062A (3885A>G), K1076Q (3927A>C), T1078A (3933A>G 3935C>T), A1087T (3960G>A), M1095V (3984A>G 3986G>T), S1096N (3988G>A 3989C>T), D1097S (3990G>A 3991A>G), H1105Q (4016T>G), I1120V (4059A>G 4061A>T), N1123S (4069A>G 4070T>C), A1124V (4072C>T), L1150I (4149C>A 4151G>A), Q1163E (4188C>G), H1175Y (4224C>T), S1195G (4284A>G 4286T>A)

| | Begin | End | Coverage | Score | Concordance | Matches | Identities | I/D/M/F* | Stop Codons |
|---|---|---|---|---|---|---|---|---|---|
| **NT** | **25** | **5403** | **99.6%** | **7554** | **70.2%** | **5379 (99.9%)** | **4583 (85.2%)** | **2/0** | |
| Codon mutations: | ATG2ACG (706T>C), AGG10AGA (731G>A), AAG11AGG (733A>G), AGT17GGT (750A>G), GCA18GCT (755A>T), GCT19GCA (758T>A), GTA20GTG (761A>G), TTA24CTA (771T>C), CAG27CAA (782G>A), ACA29ACT (788A>T), GCT30GCA (791T>A), CTG39CTA (818G>A), CTT40CTC (821T>C), AAA42AAG (827A>G), GGC44GGT (833C>T), TAT48TTT (844A>T), CTC51CTA (854C>A), GGC53GGG (860C>G), GCC54GCT (863C>T), CAA55CAG (866A>G), GCC59GCT (878C>T), GAA61GAG (884A>G), CTG62CTT (887G>T), GAC64GAT (893C>T), AAT70AAC (911T>C), TCT76TCA (929T>A), TCA77TCT (932A>T), GGA89GGG (968A>G), GTT103GTG (1010T>G), AGT120AGC (1061T>C), ACC124ACA (1073C>A), AAT125AAC (1076T>C), CTG128TTA (1083C>T 1085G>A), GTT132ATT (1095G>A), TTG133TTA (1100G>A), TCC136TCT (1109C>T), GGG138GGA (1115G>A), GCA139GCC (1118A>C), TTA143TTG (1130A>G), CCG146CCA (1139G>A), ACC147ACT (1142C>T), CTA150CTG (1151A>G), AAA154AAG (1163A>G), GGC156GGG (1169C>G), TTC159TTT (1178C>T), ACA162ACT (1187A>T), ACT168ACA (1205T>A), CCT173CCC (1220T>C), GCC178GCA (1235C>A), CAA182CAG (1247A>G), CGT187CGC (1262T>C), TCA192TCG (1277A>G), CTT204CTC (1313T>C), CAC205CAT (1316C>T), CGA206AGG (1317C>A 1319A>G), CAA208CAG (1325A>G), TTT209TTC (1328T>C), TTC210TTT (1331C>T), CAG214CAA (1343G>A), CAG218CAA (1355G>A), ATA220ATT (1361A>T), TCA222TCT (1367A>T), GTA223GTT (1370A>T), TCT225TCA (1376T>A), ACC228GCT (1383A>G 1385C>T), TCC229TCT (1388C>T), TCA230TGA (1390C>G), GCA231GCT (1394A>T), ACT232TCA (1395A>T 1397T>A), GAT233GAC (1400T>C), TCA234TCT (1403A>T), GTG235GTA (1406G>A), CTT236TTA (1407C>T 1409T>A), CCG237CCT (1412G>T), GTG239GTA (1418G>A), CAG242CCT (1426A>C 1427G>T), GAC245AAC (1434G>A), ACA246ACC (1439A>C), GTG250GTT (1451G>T), CCA251CCT (1454A>T), AGC252ACT (1456G>C 1457C>T), GAA253GAG (1460A>G), GCA254TCA (1461G>T), AGA255AAA (1465G>A), ATC257ATG (1472C>G), CGG258CGC (1475G>C), TCC259TCG (1478C>G), TCC260TCT (1481C>T), CTC262CTG (1487C>G), CGG263CGA (1490G>A), GTG264GTA (1493G>A), CCC265CCT (1496C>T), CTC267CTT (1502C>T), AGT271AGC (1514T>C), TCC275CCT (1524T>C 1526C>T), TCA276TCT (1529A>T), ATA279ATT (1538A>T), TCT281TTT (1543C>T), ACC282GCT (1545A>G 1547C>T), ACT284ACC (1553T>C), CAC286CAA (1559C>A), CTC287TTA (1560C>T 1562C>A), GGC289GGG (1568C>G), ACG291ACA (1574G>A), CGT292AGG (1575C>A 1577T>G), GTA296GTG (1589A>G), CAG298GAT (1593C>G 1595G>T), AGT301AAT (1603G>A), AGG302GGG (1605A>G), TAC303CAG (1608T>C 1610C>G), GAA304GAG (1613A>G), GAG305GAA (1616G>A), GGT307GCA (1621G>C 1622T>A), GAT309TCC (1626G>T 1627A>C 1628T>C), GGG310GGA (1631G>A), GTT311GTA (1634T>A), GTT315GTC (1646T>C), AAG317AGG (1651A>G), GCT318GCA (1655T>A), GGT320GGC (1661T>C), CTG321TTT (1662C>T 1664G>T), AAC322AAT (1667C>T), CAT325CAC (1676T>C), GTG326ATT (1677G>A 1679G>T), GAT328GAC (1685T>C), GGT331GGC (1694T>C), GAG333GAA (1700G>A), CTG340CTC (1721G>C), CCC341CCT (1724C>T), GAA342GAG (1727A>G), AAC343AAT (1730C>T), CTG347TTG (1740C>T), TCG348TCA (1745G>A), GTG349GTA (1748G>A), AGA350CGA (1749A>C), CAG351CAA (1754G>A), CAA352CAG (1757A>G), GTT353GTG (1760T>G), GCT354GCA (1763T>A), GCA355GCG (1766A>G), GCA356GCT (1769A>T), GCA359GCC (1778A>C), GCA361GCT (1784A>T), AAG363AAA (1790G>A), GGT364GGA (1793T>A), GTT365ATT (1794G>A), TGC366GGG (1797T>G 1799C>G), GTC368GTT (1805C>T), AAA370AAC (1811A>C), CAT371CAC (1814T>C), ATA372ATT (1817A>T), ATC373AAG (1819T>A 1820C>G), AAC374AAT (1823C>T), AAG377AAC (1832G>C), ATC380ATT (1841C>T), CAC381CGC (1843A>G), TTG382CTT (1845T>C 1847G>T), GAG383GAA (1850G>A), ATC385ATT (1856C>T), CAC388TAT (1863C>T 1865C>T), GTG392ATG (1875G>A), ATA393GTA (1878A>G), TAC397TAT (1892C>T), ATT399ATG (1898T>G), GTG400ACA (1899G>A 1900T>C 1901G>A), GCC402GTG (1906C>T 1907C>G), TCT404TCA (1913T>A), GAA409GAC (1928A>C), AAC410AAT (1931C>T), AAT413GAT (1938A>G), CTT414TTG (1941C>T 1943T>G), ACC415ACA (1946C>A), TTG416CTG (1947T>C), GCA417GCC (1952A>C), CAG418CAA (1955G>A), AGA419AGG (1958A>G), GAG422GAA (1967G>A), CTA423CTG (1970A>G), CAA426CAG (1979A>G), GAT429AAT (1986G>A), TAT431TAC (1994T>C), TTT434TAT (2002T>A), AAC437AAT (2012C>T), TAT441TAC (2024T>C), CCT442CCA (2027T>A), TTA450ATT (2049T>A 2051A>T), ACG451ACT (2054G>T), AAA453AGC (2059A>G 2060A>C), CGC455CGG (2066C>G), CTA458CAA (2074T>A), GTG461GCA (2083T>C 2084G>A), GCA462GCC (2087A>C), CAG465GAC (2094C>G 2096G>C), GCG466GAA (2098C>A 2099G>A), CGC467ATG (2100C>A 2101G>T 2102C>G), AGA469GAA (2106A>G 2107G>A), TTT470TTG (2111T>G), TAC471TAT (2114C>T), AAG472CGG (2115A>C 2116A>G), GAG473GCA (2119A>C 2120G>A), GCG474GCT (2123G>T), GCC475GTT (2125C>T 2126C>T), AGG476GTG (2127A>G 2128G>T), GCT477ACC (2130G>A 2132T>C), CAG478CAA (2135G>A), GTG479CTT (2136G>C 2138G>T), AAC481AAG (2144C>G), ACT486ACA (2159T>A), AGT487AGC (2162T>C), GTG488GGG (2164T>G), GAA490GTT (2170A>T 2171A>T), ATA491ATT (2174A>T), CCA492CCT (2177A>T), TCC493GCC (2178T>G), ACC494ACT (2183C>T), TCA495TCC (2186A>C), TTC496TTT (2189C>T), ACA498ACC (2195A>C), AAG499AAA (2198G>A), GTT500ACT (2199G>A 2200T>C), GCC501GCT (2204C>T), ATG502GTG (2205A>G), GAG503GAA (2210G>A), GGC505GGG (2216C>G), AGT506AGC (2219T>C), GAT508GAA (2225T>A), AGC509AGT (2228C>T), ATG510GAT (2229A>G 2230T>A 2231G>T), GCT513GCA (2240T>A), ATT514ATC (2243T>C), GAA517GAC (2252A>C), GCC519ACC (2256G>A), AAT520CAA (2259A>C 2261T>A), GAA521CAA (2262G>C), GTA523GTG (2270A>G), GTT528GTC (2285T>C), GAA531GAC (2294A>C), CAA534CAG (2303A>G), AAT537AAC (2312T>C), AAT538AAA (2315T>A), GGT540GCC (2320G>C 2321T>C), GCG541GCC (2324G>C), ACA542ATA (2326C>T), TTA546TTG (2339A>G), CAA548CAG (2345A>G), TTC558TTT (2375C>T), GAG564GAA (2393G>A), AGT571AGC (2414T>C), GTT573GTG (2420T>G), GAG576GAA (2429G>A), GTG577GTA (2432G>A), TCA579GCA (2436T>G), GCG581GCC (2444G>C), CTC582CTA (2447C>A), GCG584GCA (2453G>A), CTC586CTG (2459C>G), TCC590ACG (2469T>A 2471C>G), AAG591AAA (2474G>A), GGA596GGC (2489A>C), TCG597TCA (2492G>A), CAA598CAG (2495A>G), CTA600CTG (2501A>G), AGG601AAG (2503G>A), TAT602TAC (2507T>C), GCC605GCT (2516C>T), ATT608CTT (2523A>C), TAT615TAC (2546T>C), TCC617TCT (2552C>T), GGT620GGG (2561T>G), ATT622ATA (2567T>A), TCG623TCA (2570G>A), GGT624GGA (2573T>A), GTT629GTA (2588T>A), CTC636CTA (2609C>A), TTC642TTT (2627C>T), ATC643ATG (2630C>G), CTG648CTC (2645G>C), CAT653CAC (2660T>C), TCA657TCT (2672A>T), GGG664GGT (2693G>T), TGT667TGA (2702T>A), ACA669ACG (2708A>G), GGT672GGC (2717T>C), ATC673ATA (2720C>A), TTC676TTT (2729C>T), GCC682GCT (2747C>T), GGG684GGT (2753G>T), ATA691ATT (2774A>T), CAC697CAT (2792C>T), CCC698CCG (2795C>G), TTG699TTA (2798G>A), GCT700ACT (2799G>A), AGT701AGC (2804T>C), ATC712GTT (2835A>G 2837C>T), CAT713CAC (2840T>C), CTG714TTG (2841C>T), CAG715CAA (2846G>A), AAA717AAG (2852A>G), TCG720TCA (2861G>A), ACC721ACA (2864C>A), AAC722AAT (2867C>T), ATC723ATA (2870C>A), CCT726CCA (2879T>A), CCT727CCA (2882T>A), TTG729CTA (2886T>C 2888G>A), CGA730CGG (2891A>G), GCA731GCG (2894A>G), GCA733GCT (2900A>T), GGT736GGC (2909T>C), AAA738AAG (2915A>G), GAC740GAT (2921C>T), ACA742ACG (2927A>G), CCA745CCC (2936A>C), CGA746CGG (2939A>G), ACA749ACG (2948A>G), CAG751CAA (2954G>A), GTA752GTT (2957A>T), CTG753CTA (2960G>A), CAC755CAT (2966C>T), AAT757AAC (2972T>C), GTG760GTT (2981G>T), GCT761ACT (2982G>A), GGT765GGG (2996T>G), ACT766ACA (2999T>A), TTA768CTA (3003T>C), AGC770AGT (3011C>T), ATC771GTT (3012A>G 3014C>T), TCT773TCC (3020T>C), CCG774CCC (3023G>C), GGC776GGT (3029C>T), ATA777ATT (3032A>T), GAC781GAT (3044C>T), GGT782GGA (3047T>A), ACG783ACC (3050G>C), CTG784TTG (3051C>T), CCA786CCT (3059A>T), TCC787TCT (3062C>T), TTG788CTA (3063T>C 3065G>A), GGA790GGG (3071A>G), ATA792ATT (3077A>T), CGC794CGA (3083C>A), ACT800ACG (3101T>G), GGC801GGT (3104C>T), ACT802ACG (3107T>G), GTG804GTC (3113G>C), ATT807ATG (3122T>G), ATT809ATC (3128T>C), GAG810GAA (3131G>A), TTC814TTT (3143C>T), CAT815CAC (3146T>C), GGT817GGA (3152T>A), TTG819TTA (3158G>A), GCC820GCA (3161C>A), GGA822GGT (3167A>T), ACA825ACC (3176A>C), CTG826CTT (3179G>T), AAC827AAT (3182C>T), AGC829AAC (3187G>A), TCC831TCT (3194C>T), ACC832GCT (3195A>G 3197C>T), CCT833CCC (3200T>C), GAC835GAT (3206C>T), TTA837TTG (3212A>G), GTT842GTG (3227T>G), ATT843ATA (3230T>A), AAC845AAT (3236C>T), CTT846CTG (3239T>G), GAG847GAA (3242G>A), GGA849GGC (3248A>C), CGA850CGT (3251A>T), AAG851AGG (3253A>G), AGG855AAG (3265G>A), TGT856TGC (3269T>C), ACG857TCT (3270A>T 3272G>T), ACC859ACT (3278C>T), AAA864AAG (3293A>G), CTT866CTA (3299T>A), ACC868TCT (3303A>T 3305C>T), GGT870GGA (3311T>A), CTA875TTA (3324C>T), CCG876CCT (3329G>T), CAC883CAT (3350C>T), AGG885CGG (3354A>C), TTG886TTA (3359G>A), ACA889ACG (3368A>G), GTC890GTG (3371C>G), TTG891CTG (3372T>C), GTA895GTT (3386A>T), CTA901CTG (3404A>G), ACG903ACA (3410G>A), GTC904GTT (3413C>T), ACA907ACG (3422A>G), CAG909CAA (3428G>A), CTT910CTG (3431T>G), TGC912TGT (3437C>T), GAT915GAC (3446T>C), TTG916TTA (3449G>A), GTC917GTA (3452C>A), CTT918CTA (3455T>A), GGG919GGA (3458G>A), GGC920GGT (3461C>T), TCT923TCC (3470T>C), GTT924GTC (3473T>C), CCC926CCT (3479C>T), ATT927ATC (3482T>C), TAC928TAT (3485C>T), GGA929GGG (3488A>G), CAC930CAT (3491C>T), GGA934GGG (3503A>G), AAT936AGT (3508A>G), GCT937GCC (3512T>C), GTG938GTT (3515G>T), CTA941TTG (3522C>T 3524A>G), TTC942TTT (3527C>T), GAG944GAA (3533G>A), TCT948ACT (3543T>A), CGC950CGT (3551C>T), ATT953ATC (3560T>C), AAA955CAA (3564A>C), TTA956ATG (3567T>A 3569A>G), AAG958AAA (3575G>A), GAA959GAG (3578A>G), AAC960AAT (3581C>T), ACA963ACC (3590A>C), GGA968GGG (3605A>G), AAG970AAA (3611G>A), CCA972CCT (3617A>T), CAG973CAA (3620G>A), GTA974GTT (3623A>T), GTG976GTC (3629G>C), CCG977CAG (3631C>A), CTC978CCT (3634T>C 3635C>T), AGA979AGG (3638A>G), GAC980GAT (3641C>T), TCA983TCT (3650A>T), GGG985GGT (3656G>T), GCT986GCC (3659T>C), GTA987GTG (3662A>G), GTG991GTC (3674G>C), GAC996GAT (3689C>T), CGA999CGC (3698A>C), TTG1001CTG (3702T>C), TTA1003CTA (3708T>C), TGC1005TGT (3716C>T), GCA1006GCG (3719A>G), CCC1007CCA (3722C>A), TCC1009TCG (3728C>G), CAG1014CCC (3742A>C 3743G>C), GGG1015GGA (3746G>A), GCA1016GCT (3749A>T), CTG1017CTA (3752G>A), GTA1018GTG (3755A>G), ACA1023ACC (3770A>C), TGC1024TGT (3773C>T), CCA1025CCT (3776A>T), TTC1026TTT (3779C>T), TTC1033TTT (3800C>T), GGA1038GGT (3815A>T), TAC1042TAT (3827C>T), ATT1044ATA (3833T>A), GTA1046GTG (3839A>G), AGA1048AGG (3845A>G), GTG1049GTA (3848G>A), CAG1050CAA (3851G>A), ACT1051GCG (3852A>G 3854T>G), TCC1052TCA (3857C>A), ATA1055ATC (3866A>C), GGA1056GGT (3869A>T), GGA1057GGT (3872A>T), CTG1059CTC (3878G>C), AAC1060AAT (3881C>T), ATC1061GTA (3882A>G 3884C>A), ACC1062GCC (3885A>G), TTA1063TTG (3890A>G), GAT1064GAC (3893T>C), TCA1065TCT (3896A>T), GGA1073GGT (3920A>T), ATC1074ATT (3923C>T), AAG1076CAG (3927A>C), GGC1077GGA (3932C>A), ACC1078GCT (3933A>G 3935C>T), GTT1080GTG (3941T>G), GTC1081GTA (3944C>A), TCT1082TCC (3947T>C), GCT1083GCA (3950T>A), GGT1084GGA (3953T>A), GCA1087ACA (3960G>A), GCT1090GCA (3971T>A), GGT1094GGA (3983T>A), ATG1095GTT (3984A>G 3986G>T), AGC1096AAT (3988G>A 3989C>T), GAC1097AGC (3990G>A 3991A>G), AAC1098AAT (3995C>T), ATC1099ATT (3998C>T), TTT1100TTC (4001T>C), CAT1105CAG (4016T>G), GAG1108GAA (4025G>A), TTT1110TTC (4031T>C), CCT1111CCG (4034T>G), AGA1112CGT (4035A>C 4037A>T), CGC1113CGT (4040C>T), ACC1115ACT (4046C>T), GCC1117GCT (4052C>T), ATA1120GTT (4059A>G 4061A>T), CCA1122CCT (4067A>T), AAT1123AGC (4069A>G 4070T>C), GCT1124GTT (4072C>T), AGA1126CGA (4077A>C), CGA1133CGC (4100A>C), CTA1134CTT (4103A>T), GGA1135GGC (4106A>C), CTC1137CTT (4112C>T), ATA1147ATC (4142A>C), CTG1150ATA (4149C>A 4151G>A), TTC1160TTT (4181C>T), TTG1162TTA (4187G>A), CAA1163GAA (4188C>G), GCT1165GCA (4196T>A), TCA1169TCT (4208A>T), GCA1170GCT (4211A>T), AAG1174AAA (4223G>A), CAT1175TAT (4224C>T), TTG1176CTT (4227T>C 4229G>T), ACC1183ACT (4250C>T), TAC1184TAT (4253C>T), CTT1185CTG (4256T>G), ACT1187ACC (4262T>C), CCT1188CCA (4265T>A), TCA1191TCT (4274A>T), AGG1194CGG (4281A>C), AGT1195GGA (4284A>G 4286T>A), TTT1196TTC (4289T>C) | | | | | | | | |

#### Proteins

| | | | | | | | | | |
|---|---|---|---|---|---|---|---|---|---|
| Orf1 (YP_001040017.1) | 1 | 188 | 100% | 1072 | 88.7% | 188 (100%) | 159 (84.6%) | 0/0/0/0 | 2 |
| Protein mutations: | S5A (194T>G), L7F (202G>T), L11V (212C>G), T50A (329A>G), A53T (338G>A 340A>T), V59I (356G>A), N64T (372A>C), L90F (451A>T), Q108H (505G>C), R109K (507G>A 508G>A), I110V (509A>G 511T>A), V112I (515G>A), A124E (552C>A 553T>G), A125K (554G>A 555C>A), Q128R (564A>G 565A>T), Q130R (570A>G), T132K (576C>A), L133V (578C>G), E138D (595G>T), D141A (603A>C 604C>T), R146Q (618G>A 619C>A), E151* (632G>T), L155A (644C>G 645T>C), L158Q (654T>A), S162A (665T>G), L168I (683T>A 685G>A), N170S (689A>T 690A>C), L172P (696T>C), E184K (731G>A 733A>G) | | | | | | | | |
| Codon mutations: | TCA2TCC (187A>C), TCT5GCT (194T>G), TTG7TTT (202G>T), CTT11GTT (212C>G), GAG14GAA (223G>A), CAC17CAT (232C>T), AAG18AAA (235G>A), CAA19CAG (238A>G), GCA21GCT (244A>T), GAC22GAT (247C>T), TCG28TCA (265G>A), GTT29GTA (268T>A), GGC32GGT (277C>T), TCT33TCC (280T>C), ATA35ATC (286A>C), ATA36ATC (289A>C), AAC37AAT (292C>T), CCA40CCT (301A>T), ACT41ACC (304T>C), CTA42TTG (305C>T 307A>G), TTC44TTT (313C>T), AAG45AAA (316G>A), CCC48CCT (325C>T), ACA50GCA (329A>G), GGT51GGA (334T>A), GCA53ACT (338G>A 340A>T), CTA57CTT (352A>T), GTT59ATT (356G>A), AAC63AAT (370C>T), AAT64ACT (372A>C), AGC65AGT (376C>T), GTT69GTA (388T>A), TAT73TAC (400T>C), TTA74CTC (401T>C 403A>C), ACT79ACG (418T>G), GGC84GGT (433C>T), AAG86AAA (439G>A), TCT88TCA (445T>A), GGC89GGA (448C>A), TTA90TTT (451A>T), AGA94CGA (461A>C), ACC100ACT (481C>T), ACC101ACA (484C>A), AGA102AGG (487A>G), CTA105CTG (496A>G), GCT107GCA (502T>A), CAG108CAC (505G>C), AGG109AAA (507G>A 508G>A), ATT110GTA (509A>G 511T>A), GTA112ATA (515G>A), TCC113TCT (520C>T), TCC120TCT (541C>T), GCT122GCG (547T>G), GCT124GAG (552C>A 553T>G), GCA125AAA (554G>A 555C>A), CAA128CGT (564A>G 565A>T), CCA129CCT (568A>T), CAA130CGA (570A>G), ACG132AAG (576C>A), CTT133GTT (578C>G), CAA135CAG (586A>G), TTA136CTT (587T>C 589A>T), GAG138GAT (595G>T), CTT140TTA (599C>T 601T>A), GAC141GCT (603A>C 604C>T), GAG142GAA (607G>A), GCC143GCT (610C>T), AAA144AAG (613A>G), CGC146CAA (618G>A 619C>A), TTA147TTG (622A>G), AAA150AAG (631A>G), GAG151TAG (632G>T), AAA152AAG (637A>G), CTT155GCT (644C>G 645T>C), CTA158CAA (654T>A), TCC159TCA (658C>A), TCC162GCC (665T>G), AAA163AAG (670A>G), TTA164TTG (673A>G), TTG166TTA (679G>A), TTG168ATA (683T>A 685G>A), AAC170TCC (689A>T 690A>C), CTA172CCA (696T>C), GAT175GAC (706T>C), GAA184AAG (731G>A 733A>G) | | | | | | | | |
| polyprotein (YP_001040018.1) | 1 | 1199 | 100% | 7607 | 91.9% | 1199 (100%) | 1084 (90.4%) | 0/0/0/0 | 3 |

| | Begin | End | Coverage | Score | Concordance | Matches | Identities | I/D/M/F* | Stop Codons |
|---|---|---|---|---|---|---|---|---|---|
| **NT** | **25** | **5403** | **99.6%** | **7554** | **70.2%** | **5379 (99.9%)** | **4583 (85.2%)** | **2/0** | |
| Protein mutations: | M2T (706T>C), K11R (733A>G), S17G (750A>G), Y48F (844A>T), V132I (1095G>A), T228A (1383A>G 1385C>T), S230* (1390C>G), T232S (1395A>T 1397T>A), Q242P (1426A>C 1427G>T), D245N (1434G>A), S252T (1456G>C 1457C>T), A254S (1461G>T), R255K (1465G>A), I257M (1472C>G), S275P (1524T>C 1526C>T), S281F (1543C>T), T282A (1545A>G 1547C>T), H286Q (1559C>A), Q298D (1593C>G 1595G>T), S301N (1603G>A), R302G (1605A>G), Y303Q (1608T>C 1610C>G), G307A (1621G>C 1622T>A), D309S (1626G>T 1627A>C 1628T>C), K317R (1651A>G), L321F (1662C>T 1664G>T), V326I (1677G>A 1679G>T), V365I (1794G>A), C366G (1797T>G 1799C>G), K370N (1811A>C), I373K (1819T>A 1820C>G), K377N (1832G>C), H381R (1843A>G), H388Y (1863C>T 1865C>T), V392M (1875G>A), I393V (1878A>G), I399M (1898T>G), V400T (1899G>A 1900T>C 1901G>A), A402V (1906C>T 1907C>G), E409D (1928A>C), N413D (1938A>G), D429N (1986G>A), F434Y (2002T>A), L450I (2049T>A 2051A>T), K453S (2059A>G 2060A>C), L458Q (2074T>A), V461A (2083T>C 2084G>A), Q465D (2094C>G 2096G>C), A466E (2098C>A 2099G>A), R467M (2100C>A 2101G>T 2102C>G), R469E (2106A>G 2107G>A), F470L (2111T>G), K472R (2115A>C 2116A>G), E473A (2119A>C 2120G>A), A475V (2125C>T 2126C>T), R476V (2127A>G 2128G>T), A477T (2130G>A 2132T>C), V479L (2136G>C 2138G>T), N481K (2144C>G), V488G (2164T>G), E490V (2170A>T 2171A>T), S493A (2178T>G), V500T (2199G>A 2200T>C), M502V (2205A>G), D508E (2225T>A), M510D (2229A>G 2230T>A 2231G>T), E517D (2252A>C), A519T (2256G>A), N520Q (2259A>C 2261T>A), E521Q (2262G>C), E531D (2294A>C), N538K (2315T>A), G540A (2320G>C 2321T>C), T542I (2326C>T), S579A (2436T>G), S590T (2469T>A 2471C>G), R601K (2503G>A), I608L (2523A>C), I643M (2630C>G), C667* (2702T>A), A700T (2799G>A), I712V (2835A>G 2837C>T), A761T (2982G>A), I771V (3012A>G 3014C>T), I807M (3122T>G), S829N (3187G>A), T832A (3195A>G 3197C>T), K851R (3253A>G), R855K (3265G>A), T857S (3270A>T 3272G>T), T868S (3303A>T 3305C>T), N936S (3508A>G), S948T (3543T>A), K955Q (3564A>C), L956M (3567T>A 3569A>G), P977Q (3631C>A), L978P (3634T>C 3635C>T), Q1014P (3742A>C 3743G>C), T1051A (3852A>G 3854T>G), I1061V (3882A>G 3884C>A), T1062A (3885A>G), K1076Q (3927A>C), T1078A (3933A>G 3935C>T), A1087T (3960G>A), M1095V (3984A>G 3986G>T), S1096N (3988G>A 3989C>T), D1097S (3990G>A 3991A>G), H1105Q (4016T>G), I1120V (4059A>G 4061A>T), N1123S (4069A>G 4070T>C), A1124V (4072C>T), L1150I (4149C>A 4151G>A), Q1163E (4188C>G), H1175Y (4224C>T), S1195G (4284A>G 4286T>A) | | | | | | | | |
| Codon mutations: | ATG2ACG (706T>C), AGG10AGA (731G>A), AAG11AGG (733A>G), AGT17GGT (750A>G), GCA18GCT (755A>T), GCT19GCA (758T>A), GTA20GTG (761A>G), TTA24CTA (771T>C), CAG27CAA (782G>A), ACA29ACT (788A>T), GCT30GCA (791T>A), CTG39CTA (818G>A), CTT40CTC (821T>C), AAA42AAG (827A>G), GGC44GGT (833C>T), TAT48TTT (844A>T), CTC51CTA (854C>A), GGC53GGG (860C>G), GCC54GCT (863C>T), CAA55CAG (866A>G), GCC59GCT (878C>T), GAA61GAG (884A>G), CTG62CTT (887G>T), GAC64GAT (893C>T), AAT70AAC (911T>C), TCT76TCA (929T>A), TCA77TCT (932A>T), GGA89GGG (968A>G), GTT103GTG (1010T>G), AGT120AGC (1061T>C), ACC124ACA (1073C>A), AAT125AAC (1076T>C), CTG128TTA (1083C>T 1085G>A), GTT132ATT (1095G>A), TTG133TTA (1100G>A), TCC136TCT (1109C>T), GGG138GGA (1115G>A), GCA139GCC (1118A>C), TTA143TTG (1130A>G), CCG146CCA (1139G>A), ACC147ACT (1142C>T), CTA150CTG (1151A>G), AAA154AAG (1163A>G), GGC156GGG (1169C>G), TTC159TTT (1178C>T), ACA162ACT (1187A>T), ACT168ACA (1205T>A), CCT173CCC (1220T>C), GCC178GCA (1235C>A), CAA182CAG (1247A>G), CGT187CGC (1262T>C), TCA192TCG (1277A>G), CTT204CTC (1313T>C), CAC205CAT (1316C>T), CGA206AGG (1317C>A 1319A>G), CAA208CAG (1325A>G), TTT209TTC (1328T>C), TTC210TTT (1331C>T), CAG214CAA (1343G>A), CAG218CAA (1355G>A), ATA220ATT (1361A>T), TCA222TCT (1367A>T), GTA223GTT (1370A>T), TCT225TCA (1376T>A), ACC228GCT (1383A>G 1385C>T), TCC229TCT (1388C>T), TCA230TGA (1390C>G), GCA231GCT (1394A>T), ACT232TCA (1395A>T 1397T>A), GAT233GAC (1400T>C), TCA234TCT (1403A>T), GTG235GTA (1406G>A), CTT236TTA (1407C>T 1409T>A), CCG237CCT (1412G>T), GTG239GTA (1418G>A), CAG242CCT (1426A>C 1427G>T), GAC245AAC (1434G>A), ACA246ACC (1439A>C), GTG250GTT (1451G>T), CCA251CCT (1454A>T), AGC252ACT (1456G>C 1457C>T), GAA253GAG (1460A>G), GCA254TCA (1461G>T), AGA255AAA (1465G>A), ATC257ATG (1472C>G), CGG258CGC (1475G>C), TCC259TCG (1478C>G), TCC260TCT (1481C>T), CTC262CTG (1487C>G), CGG263CGA (1490G>A), GTG264GTA (1493G>A), CCC265CCT (1496C>T), CTC267CTT (1502C>T), AGT271AGC (1514T>C), TCC275CCT (1524T>C 1526C>T), TCA276TCT (1529A>T), ATA279ATT (1538A>T), TCT281TTT (1543C>T), ACC282GCT (1545A>G 1547C>T), ACT284ACC (1553T>C), CAC286CAA (1559C>A), CTC287TTA (1560C>T 1562C>A), GGC289GGG (1568C>G), ACG291ACA (1574G>A), CGT292AGG (1575C>A 1577T>G), GTA296GTG (1589A>G), CAG298GAT (1593C>G 1595G>T), AGT301AAT (1603G>A), AGG302GGG (1605A>G), TAC303CAG (1608T>C 1610C>G), GAA304GAG (1613A>G), GAG305GAA (1616G>A), GGT307GCA (1621G>C 1622T>A), GAT309TCC (1626G>T 1627A>C 1628T>C), GGG310GGA (1631G>A), GTT311GTA (1634T>A), GTT315GTC (1646T>C), AAG317AGG (1651A>G), GCT318GCA (1655T>A), GGT320GGC (1661T>C), CTG321TTT (1662C>T 1664G>T), AAC322AAT (1667C>T), CAT325CAC (1676T>C), GTG326ATT (1677G>A 1679G>T), GAT328GAC (1685T>C), GGT331GGC (1694T>C), GAG333GAA (1700G>A), CTG340CTC (1721G>C), CCC341CCT (1724C>T), GAA342GAG (1727A>G), AAC343AAT (1730C>T), CTG347TTG (1740C>T), TCG348TCA (1745G>A), GTG349GTA (1748G>A), AGA350CGA (1749A>C), CAG351CAA (1754G>A), CAA352CAG (1757A>G), GTT353GTG (1760T>G), GCT354GCA (1763T>A), GCA355GCG (1766A>G), GCA356GCT (1769A>T), GCA359GCC (1778A>C), GCA361GCT (1784A>T), AAG363AAA (1790G>A), GGT364GGA (1793T>A), GTT365ATT (1794G>A), TGC366GGG (1797T>G 1799C>G), GTC368GTT (1805C>T), AAA370AAC (1811A>C), CAT371CAC (1814T>C), ATA372ATT (1817A>T), ATC373AAG (1819T>A 1820C>G), AAC374AAT (1823C>T), AAG377AAC (1832G>C), ATC380ATT (1841C>T), CAC381CGC (1843A>G), TTG382CTT (1845T>C 1847G>T), GAG383GAA (1850G>A), ATC385ATT (1856C>T), CAC388TAT (1863C>T 1865C>T), GTG392ATG (1875G>A), ATA393GTA (1878A>G), TAC397TAT (1892C>T), ATT399ATG (1898T>G), GTG400ACA (1899G>A 1900T>C 1901G>A), GCC402GTG (1906C>T 1907C>G), TCT404TCA (1913T>A), GAA409GAC (1928A>C), AAC410AAT (1931C>T), AAT413GAT (1938A>G), CTT414TTG (1941C>T 1943T>G), ACC415ACA (1946C>A), TTG416CTG (1947T>C), GCA417GCC (1952A>C), CAG418CAA (1955G>A), AGA419AGG (1958A>G), GAG422GAA (1967G>A), CTA423CTG (1970A>G), CAA426CAG (1979A>G), GAT429AAT (1986G>A), TAT431TAC (1994T>C), TTT434TAT (2002T>A), AAC437AAT (2012C>T), TAT441TAC (2024T>C), CCT442CCA (2027T>A), TTA450ATT (2049T>A 2051A>T), ACG451ACT (2054G>T), AAA453AGC (2059A>G 2060A>C), CGC455CGG (2066C>G), CTA458CAA (2074T>A), GTG461GCA (2083T>C 2084G>A), GCA462GCC (2087A>C), CAG465GAC (2094C>G 2096G>C), GCG466GAA (2098C>A 2099G>A), CGC467ATG (2100C>A 2101G>T 2102C>G), AGA469GAA (2106A>G 2107G>A), TTT470TTG (2111T>G), TAC471TAT (2114C>T), AAG472CGG (2115A>C 2116A>G), GAG473GCA (2119A>C 2120G>A), GCG474GCT (2123G>T), GCC475GTT (2125C>T 2126C>T), AGG476GTG (2127A>G 2128G>T), GCT477ACC (2130G>A 2132T>C), CAG478CAA (2135G>A), GTG479CTT (2136G>C 2138G>T), AAC481AAG (2144C>G), ACT486ACA (2159T>A), AGT487AGC (2162T>C), GTG488GGG (2164T>G), GAA490GTT (2170A>T 2171A>T), ATA491ATT (2174A>T), CCA492CCT (2177A>T), TCC493GCC (2178T>G), ACC494ACT (2183C>T), TCA495TCC (2186A>C), TTC496TTT (2189C>T), ACA498ACC (2195A>C), AAG499AAA (2198G>A), GTT500ACT (2199G>A 2200T>C), GCC501GCT (2204C>T), ATG502GTG (2205A>G), GAG503GAA (2210G>A), GGC505GGG (2216C>G), AGT506AGC (2219T>C), GAT508GAA (2225T>A), AGC509AGT (2228C>T), ATG510GAT (2229A>G 2230T>A 2231G>T), GCT513GCA (2240T>A), ATT514ATC (2243T>C), GAA517GAC (2252A>C), GCC519ACC (2256G>A), AAT520CAA (2259A>C 2261T>A), GAA521CAA (2262G>C), GTA523GTG (2270A>G), GTT528GTC (2285T>C), GAA531GAC (2294A>C), CAA534CAG (2303A>G), AAT537AAC (2312T>C), AAT538AAA (2315T>A), GGT540GCC (2320G>C 2321T>C), GCG541GCC (2324G>C), ACA542ATA (2326C>T), TTA546TTG (2339A>G), CAA548CAG (2345A>G), TTC558TTT (2375C>T), GAG564GAA (2393G>A), AGT571AGC (2414T>C), GTT573GTG (2420T>G), GAG576GAA (2429G>A), GTG577GTA (2432G>A), TCA579GCA (2436T>G), GCG581GCC (2444G>C), CTC582CTA (2447C>A), GCG584GCA (2453G>A), CTC586CTG (2459C>G), TCC590ACG (2469T>A 2471C>G), AAG591AAA (2474G>A), GGA596GGC (2489A>C), TCG597TCA (2492G>A), CAA598CAG (2495A>G), CTA600CTG (2501A>G), AGG601AAG (2503G>A), TAT602TAC (2507T>C), GCC605GCT (2516C>T), ATT608CTT (2523A>C), TAT615TAC (2546T>C), TCC617TCT (2552C>T), GGT620GGG (2561T>G), ATT622ATA (2567T>A), TCG623TCA (2570G>A), GGT624GGA (2573T>A), GTT629GTA (2588T>A), CTC636CTA (2609C>A), TTC642TTT (2627C>T), ATC643ATG (2630C>G), CTG648CTC (2645G>C), CAT653CAC (2660T>C), TCA657TCT (2672A>T), GGG664GGT (2693G>T), TGT667TGA (2702T>A), ACA669ACG (2708A>G), GGT672GGC (2717T>C), ATC673ATA (2720C>A), TTC676TTT (2729C>T), GCC682GCT (2747C>T), GGG684GGT (2753G>T), ATA691ATT (2774A>T), CAC697CAT (2792C>T), CCC698CCG (2795C>G), TTG699TTA (2798G>A), GCT700ACT (2799G>A), AGT701AGC (2804T>C), ATC712GTT (2835A>G 2837C>T), CAT713CAC (2840T>C), CTG714TTG (2841C>T), CAG715CAA (2846G>A), AAA717AAG (2852A>G), TCG720TCA (2861G>A), ACC721ACA (2864C>A), AAC722AAT (2867C>T), ATC723ATA (2870C>A), CCT726CCA (2879T>A), CCT727CCA (2882T>A), TTG729CTA (2886T>C 2888G>A), CGA730CGG (2891A>G), GCA731GCG (2894A>G), GCA733GCT (2900A>T), GGT736GGC (2909T>C), AAA738AAG (2915A>G), GAC740GAT (2921C>T), ACA742ACG (2927A>G), CCA745CCC (2936A>C), CGA746CGG (2939A>G), ACA749ACG (2948A>G), CAG751CAA (2954G>A), GTA752GTT (2957A>T), CTG753CTA (2960G>A), CAC755CAT (2966C>T), AAT757AAC (2972T>C), GTG760GTT (2981G>T), GCT761ACT (2982G>A), GGT765GGG (2996T>G), ACT766ACA (2999T>A), TTA768CTA (3003T>C), AGC770AGT (3011C>T), ATC771GTT (3012A>G 3014C>T), TCT773TCC (3020T>C), CCG774CCC (3023G>C), GGC776GGT (3029C>T), ATA777ATT (3032A>T), GAC781GAT (3044C>T), GGT782GGA (3047T>A), ACG783ACC (3050G>C), CTG784TTG (3051C>T), CCA786CCT (3059A>T), TCC787TCT (3062C>T), TTG788CTA (3063T>C 3065G>A), GGA790GGG (3071A>G), ATA792ATT (3077A>T), CGC794CGA (3083C>A), ACT800ACG (3101T>G), GGC801GGT (3104C>T), ACT802ACG (3107T>G), GTG804GTC (3113G>C), ATT807ATG (3122T>G), ATT809ATC (3128T>C), GAG810GAA (3131G>A), TTC814TTT (3143C>T), CAT815CAC (3146T>C), GGT817GGA (3152T>A), TTG819TTA (3158G>A), GCC820GCA (3161C>A), GGA822GGT (3167A>T), ACA825ACC (3176A>C), CTG826CTT (3179G>T), AAC827AAT (3182C>T), AGC829AAC (3187G>A), TCC831TCT (3194C>T), ACC832GCT (3195A>G 3197C>T), CCT833CCC (3200T>C), GAC835GAT (3206C>T), TTA837TTG (3212A>G), GTT842GTG (3227T>G), ATT843ATA (3230T>A), AAC845AAT (3236C>T), CTT846CTG (3239T>G), GAG847GAA (3242G>A), GGA849GGC (3248A>C), CGA850CGT (3251A>T), AAG851AGG (3253A>G), AGG855AAG (3265G>A), TGT856TGC (3269T>C), ACG857TCT (3270A>T 3272G>T), ACC859ACT (3278C>T), AAA864AAG (3293A>G), CTT866CTA (3299T>A), ACC868TCT (3303A>T 3305C>T), GGT870GGA (3311T>A), CTA875TTA (3324C>T), CCG876CCT (3329G>T), CAC883CAT (3350C>T), AGG885CGG (3354A>C), TTG886TTA (3359G>A), ACA889ACG (3368A>G), GTC890GTG (3371C>G), TTG891CTG (3372T>C), GTA895GTT (3386A>T), CTA901CTG (3404A>G), ACG903ACA (3410G>A), GTC904GTT (3413C>T), ACA907ACG (3422A>G), CAG909CAA (3428G>A), CTT910CTG (3431T>G), TGC912TGT (3437C>T), GAT915GAC (3446T>C), TTG916TTA (3449G>A), GTC917GTA (3452C>A), CTT918CTA (3455T>A), GGG919GGA (3458G>A), GGC920GGT (3461C>T), TCT923TCC (3470T>C), GTT924GTC (3473T>C), CCC926CCT (3479C>T), ATT927ATC (3482T>C), TAC928TAT (3485C>T), GGA929GGG (3488A>G), CAC930CAT (3491C>T), GGA934GGG (3503A>G), AAT936AGT (3508A>G), GCT937GCC (3512T>C), GTG938GTT (3515G>T), CTA941TTG (3522C>T 3524A>G), TTC942TTT (3527C>T), GAG944GAA (3533G>A), TCT948ACT (3543T>A), CGC950CGT (3551C>T), ATT953ATC (3560T>C), AAA955CAA (3564A>C), TTA956ATG (3567T>A 3569A>G), AAG958AAA (3575G>A), GAA959GAG (3578A>G), AAC960AAT (3581C>T), ACA963ACC (3590A>C), GGA968GGG (3605A>G), AAG970AAA (3611G>A), CCA972CCT (3617A>T), CAG973CAA (3620G>A), GTA974GTT (3623A>T), GTG976GTC (3629G>C), CCG977CAG (3631C>A), CTC978CCT (3634T>C 3635C>T), AGA979AGG (3638A>G), GAC980GAT (3641C>T), TCA983TCT (3650A>T), GGG985GGT (3656G>T), GCT986GCC (3659T>C), GTA987GTG (3662A>G), GTG991GTC (3674G>C), GAC996GAT (3689C>T), CGA999CGC (3698A>C), TTG1001CTG (3702T>C), TTA1003CTA (3708T>C), TGC1005TGT (3716C>T), GCA1006GCG (3719A>G), CCC1007CCA (3722C>A), TCC1009TCG (3728C>G), CAG1014CCC (3742A>C 3743G>C), GGG1015GGA (3746G>A), GCA1016GCT (3749A>T), CTG1017CTA (3752G>A), GTA1018GTG (3755A>G), ACA1023ACC (3770A>C), TGC1024TGT (3773C>T), CCA1025CCT (3776A>T), TTC1026TTT (3779C>T), TTC1033TTT (3800C>T), GGA1038GGT (3815A>T), TAC1042TAT (3827C>T), ATT1044ATA (3833T>A), GTA1046GTG (3839A>G), AGA1048AGG (3845A>G), GTG1049GTA (3848G>A), CAG1050CAA (3851G>A), ACT1051GCG (3852A>G 3854T>G), TCC1052TCA (3857C>A), ATA1055ATC (3866A>C), GGA1056GGT (3869A>T), GGA1057GGT (3872A>T), CTG1059CTC (3878G>C), AAC1060AAT (3881C>T), ATC1061GTA (3882A>G 3884C>A), ACC1062GCC (3885A>G), TTA1063TTG (3890A>G), GAT1064GAC (3893T>C), TCA1065TCT (3896A>T), GGA1073GGT (3920A>T), ATC1074ATT (3923C>T), AAG1076CAG (3927A>C), GGC1077GGA (3932C>A), ACC1078GCT (3933A>G 3935C>T), GTT1080GTG (3941T>G), GTC1081GTA (3944C>A), TCT1082TCC (3947T>C), GCT1083GCA (3950T>A), GGT1084GGA (3953T>A), GCA1087ACA (3960G>A), GCT1090GCA (3971T>A), GGT1094GGA (3983T>A), ATG1095GTT (3984A>G 3986G>T), AGC1096AAT (3988G>A 3989C>T), GAC1097AGC (3990G>A 3991A>G), AAC1098AAT (3995C>T), ATC1099ATT (3998C>T), TTT1100TTC (4001T>C), CAT1105CAG (4016T>G), GAG1108GAA (4025G>A), TTT1110TTC (4031T>C), CCT1111CCG (4034T>G), AGA1112CGT (4035A>C 4037A>T), CGC1113CGT (4040C>T), ACC1115ACT (4046C>T), GCC1117GCT (4052C>T), ATA1120GTT (4059A>G 4061A>T), CCA1122CCT (4067A>T), AAT1123AGC (4069A>G 4070T>C), GCT1124GTT (4072C>T), AGA1126CGA (4077A>C), CGA1133CGC (4100A>C), CTA1134CTT (4103A>T), GGA1135GGC (4106A>C), CTC1137CTT (4112C>T), ATA1147ATC (4142A>C), CTG1150ATA (4149C>A 4151G>A), TTC1160TTT (4181C>T), TTG1162TTA (4187G>A), CAA1163GAA (4188C>G), GCT1165GCA (4196T>A), TCA1169TCT (4208A>T), GCA1170GCT (4211A>T), AAG1174AAA (4223G>A), CAT1175TAT (4224C>T), TTG1176CTT (4227T>C 4229G>T), ACC1183ACT (4250C>T), TAC1184TAT (4253C>T), CTT1185CTG (4256T>G), ACT1187ACC (4262T>C), CCT1188CCA (4265T>A), TCA1191TCT (4274A>T), AGG1194CGG (4281A>C), AGT1195GGA (4284A>G 4286T>A), TTT1196TTC (4289T>C) | | | | | | | | |

*: Inserts / Deletes / Misaligned / Frameshifts

#### Analysis details

This analysis was performed with panviral2.64

### NGS Details (UN59): Torradovirus marchitezum (segment RNA 1)

#### Assembly

| | |
|---|---|
| **Coverage Length** | 5567 (1 contig(s)) |
| **Depth Of Coverage** | 1430.4 |
| **Number Of Reads** | 59802 |
| **Reads Per Million** | 1300.04 rpm (after QC) |
| **Ambiguities** | 0 |
| **Assembly Method** | de novo + reference guided assembly |
| **Consensus Caller** | Bcf Tools |

#### Coverage Map

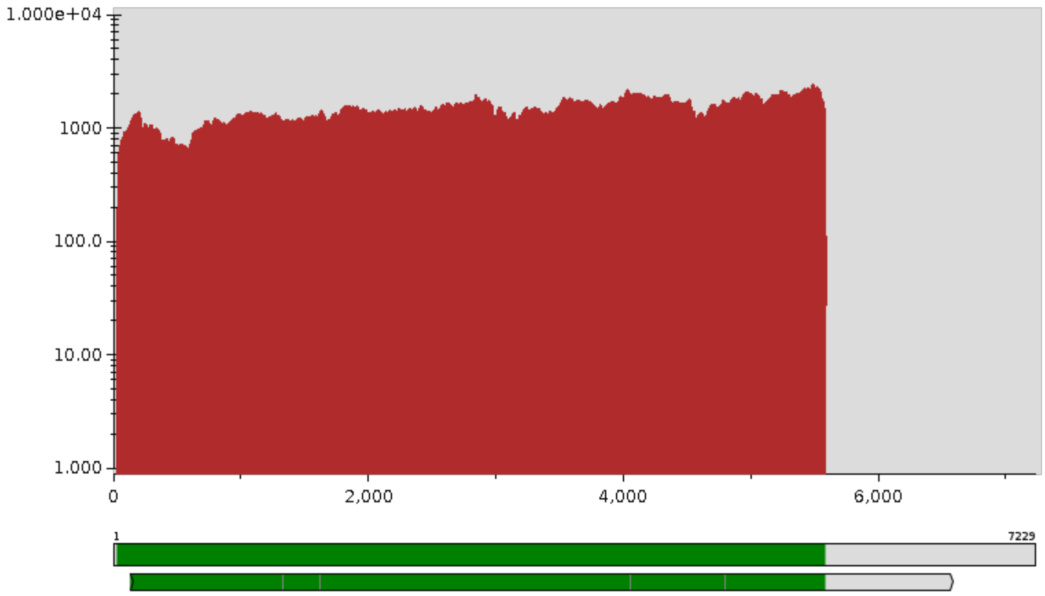

#### Assignment

| | |
|---|---|
| **Type** | Torradovirus marchitezum (Taxonomy ID: 3048376) |
| **Reference Genome** | NC_010987.1 |
| **NT Identity (%)** | 66.5281 |
| **AA Identity (%)** | 69.9448 |
| **Number Of Stop Codons** | 0 |
| **Number Of CDS** | 1 |

#### Alignment

| | |
|---|---|
| **Alignment Score** | 3552.0 (NT) + 9297.0 (AA) = 12849.0 |
| **Concordance (%)** | 54.4311 |

| Alignment Method | Global, seeded, nucleotide + amino acids (AGA) |
|---|---|

#### Genome Region

Sequence starts at position 23 and ends at position 5589 relative to NC_010987.1 reference sequence.

#### Alignment Detailed Statistics

| | Begin | End | Coverage | Score | Concordance | Matches | Identities | I/D/M/F* | Stop Codons |
|---|---|---|---|---|---|---|---|---|---|
| NT | 23 | 5589 | 77.0% | 3552 | 32.4% | 5524 (99.1%) | 3681 (66.0%) | 9/43 | |

| | Begin | End | Coverage | Score | Concordance | Matches | Identities | I/D/M/F* | Stop Codons |
|---|---|---|---|---|---|---|---|---|---|
| **NT** | **23** | **5589** | **77.0%** | **3552** | **32.4%** | **5524 (99.1%)** | **3681 (66.0%)** | **9/43** | |

Mutations:

32T>C, 37G>A, 40C>T, 59A>C, 61T>C, 62T>A, 63G>C, 66A>C, 67T>A, 68T>A, 70G>A, 71A>C, 74C>T, 75T>G, 76T>C, 77G>T, 79A>T, 81A>C, 82C>T, 85T>C, 87C>A, 89_99delAACATCTTCTT, 108A>T, 112G>C, 114C>T, 115C>T, 116A>C, 118T>C, 119A>T, 122T>G, 125A>T, 128C>A, 129C>A, 130T>A, 133_134delAT, 136T>A, 146T>A, 160T>G, 161C>G, 162T>A, 163C>G, 164C>T, 165_188delAAGGTTGCTGGTGTTGATGACGCC, 189A>C, 190G>C, 192T>C, 195T>G, 200C>T, 201A>T, 203C>A, 204A>C, 217G>C, 218T>A, 222C>G, 227A>T, 228A>T, 229A>C, 234A>T, 239C>T, 240A>T, 241A>C, 242A>T, 243T>C, 251A>G, 252G>A, 254G>T, 255G>C, 257T>C, 258A>T, 259G>C, 260T>C, 264A>G, 268C>A, 269C>T, 271C>A, 272T>A, 273A>G, 275G>T, 283A>C, 284G>C, 288T>G, 289C>T, 296T>C, 297C>A, 299T>A, 306C>G, 308G>C, 314A>G, 315T>C, 317T>A, 320G>T, 322C>T, 323T>G, 325A>C, 326T>A, 328T>C, 329G>A, 330T>G, 332C>A, 338T>A, 343T>C, 344G>C, 345G>T, 347C>T, 350T>A, 353T>C, 355G>A, 357T>G, 359A>T, 360C>G, 362T>A, 363A>G, 365T>A, 367A>G, 374G>A, 377C>T, 380A>G, 385G>C, 386T>A, 387G>A, 388C>A, 390C>A, 392T>A, 395A>G, 398T>A, 401A>C, 404T>C, 405C>A, 407T>C, 410A>G, 412C>G, 413G>T, 416C>T, 417T>G, 418C>A, 422C>T, 423T>G, 425G>T, 426T>A, 427C>A, 428C>A, 431A>T, 437C>T, 441T>A, 446C>A, 447A>G, 448A>T, 449G>A, 453A>C, 455G>T, 458T>A, 459T>C, 461A>T, 465T>A, 468T>G, 471G>T, 472A>T, 473G>A, 477T>G, 479G>C, 480T>A, 481C>A, 482C>G, 483A>G, 485T>C, 489G>T, 491T>C, 493A>T, 494A>C, 495T>C, 502T>A, 503T>C, 505C>A, 506T>A, 510G>T, 512G>A, 516A>G, 518T>A, 522A>G, 523G>T, 527A>G, 528G>T, 529A>C, 530T>A, 531G>T, 542T>G, 543G>T, 545A>T, 546T>A, 547A>T, 548T>A, 550G>C, 555C>T, 557T>A, 560A>T, 567A>T, 569A>G, 570A>G, 572A>C, 573T>A, 575G>T, 578A>G, 579G>C, 581G>A, 582C>T, 584T>A, 587A>G, 589T>A, 593T>A, 596T>C, 600A>G, 601G>T, 602T>A, 611T>A, 612G>A, 613A>C, 615G>C, 616G>C, 618G>T, 621T>A, 622C>A, 626C>T, 627A>G, 628C>T, 629T>A, 632C>T, 633A>G, 634A>T, 635A>T, 637A>C, 639A>T, 641C>G, 646T>C, 647G>A, 648A>G, 650A>T, 653C>T, 659C>T, 660A>T, 662G>T, 665T>C, 668A>T, 669A>T, 670G>C, 671T>A, 672C>T, 674A>G, 677C>T, 697A>C, 702G>T, 704C>T, 706A>G, 707A>G, 708C>A, 709G>A, 710A>G, 713C>T, 714A>T, 717A>C, 718T>A, 720G>C, 725C>G, 731T>A, 732A>G, 733G>C, 734T>A, 735A>G, 737G>A, 746G>T, 750T>A, 751C>G, 752A>T, 753G>T, 755T>A, 756T>A, 758A>G, 770G>T, 771T>A, 772C>G, 773T>C, 776G>A, 777T>C, 779G>T, 780A>T, 782G>A, 786A>T, 789A>T, 791A>G, 803G>T, 806T>C, 809G>A, 812G>T, 815G>A, 818G>A, 821C>T, 822A>G, 827A>T, 830G>T, 836T>G, 848C>T, 849T>C, 851G>T, 852A>T, 853G>C, 854C>A, 861T>C, 866C>A, 872T>C, 873C>T, 875C>G, 878T>C, 881G>T, 882A>G, 884A>T, 896T>G, 899G>A, 911A>G, 914A>G, 918A>C, 919C>A, 920T>G, 921C>T, 923T>G, 927C>A, 929T>G, 932T>C, 938A>G, 944C>T, 947C>A, 950C>T, 953G>A, 956T>C, 974A>G, 978A>G, 980A>G, 983C>T, 984G>A, 988C>T, 989C>T, 995A>C, 998C>T, 1010G>A, 1019T>G, 1024G>A, 1028A>T, 1031G>C, 1032C>T, 1034G>A, 1040T>C, 1046G>T, 1050G>C, 1059C>A, 1060A>C, 1064G>A, 1068G>T, 1069A>C, 1070T>A, 1072C>A, 1073A>G, 1076G>T, 1077T>G, 1078G>C, 1082T>A, 1085T>A, 1086G>T, 1088A>T, 1090C>T, 1100G>A, 1103C>T, 1109C>A, 1113A>C, 1114A>C, 1115G>T, 1118T>A, 1121C>T, 1124T>A, 1126C>T, 1127C>T, 1130A>G, 1134A>C, 1136C>A, 1140C>T, 1142G>A, 1144G>A, 1145G>A, 1148A>G, 1149G>C, 1151A>G, 1154T>C, 1155C>G, 1160G>A, 1163C>A, 1178A>C, 1193G>A, 1199C>T, 1200A>T, 1215A>T, 1216G>C, 1226T>A, 1227G>A, 1232G>A, 1233T>C, 1238A>T, 1242G>T, 1244T>G, 1245T>G, 1246A>T, 1253A>G, 1254A>G, 1255G>C, 1256T>C, 1257T>A, 1259T>C, 1260A>C, 1261G>A, 1262G>A, 1265C>T, 1266A>T, 1267T>C, 1268G>T, 1271C>T, 1280G>A, 1284C>A, 1286T>G, 1292A>G, 1298T>C, 1304A>G, 1313C>T, 1316G>A, 1317T>G, 1319C>T, 1325G>A, 1337T>A, 1340A>G, 1342C>G, 1350T>C, 1352A>T, 1355T>C, 1367G>T, 1368T>C, 1370G>C, 1374A>G, 1375G>C, 1385G>A, 1391A>G, 1394C>G, 1397T>C, 1400G>C, 1401C>T, 1403A>G, 1409A>G, 1418G>A, 1421G>A, 1445T>A, 1448C>T, 1457T>A, 1458A>G, 1460C>G, 1463G>T, 1472C>T, 1478A>T, 1481A>T, 1482T>C, 1484G>T, 1487A>G, 1496T>A, 1505C>T, 1508A>G, 1511T>C, 1514T>A, 1517A>G, 1529T>C, 1538A>T, 1541A>C, 1544A>G, 1547A>T, 1553C>T, 1554T>C, 1556A>T, 1565C>T, 1571T>C, 1577C>T, 1583C>A, 1589T>C, 1592C>T, 1595C>A, 1598C>A, 1604T>C, 1607A>C, 1616A>C, 1623C>T, 1628C>A, 1631A>T, 1637T>A, 1638G>A, 1640C>T, 1643A>G, 1649T>G, 1652T>A, 1655A>G, 1661A>T, 1667T>A, 1673G>A, 1674C>A, 1679A>G, 1682G>A, 1685C>T, 1688C>T, 1689C>T, 1691G>A, 1694G>T, 1700T>C, 1702T>G, 1703G>A, 1708A>T, 1715G>T, 1719T>G, 1720C>A, 1721C>T, 1722A>C, 1724T>A, 1725G>A, 1727A>T, 1729A>C, 1733T>A, 1736G>A, 1738G>C, 1739A>C, 1745C>A, 1746A>G, 1747G>T, 1748A>T, 1752A>G, 1754C>T, 1755A>G, 1757A>G, 1763G>A, 1781T>C, 1787T>A, 1790C>T, 1793G>A, 1794A>G, 1795A>C, 1799G>A, 1800A>T, 1806A>G, 1807G>A, 1811T>C, 1814T>C, 1818G>C, 1820G>A, 1829A>G, 1832C>A, 1835A>G, 1838T>C, 1840A>G, 1844A>G, 1847T>C, 1848G>T, 1849C>A, 1850A>T, 1853A>G, 1856A>G, 1861A>G, 1862G>C, 1866T>C, 1868G>T, 1870T>A, 1872G>A, 1874A>T, 1875G>A, 1878G>A, 1879A>C, 1883T>A, 1886C>A, 1887A>T, 1889A>G, 1890A>G, 1891G>C, 1893C>G, 1894C>A, 1895T>C, 1899A>G, 1901A>T, 1910T>C, 1911C>A, 1913A>T, 1917T>G, 1921C>T, 1922A>T, 1928T>A, 1932A>C, 1934A>G, 1936C>A, 1937G>T, 1941G>T, 1943G>C, 1944G>C, 1946T>A, 1948G>A, 1949G>T, 1952T>C, 1953_1955delAGA, 1958G>A, 1960G>A, 1961C>A, 1962A>G, 1963C>T, 1964C>T, 1965A>T, 1966A>T, 1969A>C, 1970A>T, 1971G>C, 1972A>C, 1976A>G, 1980A>G, 1981T>C, 1983T>A, 1984T>C, 1985G>T, 1986G>T, 1987T>A, 1988G>T, 1989C>G, 1990C>A, 1991C>G, 1992A>C, 1993C>A, 1994T>G, 1995G>A, 1996T>C, 1997T>A, 1998G>A, 2000G>T, 2002G>C, 2003T>A, 2007A>G, 2008A>C, 2012T>A, 2014G>A, 2016G>A, 2018G>T, 2019A>G, 2021C>T, 2024T>C, 2025G>T, 2027A>C, 2028A>C, 2029G>A, 2030A>G, 2033G>A, 2039G>A, 2042A>T, 2048A>C, 2049T>C, 2051G>C, 2060A>T, 2066G>A, 2069C>G, 2073T>A, 2076G>A, 2079C>A, 2080C>A, 2081T>A, 2082G>T, 2083A>T, 2084G>T, 2087A>G, 2088C>T, 2093T>C, 2096T>A, 2104A>T, 2105T>C, 2106C>T, 2107A>C, 2108T>C, 2117C>T, 2120G>C, 2127T>G, 2128C>T, 2129T>G, 2132T>A, 2133G>T, 2134A>C, 2136G>T, 2137A>C, 2141A>T, 2142C>G, 2143A>C, 2144T>C, 2148G>C, 2152T>A, 2156C>T, 2157A>G, 2159C>A, 2161A>T, 2162A>G, 2166A>G, 2167G>T, 2169G>A, 2170T>A, 2171C>T, 2172A>G, 2174C>T, 2175A>T, 2181T>G, 2182A>T, 2183T>C, 2185G>A, 2193G>A, 2194A>G, 2196G>C, 2199G>C, 2201A>G, 2208C>G, 2209G>C, 2210A>T, 2219A>G, 2237C>T, 2241A>G, 2243A>C, 2249C>T, 2252T>C, 2268G>A, 2269C>G, 2270A>C, 2271T>A, 2279C>T, 2285G>A, 2286A>G, 2289A>T, 2290A>T, 2291A>G, 2294A>G, 2296C>A, 2297A>T, 2300G>C, 2303G>A, 2304A>C, 2305C>G, 2306A>C, 2307G>A, 2309G>T, 2310A>C, 2312G>T, 2313A>G, 2314G>A, 2316A>C, 2317C>A, 2318T>A, 2321A>G, 2322T>A, 2323T>A, 2324T>A, 2327A>G, 2328C>T, 2331T>A, 2332C>A, 2333T>G, 2334T>G, 2335G>C, 2336C>G, 2337A>C, 2340G>C, 2342T>G, 2345A>T, 2350T>C, 2351T>A, 2354C>T, 2357C>A, 2360C>T, 2362T>C, 2363A>G, 2369C>T, 2372G>A, 2381A>G, 2385T>A, 2386T>G, 2387T>A, 2390T>A, 2396T>C, 2397T>C, 2399A>G, 2405A>T, 2408A>T, 2415A>C, 2417A>C, 2426C>T, 2428A>G, 2429C>G, 2435A>T, 2436T>G, 2438_2439insGGT, 2440A>C, 2441A>T, 2444G>C, 2445C>A, 2446C>A, 2447A>G, 2451G>A, 2452A>G, 2454T>A, 2455C>A, 2456A>T, 2462C>T, 2464A>T, 2465A>C, 2466A>G, 2467T>A, 2468T>A, 2472T>A, 2473A>T, 2474C>G, 2475G>A, 2476A>T, 2478A>G, 2479A>G, 2480A>T, 2484G>A, 2485T>A, 2486T>G, 2489A>C, 2490T>A, 2492T>A, 2495C>G, 2497A>T, 2498G>C, 2499G>C, 2501T>G, 2503C>T, 2504A>G, 2508T>A, 2509G>A, 2510G>T, 2511A>G, 2512G>A, 2514C>T, 2516T>G, 2521A>T, 2522T>C, 2523G>A, 2524C>T, 2525A>C, 2526C>T, 2530G>A, 2531A>G, 2537C>T, 2546A>C, 2549T>A, 2556A>T, 2558A>G, 2561G>A, 2564T>A, 2573A>C, 2574G>A, 2575T>C, 2576T>C, 2577T>A, 2579T>G, 2580G>A, 2582A>T, 2584G>C, 2586A>G, 2591A>T, 2594A>T, 2597T>C, 2598T>A, 2600G>T, 2601G>A, 2603G>T, 2607T>C, 2608G>T, 2609G>C, 2610C>A, 2613A>G, 2614G>C, 2615T>A, 2616A>G, 2617A>C, 2618C>A, 2619T>A, 2623T>C, 2627C>T, 2628A>T, 2629A>T, 2630T>A, 2633A>T, 2634T>G, 2635C>A, 2636G>T, 2645A>G, 2648T>C, 2651G>A, 2660A>G, 2672A>C, 2678T>C, 2684G>A, 2687C>T, 2690G>A, 2693T>G, 2696T>G, 2698A>C, 2699G>T, 2699_2700insCAT, 2700C>T, 2703A>G, 2704G>C, 2708T>A, 2709C>T, 2710T>C, 2711T>C, 2712C>T, 2714C>G, 2715G>C, 2717T>A, 2720T>G, 2724A>G, 2725G>C, 2726T>G, 2727T>G, 2729T>A, 2730C>G, 2732A>C, 2733T>A, 2734C>A, 2735A>C, 2736T>A, 2737T>A, 2738G>C, 2739G>C, 2740C>A, 2741A>T, 2742C>G, 2743A>T, 2744A>G, 2747T>C, 2747_2748insCGA, 2753T>A, 2755C>A, 2756T>A, 2757G>C, 2759A>G, 2771A>T, 2775C>G, 2777G>A, 2780C>T, 2783C>G, 2784A>C, 2786G>C, 2792T>C, 2798G>T, 2801G>T, 2804A>G, 2808C>A, 2812G>C, 2822G>C, 2825C>T, 2828T>C, 2834C>T, 2837C>G, 2840C>T, 2843G>T, 2858A>T, 2859A>C, 2861G>C, 2864C>T, 2867A>G, 2870T>A, 2876A>C, 2879C>T, 2880C>A, 2881A>G, 2882C>A, 2885T>C, 2886G>A, 2887C>T, 2888T>G, 2889C>A, 2891A>G, 2897T>C, 2903A>T, 2907A>T, 2909A>C, 2915C>T, 2920C>A, 2921C>G, 2922A>C, 2927T>C, 2928C>A, 2929G>T, 2930A>T, 2932C>G, 2936A>T, 2939T>G, 2942A>G, 2944A>T, 2945C>T, 2948C>T, 2953G>C, 2954A>C, 2955G>C, 2964A>T, 2966T>G, 2969T>A, 2975C>T, 2981C>T, 2982A>G, 2984C>G, 2988C>T, 2994C>A, 2996C>G, 2999T>A, 3002T>G, 3017A>C, 3020G>A, 3021C>T, 3023T>A, 3026C>T, 3029C>A, 3030C>A, 3032T>A, 3038C>T, 3044A>T, 3045A>C, 3047A>C, 3050G>A, 3056G>T, 3062C>T, 3064T>A, 3066A>T, 3068A>C, 3078C>A, 3079A>T, 3080A>G, 3083A>C, 3090T>G, 3091G>T, 3092T>G, 3095A>G, 3101T>A, 3104G>T, 3107T>A, 3109C>A, 3110A>G, 3116A>C, 3117C>A, 3118G>A, 3119T>G, 3120C>A, 3121T>A, 3122T>A, 3123G>T, 3124G>C, 3131A>G, 3135T>G, 3137T>G, 3140C>T, 3143G>A, 3144A>T, 3145T>A, 3146G>T, 3167A>C, 3173A>C, 3175T>A, 3176T>A, 3182G>C, 3183G>A, 3184A>C, 3190A>G, 3192C>T, 3194G>A, 3195A>G, 3197C>T, 3200C>G, 3201T>A, 3202C>A, 3203C>T, 3204A>G, 3205G>C, 3206T>A, 3210A>G, 3211A>C, 3212G>T, 3213A>C, 3216T>A, 3221T>A, 3225C>G, 3226A>T, 3227A>T, 3228T>G, 3230T>A, 3236A>G, 3245A>T, 3249G>A, 3251G>T, 3254A>G, 3260T>A, 3263C>T, 3269C>T, 3272G>T, 3273T>C, 3275G>T, 3278A>C, 3279C>T, 3281G>A, 3284A>G, 3289G>C, 3293T>G, 3299T>C, 3305A>G, 3308T>G, 3309G>T, 3310T>G, 3311G>C, 3320C>A, 3323C>A, 3326C>T, 3335A>T, 3343G>A, 3344C>T, 3345G>A, 3346T>C, 3347A>C, 3350A>T, 3353T>A, 3356A>G, 3357C>A, 3358G>A, 3359T>G, 3362A>T, 3364T>A, 3365T>C, 3366A>G, 3368C>G, 3369T>G, 3377T>C, 3380T>A, 3386T>A, 3387A>G, 3389C>G, 3391A>T, 3392C>T, 3395G>A, 3398A>T, 3402C>T, 3404A>G, 3407G>A, 3416A>C, 3419T>G, 3422G>A, 3425G>C, 3426G>A, 3428C>T, 3433C>T, 3434A>G, 3437A>T, 3452T>A, 3453A>C, 3455T>G, 3456C>T, 3462T>C, 3463C>A, 3464A>C, 3465A>C, 3467T>G, 3470A>G, 3471A>T, 3472G>C, 3475G>A, 3476A>G, 3479A>G, 3480A>G, 3481C>A, 3483C>A, 3484A>G, 3485G>A, 3486C>T, 3488T>C, 3497A>G, 3500C>A, 3503A>G, 3507T>G, 3508A>C, 3509C>T, 3511A>T, 3513C>T, 3515T>G, 3518A>C, 3521A>T, 3524T>A, 3527A>G, 3528C>T, 3529A>T, 3533G>A, 3534T>A, 3539C>T, 3542C>T, 3548T>G, 3551C>A, 3557G>A, 3560C>A, 3563T>C, 3566G>T, 3567C>A, 3568G>A, 3569G>A, 3570A>C, 3571A>G, 3572A>C, 3575C>T, 3577A>C, 3578A>C, 3579T>C, 3587A>G, 3590A>T, 3593C>T, 3596A>G, 3597C>G, 3598C>A, 3602A>T, 3603G>C, 3608A>T, 3611A>C, 3620T>C, 3624A>T, 3626A>T, 3628T>C, 3629T>C, 3630C>A, 3632T>A, 3641A>G, 3644T>A, 3647T>A, 3650T>G, 3656C>T, 3671G>T, 3674A>C, 3675A>G, 3676T>A, 3680A>G, 3683C>T, 3697G>C, 3698T>C, 3699G>A, 3700G>A, 3701A>C, 3707T>G, 3710T>G, 3711G>C, 3712A>C, 3713C>A, 3714A>G, 3719C>A, 3722A>G, 3725A>G, 3728A>C, 3731C>T, 3734A>G, 3735A>G, 3736A>G, 3740C>T, 3743G>A, 3744T>G, 3745G>C, 3746C>A, 3749G>C, 3750C>A, 3752G>T, 3755A>T, 3759A>G, 3761C>G, 3762A>C, 3764G>T, 3765C>A, 3766G>A, 3767T>G, 3769G>T, 3770C>G, 3774A>C, 3775A>G, 3776G>A, 3777A>G, 3779C>G, 3782T>C, 3788A>G, 3789C>A, 3790G>A, 3791T>A, 3794A>T, 3795T>G, 3796G>C, 3797T>A, 3798G>A, 3799A>T, 3800A>T, 3803G>A, 3804G>A, 3827A>G, 3838G>T, 3839T>C, 3845C>T, 3847A>T, 3853G>C, 3854C>A, 3860G>A, 3861C>T, 3869A>G, 3872T>A, 3875C>T, 3878A>T, 3884C>T, 3887C>T, 3894A>T, 3899G>A, 3900A>C, 3902G>T, 3905G>T, 3906A>T, 3907T>C, 3908G>T, 3914G>A, 3917A>G, 3927C>G, 3930T>G, 3933A>C, 3934C>A, 3935T>A, 3936G>C, 3937G>C, 3938C>A, 3941A>C, 3942G>T, 3943G>C, 3944A>G, 3945G>C, 3950A>G, 3953G>T, 3954G>C, 3955A>T, 3956A>T, 3965A>G, 3966G>A, 3971T>A, 3977A>C, 3983A>T, 3996C>T, 3998A>G, 4001T>C, 4002G>T, 4003T>C, 4008A>G, 4010T>A, 4014A>C, 4019C>A, 4021T>C, 4022C>A, 4023C>T, 4024A>T, 4028G>T, 4031T>C, 4034G>T, 4040C>A, 4046A>G, 4049C>T, 4052C>G, 4064C>G, 4065C>T, 4067C>G, 4079A>G, 4088C>G, 4094A>G, 4101A>C, 4103G>C, 4106C>G, 4109T>C, 4112G>A, 4115A>C, 4116C>T, 4118T>G, 4130C>T, 4137T>C, 4142C>G, 4143A>C, 4148G>A, 4151T>C, 4154T>C, 4160T>C, 4163C>T, 4166A>T, 4169C>T, 4170A>T, 4171C>G, 4172A>T, 4182T>G, 4183T>C, 4184G>A, 4190T>C, 4192C>T, 4193T>A, 4194A>C, 4196A>C, 4199A>T, 4205A>G, 4208T>A, 4214T>C, 4215G>A, 4220T>A, 4223A>G, 4226T>C, 4227T>C, 4228C>A, 4239T>G, 4242T>C, 4247A>T, 4249A>G, 4251G>A, 4253G>C, 4259A>G, 4260G>T, 4262T>A, 4265G>A, 4268A>T, 4271C>T, 4273T>A, 4280T>C, 4286T>C, 4292T>C, 4295C>G, 4316T>C, 4317G>A, 4319T>C, 4325T>A, 4331G>A, 4344T>C, 4346G>A, 4354A>C, 4355A>T, 4357A>T, 4358C>T, 4361C>T, 4364A>T, 4365T>G, 4366C>A, 4367A>T, 4368A>G, 4376C>G, 4385T>G, 4388C>A, 4395C>A, 4397A>G, 4400T>C, 4401G>A, 4404C>T, 4406T>A, 4418A>C, 4424T>C, 4430C>T, 4433A>C, 4436G>C, 4439C>T, 4443G>A, 4445T>C, 4448A>G, 4449G>T, 4455A>C, 4457G>C, 4458T>C, 4460G>A, 4463T>G, 4466G>T, 4469C>A, 4475A>C, 4478A>T, 4481A>T, 4487A>T, 4488C>T, 4490A>G, 4493G>T, 4499C>A, 4502C>T, 4508T>A, 4509C>T, 4511C>G, 4514C>A, 4530C>A, 4532A>G, 4535T>C, 4538C>G, 4548T>C, 4553A>G, 4556A>G, 4562C>A, 4566A>G, 4568T>C, 4569G>A, 4570C>A, 4571T>A, 4576G>A, 4577G>A, 4578G>T, 4581A>G, 4583A>G, 4586A>C, 4587G>A, 4592A>G, 4597G>A, 4601T>C, 4611C>T, 4613A>G, 4622T>A, 4625T>A, 4631A>T, 4637C>T, 4640T>C, 4641C>T, 4643T>G, 4649T>G, 4652A>T, 4656T>A, 4658A>G, 4662C>T, 4664C>G, 4665C>A, 4666A>G, 4673T>C, 4674T>G, 4675C>A, 4679G>A, 4683A>T, 4684A>C, 4685T>C, 4688A>T, 4692C>A, 4693A>G, 4694T>A, 4697G>A, 4698T>C, 4700A>G, 4704C>G, 4705T>C, 4706G>T, 4709C>T, 4712T>C, 4715A>C, 4716A>G, 4731C>T, 4733T>G, 4739A>G, 4742G>A, 4751T>A, 4752A>G, 4754T>G, 4755G>A, 4758G>A, 4759T>C, 4761C>A, 4768A>T, 4769A>G, 4784A>C, 4787T>C, 4793G>C, 4799A>G, 4805T>C, 4808A>G, 4817T>A, 4818C>T, 4820T>G, 4826C>T, 4829C>A, 4835T>A, 4838T>A, 4844G>A, 4854A>G, 4868G>T, 4871T>C, 4877A>T, 4880A>G, 4883T>A, 4886C>A, 4887C>G, 4888C>T, 4889A>G, 4892T>A, 4901C>T, 4902A>G, 4903C>T, 4904T>A, 4907G>A, 4910C>T, 4914A>T, 4915T>A, 4916G>T, 4923G>A, 4925G>T, 4934A>C, 4937A>C, 4938C>A, 4940T>A, 4946T>C, 4955C>T, 4958T>G, 4960G>A, 4961T>G, 4963T>C, 4964G>C, 4965G>A, 4973C>A, 4975A>G, 4976G>T, 4977C>T, 4982C>A, 4983T>A, 4984T>A, 4985A>T, 4993A>T, 4994G>T, 4995A>G, 4996C>G, 4997G>A, 5000G>T, 5001T>C, 5003G>C, 5004G>A, 5006T>G, 5007G>C, 5009A>G, 5012G>A, 5013G>A, 5014G>A, 5015T>C, 5016T>C, 5019A>C, 5020G>C, 5021A>C, 5022G>A, 5023C>T, 5024T>C, 5025G>A, 5031T>A, 5033T>G, 5036T>C, 5037T>A, 5039C>T, 5042T>C, 5044T>A, 5045G>T, 5049G>C, 5051G>T, 5055C>A, 5056A>G, 5057G>T, 5058G>C, 5064A>T, 5066T>G, 5067G>A, 5070A>G, 5075A>C, 5081G>A, 5085T>G, 5086C>A, 5087A>G, 5091G>A, 5092C>T, 5093C>T, 5094C>G, 5095T>G, 5097A>G, 5098C>T, 5099A>T, 5100A>T, 5102T>G, 5103T>A, 5104G>T, 5105T>G, 5108A>G, 5109A>C, 5111G>A, 5112G>T, 5115A>G, 5117A>T, 5120A>T, 5121G>A, 5122C>A, 5123T>A, 5125T>A, 5127T>A, 5129C>A, 5132T>C, 5135G>C, 5138A>T, 5141G>A, 5142G>C, 5143A>C, 5144C>T, 5147A>T, 5150T>G, 5153G>A, 5159C>T, 5160_5162delACA, 5172C>G, 5173G>A, 5174T>G, 5176A>G, 5178C>G, 5180G>C, 5181A>C, 5184T>G, 5185G>A, 5186G>A, 5192A>G, 5195A>T, 5196A>G, 5198C>T, 5199A>G, 5201T>A, 5202G>T, 5203G>T, 5207T>C, 5211T>A, 5212C>G, 5213A>T, 5216A>T, 5219T>G, 5226G>T, 5228C>A, 5229C>A, 5232T>C, 5234C>A, 5235A>C, 5236C>A, 5237G>A, 5241G>T, 5242A>C, 5243T>A, 5244G>T, 5245A>C, 5249G>T, 5250T>A, 5255T>C, 5256A>C, 5258T>G, 5264C>T, 5267T>C, 5270A>T, 5273T>A, 5276A>C, 5277A>C, 5278A>G, 5279G>C, 5281A>G, 5282G>A, 5283T>A, 5285C>A, 5287A>C, 5288T>A, 5294T>G, 5300T>G, 5303A>T, 5309C>T, 5318C>T, 5333G>A, 5339G>C, 5346A>T, 5349T>G, 5354T>A, 5357A>C, 5358A>C, 5359T>A, 5364G>C, 5365C>A, 5366C>T, 5370A>C, 5371A>G, 5372G>C, 5375T>G, 5376G>A, 5377T>G, 5378G>T, 5379T>C, 5383C>A, 5386G>T, 5387A>T, 5390A>T, 5391C>G, 5395T>G, 5396A>G, 5399A>T, 5403A>T, 5404A>C, 5409G>A, 5410T>C, 5411T>C, 5417G>T, 5423A>C, 5429A>T, 5435T>A, 5436G>A, 5438C>A, 5440G>C, 5441C>T, 5447T>C, 5456G>A, 5457C>T, 5460A>T, 5462G>T, 5465C>A, 5468A>G, 5471A>C, 5474A>T, 5475A>C, 5476G>A, 5477G>T, 5483T>C, 5484T>A, 5486T>G, 5488T>C, 5489G>C, 5490G>A, 5491A>C, 5492G>A, 5496A>C, 5498A>T, 5501A>T, 5504T>G, 5505A>C, 5507A>C, 5508T>C, 5510G>A, 5513C>A, 5516A>C, 5519T>G, 5521G>C, 5522A>T, 5523A>G, 5524A>G, 5525T>G, 5531T>G, 5534C>T, 5537C>G, 5540T>C, 5546T>C, 5549T>C, 5555C>T, 5560T>C, 5564G>A, 5567T>C, 5570T>G, 5573T>C, 5574C>A, 5576A>G

#### CDS

| | Begin | End | Coverage | Score | Concordance | Matches | Identities | I/D/M/F* | Stop Codons |
|---|---|---|---|---|---|---|---|---|---|
| **NT** | **23** | **5589** | **77.0%** | **3552** | **32.4%** | **5524 (99.1%)** | **3681 (66.0%)** | **9/43** | |
| ToMarV_RNA1gp1 | 1 | 1817 | 84.4% | 9297 | 73.5% | 1807 (99.3%) | 1266 (69.6%) | 3/10/0/0 | 0 |
| Protein mutations: | F7W (160T>G 161C>G), K9_A16del (165_188delAAGGTTGCTGGTGTTGATGACGCC), S17P (189A>C 190G>C), S18P (192T>C), S19A (195T>G), T21S (201A>T 203C>A), T22P (204A>C), G26A (217G>C 218T>A), L28V (222C>G), N30S (228A>T 229A>C), I32F (234A>T), K34S (240A>T 241A>C 242A>T), F35L (243T>C), A38T (252G>A 254G>T), V39L (255G>C 257T>C), I42V (264A>G), T43N (268C>A 269C>T), T44K (271C>A 272T>A), M45V (273A>G 275G>T), K48T (283A>C 284G>C), S50V (288T>G 289C>T), L53I (297C>A 299T>A), L56V (306C>G 308G>C), S59P (315T>C 317T>A), T61M (322C>T 323T>G), D62A (325A>C 326T>A), V63A (328T>C 329G>A), S64A (330T>G 332C>A), V68A (343T>C 344G>C), A69S (345G>T 347C>T), S72N (355G>A), S73A (357T>G 359A>T), L74V (360C>G 362T>A), N75E (363A>G 365T>A), K76R (367A>G), G82A (385G>C 386T>A), A83K (387G>A 388C>A), L84I (390C>A 392T>A), I85M (395A>G), D86E (398T>A), L89I (405C>A 407T>C), A91G (412C>G 413G>T), S93D (417T>G 418C>A), L95V (423T>G 425G>T), S96K (426T>A 427C>A 428C>A), S101T (441T>A), K103V (447A>G 448A>T 449G>A), M105L (453A>C 455G>T), L109M (465T>A), F110V (468T>G), E111L (471G>T 472A>T 473G>A), L113V (477T>G 479G>C), S114K (480T>A 481C>A 482C>G), N115D (483A>G 485T>C), V117F (489G>T 491T>C), K118I (493A>T 494A>C), F121Y (502T>A 503T>C), A122E (505C>A 506T>A), V124L (510G>T 512G>A), N126E (516A>G 518T>A), S128V (522A>G 523G>T), D130S (528G>T 529A>C 530T>A), V131L (531G>T), V135F (543G>T 545A>T), Y136I (546T>A 547A>T 548T>A), G137A (550G>C), I143L (567A>T 569A>G), T144A (570A>G 572A>C), L145I (573T>A 575G>T), V147L (579G>C 581G>A), F150Y (589T>A), S154V (600A>G 601G>T 602T>A), E158T (612G>A 613A>C), G159P (615G>C 616G>C), V160F (618G>T), S161N (621T>A 622C>A), T163V (627A>G 628C>T 629T>A), K165V (633A>G 634A>T 635A>T), N166T (637A>C), I167L (639A>T 641C>G), L169S (646T>C 647G>A), I170V (648A>G 650A>T), M174F (660A>T 662G>T), K186T (697A>C), A188S (702G>T 704C>T), E189G (706A>G 707A>G), R190K (708C>A 709G>A 710A>G), T192S (714A>T), M193Q (717A>C 718T>A), V194L (720G>C), S198A (732A>G 733G>C 734T>A), M199V (735A>G 737G>A), V205L (753G>T 755T>A), S206T (756T>A 758A>G), M214L (780A>T 782G>A), T216S (786A>T), I217L (789A>T 791A>G), I228V (822A>G), T248A (882A>G 884A>T), T260Q (918A>C 919C>A 920T>G), I280V (978A>G 980A>G), G282S (984G>A), T283I (988C>T 989C>T), S295N (1024G>A), E304Q (1050G>C), Q307T (1059C>A 1060A>C), D310S (1068G>T 1069A>C 1070T>A), A311E (1072C>A 1073A>G), C313A (1077T>G 1078G>C), D314E (1082T>A), A316S (1086G>T 1088A>T), T317M (1090C>T), K325P (1113A>C 1114A>C 1115G>T), T329I (1126C>T 1127C>T), N332Q (1134A>C 1136C>A), R335K (1144G>A 1145G>A), E337Q (1149G>C 1151A>G), L339V (1155C>G), M354L (1200A>T), A363T (1227G>A), E366D (1238A>T), V368L (1242G>T 1244T>G), Y369V (1245T>G 1246A>T), S372A (1254A>G 1255G>C 1256T>C), C373S (1257T>A 1259T>C), R374Q (1260A>C 1261G>A 1262G>A), M376S (1266A>T 1267T>C 1268G>T), S393A (1317T>G 1319C>T), S401C (1342C>G), R412A (1374A>G 1375G>C), I440V (1458A>G 1460C>G), A500T (1638G>A 1640C>T), M521R (1702T>G 1703G>A), E523V (1708A>T), E525D (1715G>T), S527D (1719T>G 1720C>A 1721C>T), T528P (1722A>C 1724T>A), E529N (1725G>A 1727A>T), D530A (1729A>C), G533A (1738G>C 1739A>C), R536V (1746A>G 1747G>T 1748A>T), N538D (1752A>G 1754C>T), I539V (1755A>G 1757A>G), N552A (1794A>G 1795A>C), T554S (1800A>T), S556D (1806A>G 1807G>A), E560Q (1818G>C 1820G>A), Q567R (1840A>G), A570Y (1848G>T 1849C>A 1850A>T), K574S (1861A>G 1862G>C), F577Y (1870T>A), E578N (1872G>A 1874A>T), A579T (1875G>A), E580T (1878G>A 1879A>C), I583L (1887A>T 1889A>G), S584A (1890A>G 1891G>C), P585D (1893C>G 1894C>A 1895T>C), I587V (1899A>G 1901A>T), Q591N (1911C>A 1913A>T), S593A (1917T>G), A594V (1921C>T 1922A>T), D596E (1928T>A), I598L (1932A>C 1934A>G), A599D (1936C>A 1937G>T), V601F (1941G>T 1943G>C), D602Q (1944G>C 1946T>A), G603D (1948G>A 1949G>T), R605del (1953_1955delAGA), R607Q (1960G>A 1961C>A), T608V (1962A>G 1963C>T 1964C>T), K609L (1965A>T 1966A>T), Q610P (1969A>C 1970A>T), D611P (1971G>C 1972A>C), I614A (1980A>G 1981T>C), L615T (1983T>A 1984T>C 1985G>T), V616Y (1986G>T 1987T>A 1988G>T), P617E (1989C>G 1990C>A 1991C>G), T618Q (1992A>C 1993C>A 1994T>G), V619T (1995G>A 1996T>C 1997T>A), V620I (1998G>A 2000G>T), S621T (2002G>C 2003T>A), N623A (2007A>G 2008A>C), D624E (2012T>A), R625K (2014G>A), V626I (2016G>A 2018G>T), I627V (2019A>G 2021C>T), A629S (2025G>T 2027A>C), R630Q (2028A>C 2029G>A 2030A>G), L645M (2073T>A), D646N (2076G>A), P647K (2079C>A 2080C>A 2081T>A), E648F (2082G>T 2083A>T 2084G>T), L650F (2088C>T), H655L (2104A>T 2105T>C), H656S (2106C>T 2107A>C 2108T>C), M660I (2120G>C), S663V (2127T>G 2128C>T 2129T>G), D665S (2133G>T 2134A>C), D666S (2136G>T 2137A>C), E667D (2141A>T), H668A (2142C>G 2143A>C 2144T>C), E670Q (2148G>C), I671N (2152T>A), N673E (2157A>G 2159C>A), K674M (2161A>T 2162A>G), R676V (2166A>G 2167G>T), V677N (2169G>A 2170T>A 2171C>T), N678D (2172A>G 2174C>T), M679L (2175A>T), Y681V (2181T>G 2182A>T 2183T>C), R682K (2185G>A), D685S (2193G>A 2194A>G), A686P (2196G>C), E687Q (2199G>C 2201A>G), R690A (2208C>G 2209G>C 2210A>T), I701V (2241A>G 2243A>C), A710S (2268G>A 2269C>G 2270A>C), S711T (2271T>A), K716E (2286A>G), K717L (2289A>T 2290A>T 2291A>G), A719D (2296C>A 2297A>T), T722R (2304A>C 2305C>G 2306A>C), E723N (2307G>A 2309G>T), M724L (2310A>C 2312G>T), R725E (2313A>G 2314G>A), T726Q (2316A>C 2317C>A 2318T>A), F728K (2322T>A 2323T>A 2324T>A), H730Y (2328C>T), S731K (2331T>A 2332C>A 2333T>G), C732A (2334T>G 2335G>C 2336C>G), I733L (2337A>C), D734Q (2340G>C 2342T>G), V737A (2350T>C 2351T>A), I741T (2362T>C 2363A>G), F749R (2385T>A 2386T>G 2387T>A), N763R (2428A>G 2429C>G), S766A (2436T>G), S766_K767insG (2438_2439insGGT), K767T (2440A>C 2441A>T), E768D (2444G>C), P769K (2445C>A 2446C>A 2447A>G), E771R (2451G>A 2452A>G), S772N (2454T>A 2455C>A 2456A>T), K775I (2464A>T 2465A>C), I776E (2466A>G 2467T>A 2468T>A), Y778M (2472T>A 2473A>T 2474C>G), H779I (2475C>A 2476A>T), K780G (2478A>G 2479A>G 2480A>T), V782K (2484G>A 2485T>A 2486T>G), E783D (2489A>C), Y784K (2490T>A 2492T>A), I785M (2495C>G), K786I (2497A>T 2498G>C), D787Q (2499G>C 2501T>G), T788M (2503C>T 2504A>G), W790N (2508T>A 2509G>A 2510G>T), S791D (2511A>G 2512G>A), Y794F (2521A>T 2522T>C), A795I (2523G>A 2524C>T 2525A>C), R797K (2530G>A 2531A>G), I806L (2556A>T 2558A>G), M807I (2561G>A), V812T (2574G>A 2575T>C 2576T>C), F813M (2577T>A 2579T>G), G814S (2580G>A 2582A>T), G815A (2584G>C), I816V (2586A>G), L820I (2598T>A 2600G>T), A821T (2601G>A 2603G>T), W823L (2607T>C 2608G>T 2609G>C), Q824K (2610C>A), S825A (2613A>G 2614G>C 2615T>A), N826A (2616A>G 2617A>C 2618C>A), S827T (2619T>A), F828S (2623T>C), N830L (2628A>T 2629A>T 2630T>A), S832D (2634T>G 2635C>A 2636G>T), N852K (2696T>G), K853T (2698A>C 2699G>T), K853_L854insH (2699_2700insCAT), L854F (2700C>T), R855A (2703A>G 2704G>C), N856K (2708T>A), L857S (2709C>T 2710T>C 2711T>C), V859L (2715G>C 2717T>A), S862A (2724A>G 2725G>C 2726T>G), S863A (2727T>G 2729T>A), Q864D (2730C>G 2732A>C), S865N (2733T>A 2734C>A 2735A>C), L866N (2736T>A 2737T>A 2738G>C), A867H (2739G>C 2740C>A 2741A>T), Q868V (2742C>G 2743A>T 2744A>G), D869_W870insR (2747_2748insCGA), A872E (2755C>A 2756T>A), E873Q (2757G>C 2759A>G), Q879E (2775C>G 2777G>A), K882H (2784A>C 2786G>C), L890I (2808C>A), G891A (2812G>C), M907L (2859A>C 2861G>C), H914R (2880C>A 2881A>G 2882C>A), A916M (2886G>A 2887C>T 2888T>G), T923S (2907A>T 2909A>C), A927E (2920C>A 2921C>G), N928H (2922A>C), R930I (2928C>A 2929G>T 2930A>T), S931C (2932C>G), K932N (2936A>T), Y935F (2944A>T 2945C>T), R938T (2953G>C 2954A>C), E939Q (2955G>C), I942L (2964A>T 2966T>G), I948V (2982A>G 2984C>G), P961S (3021C>T 3023T>A), E972D (3056G>T), F975Y (3064T>A), T976S (3066A>T 3068A>C), Q980M (3078C>A 3079A>T 3080A>G), C984V (3090T>G 3091G>T 3092T>G), T990K (3109C>A 3110A>G), R993K (3117C>A 3118G>A 3119T>G), L994K (3120C>A 3121T>A 3122T>A), G995S (3123G>T 3124G>C), F999V (3135T>G 3137T>G), M1002Y (3144A>T 3145T>A 3146G>T), V1012E (3175T>A 3176T>A), E1015T (3183G>A 3184A>C), E1017G (3190A>G), N1019D (3195A>G 3197C>T), S1021N (3201T>A 3202C>A 3203C>T), S1022A (3204A>G 3205G>C 3206T>A), K1024A (3210A>G 3211A>C 3212G>T), I1025L (3213A>C), C1026S (3216T>A), Q1029V (3225C>G 3226A>T 3227A>T), S1030A (3228T>G 3230T>A), G1037S (3249G>A 3251G>T), S1050T (3289G>C), V1057C (3309G>T 3310T>G 3311G>C), G1068D (3343G>A 3344C>T), V1069T (3345G>A 3346T>C 3347A>C), R1073K (3357C>A 3358G>A 3359T>G), F1075Y (3364T>A 3365T>C), I1076V (3366A>G 3368C>G), S1077A (3369T>G), I1083V (3387A>G 3389C>G), Y1084F (3391A>T 3392C>T), E1086D (3398A>T), V1096I (3426G>A 3428C>T), A1098V (3433C>T 3434A>G), D1104E (3452T>A), I1105L (3453A>C 3455T>G), S1108H (3462T>C 3463C>A 3464A>C), I1109L (3465A>C 3467T>G), R1112K (3475G>A 3476A>G), T1114E (3480A>G 3481C>A), Q1115R (3483C>A 3484A>G 3485G>A), L1116F (3486C>T 3488T>C), N1120K (3500C>A), Y1123A (3507T>G 3508A>C 3509C>T), Y1124F (3511A>T), Q1130L (3528C>T 3529A>T), L1132M (3534T>A), R1143K (3567C>A 3568G>A 3569G>A), K1144R (3570A>C 3571A>G 3572A>C), Q1146P (3577A>C 3578A>C), P1153D (3597C>G 3598C>A), E1155Q (3603G>C), T1162S (3624A>T 3626A>T), V1163A (3628T>C 3629T>C), H1164K (3630C>A 3632T>A), D1169E (3647T>A), M1179E (3675A>G 3676T>A), C1186S (3697G>C 3698T>C), G1187N (3699G>A 3700G>A 3701A>C), D1189E (3707T>G), D1190E (3710T>G), D1191P (3711G>C 3712A>C 3713C>A), I1192V (3714A>G), N1199G (3735A>G 3736A>G), C1202A (3744T>G 3745G>C 3746C>A), L1204I (3750C>A 3752G>T), I1207V (3759A>G 3761C>G), M1208L (3762A>C 3764G>T), R1209K (3765C>A 3766G>A 3767T>G), S1210M (3769G>T 3770C>G), K1212R (3774A>C 3775A>G 3776G>A), I1213V (3777A>G 3779C>G), R1217K (3789C>A 3790G>A 3791T>A), C1219A (3795T>G 3796G>C 3797T>A), E1220I (3798G>A 3799A>T 3800A>T), D1222N (3804G>A), C1233F (3838G>T 3839T>C), Y1236F (3847A>T), G1238A (3853G>C 3854C>A), M1252L (3894A>T), M1254L (3900A>C 3902G>T), M1256S (3906A>T 3907T>C 3908G>T), Q1263E (3927C>G), S1264A (3930T>G), T1265Q (3933A>C 3934C>A 3935T>A), G1266P (3936G>C 3937G>C 3938C>A), E1267D (3941A>C), G1268S (3942G>T 3943G>C 3944A>G), E1269Q (3945G>C), E1272L (3954G>C 3955A>T 3956A>T), D1276N (3966G>A), E1281D (3983A>T), V1288S (4002G>T 4003T>C), I1290V (4008A>G 4010T>A), K1292Q (4014A>C), I1294T (4021T>C 4022C>A), H1295F (4023C>T 4024A>T), T1344C (4170A>T 4171C>G 4172A>T), L1348A (4182T>G 4183T>C 4184G>A), A1351V (4192C>T 4193T>A), I1352L (4194A>C 4196A>C), D1359N (4215G>A), S1363H (4227T>C 4228C>A), L1367V (4239T>G), N1370S (4249A>G), G1371S (4251G>A 4253G>C), A1374S (4260G>T 4262T>A), V1378E (4273T>A), V1393I (4317G>A 4319T>C), E1405A (4354A>C 4355A>T), Y1406F (4357A>T 4358C>T), S1409D (4365T>G 4366C>A 4367A>T), K1410E (4368A>G), D1415E (4385T>G), A1421T (4401G>A), D1435N (4443G>A 4445T>C), V1437L (4449G>T), I1476V (4566A>G 4568T>C), A1477K (4569G>A 4570C>A 4571T>A), R1479K (4576G>A 4577G>A), A1480S (4578G>T), I1481V (4581A>G 4583A>G), V1483I (4587G>A), S1486N (4597G>A), L1506M (4656T>A 4658A>G), Q1509R (4665C>A 4666A>G), S1512D (4674T>G 4675C>A), N1515S (4683A>T 4684A>C 4685T>C), H1518R (4692C>A 4693A>G 4694T>A), L1522A (4704C>G 4705T>C 4706G>T), I1526V (4716A>G), N1538E (4752A>G 4754T>G), E1539K (4755G>A), V1540T (4758G>A 4759T>C), Q1541K (4761C>A), Q1543L (4768A>T 4769A>G), H1563Q (4829C>A), I1572V (4854A>G), M1576I (4868G>T), P1583V (4887C>G 4888C>T 4889A>G), T1588V (4902A>G 4903C>T 4904T>A), M1592Y (4914A>T 4915T>A 4916G>T), V1595I (4923G>A 4925G>T), R1607Q (4960G>A 4961T>G), V1608A (4963T>C 4964G>C), A1609T (4965G>A), D1611E (4973C>A), K1612S (4975A>G 4976G>T), L1613F (4977C>T), L1615N (4983T>A 4984T>A 4985A>T), Q1618L (4993A>T 4994G>T), T1619G (4995A>G 4996C>G 4997G>A), D1622K (5004G>A 5006T>G), E1623Q (5007G>C 5009A>G), G1625N (5013G>A 5014G>A 5015T>C), F1626L (5016T>C), R1627P (5019A>C 5020G>C 5021A>C), A1628I (5022G>A 5023C>T 5024T>C), V1629M (5025G>A), F1631M (5031T>A 5033T>G), S1633T (5037T>A 5039C>T), L1635Y (5044T>A 5045G>T), V1637L (5049G>C 5051G>T), Q1639S (5055C>A 5056A>G 5057G>T), E1640Q (5058G>C), I1642L (5064A>T 5066T>G), A1643T (5067G>A), K1644E (5070A>G), S1649E (5085T>G 5086C>A 5087A>G), A1651I (5091G>A 5092C>T 5093C>T), L1652G (5094C>G 5095T>G), T1653V (5097A>G 5098C>T 5099A>T), I1654L (5100A>T 5102T>G), C1655M (5103T>A 5104G>T 5105T>G), M1657L (5109A>C 5111G>A), A1658S (5112G>T), K1659D (5115A>G 5117A>T), A1661K (5121G>A 5122C>A 5123T>A), I1662N (5125T>A), S1663T (5127T>A 5129C>A), D1668P (5142G>C 5143A>C 5144C>T), T1674del (5160_5162delACA), R1678E (5172C>G 5173G>A 5174T>G), D1679G (5176A>G), L1680V (5178C>G 5180G>C), N1681H (5181A>C), W1682E (5184T>G 5185G>A 5186G>A), N1686D (5196A>G 5198C>T), I1687V (5199A>G 5201T>A), G1688F (5202G>T 5203G>T), A1696S (5226G>T 5228C>A), H1697N (5229C>A), Y1698Q (5232T>C 5234C>A), T1699Q (5235A>C 5236C>A 5237G>A), D1701S (5241G>T 5242A>C 5243T>A), E1702S (5244G>T 5245A>C), Y1704N (5250T>A), I1706L (5256A>C 5258T>G), K1713R (5277A>C 5278A>G 5279G>C), K1714R (5281A>G 5282G>A), S1715T (5283T>A 5285C>A), N1716T (5287A>C 5288T>A), M1731I (5333G>A), M1736L (5346A>T), S1737A (5349T>G), M1740Q (5358A>C 5359T>A), A1742H (5364G>C 5365C>A 5366C>T), K1744R (5370A>C 5371A>G 5372G>C), N1745K (5375T>G), V1746S (5376G>A 5377T>G 5378G>T), A1748E (5383C>A), G1749V (5386G>T 5387A>T), H1751D (5391C>G), V1752G (5395T>G 5396A>G), N1755S (5403A>T 5404A>C), V1757T (5409G>A 5410T>C 5411T>C), V1766I (5436G>A 5438C>A), G1767A (5440G>C 5441C>T), M1774F (5460A>T 5462G>T), R1779H (5475A>C 5476G>A 5477G>T), C1782R (5484T>A 5486T>G), V1783A (5488T>C 5489G>C), E1784T (5490G>A 5491A>C 5492G>A), I1786L (5496A>C 5498A>T), I1793M (5519T>G), G1794A (5521G>C 5522A>T), N1795G (5523A>G 5524A>G 5525T>G), V1807A (5560T>C), F1810L (5570T>G), Q1812K (5574C>A 5576A>G) | | | | | | | | |

| | Begin | End | Coverage | Score | Concordance | Matches | Identities | I/D/M/F* | Stop Codons |
|---|---|---|---|---|---|---|---|---|---|
| **NT** | **23** | **5589** | **77.0%** | **3552** | **32.4%** | **5524 (99.1%)** | **3681 (66.0%)** | **9/43** | |

Codon mutations:

GCT2GCA (146T>A), TTC7TGG (160T>G 161C>G), TCC8AGT (162T>A 163C>G 164C>T), AAG9_GCC16del (165_188delAAGGTTGCTGGTGTTGATGACGCC), AGT17CCT (189A>C 190G>C), TCT18CCT (192T>C), TCT19GCT (195T>G), TCC20TCT (200C>T), ACC21TCA (201A>T 203C>A), ACT22CCT (204A>C), GGT26GCA (217G>C 218T>A), CTT28GTT (222C>G), TCA29TCT (227A>T), AAT30TCT (228A>T 229A>C), ATT32TTT (234A>T), TCC33TCT (239C>T), AAA34TCT (240A>T 241A>C 242A>T), TTT35CTT (243T>C), AAA37AAG (251A>G), GCG38ACT (252G>A 254G>T), GTT39CTC (255G>C 257T>C), AGT40TCC (258A>T 259G>C 260T>C), ATT42GTT (264A>G), ACC43AAT (268C>A 269C>T), ACT44AAA (271C>A 272T>A), ATG45GTT (273A>G 275G>T), AAG48ACC (283A>C 284G>C), TCA50GTA (288T>G 289C>T), CAT52CAC (296T>C), CTT53ATA (297C>A 299T>A), CTG56GTC (306C>G 308G>C), CCA58CCG (314A>G), TCT59CCA (315T>C 317T>A), GTG60GTT (320G>T), ACT61ATG (322C>T 323T>G), GAT62GCA (325A>C 326T>A), GTG63GCA (328T>C 329G>A), TCC64GCA (330T>G 332C>A), TCT66TCA (338T>A), GTG68GCC (343T>C 344G>C), GCC69TCT (345G>T 347C>T), ACT70ACA (350T>A), TGT71TGC (353T>C), AGT72AAT (355G>A), TCA73GCT (357T>G 359A>T), CTT74GTA (360C>G 362T>A), AAT75GAA (363A>G 365T>A), AAA76AGA (367A>G), TTG78TTA (374G>A), GAC79GAT (377C>T), AAA80AAG (380A>G), GGT82GCA (385G>C 386T>A), GCA83AAA (387G>A 388C>A), CTT84ATA (390C>A 392T>A), ATA85ATG (395A>G), GAT86GAA (398T>A), CCA87CCC (401A>C), TTT88TTC (404T>C), CTT89ATC (405C>A 407T>C), AAA90AAG (410A>G), GCG91GGT (412C>G 413G>T), TAC92TAT (416C>T), TCT93GAT (417T>G 418C>A), TTC94TTT (422C>T), TTG95GTT (423T>G 425G>T), TCC96AAA (426T>A 427C>A 428C>A), ACA97ACT (431A>T), TAC99TAT (437C>T), TCT101ACT (441T>A), ATC102ATA (446C>A), AAG103GTA (447A>G 448A>T 449G>A), ATG105CTT (453A>C 455G>T), GTT106GTA (458T>A), TTA107CTT (459T>C 461A>T), TTG109ATG (465T>A), TTT110GTT (468T>G), GAG111TTA (471G>T 472A>T 473G>A), TTG113GTC (477T>G 479G>C), TCC114AAG (480T>A 481C>A 482C>G), AAT115GAC (483A>G 485T>C), GTT117TTC (489G>T 491T>C), AAA118ATC (493A>T 494A>C), TTA119CTA (495T>C), TTT121TAC (502T>A 503T>C), GCT122GAA (505C>A 506T>A), GTG124TTA (510G>T 512G>A), AAT126GAA (516A>G 518T>A), AGT128GTT (522A>G 523G>T), GAA129GAG (527A>G), GAT130TCA (528G>T 529A>C 530T>A), GTA131TTA (531G>T), GTT134GTG (542T>G), GTA135TTT (543G>T 545A>T), TAT136ATA (546T>A 547A>T 548T>A), GGT137GCT (550G>C), CTT139TTA (555C>T 557T>A), GTA140GTT (560A>T), ATA143TTG (567A>T 569A>G), ACA144GCC (570A>G 572A>C), TTG145ATT (573T>A 575G>T), TTA146TTG (578A>G), GTG147CTA (579G>C 581G>A), CTT148TTA (582C>T 584T>A), CTA149CTG (587A>G), TTT150TAT (589T>A), GTT151GTA (593T>A), TGT152TGC (596T>C), AGT154GTA (600A>G 601G>T 602T>A), GTT157GTA (611T>A), GAG158ACG (612G>A 613A>C), GGT159CCT (615G>C 616G>C), GTT160TTT (618G>T), TCT161AAT (621T>A 622C>A), AAC162AAT (626C>T), ACT163GTA (627A>G 628C>T 629T>A), GTC164GTT (632C>T), AAA165GTT (633A>G 634A>T 635A>T), AAT166ACT (637A>C), ATC167TTG (639A>T 641C>G), TTG169TCA (646T>C 647G>A), ATA170GTT (648A>G 650A>T), GTC171GTT (653C>T), AAC173AAT (659C>T), ATG174TTT (660A>T 662G>T), TTT175TTC (665T>C), TCA176TCT (668A>T), AGT177TCA (669A>T 670G>C 671T>A), CTA178TTG (672C>T 674A>G), TAC179TAT (677C>T), AAA186ACA (697A>C), GCC188TCT (702G>T 704C>T), GAA189GGG (706A>G 707A>G), CGA190AAG (708C>A 709G>A 710A>G), TTC191TTT (713C>T), ACC192TCC (714A>T), ATG193CAG (717A>C 718T>A), GTG194CTG (720G>C), GCC195GCG (725C>G), GCT197GCA (731T>A), AGT198GCA (732A>G 733G>C 734T>A), ATG199GTA (735A>G 737G>A), GGG202GGT (746G>T), TCA204AGT (750T>A 751C>G 752A>T), GTT205TTA (753G>T 755T>A), TCA206ACG (756T>A 758A>G), ACG210ACT (770G>T), TCT211AGC (771T>A 772C>G 773T>C), CAG212CAA (776G>A), TTG213CTT (777T>C 779G>T), ATG214TTA (780A>T 782G>A), ACA216TCA (786A>T), ATA217TTG (789A>T 791A>G), GGG221GGT (803G>T), ATT222ATC (806T>C), TCG223TCA (809G>A), ACG224ACT (812G>T), TTG225TTA (815G>A), GTG226GTA (818G>A), TTC227TTT (821C>T), ATT228GTT (822A>G), GCA229GCT (827A>T), GTG230GTT (830G>T), GGT232GGG (836T>G), GGC236GGT (848C>T), TTG237CTT (849T>C 851G>T), AGC238TCA (852A>T 853G>C 854C>A), TTG241CTG (861T>C), TCC242TCA (866C>A), ATT244ATC (872T>C), CTC245TTG (873C>T 875C>G), TAT246TAC (878T>C), TCG247TCT (881G>T), ACA248GCT (882A>G 884A>T), GCT252GCG (896T>G), CAG253CAA (899G>A), CAA257CAG (911A>G), CTA258CTG (914A>G), ACT260CAG (918A>C 919C>A 920T>G), CTT261TTG (921C>T 923T>G), CGT263AGG (927C>A 929T>G), AAT264AAC (932T>C), AAA266AAG (938A>G), TGC268TGT (944C>T), ACC269ACA (947C>A), TCC270TCT (950C>T), CAG271CAA (953G>A), GCT272GCC (956T>C), GAA278GAG (974A>G), ATA280GTG (978A>G 980A>G), GTC281GTT (983C>T), GGC282AGC (984G>A), ACC283ATT (988C>T 989C>T), GGA285GGC (995A>C), TTC286TTT (998C>T), GTG290GTA (1010G>A), GCT293GCG (1019T>G), AGT295AAT (1024G>A), GCA296GCT (1028A>T), ACG297ACC (1031G>C), CTG298TTA (1032C>T 1034G>A), ACT300ACC (1040T>C), CTG302CTT (1046G>T), GAG304CAG (1050G>C), CAG307ACG (1059C>A 1060A>C), GAG308GAA (1064G>A), GAT310TCA (1068G>T 1069A>C 1070T>A), GCA311GAG (1072C>A 1073A>G), GTG312GTT (1076G>T), TGT313GCT (1077T>G 1078G>C), GAT314GAA (1082T>A), CCT315CCA (1085T>A), GCA316TCT (1086G>T 1088A>T), ACG317ATG (1090C>T), GAG320GAA (1100G>A), AAC321AAT (1103C>T), GCC323GCA (1109C>A), AAG325CCT (1113A>C 1114A>C 1115G>T), GCT326GCA (1118T>A), TTC327TTT (1121C>T), CCT328CCA (1124T>A), ACC329ATT (1126C>T 1127C>T), AAA330AAG (1130A>G), AAC332CAA (1134A>C 1136C>A), CTG334TTA (1140C>T 1142G>A), AGG335AAA (1144G>A 1145G>A), GAA336GAG (1148A>G), GAA337CAG (1149G>C 1151A>G), GCT338GCC (1154T>C), CTC339GTC (1155C>G), AAG340AAA (1160G>A), ATC341ATA (1163C>A), GCA346GCC (1178A>C), GCG351GCA (1193G>A), TTC353TTT (1199C>T), ATG354TTG (1200A>T), AGT359TCT (1215A>T 1216G>C), ATT362ATA (1226T>A), GCG363ACG (1227G>A), CAG364CAA (1232G>A), TTA365CTA (1233T>C), GAA366GAT (1238A>T), GTT368TTG (1242G>T 1244T>G), TAT369GTT (1245T>G 1246A>T), GAA371GAG (1253A>G), AGT372GCC (1254A>G 1255G>C 1256T>C), TGT373AGC (1257T>A 1259T>C), AGG374CAA (1260A>C 1261G>A 1262G>A), CAC375CAT (1265C>T), ATG376TCT (1266A>T 1267T>C 1268G>T), GGC377GGT (1271C>T), CAG380CAA (1280G>A), CGT382AGG (1284C>A 1286T>G), GAA384GAG (1292A>G), TTT386TTC (1298T>C), GTA388GTG (1304A>G), TTC391TTT (1313C>T), GGG392GGA (1316G>A), TCC393GCT (1317T>G 1319C>T), GGG395GGA (1325G>A), TCT399TCA (1337T>A), ACA400ACG (1340A>G), TCC401TGC (1342C>G), TTA404CTT (1350T>C 1352A>T), TTT405TTC (1355T>C), GTG409GTT (1367G>T), TTG410CTC (1368T>C 1370G>C), AGA412GCA (1374A>G 1375G>C), GAG415GAA (1385G>A), AAA417AAG (1391A>G), CTC418CTG (1394C>G), AAT419AAC (1397T>C), CGG420CGC (1400G>C), CTA421TTG (1401C>T 1403A>G), GCA423GCG (1409A>G), AAG426AAA (1418G>A), AGG427AGA (1421G>A), GCT435GCA (1445T>A), CAC436CAT (1448C>T), GCT439GCA (1457T>A), ATC440GTG (1458A>G 1460C>G), CTG441CTT (1463G>T), GAC444GAT (1472C>T), GGA446GGT (1478A>T), GCA447GCT (1481A>T), TTG448CTT (1482T>C 1484G>T), CGA449CGG (1487A>G), GCT452GCA (1496T>A), TGC455TGT (1505C>T), CAA456CAG (1508A>G), GAT457GAC (1511T>C), ATT458ATA (1514T>A), AAA459AAG (1517A>G), GAT463GAC (1529T>C), TCA466TCT (1538A>T), ACA467ACC (1541A>C), CAA468CAG (1544A>G), CCA469CCT (1547A>T), CCC471CCT (1553C>T), TTA472CTT (1554T>C 1556A>T), GCC475GCT (1565C>T), GTT477GTC (1571T>C), GAC479GAT (1577C>T), GGC481GGA (1583C>A), CAT483CAC (1589T>C), TTC484TTT (1592C>T), ACC485ACA (1595C>A), TCC486TCA (1598C>A), TAT488TAC (1604T>C), ATA489ATC (1607A>C), ACA492ACC (1616A>C), CTG495TTG (1623C>T), ATC496ATA (1628C>A), TCA497TCT (1631A>T), CCT499CCA (1637T>A), GCC500ACT (1638G>A 1640C>T), CAA501CAG (1643A>G), GGT503GGG (1649T>G), CTT504CTA (1652T>A), ACA505ACG (1655A>G), CCA507CCT (1661A>T), GCT509GCA (1667T>A), GAG511GAA (1673G>A), CGG512AGG (1674C>A), AGA513AGG (1679A>G), AGG514AGA (1682G>A), GAC515GAT (1685C>T), ATC516ATT (1688C>T), CTG517TTA (1689C>T 1691G>A), GTG518GTT (1694G>T), TGT520TGC (1700T>C), ATG521AGA (1702T>G 1703G>A), GAG523GTG (1708A>T), GAG525GAT (1715G>T), TCC527GAT (1719T>G 1720C>A 1721C>T), ACT528CCA (1722A>C 1724T>A), GAA529AAT (1725G>A 1727A>T), GAT530GCT (1729A>C), CCT531CCA (1733T>A), ACG532ACA (1736G>A), GGA533GCC (1738G>C 1739A>C), CTC535CTA (1745C>A), AGA536GTT (1746A>G 1747G>T 1748A>T), AAC538GAT (1752A>G 1754C>T), ATA539GTG (1755A>G 1757A>G), GAG541GAA (1763G>A), GCT547GCC (1781T>C), ACT549ACA (1787T>A), CAC550CAT (1790C>T), AGG551AGA (1793G>A), AAT552GCT (1794A>G 1795A>C), TTG553TTA (1799G>A), ACC554TCC (1800A>T), AGT556GAT (1806A>G 1807G>A), GAT557GAC (1811T>C), CTT558CTC (1814T>C), GAG560CAA (1818G>C 1820G>A), GTA563GTG (1829A>G), GCC564GCA (1832C>A), AAA565AAG (1835A>G), TGT566TGC (1838T>C), CAG567CGG (1840A>G), GTA568GTG (1844A>G), CAT569CAC (1847T>C), GCA570TAT (1848G>T 1849C>A 1850A>T), GAA571GAG (1853A>G), GTA572GTG (1856A>G), AAG574AGC (1861A>G 1862G>C), TTG576CTT (1866T>C 1868G>T), TTT577TAT (1870T>A), GAA578AAT (1872G>A 1874A>T), GCT579ACT (1875G>A), GAA580ACA (1878G>A 1879A>C), TCT581TCA (1883T>A), GGC582GGA (1886C>A), ATA583TTG (1887A>T 1889A>G), AGT584GCT (1890A>G 1891G>C), CCT585GAC (1893C>G 1894C>A 1895T>C), ATA587GTT (1899A>G 1901A>T), GTT590GTC (1910T>C), CAA591AAT (1911C>A 1913A>T), TCT593GCT (1917T>G), GCA594GTT (1921C>T 1922A>T), GAT596GAA (1928T>A), ATA598CTG (1932A>C 1934A>G), GCG599GAT (1936C>A 1937G>T), GTG601TTC (1941G>T 1943G>C), GAT602CAA (1944G>C 1946T>A), GGG603GAT (1948G>A 1949G>T), GCT604GCC (1952T>C), AGA605del (1953_1955delAGA), TTG606TTA (1958G>A), CGC607CAA (1960G>A 1961C>A), ACC608GTT (1962A>G 1963C>T 1964C>T), AAG609TTG (1965A>T 1966A>T), CAA610CCT (1969A>C 1970A>T), GAT611CCT (1971G>C 1972A>C), GAA612GAG (1976A>G), ATA614GCA (1980A>G 1981T>C), TTG615ACT (1983T>A 1984T>C 1985G>T), GTG616TAT (1986G>T 1987T>A 1988G>T), CCC617GAG (1989C>G 1990C>A 1991C>G), ACT618CAG (1992A>C 1993C>A 1994T>G), GTT619ACA (1995G>A 1996T>C 1997T>A), GTG620ATT (1998G>A 2000G>T), AGT621ACA (2002G>C 2003T>A), AAT623GCT (2007A>G 2008A>C), GAT624GAA (2012T>A), AGA625AAA (2014G>A), GTG626ATT (2016G>A 2018G>T), ATC627GTT (2019A>G 2021C>T), TAT628TAC (2024T>C), GCA629TCC (2025G>T 2027A>C), AGA630CAG (2028A>C 2029G>A 2030A>G), GAG631GAA (2033G>A), ACG633ACA (2039G>A), GTA634GTT (2042A>T), GCA636GCC (2048A>C), TTG637CTC (2049T>C 2051G>C), GCA640GCT (2060A>T), CAG642CAA (2066G>A), GGC643GGG (2069C>G), TTG645ATG (2073T>A), GAC646AAC (2076G>A), CCT647AAA (2079C>A 2080C>A 2081T>A), GAG648TTT (2082G>T 2083A>T 2084G>T), GAA649GAG (2087A>G), CTT650TTT (2088C>T), TTT651TTC (2093T>C), CCT652CCA (2096T>A), CAT655CTC (2104A>T 2105T>C), CAT656TCC (2106C>T 2107A>C 2108T>C), GCC659GCT (2117C>T), ATG660ATC (2120G>C), TCT663GTG (2127T>G 2128C>T 2129T>G), CTT664CTA (2132T>A), GAT665TCT (2133G>T 2134A>C), GAT666TCT (2136G>T 2137A>C), GAA667GAT (2141A>T), CAT668GCC (2142C>G 2143A>C 2144T>C), GAG670CAG (2148G>C), ATT671AAT (2152T>A), TTC672TTT (2156C>T), AAC673GAA (2157A>G 2159C>A), AAA674ATG (2161A>T 2162A>G), AGA676GTA (2166A>G 2167G>T), GTC677AAT (2169G>A 2170T>A 2171C>T), AAC678GAT (2172A>G 2174C>T), ATG679TTG (2175A>T), TAT681GTC (2181T>G 2182A>T 2183T>C), AGA682AAA (2185G>A), GAT685AGT (2193G>A 2194A>G), GCT686CCT (2196G>C), GAA687CAG (2199G>C 2201A>G), CGA690GCT (2208C>G 2209G>C 2210A>T), GTA693GTG (2219A>G), GAC699GAT (2237C>T), ATA701GTC (2241A>G 2243A>C), CAC703CAT (2249C>T), TTT704TTC (2252T>C), GCA710AGC (2268G>A 2269C>G 2270A>C), TCA711ACA (2271T>A), ATC713ATT (2279C>T), GAG715GAA (2285G>A), AAG716GAG (2286A>G), AAA717TTG (2289A>T 2290A>T 2291A>G), CTA718CTG (2294A>G), GCA719GAT (2296C>A 2297A>T), GTG720GTC (2300G>C), CAG721CAA (2303G>A), ACA722CGC (2304A>C 2305C>G 2306A>C), GAG723AAT (2307G>A 2309G>T), ATG724CTT (2310A>C 2312G>T), AGG725GAG (2313A>G 2314G>A), ACT726CAA (2316A>C 2317C>A 2318T>A), GGA727GGG (2321A>G), TTT728AAA (2322T>A 2323T>A 2324T>A), GCA729GCG (2327A>G), CAT730TAT (2328C>T), TCT731AAG (2331T>A 2332C>A 2333T>G), TGC732GCG (2334T>G 2335G>C 2336C>G), ATT733CTT (2337A>C), GAT734CAG (2340G>C 2342T>G), GCA735GCT (2345A>T), GTT737GCA (2350T>C 2351T>A), GAC738GAT (2354C>T), ACC739ACA (2357C>A), CTC740CTT (2360C>T), ATA741ACG (2362T>C 2363A>G), ATC743ATT (2369C>T), GAG744GAA (2372G>A), CCA747CCG (2381A>G), TTT749AGA (2385T>A 2386T>G 2387T>A), GTT750GTA (2390T>A), TTT752TTC (2396T>C), TTA753CTG (2397T>C 2399A>G), ACA755ACT (2405A>T), GCA756GCT (2408A>T), AGA759CGC (2415A>C 2417A>C), TGC762TGT (2426C>T), AAC763AGG (2428A>G 2429C>G), GTA765GTT (2435A>T), TCA766GCA (2436T>G), TCA766_AAA767insGGT (2438_2439insGGT), AAA767ACT (2440A>C 2441A>T), GAG768GAC (2444G>C), CCA769AAG (2445C>A 2446C>A 2447A>G), GAA771AGA (2451G>A 2452A>G), TCA772AAT (2454T>A 2455C>A 2456A>T), ATC774ATT (2462C>T), AAA775ATC (2464A>T 2465A>C), ATT776GAA (2466A>G 2467T>A 2468T>A), TAC778ATG (2472T>A 2473A>T 2474C>G), CAT779ATT (2475C>A 2476A>T), AAA780GGT (2478A>G 2479A>G 2480A>T), GTT782AAG (2484G>A 2485T>A 2486T>G), GAA783GAC (2489A>C), TAT784AAA (2490T>A 2492T>A), ATC785ATG (2495C>G), AAG786ATC (2497A>T 2498G>C), GAT787CAG (2499G>C 2501T>G), ACA788ATG (2503C>T 2504A>G), TGG790AAT (2508T>A 2509G>A 2510G>T), AGC791GAC (2511A>G 2512G>A), CTT792TTG (2514C>T 2516T>G), TAT794TTC (2521A>T 2522T>C), GCA795ATC (2523G>A 2524C>T 2525A>C), CTG796TTG (2526C>T), AGA797AAG (2530G>A 2531A>G), CTC799CTT (2537C>T), GCA802GCC (2546A>C), GGT803GGA (2549T>A), ATA806TTG (2556A>T 2558A>G), ATG807ATA (2561G>A), GCT808GCA (2564T>A), GGA811GGC (2573A>C), GTT812ACC (2574G>A 2575T>C 2576T>C), TTT813ATG (2577T>A 2579T>G), GGA814AGT (2580G>A 2582A>T), GGC815GCC (2584G>C), ATT816GTT (2586A>G), ACA817ACT (2591A>T), GCA818GCT (2594A>T), TTT819TTC (2597T>C), TTG820ATT (2598T>A 2600G>T), GCG821ACT (2601G>A 2603G>T), TGG823CTC (2607T>C 2608G>T 2609G>C), CAG824AAG (2610C>A), AGT825GCA (2613A>G 2614G>C 2615T>A), AAC826GCA (2616A>G 2617A>C 2618C>A), TCT827ACT (2619T>A), TTC828TCC (2623T>C), CCC829CCT (2627C>T), AAT830TTA (2628A>T 2629A>T 2630T>A), GCA831GCT (2633A>T), TCG832GAT (2634T>G 2635C>A 2636G>T), GGA835GGG (2645A>G), GGT836GGC (2648T>C), GTG837GTA (2651G>A), GAA840GAG (2660A>G), ATA844ATC (2672A>C), AGT846AGC (2678T>C), AAG848AAA (2684G>A), AAC849AAT (2687C>T), AAG850AAA (2690G>A), GGT851GGG (2693T>G), AAT852AAG (2696T>G), AAG853ACT (2698A>C 2699G>T), AAG853_CTC854insCAT (2699_2700insCAT), CTC854TTC (2700C>T), AGA855GCA (2703A>G 2704G>C), AAT856AAA (2708T>A), CTT857TCC (2709C>T 2710T>C 2711T>C), CTC858TTG (2712C>T 2714C>G), GTT859CTA (2715G>C 2717T>A), GGT860GGG (2720T>G), AGT862GCG (2724A>G 2725G>C 2726T>G), TCT863GCA (2727T>G 2729T>A), CAA864GAC (2730C>G 2732A>C), TCA865AAC (2733T>A 2734C>A 2735A>C), TTG866AAC (2736T>A 2737T>A 2738G>C), GCA867CAT (2739G>C 2740C>A 2741A>T), CAA868GTG (2742C>G 2743A>T 2744A>G), GAT869GAC (2747T>C), GAT869_TGG870insCGA (2747_2748insCGA), GCT871GCA (2753T>A), GCT872GAA (2755C>A 2756T>A), GAA873CAG (2757G>C 2759A>G), GTA877GTT (2771A>T), CAG879GAA (2775C>G 2777G>A), AGC880AGT (2780C>T), CTC881CTG (2783C>G), AAG882CAC (2784A>C 2786G>C), AAT884AAC (2792T>C), GTG886GTT (2798G>T), GTG887GTT (2801G>T), TTA888TTG (2804A>G), CTT890ATT (2808C>A), GGT891GCT (2812G>C), GTG894GTC (2822G>C), TAC895TAT (2825C>T), TTT896TTC (2828T>C), GGC898GGT (2834C>T), ACC899ACG (2837C>G), TAC900TAT (2840C>T), GTG901GTT (2843G>T), ATA906ATT (2858A>T), ATG907CTC (2859A>C 2861G>C), ACC908ACT (2864C>T), GTA909GTG (2867A>G), GCT910GCA (2870T>A), GCA912GCC (2876A>C), TTC913TTT (2879C>T), CAC914AGA (2880C>A 2881A>G 2882C>A), AAT915AAC (2885T>C), GCT916ATG (2886G>A 2887C>T 2888T>G), CGA917AGG (2889C>A 2891A>G), GGT919GGC (2897T>C), CCA921CCT (2903A>T), ACA923TCC (2907A>T 2909A>C), ATC925ATT (2915C>T), GCC927GAG (2920C>A 2921C>G), AAT928CAT (2922A>C), TCT929TCC (2927T>C), CGA930ATT (2928C>A 2929G>T 2930A>T), TCT931TGT (2932C>G), AAA932AAT (2936A>T), GTT933GTG (2939T>G), CAA934CAG (2942A>G), TAC935TTT (2944A>T 2945C>T), AAC936AAT (2948C>T), AGA938ACC (2953G>C 2954A>C), GAA939CAA (2955G>C), ATT942TTG (2964A>T 2966T>G), ATT943ATA (2969T>A), GGC945GGT (2975C>T), GAC947GAT (2981C>T), ATC948GTG (2982A>G 2984C>G), CTG950TTG (2988C>T), CGC952AGG (2994C>A 2996C>G), GTT953GTA (2999T>A), GGT954GGG (3002T>G), CCA959CCC (3017A>C), AAG960AAA (3020G>A), CCT961TCA (3021C>T 3023T>A), GAC962GAT (3026C>T), ATC963ATA (3029C>A), CGT964AGA (3030C>A 3032T>A), CAC966CAT (3038C>T), CCA968CCT (3044A>T), AGA969CGC (3045A>C 3047A>C), AGG970AGA (3050G>A), GAG972GAT (3056G>T), TGC974TGT (3062C>T), TTC975TAC (3064T>A), ACA976TCC (3066A>T 3068A>C), CAA980ATG

(3078C>A 3079A>T 3080A>G), GGA981GGC (3083A>C), TGT984GTG (3090T>G 3091G>T 3092T>G), AGA985AGG (3095A>G), GTT987GTA (3101T>A), GCG988GCT (3104G>T), TCT989TCA (3107T>A), ACA990AAG (3109C>A 3110A>G), CCA992CCC (3116A>C), CGT993AAG (3117C>A 3118G>A 3119T>G), CTT994AAA (3120C>A 3121T>A 3122T>A), GGC995TCC (3123G>T 3124G>C), TTA997TTG (3131A>G), TTT999GTG (3135T>G 3137T>G), CTC1000CTT (3140C>T), AAG1001AAA (3143G>A), ATG1002TAT (3144A>T 3145T>A 3146G>T), GGA1009GGC (3167A>C), ACA1011ACC (3173A>C), GTT1012GAA (3175T>A 3176T>A), GTG1014GTC (3182G>C), GAA1015ACA (3183G>A 3184A>C), GAA1017GGA (3190A>G), CTG1018TTA (3192C>T 3194G>A), AAC1019GAT (3195A>G 3197C>T), TCC1020TCG (3200C>G), TCC1021AAT (3201T>A 3202C>A 3203C>T), AGT1022GCA (3204A>G 3205G>C 3206T>A), AAG1024GCT (3210A>G 3211A>C 3212G>T), ATT1025CTT (3213A>C), TGC1026AGC (3216T>A), TCT1027TCA (3221T>A), CAA1029GTT (3225C>G 3226A>T 3227A>T), TCT1030GCA (3228T>G 3230T>A), GAA1032GAG (3236A>G), ATA1035ATT (3245A>T), GGG1037AGT (3249G>A 3251G>T), GAA1038GAG (3254A>G), GGT1040GGA (3260T>A), GAC1041GAT (3263C>T), GGC1043GGT (3269C>T), ACG1044ACT (3272G>T), TTG1045CTT (3273T>C 3275G>T), CTA1046CTC (3278A>C), CTG1047TTA (3279C>T 3281G>A), TTA1048TTG (3284A>G), AGT1050ACT (3289G>C), GTT1051GTG (3293T>G), AAT1053AAC (3299T>C), CAA1055CAG (3305A>G), CCT1056CCG (3308T>G), GTG1057TGC (3309G>T 3310T>G 3311G>C), GGC1060GGA (3320C>A), ATC1061ATA (3323C>A), CAC1062CAT (3326C>T), TCA1065TCT (3335A>T), GGC1068GAT (3343G>A 3344C>T), GTA1069ACC (3345G>A 3346T>C 3347A>C), GCA1070GCT (3350A>T), GCT1071GCA (3353T>A), GAA1072GAG (3356A>G), CGT1073AAG (3357C>A 3358G>A 3359T>G), GGA1074GGT (3362A>T), TTT1075TAC (3364T>A 3365T>C), ATC1076GTG (3366A>G 3368C>G), TCT1077GCT (3369T>G), AAT1079AAC (3377T>C), GCT1080GCA (3380T>A), GCT1082GCA (3386T>A), ATC1083GTG (3387A>G 3389C>G), TAC1084TTT (3391A>T 3392C>T), AGG1085AGA (3395G>A), GAA1086GAT (3398A>T), CTA1088TTG (3402C>T 3404A>G), GAG1089GAA (3407G>A), CCA1092CCC (3416A>C), ACT1093ACG (3419T>G), GGG1094GGA (3422G>A), CCG1095CCC (3425G>C), GTC1096ATT (3426G>A 3428C>T), GCA1098GTG (3433C>T 3434A>G), GCA1099GCT (3437A>T), GAT1104GAA (3452T>A), ATT1105CTG (3453A>C 3455T>G), CTG1106TTG (3456C>T), TCA1108CAC (3462T>C 3463C>A 3464A>C), ATT1109CTG (3465A>C 3467T>G), AGA1110AGG (3470A>G), AGC1111TCC (3471A>T 3472G>C), AGA1112AAG (3475G>A 3476A>G), GAA1113GAG (3479A>G), ACA1114GAA (3480A>G 3481C>A), CAG1115AGA (3483C>A 3484A>G 3485G>A), CTT1116TTC (3486C>T 3488T>C), GAA1119GAG (3497A>G), AAC1120AAA (3500C>A), CAA1121CAG (3503A>G), TAC1123GCT (3507T>G 3508A>C 3509C>T), TAC1124TTC (3511A>T), CTT1125TTG (3513C>T 3515T>G), GGA1126GGC (3518A>C), ACA1127ACT (3521A>T), GTT1128GTA (3524T>A), CCA1129CCG (3527A>G), CAG1130TTG (3528C>T 3529A>T), GAG1131GAA (3533G>A), TTG1132ATG (3534T>A), GCC1133GCT (3539C>T), GCC1134GCT (3542C>T), GTT1136GTG (3548T>G), CCC1137CCA (3551C>A), AAG1139AAA (3557G>A), ACC1140ACA (3560C>A), ACT1141ACC (3563T>C), CTG1142CTT (3566G>T), CGG1143AAA (3567C>A 3568G>A 3569G>A), AAA1144CGC (3570A>C 3571A>G 3572A>C), AGC1145AGT (3575C>T), CAA1146CCC (3577A>C 3578A>C), TTG1147CTG (3579T>C), GAA1149GAG (3587A>G), GCA1150GCT (3590A>T), TTC1151TTT (3593C>T), GGA1152GGG (3596A>G), CCT1153GAT (3597C>G 3598C>A), GCA1154GCT (3602A>T), GAG1155CAG (3603G>C), ACA1156ACT (3608A>T), GCA1157GCC (3611A>C), ATT1160ATC (3620T>C), ACA1162TCT (3624A>T 3626A>T), GTT1163GCC (3628T>C 3629T>C), CAT1164AAA (3630C>A 3632T>A), AGA1167AGG (3641A>G), GGT1168GGA (3644T>A), GAT1169GAA (3647T>A), GGT1170GGG (3650T>G), GAC1172GAT (3656C>T), GGG1177GGT (3671G>T), GTA1178GTC (3674A>C), ATG1179GAG (3675A>G 3676T>A), AAA1180AAG (3680A>G), TAC1181TAT (3683C>T), TGT1186TCC (3697G>C 3698T>C), GGA1187AAC (3699G>A 3700G>A 3701A>C), GAT1189GAG (3707T>G), GAT1190GAG (3710T>G), GAC1191CCA (3711G>C 3712A>C 3713C>A), ATT1192GTT (3714A>G), GCC1193GCA (3719C>A), AAA1194AAG (3722A>G), CTA1195CTG (3725A>G), GCA1196GCC (3728A>C), TTC1197TTT (3731C>T), GAA1198GAG (3734A>G), AAT1199GGT (3735A>G 3736A>G), CTC1200CTT (3740C>T), AAG1201AAA (3743G>A), TGC1202GCA (3744T>G 3745G>C 3746C>A), TCG1203TCC (3749G>C), CTG1204ATT (3750C>A 3752G>T), CTA1205CTT (3755A>T), ATC1207GTG (3759A>G 3761C>G), ATG1208CTT (3762A>C 3764G>T), CGT1209AAG (3765C>A 3766G>A 3767T>G), AGC1210ATG (3769G>T 3770C>G), AAG1212CGA (3774A>C 3775A>G 3776G>A), ATC1213GTG (3777A>G 3779C>G), CCT1214CCC (3782T>C), GGA1216GGG (3788A>G), CGT1217AAA (3789C>A 3790G>A 3791T>A), CCA1218CCT (3794A>T), TGT1219GCA (3795T>G 3796G>C 3797T>A), GAA1220ATT (3798G>A 3799A>T 3800A>T), AGG1221AGA (3803G>A), GAT1222AAT (3804G>A), GGA1229GGG (3827A>G), TGT1233TTC (3838G>T 3839T>C), TAC1235TAT (3845C>T), TAT1236TTT (3847A>T), GGC1238GCA (3853G>C 3854C>A), GAG1240GAA (3860G>A), CTG1241TTG (3861C>T), ACA1243ACG (3869A>G), TCT1244TCA (3872T>A), TGC1245TGT (3875C>T), GGA1246GGT (3878A>T), CCC1248CCT (3884C>T), TTC1249TTT (3887C>T), ATG1252TTG (3894A>T), GGG1253GGA (3899G>A), ATG1254CTT (3900A>C 3902G>T), GGG1255GGT (3905G>T), ATG1256TCT (3906A>T 3907T>C 3908G>T), AAG1258AAA (3914G>A), AGA1259AGG (3917A>G), CAA1263GAA (3927C>G), TCC1264GCC (3930T>G), ACT1265CAA (3933A>C 3934C>A 3935T>A), GGC1266CCA (3936G>C 3937G>C 3938C>A), GAA1267GAC (3941A>C), GGA1268TCG (3942G>T 3943G>C 3944A>G), GAG1269CAG (3945G>C), AGA1270AGG (3950A>G), GTG1271GTT (3953G>T), GAA1272CTT (3954G>C 3955A>T 3956A>T), AGA1275AGG (3965A>G), GAC1276AAC (3966G>A), ACT1277ACA (3971T>A), GTA1279GTC (3977A>C), GAA1281GAT (3983A>T), CTA1286TTG (3996C>T 3998A>G), GAT1287GAC (4001T>C), GTG1288TCG (4002G>T 4003T>C), ATT1290GTA (4008A>G 4010T>A), AAA1292CAA (4014A>C), GGC1293GGA (4019C>A), ATC1294ACA (4021T>C 4022C>A), CAT1295TTT (4023C>T 4024A>T), GTG1296GTT (4028G>T), GAT1297GAC (4031T>C), CTG1298CTT (4034G>T), ACC1300ACA (4040C>A), CAA1302CAG (4046A>G), TGC1303TGT (4049C>T), GCC1304GCG (4052C>G), CGC1308CGG (4064C>G), CTC1309TTG (4065C>T 4067C>G), AAA1313AAG (4079A>G), GGC1316GGG (4088C>G), AGA1318AGG (4094A>G), AGG1321CGC (4101A>C 4103G>C), CTC1322CTG (4106C>G), TTT1323TTC (4109T>C), GAG1324GAA (4112G>A), ATA1325ATC (4115A>C), CTT1326TTG (4116C>T 4118T>G), TAC1330TAT (4130C>T), TTG1333CTG (4137T>C), GTC1334GTG (4142C>G), AGG1335CGG (4143A>C), AAG1336AAA (4148G>A), TAT1337TAC (4151T>C), TTT1338TTC (4154T>C), GAT1340GAC (4160T>C), TTC1341TTT (4163C>T), TCA1342TCT (4166A>T), GCC1343GCT (4169C>T), ACA1344TGT (4170A>T 4171C>G 4172A>T), TTG1348GCA (4182T>G 4183T>C 4184G>A), AAT1350AAC (4190T>C), GCT1351GTA (4192C>T 4193T>A), ATA1352CTC (4194A>C 4196A>C), CCA1353CCT (4199A>T), AAA1355AAG (4205A>G), GTT1356GTA (4208T>A), ATT1358ATC (4214T>C), GAT1359AAT (4215G>A), CCT1360CCA (4220T>A), ACA1361ACG (4223A>G), AGT1362AGC (4226T>C), TCT1363CAT (4227T>C 4228C>A), TTG1367GTG (4239T>G), TTG1368CTG (4242T>C), GCA1369GCT (4247A>T), AAT1370AGT (4249A>G), GGG1371AGC (4251G>A 4253G>C), AGA1373AGG (4259A>G), GCT1374TCA (4260G>T 4262T>A), GTG1375GTA (4265G>A), TCA1376TCT (4268A>T), GAC1377GAT (4271C>T), GTG1378GAG (4273T>A), TTT1380TTC (4280T>C), GCT1382GCC (4286T>C), TAT1384TAC (4292T>C), TCC1385TCG (4295C>G), CCT1392CCC (4316T>C), GTT1393ATC (4317G>A 4319T>C), GCT1395GCA (4325T>A), CAG1397CAA (4331G>A), TTG1402CTA (4344T>C 4346G>A), GAA1405GCT (4354A>C 4355A>T), TAC1406TTT (4357A>T 4358C>T), TAC1407TAT (4361C>T), GGA1408GGT (4364A>T), TCA1409GAT (4365T>G 4366C>A 4367A>T), AAG1410GAG (4368A>G), GGC1412GGG (4376C>G), GAT1415GAG (4385T>G), TCC1416TCA (4388C>A), CGA1419AGG (4395C>A 4397A>G), CAT1420CAC (4400T>C), GCA1421ACA (4401G>A), CTT1422TTA (4404C>T 4406T>A), GCA1426GCC (4418A>C), TGT1428TGC (4424T>C), TAC1430TAT (4430C>T), ACA1431ACC (4433A>C), CTG1432CTC (4436G>C), TGC1433TGT (4439C>T), GAT1435AAC (4443G>A 4445T>C), AAA1436AAG (4448A>G), GTG1437TTG (4449G>T), AGG1439CGC (4455A>C 4457G>C), TTG1440CTA (4458T>C 4460G>A), GTT1441GTG (4463T>G), GGG1442GGT (4466G>T), GGC1443GGA (4469C>A), CCA1445CCC (4475A>C), TCA1446TCT (4478A>T), GGA1447GGT (4481A>T), GCA1449GCT (4487A>T), CTA1450TTG (4488C>T 4490A>G), ACG1451ACT (4493G>T), ATC1453ATA (4499C>A), TTC1454TTT (4502C>T), TCT1456TCA (4508T>A), CTC1457TTG (4509C>T 4511C>G), CTC1458CTA (4514C>A), CGA1464AGG (4530C>A 4532A>G), TAT1465TAC (4535T>C), GCC1466GCG (4538C>G), TTG1470CTG (4548T>C), TTA1471TTG (4553A>G), AGA1472AGG (4556A>G), CCC1474CCA (4562C>A), ATT1476GTC (4566A>G 4568T>C), GCT1477AAA (4569G>A 4570C>A 4571T>A), AGG1479AAA (4576G>A 4577G>A), GCT1480TCT (4578G>T), ATA1481GTG (4581A>G 4583A>G), GGA1482GGC (4586A>C), GTT1483ATT (4587G>A), AAA1484AAG (4592A>G), AGT1486AAT (4597G>A), GAT1487GAC (4601T>C), CTA1491TTG (4611C>T 4613A>G), GCT1494GCA (4622T>A), GTT1495GTA (4625T>A), GGA1497GGT (4631A>T), GAC1499GAT (4637C>T), AAT1500AAC (4640T>C), CTT1501TTG (4641C>T 4643T>G), GCT1503GCG (4649T>G), GTA1504GTT (4652A>T), TTA1506ATG (4656T>A 4658A>G), CTC1508TTG (4662C>T 4664C>G), CAG1509AGG (4665C>A 4666A>G), TAT1511TAC (4673T>C), TCT1512GAT (4674T>G 4675C>A), CTG1513CTA (4679G>A), AAT1515TCC (4683A>T 4684A>C 4685T>C), ATA1516ATT (4688A>T), CAT1518AGA (4692C>A 4693A>G 4694T>A), GAG1519GAA (4697G>A), TTA1520CTG (4698T>C 4700A>G), CTG1522GCT (4704C>G 4705T>C 4706G>T), GTC1523GTT (4709C>T), AAT1524AAC (4712T>C), GTA1525GTC (4715A>C), ATC1526GTC (4716A>G), CTT1531TTG (4731C>T 4733T>G), AAA1533AAG (4739A>G), TCG1534TCA (4742G>A), GTT1537GTA (4751T>A), AAT1538GAG (4752A>G 4754T>G), GAG1539AAG (4755G>A), GTA1540ACA (4758G>A 4759T>C), CAA1541AAA (4761C>A), CAA1543CTG (4768A>T 4769A>G), CTA1548CTC (4784A>C), ACT1549ACC (4787T>C), CTG1551CTC (4793G>C), AGA1553AGG (4799A>G), TTT1555TTC (4805T>C), AAA1556AAG (4808A>G), GCT1559GCA (4817T>A), CTT1560TTG (4818C>T 4820T>G), TAC1562TAT (4826C>T), CAC1563CAA (4829C>A), GCT1565GCA (4835T>A), CCT1566CCA (4838T>A), AAG1568AAA (4844G>A), ATC1572GTC (4854A>G), ATG1576ATT (4868G>T), TAT1577TAC (4871T>C), ATA1579ATT (4877A>T), AGA1580AGG (4880A>G), CCT1581CCA (4883T>A), GCC1582GCA (4886C>A), CCA1583GTG (4887C>G 4888C>T 4889A>G), GGT1584GGA (4892T>A), GAC1587GAT (4901C>T), ACT1588GTA (4902A>G 4903C>T 4904T>A), CAG1589CAA (4907G>A), GCC1590GCT (4910C>T), ATG1592TAT (4914A>T 4915T>A 4916G>T), GTG1595ATT (4923G>A 4925G>T), GGA1598GGC (4934A>C), ATA1599ATC (4937A>C), CGT1600AGA (4938C>A 4940T>A), GCT1602GCC (4946T>C), CAC1605CAT (4955C>T), GGT1606GGG (4958T>G), CGT1607CAG (4960G>A 4961T>G), GTG1608GCC (4963T>C 4964G>C), GCT1609ACT (4965G>A), GAC1611GAA (4973C>A), AAG1612AGT (4975A>G 4976G>T), CTT1613TTT (4977C>T), GTC1614GTA (4982C>A), TTA1615AAT (4983T>A 4984T>A 4985A>T), CAG1618CTT (4993A>T 4994G>T), ACG1619GGA (4995A>G 4996C>G 4997G>A), GCG1620GCT (5000G>T), TTG1621CTC (5001T>C 5003G>C), GAT1622AAG (5004G>A 5006T>G), GAA1623CAG (5007G>C 5009A>G), AGG1624AGA (5012G>A), GGT1625AAC (5013G>A 5014G>A 5015T>C), TTC1626CTC (5016T>C), AGA1627CCC (5019A>C 5020G>C 5021A>C), GCT1628ATC (5022G>A 5023C>T 5024T>C), GTG1629ATG (5025G>A), TTT1631ATG (5031T>A 5033T>G), CCT1632CCC (5036T>C), TCC1633ACT (5037T>A 5039C>T), TAT1634TAC (5042T>C), TTG1635TAT (5044T>A 5045G>T), GTG1637CTT (5049G>C 5051G>T), CAG1639AGT (5055C>A 5056A>G 5057G>T), GAA1640CAA (5058G>C), ATT1642TTG (5064A>T 5066T>G), GCA1643ACA (5067G>A), AAG1644GAG (5070A>G), GTA1645GTC (5075A>C), GGG1647GGA (5081G>A), TCA1649GAG (5085T>G 5086C>A 5087A>G), GCC1651ATT (5091G>A 5092C>T 5093C>T), CTG1652GGG (5094C>G 5095T>G), ACA1653GTT (5097A>G 5098C>T 5099A>T), ATT1654TTG (5100A>T 5102T>G), TGT1655ATG (5103T>A 5104G>T 5105T>G), GAA1656GAG (5108A>G), ATG1657CTA (5109A>C 5111G>A), GCA1658TCA (5112G>T), AAA1659GAT (5115A>G 5117A>T), GCA1660GCT (5120A>T), GCT1661AAA (5121G>A 5122C>A 5123T>A), ATT1662AAT (5125T>A), TCC1663ACA (5127T>A 5129C>A), TAT1664TAC (5132T>C), ACG1665ACC (5135G>C), CCA1666CCT (5138A>T), TTG1667TTA (5141G>A), GAC1668CCT (5142G>C 5143A>C 5144C>T), GCA1669GCT (5147A>T), GGT1670GGG (5150T>G), GAG1671GAA (5153G>A), ATC1673ATT (5159C>T), ACA1674del (5160_5162delACA), CGT1678GAG (5172C>G 5173G>A 5174T>G), GAT1679GGT (5176A>G), CTG1680GTC (5178C>G 5180G>C), AAT1681CAT (5181A>C), TGG1682GAA (5184T>G 5185G>A 5186G>A), GCA1684GCG (5192A>G), CCA1685CCT (5195A>T), AAC1686GAT (5196A>G 5198C>T), ATT1687GTA (5199A>G 5201T>A), GGT1688TTT (5202G>T 5203G>T), TTT1689TTC (5207T>C), TCA1691AGT (5211T>A 5212C>G 5213A>T), GCA1692GCT (5216A>T), CGT1693CGG (5219T>G), GCC1696TCA (5226G>T 5228C>A), CAC1697AAC (5229C>A), TAC1698CAA (5232T>C 5234C>A), ACG1699CAA (5235A>C 5236C>A 5237G>A), GAT1701TCA (5241G>T 5242A>C 5243T>A), GAA1702TCA (5244G>T 5245A>C), GGG1703GGT (5249G>T), TAC1704AAC (5250T>A), ATT1705ATC (5255T>C), ATT1706CTG (5256A>C 5258T>G), AAC1708AAT (5264C>T), TGT1709TGC (5267T>C), ACA1710ACT (5270A>T), GGT1711GGA (5273T>A), GCA1712GCC (5276A>C), AAG1713CGC (5277A>C 5278A>G 5279G>C), AAG1714AGA (5281A>G 5282G>A), TCC1715ACA (5283T>A 5285C>A), AAT1716ACA (5287A>C 5288T>A), GTT1718GTG (5294T>G), GGT1720GGG (5300T>G), CCA1721CCT (5303A>T), AAC1723AAT (5309C>T), GAC1726GAT (5318C>T), ATG1731ATA (5333G>A), CCG1733CCC (5339G>C), ATG1736TTG (5346A>T), TCA1737GCA (5349T>G), GCT1738GCA (5354T>A), ATA1739ATC (5357A>C), ATG1740CAG (5358A>C 5359T>A), GCC1742CAT (5364G>C 5365C>A 5366C>T), AAG1744CGC (5370A>C 5371A>G 5372G>C), AAT1745AAG (5375T>G), GTG1746AGT (5376G>A 5377T>G 5378G>T), TTG1747CTG (5379T>C), GCA1748GAA (5383C>A), GGA1749GTT (5386G>T 5387A>T), GGA1750GGT (5390A>T), CAT1751GAT (5391C>G), GTA1752GGG (5395T>G 5396A>G), GCA1753GCT (5399A>T), AAT1755TCT (5403A>T 5404A>C), GTT1757ACC (5409G>A 5410T>C 5411T>C), GTG1759GTT (5417G>T), GGA1761GGC (5423A>C), GGA1763GGT (5429A>T), GCT1765GCA (5435T>A), GTC1766ATA (5436G>A 5438C>A), GGC1767GCT (5440G>C 5441C>T), ATT1769ATC (5447T>C), GCG1772GCA (5456G>A), CTG1773TTG (5457C>T), ATG1774TTT (5460A>T 5462G>T), GCC1775GCA (5465C>A), TTA1776TTG (5468A>G), GCA1777GCC (5471A>C), ACA1778ACT (5474A>T), AGG1779CAT (5475A>C 5476G>A 5477G>T), TAT1781TAC (5483T>C), TGT1782AGG (5484T>A 5486T>G), GTG1783GCC (5488T>C 5489G>C), GAG1784ACA (5490G>A 5491A>C 5492G>A), ATA1786CTT (5496A>C 5498A>T), ATA1787ATT (5501A>T), GTT1788GTG (5504T>G), AGA1789CGC (5505A>C 5507A>C), TTG1790CTA (5508T>C 5510G>A), CGC1791CGA (5513C>A), ACA1792ACC (5516A>C), ATT1793ATG (5519T>G), GGA1794GCT (5521G>C 5522A>T), AAT1795GGG (5523A>G 5524A>G 5525T>G), CTT1797CTG (5531T>G), GAC1798GAT (5534C>T), CTC1799CTG (5537C>G), AAT1800AAC (5540T>C), TAT1802TAC (5546T>C), CCT1803CCC (5549T>C), GGC1805GGT (5555C>T), GTG1807GCG (5560T>C), CAG1808CAA (5564G>A), TAT1809TAC (5567T>C), TTT1810TTG (5570T>G), CTT1811CTC (5573T>C), CAA1812AAG (5574C>A 5576A>G)

#### Proteins

| | | | | | | | | | |
|---|---|---|---|---|---|---|---|---|---|
| polyprotein (YP_001976147.1) | 1 | 1817 | 84.4% | 9297 | 73.5% | 1807 (99.3%) | 1266 (69.6%) | 3/10/0/0 | 0 |

| | Begin | End | Coverage | Score | Concordance | Matches | Identities | I/D/M/F* | Stop Codons |
|---|---|---|---|---|---|---|---|---|---|
| **NT** | **23** | **5589** | **77.0%** | **3552** | **32.4%** | **5524 (99.1%)** | **3681 (66.0%)** | **9/43** | |

Protein mutations:

F7W (160T>G 161C>G), K9_A16del (165_188delAAGGTTGCTGGTGTTGATGACGCC), S17P (189A>C 190G>C), S18P (192T>C), S19A (195T>G), T21S (201A>T 203C>A), T22P (204A>C), G26A (217G>C 218T>A), L28V (222C>G), N30S (228A>T 229A>C), I32F (234A>T), K34S (240A>T 241A>C 242A>T), F35L (243T>C), A38T (252G>A 254G>T), V39L (255G>C 257T>C), I42V (264A>G), T43N (268C>A 269C>T), T44K (271C>A 272T>A), M45V (273A>G 275G>T), K48T (283A>C 284G>C), S50V (288T>G 289C>T), L53I (297C>A 299T>A), L56V (306C>G 308G>C), S59P (315T>C 317T>A), T61M (322C>T 323T>G), D62A (325A>C 326T>A), V63A (328T>C 329G>A), S64A (330T>G 332C>A), V68A (343T>C 344G>C), A69S (345G>T 347C>T), S72N (355G>A), S73A (357T>G 359A>T), L74V (360C>G 362T>A), N75E (363A>G 365T>A), K76R (367A>G), G82A (385G>C 386T>A), A83K (387G>A 388C>A), L84I (390C>A 392T>A), I85M (395A>G), D86E (398T>A), L89I (405C>A 407T>C), A91G (412C>G 413G>T), S93D (417T>G 418C>A), L95V (423T>G 425G>T), S96K (426T>A 427C>A 428C>A), S101T (441T>A), K103V (447A>G 448A>T 449G>A), M105L (453A>C 455G>T), L109M (465T>A), F110V (468T>G), E111L (471G>T 472A>T 473G>A), L113V (477T>G 479G>C), S114K (480T>A 481C>A 482C>G), N115D (483A>G 485T>C), V117F (489G>T 491T>C), K118I (493A>T 494A>C), F121Y (502T>A 503T>C), A122E (505C>A 506T>A), V124L (510G>T 512G>A), N126E (516A>G 518T>A), S128V (522A>G 523G>T), D130S (528G>T 529A>C 530T>A), V131L (531G>T), V135F (543G>T 545A>T), Y136I (546T>A 547A>T 548T>A), G137A (550G>C), I143L (567A>T 569A>G), T144A (570A>G 572A>C), L145I (573T>A 575G>T), V147L (579G>C 581G>A), F150Y (589T>A), S154V (600A>G 601G>T 602T>A), E158T (612G>A 613A>C), G159P (615G>C 616G>C), V160F (618G>T), S161N (621T>A 622C>A), T163V (627A>G 628C>T 629T>A), K165V (633A>G 634A>T 635A>T), N166T (637A>C), I167L (639A>T 641C>G), L169S (646T>C 647G>A), I170V (648A>G 650A>T), M174F (660A>T 662G>T), K186T (697A>C), A188S (702G>T 704C>T), E189G (706A>G 707A>G), R190K (708C>A 709G>A 710A>G), T192S (714A>T), M193Q (717A>C 718T>A), V194L (720G>C), S198A (732A>G 733G>C 734T>A), M199V (735A>G 737G>A), V205L (753G>T 755T>A), S206T (756T>A 758A>G), M214L (780A>T 782G>A), T216S (786A>T), I217L (789A>T 791A>G), I228V (822A>G), T248A (882A>G 884A>T), T260Q (918A>C 919C>A 920T>G), I280V (978A>G 980A>G), G282S (984G>A), T283I (988C>T 989C>T), S295N (1024G>A), E304Q (1050G>C), Q307T (1059C>A 1060A>C), D310S (1068G>T 1069A>C 1070T>A), A311E (1072C>A 1073A>G), C313A (1077T>G 1078G>C), D314E (1082T>A), A316S (1086G>T 1088A>T), T317M (1090C>T), K325P (1113A>C 1114A>C 1115G>T), T329I (1126C>T 1127C>T), N332Q (1134A>C 1136C>A), R335K (1144G>A 1145G>A), E337Q (1149G>C 1151A>G), L339V (1155C>G), M354L (1200A>T), A363T (1227G>A), E366D (1238A>T), V368L (1242G>T 1244T>G), Y369V (1245T>G 1246A>T), S372A (1254A>G 1255G>C 1256T>C), C373S (1257T>A 1259T>C), R374Q (1260A>C 1261G>A 1262G>A), M376S (1266A>T 1267T>C 1268G>T), S393A (1317T>G 1319C>T), S401C (1342C>G), R412A (1374A>G 1375G>C), I440V (1458A>G 1460C>G), A500T (1638G>A 1640C>T), M521R (1702T>G 1703G>A), E523V (1708A>T), E525D (1715G>T), S527D (1719T>G 1720C>A 1721C>T), T528P (1722A>C 1724T>A), E529N (1725G>A 1727A>T), D530A (1729A>C), G533A (1738G>C 1739A>C), R536V (1746A>G 1747G>T 1748A>T), N538D (1752A>G 1754C>T), I539V (1755A>G 1757A>G), N552A (1794A>G 1795A>C), T554S (1800A>T), S556D (1806A>G 1807G>A), E560Q (1818G>C 1820G>A), Q567R (1840A>G), A570Y (1848G>T 1849C>A 1850A>T), K574S (1861A>G 1862G>C), F577Y (1870T>A), E578N (1872G>A 1874A>T), A579T (1875G>A), E580T (1878G>A 1879A>C), I583L (1887A>T 1889A>G), S584A (1890A>G 1891G>C), P585D (1893C>G 1894C>A 1895T>C), I587V (1899A>G 1901A>T), Q591N (1911C>A 1913A>T), S593A (1917T>G), A594V (1921C>T 1922A>T), D596E (1928T>A), I598L (1932A>C 1934A>G), A599D (1936C>A 1937G>T), V601F (1941G>T 1943G>C), D602Q (1944G>C 1946T>A), G603D (1948G>A 1949G>T), R605del (1953_1955delAGA), R607Q (1960G>A 1961C>A), T608V (1962A>G 1963C>T 1964C>T), K609L (1965A>T 1966A>T), Q610P (1969A>C 1970A>T), D611P (1971G>C 1972A>C), I614A (1980A>G 1981T>C), L615T (1983T>A 1984T>C 1985G>T), V616Y (1986G>T 1987T>A 1988G>T), P617E (1989C>G 1990C>A 1991C>G), T618Q (1992A>C 1993C>A 1994T>G), V619T (1995G>A 1996T>C 1997T>A), V620I (1998G>A 2000G>T), S621T (2002G>C 2003T>A), N623A (2007A>G 2008A>C), D624E (2012T>A), R625K (2014G>A), V626I (2016G>A 2018G>T), I627V (2019A>G 2021C>T), A629S (2025G>T 2027A>C), R630Q (2028A>C 2029G>A 2030A>G), L645M (2073T>A), D646N (2076G>A), P647K (2079C>A 2080C>A 2081T>A), E648F (2082G>T 2083A>T 2084G>T), L650F (2088C>T), H655L (2104A>T 2105T>C), H656S (2106C>T 2107A>C 2108T>C), M660I (2120G>C), S663V (2127T>G 2128C>T 2129T>G), D665S (2133G>T 2134A>C), D666S (2136G>T 2137A>C), E667D (2141A>T), H668A (2142C>G 2143A>C 2144T>C), E670Q (2148G>C), I671N (2152T>A), N673E (2157A>G 2159C>A), K674M (2161A>T 2162A>G), R676V (2166A>G 2167G>T), V677N (2169G>A 2170T>A 2171C>T), N678D (2172A>G 2174C>T), M679L (2175A>T), Y681V (2181T>G 2182A>T 2183T>C), R682K (2185G>A), D685S (2193G>A 2194A>G), A686P (2196G>C), E687Q (2199G>C 2201A>G), R690A (2208C>G 2209G>C 2210A>T), I701V (2241A>G 2243A>C), A710S (2268G>A 2269C>G 2270A>C), S711T (2271T>A), K716E (2286A>G), K717L (2289A>T 2290A>T 2291A>G), A719D (2296C>A 2297A>T), T722R (2304A>C 2305C>G 2306A>C), E723N (2307G>A 2309G>T), M724L (2310A>C 2312G>T), R725E (2313A>G 2314G>A), T726Q (2316A>C 2317C>A 2318T>A), F728K (2322T>A 2323T>A 2324T>A), H730Y (2328C>T), S731K (2331T>A 2332C>A 2333T>G), C732A (2334T>G 2335G>C 2336C>G), I733L (2337A>C), D734Q (2340G>C 2342T>G), V737A (2350T>C 2351T>A), I741T (2362T>C 2363A>G), F749R (2385T>A 2386T>G 2387T>A), N763R (2428A>G 2429C>G), S766A (2436T>G), S766_K767insG (2438_2439insGGT), K767T (2440A>C 2441A>T), E768D (2444G>C), P769K (2445C>A 2446C>A 2447A>G), E771R (2451G>A 2452A>G), S772N (2454T>A 2455C>A 2456A>T), K775I (2464A>T 2465A>C), I776E (2466A>G 2467T>A 2468T>A), Y778M (2472T>A 2473A>T 2474C>G), H779I (2475C>A 2476A>T), K780G (2478A>G 2479A>G 2480A>T), V782K (2484G>A 2485T>A 2486T>G), E783D (2489A>C), Y784K (2490T>A 2492T>A), I785M (2495C>G), K786I (2497A>T 2498G>C), D787Q (2499G>C 2501T>G), T788M (2503C>T 2504A>G), W790N (2508T>A 2509G>A 2510G>T), S791D (2511A>G 2512G>A), Y794F (2521A>T 2522T>C), A795I (2523G>A 2524C>T 2525A>C), R797K (2530G>A 2531A>G), I806L (2556A>T 2558A>G), M807I (2561G>A), V812T (2574G>A 2575T>C 2576T>C), F813M (2577T>A 2579T>G), G814S (2580G>A 2582A>T), G815A (2584G>C), I816V (2586A>G), L820I (2598T>A 2600G>T), A821T (2601G>A 2603G>T), W823L (2607T>C 2608G>T 2609G>C), Q824K (2610C>A), S825A (2613A>G 2614G>C 2615T>A), N826A (2616A>G 2617A>C 2618C>A), S827T (2619T>A), F828S (2623T>C), N830L (2628A>T 2629A>T 2630T>A), S832D (2634T>G 2635C>A 2636G>T), N852K (2696T>G), K853T (2698A>C 2699G>T), K853_L854insH (2699_2700insCAT), L854F (2700C>T), R855A (2703A>G 2704G>C), N856K (2708T>A), L857S (2709C>T 2710T>C 2711T>C), V859L (2715G>C 2717T>A), S862A (2724A>G 2725G>C 2726T>G), S863A (2727T>G 2729T>A), Q864D (2730C>G 2732A>C), S865N (2733T>A 2734C>A 2735A>C), L866N (2736T>A 2737T>A 2738G>C), A867H (2739G>C 2740C>A 2741A>T), Q868V (2742C>G 2743A>T 2744A>G), D869_W870insR (2747_2748insCGA), A872E (2755C>A 2756T>A), E873Q (2757G>C 2759A>G), Q879E (2775C>G 2777G>A), K882H (2784A>C 2786G>C), L890I (2808C>A), G891A (2812G>C), M907L (2859A>C 2861G>C), H914R (2880C>A 2881A>G 2882C>A), A916M (2886G>A 2887C>T 2888T>G), T923S (2907A>T 2909A>C), A927E (2920C>A 2921C>G), N928H (2922A>C), R930I (2928C>A 2929G>T 2930A>T), S931C (2932C>G), K932N (2936A>T), Y935F (2944A>T 2945C>T), R938T (2953G>C 2954A>C), E939Q (2955G>C), I942L (2964A>T 2966T>G), I948V (2982A>G 2984C>G), P961S (3021C>T 3023T>A), E972D (3056G>T), F975Y (3064T>A), T976S (3066A>T 3068A>C), Q980M (3078C>A 3079A>T 3080A>G), C984V (3090T>G 3091G>T 3092T>G), T990K (3109C>A 3110A>G), R993K (3117C>A 3118G>A 3119T>G), L994K (3120C>A 3121T>A 3122T>A), G995S (3123G>T 3124G>C), F999V (3135T>G 3137T>G), M1002Y (3144A>T 3145T>A 3146G>T), V1012E (3175T>A 3176T>A), E1015T (3183G>A 3184A>C), E1017G (3190A>G), N1019D (3195A>G 3197C>T), S1021N (3201T>A 3202C>A 3203C>T), S1022A (3204A>G 3205G>C 3206T>A), K1024A (3210A>G 3211A>C 3212G>T), I1025L (3213A>C), C1026S (3216T>A), Q1029V (3225C>G 3226A>T 3227A>T), S1030A (3228T>G 3230T>A), G1037S (3249G>A 3251G>T), S1050T (3289G>C), V1057C (3309G>T 3310T>G 3311G>C), G1068D (3343G>A 3344C>T), V1069T (3345G>A 3346T>C 3347A>C), R1073K (3357C>A 3358G>A 3359T>G), F1075Y (3364T>A 3365T>C), I1076V (3366A>G 3368C>G), S1077A (3369T>G), I1083V (3387A>G 3389C>G), Y1084F (3391A>T 3392C>T), E1086D (3398A>T), V1096I (3426G>A 3428C>T), A1098V (3433C>T 3434A>G), D1104E (3452T>A), I1105L (3453A>C 3455T>G), S1108H (3462T>C 3463C>A 3464A>C), I1109L (3465A>C 3467T>G), R1112K (3475G>A 3476A>G), T1114E (3480A>G 3481C>A), Q1115R (3483C>A 3484A>G 3485G>A), L1116F (3486C>T 3488T>C), N1120K (3500C>A), Y1123A (3507T>G 3508A>C 3509C>T), Y1124F (3511A>T), Q1130L (3528C>T 3529A>T), L1132M (3534T>A), R1143K (3567C>A 3568G>A 3569G>A), K1144R (3570A>C 3571A>G 3572A>C), Q1146P (3577A>C 3578A>C), P1153D (3597C>G 3598C>A), E1155Q (3603G>C), T1162S (3624A>T 3626A>T), V1163A (3628T>C 3629T>C), H1164K (3630C>A 3632T>A), D1169E (3647T>A), M1179E (3675A>G 3676T>A), C1186S (3697G>C 3698T>C), G1187N (3699G>A 3700G>A 3701A>C), D1189E (3707T>G), D1190E (3710T>G), D1191P (3711G>C 3712A>C 3713C>A), I1192V (3714A>G), N1199G (3735A>G 3736A>G), C1202A (3744T>G 3745G>C 3746C>A), L1204I (3750C>A 3752G>T), I1207V (3759A>G 3761C>G), M1208L (3762A>C 3764G>T), R1209K (3765C>A 3766G>A 3767T>G), S1210M (3769G>T 3770C>G), K1212R (3774A>C 3775A>G 3776G>A), I1213V (3777A>G 3779C>G), R1217K (3789C>A 3790G>A 3791T>A), C1219A (3795T>G 3796G>C 3797T>A), E1220I (3798G>A 3799A>T 3800A>T), D1222N (3804G>A), C1233F (3838G>T 3839T>C), Y1236F (3847A>T), G1238A (3853G>C 3854C>A), M1252L (3894A>T), M1254L (3900A>C 3902G>T), M1256S (3906A>T 3907T>C 3908G>T), Q1263E (3927C>G), S1264A (3930T>G), T1265Q (3933A>C 3934C>A 3935T>A), G1266P (3936G>C 3937G>C 3938C>A), E1267D (3941A>C), G1268S (3942G>T 3943G>C 3944A>G), E1269Q (3945G>C), E1272L (3954G>C 3955A>T 3956A>T), D1276N (3966G>A), E1281D (3983A>T), V1288S (4002G>T 4003T>C), I1290V (4008A>G 4010T>A), K1292Q (4014A>C), I1294T (4021T>C 4022C>A), H1295F (4023C>T 4024A>T), T1344C (4170A>T 4171C>G 4172A>T), L1348A (4182T>G 4183T>C 4184G>A), A1351V (4192C>T 4193T>A), I1352L (4194A>C 4196A>C), D1359N (4215G>A), S1363H (4227T>C 4228C>A), L1367V (4239T>G), N1370S (4249A>G), G1371S (4251G>A 4253G>C), A1374S (4260G>T 4262T>A), V1378E (4273T>A), V1393I (4317G>A 4319T>C), E1405A (4354A>C 4355A>T), Y1406F (4357A>T 4358C>T), S1409D (4365T>G 4366C>A 4367A>T), K1410E (4368A>G), D1415E (4385T>G), A1421T (4401G>A), D1435N (4443G>A 4445T>C), V1437L (4449G>T), I1476V (4566A>G 4568T>C), A1477K (4569G>A 4570C>A 4571T>A), R1479K (4576G>A 4577G>A), A1480S (4578G>T), I1481V (4581A>G 4583A>G), V1483I (4587G>A), S1486N (4597G>A), L1506M (4656T>A 4658A>G), Q1509R (4665C>A 4666A>G), S1512D (4674T>G 4675C>A), N1515S (4683A>T 4684A>C 4685T>C), H1518R (4692C>A 4693A>G 4694T>A), L1522A (4704C>G 4705T>C 4706G>T), I1526V (4716A>G), N1538E (4752A>G 4754T>G), E1539K (4755G>A), V1540T (4758G>A 4759T>C), Q1541K (4761C>A), Q1543L (4768A>T 4769A>G), H1563Q (4829C>A), I1572V (4854A>G), M1576I (4868G>T), P1583V (4887C>G 4888C>T 4889A>G), T1588V (4902A>G 4903C>T 4904T>A), M1592Y (4914A>T 4915T>A 4916G>T), V1595I (4923G>A 4925G>T), R1607Q (4960G>A 4961T>G), V1608A (4963T>C 4964G>C), A1609T (4965G>A), D1611E (4973C>A), K1612S (4975A>G 4976G>T), L1613F (4977C>T), L1615N (4983T>A 4984T>A 4985A>T), Q1618L (4993A>T 4994G>T), T1619G (4995A>G 4996C>G 4997G>A), D1622K (5004G>A 5006T>G), E1623Q (5007G>C 5009A>G), G1625N (5013G>A 5014G>A 5015T>C), F1626L (5016T>C), R1627P (5019A>C 5020G>C 5021A>C), A1628I (5022G>A 5023C>T 5024T>C), V1629M (5025G>A), F1631M (5031T>A 5033T>G), S1633T (5037T>A 5039C>T), L1635Y (5044T>A 5045G>T), V1637L (5049G>C 5051G>T), Q1639S (5055C>A 5056A>G 5057G>T), E1640Q (5058G>C), I1642L (5064A>T 5066T>G), A1643T (5067G>A), K1644E (5070A>G), S1649E (5085T>G 5086C>A 5087A>G), A1651I (5091G>A 5092C>T 5093C>T), L1652G (5094C>G 5095T>G), T1653V (5097A>G 5098C>T 5099A>T), I1654L (5100A>T 5102T>G), C1655M (5103T>A 5104G>T 5105T>G), M1657L (5109A>C 5111G>A), A1658S (5112G>T), K1659D (5115A>G 5117A>T), A1661K (5121G>A 5122C>A 5123T>A), I1662N (5125T>A), S1663T (5127T>A 5129C>A), D1668P (5142G>C 5143A>C 5144C>T), T1674del (5160_5162delACA), R1678E (5172C>G 5173G>A 5174T>G), D1679G (5176A>G), L1680V (5178C>G 5180G>C), N1681H (5181A>C), W1682E (5184T>G 5185G>A 5186G>A), N1686D (5196A>G 5198C>T), I1687V (5199A>G 5201T>A), G1688F (5202G>T 5203G>T), A1696S (5226G>T 5228C>A), H1697N (5229C>A), Y1698Q (5232T>C 5234C>A), T1699Q (5235A>C 5236C>A 5237G>A), D1701S (5241G>T 5242A>C 5243T>A), E1702S (5244G>T 5245A>C), Y1704N (5250T>A), I1706L (5256A>C 5258T>G), K1713R (5277A>C 5278A>G 5279G>C), K1714R (5281A>G 5282G>A), S1715T (5283T>A 5285C>A), N1716T (5287A>C 5288T>A), M1731I (5333G>A), M1736L (5346A>T), S1737A (5349T>G), M1740Q (5358A>C 5359T>A), A1742H (5364G>C 5365C>A 5366C>T), K1744R (5370A>C 5371A>G 5372G>C), N1745K (5375T>G), V1746S (5376G>A 5377T>G 5378G>T), A1748E (5383C>A), G1749V (5386G>T 5387A>T), H1751D (5391C>G), V1752G (5395T>G 5396A>G), N1755S (5403A>T 5404A>C), V1757T (5409G>A 5410T>C 5411T>C), V1766I (5436G>A 5438C>A), G1767A (5440G>C 5441C>T), M1774F (5460A>T 5462G>T), R1779H (5475A>C 5476G>A 5477G>T), C1782R (5484T>A 5486T>G), V1783A (5488T>C 5489G>C), E1784T (5490G>A 5491A>C 5492G>A), I1786L (5496A>C 5498A>T), I1793M (5519T>G), G1794A (5521G>C 5522A>T), N1795G (5523A>G 5524A>G 5525T>G), V1807A (5560T>C), F1810L (5570T>G), Q1812K (5574C>A 5576A>G)

| | Begin | End | Coverage | Score | Concordance | Matches | Identities | I/D/M/F* | Stop Codons |
|---|---|---|---|---|---|---|---|---|---|
| **NT** | **23** | **5589** | **77.0%** | **3552** | **32.4%** | **5524 (99.1%)** | **3681 (66.0%)** | **9/43** | |

Codon mutations:

GCT2GCA (146T>A), TTC7TGG (160T>G 161C>G), TCC8AGT (162T>A 163C>G 164C>T), AAG9_GCC16del (165_188delAAGGTTGCTGGTGTTGATGACGCC), AGT17CCT (189A>C 190G>C), TCT18CCT (192T>C), TCT19GCT (195T>G), TCC20TCT (200C>T), ACC21TCA (201A>T 203C>A), ACT22CCT (204A>C), GGT26GCA (217G>C 218T>A), CTT28GTT (222C>G), TCA29TCT (227A>T), AAT30TCT (228A>T 229A>C), ATT32TTT (234A>T), TCC33TCT (239C>T), AAA34TCT (240A>T 241A>C 242A>T), TTT35CTT (243T>C), AAA37AAG (251A>G), GCG38ACT (252G>A 254G>T), GTT39CTC (255G>C 257T>C), AGT40TCC (258A>T 259G>C 260T>C), ATT42GTT (264A>G), ACC43AAT (268C>A 269C>T), ACT44AAA (271C>A 272T>A), ATG45GTT (273A>G 275G>T), AAG48ACC (283A>C 284G>C), TCA50GTA (288T>G 289C>T), CAT52CAC (296T>C), CTT53ATA (297C>A 299T>A), CTG56GTC (306C>G 308G>C), CCA58CCG (314A>G), TCT59CCA (315T>C 317T>A), GTG60GTT (320G>T), ACT61ATG (322C>T 323T>G), GAT62GCA (325A>C 326T>A), GTG63GCA (328T>C 329G>A), TCC64GCA (330T>G 332C>A), TCT66TCA (338T>A), GTG68GCC (343T>C 344G>C), GCC69TCT (345G>T 347C>T), ACT70ACA (350T>A), TGT71TGC (353T>C), AGT72AAT (355G>A), TCA73GCT (357T>G 359A>T), CTT74GTA (360C>G 362T>A), AAT75GAA (363A>G 365T>A), AAA76AGA (367A>G), TTG78TTA (374G>A), GAC79GAT (377C>T), AAA80AAG (380A>G), GGT82GCA (385G>C 386T>A), GCA83AAA (387G>A 388C>A), CTT84ATA (390C>A 392T>A), ATA85ATG (395A>G), GAT86GAA (398T>A), CCA87CCC (401A>C), TTT88TTC (404T>C), CTT89ATC (405C>A 407T>C), AAA90AAG (410A>G), GCG91GGT (412C>G 413G>T), TAC92TAT (416C>T), TCT93GAT (417T>G 418C>A), TTC94TTT (422C>T), TTG95GTT (423T>G 425G>T), TCC96AAA (426T>A 427C>A 428C>A), ACA97ACT (431A>T), TAC99TAT (437C>T), TCT101ACT (441T>A), ATC102ATA (446C>A), AAG103GTA (447A>G 448A>T 449G>A), ATG105CTT (453A>C 455G>T), GTT106GTA (458T>A), TTA107CTT (459T>C 461A>T), TTG109ATG (465T>A), TTT110GTT (468T>G), GAG111TTA (471G>T 472A>T 473G>A), TTG113GTC (477T>G 479G>C), TCC114AAG (480T>A 481C>A 482C>G), AAT115GAC (483A>G 485T>C), GTT117TTC (489G>T 491T>C), AAA118ATC (493A>T 494A>C), TTA119CTA (495T>C), TTT121TAC (502T>A 503T>C), GCT122GAA (505C>A 506T>A), GTG124TTA (510G>T 512G>A), AAT126GAA (516A>G 518T>A), AGT128GTT (522A>G 523G>T), GAA129GAG (527A>G), GAT130TCA (528G>T 529A>C 530T>A), GTA131TTA (531G>T), GTT134GTG (542T>G), GTA135TTT (543G>T 545A>T), TAT136ATA (546T>A 547A>T 548T>A), GGT137GCT (550G>C), CTT139TTA (555C>T 557T>A), GTA140GTT (560A>T), ATA143TTG (567A>T 569A>G), ACA144GCC (570A>G 572A>C), TTG145ATT (573T>A 575G>T), TTA146TTG (578A>G), GTG147CTA (579G>C 581G>A), CTT148TTA (582C>T 584T>A), CTA149CTG (587A>G), TTT150TAT (589T>A), GTT151GTA (593T>A), TGT152TGC (596T>C), AGT154GTA (600A>G 601G>T 602T>A), GTT157GTA (611T>A), GAG158ACG (612G>A 613A>C), GGT159CCT (615G>C 616G>C), GTT160TTT (618G>T), TCT161AAT (621T>A 622C>A), AAC162AAT (626C>T), ACT163GTA (627A>G 628C>T 629T>A), GTC164GTT (632C>T), AAA165GTT (633A>G 634A>T 635A>T), AAT166ACT (637A>C), ATC167TTG (639A>T 641C>G), TTG169TCA (646T>C 647G>A), ATA170GTT (648A>G 650A>T), GTC171GTT (653C>T), AAC173AAT (659C>T), ATG174TTT (660A>T 662G>T), TTT175TTC (665T>C), TCA176TCT (668A>T), AGT177TCA (669A>T 670G>C 671T>A), CTA178TTG (672C>T 674A>G), TAC179TAT (677C>T), AAA186ACA (697A>C), GCC188TCT (702G>T 704C>T), GAA189GGG (706A>G 707A>G), CGA190AAG (708C>A 709G>A 710A>G), TTC191TTT (713C>T), ACC192TCC (714A>T), ATG193CAG (717A>C 718T>A), GTG194CTG (720G>C), GCC195GCG (725C>G), GCT197GCA (731T>A), AGT198GCA (732A>G 733G>C 734T>A), ATG199GTA (735A>G 737G>A), GGG202GGT (746G>T), TCA204AGT (750T>A 751C>G 752A>T), GTT205TTA (753G>T 755T>A), TCA206ACG (756T>A 758A>G), ACG210ACT (770G>T), TCT211AGC (771T>A 772C>G 773T>C), CAG212CAA (776G>A), TTG213CTT (777T>C 779G>T), ATG214TTA (780A>T 782G>A), ACA216TCA (786A>T), ATA217TTG (789A>T 791A>G), GGG221GGT (803G>T), ATT222ATC (806T>C), TCG223TCA (809G>A), ACG224ACT (812G>T), TTG225TTA (815G>A), GTG226GTA (818G>A), TTC227TTT (821C>T), ATT228GTT (822A>G), GCA229GCT (827A>T), GTG230GTT (830G>T), GGT232GGG (836T>G), GGC236GGT (848C>T), TTG237CTT (849T>C 851G>T), AGC238TCA (852A>T 853G>C 854C>A), TTG241CTG (861T>C), TCC242TCA (866C>A), ATT244ATC (872T>C), CTC245TTG (873C>T 875C>G), TAT246TAC (878T>C), TCG247TCT (881G>T), ACA248GCT (882A>G 884A>T), GCT252GCG (896T>G), CAG253CAA (899G>A), CAA257CAG (911A>G), CTA258CTG (914A>G), ACT260CAG (918A>C 919C>A 920T>G), CTT261TTG (921C>T 923T>G), CGT263AGG (927C>A 929T>G), AAT264AAC (932T>C), AAA266AAG (938A>G), TGC268TGT (944C>T), ACC269ACA (947C>A), TCC270TCT (950C>T), CAG271CAA (953G>A), GCT272GCC (956T>C), GAA278GAG (974A>G), ATA280GTG (978A>G 980A>G), GTC281GTT (983C>T), GGC282AGC (984G>A), ACC283ATT (988C>T 989C>T), GGA285GGC (995A>C), TTC286TTT (998C>T), GTG290GTA (1010G>A), GCT293GCG (1019T>G), AGT295AAT (1024G>A), GCA296GCT (1028A>T), ACG297ACC (1031G>C), CTG298TTA (1032C>T 1034G>A), ACT300ACC (1040T>C), CTG302CTT (1046G>T), GAG304CAG (1050G>C), CAG307ACG (1059C>A 1060A>C), GAG308GAA (1064G>A), GAT310TCA (1068G>T 1069A>C 1070T>A), GCA311GAG (1072C>A 1073A>G), GTG312GTT (1076G>T), TGT313GCT (1077T>G 1078G>C), GAT314GAA (1082T>A), CCT315CCA (1085T>A), GCA316TCT (1086G>T 1088A>T), ACG317ATG (1090C>T), GAG320GAA (1100G>A), AAC321AAT (1103C>T), GCC323GCA (1109C>A), AAG325CCT (1113A>C 1114A>C 1115G>T), GCT326GCA (1118T>A), TTC327TTT (1121C>T), CCT328CCA (1124T>A), ACC329ATT (1126C>T 1127C>T), AAA330AAG (1130A>G), AAC332CAA (1134A>C 1136C>A), CTG334TTA (1140C>T 1142G>A), AGG335AAA (1144G>A 1145G>A), GAA336GAG (1148A>G), GAA337CAG (1149G>C 1151A>G), GCT338GCC (1154T>C), CTC339GTC (1155C>G), AAG340AAA (1160G>A), ATC341ATA (1163C>A), GCA346GCC (1178A>C), GCG351GCA (1193G>A), TTC353TTT (1199C>T), ATG354TTG (1200A>T), AGT359TCT (1215A>T 1216G>C), ATT362ATA (1226T>A), GCG363ACG (1227G>A), CAG364CAA (1232G>A), TTA365CTA (1233T>C), GAA366GAT (1238A>T), GTT368TTG (1242G>T 1244T>G), TAT369GTT (1245T>G 1246A>T), GAA371GAG (1253A>G), AGT372GCC (1254A>G 1255G>C 1256T>C), TGT373AGC (1257T>A 1259T>C), AGG374CAA (1260A>C 1261G>A 1262G>A), CAC375CAT (1265C>T), ATG376TCT (1266A>T 1267T>C 1268G>T), GGC377GGT (1271C>T), CAG380CAA (1280G>A), CGT382AGG (1284C>A 1286T>G), GAA384GAG (1292A>G), TTT386TTC (1298T>C), GTA388GTG (1304A>G), TTC391TTT (1313C>T), GGG392GGA (1316G>A), TCC393GCT (1317T>G 1319C>T), GGG395GGA (1325G>A), TCT399TCA (1337T>A), ACA400ACG (1340A>G), TCC401TGC (1342C>G), TTA404CTT (1350T>C 1352A>T), TTT405TTC (1355T>C), GTG409GTT (1367G>T), TTG410CTC (1368T>C 1370G>C), AGA412GCA (1374A>G 1375G>C), GAG415GAA (1385G>A), AAA417AAG (1391A>G), CTC418CTG (1394C>G), AAT419AAC (1397T>C), CGG420CGC (1400G>C), CTA421TTG (1401C>T 1403A>G), GCA423GCG (1409A>G), AAG426AAA (1418G>A), AGG427AGA (1421G>A), GCT435GCA (1445T>A), CAC436CAT (1448C>T), GCT439GCA (1457T>A), ATC440GTG (1458A>G 1460C>G), CTG441CTT (1463G>T), GAC444GAT (1472C>T), GGA446GGT (1478A>T), GCA447GCT (1481A>T), TTG448CTT (1482T>C 1484G>T), CGA449CGG (1487A>G), GCT452GCA (1496T>A), TGC455TGT (1505C>T), CAA456CAG (1508A>G), GAT457GAC (1511T>C), ATT458ATA (1514T>A), AAA459AAG (1517A>G), GAT463GAC (1529T>C), TCA466TCT (1538A>T), ACA467ACC (1541A>C), CAA468CAG (1544A>G), CCA469CCT (1547A>T), CCC471CCT (1553C>T), TTA472CTT (1554T>C 1556A>T), GCC475GCT (1565C>T), GTT477GTC (1571T>C), GAC479GAT (1577C>T), GGC481GGA (1583C>A), CAT483CAC (1589T>C), TTC484TTT (1592C>T), ACC485ACA (1595C>A), TCC486TCA (1598C>A), TAT488TAC (1604T>C), ATA489ATC (1607A>C), ACA492ACC (1616A>C), CTG495TTG (1623C>T), ATC496ATA (1628C>A), TCA497TCT (1631A>T), CCT499CCA (1637T>A), GCC500ACT (1638G>A 1640C>T), CAA501CAG (1643A>G), GGT503GGG (1649T>G), CTT504CTA (1652T>A), ACA505ACG (1655A>G), CCA507CCT (1661A>T), GCT509GCA (1667T>A), GAG511GAA (1673G>A), CGG512AGG (1674C>A), AGA513AGG (1679A>G), AGG514AGA (1682G>A), GAC515GAT (1685C>T), ATC516ATT (1688C>T), CTG517TTA (1689C>T 1691G>A), GTG518GTT (1694G>T), TGT520TGC (1700T>C), ATG521AGA (1702T>G 1703G>A), GAG523GTG (1708A>T), GAG525GAT (1715G>T), TCC527GAT (1719T>G 1720C>A 1721C>T), ACT528CCA (1722A>C 1724T>A), GAA529AAT (1725G>A 1727A>T), GAT530GCT (1729A>C), CCT531CCA (1733T>A), ACG532ACA (1736G>A), GGA533GCC (1738G>C 1739A>C), CTC535CTA (1745C>A), AGA536GTT (1746A>G 1747G>T 1748A>T), AAC538GAT (1752A>G 1754C>T), ATA539GTG (1755A>G 1757A>G), GAG541GAA (1763G>A), GCT547GCC (1781T>C), ACT549ACA (1787T>A), CAC550CAT (1790C>T), AGG551AGA (1793G>A), AAT552GCT (1794A>G 1795A>C), TTG553TTA (1799G>A), ACC554TCC (1800A>T), AGT556GAT (1806A>G 1807G>A), GAT557GAC (1811T>C), CTT558CTC (1814T>C), GAG560CAA (1818G>C 1820G>A), GTA563GTG (1829A>G), GCC564GCA (1832C>A), AAA565AAG (1835A>G), TGT566TGC (1838T>C), CAG567CGG (1840A>G), GTA568GTG (1844A>G), CAT569CAC (1847T>C), GCA570TAT (1848G>T 1849C>A 1850A>T), GAA571GAG (1853A>G), GTA572GTG (1856A>G), AAG574AGC (1861A>G 1862G>C), TTG576CTT (1866T>C 1868G>T), TTT577TAT (1870T>A), GAA578AAT (1872G>A 1874A>T), GCT579ACT (1875G>A), GAA580ACA (1878G>A 1879A>C), TCT581TCA (1883T>A), GGC582GGA (1886C>A), ATA583TTG (1887A>T 1889A>G), AGT584GCT (1890A>G 1891G>C), CCT585GAC (1893C>G 1894C>A 1895T>C), ATA587GTT (1899A>G 1901A>T), GTT590GTC (1910T>C), CAA591AAT (1911C>A 1913A>T), TCT593GCT (1917T>G), GCA594GTT (1921C>T 1922A>T), GAT596GAA (1928T>A), ATA598CTG (1932A>C 1934A>G), GCG599GAT (1936C>A 1937G>T), GTG601TTC (1941G>T 1943G>C), GAT602CAA (1944G>C 1946T>A), GGG603GAT (1948G>A 1949G>T), GCT604GCC (1952T>C), AGA605del (1953_1955delAGA), TTG606TTA (1958G>A), CGC607CAA (1960G>A 1961C>A), ACC608GTT (1962A>G 1963C>T 1964C>T), AAG609TTG (1965A>T 1966A>T), CAA610CCT (1969A>C 1970A>T), GAT611CCT (1971G>C 1972A>C), GAA612GAG (1976A>G), ATA614GCA (1980A>G 1981T>C), TTG615ACT (1983T>A 1984T>C 1985G>T), GTG616TAT (1986G>T 1987T>A 1988G>T), CCC617GAG (1989C>G 1990C>A 1991C>G), ACT618CAG (1992A>C 1993C>A 1994T>G), GTT619ACA (1995G>A 1996T>C 1997T>A), GTG620ATT (1998G>A 2000G>T), AGT621ACA (2002G>C 2003T>A), AAT623GCT (2007A>G 2008A>C), GAT624GAA (2012T>A), AGA625AAA (2014G>A), GTG626ATT (2016G>A 2018G>T), ATC627GTT (2019A>G 2021C>T), TAT628TAC (2024T>C), GCA629TCC (2025G>T 2027A>C), AGA630CAG (2028A>C 2029G>A 2030A>G), GAG631GAA (2033G>A), ACG633ACA (2039G>A), GTA634GTT (2042A>T), GCA636GCC (2048A>C), TTG637CTC (2049T>C 2051G>C), GCA640GCT (2060A>T), CAG642CAA (2066G>A), GGC643GGG (2069C>G), TTG645ATG (2073T>A), GAC646AAC (2076G>A), CCT647AAA (2079C>A 2080C>A 2081T>A), GAG648TTT (2082G>T 2083A>T 2084G>T), GAA649GAG (2087A>G), CTT650TTT (2088C>T), TTT651TTC (2093T>C), CCT652CCA (2096T>A), CAT655CTC (2104A>T 2105T>C), CAT656TCC (2106C>T 2107A>C 2108T>C), GCC659GCT (2117C>T), ATG660ATC (2120G>C), TCT663GTG (2127T>G 2128C>T 2129T>G), CTT664CTA (2132T>A), GAT665TCT (2133G>T 2134A>C), GAT666TCT (2136G>T 2137A>C), GAA667GAT (2141A>T), CAT668GCC (2142C>G 2143A>C 2144T>C), GAG670CAG (2148G>C), ATT671AAT (2152T>A), TTC672TTT (2156C>T), AAC673GAA (2157A>G 2159C>A), AAA674ATG (2161A>T 2162A>G), AGA676GTA (2166A>G 2167G>T), GTC677AAT (2169G>A 2170T>A 2171C>T), AAC678GAT (2172A>G 2174C>T), ATG679TTG (2175A>T), TAT681GTC (2181T>G 2182A>T 2183T>C), AGA682AAA (2185G>A), GAT685AGT (2193G>A 2194A>G), GCT686CCT (2196G>C), GAA687CAG (2199G>C 2201A>G), CGA690GCT (2208C>G 2209G>C 2210A>T), GTA693GTG (2219A>G), GAC699GAT (2237C>T), ATA701GTC (2241A>G 2243A>C), CAC703CAT (2249C>T), TTT704TTC (2252T>C), GCA710AGC (2268G>A 2269C>G 2270A>C), TCA711ACA (2271T>A), ATC713ATT (2279C>T), GAG715GAA (2285G>A), AAG716GAG (2286A>G), AAA717TTG (2289A>T 2290A>T 2291A>G), CTA718CTG (2294A>G), GCA719GAT (2296C>A 2297A>T), GTG720GTC (2300G>C), CAG721CAA (2303G>A), ACA722CGC (2304A>C 2305C>G 2306A>C), GAG723AAT (2307G>A 2309G>T), ATG724CTT (2310A>C 2312G>T), AGG725GAG (2313A>G 2314G>A), ACT726CAA (2316A>C 2317C>A 2318T>A), GGA727GGG (2321A>G), TTT728AAA (2322T>A 2323T>A 2324T>A), GCA729GCG (2327A>G), CAT730TAT (2328C>T), TCT731AAG (2331T>A 2332C>A 2333T>G), TGC732GCG (2334T>G 2335G>C 2336C>G), ATT733CTT (2337A>C), GAT734CAG (2340G>C 2342T>G), GCA735GCT (2345A>T), GTT737GCA (2350T>C 2351T>A), GAC738GAT (2354C>T), ACC739ACA (2357C>A), CTC740CTT (2360C>T), ATA741ACG (2362T>C 2363A>G), ATC743ATT (2369C>T), GAG744GAA (2372G>A), CCA747CCG (2381A>G), TTT749AGA (2385T>A 2386T>G 2387T>A), GTT750GTA (2390T>A), TTT752TTC (2396T>C), TTA753CTG (2397T>C 2399A>G), ACA755ACT (2405A>T), GCA756GCT (2408A>T), AGA759CGC (2415A>C 2417A>C), TGC762TGT (2426C>T), AAC763AGG (2428A>G 2429C>G), GTA765GTT (2435A>T), TCA766GCA (2436T>G), TCA766_AAA767insGGT (2438_2439insGGT), AAA767ACT (2440A>C 2441A>T), GAG768GAC (2444G>C), CCA769AAG (2445C>A 2446C>A 2447A>G), GAA771AGA (2451G>A 2452A>G), TCA772AAT (2454T>A 2455C>A 2456A>T), ATC774ATT (2462C>T), AAA775ATC (2464A>T 2465A>C), ATT776GAA (2466A>G 2467T>A 2468T>A), TAC778ATG (2472T>A 2473A>T 2474C>G), CAT779ATT (2475C>A 2476A>T), AAA780GGT (2478A>G 2479A>G 2480A>T), GTT782AAG (2484G>A 2485T>A 2486T>G), GAA783GAC (2489A>C), TAT784AAA (2490T>A 2492T>A), ATC785ATG (2495C>G), AAG786ATC (2497A>T 2498G>C), GAT787CAG (2499G>C 2501T>G), ACA788ATG (2503C>T 2504A>G), TGG790AAT (2508T>A 2509G>A 2510G>T), AGC791GAC (2511A>G 2512G>A), CTT792TTG (2514C>T 2516T>G), TAT794TTC (2521A>T 2522T>C), GCA795ATC (2523G>A 2524C>T 2525A>C), CTG796TTG (2526C>T), AGA797AAG (2530G>A 2531A>G), CTC799CTT (2537C>T), GCA802GCC (2546A>C), GGT803GGA (2549T>A), ATA806TTG (2556A>T 2558A>G), ATG807ATA (2561G>A), GCT808GCA (2564T>A), GGA811GGC (2573A>C), GTT812ACC (2574G>A 2575T>C 2576T>C), TTT813ATG (2577T>A 2579T>G), GGA814AGT (2580G>A 2582A>T), GGC815GCC (2584G>C), ATT816GTT (2586A>G), ACA817ACT (2591A>T), GCA818GCT (2594A>T), TTT819TTC (2597T>C), TTG820ATT (2598T>A 2600G>T), GCG821ACT (2601G>A 2603G>T), TGG823CTC (2607T>C 2608G>T 2609G>C), CAG824AAG (2610C>A), AGT825GCA (2613A>G 2614G>C 2615T>A), AAC826GCA (2616A>G 2617A>C 2618C>A), TCT827ACT (2619T>A), TTC828TCC (2623T>C), CCC829CCT (2627C>T), AAT830TTA (2628A>T 2629A>T 2630T>A), GCA831GCT (2633A>T), TCG832GAT (2634T>G 2635C>A 2636G>T), GGA835GGG (2645A>G), GGT836GGC (2648T>C), GTG837GTA (2651G>A), GAA840GAG (2660A>G), ATA844ATC (2672A>C), AGT846AGC (2678T>C), AAG848AAA (2684G>A), AAC849AAT (2687C>T), AAG850AAA (2690G>A), GGT851GGG (2693T>G), AAT852AAG (2696T>G), AAG853ACT (2698A>C 2699G>T), AAG853_CTC854insCAT (2699_2700insCAT), CTC854TTC (2700C>T), AGA855GCA (2703A>G 2704G>C), AAT856AAA (2708T>A), CTT857TCC (2709C>T 2710T>C 2711T>C), CTC858TTG (2712C>T 2714C>G), GTT859CTA (2715G>C 2717T>A), GGT860GGG (2720T>G), AGT862GCG (2724A>G 2725G>C 2726T>G), TCT863GCA (2727T>G 2729T>A), CAA864GAC (2730C>G 2732A>C), TCA865AAC (2733T>A 2734C>A 2735A>C), TTG866AAC (2736T>A 2737T>A 2738G>C), GCA867CAT (2739G>C 2740C>A 2741A>T), CAA868GTG (2742C>G 2743A>T 2744A>G), GAT869GAC (2747T>C), GAT869_TGG870insCGA (2747_2748insCGA), GCT871GCA (2753T>A), GCT872GAA (2755C>A 2756T>A), GAA873CAG (2757G>C 2759A>G), GTA877GTT (2771A>T), CAG879GAA (2775C>G 2777G>A), AGC880AGT (2780C>T), CTC881CTG (2783C>G), AAG882CAC (2784A>C 2786G>C), AAT884AAC (2792T>C), GTG886GTT (2798G>T), GTG887GTT (2801G>T), TTA888TTG (2804A>G), CTT890ATT (2808C>A), GGT891GCT (2812G>C), GTG894GTC (2822G>C), TAC895TAT (2825C>T), TTT896TTC (2828T>C), GGC898GGT (2834C>T), ACC899ACG (2837C>G), TAC900TAT (2840C>T), GTG901GTT (2843G>T), ATA906ATT (2858A>T), ATG907CTC (2859A>C 2861G>C), ACC908ACT (2864C>T), GTA909GTG (2867A>G), GCT910GCA (2870T>A), GCA912GCC (2876A>C), TTC913TTT (2879C>T), CAC914AGA (2880C>A 2881A>G 2882C>A), AAT915AAC (2885T>C), GCT916ATG (2886G>A 2887C>T 2888T>G), CGA917AGG (2889C>A 2891A>G), GGT919GGC (2897T>C), CCA921CCT (2903A>T), ACA923TCC (2907A>T 2909A>C), ATC925ATT (2915C>T), GCC927GAG (2920C>A 2921C>G), AAT928CAT (2922A>C), TCT929TCC (2927T>C), CGA930ATT (2928C>A 2929G>T 2930A>T), TCT931TGT (2932C>G), AAA932AAT (2936A>T), GTT933GTG (2939T>G), CAA934CAG (2942A>G), TAC935TTT (2944A>T 2945C>T), AAC936AAT (2948C>T), AGA938ACC (2953G>C 2954A>C), GAA939CAA (2955G>C), ATT942TTG (2964A>T 2966T>G), ATT943ATA (2969T>A), GGC945GGT (2975C>T), GAC947GAT (2981C>T), ATC948GTG (2982A>G 2984C>G), CTG950TTG (2988C>T), CGC952AGG (2994C>A 2996C>G), GTT953GTA (2999T>A), GGT954GGG (3002T>G), CCA959CCC (3017A>C), AAG960AAA (3020G>A), CCT961TCA (3021C>T 3023T>A), GAC962GAT (3026C>T), ATC963ATA (3029C>A), CGT964AGA (3030C>A 3032T>A), CAC966CAT (3038C>T), CCA968CCT (3044A>T), AGA969CGC (3045A>C 3047A>C), AGG970AGA (3050G>A), GAG972GAT (3056G>T), TGC974TGT (3062C>T), TTC975TAC (3064T>A), ACA976TCC (3066A>T 3068A>C), CAA980ATG

| | | | | | | | | | |
|---|---|---|---|---|---|---|---|---|---|
| | (3078C>A 3079A>T 3080A>G), GGA981GGC (3083A>C), TGT984GTG (3090T>G 3091G>T 3092T>G), AGA985AGG (3095A>G), GTT987GTA (3101T>A), GCG988GCT (3104G>T), TCT989TCA (3107T>A), ACA990AAG (3109C>A 3110A>G), CCA992CCC (3116A>C), CGT993AAG (3117C>A 3118G>A 3119T>G), CTT994AAA (3120C>A 3121T>A 3122T>A), GGC995TCC (3123G>T 3124G>C), TTA997TTG (3131A>G), TTT999GTG (3135T>G 3137T>G), CTC1000CTT (3140C>T), AAG1001AAA (3143G>A), ATG1002TAT (3144A>T 3145T>A 3146G>T), GGA1009GGC (3167A>C), ACA1011ACC (3173A>C), GTT1012GAA (3175T>A 3176T>A), GTG1014GTC (3182G>C), GAA1015ACA (3183G>A 3184A>C), GAA1017GGA (3190A>G), CTG1018TTA (3192C>T 3194G>A), AAC1019GAT (3195A>G 3197C>T), TCC1020TCG (3200C>G), TCC1021AAT (3201T>A 3202C>A 3203C>T), AGT1022GCA (3204A>G 3205G>C 3206T>A), AAG1024GCT (3210A>G 3211A>C 3212G>T), ATT1025CTT (3213A>C), TGC1026AGC (3216T>A), TCT1027TCA (3221T>A), CAA1029GTT (3225C>G 3226A>T 3227A>T), TCT1030GCA (3228T>G 3230T>A), GAA1032GAG (3236A>G), ATA1035ATT (3245A>T), GGG1037AGT (3249G>A 3251G>T), GAA1038GAG (3254A>G), GGT1040GGA (3260T>A), GAC1041GAT (3263C>T), GGC1043GGT (3269C>T), ACG1044ACT (3272G>T), TTG1045CTT (3273T>C 3275G>T), CTA1046CTC (3278A>C), CTG1047TTA (3279C>T 3281G>A), TTA1048TTG (3284A>G), AGT1050ACT (3289G>C), GTT1051GTG (3293T>G), AAT1053AAC (3299T>C), CAA1055CAG (3305A>G), CCT1056CCG (3308T>G), GTG1057TGC (3309G>T 3310T>G 3311G>C), GGC1060GGA (3320C>A), ATC1061ATA (3323C>A), CAC1062CAT (3326C>T), TCA1065TCT (3335A>T), GGC1068GAT (3343G>A 3344C>T), GTA1069ACC (3345G>A 3346T>C 3347A>C), GCA1070GCT (3350A>T), GCT1071GCA (3353T>A), GAA1072GAG (3356A>G), CGT1073AAG (3357C>A 3358G>A 3359T>G), GGA1074GGT (3362A>T), TTT1075TAC (3364T>A 3365T>C), ATC1076GTG (3366A>G 3368C>G), TCT1077GCT (3369T>G), AAT1079AAC (3377T>C), GCT1080GCA (3380T>A), GCT1082GCA (3386T>A), ATC1083GTG (3387A>G 3389C>G), TAC1084TTT (3391A>T 3392C>T), AGG1085AGA (3395G>A), GAA1086GAT (3398A>T), CTA1088TTG (3402C>T 3404A>G), GAG1089GAA (3407G>A), CCA1092CCC (3416A>C), ACT1093ACG (3419T>G), GGG1094GGA (3422G>A), CCG1095CCC (3425G>C), GTC1096ATT (3426G>A 3428C>T), GCA1098GTG (3433C>T 3434A>G), GCA1099GCT (3437A>T), GAT1104GAA (3452T>A), ATT1105CTG (3453A>C 3455T>G), CTG1106TTG (3456C>T), TCA1108CAC (3462T>C 3463C>A 3464A>C), ATT1109CTG (3465A>C 3467T>G), AGA1110AGG (3470A>G), AGC1111TCC (3471A>T 3472G>C), AGA1112AAG (3475G>A 3476A>G), GAA1113GAG (3479A>G), ACA1114GAA (3480A>G 3481C>A), CAG1115AGA (3483C>A 3484A>G 3485G>A), CTT1116TTC (3486C>T 3488T>C), GAA1119GAG (3497A>G), AAC1120AAA (3500C>A), CAA1121CAG (3503A>G), TAC1123GCT (3507T>G 3508A>C 3509C>T), TAC1124TTC (3511A>T), CTT1125TTG (3513C>T 3515T>G), GGA1126GGC (3518A>C), ACA1127ACT (3521A>T), GTT1128GTA (3524T>A), CCA1129CCG (3527A>G), CAG1130TTG (3528C>T 3529A>T), GAG1131GAA (3533G>A), TTG1132ATG (3534T>A), GCC1133GCT (3539C>T), GCC1134GCT (3542C>T), GTT1136GTG (3548T>G), CCC1137CCA (3551C>A), AAG1139AAA (3557G>A), ACC1140ACA (3560C>A), ACT1141ACC (3563T>C), CTG1142CTT (3566G>T), CGG1143AAA (3567C>A 3568G>A 3569G>A), AAA1144CGC (3570A>C 3571A>G 3572A>C), AGC1145AGT (3575C>T), CAA1146CCC (3577A>C 3578A>C), TTG1147CTG (3579T>C), GAA1149GAG (3587A>G), GCA1150GCT (3590A>T), TTC1151TTT (3593C>T), GGA1152GGG (3596A>G), CCT1153GAT (3597C>G 3598C>A), GCA1154GCT (3602A>T), GAG1155CAG (3603G>C), ACA1156ACT (3608A>T), GCA1157GCC (3611A>C), ATT1160ATC (3620T>C), ACA1162TCT (3624A>T 3626A>T), GTT1163GCC (3628T>C 3629T>C), CAT1164AAA (3630C>A 3632T>A), AGA1167AGG (3641A>G), GGT1168GGA (3644T>A), GAT1169GAA (3647T>A), GGT1170GGG (3650T>G), GAC1172GAT (3656C>T), GGG1177GGT (3671G>T), GTA1178GTC (3674A>C), ATG1179GAG (3675A>G 3676T>A), AAA1180AAG (3680A>G), TAC1181TAT (3683C>T), TGT1186TCC (3697G>C 3698T>C), GGA1187AAC (3699G>A 3700G>A 3701A>C), GAT1189GAG (3707T>G), GAT1190GAG (3710T>G), GAC1191CCA (3711G>C 3712A>C 3713C>A), ATT1192GTT (3714A>G), GCC1193GCA (3719C>A), AAA1194AAG (3722A>G), CTA1195CTG (3725A>G), GCA1196GCC (3728A>C), TTC1197TTT (3731C>T), GAA1198GAG (3734A>G), AAT1199GGT (3735A>G 3736A>G), CTC1200CTT (3740C>T), AAG1201AAA (3743G>A), TGC1202GCA (3744T>G 3745G>C 3746C>A), TCG1203TCC (3749G>C), CTG1204ATT (3750C>A 3752G>T), CTA1205CTT (3755A>T), ATC1207GTG (3759A>G 3761C>G), ATG1208CTT (3762A>C 3764G>T), CGT1209AAG (3765C>A 3766G>A 3767T>G), AGC1210ATG (3769G>T 3770C>G), AAG1212CGA (3774A>C 3775A>G 3776G>A), ATC1213GTG (3777A>G 3779C>G), CCT1214CCC (3782T>C), GGA1216GGG (3788A>G), CGT1217AAA (3789C>A 3790G>A 3791T>A), CCA1218CCT (3794A>T), TGT1219GCA (3795T>G 3796G>C 3797T>A), GAA1220ATT (3798G>A 3799A>T 3800A>T), AGG1221AGA (3803G>A), GAT1222AAT (3804G>A), GGA1229GGG (3827A>G), TGT1233TTC (3838G>T 3839T>C), TAC1235TAT (3845C>T), TAT1236TTT (3847A>T), GGC1238GCA (3853G>C 3854C>A), GAG1240GAA (3860G>A), CTG1241TTG (3861C>T), ACA1243ACG (3869A>G), TCT1244TCA (3872T>A), TGC1245TGT (3875C>T), GGA1246GGT (3878A>T), CCC1248CCT (3884C>T), TTC1249TTT (3887C>T), ATG1252TTG (3894A>T), GGG1253GGA (3899G>A), ATG1254CTT (3900A>C 3902G>T), GGG1255GGT (3905G>T), ATG1256TCT (3906A>T 3907T>C 3908G>T), AAG1258AAA (3914G>A), AGA1259AGG (3917A>G), CAA1263GAA (3927C>G), TCC1264GCC (3930T>G), ACT1265CAA (3933A>C 3934C>A 3935T>A), GGC1266CCA (3936G>C 3937G>C 3938C>A), GAA1267GAC (3941A>C), GGA1268TCG (3942G>T 3943G>C 3944A>G), GAG1269CAG (3945G>C), AGA1270AGG (3950A>G), GTG1271GTT (3953G>T), GAA1272CTT (3954G>C 3955A>T 3956A>T), AGA1275AGG (3965A>G), GAC1276AAC (3966G>A), ACT1277ACA (3971T>A), GTA1279GTC (3977A>C), GAA1281GAT (3983A>T), CTA1286TTG (3996C>T 3998A>G), GAT1287GAC (4001T>C), GTG1288TCG (4002G>T 4003T>C), ATT1290GTA (4008A>G 4010T>A), AAA1292CAA (4014A>C), GGC1293GGA (4019C>A), ATC1294ACA (4021T>C 4022C>A), CAT1295TTT (4023C>T 4024A>T), GTG1296GTT (4028G>T), GAT1297GAC (4031T>C), CTG1298CTT (4034G>T), ACC1300ACA (4040C>A), CAA1302CAG (4046A>G), TGC1303TGT (4049C>T), GCC1304GCG (4052C>G), CGC1308CGG (4064C>G), CTC1309TTG (4065C>T 4067C>G), AAA1313AAG (4079A>G), GGC1316GGG (4088C>G), AGA1318AGG (4094A>G), AGG1321CGC (4101A>C 4103G>C), CTC1322CTG (4106C>G), TTT1323TTC (4109T>C), GAG1324GAA (4112G>A), ATA1325ATC (4115A>C), CTT1326TTG (4116C>T 4118T>G), TAC1330TAT (4130C>T), TTG1333CTG (4137T>C), GTC1334GTG (4142C>G), AGG1335CGG (4143A>C), AAG1336AAA (4148G>A), TAT1337TAC (4151T>C), TTT1338TTC (4154T>C), GAT1340GAC (4160T>C), TTC1341TTT (4163C>T), TCA1342TCT (4166A>T), GCC1343GCT (4169C>T), ACA1344TGT (4170A>T 4171C>G 4172A>T), TTG1348GCA (4182T>G 4183T>C 4184G>A), AAT1350AAC (4190T>C), GCT1351GTA (4192C>T 4193T>A), ATA1352CTC (4194A>C 4196A>C), CCA1353CCT (4199A>T), AAA1355AAG (4205A>G), GTT1356GTA (4208T>A), ATT1358ATC (4214T>C), GAT1359AAT (4215G>A), CCT1360CCA (4220T>A), ACA1361ACG (4223A>G), AGT1362AGC (4226T>C), TCT1363CAT (4227T>C 4228C>A), TTG1367GTG (4239T>G), TTG1368CTG (4242T>C), GCA1369GCT (4247A>T), AAT1370AGT (4249A>G), GGG1371AGC (4251G>A 4253G>C), AGA1373AGG (4259A>G), GCT1374TCA (4260G>T 4262T>A), GTG1375GTA (4265G>A), TCA1376TCT (4268A>T), GAC1377GAT (4271C>T), GTG1378GAG (4273T>A), TTT1380TTC (4280T>C), GCT1382GCC (4286T>C), TAT1384TAC (4292T>C), TCC1385TCG (4295C>G), CCT1392CCC (4316T>C), GTT1393ATC (4317G>A 4319T>C), GCT1395GCA (4325T>A), CAG1397CAA (4331G>A), TTG1402CTA (4344T>C 4346G>A), GAA1405GCT (4354A>C 4355A>T), TAC1406TTT (4357A>T 4358C>T), TAC1407TAT (4361C>T), GGA1408GGT (4364A>T), TCA1409GAT (4365T>G 4366C>A 4367A>T), AAG1410GAG (4368A>G), GGC1412GGG (4376C>G), GAT1415GAG (4385T>G), TCC1416TCA (4388C>A), CGA1419AGG (4395C>A 4397A>G), CAT1420CAC (4400T>C), GCA1421ACA (4401G>A), CTT1422TTA (4404C>T 4406T>A), GCA1426GCC (4418A>C), TGT1428TGC (4424T>C), TAC1430TAT (4430C>T), ACA1431ACC (4433A>C), CTG1432CTC (4436G>C), TGC1433TGT (4439C>T), GAT1435AAC (4443G>A 4445T>C), AAA1436AAG (4448A>G), GTG1437TTG (4449G>T), AGG1439CGC (4455A>C 4457G>C), TTG1440CTA (4458T>C 4460G>A), GTT1441GTG (4463T>G), GGG1442GGT (4466G>T), GGC1443GGA (4469C>A), CCA1445CCC (4475A>C), TCA1446TCT (4478A>T), GGA1447GGT (4481A>T), GCA1449GCT (4487A>T), CTA1450TTG (4488C>T 4490A>G), ACG1451ACT (4493G>T), ATC1453ATA (4499C>A), TTC1454TTT (4502C>T), TCT1456TCA (4508T>A), CTC1457TTG (4509C>T 4511C>G), CTC1458CTA (4514C>A), CGA1464AGG (4530C>A 4532A>G), TAT1465TAC (4535T>C), GCC1466GCG (4538C>G), TTG1470CTG (4548T>C), TTA1471TTG (4553A>G), AGA1472AGG (4556A>G), CCC1474CCA (4562C>A), ATT1476GTC (4566A>G 4568T>C), GCT1477AAA (4569G>A 4570C>A 4571T>A), AGG1479AAA (4576G>A 4577G>A), GCT1480TCT (4578G>T), ATA1481GTG (4581A>G 4583A>G), GGA1482GGC (4586A>C), GTT1483ATT (4587G>A), AAA1484AAG (4592A>G), AGT1486AAT (4597G>A), GAT1487GAC (4601T>C), CTA1491TTG (4611C>T 4613A>G), GCT1494GCA (4622T>A), GTT1495GTA (4625T>A), GGA1497GGT (4631A>T), GAC1499GAT (4637C>T), AAT1500AAC (4640T>C), CTT1501TTG (4641C>T 4643T>G), GCT1503GCG (4649T>G), GTA1504GTT (4652A>T), TTA1506ATG (4656T>A 4658A>G), CTC1508TTG (4662C>T 4664C>G), CAG1509AGG (4665C>A 4666A>G), TAT1511TAC (4673T>C), TCT1512GAT (4674T>G 4675C>A), CTG1513CTA (4679G>A), AAT1515TCC (4683A>T 4684A>C 4685T>C), ATA1516ATT (4688A>T), CAT1518AGA (4692C>A 4693A>G 4694T>A), GAG1519GAA (4697G>A), TTA1520CTG (4698T>C 4700A>G), CTG1522GCT (4704C>G 4705T>C 4706G>T), GTC1523GTT (4709C>T), AAT1524AAC (4712T>C), GTA1525GTC (4715A>C), ATC1526GTC (4716A>G), CTT1531TTG (4731C>T 4733T>G), AAA1533AAG (4739A>G), TCG1534TCA (4742G>A), GTT1537GTA (4751T>A), AAT1538GAG (4752A>G 4754T>G), GAG1539AAG (4755G>A), GTA1540ACA (4758G>A 4759T>C), CAA1541AAA (4761C>A), CAA1543CTG (4768A>T 4769A>G), CTA1548CTC (4784A>C), ACT1549ACC (4787T>C), CTG1551CTC (4793G>C), AGA1553AGG (4799A>G), TTT1555TTC (4805T>C), AAA1556AAG (4808A>G), GCT1559GCA (4817T>A), CTT1560TTG (4818C>T 4820T>G), TAC1562TAT (4826C>T), CAC1563CAA (4829C>A), GCT1565GCA (4835T>A), CCT1566CCA (4838T>A), AAG1568AAA (4844G>A), ATC1572GTC (4854A>G), ATG1576ATT (4868G>T), TAT1577TAC (4871T>C), ATA1579ATT (4877A>T), AGA1580AGG (4880A>G), CCT1581CCA (4883T>A), GCC1582GCA (4886C>A), CCA1583GTG (4887C>G 4888C>T 4889A>G), GGT1584GGA (4892T>A), GAC1587GAT (4901C>T), ACT1588GTA (4902A>G 4903C>T 4904T>A), CAG1589CAA (4907G>A), GCC1590GCT (4910C>T), ATG1592TAT (4914A>T 4915T>A 4916G>T), GTG1595ATT (4923G>A 4925G>T), GGA1598GGC (4934A>C), ATA1599ATC (4937A>C), CGT1600AGA (4938C>A 4940T>A), GCT1602GCC (4946T>C), CAC1605CAT (4955C>T), GGT1606GGG (4958T>G), CGT1607CAG (4960G>A 4961T>G), GTG1608GCC (4963T>C 4964G>C), GCT1609ACT (4965G>A), GAC1611GAA (4973C>A), AAG1612AGT (4975A>G 4976G>T), CTT1613TTT (4977C>T), GTC1614GTA (4982C>A), TTA1615AAT (4983T>A 4984T>A 4985A>T), CAG1618CTT (4993A>T 4994G>T), ACG1619GGA (4995A>G 4996C>G 4997G>A), GCG1620GCT (5000G>T), TTG1621CTC (5001T>C 5003G>C), GAT1622AAG (5004G>A 5006T>G), GAA1623CAG (5007G>C 5009A>G), AGG1624AGA (5012G>A), GGT1625AAC (5013G>A 5014G>A 5015T>C), TTC1626CTC (5016T>C), AGA1627CCC (5019A>C 5020G>C 5021A>C), GCT1628ATC (5022G>A 5023C>T 5024T>C), GTG1629ATG (5025G>A), TTT1631ATG (5031T>A 5033T>G), CCT1632CCC (5036T>C), TCC1633ACT (5037T>A 5039C>T), TAT1634TAC (5042T>C), TTG1635TAT (5044T>A 5045G>T), GTG1637CTT (5049G>C 5051G>T), CAG1639AGT (5055C>A 5056A>G 5057G>T), GAA1640CAA (5058G>C), ATT1642TTG (5064A>T 5066T>G), GCA1643ACA (5067G>A), AAG1644GAG (5070A>G), GTA1645GTC (5075A>C), GGG1647GGA (5081G>A), TCA1649GAG (5085T>G 5086C>A 5087A>G), GCC1651ATT (5091G>A 5092C>T 5093C>T), CTG1652GGG (5094C>G 5095T>G), ACA1653GTT (5097A>G 5098C>T 5099A>T), ATT1654TTG (5100A>T 5102T>G), TGT1655ATG (5103T>A 5104G>T 5105T>G), GAA1656GAG (5108A>G), ATG1657CTA (5109A>C 5111G>A), GCA1658TCA (5112G>T), AAA1659GAT (5115A>G 5117A>T), GCA1660GCT (5120A>T), GCT1661AAA (5121G>A 5122C>A 5123T>A), ATT1662AAT (5125T>A), TCC1663ACA (5127T>A 5129C>A), TAT1664TAC (5132T>C), ACG1665ACC (5135G>C), CCA1666CCT (5138A>T), TTG1667TTA (5141G>A), GAC1668CCT (5142G>C 5143A>C 5144C>T), GCA1669GCT (5147A>T), GGT1670GGG (5150T>G), GAG1671GAA (5153G>A), ATC1673ATT (5159C>T), ACA1674del (5160_5162delACA), CGT1678GAG (5172C>G 5173G>A 5174T>G), GAT1679GGT (5176A>G), CTG1680GTC (5178C>G 5180G>C), AAT1681CAT (5181A>C), TGG1682GAA (5184T>G 5185G>A 5186G>A), GCA1684GCG (5192A>G), CCA1685CCT (5195A>T), AAC1686GAT (5196A>G 5198C>T), ATT1687GTA (5199A>G 5201T>A), GGT1688TTT (5202G>T 5203G>T), TTT1689TTC (5207T>C), TCA1691AGT (5211T>A 5212C>G 5213A>T), GCA1692GCT (5216A>T), CGT1693CGG (5219T>G), GCC1696TCA (5226G>T 5228C>A), CAC1697AAC (5229C>A), TAC1698CAA (5232T>C 5234C>A), ACG1699CAA (5235A>C 5236C>A 5237G>A), GAT1701TCA (5241G>T 5242A>C 5243T>A), GAA1702TCA (5244G>T 5245A>C), GGG1703GGT (5249G>T), TAC1704AAC (5250T>A), ATT1705ATC (5255T>C), ATT1706CTG (5256A>C 5258T>G), AAC1708AAT (5264C>T), TGT1709TGC (5267T>C), ACA1710ACT (5270A>T), GGT1711GGA (5273T>A), GCA1712GCC (5276A>C), AAG1713CGC (5277A>C 5278A>G 5279G>C), AAG1714AGA (5281A>G 5282G>A), TCC1715ACA (5283T>A 5285C>A), AAT1716ACA (5287A>C 5288T>A), GTT1718GTG (5294T>G), GGT1720GGG (5300T>G), CCA1721CCT (5303A>T), AAC1723AAT (5309C>T), GAC1726GAT (5318C>T), ATG1731ATA (5333G>A), CCG1733CCC (5339G>C), ATG1736TTG (5346A>T), TCA1737GCA (5349T>G), GCT1738GCA (5354T>A), ATA1739ATC (5357A>C), ATG1740CAG (5358A>C 5359T>A), GCC1742CAT (5364G>C 5365C>A 5366C>T), AAG1744CGC (5370A>C 5371A>G 5372G>C), AAT1745AAG (5375T>G), GTG1746AGT (5376G>A 5377T>G 5378G>T), TTG1747CTG (5379T>C), GCA1748GAA (5383C>A), GGA1749GTT (5386G>T 5387A>T), GGA1750GGT (5390A>T), CAT1751GAT (5391C>G), GTA1752GGG (5395T>G 5396A>G), GCA1753GCT (5399A>T), AAT1755TCT (5403A>T 5404A>C), GTT1757ACC (5409G>A 5410T>C 5411T>C), GTG1759GTT (5417G>T), GGA1761GGC (5423A>C), GGA1763GGT (5429A>T), GCT1765GCA (5435T>A), GTC1766ATA (5436G>A 5438C>A), GGC1767GCT (5440G>C 5441C>T), ATT1769ATC (5447T>C), GCG1772GCA (5456G>A), CTG1773TTG (5457C>T), ATG1774TTT (5460A>T 5462G>T), GCC1775GCA (5465C>A), TTA1776TTG (5468A>G), GCA1777GCC (5471A>C), ACA1778ACT (5474A>T), AGG1779CAT (5475A>C 5476G>A 5477G>T), TAT1781TAC (5483T>C), TGT1782AGG (5484T>A 5486T>G), GTG1783GCC (5488T>C 5489G>C), GAG1784ACA (5490G>A 5491A>C 5492G>A), ATA1786CTT (5496A>C 5498A>T), ATA1787ATT (5501A>T), GTT1788GTG (5504T>G), AGA1789CGC (5505A>C 5507A>C), TTG1790CTA (5508T>C 5510G>A), CGC1791CGA (5513C>A), ACA1792ACC (5516A>C), ATT1793ATG (5519T>G), GGA1794GCT (5521G>C 5522A>T), AAT1795GGG (5523A>G 5524A>G 5525T>G), CTT1797CTG (5531T>G), GAC1798GAT (5534C>T), CTC1799CTG (5537C>G), AAT1800AAC (5540T>C), TAT1802TAC (5546T>C), CCT1803CCC (5549T>C), GGC1805GGT (5555C>T), GTG1807GCG (5560T>C), CAG1808CAA (5564G>A), TAT1809TAC (5567T>C), TTT1810TTG (5570T>G), CTT1811CTC (5573T>C), CAA1812AAG (5574C>A 5576A>G) | | | | | | | | |
| helicase (YP_001976154.1) | 1 | 98 | 100% | 654 | 97.5% | 98 (100%) | 95 (96.9%) | 0/0/0/0 | 0 |
| Protein mutations: | S5C (1342C>G), R16A (1374A>G 1375G>C), I44V (1458A>G 1460C>G) | | | | | | | | |
| Codon mutations: | TCT3TCA (1337T>A), ACA4ACG (1340A>G), TCC5TGC (1342C>G), TTA8CTT (1350T>C 1352A>T), TTT9TTC (1355T>C), GTG13GTT (1367G>T), TTG14CTC (1368T>C 1370G>C), AGA16GCA (1374A>G 1375G>C), GAG19GAA (1385G>A), AAA21AAG (1391A>G), CTC22CTG (1394C>G), AAT23AAC (1397T>C), CGG24CGC (1400G>C), CTA25TTG (1401C>T 1403A>G), GCA27GCG (1409A>G), AAG30AAA (1418G>A), AGG31AGA (1421G>A), GCT39GCA (1445T>A), CAC40CAT (1448C>T), GCT43GCA (1457T>A), ATC44GTG (1458A>G 1460C>G), CTG45CTT (1463G>T), GAC48GAT (1472C>T), GGA50GGT (1478A>T), GCA51GCT (1481A>T), TTG52CTT (1482T>C 1484G>T), CGA53CGG (1487A>G), GCT56GCA (1496T>A), TGC59TGT (1505C>T), CAA60CAG (1508A>G), GAT61GAC (1511T>C), ATT62ATA (1514T>A), AAA63AAG (1517A>G), GAT67GAC (1529T>C), TCA70TCT (1538A>T), ACA71ACC (1541A>C), CAA72CAG (1544A>G), CCA73CCT (1547A>T), CCC75CCT (1553C>T), TTA76CTT (1554T>C 1556A>T), GCC79GCT (1565C>T), GTT81GTC (1571T>C), GAC83GAT (1577C>T), GGC85GGA (1583C>A), CAT87CAC (1589T>C), TTC88TTT (1592C>T), ACC89ACA (1595C>A), TCC90TCA (1598C>A), TAT92TAC (1604T>C), ATA93ATC (1607A>C), ACA96ACC (1616A>C) | | | | | | | | |

| | Begin | End | Coverage | Score | Concordance | Matches | Identities | I/D/M/F* | Stop Codons |
|---|---|---|---|---|---|---|---|---|---|
| **NT** | **23** | **5589** | **77.0%** | **3552** | **32.4%** | **5524 (99.1%)** | **3681 (66.0%)** | **9/43** | |
| RNA-dependent RNA polymerase (YP_001976153.1) | 1 | 249 | 100% | 1552 | 88.4% | 249 (100%) | 210 (84.3%) | 0/0/0/0 | 0 |
| Protein mutations: | T40C (4170A>T 4171C>G 4172A>T), L44A (4182T>G 4183T>C 4184G>A), A47V (4192C>T 4193T>A), I48L (4194A>C 4196A>C), D55N (4215G>A), S59H (4227T>C 4228C>A), L63V (4239T>G), N66S (4249A>G), G67S (4251G>A 4253G>C), A70S (4260G>T 4262T>A), V74E (4273T>A), V89I (4317G>A 4319T>C), E101A (4354A>C 4355A>T), Y102F (4357A>T 4358C>T), S105D (4365T>G 4366C>A 4367A>T), K106E (4368A>G), D111E (4385T>G), A117T (4401G>A), D131N (4443G>A 4445T>C), V133L (4449G>T), I172V (4566A>G 4568T>C), A173K (4569G>A 4570C>A 4571T>A), R175K (4576G>A 4577G>A), A176S (4578G>T), I177V (4581A>G 4583A>G), V179I (4587G>A), S182N (4597G>A), L202M (4656T>A 4658A>G), Q205R (4665C>A 4666A>G), S208D (4674T>G 4675C>A), N211S (4683A>T 4684A>C 4685T>C), H214R (4692C>A 4693A>G 4694T>A), L218A (4704C>G 4705T>C 4706G>T), I222V (4716A>G), N234E (4752A>G 4754T>G), E235K (4755G>A), V236T (4758G>A 4759T>C), Q237K (4761C>A), Q239L (4768A>T 4769A>G) | | | | | | | | |
| Codon mutations: | CGC4CGG (4064C>G), CTC5TTG (4065C>T 4067C>G), AAA9AAG (4079A>G), GGC12GGG (4088C>G), AGA14AGG (4094A>G), AGG17CGC (4101A>C 4103G>C), CTC18CTG (4106C>G), TTT19TTC (4109T>C), GAG20GAA (4112G>A), ATA21ATC (4115A>C), CTT22TTG (4116C>T 4118T>G), TAC26TAT (4130C>T), TTG29CTG (4137T>C), GTC30GTG (4142C>G), AGG31CGG (4143A>C), AAG32AAA (4148G>A), TAT33TAC (4151T>C), TTT34TTC (4154T>C), GAT36GAC (4160T>C), TTC37TTT (4163C>T), TCA38TCT (4166A>T), GCC39GCT (4169C>T), ACA40TGT (4170A>T 4171C>G 4172A>T), TTG44GCA (4182T>G 4183T>C 4184G>A), AAT46AAC (4190T>C), GCT47GTA (4192C>T 4193T>A), ATA48CTC (4194A>C 4196A>C), CCA49CCT (4199A>T), AAA51AAG (4205A>G), GTT52GTA (4208T>A), ATT54ATC (4214T>C), GAT55AAT (4215G>A), CCT56CCA (4220T>A), ACA57ACG (4223A>G), AGT58AGC (4226T>C), TCT59CAT (4227T>C 4228C>A), TTG63GTG (4239T>G), TTG64CTG (4242T>C), GCA65GCT (4247A>T), AAT66AGT (4249A>G), GGG67AGC (4251G>A 4253G>C), AGA69AGG (4259A>G), GCT70TCA (4260G>T 4262T>A), GTG71GTA (4265G>A), TCA72TCT (4268A>T), GAC73GAT (4271C>T), GTG74GAG (4273T>A), TTT76TTC (4280T>C), GCT78GCC (4286T>C), TAT80TAC (4292T>C), TCC81TCG (4295C>G), CCT88CCC (4316T>C), GTT89ATC (4317G>A 4319T>C), GCT91GCA (4325T>A), CAG93CAA (4331G>A), TTG98CTA (4344T>C 4346G>A), GAA101GCT (4354A>C 4355A>T), TAC102TTT (4357A>T 4358C>T), TAC103TAT (4361C>T), GGA104GGT (4364A>T), TCA105GAT (4365T>G 4366C>A 4367A>T), AAG106GAG (4368A>G), GGC108GGG (4376C>G), GAT111GAG (4385T>G), TCC112TCA (4388C>A), CGA115AGG (4395C>A 4397A>G), CAT116CAC (4400T>C), GCA117ACA (4401G>A), CTT118TTA (4404C>T 4406T>A), GCA122GCC (4418A>C), TGT124TGC (4424T>C), TAC126TAT (4430C>T), ACA127ACC (4433A>C), CTG128CTC (4436G>C), TGC129TGT (4439C>T), GAT131AAC (4443G>A 4445T>C), AAA132AAG (4448A>G), GTG133TTG (4449G>T), AGG135CGC (4455A>C 4457G>C), TTG136CTA (4458T>C 4460G>A), GTT137GTG (4463T>G), GGG138GGT (4466G>T), GGC139GGA (4469C>A), CCA141CCC (4475A>C), TCA142TCT (4478A>T), GGA143GGT (4481A>T), GCA145GCT (4487A>T), CTA146TTG (4488C>T 4490A>G), ACG147ACT (4493G>T), ATC149ATA (4499C>A), TTC150TTT (4502C>T), TCT152TCA (4508T>A), CTC153TTG (4509C>T 4511C>G), CTC154CTA (4514C>A), CGA160AGG (4530C>A 4532A>G), TAT161TAC (4535T>C), GCC162GCG (4538C>G), TTG166CTG (4548T>C), TTA167TTG (4553A>G), AGA168AGG (4556A>G), CCC170CCA (4562C>A), ATT172GTC (4566A>G 4568T>C), GCT173AAA (4569G>A 4570C>A 4571T>A), AGG175AAA (4576G>A 4577G>A), GCT176TCT (4578G>T), ATA177GTG (4581A>G 4583A>G), GGA178GGC (4586A>C), GTT179ATT (4587G>A), AAA180AAG (4592A>G), AGT182AAT (4597G>A), GAT183GAC (4601T>C), CTA187TTG (4611C>T 4613A>G), GCT190GCA (4622T>A), GTT191GTA (4625T>A), GGA193GGT (4631A>T), GAC195GAT (4637C>T), AAT196AAC (4640T>C), CTT197TTG (4641C>T 4643T>G), GCT199GCG (4649T>G), GTA200GTT (4652A>T), TTA202ATG (4656T>A 4658A>G), CTC204TTG (4662C>T 4664C>G), CAG205AGG (4665C>A 4666A>G), TAT207TAC (4673T>C), TCT208GAT (4674T>G 4675C>A), CTG209CTA (4679G>A), AAT211TCC (4683A>T 4684A>C 4685T>C), ATA212ATT (4688A>T), CAT214AGA (4692C>A 4693A>G 4694T>A), GAG215GAA (4697G>A), TTA216CTG (4698T>C 4700A>G), CTG218GCT (4704C>G 4705T>C 4706G>T), GTC219GTT (4709C>T), AAT220AAC (4712T>C), GTA221GTC (4715A>C), ATC222GTC (4716A>G), CTT227TTG (4731C>T 4733T>G), AAA229AAG (4739A>G), TCG230TCA (4742G>A), GTT233GTA (4751T>A), AAT234GAG (4752A>G 4754T>G), GAG235AAG (4755G>A), GTA236ACA (4758G>A 4759T>C), CAA237AAA (4761C>A), CAA239CTG (4768A>T 4769A>G), CTA244CTC (4784A>C), ACT245ACC (4787T>C), CTG247CTC (4793G>C), AGA249AGG (4799A>G) | | | | | | | | |

*: Inserts / Deletes / Misaligned / Frameshifts

#### Analysis details

This analysis was performed with panviral2.64

### NGS Details (UN59): Torradovirus marchitezum (segment RNA 2)

#### Assembly

| | |
|---|---|
| **Coverage Length** | 4656 (1 contig(s)) |
| **Depth Of Coverage** | 3539.8 |
| **Number Of Reads** | 128651 |
| **Reads Per Million** | 2796.76 rpm (after QC) |
| **Ambiguities** | 0 |
| **Assembly Method** | de novo + reference guided assembly |
| **Consensus Caller** | Bcf Tools |

#### Coverage Map

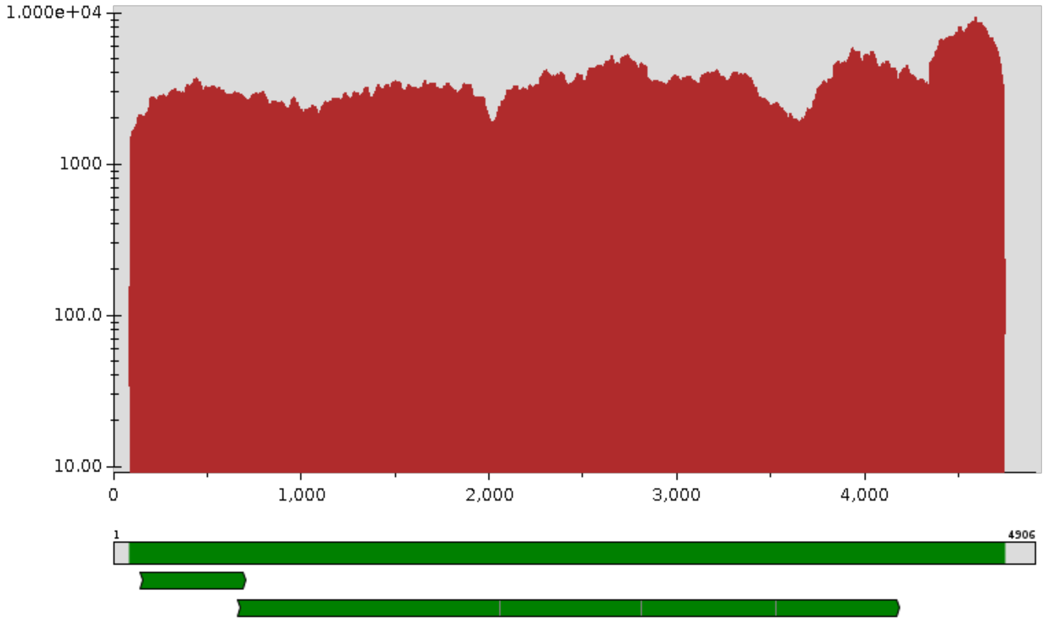

#### Assignment

| | |
|---|---|
| **Type** | Torradovirus marchitezum (Taxonomy ID: 3048376) |
| **Reference Genome** | NC_010988.1 |
| **NT Identity (%)** | 64.4751 |
| **AA Identity (%)** | 67.9739 |
| **Number Of Stop Codons** | 2 |
| **Number Of CDS** | 2 |

#### Alignment

| | |
|---|---|
| **Alignment Score** | 2600.0 (NT) + 6793.0 (AA) = 9393.0 |
| **Concordance (%)** | 50.8032 |

| **Alignment Method** | Global, seeded, nucleotide + amino acids (AGA) |
| --- | --- |

#### Genome Region

Sequence starts at position 89 and ends at position 4744 relative to NC_010988.1 reference sequence.

#### Alignment Detailed Statistics

| | Begin | End | Coverage | Score | Concordance | Matches | Identities | I/D/M/F* | Stop Codons |
| --- | --- | --- | --- | --- | --- | --- | --- | --- | --- |
| **NT** | **89** | **4744** | **94.9%** | **2600** | **28.6%** | **4608 (96.7%)** | **3040 (63.8%)** | **107/48** | |
| Mutations: | 101G>T, 106C>A, 109A>C, 110T>C, 111T>C, 118T>C, 120C>A, 121T>A, 122T>A, 124G>C, 128T>A, 129G>A, 130T>A, 132G>T, 133_134insCACTCT, 135C>T, 137T>G, 144A>T, 147C>T, 150T>C, 151G>A, 153G>T, 154C>A, 155G>A, 156T>G, 157T>C, 159G>C, 162T>C, 165A>C, 166T>C, 167C>A, 168T>A, 169G>A, 174C>G, 177A>T, 182C>A, 183T>A, 186T>C, 190C>A, 198T>A, 201C>A, 202A>G, 204C>A, 207G>T, 209A>C, 210T>C, 214A>G, 220T>A, 221C>G, 229G>A, 230T>C, 234C>T, 235A>G, 236C>T, 237C>T, 241T>A, 243G>T, 246A>T, 261C>T, 262C>T, 264T>A, 267C>T, 273A>G, 274G>A, 277G>T, 278T>G, 282T>C, 291T>A, 298G>A, 300C>T, 303A>G, 306T>G, 307T>C, 309A>T, 310A>T, 312C>T, 315T>G, 316T>A, 317C>G, 324G>A, 333G>T, 336A>C, 340T>G, 342A>T, 348T>A, 351A>C, 354C>T, 357T>C, 358T>C, 360G>A, 361T>C, 363G>A, 374A>C, 375T>G, 379C>G, 380C>A, 381C>T, 384T>C, 387G>A, 388A>G, 389A>C, 390C>A, 402T>A, 405T>C, 408A>G, 409C>A, 414C>G, 416A>G, 417T>A, 423A>C, 424G>A, 426G>T, 427C>A, 428G>A, 429T>A, 430C>T, 432T>G, 441C>A, 442A>C, 444A>G, 448G>T, 450C>G, 451C>T, 453A>G, 456G>A, 458A>C, 459A>C, 462C>T, 465G>A, 468T>G, 469T>C, 470C>A, 477A>T, 478G>T, 480A>T, 481T>A, 483A>C, 489A>T, 492T>A, 495T>C, 496C>A, 497C>A, 498T>C, 499T>C, 501T>C, 508_525delGACAGTAAAGGCAAGAGT, 527A>C, 529G>T, 532G>T, 533A>C, 534G>A, 535C>G, 538C>A, 540C>T, 541A>G, 544_549delCGAGAG, 550G>A, 551A>G, 553C>T, 555C>A, 557T>G, 558T>C, 559A>G, 560A>G, 564A>G, 565G>A, 566T>A, 567T>C, 568G>A, 572T>C, 573T>A, 576C>T, 579A>G, 580C>G, 581G>A, 582T>A, 584A>C, 585G>C, 586T>C, 588A>T, 591G>A, 592A>G, 593G>T, 594A>T, 595T>G, 596T>C, 597T>C, 598C>A, 599A>G, 600G>A, 606G>A, 607T>C, 610G>C, 611C>A, 613A>G, 614G>A, 615T>A, 616C>G, 617A>C, 619A>C, 620A>G, 621A>G, 629A>T, 631C>G, 632A>T, 633G>C, 639A>C, 640C>A, 641A>G, 642A>G, 643C>T, 645T>G, 649C>A, 651C>T, 652T>C, 653C>A, 654C>A, 655A>G, 656A>G, 661G>C, 662T>C, 663T>A, 666T>C, 669T>C, 670A>G, 672T>G, 675C>T, 676A>G, 681C>T, 682T>G, 687T>C, 693T>G, 694G>A, 695A>G, 696A>T, 697A>G, 698G>A, 702G>C, 706C>A, 710A>G, 712T>C, 714A>G, 716C>T, 717A>C, 718G>A, 719_721delCAG, 722G>A, 724G>A, 729C>T, 730C>A, 733C>T, 736G>C, 738G>A, 739G>A, 745A>T, 748A>T, 751A>G, 754A>C, 760T>G, 766G>T, 769G>A, 771C>A, 772C>A, 775A>T, 784T>C, 785T>C, 787G>T, 790C>T, 792G>A, 796G>A, 797C>T, 799T>G, 802C>A, 811C>A, 814C>T, 817G>C, 818G>A, 820T>A, 821C>T, 823T>G, 828G>C, 829T>A, 835A>G, 841G>A, 844A>G, 853G>T, 854T>C, 856G>T, 858T>A, 862C>T, 874A>T, 875T>A, 876C>G, 877A>T, 880G>A, 881A>G, 883C>G, 884A>G, 892T>A, 899A>G, 901A>T, 904C>T, 908C>A, 909A>G, 910T>G, 912A>C, 913G>C, 914T>A, 916T>A, 917G>A, 918T>C, 919T>A, 922G>A, 925T>A, 926A>C, 927A>G, 928G>T, 931C>T, 936C>G, 937A>G, 940C>T, 943G>A, 949T>C, 953C>T, 955G>A, 956T>C, 958T>C, 961T>G, 965C>T, 967T>G, 970C>T, 973C>T, 979C>T, 982A>C, 987T>C, 988T>A, 993G>A, 997G>A, 1003A>T, 1004A>G, 1005G>C, 1006G>C, 1009T>C, 1012G>A, 1015A>C, 1018T>C, 1019C>T, 1021T>G, 1024T>C, 1027A>T, 1030T>C, 1033T>C, 1040C>A, 1041T>C, 1042C>T, 1043T>G, 1044C>A, 1045A>T, 1048G>T, 1057A>G, 1063G>A, 1065C>T, 1066A>T, 1067C>T, 1071C>A, 1075G>T, 1076G>T, 1078C>T, 1080A>G, 1081T>A, 1087G>A, 1090G>A, 1093A>C, 1099C>A, 1105C>T, 1108C>T, 1111C>T, 1127C>T, 1129C>G, 1132A>G, 1135A>G, 1138G>C, 1156C>A, 1162G>A, 1165C>T, 1167A>T, 1168A>C, 1174C>G, 1177G>A, 1180A>T, 1183G>A, 1189G>T, 1195C>T, 1198G>A, 1204C>A, 1207A>T, 1223G>A, 1224A>G, 1225C>T, 1226A>C, 1228A>T, 1229A>C, 1231G>T, 1237A>T, 1240T>A, 1244T>A, 1247C>G, 1249G>C, 1252A>T, 1255A>C, 1258A>T, 1262T>A, 1263C>G, 1270C>T, 1273A>T, 1276A>G, 1279A>T, 1282C>T, 1285T>C, 1286C>A, 1288T>A, 1291G>T, 1294A>G, 1306T>C, 1309A>G, 1315G>A, 1318T>A, 1321T>A, 1327T>C, 1330A>T, 1333T>A, 1339T>G, 1342T>C, 1345G>C, 1347G>A, 1351_1352insGTA, 1354A>G, 1357T>A, 1359C>G, 1363A>G, 1364A>C, 1367_1369delGCT, 1370C>T, 1372T>G, 1375A>G, 1380A>G, 1382G>C, 1384C>T, 1390T>G, 1393A>T, 1395A>C, 1396A>C, 1397C>G, 1398A>C, 1399G>A, 1405G>T, 1407G>A, 1408C>T, 1411A>C, 1418G>A, 1420C>T, 1423C>G, 1424A>C, 1425T>C, 1427A>C, 1429T>G, 1432A>G, 1433A>G, 1434G>C, 1435C>A, 1437C>A, 1442A>G, 1444C>G, 1447G>C, 1448A>T, 1449G>C, 1456T>A, 1462A>G, 1471T>C, 1474C>A, 1477A>G, 1480G>A, 1488A>G, 1492C>A, 1496C>G, 1498G>A, 1501C>A, 1508C>T, 1513G>T, 1513_1514insGACTTT, 1517G>T, 1518T>C, 1521G>C, 1523A>T, 1525A>T, 1526T>A, 1527C>G, 1529T>G, 1531C>G, 1533T>A, 1534A>T, 1536T>A, 1537C>G, 1538A>G, 1540C>A, 1541A>G, 1542C>T, 1543A>G, 1544C>A, 1546T>A, 1549A>G, 1555A>T, 1557T>G, 1559C>A, 1560A>T, 1565_1567delGAG, 1574T>A, 1575G>C, 1576T>G, 1577G>T, 1578G>C, 1579G>T, 1580G>C, 1581G>T, 1583T>C, 1591A>T, 1594A>C, 1595_1600delCCCAAG, 1603A>G, 1605A>T, 1606G>A, 1607G>T, 1608G>C, 1609C>A, 1615C>A, 1616T>G, 1617T>G, 1618C>A, 1620A>C, 1621C>A, 1622A>C, 1624G>C, 1625G>C, 1626G>A, 1627T>A, 1629C>A, 1630C>A, 1631A>G, 1633T>C, 1639T>C, 1644T>A, 1645G>T, 1651G>T, 1652G>C, 1654G>A, 1655T>A, 1656C>G, 1657G>T, 1660C>T, 1662C>A, 1664C>G, 1666C>T, 1667A>T, 1671A>C, 1672T>C, 1679G>A, 1680A>G, 1682G>A, 1684T>C, 1692A>C, 1693A>C, 1695G>T, 1696A>T, 1700A>G, 1702A>G, 1703G>C, 1705T>C, 1706C>G, 1711G>A, 1712G>A, 1714A>T, 1715C>G, 1717A>T, 1719T>A, 1721A>G, 1723C>T, 1726C>T, 1727T>G, 1729T>G, 1730A>C, 1731G>A, 1732T>G, 1733G>A, 1740A>G, 1742A>G, 1743G>C, 1744A>T, 1747A>T, 1750C>T, 1751G>C, 1753T>C, 1754A>T, 1755T>G, 1759G>C, 1764G>A, 1769G>A, 1773C>A, 1774A>G, 1776A>T, 1778C>A, 1781C>G, 1783T>C, 1784C>T, 1789T>C, 1790A>T, 1791A>G, 1795C>A, 1799A>T, 1800G>T, 1801G>T, 1804G>A, 1807C>T, 1810A>C, 1813C>T, 1818A>C, 1828A>T, 1831A>T, 1834A>T, 1835C>T, 1837T>G, 1839A>C, 1840A>G, 1843T>A, 1849T>A, 1858T>G, 1859A>G, 1861C>T, 1862C>G, 1863A>G, 1864A>G, 1865G>A, 1866G>T, 1867C>G, 1871A>G, 1872C>A, 1873T>A, 1874C>A, 1876C>A, 1879T>A, 1880T>C, 1881T>C, 1882C>A, 1891T>C, 1892G>T, 1895A>T, 1898T>A, 1899C>G, 1900T>C, 1903C>G, 1907G>C, 1915T>G, 1918G>T, 1924C>A, 1930T>C, 1931C>A, 1932A>T, 1933A>G, 1936A>G, 1939T>C, 1943T>C, 1944C>A, 1945C>A, 1948C>T, 1953T>G, 1954T>G, 1955C>T, 1957A>T, 1958G>T, 1961C>G, 1963G>T, 1964A>T, 1965C>T, 1966T>A, 1968A>C, 1969C>A, 1972T>C, 1975A>G, 1981G>T, 1982A>G, 1984T>C, 1991T>G, 1992G>T, 1993C>G, 2002A>T, 2003T>C, 2005A>T, 2008C>A, 2009T>C, 2012A>G, 2013A>C, 2017G>T, 2018C>A, 2020T>G, 2020_2021insGTGAAA, 2023G>T, 2024T>C, 2025T>C, 2026G>C, 2027A>T, 2029T>G, 2030G>A, 2033G>A, 2034A>G, 2035G>T, 2038G>T, 2040A>T, 2043T>C, 2044G>A, 2045G>T, 2046C>G, 2050_2051insTTGAAT, 2051A>C, 2053A>C, 2054A>T, 2055G>T, 2057T>G, 2058C>G, 2059T>A, 2060G>A, 2063A>C, 2064G>C, 2066C>A, 2067A>G, 2068A>C, 2069G>A, 2070C>T, 2071A>G, 2072C>A, 2073A>G, 2074G>T, 2078A>G, 2081G>C, 2082C>A, 2085G>C, 2086G>T, 2089C>A, 2091T>A, 2092A>T, 2095T>C, 2095_2096insAGC, 2096G>A, 2098A>G, 2099A>T, 2101T>G, 2104G>A, 2105C>A, 2112A>G, 2113C>A, 2115G>A, 2116A>G, 2120A>C, 2121C>T, 2122T>G, 2128C>A, 2134G>T, 2135T>A, 2140T>C, 2143C>A, 2144A>G, 2146G>A, 2147A>C, 2149G>T, 2152A>G, 2152_2153insAGT, 2155T>G, 2156C>G, 2157C>G, 2158C>T, 2160C>T, 2162A>C, 2164G>A, 2165C>A, 2166C>G, 2168G>A, 2170T>G, 2171A>G, 2174A>G, 2176A>G, 2177G>T, 2178A>C, 2184T>A, 2185T>G, 2188A>T, 2191G>A, 2193G>A, 2194A>T, 2195G>C, 2197A>G, 2202A>G, 2203A>G, 2204G>A, 2209T>A, 2210C>G, 2212A>G, 2216G>A, 2217G>A, 2221T>C, 2222G>A, 2225A>T, 2227T>C, 2233C>T, 2239A>G, 2240C>G, 2241C>A, 2242A>G, 2249G>A, 2250A>T, 2251C>G, 2252A>G, 2253C>G, 2254A>T, 2255T>G, 2256C>A, 2257C>T, 2258A>G, 2259T>C, 2260G>C, 2261G>C, 2263A>G, 2264G>T, 2266T>A, 2267G>A, 2269A>T, 2275T>G, 2276A>T, 2281G>A, 2284A>G, 2287G>A, 2289T>C, 2290G>A, 2296A>G, 2302C>T, 2308T>G, 2311C>T, 2323A>G, 2324G>A, 2335C>T, 2336G>C, 2338T>G, 2342G>A, 2344G>A, 2348T>A, 2349C>G, 2351G>A, 2352C>A, 2353T>C, 2365G>A, 2368A>G, 2371A>T, 2372A>C, 2374G>T, 2375A>G, 2376G>C, 2377C>A, 2378T>C, 2380G>A, 2381G>T, 2383T>C, 2384T>C, 2386G>C, 2389C>A, 2390G>A, 2391C>T, 2392T>G, 2395C>A, 2396C>T, 2398A>G, 2401C>T, 2402T>A, 2403C>G, 2404A>C, 2407A>G, 2409G>A, 2410C>G, 2413A>G, 2417A>G, 2422A>C, 2428T>G, 2429G>T, 2431G>C, 2440T>C, 2446T>C, 2449C>T, 2452T>C, 2455A>T, 2458T>A, 2459C>A, 2460T>A, 2461A>C, 2462A>C, 2464C>T, 2469A>T, 2470C>T, 2474A>G, 2476A>C, 2482T>G, 2485T>C, 2486G>A, 2495G>A, 2497T>G, 2503T>G, 2504A>G, 2506C>G, 2509T>G, 2512T>A, 2515A>G, 2518G>A, 2521T>C, 2536A>G, 2539T>C, 2544T>C, 2545G>A, 2552C>A, 2556A>C, 2557G>C, 2572C>T, 2575C>A, 2581A>C, 2582T>A, 2584G>T, 2596G>A, 2600A>C, 2601C>A, 2602A>T, 2603T>C, 2608T>G, 2618C>G, 2624A>T, 2625A>C, 2631A>G, 2633G>A, 2639T>G, 2640G>T, 2641C>G, 2644T>C, 2645A>T, 2647C>A, 2650A>T, 2651A>C, 2652C>A, 2656G>T, 2662C>T, 2664A>C, 2665G>C, 2667A>T, 2668C>T, 2671G>A, 2674T>A, 2675C>T, 2681G>A, 2682A>C, 2683A>T, 2684G>T, 2685G>C, 2686T>A, 2687A>C, 2688C>T, 2689A>G, 2692A>T, 2694C>T, 2695C>T, 2696A>C, 2701T>A, 2702T>C, 2704A>T, 2707T>C, 2713G>T, 2728A>C, 2731C>T, 2737A>G, 2738T>G, 2740T>A, 2746A>C, 2752G>A, 2756A>G, 2758T>G, 2759A>T, 2760G>C, 2761T>A, 2762G>A, 2767A>T, 2770C>T, 2773T>C, 2774T>A, 2776G>A, 2780T>C, 2783C>T, 2784A>C, 2785G>C, 2791A>G, 2795T>C, 2797A>C, 2803C>A, 2810C>A, 2812T>G, 2822C>A, 2824A>G, 2825A>T, 2829T>G, 2832C>G, 2833A>G, 2836G>A, 2842A>G, 2844A>T, 2848C>A, 2852A>C, 2853A>G, 2855G>C, 2857G>A, 2860G>C, 2861C>A, 2862A>C, 2863G>A, 2866A>C, 2868A>T, 2869T>C, 2873C>A, 2874C>G, 2875A>T, 2876A>C, 2878G>C, 2881T>C, 2884A>T, 2887T>A, 2896T>G, 2897T>C, 2899G>T, 2900T>C, 2902A>G, 2905T>C, 2908T>C, 2915G>A, 2917G>T, 2918A>G, 2919C>T, 2921T>G, 2923A>T, 2924T>G, 2926C>A, 2927C>G, 2929A>C, 2941A>T, 2950C>T, 2951A>G, 2953A>C, 2956T>C, 2959A>C, 2971A>T, 2977G>A, 2986C>T, 2988C>T, 2992G>A, 2998G>A, 3001T>A, 3002T>C, 3010T>A, 3016A>T, 3019C>A, 3022G>A, 3031G>A, 3039C>G, 3040T>A, 3043A>C, 3046T>C, 3059A>G, 3061T>G, 3064C>T, 3065A>G, 3076C>T, 3077T>C, 3088A>T, 3091T>G, 3097G>T, 3100A>T, 3103T>A, 3106A>G, 3107C>T, 3109T>G, 3112A>C, 3115T>G, 3119A>G, 3120A>C, 3121C>A, 3124C>A, 3125A>G, 3127A>G, 3131A>T, 3135A>C, 3136T>A, 3139T>G, 3151T>G, 3160A>C, 3169A>G, 3172C>T, 3176C>T, 3178T>G, 3187T>G, 3192A>G, 3193A>G, 3198G>A, 3199G>A, 3203C>A, 3204G>A, 3205G>A, 3208T>C, 3211T>A, 3212A>C, 3214T>G, 3217A>C, 3229A>G, 3230A>T, 3231A>G, 3232A>C, 3235T>C, 3236T>C, 3238G>C, 3241T>G, 3244C>T, 3247A>C, 3250G>A, 3253A>C, 3256G>A, 3262C>A, 3265G>A, 3275C>A, 3280C>T, 3281T>C, 3282T>A, 3283T>A, 3286C>T, 3290T>A, 3295C>T, 3296T>C, 3301T>C, 3307G>T, 3310A>T, 3322C>G, 3331G>A, 3335A>T, 3337A>G, 3340T>C, 3347T>A, 3349C>T, 3355T>A, 3358C>A, 3367G>A, 3373G>A, 3374T>C, 3377C>A, 3383A>G, 3388T>C, 3391T>G, 3392T>C, 3394G>T, 3397T>G, 3400C>T, 3401A>T, 3403A>T, 3406A>G, 3409T>A, 3412G>T, 3415A>G, 3421A>T, 3427G>T, 3436A>C, 3439A>G, 3442C>G, 3444A>G, 3448C>T, 3449T>G, 3451A>T, 3454T>G, 3460C>T, 3463T>G, 3466C>T, 3476G>A, 3478T>C, 3479T>C, 3481A>G, 3482T>A, 3483C>A, 3484A>T, 3490T>C, 3497A>C, 3502A>G, 3505G>A, 3506T>C, 3514G>A, 3517C>A, 3520C>T, 3521A>G, 3526G>T, 3527G>A, 3528A>T, 3529G>C, 3531A>T, 3532C>T, 3533C>G, 3535G>C, 3536G>C, 3544C>G, 3548_3553delGATCCG, 3554C>A, 3555C>G, 3561A>C, 3562G>A, 3563G>T, 3569A>G, 3570G>T, 3572A>T, 3573T>C, 3574T>C, 3575C>T, 3577G>C, 3578T>C, 3579C>A, 3580C>G, 3583A>C, 3586A>G, 3589G>A, 3592A>G, 3595T>C, 3596T>A, 3598C>A, 3604T>A, 3607T>A, 3610A>G, 3619C>T, 3622C>T, 3623A>G, 3624T>C, 3625G>T, 3627G>C, 3628T>A, 3638G>A, 3639A>C, 3640C>A, 3643A>C, 3650T>G, 3652G>A, 3655A>T, 3656A>T, 3658T>G, 3661T>A, 3667A>C, 3670A>T, 3676C>A, 3677A>G, 3679A>G, 3680A>C, 3682G>C, 3688A>T, 3690G>C, 3691G>A, 3692C>T, 3693A>C, 3700T>A, 3701G>A, 3706G>A, 3715C>A, 3721T>C, 3727T>C, 3733T>C, 3740A>T, 3742A>C, 3745A>T, 3760A>T, 3763T>A, 3770A>G, 3772T>A, 3776T>A, 3779A>T, 3784G>A, 3787A>G, 3791A>C, 3794G>A, 3795T>C, 3796A>T, 3799A>G, 3800T>A, 3801C>G, 3803A>T, 3804G>C, 3805T>C, 3806C>A, 3807C>G, 3808T>C, 3810A>G, 3812G>A, 3814T>A, 3817A>T, 3821G>A, 3823A>C, 3824C>T, 3826A>G, 3829T>C, 3830G>A, 3835T>A, 3836T>C, 3841T>C, 3842G>A, 3844C>A, 3850C>G, 3859T>C, 3861C>A, 3863G>T, 3864C>T, 3868G>T, 3869C>G, 3871T>A, 3872A>T, 3873A>C, 3875A>G, 3876A>C, 3877A>G, 3881A>G, 3882A>C, 3883T>A, 3884T>C, 3890G>T, 3901T>C, 3904A>T, 3907C>A, 3910A>T, 3925A>C, 3928C>T, 3931T>G, 3934G>T, 3935G>A, 3937A>C, 3940A>C, 3943T>C, 3945C>T, 3946G>C, 3952A>T, 3955C>T, 3956A>G, 3957C>T, 3958T>A, 3971G>C, 3976T>C, 3979C>T, 3982A>T, 3983A>C, 3985G>C, 3988G>T, 3989C>T, 3990A>T, 3991T>C, 3996G>A, 3998A>G, 3999T>C, 4000G>C, 4002G>A, 4003G>A, 4004G>T, 4005A>C, 4006A>C, 4008T>A, 4009C>T, 4010T>G, 4011C>A, 4012A>T, 4014G>A, 4015C>G, 4016A>G, 4018G>A, 4019C>A, 4020A>G, 4021A>T, 4024C>A, 4025C>A, 4027C>A, 4030C>T, 4031A>G, 4033G>T, 4036A>T, 4037C>T, 4039A>G, 4045T>C, 4047G>A, 4051T>A, 4054A>C, 4055A>T, 4056A>T, 4057T>G, 4058C>T, 4061A>C, 4063C>A, 4066A>C, 4070T>C, 4072G>T, 4075T>A, 4078T>A, 4079T>G, 4080C>T, 4081C>T, 4083C>A, 4084C>A, 4085A>C, 4087A>T, 4088G>A, 4090G>T, 4096C>A, 4102G>A, 4105A>C, 4106C>A, 4108T>A, 4109A>G, 4111A>C, 4112T>A, 4113C>A, 4114T>G, 4117T>A, 4120A>T, 4122T>C, 4129C>T, 4132G>A, 4144C>T, 4150T>C, 4153T>A, 4154T>C, 4155C>T, 4159C>A, 4163C>A, 4168A>G, 4171A>G, 4177T>C, 4180C>A, 4181A>G, 4189A>G, 4192T>G, 4195C>T, 4198C>T, 4210C>A, 4213A>T, 4226C>T, 4228A>G, 4231T>C, 4242T>A, 4247A>G, 4248A>C, 4259C>T, 4261A>G, 4267T>A, 4274T>C, 4276G>A, 4279A>T, 4280A>T, 4281A>T, 4283A>C, 4284C>T, 4285A>T, 4286C>T, 4287T>G, 4289A>T, 4291C>A, 4291_4292insCGTA, 4294C>T, 4304T>G, 4306G>A, 4309G>A, 4312G>T, 4312_4313insAGTAAATAAATAAAGT, 4317G>T, 4320G>T, 4324T>G, 4325T>G, 4330G>C, 4331C>A, 4333A>G, 4336A>T, 4339A>G, 4340G>T, 4342G>T, 4343A>T, 4345A>T, 4349A>G, 4351G>A, 4363T>C, 4364C>T, 4365T>C, 4370A>T, 4372A>G, 4373G>A, 4374A>G, 4384C>T, 4385C>G, 4386T>A, 4391T>G, 4393C>T, 4394A>G, 4398A>T, 4399G>T, 4400A>G, 4404A>G, 4415T>C, 4420delA, 4422A>G, 4426G>A, 4437C>T, 4441T>C, 4449A>T, 4451T>G, 4477_4478insTCCTCCACA, 4479A>C, 4480T>A, 4482T>A, 4486G>T, 4488T>A, 4489T>A, 4490A>C, 4491A>G, 4492C>G, 4493A>T, 4497C>G, 4499C>G, 4500G>T, 4501T>G, 4502T>A, 4506A>T, 4510T>A, 4510_4511insTCACTTA, 4518A>G, 4519G>A, 4520A>C, 4521C>A, 4523T>C, 4525A>G, 4526T>G, 4532C>G, 4534A>T, 4535A>T, 4539A>C, 4540_4541insAGTTAACGCGAACACAGGTTGTTTCCCTATAGAAACTA, 4542C>A, 4544T>C, 4548T>G, 4552C>G, 4553A>C, 4554T>A, 4556T>C, 4557A>T, 4559G>A, 4562A>G, 4563_4564delTA, 4566A>G, 4568G>A, 4569A>G, 4570G>C, 4581C>T, 4594T>C, 4595T>A, 4602C>G, 4606A>T, 4613T>C, 4614C>T, 4616C>T, 4618A>G, 4626T>C, 4628G>A, 4630G>A, 4632C>T, 4637T>A, 4639G>A, 4642A>T, 4643G>T, 4646C>A, 4654C>T, 4675G>C, 4680A>G, 4684T>A, 4685T>A, 4688C>T, 4689C>G, 4704G>C, 4706G>A, 4708T>A, 4722T>G | | | | | | | | |

**CDS**

| | Begin | End | Coverage | Score | Concordance | Matches | Identities | I/D/M/F* | Stop Codons |
|---|---|---|---|---|---|---|---|---|---|
| **NT** | **89** | **4744** | **94.9%** | **2600** | **28.6%** | **4608 (96.7%)** | **3040 (63.8%)** | **107/48** | |
| ToMarV_RNA2gp1 | 1 | 191 | 100% | 784 | 66.7% | 183 (95.8%) | 116 (60.7%) | 0/8/0/0 | 1 |
| Protein mutations: | G5S (151G>A 153G>T), R6K (154C>A 155G>A 156T>G), S10Q (166T>C 167C>A 168T>A), V11I (169G>A), D12E (174C>G), E13D (177A>T), A15E (182C>A 183T>A), H18N (190C>A), T22A (202A>G 204C>A), N24T (209A>C 210T>C), I26V (214A>G), V31T (229G>A 230T>C), T33V (235A>G 236C>T 237C>T), L35I (241T>A 243G>T), V46I (274G>A), V47C (277G>T 278T>G), V54I (298G>A 300C>T), T58S (310A>T 312C>T), L68V (340T>G 342A>T), N79T (374A>C 375T>G), P81D (379C>G 380C>A 381C>T), N84A (388A>G 389A>C 390C>A), L91I (409C>A), H93R (416A>G 417T>A), V96I (424G>A 426G>T), R97K (427C>A 428G>A 429T>A), V104L (448G>T 450C>G), E107A (458A>C 459A>C), S111H (469T>C 470C>A), A114S (478G>T 480A>T), S115T (481T>A 483A>C), P120N (496C>A 497C>A 498T>C), S121P (499T>C 501T>C), D124_S129del (508_525delGACAGTAAAGGCAAGAGT), K130T (527A>C), V131L (529G>T), E132S (532G>T 533A>C 534G>A), Q133E (535C>G), P134T (538C>A 540C>T), T135A (541A>G), R136_E137del (544_549delCGAGAG), D138S (550G>A 551A>G), I140S (557T>G 558T>C), K141G (559A>G 560A>G), V143N (565G>A 566T>A 567T>C), E144K (568G>A), V145A (572T>C 573T>A), R148E (580C>G 581G>A 582T>A), E149A (584A>C 585G>C), R152V (592A>G 593G>T 594A>T), F153A (595T>G 596T>C 597T>C), Q154R (598C>A 599A>G 600G>A), A158Q (610G>C 611C>A), S159E (613A>G 614G>A 615T>A), Q160A (616C>G 617A>C), K161R (619A>C 620A>G 621A>G), N164I (629A>T), Q165V (631C>G 632A>T 633G>C), Q168R (640C>A 641A>G 642A>G), L171I (649C>A 651C>T), S172Q (652T>C 653C>A 654C>A), N173G (655A>G 656A>G), V175P (661G>C 662T>C 663T>A), N178E (670A>G 672T>G), I180V (676A>G), S182A (682T>G), E186S (694G>A 695A>G 696A>T), S187D (697A>G 698G>A), Q190K (706C>A) | | | | | | | | |
| Codon mutations: | TCA2TCT (144A>T), TTC3TTT (147C>T), ATT4ATC (150T>C), GGG5AGT (151G>A 153G>T), CGT6AAG (154C>A 155G>A 156T>G), TTG7CTC (157T>C 159G>C), AAT8AAC (162T>C), ACA9ACC (165A>C), TCT10CAA (166T>C 167C>A 168T>A), GTA11ATA (169G>A), GAC12GAG (174C>G), GAA13GAT (177A>T), GCT15GAA (182C>A 183T>A), TTT16TTC (186T>C), CAT18AAT (190C>A), GTT20GTA (198T>A), GCC21GCA (201C>A), ACC22GCA (202A>G 204C>A), TCG23TCT (207G>T), AAT24ACC (209A>C 210T>C), ATT26GTT (214A>G), TCT28AGT (220T>A 221C>G), GTT31ACT (229G>A 230T>C), GGC32GGT (234C>T), ACC33GTT (235A>G 236C>T 237C>T), TTG35ATT (241T>A 243G>T), ATA36ATT (246A>T), ACC41ACT (261C>T), CTT42TTA (262C>T 264T>A), GAC43GAT (267C>T), AAA45AAG (273A>G), GTT46ATT (274G>A), GTT47TGT (277G>T 278T>G), CCT48CCC (282T>C), GGT51GGA (291T>A), GTC54ATT (298G>A 300C>T), TCA55TCG (303A>G), GTT56GTG (306T>G), TTA57CTT (307T>C 309A>T), ACC58TCT (310A>T 312C>T), GTT59GTG (315T>G), TCT60AGT (316T>A 317C>G), GAG62GAA (324G>A), ACG65ACT (333G>T), CCA66CCC (336A>C), TTA68GTT (340T>G 342A>T), CCT70CCA (348T>A), GGA71GGC (351A>C), CAC72CAT (354C>T), TAT73TAC (357T>C), TTG74CTA (358T>C 360G>A), TTG75CTA (361T>C 363G>A), AAT79ACG (374A>C 375T>G), CCC81GAT (379C>G 380C>A 381C>T), ATT82ATC (384T>C), AAG83AAA (387G>A), AAC84GCA (388A>G 389A>C 390C>A), TCT88TCA (402T>A), GGT89GGC (405T>C), TTA90TTG (408A>G), CTT91ATT (409C>A), GTC92GTG (414C>G), CAT93CGA (416A>G 417T>A), TCA95TCC (423A>C), GTG96ATT (424G>A 426G>T), CGT97AAA (427C>A 428G>A 429T>A), CTT98TTG (430C>T 432T>G), ACC101ACA (441C>A), AGA102CGG (442A>C 444A>G), GTC104TTG (448G>T 450C>G), CTA105TTG (451C>T 453A>G), GAG106GAA (456G>A), GAA107GCC (458A>C 459A>C), AAC108AAT (462C>T), AAG109AAA (465G>A), GTT110GTG (468T>G), TCT111CAT (469T>C 470C>A), TCA113TCT (477A>T), GCA114TCT (478G>T 480A>T), TCA115ACC (481T>A 483A>C), TCA117TCT (489A>T), TCT118TCA (492T>A), TCT119TCC (495T>C), CCT120AAC (496C>A 497C>A 498T>C), TCT121CCC (499T>C 501T>C), GAC124_AGT129del (508_525delGACAGTAAAGGCAAGAGT), AAA130ACA (527A>C), GTA131TTA (529G>T), GAG132TCA (532G>T 533A>C 534G>A), CAA133GAA (535C>G), CCC134ACT (538C>A 540C>T), ACA135GCA (541A>G), CGA136_GAG137del (544_549delCGAGAG), GAT138AGT (550G>A 551A>G), CTC139TTA (553C>T 555C>A), ATT140AGC (557T>G 558T>C), AAA141GGA (559A>G 560A>G), GAA142GAG (564A>G), GTT143AAC (565G>A 566T>A 567T>C), GAG144AAG (568G>A), GTT145GCA (572T>C 573T>A), CTC146CTT (576C>T), AAA147AAG (579A>G), CGT148GAA (580C>G 581G>A 582T>A), GAG149GCC (584A>C 585G>C), TTA150CTT (586T>C 588A>T), GAG151GAA (591G>A), AGA152GTT (592A>G 593G>T 594A>T), TTT153GCC (595T>G 596T>C 597T>C), CAG154AGA (598C>A 599A>G 600G>A), GAG156GAA (606G>A), TTG157CTG (607T>C), GCA158CAA (610G>C 611C>A), AGT159GAA (613A>G 614G>A 615T>A), CAA160GCA (616C>G 617A>C), AAA161CGG (619A>C 620A>G 621A>G), AAT164ATT (629A>T), CAG165GTC (631C>G 632A>T 633G>C), CTA167CTC (639A>C), CAA168AGG (640C>A 641A>G 642A>G), CTT169TTG (643C>T 645T>G), CTC171ATT (649C>A 651C>T), TCC172CAA (652T>C 653C>A 654C>A), AAT173GGT (655A>G 656A>G), GTT175CCA (661G>C 662T>C 663T>A), AGT176AGC (666T>C), AAT177AAC (669T>C), AAT178GAG (670A>G 672T>G), GAC179GAT (675C>T), ATC180GTC (676A>G), TTC181TTT (681C>T), TCA182GCA (682T>G), GGT183GGC (687T>C), ACT185ACG (693T>G), GAA186AGT (694G>A 695A>G 696A>T), AGT187GAT (697A>G 698G>A), GGG188GGC (702G>C), CAG190AAG (706C>A), TAA191TGA (710A>G) | | | | | | | | |
| ToMarV_RNA2gp2 | 1 | 1192 | 100% | 6009 | 73.0% | 1185 (98.7%) | 820 (68.3%) | 9/7/0/0 | 1 |
| Protein mutations: | M1R (672T>G), T2M (675C>T 676A>G), S4L (681C>T 682T>G), V6A (687T>C), L8R (693T>G 694G>A), K9V (695A>G 696A>T 697A>G), V10M (698G>A), G11A (702G>C), N14D (710A>G 712T>C), H15R (714A>G), Q16S (716C>T 717A>C 718G>A), Q17del (719_721delCAG), E18K (722G>A 724G>A), A20V (729C>T 730C>A), E22D (736G>C), R23K (738G>A 739G>A), A34E (771C>A 772C>A), R41K (792G>A), V50I (818G>A 820T>A), G53A (828G>C 829T>A), E61D (853G>T), F63Y (858T>A), I71V (881A>G 883C>G), T72A (884A>G), T77A (899A>G 901A>T), H80R (908C>A 909A>G 910T>G), K81T (912A>C 913G>C), F82I (914T>A 916T>A), V83T (917G>A 918T>C 919T>A), K86R (926A>C 927A>G 928G>T), A89G (936C>G 937A>G), Y96H (956T>C 958T>C), V106A (987T>C 988T>A), S108N (993G>A), R112A (1004A>G 1005G>C 1006G>C), L124T (1040C>A 1041T>C 1042C>T), S125D (1043T>G 1044C>A 1045A>T), T132I (1065C>T 1066A>T), S134Y (1071C>A), L135F (1075G>T), A136S (1076G>T 1078C>T), D137G (1080A>G 1081T>A), Q166L (1167A>T 1168A>C), D185S (1223G>A 1224A>G 1225C>T), T186P (1226A>C 1228A>T), S192T (1244T>A), Q193D (1247C>G 1249G>C), Q194H (1252A>T), E195D (1255A>C), S226N (1347G>A), G227_Q228insV (1351_1352insGTA), S229R (1357T>A), T230S (1359C>G), M232L (1364A>C), A233del (1367_1369delGCT), K237R (1380A>G), A238P (1382G>C 1384C>T), E242A (1395A>C 1396A>C), Q243A (1397C>G 1398A>C 1399G>A), G246D (1407G>A 1408C>T), D250N (1418G>A 1420C>T), M252P (1424A>C 1425T>C), N253Q (1427A>C 1429T>G), S255A (1433A>G 1434G>C 1435C>A), T256N (1437C>A), I258V (1442A>G 1444C>G), K273R (1488A>G), P276A (1496C>G 1498G>A), P281_A282insDF (1513_1514insGACTTT), V283S (1517G>T 1518T>C), S284T (1521G>C), T285S (1523A>T 1525A>T), S286R (1526T>A 1527C>G), S287A (1529T>G 1531C>G), L288Y (1533T>A 1534A>T), L289Q (1536T>A 1537C>G), S290G (1538A>G 1540C>A), T291V (1541A>G 1542C>T 1543A>G), V296G (1557T>G), H297I (1559C>A 1560A>T), E299del (1565_1567delGAG), C302T (1574T>A 1575G>C 1576T>G), G303S (1577G>T 1578G>C 1579G>T), G304L (1580G>C 1581G>T), P309_K310del (1595_1600delCCCAAG), K312I (1605A>T 1606G>A), G313S (1607G>T 1608G>C 1609C>A), F316G (1616T>G 1617T>G 1618C>A), N317T (1620A>C 1621C>A), M318L (1622A>C 1624G>C), G319Q (1625G>C 1626G>A 1627T>A), A320E (1629C>A 1630C>A), I321V (1631A>G 1633T>C), L325Y (1644T>A 1645G>T), M327I (1651G>T), E328Q (1652G>C 1654G>A), A331E (1662C>A), H332D (1664C>G 1666C>T), I333F (1667A>T), D334A (1671A>C 1672T>C), D337S (1679G>A 1680A>G), D338N (1682G>A 1684T>C), E341A (1692A>C 1693A>C), R342L (1695G>T 1696A>T), I344V (1700A>G 1702A>G), A345P (1703G>C 1705T>C), Q346E (1706C>G), V348I (1712G>A 1714A>T), Q349D (1715C>G 1717A>T), F350Y (1719T>A), I351V (1721A>G 1723C>T), F353V (1727T>G 1729T>G), S354Q (1730A>C 1731G>A 1732T>G), E355K (1733G>A), K357R (1740A>G), R358A (1742A>G 1743G>C 1744A>T), V361L (1751G>C 1753T>C), I362C (1754A>T 1755T>G), R365K (1764G>A), V367I (1769G>A), A368E (1773C>A 1774A>G), K369M (1776A>T), H371D (1781C>G 1783T>C), N374C (1790A>T 1791A>G), R377F (1799A>T 1800G>T 1801G>T), D383A (1818A>C), E387D (1831A>T), E390A (1839A>C 1840A>G), N396K (1858T>G), I397V (1859A>G 1861C>T), Q398G (1862C>G 1863A>G 1864A>G), G399M (1865G>A 1866G>T 1867C>G), T401E (1871A>G 1872C>A 1873T>A), F404P (1880T>C 1881T>C 1882C>A), A408S (1892G>T), M409L (1895A>T), E413Q (1907G>C), Q421M (1931C>A 1932A>T 1933A>G), S425Q (1943T>C 1944C>A 1945C>A), I428R (1953T>G 1954T>G), Q429Y (1955C>T 1957A>T), G430W (1958G>T), Q431D (1961C>G 1963G>T), T432L (1964A>T 1965C>T 1966T>A), N433T (1968A>C 1969C>A), N438D (1982A>G 1984T>C), C441V (1991T>G 1992G>T 1993C>G), K448A (2012A>G 2013A>C), E449D (2017G>T), R450_Q451insVK (2020_2021insGTGAAA), Q451H (2023G>T), L452P (2024T>C 2025T>C 2026G>C), I453L (2027A>T 2029T>G), V454M (2030G>A), E455S (2033G>A 2034A>G 2035G>T), E456D (2038G>T), Q457L (2040A>T), V458A (2043T>C 2044G>A), A459C (2045G>T 2046C>G), Q460_R461insLN (2050_2051insTTGAAT), R462L (2054A>T 2055G>T), S463G (2057T>G 2058C>G 2059T>A), E464K (2060G>A), R465P (2063A>C 2064G>C), Q466S (2066C>A 2067A>G 2068A>C), A467M (2069G>A 2070C>T 2071A>G), Q468S (2072C>A 2073A>G 2074G>T), T470A (2078A>G), A471Q (2081G>C 2082C>A), R472T (2085G>C 2086G>T), I474N (2091T>A 2092A>T), A475_E476insS (2095_2096insAGC), E476K (2096G>A 2098A>G), S477W (2099A>T 2101T>G), P479T (2105C>A), D481G (2112A>G 2113C>A), R482K (2115G>A 2116A>G), T484L (2120A>C 2121C>T 2122T>G), S489T (2135T>A), T492A (2144A>G 2146G>A), M493L (2147A>C 2149G>T), E494_D495insS (2152_2153insAGT), D495E (2155T>G), P496G (2156C>G 2157C>G 2158C>T), T497I (2160C>T), K498Q (2162A>C 2164G>A), P499R (2165C>A 2166C>G), D500K (2168G>A 2170T>G), K501E (2171A>G), I502V (2174A>G 2176A>G), E503S (2177G>T 2178A>C), V505E (2184T>A 2185T>G), G508D (2193G>A 2194A>T), A509P (2195G>C 2197A>G), E511G (2202A>G 2203A>G), E512K (2204G>A), Q514E (2210C>G 2212A>G), G516N (2216G>A 2217G>A), V518I (2222G>A), I519F (2225A>T 2227T>C), P524E (2240C>G 2241C>A 2242A>G), D527M (2249G>A 2250A>T 2251C>G), T528G (2252A>G 2253C>G 2254A>T), S529D (2255T>G 2256C>A 2257C>T), M530A (2258A>G 2259T>C 2260G>C), A531P (2261G>C 2263A>G), V532L (2264G>T 2266T>A), E533N (2267G>A 2269A>T), D535E (2275T>G), M536L (2276A>T), V540A (2289T>C 2290G>A), D546E (2308T>G), V552I (2324G>A), V556L (2336G>C 2338T>G), E558K (2342G>A 2344G>A), A561N (2351G>A 2352C>A 2353T>C), M568L (2372A>C 2374G>T), S569A (2375A>G 2376G>C 2377C>A), A571S (2381G>T 2383T>C), A574M (2390G>A 2391C>T 2392T>G), S580K (2409G>A 2410C>G), T583A (2417A>G), A587S (2429G>T 2431G>C), L597N (2459C>A 2460T>A 2461A>C), I598L (2462A>C 2464C>T), Y600F (2469A>T 2470C>T), I602V (2474A>G 2476A>C), V606I (2486G>A), V609M (2495G>A 2497T>G), I612V (2504A>G 2506C>G), V625A (2544T>C 2545G>A), K629T (2556A>C 2557G>C), L638I (2582T>A 2584G>T), T644H (2600A>C 2601C>A 2602A>T), Q650E (2618C>G), N652S (2624A>T 2625A>C), E654G (2631A>G), E655K (2633G>A), C657V (2639T>G 2640G>T 2641C>G), T659S (2645A>T 2647C>A), T661Q (2651A>C 2652C>A), K665T (2664A>C 2665G>C), Y666F (2667A>T 2668C>T), E671T (2681G>A 2682A>C 2683A>T), G672S (2684G>T 2685G>C 2686T>A), T673L (2687A>C 2688C>T 2689A>G), A675V (2694C>T 2695C>T), T676P (2696A>C), D677E (2701T>A), S690A (2738T>G 2740T>A), N696E (2756A>G 2758T>G), V698I (2762G>A), L702I (2774T>A 2776G>A), Q705S (2783C>T 2784A>C 2785G>C), L714M (2810C>A 2812T>G), M719L (2825A>T), I720R (2829T>G), A721G (2832C>G 2833A>G), Y725F (2844A>T), K728R (2852A>C 2853A>G), A729P (2855G>C 2857G>A), Q731T (2861C>A 2862A>C 2863G>A), Y733F (2868A>T 2869T>C), P735S (2873C>A 2874C>G 2875A>T), G749S (2915G>A 2917G>T), T750V (2918A>G 2919C>T), S751A (2921T>G 2923A>T), S752A (2924T>G 2926C>A), P753A (2927C>G 2929A>C), I761V (2951A>G 2953A>C), T773I (2988C>T), T790R (3039C>G 3040T>A), I797V (3059A>G 3061T>G), I799V (3065A>G), N817A (3119A>G 3120A>C 3121C>A), K819E (3125A>G 3127A>G), T821S (3131A>T), N822T (3135A>C 3136T>A), F827L (3151T>G), K841R (3192A>G 3193A>G), R843Q (3198G>A 3199G>A), R845K (3203C>A 3204G>A 3205G>A), I848L (3212A>C 3214T>G), K854C (3230A>T 3231A>G 3232A>C), Q869K (3275C>A), F871Q (3281T>C 3282T>A 3283T>A), L874M (3290T>A), I889L (3335A>T 3337A>G), S893T (3347T>A 3349C>T), C902R (3374T>C), L903I (3377C>A), N905D (3383A>G), T911S (3401A>T 3403A>T), K925R (3444A>G), S927A (3449T>G 3451A>T), D936N (3476G>A 3478T>C), S938N (3482T>A 3483C>A 3484A>T), I943L (3497A>C), D949E (3517C>A), I951V (3521A>G), E952D (3526G>T), E953I (3527G>A 3528A>T 3529G>C), Y954F (3531A>T 3532C>T), Q955D (3533C>G 3535G>C), E956Q (3536G>C), D960_P961del (3548_3553delGATCCG), P962S (3554C>A 3555C>G), K964T (3561A>C 3562G>A), A965S (3563G>T), S967V (3569A>G 3570G>T), I968S (3572A>T 3573T>C 3574T>C), L969F (3575C>T 3577G>C), S970Q (3578T>C 3579C>A 3580C>G), S976T (3596T>A 3598C>A), M985A (3623A>G 3624T>C 3625G>T), G986A (3627G>C 3628T>A), D990T (3638G>A 3639A>C 3640C>A), E991D (3643A>C), L994V (3650T>G 3652G>A), I996L (3656A>T 3658T>G), I1003V (3677A>G 3679A>G), E1006D (3688A>T), G1007A (3690G>C 3691G>A), H1008S (3692C>T 3693A>C), V1011I (3701G>A), T1024S (3740A>T 3742A>C), N1034E (3770A>G 3772T>A), S1036T (3776T>A), I1037F (3779A>T), I1039M (3787A>G), V1042T (3794G>A 3795T>C 3796A>T), P1046S (3806C>A 3807C>G 3808T>C), N1047S (3810A>G), V1048I (3812G>A 3814T>A), V1051I (3821G>A 3823A>C), V1054I (3830G>A), F1056L (3836T>C), A1058T (3842G>A 3844C>A), P1064Q (3861C>A), A1065F (3863G>T 3864C>T), L1067V (3869C>G 3871T>A), N1068S (3872A>T 3873A>C), K1069A (3875A>G 3876A>C 3877A>G), N1071A (3881A>G 3882A>C 3883T>A), Y1072H (3884T>C), V1074L (3890G>T), A1089T (3935G>A 3937A>C), T1092I (3945C>T 3946G>C), T1096V (3956A>G 3957C>T 3958T>A), E1101Q (3971G>C), H1107F (3989C>T 3990A>T 3991T>C), R1109K (3996G>A), M1110A (3998A>G 3999T>C 4000G>C), R1111K (4002G>A 4003G>A), E1112S (4004G>T 4005A>C 4006A>C), F1113Y (4008T>A 4009C>T), S1114D (4010T>G 4011C>A 4012A>T), S1115K (4014G>A 4015C>G), K1116E (4016A>G 4018G>A), Q1117S (4019C>A 4020A>G 4021A>T), M1121V (4031A>G 4033G>T), R1126K (4047G>A), N1129L (4055A>T 4056A>T 4057T>G), I1131L (4061A>C 4063C>A), S1137V (4079T>G 4080C>T 4081C>T), A1138E (4083C>A 4084C>A), I1139L (4085A>C 4087A>T), V1140I (4088G>A 4090G>T), L1146I (4106C>A 4108T>A), I1147V (4109A>G 4111A>C), S1148K (4112T>A 4113C>A 4114T>G), L1151P (4122T>C), S1162L (4154T>C 4155C>T), H1165N (4163C>A), N1171D (4181A>G), I1191N (4242T>A) | | | | | | | | |

| | Begin | End | Coverage | Score | Concordance | Matches | Identities | I/D/M/F* | Stop Codons |
|---|---|---|---|---|---|---|---|---|---|
| **NT** | **89** | **4744** | **94.9%** | **2600** | **28.6%** | **4608 (96.7%)** | **3040 (63.8%)** | **107/48** | |

Codon mutations:

ATG1AGG (672T>G), ACA2ATG (675C>T 676A>G), TCT4TTG (681C>T 682T>G), GTT6GCT (687T>C), CTG8CGA (693T>G 694G>A), AAA9GTG (695A>G 696A>T 697A>G), GTG10ATG (698G>A), GGC11GCC (702G>C), CCC12CCA (706C>A), AAT14GAC (710A>G 712T>C), CAC15CGC (714A>G), CAG16TCA (716C>T 717A>C 718G>A), CAG17del (719_721delCAG), GAG18AAA (722G>A 724G>A), GCC20GTA (729C>T 730C>A), TCC21TCT (733C>T), GAG22GAC (736G>C), AGG23AAA (738G>A 739G>A), ATA25ATT (745A>T), TCA26TCT (748A>T), CAA27CAG (751A>G), ATA28ATC (754A>C), GCT30GCG (760T>G), GTG32GTT (766G>T), GAG33GAA (769G>A), GCC34GAA (771C>A 772C>A), GGA35GGT (775A>T), AAT38AAC (784T>C), TTG39CTT (785T>C 787G>T), CTC40CTT (790C>T), AGA41AAA (792G>A), AAG42AAA (796G>A), CTT43TTG (797C>T 799T>G), GGC44GGA (802C>A), TCC47TCA (811C>A), TAC48TAT (814C>T), GGG49GGC (817G>C), GTT50ATA (818G>A 820T>A), CTT51TTG (821C>T 823T>G), GGT53GCA (828G>C 829T>A), CAA55CAG (835A>G), AAG57AAA (841G>A), AGA58AGG (844A>G), GAG61GAT (853G>T), TTG62CTT (854T>C 856G>T), TTT63TAT (858T>A), GAC64GAT (862C>T), GTA68GTT (874A>T), TCA69AGT (875T>A 876C>G 877A>T), AAG70AAA (880G>A), ATC71GTG (881A>G 883C>G), ACA72GCA (884A>G), CTT74CTA (892T>A), ACA77GCT (899A>G 901A>T), TTC78TTT (904C>T), CAT80AGG (908C>A 909A>G 910T>G), AAG81ACC (912A>C 913G>C), TTT82ATA (914T>A 916T>A), GTT83ACA (917G>A 918T>C 919T>A), GAG84GAA (922G>A), TCT85TCA (925T>A), AAG86CGT (926A>C 927A>G 928G>T), GAC87GAT (931C>T), GCA89GGG (936C>G 937A>G), AAC90AAT (940C>T), TTG91TTA (943G>A), TTT93TTC (949T>C), CTG95TTA (953C>T 955G>A), TAT96CAC (956T>C 958T>C), GGT97GGG (961T>G), CTT99TTG (965C>T 967T>G), TTC100TTT (970C>T), TTC101TTT (973C>T), GTC103GTT (979C>T), CCA104CCC (982A>C), GTT106GCA (987T>C 988T>A), AGT108AAT (993G>A), GGG109GGA (997G>A), GGA111GGT (1003A>T), AGG112GCC (1004A>G 1005G>C 1006G>C), GTT113GTC (1009T>C), AAG114AAA (1012G>A), ATA115ATC (1015A>C), AGT116AGC (1018T>C), CTT117TTG (1019C>T 1021T>G), TGT118TGC (1024T>C), TCA119TCT (1027A>T), AGT120AGC (1030T>C), AAT121AAC (1033T>C), CTC124ACT (1040C>A 1041T>C 1042C>T), TCA125GAT (1043T>G 1044C>A 1045A>T), CCG126CCT (1048G>T), CAA129CAG (1057A>G), AAG131AAA (1063G>A), ACA132ATT (1065C>T 1066A>T), CTG133TTG (1067C>T), TCC134TAC (1071C>A), TTG135TTT (1075G>T), GCC136TCT (1076G>T 1078C>T), GAT137GGA (1080A>G 1081T>A), GCG139GCA (1087G>A), CAG140CAA (1090G>A), GCA141GCC (1093A>C), CTC143CTA (1099C>A), AGC145AGT (1105C>T), CCC146CCT (1108C>T), AGC147AGT (1111C>T), CTC153TTG (1127C>T 1129C>G), AAA154AAG (1132A>G), AGA155AGG (1135A>G), GGG156GGC (1138G>C), ACC162ACA (1156C>A), GAG164GAA (1162G>A), TGC165TGT (1165C>T), CAA166CTC (1167A>T 1168A>C), ACC168ACG (1174C>G), AGG169AGA (1177G>A), GCA170GCT (1180A>T), CAG171CAA (1183G>A), CCG173CCT (1189G>T), TCC175TCT (1195C>T), GTG176GTA (1198G>A), GCC178GCA (1204C>A), ATA179ATT (1207A>T), GAC185AGT (1223G>A 1224A>G 1225C>T), ACA186CCT (1226A>C 1228A>T), AGG187CGT (1229A>C 1231G>T), GCA189GCT (1237A>T), GTT190GTA (1240T>A), TCA192ACA (1244T>A), CAG193GAC (1247C>G 1249G>C), CAA194CAT (1252A>T), GAA195GAC (1255A>C), ACA196ACT (1258A>T), TCT198AGT (1262T>A 1263C>G), GCC200GCT (1270C>T), ATA201ATT (1273A>T), GAA202GAG (1276A>G), GCA203GCT (1279A>T), CTC204CTT (1282C>T), AAT205AAC (1285T>C), CGT206AGA (1286C>A 1288T>A), CCG207CCT (1291G>T), CAA208CAG (1294A>G), GAT212GAC (1306T>C), AGA213AGG (1309A>G), GAG215GAA (1315G>A), GCT216GCA (1318T>A), GCT217GCA (1321T>A), TAT219TAC (1327T>C), ATA220ATT (1330A>T), GCT221GCA (1333T>A), GTT223GTG (1339T>G), TAT224TAC (1342T>C), TCG225TCC (1345G>C), AGT226AAT (1347G>A), GGG227_CAA228insGTA (1351_1352insGTA), CAA228CAG (1354A>G), AGT229AGA (1357T>A), ACT230AGT (1359C>G), CAA231CAG (1363A>G), ATG232CTG (1364A>C), GCT233del (1367_1369delGCT), CTT234TTG (1370C>T 1372T>G), GAA235GAG (1375A>G), AAA237AGA (1380A>G), GCC238CCT (1382G>C 1384C>T), GTT240GTG (1390T>G), GGA241GGT (1393A>T), GAA242GCC (1395A>C 1396A>C), CAG243GCA (1397C>G 1398A>C 1399G>A), GGG245GGT (1405G>T), GGC246GAT (1407G>A 1408C>T), ACA247ACC (1411A>C), GAC250AAT (1418G>A 1420C>T), GTC251GTG (1423C>G), ATG252CCG (1424A>C 1425T>C), AAT253CAG (1427A>C 1429T>G), GAA254GAG (1432A>G), AGC255GCA (1433A>G 1434G>C 1435C>A), ACT256AAT (1437C>A), ATC258GTG (1442A>G 1444C>G), CGG259CGC (1447G>C), AGT260TCT (1448A>T 1449G>C), TCT262TCA (1456T>A), AGA264AGG (1462A>G), ACT267ACC (1471T>C), CTC268CTA (1474C>A), AAA269AAG (1477A>G), GTG270GTA (1480G>A), AAG273AGG (1488A>G), CGC274CGA (1492C>A), CCG276GCA (1496C>G 1498G>A), TCC277TCA (1501C>A), CTG280TTG (1508C>T), CCG281CCT (1513G>T), CCG281_GCT282insGACTTT (1513_1514insGACTTT), GTA283TCA (1517G>T 1518T>C), AGT284ACT (1521G>C), ACA285TCT (1523A>T 1525A>T), TCA286AGA (1526T>A 1527C>G), TCC287GCG (1529T>G 1531C>G), TTA288TAT (1533T>A 1534A>T), CTC289CAG (1536T>A 1537C>G), AGC290GGA (1538A>G 1540C>A), ACA291GTG (1541A>G 1542C>T 1543A>G), CGT292AGA (1544C>A 1546T>A), GAA293GAG (1549A>G), ACA295ACT (1555A>T), GTT296GGT (1557T>G), CAT297ATT (1559C>A 1560A>T), GAG299del (1565_1567delGAG), TGT302ACG (1574T>A 1575G>C 1576T>G), GGG303TCT (1577G>T 1578G>C 1579G>T), GGA304CTA (1580G>C 1581G>T), TTG305CTG (1583T>C), CCA307CCT (1591A>T), GCA308GCC (1594A>C), CCC309_AAG310del (1595_1600delCCCAAG), AAA311AAG (1603A>G), AAG312ATA (1605A>T 1606G>A), GGC313TCA (1607G>T 1608G>C 1609C>A), GCC315GCA (1615C>A), TTC316GGA (1616T>G 1617T>G 1618C>A), AAC317ACA (1620A>C 1621C>A), ATG318CTC (1622A>C 1624G>C), GGT319CAA (1625G>C 1626G>A 1627T>A), GCC320GAA (1629C>A 1630C>A), ATT321GTC (1631A>G 1633T>C), GAT323GAC (1639T>C), TTG325TAT (1644T>A 1645G>T), ATG327ATT (1651G>T), GAG328CAA (1652G>C 1654G>A), TCG329AGT (1655T>A 1656C>G 1657G>T), TTC330TTT (1660C>T), GCA331GAA (1662C>A), CAC332GAT (1664C>G 1666C>T), ATT333TTT (1667A>T), GAT334GCC (1671A>C 1672T>C), GAT337AGT (1679G>A 1680A>G), GAT338AAC (1682G>A 1684T>C), GAA341GCC (1692A>C 1693A>C), CGA342CTT (1695G>T 1696A>T), ATA344GTG (1700A>G 1702A>G), GCT345CCC (1703G>C 1705T>C), CAG346GAG (1706C>G), CAG347CAA (1711G>A), GTA348ATT (1712G>A 1714A>T), CAA349GAT (1715C>G 1717A>T), TTT350TAT (1719T>A), ATC351GTT (1721A>G 1723C>T), CTC352CTT (1726C>T), TTT353GTG (1727T>G 1729T>G), AGT354CAG (1730A>C 1731G>A 1732T>G), GAG355AAG (1733G>A), AAA357AGA (1740A>G), AGA358GCT (1742A>G 1743G>C 1744A>T), GGA359GGT (1747A>T), AAC360AAT (1750C>T), GTT361CTC (1751G>C 1753T>C), ATT362TGT (1754A>T 1755T>G), GTG363GTC (1759G>C), AGA365AAA (1764G>A), GTT367ATT (1769G>A), GCA368GAG (1773C>A 1774A>G), AAG369ATG (1776A>T), CGG370AGG (1778C>A), CAT371GAC (1781C>G 1783T>C), CTG372TTG (1784C>T), CAT373CAC (1789T>C), AAT374TGT (1790A>T 1791A>G), ATC375ATA (1795C>A), AGG377TTT (1799A>T 1800G>T 1801G>T), GAG378GAA (1804G>A), CAC379CAT (1807C>T), ATA380ATC (1810A>C), ACC381ACT (1813C>T), GAC383GCC (1818A>C), GTA386GTT (1828A>T), GAA387GAT (1831A>T), ATA388ATT (1834A>T), CTT389TTG (1835C>T 1837T>G), GAA390GCG (1839A>C 1840A>G), GGT391GGA (1843T>A), GGT393GGA (1849T>A), AAT396AAG (1858T>G), ATC397GTT (1859A>G 1861C>T), CAA398GGG (1862C>G 1863A>G 1864A>G), GGC399ATG (1865G>A 1866G>T 1867C>G), ACT401GAA (1871A>G 1872C>A 1873T>A), CGC402AGA (1874C>A 1876C>A), ACT403ACA (1879T>A), TTC404CCA (1880T>C 1881T>C 1882C>A), TAT407TAC (1891T>C), GCT408TCT (1892G>T), ATG409TTG (1895A>T), TCT410AGC (1898T>A 1899C>G 1900T>C), CTC411CTG (1903C>G), GAG413CAG (1907G>C), GTT415GTG (1915T>G), GTG416GTT (1918G>T), CTC418CTA (1924C>A), CAT420CAC (1930T>C), CAA421ATG (1931C>A 1932A>T 1933A>G), AGA422AGG (1936A>G), GAT423GAC (1939T>C), TCC425CAA (1943T>C 1944C>A 1945C>A), TTC426TTT (1948C>T), ATT428AGG (1953T>G 1954T>G), CAA429TAT (1955C>T 1957A>T), GGG430TGG (1958G>T), CAG431GAT (1961C>G 1963G>T), ACT432TTA (1964A>T 1965C>T 1966T>A), AAC433ACA (1968A>C 1969C>A), AAT434AAC (1972T>C), CCA435CCG (1975A>G), CCG437CCT (1981G>T), AAT438GAC (1982A>G 1984T>C), TGC441GTG (1991T>G 1992G>T 1993C>G), GGA444GGT (2002A>T), TTA445CTT (2003T>C 2005A>T), ACC446ACA (2008C>A), TTG447CTG (2009T>C), AAA448GCA (2012A>G 2013A>C), GAG449GAT (2017G>T), CGT450AGG (2018C>A 2020T>G), CGT450_CAG451insGTGAAA (2020_2021insGTGAAA), CAG451CAT (2023G>T), TTG452CCC (2024T>C 2025T>C 2026G>C), ATT453TTG (2027A>T 2029T>G), GTG454ATG (2030G>A), GAG455AGT (2033G>A 2034A>G 2035G>T), GAG456GAT (2038G>T), CAA457CTA (2040A>T), GTG458GCA (2043T>C 2044G>A), GCC459TGC (2045G>T 2046C>G), CAA460_AGA461insTTGAAT (2050_2051insTTGAAT), AGA461CGC (2051A>C 2053A>C), AGG462TTG (2054A>T 2055G>T), TCT463GGA (2057T>G 2058C>G 2059T>A), GAA464AAA (2060G>A), AGA465CCA (2063A>C 2064G>C), CAA466AGC (2066C>A 2067A>G 2068A>C), GCA467ATG (2069G>A 2070C>T 2071A>G), CAG468AGT (2072C>A 2073A>G 2074G>T), ACA470GCA (2078A>G), GCA471CAA (2081G>C 2082C>A), AGG472ACT (2085G>C 2086G>T), GGC473GGA (2089C>A), ATA474AAT (2091T>A 2092A>T), GCT475GCC (2095T>C), GCT475_GAA476insAGC (2095_2096insAGC), GAA476AAG (2096G>A 2098A>G), AGT477TGG (2099A>T 2101T>G), CAG478CAA (2104G>A), CCT479ACT (2105C>A), GAC481GGA (2112A>G 2113C>A), AGA482AAG (2115G>A 2116A>G), ACT484CTG (2120A>C 2121C>T 2122T>G), TCC486TCA (2128C>A), GTG488GTT (2134G>T), TCA489ACA (2135T>A), ACT490ACC (2140T>C), ACC491ACA (2143C>A), ACG492GCA (2144A>G 2146G>A), ATG493CTT (2147A>C 2149G>T), GAA494GAG (2152A>G), GAA494_GAT495insAGT (2152_2153insAGT), GAT495GAG (2155T>G), CCC496GGT (2156C>G 2157C>G 2158C>T), ACC497ATC (2160C>T), AAG498CAA (2162A>C 2164G>A), CCA499AGA (2165C>A 2166C>G), GAT500AAG (2168G>A 2170T>G), AAG501GAG (2171A>G), ATA502GTG (2174A>G 2176A>G), GAA503TCA (2177G>T 2178A>C), GTT505GAG (2184T>A 2185T>G), GCA506GCT (2188A>T), GAG507GAA (2191G>A), GGA508GAT (2193G>A 2194A>T), GCA509CCG (2195G>C 2197A>G), GAA511GGG (2202A>G 2203A>G), GAA512AAA (2204G>A), ACT513ACA (2209T>A), CAA514GAG (2210C>G 2212A>G), GGT516AAT (2216G>A 2217G>A), GAT517GAC (2221T>C), GTT518ATT (2222G>A), ATT519TTC (2225A>T 2227T>C), GAC521GAT (2233C>T), GGA523GGG (2239A>G), CCA524GAG (2240C>G 2241C>A 2242A>G), GAC527ATG (2249G>A 2250A>T 2251C>G), ACA528GGT (2252A>G 2253C>G 2254A>T), TCC529GAT (2255T>G 2256C>A 2257C>T), ATG530GCC (2258A>G 2259T>C 2260G>C), GCA531CCG (2261G>C 2263A>G), GTT532TTA (2264G>T 2266T>A), GAA533AAT (2267G>A 2269A>T), GAT535GAG (2275T>G), ATG536TTG (2276A>T), CAG537CAA (2281G>A), CAA538CAG (2284A>G), CCG539CCA (2287G>A), GTG540GCA (2289T>C 2290G>A), GTA542GTG (2296A>G), AGC544AGT (2302C>T), GAT546GAG (2308T>G), TTC547TTT (2311C>T), GGA551GGG (2323A>G), GTT552ATT (2324G>A), TTC555TTT (2335C>T), GTT556CTG (2336G>C 2338T>G), GAG558AAA (2342G>A 2344G>A), TCC560AGC (2348T>A 2349C>G), GCT561AAC (2351G>A 2352C>A 2353T>C), GAG565GAA (2365G>A), CAA566CAG (2368A>G), GTA567GTT (2371A>T), ATG568CTT (2372A>C 2374G>T), AGC569GCA (2375A>G 2376G>C 2377C>A), TTG570CTA (2378T>C 2380G>A), GCT571TCC (2381G>T 2383T>C), TTG572CTC (2384T>C 2386G>C), CCC573CCA (2389C>A), GCT574ATG (2390G>A 2391C>T 2392T>G), GCC575GCA (2395C>A), CTA576TTG (2396C>T 2398A>G), TTC577TTT (2401C>T), TCA578AGC (2402T>A 2403C>G 2404A>C), AAA579AAG (2407A>G), AGC580AAG (2409G>A 2410C>G), AAA581AAG (2413A>G), ACT583GCT (2417A>G), TCA584TCC (2422A>C), GGT586GGG (2428T>G), GCG587TCC (2429G>T 2431G>C), CTT590CTC (2440T>C), TAT592TAC (2446T>C), TAC593TAT (2449C>T), GAT594GAC (2452T>C), GCA595GCT (2455A>T), GCT596GCA (2458T>A), CTA597AAC (2459C>A 2460T>A 2461A>C), ATC598CTT (2462A>C 2464C>T), TAC600TTT (2469A>T 2470C>T), ATA602GTC (2474A>G 2476A>C), CTT604CTG (2482T>G), TAT605TAC (2485T>C), GTT606ATT (2486G>A), GTT609ATG (2495G>A 2497T>G), GCT611GCG (2503T>G), ATC612GTG (2504A>G 2506C>G), TCT613TCG (2509T>G), GGT614GGA (2512T>A), CAA615CAG (2515A>G), CTG616CTA (2518G>A), GCT617GCC (2521T>C), GAA622GAG (2536A>G), TGT623TGC (2539T>C), GTG625GCA (2544T>C 2545G>A), CGA628AGA (2552C>A), AAG629ACC (2556A>C 2557G>C), AAC634AAT (2572C>T), ATC635ATA (2575C>A), ACA637ACC (2581A>C), TTG638ATT (2582T>A 2584G>T), AAG642AAA (2596G>A), ACA644CAT (2600A>C 2601C>A 2602A>T), TTG645CTG (2603T>C), GTT646GTG (2608T>G), CAA650GAA (2618C>G), AAC652TCC (2624A>T 2625A>C), GAG654GGG (2631A>G), GAG655AAG (2633G>A), TGC657GTG (2639T>G 2640G>T 2641C>G), TTT658TTC (2644T>C), ACC659TCA (2645A>T 2647C>A), CCA660CCT (2650A>T), ACA661CAA (2651A>C 2652C>A), GGG662GGT (2656G>T), GGC664GGT (2662C>T), AAG665ACC (2664A>C 2665G>C), TAC666TTT (2667A>T 2668C>T), GTG667GTA (2671G>A), CCT668CCA (2674T>A), CTG669TTG (2675C>T), GAA671ACT (2681G>A 2682A>C 2683A>T), GGT672TCA (2684G>T 2685G>C 2686T>A), ACA673CTG (2687A>C 2688C>T 2689A>G), GGA674GGT (2692A>T), GCC675GTT (2694C>T 2695C>T), ACT676CCT (2696A>C), GAT677GAA (2701T>A), TTA678CTT (2702T>C 2704A>T), GGT679GGC (2707T>C), GTG681GTT (2713G>T), ACA686ACC (2728A>C), CAC687CAT (2731C>T), TTA689TTG (2737A>G), TCT690GCA (2738T>G 2740T>A), GCA692GCC (2746A>C), GAG694GAA (2752G>A), AAT696GAG (2756A>G 2758T>G), AGT697TCA (2759A>T 2760G>C 2761T>A), GTA698ATA (2762G>A), CCA699CCT (2767A>T), TGC700TGT (2770C>T), CAT701CAC (2773T>C), TTG702ATA (2774T>A 2776G>A), TTA704CTA (2780T>C), CAG705TCC (2783C>T 2784A>C 2785G>C), AAA707AAG (2791A>G), TTA709CTC (2795T>C 2797A>C), ACC711ACA (2803C>A), CTT714ATG (2810C>A 2812T>G), CGA718AGG (2822C>A 2824A>G), ATG719TTG (2825A>T), ATA720AGA (2829T>G), GCA721GGG (2832C>G 2833A>G), CAG722CAA (2836G>A), CAA724CAG (2842A>G), TAT725TTT (2844A>T), GGC726GGA (2848C>A), AAG728CGG (2852A>C 2853A>G), GCG729CCA (2855G>C 2857G>A), GGG730GGC (2860G>C), CAG731ACA (2861C>A 2862A>C 2863G>A), ACA732ACC (2866A>C), TAT733TTC (2868A>T 2869T>C), CCA735AGT (2873C>A 2874C>G 2875A>T), AGG736CGC (2876A>C 2878G>C), TTT737TTC (2881T>C), CCA738CCT (2884A>T), ACT739ACA (2887T>A), GTT742GTG (2896T>G), TTG743CTT (2897T>C 2899G>T), TTA744CTG (2900T>C 2902A>G), CAT745CAC (2905T>C), TAT746TAC (2908T>C), GGG749AGT (2915G>A 2917G>T), ACA750GTA (2918A>G 2919C>T), TCA751GCT (2921T>G 2923A>T), TCC752GCA (2924T>G 2926C>A), CCA753GCC (2927C>G 2929A>C), ACA757ACT (2941A>T), AGC760AGT (2950C>T), ATA761GTC (2951A>G 2953A>C), TTT762TTC (2956T>C), TCA763TCC (2959A>C), ATA767ATT (2971A>T), GAG769GAA (2977G>A), GGC772GGT (2986C>T), ACC773ATC (2988C>T), TTG774TTA (2992G>A), CCG776CCA (2998G>A), TCT777TCA (3001T>A), TTG778CTG (3002T>C), GGT780GGA (3010T>A), ATA782ATT (3016A>T), GCC783GCA (3019C>A), AGG784AGA (3022G>A), AAG787AAA (3031G>A), ACT790AGA (3039C>G 3040T>A), GGA791GGC (3043A>C), ACT792ACC (3046T>C), ATT797GTG (3059A>G 3061T>G), TGC798TGT (3064C>T), ATT799GTT (3065A>G), ACC802ACT (3076C>T), TTG803CTG (3077T>C), TCA806TCT (3088A>T), GGT807GGG (3091T>G), CTG809CTT (3097G>T), GCA810GCT (3100A>T), ATT811ATA (3103T>A), GGA812GGG (3106A>G), CTT813TTG (3107C>T 3109T>G), GGA814GGC (3112A>C), ACT815ACG (3115T>G), AAC817GCA (3119A>G 3120A>C 3121C>A), ACC818ACA (3124C>A), AAA819GAG (3125A>G 3127A>G), ACC821TCC (3131A>T), AAT822ACA (3135A>C 3136T>A), GCT823GCG (3139T>G), TTT827TTG (3151T>G), CCA830CCC (3160A>C), GTA833GTG (3169A>G), TGC834TGT (3172C>T), CTT836TTG (3176C>T 3178T>G), GGT839GGG (3187T>G), AAA841AGG (3192A>G 3193A>G), CGG843CAA (3198G>A 3199G>A), CGG845AAA (3203C>A 3204G>A 3205G>A), TGT846TGC (3208T>C), TCT847TCA (3211T>A), ATT848CTG (3212A>C 3214T>G), ACA849ACC (3217A>C), GGA853GGG (3229A>G), AAA854TGC (3230A>T 3231A>G 3232A>C), AAT855AAC (3235T>C), TTG856CTC (3236T>C 3238G>C), CTT857CTG (3241T>G), TCC858TCT (3244C>T), ACA859ACC (3247A>C), GGG860GGA (3250G>A), CGA861CGC (3253A>C), AAG862AAA (3256G>A), TCC864TCA (3262C>A), TTG865TTA (3265G>A), CAG869AAG (3275C>A), CAC870CAT (3280C>T), TTT871CAA (3281T>C 3282T>A 3283T>A), TCC872TCT (3286C>T), TTG874ATG (3290T>A), CGC875CGT (3295C>T), TTG876CTG (3296T>C), TTT877TTC (3301T>C), ACG879ACT (3307G>T), GTA880GTT (3310A>T), CTC884CTG (3322C>G), ACG887ACA (3331G>A), ATA889TTG (3335A>T 3337A>G), CAT890CAC (3340T>C), TCC893ACT (3347T>A 3349C>T), GGT895GGA (3355T>A), GTC896GTA (3358C>A), CAG899CAA (3367G>A), AAG901AAA (3373G>A), TGC902CGC (3374T>C), CTT903ATT (3377C>A), AAT905GAT (3383A>G), CTT906CTC (3388T>C), ACT907ACG (3391T>G), TTG908CTT (3392T>C 3394G>T), GGT909GGG (3397T>G), GGC910GGT (3400C>T), ACA911TCT (3401A>T 3403A>T), GTA912GTG (3406A>G), TCT913TCA (3409T>A), GTG914GTT (3412G>T), AAA915AAG (3415A>G), ATA917ATT (3421A>T), GGG919GGT (3427G>T), ACA922ACC (3436A>C), AAA923AAG (3439A>G), GGC924GGG (3442C>G), AAA925AGA (3444A>G), AGC926AGT (3448C>T), TCA927GCT (3449T>G 3451A>T), GTT928GTG (3454T>G), TTC930TTT (3460C>T),

| | |
|---|---|
| | CTT931CTG (3463T>G), TTC932TTT (3466C>T), GAT936AAC (3476G>A 3478T>C), TTA937CTG (3479T>C 3481A>G), TCA938AAT (3482T>A 3483C>A 3484A>T), CGT940CGC (3490T>C), ATT943CTT (3497A>C), GAA944GAG (3502A>G), AAG945AAA (3505G>A), TTG946CTG (3506T>C), AAG948AAA (3514G>A), GAC949GAA (3517C>A), AAC950AAT (3520C>T), ATT951GTT (3521A>G), GAG952GAT (3526G>T), GAG953ATC (3527G>A 3528A>T 3529G>C), TAC954TTT (3531A>T 3532C>T), CAG955GAC (3533C>G 3535G>C), GAG956CAG (3536G>C), GGC958GGG (3544C>G), GAT960_CCG961del (3548_3553delGATCCG), CCC962AGC (3554C>A 3555C>G), AAG964ACA (3561A>C 3562G>A), GCT965TCT (3563G>T), AGT967GTT (3569A>G 3570G>T), ATT968TCC (3572A>T 3573T>C 3574T>C), CTG969TTC (3575C>T 3577G>C), TCC970CAG (3578T>C 3579C>A 3580C>G), ATA971ATC (3583A>C), AGA972AGG (3586A>G), GAG973GAA (3589G>A), AAA974AAG (3592A>G), TTT975TTC (3595T>C), TCC976ACA (3596T>A 3598C>A), GGT978GGA (3604T>A), GCT979GCA (3607T>A), GTA980GTG (3610A>G), TTC983TTT (3619C>T), TGC984TGT (3622C>T), ATG985GCT (3623A>G 3624T>C 3625G>T), GGT986GCA (3627G>C 3628T>A), GAC990ACA (3638G>A 3639A>C 3640C>A), GAA991GAC (3643A>C), TTG994GTA (3650T>G 3652G>A), GTA995GTT (3655A>T), ATT996TTG (3656A>T 3658T>G), CCT997CCA (3661T>A), GCA999GCC (3667A>C), CCA1000CCT (3670A>T), TCC1002TCA (3676C>A), ATA1003GTG (3677A>G 3679A>G), AGG1004CGC (3680A>C 3682G>C), GAA1006GAT (3688A>T), GGG1007GCA (3690G>C 3691G>A), CAC1008TCC (3692C>T 3693A>C), CCT1010CCA (3700T>A), GTT1011ATT (3701G>A), AAG1012AAA (3706G>A), ATC1015ATA (3715C>A), TGT1017TGC (3721T>C), TTT1019TTC (3727T>C), GAT1021GAC (3733T>C), ACA1024TCC (3740A>T 3742A>C), TCA1025TCT (3745A>T), TCA1030TCT (3760A>T), GGT1031GGA (3763T>A), AAT1034GAA (3770A>G 3772T>A), TCA1036ACA (3776T>A), ATT1037TTT (3779A>T), GTG1038GTA (3784G>A), ATA1039ATG (3787A>G), AGA1041CGA (3791A>C), GTA1042ACT (3794G>A 3795T>C 3796A>T), CAA1043CAG (3799A>G), TCC1044AGC (3800T>A 3801C>G), AGT1045TCC (3803A>T 3804G>C 3805T>C), CCT1046AGC (3806C>A 3807C>G 3808T>C), AAT1047AGT (3810A>G), GTT1048ATA (3812G>A 3814T>A), GGA1049GGT (3817A>T), GTA1051ATC (3821G>A 3823A>C), CTA1052TTG (3824C>T 3826A>G), AAT1053AAC (3829T>C), GTT1054ATT (3830G>A), GCT1055GCA (3835T>A), TTT1056CTT (3836T>C), GAT1057GAC (3841T>C), GCC1058ACA (3842G>A 3844C>A), GGC1060GGG (3850C>G), TTT1063TTC (3859T>C), CCA1064CAA (3861C>A), GCT1065TTT (3863G>T 3864C>T), GGG1066GGT (3868G>T), CTT1067GTA (3869C>G 3871T>A), AAT1068TCT (3872A>T 3873A>C), AAA1069GCG (3875A>G 3876A>C 3877A>G), AAT1071GCA (3881A>G 3882A>C 3883T>A), TAT1072CAT (3884T>C), GTA1074TTA (3890G>T), GGT1077GGC (3901T>C), GGA1078GGT (3904A>T), GGC1079GGA (3907C>A), ACA1080ACT (3910A>T), TCA1085TCC (3925A>C), TAC1086TAT (3928C>T), GGT1087GGG (3931T>G), GTG1088GTT (3934G>T), GCA1089ACC (3935G>A 3937A>C), ACA1090ACC (3940A>C), AAT1091AAC (3943T>C), ACG1092ATC (3945C>T 3946G>C), TCA1094TCT (3952A>T), TTC1095TTT (3955C>T), ACT1096GTA (3956A>G 3957C>T 3958T>A), GAG1101CAG (3971G>C), TTT1102TTC (3976T>C), TTC1103TTT (3979C>T), CCA1104CCT (3982A>T), AGG1105CGC (3983A>C 3985G>C), CGG1106CGT (3988G>T), CAT1107TTC (3989C>T 3990A>T 3991T>C), AGG1109AAG (3996G>A), ATG1110GCC (3998A>G 3999T>C 4000G>C), AGG1111AAA (4002G>A 4003G>A), GAA1112TCC (4004G>T 4005A>C 4006A>C), TTC1113TAT (4008T>A 4009C>T), TCA1114GAT (4010T>G 4011C>A 4012A>T), AGC1115AAG (4014G>A 4015C>G), AAG1116GAA (4016A>G 4018G>A), CAA1117AGT (4019C>A 4020A>G 4021A>T), TCC1118TCA (4024C>A), CGC1119AGA (4025C>A 4027C>A), ATC1120ATT (4030C>T), ATG1121GTT (4031A>G 4033G>T), TCA1122TCT (4036A>T), CTA1123TTG (4037C>T 4039A>G), GAT1125GAC (4045T>C), AGG1126AAG (4047G>A), CTT1127CTA (4051T>A), GGA1128GGC (4054A>C), AAT1129TTG (4055A>T 4056A>T 4057T>G), CTG1130TTG (4058C>T), ATC1131CTA (4061A>C 4063C>A), ATA1132ATC (4066A>C), TTG1134CTT (4070T>C 4072G>T), CCT1135CCA (4075T>A), CCT1136CCA (4078T>A), TCC1137GTT (4079T>G 4080C>T 4081C>T), GCC1138GAA (4083C>A 4084C>A), ATA1139CTT (4085A>C 4087A>T), GTG1140ATT (4088G>A 4090G>T), TCC1142TCA (4096C>A), GAG1144GAA (4102G>A), ATA1145ATC (4105A>C), CTT1146ATA (4106C>A 4108T>A), ATA1147GTC (4109A>G 4111A>C), TCT1148AAG (4112T>A 4113C>A 4114T>G), CCT1149CCA (4117T>A), GGA1150GGT (4120A>T), CTT1151CCT (4122T>C), TTC1153TTT (4129C>T), AAG1154AAA (4132G>A), GCC1158GCT (4144C>T), CCT1160CCC (4150T>C), CCT1161CCA (4153T>A), TCT1162CTT (4154T>C 4155C>T), GCC1163GCA (4159C>A), CAT1165AAT (4163C>A), GAA1166GAG (4168A>G), AAA1167AAG (4171A>G), CTT1169CTC (4177T>C), GGC1170GGA (4180C>A), AAT1171GAT (4181A>G), CAA1173CAG (4189A>G), ACT1174ACG (4192T>G), CAC1175CAT (4195C>T), ACC1176ACT (4198C>T), ACC1180ACA (4210C>A), TCA1181TCT (4213A>T), CTA1186TTG (4226C>T 4228A>G), CGT1187CGC (4231T>C), ATT1191AAT (4242T>A) |

#### Proteins

| | | | | | | | | | |
|---|---|---|---|---|---|---|---|---|---|
| hypothetical protein (YP_001976148.1) | 1 | 191 | 100% | 784 | 66.7% | 183 (95.8%) | 116 (60.7%) | 0/8/0/0 | 1 |
| Protein mutations: | G5S (151G>A 153G>T), R6K (154C>A 155G>A 156T>G), S10Q (166T>C 167C>A 168T>A), V11I (169G>A), D12E (174C>G), E13D (177A>T), A15E (182C>A 183T>A), H18N (190C>A), T22A (202A>G 204C>A), N24T (209A>C 210T>C), I26V (214A>G), V31T (229G>A 230T>C), T33V (235A>G 236C>T 237C>T), L35I (241T>A 243G>T), V46I (274G>A), V47C (277G>T 278T>G), V54I (298G>A 300C>T), T58S (310A>T 312C>T), L68V (340T>G 342A>T), N79T (374A>C 375T>G), P81D (379C>G 380C>A 381C>T), N84A (388A>G 389A>C 390C>A), L91I (409C>A), H93R (416A>G 417T>A), V96I (424G>A 426G>T), R97K (427C>A 428G>A 429T>A), V104L (448G>T 450C>G), E107A (458A>C 459A>C), S111H (469T>C 470C>A), A114S (478G>T 480A>T), S115T (481T>A 483A>C), P120N (496C>A 497C>A 498T>C), S121P (499T>C 501T>C), D124_S129del (508_525delGACAGTAAAGGCAAGAGT), K130T (527A>C), V131L (529G>T), E132S (532G>T 533A>C 534G>A), Q133E (535C>G), P134T (538C>A 540C>T), T135A (541A>G), R136_E137del (544_549delCGAGAG), D138S (550G>A 551A>G), I140S (557T>G 558T>C), K141G (559A>G 560A>G), V143N (565G>A 566T>A 567T>C), E144K (568G>A), V145A (572T>C 573T>A), R148E (580C>G 581G>A 582T>A), E149A (584A>C 585G>C), R152V (592A>G 593G>T 594A>T), F153A (595T>G 596T>C 597T>C), Q154R (598C>A 599A>G 600G>A), A158Q (610G>C 611C>A), S159E (613A>G 614G>A 615T>A), Q160A (616C>G 617A>C), K161R (619A>C 620A>G 621A>G), N164I (629A>T), Q165V (631C>G 632A>T 633G>C), Q168R (640C>A 641A>G 642A>G), L171I (649C>A 651C>T), S172Q (652T>C 653C>A 654C>A), N173G (655A>G 656A>G), V175P (661G>C 662T>C 663T>A), N178E (670A>G 672T>G), I180V (676A>G), S182A (682T>G), E186S (694G>A 695A>G 696A>T), S187D (697A>G 698G>A), Q190K (706C>A) | | | | | | | | |
| Codon mutations: | TCA2TCT (144A>T), TTC3TTT (147C>T), ATT4ATC (150T>C), GGG5AGT (151G>A 153G>T), CGT6AAG (154C>A 155G>A 156T>G), TTG7CTC (157T>C 159G>C), AAT8AAC (162T>C), ACA9ACC (165A>C), TCT10CAA (166T>C 167C>A 168T>A), GTA11ATA (169G>A), GAC12GAG (174C>G), GAA13GAT (177A>T), GCT15GAA (182C>A 183T>A), TTT16TTC (186T>C), CAT18AAT (190C>A), GTT20GTA (198T>A), GCC21GCA (201C>A), ACC22GCA (202A>G 204C>A), TCG23TCT (207G>T), AAT24ACC (209A>C 210T>C), ATT26GTT (214A>G), TCT28AGT (220T>A 221C>G), GTT31ACT (229G>A 230T>C), GGC32GGT (234C>T), ACC33GTT (235A>G 236C>T 237C>T), TTG35ATT (241T>A 243G>T), ATA36ATT (246A>T), ACC41ACT (261C>T), CTT42TTA (262C>T 264T>A), GAC43GAT (267C>T), AAA45AAG (273A>G), GTT46ATT (274G>A), GTT47TGT (277G>T 278T>G), CCT48CCC (282T>C), GGT51GGA (291T>A), GTC54ATT (298G>A 300C>T), TCA55TCG (303A>G), GTT56GTG (306T>G), TTA57CTT (307T>C 309A>T), ACC58TCT (310A>T 312C>T), GTT59GTG (315T>G), TCT60AGT (316T>A 317C>G), GAG62GAA (324G>A), ACG65ACT (333G>T), CCA66CCC (336A>C), TTA68GTT (340T>G 342A>T), CCT70CCA (348T>A), GGA71GGC (351A>C), CAC72CAT (354C>T), TAT73TAC (357T>C), TTG74CTA (358T>C 360G>A), TTG75CTA (361T>C 363G>A), AAT79ACG (374A>C 375T>G), CCC81GAT (379C>G 380C>A 381C>T), ATT82ATC (384T>C), AAG83AAA (387G>A), AAC84GCA (388A>G 389A>C 390C>A), TCT88TCA (402T>A), GGT89GGC (405T>C), TTA90TTG (408A>G), CTT91ATT (409C>A), GTC92GTG (414C>G), CAT93CGA (416A>G 417T>A), TCA95TCC (423A>C), GTG96ATT (424G>A 426G>T), CGT97AAA (427C>A 428G>A 429T>A), CTT98TTG (430C>T 432T>G), ACC101ACA (441C>A), AGA102CGG (442A>C 444A>G), GTC104TTG (448G>T 450C>G), CTA105TTG (451C>T 453A>G), GAG106GAA (456G>A), GAA107GCC (458A>C 459A>C), AAC108AAT (462C>T), AAG109AAA (465G>A), GTT110GTG (468T>G), TCT111CAT (469T>C 470C>A), TCA113TCT (477A>T), GCA114TCT (478G>T 480A>T), TCA115ACC (481T>A 483A>C), TCA117TCT (489A>T), TCT118TCA (492T>A), TCT119TCC (495T>C), CCT120AAC (496C>A 497C>A 498T>C), TCT121CCC (499T>C 501T>C), GAC124_AGT129del (508_525delGACAGTAAAGGCAAGAGT), AAA130ACA (527A>C), GTA131TTA (529G>T), GAG132TCA (532G>T 533A>C 534G>A), CAA133GAA (535C>G), CCC134ACT (538C>A 540C>T), ACA135GCA (541A>G), CGA136_GAG137del (544_549delCGAGAG), GAT138AGT (550G>A 551A>G), CTC139TTA (553C>T 555C>A), ATT140AGC (557T>G 558T>C), AAA141GGA (559A>G 560A>G), GAA142GAG (564A>G), GTT143AAC (565G>A 566T>A 567T>C), GAG144AAG (568G>A), GTT145GCA (572T>C 573T>A), CTC146CTT (576C>T), AAA147AAG (579A>G), CGT148GAA (580C>G 581G>A 582T>A), GAG149GCC (584A>C 585G>C), TTA150CTT (586T>C 588A>T), GAG151GAA (591G>A), AGA152GTT (592A>G 593G>T 594A>T), TTT153GCC (595T>G 596T>C 597T>C), CAG154AGA (598C>A 599A>G 600G>A), GAG156GAA (606G>A), TTG157CTG (607T>C), GCA158CAA (610G>C 611C>A), AGT159GAA (613A>G 614G>A 615T>A), CAA160GCA (616C>G 617A>C), AAA161CGG (619A>C 620A>G 621A>G), AAT164ATT (629A>T), CAG165GTC (631C>G 632A>T 633G>C), CTA167CTC (639A>C), CAA168AGG (640C>A 641A>G 642A>G), CTT169TTG (643C>T 645T>G), CTC171ATT (649C>A 651C>T), TCC172CAA (652T>C 653C>A 654C>A), AAT173GGT (655A>G 656A>G), GTT175CCA (661G>C 662T>C 663T>A), AGT176AGC (666T>C), AAT177AAC (669T>C), AAT178GAG (670A>G 672T>G), GAC179GAT (675C>T), ATC180GTC (676A>G), TTC181TTT (681C>T), TCA182GCA (682T>G), GGT183GGC (687T>C), ACT185ACG (693T>G), GAA186AGT (694G>A 695A>G 696A>T), AGT187GAT (697A>G 698G>A), GGG188GGC (702G>C), CAG190AAG (706C>A), TAA191TGA (710A>G) | | | | | | | | |
| polyprotein (YP_001976149.1) | 1 | 1192 | 100% | 6009 | 73.0% | 1185 (98.7%) | 820 (68.3%) | 9/7/0/0 | 1 |

| | Begin | End | Coverage | Score | Concordance | Matches | Identities | I/D/M/F* | Stop Codons |
|---|---|---|---|---|---|---|---|---|---|
| **NT** | **89** | **4744** | **94.9%** | **2600** | **28.6%** | **4608 (96.7%)** | **3040 (63.8%)** | **107/48** | |
| Protein mutations: | M1R (672T>G), T2M (675C>T 676A>G), S4L (681C>T 682T>G), V6A (687T>C), L8R (693T>G 694G>A), K9V (695A>G 696A>T 697A>G), V10M (698G>A), G11A (702G>C), N14D (710A>G 712T>C), H15R (714A>G), Q16S (716C>T 717A>C 718G>A), Q17del (719_721delCAG), E18K (722G>A 724G>A), A20V (729C>T 730C>A), E22D (736G>C), R23K (738G>A 739G>A), A34E (771C>A 772C>A), R41K (792G>A), V50I (818G>A 820T>A), G53A (828G>C 829T>A), E61D (853G>T), F63Y (858T>A), I71V (881A>G 883C>G), T72A (884A>G), T77A (899A>G 901A>T), H80R (908C>A 909A>G 910T>G), K81T (912A>C 913G>C), F82I (914T>A 916T>A), V83T (917G>A 918T>C 919T>A), K86R (926A>C 927A>G 928G>T), A89G (936C>G 937A>G), Y96H (956T>C 958T>C), V106A (987T>C 988T>A), S108N (993G>A), R112A (1004A>G 1005G>C 1006G>C), L124T (1040C>A 1041T>C 1042C>T), S125D (1043T>G 1044C>A 1045A>T), T132I (1065C>T 1066A>T), S134Y (1071C>A), L135F (1075G>T), A136S (1076G>T 1078C>T), D137G (1080A>G 1081T>A), Q166L (1167A>T 1168A>C), D185S (1223G>A 1224A>G 1225C>T), T186P (1226A>C 1228A>T), S192T (1244T>A), Q193D (1247C>G 1249G>C), Q194H (1252A>T), E195D (1255A>C), S226N (1347G>A), G227_Q228insV (1351_1352insGTA), S229R (1357T>A), T230S (1359C>G), M232L (1364A>C), A233del (1367_1369delGCT), K237R (1380A>G), A238P (1382G>C 1384C>T), E242A (1395A>C 1396A>C), Q243A (1397C>G 1398A>C 1399G>A), G246D (1407G>A 1408C>T), D250N (1418G>A 1420C>T), M252P (1424A>C 1425T>C), N253Q (1427A>C 1429T>G), S255A (1433A>G 1434G>C 1435C>A), T256N (1437C>A), I258V (1442A>G 1444C>G), K273R (1488A>G), P276A (1496C>G 1498G>A), P281_A282insDF (1513_1514insGACTTT), V283S (1517G>T 1518T>C), S284T (1521G>C), T285S (1523A>T 1525A>T), S286R (1526T>A 1527C>G), S287A (1529T>G 1531C>G), L288Y (1533T>A 1534A>T), L289Q (1536T>A 1537C>G), S290G (1538A>G 1540C>A), T291V (1541A>G 1542C>T 1543A>G), V296G (1557T>G), H297I (1559C>A 1560A>T), E299del (1565_1567delGAG), C302T (1574T>A 1575G>C 1576T>G), G303S (1577G>T 1578G>C 1579G>T), G304L (1580G>C 1581G>T), P309_K310del (1595_1600delCCCAAG), K312I (1605A>T 1606G>A), G313S (1607G>T 1608G>C 1609C>A), F316G (1616T>G 1617T>G 1618C>A), N317T (1620A>C 1621C>A), M318L (1622A>C 1624G>C), G319Q (1625G>C 1626G>A 1627T>A), A320E (1629C>A 1630C>A), I321V (1631A>G 1633T>C), L325Y (1644T>A 1645G>T), M327I (1651G>T), E328Q (1652G>C 1654G>A), A331E (1662C>A), H332D (1664C>G 1666C>T), I333F (1667A>T), D334A (1671A>C 1672T>C), D337S (1679G>A 1680A>G), D338N (1682G>A 1684T>C), E341A (1692A>C 1693A>C), R342L (1695G>T 1696A>T), I344V (1700A>G 1702A>G), A345P (1703G>C 1705T>C), Q346E (1706C>G), V348I (1712G>A 1714A>T), Q349D (1715C>G 1717A>T), F350Y (1719T>A), I351V (1721A>G 1723C>T), F353V (1727T>G 1729T>G), S354Q (1730A>C 1731G>A 1732T>G), E355K (1733G>A), K357R (1740A>G), R358A (1742A>G 1743G>C 1744A>T), V361L (1751G>C 1753T>C), I362C (1754A>T 1755T>G), R365K (1764G>A), V367I (1769G>A), A368E (1773C>A 1774A>G), K369M (1776A>T), H371D (1781C>G 1783T>C), N374C (1790A>T 1791A>G), R377F (1799A>T 1800G>T 1801G>T), D383A (1818A>C), E387D (1831A>T), E390A (1839A>C 1840A>G), N396K (1858T>G), I397V (1859A>G 1861C>T), Q398G (1862C>G 1863A>G 1864A>G), G399M (1865G>A 1866G>T 1867C>G), T401E (1871A>G 1872C>A 1873T>A), F404P (1880T>C 1881T>C 1882C>A), A408S (1892G>T), M409L (1895A>T), E413Q (1907G>C), Q421M (1931C>A 1932A>T 1933A>G), S425Q (1943T>C 1944C>A 1945C>A), I428R (1953T>G 1954T>G), Q429Y (1955C>T 1957A>T), G430W (1958G>T), Q431D (1961C>G 1963G>T), T432L (1964A>T 1965C>T 1966T>A), N433T (1968A>C 1969C>A), N438D (1982A>G 1984T>C), C441V (1991T>G 1992G>T 1993C>G), K448A (2012A>G 2013A>C), E449D (2017G>T), R450_Q451insVK (2020_2021insGTGAAA), Q451H (2023G>T), L452P (2024T>C 2025T>C 2026G>C), I453L (2027A>T 2029T>G), V454M (2030G>A), E455S (2033G>A 2034A>G 2035G>T), E456D (2038G>T), Q457L (2040A>T), V458A (2043T>C 2044G>A), A459C (2045G>T 2046C>G), Q460_R461insLN (2050_2051insTTGAAT), R462L (2054A>T 2055G>T), S463G (2057T>G 2058C>G 2059T>A), E464K (2060G>A), R465P (2063A>C 2064G>C), Q466S (2066C>A 2067A>G 2068A>C), A467M (2069G>A 2070C>T 2071A>G), Q468S (2072C>A 2073A>G 2074G>T), T470A (2078A>G), A471Q (2081G>C 2082C>A), R472T (2085G>C 2086G>T), I474N (2091T>A 2092A>T), A475_E476insS (2095_2096insAGC), E476K (2096G>A 2098A>G), S477W (2099A>T 2101T>G), P479T (2105C>A), D481G (2112A>G 2113C>A), R482K (2115G>A 2116A>G), T484L (2120A>C 2121C>T 2122T>G), S489T (2135T>A), T492A (2144A>G 2146G>A), M493L (2147A>C 2149G>T), E494_D495insS (2152_2153insAGT), D495E (2155T>G), P496G (2156C>G 2157C>G 2158C>T), T497I (2160C>T), K498Q (2162A>C 2164G>A), P499R (2165C>A 2166C>G), D500K (2168G>A 2170T>G), K501E (2171A>G), I502V (2174A>G 2176A>G), E503S (2177G>T 2178A>C), V505E (2184T>A 2185T>G), G508D (2193G>A 2194A>T), A509P (2195G>C 2197A>G), E511G (2202A>G 2203A>G), E512K (2204G>A), Q514E (2210C>G 2212A>G), G516N (2216G>A 2217G>A), V518I (2222G>A), I519F (2225A>T 2227T>C), P524E (2240C>G 2241C>A 2242A>G), D527M (2249G>A 2250A>T 2251C>G), T528G (2252A>G 2253C>G 2254A>T), S529D (2255T>G 2256C>A 2257C>T), M530A (2258A>G 2259T>C 2260G>C), A531P (2261G>C 2263A>G), V532L (2264G>T 2266T>A), E533N (2267G>A 2269A>T), D535E (2275T>G), M536L (2276A>T), V540A (2289T>C 2290G>A), D546E (2308T>G), V552I (2324G>A), V556L (2336G>C 2338T>G), E558K (2342G>A 2344G>A), A561N (2351G>A 2352C>A 2353T>C), M568L (2372A>C 2374G>T), S569A (2375A>G 2376G>C 2377C>A), A571S (2381G>T 2383T>C), A574M (2390G>A 2391C>T 2392T>G), S580K (2409G>A 2410C>G), T583A (2417A>G), A587S (2429G>T 2431G>C), L597N (2459C>A 2460T>A 2461A>C), I598L (2462A>C 2464C>T), Y600F (2469A>T 2470C>T), I602V (2474A>G 2476A>C), V606I (2486G>A), V609M (2495G>A 2497T>G), I612V (2504A>G 2506C>G), V625A (2544T>C 2545G>A), K629T (2556A>C 2557G>C), L638I (2582T>A 2584G>T), T644H (2600A>C 2601C>A 2602A>T), Q650E (2618C>G), N652S (2624A>T 2625A>C), E654G (2631A>G), E655K (2633G>A), C657V (2639T>G 2640G>T 2641C>G), T659S (2645A>T 2647C>A), T661Q (2651A>C 2652C>A), K665T (2664A>C 2665G>C), Y666F (2667A>T 2668C>T), E671T (2681G>A 2682A>C 2683A>T), G672S (2684G>T 2685G>C 2686T>A), T673L (2687A>C 2688C>T 2689A>G), A675V (2694C>T 2695C>T), T676P (2696A>C), D677E (2701T>A), S690A (2738T>G 2740T>A), N696E (2756A>G 2758T>G), V698I (2762G>A), L702I (2774T>A 2776G>A), Q705S (2783C>T 2784A>C 2785G>C), L714M (2810C>A 2812T>G), M719L (2825A>T), I720R (2829T>G), A721G (2832C>G 2833A>G), Y725F (2844A>T), K728R (2852A>C 2853A>G), A729P (2855G>C 2857G>A), Q731T (2861C>A 2862A>C 2863G>A), Y733F (2868A>T 2869T>C), P735S (2873C>A 2874C>G 2875A>T), G749S (2915G>A 2917G>T), T750V (2918A>G 2919C>T), S751A (2921T>G 2923A>T), S752A (2924T>G 2926C>A), P753A (2927C>G 2929A>C), I761V (2951A>G 2953A>C), T773I (2988C>T), T790R (3039C>G 3040T>A), I797V (3059A>G 3061T>G), I799V (3065A>G), N817A (3119A>G 3120A>C 3121C>A), K819E (3125A>G 3127A>G), T821S (3131A>T), N822T (3135A>C 3136T>A), F827L (3151T>G), K841R (3192A>G 3193A>G), R843Q (3198G>A 3199G>A), R845K (3203C>A 3204G>A 3205G>A), I848L (3212A>C 3214T>G), K854C (3230A>T 3231A>G 3232A>C), Q869K (3275C>A), F871Q (3281T>C 3282T>A 3283T>A), L874M (3290T>A), I889L (3335A>T 3337A>G), S893T (3347T>A 3349C>T), C902R (3374T>C), L903I (3377C>A), N905D (3383A>G), T911S (3401A>T 3403A>T), K925R (3444A>G), S927A (3449T>G 3451A>T), D936N (3476G>A 3478T>C), S938N (3482T>A 3483C>A 3484A>T), I943L (3497A>C), D949E (3517C>A), I951V (3521A>G), E952D (3526G>T), E953I (3527G>A 3528A>T 3529G>C), Y954F (3531A>T 3532C>T), Q955D (3533C>G 3535G>C), E956Q (3536G>C), D960_P961del (3548_3553delGATCCG), P962S (3554C>A 3555C>G), K964T (3561A>C 3562G>A), A965S (3563G>T), S967V (3569A>G 3570G>T), I968S (3572A>T 3573T>C 3574T>C), L969F (3575C>T 3577G>C), S970Q (3578T>C 3579C>A 3580C>G), S976T (3596T>A 3598C>A), M985A (3623A>G 3624T>C 3625G>T), G986A (3627G>C 3628T>A), D990T (3638G>A 3639A>C 3640C>A), E991D (3643A>C), L994V (3650T>G 3652G>A), I996L (3656A>T 3658T>G), I1003V (3677A>G 3679A>G), E1006D (3688A>T), G1007A (3690G>C 3691G>A), H1008S (3692C>T 3693A>C), V1011I (3701G>A), T1024S (3740A>T 3742A>C), N1034E (3770A>G 3772T>A), S1036T (3776T>A), I1037F (3779A>T), I1039M (3787A>G), V1042T (3794G>A 3795T>C 3796A>T), P1046S (3806C>A 3807C>G 3808T>C), N1047S (3810A>G), V1048I (3812G>A 3814T>A), V1051I (3821G>A 3823A>C), V1054I (3830G>A), F1056L (3836T>C), A1058T (3842G>A 3844C>A), P1064Q (3861C>A), A1065F (3863G>T 3864C>T), L1067V (3869C>G 3871T>A), N1068S (3872A>T 3873A>C), K1069A (3875A>G 3876A>C 3877A>G), N1071A (3881A>G 3882A>C 3883T>A), Y1072H (3884T>C), V1074L (3890G>T), A1089T (3935G>A 3937A>C), T1092I (3945C>T 3946G>C), T1096V (3956A>G 3957C>T 3958T>A), E1101Q (3971G>C), H1107F (3989C>T 3990A>T 3991T>C), R1109K (3996G>A), M1110A (3998A>G 3999T>C 4000G>C), R1111K (4002G>A 4003G>A), E1112S (4004G>T 4005A>C 4006A>C), F1113Y (4008T>A 4009C>T), S1114D (4010T>G 4011C>A 4012A>T), S1115K (4014G>A 4015C>G), K1116E (4016A>G 4018G>A), Q1117S (4019C>A 4020A>G 4021A>T), M1121V (4031A>G 4033G>T), R1126K (4047G>A), N1129L (4055A>T 4056A>T 4057T>G), I1131L (4061A>C 4063C>A), S1137V (4079T>G 4080C>T 4081C>T), A1138E (4083C>A 4084C>A), I1139L (4085A>C 4087A>T), V1140I (4088G>A 4090G>T), L1146I (4106C>A 4108T>A), I1147V (4109A>G 4111A>C), S1148K (4112T>A 4113C>A 4114T>G), L1151P (4122T>C), S1162L (4154T>C 4155C>T), H1165N (4163C>A), N1171D (4181A>G), I1191N (4242T>A) | | | | | | | | |

| | Begin | End | Coverage | Score | Concordance | Matches | Identities | I/D/M/F* | Stop Codons |
|---|---|---|---|---|---|---|---|---|---|
| NT | 89 | 4744 | 94.9% | 2600 | 28.6% | 4608 (96.7%) | 3040 (63.8%) | 107/48 | |

Codon mutations:

ATG1AGG (672T>G), ACA2ATG (675C>T 676A>G), TCT4TTG (681C>T 682T>G), GTT6GCT (687T>C), CTG8CGA (693T>G 694G>A), AAA9GTG (695A>G 696A>T 697A>G), GTG10ATG (698G>A), GGC11GCC (702G>C), CCC12CCA (706C>A), AAT14GAC (710A>G 712T>C), CAC15CGC (714A>G), CAG16TCA (716C>T 717A>C 718G>A), CAG17del (719_721delCAG), GAG18AAA (722G>A 724G>A), GCC20GTA (729C>T 730C>A), TCC21TCT (733C>T), GAG22GAC (736G>C), AGG23AAA (738G>A 739G>A), ATA25ATT (745A>T), TCA26TCT (748A>T), CAA27CAG (751A>G), ATA28ATC (754A>C), GCT30GCG (760T>G), GTG32GTT (766G>T), GAG33GAA (769G>A), GCC34GAA (771C>A 772C>A), GGA35GGT (775A>T), AAT38AAC (784T>C), TTG39CTT (785T>C 787G>T), CTC40CTT (790C>T), AGA41AAA (792G>A), AAG42AAA (796G>A), CTT43TTG (797C>T 799T>G), GGC44GGA (802C>A), TCC47TCA (811C>A), TAC48TAT (814C>T), GGG49GGC (817G>C), GTT50ATA (818G>A 820T>A), CTT51TTG (821C>T 823T>G), GGT53GCA (828G>C 829T>A), CAA55CAG (835A>G), AAG57AAA (841G>A), AGA58AGG (844A>G), GAG61GAT (853G>T), TTG62CTT (854T>C 856G>T), TTT63TAT (858T>A), GAC64GAT (862C>T), GTA68GTT (874A>T), TCA69AGT (875T>A 876C>G 877A>T), AAG70AAA (880G>A), ATC71GTG (881A>G 883C>G), ACA72GCA (884A>G), CTT74CTA (892T>A), ACA77GCT (899A>G 901A>T), TTC78TTT (904C>T), CAT80AGG (908C>A 909A>G 910T>G), AAG81ACC (912A>C 913G>C), TTT82ATA (914T>A 916T>A), GTT83ACA (917G>A 918T>C 919T>A), GAG84GAA (922G>A), TCT85TCA (925T>A), AAG86CGT (926A>C 927A>G 928G>T), GAC87GAT (931C>T), GCA89GGG (936C>G 937A>G), AAC90AAT (940C>T), TTG91TTA (943G>A), TTT93TTC (949T>C), CTG95TTA (953C>T 955G>A), TAT96CAC (956T>C 958T>C), GGT97GGG (961T>G), CTT99TTG (965C>T 967T>G), TTC100TTT (970C>T), TTC101TTT (973C>T), GTC103GTT (979C>T), CCA104CCC (982A>C), GTT106GCA (987T>C 988T>A), AGT108AAT (993G>A), GGG109GGA (997G>A), GGA111GGT (1003A>T), AGG112GCC (1004A>G 1005G>C 1006G>C), GTT113GTC (1009T>C), AAG114AAA (1012G>A), ATA115ATC (1015A>C), AGT116AGC (1018T>C), CTT117TTG (1019C>T 1021T>G), TGT118TGC (1024T>C), TCA119TCT (1027A>T), AGT120AGC (1030T>C), AAT121AAC (1033T>C), CTC124ACT (1040C>A 1041T>C 1042C>T), TCA125GAT (1043T>G 1044C>A 1045A>T), CCG126CCT (1048G>T), CAA129CAG (1057A>G), AAG131AAA (1063G>A), ACA132ATT (1065C>T 1066A>T), CTG133TTG (1067C>T), TCC134TAC (1071C>A), TTG135TTT (1075G>T), GCC136TCT (1076G>T 1078C>T), GAT137GGA (1080A>G 1081T>A), GCG139GCA (1087G>A), CAG140CAA (1090G>A), GCA141GCC (1093A>C), CTC143CTA (1099C>A), AGC145AGT (1105C>T), CCC146CCT (1108C>T), AGC147AGT (1111C>T), CTC153TTG (1127C>T 1129C>G), AAA154AAG (1132A>G), AGA155AGG (1135A>G), GGG156GGC (1138G>C), ACC162ACA (1156C>A), GAG164GAA (1162G>A), TGC165TGT (1165C>T), CAA166CTC (1167A>T 1168A>C), ACC168ACG (1174C>G), AGG169AGA (1177G>A), GCA170GCT (1180A>T), CAG171CAA (1183G>A), CCG173CCT (1189G>T), TCC175TCT (1195C>T), GTG176GTA (1198G>A), GCC178GCA (1204C>A), ATA179ATT (1207A>T), GAC185AGT (1223G>A 1224A>G 1225C>T), ACA186CCT (1226A>C 1228A>T), AGG187CGT (1229A>C 1231G>T), GCA189GCT (1237A>T), GTT190GTA (1240T>A), TCA192ACA (1244T>A), CAG193GAC (1247C>G 1249G>C), CAA194CAT (1252A>T), GAA195GAC (1255A>C), ACA196ACT (1258A>T), TCT198AGT (1262T>A 1263C>G), GCC200GCT (1270C>T), ATA201ATT (1273A>T), GAA202GAG (1276A>G), GCA203GCT (1279A>T), CTC204CTT (1282C>T), AAT205AAC (1285T>C), CGT206AGA (1286C>A 1288T>A), CCG207CCT (1291G>T), CAA208CAG (1294A>G), GAT212GAC (1306T>C), AGA213AGG (1309A>G), GAG215GAA (1315G>A), GCT216GCA (1318T>A), GCT217GCA (1321T>A), TAT219TAC (1327T>C), ATA220ATT (1330A>T), GCT221GCA (1333T>A), GTT223GTG (1339T>G), TAT224TAC (1342T>C), TCG225TCC (1345G>C), AGT226AAT (1347G>A), GGG227_CAA228insGTA (1351_1352insGTA), CAA228CAG (1354A>G), AGT229AGA (1357T>A), ACT230AGT (1359C>G), CAA231CAG (1363A>G), ATG232CTG (1364A>C), GCT233del (1367_1369delGCT), CTT234TTG (1370C>T 1372T>G), GAA235GAG (1375A>G), AAA237AGA (1380A>G), GCC238CCT (1382G>C 1384C>T), GTT240GTG (1390T>G), GGA241GGT (1393A>T), GAA242GCC (1395A>C 1396A>C), CAG243GCA (1397C>G 1398A>C 1399G>A), GGG245GGT (1405G>T), GGC246GAT (1407G>A 1408C>T), ACA247ACC (1411A>C), GAC250AAT (1418G>A 1420C>T), GTC251GTG (1423C>G), ATG252CCG (1424A>C 1425T>C), AAT253CAG (1427A>C 1429T>G), GAA254GAG (1432A>G), AGC255GCA (1433A>G 1434G>C 1435C>A), ACT256AAT (1437C>A), ATC258GTG (1442A>G 1444C>G), CGG259CGC (1447G>C), AGT260TCT (1448A>T 1449G>C), TCT262TCA (1456T>A), AGA264AGG (1462A>G), ACT267ACC (1471T>C), CTC268CTA (1474C>A), AAA269AAG (1477A>G), GTG270GTA (1480G>A), AAG273AGG (1488A>G), CGC274CGA (1492C>A), CCG276GCA (1496C>G 1498G>A), TCC277TCA (1501C>A), CTG280TTG (1508C>T), CCG281CCT (1513G>T), CCG281_GCT282insGACTTT (1513_1514insGACTTT), GTA283TCA (1517G>T 1518T>C), AGT284ACT (1521G>C), ACA285TCT (1523A>T 1525A>T), TCA286AGA (1526T>A 1527C>G), TCC287GCG (1529T>G 1531C>G), TTA288TAT (1533T>A 1534A>T), CTC289CAG (1536T>A 1537C>G), AGC290GGA (1538A>G 1540C>A), ACA291GTG (1541A>G 1542C>T 1543A>G), CGT292AGA (1544C>A 1546T>A), GAA293GAG (1549A>G), ACA295ACT (1555A>T), GTT296GGT (1557T>G), CAT297ATT (1559C>A 1560A>T), GAG299del (1565_1567delGAG), TGT302ACG (1574T>A 1575G>C 1576T>G), GGG303TCT (1577G>T 1578G>C 1579G>T), GGA304CTA (1580G>C 1581G>T), TTG305CTG (1583T>C), CCA307CCT (1591A>T), GCA308GCC (1594A>C), CCC309_AAG310del (1595_1600delCCCAAG), AAA311AAG (1603A>G), AAG312ATA (1605A>T 1606G>A), GGC313TCA (1607G>T 1608G>C 1609C>A), GCC315GCA (1615C>A), TTC316GGA (1616T>G 1617T>G 1618C>A), AAC317ACA (1620A>C 1621C>A), ATG318CTC (1622A>C 1624G>C), GGT319CAA (1625G>C 1626G>A 1627T>A), GCC320GAA (1629C>A 1630C>A), ATT321GTC (1631A>G 1633T>C), GAT323GAC (1639T>C), TTG325TAT (1644T>A 1645G>T), ATG327ATT (1651G>T), GAG328CAA (1652G>C 1654G>A), TCG329AGT (1655T>A 1656C>G 1657G>T), TTC330TTT (1660C>T), GCA331GAA (1662C>A), CAC332GAT (1664C>G 1666C>T), ATT333TTT (1667A>T), GAT334GCC (1671A>C 1672T>C), GAT337AGT (1679G>A 1680A>G), GAT338AAC (1682G>A 1684T>C), GAA341GCC (1692A>C 1693A>C), CGA342CTT (1695G>T 1696A>T), ATA344GTG (1700A>G 1702A>G), GCT345CCC (1703G>C 1705T>C), CAG346GAG (1706C>G), CAG347CAA (1711G>A), GTA348ATT (1712G>A 1714A>T), CAA349GAT (1715C>G 1717A>T), TTT350TAT (1719T>A), ATC351GTT (1721A>G 1723C>T), CTC352CTT (1726C>T), TTT353GTG (1727T>G 1729T>G), AGT354CAG (1730A>C 1731G>A 1732T>G), GAG355AAG (1733G>A), AAA357AGA (1740A>G), AGA358GCT (1742A>G 1743G>C 1744A>T), GGA359GGT (1747A>T), AAC360AAT (1750C>T), GTT361CTC (1751G>C 1753T>C), ATT362TGT (1754A>T 1755T>G), GTG363GTC (1759G>C), AGA365AAA (1764G>A), GTT367ATT (1769G>A), GCA368GAG (1773C>A 1774A>G), AAG369ATG (1776A>T), CGG370AGG (1778C>A), CAT371GAC (1781C>G 1783T>C), CTG372TTG (1784C>T), CAT373CAC (1789T>C), AAT374TGT (1790A>T 1791A>G), ATC375ATA (1795C>A), AGG377TTT (1799A>T 1800G>T 1801G>T), GAG378GAA (1804G>A), CAC379CAT (1807C>T), ATA380ATC (1810A>C), ACC381ACT (1813C>T), GAC383GCC (1818A>C), GTA386GTT (1828A>T), GAA387GAT (1831A>T), ATA388ATT (1834A>T), CTT389TTG (1835C>T 1837T>G), GAA390GCG (1839A>C 1840A>G), GGT391GGA (1843T>A), GGT393GGA (1849T>A), AAT396AAG (1858T>G), ATC397GTT (1859A>G 1861C>T), CAA398GGG (1862C>G 1863A>G 1864A>G), GGC399ATG (1865G>A 1866G>T 1867C>G), ACT401GAA (1871A>G 1872C>A 1873T>A), CGC402AGA (1874C>A 1876C>A), ACT403ACA (1879T>A), TTC404CCA (1880T>C 1881T>C 1882C>A), TAT407TAC (1891T>C), GCT408TCT (1892G>T), ATG409TTG (1895A>T), TCT410AGC (1898T>A 1899C>G 1900T>C), CTC411CTG (1903C>G), GAG413CAG (1907G>C), GTT415GTG (1915T>G), GTG416GTT (1918G>T), CTC418CTA (1924C>A), CAT420CAC (1930T>C), CAA421ATG (1931C>A 1932A>T 1933A>G), AGA422AGG (1936A>G), GAT423GAC (1939T>C), TCC425CAA (1943T>C 1944C>A 1945C>A), TTC426TTT (1948C>T), ATT428AGG (1953T>G 1954T>G), CAA429TAT (1955C>T 1957A>T), GGG430TGG (1958G>T), CAG431GAT (1961C>G 1963G>T), ACT432TTA (1964A>T 1965C>T 1966T>A), AAC433ACA (1968A>C 1969C>A), AAT434AAC (1972T>C), CCA435CCG (1975A>G), CCG437CCT (1981G>T), AAT438GAC (1982A>G 1984T>C), TGC441GTG (1991T>G 1992G>T 1993C>G), GGA444GGT (2002A>T), TTA445CTT (2003T>C 2005A>T), ACC446ACA (2008C>A), TTG447CTG (2009T>C), AAA448GCA (2012A>G 2013A>C), GAG449GAT (2017G>T), CGT450AGG (2018C>A 2020T>G), CGT450_CAG451insGTGAAA (2020_2021insGTGAAA), CAG451CAT (2023G>T), TTG452CCC (2024T>C 2025T>C 2026G>C), ATT453TTG (2027A>T 2029T>G), GTG454ATG (2030G>A), GAG455AGT (2033G>A 2034A>G 2035G>T), GAG456GAT (2038G>T), CAA457CTA (2040A>T), GTG458GCA (2043T>C 2044G>A), GCC459TGC (2045G>T 2046C>G), CAA460_AGA461insTTGAAT (2050_2051insTTGAAT), AGA461CGC (2051A>C 2053A>C), AGG462TTG (2054A>T 2055G>T), TCT463GGA (2057T>G 2058C>G 2059T>A), GAA464AAA (2060G>A), AGA465CCA (2063A>C 2064G>C), CAA466AGC (2066C>A 2067A>G 2068A>C), GCA467ATG (2069G>A 2070C>T 2071A>G), CAG468AGT (2072C>A 2073A>G 2074G>T), ACA470GCA (2078A>G), GCA471CAA (2081G>C 2082C>A), AGG472ACT (2085G>C 2086G>T), GGC473GGA (2089C>A), ATA474AAT (2091T>A 2092A>T), GCT475GCC (2095T>C), GCT475_GAA476insAGC (2095_2096insAGC), GAA476AAG (2096G>A 2098A>G), AGT477TGG (2099A>T 2101T>G), CAG478CAA (2104G>A), CCT479ACT (2105C>A), GAC481GGA (2112A>G 2113C>A), AGA482AAG (2115G>A 2116A>G), ACT484CTG (2120A>C 2121C>T 2122T>G), TCC486TCA (2128C>A), GTG488GTT (2134G>T), TCA489ACA (2135T>A), ACT490ACC (2140T>C), ACC491ACA (2143C>A), ACG492GCA (2144A>G 2146G>A), ATG493CTT (2147A>C 2149G>T), GAA494GAG (2152A>G), GAA494_GAT495insAGT (2152_2153insAGT), GAT495GAG (2155T>G), CCC496GGT (2156C>G 2157C>G 2158C>T), ACC497ATC (2160C>T), AAG498CAA (2162A>C 2164G>A), CCA499AGA (2165C>A 2166C>G), GAT500AAG (2168G>A 2170T>G), AAG501GAG (2171A>G), ATA502GTG (2174A>G 2176A>G), GAA503TCA (2177G>T 2178A>C), GTT505GAG (2184T>A 2185T>G), GCA506GCT (2188A>T), GAG507GAA (2191G>A), GGA508GAT (2193G>A 2194A>T), GCA509CCG (2195G>C 2197A>G), GAA511GGG (2202A>G 2203A>G), GAA512AAA (2204G>A), ACT513ACA (2209T>A), CAA514GAG (2210C>G 2212A>G), GGT516AAT (2216G>A 2217G>A), GAT517GAC (2221T>C), GTT518ATT (2222G>A), ATT519TTC (2225A>T 2227T>C), GAC521GAT (2233C>T), GGA523GGG (2239A>G), CCA524GAG (2240C>G 2241C>A 2242A>G), GAC527ATG (2249G>A 2250A>T 2251C>G), ACA528GGT (2252A>G 2253C>G 2254A>T), TCC529GAT (2255T>G 2256C>A 2257C>T), ATG530GCC (2258A>G 2259T>C 2260G>C), GCA531CCG (2261G>C 2263A>G), GTT532TTA (2264G>T 2266T>A), GAA533AAT (2267G>A 2269A>T), GAT535GAG (2275T>G), ATG536TTG (2276A>T), CAG537CAA (2281G>A), CAA538CAG (2284A>G), CCG539CCA (2287G>A), GTG540GCA (2289T>C 2290G>A), GTA542GTG (2296A>G), AGC544AGT (2302C>T), GAT546GAG (2308T>G), TTC547TTT (2311C>T), GGA551GGG (2323A>G), GTT552ATT (2324G>A), TTC555TTT (2335C>T), GTT556CTG (2336G>C 2338T>G), GAG558AAA (2342G>A 2344G>A), TCC560AGC (2348T>A 2349C>G), GCT561AAC (2351G>A 2352C>A 2353T>C), GAG565GAA (2365G>A), CAA566CAG (2368A>G), GTA567GTT (2371A>T), ATG568CTT (2372A>C 2374G>T), AGC569GCA (2375A>G 2376G>C 2377C>A), TTG570CTA (2378T>C 2380G>A), GCT571TCC (2381G>T 2383T>C), TTG572CTC (2384T>C 2386G>C), CCC573CCA (2389C>A), GCT574ATG (2390G>A 2391C>T 2392T>G), GCC575GCA (2395C>A), CTA576TTG (2396C>T 2398A>G), TTC577TTT (2401C>T), TCA578AGC (2402T>A 2403C>G 2404A>C), AAA579AAG (2407A>G), AGC580AAG (2409G>A 2410C>G), AAA581AAG (2413A>G), ACT583GCT (2417A>G), TCA584TCC (2422A>C), GGT586GGG (2428T>G), GCG587TCC (2429G>T 2431G>C), CTT590CTC (2440T>C), TAT592TAC (2446T>C), TAC593TAT (2449C>T), GAT594GAC (2452T>C), GCA595GCT (2455A>T), GCT596GCA (2458T>A), CTA597AAC (2459C>A 2460T>A 2461A>C), ATC598CTT (2462A>C 2464C>T), TAC600TTT (2469A>T 2470C>T), ATA602GTC (2474A>G 2476A>C), CTT604CTG (2482T>G), TAT605TAC (2485T>C), GTT606ATT (2486G>A), GTT609ATG (2495G>A 2497T>G), GCT611GCG (2503T>G), ATC612GTG (2504A>G 2506C>G), TCT613TCG (2509T>G), GGT614GGA (2512T>A), CAA615CAG (2515A>G), CTG616CTA (2518G>A), GCT617GCC (2521T>C), GAA622GAG (2536A>G), TGT623TGC (2539T>C), GTG625GCA (2544T>C 2545G>A), CGA628AGA (2552C>A), AAG629ACC (2556A>C 2557G>C), AAC634AAT (2572C>T), ATC635ATA (2575C>A), ACA637ACC (2581A>C), TTG638ATT (2582T>A 2584G>T), AAG642AAA (2596G>A), ACA644CAT (2600A>C 2601C>A 2602A>T), TTG645CTG (2603T>C), GTT646GTG (2608T>G), CAA650GAA (2618C>G), AAC652TCC (2624A>T 2625A>C), GAG654GGG (2631A>G), GAG655AAG (2633G>A), TGC657GTG (2639T>G 2640G>T 2641C>G), TTT658TTC (2644T>C), ACC659TCA (2645A>T 2647C>A), CCA660CCT (2650A>T), ACA661CAA (2651A>C 2652C>A), GGG662GGT (2656G>T), GGC664GGT (2662C>T), AAG665ACC (2664A>C 2665G>C), TAC666TTT (2667A>T 2668C>T), GTG667GTA (2671G>A), CCT668CCA (2674T>A), CTG669TTG (2675C>T), GAA671ACT (2681G>A 2682A>C 2683A>T), GGT672TCA (2684G>T 2685G>C 2686T>A), ACA673CTG (2687A>C 2688C>T 2689A>G), GGA674GGT (2692A>T), GCC675GTT (2694C>T 2695C>T), ACT676CCT (2696A>C), GAT677GAA (2701T>A), TTA678CTT (2702T>C 2704A>T), GGT679GGC (2707T>C), GTG681GTT (2713G>T), ACA686ACC (2728A>C), CAC687CAT (2731C>T), TTA689TTG (2737A>G), TCT690GCA (2738T>G 2740T>A), GCA692GCC (2746A>C), GAG694GAA (2752G>A), AAT696GAG (2756A>G 2758T>G), AGT697TCA (2759A>T 2760G>C 2761T>A), GTA698ATA (2762G>A), CCA699CCT (2767A>T), TGC700TGT (2770C>T), CAT701CAC (2773T>C), TTG702ATA (2774T>A 2776G>A), TTA704CTA (2780T>C), CAG705TCC (2783C>T 2784A>C 2785G>C), AAA707AAG (2791A>G), TTA709CTC (2795T>C 2797A>C), ACC711ACA (2803C>A), CTT714ATG (2810C>A 2812T>G), CGA718AGG (2822C>A 2824A>G), ATG719TTG (2825A>T), ATA720AGA (2829T>G), GCA721GGG (2832C>G 2833A>G), CAG722CAA (2836G>A), CAA724CAG (2842A>G), TAT725TTT (2844A>T), GGC726GGA (2848C>A), AAG728CGG (2852A>C 2853A>G), GCG729CCA (2855G>C 2857G>A), GGG730GGC (2860G>C), CAG731ACA (2861C>A 2862A>C 2863G>A), ACA732ACC (2866A>C), TAT733TTC (2868A>T 2869T>C), CCA735AGT (2873C>A 2874C>G 2875A>T), AGG736CGC (2876A>C 2878G>C), TTT737TTC (2881T>C), CCA738CCT (2884A>T), ACT739ACA (2887T>A), GTT742GTG (2896T>G), TTG743CTT (2897T>C 2899G>T), TTA744CTG (2900T>C 2902A>G), CAT745CAC (2905T>C), TAT746TAC (2908T>C), GGG749AGT (2915G>A 2917G>T), ACA750GTA (2918A>G 2919C>T), TCA751GCT (2921T>G 2923A>T), TCC752GCA (2924T>G 2926C>A), CCA753GCC (2927C>G 2929A>C), ACA757ACT (2941A>T), AGC760AGT (2950C>T), ATA761GTC (2951A>G 2953A>C), TTT762TTC (2956T>C), TCA763TCC (2959A>C), ATA767ATT (2971A>T), GAG769GAA (2977G>A), GGC772GGT (2986C>T), ACC773ATC (2988C>T), TTG774TTA (2992G>A), CCG776CCA (2998G>A), TCT777TCA (3001T>A), TTG778CTG (3002T>C), GGT780GGA (3010T>A), ATA782ATT (3016A>T), GCC783GCA (3019C>A), AGG784AGA (3022G>A), AAG787AAA (3031G>A), ACT790AGA (3039C>G 3040T>A), GGA791GGC (3043A>C), ACT792ACC (3046T>C), ATT797GTG (3059A>G 3061T>G), TGC798TGT (3064C>T), ATT799GTT (3065A>G), ACC802ACT (3076C>T), TTG803CTG (3077T>C), TCA806TCT (3088A>T), GGT807GGG (3091T>G), CTG809CTT (3097G>T), GCA810GCT (3100A>T), ATT811ATA (3103T>A), GGA812GGG (3106A>G), CTT813TTG (3107C>T 3109T>G), GGA814GGC (3112A>C), ACT815ACG (3115T>G), AAC817GCA (3119A>G 3120A>C 3121C>A), ACC818ACA (3124C>A), AAA819GAG (3125A>G 3127A>G), ACC821TCC (3131A>T), AAT822ACA (3135A>C 3136T>A), GCT823GCG (3139T>G), TTT827TTG (3151T>G), CCA830CCC (3160A>C), GTA833GTG (3169A>G), TGC834TGT (3172C>T), CTT836TTG (3176C>T 3178T>G), GGT839GGG (3187T>G), AAA841AGG (3192A>G 3193A>G), CGG843CAA (3198G>A 3199G>A), CGG845AAA (3203C>A 3204G>A 3205G>A), TGT846TGC (3208T>C), TCT847TCA (3211T>A), ATT848CTG (3212A>C 3214T>G), ACA849ACC (3217A>C), GGA853GGG (3229A>G), AAA854TGC (3230A>T 3231A>G 3232A>C), AAT855AAC (3235T>C), TTG856CTC (3236T>C 3238G>C), CTT857CTG (3241T>G), TCC858TCT (3244C>T), ACA859ACC (3247A>C), GGG860GGA (3250G>A), CGA861CGC (3253A>C), AAG862AAA (3256G>A), TCC864TCA (3262C>A), TTG865TTA (3265G>A), CAG869AAG (3275C>A), CAC870CAT (3280C>T), TTT871CAA (3281T>C 3282T>A 3283T>A), TCC872TCT (3286C>T), TTG874ATG (3290T>A), CGC875CGT (3295C>T), TTG876CTG (3296T>C), TTT877TTC (3301T>C), ACG879ACT (3307G>T), GTA880GTT (3310A>T), CTC884CTG (3322C>G), ACG887ACA (3331G>A), ATA889TTG (3335A>T 3337A>G), CAT890CAC (3340T>C), TCC893ACT (3347T>A 3349C>T), GGT895GGA (3355T>A), GTC896GTA (3358C>A), CAG899CAA (3367G>A), AAG901AAA (3373G>A), TGC902CGC (3374T>C), CTT903ATT (3377C>A), AAT905GAT (3383A>G), CTT906CTC (3388T>C), ACT907ACG (3391T>G), TTG908CTT (3392T>C 3394G>T), GGT909GGG (3397T>G), GGC910GGT (3400C>T), ACA911TCT (3401A>T 3403A>T), GTA912GTG (3406A>G), TCT913TCA (3409T>A), GTG914GTT (3412G>T), AAA915AAG (3415A>G), ATA917ATT (3421A>T), GGG919GGT (3427G>T), ACA922ACC (3436A>C), AAA923AAG (3439A>G), GGC924GGG (3442C>G), AAA925AGA (3444A>G), AGC926AGT (3448C>T), TCA927GCT (3449T>G 3451A>T), GTT928GTG (3454T>G), TTC930TTT (3460C>T),

| | | | | | | | | | |
|---|---|---|---|---|---|---|---|---|---|
| | CTT931CTG (3463T>G), TTC932TTT (3466C>T), GAT936AAC (3476G>A 3478T>C), TTA937CTG (3479T>C 3481A>G), TCA938AAT (3482T>A 3483C>A 3484A>T), CGT940CGC (3490T>C), ATT943CTT (3497A>C), GAA944GAG (3502A>G), AAG945AAA (3505G>A), TTG946CTG (3506T>C), AAG948AAA (3514G>A), GAC949GAA (3517C>A), AAC950AAT (3520C>T), ATT951GTT (3521A>G), GAG952GAT (3526G>T), GAG953ATC (3527G>A 3528A>T 3529G>C), TAC954TTT (3531A>T 3532C>T), CAG955GAC (3533C>G 3535G>C), GAG956CAG (3536G>C), GGC958GGG (3544C>G), GAT960_CCG961del (3548_3553delGATCCG), CCC962AGC (3554C>A 3555C>G), AAG964ACA (3561A>C 3562G>A), GCT965TCT (3563G>T), AGT967GTT (3569A>G 3570G>T), ATT968TCC (3572A>T 3573T>C 3574T>C), CTG969TTC (3575C>T 3577G>C), TCC970CAG (3578T>C 3579C>A 3580C>G), ATA971ATC (3583A>C), AGA972AGG (3586A>G), GAG973GAA (3589G>A), AAA974AAG (3592A>G), TTT975TTC (3595T>C), TCC976ACA (3596T>A 3598C>A), GGT978GGA (3604T>A), GCT979GCA (3607T>A), GTA980GTG (3610A>G), TTC983TTT (3619C>T), TGC984TGT (3622C>T), ATG985GCT (3623A>G 3624T>C 3625G>T), GGT986GCA (3627G>C 3628T>A), GAC990ACA (3638G>A 3639A>C 3640C>A), GAA991GAC (3643A>C), TTG994GTA (3650T>G 3652G>A), GTA995GTT (3655A>T), ATT996TTG (3656A>T 3658T>G), CCT997CCA (3661T>A), GCA999GCC (3667A>C), CCA1000CCT (3670A>T), TCC1002TCA (3676C>A), ATA1003GTG (3677A>G 3679A>G), AGG1004CGC (3680A>C 3682G>C), GAA1006GAT (3688A>T), GGG1007GCA (3690G>C 3691G>A), CAC1008TCC (3692C>T 3693A>C), CCT1010CCA (3700T>A), GTT1011ATT (3701G>A), AAG1012AAA (3706G>A), ATC1015ATA (3715C>A), TGT1017TGC (3721T>C), TTT1019TTC (3727T>C), GAT1021GAC (3733T>C), ACA1024TCC (3740A>T 3742A>C), TCA1025TCT (3745A>T), TCA1030TCT (3760A>T), GGT1031GGA (3763T>A), AAT1034GAA (3770A>G 3772T>A), TCA1036ACA (3776T>A), ATT1037TTT (3779A>T), GTG1038GTA (3784G>A), ATA1039ATG (3787A>G), AGA1041CGA (3791A>C), GTA1042ACT (3794G>A 3795T>C 3796A>T), CAA1043CAG (3799A>G), TCC1044AGC (3800T>A 3801C>G), AGT1045TCC (3803A>T 3804G>C 3805T>C), CCT1046AGC (3806C>A 3807C>G 3808T>C), AAT1047AGT (3810A>G), GTT1048ATA (3812G>A 3814T>A), GGA1049GGT (3817A>T), GTA1051ATC (3821G>A 3823A>C), CTA1052TTG (3824C>T 3826A>G), AAT1053AAC (3829T>C), GTT1054ATT (3830G>A), GCT1055GCA (3835T>A), TTT1056CTT (3836T>C), GAT1057GAC (3841T>C), GCC1058ACA (3842G>A 3844C>A), GGC1060GGG (3850C>G), TTT1063TTC (3859T>C), CCA1064CAA (3861C>A), GCT1065TTT (3863G>T 3864C>T), GGG1066GGT (3868G>T), CTT1067GTA (3869C>G 3871T>A), AAT1068TCT (3872A>T 3873A>C), AAA1069GCG (3875A>G 3876A>C 3877A>G), AAT1071GCA (3881A>G 3882A>C 3883T>A), TAT1072CAT (3884T>C), GTA1074TTA (3890G>T), GGT1077GGC (3901T>C), GGA1078GGT (3904A>T), GGC1079GGA (3907C>A), ACA1080ACT (3910A>T), TCA1085TCC (3925A>C), TAC1086TAT (3928C>T), GGT1087GGG (3931T>G), GTG1088GTT (3934G>T), GCA1089ACC (3935G>A 3937A>C), ACA1090ACC (3940A>C), AAT1091AAC (3943T>C), ACG1092ATC (3945C>T 3946G>C), TCA1094TCT (3952A>T), TTC1095TTT (3955C>T), ACT1096GTA (3956A>G 3957C>T 3958T>A), GAG1101CAG (3971G>C), TTT1102TTC (3976T>C), TTC1103TTT (3979C>T), CCA1104CCT (3982A>T), AGG1105CGC (3983A>C 3985G>C), CGG1106CGT (3988G>T), CAT1107TTC (3989C>T 3990A>T 3991T>C), AGG1109AAG (3996G>A), ATG1110GCC (3998A>G 3999T>C 4000G>C), AGG1111AAA (4002G>A 4003G>A), GAA1112TCC (4004G>T 4005A>C 4006A>C), TTC1113TAT (4008T>A 4009C>T), TCA1114GAT (4010T>G 4011C>A 4012A>T), AGC1115AAG (4014G>A 4015C>G), AAG1116GAA (4016A>G 4018G>A), CAA1117AGT (4019C>A 4020A>G 4021A>T), TCC1118TCA (4024C>A), CGC1119AGA (4025C>A 4027C>A), ATC1120ATT (4030C>T), ATG1121GTT (4031A>G 4033G>T), TCA1122TCT (4036A>T), CTA1123TTG (4037C>T 4039A>G), GAT1125GAC (4045T>C), AGG1126AAG (4047G>A), CTT1127CTA (4051T>A), GGA1128GGC (4054A>C), AAT1129TTG (4055A>T 4056A>T 4057T>G), CTG1130TTG (4058C>T), ATC1131CTA (4061A>C 4063C>A), ATA1132ATC (4066A>C), TTG1134CTT (4070T>C 4072G>T), CCT1135CCA (4075T>A), CCT1136CCA (4078T>A), TCC1137GTT (4079T>G 4080C>T 4081C>T), GCC1138GAA (4083C>A 4084C>A), ATA1139CTT (4085A>C 4087A>T), GTG1140ATT (4088G>A 4090G>T), TCC1142TCA (4096C>A), GAG1144GAA (4102G>A), ATA1145ATC (4105A>C), CTT1146ATA (4106C>A 4108T>A), ATA1147GTC (4109A>G 4111A>C), TCT1148AAG (4112T>A 4113C>A 4114T>G), CCT1149CCA (4117T>A), GGA1150GGT (4120A>T), CTT1151CCT (4122T>C), TTC1153TTT (4129C>T), AAG1154AAA (4132G>A), GCC1158GCT (4144C>T), CCT1160CCC (4150T>C), CCT1161CCA (4153T>A), TCT1162CTT (4154T>C 4155C>T), GCC1163GCA (4159C>A), CAT1165AAT (4163C>A), GAA1166GAG (4168A>G), AAA1167AAG (4171A>G), CTT1169CTC (4177T>C), GGC1170GGA (4180C>A), AAT1171GAT (4181A>G), CAA1173CAG (4189A>G), ACT1174ACG (4192T>G), CAC1175CAT (4195C>T), ACC1176ACT (4198C>T), ACC1180ACA (4210C>A), TCA1181TCT (4213A>T), CTA1186TTG (4226C>T 4228A>G), CGT1187CGC (4231T>C), ATT1191AAT (4242T>A) | | | | | | | | |
| Coat protein A (YP_001976155.1) | 1 | 254 | 100% | 1189 | 70.1% | 254 (99.2%) | 165 (64.5%) | 2/0/0/0 | 0 |
| Protein mutations: | T2A (2078A>G), A3Q (2081G>C 2082C>A), R4T (2085G>C 2086G>T), I6N (2091T>A 2092A>T), A7_E8insS (2095_2096insAGC), E8K (2096G>A 2098A>G), S9W (2099A>T 2101T>G), P11T (2105C>A), D13G (2112A>G 2113C>A), R14K (2115G>A 2116A>G), T16L (2120A>C 2121C>T 2122T>G), S21T (2135T>A), T24A (2144A>G 2146G>A), M25L (2147A>C 2149G>T), E26_D27insS (2152_2153insAGT), D27E (2155T>G), P28G (2156C>G 2157C>G 2158C>T), T29I (2160C>T), K30Q (2162A>C 2164G>A), P31R (2165C>A 2166C>G), D32K (2168G>A 2170T>G), K33E (2171A>G), I34V (2174A>G 2176A>G), E35S (2177G>T 2178A>C), V37E (2184T>A 2185T>G), G40D (2193G>A 2194A>T), A41P (2195G>C 2197A>G), E43G (2202A>G 2203A>G), E44K (2204G>A), Q46E (2210C>G 2212A>G), G48N (2216G>A 2217G>A), V50I (2222G>A), I51F (2225A>T 2227T>C), P56E (2240C>G 2241C>A 2242A>G), D59M (2249G>A 2250A>T 2251C>G), T60G (2252A>G 2253C>G 2254A>T), S61D (2255T>G 2256C>A 2257C>T), M62A (2258A>G 2259T>C 2260G>C), A63P (2261G>C 2263A>G), V64L (2264G>T 2266T>A), E65N (2267G>A 2269A>T), D67E (2275T>G), M68L (2276A>T), V72A (2289T>C 2290G>A), D78E (2308T>G), V84I (2324G>A), V88L (2336G>C 2338T>G), E90K (2342G>A 2344G>A), A93N (2351G>A 2352C>A 2353T>C), M100L (2372A>C 2374G>T), S101A (2375A>G 2376G>C 2377C>A), A103S (2381G>T 2383T>C), A106M (2390G>A 2391C>T 2392T>G), S112K (2409G>A 2410C>G), T115A (2417A>G), A119S (2429G>T 2431G>C), L129N (2459C>A 2460T>A 2461A>C), I130L (2462A>C 2464C>T), Y132F (2469A>T 2470C>T), I134V (2474A>G 2476A>C), V138I (2486G>A), V141M (2495G>A 2497T>G), I144V (2504A>G 2506C>G), V157A (2544T>C 2545G>A), K161T (2556A>C 2557G>C), L170I (2582T>A 2584G>T), T176H (2600A>C 2601C>A 2602A>T), Q182E (2618C>G), N184S (2624A>T 2625A>C), E186G (2631A>G), E187K (2633G>A), C189V (2639T>G 2640G>T 2641C>G), T191S (2645A>T 2647C>A), T193Q (2651A>C 2652C>A), K197T (2664A>C 2665G>C), Y198F (2667A>T 2668C>T), E203T (2681G>A 2682A>C 2683A>T), G204S (2684G>T 2685G>C 2686T>A), T205L (2687A>C 2688C>T 2689A>G), A207V (2694C>T 2695C>T), T208P (2696A>C), D209E (2701T>A), S222A (2738T>G 2740T>A), N228E (2756A>G 2758T>G), V230I (2762G>A), L234I (2774T>A 2776G>A), Q237S (2783C>T 2784A>C 2785G>C), L246M (2810C>A 2812T>G), M251L (2825A>T), I252R (2829T>G), A253G (2832C>G 2833A>G) | | | | | | | | |
| Codon mutations: | ACA2GCA (2078A>G), GCA3CAA (2081G>C 2082C>A), AGG4ACT (2085G>C 2086G>T), GGC5GGA (2089C>A), ATA6AAT (2091T>A 2092A>T), GCT7GCC (2095T>C), GCT7_GAA8insAGC (2095_2096insAGC), GAA8AAG (2096G>A 2098A>G), AGT9TGG (2099A>T 2101T>G), CAG10CAA (2104G>A), CCT11ACT (2105C>A), GAC13GGA (2112A>G 2113C>A), AGA14AAG (2115G>A 2116A>G), ACT16CTG (2120A>C 2121C>T 2122T>G), TCC18TCA (2128C>A), GTG20GTT (2134G>T), TCA21ACA (2135T>A), ACT22ACC (2140T>C), ACC23ACA (2143C>A), ACG24GCA (2144A>G 2146G>A), ATG25CTT (2147A>C 2149G>T), GAA26GAG (2152A>G), GAA26_GAT27insAGT (2152_2153insAGT), GAT27GAG (2155T>G), CCC28GGT (2156C>G 2157C>G 2158C>T), ACC29ATC (2160C>T), AAG30CAA (2162A>C 2164G>A), CCA31AGA (2165C>A 2166C>G), GAT32AAG (2168G>A 2170T>G), AAG33GAG (2171A>G), ATA34GTG (2174A>G 2176A>G), GAA35TCA (2177G>T 2178A>C), GTT37GAG (2184T>A 2185T>G), GCA38GCT (2188A>T), GAG39GAA (2191G>A), GGA40GAT (2193G>A 2194A>T), GCA41CCG (2195G>C 2197A>G), GAA43GGG (2202A>G 2203A>G), GAA44AAA (2204G>A), ACT45ACA (2209T>A), CAA46GAG (2210C>G 2212A>G), GGT48AAT (2216G>A 2217G>A), GAT49GAC (2221T>C), GTT50ATT (2222G>A), ATT51TTC (2225A>T 2227T>C), GAC53GAT (2233C>T), GGA55GGG (2239A>G), CCA56GAG (2240C>G 2241C>A 2242A>G), GAC59ATG (2249G>A 2250A>T 2251C>G), ACA60GGT (2252A>G 2253C>G 2254A>T), TCC61GAT (2255T>G 2256C>A 2257C>T), ATG62GCC (2258A>G 2259T>C 2260G>C), GCA63CCG (2261G>C 2263A>G), GTT64TTA (2264G>T 2266T>A), GAA65AAT (2267G>A 2269A>T), GAT67GAG (2275T>G), ATG68TTG (2276A>T), CAG69CAA (2281G>A), CAA70CAG (2284A>G), CCG71CCA (2287G>A), GTG72GCA (2289T>C 2290G>A), GTA74GTG (2296A>G), AGC76AGT (2302C>T), GAT78GAG (2308T>G), TTC79TTT (2311C>T), GGA83GGG (2323A>G), GTT84ATT (2324G>A), TTC87TTT (2335C>T), GTT88CTG (2336G>C 2338T>G), GAG90AAA (2342G>A 2344G>A), TCC92AGC (2348T>A 2349C>G), GCT93AAC (2351G>A 2352C>A 2353T>C), GAG97GAA (2365G>A), CAA98CAG (2368A>G), GTA99GTT (2371A>T), ATG100CTT (2372A>C 2374G>T), AGC101GCA (2375A>G 2376G>C 2377C>A), TTG102CTA (2378T>C 2380G>A), GCT103TCC (2381G>T 2383T>C), TTG104CTC (2384T>C 2386G>C), CCC105CCA (2389C>A), GCT106ATG (2390G>A 2391C>T 2392T>G), GCC107GCA (2395C>A), CTA108TTG (2396C>T 2398A>G), TTC109TTT (2401C>T), TCA110AGC (2402T>A 2403C>G 2404A>C), AAA111AAG (2407A>G), AGC112AAG (2409G>A 2410C>G), AAA113AAG (2413A>G), ACT115GCT (2417A>G), TCA116TCC (2422A>C), GGT118GGG (2428T>G), GCG119TCC (2429G>T 2431G>C), CTT122CTC (2440T>C), TAT124TAC (2446T>C), TAC125TAT (2449C>T), GAT126GAC (2452T>C), GCA127GCT (2455A>T), GCT128GCA (2458T>A), CTA129AAC (2459C>A 2460T>A 2461A>C), ATC130CTT (2462A>C 2464C>T), TAC132TTT (2469A>T 2470C>T), ATA134GTC (2474A>G 2476A>C), CTT136CTG (2482T>G), TAT137TAC (2485T>C), GTT138ATT (2486G>A), GTT141ATG (2495G>A 2497T>G), GCT143GCG (2503T>G), ATC144GTG (2504A>G 2506C>G), TCT145TCG (2509T>G), GGT146GGA (2512T>A), CAA147CAG (2515A>G), CTG148CTA (2518G>A), GCT149GCC (2521T>C), GAA154GAG (2536A>G), TGT155TGC (2539T>C), GTG157GCA (2544T>C 2545G>A), CGA160AGA (2552C>A), AAG161ACC (2556A>C 2557G>C), AAC166AAT (2572C>T), ATC167ATA (2575C>A), ACA169ACC (2581A>C), TTG170ATT (2582T>A 2584G>T), AAG174AAA (2596G>A), ACA176CAT (2600A>C 2601C>A 2602A>T), TTG177CTG (2603T>C), GTT178GTG (2608T>G), CAA182GAA (2618C>G), AAC184TCC (2624A>T 2625A>C), GAG186GGG (2631A>G), GAG187AAG (2633G>A), TGC189GTG (2639T>G 2640G>T 2641C>G), TTT190TTC (2644T>C), ACC191TCA (2645A>T 2647C>A), CCA192CCT (2650A>T), ACA193CAA (2651A>C 2652C>A), GGG194GGT (2656G>T), GGC196GGT (2662C>T), AAG197ACC (2664A>C 2665G>C), TAC198TTT (2667A>T 2668C>T), GTG199GTA (2671G>A), CCT200CCA (2674T>A), CTG201TTG (2675C>T), GAA203ACT (2681G>A 2682A>C 2683A>T), GGT204TCA (2684G>T 2685G>C 2686T>A), ACA205CTG (2687A>C 2688C>T 2689A>G), GGA206GGT (2692A>T), GCC207GTT (2694C>T 2695C>T), ACT208CCT (2696A>C), GAT209GAA (2701T>A), TTA210CTT (2702T>C 2704A>T), GGT211GGC (2707T>C), GTG213GTT (2713G>T), ACA218ACC (2728A>C), CAC219CAT (2731C>T), TTA221TTG (2737A>G), TCT222GCA (2738T>G 2740T>A), GCA224GCC (2746A>C), GAG226GAA (2752G>A), AAT228GAG (2756A>G 2758T>G), AGT229TCA (2759A>T 2760G>C 2761T>A), GTA230ATA (2762G>A), CCA231CCT (2767A>T), TGC232TGT (2770C>T), CAT233CAC (2773T>C), TTG234ATA (2774T>A 2776G>A), TTA236CTA (2780T>C), CAG237TCC (2783C>T 2784A>C 2785G>C), AAA239AAG (2791A>G), TTA241CTC (2795T>C 2797A>C), ACC243ACA (2803C>A), CTT246ATG (2810C>A 2812T>G), CGA250AGG (2822C>A 2824A>G), ATG251TTG (2825A>T), ATA252AGA (2829T>G), GCA253GGG (2832C>G 2833A>G), CAG254CAA (2836G>A) | | | | | | | | |
| Coat protein B (YP_001976156.1) | 1 | 244 | 100% | 1420 | 83.1% | 242 (99.2%) | 192 (78.7%) | 0/2/0/0 | 0 |
| Protein mutations: | Y3F (2844A>T), K6R (2852A>C 2853A>G), A7P (2855G>C 2857G>A), Q9T (2861C>A 2862A>C 2863G>A), Y11F (2868A>T 2869T>C), P13S (2873C>A 2874C>G 2875A>T), G27S (2915G>A 2917G>T), T28V (2918A>G 2919C>T), S29A (2921T>G 2923A>T), S30A (2924T>G 2926C>A), P31A (2927C>G 2929A>C), I39V (2951A>G 2953A>C), T51I (2988C>T), T68R (3039C>G 3040T>A), I75V (3059A>G 3061T>G), I77V (3065A>G), N95A (3119A>G 3120A>C 3121C>A), K97E (3125A>G 3127A>G), T99S (3131A>T), N100T (3135A>C 3136T>A), F105L (3151T>G), K119R (3192A>G 3193A>G), R121Q (3198G>A 3199G>A), R123K (3203C>A 3204G>A 3205G>A), I126L (3212A>C 3214T>G), K132C (3230A>T 3231A>G 3232A>C), Q147K (3275C>A), F149Q (3281T>C 3282T>A 3283T>A), L152M (3290T>A), I167L (3335A>T 3337A>G), S171T (3347T>A 3349C>T), C180R (3374T>C), L181I (3377C>A), N183D (3383A>G), T189S (3401A>T 3403A>T), K203R (3444A>G), S205A (3449T>G 3451A>T), D214N (3476G>A 3478T>C), S216N (3482T>A 3483C>A 3484A>T), I221L (3497A>C), D227E (3517C>A), I229V (3521A>G), E230D (3526G>T), E231I (3527G>A 3528A>T 3529G>C), Y232F (3531A>T 3532C>T), Q233D (3533C>G 3535G>C), E234Q (3536G>C), D238_P239del (3548_3553delGATCCG), P240S (3554C>A 3555C>G), K242T (3561A>C 3562G>A), A243S (3563G>T) | | | | | | | | |

| | Begin | End | Coverage | Score | Concordance | Matches | Identities | I/D/M/F* | Stop Codons |
|---|---|---|---|---|---|---|---|---|---|
| **NT** | **89** | **4744** | **94.9%** | **2600** | **28.6%** | **4608 (96.7%)** | **3040 (63.8%)** | **107/48** | |
| Codon mutations: | CAA2CAG (2842A>G), TAT3TTT (2844A>T), GGC4GGA (2848C>A), AAG6CGG (2852A>C 2853A>G), GCG7CCA (2855G>C 2857G>A), GGG8GGC (2860G>C), CAG9ACA (2861C>A 2862A>C 2863G>A), ACA10ACC (2866A>C), TAT11TTC (2868A>T 2869T>C), CCA13AGT (2873C>A 2874C>G 2875A>T), AGG14CGC (2876A>C 2878G>C), TTT15TTC (2881T>C), CCA16CCT (2884A>T), ACT17ACA (2887T>A), GTT20GTG (2896T>G), TTG21CTT (2897T>C 2899G>T), TTA22CTG (2900T>C 2902A>G), CAT23CAC (2905T>C), TAT24TAC (2908T>C), GGG27AGT (2915G>A 2917G>T), ACA28GTA (2918A>G 2919C>T), TCA29GCT (2921T>G 2923A>T), TCC30GCA (2924T>G 2926C>A), CCA31GCC (2927C>G 2929A>C), ACA35ACT (2941A>T), AGC38AGT (2950C>T), ATA39GTC (2951A>G 2953A>C), TTT40TTC (2956T>C), TCA41TCC (2959A>C), ATA45ATT (2971A>T), GAG47GAA (2977G>A), GGC50GGT (2986C>T), ACC51ATC (2988C>T), TTG52TTA (2992G>A), CCG54CCA (2998G>A), TCT55TCA (3001T>A), TTG56CTG (3002T>C), GGT58GGA (3010T>A), ATA60ATT (3016A>T), GCC61GCA (3019C>A), AGG62AGA (3022G>A), AAG65AAA (3031G>A), ACT68AGA (3039C>G 3040T>A), GGA69GGC (3043A>C), ACT70ACC (3046T>C), ATT75GTG (3059A>G 3061T>G), TGC76TGT (3064C>T), ATT77GTT (3065A>G), ACC80ACT (3076C>T), TTG81CTG (3077T>C), TCA84TCT (3088A>T), GGT85GGG (3091T>G), CTG87CTT (3097G>T), GCA88GCT (3100A>T), ATT89ATA (3103T>A), GGA90GGG (3106A>G), CTT91TTG (3107C>T 3109T>G), GGA92GGC (3112A>C), ACT93ACG (3115T>G), AAC95GCA (3119A>G 3120A>C 3121C>A), ACC96ACA (3124C>A), AAA97GAG (3125A>G 3127A>G), ACC99TCC (3131A>T), AAT100ACA (3135A>C 3136T>A), GCT101GCG (3139T>G), TTT105TTG (3151T>G), CCA108CCC (3160A>C), GTA111GTG (3169A>G), TGC112TGT (3172C>T), CTT114TTG (3176C>T 3178T>G), GGT117GGG (3187T>G), AAA119AGG (3192A>G 3193A>G), CGG121CAA (3198G>A 3199G>A), CGG123AAA (3203C>A 3204G>A 3205G>A), TGT124TGC (3208T>C), TCT125TCA (3211T>A), ATT126CTG (3212A>C 3214T>G), ACA127ACC (3217A>C), GGA131GGG (3229A>G), AAA132TGC (3230A>T 3231A>G 3232A>C), AAT133AAC (3235T>C), TTG134CTC (3236T>C 3238G>C), CTT135CTG (3241T>G), TCC136TCT (3244C>T), ACA137ACC (3247A>C), GGG138GGA (3250G>A), CGA139CGC (3253A>C), AAG140AAA (3256G>A), TCC142TCA (3262C>A), TTG143TTA (3265G>A), CAG147AAG (3275C>A), CAC148CAT (3280C>T), TTT149CAA (3281T>C 3282T>A 3283T>A), TCC150TCT (3286C>T), TTG152ATG (3290T>A), CGC153CGT (3295C>T), TTG154CTG (3296T>C), TTT155TTC (3301T>C), ACG157ACT (3307G>T), GTA158GTT (3310A>T), CTC162CTG (3322C>G), ACG165ACA (3331G>A), ATA167TTG (3335A>T 3337A>G), CAT168CAC (3340T>C), TCC171ACT (3347T>A 3349C>T), GGT173GGA (3355T>A), GTC174GTA (3358C>A), CAG177CAA (3367G>A), AAG179AAA (3373G>A), TGC180CGC (3374T>C), CTT181ATT (3377C>A), AAT183GAT (3383A>G), CTT184CTC (3388T>C), ACT185ACG (3391T>G), TTG186CTT (3392T>C 3394G>T), GGT187GGG (3397T>G), GGC188GGT (3400C>T), ACA189TCT (3401A>T 3403A>T), GTA190GTG (3406A>G), TCT191TCA (3409T>A), GTG192GTT (3412G>T), AAA193AAG (3415A>G), ATA195ATT (3421A>T), GGG197GGT (3427G>T), ACA200ACC (3436A>C), AAA201AAG (3439A>G), GGC202GGG (3442C>G), AAA203AGA (3444A>G), AGC204AGT (3448C>T), TCA205GCT (3449T>G 3451A>T), GTT206GTG (3454T>G), TTC208TTT (3460C>T), CTT209CTG (3463T>G), TTC210TTT (3466C>T), GAT214AAC (3476G>A 3478T>C), TTA215CTG (3479T>C 3481A>G), TCA216AAT (3482T>A 3483C>A 3484A>T), CGT218CGC (3490T>C), ATT221CTT (3497A>C), GAA222GAG (3502A>G), AAG223AAA (3505G>A), TTG224CTG (3506T>C), AAG226AAA (3514G>A), GAC227GAA (3517C>A), AAC228AAT (3520C>T), ATT229GTT (3521A>G), GAG230GAT (3526G>T), GAG231ATC (3527G>A 3528A>T 3529G>C), TAC232TTT (3531A>T 3532C>T), CAG233GAC (3533C>G 3535G>C), GAG234CAG (3536G>C), GGC236GGG (3544C>G), GAT238_CCG239del (3548_3553delGATCCG), CCC240AGC (3554C>A 3555C>G), AAG242ACA (3561A>C 3562G>A), GCT243TCT (3563G>T) | | | | | | | | |
| Coat protein C (YP_001976157.1) | 1 | 225 | 100% | 1273 | 77.2% | 225 (100%) | 158 (70.2%) | 0/0/0/0 | 0 |
| Protein mutations: | S1V (3569A>G 3570G>T), I2S (3572A>T 3573T>C 3574T>C), L3F (3575C>T 3577G>C), S4Q (3578T>C 3579C>A 3580C>G), S10T (3596T>A 3598C>A), M19A (3623A>G 3624T>C 3625G>T), G20A (3627G>C 3628T>A), D24T (3638G>A 3639A>C 3640C>A), E25D (3643A>C), L28V (3650T>G 3652G>A), I30L (3656A>T 3658T>G), I37V (3677A>G 3679A>G), E40D (3688A>T), G41A (3690G>C 3691G>A), H42S (3692C>T 3693A>C), V45I (3701G>A), T58S (3740A>T 3742A>C), N68E (3770A>G 3772T>A), S70T (3776T>A), I71F (3779A>T), I73M (3787A>G), V76T (3794G>A 3795T>C 3796A>T), P80S (3806C>A 3807C>G 3808T>C), N81S (3810A>G), V82I (3812G>A 3814T>A), V85I (3821G>A 3823A>C), V88I (3830G>A), F90L (3836T>C), A92T (3842G>A 3844C>A), P98Q (3861C>A), A99F (3863G>T 3864C>T), L101V (3869C>G 3871T>A), N102S (3872A>T 3873A>C), K103A (3875A>G 3876A>C 3877A>G), N105A (3881A>G 3882A>C 3883T>A), Y106H (3884T>C), V108L (3890G>T), A123T (3935G>A 3937A>C), T126I (3945C>T 3946G>C), T130V (3956A>G 3957C>T 3958T>A), E135Q (3971G>C), H141F (3989C>T 3990A>T 3991T>C), R143K (3996G>A), M144A (3998A>G 3999T>C 4000G>C), R145K (4002G>A 4003G>A), E146S (4004G>T 4005A>C 4006A>C), F147Y (4008T>A 4009C>T), S148D (4010T>G 4011C>A 4012A>T), S149K (4014G>A 4015C>G), K150E (4016A>G 4018G>A), Q151S (4019C>A 4020A>G 4021A>T), M155V (4031A>G 4033G>T), R160K (4047G>A), N163L (4055A>T 4056A>T 4057T>G), I165L (4061A>C 4063C>A), S171V (4079T>G 4080C>T 4081C>T), A172E (4083C>A 4084C>A), I173L (4085A>C 4087A>T), V174I (4088G>A 4090G>T), L180I (4106C>A 4108T>A), I181V (4109A>G 4111A>C), S182K (4112T>A 4113C>A 4114T>G), L185P (4122T>C), S196L (4154T>C 4155C>T), H199N (4163C>A), N205D (4181A>G), I225N (4242T>A) | | | | | | | | |
| Codon mutations: | AGT1GTT (3569A>G 3570G>T), ATT2TCC (3572A>T 3573T>C 3574T>C), CTG3TTC (3575C>T 3577G>C), TCC4CAG (3578T>C 3579C>A 3580C>G), ATA5ATC (3583A>C), AGA6AGG (3586A>G), GAG7GAA (3589G>A), AAA8AAG (3592A>G), TTT9TTC (3595T>C), TCC10ACA (3596T>A 3598C>A), GGT12GGA (3604T>A), GCT13GCA (3607T>A), GTA14GTG (3610A>G), TTC17TTT (3619C>T), TGC18TGT (3622C>T), ATG19GCT (3623A>G 3624T>C 3625G>T), GGT20GCA (3627G>C 3628T>A), GAC24ACA (3638G>A 3639A>C 3640C>A), GAA25GAC (3643A>C), TTG28GTA (3650T>G 3652G>A), GTA29GTT (3655A>T), ATT30TTG (3656A>T 3658T>G), CCT31CCA (3661T>A), GCA33GCC (3667A>C), CCA34CCT (3670A>T), TCC36TCA (3676C>A), ATA37GTG (3677A>G 3679A>G), AGG38CGC (3680A>C 3682G>C), GAA40GAT (3688A>T), GGG41GCA (3690G>C 3691G>A), CAC42TCC (3692C>T 3693A>C), CCT44CCA (3700T>A), GTT45ATT (3701G>A), AAG46AAA (3706G>A), ATC49ATA (3715C>A), TGT51TGC (3721T>C), TTT53TTC (3727T>C), GAT55GAC (3733T>C), ACA58TCC (3740A>T 3742A>C), TCA59TCT (3745A>T), TCA64TCT (3760A>T), GGT65GGA (3763T>A), AAT68GAA (3770A>G 3772T>A), TCA70ACA (3776T>A), ATT71TTT (3779A>T), GTG72GTA (3784G>A), ATA73ATG (3787A>G), AGA75CGA (3791A>C), GTA76ACT (3794G>A 3795T>C 3796A>T), CAA77CAG (3799A>G), TCC78AGC (3800T>A 3801C>G), AGT79TCC (3803A>T 3804G>C 3805T>C), CCT80AGC (3806C>A 3807C>G 3808T>C), AAT81AGT (3810A>G), GTT82ATA (3812G>A 3814T>A), GGA83GGT (3817A>T), GTA85ATC (3821G>A 3823A>C), CTA86TTG (3824C>T 3826A>G), AAT87AAC (3829T>C), GTT88ATT (3830G>A), GCT89GCA (3835T>A), TTT90CTT (3836T>C), GAT91GAC (3841T>C), GCC92ACA (3842G>A 3844C>A), GGC94GGG (3850C>G), TTT97TTC (3859T>C), CCA98CAA (3861C>A), GCT99TTT (3863G>T 3864C>T), GGG100GGT (3868G>T), CTT101GTA (3869C>G 3871T>A), AAT102TCT (3872A>T 3873A>C), AAA103GCG (3875A>G 3876A>C 3877A>G), AAT105GCA (3881A>G 3882A>C 3883T>A), TAT106CAT (3884T>C), GTA108TTA (3890G>T), GGT111GGC (3901T>C), GGA112GGT (3904A>T), GGC113GGA (3907C>A), ACA114ACT (3910A>T), TCA119TCC (3925A>C), TAC120TAT (3928C>T), GGT121GGG (3931T>G), GTG122GTT (3934G>T), GCA123ACC (3935G>A 3937A>C), ACA124ACC (3940A>C), AAT125AAC (3943T>C), ACG126ATC (3945C>T 3946G>C), TCA128TCT (3952A>T), TTC129TTT (3955C>T), ACT130GTA (3956A>G 3957C>T 3958T>A), GAG135CAG (3971G>C), TTT136TTC (3976T>C), TTC137TTT (3979C>T), CCA138CCT (3982A>T), AGG139CGC (3983A>C 3985G>C), CGG140CGT (3988G>T), CAT141TTC (3989C>T 3990A>T 3991T>C), AGG143AAG (3996G>A), ATG144GCC (3998A>G 3999T>C 4000G>C), AGG145AAA (4002G>A 4003G>A), GAA146TCC (4004G>T 4005A>C 4006A>C), TTC147TAT (4008T>A 4009C>T), TCA148GAT (4010T>G 4011C>A 4012A>T), AGC149AAG (4014G>A 4015C>G), AAG150GAA (4016A>G 4018G>A), CAA151AGT (4019C>A 4020A>G 4021A>T), TCC152TCA (4024C>A), CGC153AGA (4025C>A 4027C>A), ATC154ATT (4030C>T), ATG155GTT (4031A>G 4033G>T), TCA156TCT (4036A>T), CTA157TTG (4037C>T 4039A>G), GAT159GAC (4045T>C), AGG160AAG (4047G>A), CTT161CTA (4051T>A), GGA162GGC (4054A>C), AAT163TTG (4055A>T 4056A>T 4057T>G), CTG164TTG (4058C>T), ATC165CTA (4061A>C 4063C>A), ATA166ATC (4066A>C), TTG168CTT (4070T>C 4072G>T), CCT169CCA (4075T>A), CCT170CCA (4078T>A), TCC171GTT (4079T>G 4080C>T 4081C>T), GCC172GAA (4083C>A 4084C>A), ATA173CTT (4085A>C 4087A>T), GTG174ATT (4088G>A 4090G>T), TCC176TCA (4096C>A), GAG178GAA (4102G>A), ATA179ATC (4105A>C), CTT180ATA (4106C>A 4108T>A), ATA181GTC (4109A>G 4111A>C), TCT182AAG (4112T>A 4113C>A 4114T>G), CCT183CCA (4117T>A), GGA184GGT (4120A>T), CTT185CCT (4122T>C), TTC187TTT (4129C>T), AAG188AAA (4132G>A), GCC192GCT (4144C>T), CCT194CCC (4150T>C), CCT195CCA (4153T>A), TCT196CTT (4154T>C 4155C>T), GCC197GCA (4159C>A), CAT199AAT (4163C>A), GAA200GAG (4168A>G), AAA201AAG (4171A>G), CTT203CTC (4177T>C), GGC204GGA (4180C>A), AAT205GAT (4181A>G), CAA207CAG (4189A>G), ACT208ACG (4192T>G), CAC209CAT (4195C>T), ACC210ACT (4198C>T), ACC214ACA (4210C>A), TCA215TCT (4213A>T), CTA220TTG (4226C>T 4228A>G), CGT221CGC (4231T>C), ATT225AAT (4242T>A) | | | | | | | | |

*: Inserts / Deletes / Misaligned / Frameshifts

#### Analysis details

This analysis was performed with panviral2.64

### NGS Details (UN59): Bracoviriform glomeratae (segment NC_043292.1)

#### Assembly

| | |
|---|---|
| **Coverage Length** | 308 (1 contig(s)) |
| **Depth Of Coverage** | 4640.4 |
| **Number Of Reads** | 12058 |
| **Reads Per Million** | 262.13 rpm (after QC) |
| **Ambiguities** | 0 |
| **Assembly Method** | de novo + reference guided assembly |
| **Consensus Caller** | Bcf Tools |

#### Coverage Map

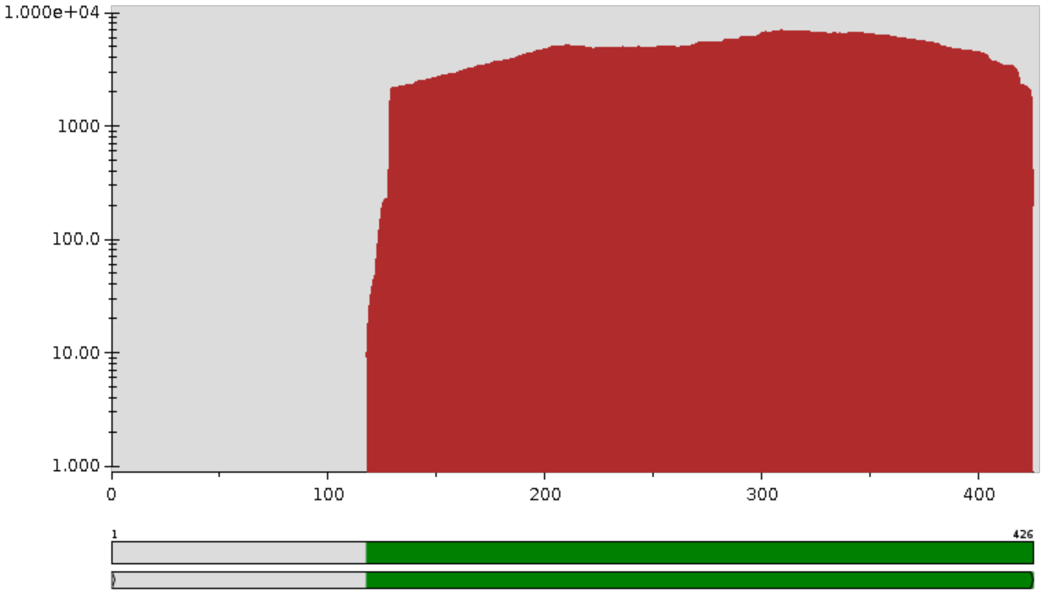

#### Assignment

| | |
|---|---|
| **Type** | Bracoviriform glomeratae (Taxonomy ID: 257816) |
| **Reference Genome** | NC_043292.1 |
| **NT Identity (%)** | 72.7273 |
| **AA Identity (%)** | 84.3137 |
| **Number Of Stop Codons** | 1 |
| **Number Of CDS** | 1 |

#### Alignment

| | |
|---|---|
| **Alignment Score** | 280.0 (NT) + 559.0 (AA) = 839.0 |
| **Concordance (%)** | 66.3241 |

| Alignment Method | Global, seeded, nucleotide + amino acids (AGA) |
|---|---|

#### Genome Region

Sequence starts at position 119 and ends at position 426 relative to NC_043292.1 reference sequence.

#### Alignment Detailed Statistics

| | Begin | End | Coverage | Score | Concordance | Matches | Identities | I/D/M/F* | Stop Codons |
|---|---|---|---|---|---|---|---|---|---|
| **NT** | **119** | **426** | **72.3%** | **280** | **45.5%** | **308 (100%)** | **224 (72.7%)** | **0/0** | |
| Mutations: | 126G>T, 129C>A, 132A>G, 136A>G, 137C>G, 138G>C, 150A>C, 153A>G, 155C>G, 156G>A, 159C>A, 166C>A, 168T>G, 169T>C, 171T>C, 172C>A, 174C>G, 177A>G, 183T>A, 184C>A, 189T>C, 198A>G, 201A>C, 207C>T, 214A>G, 219C>T, 222T>C, 223C>A, 225T>G, 228G>T, 235C>A, 237T>G, 243A>T, 246C>G, 255C>T, 261A>G, 262T>C, 264A>G, 265G>A, 273A>G, 278T>C, 279T>C, 284A>G, 285T>A, 288T>G, 289T>C, 300C>T, 301C>T, 303C>G, 309C>T, 312T>G, 315C>T, 322G>T, 324T>C, 329T>C, 333T>C, 336C>T, 339A>G, 345A>C, 347A>G, 348A>G, 349C>A, 351T>A, 360C>T, 363C>T, 366C>T, 375C>T, 384A>C, 385T>C, 387G>T, 390A>G, 391C>A, 392A>G, 393T>A, 394A>C, 399C>A, 400C>A, 402T>G, 404T>C, 405G>T, 406A>C, 414T>A, 416A>T, 420A>G | | | | | | | | |

##### CDS

| | Begin | End | Coverage | Score | Concordance | Matches | Identities | I/D/M/F* | Stop Codons |
|---|---|---|---|---|---|---|---|---|---|
| FK954_p501 | 41 | 142 | 71.8% | 559 | 85.1% | 102 (100%) | 86 (84.3%) | 0/0/0/0 | 1 |
| Protein mutations: | L42F (126G>T), T46G (136A>G 137C>G 138G>C), A52G (155C>G 156G>A), Y57H (169T>C 171T>C), T72A (214A>G), V89I (265G>A), I93T (278T>C 279T>C), D95G (284A>G 285T>A), A108S (322G>T 324T>C), I110T (329T>C), K116R (347A>G 348A>G), H131R (391C>A 392A>G 393T>A), K132Q (394A>C), M135T (404T>C 405G>T), I136L (406A>C), Y139F (416A>T) | | | | | | | | |
| Codon mutations: | TTG42TTT (126G>T), GGC43GGA (129C>A), AAA44AAG (132A>G), ACG46GGC (136A>G 137C>G 138G>C), GGA50GGC (150A>C), AAA51AAG (153A>G), GCG52GGA (155C>G 156G>A), GGC53GGA (159C>A), CGT56AGG (166C>A 168T>G), TAT57CAC (169T>C 171T>C), CGC58AGG (172C>A 174C>G), AAA59AAG (177A>G), CTT61CTA (183T>A), CGA62AGA (184C>A), GAT63GAC (189T>C), CAA66CAG (198A>G), GGA67GGC (201A>C), ACC69ACT (207C>T), ACT72GCT (214A>G), ATC73ATT (219C>T), CGT74CGC (222T>C), CGT75AGG (223C>A 225T>G), CTG76CTT (228G>T), CGT79AGG (235C>A 237T>G), GGA81GGT (243A>T), GTC82GTG (246C>G), ATC85ATT (255C>T), GGA87GGG (261A>G), TTA88CTG (262T>C 264A>G), GTC89ATC (265G>A), GAA91GAG (273A>G), ATT93ACC (278T>C 279T>C), GAT95GGA (284A>G 285T>A), GTT96GTG (288T>G), TTG97CTG (289T>C), TTC100TTT (300C>T), CTC101TTG (301C>T 303C>G), AAC103AAT (309C>T), GTT104GTG (312T>G), ATC105ATT (315C>T), GCT108TCC (322G>T 324T>C), ATC110ACC (329T>C), TAT111TAC (333T>C), ACC112ACT (336C>T), GAA113GAG (339A>G), GCA115GCC (345A>C), AAA116AGG (347A>G 348A>G), CGT117AGA (349C>A 351T>A), GTC120GTT (360C>T), ACC121ACT (363C>T), GCC122GCT (366C>T), GTC125GTT (375C>T), GCA128GCC (384A>C), TTG129CTT (385T>C 387G>T), AAA130AAG (390A>G), CAT131AGA (391C>A 392A>G 393T>A), AAA132CAA (394A>C), GGC133GGA (399C>A), CGT134AGG (400C>A 402T>G), ATG135ACT (404T>C 405G>T), ATC136CTC (406A>C), GGT138GGA (414T>A), TAT139TTT (416A>T), GGA140GGG (420A>G) | | | | | | | | |

##### Proteins

| | Begin | End | Coverage | Score | Concordance | Matches | Identities | I/D/M/F* | Stop Codons |
|---|---|---|---|---|---|---|---|---|---|
| putative histone 4 (YP_009665791.1) | 41 | 142 | 71.8% | 559 | 85.1% | 102 (100%) | 86 (84.3%) | 0/0/0/0 | 1 |
| Protein mutations: | L42F (126G>T), T46G (136A>G 137C>G 138G>C), A52G (155C>G 156G>A), Y57H (169T>C 171T>C), T72A (214A>G), V89I (265G>A), I93T (278T>C 279T>C), D95G (284A>G 285T>A), A108S (322G>T 324T>C), I110T (329T>C), K116R (347A>G 348A>G), H131R (391C>A 392A>G 393T>A), K132Q (394A>C), M135T (404T>C 405G>T), I136L (406A>C), Y139F (416A>T) | | | | | | | | |
| Codon mutations: | TTG42TTT (126G>T), GGC43GGA (129C>A), AAA44AAG (132A>G), ACG46GGC (136A>G 137C>G 138G>C), GGA50GGC (150A>C), AAA51AAG (153A>G), GCG52GGA (155C>G 156G>A), GGC53GGA (159C>A), CGT56AGG (166C>A 168T>G), TAT57CAC (169T>C 171T>C), CGC58AGG (172C>A 174C>G), AAA59AAG (177A>G), CTT61CTA (183T>A), CGA62AGA (184C>A), GAT63GAC (189T>C), CAA66CAG (198A>G), GGA67GGC (201A>C), ACC69ACT (207C>T), ACT72GCT (214A>G), ATC73ATT (219C>T), CGT74CGC (222T>C), CGT75AGG (223C>A 225T>G), CTG76CTT (228G>T), CGT79AGG (235C>A 237T>G), GGA81GGT (243A>T), GTC82GTG (246C>G), ATC85ATT (255C>T), GGA87GGG (261A>G), TTA88CTG (262T>C 264A>G), GTC89ATC (265G>A), GAA91GAG (273A>G), ATT93ACC (278T>C 279T>C), GAT95GGA (284A>G 285T>A), GTT96GTG (288T>G), TTG97CTG (289T>C), TTC100TTT (300C>T), CTC101TTG (301C>T 303C>G), AAC103AAT (309C>T), GTT104GTG (312T>G), ATC105ATT (315C>T), GCT108TCC (322G>T 324T>C), ATC110ACC (329T>C), TAT111TAC (333T>C), ACC112ACT (336C>T), GAA113GAG (339A>G), GCA115GCC (345A>C), AAA116AGG (347A>G 348A>G), CGT117AGA (349C>A 351T>A), GTC120GTT (360C>T), ACC121ACT (363C>T), GCC122GCT (366C>T), GTC125GTT (375C>T), GCA128GCC (384A>C), TTG129CTT (385T>C 387G>T), AAA130AAG (390A>G), CAT131AGA (391C>A 392A>G 393T>A), AAA132CAA (394A>C), GGC133GGA (399C>A), CGT134AGG (400C>A 402T>G), ATG135ACT (404T>C 405G>T), ATC136CTC (406A>C), GGT138GGA (414T>A), TAT139TTT (416A>T), GGA140GGG (420A>G) | | | | | | | | |

*: Inserts / Deletes / Misaligned / Frameshifts

#### Analysis details

This analysis was performed with panviral2.64

### NGS Details (UN59): Diachasmimorpha longicaudata entomopoxvirus (segment NC_043455.1)

#### Assembly

| | |
|---|---|
| **Coverage Length** | 1121 (1 contig(s)) |
| **Depth Of Coverage** | 876.9 |
| **Number Of Reads** | 7928 |
| **Reads Per Million** | 172.35 rpm (after QC) |
| **Ambiguities** | 0 |
| **Assembly Method** | de novo + reference guided assembly |
| **Consensus Caller** | Bcf Tools |

#### Coverage Map

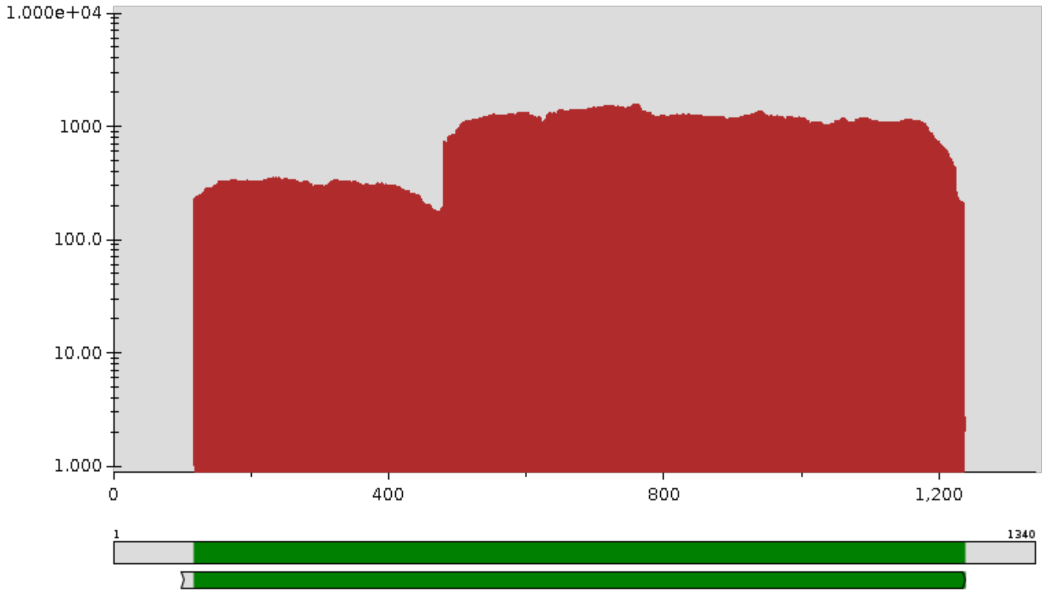

#### Assignment

| | |
|---|---|
| **Type** | Diachasmimorpha longicaudata entomopoxvirus (Taxonomy ID: 109981) |
| **Reference Genome** | NC_043455.1 |
| **NT Identity (%)** | 59.1436 |
| **AA Identity (%)** | 55.3476 |
| **Number Of Stop Codons** | 0 |
| **Number Of CDS** | 1 |

#### Alignment

| | |
|---|---|
| **Alignment Score** | 392.0 (NT) + 1456.0 (AA) = 1848.0 |
| **Concordance (%)** | 39.4535 |

| Alignment Method | Global, seeded, nucleotide + amino acids (AGA) |
|---|---|

#### Genome Region

Sequence starts at position 118 and ends at position 1238 relative to NC_043455.1 reference sequence.

#### Alignment Detailed Statistics

| | Begin | End | Coverage | Score | Concordance | Matches | Identities | I/D/M/F* | Stop Codons |
|---|---|---|---|---|---|---|---|---|---|
| **NT** | **118** | **1238** | **83.7%** | **392** | **17.6%** | **1118 (99.5%)** | **663 (59.0%)** | **3/3** | |

Mutations:

119G>A, 120A>G, 121A>T, 126A>C, 127C>T, 128C>G, 130T>G, 136C>G, 137C>A, 139C>T, 142G>A, 143A>G, 145A>T, 146A>G, 148T>C, 149A>C, 151C>T, 154A>T, 156A>G, 157G>A, 160T>G, 163A>T, 166T>C, 167T>C, 168C>A, 169T>G, 171G>A, 172T>C, 175A>G, 178T>C, 187T>A, 190A>T, 191A>G, 193T>C, 203A>C, 204A>G, 208C>T, 209A>G, 211T>A, 214T>G, 217A>T, 218T>A, 219G>T, 223T>A, 226T>G, 230A>C, 231A>G, 232A>T, 238G>C, 241A>T, 243T>C, 244C>T, 247A>G, 256G>C, 259A>T, 262A>C, 271A>C, 272G>T, 274A>C, 276C>T, 277A>G, 278T>A, 279A>T, 283A>T, 284A>C, 286C>T, 288G>C, 289T>C, 294T>G, 295A>C, 298G>A, 299C>G, 300A>T, 301A>T, 302A>G, 304A>G, 310T>C, 311T>A, 312C>A, 313T>A, 320A>G, 322T>G, 323A>G, 325C>A, 331T>A, 337C>A, 341A>T, 343C>T, 346A>T, 352T>G, 355A>G, 356T>C, 358G>T, 361T>A, 362C>G, 363T>C, 364C>T, 368G>A, 370T>G, 371C>G, 374C>A, 375G>A, 376T>G, 379A>G, 380T>A, 382G>T, 383C>T, 384A>T, 385A>G, 386A>G, 394G>A, 395A>G, 400T>C, 401C>A, 403A>T, 404_406delTAT, 409T>C, 410T>A, 412T>A, 413A>C, 415A>G, 416T>G, 417G>C, 418T>A, 421A>T, 423T>C, 424T>G, 438C>A, 442T>C, 443A>G, 445C>G, 446A>G, 447A>G, 448G>T, 452A>G, 453G>A, 454C>T, 455C>A, 456A>T, 457G>C, 458G>A, 459A>G, 462C>A, 463T>G, 467A>G, 469_470insCAT, 470A>G, 471A>G, 474C>T, 475C>A, 478G>A, 481T>A, 482T>G, 485A>G, 490A>T, 496T>A, 499A>C, 502A>T, 503A>G, 505G>C, 506A>T, 511T>C, 512C>A, 514A>G, 517C>G, 518A>C, 519C>G, 521C>A, 523A>G, 524A>C, 526A>G, 527A>T, 528G>C, 530A>C, 532C>T, 533G>C, 534A>G, 535T>C, 536A>C, 538C>T, 539A>G, 541A>C, 542G>C, 543C>A, 544A>C, 547T>C, 550A>G, 553A>G, 554G>T, 559A>T, 560A>T, 562A>G, 574T>C, 577G>A, 581T>C, 583G>C, 584A>T, 585T>C, 586A>T, 587G>A, 588A>G, 589T>A, 590A>G, 591A>G, 595T>C, 596T>A, 597T>A, 602A>C, 604A>G, 607A>T, 608C>T, 610A>T, 616T>A, 620G>C, 622A>G, 623T>C, 628T>G, 629G>C, 630A>C, 632A>C, 633G>C, 634T>A, 635C>A, 637T>A, 638G>A, 646T>G, 647A>G, 648T>G, 650T>G, 652A>T, 655A>C, 658A>T, 661A>T, 664C>T, 665G>A, 667T>G, 671T>C, 674A>G, 675G>A, 678T>C, 680A>C, 682C>T, 683A>G, 685T>A, 687C>T, 688A>T, 689T>A, 692C>A, 693A>G, 695G>A, 696T>A, 697C>G, 700T>C, 705G>A, 706A>C, 707G>A, 709T>A, 712C>T, 715A>G, 717A>G, 718A>G, 724G>T, 727A>G, 730A>G, 731A>C, 732A>G, 735C>A, 739A>G, 740T>C, 742G>C, 745A>T, 748C>T, 754C>T, 757T>C, 759G>A, 765A>T, 770A>G, 772A>C, 778A>C, 779A>G, 781G>T, 784A>G, 785A>G, 787T>A, 790T>A, 792T>G, 793T>G, 796A>G, 797G>C, 798C>T, 799A>T, 810T>G, 823C>G, 824C>A, 825A>C, 826C>T, 829A>G, 830A>G, 831G>C, 833C>A, 838T>C, 841A>G, 843C>G, 844A>T, 845C>G, 847A>C, 850A>C, 854T>G, 855G>T, 856T>G, 859T>C, 862A>C, 863C>A, 865T>G, 866A>C, 868A>T, 869C>A, 878G>T, 879T>G, 880A>G, 885T>C, 886G>T, 889A>T, 890T>A, 891G>A, 892T>G, 893T>A, 895A>G, 896A>C, 897C>G, 898A>C, 900A>G, 901T>C, 902A>C, 904A>T, 905A>G, 908T>C, 909T>A, 915C>T, 919A>T, 920A>G, 921G>C, 922C>A, 924T>C, 925C>T, 934T>C, 938T>G, 939C>A, 943A>G, 944C>A, 946A>C, 947G>A, 948A>C, 949A>T, 950C>A, 952T>A, 956T>A, 958T>C, 965A>C, 966A>G, 970A>G, 973T>C, 974A>C, 977G>T, 978A>C, 982T>G, 983A>T, 984A>C, 986A>T, 989A>C, 991A>T, 992A>G, 994A>G, 997T>C, 998C>A, 1000T>C, 1001T>A, 1007A>G, 1012A>C, 1013T>C, 1015A>T, 1018A>T, 1019A>C, 1021A>T, 1024C>T, 1033T>A, 1036A>G, 1040A>G, 1042A>C, 1045T>C, 1046T>C, 1048A>T, 1051T>C, 1054C>T, 1057T>C, 1063T>C, 1064T>C, 1070C>A, 1071A>C, 1073A>C, 1075T>A, 1076A>C, 1077G>C, 1087T>C, 1088A>C, 1090A>G, 1103A>C, 1105A>T, 1111T>A, 1112A>C, 1120C>A, 1121C>A, 1123A>G, 1126A>G, 1129G>T, 1130A>G, 1141T>C, 1145A>G, 1147A>C, 1150T>C, 1151G>A, 1153A>G, 1156C>T, 1159A>T, 1160T>G, 1161T>A, 1163C>A, 1164C>G, 1165A>G, 1167A>T, 1172G>T, 1173C>T, 1177A>C, 1178T>A, 1181G>C, 1183A>G, 1185C>A, 1186G>A, 1192T>C, 1195T>C, 1196A>G, 1197C>T, 1199A>G, 1200A>T, 1204A>C, 1207T>G, 1210A>G, 1211A>C, 1213G>C, 1218A>C, 1219G>C, 1223A>G, 1227T>C, 1232T>C, 1234G>C, 1235A>C, 1237A>G

**CDS**

| | Begin | End | Coverage | Score | Concordance | Matches | Identities | I/D/M/F* | Stop Codons |
|---|---|---|---|---|---|---|---|---|---|
| FLA14_p101 | 7 | 380 | 98.4% | 1456 | 58.9% | 373 (99.5%) | 207 (55.2%) | 1/1/0/0 | 0 |

Protein mutations:

E7S (119G>A 120A>G 121A>T), D9A (126A>C 127C>T), H10E (128C>G 130T>G), L13I (137C>A 139C>T), K15D (143A>G 145A>T), N16D (146A>G 148T>C), I17L (149A>C 151C>T), K19R (156A>G 157G>A), S23Q (167T>C 168C>A 169T>G), C24Y (171G>A 172T>C), T31A (191A>G 193T>C), K35R (203A>C 204A>G), I37V (209A>G 211T>A), F38L (214T>G), C40I (218T>A 219G>T), K44R (230A>C 231A>G 232A>T), V48A (243T>C 244C>T), A58S (272G>T 274A>C), T59M (276C>T 277A>G), Y60I (278T>A 279A>T), I62L (284A>C 286C>T), S63T (288G>C 289T>C), L65C (294T>G 295A>C), Q67V (299C>G 300A>T 301A>T), I68V (302A>G 304A>G), S71K (311T>A 312C>A 313T>A), N74E (320A>G 322T>G), I75V (323A>G 325C>A), T81S (341A>T 343C>T), L88A (362C>G 363T>C 364C>T), A90T (368G>A 370T>G), Q91E (371C>G), R92K (374C>A 375G>A 376T>G), L94I (380T>A 382G>T), Q95L (383C>T 384A>T 385A>G), T96A (386A>G), N99D (395A>G), L101I (401C>A 403A>T), Y102del (404_406delTAT), F104I (410T>A 412T>A), K105Q (413A>C 415A>G), C106A (416T>G 417G>C 418T>A), Q107H (421A>T), V108A (423T>C 424T>G), T113K (438C>A), I115V (443A>G 445C>G), K116G (446A>G 447A>G 448G>T), S118D (452A>G 453G>A 454C>T), Q119I (455C>A 456A>T 457G>C), E120R (458G>A 459A>G), T121K (462C>A 463T>G), K123E (467A>G), K123_K124insH (469_470insCAT), K124G (470A>G 471A>G), A125V (474C>T 475C>A), L128V (482T>G), I129V (485A>G), M135V (503A>G 505G>C), I136F (506A>T), L138M (512C>A 514A>G), T140R (518A>C 519C>G), K142Q (524A>C 526A>G), I144L (530A>C 532C>T), D145R (533G>C 534A>G 535T>C), T146P (536A>C 538C>T), K147D (539A>G 541A>C), A148H (542G>C 543C>A 544A>C), I151M (553A>G), V152F (554G>T), I154L (560A>T 562A>G), I162S (584A>T 585T>C 586A>T), D163R (587G>A 588A>G 589T>A), N164G (590A>G 591A>G), L166K (596T>A 597T>A), K168Q (602A>C 604A>G), Q170Y (608C>T 610A>T), E174Q (620G>C 622A>G), F175L (623T>C), F176L (628T>G), E177P (629G>C 630A>C), S178P (632A>C 633G>C 634T>A), H179K (635C>A 637T>A), V180I (638G>A), I183G (647A>G 648T>G), L184V (650T>G 652A>T), L185F (655A>C), V189M (665G>A 667T>G), S191P (671T>C), R192E (674A>G 675G>A), V193A (678T>C), I194L (680A>C 682C>T), N195E (683A>G 685T>A), T196I (687C>T 688A>T), S197T (689T>A), Q198R (692C>A 693A>G), V199K (695G>A 696T>A 697C>G), R202N (705G>A 706A>C), D203K (707G>A 709T>A), K206R (717A>G 718A>G), N211R (731A>C 732A>G), A212D (735C>A), R220K (759G>A), Y222F (765A>T), I224V (770A>G 772A>C), K227D (779A>G 781G>T), N229E (785A>G 787T>A), D230E (790T>A), F231W (792T>G 793T>G), A233L (797G>C 798C>T 799A>T), F237C (810T>G), D241E (823C>G), H242T (824C>A 825A>C 826C>T), S244A (830A>G 831G>C), L245I (833C>A), T248S (843C>G 844A>T), L249V (845C>G 847A>C), C252V (854T>G 855G>T 856T>G), Q257K (869C>A), V260W (878G>T 879T>G 880A>G), M262T (885T>C 886G>T), E263D (889A>T), C264K (890T>A 891G>A 892T>G), L265M (893T>A 895A>G), T266R (896A>C 897C>G 898A>C), N267S (900A>G 901T>C), N269D (905A>G), F270H (908T>C 909T>A), A272V (915C>T), S274A (920A>G 921G>C 922C>A), I275T (924T>C 925C>T), S280D (938T>G 939C>A), Q282N (944C>A 946A>C), E283T (947G>A 948A>C 949A>T), F286I (956T>A 958T>C), K289R (965A>C 966A>G), D293S (977G>T 978A>C), K295S (983A>T 984A>C), T296S (986A>T), I298V (992A>G 994A>G), L300I (998C>A 1000T>C), S301T (1001T>A), N303D (1007A>G), I314V (1040A>G 1042A>C), H324T (1070C>A 1071A>C), N325Q (1073A>C 1075T>A), R326P (1076A>C 1077G>C), I330L (1088A>C 1090A>G), I344V (1130A>G), I349V (1145A>G 1147A>C), E351K (1151G>A 1153A>G), E353D (1159A>T), L354E (1160T>G 1161T>A), P355R (1163C>A 1164C>G 1165A>G), K356M (1167A>T), A358F (1172G>T 1173C>T), E359D (1177A>C), L360I (1178T>A), E361Q (1181G>C 1183A>G), T362K (1185C>A 1186G>A), T366V (1196A>G 1197C>T), K367V (1199A>G 1200A>T), D369E (1207T>G), M371L (1211A>C 1213G>C), E373A (1218A>C 1219G>C), I375V (1223A>G), V376A (1227T>C), I379L (1235A>C 1237A>G)

| | Begin | End | Coverage | Score | Concordance | Matches | Identities | I/D/M/F* | Stop Codons |
|---|---|---|---|---|---|---|---|---|---|
| **NT** | **118** | **1238** | **83.7%** | **392** | **17.6%** | **1118 (99.5%)** | **663 (59.0%)** | **3/3** | |
| Codon mutations: | GAA7AGT (119G>A 120A>G 121A>T), GAC9GCT (126A>C 127C>T), CAT10GAG (128C>G 130T>G), GGC12GGG (136C>G), CTC13ATT (137C>A 139C>T), AAG14AAA (142G>A), AAA15GAT (143A>G 145A>T), AAT16GAC (146A>G 148T>C), ATC17CTT (149A>C 151C>T), CTA18CTT (154A>T), AAG19AGA (156A>G 157G>A), GGT20GGG (160T>G), ATA21ATT (163A>T), TAT22TAC (166T>C), TCT23CAG (167T>C 168C>A 169T>G), TGT24TAC (171G>A 172T>C), GGA25GGG (175A>G), TTT26TTC (178T>C), CCT29CCA (187T>A), TCA30TCT (190A>T), ACT31GCC (191A>G 193T>C), AAA35CGA (203A>C 204A>G), GCC36GCT (208C>T), ATT37GTA (209A>G 211T>A), TTT38TTG (214T>G), CCA39CCT (217A>T), TGT40ATT (218T>A 219G>T), ATT41ATA (223T>A), TCT42TCG (226T>G), AAA44CGT (230A>C 231A>G 232A>T), GTG46GTC (238G>C), ATA47ATT (241A>T), GTC48GCT (243T>C 244C>T), CAA49CAG (247A>G), TCG52TCC (256G>C), GGA53GGT (259A>T), ACA54ACC (262A>C), ACA57ACC (271A>C), GCA58TCC (272G>T 274A>C), ACA59ATG (276C>T 277A>G), TAT60ATT (278T>A 279A>T), GCA61GCT (283A>T), ATC62CTT (284A>C 286C>T), AGT63ACC (288G>C 289T>C), TTA65TGC (294T>G 295A>C), CAG66CAA (298G>A), CAA67GTT (299C>G 300A>T 301A>T), ATA68GTG (302A>G 304A>G), ACT70ACC (310T>C), TCT71AAA (311T>A 312C>A 313T>A), AAT74GAG (320A>G 322T>G), ATC75GTA (323A>G 325C>A), GCT77GCA (331T>A), ATC79ATA (337C>A), ACC81TCT (341A>T 343C>T), CCA82CCT (346A>T), CGT84CGG (352T>G), GAA85GAG (355A>G), TTG86CTT (356T>C 358G>T), GCT87GCA (361T>A), CTC88GCT (362C>G 363T>C 364C>T), GCT90ACG (368G>A 370T>G), CAA91GAA (371C>G), CGT92AAG (374C>A 375G>A 376T>G), GTA93GTG (379A>G), TTG94ATT (380T>A 382G>T), CAA95TTG (383C>T 384A>T 385A>G), ACA96GCA (386A>G), GGG98GGA (394G>A), AAT99GAT (395A>G), TAT100TAC (400T>C), CTA101ATT (401C>A 403A>T), TAT102del (404_406delTAT), AAT103AAC (409T>C), TTT104ATA (410T>A 412T>A), AAA105CAG (413A>C 415A>G), TGT106GCA (416T>G 417G>C 418T>A), CAA107CAT (421A>T), GTT108GCG (423T>C 424T>G), ACA113AAA (438C>A), AGT114AGC (442T>C), ATC115GTG (443A>G 445C>G), AAG116GGT (446A>G 447A>G 448G>T), AGC118GAT (452A>G 453G>A 454C>T), CAG119ATC (455C>A 456A>T 457G>C), GAA120AGA (458G>A 459A>G), ACT121AAG (462C>A 463T>G), AAG123GAG (467A>G), AAG123_AAA124insCAT (469_470insCAT), AAA124GGA (470A>G 471A>G), GCC125GTA (474C>T 475C>A), CAG126CAA (478G>A), GTT127GTA (481T>A), TTG128GTG (482T>G), ATT129GTT (485A>G), GGA130GGT (490A>T), CCT132CCA (496T>A), GGA133GGC (499A>C), CGA134CGT (502A>T), ATG135GTC (503A>G 505G>C), ATT136TTT (506A>T), GAT137GAC (511T>C), CTA138ATG (512C>A 514A>G), CTC139CTG (517C>G), ACC140CGC (518A>C 519C>G), CGA141AGG (521C>A 523A>G), AAA142CAG (524A>C 526A>G), AGT143TCT (527A>T 528G>C), ATC144CTT (530A>C 532C>T), GAT145CGC (533G>C 534A>G 535T>C), ACC146CCT (536A>C 538C>T), AAA147GAC (539A>G 541A>C), GCA148CAC (542G>C 543C>A 544A>C), ATT149ATC (547T>C), AAA150AAG (550A>G), ATA151ATG (553A>G), GTT152TTT (554G>T), GTA153GTT (559A>T), ATA154TTG (560A>T 562A>G), GAT158GAC (574T>C), GAG159GAA (577G>A), TTG161CTC (581T>C 583G>C), ATA162TCT (584A>T 585T>C 586A>T), GAT163AGA (587G>A 588A>G 589T>A), AAT164GGT (590A>G 591A>G), TTT165TTC (595T>C), TTG166AAG (596T>A 597T>A), AAA168CAG (602A>C 604A>G), ATA169ATT (607A>T), CAA170TAT (608C>T 610A>T), ATT172ATA (616T>A), GAA174CAG (620G>C 622A>G), TTT175CTT (623T>C), TTT176TTG (628T>G), GAA177CCA (629G>C 630A>C), AGT178CCA (632A>C 633G>C 634T>A), CAT179AAA (635C>A 637T>A), GTC180ATC (638G>A), GTT182GTG (646T>G), ATT183GGT (647A>G 648T>G), TTA184GTT (650T>G 652A>T), TTA185TTC (655A>C), TCA186TCT (658A>T), GCA187GCT (661A>T), ACC188ACT (664C>T), GTT189ATG (665G>A 667T>G), TCA191CCA (671T>C), AGG192GAG (674A>G 675G>A), GTT193GCT (678T>C), ATC194CTT (680A>C 682C>T), AAT195GAA (683A>G 685T>A), ACA196ATT (687C>T 688A>T), TCT197ACT (689T>A), CAG198AGG (692C>A 693A>G), GTC199AAG (695G>A 696T>A 697C>G), TTT200TTC (700T>C), AGA202AAC (705G>A 706A>C), GAT203AAA (707G>A 709T>A), CCC204CCT (712C>T), GTA205GTG (715A>G), AAA206AGG (717A>G 718A>G), CTG208CTT (724G>T), GTA209GTG (727A>G), AAA210AAG (730A>G), AAT211CGT (731A>C 732A>G), GCT212GAT (735C>A), GAA213GAG (739A>G), TTG214CTC (740T>C 742G>C), ACA215ACT (745A>T), CTC216CTT (748C>T), GGC218GGT (754C>T), ATT219ATC (757T>C), AGG220AAG (759G>A), TAT222TTT (765A>T), ATA224GTC (770A>G 772A>C), GTA226GTC (778A>C), AAG227GAT (779A>G 781G>T), AAA228AAG (784A>G), AAT229GAA (785A>G 787T>A), GAT230GAA (790T>A), TTT231TGG (792T>G 793T>G), AAA232AAG (796A>G), GCA233CTT (797G>C 798C>T 799A>T), TTT237TGT (810T>G), GAC241GAG (823C>G), CAC242ACT (824C>A 825A>C 826C>T), TTA243TTG (829A>G), AGC244GCC (830A>G 831G>C), CTC245ATC (833C>A), ACT246ACC (838T>C), CAA247CAG (841A>G), ACA248AGT (843C>G 844A>T), CTA249GTC (845C>G 847A>C), ATA250ATC (850A>C), TGT252GTG (854T>G 855G>T 856T>G), AAT253AAC (859T>C), ACA254ACC (862A>C), CGT255AGG (863C>A 865T>G), AGA256CGT (866A>C 868A>T), CAA257AAA (869C>A), GTA260TGG (878G>T 879T>G 880A>G), ATG262ACT (885T>C 886G>T), GAA263GAT (889A>T), TGT264AAG (890T>A 891G>A 892T>G), TTA265ATG (893T>A 895A>G), ACA266CGC (896A>C 897C>G 898A>C), AAT267AGC (900A>G 901T>C), AGA268CGT (902A>C 904A>T), AAT269GAT (905A>G), TTC270CAC (908T>C 909T>A), GCA272GTA (915C>T), TCA273TCT (919A>T), AGC274GCA (920A>G 921G>C 922C>A), ATC275ACT (924T>C 925C>T), GAT278GAC (934T>C), TCC280GAC (938T>G 939C>A), CAA281CAG (943A>G), CAA282AAC (944C>A 946A>C), GAA283ACT (947G>A 948A>C 949A>T), CGT284AGA (950C>A 952T>A), TTT286ATC (956T>A 958T>C), AAA289CGA (965A>C 966A>G), GAA290GAG (970A>G), TTT291TTC (973T>C), AGA292CGA (974A>C), GAT293TCT (977G>T 978A>C), GGT294GGG (982T>G), AAA295TCA (983A>T 984A>C), ACT296TCT (986A>T), AGA297CGT (989A>C 991A>T), ATA298GTG (992A>G 994A>G), CTT299CTC (997T>C), CTT300ATC (998C>A 1000T>C), TCA301ACA (1001T>A), AAT303GAT (1007A>G), CTA304CTC (1012A>C), TTA305CTT (1013T>C 1015A>T), GCA306GCT (1018A>T), AGA307CGT (1019A>C 1021A>T), GGC308GGT (1024C>T), GTT311GTA (1033T>A), CAA312CAG (1036A>G), ATA314GTC (1040A>G 1042A>C), TCT315TCC (1045T>C), TTA316CTT (1046T>C 1048A>T), GTT317GTC (1051T>C), ATC318ATT (1054C>T), AAT319AAC (1057T>C), GAT321GAC (1063T>C), TTG322CTG (1064T>C), CAT324ACT (1070C>A 1071A>C), AAT325CAA (1073A>C 1075T>A), AGA326CCA (1076A>C 1077G>C), TAT329TAC (1087T>C), ATA330CTG (1088A>C 1090A>G), AGA335CGT (1103A>C 1105A>T), GGT337GGA (1111T>A), AGA338CGA (1112A>C), GGC340GGA (1120C>A), CGA341AGG (1121C>A 1123A>G), AAA342AAG (1126A>G), GGG343GGT (1129G>T), ATT344GTT (1130A>G), AAT347AAC (1141T>C), ATA349GTC (1145A>G 1147A>C), ACT350ACC (1150T>C), GAA351AAG (1151G>A 1153A>G), GAC352GAT (1156C>T), GAA353GAT (1159A>T), TTA354GAA (1160T>G 1161T>A), CCA355AGG (1163C>A 1164C>G 1165A>G), AAG356ATG (1167A>T), GCT358TTT (1172G>T 1173C>T), GAA359GAC (1177A>C), TTA360ATA (1178T>A), GAA361CAG (1181G>C 1183A>G), ACG362AAA (1185C>A 1186G>A), TAT364TAC (1192T>C), AAT365AAC (1195T>C), ACC366GTC (1196A>G 1197C>T), AAA367GTA (1199A>G 1200A>T), ATA368ATC (1204A>C), GAT369GAG (1207T>G), GAA370GAG (1210A>G), ATG371CTC (1211A>C 1213G>C), GAG373GCC (1218A>C 1219G>C), ATT375GTT (1223A>G), GTT376GCT (1227T>C), TTG378CTC (1232T>C 1234G>C), ATA379CTG (1235A>C 1237A>G) | | | | | | | | |

#### Proteins

| | | | | | | | | | |
|---|---|---|---|---|---|---|---|---|---|
| hypothetical protein (YP_009666148.1) | 7 | 380 | 98.4% | 1456 | 58.9% | 373 (99.5%) | 207 (55.2%) | 1/1/0/0 | 0 |
| Protein mutations: | E7S (119G>A 120A>G 121A>T), D9A (126A>C 127C>T), H10E (128C>G 130T>G), L13I (137C>A 139C>T), K15D (143A>G 145A>T), N16D (146A>G 148T>C), I17L (149A>C 151C>T), K19R (156A>G 157G>A), S23Q (167T>C 168C>A 169T>G), C24Y (171G>A 172T>C), T31A (191A>G 193T>C), K35R (203A>C 204A>G), I37V (209A>G 211T>A), F38L (214T>G), C40I (218T>A 219G>T), K44R (230A>C 231A>G 232A>T), V48A (243T>C 244C>T), A58S (272G>T 274A>C), T59M (276C>T 277A>G), Y60I (278T>A 279A>T), I62L (284A>C 286C>T), S63T (288G>C 289T>C), L65C (294T>G 295A>C), Q67V (299C>G 300A>T 301A>T), I68V (302A>G 304A>G), S71K (311T>A 312C>A 313T>A), N74E (320A>G 322T>G), I75V (323A>G 325C>A), T81S (341A>T 343C>T), L88A (362C>G 363T>C 364C>T), A90T (368G>A 370T>G), Q91E (371C>G), R92K (374C>A 375G>A 376T>G), L94I (380T>A 382G>T), Q95L (383C>T 384A>T 385A>G), T96A (386A>G), N99D (395A>G), L101I (401C>A 403A>T), Y102del (404_406delTAT), F104I (410T>A 412T>A), K105Q (413A>C 415A>G), C106A (416T>G 417G>C 418T>A), Q107H (421A>T), V108A (423T>C 424T>G), T113K (438C>A), I115V (443A>G 445C>G), K116G (446A>G 447A>G 448G>T), S118D (452A>G 453G>A 454C>T), Q119I (455C>A 456A>T 457G>C), E120R (458G>A 459A>G), T121K (462C>A 463T>G), K123E (467A>G), K123_K124insH (469_470insCAT), K124G (470A>G 471A>G), A125V (474C>T 475C>A), L128V (482T>G), I129V (485A>G), M135V (503A>G 505G>C), I136F (506A>T), L138M (512C>A 514A>G), T140R (518A>C 519C>G), K142Q (524A>C 526A>G), I144L (530A>C 532C>T), D145R (533G>C 534A>G 535T>C), T146P (536A>C 538C>T), K147D (539A>G 541A>C), A148H (542G>C 543C>A 544A>C), I151M (553A>G), V152F (554G>T), I154L (560A>T 562A>G), I162S (584A>T 585T>C 586A>T), D163R (587G>A 588A>G 589T>A), N164G (590A>G 591A>G), L166K (596T>A 597T>A), K168Q (602A>C 604A>G), Q170Y (608C>T 610A>T), E174Q (620G>C 622A>G), F175L (623T>C), F176L (628T>G), E177P (629G>C 630A>C), S178P (632A>C 633G>C 634T>A), H179K (635C>A 637T>A), V180I (638G>A), I183G (647A>G 648T>G), L184V (650T>G 652A>T), L185F (655A>C), V189M (665G>A 667T>G), S191P (671T>C), R192E (674A>G 675G>A), V193A (678T>C), I194L (680A>C 682C>T), N195E (683A>G 685T>A), T196I (687C>T 688A>T), S197T (689T>A), Q198R (692C>A 693A>G), V199K (695G>A 696T>A 697C>G), R202N (705G>A 706A>C), D203K (707G>A 709T>A), K206R (717A>G 718A>G), N211R (731A>C 732A>G), A212D (735C>A), R220K (759G>A), Y222F (765A>T), I224V (770A>G 772A>C), K227D (779A>G 781G>T), N229E (785A>G 787T>A), D230E (790T>A), F231W (792T>G 793T>G), A233L (797G>C 798C>T 799A>T), F237C (810T>G), D241E (823C>G), H242T (824C>A 825A>C 826C>T), S244A (830A>G 831G>C), L245I (833C>A), T248S (843C>G 844A>T), L249V (845C>G 847A>C), C252V (854T>G 855G>T 856T>G), Q257K (869C>A), V260W (878G>T 879T>G 880A>G), M262T (885T>C 886G>T), E263D (889A>T), C264K (890T>A 891G>A 892T>G), L265M (893T>A 895A>G), T266R (896A>C 897C>G 898A>C), N267S (900A>G 901T>C), N269D (905A>G), F270H (908T>C 909T>A), A272V (915C>T), S274A (920A>G 921G>C 922C>A), I275T (924T>C 925C>T), S280D (938T>G 939C>A), Q282N (944C>A 946A>C), E283T (947G>A 948A>C 949A>T), F286I (956T>A 958T>C), K289R (965A>C 966A>G), D293S (977G>T 978A>C), K295S (983A>T 984A>C), T296S (986A>T), I298V (992A>G 994A>G), L300I (998C>A 1000T>C), S301T (1001T>A), N303D (1007A>G), I314V (1040A>G 1042A>C), H324T (1070C>A 1071A>C), N325Q (1073A>C 1075T>A), R326P (1076A>C 1077G>C), I330L (1088A>C 1090A>G), I344V (1130A>G), I349V (1145A>G 1147A>C), E351K (1151G>A 1153A>G), E353D (1159A>T), L354E (1160T>G 1161T>A), P355R (1163C>A 1164C>G 1165A>G), K356M (1167A>T), A358F (1172G>T 1173C>T), E359D (1177A>C), L360I (1178T>A), E361Q (1181G>C 1183A>G), T362K (1185C>A 1186G>A), T366V (1196A>G 1197C>T), K367V (1199A>G 1200A>T), D369E (1207T>G), M371L (1211A>C 1213G>C), E373A (1218A>C 1219G>C), I375V (1223A>G), V376A (1227T>C), I379L (1235A>C 1237A>G) | | | | | | | | |

| | Begin | End | Coverage | Score | Concordance | Matches | Identities | I/D/M/F* | Stop Codons |
|---|---|---|---|---|---|---|---|---|---|
| **NT** | **118** | **1238** | **83.7%** | **392** | **17.6%** | **1118 (99.5%)** | **663 (59.0%)** | **3/3** | |

Codon mutations: GAA7AGT (119G>A 120A>G 121A>T), GAC9GCT (126A>C 127C>T), CAT10GAG (128C>G 130T>G), GGC12GGG (136C>G), CTC13ATT (137C>A 139C>T), AAG14AAA (142G>A), AAA15GAT (143A>G 145A>T), AAT16GAC (146A>G 148T>C), ATC17CTT (149A>C 151C>T), CTA18CTT (154A>T), AAG19AGA (156A>G 157G>A), GGT20GGG (160T>G), ATA21ATT (163A>T), TAT22TAC (166T>C), TCT23CAG (167T>C 168C>A 169T>G), TGT24TAC (171G>A 172T>C), GGA25GGG (175A>G), TTT26TTC (178T>C), CCT29CCA (187T>A), TCA30TCT (190A>T), ACT31GCC (191A>G 193T>C), AAA35CGA (203A>C 204A>G), GCC36GCT (208C>T), ATT37GTA (209A>G 211T>A), TTT38TTG (214T>G), CCA39CCT (217A>T), TGT40ATT (218T>A 219G>T), ATT41ATA (223T>A), TCT42TCG (226T>G), AAA44CGT (230A>C 231A>G 232A>T), GTG46GTC (238G>C), ATA47ATT (241A>T), GTC48GCT (243T>C 244C>T), CAA49CAG (247A>G), TCG52TCC (256G>C), GGA53GGT (259A>T), ACA54ACC (262A>C), ACA57ACC (271A>C), GCA58TCC (272G>T 274A>C), ACA59ATG (276C>T 277A>G), TAT60ATT (278T>A 279A>T), GCA61GCT (283A>T), ATC62CTT (284A>C 286C>T), AGT63ACC (288G>C 289T>C), TTA65TGC (294T>G 295A>C), CAG66CAA (298G>A), CAA67GTT (299C>G 300A>T 301A>T), ATA68GTG (302A>G 304A>G), ACT70ACC (310T>C), TCT71AAA (311T>A 312C>A 313T>A), AAT74GAG (320A>G 322T>G), ATC75GTA (323A>G 325C>A), GCT77GCA (331T>A), ATC79ATA (337C>A), ACC81TCT (341A>T 343C>T), CCA82CCT (346A>T), CGT84CGG (352T>G), GAA85GAG (355A>G), TTG86CTT (356T>C 358G>T), GCT87GCA (361T>A), CTC88GCT (362C>G 363T>C 364C>T), GCT90ACG (368G>A 370T>G), CAA91GAA (371C>G), CGT92AAG (374C>A 375G>A 376T>G), GTA93GTG (379A>G), TTG94ATT (380T>A 382G>T), CAA95TTG (383C>T 384A>T 385A>G), ACA96GCA (386A>G), GGG98GGA (394G>A), AAT99GAT (395A>G), TAT100TAC (400T>C), CTA101ATT (401C>A 403A>T), TAT102del (404_406delTAT), AAT103AAC (409T>C), TTT104ATA (410T>A 412T>A), AAA105CAG (413A>C 415A>G), TGT106GCA (416T>G 417G>C 418T>A), CAA107CAT (421A>T), GTT108GCG (423T>C 424T>G), ACA113AAA (438C>A), AGT114AGC (442T>C), ATC115GTG (443A>G 445C>G), AAG116GGT (446A>G 447A>G 448G>T), AGC118GAT (452A>G 453G>A 454C>T), CAG119ATC (455C>A 456A>T 457G>C), GAA120AGA (458G>A 459A>G), ACT121AAG (462C>A 463T>G), AAG123GAG (467A>G), AAG123_AAA124insCAT (469_470insCAT), AAA124GGA (470A>G 471A>G), GCC125GTA (474C>T 475C>A), CAG126CAA (478G>A), GTT127GTA (481T>A), TTG128GTG (482T>G), ATT129GTT (485A>G), GGA130GGT (490A>T), CCT132CCA (496T>A), GGA133GGC (499A>C), CGA134CGT (502A>T), ATG135GTC (503A>G 505G>C), ATT136TTT (506A>T), GAT137GAC (511T>C), CTA138ATG (512C>A 514A>G), CTC139CTG (517C>G), ACC140CGC (518A>C 519C>G), CGA141AGG (521C>A 523A>G), AAA142CAG (524A>C 526A>G), AGT143TCT (527A>T 528G>C), ATC144CTT (530A>C 532C>T), GAT145CGC (533G>C 534A>G 535T>C), ACC146CCT (536A>C 538C>T), AAA147GAC (539A>G 541A>C), GCA148CAC (542G>C 543C>A 544A>C), ATT149ATC (547T>C), AAA150AAG (550A>G), ATA151ATG (553A>G), GTT152TTT (554G>T), GTA153GTT (559A>T), ATA154TTG (560A>T 562A>G), GAT158GAC (574T>C), GAG159GAA (577G>A), TTG161CTC (581T>C 583G>C), ATA162TCT (584A>T 585T>C 586A>T), GAT163AGA (587G>A 588A>G 589T>A), AAT164GGT (590A>G 591A>G), TTT165TTC (595T>C), TTG166AAG (596T>A 597T>A), AAA168CAG (602A>C 604A>G), ATA169ATT (607A>T), CAA170TAT (608C>T 610A>T), ATT172ATA (616T>A), GAA174CAG (620G>C 622A>G), TTT175CTT (623T>C), TTT176TTG (628T>G), GAA177CCA (629G>C 630A>C), AGT178CCA (632A>C 633G>C 634T>A), CAT179AAA (635C>A 637T>A), GTC180ATC (638G>A), GTT182GTG (646T>G), ATT183GGT (647A>G 648T>G), TTA184GTT (650T>G 652A>T), TTA185TTC (655A>C), TCA186TCT (658A>T), GCA187GCT (661A>T), ACC188ACT (664C>T), GTT189ATG (665G>A 667T>G), TCA191CCA (671T>C), AGG192GAG (674A>G 675G>A), GTT193GCT (678T>C), ATC194CTT (680A>C 682C>T), AAT195GAA (683A>G 685T>A), ACA196ATT (687C>T 688A>T), TCT197ACT (689T>A), CAG198AGG (692C>A 693A>G), GTC199AAG (695G>A 696T>A 697C>G), TTT200TTC (700T>C), AGA202AAC (705G>A 706A>C), GAT203AAA (707G>A 709T>A), CCC204CCT (712C>T), GTA205GTG (715A>G), AAA206AGG (717A>G 718A>G), CTG208CTT (724G>T), GTA209GTG (727A>G), AAA210AAG (730A>G), AAT211CGT (731A>C 732A>G), GCT212GAT (735C>A), GAA213GAG (739A>G), TTG214CTC (740T>C 742G>C), ACA215ACT (745A>T), CTC216CTT (748C>T), GGC218GGT (754C>T), ATT219ATC (757T>C), AGG220AAG (759G>A), TAT222TTT (765A>T), ATA224GTC (770A>G 772A>C), GTA226GTC (778A>C), AAG227GAT (779A>G 781G>T), AAA228AAG (784A>G), AAT229GAA (785A>G 787T>A), GAT230GAA (790T>A), TTT231TGG (792T>G 793T>G), AAA232AAG (796A>G), GCA233CTT (797G>C 798C>T 799A>T), TTT237TGT (810T>G), GAC241GAG (823C>G), CAC242ACT (824C>A 825A>C 826C>T), TTA243TTG (829A>G), AGC244GCC (830A>G 831G>C), CTC245ATC (833C>A), ACT246ACC (838T>C), CAA247CAG (841A>G), ACA248AGT (843C>G 844A>T), CTA249GTC (845C>G 847A>C), ATA250ATC (850A>C), TGT252GTG (854T>G 855G>T 856T>G), AAT253AAC (859T>C), ACA254ACC (862A>C), CGT255AGG (863C>A 865T>G), AGA256CGT (866A>C 868A>T), CAA257AAA (869C>A), GTA260TGG (878G>T 879T>G 880A>G), ATG262ACT (885T>C 886G>T), GAA263GAT (889A>T), TGT264AAG (890T>A 891G>A 892T>G), TTA265ATG (893T>A 895A>G), ACA266CGC (896A>C 897C>G 898A>C), AAT267AGC (900A>G 901T>C), AGA268CGT (902A>C 904A>T), AAT269GAT (905A>G), TTC270CAC (908T>C 909T>A), GCA272GTA (915C>T), TCA273TCT (919A>T), AGC274GCA (920A>G 921G>C 922C>A), ATC275ACT (924T>C 925C>T), GAT278GAC (934T>C), TCC280GAC (938T>G 939C>A), CAA281CAG (943A>G), CAA282AAC (944C>A 946A>C), GAA283ACT (947G>A 948A>C 949A>T), CGT284AGA (950C>A 952T>A), TTT286ATC (956T>A 958T>C), AAA289CGA (965A>C 966A>G), GAA290GAG (970A>G), TTT291TTC (973T>C), AGA292CGA (974A>C), GAT293TCT (977G>T 978A>C), GGT294GGG (982T>G), AAA295TCA (983A>T 984A>C), ACT296TCT (986A>T), AGA297CGT (989A>C 991A>T), ATA298GTG (992A>G 994A>G), CTT299CTC (997T>C), CTT300ATC (998C>A 1000T>C), TCA301ACA (1001T>A), AAT303GAT (1007A>G), CTA304CTC (1012A>C), TTA305CTT (1013T>C 1015A>T), GCA306GCT (1018A>T), AGA307CGT (1019A>C 1021A>T), GGC308GGT (1024C>T), GTT311GTA (1033T>A), CAA312CAG (1036A>G), ATA314GTC (1040A>G 1042A>C), TCT315TCC (1045T>C), TTA316CTT (1046T>C 1048A>T), GTT317GTC (1051T>C), ATC318ATT (1054C>T), AAT319AAC (1057T>C), GAT321GAC (1063T>C), TTG322CTG (1064T>C), CAT324ACT (1070C>A 1071A>C), AAT325CAA (1073A>C 1075T>A), AGA326CCA (1076A>C 1077G>C), TAT329TAC (1087T>C), ATA330CTG (1088A>C 1090A>G), AGA335CGT (1103A>C 1105A>T), GGT337GGA (1111T>A), AGA338CGA (1112A>C), GGC340GGA (1120C>A), CGA341AGG (1121C>A 1123A>G), AAA342AAG (1126A>G), GGG343GGT (1129G>T), ATT344GTT (1130A>G), AAT347AAC (1141T>C), ATA349GTC (1145A>G 1147A>C), ACT350ACC (1150T>C), GAA351AAG (1151G>A 1153A>G), GAC352GAT (1156C>T), GAA353GAT (1159A>T), TTA354GAA (1160T>G 1161T>A), CCA355AGG (1163C>A 1164C>G 1165A>G), AAG356ATG (1167A>T), GCT358TTT (1172G>T 1173C>T), GAA359GAC (1177A>C), TTA360ATA (1178T>A), GAA361CAG (1181G>C 1183A>G), ACG362AAA (1185C>A 1186G>A), TAT364TAC (1192T>C), AAT365AAC (1195T>C), ACC366GTC (1196A>G 1197C>T), AAA367GTA (1199A>G 1200A>T), ATA368ATC (1204A>C), GAT369GAG (1207T>G), GAA370GAG (1210A>G), ATG371CTC (1211A>C 1213G>C), GAG373GCC (1218A>C 1219G>C), ATT375GTT (1223A>G), GTT376GCT (1227T>C), TTG378CTC (1232T>C 1234G>C), ATA379CTG (1235A>C 1237A>G)

*: Inserts / Deletes / Misaligned / Frameshifts

#### Analysis details

This analysis was performed with panviral2.64

### NGS Details (UN59): Potato virus Y

#### Assembly

| | |
|---|---|
| **Coverage Length** | 8670 (6 contig(s)) |
| **Depth Of Coverage** | 11.7 |
| **Number Of Reads** | 757 |
| **Reads Per Million** | 16.46 rpm (after QC) |
| **Ambiguities** | 0 |
| **Assembly Method** | de novo + reference guided assembly |
| **Consensus Caller** | Bcf Tools |

#### Coverage Map

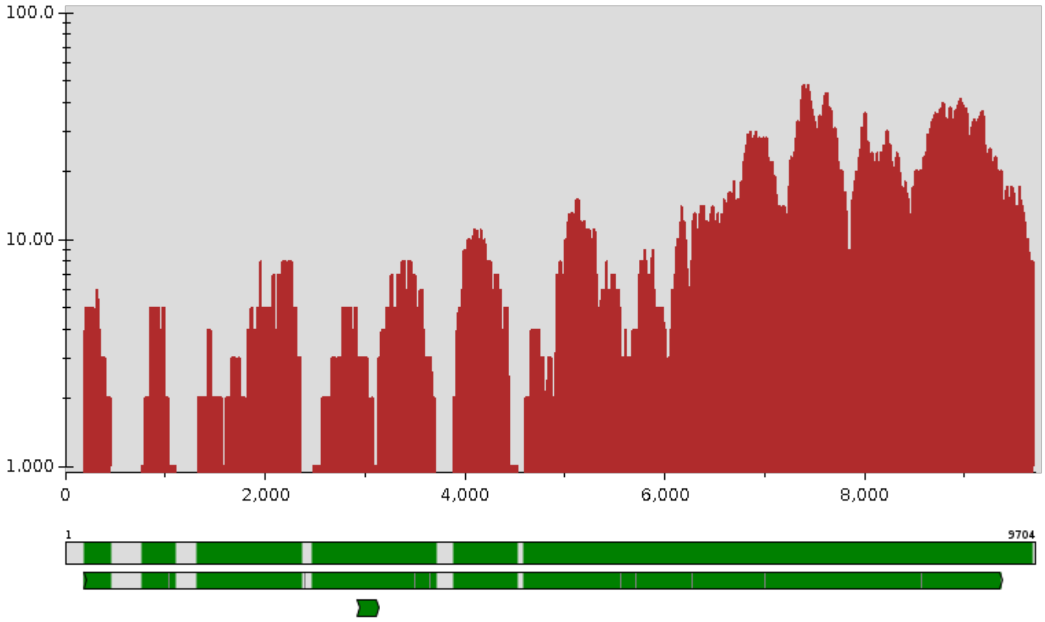

#### Assignment

| | |
|---|---|
| **Type** | Potato virus Y (Taxonomy ID: 12216) |
| **Reference Genome** | NC_001616.1 |
| **NT Identity (%)** | 83.3353 |
| **AA Identity (%)** | 90.1748 |
| **Number Of Stop Codons** | 4 |
| **Number Of CDS** | 2 |

#### Alignment

| | |
|---|---|
| **Alignment Score** | 11427.0 (NT) + 17126.0 (AA) = 28553.0 |
| **Concordance (%)** | 78.8456 |

| Alignment Method | Global, seeded, nucleotide + amino acids (AGA) |
|---|---|

#### Genome Region

Sequence starts at position 183 and ends at position 9687 relative to NC_001616.1 reference sequence.

#### Alignment Detailed Statistics

| | Begin | End | Coverage | Score | Concordance | Matches | Identities | I/D/M/F* | Stop Codons |
|---|---|---|---|---|---|---|---|---|---|
| **NT** | **183** | **9687** | **89.3%** | **11427** | **66.3%** | **8654 (99.8%)** | **7216 (83.2%)** | **5/16** | |

Mutations:

183C>T, 193T>C, 194T>C, 196C>A, 202A>G, 208C>T, 209T>C, 210G>T, 211T>G, 220G>A, 221T>A, 223T>G, 226A>G, 235A>T, 241C>T, 247A>T, 251T>C, 255G>T, 256C>T, 258A>G, 262T>C, 263A>G, 265T>G, 267T>C, 268G>A, 273A>T, 277A>T, 280A>G, 283G>T, 286G>A, 287G>A, 289T>C, 293G>A, 294T>C, 301T>C, 304C>T, 305G>A, 306C>T, 309A>G, 311C>T, 316A>T, 317A>G, 318C>T, 319A>G, 322A>G, 331A>T, 339T>A, 340C>A, 342A>G, 347A>G, 348A>C, 349A>G, 350T>C, 355T>A, 356A>G, 357C>T, 364T>C, 365G>A, 366T>C, 367G>A, 368C>T, 369T>C, 373A>G, 376C>T, 384T>G, 388G>A, 391C>T, 394A>G, 397C>T, 400G>A, 406T>C, 408C>T, 409C>T, 412G>A, 415A>T, 416A>G, 417T>A, 424T>C, 427G>A, 433A>G, 436G>C, 437G>A, 440A>G, 441G>A, 442G>A, 443A>C, 444A>G, 445G>A, 448T>A, 454A>G, 765T>A, 770T>C, 773C>A, 774G>A, 775A>G, 776C>A, 790C>T, 801T>A, 813G>C, 814A>G, 816T>A, 818T>C, 823A>G, 825G>A, 826T>A, 833C>A, 835G>A, 841C>T, 853T>C, 859C>T, 860C>T, 862C>T, 863A>G, 865T>C, 867T>C, 869A>G, 871C>T, 872A>T, 874A>G, 879G>A, 886T>C, 896A>G, 898C>T, 903A>G, 906C>A, 907A>C, 909A>C, 910A>C, 912G>A, 913C>T, 922C>A, 931T>A, 938T>C, 939C>T, 940A>T, 943A>G, 952C>T, 967A>G, 973A>G, 976G>C, 980T>A, 982G>C, 991A>G, 994T>C, 999G>A, 1003T>C, 1007C>A, 1008A>G, 1009G>A, 1010A>G, 1012T>G, 1013A>G, 1018A>G, 1019A>C, 1021C>G, 1030C>T, 1039G>A, 1042T>C, 1045C>A, 1048C>A, 1051T>C, 1060G>A, 1063T>A, 1064C>T, 1069C>T, 1083G>A, 1092A>G, 1093T>C, 1097T>G, 1099G>A, 1105C>T, 1318T>A, 1321G>A, 1327G>A, 1329T>C, 1333G>T, 1336A>G, 1343C>T, 1345A>G, 1354G>A, 1357G>T, 1360C>T, 1367C>T, 1369C>G, 1373C>A, 1378C>T, 1384G>A, 1385A>G, 1393A>G, 1396C>T, 1405G>T, 1408A>G, 1411G>A, 1415G>T, 1417A>T, 1420G>C, 1429T>C, 1432A>G, 1436G>A, 1438C>T, 1439T>C, 1441A>G, 1444T>C, 1453C>T, 1456G>A, 1468A>G, 1492A>G, 1499T>C, 1501G>A, 1504T>C, 1507G>A, 1510C>T, 1513A>G, 1516A>G, 1519A>T, 1522G>A, 1525C>T, 1528G>A, 1534C>T, 1540T>A, 1546C>T, 1555A>G, 1561T>C, 1564A>C, 1565_1568delTCTT, 1573C>T, 1576A>G, 1583T>C, 1588T>A, 1594G>A, 1600C>T, 1618G>A, 1621C>T, 1624C>T, 1631C>T, 1639A>G, 1651C>T, 1654C>G, 1675T>C, 1693A>G, 1702T>C, 1707A>G, 1718G>A, 1726A>G, 1729C>T, 1732A>C, 1738T>C, 1742A>G, 1744C>T, 1747C>T, 1753T>C, 1758G>A, 1768G>A, 1786C>T, 1789A>G, 1790G>A, 1795C>T, 1805T>C, 1807A>G, 1813G>A, 1819G>A, 1825G>A, 1837C>T, 1840T>C, 1849A>G, 1858C>G, 1864A>G, 1870C>T, 1876T>C, 1879G>T, 1882A>G, 1888T>A, 1891T>C, 1906T>C, 1915A>T, 1930T>C, 1936C>T, 1937A>G, 1939T>C, 1945A>G, 1951C>T, 1960A>T, 1978A>G, 1981T>A, 1984C>T, 1987T>C, 1996C>T, 2004T>A, 2020A>G, 2023G>A, 2026T>C, 2029G>T, 2032G>A, 2059T>C, 2068T>G, 2071C>T, 2072G>A, 2074G>A, 2087C>T, 2089G>A, 2101C>T, 2104G>A, 2107G>A, 2116G>A, 2122C>T, 2137C>T, 2158T>C, 2164C>T, 2170A>G, 2182T>C, 2188G>A, 2191C>A, 2208G>A, 2218T>C, 2224C>T, 2227A>T, 2242G>A, 2245C>T, 2254G>A, 2257T>C, 2260C>T, 2266C>G, 2272A>G, 2275G>A, 2293A>G, 2299C>T, 2302G>A, 2305G>A, 2314A>T, 2317T>C, 2323T>C, 2326A>G, 2338C>T, 2341G>A, 2347A>G, 2350T>C, 2356G>A, 2365T>C, 2476T>C, 2482G>A, 2491G>A, 2497G>T, 2503G>A, 2518A>G, 2521T>C, 2523A>G, 2533A>G, 2539G>A, 2545A>G, 2551G>A, 2554T>A, 2558C>T, 2561T>C, 2563G>C, 2569A>G, 2572A>G, 2584T>C, 2587C>T, 2593G>C, 2599T>C, 2605T>C, 2608T>C, 2620T>C, 2629G>A, 2632G>A, 2634G>A, 2638G>A, 2644T>A, 2653A>G, 2664C>T, 2665T>C, 2678C>T, 2683A>G, 2689T>C, 2698A>C, 2707A>G, 2710G>A, 2716A>G, 2719A>G, 2722C>T, 2723G>A, 2728A>G, 2734G>A, 2737T>A, 2747G>A, 2749T>A, 2752A>T, 2755T>A, 2761C>T, 2764C>T, 2779T>C, 2785G>T, 2791C>T, 2795C>A, 2806C>T, 2816T>C, 2818G>C, 2822G>A, 2824G>A, 2827G>A, 2836T>C, 2839G>A, 2842T>C, 2863C>A, 2864C>T, 2882C>A, 2885A>C, 2886G>A, 2893C>T, 2895C>T, 2902C>T, 2905A>G, 2911T>C, 2923T>C, 2931A>G, 2940A>G, 2947T>C, 2962T>C, 2971_2972insA, 2989T>C, 2991C>T, 2994A>G, 2999G>A, 3000A>G, 3016delC, 3021G>A, 3024A>T, 3025C>T, 3036C>T, 3039A>G, 3040A>C, 3042A>C, 3043G>A, 3052G>A, 3061A>G, 3082G>A, 3091C>T, 3099C>T, 3100C>A, 3104G>A, 3105T>G, 3106G>A, 3109C>T, 3112A>G, 3118T>C, 3121C>T, 3127A>G, 3132G>A, 3137C>A, 3138G>A, 3139A>G, 3151T>C, 3152T>A, 3154C>T, 3165G>A, 3169A>G, 3199C>T, 3203A>C, 3208T>C, 3214A>G, 3217C>T, 3218T>C, 3223C>A, 3230G>A, 3235C>T, 3257T>C, 3262C>T, 3265C>T, 3266G>A, 3268A>G, 3271G>A, 3275G>A, 3284G>A, 3301G>A, 3315G>A, 3316A>G, 3318A>C, 3322C>T, 3325G>A, 3328G>A, 3337A>T, 3349G>A, 3355C>T, 3358C>T, 3359A>G, 3367A>G, 3370T>C, 3377T>C, 3379A>G, 3394A>G, 3397C>T, 3400T>G, 3406A>G, 3418C>T, 3421T>A, 3427T>C, 3433G>A, 3436A>T, 3439A>G, 3442C>T, 3445T>C, 3457G>A, 3460T>C, 3463T>A, 3469G>T, 3472G>A, 3496C>T, 3499G>A, 3505C>T, 3513T>G, 3520A>G, 3529A>G, 3532A>G, 3545A>G, 3548G>A, 3551T>C, 3554G>A, 3556C>T, 3583T>C, 3586T>C, 3589C>T, 3595C>T, 3607T>C, 3613T>C, 3616G>A, 3619T>C, 3623C>G, 3628C>T, 3632C>A, 3637T>C, 3638T>C, 3643A>G, 3661A>G, 3670G>A, 3676G>C, 3679T>A, 3688G>A, 3694T>A, 3697G>A, 3698A>G, 3704G>T, 3706T>A, 3709T>A, 3883C>T, 3886A>G, 3892A>G, 3895C>T, 3898T>C, 3904A>G, 3905C>A, 3907G>A, 3913T>C, 3928G>A, 3931A>T, 3937G>A, 3955T>C, 3958G>A, 3961C>T, 3964A>G, 3967T>C, 3976A>G, 3979T>C, 3985A>T, 3997A>T, 4000G>A, 4006C>T, 4009T>G, 4012C>T, 4015A>G, 4024T>C, 4030A>G, 4033A>T, 4042G>A, 4052C>T, 4054G>A, 4057T>C, 4063T>A, 4066A>G, 4073A>G, 4078T>C, 4081C>T, 4084T>C, 4096C>T, 4108C>T, 4111A>T, 4117G>A, 4118C>T, 4120A>G, 4126C>T, 4128T>A, 4129C>T, 4132C>T, 4135T>C, 4147A>G, 4148T>C, 4151G>T, 4162C>T, 4166G>A, 4171A>C, 4180G>A, 4190C>T, 4198T>C, 4201C>T, 4210G>T, 4213C>T, 4219T>C, 4220C>T, 4223C>A, 4225G>A, 4231T>G, 4237T>C, 4252A>G, 4253T>C, 4255A>G, 4258A>G, 4270T>C, 4282A>G, 4285G>A, 4291A>G, 4303G>A, 4312C>T, 4319A>G, 4321A>G, 4324G>T, 4327G>A, 4336G>T, 4339T>C, 4348A>T, 4360A>G, 4366T>C, 4369T>A, 4375T>C, 4387T>C, 4395T>A, 4406G>A, 4444C>A, 4450C>T, 4456C>T, 4465C>T, 4475A>G, 4477G>T, 4480C>T, 4481A>T, 4495C>T, 4507G>A, 4510C>T, 4513T>G, 4514T>C, 4517C>T, 4528C>T, 4534A>G, 4591G>A, 4597G>A, 4600C>T, 4606C>T, 4609G>A, 4615G>A, 4618C>T, 4625T>C, 4627G>T, 4648C>T, 4651T>C, 4660G>A, 4663C>T, 4669T>A, 4679G>A, 4681G>C, 4684T>C, 4687T>G, 4699G>A, 4702A>G, 4708A>G, 4711G>A, 4712C>T, 4738A>G, 4747A>G, 4748T>C, 4765T>C, 4796G>A, 4798C>T, 4803C>G, 4804T>C, 4807G>A, 4810G>T, 4813T>G, 4816T>C, 4831T>C, 4834C>T, 4840A>T, 4852A>G, 4855C>T, 4867T>C, 4870G>C, 4873T>A, 4903A>T, 4909G>A, 4915T>C, 4918A>G, 4919T>C, 4921G>A, 4924T>C, 4927C>G, 4933T>C, 4936C>A, 4939G>A, 4942T>C, 4945C>T, 4963A>G, 4972T>C, 4990T>C, 5002A>G, 5023T>C, 5024T>C, 5026G>A, 5029C>T, 5032T>C, 5038T>C, 5044A>T, 5053A>T, 5056G>A, 5057A>G, 5060A>G, 5071G>T, 5090C>A, 5095A>G, 5098G>T, 5100C>T, 5104A>G, 5107A>T, 5110T>C, 5113A>G, 5116G>A, 5119A>G, 5120G>T, 5122C>T, 5126A>G, 5137T>C, 5140C>T, 5143A>G, 5146G>A, 5161C>A, 5167A>G, 5173T>G, 5182G>A, 5183G>A, 5188C>T, 5191G>A, 5197A>G, 5200C>T, 5206C>T, 5209A>G, 5230G>A, 5236C>T, 5239C>T, 5245C>T, 5263A>T, 5269C>T, 5272C>T, 5275C>A, 5278C>T, 5281A>C, 5284A>G, 5287T>C, 5288C>T, 5293A>T, 5294T>C, 5296G>A, 5302G>A, 5311T>A, 5314A>G, 5323A>T, 5344A>G, 5347T>C, 5353C>T, 5365C>T, 5386T>C, 5395C>A, 5422A>G, 5425T>C, 5431C>T, 5437G>A, 5452T>C, 5458G>A, 5467T>C, 5470A>G, 5471C>T, 5473A>G, 5482C>T, 5485A>T, 5488T>C, 5489T>C, 5491G>A, 5497T>C, 5506T>C, 5509G>A, 5512G>A, 5516T>C, 5530T>C, 5533G>A, 5536G>T, 5539G>A, 5548T>G, 5554C>T, 5560T>A, 5561G>A, 5563G>A, 5566G>C, 5569A>G, 5584C>G, 5593G>A, 5596G>A, 5597A>G, 5605C>A, 5617G>T, 5619C>T, 5620T>G, 5629G>A, 5630A>C, 5641C>T, 5650A>T, 5662A>T, 5665A>G, 5689A>G, 5693G>A, 5698G>A, 5713A>G, 5716G>C, 5719A>G, 5743C>A, 5755C>A, 5758T>C, 5764T>C, 5770A>G, 5779C>T, 5782T>C, 5791C>T, 5803C>T, 5836G>A, 5839A>G, 5845A>G, 5848A>G, 5857T>C, 5860C>T, 5863A>T, 5869T>A, 5875T>C, 5884C>T, 5893C>T, 5894A>G, 5896C>G, 5899C>T, 5908G>A, 5911T>C, 5923G>A, 5938A>G, 5941C>T, 5944T>C, 5947T>C, 5959G>A, 5960C>G, 5961G>C, 5962G>C, 5974A>G, 5986T>A, 5989C>T, 5990A>G, 5998T>C, 6008A>C, 6010A>T, 6019A>T, 6022G>T, 6025A>T, 6030A>C, 6049C>A, 6058G>A, 6061A>G, 6064C>A, 6070T>C, 6072G>A, 6073T>C, 6074A>C, 6079G>A, 6081C>A, 6085A>T, 6091A>T, 6094C>T, 6111G>C, 6112T>C, 6118A>G, 6124G>A, 6127G>A, 6128A>G, 6130T>C, 6133T>C, 6136A>G, 6148C>T, 6157A>G, 6158G>A, 6163T>C, 6169A>G, 6170A>T, 6172A>G, 6178C>T, 6187A>G, 6190T>C, 6196G>A, 6203C>T, 6205C>T, 6208A>G, 6209C>T, 6211A>G, 6217G>A, 6229T>A, 6230G>A, 6238C>T, 6247G>A, 6253A>T, 6257G>A, 6258C>A, 6274T>C, 6282A>G, 6286G>A, 6298C>T, 6301G>A, 6304A>G, 6307C>T, 6313C>T, 6322C>T, 6328A>T, 6331T>G, 6334T>C, 6337G>A, 6338C>T, 6358T>C, 6361G>A, 6362G>A, 6367A>T, 6370G>A, 6376C>T, 6379T>G, 6385A>C, 6391A>T, 6394A>G, 6397C>T, 6401G>A, 6405C>T, 6412C>T, 6418A>G, 6421T>C, 6429A>T, 6436T>A, 6445G>A, 6450A>G, 6454C>A, 6460C>T, 6463T>A, 6469C>T, 6472G>A, 6475G>A, 6481T>C, 6485C>A, 6487C>T, 6490T>C, 6496C>T, 6502G>A, 6514T>C, 6517G>A, 6527C>A, 6532C>A, 6541G>A, 6544A>G, 6556C>T, 6559T>C, 6565G>A, 6569T>C, 6571G>A, 6586T>A, 6588C>T, 6598A>G, 6601A>G, 6604T>C, 6607T>C, 6616A>G, 6622C>T, 6631G>A, 6634G>A, 6635T>C, 6640T>A, 6646G>C, 6647A>G, 6652C>T, 6664C>T, 6670T>G, 6677A>G, 6679A>G, 6685C>T, 6691A>T, 6694C>T, 6715A>G, 6716G>A, 6719A>G, 6724A>G, 6734C>T, 6742G>A, 6745G>A, 6754C>T, 6760A>G, 6764A>T, 6766A>G, 6769C>T, 6775T>A, 6782C>T, 6787A>T, 6790C>T, 6795C>T, 6796A>G, 6799C>A, 6802C>A, 6808C>T, 6811C>T, 6814C>T, 6823C>T, 6829A>G, 6843A>G, 6847C>T, 6853A>C, 6857A>G, 6862G>A, 6871A>G, 6875G>A, 6876T>C, 6877C>A, 6880G>A, 6883T>A, 6889T>G, 6901C>T, 6916C>T, 6919G>A, 6925A>G, 6928T>C, 6934C>A, 6949G>C, 6955C>T, 6958A>G, 6976A>G, 6980C>T, 6985C>T, 6994T>C, 6997A>T, 7000G>T, 7001G>A, 7003G>A, 7012T>C, 7015G>A, 7018T>C, 7032T>A, 7039C>T, 7040T>C, 7042G>A, 7048A>G, 7060T>A, 7063C>A, 7066A>G, 7078C>T, 7081A>G, 7082T>C, 7084A>G, 7087A>G, 7090C>A, 7096T>C, 7099A>G, 7102T>C, 7111G>A, 7117A>G, 7125C>A, 7129A>G, 7133C>T, 7135G>A, 7141G>A, 7144T>C, 7145G>T, 7147A>G, 7150G>A, 7156G>A, 7159A>T, 7160_7165delGAGGCA, 7171C>T, 7189G>T, 7198A>G, 7201C>T, 7205C>T, 7210T>C, 7216T>A, 7219G>A, 7225C>A, 7228G>A, 7240G>A, 7243T>C, 7249A>G, 7252T>G, 7258T>A, 7261T>C, 7264T>A, 7267C>G, 7270G>A, 7273_7274insT, 7274C>G, 7275G>T, 7281A>T, 7284T>C, 7296C>T, 7311C>T, 7314C>T, 7317G>A, 7321T>A, 7322delG, 7324G>A, 7327C>T, 7330C>T, 7333C>T, 7334A>C, 7342T>C, 7343G>A, 7344C>A, 7345A>C, 7354T>C, 7363A>G, 7369T>C, 7372C>T, 7378G>T, 7381C>T, 7396A>T, 7399C>T, 7405C>T, 7407G>T, 7408T>G, 7414G>T, 7415T>G, 7420A>G, 7429C>T, 7432T>C, 7441T>C, 7459G>A, 7462A>G, 7468A>T, 7471C>A, 7477A>G, 7480C>T, 7486G>T, 7495T>C, 7498A>G, 7501T>C, 7503T>C, 7504G>A, 7513T>A, 7519C>T, 7522A>G, 7531G>A, 7537G>A, 7540C>T, 7546T>C, 7570G>A, 7573G>A, 7579G>A, 7588T>C, 7595C>T, 7604T>C, 7606G>A, 7615T>A, 7624T>C, 7633C>T, 7636C>T, 7639T>C, 7648T>C, 7666A>G, 7672T>C, 7678A>T, 7684G>A, 7690T>C, 7696T>C, 7702T>G, 7709G>A, 7713A>G, 7714A>G, 7715C>T, 7726T>C, 7735G>A, 7744A>G, 7747C>T, 7762C>T, 7768G>A, 7771T>C, 7780A>G, 7783A>C, 7786T>C, 7793C>T, 7795A>G, 7798C>T, 7804T>C, 7812C>T, 7834A>G, 7837C>T, 7844G>T, 7846G>A, 7855G>A, 7861G>A, 7870A>G, 7873C>T, 7876T>A, 7879G>A, 7882T>A, 7883G>A, 7894A>T, 7897T>C, 7921C>T, 7927G>A, 7936T>C, 7939T>C, 7942T>C, 7966G>A, 7969T>C, 7975T>C, 7987C>T, 7990T>C, 8005T>C, 8032A>G, 8038C>T, 8041C>T, 8043G>A, 8080G>A, 8089G>A, 8095G>A, 8098G>A, 8107C>T, 8110T>C, 8113C>T, 8116T>C, 8128A>G, 8146T>C, 8151A>G, 8152C>T, 8164G>A, 8178G>A, 8182G>A, 8191G>A, 8197C>T, 8203C>T, 8208G>A, 8213C>T, 8218A>G, 8221C>T, 8224G>A, 8239A>G, 8260T>C, 8263A>G, 8281T>C, 8284A>G, 8294C>G, 8296A>G, 8299G>A, 8302C>T, 8314G>A, 8317T>C, 8320C>T, 8326T>A, 8335G>A, 8338C>T, 8349C>T, 8353A>G, 8362C>T, 8365A>G, 8368C>T, 8380C>T, 8383A>G, 8389A>G, 8404T>C, 8408G>T, 8413A>G, 8419G>T, 8434T>C, 8440T>C, 8452G>A, 8458A>G, 8460G>A, 8464A>G, 8467G>A, 8480G>A, 8482T>A, 8497G>A, 8498C>T, 8500A>G, 8502G>A, 8503A>G, 8506C>T, 8515A>T, 8524C>T, 8527A>C, 8528T>C, 8530A>G, 8539G>C, 8546C>T, 8547T>G, 8552T>A, 8563A>G, 8574C>G, 8585A>G, 8598G>A, 8602C>T, 8603A>T, 8604A>C, 8605C>T, 8620A>G, 8622C>A, 8625A>G, 8626G>A, 8629G>A, 8632C>T, 8638C>T, 8641G>A, 8647C>T, 8648C>T, 8649C>T, 8650G>C, 8653C>T, 8656A>G, 8658G>A, 8664A>C, 8665T>G, 8674T>G, 8679C>T, 8680A>T, 8683C>A, 8692G>A, 8695A>T, 8707G>A, 8708A>C, 8716G>A, 8722C>T, 8725G>A, 8731A>G, 8737A>G, 8743C>T, 8745C>A, 8746A>G, 8749C>T, 8752G>A, 8755A>T, 8770C>T, 8771T>C, 8776A>G, 8779T>C, 8782G>A, 8791T>C, 8797A>G, 8800A>G, 8803A>G, 8809T>C, 8812T>C, 8818T>C, 8824G>A, 8827A>G, 8848G>A, 8854T>C, 8856A>G, 8857G>A, 8863G>A, 8865G>A, 8867A>C, 8869G>T, 8872A>G, 8883G>A, 8888A>T, 8929C>T, 8954G>A, 8959C>T, 8962A>T, 8969G>A, 8983G>A, 8984A>G, 8986T>C, 8988A>T, 8998G>A, 9004G>A, 9007G>A, 9016C>T, 9037C>A, 9085A>T, 9097C>T, 9131A>G, 9136A>T, 9145G>T, 9149A>G, 9150T>G, 9151G>A, 9152G>A, 9157A>G, 9160G>T, 9163T>C, 9184G>A, 9196A>G, 9214G>A, 9217A>G, 9220G>A, 9244A>T, 9247G>A, 9256C>T, 9260C>T, 9262T>C, 9274G>A, 9280C>T, 9289C>T, 9304G>A, 9334C>T, 9364C>G, 9377_9378insT, 9387C>T, 9404G>A, 9411C>T, 9412A>G, 9414A>T, 9432C>T, 9437C>T, 9454A>G, 9458T>C, 9467G>A, 9470_9471insT, 9482G>A, 9483C>T, 9498T>C, 9516T>G, 9517delA, 9533G>A, 9534G>T, 9548A>T, 9551C>T, 9553_9554delTA, 9556A>T, 9557T>A, 9558A>T, 9574T>A, 9580_9581insT, 9582T>C, 9583G>A, 9590C>T, 9591T>C, 9607C>T, 9610T>C, 9615delC, 9620A>G, 9648A>G

##### CDS

| | | | | | | | | | |
|---|---|---|---|---|---|---|---|---|---|
| PVYgp1 | 1 | 3064 | 90.9% | 17031 | 88.8% | 2782 (99.8%) | 2525 (90.6%) | 2/3/5/5 | 2 |

| | Begin | End | Coverage | Score | Concordance | Matches | Identities | I/D/M/F* | Stop Codons |
|---|---|---|---|---|---|---|---|---|---|
| **NT** | **183** | **9687** | **89.3%** | **11427** | **66.3%** | **8654 (99.8%)** | **7216 (83.2%)** | **5/16** | |
| Protein mutations: | Y4Q (194T>C 196C>A), C9L (209T>C 210G>T 211T>G), F13M (221T>A 223T>G), S23P (251T>C), C24F (255G>T 256C>T), E25G (258A>G), I27V (263A>G 265T>G), V28A (267T>C 268G>A), E30V (273A>T), A35T (287G>A 289T>C), V37T (293G>A 294T>C), A41I (305G>A 306C>T), D42G (309A>G), E44D (316A>T), T45V (317A>G 318C>T 319A>G), L52Q (339T>A 340C>A), K53R (342A>G), K55A (347A>G 348A>C 349A>G), Y56H (350T>C), T58V (356A>G 357C>T), V61T (365G>A 366T>C 367G>A), L62S (368C>T 369T>C), F67C (384T>G), A75V (408C>T 409C>T), M78E (416A>G 417T>A), E85K (437G>A), R86E (440A>G 441G>A 442G>A), K87R (443A>C 444A>G 445G>A), D88E (448T>A), M194K (765T>A), R197K (773C>A 774G>A 775A>G), M206K (801T>A), G210A (813G>C 814A>G), L211H (816T>A), R214Q (825G>A 826T>A), R226C (860C>T 862C>T), T227A (863A>G 865T>C), I228T (867T>C), N229D (869A>G 871C>T), I230L (872A>T 874A>G), R232K (879G>A), I238V (896A>G 898C>T), N240S (903A>G), T241N (906C>A 907A>C), K242T (909A>C 910A>C), S243N (912G>A 913C>T), S252L (938T>C 939C>T 940A>T), L266I (980T>A 982G>C), R272K (999G>A), Q275R (1007C>A 1008A>G 1009G>A), S276G (1010A>G 1012T>G), I277V (1013A>G), N279Q (1019A>C 1021C>G), D288E (1048C>A), R300Q (1083G>A), Y303C (1092A>G 1093T>C), S305A (1097T>G 1099G>A), I382T (1329T>C), L397I (1373C>A), I401V (1385A>G), E407D (1405G>T), A411S (1415G>T 1417A>T), V418I (1436G>A 1438C>T), S461del (1565_1568delTCTT), E508G (1707A>G), A512T (1718G>A), I520V (1742A>G 1744C>T), S525N (1758G>A), V536I (1790G>A), I585V (1937A>G 1939T>C), F607Y (2004T>A), I628M (2068T>G), V630I (2072G>A 2074G>A), R675K (2208G>A), D694E (2266C>G), K780R (2523A>G), R817K (2634G>A), A827V (2664C>T 2665T>C), V847I (2723G>A), A855T (2747G>A 2749T>A), H871N (2795C>A), V880I (2822G>A 2824G>A), P900T (2882C>A), S901H (2885A>C 2886G>A), T904M (2895C>T), N916S (2931A>G), N919S (2940A>G), E929_N930insX (2971_2972insA), T936I (2991C>T), H937R (2994A>G), E939R (2999G>A 3000A>G), R946Q (3021G>A), Y947F (3024A>T 3025C>T), T951I (3036C>T), E952G (3039A>G 3040A>C), K953T (3042A>C 3043G>A), A972V (3099C>T 3100C>A), V974R (3104G>A 3105T>G 3106G>A), S983N (3132G>A), R985K (3137C>A 3138G>A 3139A>G), F990I (3152T>A 3154C>T), C994Y (3165G>A), F1012L (3218T>C), V1016I (3230G>A), V1028M (3266G>A 3268A>G), V1031M (3275G>A), A1034T (3284G>A), R1044K (3315G>A 3316A>G), E1045A (3318A>C), M1059V (3359A>G), D1072E (3400T>G), V1110G (3513T>G), M1121V (3545A>G), A1122T (3548G>A), V1124I (3554G>A 3556C>T), L1147V (3623C>G), L1150M (3632C>A), Y1152H (3638T>C), K1164N (3676G>C), N1165K (3679T>A), N1170K (3694T>A), I1172V (3698A>G), D1174* (3704G>T 3706T>A), F1175L (3709T>A), I1297V (4073A>G), F1315Y (4128T>A 4129C>T), A1323S (4151G>T), V1328I (4166G>A), L1347I (4223C>A 4225G>A), I1379V (4319A>G 4321A>G), F1404Y (4395T>A), V1408I (4406G>A), M1431V (4475A>G 4477G>T), T1433S (4481A>T), C1444R (4514T>C), V1499I (4679G>A 4681G>C), V1538I (4796G>A 4798C>T), T1540S (4803C>G 4804T>C), S1625G (5057A>G), T1626A (5060A>G), L1636I (5090C>A), A1639V (5100C>T), E1641D (5107A>T), V1646F (5120G>T 5122C>T), I1648V (5126A>G), V1667I (5183G>A), A1793T (5561G>A 5563G>A), I1805V (5597A>G), N1807K (5605C>A), A1812V (5619C>T 5620T>G), I1816L (5630A>C), V1837I (5693G>A), I1904V (5894A>G 5896C>G), R1926A (5960C>G 5961G>C 5962G>C), I1936V (5990A>G), E1945D (6019A>T), K1949T (6030A>C), D1955E (6049C>A), M1958I (6058G>A), S1963N (6072G>A 6073T>C), N1964H (6074A>C), T1966N (6081C>A), C1976S (6111G>C 6112T>C), I1982V (6128A>G 6130T>C), V1992I (6158G>A), T1996S (6170A>T 6172A>G), L2007F (6203C>T 6205C>T), V2016I (6230G>A), A2025K (6257G>A 6258C>A), K2033R (6282A>G), A2060T (6362G>A), V2073I (6401G>A), A2074V (6405C>T), Y2082F (6429A>T), Q2089R (6450A>G), H2101N (6485C>A 6487C>T), L2115I (6527C>A), T2135I (6588C>T), Y2151H (6635T>C), I2155V (6647A>G), I2165V (6677A>G 6679A>G), D2178N (6716G>A), N2179D (6719A>G), I2194L (6764A>T 6766A>G), A2204V (6795C>T 6796A>G), H2205Q (6799C>A), K2220R (6843A>G), N2225D (6857A>G), V2231T (6875G>A 6876T>C 6877C>A), D2250E (6934C>A), V2273I (7001G>A 7003G>A), F2283Y (7032T>A), T2314K (7125C>A), A2321S (7145G>T 7147A>G), E2326_A2327del (7160_7165delGAGGCA), D2344E (7216T>A), D2358E (7258T>A), D2363_R2364insX (7273_7274insT), R2364V (7274C>G 7275G>T), H2366L (7281A>T), L2367S (7284T>C), S2371L (7296C>T), S2376F (7311C>T), T2377I (7314C>T), C2378Y (7317G>A), N2379K (7321T>A), K2384Q (7334A>C), A2387N (7343G>A 7344C>A 7345A>C), S2408M (7407G>T 7408T>G), C2411G (7415T>G), L2440S (7503T>C 7504G>A), D2509N (7709G>A), K2510R (7713A>G 7714A>G), T2543I (7812C>T), V2554L (7844G>T 7846G>A), V2567I (7883G>A), S2620N (8043G>A), N2656S (8151A>G 8152C>T), R2665K (8178G>A), R2675K (8208G>A), P2704A (8294C>G 8296A>G), S2722F (8349C>T), A2742S (8408G>T), M2756I (8452G>A), R2759K (8460G>A), A2766T (8480G>A 8482T>A), R2773K (8502G>A 8503A>G), E2777D (8515A>T), E2785D (8539G>C), L2788C (8546C>T 8547T>G), S2790T (8552T>A), A2797G (8574C>G), I2801V (8585A>G), G2805E (8598G>A), N2807S (8603A>T 8604A>C 8605C>T), P2813Q (8622C>A), E2814G (8625A>G 8626G>A), P2822F (8648C>T 8649C>T 8650G>C), G2825E (8658G>A), D2827A (8664A>C 8665T>G), A2832V (8679C>T 8680A>T), T2854K (8745C>A 8746A>G), E2891G (8856A>G 8857G>A), R2894Q (8865G>A), M2895L (8867A>C 8869G>T), G2900E (8883G>A), T2902S (8888A>T), V2924I (8954G>A), V2929I (8969G>A), N2934D (8984A>G 8986T>C), E2935V (8988A>T), I2983V (9131A>G), M2989G (9149A>G 9150T>G 9151G>A), G2990S (9152G>A), P3026S (9260C>T 9262T>C) | | | | | | | | |

| | Begin | End | Coverage | Score | Concordance | Matches | Identities | I/D/M/F* | Stop Codons |
|---|---|---|---|---|---|---|---|---|---|
| **NT** | **183** | **9687** | **89.3%** | **11427** | **66.3%** | **8654 (99.8%)** | **7216 (83.2%)** | **5/16** | |

Codon mutations:

ACT3ACC (193T>C), TAC4CAA (194T>C 196C>A), TCA6TCG (202A>G), ATC8ATT (208C>T), TGT9CTG (209T>C 210G>T 211T>G), TCG12TCA (220G>A), TTT13ATG (221T>A 223T>G), GAA14GAG (226A>G), CTA17CTT (235A>T), TAC19TAT (241C>T), CCA21CCT (247A>T), TCT23CCT (251T>C), TGC24TTT (255G>T 256C>T), GAG25GGG (258A>G), CAT26CAC (262T>C), ATT27GTG (263A>G 265T>G), GTG28GCA (267T>C 268G>A), GAA30GTA (273A>T), CGA31CGT (277A>T), GAA32GAG (280A>G), GTG33GTT (283G>T), CCG34CCA (286G>A), GCT35ACC (287G>A 289T>C), GTT37ACT (293G>A 294T>C), CCT39CCC (301T>C), TTC40TTT (304C>T), GCA41ATA (305G>A 306C>T), GAT42GGT (309A>G), CTG43TTG (311C>T), GAA44GAT (316A>T), ACA45GTG (317A>G 318C>T 319A>G), CAA46CAG (322A>G), GCA49GCT (331A>T), CTC52CAA (339T>A 340C>A), AAG53AGG (342A>G), AAA55GCG (347A>G 348A>C 349A>G), TAT56CAT (350T>C), GCT57GCA (355T>A), ACT58GTT (356A>G 357C>T), CGT60CGC (364T>C), GTG61ACA (365G>A 366T>C 367G>A), CTC62TCC (368C>T 369T>C), AAA63AAG (373A>G), AAC64AAT (376C>T), TTT67TGT (384T>G), ACG68ACA (388G>A), TAC69TAT (391C>T), CGA70CGG (394A>G), TAC71TAT (397C>T), AAG72AAA (400G>A), GAT74GAC (406T>C), GCC75GTT (408C>T 409C>T), CAG76CAA (412G>A), ATA77ATT (415A>T), ATG78GAG (416A>G 417T>A), ATT80ATC (424T>C), CAG81CAA (427G>A), AAA83AAG (433A>G), CTG84CTC (436G>C), GAG85AAG (437G>A), AGG86GAA (440A>G 441G>A 442G>A), AAG87CGA (443A>C 444A>G 445G>A), GAT88GAA (448T>A), GAA90GAG (454A>G), ATG194AAG (765T>A), TTG196CTG (770T>C), CGA197AAG (773C>A 774G>A 775A>G), CGG198AGG (776C>A), TTC202TTT (790C>T), ATG206AAG (801T>A), GGA210GCG (813G>C 814A>G), CTT211CAT (816T>A), TTG212CTG (818T>C), CAA213CAG (823A>G), CGT214CAA (825G>A 826T>A), CGG217AGA (833C>A 835G>A), GAC219GAT (841C>T), AAT223AAC (853T>C), GTC225GTT (859C>T), CGC226TGT (860C>T 862C>T), ACT227GCC (863A>G 865T>C), ATC228ACC (867T>C), AAC229GAT (869A>G 871C>T), ATA230TTG (872A>T 874A>G), AGG232AAG (879G>A), GAT234GAC (886T>C), ATC238GTT (896A>G 898C>T), AAC240AGC (903A>G), ACA241AAC (906C>A 907A>C), AAA242ACC (909A>C 910A>C), AGC243AAT (912G>A 913C>T), GGC246GGA (922C>A), GGT249GGA (931T>A), TCA252CTT (938T>C 939C>T 940A>T), GGA253GGG (943A>G), TTC256TTT (952C>T), TCA261TCG (967A>G), GAA263GAG (973A>G), GGG264GGC (976G>C), TTG266ATC (980T>A 982G>C), GCA269GCG (991A>G), CGT270CGC (994T>C), AGA272AAA (999G>A), GTT273GTC (1003T>C), CAG275AGA (1007C>A 1008A>G 1009G>A), AGT276GGG (1010A>G 1012T>G), ATT277GTT (1013A>G), TTA278TTG (1018A>G), AAC279CAG (1019A>C 1021C>G), ATC282ATT (1030C>T), TCG285TCA (1039G>A), AAT286AAC (1042T>C), GCC287GCA (1045C>A), GAC288GAA (1048C>A), AAT289AAC (1051T>C), AAG292AAA (1060G>A), GGT293GGA (1063T>A), CTG294TTG (1064C>T), GAC295GAT (1069C>T), CGA300CAA (1083G>A), TAT303TGC (1092A>G 1093T>C), TCG305GCA (1097T>G 1099G>A), CAC307CAT (1105C>T), AAT378..A (1318T>A), AAG379AAA (1321G>A), TTG381TTA (1327G>A), ATA382ACA (1329T>C), GCG383GCT (1333G>T), TTA384TTG (1336A>G), CTA387TTG (1343C>T 1345A>G), CCG390CCA (1354G>A), GTG391GTT (1357G>T), GAC392GAT (1360C>T), CTC395TTG (1367C>T 1369C>G), CTT397ATT (1373C>A), TTC398TTT (1378C>T), GAG400GAA (1384G>A), ATA401GTA (1385A>G), AAA403AAG (1393A>G), TCC404TCT (1396C>T), GAG407GAT (1405G>T), AAA408AAG (1408A>G), CAG409CAA (1411G>A), GCA411TCT (1415G>T 1417A>T), CCG412CCC (1420G>C), AAT415AAC (1429T>C), TTA416TTG (1432A>G), GTC418ATT (1436G>A 1438C>T), TTA419CTG (1439T>C 1441A>G), AAT420AAC (1444T>C), TTC423TTT (1453C>T), CTG424CTA (1456G>A), GAA428GAG (1468A>G), GTA436GTG (1492A>G), TTG439CTA (1499T>C 1501G>A), AGT440AGC (1504T>C), TTG441TTA (1507G>A), CTC442CTT (1510C>T), GAA443GAG (1513A>G), TTA444TTG (1516A>G), GCA445GCT (1519A>T), AGG446AGA (1522G>A), TTC447TTT (1525C>T), CAG448CAA (1528G>A), AAC450AAT (1534C>T), ACT452ACA (1540T>A), AAC454AAT (1546C>T), AAA457AAG (1555A>G), GAT459GAC (1561T>C), ATA460ATC (1564A>C), TCT461del (1565_1568delTCTT), TTC462-TC (1565_1568delTCTT), TTC463TTT (1573C>T), AGA464AGG (1576A>G), TTA467CTA (1583T>C), TCT468TCA (1588T>A), AAG470AAA (1594G>A), AAC472AAT (1600C>T), TCG478TCA (1618G>A), TGC479TGT (1621C>T), GAC480GAT (1624C>T), CTG483TTG (1631C>T), AAA485AAG (1639A>G), TTC489TTT (1651C>T), CTC490CTG (1654C>G), CAT497CAC (1675T>C), TCA503TCG (1693A>G), TTT506TTC (1702T>C), GAA508GGA (1707A>G), GCA512ACA (1718G>A), GGA514GGG (1726A>G), TAC515TAT (1729C>T), TCA516TCC (1732A>C), TAT518TAC (1738T>C), ATC520GTT (1742A>G 1744C>T), CGC521CGT (1747C>T), CAT523CAC (1753T>C), AGT525AAT (1758G>A), AGG528AGA (1768G>A), AAC534AAT (1786C>T), TTA535TTG (1789A>G), GTT536ATT (1790G>A), GTC537GTT (1795C>T), TTA541CTG (1805T>C 1807A>G), GAG543GAA (1813G>A), AGG545AGA (1819G>A), AAG547AAA (1825G>A), GAC551GAT (1837C>T), TAT552TAC (1840T>C), CAA555CAG (1849A>G), GTC558GTG (1858C>G), AAA560AAG (1864A>G), TGC562TGT (1870C>T), AGT564AGC (1876T>C), TCG565TCT (1879G>T), AAA566AAG (1882A>G), GGT568GGA (1888T>A), AAT569AAC (1891T>C), TGT574TGC (1906T>C), ACA577ACT (1915A>T), GGT582GGC (1930T>C), GCC584GCT (1936C>T), ATT585GTC (1937A>G 1939T>C), TCA587TCG (1945A>G), TTC589TTT (1951C>T), CCA592CCT (1960A>T), GTA598GTG (1978A>G), ATT599ATA (1981T>A), GGC600GGT (1984C>T), AAT601AAC (1987T>C), GAC604GAT (1996C>T), TTT607TAT (2004T>A), AAA612AAG (2020A>G), GGG613GGA (2023G>A), GAT614GAC (2026T>C), TCG615TCT (2029G>T), GAG616GAA (2032G>A), TAT625TAC (2059T>C), ATT628ATG (2068T>G), AAC629AAT (2071C>T), GTG630ATA (2072G>A 2074G>A), CTG635TTA (2087C>T 2089G>A), AGC639AGT (2101C>T), GAG640GAA (2104G>A), GAG641GAA (2107G>A), AAG644AAA (2116G>A), TTC646TTT (2122C>T), CGC651CGT (2137C>T), CTT658CTC (2158T>C), ACC660ACT (2164C>T), CCA662CCG (2170A>G), GAT666GAC (2182T>C), GCG668GCA (2188G>A), ACC669ACA (2191C>A), AGA675AAA (2208G>A), TAT678TAC (2218T>C), GAC680GAT (2224C>T), GTA681GTT (2227A>T), TTG686TTA (2242G>A), CCC687CCT (2245C>T), TTG690TTA (2254G>A), GTT691GTC (2257T>C), GAC692GAT (2260C>T), GAC694GAG (2266C>G), CAA696CAG (2272A>G), ACG697ACA (2275G>A), TCA703TCG (2293A>G), GGC705GGT (2299C>T), TCG706TCA (2302G>A), CAG707CAA (2305G>A), GGA710GGT (2314A>T), TAT711TAC (2317T>C), ATT713ATC (2323T>C), CTA714CTG (2326A>G), AGC718AGT (2338C>T), GTG719GTA (2341G>A), CAA721CAG (2347A>G), CTT722CTC (2350T>C), TTG724TTA (2356G>A), AAT727AAC (2365T>C), TCT764.CC (2476T>C), TCG766TCA (2482G>A), CTG769CTA (2491G>A), CTG771CTT (2497G>T), TTG773TTA (2503G>A), AGA778AGG (2518A>G), CCT779CCC (2521T>C), AAG780AGG (2523A>G), AGA783AGG (2533A>G), TTG785TTA (2539G>A), TTA787TTG (2545A>G), GAG789GAA (2551G>A), CCT790CCA (2554T>A), CTG792TTG (2558C>T), TTG793CTC (2561T>C 2563G>C), CTA795CTG (2569A>G), TCA796TCG (2572A>G), CCT800CCC (2584T>C), GGC801GGT (2587C>T), CTG803CTC (2593G>C), GCT805GCC (2599T>C), TAT807TAC (2605T>C), AAT808AAC (2608T>C), TTT812TTC (2620T>C), GCG815GCA (2629G>A), GTG816GTA (2632G>A), AGG817AAG (2634G>A), TTG818TTA (2638G>A), ATT820ATA (2644T>A), AAA823AAG (2653A>G), GCT827GTC (2664C>T 2665T>C), CTA832TTA (2678C>T), CTA833CTG (2683A>G), GCT835GCC (2689T>C), CTA838CTC (2698A>C), TCA841TCG (2707A>G), GCG842GCA (2710G>A), GAA844GAG (2716A>G), ACA845ACG (2719A>G), CTC846CTT (2722C>T), GTC847ATC (2723G>A), GCA848GCG (2728A>G), AGG850AGA (2734G>A), ATT851ATA (2737T>A), GCT855ACA (2747G>A 2749T>A), GCA856GCT (2752A>T), GCT857GCA (2755T>A), GAC859GAT (2761C>T), CTC860CTT (2764C>T), TGT865TGC (2779T>C), GGG867GGT (2785G>T), AAC869AAT (2791C>T), CAT871AAT (2795C>A), TAC874TAT (2806C>T), TTG878CTC (2816T>C 2818G>C), GTG880ATA (2822G>A 2824G>A), TTG881TTA (2827G>A), GTT884GTC (2836T>C), AAG885AAA (2839G>A), AAT886AAC (2842T>C), ACC893ACA (2863C>A), CTA894TTA (2864C>T), CCA900ACA (2882C>A), AGT901CAT (2885A>C 2886G>A), AAC903AAT (2893C>T), ACG904ATG (2895C>T), GTC906GTT (2902C>T), GTA907GTG (2905A>G), ATT909ATC (2911T>C), AAT913AAC (2923T>C), AAT916AGT (2931A>G), AAC919AGC (2940A>G), GCT921GCC (2947T>C), ACT926ACC (2962T>C), GAA929_AAT930insA-- (2971_2972insA), GGT935GGC (2989T>C), ACT936ATT (2991C>T), CAT937CGT (2994A>G), GAG939AGG (2999G>A 3000A>G), TCC944TC- (3016delC), CGG946CAG (3021G>A), TAC947TTT (3024A>T 3025C>T), ACA951ATA (3036C>T), GAA952GGC (3039A>G 3040A>C), AAG953ACA (3042A>C 3043G>A), TTG956TTA (3052G>A), TTA959TTG (3061A>G), GCG966GCA (3082G>A), GGC969GGT (3091C>T), GCC972GTA (3099C>T 3100C>A), GTG974AGA (3104G>A 3105T>G 3106G>A), GTC975GTT (3109C>T), AAA976AAG (3112A>G), ACT978ACC (3118T>C), GCC979GCT (3121C>T), GGA981GGG (3127A>G), AGC983AAC (3132G>A), CGA985AAG (3137C>A 3138G>A 3139A>G), TAT989TAC (3151T>C), TTC990ATT (3152T>A 3154C>T), TGT994TAT (3165G>A), GTA995GTG (3169A>G), ATC1005ATT (3199C>T), AGG1007CGG (3203A>C), CGT1008CGC (3208T>C), CCA1010CCG (3214A>G), ACC1011ACT (3217C>T), TTT1012CTT (3218T>C), GTC1013GTA (3223C>A), GTT1016ATT (3230G>A), AAC1017AAT (3235C>T), TTA1025CTA (3257T>C), ACC1026ACT (3262C>T), AGC1027AGT (3265C>T), GTA1028ATG (3266G>A 3268A>G), GTG1029GTA (3271G>A), GTG1031ATG (3275G>A), GCA1034ACA (3284G>A), CAG1039CAA (3301G>A), AGA1044AAG (3315G>A 3316A>G), GAA1045GCA (3318A>C), ATC1046ATT (3322C>T), GAG1047GAA (3325G>A), TTG1048TTA (3328G>A), ATA1051ATT (3337A>T), GAG1055GAA (3349G>A), GTC1057GTT (3355C>T), TGC1058TGT (3358C>T), ATG1059GTG (3359A>G), CTA1061CTG (3367A>G), TAT1062TAC (3370T>C), TTA1065CTG (3377T>C 3379A>G), GAA1070GAG (3394A>G), CGC1071CGT (3397C>T), GAT1072GAG (3400T>G), ACA1074ACG (3406A>G), TAC1078TAT (3418C>T), ATT1079ATA (3421T>A), TAT1081TAC (3427T>C), AAG1083AAA (3433G>A), TCA1084TCT (3436A>T), GTA1085GTG (3439A>G), AAC1086AAT (3442C>T), CCT1087CCC (3445T>C), CAG1091CAA (3457G>A), TTT1092TTC (3460T>C), GCT1093GCA (3463T>A), GCG1095GCT (3469G>T), CAG1096CAA (3472G>A), CAC1104CAT (3496C>T), CAG1105CAA (3499G>A), TCC1107TCT (3505C>T), GTT1110GGT (3513T>G), AAA1112AAG (3520A>G), GAA1115GAG (3529A>G), CAA1116CAG (3532A>G), ATG1121GTG (3545A>G), GCT1122ACT (3548G>A), TTA1123CTA (3551T>C), GTC1124ATT (3554G>A 3556C>T), AGT1133AGC (3583T>C), GAT1134GAC (3586T>C), TGC1135TGT (3589C>T), TTC1137TTT (3595C>T), AAT1141AAC (3607T>C), TTT1143TTC (3613T>C), AAG1144AAA (3616G>A), GGT1145GGC (3619T>C), CTT1147GTT (3623C>G), TCC1148TCT (3628C>T), CTG1150ATG (3632C>A), GAT1151GAC (3637T>C), TAT1152CAT (3638T>C), GAA1153GAG (3643A>G), TTA1159TTG (3661A>G), GTG1162GTA (3670G>A), AAG1164AAC (3676G>C), AAT1165AAA (3679T>A), GAG1168GAA (3688G>A), AAT1170AAA (3694T>A), GAG1171GAA (3697G>A), ATT1172GTT (3698A>G), GAT1174TAA (3704G>T 3706T>A), TTT1175TTA (3709T>A), AGC1233..T (3883C>T), GAA1234GAG (3886A>G), CTA1236CTG (3892A>G), GAC1237GAT (3895C>T), TTT1238TTC (3898T>C), GTA1240GTG (3904A>G), CGG1241AGA (3905C>A 3907G>A), GCT1243GCC (3913T>C), AAG1248AAA (3928G>A), TCA1249TCT (3931A>T), GGG1251GGA (3937G>A), AGT1257AGC (3955T>C), GTG1258GTA (3958G>A), GCC1259GCT (3961C>T), GGA1260GGG (3964A>G), TCT1261TCC (3967T>C), TTA1264TTG (3976A>G), ATT1265ATC (3979T>C), CCA1267CCT (3985A>T), CTA1271CTT (3997A>T), GCG1272GCA (4000G>A), AAC1274AAT (4006C>T), GTT1275GTG (4009T>G), TTC1276TTT (4012C>T), AAA1277AAG (4015A>G), TCT1280TCC (4024T>C), GAA1282GAG (4030A>G), CCA1283CCT (4033A>T), AAG1286AAA (4042G>A), CTG1290TTA (4052C>T 4054G>A), CGT1291CGC (4057T>C), CGT1293CGA (4063T>A), GGA1294GGG (4066A>G), ATA1297GTA (4073A>G), TTT1298TTC (4078T>C), GGC1299GGT (4081C>T), TCT1300TCC (4084T>C), TCC1304TCT (4096C>T), AGC1308AGT (4108C>T), GGA1309GGT (4111A>T), GCG1311GCA (4117G>A), CTA1312TTG (4118C>T 4120A>G), TAC1314TAT (4126C>T), TTC1315TAT (4128T>A 4129C>T), GCC1316GCT (4132C>T), AAT1317AAC (4135T>C), CAA1321CAG (4147A>G), TTA1322CTA (4148T>C), GCT1323TCT (4151G>T), AAC1326AAT (4162C>T), GTA1328ATA (4166G>A), ATA1329ATC (4171A>C), GAG1332GAA (4180G>A), CTG1336TTG (4190C>T), CCT1338CCC (4198T>C), TCC1339TCT (4201C>T), GCG1342GCT (4210G>T), TTC1343TTT (4213C>T), AGT1345AGC (4219T>C), CTG1346TTG (4220C>T), CTG1347ATA (4223C>A 4225G>A), GTT1349GTG (4231T>G), CAT1351CAC (4237T>C), GTA1356GTG (4252A>G), TTA1357CTG (4253T>C 4255A>G), AAA1358AAG (4258A>G), ACT1362ACC (4270T>C), AGA1366AGG (4282A>G), GAG1367GAA (4285G>A), GAA1369GAG (4291A>G), CAG1373CAA (4303G>A), GTC1376GTT (4312C>T), ATA1379GTG (4319A>G 4321A>G), GTG1380GTT (4324G>T), GAG1381GAA (4327G>A), CTG1384CTT (4336G>T), TCT1385TCC (4339T>C), TCA1388TCT (4348A>T), GCA1392GCG (4360A>G), GGT1394GGC (4366T>C), TCT1395TCA (4369T>A), ACT1397ACC (4375T>C), GTT1401GTC (4387T>C), TTT1404TAT (4395T>A), GTA1408ATA (4406G>A), ACC1420ACA (4444C>A), GCC1422GCT (4450C>T), CTC1424CTT (4456C>T), GAC1427GAT (4465C>T), ATG1431GTT (4475A>G 4477G>T), GTC1432GTT (4480C>T), ACA1433TCA (4481A>T), GGC1437GGT (4495C>T), AAG1441AAA (4507G>A), CAC1442CAT (4510C>T), GGT1443GGG (4513T>G), TGC1444CGC (4514T>C), CTA1445TTA (4517C>T), GTC1448GTT (4528C>T), AAA1450AAG (4534A>G), GTG1469GTA (4591G>A), TTG1471TTA (4597G>A), GAC1472GAT (4600C>T), GAC1474GAT (4606C>T), GTG1475GTA (4609G>A), GTG1477GTA (4615G>A), GAC1478GAT (4618C>T), TTG1481CTT (4625T>C 4627G>T), GAC1488GAT (4648C>T), ATT1489ATC (4651T>C), AGG1492AGA (4660G>A), AGC1493AGT (4663C>T), GCT1495GCA (4669T>A), GTG1499ATC (4679G>A 4681G>C), AGT1500AGC (4684T>C), GTT1501GTG (4687T>G), GAG1505GAA (4699G>A), AGA1506AGG (4702A>G), CAA1508CAG (4708A>G), AGG1509AGA (4711G>A), CTG1510TTG (4712C>T), AAA1518AAG (4738A>G), GCA1521GCG (4747A>G), TTG1522CTG (4748T>C), ACT1527ACC (4765T>C), GTC1538ATT (4796G>A 4798C>T), ACT1540AGC (4803C>G 4804T>C), GAG1541GAA (4807G>A), GCG1542GCT (4810G>T), GCT1543GCG (4813T>G), CTT1544CTC (4816T>C), TAT1549TAC (4831T>C), AAC1550AAT (4834C>T), CCA1552CCT (4840A>T), GGA1556GGG (4852A>G), GGC1557GGT (4855C>T), AGT1561AGC (4867T>C), CTG1562CTC (4870G>C), ATT1563ATA (4873T>A), ACA1573ACT (4903A>T), CAG1575CAA (4909G>A), TTT1577TTC (4915T>C), GAA1578GAG (4918A>G), TTG1579CTA (4919T>C 4921G>A), AGT1580AGC (4924T>C), CCC1581CCG (4927C>G), TTT1583TTC (4933T>C), ATC1584ATA (4936C>A), CAG1585CAA (4939G>A), AAT1586AAC (4942T>C), TTC1587TTT (4945C>T), TCA1593TCG (4963A>G), CCT1596CCC (4972T>C), CTT1602CTC (4990T>C), AAA1606AAG (5002A>G), CCT1613CCC (5023T>C), TTG1614CTA (5024T>C 5026G>A), TGC1615TGT (5029C>T), GAT1616GAC (5032T>C), TCT1618TCC (5038T>C), CCA1620CCT (5044A>T), GCA1623GCT (5053A>T), TCG1624TCA (5056G>A), AGT1625GGT (5057A>G), ACT1626GCT (5060A>G), TCG1629TCT (5071G>T), CTT1636ATT (5090C>A), GGA1637GGG (5095A>G), GTG1638GTT (5098G>T), GCC1639GTC (5100C>T), TTA1640TTG (5104A>G), GAA1641GAT (5107A>T), ATT1642ATC (5110T>C), CCA1643CCG (5113A>G), AAG1644AAA (5116G>A), CAA1645CAG (5119A>G), GTC1646TTT (5120G>T 5122C>T), ATT1648GTT (5126A>G), CAT1651CAC (5137T>C), ATC1652ATT (5140C>T), AAA1653AAG (5143A>G), GAG1654GAA (5146G>A), CTC1659CTA (5161C>A), GAA1661GAG (5167A>G), CTT1663CTG (5173T>G), ACG1666ACA (5182G>A), GTT1667ATT (5183G>A), GTC1668GTT (5188C>T), AAG1669AAA (5191G>A), AAA1671AAG (5197A>G), GAC1672GAT (5200C>T), TGC1674TGT (5206C>T), TTA1675TTG (5209A>G), TCG1682TCA (5230G>A), ATC1684ATT (5236C>T), AGC1685AGT (5239C>T), ATC1687ATT (5245C>T), ACA1693ACT (5263A>T), CTC1695CTT (5269C>T), TTC1696TTT (5272C>T), GCC1697GCA (5275C>A), ATC1698ATT (5278C>T), CCA1699CCC (5281A>C), AGA1700AGG (5284A>G), ACT1701ACC (5287T>C), CTA1702TTA (5288C>T), ATA1703ATT (5293A>T), TTG1704CTA (5294T>C 5296G>A), GAG1706GAA (5302G>A), CTT1709CTA (5311T>A), GAA1710GAG (5314A>G), CGA1713CGT (5323A>T), AGA1720AGG (5344A>G), AGT1721AGC (5347T>C), ATC1723ATT (5353C>T), TGC1727TGT (5365C>T), GTT1734GTC (5386T>C), ACC1737ACA (5395C>A), AAA1746AAG (5422A>G), GAT1747GAC (5425T>C), ACC1749ACT (5431C>T), GAG1751GAA (5437G>A), CTT1756CTC (5452T>C), AAG1758AAA (5458G>A), AGT1761AGC (5467T>C), CAA1762CAG (5470A>G), CTA1763TTG (5471C>T 5473A>G), TTC1766TTT (5482C>T), TCA1767TCT (5485A>T), AAT1768AAC (5488T>C), TTG1769CTA (5489T>C 5491G>A), GGT1771GGC (5497T>C), TGT1774TGC (5506T>C), GAG1775GAA (5509G>A), GAG1776GAA (5512G>A), TTA1778CTA (5516T>C), TAT1782TAC (5530T>C), GAG1783GAA (5533G>A), TCG1784TCT (5536G>T), TTG1785TTA (5539G>A), GTT1788GTG (5548T>G), CAC1790CAT (5554C>T), GCT1792GCA (5560T>A), GCG1793ACA (5561G>A 5563G>A), ACG1794ACC (5566G>C), TCA1795TCG (5569A>G), CTC1800CTG (5584C>G), AAG1803AAA (5593G>A), GGG1804GGA (5596G>A), ATT1805GTT (5597A>G), AAC1807AAA (5605C>A), GTG1811GTT (5617G>T), GCT1812GTG (5619C>T 5620T>G), TTG1815TTA (5629G>A), ATC1816CTC (5630A>C), GGC1819GGT (5641C>T), GCA1822GCT (5650A>T), ATA1826ATT (5662A>T), GGA1827GGG (5665A>G), CAA1835CAG (5689A>G), GTT1837ATT (5693G>A), GAG1838GAA

| | | | | | | | | | |
|---|---|---|---|---|---|---|---|---|---|
| | (5698G>A), CAA1843CAG (5713A>G), GGG1844GGC (5716G>C), AAA1845AAG (5719A>G), GCC1853GCA (5743C>A), CGC1857CGA (5755C>A), CAT1858CAC (5758T>C), CGT1860CGC (5764T>C), AAA1862AAG (5770A>G), GGC1865GGT (5779C>T), TTT1866TTC (5782T>C), GAC1869GAT (5791C>T), GAC1873GAT (5803C>T), AGG1884AGA (5836G>A), AAA1885AAG (5839A>G), GGA1887GGG (5845A>G), AAA1888AAG (5848A>G), GGT1891GGC (5857T>C), ACC1892ACT (5860C>T), ACA1893ACT (5863A>T), GGT1895GGA (5869T>A), GGT1897GGC (5875T>C), AGC1900AGT (5884C>T), TTC1903TTT (5893C>T), ATC1904GTG (5894A>G 5896C>G), AAC1905AAT (5899C>T), GGG1908GGA (5908G>A), TTT1909TTC (5911T>C), GAG1913GAA (5923G>A), CAA1918CAG (5938A>G), TTC1919TTT (5941C>T), GTT1920GTC (5944T>C), GAT1921GAC (5947T>C), GGG1925GGA (5959G>A), CGG1926GCC (5960C>G 5961G>C 5962G>C), GAA1930GAG (5974A>G), GCT1934GCA (5986T>A), GAC1935GAT (5989C>T), ATT1936GTT (5990A>G), GAT1938GAC (5998T>C), AGA1942CGT (6008A>C 6010A>T), GAA1945GAT (6019A>T), GTG1946GTT (6022G>T), CGA1947CGT (6025A>T), AAA1949ACA (6030A>C), GAC1955GAA (6049C>A), ATG1958ATA (6058G>A), CAA1959CAG (6061A>G), GCC1960GCA (6064C>A), GGT1962GGC (6070T>C), AGT1963AAC (6072G>A 6073T>C), AAC1964CAC (6074A>C), ACG1965ACA (6079G>A), ACC1966AAC (6081C>A), ATA1967ATT (6085A>T), GCA1969GCT (6091A>T), TAC1970TAT (6094C>T), TGT1976TCC (6111G>C 6112T>C), AAA1978AAG (6118A>G), TTG1980TTA (6124G>A), AAG1981AAA (6127G>A), ATT1982GTC (6128A>G 6130T>C), GAT1983GAC (6133T>C), TTA1984TTG (6136A>G), AAC1988AAT (6148C>T), AAA1991AAG (6157A>G), GTT1992ATT (6158G>A), TGT1993TGC (6163T>C), AAA1995AAG (6169A>G), ACA1996TCG (6170A>T 6172A>G), GGC1998GGT (6178C>T), AAA2001AAG (6187A>G), TTT2002TTC (6190T>C), GAG2004GAA (6196G>A), CTC2007TTT (6203C>T 6205C>T), GAA2008GAG (6208A>G), CTA2009TTG (6209C>T 6211A>G), CAG2011CAA (6217G>A), GCT2015GCA (6229T>A), GTA2016ATA (6230G>A), GTC2018GTT (6238C>T), AAG2021AAA (6247G>A), ATA2023ATT (6253A>T), GCA2025AAA (6257G>A 6258C>A), CAT2030CAC (6274T>C), AAA2033AGA (6282A>G), TCG2034TCA (6286G>A), GGC2038GGT (6298C>T), TTG2039TTA (6301G>A), AGA2040AGG (6304A>G), GAC2041GAT (6307C>T), AAC2043AAT (6313C>T), GCC2046GCT (6322C>T), ACA2048ACT (6328A>T), GTT2049GTG (6331T>G), TGT2050TGC (6334T>C), AGG2051AGA (6337G>A), CTG2052TTG (6338C>T), TAT2058TAC (6358T>C), GGG2059GGA (6361G>A), GCA2060ACA (6362G>A), TCA2061TCT (6367A>T), GAG2062GAA (6370G>A), TAC2064TAT (6376C>T), GGT2065GGG (6379T>G), GGA2067GGC (6385A>C), GGA2069GGT (6391A>T), GCA2070GCG (6394A>G), TAC2071TAT (6397C>T), GTA2073ATA (6401G>A), GCG2074GTG (6405C>T), CAC2076CAT (6412C>T), TTA2078TTG (6418A>G), TTT2079TTC (6421T>C), TAC2082TTC (6429A>T), GGT2084GGA (6436T>A), GAG2087GAA (6445G>A), CAA2089CGA (6450A>G), TCC2090TCA (6454C>A), CAC2092CAT (6460C>T), GGT2093GGA (6463T>A), TTC2095TTT (6469C>T), AGG2096AGA (6472G>A), GTG2097GTA (6475G>A), AAT2099AAC (6481T>C), CAC2101AAT (6485C>A 6487C>T), AGT2102AGC (6490T>C), AGC2104AGT (6496C>T), CTG2106CTA (6502G>A), GGT2110GGC (6514T>C), AGG2111AGA (6517G>A), CTC2115ATC (6527C>A), ATC2116ATA (6532C>A), CCG2119CCA (6541G>A), AAA2120AAG (6544A>G), GTC2124GTT (6556C>T), TTT2125TTC (6559T>C), CAG2127CAA (6565G>A), TTG2129CTA (6569T>C 6571G>A), CCT2134CCA (6586T>A), ACA2135ATA (6588C>T), GAA2138GAG (6598A>G), AGA2139AGG (6601A>G), ATT2140ATC (6604T>C), TGT2141TGC (6607T>C), GGA2144GGG (6616A>G), AAC2146AAT (6622C>T), GAG2149GAA (6631G>A), AAG2150AAA (6634G>A), TAT2151CAT (6635T>C), GCT2152GCA (6640T>A), TCG2154TCC (6646G>C), ATC2155GTC (6647A>G), ATC2156ATT (6652C>T), AGC2160AGT (6664C>T), ACT2162ACG (6670T>G), ATA2165GTG (6677A>G 6679A>G), GGC2167GGT (6685C>T), ACA2169ACT (6691A>T), TTC2170TTT (6694C>T), ACA2177ACG (6715A>G), GAT2178AAT (6716G>A), AAT2179GAT (6719A>G), GGA2180GGG (6724A>G), CTA2184TTA (6734C>T), GTG2186GTA (6742G>A), GTG2187GTA (6745G>A), GCC2190GCT (6754C>T), GGA2192GGG (6760A>G), ATA2194TTG (6764A>T 6766A>G), GTC2195GTT (6769C>T), ATT2197ATA (6775T>A), CTG2200TTG (6782C>T), GCA2201GCT (6787A>T), AAC2202AAT (6790C>T), GCA2204GTG (6795C>T 6796A>G), CAC2205CAA (6799C>A), ACC2206ACA (6802C>A), AAC2208AAT (6808C>T), TAC2209TAT (6811C>T), TAC2210TAT (6814C>T), TTC2213TTT (6823C>T), GAA2215GAG (6829A>G), AAG2220AGG (6843A>G), TAC2221TAT (6847C>T), CGA2223CGC (6853A>C), AAT2225GAT (6857A>G), GAG2226GAA (6862G>A), GAA2229GAG (6871A>G), GTC2231ACA (6875G>A 6876T>C 6877C>A), AAG2232AAA (6880G>A), TCT2233TCA (6883T>A), GTT2235GTG (6889T>G), GAC2239GAT (6901C>T), GGC2244GGT (6916C>T), CCG2245CCA (6919G>A), AAA2247AAG (6925A>G), CTT2248CTC (6928T>C), GAC2250GAA (6934C>A), GGG2255GGC (6949G>C), TTC2257TTT (6955C>T), AAA2258AAG (6958A>G), CAA2264CAG (6976A>G), CTA2266TTA (6980C>T), ATC2267ATT (6985C>T), GAT2270GAC (6994T>C), GTA2271GTT (6997A>T), GTG2272GTT (7000G>T), GTG2273ATA (7001G>A 7003G>A), GCT2276GCC (7012T>C), AAG2277AAA (7015G>A), CAT2278CAC (7018T>C), TTT2283TAT (7032T>A), GCC2285GCT (7039C>T), TTG2286CTA (7040T>C 7042G>A), GGA2288GGG (7048A>G), GCT2292GCA (7060T>A), GTC2293GTA (7063C>A), GCA2294GCG (7066A>G), AGC2298AGT (7078C>T), CAA2299CAG (7081A>G), TTA2300CTG (7082T>C 7084A>G), GTA2301GTG (7087A>G), ACC2302ACA (7090C>A), CAT2304CAC (7096T>C), GTA2305GTG (7099A>G), GTT2306GTC (7102T>C), GAG2309GAA (7111G>A), CGA2311CGG (7117A>G), ACA2314AAA (7125C>A), GAA2315GAG (7129A>G), CTG2317TTA (7133C>T 7135G>A), GTG2319GTA (7141G>A), GAT2320GAC (7144T>C), GCA2321TCG (7145G>T 7147A>G), GAG2322GAA (7150G>A), GAG2324GAA (7156G>A), GCA2325GCT (7159A>T), GAG2326_GCA2327del (7160_7165delGAGGCA), TTC2329TTT (7171C>T), GCG2335GCT (7189G>T), AAA2338AAG (7198A>G), AGC2339AGT (7201C>T), CTA2341TTA (7205C>T), AAT2342AAC (7210T>C), GAT2344GAA (7216T>A), GCG2345GCA (7219G>A), ATC2347ATA (7225C>A), AAG2348AAA (7228G>A), AAG2352AAA (7240G>A), TAT2353TAC (7243T>C), AAA2355AAG (7249A>G), CCT2356CCG (7252T>G), GAT2358GAA (7258T>A), GTT2359GTC (7261T>C), GGT2360GGA (7264T>A), GTC2361GTG (7267C>G), GTG2362GTA (7270G>A), GAT2363_CGG2364insT-- (7273_7274insT), CGG2364GTG (7274C>G 7275G>T), CAT2366CTT (7281A>T), TTG2367TCG (7284T>C), TCA2371TTA (7296C>T), TCT2376TTT (7311C>T), ACC2377ATC (7314C>T), TGC2378TAC (7317G>A), AAT2379AAA (7321T>A), GTG2380-TA (7322delG 7324G>A), CAC2381CAT (7327C>T), GGC2382GGT (7330C>T), TTC2383TTT (7333C>T), AAG2384CAG (7334A>C), TGT2386TGC (7342T>C), GCA2387AAC (7343G>A 7344C>A 7345A>C), ACT2390ACC (7354T>C), CAA2393CAG (7363A>G), ATT2395ATC (7369T>C), TTC2396TTT (7372C>T), GCG2398GCT (7378G>T), CTC2399CTT (7381C>T), GCA2404GCT (7396A>T), GTC2405GTT (7399C>T), GCC2407GCT (7405C>T), AGT2408ATG (7407G>T 7408T>G), GGG2410GGT (7414G>T), TGC2411GGC (7415T>G), AAA2412AAG (7420A>G), GAC2415GAT (7429C>T), TAT2416TAC (7432T>C), CAT2419CAC (7441T>C), AAG2425AAA (7459G>A), GAA2426GAG (7462A>G), ATA2428ATT (7468A>T), GTC2429GTA (7471C>A), CAA2431CAG (7477A>G), AGC2432AGT (7480C>T), CTG2434CTT (7486G>T), TAT2437TAC (7495T>C), AAA2438AAG (7498A>G), GGT2439GGC (7501T>C), TTG2440TCA (7503T>C 7504G>A), ATT2443ATA (7513T>A), AAC2445AAT (7519C>T), GGA2446GGG (7522A>G), AAG2449AAA (7531G>A), GAG2451GAA (7537G>A), CTC2452CTT (7540C>T), TGT2454TGC (7546T>C), AAG2462AAA (7570G>A), ACG2463ACA (7573G>A), ACG2465ACA (7579G>A), GCT2468GCC (7588T>C), CTA2471TTA (7595C>T), TTG2474CTA (7604T>C 7606G>A), GGT2477GGA (7615T>A), TGT2480TGC (7624T>C), GAC2483GAT (7633C>T), TTC2484TTT (7636C>T), AAT2485AAC (7639T>C), TTT2488TTC (7648T>C), GAA2494GAG (7666A>G), TGT2496TGC (7672T>C), ACA2498ACT (7678A>T), GGG2500GGA (7684G>A), ACT2502ACC (7690T>C), TTT2504TTC (7696T>C), GGT2506GGG (7702T>G), GAT2509AAT (7709G>A), AAA2510AGG (7713A>G 7714A>G), CTG2511TTG (7715C>T), CGT2514CGC (7726T>C), GAG2517GAA (7735G>A), GTA2520GTG (7744A>G), TAC2521TAT (7747C>T), GGC2526GGT (7762C>T), CAG2528CAA (7768G>A), TTT2529TTC (7771T>C), TCA2532TCG (7780A>G), CTA2533CTC (7783A>C), ACT2534ACC (7786T>C), CTA2537TTG (7793C>T 7795A>G), ATC2538ATT (7798C>T), GCT2540GCC (7804T>C), ACC2543ATC (7812C>T), GAA2550GAG (7834A>G), GAC2551GAT (7837C>T), GTG2554TTA (7844G>T 7846G>A), CAG2557CAA (7855G>A), CTG2559CTA (7861G>A), TTA2562TTG (7870A>G), TAC2563TAT (7873C>T), ACT2564ACA (7876T>A), GAG2565GAA (7879G>A), ATT2566ATA (7882T>A), GTT2567ATT (7883G>A), CCA2570CCT (7894A>T), ATT2571ATC (7897T>C), GTC2579GTT (7921C>T), AAG2581AAA (7927G>A), GGT2584GGC (7936T>C), AAT2585AAC (7939T>C), AAT2586AAC (7942T>C), GTG2594GTA (7966G>A), GAT2595GAC (7969T>C), TCT2597TCC (7975T>C), GTC2601GTT (7987C>T), CTT2602CTC (7990T>C), GCT2607GCC (8005T>C), GAA2616GAG (8032A>G), ATC2618ATT (8038C>T), GAC2619GAT (8041C>T), AGC2620AAC (8043G>A), TTG2632TTA (8080G>A), GTG2635GTA (8089G>A), CCG2637CCA (8095G>A), GAG2638GAA (8098G>A), AGC2641AGT (8107C>T), ATT2642ATC (8110T>C), CTC2643CTT (8113C>T), GAT2644GAC (8116T>C), CAA2648CAG (8128A>G), GGT2654GGC (8146T>C), AAC2656AGT (8151A>G 8152C>T), TCG2660TCA (8164G>A), AGG2665AAG (8178G>A), AAG2666AAA (8182G>A), TTG2669TTA (8191G>A), TTC2671TTT (8197C>T), TCC2673TCT (8203C>T), AGA2675AAA (8208G>A), CTG2677TTG (8213C>T), CTA2678CTG (8218A>G), ATC2679ATT (8221C>T), GAG2680GAA (8224G>A), CCA2685CCG (8239A>G), ATT2692ATC (8260T>C), GTA2693GTG (8263A>G), GAT2699GAC (8281T>C), AGA2700AGG (8284A>G), CCA2704GCG (8294C>G 8296A>G), GAG2705GAA (8299G>A), CAC2706CAT (8302C>T), GCG2710GCA (8314G>A), ATT2711ATC (8317T>C), TGC2712TGT (8320C>T), GCT2714GCA (8326T>A), GAG2717GAA (8335G>A), TCC2718TCT (8338C>T), TCT2722TTT (8349C>T), GAA2723GAG (8353A>G), CAC2726CAT (8362C>T), CAA2727CAG (8365A>G), ATC2728ATT (8368C>T), TAC2732TAT (8380C>T), TCA2733TCG (8383A>G), TTA2735TTG (8389A>G), CCT2740CCC (8404T>C), GCA2742TCA (8408G>T), ACA2743ACG (8413A>G), GCG2745GCT (8419G>T), GCT2750GCC (8434T>C), TAT2752TAC (8440T>C), ATG2756ATA (8452G>A), CTA2758CTG (8458A>G), AGG2759AAG (8460G>A), AAA2760AAG (8464A>G), CTG2761CTA (8467G>A), GCT2766ACA (8480G>A 8482T>A), GAG2771GAA (8497G>A), CTA2772TTG (8498C>T 8500A>G), AGA2773AAG (8502G>A 8503A>G), GCC2774GCT (8506C>T), GAA2777GAT (8515A>T), GTC2780GTT (8524C>T), GCA2781GCC (8527A>C), TTA2782CTG (8528T>C 8530A>G), GAG2785GAC (8539G>C), CTT2788TGT (8546C>T 8547T>G), TCT2790ACT (8552T>A), GTA2793GTG (8563A>G), GCA2797GGA (8574C>G), ATT2801GTT (8585A>G), GGA2805GAA (8598G>A), AGC2806AGT (8602C>T), AAC2807TCT (8603A>T 8604A>C 8605C>T), AAA2812AAG (8620A>G), CCA2813CAA (8622C>A), GAG2814GGA (8625A>G 8626G>A), CAG2815CAA (8629G>A), GGC2816GGT (8632C>T), ATC2818ATT (8638C>T), CAG2819CAA (8641G>A), AAC2821AAT (8647C>T), CCG2822TTC (8648C>T 8649C>T 8650G>C), AAC2823AAT (8653C>T), AAA2824AAG (8656A>G), GGA2825GAA (8658G>A), GAT2827GCG (8664A>C 8665T>G), GTT2830GTG (8674T>G), GCA2832GTT (8679C>T 8680A>T), GGC2833GGA (8683C>A), GGG2836GGA (8692G>A), ACA2837ACT (8695A>T), CCG2841CCA (8707G>A), AGA2842CGA (8708A>C), AAG2844AAA (8716G>A), ATC2846ATT (8722C>T), ACG2847ACA (8725G>A), AAA2849AAG (8731A>G), AGA2851AGG (8737A>G), CCC2853CCT (8743C>T), ACA2854AAG (8745C>A 8746A>G), AGC2855AGT (8749C>T), AAG2856AAA (8752G>A), GGA2857GGT (8755A>T), AAC2862AAT (8770C>T), TTA2863CTA (8771T>C), GAA2864GAG (8776A>G), CAT2865CAC (8779T>C), TTG2866TTA (8782G>A), TAT2869TAC (8791T>C), CCA2871CCG (8797A>G), CAA2872CAG (8800A>G), CAA2873CAG (8803A>G), GAT2875GAC (8809T>C), ATT2876ATC (8812T>C), AAT2878AAC (8818T>C), CGG2880CGA (8824G>A), GCA2881GCG (8827A>G), ACG2888ACA (8848G>A), TAT2890TAC (8854T>C), GAG2891GGA (8856A>G 8857G>A), GTG2893GTA (8863G>A), CGG2894CAG (8865G>A), ATG2895CTT (8867A>C 8869G>T), GCA2896GCG (8872A>G), GGA2900GAA (8883G>A), ACT2902TCT (8888A>T), TGC2915TGT (8929C>T), GTC2924ATC (8954G>A), AAC2925AAT (8959C>T), GGA2926GGT (8962A>T), GTT2929ATT (8969G>A), GGG2933GGA (8983G>A), AAT2934GAC (8984A>G 8986T>C), GAA2935GTA (8988A>T), GAG2938GAA (8998G>A), CCG2940CCA (9004G>A), TTG2941TTA (9007G>A), ATC2944ATT (9016C>T), ACC2951ACA (9037C>A), ATA2967ATT (9085A>T), AAC2971AAT (9097C>T), ATT2983GTT (9131A>G), CGA2984CGT (9136A>T), CGG2987CGT (9145G>T), ATG2989GGA (9149A>G 9150T>G 9151G>A), GGT2990AGT (9152G>A), TTA2991TTG (9157A>G), GCG2992GCT (9160G>T), CGT2993CGC (9163T>C), GAG3000GAA (9184G>A), CGA3004CGG (9196A>G), AGG3010AGA (9214G>A), GAA3011GAG (9217A>G), GCG3012GCA (9220G>A), GCA3020GCT (9244A>T), TTG3021TTA (9247G>A), GCC3024GCT (9256C>T), CCT3026TCC (9260C>T 9262T>C), GGG3030GGA (9274G>A), GAC3032GAT (9280C>T), ATC3035ATT (9289C>T), GAG3040GAA (9304G>A), GTC3050GTT (9334C>T), GTC3060GTG (9364C>G) | | | | | | | | |
| PVYgp2 | 1 | 75 | 100% | 95 | 18.1% | 75 (98.7%) | 54 (71.1%) | 1/0/3/2 | 2 |
| Protein mutations: | R17_K17insX (2971_2972insA), Y23H (2989T>C 2991C>T), R26K (2999G>A 3000A>G), H35Y (3025C>T), K40H (3040A>C 3042A>C), G41S (3043G>A), E44K (3052G>A), I47V (3061A>G), V54I (3082G>A), P57S (3091C>T), P60T (3100C>A), G61E (3104G>A 3105T>G), G62S (3106G>A), Q63* (3109C>T), R64G (3112A>G), C66R (3118T>C), L67F (3121C>T), I69V (3127A>G), A72E (3137C>A 3138G>A), I73V (3139A>G) | | | | | | | | |
| Codon mutations: | TTA2CTA (2923T>C), AAA4AAG (2931A>G), GAA7GAG (2940A>G), TTG10CTG (2947T>C), TTG15CTG (2962T>C), AGA17_AAA17insAA- (2971_2972insA), TAC23CAT (2989T>C 2991C>T), TCA24TCG (2994A>G), AGA26AAG (2999G>A 3000A>G), CAC32-AC (3016delC), TCG33TCA (3021G>A), GTA34GTT (3024A>T), CAT35TAT (3025C>T), CAC38CAT (3036C>T), AGA39AGG (3039A>G), AAA40CAC (3040A>C 3042A>C), GGC41AGC (3043G>A), GAA44AAA (3052G>A), ATA47GTA (3061A>G), GTT54ATT (3082G>A), CCG57TCG (3091C>T), CGC59CGT (3099C>T), CCA60ACA (3100C>A), GGT61GAG (3104G>A 3105T>G), GGT62AGT (3106G>A), CAA63TAA (3109C>T), AGG64GGG (3112A>G), TGC66CGC (3118T>C), CTC67TTC (3121C>T), ATT69GTT (3127A>G), GAG70GAA (3132G>A), GCG72GAA (3137C>A 3138G>A), ATT73GTT (3139A>G) | | | | | | | | |

#### Proteins

| | | | | | | | | | |
|---|---|---|---|---|---|---|---|---|---|
| polyprotein (NP_056759.1) | 1 | 3064 | 90.9% | 17031 | 88.8% | 2782 (99.8%) | 2525 (90.6%) | 2/3/5/5 | 2 |

| | Begin | End | Coverage | Score | Concordance | Matches | Identities | I/D/M/F* | Stop Codons |
|---|---|---|---|---|---|---|---|---|---|
| **NT** | **183** | **9687** | **89.3%** | **11427** | **66.3%** | **8654 (99.8%)** | **7216 (83.2%)** | **5/16** | |
| Protein mutations: | Y4Q (194T>C 196C>A), C9L (209T>C 210G>T 211T>G), F13M (221T>A 223T>G), S23P (251T>C), C24F (255G>T 256C>T), E25G (258A>G), I27V (263A>G 265T>G), V28A (267T>C 268G>A), E30V (273A>T), A35T (287G>A 289T>C), V37T (293G>A 294T>C), A41I (305G>A 306C>T), D42G (309A>G), E44D (316A>T), T45V (317A>G 318C>T 319A>G), L52Q (339T>A 340C>A), K53R (342A>G), K55A (347A>G 348A>C 349A>G), Y56H (350T>C), T58V (356A>G 357C>T), V61T (365G>A 366T>C 367G>A), L62S (368C>T 369T>C), F67C (384T>G), A75V (408C>T 409C>T), M78E (416A>G 417T>A), E85K (437G>A), R86E (440A>G 441G>A 442G>A), K87R (443A>C 444A>G 445G>A), D88E (448T>A), M194K (765T>A), R197K (773C>A 774G>A 775A>G), M206K (801T>A), G210A (813G>C 814A>G), L211H (816T>A), R214Q (825G>A 826T>A), R226C (860C>T 862C>T), T227A (863A>G 865T>C), I228T (867T>C), N229D (869A>G 871C>T), I230L (872A>T 874A>G), R232K (879G>A), I238V (896A>G 898C>T), N240S (903A>G), T241N (906C>A 907A>C), K242T (909A>C 910A>C), S243N (912G>A 913C>T), S252L (938T>C 939C>T 940A>T), L266I (980T>A 982G>C), R272K (999G>A), Q275R (1007C>A 1008A>G 1009G>A), S276G (1010A>G 1012T>G), I277V (1013A>G), N279Q (1019A>C 1021C>G), D288E (1048C>A), R300Q (1083G>A), Y303C (1092A>G 1093T>C), S305A (1097T>G 1099G>A), I382T (1329T>C), L397I (1373C>A), I401V (1385A>G), E407D (1405G>T), A411S (1415G>T 1417A>T), V418I (1436G>A 1438C>T), S461del (1565_1568delTCTT), E508G (1707A>G), A512T (1718G>A), I520V (1742A>G 1744C>T), S525N (1758G>A), V536I (1790G>A), I585V (1937A>G 1939T>C), F607Y (2004T>A), I628M (2068T>G), V630I (2072G>A 2074G>A), R675K (2208G>A), D694E (2266C>G), K780R (2523A>G), R817K (2634G>A), A827V (2664C>T 2665T>C), V847I (2723G>A), A855T (2747G>A 2749T>A), H871N (2795C>A), V880I (2822G>A 2824G>A), P900T (2882C>A), S901H (2885A>C 2886G>A), T904M (2895C>T), N916S (2931A>G), N919S (2940A>G), E929_N930insX (2971_2972insA), T936I (2991C>T), H937R (2994A>G), E939R (2999G>A 3000A>G), R946Q (3021G>A), Y947F (3024A>T 3025C>T), T951I (3036C>T), E952G (3039A>G 3040A>C), K953T (3042A>C 3043G>A), A972V (3099C>T 3100C>A), V974R (3104G>A 3105T>G 3106G>A), S983N (3132G>A), R985K (3137C>A 3138G>A 3139A>G), F990I (3152T>A 3154C>T), C994Y (3165G>A), F1012L (3218T>C), V1016I (3230G>A), V1028M (3266G>A 3268A>G), V1031M (3275G>A), A1034T (3284G>A), R1044K (3315G>A 3316A>G), E1045A (3318A>C), M1059V (3359A>G), D1072E (3400T>G), V1110G (3513T>G), M1121V (3545A>G), A1122T (3548G>A), V1124I (3554G>A 3556C>T), L1147V (3623C>G), L1150M (3632C>A), Y1152H (3638T>C), K1164N (3676G>C), N1165K (3679T>A), N1170K (3694T>A), I1172V (3698A>G), D1174* (3704G>T 3706T>A), F1175L (3709T>A), I1297V (4073A>G), F1315Y (4128T>A 4129C>T), A1323S (4151G>T), V1328I (4166G>A), L1347I (4223C>A 4225G>A), I1379V (4319A>G 4321A>G), F1404Y (4395T>A), V1408I (4406G>A), M1431V (4475A>G 4477G>T), T1433S (4481A>T), C1444R (4514T>C), V1499I (4679G>A 4681G>C), V1538I (4796G>A 4798C>T), T1540S (4803C>G 4804T>C), S1625G (5057A>G), T1626A (5060A>G), L1636I (5090C>A), A1639V (5100C>T), E1641D (5107A>T), V1646F (5120G>T 5122C>T), I1648V (5126A>G), V1667I (5183G>A), A1793T (5561G>A 5563G>A), I1805V (5597A>G), N1807K (5605C>A), A1812V (5619C>T 5620T>G), I1816L (5630A>C), V1837I (5693G>A), I1904V (5894A>G 5896C>G), R1926A (5960C>G 5961G>C 5962G>C), I1936V (5990A>G), E1945D (6019A>T), K1949T (6030A>C), D1955E (6049C>A), M1958I (6058G>A), S1963N (6072G>A 6073T>C), N1964H (6074A>C), T1966N (6081C>A), C1976S (6111G>C 6112T>C), I1982V (6128A>G 6130T>C), V1992I (6158G>A), T1996S (6170A>T 6172A>G), L2007F (6203C>T 6205C>T), V2016I (6230G>A), A2025K (6257G>A 6258C>A), K2033R (6282A>G), A2060T (6362G>A), V2073I (6401G>A), A2074V (6405C>T), Y2082F (6429A>T), Q2089R (6450A>G), H2101N (6485C>A 6487C>T), L2115I (6527C>A), T2135I (6588C>T), Y2151H (6635T>C), I2155V (6647A>G), I2165V (6677A>G 6679A>G), D2178N (6716G>A), N2179D (6719A>G), I2194L (6764A>T 6766A>G), A2204V (6795C>T 6796A>G), H2205Q (6799C>A), K2220R (6843A>G), N2225D (6857A>G), V2231T (6875G>A 6876T>C 6877C>A), D2250E (6934C>A), V2273I (7001G>A 7003G>A), F2283Y (7032T>A), T2314K (7125C>A), A2321S (7145G>T 7147A>G), E2326_A2327del (7160_7165delGAGGCA), D2344E (7216T>A), D2358E (7258T>A), D2363_R2364insX (7273_7274insT), R2364V (7274C>G 7275G>T), H2366L (7281A>T), L2367S (7284T>C), S2371L (7296C>T), S2376F (7311C>T), T2377I (7314C>T), C2378Y (7317G>A), N2379K (7321T>A), K2384Q (7334A>C), A2387N (7343G>A 7344C>A 7345A>C), S2408M (7407G>T 7408T>G), C2411G (7415T>G), L2440S (7503T>C 7504G>A), D2509N (7709G>A), K2510R (7713A>G 7714A>G), T2543I (7812C>T), V2554L (7844G>T 7846G>A), V2567I (7883G>A), S2620N (8043G>A), N2656S (8151A>G 8152C>T), R2665K (8178G>A), R2675K (8208G>A), P2704A (8294C>G 8296A>G), S2722F (8349C>T), A2742S (8408G>T), M2756I (8452G>A), R2759K (8460G>A), A2766T (8480G>A 8482T>A), R2773K (8502G>A 8503A>G), E2777D (8515A>T), E2785D (8539G>C), L2788C (8546C>T 8547T>G), S2790T (8552T>A), A2797G (8574C>G), I2801V (8585A>G), G2805E (8598G>A), N2807S (8603A>T 8604A>C 8605C>T), P2813Q (8622C>A), E2814G (8625A>G 8626G>A), P2822F (8648C>T 8649C>T 8650G>C), G2825E (8658G>A), D2827A (8664A>C 8665T>G), A2832V (8679C>T 8680A>T), T2854K (8745C>A 8746A>G), E2891G (8856A>G 8857G>A), R2894Q (8865G>A), M2895L (8867A>C 8869G>T), G2900E (8883G>A), T2902S (8888A>T), V2924I (8954G>A), V2929I (8969G>A), N2934D (8984A>G 8986T>C), E2935V (8988A>T), I2983V (9131A>G), M2989G (9149A>G 9150T>G 9151G>A), G2990S (9152G>A), P3026S (9260C>T 9262T>C) | | | | | | | | |

| | Begin | End | Coverage | Score | Concordance | Matches | Identities | I/D/M/F* | Stop Codons |
|---|---|---|---|---|---|---|---|---|---|
| NT | 183 | 9687 | 89.3% | 11427 | 66.3% | 8654 (99.8%) | 7216 (83.2%) | 5/16 | |

Codon mutations:

ACT3ACC (193T>C), TAC4CAA (194T>C 196C>A), TCA6TCG (202A>G), ATC8ATT (208C>T), TGT9CTG (209T>C 210G>T 211T>G), TCG12TCA (220G>A), TTT13ATG (221T>A 223T>G), GAA14GAG (226A>G), CTA17CTT (235A>T), TAC19TAT (241C>T), CCA21CCT (247A>T), TCT23CCT (251T>C), TGC24TTT (255G>T 256C>T), GAG25GGG (258A>G), CAT26CAC (262T>C), ATT27GTG (263A>G 265T>G), GTG28GCA (267T>C 268G>A), GAA30GTA (273A>T), CGA31CGT (277A>T), GAA32GAG (280A>G), GTG33GTT (283G>T), CCG34CCA (286G>A), GCT35ACC (287G>A 289T>C), GTT37ACT (293G>A 294T>C), CCT39CCC (301T>C), TTC40TTT (304C>T), GCA41ATA (305G>A 306C>T), GAT42GGT (309A>G), CTG43TTG (311C>T), GAA44GAT (316A>T), ACA45GTG (317A>G 318C>T 319A>G), CAA46CAG (322A>G), GCA49GCT (331A>T), CTC52CAA (339T>A 340C>A), AAG53AGG (342A>G), AAA55GCG (347A>G 348A>C 349A>G), TAT56CAT (350T>C), GCT57GCA (355T>A), ACT58GTT (356A>G 357C>T), CGT60CGC (364T>C), GTG61ACA (365G>A 366T>C 367G>A), CTC62TCC (368C>T 369T>C), AAA63AAG (373A>G), AAC64AAT (376C>T), TTT67TGT (384T>G), ACG68ACA (388G>A), TAC69TAT (391C>T), CGA70CGG (394A>G), TAC71TAT (397C>T), AAG72AAA (400G>A), GAT74GAC (406T>C), GCC75GTT (408C>T 409C>T), CAG76CAA (412G>A), ATA77ATT (415A>T), ATG78GAG (416A>G 417T>A), ATT80ATC (424T>C), CAG81CAA (427G>A), AAA83AAG (433A>G), CTG84CTC (436G>C), GAG85AAG (437G>A), AGG86GAA (440A>G 441G>A 442G>A), AAG87CGA (443A>C 444A>G 445G>A), GAT88GAA (448T>A), GAA90GAG (454A>G), ATG194AAG (765T>A), TTG196CTG (770T>C), CGA197AAG (773C>A 774G>A 775A>G), CGG198AGG (776C>A), TTC202TTT (790C>T), ATG206AAG (801T>A), GGA210GCG (813G>C 814A>G), CTT211CAT (816T>A), TTG212CTG (818T>C), CAA213CAG (823A>G), CGT214CAA (825G>A 826T>A), CGG217AGA (833C>A 835G>A), GAC219GAT (841C>T), AAT223AAC (853T>C), GTC225GTT (859C>T), CGC226TGT (860C>T 862C>T), ACT227GCC (863A>G 865T>C), ATC228ACC (867T>C), AAC229GAT (869A>G 871C>T), ATA230TTG (872A>T 874A>G), AGG232AAG (879G>A), GAT234GAC (886T>C), ATC238GTT (896A>G 898C>T), AAC240AGC (903A>G), ACA241AAC (906C>A 907A>C), AAA242ACC (909A>C 910A>C), AGC243AAT (912G>A 913C>T), GGC246GGA (922C>A), GGT249GGA (931T>A), TCA252CTT (938T>C 939C>T 940A>T), GGA253GGG (943A>G), TTC256TTT (952C>T), TCA261TCG (967A>G), GAA263GAG (973A>G), GGG264GGC (976G>C), TTG266ATC (980T>A 982G>C), GCA269GCG (991A>G), CGT270CGC (994T>C), AGA272AAA (999G>A), GTT273GTC (1003T>C), CAG275AGA (1007C>A 1008A>G 1009G>A), AGT276GGG (1010A>G 1012T>G), ATT277GTT (1013A>G), TTA278TTG (1018A>G), AAC279CAG (1019A>C 1021C>G), ATC282ATT (1030C>T), TCG285TCA (1039G>A), AAT286AAC (1042T>C), GCC287GCA (1045C>A), GAC288GAA (1048C>A), AAT289AAC (1051T>C), AAG292AAA (1060G>A), GGT293GGA (1063T>A), CTG294TTG (1064C>T), GAC295GAT (1069C>T), CGA300CAA (1083G>A), TAT303TGC (1092A>G 1093T>C), TCG305GCA (1097T>G 1099G>A), CAC307CAT (1105C>T), AAT378..A (1318T>A), AAG379AAA (1321G>A), TTG381TTA (1327G>A), ATA382ACA (1329T>C), GCG383GCT (1333G>T), TTA384TTG (1336A>G), CTA387TTG (1343C>T 1345A>G), CCG390CCA (1354G>A), GTG391GTT (1357G>T), GAC392GAT (1360C>T), CTC395TTG (1367C>T 1369C>G), CTT397ATT (1373C>A), TTC398TTT (1378C>T), GAG400GAA (1384G>A), ATA401GTA (1385A>G), AAA403AAG (1393A>G), TCC404TCT (1396C>T), GAG407GAT (1405G>T), AAA408AAG (1408A>G), CAG409CAA (1411G>A), GCA411TCT (1415G>T 1417A>T), CCG412CCC (1420G>C), AAT415AAC (1429T>C), TTA416TTG (1432A>G), GTC418ATT (1436G>A 1438C>T), TTA419CTG (1439T>C 1441A>G), AAT420AAC (1444T>C), TTC423TTT (1453C>T), CTG424CTA (1456G>A), GAA428GAG (1468A>G), GTA436GTG (1492A>G), TTG439CTA (1499T>C 1501G>A), AGT440AGC (1504T>C), TTG441TTA (1507G>A), CTC442CTT (1510C>T), GAA443GAG (1513A>G), TTA444TTG (1516A>G), GCA445GCT (1519A>T), AGG446AGA (1522G>A), TTC447TTT (1525C>T), CAG448CAA (1528G>A), AAC450AAT (1534C>T), ACT452ACA (1540T>A), AAC454AAT (1546C>T), AAA457AAG (1555A>G), GAT459GAC (1561T>C), ATA460ATC (1564A>C), TCT461del (1565_1568delTCTT), TTC462-TC (1565_1568delTCTT), TTC463TTT (1573C>T), AGA464AGG (1576A>G), TTA467CTA (1583T>C), TCT468TCA (1588T>A), AAG470AAA (1594G>A), AAC472AAT (1600C>T), TCG478TCA (1618G>A), TGC479TGT (1621C>T), GAC480GAT (1624C>T), CTG483TTG (1631C>T), AAA485AAG (1639A>G), TTC489TTT (1651C>T), CTC490CTG (1654C>G), CAT497CAC (1675T>C), TCA503TCG (1693A>G), TTT506TTC (1702T>C), GAA508GGA (1707A>G), GCA512ACA (1718G>A), GGA514GGG (1726A>G), TAC515TAT (1729C>T), TCA516TCC (1732A>C), TAT518TAC (1738T>C), ATC520GTT (1742A>G 1744C>T), CGC521CGT (1747C>T), CAT523CAC (1753T>C), AGT525AAT (1758G>A), AGG528AGA (1768G>A), AAC534AAT (1786C>T), TTA535TTG (1789A>G), GTT536ATT (1790G>A), GTC537GTT (1795C>T), TTA541CTG (1805T>C 1807A>G), GAG543GAA (1813G>A), AGG545AGA (1819G>A), AAG547AAA (1825G>A), GAC551GAT (1837C>T), TAT552TAC (1840T>C), CAA555CAG (1849A>G), GTC558GTG (1858C>G), AAA560AAG (1864A>G), TGC562TGT (1870C>T), AGT564AGC (1876T>C), TCG565TCT (1879G>T), AAA566AAG (1882A>G), GGT568GGA (1888T>A), AAT569AAC (1891T>C), TGT574TGC (1906T>C), ACA577ACT (1915A>T), GGT582GGC (1930T>C), GCC584GCT (1936C>T), ATT585GTC (1937A>G 1939T>C), TCA587TCG (1945A>G), TTC589TTT (1951C>T), CCA592CCT (1960A>T), GTA598GTG (1978A>G), ATT599ATA (1981T>A), GGC600GGT (1984C>T), AAT601AAC (1987T>C), GAC604GAT (1996C>T), TTT607TAT (2004T>A), AAA612AAG (2020A>G), GGG613GGA (2023G>A), GAT614GAC (2026T>C), TCG615TCT (2029G>T), GAG616GAA (2032G>A), TAT625TAC (2059T>C), ATT628ATG (2068T>G), AAC629AAT (2071C>T), GTG630ATA (2072G>A 2074G>A), CTG635TTA (2087C>T 2089G>A), AGC639AGT (2101C>T), GAG640GAA (2104G>A), GAG641GAA (2107G>A), AAG644AAA (2116G>A), TTC646TTT (2122C>T), CGC651CGT (2137C>T), CTT658CTC (2158T>C), ACC660ACT (2164C>T), CCA662CCG (2170A>G), GAT666GAC (2182T>C), GCG668GCA (2188G>A), ACC669ACA (2191C>A), AGA675AAA (2208G>A), TAT678TAC (2218T>C), GAC680GAT (2224C>T), GTA681GTT (2227A>T), TTG686TTA (2242G>A), CCC687CCT (2245C>T), TTG690TTA (2254G>A), GTT691GTC (2257T>C), GAC692GAT (2260C>T), GAC694GAG (2266C>G), CAA696CAG (2272A>G), ACG697ACA (2275G>A), TCA703TCG (2293A>G), GGC705GGT (2299C>T), TCG706TCA (2302G>A), CAG707CAA (2305G>A), GGA710GGT (2314A>T), TAT711TAC (2317T>C), ATT713ATC (2323T>C), CTA714CTG (2326A>G), AGC718AGT (2338C>T), GTG719GTA (2341G>A), CAA721CAG (2347A>G), CTT722CTC (2350T>C), TTG724TTA (2356G>A), AAT727AAC (2365T>C), TCT764.CC (2476T>C), TCG766TCA (2482G>A), CTG769CTA (2491G>A), CTG771CTT (2497G>T), TTG773TTA (2503G>A), AGA778AGG (2518A>G), CCT779CCC (2521T>C), AAG780AGG (2523A>G), AGA783AGG (2533A>G), TTG785TTA (2539G>A), TTA787TTG (2545A>G), GAG789GAA (2551G>A), CCT790CCA (2554T>A), CTG792TTG (2558C>T), TTG793CTC (2561T>C 2563G>C), CTA795CTG (2569A>G), TCA796TCG (2572A>G), CCT800CCC (2584T>C), GGC801GGT (2587C>T), CTG803CTC (2593G>C), GCT805GCC (2599T>C), TAT807TAC (2605T>C), AAT808AAC (2608T>C), TTT812TTC (2620T>C), GCG815GCA (2629G>A), GTG816GTA (2632G>A), AGG817AAG (2634G>A), TTG818TTA (2638G>A), ATT820ATA (2644T>A), AAA823AAG (2653A>G), GCT827GTC (2664C>T 2665T>C), CTA832TTA (2678C>T), CTA833CTG (2683A>G), GCT835GCC (2689T>C), CTA838CTC (2698A>C), TCA841TCG (2707A>G), GCG842GCA (2710G>A), GAA844GAG (2716A>G), ACA845ACG (2719A>G), CTC846CTT (2722C>T), GTC847ATC (2723G>A), GCA848GCG (2728A>G), AGG850AGA (2734G>A), ATT851ATA (2737T>A), GCT855ACA (2747G>A 2749T>A), GCA856GCT (2752A>T), GCT857GCA (2755T>A), GAC859GAT (2761C>T), CTC860CTT (2764C>T), TGT865TGC (2779T>C), GGG867GGT (2785G>T), AAC869AAT (2791C>T), CAT871AAT (2795C>A), TAC874TAT (2806C>T), TTG878CTC (2816T>C 2818G>C), GTG880ATA (2822G>A 2824G>A), TTG881TTA (2827G>A), GTT884GTC (2836T>C), AAG885AAA (2839G>A), AAT886AAC (2842T>C), ACC893ACA (2863C>A), CTA894TTA (2864C>T), CCA900ACA (2882C>A), AGT901CAT (2885A>C 2886G>A), AAC903AAT (2893C>T), ACG904ATG (2895C>T), GTC906GTT (2902C>T), GTA907GTG (2905A>G), ATT909ATC (2911T>C), AAT913AAC (2923T>C), AAT916AGT (2931A>G), AAC919AGC (2940A>G), GCT921GCC (2947T>C), ACT926ACC (2962T>C), GAA929_AAT930insA-- (2971_2972insA), GGT935GGC (2989T>C), ACT936ATT (2991C>T), CAT937CGT (2994A>G), GAG939AGG (2999G>A 3000A>G), TCC944TC- (3016delC), CGG946CAG (3021G>A), TAC947TTT (3024A>T 3025C>T), ACA951ATA (3036C>T), GAA952GGC (3039A>G 3040A>C), AAG953ACA (3042A>C 3043G>A), TTG956TTA (3052G>A), TTA959TTG (3061A>G), GCG966GCA (3082G>A), GGC969GGT (3091C>T), GCC972GTA (3099C>T 3100C>A), GTG974AGA (3104G>A 3105T>G 3106G>A), GTC975GTT (3109C>T), AAA976AAG (3112A>G), ACT978ACC (3118T>C), GCC979GCT (3121C>T), GGA981GGG (3127A>G), AGC983AAC (3132G>A), CGA985AAG (3137C>A 3138G>A 3139A>G), TAT989TAC (3151T>C), TTC990ATT (3152T>A 3154C>T), TGT994TAT (3165G>A), GTA995GTG (3169A>G), ATC1005ATT (3199C>T), AGG1007CGG (3203A>C), CGT1008CGC (3208T>C), CCA1010CCG (3214A>G), ACC1011ACT (3217C>T), TTT1012CTT (3218T>C), GTC1013GTA (3223C>A), GTT1016ATT (3230G>A), AAC1017AAT (3235C>T), TTA1025CTA (3257T>C), ACC1026ACT (3262C>T), AGC1027AGT (3265C>T), GTA1028ATG (3266G>A 3268A>G), GTG1029GTA (3271G>A), GTG1031ATG (3275G>A), GCA1034ACA (3284G>A), CAG1039CAA (3301G>A), AGA1044AAG (3315G>A 3316A>G), GAA1045GCA (3318A>C), ATC1046ATT (3322C>T), GAG1047GAA (3325G>A), TTG1048TTA (3328G>A), ATA1051ATT (3337A>T), GAG1055GAA (3349G>A), GTC1057GTT (3355C>T), TGC1058TGT (3358C>T), ATG1059GTG (3359A>G), CTA1061CTG (3367A>G), TAT1062TAC (3370T>C), TTA1065CTG (3377T>C 3379A>G), GAA1070GAG (3394A>G), CGC1071CGT (3397C>T), GAT1072GAG (3400T>G), ACA1074ACG (3406A>G), TAC1078TAT (3418C>T), ATT1079ATA (3421T>A), TAT1081TAC (3427T>C), AAG1083AAA (3433G>A), TCA1084TCT (3436A>T), GTA1085GTG (3439A>G), AAC1086AAT (3442C>T), CCT1087CCC (3445T>C), CAG1091CAA (3457G>A), TTT1092TTC (3460T>C), GCT1093GCA (3463T>A), GCG1095GCT (3469G>T), CAG1096CAA (3472G>A), CAC1104CAT (3496C>T), CAG1105CAA (3499G>A), TCC1107TCT (3505C>T), GTT1110GGT (3513T>G), AAA1112AAG (3520A>G), GAA1115GAG (3529A>G), CAA1116CAG (3532A>G), ATG1121GTG (3545A>G), GCT1122ACT (3548G>A), TTA1123CTA (3551T>C), GTC1124ATT (3554G>A 3556C>T), AGT1133AGC (3583T>C), GAT1134GAC (3586T>C), TGC1135TGT (3589C>T), TTC1137TTT (3595C>T), AAT1141AAC (3607T>C), TTT1143TTC (3613T>C), AAG1144AAA (3616G>A), GGT1145GGC (3619T>C), CTT1147GTT (3623C>G), TCC1148TCT (3628C>T), CTG1150ATG (3632C>A), GAT1151GAC (3637T>C), TAT1152CAT (3638T>C), GAA1153GAG (3643A>G), TTA1159TTG (3661A>G), GTG1162GTA (3670G>A), AAG1164AAC (3676G>C), AAT1165AAA (3679T>A), GAG1168GAA (3688G>A), AAT1170AAA (3694T>A), GAG1171GAA (3697G>A), ATT1172GTT (3698A>G), GAT1174TAA (3704G>T 3706T>A), TTT1175TTA (3709T>A), AGC1233..T (3883C>T), GAA1234GAG (3886A>G), CTA1236CTG (3892A>G), GAC1237GAT (3895C>T), TTT1238TTC (3898T>C), GTA1240GTG (3904A>G), CGG1241AGA (3905C>A 3907G>A), GCT1243GCC (3913T>C), AAG1248AAA (3928G>A), TCA1249TCT (3931A>T), GGG1251GGA (3937G>A), AGT1257AGC (3955T>C), GTG1258GTA (3958G>A), GCC1259GCT (3961C>T), GGA1260GGG (3964A>G), TCT1261TCC (3967T>C), TTA1264TTG (3976A>G), ATT1265ATC (3979T>C), CCA1267CCT (3985A>T), CTA1271CTT (3997A>T), GCG1272GCA (4000G>A), AAC1274AAT (4006C>T), GTT1275GTG (4009T>G), TTC1276TTT (4012C>T), AAA1277AAG (4015A>G), TCT1280TCC (4024T>C), GAA1282GAG (4030A>G), CCA1283CCT (4033A>T), AAG1286AAA (4042G>A), CTG1290TTA (4052C>T 4054G>A), CGT1291CGC (4057T>C), CGT1293CGA (4063T>A), GGA1294GGG (4066A>G), ATA1297GTA (4073A>G), TTT1298TTC (4078T>C), GGC1299GGT (4081C>T), TCT1300TCC (4084T>C), TCC1304TCT (4096C>T), AGC1308AGT (4108C>T), GGA1309GGT (4111A>T), GCG1311GCA (4117G>A), CTA1312TTG (4118C>T 4120A>G), TAC1314TAT (4126C>T), TTC1315TAT (4128T>A 4129C>T), GCC1316GCT (4132C>T), AAT1317AAC (4135T>C), CAA1321CAG (4147A>G), TTA1322CTA (4148T>C), GCT1323TCT (4151G>T), AAC1326AAT (4162C>T), GTA1328ATA (4166G>A), ATA1329ATC (4171A>C), GAG1332GAA (4180G>A), CTG1336TTG (4190C>T), CCT1338CCC (4198T>C), TCC1339TCT (4201C>T), GCG1342GCT (4210G>T), TTC1343TTT (4213C>T), AGT1345AGC (4219T>C), CTG1346TTG (4220C>T), CTG1347ATA (4223C>A 4225G>A), GTT1349GTG (4231T>G), CAT1351CAC (4237T>C), GTA1356GTG (4252A>G), TTA1357CTG (4253T>C 4255A>G), AAA1358AAG (4258A>G), ACT1362ACC (4270T>C), AGA1366AGG (4282A>G), GAG1367GAA (4285G>A), GAA1369GAG (4291A>G), CAG1373CAA (4303G>A), GTC1376GTT (4312C>T), ATA1379GTG (4319A>G 4321A>G), GTG1380GTT (4324G>T), GAG1381GAA (4327G>A), CTG1384CTT (4336G>T), TCT1385TCC (4339T>C), TCA1388TCT (4348A>T), GCA1392GCG (4360A>G), GGT1394GGC (4366T>C), TCT1395TCA (4369T>A), ACT1397ACC (4375T>C), GTT1401GTC (4387T>C), TTT1404TAT (4395T>A), GTA1408ATA (4406G>A), ACC1420ACA (4444C>A), GCC1422GCT (4450C>T), CTC1424CTT (4456C>T), GAC1427GAT (4465C>T), ATG1431GTT (4475A>G 4477G>T), GTC1432GTT (4480C>T), ACA1433TCA (4481A>T), GGC1437GGT (4495C>T), AAG1441AAA (4507G>A), CAC1442CAT (4510C>T), GGT1443GGG (4513T>G), TGC1444CGC (4514T>C), CTA1445TTA (4517C>T), GTC1448GTT (4528C>T), AAA1450AAG (4534A>G), GTG1469GTA (4591G>A), TTG1471TTA (4597G>A), GAC1472GAT (4600C>T), GAC1474GAT (4606C>T), GTG1475GTA (4609G>A), GTG1477GTA (4615G>A), GAC1478GAT (4618C>T), TTG1481CTT (4625T>C 4627G>T), GAC1488GAT (4648C>T), ATT1489ATC (4651T>C), AGG1492AGA (4660G>A), AGC1493AGT (4663C>T), GCT1495GCA (4669T>A), GTG1499ATC (4679G>A 4681G>C), AGT1500AGC (4684T>C), GTT1501GTG (4687T>G), GAG1505GAA (4699G>A), AGA1506AGG (4702A>G), CAA1508CAG (4708A>G), AGG1509AGA (4711G>A), CTG1510TTG (4712C>T), AAA1518AAG (4738A>G), GCA1521GCG (4747A>G), TTG1522CTG (4748T>C), ACT1527ACC (4765T>C), GTC1538ATT (4796G>A 4798C>T), ACT1540AGC (4803C>G 4804T>C), GAG1541GAA (4807G>A), GCG1542GCT (4810G>T), GCT1543GCG (4813T>G), CTT1544CTC (4816T>C), TAT1549TAC (4831T>C), AAC1550AAT (4834C>T), CCA1552CCT (4840A>T), GGA1556GGG (4852A>G), GGC1557GGT (4855C>T), AGT1561AGC (4867T>C), CTG1562CTC (4870G>C), ATT1563ATA (4873T>A), ACA1573ACT (4903A>T), CAG1575CAA (4909G>A), TTT1577TTC (4915T>C), GAA1578GAG (4918A>G), TTG1579CTA (4919T>C 4921G>A), AGT1580AGC (4924T>C), CCC1581CCG (4927C>G), TTT1583TTC (4933T>C), ATC1584ATA (4936C>A), CAG1585CAA (4939G>A), AAT1586AAC (4942T>C), TTC1587TTT (4945C>T), TCA1593TCG (4963A>G), CCT1596CCC (4972T>C), CTT1602CTC (4990T>C), AAA1606AAG (5002A>G), CCT1613CCC (5023T>C), TTG1614CTA (5024T>C 5026G>A), TGC1615TGT (5029C>T), GAT1616GAC (5032T>C), TCT1618TCC (5038T>C), CCA1620CCT (5044A>T), GCA1623GCT (5053A>T), TCG1624TCA (5056G>A), AGT1625GGT (5057A>G), ACT1626GCT (5060A>G), TCG1629TCT (5071G>T), CTT1636ATT (5090C>A), GGA1637GGG (5095A>G), GTG1638GTT (5098G>T), GCC1639GTC (5100C>T), TTA1640TTG (5104A>G), GAA1641GAT (5107A>T), ATT1642ATC (5110T>C), CCA1643CCG (5113A>G), AAG1644AAA (5116G>A), CAA1645CAG (5119A>G), GTC1646TTT (5120G>T 5122C>T), ATT1648GTT (5126A>G), CAT1651CAC (5137T>C), ATC1652ATT (5140C>T), AAA1653AAG (5143A>G), GAG1654GAA (5146G>A), CTC1659CTA (5161C>A), GAA1661GAG (5167A>G), CTT1663CTG (5173T>G), ACG1666ACA (5182G>A), GTT1667ATT (5183G>A), GTC1668GTT (5188C>T), AAG1669AAA (5191G>A), AAA1671AAG (5197A>G), GAC1672GAT (5200C>T), TGC1674TGT (5206C>T), TTA1675TTG (5209A>G), TCG1682TCA (5230G>A), ATC1684ATT (5236C>T), AGC1685AGT (5239C>T), ATC1687ATT (5245C>T), ACA1693ACT (5263A>T), CTC1695CTT (5269C>T), TTC1696TTT (5272C>T), GCC1697GCA (5275C>A), ATC1698ATT (5278C>T), CCA1699CCC (5281A>C), AGA1700AGG (5284A>G), ACT1701ACC (5287T>C), CTA1702TTA (5288C>T), ATA1703ATT (5293A>T), TTG1704CTA (5294T>C 5296G>A), GAG1706GAA (5302G>A), CTT1709CTA (5311T>A), GAA1710GAG (5314A>G), CGA1713CGT (5323A>T), AGA1720AGG (5344A>G), AGT1721AGC (5347T>C), ATC1723ATT (5353C>T), TGC1727TGT (5365C>T), GTT1734GTC (5386T>C), ACC1737ACA (5395C>A), AAA1746AAG (5422A>G), GAT1747GAC (5425T>C), ACC1749ACT (5431C>T), GAG1751GAA (5437G>A), CTT1756CTC (5452T>C), AAG1758AAA (5458G>A), AGT1761AGC (5467T>C), CAA1762CAG (5470A>G), CTA1763TTG (5471C>T 5473A>G), TTC1766TTT (5482C>T), TCA1767TCT (5485A>T), AAT1768AAC (5488T>C), TTG1769CTA (5489T>C 5491G>A), GGT1771GGC (5497T>C), TGT1774TGC (5506T>C), GAG1775GAA (5509G>A), GAG1776GAA (5512G>A), TTA1778CTA (5516T>C), TAT1782TAC (5530T>C), GAG1783GAA (5533G>A), TCG1784TCT (5536G>T), TTG1785TTA (5539G>A), GTT1788GTG (5548T>G), CAC1790CAT (5554C>T), GCT1792GCA (5560T>A), GCG1793ACA (5561G>A 5563G>A), ACG1794ACC (5566G>C), TCA1795TCG (5569A>G), CTC1800CTG (5584C>G), AAG1803AAA (5593G>A), GGG1804GGA (5596G>A), ATT1805GTT (5597A>G), AAC1807AAA (5605C>A), GTG1811GTT (5617G>T), GCT1812GTG (5619C>T 5620T>G), TTG1815TTA (5629G>A), ATC1816CTC (5630A>C), GGC1819GGT (5641C>T), GCA1822GCT (5650A>T), ATA1826ATT (5662A>T), GGA1827GGG (5665A>G), CAA1835CAG (5689A>G), GTT1837ATT (5693G>A), GAG1838GAA

| | | | | | | | | | |
|---|---|---|---|---|---|---|---|---|---|
| | (5698G>A), CAA1843CAG (5713A>G), GGG1844GGC (5716G>C), AAA1845AAG (5719A>G), GCC1853GCA (5743C>A), CGC1857CGA (5755C>A), CAT1858CAC (5758T>C), CGT1860CGC (5764T>C), AAA1862AAG (5770A>G), GGC1865GGT (5779C>T), TTT1866TTC (5782T>C), GAC1869GAT (5791C>T), GAC1873GAT (5803C>T), AGG1884AGA (5836G>A), AAA1885AAG (5839A>G), GGA1887GGG (5845A>G), AAA1888AAG (5848A>G), GGT1891GGC (5857T>C), ACC1892ACT (5860C>T), ACA1893ACT (5863A>T), GGT1895GGA (5869T>A), GGT1897GGC (5875T>C), AGC1900AGT (5884C>T), TTC1903TTT (5893C>T), ATC1904GTG (5894A>G 5896C>G), AAC1905AAT (5899C>T), GGG1908GGA (5908G>A), TTT1909TTC (5911T>C), GAG1913GAA (5923G>A), CAA1918CAG (5938A>G), TTC1919TTT (5941C>T), GTT1920GTC (5944T>C), GAT1921GAC (5947T>C), GGG1925GGA (5959G>A), CGG1926GCC (5960C>G 5961G>C 5962G>C), GAA1930GAG (5974A>G), GCT1934GCA (5986T>A), GAC1935GAT (5989C>T), ATT1936GTT (5990A>G), GAT1938GAC (5998T>C), AGA1942CGT (6008A>C 6010A>T), GAA1945GAT (6019A>T), GTG1946GTT (6022G>T), CGA1947CGT (6025A>T), AAA1949ACA (6030A>C), GAC1955GAA (6049C>A), ATG1958ATA (6058G>A), CAA1959CAG (6061A>G), GCC1960GCA (6064C>A), GGT1962GGC (6070T>C), AGT1963AAC (6072G>A 6073T>C), AAC1964CAC (6074A>C), ACG1965ACA (6079G>A), ACC1966AAC (6081C>A), ATA1967ATT (6085A>T), GCA1969GCT (6091A>T), TAC1970TAT (6094C>T), TGT1976TCC (6111G>C 6112T>C), AAA1978AAG (6118A>G), TTG1980TTA (6124G>A), AAG1981AAA (6127G>A), ATT1982GTC (6128A>G 6130T>C), GAT1983GAC (6133T>C), TTA1984TTG (6136A>G), AAC1988AAT (6148C>T), AAA1991AAG (6157A>G), GTT1992ATT (6158G>A), TGT1993TGC (6163T>C), AAA1995AAG (6169A>G), ACA1996TCG (6170A>T 6172A>G), GGC1998GGT (6178C>T), AAA2001AAG (6187A>G), TTT2002TTC (6190T>C), GAG2004GAA (6196G>A), CTC2007TTT (6203C>T 6205C>T), GAA2008GAG (6208A>G), CTA2009TTG (6209C>T 6211A>G), CAG2011CAA (6217G>A), GCT2015GCA (6229T>A), GTA2016ATA (6230G>A), GTC2018GTT (6238C>T), AAG2021AAA (6247G>A), ATA2023ATT (6253A>T), GCA2025AAA (6257G>A 6258C>A), CAT2030CAC (6274T>C), AAA2033AGA (6282A>G), TCG2034TCA (6286G>A), GGC2038GGT (6298C>T), TTG2039TTA (6301G>A), AGA2040AGG (6304A>G), GAC2041GAT (6307C>T), AAC2043AAT (6313C>T), GCC2046GCT (6322C>T), ACA2048ACT (6328A>T), GTT2049GTG (6331T>G), TGT2050TGC (6334T>C), AGG2051AGA (6337G>A), CTG2052TTG (6338C>T), TAT2058TAC (6358T>C), GGG2059GGA (6361G>A), GCA2060ACA (6362G>A), TCA2061TCT (6367A>T), GAG2062GAA (6370G>A), TAC2064TAT (6376C>T), GGT2065GGG (6379T>G), GGA2067GGC (6385A>C), GGA2069GGT (6391A>T), GCA2070GCG (6394A>G), TAC2071TAT (6397C>T), GTA2073ATA (6401G>A), GCG2074GTG (6405C>T), CAC2076CAT (6412C>T), TTA2078TTG (6418A>G), TTT2079TTC (6421T>C), TAC2082TTC (6429A>T), GGT2084GGA (6436T>A), GAG2087GAA (6445G>A), CAA2089CGA (6450A>G), TCC2090TCA (6454C>A), CAC2092CAT (6460C>T), GGT2093GGA (6463T>A), TTC2095TTT (6469C>T), AGG2096AGA (6472G>A), GTG2097GTA (6475G>A), AAT2099AAC (6481T>C), CAC2101AAT (6485C>A 6487C>T), AGT2102AGC (6490T>C), AGC2104AGT (6496C>T), CTG2106CTA (6502G>A), GGT2110GGC (6514T>C), AGG2111AGA (6517G>A), CTC2115ATC (6527C>A), ATC2116ATA (6532C>A), CCG2119CCA (6541G>A), AAA2120AAG (6544A>G), GTC2124GTT (6556C>T), TTT2125TTC (6559T>C), CAG2127CAA (6565G>A), TTG2129CTA (6569T>C 6571G>A), CCT2134CCA (6586T>A), ACA2135ATA (6588C>T), GAA2138GAG (6598A>G), AGA2139AGG (6601A>G), ATT2140ATC (6604T>C), TGT2141TGC (6607T>C), GGA2144GGG (6616A>G), AAC2146AAT (6622C>T), GAG2149GAA (6631G>A), AAG2150AAA (6634G>A), TAT2151CAT (6635T>C), GCT2152GCA (6640T>A), TCG2154TCC (6646G>C), ATC2155GTC (6647A>G), ATC2156ATT (6652C>T), AGC2160AGT (6664C>T), ACT2162ACG (6670T>G), ATA2165GTG (6677A>G 6679A>G), GGC2167GGT (6685C>T), ACA2169ACT (6691A>T), TTC2170TTT (6694C>T), ACA2177ACG (6715A>G), GAT2178AAT (6716G>A), AAT2179GAT (6719A>G), GGA2180GGG (6724A>G), CTA2184TTA (6734C>T), GTG2186GTA (6742G>A), GTG2187GTA (6745G>A), GCC2190GCT (6754C>T), GGA2192GGG (6760A>G), ATA2194TTG (6764A>T 6766A>G), GTC2195GTT (6769C>T), ATT2197ATA (6775T>A), CTG2200TTG (6782C>T), GCA2201GCT (6787A>T), AAC2202AAT (6790C>T), GCA2204GTG (6795C>T 6796A>G), CAC2205CAA (6799C>A), ACC2206ACA (6802C>A), AAC2208AAT (6808C>T), TAC2209TAT (6811C>T), TAC2210TAT (6814C>T), TTC2213TTT (6823C>T), GAA2215GAG (6829A>G), AAG2220AGG (6843A>G), TAC2221TAT (6847C>T), CGA2223CGC (6853A>C), AAT2225GAT (6857A>G), GAG2226GAA (6862G>A), GAA2229GAG (6871A>G), GTC2231ACA (6875G>A 6876T>C 6877C>A), AAG2232AAA (6880G>A), TCT2233TCA (6883T>A), GTT2235GTG (6889T>G), GAC2239GAT (6901C>T), GGC2244GGT (6916C>T), CCG2245CCA (6919G>A), AAA2247AAG (6925A>G), CTT2248CTC (6928T>C), GAC2250GAA (6934C>A), GGG2255GGC (6949G>C), TTC2257TTT (6955C>T), AAA2258AAG (6958A>G), CAA2264CAG (6976A>G), CTA2266TTA (6980C>T), ATC2267ATT (6985C>T), GAT2270GAC (6994T>C), GTA2271GTT (6997A>T), GTG2272GTT (7000G>T), GTG2273ATA (7001G>A 7003G>A), GCT2276GCC (7012T>C), AAG2277AAA (7015G>A), CAT2278CAC (7018T>C), TTT2283TAT (7032T>A), GCC2285GCT (7039C>T), TTG2286CTA (7040T>C 7042G>A), GGA2288GGG (7048A>G), GCT2292GCA (7060T>A), GTC2293GTA (7063C>A), GCA2294GCG (7066A>G), AGC2298AGT (7078C>T), CAA2299CAG (7081A>G), TTA2300CTG (7082T>C 7084A>G), GTA2301GTG (7087A>G), ACC2302ACA (7090C>A), CAT2304CAC (7096T>C), GTA2305GTG (7099A>G), GTT2306GTC (7102T>C), GAG2309GAA (7111G>A), CGA2311CGG (7117A>G), ACA2314AAA (7125C>A), GAA2315GAG (7129A>G), CTG2317TTA (7133C>T 7135G>A), GTG2319GTA (7141G>A), GAT2320GAC (7144T>C), GCA2321TCG (7145G>T 7147A>G), GAG2322GAA (7150G>A), GAG2324GAA (7156G>A), GCA2325GCT (7159A>T), GAG2326_GCA2327del (7160_7165delGAGGCA), TTC2329TTT (7171C>T), GCG2335GCT (7189G>T), AAA2338AAG (7198A>G), AGC2339AGT (7201C>T), CTA2341TTA (7205C>T), AAT2342AAC (7210T>C), GAT2344GAA (7216T>A), GCG2345GCA (7219G>A), ATC2347ATA (7225C>A), AAG2348AAA (7228G>A), AAG2352AAA (7240G>A), TAT2353TAC (7243T>C), AAA2355AAG (7249A>G), CCT2356CCG (7252T>G), GAT2358GAA (7258T>A), GTT2359GTC (7261T>C), GGT2360GGA (7264T>A), GTC2361GTG (7267C>G), GTG2362GTA (7270G>A), GAT2363_CGG2364insT-- (7273_7274insT), CGG2364GTG (7274C>G 7275G>T), CAT2366CTT (7281A>T), TTG2367TCG (7284T>C), TCA2371TTA (7296C>T), TCT2376TTT (7311C>T), ACC2377ATC (7314C>T), TGC2378TAC (7317G>A), AAT2379AAA (7321T>A), GTG2380-TA (7322delG 7324G>A), CAC2381CAT (7327C>T), GGC2382GGT (7330C>T), TTC2383TTT (7333C>T), AAG2384CAG (7334A>C), TGT2386TGC (7342T>C), GCA2387AAC (7343G>A 7344C>A 7345A>C), ACT2390ACC (7354T>C), CAA2393CAG (7363A>G), ATT2395ATC (7369T>C), TTC2396TTT (7372C>T), GCG2398GCT (7378G>T), CTC2399CTT (7381C>T), GCA2404GCT (7396A>T), GTC2405GTT (7399C>T), GCC2407GCT (7405C>T), AGT2408ATG (7407G>T 7408T>G), GGG2410GGT (7414G>T), TGC2411GGC (7415T>G), AAA2412AAG (7420A>G), GAC2415GAT (7429C>T), TAT2416TAC (7432T>C), CAT2419CAC (7441T>C), AAG2425AAA (7459G>A), GAA2426GAG (7462A>G), ATA2428ATT (7468A>T), GTC2429GTA (7471C>A), CAA2431CAG (7477A>G), AGC2432AGT (7480C>T), CTG2434CTT (7486G>T), TAT2437TAC (7495T>C), AAA2438AAG (7498A>G), GGT2439GGC (7501T>C), TTG2440TCA (7503T>C 7504G>A), ATT2443ATA (7513T>A), AAC2445AAT (7519C>T), GGA2446GGG (7522A>G), AAG2449AAA (7531G>A), GAG2451GAA (7537G>A), CTC2452CTT (7540C>T), TGT2454TGC (7546T>C), AAG2462AAA (7570G>A), ACG2463ACA (7573G>A), ACG2465ACA (7579G>A), GCT2468GCC (7588T>C), CTA2471TTA (7595C>T), TTG2474CTA (7604T>C 7606G>A), GGT2477GGA (7615T>A), TGT2480TGC (7624T>C), GAC2483GAT (7633C>T), TTC2484TTT (7636C>T), AAT2485AAC (7639T>C), TTT2488TTC (7648T>C), GAA2494GAG (7666A>G), TGT2496TGC (7672T>C), ACA2498ACT (7678A>T), GGG2500GGA (7684G>A), ACT2502ACC (7690T>C), TTT2504TTC (7696T>C), GGT2506GGG (7702T>G), GAT2509AAT (7709G>A), AAA2510AGG (7713A>G 7714A>G), CTG2511TTG (7715C>T), CGT2514CGC (7726T>C), GAG2517GAA (7735G>A), GTA2520GTG (7744A>G), TAC2521TAT (7747C>T), GGC2526GGT (7762C>T), CAG2528CAA (7768G>A), TTT2529TTC (7771T>C), TCA2532TCG (7780A>G), CTA2533CTC (7783A>C), ACT2534ACC (7786T>C), CTA2537TTG (7793C>T 7795A>G), ATC2538ATT (7798C>T), GCT2540GCC (7804T>C), ACC2543ATC (7812C>T), GAA2550GAG (7834A>G), GAC2551GAT (7837C>T), GTG2554TTA (7844G>T 7846G>A), CAG2557CAA (7855G>A), CTG2559CTA (7861G>A), TTA2562TTG (7870A>G), TAC2563TAT (7873C>T), ACT2564ACA (7876T>A), GAG2565GAA (7879G>A), ATT2566ATA (7882T>A), GTT2567ATT (7883G>A), CCA2570CCT (7894A>T), ATT2571ATC (7897T>C), GTC2579GTT (7921C>T), AAG2581AAA (7927G>A), GGT2584GGC (7936T>C), AAT2585AAC (7939T>C), AAT2586AAC (7942T>C), GTG2594GTA (7966G>A), GAT2595GAC (7969T>C), TCT2597TCC (7975T>C), GTC2601GTT (7987C>T), CTT2602CTC (7990T>C), GCT2607GCC (8005T>C), GAA2616GAG (8032A>G), ATC2618ATT (8038C>T), GAC2619GAT (8041C>T), AGC2620AAC (8043G>A), TTG2632TTA (8080G>A), GTG2635GTA (8089G>A), CCG2637CCA (8095G>A), GAG2638GAA (8098G>A), AGC2641AGT (8107C>T), ATT2642ATC (8110T>C), CTC2643CTT (8113C>T), GAT2644GAC (8116T>C), CAA2648CAG (8128A>G), GGT2654GGC (8146T>C), AAC2656AGT (8151A>G 8152C>T), TCG2660TCA (8164G>A), AGG2665AAG (8178G>A), AAG2666AAA (8182G>A), TTG2669TTA (8191G>A), TTC2671TTT (8197C>T), TCC2673TCT (8203C>T), AGA2675AAA (8208G>A), CTG2677TTG (8213C>T), CTA2678CTG (8218A>G), ATC2679ATT (8221C>T), GAG2680GAA (8224G>A), CCA2685CCG (8239A>G), ATT2692ATC (8260T>C), GTA2693GTG (8263A>G), GAT2699GAC (8281T>C), AGA2700AGG (8284A>G), CCA2704GCG (8294C>G 8296A>G), GAG2705GAA (8299G>A), CAC2706CAT (8302C>T), GCG2710GCA (8314G>A), ATT2711ATC (8317T>C), TGC2712TGT (8320C>T), GCT2714GCA (8326T>A), GAG2717GAA (8335G>A), TCC2718TCT (8338C>T), TCT2722TTT (8349C>T), GAA2723GAG (8353A>G), CAC2726CAT (8362C>T), CAA2727CAG (8365A>G), ATC2728ATT (8368C>T), TAC2732TAT (8380C>T), TCA2733TCG (8383A>G), TTA2735TTG (8389A>G), CCT2740CCC (8404T>C), GCA2742TCA (8408G>T), ACA2743ACG (8413A>G), GCG2745GCT (8419G>T), GCT2750GCC (8434T>C), TAT2752TAC (8440T>C), ATG2756ATA (8452G>A), CTA2758CTG (8458A>G), AGG2759AAG (8460G>A), AAA2760AAG (8464A>G), CTG2761CTA (8467G>A), GCT2766ACA (8480G>A 8482T>A), GAG2771GAA (8497G>A), CTA2772TTG (8498C>T 8500A>G), AGA2773AAG (8502G>A 8503A>G), GCC2774GCT (8506C>T), GAA2777GAT (8515A>T), GTC2780GTT (8524C>T), GCA2781GCC (8527A>C), TTA2782CTG (8528T>C 8530A>G), GAG2785GAC (8539G>C), CTT2788TGT (8546C>T 8547T>G), TCT2790ACT (8552T>A), GTA2793GTG (8563A>G), GCA2797GGA (8574C>G), ATT2801GTT (8585A>G), GGA2805GAA (8598G>A), AGC2806AGT (8602C>T), AAC2807TCT (8603A>T 8604A>C 8605C>T), AAA2812AAG (8620A>G), CCA2813CAA (8622C>A), GAG2814GGA (8625A>G 8626G>A), CAG2815CAA (8629G>A), GGC2816GGT (8632C>T), ATC2818ATT (8638C>T), CAG2819CAA (8641G>A), AAC2821AAT (8647C>T), CCG2822TTC (8648C>T 8649C>T 8650G>C), AAC2823AAT (8653C>T), AAA2824AAG (8656A>G), GGA2825GAA (8658G>A), GAT2827GCG (8664A>C 8665T>G), GTT2830GTG (8674T>G), GCA2832GTT (8679C>T 8680A>T), GGC2833GGA (8683C>A), GGG2836GGA (8692G>A), ACA2837ACT (8695A>T), CCG2841CCA (8707G>A), AGA2842CGA (8708A>C), AAG2844AAA (8716G>A), ATC2846ATT (8722C>T), ACG2847ACA (8725G>A), AAA2849AAG (8731A>G), AGA2851AGG (8737A>G), CCC2853CCT (8743C>T), ACA2854AAG (8745C>A 8746A>G), AGC2855AGT (8749C>T), AAG2856AAA (8752G>A), GGA2857GGT (8755A>T), AAC2862AAT (8770C>T), TTA2863CTA (8771T>C), GAA2864GAG (8776A>G), CAT2865CAC (8779T>C), TTG2866TTA (8782G>A), TAT2869TAC (8791T>C), CCA2871CCG (8797A>G), CAA2872CAG (8800A>G), CAA2873CAG (8803A>G), GAT2875GAC (8809T>C), ATT2876ATC (8812T>C), AAT2878AAC (8818T>C), CGG2880CGA (8824G>A), GCA2881GCG (8827A>G), ACG2888ACA (8848G>A), TAT2890TAC (8854T>C), GAG2891GGA (8856A>G 8857G>A), GTG2893GTA (8863G>A), CGG2894CAG (8865G>A), ATG2895CTT (8867A>C 8869G>T), GCA2896GCG (8872A>G), GGA2900GAA (8883G>A), ACT2902TCT (8888A>T), TGC2915TGT (8929C>T), GTC2924ATC (8954G>A), AAC2925AAT (8959C>T), GGA2926GGT (8962A>T), GTT2929ATT (8969G>A), GGG2933GGA (8983G>A), AAT2934GAC (8984A>G 8986T>C), GAA2935GTA (8988A>T), GAG2938GAA (8998G>A), CCG2940CCA (9004G>A), TTG2941TTA (9007G>A), ATC2944ATT (9016C>T), ACC2951ACA (9037C>A), ATA2967ATT (9085A>T), AAC2971AAT (9097C>T), ATT2983GTT (9131A>G), CGA2984CGT (9136A>T), CGG2987CGT (9145G>T), ATG2989GGA (9149A>G 9150T>G 9151G>A), GGT2990AGT (9152G>A), TTA2991TTG (9157A>G), GCG2992GCT (9160G>T), CGT2993CGC (9163T>C), GAG3000GAA (9184G>A), CGA3004CGG (9196A>G), AGG3010AGA (9214G>A), GAA3011GAG (9217A>G), GCG3012GCA (9220G>A), GCA3020GCT (9244A>T), TTG3021TTA (9247G>A), GCC3024GCT (9256C>T), CCT3026TCC (9260C>T 9262T>C), GGG3030GGA (9274G>A), GAC3032GAT (9280C>T), ATC3035ATT (9289C>T), GAG3040GAA (9304G>A), GTC3050GTT (9334C>T), GTC3060GTG (9364C>G) | | | | | | | | |
| P1 protein (NP_734243.1) | 1 | 284 | 64.1% | 917 | 72.8% | 182 (100%) | 129 (70.9%) | 0/0/0/0 | 0 |
| Protein mutations: | Y4Q (194T>C 196C>A), C9L (209T>C 210G>T 211T>G), F13M (221T>A 223T>G), S23P (251T>C), C24F (255G>T 256C>T), E25G (258A>G), I27V (263A>G 265T>G), V28A (267T>C 268G>A), E30V (273A>T), A35T (287G>A 289T>C), V37T (293G>A 294T>C), A41I (305G>A 306C>T), D42G (309A>G), E44D (316A>T), T45V (317A>G 318C>T 319A>G), L52Q (339T>A 340C>A), K53R (342A>G), K55A (347A>G 348A>C 349A>G), Y56H (350T>C), T58V (356A>G 357C>T), V61T (365G>A 366T>C 367G>A), L62S (368C>T 369T>C), F67C (384T>G), A75V (408C>T 409C>T), M78E (416A>G 417T>A), E85K (437G>A), R86E (440A>G 441G>A 442G>A), K87R (443A>C 444A>G 445G>A), D88E (448T>A), M194K (765T>A), R197K (773C>A 774G>A 775A>G), M206K (801T>A), G210A (813G>C 814A>G), L211H (816T>A), R214Q (825G>A 826T>A), R226C (860C>T 862C>T), T227A (863A>G 865T>C), I228T (867T>C), N229D (869A>G 871C>T), I230L (872A>T 874A>G), R232K (879G>A), I238V (896A>G 898C>T), N240S (903A>G), T241N (906C>A 907A>C), K242T (909A>C 910A>C), S243N (912G>A 913C>T), S252L (938T>C 939C>T 940A>T), L266I (980T>A 982G>C), R272K (999G>A), Q275R (1007C>A 1008A>G 1009G>A), S276G (1010A>G 1012T>G), I277V (1013A>G), N279Q (1019A>C 1021C>G) | | | | | | | | |
| Codon mutations: | ACT3ACC (193T>C), TAC4CAA (194T>C 196C>A), TCA6TCG (202A>G), ATC8ATT (208C>T), TGT9CTG (209T>C 210G>T 211T>G), TCG12TCA (220G>A), TTT13ATG (221T>A 223T>G), GAA14GAG (226A>G), CTA17CTT (235A>T), TAC19TAT (241C>T), CCA21CCT (247A>T), TCT23CCT (251T>C), TGC24TTT (255G>T 256C>T), GAG25GGG (258A>G), CAT26CAC (262T>C), ATT27GTG (263A>G 265T>G), GTG28GCA (267T>C 268G>A), GAA30GTA (273A>T), CGA31CGT (277A>T), GAA32GAG (280A>G), GTG33GTT (283G>T), CCG34CCA (286G>A), GCT35ACC (287G>A 289T>C), GTT37ACT (293G>A 294T>C), CCT39CCC (301T>C), TTC40TTT (304C>T), GCA41ATA (305G>A 306C>T), GAT42GGT (309A>G), CTG43TTG (311C>T), GAA44GAT (316A>T), ACA45GTG (317A>G 318C>T 319A>G), CAA46CAG (322A>G), GCA49GCT (331A>T), CTC52CAA (339T>A 340C>A), AAG53AGG (342A>G), AAA55GCG (347A>G 348A>C 349A>G), TAT56CAT (350T>C), GCT57GCA (355T>A), ACT58GTT (356A>G 357C>T), CGT60CGC (364T>C), GTG61ACA (365G>A 366T>C 367G>A), CTC62TCC (368C>T 369T>C), AAA63AAG (373A>G), AAC64AAT (376C>T), TTT67TGT (384T>G), ACG68ACA (388G>A), TAC69TAT (391C>T), CGA70CGG (394A>G), TAC71TAT (397C>T), AAG72AAA (400G>A), GAT74GAC (406T>C), GCC75GTT (408C>T 409C>T), CAG76CAA (412G>A), ATA77ATT (415A>T), ATG78GAG (416A>G 417T>A), ATT80ATC (424T>C), CAG81CAA (427G>A), AAA83AAG (433A>G), CTG84CTC (436G>C), GAG85AAG (437G>A), AGG86GAA (440A>G 441G>A 442G>A), AAG87CGA (443A>C 444A>G 445G>A), GAT88GAA (448T>A), GAA90GAG (454A>G), ATG194AAG (765T>A), TTG196CTG (770T>C), CGA197AAG (773C>A 774G>A 775A>G), CGG198AGG (776C>A), TTC202TTT (790C>T), ATG206AAG (801T>A), GGA210GCG (813G>C 814A>G), CTT211CAT (816T>A), TTG212CTG (818T>C), CAA213CAG (823A>G), CGT214CAA (825G>A 826T>A), CGG217AGA (833C>A 835G>A), GAC219GAT (841C>T), AAT223AAC (853T>C), GTC225GTT (859C>T), CGC226TGT (860C>T 862C>T), ACT227GCC (863A>G 865T>C), ATC228ACC (867T>C), AAC229GAT (869A>G 871C>T), ATA230TTG (872A>T 874A>G), AGG232AAG (879G>A), GAT234GAC (886T>C), ATC238GTT (896A>G 898C>T), AAC240AGC (903A>G), ACA241AAC (906C>A 907A>C), AAA242ACC (909A>C 910A>C), AGC243AAT (912G>A 913C>T), GGC246GGA (922C>A), GGT249GGA (931T>A), TCA252CTT (938T>C 939C>T 940A>T), GGA253GGG (943A>G), TTC256TTT (952C>T), TCA261TCG (967A>G), GAA263GAG (973A>G), GGG264GGC (976G>C), TTG266ATC (980T>A 982G>C), GCA269GCG (991A>G), CGT270CGC (994T>C), AGA272AAA (999G>A), GTT273GTC (1003T>C), CAG275AGA (1007C>A 1008A>G 1009G>A), AGT276GGG (1010A>G 1012T>G), ATT277GTT (1013A>G), TTA278TTG (1018A>G), AAC279CAG (1019A>C 1021C>G), ATC282ATT (1030C>T) | | | | | | | | |
| HC-Pro protein (NP_734244.1) | 1 | 444 | 82.2% | 2446 | 90.7% | 374 (99.7%) | 351 (93.6%) | 0/1/1/1 | 0 |

| | Begin | End | Coverage | Score | Concordance | Matches | Identities | I/D/M/F* | Stop Codons |
|---|---|---|---|---|---|---|---|---|---|
| **NT** | **183** | **9687** | **89.3%** | **11427** | **66.3%** | **8654 (99.8%)** | **7216 (83.2%)** | **5/16** | |
| Protein mutations: | D4E (1048C>A), R16Q (1083G>A), Y19C (1092A>G 1093T>C), S21A (1097T>G 1099G>A), I98T (1329T>C), L113I (1373C>A), I117V (1385A>G), E123D (1405G>T), A127S (1415G>T 1417A>T), V134I (1436G>A 1438C>T), S177del (1565_1568delTCTT), E224G (1707A>G), A228T (1718G>A), I236V (1742A>G 1744C>T), S241N (1758G>A), V252I (1790G>A), I301V (1937A>G 1939T>C), F323Y (2004T>A), I344M (2068T>G), V346I (2072G>A 2074G>A), R391K (2208G>A), D410E (2266C>G) | | | | | | | | |
| Codon mutations: | TCG1TCA (1039G>A), AAT2AAC (1042T>C), GCC3GCA (1045C>A), GAC4GAA (1048C>A), AAT5AAC (1051T>C), AAG8AAA (1060G>A), GGT9GGA (1063T>A), CTG10TTG (1064C>T), GAC11GAT (1069C>T), CGA16CAA (1083G>A), TAT19TGC (1092A>G 1093T>C), TCG21GCA (1097T>G 1099G>A), CAC23CAT (1105C>T), AAT94..A (1318T>A), AAG95AAA (1321G>A), TTG97TTA (1327G>A), ATA98ACA (1329T>C), GCG99GCT (1333G>T), TTA100TTG (1336A>G), CTA103TTG (1343C>T 1345A>G), CCG106CCA (1354G>A), GTG107GTT (1357G>T), GAC108GAT (1360C>T), CTC111TTG (1367C>T 1369C>G), CTT113ATT (1373C>A), TTC114TTT (1378C>T), GAG116GAA (1384G>A), ATA117GTA (1385A>G), AAA119AAG (1393A>G), TCC120TCT (1396C>T), GAG123GAT (1405G>T), AAA124AAG (1408A>G), CAG125CAA (1411G>A), GCA127TCT (1415G>T 1417A>T), CCG128CCC (1420G>C), AAT131AAC (1429T>C), TTA132TTG (1432A>G), GTC134ATT (1436G>A 1438C>T), TTA135CTG (1439T>C 1441A>G), AAT136AAC (1444T>C), TTC139TTT (1453C>T), CTG140CTA (1456G>A), GAA144GAG (1468A>G), GTA152GTG (1492A>G), TTG155CTA (1499T>C 1501G>A), AGT156AGC (1504T>C), TTG157TTA (1507G>A), CTC158CTT (1510C>T), GAA159GAG (1513A>G), TTA160TTG (1516A>G), GCA161GCT (1519A>T), AGG162AGA (1522G>A), TTC163TTT (1525C>T), CAG164CAA (1528G>A), AAC166AAT (1534C>T), ACT168ACA (1540T>A), AAC170AAT (1546C>T), AAA173AAG (1555A>G), GAT175GAC (1561T>C), ATA176ATC (1564A>C), TCT177del (1565_1568delTCTT), TTC178-TC (1565_1568delTCTT), TTC179TTT (1573C>T), AGA180AGG (1576A>G), TTA183CTA (1583T>C), TCT184TCA (1588T>A), AAG186AAA (1594G>A), AAC188AAT (1600C>T), TCG194TCA (1618G>A), TGC195TGT (1621C>T), GAC196GAT (1624C>T), CTG199TTG (1631C>T), AAA201AAG (1639A>G), TTC205TTT (1651C>T), CTC206CTG (1654C>G), CAT213CAC (1675T>C), TCA219TCG (1693A>G), TTT222TTC (1702T>C), GAA224GGA (1707A>G), GCA228ACA (1718G>A), GGA230GGG (1726A>G), TAC231TAT (1729C>T), TCA232TCC (1732A>C), TAT234TAC (1738T>C), ATC236GTT (1742A>G 1744C>T), CGC237CGT (1747C>T), CAT239CAC (1753T>C), AGT241AAT (1758G>A), AGG244AGA (1768G>A), AAC250AAT (1786C>T), TTA251TTG (1789A>G), GTT252ATT (1790G>A), GTC253GTT (1795C>T), TTA257CTG (1805T>C 1807A>G), GAG259GAA (1813G>A), AGG261AGA (1819G>A), AAG263AAA (1825G>A), GAC267GAT (1837C>T), TAT268TAC (1840T>C), CAA271CAG (1849A>G), GTC274GTG (1858C>G), AAA276AAG (1864A>G), TGC278TGT (1870C>T), AGT280AGC (1876T>C), TCG281TCT (1879G>T), AAA282AAG (1882A>G), GGT284GGA (1888T>A), AAT285AAC (1891T>C), TGT290TGC (1906T>C), ACA293ACT (1915A>T), GGT298GGC (1930T>C), GCC300GCT (1936C>T), ATT301GTC (1937A>G 1939T>C), TCA303TCG (1945A>G), TTC305TTT (1951C>T), CCA308CCT (1960A>T), GTA314GTG (1978A>G), ATT315ATA (1981T>A), GGC316GGT (1984C>T), AAT317AAC (1987T>C), GAC320GAT (1996C>T), TTT323TAT (2004T>A), AAA328AAG (2020A>G), GGG329GGA (2023G>A), GAT330GAC (2026T>C), TCG331TCT (2029G>T), GAG332GAA (2032G>A), TAT341TAC (2059T>C), ATT344ATG (2068T>G), AAC345AAT (2071C>T), GTG346ATA (2072G>A 2074G>A), CTG351TTA (2087C>T 2089G>A), AGC355AGT (2101C>T), GAG356GAA (2104G>A), GAG357GAA (2107G>A), AAG360AAA (2116G>A), TTC362TTT (2122C>T), CGC367CGT (2137C>T), CTT374CTC (2158T>C), ACC376ACT (2164C>T), CCA378CCG (2170A>G), GAT382GAC (2182T>C), GCG384GCA (2188G>A), ACC385ACA (2191C>A), AGA391AAA (2208G>A), TAT394TAC (2218T>C), GAC396GAT (2224C>T), GTA397GTT (2227A>T), TTG402TTA (2242G>A), CCC403CCT (2245C>T), TTG406TTA (2254G>A), GTT407GTC (2257T>C), GAC408GAT (2260C>T), GAC410GAG (2266C>G), CAA412CAG (2272A>G), ACG413ACA (2275G>A), TCA419TCG (2293A>G), GGC421GGT (2299C>T), TCG422TCA (2302G>A), CAG423CAA (2305G>A), GGA426GGT (2314A>T), TAT427TAC (2317T>C), ATT429ATC (2323T>C), CTA430CTG (2326A>G), AGC434AGT (2338C>T), GTG435GTA (2341G>A), CAA437CAG (2347A>G), CTT438CTC (2350T>C), TTG440TTA (2356G>A), AAT443AAC (2365T>C) | | | | | | | | |
| P3 protein (NP_734251.1) | 25 | 365 | 93.4% | 1779 | 78.7% | 341 (99.7%) | 305 (89.2%) | 1/0/2/2 | 0 |
| Protein mutations: | K40R (2523A>G), R77K (2634G>A), A87V (2664C>T 2665T>C), V107I (2723G>A), A115T (2747G>A 2749T>A), H131N (2795C>A), V140I (2822G>A 2824G>A), P160T (2882C>A), S161H (2885A>C 2886G>A), T164M (2895C>T), N176S (2931A>G), N179S (2940A>G), E189_N190insX (2971_2972insA), T196I (2991C>T), H197R (2994A>G), E199R (2999G>A 3000A>G), R206Q (3021G>A), Y207F (3024A>T 3025C>T), T211I (3036C>T), E212G (3039A>G 3040A>C), K213T (3042A>C 3043G>A), A232V (3099C>T 3100C>A), V234R (3104G>A 3105T>G 3106G>A), S243N (3132G>A), R245K (3137C>A 3138G>A 3139A>G), F250I (3152T>A 3154C>T), C254Y (3165G>A), F272L (3218T>C), V276I (3230G>A), V288M (3266G>A 3268A>G), V291M (3275G>A), A294T (3284G>A), R304K (3315G>A 3316A>G), E305A (3318A>C), M319V (3359A>G), D332E (3400T>G) | | | | | | | | |
| Codon mutations: | TCT24.CC (2476T>C), TCG26TCA (2482G>A), CTG29CTA (2491G>A), CTG31CTT (2497G>T), TTG33TTA (2503G>A), AGA38AGG (2518A>G), CCT39CCC (2521T>C), AAG40AGG (2523A>G), AGA43AGG (2533A>G), TTG45TTA (2539G>A), TTA47TTG (2545A>G), GAG49GAA (2551G>A), CCT50CCA (2554T>A), CTG52TTG (2558C>T), TTG53CTC (2561T>C 2563G>C), CTA55CTG (2569A>G), TCA56TCG (2572A>G), CCT60CCC (2584T>C), GGC61GGT (2587C>T), CTG63CTC (2593G>C), GCT65GCC (2599T>C), TAT67TAC (2605T>C), AAT68AAC (2608T>C), TTT72TTC (2620T>C), GCG75GCA (2629G>A), GTG76GTA (2632G>A), AGG77AAG (2634G>A), TTG78TTA (2638G>A), ATT80ATA (2644T>A), AAA83AAG (2653A>G), GCT87GTC (2664C>T 2665T>C), CTA92TTA (2678C>T), CTA93CTG (2683A>G), GCT95GCC (2689T>C), CTA98CTC (2698A>C), TCA101TCG (2707A>G), GCG102GCA (2710G>A), GAA104GAG (2716A>G), ACA105ACG (2719A>G), CTC106CTT (2722C>T), GTC107ATC (2723G>A), GCA108GCG (2728A>G), AGG110AGA (2734G>A), ATT111ATA (2737T>A), GCT115ACA (2747G>A 2749T>A), GCA116GCT (2752A>T), GCT117GCA (2755T>A), GAC119GAT (2761C>T), CTC120CTT (2764C>T), TGT125TGC (2779T>C), GGG127GGT (2785G>T), AAC129AAT (2791C>T), CAT131AAT (2795C>A), TAC134TAT (2806C>T), TTG138CTC (2816T>C 2818G>C), GTG140ATA (2822G>A 2824G>A), TTG141TTA (2827G>A), GTT144GTC (2836T>C), AAG145AAA (2839G>A), AAT146AAC (2842T>C), ACC153ACA (2863C>A), CTA154TTA (2864C>T), CCA160ACA (2882C>A), AGT161CAT (2885A>C 2886G>A), AAC163AAT (2893C>T), ACG164ATG (2895C>T), GTC166GTT (2902C>T), GTA167GTG (2905A>G), ATT169ATC (2911T>C), AAT173AAC (2923T>C), AAT176AGT (2931A>G), AAC179AGC (2940A>G), GCT181GCC (2947T>C), ACT186ACC (2962T>C), GAA189_AAT190insA-- (2971_2972insA), GGT195GGC (2989T>C), ACT196ATT (2991C>T), CAT197CGT (2994A>G), GAG199AGG (2999G>A 3000A>G), TCC204TC- (3016delC), CGG206CAG (3021G>A), TAC207TTT (3024A>T 3025C>T), ACA211ATA (3036C>T), GAA212GGC (3039A>G 3040A>C), AAG213ACA (3042A>C 3043G>A), TTG216TTA (3052G>A), TTA219TTG (3061A>G), GCG226GCA (3082G>A), GGC229GGT (3091C>T), GCC232GTA (3099C>T 3100C>A), GTG234AGA (3104G>A 3105T>G 3106G>A), GTC235GTT (3109C>T), AAA236AAG (3112A>G), ACT238ACC (3118T>C), GCC239GCT (3121C>T), GGA241GGG (3127A>G), AGC243AAC (3132G>A), CGA245AAG (3137C>A 3138G>A 3139A>G), TAT249TAC (3151T>C), TTC250ATT (3152T>A 3154C>T), TGT254TAT (3165G>A), GTA255GTG (3169A>G), ATC265ATT (3199C>T), AGG267CGG (3203A>C), CGT268CGC (3208T>C), CCA270CCG (3214A>G), ACC271ACT (3217C>T), TTT272CTT (3218T>C), GTC273GTA (3223C>A), GTT276ATT (3230G>A), AAC277AAT (3235C>T), TTA285CTA (3257T>C), ACC286ACT (3262C>T), AGC287AGT (3265C>T), GTA288ATG (3266G>A 3268A>G), GTG289GTA (3271G>A), GTG291ATG (3275G>A), GCA294ACA (3284G>A), CAG299CAA (3301G>A), AGA304AAG (3315G>A 3316A>G), GAA305GCA (3318A>C), ATC306ATT (3322C>T), GAG307GAA (3325G>A), TTG308TTA (3328G>A), ATA311ATT (3337A>T), GAG315GAA (3349G>A), GTC317GTT (3355C>T), TGC318TGT (3358C>T), ATG319GTG (3359A>G), CTA321CTG (3367A>G), TAT322TAC (3370T>C), TTA325CTG (3377T>C 3379A>G), GAA330GAG (3394A>G), CGC331CGT (3397C>T), GAT332GAG (3400T>G), ACA334ACG (3406A>G), TAC338TAT (3418C>T), ATT339ATA (3421T>A), TAT341TAC (3427T>C), AAG343AAA (3433G>A), TCA344TCT (3436A>T), GTA345GTG (3439A>G), AAC346AAT (3442C>T), CCT347CCC (3445T>C), CAG351CAA (3457G>A), TTT352TTC (3460T>C), GCT353GCA (3463T>A), GCG355GCT (3469G>T), CAG356CAA (3472G>A), CAC364CAT (3496C>T), CAG365CAA (3499G>A) | | | | | | | | |
| 6K1 protein (NP_734245.1) | 1 | 52 | 100% | 305 | 90.5% | 52 (100%) | 45 (86.5%) | 0/0/0/0 | 0 |
| Protein mutations: | V5G (3513T>G), M16V (3545A>G), A17T (3548G>A), V19I (3554G>A 3556C>T), L42V (3623C>G), L45M (3632C>A), Y47H (3638T>C) | | | | | | | | |
| Codon mutations: | TCC2TCT (3505C>T), GTT5GGT (3513T>G), AAA7AAG (3520A>G), GAA10GAG (3529A>G), CAA11CAG (3532A>G), ATG16GTG (3545A>G), GCT17ACT (3548G>A), TTA18CTA (3551T>C), GTC19ATT (3554G>A 3556C>T), AGT28AGC (3583T>C), GAT29GAC (3586T>C), TGC30TGT (3589C>T), TTC32TTT (3595C>T), AAT36AAC (3607T>C), TTT38TTC (3613T>C), AAG39AAA (3616G>A), GGT40GGC (3619T>C), CTT42GTT (3623C>G), TCC43TCT (3628C>T), CTG45ATG (3632C>A), GAT46GAC (3637T>C), TAT47CAT (3638T>C), GAA48GAG (3643A>G) | | | | | | | | |
| CI protein (NP_734246.1) | 1 | 634 | 88.6% | 3611 | 96.1% | 562 (100%) | 532 (94.7%) | 0/0/0/0 | 1 |
| Protein mutations: | K7N (3676G>C), N8K (3679T>A), N13K (3694T>A), I15V (3698A>G), D17* (3704G>T 3706T>A), F18L (3709T>A), I140V (4073A>G), F158Y (4128T>A 4129C>T), A166S (4151G>T), V171I (4166G>A), L190I (4223C>A 4225G>A), I222V (4319A>G 4321A>G), F247Y (4395T>A), V251I (4406G>A), M274V (4475A>G 4477G>T), T276S (4481A>T), C287R (4514T>C), V342I (4679G>A 4681G>C), V381I (4796G>A 4798C>T), T383S (4803C>G 4804T>C), S468G (5057A>G), T469A (5060A>G), L479I (5090C>A), A482V (5100C>T), E484D (5107A>T), V489F (5120G>T 5122C>T), I491V (5126A>G), V510I (5183G>A) | | | | | | | | |

| | Begin | End | Coverage | Score | Concordance | Matches | Identities | I/D/M/F* | Stop Codons |
|---|---|---|---|---|---|---|---|---|---|
| **NT** | **183** | **9687** | **89.3%** | **11427** | **66.3%** | **8654 (99.8%)** | **7216 (83.2%)** | **5/16** | |
| Codon mutations: | TTA2TTG (3661A>G), GTG5GTA (3670G>A), AAG7AAC (3676G>C), AAT8AAA (3679T>A), GAG11GAA (3688G>A), AAT13AAA (3694T>A), GAG14GAA (3697G>A), ATT15GTT (3698A>G), GAT17TAA (3704G>T 3706T>A), TTT18TTA (3709T>A), AGC76..T (3883C>T), GAA77GAG (3886A>G), CTA79CTG (3892A>G), GAC80GAT (3895C>T), TTT81TTC (3898T>C), GTA83GTG (3904A>G), CGG84AGA (3905C>A 3907G>A), GCT86GCC (3913T>C), AAG91AAA (3928G>A), TCA92TCT (3931A>T), GGG94GGA (3937G>A), AGT100AGC (3955T>C), GTG101GTA (3958G>A), GCC102GCT (3961C>T), GGA103GGG (3964A>G), TCT104TCC (3967T>C), TTA107TTG (3976A>G), ATT108ATC (3979T>C), CCA110CCT (3985A>T), CTA114CTT (3997A>T), GCG115GCA (4000G>A), AAC117AAT (4006C>T), GTT118GTG (4009T>G), TTC119TTT (4012C>T), AAA120AAG (4015A>G), TCT123TCC (4024T>C), GAA125GAG (4030A>G), CCA126CCT (4033A>T), AAG129AAA (4042G>A), CTG133TTA (4052C>T 4054G>A), CGT134CGC (4057T>C), CGT136CGA (4063T>A), GGA137GGG (4066A>G), ATA140GTA (4073A>G), TTT141TTC (4078T>C), GGC142GGT (4081C>T), TCT143TCC (4084T>C), TCC147TCT (4096C>T), AGC151AGT (4108C>T), GGA152GGT (4111A>T), GCG154GCA (4117G>A), CTA155TTG (4118C>T 4120A>G), TAC157TAT (4126C>T), TTC158TAT (4128T>A 4129C>T), GCC159GCT (4132C>T), AAT160AAC (4135T>C), CAA164CAG (4147A>G), TTA165CTA (4148T>C), GCT166TCT (4151G>T), AAC169AAT (4162C>T), GTA171ATA (4166G>A), ATA172ATC (4171A>C), GAG175GAA (4180G>A), CTG179TTG (4190C>T), CCT181CCC (4198T>C), TCC182TCT (4201C>T), GCG185GCT (4210G>T), TTC186TTT (4213C>T), AGT188AGC (4219T>C), CTG189TTG (4220C>T), CTG190ATA (4223C>A 4225G>A), GTT192GTG (4231T>G), CAT194CAC (4237T>C), GTA199GTG (4252A>G), TTA200CTG (4253T>C 4255A>G), AAA201AAG (4258A>G), ACT205ACC (4270T>C), AGA209AGG (4282A>G), GAG210GAA (4285G>A), GAA212GAG (4291A>G), CAG216CAA (4303G>A), GTC219GTT (4312C>T), ATA222GTG (4319A>G 4321A>G), GTG223GTT (4324G>T), GAG224GAA (4327G>A), CTG227CTT (4336G>T), TCT228TCC (4339T>C), TCA231TCT (4348A>T), GCA235GCG (4360A>G), GGT237GGC (4366T>C), TCT238TCA (4369T>A), ACT240ACC (4375T>C), GTT244GTC (4387T>C), TTT247TAT (4395T>A), GTA251ATA (4406G>A), ACC263ACA (4444C>A), GCC265GCT (4450C>T), CTC267CTT (4456C>T), GAC270GAT (4465C>T), ATG274GTT (4475A>G 4477G>T), GTC275GTT (4480C>T), ACA276TCA (4481A>T), GGC280GGT (4495C>T), AAG284AAA (4507G>A), CAC285CAT (4510C>T), GGT286GGG (4513T>G), TGC287CGC (4514T>C), CTA288TTA (4517C>T), GTC291GTT (4528C>T), AAA293AAG (4534A>G), GTG312GTA (4591G>A), TTG314TTA (4597G>A), GAC315GAT (4600C>T), GAC317GAT (4606C>T), GTG318GTA (4609G>A), GTG320GTA (4615G>A), GAC321GAT (4618C>T), TTG324CTT (4625T>C 4627G>T), GAC331GAT (4648C>T), ATT332ATC (4651T>C), AGG335AGA (4660G>A), AGC336AGT (4663C>T), GCT338GCA (4669T>A), GTG342ATC (4679G>A 4681G>C), AGT343AGC (4684T>C), GTT344GTG (4687T>G), GAG348GAA (4699G>A), AGA349AGG (4702A>G), CAA351CAG (4708A>G), AGG352AGA (4711G>A), CTG353TTG (4712C>T), AAA361AAG (4738A>G), GCA364GCG (4747A>G), TTG365CTG (4748T>C), ACT370ACC (4765T>C), GTC381ATT (4796G>A 4798C>T), ACT383AGC (4803C>G 4804T>C), GAG384GAA (4807G>A), GCG385GCT (4810G>T), GCT386GCG (4813T>G), CTT387CTC (4816T>C), TAT392TAC (4831T>C), AAC393AAT (4834C>T), CCA395CCT (4840A>T), GGA399GGG (4852A>G), GGC400GGT (4855C>T), AGT404AGC (4867T>C), CTG405CTC (4870G>C), ATT406ATA (4873T>A), ACA416ACT (4903A>T), CAG418CAA (4909G>A), TTT420TTC (4915T>C), GAA421GAG (4918A>G), TTG422CTA (4919T>C 4921G>A), AGT423AGC (4924T>C), CCC424CCG (4927C>G), TTT426TTC (4933T>C), ATC427ATA (4936C>A), CAG428CAA (4939G>A), AAT429AAC (4942T>C), TTC430TTT (4945C>T), TCA436TCG (4963A>G), CCT439CCC (4972T>C), CTT445CTC (4990T>C), AAA449AAG (5002A>G), CCT456CCC (5023T>C), TTG457CTA (5024T>C 5026G>A), TGC458TGT (5029C>T), GAT459GAC (5032T>C), TCT461TCC (5038T>C), CCA463CCT (5044A>T), GCA466GCT (5053A>T), TCG467TCA (5056G>A), AGT468GGT (5057A>G), ACT469GCT (5060A>G), TCG472TCT (5071G>T), CTT479ATT (5090C>A), GGA480GGG (5095A>G), GTG481GTT (5098G>T), GCC482GTC (5100C>T), TTA483TTG (5104A>G), GAA484GAT (5107A>T), ATT485ATC (5110T>C), CCA486CCG (5113A>G), AAG487AAA (5116G>A), CAA488CAG (5119A>G), GTC489TTT (5120G>T 5122C>T), ATT491GTT (5126A>G), CAT494CAC (5137T>C), ATC495ATT (5140C>T), AAA496AAG (5143A>G), GAG497GAA (5146G>A), CTC502CTA (5161C>A), GAA504GAG (5167A>G), CTT506CTG (5173T>G), ACG509ACA (5182G>A), GTT510ATT (5183G>A), GTC511GTT (5188C>T), AAG512AAA (5191G>A), AAA514AAG (5197A>G), GAC515GAT (5200C>T), TGC517TGT (5206C>T), TTA518TTG (5209A>G), TCG525TCA (5230G>A), ATC527ATT (5236C>T), AGC528AGT (5239C>T), ATC530ATT (5245C>T), ACA536ACT (5263A>T), CTC538CTT (5269C>T), TTC539TTT (5272C>T), GCC540GCA (5275C>A), ATC541ATT (5278C>T), CCA542CCC (5281A>C), AGA543AGG (5284A>G), ACT544ACC (5287T>C), CTA545TTA (5288C>T), ATA546ATT (5293A>T), TTG547CTA (5294T>C 5296G>A), GAG549GAA (5302G>A), CTT552CTA (5311T>A), GAA553GAG (5314A>G), CGA556CGT (5323A>T), AGA563AGG (5344A>G), AGT564AGC (5347T>C), ATC566ATT (5353C>T), TGC570TGT (5365C>T), GTT577GTC (5386T>C), ACC580ACA (5395C>A), AAA589AAG (5422A>G), GAT590GAC (5425T>C), ACC592ACT (5431C>T), GAG594GAA (5437G>A), CTT599CTC (5452T>C), AAG601AAA (5458G>A), AGT604AGC (5467T>C), CAA605CAG (5470A>G), CTA606TTG (5471C>T 5473A>G), TTC609TTT (5482C>T), TCA610TCT (5485A>T), AAT611AAC (5488T>C), TTG612CTA (5489T>C 5491G>A), GGT614GGC (5497T>C), TGT617TGC (5506T>C), GAG618GAA (5509G>A), GAG619GAA (5512G>A), TTA621CTA (5516T>C), TAT625TAC (5530T>C), GAG626GAA (5533G>A), TCG627TCT (5536G>T), TTG628TTA (5539G>A), GTT631GTG (5548T>G), CAC633CAT (5554C>T) | | | | | | | | |
| 6K2 protein (NP_734247.1) | 1 | 52 | 100% | 303 | 93.5% | 52 (100%) | 46 (88.5%) | 0/0/0/0 | 0 |
| Protein mutations: | A2T (5561G>A 5563G>A), I14V (5597A>G), N16K (5605C>A), A21V (5619C>T 5620T>G), I25L (5630A>C), V46I (5693G>A) | | | | | | | | |
| Codon mutations: | GCT1GCA (5560T>A), GCG2ACA (5561G>A 5563G>A), ACG3ACC (5566G>C), TCA4TCG (5569A>G), CTC9CTG (5584C>G), AAG12AAA (5593G>A), GGG13GGA (5596G>A), ATT14GTT (5597A>G), AAC16AAA (5605C>A), GTG20GTT (5617G>T), GCT21GTG (5619C>T 5620T>G), TTG24TTA (5629G>A), ATC25CTC (5630A>C), GGC28GGT (5641C>T), GCA31GCT (5650A>T), ATA35ATT (5662A>T), GGA36GGG (5665A>G), CAA44CAG (5689A>G), GTT46ATT (5693G>A), GAG47GAA (5698G>A), CAA52CAG (5713A>G) | | | | | | | | |
| NIa-VPg protein (NP_734252.1) | 1 | 188 | 100% | 1230 | 93.5% | 188 (100%) | 171 (91.0%) | 0/0/0/0 | 0 |
| Protein mutations: | I61V (5894A>G 5896C>G), R83A (5960C>G 5961G>C 5962G>C), I93V (5990A>G), E102D (6019A>T), K106T (6030A>C), D112E (6049C>A), M115I (6058G>A), S120N (6072G>A 6073T>C), N121H (6074A>C), T123N (6081C>A), C133S (6111G>C 6112T>C), I139V (6128A>G 6130T>C), V149I (6158G>A), T153S (6170A>T 6172A>G), L164F (6203C>T 6205C>T), V173I (6230G>A), A182K (6257G>A 6258C>A) | | | | | | | | |
| Codon mutations: | GGG1GGC (5716G>C), AAA2AAG (5719A>G), GCC10GCA (5743C>A), CGC14CGA (5755C>A), CAT15CAC (5758T>C), CGT17CGC (5764T>C), AAA19AAG (5770A>G), GGC22GGT (5779C>T), TTT23TTC (5782T>C), GAC26GAT (5791C>T), GAC30GAT (5803C>T), AGG41AGA (5836G>A), AAA42AAG (5839A>G), GGA44GGG (5845A>G), AAA45AAG (5848A>G), GGT48GGC (5857T>C), ACC49ACT (5860C>T), ACA50ACT (5863A>T), GGT52GGA (5869T>A), GGT54GGC (5875T>C), AGC57AGT (5884C>T), TTC60TTT (5893C>T), ATC61GTG (5894A>G 5896C>G), AAC62AAT (5899C>T), GGG65GGA (5908G>A), TTT66TTC (5911T>C), GAG70GAA (5923G>A), CAA75CAG (5938A>G), TTC76TTT (5941C>T), GTT77GTC (5944T>C), GAT78GAC (5947T>C), GGG82GGA (5959G>A), CGG83GCC (5960C>G 5961G>C 5962G>C), GAA87GAG (5974A>G), GCT91GCA (5986T>A), GAC92GAT (5989C>T), ATT93GTT (5990A>G), GAT95GAC (5998T>C), AGA99CGT (6008A>C 6010A>T), GAA102GAT (6019A>T), GTG103GTT (6022G>T), CGA104CGT (6025A>T), AAA106ACA (6030A>C), GAC112GAA (6049C>A), ATG115ATA (6058G>A), CAA116CAG (6061A>G), GCC117GCA (6064C>A), GGT119GGC (6070T>C), AGT120AAC (6072G>A 6073T>C), AAC121CAC (6074A>C), ACG122ACA (6079G>A), ACC123AAC (6081C>A), ATA124ATT (6085A>T), GCA126GCT (6091A>T), TAC127TAT (6094C>T), TGT133TCC (6111G>C 6112T>C), AAA135AAG (6118A>G), TTG137TTA (6124G>A), AAG138AAA (6127G>A), ATT139GTC (6128A>G 6130T>C), GAT140GAC (6133T>C), TTA141TTG (6136A>G), AAC145AAT (6148C>T), AAA148AAG (6157A>G), GTT149ATT (6158G>A), TGT150TGC (6163T>C), AAA152AAG (6169A>G), ACA153TCG (6170A>T 6172A>G), GGC155GGT (6178C>T), AAA158AAG (6187A>G), TTT159TTC (6190T>C), GAG161GAA (6196G>A), CTC164TTT (6203C>T 6205C>T), GAA165GAG (6208A>G), CTA166TTG (6209C>T 6211A>G), CAG168CAA (6217G>A), GCT172GCA (6229T>A), GTA173ATA (6230G>A), GTC175GTT (6238C>T), AAG178AAA (6247G>A), ATA180ATT (6253A>T), GCA182AAA (6257G>A 6258C>A), CAT187CAC (6274T>C) | | | | | | | | |
| NIa-Pro protein (NP_734248.1) | 1 | 244 | 100% | 1651 | 93.5% | 244 (100%) | 222 (91.0%) | 0/0/0/0 | 0 |
| Protein mutations: | K2R (6282A>G), A29T (6362G>A), V42I (6401G>A), A43V (6405C>T), Y51F (6429A>T), Q58R (6450A>G), H70N (6485C>A 6487C>T), L84I (6527C>A), T104I (6588C>T), Y120H (6635T>C), I124V (6647A>G), I134V (6677A>G 6679A>G), D147N (6716G>A), N148D (6719A>G), I163L (6764A>T 6766A>G), A173V (6795C>T 6796A>G), H174Q (6799C>A), K189R (6843A>G), N194D (6857A>G), V200T (6875G>A 6876T>C 6877C>A), D219E (6934C>A), V242I (7001G>A 7003G>A) | | | | | | | | |
| Codon mutations: | AAA2AGA (6282A>G), TCG3TCA (6286G>A), GGC7GGT (6298C>T), TTG8TTA (6301G>A), AGA9AGG (6304A>G), GAC10GAT (6307C>T), AAC12AAT (6313C>T), GCC15GCT (6322C>T), ACA17ACT (6328A>T), GTT18GTG (6331T>G), TGT19TGC (6334T>C), AGG20AGA (6337G>A), CTG21TTG (6338C>T), TAT27TAC (6358T>C), GGG28GGA (6361G>A), GCA29ACA (6362G>A), TCA30TCT (6367A>T), GAG31GAA (6370G>A), TAC33TAT (6376C>T), GGT34GGG (6379T>G), GGA36GGC (6385A>C), GGA38GGT (6391A>T), GCA39GCG (6394A>G), TAC40TAT (6397C>T), GTA42ATA (6401G>A), GCG43GTG (6405C>T), CAC45CAT (6412C>T), TTA47TTG (6418A>G), TTT48TTC (6421T>C), TAC51TTC (6429A>T), GGT53GGA (6436T>A), GAG56GAA (6445G>A), CAA58CGA (6450A>G), TCC59TCA (6454C>A), CAC61CAT (6460C>T), GGT62GGA (6463T>A), TTC64TTT (6469C>T), AGG65AGA (6472G>A), GTG66GTA (6475G>A), AAT68AAC (6481T>C), CAC70AAT (6485C>A 6487C>T), AGT71AGC (6490T>C), AGC73AGT (6496C>T), CTG75CTA (6502G>A), GGT79GGC (6514T>C), AGG80AGA (6517G>A), CTC84ATC (6527C>A), ATC85ATA (6532C>A), CCG88CCA (6541G>A), AAA89AAG (6544A>G), GTC93GTT (6556C>T), TTT94TTC (6559T>C), CAG96CAA (6565G>A), TTG98CTA (6569T>C 6571G>A), CCT103CCA (6586T>A), ACA104ATA (6588C>T), GAA107GAG (6598A>G), AGA108AGG (6601A>G), ATT109ATC (6604T>C), TGT110TGC (6607T>C), GGA113GGG (6616A>G), AAC115AAT (6622C>T), GAG118GAA (6631G>A), AAG119AAA (6634G>A), TAT120CAT (6635T>C), GCT121GCA (6640T>A), TCG123TCC (6646G>C), ATC124GTC (6647A>G), ATC125ATT (6652C>T), AGC129AGT (6664C>T), ACT131ACG (6670T>G), ATA134GTG (6677A>G 6679A>G), GGC136GGT (6685C>T), ACA138ACT (6691A>T), TTC139TTT (6694C>T), ACA146ACG (6715A>G), GAT147AAT (6716G>A), AAT148GAT (6719A>G), GGA149GGG (6724A>G), CTA153TTA (6734C>T), GTG155GTA (6742G>A), GTG156GTA (6745G>A), GCC159GCT (6754C>T), GGA161GGG (6760A>G), ATA163TTG (6764A>T 6766A>G), GTC164GTT (6769C>T), ATT166ATA (6775T>A), CTG169TTG (6782C>T), GCA170GCT (6787A>T), AAC171AAT (6790C>T), GCA173GTG (6795C>T 6796A>G), CAC174CAA (6799C>A), ACC175ACA (6802C>A), AAC177AAT (6808C>T), TAC178TAT (6811C>T), TAC179TAT (6814C>T), TTC182TTT (6823C>T), GAA184GAG (6829A>G), AAG189AGG (6843A>G), TAC190TAT (6847C>T), CGA192CGC (6853A>C), AAT194GAT (6857A>G), GAG195GAA (6862G>A), GAA198GAG (6871A>G), GTC200ACA (6875G>A 6876T>C 6877C>A), AAG201AAA (6880G>A), TCT202TCA (6883T>A), GTT204GTG (6889T>G), GAC208GAT (6901C>T), GGC213GGT (6916C>T), CCG214CCA (6919G>A), AAA216AAG (6925A>G), CTT217CTC (6928T>C), GAC219GAA (6934C>A), GGG224GGC (6949G>C), TTC226TTT (6955C>T), AAA227AAG (6958A>G), CAA233CAG (6976A>G), CTA235TTA (6980C>T), ATC236ATT (6985C>T), GAT239GAC (6994T>C), GTA240GTT (6997A>T), GTG241GTT (7000G>T), GTG242ATA (7001G>A 7003G>A) | | | | | | | | |
| NIb protein (NP_734249.1) | 1 | 521 | 100% | 3129 | 86.1% | 519 (99.4%) | 480 (92.0%) | 1/2/2/2 | 0 |
| Protein mutations: | F8Y (7032T>A), T39K (7125C>A), A46S (7145G>T 7147A>G), E51_A52del (7160_7165delGAGGCA), D69E (7216T>A), D83E (7258T>A), D88_R89insX (7273_7274insT), R89V (7274C>G 7275G>T), H91L (7281A>T), L92S (7284T>C), S96L (7296C>T), S101F (7311C>T), T102I (7314C>T), C103Y (7317G>A), N104K (7321T>A), K109Q (7334A>C), A112N (7343G>A 7344C>A 7345A>C), S133M (7407G>T 7408T>G), C136G (7415T>G), L165S (7503T>C 7504G>A), D234N (7709G>A), K235R (7713A>G 7714A>G), T268I (7812C>T), V279L (7844G>T 7846G>A), V292I (7883G>A), S345N (8043G>A), N381S (8151A>G 8152C>T), R390K (8178G>A), R400K (8208G>A), P429A (8294C>G 8296A>G), S447F (8349C>T), A467S (8408G>T), M481I (8452G>A), R484K (8460G>A), A491T (8480G>A 8482T>A), R498K (8502G>A 8503A>G), E502D (8515A>T), E510D (8539G>C), L513C (8546C>T 8547T>G), S515T (8552T>A) | | | | | | | | |

| | Begin | End | Coverage | Score | Concordance | Matches | Identities | I/D/M/F* | Stop Codons |
|---|---|---|---|---|---|---|---|---|---|
| **NT** | **183** | **9687** | **89.3%** | **11427** | **66.3%** | **8654 (99.8%)** | **7216 (83.2%)** | **5/16** | |
| Codon mutations: | GCT1GCC (7012T>C), AAG2AAA (7015G>A), CAT3CAC (7018T>C), TTT8TAT (7032T>A), GCC10GCT (7039C>T), TTG11CTA (7040T>C 7042G>A), GGA13GGG (7048A>G), GCT17GCA (7060T>A), GTC18GTA (7063C>A), GCA19GCG (7066A>G), AGC23AGT (7078C>T), CAA24CAG (7081A>G), TTA25CTG (7082T>C 7084A>G), GTA26GTG (7087A>G), ACC27ACA (7090C>A), CAT29CAC (7096T>C), GTA30GTG (7099A>G), GTT31GTC (7102T>C), GAG34GAA (7111G>A), CGA36CGG (7117A>G), ACA39AAA (7125C>A), GAA40GAG (7129A>G), CTG42TTA (7133C>T 7135G>A), GTG44GTA (7141G>A), GAT45GAC (7144T>C), GCA46TCG (7145G>T 7147A>G), GAG47GAA (7150G>A), GAG49GAA (7156G>A), GCA50GCT (7159A>T), GAG51_GCA52del (7160_7165delGAGGCA), TTC54TTT (7171C>T), GCG60GCT (7189G>T), AAA63AAG (7198A>G), AGC64AGT (7201C>T), CTA66TTA (7205C>T), AAT67AAC (7210T>C), GAT69GAA (7216T>A), GCG70GCA (7219G>A), ATC72ATA (7225C>A), AAG73AAA (7228G>A), AAG77AAA (7240G>A), TAT78TAC (7243T>C), AAA80AAG (7249A>G), CCT81CCG (7252T>G), GAT83GAA (7258T>A), GTT84GTC (7261T>C), GGT85GGA (7264T>A), GTC86GTG (7267C>G), GTG87GTA (7270G>A), GAT88_CGG89insT-- (7273_7274insT), CGG89GTG (7274C>G 7275G>T), CAT91CTT (7281A>T), TTG92TCG (7284T>C), TCA96TTA (7296C>T), TCT101TTT (7311C>T), ACC102ATC (7314C>T), TGC103TAC (7317G>A), AAT104AAA (7321T>A), GTG105-TA (7322delG 7324G>A), CAC106CAT (7327C>T), GGC107GGT (7330C>T), TTC108TTT (7333C>T), AAG109CAG (7334A>C), TGT111TGC (7342T>C), GCA112AAC (7343G>A 7344C>A 7345A>C), ACT115ACC (7354T>C), CAA118CAG (7363A>G), ATT120ATC (7369T>C), TTC121TTT (7372C>T), GCG123GCT (7378G>T), CTC124CTT (7381C>T), GCA129GCT (7396A>T), GTC130GTT (7399C>T), GCC132GCT (7405C>T), AGT133ATG (7407G>T 7408T>G), GGG135GGT (7414G>T), TGC136GGC (7415T>G), AAA137AAG (7420A>G), GAC140GAT (7429C>T), TAT141TAC (7432T>C), CAT144CAC (7441T>C), AAG150AAA (7459G>A), GAA151GAG (7462A>G), ATA153ATT (7468A>T), GTC154GTA (7471C>A), CAA156CAG (7477A>G), AGC157AGT (7480C>T), CTG159CTT (7486G>T), TAT162TAC (7495T>C), AAA163AAG (7498A>G), GGT164GGC (7501T>C), TTG165TCA (7503T>C 7504G>A), ATT168ATA (7513T>A), AAC170AAT (7519C>T), GGA171GGG (7522A>G), AAG174AAA (7531G>A), GAG176GAA (7537G>A), CTC177CTT (7540C>T), TGT179TGC (7546T>C), AAG187AAA (7570G>A), ACG188ACA (7573G>A), ACG190ACA (7579G>A), GCT193GCC (7588T>C), CTA196TTA (7595C>T), TTG199CTA (7604T>C 7606G>A), GGT202GGA (7615T>A), TGT205TGC (7624T>C), GAC208GAT (7633C>T), TTC209TTT (7636C>T), AAT210AAC (7639T>C), TTT213TTC (7648T>C), GAA219GAG (7666A>G), TGT221TGC (7672T>C), ACA223ACT (7678A>T), GGG225GGA (7684G>A), ACT227ACC (7690T>C), TTT229TTC (7696T>C), GGT231GGG (7702T>G), GAT234AAT (7709G>A), AAA235AGG (7713A>G 7714A>G), CTG236TTG (7715C>T), CGT239CGC (7726T>C), GAG242GAA (7735G>A), GTA245GTG (7744A>G), TAC246TAT (7747C>T), GGC251GGT (7762C>T), CAG253CAA (7768G>A), TTT254TTC (7771T>C), TCA257TCG (7780A>G), CTA258CTC (7783A>C), ACT259ACC (7786T>C), CTA262TTG (7793C>T 7795A>G), ATC263ATT (7798C>T), GCT265GCC (7804T>C), ACC268ATC (7812C>T), GAA275GAG (7834A>G), GAC276GAT (7837C>T), GTG279TTA (7844G>T 7846G>A), CAG282CAA (7855G>A), CTG284CTA (7861G>A), TTA287TTG (7870A>G), TAC288TAT (7873C>T), ACT289ACA (7876T>A), GAG290GAA (7879G>A), ATT291ATA (7882T>A), GTT292ATT (7883G>A), CCA295CCT (7894A>T), ATT296ATC (7897T>C), GTC304GTT (7921C>T), AAG306AAA (7927G>A), GGT309GGC (7936T>C), AAT310AAC (7939T>C), AAT311AAC (7942T>C), GTG319GTA (7966G>A), GAT320GAC (7969T>C), TCT322TCC (7975T>C), GTC326GTT (7987C>T), CTT327CTC (7990T>C), GCT332GCC (8005T>C), GAA341GAG (8032A>G), ATC343ATT (8038C>T), GAC344GAT (8041C>T), AGC345AAC (8043G>A), TTG357TTA (8080G>A), GTG360GTA (8089G>A), CCG362CCA (8095G>A), GAG363GAA (8098G>A), AGC366AGT (8107C>T), ATT367ATC (8110T>C), CTC368CTT (8113C>T), GAT369GAC (8116T>C), CAA373CAG (8128A>G), GGT379GGC (8146T>C), AAC381AGT (8151A>G 8152C>T), TCG385TCA (8164G>A), AGG390AAG (8178G>A), AAG391AAA (8182G>A), TTG394TTA (8191G>A), TTC396TTT (8197C>T), TCC398TCT (8203C>T), AGA400AAA (8208G>A), CTG402TTG (8213C>T), CTA403CTG (8218A>G), ATC404ATT (8221C>T), GAG405GAA (8224G>A), CCA410CCG (8239A>G), ATT417ATC (8260T>C), GTA418GTG (8263A>G), GAT424GAC (8281T>C), AGA425AGG (8284A>G), CCA429GCG (8294C>G 8296A>G), GAG430GAA (8299G>A), CAC431CAT (8302C>T), GCG435GCA (8314G>A), ATT436ATC (8317T>C), TGC437TGT (8320C>T), GCT439GCA (8326T>A), GAG442GAA (8335G>A), TCC443TCT (8338C>T), TCT447TTT (8349C>T), GAA448GAG (8353A>G), CAC451CAT (8362C>T), CAA452CAG (8365A>G), ATC453ATT (8368C>T), TAC457TAT (8380C>T), TCA458TCG (8383A>G), TTA460TTG (8389A>G), CCT465CCC (8404T>C), GCA467TCA (8408G>T), ACA468ACG (8413A>G), GCG470GCT (8419G>T), GCT475GCC (8434T>C), TAT477TAC (8440T>C), ATG481ATA (8452G>A), CTA483CTG (8458A>G), AGG484AAG (8460G>A), AAA485AAG (8464A>G), CTG486CTA (8467G>A), GCT491ACA (8480G>A 8482T>A), GAG496GAA (8497G>A), CTA497TTG (8498C>T 8500A>G), AGA498AAG (8502G>A 8503A>G), GCC499GCT (8506C>T), GAA502GAT (8515A>T), GTC505GTT (8524C>T), GCA506GCC (8527A>C), TTA507CTG (8528T>C 8530A>G), GAG510GAC (8539G>C), CTT513TGT (8546C>T 8547T>G), TCT515ACT (8552T>A), GTA518GTG (8563A>G) | | | | | | | | |
| coat protein (NP_734250.1) | 1 | 267 | 100% | 1659 | 91.0% | 267 (100%) | 243 (91.0%) | 0/0/0/0 | 0 |
| Protein mutations: | A1G (8574C>G), I5V (8585A>G), G9E (8598G>A), N11S (8603A>T 8604A>C 8605C>T), P17Q (8622C>A), E18G (8625A>G 8626G>A), P26F (8648C>T 8649C>T 8650G>C), G29E (8658G>A), D31A (8664A>C 8665T>G), A36V (8679C>T 8680A>T), T58K (8745C>A 8746A>G), E95G (8856A>G 8857G>A), R98Q (8865G>A), M99L (8867A>C 8869G>T), G104E (8883G>A), T106S (8888A>T), V128I (8954G>A), V133I (8969G>A), N138D (8984A>G 8986T>C), E139V (8988A>T), I187V (9131A>G), M193G (9149A>G 9150T>G 9151G>A), G194S (9152G>A), P230S (9260C>T 9262T>C) | | | | | | | | |
| Codon mutations: | GCA1GGA (8574C>G), ATT5GTT (8585A>G), GGA9GAA (8598G>A), AGC10AGT (8602C>T), AAC11TCT (8603A>T 8604A>C 8605C>T), AAA16AAG (8620A>G), CCA17CAA (8622C>A), GAG18GGA (8625A>G 8626G>A), CAG19CAA (8629G>A), GGC20GGT (8632C>T), ATC22ATT (8638C>T), CAG23CAA (8641G>A), AAC25AAT (8647C>T), CCG26TTC (8648C>T 8649C>T 8650G>C), AAC27AAT (8653C>T), AAA28AAG (8656A>G), GGA29GAA (8658G>A), GAT31GCG (8664A>C 8665T>G), GTT34GTG (8674T>G), GCA36GTT (8679C>T 8680A>T), GGC37GGA (8683C>A), GGG40GGA (8692G>A), ACA41ACT (8695A>T), CCG45CCA (8707G>A), AGA46CGA (8708A>C), AAG48AAA (8716G>A), ATC50ATT (8722C>T), ACG51ACA (8725G>A), AAA53AAG (8731A>G), AGA55AGG (8737A>G), CCC57CCT (8743C>T), ACA58AAG (8745C>A 8746A>G), AGC59AGT (8749C>T), AAG60AAA (8752G>A), GGA61GGT (8755A>T), AAC66AAT (8770C>T), TTA67CTA (8771T>C), GAA68GAG (8776A>G), CAT69CAC (8779T>C), TTG70TTA (8782G>A), TAT73TAC (8791T>C), CCA75CCG (8797A>G), CAA76CAG (8800A>G), CAA77CAG (8803A>G), GAT79GAC (8809T>C), ATT80ATC (8812T>C), AAT82AAC (8818T>C), CGG84CGA (8824G>A), GCA85GCG (8827A>G), ACG92ACA (8848G>A), TAT94TAC (8854T>C), GAG95GGA (8856A>G 8857G>A), GTG97GTA (8863G>A), CGG98CAG (8865G>A), ATG99CTT (8867A>C 8869G>T), GCA100GCG (8872A>G), GGA104GAA (8883G>A), ACT106TCT (8888A>T), TGC119TGT (8929C>T), GTC128ATC (8954G>A), AAC129AAT (8959C>T), GGA130GGT (8962A>T), GTT133ATT (8969G>A), GGG137GGA (8983G>A), AAT138GAC (8984A>G 8986T>C), GAA139GTA (8988A>T), GAG142GAA (8998G>A), CCG144CCA (9004G>A), TTG145TTA (9007G>A), ATC148ATT (9016C>T), ACC155ACA (9037C>A), ATA171ATT (9085A>T), AAC175AAT (9097C>T), ATT187GTT (9131A>G), CGA188CGT (9136A>T), CGG191CGT (9145G>T), ATG193GGA (9149A>G 9150T>G 9151G>A), GGT194AGT (9152G>A), TTA195TTG (9157A>G), GCG196GCT (9160G>T), CGT197CGC (9163T>C), GAG204GAA (9184G>A), CGA208CGG (9196A>G), AGG214AGA (9214G>A), GAA215GAG (9217A>G), GCG216GCA (9220G>A), GCA224GCT (9244A>T), TTG225TTA (9247G>A), GCC228GCT (9256C>T), CCT230TCC (9260C>T 9262T>C), GGG234GGA (9274G>A), GAC236GAT (9280C>T), ATC239ATT (9289C>T), GAG244GAA (9304G>A), GTC254GTT (9334C>T), GTC264GTG (9364C>G) | | | | | | | | |
| PIPO (YP_006393460.1) | 1 | 75 | 100% | 95 | 18.1% | 75 (98.7%) | 54 (71.1%) | 1/0/3/2 | 2 |
| Protein mutations: | R17_K17insX (2971_2972insA), Y23H (2989T>C 2991C>T), R26K (2999G>A 3000A>G), H35Y (3025C>T), K40H (3040A>C 3042A>C), G41S (3043G>A), E44K (3052G>A), I47V (3061A>G), V54I (3082G>A), P57S (3091C>T), P60T (3100C>A), G61E (3104G>A 3105T>G), G62S (3106G>A), Q63* (3109C>T), R64G (3112A>G), C66R (3118T>C), L67F (3121C>T), I69V (3127A>G), A72E (3137C>A 3138G>A), I73V (3139A>G) | | | | | | | | |
| Codon mutations: | TTA2CTA (2923T>C), AAA4AAG (2931A>G), GAA7GAG (2940A>G), TTG10CTG (2947T>C), TTG15CTG (2962T>C), AGA17_AAA17insAA- (2971_2972insA), TAC23CAT (2989T>C 2991C>T), TCA24TCG (2994A>G), AGA26AAG (2999G>A 3000A>G), CAC32-AC (3016delC), TCG33TCA (3021G>A), GTA34GTT (3024A>T), CAT35TAT (3025C>T), CAC38CAT (3036C>T), AGA39AGG (3039A>G), AAA40CAC (3040A>C 3042A>C), GGC41AGC (3043G>A), GAA44AAA (3052G>A), ATA47GTA (3061A>G), GTT54ATT (3082G>A), CCG57TCG (3091C>T), CGC59CGT (3099C>T), CCA60ACA (3100C>A), GGT61GAG (3104G>A 3105T>G), GGT62AGT (3106G>A), CAA63TAA (3109C>T), AGG64GGG (3112A>G), TGC66CGC (3118T>C), CTC67TTC (3121C>T), ATT69GTT (3127A>G), GAG70GAA (3132G>A), GCG72GAA (3137C>A 3138G>A), ATT73GTT (3139A>G) | | | | | | | | |

*: Inserts / Deletes / Misaligned / Frameshifts

#### Analysis details

This analysis was performed with panviral2.64

### NGS Details (UN59): Duamitovirus soch1

#### Assembly

| | |
|---|---|
| **Coverage Length** | 2273 (1 contig(s)) |
| **Depth Of Coverage** | 39.0 |
| **Number Of Reads** | 734 |
| **Reads Per Million** | 15.96 rpm (after QC) |
| **Ambiguities** | 0 |
| **Assembly Method** | de novo + reference guided assembly |
| **Consensus Caller** | Bcf Tools |

#### Coverage Map

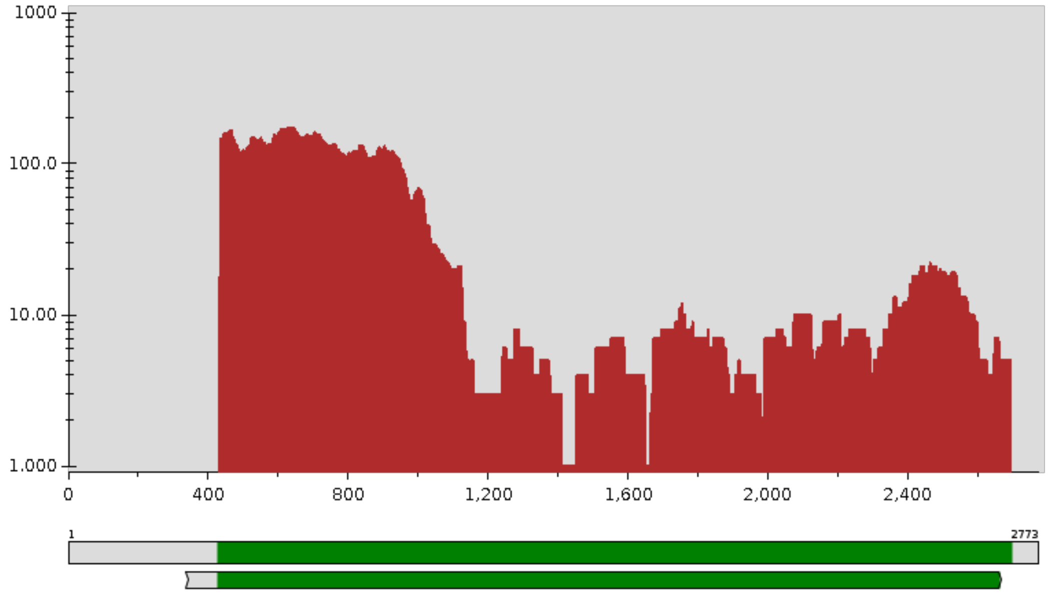

#### Assignment

| | |
|---|---|
| **Type** | Duamitovirus soch1 (Taxonomy ID: 2955838) |
| **Reference Genome** | NC_076524.1 |
| **NT Identity (%)** | 67.5497 |
| **AA Identity (%)** | 68.0108 |
| **Number Of Stop Codons** | 4 |
| **Number Of CDS** | 1 |

#### Alignment

| | |
|---|---|
| **Alignment Score** | 1546.0 (NT) + 3373.0 (AA) = 4919.0 |
| **Concordance (%)** | 51.6287 |

| **Alignment Method** | Global, seeded, nucleotide + amino acids (AGA) |
|---|---|

#### Genome Region

Sequence starts at position 428 and ends at position 2700 relative to NC_076524.1 reference sequence.

#### Alignment Detailed Statistics

| | Begin | End | Coverage | Score | Concordance | Matches | Identities | I/D/M/F* | Stop Codons |
|---|---|---|---|---|---|---|---|---|---|
| **NT** | **428** | **2700** | **82.0%** | **1546** | **34.4%** | **2264 (99.6%)** | **1530 (67.3%)** | **1/9** | |
| Mutations: | 437A>G, 441A>C, 443C>T, 447C>T, 459C>T, 466C>T, 470C>T, 471G>A, 472C>T, 473T>G, 477T>A, 480A>C, 481C>A, 482A>T, 485A>G, 489A>G, 494T>C, 497T>A, 500T>G, 503A>C, 506T>C, 509C>T, 512T>C, 513G>A, 516T>A, 517C>A, 518T>A, 520A>C, 524G>T, 525A>G, 526C>T, 527T>G, 529C>A, 530C>A, 531T>C, 533A>G, 534C>A, 535G>A, 536C>A, 539G>A, 540A>C, 541A>G, 542G>T, 545G>A, 548C>T, 550A>T, 556T>A, 557C>T, 560A>T, 569C>T, 575G>A, 581C>T, 582G>A, 587A>T, 589G>C, 591T>C, 593A>T, 599A>T, 602C>T, 612T>A, 613C>G, 614C>T, 616T>A, 618C>T, 619G>C, 620A>T, 624G>A, 626A>T, 629A>C, 630C>A, 632A>C, 633C>A, 635T>G, 638G>T, 641G>T, 644T>A, 647G>T, 651C>T, 653C>G, 656A>C, 657A>C, 661C>G, 662G>T, 665T>A, 668C>T, 672C>A, 673G>A, 674T>G, 680C>A, 683C>T, 684A>G, 685A>C, 686A>C, 692G>A, 695T>A, 699G>C, 700C>A, 701A>C, 702A>G, 705C>A, 707T>G, 708A>G, 709G>C, 710G>A, 713A>G, 714G>T, 715A>C, 719A>T, 722C>T, 737T>C, 746C>T, 752A>T, 761G>A, 765G>T, 767T>A, 770G>A, 771G>T, 772T>A, 773T>G, 774T>A, 776G>T, 777A>G, 778A>C, 779A>G, 782T>G, 796G>A, 797T>G, 803A>C, 809T>C, 810G>C, 815C>A, 818A>T, 821T>G, 824C>T, 827A>T, 831C>T, 833T>A, 841G>A, 842G>T, 850T>G, 851A>T, 857G>T, 860G>T, 862A>G, 869T>G, 875G>T, 880T>G, 881C>A, 885C>A, 887T>G, 892A>G, 905A>T, 908C>A, 909T>C, 910T>A, 916G>A, 918G>T, 919G>C, 923T>A, 926A>T, 927A>C, 929G>T, 934T>A, 935T>G, 938G>T, 942T>A, 943C>A, 948C>A, 971C>A, 972C>T, 975A>C, 976A>T, 977T>C, 983T>C, 990A>C, 992G>C, 993T>C, 994C>G, 995A>G, 997A>T, 998C>T, 999G>C, 1000G>A, 1001T>A, 1003T>A, 1004A>T, 1005G>T, 1007C>G, 1010A>G, 1016C>T, 1017A>G, 1018A>T, 1019A>T, 1022T>A, 1025T>C, 1026C>G, 1027A>G, 1030A>C, 1031G>A, 1034T>G, 1035A>G, 1036G>C, 1044T>G, 1046C>T, 1049G>A, 1050T>G, 1051T>A, 1052T>A, 1054C>A, 1055A>C, 1061T>C, 1067T>G, 1073G>A, 1076A>G, 1083C>A, 1088G>T, 1091C>G, 1099G>T, 1104A>G, 1105C>T, 1106A>G, 1115T>G, 1121A>C, 1122T>A, 1124C>T, 1127G>A, 1130C>T, 1133C>G, 1142A>C, 1145C>T, 1148G>T, 1151G>T, 1152T>C, 1154G>T, 1160C>T, 1161A>C, 1164T>A, 1165C>T, 1166T>G, 1169C>T, 1172C>A, 1174T>A, 1175T>C, 1178T>A, 1181A>T, 1184C>T, 1185A>T, 1186G>C, 1187G>T, 1188G>A, 1189C>A, 1191G>A, 1192C>A, 1193A>T, 1196T>C, 1197_1199delGAA, 1205A>T, 1211C>T, 1212T>G, 1213T>A, 1217T>C, 1220C>T, 1223A>T, 1224C>T, 1225A>T, 1226C>A, 1232G>A, 1236A>C, 1237G>A, 1238T>A, 1241C>A, 1247A>C, 1248T>C, 1249G>T, 1250G>T, 1253T>C, 1256A>T, 1258C>G, 1265G>A, 1266A>G, 1268T>G, 1269C>G, 1271T>C, 1274T>C, 1276G>A, 1280G>A, 1283G>T, 1292T>G, 1301G>A, 1302C>T, 1304T>G, 1306G>C, 1308G>C, 1309C>A, 1310A>G, 1312T>C, 1313G>C, 1316A>T, 1319G>A, 1322C>T, 1328A>G, 1332A>C, 1334A>G, 1338G>A, 1340A>C, 1345T>C, 1346A>T, 1349G>A, 1355C>T, 1357A>C, 1358C>T, 1376C>G, 1379C>T, 1382T>C, 1385T>C, 1388C>T, 1389G>A, 1390A>T, 1403A>T, 1406C>A, 1409G>A, 1412T>A, 1414A>G, 1415A>T, 1416A>C, 1418A>T, 1424A>T, 1427C>T, 1430G>T, 1433T>A, 1437G>A, 1445G>A, 1446A>C, 1447A>G, 1448A>T, 1451T>G, 1455T>C, 1456T>A, 1458G>T, 1459C>A, 1461T>C, 1465A>G, 1466G>A, 1468A>T, 1469A>T, 1472G>A, 1474A>G, 1475C>A, 1476T>C, 1477T>C, 1478T>G, 1481A>G, 1482A>T, 1483A>C, 1490T>C, 1502A>C, 1505G>A, 1514G>A, 1515G>A, 1526G>A, 1535C>T, 1538G>A, 1541T>C, 1545C>T, 1547T>A, 1550G>C, 1551T>G, 1552C>A, 1553T>A, 1559A>T, 1565G>T, 1566C>A, 1567C>G, 1571C>T, 1574G>A, 1580C>T, 1589T>C, 1593_1597delAGCTC, 1598T>A, 1613G>T, 1614A>T, 1616C>T, 1619T>C, 1622T>A, 1623A>T, 1624A>C, 1625C>A, 1626A>C, 1627A>C, 1628A>T, 1631G>T, 1632T>A, 1634A>G, 1635A>G, 1636C>T, 1637A>G, 1642G>A, 1643A>G, 1644G>A, 1646G>T, 1647T>A, 1648A>G, 1652G>A, 1653A>G, 1664G>T, 1670G>A, 1673G>A, 1676T>G, 1679G>A, 1682C>T, 1684T>A, 1686C>T, 1694T>A, 1698G>T, 1700A>C, 1703T>A, 1706C>T, 1707T>G, 1710T>C, 1712G>T, 1715T>C, 1721C>T, 1727A>G, 1734T>C, 1739A>T, 1744G>A, 1745C>A, 1746A>T, 1748A>T, 1754T>C, 1758G>A, 1759G>C, 1760A>G, 1761T>A, 1765A>C, 1766C>T, 1767T>A, 1769T>C, 1772A>T, 1777G>C, 1778G>T, 1787T>C, 1788T>C, 1790A>G, 1791C>T, 1819G>C, 1820T>A, 1821A>G, 1823A>G, 1829C>G, 1830A>C, 1831A>G, 1832T>G, 1833G>C, 1840A>G, 1847A>G, 1848T>C, 1856A>G, 1859C>T, 1860A>G, 1861A>G, 1865C>T, 1867A>C, 1871T>C, 1874G>T, 1875A>T, 1878C>G, 1880A>T, 1886T>G, 1887T>A, 1888T>G, 1889G>A, 1895T>C, 1898T>G, 1901T>C, 1910T>C, 1919G>T, 1925G>A, 1934T>C, 1937A>G, 1938T>G, 1939C>A, 1940A>C, 1950G>T, 1952T>C, 1953T>C, 1955A>T, 1964T>C, 1967C>T, 1976T>A, 1980T>A, 1981T>C, 1982A>T, 1985T>C, 1989C>A, 1991T>G, 1994T>A, 2000T>G, 2003A>T, 2006T>G, 2009C>T, 2010G>C, 2011C>A, 2012C>A, 2020T>C, 2028G>T, 2029A>C, 2030T>G, 2034A>T, 2035A>C, 2036C>A, 2038T>C, 2042A>G, 2043G>A, 2050A>C, 2051A>C, 2053G>T, 2054A>T, 2057G>T, 2058G>T, 2063T>G, 2069G>A, 2071A>T, 2074A>G, 2075G>A, 2078A>T, 2082G>T, 2084G>T, 2087C>T, 2090A>G, 2091T>A, 2093C>G, 2094A>T, 2099C>T, 2103A>T, 2104G>C, 2105C>T, 2108G>A, 2117G>A, 2118C>A, 2120T>A, 2123A>G, 2129T>G, 2133C>G, 2134A>C, 2135T>A, 2136C>A, 2137T>G, 2138T>C, 2141G>A, 2145G>C, 2146T>A, 2147T>C, 2148C>A, 2156C>T, 2157T>C, 2160C>T, 2166T>C, 2167A>T, 2173G>A, 2174G>T, 2183A>G, 2186G>A, 2187G>T, 2188G>T, 2191T>A, 2192G>A, 2195C>T, 2204C>T, 2206A>T, 2207_2208insT, 2213A>G, 2216G>A, 2223G>A, 2226A>G, 2227G>A, 2231C>T, 2236T>G, 2237T>G, 2240A>G, 2248A>C, 2250G>C, 2251T>C, 2259T>A, 2263A>G, 2264G>A, 2265A>C, 2276G>T, 2279T>G, 2280C>T, 2288C>T, 2297A>G, 2298G>A, 2299T>G, 2300T>A, 2306A>C, 2309A>T, 2313C>T, 2315T>C, 2317C>T, 2322C>T, 2325C>G, 2326G>T, 2328C>A, 2330A>T, 2336A>T, 2337G>A, 2346T>C, 2348A>T, 2349A>C, 2350A>G, 2355C>G, 2357C>T, 2358A>T, 2359A>C, 2360C>T, 2362G>A, 2369T>C, 2372G>T, 2377G>T, 2378T>G, 2382A>T, 2383A>C, 2384C>T, 2385C>T, 2386C>T, 2387A>T, 2390T>A, 2396C>T, 2399T>C, 2400A>G, 2402G>A, 2404G>A, 2405T>A, 2406C>T, 2408C>G, 2411C>G, 2414C>T, 2415T>G, 2417C>T, 2420T>C, 2423A>T, 2425G>A, 2430A>T, 2431C>A, 2432A>G, 2435C>A, 2441A>G, 2442A>C, 2444G>A, 2445C>T, 2446T>C, 2447C>G, 2448G>A, 2450A>T, 2451A>G, 2452C>A, 2453T>A, 2456T>C, 2457T>C, 2459T>G, 2462T>A, 2465A>G, 2466A>G, 2477C>T, 2478C>T, 2489A>G, 2492T>C, 2495C>T, 2504T>A, 2510T>C, 2514A>G, 2518C>T, 2519C>G, 2524C>T, 2525C>T, 2528T>A, 2532C>A, 2533A>G, 2536G>A, 2537G>T, 2543T>G, 2551A>G, 2552C>T, 2555T>C, 2556G>T, 2557T>C, 2561T>C, 2568A>G, 2571A>C, 2572A>G, 2582A>T, 2585G>A, 2597G>A, 2598A>G, 2599A>C, 2600C>T, 2601G>A, 2609T>A, 2613T>G, 2615T>A, 2618T>A, 2622G>T, 2624A>T, 2626A>T, 2627C>A, 2629A>C, 2631A>T, 2634A>C, 2635C>T, 2638G>T, 2639A>G, 2644A>G, 2646G>T, 2647A>G, 2648A>T, 2649G>A, 2650T>G, 2652G>T, 2653G>C, 2654C>A, 2656T>A, 2657A>T, 2658A>G, 2659G>A, 2660A>C, 2661C>T, 2664T>A, 2665A>G, 2667delG, 2674T>A, 2678C>T, 2681G>A, 2686T>A, 2689T>C, 2690G>C, 2694A>G, 2695T>A | | | | | | | | |

##### CDS

| | Begin | End | Coverage | Score | Concordance | Matches | Identities | I/D/M/F* | Stop Codons |
|---|---|---|---|---|---|---|---|---|---|
| RdRp | 32 | 777 | 95.9% | 3373 | 64.7% | 743 (99.6%) | 506 (67.8%) | 1/2/2/2 | 4 |
| Protein mutations: | I36L (441A>C 443C>T), L38F (447C>T), A44V (466C>T), A46M (471G>A 472C>T 473T>G), L48M (477T>A), T49H (480A>C 481C>A 482A>T), I52V (489A>G), V60I (513G>A), S61K (516T>A 517C>A 518T>A), K62T (520A>C), T64V (525A>G 526C>T 527T>G), T65K (529C>A 530C>A), R67K (534C>A 535G>A 536C>A), K69R (540A>C 541A>G 542G>T), Y72F (550A>T), F74Y (556T>A 557C>T), A83T (582G>A), C85S (589G>C), F94Y (616T>A), R95S (618C>T 619G>C 620A>T), E97N (624G>A 626A>T), E98D (629A>C), P99T (630C>A 632A>C), L100M (633C>A 635T>G), S109C (661C>G 662G>T), R113K (672C>A 673G>A 674T>G), K117A (684A>G 685A>C 686A>C), A122H (699G>C 700C>A 701A>C), I123V (702A>G), R125A (708A>G 709G>C 710G>A), D127S (714G>T 715A>C), E128D (719A>T), A144S (765G>T 767T>A), V146* (771G>T 772T>A 773T>G), L147I (774T>A 776G>T), K148A (777A>G 778A>C 779A>G), S154K (796G>A 797T>G), E159Q (810G>C), P166S (831C>T 833T>A), G169D (841G>A 842G>T), L172R (850T>G 851A>T), E176G (862A>G), I178M (869T>G), F182* (880T>G 881C>A), H184K (885C>A 887T>G), Q186R (892A>G), F192H (909T>C 910T>A), R194Q (916G>A), G195S (918G>T 919G>C), M198L (927A>C 929G>T), I200K (934T>A 935T>G), S203N (942T>A 943C>A), Q205K (948C>A), N214L (975A>C 976A>T 977T>C), S220R (993T>C 994C>G 995A>G), Y221F (997A>T 998C>T), G222Q (999G>C 1000G>A 1001T>A), L223Y (1003T>A 1004A>T), A224S (1005G>T 1007C>G), K228V (1017A>G 1018A>T 1019A>T), Q231G (1026C>G 1027A>G), E232A (1030A>C 1031G>A), S234A (1035A>G 1036G>C), F237V (1044T>G 1046C>T), F239E (1050T>G 1051T>A 1052T>A), P240H (1054C>A 1055A>C), L250I (1083C>A), W255L (1099G>T), T257V (1104A>G 1105C>T 1106A>G), I260M (1115T>G), F263I (1122T>A 1124C>T), M272I (1151G>T), K276Q (1161A>C), S277M (1164T>A 1165C>T 1166T>G), F279L (1172C>A), F280Y (1174T>A 1175T>C), L282F (1181A>T), R284S (1185A>T 1186G>C 1187G>T), A285N (1188G>A 1189C>A), A286N (1191G>A 1192C>A 1193A>T), E288del (1197_1199delGAA), L290F (1205A>T), L293E (1212T>G 1213T>A), E296D (1223A>T), H297L (1224C>T 1225A>T 1226C>A), S301Q (1236A>C 1237G>A 1238T>A), W305L (1248T>C 1249G>T 1250G>T), T308S (1258C>G), S311G (1266A>G 1268T>G), P312A (1269C>G 1271T>C), R314Q (1276G>A), G324A (1306G>C), A325Q (1308G>C 1309C>A 1310A>G), V326A (1312T>C 1313G>C), V335I (1338G>A 1340A>C), V337A (1345T>C 1346A>T), M338I (1349G>A), Y341S (1357A>C 1358C>T), D352I (1389G>A 1390A>T), K360S (1414A>G 1415A>T), I361L (1416A>C 1418A>T), D368N (1437G>A), K371R (1446A>C 1447A>G 1448A>T), L374Q (1455T>C 1456T>A), A375Y (1458G>T 1459C>A), K377R (1465A>G 1466G>A), Q378L (1468A>T 1469A>T), N380R (1474A>G 1475C>A), F381P (1476T>C 1477T>C 1478T>G), N383S (1482A>T 1483A>C), L389F (1502A>C), D394N (1515G>A), M401I (1538G>A), M405I (1550G>C), S406E (1551T>G 1552C>A 1553T>A), P411S (1566C>A 1567C>G), S420del (1593_1597delAGCTC), I427F (1614A>T 1616C>T), N430S (1623A>T 1624A>C 1625C>A), K431P (1626A>C 1627A>C 1628A>T), L433M (1632T>A 1634A>G), T434V (1635A>G 1636C>T 1637A>G), R436K (1642G>A 1643A>G), V437I (1644G>A 1646G>T), Y438S (1647T>A 1648A>G), I440V (1653A>G), F450Y (1684T>A), H451Y (1686C>T), A455S (1698G>T 1700A>C), S458A (1707T>G), I464M (1727A>G), S470K (1744G>A 1745C>A), K471Y (1746A>T 1748A>T), G475T (1758G>A 1759G>C 1760A>G), C476S (1761T>A), N477T (1765A>C 1766C>T), S478T (1767T>A 1769T>C), G481A (1777G>C 1778G>T), S495T (1819G>C 1820T>A), K496E (1821A>G 1823A>G), N499R (1830A>C 1831A>G 1832T>G), E500Q (1833G>C), K502R (1840A>G), N509G (1860A>G 1861A>G), K511T (1867A>C), I514L (1875A>T), P515A (1878C>G 1880A>T), L518R (1887T>A 1888T>G 1889G>A), D521E (1898T>G), S535D (1938T>G 1939C>A 1940A>C), D539Y (1950G>T 1952T>C), L549T (1980T>A 1981T>C 1982A>T), A559Q (2010G>C 2011C>A 2012C>A), I562T (2020T>C), D565S (2028G>T 2029A>C 2030T>G), N567S (2034A>T 2035A>C 2036C>A), I568T (2038T>C), V570I (2043G>A), Q572P (2050A>C 2051A>C), R573I (2053G>T 2054A>T), L574F (2057G>T), G575W (2058G>T), Y579F (2071A>T), K580R (2074A>G 2075G>A), A583S (2082G>T 2084G>T), F586M (2091T>A 2093C>G), T587S (2094A>T), H600A (2133C>G 2134A>C 2135T>A), L601S (2136C>A 2137T>G 2138T>C), V604H (2145G>C 2146T>A 2147T>C), Y611L (2166T>C 2167A>T), W613Y (2173G>A 2174G>T), G618F (2187G>T 2188G>T), M619K (2191T>A 2192G>A), Y624F (2206A>T), Y624_L625insX (2207_2208insT), V630I (2223G>A), S631D (2226A>G 2227G>A), L634R (2236T>G 2237T>G), K638T (2248A>C), V639P (2250G>C 2251T>C), L642I (2259T>A), Q643R (2263A>G 2264G>A), I644L (2265A>C), K647N (2276G>T), D648E (2279T>G), V655R (2298G>A 2299T>G 2300T>A), R660C (2313C>T 2315T>C), T661I (2317C>T), R664V (2325C>G 2326G>T), Q665N (2328C>A 2330A>T), E668K (2337G>A), K672R (2349A>C 2350A>G), L674V (2355C>G 2357C>T), N675S (2358A>T 2359A>C 2360C>T), W676* (2362G>A), C681L (2377G>T 2378T>G), N683S (2382A>T 2383A>C 2384C>T), P684F (2385C>T 2386C>T 2387A>T), D685E (2390T>A), K689E (2400A>G 2402G>A), S690K (2404G>A 2405T>A), I692M (2411C>G), S694A (2415T>G 2417C>T), C697Y (2425G>A), T699* (2430A>T 2431C>A 2432A>G), L704S (2445C>T 2446T>C 2447C>G), E705N (2448G>A 2450A>T), T706E (2451A>G 2452C>A 2453T>A), F708L (2457T>C 2459T>G), D709E (2462T>A), I711V (2466A>G), N727D (2514A>G), P728L (2518C>T 2519C>G), T730I (2524C>T 2525C>T), Q733R (2532C>A 2533A>G), W734Y (2536G>A 2537G>T), F736L (2543T>G), N739S (2551A>G 2552C>T), V741S (2556G>T 2557T>C), N745D (2568A>G), K746R (2571A>C 2572A>G), L749F (2582A>T), N755A (2598A>G 2599A>C 2600C>T), D756N (2601G>A), S760A (2613T>G 2615T>A), E763Y (2622G>T 2624A>T), Y764L (2626A>T 2627C>A), Q765P (2629A>C), T766S (2631A>T), T767L (2634A>C 2635C>T), G768V (2638G>T 2639A>G), E770G (2644A>G), E771C (2646G>T 2647A>G 2648A>T), V772R (2649G>A 2650T>G), G773S (2652G>T 2653G>C 2654C>A), L774Y (2656T>A 2657A>T), R775D (2658A>G 2659G>A 2660A>C), L776F (2661C>T), *777R (2664T>A 2665A>G) | | | | | | | | |

| | Begin | End | Coverage | Score | Concordance | Matches | Identities | I/D/M/F* | Stop Codons |
|---|---|---|---|---|---|---|---|---|---|
| **NT** | **428** | **2700** | **82.0%** | **1546** | **34.4%** | **2264 (99.6%)** | **1530 (67.3%)** | **1/9** | |
| Codon mutations: | AAA34AAG (437A>G), ATC36CTT (441A>C 443C>T), CTT38TTT (447C>T), CTA42TTA (459C>T), GCC44GTC (466C>T), TCC45TCT (470C>T), GCT46ATG (471G>A 472C>T 473T>G), TTG48ATG (477T>A), ACA49CAT (480A>C 481C>A 482A>T), AAA50AAG (485A>G), ATC52GTC (489A>G), GCT53GCC (494T>C), CTT54CTA (497T>A), GCT55GCG (500T>G), GGA56GGC (503A>C), TGT57TGC (506T>C), GAC58GAT (509C>T), TTT59TTC (512T>C), GTA60ATA (513G>A), TCT61AAA (516T>A 517C>A 518T>A), AAA62ACA (520A>C), GTG63GTT (524G>T), ACT64GTG (525A>G 526C>T 527T>G), ACC65AAA (529C>A 530C>A), TTA66CTG (531T>C 533A>G), CGC67AAA (534C>A 535G>A 536C>A), CGG68CGA (539G>A), AAG69CGT (540A>C 541A>G 542G>T), TCG70TCA (545G>A), GGC71GGT (548C>T), TAT72TTT (550A>T), TTC74TAT (556T>A 557C>T), ACA75ACT (560A>T), TAC78TAT (569C>T), AAG80AAA (575G>A), TGC82TGT (581C>T), GCA83ACA (582G>A), GTA84GTT (587A>T), TGC85TCC (589G>C), TTA86CTT (591T>C 593A>T), CGA88CGT (599A>T), TAC89TAT (602C>T), TCC93AGT (612T>A 613C>G 614C>T), TTT94TAT (616T>A), CGA95TCT (618C>T 619G>C 620A>T), GAA97AAT (624G>A 626A>T), GAA98GAC (629A>C), CCA99ACC (630C>A 632A>C), CTT100ATG (633C>A 635T>G), TCG101TCT (638G>T), GTG102GTT (641G>T), CCT103CCA (644T>A), GTG104GTT (647G>T), CTC106TTG (651C>T 653C>G), ACA107ACC (656A>C), AGA108CGA (657A>C), TCG109TGT (661C>G 662G>T), GGT110GGA (665T>A), ATC111ATT (668C>T), CGT113AAG (672C>A 673G>A 674T>G), ATC115ATA (680C>A), CCC116CCT (683C>T), AAA117GCC (684A>G 685A>C 686A>C), TTG119TTA (692G>A), CGT120CGA (695T>A), GCA122CAC (699G>C 700C>A 701A>C), ATA123GTA (702A>G), CGT124AGG (705C>A 707T>G), AGG125GCA (708A>G 709G>C 710G>A), AAA126AAG (713A>G), GAT127TCT (714G>T 715A>C), GAA128GAT (719A>T), CAC129CAT (722C>T), GTT134GTC (737T>C), TAC137TAT (746C>T), TCA139TCT (752A>T), GGG142GGA (761G>A), GCT144TCA (765G>T 767T>A), AAG145AAA (770G>A), GTT146TAG (771G>T 772T>A 773T>G), TTG147ATT (774T>A 776G>T), AAA148GCG (777A>G 778A>C 779A>G), GTT149GTG (782T>G), AGT154AAG (796G>A 797T>G), GCA156GCC (803A>C), TTT158TTC (809T>C), GAA159CAA (810G>C), TCC160TCA (815C>A), ATA161ATT (818A>T), ACT162ACG (821T>G), ACC163ACT (824C>T), CCA164CCT (827A>T), CCT166TCA (831C>T 833T>A), GGG169GAT (841G>A 842G>T), CTA172CGT (850T>G 851A>T), GTG174GTT (857G>T), CTG175CTT (860G>T), GAA176GGA (862A>G), ATT178ATG (869T>G), ACG180ACT (875G>T), TTC182TGA (880T>G 881C>A), CAT184AAG (885C>A 887T>G), CAA186CGA (892A>G), CTA190CTT (905A>T), CCC191CCA (908C>A), TTT192CAT (909T>C 910T>A), CGA194CAA (916G>A), GGC195TCC (918G>T 919G>C), ATT196ATA (923T>A), CCA197CCT (926A>T), ATG198CTT (927A>C 929G>T), ATT200AAG (934T>A 935T>G), GGG201GGT (938G>T), TCT203AAT (942T>A 943C>A), CAA205AAA (948C>A), CCC212CCA (971C>A), CTA213TTA (972C>T), AAT214CTC (975A>C 976A>T 977T>C), AAT216AAC (983T>C), AGG219CGC (990A>C 992G>C), TCA220CGG (993T>C 994C>G 995A>G), TAC221TTT (997A>T 998C>T), GGT222CAA (999G>C 1000G>A 1001T>A), TTA223TAT (1003T>A 1004A>T), GCC224TCG (1005G>T 1007C>G), GTA225GTG (1010A>G), GAC227GAT (1016C>T), AAA228GTT (1017A>G 1018A>T 1019A>T), GTT229GTA (1022T>A), GAT230GAC (1025T>C), CAA231GGA (1026C>G 1027A>G), GAG232GCA (1030A>C 1031G>A), GCT233GCG (1034T>G), AGT234GCT (1035A>G 1036G>C), TTC237GTT (1044T>G 1046C>T), AAG238AAA (1049G>A), TTT239GAA (1050T>G 1051T>A 1052T>A), CCA240CAC (1054C>A 1055A>C), ATT242ATC (1061T>C), GTT244GTG (1067T>G), TTG246TTA (1073G>A), AAA247AAG (1076A>G), CTA250ATA (1083C>A), GCG251GCT (1088G>T), GCC252GCG (1091C>G), TGG255TTG (1099G>T), ACA257GTG (1104A>G 1105C>T 1106A>G), ATT260ATG (1115T>G), TCA262TCC (1121A>C), TTC263ATT (1122T>A 1124C>T), CAG264CAA (1127G>A), GAC265GAT (1130C>T), GGC266GGG (1133C>G), TCA269TCC (1142A>C), CCC270CCT (1145C>T), GGG271GGT (1148G>T), ATG272ATT (1151G>T), TTG273CTT (1152T>C 1154G>T), TCC275TCT (1160C>T), AAA276CAA (1161A>C), TCT277ATG (1164T>A 1165C>T 1166T>G), GTC278GTT (1169C>T), TTC279TTA (1172C>A), TTT280TAC (1174T>A 1175T>C), CCT281CCA (1178T>A), TTA282TTT (1181A>T), GAC283GAT (1184C>T), AGG284TCT (1185A>T 1186G>C 1187G>T), GCT285AAT (1188G>A 1189C>A), GCA286AAT (1191G>A 1192C>A 1193A>T), ACT287ACC (1196T>C), GAA288del (1197_1199delGAA), TTA290TTT (1205A>T), AAC292AAT (1211C>T), TTA293GAA (1212T>G 1213T>A), TCT294TCC (1217T>C), CTC295CTT (1220C>T), GAA296GAT (1223A>T), CAC297TTA (1224C>T 1225A>T 1226C>A), GAG299GAA (1232G>A), AGT301CAA (1236A>C 1237G>A 1238T>A), GTC302GTA (1241C>A), CCA304CCC (1247A>C), TGG305CTT (1248T>C 1249G>T 1250G>T), TTT306TTC (1253T>C), TCA307TCT (1256A>T), ACC308AGC (1258C>G), TTG310TTA (1265G>A), AGT311GGG (1266A>G 1268T>G), CCT312GCC (1269C>G 1271T>C), TAT313TAC (1274T>C), CGA314CAA (1276G>A), GGG315GGA (1280G>A), GTG316GTT (1283G>T), GTT319GTG (1292T>G), AGG322AGA (1301G>A), CTT323TTG (1302C>T 1304T>G), GGT324GCT (1306G>C), GCA325CAG (1308G>C 1309C>A 1310A>G), GTG326GCC (1312T>C 1313G>C), ATA327ATT (1316A>T), GAG328GAA (1319G>A), GGC329GGT (1322C>T), GGA331GGG (1328A>G), AGA333CGG (1332A>C 1334A>G), GTA335ATC (1338G>A 1340A>C), GTA337GCT (1345T>C 1346A>T), ATG338ATA (1349G>A), AAC340AAT (1355C>T), TAC341TCT (1357A>C 1358C>T), CTC347CTG (1376C>G), CAC348CAT (1379C>T), CCT349CCC (1382T>C), GTT350GTC (1385T>C), CAC351CAT (1388C>T), GAT352ATT (1389G>A 1390A>T), ACA356ACT (1403A>T), GTC357GTA (1406C>A), CTG358CTA (1409G>A), TCT359TCA (1412T>A), AAA360AGT (1414A>G 1415A>T), ATA361CTT (1416A>C 1418A>T), ACA363ACT (1424A>T), GAC364GAT (1427C>T), GGG365GGT (1430G>T), ACT366ACA (1433T>A), GAC368AAC (1437G>A), GAG370GAA (1445G>A), AAA371CGT (1446A>C 1447A>G 1448A>T), CCT372CCG (1451T>G), TTG374CAG (1455T>C 1456T>A), GCT375TAT (1458G>T 1459C>A), TTA376CTA (1461T>C), AAG377AGA (1465A>G 1466G>A), CAA378CTT (1468A>T 1469A>T), AAG379AAA (1472G>A), AAC380AGA (1474A>G 1475C>A), TTT381CCG (1476T>C 1477T>C 1478T>G), AAA382AAG (1481A>G), AAT383TCT (1482A>T 1483A>C), TAT385TAC (1490T>C), TTA389TTC (1502A>C), AAG390AAA (1505G>A), ACG393ACA (1514G>A), GAC394AAC (1515G>A), CCG397CCA (1526G>A), GTC400GTT (1535C>T), ATG401ATA (1538G>A), TAT402TAC (1541T>C), CTT404TTA (1545C>T 1547T>A), ATG405ATC (1550G>C), TCT406GAA (1551T>G 1552C>A 1553T>A), ATA408ATT (1559A>T), GGG410GGT (1565G>T), CCT411AGT (1566C>A 1567C>G), ACC412ACT (1571C>T), TTG413TTA (1574G>A), TCC415TCT (1580C>T), GTT418GTC (1589T>C), AGC420del (1593_1597delAGCTC), TCT421--A (1593_1597delAGCTC 1598T>A), ACG426ACT (1613G>T), ATC427TTT (1614A>T 1616C>T), CTT428CTC (1619T>C), GTT429GTA (1622T>A), AAC430TCA (1623A>T 1624A>C 1625C>A), AAA431CCT (1626A>C 1627A>C 1628A>T), CCG432CCT (1631G>T), TTA433ATG (1632T>A 1634A>G), ACA434GTG (1635A>G 1636C>T 1637A>G), AGA436AAG (1642G>A 1643A>G), GTG437ATT (1644G>A 1646G>T), TAT438AGT (1647T>A 1648A>G), GAG439GAA (1652G>A), ATA440GTA (1653A>G), CTG443CTT (1664G>T), GGG445GGA (1670G>A), CAG446CAA (1673G>A), CCT447CCG (1676T>G), TTG448TTA (1679G>A), GGC449GGT (1682C>T), TTC450TAC (1684T>A), CAC451TAC (1686C>T), TCT453TCA (1694T>A), GCA455TCC (1698G>T 1700A>C), CTT456CTA (1703T>A), TTC457TTT (1706C>T), TCG458GCG (1707T>G), TTG459CTT (1710T>C 1712G>T), TCT460TCC (1715T>C), CAC462CAT (1721C>T), ATA464ATG (1727A>G), TTA467CTA (1734T>C), GCA468GCT (1739A>T), AGC470AAA (1744G>A 1745C>A), AAA471TAT (1746A>T 1748A>T), TAT473TAC (1754T>C), GGA475ACG (1758G>A 1759G>C 1760A>G), TGT476AGT (1761T>A), AAC477ACT (1765A>C 1766C>T), TCT478ACC (1767T>A 1769T>C), CCA479CCT (1772A>T), GGG481GCT (1777G>C 1778G>T), GCT484GCC (1787T>C), TTA485CTG (1788T>C 1790A>G), CTA486TTA (1791C>T), AGT495ACA (1819G>C 1820T>A), AAA496GAG (1821A>G 1823A>G), GCC498GCG (1829C>G), AAT499CGG (1830A>C 1831A>G 1832T>G), GAG500CAG (1833G>C), AAG502AGG (1840A>G), TTA504TTG (1847A>G), TTG505CTG (1848T>C), AGA507AGG (1856A>G), CTC508CTT (1859C>T), AAT509GGT (1860A>G 1861A>G), GTC510GTT (1865C>T), AAA511ACA (1867A>C), ATT512ATC (1871T>C), TCG513TCT (1874G>T), ATA514TTA (1875A>T), CCA515GCT (1878C>G 1880A>T), TCT517TCG (1886T>G), TTG518AGA (1887T>A 1888T>G 1889G>A), TCT520TCC (1895T>C), GAT521GAG (1898T>G), AAT522AAC (1901T>C), ATT525ATC (1910T>C), GCG528GCT (1919G>T), AGG530AGA (1925G>A), ACT533ACC (1934T>C), AAA534AAG (1937A>G), TCA535GAC (1938T>G 1939C>A 1940A>C), GAT539TAC (1950G>T 1952T>C), TTA540CTT (1953T>C 1955A>T), ATT543ATC (1964T>C), TCC544TCT (1967C>T), GCT547GCA (1976T>A), TTA549ACT (1980T>A 1981T>C 1982A>T), TCT550TCC (1985T>C), CGT552AGG (1989C>A 1991T>G), ACT553ACA (1994T>A), GTT555GTG (2000T>G), GGA556GGT (2003A>T), CTT557CTG (2006T>G), TGC558TGT (2009C>T), GCC559CAA (2010G>C 2011C>A 2012C>A), ATC562ACC (2020T>C), GAT565TCG (2028G>T 2029A>C 2030T>G), AAC567TCA (2034A>T 2035A>C 2036C>A), ATA568ACA (2038T>C), TCA569TCG (2042A>G), GTT570ATT (2043G>A), CAA572CCC (2050A>C 2051A>C), AGA573ATT (2053G>T 2054A>T), TTG574TTT (2057G>T), GGG575TGG (2058G>T), GGT576GGG (2063T>G), GGG578GGA (2069G>A), TAC579TTC (2071A>T), AAG580AGA (2074A>G 2075G>A), GTA581GTT (2078A>T), GCG583TCT (2082G>T 2084G>T), CGC584CGT (2087C>T), TTA585TTG (2090A>G), TTC586ATG (2091T>A 2093C>G), ACT587TCT (2094A>T), ACC588ACT (2099C>T), AGC590TCT (2103A>T 2104G>C 2105C>T), AAG591AAA (2108G>A), GAG594GAA (2117G>A), CGT595AGA (2118C>A 2120T>A), CTA596CTG (2123A>G), GCT598GCG (2129T>G), CAT600GCA (2133C>G 2134A>C 2135T>A), CTT601AGC (2136C>A 2137T>G 2138T>C), AAG602AAA (2141G>A), GTT604CAC (2145G>C 2146T>A 2147T>C), CGG605AGG (2148C>A), CAC607CAT (2156C>T), TTA608CTA (2157T>C), CTA609TTA (2160C>T), TAC611CTC (2166T>C 2167A>T), TGG613TAT (2173G>A 2174G>T), GGA616GGG (2183A>G), AGG617AGA (2186G>A), GGT618TTT (2187G>T 2188G>T), ATG619AAA (2191T>A 2192G>A), CCC620CCT (2195C>T), CCC623CCT (2204C>T), TAT624TTT (2206A>T), TAT624_TTG625insT-- (2207_2208insT), AAA626AAG (2213A>G), GGG627GGA (2216G>A), GTA630ATA (2223G>A), AGT631GAT (2226A>G 2227G>A), TAC632TAT (2231C>T), CTT634CGG (2236T>G 2237T>G), AAA635AAG (2240A>G), AAA638ACA (2248A>C), GTT639CCT (2250G>C 2251T>C), TTA642ATA (2259T>A), CAG643CGA (2263A>G 2264G>A), ATC644CTC (2265A>C), AAG647AAT (2276G>T), GAT648GAG (2279T>G), CTG649TTG (2280C>T), TTC651TTT (2288C>T), GAA654GAG (2297A>G), GTT655AGA (2298G>A 2299T>G 2300T>A), ATA657ATC (2306A>C), CTA658CTT (2309A>T), CGT660TGC (2313C>T 2315T>C), ACT661ATT (2317C>T), CTA663TTA (2322C>T), CGC664GTC (2325C>G 2326G>T), CAA665AAT (2328C>A 2330A>T), GTA667GTT (2336A>T), GAA668AAA (2337G>A), TTA671CTT (2346T>C 2348A>T), AAG672CGG (2349A>C 2350A>G), CTC674GTT (2355C>G 2357C>T), AAC675TCT (2358A>T 2359A>C 2360C>T), TGG676TAG (2362G>A), CAT678CAC (2369T>C), ACG679ACT (2372G>T), TGT681TTG (2377G>T 2378T>G), AAC683TCT (2382A>T 2383A>C 2384C>T), CCA684TTT (2385C>T 2386C>T 2387A>T), GAT685GAA (2390T>A), TCC687TCT (2396C>T), ATT688ATC (2399T>C), AAG689GAA (2400A>G 2402G>A), AGT690AAA (2404G>A 2405T>A), CTC691TTG (2406C>T 2408C>G), ATC692ATG (2411C>G), GAC693GAT (2414C>T), TCC694GCT (2415T>G 2417C>T), CCT695CCC (2420T>C), ATA696ATT (2423A>T), TGC697TAC (2425G>A), ACA699TAG (2430A>T 2431C>A 2432A>G), TCC700TCA (2435C>A), AAA702AAG (2441A>G), AGG703CGA (2442A>C 2444G>A), CTC704TCG (2445C>T 2446T>C 2447C>G), GAA705AAT (2448G>A 2450A>T), ACT706GAA (2451A>G 2452C>A 2453T>A), GAT707GAC (2456T>C), TTT708CTG (2457T>C 2459T>G), GAT709GAA (2462T>A), TTA710TTG (2465A>G), ATC711GTC (2466A>G), GGC714GGT (2477C>T), CTA715TTA (2478C>T), AAA718AAG (2489A>G), TGT719TGC (2492T>C), TAC720TAT (2495C>T), GGT723GGA (2504T>A), GGT725GGC (2510T>C), AAT727GAT (2514A>G), CCC728CTG (2518C>T 2519C>G), ACC730ATT (2524C>T 2525C>T), ACT731ACA (2528T>A), CAG733AGG (2532C>A 2533A>G), TGG734TAT (2536G>A 2537G>T), TTT736TTG (2543T>G), AAC739AGT (2551A>G 2552C>T), ACT740ACC (2555T>C), GTT741TCT (2556G>T 2557T>C), ATT742ATC (2561T>C), AAT745GAT (2568A>G), AAA746CGA (2571A>C 2572A>G), TTA749TTT (2582A>T), GGG750GGA (2585G>A), GGG754GGA (2597G>A), AAC755GCT (2598A>G 2599A>C 2600C>T), GAT756AAT (2601G>A), ATT758ATA (2609T>A), TCT760GCA (2613T>G 2615T>A), CCT761CCA (2618T>A), GAA763TAT (2622G>T 2624A>T), TAC764TTA (2626A>T 2627C>A), CAA765CCA (2629A>C), ACT766TCT (2631A>T), ACA767CTA (2634A>C 2635C>T), GGA768GTG (2638G>T 2639A>G), GAG770GGG (2644A>G), GAA771TGT (2646G>T 2647A>G 2648A>T), GTG772AGG (2649G>A 2650T>G), GGC773TCA (2652G>T 2653G>C 2654C>A), TTA774TAT (2656T>A 2657A>T), AGA775GAC (2658A>G 2659G>A 2660A>C), CTT776TTT (2661C>T), TAG777AGG (2664T>A 2665A>G) | | | | | | | | |

#### Proteins

| | Begin | End | Coverage | Score | Concordance | Matches | Identities | I/D/M/F* | Stop Codons |
|---|---|---|---|---|---|---|---|---|---|
| RNA-dependent RNA polymerase (YP_010798874.1) | 32 | 777 | 95.9% | 3373 | 64.7% | 743 (99.6%) | 506 (67.8%) | 1/2/2/2 | 4 |

| | Begin | End | Coverage | Score | Concordance | Matches | Identities | I/D/M/F* | Stop Codons |
|---|---|---|---|---|---|---|---|---|---|
| **NT** | **428** | **2700** | **82.0%** | **1546** | **34.4%** | **2264 (99.6%)** | **1530 (67.3%)** | **1/9** | |
| Protein mutations: | I36L (441A>C 443C>T), L38F (447C>T), A44V (466C>T), A46M (471G>A 472C>T 473T>G), L48M (477T>A), T49H (480A>C 481C>A 482A>T), I52V (489A>G), V60I (513G>A), S61K (516T>A 517C>A 518T>A), K62T (520A>C), T64V (525A>G 526C>T 527T>G), T65K (529C>A 530C>A), R67K (534C>A 535G>A 536C>A), K69R (540A>C 541A>G 542G>T), Y72F (550A>T), F74Y (556T>A 557C>T), A83T (582G>A), C85S (589G>C), F94Y (616T>A), R95S (618C>T 619G>C 620A>T), E97N (624G>A 626A>T), E98D (629A>C), P99T (630C>A 632A>C), L100M (633C>A 635T>G), S109C (661C>G 662G>T), R113K (672C>A 673G>A 674T>G), K117A (684A>G 685A>C 686A>C), A122H (699G>C 700C>A 701A>C), I123V (702A>G), R125A (708A>G 709G>C 710G>A), D127S (714G>T 715A>C), E128D (719A>T), A144S (765G>T 767T>A), V146* (771G>T 772T>A 773T>G), L147I (774T>A 776G>T), K148A (777A>G 778A>C 779A>G), S154K (796G>A 797T>G), E159Q (810G>C), P166S (831C>T 833T>A), G169D (841G>A 842G>T), L172R (850T>G 851A>T), E176G (862A>G), I178M (869T>G), F182* (880T>G 881C>A), H184K (885C>A 887T>G), Q186R (892A>G), F192H (909T>C 910T>A), R194Q (916G>A), G195S (918G>T 919G>C), M198L (927A>C 929G>T), I200K (934T>A 935T>G), S203N (942T>A 943C>A), Q205K (948C>A), N214L (975A>C 976A>T 977T>C), S220R (993T>C 994C>G 995A>G), Y221F (997A>T 998C>T), G222Q (999G>C 1000G>A 1001T>A), L223Y (1003T>A 1004A>T), A224S (1005G>T 1007C>G), K228V (1017A>G 1018A>T 1019A>T), Q231G (1026C>G 1027A>G), E232A (1030A>C 1031G>A), S234A (1035A>G 1036G>C), F237V (1044T>G 1046C>T), F239E (1050T>G 1051T>A 1052T>A), P240H (1054C>A 1055A>C), L250I (1083C>A), W255L (1099G>T), T257V (1104A>G 1105C>T 1106A>G), I260M (1115T>G), F263I (1122T>A 1124C>T), M272I (1151G>T), K276Q (1161A>C), S277M (1164T>A 1165C>T 1166T>G), F279L (1172C>A), F280Y (1174T>A 1175T>C), L282F (1181A>T), R284S (1185A>T 1186G>C 1187G>T), A285N (1188G>A 1189C>A), A286N (1191G>A 1192C>A 1193A>T), E288del (1197_1199delGAA), L290F (1205A>T), L293E (1212T>G 1213T>A), E296D (1223A>T), H297L (1224C>T 1225A>T 1226C>A), S301Q (1236A>C 1237G>A 1238T>A), W305L (1248T>C 1249G>T 1250G>T), T308S (1258C>G), S311G (1266A>G 1268T>G), P312A (1269C>G 1271T>C), R314Q (1276G>A), G324A (1306G>C), A325Q (1308G>C 1309C>A 1310A>G), V326A (1312T>C 1313G>C), V335I (1338G>A 1340A>C), V337A (1345T>C 1346A>T), M338I (1349G>A), Y341S (1357A>C 1358C>T), D352I (1389G>A 1390A>T), K360S (1414A>G 1415A>T), I361L (1416A>C 1418A>T), D368N (1437G>A), K371R (1446A>C 1447A>G 1448A>T), L374Q (1455T>C 1456T>A), A375Y (1458G>T 1459C>A), K377R (1465A>G 1466G>A), Q378L (1468A>T 1469A>T), N380R (1474A>G 1475C>A), F381P (1476T>C 1477T>C 1478T>G), N383S (1482A>T 1483A>C), L389F (1502A>C), D394N (1515G>A), M401I (1538G>A), M405I (1550G>C), S406E (1551T>G 1552C>A 1553T>A), P411S (1566C>A 1567C>G), S420del (1593_1597delAGCTC), I427F (1614A>T 1616C>T), N430S (1623A>T 1624A>C 1625C>A), K431P (1626A>C 1627A>C 1628A>T), L433M (1632T>A 1634A>G), T434V (1635A>G 1636C>T 1637A>G), R436K (1642G>A 1643A>G), V437I (1644G>A 1646G>T), Y438S (1647T>A 1648A>G), I440V (1653A>G), F450Y (1684T>A), H451Y (1686C>T), A455S (1698G>T 1700A>C), S458A (1707T>G), I464M (1727A>G), S470K (1744G>A 1745C>A), K471Y (1746A>T 1748A>T), G475T (1758G>A 1759G>C 1760A>G), C476S (1761T>A), N477T (1765A>C 1766C>T), S478T (1767T>A 1769T>C), G481A (1777G>C 1778G>T), S495T (1819G>C 1820T>A), K496E (1821A>G 1823A>G), N499R (1830A>C 1831A>G 1832T>G), E500Q (1833G>C), K502R (1840A>G), N509G (1860A>G 1861A>G), K511T (1867A>C), I514L (1875A>T), P515A (1878C>G 1880A>T), L518R (1887T>A 1888T>G 1889G>A), D521E (1898T>G), S535D (1938T>G 1939C>A 1940A>C), D539Y (1950G>T 1952T>C), L549T (1980T>A 1981T>C 1982A>T), A559Q (2010G>C 2011C>A 2012C>A), I562T (2020T>C), D565S (2028G>T 2029A>C 2030T>G), N567S (2034A>T 2035A>C 2036C>A), I568T (2038T>C), V570I (2043G>A), Q572P (2050A>C 2051A>C), R573I (2053G>T 2054A>T), L574F (2057G>T), G575W (2058G>T), Y579F (2071A>T), K580R (2074A>G 2075G>A), A583S (2082G>T 2084G>T), F586M (2091T>A 2093C>G), T587S (2094A>T), H600A (2133C>G 2134A>C 2135T>A), L601S (2136C>A 2137T>G 2138T>C), V604H (2145G>C 2146T>A 2147T>C), Y611L (2166T>C 2167A>T), W613Y (2173G>A 2174G>T), G618F (2187G>T 2188G>T), M619K (2191T>A 2192G>A), Y624F (2206A>T), Y624_L625insX (2207_2208insT), V630I (2223G>A), S631D (2226A>G 2227G>A), L634R (2236T>G 2237T>G), K638T (2248A>C), V639P (2250G>C 2251T>C), L642I (2259T>A), Q643R (2263A>G 2264G>A), I644L (2265A>C), K647N (2276G>T), D648E (2279T>G), V655R (2298G>A 2299T>G 2300T>A), R660C (2313C>T 2315T>C), T661I (2317C>T), R664V (2325C>G 2326G>T), Q665N (2328C>A 2330A>T), E668K (2337G>A), K672R (2349A>C 2350A>G), L674V (2355C>G 2357C>T), N675S (2358A>T 2359A>C 2360C>T), W676* (2362G>A), C681L (2377G>T 2378T>G), N683S (2382A>T 2383A>C 2384C>T), P684F (2385C>T 2386C>T 2387A>T), D685E (2390T>A), K689E (2400A>G 2402G>A), S690K (2404G>A 2405T>A), I692M (2411C>G), S694A (2415T>G 2417C>T), C697Y (2425G>A), T699* (2430A>T 2431C>A 2432A>G), L704S (2445C>T 2446T>C 2447C>G), E705N (2448G>A 2450A>T), T706E (2451A>G 2452C>A 2453T>A), F708L (2457T>C 2459T>G), D709E (2462T>A), I711V (2466A>G), N727D (2514A>G), P728L (2518C>T 2519C>G), T730I (2524C>T 2525C>T), Q733R (2532C>A 2533A>G), W734Y (2536G>A 2537G>T), F736L (2543T>G), N739S (2551A>G 2552C>T), V741S (2556G>T 2557T>C), N745D (2568A>G), K746R (2571A>C 2572A>G), L749F (2582A>T), N755A (2598A>G 2599A>C 2600C>T), D756N (2601G>A), S760A (2613T>G 2615T>A), E763Y (2622G>T 2624A>T), Y764L (2626A>T 2627C>A), Q765P (2629A>C), T766S (2631A>T), T767L (2634A>C 2635C>T), G768V (2638G>T 2639A>G), E770G (2644A>G), E771C (2646G>T 2647A>G 2648A>T), V772R (2649G>A 2650T>G), G773S (2652G>T 2653G>C 2654C>A), L774Y (2656T>A 2657A>T), R775D (2658A>G 2659G>A 2660A>C), L776F (2661C>T), *777R (2664T>A 2665A>G) | | | | | | | | |

| | Begin | End | Coverage | Score | Concordance | Matches | Identities | I/D/M/F* | Stop Codons |
|---|---|---|---|---|---|---|---|---|---|
| **NT** | **428** | **2700** | **82.0%** | **1546** | **34.4%** | **2264 (99.6%)** | **1530 (67.3%)** | **1/9** | |

Codon mutations:

AAA34AAG (437A>G), ATC36CTT (441A>C 443C>T), CTT38TTT (447C>T), CTA42TTA (459C>T), GCC44GTC (466C>T), TCC45TCT (470C>T), GCT46ATG (471G>A 472C>T 473T>G), TTG48ATG (477T>A), ACA49CAT (480A>C 481C>A 482A>T), AAA50AAG (485A>G), ATC52GTC (489A>G), GCT53GCC (494T>C), CTT54CTA (497T>A), GCT55GCG (500T>G), GGA56GGC (503A>C), TGT57TGC (506T>C), GAC58GAT (509C>T), TTT59TTC (512T>C), GTA60ATA (513G>A), TCT61AAA (516T>A 517C>A 518T>A), AAA62ACA (520A>C), GTG63GTT (524G>T), ACT64GTG (525A>G 526C>T 527T>G), ACC65AAA (529C>A 530C>A), TTA66CTG (531T>C 533A>G), CGC67AAA (534C>A 535G>A 536C>A), CGG68CGA (539G>A), AAG69CGT (540A>C 541A>G 542G>T), TCG70TCA (545G>A), GGC71GGT (548C>T), TAT72TTT (550A>T), TTC74TAT (556T>A 557C>T), ACA75ACT (560A>T), TAC78TAT (569C>T), AAG80AAA (575G>A), TGC82TGT (581C>T), GCA83ACA (582G>A), GTA84GTT (587A>T), TGC85TCC (589G>C), TTA86CTT (591T>C 593A>T), CGA88CGT (599A>T), TAC89TAT (602C>T), TCC93AGT (612T>A 613C>G 614C>T), TTT94TAT (616T>A), CGA95TCT (618C>T 619G>C 620A>T), GAA97AAT (624G>A 626A>T), GAA98GAC (629A>C), CCA99ACC (630C>A 632A>C), CTT100ATG (633C>A 635T>G), TCG101TCT (638G>T), GTG102GTT (641G>T), CCT103CCA (644T>A), GTG104GTT (647G>T), CTC106TTG (651C>T 653C>G), ACA107ACC (656A>C), AGA108CGA (657A>C), TCG109TGT (661C>G 662G>T), GGT110GGA (665T>A), ATC111ATT (668C>T), CGT113AAG (672C>A 673G>A 674T>G), ATC115ATA (680C>A), CCC116CCT (683C>T), AAA117GCC (684A>G 685A>C 686A>C), TTG119TTA (692G>A), CGT120CGA (695T>A), GCA122CAC (699G>C 700C>A 701A>C), ATA123GTA (702A>G), CGT124AGG (705C>A 707T>G), AGG125GCA (708A>G 709G>C 710G>A), AAA126AAG (713A>G), GAT127TCT (714G>T 715A>C), GAA128GAT (719A>T), CAC129CAT (722C>T), GTT134GTC (737T>C), TAC137TAT (746C>T), TCA139TCT (752A>T), GGG142GGA (761G>A), GCT144TCA (765G>T 767T>A), AAG145AAA (770G>A), GTT146TAG (771G>T 772T>A 773T>G), TTG147ATT (774T>A 776G>T), AAA148GCG (777A>G 778A>C 779A>G), GTT149GTG (782T>G), AGT154AAG (796G>A 797T>G), GCA156GCC (803A>C), TTT158TTC (809T>C), GAA159CAA (810G>C), TCC160TCA (815C>A), ATA161ATT (818A>T), ACT162ACG (821T>G), ACC163ACT (824C>T), CCA164CCT (827A>T), CCT166TCA (831C>T 833T>A), GGG169GAT (841G>A 842G>T), CTA172CGT (850T>G 851A>T), GTG174GTT (857G>T), CTG175CTT (860G>T), GAA176GGA (862A>G), ATT178ATG (869T>G), ACG180ACT (875G>T), TTC182TGA (880T>G 881C>A), CAT184AAG (885C>A 887T>G), CAA186CGA (892A>G), CTA190CTT (905A>T), CCC191CCA (908C>A), TTT192CAT (909T>C 910T>A), CGA194CAA (916G>A), GGC195TCC (918G>T 919G>C), ATT196ATA (923T>A), CCA197CCT (926A>T), ATG198CTT (927A>C 929G>T), ATT200AAG (934T>A 935T>G), GGG201GGT (938G>T), TCT203AAT (942T>A 943C>A), CAA205AAA (948C>A), CCC212CCA (971C>A), CTA213TTA (972C>T), AAT214CTC (975A>C 976A>T 977T>C), AAT216AAC (983T>C), AGG219CGC (990A>C 992G>C), TCA220CGG (993T>C 994C>G 995A>G), TAC221TTT (997A>T 998C>T), GGT222CAA (999G>C 1000G>A 1001T>A), TTA223TAT (1003T>A 1004A>T), GCC224TCG (1005G>T 1007C>G), GTA225GTG (1010A>G), GAC227GAT (1016C>T), AAA228GTT (1017A>G 1018A>T 1019A>T), GTT229GTA (1022T>A), GAT230GAC (1025T>C), CAA231GGA (1026C>G 1027A>G), GAG232GCA (1030A>C 1031G>A), GCT233GCG (1034T>G), AGT234GCT (1035A>G 1036G>C), TTC237GTT (1044T>G 1046C>T), AAG238AAA (1049G>A), TTT239GAA (1050T>G 1051T>A 1052T>A), CCA240CAC (1054C>A 1055A>C), ATT242ATC (1061T>C), GTT244GTG (1067T>G), TTG246TTA (1073G>A), AAA247AAG (1076A>G), CTA250ATA (1083C>A), GCG251GCT (1088G>T), GCC252GCG (1091C>G), TGG255TTG (1099G>T), ACA257GTG (1104A>G 1105C>T 1106A>G), ATT260ATG (1115T>G), TCA262TCC (1121A>C), TTC263ATT (1122T>A 1124C>T), CAG264CAA (1127G>A), GAC265GAT (1130C>T), GGC266GGG (1133C>G), TCA269TCC (1142A>C), CCC270CCT (1145C>T), GGG271GGT (1148G>T), ATG272ATT (1151G>T), TTG273CTT (1152T>C 1154G>T), TCC275TCT (1160C>T), AAA276CAA (1161A>C), TCT277ATG (1164T>A 1165C>T 1166T>G), GTC278GTT (1169C>T), TTC279TTA (1172C>A), TTT280TAC (1174T>A 1175T>C), CCT281CCA (1178T>A), TTA282TTT (1181A>T), GAC283GAT (1184C>T), AGG284TCT (1185A>T 1186G>C 1187G>T), GCT285AAT (1188G>A 1189C>A), GCA286AAT (1191G>A 1192C>A 1193A>T), ACT287ACC (1196T>C), GAA288del (1197_1199delGAA), TTA290TTT (1205A>T), AAC292AAT (1211C>T), TTA293GAA (1212T>G 1213T>A), TCT294TCC (1217T>C), CTC295CTT (1220C>T), GAA296GAT (1223A>T), CAC297TTA (1224C>T 1225A>T 1226C>A), GAG299GAA (1232G>A), AGT301CAA (1236A>C 1237G>A 1238T>A), GTC302GTA (1241C>A), CCA304CCC (1247A>C), TGG305CTT (1248T>C 1249G>T 1250G>T), TTT306TTC (1253T>C), TCA307TCT (1256A>T), ACC308AGC (1258C>G), TTG310TTA (1265G>A), AGT311GGG (1266A>G 1268T>G), CCT312GCC (1269C>G 1271T>C), TAT313TAC (1274T>C), CGA314CAA (1276G>A), GGG315GGA (1280G>A), GTG316GTT (1283G>T), GTT319GTG (1292T>G), AGG322AGA (1301G>A), CTT323TTG (1302C>T 1304T>G), GGT324GCT (1306G>C), GCA325CAG (1308G>C 1309C>A 1310A>G), GTG326GCC (1312T>C 1313G>C), ATA327ATT (1316A>T), GAG328GAA (1319G>A), GGC329GGT (1322C>T), GGA331GGG (1328A>G), AGA333CGG (1332A>C 1334A>G), GTA335ATC (1338G>A 1340A>C), GTA337GCT (1345T>C 1346A>T), ATG338ATA (1349G>A), AAC340AAT (1355C>T), TAC341TCT (1357A>C 1358C>T), CTC347CTG (1376C>G), CAC348CAT (1379C>T), CCT349CCC (1382T>C), GTT350GTC (1385T>C), CAC351CAT (1388C>T), GAT352ATT (1389G>A 1390A>T), ACA356ACT (1403A>T), GTC357GTA (1406C>A), CTG358CTA (1409G>A), TCT359TCA (1412T>A), AAA360AGT (1414A>G 1415A>T), ATA361CTT (1416A>C 1418A>T), ACA363ACT (1424A>T), GAC364GAT (1427C>T), GGG365GGT (1430G>T), ACT366ACA (1433T>A), GAC368AAC (1437G>A), GAG370GAA (1445G>A), AAA371CGT (1446A>C 1447A>G 1448A>T), CCT372CCG (1451T>G), TTG374CAG (1455T>C 1456T>A), GCT375TAT (1458G>T 1459C>A), TTA376CTA (1461T>C), AAG377AGA (1465A>G 1466G>A), CAA378CTT (1468A>T 1469A>T), AAG379AAA (1472G>A), AAC380AGA (1474A>G 1475C>A), TTT381CCG (1476T>C 1477T>C 1478T>G), AAA382AAG (1481A>G), AAT383TCT (1482A>T 1483A>C), TAT385TAC (1490T>C), TTA389TTC (1502A>C), AAG390AAA (1505G>A), ACG393ACA (1514G>A), GAC394AAC (1515G>A), CCG397CCA (1526G>A), GTC400GTT (1535C>T), ATG401ATA (1538G>A), TAT402TAC (1541T>C), CTT404TTA (1545C>T 1547T>A), ATG405ATC (1550G>C), TCT406GAA (1551T>G 1552C>A 1553T>A), ATA408ATT (1559A>T), GGG410GGT (1565G>T), CCT411AGT (1566C>A 1567C>G), ACC412ACT (1571C>T), TTG413TTA (1574G>A), TCC415TCT (1580C>T), GTT418GTC (1589T>C), AGC420del (1593_1597delAGCTC), TCT421--A (1593_1597delAGCTC 1598T>A), ACG426ACT (1613G>T), ATC427TTT (1614A>T 1616C>T), CTT428CTC (1619T>C), GTT429GTA (1622T>A), AAC430TCA (1623A>T 1624A>C 1625C>A), AAA431CCT (1626A>C 1627A>C 1628A>T), CCG432CCT (1631G>T), TTA433ATG (1632T>A 1634A>G), ACA434GTG (1635A>G 1636C>T 1637A>G), AGA436AAG (1642G>A 1643A>G), GTG437ATT (1644G>A 1646G>T), TAT438AGT (1647T>A 1648A>G), GAG439GAA (1652G>A), ATA440GTA (1653A>G), CTG443CTT (1664G>T), GGG445GGA (1670G>A), CAG446CAA (1673G>A), CCT447CCG (1676T>G), TTG448TTA (1679G>A), GGC449GGT (1682C>T), TTC450TAC (1684T>A), CAC451TAC (1686C>T), TCT453TCA (1694T>A), GCA455TCC (1698G>T 1700A>C), CTT456CTA (1703T>A), TTC457TTT (1706C>T), TCG458GCG (1707T>G), TTG459CTT (1710T>C 1712G>T), TCT460TCC (1715T>C), CAC462CAT (1721C>T), ATA464ATG (1727A>G), TTA467CTA (1734T>C), GCA468GCT (1739A>T), AGC470AAA (1744G>A 1745C>A), AAA471TAT (1746A>T 1748A>T), TAT473TAC (1754T>C), GGA475ACG (1758G>A 1759G>C 1760A>G), TGT476AGT (1761T>A), AAC477ACT (1765A>C 1766C>T), TCT478ACC (1767T>A 1769T>C), CCA479CCT (1772A>T), GGG481GCT (1777G>C 1778G>T), GCT484GCC (1787T>C), TTA485CTG (1788T>C 1790A>G), CTA486TTA (1791C>T), AGT495ACA (1819G>C 1820T>A), AAA496GAG (1821A>G 1823A>G), GCC498GCG (1829C>G), AAT499CGG (1830A>C 1831A>G 1832T>G), GAG500CAG (1833G>C), AAG502AGG (1840A>G), TTA504TTG (1847A>G), TTG505CTG (1848T>C), AGA507AGG (1856A>G), CTC508CTT (1859C>T), AAT509GGT (1860A>G 1861A>G), GTC510GTT (1865C>T), AAA511ACA (1867A>C), ATT512ATC (1871T>C), TCG513TCT (1874G>T), ATA514TTA (1875A>T), CCA515GCT (1878C>G 1880A>T), TCT517TCG (1886T>G), TTG518AGA (1887T>A 1888T>G 1889G>A), TCT520TCC (1895T>C), GAT521GAG (1898T>G), AAT522AAC (1901T>C), ATT525ATC (1910T>C), GCG528GCT (1919G>T), AGG530AGA (1925G>A), ACT533ACC (1934T>C), AAA534AAG (1937A>G), TCA535GAC (1938T>G 1939C>A 1940A>C), GAT539TAC (1950G>T 1952T>C), TTA540CTT (1953T>C 1955A>T), ATT543ATC (1964T>C), TCC544TCT (1967C>T), GCT547GCA (1976T>A), TTA549ACT (1980T>A 1981T>C 1982A>T), TCT550TCC (1985T>C), CGT552AGG (1989C>A 1991T>G), ACT553ACA (1994T>A), GTT555GTG (2000T>G), GGA556GGT (2003A>T), CTT557CTG (2006T>G), TGC558TGT (2009C>T), GCC559CAA (2010G>C 2011C>A 2012C>A), ATC562ACC (2020T>C), GAT565TCG (2028G>T 2029A>C 2030T>G), AAC567TCA (2034A>T 2035A>C 2036C>A), ATA568ACA (2038T>C), TCA569TCG (2042A>G), GTT570ATT (2043G>A), CAA572CCC (2050A>C 2051A>C), AGA573ATT (2053G>T 2054A>T), TTG574TTT (2057G>T), GGG575TGG (2058G>T), GGT576GGG (2063T>G), GGG578GGA (2069G>A), TAC579TTC (2071A>T), AAG580AGA (2074A>G 2075G>A), GTA581GTT (2078A>T), GCG583TCT (2082G>T 2084G>T), CGC584CGT (2087C>T), TTA585TTG (2090A>G), TTC586ATG (2091T>A 2093C>G), ACT587TCT (2094A>T), ACC588ACT (2099C>T), AGC590TCT (2103A>T 2104G>C 2105C>T), AAG591AAA (2108G>A), GAG594GAA (2117G>A), CGT595AGA (2118C>A 2120T>A), CTA596CTG (2123A>G), GCT598GCG (2129T>G), CAT600GCA (2133C>G 2134A>C 2135T>A), CTT601AGC (2136C>A 2137T>G 2138T>C), AAG602AAA (2141G>A), GTT604CAC (2145G>C 2146T>A 2147T>C), CGG605AGG (2148C>A), CAC607CAT (2156C>T), TTA608CTA (2157T>C), CTA609TTA (2160C>T), TAC611CTC (2166T>C 2167A>T), TGG613TAT (2173G>A 2174G>T), GGA616GGG (2183A>G), AGG617AGA (2186G>A), GGT618TTT (2187G>T 2188G>T), ATG619AAA (2191T>A 2192G>A), CCC620CCT (2195C>T), CCC623CCT (2204C>T), TAT624TTT (2206A>T), TAT624_TTG625insT-- (2207_2208insT), AAA626AAG (2213A>G), GGG627GGA (2216G>A), GTA630ATA (2223G>A), AGT631GAT (2226A>G 2227G>A), TAC632TAT (2231C>T), CTT634CGG (2236T>G 2237T>G), AAA635AAG (2240A>G), AAA638ACA (2248A>C), GTT639CCT (2250G>C 2251T>C), TTA642ATA (2259T>A), CAG643CGA (2263A>G 2264G>A), ATC644CTC (2265A>C), AAG647AAT (2276G>T), GAT648GAG (2279T>G), CTG649TTG (2280C>T), TTC651TTT (2288C>T), GAA654GAG (2297A>G), GTT655AGA (2298G>A 2299T>G 2300T>A), ATA657ATC (2306A>C), CTA658CTT (2309A>T), CGT660TGC (2313C>T 2315T>C), ACT661ATT (2317C>T), CTA663TTA (2322C>T), CGC664GTC (2325C>G 2326G>T), CAA665AAT (2328C>A 2330A>T), GTA667GTT (2336A>T), GAA668AAA (2337G>A), TTA671CTT (2346T>C 2348A>T), AAG672CGG (2349A>C 2350A>G), CTC674GTT (2355C>G 2357C>T), AAC675TCT (2358A>T 2359A>C 2360C>T), TGG676TAG (2362G>A), CAT678CAC (2369T>C), ACG679ACT (2372G>T), TGT681TTG (2377G>T 2378T>G), AAC683TCT (2382A>T 2383A>C 2384C>T), CCA684TTT (2385C>T 2386C>T 2387A>T), GAT685GAA (2390T>A), TCC687TCT (2396C>T), ATT688ATC (2399T>C), AAG689GAA (2400A>G 2402G>A), AGT690AAA (2404G>A 2405T>A), CTC691TTG (2406C>T 2408C>G), ATC692ATG (2411C>G), GAC693GAT (2414C>T), TCC694GCT (2415T>G 2417C>T), CCT695CCC (2420T>C), ATA696ATT (2423A>T), TGC697TAC (2425G>A), ACA699TAG (2430A>T 2431C>A 2432A>G), TCC700TCA (2435C>A), AAA702AAG (2441A>G), AGG703CGA (2442A>C 2444G>A), CTC704TCG (2445C>T 2446T>C 2447C>G), GAA705AAT (2448G>A 2450A>T), ACT706GAA (2451A>G 2452C>A 2453T>A), GAT707GAC (2456T>C), TTT708CTG (2457T>C 2459T>G), GAT709GAA (2462T>A), TTA710TTG (2465A>G), ATC711GTC (2466A>G), GGC714GGT (2477C>T), CTA715TTA (2478C>T), AAA718AAG (2489A>G), TGT719TGC (2492T>C), TAC720TAT (2495C>T), GGT723GGA (2504T>A), GGT725GGC (2510T>C), AAT727GAT (2514A>G), CCC728CTG (2518C>T 2519C>G), ACC730ATT (2524C>T 2525C>T), ACT731ACA (2528T>A), CAG733AGG (2532C>A 2533A>G), TGG734TAT (2536G>A 2537G>T), TTT736TTG (2543T>G), AAC739AGT (2551A>G 2552C>T), ACT740ACC (2555T>C), GTT741TCT (2556G>T 2557T>C), ATT742ATC (2561T>C), AAT745GAT (2568A>G), AAA746CGA (2571A>C 2572A>G), TTA749TTT (2582A>T), GGG750GGA (2585G>A), GGG754GGA (2597G>A), AAC755GCT (2598A>G 2599A>C 2600C>T), GAT756AAT (2601G>A), ATT758ATA (2609T>A), TCT760GCA (2613T>G 2615T>A), CCT761CCA (2618T>A), GAA763TAT (2622G>T 2624A>T), TAC764TTA (2626A>T 2627C>A), CAA765CCA (2629A>C), ACT766TCT (2631A>T), ACA767CTA (2634A>C 2635C>T), GGA768GTG (2638G>T 2639A>G), GAG770GGG (2644A>G), GAA771TGT (2646G>T 2647A>G 2648A>T), GTG772AGG (2649G>A 2650T>G), GGC773TCA (2652G>T 2653G>C 2654C>A), TTA774TAT (2656T>A 2657A>T), AGA775GAC (2658A>G 2659G>A 2660A>C), CTT776TTT (2661C>T), TAG777AGG (2664T>A 2665A>G)

*: Inserts / Deletes / Misaligned / Frameshifts

#### Analysis details

This analysis was performed with panviral2.64

### NGS Details (UN59): Solendovirus venanicotianae

#### Assembly

| | |
|---|---|
| **Coverage Length** | 2286 (5 contig(s)) |
| **Depth Of Coverage** | 14.6 |
| **Number Of Reads** | 285 |
| **Reads Per Million** | 6.20 rpm (after QC) |
| **Ambiguities** | 0 |
| **Assembly Method** | de novo + reference guided assembly |
| **Consensus Caller** | Bcf Tools |

#### Coverage Map

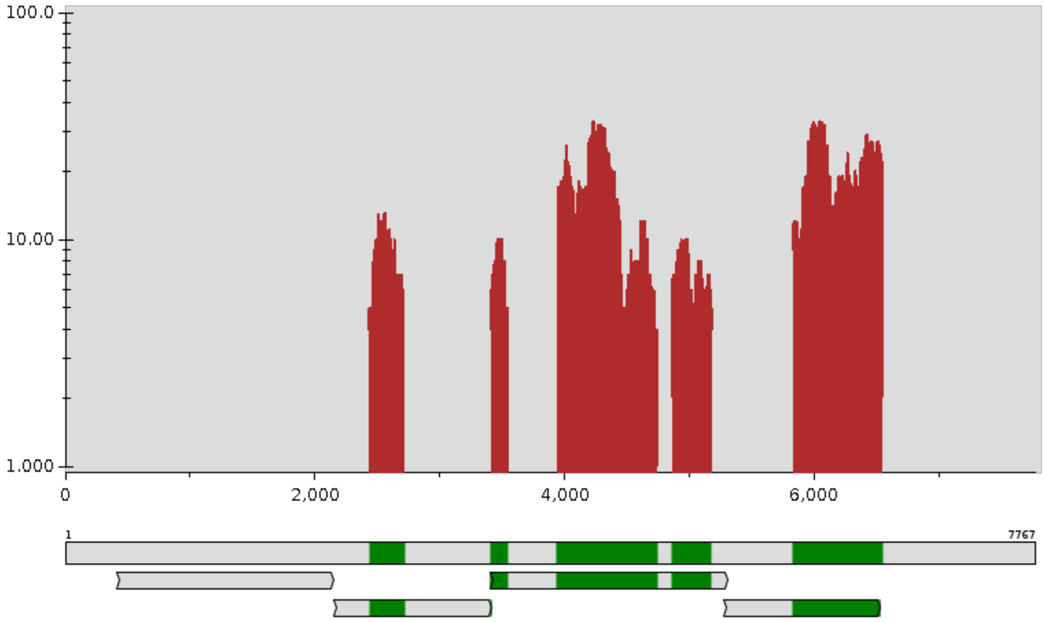

#### Assignment

| | |
|---|---|
| **Type** | Solendovirus venanicotianae (Taxonomy ID: 3048371) |
| **Reference Genome** | NC_003378.1 |
| **NT Identity (%)** | 79.8494 |
| **AA Identity (%)** | 76.8097 |
| **Number Of Stop Codons** | 4 |
| **Number Of CDS** | 4 |

#### Alignment

| | |
|---|---|
| **Alignment Score** | 2623.0 (NT) + 3754.0 (AA) = 6377.0 |
| **Concordance (%)** | 68.6585 |

| Alignment Method | Global, seeded, nucleotide + amino acids (AGA) |
|---|---|

#### Genome Region

Sequence starts at position 2433 and ends at position 6550 relative to NC_003378.1 reference sequence.

#### Alignment Detailed Statistics

| | Begin | End | Coverage | Score | Concordance | Matches | Identities | I/D/M/F* | Stop Codons |
|---|---|---|---|---|---|---|---|---|---|
| **NT** | **2433** | **6550** | **29.4%** | **2623** | **59.0%** | **2257 (98.7%)** | **1803 (78.8%)** | **1/29** | |
| Mutations: | 2442A>C, 2443C>A, 2446G>A, 2448T>G, 2454G>A, 2455G>A, 2456C>T, 2457A>G, 2463C>A, 2464C>G, 2465C>A, 2472G>A, 2482T>C, 2493T>A, 2511A>T, 2514T>C, 2535C>A, 2551C>T, 2571C>A, 2590A>G, 2598T>C, 2602A>G, 2604A>T, 2622T>C, 2628G>A, 2637T>C, 2643A>G, 2646C>T, 2648C>T, 2652C>T, 2665C>A, 2668A>G, 2670T>A, 2676A>T, 2687A>G, 2692T>C, 2709G>A, 2717G>A, 3406G>A, 3409G>T, 3410A>T, 3419A>T, 3422C>T, 3431A>C, 3434A>G, 3437C>A, 3443A>T, 3452T>C, 3456A>G, 3464C>T, 3467T>C, 3470A>T, 3500T>C, 3506T>C, 3518T>C, 3525A>G, 3526C>A, 3530T>A, 3536G>A, 3537G>A, 3547A>C, 3550C>T, 3936T>C, 3942A>G, 3952A>G, 3956T>A, 3957A>G, 3962T>A, 3963C>T, 3966C>A, 3977G>T, 3989A>T, 3991A>C, 4010T>A, 4011G>A, 4013A>T, 4014G>A, 4016G>A, 4017A>G, 4025A>T, 4028A>C, 4040T>C, 4041C>T, 4045A>G, 4046A>T, 4049T>C, 4051A>G, 4052C>T, 4056A>G, 4061T>C, 4062T>C, 4063T>A, 4066C>A, 4070A>G, 4073T>C, 4079A>G, 4086A>G, 4088A>T, 4091T>A, 4097G>A, 4100T>A, 4103T>A, 4104A>G, 4105A>G, 4115A>G, 4116G>A, 4121T>C, 4122A>G, 4124T>A, 4125A>G, 4127C>T, 4133T>C, 4161C>T, 4181A>T, 4190G>A, 4196C>T, 4199A>T, 4202C>T, 4212T>C, 4241T>C, 4250C>A, 4253T>C, 4265G>A, 4275A>G, 4286T>C, 4292C>T, 4298T>C, 4307C>T, 4313G>A, 4325T>C, 4328T>C, 4329T>C, 4331G>A, 4334A>G, 4340G>A, 4343T>A, 4355A>G, 4362G>A, 4364T>A, 4388T>C, 4400T>A, 4406A>T, 4409T>C, 4415T>C, 4421T>C, 4423C>T, 4436C>T, 4442T>C, 4448G>T, 4449G>A, 4451C>T, 4454C>T, 4457T>C, 4463T>C, 4475T>C, 4484A>T, 4487T>C, 4493T>C, 4500C>T, 4503C>T, 4508T>C, 4513G>A, 4540T>C, 4545A>C, 4550C>T, 4551G>A, 4552C>T, 4554C>A, 4559T>A, 4560A>G, 4568C>A, 4571T>A, 4574T>A, 4581T>C, 4591C>A, 4592C>G, 4598A>T, 4601A>G, 4604T>A, 4605A>T, 4607G>T, 4626T>C, 4643T>C, 4653A>G, 4660T>C, 4665A>C, 4666C>A, 4667T>A, 4679A>G, 4687T>A, 4691T>C, 4692C>T, 4693T>C, 4694T>A, 4697T>A, 4701A>G, 4703C>A, 4704A>T, 4706T>A, 4721A>G, 4722T>C, 4725C>T, 4734C>T, 4740A>T, 4868G>A, 4869A>T, 4872T>C, 4874G>T, 4897A>G, 4898C>T, 4907T>C, 4913C>T, 4919A>T, 4924C>T, 4925C>A, 4931T>A, 4938A>G, 4944T>G, 4946A>G, 4950A>G, 4953T>C, 4955T>A, 4970T>A, 4971C>T, 4973T>A, 4976G>A, 4979T>C, 4982A>G, 4986A>G, 4988T>A, 5011G>A, 5012C>T, 5018C>T, 5021T>C, 5027A>T, 5028T>A, 5030T>A, 5032A>T, 5033T>C, 5034A>T, 5037G>A, 5039A>C, 5040C>T, 5043C>G, 5046G>A, 5047A>T, 5053G>A, 5055G>A, 5076T>C, 5077T>A, 5079G>T, 5094C>T, 5117A>T, 5120T>G, 5123C>T, 5125C>A, 5128G>A, 5135T>A, 5136G>A, 5147C>T, 5150T>C, 5156A>G, 5174T>A, 5180A>G, 5833G>C, 5841T>C, 5844T>A, 5850C>A, 5851C>A, 5853T>A, 5857A>C, 5858C>A, 5859A>G, 5865C>T, 5868T>C, 5872T>C, 5874G>T, 5877A>T, 5880T>A, 5881T>C, 5882T>A, 5885C>T, 5886A>C, 5889T>A, 5892A>T, 5894G>A, 5897C>T, 5898A>C, 5901A>T, 5902T>C, 5907A>T, 5908G>A, 5909C>G, 5910C>A, 5913C>A, 5916C>T, 5921A>G, 5922C>T, 5925C>T, 5935G>A, 5936A>G, 5940T>C, 5941G>A, 5950C>T, 5952G>C, 5953A>G, 5954C>A, 5955A>T, 5959C>G, 5964G>A, 5968G>T, 5970C>A, 5972A>C, 5979T>A, 5991G>A, 5997C>T, 5999T>C, 6000G>A, 6001C>A, 6002A>C, 6009G>A, 6012G>A, 6015A>T, 6018T>A, 6024G>C, 6026C>A, 6027A>C, 6033C>T, 6036C>T, 6037G>A, 6040C>A, 6041G>A, 6042A>G, 6045C>T, 6049T>A, 6051G>A, 6054T>A, 6055A>C, 6063G>T, 6067C>T, 6071A>T, 6075A>T, 6078G>A, 6081A>T, 6084G>A, 6087T>A, 6088A>G, 6090C>G, 6105A>G, 6108T>A, 6114C>A, 6120G>A, 6121G>A, 6123A>T, 6127A>T, 6129C>A, 6139A>G, 6141G>C, 6148G>C, 6154G>C, 6155A>C, 6163G>C, 6164A>C, 6169G>A, 6171A>T, 6172A>G, 6174T>G, 6177T>A, 6181G>A, 6182A>G, 6183A>T, 6186C>T, 6188A>G, 6189T>C, 6201G>A, 6204A>T, 6211A>C, 6219T>G, 6223C>A, 6225A>C, 6232C>T, 6246T>C, 6247C>A, 6248T>C, 6253C>G, 6254A>G, 6258T>C, 6259G>C, 6264T>G, 6269G>A, 6270C>T, 6271T>C, 6277_6283delTCAAGGG, 6285A>C, 6290C>T, 6298_6315delTATTCAAATTGCAGAGAT, 6318A>C, 6319_6321delAGA, 6325G>A, 6326C>G, 6340C>G, 6341T>C, 6344G>T, 6345A>G, 6352G>A, 6354C>A, 6355T>A, 6357A>G, 6359G>C, 6360T>A, 6361C>T, 6363T>G, 6369A>T, 6375A>G, 6376C>A, 6380C>A, 6381T>A, 6382A>C, 6384A>T, 6386C>T, 6387C>T, 6401G>A, 6402G>A, 6404G>A, 6406delA, 6407A>G, 6408T>A, 6411C>T, 6414C>T, 6417C>G, 6418C>G, 6419C>A, 6420A>T, 6423T>A, 6424C>T, 6426G>A, 6427T>C, 6446T>A, 6451A>G, 6453T>G, 6456C>A, 6462T>C, 6471C>T, 6477C>T, 6480A>G, 6489A>C, 6502A>G, 6504T>A, 6505G>A, 6507T>C, 6513C>T, 6522G>A, 6525A>C, 6526T>G, 6530T>A, 6531C>A, 6532_6533insA, 6537G>T, 6542A>T | | | | | | | | |
| **CDS** | | | | | | | | | |
| Tvvgp2 | 94 | 420 | 23.6% | 549 | 83.6% | 99 (100%) | 82 (82.8%) | 0/0/0/0 | 1 |
| Protein mutations: | Q97K (2443C>A), D98K (2446G>A 2448T>G), A101M (2455G>A 2456C>T 2457A>G), N103K (2463C>A), P104E (2464C>G 2465C>A), I146V (2590A>G), I150V (2602A>G 2604A>T), I163M (2643A>G), A165V (2648C>T), L171I (2665C>A), N172E (2668A>G 2670T>A), R174S (2676A>T), K178R (2687A>G), G188E (2717G>A), E418K (3406G>A), E419L (3409G>T 3410A>T) | | | | | | | | |
| Codon mutations: | ATA96ATC (2442A>C), CAA97AAA (2443C>A), GAT98AAG (2446G>A 2448T>G), AAG100AAA (2454G>A), GCA101ATG (2455G>A 2456C>T 2457A>G), AAC103AAA (2463C>A), CCA104GAA (2464C>G 2465C>A), AAG106AAA (2472G>A), TTA110CTA (2482T>C), ACT113ACA (2493T>A), GCA119GCT (2511A>T), TGT120TGC (2514T>C), ACC127ACA (2535C>A), CTA133TTA (2551C>T), ATC139ATA (2571C>A), ATA146GTA (2590A>G), AGT148AGC (2598T>C), ATA150GTT (2602A>G 2604A>T), TAT156TAC (2622T>C), AAG158AAA (2628G>A), TTT161TTC (2637T>C), ATA163ATG (2643A>G), AGC164AGT (2646C>T), GCC165GTC (2648C>T), AAC166AAT (2652C>T), CTA171ATA (2665C>A), AAT172GAA (2668A>G 2670T>A), AGA174AGT (2676A>T), AAA178AGA (2687A>G), TTA180CTA (2692T>C), AAG185AAA (2709G>A), GGA188GAA (2717G>A), GAA418AAA (3406G>A), GAA419TTA (3409G>T 3410A>T) | | | | | | | | |
| Tvvgp3 | 1 | 591 | 66.8% | 2399 | 80.4% | 425 (100%) | 342 (80.5%) | 0/0/0/0 | 3 |
| Protein mutations: | N16D (3456A>G), T39E (3525A>G 3526C>A), D40E (3530T>A), W42* (3536G>A), E43K (3537G>A), K46T (3547A>C), T47I (3550C>T), K178E (3942A>G), N181S (3952A>G), D182E (3956T>A), N183D (3957A>G), Q186K (3966C>A), E189D (3977G>T), K194T (3991A>C), E201N (4011G>A 4013A>T), E202K (4014G>A 4016G>A), N203D (4017A>G), K212S (4045A>G 4046A>T), N214S (4051A>G 4052C>T), N216D (4056A>G), L218Q (4062T>C 4063T>A), T219K (4066C>A), I223M (4079A>G), K226D (4086A>G 4088A>T), D227E (4091T>A), D230E (4100T>A), N231K (4103T>A), K232G (4104A>G 4105A>G), E236K (4116G>A), N238E (4122A>G 4124T>A), S239G (4125A>G 4127C>T), H251Y (4161C>T), I289V (4275A>G), D311E (4343T>A), A318T (4362G>A 4364T>A), A338V (4423C>T), M346I (4448G>T), D347N (4449G>A 4451C>T), R368K (4513G>A), L377S (4540T>C), K379Q (4545A>C), A381I (4551G>A 4552C>T), H382N (4554C>A), I384V (4560A>G), N386K (4568C>A), T394K (4591C>A 4592C>G), M399F (4605A>T 4607G>T), I415V (4653A>G), M417T (4660T>C), T419Q (4665A>C 4666C>A 4667T>A), I426N (4687T>A), L428S (4692C>T 4693T>C 4694T>A), D429E (4697T>A), N431E (4701A>G 4703C>A), I432L (4704A>T 4706T>A), Q439* (4725C>T), I444L (4740A>T), I487L (4869A>T), Y496C (4897A>G 4898C>T), K503N (4919A>T), T505I (4924C>T 4925C>A), T510A (4938A>G), S512A (4944T>G 4946A>G), N514D (4950A>G), Y515Q (4953T>C 4955T>A), N526E (4986A>G 4988T>A), C534Y (5011G>A 5012C>T), S540T (5028T>A 5030T>A), Y541F (5032A>T 5033T>C), T542S (5034A>T), E543N (5037G>A 5039A>C), P544S (5040C>T), Q545E (5043C>G), E546I (5046G>A 5047A>T), W548* (5053G>A), E549K (5055G>A), F556H (5076T>C 5077T>A), A557S (5079G>T), T572N (5125C>A), R573K (5128G>A), V576I (5136G>A) | | | | | | | | |
| Codon mutations: | ATA3ATT (3419A>T), TAC4TAT (3422C>T), GCA7GCC (3431A>C), AAA8AAG (3434A>G), ATC9ATA (3437C>A), GTA11GTT (3443A>T), TAT14TAC (3452T>C), AAT16GAT (3456A>G), TAC18TAT (3464C>T), TAT19TAC (3467T>C), ACA20ACT (3470A>T), AAT30AAC (3500T>C), TGT32TGC (3506T>C), TGT36TGC (3518T>C), ACA39GAA (3525A>G 3526C>A), GAT40GAA (3530T>A), TGG42TGA (3536G>A), GAA43AAA (3537G>A), AAA46ACA (3547A>C), ACA47ATA (3550C>T), TTA176CTA (3936T>C), AAA178GAA (3942A>G), AAT181AGT (3952A>G), GAT182GAA (3956T>A), AAC183GAC (3957A>G), CCT184CCA (3962T>A), CTA185TTA (3963C>T), CAA186AAA (3966C>A), GAG189GAT (3977G>T), ACA193ACT (3989A>T), AAA194ACA (3991A>C), ATT200ATA (4010T>A), GAA201AAT (4011G>A 4013A>T), GAG202AAA (4014G>A 4016G>A), AAT203GAT (4017A>G), ATA205ATT (4025A>T), ATA206ATC (4028A>C), CCT210CCC (4040T>C), CTA211TTA (4041C>T), AAA212AGT (4045A>G 4046A>T), TAT213TAC (4049T>C), AAC214AGT (4051A>G 4052C>T), AAT216GAT (4056A>G), GAT217GAC (4061T>C), TTA218CAA (4062T>C 4063T>A), ACA219AAA (4066C>A), GAA220GAG (4070A>G), TTT221TTC (4073T>C), ATA223ATG (4079A>G), AAA226GAT (4086A>G 4088A>T), GAT227GAA (4091T>A), TTG229TTA (4097G>A), GAT230GAA (4100T>A), AAT231AAA (4103T>A), AAA232GGA (4104A>G 4105A>G), CAA235CAG (4115A>G), GAA236AAA (4116G>A), AGT237AGC (4121T>C), AAT238GAA (4122A>G 4124T>A), AGC239GGT (4125A>G 4127C>T), CAT241CAC (4133T>C), CAT251TAT (4161C>T), GGA257GGT (4181A>T), AGG260AGA (4190G>A), GTC262GTT (4196C>T), ATA263ATT (4199A>T), GAC264GAT (4202C>T), TTA268CTA (4212T>C), TAT277TAC (4241T>C), CCC280CCA (4250C>A), AAT281AAC (4253T>C), AAG285AAA (4265G>A), ATA289GTA (4275A>G), TAT292TAC (4286T>C), TAC294TAT (4292C>T), AGT296AGC (4298T>C), GAC299GAT (4307C>T), AAG301AAA (4313G>A), TAT305TAC (4325T>C), CAT306CAC (4328T>C), TTG307CTA (4329T>C 4331G>A), AAA308AAG (4334A>G), GAG310GAA (4340G>A), GAT311GAA (4343T>A), AAA315AAG (4355A>G), GCT318ACA (4362G>A 4364T>A), TAT326TAC (4388T>C), GTT330GTA (4400T>A), CCA332CCT (4406A>T), TTT333TTC (4409T>C), TAT335TAC (4415T>C), AAT337AAC (4421T>C), GCA338GTA (4423C>T), TAC342TAT (4436C>T), CAT344CAC (4442T>C), ATG346ATT (4448G>T), GAC347AAT (4449G>A 4451C>T), AAC348AAT (4454C>T), TAT349TAC (4457T>C), AAT351AAC (4463T>C), AAT355AAC (4475T>C), GTA358GTT (4484A>T), TAT359TAC (4487T>C), GAT361GAC (4493T>C), CTA364TTA (4500C>T), CTA365TTA (4503C>T), TAT366TAC (4508T>C), AGA368AAA (4513G>A), TTA377TCA (4540T>C), AAA379CAA (4545A>C), TTC380TTT (4550C>T), GCA381ATA (4551G>A 4552C>T), CAT382AAT (4554C>A), ATT383ATA (4559T>A), ATA384GTA (4560A>G), AAC386AAA (4568C>A), TCT387TCA (4571T>A), GGT388GGA (4574T>A), TTA391CTA (4581T>C), ACC394AAG (4591C>A 4592C>G), GCA396GCT (4598A>T), GAA397GAG (4601A>G), ATT398ATA (4604T>A), ATG399TTT (4605A>T 4607G>T), TTA406CTA (4626T>C), GAT411GAC (4643T>C), ATA415GTA (4653A>G), ATG417ACG (4660T>C), ACT419CAA (4665A>C 4666C>A 4667T>A), CAA423CAG (4679A>G), ATC426AAC (4687T>A), AAT427AAC (4691T>C), CTT428TCA (4692C>T 4693T>C 4694T>A), GAT429GAA (4697T>A), AAC431GAA (4701A>G 4703C>A), ATT432TTA (4704A>T 4706T>A), AAA437AAG (4721A>G), TTA438CTA (4722T>C), CAA439TAA (4725C>T), CTA442TTA (4734C>T), ATA444TTA (4740A>T), AAG486AAA (4868G>A), ATA487TTA (4869A>T), TTG488CTT (4872T>C 4874G>T), TAC496TGT (4897A>G 4898C>T), GAT499GAC (4907T>C), AAC501AAT (4913C>T), AAA503AAT (4919A>T), ACC505ATA (4924C>T 4925C>A), ATT507ATA (4931T>A), ACA510GCA (4938A>G), TCA512GCG (4944T>G 4946A>G), AAT514GAT (4950A>G), TAT515CAA (4953T>C 4955T>A), GTT520GTA (4970T>A), CTT521TTA (4971C>T 4973T>A), AAG522AAA (4976G>A), TAT523TAC (4979T>C), AGA524AGG (4982A>G), AAT526GAA (4986A>G 4988T>A), TGC534TAT (5011G>A 5012C>T), TAC536TAT (5018C>T), TAT537TAC (5021T>C), GGA539GGT (5027A>T), TCT540ACA (5028T>A 5030T>A), TAT541TTC (5032A>T 5033T>C), ACA542TCA (5034A>T), GAA543AAC (5037G>A 5039A>C), CCA544TCA (5040C>T), CAA545GAA (5043C>G), GAA546ATA (5046G>A 5047A>T), TGG548TAG (5053G>A), GAA549AAA (5055G>A), TTT556CAT (5076T>C 5077T>A), GCA557TCA (5079G>T), CTA562TTA (5094C>T), ATA569ATT (5117A>T), GTT570GTG (5120T>G), TAC571TAT (5123C>T), ACC572AAC (5125C>A), AGA573AAA (5128G>A), ATT575ATA (5135T>A), GTA576ATA (5136G>A), GAC579GAT (5147C>T), AAT580AAC (5150T>C), CAA582CAG (5156A>G), ACT588ACA (5174T>A), AAA590AAG (5180A>G) | | | | | | | | |
| Tvvgp4 | 187 | 418 | 55.3% | 806 | 54.0% | 222 (96.1%) | 149 (64.5%) | 0/9/2/2 | 0 |

| | Begin | End | Coverage | Score | Concordance | Matches | Identities | I/D/M/F* | Stop Codons |
|---|---|---|---|---|---|---|---|---|---|
| **NT** | **2433** | **6550** | **29.4%** | **2623** | **59.0%** | **2257 (98.7%)** | **1803 (78.8%)** | **1/29** | |
| Protein mutations: | E187Q (5833G>C), H192Q (5850C>A), H193K (5851C>A 5853T>A), T195Q (5857A>C 5858C>A 5859A>G), L203Q (5881T>C 5882T>A), P204L (5885C>T 5886A>C), N205K (5889T>A), S207N (5894G>A), A208V (5897C>T 5898A>C), K209N (5901A>T), A212R (5908G>A 5909C>G 5910C>A), N216S (5921A>G 5922C>T), D221S (5935G>A 5936A>G), V223I (5941G>A), Q226Y (5950C>T 5952G>C), T227D (5953A>G 5954C>A 5955A>T), Q229E (5959C>G), A232S (5968G>T 5970C>A), N233T (5972A>C), M242T (5999T>C 6000G>A), Q243T (6001C>A 6002A>C), T251N (6026C>A 6027A>C), V255I (6037G>A), R256K (6040C>A 6041G>A 6042A>G), S259T (6049T>A 6051G>A), T261P (6055A>C), E263D (6063G>T), Y266F (6071A>T), E267D (6075A>T), I272V (6088A>G 6090C>G), E283N (6121G>A 6123A>T), I285L (6127A>T 6129C>A), K289D (6139A>G 6141G>C), E292Q (6148G>C), E294P (6154G>C 6155A>C), E297P (6163G>C 6164A>C), V299I (6169G>A 6171A>T), N300E (6172A>G 6174T>G), E303S (6181G>A 6182A>G 6183A>T), Y305C (6188A>G 6189T>C), I313L (6211A>C), L317I (6223C>A 6225A>C), L325T (6247C>A 6248T>C), Q327G (6253C>G 6254A>G), A329P (6259G>C), I330M (6264T>G), S332N (6269G>A 6270C>T), Y333H (6271T>C), S335_R336del (6277_6283delTCAAGGG), T339I (6290C>T), Y342_D347del (6298_6315delTATTCAAATTGCAGAGAT), R349del (6319_6321delAGA), A351R (6325G>A 6326C>G), L356A (6340C>G 6341T>C), R357M (6344G>T 6345A>G), V360I (6352G>A 6354C>A), L361M (6355T>A 6357A>G), S362T (6359G>C 6360T>A), K365N (6369A>T), Q368K (6376C>A), P369Q (6380C>A 6381T>A), T370P (6382A>C 6384A>T), T371I (6386C>T 6387C>T), R376K (6401G>A 6402G>A), R377K (6404G>A), P382D (6418C>G 6419C>A 6420A>T), D383E (6423T>A), L391Q (6446T>A), S393G (6451A>G 6453T>G), H394Q (6456C>A), K405N (6489A>C), I410V (6502A>G 6504T>A), V411I (6505G>A 6507T>C), E417D (6525A>C), *418E (6526T>G) | | | | | | | | |
| Codon mutations: | GAA187CAA (5833G>C), TAT189TAC (5841T>C), ATT190ATA (5844T>A), CAC192CAA (5850C>A), CAT193AAA (5851C>A 5853T>A), ACA195CAG (5857A>C 5858C>A 5859A>G), TAC197TAT (5865C>T), AAT198AAC (5868T>C), TTG200CTT (5872T>C 5874G>T), ATA201ATT (5877A>T), GCT202GCA (5880T>A), TTA203CAA (5881T>C 5882T>A), CCA204CTC (5885C>T 5886A>C), AAT205AAA (5889T>A), ACA206ACT (5892A>T), AGT207AAT (5894G>A), GCA208GTC (5897C>T 5898A>C), AAA209AAT (5901A>T), TTA210CTA (5902T>C), GTA211GTT (5907A>T), GCC212AGA (5908G>A 5909C>G 5910C>A), ACC213ACA (5913C>A), TGC214TGT (5916C>T), AAC216AGT (5921A>G 5922C>T), TAC217TAT (5925C>T), GAT221AGT (5935G>A 5936A>G), ACT222ACC (5940T>C), GTA223ATA (5941G>A), CAG226TAC (5950C>T 5952G>C), ACA227GAT (5953A>G 5954C>A 5955A>T), CAA229GAA (5959C>G), GAG230GAA (5964G>A), GCC232TCA (5968G>T 5970C>A), AAT233ACT (5972A>C), CCT235CCA (5979T>A), AAG239AAA (5991G>A), TTC241TTT (5997C>T), ATG242ACA (5999T>C 6000G>A), CAA243ACA (6001C>A 6002A>C), AAG245AAA (6009G>A), AGG246AGA (6012G>A), ATA247ATT (6015A>T), ACT248ACA (6018T>A), GGG250GGC (6024G>C), ACA251AAC (6026C>A 6027A>C), TTC253TTT (6033C>T), TAC254TAT (6036C>T), GTA255ATA (6037G>A), CGA256AAG (6040C>A 6041G>A 6042A>G), TTC257TTT (6045C>T), TCG259ACA (6049T>A 6051G>A), GCT260GCA (6054T>A), ACA261CCA (6055A>C), GAG263GAT (6063G>T), CTA265TTA (6067C>T), TAT266TTT (6071A>T), GAA267GAT (6075A>T), GAG268GAA (6078G>A), ATA269ATT (6081A>T), AAG270AAA (6084G>A), CCT271CCA (6087T>A), ATC272GTG (6088A>G 6090C>G), AAA277AAG (6105A>G), ATT278ATA (6108T>A), CTC280CTA (6114C>A), AGG282AGA (6120G>A), GAA283AAT (6121G>A 6123A>T), ATC285TTA (6127A>T 6129C>A), AAG289GAC (6139A>G 6141G>C), GAA292CAA (6148G>C), GAA294CCA (6154G>C 6155A>C), GAA297CCA (6163G>C 6164A>C), GTA299ATT (6169G>A 6171A>T), AAT300GAG (6172A>G 6174T>G), ATT301ATA (6177T>A), GAA303AGT (6181G>A 6182A>G 6183A>T), TTC304TTT (6186C>T), TAT305TGC (6188A>G 6189T>C), AGG309AGA (6201G>A), ATA310ATT (6204A>T), ATA313CTA (6211A>C), ACT315ACG (6219T>G), CTA317ATC (6223C>A 6225A>C), CTA320TTA (6232C>T), TAT324TAC (6246T>C), CTA325ACA (6247C>A 6248T>C), CAG327GGG (6253C>G 6254A>G), AAT328AAC (6258T>C), GCA329CCA (6259G>C), ATT330ATG (6264T>G), AGC332AAT (6269G>A 6270C>T), TAT333CAT (6271T>C), TCA335_AGG336del (6277_6283delTCAAGGG), GAA337-AC (6277_6283delTCAAGGG 6285A>C), ACA339ATA (6290C>T), TAT342_GAT347del (6298_6315delTATTCAAATTGCAGAGAT), ATA348ATC (6318A>C), AGA349del (6319_6321delAGA), GCA351AGA (6325G>A 6326C>G), CTA356GCA (6340C>G 6341T>C), AGA357ATG (6344G>T 6345A>G), GTC360ATA (6352G>A 6354C>A), TTA361ATG (6355T>A 6357A>G), AGT362ACA (6359G>C 6360T>A), CTT363TTG (6361C>T 6363T>G), AAA365AAT (6369A>T), GAA367GAG (6375A>G), CAA368AAA (6376C>A), CCT369CAA (6380C>A 6381T>A), ACA370CCT (6382A>C 6384A>T), ACC371ATT (6386C>T 6387C>T), AGG376AAA (6401G>A 6402G>A), AGG377AAG (6404G>A), AAT378-GA (6406delA 6407A>G 6408T>A), TTC379TTT (6411C>T), ATC380ATT (6414C>T), TCC381TCG (6417C>G), CCA382GAT (6418C>G 6419C>A 6420A>T), GAT383GAA (6423T>A), CTG384TTA (6424C>T 6426G>A), TTA385CTA (6427T>C), CTA391CAA (6446T>A), AGT393GGG (6451A>G 6453T>G), CAC394CAA (6456C>A), TAT396TAC (6462T>C), CAC399CAT (6471C>T), TGC401TGT (6477C>T), TCA402TCG (6480A>G), AAA405AAC (6489A>C), ATT410GTA (6502A>G 6504T>A), GTT411ATC (6505G>A 6507T>C), GAC413GAT (6513C>T), CTG416CTA (6522G>A), GAA417GAC (6525A>C), TAA418GAA (6526T>G) | | | | | | | | |

#### Proteins

| | | | | | | | | | |
|---|---|---|---|---|---|---|---|---|---|
| putative cell-to-cell movement protein (NP_569140.1) | 94 | 420 | 23.6% | 549 | 83.6% | 99 (100%) | 82 (82.8%) | 0/0/0/0 | 1 |
| Protein mutations: | Q97K (2443C>A), D98K (2446G>A 2448T>G), A101M (2455G>A 2456C>T 2457A>G), N103K (2463C>A), P104E (2464C>G 2465C>A), I146V (2590A>G), I150V (2602A>G 2604A>T), I163M (2643A>G), A165V (2648C>T), L171I (2665C>A), N172E (2668A>G 2670T>A), R174S (2676A>T), K178R (2687A>G), G188E (2717G>A), E418K (3406G>A), E419L (3409G>T 3410A>T) | | | | | | | | |
| Codon mutations: | ATA96ATC (2442A>C), CAA97AAA (2443C>A), GAT98AAG (2446G>A 2448T>G), AAG100AAA (2454G>A), GCA101ATG (2455G>A 2456C>T 2457A>G), AAC103AAA (2463C>A), CCA104GAA (2464C>G 2465C>A), AAG106AAA (2472G>A), TTA110CTA (2482T>C), ACT113ACA (2493T>A), GCA119GCT (2511A>T), TGT120TGC (2514T>C), ACC127ACA (2535C>A), CTA133TTA (2551C>T), ATC139ATA (2571C>A), ATA146GTA (2590A>G), AGT148AGC (2598T>C), ATA150GTT (2602A>G 2604A>T), TAT156TAC (2622T>C), AAG158AAA (2628G>A), TTT161TTC (2637T>C), ATA163ATG (2643A>G), AGC164AGT (2646C>T), GCC165GTC (2648C>T), AAC166AAT (2652C>T), CTA171ATA (2665C>A), AAT172GAA (2668A>G 2670T>A), AGA174AGT (2676A>T), AAA178AGA (2687A>G), TTA180CTA (2692T>C), AAG185AAA (2709G>A), GGA188GAA (2717G>A), GAA418AAA (3406G>A), GAA419TTA (3409G>T 3410A>T) | | | | | | | | |
| polyprotein (NP_569141.1) | 1 | 591 | 66.8% | 2399 | 80.4% | 425 (100%) | 342 (80.5%) | 0/0/0/0 | 3 |
| Protein mutations: | N16D (3456A>G), T39E (3525A>G 3526C>A), D40E (3530T>A), W42* (3536G>A), E43K (3537G>A), K46T (3547A>C), T47I (3550C>T), K178E (3942A>G), N181S (3952A>G), D182E (3956T>A), N183D (3957A>G), Q186K (3966C>A), E189D (3977G>T), K194T (3991A>C), E201N (4011G>A 4013A>T), E202K (4014G>A 4016G>A), N203D (4017A>G), K212S (4045A>G 4046A>T), N214S (4051A>G 4052C>T), N216D (4056A>G), L218Q (4062T>C 4063T>A), T219K (4066C>A), I223M (4079A>G), K226D (4086A>G 4088A>T), D227E (4091T>A), D230E (4100T>A), N231K (4103T>A), K232G (4104A>G 4105A>G), E236K (4116G>A), N238E (4122A>G 4124T>A), S239G (4125A>G 4127C>T), H251Y (4161C>T), I289V (4275A>G), D311E (4343T>A), A318T (4362G>A 4364T>A), A338V (4423C>T), M346I (4448G>T), D347N (4449G>A 4451C>T), R368K (4513G>A), L377S (4540T>C), K379Q (4545A>C), A381I (4551G>A 4552C>T), H382N (4554C>A), I384V (4560A>G), N386K (4568C>A), T394K (4591C>A 4592C>G), M399F (4605A>T 4607G>T), I415V (4653A>G), M417T (4660T>C), T419Q (4665A>C 4666C>A 4667T>A), I426N (4687T>A), L428S (4692C>T 4693T>C 4694T>A), D429E (4697T>A), N431E (4701A>G 4703C>A), I432L (4704A>T 4706T>A), Q439* (4725C>T), I444L (4740A>T), I487L (4869A>T), Y496C (4897A>G 4898C>T), K503N (4919A>T), T505I (4924C>T 4925C>A), T510A (4938A>G), S512A (4944T>G 4946A>G), N514D (4950A>G), Y515Q (4953T>C 4955T>A), N526E (4986A>G 4988T>A), C534Y (5011G>A 5012C>T), S540T (5028T>A 5030T>A), Y541F (5032A>T 5033T>C), T542S (5034A>T), E543N (5037G>A 5039A>C), P544S (5040C>T), Q545E (5043C>G), E546I (5046G>A 5047A>T), W548* (5053G>A), E549K (5055G>A), F556H (5076T>C 5077T>A), A557S (5079G>T), T572N (5125C>A), R573K (5128G>A), V576I (5136G>A) | | | | | | | | |
| Codon mutations: | ATA3ATT (3419A>T), TAC4TAT (3422C>T), GCA7GCC (3431A>C), AAA8AAG (3434A>G), ATC9ATA (3437C>A), GTA11GTT (3443A>T), TAT14TAC (3452T>C), AAT16GAT (3456A>G), TAC18TAT (3464C>T), TAT19TAC (3467T>C), ACA20ACT (3470A>T), AAT30AAC (3500T>C), TGT32TGC (3506T>C), TGT36TGC (3518T>C), ACA39GAA (3525A>G 3526C>A), GAT40GAA (3530T>A), TGG42TGA (3536G>A), GAA43AAA (3537G>A), AAA46ACA (3547A>C), ACA47ATA (3550C>T), TTA176CTA (3936T>C), AAA178GAA (3942A>G), AAT181AGT (3952A>G), GAT182GAA (3956T>A), AAC183GAC (3957A>G), CCT184CCA (3962T>A), CTA185TTA (3963C>T), CAA186AAA (3966C>A), GAG189GAT (3977G>T), ACA193ACT (3989A>T), AAA194ACA (3991A>C), ATT200ATA (4010T>A), GAA201AAT (4011G>A 4013A>T), GAG202AAA (4014G>A 4016G>A), AAT203GAT (4017A>G), ATA205ATT (4025A>T), ATA206ATC (4028A>C), CCT210CCC (4040T>C), CTA211TTA (4041C>T), AAA212AGT (4045A>G 4046A>T), TAT213TAC (4049T>C), AAC214AGT (4051A>G 4052C>T), AAT216GAT (4056A>G), GAT217GAC (4061T>C), TTA218CAA (4062T>C 4063T>A), ACA219AAA (4066C>A), GAA220GAG (4070A>G), TTT221TTC (4073T>C), ATA223ATG (4079A>G), AAA226GAT (4086A>G 4088A>T), GAT227GAA (4091T>A), TTG229TTA (4097G>A), GAT230GAA (4100T>A), AAT231AAA (4103T>A), AAA232GGA (4104A>G 4105A>G), CAA235CAG (4115A>G), GAA236AAA (4116G>A), AGT237AGC (4121T>C), AAT238GAA (4122A>G 4124T>A), AGC239GGT (4125A>G 4127C>T), CAT241CAC (4133T>C), CAT251TAT (4161C>T), GGA257GGT (4181A>T), AGG260AGA (4190G>A), GTC262GTT (4196C>T), ATA263ATT (4199A>T), GAC264GAT (4202C>T), TTA268CTA (4212T>C), TAT277TAC (4241T>C), CCC280CCA (4250C>A), AAT281AAC (4253T>C), AAG285AAA (4265G>A), ATA289GTA (4275A>G), TAT292TAC (4286T>C), TAC294TAT (4292C>T), AGT296AGC (4298T>C), GAC299GAT (4307C>T), AAG301AAA (4313G>A), TAT305TAC (4325T>C), CAT306CAC (4328T>C), TTG307CTA (4329T>C 4331G>A), AAA308AAG (4334A>G), GAG310GAA (4340G>A), GAT311GAA (4343T>A), AAA315AAG (4355A>G), GCT318ACA (4362G>A 4364T>A), TAT326TAC (4388T>C), GTT330GTA (4400T>A), CCA332CCT (4406A>T), TTT333TTC (4409T>C), TAT335TAC (4415T>C), AAT337AAC (4421T>C), GCA338GTA (4423C>T), TAC342TAT (4436C>T), CAT344CAC (4442T>C), ATG346ATT (4448G>T), GAC347AAT (4449G>A 4451C>T), AAC348AAT (4454C>T), TAT349TAC (4457T>C), AAT351AAC (4463T>C), AAT355AAC (4475T>C), GTA358GTT (4484A>T), TAT359TAC (4487T>C), GAT361GAC (4493T>C), CTA364TTA (4500C>T), CTA365TTA (4503C>T), TAT366TAC (4508T>C), AGA368AAA (4513G>A), TTA377TCA (4540T>C), AAA379CAA (4545A>C), TTC380TTT (4550C>T), GCA381ATA (4551G>A 4552C>T), CAT382AAT (4554C>A), ATT383ATA (4559T>A), ATA384GTA (4560A>G), AAC386AAA (4568C>A), TCT387TCA (4571T>A), GGT388GGA (4574T>A), TTA391CTA (4581T>C), ACC394AAG (4591C>A 4592C>G), GCA396GCT (4598A>T), GAA397GAG (4601A>G), ATT398ATA (4604T>A), ATG399TTT (4605A>T 4607G>T), TTA406CTA (4626T>C), GAT411GAC (4643T>C), ATA415GTA (4653A>G), ATG417ACG (4660T>C), ACT419CAA (4665A>C 4666C>A 4667T>A), CAA423CAG (4679A>G), ATC426AAC (4687T>A), AAT427AAC (4691T>C), CTT428TCA (4692C>T 4693T>C 4694T>A), GAT429GAA (4697T>A), AAC431GAA (4701A>G 4703C>A), ATT432TTA (4704A>T 4706T>A), AAA437AAG (4721A>G), TTA438CTA (4722T>C), CAA439TAA (4725C>T), CTA442TTA (4734C>T), ATA444TTA (4740A>T), AAG486AAA (4868G>A), ATA487TTA (4869A>T), TTG488CTT (4872T>C 4874G>T), TAC496TGT (4897A>G 4898C>T), GAT499GAC (4907T>C), AAC501AAT (4913C>T), AAA503AAT (4919A>T), ACC505ATA (4924C>T 4925C>A), ATT507ATA (4931T>A), ACA510GCA (4938A>G), TCA512GCG (4944T>G 4946A>G), AAT514GAT (4950A>G), TAT515CAA (4953T>C 4955T>A), GTT520GTA (4970T>A), CTT521TTA (4971C>T 4973T>A), AAG522AAA (4976G>A), TAT523TAC (4979T>C), AGA524AGG (4982A>G), AAT526GAA (4986A>G 4988T>A), TGC534TAT (5011G>A 5012C>T), TAC536TAT (5018C>T), TAT537TAC (5021T>C), GGA539GGT (5027A>T), TCT540ACA (5028T>A 5030T>A), TAT541TTC (5032A>T 5033T>C), ACA542TCA (5034A>T), GAA543AAC (5037G>A 5039A>C), CCA544TCA (5040C>T), CAA545GAA (5043C>G), GAA546ATA (5046G>A 5047A>T), TGG548TAG (5053G>A), GAA549AAA (5055G>A), TTT556CAT (5076T>C 5077T>A), GCA557TCA (5079G>T), CTA562TTA (5094C>T), ATA569ATT (5117A>T), GTT570GTG (5120T>G), TAC571TAT (5123C>T), ACC572AAC (5125C>A), AGA573AAA (5128G>A), ATT575ATA (5135T>A), GTA576ATA (5136G>A), GAC579GAT (5147C>T), AAT580AAC (5150T>C), CAA582CAG (5156A>G), ACT588ACA (5174T>A), AAA590AAG (5180A>G) | | | | | | | | |
| putative transactivator factor (NP_569142.1) | 187 | 418 | 55.3% | 806 | 54.0% | 222 (96.1%) | 149 (64.5%) | 0/9/2/2 | 0 |

| | Begin | End | Coverage | Score | Concordance | Matches | Identities | I/D/M/F* | Stop Codons |
|---|---|---|---|---|---|---|---|---|---|
| **NT** | **2433** | **6550** | **29.4%** | **2623** | **59.0%** | **2257 (98.7%)** | **1803 (78.8%)** | **1/29** | |

| | |
|---|---|
| Protein mutations: | E187Q (5833G>C), H192Q (5850C>A), H193K (5851C>A 5853T>A), T195Q (5857A>C 5858C>A 5859A>G), L203Q (5881T>C 5882T>A), P204L (5885C>T 5886A>C), N205K (5889T>A), S207N (5894G>A), A208V (5897C>T 5898A>C), K209N (5901A>T), A212R (5908G>A 5909C>G 5910C>A), N216S (5921A>G 5922C>T), D221S (5935G>A 5936A>G), V223I (5941G>A), Q226Y (5950C>T 5952G>C), T227D (5953A>G 5954C>A 5955A>T), Q229E (5959C>G), A232S (5968G>T 5970C>A), N233T (5972A>C), M242T (5999T>C 6000G>A), Q243T (6001C>A 6002A>C), T251N (6026C>A 6027A>C), V255I (6037G>A), R256K (6040C>A 6041G>A 6042A>G), S259T (6049T>A 6051G>A), T261P (6055A>C), E263D (6063G>T), Y266F (6071A>T), E267D (6075A>T), I272V (6088A>G 6090C>G), E283N (6121G>A 6123A>T), I285L (6127A>T 6129C>A), K289D (6139A>G 6141G>C), E292Q (6148G>C), E294P (6154G>C 6155A>C), E297P (6163G>C 6164A>C), V299I (6169G>A 6171A>T), N300E (6172A>G 6174T>G), E303S (6181G>A 6182A>G 6183A>T), Y305C (6188A>G 6189T>C), I313L (6211A>C), L317I (6223C>A 6225A>C), L325T (6247C>A 6248T>C), Q327G (6253C>G 6254A>G), A329P (6259G>C), I330M (6264T>G), S332N (6269G>A 6270C>T), Y333H (6271T>C), S335_R336del (6277_6283delTCAAGGG), T339I (6290C>T), Y342_D347del (6298_6315delTATTCAAATTGCAGAGAT), R349del (6319_6321delAGA), A351R (6325G>A 6326C>G), L356A (6340C>G 6341T>C), R357M (6344G>T 6345A>G), V360I (6352G>A 6354C>A), L361M (6355T>A 6357A>G), S362T (6359G>C 6360T>A), K365N (6369A>T), Q368K (6376C>A), P369Q (6380C>A 6381T>A), T370P (6382A>C 6384A>T), T371I (6386C>T 6387C>T), R376K (6401G>A 6402G>A), R377K (6404G>A), P382D (6418C>G 6419C>A 6420A>T), D383E (6423T>A), L391Q (6446T>A), S393G (6451A>G 6453T>G), H394Q (6456C>A), K405N (6489A>C), I410V (6502A>G 6504T>A), V411I (6505G>A 6507T>C), E417D (6525A>C), *418E (6526T>G) |
| Codon mutations: | GAA187CAA (5833G>C), TAT189TAC (5841T>C), ATT190ATA (5844T>A), CAC192CAA (5850C>A), CAT193AAA (5851C>A 5853T>A), ACA195CAG (5857A>C 5858C>A 5859A>G), TAC197TAT (5865C>T), AAT198AAC (5868T>C), TTG200CTT (5872T>C 5874G>T), ATA201ATT (5877A>T), GCT202GCA (5880T>A), TTA203CAA (5881T>C 5882T>A), CCA204CTC (5885C>T 5886A>C), AAT205AAA (5889T>A), ACA206ACT (5892A>T), AGT207AAT (5894G>A), GCA208GTC (5897C>T 5898A>C), AAA209AAT (5901A>T), TTA210CTA (5902T>C), GTA211GTT (5907A>T), GCC212AGA (5908G>A 5909C>G 5910C>A), ACC213ACA (5913C>A), TGC214TGT (5916C>T), AAC216AGT (5921A>G 5922C>T), TAC217TAT (5925C>T), GAT221AGT (5935G>A 5936A>G), ACT222ACC (5940T>C), GTA223ATA (5941G>A), CAG226TAC (5950C>T 5952G>C), ACA227GAT (5953A>G 5954C>A 5955A>T), CAA229GAA (5959C>G), GAG230GAA (5964G>A), GCC232TCA (5968G>T 5970C>A), AAT233ACT (5972A>C), CCT235CCA (5979T>A), AAG239AAA (5991G>A), TTC241TTT (5997C>T), ATG242ACA (5999T>C 6000G>A), CAA243ACA (6001C>A 6002A>C), AAG245AAA (6009G>A), AGG246AGA (6012G>A), ATA247ATT (6015A>T), ACT248ACA (6018T>A), GGG250GGC (6024G>C), ACA251AAC (6026C>A 6027A>C), TTC253TTT (6033C>T), TAC254TAT (6036C>T), GTA255ATA (6037G>A), CGA256AAG (6040C>A 6041G>A 6042A>G), TTC257TTT (6045C>T), TCG259ACA (6049T>A 6051G>A), GCT260GCA (6054T>A), ACA261CCA (6055A>C), GAG263GAT (6063G>T), CTA265TTA (6067C>T), TAT266TTT (6071A>T), GAA267GAT (6075A>T), GAG268GAA (6078G>A), ATA269ATT (6081A>T), AAG270AAA (6084G>A), CCT271CCA (6087T>A), ATC272GTG (6088A>G 6090C>G), AAA277AAG (6105A>G), ATT278ATA (6108T>A), CTC280CTA (6114C>A), AGG282AGA (6120G>A), GAA283AAT (6121G>A 6123A>T), ATC285TTA (6127A>T 6129C>A), AAG289GAC (6139A>G 6141G>C), GAA292CAA (6148G>C), GAA294CCA (6154G>C 6155A>C), GAA297CCA (6163G>C 6164A>C), GTA299ATT (6169G>A 6171A>T), AAT300GAG (6172A>G 6174T>G), ATT301ATA (6177T>A), GAA303AGT (6181G>A 6182A>G 6183A>T), TTC304TTT (6186C>T), TAT305TGC (6188A>G 6189T>C), AGG309AGA (6201G>A), ATA310ATT (6204A>T), ATA313CTA (6211A>C), ACT315ACG (6219T>G), CTA317ATC (6223C>A 6225A>C), CTA320TTA (6232C>T), TAT324TAC (6246T>C), CTA325ACA (6247C>A 6248T>C), CAG327GGG (6253C>G 6254A>G), AAT328AAC (6258T>C), GCA329CCA (6259G>C), ATT330ATG (6264T>G), AGC332AAT (6269G>A 6270C>T), TAT333CAT (6271T>C), TCA335_AGG336del (6277_6283delTCAAGGG), GAA337-AC (6277_6283delTCAAGGG 6285A>C), ACA339ATA (6290C>T), TAT342_GAT347del (6298_6315delTATTCAAATTGCAGAGAT), ATA348ATC (6318A>C), AGA349del (6319_6321delAGA), GCA351AGA (6325G>A 6326C>G), CTA356GCA (6340C>G 6341T>C), AGA357ATG (6344G>T 6345A>G), GTC360ATA (6352G>A 6354C>A), TTA361ATG (6355T>A 6357A>G), AGT362ACA (6359G>C 6360T>A), CTT363TTG (6361C>T 6363T>G), AAA365AAT (6369A>T), GAA367GAG (6375A>G), CAA368AAA (6376C>A), CCT369CAA (6380C>A 6381T>A), ACA370CCT (6382A>C 6384A>T), ACC371ATT (6386C>T 6387C>T), AGG376AAA (6401G>A 6402G>A), AGG377AAG (6404G>A), AAT378-GA (6406delA 6407A>G 6408T>A), TTC379TTT (6411C>T), ATC380ATT (6414C>T), TCC381TCG (6417C>G), CCA382GAT (6418C>G 6419C>A 6420A>T), GAT383GAA (6423T>A), CTG384TTA (6424C>T 6426G>A), TTA385CTA (6427T>C), CTA391CAA (6446T>A), AGT393GGG (6451A>G 6453T>G), CAC394CAA (6456C>A), TAT396TAC (6462T>C), CAC399CAT (6471C>T), TGC401TGT (6477C>T), TCA402TCG (6480A>G), AAA405AAC (6489A>C), ATT410GTA (6502A>G 6504T>A), GTT411ATC (6505G>A 6507T>C), GAC413GAT (6513C>T), CTG416CTA (6522G>A), GAA417GAC (6525A>C), TAA418GAA (6526T>G) |

*: Inserts / Deletes / Misaligned / Frameshifts

#### Analysis details

This analysis was performed with panviral2.64

### NGS Details (UN59): Tomato chocolate spot virus (segment RNA 1)

#### Assembly

| | |
|---|---|
| **Coverage Length** | 464 (2 contig(s)) |
| **Depth Of Coverage** | 1311.6 |
| **Number Of Reads** | 5597 |
| **Reads Per Million** | 121.67 rpm (after QC) |
| **Ambiguities** | 0 |
| **Assembly Method** | de novo + reference guided assembly |
| **Consensus Caller** | Bcf Tools |

#### Coverage Map

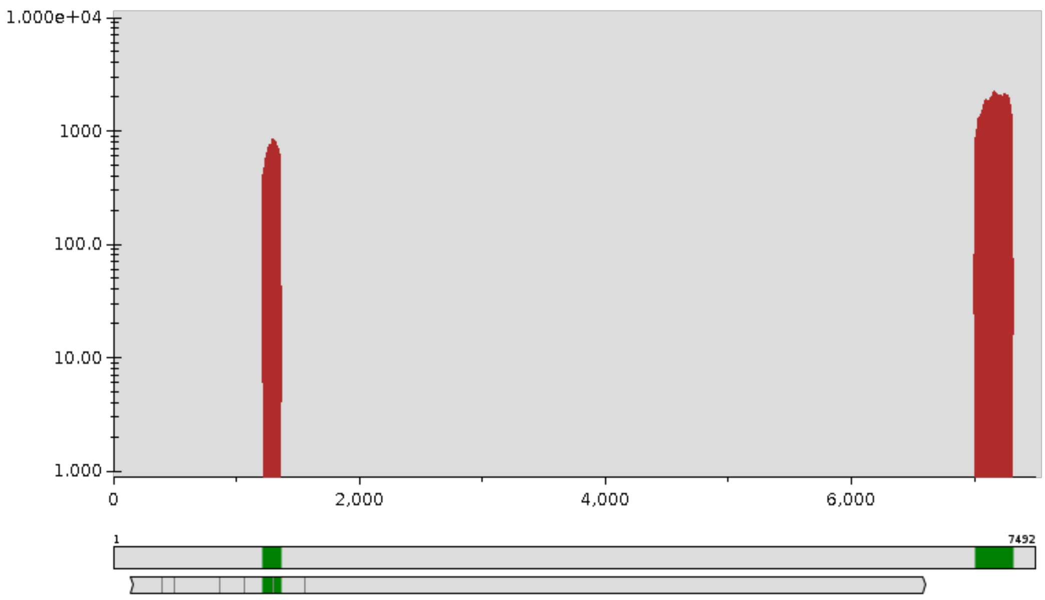

#### Assignment

| | |
|---|---|
| **Type** | Tomato chocolate spot virus (Taxonomy ID: 661101) |
| **Reference Genome** | NC_013075.1 |
| **NT Identity (%)** | 79.5259 |
| **AA Identity (%)** | 80.3922 |
| **Number Of Stop Codons** | 0 |
| **Number Of CDS** | 1 |

#### Alignment

| | |
|---|---|
| **Alignment Score** | 548.0 (NT) + 310.0 (AA) = 858.0 |
| **Concordance (%)** | 66.5632 |

| Alignment Method | Global, seeded, nucleotide + amino acids (AGA) |
|---|---|

#### Genome Region

Sequence starts at position 1213 and ends at position 7314 relative to NC_013075.1 reference sequence.

#### Alignment Detailed Statistics

| | Begin | End | Coverage | Score | Concordance | Matches | Identities | I/D/M/F* | Stop Codons |
|---|---|---|---|---|---|---|---|---|---|
| **NT** | **1213** | **7314** | **6.2%** | **548** | **59.1%** | **464 (100%)** | **369 (79.5%)** | **0/0** | |
| Mutations: | 1222G>A, 1228A>G, 1229A>T, 1230T>G, 1233G>A, 1234C>T, 1236T>G, 1239A>T, 1243G>C, 1245G>C, 1246T>A, 1247A>T, 1248T>A, 1251T>A, 1254G>A, 1256G>A, 1260C>T, 1261A>C, 1262G>A, 1266C>T, 1268T>C, 1269G>A, 1275C>G, 1278A>T, 1281A>G, 1287G>A, 1293G>A, 1305G>T, 1317A>T, 1319C>A, 1320T>G, 1326C>A, 1329T>C, 1335G>A, 1338A>C, 1341A>G, 1342T>A, 1343C>G, 1348A>C, 1351T>C, 7013A>T, 7014T>A, 7018C>G, 7019A>G, 7020A>T, 7023C>A, 7042G>T, 7044A>T, 7062G>A, 7070T>C, 7071C>T, 7084A>G, 7087T>C, 7098C>T, 7099T>C, 7107G>A, 7116C>T, 7118T>A, 7121G>A, 7122T>C, 7123C>T, 7124T>G, 7137A>T, 7138G>A, 7139A>G, 7140C>T, 7148A>G, 7151A>T, 7159A>T, 7162T>C, 7165G>A, 7171T>G, 7173C>T, 7177A>C, 7185G>A, 7199C>T, 7208A>T, 7209T>A, 7210C>T, 7211A>G, 7213C>T, 7214A>T, 7229T>C, 7245T>C, 7250G>A, 7253T>A, 7254G>A, 7256T>C, 7257G>A, 7258G>T, 7259C>G, 7274G>C, 7275T>C, 7292T>G, 7303T>A | | | | | | | | |

##### CDS

| | Begin | End | Coverage | Score | Concordance | Matches | Identities | I/D/M/F* | Stop Codons |
|---|---|---|---|---|---|---|---|---|---|
| ToChSV_s1_gp1 | 359 | 409 | 2.4% | 310 | 83.6% | 51 (100%) | 41 (80.4%) | 0/0/0/0 | 0 |
| Protein mutations: | A362T (1222G>A), N364V (1228A>G 1229A>T 1230T>G), E367D (1239A>T), V369L (1243G>C 1245G>C), Y370I (1246T>A 1247A>T 1248T>A), S373N (1256G>A), R375Q (1261A>C 1262G>A), M377T (1268T>C 1269G>A), A394E (1319C>A 1320T>G) | | | | | | | | |
| Codon mutations: | GCA362ACA (1222G>A), AAT364GTG (1228A>G 1229A>T 1230T>G), CAG365CAA (1233G>A), CTT366TTG (1234C>T 1236T>G), GAA367GAT (1239A>T), GTG369CTC (1243G>C 1245G>C), TAT370ATA (1246T>A 1247A>T 1248T>A), GCT371GCA (1251T>A), GAG372GAA (1254G>A), AGC373AAC (1256G>A), TGC374TGT (1260C>T), AGA375CAA (1261A>C 1262G>A), CAC376CAT (1266C>T), ATG377ACA (1268T>C 1269G>A), GTC379GTG (1275C>G), GGA380GGT (1278A>T), CAA381CAG (1281A>G), AGG383AGA (1287G>A), GAG385GAA (1293G>A), GTG389GTT (1305G>T), GGA393GGT (1317A>T), GCT394GAG (1319C>A 1320T>G), GGC396GGA (1326C>A), TGT397TGC (1329T>C), AAG399AAA (1335G>A), TCA400TCC (1338A>C), ACA401ACG (1341A>G), TCC402AGC (1342T>A 1343C>G), AGA404CGA (1348A>C), TTG405CTG (1351T>C) | | | | | | | | |

##### Proteins

| | Begin | End | Coverage | Score | Concordance | Matches | Identities | I/D/M/F* | Stop Codons |
|---|---|---|---|---|---|---|---|---|---|
| polyprotein (YP_003097229.1) | 359 | 409 | 2.4% | 310 | 83.6% | 51 (100%) | 41 (80.4%) | 0/0/0/0 | 0 |
| Protein mutations: | A362T (1222G>A), N364V (1228A>G 1229A>T 1230T>G), E367D (1239A>T), V369L (1243G>C 1245G>C), Y370I (1246T>A 1247A>T 1248T>A), S373N (1256G>A), R375Q (1261A>C 1262G>A), M377T (1268T>C 1269G>A), A394E (1319C>A 1320T>G) | | | | | | | | |
| Codon mutations: | GCA362ACA (1222G>A), AAT364GTG (1228A>G 1229A>T 1230T>G), CAG365CAA (1233G>A), CTT366TTG (1234C>T 1236T>G), GAA367GAT (1239A>T), GTG369CTC (1243G>C 1245G>C), TAT370ATA (1246T>A 1247A>T 1248T>A), GCT371GCA (1251T>A), GAG372GAA (1254G>A), AGC373AAC (1256G>A), TGC374TGT (1260C>T), AGA375CAA (1261A>C 1262G>A), CAC376CAT (1266C>T), ATG377ACA (1268T>C 1269G>A), GTC379GTG (1275C>G), GGA380GGT (1278A>T), CAA381CAG (1281A>G), AGG383AGA (1287G>A), GAG385GAA (1293G>A), GTG389GTT (1305G>T), GGA393GGT (1317A>T), GCT394GAG (1319C>A 1320T>G), GGC396GGA (1326C>A), TGT397TGC (1329T>C), AAG399AAA (1335G>A), TCA400TCC (1338A>C), ACA401ACG (1341A>G), TCC402AGC (1342T>A 1343C>G), AGA404CGA (1348A>C), TTG405CTG (1351T>C) | | | | | | | | |
| RNA-dependent RNA polymerase (YP_003097234.1) | 1 | 20 | 23.8% | 141 | 91.0% | 20 (100%) | 18 (90.0%) | 0/0/0/0 | 0 |
| Protein mutations: | A5E (1319C>A 1320T>G) | | | | | | | | |
| Codon mutations: | GGA4GGT (1317A>T), GCT5GAG (1319C>A 1320T>G), GGC7GGA (1326C>A), TGT8TGC (1329T>C), AAG10AAA (1335G>A), TCA11TCC (1338A>C), ACA12ACG (1341A>G), TCC13AGC (1342T>A 1343C>G), AGA15CGA (1348A>C), TTG16CTG (1351T>C) | | | | | | | | |

*: Inserts / Deletes / Misaligned / Frameshifts

#### Analysis details

This analysis was performed with panviral2.64

### NGS Details (UN59): Tomato chocolate spot virus (segment RNA2)

#### Assembly

| | |
|---|---|
| **Coverage Length** | 810 (2 contig(s)) |
| **Depth Of Coverage** | 1360.0 |
| **Number Of Reads** | 9765 |
| **Reads Per Million** | 212.28 rpm (after QC) |
| **Ambiguities** | 0 |
| **Assembly Method** | de novo + reference guided assembly |
| **Consensus Caller** | Bcf Tools |

#### Coverage Map

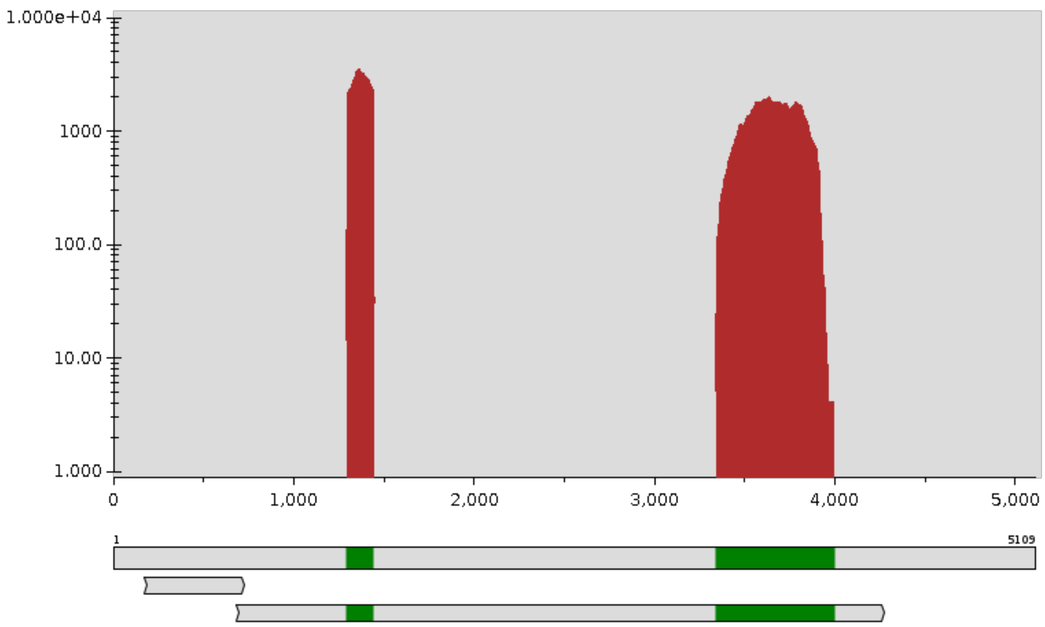

#### Assignment

| | |
|---|---|
| **Type** | Tomato chocolate spot virus (Taxonomy ID: 661101) |
| **Reference Genome** | NC_013076.1 |
| **NT Identity (%)** | 69.7385 |
| **AA Identity (%)** | 75.3731 |
| **Number Of Stop Codons** | 0 |
| **Number Of CDS** | 2 |

#### Alignment

| | |
|---|---|
| **Alignment Score** | 609.0 (NT) + 1363.0 (AA) = 1972.0 |
| **Concordance (%)** | 59.8301 |

| Alignment Method | Global, seeded, nucleotide + amino acids (AGA) |
|---|---|

#### Genome Region

Sequence starts at position 1293 and ends at position 3998 relative to NC_013076.1 reference sequence.

#### Alignment Detailed Statistics

| | Begin | End | Coverage | Score | Concordance | Matches | Identities | I/D/M/F* | Stop Codons |
|---|---|---|---|---|---|---|---|---|---|
| **NT** | **1293** | **3998** | **15.9%** | **609** | **38.5%** | **803 (99.1%)** | **560 (69.1%)** | **0/7** | |

| | |
|---|---|
| Mutations: | 1306T>A, 1309A>G, 1312T>C, 1315C>T, 1316C>T, 1317A>C, 1318A>T, 1327G>A, 1333T>A, 1336C>A, 1351A>T, 1354A>C, 1360T>G, 1366G>A, 1367C>G, 1369A>G, 1372T>G, 1373T>G, 1375C>G, 1376C>T, 1377A>C, 1378G>T, 1379A>C, 1380T>A, 1381G>A, 1385T>C, 1387G>A, 1394C>A, 1396T>A, 1397G>C, 1405T>A, 1408A>T, 1410A>C, 1411G>A, 1412A>C, 1413G>T, 1414G>T, 1423T>C, 1426A>T, 1429A>C, 1433G>A, 3355T>G, 3358G>T, 3361C>A, 3367A>C, 3368G>T, 3370T>G, 3376T>G, 3379C>T, 3380T>A, 3382A>T, 3388T>A, 3391C>A, 3394G>A, 3400G>A, 3401C>T, 3403T>G, 3407T>C, 3410C>A, 3416A>G, 3419T>C, 3421G>C, 3422G>A, 3423T>C, 3424C>G, 3427A>T, 3430T>G, 3434A>T, 3436G>T, 3439T>G, 3442T>A, 3445G>T, 3454C>T, 3460A>T, 3463C>T, 3469A>C, 3472A>G, 3475A>G, 3477A>G, 3478G>A, 3484G>T, 3490C>T, 3496T>G, 3499C>T, 3505G>A, 3509G>A, 3511T>C, 3512A>C, 3516C>A, 3517C>T, 3530A>C, 3535A>G, 3538G>A, 3541C>G, 3544G>A, 3547G>A, 3550T>A, 3553C>T, 3556G>T, 3559A>T, 3560C>A, 3561A>T, 3562A>C, 3564A>T, 3568T>C, 3569G>C, 3570C>A, 3571C>G, 3574G>A, 3577T>G, 3581_3586delCAGCCA, 3587G>A, 3588T>G, 3593C>A, 3594A>C, 3596A>T, 3597G>C, 3598C>T, 3601G>A, 3604G>T, 3605A>T, 3606A>C, 3607T>C, 3610G>C, 3611T>C, 3612C>A, 3613C>G, 3616T>C, 3622G>A, 3625A>G, 3628T>C, 3629T>A, 3637T>A, 3643C>G, 3646G>A, 3652C>T, 3655C>T, 3656A>G, 3657T>C, 3658G>T, 3660G>C, 3661C>A, 3671G>A, 3672A>C, 3676A>C, 3679G>A, 3683C>G, 3685T>A, 3688A>T, 3689A>T, 3691A>G, 3700T>C, 3703C>T, 3721A>T, 3724T>A, 3725C>T, 3726A>C, 3727G>C, 3731G>C, 3733T>A, 3734G>A, 3739G>A, 3751C>T, 3754T>C, 3757T>A, 3763C>A, 3766T>C, 3773A>T, 3775A>C, 3778C>T, 3787C>T, 3793A>T, 3796C>A, 3799C>T, 3800C>T, 3802C>G, 3803A>G, 3805T>A, 3809T>A, 3811C>A, 3812A>T, 3814A>T, 3817T>A, 3818G>A, 3827G>A, 3828T>C, 3833T>A, 3834C>G, 3836A>T, 3837G>C, 3839C>A, 3840C>G, 3841T>C, 3843A>G, 3845G>A, 3847C>A, 3850A>T, 3853C>T, 3856T>C, 3857C>T, 3862T>C, 3863G>A, 3865A>T, 3868T>A, 3875T>A, 3877C>A, 3880T>A, 3883T>G, 3889A>T, 3892T>C, 3894C>A, 3896G>T, 3897C>T, 3898A>T, 3901C>T, 3902C>G, 3904G>A, 3905A>T, 3906G>C, 3908A>G, 3909A>C, 3914A>G, 3915A>C, 3916T>A, 3917T>C, 3922A>G, 3923G>T, 3925T>A, 3934A>C, 3937A>T, 3941T>A, 3943C>T, 3946G>A, 3950delA, 3951A>G, 3952C>A, 3964T>G, 3970A>C, 3973A>C, 3979A>C, 3982T>C, 3989A>G, 3990C>T, 3991G>A, 3994T>G |

##### CDS

| | Begin | End | Coverage | Score | Concordance | Matches | Identities | I/D/M/F* | Stop Codons |
|---|---|---|---|---|---|---|---|---|---|
| ToChV_s2_gp2 | 204 | 1105 | 22.5% | 1363 | 72.9% | 268 (99.3%) | 202 (74.8%) | 0/2/1/1 | 0 |

| | |
|---|---|
| Protein mutations: | Q211S (1316C>T 1317A>C 1318A>T), Q228E (1367C>G 1369A>G), S229R (1372T>G), S230A (1373T>G 1375C>G), Q231S (1376C>T 1377A>C 1378G>T), M232Q (1379A>C 1380T>A 1381G>A), A238P (1397G>C), E242A (1410A>C 1411G>A), R243L (1412A>C 1413G>T 1414G>T), D250N (1433G>A), V895L (3368G>T 3370T>G), S899T (3380T>A 3382A>T), C908R (3407T>C), L909I (3410C>A), N911D (3416A>G), V913T (3422G>A 3423T>C 3424C>G), T917S (3434A>T 3436G>T), K931R (3477A>G 3478G>A), D942N (3509G>A 3511T>C), M943L (3512A>C), T944N (3516C>A 3517C>T), I949L (3530A>C), D955E (3550T>A), E958D (3559A>T), Q959I (3560C>A 3561A>T 3562A>C), Y960F (3564A>T), A962Q (3569G>C 3570C>A 3571C>G), Q966_P967del (3581_3586delCAGCCA), V968S (3587G>A 3588T>G), Q970T (3593C>A 3594A>C), N974S (3605A>T 3606A>C 3607T>C), L975F (3610G>C), S976Q (3611T>C 3612C>A 3613C>G), S982T (3629T>A), M991A (3656A>G 3657T>C 3658G>T), G992A (3660G>C 3661C>A), E996T (3671G>A 3672A>C), E997D (3676A>C), L1000V (3683C>G 3685T>A), I1002L (3689A>T 3691A>G), E1012D (3721A>T), Q1014S (3725C>T 3726A>C 3727G>C), A1016P (3731G>C 3733T>A), V1017I (3734G>A), T1030S (3773A>T 3775A>C), N1040E (3803A>G 3805T>A), S1042T (3809T>A 3811C>A), I1043F (3812A>T 3814A>T), V1045M (3818G>A), V1048T (3827G>A 3828T>C), P1052S (3839C>A 3840C>G 3841T>C), N1053S (3843A>G), V1054I (3845G>A 3847C>A), V1060I (3863G>A 3865A>T), S1064T (3875T>A 3877C>A), P1070Q (3894C>A), A1071F (3896G>T 3897C>T 3898A>T), L1073V (3902C>G 3904G>A), K1075A (3908A>G 3909A>C), N1077A (3914A>G 3915A>C 3916T>A), Y1078H (3917T>C), V1080L (3923G>T 3925T>A), S1086T (3941T>A 3943C>T), T1102V (3989A>G 3990C>T 3991G>A) |
| Codon mutations: | CCT207CCA (1306T>A), CAA208CAG (1309A>G), TTT209TTC (1312T>C), TTC210TTT (1315C>T), CAA211TCT (1316C>T 1317A>C 1318A>T), CAG214CAA (1327G>A), GCT216GCA (1333T>A), GCC217GCA (1336C>A), TCA222TCT (1351A>T), GTA223GTC (1354A>C), TCT225TCG (1360T>G), GGG227GGA (1366G>A), CAA228GAG (1367C>G 1369A>G), AGT229AGG (1372T>G), TCC230GCG (1373T>G 1375C>G), CAG231TCT (1376C>T 1377A>C 1378G>T), ATG232CAA (1379A>C 1380T>A 1381G>A), TTG234CTA (1385T>C 1387G>A), CGT237AGA (1394C>A 1396T>A), GCT238CCT (1397G>C), GTT240GTA (1405T>A), GGA241GGT (1408A>T), GAG242GCA (1410A>C 1411G>A), AGG243CTT (1412A>C 1413G>T 1414G>T), AAT246AAC (1423T>C), ACA247ACT (1426A>T), CGA248CGC (1429A>C), GAT250AAT (1433G>A), CTT890CTG (3355T>G), GTG891GTT (3358G>T), TCC892TCA (3361C>A), TCA894TCC (3367A>C), GTT895TTG (3368G>T 3370T>G), CTT897CTG (3376T>G), GAC898GAT (3379C>T), TCA899ACT (3380T>A 3382A>T), GGT901GGA (3388T>A), GTC902GTA (3391C>A), ACG903ACA (3394G>A), CAG905CAA (3400G>A), CTT906TTG (3401C>T 3403T>G), TGC908CGC (3407T>C), CTT909ATT (3410C>A), AAT911GAT (3416A>G), TTG912CTC (3419T>C 3421G>C), GTC913ACG (3422G>A 3423T>C 3424C>G), CTA914CTT (3427A>T), GGT915GGG (3430T>G), ACG917TCT (3434A>T 3436G>T), GTT918GTG (3439T>G), TCT919TCA (3442T>A), GTG920GTT (3445G>T), ATC923ATT (3454C>T), GGA925GGT (3460A>T), CAC926CAT (3463C>T), ACA928ACC (3469A>C), AAA929AAG (3472A>G), GGA930GGG (3475A>G), AAG931AGA (3477A>G 3478G>A), GCG933GCT (3484G>T), GAC935GAT (3490C>T), CTT937CTG (3496T>G), TTC938TTT (3499C>T), GAG940GAA (3505G>A), GAT942AAC (3509G>A 3511T>C), ATG943CTG (3512A>C), ACC944AAT (3516C>A 3517C>T), ATT949CTT (3530A>C), GAA950GAG (3535A>G), AAG951AAA (3538G>A), CTC952CTG (3541C>G), AGG953AGA (3544G>A), AAG954AAA (3547G>A), GAT955GAA (3550T>A), AAC956AAT (3553C>T), GTG957GTT (3556G>T), GAA958GAT (3559A>T), CAA959ATC (3560C>A 3561A>T 3562A>C), TAT960TTT (3564A>T), GAT961GAC (3568T>C), GCC962CAG (3569G>C 3570C>A 3571C>G), AAG963AAA (3574G>A), GGT964GGG (3577T>G), CAG966_CCA967del (3581_3586delCAGCCA), GTC968AGC (3587G>A 3588T>G), CAA970ACA (3593C>A 3594A>C), AGC971TCT (3596A>T 3597G>C 3598C>T), CAG972CAA (3601G>A), GTG973GTT (3604G>T), AAT974TCC (3605A>T 3606A>C 3607T>C), TTG975TTC (3610G>C), TCC976CAG (3611T>C 3612C>A 3613C>G), ATT977ATC (3616T>C), GAG979GAA (3622G>A), AAA980AAG (3625A>G), TTT981TTC (3628T>C), TCA982ACA (3629T>A), GGT984GGA (3637T>A), GTC986GTG (3643C>G), CAG987CAA (3646G>A), TTC989TTT (3652C>T), TGC990TGT (3655C>T), ATG991GCT (3656A>G 3657T>C 3658G>T), GGC992GCA (3660G>C 3661C>A), GAA996ACA (3671G>A 3672A>C), GAA997GAC (3676A>C), AGG998AGA (3679G>A), CTT1000GTA (3683C>G 3685T>A), GTA1001GTT (3688A>T), ATA1002TTG (3689A>T 3691A>G), GCT1005GCC (3700T>C), CCC1006CCT (3703C>T), GAA1012GAT (3721A>T), GCT1013GCA (3724T>A), CAG1014TCC (3725C>T 3726A>C 3727G>C), GCT1016CCA (3731G>C 3733T>A), GTT1017ATT (3734G>A), AAG1018AAA (3739G>A), ACC1022ACT (3751C>T), TGT1023TGC (3754T>C), CCT1024CCA (3757T>A), ATC1026ATA (3763C>A), GAT1027GAC (3766T>C), ACA1030TCC (3773A>T 3775A>C), TCC1031TCT (3778C>T), TAC1034TAT (3787C>T), TCA1036TCT (3793A>T), GGC1037GGA (3796C>A), AGC1038AGT (3799C>T), CTC1039TTG (3800C>T 3802C>G), AAT1040GAA (3803A>G 3805T>A), TCC1042ACA (3809T>A 3811C>A), ATA1043TTT (3812A>T 3814A>T), GTT1044GTA (3817T>A), GTG1045ATG (3818G>A), GTT1048ACT (3827G>A 3828T>C), TCC1050AGC (3833T>A 3834C>G), AGC1051TCC (3836A>T 3837G>C), CCT1052AGC (3839C>A 3840C>G 3841T>C), AAT1053AGT (3843A>G), GTC1054ATA (3845G>A 3847C>A), GGA1055GGT (3850A>T), GGC1056GGT (3853C>T), ATT1057ATC (3856T>C), CTG1058TTG (3857C>T), AAT1059AAC (3862T>C), GTA1060ATT (3863G>A 3865A>T), GCT1061GCA (3868T>A), TCC1064ACA (3875T>A 3877C>A), TCT1065TCA (3880T>A), GGT1066GGG (3883T>G), CCA1068CCT (3889A>T), TTT1069TTC (3892T>C), CCA1070CAA (3894C>A), GCA1071TTT (3896G>T 3897C>T 3898A>T), GGC1072GGT (3901C>T), CTG1073GTA (3902C>G 3904G>A), AGT1074TCT (3905A>T 3906G>C), AAG1075GCG (3908A>G 3909A>C), AAT1077GCA (3914A>G 3915A>C 3916T>A), TAT1078CAT (3917T>C), GTA1079GTG (3922A>G), GTT1080TTA (3923G>T 3925T>A), GGA1083GGC (3934A>C), GGA1084GGT (3937A>T), TCC1086ACT (3941T>A 3943C>T), AAG1087AAA (3946G>A), AAC1089-GA (3950delA 3951A>G 3952C>A), GGT1093GGG (3964T>G), ACA1095ACC (3970A>C), ACA1096ACC (3973A>C), ATA1098ATC (3979A>C), TTT1099TTC (3982T>C), ACG1102GTA (3989A>G 3990C>T 3991G>A), GTT1103GTG (3994T>G) |

##### Proteins

| | Begin | End | Coverage | Score | Concordance | Matches | Identities | I/D/M/F* | Stop Codons |
|---|---|---|---|---|---|---|---|---|---|
| polyprotein (YP_003097231.1) | 204 | 1105 | 22.5% | 1363 | 72.9% | 268 (99.3%) | 202 (74.8%) | 0/2/1/1 | 0 |

| | |
|---|---|
| Protein mutations: | Q211S (1316C>T 1317A>C 1318A>T), Q228E (1367C>G 1369A>G), S229R (1372T>G), S230A (1373T>G 1375C>G), Q231S (1376C>T 1377A>C 1378G>T), M232Q (1379A>C 1380T>A 1381G>A), A238P (1397G>C), E242A (1410A>C 1411G>A), R243L (1412A>C 1413G>T 1414G>T), D250N (1433G>A), V895L (3368G>T 3370T>G), S899T (3380T>A 3382A>T), C908R (3407T>C), L909I (3410C>A), N911D (3416A>G), V913T (3422G>A 3423T>C 3424C>G), T917S (3434A>T 3436G>T), K931R (3477A>G 3478G>A), D942N (3509G>A 3511T>C), M943L (3512A>C), T944N (3516C>A 3517C>T), I949L (3530A>C), D955E (3550T>A), E958D (3559A>T), Q959I (3560C>A 3561A>T 3562A>C), Y960F (3564A>T), A962Q (3569G>C 3570C>A 3571C>G), Q966_P967del (3581_3586delCAGCCA), V968S (3587G>A 3588T>G), Q970T (3593C>A 3594A>C), N974S (3605A>T 3606A>C 3607T>C), L975F (3610G>C), S976Q (3611T>C 3612C>A 3613C>G), S982T (3629T>A), M991A (3656A>G 3657T>C 3658G>T), G992A (3660G>C 3661C>A), E996T (3671G>A 3672A>C), E997D (3676A>C), L1000V (3683C>G 3685T>A), I1002L (3689A>T 3691A>G), E1012D (3721A>T), Q1014S (3725C>T 3726A>C 3727G>C), A1016P (3731G>C 3733T>A), V1017I (3734G>A), T1030S (3773A>T 3775A>C), N1040E (3803A>G 3805T>A), S1042T (3809T>A 3811C>A), I1043F (3812A>T 3814A>T), V1045M (3818G>A), V1048T (3827G>A 3828T>C), P1052S (3839C>A 3840C>G 3841T>C), N1053S (3843A>G), V1054I (3845G>A 3847C>A), V1060I (3863G>A 3865A>T), S1064T (3875T>A 3877C>A), P1070Q (3894C>A), A1071F (3896G>T 3897C>T 3898A>T), L1073V (3902C>G 3904G>A), K1075A (3908A>G 3909A>C), N1077A (3914A>G 3915A>C 3916T>A), Y1078H (3917T>C), V1080L (3923G>T 3925T>A), S1086T (3941T>A 3943C>T), T1102V (3989A>G 3990C>T 3991G>A) |

| | Begin | End | Coverage | Score | Concordance | Matches | Identities | I/D/M/F* | Stop Codons |
|---|---|---|---|---|---|---|---|---|---|
| **NT** | **1293** | **3998** | **15.9%** | **609** | **38.5%** | **803 (99.1%)** | **560 (69.1%)** | **0/7** | |
| Codon mutations: | CCT207CCA (1306T>A), CAA208CAG (1309A>G), TTT209TTC (1312T>C), TTC210TTT (1315C>T), CAA211TCT (1316C>T 1317A>C 1318A>T), CAG214CAA (1327G>A), GCT216GCA (1333T>A), GCC217GCA (1336C>A), TCA222TCT (1351A>T), GTA223GTC (1354A>C), TCT225TCG (1360T>G), GGG227GGA (1366G>A), CAA228GAG (1367C>G 1369A>G), AGT229AGG (1372T>G), TCC230GCG (1373T>G 1375C>G), CAG231TCT (1376C>T 1377A>C 1378G>T), ATG232CAA (1379A>C 1380T>A 1381G>A), TTG234CTA (1385T>C 1387G>A), CGT237AGA (1394C>A 1396T>A), GCT238CCT (1397G>C), GTT240GTA (1405T>A), GGA241GGT (1408A>T), GAG242GCA (1410A>C 1411G>A), AGG243CTT (1412A>C 1413G>T 1414G>T), AAT246AAC (1423T>C), ACA247ACT (1426A>T), CGA248CGC (1429A>C), GAT250AAT (1433G>A), CTT890CTG (3355T>G), GTG891GTT (3358G>T), TCC892TCA (3361C>A), TCA894TCC (3367A>C), GTT895TTG (3368G>T 3370T>G), CTT897CTG (3376T>G), GAC898GAT (3379C>T), TCA899ACT (3380T>A 3382A>T), GGT901GGA (3388T>A), GTC902GTA (3391C>A), ACG903ACA (3394G>A), CAG905CAA (3400G>A), CTT906TTG (3401C>T 3403T>G), TGC908CGC (3407T>C), CTT909ATT (3410C>A), AAT911GAT (3416A>G), TTG912CTC (3419T>C 3421G>C), GTC913ACG (3422G>A 3423T>C 3424C>G), CTA914CTT (3427A>T), GGT915GGG (3430T>G), ACG917TCT (3434A>T 3436G>T), GTT918GTG (3439T>G), TCT919TCA (3442T>A), GTG920GTT (3445G>T), ATC923ATT (3454C>T), GGA925GGT (3460A>T), CAC926CAT (3463C>T), ACA928ACC (3469A>C), AAA929AAG (3472A>G), GGA930GGG (3475A>G), AAG931AGA (3477A>G 3478G>A), GCG933GCT (3484G>T), GAC935GAT (3490C>T), CTT937CTG (3496T>G), TTC938TTT (3499C>T), GAG940GAA (3505G>A), GAT942AAC (3509G>A 3511T>C), ATG943CTG (3512A>C), ACC944AAT (3516C>A 3517C>T), ATT949CTT (3530A>C), GAA950GAG (3535A>G), AAG951AAA (3538G>A), CTC952CTG (3541C>G), AGG953AGA (3544G>A), AAG954AAA (3547G>A), GAT955GAA (3550T>A), AAC956AAT (3553C>T), GTG957GTT (3556G>T), GAA958GAT (3559A>T), CAA959ATC (3560C>A 3561A>T 3562A>C), TAT960TTT (3564A>T), GAT961GAC (3568T>C), GCC962CAG (3569G>C 3570C>A 3571C>G), AAG963AAA (3574G>A), GGT964GGG (3577T>G), CAG966_CCA967del (3581_3586delCAGCCA), GTC968AGC (3587G>A 3588T>G), CAA970ACA (3593C>A 3594A>C), AGC971TCT (3596A>T 3597G>C 3598C>T), CAG972CAA (3601G>A), GTG973GTT (3604G>T), AAT974TCC (3605A>T 3606A>C 3607T>C), TTG975TTC (3610G>C), TCC976CAG (3611T>C 3612C>A 3613C>G), ATT977ATC (3616T>C), GAG979GAA (3622G>A), AAA980AAG (3625A>G), TTT981TTC (3628T>C), TCA982ACA (3629T>A), GGT984GGA (3637T>A), GTC986GTG (3643C>G), CAG987CAA (3646G>A), TTC989TTT (3652C>T), TGC990TGT (3655C>T), ATG991GCT (3656A>G 3657T>C 3658G>T), GGC992GCA (3660G>C 3661C>A), GAA996ACA (3671G>A 3672A>C), GAA997GAC (3676A>C), AGG998AGA (3679G>A), CTT1000GTA (3683C>G 3685T>A), GTA1001GTT (3688A>T), ATA1002TTG (3689A>T 3691A>G), GCT1005GCC (3700T>C), CCC1006CCT (3703C>T), GAA1012GAT (3721A>T), GCT1013GCA (3724T>A), CAG1014TCC (3725C>T 3726A>C 3727G>C), GCT1016CCA (3731G>C 3733T>A), GTT1017ATT (3734G>A), AAG1018AAA (3739G>A), ACC1022ACT (3751C>T), TGT1023TGC (3754T>C), CCT1024CCA (3757T>A), ATC1026ATA (3763C>A), GAT1027GAC (3766T>C), ACA1030TCC (3773A>T 3775A>C), TCC1031TCT (3778C>T), TAC1034TAT (3787C>T), TCA1036TCT (3793A>T), GGC1037GGA (3796C>A), AGC1038AGT (3799C>T), CTC1039TTG (3800C>T 3802C>G), AAT1040GAA (3803A>G 3805T>A), TCC1042ACA (3809T>A 3811C>A), ATA1043TTT (3812A>T 3814A>T), GTT1044GTA (3817T>A), GTG1045ATG (3818G>A), GTT1048ACT (3827G>A 3828T>C), TCC1050AGC (3833T>A 3834C>G), AGC1051TCC (3836A>T 3837G>C), CCT1052AGC (3839C>A 3840C>G 3841T>C), AAT1053AGT (3843A>G), GTC1054ATA (3845G>A 3847C>A), GGA1055GGT (3850A>T), GGC1056GGT (3853C>T), ATT1057ATC (3856T>C), CTG1058TTG (3857C>T), AAT1059AAC (3862T>C), GTA1060ATT (3863G>A 3865A>T), GCT1061GCA (3868T>A), TCC1064ACA (3875T>A 3877C>A), TCT1065TCA (3880T>A), GGT1066GGG (3883T>G), CCA1068CCT (3889A>T), TTT1069TTC (3892T>C), CCA1070CAA (3894C>A), GCA1071TTT (3896G>T 3897C>T 3898A>T), GGC1072GGT (3901C>T), CTG1073GTA (3902C>G 3904G>A), AGT1074TCT (3905A>T 3906G>C), AAG1075GCG (3908A>G 3909A>C), AAT1077GCA (3914A>G 3915A>C 3916T>A), TAT1078CAT (3917T>C), GTA1079GTG (3922A>G), GTT1080TTA (3923G>T 3925T>A), GGA1083GGC (3934A>C), GGA1084GGT (3937A>T), TCC1086ACT (3941T>A 3943C>T), AAG1087AAA (3946G>A), AAC1089-GA (3950delA 3951A>G 3952C>A), GGT1093GGG (3964T>G), ACA1095ACC (3970A>C), ACA1096ACC (3973A>C), ATA1098ATC (3979A>C), TTT1099TTC (3982T>C), ACG1102GTA (3989A>G 3990C>T 3991G>A), GTT1103GTG (3994T>G) | | | | | | | | |

*: Inserts / Deletes / Misaligned / Frameshifts

#### Analysis details

This analysis was performed with panviral2.64

### NGS Details (UN59): Tomato chocolate spot virus (segment RNA 1)

#### Assembly

| | |
|---|---|
| **Coverage Length** | 1210 (2 contig(s)) |
| **Depth Of Coverage** | 7250.8 |
| **Number Of Reads** | 56205 |
| **Reads Per Million** | 1221.85 rpm (after QC) |
| **Ambiguities** | 0 |
| **Assembly Method** | de novo + reference guided assembly |
| **Consensus Caller** | Bcf Tools |

#### Coverage Map

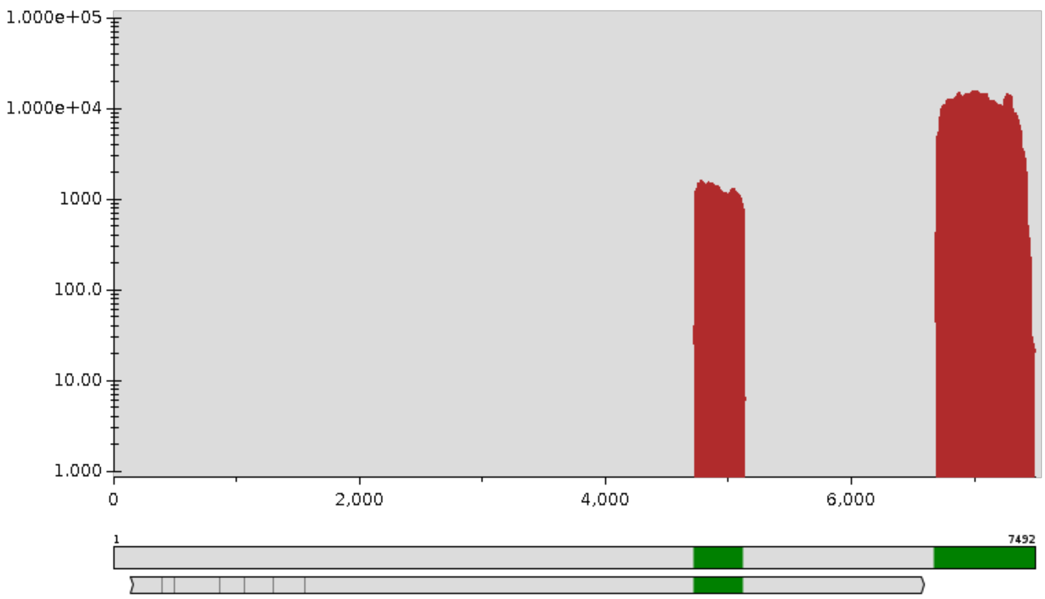

#### Assignment

| | |
|---|---|
| **Type** | Tomato chocolate spot virus (Taxonomy ID: 661101) |
| **Reference Genome** | NC_013075.1 |
| **NT Identity (%)** | 67.5094 |
| **AA Identity (%)** | 66.1765 |
| **Number Of Stop Codons** | 0 |
| **Number Of CDS** | 1 |

#### Alignment

| | |
|---|---|
| **Alignment Score** | 327.0 (NT) + 683.0 (AA) = 1010.0 |
| **Concordance (%)** | 37.9699 |

| Alignment Method | Global, seeded, nucleotide + amino acids (AGA) |
|---|---|

#### Genome Region

Sequence starts at position 4726 and ends at position 7492 relative to NC_013075.1 reference sequence.

#### Alignment Detailed Statistics

| | Begin | End | Coverage | Score | Concordance | Matches | Identities | I/D/M/F* | Stop Codons |
|---|---|---|---|---|---|---|---|---|---|
| **NT** | **4726** | **7492** | **16.2%** | **327** | **19.2%** | **1049 (85.4%)** | **721 (58.7%)** | **19/161** | |

Mutations: 4735T>C, 4737G>T, 4740C>T, 4744T>A, 4745C>A, 4746A>C, 4750G>A, 4755G>C, 4761G>T, 4764G>C, 4765C>A, 4767G>A, 4770T>C, 4785A>G, 4786C>T, 4791T>G, 4795C>T, 4806G>C, 4813A>C, 4815A>C, 4818T>C, 4822C>T, 4827C>T, 4833T>A, 4834A>C, 4836G>A, 4839A>C, 4842G>C, 4845T>C, 4854A>T, 4860A>T, 4861A>G, 4863A>G, 4881A>C, 4882C>A, 4891C>G, 4892C>T, 4893C>T, 4896T>G, 4902T>A, 4906G>A, 4907C>T, 4908A>C, 4909C>A, 4910A>C, 4911G>T, 4914A>C, 4918A>T, 4919T>A, 4920G>T, 4929T>G, 4935T>G, 4938C>T, 4939A>G, 4950C>T, 4953T>C, 4956T>G, 4962T>A, 4965A>G, 4968T>C, 4970C>T, 4971A>T, 4977G>C, 4979A>C, 4984G>A, 4985T>C, 4986G>A, 4988C>A, 4989A>C, 4990G>A, 4995G>C, 4999G>A, 5005C>A, 5007G>A, 5009A>T, 5010T>G, 5011G>C, 5013A>G, 5016T>A, 5019A>C, 5022T>G, 5024G>C, 5025T>A, 5026G>A, 5027C>T, 5029A>G, 5030C>T, 5031A>G, 5032A>C, 5033T>A, 5037C>T, 5040T>A, 5043C>T, 5046C>T, 5047C>G, 5048T>C, 5049G>T, 5053G>C, 5054T>A, 5055C>A, 5059C>A, 5060A>G, 5061A>T, 5064A>G, 5068A>C, 5070T>A, 5071G>C, 5072C>A, 5073C>A, 5074A>G, 5075A>C, 5076G>T, 5086A>G, 5088T>C, 5089G>A, 5090C>G, 5092G>A, 5094T>C, 5095G>A, 5097G>A, 5098A>G, 5099T>G, 5101A>G, 5103A>T, 5104A>C, 5106C>G, 5107T>C, 5108G>T, 5109C>G, 5113A>C, 5116G>A, 5117C>G, 5118A>C, 5120G>A, 5121A>G, 6706A>G, 6707T>A, 6721G>C, 6722G>C, 6729T>C, 6731G>A, 6735G>A, 6736T>G, 6740C>A, 6741G>A, 6753G>T, 6757T>G, 6758_6759insT, 6759_6760insG, 6772C>T, 6777T>C, 6795A>G, 6806C>T, 6814C>T, 6815_6834delAATTGTTTGGTGTTGTCAAC, 6837G>T, 6840G>A, 6842C>A, 6843T>C, 6845T>G, 6847_6850delGCCA, 6853A>G, 6855A>T, 6857C>A, 6859C>T, 6860C>G, 6861C>G, 6862C>T, 6863T>C, 6867_6868delGC, 6870T>G, 6872A>G, 6873T>A, 6878A>G, 6879A>T, 6881G>A, 6885A>C, 6886G>T, 6887A>T, 6889_6894delGTGGAG, 6897G>A, 6903C>A, 6906G>C, 6908A>C, 6909A>G, 6911C>G, 6912A>G, 6914A>G, 6915A>T, 6918A>T, 6919_6921delGAG, 6923C>T, 6924T>C, 6926T>C, 6930_6937delGGTCGGGC, 6942G>C, 6944_6946delTGG, 6949G>A, 6950C>G, 6951C>T, 6953G>C, 6955A>G, 6956T>G, 6958A>T, 6960A>T, 6961G>T, 6962C>T, 6964T>C, 6965T>C, 6967G>A, 6969G>A, 6970T>G, 6972T>A, 6973G>A, 6976T>A, 6978T>A, 6982_6985delCATC, 6987G>T, 6989T>A, 6991T>C, 6996A>C, 7001_7012delCTCGTTTGGAGT, 7014T>G, 7016A>G, 7018C>G, 7020A>T, 7022delT, 7026G>T, 7030_7035delTGGTGT, 7040A>C, 7041C>G, 7042G>A, 7047_7067delGCAATGACACCCGGGGTGTGC, 7073C>G, 7077T>C, 7079A>T, 7080A>G, 7084_7088delAAGTA, 7091G>A, 7093T>C, 7097T>C, 7098C>T, 7099T>G, 7101C>A, 7103A>T, 7104T>A, 7105A>G, 7108A>G, 7109A>T, 7110C>G, 7111T>C, 7115_7123delGCGTTGGTC, 7125C>G, 7128C>G, 7130C>T, 7131A>G, 7132A>T, 7134delG, 7136G>C, 7138G>C, 7142C>A, 7143G>C, 7146T>C, 7147G>C, 7148_7161delAAAATCGATAAATT, 7165G>C, 7169C>G, 7172G>T, 7173delC, 7177A>C, 7180_7183delCTGC, 7186T>G, 7188G>T, 7190T>C, 7193G>A, 7195_7218delGCAACAGTGAAGAATCATCACACC, 7220C>G, 7223C>A, 7224T>C, 7225T>C, 7227C>T, 7228C>G, 7230A>C, 7231G>T, 7234T>G, 7235C>G, 7238C>T, 7241G>C, 7244G>A, 7245T>A, 7247_7248delTC, 7251A>T, 7259_7260delCT, 7262C>G, 7264T>A, 7266G>A, 7270A>C, 7273_7274delAG, 7288G>C, 7289T>C, 7291T>G, 7292T>G, 7293G>A, 7294C>T, 7296_7297insGCTTCTCCTAGTTAA, 7298G>A, 7299A>C, 7300C>G, 7302A>G, 7303T>C, 7304G>A, 7308C>T, 7310T>A, 7311A>T, 7314G>C, 7315T>C, 7316T>A, 7317T>G, 7318G>T, 7319G>A, 7321T>A, 7325C>A, 7327T>A, 7328A>T, 7330A>T, 7335_7336insT, 7342T>G, 7346A>T, 7348A>T, 7350G>C, 7351G>T, 7352_7353insC, 7358C>G, 7367C>T, 7369G>A, 7372G>T, 7373delA, 7375T>C, 7376G>C, 7378A>C, 7381C>G, 7385C>T, 7387T>G, 7388T>C, 7389C>A, 7390A>T, 7392T>C, 7393_7396delTATT, 7407G>T, 7410delG, 7425A>C, 7428delC, 7430T>C, 7431C>A, 7447A>C, 7453A>G, 7454G>A, 7455A>T, 7460C>G

##### CDS

| | Begin | End | Coverage | Score | Concordance | Matches | Identities | I/D/M/F* | Stop Codons |
|---|---|---|---|---|---|---|---|---|---|
| ToChSV_s1_gp1 | 1530 | 1665 | 6.3% | 683 | 71.1% | 136 (100%) | 90 (66.2%) | 0/0/0/0 | 0 |

Protein mutations: S1536N (4744T>A 4745C>A 4746A>C), D1538N (4750G>A), E1541D (4761G>T), Q1543K (4765C>A 4767G>A), H1565Q (4833T>A), M1566L (4834A>C 4836G>A), I1575V (4861A>G 4863A>G), P1585V (4891C>G 4892C>T 4893C>T), A1590I (4906G>A 4907C>T 4908A>C), Q1591T (4909C>A 4910A>C 4911G>T), M1594Y (4918A>T 4919T>A 4920G>T), I1601V (4939A>G), H1606Q (4956T>G), A1611V (4970C>T 4971A>T), E1613D (4977G>C), K1614T (4979A>C), V1616T (4984G>A 4985T>C 4986G>A), A1617D (4988C>A 4989A>C), D1618N (4990G>A), E1621K (4999G>A), L1623I (5005C>A 5007G>A), N1624M (5009A>T 5010T>G), E1625Q (5011G>C 5013A>G), F1628L (5022T>G), R1629P (5024G>C 5025T>A), A1630I (5026G>A 5027C>T), T1631V (5029A>G 5030C>T 5031A>G), I1632H (5032A>C 5033T>A), L1637A (5047C>G 5048T>C 5049G>T), V1639Q (5053G>C 5054T>A 5055C>A), Q1641S (5059C>A 5060A>G 5061A>T), I1644L (5068A>C 5070T>A), A1645Q (5071G>C 5072C>A 5073C>A), K1646A (5074A>G 5075A>C 5076G>T), N1650D (5086A>G 5088T>C), A1651S (5089G>A 5090C>G), D1652N (5092G>A 5094T>C), V1653I (5095G>A 5097G>A), M1654G (5098A>G 5099T>G), T1655A (5101A>G 5103A>T), I1656L (5104A>C 5106C>G), C1657L (5107T>C 5108G>T 5109C>G), M1659L (5113A>C), A1660S (5116G>A 5117C>G 5118A>C), R1661K (5120G>A 5121A>G)

Codon mutations: TTG1533CTT (4735T>C 4737G>T), GAC1534GAT (4740C>T), TCA1536AAC (4744T>A 4745C>A 4746A>C), GAT1538AAT (4750G>A), GTG1539GTC (4755G>C), GAG1541GAT (4761G>T), GTG1542GTC (4764G>C), CAG1543AAA (4765C>A 4767G>A), TTT1544TTC (4770T>C), GAA1549GAG (4785A>G), CTG1550TTG (4786C>T), ACT1551ACG (4791T>G), CTG1553TTG (4795C>T), GGG1556GGC (4806G>C), AGA1559CGC (4813A>C 4815A>C), CAT1560CAC (4818T>C), CTG1562TTG (4822C>T), GGC1563GGT (4827C>T), CAT1565CAA (4833T>A), ATG1566CTA (4834A>C 4836G>A), GCA1567GCC (4839A>C), CCG1568CCC (4842G>C), CTT1569CTC (4845T>C), GTA1572GTT (4854A>T), ATA1574ATT (4860A>T), ATA1575GTG (4861A>G 4863A>G), ATA1581ATC (4881A>C), CGA1582AGA (4882C>A), CCC1585GTT (4891C>G 4892C>T 4893C>T), GGT1586GGG (4896T>G), CCT1588CCA (4902T>A), GCA1590ATC (4906G>A 4907C>T 4908A>C), CAG1591ACT (4909C>A 4910A>C 4911G>T), GCA1592GCC (4914A>C), ATG1594TAT (4918A>T 4919T>A 4920G>T), GTT1597GTG (4929T>G), ACT1599ACG (4935T>G), GGC1600GGT (4938C>T), ATA1601GTA (4939A>G), GCC1604GCT (4950C>T), TTT1605TTC (4953T>C), CAT1606CAG (4956T>G), GGT1608GGA (4962T>A), CAA1609CAG (4965A>G), GTT1610GTC (4968T>C), GCA1611GTT (4970C>T 4971A>T), GAG1613GAC (4977G>C), AAA1614ACA (4979A>C), GTG1616ACA (4984G>A 4985T>C 4986G>A), GCA1617GAC (4988C>A 4989A>C), GAT1618AAT (4990G>A), GTG1619GTC (4995G>C), GAA1621AAA (4999G>A), CTG1623ATA (5005C>A 5007G>A), AAT1624ATG (5009A>T 5010T>G), GAA1625CAG (5011G>C 5013A>G), CGT1626CGA (5016T>A), GGA1627GGC (5019A>C), TTT1628TTG (5022T>G), CGT1629CCA (5024G>C 5025T>A), GCC1630ATC (5026G>A 5027C>T), ACA1631GTG (5029A>G 5030C>T 5031A>G), ATT1632CAT (5032A>C 5033T>A), TTC1633TTT (5037C>T), CCT1634CCA (5040T>A), TCC1635TCT (5043C>T), TAC1636TAT (5046C>T), CTG1637GCT (5047C>G 5048T>C 5049G>T), GTC1639CAA (5053G>C 5054T>A 5055C>A), CAA1641AGT (5059C>A 5060A>G 5061A>T), GAA1642GAG (5064A>G), ATT1644CTA (5068A>C 5070T>A), GCC1645CAA (5071G>C 5072C>A 5073C>A), AAG1646GCT (5074A>G 5075A>C 5076G>T), AAT1650GAC (5086A>G 5088T>C), GCT1651AGT (5089G>A 5090C>G), GAT1652AAC (5092G>A 5094T>C), GTG1653ATA (5095G>A 5097G>A), ATG1654GGG (5098A>G 5099T>G), ACA1655GCT (5101A>G 5103A>T), ATC1656CTG (5104A>C 5106C>G), TGC1657CTG (5107T>C 5108G>T 5109C>G), ATG1659CTG (5113A>C), GCA1660AGC (5116G>A 5117C>G 5118A>C), AGA1661AAG (5120G>A 5121A>G)

##### Proteins

| | Begin | End | Coverage | Score | Concordance | Matches | Identities | I/D/M/F* | Stop Codons |
|---|---|---|---|---|---|---|---|---|---|
| polyprotein (YP_003097229.1) | 1530 | 1665 | 6.3% | 683 | 71.1% | 136 (100%) | 90 (66.2%) | 0/0/0/0 | 0 |

Protein mutations: S1536N (4744T>A 4745C>A 4746A>C), D1538N (4750G>A), E1541D (4761G>T), Q1543K (4765C>A 4767G>A), H1565Q (4833T>A), M1566L (4834A>C 4836G>A), I1575V (4861A>G 4863A>G), P1585V (4891C>G 4892C>T 4893C>T), A1590I (4906G>A 4907C>T 4908A>C), Q1591T (4909C>A 4910A>C 4911G>T), M1594Y (4918A>T 4919T>A 4920G>T), I1601V (4939A>G), H1606Q (4956T>G), A1611V (4970C>T 4971A>T), E1613D (4977G>C), K1614T (4979A>C), V1616T (4984G>A 4985T>C 4986G>A), A1617D (4988C>A 4989A>C), D1618N (4990G>A), E1621K (4999G>A), L1623I (5005C>A 5007G>A), N1624M (5009A>T 5010T>G), E1625Q (5011G>C 5013A>G), F1628L (5022T>G), R1629P (5024G>C 5025T>A), A1630I (5026G>A 5027C>T), T1631V (5029A>G 5030C>T 5031A>G), I1632H (5032A>C 5033T>A), L1637A (5047C>G 5048T>C 5049G>T), V1639Q (5053G>C 5054T>A 5055C>A), Q1641S (5059C>A 5060A>G 5061A>T), I1644L (5068A>C 5070T>A), A1645Q (5071G>C 5072C>A 5073C>A), K1646A (5074A>G 5075A>C 5076G>T), N1650D (5086A>G 5088T>C), A1651S (5089G>A 5090C>G), D1652N (5092G>A 5094T>C), V1653I (5095G>A 5097G>A), M1654G (5098A>G 5099T>G), T1655A (5101A>G 5103A>T), I1656L (5104A>C 5106C>G), C1657L (5107T>C 5108G>T 5109C>G), M1659L (5113A>C), A1660S (5116G>A 5117C>G 5118A>C), R1661K (5120G>A 5121A>G)

Codon mutations: TTG1533CTT (4735T>C 4737G>T), GAC1534GAT (4740C>T), TCA1536AAC (4744T>A 4745C>A 4746A>C), GAT1538AAT (4750G>A), GTG1539GTC (4755G>C), GAG1541GAT (4761G>T), GTG1542GTC (4764G>C), CAG1543AAA (4765C>A 4767G>A), TTT1544TTC (4770T>C), GAA1549GAG (4785A>G), CTG1550TTG (4786C>T), ACT1551ACG (4791T>G), CTG1553TTG (4795C>T), GGG1556GGC (4806G>C), AGA1559CGC (4813A>C 4815A>C), CAT1560CAC (4818T>C), CTG1562TTG (4822C>T), GGC1563GGT (4827C>T), CAT1565CAA (4833T>A), ATG1566CTA (4834A>C 4836G>A), GCA1567GCC (4839A>C), CCG1568CCC (4842G>C), CTT1569CTC (4845T>C), GTA1572GTT (4854A>T), ATA1574ATT (4860A>T), ATA1575GTG (4861A>G 4863A>G), ATA1581ATC (4881A>C), CGA1582AGA (4882C>A), CCC1585GTT (4891C>G 4892C>T 4893C>T), GGT1586GGG (4896T>G), CCT1588CCA (4902T>A), GCA1590ATC (4906G>A 4907C>T 4908A>C), CAG1591ACT (4909C>A 4910A>C 4911G>T), GCA1592GCC (4914A>C), ATG1594TAT (4918A>T 4919T>A 4920G>T), GTT1597GTG (4929T>G), ACT1599ACG (4935T>G), GGC1600GGT (4938C>T), ATA1601GTA (4939A>G), GCC1604GCT (4950C>T), TTT1605TTC (4953T>C), CAT1606CAG (4956T>G), GGT1608GGA (4962T>A), CAA1609CAG (4965A>G), GTT1610GTC (4968T>C), GCA1611GTT (4970C>T 4971A>T), GAG1613GAC (4977G>C), AAA1614ACA (4979A>C), GTG1616ACA (4984G>A 4985T>C 4986G>A), GCA1617GAC (4988C>A 4989A>C), GAT1618AAT (4990G>A), GTG1619GTC (4995G>C), GAA1621AAA (4999G>A), CTG1623ATA (5005C>A 5007G>A), AAT1624ATG (5009A>T 5010T>G), GAA1625CAG (5011G>C 5013A>G), CGT1626CGA (5016T>A), GGA1627GGC (5019A>C), TTT1628TTG (5022T>G), CGT1629CCA (5024G>C 5025T>A), GCC1630ATC (5026G>A 5027C>T), ACA1631GTG (5029A>G 5030C>T 5031A>G), ATT1632CAT (5032A>C 5033T>A), TTC1633TTT (5037C>T), CCT1634CCA (5040T>A), TCC1635TCT (5043C>T), TAC1636TAT (5046C>T), CTG1637GCT (5047C>G 5048T>C 5049G>T), GTC1639CAA (5053G>C 5054T>A 5055C>A), CAA1641AGT (5059C>A 5060A>G 5061A>T), GAA1642GAG (5064A>G), ATT1644CTA (5068A>C 5070T>A), GCC1645CAA (5071G>C 5072C>A 5073C>A), AAG1646GCT (5074A>G 5075A>C 5076G>T), AAT1650GAC (5086A>G 5088T>C), GCT1651AGT (5089G>A 5090C>G), GAT1652AAC (5092G>A 5094T>C), GTG1653ATA (5095G>A 5097G>A), ATG1654GGG (5098A>G 5099T>G), ACA1655GCT (5101A>G 5103A>T), ATC1656CTG (5104A>C 5106C>G), TGC1657CTG (5107T>C 5108G>T 5109C>G), ATG1659CTG (5113A>C), GCA1660AGC (5116G>A 5117C>G 5118A>C), AGA1661AAG (5120G>A 5121A>G)

*: Inserts / Deletes / Misaligned / Frameshifts

#### Analysis details

This analysis was performed with panviral2.64

### NGS Details (UN59): Lausannevirus

#### Assembly

| | |
|---|---|
| **Coverage Length** | 176 (1 contig(s)) |
| **Depth Of Coverage** | 9728.0 |
| **Number Of Reads** | 18309 |
| **Reads Per Million** | 398.02 rpm (after QC) |
| **Ambiguities** | 0 |
| **Assembly Method** | de novo + reference guided assembly |
| **Consensus Caller** | Bcf Tools |

#### Coverage Map

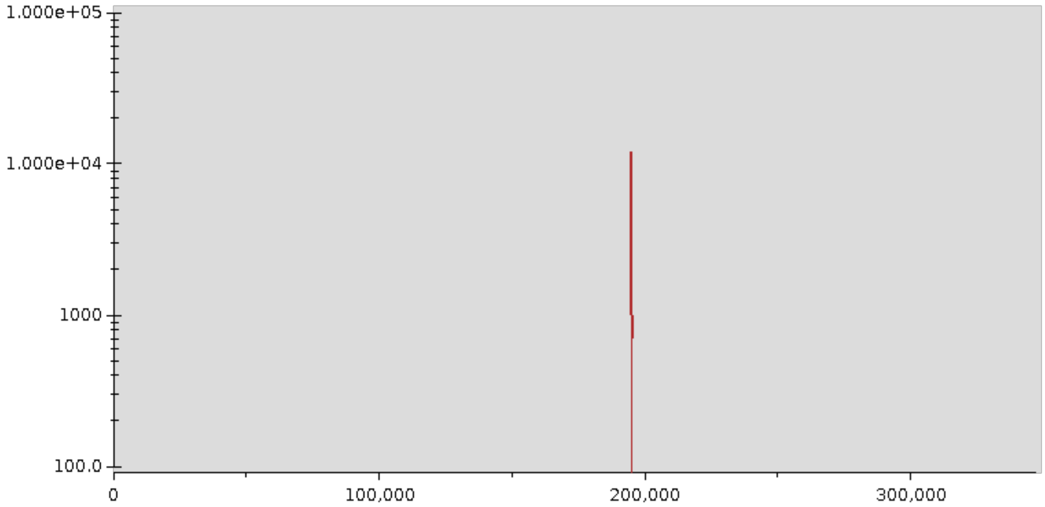

#### Assignment

| | |
|---|---|
| **Type** | Lausannevirus (Taxonomy ID: 999883) |
| **Reference Genome** | NC_015326.1 |
| **NT Identity (%)** | 81.8182 |
| **AA Identity (%)** | 94.9153 |
| **Number Of Stop Codons** | 0 |
| **Number Of CDS** | 444 |

#### Alignment

| | |
|---|---|
| **Alignment Score** | 224.0 (NT) + 364.0 (AA) = 588.0 |
| **Concordance (%)** | 80.7692 |
| **Alignment Method** | Local, heuristic, nucleotide (BLASTN) |

#### Genome Region

Sequence starts at position 194923 and ends at position 195098 relative to NC_015326.1 reference sequence.

#### Alignment Detailed Statistics

| | Begin | End | Coverage | Score | Concordance | Matches | Identities | I/D/M/F* | Stop Codons |
|---|---|---|---|---|---|---|---|---|---|
| **NT** | **194923** | **195098** | **0.1%** | **224** | **63.6%** | **176 (100%)** | **144 (81.8%)** | **0/0** | |

Mutations: 194940A>C, 194946G>T, 194952T>C, 194955T>C, 194958A>G, 194967A>G, 194970G>C, 194979T>G, 194981T>G, 194982C>G, 194988T>C, 195003T>A, 195012A>C, 195015T>C, 195017G>T, 195018T>C, 195021T>C, 195033A>G, 195036G>C, 195042T>C, 195048T>C, 195054T>C, 195057T>A, 195058T>G, 195059G>C, 195060C>T, 195063G>C, 195066G>A, 195069C>G, 195072A>G, 195081C>A, 195087T>C

*: Inserts / Deletes / Misaligned / Frameshifts

#### Analysis details

This analysis was performed with panviral2.64

### NGS Details (UN59): Makelovirus prm1

#### Assembly

| | |
|---|---|
| **Coverage Length** | 1014 (1 contig(s)) |
| **Depth Of Coverage** | 557.0 |
| **Number Of Reads** | 4406 |
| **Reads Per Million** | 95.78 rpm (after QC) |
| **Ambiguities** | 0 |
| **Assembly Method** | de novo + reference guided assembly |
| **Consensus Caller** | Bcf Tools |

#### Coverage Map

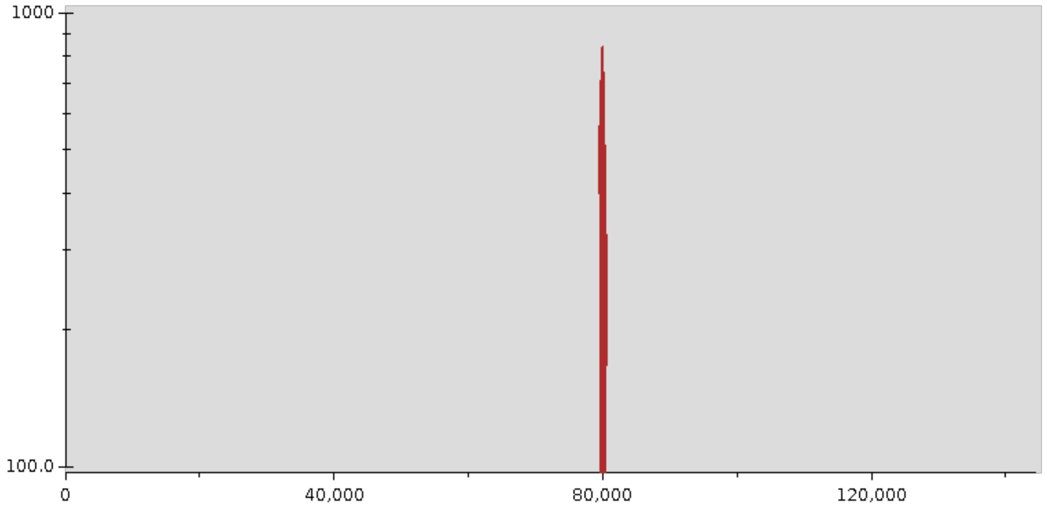

#### Assignment

| | |
|---|---|
| **Type** | Makelovirus prm1 (Taxonomy ID: 2956181) |
| **Reference Genome** | NC_055761.1 |
| **NT Identity (%)** | 76.8245 |
| **AA Identity (%)** | 87.574 |
| **Number Of Stop Codons** | 0 |
| **Number Of CDS** | 190 |

#### Alignment

| | |
|---|---|
| **Alignment Score** | 1088.0 (NT) + 2258.0 (AA) = 3346.0 |
| **Concordance (%)** | 74.4548 |
| **Alignment Method** | Local, heuristic, nucleotide (BLASTN) |

#### Genome Region

Sequence starts at position 79506 and ends at position 80519 relative to NC_055761.1 reference sequence.

#### Alignment Detailed Statistics

| | Begin | End | Coverage | Score | Concordance | Matches | Identities | I/D/M/F* | Stop Codons |
|---|---|---|---|---|---|---|---|---|---|
| **NT** | **79506** | **80519** | **0.7%** | **1088** | **53.6%** | **1014 (100%)** | **779 (76.8%)** | **0/0** | |
| Mutations: | 79519A>G, 79520A>T, 79522A>G, 79523A>C, 79529C>T, 79530G>A, 79532G>C, 79536G>A, 79541G>T, 79542T>A, 79543C>G, 79544T>C, 79547A>T, 79548A>G, 79550C>A, 79563G>A, 79565C>T, 79568T>A, 79574C>T, 79580G>T, 79581C>T, 79592A>T, 79595T>C, 79596C>T, 79598G>A, 79602G>A, 79604A>G, 79610C>T, 79611G>T, 79612T>C, 79613C>T, 79614T>G, 79615G>T, 79616C>A, 79619C>T, 79624C>T, 79628G>C, 79632G>A, 79643C>T, 79646T>A, 79649G>A, 79655C>T, 79658C>T, 79664C>T, 79673A>T, 79676C>T, 79677G>T, 79679T>A, 79682T>G, 79688G>A, 79689A>C, 79691G>T, 79694T>C, 79703C>T, 79712T>C, 79718T>C, 79719G>A, 79722G>A, 79724A>T, 79728T>A, 79733C>T, 79734A>G, 79735A>C, 79736C>A, 79739A>T, 79742T>A, 79746C>T, 79748T>A, 79751C>T, 79754C>T, 79760C>A, 79763T>C, 79772T>G, 79779C>G, 79791C>T, 79793C>A, 79799T>C, 79811C>T, 79812C>G, 79817G>A, 79818G>A, 79820C>T, 79823A>T, 79824T>C, 79826T>A, 79833C>T, 79835G>A, 79836A>C, 79838C>T, 79842A>G, 79844C>A, 79845T>G, 79846T>C, 79847C>T, 79850C>T, 79871A>G, 79874C>T, 79875T>A, 79876C>G, 79879A>T, 79880T>C, 79895T>A, 79898C>T, 79904C>T, 79905T>G, 79906G>C, 79907C>T, 79916C>T, 79919T>A, 79922A>T, 79931T>A, 79937A>T, 79938T>A, 79940T>C, 79946C>T, 79950C>T, 79953G>A, 79958T>C, 79961T>A, 79962T>A, 79973T>A, 79974T>A, 79975C>G, 79976G>T, 79979C>T, 79985C>T, 79994C>T, 79997C>A, 80000C>A, 80018C>T, 80025C>A, 80030C>A, 80039A>T, 80042A>G, 80054G>T, 80063C>A, 80066C>T, 80069T>C, 80073C>T, 80075T>A, 80078T>C, 80090C>A, 80096T>C, 80099T>C, 80105G>A, 80114A>T, 80120C>T, 80126T>C, 80127C>T, 80132T>A, 80139T>A, 80140C>G, 80141G>T, 80142C>T, 80145G>A, 80147T>C, 80148C>A, 80150T>G, 80159C>A, 80162G>A, 80165C>T, 80168G>A, 80171C>T, 80172C>G, 80173A>C, 80174G>T, 80177C>T, 80178T>G, 80180T>A, 80188A>G, 80189G>A, 80201A>G, 80207G>A, 80210C>T, 80213C>T, 80216C>T, 80222G>A, 80225T>C, 80237C>T, 80240C>T, 80243T>C, 80246C>A, 80247C>T, 80261C>T, 80265T>A, 80266C>G, 80267C>T, 80279C>T, 80286C>T, 80288G>A, 80291T>C, 80300T>A, 80306A>T, 80315T>A, 80318C>A, 80333T>C, 80336A>T, 80337C>T, 80339T>A, 80343G>A, 80346T>A, 80347C>G, 80351C>T, 80357A>T, 80366G>A, 80369C>T, 80378C>T, 80387G>A, 80390C>T, 80391A>G, 80393C>A, 80394A>G, 80396G>T, 80399T>C, 80400G>A, 80409A>C, 80410A>G, 80411A>T, 80414C>A, 80415C>A, 80417C>T, 80433G>A, 80435T>C, 80436C>A, 80438A>C, 80447C>T, 80453G>T, 80456G>T, 80462G>A, 80465A>T, 80471C>T, 80474G>A, 80486C>T, 80498G>A, 80504G>A | | | | | | | | |

*: Inserts / Deletes / Misaligned / Frameshifts

#### Analysis details

This analysis was performed with panviral2.64

### NGS Details (UN59): Lausannevirus

#### Assembly

| | |
|---|---|
| **Coverage Length** | 129 (1 contig(s)) |
| **Depth Of Coverage** | 709.4 |
| **Number Of Reads** | 1051 |
| **Reads Per Million** | 22.85 rpm (after QC) |
| **Ambiguities** | 0 |
| **Assembly Method** | de novo + reference guided assembly |
| **Consensus Caller** | Bcf Tools |

#### Coverage Map

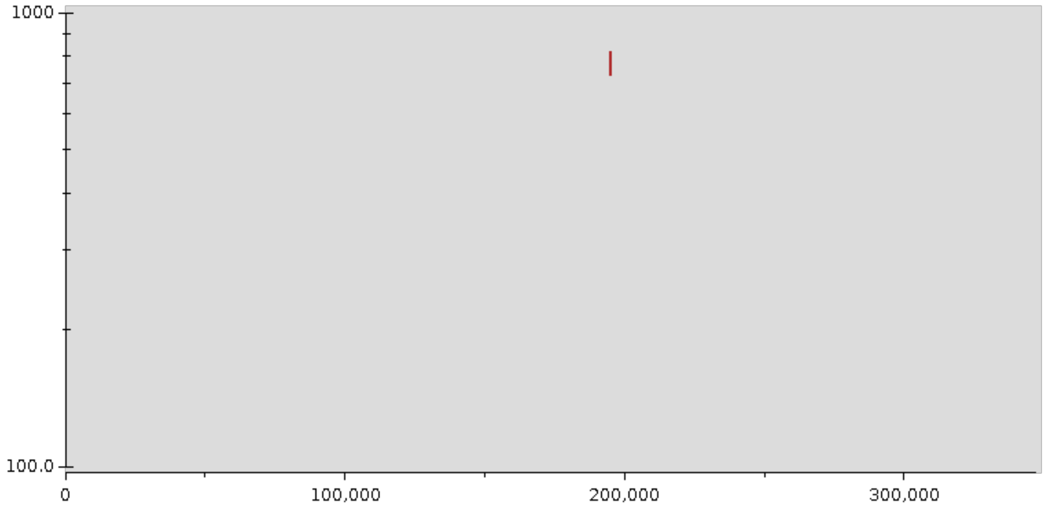

#### Assignment

| | |
|---|---|
| **Type** | Lausannevirus (Taxonomy ID: 999883) |
| **Reference Genome** | NC_015326.1 |
| **NT Identity (%)** | 79.845 |
| **AA Identity (%)** | 95.3488 |
| **Number Of Stop Codons** | 1 |
| **Number Of CDS** | 444 |

#### Alignment

| | |
|---|---|
| **Alignment Score** | 154.0 (NT) + 273.0 (AA) = 427.0 |
| **Concordance (%)** | 77.9197 |
| **Alignment Method** | Local, heuristic, nucleotide (BLASTN) |

#### Genome Region

Sequence starts at position 194922 and ends at position 195050 relative to NC_015326.1 reference sequence.

#### Alignment Detailed Statistics

| | Begin | End | Coverage | Score | Concordance | Matches | Identities | I/D/M/F* | Stop Codons |
|---|---|---|---|---|---|---|---|---|---|
| NT | 194922 | 195050 | 0.1% | 154 | 59.7% | 129 (100%) | 103 (79.8%) | 0/0 | |

Mutations: 194937G>A, 194940A>T, 194941A>C, 194942G>A, 194943A>G, 194946G>T, 194953T>A, 194955T>C, 194967A>G, 194973A>C, 194976A>G, 194979T>C, 194982C>T, 194988T>C, 194991G>A, 194994C>T, 195000A>C, 195003T>G, 195006G>A, 195012A>C, 195014G>A, 195015T>C, 195017G>T, 195018T>C, 195036G>A, 195039T>C

*: Inserts / Deletes / Misaligned / Frameshifts

#### Analysis details

This analysis was performed with panviral2.64

### NGS Details (UN59): Betabaculovirus disaccharalis

#### Assembly

| | |
|---|---|
| **Coverage Length** | 137 (1 contig(s)) |
| **Depth Of Coverage** | 510.6 |
| **Number Of Reads** | 905 |
| **Reads Per Million** | 19.67 rpm (after QC) |
| **Ambiguities** | 0 |
| **Assembly Method** | de novo + reference guided assembly |
| **Consensus Caller** | Bcf Tools |

#### Coverage Map

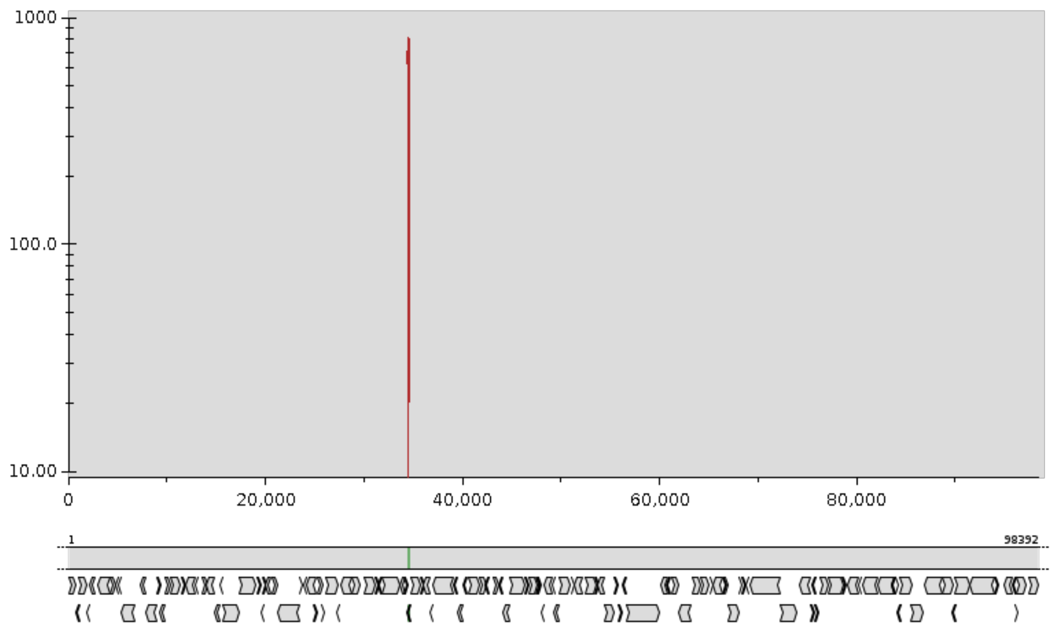

#### Assignment

| | |
|---|---|
| **Type** | Betabaculovirus disaccharalis (Taxonomy ID: 3047795) |
| **Reference Genome** | NC_028491.1 |
| **NT Identity (%)** | 83.9416 |
| **AA Identity (%)** | 91.3043 |
| **Number Of Stop Codons** | 0 |
| **Number Of CDS** | 125 |

#### Alignment

| | |
|---|---|
| **Alignment Score** | 186.0 (NT) + 285.0 (AA) = 471.0 |
| **Concordance (%)** | 81.2069 |

| Alignment Method | Local, heuristic, nucleotide (BLASTN) |
|---|---|

#### Genome Region

Sequence starts at position 34471 and ends at position 34607 relative to NC_028491.1 reference sequence.

#### Alignment Detailed Statistics

| | Begin | End | Coverage | Score | Concordance | Matches | Identities | I/D/M/F* | Stop Codons |
|---|---|---|---|---|---|---|---|---|---|
| **NT** | **34471** | **34607** | **0.1%** | **186** | **67.9%** | **137 (100%)** | **115 (83.9%)** | **0/0** | |
| Mutations: | 34482G>T, 34485T>C, 34488A>C, 34490G>A, 34513C>G, 34514T>A, 34521T>C, 34524A>G, 34533A>G, 34538A>G, 34542T>A, 34545T>C, 34547G>T, 34549T>C, 34554T>C, 34561T>G, 34569G>A, 34575T>G, 34578T>C, 34587T>C, 34593C>T, 34596C>G | | | | | | | | |
| **CDS** | | | | | | | | | |
| v-ubq | 57 | 102 | 38.7% | 285 | 86.9% | 46 (100%) | 42 (91.3%) | 0/0/0/0 | 0 |
| Protein mutations: | Q72P (34561T>G), E76G (34549T>C), S80P (34538A>G) | | | | | | | | |
| Codon mutations: | CCG60CCC (34596C>G), CCG61CCA (34593C>T), CAA63CAG (34587T>C), TTA66TTG (34578T>C), ATA67ATC (34575T>G), GCC69GCT (34569G>A), CAA72CCA (34561T>G), GAA74GAG (34554T>C), GAG76GGG (34549T>C), CGA77AGG (34545T>C 34547G>T), ACA78ACT (34542T>A), TCA80CCA (34538A>G), GAT81GAC (34533A>G), ATT84ATC (34524A>G), CAA85CAG (34521T>C), AGC88TCC (34513C>G 34514T>A), CTT96TTG (34488A>C 34490G>A), AGA97AGG (34485T>C), GGC98GGA (34482G>T) | | | | | | | | |
| **Proteins** | | | | | | | | | |
| ubiquitin-like protein (YP_009182246.1) | 57 | 102 | 38.7% | 285 | 86.9% | 46 (100%) | 42 (91.3%) | 0/0/0/0 | 0 |
| Protein mutations: | Q72P (34561T>G), E76G (34549T>C), S80P (34538A>G) | | | | | | | | |
| Codon mutations: | CCG60CCC (34596C>G), CCG61CCA (34593C>T), CAA63CAG (34587T>C), TTA66TTG (34578T>C), ATA67ATC (34575T>G), GCC69GCT (34569G>A), CAA72CCA (34561T>G), GAA74GAG (34554T>C), GAG76GGG (34549T>C), CGA77AGG (34545T>C 34547G>T), ACA78ACT (34542T>A), TCA80CCA (34538A>G), GAT81GAC (34533A>G), ATT84ATC (34524A>G), CAA85CAG (34521T>C), AGC88TCC (34513C>G 34514T>A), CTT96TTG (34488A>C 34490G>A), AGA97AGG (34485T>C), GGC98GGA (34482G>T) | | | | | | | | |

*: Inserts / Deletes / Misaligned / Frameshifts

#### Analysis details

This analysis was performed with panviral2.64

### NGS Details (UN59): Betabaculovirus disaccharalis

#### Assembly

| | |
|---|---|
| **Coverage Length** | 134 (1 contig(s)) |
| **Depth Of Coverage** | 486.0 |
| **Number Of Reads** | 820 |
| **Reads Per Million** | 17.83 rpm (after QC) |
| **Ambiguities** | 0 |
| **Assembly Method** | de novo + reference guided assembly |
| **Consensus Caller** | Bcf Tools |

#### Coverage Map

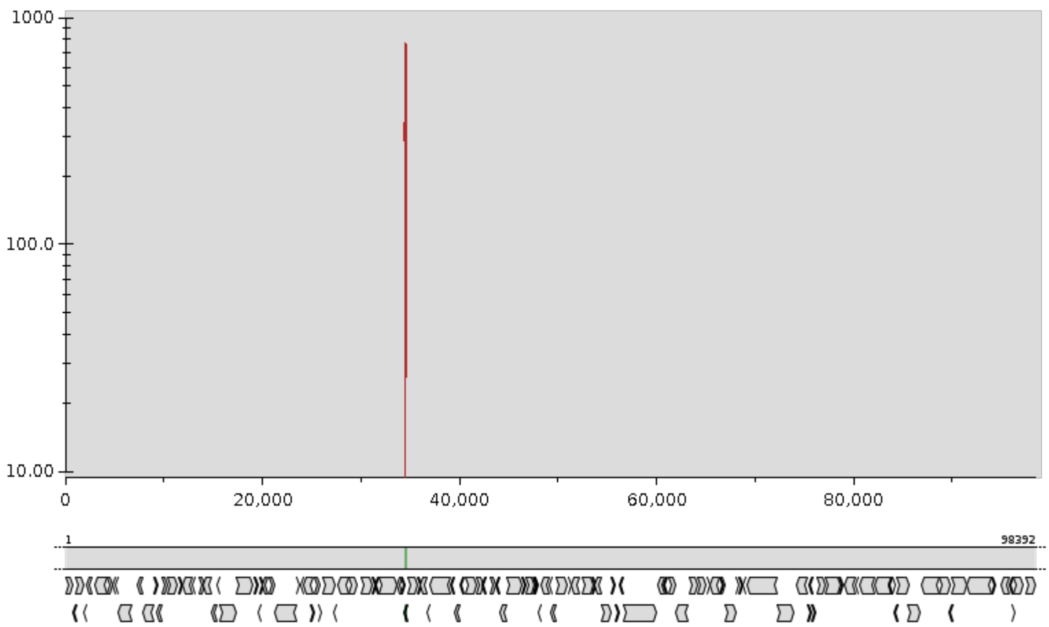

#### Assignment

| | |
|---|---|
| **Type** | Betabaculovirus disaccharalis (Taxonomy ID: 3047795) |
| **Reference Genome** | NC_028491.1 |
| **NT Identity (%)** | 82.8358 |
| **AA Identity (%)** | 88.8889 |
| **Number Of Stop Codons** | 0 |
| **Number Of CDS** | 125 |

#### Alignment

| | |
|---|---|
| **Alignment Score** | 176.0 (NT) + 279.0 (AA) = 455.0 |
| **Concordance (%)** | 80.3887 |

| Alignment Method | Local, heuristic, nucleotide (BLASTN) |
|---|---|

#### Genome Region

Sequence starts at position 34474 and ends at position 34607 relative to NC_028491.1 reference sequence.

#### Alignment Detailed Statistics

| | Begin | End | Coverage | Score | Concordance | Matches | Identities | I/D/M/F* | Stop Codons |
|---|---|---|---|---|---|---|---|---|---|
| **NT** | **34474** | **34607** | **0.1%** | **176** | **65.7%** | **134 (100%)** | **111 (82.8%)** | **0/0** | |
| Mutations: | 34482G>T, 34485T>C, 34488A>C, 34490G>A, 34510G>T, 34513C>G, 34514T>A, 34518C>A, 34521T>C, 34524A>G, 34533A>G, 34538A>C, 34542T>A, 34545T>C, 34547G>T, 34549T>C, 34554T>C, 34569G>A, 34575T>G, 34578T>C, 34587T>C, 34593C>T, 34596C>G | | | | | | | | |
| **CDS** | | | | | | | | | |
| v-ubq | 57 | 101 | 37.8% | 279 | 90.6% | 45 (100%) | 40 (88.9%) | 0/0/0/0 | 0 |
| Protein mutations: | E76G (34549T>C), S80A (34538A>C), K86N (34518C>A), T89N (34510G>T) | | | | | | | | |
| Codon mutations: | CCG60CCC (34596C>G), CCG61CCA (34593C>T), CAA63CAG (34587T>C), TTA66TTG (34578T>C), ATA67ATC (34575T>G), GCC69GCT (34569G>A), GAA74GAG (34554T>C), GAG76GGG (34549T>C), CGA77AGG (34545T>C 34547G>T), ACA78ACT (34542T>A), TCA80GCA (34538A>C), GAT81GAC (34533A>G), ATT84ATC (34524A>G), CAA85CAG (34521T>C), AAG86AAT (34518C>A), AGC88TCC (34513C>G 34514T>A), ACT89AAT (34510G>T), CTT96TTG (34488A>C 34490G>A), AGA97AGG (34485T>C), GGC98GGA (34482G>T) | | | | | | | | |
| **Proteins** | | | | | | | | | |
| ubiquitin-like protein (YP_009182246.1) | 57 | 101 | 37.8% | 279 | 90.6% | 45 (100%) | 40 (88.9%) | 0/0/0/0 | 0 |
| Protein mutations: | E76G (34549T>C), S80A (34538A>C), K86N (34518C>A), T89N (34510G>T) | | | | | | | | |
| Codon mutations: | CCG60CCC (34596C>G), CCG61CCA (34593C>T), CAA63CAG (34587T>C), TTA66TTG (34578T>C), ATA67ATC (34575T>G), GCC69GCT (34569G>A), GAA74GAG (34554T>C), GAG76GGG (34549T>C), CGA77AGG (34545T>C 34547G>T), ACA78ACT (34542T>A), TCA80GCA (34538A>C), GAT81GAC (34533A>G), ATT84ATC (34524A>G), CAA85CAG (34521T>C), AAG86AAT (34518C>A), AGC88TCC (34513C>G 34514T>A), ACT89AAT (34510G>T), CTT96TTG (34488A>C 34490G>A), AGA97AGG (34485T>C), GGC98GGA (34482G>T) | | | | | | | | |

*: Inserts / Deletes / Misaligned / Frameshifts

#### Analysis details

This analysis was performed with panviral2.64

### NGS Details (UN59): Cladosporium fulvum T-1 virus

#### Assembly

| | |
|---|---|
| **Coverage Length** | 1462 (4 contig(s)) |
| **Depth Of Coverage** | 42.1 |
| **Number Of Reads** | 529 |
| **Reads Per Million** | 11.50 rpm (after QC) |
| **Ambiguities** | 0 |
| **Assembly Method** | de novo + reference guided assembly |
| **Consensus Caller** | Bcf Tools |

#### Coverage Map

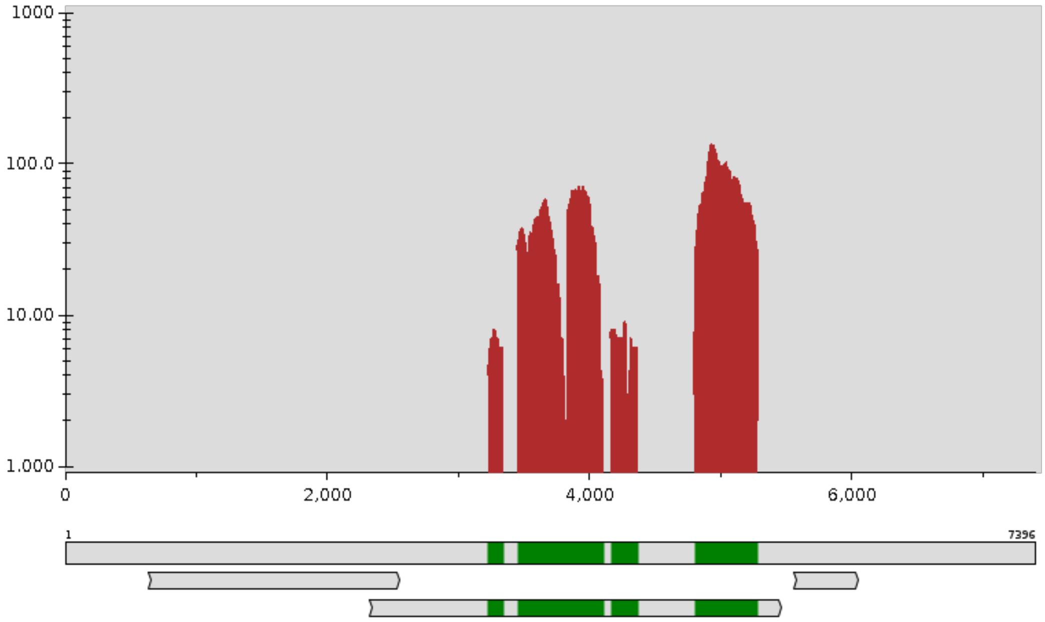

#### Assignment

| | |
|---|---|
| **Type** | Cladosporium fulvum T-1 virus (Taxonomy ID: 2052899) |
| **Reference Genome** | NC_043491.1 |
| **NT Identity (%)** | 54.095 |
| **AA Identity (%)** | 47.205 |
| **Number Of Stop Codons** | 3 |
| **Number Of CDS** | 3 |

#### Alignment

| | |
|---|---|
| **Alignment Score** | 193.0 (NT) + 1658.0 (AA) = 1851.0 |
| **Concordance (%)** | 29.7924 |

| | |
|---|---|
| **Alignment Method** | Global, seeded, nucleotide + amino acids (AGA) |

#### Genome Region

Sequence starts at position 3231 and ends at position 5281 relative to NC_043491.1 reference sequence.

#### Alignment Detailed Statistics

| | Begin | End | Coverage | Score | Concordance | Matches | Identities | I/D/M/F* | Stop Codons |
|---|---|---|---|---|---|---|---|---|---|
| **NT** | **3231** | **5281** | **19.8%** | **193** | **6.8%** | **1441 (97.8%)** | **786 (53.3%)** | **12/21** | |
| Mutations: | 3231C>A, 3232C>G, 3233T>C, 3235A>G, 3238T>A, 3242C>A, 3244C>G, 3245C>T, 3246A>T, 3251T>C, 3252G>C, 3253_3254insAAGAGGGAGGTG, 3256T>C, 3266A>G, 3268C>G, 3269A>C, 3271T>C, 3272C>G, 3274G>A, 3280G>T, 3281A>G, 3282A>C, 3283A>C, 3284G>A, 3285A>C, 3286G>A, 3289T>G, 3293T>G, 3294G>C, 3297G>C, 3298A>T, 3299C>G, 3300C>T, 3301C>G, 3303T>C, 3307T>C, 3309A>G, 3310A>C, 3314T>G, 3316T>A, 3317G>C, 3318A>C, 3319G>T, 3320A>C, 3321A>C, 3322A>G, 3328A>T, 3329C>G, 3331G>A, 3332A>G, 3333C>A, 3334C>A, 3335C>T, 3338C>A, 3340A>G, 3443C>A, 3445C>G, 3446G>T, 3447T>G, 3448A>C, 3449C>A, 3450A>T, 3451A>G, 3454C>T, 3457C>T, 3460A>G, 3461A>G, 3462A>C, 3464T>C, 3470G>A, 3473A>G, 3475C>A, 3478G>T, 3479A>G, 3481C>G, 3488C>A, 3489G>A, 3490A>G, 3493T>C, 3496G>A, 3497C>G, 3499A>T, 3502C>T, 3503A>T, 3504A>T, 3505C>G, 3508C>G, 3511A>T, 3514A>T, 3515G>C, 3516C>T, 3517A>G, 3518C>T, 3519A>T, 3520A>G, 3525G>A, 3526A>G, 3529A>G, 3531C>G, 3534G>A, 3536T>G, 3538A>G, 3539G>T, 3540A>G, 3546A>T, 3548A>T, 3550G>T, 3556T>A, 3559C>T, 3562A>T, 3563C>A, 3565A>G, 3566G>T, 3567A>C, 3568C>A, 3570C>G, 3573T>A, 3574C>T, 3578G>C, 3579C>A, 3580T>G, 3581A>G, 3583C>T, 3584C>A, 3586A>G, 3589G>A, 3598A>G, 3601A>T, 3603A>T, 3604A>G, 3605T>C, 3606G>C, 3607G>T, 3610A>G, 3613C>T, 3614G>A, 3619C>T, 3620A>G, 3621G>T, 3622G>A, 3625A>T, 3628A>G, 3631C>T, 3634A>G, 3635C>T, 3636T>A, 3637C>A, 3640C>T, 3647T>C, 3649G>T, 3652C>A, 3658A>G, 3659A>T, 3661G>T, 3664A>G, 3670C>T, 3673C>T, 3676A>C, 3679C>G, 3682A>T, 3683T>A, 3687G>T, 3688C>T, 3689C>T, 3690A>G, 3692G>A, 3694C>T, 3695C>T, 3697T>A, 3698G>A, 3700C>G, 3703C>T, 3706A>C, 3707A>G, 3708C>T, 3712T>G, 3713A>T, 3714G>A, 3715A>T, 3719C>T, 3720T>A, 3721A>T, 3724C>T, 3727C>T, 3728G>A, 3729T>A, 3732G>T, 3733C>T, 3736C>A, 3739T>A, 3741C>T, 3742T>G, 3744A>T, 3748G>A, 3751C>T, 3757A>T, 3758C>G, 3760G>C, 3761G>A, 3763C>A, 3766C>T, 3768C>G, 3769A>T, 3771A>G, 3774G>A, 3775A>T, 3776T>A, 3778C>G, 3779C>T, 3781C>G, 3782C>G, 3784G>T, 3791A>G, 3792C>T, 3793C>G, 3796G>C, 3799A>T, 3800G>T, 3802T>G, 3803C>A, 3804A>G, 3805A>T, 3806G>A, 3807A>T, 3808T>G, 3814C>G, 3815G>C, 3818C>A, 3820A>T, 3821C>T, 3823C>G, 3825C>G, 3827A>G, 3829G>A, 3831C>A, 3833G>C, 3834G>A, 3836T>C, 3839A>T, 3841G>T, 3842A>G, 3843C>T, 3844A>T, 3845G>A, 3846C>A, 3848C>A, 3849C>T, 3850C>G, 3856A>G, 3859C>T, 3862A>G, 3865C>T, 3866C>G, 3867A>C, 3869A>C, 3871G>A, 3877A>G, 3878G>A, 3880C>A, 3886T>C, 3887T>C, 3892C>G, 3893T>C, 3894T>A, 3896A>C, 3898C>A, 3899A>G, 3901C>T, 3905A>G, 3906C>A, 3909C>A, 3910A>C, 3911G>C, 3912G>A, 3913G>A, 3914A>G, 3918C>G, 3922C>G, 3925C>T, 3929G>A, 3930C>A, 3931A>G, 3934G>A, 3935A>G, 3936C>T, 3937A>G, 3940G>A, 3941T>G, 3943A>C, 3946C>T, 3947A>G, 3948G>T, 3949A>T, 3952A>T, 3957C>A, 3960A>C, 3964G>C, 3967G>A, 3970A>T, 3973C>G, 3980G>C, 3982A>G, 3983C>A, 3984A>G, 3988A>T, 3992C>T, 3994T>G, 3997A>G, 4000C>G, 4003C>A, 4006C>T, 4009C>T, 4010A>T, 4012C>T, 4025A>G, 4026A>C, 4027G>T, 4029A>G, 4030C>A, 4033T>C, 4036A>G, 4041C>A, 4052A>T, 4057G>T, 4058A>G, 4059T>A, 4060G>T, 4061C>T, 4063T>G, 4064A>C, 4065C>T, 4066A>G, 4068G>A, 4069A>G, 4072A>G, 4073G>A, 4075C>G, 4081C>G, 4085A>G, 4086A>T, 4091G>T, 4092G>C, 4093A>G, 4094A>G, 4097G>A, 4100C>T, 4101A>G, 4102G>C, 4103A>G, 4104C>A, 4165T>C, 4166G>C, 4167G>A, 4168T>C, 4170G>C, 4175G>C, 4176A>C, 4178G>T, 4179T>A, 4181C>G, 4183C>A, 4184A>G, 4186C>G, 4187G>C, 4192C>G, 4195C>T, 4198G>C, 4201T>A, 4204T>C, 4205A>T, 4207G>C, 4210A>C, 4211A>T, 4216C>A, 4218C>G, 4219A>G, 4220T>G, 4221G>T, 4222T>C, 4225A>T, 4227C>T, 4228A>G, 4231G>A, 4232_4237delACACAC, 4240T>A, 4244A>G, 4248G>A, 4252C>T, 4255A>G, 4261C>T, 4264T>C, 4265T>G, 4267T>G, 4268T>A, 4269C>G, 4273G>C, 4276A>G, 4277A>C, 4279G>A, 4281C>A, 4283A>G, 4284C>A, 4285A>G, 4286G>A, 4288G>A, 4293A>G, 4294G>A, 4295A>C, 4296A>G, 4297C>G, 4301G>A, 4302A>C, 4303C>A, 4304A>G, 4309T>A, 4312C>G, 4319C>A, 4321T>G, 4322C>A, 4323T>C, 4330T>C, 4333T>C, 4334G>C, 4335C>A, 4337G>T, 4338C>G, 4340A>C, 4344A>G, 4345A>C, 4346C>A, 4347A>C, 4348T>C, 4355G>C, 4356T>A, 4357G>C, 4361G>T, 4363C>G, 4364G>C, 4365A>T, 4366G>T, 4804A>G, 4806T>G, 4807C>G, 4808C>A, 4809C>A, 4810A>T, 4811C>G, 4812A>G, 4813G>T, 4818G>A, 4820C>A, 4821T>G, 4822T>G, 4823A>G, 4825G>C, 4826G>A, 4828G>T, 4829C>G, 4830T>C, 4832C>G, 4833G>A, 4834C>A, 4836A>T, 4837C>T, 4838A>G, 4840C>A, 4841A>G, 4842A>C, 4843G>C, 4844A>C, 4849C>T, 4850G>C, 4851T>C, 4853C>A, 4855C>T, 4865A>G, 4866A>T, 4870A>G, 4872C>T, 4873T>A, 4875C>A, 4878G>A, 4879G>C, 4882C>A, 4883G>A, 4884C>G, 4886A>C, 4887A>C, 4889T>G, 4890A>G, 4894T>A, 4895C>T, 4896A>T, 4897C>A, 4903G>A, 4904T>G, 4905T>A, 4908G>T, 4909G>T, 4910A>G, 4911C>A, 4913C>A, 4914C>T, 4915A>T, 4918A>T, 4921G>T, 4922A>T, 4923A>G, 4924A>G, 4925C>A, 4926C>A, 4927A>G, 4933C>A, 4939T>A, 4941C>A, 4942G>T, 4951C>T, 4952A>G, 4955A>G, 4956C>T, 4958A>G, 4959A>G, 4961C>T, 4963C>G, 4966G>A, 4968G>A, 4970T>A, 4972A>T, 4974A>G, 4976_4987delGATCGAGTCACA, 4988G>A, 4993A>G, 4994_4996delGCC, 4997T>C, 5002C>T, 5003A>T, 5004T>C, 5005G>T, 5009C>T, 5010T>G, 5011A>G, 5014C>G, 5017G>A, 5020C>T, 5024A>C, 5027C>A, 5029C>G, 5032A>C, 5037A>C, 5038T>A, 5041A>C, 5048A>C, 5050T>A, 5053T>A, 5055C>T, 5056A>C, 5057T>A, 5058C>A, 5059A>G, 5060G>A, 5061A>C, 5062A>T, 5064T>C, 5065A>T, 5067A>T, 5070C>G, 5074A>T, 5077G>A, 5078C>A, 5080G>T, 5081C>T, 5082T>A, 5083C>T, 5085G>C, 5086A>C, 5087T>A, 5089C>A, 5092C>G, 5093G>T, 5094T>A, 5095A>T, 5096C>A, 5098G>T, 5099G>C, 5100A>G, 5101C>T, 5102A>G, 5103G>A, 5108A>G, 5111C>A, 5114T>C, 5115A>T, 5119C>T, 5122A>T, 5123T>A, 5125C>T, 5128G>C, 5129G>T, 5130A>T, 5131A>G, 5132G>T, 5133T>C, 5134A>C, 5135T>A, 5141A>T, 5143A>T, 5146C>T, 5151A>G, 5154A>C, 5155G>T, 5156C>T, 5157T>A, 5158C>A, 5167A>G, 5168A>C, 5172A>T, 5173C>T, 5179G>A, 5180A>T, 5182G>A, 5183C>T, 5185C>T, 5186A>T, 5187T>A, 5189G>A, 5190G>A, 5191A>G, 5192A>G, 5193C>G, 5194G>A, 5195A>C, 5200A>C, 5202T>C, 5206G>A, 5207C>G, 5208A>T, 5209C>T, 5213T>C, 5215G>A, 5216T>A, 5217C>G, 5218A>C, 5221A>T, 5226A>T, 5227C>T, 5230T>C, 5233A>T, 5234G>C, 5238C>T, 5239G>A, 5245G>A, 5248A>G, 5249A>G, 5251G>A, 5255A>C, 5262A>T, 5263C>T, 5269A>C, 5276C>G | | | | | | | | |

| **CDS** | | | | | | | | | |
|---|---|---|---|---|---|---|---|---|---|
| homologue_of_retroviral_POL_genes | 303 | 985 | 46.5% | 1658 | 48.7% | 479 (97.8%) | 228 (46.5%) | 4/7/0/0 | 3 |

| | Begin | End | Coverage | Score | Concordance | Matches | Identities | I/D/M/F* | Stop Codons |
|---|---|---|---|---|---|---|---|---|---|
| **NT** | **3231** | **5281** | **19.8%** | **193** | **6.8%** | **1441 (97.8%)** | **786 (53.3%)** | **12/21** | |
| Protein mutations: | H306K (3242C>A 3244C>G), Q307L (3245C>T 3246A>T), W309P (3251T>C 3252G>C), W309_D310insKREV (3253_3254insAAGAGGGAGGTG), N314E (3266A>G 3268C>G), I315L (3269A>C 3271T>C), Q316E (3272C>G 3274G>A), K319A (3281A>G 3282A>C 3283A>C), E320T (3284G>A 3285A>C 3286G>A), W323A (3293T>G 3294G>C), G324A (3297G>C 3298A>T), P325V (3299C>G 3300C>T 3301C>G), L326P (3303T>C), Q328R (3309A>G 3310A>C), S330A (3314T>G 3316T>A), E331P (3317G>C 3318A>C 3319G>T), K332P (3320A>C 3321A>C 3322A>G), Q335E (3329C>G 3331G>A), T336E (3332A>G 3333C>A 3334C>A), L373M (3443C>A 3445C>G), V374C (3446G>T 3447T>G 3448A>C), Q375M (3449C>A 3450A>T 3451A>G), K379A (3461A>G 3462A>C), E382K (3470G>A), I383V (3473A>G 3475C>A), I385V (3479A>G 3481C>G), R388K (3488C>A 3489G>A 3490A>G), L391V (3497C>G 3499A>T), N393L (3503A>T 3504A>T 3505C>G), I394M (3508C>G), E395D (3511A>T), E396D (3514A>T), A397L (3515G>C 3516C>T 3517A>G), Q398L (3518C>T 3519A>T 3520A>G), R400K (3525G>A 3526A>G), T402S (3531C>G), G403E (3534G>A), S404A (3536T>G 3538A>G), D405C (3539G>T 3540A>G), Y407F (3546A>T), T408S (3548A>T 3550G>T), D414S (3566G>T 3567A>C 3568C>A), A415G (3570C>G), F416Y (3573T>A 3574C>T), A418Q (3578G>C 3579C>A 3580T>G), I419V (3581A>G 3583C>T), M421I (3589G>A), E425D (3601A>T), E426V (3603A>T 3604A>G), W427P (3605T>C 3606G>C 3607G>T), A430T (3614G>A), R432V (3620A>G 3621G>T 3622G>A), L437* (3635C>T 3636T>A 3637C>A), M445F (3659A>T 3661G>T), S453T (3683T>A), C454F (3687G>T 3688C>T), Q455W (3689C>T 3690A>G), D456N (3692G>A 3694C>T), V458M (3698G>A 3700C>G), E460D (3706A>C), T461V (3707A>G 3708C>T), R463Y (3713A>T 3714G>A 3715A>T), L465Y (3719C>T 3720T>A 3721A>T), V468K (3728G>A 3729T>A), C469F (3732G>T 3733C>T), A472V (3741C>T 3742T>G), Y473F (3744A>T), M474I (3748G>A), L478V (3758C>G 3760G>C), V479I (3761G>A 3763C>A), T481S (3768C>G 3769A>T), K482R (3771A>G), G483D (3774G>A 3775A>T), S484T (3776T>A 3778C>G), Q486D (3782C>G 3784G>T), T489V (3791A>G 3792C>T 3793C>G), K490N (3796G>C), Q491H (3799A>T), V492L (3800G>T 3802T>G), Q493S (3803C>A 3804A>G 3805A>T), D494M (3806G>A 3807A>T 3808T>G), F496L (3814C>G), E497Q (3815G>C), R498S (3818C>A 3820A>T), T500R (3825C>G), K501E (3827A>G 3829G>A), S502Y (3831C>A), G503Q (3833G>C 3834G>A), F504L (3836T>C), K505Y (3839A>T 3841G>T), T506V (3842A>G 3843C>T 3844A>T), A507K (3845G>A 3846C>A), P508M (3848C>A 3849C>T 3850C>G), H514A (3866C>G 3867A>C), K515Q (3869A>C 3871G>A), V518I (3878G>A 3880C>A), F523H (3893T>C 3894T>A), I524L (3896A>C 3898C>A), I525V (3899A>G 3901C>T), T527E (3905A>G 3906C>A), T528N (3909C>A 3910A>C), G529Q (3911G>C 3912G>A 3913G>A), I530V (3914A>G), T531R (3918C>G), I532M (3922C>G), A535K (3929G>A 3930C>A 3931A>G), T537V (3935A>G 3936C>T 3937A>G), S539A (3941T>G 3943A>C), R541V (3947A>G 3948G>T 3949A>T), E542D (3952A>T), P544Q (3957C>A), E545A (3960A>C), V552L (3980G>C 3982A>G), Q553R (3983C>A 3984A>G), N562Y (4010A>T 4012C>T), K567A (4025A>G 4026A>C 4027G>T), D568G (4029A>G 4030C>A), T572K (4041C>A), M576L (4052A>T), M578D (4058A>G 4059T>A 4060G>T), T580L (4064A>C 4065C>T 4066A>G), R581K (4068G>A 4069A>G), D583K (4073G>A 4075C>G), N585K (4081C>G), K587V (4085A>G 4086A>T), G589S (4091G>T 4092G>C 4093A>G), K590E (4094A>G), E591K (4097G>A), Q592C (4100C>T 4101A>G 4102G>C), G614H (4166G>C 4167G>A 4168T>C), S615T (4170G>C), E617P (4175G>C 4176A>C), V618Y (4178G>T 4179T>A), H619E (4181C>G 4183C>A), I620V (4184A>G 4186C>G), E621Q (4187G>C), M627F (4205A>T 4207G>C), I629L (4211A>T), A631G (4218C>G 4219A>G), C632V (4220T>G 4221G>T 4222T>C), T634M (4227C>T 4228A>G), T636_H637del (4232_4237delACACAC), D638E (4240T>A), K640E (4244A>G), R641K (4248G>A), Y647E (4265T>G 4267T>G), M651L (4277A>C 4279G>A), T652N (4281C>A), T653E (4283A>G 4284C>A 4285A>G), A654T (4286G>A 4288G>A), Q656R (4293A>G 4294G>A), N657R (4295A>C 4296A>G 4297C>G), D659T (4301G>A 4302A>C 4303C>A), I660V (4304A>G), H661Q (4309T>A), D662E (4312C>G), L665M (4319C>A 4321T>G), L666T (4322C>A 4323T>C), A670H (4334G>C 4335C>A), A671C (4337G>T 4338C>G), M672L (4340A>C), Q673R (4344A>G 4345A>C), H674T (4346C>A 4347A>C 4348T>C), V677H (4355G>C 4356T>A 4357G>C), V679L (4361G>T 4363C>G), E680L (4364G>C 4365A>T 4366G>T), F827W (4806T>G 4807C>G), P828N (4808C>A 4809C>A 4810A>T), Q829G (4811C>G 4812A>G 4813G>T), R831K (4818G>A), L832R (4820C>A 4821T>G 4822T>G), K833D (4823A>G 4825G>C), V834I (4826G>A 4828G>T), L835A (4829C>G 4830T>C), R836E (4832C>G 4833G>A 4834C>A), Y837F (4836A>T 4837C>T), I838V (4838A>G 4840C>A), K839A (4841A>G 4842A>C 4843G>C), K840Q (4844A>C), V842P (4850G>C 4851T>C), H843N (4853C>A 4855C>T), N847V (4865A>G 4866A>T), A849V (4872C>T 4873T>A), A850E (4875C>A), R851H (4878G>A 4879G>C), H852Q (4882C>A), A853R (4883G>A 4884C>G), K854P (4886A>C 4887A>C), Y855G (4889T>G 4890A>G), H857L (4895C>T 4896A>T 4897C>A), F860D (4904T>G 4905T>A), R861I (4908G>T 4909G>T), T862E (4910A>G 4911C>A), P863I (4913C>A 4914C>T 4915A>T), K866W (4922A>T 4923A>G 4924A>G), P867K (4925C>A 4926C>A 4927A>G), D869E (4933C>A), T872N (4941C>A 4942G>T), I876V (4952A>G), T877V (4955A>G 4956C>T), K878G (4958A>G 4959A>G), R881K (4968G>A), S882T (4970T>A 4972A>T), K883R (4974A>G), D884_T887del (4976_4987delGATCGAGTCACA), G888R (4988G>A), A890del (4994_4996delGCC), Y891H (4997T>C), M893S (5003A>T 5004T>C 5005G>T), L895W (5009C>T 5010T>G 5011A>G), M897I (5017G>A), L901M (5027C>A 5029C>G), Y904S (5037A>C 5038T>A), I908L (5048A>C 5050T>A), A910V (5055C>T 5056A>C), S911K (5057T>A 5058C>A 5059A>G), E912T (5060G>A 5061A>C 5062A>T), I913T (5064T>C 5065A>T), Y914F (5067A>T), T915S (5070C>G), Q918N (5078C>A 5080G>T), L919Y (5081C>T 5082T>A 5083C>T), G920A (5085G>C 5086A>C), Y921K (5087T>A 5089C>A), V923Y (5093G>T 5094T>A 5095A>T), L924I (5096C>A 5098G>T), D925R (5099G>C 5100A>G 5101C>T), R926E (5102A>G 5103G>A), I928V (5108A>G), Y930L (5114T>C 5115A>T), F933I (5123T>A 5125C>T), E935L (5129G>T 5130A>T 5131A>G), V936S (5132G>T 5133T>C 5134A>C), F937I (5135T>A), T939S (5141A>T 5143A>T), D942G (5151A>G), K943T (5154A>C 5155G>T), L944* (5156C>T 5157T>A 5158C>A), N948H (5168A>C), Y949F (5172A>T 5173C>T), T952S (5180A>T 5182G>A), L953F (5183C>T 5185C>T), M954* (5186A>T 5187T>A), G955K (5189G>A 5190G>A 5191A>G), T956G (5192A>G 5193C>G 5194G>A), I957L (5195A>C), I959T (5202T>C), H961V (5207C>G 5208A>T 5209C>T), Y967F (5226A>T 5227C>T), E970Q (5234G>C), T971I (5238C>T 5239G>A), T975A (5249A>G 5251G>A), N979I (5262A>T 5263C>T), Q984E (5276C>G) | | | | | | | | |

| | Begin | End | Coverage | Score | Concordance | Matches | Identities | I/D/M/F* | Stop Codons |
|---|---|---|---|---|---|---|---|---|---|
| **NT** | **3231** | **5281** | **19.8%** | **193** | **6.8%** | **1441 (97.8%)** | **786 (53.3%)** | **12/21** | |
| Codon mutations: | GCC302.AG (3231C>A 3232C>G), TTA303CTG (3233T>C 3235A>G), CCT304CCA (3238T>A), CAC306AAG (3242C>A 3244C>G), CAA307TTA (3245C>T 3246A>T), TGG309CCG (3251T>C 3252G>C), TGG309_GAT310insAAGAGGGAGGTG (3253_3254insAAGAGGGAGGTG), GAT310GAC (3256T>C), AAC314GAG (3266A>G 3268C>G), ATT315CTC (3269A>C 3271T>C), CAG316GAA (3272C>G 3274G>A), GGG318GGT (3280G>T), AAA319GCC (3281A>G 3282A>C 3283A>C), GAG320ACA (3284G>A 3285A>C 3286G>A), CCT321CCG (3289T>G), TGG323GCG (3293T>G 3294G>C), GGA324GCT (3297G>C 3298A>T), CCC325GTG (3299C>G 3300C>T 3301C>G), CTA326CCA (3303T>C), TAT327TAC (3307T>C), CAA328CGC (3309A>G 3310A>C), TCT330GCA (3314T>G 3316T>A), GAG331CCT (3317G>C 3318A>C 3319G>T), AAA332CCG (3320A>C 3321A>C 3322A>G), CTA334CTT (3328A>T), CAG335GAA (3329C>G 3331G>A), ACC336GAA (3332A>G 3333C>A 3334C>A), CTA337TTA (3335C>T), CGA338AGG (3338C>A 3340A>G), CTC373ATG (3443C>A 3445C>G), GTA374TGC (3446G>T 3447T>G 3448A>C), CAA375ATG (3449C>A 3450A>T 3451A>G), GAC376GAT (3454C>T), TAC377TAT (3457C>T), CGA378CGG (3460A>G), AAG379GCG (3461A>G 3462A>C), TTG380CTG (3464T>C), GAG382AAG (3470G>A), ATC383GTA (3473A>G 3475C>A), ACG384ACT (3478G>T), ATC385GTG (3479A>G 3481C>G), CGA388AAG (3488C>A 3489G>A 3490A>G), TAT389TAC (3493T>C), CCG390CCA (3496G>A), CTA391GTT (3497C>G 3499A>T), CCC392CCT (3502C>T), AAC393TTG (3503A>T 3504A>T 3505C>G), ATC394ATG (3508C>G), GAA395GAT (3511A>T), GAA396GAT (3514A>T), GCA397CTG (3515G>C 3516C>T 3517A>G), CAA398TTG (3518C>T 3519A>T 3520A>G), AGA400AAG (3525G>A 3526A>G), TTA401TTG (3529A>G), ACC402AGC (3531C>G), GGA403GAA (3534G>A), TCA404GCG (3536T>G 3538A>G), GAC405TGC (3539G>T 3540A>G), TAC407TTC (3546A>T), ACG408TCT (3548A>T 3550G>T), ATT410ATA (3556T>A), GAC411GAT (3559C>T), CTA412CTT (3562A>T), CGA413AGG (3563C>A 3565A>G), GAC414TCA (3566G>T 3567A>C 3568C>A), GCC415GGC (3570C>G), TTC416TAT (3573T>A 3574C>T), GCT418CAG (3578G>C 3579C>A 3580T>G), ATC419GTT (3581A>G 3583C>T), CGA420AGG (3584C>A 3586A>G), ATG421ATA (3589G>A), GGA424GGG (3598A>G), GAA425GAT (3601A>T), GAA426GTG (3603A>T 3604A>G), TGG427CCT (3605T>C 3606G>C 3607G>T), AAA428AAG (3610A>G), ACC429ACT (3613C>T), GCT430ACT (3614G>A), TTC431TTT (3619C>T), AGG432GTA (3620A>G 3621G>T 3622G>A), ACA433ACT (3625A>T), AGA434AGG (3628A>G), TAC435TAT (3631C>T), GGA436GGG (3634A>G), CTC437TAA (3635C>T 3636T>A 3637C>A), TAC438TAT (3640C>T), TTG441CTT (3647T>C 3649G>T), GTC442GTA (3652C>A), CCA444CCG (3658A>G), ATG445TTT (3659A>T 3661G>T), GGA446GGG (3664A>G), ACC448ACT (3670C>T), AAC449AAT (3673C>T), GCA450GCC (3676A>C), CCC451CCG (3679C>G), GCA452GCT (3682A>T), TCC453ACC (3683T>A), TGC454TTT (3687G>T 3688C>T), CAG455TGG (3689C>T 3690A>G), GAC456AAT (3692G>A 3694C>T), CTT457TTA (3695C>T 3697T>A), GTC458ATG (3698G>A 3700C>G), AAC459AAT (3703C>T), GAA460GAC (3706A>C), ACA461GTA (3707A>G 3708C>T), CTT462CTG (3712T>G), AGA463TAT (3713A>T 3714G>A 3715A>T), CTA465TAT (3719C>T 3720T>A 3721A>T), CTC466CTT (3724C>T), GAC467GAT (3727C>T), GTG468AAG (3728G>A 3729T>A), TGC469TTT (3732G>T 3733C>T), GTC470GTA (3736C>A), GTT471GTA (3739T>A), GCT472GTG (3741C>T 3742T>G), TAC473TTC (3744A>T), ATG474ATA (3748G>A), GAC475GAT (3751C>T), ATA477ATT (3757A>T), CTG478GTC (3758C>G 3760G>C), GTC479ATA (3761G>A 3763C>A), TAC480TAT (3766C>T), ACA481AGT (3768C>G 3769A>T), AAA482AGA (3771A>G), GGA483GAT (3774G>A 3775A>T), TCC484ACG (3776T>A 3778C>G), CTC485TTG (3779C>T 3781C>G), CAG486GAT (3782C>G 3784G>T), ACC489GTG (3791A>G 3792C>T 3793C>G), AAG490AAC (3796G>C), CAA491CAT (3799A>T), GTT492TTG (3800G>T 3802T>G), CAA493AGT (3803C>A 3804A>G 3805A>T), GAT494ATG (3806G>A 3807A>T 3808T>G), TTC496TTG (3814C>G), GAA497CAA (3815G>C), CGA498AGT (3818C>A 3820A>T), CTC499TTG (3821C>T 3823C>G), ACG500AGG (3825C>G), AAG501GAA (3827A>G 3829G>A), TCC502TAC (3831C>A), GGA503CAA (3833G>C 3834G>A), TTC504CTC (3836T>C), AAG505TAT (3839A>T 3841G>T), ACA506GTT (3842A>G 3843C>T 3844A>T), GCA507AAA (3845G>A 3846C>A), CCC508ATG (3848C>A 3849C>T 3850C>G), AAA510AAG (3856A>G), TGC511TGT (3859C>T), GAA512GAG (3862A>G), TTC513TTT (3865C>T), CAC514GCC (3866C>G 3867A>C), AAG515CAA (3869A>C 3871G>A), GAA517GAG (3877A>G), GTC518ATA (3878G>A 3880C>A), TTT520TTC (3886T>C), TTA521CTA (3887T>C), GGC522GGG (3892C>G), TTT523CAT (3893T>C 3894T>A), ATC524CTA (3896A>C 3898C>A), ATC525GTT (3899A>G 3901C>T), ACA527GAA (3905A>G 3906C>A), ACA528AAC (3909C>A 3910A>C), GGG529CAA (3911G>C 3912G>A 3913G>A), ATA530GTA (3914A>G), ACG531AGG (3918C>G), ATC532ATG (3922C>G), GAC533GAT (3925C>T), GCA535AAG (3929G>A 3930C>A 3931A>G), AAG536AAA (3934G>A), ACA537GTG (3935A>G 3936C>T 3937A>G), CAG538CAA (3940G>A), TCA539GCC (3941T>G 3943A>C), ATC540ATT (3946C>T), AGA541GTT (3947A>G 3948G>T 3949A>T), GAA542GAT (3952A>T), CCA544CAA (3957C>A), GAA545GCA (3960A>C), CCG546CCC (3964G>C), AAG547AAA (3967G>A), ACA548ACT (3970A>T), GTC549GTG (3973C>G), GTA552CTG (3980G>C 3982A>G), CAG553AGG (3983C>A 3984A>G), TCA554TCT (3988A>T), CTT556TTG (3992C>T 3994T>G), GGA557GGG (3997A>G), CTC558CTG (4000C>G), GCC559GCA (4003C>A), AAC560AAT (4006C>T), TAC561TAT (4009C>T), AAC562TAT (4010A>T 4012C>T), AAG567GCT (4025A>G 4026A>C 4027G>T), GAC568GGA (4029A>G 4030C>A), TAT569TAC (4033T>C), TCA570TCG (4036A>G), ACA572AAA (4041C>A), ATG576TTG (4052A>T), ACG577ACT (4057G>T), ATG578GAT (4058A>G 4059T>A 4060G>T), CTT579TTG (4061C>T 4063T>G), ACA580CTG (4064A>C 4065C>T 4066A>G), AGA581AAG (4068G>A 4069A>G), AAA582AAG (4072A>G), GAC583AAG (4073G>A 4075C>G), AAC585AAG (4081C>G), AAA587GTA (4085A>G 4086A>T), GGA589TCG (4091G>T 4092G>C 4093A>G), AAA590GAA (4094A>G), GAA591AAA (4097G>A), CAG592TGC (4100C>T 4101A>G 4102G>C), ACC593GA. (4103A>G 4104C>A), GAT613GAC (4165T>C), GGT614CAC (4166G>C 4167G>A 4168T>C), AGC615ACC (4170G>C), GAA617CCA (4175G>C 4176A>C), GTC618TAC (4178G>T 4179T>A), CAC619GAA (4181C>G 4183C>A), ATC620GTG (4184A>G 4186C>G), GAG621CAG (4187G>C), ACC622ACG (4192C>G), GAC623GAT (4195C>T), GCG624GCC (4198G>C), TCT625TCA (4201T>A), GAT626GAC (4204T>C), ATG627TTC (4205A>T 4207G>C), GCA628GCC (4210A>C), ATA629TTA (4211A>T), GGC630GGA (4216C>A), GCA631GGG (4218C>G 4219A>G), TGT632GTC (4220T>G 4221G>T 4222T>C), CTA633CTT (4225A>T), ACA634ATG (4227C>T 4228A>G), CAG635CAA (4231G>A), ACA636_CAC637del (4232_4237delACACAC), GAT638GAA (4240T>A), AAA640GAA (4244A>G), AGA641AAA (4248G>A), CAC642CAT (4252C>T), CCA643CCG (4255A>G), GCC645GCT (4261C>T), TAT646TAC (4264T>C), TAT647GAG (4265T>G 4267T>G), TCC648AGC (4268T>A 4269C>G), CGG649CGC (4273G>C), AAA650AAG (4276A>G), ATG651CTA (4277A>C 4279G>A), ACC652AAC (4281C>A), ACA653GAG (4283A>G 4284C>A 4285A>G), GCG654ACA (4286G>A 4288G>A), CAG656CGA (4293A>G 4294G>A), AAC657CGG (4295A>C 4296A>G 4297C>G), GAC659ACA (4301G>A 4302A>C 4303C>A), ATC660GTC (4304A>G), CAT661CAA (4309T>A), GAC662GAG (4312C>G), CTT665ATG (4319C>A 4321T>G), CTA666ACA (4322C>A 4323T>C), ATT668ATC (4330T>C), GTT669GTC (4333T>C), GCC670CAC (4334G>C 4335C>A), GCC671TGC (4337G>T 4338C>G), ATG672CTG (4340A>C), CAA673CGC (4344A>G 4345A>C), CAT674ACC (4346C>A 4347A>C 4348T>C), GTG677CAC (4355G>C 4356T>A 4357G>C), GTC679TTG (4361G>T 4363C>G), GAG680CTT (4364G>C 4365A>T 4366G>T), TCA826TCG (4804A>G), TTC827TGG (4806T>G 4807C>G), CCA828AAT (4808C>A 4809C>A 4810A>T), CAG829GGT (4811C>G 4812A>G 4813G>T), AGA831AAA (4818G>A), CTT832AGG (4820C>A 4821T>G 4822T>G), AAG833GAC (4823A>G 4825G>C), GTG834ATT (4826G>A 4828G>T), CTA835GCA (4829C>G 4830T>C), CGC836GAA (4832C>G 4833G>A 4834C>A), TAC837TTT (4836A>T 4837C>T), ATC838GTA (4838A>G 4840C>A), AAG839GCC (4841A>G 4842A>C 4843G>C), AAA840CAA (4844A>C), TGC841TGT (4849C>T), GTA842CCA (4850G>C 4851T>C), CAC843AAT (4853C>A 4855C>T), AAC847GTC (4865A>G 4866A>T), AAA848AAG (4870A>G), GCT849GTA (4872C>T 4873T>A), GCA850GAA (4875C>A), CGG851CAC (4878G>A 4879G>C), CAC852CAA (4882C>A), GCG853AGG (4883G>A 4884C>G), AAA854CCA (4886A>C 4887A>C), TAC855GGC (4889T>G 4890A>G), GGT856GGA (4894T>A), CAC857TTA (4895C>T 4896A>T 4897C>A), CAG859CAA (4903G>A), TTC860GAC (4904T>G 4905T>A), AGG861ATT (4908G>T 4909G>T), ACA862GAA (4910A>G 4911C>A), CCA863ATT (4913C>A 4914C>T 4915A>T), CCA864CCT (4918A>T), ACG865ACT (4921G>T), AAA866TGG (4922A>T 4923A>G 4924A>G), CCA867AAG (4925C>A 4926C>A 4927A>G), GAC869GAA (4933C>A), GTT871GTA (4939T>A), ACG872AAT (4941C>A 4942G>T), TTC875TTT (4951C>T), ATT876GTT (4952A>G), ACG877GTG (4955A>G 4956C>T), AAA878GGA (4958A>G 4959A>G), CTC879TTG (4961C>T 4963C>G), CCG880CCA (4966G>A), AGG881AAG (4968G>A), TCA882ACT (4970T>A 4972A>T), AAG883AGG (4974A>G), GAT884_ACA887del (4976_4987delGATCGAGTCACA), GGA888AGA (4988G>A), CAA889CAG (4993A>G), GCC890del (4994_4996delGCC), TAT891CAT (4997T>C), GAC892GAT (5002C>T), ATG893TCT (5003A>T 5004T>C 5005G>T), CTA895TGG (5009C>T 5010T>G 5011A>G), GTC896GTG (5014C>G), ATG897ATA (5017G>A), GTC898GTT (5020C>T), AGA900CGA (5024A>C), CTC901ATG (5027C>A 5029C>G), ACA902ACC (5032A>C), TAT904TCA (5037A>C 5038T>A), GCA905GCC (5041A>C), ATT908CTA (5048A>C 5050T>A), CCT909CCA (5053T>A), GCA910GTC (5055C>T 5056A>C), TCA911AAG (5057T>A 5058C>A 5059A>G), GAA912ACT (5060G>A 5061A>C 5062A>T), ATA913ACT (5064T>C 5065A>T), TAC914TTC (5067A>T), ACT915AGT (5070C>G), GCA916GCT (5074A>T), GAG917GAA (5077G>A), CAG918AAT (5078C>A 5080G>T), CTC919TAT (5081C>T 5082T>A 5083C>T), GGA920GCC (5085G>C 5086A>C), TAC921AAA (5087T>A 5089C>A), CTC922CTG (5092C>G), GTA923TAT (5093G>T 5094T>A 5095A>T), CTG924ATT (5096C>A 5098G>T), GAC925CGT (5099G>C 5100A>G 5101C>T), AGA926GAA (5102A>G 5103G>A), ATC928GTC (5108A>G), CGA929AGA (5111C>A), TAT930CTT (5114T>C 5115A>T), CAC931CAT (5119C>T), GGA932GGT (5122A>T), TTC933ATT (5123T>A 5125C>T), CCG934CCC (5128G>C), GAA935TTG (5129G>T 5130A>T 5131A>G), GTA936TCC (5132G>T 5133T>C 5134A>C), TTC937ATC (5135T>A), ACA939TCT (5141A>T 5143A>T), GAC940GAT (5146C>T), GAC942GGC (5151A>G), AAG943ACT (5154A>C 5155G>T), CTC944TAA (5156C>T 5157T>A 5158C>A), TCA947TCG (5167A>G), AAC948CAC (5168A>C), TAC949TTT (5172A>T 5173C>T), AAG951AAA (5179G>A), ACG952TCA (5180A>T 5182G>A), CTC953TTT (5183C>T 5185C>T), ATG954TAG (5186A>T 5187T>A), GGA955AAG (5189G>A 5190G>A 5191A>G), ACG956GGA (5192A>G 5193C>G 5194G>A), ATT957CTT (5195A>C), GGA958GGC (5200A>C), ATC959ACC (5202T>C), AAG960AAA (5206G>A), CAC961GTT (5207C>G 5208A>T 5209C>T), TTG963CTA (5213T>C 5215G>A), TCA964AGC (5216T>A 5217C>G 5218A>C), ACA965ACT (5221A>T), TAC967TTT (5226A>T 5227C>T), CAT968CAC (5230T>C), CCA969CCT (5233A>T), GAG970CAG (5234G>C), ACG971ATA (5238C>T 5239G>A), GGG973GGA (5245G>A), CAA974CAG (5248A>G), ACG975GCA (5249A>G 5251G>A), AGA977CGA (5255A>C), AAC979ATT (5262A>T 5263C>T), ACA981ACC (5269A>C), CAA984GAA (5276C>G) | | | | | | | | |

#### Proteins

| | Begin | End | Coverage | Score | Concordance | Matches | Identities | I/D/M/F* | Stop Codons |
|---|---|---|---|---|---|---|---|---|---|
| Reverse Transcriptase (YP_009666308.1) | 303 | 985 | 46.5% | 1658 | 48.7% | 479 (97.8%) | 228 (46.5%) | 4/7/0/0 | 3 |

| | Begin | End | Coverage | Score | Concordance | Matches | Identities | I/D/M/F* | Stop Codons |
|---|---|---|---|---|---|---|---|---|---|
| **NT** | **3231** | **5281** | **19.8%** | **193** | **6.8%** | **1441 (97.8%)** | **786 (53.3%)** | **12/21** | |
| Protein mutations: | H306K (3242C>A 3244C>G), Q307L (3245C>T 3246A>T), W309P (3251T>C 3252G>C), W309_D310insKREV (3253_3254insAAGAGGGAGGTG), N314E (3266A>G 3268C>G), I315L (3269A>C 3271T>C), Q316E (3272C>G 3274G>A), K319A (3281A>G 3282A>C 3283A>C), E320T (3284G>A 3285A>C 3286G>A), W323A (3293T>G 3294G>C), G324A (3297G>C 3298A>T), P325V (3299C>G 3300C>T 3301C>G), L326P (3303T>C), Q328R (3309A>G 3310A>C), S330A (3314T>G 3316T>A), E331P (3317G>C 3318A>C 3319G>T), K332P (3320A>C 3321A>C 3322A>G), Q335E (3329C>G 3331G>A), T336E (3332A>G 3333C>A 3334C>A), L373M (3443C>A 3445C>G), V374C (3446G>T 3447T>G 3448A>C), Q375M (3449C>A 3450A>T 3451A>G), K379A (3461A>G 3462A>C), E382K (3470G>A), I383V (3473A>G 3475C>A), I385V (3479A>G 3481C>G), R388K (3488C>A 3489G>A 3490A>G), L391V (3497C>G 3499A>T), N393L (3503A>T 3504A>T 3505C>G), I394M (3508C>G), E395D (3511A>T), E396D (3514A>T), A397L (3515G>C 3516C>T 3517A>G), Q398L (3518C>T 3519A>T 3520A>G), R400K (3525G>A 3526A>G), T402S (3531C>G), G403E (3534G>A), S404A (3536T>G 3538A>G), D405C (3539G>T 3540A>G), Y407F (3546A>T), T408S (3548A>T 3550G>T), D414S (3566G>T 3567A>C 3568C>A), A415G (3570C>G), F416Y (3573T>A 3574C>T), A418Q (3578G>C 3579C>A 3580T>G), I419V (3581A>G 3583C>T), M421I (3589G>A), E425D (3601A>T), E426V (3603A>T 3604A>G), W427P (3605T>C 3606G>C 3607G>T), A430T (3614G>A), R432V (3620A>G 3621G>T 3622G>A), L437* (3635C>T 3636T>A 3637C>A), M445F (3659A>T 3661G>T), S453T (3683T>A), C454F (3687G>T 3688C>T), Q455W (3689C>T 3690A>G), D456N (3692G>A 3694C>T), V458M (3698G>A 3700C>G), E460D (3706A>C), T461V (3707A>G 3708C>T), R463Y (3713A>T 3714G>A 3715A>T), L465Y (3719C>T 3720T>A 3721A>T), V468K (3728G>A 3729T>A), C469F (3732G>T 3733C>T), A472V (3741C>T 3742T>G), Y473F (3744A>T), M474I (3748G>A), L478V (3758C>G 3760G>C), V479I (3761G>A 3763C>A), T481S (3768C>G 3769A>T), K482R (3771A>G), G483D (3774G>A 3775A>T), S484T (3776T>A 3778C>G), Q486D (3782C>G 3784G>T), T489V (3791A>G 3792C>T 3793C>G), K490N (3796G>C), Q491H (3799A>T), V492L (3800G>T 3802T>G), Q493S (3803C>A 3804A>G 3805A>T), D494M (3806G>A 3807A>T 3808T>G), F496L (3814C>G), E497Q (3815G>C), R498S (3818C>A 3820A>T), T500R (3825C>G), K501E (3827A>G 3829G>A), S502Y (3831C>A), G503Q (3833G>C 3834G>A), F504L (3836T>C), K505Y (3839A>T 3841G>T), T506V (3842A>G 3843C>T 3844A>T), A507K (3845G>A 3846C>A), P508M (3848C>A 3849C>T 3850C>G), H514A (3866C>G 3867A>C), K515Q (3869A>C 3871G>A), V518I (3878G>A 3880C>A), F523H (3893T>C 3894T>A), I524L (3896A>C 3898C>A), I525V (3899A>G 3901C>T), T527E (3905A>G 3906C>A), T528N (3909C>A 3910A>C), G529Q (3911G>C 3912G>A 3913G>A), I530V (3914A>G), T531R (3918C>G), I532M (3922C>G), A535K (3929G>A 3930C>A 3931A>G), T537V (3935A>G 3936C>T 3937A>G), S539A (3941T>G 3943A>C), R541V (3947A>G 3948G>T 3949A>T), E542D (3952A>T), P544Q (3957C>A), E545A (3960A>C), V552L (3980G>C 3982A>G), Q553R (3983C>A 3984A>G), N562Y (4010A>T 4012C>T), K567A (4025A>G 4026A>C 4027G>T), D568G (4029A>G 4030C>A), T572K (4041C>A), M576L (4052A>T), M578D (4058A>G 4059T>A 4060G>T), T580L (4064A>C 4065C>T 4066A>G), R581K (4068G>A 4069A>G), D583K (4073G>A 4075C>G), N585K (4081C>G), K587V (4085A>G 4086A>T), G589S (4091G>T 4092G>C 4093A>G), K590E (4094A>G), E591K (4097G>A), Q592C (4100C>T 4101A>G 4102G>C), G614H (4166G>C 4167G>A 4168T>C), S615T (4170G>C), E617P (4175G>C 4176A>C), V618Y (4178G>T 4179T>A), H619E (4181C>G 4183C>A), I620V (4184A>G 4186C>G), E621Q (4187G>C), M627F (4205A>T 4207G>C), I629L (4211A>T), A631G (4218C>G 4219A>G), C632V (4220T>G 4221G>T 4222T>C), T634M (4227C>T 4228A>G), T636_H637del (4232_4237delACACAC), D638E (4240T>A), K640E (4244A>G), R641K (4248G>A), Y647E (4265T>G 4267T>G), M651L (4277A>C 4279G>A), T652N (4281C>A), T653E (4283A>G 4284C>A 4285A>G), A654T (4286G>A 4288G>A), Q656R (4293A>G 4294G>A), N657R (4295A>C 4296A>G 4297C>G), D659T (4301G>A 4302A>C 4303C>A), I660V (4304A>G), H661Q (4309T>A), D662E (4312C>G), L665M (4319C>A 4321T>G), L666T (4322C>A 4323T>C), A670H (4334G>C 4335C>A), A671C (4337G>T 4338C>G), M672L (4340A>C), Q673R (4344A>G 4345A>C), H674T (4346C>A 4347A>C 4348T>C), V677H (4355G>C 4356T>A 4357G>C), V679L (4361G>T 4363C>G), E680L (4364G>C 4365A>T 4366G>T), F827W (4806T>G 4807C>G), P828N (4808C>A 4809C>A 4810A>T), Q829G (4811C>G 4812A>G 4813G>T), R831K (4818G>A), L832R (4820C>A 4821T>G 4822T>G), K833D (4823A>G 4825G>C), V834I (4826G>A 4828G>T), L835A (4829C>G 4830T>C), R836E (4832C>G 4833G>A 4834C>A), Y837F (4836A>T 4837C>T), I838V (4838A>G 4840C>A), K839A (4841A>G 4842A>C 4843G>C), K840Q (4844A>C), V842P (4850G>C 4851T>C), H843N (4853C>A 4855C>T), N847V (4865A>G 4866A>T), A849V (4872C>T 4873T>A), A850E (4875C>A), R851H (4878G>A 4879G>C), H852Q (4882C>A), A853R (4883G>A 4884C>G), K854P (4886A>C 4887A>C), Y855G (4889T>G 4890A>G), H857L (4895C>T 4896A>T 4897C>A), F860D (4904T>G 4905T>A), R861I (4908G>T 4909G>T), T862E (4910A>G 4911C>A), P863I (4913C>A 4914C>T 4915A>T), K866W (4922A>T 4923A>G 4924A>G), P867K (4925C>A 4926C>A 4927A>G), D869E (4933C>A), T872N (4941C>A 4942G>T), I876V (4952A>G), T877V (4955A>G 4956C>T), K878G (4958A>G 4959A>G), R881K (4968G>A), S882T (4970T>A 4972A>T), K883R (4974A>G), D884_T887del (4976_4987delGATCGAGTCACA), G888R (4988G>A), A890del (4994_4996delGCC), Y891H (4997T>C), M893S (5003A>T 5004T>C 5005G>T), L895W (5009C>T 5010T>G 5011A>G), M897I (5017G>A), L901M (5027C>A 5029C>G), Y904S (5037A>C 5038T>A), I908L (5048A>C 5050T>A), A910V (5055C>T 5056A>C), S911K (5057T>A 5058C>A 5059A>G), E912T (5060G>A 5061A>C 5062A>T), I913T (5064T>C 5065A>T), Y914F (5067A>T), T915S (5070C>G), Q918N (5078C>A 5080G>T), L919Y (5081C>T 5082T>A 5083C>T), G920A (5085G>C 5086A>C), Y921K (5087T>A 5089C>A), V923Y (5093G>T 5094T>A 5095A>T), L924I (5096C>A 5098G>T), D925R (5099G>C 5100A>G 5101C>T), R926E (5102A>G 5103G>A), I928V (5108A>G), Y930L (5114T>C 5115A>T), F933I (5123T>A 5125C>T), E935L (5129G>T 5130A>T 5131A>G), V936S (5132G>T 5133T>C 5134A>C), F937I (5135T>A), T939S (5141A>T 5143A>T), D942G (5151A>G), K943T (5154A>C 5155G>T), L944* (5156C>T 5157T>A 5158C>A), N948H (5168A>C), Y949F (5172A>T 5173C>T), T952S (5180A>T 5182G>A), L953F (5183C>T 5185C>T), M954* (5186A>T 5187T>A), G955K (5189G>A 5190G>A 5191A>G), T956G (5192A>G 5193C>G 5194G>A), I957L (5195A>C), I959T (5202T>C), H961V (5207C>G 5208A>T 5209C>T), Y967F (5226A>T 5227C>T), E970Q (5234G>C), T971I (5238C>T 5239G>A), T975A (5249A>G 5251G>A), N979I (5262A>T 5263C>T), Q984E (5276C>G) | | | | | | | | |

| | Begin | End | Coverage | Score | Concordance | Matches | Identities | I/D/M/F* | Stop Codons |
|---|---|---|---|---|---|---|---|---|---|
| **NT** | **3231** | **5281** | **19.8%** | **193** | **6.8%** | **1441 (97.8%)** | **786 (53.3%)** | **12/21** | |

Codon mutations:

GCC302.AG (3231C>A 3232C>G), TTA303CTG (3233T>C 3235A>G), CCT304CCA (3238T>A), CAC306AAG (3242C>A 3244C>G), CAA307TTA (3245C>T 3246A>T), TGG309CCG (3251T>C 3252G>C), TGG309_GAT310insAAGAGGGAGGTG (3253_3254insAAGAGGGAGGTG), GAT310GAC (3256T>C), AAC314GAG (3266A>G 3268C>G), ATT315CTC (3269A>C 3271T>C), CAG316GAA (3272C>G 3274G>A), GGG318GGT (3280G>T), AAA319GCC (3281A>G 3282A>C 3283A>C), GAG320ACA (3284G>A 3285A>C 3286G>A), CCT321CCG (3289T>G), TGG323GCG (3293T>G 3294G>C), GGA324GCT (3297G>C 3298A>T), CCC325GTG (3299C>G 3300C>T 3301C>G), CTA326CCA (3303T>C), TAT327TAC (3307T>C), CAA328CGC (3309A>G 3310A>C), TCT330GCA (3314T>G 3316T>A), GAG331CCT (3317G>C 3318A>C 3319G>T), AAA332CCG (3320A>C 3321A>C 3322A>G), CTA334CTT (3328A>T), CAG335GAA (3329C>G 3331G>A), ACC336GAA (3332A>G 3333C>A 3334C>A), CTA337TTA (3335C>T), CGA338AGG (3338C>A 3340A>G), CTC373ATG (3443C>A 3445C>G), GTA374TGC (3446G>T 3447T>G 3448A>C), CAA375ATG (3449C>A 3450A>T 3451A>G), GAC376GAT (3454C>T), TAC377TAT (3457C>T), CGA378CGG (3460A>G), AAG379GCG (3461A>G 3462A>C), TTG380CTG (3464T>C), GAG382AAG (3470G>A), ATC383GTA (3473A>G 3475C>A), ACG384ACT (3478G>T), ATC385GTG (3479A>G 3481C>G), CGA388AAG (3488C>A 3489G>A 3490A>G), TAT389TAC (3493T>C), CCG390CCA (3496G>A), CTA391GTT (3497C>G 3499A>T), CCC392CCT (3502C>T), AAC393TTG (3503A>T 3504A>T 3505C>G), ATC394ATG (3508C>G), GAA395GAT (3511A>T), GAA396GAT (3514A>T), GCA397CTG (3515G>C 3516C>T 3517A>G), CAA398TTG (3518C>T 3519A>T 3520A>G), AGA400AAG (3525G>A 3526A>G), TTA401TTG (3529A>G), ACC402AGC (3531C>G), GGA403GAA (3534G>A), TCA404GCG (3536T>G 3538A>G), GAC405TGC (3539G>T 3540A>G), TAC407TTC (3546A>T), ACG408TCT (3548A>T 3550G>T), ATT410ATA (3556T>A), GAC411GAT (3559C>T), CTA412CTT (3562A>T), CGA413AGG (3563C>A 3565A>G), GAC414TCA (3566G>T 3567A>C 3568C>A), GCC415GGC (3570C>G), TTC416TAT (3573T>A 3574C>T), GCT418CAG (3578G>C 3579C>A 3580T>G), ATC419GTT (3581A>G 3583C>T), CGA420AGG (3584C>A 3586A>G), ATG421ATA (3589G>A), GGA424GGG (3598A>G), GAA425GAT (3601A>T), GAA426GTG (3603A>T 3604A>G), TGG427CCT (3605T>C 3606G>C 3607G>T), AAA428AAG (3610A>G), ACC429ACT (3613C>T), GCT430ACT (3614G>A), TTC431TTT (3619C>T), AGG432GTA (3620A>G 3621G>T 3622G>A), ACA433ACT (3625A>T), AGA434AGG (3628A>G), TAC435TAT (3631C>T), GGA436GGG (3634A>G), CTC437TAA (3635C>T 3636T>A 3637C>A), TAC438TAT (3640C>T), TTG441CTT (3647T>C 3649G>T), GTC442GTA (3652C>A), CCA444CCG (3658A>G), ATG445TTT (3659A>T 3661G>T), GGA446GGG (3664A>G), ACC448ACT (3670C>T), AAC449AAT (3673C>T), GCA450GCC (3676A>C), CCC451CCG (3679C>G), GCA452GCT (3682A>T), TCC453ACC (3683T>A), TGC454TTT (3687G>T 3688C>T), CAG455TGG (3689C>T 3690A>G), GAC456AAT (3692G>A 3694C>T), CTT457TTA (3695C>T 3697T>A), GTC458ATG (3698G>A 3700C>G), AAC459AAT (3703C>T), GAA460GAC (3706A>C), ACA461GTA (3707A>G 3708C>T), CTT462CTG (3712T>G), AGA463TAT (3713A>T 3714G>A 3715A>T), CTA465TAT (3719C>T 3720T>A 3721A>T), CTC466CTT (3724C>T), GAC467GAT (3727C>T), GTG468AAG (3728G>A 3729T>A), TGC469TTT (3732G>T 3733C>T), GTC470GTA (3736C>A), GTT471GTA (3739T>A), GCT472GTG (3741C>T 3742T>G), TAC473TTC (3744A>T), ATG474ATA (3748G>A), GAC475GAT (3751C>T), ATA477ATT (3757A>T), CTG478GTC (3758C>G 3760G>C), GTC479ATA (3761G>A 3763C>A), TAC480TAT (3766C>T), ACA481AGT (3768C>G 3769A>T), AAA482AGA (3771A>G), GGA483GAT (3774G>A 3775A>T), TCC484ACG (3776T>A 3778C>G), CTC485TTG (3779C>T 3781C>G), CAG486GAT (3782C>G 3784G>T), ACC489GTG (3791A>G 3792C>T 3793C>G), AAG490AAC (3796G>C), CAA491CAT (3799A>T), GTT492TTG (3800G>T 3802T>G), CAA493AGT (3803C>A 3804A>G 3805A>T), GAT494ATG (3806G>A 3807A>T 3808T>G), TTC496TTG (3814C>G), GAA497CAA (3815G>C), CGA498AGT (3818C>A 3820A>T), CTC499TTG (3821C>T 3823C>G), ACG500AGG (3825C>G), AAG501GAA (3827A>G 3829G>A), TCC502TAC (3831C>A), GGA503CAA (3833G>C 3834G>A), TTC504CTC (3836T>C), AAG505TAT (3839A>T 3841G>T), ACA506GTT (3842A>G 3843C>T 3844A>T), GCA507AAA (3845G>A 3846C>A), CCC508ATG (3848C>A 3849C>T 3850C>G), AAA510AAG (3856A>G), TGC511TGT (3859C>T), GAA512GAG (3862A>G), TTC513TTT (3865C>T), CAC514GCC (3866C>G 3867A>C), AAG515CAA (3869A>C 3871G>A), GAA517GAG (3877A>G), GTC518ATA (3878G>A 3880C>A), TTT520TTC (3886T>C), TTA521CTA (3887T>C), GGC522GGG (3892C>G), TTT523CAT (3893T>C 3894T>A), ATC524CTA (3896A>C 3898C>A), ATC525GTT (3899A>G 3901C>T), ACA527GAA (3905A>G 3906C>A), ACA528AAC (3909C>A 3910A>C), GGG529CAA (3911G>C 3912G>A 3913G>A), ATA530GTA (3914A>G), ACG531AGG (3918C>G), ATC532ATG (3922C>G), GAC533GAT (3925C>T), GCA535AAG (3929G>A 3930C>A 3931A>G), AAG536AAA (3934G>A), ACA537GTG (3935A>G 3936C>T 3937A>G), CAG538CAA (3940G>A), TCA539GCC (3941T>G 3943A>C), ATC540ATT (3946C>T), AGA541GTT (3947A>G 3948G>T 3949A>T), GAA542GAT (3952A>T), CCA544CAA (3957C>A), GAA545GCA (3960A>C), CCG546CCC (3964G>C), AAG547AAA (3967G>A), ACA548ACT (3970A>T), GTC549GTG (3973C>G), GTA552CTG (3980G>C 3982A>G), CAG553AGG (3983C>A 3984A>G), TCA554TCT (3988A>T), CTT556TTG (3992C>T 3994T>G), GGA557GGG (3997A>G), CTC558CTG (4000C>G), GCC559GCA (4003C>A), AAC560AAT (4006C>T), TAC561TAT (4009C>T), AAC562TAT (4010A>T 4012C>T), AAG567GCT (4025A>G 4026A>C 4027G>T), GAC568GGA (4029A>G 4030C>A), TAT569TAC (4033T>C), TCA570TCG (4036A>G), ACA572AAA (4041C>A), ATG576TTG (4052A>T), ACG577ACT (4057G>T), ATG578GAT (4058A>G 4059T>A 4060G>T), CTT579TTG (4061C>T 4063T>G), ACA580CTG (4064A>C 4065C>T 4066A>G), AGA581AAG (4068G>A 4069A>G), AAA582AAG (4072A>G), GAC583AAG (4073G>A 4075C>G), AAC585AAG (4081C>G), AAA587GTA (4085A>G 4086A>T), GGA589TCG (4091G>T 4092G>C 4093A>G), AAA590GAA (4094A>G), GAA591AAA (4097G>A), CAG592TGC (4100C>T 4101A>G 4102G>C), ACC593GA. (4103A>G 4104C>A), GAT613GAC (4165T>C), GGT614CAC (4166G>C 4167G>A 4168T>C), AGC615ACC (4170G>C), GAA617CCA (4175G>C 4176A>C), GTC618TAC (4178G>T 4179T>A), CAC619GAA (4181C>G 4183C>A), ATC620GTG (4184A>G 4186C>G), GAG621CAG (4187G>C), ACC622ACG (4192C>G), GAC623GAT (4195C>T), GCG624GCC (4198G>C), TCT625TCA (4201T>A), GAT626GAC (4204T>C), ATG627TTC (4205A>T 4207G>C), GCA628GCC (4210A>C), ATA629TTA (4211A>T), GGC630GGA (4216C>A), GCA631GGG (4218C>G 4219A>G), TGT632GTC (4220T>G 4221G>T 4222T>C), CTA633CTT (4225A>T), ACA634ATG (4227C>T 4228A>G), CAG635CAA (4231G>A), ACA636_CAC637del (4232_4237delACACAC), GAT638GAA (4240T>A), AAA640GAA (4244A>G), AGA641AAA (4248G>A), CAC642CAT (4252C>T), CCA643CCG (4255A>G), GCC645GCT (4261C>T), TAT646TAC (4264T>C), TAT647GAG (4265T>G 4267T>G), TCC648AGC (4268T>A 4269C>G), CGG649CGC (4273G>C), AAA650AAG (4276A>G), ATG651CTA (4277A>C 4279G>A), ACC652AAC (4281C>A), ACA653GAG (4283A>G 4284C>A 4285A>G), GCG654ACA (4286G>A 4288G>A), CAG656CGA (4293A>G 4294G>A), AAC657CGG (4295A>C 4296A>G 4297C>G), GAC659ACA (4301G>A 4302A>C 4303C>A), ATC660GTC (4304A>G), CAT661CAA (4309T>A), GAC662GAG (4312C>G), CTT665ATG (4319C>A 4321T>G), CTA666ACA (4322C>A 4323T>C), ATT668ATC (4330T>C), GTT669GTC (4333T>C), GCC670CAC (4334G>C 4335C>A), GCC671TGC (4337G>T 4338C>G), ATG672CTG (4340A>C), CAA673CGC (4344A>G 4345A>C), CAT674ACC (4346C>A 4347A>C 4348T>C), GTG677CAC (4355G>C 4356T>A 4357G>C), GTC679TTG (4361G>T 4363C>G), GAG680CTT (4364G>C 4365A>T 4366G>T), TCA826TCG (4804A>G), TTC827TGG (4806T>G 4807C>G), CCA828AAT (4808C>A 4809C>A 4810A>T), CAG829GGT (4811C>G 4812A>G 4813G>T), AGA831AAA (4818G>A), CTT832AGG (4820C>A 4821T>G 4822T>G), AAG833GAC (4823A>G 4825G>C), GTG834ATT (4826G>A 4828G>T), CTA835GCA (4829C>G 4830T>C), CGC836GAA (4832C>G 4833G>A 4834C>A), TAC837TTT (4836A>T 4837C>T), ATC838GTA (4838A>G 4840C>A), AAG839GCC (4841A>G 4842A>C 4843G>C), AAA840CAA (4844A>C), TGC841TGT (4849C>T), GTA842CCA (4850G>C 4851T>C), CAC843AAT (4853C>A 4855C>T), AAC847GTC (4865A>G 4866A>T), AAA848AAG (4870A>G), GCT849GTA (4872C>T 4873T>A), GCA850GAA (4875C>A), CGG851CAC (4878G>A 4879G>C), CAC852CAA (4882C>A), GCG853AGG (4883G>A 4884C>G), AAA854CCA (4886A>C 4887A>C), TAC855GGC (4889T>G 4890A>G), GGT856GGA (4894T>A), CAC857TTA (4895C>T 4896A>T 4897C>A), CAG859CAA (4903G>A), TTC860GAC (4904T>G 4905T>A), AGG861ATT (4908G>T 4909G>T), ACA862GAA (4910A>G 4911C>A), CCA863ATT (4913C>A 4914C>T 4915A>T), CCA864CCT (4918A>T), ACG865ACT (4921G>T), AAA866TGG (4922A>T 4923A>G 4924A>G), CCA867AAG (4925C>A 4926C>A 4927A>G), GAC869GAA (4933C>A), GTT871GTA (4939T>A), ACG872AAT (4941C>A 4942G>T), TTC875TTT (4951C>T), ATT876GTT (4952A>G), ACG877GTG (4955A>G 4956C>T), AAA878GGA (4958A>G 4959A>G), CTC879TTG (4961C>T 4963C>G), CCG880CCA (4966G>A), AGG881AAG (4968G>A), TCA882ACT (4970T>A 4972A>T), AAG883AGG (4974A>G), GAT884_ACA887del (4976_4987delGATCGAGTCACA), GGA888AGA (4988G>A), CAA889CAG (4993A>G), GCC890del (4994_4996delGCC), TAT891CAT (4997T>C), GAC892GAT (5002C>T), ATG893TCT (5003A>T 5004T>C 5005G>T), CTA895TGG (5009C>T 5010T>G 5011A>G), GTC896GTG (5014C>G), ATG897ATA (5017G>A), GTC898GTT (5020C>T), AGA900CGA (5024A>C), CTC901ATG (5027C>A 5029C>G), ACA902ACC (5032A>C), TAT904TCA (5037A>C 5038T>A), GCA905GCC (5041A>C), ATT908CTA (5048A>C 5050T>A), CCT909CCA (5053T>A), GCA910GTC (5055C>T 5056A>C), TCA911AAG (5057T>A 5058C>A 5059A>G), GAA912ACT (5060G>A 5061A>C 5062A>T), ATA913ACT (5064T>C 5065A>T), TAC914TTC (5067A>T), ACT915AGT (5070C>G), GCA916GCT (5074A>T), GAG917GAA (5077G>A), CAG918AAT (5078C>A 5080G>T), CTC919TAT (5081C>T 5082T>A 5083C>T), GGA920GCC (5085G>C 5086A>C), TAC921AAA (5087T>A 5089C>A), CTC922CTG (5092C>G), GTA923TAT (5093G>T 5094T>A 5095A>T), CTG924ATT (5096C>A 5098G>T), GAC925CGT (5099G>C 5100A>G 5101C>T), AGA926GAA (5102A>G 5103G>A), ATC928GTC (5108A>G), CGA929AGA (5111C>A), TAT930CTT (5114T>C 5115A>T), CAC931CAT (5119C>T), GGA932GGT (5122A>T), TTC933ATT (5123T>A 5125C>T), CCG934CCC (5128G>C), GAA935TTG (5129G>T 5130A>T 5131A>G), GTA936TCC (5132G>T 5133T>C 5134A>C), TTC937ATC (5135T>A), ACA939TCT (5141A>T 5143A>T), GAC940GAT (5146C>T), GAC942GGC (5151A>G), AAG943ACT (5154A>C 5155G>T), CTC944TAA (5156C>T 5157T>A 5158C>A), TCA947TCG (5167A>G), AAC948CAC (5168A>C), TAC949TTT (5172A>T 5173C>T), AAG951AAA (5179G>A), ACG952TCA (5180A>T 5182G>A), CTC953TTT (5183C>T 5185C>T), ATG954TAG (5186A>T 5187T>A), GGA955AAG (5189G>A 5190G>A 5191A>G), ACG956GGA (5192A>G 5193C>G 5194G>A), ATT957CTT (5195A>C), GGA958GGC (5200A>C), ATC959ACC (5202T>C), AAG960AAA (5206G>A), CAC961GTT (5207C>G 5208A>T 5209C>T), TTG963CTA (5213T>C 5215G>A), TCA964AGC (5216T>A 5217C>G 5218A>C), ACA965ACT (5221A>T), TAC967TTT (5226A>T 5227C>T), CAT968CAC (5230T>C), CCA969CCT (5233A>T), GAG970CAG (5234G>C), ACG971ATA (5238C>T 5239G>A), GGG973GGA (5245G>A), CAA974CAG (5248A>G), ACG975GCA (5249A>G 5251G>A), AGA977CGA (5255A>C), AAC979ATT (5262A>T 5263C>T), ACA981ACC (5269A>C), CAA984GAA (5276C>G)

*: Inserts / Deletes / Misaligned / Frameshifts

#### Analysis details

This analysis was performed with panviral2.64

### NGS Details (UN59): Epiphyllum badnavirus 1

#### Assembly

| | |
|---|---|
| **Coverage Length** | 414 (1 contig(s)) |
| **Depth Of Coverage** | 143.5 |
| **Number Of Reads** | 512 |
| **Reads Per Million** | 11.13 rpm (after QC) |
| **Ambiguities** | 0 |
| **Assembly Method** | de novo + reference guided assembly |
| **Consensus Caller** | Bcf Tools |

#### Coverage Map

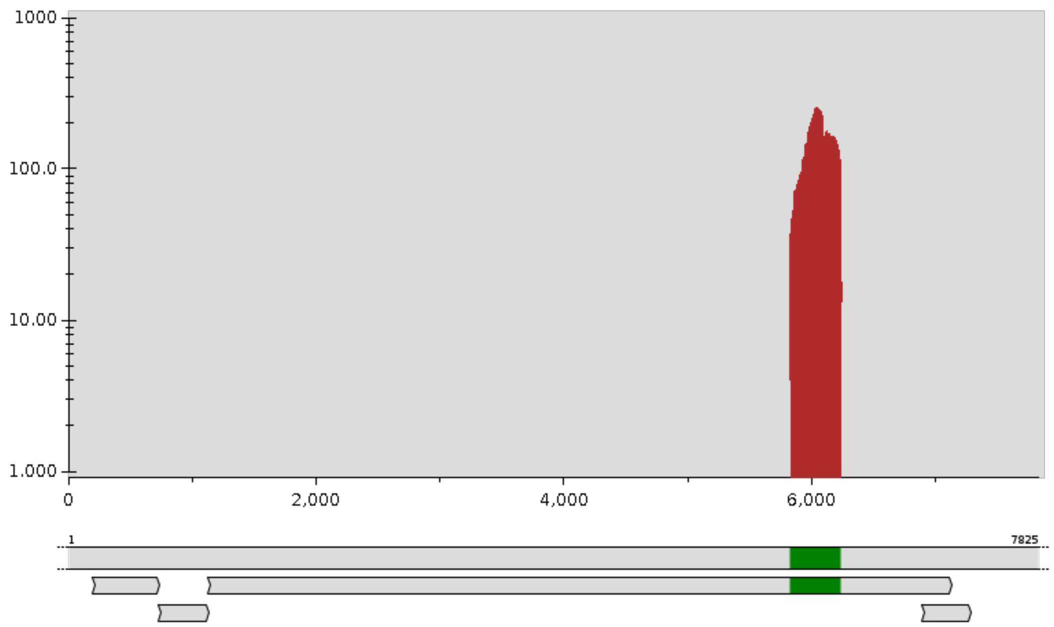

#### Assignment

| | |
|---|---|
| **Type** | Epiphyllum badnavirus 1 (Taxonomy ID: 2518008) |
| **Reference Genome** | NC_076247.1 |
| **NT Identity (%)** | 52.518 |
| **AA Identity (%)** | 46.7626 |
| **Number Of Stop Codons** | 0 |
| **Number Of CDS** | 4 |

#### Alignment

| | |
|---|---|
| **Alignment Score** | 36.0 (NT) + 441.0 (AA) = 477.0 |
| **Concordance (%)** | 26.5295 |

| Alignment Method | Global, seeded, nucleotide + amino acids (AGA) |
|---|---|

#### Genome Region

Sequence starts at position 5825 and ends at position 6238 relative to NC_076247.1 reference sequence.

#### Alignment Detailed Statistics

| | Begin | End | Coverage | Score | Concordance | Matches | Identities | I/D/M/F* | Stop Codons |
|---|---|---|---|---|---|---|---|---|---|
| **NT** | **5825** | **6238** | **5.3%** | **36** | **4.3%** | **414 (99.3%)** | **219 (52.5%)** | **3/0** | |
| Mutations: | 5835C>T, 5836A>T, 5837G>C, 5841C>G, 5847C>T, 5851T>A, 5853C>T, 5856T>C, 5857C>T, 5861A>G, 5862G>A, 5863A>T, 5864G>C, 5865C>G, 5868A>T, 5870T>A, 5871C>T, 5874C>T, 5878A>T, 5880T>G, 5882T>G, 5883G>A, 5884A>G, 5886G>T, 5887G>C, 5888A>G, 5889C>A, 5892A>G, 5893G>T, 5894A>C, 5896T>G, 5897C>A, 5898C>T, 5904T>G, 5905T>A, 5906G>A, 5910C>T, 5913C>G, 5917C>A, 5918T>G, 5924C>G, 5925A>C, 5926G>T, 5931A>C, 5933T>A, 5937C>T, 5940A>G, 5942G>T, 5943G>T, 5944T>G, 5946A>G, 5953C>T, 5955A>T, 5958C>T, 5961A>G, 5966A>C, 5967G>T, 5981T>C, 5982G>T, 5986C>A, 5987A>T, 5988A>G, 5989C>G, 5990G>A, 5991A>G, 5992A>T, 5993A>T, 5998G>A, 6001A>C, 6002A>G, 6003C>A, 6004T>G, 6005G>T, 6006T>G, 6011G>A, 6012_6013insTCT, 6013G>T, 6015C>T, 6018G>T, 6019T>G, 6020C>A, 6021A>T, 6022G>C, 6023G>A, 6024C>G, 6030A>G, 6031G>A, 6032C>T, 6033A>T, 6036T>G, 6038A>T, 6042T>A, 6048C>T, 6052C>T, 6057C>T, 6059T>A, 6060C>T, 6061A>T, 6062G>C, 6063T>C, 6066C>G, 6068C>G, 6072A>G, 6073G>A, 6074A>G, 6078C>T, 6083C>T, 6084A>G, 6085C>A, 6090C>T, 6091C>T, 6095A>G, 6096G>A, 6098C>T, 6099C>T, 6100A>G, 6103C>T, 6109G>A, 6110T>G, 6111T>G, 6113G>T, 6114T>G, 6115G>C, 6116A>G, 6118A>G, 6119G>A, 6120A>T, 6124G>A, 6125G>A, 6127C>T, 6130A>T, 6131T>A, 6132C>T, 6133C>G, 6134T>C, 6136T>A, 6137C>A, 6138T>G, 6139C>T, 6140C>T, 6141T>C, 6142A>T, 6144G>A, 6148A>T, 6149T>G, 6150G>T, 6151A>G, 6154A>G, 6156A>T, 6157G>T, 6158C>G, 6159T>G, 6160G>C, 6161C>T, 6165A>G, 6166G>T, 6167A>C, 6168G>A, 6169A>G, 6171T>A, 6172G>T, 6173A>C, 6177C>T, 6180G>A, 6183A>T, 6184A>C, 6185T>A, 6186C>T, 6187A>G, 6188C>T, 6189C>G, 6190C>G, 6192A>G, 6193G>A, 6194G>C, 6195A>T, 6199C>G, 6201G>T, 6202C>G, 6203A>G, 6204G>C, 6205C>A, 6207G>T, 6209A>T, 6211T>G, 6213G>T, 6214C>G, 6216A>T, 6219C>G, 6220C>G, 6221A>C, 6224T>A, 6225C>A, 6226A>G, 6228C>G, 6229A>G | | | | | | | | |

##### CDS

| | Begin | End | Coverage | Score | Concordance | Matches | Identities | I/D/M/F* | Stop Codons |
|---|---|---|---|---|---|---|---|---|---|
| QKM20_gp3 | 1567 | 1704 | 6.9% | 441 | 45.0% | 138 (99.3%) | 65 (46.8%) | 1/0/0/0 | 0 |
| Protein mutations: | F1575I (5851T>A 5853C>T), K1578R (5861A>G 5862G>A), F1581Y (5870T>A 5871C>T), I1584L (5878A>T 5880T>G), M1585R (5882T>G 5883G>A), M1586V (5884A>G 5886G>T), D1587R (5887G>C 5888A>G 5889C>A), E1589S (5893G>T 5894A>C), S1590D (5896T>G 5897C>A 5898C>T), W1593K (5905T>A 5906G>A), L1597R (5917C>A 5918T>G), P1599R (5924C>G 5925A>C), D1600Y (5926G>T), L1602H (5933T>A), W1605F (5942G>T 5943G>T), L1606V (5944T>G 5946A>G), P1609S (5953C>T 5955A>T), K1613T (5966A>C 5967G>T), V1618A (5981T>C 5982G>T), Q1620M (5986C>A 5987A>T 5988A>G), R1621E (5989C>G 5990G>A 5991A>G), K1622L (5992A>T 5993A>T), D1624N (5998G>A), N1625R (6001A>C 6002A>G 6003C>A), C1626V (6004T>G 6005G>T 6006T>G), R1628K (6011G>A), R1628_D1629insS (6012_6013insTCT), D1629Y (6013G>T 6015C>T), S1631D (6019T>G 6020C>A 6021A>T), G1632Q (6022G>C 6023G>A 6024C>G), A1635I (6031G>A 6032C>T 6033A>T), Y1637F (6038A>T), F1644Y (6059T>A 6060C>T), N1646K (6066C>G), T1647S (6068C>G), D1649S (6073G>A 6074A>G), A1652V (6083C>T 6084A>G), Q1653K (6085C>A), K1656R (6095A>G 6096G>A), S1657F (6098C>T 6099C>T), M1658V (6100A>G), V1661R (6109G>A 6110T>G 6111T>G), C1662L (6113G>T 6114T>G), E1663R (6115G>C 6116A>G), R1664D (6118A>G 6119G>A 6120A>T), G1666K (6124G>A 6125G>A), I1668Y (6130A>T 6131T>A 6132C>T), L1669A (6133C>G 6134T>C), S1670K (6136T>A 6137C>A 6138T>G), P1671F (6139C>T 6140C>T 6141T>C), T1672S (6142A>T 6144G>A), M1674C (6148A>T 6149T>G 6150G>T), K1675E (6151A>G), I1676V (6154A>G 6156A>T), A1677W (6157G>T 6158C>G 6159T>G), A1678L (6160G>C 6161C>T), E1680S (6166G>T 6167A>C 6168G>A), I1681V (6169A>G 6171T>A), D1682S (6172G>T 6173A>C), I1686H (6184A>C 6185T>A 6186C>T), T1687V (6187A>G 6188C>T 6189C>G), L1688V (6190C>G 6192A>G), G1689T (6193G>A 6194G>C 6195A>T), Q1691D (6199C>G 6201G>T), Q1692G (6202C>G 6203A>G 6204G>C), L1693I (6205C>A 6207G>T), K1694M (6209A>T), L1695V (6211T>G 6213G>T), Q1696D (6214C>G 6216A>T), H1698A (6220C>G 6221A>C), I1699K (6224T>A 6225C>A), I1700V (6226A>G 6228C>G), T1701A (6229A>G) | | | | | | | | |
| Codon mutations: | GCC1569GCT (5835C>T), AGT1570TCT (5836A>T 5837G>C), GTC1571GTG (5841C>G), TCC1573TCT (5847C>T), TTC1575ATT (5851T>A 5853C>T), GAT1576GAC (5856T>C), CTG1577TTG (5857C>T), AAG1578AGA (5861A>G 5862G>A), AGC1579TCG (5863A>T 5864G>C 5865C>G), GGA1580GGT (5868A>T), TTC1581TAT (5870T>A 5871C>T), CAC1582CAT (5874C>T), ATT1584TTG (5878A>T 5880T>G), ATG1585AGA (5882T>G 5883G>A), ATG1586GTT (5884A>G 5886G>T), GAC1587CGA (5887G>C 5888A>G 5889C>A), GAA1588GAG (5892A>G), GAA1589TCA (5893G>T 5894A>C), TCC1590GAT (5896T>G 5897C>A 5898C>T), CCT1592CCG (5904T>G), TGG1593AAG (5905T>A 5906G>A), ACC1594ACT (5910C>T), GCC1595GCG (5913C>G), CTG1597AGG (5917C>A 5918T>G), CCA1599CGC (5924C>G 5925A>C), GAT1600TAT (5926G>T), GGA1601GGC (5931A>C), CTT1602CAT (5933T>A), TAC1603TAT (5937C>T), GAA1604GAG (5940A>G), TGG1605TTT (5942G>T 5943G>T), TTA1606GTG (5944T>G 5946A>G), CCA1609TCT (5953C>T 5955A>T), TTC1610TTT (5958C>T), GGA1611GGG (5961A>G), AAG1613ACT (5966A>C 5967G>T), GTG1618GCT (5981T>C 5982G>T), CAA1620ATG (5986C>A 5987A>T 5988A>G), CGA1621GAG (5989C>G 5990G>A 5991A>G), AAG1622TTG (5992A>T 5993A>T), GAC1624AAC (5998G>A), AAC1625CGA (6001A>C 6002A>G 6003C>A), TGT1626GTG (6004T>G 6005G>T 6006T>G), AGA1628AAA (6011G>A), AGA1628_GAC1629insTCT (6012_6013insTCT), GAC1629TAT (6013G>T 6015C>T), CTG1630CTT (6018G>T), TCA1631GAT (6019T>G 6020C>A 6021A>T), GGC1632CAG (6022G>C 6023G>A 6024C>G), GTA1634GTG (6030A>G), GCA1635ATT (6031G>A 6032C>T 6033A>T), GTT1636GTG (6036T>G), TAC1637TTC (6038A>T), ATT1638ATA (6042T>A), GAC1640GAT (6048C>T), CTG1642TTG (6052C>T), GTC1643GTT (6057C>T), TTC1644TAT (6059T>A 6060C>T), AGT1645TCC (6061A>T 6062G>C 6063T>C), AAC1646AAG (6066C>G), ACT1647AGT (6068C>G), GAA1648GAG (6072A>G), GAT1649AGT (6073G>A 6074A>G), GAC1650GAT (6078C>T), GCA1652GTG (6083C>T 6084A>G), CAG1653AAG (6085C>A), CAC1654CAT (6090C>T), CTG1655TTG (6091C>T), AAG1656AGA (6095A>G 6096G>A), TCC1657TTT (6098C>T 6099C>T), ATG1658GTG (6100A>G), CTG1659TTG (6103C>T), GTT1661AGG (6109G>A 6110T>G 6111T>G), TGT1662TTG (6113G>T 6114T>G), GAG1663CGG (6115G>C 6116A>G), AGA1664GAT (6118A>G 6119G>A 6120A>T), GGG1666AAG (6124G>A 6125G>A), CTG1667TTG (6127C>T), ATC1668TAT (6130A>T 6131T>A 6132C>T), CTC1669GCC (6133C>G 6134T>C), TCT1670AAG (6136T>A 6137C>A 6138T>G), CCT1671TTC (6139C>T 6140C>T 6141T>C), ACG1672TCA (6142A>T 6144G>A), ATG1674TGT (6148A>T 6149T>G 6150G>T), AAG1675GAG (6151A>G), ATA1676GTT (6154A>G 6156A>T), GCT1677TGG (6157G>T 6158C>G 6159T>G), GCA1678CTA (6160G>C 6161C>T), CAA1679CAG (6165A>G), GAG1680TCA (6166G>T 6167A>C 6168G>A), ATT1681GTA (6169A>G 6171T>A), GAC1682TCC (6172G>T 6173A>C), TTC1683TTT (6177C>T), CTG1684CTA (6180G>A), GGA1685GGT (6183A>T), ATC1686CAT (6184A>C 6185T>A 6186C>T), ACC1687GTG (6187A>G 6188C>T 6189C>G), CTA1688GTG (6190C>G 6192A>G), GGA1689ACT (6193G>A 6194G>C 6195A>T), CAG1691GAT (6199C>G 6201G>T), CAG1692GGC (6202C>G 6203A>G 6204G>C), CTG1693ATT (6205C>A 6207G>T), AAG1694ATG (6209A>T), TTG1695GTT (6211T>G 6213G>T), CAA1696GAT (6214C>G 6216A>T), CCC1697CCG (6219C>G), CAC1698GCC (6220C>G 6221A>C), ATC1699AAA (6224T>A 6225C>A), ATC1700GTG (6226A>G 6228C>G), ACC1701GCC (6229A>G) | | | | | | | | |

##### Proteins

| | Begin | End | Coverage | Score | Concordance | Matches | Identities | I/D/M/F* | Stop Codons |
|---|---|---|---|---|---|---|---|---|---|
| polyprotein (YP_010797894.1) | 1567 | 1704 | 6.9% | 441 | 45.0% | 138 (99.3%) | 65 (46.8%) | 1/0/0/0 | 0 |
| Protein mutations: | F1575I (5851T>A 5853C>T), K1578R (5861A>G 5862G>A), F1581Y (5870T>A 5871C>T), I1584L (5878A>T 5880T>G), M1585R (5882T>G 5883G>A), M1586V (5884A>G 5886G>T), D1587R (5887G>C 5888A>G 5889C>A), E1589S (5893G>T 5894A>C), S1590D (5896T>G 5897C>A 5898C>T), W1593K (5905T>A 5906G>A), L1597R (5917C>A 5918T>G), P1599R (5924C>G 5925A>C), D1600Y (5926G>T), L1602H (5933T>A), W1605F (5942G>T 5943G>T), L1606V (5944T>G 5946A>G), P1609S (5953C>T 5955A>T), K1613T (5966A>C 5967G>T), V1618A (5981T>C 5982G>T), Q1620M (5986C>A 5987A>T 5988A>G), R1621E (5989C>G 5990G>A 5991A>G), K1622L (5992A>T 5993A>T), D1624N (5998G>A), N1625R (6001A>C 6002A>G 6003C>A), C1626V (6004T>G 6005G>T 6006T>G), R1628K (6011G>A), R1628_D1629insS (6012_6013insTCT), D1629Y (6013G>T 6015C>T), S1631D (6019T>G 6020C>A 6021A>T), G1632Q (6022G>C 6023G>A 6024C>G), A1635I (6031G>A 6032C>T 6033A>T), Y1637F (6038A>T), F1644Y (6059T>A 6060C>T), N1646K (6066C>G), T1647S (6068C>G), D1649S (6073G>A 6074A>G), A1652V (6083C>T 6084A>G), Q1653K (6085C>A), K1656R (6095A>G 6096G>A), S1657F (6098C>T 6099C>T), M1658V (6100A>G), V1661R (6109G>A 6110T>G 6111T>G), C1662L (6113G>T 6114T>G), E1663R (6115G>C 6116A>G), R1664D (6118A>G 6119G>A 6120A>T), G1666K (6124G>A 6125G>A), I1668Y (6130A>T 6131T>A 6132C>T), L1669A (6133C>G 6134T>C), S1670K (6136T>A 6137C>A 6138T>G), P1671F (6139C>T 6140C>T 6141T>C), T1672S (6142A>T 6144G>A), M1674C (6148A>T 6149T>G 6150G>T), K1675E (6151A>G), I1676V (6154A>G 6156A>T), A1677W (6157G>T 6158C>G 6159T>G), A1678L (6160G>C 6161C>T), E1680S (6166G>T 6167A>C 6168G>A), I1681V (6169A>G 6171T>A), D1682S (6172G>T 6173A>C), I1686H (6184A>C 6185T>A 6186C>T), T1687V (6187A>G 6188C>T 6189C>G), L1688V (6190C>G 6192A>G), G1689T (6193G>A 6194G>C 6195A>T), Q1691D (6199C>G 6201G>T), Q1692G (6202C>G 6203A>G 6204G>C), L1693I (6205C>A 6207G>T), K1694M (6209A>T), L1695V (6211T>G 6213G>T), Q1696D (6214C>G 6216A>T), H1698A (6220C>G 6221A>C), I1699K (6224T>A 6225C>A), I1700V (6226A>G 6228C>G), T1701A (6229A>G) | | | | | | | | |

| | Begin | End | Coverage | Score | Concordance | Matches | Identities | I/D/M/F* | Stop Codons |
|---|---|---|---|---|---|---|---|---|---|
| **NT** | **5825** | **6238** | **5.3%** | **36** | **4.3%** | **414 (99.3%)** | **219 (52.5%)** | **3/0** | |
| Codon mutations: | GCC1569GCT (5835C>T), AGT1570TCT (5836A>T 5837G>C), GTC1571GTG (5841C>G), TCC1573TCT (5847C>T), TTC1575ATT (5851T>A 5853C>T), GAT1576GAC (5856T>C), CTG1577TTG (5857C>T), AAG1578AGA (5861A>G 5862G>A), AGC1579TCG (5863A>T 5864G>C 5865C>G), GGA1580GGT (5868A>T), TTC1581TAT (5870T>A 5871C>T), CAC1582CAT (5874C>T), ATT1584TTG (5878A>T 5880T>G), ATG1585AGA (5882T>G 5883G>A), ATG1586GTT (5884A>G 5886G>T), GAC1587CGA (5887G>C 5888A>G 5889C>A), GAA1588GAG (5892A>G), GAA1589TCA (5893G>T 5894A>C), TCC1590GAT (5896T>G 5897C>A 5898C>T), CCT1592CCG (5904T>G), TGG1593AAG (5905T>A 5906G>A), ACC1594ACT (5910C>T), GCC1595GCG (5913C>G), CTG1597AGG (5917C>A 5918T>G), CCA1599CGC (5924C>G 5925A>C), GAT1600TAT (5926G>T), GGA1601GGC (5931A>C), CTT1602CAT (5933T>A), TAC1603TAT (5937C>T), GAA1604GAG (5940A>G), TGG1605TTT (5942G>T 5943G>T), TTA1606GTG (5944T>G 5946A>G), CCA1609TCT (5953C>T 5955A>T), TTC1610TTT (5958C>T), GGA1611GGG (5961A>G), AAG1613ACT (5966A>C 5967G>T), GTG1618GCT (5981T>C 5982G>T), CAA1620ATG (5986C>A 5987A>T 5988A>G), CGA1621GAG (5989C>G 5990G>A 5991A>G), AAG1622TTG (5992A>T 5993A>T), GAC1624AAC (5998G>A), AAC1625CGA (6001A>C 6002A>G 6003C>A), TGT1626GTG (6004T>G 6005G>T 6006T>G), AGA1628AAA (6011G>A), AGA1628_GAC1629insTCT (6012_6013insTCT), GAC1629TAT (6013G>T 6015C>T), CTG1630CTT (6018G>T), TCA1631GAT (6019T>G 6020C>A 6021A>T), GGC1632CAG (6022G>C 6023G>A 6024C>G), GTA1634GTG (6030A>G), GCA1635ATT (6031G>A 6032C>T 6033A>T), GTT1636GTG (6036T>G), TAC1637TTC (6038A>T), ATT1638ATA (6042T>A), GAC1640GAT (6048C>T), CTG1642TTG (6052C>T), GTC1643GTT (6057C>T), TTC1644TAT (6059T>A 6060C>T), AGT1645TCC (6061A>T 6062G>C 6063T>C), AAC1646AAG (6066C>G), ACT1647AGT (6068C>G), GAA1648GAG (6072A>G), GAT1649AGT (6073G>A 6074A>G), GAC1650GAT (6078C>T), GCA1652GTG (6083C>T 6084A>G), CAG1653AAG (6085C>A), CAC1654CAT (6090C>T), CTG1655TTG (6091C>T), AAG1656AGA (6095A>G 6096G>A), TCC1657TTT (6098C>T 6099C>T), ATG1658GTG (6100A>G), CTG1659TTG (6103C>T), GTT1661AGG (6109G>A 6110T>G 6111T>G), TGT1662TTG (6113G>T 6114T>G), GAG1663CGG (6115G>C 6116A>G), AGA1664GAT (6118A>G 6119G>A 6120A>T), GGG1666AAG (6124G>A 6125G>A), CTG1667TTG (6127C>T), ATC1668TAT (6130A>T 6131T>A 6132C>T), CTC1669GCC (6133C>G 6134T>C), TCT1670AAG (6136T>A 6137C>A 6138T>G), CCT1671TTC (6139C>T 6140C>T 6141T>C), ACG1672TCA (6142A>T 6144G>A), ATG1674TGT (6148A>T 6149T>G 6150G>T), AAG1675GAG (6151A>G), ATA1676GTT (6154A>G 6156A>T), GCT1677TGG (6157G>T 6158C>G 6159T>G), GCA1678CTA (6160G>C 6161C>T), CAA1679CAG (6165A>G), GAG1680TCA (6166G>T 6167A>C 6168G>A), ATT1681GTA (6169A>G 6171T>A), GAC1682TCC (6172G>T 6173A>C), TTC1683TTT (6177C>T), CTG1684CTA (6180G>A), GGA1685GGT (6183A>T), ATC1686CAT (6184A>C 6185T>A 6186C>T), ACC1687GTG (6187A>G 6188C>T 6189C>G), CTA1688GTG (6190C>G 6192A>G), GGA1689ACT (6193G>A 6194G>C 6195A>T), CAG1691GAT (6199C>G 6201G>T), CAG1692GGC (6202C>G 6203A>G 6204G>C), CTG1693ATT (6205C>A 6207G>T), AAG1694ATG (6209A>T), TTG1695GTT (6211T>G 6213G>T), CAA1696GAT (6214C>G 6216A>T), CCC1697CCG (6219C>G), CAC1698GCC (6220C>G 6221A>C), ATC1699AAA (6224T>A 6225C>A), ATC1700GTG (6226A>G 6228C>G), ACC1701GCC (6229A>G) | | | | | | | | |

*: Inserts / Deletes / Misaligned / Frameshifts

#### Analysis details

This analysis was performed with panviral2.64

### NGS Details (UN59): Human gammaherpesvirus 8

#### Assembly

| | |
|---|---|
| **Coverage Length** | 135 (1 contig(s)) |
| **Depth Of Coverage** | 265.0 |
| **Number Of Reads** | 421 |
| **Reads Per Million** | 9.15 rpm (after QC) |
| **Ambiguities** | 0 |
| **Assembly Method** | de novo + reference guided assembly |
| **Consensus Caller** | Bcf Tools |

#### Coverage Map

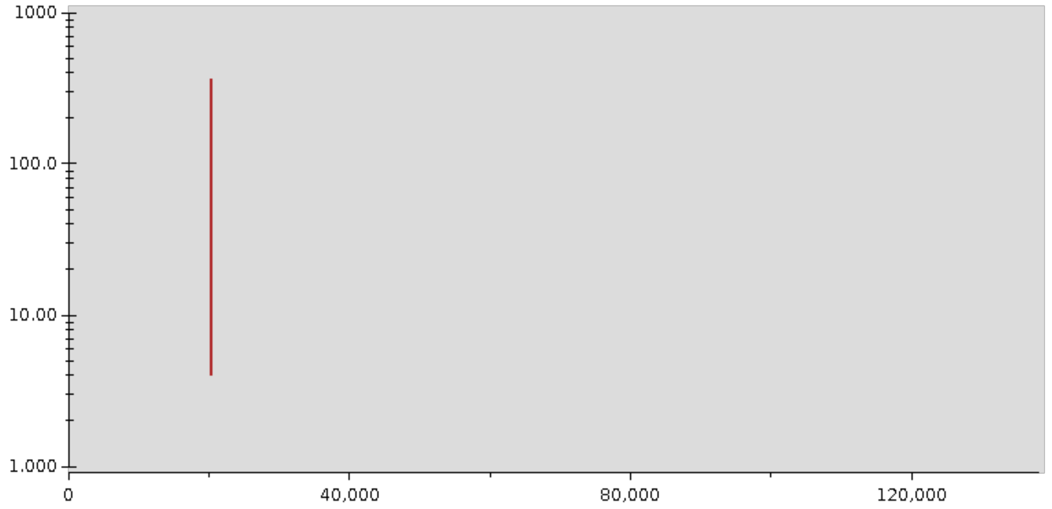

#### Assignment

| | |
|---|---|
| **Type** | Human gammaherpesvirus 8 (Taxonomy ID: 37296) |
| **Subtype** | Could not assign |
| **Reference Genome** | NC_009333.1 |
| **NT Identity (%)** | 83.7037 |
| **AA Identity (%)** | 84.4444 |
| **Number Of Stop Codons** | 0 |
| **Number Of CDS** | 86 |

#### Alignment

| | |
|---|---|
| **Alignment Score** | 182.0 (NT) + 287.0 (AA) = 469.0 |
| **Concordance (%)** | 78.2972 |
| **Alignment Method** | Local, heuristic, nucleotide (BLASTN) |

#### Genome Region

Sequence starts at position 20273 and ends at position 20407 relative to NC_009333.1 reference sequence.

#### Alignment Detailed Statistics

| | Begin | End | Coverage | Score | Concordance | Matches | Identities | I/D/M/F* | Stop Codons |
|---|---|---|---|---|---|---|---|---|---|
| NT | 20273 | 20407 | 0.1% | 182 | 67.4% | 135 (100%) | 113 (83.7%) | 0/0 | |

Mutations: 20293A>T, 20296A>C, 20311G>A, 20314T>G, 20315C>G, 20317C>A, 20320C>T, 20322T>G, 20331G>T, 20341A>C, 20347A>C, 20350G>A, 20352C>T, 20353A>G, 20369C>G, 20370A>C, 20371T>A, 20376A>T, 20383G>A, 20386C>T, 20389A>G, 20392C>T

*: Inserts / Deletes / Misaligned / Frameshifts

#### Analysis details

This analysis was performed with panviral2.64

### NGS Details (UN59): Tomato necrotic dwarf virus (segment RNA1)

#### Assembly

| | |
|---|---|
| **Coverage Length** | 303 (1 contig(s)) |
| **Depth Of Coverage** | 176.1 |
| **Number Of Reads** | 407 |
| **Reads Per Million** | 8.85 rpm (after QC) |
| **Ambiguities** | 0 |
| **Assembly Method** | de novo + reference guided assembly |
| **Consensus Caller** | Bcf Tools |

#### Coverage Map

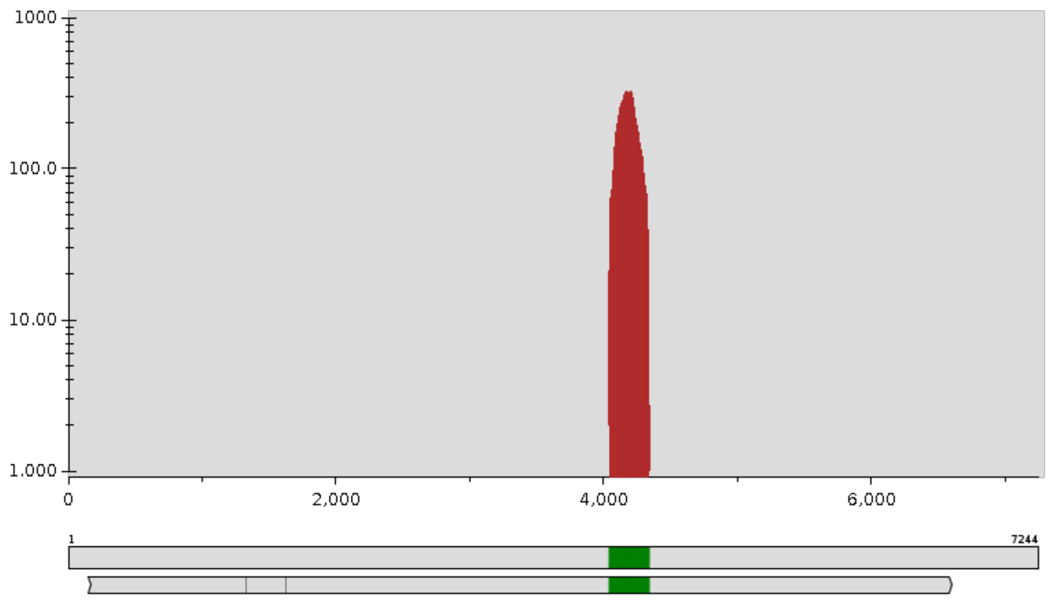

#### Assignment

| | |
|---|---|
| **Type** | Tomato necrotic dwarf virus (Taxonomy ID: 1481465) |
| **Reference Genome** | NC_027926.1 |
| **NT Identity (%)** | 75.2475 |
| **AA Identity (%)** | 86.1386 |
| **Number Of Stop Codons** | 0 |
| **Number Of CDS** | 1 |

#### Alignment

| | |
|---|---|
| **Alignment Score** | 306.0 (NT) + 643.0 (AA) = 949.0 |
| **Concordance (%)** | 71.9484 |

| Alignment Method | Global, seeded, nucleotide + amino acids (AGA) |
|---|---|

#### Genome Region

Sequence starts at position 4040 and ends at position 4342 relative to NC_027926.1 reference sequence.

#### Alignment Detailed Statistics

| | Begin | End | Coverage | Score | Concordance | Matches | Identities | I/D/M/F* | Stop Codons |
|---|---|---|---|---|---|---|---|---|---|
| **NT** | **4040** | **4342** | **4.2%** | **306** | **50.5%** | **303 (100%)** | **228 (75.2%)** | **0/0** | |
| Mutations: | 4050A>C, 4053A>G, 4059C>G, 4062G>A, 4068G>A, 4071C>G, 4072C>T, 4074T>G, 4077T>A, 4083A>G, 4089T>C, 4090T>C, 4104A>G, 4107A>C, 4108A>C, 4110A>C, 4111T>C, 4113A>G, 4116T>C, 4119G>A, 4122A>C, 4137C>T, 4149C>G, 4150A>C, 4161T>C, 4173C>T, 4176C>T, 4177A>T, 4178C>G, 4179C>T, 4188A>T, 4189T>G, 4190T>C, 4197T>C, 4200C>A, 4201A>C, 4206A>T, 4209T>C, 4215T>A, 4218A>T, 4221A>C, 4222G>A, 4230C>G, 4233T>C, 4234T>C, 4235C>A, 4243T>A, 4245T>A, 4246C>G, 4248T>G, 4251T>G, 4254C>T, 4256A>G, 4257C>T, 4259A>G, 4263C>T, 4267G>T, 4269T>A, 4279A>G, 4280C>A, 4281T>G, 4284G>A, 4287T>C, 4293T>C, 4302A>G, 4305T>C, 4311C>T, 4314G>A, 4315C>A, 4317C>A, 4323A>C, 4324G>A, 4326G>C, 4329C>T, 4332T>A | | | | | | | | |
| **CDS** | | | | | | | | | |
| APL33_sRA11gp1 | 1298 | 1398 | 4.7% | 643 | 88.3% | 101 (100%) | 87 (86.1%) | 0/0/0/0 | 0 |
| Protein mutations: | Y1314H (4090T>C), T1343C (4177A>T 4178C>G 4179C>T), L1347A (4189T>G 4190T>C), I1351L (4201A>C), D1358N (4222G>A), S1362H (4234T>C 4235C>A), S1365T (4243T>A 4245T>A), L1366V (4246C>G 4248T>G), N1369S (4256A>G 4257C>T), N1370S (4259A>G), A1373S (4267G>T 4269T>A), T1377E (4279A>G 4280C>A 4281T>G), V1392I (4324G>A 4326G>C) | | | | | | | | |
| Codon mutations: | ACA1300ACC (4050A>C), CAA1301CAG (4053A>G), GCC1303GCG (4059C>G), AAG1304AAA (4062G>A), GAG1306GAA (4068G>A), CGC1307CGG (4071C>G), CTT1308TTG (4072C>T 4074T>G), CCT1309CCA (4077T>A), GAA1311GAG (4083A>G), ATT1313ATC (4089T>C), TAT1314CAT (4090T>C), AAA1318AAG (4104A>G), ACA1319ACC (4107A>C), AGA1320CGC (4108A>C 4110A>C), TTA1321CTG (4111T>C 4113A>G), TTT1322TTC (4116T>C), GAG1323GAA (4119G>A), ATA1324ATC (4122A>C), TAC1329TAT (4137C>T), GTC1333GTG (4149C>G), AGG1334CGG (4150A>C), TTT1337TTC (4161T>C), TCC1341TCT (4173C>T), GCC1342GCT (4176C>T), ACC1343TGT (4177A>T 4178C>G 4179C>T), GCA1346GCT (4188A>T), TTA1347GCA (4189T>G 4190T>C), AAT1349AAC (4197T>C), GTC1350GTA (4200C>A), ATC1351CTC (4201A>C), CCA1352CCT (4206A>T), TGT1353TGC (4209T>C), GTT1355GTA (4215T>A), GGA1356GGT (4218A>T), ATA1357ATC (4221A>C), GAT1358AAT (4222G>A), ACC1360ACG (4230C>G), AGT1361AGC (4233T>C), TCT1362CAT (4234T>C 4235C>A), TCT1365ACA (4243T>A 4245T>A), CTT1366GTG (4246C>G 4248T>G), CTT1367CTG (4251T>G), GCC1368GCT (4254C>T), AAC1369AGT (4256A>G 4257C>T), AAC1370AGC (4259A>G), TTC1371TTT (4263C>T), GCT1373TCA (4267G>T 4269T>A), ACT1377GAG (4279A>G 4280C>A 4281T>G), GGG1378GGA (4284G>A), TTT1379TTC (4287T>C), GCT1381GCC (4293T>C), TCA1384TCG (4302A>G), AGT1385AGC (4305T>C), GAC1387GAT (4311C>T), GGG1388GGA (4314G>A), CGC1389AGA (4315C>A 4317C>A), CCA1391CCC (4323A>C), GTG1392ATC (4324G>A 4326G>C), TTC1393TTT (4329C>T), GCT1394GCA (4332T>A) | | | | | | | | |
| **Proteins** | | | | | | | | | |
| polyprotein (YP_009165993.1) | 1298 | 1398 | 4.7% | 643 | 88.3% | 101 (100%) | 87 (86.1%) | 0/0/0/0 | 0 |
| Protein mutations: | Y1314H (4090T>C), T1343C (4177A>T 4178C>G 4179C>T), L1347A (4189T>G 4190T>C), I1351L (4201A>C), D1358N (4222G>A), S1362H (4234T>C 4235C>A), S1365T (4243T>A 4245T>A), L1366V (4246C>G 4248T>G), N1369S (4256A>G 4257C>T), N1370S (4259A>G), A1373S (4267G>T 4269T>A), T1377E (4279A>G 4280C>A 4281T>G), V1392I (4324G>A 4326G>C) | | | | | | | | |
| Codon mutations: | ACA1300ACC (4050A>C), CAA1301CAG (4053A>G), GCC1303GCG (4059C>G), AAG1304AAA (4062G>A), GAG1306GAA (4068G>A), CGC1307CGG (4071C>G), CTT1308TTG (4072C>T 4074T>G), CCT1309CCA (4077T>A), GAA1311GAG (4083A>G), ATT1313ATC (4089T>C), TAT1314CAT (4090T>C), AAA1318AAG (4104A>G), ACA1319ACC (4107A>C), AGA1320CGC (4108A>C 4110A>C), TTA1321CTG (4111T>C 4113A>G), TTT1322TTC (4116T>C), GAG1323GAA (4119G>A), ATA1324ATC (4122A>C), TAC1329TAT (4137C>T), GTC1333GTG (4149C>G), AGG1334CGG (4150A>C), TTT1337TTC (4161T>C), TCC1341TCT (4173C>T), GCC1342GCT (4176C>T), ACC1343TGT (4177A>T 4178C>G 4179C>T), GCA1346GCT (4188A>T), TTA1347GCA (4189T>G 4190T>C), AAT1349AAC (4197T>C), GTC1350GTA (4200C>A), ATC1351CTC (4201A>C), CCA1352CCT (4206A>T), TGT1353TGC (4209T>C), GTT1355GTA (4215T>A), GGA1356GGT (4218A>T), ATA1357ATC (4221A>C), GAT1358AAT (4222G>A), ACC1360ACG (4230C>G), AGT1361AGC (4233T>C), TCT1362CAT (4234T>C 4235C>A), TCT1365ACA (4243T>A 4245T>A), CTT1366GTG (4246C>G 4248T>G), CTT1367CTG (4251T>G), GCC1368GCT (4254C>T), AAC1369AGT (4256A>G 4257C>T), AAC1370AGC (4259A>G), TTC1371TTT (4263C>T), GCT1373TCA (4267G>T 4269T>A), ACT1377GAG (4279A>G 4280C>A 4281T>G), GGG1378GGA (4284G>A), TTT1379TTC (4287T>C), GCT1381GCC (4293T>C), TCA1384TCG (4302A>G), AGT1385AGC (4305T>C), GAC1387GAT (4311C>T), GGG1388GGA (4314G>A), CGC1389AGA (4315C>A 4317C>A), CCA1391CCC (4323A>C), GTG1392ATC (4324G>A 4326G>C), TTC1393TTT (4329C>T), GCT1394GCA (4332T>A) | | | | | | | | |

*: Inserts / Deletes / Misaligned / Frameshifts

#### Analysis details

This analysis was performed with panviral2.64

### NGS Details (UN59): Badnavirus maculasmallanthi

#### Assembly

| | |
|---|---|
| **Coverage Length** | 296 (1 contig(s)) |
| **Depth Of Coverage** | 121.8 |
| **Number Of Reads** | 339 |
| **Reads Per Million** | 7.37 rpm (after QC) |
| **Ambiguities** | 0 |
| **Assembly Method** | de novo + reference guided assembly |
| **Consensus Caller** | Bcf Tools |

#### Coverage Map

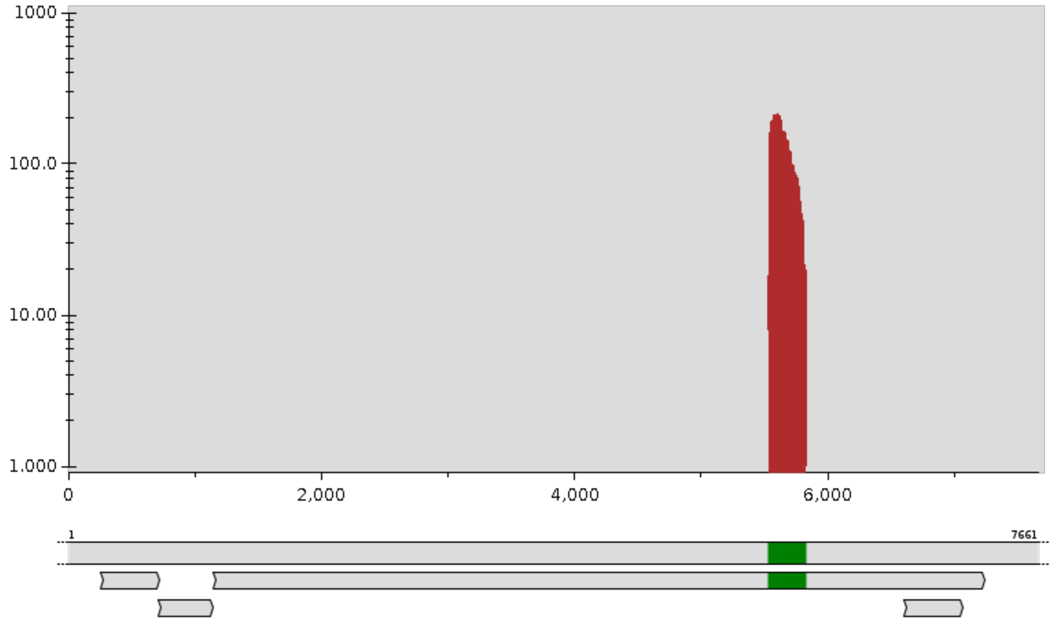

#### Assignment

| | |
|---|---|
| **Type** | Badnavirus maculasmallanthi (Taxonomy ID: 3048453) |
| **Reference Genome** | NC_026472.1 |
| **NT Identity (%)** | 56.5217 |
| **AA Identity (%)** | 53.5354 |
| **Number Of Stop Codons** | 0 |
| **Number Of CDS** | 4 |

#### Alignment

| | |
|---|---|
| **Alignment Score** | 72.0 (NT) + 371.0 (AA) = 443.0 |
| **Concordance (%)** | 34.5824 |

| Alignment Method | Global, seeded, nucleotide + amino acids (AGA) |
|---|---|

#### Genome Region

Sequence starts at position 5534 and ends at position 5829 relative to NC_026472.1 reference sequence.

#### Alignment Detailed Statistics

| | Begin | End | Coverage | Score | Concordance | Matches | Identities | I/D/M/F* | Stop Codons |
|---|---|---|---|---|---|---|---|---|---|
| **NT** | **5534** | **5829** | **3.9%** | **72** | **12.2%** | **296 (99.0%)** | **169 (56.5%)** | **3/0** | |
| Mutations: | 5544A>T, 5545A>G, 5547A>C, 5548A>C, 5549G>A, 5550A>G, 5551A>T, 5554G>C, 5555G>A, 5557A>G, 5558A>G, 5559T>A, 5562A>T, 5563A>T, 5564A>C, 5565G>T, 5566A>G, 5568A>G, 5574A>T, 5578T>A, 5584C>T, 5588A>G, 5589G>A, 5592T>G, 5595A>T, 5597T>A, 5604A>G, 5605G>T, 5607T>G, 5608G>A, 5609C>G, 5610C>A, 5611A>G, 5613G>T, 5614G>C, 5615A>G, 5616A>G, 5619A>G, 5620G>T, 5621A>C, 5623T>G, 5624C>A, 5625C>T, 5628A>C, 5631C>G, 5632T>A, 5633G>A, 5637G>T, 5640C>G, 5643T>C, 5644T>A, 5645T>G, 5646A>G, 5648T>C, 5649C>T, 5651C>G, 5652A>C, 5653G>T, 5654G>A, 5655A>T, 5658A>T, 5659T>C, 5660T>A, 5661A>T, 5664C>T, 5667A>G, 5669G>T, 5670G>T, 5671C>G, 5673T>G, 5680C>T, 5682A>T, 5685C>T, 5688A>G, 5691C>T, 5693A>C, 5694A>C, 5700A>C, 5703C>T, 5707A>G, 5708T>C, 5712T>C, 5713C>A, 5714A>T, 5716A>G, 5717G>A, 5718A>G, 5719A>T, 5720A>T, 5721A>G, 5725G>A, 5728A>C, 5729A>G, 5730A>G, 5731T>G, 5732G>T, 5733C>G, 5736C>T, 5739A>G, 5740G>T, 5741A>C, 5742_5743insTAT, 5743A>T, 5744C>T, 5745A>G, 5748A>T, 5749G>C, 5755A>G, 5757C>G, 5758G>A, 5759C>T, 5765A>T, 5766C>T, 5769C>A, 5772C>T, 5779C>T, 5784A>G, 5787C>T, 5793C>A, 5799A>G, 5800G>A, 5801C>G, 5802G>T, 5813G>A, 5820A>G, 5822A>G, 5823G>A, 5827A>G | | | | | | | | |

##### CDS

| | Begin | End | Coverage | Score | Concordance | Matches | Identities | I/D/M/F* | Stop Codons |
|---|---|---|---|---|---|---|---|---|---|
| UF61_gp3 | 1465 | 1562 | 4.8% | 371 | 53.2% | 98 (99.0%) | 53 (53.5%) | 1/0/0/0 | 0 |
| Protein mutations: | K1468D (5545A>G 5547A>C), R1469Q (5548A>C 5549G>A 5550A>G), I1470L (5551A>T), G1471Q (5554G>C 5555G>A), N1472G (5557A>G 5558A>G 5559T>A), K1474S (5563A>T 5564A>C 5565G>T), I1475V (5566A>G 5568A>G), F1479I (5578T>A), K1482R (5588A>G 5589G>A), F1485Y (5597T>A), V1488L (5605G>T 5607T>G), A1489R (5608G>A 5609C>G 5610C>A), M1490V (5611A>G 5613G>T), E1491R (5614G>C 5615A>G 5616A>G), E1493S (5620G>T 5621A>C), S1494D (5623T>G 5624C>A 5625C>T), W1497K (5632T>A 5633G>A), L1501R (5644T>A 5645T>G 5646A>G), I1502T (5648T>C 5649C>T), P1503R (5651C>G 5652A>C), G1504Y (5653G>T 5654G>A 5655A>T), L1506H (5659T>C 5660T>A 5661A>T), W1509F (5669G>T 5670G>T), L1510V (5671C>G 5673T>G), P1513S (5680C>T 5682A>T), K1517T (5693A>C 5694A>C), I1522A (5707A>G 5708T>C), Q1524M (5713C>A 5714A>T), R1525E (5716A>G 5717G>A 5718A>G), K1526L (5719A>T 5720A>T 5721A>G), D1528N (5725G>A), K1529R (5728A>C 5729A>G 5730A>G), C1530V (5731T>G 5732G>T 5733C>G), D1533S (5740G>T 5741A>C), D1533_T1534insY (5742_5743insTAT), T1534L (5743A>T 5744C>T 5745A>G), E1535D (5748A>T), E1536Q (5749G>C), I1538V (5755A>G 5757C>G), A1539I (5758G>A 5759C>T), Y1541F (5765A>T 5766C>T), N1550K (5793C>A), A1553S (5800G>A 5801C>G 5802G>T), R1557K (5813G>A), K1560R (5822A>G 5823G>A), M1562V (5827A>G) | | | | | | | | |
| Codon mutations: | CTA1467CTT (5544A>T), AAA1468GAC (5545A>G 5547A>C), AGA1469CAG (5548A>C 5549G>A 5550A>G), ATA1470TTA (5551A>T), GGG1471CAG (5554G>C 5555G>A), AAT1472GGA (5557A>G 5558A>G 5559T>A), GCA1473GCT (5562A>T), AAG1474TCT (5563A>T 5564A>C 5565G>T), ATA1475GTG (5566A>G 5568A>G), TCA1477TCT (5574A>T), TTT1479ATT (5578T>A), CTG1481TTG (5584C>T), AAG1482AGA (5588A>G 5589G>A), TCT1483TCG (5592T>G), GGA1484GGT (5595A>T), TTT1485TAT (5597T>A), CAA1487CAG (5604A>G), GTT1488TTG (5605G>T 5607T>G), GCC1489AGA (5608G>A 5609C>G 5610C>A), ATG1490GTT (5611A>G 5613G>T), GAA1491CGG (5614G>C 5615A>G 5616A>G), GAA1492GAG (5619A>G), GAA1493TCA (5620G>T 5621A>C), TCC1494GAT (5623T>G 5624C>A 5625C>T), ATA1495ATC (5628A>C), CCC1496CCG (5631C>G), TGG1497AAG (5632T>A 5633G>A), ACG1498ACT (5637G>T), GCC1499GCG (5640C>G), TTT1500TTC (5643T>C), TTA1501AGG (5644T>A 5645T>G 5646A>G), ATC1502ACT (5648T>C 5649C>T), CCA1503CGC (5651C>G 5652A>C), GGA1504TAT (5653G>T 5654G>A 5655A>T), GGA1505GGT (5658A>T), TTA1506CAT (5659T>C 5660T>A 5661A>T), TAC1507TAT (5664C>T), GAA1508GAG (5667A>G), TGG1509TTT (5669G>T 5670G>T), CTT1510GTG (5671C>G 5673T>G), CCA1513TCT (5680C>T 5682A>T), TTC1514TTT (5685C>T), GGA1515GGG (5688A>G), CTC1516CTT (5691C>T), AAA1517ACC (5693A>C 5694A>C), GCA1519GCC (5700A>C), CCC1520CCT (5703C>T), ATT1522GCT (5707A>G 5708T>C), TTT1523TTC (5712T>C), CAG1524ATG (5713C>A 5714A>T), AGA1525GAG (5716A>G 5717G>A 5718A>G), AAA1526TTG (5719A>T 5720A>T 5721A>G), GAT1528AAT (5725G>A), AAA1529CGG (5728A>C 5729A>G 5730A>G), TGC1530GTG (5731T>G 5732G>T 5733C>G), TTC1531TTT (5736C>T), AAA1532AAG (5739A>G), GAC1533TCC (5740G>T 5741A>C), GAC1533_ACA1534insTAT (5742_5743insTAT), ACA1534TTG (5743A>T 5744C>T 5745A>G), GAA1535GAT (5748A>T), GAG1536CAG (5749G>C), ATC1538GTG (5755A>G 5757C>G), GCC1539ATC (5758G>A 5759C>T), TAC1541TTT (5765A>T 5766C>T), ATC1542ATA (5769C>A), GAC1543GAT (5772C>T), CTG1546TTG (5779C>T), GTA1547GTG (5784A>G), TAC1548TAT (5787C>T), AAC1550AAA (5793C>A), GAA1552GAG (5799A>G), GCG1553AGT (5800G>A 5801C>G 5802G>T), AGG1557AAG (5813G>A), TTA1559TTG (5820A>G), AAG1560AGA (5822A>G 5823G>A), ATG1562GTG (5827A>G) | | | | | | | | |

##### Proteins

| | Begin | End | Coverage | Score | Concordance | Matches | Identities | I/D/M/F* | Stop Codons |
|---|---|---|---|---|---|---|---|---|---|
| ORF3 (YP_009121747.1) | 1465 | 1562 | 4.8% | 371 | 53.2% | 98 (99.0%) | 53 (53.5%) | 1/0/0/0 | 0 |
| Protein mutations: | K1468D (5545A>G 5547A>C), R1469Q (5548A>C 5549G>A 5550A>G), I1470L (5551A>T), G1471Q (5554G>C 5555G>A), N1472G (5557A>G 5558A>G 5559T>A), K1474S (5563A>T 5564A>C 5565G>T), I1475V (5566A>G 5568A>G), F1479I (5578T>A), K1482R (5588A>G 5589G>A), F1485Y (5597T>A), V1488L (5605G>T 5607T>G), A1489R (5608G>A 5609C>G 5610C>A), M1490V (5611A>G 5613G>T), E1491R (5614G>C 5615A>G 5616A>G), E1493S (5620G>T 5621A>C), S1494D (5623T>G 5624C>A 5625C>T), W1497K (5632T>A 5633G>A), L1501R (5644T>A 5645T>G 5646A>G), I1502T (5648T>C 5649C>T), P1503R (5651C>G 5652A>C), G1504Y (5653G>T 5654G>A 5655A>T), L1506H (5659T>C 5660T>A 5661A>T), W1509F (5669G>T 5670G>T), L1510V (5671C>G 5673T>G), P1513S (5680C>T 5682A>T), K1517T (5693A>C 5694A>C), I1522A (5707A>G 5708T>C), Q1524M (5713C>A 5714A>T), R1525E (5716A>G 5717G>A 5718A>G), K1526L (5719A>T 5720A>T 5721A>G), D1528N (5725G>A), K1529R (5728A>C 5729A>G 5730A>G), C1530V (5731T>G 5732G>T 5733C>G), D1533S (5740G>T 5741A>C), D1533_T1534insY (5742_5743insTAT), T1534L (5743A>T 5744C>T 5745A>G), E1535D (5748A>T), E1536Q (5749G>C), I1538V (5755A>G 5757C>G), A1539I (5758G>A 5759C>T), Y1541F (5765A>T 5766C>T), N1550K (5793C>A), A1553S (5800G>A 5801C>G 5802G>T), R1557K (5813G>A), K1560R (5822A>G 5823G>A), M1562V (5827A>G) | | | | | | | | |
| Codon mutations: | CTA1467CTT (5544A>T), AAA1468GAC (5545A>G 5547A>C), AGA1469CAG (5548A>C 5549G>A 5550A>G), ATA1470TTA (5551A>T), GGG1471CAG (5554G>C 5555G>A), AAT1472GGA (5557A>G 5558A>G 5559T>A), GCA1473GCT (5562A>T), AAG1474TCT (5563A>T 5564A>C 5565G>T), ATA1475GTG (5566A>G 5568A>G), TCA1477TCT (5574A>T), TTT1479ATT (5578T>A), CTG1481TTG (5584C>T), AAG1482AGA (5588A>G 5589G>A), TCT1483TCG (5592T>G), GGA1484GGT (5595A>T), TTT1485TAT (5597T>A), CAA1487CAG (5604A>G), GTT1488TTG (5605G>T 5607T>G), GCC1489AGA (5608G>A 5609C>G 5610C>A), ATG1490GTT (5611A>G 5613G>T), GAA1491CGG (5614G>C 5615A>G 5616A>G), GAA1492GAG (5619A>G), GAA1493TCA (5620G>T 5621A>C), TCC1494GAT (5623T>G 5624C>A 5625C>T), ATA1495ATC (5628A>C), CCC1496CCG (5631C>G), TGG1497AAG (5632T>A 5633G>A), ACG1498ACT (5637G>T), GCC1499GCG (5640C>G), TTT1500TTC (5643T>C), TTA1501AGG (5644T>A 5645T>G 5646A>G), ATC1502ACT (5648T>C 5649C>T), CCA1503CGC (5651C>G 5652A>C), GGA1504TAT (5653G>T 5654G>A 5655A>T), GGA1505GGT (5658A>T), TTA1506CAT (5659T>C 5660T>A 5661A>T), TAC1507TAT (5664C>T), GAA1508GAG (5667A>G), TGG1509TTT (5669G>T 5670G>T), CTT1510GTG (5671C>G 5673T>G), CCA1513TCT (5680C>T 5682A>T), TTC1514TTT (5685C>T), GGA1515GGG (5688A>G), CTC1516CTT (5691C>T), AAA1517ACC (5693A>C 5694A>C), GCA1519GCC (5700A>C), CCC1520CCT (5703C>T), ATT1522GCT (5707A>G 5708T>C), TTT1523TTC (5712T>C), CAG1524ATG (5713C>A 5714A>T), AGA1525GAG (5716A>G 5717G>A 5718A>G), AAA1526TTG (5719A>T 5720A>T 5721A>G), GAT1528AAT (5725G>A), AAA1529CGG (5728A>C 5729A>G 5730A>G), TGC1530GTG (5731T>G 5732G>T 5733C>G), TTC1531TTT (5736C>T), AAA1532AAG (5739A>G), GAC1533TCC (5740G>T 5741A>C), GAC1533_ACA1534insTAT (5742_5743insTAT), ACA1534TTG (5743A>T 5744C>T 5745A>G), GAA1535GAT (5748A>T), GAG1536CAG (5749G>C), ATC1538GTG (5755A>G 5757C>G), GCC1539ATC (5758G>A 5759C>T), TAC1541TTT (5765A>T 5766C>T), ATC1542ATA (5769C>A), GAC1543GAT (5772C>T), CTG1546TTG (5779C>T), GTA1547GTG (5784A>G), TAC1548TAT (5787C>T), AAC1550AAA (5793C>A), GAA1552GAG (5799A>G), GCG1553AGT (5800G>A 5801C>G 5802G>T), AGG1557AAG (5813G>A), TTA1559TTG (5820A>G), AAG1560AGA (5822A>G 5823G>A), ATG1562GTG (5827A>G) | | | | | | | | |

*: Inserts / Deletes / Misaligned / Frameshifts

#### Analysis details

This analysis was performed with panviral2.64

### NGS Details (UN59): Cassava brown streak virus

#### Assembly

| | |
|---|---|
| **Coverage Length** | 530 (1 contig(s)) |
| **Depth Of Coverage** | 61.7 |
| **Number Of Reads** | 273 |
| **Reads Per Million** | 5.93 rpm (after QC) |
| **Ambiguities** | 0 |
| **Assembly Method** | de novo + reference guided assembly |
| **Consensus Caller** | Bcf Tools |

#### Coverage Map

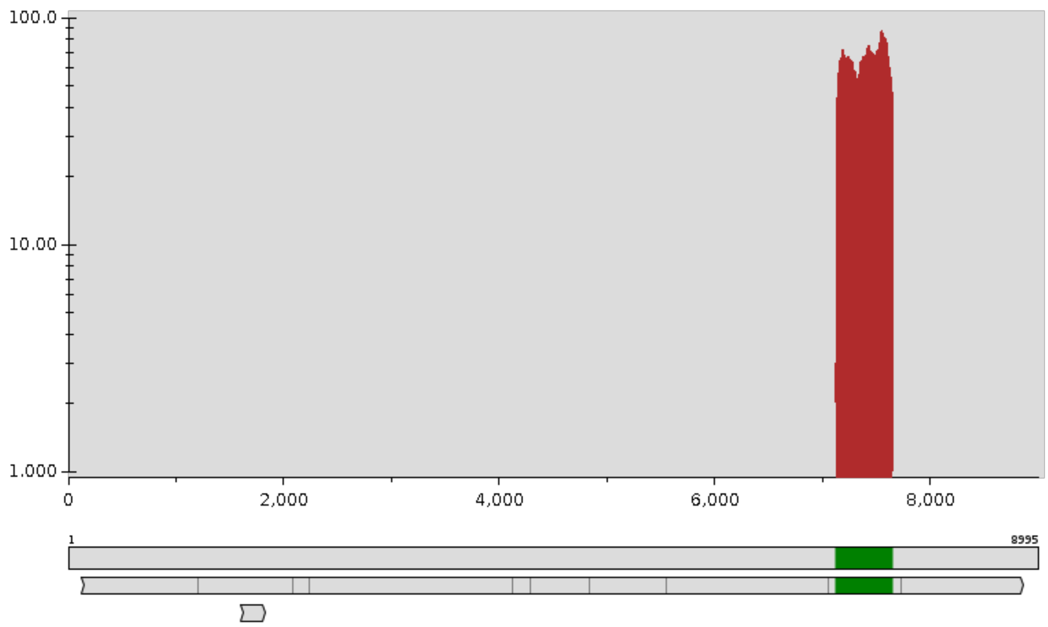

#### Assignment

| | |
|---|---|
| **Type** | Cassava brown streak virus (Taxonomy ID: 137758) |
| **Reference Genome** | NC_012698.2 |
| **NT Identity (%)** | 61.1321 |
| **AA Identity (%)** | 58.1921 |
| **Number Of Stop Codons** | 0 |
| **Number Of CDS** | 2 |

#### Alignment

| | |
|---|---|
| **Alignment Score** | 236.0 (NT) + 748.0 (AA) = 984.0 |
| **Concordance (%)** | 43.0258 |

| Alignment Method | Global, seeded, nucleotide + amino acids (AGA) |
|---|---|

#### Genome Region

Sequence starts at position 7121 and ends at position 7650 relative to NC_012698.2 reference sequence.

#### Alignment Detailed Statistics

| | Begin | End | Coverage | Score | Concordance | Matches | Identities | I/D/M/F* | Stop Codons |
|---|---|---|---|---|---|---|---|---|---|
| **NT** | **7121** | **7650** | **5.9%** | **236** | **22.3%** | **530 (100%)** | **324 (61.1%)** | **0/0** | |
| Mutations: | 7130A>G, 7133A>G, 7134A>G, 7136T>G, 7142C>T, 7145A>T, 7148A>C, 7151G>A, 7157A>T, 7158C>A, 7163A>G, 7167A>G, 7175A>T, 7177A>G, 7178A>G, 7179C>G, 7180A>C, 7184A>T, 7185T>C, 7190T>C, 7192C>A, 7193C>G, 7194A>T, 7202A>C, 7203A>T, 7205C>T, 7206A>C, 7207T>A, 7208C>G, 7211T>C, 7213G>T, 7214G>T, 7217A>G, 7218G>C, 7229A>T, 7234C>T, 7235A>T, 7241A>G, 7242A>G, 7243C>A, 7244A>G, 7245G>C, 7246T>C, 7250G>A, 7253A>T, 7256T>A, 7257A>T, 7258T>C, 7270T>C, 7271A>T, 7272C>A, 7273G>A, 7274T>A, 7275G>A, 7277A>T, 7280G>T, 7281T>G, 7283T>C, 7284G>A, 7286G>A, 7287T>G, 7288T>A, 7290A>G, 7292T>G, 7293G>A, 7294G>A, 7295A>T, 7298G>A, 7304T>G, 7307C>A, 7310A>T, 7313A>G, 7319A>T, 7320A>T, 7323T>C, 7325G>C, 7331T>C, 7332G>A, 7334T>C, 7337T>C, 7340C>T, 7343T>G, 7355A>G, 7358A>T, 7361C>T, 7364T>C, 7374A>T, 7377G>C, 7380G>A, 7381G>A, 7385T>C, 7388A>T, 7390T>A, 7391A>T, 7397A>T, 7398T>C, 7400A>T, 7401T>A, 7403T>C, 7406G>C, 7409G>A, 7410G>T, 7413G>A, 7414A>T, 7416C>G, 7418G>T, 7422C>G, 7424A>G, 7425A>G, 7432T>C, 7433G>A, 7436T>A, 7437A>T, 7438G>A, 7442T>C, 7443C>A, 7445C>G, 7448C>T, 7449G>A, 7451G>C, 7455G>T, 7457T>A, 7458T>C, 7462T>C, 7463A>T, 7464A>C, 7465A>T, 7467A>G, 7468A>G, 7469A>G, 7470G>C, 7471T>C, 7472T>A, 7473G>A, 7474G>A, 7476G>A, 7477A>C, 7481T>G, 7484T>G, 7487A>G, 7489T>C, 7490C>T, 7494A>G, 7495A>T, 7496G>T, 7499T>A, 7500G>A, 7501T>A, 7502G>A, 7503C>A, 7504T>C, 7505A>G, 7506A>C, 7507G>T, 7508A>G, 7511T>A, 7512G>A, 7517T>A, 7520A>G, 7521A>C, 7522T>C, 7523G>A, 7524C>G, 7526A>T, 7527C>A, 7529C>G, 7535A>C, 7539T>A, 7540C>A, 7541A>T, 7544T>C, 7547G>A, 7553C>T, 7556A>G, 7559T>A, 7562C>T, 7568A>T, 7570T>A, 7572A>G, 7573A>G, 7576G>A, 7577G>T, 7578A>G, 7579G>A, 7580A>C, 7581A>C, 7586A>T, 7588T>A, 7589T>C, 7599A>C, 7600T>C, 7601G>C, 7602G>A, 7603C>A, 7604C>G, 7607G>A, 7610G>A, 7616T>C, 7618T>A, 7622A>T, 7628T>C, 7631A>C, 7632T>G, 7633T>G, 7635C>A, 7636G>A | | | | | | | | |

##### CDS

| | Begin | End | Coverage | Score | Concordance | Matches | Identities | I/D/M/F* | Stop Codons |
|---|---|---|---|---|---|---|---|---|---|
| CBSV_gp1 | 2333 | 2509 | 6.1% | 748 | 60.3% | 177 (100%) | 103 (58.2%) | 0/0/0/0 | 0 |
| Protein mutations: | I2337V (7134A>G 7136T>G), Q2345K (7158C>A), K2348E (7167A>G), K2351R (7177A>G 7178A>G), Q2352A (7179C>G 7180A>C), F2354L (7185T>C), P2356Q (7192C>A 7193C>G), T2357S (7194A>T), I2360F (7203A>T 7205C>T), I2361Q (7206A>C 7207T>A 7208C>G), R2363L (7213G>T 7214G>T), V2365L (7218G>C), P2370L (7234C>T 7235A>T), T2373E (7242A>G 7243C>A 7244A>G), V2374P (7245G>C 7246T>C), E2376D (7253A>T), I2378S (7257A>T 7258T>C), V2382A (7270T>C 7271A>T), R2383K (7272C>A 7273G>A 7274T>A), V2384I (7275G>A 7277A>T), S2386A (7281T>G 7283T>C), E2387K (7284G>A 7286G>A), L2388E (7287T>G 7288T>A), I2389V (7290A>G 7292T>G), G2390N (7293G>A 7294G>A 7295A>T), S2399C (7320A>T), D2403N (7332G>A 7334T>C), N2406K (7343T>G), M2417L (7374A>T), E2418Q (7377G>C), G2419K (7380G>A 7381G>A), L2422H (7390T>A 7391A>T), Y2426N (7401T>A 7403T>C), K2427N (7406G>C), V2429L (7410G>T), E2430M (7413G>A 7414A>T), P2431A (7416C>G 7418G>T), Q2433E (7422C>G 7424A>G), N2434D (7425A>G), M2436T (7432T>C 7433G>A), S2438Y (7437A>T 7438G>A), L2440M (7443C>A 7445C>G), V2442I (7449G>A 7451G>C), A2444S (7455G>T 7457T>A), F2445L (7458T>C), V2446A (7462T>C 7463A>T), N2447L (7464A>C 7465A>T), K2448G (7467A>G 7468A>G 7469A>G), V2449P (7470G>C 7471T>C 7472T>A), G2450N (7473G>A 7474G>A), D2451T (7476G>A 7477A>C), D2452E (7481T>G), I2454M (7487A>G), I2455T (7489T>C 7490C>T), K2457V (7494A>G 7495A>T 7496G>T), V2459K (7500G>A 7501T>A 7502G>A), L2460T (7503C>A 7504T>C 7505A>G), R2461L (7506A>C 7507G>T 7508A>G), E2463K (7512G>A), M2466P (7521A>C 7522T>C 7523G>A), P2467A (7524C>G 7526A>T), S2472N (7539T>A 7540C>A 7541A>T), L2482H (7570T>A), N2483G (7572A>G 7573A>G), W2484Y (7576G>A 7577G>T), R2485D (7578A>G 7579G>A 7580A>C), K2486Q (7581A>C), F2488Y (7588T>A 7589T>C), M2492P (7599A>C 7600T>C 7601G>C), A2493K (7602G>A 7603C>A 7604C>G), M2498K (7618T>A), F2503G (7632T>G 7633T>G), R2504K (7635C>A 7636G>A) | | | | | | | | |
| Codon mutations: | GCA2335GCG (7130A>G), CCA2336CCG (7133A>G), ATT2337GTG (7134A>G 7136T>G), TTC2339TTT (7142C>T), GTA2340GTT (7145A>T), ACA2341ACC (7148A>C), GGG2342GGA (7151G>A), GCA2344GCT (7157A>T), CAA2345AAA (7158C>A), AAA2346AAG (7163A>G), AAG2348GAG (7167A>G), GTA2350GTT (7175A>T), AAA2351AGG (7177A>G 7178A>G), CAA2352GCA (7179C>G 7180A>C), ATA2353ATT (7184A>T), TTT2354CTT (7185T>C), GGT2355GGC (7190T>C), CCC2356CAG (7192C>A 7193C>G), ACT2357TCT (7194A>T), CCA2359CCC (7202A>C), ATC2360TTT (7203A>T 7205C>T), ATC2361CAG (7206A>C 7207T>A 7208C>G), TCT2362TCC (7211T>C), CGG2363CTT (7213G>T 7214G>T), AAA2364AAG (7217A>G), GTT2365CTT (7218G>C), CCA2368CCT (7229A>T), CCA2370CTT (7234C>T 7235A>T), GGA2372GGG (7241A>G), ACA2373GAG (7242A>G 7243C>A 7244A>G), GTT2374CCT (7245G>C 7246T>C), GAG2375GAA (7250G>A), GAA2376GAT (7253A>T), ATT2377ATA (7256T>A), ATC2378TCC (7257A>T 7258T>C), GTA2382GCT (7270T>C 7271A>T), CGT2383AAA (7272C>A 7273G>A 7274T>A), GTA2384ATT (7275G>A 7277A>T), GCG2385GCT (7280G>T), TCT2386GCC (7281T>G 7283T>C), GAG2387AAA (7284G>A 7286G>A), TTG2388GAG (7287T>G 7288T>A), ATT2389GTG (7290A>G 7292T>G), GGA2390AAT (7293G>A 7294G>A 7295A>T), GGG2391GGA (7298G>A), GTT2393GTG (7304T>G), CTC2394CTA (7307C>A), GTA2395GTT (7310A>T), GAA2396GAG (7313A>G), ACA2398ACT (7319A>T), AGT2399TGT (7320A>T), TTG2400CTC (7323T>C 7325G>C), TTT2402TTC (7331T>C), GAT2403AAC (7332G>A 7334T>C), GCT2404GCC (7337T>C), CTC2405CTT (7340C>T), AAT2406AAG (7343T>G), GGA2410GGG (7355A>G), CCA2411CCT (7358A>T), TAC2412TAT (7361C>T), ATT2413ATC (7364T>C), ATG2417TTG (7374A>T), GAA2418CAA (7377G>C), GGG2419AAG (7380G>A 7381G>A), ATT2420ATC (7385T>C), GGA2421GGT (7388A>T), CTA2422CAT (7390T>A 7391A>T), GGA2424GGT (7397A>T), TTA2425CTT (7398T>C 7400A>T), TAT2426AAC (7401T>A 7403T>C), AAG2427AAC (7406G>C), TTG2428TTA (7409G>A), GTG2429TTG (7410G>T), GAG2430ATG (7413G>A 7414A>T), CCG2431GCT (7416C>G 7418G>T), CAA2433GAG (7422C>G 7424A>G), AAT2434GAT (7425A>G), ATG2436ACA (7432T>C 7433G>A), GCT2437GCA (7436T>A), AGT2438TAT (7437A>T 7438G>A), GCT2439GCC (7442T>C), CTC2440ATG (7443C>A 7445C>G), TGC2441TGT (7448C>T), GTG2442ATC (7449G>A 7451G>C), GCT2444TCA (7455G>T 7457T>A), TTT2445CTT (7458T>C), GTA2446GCT (7462T>C 7463A>T), AAT2447CTT (7464A>C 7465A>T), AAA2448GGG (7467A>G 7468A>G 7469A>G), GTT2449CCA (7470G>C 7471T>C 7472T>A), GGT2450AAT (7473G>A 7474G>A), GAT2451ACT (7476G>A 7477A>C), GAT2452GAG (7481T>G), CCT2453CCG (7484T>G), ATA2454ATG (7487A>G), ATC2455ACT (7489T>C 7490C>T), AAG2457GTT (7494A>G 7495A>T 7496G>T), GGT2458GGA (7499T>A), GTG2459AAA (7500G>A 7501T>A 7502G>A), CTA2460ACG (7503C>A 7504T>C 7505A>G), AGA2461CTG (7506A>C 7507G>T 7508A>G), GGT2462GGA (7511T>A), GAG2463AAG (7512G>A), ATT2464ATA (7517T>A), GTA2465GTG (7520A>G), ATG2466CCA (7521A>C 7522T>C 7523G>A), CCA2467GCT (7524C>G 7526A>T), CGC2468AGG (7527C>A 7529C>G), CCA2470CCC (7535A>C), TCA2472AAT (7539T>A 7540C>A 7541A>T), TTT2473TTC (7544T>C), GGG2474GGA (7547G>A), GAC2476GAT (7553C>T), CCA2477CCG (7556A>G), ATT2478ATA (7559T>A), TTC2479TTT (7562C>T), CCA2481CCT (7568A>T), CTT2482CAT (7570T>A), AAC2483GGC (7572A>G 7573A>G), TGG2484TAT (7576G>A 7577G>T), AGA2485GAC (7578A>G 7579G>A 7580A>C), AAG2486CAG (7581A>C), ACA2487ACT (7586A>T), TTT2488TAC (7588T>A 7589T>C), ATG2492CCC (7599A>C 7600T>C 7601G>C), GCC2493AAG (7602G>A 7603C>A 7604C>G), GAG2494GAA (7607G>A), GAG2495GAA (7610G>A), AAT2497AAC (7616T>C), ATG2498AAG (7618T>A), ATA2499ATT (7622A>T), CAT2501CAC (7628T>C), CGA2502CGC (7631A>C), TTT2503GGT (7632T>G 7633T>G), CGA2504AAA (7635C>A 7636G>A) | | | | | | | | |

##### Proteins

| | Begin | End | Coverage | Score | Concordance | Matches | Identities | I/D/M/F* | Stop Codons |
|---|---|---|---|---|---|---|---|---|---|
| polyprotein (YP_007027011.1) | 2333 | 2509 | 6.1% | 748 | 60.3% | 177 (100%) | 103 (58.2%) | 0/0/0/0 | 0 |
| Protein mutations: | I2337V (7134A>G 7136T>G), Q2345K (7158C>A), K2348E (7167A>G), K2351R (7177A>G 7178A>G), Q2352A (7179C>G 7180A>C), F2354L (7185T>C), P2356Q (7192C>A 7193C>G), T2357S (7194A>T), I2360F (7203A>T 7205C>T), I2361Q (7206A>C 7207T>A 7208C>G), R2363L (7213G>T 7214G>T), V2365L (7218G>C), P2370L (7234C>T 7235A>T), T2373E (7242A>G 7243C>A 7244A>G), V2374P (7245G>C 7246T>C), E2376D (7253A>T), I2378S (7257A>T 7258T>C), V2382A (7270T>C 7271A>T), R2383K (7272C>A 7273G>A 7274T>A), V2384I (7275G>A 7277A>T), S2386A (7281T>G 7283T>C), E2387K (7284G>A 7286G>A), L2388E (7287T>G 7288T>A), I2389V (7290A>G 7292T>G), G2390N (7293G>A 7294G>A 7295A>T), S2399C (7320A>T), D2403N (7332G>A 7334T>C), N2406K (7343T>G), M2417L (7374A>T), E2418Q (7377G>C), G2419K (7380G>A 7381G>A), L2422H (7390T>A 7391A>T), Y2426N (7401T>A 7403T>C), K2427N (7406G>C), V2429L (7410G>T), E2430M (7413G>A 7414A>T), P2431A (7416C>G 7418G>T), Q2433E (7422C>G 7424A>G), N2434D (7425A>G), M2436T (7432T>C 7433G>A), S2438Y (7437A>T 7438G>A), L2440M (7443C>A 7445C>G), V2442I (7449G>A 7451G>C), A2444S (7455G>T 7457T>A), F2445L (7458T>C), V2446A (7462T>C 7463A>T), N2447L (7464A>C 7465A>T), K2448G (7467A>G 7468A>G 7469A>G), V2449P (7470G>C 7471T>C 7472T>A), G2450N (7473G>A 7474G>A), D2451T (7476G>A 7477A>C), D2452E (7481T>G), I2454M (7487A>G), I2455T (7489T>C 7490C>T), K2457V (7494A>G 7495A>T 7496G>T), V2459K (7500G>A 7501T>A 7502G>A), L2460T (7503C>A 7504T>C 7505A>G), R2461L (7506A>C 7507G>T 7508A>G), E2463K (7512G>A), M2466P (7521A>C 7522T>C 7523G>A), P2467A (7524C>G 7526A>T), S2472N (7539T>A 7540C>A 7541A>T), L2482H (7570T>A), N2483G (7572A>G 7573A>G), W2484Y (7576G>A 7577G>T), R2485D (7578A>G 7579G>A 7580A>C), K2486Q (7581A>C), F2488Y (7588T>A 7589T>C), M2492P (7599A>C 7600T>C 7601G>C), A2493K (7602G>A 7603C>A 7604C>G), M2498K (7618T>A), F2503G (7632T>G 7633T>G), R2504K (7635C>A 7636G>A) | | | | | | | | |

| | Begin | End | Coverage | Score | Concordance | Matches | Identities | I/D/M/F* | Stop Codons |
|---|---|---|---|---|---|---|---|---|---|
| **NT** | **7121** | **7650** | **5.9%** | **236** | **22.3%** | **530 (100%)** | **324 (61.1%)** | **0/0** | |
| Codon mutations: | GCA2335GCG (7130A>G), CCA2336CCG (7133A>G), ATT2337GTG (7134A>G 7136T>G), TTC2339TTT (7142C>T), GTA2340GTT (7145A>T), ACA2341ACC (7148A>C), GGG2342GGA (7151G>A), GCA2344GCT (7157A>T), CAA2345AAA (7158C>A), AAA2346AAG (7163A>G), AAG2348GAG (7167A>G), GTA2350GTT (7175A>T), AAA2351AGG (7177A>G 7178A>G), CAA2352GCA (7179C>G 7180A>C), ATA2353ATT (7184A>T), TTT2354CTT (7185T>C), GGT2355GGC (7190T>C), CCC2356CAG (7192C>A 7193C>G), ACT2357TCT (7194A>T), CCA2359CCC (7202A>C), ATC2360TTT (7203A>T 7205C>T), ATC2361CAG (7206A>C 7207T>A 7208C>G), TCT2362TCC (7211T>C), CGG2363CTT (7213G>T 7214G>T), AAA2364AAG (7217A>G), GTT2365CTT (7218G>C), CCA2368CCT (7229A>T), CCA2370CTT (7234C>T 7235A>T), GGA2372GGG (7241A>G), ACA2373GAG (7242A>G 7243C>A 7244A>G), GTT2374CCT (7245G>C 7246T>C), GAG2375GAA (7250G>A), GAA2376GAT (7253A>T), ATT2377ATA (7256T>A), ATC2378TCC (7257A>T 7258T>C), GTA2382GCT (7270T>C 7271A>T), CGT2383AAA (7272C>A 7273G>A 7274T>A), GTA2384ATT (7275G>A 7277A>T), GCG2385GCT (7280G>T), TCT2386GCC (7281T>G 7283T>C), GAG2387AAA (7284G>A 7286G>A), TTG2388GAG (7287T>G 7288T>A), ATT2389GTG (7290A>G 7292T>G), GGA2390AAT (7293G>A 7294G>A 7295A>T), GGG2391GGA (7298G>A), GTT2393GTG (7304T>G), CTC2394CTA (7307C>A), GTA2395GTT (7310A>T), GAA2396GAG (7313A>G), ACA2398ACT (7319A>T), AGT2399TGT (7320A>T), TTG2400CTC (7323T>C 7325G>C), TTT2402TTC (7331T>C), GAT2403AAC (7332G>A 7334T>C), GCT2404GCC (7337T>C), CTC2405CTT (7340C>T), AAT2406AAG (7343T>G), GGA2410GGG (7355A>G), CCA2411CCT (7358A>T), TAC2412TAT (7361C>T), ATT2413ATC (7364T>C), ATG2417TTG (7374A>T), GAA2418CAA (7377G>C), GGG2419AAG (7380G>A 7381G>A), ATT2420ATC (7385T>C), GGA2421GGT (7388A>T), CTA2422CAT (7390T>A 7391A>T), GGA2424GGT (7397A>T), TTA2425CTT (7398T>C 7400A>T), TAT2426AAC (7401T>A 7403T>C), AAG2427AAC (7406G>C), TTG2428TTA (7409G>A), GTG2429TTG (7410G>T), GAG2430ATG (7413G>A 7414A>T), CCG2431GCT (7416C>G 7418G>T), CAA2433GAG (7422C>G 7424A>G), AAT2434GAT (7425A>G), ATG2436ACA (7432T>C 7433G>A), GCT2437GCA (7436T>A), AGT2438TAT (7437A>T 7438G>A), GCT2439GCC (7442T>C), CTC2440ATG (7443C>A 7445C>G), TGC2441TGT (7448C>T), GTG2442ATC (7449G>A 7451G>C), GCT2444TCA (7455G>T 7457T>A), TTT2445CTT (7458T>C), GTA2446GCT (7462T>C 7463A>T), AAT2447CTT (7464A>C 7465A>T), AAA2448GGG (7467A>G 7468A>G 7469A>G), GTT2449CCA (7470G>C 7471T>C 7472T>A), GGT2450AAT (7473G>A 7474G>A), GAT2451ACT (7476G>A 7477A>C), GAT2452GAG (7481T>G), CCT2453CCG (7484T>G), ATA2454ATG (7487A>G), ATC2455ACT (7489T>C 7490C>T), AAG2457GTT (7494A>G 7495A>T 7496G>T), GGT2458GGA (7499T>A), GTG2459AAA (7500G>A 7501T>A 7502G>A), CTA2460ACG (7503C>A 7504T>C 7505A>G), AGA2461CTG (7506A>C 7507G>T 7508A>G), GGT2462GGA (7511T>A), GAG2463AAG (7512G>A), ATT2464ATA (7517T>A), GTA2465GTG (7520A>G), ATG2466CCA (7521A>C 7522T>C 7523G>A), CCA2467GCT (7524C>G 7526A>T), CGC2468AGG (7527C>A 7529C>G), CCA2470CCC (7535A>C), TCA2472AAT (7539T>A 7540C>A 7541A>T), TTT2473TTC (7544T>C), GGG2474GGA (7547G>A), GAC2476GAT (7553C>T), CCA2477CCG (7556A>G), ATT2478ATA (7559T>A), TTC2479TTT (7562C>T), CCA2481CCT (7568A>T), CTT2482CAT (7570T>A), AAC2483GGC (7572A>G 7573A>G), TGG2484TAT (7576G>A 7577G>T), AGA2485GAC (7578A>G 7579G>A 7580A>C), AAG2486CAG (7581A>C), ACA2487ACT (7586A>T), TTT2488TAC (7588T>A 7589T>C), ATG2492CCC (7599A>C 7600T>C 7601G>C), GCC2493AAG (7602G>A 7603C>A 7604C>G), GAG2494GAA (7607G>A), GAG2495GAA (7610G>A), AAT2497AAC (7616T>C), ATG2498AAG (7618T>A), ATA2499ATT (7622A>T), CAT2501CAC (7628T>C), CGA2502CGC (7631A>C), TTT2503GGT (7632T>G 7633T>G), CGA2504AAA (7635C>A 7636G>A) | | | | | | | | |
| HAM1-like protein (YP_007032446.1) | 25 | 201 | 78.3% | 748 | 60.3% | 177 (100%) | 103 (58.2%) | 0/0/0/0 | 0 |
| Protein mutations: | I29V (7134A>G 7136T>G), Q37K (7158C>A), K40E (7167A>G), K43R (7177A>G 7178A>G), Q44A (7179C>G 7180A>C), F46L (7185T>C), P48Q (7192C>A 7193C>G), T49S (7194A>T), I52F (7203A>T 7205C>T), I53Q (7206A>C 7207T>A 7208C>G), R55L (7213G>T 7214G>T), V57L (7218G>C), P62L (7234C>T 7235A>T), T65E (7242A>G 7243C>A 7244A>G), V66P (7245G>C 7246T>C), E68D (7253A>T), I70S (7257A>T 7258T>C), V74A (7270T>C 7271A>T), R75K (7272C>A 7273G>A 7274T>A), V76I (7275G>A 7277A>T), S78A (7281T>G 7283T>C), E79K (7284G>A 7286G>A), L80E (7287T>G 7288T>A), I81V (7290A>G 7292T>G), G82N (7293G>A 7294G>A 7295A>T), S91C (7320A>T), D95N (7332G>A 7334T>C), N98K (7343T>G), M109L (7374A>T), E110Q (7377G>C), G111K (7380G>A 7381G>A), L114H (7390T>A 7391A>T), Y118N (7401T>A 7403T>C), K119N (7406G>C), V121L (7410G>T), E122M (7413G>A 7414A>T), P123A (7416C>G 7418G>T), Q125E (7422C>G 7424A>G), N126D (7425A>G), M128T (7432T>C 7433G>A), S130Y (7437A>T 7438G>A), L132M (7443C>A 7445C>G), V134I (7449G>A 7451G>C), A136S (7455G>T 7457T>A), F137L (7458T>C), V138A (7462T>C 7463A>T), N139L (7464A>C 7465A>T), K140G (7467A>G 7468A>G 7469A>G), V141P (7470G>C 7471T>C 7472T>A), G142N (7473G>A 7474G>A), D143T (7476G>A 7477A>C), D144E (7481T>G), I146M (7487A>G), I147T (7489T>C 7490C>T), K149V (7494A>G 7495A>T 7496G>T), V151K (7500G>A 7501T>A 7502G>A), L152T (7503C>A 7504T>C 7505A>G), R153L (7506A>C 7507G>T 7508A>G), E155K (7512G>A), M158P (7521A>C 7522T>C 7523G>A), P159A (7524C>G 7526A>T), S164N (7539T>A 7540C>A 7541A>T), L174H (7570T>A), N175G (7572A>G 7573A>G), W176Y (7576G>A 7577G>T), R177D (7578A>G 7579G>A 7580A>C), K178Q (7581A>C), F180Y (7588T>A 7589T>C), M184P (7599A>C 7600T>C 7601G>C), A185K (7602G>A 7603C>A 7604C>G), M190K (7618T>A), F195G (7632T>G 7633T>G), R196K (7635C>A 7636G>A) | | | | | | | | |
| Codon mutations: | GCA27GCG (7130A>G), CCA28CCG (7133A>G), ATT29GTG (7134A>G 7136T>G), TTC31TTT (7142C>T), GTA32GTT (7145A>T), ACA33ACC (7148A>C), GGG34GGA (7151G>A), GCA36GCT (7157A>T), CAA37AAA (7158C>A), AAA38AAG (7163A>G), AAG40GAG (7167A>G), GTA42GTT (7175A>T), AAA43AGG (7177A>G 7178A>G), CAA44GCA (7179C>G 7180A>C), ATA45ATT (7184A>T), TTT46CTT (7185T>C), GGT47GGC (7190T>C), CCC48CAG (7192C>A 7193C>G), ACT49TCT (7194A>T), CCA51CCC (7202A>C), ATC52TTT (7203A>T 7205C>T), ATC53CAG (7206A>C 7207T>A 7208C>G), TCT54TCC (7211T>C), CGG55CTT (7213G>T 7214G>T), AAA56AAG (7217A>G), GTT57CTT (7218G>C), CCA60CCT (7229A>T), CCA62CTT (7234C>T 7235A>T), GGA64GGG (7241A>G), ACA65GAG (7242A>G 7243C>A 7244A>G), GTT66CCT (7245G>C 7246T>C), GAG67GAA (7250G>A), GAA68GAT (7253A>T), ATT69ATA (7256T>A), ATC70TCC (7257A>T 7258T>C), GTA74GCT (7270T>C 7271A>T), CGT75AAA (7272C>A 7273G>A 7274T>A), GTA76ATT (7275G>A 7277A>T), GCG77GCT (7280G>T), TCT78GCC (7281T>G 7283T>C), GAG79AAA (7284G>A 7286G>A), TTG80GAG (7287T>G 7288T>A), ATT81GTG (7290A>G 7292T>G), GGA82AAT (7293G>A 7294G>A 7295A>T), GGG83GGA (7298G>A), GTT85GTG (7304T>G), CTC86CTA (7307C>A), GTA87GTT (7310A>T), GAA88GAG (7313A>G), ACA90ACT (7319A>T), AGT91TGT (7320A>T), TTG92CTC (7323T>C 7325G>C), TTT94TTC (7331T>C), GAT95AAC (7332G>A 7334T>C), GCT96GCC (7337T>C), CTC97CTT (7340C>T), AAT98AAG (7343T>G), GGA102GGG (7355A>G), CCA103CCT (7358A>T), TAC104TAT (7361C>T), ATT105ATC (7364T>C), ATG109TTG (7374A>T), GAA110CAA (7377G>C), GGG111AAG (7380G>A 7381G>A), ATT112ATC (7385T>C), GGA113GGT (7388A>T), CTA114CAT (7390T>A 7391A>T), GGA116GGT (7397A>T), TTA117CTT (7398T>C 7400A>T), TAT118AAC (7401T>A 7403T>C), AAG119AAC (7406G>C), TTG120TTA (7409G>A), GTG121TTG (7410G>T), GAG122ATG (7413G>A 7414A>T), CCG123GCT (7416C>G 7418G>T), CAA125GAG (7422C>G 7424A>G), AAT126GAT (7425A>G), ATG128ACA (7432T>C 7433G>A), GCT129GCA (7436T>A), AGT130TAT (7437A>T 7438G>A), GCT131GCC (7442T>C), CTC132ATG (7443C>A 7445C>G), TGC133TGT (7448C>T), GTG134ATC (7449G>A 7451G>C), GCT136TCA (7455G>T 7457T>A), TTT137CTT (7458T>C), GTA138GCT (7462T>C 7463A>T), AAT139CTT (7464A>C 7465A>T), AAA140GGG (7467A>G 7468A>G 7469A>G), GTT141CCA (7470G>C 7471T>C 7472T>A), GGT142AAT (7473G>A 7474G>A), GAT143ACT (7476G>A 7477A>C), GAT144GAG (7481T>G), CCT145CCG (7484T>G), ATA146ATG (7487A>G), ATC147ACT (7489T>C 7490C>T), AAG149GTT (7494A>G 7495A>T 7496G>T), GGT150GGA (7499T>A), GTG151AAA (7500G>A 7501T>A 7502G>A), CTA152ACG (7503C>A 7504T>C 7505A>G), AGA153CTG (7506A>C 7507G>T 7508A>G), GGT154GGA (7511T>A), GAG155AAG (7512G>A), ATT156ATA (7517T>A), GTA157GTG (7520A>G), ATG158CCA (7521A>C 7522T>C 7523G>A), CCA159GCT (7524C>G 7526A>T), CGC160AGG (7527C>A 7529C>G), CCA162CCC (7535A>C), TCA164AAT (7539T>A 7540C>A 7541A>T), TTT165TTC (7544T>C), GGG166GGA (7547G>A), GAC168GAT (7553C>T), CCA169CCG (7556A>G), ATT170ATA (7559T>A), TTC171TTT (7562C>T), CCA173CCT (7568A>T), CTT174CAT (7570T>A), AAC175GGC (7572A>G 7573A>G), TGG176TAT (7576G>A 7577G>T), AGA177GAC (7578A>G 7579G>A 7580A>C), AAG178CAG (7581A>C), ACA179ACT (7586A>T), TTT180TAC (7588T>A 7589T>C), ATG184CCC (7599A>C 7600T>C 7601G>C), GCC185AAG (7602G>A 7603C>A 7604C>G), GAG186GAA (7607G>A), GAG187GAA (7610G>A), AAT189AAC (7616T>C), ATG190AAG (7618T>A), ATA191ATT (7622A>T), CAT193CAC (7628T>C), CGA194CGC (7631A>C), TTT195GGT (7632T>G 7633T>G), CGA196AAA (7635C>A 7636G>A) | | | | | | | | |

*: Inserts / Deletes / Misaligned / Frameshifts

#### Analysis details

This analysis was performed with panviral2.64

### NGS Details (UN59): Caulimovirus venafragariae

#### Assembly

| | |
|---|---|
| **Coverage Length** | 1219 (3 contig(s)) |
| **Depth Of Coverage** | 12.6 |
| **Number Of Reads** | 134 |
| **Reads Per Million** | 2.91 rpm (after QC) |
| **Ambiguities** | 0 |
| **Assembly Method** | de novo + reference guided assembly |
| **Consensus Caller** | Bcf Tools |

#### Coverage Map

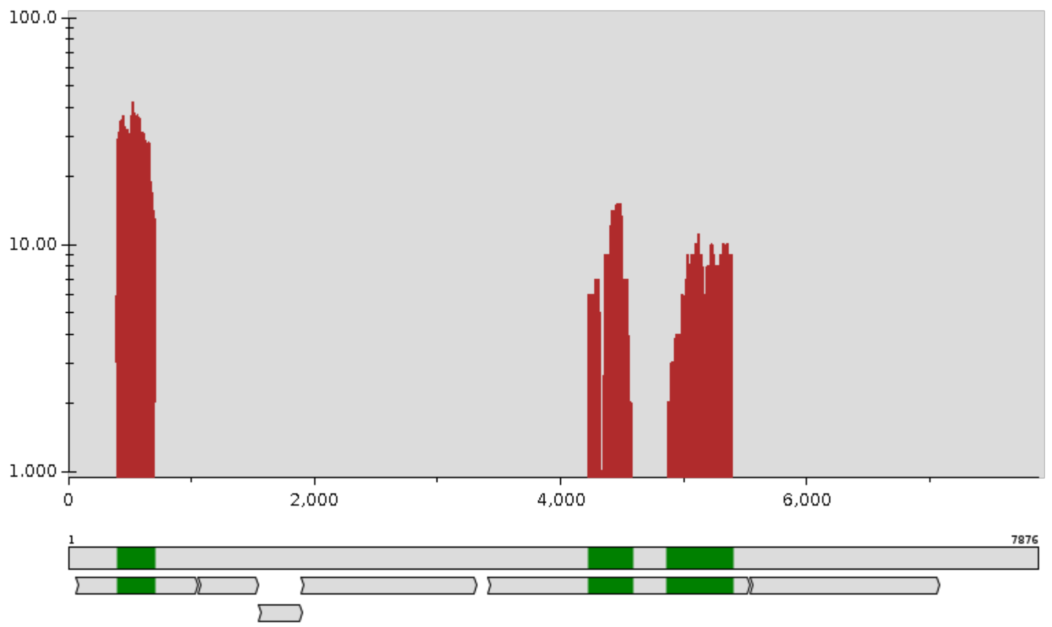

#### Assignment

| | |
|---|---|
| **Type** | Caulimovirus venafragariae (Taxonomy ID: 3048344) |
| **Reference Genome** | NC_001725.1 |
| **NT Identity (%)** | 56.6445 |
| **AA Identity (%)** | 48.5 |
| **Number Of Stop Codons** | 0 |
| **Number Of CDS** | 6 |

#### Alignment

| | |
|---|---|
| **Alignment Score** | 269.0 (NT) + 1436.0 (AA) = 1705.0 |
| **Concordance (%)** | 33.7423 |

| Alignment Method | Global, seeded, nucleotide + amino acids (AGA) |
|---|---|

#### Genome Region

Sequence starts at position 395 and ends at position 5402 relative to NC_001725.1 reference sequence.

#### Alignment Detailed Statistics

| | Begin | End | Coverage | Score | Concordance | Matches | Identities | I/D/M/F* | Stop Codons |
|---|---|---|---|---|---|---|---|---|---|
| **NT** | **395** | **5402** | **15.5%** | **269** | **11.4%** | **1201 (98.3%)** | **682 (55.8%)** | **3/18** | |

Mutations: 403A>C, 405A>T, 411A>G, 412G>A, 416T>A, 418A>G, 424A>T, 426A>G, 430T>G, 431C>G, 432T>A, 433G>A, 435T>A, 436A>C, 441T>C, 445A>G, 447C>G, 451A>T, 452G>T, 456C>T, 459T>C, 461G>A, 462G>A, 463A>G, 464C>A, 468A>T, 472G>A, 474C>T, 476C>G, 483T>A, 485G>T, 486T>A, 487G>A, 490G>A, 491C>A, 492A>C, 495C>T, 496C>T, 497T>C, 501C>T, 502C>G, 503G>A, 504G>A, 508A>T, 510G>T, 512A>T, 513A>T, 516C>T, 517G>A, 521A>G, 525C>A, 527C>G, 529G>A, 530T>A, 531C>T, 532T>C, 534C>T, 537C>A, 538G>A, 539G>T, 544A>G, 552C>T, 553C>T, 555T>A, 556T>G, 558C>A, 562G>A, 563G>C, 564G>T, 567A>G, 568C>T, 570T>A, 571A>C, 573C>T, 579C>A, 582C>T, 583A>T, 585C>T, 588A>C, 590A>G, 592A>T, 593T>A, 594C>T, 596C>G, 598G>A, 600T>A, 601A>T, 602G>C, 603T>A, 604C>T, 606T>A, 608G>A, 612C>T, 613C>G, 617C>A, 618G>C, 619A>T, 624C>T, 625A>G, 627G>A, 628A>G, 630T>A, 633A>G, 635C>G, 636A>T, 637C>T, 639A>G, 640_642delGCT, 648C>T, 648_649insCAG, 649T>G, 651T>C, 652G>A, 656A>G, 658G>A, 663A>T, 666G>A, 667A>T, 669G>C, 670C>A, 672T>G, 673G>C, 674A>C, 675G>A, 678A>T, 685C>A, 686C>T, 687C>T, 688T>A, 689A>T, 690T>G, 692C>G, 693T>C, 696A>T, 698C>A, 699C>T, 4222T>A, 4223T>A, 4227C>T, 4228A>C, 4230T>C, 4232A>G, 4235A>T, 4236A>C, 4242T>A, 4244A>G, 4245G>A, 4246T>G, 4247G>T, 4259A>T, 4260G>A, 4264T>A, 4265C>A, 4266A>T, 4267C>G, 4268C>A, 4269T>G, 4275C>T, 4276C>A, 4277G>A, 4279G>A, 4280A>C, 4281A>T, 4284G>A, 4292C>T, 4299C>T, 4300G>C, 4306C>T, 4309C>T, 4311C>A, 4314G>T, 4315C>A, 4317T>G, 4318G>A, 4319G>A, 4320T>A, 4321A>T, 4323C>A, 4326C>A, 4330C>G, 4331C>A, 4332T>A, 4335C>T, 4338G>A, 4348T>A, 4349C>G, 4350C>T, 4351T>A, 4352C>G, 4353A>T, 4356A>T, 4359C>A, 4362C>T, 4371G>A, 4374T>C, 4380C>G, 4383G>A, 4384A>C, 4385T>A, 4386C>A, 4395C>A, 4401A>T, 4402A>C, 4410A>T, 4413T>A, 4423A>G, 4426T>C, 4428A>T, 4432G>A, 4434C>A, 4435C>A, 4437T>C, 4438A>T, 4439C>G, 4440A>T, 4442A>T, 4443G>T, 4444G>T, 4445G>T, 4446A>T, 4449T>C, 4452C>A, 4456C>T, 4458A>T, 4459C>A, 4461T>C, 4467C>T, 4470G>A, 4471G>A, 4472A>C, 4473G>T, 4474C>A, 4476A>T, 4479G>T, 4480C>A, 4482T>A, 4483C>A, 4485A>T, 4486A>C, 4487G>T, 4489A>G, 4492G>A, 4493G>A, 4496G>T, 4497T>A, 4500G>A, 4505T>A, 4506T>C, 4508A>T, 4509C>T, 4512T>C, 4513T>A, 4514C>A, 4515C>A, 4521C>T, 4528T>A, 4529C>G, 4543G>A, 4546C>A, 4548C>A, 4551T>C, 4552G>T, 4554T>A, 4555C>G, 4556C>A, 4557A>G, 4561A>T, 4568A>C, 4569G>T, 4575C>A, 4578T>A, 4859T>C, 4861G>A, 4865G>A, 4866A>T, 4869A>C, 4871A>T, 4873_4875delCTC, 4876A>G, 4879G>T, 4881C>A, 4884G>A, 4886G>C, 4890T>C, 4893C>A, 4894G>T, 4895G>C, 4897C>A, 4898T>A, 4900C>A, 4902C>G, 4903T>A, 4905A>C, 4906G>T, 4907T>G, 4910C>A, 4911T>A, 4915C>T, 4921C>A, 4926T>A, 4928G>A, 4929T>C, 4930G>A, 4931A>C, 4932T>A, 4934G>A, 4935G>A, 4936A>C, 4938T>A, 4940C>A, 4941C>A, 4942T>A, 4944A>C, 4950A>G, 4954T>C, 4956A>T, 4959C>A, 4960C>T, 4961T>G, 4962C>T, 4963C>T, 4971T>C, 4972A>G, 4973T>C, 4974C>A, 4975T>A, 4976C>G, 4977C>T, 4979C>A, 4980T>A, 4982A>T, 4984T>A, 4986T>A, 4990A>G, 4992A>C, 4993A>C, 5001C>A, 5002C>A, 5003T>A, 5004T>G, 5007G>A, 5008T>A, 5009C>A, 5010A>T, 5012C>T, 5019G>A, 5020G>A, 5021T>G, 5022T>G, 5023A>C, 5024A>T, 5025A>T, 5026C>A, 5029A>C, 5030A>G, 5031G>A, 5035G>T, 5036A>C, 5037G>C, 5039T>A, 5040A>C, 5041A>C, 5042C>G, 5043T>A, 5044T>A, 5045G>T, 5047T>G, 5048C>G, 5049T>A, 5055T>A, 5058G>T, 5059A>G, 5062G>C, 5064C>T, 5067G>C, 5068G>A, 5071A>T, 5072C>G, 5076C>T, 5077A>C, 5078G>A, 5081A>G, 5082A>C, 5083A>C, 5088G>A, 5089A>C, 5090G>A, 5091T>A, 5092C>G, 5093T>A, 5095G>T, 5096T>G, 5097A>T, 5098A>T, 5099A>C, 5100A>C, 5102C>A, 5103T>A, 5110G>A, 5112C>G, 5115T>A, 5116T>A, 5117A>G, 5118C>A, 5119A>C, 5120A>T, 5124T>A, 5125T>G, 5126C>A, 5130A>G, 5133G>A, 5136T>C, 5139G>C, 5140C>T, 5141C>T, 5142T>A, 5143A>G, 5145C>T, 5146A>C, 5148C>A, 5149G>C, 5152T>A, 5153G>C, 5154C>A, 5160A>T, 5167G>T, 5182A>T, 5184C>A, 5185C>T, 5187C>A, 5188A>C, 5191G>A, 5193A>T, 5194A>G, 5196A>C, 5199T>A, 5200C>A, 5201C>A, 5202A>T, 5206_5217delGGTAAAGAGGTA, 5220A>T, 5223C>T, 5230G>A, 5233T>A, 5234C>G, 5235A>T, 5238A>C, 5241C>A, 5247A>T, 5248C>G, 5249C>A, 5251G>A, 5253T>C, 5256G>A, 5258A>C, 5259G>T, 5262T>A, 5266C>T, 5267A>C, 5268T>A, 5270G>C, 5271T>A, 5284A>T, 5286C>A, 5287C>T, 5289T>G, 5290T>G, 5295T>A, 5296A>G, 5298T>A, 5300A>G, 5301G>A, 5303C>G, 5304A>C, 5307C>A, 5308A>C, 5309A>G, 5311G>A, 5312C>A, 5313T>A, 5317A>T, 5318G>C, 5319A>T, 5324A>T, 5325T>C, 5326A>C, 5328C>A, 5331T>A, 5334T>A, 5335T>A, 5337T>G, 5338A>C, 5343T>C, 5344C>A, 5346T>C, 5347G>A, 5349A>C, 5352G>A, 5353A>T, 5355T>A, 5358T>C, 5364T>A, 5365A>C, 5367T>G, 5369C>T, 5370T>A, 5371G>T, 5373T>A, 5374T>G, 5375A>G, 5379T>G, 5380G>A, 5382G>A, 5383A>T, 5384G>T, 5385A>T, 5387C>A, 5388A>T, 5391T>A, 5392A>C, 5395G>C, 5398G>T, 5399G>C

##### CDS

| | Begin | End | Coverage | Score | Concordance | Matches | Identities | I/D/M/F* | Stop Codons |
|---|---|---|---|---|---|---|---|---|---|
| ORF_I | 111 | 212 | 31.0% | 332 | 49.3% | 101 (98.1%) | 44 (42.7%) | 1/1/0/0 | 0 |

Protein mutations: K113H (403A>C 405A>T), I115M (411A>G), D116N (412G>A), F117Y (416T>A), I118V (418A>G), I120L (424A>T 426A>G), S122G (430T>G 431C>G 432T>A), V123I (433G>A 435T>A), K124Q (436A>C), I127V (445A>G 447C>G), S129F (451A>T 452G>T), R132K (461G>A 462G>A), T133E (463A>G 464C>A), D136N (472G>A 474C>T), S137C (476C>G), S140I (485G>T 486T>A), V141I (487G>A), A142N (490G>A 491C>A 492A>C), L144S (496C>T 497T>C), R146E (502C>G 503G>A 504G>A), M148F (508A>T 510G>T), K149I (512A>T 513A>T), A151T (517G>A), K152R (521A>G), D153E (525C>A), A154G (527C>G), V155N (529G>A 530T>A 531C>T), F156L (532T>C 534C>T), G158I (538G>A 539G>T), K160E (544A>G), S164A (556T>G 558C>A), G166T (562G>A 563G>C 564G>T), I169L (571A>C 573C>T), N173Y (583A>T 585C>T), K175R (590A>G), I176Y (592A>T 593T>A 594C>T), S177C (596C>G), V178I (598G>A 600T>A), R181K (608G>A), P183A (613C>G), T184N (617C>A 618G>C), I185F (619A>T), K187E (625A>G 627G>A), T188A (628A>G 630T>A), T190S (635C>G 636A>T), A192del (640_642delGCT), F194_F195insQ (648_649insCAG), F195V (649T>G 651T>C), E196K (652G>A), K197R (656A>G), E198K (658G>A), E199D (663A>T), M201F (667A>T 669G>C), H202K (670C>A 672T>G), E203P (673G>C 674A>C 675G>A), P207I (685C>A 686C>T 687C>T), Y208M (688T>A 689A>T 690T>G), T209S (692C>G 693T>C), S211Y (698C>A 699C>T)

Codon mutations: AAA113CAT (403A>C 405A>T), ATA115ATG (411A>G), GAT116AAT (412G>A), TTT117TAT (416T>A), ATT118GTT (418A>G), ATA120TTG (424A>T 426A>G), TCT122GGA (430T>G 431C>G 432T>A), GTT123ATA (433G>A 435T>A), AAA124CAA (436A>C), ATT125ATC (441T>C), ATC127GTG (445A>G 447C>G), AGC129TTC (451A>T 452G>T), ACC130ACT (456C>T), TTT131TTC (459T>C), AGG132AAA (461G>A 462G>A), ACA133GAA (463A>G 464C>A), GGA134GGT (468A>T), GAC136AAT (472G>A 474C>T), TCT137TGT (476C>G), ATT139ATA (483T>A), AGT140ATA (485G>T 486T>A), GTA141ATA (487G>A), GCA142AAC (490G>A 491C>A 492A>C), CTC143CTT (495C>T), CTT144TCT (496C>T 497T>C), GAC145GAT (501C>T), CGG146GAA (502C>G 503G>A 504G>A), ATG148TTT (508A>T 510G>T), AAA149ATT (512A>T 513A>T), AAC150AAT (516C>T), GCT151ACT (517G>A), AAA152AGA (521A>G), GAC153GAA (525C>A), GCA154GGA (527C>G), GTC155AAT (529G>A 530T>A 531C>T), TTC156CTT (532T>C 534C>T), GGC157GGA (537C>A), GGA158ATA (538G>A 539G>T), AAA160GAA (544A>G), AAC162AAT (552C>T), CTT163TTA (553C>T 555T>A), TCC164GCA (556T>G 558C>A), GGG166ACT (562G>A 563G>C 564G>T), AAA167AAG (567A>G), CTT168TTA (568C>T 570T>A), ATC169CTT (571A>C 573C>T), ACC171ACA (579C>A), TAC172TAT (582C>T), AAC173TAT (583A>T 585C>T), CCA174CCC (588A>C), AAA175AGA (590A>G), ATC176TAT (592A>T 593T>A 594C>T), TCT177TGT (596C>G), GTT178ATA (598G>A 600T>A), AGT179TCA (601A>T 602G>C 603T>A), CTT180TTA (604C>T 606T>A), AGA181AAA (608G>A), GAC182GAT (612C>T), CCT183GCT (613C>G), ACG184AAC (617C>A 618G>C), ATT185TTT (619A>T), AAC186AAT (624C>T), AAG187GAA (625A>G 627G>A), ACT188GCA (628A>G 630T>A), TTA189TTG (633A>G), ACA190AGT (635C>G 636A>T), CTA191TTG (637C>T 639A>G), GCT192del (640_642delGCT), TTC194TTT (648C>T), TTC194_TTT195insCAG (648_649insCAG), TTT195GTC (649T>G 651T>C), GAA196AAA (652G>A), AAA197AGA (656A>G), GAA198AAA (658G>A), GAA199GAT (663A>T), TTG200TTA (666G>A), ATG201TTC (667A>T 669G>C), CAT202AAG (670C>A 672T>G), GAG203CCA (673G>C 674A>C 675G>A), GGA204GGT (678A>T), CCC207ATT (685C>A 686C>T 687C>T), TAT208ATG (688T>A 689A>T 690T>G), ACT209AGC (692C>G 693T>C), ATA210ATT (696A>T), TCC211TAT (698C>A 699C>T)

| | Begin | End | Coverage | Score | Concordance | Matches | Identities | I/D/M/F* | Stop Codons |
|---|---|---|---|---|---|---|---|---|---|
| ORF_V | 273 | 666 | 42.7% | 1104 | 53.9% | 298 (98.3%) | 150 (49.5%) | 0/5/0/0 | 0 |

Protein mutations: L273K (4222T>A 4223T>A), T275P (4228A>C 4230T>C), N276S (4232A>G), K277I (4235A>T 4236A>C), K280R (4244A>G 4245G>A), C281V (4246T>G 4247G>T), Q285L (4259A>T 4260G>A), S287N (4264T>A 4265C>A 4266A>T), P288E (4267C>G 4268C>A 4269T>G), R291K (4276C>A 4277G>A), E292T (4279G>A 4280A>C 4281A>T), T296I (4292C>T), E299Q (4300G>C), K303N (4314G>T), L304M (4315C>A 4317T>G), G305K (4318G>A 4319G>A 4320T>A), I306L (4321A>T 4323C>A), P309E (4330C>G 4331C>A 4332T>A), I327Q (4384A>C 4385T>A 4386C>A), K340E (4423A>G), D343K (4432G>A 4434C>A), H344N (4435C>A 4437T>C), T345C (4438A>T 4439C>G 4440A>T), K346I (4442A>T 4443G>T), G347F (4444G>T 4445G>T 4446A>T), L351F (4456C>T 4458A>T), L352I (4459C>A 4461T>C), E356T (4471G>A 4472A>C 4473G>T), Q357N (4474C>A 4476A>T), L359I (4480C>A 4482T>A), Q360N (4483C>A 4485A>T), R361L (4486A>C 4487G>T), I362V (4489A>G), G363K (4492G>A 4493G>A), G364V (4496G>T 4497T>A), F367Y (4505T>A 4506T>C), Y368F (4508A>T 4509C>T), S370K (4513T>A 4514C>A 4515C>A), V380I (4543G>A), A383S (4552G>T 4554T>A), P384E (4555C>G 4556C>A 4557A>G), T386S (4561A>T), Q388P (4568A>C 4569G>T), E486K (4861G>A), R487N (4865G>A 4866A>T), N489I (4871A>T), L490del (4873_4875delCTC), K491E (4876A>G), V492L (4879G>T 4881C>A), S494T (4886G>C), G497S (4894G>T 4895G>C), L498K (4897C>A 4898T>A), H499K (4900C>A 4902C>G), L500I (4903T>A 4905A>C), V501C (4906G>T 4907T>G), A502E (4910C>A 4911T>A), P504S (4915C>T), Q506K (4921C>A), S508N (4928G>A 4929T>C), D509T (4930G>A 4931A>C 4932T>A), R510K (4934G>A 4935G>A), N511Q (4936A>C 4938T>A), A512E (4940C>A 4941C>A), L513I (4942T>A 4944A>C), L519C (4960C>T 4961T>G 4962C>T), I523A (4972A>G 4973T>C 4974C>A), A525E (4979C>A 4980T>A), Y526F (4982A>T), F527I (4984T>A 4986T>A), K529D (4990A>G 4992A>C), I530L (4993A>C), N532K (5001C>A), L533K (5002C>A 5003T>A 5004T>G), S535N (5008T>A 5009C>A 5010A>T), P536L (5012C>T), V539R (5020G>A 5021T>G 5022T>G), K540L (5023A>C 5024A>T 5025A>T), L541I (5026C>A), K542R (5029A>C 5030A>G 5031G>A), E544S (5035G>T 5036A>C 5037G>C), I545N (5039T>A 5040A>C), T546R (5041A>C 5042C>G 5043T>A), W547M (5044T>A 5045G>T), S548G (5047T>G 5048C>G 5049T>A), E551D (5058G>T), K552E (5059A>G), D553H (5062G>C 5064C>T), E555K (5068G>A), T556C (5071A>T 5072C>G), R558Q (5077A>C 5078G>A), K559S (5081A>G 5082A>C), I560L (5083A>C), S562Q (5089A>C 5090G>A 5091T>A), L563E (5092C>G 5093T>A), V564C (5095G>T 5096T>G 5097A>T), K565S (5098A>T 5099A>C 5100A>C), T566K (5102C>A 5103T>A), D569K (5110G>A 5112C>G), Y571R (5116T>A 5117A>G 5118C>A), N572L (5119A>C 5120A>T), S574E (5125T>G 5126C>A), K578N (5139G>C), P579L (5140C>T 5141C>T 5142T>A), I580V (5143A>G 5145C>T), I581L (5146A>C 5148C>A), E582Q (5149G>C), C583T (5152T>A 5153G>C 5154C>A), D588Y (5167G>T), I593L (5182A>T 5184C>A), K595Q (5188A>C), A596T (5191G>A 5193A>T), K597D (5194A>G 5196A>C), P599N (5200C>A 5201C>A 5202A>T), G601_V604del (5206_5217delGGTAAAGAGGTA), A609T (5230G>A), K614N (5247A>T), P615E (5248C>G 5249C>A), A616T (5251G>A 5253T>C), K618T (5258A>C 5259G>T), N619K (5262T>A), H621S (5266C>T 5267A>C 5268T>A), S622T (5270G>C 5271T>A), I627L (5284A>T 5286C>A), S629A (5290T>G), I631V (5296A>G 5298T>A), K632R (5300A>G 5301G>A), A633G (5303C>G 5304A>C), K635R (5308A>C 5309A>G), A636K (5311G>A 5312C>A 5313T>A), R638S (5317A>T 5318G>C 5319A>T), Y640F (5324A>T 5325T>C), I641L (5326A>C 5328C>A), Y644K (5335T>A 5337T>G), K645Q (5338A>C), L647I (5344C>A 5346T>C), V648I (5347G>A 5349A>C), T650S (5353A>T 5355T>A), N654Q (5365A>C 5367T>G), A655V (5369C>T 5370T>A), A656S (5371G>T 5373T>A), Y657G (5374T>G 5375A>G), F658L (5379T>G), V659I (5380G>A 5382G>A), R660F (5383A>T 5384G>T 5385A>T), T661N (5387C>A 5388A>T), N662K (5391T>A), I663L (5392A>C), A664P (5395G>C), G665S (5398G>T 5399G>C)

| | Begin | End | Coverage | Score | Concordance | Matches | Identities | I/D/M/F* | Stop Codons |
|---|---|---|---|---|---|---|---|---|---|
| **NT** | **395** | **5402** | **15.5%** | **269** | **11.4%** | **1201 (98.3%)** | **682 (55.8%)** | **3/18** | |
| Codon mutations: | TTA273AAA (4222T>A 4223T>A), GAC274GAT (4227C>T), ACT275CCC (4228A>C 4230T>C), AAC276AGC (4232A>G), AAA277ATC (4235A>T 4236A>C), ATT279ATA (4242T>A), AAG280AGA (4244A>G 4245G>A), TGT281GTT (4246T>G 4247G>T), CAG285CTA (4259A>T 4260G>A), TCA287AAT (4264T>A 4265C>A 4266A>T), CCT288GAG (4267C>G 4268C>A 4269T>G), GAC290GAT (4275C>T), CGA291AAA (4276C>A 4277G>A), GAA292ACT (4279G>A 4280A>C 4281A>T), GAG293GAA (4284G>A), ACT296ATT (4292C>T), ATC298ATT (4299C>T), GAA299CAA (4300G>C), CTA301TTA (4306C>T), CTC302TTA (4309C>T 4311C>A), AAG303AAT (4314G>T), CTT304ATG (4315C>A 4317T>G), GGT305AAA (4318G>A 4319G>A 4320T>A), ATC306TTA (4321A>T 4323C>A), ATC307ATA (4326C>A), CCT309GAA (4330C>G 4331C>A 4332T>A), AGC310AGT (4335C>T), AAG311AAA (4338G>A), TCC315AGT (4348T>A 4349C>G 4350C>T), TCA316AGT (4351T>A 4352C>G 4353A>T), CCA317CCT (4356A>T), GCC318GCA (4359C>A), TTC319TTT (4362C>T), AGG322AGA (4371G>A), AAT323AAC (4374T>C), GCC325GCG (4380C>G), GAG326GAA (4383G>A), ATC327CAA (4384A>C 4385T>A 4386C>A), GGC330GGA (4395C>A), GCA332GCT (4401A>T), AGA333CGA (4402A>C), GTA335GTT (4410A>T), ATT336ATA (4413T>A), AAG340GAG (4423A>G), TTA341CTT (4426T>C 4428A>T), GAC343AAA (4432G>A 4434C>A), CAT344AAC (4435C>A 4437T>C), ACA345TGT (4438A>T 4439C>G 4440A>T), AAG346ATT (4442A>T 4443G>T), GGA347TTT (4444G>T 4445G>T 4446A>T), GAT348GAC (4449T>C), GGC349GGA (4452C>A), CTA351TTT (4456C>T 4458A>T), CTT352ATC (4459C>A 4461T>C), AAC354AAT (4467C>T), AAG355AAA (4470G>A), GAG356ACT (4471G>A 4472A>C 4473G>T), CAA357AAT (4474C>A 4476A>T), CTG358CTT (4479G>T), CTT359ATA (4480C>A 4482T>A), CAA360AAT (4483C>A 4485A>T), AGA361CTA (4486A>C 4487G>T), ATC362GTC (4489A>G), GGA363AAA (4492G>A 4493G>A), GGT364GTA (4496G>T 4497T>A), AAG365AAA (4500G>A), TTT367TAC (4505T>A 4506T>C), TAC368TTT (4508A>T 4509C>T), TCT369TCC (4512T>C), TCC370AAA (4513T>A 4514C>A 4515C>A), GAC372GAT (4521C>T), TCT375AGT (4528T>A 4529C>G), GTA380ATA (4543G>A), CGC381AGA (4546C>A 4548C>A), CTT382CTC (4551T>C), GCT383TCA (4552G>T 4554T>A), CCA384GAG (4555C>G 4556C>A 4557A>G), ACA386TCA (4561A>T), CAG388CCT (4568A>C 4569G>T), ACC390ACA (4575C>A), GCT391GCA (4578T>A), ATA485.CA (4859T>C), GAA486AAA (4861G>A), AGA487AAT (4865G>A 4866A>T), GGA488GGC (4869A>C), AAT489ATT (4871A>T), CTC490del (4873_4875delCTC), AAA491GAA (4876A>G), GTC492TTA (4879G>T 4881C>A), CAG493CAA (4884G>A), AGT494ACT (4886G>C), CAT495CAC (4890T>C), ATC496ATA (4893C>A), GGA497TCA (4894G>T 4895G>C), CTA498AAA (4897C>A 4898T>A), CAC499AAG (4900C>A 4902C>G), TTA500ATC (4903T>A 4905A>C), GTT501TGT (4906G>T 4907T>G), GCT502GAA (4910C>A 4911T>A), CCA504TCA (4915C>T), CAA506AAA (4921C>A), CTT507CTA (4926T>A), AGT508AAC (4928G>A 4929T>C), GAT509ACA (4930G>A 4931A>C 4932T>A), AGG510AAA (4934G>A 4935G>A), AAT511CAA (4936A>C 4938T>A), GCC512GAA (4940C>A 4941C>A), TTA513ATC (4942T>A 4944A>C), AGA515AGG (4950A>G), TTA517CTT (4954T>C 4956A>T), GGC518GGA (4959C>A), CTC519TGT (4960C>T 4961T>G 4962C>T), CTA520TTA (4963C>T), TAT522TAC (4971T>C), ATC523GCA (4972A>G 4973T>C 4974C>A), TCC524AGT (4975T>A 4976C>G 4977C>T), GCT525GAA (4979C>A 4980T>A), TAT526TTT (4982A>T), TTT527ATA (4984T>A 4986T>A), AAA529GAC (4990A>G 4992A>C), ATA530CTA (4993A>C), AAC532AAA (5001C>A), CTT533AAG (5002C>A 5003T>A 5004T>G), AGG534AGA (5007G>A), TCA535AAT (5008T>A 5009C>A 5010A>T), CCT536CTT (5012C>T), CAG538CAA (5019G>A), GTT539AGG (5020G>A 5021T>G 5022T>G), AAA540CTT (5023A>C 5024A>T 5025A>T), CTT541ATT (5026C>A), AAG542CGA (5029A>C 5030A>G 5031G>A), GAG544TCC (5035G>T 5036A>C 5037G>C), ATA545AAC (5039T>A 5040A>C), ACT546CGA (5041A>C 5042C>G 5043T>A), TGG547ATG (5044T>A 5045G>T), TCT548GGA (5047T>G 5048C>G 5049T>A), ACT550ACA (5055T>A), GAG551GAT (5058G>T), AAA552GAA (5059A>G), GAC553CAT (5062G>C 5064C>T), ACG554ACC (5067G>C), GAA555AAA (5068G>A), ACT556TGT (5071A>T 5072C>G), GTC557GTT (5076C>T), AGA558CAA (5077A>C 5078G>A), AAA559AGC (5081A>G 5082A>C), ATA560CTA (5083A>C), AAG561AAA (5088G>A), AGT562CAA (5089A>C 5090G>A 5091T>A), CTA563GAA (5092C>G 5093T>A), GTA564TGT (5095G>T 5096T>G 5097A>T), AAA565TCC (5098A>T 5099A>C 5100A>C), ACT566AAA (5102C>A 5103T>A), GAC569AAG (5110G>A 5112C>G), CTT570CTA (5115T>A), TAC571AGA (5116T>A 5117A>G 5118C>A), AAT572CTT (5119A>C 5120A>T), CCT573CCA (5124T>A), TCA574GAA (5125T>G 5126C>A), CCA575CCG (5130A>G), GAG576GAA (5133G>A), GAT577GAC (5136T>C), AAG578AAC (5139G>C), CCT579TTA (5140C>T 5141C>T 5142T>A), ATC580GTT (5143A>G 5145C>T), ATC581CTA (5146A>C 5148C>A), GAA582CAA (5149G>C), TGC583ACA (5152T>A 5153G>C 5154C>A), GCA585GCT (5160A>T), GAC588TAC (5167G>T), ATC593TTA (5182A>T 5184C>A), CTC594TTA (5185C>T 5187C>A), AAA595CAA (5188A>C), GCA596ACT (5191G>A 5193A>T), AAA597GAC (5194A>G 5196A>C), CTT598CTA (5199T>A), CCA599AAT (5200C>A 5201C>A 5202A>T), GGT601_GTA604del (5206_5217delGGTAAAGAGGTA), ATA605ATT (5220A>T), TGC606TGT (5223C>T), GCT609ACT (5230G>A), TCA610AGT (5233T>A 5234C>G 5235A>T), GGA611GGC (5238A>C), ACC612ACA (5241C>A), AAA614AAT (5247A>T), CCA615GAA (5248C>G 5249C>A), GCT616ACC (5251G>A 5253T>C), GAG617GAA (5256G>A), AAG618ACT (5258A>C 5259G>T), AAT619AAA (5262T>A), CAT621TCA (5266C>T 5267A>C 5268T>A), AGT622ACA (5270G>C 5271T>A), ATC627TTA (5284A>T 5286C>A), CTT628TTG (5287C>T 5289T>G), TCC629GCC (5290T>G), ATT630ATA (5295T>A), ATT631GTA (5296A>G 5298T>A), AAG632AGA (5300A>G 5301G>A), GCA633GGC (5303C>G 5304A>C), ATC634ATA (5307C>A), AAA635CGA (5308A>C 5309A>G), GCT636AAA (5311G>A 5312C>A 5313T>A), AGA638TCT (5317A>T 5318G>C 5319A>T), TAT640TTC (5324A>T 5325T>C), ATC641CTA (5326A>C 5328C>A), CTT642CTA (5331T>A), CCT643CCA (5334T>A), TAT644AAG (5335T>A 5337T>G), AAA645CAA (5338A>C), TTT646TTC (5343T>C), CTT647ATC (5344C>A 5346T>C), GTA648ATC (5347G>A 5349A>C), AGG649AGA (5352G>A), ACT650TCA (5353A>T 5355T>A), GAT651GAC (5358T>C), ACT653ACA (5364T>A), AAT654CAG (5365A>C 5367T>G), GCT655GTA (5369C>T 5370T>A), GCT656TCA (5371G>T 5373T>A), TAT657GGT (5374T>G 5375A>G), TTT658TTG (5379T>G), GTG659ATA (5380G>A 5382G>A), AGA660TTT (5383A>T 5384G>T 5385A>T), ACA661AAT (5387C>A 5388A>T), AAT662AAA (5391T>A), ATT663CTT (5392A>C), GCA664CCA (5395G>C), GGT665TCT (5398G>T 5399G>C) | | | | | | | | |

#### Proteins

| | Begin | End | Coverage | Score | Concordance | Matches | Identities | I/D/M/F* | Stop Codons |
|---|---|---|---|---|---|---|---|---|---|
| hypothetical protein (NP_043929.1) | 111 | 212 | 31.0% | 332 | 49.3% | 101 (98.1%) | 44 (42.7%) | 1/1/0/0 | 0 |
| Protein mutations: | K113H (403A>C 405A>T), I115M (411A>G), D116N (412G>A), F117Y (416T>A), I118V (418A>G), I120L (424A>T 426A>G), S122G (430T>G 431C>G 432T>A), V123I (433G>A 435T>A), K124Q (436A>C), I127V (445A>G 447C>G), S129F (451A>T 452G>T), R132K (461G>A 462G>A), T133E (463A>G 464C>A), D136N (472G>A 474C>T), S137C (476C>G), S140I (485G>T 486T>A), V141I (487G>A), A142N (490G>A 491C>A 492A>C), L144S (496C>T 497T>C), R146E (502C>G 503G>A 504G>A), M148F (508A>T 510G>T), K149I (512A>T 513A>T), A151T (517G>A), K152R (521A>G), D153E (525C>A), A154G (527C>G), V155N (529G>A 530T>A 531C>T), F156L (532T>C 534C>T), G158I (538G>A 539G>T), K160E (544A>G), S164A (556T>G 558C>A), G166T (562G>A 563G>C 564G>T), I169L (571A>C 573C>T), N173Y (583A>T 585C>T), K175R (590A>G), I176Y (592A>T 593T>A 594C>T), S177C (596C>G), V178I (598G>A 600T>A), R181K (608G>A), P183A (613C>G), T184N (617C>A 618G>C), I185F (619A>T), K187E (625A>G 627G>A), T188A (628A>G 630T>A), T190S (635C>G 636A>T), A192del (640_642delGCT), F194_F195insQ (648_649insCAG), F195V (649T>G 651T>C), E196K (652G>A), K197R (656A>G), E198K (658G>A), E199D (663A>T), M201F (667A>T 669G>C), H202K (670C>A 672T>G), E203P (673G>C 674A>C 675G>A), P207I (685C>A 686C>T 687C>T), Y208M (688T>A 689A>T 690T>G), T209S (692C>G 693T>C), S211Y (698C>A 699C>T) | | | | | | | | |
| Codon mutations: | AAA113CAT (403A>C 405A>T), ATA115ATG (411A>G), GAT116AAT (412G>A), TTT117TAT (416T>A), ATT118GTT (418A>G), ATA120TTG (424A>T 426A>G), TCT122GGA (430T>G 431C>G 432T>A), GTT123ATA (433G>A 435T>A), AAA124CAA (436A>C), ATT125ATC (441T>C), ATC127GTG (445A>G 447C>G), AGC129TTC (451A>T 452G>T), ACC130ACT (456C>T), TTT131TTC (459T>C), AGG132AAA (461G>A 462G>A), ACA133GAA (463A>G 464C>A), GGA134GGT (468A>T), GAC136AAT (472G>A 474C>T), TCT137TGT (476C>G), ATT139ATA (483T>A), AGT140ATA (485G>T 486T>A), GTA141ATA (487G>A), GCA142AAC (490G>A 491C>A 492A>C), CTC143CTT (495C>T), CTT144TCT (496C>T 497T>C), GAC145GAT (501C>T), CGG146GAA (502C>G 503G>A 504G>A), ATG148TTT (508A>T 510G>T), AAA149ATT (512A>T 513A>T), AAC150AAT (516C>T), GCT151ACT (517G>A), AAA152AGA (521A>G), GAC153GAA (525C>A), GCA154GGA (527C>G), GTC155AAT (529G>A 530T>A 531C>T), TTC156CTT (532T>C 534C>T), GGC157GGA (537C>A), GGA158ATA (538G>A 539G>T), AAA160GAA (544A>G), AAC162AAT (552C>T), CTT163TTA (553C>T 555T>A), TCC164GCA (556T>G 558C>A), GGG166ACT (562G>A 563G>C 564G>T), AAA167AAG (567A>G), CTT168TTA (568C>T 570T>A), ATC169CTT (571A>C 573C>T), ACC171ACA (579C>A), TAC172TAT (582C>T), AAC173TAT (583A>T 585C>T), CCA174CCC (588A>C), AAA175AGA (590A>G), ATC176TAT (592A>T 593T>A 594C>T), TCT177TGT (596C>G), GTT178ATA (598G>A 600T>A), AGT179TCA (601A>T 602G>C 603T>A), CTT180TTA (604C>T 606T>A), AGA181AAA (608G>A), GAC182GAT (612C>T), CCT183GCT (613C>G), ACG184AAC (617C>A 618G>C), ATT185TTT (619A>T), AAC186AAT (624C>T), AAG187GAA (625A>G 627G>A), ACT188GCA (628A>G 630T>A), TTA189TTG (633A>G), ACA190AGT (635C>G 636A>T), CTA191TTG (637C>T 639A>G), GCT192del (640_642delGCT), TTC194TTT (648C>T), TTC194_TTT195insCAG (648_649insCAG), TTT195GTC (649T>G 651T>C), GAA196AAA (652G>A), AAA197AGA (656A>G), GAA198AAA (658G>A), GAA199GAT (663A>T), TTG200TTA (666G>A), ATG201TTC (667A>T 669G>C), CAT202AAG (670C>A 672T>G), GAG203CCA (673G>C 674A>C 675G>A), GGA204GGT (678A>T), CCC207ATT (685C>A 686C>T 687C>T), TAT208ATG (688T>A 689A>T 690T>G), ACT209AGC (692C>G 693T>C), ATA210ATT (696A>T), TCC211TAT (698C>A 699C>T) | | | | | | | | |
| hypothetical protein (NP_043933.1) | 273 | 666 | 42.7% | 1104 | 53.9% | 298 (98.3%) | 150 (49.5%) | 0/5/0/0 | 0 |
| Protein mutations: | L273K (4222T>A 4223T>A), T275P (4228A>C 4230T>C), N276S (4232A>G), K277I (4235A>T 4236A>C), K280R (4244A>G 4245G>A), C281V (4246T>G 4247G>T), Q285L (4259A>T 4260G>A), S287N (4264T>A 4265C>A 4266A>T), P288E (4267C>G 4268C>A 4269T>G), R291K (4276C>A 4277G>A), E292T (4279G>A 4280A>C 4281A>T), T296I (4292C>T), E299Q (4300G>C), K303N (4314G>T), L304M (4315C>A 4317T>G), G305K (4318G>A 4319G>A 4320T>A), I306L (4321A>T 4323C>A), P309E (4330C>G 4331C>A 4332T>A), I327Q (4384A>C 4385T>A 4386C>A), K340E (4423A>G), D343K (4432G>A 4434C>A), H344N (4435C>A 4437T>C), T345C (4438A>T 4439C>G 4440A>T), K346I (4442A>T 4443G>T), G347F (4444G>T 4445G>T 4446A>T), L351F (4456C>T 4458A>T), L352I (4459C>A 4461T>C), E356T (4471G>A 4472A>C 4473G>T), Q357N (4474C>A 4476A>T), L359I (4480C>A 4482T>A), Q360N (4483C>A 4485A>T), R361L (4486A>C 4487G>T), I362V (4489A>G), G363K (4492G>A 4493G>A), G364V (4496G>T 4497T>A), F367Y (4505T>A 4506T>C), Y368F (4508A>T 4509C>T), S370K (4513T>A 4514C>A 4515C>A), V380I (4543G>A), A383S (4552G>T 4554T>A), P384E (4555C>G 4556C>A 4557A>G), T386S (4561A>T), Q388P (4568A>C 4569G>T), E486K (4861G>A), R487N (4865G>A 4866A>T), N489I (4871A>T), L490del (4873_4875delCTC), K491E (4876A>G), V492L (4879G>T 4881C>A), S494T (4886G>C), G497S (4894G>T 4895G>C), L498K (4897C>A 4898T>A), H499K (4900C>A 4902C>G), L500I (4903T>A 4905A>C), V501C (4906G>T 4907T>G), A502E (4910C>A 4911T>A), P504S (4915C>T), Q506K (4921C>A), S508N (4928G>A 4929T>C), D509T (4930G>A 4931A>C 4932T>A), R510K (4934G>A 4935G>A), N511Q (4936A>C 4938T>A), A512E (4940C>A 4941C>A), L513I (4942T>A 4944A>C), L519C (4960C>T 4961T>G 4962C>T), I523A (4972A>G 4973T>C 4974C>A), A525E (4979C>A 4980T>A), Y526F (4982A>T), F527I (4984T>A 4986T>A), K529D (4990A>G 4992A>C), I530L (4993A>C), N532K (5001C>A), L533K (5002C>A 5003T>A 5004T>G), S535N (5008T>A 5009C>A 5010A>T), P536L (5012C>T), V539R (5020G>A 5021T>G 5022T>G), K540L (5023A>C 5024A>T 5025A>T), L541I (5026C>A), K542R (5029A>C 5030A>G 5031G>A), E544S (5035G>T 5036A>C 5037G>C), I545N (5039T>A 5040A>C), T546R (5041A>C 5042C>G 5043T>A), W547M (5044T>A 5045G>T), S548G (5047T>G 5048C>G 5049T>A), E551D (5058G>T), K552E (5059A>G), D553H (5062G>C 5064C>T), E555K (5068G>A), T556C (5071A>T 5072C>G), R558Q (5077A>C 5078G>A), K559S (5081A>G 5082A>C), I560L (5083A>C), S562Q (5089A>C 5090G>A 5091T>A), L563E (5092C>G 5093T>A), V564C (5095G>T 5096T>G 5097A>T), K565S (5098A>T 5099A>C 5100A>C), T566K (5102C>A 5103T>A), D569K (5110G>A 5112C>G), Y571R (5116T>A 5117A>G 5118C>A), N572L (5119A>C 5120A>T), S574E (5125T>G 5126C>A), K578N (5139G>C), P579L (5140C>T 5141C>T 5142T>A), I580V (5143A>G 5145C>T), I581L (5146A>C 5148C>A), E582Q (5149G>C), C583T (5152T>A 5153G>C 5154C>A), D588Y (5167G>T), I593L (5182A>T 5184C>A), K595Q (5188A>C), A596T (5191G>A 5193A>T), K597D (5194A>G 5196A>C), P599N (5200C>A 5201C>A 5202A>T), G601_V604del (5206_5217delGGTAAAGAGGTA), A609T (5230G>A), K614N (5247A>T), P615E (5248C>G 5249C>A), A616T (5251G>A 5253T>C), K618T (5258A>C 5259G>T), N619K (5262T>A), H621S (5266C>T 5267A>C 5268T>A), S622T (5270G>C 5271T>A), I627L (5284A>T 5286C>A), S629A (5290T>G), I631V (5296A>G 5298T>A), K632R (5300A>G 5301G>A), A633G (5303C>G 5304A>C), K635R (5308A>C 5309A>G), A636K (5311G>A 5312C>A 5313T>A), R638S (5317A>T 5318G>C 5319A>T), Y640F (5324A>T 5325T>C), I641L (5326A>C 5328C>A), Y644K (5335T>A 5337T>G), K645Q (5338A>C), L647I (5344C>A 5346T>C), V648I (5347G>A 5349A>C), T650S (5353A>T 5355T>A), N654Q (5365A>C 5367T>G), A655V (5369C>T 5370T>A), A656S (5371G>T 5373T>A), Y657G (5374T>G 5375A>G), F658L (5379T>G), V659I (5380G>A 5382G>A), R660F (5383A>T 5384G>T 5385A>T), T661N (5387C>A 5388A>T), N662K (5391T>A), I663L (5392A>C), A664P (5395G>C), G665S (5398G>T 5399G>C) | | | | | | | | |

| | Begin | End | Coverage | Score | Concordance | Matches | Identities | I/D/M/F* | Stop Codons |
|---|---|---|---|---|---|---|---|---|---|
| **NT** | **395** | **5402** | **15.5%** | **269** | **11.4%** | **1201 (98.3%)** | **682 (55.8%)** | **3/18** | |
| Codon mutations: | TTA273AAA (4222T>A 4223T>A), GAC274GAT (4227C>T), ACT275CCC (4228A>C 4230T>C), AAC276AGC (4232A>G), AAA277ATC (4235A>T 4236A>C), ATT279ATA (4242T>A), AAG280AGA (4244A>G 4245G>A), TGT281GTT (4246T>G 4247G>T), CAG285CTA (4259A>T 4260G>A), TCA287AAT (4264T>A 4265C>A 4266A>T), CCT288GAG (4267C>G 4268C>A 4269T>G), GAC290GAT (4275C>T), CGA291AAA (4276C>A 4277G>A), GAA292ACT (4279G>A 4280A>C 4281A>T), GAG293GAA (4284G>A), ACT296ATT (4292C>T), ATC298ATT (4299C>T), GAA299CAA (4300G>C), CTA301TTA (4306C>T), CTC302TTA (4309C>T 4311C>A), AAG303AAT (4314G>T), CTT304ATG (4315C>A 4317T>G), GGT305AAA (4318G>A 4319G>A 4320T>A), ATC306TTA (4321A>T 4323C>A), ATC307ATA (4326C>A), CCT309GAA (4330C>G 4331C>A 4332T>A), AGC310AGT (4335C>T), AAG311AAA (4338G>A), TCC315AGT (4348T>A 4349C>G 4350C>T), TCA316AGT (4351T>A 4352C>G 4353A>T), CCA317CCT (4356A>T), GCC318GCA (4359C>A), TTC319TTT (4362C>T), AGG322AGA (4371G>A), AAT323AAC (4374T>C), GCC325GCG (4380C>G), GAG326GAA (4383G>A), ATC327CAA (4384A>C 4385T>A 4386C>A), GGC330GGA (4395C>A), GCA332GCT (4401A>T), AGA333CGA (4402A>C), GTA335GTT (4410A>T), ATT336ATA (4413T>A), AAG340GAG (4423A>G), TTA341CTT (4426T>C 4428A>T), GAC343AAA (4432G>A 4434C>A), CAT344AAC (4435C>A 4437T>C), ACA345TGT (4438A>T 4439C>G 4440A>T), AAG346ATT (4442A>T 4443G>T), GGA347TTT (4444G>T 4445G>T 4446A>T), GAT348GAC (4449T>C), GGC349GGA (4452C>A), CTA351TTT (4456C>T 4458A>T), CTT352ATC (4459C>A 4461T>C), AAC354AAT (4467C>T), AAG355AAA (4470G>A), GAG356ACT (4471G>A 4472A>C 4473G>T), CAA357AAT (4474C>A 4476A>T), CTG358CTT (4479G>T), CTT359ATA (4480C>A 4482T>A), CAA360AAT (4483C>A 4485A>T), AGA361CTA (4486A>C 4487G>T), ATC362GTC (4489A>G), GGA363AAA (4492G>A 4493G>A), GGT364GTA (4496G>T 4497T>A), AAG365AAA (4500G>A), TTT367TAC (4505T>A 4506T>C), TAC368TTT (4508A>T 4509C>T), TCT369TCC (4512T>C), TCC370AAA (4513T>A 4514C>A 4515C>A), GAC372GAT (4521C>T), TCT375AGT (4528T>A 4529C>G), GTA380ATA (4543G>A), CGC381AGA (4546C>A 4548C>A), CTT382CTC (4551T>C), GCT383TCA (4552G>T 4554T>A), CCA384GAG (4555C>G 4556C>A 4557A>G), ACA386TCA (4561A>T), CAG388CCT (4568A>C 4569G>T), ACC390ACA (4575C>A), GCT391GCA (4578T>A), ATA485.CA (4859T>C), GAA486AAA (4861G>A), AGA487AAT (4865G>A 4866A>T), GGA488GGC (4869A>C), AAT489ATT (4871A>T), CTC490del (4873_4875delCTC), AAA491GAA (4876A>G), GTC492TTA (4879G>T 4881C>A), CAG493CAA (4884G>A), AGT494ACT (4886G>C), CAT495CAC (4890T>C), ATC496ATA (4893C>A), GGA497TCA (4894G>T 4895G>C), CTA498AAA (4897C>A 4898T>A), CAC499AAG (4900C>A 4902C>G), TTA500ATC (4903T>A 4905A>C), GTT501TGT (4906G>T 4907T>G), GCT502GAA (4910C>A 4911T>A), CCA504TCA (4915C>T), CAA506AAA (4921C>A), CTT507CTA (4926T>A), AGT508AAC (4928G>A 4929T>C), GAT509ACA (4930G>A 4931A>C 4932T>A), AGG510AAA (4934G>A 4935G>A), AAT511CAA (4936A>C 4938T>A), GCC512GAA (4940C>A 4941C>A), TTA513ATC (4942T>A 4944A>C), AGA515AGG (4950A>G), TTA517CTT (4954T>C 4956A>T), GGC518GGA (4959C>A), CTC519TGT (4960C>T 4961T>G 4962C>T), CTA520TTA (4963C>T), TAT522TAC (4971T>C), ATC523GCA (4972A>G 4973T>C 4974C>A), TCC524AGT (4975T>A 4976C>G 4977C>T), GCT525GAA (4979C>A 4980T>A), TAT526TTT (4982A>T), TTT527ATA (4984T>A 4986T>A), AAA529GAC (4990A>G 4992A>C), ATA530CTA (4993A>C), AAC532AAA (5001C>A), CTT533AAG (5002C>A 5003T>A 5004T>G), AGG534AGA (5007G>A), TCA535AAT (5008T>A 5009C>A 5010A>T), CCT536CTT (5012C>T), CAG538CAA (5019G>A), GTT539AGG (5020G>A 5021T>G 5022T>G), AAA540CTT (5023A>C 5024A>T 5025A>T), CTT541ATT (5026C>A), AAG542CGA (5029A>C 5030A>G 5031G>A), GAG544TCC (5035G>T 5036A>C 5037G>C), ATA545AAC (5039T>A 5040A>C), ACT546CGA (5041A>C 5042C>G 5043T>A), TGG547ATG (5044T>A 5045G>T), TCT548GGA (5047T>G 5048C>G 5049T>A), ACT550ACA (5055T>A), GAG551GAT (5058G>T), AAA552GAA (5059A>G), GAC553CAT (5062G>C 5064C>T), ACG554ACC (5067G>C), GAA555AAA (5068G>A), ACT556TGT (5071A>T 5072C>G), GTC557GTT (5076C>T), AGA558CAA (5077A>C 5078G>A), AAA559AGC (5081A>G 5082A>C), ATA560CTA (5083A>C), AAG561AAA (5088G>A), AGT562CAA (5089A>C 5090G>A 5091T>A), CTA563GAA (5092C>G 5093T>A), GTA564TGT (5095G>T 5096T>G 5097A>T), AAA565TCC (5098A>T 5099A>C 5100A>C), ACT566AAA (5102C>A 5103T>A), GAC569AAG (5110G>A 5112C>G), CTT570CTA (5115T>A), TAC571AGA (5116T>A 5117A>G 5118C>A), AAT572CTT (5119A>C 5120A>T), CCT573CCA (5124T>A), TCA574GAA (5125T>G 5126C>A), CCA575CCG (5130A>G), GAG576GAA (5133G>A), GAT577GAC (5136T>C), AAG578AAC (5139G>C), CCT579TTA (5140C>T 5141C>T 5142T>A), ATC580GTT (5143A>G 5145C>T), ATC581CTA (5146A>C 5148C>A), GAA582CAA (5149G>C), TGC583ACA (5152T>A 5153G>C 5154C>A), GCA585GCT (5160A>T), GAC588TAC (5167G>T), ATC593TTA (5182A>T 5184C>A), CTC594TTA (5185C>T 5187C>A), AAA595CAA (5188A>C), GCA596ACT (5191G>A 5193A>T), AAA597GAC (5194A>G 5196A>C), CTT598CTA (5199T>A), CCA599AAT (5200C>A 5201C>A 5202A>T), GGT601_GTA604del (5206_5217delGGTAAAGAGGTA), ATA605ATT (5220A>T), TGC606TGT (5223C>T), GCT609ACT (5230G>A), TCA610AGT (5233T>A 5234C>G 5235A>T), GGA611GGC (5238A>C), ACC612ACA (5241C>A), AAA614AAT (5247A>T), CCA615GAA (5248C>G 5249C>A), GCT616ACC (5251G>A 5253T>C), GAG617GAA (5256G>A), AAG618ACT (5258A>C 5259G>T), AAT619AAA (5262T>A), CAT621TCA (5266C>T 5267A>C 5268T>A), AGT622ACA (5270G>C 5271T>A), ATC627TTA (5284A>T 5286C>A), CTT628TTG (5287C>T 5289T>G), TCC629GCC (5290T>G), ATT630ATA (5295T>A), ATT631GTA (5296A>G 5298T>A), AAG632AGA (5300A>G 5301G>A), GCA633GGC (5303C>G 5304A>C), ATC634ATA (5307C>A), AAA635CGA (5308A>C 5309A>G), GCT636AAA (5311G>A 5312C>A 5313T>A), AGA638TCT (5317A>T 5318G>C 5319A>T), TAT640TTC (5324A>T 5325T>C), ATC641CTA (5326A>C 5328C>A), CTT642CTA (5331T>A), CCT643CCA (5334T>A), TAT644AAG (5335T>A 5337T>G), AAA645CAA (5338A>C), TTT646TTC (5343T>C), CTT647ATC (5344C>A 5346T>C), GTA648ATC (5347G>A 5349A>C), AGG649AGA (5352G>A), ACT650TCA (5353A>T 5355T>A), GAT651GAC (5358T>C), ACT653ACA (5364T>A), AAT654CAG (5365A>C 5367T>G), GCT655GTA (5369C>T 5370T>A), GCT656TCA (5371G>T 5373T>A), TAT657GGT (5374T>G 5375A>G), TTT658TTG (5379T>G), GTG659ATA (5380G>A 5382G>A), AGA660TTT (5383A>T 5384G>T 5385A>T), ACA661AAT (5387C>A 5388A>T), AAT662AAA (5391T>A), ATT663CTT (5392A>C), GCA664CCA (5395G>C), GGT665TCT (5398G>T 5399G>C) | | | | | | | | |

*: Inserts / Deletes / Misaligned / Frameshifts

#### Analysis details

This analysis was performed with panviral2.64

### NGS Details (UN59): Dioscovirus dioscoreae

#### Assembly

| | |
|---|---|
| **Coverage Length** | 1200 (2 contig(s)) |
| **Depth Of Coverage** | 8.0 |
| **Number Of Reads** | 81 |
| **Reads Per Million** | 1.76 rpm (after QC) |
| **Ambiguities** | 0 |
| **Assembly Method** | de novo + reference guided assembly |
| **Consensus Caller** | Bcf Tools |

#### Coverage Map

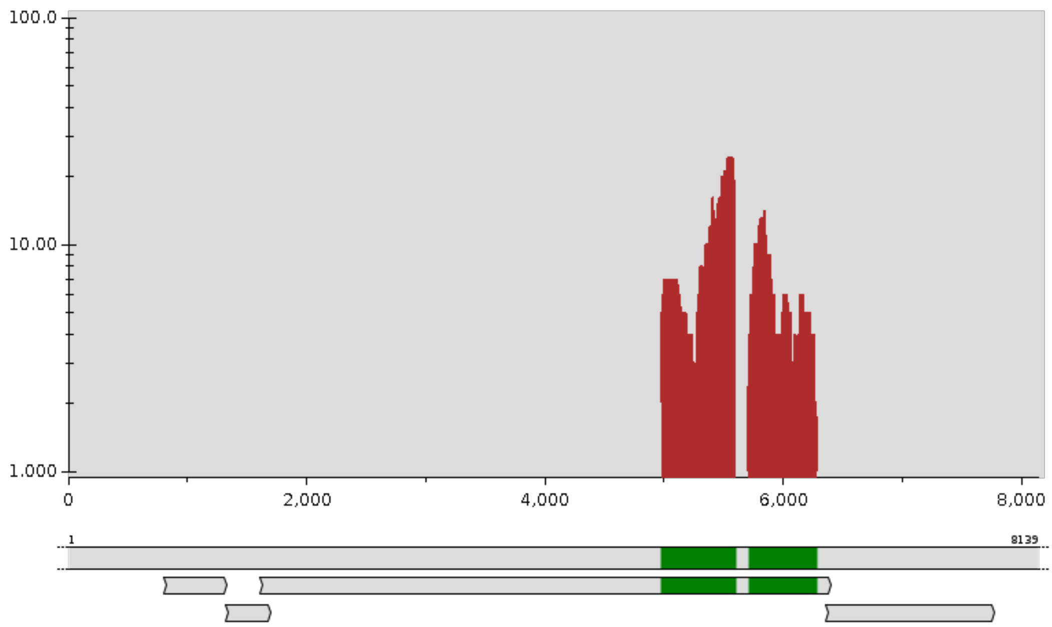

#### Assignment

| | |
|---|---|
| **Type** | Dioscovirus dioscoreae (Taxonomy ID: 3052184) |
| **Reference Genome** | NC_040712.1 |
| **NT Identity (%)** | 61.9917 |
| **AA Identity (%)** | 52.2388 |
| **Number Of Stop Codons** | 1 |
| **Number Of CDS** | 4 |

#### Alignment

| | |
|---|---|
| **Alignment Score** | 547.0 (NT) + 1411.0 (AA) = 1958.0 |
| **Concordance (%)** | 39.0292 |

| Alignment Method | Global, seeded, nucleotide + amino acids (AGA) |
|---|---|

#### Genome Region

Sequence starts at position 4979 and ends at position 6282 relative to NC_040712.1 reference sequence.

#### Alignment Detailed Statistics

| | Begin | End | Coverage | Score | Concordance | Matches | Identities | I/D/M/F* | Stop Codons |
|---|---|---|---|---|---|---|---|---|---|
| **NT** | **4979** | **6282** | **14.7%** | **547** | **23.1%** | **1196 (98.9%)** | **747 (61.8%)** | **9/4** | |
| Mutations: | 4983C>A, 4986C>A, 4988T>A, 4989A>T, 4990A>T, 4992T>C, 4994G>A, 4997A>T, 5001A>T, 5003C>A, 5004C>G, 5009T>A, 5012T>A, 5015G>A, 5016A>G, 5018T>A, 5019G>A, 5021C>A, 5024A>C, 5025T>C, 5029A>T, 5030G>A, 5031T>C, 5038A>G, 5040A>G, 5041A>C, 5042G>C, 5046C>A, 5047A>G, 5048G>A, 5051T>A, 5053G>T, 5054A>G, 5057C>T, 5063G>A, 5066C>T, 5069C>A, 5078T>A, 5081T>A, 5082C>T, 5083T>A, 5091C>A, 5093G>A, 5094G>A, 5095A>T, 5096C>A, 5097A>G, 5098A>C, 5099G>A, 5100C>G, 5102T>A, 5105T>A, 5106A>G, 5107T>A, 5109C>G, 5112T>A, 5115C>A, 5116C>A, 5117A>T, 5118G>C, 5121A>G, 5122T>A, 5123G>C, 5124G>A, 5125C>T, 5126G>A, 5127A>G, 5132A>T, 5134A>T, 5136G>C, 5138A>G, 5140A>T, 5141G>A, 5144T>A, 5147T>A, 5149C>A, 5157C>T, 5159G>A, 5161C>A, 5162A>T, 5163C>T, 5165C>A, 5166A>G, 5167A>G, 5168G>A, 5171T>A, 5174C>A, 5178C>A, 5179C>G, 5180T>A, 5183C>A, 5184C>A, 5185A>C, 5187A>T, 5188G>C, 5190C>A, 5192C>A, 5195T>C, 5198G>A, 5199A>T, 5207T>A, 5213T>G, 5216C>A, 5219C>A, 5220delA, 5222A>C, 5226T>A, 5227C>A, 5228T>C, 5232C>A, 5233A>T, 5237A>C, 5243T>A, 5247A>G, 5248G>C, 5249T>C, 5250C>A, 5252T>A, 5258A>T, 5265T>A, 5266A>G, 5268A>C, 5270G>A, 5273_5274insGAA, 5276A>T, 5288A>T, 5289T>A, 5290A>G, 5291C>A, 5293A>C, 5297C>T, 5298C>G, 5299A>C, 5303C>T, 5305C>A, 5307C>T, 5309C>A, 5310C>T, 5313A>G, 5314G>A, 5317T>A, 5318T>A, 5322T>A, 5324T>A, 5325C>T, 5327C>G, 5328C>A, 5330A>T, 5331C>T, 5332T>G, 5335A>G, 5340A>C, 5344A>G, 5345C>T, 5348G>A, 5350T>A, 5351T>A, 5352G>A, 5356A>T, 5357C>T, 5360C>T, 5361A>G, 5370C>T, 5371T>G, 5372A>T, 5375G>A, 5378A>G, 5383T>A, 5385C>T, 5387C>G, 5390G>A, 5391A>G, 5393T>A, 5395T>A, 5396G>A, 5400G>C, 5402T>C, 5406C>G, 5408G>C, 5429T>C, 5430G>A, 5431T>C, 5432C>T, 5435C>T, 5437C>T, 5438A>T, 5439C>G, 5444T>A, 5457A>T, 5461T>C, 5462A>C, 5471C>T, 5474A>T, 5475C>T, 5477C>A, 5482A>C, 5483T>A, 5486A>G, 5492A>T, 5493G>A, 5498C>T, 5502C>A, 5511G>A, 5514A>G, 5519A>G, 5523A>G, 5524A>G, 5526A>G, 5528A>C, 5532T>A, 5533C>A, 5535G>A, 5537A>C, 5543C>T, 5545G>T, 5546T>A, 5549T>C, 5553A>G, 5558C>T, 5567G>A, 5568A>G, 5574T>A, 5575C>G, 5576T>C, 5577G>A, 5580A>G, 5581G>A, 5589C>G, 5595G>T, 5598C>G, 5599A>G, 5604T>C, 5606A>T, 5711A>T, 5714A>G, 5717A>G, 5718G>A, 5720T>A, 5721C>A, 5726C>A, 5739T>G, 5740T>C, 5741A>T, 5748A>G, 5756A>T, 5757T>A, 5759T>G, 5762T>A, 5763_5765delGAA, 5768C>T, 5769C>A, 5777A>T, 5779C>A, 5781T>A, 5782T>C, 5786G>A, 5788G>A, 5790C>T, 5795G>A, 5798G>A, 5807G>A, 5808A>T, 5811T>G, 5813A>G, 5816C>T, 5822C>T, 5828T>C, 5830A>T, 5834C>A, 5836C>A, 5843G>A, 5844A>G, 5846T>A, 5850T>A, 5851A>T, 5853A>G, 5855T>A, 5856A>G, 5859A>C, 5860T>C, 5864T>A, 5868A>G, 5869A>C, 5870T>A, 5873G>A, 5874T>A, 5875G>T, 5876T>A, 5877T>G, 5878C>G, 5879C>A, 5882C>T, 5889G>C, 5893G>A, 5895A>C, 5897A>C, 5903T>C, 5904T>A, 5906T>A, 5907C>G, 5910G>A, 5912C>T, 5913T>A, 5914G>T, 5915G>T, 5918G>A, 5919A>C, 5924T>A, 5925A>C, 5926G>A, 5928A>C, 5933C>T, 5940G>A, 5941T>A, 5945T>A, 5947C>A, 5948C>A, 5951A>C, 5952T>A, 5957C>A, 5958C>G, 5959C>A, 5960T>C, 5963G>A, 5965T>G, 5966T>C, 5967T>A, 5973A>C, 5974A>T, 5976C>G, 5977C>A, 5979C>T, 5980A>C, 5982A>G, 5983G>A, 5988A>C, 5990T>A, 5999G>A, 6005T>C, 6009T>A, 6010C>G, 6011T>C, 6014A>C, 6015G>C, 6026G>A, 6028G>C, 6029A>T, 6030A>G, 6032T>A, 6039T>G, 6040G>C, 6041G>A, 6047C>A, 6048T>A, 6049C>A, 6050T>C, 6051C>A, 6054C>T, 6056A>C, 6057G>A, 6058A>G, 6060T>C, 6062A>T, 6063C>A, 6064C>A, 6065T>A, 6066T>A, 6067C>A, 6069A>G, 6070C>A, 6082C>G, 6083A>T, 6084C>G, 6085G>C, 6086G>T, 6089T>C, 6090T>C, 6091G>A, 6092T>A, 6100G>A, 6101C>A, 6104T>C, 6107_6108insGAAAAG, 6108T>G, 6109C>G, 6111G>A, 6112C>A, 6116C>G, 6117T>A, 6118C>G, 6120G>A, 6121C>G, 6122A>T, 6130C>T, 6131T>C, 6135A>G, 6138A>T, 6140G>A, 6144T>A, 6145G>T, 6146T>A, 6154T>G, 6156T>C, 6159G>A, 6163A>G, 6164G>T, 6167T>C, 6171A>T, 6175T>A, 6177T>A, 6179A>T, 6181A>T, 6182T>A, 6183G>A, 6185G>C, 6190A>G, 6191A>T, 6195T>A, 6197T>A, 6198A>G, 6199C>T, 6200C>A, 6201C>G, 6203T>A, 6206G>A, 6212C>T, 6215T>C, 6216A>G, 6217T>A, 6218T>A, 6221A>T, 6224T>A, 6230T>A, 6234T>C, 6237A>C, 6239C>A, 6243C>A, 6245T>A, 6246A>G, 6248T>C, 6249A>C, 6251T>G, 6257A>T, 6258T>A, 6259T>G, 6260G>T, 6262A>G, 6265T>A, 6268A>G, 6269T>A, 6270C>A, 6272T>A, 6276G>C | | | | | | | | |

##### CDS

| | Begin | End | Coverage | Score | Concordance | Matches | Identities | I/D/M/F* | Stop Codons |
|---|---|---|---|---|---|---|---|---|---|
| EXK67_gp3 | 1124 | 1558 | 25.1% | 1411 | 49.9% | 399 (99.0%) | 210 (52.1%) | 3/1/1/1 | 1 |
| Protein mutations: | L1125I (4983C>A), H1126K (4986C>A 4988T>A), K1127L (4989A>T 4990A>T), E1129D (4997A>T), I1131L (5001A>T 5003C>A), Q1132E (5004C>G), N1136E (5016A>G 5018T>A), D1137K (5019G>A 5021C>A), K1140I (5029A>T 5030G>A), Y1141H (5031T>C), E1143G (5038A>G), K1144A (5040A>G 5041A>C 5042G>C), Q1146R (5046C>A 5047A>G 5048G>A), R1148M (5053G>T 5054A>G), D1157E (5081T>A), L1158Y (5082C>T 5083T>A), Q1161K (5091C>A 5093G>A), D1162I (5094G>A 5095A>T 5096C>A), K1163A (5097A>G 5098A>C 5099G>A), P1164A (5100C>G 5102T>A), I1166E (5106A>G 5107T>A), P1167A (5109C>G), S1168T (5112T>A), P1169N (5115C>A 5116C>A 5117A>T), E1170Q (5118G>C), M1171D (5121A>G 5122T>A 5123G>C), A1172I (5124G>A 5125C>T 5126G>A), K1173E (5127A>G), E1174D (5132A>T), Y1175F (5134A>T), E1176Q (5136G>C 5138A>G), K1177I (5140A>T 5141G>A), H1178Q (5144T>A), T1180K (5149C>A), A1184D (5161C>A 5162A>T), K1186G (5166A>G 5167A>G 5168G>A), P1190R (5178C>A 5179C>G 5180T>A), Q1192T (5184C>A 5185A>C), T1197S (5199A>T), I1201M (5213T>G), N1203K (5219C>A), S1206N (5226T>A 5227C>A 5228T>C), Q1208I (5232C>A 5233A>T), S1213A (5247A>G 5248G>C 5249T>C), Y1219S (5265T>A 5266A>G), K1220Q (5268A>C 5270G>A), R1221_L1222insE (5273_5274insGAA), Y1227R (5289T>A 5290A>G 5291C>A), K1228T (5293A>C), Q1230A (5298C>G 5299A>C), T1232K (5305C>A), P1234S (5310C>T), S1235D (5313A>G 5314G>A), I1236K (5317T>A 5318T>A), Y1238K (5322T>A 5324T>A), L1240I (5328C>A 5330A>T), L1241C (5331C>T 5332T>G), K1242R (5335A>G), K1244Q (5340A>C), D1245G (5344A>G 5345C>T), I1247K (5350T>A 5351T>A), V1248I (5352G>A), Y1249F (5356A>T 5357C>T), K1251E (5361A>G), L1254C (5370C>T 5371T>G 5372A>T), F1258Y (5383T>A), H1259* (5385C>T 5387C>G), I1261V (5391A>G 5393T>A), M1262K (5395T>A 5396G>A), D1264H (5400G>C 5402T>C), Q1266D (5406C>G 5408G>C), V1274T (5430G>A 5431T>C 5432C>T), P1276L (5437C>T 5438A>T), Q1277E (5439C>G), I1283L (5457A>T), V1284A (5461T>C 5462A>C), N1291T (5482A>C 5483T>A), V1295I (5493G>A), D1301N (5511G>A), N1302D (5514A>G), I1303M (5519A>G), K1305G (5523A>G 5524A>G), K1306D (5526A>G 5528A>C), S1308K (5532T>A 5533C>A), E1309N (5535G>A 5537A>C), C1312L (5545G>T 5546T>A), I1315V (5553A>G), I1320V (5568A>G), E1323K (5577G>A), S1324D (5580A>G 5581G>A), Q1327E (5589C>G), V1329L (5595G>T), Q1330G (5598C>G 5599A>G), V1370I (5718G>A 5720T>A), Q1371K (5721C>A), L1377A (5739T>G 5740T>C 5741A>T), I1380V (5748A>G), E1382D (5756A>T), F1383M (5757T>A 5759T>G), E1385del (5763_5765delGAA), Q1387K (5769C>A), E1389D (5777A>T), T1390K (5779C>A), L1391T (5781T>A 5782T>C), G1393D (5788G>A), I1400L (5808A>T), L1401V (5811T>G 5813A>G), Y1407F (5830A>T), P1409Q (5836C>A), S1412G (5844A>G 5846T>A), Y1414I (5850T>A 5851A>T), T1415A (5853A>G 5855T>A), R1416G (5856A>G), I1417P (5859A>C 5860T>C), F1418L (5864T>A), N1420A (5868A>G 5869A>C 5870T>A), C1422I (5874T>A 5875G>T 5876T>A), S1423G (5877T>G 5878C>G 5879C>A), E1427Q (5889G>C), R1428K (5893G>A), K1429H (5895A>C 5897A>C), S1432T (5904T>A 5906T>A), Q1433E (5907C>G), D1434N (5910G>A 5912C>T), W1435I (5913T>A 5914G>T 5915G>T), M1437L (5919A>C), R1439Q (5925A>C 5926G>A), K1440Q (5928A>C), V1444K (5940G>A 5941T>A), T1446K (5947C>A 5948C>A), K1447N (5951A>C), L1448I (5952T>A), P1450D (5958C>G 5959C>A 5960T>C), I1452S (5965T>G 5966T>C), L1453I (5967T>A), K1455L (5973A>C 5974A>T), P1456E (5976C>G 5977C>A), Q1457S (5979C>T 5980A>C), S1458D (5982A>G 5983G>A), I1460L (5988A>C 5990T>A), E1469Q (6015G>C), G1473A (6028G>C 6029A>T), I1474V (6030A>G 6032T>A), W1477A (6039T>G 6040G>C 6041G>A), S1480N (6048T>A 6049C>A 6050T>C), Q1481K (6051C>A), Q1482Y (6054C>T 6056A>C), D1483S (6057G>A 6058A>G), S1484P (6060T>C 6062A>T), P1485K (6063C>A 6064C>A 6065T>A), S1486K (6066T>A 6067C>A), T1487E (6069A>G 6070C>A), S1491C (6082C>G 6083A>T), R1492A (6084C>G 6085G>C 6086G>T), C1494Q (6090T>C 6091G>A 6092T>A), S1497K (6100G>A 6101C>A), K1499_S1500insEK (6107_6108insGAAAAG), S1500G (6108T>G 6109C>G), A1501N (6111G>A 6112C>A), I1502M (6116C>G), A1504S (6120G>A 6121C>G 6122A>T), A1507V (6130C>T 6131T>C), I1509V (6135A>G), M1510L (6138A>T 6140G>A), C1512I (6144T>A 6145G>T 6146T>A), V1515G (6154T>G), D1517N (6159G>A), K1518S (6163A>G 6164G>T), I1521L (6171A>T), F1522Y (6175T>A), L1523I (6177T>A 6179A>T), Y1524L (6181A>T 6182T>A), E1525N (6183G>A 6185G>C), K1527S (6190A>G 6191A>T), F1529I (6195T>A 6197T>A), T1530V (6198A>G 6199C>T 6200C>A), L1531V (6201C>G 6203T>A), I1536E (6216A>G 6217T>A 6218T>A), N1540K (6230T>A), Y1542H (6234T>C), N1543Q (6237A>C 6239C>A), L1545I (6243C>A 6245T>A), N1546D (6246A>G 6248T>C), N1547Q (6249A>C 6251T>G), K1549N (6257A>T), L1550S (6258T>A 6259T>G 6260G>T), N1551S (6262A>G), I1552K (6265T>A), N1553R (6268A>G 6269T>A), V1556L (6276G>C) | | | | | | | | |

| | Begin | End | Coverage | Score | Concordance | Matches | Identities | I/D/M/F* | Stop Codons |
|---|---|---|---|---|---|---|---|---|---|
| **NT** | **4979** | **6282** | **14.7%** | **547** | **23.1%** | **1196 (98.9%)** | **747 (61.8%)** | **9/4** | |

Codon mutations:

CTA1125ATA (4983C>A), CAT1126AAA (4986C>A 4988T>A), AAA1127TTA (4989A>T 4990A>T), TTG1128CTA (4992T>C 4994G>A), GAA1129GAT (4997A>T), ATC1131TTA (5001A>T 5003C>A), CAA1132GAA (5004C>G), ATT1133ATA (5009T>A), ATT1134ATA (5012T>A), GGG1135GGA (5015G>A), AAT1136GAA (5016A>G 5018T>A), GAC1137AAA (5019G>A 5021C>A), CCA1138CCC (5024A>C), TTA1139CTA (5025T>C), AAG1140ATA (5029A>T 5030G>A), TAT1141CAT (5031T>C), GAA1143GGA (5038A>G), AAG1144GCC (5040A>G 5041A>C 5042G>C), CAG1146AGA (5046C>A 5047A>G 5048G>A), ATT1147ATA (5051T>A), AGA1148ATG (5053G>T 5054A>G), TGC1149TGT (5057C>T), CTG1151CTA (5063G>A), GAC1152GAT (5066C>T), ATC1153ATA (5069C>A), CCT1156CCA (5078T>A), GAT1157GAA (5081T>A), CTT1158TAT (5082C>T 5083T>A), CAG1161AAA (5091C>A 5093G>A), GAC1162ATA (5094G>A 5095A>T 5096C>A), AAG1163GCA (5097A>G 5098A>C 5099G>A), CCT1164GCA (5100C>G 5102T>A), ATT1165ATA (5105T>A), ATA1166GAA (5106A>G 5107T>A), CCT1167GCT (5109C>G), TCA1168ACA (5112T>A), CCA1169AAT (5115C>A 5116C>A 5117A>T), GAA1170CAA (5118G>C), ATG1171GAC (5121A>G 5122T>A 5123G>C), GCG1172ATA (5124G>A 5125C>T 5126G>A), AAA1173GAA (5127A>G), GAA1174GAT (5132A>T), TAT1175TTT (5134A>T), GAA1176CAG (5136G>C 5138A>G), AAG1177ATA (5140A>T 5141G>A), CAT1178CAA (5144T>A), ATT1179ATA (5147T>A), ACA1180AAA (5149C>A), CTG1183TTA (5157C>T 5159G>A), GCA1184GAT (5161C>A 5162A>T), CTC1185TTA (5163C>T 5165C>A), AAG1186GGA (5166A>G 5167A>G 5168G>A), GTT1187GTA (5171T>A), ATC1188ATA (5174C>A), CCT1190AGA (5178C>A 5179C>G 5180T>A), TCC1191TCA (5183C>A), CAA1192ACA (5184C>A 5185A>C), AGT1193TCT (5187A>T 5188G>C), CGC1194AGA (5190C>A 5192C>A), CAT1195CAC (5195T>C), AGG1196AGA (5198G>A), ACA1197TCA (5199A>T), GCT1199GCA (5207T>A), ATT1201ATG (5213T>G), GTC1202GTA (5216C>A), AAC1203AAA (5219C>A), AAA1204-AC (5220delA 5222A>C), TCT1206AAC (5226T>A 5227C>A 5228T>C), CAA1208ATA (5232C>A 5233A>T), GTA1209GTC (5237A>C), GGT1211GGA (5243T>A), AGT1213GCC (5247A>G 5248G>C 5249T>C), CGT1214AGA (5250C>A 5252T>A), GTA1216GTT (5258A>T), TAT1219AGT (5265T>A 5266A>G), AAG1220CAA (5268A>C 5270G>A), CGA1221_CTA1222insGAA (5273_5274insGAA), CTA1222CTT (5276A>T), ACA1226ACT (5288A>T), TAC1227AGA (5289T>A 5290A>G 5291C>A), AAA1228ACA (5293A>C), GAC1229GAT (5297C>T), CAA1230GCA (5298C>G 5299A>C), TAC1231TAT (5303C>T), ACA1232AAA (5305C>A), CTC1233TTA (5307C>T 5309C>A), CCA1234TCA (5310C>T), AGT1235GAT (5313A>G 5314G>A), ATT1236AAA (5317T>A 5318T>A), TAT1238AAA (5322T>A 5324T>A), CTC1239TTG (5325C>T 5327C>G), CTA1240ATT (5328C>A 5330A>T), CTT1241TGT (5331C>T 5332T>G), AAA1242AGA (5335A>G), AAA1244CAA (5340A>C), GAC1245GGT (5344A>G 5345C>T), AAG1246AAA (5348G>A), ATT1247AAA (5350T>A 5351T>A), GTC1248ATC (5352G>A), TAC1249TTT (5356A>T 5357C>T), AGC1250AGT (5360C>T), AAA1251GAA (5361A>G), CTA1254TGT (5370C>T 5371T>G 5372A>T), AAG1255AAA (5375G>A), TCA1256TCG (5378A>G), TTC1258TAC (5383T>A), CAC1259TAG (5385C>T 5387C>G), CAG1260CAA (5390G>A), ATT1261GTA (5391A>G 5393T>A), ATG1262AAA (5395T>A 5396G>A), GAT1264CAC (5400G>C 5402T>C), CAG1266GAC (5406C>G 5408G>C), TTT1273TTC (5429T>C), GTC1274ACT (5430G>A 5431T>C 5432C>T), TGC1275TGT (5435C>T), CCA1276CTT (5437C>T 5438A>T), CAG1277GAG (5439C>G), GGT1278GGA (5444T>A), ATA1283TTA (5457A>T), GTA1284GCC (5461T>C 5462A>C), TTC1287TTT (5471C>T), GGA1288GGT (5474A>T), CTC1289TTA (5475C>T 5477C>A), AAT1291ACA (5482A>C 5483T>A), GCA1292GCG (5486A>G), TCA1294TCT (5492A>T), GTA1295ATA (5493G>A), TTC1296TTT (5498C>T), CGA1298AGA (5502C>A), GAT1301AAT (5511G>A), AAT1302GAT (5514A>G), ATA1303ATG (5519A>G), AAA1305GGA (5523A>G 5524A>G), AAA1306GAC (5526A>G 5528A>C), TCA1308AAA (5532T>A 5533C>A), GAA1309AAC (5535G>A 5537A>C), GTC1311GTT (5543C>T), TGT1312TTA (5545G>T 5546T>A), GTT1313GTC (5549T>C), ATA1315GTA (5553A>G), GAC1316GAT (5558C>T), TTG1319TTA (5567G>A), ATA1320GTA (5568A>G), TCT1322AGC (5574T>A 5575C>G 5576T>C), GAA1323AAA (5577G>A), AGT1324GAT (5580A>G 5581G>A), CAA1327GAA (5589C>G), GTA1329TTA (5595G>T), CAA1330GGA (5598C>G 5599A>G), TTA1332CTT (5604T>C 5606A>T), GAA1367..T (5711A>T), GGA1368GGG (5714A>G), AAA1369AAG (5717A>G), GTT1370ATA (5718G>A 5720T>A), CAA1371AAA (5721C>A), CTC1372CTA (5726C>A), TTA1377GCT (5739T>G 5740T>C 5741A>T), ATA1380GTA (5748A>G), GAA1382GAT (5756A>T), TTT1383ATG (5757T>A 5759T>G), CCT1384CCA (5762T>A), GAA1385del (5763_5765delGAA), GAC1386GAT (5768C>T), CAA1387AAA (5769C>A), GAA1389GAT (5777A>T), ACA1390AAA (5779C>A), TTA1391ACA (5781T>A 5782T>C), AAG1392AAA (5786G>A), GGT1393GAT (5788G>A), CTA1394TTA (5790C>T), CAG1395CAA (5795G>A), AAG1396AAA (5798G>A), GGG1399GGA (5807G>A), ATA1400TTA (5808A>T), TTA1401GTG (5811T>G 5813A>G), AAC1402AAT (5816C>T), GCC1404GCT (5822C>T), AAT1406AAC (5828T>C), TAT1407TTT (5830A>T), ATC1408ATA (5834C>A), CCA1409CAA (5836C>A), TTG1411TTA (5843G>A), AGT1412GGA (5844A>G 5846T>A), TAT1414ATT (5850T>A 5851A>T), ACT1415GCA (5853A>G 5855T>A), AGA1416GGA (5856A>G), ATA1417CCA (5859A>C 5860T>C), TTT1418TTA (5864T>A), AAT1420GCA (5868A>G 5869A>C 5870T>A), AAG1421AAA (5873G>A), TGT1422ATA (5874T>A 5875G>T 5876T>A), TCC1423GGA (5877T>G 5878C>G 5879C>A), AGC1424AGT (5882C>T), GAA1427CAA (5889G>C), AGA1428AAA (5893G>A), AAA1429CAC (5895A>C 5897A>C), AAT1431AAC (5903T>C), TCT1432ACA (5904T>A 5906T>A), CAA1433GAA (5907C>G), GAC1434AAT (5910G>A 5912C>T), TGG1435ATT (5913T>A 5914G>T 5915G>T), AAG1436AAA (5918G>A), ATG1437CTG (5919A>C), GTT1438GTA (5924T>A), AGA1439CAA (5925A>C 5926G>A), AAA1440CAA (5928A>C), ATC1441ATT (5933C>T), GTA1444AAA (5940G>A 5941T>A), ATT1445ATA (5945T>A), ACC1446AAA (5947C>A 5948C>A), AAA1447AAC (5951A>C), TTA1448ATA (5952T>A), CCC1449CCA (5957C>A), CCT1450GAC (5958C>G 5959C>A 5960T>C), TTG1451TTA (5963G>A), ATT1452AGC (5965T>G 5966T>C), TTA1453ATA (5967T>A), AAA1455CTA (5973A>C 5974A>T), CCA1456GAA (5976C>G 5977C>A), CAA1457TCA (5979C>T 5980A>C), AGT1458GAT (5982A>G 5983G>A), ATT1460CTA (5988A>C 5990T>A), GAG1463GAA (5999G>A), GAT1465GAC (6005T>C), TCT1467AGC (6009T>A 6010C>G 6011T>C), CTA1468CTC (6014A>C), GAA1469CAA (6015G>C), GGG1472GGA (6026G>A), GGA1473GCT (6028G>C 6029A>T), ATT1474GTA (6030A>G 6032T>A), TGG1477GCA (6039T>G 6040G>C 6041G>A), CCC1479CCA (6047C>A), TCT1480AAC (6048T>A 6049C>A 6050T>C), CAA1481AAA (6051C>A), CAA1482TAC (6054C>T 6056A>C), GAT1483AGT (6057G>A 6058A>G), TCA1484CCT (6060T>C 6062A>T), CCT1485AAA (6063C>A 6064C>A 6065T>A), TCA1486AAA (6066T>A 6067C>A), ACA1487GAA (6069A>G 6070C>A), TCA1491TGT (6082C>G 6083A>T), CGG1492GCT (6084C>G 6085G>C 6086G>T), TAT1493TAC (6089T>C), TGT1494CAA (6090T>C 6091G>A 6092T>A), AGC1497AAA (6100G>A 6101C>A), TAT1498TAC (6104T>C), AAA1499_TCT1500insGAAAAG (6107_6108insGAAAAG), TCT1500GGT (6108T>G 6109C>G), GCC1501AAC (6111G>A 6112C>A), ATC1502ATG (6116C>G), TCT1503AGT (6117T>A 6118C>G), GCA1504AGT (6120G>A 6121C>G 6122A>T), GCT1507GTC (6130C>T 6131T>C), ATA1509GTA (6135A>G), ATG1510TTA (6138A>T 6140G>A), TGT1512ATA (6144T>A 6145G>T 6146T>A), GTA1515GGA (6154T>G), TTA1516CTA (6156T>C), GAT1517AAT (6159G>A), AAG1518AGT (6163A>G 6164G>T), TTT1519TTC (6167T>C), ATA1521TTA (6171A>T), TTT1522TAT (6175T>A), TTA1523ATT (6177T>A 6179A>T), TAT1524TTA (6181A>T 6182T>A), GAG1525AAC (6183G>A 6185G>C), AAA1527AGT (6190A>G 6191A>T), TTT1529ATA (6195T>A 6197T>A), ACC1530GTA (6198A>G 6199C>T 6200C>A), CTT1531GTA (6201C>G 6203T>A), AGG1532AGA (6206G>A), GAC1534GAT (6212C>T), TGT1535TGC (6215T>C), ATT1536GAA (6216A>G 6217T>A 6218T>A), GCA1537GCT (6221A>T), ATT1538ATA (6224T>A), AAT1540AAA (6230T>A), TAT1542CAT (6234T>C), AAC1543CAA (6237A>C 6239C>A), CTT1545ATA (6243C>A 6245T>A), AAT1546GAC (6246A>G 6248T>C), AAT1547CAG (6249A>C 6251T>G), AAA1549AAT (6257A>T), TTG1550AGT (6258T>A 6259T>G 6260G>T), AAT1551AGT (6262A>G), ATA1552AAA (6265T>A), AAT1553AGA (6268A>G 6269T>A), CGT1554AGA (6270C>A 6272T>A), GTA1556CTA (6276G>C)

#### Proteins

| | Begin | End | Coverage | Score | Concordance | Matches | Identities | I/D/M/F* | Stop Codons |
|---|---|---|---|---|---|---|---|---|---|
| ORF3 (YP_009553219.1) | 1124 | 1558 | 25.1% | 1411 | 49.9% | 399 (99.0%) | 210 (52.1%) | 3/1/1/1 | 1 |

Protein mutations:

L1125I (4983C>A), H1126K (4986C>A 4988T>A), K1127L (4989A>T 4990A>T), E1129D (4997A>T), I1131L (5001A>T 5003C>A), Q1132E (5004C>G), N1136E (5016A>G 5018T>A), D1137K (5019G>A 5021C>A), K1140I (5029A>T 5030G>A), Y1141H (5031T>C), E1143G (5038A>G), K1144A (5040A>G 5041A>C 5042G>C), Q1146R (5046C>A 5047A>G 5048G>A), R1148M (5053G>T 5054A>G), D1157E (5081T>A), L1158Y (5082C>T 5083T>A), Q1161K (5091C>A 5093G>A), D1162I (5094G>A 5095A>T 5096C>A), K1163A (5097A>G 5098A>C 5099G>A), P1164A (5100C>G 5102T>A), I1166E (5106A>G 5107T>A), P1167A (5109C>G), S1168T (5112T>A), P1169N (5115C>A 5116C>A 5117A>T), E1170Q (5118G>C), M1171D (5121A>G 5122T>A 5123G>C), A1172I (5124G>A 5125C>T 5126G>A), K1173E (5127A>G), E1174D (5132A>T), Y1175F (5134A>T), E1176Q (5136G>C 5138A>G), K1177I (5140A>T 5141G>A), H1178Q (5144T>A), T1180K (5149C>A), A1184D (5161C>A 5162A>T), K1186G (5166A>G 5167A>G 5168G>A), P1190R (5178C>A 5179C>G 5180T>A), Q1192T (5184C>A 5185A>C), T1197S (5199A>T), I1201M (5213T>G), N1203K (5219C>A), S1206N (5226T>A 5227C>A 5228T>C), Q1208I (5232C>A 5233A>T), S1213A (5247A>G 5248G>C 5249T>C), Y1219S (5265T>A 5266A>G), K1220Q (5268A>C 5270G>A), R1221_L1222insE (5273_5274insGAA), Y1227R (5289T>A 5290A>G 5291C>A), K1228T (5293A>C), Q1230A (5298C>G 5299A>C), T1232K (5305C>A), P1234S (5310C>T), S1235D (5313A>G 5314G>A), I1236K (5317T>A 5318T>A), Y1238K (5322T>A 5324T>A), L1240I (5328C>A 5330A>T), L1241C (5331C>T 5332T>G), K1242R (5335A>G), K1244Q (5340A>C), D1245G (5344A>G 5345C>T), I1247K (5350T>A 5351T>A), V1248I (5352G>A), Y1249F (5356A>T 5357C>T), K1251E (5361A>G), L1254C (5370C>T 5371T>G 5372A>T), F1258Y (5383T>A), H1259* (5385C>T 5387C>G), I1261V (5391A>G 5393T>A), M1262K (5395T>A 5396G>A), D1264H (5400G>C 5402T>C), Q1266D (5406C>G 5408G>C), V1274T (5430G>A 5431T>C 5432C>T), P1276L (5437C>T 5438A>T), Q1277E (5439C>G), I1283L (5457A>T), V1284A (5461T>C 5462A>C), N1291T (5482A>C 5483T>A), V1295I (5493G>A), D1301N (5511G>A), N1302D (5514A>G), I1303M (5519A>G), K1305G (5523A>G 5524A>G), K1306D (5526A>G 5528A>C), S1308K (5532T>A 5533C>A), E1309N (5535G>A 5537A>C), C1312L (5545G>T 5546T>A), I1315V (5553A>G), I1320V (5568A>G), E1323K (5577G>A), S1324D (5580A>G 5581G>A), Q1327E (5589C>G), V1329L (5595G>T), Q1330G (5598C>G 5599A>G), V1370I (5718G>A 5720T>A), Q1371K (5721C>A), L1377A (5739T>G 5740T>C 5741A>T), I1380V (5748A>G), E1382D (5756A>T), F1383M (5757T>A 5759T>G), E1385del (5763_5765delGAA), Q1387K (5769C>A), E1389D (5777A>T), T1390K (5779C>A), L1391T (5781T>A 5782T>C), G1393D (5788G>A), I1400L (5808A>T), L1401V (5811T>G 5813A>G), Y1407F (5830A>T), P1409Q (5836C>A), S1412G (5844A>G 5846T>A), Y1414I (5850T>A 5851A>T), T1415A (5853A>G 5855T>A), R1416G (5856A>G), I1417P (5859A>C 5860T>C), F1418L (5864T>A), N1420A (5868A>G 5869A>C 5870T>A), C1422I (5874T>A 5875G>T 5876T>A), S1423G (5877T>G 5878C>G 5879C>A), E1427Q (5889G>C), R1428K (5893G>A), K1429H (5895A>C 5897A>C), S1432T (5904T>A 5906T>A), Q1433E (5907C>G), D1434N (5910G>A 5912C>T), W1435I (5913T>A 5914G>T 5915G>T), M1437L (5919A>C), R1439Q (5925A>C 5926G>A), K1440Q (5928A>C), V1444K (5940G>A 5941T>A), T1446K (5947C>A 5948C>A), K1447N (5951A>C), L1448I (5952T>A), P1450D (5958C>G 5959C>A 5960T>C), I1452S (5965T>G 5966T>C), L1453I (5967T>A), K1455L (5973A>C 5974A>T), P1456E (5976C>G 5977C>A), Q1457S (5979C>T 5980A>C), S1458D (5982A>G 5983G>A), I1460L (5988A>C 5990T>A), E1469Q (6015G>C), G1473A (6028G>C 6029A>T), I1474V (6030A>G 6032T>A), W1477A (6039T>G 6040G>C 6041G>A), S1480N (6048T>A 6049C>A 6050T>C), Q1481K (6051C>A), Q1482Y (6054C>T 6056A>C), D1483S (6057G>A 6058A>G), S1484P (6060T>C 6062A>T), P1485K (6063C>A 6064C>A 6065T>A), S1486K (6066T>A 6067C>A), T1487E (6069A>G 6070C>A), S1491C (6082C>G 6083A>T), R1492A (6084C>G 6085G>C 6086G>T), C1494Q (6090T>C 6091G>A 6092T>A), S1497K (6100G>A 6101C>A), K1499_S1500insEK (6107_6108insGAAAAG), S1500G (6108T>G 6109C>G), A1501N (6111G>A 6112C>A), I1502M (6116C>G), A1504S (6120G>A 6121C>G 6122A>T), A1507V (6130C>T 6131T>C), I1509V (6135A>G), M1510L (6138A>T 6140G>A), C1512I (6144T>A 6145G>T 6146T>A), V1515G (6154T>G), D1517N (6159G>A), K1518S (6163A>G 6164G>T), I1521L (6171A>T), F1522Y (6175T>A), L1523I (6177T>A 6179A>T), Y1524L (6181A>T 6182T>A), E1525N (6183G>A 6185G>C), K1527S (6190A>G 6191A>T), F1529I (6195T>A 6197T>A), T1530V (6198A>G 6199C>T 6200C>A), L1531V (6201C>G 6203T>A), I1536E (6216A>G 6217T>A 6218T>A), N1540K (6230T>A), Y1542H (6234T>C), N1543Q (6237A>C 6239C>A), L1545I (6243C>A 6245T>A), N1546D (6246A>G 6248T>C), N1547Q (6249A>C 6251T>G), K1549N (6257A>T), L1550S (6258T>A 6259T>G 6260G>T), N1551S (6262A>G), I1552K (6265T>A), N1553R (6268A>G 6269T>A), V1556L (6276G>C)

| | Begin | End | Coverage | Score | Concordance | Matches | Identities | I/D/M/F* | Stop Codons |
|---|---|---|---|---|---|---|---|---|---|
| **NT** | **4979** | **6282** | **14.7%** | **547** | **23.1%** | **1196 (98.9%)** | **747 (61.8%)** | **9/4** | |

Codon mutations: CTA1125ATA (4983C>A), CAT1126AAA (4986C>A 4988T>A), AAA1127TTA (4989A>T 4990A>T), TTG1128CTA (4992T>C 4994G>A), GAA1129GAT (4997A>T), ATC1131TTA (5001A>T 5003C>A), CAA1132GAA (5004C>G), ATT1133ATA (5009T>A), ATT1134ATA (5012T>A), GGG1135GGA (5015G>A), AAT1136GAA (5016A>G 5018T>A), GAC1137AAA (5019G>A 5021C>A), CCA1138CCC (5024A>C), TTA1139CTA (5025T>C), AAG1140ATA (5029A>T 5030G>A), TAT1141CAT (5031T>C), GAA1143GGA (5038A>G), AAG1144GCC (5040A>G 5041A>C 5042G>C), CAG1146AGA (5046C>A 5047A>G 5048G>A), ATT1147ATA (5051T>A), AGA1148ATG (5053G>T 5054A>G), TGC1149TGT (5057C>T), CTG1151CTA (5063G>A), GAC1152GAT (5066C>T), ATC1153ATA (5069C>A), CCT1156CCA (5078T>A), GAT1157GAA (5081T>A), CTT1158TAT (5082C>T 5083T>A), CAG1161AAA (5091C>A 5093G>A), GAC1162ATA (5094G>A 5095A>T 5096C>A), AAG1163GCA (5097A>G 5098A>C 5099G>A), CCT1164GCA (5100C>G 5102T>A), ATT1165ATA (5105T>A), ATA1166GAA (5106A>G 5107T>A), CCT1167GCT (5109C>G), TCA1168ACA (5112T>A), CCA1169AAT (5115C>A 5116C>A 5117A>T), GAA1170CAA (5118G>C), ATG1171GAC (5121A>G 5122T>A 5123G>C), GCG1172ATA (5124G>A 5125C>T 5126G>A), AAA1173GAA (5127A>G), GAA1174GAT (5132A>T), TAT1175TTT (5134A>T), GAA1176CAG (5136G>C 5138A>G), AAG1177ATA (5140A>T 5141G>A), CAT1178CAA (5144T>A), ATT1179ATA (5147T>A), ACA1180AAA (5149C>A), CTG1183TTA (5157C>T 5159G>A), GCA1184GAT (5161C>A 5162A>T), CTC1185TTA (5163C>T 5165C>A), AAG1186GGA (5166A>G 5167A>G 5168G>A), GTT1187GTA (5171T>A), ATC1188ATA (5174C>A), CCT1190AGA (5178C>A 5179C>G 5180T>A), TCC1191TCA (5183C>A), CAA1192ACA (5184C>A 5185A>C), AGT1193TCT (5187A>T 5188G>C), CGC1194AGA (5190C>A 5192C>A), CAT1195CAC (5195T>C), AGG1196AGA (5198G>A), ACA1197TCA (5199A>T), GCT1199GCA (5207T>A), ATT1201ATG (5213T>G), GTC1202GTA (5216C>A), AAC1203AAA (5219C>A), AAA1204-AC (5220delA 5222A>C), TCT1206AAC (5226T>A 5227C>A 5228T>C), CAA1208ATA (5232C>A 5233A>T), GTA1209GTC (5237A>C), GGT1211GGA (5243T>A), AGT1213GCC (5247A>G 5248G>C 5249T>C), CGT1214AGA (5250C>A 5252T>A), GTA1216GTT (5258A>T), TAT1219AGT (5265T>A 5266A>G), AAG1220CAA (5268A>C 5270G>A), CGA1221_CTA1222insGAA (5273_5274insGAA), CTA1222CTT (5276A>T), ACA1226ACT (5288A>T), TAC1227AGA (5289T>A 5290A>G 5291C>A), AAA1228ACA (5293A>C), GAC1229GAT (5297C>T), CAA1230GCA (5298C>G 5299A>C), TAC1231TAT (5303C>T), ACA1232AAA (5305C>A), CTC1233TTA (5307C>T 5309C>A), CCA1234TCA (5310C>T), AGT1235GAT (5313A>G 5314G>A), ATT1236AAA (5317T>A 5318T>A), TAT1238AAA (5322T>A 5324T>A), CTC1239TTG (5325C>T 5327C>G), CTA1240ATT (5328C>A 5330A>T), CTT1241TGT (5331C>T 5332T>G), AAA1242AGA (5335A>G), AAA1244CAA (5340A>C), GAC1245GGT (5344A>G 5345C>T), AAG1246AAA (5348G>A), ATT1247AAA (5350T>A 5351T>A), GTC1248ATC (5352G>A), TAC1249TTT (5356A>T 5357C>T), AGC1250AGT (5360C>T), AAA1251GAA (5361A>G), CTA1254TGT (5370C>T 5371T>G 5372A>T), AAG1255AAA (5375G>A), TCA1256TCG (5378A>G), TTC1258TAC (5383T>A), CAC1259TAG (5385C>T 5387C>G), CAG1260CAA (5390G>A), ATT1261GTA (5391A>G 5393T>A), ATG1262AAA (5395T>A 5396G>A), GAT1264CAC (5400G>C 5402T>C), CAG1266GAC (5406C>G 5408G>C), TTT1273TTC (5429T>C), GTC1274ACT (5430G>A 5431T>C 5432C>T), TGC1275TGT (5435C>T), CCA1276CTT (5437C>T 5438A>T), CAG1277GAG (5439C>G), GGT1278GGA (5444T>A), ATA1283TTA (5457A>T), GTA1284GCC (5461T>C 5462A>C), TTC1287TTT (5471C>T), GGA1288GGT (5474A>T), CTC1289TTA (5475C>T 5477C>A), AAT1291ACA (5482A>C 5483T>A), GCA1292GCG (5486A>G), TCA1294TCT (5492A>T), GTA1295ATA (5493G>A), TTC1296TTT (5498C>T), CGA1298AGA (5502C>A), GAT1301AAT (5511G>A), AAT1302GAT (5514A>G), ATA1303ATG (5519A>G), AAA1305GGA (5523A>G 5524A>G), AAA1306GAC (5526A>G 5528A>C), TCA1308AAA (5532T>A 5533C>A), GAA1309AAC (5535G>A 5537A>C), GTC1311GTT (5543C>T), TGT1312TTA (5545G>T 5546T>A), GTT1313GTC (5549T>C), ATA1315GTA (5553A>G), GAC1316GAT (5558C>T), TTG1319TTA (5567G>A), ATA1320GTA (5568A>G), TCT1322AGC (5574T>A 5575C>G 5576T>C), GAA1323AAA (5577G>A), AGT1324GAT (5580A>G 5581G>A), CAA1327GAA (5589C>G), GTA1329TTA (5595G>T), CAA1330GGA (5598C>G 5599A>G), TTA1332CTT (5604T>C 5606A>T), GAA1367..T (5711A>T), GGA1368GGG (5714A>G), AAA1369AAG (5717A>G), GTT1370ATA (5718G>A 5720T>A), CAA1371AAA (5721C>A), CTC1372CTA (5726C>A), TTA1377GCT (5739T>G 5740T>C 5741A>T), ATA1380GTA (5748A>G), GAA1382GAT (5756A>T), TTT1383ATG (5757T>A 5759T>G), CCT1384CCA (5762T>A), GAA1385del (5763_5765delGAA), GAC1386GAT (5768C>T), CAA1387AAA (5769C>A), GAA1389GAT (5777A>T), ACA1390AAA (5779C>A), TTA1391ACA (5781T>A 5782T>C), AAG1392AAA (5786G>A), GGT1393GAT (5788G>A), CTA1394TTA (5790C>T), CAG1395CAA (5795G>A), AAG1396AAA (5798G>A), GGG1399GGA (5807G>A), ATA1400TTA (5808A>T), TTA1401GTG (5811T>G 5813A>G), AAC1402AAT (5816C>T), GCC1404GCT (5822C>T), AAT1406AAC (5828T>C), TAT1407TTT (5830A>T), ATC1408ATA (5834C>A), CCA1409CAA (5836C>A), TTG1411TTA (5843G>A), AGT1412GGA (5844A>G 5846T>A), TAT1414ATT (5850T>A 5851A>T), ACT1415GCA (5853A>G 5855T>A), AGA1416GGA (5856A>G), ATA1417CCA (5859A>C 5860T>C), TTT1418TTA (5864T>A), AAT1420GCA (5868A>G 5869A>C 5870T>A), AAG1421AAA (5873G>A), TGT1422ATA (5874T>A 5875G>T 5876T>A), TCC1423GGA (5877T>G 5878C>G 5879C>A), AGC1424AGT (5882C>T), GAA1427CAA (5889G>C), AGA1428AAA (5893G>A), AAA1429CAC (5895A>C 5897A>C), AAT1431AAC (5903T>C), TCT1432ACA (5904T>A 5906T>A), CAA1433GAA (5907C>G), GAC1434AAT (5910G>A 5912C>T), TGG1435ATT (5913T>A 5914G>T 5915G>T), AAG1436AAA (5918G>A), ATG1437CTG (5919A>C), GTT1438GTA (5924T>A), AGA1439CAA (5925A>C 5926G>A), AAA1440CAA (5928A>C), ATC1441ATT (5933C>T), GTA1444AAA (5940G>A 5941T>A), ATT1445ATA (5945T>A), ACC1446AAA (5947C>A 5948C>A), AAA1447AAC (5951A>C), TTA1448ATA (5952T>A), CCC1449CCA (5957C>A), CCT1450GAC (5958C>G 5959C>A 5960T>C), TTG1451TTA (5963G>A), ATT1452AGC (5965T>G 5966T>C), TTA1453ATA (5967T>A), AAA1455CTA (5973A>C 5974A>T), CCA1456GAA (5976C>G 5977C>A), CAA1457TCA (5979C>T 5980A>C), AGT1458GAT (5982A>G 5983G>A), ATT1460CTA (5988A>C 5990T>A), GAG1463GAA (5999G>A), GAT1465GAC (6005T>C), TCT1467AGC (6009T>A 6010C>G 6011T>C), CTA1468CTC (6014A>C), GAA1469CAA (6015G>C), GGG1472GGA (6026G>A), GGA1473GCT (6028G>C 6029A>T), ATT1474GTA (6030A>G 6032T>A), TGG1477GCA (6039T>G 6040G>C 6041G>A), CCC1479CCA (6047C>A), TCT1480AAC (6048T>A 6049C>A 6050T>C), CAA1481AAA (6051C>A), CAA1482TAC (6054C>T 6056A>C), GAT1483AGT (6057G>A 6058A>G), TCA1484CCT (6060T>C 6062A>T), CCT1485AAA (6063C>A 6064C>A 6065T>A), TCA1486AAA (6066T>A 6067C>A), ACA1487GAA (6069A>G 6070C>A), TCA1491TGT (6082C>G 6083A>T), CGG1492GCT (6084C>G 6085G>C 6086G>T), TAT1493TAC (6089T>C), TGT1494CAA (6090T>C 6091G>A 6092T>A), AGC1497AAA (6100G>A 6101C>A), TAT1498TAC (6104T>C), AAA1499_TCT1500insGAAAAG (6107_6108insGAAAAG), TCT1500GGT (6108T>G 6109C>G), GCC1501AAC (6111G>A 6112C>A), ATC1502ATG (6116C>G), TCT1503AGT (6117T>A 6118C>G), GCA1504AGT (6120G>A 6121C>G 6122A>T), GCT1507GTC (6130C>T 6131T>C), ATA1509GTA (6135A>G), ATG1510TTA (6138A>T 6140G>A), TGT1512ATA (6144T>A 6145G>T 6146T>A), GTA1515GGA (6154T>G), TTA1516CTA (6156T>C), GAT1517AAT (6159G>A), AAG1518AGT (6163A>G 6164G>T), TTT1519TTC (6167T>C), ATA1521TTA (6171A>T), TTT1522TAT (6175T>A), TTA1523ATT (6177T>A 6179A>T), TAT1524TTA (6181A>T 6182T>A), GAG1525AAC (6183G>A 6185G>C), AAA1527AGT (6190A>G 6191A>T), TTT1529ATA (6195T>A 6197T>A), ACC1530GTA (6198A>G 6199C>T 6200C>A), CTT1531GTA (6201C>G 6203T>A), AGG1532AGA (6206G>A), GAC1534GAT (6212C>T), TGT1535TGC (6215T>C), ATT1536GAA (6216A>G 6217T>A 6218T>A), GCA1537GCT (6221A>T), ATT1538ATA (6224T>A), AAT1540AAA (6230T>A), TAT1542CAT (6234T>C), AAC1543CAA (6237A>C 6239C>A), CTT1545ATA (6243C>A 6245T>A), AAT1546GAC (6246A>G 6248T>C), AAT1547CAG (6249A>C 6251T>G), AAA1549AAT (6257A>T), TTG1550AGT (6258T>A 6259T>G 6260G>T), AAT1551AGT (6262A>G), ATA1552AAA (6265T>A), AAT1553AGA (6268A>G 6269T>A), CGT1554AGA (6270C>A 6272T>A), GTA1556CTA (6276G>C)

*: Inserts / Deletes / Misaligned / Frameshifts

#### Analysis details

This analysis was performed with panviral2.64

### NGS Details (UN59): Badnavirus occultipomeae

#### Assembly

| | |
|---|---|
| **Coverage Length** | 255 (1 contig(s)) |
| **Depth Of Coverage** | 30.9 |
| **Number Of Reads** | 80 |
| **Reads Per Million** | 1.74 rpm (after QC) |
| **Ambiguities** | 0 |
| **Assembly Method** | de novo + reference guided assembly |
| **Consensus Caller** | Bcf Tools |

#### Coverage Map

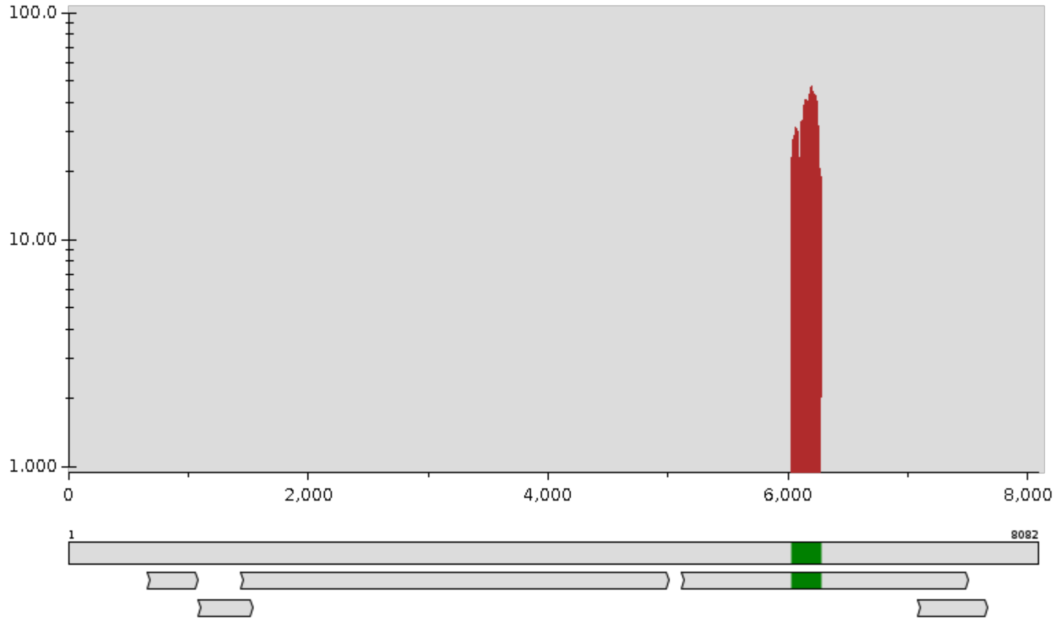

#### Assignment

| | |
|---|---|
| **Type** | Badnavirus occultipomeae (Taxonomy ID: 3048353) |
| **Reference Genome** | NC_015655.1 |
| **NT Identity (%)** | 58.1395 |
| **AA Identity (%)** | 50.0 |
| **Number Of Stop Codons** | 0 |
| **Number Of CDS** | 5 |

#### Alignment

| | |
|---|---|
| **Alignment Score** | 78.0 (NT) + 347.0 (AA) = 425.0 |
| **Concordance (%)** | 37.7107 |

| **Alignment Method** | Global, seeded, nucleotide + amino acids (AGA) |
|---|---|

#### Genome Region

Sequence starts at position 6025 and ends at position 6279 relative to NC_015655.1 reference sequence.

#### Alignment Detailed Statistics

| | Begin | End | Coverage | Score | Concordance | Matches | Identities | I/D/M/F* | Stop Codons |
|---|---|---|---|---|---|---|---|---|---|
| **NT** | **6025** | **6279** | **3.2%** | **78** | **15.3%** | **255 (98.8%)** | **150 (58.1%)** | **3/0** | |
| Mutations: | 6033A>T, 6036C>T, 6042A>G, 6043T>A, 6051C>T, 6052A>C, 6053A>G, 6054A>T, 6055G>T, 6057A>T, 6060C>G, 6062T>A, 6066T>C, 6070A>C, 6072A>T, 6075A>G, 6076A>G, 6079G>A, 6081G>A, 6084A>G, 6087G>T, 6088T>G, 6089C>A, 6090C>T, 6091A>G, 6092A>T, 6093A>T, 6097T>A, 6098G>A, 6099G>A, 6101C>T, 6102T>G, 6114T>G, 6115C>A, 6116C>G, 6117A>G, 6118G>T, 6120A>T, 6125T>A, 6126C>T, 6128A>T, 6129T>C, 6130G>A, 6135T>C, 6136G>C, 6137A>T, 6138A>T, 6145C>T, 6147C>T, 6153A>T, 6154C>T, 6156G>A, 6158T>C, 6159G>T, 6163G>T, 6165A>C, 6168A>T, 6169G>T, 6173A>T, 6174T>G, 6178C>A, 6179A>T, 6181A>G, 6182G>A, 6183A>T, 6184A>T, 6185A>T, 6190G>A, 6192T>C, 6194A>G, 6195T>G, 6197C>T, 6198A>T, 6201T>C, 6203G>A, 6205G>C, 6206G>C, 6207A>T, 6207_6208insTTC, 6208A>C, 6209C>T, 6210A>C, 6213T>C, 6214G>A, 6215C>T, 6216A>G, 6220A>G, 6222C>G, 6224C>T, 6225A>G, 6228A>C, 6230A>T, 6240C>T, 6241A>G, 6243A>G, 6244T>C, 6247G>A, 6251T>A, 6255T>C, 6256G>C, 6259A>G, 6260A>G, 6261T>A, 6262G>A, 6263A>G, 6264A>G | | | | | | | | |

##### CDS

| | Begin | End | Coverage | Score | Concordance | Matches | Identities | I/D/M/F* | Stop Codons |
|---|---|---|---|---|---|---|---|---|---|
| SPBVa_gp4 | 304 | 388 | 10.7% | 347 | 56.2% | 85 (98.8%) | 43 (50.0%) | 1/0/0/0 | 0 |
| Protein mutations: | F310I (6043T>A), K313R (6052A>C 6053A>G 6054A>T), A314S (6055G>T 6057A>T), F316Y (6062T>A), I319L (6070A>C 6072A>T), M321V (6076A>G), E322K (6079G>A 6081G>A), K324N (6087G>T), S325D (6088T>G 6089C>A 6090C>T), K326V (6091A>G 6092A>T 6093A>T), W328K (6097T>A 6098G>A 6099G>A), T329M (6101C>T 6102T>G), P334R (6115C>A 6116C>G 6117A>G), E335Y (6118G>T 6120A>T), L337H (6125T>A 6126C>T), Y338F (6128A>T 6129T>C), E339K (6130G>A), E341L (6136G>C 6137A>T 6138A>T), P344S (6145C>T 6147C>T), M348T (6158T>C 6159G>T), A350S (6163G>T 6165A>C), A352S (6169G>T), D353V (6173A>T 6174T>G), Q355M (6178C>A 6179A>T), R356D (6181A>G 6182G>A 6183A>T), K357L (6184A>T 6185A>T), D359N (6190G>A 6192T>C), N360R (6194A>G 6195T>G), A361V (6197C>T 6198A>T), R363K (6203G>A), G364P (6205G>C 6206G>C 6207A>T), G364_T365insF (6207_6208insTTC), T365L (6208A>C 6209C>T 6210A>C), A367M (6214G>A 6215C>T 6216A>G), I369V (6220A>G 6222C>G), A370V (6224C>T 6225A>G), Y372F (6230A>T), I376V (6241A>G 6243A>G), V378I (6247G>A), F379Y (6251T>A), E381Q (6256G>C), N382G (6259A>G 6260A>G 6261T>A), E383R (6262G>A 6263A>G 6264A>G) | | | | | | | | |
| Codon mutations: | ATA306ATT (6033A>T), TTC307TTT (6036C>T), AAA309AAG (6042A>G), TTT310ATT (6043T>A), CTC312CTT (6051C>T), AAA313CGT (6052A>C 6053A>G 6054A>T), GCA314TCT (6055G>T 6057A>T), GGC315GGG (6060C>G), TTT316TAT (6062T>A), CAT317CAC (6066T>C), ATA319CTT (6070A>C 6072A>T), AGA320AGG (6075A>G), ATG321GTG (6076A>G), GAG322AAA (6079G>A 6081G>A), GAA323GAG (6084A>G), AAG324AAT (6087G>T), TCC325GAT (6088T>G 6089C>A 6090C>T), AAA326GTT (6091A>G 6092A>T 6093A>T), TGG328AAA (6097T>A 6098G>A 6099G>A), ACT329ATG (6101C>T 6102T>G), ACT333ACG (6114T>G), CCA334AGG (6115C>A 6116C>G 6117A>G), GAA335TAT (6118G>T 6120A>T), CTC337CAT (6125T>A 6126C>T), TAT338TTC (6128A>T 6129T>C), GAA339AAA (6130G>A), TTT340TTC (6135T>C), GAA341CTT (6136G>C 6137A>T 6138A>T), CCC344TCT (6145C>T 6147C>T), GGA346GGT (6153A>T), CTG347TTA (6154C>T 6156G>A), ATG348ACT (6158T>C 6159G>T), GCA350TCC (6163G>T 6165A>C), CCA351CCT (6168A>T), GCA352TCA (6169G>T), GAT353GTG (6173A>T 6174T>G), CAG355ATG (6178C>A 6179A>T), AGA356GAT (6181A>G 6182G>A 6183A>T), AAG357TTG (6184A>T 6185A>T), GAT359AAC (6190G>A 6192T>C), AAT360AGG (6194A>G 6195T>G), GCA361GTT (6197C>T 6198A>T), TTT362TTC (6201T>C), AGG363AAG (6203G>A), GGA364CCT (6205G>C 6206G>C 6207A>T), GGA364_ACA365insTTC (6207_6208insTTC), ACA365CTC (6208A>C 6209C>T 6210A>C), GAT366GAC (6213T>C), GCA367ATG (6214G>A 6215C>T 6216A>G), ATC369GTG (6220A>G 6222C>G), GCA370GTG (6224C>T 6225A>G), GTA371GTC (6228A>C), TAT372TTT (6230A>T), GAC375GAT (6240C>T), ATA376GTG (6241A>G 6243A>G), TTG377CTG (6244T>C), GTA378ATA (6247G>A), TTC379TAC (6251T>A), TCT380TCC (6255T>C), GAA381CAA (6256G>C), AAT382GGA (6259A>G 6260A>G 6261T>A), GAA383AGG (6262G>A 6263A>G 6264A>G) | | | | | | | | |

##### Proteins

| | Begin | End | Coverage | Score | Concordance | Matches | Identities | I/D/M/F* | Stop Codons |
|---|---|---|---|---|---|---|---|---|---|
| RNaseH/reverse transcriptase (YP_004581513.1) | 304 | 388 | 10.7% | 347 | 56.2% | 85 (98.8%) | 43 (50.0%) | 1/0/0/0 | 0 |
| Protein mutations: | F310I (6043T>A), K313R (6052A>C 6053A>G 6054A>T), A314S (6055G>T 6057A>T), F316Y (6062T>A), I319L (6070A>C 6072A>T), M321V (6076A>G), E322K (6079G>A 6081G>A), K324N (6087G>T), S325D (6088T>G 6089C>A 6090C>T), K326V (6091A>G 6092A>T 6093A>T), W328K (6097T>A 6098G>A 6099G>A), T329M (6101C>T 6102T>G), P334R (6115C>A 6116C>G 6117A>G), E335Y (6118G>T 6120A>T), L337H (6125T>A 6126C>T), Y338F (6128A>T 6129T>C), E339K (6130G>A), E341L (6136G>C 6137A>T 6138A>T), P344S (6145C>T 6147C>T), M348T (6158T>C 6159G>T), A350S (6163G>T 6165A>C), A352S (6169G>T), D353V (6173A>T 6174T>G), Q355M (6178C>A 6179A>T), R356D (6181A>G 6182G>A 6183A>T), K357L (6184A>T 6185A>T), D359N (6190G>A 6192T>C), N360R (6194A>G 6195T>G), A361V (6197C>T 6198A>T), R363K (6203G>A), G364P (6205G>C 6206G>C 6207A>T), G364_T365insF (6207_6208insTTC), T365L (6208A>C 6209C>T 6210A>C), A367M (6214G>A 6215C>T 6216A>G), I369V (6220A>G 6222C>G), A370V (6224C>T 6225A>G), Y372F (6230A>T), I376V (6241A>G 6243A>G), V378I (6247G>A), F379Y (6251T>A), E381Q (6256G>C), N382G (6259A>G 6260A>G 6261T>A), E383R (6262G>A 6263A>G 6264A>G) | | | | | | | | |
| Codon mutations: | ATA306ATT (6033A>T), TTC307TTT (6036C>T), AAA309AAG (6042A>G), TTT310ATT (6043T>A), CTC312CTT (6051C>T), AAA313CGT (6052A>C 6053A>G 6054A>T), GCA314TCT (6055G>T 6057A>T), GGC315GGG (6060C>G), TTT316TAT (6062T>A), CAT317CAC (6066T>C), ATA319CTT (6070A>C 6072A>T), AGA320AGG (6075A>G), ATG321GTG (6076A>G), GAG322AAA (6079G>A 6081G>A), GAA323GAG (6084A>G), AAG324AAT (6087G>T), TCC325GAT (6088T>G 6089C>A 6090C>T), AAA326GTT (6091A>G 6092A>T 6093A>T), TGG328AAA (6097T>A 6098G>A 6099G>A), ACT329ATG (6101C>T 6102T>G), ACT333ACG (6114T>G), CCA334AGG (6115C>A 6116C>G 6117A>G), GAA335TAT (6118G>T 6120A>T), CTC337CAT (6125T>A 6126C>T), TAT338TTC (6128A>T 6129T>C), GAA339AAA (6130G>A), TTT340TTC (6135T>C), GAA341CTT (6136G>C 6137A>T 6138A>T), CCC344TCT (6145C>T 6147C>T), GGA346GGT (6153A>T), CTG347TTA (6154C>T 6156G>A), ATG348ACT (6158T>C 6159G>T), GCA350TCC (6163G>T 6165A>C), CCA351CCT (6168A>T), GCA352TCA (6169G>T), GAT353GTG (6173A>T 6174T>G), CAG355ATG (6178C>A 6179A>T), AGA356GAT (6181A>G 6182G>A 6183A>T), AAG357TTG (6184A>T 6185A>T), GAT359AAC (6190G>A 6192T>C), AAT360AGG (6194A>G 6195T>G), GCA361GTT (6197C>T 6198A>T), TTT362TTC (6201T>C), AGG363AAG (6203G>A), GGA364CCT (6205G>C 6206G>C 6207A>T), GGA364_ACA365insTTC (6207_6208insTTC), ACA365CTC (6208A>C 6209C>T 6210A>C), GAT366GAC (6213T>C), GCA367ATG (6214G>A 6215C>T 6216A>G), ATC369GTG (6220A>G 6222C>G), GCA370GTG (6224C>T 6225A>G), GTA371GTC (6228A>C), TAT372TTT (6230A>T), GAC375GAT (6240C>T), ATA376GTG (6241A>G 6243A>G), TTG377CTG (6244T>C), GTA378ATA (6247G>A), TTC379TAC (6251T>A), TCT380TCC (6255T>C), GAA381CAA (6256G>C), AAT382GGA (6259A>G 6260A>G 6261T>A), GAA383AGG (6262G>A 6263A>G 6264A>G) | | | | | | | | |

*: Inserts / Deletes / Misaligned / Frameshifts

#### Analysis details

This analysis was performed with panviral2.64

### NGS Details (UN59): Badnavirus venabougainvilleae

#### Assembly

| | |
|---|---|
| **Coverage Length** | 285 (1 contig(s)) |
| **Depth Of Coverage** | 25.3 |
| **Number Of Reads** | 73 |
| **Reads Per Million** | 1.59 rpm (after QC) |
| **Ambiguities** | 0 |
| **Assembly Method** | de novo + reference guided assembly |
| **Consensus Caller** | Bcf Tools |

#### Coverage Map

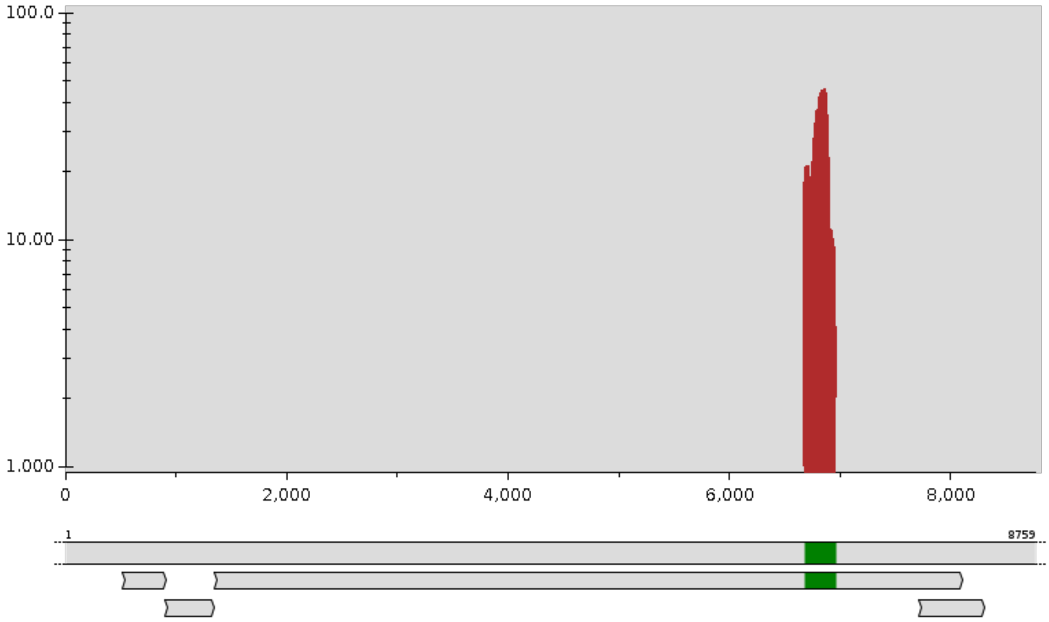

#### Assignment

| | |
|---|---|
| **Type** | Badnavirus venabougainvilleae (Taxonomy ID: 3047697) |
| **Reference Genome** | NC_011592.1 |
| **NT Identity (%)** | 55.2083 |
| **AA Identity (%)** | 46.875 |
| **Number Of Stop Codons** | 1 |
| **Number Of CDS** | 4 |

#### Alignment

| | |
|---|---|
| **Alignment Score** | 54.0 (NT) + 333.0 (AA) = 387.0 |
| **Concordance (%)** | 30.5687 |

| Alignment Method | Global, seeded, nucleotide + amino acids (AGA) |
|---|---|

#### Genome Region

Sequence starts at position 6678 and ends at position 6962 relative to NC_011592.1 reference sequence.

#### Alignment Detailed Statistics

| | Begin | End | Coverage | Score | Concordance | Matches | Identities | I/D/M/F* | Stop Codons |
|---|---|---|---|---|---|---|---|---|---|
| **NT** | **6678** | **6962** | **3.3%** | **54** | **9.5%** | **285 (99.0%)** | **159 (55.2%)** | **3/0** | |
| Mutations: | 6681A>T, 6682A>T, 6683T>G, 6686A>T, 6693A>G, 6694T>A, 6696T>A, 6702G>T, 6703A>C, 6704A>G, 6705G>T, 6706A>T, 6707G>C, 6708T>G, 6711A>T, 6713T>A, 6717T>C, 6721G>T, 6723T>A, 6724G>A, 6725C>A, 6726T>G, 6729G>T, 6730G>A, 6731A>G, 6732T>G, 6733C>G, 6734C>A, 6735A>G, 6736G>A, 6737A>G, 6738A>T, 6739T>G, 6740C>A, 6741C>T, 6744A>T, 6747A>T, 6748T>A, 6749G>A, 6753G>A, 6756C>G, 6761T>G, 6762T>G, 6764C>T, 6765T>G, 6766C>A, 6767C>G, 6768A>G, 6769C>T, 6771A>T, 6776T>A, 6777G>T, 6785G>T, 6786G>T, 6789A>T, 6790G>T, 6796C>T, 6798A>C, 6805C>T, 6809A>C, 6810G>T, 6813T>C, 6816A>T, 6818C>T, 6819A>G, 6821C>T, 6823A>G, 6828C>T, 6829C>A, 6830A>T, 6831A>G, 6832A>G, 6833G>A, 6834G>T, 6835A>C, 6836A>T, 6837G>T, 6841G>A, 6843C>T, 6845A>G, 6846C>G, 6847T>A, 6848G>T, 6849C>A, 6851T>A, 6852T>C, 6853T>A, 6854C>A, 6857G>C, 6858A>T, 6862A>C, 6863G>T, 6864C>T, 6867T>C, 6867_6868insATG, 6869T>A, 6870C>T, 6873T>G, 6875C>T, 6876T>G, 6879A>G, 6880T>C, 6881A>T, 6882C>T, 6894A>T, 6900G>C, 6902T>A, 6907G>C, 6909G>A, 6910A>G, 6913G>T, 6915A>G, 6916A>C, 6917G>A, 6921C>A, 6924T>C, 6925G>A, 6926A>T, 6928A>G, 6930G>A, 6934C>T, 6936C>G, 6940G>A, 6941T>C, 6946C>T, 6948T>G, 6951A>G | | | | | | | | |

##### CDS

| | Begin | End | Coverage | Score | Concordance | Matches | Identities | I/D/M/F* | Stop Codons |
|---|---|---|---|---|---|---|---|---|---|
| BSCVV_gp3 | 1779 | 1873 | 4.2% | 333 | 47.1% | 95 (99.0%) | 45 (46.9%) | 1/0/0/0 | 1 |
| Protein mutations: | K1779N (6681A>T), I1780C (6682A>T 6683T>G), Y1781F (6686A>T), F1784I (6694T>A 6696T>A), K1787R (6703A>C 6704A>G 6705G>T), F1790Y (6713T>A), V1793L (6721G>T 6723T>A), A1794K (6724G>A 6725C>A 6726T>G), M1795I (6729G>T), D1796R (6730G>A 6731A>G 6732T>G), P1797E (6733C>G 6734C>A 6735A>G), E1798S (6736G>A 6737A>G 6738A>T), S1799D (6739T>G 6740C>A 6741C>T), W1802K (6748T>A 6749G>A), L1806R (6761T>G 6762T>G), T1807M (6764C>T 6765T>G), P1808R (6766C>A 6767C>G 6768A>G), Q1809Y (6769C>T 6771A>T), L1811Y (6776T>A 6777G>T), W1814F (6785G>T 6786G>T), V1816F (6790G>T), P1818S (6796C>T 6798A>C), K1822T (6809A>C 6810G>T), P1825L (6818C>T 6819A>G), A1826V (6821C>T), I1827V (6823A>G), Q1829M (6829C>A 6830A>T 6831A>G), R1830D (6832A>G 6833G>A 6834G>T), K1831L (6835A>C 6836A>T 6837G>T), D1833N (6841G>A 6843C>T), N1834R (6845A>G 6846C>G), C1835I (6847T>A 6848G>T 6849C>A), F1836Y (6851T>A 6852T>C), S1837K (6853T>A 6854C>A), R1838P (6857G>C 6858A>T), S1840L (6862A>C 6863G>T 6864C>T), D1841_F1842insM (6867_6868insATG), F1842Y (6869T>A 6870C>T), I1843M (6873T>G), A1844V (6875C>T 6876T>G), Y1846L (6880T>C 6881A>T 6882C>T), F1853Y (6902T>A), E1855Q (6907G>C 6909G>A), S1856G (6910A>G), E1857* (6913G>T 6915A>G), R1858Q (6916A>C 6917G>A), D1859E (6921C>A), E1861M (6925G>A 6926A>T), K1862E (6928A>G 6930G>A), V1866T (6940G>A 6941T>C) | | | | | | | | |
| Codon mutations: | AAA1779AAT (6681A>T), ATC1780TGC (6682A>T 6683T>G), TAC1781TTC (6686A>T), AAA1783AAG (6693A>G), TTT1784ATA (6694T>A 6696T>A), CTG1786CTT (6702G>T), AAG1787CGT (6703A>C 6704A>G 6705G>T), AGT1788TCG (6706A>T 6707G>C 6708T>G), GGA1789GGT (6711A>T), TTT1790TAT (6713T>A), CAT1791CAC (6717T>C), GTT1793TTA (6721G>T 6723T>A), GCT1794AAG (6724G>A 6725C>A 6726T>G), ATG1795ATT (6729G>T), GAT1796AGG (6730G>A 6731A>G 6732T>G), CCA1797GAG (6733C>G 6734C>A 6735A>G), GAA1798AGT (6736G>A 6737A>G 6738A>T), TCC1799GAT (6739T>G 6740C>A 6741C>T), ATA1800ATT (6744A>T), CCA1801CCT (6747A>T), TGG1802AAG (6748T>A 6749G>A), ACG1803ACA (6753G>A), GCC1804GCG (6756C>G), CTT1806CGG (6761T>G 6762T>G), ACT1807ATG (6764C>T 6765T>G), CCA1808AGG (6766C>A 6767C>G 6768A>G), CAA1809TAT (6769C>T 6771A>T), TTG1811TAT (6776T>A 6777G>T), TGG1814TTT (6785G>T 6786G>T), CTA1815CTT (6789A>T), GTC1816TTC (6790G>T), CCA1818TCC (6796C>T 6798A>C), CTA1821TTA (6805C>T), AAG1822ACT (6809A>C 6810G>T), AAT1823AAC (6813T>C), GCA1824GCT (6816A>T), CCA1825CTG (6818C>T 6819A>G), GCT1826GTT (6821C>T), ATC1827GTC (6823A>G), TTC1828TTT (6828C>T), CAA1829ATG (6829C>A 6830A>T 6831A>G), AGG1830GAT (6832A>G 6833G>A 6834G>T), AAG1831CTT (6835A>C 6836A>T 6837G>T), GAC1833AAT (6841G>A 6843C>T), AAC1834AGG (6845A>G 6846C>G), TGC1835ATA (6847T>A 6848G>T 6849C>A), TTT1836TAC (6851T>A 6852T>C), TCA1837AAA (6853T>A 6854C>A), CGA1838CCT (6857G>C 6858A>T), AGC1840CTT (6862A>C 6863G>T 6864C>T), GAT1841GAC (6867T>C), GAT1841_TTC1842insATG (6867_6868insATG), TTC1842TAT (6869T>A 6870C>T), ATT1843ATG (6873T>G), GCT1844GTG (6875C>T 6876T>G), GTA1845GTG (6879A>G), TAC1846CTT (6880T>C 6881A>T 6882C>T), ATA1850ATT (6894A>T), GTG1852GTC (6900G>C), TTT1853TAT (6902T>A), GAG1855CAA (6907G>C 6909G>A), AGT1856GGT (6910A>G), GAA1857TAG (6913G>T 6915A>G), AGA1858CAA (6916A>C 6917G>A), GAC1859GAA (6921C>A), CAT1860CAC (6924T>C), GAG1861ATG (6925G>A 6926A>T), AAG1862GAA (6928A>G 6930G>A), CTC1864TTG (6934C>T 6936C>G), GTT1866ACT (6940G>A 6941T>C), CTT1868TTG (6946C>T 6948T>G), CAA1869CAG (6951A>G) | | | | | | | | |

##### Proteins

| | Begin | End | Coverage | Score | Concordance | Matches | Identities | I/D/M/F* | Stop Codons |
|---|---|---|---|---|---|---|---|---|---|
| polyprotein (YP_002321513.1) | 1779 | 1873 | 4.2% | 333 | 47.1% | 95 (99.0%) | 45 (46.9%) | 1/0/0/0 | 1 |
| Protein mutations: | K1779N (6681A>T), I1780C (6682A>T 6683T>G), Y1781F (6686A>T), F1784I (6694T>A 6696T>A), K1787R (6703A>C 6704A>G 6705G>T), F1790Y (6713T>A), V1793L (6721G>T 6723T>A), A1794K (6724G>A 6725C>A 6726T>G), M1795I (6729G>T), D1796R (6730G>A 6731A>G 6732T>G), P1797E (6733C>G 6734C>A 6735A>G), E1798S (6736G>A 6737A>G 6738A>T), S1799D (6739T>G 6740C>A 6741C>T), W1802K (6748T>A 6749G>A), L1806R (6761T>G 6762T>G), T1807M (6764C>T 6765T>G), P1808R (6766C>A 6767C>G 6768A>G), Q1809Y (6769C>T 6771A>T), L1811Y (6776T>A 6777G>T), W1814F (6785G>T 6786G>T), V1816F (6790G>T), P1818S (6796C>T 6798A>C), K1822T (6809A>C 6810G>T), P1825L (6818C>T 6819A>G), A1826V (6821C>T), I1827V (6823A>G), Q1829M (6829C>A 6830A>T 6831A>G), R1830D (6832A>G 6833G>A 6834G>T), K1831L (6835A>C 6836A>T 6837G>T), D1833N (6841G>A 6843C>T), N1834R (6845A>G 6846C>G), C1835I (6847T>A 6848G>T 6849C>A), F1836Y (6851T>A 6852T>C), S1837K (6853T>A 6854C>A), R1838P (6857G>C 6858A>T), S1840L (6862A>C 6863G>T 6864C>T), D1841_F1842insM (6867_6868insATG), F1842Y (6869T>A 6870C>T), I1843M (6873T>G), A1844V (6875C>T 6876T>G), Y1846L (6880T>C 6881A>T 6882C>T), F1853Y (6902T>A), E1855Q (6907G>C 6909G>A), S1856G (6910A>G), E1857* (6913G>T 6915A>G), R1858Q (6916A>C 6917G>A), D1859E (6921C>A), E1861M (6925G>A 6926A>T), K1862E (6928A>G 6930G>A), V1866T (6940G>A 6941T>C) | | | | | | | | |
| Codon mutations: | AAA1779AAT (6681A>T), ATC1780TGC (6682A>T 6683T>G), TAC1781TTC (6686A>T), AAA1783AAG (6693A>G), TTT1784ATA (6694T>A 6696T>A), CTG1786CTT (6702G>T), AAG1787CGT (6703A>C 6704A>G 6705G>T), AGT1788TCG (6706A>T 6707G>C 6708T>G), GGA1789GGT (6711A>T), TTT1790TAT (6713T>A), CAT1791CAC (6717T>C), GTT1793TTA (6721G>T 6723T>A), GCT1794AAG (6724G>A 6725C>A 6726T>G), ATG1795ATT (6729G>T), GAT1796AGG (6730G>A 6731A>G 6732T>G), CCA1797GAG (6733C>G 6734C>A 6735A>G), GAA1798AGT (6736G>A 6737A>G 6738A>T), TCC1799GAT (6739T>G 6740C>A 6741C>T), ATA1800ATT (6744A>T), CCA1801CCT (6747A>T), TGG1802AAG (6748T>A 6749G>A), ACG1803ACA (6753G>A), GCC1804GCG (6756C>G), CTT1806CGG (6761T>G 6762T>G), ACT1807ATG (6764C>T 6765T>G), CCA1808AGG (6766C>A 6767C>G 6768A>G), CAA1809TAT (6769C>T 6771A>T), TTG1811TAT (6776T>A 6777G>T), TGG1814TTT (6785G>T 6786G>T), CTA1815CTT (6789A>T), GTC1816TTC (6790G>T), CCA1818TCC (6796C>T 6798A>C), CTA1821TTA (6805C>T), AAG1822ACT (6809A>C 6810G>T), AAT1823AAC (6813T>C), GCA1824GCT (6816A>T), CCA1825CTG (6818C>T 6819A>G), GCT1826GTT (6821C>T), ATC1827GTC (6823A>G), TTC1828TTT (6828C>T), CAA1829ATG (6829C>A 6830A>T 6831A>G), AGG1830GAT (6832A>G 6833G>A 6834G>T), AAG1831CTT (6835A>C 6836A>T 6837G>T), GAC1833AAT (6841G>A 6843C>T), AAC1834AGG (6845A>G 6846C>G), TGC1835ATA (6847T>A 6848G>T 6849C>A), TTT1836TAC (6851T>A 6852T>C), TCA1837AAA (6853T>A 6854C>A), CGA1838CCT (6857G>C 6858A>T), AGC1840CTT (6862A>C 6863G>T 6864C>T), GAT1841GAC (6867T>C), GAT1841_TTC1842insATG (6867_6868insATG), TTC1842TAT (6869T>A 6870C>T), ATT1843ATG (6873T>G), GCT1844GTG (6875C>T 6876T>G), GTA1845GTG (6879A>G), TAC1846CTT (6880T>C 6881A>T 6882C>T), ATA1850ATT (6894A>T), GTG1852GTC (6900G>C), TTT1853TAT (6902T>A), GAG1855CAA (6907G>C 6909G>A), AGT1856GGT (6910A>G), GAA1857TAG (6913G>T 6915A>G), AGA1858CAA (6916A>C 6917G>A), GAC1859GAA (6921C>A), CAT1860CAC (6924T>C), GAG1861ATG (6925G>A 6926A>T), AAG1862GAA (6928A>G 6930G>A), CTC1864TTG (6934C>T 6936C>G), GTT1866ACT (6940G>A 6941T>C), CTT1868TTG (6946C>T 6948T>G), CAA1869CAG (6951A>G) | | | | | | | | |

*: Inserts / Deletes / Misaligned / Frameshifts

#### Analysis details

This analysis was performed with panviral2.64

### NGS Details (UN59): Badnavirus occultipomeae

#### Assembly

| | |
|---|---|
| **Coverage Length** | 301 (1 contig(s)) |
| **Depth Of Coverage** | 18.2 |
| **Number Of Reads** | 50 |
| **Reads Per Million** | 1.09 rpm (after QC) |
| **Ambiguities** | 0 |
| **Assembly Method** | de novo + reference guided assembly |
| **Consensus Caller** | Bcf Tools |

#### Coverage Map

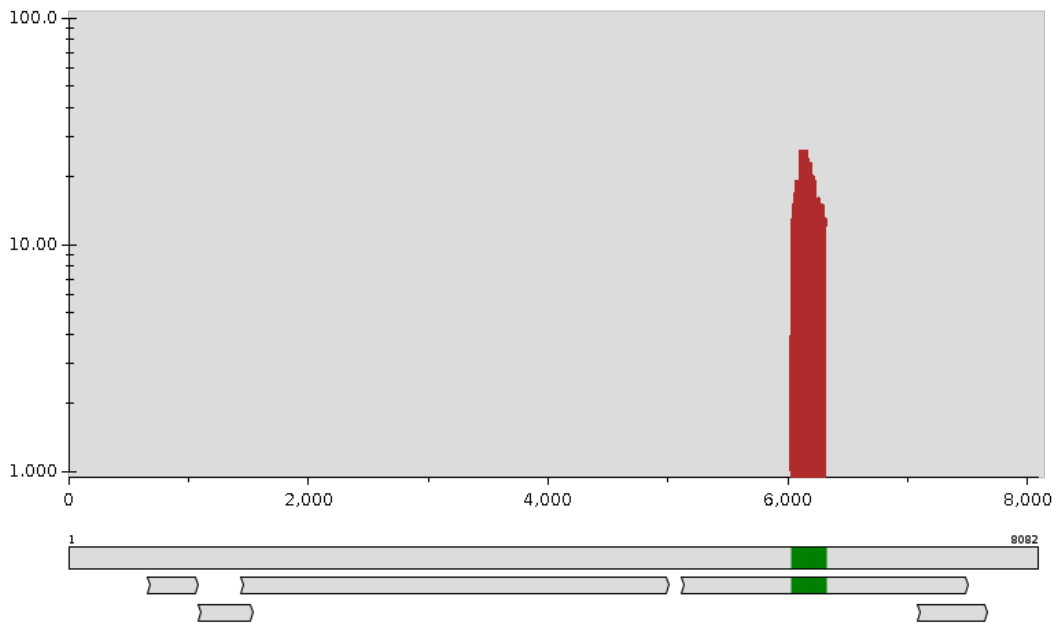

#### Assignment

| | |
|---|---|
| **Type** | Badnavirus occultipomeae (Taxonomy ID: 3048353) |
| **Reference Genome** | NC_015655.1 |
| **NT Identity (%)** | 57.7558 |
| **AA Identity (%)** | 49.0196 |
| **Number Of Stop Codons** | 2 |
| **Number Of CDS** | 5 |

#### Alignment

| | |
|---|---|
| **Alignment Score** | 78.0 (NT) + 196.0 (AA) = 274.0 |
| **Concordance (%)** | 24.2525 |

| Alignment Method | Global, seeded, nucleotide + amino acids (AGA) |
|---|---|

#### Genome Region

Sequence starts at position 6025 and ends at position 6325 relative to NC_015655.1 reference sequence.

#### Alignment Detailed Statistics

| | Begin | End | Coverage | Score | Concordance | Matches | Identities | I/D/M/F* | Stop Codons |
|---|---|---|---|---|---|---|---|---|---|
| **NT** | **6025** | **6325** | **3.7%** | **78** | **13.2%** | **300 (98.7%)** | **175 (57.6%)** | **3/1** | |
| Mutations: | 6031A>G, 6036C>T, 6039A>T, 6042A>G, 6043T>A, 6045T>A, 6048T>C, 6051C>G, 6053A>G, 6060C>T, 6062T>A, 6069G>A, 6072A>T, 6079G>C, 6081G>T, 6084A>G, 6086A>T, 6087G>A, 6088T>G, 6089C>A, 6092A>T, 6094C>T, 6095C>A, 6096A>C, 6097T>A, 6098G>A, 6109T>A, 6114T>A, 6116C>A, 6117A>T, 6118G>C, 6119A>T, 6120A>G, 6125T>A, 6126C>T, 6135T>C, 6136G>A, 6138A>G, 6147C>T, 6153A>T, 6154C>T, 6156G>A, 6158T>C, 6159G>C, 6162T>C, 6165A>C, 6168A>T, 6171A>C, 6172G>A, 6173A>C, 6174T>C, 6178C>T, 6183A>T, 6184A>T, 6185A>T, 6186G>A, 6190G>A, 6191A>G, 6192T>C, 6193A>C, 6197C>T, 6198A>G, 6202delA, 6203G>T, 6204G>A, 6208A>T, 6209C>T, 6210_6211insCTT, 6217T>G, 6220A>T, 6221T>G, 6222C>T, 6224C>T, 6225A>T, 6226G>A, 6227T>G, 6228A>T, 6230A>T, 6232A>T, 6235G>A, 6240C>T, 6243A>T, 6244T>C, 6249A>G, 6251T>A, 6252C>T, 6253T>A, 6254C>G, 6256G>C, 6257A>C, 6258A>T, 6262G>A, 6263A>T, 6264A>C, 6265G>T, 6267A>G, 6268G>A, 6269A>C, 6270G>T, 6275A>T, 6276G>A, 6277G>T, 6278A>G, 6279T>G, 6282T>G, 6283C>T, 6285C>G, 6286T>G, 6287T>A, 6288A>G, 6291T>G, 6294G>C, 6295G>T, 6296C>T, 6297C>G, 6300A>T, 6304G>C, 6306G>T, 6309A>C, 6315A>G, 6316G>A, 6317G>A, 6319C>T, 6320T>G, 6321T>G, 6322A>T, 6325C>T | | | | | | | | |

##### CDS

| | Begin | End | Coverage | Score | Concordance | Matches | Identities | I/D/M/F* | Stop Codons |
|---|---|---|---|---|---|---|---|---|---|
| SPBVa_gp4 | 304 | 404 | 12.7% | 196 | 27.3% | 101 (99.0%) | 50 (49.0%) | 1/0/1/1 | 2 |
| Protein mutations: | I306V (6031A>G), F310I (6043T>A 6045T>A), K313R (6053A>G), F316Y (6062T>A), E322H (6079G>C 6081G>T), K324I (6086A>T 6087G>A), S325D (6088T>G 6089C>A), K326I (6092A>T), P327Y (6094C>T 6095C>A 6096A>C), W328K (6097T>A 6098G>A), W332R (6109T>A), P334H (6116C>A 6117A>T), E335L (6118G>C 6119A>T 6120A>G), L337H (6125T>A 6126C>T), E341K (6136G>A 6138A>G), M348T (6158T>C 6159G>C), D353T (6172G>A 6173A>C 6174T>C), Q355* (6178C>T), R356S (6183A>T), K357L (6184A>T 6185A>T 6186G>A), D359S (6190G>A 6191A>G 6192T>C), N360H (6193A>C), A361V (6197C>T 6198A>G), T365L (6208A>T 6209C>T), T365_D366insL (6210_6211insCTT), F368V (6217T>G), I369C (6220A>T 6221T>G 6222C>T), A370V (6224C>T 6225A>T), V371S (6226G>A 6227T>G 6228A>T), Y372F (6230A>T), I373F (6232A>T), D374N (6235G>A), F379Y (6251T>A 6252C>T), E381P (6256G>C 6257A>C 6258A>T), E383I (6262G>A 6263A>T 6264A>C), E384* (6265G>T 6267A>G), E385T (6268G>A 6269A>C 6270G>T), E387V (6275A>T 6276G>A), D388W (6277G>T 6278A>G 6279T>G), H389Q (6282T>G), L391E (6286T>G 6287T>A 6288A>G), N392K (6291T>G), L393F (6294G>C), A394L (6295G>T 6296C>T 6297C>G), Q395H (6300A>T), V397L (6304G>C 6306G>T), R398S (6309A>C), G401K (6316G>A 6317G>A), L402W (6319C>T 6320T>G 6321T>G), I403F (6322A>T) | | | | | | | | |
| Codon mutations: | ATA306GTA (6031A>G), TTC307TTT (6036C>T), TCA308TCT (6039A>T), AAA309AAG (6042A>G), TTT310ATA (6043T>A 6045T>A), GAT311GAC (6048T>C), CTC312CTG (6051C>G), AAA313AGA (6053A>G), GGC315GGT (6060C>T), TTT316TAT (6062T>A), CAG318CAA (6069G>A), ATA319ATT (6072A>T), GAG322CAT (6079G>C 6081G>T), GAA323GAG (6084A>G), AAG324ATA (6086A>T 6087G>A), TCC325GAC (6088T>G 6089C>A), AAA326ATA (6092A>T), CCA327TAC (6094C>T 6095C>A 6096A>C), TGG328AAG (6097T>A 6098G>A), TGG332AGG (6109T>A), ACT333ACA (6114T>A), CCA334CAT (6116C>A 6117A>T), GAA335CTG (6118G>C 6119A>T 6120A>G), CTC337CAT (6125T>A 6126C>T), TTT340TTC (6135T>C), GAA341AAG (6136G>A 6138A>G), CCC344CCT (6147C>T), GGA346GGT (6153A>T), CTG347TTA (6154C>T 6156G>A), ATG348ACC (6158T>C 6159G>C), AAT349AAC (6162T>C), GCA350GCC (6165A>C), CCA351CCT (6168A>T), GCA352GCC (6171A>C), GAT353ACC (6172G>A 6173A>C 6174T>C), CAG355TAG (6178C>T), AGA356AGT (6183A>T), AAG357TTA (6184A>T 6185A>T 6186G>A), GAT359AGC (6190G>A 6191A>G 6192T>C), AAT360CAT (6193A>C), GCA361GTG (6197C>T 6198A>G), AGG363-TA (6202delA 6203G>T 6204G>A), ACA365TTA (6208A>T 6209C>T), ACA365_GAT366insCTT (6210_6211insCTT), TTT368GTT (6217T>G), ATC369TGT (6220A>T 6221T>G 6222C>T), GCA370GTT (6224C>T 6225A>T), GTA371AGT (6226G>A 6227T>G 6228A>T), TAT372TTT (6230A>T), ATT373TTT (6232A>T), GAT374AAT (6235G>A), GAC375GAT (6240C>T), ATA376ATT (6243A>T), TTG377CTG (6244T>C), GTA378GTG (6249A>G), TTC379TAT (6251T>A 6252C>T), TCT380AGT (6253T>A 6254C>G), GAA381CCT (6256G>C 6257A>C 6258A>T), GAA383ATC (6262G>A 6263A>T 6264A>C), GAA384TAG (6265G>T 6267A>G), GAG385ACT (6268G>A 6269A>C 6270G>T), GAG387GTA (6275A>T 6276G>A), GAT388TGG (6277G>T 6278A>G 6279T>G), CAT389CAG (6282T>G), CTC390TTG (6283C>T 6285C>G), TTA391GAG (6286T>G 6287T>A 6288A>G), AAT392AAG (6291T>G), TTG393TTC (6294G>C), GCC394TTG (6295G>T 6296C>T 6297C>G), CAA395CAT (6300A>T), GTG397CTT (6304G>C 6306G>T), AGA398AGC (6309A>C), GAA400GAG (6315A>G), GGG401AAG (6316G>A 6317G>A), CTT402TGG (6319C>T 6320T>G 6321T>G), ATT403TTT (6322A>T), CTA404T.. (6325C>T) | | | | | | | | |

##### Proteins

| | Begin | End | Coverage | Score | Concordance | Matches | Identities | I/D/M/F* | Stop Codons |
|---|---|---|---|---|---|---|---|---|---|
| RNaseH/reverse transcriptase (YP_004581513.1) | 304 | 404 | 12.7% | 196 | 27.3% | 101 (99.0%) | 50 (49.0%) | 1/0/1/1 | 2 |
| Protein mutations: | I306V (6031A>G), F310I (6043T>A 6045T>A), K313R (6053A>G), F316Y (6062T>A), E322H (6079G>C 6081G>T), K324I (6086A>T 6087G>A), S325D (6088T>G 6089C>A), K326I (6092A>T), P327Y (6094C>T 6095C>A 6096A>C), W328K (6097T>A 6098G>A), W332R (6109T>A), P334H (6116C>A 6117A>T), E335L (6118G>C 6119A>T 6120A>G), L337H (6125T>A 6126C>T), E341K (6136G>A 6138A>G), M348T (6158T>C 6159G>C), D353T (6172G>A 6173A>C 6174T>C), Q355* (6178C>T), R356S (6183A>T), K357L (6184A>T 6185A>T 6186G>A), D359S (6190G>A 6191A>G 6192T>C), N360H (6193A>C), A361V (6197C>T 6198A>G), T365L (6208A>T 6209C>T), T365_D366insL (6210_6211insCTT), F368V (6217T>G), I369C (6220A>T 6221T>G 6222C>T), A370V (6224C>T 6225A>T), V371S (6226G>A 6227T>G 6228A>T), Y372F (6230A>T), I373F (6232A>T), D374N (6235G>A), F379Y (6251T>A 6252C>T), E381P (6256G>C 6257A>C 6258A>T), E383I (6262G>A 6263A>T 6264A>C), E384* (6265G>T 6267A>G), E385T (6268G>A 6269A>C 6270G>T), E387V (6275A>T 6276G>A), D388W (6277G>T 6278A>G 6279T>G), H389Q (6282T>G), L391E (6286T>G 6287T>A 6288A>G), N392K (6291T>G), L393F (6294G>C), A394L (6295G>T 6296C>T 6297C>G), Q395H (6300A>T), V397L (6304G>C 6306G>T), R398S (6309A>C), G401K (6316G>A 6317G>A), L402W (6319C>T 6320T>G 6321T>G), I403F (6322A>T) | | | | | | | | |
| Codon mutations: | ATA306GTA (6031A>G), TTC307TTT (6036C>T), TCA308TCT (6039A>T), AAA309AAG (6042A>G), TTT310ATA (6043T>A 6045T>A), GAT311GAC (6048T>C), CTC312CTG (6051C>G), AAA313AGA (6053A>G), GGC315GGT (6060C>T), TTT316TAT (6062T>A), CAG318CAA (6069G>A), ATA319ATT (6072A>T), GAG322CAT (6079G>C 6081G>T), GAA323GAG (6084A>G), AAG324ATA (6086A>T 6087G>A), TCC325GAC (6088T>G 6089C>A), AAA326ATA (6092A>T), CCA327TAC (6094C>T 6095C>A 6096A>C), TGG328AAG (6097T>A 6098G>A), TGG332AGG (6109T>A), ACT333ACA (6114T>A), CCA334CAT (6116C>A 6117A>T), GAA335CTG (6118G>C 6119A>T 6120A>G), CTC337CAT (6125T>A 6126C>T), TTT340TTC (6135T>C), GAA341AAG (6136G>A 6138A>G), CCC344CCT (6147C>T), GGA346GGT (6153A>T), CTG347TTA (6154C>T 6156G>A), ATG348ACC (6158T>C 6159G>C), AAT349AAC (6162T>C), GCA350GCC (6165A>C), CCA351CCT (6168A>T), GCA352GCC (6171A>C), GAT353ACC (6172G>A 6173A>C 6174T>C), CAG355TAG (6178C>T), AGA356AGT (6183A>T), AAG357TTA (6184A>T 6185A>T 6186G>A), GAT359AGC (6190G>A 6191A>G 6192T>C), AAT360CAT (6193A>C), GCA361GTG (6197C>T 6198A>G), AGG363-TA (6202delA 6203G>T 6204G>A), ACA365TTA (6208A>T 6209C>T), ACA365_GAT366insCTT (6210_6211insCTT), TTT368GTT (6217T>G), ATC369TGT (6220A>T 6221T>G 6222C>T), GCA370GTT (6224C>T 6225A>T), GTA371AGT (6226G>A 6227T>G 6228A>T), TAT372TTT (6230A>T), ATT373TTT (6232A>T), GAT374AAT (6235G>A), GAC375GAT (6240C>T), ATA376ATT (6243A>T), TTG377CTG (6244T>C), GTA378GTG (6249A>G), TTC379TAT (6251T>A 6252C>T), TCT380AGT (6253T>A 6254C>G), GAA381CCT (6256G>C 6257A>C 6258A>T), GAA383ATC (6262G>A 6263A>T 6264A>C), GAA384TAG (6265G>T 6267A>G), GAG385ACT (6268G>A 6269A>C 6270G>T), GAG387GTA (6275A>T 6276G>A), GAT388TGG (6277G>T 6278A>G 6279T>G), CAT389CAG (6282T>G), CTC390TTG (6283C>T 6285C>G), TTA391GAG (6286T>G 6287T>A 6288A>G), AAT392AAG (6291T>G), TTG393TTC (6294G>C), GCC394TTG (6295G>T 6296C>T 6297C>G), CAA395CAT (6300A>T), GTG397CTT (6304G>C 6306G>T), AGA398AGC (6309A>C), GAA400GAG (6315A>G), GGG401AAG (6316G>A 6317G>A), CTT402TGG (6319C>T 6320T>G 6321T>G), ATT403TTT (6322A>T), CTA404T.. (6325C>T) | | | | | | | | |

*: Inserts / Deletes / Misaligned / Frameshifts

#### Analysis details

This analysis was performed with panviral2.64

### NGS Details (UN59): Badnavirus venaribis

#### Assembly

| | |
|---|---|
| **Coverage Length** | 441 (1 contig(s)) |
| **Depth Of Coverage** | 9.7 |
| **Number Of Reads** | 36 |
| **Reads Per Million** | 0.78 rpm (after QC) |
| **Ambiguities** | 0 |
| **Assembly Method** | de novo + reference guided assembly |
| **Consensus Caller** | Bcf Tools |

#### Coverage Map

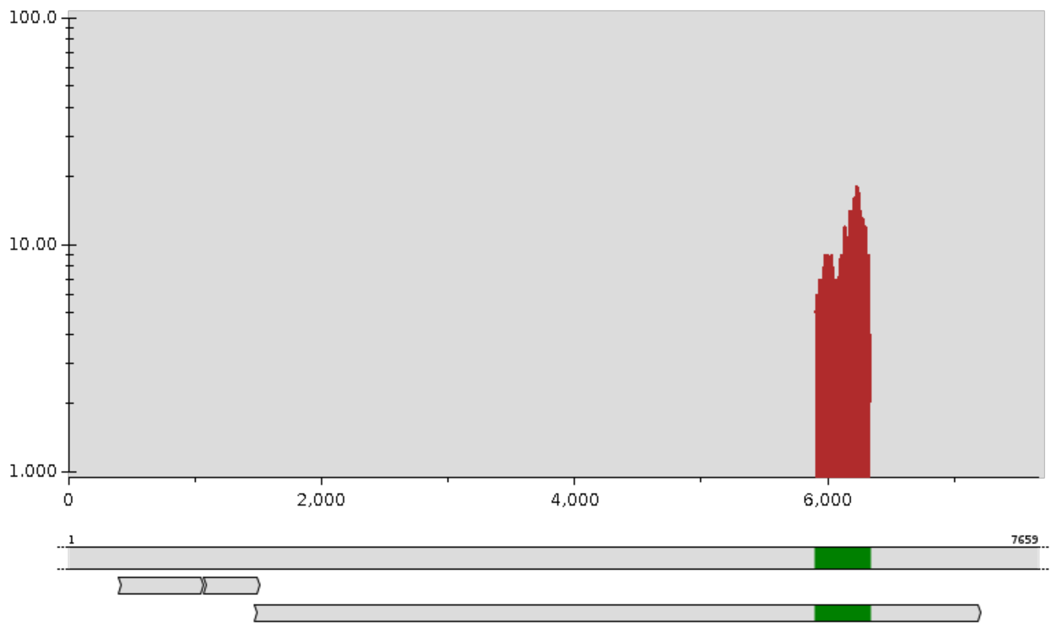

#### Assignment

| | |
|---|---|
| **Type** | Badnavirus venaribis (Taxonomy ID: 3047843) |
| **Reference Genome** | NC_018105.1 |
| **NT Identity (%)** | 52.9279 |
| **AA Identity (%)** | 45.9459 |
| **Number Of Stop Codons** | 0 |
| **Number Of CDS** | 3 |

#### Alignment

| | |
|---|---|
| **Alignment Score** | 37.0 (NT) + 494.0 (AA) = 531.0 |
| **Concordance (%)** | 28.2297 |

| Alignment Method | Global, seeded, nucleotide + amino acids (AGA) |
|---|---|

#### Genome Region

Sequence starts at position 5902 and ends at position 6342 relative to NC_018105.1 reference sequence.

#### Alignment Detailed Statistics

| | Begin | End | Coverage | Score | Concordance | Matches | Identities | I/D/M/F* | Stop Codons |
|---|---|---|---|---|---|---|---|---|---|
| **NT** | **5902** | **6342** | **5.8%** | **37** | **4.3%** | **438 (98.0%)** | **235 (52.6%)** | **6/3** | |
| Mutations: | 5902G>T, 5903T>G, 5904C>T, 5905T>G, 5907C>A, 5913T>C, 5914C>A, 5916C>G, 5918C>A, 5919T>A, 5920C>T, 5922G>A, 5927A>C, 5934G>T, 5935C>G, 5936A>T, 5943C>T, 5944C>A, 5948A>T, 5949C>T, 5950A>C, 5953T>A, 5955G>T, 5958T>A, 5960G>T, 5961C>T, 5964C>T, 5967C>G, 5968T>G, 5969C>A, 5970G>A, 5971A>T, 5973C>A, 5974G>T, 5978C>A, 5979A>T, 5981C>A, 5982T>G, 5983G>T, 5985T>G, 5986G>C, 5987G>A, 5988C>T, 5989A>G, 5990A>G, 5991C>A, 5997A>G, 5998A>T, 5999T>A, 6000C>T, 6002A>T, 6003C>T, 6004A>T, 6005G>C, 6006C>T, 6009A>G, 6010T>C, 6012C>A, 6020A>G, 6021G>A, 6022G>T, 6029T>A, 6037G>A, 6039C>A, 6040C>A, 6041T>G, 6046G>T, 6047A>T, 6048A>T, 6054C>T, 6055T>A, 6056C>A, 6060C>A, 6064T>A, 6065G>A, 6066G>A, 6069A>G, 6070G>A, 6072G>A, 6077T>G, 6078C>A, 6081G>C, 6083C>A, 6084C>T, 6085G>C, 6086T>A, 6087A>G, 6090A>G, 6091T>C, 6092T>A, 6099A>G, 6101G>T, 6102G>C, 6105G>A, 6108A>G, 6114T>C, 6117C>T, 6120T>C, 6121A>C, 6124G>A, 6126T>C, 6129C>T, 6132T>C, 6135A>C, 6136G>T, 6138G>C, 6139G>A, 6140T>C, 6141T>A, 6144C>T, 6150G>C, 6151A>C, 6152A>T, 6153A>C, 6157G>A, 6160A>C, 6162C>A, 6163T>A, 6164G>T, 6165C>A, 6168_6169insAATGCT, 6171C>T, 6172_6174delAAG, 6177G>T, 6181G>A, 6183G>A, 6186C>T, 6187G>A, 6190G>C, 6191C>T, 6197A>T, 6198T>C, 6199A>T, 6201C>T, 6204C>T, 6207C>T, 6210C>A, 6213C>T, 6214G>A, 6216C>A, 6218T>A, 6221G>A, 6224A>G, 6225T>C, 6226T>A, 6227C>A, 6228C>T, 6229C>T, 6230T>G, 6232C>T, 6233A>C, 6234A>T, 6237A>T, 6241G>C, 6242A>T, 6243G>C, 6244A>T, 6245G>C, 6246C>A, 6250T>C, 6252A>G, 6254G>A, 6255G>A, 6260T>C, 6261G>T, 6262C>T, 6264G>C, 6265G>C, 6268G>A, 6271T>C, 6272G>A, 6273C>T, 6279G>C, 6284G>T, 6285G>T, 6286T>C, 6288G>T, 6289G>T, 6291G>T, 6294G>C, 6295T>A, 6296C>A, 6297A>G, 6298C>A, 6299C>A, 6303A>C, 6306A>G, 6307A>T, 6308T>G, 6309G>T, 6310A>G, 6311A>C, 6312G>T, 6313G>T, 6317C>G, 6318T>G, 6324A>T, 6325A>C, 6326C>A, 6327A>G, 6328G>A, 6330A>T, 6333A>C, 6335T>A, 6342G>T | | | | | | | | |

##### CDS

| | Begin | End | Coverage | Score | Concordance | Matches | Identities | I/D/M/F* | Stop Codons |
|---|---|---|---|---|---|---|---|---|---|
| B227_gp3 | 1479 | 1625 | 7.7% | 494 | 48.6% | 146 (98.0%) | 68 (45.6%) | 2/1/0/0 | 0 |
| Protein mutations: | V1479C (5902G>T 5903T>G 5904C>T), F1480V (5905T>G 5907C>A), A1484E (5918C>A 5919T>A), N1487T (5927A>C), H1490V (5935C>G 5936A>T), Q1493K (5944C>A), Y1494F (5948A>T 5949C>T), T1495P (5950A>C), L1496I (5953T>A 5955G>T), G1498V (5960G>T 5961C>T), N1500K (5967C>G), S1501E (5968T>G 5969C>A 5970G>A), I1502L (5971A>T 5973C>A), V1503L (5974G>T), A1504D (5978C>A 5979A>T), A1505E (5981C>A 5982T>G), V1506L (5983G>T 5985T>G), G1507H (5986G>C 5987G>A 5988C>T), N1508G (5989A>G 5990A>G 5991C>A), I1511Y (5998A>T 5999T>A 6000C>T), Y1512F (6002A>T 6003C>T), F1515L (6010T>C 6012C>A), K1518R (6020A>G 6021G>A), A1519S (6022G>T), F1521Y (6029T>A), V1524I (6037G>A 6039C>A), L1525R (6040C>A 6041T>G), E1527F (6046G>T 6047A>T 6048A>T), S1530N (6055T>A 6056C>A), W1533K (6064T>A 6065G>A 6066G>A), A1535T (6070G>A 6072G>A), I1537R (6077T>G 6078C>A), P1539H (6083C>A 6084C>T), V1540Q (6085G>C 6086T>A 6087A>G), F1542H (6091T>C 6092T>A), W1545F (6101G>T 6102G>C), I1552L (6121A>C), A1553T (6124G>A 6126T>C), A1557S (6136G>T 6138G>C), V1558T (6139G>A 6140T>C 6141T>A), R1561S (6150G>C), K1562L (6151A>C 6152A>T 6153A>C), D1564N (6157G>A), N1565Q (6160A>C 6162C>A), C1566I (6163T>A 6164G>T 6165C>A), F1567_H1568insNA (6168_6169insAATGCT), K1569del (6172_6174delAAG), E1572K (6181G>A 6183G>A), V1574I (6187G>A), A1575L (6190G>C 6191C>T), Y1577F (6197A>T 6198T>C), I1578F (6199A>T 6201C>T), V1583I (6214G>A 6216C>A), F1584Y (6218T>A), S1585N (6221G>A), N1586S (6224A>G 6225T>C), S1587N (6226T>A 6227C>A 6228C>T), L1588W (6229C>T 6230T>G), Q1589S (6232C>T 6233A>C 6234A>T), E1590D (6237A>T), E1592L (6241G>C 6242A>T 6243G>C), R1596Q (6254G>A 6255G>A), M1598T (6260T>C 6261G>T), L1599F (6262C>T 6264G>C), E1600Q (6265G>C), V1601I (6268G>A), C1602H (6271T>C 6272G>A 6273C>T), K1604N (6279G>C), G1606V (6284G>T 6285G>T), V1608F (6289G>T 6291G>T), S1610K (6295T>A 6296C>A 6297A>G), P1611K (6298C>A 6299C>A), M1614C (6307A>T 6308T>G 6309G>T), K1615A (6310A>G 6311A>C 6312G>T), V1616F (6313G>T), A1617G (6317C>G 6318T>G), T1620Q (6325A>C 6326C>A 6327A>G), V1621I (6328G>A 6330A>T), E1622D (6333A>C), F1623Y (6335T>A) | | | | | | | | |
| Codon mutations: | GTC1479TGT (5902G>T 5903T>G 5904C>T), TTC1480GTA (5905T>G 5907C>A), TAT1482TAC (5913T>C), CGC1483AGG (5914C>A 5916C>G), GCT1484GAA (5918C>A 5919T>A), CTG1485TTA (5920C>T 5922G>A), AAC1487ACC (5927A>C), ACG1489ACT (5934G>T), CAC1490GTC (5935C>G 5936A>T), GAC1492GAT (5943C>T), CAG1493AAG (5944C>A), TAC1494TTT (5948A>T 5949C>T), ACA1495CCA (5950A>C), TTG1496ATT (5953T>A 5955G>T), CCT1497CCA (5958T>A), GGC1498GTT (5960G>T 5961C>T), ATC1499ATT (5964C>T), AAC1500AAG (5967C>G), TCG1501GAA (5968T>G 5969C>A 5970G>A), ATC1502TTA (5971A>T 5973C>A), GTA1503TTA (5974G>T), GCA1504GAT (5978C>A 5979A>T), GCT1505GAG (5981C>A 5982T>G), GTT1506TTG (5983G>T 5985T>G), GGC1507CAT (5986G>C 5987G>A 5988C>T), AAC1508GGA (5989A>G 5990A>G 5991C>A), AAA1510AAG (5997A>G), ATC1511TAT (5998A>T 5999T>A 6000C>T), TAC1512TTT (6002A>T 6003C>T), AGC1513TCT (6004A>T 6005G>C 6006C>T), AAA1514AAG (6009A>G), TTC1515CTA (6010T>C 6012C>A), AAG1518AGA (6020A>G 6021G>A), GCT1519TCT (6022G>T), TTC1521TAC (6029T>A), GTC1524ATA (6037G>A 6039C>A), CTA1525AGA (6040C>A 6041T>G), GAA1527TTT (6046G>T 6047A>T 6048A>T), TCC1529TCT (6054C>T), TCT1530AAT (6055T>A 6056C>A), ATC1531ATA (6060C>A), TGG1533AAA (6064T>A 6065G>A 6066G>A), ACA1534ACG (6069A>G), GCG1535ACA (6070G>A 6072G>A), ATC1537AGA (6077T>G 6078C>A), ACG1538ACC (6081G>C), CCC1539CAT (6083C>A 6084C>T), GTA1540CAG (6085G>C 6086T>A 6087A>G), GGA1541GGG (6090A>G), TTC1542CAC (6091T>C 6092T>A), GAA1544GAG (6099A>G), TGG1545TTC (6101G>T 6102G>C), TTG1546TTA (6105G>A), GTA1547GTG (6108A>G), CCT1549CCC (6114T>C), TTC1550TTT (6117C>T), GGT1551GGC (6120T>C), ATT1552CTT (6121A>C), GCT1553ACC (6124G>A 6126T>C), AAC1554AAT (6129C>T), GCT1555GCC (6132T>C), CCA1556CCC (6135A>C), GCG1557TCC (6136G>T 6138G>C), GTT1558ACA (6139G>A 6140T>C 6141T>A), TTC1559TTT (6144C>T), AGG1561AGC (6150G>C), AAA1562CTC (6151A>C 6152A>T 6153A>C), GAT1564AAT (6157G>A), AAC1565CAA (6160A>C 6162C>A), TGC1566ATA (6163T>A 6164G>T 6165C>A), TTC1567_CAC1568insAATGCT (6168_6169insAATGCT), CAC1568CAT (6171C>T), AAG1569del (6172_6174delAAG), CTG1570CTT (6177G>T), GAG1572AAA (6181G>A 6183G>A), TTC1573TTT (6186C>T), GTC1574ATC (6187G>A), GCG1575CTG (6190G>C 6191C>T), TAT1577TTC (6197A>T 6198T>C), ATC1578TTT (6199A>T 6201C>T), GAC1579GAT (6204C>T), GAC1580GAT (6207C>T), ATC1581ATA (6210C>A), CTC1582CTT (6213C>T), GTC1583ATA (6214G>A 6216C>A), TTC1584TAC (6218T>A), AGC1585AAC (6221G>A), AAT1586AGC (6224A>G 6225T>C), TCC1587AAT (6226T>A 6227C>A 6228C>T), CTG1588TGG (6229C>T 6230T>G), CAA1589TCT (6232C>T 6233A>C 6234A>T), GAA1590GAT (6237A>T), GAG1592CTC (6241G>C 6242A>T 6243G>C), AGC1593TCA (6244A>T 6245G>C 6246C>A), TTA1595CTG (6250T>C 6252A>G), CGG1596CAA (6254G>A 6255G>A), ATG1598ACT (6260T>C 6261G>T), CTG1599TTC (6262C>T 6264G>C), GAA1600CAA (6265G>C), GTT1601ATT (6268G>A), TGC1602CAT (6271T>C 6272G>A 6273C>T), AAG1604AAC (6279G>C), GGG1606GTT (6284G>T 6285G>T), TTG1607CTT (6286T>C 6288G>T), GTG1608TTT (6289G>T 6291G>T), CTG1609CTC (6294G>C), TCA1610AAG (6295T>A 6296C>A 6297A>G), CCA1611AAA (6298C>A 6299C>A), ACA1612ACC (6303A>C), AAA1613AAG (6306A>G), ATG1614TGT (6307A>T 6308T>G 6309G>T), AAG1615GCT (6310A>G 6311A>C 6312G>T), GTT1616TTT (6313G>T), GCT1617GGG (6317C>G 6318T>G), ACA1619ACT (6324A>T), ACA1620CAG (6325A>C 6326C>A 6327A>G), GTA1621ATT (6328G>A 6330A>T), GAA1622GAC (6333A>C), TTC1623TAC (6335T>A), GGG1625GGT (6342G>T) | | | | | | | | |

##### Proteins

| | Begin | End | Coverage | Score | Concordance | Matches | Identities | I/D/M/F* | Stop Codons |
|---|---|---|---|---|---|---|---|---|---|
| polyprotein (YP_006495799.1) | 1479 | 1625 | 7.7% | 494 | 48.6% | 146 (98.0%) | 68 (45.6%) | 2/1/0/0 | 0 |
| Protein mutations: | V1479C (5902G>T 5903T>G 5904C>T), F1480V (5905T>G 5907C>A), A1484E (5918C>A 5919T>A), N1487T (5927A>C), H1490V (5935C>G 5936A>T), Q1493K (5944C>A), Y1494F (5948A>T 5949C>T), T1495P (5950A>C), L1496I (5953T>A 5955G>T), G1498V (5960G>T 5961C>T), N1500K (5967C>G), S1501E (5968T>G 5969C>A 5970G>A), I1502L (5971A>T 5973C>A), V1503L (5974G>T), A1504D (5978C>A 5979A>T), A1505E (5981C>A 5982T>G), V1506L (5983G>T 5985T>G), G1507H (5986G>C 5987G>A 5988C>T), N1508G (5989A>G 5990A>G 5991C>A), I1511Y (5998A>T 5999T>A 6000C>T), Y1512F (6002A>T 6003C>T), F1515L (6010T>C 6012C>A), K1518R (6020A>G 6021G>A), A1519S (6022G>T), F1521Y (6029T>A), V1524I (6037G>A 6039C>A), L1525R (6040C>A 6041T>G), E1527F (6046G>T 6047A>T 6048A>T), S1530N (6055T>A 6056C>A), W1533K (6064T>A 6065G>A 6066G>A), A1535T (6070G>A 6072G>A), I1537R (6077T>G 6078C>A), P1539H (6083C>A 6084C>T), V1540Q (6085G>C 6086T>A 6087A>G), F1542H (6091T>C 6092T>A), W1545F (6101G>T 6102G>C), I1552L (6121A>C), A1553T (6124G>A 6126T>C), A1557S (6136G>T 6138G>C), V1558T (6139G>A 6140T>C 6141T>A), R1561S (6150G>C), K1562L (6151A>C 6152A>T 6153A>C), D1564N (6157G>A), N1565Q (6160A>C 6162C>A), C1566I (6163T>A 6164G>T 6165C>A), F1567_H1568insNA (6168_6169insAATGCT), K1569del (6172_6174delAAG), E1572K (6181G>A 6183G>A), V1574I (6187G>A), A1575L (6190G>C 6191C>T), Y1577F (6197A>T 6198T>C), I1578F (6199A>T 6201C>T), V1583I (6214G>A 6216C>A), F1584Y (6218T>A), S1585N (6221G>A), N1586S (6224A>G 6225T>C), S1587N (6226T>A 6227C>A 6228C>T), L1588W (6229C>T 6230T>G), Q1589S (6232C>T 6233A>C 6234A>T), E1590D (6237A>T), E1592L (6241G>C 6242A>T 6243G>C), R1596Q (6254G>A 6255G>A), M1598T (6260T>C 6261G>T), L1599F (6262C>T 6264G>C), E1600Q (6265G>C), V1601I (6268G>A), C1602H (6271T>C 6272G>A 6273C>T), K1604N (6279G>C), G1606V (6284G>T 6285G>T), V1608F (6289G>T 6291G>T), S1610K (6295T>A 6296C>A 6297A>G), P1611K (6298C>A 6299C>A), M1614C (6307A>T 6308T>G 6309G>T), K1615A (6310A>G 6311A>C 6312G>T), V1616F (6313G>T), A1617G (6317C>G 6318T>G), T1620Q (6325A>C 6326C>A 6327A>G), V1621I (6328G>A 6330A>T), E1622D (6333A>C), F1623Y (6335T>A) | | | | | | | | |

| | Begin | End | Coverage | Score | Concordance | Matches | Identities | I/D/M/F* | Stop Codons |
|---|---|---|---|---|---|---|---|---|---|
| **NT** | **5902** | **6342** | **5.8%** | **37** | **4.3%** | **438 (98.0%)** | **235 (52.6%)** | **6/3** | |
| Codon mutations: | GTC1479TGT (5902G>T 5903T>G 5904C>T), TTC1480GTA (5905T>G 5907C>A), TAT1482TAC (5913T>C), CGC1483AGG (5914C>A 5916C>G), GCT1484GAA (5918C>A 5919T>A), CTG1485TTA (5920C>T 5922G>A), AAC1487ACC (5927A>C), ACG1489ACT (5934G>T), CAC1490GTC (5935C>G 5936A>T), GAC1492GAT (5943C>T), CAG1493AAG (5944C>A), TAC1494TTT (5948A>T 5949C>T), ACA1495CCA (5950A>C), TTG1496ATT (5953T>A 5955G>T), CCT1497CCA (5958T>A), GGC1498GTT (5960G>T 5961C>T), ATC1499ATT (5964C>T), AAC1500AAG (5967C>G), TCG1501GAA (5968T>G 5969C>A 5970G>A), ATC1502TTA (5971A>T 5973C>A), GTA1503TTA (5974G>T), GCA1504GAT (5978C>A 5979A>T), GCT1505GAG (5981C>A 5982T>G), GTT1506TTG (5983G>T 5985T>G), GGC1507CAT (5986G>C 5987G>A 5988C>T), AAC1508GGA (5989A>G 5990A>G 5991C>A), AAA1510AAG (5997A>G), ATC1511TAT (5998A>T 5999T>A 6000C>T), TAC1512TTT (6002A>T 6003C>T), AGC1513TCT (6004A>T 6005G>C 6006C>T), AAA1514AAG (6009A>G), TTC1515CTA (6010T>C 6012C>A), AAG1518AGA (6020A>G 6021G>A), GCT1519TCT (6022G>T), TTC1521TAC (6029T>A), GTC1524ATA (6037G>A 6039C>A), CTA1525AGA (6040C>A 6041T>G), GAA1527TTT (6046G>T 6047A>T 6048A>T), TCC1529TCT (6054C>T), TCT1530AAT (6055T>A 6056C>A), ATC1531ATA (6060C>A), TGG1533AAA (6064T>A 6065G>A 6066G>A), ACA1534ACG (6069A>G), GCG1535ACA (6070G>A 6072G>A), ATC1537AGA (6077T>G 6078C>A), ACG1538ACC (6081G>C), CCC1539CAT (6083C>A 6084C>T), GTA1540CAG (6085G>C 6086T>A 6087A>G), GGA1541GGG (6090A>G), TTC1542CAC (6091T>C 6092T>A), GAA1544GAG (6099A>G), TGG1545TTC (6101G>T 6102G>C), TTG1546TTA (6105G>A), GTA1547GTG (6108A>G), CCT1549CCC (6114T>C), TTC1550TTT (6117C>T), GGT1551GGC (6120T>C), ATT1552CTT (6121A>C), GCT1553ACC (6124G>A 6126T>C), AAC1554AAT (6129C>T), GCT1555GCC (6132T>C), CCA1556CCC (6135A>C), GCG1557TCC (6136G>T 6138G>C), GTT1558ACA (6139G>A 6140T>C 6141T>A), TTC1559TTT (6144C>T), AGG1561AGC (6150G>C), AAA1562CTC (6151A>C 6152A>T 6153A>C), GAT1564AAT (6157G>A), AAC1565CAA (6160A>C 6162C>A), TGC1566ATA (6163T>A 6164G>T 6165C>A), TTC1567_CAC1568insAATGCT (6168_6169insAATGCT), CAC1568CAT (6171C>T), AAG1569del (6172_6174delAAG), CTG1570CTT (6177G>T), GAG1572AAA (6181G>A 6183G>A), TTC1573TTT (6186C>T), GTC1574ATC (6187G>A), GCG1575CTG (6190G>C 6191C>T), TAT1577TTC (6197A>T 6198T>C), ATC1578TTT (6199A>T 6201C>T), GAC1579GAT (6204C>T), GAC1580GAT (6207C>T), ATC1581ATA (6210C>A), CTC1582CTT (6213C>T), GTC1583ATA (6214G>A 6216C>A), TTC1584TAC (6218T>A), AGC1585AAC (6221G>A), AAT1586AGC (6224A>G 6225T>C), TCC1587AAT (6226T>A 6227C>A 6228C>T), CTG1588TGG (6229C>T 6230T>G), CAA1589TCT (6232C>T 6233A>C 6234A>T), GAA1590GAT (6237A>T), GAG1592CTC (6241G>C 6242A>T 6243G>C), AGC1593TCA (6244A>T 6245G>C 6246C>A), TTA1595CTG (6250T>C 6252A>G), CGG1596CAA (6254G>A 6255G>A), ATG1598ACT (6260T>C 6261G>T), CTG1599TTC (6262C>T 6264G>C), GAA1600CAA (6265G>C), GTT1601ATT (6268G>A), TGC1602CAT (6271T>C 6272G>A 6273C>T), AAG1604AAC (6279G>C), GGG1606GTT (6284G>T 6285G>T), TTG1607CTT (6286T>C 6288G>T), GTG1608TTT (6289G>T 6291G>T), CTG1609CTC (6294G>C), TCA1610AAG (6295T>A 6296C>A 6297A>G), CCA1611AAA (6298C>A 6299C>A), ACA1612ACC (6303A>C), AAA1613AAG (6306A>G), ATG1614TGT (6307A>T 6308T>G 6309G>T), AAG1615GCT (6310A>G 6311A>C 6312G>T), GTT1616TTT (6313G>T), GCT1617GGG (6317C>G 6318T>G), ACA1619ACT (6324A>T), ACA1620CAG (6325A>C 6326C>A 6327A>G), GTA1621ATT (6328G>A 6330A>T), GAA1622GAC (6333A>C), TTC1623TAC (6335T>A), GGG1625GGT (6342G>T) | | | | | | | | |

*: Inserts / Deletes / Misaligned / Frameshifts

#### Analysis details

This analysis was performed with panviral2.64

### NGS Details (UN59): Dioscorea bacilliform virus

#### Assembly

| | |
|---|---|
| **Coverage Length** | 278 (1 contig(s)) |
| **Depth Of Coverage** | 14.2 |
| **Number Of Reads** | 35 |
| **Reads Per Million** | 0.76 rpm (after QC) |
| **Ambiguities** | 0 |
| **Assembly Method** | de novo + reference guided assembly |
| **Consensus Caller** | Bcf Tools |

#### Coverage Map

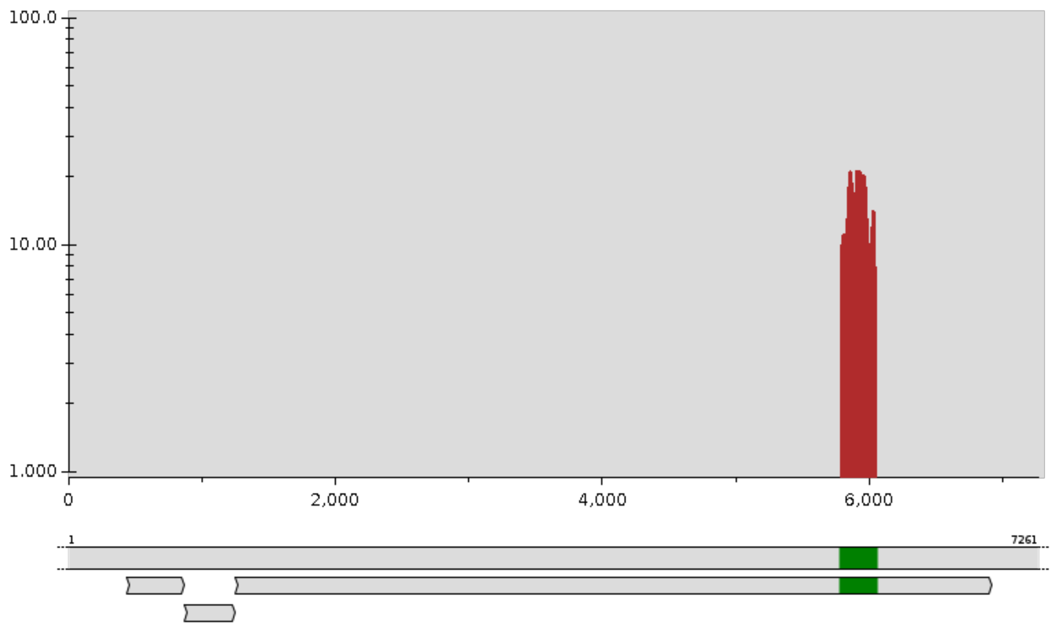

#### Assignment

| | |
|---|---|
| **Type** | Dioscorea bacilliform virus (Taxonomy ID: 52996) |
| **Reference Genome** | NC_009010.1 |
| **NT Identity (%)** | 55.1601 |
| **AA Identity (%)** | 45.1613 |
| **Number Of Stop Codons** | 1 |
| **Number Of CDS** | 3 |

#### Alignment

| | |
|---|---|
| **Alignment Score** | 52.0 (NT) + 293.0 (AA) = 345.0 |
| **Concordance (%)** | 28.6545 |

| Alignment Method | Global, seeded, nucleotide + amino acids (AGA) |
|---|---|

#### Genome Region

Sequence starts at position 5783 and ends at position 6060 relative to NC_009010.1 reference sequence.

#### Alignment Detailed Statistics

| | Begin | End | Coverage | Score | Concordance | Matches | Identities | I/D/M/F* | Stop Codons |
|---|---|---|---|---|---|---|---|---|---|
| **NT** | **5783** | **6060** | **3.8%** | **52** | **9.4%** | **278 (98.9%)** | **155 (55.2%)** | **3/0** | |
| Mutations: | 5790G>A, 5793G>A, 5796G>C, 5800C>A, 5801T>G, 5803G>A, 5805G>T, 5807C>A, 5808A>T, 5809G>T, 5810G>T, 5811A>T, 5816T>A, 5817T>C, 5819A>T, 5820C>T, 5825G>A, 5826G>T, 5827T>G, 5832G>T, 5838A>C, 5841C>T, 5844A>C, 5848A>T, 5849A>C, 5850G>A, 5857C>A, 5859A>T, 5862A>C, 5863G>A, 5864T>C, 5865G>A, 5868C>T, 5871G>A, 5872A>G, 5874G>T, 5875A>C, 5876A>T, 5881G>A, 5884A>T, 5886C>T, 5892C>T, 5894G>A, 5895A>G, 5897G>C, 5898T>C, 5898_5899insTTT, 5899A>C, 5900C>T, 5902G>A, 5903A>G, 5904A>G, 5905G>A, 5908T>C, 5909T>A, 5911A>G, 5913T>A, 5914G>C, 5915C>T, 5919A>T, 5921A>T, 5923A>T, 5931T>C, 5937G>T, 5938G>A, 5940G>C, 5942T>A, 5943T>C, 5944T>A, 5945C>G, 5946T>C, 5947G>A, 5948A>G, 5952A>T, 5953G>C, 5954A>T, 5965T>C, 5970G>A, 5973T>C, 5974C>T, 5976G>A, 5980A>T, 5981T>C, 5982A>T, 5983C>G, 5985G>T, 5986C>T, 5989C>G, 5992A>G, 5994C>T, 5995T>A, 5996G>T, 5997T>G, 6003A>C, 6007G>C, 6008G>A, 6012T>A, 6013G>T, 6016C>G, 6017T>C, 6020G>A, 6021C>G, 6023C>A, 6024A>G, 6025A>T, 6027C>T, 6031A>T, 6032T>G, 6033G>C, 6034A>T, 6035A>T, 6036G>C, 6037A>T, 6039A>T, 6043A>G, 6044C>T, 6045T>G, 6047A>G, 6050C>G, 6051A>G, 6054C>T, 6057G>A, 6059T>A, 6060C>T | | | | | | | | |

##### CDS

| | Begin | End | Coverage | Score | Concordance | Matches | Identities | I/D/M/F* | Stop Codons |
|---|---|---|---|---|---|---|---|---|---|
| DBV_gp3 | 1514 | 1605 | 4.9% | 293 | 44.9% | 92 (98.9%) | 42 (45.2%) | 1/0/0/0 | 1 |
| Protein mutations: | W1515* (5790G>A), L1519R (5800C>A 5801T>G), A1520T (5803G>A 5805G>T), P1521H (5807C>A 5808A>T), G1522F (5809G>T 5810G>T 5811A>T), L1524H (5816T>A 5817T>C), Y1525F (5819A>T 5820C>T), W1527Y (5825G>A 5826G>T), L1528V (5827T>G), K1535S (5848A>T 5849A>C 5850G>A), P1538T (5857C>A 5859A>T), V1540T (5863G>A 5864T>C 5865G>A), R1543G (5872A>G 5874G>T), K1544L (5875A>C 5876A>T), D1546N (5881G>A), N1547Y (5884A>T 5886C>T), R1550Q (5894G>A 5895A>G), G1551A (5897G>C 5898T>C), G1551_T1552insF (5898_5899insTTT), T1552L (5899A>C 5900C>T), E1553R (5902G>A 5903A>G 5904A>G), E1554K (5905G>A), F1555H (5908T>C 5909T>A), I1556V (5911A>G 5913T>A), A1557L (5914G>C 5915C>T), Y1559F (5921A>T), I1560F (5923A>T), V1565I (5938G>A 5940G>C), F1566Y (5942T>A 5943T>C), E1568R (5947G>A 5948A>G), E1570L (5953G>C 5954A>T), I1579S (5980A>T 5981T>C 5982A>T), L1580V (5983C>G 5985G>T), L1581F (5986C>T), Q1582E (5989C>G), I1583V (5992A>G 5994C>T), C1584M (5995T>A 5996G>T 5997T>G), Q1586H (6003A>C), G1588Q (6007G>C 6008G>A), V1590F (6013G>T), L1591A (6016C>G 6017T>C), S1592K (6020G>A 6021C>G), P1593Q (6023C>A 6024A>G), T1594S (6025A>T 6027C>T), M1596C (6031A>T 6032T>G 6033G>C), K1597F (6034A>T 6035A>T 6036G>C), I1598F (6037A>T 6039A>T), T1600V (6043A>G 6044C>T 6045T>G), K1601R (6047A>G), T1602R (6050C>G 6051A>G), F1605Y (6059T>A 6060C>T) | | | | | | | | |
| Codon mutations: | TGG1515TGA (5790G>A), ACG1516ACA (5793G>A), GCG1517GCC (5796G>C), CTA1519AGA (5800C>A 5801T>G), GCG1520ACT (5803G>A 5805G>T), CCA1521CAT (5807C>A 5808A>T), GGA1522TTT (5809G>T 5810G>T 5811A>T), CTT1524CAC (5816T>A 5817T>C), TAC1525TTT (5819A>T 5820C>T), TGG1527TAT (5825G>A 5826G>T), TTG1528GTG (5827T>G), GTG1529GTT (5832G>T), CCA1531CCC (5838A>C), TTC1532TTT (5841C>T), GGA1533GGC (5844A>C), AAG1535TCA (5848A>T 5849A>C 5850G>A), CCA1538ACT (5857C>A 5859A>T), GCA1539GCC (5862A>C), GTG1540ACA (5863G>A 5864T>C 5865G>A), TTC1541TTT (5868C>T), CAG1542CAA (5871G>A), AGG1543GGT (5872A>G 5874G>T), AAG1544CTG (5875A>C 5876A>T), GAC1546AAC (5881G>A), AAC1547TAT (5884A>T 5886C>T), TTC1549TTT (5892C>T), CGA1550CAG (5894G>A 5895A>G), GGT1551GCC (5897G>C 5898T>C), GGT1551_ACT1552insTTT (5898_5899insTTT), ACT1552CTT (5899A>C 5900C>T), GAA1553AGG (5902G>A 5903A>G 5904A>G), GAA1554AAA (5905G>A), TTT1555CAT (5908T>C 5909T>A), ATT1556GTA (5911A>G 5913T>A), GCT1557CTT (5914G>C 5915C>T), GTA1558GTT (5919A>T), TAC1559TTC (5921A>T), ATT1560TTT (5923A>T), GAT1562GAC (5931T>C), CTG1564CTT (5937G>T), GTG1565ATC (5938G>A 5940G>C), TTT1566TAC (5942T>A 5943T>C), TCT1567AGC (5944T>A 5945C>G 5946T>C), GAA1568AGA (5947G>A 5948A>G), ACA1569ACT (5952A>T), GAA1570CTA (5953G>C 5954A>T), TTG1574CTG (5965T>C), AAG1575AAA (5970G>A), CAT1576CAC (5973T>C), CTG1577TTA (5974C>T 5976G>A), ATA1579TCT (5980A>T 5981T>C 5982A>T), CTG1580GTT (5983C>G 5985G>T), CTT1581TTT (5986C>T), CAA1582GAA (5989C>G), ATC1583GTT (5992A>G 5994C>T), TGT1584ATG (5995T>A 5996G>T 5997T>G), CAA1586CAC (6003A>C), GGG1588CAG (6007G>C 6008G>A), CTT1589CTA (6012T>A), GTT1590TTT (6013G>T), CTT1591GCT (6016C>G 6017T>C), AGC1592AAG (6020G>A 6021C>G), CCA1593CAG (6023C>A 6024A>G), ACC1594TCT (6025A>T 6027C>T), ATG1596TGC (6031A>T 6032T>G 6033G>C), AAG1597TTC (6034A>T 6035A>T 6036G>C), ATA1598TTT (6037A>T 6039A>T), ACT1600GTG (6043A>G 6044C>T 6045T>G), AAA1601AGA (6047A>G), ACA1602AGG (6050C>G 6051A>G), ATC1603ATT (6054C>T), GAG1604GAA (6057G>A), TTC1605TAT (6059T>A 6060C>T) | | | | | | | | |

##### Proteins

| | Begin | End | Coverage | Score | Concordance | Matches | Identities | I/D/M/F* | Stop Codons |
|---|---|---|---|---|---|---|---|---|---|
| ORF3 protein (YP_001036293.1) | 1514 | 1605 | 4.9% | 293 | 44.9% | 92 (98.9%) | 42 (45.2%) | 1/0/0/0 | 1 |
| Protein mutations: | W1515* (5790G>A), L1519R (5800C>A 5801T>G), A1520T (5803G>A 5805G>T), P1521H (5807C>A 5808A>T), G1522F (5809G>T 5810G>T 5811A>T), L1524H (5816T>A 5817T>C), Y1525F (5819A>T 5820C>T), W1527Y (5825G>A 5826G>T), L1528V (5827T>G), K1535S (5848A>T 5849A>C 5850G>A), P1538T (5857C>A 5859A>T), V1540T (5863G>A 5864T>C 5865G>A), R1543G (5872A>G 5874G>T), K1544L (5875A>C 5876A>T), D1546N (5881G>A), N1547Y (5884A>T 5886C>T), R1550Q (5894G>A 5895A>G), G1551A (5897G>C 5898T>C), G1551_T1552insF (5898_5899insTTT), T1552L (5899A>C 5900C>T), E1553R (5902G>A 5903A>G 5904A>G), E1554K (5905G>A), F1555H (5908T>C 5909T>A), I1556V (5911A>G 5913T>A), A1557L (5914G>C 5915C>T), Y1559F (5921A>T), I1560F (5923A>T), V1565I (5938G>A 5940G>C), F1566Y (5942T>A 5943T>C), E1568R (5947G>A 5948A>G), E1570L (5953G>C 5954A>T), I1579S (5980A>T 5981T>C 5982A>T), L1580V (5983C>G 5985G>T), L1581F (5986C>T), Q1582E (5989C>G), I1583V (5992A>G 5994C>T), C1584M (5995T>A 5996G>T 5997T>G), Q1586H (6003A>C), G1588Q (6007G>C 6008G>A), V1590F (6013G>T), L1591A (6016C>G 6017T>C), S1592K (6020G>A 6021C>G), P1593Q (6023C>A 6024A>G), T1594S (6025A>T 6027C>T), M1596C (6031A>T 6032T>G 6033G>C), K1597F (6034A>T 6035A>T 6036G>C), I1598F (6037A>T 6039A>T), T1600V (6043A>G 6044C>T 6045T>G), K1601R (6047A>G), T1602R (6050C>G 6051A>G), F1605Y (6059T>A 6060C>T) | | | | | | | | |
| Codon mutations: | TGG1515TGA (5790G>A), ACG1516ACA (5793G>A), GCG1517GCC (5796G>C), CTA1519AGA (5800C>A 5801T>G), GCG1520ACT (5803G>A 5805G>T), CCA1521CAT (5807C>A 5808A>T), GGA1522TTT (5809G>T 5810G>T 5811A>T), CTT1524CAC (5816T>A 5817T>C), TAC1525TTT (5819A>T 5820C>T), TGG1527TAT (5825G>A 5826G>T), TTG1528GTG (5827T>G), GTG1529GTT (5832G>T), CCA1531CCC (5838A>C), TTC1532TTT (5841C>T), GGA1533GGC (5844A>C), AAG1535TCA (5848A>T 5849A>C 5850G>A), CCA1538ACT (5857C>A 5859A>T), GCA1539GCC (5862A>C), GTG1540ACA (5863G>A 5864T>C 5865G>A), TTC1541TTT (5868C>T), CAG1542CAA (5871G>A), AGG1543GGT (5872A>G 5874G>T), AAG1544CTG (5875A>C 5876A>T), GAC1546AAC (5881G>A), AAC1547TAT (5884A>T 5886C>T), TTC1549TTT (5892C>T), CGA1550CAG (5894G>A 5895A>G), GGT1551GCC (5897G>C 5898T>C), GGT1551_ACT1552insTTT (5898_5899insTTT), ACT1552CTT (5899A>C 5900C>T), GAA1553AGG (5902G>A 5903A>G 5904A>G), GAA1554AAA (5905G>A), TTT1555CAT (5908T>C 5909T>A), ATT1556GTA (5911A>G 5913T>A), GCT1557CTT (5914G>C 5915C>T), GTA1558GTT (5919A>T), TAC1559TTC (5921A>T), ATT1560TTT (5923A>T), GAT1562GAC (5931T>C), CTG1564CTT (5937G>T), GTG1565ATC (5938G>A 5940G>C), TTT1566TAC (5942T>A 5943T>C), TCT1567AGC (5944T>A 5945C>G 5946T>C), GAA1568AGA (5947G>A 5948A>G), ACA1569ACT (5952A>T), GAA1570CTA (5953G>C 5954A>T), TTG1574CTG (5965T>C), AAG1575AAA (5970G>A), CAT1576CAC (5973T>C), CTG1577TTA (5974C>T 5976G>A), ATA1579TCT (5980A>T 5981T>C 5982A>T), CTG1580GTT (5983C>G 5985G>T), CTT1581TTT (5986C>T), CAA1582GAA (5989C>G), ATC1583GTT (5992A>G 5994C>T), TGT1584ATG (5995T>A 5996G>T 5997T>G), CAA1586CAC (6003A>C), GGG1588CAG (6007G>C 6008G>A), CTT1589CTA (6012T>A), GTT1590TTT (6013G>T), CTT1591GCT (6016C>G 6017T>C), AGC1592AAG (6020G>A 6021C>G), CCA1593CAG (6023C>A 6024A>G), ACC1594TCT (6025A>T 6027C>T), ATG1596TGC (6031A>T 6032T>G 6033G>C), AAG1597TTC (6034A>T 6035A>T 6036G>C), ATA1598TTT (6037A>T 6039A>T), ACT1600GTG (6043A>G 6044C>T 6045T>G), AAA1601AGA (6047A>G), ACA1602AGG (6050C>G 6051A>G), ATC1603ATT (6054C>T), GAG1604GAA (6057G>A), TTC1605TAT (6059T>A 6060C>T) | | | | | | | | |

*: Inserts / Deletes / Misaligned / Frameshifts

#### Analysis details

This analysis was performed with panviral2.64

### NGS Details (UN59): Errantivirus

#### Assembly

| | |
|---|---|
| **Coverage Length** | 498 (10 contig(s)) |
| **Depth Of Coverage** | 8.5 |
| **Number Of Reads** | 32 |
| **Reads Per Million** | 0.70 rpm (after QC) |
| **Ambiguities** | 0 |
| **Assembly Method** | de novo + reference guided assembly |
| **Consensus Caller** | Bcf Tools |

#### Coverage Map

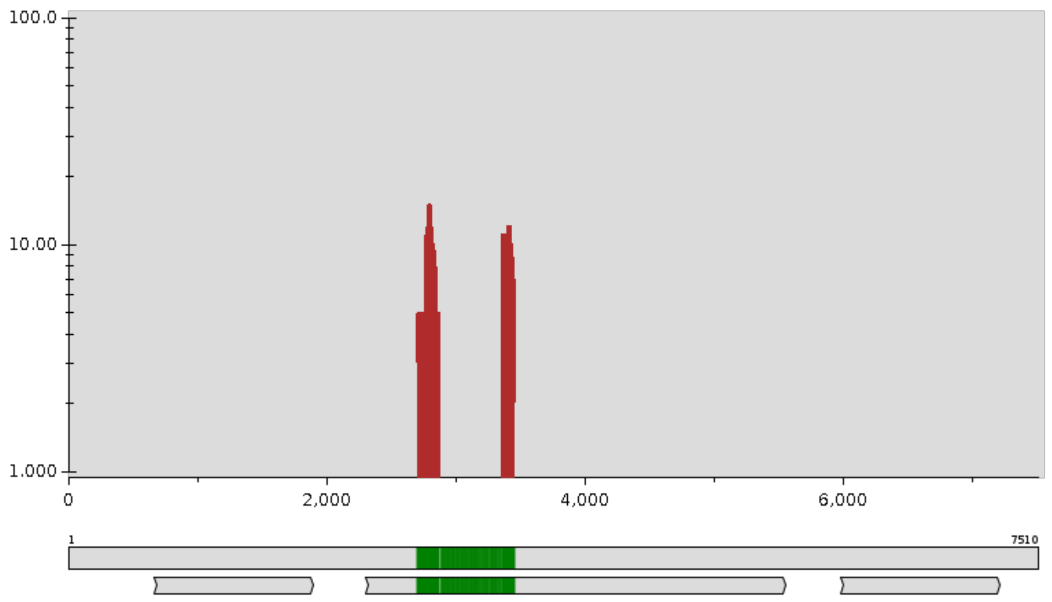

#### Assignment

| | |
|---|---|
| **Type** | Errantivirus (Taxonomy ID: 186666) |
| **Reference Genome** | NC_038512.1 |
| **NT Identity (%)** | 31.4168 |
| **AA Identity (%)** | 30.3279 |
| **Number Of Stop Codons** | 1 |
| **Number Of CDS** | 3 |

#### Alignment

| | |
|---|---|
| **Alignment Score** | -391.0 (NT) + -161.0 (AA) = -552.0 |
| **Concordance (%)** | -36.7318 |

| Alignment Method | Global, seeded, nucleotide + amino acids (AGA) |
|---|---|

#### Genome Region

Sequence starts at position 2698 and ends at position 3462 relative to NC_038512.1 reference sequence.

#### Alignment Detailed Statistics

| | Begin | End | Coverage | Score | Concordance | Matches | Identities | I/D/M/F* | Stop Codons |
|---|---|---|---|---|---|---|---|---|---|
| **NT** | **2698** | **3462** | **6.6%** | **-391** | **-41.1%** | **486 (97.4%)** | **153 (30.7%)** | **1/12** | |
| Mutations: | 2699A>T, 2700G>T, 2701A>G, 2702A>C, 2703C>A, 2705A>T, 2709A>G, 2710C>T, 2712A>T, 2713A>G, 2715T>A, 2716A>G, 2719C>T, 2723T>A, 2725C>T, 2727C>A, 2728C>G, 2730A>G, 2732A>C, 2733G>C, 2737C>T, 2738C>A, 2740G>A, 2743C>T, 2745C>T, 2746C>T, 2748T>C, 2750A>G, 2753C>A, 2755T>A, 2756C>A, 2757G>A, 2758C>A, 2759C>G, 2761G>T, 2763A>T, 2764A>T, 2765G>A, 2767T>G, 2769G>A, 2773C>G, 2777A>C, 2779C>G, 2781C>A, 2782G>A, 2783A>G, 2785A>T, 2786A>T, 2788G>A, 2791G>A, 2794C>G, 2795C>A, 2806A>T, 2807A>C, 2809A>G, 2812A>T, 2813T>A, 2814C>G, 2816G>A, 2818C>A, 2821T>C, 2822G>C, 2824A>G, 2828A>G, 2830C>G, 2830_2831insT, 2831T>G, 2833A>T, 2836C>G, 2837A>G, 2839A>G, 2841G>T, 2842G>A, 2843G>T, 2849C>A, 2850C>G, 2851C>A, 2857A>G, 2859T>A, 2860C>A, 2863C>T, 2865C>G, 2869T>C, 2870_2881delGGGAAACAAAAG, 2885C>A, 2887T>G, 2888C>A, 2890C>G, 2891G>T, 2892T>G, 2893A>T, 2896T>A, 2901T>A, 2903C>A, 2905T>G, 2906A>G, 2908G>T, 2915G>A, 2919A>T, 2920G>C, 2924A>G, 2926C>G, 2927G>A, 2929T>G, 2932C>T, 2935A>G, 2937A>T, 2938C>T, 2941G>A, 2944A>T, 2949A>T, 2950C>T, 2953A>T, 2954A>G, 2955G>A, 2959C>G, 2960G>T, 2963C>T, 2968C>T, 2969A>G, 2974A>G, 2975G>A, 2976G>A, 2978A>G, 2979A>G, 2980G>T, 2981T>G, 2982G>C, 2983C>T, 2984C>G, 2985A>T, 2986A>T, 2987T>A, 2988A>T, 2989C>T, 2993A>T, 2995C>A, 2997C>A, 2998C>G, 2999T>C, 3001A>T, 3008G>A, 3009C>G, 3011A>T, 3012G>C, 3016G>C, 3018T>A, 3020T>C, 3025G>A, 3026G>A, 3028G>T, 3029G>A, 3030A>G, 3035G>A, 3037C>G, 3038C>G, 3039C>A, 3040T>G, 3041C>G, 3043A>G, 3047A>G, 3050T>C, 3052G>C, 3058C>T, 3061G>T, 3066A>G, 3067C>G, 3068G>A, 3069T>C, 3071G>C, 3073A>C, 3074C>T, 3076C>T, 3079G>C, 3084T>A, 3092C>T, 3094T>A, 3095C>G, 3096G>T, 3097A>C, 3104A>T, 3106G>T, 3109A>G, 3110T>C, 3112A>C, 3114A>C, 3115A>T, 3118C>T, 3119T>G, 3124A>G, 3127T>C, 3130T>A, 3137A>G, 3139A>G, 3140G>C, 3142T>G, 3146G>A, 3148C>T, 3149A>C, 3154C>G, 3155C>T, 3157A>C, 3159G>A, 3161G>C, 3162G>C, 3164C>T, 3165T>A, 3166C>T, 3168A>T, 3169A>C, 3170A>T, 3171A>G, 3172T>A, 3176A>T, 3178C>T, 3179T>G, 3180G>T, 3181T>A, 3182C>T, 3184C>G, 3189A>T, 3190C>T, 3191C>T, 3196C>T, 3202T>C, 3203A>C, 3205T>G, 3206G>A, 3208C>G, 3215A>T, 3218T>A, 3219C>A, 3220C>T, 3221C>T, 3223A>G, 3224C>G, 3226G>A, 3227G>A, 3232C>T, 3233C>T, 3236G>T, 3237A>T, 3239A>C, 3241C>T, 3242C>T, 3245G>A, 3246A>G, 3248C>A, 3261G>C, 3268A>T, 3271A>C, 3273G>A, 3274T>C, 3277C>A, 3280C>G, 3281A>T, 3283A>T, 3284A>G, 3286T>C, 3287C>A, 3289A>G, 3292G>A, 3293G>A, 3294A>G, 3300C>G, 3301C>T, 3303A>C, 3304A>T, 3307C>T, 3308T>G, 3309T>G, 3311A>T, 3313G>A, 3317G>A, 3318A>G, 3319A>G, 3320A>G, 3321C>T, 3322T>G, 3327A>T, 3331T>A, 3338A>T, 3340C>G, 3343A>T, 3345G>C, 3346C>T, 3347A>T, 3353G>A, 3354G>T, 3359A>C, 3360A>G, 3361G>A, 3362C>G, 3363C>T, 3364T>G, 3365A>G, 3367C>T, 3368C>A, 3371G>C, 3373T>A, 3380T>G, 3381C>A, 3382C>G, 3385T>A, 3386A>G, 3388T>C, 3389C>A, 3394A>C, 3396A>G, 3397T>G, 3399T>C, 3400G>C, 3402T>A, 3403T>A, 3406A>C, 3408A>C, 3409G>C, 3410A>T, 3412C>T, 3415T>A, 3417A>C, 3418G>T, 3426A>G, 3428C>A, 3429A>G, 3430A>T, 3433T>C, 3436A>G, 3439C>T, 3440C>T, 3442T>G, 3443C>G, 3444T>C, 3445C>T, 3448T>G | | | | | | | | |

##### CDS

| | Begin | End | Coverage | Score | Concordance | Matches | Identities | I/D/M/F* | Stop Codons |
|---|---|---|---|---|---|---|---|---|---|
| D1R33_gp2 | 132 | 386 | 11.5% | -161 | -10.8% | 121 (96.0%) | 37 (29.4%) | 1/4/14/1 | 1 |
| Protein mutations: | R132L (2699A>T 2700G>T 2701A>G), T133Q (2702A>C 2703C>A), T134S (2705A>T), D135G (2709A>G 2710C>T), E136V (2712A>T 2713A>G), V137E (2715T>A 2716A>G), Y140N (2723T>A 2725C>T), T141K (2727C>A 2728C>G), K142R (2730A>G), S143P (2732A>C 2733G>C), P147L (2745C>T 2746C>T), F148S (2748T>C), I149V (2750A>G), H150K (2753C>A 2755T>A), R151K (2756C>A 2757G>A 2758C>A), Q152D (2759C>G 2761G>T), E153V (2763A>T 2764A>T), V154M (2765G>A 2767T>G), R155K (2769G>A), D156E (2773C>G), I158L (2777A>C 2779C>G), T159K (2781C>A 2782G>A), K160D (2783A>G 2785A>T), M161L (2786A>T 2788G>A), D163E (2794C>G), Q164K (2795C>A), S170R (2813T>A 2814C>G), D171K (2816G>A 2818C>A), A173P (2822G>C 2824A>G), S175G (2828A>G 2830C>G), S175_S176insX (2830_2831insT), S176A (2831T>G 2833A>T), I178V (2837A>G 2839A>G), W179L (2841G>T 2842G>A), V180F (2843G>T), P182R (2849C>A 2850C>G 2851C>A), I185K (2859T>A 2860C>A), A187G (2865C>G), G189_K192del (2870_2881delGGGAAACAAAAG), V196C (2891G>T 2892T>G 2893A>T), K225G (2978A>G 2979A>G 2980G>T), C226A (2981T>G 2982G>C 2983C>T), Q227V (2984C>G 2985A>T 2986A>T), Y228I (2987T>A 2988A>T 2989C>T), P245E (3038C>G 3039C>A 3040T>G), R264V (3095C>G 3096G>T 3097A>C), L287Y (3164C>T 3165T>A 3166C>T), N289* (3170A>T 3171A>G 3172T>A), C292V (3179T>G 3180G>T 3181T>A), S305N (3218T>A 3219C>A 3220C>T), E338R (3317G>A 3318A>G 3319A>G), T339V (3320A>G 3321C>T 3322T>G), G350I (3353G>A 3354G>T), K352R (3359A>C 3360A>G 3361G>A), P353V (3362C>G 3363C>T 3364T>G), N354D (3365A>G 3367C>T), P355T (3368C>A), D356Q (3371G>C 3373T>A), S359E (3380T>G 3381C>A 3382C>G), I361V (3386A>G 3388T>C), Q362K (3389C>A), K363N (3394A>C), Y364W (3396A>G 3397T>G), L365P (3399T>C 3400G>C), I366K (3402T>A 3403T>A), K368T (3408A>C 3409G>C), T369S (3410A>T 3412C>T), K371T (3417A>C 3418G>T), K374R (3426A>G), Q375S (3428C>A 3429A>G 3430A>T), L380A (3443C>G 3444T>C 3445C>T) | | | | | | | | |
| Codon mutations: | AGA132TTG (2699A>T 2700G>T 2701A>G), ACA133CAA (2702A>C 2703C>A), ACG134TCG (2705A>T), GAC135GGT (2709A>G 2710C>T), GAA136GTG (2712A>T 2713A>G), GTA137GAG (2715T>A 2716A>G), CCC138CCT (2719C>T), TAC140AAT (2723T>A 2725C>T), ACC141AAG (2727C>A 2728C>G), AAA142AGA (2730A>G), AGT143CCT (2732A>C 2733G>C), TAC144TAT (2737C>T), CGG145AGA (2738C>A 2740G>A), TAC146TAT (2743C>T), CCC147CTT (2745C>T 2746C>T), TTC148TCC (2748T>C), ATC149GTC (2750A>G), CAT150AAA (2753C>A 2755T>A), CGC151AAA (2756C>A 2757G>A 2758C>A), CAG152GAT (2759C>G 2761G>T), GAA153GTT (2763A>T 2764A>T), GTT154ATG (2765G>A 2767T>G), AGG155AAG (2769G>A), GAC156GAG (2773C>G), ATC158CTG (2777A>C 2779C>G), ACG159AAA (2781C>A 2782G>A), AAA160GAT (2783A>G 2785A>T), ATG161TTA (2786A>T 2788G>A), TTG162TTA (2791G>A), GAC163GAG (2794C>G), CAA164AAA (2795C>A), ATA167ATT (2806A>T), AGA168CGG (2807A>C 2809A>G), CCA169CCT (2812A>T), TCA170AGA (2813T>A 2814C>G), GAC171AAA (2816G>A 2818C>A), TCT172TCC (2821T>C), GCA173CCG (2822G>C 2824A>G), AGC175GGG (2828A>G 2830C>G), AGC175_TCA176insT-- (2830_2831insT), TCA176GCT (2831T>G 2833A>T), CCC177CCG (2836C>G), ATA178GTG (2837A>G 2839A>G), TGG179TTA (2841G>T 2842G>A), GTT180TTT (2843G>T), CCC182AGA (2849C>A 2850C>G 2851C>A), AAA184AAG (2857A>G), ATC185AAA (2859T>A 2860C>A), GAC186GAT (2863C>T), GCT187GGT (2865C>G), TCT188TCC (2869T>C), GGG189_AAG192del (2870_2881delGGGAAACAAAAG), CGT194A.G (2885C>A 2887T>G), CTC195A.G (2888C>A 2890C>G), GTA196TGT (2891G>T 2892T>G 2893A>T), GTT197..A (2896T>A), TTC199.A. (2901T>A), CGT200A.G (2903C>A 2905T>G), AAG201G.T (2906A>G 2908G>T), GAG204A.. (2915G>A), AAG205.TC (2919A>T 2920G>C), ATC207G.G (2924A>G 2926C>G), GAT208A.G (2927G>A 2929T>G), GAC209..T (2932C>T), AAA210..G (2935A>G), TAC211.TT (2937A>T 2938C>T), CCG212..A (2941G>A), ATA213..T (2944A>T), AAC215.TT (2949A>T 2950C>T), ATA216..T (2953A>T), AGT217GA. (2954A>G 2955G>A), GAC218..G (2959C>G), GTA219T.. (2960G>T), CTT220T.. (2963C>T), GAC221..T (2968C>T), AAG222G.. (2969A>G), TTA223..G (2974A>G), GGT224AA. (2975G>A 2976G>A), AAG225GGT (2978A>G 2979A>G 2980G>T), TGC226GCT (2981T>G 2982G>C 2983C>T), CAA227GTT (2984C>G 2985A>T 2986A>T), TAC228ATT (2987T>A 2988A>T 2989C>T), ACC230T.A (2993A>T 2995C>A), ACC231.AG (2997C>A 2998C>G), TTA232C.T (2999T>C 3001A>T), GCA235AG. (3008G>A 3009C>G), AGT236TC. (3011A>T 3012G>C), GGG237..C (3016G>C), TTT238.A. (3018T>A), TAT239C.. (3020T>C), CAG240..A (3025G>A), GTG241A.T (3026G>A 3028G>T), GAG242AG. (3029G>A 3030A>G), GAC244A.G (3035G>A 3037C>G), CCT245GAG (3038C>G 3039C>A 3040T>G), CAA246G.G (3041C>G 3043A>G), ATA248G.. (3047A>G), TCG249C.C (3050T>C 3052G>C), ACC251..T (3058C>T), GCG252..T (3061G>T), AAC254.GG (3066A>G 3067C>G), GTA255AC. (3068G>A 3069T>C), GAA256C.C (3071G>C 3073A>C), CAC257T.T (3074C>T 3076C>T), GGG258..C (3079G>C), TTT260.A. (3084T>A), CTT263T.A (3092C>T 3094T>A), CGA264GTC (3095C>G 3096G>T 3097A>C), ATG267T.T (3104A>T 3106G>T), GGA268..G (3109A>G), TTA269C.C (3110T>C 3112A>C), AAA270.CT (3114A>C 3115A>T), AAC271..T (3118C>T), TCA272G.. (3119T>G), CCA273..G (3124A>G), TCT274..C (3127T>C), ACT275..A (3130T>A), AGA278G.G (3137A>G 3139A>G), GTT279C.G (3140G>C 3142T>G), GAC281A.T (3146G>A 3148C>T), AAT282C.. (3149A>C), GTC283..G (3154C>G), CTA284T.C (3155C>T 3157A>C), AGA285.A. (3159G>A), GGT286CC. (3161G>C 3162G>C), CTC287TAT (3164C>T 3165T>A 3166C>T), CAA288.TC (3168A>T 3169A>C), AAT289TGA (3170A>T 3171A>G 3172T>A), ATC291T.T (3176A>T 3178C>T), TGT292GTA (3179T>G 3180G>T 3181T>A), CTC293T.G (3182C>T 3184C>G), TAC295.TT (3189A>T 3190C>T), CTT296T.. (3191C>T), GAC297..T (3196C>T), ATT299..C (3202T>C), ATT300C.G (3203A>C 3205T>G), GTC301A.G (3206G>A 3208C>G), ACT304T.. (3215A>T), TCC305AAT (3218T>A 3219C>A 3220C>T), CTA306T.G (3221C>T 3223A>G), CAG307G.A (3224C>G 3226G>A), GAA308A.. (3227G>A), CAC309..T (3232C>T), CTG310T.. (3233C>T), GAG311TT. (3236G>T 3237A>T), AAC312C.T (3239A>C 3241C>T), CTG313T.. (3242C>T), GAA314AG. (3245G>A 3246A>G), CGA315A.. (3248C>A), AGA319.C. (3261G>C), AGA321..T (3268A>T), GAA322..C (3271A>C), AGT323.AC (3273G>A 3274T>C), AAC324..A (3277C>A), TTC325..G (3280C>G), AAA326T.T (3281A>T 3283A>T), ATT327G.C (3284A>G 3286T>C), CAA328A.G (3287C>A 3289A>G), ATG329..A (3292G>A), GAC330AG. (3293G>A 3294A>G), TCC332.GT (3300C>G 3301C>T), GAA333.CT (3303A>C 3304A>T), TTC334..T (3307C>T), TTG335GG. (3308T>G 3309T>G), AAG336T.A (3311A>T 3313G>A), GAA338AGG (3317G>A 3318A>G 3319A>G), ACT339GTG (3320A>G 3321C>T 3322T>G), TAT341.T. (3327A>T), CTT342..A (3331T>A), ATC345T.G (3338A>T 3340C>G), ATA346..T (3343A>T), AGC347.CT (3345G>C 3346C>T), AGG348T.. (3347A>T), GGT350ATT (3353G>A 3354G>T), AAG352CGA (3359A>C 3360A>G 3361G>A), CCT353GTG (3362C>G 3363C>T 3364T>G), AAC354GAT (3365A>G 3367C>T), CCT355ACT (3368C>A), GAT356CAA (3371G>C 3373T>A), TCC359GAG (3380T>G 3381C>A 3382C>G), GCT360GCA (3385T>A), ATT361GTC (3386A>G 3388T>C), CAA362AAA (3389C>A), AAA363AAC (3394A>C), TAT364TGG (3396A>G 3397T>G), CTG365CCC (3399T>C 3400G>C), ATT366AAA (3402T>A 3403T>A), CCA367CCC (3406A>C), AAG368ACC (3408A>C 3409G>C), ACC369TCT (3410A>T 3412C>T), CCT370CCA (3415T>A), AAG371ACT (3417A>C 3418G>T), AAA374AGA (3426A>G), CAA375AGT (3428C>A 3429A>G 3430A>T), TTT376TTC (3433T>C), TTA377TTG (3436A>G), GGC378GGT (3439C>T), CTT379TTG (3440C>T 3442T>G), CTC380GCT (3443C>G 3444T>C 3445C>T), GGT381GGG (3448T>G) | | | | | | | | |

##### Proteins

| | Begin | End | Coverage | Score | Concordance | Matches | Identities | I/D/M/F* | Stop Codons |
|---|---|---|---|---|---|---|---|---|---|
| ORF B (YP_009507248.1) | 132 | 386 | 11.5% | -161 | -10.8% | 121 (96.0%) | 37 (29.4%) | 1/4/14/1 | 1 |
| Protein mutations: | R132L (2699A>T 2700G>T 2701A>G), T133Q (2702A>C 2703C>A), T134S (2705A>T), D135G (2709A>G 2710C>T), E136V (2712A>T 2713A>G), V137E (2715T>A 2716A>G), Y140N (2723T>A 2725C>T), T141K (2727C>A 2728C>G), K142R (2730A>G), S143P (2732A>C 2733G>C), P147L (2745C>T 2746C>T), F148S (2748T>C), I149V (2750A>G), H150K (2753C>A 2755T>A), R151K (2756C>A 2757G>A 2758C>A), Q152D (2759C>G 2761G>T), E153V (2763A>T 2764A>T), V154M (2765G>A 2767T>G), R155K (2769G>A), D156E (2773C>G), I158L (2777A>C 2779C>G), T159K (2781C>A 2782G>A), K160D (2783A>G 2785A>T), M161L (2786A>T 2788G>A), D163E (2794C>G), Q164K (2795C>A), S170R (2813T>A 2814C>G), D171K (2816G>A 2818C>A), A173P (2822G>C 2824A>G), S175G (2828A>G 2830C>G), S175_S176insX (2830_2831insT), S176A (2831T>G 2833A>T), I178V (2837A>G 2839A>G), W179L (2841G>T 2842G>A), V180F (2843G>T), P182R (2849C>A 2850C>G 2851C>A), I185K (2859T>A 2860C>A), A187G (2865C>G), G189_K192del (2870_2881delGGGAAACAAAAG), V196C (2891G>T 2892T>G 2893A>T), K225G (2978A>G 2979A>G 2980G>T), C226A (2981T>G 2982G>C 2983C>T), Q227V (2984C>G 2985A>T 2986A>T), Y228I (2987T>A 2988A>T 2989C>T), P245E (3038C>G 3039C>A 3040T>G), R264V (3095C>G 3096G>T 3097A>C), L287Y (3164C>T 3165T>A 3166C>T), N289* (3170A>T 3171A>G 3172T>A), C292V (3179T>G 3180G>T 3181T>A), S305N (3218T>A 3219C>A 3220C>T), E338R (3317G>A 3318A>G 3319A>G), T339V (3320A>G 3321C>T 3322T>G), G350I (3353G>A 3354G>T), K352R (3359A>C 3360A>G 3361G>A), P353V (3362C>G 3363C>T 3364T>G), N354D (3365A>G 3367C>T), P355T (3368C>A), D356Q (3371G>C 3373T>A), S359E (3380T>G 3381C>A 3382C>G), I361V (3386A>G 3388T>C), Q362K (3389C>A), K363N (3394A>C), Y364W (3396A>G 3397T>G), L365P (3399T>C 3400G>C), I366K (3402T>A 3403T>A), K368T (3408A>C 3409G>C), T369S (3410A>T 3412C>T), K371T (3417A>C 3418G>T), K374R (3426A>G), Q375S (3428C>A 3429A>G 3430A>T), L380A (3443C>G 3444T>C 3445C>T) | | | | | | | | |

| | Begin | End | Coverage | Score | Concordance | Matches | Identities | I/D/M/F* | Stop Codons |
|---|---|---|---|---|---|---|---|---|---|
| **NT** | **2698** | **3462** | **6.6%** | **-391** | **-41.1%** | **486 (97.4%)** | **153 (30.7%)** | **1/12** | |
| Codon mutations: | AGA132TTG (2699A>T 2700G>T 2701A>G), ACA133CAA (2702A>C 2703C>A), ACG134TCG (2705A>T), GAC135GGT (2709A>G 2710C>T), GAA136GTG (2712A>T 2713A>G), GTA137GAG (2715T>A 2716A>G), CCC138CCT (2719C>T), TAC140AAT (2723T>A 2725C>T), ACC141AAG (2727C>A 2728C>G), AAA142AGA (2730A>G), AGT143CCT (2732A>C 2733G>C), TAC144TAT (2737C>T), CGG145AGA (2738C>A 2740G>A), TAC146TAT (2743C>T), CCC147CTT (2745C>T 2746C>T), TTC148TCC (2748T>C), ATC149GTC (2750A>G), CAT150AAA (2753C>A 2755T>A), CGC151AAA (2756C>A 2757G>A 2758C>A), CAG152GAT (2759C>G 2761G>T), GAA153GTT (2763A>T 2764A>T), GTT154ATG (2765G>A 2767T>G), AGG155AAG (2769G>A), GAC156GAG (2773C>G), ATC158CTG (2777A>C 2779C>G), ACG159AAA (2781C>A 2782G>A), AAA160GAT (2783A>G 2785A>T), ATG161TTA (2786A>T 2788G>A), TTG162TTA (2791G>A), GAC163GAG (2794C>G), CAA164AAA (2795C>A), ATA167ATT (2806A>T), AGA168CGG (2807A>C 2809A>G), CCA169CCT (2812A>T), TCA170AGA (2813T>A 2814C>G), GAC171AAA (2816G>A 2818C>A), TCT172TCC (2821T>C), GCA173CCG (2822G>C 2824A>G), AGC175GGG (2828A>G 2830C>G), AGC175_TCA176insT-- (2830_2831insT), TCA176GCT (2831T>G 2833A>T), CCC177CCG (2836C>G), ATA178GTG (2837A>G 2839A>G), TGG179TTA (2841G>T 2842G>A), GTT180TTT (2843G>T), CCC182AGA (2849C>A 2850C>G 2851C>A), AAA184AAG (2857A>G), ATC185AAA (2859T>A 2860C>A), GAC186GAT (2863C>T), GCT187GGT (2865C>G), TCT188TCC (2869T>C), GGG189_AAG192del (2870_2881delGGGAAACAAAAG), CGT194A.G (2885C>A 2887T>G), CTC195A.G (2888C>A 2890C>G), GTA196TGT (2891G>T 2892T>G 2893A>T), GTT197..A (2896T>A), TTC199.A. (2901T>A), CGT200A.G (2903C>A 2905T>G), AAG201G.T (2906A>G 2908G>T), GAG204A.. (2915G>A), AAG205.TC (2919A>T 2920G>C), ATC207G.G (2924A>G 2926C>G), GAT208A.G (2927G>A 2929T>G), GAC209..T (2932C>T), AAA210..G (2935A>G), TAC211.TT (2937A>T 2938C>T), CCG212..A (2941G>A), ATA213..T (2944A>T), AAC215.TT (2949A>T 2950C>T), ATA216..T (2953A>T), AGT217GA. (2954A>G 2955G>A), GAC218..G (2959C>G), GTA219T.. (2960G>T), CTT220T.. (2963C>T), GAC221..T (2968C>T), AAG222G.. (2969A>G), TTA223..G (2974A>G), GGT224AA. (2975G>A 2976G>A), AAG225GGT (2978A>G 2979A>G 2980G>T), TGC226GCT (2981T>G 2982G>C 2983C>T), CAA227GTT (2984C>G 2985A>T 2986A>T), TAC228ATT (2987T>A 2988A>T 2989C>T), ACC230T.A (2993A>T 2995C>A), ACC231.AG (2997C>A 2998C>G), TTA232C.T (2999T>C 3001A>T), GCA235AG. (3008G>A 3009C>G), AGT236TC. (3011A>T 3012G>C), GGG237..C (3016G>C), TTT238.A. (3018T>A), TAT239C.. (3020T>C), CAG240..A (3025G>A), GTG241A.T (3026G>A 3028G>T), GAG242AG. (3029G>A 3030A>G), GAC244A.G (3035G>A 3037C>G), CCT245GAG (3038C>G 3039C>A 3040T>G), CAA246G.G (3041C>G 3043A>G), ATA248G.. (3047A>G), TCG249C.C (3050T>C 3052G>C), ACC251..T (3058C>T), GCG252..T (3061G>T), AAC254.GG (3066A>G 3067C>G), GTA255AC. (3068G>A 3069T>C), GAA256C.C (3071G>C 3073A>C), CAC257T.T (3074C>T 3076C>T), GGG258..C (3079G>C), TTT260.A. (3084T>A), CTT263T.A (3092C>T 3094T>A), CGA264GTC (3095C>G 3096G>T 3097A>C), ATG267T.T (3104A>T 3106G>T), GGA268..G (3109A>G), TTA269C.C (3110T>C 3112A>C), AAA270.CT (3114A>C 3115A>T), AAC271..T (3118C>T), TCA272G.. (3119T>G), CCA273..G (3124A>G), TCT274..C (3127T>C), ACT275..A (3130T>A), AGA278G.G (3137A>G 3139A>G), GTT279C.G (3140G>C 3142T>G), GAC281A.T (3146G>A 3148C>T), AAT282C.. (3149A>C), GTC283..G (3154C>G), CTA284T.C (3155C>T 3157A>C), AGA285.A. (3159G>A), GGT286CC. (3161G>C 3162G>C), CTC287TAT (3164C>T 3165T>A 3166C>T), CAA288.TC (3168A>T 3169A>C), AAT289TGA (3170A>T 3171A>G 3172T>A), ATC291T.T (3176A>T 3178C>T), TGT292GTA (3179T>G 3180G>T 3181T>A), CTC293T.G (3182C>T 3184C>G), TAC295.TT (3189A>T 3190C>T), CTT296T.. (3191C>T), GAC297..T (3196C>T), ATT299..C (3202T>C), ATT300C.G (3203A>C 3205T>G), GTC301A.G (3206G>A 3208C>G), ACT304T.. (3215A>T), TCC305AAT (3218T>A 3219C>A 3220C>T), CTA306T.G (3221C>T 3223A>G), CAG307G.A (3224C>G 3226G>A), GAA308A.. (3227G>A), CAC309..T (3232C>T), CTG310T.. (3233C>T), GAG311TT. (3236G>T 3237A>T), AAC312C.T (3239A>C 3241C>T), CTG313T.. (3242C>T), GAA314AG. (3245G>A 3246A>G), CGA315A.. (3248C>A), AGA319.C. (3261G>C), AGA321..T (3268A>T), GAA322..C (3271A>C), AGT323.AC (3273G>A 3274T>C), AAC324..A (3277C>A), TTC325..G (3280C>G), AAA326T.T (3281A>T 3283A>T), ATT327G.C (3284A>G 3286T>C), CAA328A.G (3287C>A 3289A>G), ATG329..A (3292G>A), GAC330AG. (3293G>A 3294A>G), TCC332.GT (3300C>G 3301C>T), GAA333.CT (3303A>C 3304A>T), TTC334..T (3307C>T), TTG335GG. (3308T>G 3309T>G), AAG336T.A (3311A>T 3313G>A), GAA338AGG (3317G>A 3318A>G 3319A>G), ACT339GTG (3320A>G 3321C>T 3322T>G), TAT341.T. (3327A>T), CTT342..A (3331T>A), ATC345T.G (3338A>T 3340C>G), ATA346..T (3343A>T), AGC347.CT (3345G>C 3346C>T), AGG348T.. (3347A>T), GGT350ATT (3353G>A 3354G>T), AAG352CGA (3359A>C 3360A>G 3361G>A), CCT353GTG (3362C>G 3363C>T 3364T>G), AAC354GAT (3365A>G 3367C>T), CCT355ACT (3368C>A), GAT356CAA (3371G>C 3373T>A), TCC359GAG (3380T>G 3381C>A 3382C>G), GCT360GCA (3385T>A), ATT361GTC (3386A>G 3388T>C), CAA362AAA (3389C>A), AAA363AAC (3394A>C), TAT364TGG (3396A>G 3397T>G), CTG365CCC (3399T>C 3400G>C), ATT366AAA (3402T>A 3403T>A), CCA367CCC (3406A>C), AAG368ACC (3408A>C 3409G>C), ACC369TCT (3410A>T 3412C>T), CCT370CCA (3415T>A), AAG371ACT (3417A>C 3418G>T), AAA374AGA (3426A>G), CAA375AGT (3428C>A 3429A>G 3430A>T), TTT376TTC (3433T>C), TTA377TTG (3436A>G), GGC378GGT (3439C>T), CTT379TTG (3440C>T 3442T>G), CTC380GCT (3443C>G 3444T>C 3445C>T), GGT381GGG (3448T>G) | | | | | | | | |

*: Inserts / Deletes / Misaligned / Frameshifts

#### Analysis details

This analysis was performed with panviral2.64

### NGS Details (UN59): Pinus nigra virus 1

#### Assembly

| | |
|---|---|
| **Coverage Length** | 535 (1 contig(s)) |
| **Depth Of Coverage** | 6.5 |
| **Number Of Reads** | 27 |
| **Reads Per Million** | 0.59 rpm (after QC) |
| **Ambiguities** | 0 |
| **Assembly Method** | de novo + reference guided assembly |
| **Consensus Caller** | Bcf Tools |

#### Coverage Map

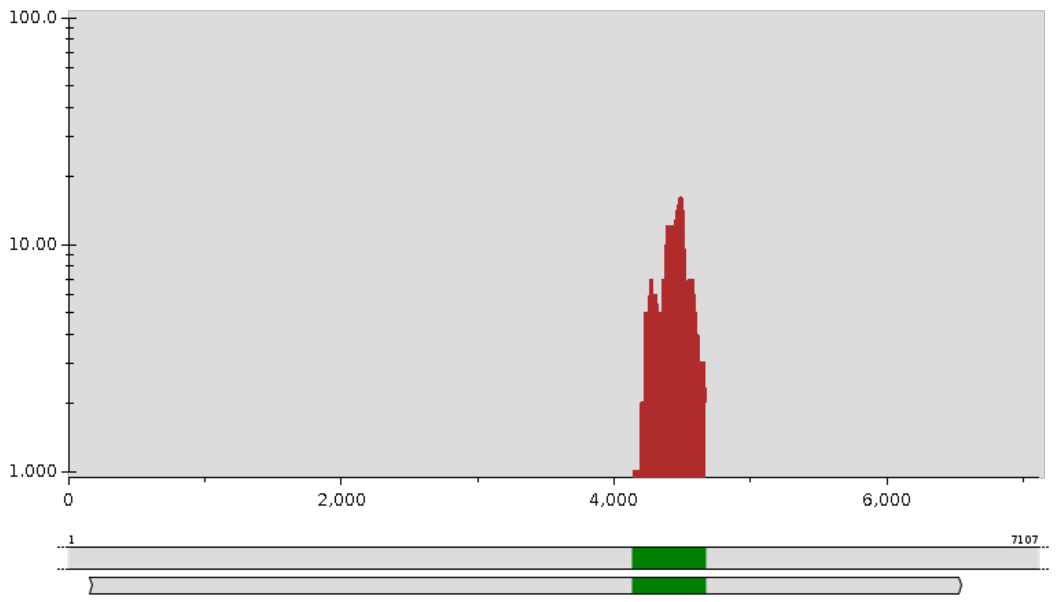

#### Assignment

| | |
|---|---|
| **Type** | Pinus nigra virus 1 (Taxonomy ID: 2267679) |
| **Reference Genome** | NC_040841.1 |
| **NT Identity (%)** | 56.6485 |
| **AA Identity (%)** | 43.4783 |
| **Number Of Stop Codons** | 1 |
| **Number Of CDS** | 1 |

#### Alignment

| | |
|---|---|
| **Alignment Score** | 124.0 (NT) + 313.0 (AA) = 437.0 |
| **Concordance (%)** | 22.1215 |

| Alignment Method | Global, seeded, nucleotide + amino acids (AGA) |
|---|---|

#### Genome Region

Sequence starts at position 4141 and ends at position 4675 relative to NC_040841.1 reference sequence.

#### Alignment Detailed Statistics

| | Begin | End | Coverage | Score | Concordance | Matches | Identities | I/D/M/F* | Stop Codons |
|---|---|---|---|---|---|---|---|---|---|
| **NT** | **4141** | **4675** | **7.5%** | **124** | **11.8%** | **533 (96.7%)** | **311 (56.4%)** | **16/2** | |
| Mutations: | 4141G>A, 4147G>A, 4151G>C, 4152C>A, 4153C>A, 4154T>A, 4167G>A, 4168G>A, 4175T>A, 4177T>A, 4180C>A, 4184G>T, 4186T>A, 4187A>G, 4190T>A, 4192T>G, 4193G>A, 4195T>A, 4199A>T, 4202G>A, 4204T>A, 4205A>G, 4206C>T, 4210C>T, 4223T>C, 4226A>G, 4228T>A, 4229C>T, 4230A>G, 4232A>T, 4233G>C, 4234T>A, 4235C>G, 4236C>A, 4238G>A, 4239A>C, 4240T>A, 4241G>T, 4242T>A, 4243A>C, 4246G>A, 4247A>G, 4249T>A, 4253T>C, 4255G>T, 4257A>C, 4258G>A, 4261C>T, 4262T>C, 4264A>T, 4265G>A, 4268A>G, 4269T>A, 4270T>A, 4274C>G, 4275T>C, 4276T>A, 4279A>T, 4280G>C, 4281A>T, 4282A>C, 4283G>T, 4284T>G, 4285C>T, 4286A>G, 4287A>T, 4288G>C, 4291A>C, 4292C>A, 4294T>C, 4296G>A, 4297A>C, 4300T>A, 4306G>A, 4307T>A, 4308C>G, 4309A>C, 4310G>C, 4314G>A, 4319A>G, 4320T>C, 4321G>T, 4322T>G, 4323C>A, 4324C>A, 4325T>A, 4328T>A, 4330T>C, 4331C>A, 4333G>A, 4333_4334insA, 4334G>A, 4337A>G, 4339T>A, 4342T>A, 4345T>A, 4348T>C, 4349T>C, 4351G>T, 4357T>G, 4359A>T, 4361G>A, 4364G>T, 4369T>A, 4370G>A, 4371G>A, 4373T>G, 4374C>G, 4375T>G, 4378T>A, 4379C>G, 4381A>G, 4382A>T, 4384G>A, 4388C>A, 4390C>T, 4394G>C, 4396G>C, 4398T>A, 4400_4401delAC, 4406T>A, 4409A>T, 4410C>T, 4412C>G, 4414G>A, 4429G>A, 4431A>C, 4432G>A, 4433G>A, 4434A>C, 4438G>A, 4439A>C, 4447C>A, 4456C>T, 4457A>C, 4463G>T, 4464T>G, 4466C>T, 4468G>A, 4473A>T, 4475T>G, 4476T>C, 4477G>A, 4478C>G, 4479A>G, 4480C>A, 4481A>G, 4485A>T, 4486T>C, 4488T>C, 4489T>A, 4493A>G, 4495C>A, 4496C>T, 4498C>A, 4499T>G, 4502G>A, 4509C>G, 4511G>A, 4512C>A, 4515C>T, 4516_4517insCTACAAAAACTC, 4519A>C, 4524G>A, 4526C>T, 4527A>C, 4528C>A, 4531C>T, 4532A>C, 4533A>G, 4534T>A, 4535G>T, 4536G>T, 4537A>G, 4538G>T, 4544T>A, 4546A>C, 4547G>A, 4553G>C, 4554C>A, 4558A>T, 4559G>C, 4560C>G, 4561A>T, 4562G>T, 4563C>G, 4564G>T, 4568A>C, 4570G>A, 4573G>A, 4576T>C, 4589C>G, 4590A>T, 4591G>T, 4592C>A, 4594T>G, 4607C>A, 4608A>G, 4609A>G, 4611C>T, 4617G>A, 4618_4619insCCA, 4621T>A, 4623G>A, 4625C>A, 4626T>A, 4627A>T, 4628T>C, 4631A>G, 4633T>A, 4642T>A, 4643G>A, 4647C>T, 4648T>C, 4651A>C, 4652C>G, 4653T>A, 4655G>T, 4658T>C, 4659T>A, 4664T>G, 4665C>G, 4669T>A, 4670G>T, 4672A>G, 4673T>C | | | | | | | | |

##### CDS

| | Begin | End | Coverage | Score | Concordance | Matches | Identities | I/D/M/F* | Stop Codons |
|---|---|---|---|---|---|---|---|---|---|
| EXL67_gp1 | 1330 | 1507 | 8.4% | 313 | 24.9% | 178 (96.7%) | 80 (43.5%) | 6/0/2/2 | 1 |
| Protein mutations: | A1333Q (4151G>C 4152C>A 4153C>A), F1334I (4154T>A), R1338K (4167G>A 4168G>A), Y1341K (4175T>A 4177T>A), A1344S (4184G>T 4186T>A), K1345E (4187A>G), Y1346K (4190T>A 4192T>G), D1347K (4193G>A 4195T>A), I1349L (4199A>T), V1350I (4202G>A 4204T>A), T1351V (4205A>G 4206C>T), I1358V (4226A>G 4228T>A), H1359C (4229C>T 4230A>G), P1361E (4235C>G 4236C>A), D1362T (4238G>A 4239A>C 4240T>A), V1363Y (4241G>T 4242T>A 4243A>C), N1365E (4247A>G 4249T>A), K1368T (4257A>C 4258G>A), E1371K (4265G>A), I1372E (4268A>G 4269T>A 4270T>A), L1374A (4274C>G 4275T>C 4276T>A), E1375D (4279A>T), E1376L (4280G>C 4281A>T 4282A>C), V1377C (4283G>T 4284T>G 4285C>T), K1378V (4286A>G 4287A>T 4288G>C), K1379N (4291A>C), H1380N (4292C>A 4294T>C), G1381D (4296G>A 4297A>C), E1386Q (4310G>C), R1387K (4314G>A), M1389A (4319A>G 4320T>C 4321G>T), S1390E (4322T>G 4323C>A 4324C>A), L1391I (4325T>A), F1392I (4328T>A 4330T>C), Q1393K (4331C>A 4333G>A), Q1393_D1394insX (4333_4334insA), D1394N (4334G>A), N1395E (4337A>G 4339T>A), D1397E (4345T>A), I1401M (4357T>G), N1402I (4359A>T), V1403I (4361G>A), A1404S (4364G>T), N1405K (4369T>A), G1406N (4370G>A 4371G>A), S1407G (4373T>G 4374C>G 4375T>G), Q1409E (4379C>G 4381A>G), M1410L (4382A>T 4384G>A), P1412T (4388C>A 4390C>T), V1414L (4394G>C 4396G>C), L1415Q (4398T>A), L1418I (4406T>A), T1419L (4409A>T 4410C>T), Q1420E (4412C>G 4414G>A), K1426T (4431A>C 4432G>A), D1427T (4433G>A 4434A>C), K1429Q (4439A>C), I1435L (4457A>C), V1437C (4463G>T 4464T>G), Y1440F (4473A>T), L1441A (4475T>G 4476T>C 4477G>A), H1442G (4478C>G 4479A>G 4480C>A), K1443E (4481A>G), Y1444F (4485A>T 4486T>C), I1445T (4488T>C 4489T>A), N1447E (4493A>G 4495C>A), S1449A (4499T>G), E1450K (4502G>A), T1452R (4509C>G), A1453N (4511G>A 4512C>A), P1454L (4515C>T), P1454_I1455insLQKL (4516_4517insCTACAAAAACTC), R1457K (4524G>A), H1458S (4526C>T 4527A>C 4528C>A), N1460R (4532A>C 4533A>G 4534T>A), G1461L (4535G>T 4536G>T 4537A>G), G1462* (4538G>T), S1464T (4544T>A 4546A>C), D1465N (4547G>A), A1467H (4553G>C 4554C>A), A1469R (4559G>C 4560C>G 4561A>T), A1470C (4562G>T 4563C>G 4564G>T), K1472Q (4568A>C 4570G>A), Q1479V (4589C>G 4590A>T 4591G>T), H1480K (4592C>A 4594T>G), Q1485R (4607C>A 4608A>G 4609A>G), P1486L (4611C>T), G1488D (4617G>A), G1488_D1489insP (4618_4619insCCA), D1489E (4621T>A), G1490D (4623G>A), L1491N (4625C>A 4626T>A 4627A>T), I1493V (4631A>G 4633T>A), D1497N (4643G>A), A1498V (4647C>T 4648T>C), L1500D (4652C>G 4653T>A), D1501Y (4655G>T), F1502H (4658T>C 4659T>A), S1504G (4664T>G 4665C>G), V1506L (4670G>T 4672A>G) | | | | | | | | |
| Codon mutations: | AAG1329..A (4141G>A), GCG1331GCA (4147G>A), GCC1333CAA (4151G>C 4152C>A 4153C>A), TTC1334ATC (4154T>A), AGG1338AAA (4167G>A 4168G>A), TAT1341AAA (4175T>A 4177T>A), ATC1342ATA (4180C>A), GCT1344TCA (4184G>T 4186T>A), AAA1345GAA (4187A>G), TAT1346AAG (4190T>A 4192T>G), GAT1347AAA (4193G>A 4195T>A), ATA1349TTA (4199A>T), GTT1350ATA (4202G>A 4204T>A), ACA1351GTA (4205A>G 4206C>T), TAC1352TAT (4210C>T), TTA1357CTA (4223T>C), ATT1358GTA (4226A>G 4228T>A), CAT1359TGT (4229C>T 4230A>G), AGT1360TCA (4232A>T 4233G>C 4234T>A), CCA1361GAA (4235C>G 4236C>A), GAT1362ACA (4238G>A 4239A>C 4240T>A), GTA1363TAC (4241G>T 4242T>A 4243A>C), CAG1364CAA (4246G>A), AAT1365GAA (4247A>G 4249T>A), TTG1367CTT (4253T>C 4255G>T), AAG1368ACA (4257A>C 4258G>A), CAC1369CAT (4261C>T), TTA1370CTT (4262T>C 4264A>T), GAA1371AAA (4265G>A), ATT1372GAA (4268A>G 4269T>A 4270T>A), CTT1374GCA (4274C>G 4275T>C 4276T>A), GAA1375GAT (4279A>T), GAA1376CTC (4280G>C 4281A>T 4282A>C), GTC1377TGT (4283G>T 4284T>G 4285C>T), AAG1378GTC (4286A>G 4287A>T 4288G>C), AAA1379AAC (4291A>C), CAT1380AAC (4292C>A 4294T>C), GGA1381GAC (4296G>A 4297A>C), ATT1382ATA (4300T>A), TTG1384TTA (4306G>A), TCA1385AGC (4307T>A 4308C>G 4309A>C), GAA1386CAA (4310G>C), AGA1387AAA (4314G>A), ATG1389GCT (4319A>G 4320T>C 4321G>T), TCC1390GAA (4322T>G 4323C>A 4324C>A), TTA1391ATA (4325T>A), TTT1392ATC (4328T>A 4330T>C), CAG1393AAA (4331C>A 4333G>A), CAG1393_GAT1394insA-- (4333_4334insA), GAT1394AAT (4334G>A), AAT1395GAA (4337A>G 4339T>A), ATT1396ATA (4342T>A), GAT1397GAA (4345T>A), TTT1398TTC (4348T>C), TTG1399CTT (4349T>C 4351G>T), ATT1401ATG (4357T>G), AAT1402ATT (4359A>T), GTT1403ATT (4361G>A), GCA1404TCA (4364G>T), AAT1405AAA (4369T>A), GGT1406AAT (4370G>A 4371G>A), TCT1407GGG (4373T>G 4374C>G 4375T>G), ATT1408ATA (4378T>A), CAA1409GAG (4379C>G 4381A>G), ATG1410TTA (4382A>T 4384G>A), CCC1412ACT (4388C>A 4390C>T), GTG1414CTC (4394G>C 4396G>C), CTA1415CAA (4398T>A), ACA1416--A (4400_4401delAC), TTA1418ATA (4406T>A), ACA1419TTA (4409A>T 4410C>T), CAG1420GAA (4412C>G 4414G>A), TTG1425TTA (4429G>A), AAG1426ACA (4431A>C 4432G>A), GAC1427ACC (4433G>A 4434A>C), AAG1428AAA (4438G>A), AAA1429CAA (4439A>C), ATC1431ATA (4447C>A), TTC1434TTT (4456C>T), ATT1435CTT (4457A>C), GTT1437TGT (4463G>T 4464T>G), CTG1438TTA (4466C>T 4468G>A), TAT1440TTT (4473A>T), TTG1441GCA (4475T>G 4476T>C 4477G>A), CAC1442GGA (4478C>G 4479A>G 4480C>A), AAA1443GAA (4481A>G), TAT1444TTC (4485A>T 4486T>C), ATT1445ACA (4488T>C 4489T>A), AAC1447GAA (4493A>G 4495C>A), CTC1448TTA (4496C>T 4498C>A), TCA1449GCA (4499T>G), GAA1450AAA (4502G>A), ACA1452AGA (4509C>G), GCT1453AAT (4511G>A 4512C>A), CCT1454CTT (4515C>T), CCT1454_ATA1455insCTACAAAAACTC (4516_4517insCTACAAAAACTC), ATA1455ATC (4519A>C), AGA1457AAA (4524G>A), CAC1458TCA (4526C>T 4527A>C 4528C>A), AAC1459AAT (4531C>T), AAT1460CGA (4532A>C 4533A>G 4534T>A), GGA1461TTG (4535G>T 4536G>T 4537A>G), GGA1462TGA (4538G>T), TCA1464ACC (4544T>A 4546A>C), GAT1465AAT (4547G>A), GCT1467CAT (4553G>C 4554C>A), ACA1468ACT (4558A>T), GCA1469CGT (4559G>C 4560C>G 4561A>T), GCG1470TGT (4562G>T 4563C>G 4564G>T), AAG1472CAA (4568A>C 4570G>A), AAG1473AAA (4573G>A), CTT1474CTC (4576T>C), CAG1479GTT (4589C>G 4590A>T 4591G>T), CAT1480AAG (4592C>A 4594T>G), CAA1485AGG (4607C>A 4608A>G 4609A>G), CCA1486CTA (4611C>T), GGT1488GAT (4617G>A), GGT1488_GAT1489insCCA (4618_4619insCCA), GAT1489GAA (4621T>A), GGC1490GAC (4623G>A), CTA1491AAT (4625C>A 4626T>A 4627A>T), TTG1492CTG (4628T>C), ATT1493GTA (4631A>G 4633T>A), ACT1496ACA (4642T>A), GAT1497AAT (4643G>A), GCT1498GTC (4647C>T 4648T>C), TCA1499TCC (4651A>C), CTT1500GAT (4652C>G 4653T>A), GAT1501TAT (4655G>T), TTT1502CAT (4658T>C 4659T>A), TCA1504GGA (4664T>G 4665C>G), GCT1505GCA (4669T>A), GTA1506TTG (4670G>T 4672A>G), TTA1507CTA (4673T>C) | | | | | | | | |

##### Proteins

| | Begin | End | Coverage | Score | Concordance | Matches | Identities | I/D/M/F* | Stop Codons |
|---|---|---|---|---|---|---|---|---|---|
| polyprotein (YP_009553669.1) | 1330 | 1507 | 8.4% | 313 | 24.9% | 178 (96.7%) | 80 (43.5%) | 6/0/2/2 | 1 |
| Protein mutations: | A1333Q (4151G>C 4152C>A 4153C>A), F1334I (4154T>A), R1338K (4167G>A 4168G>A), Y1341K (4175T>A 4177T>A), A1344S (4184G>T 4186T>A), K1345E (4187A>G), Y1346K (4190T>A 4192T>G), D1347K (4193G>A 4195T>A), I1349L (4199A>T), V1350I (4202G>A 4204T>A), T1351V (4205A>G 4206C>T), I1358V (4226A>G 4228T>A), H1359C (4229C>T 4230A>G), P1361E (4235C>G 4236C>A), D1362T (4238G>A 4239A>C 4240T>A), V1363Y (4241G>T 4242T>A 4243A>C), N1365E (4247A>G 4249T>A), K1368T (4257A>C 4258G>A), E1371K (4265G>A), I1372E (4268A>G 4269T>A 4270T>A), L1374A (4274C>G 4275T>C 4276T>A), E1375D (4279A>T), E1376L (4280G>C 4281A>T 4282A>C), V1377C (4283G>T 4284T>G 4285C>T), K1378V (4286A>G 4287A>T 4288G>C), K1379N (4291A>C), H1380N (4292C>A 4294T>C), G1381D (4296G>A 4297A>C), E1386Q (4310G>C), R1387K (4314G>A), M1389A (4319A>G 4320T>C 4321G>T), S1390E (4322T>G 4323C>A 4324C>A), L1391I (4325T>A), F1392I (4328T>A 4330T>C), Q1393K (4331C>A 4333G>A), Q1393_D1394insX (4333_4334insA), D1394N (4334G>A), N1395E (4337A>G 4339T>A), D1397E (4345T>A), I1401M (4357T>G), N1402I (4359A>T), V1403I (4361G>A), A1404S (4364G>T), N1405K (4369T>A), G1406N (4370G>A 4371G>A), S1407G (4373T>G 4374C>G 4375T>G), Q1409E (4379C>G 4381A>G), M1410L (4382A>T 4384G>A), P1412T (4388C>A 4390C>T), V1414L (4394G>C 4396G>C), L1415Q (4398T>A), L1418I (4406T>A), T1419L (4409A>T 4410C>T), Q1420E (4412C>G 4414G>A), K1426T (4431A>C 4432G>A), D1427T (4433G>A 4434A>C), K1429Q (4439A>C), I1435L (4457A>C), V1437C (4463G>T 4464T>G), Y1440F (4473A>T), L1441A (4475T>G 4476T>C 4477G>A), H1442G (4478C>G 4479A>G 4480C>A), K1443E (4481A>G), Y1444F (4485A>T 4486T>C), I1445T (4488T>C 4489T>A), N1447E (4493A>G 4495C>A), S1449A (4499T>G), E1450K (4502G>A), T1452R (4509C>G), A1453N (4511G>A 4512C>A), P1454L (4515C>T), P1454_I1455insLQKL (4516_4517insCTACAAAAACTC), R1457K (4524G>A), H1458S (4526C>T 4527A>C 4528C>A), N1460R (4532A>C 4533A>G 4534T>A), G1461L (4535G>T 4536G>T 4537A>G), G1462* (4538G>T), S1464T (4544T>A 4546A>C), D1465N (4547G>A), A1467H (4553G>C 4554C>A), A1469R (4559G>C 4560C>G 4561A>T), A1470C (4562G>T 4563C>G 4564G>T), K1472Q (4568A>C 4570G>A), Q1479V (4589C>G 4590A>T 4591G>T), H1480K (4592C>A 4594T>G), Q1485R (4607C>A 4608A>G 4609A>G), P1486L (4611C>T), G1488D (4617G>A), G1488_D1489insP (4618_4619insCCA), D1489E (4621T>A), G1490D (4623G>A), L1491N (4625C>A 4626T>A 4627A>T), I1493V (4631A>G 4633T>A), D1497N (4643G>A), A1498V (4647C>T 4648T>C), L1500D (4652C>G 4653T>A), D1501Y (4655G>T), F1502H (4658T>C 4659T>A), S1504G (4664T>G 4665C>G), V1506L (4670G>T 4672A>G) | | | | | | | | |

| | Begin | End | Coverage | Score | Concordance | Matches | Identities | I/D/M/F* | Stop Codons |
|---|---|---|---|---|---|---|---|---|---|
| **NT** | **4141** | **4675** | **7.5%** | **124** | **11.8%** | **533 (96.7%)** | **311 (56.4%)** | **16/2** | |
| Codon mutations: | AAG1329..A (4141G>A), GCG1331GCA (4147G>A), GCC1333CAA (4151G>C 4152C>A 4153C>A), TTC1334ATC (4154T>A), AGG1338AAA (4167G>A 4168G>A), TAT1341AAA (4175T>A 4177T>A), ATC1342ATA (4180C>A), GCT1344TCA (4184G>T 4186T>A), AAA1345GAA (4187A>G), TAT1346AAG (4190T>A 4192T>G), GAT1347AAA (4193G>A 4195T>A), ATA1349TTA (4199A>T), GTT1350ATA (4202G>A 4204T>A), ACA1351GTA (4205A>G 4206C>T), TAC1352TAT (4210C>T), TTA1357CTA (4223T>C), ATT1358GTA (4226A>G 4228T>A), CAT1359TGT (4229C>T 4230A>G), AGT1360TCA (4232A>T 4233G>C 4234T>A), CCA1361GAA (4235C>G 4236C>A), GAT1362ACA (4238G>A 4239A>C 4240T>A), GTA1363TAC (4241G>T 4242T>A 4243A>C), CAG1364CAA (4246G>A), AAT1365GAA (4247A>G 4249T>A), TTG1367CTT (4253T>C 4255G>T), AAG1368ACA (4257A>C 4258G>A), CAC1369CAT (4261C>T), TTA1370CTT (4262T>C 4264A>T), GAA1371AAA (4265G>A), ATT1372GAA (4268A>G 4269T>A 4270T>A), CTT1374GCA (4274C>G 4275T>C 4276T>A), GAA1375GAT (4279A>T), GAA1376CTC (4280G>C 4281A>T 4282A>C), GTC1377TGT (4283G>T 4284T>G 4285C>T), AAG1378GTC (4286A>G 4287A>T 4288G>C), AAA1379AAC (4291A>C), CAT1380AAC (4292C>A 4294T>C), GGA1381GAC (4296G>A 4297A>C), ATT1382ATA (4300T>A), TTG1384TTA (4306G>A), TCA1385AGC (4307T>A 4308C>G 4309A>C), GAA1386CAA (4310G>C), AGA1387AAA (4314G>A), ATG1389GCT (4319A>G 4320T>C 4321G>T), TCC1390GAA (4322T>G 4323C>A 4324C>A), TTA1391ATA (4325T>A), TTT1392ATC (4328T>A 4330T>C), CAG1393AAA (4331C>A 4333G>A), CAG1393_GAT1394insA-- (4333_4334insA), GAT1394AAT (4334G>A), AAT1395GAA (4337A>G 4339T>A), ATT1396ATA (4342T>A), GAT1397GAA (4345T>A), TTT1398TTC (4348T>C), TTG1399CTT (4349T>C 4351G>T), ATT1401ATG (4357T>G), AAT1402ATT (4359A>T), GTT1403ATT (4361G>A), GCA1404TCA (4364G>T), AAT1405AAA (4369T>A), GGT1406AAT (4370G>A 4371G>A), TCT1407GGG (4373T>G 4374C>G 4375T>G), ATT1408ATA (4378T>A), CAA1409GAG (4379C>G 4381A>G), ATG1410TTA (4382A>T 4384G>A), CCC1412ACT (4388C>A 4390C>T), GTG1414CTC (4394G>C 4396G>C), CTA1415CAA (4398T>A), ACA1416--A (4400_4401delAC), TTA1418ATA (4406T>A), ACA1419TTA (4409A>T 4410C>T), CAG1420GAA (4412C>G 4414G>A), TTG1425TTA (4429G>A), AAG1426ACA (4431A>C 4432G>A), GAC1427ACC (4433G>A 4434A>C), AAG1428AAA (4438G>A), AAA1429CAA (4439A>C), ATC1431ATA (4447C>A), TTC1434TTT (4456C>T), ATT1435CTT (4457A>C), GTT1437TGT (4463G>T 4464T>G), CTG1438TTA (4466C>T 4468G>A), TAT1440TTT (4473A>T), TTG1441GCA (4475T>G 4476T>C 4477G>A), CAC1442GGA (4478C>G 4479A>G 4480C>A), AAA1443GAA (4481A>G), TAT1444TTC (4485A>T 4486T>C), ATT1445ACA (4488T>C 4489T>A), AAC1447GAA (4493A>G 4495C>A), CTC1448TTA (4496C>T 4498C>A), TCA1449GCA (4499T>G), GAA1450AAA (4502G>A), ACA1452AGA (4509C>G), GCT1453AAT (4511G>A 4512C>A), CCT1454CTT (4515C>T), CCT1454_ATA1455insCTACAAAAACTC (4516_4517insCTACAAAAACTC), ATA1455ATC (4519A>C), AGA1457AAA (4524G>A), CAC1458TCA (4526C>T 4527A>C 4528C>A), AAC1459AAT (4531C>T), AAT1460CGA (4532A>C 4533A>G 4534T>A), GGA1461TTG (4535G>T 4536G>T 4537A>G), GGA1462TGA (4538G>T), TCA1464ACC (4544T>A 4546A>C), GAT1465AAT (4547G>A), GCT1467CAT (4553G>C 4554C>A), ACA1468ACT (4558A>T), GCA1469CGT (4559G>C 4560C>G 4561A>T), GCG1470TGT (4562G>T 4563C>G 4564G>T), AAG1472CAA (4568A>C 4570G>A), AAG1473AAA (4573G>A), CTT1474CTC (4576T>C), CAG1479GTT (4589C>G 4590A>T 4591G>T), CAT1480AAG (4592C>A 4594T>G), CAA1485AGG (4607C>A 4608A>G 4609A>G), CCA1486CTA (4611C>T), GGT1488GAT (4617G>A), GGT1488_GAT1489insCCA (4618_4619insCCA), GAT1489GAA (4621T>A), GGC1490GAC (4623G>A), CTA1491AAT (4625C>A 4626T>A 4627A>T), TTG1492CTG (4628T>C), ATT1493GTA (4631A>G 4633T>A), ACT1496ACA (4642T>A), GAT1497AAT (4643G>A), GCT1498GTC (4647C>T 4648T>C), TCA1499TCC (4651A>C), CTT1500GAT (4652C>G 4653T>A), GAT1501TAT (4655G>T), TTT1502CAT (4658T>C 4659T>A), TCA1504GGA (4664T>G 4665C>G), GCT1505GCA (4669T>A), GTA1506TTG (4670G>T 4672A>G), TTA1507CTA (4673T>C) | | | | | | | | |

*: Inserts / Deletes / Misaligned / Frameshifts

#### Analysis details

This analysis was performed with panviral2.64

### NGS Details (UN59): Duamitovirus peex1

#### Assembly

| | |
|---|---|
| **Coverage Length** | 388 (1 contig(s)) |
| **Depth Of Coverage** | 6.5 |
| **Number Of Reads** | 26 |
| **Reads Per Million** | 0.57 rpm (after QC) |
| **Ambiguities** | 0 |
| **Assembly Method** | de novo + reference guided assembly |
| **Consensus Caller** | Bcf Tools |

#### Coverage Map

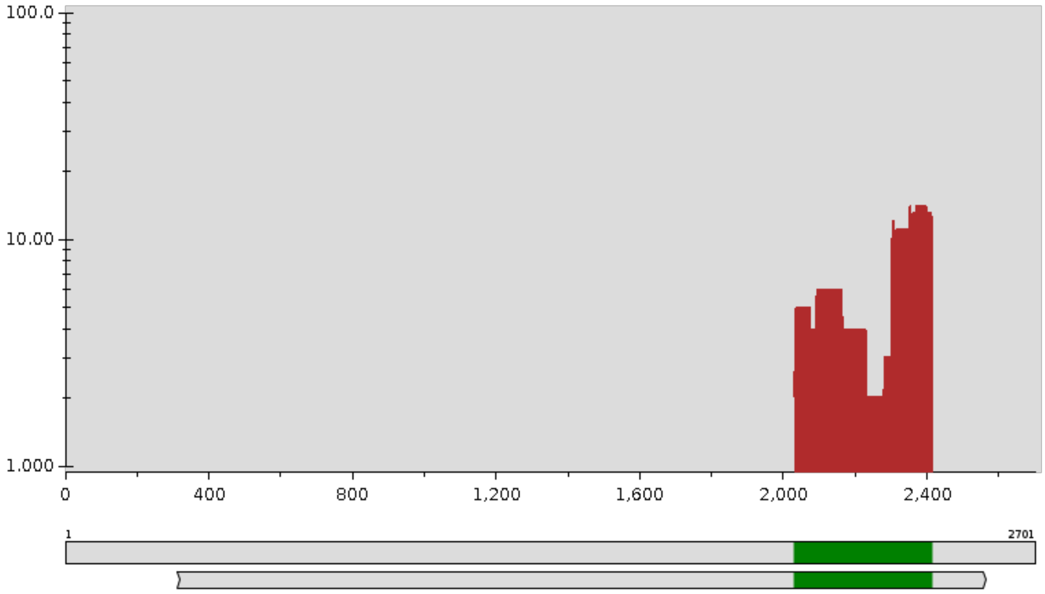

#### Assignment

| | |
|---|---|
| **Type** | Duamitovirus peex1 (Taxonomy ID: 2955799) |
| **Reference Genome** | NC_076525.1 |
| **NT Identity (%)** | 71.9072 |
| **AA Identity (%)** | 65.8915 |
| **Number Of Stop Codons** | 0 |
| **Number Of CDS** | 1 |

#### Alignment

| | |
|---|---|
| **Alignment Score** | 340.0 (NT) + 633.0 (AA) = 973.0 |
| **Concordance (%)** | 56.5041 |

| Alignment Method | Global, seeded, nucleotide + amino acids (AGA) |
|---|---|

#### Genome Region

Sequence starts at position 2030 and ends at position 2417 relative to NC_076525.1 reference sequence.

#### Alignment Detailed Statistics

| | Begin | End | Coverage | Score | Concordance | Matches | Identities | I/D/M/F* | Stop Codons |
|---|---|---|---|---|---|---|---|---|---|
| **NT** | **2030** | **2417** | **14.4%** | **340** | **43.8%** | **388 (100%)** | **279 (71.9%)** | **0/0** | |
| Mutations: | 2036T>G, 2040A>T, 2042G>T, 2051T>A, 2052A>C, 2053A>C, 2057C>A, 2066G>A, 2067A>C, 2069T>A, 2075T>G, 2077T>C, 2088T>C, 2090T>A, 2091T>A, 2095T>G, 2096A>G, 2098C>T, 2102G>T, 2104G>T, 2105G>A, 2109T>C, 2119G>T, 2120G>T, 2133G>T, 2134G>T, 2135A>T, 2136A>G, 2137A>G, 2138G>A, 2139C>A, 2145G>A, 2156A>G, 2158G>A, 2159A>G, 2161G>C, 2162T>G, 2165G>A, 2171C>A, 2173C>A, 2180T>G, 2184A>G, 2188A>C, 2189G>T, 2190T>G, 2195A>G, 2202G>C, 2210G>A, 2211T>A, 2213G>C, 2214A>T, 2216A>T, 2223C>G, 2225A>T, 2226C>T, 2231G>A, 2234A>T, 2240T>A, 2245A>G, 2246T>G, 2249G>A, 2255A>T, 2258A>G, 2261C>G, 2268G>A, 2270A>C, 2272G>T, 2274C>A, 2283G>A, 2292C>T, 2294T>G, 2295C>A, 2296G>C, 2297A>T, 2306A>T, 2311A>G, 2312T>C, 2315C>T, 2321G>T, 2325A>G, 2326A>C, 2327T>A, 2330C>A, 2331C>G, 2333G>T, 2339G>A, 2345A>T, 2348C>T, 2351A>G, 2352C>T, 2354T>C, 2363G>T, 2373G>A, 2374C>G, 2375G>T, 2377C>A, 2378T>A, 2379T>A, 2388C>A, 2390T>A, 2393T>A, 2394C>A, 2396G>C, 2398C>A, 2399T>A, 2406C>G, 2411A>G, 2412G>A, 2414G>C | | | | | | | | |

##### CDS

| | Begin | End | Coverage | Score | Concordance | Matches | Identities | I/D/M/F* | Stop Codons |
|---|---|---|---|---|---|---|---|---|---|
| RdRp | 574 | 702 | 17.2% | 633 | 66.6% | 129 (100%) | 85 (65.9%) | 0/0/0/0 | 0 |
| Protein mutations: | T577S (2040A>T 2042G>T), K581P (2052A>C 2053A>C), I586L (2067A>C 2069T>A), V589A (2077T>C), S593P (2088T>C 2090T>A), Y594N (2091T>A), I595R (2095T>G 2096A>G), S596L (2098C>T), Q597H (2102G>T), R598L (2104G>T 2105G>A), S600P (2109T>C), W603F (2119G>T 2120G>T), G608F (2133G>T 2134G>T 2135A>T), K609G (2136A>G 2137A>G 2138G>A), P610T (2139C>A), D612N (2145G>A), R616K (2158G>A 2159A>G), G617A (2161G>C 2162T>G), A621D (2173C>A), K625E (2184A>G), E626A (2188A>C 2189G>T), L627V (2190T>G), E631Q (2202G>C), L634I (2211T>A 2213G>C), I635F (2214A>T 2216A>T), Q638D (2223C>G 2225A>T), L641F (2234A>T), H645R (2245A>G 2246T>G), V653I (2268G>A 2270A>C), R654L (2272G>T), H655N (2274C>A), E658K (2283G>A), R662T (2295C>A 2296G>C 2297A>T), Y667C (2311A>G 2312T>C), N672A (2325A>G 2326A>C 2327T>A), P674A (2331C>G 2333G>T), L681F (2352C>T 2354T>C), A688S (2373G>A 2374C>G 2375G>T), T689K (2377C>A 2378T>A), S690T (2379T>A), Q695N (2394C>A 2396G>C), T696K (2398C>A 2399T>A), H699D (2406C>G), V701I (2412G>A 2414G>C) | | | | | | | | |
| Codon mutations: | CTT575CTG (2036T>G), ACG577TCT (2040A>T 2042G>T), TCT580TCA (2051T>A), AAA581CCA (2052A>C 2053A>C), CGC582CGA (2057C>A), AGG585AGA (2066G>A), ATT586CTA (2067A>C 2069T>A), GCT588GCG (2075T>G), GTT589GCT (2077T>C), TCT593CCA (2088T>C 2090T>A), TAC594AAC (2091T>A), ATA595AGG (2095T>G 2096A>G), TCA596TTA (2098C>T), CAG597CAT (2102G>T), CGG598CTA (2104G>T 2105G>A), TCT600CCT (2109T>C), TGG603TTT (2119G>T 2120G>T), GGA608TTT (2133G>T 2134G>T 2135A>T), AAG609GGA (2136A>G 2137A>G 2138G>A), CCC610ACC (2139C>A), GAT612AAT (2145G>A), TTA615TTG (2156A>G), AGA616AAG (2158G>A 2159A>G), GGT617GCG (2161G>C 2162T>G), AAG618AAA (2165G>A), ATC620ATA (2171C>A), GCT621GAT (2173C>A), CTT623CTG (2180T>G), AAG625GAG (2184A>G), GAG626GCT (2188A>C 2189G>T), TTA627GTA (2190T>G), AAA628AAG (2195A>G), GAG631CAG (2202G>C), CAG633CAA (2210G>A), TTG634ATC (2211T>A 2213G>C), ATA635TTT (2214A>T 2216A>T), CAA638GAT (2223C>G 2225A>T), CTG639TTG (2226C>T), GTG640GTA (2231G>A), TTA641TTT (2234A>T), GGT643GGA (2240T>A), CAT645CGG (2245A>G 2246T>G), GAG646GAA (2249G>A), CTA648CTT (2255A>T), GAA649GAG (2258A>G), CGC650CGG (2261C>G), GTA653ATC (2268G>A 2270A>C), CGT654CTT (2272G>T), CAT655AAT (2274C>A), GAA658AAA (2283G>A), CTT661TTG (2292C>T 2294T>G), CGA662ACT (2295C>A 2296G>C 2297A>T), TCA665TCT (2306A>T), TAT667TGC (2311A>G 2312T>C), CAC668CAT (2315C>T), GTG670GTT (2321G>T), AAT672GCA (2325A>G 2326A>C 2327T>A), TCC673TCA (2330C>A), CCG674GCT (2331C>G 2333G>T), GTG676GTA (2339G>A), ATA678ATT (2345A>T), GAC679GAT (2348C>T), CAA680CAG (2351A>G), CTT681TTC (2352C>T 2354T>C), GTG684GTT (2363G>T), GCG688AGT (2373G>A 2374C>G 2375G>T), ACT689AAA (2377C>A 2378T>A), TCT690ACT (2379T>A), CGT693AGA (2388C>A 2390T>A), ACT694ACA (2393T>A), CAG695AAC (2394C>A 2396G>C), ACT696AAA (2398C>A 2399T>A), CAT699GAT (2406C>G), TTA700TTG (2411A>G), GTG701ATC (2412G>A 2414G>C) | | | | | | | | |

##### Proteins

| | Begin | End | Coverage | Score | Concordance | Matches | Identities | I/D/M/F* | Stop Codons |
|---|---|---|---|---|---|---|---|---|---|
| RNA-dependent RNA polymerase (YP_010798875.1) | 574 | 702 | 17.2% | 633 | 66.6% | 129 (100%) | 85 (65.9%) | 0/0/0/0 | 0 |
| Protein mutations: | T577S (2040A>T 2042G>T), K581P (2052A>C 2053A>C), I586L (2067A>C 2069T>A), V589A (2077T>C), S593P (2088T>C 2090T>A), Y594N (2091T>A), I595R (2095T>G 2096A>G), S596L (2098C>T), Q597H (2102G>T), R598L (2104G>T 2105G>A), S600P (2109T>C), W603F (2119G>T 2120G>T), G608F (2133G>T 2134G>T 2135A>T), K609G (2136A>G 2137A>G 2138G>A), P610T (2139C>A), D612N (2145G>A), R616K (2158G>A 2159A>G), G617A (2161G>C 2162T>G), A621D (2173C>A), K625E (2184A>G), E626A (2188A>C 2189G>T), L627V (2190T>G), E631Q (2202G>C), L634I (2211T>A 2213G>C), I635F (2214A>T 2216A>T), Q638D (2223C>G 2225A>T), L641F (2234A>T), H645R (2245A>G 2246T>G), V653I (2268G>A 2270A>C), R654L (2272G>T), H655N (2274C>A), E658K (2283G>A), R662T (2295C>A 2296G>C 2297A>T), Y667C (2311A>G 2312T>C), N672A (2325A>G 2326A>C 2327T>A), P674A (2331C>G 2333G>T), L681F (2352C>T 2354T>C), A688S (2373G>A 2374C>G 2375G>T), T689K (2377C>A 2378T>A), S690T (2379T>A), Q695N (2394C>A 2396G>C), T696K (2398C>A 2399T>A), H699D (2406C>G), V701I (2412G>A 2414G>C) | | | | | | | | |
| Codon mutations: | CTT575CTG (2036T>G), ACG577TCT (2040A>T 2042G>T), TCT580TCA (2051T>A), AAA581CCA (2052A>C 2053A>C), CGC582CGA (2057C>A), AGG585AGA (2066G>A), ATT586CTA (2067A>C 2069T>A), GCT588GCG (2075T>G), GTT589GCT (2077T>C), TCT593CCA (2088T>C 2090T>A), TAC594AAC (2091T>A), ATA595AGG (2095T>G 2096A>G), TCA596TTA (2098C>T), CAG597CAT (2102G>T), CGG598CTA (2104G>T 2105G>A), TCT600CCT (2109T>C), TGG603TTT (2119G>T 2120G>T), GGA608TTT (2133G>T 2134G>T 2135A>T), AAG609GGA (2136A>G 2137A>G 2138G>A), CCC610ACC (2139C>A), GAT612AAT (2145G>A), TTA615TTG (2156A>G), AGA616AAG (2158G>A 2159A>G), GGT617GCG (2161G>C 2162T>G), AAG618AAA (2165G>A), ATC620ATA (2171C>A), GCT621GAT (2173C>A), CTT623CTG (2180T>G), AAG625GAG (2184A>G), GAG626GCT (2188A>C 2189G>T), TTA627GTA (2190T>G), AAA628AAG (2195A>G), GAG631CAG (2202G>C), CAG633CAA (2210G>A), TTG634ATC (2211T>A 2213G>C), ATA635TTT (2214A>T 2216A>T), CAA638GAT (2223C>G 2225A>T), CTG639TTG (2226C>T), GTG640GTA (2231G>A), TTA641TTT (2234A>T), GGT643GGA (2240T>A), CAT645CGG (2245A>G 2246T>G), GAG646GAA (2249G>A), CTA648CTT (2255A>T), GAA649GAG (2258A>G), CGC650CGG (2261C>G), GTA653ATC (2268G>A 2270A>C), CGT654CTT (2272G>T), CAT655AAT (2274C>A), GAA658AAA (2283G>A), CTT661TTG (2292C>T 2294T>G), CGA662ACT (2295C>A 2296G>C 2297A>T), TCA665TCT (2306A>T), TAT667TGC (2311A>G 2312T>C), CAC668CAT (2315C>T), GTG670GTT (2321G>T), AAT672GCA (2325A>G 2326A>C 2327T>A), TCC673TCA (2330C>A), CCG674GCT (2331C>G 2333G>T), GTG676GTA (2339G>A), ATA678ATT (2345A>T), GAC679GAT (2348C>T), CAA680CAG (2351A>G), CTT681TTC (2352C>T 2354T>C), GTG684GTT (2363G>T), GCG688AGT (2373G>A 2374C>G 2375G>T), ACT689AAA (2377C>A 2378T>A), TCT690ACT (2379T>A), CGT693AGA (2388C>A 2390T>A), ACT694ACA (2393T>A), CAG695AAC (2394C>A 2396G>C), ACT696AAA (2398C>A 2399T>A), CAT699GAT (2406C>G), TTA700TTG (2411A>G), GTG701ATC (2412G>A 2414G>C) | | | | | | | | |

*: Inserts / Deletes / Misaligned / Frameshifts

#### Analysis details

This analysis was performed with panviral2.64

### NGS Details (UN59): Caulimovirus latensarmoraciae

#### Assembly

| | |
|---|---|
| **Coverage Length** | 342 (1 contig(s)) |
| **Depth Of Coverage** | 9.8 |
| **Number Of Reads** | 26 |
| **Reads Per Million** | 0.57 rpm (after QC) |
| **Ambiguities** | 0 |
| **Assembly Method** | de novo + reference guided assembly |
| **Consensus Caller** | Bcf Tools |

#### Coverage Map

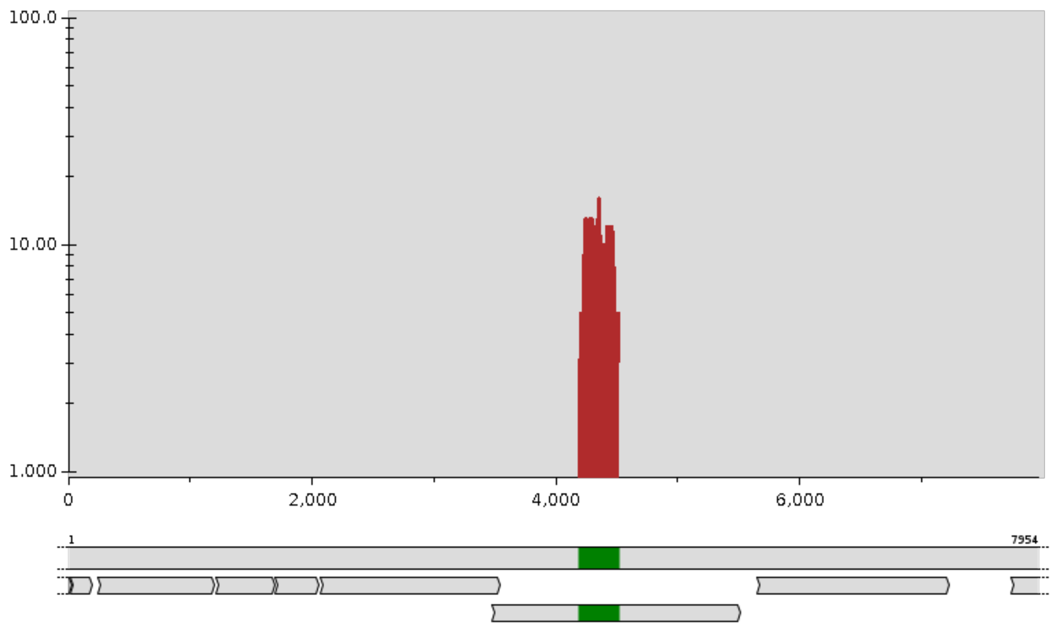

#### Assignment

| | |
|---|---|
| **Type** | Caulimovirus latensarmoraciae (Taxonomy ID: 3047955) |
| **Reference Genome** | NC_018858.1 |
| **NT Identity (%)** | 59.9415 |
| **AA Identity (%)** | 56.1404 |
| **Number Of Stop Codons** | 0 |
| **Number Of CDS** | 8 |

#### Alignment

| | |
|---|---|
| **Alignment Score** | 136.0 (NT) + 482.0 (AA) = 618.0 |
| **Concordance (%)** | 43.338 |

| Alignment Method | Global, seeded, nucleotide + amino acids (AGA) |
|---|---|

#### Genome Region

Sequence starts at position 4182 and ends at position 4523 relative to NC_018858.1 reference sequence.

#### Alignment Detailed Statistics

| | Begin | End | Coverage | Score | Concordance | Matches | Identities | I/D/M/F* | Stop Codons |
|---|---|---|---|---|---|---|---|---|---|
| **NT** | **4182** | **4523** | **4.3%** | **136** | **19.9%** | **342 (100%)** | **205 (59.9%)** | **0/0** | |
| Mutations: | 4182A>G, 4186C>A, 4189C>G, 4190A>T, 4192C>A, 4195A>T, 4199T>A, 4200C>A, 4209C>G, 4210G>C, 4212A>T, 4213G>C, 4214G>A, 4219C>A, 4221A>G, 4222G>A, 4231C>G, 4235A>C, 4236A>T, 4240T>C, 4242G>A, 4243C>T, 4244C>G, 4245C>A, 4247A>C, 4248T>A, 4249G>A, 4252C>T, 4253C>A, 4254G>A, 4256G>A, 4257A>C, 4258A>T, 4265G>A, 4267G>A, 4269A>T, 4270G>T, 4273A>G, 4286C>T, 4288G>A, 4289G>A, 4291C>T, 4292C>A, 4294C>G, 4297G>A, 4298G>T, 4303C>A, 4306G>A, 4308C>A, 4309C>T, 4318C>T, 4326T>G, 4327G>T, 4328G>A, 4329C>G, 4330A>C, 4336G>A, 4339C>T, 4340T>A, 4342A>G, 4345C>A, 4347A>G, 4348C>A, 4352G>C, 4354A>T, 4356C>G, 4357T>C, 4360G>A, 4361A>C, 4364C>A, 4365G>A, 4366C>A, 4367A>C, 4369A>T, 4375G>A, 4376A>G, 4377A>C, 4378G>T, 4381T>A, 4387G>A, 4388G>A, 4390C>A, 4396C>T, 4401C>A, 4402A>G, 4403A>C, 4405G>T, 4409G>A, 4411T>A, 4412G>A, 4413C>A, 4414G>T, 4415A>T, 4416C>G, 4420A>C, 4421G>T, 4422G>T, 4428C>G, 4432C>T, 4433A>T, 4434A>T, 4435T>C, 4436C>A, 4438T>C, 4441C>A, 4444C>T, 4447G>A, 4448G>A, 4451G>A, 4453A>C, 4456C>T, 4457C>A, 4459T>A, 4461C>A, 4462G>T, 4465C>T, 4466A>G, 4467T>C, 4468C>A, 4470G>A, 4477G>A, 4479A>C, 4480A>C, 4481A>T, 4482T>A, 4486C>T, 4489T>A, 4490T>A, 4491C>A, 4492C>A, 4498C>T, 4505T>A, 4506C>G, 4507A>T, 4513C>T, 4520G>A, 4522T>A | | | | | | | | |

##### CDS

| | Begin | End | Coverage | Score | Concordance | Matches | Identities | I/D/M/F* | Stop Codons |
|---|---|---|---|---|---|---|---|---|---|
| D306_gp7 | 236 | 349 | 16.8% | 482 | 63.5% | 114 (100%) | 64 (56.1%) | 0/0/0/0 | 0 |
| Protein mutations: | S237R (4189C>G), I238L (4190A>T 4192C>A), K239N (4195A>T), S241K (4199T>A 4200C>A), T244S (4209C>G 4210G>C), K245I (4212A>T 4213G>C), V246I (4214G>A), K248R (4221A>G 4222G>A), K253L (4235A>C 4236A>T), S255N (4242G>A 4243C>T), P256E (4244C>G 4245C>A), M257Q (4247A>C 4248T>A 4249G>A), R259K (4253C>A 4254G>A), E260T (4256G>A 4257A>C 4258A>T), E263K (4265G>A 4267G>A), K264I (4269A>T 4270G>T), D271N (4289G>A 4291C>T), L272M (4292C>A 4294C>G), V274L (4298G>T), P277H (4308C>A 4309C>T), M283S (4326T>G 4327G>T), A284S (4328G>A 4329C>G 4330A>C), L288M (4340T>A 4342A>G), N290R (4347A>G 4348C>A), E292H (4352G>C 4354A>T), A293G (4356C>G 4357T>C), K295Q (4361A>C), R296K (4364C>A 4365G>A 4366C>A), K300A (4376A>G 4377A>C 4378G>T), V304I (4388G>A 4390C>A), A308E (4401C>A 4402A>G), M309L (4403A>C 4405G>T), D311K (4409G>A 4411T>A), A312N (4412G>A 4413C>A 4414G>T), T313C (4415A>T 4416C>G), G315F (4421G>T 4422G>T), A317G (4428C>G), N319F (4433A>T 4434A>T 4435T>C), L320I (4436C>A 4438T>C), D324N (4448G>A), E325N (4451G>A 4453A>C), L327I (4457C>A 4459T>A), T328N (4461C>A 4462G>T), I330A (4466A>G 4467T>C 4468C>A), R331K (4470G>A), K334T (4479A>C 4480A>C), I335Y (4481A>T 4482T>A), S338K (4490T>A 4491C>A 4492C>A), V348I (4520G>A 4522T>A) | | | | | | | | |
| Codon mutations: | CAA235.GA (4182A>G), GCC236GCA (4186C>A), AGC237AGG (4189C>G), ATC238TTA (4190A>T 4192C>A), AAA239AAT (4195A>T), TCG241AAG (4199T>A 4200C>A), ACG244AGC (4209C>G 4210G>C), AAG245ATC (4212A>T 4213G>C), GTT246ATT (4214G>A), ATC247ATA (4219C>A), AAG248AGA (4221A>G 4222G>A), CCC251CCG (4231C>G), AAA253CTA (4235A>C 4236A>T), TAT254TAC (4240T>C), AGC255AAT (4242G>A 4243C>T), CCA256GAA (4244C>G 4245C>A), ATG257CAA (4247A>C 4248T>A 4249G>A), GAC258GAT (4252C>T), CGA259AAA (4253C>A 4254G>A), GAA260ACT (4256G>A 4257A>C 4258A>T), GAG263AAA (4265G>A 4267G>A), AAG264ATT (4269A>T 4270G>T), CAA265CAG (4273A>G), CTG270TTA (4286C>T 4288G>A), GAC271AAT (4289G>A 4291C>T), CTC272ATG (4292C>A 4294C>G), AAG273AAA (4297G>A), GTA274TTA (4298G>T), ATC275ATA (4303C>A), AGG276AGA (4306G>A), CCC277CAT (4308C>A 4309C>T), AGC280AGT (4318C>T), ATG283AGT (4326T>G 4327G>T), GCA284AGC (4328G>A 4329C>G 4330A>C), GCG286GCA (4336G>A), TTC287TTT (4339C>T), TTA288ATG (4340T>A 4342A>G), GTC289GTA (4345C>A), AAC290AGA (4347A>G 4348C>A), GAA292CAT (4352G>C 4354A>T), GCT293GGC (4356C>G 4357T>C), GAG294GAA (4360G>A), AAA295CAA (4361A>C), CGC296AAA (4364C>A 4365G>A 4366C>A), AGA297CGT (4367A>C 4369A>T), AAG299AAA (4375G>A), AAG300GCT (4376A>G 4377A>C 4378G>T), CGT301CGA (4381T>A), GTG303GTA (4387G>A), GTC304ATA (4388G>A 4390C>A), TAC306TAT (4396C>T), GCA308GAG (4401C>A 4402A>G), ATG309CTT (4403A>C 4405G>T), GAT311AAA (4409G>A 4411T>A), GCG312AAT (4412G>A 4413C>A 4414G>T), ACT313TGT (4415A>T 4416C>G), GTA314GTC (4420A>C), GGC315TTC (4421G>T 4422G>T), GCA317GGA (4428C>G), TAC318TAT (4432C>T), AAT319TTC (4433A>T 4434A>T 4435T>C), CTT320ATC (4436C>A 4438T>C), CCC321CCA (4441C>A), AAC322AAT (4444C>T), AAG323AAA (4447G>A), GAT324AAT (4448G>A), GAA325AAC (4451G>A 4453A>C), CTC326CTT (4456C>T), CTT327ATA (4457C>A 4459T>A), ACG328AAT (4461C>A 4462G>T), CTC329CTT (4465C>T), ATC330GCA (4466A>G 4467T>C 4468C>A), AGA331AAA (4470G>A), AAG333AAA (4477G>A), AAA334ACC (4479A>C 4480A>C), ATC335TAC (4481A>T 4482T>A), TTC336TTT (4486C>T), TCT337TCA (4489T>A), TCC338AAA (4490T>A 4491C>A 4492C>A), GAC340GAT (4498C>T), TCA343AGT (4505T>A 4506C>G 4507A>T), TTC345TTT (4513C>T), GTT348ATA (4520G>A 4522T>A) | | | | | | | | |

##### Proteins

| | Begin | End | Coverage | Score | Concordance | Matches | Identities | I/D/M/F* | Stop Codons |
|---|---|---|---|---|---|---|---|---|---|
| polyprotein (YP_006907834.1) | 236 | 349 | 16.8% | 482 | 63.5% | 114 (100%) | 64 (56.1%) | 0/0/0/0 | 0 |
| Protein mutations: | S237R (4189C>G), I238L (4190A>T 4192C>A), K239N (4195A>T), S241K (4199T>A 4200C>A), T244S (4209C>G 4210G>C), K245I (4212A>T 4213G>C), V246I (4214G>A), K248R (4221A>G 4222G>A), K253L (4235A>C 4236A>T), S255N (4242G>A 4243C>T), P256E (4244C>G 4245C>A), M257Q (4247A>C 4248T>A 4249G>A), R259K (4253C>A 4254G>A), E260T (4256G>A 4257A>C 4258A>T), E263K (4265G>A 4267G>A), K264I (4269A>T 4270G>T), D271N (4289G>A 4291C>T), L272M (4292C>A 4294C>G), V274L (4298G>T), P277H (4308C>A 4309C>T), M283S (4326T>G 4327G>T), A284S (4328G>A 4329C>G 4330A>C), L288M (4340T>A 4342A>G), N290R (4347A>G 4348C>A), E292H (4352G>C 4354A>T), A293G (4356C>G 4357T>C), K295Q (4361A>C), R296K (4364C>A 4365G>A 4366C>A), K300A (4376A>G 4377A>C 4378G>T), V304I (4388G>A 4390C>A), A308E (4401C>A 4402A>G), M309L (4403A>C 4405G>T), D311K (4409G>A 4411T>A), A312N (4412G>A 4413C>A 4414G>T), T313C (4415A>T 4416C>G), G315F (4421G>T 4422G>T), A317G (4428C>G), N319F (4433A>T 4434A>T 4435T>C), L320I (4436C>A 4438T>C), D324N (4448G>A), E325N (4451G>A 4453A>C), L327I (4457C>A 4459T>A), T328N (4461C>A 4462G>T), I330A (4466A>G 4467T>C 4468C>A), R331K (4470G>A), K334T (4479A>C 4480A>C), I335Y (4481A>T 4482T>A), S338K (4490T>A 4491C>A 4492C>A), V348I (4520G>A 4522T>A) | | | | | | | | |
| Codon mutations: | CAA235.GA (4182A>G), GCC236GCA (4186C>A), AGC237AGG (4189C>G), ATC238TTA (4190A>T 4192C>A), AAA239AAT (4195A>T), TCG241AAG (4199T>A 4200C>A), ACG244AGC (4209C>G 4210G>C), AAG245ATC (4212A>T 4213G>C), GTT246ATT (4214G>A), ATC247ATA (4219C>A), AAG248AGA (4221A>G 4222G>A), CCC251CCG (4231C>G), AAA253CTA (4235A>C 4236A>T), TAT254TAC (4240T>C), AGC255AAT (4242G>A 4243C>T), CCA256GAA (4244C>G 4245C>A), ATG257CAA (4247A>C 4248T>A 4249G>A), GAC258GAT (4252C>T), CGA259AAA (4253C>A 4254G>A), GAA260ACT (4256G>A 4257A>C 4258A>T), GAG263AAA (4265G>A 4267G>A), AAG264ATT (4269A>T 4270G>T), CAA265CAG (4273A>G), CTG270TTA (4286C>T 4288G>A), GAC271AAT (4289G>A 4291C>T), CTC272ATG (4292C>A 4294C>G), AAG273AAA (4297G>A), GTA274TTA (4298G>T), ATC275ATA (4303C>A), AGG276AGA (4306G>A), CCC277CAT (4308C>A 4309C>T), AGC280AGT (4318C>T), ATG283AGT (4326T>G 4327G>T), GCA284AGC (4328G>A 4329C>G 4330A>C), GCG286GCA (4336G>A), TTC287TTT (4339C>T), TTA288ATG (4340T>A 4342A>G), GTC289GTA (4345C>A), AAC290AGA (4347A>G 4348C>A), GAA292CAT (4352G>C 4354A>T), GCT293GGC (4356C>G 4357T>C), GAG294GAA (4360G>A), AAA295CAA (4361A>C), CGC296AAA (4364C>A 4365G>A 4366C>A), AGA297CGT (4367A>C 4369A>T), AAG299AAA (4375G>A), AAG300GCT (4376A>G 4377A>C 4378G>T), CGT301CGA (4381T>A), GTG303GTA (4387G>A), GTC304ATA (4388G>A 4390C>A), TAC306TAT (4396C>T), GCA308GAG (4401C>A 4402A>G), ATG309CTT (4403A>C 4405G>T), GAT311AAA (4409G>A 4411T>A), GCG312AAT (4412G>A 4413C>A 4414G>T), ACT313TGT (4415A>T 4416C>G), GTA314GTC (4420A>C), GGC315TTC (4421G>T 4422G>T), GCA317GGA (4428C>G), TAC318TAT (4432C>T), AAT319TTC (4433A>T 4434A>T 4435T>C), CTT320ATC (4436C>A 4438T>C), CCC321CCA (4441C>A), AAC322AAT (4444C>T), AAG323AAA (4447G>A), GAT324AAT (4448G>A), GAA325AAC (4451G>A 4453A>C), CTC326CTT (4456C>T), CTT327ATA (4457C>A 4459T>A), ACG328AAT (4461C>A 4462G>T), CTC329CTT (4465C>T), ATC330GCA (4466A>G 4467T>C 4468C>A), AGA331AAA (4470G>A), AAG333AAA (4477G>A), AAA334ACC (4479A>C 4480A>C), ATC335TAC (4481A>T 4482T>A), TTC336TTT (4486C>T), TCT337TCA (4489T>A), TCC338AAA (4490T>A 4491C>A 4492C>A), GAC340GAT (4498C>T), TCA343AGT (4505T>A 4506C>G 4507A>T), TTC345TTT (4513C>T), GTT348ATA (4520G>A 4522T>A) | | | | | | | | |

*: Inserts / Deletes / Misaligned / Frameshifts

#### Analysis details

This analysis was performed with panviral2.64

### NGS Details (UN59): Badnavirus volubetulae

#### Assembly

| | |
|---|---|
| **Coverage Length** | 391 (7 contig(s)) |
| **Depth Of Coverage** | 7.7 |
| **Number Of Reads** | 25 |
| **Reads Per Million** | 0.54 rpm (after QC) |
| **Ambiguities** | 0 |
| **Assembly Method** | de novo + reference guided assembly |
| **Consensus Caller** | Bcf Tools |

#### Coverage Map

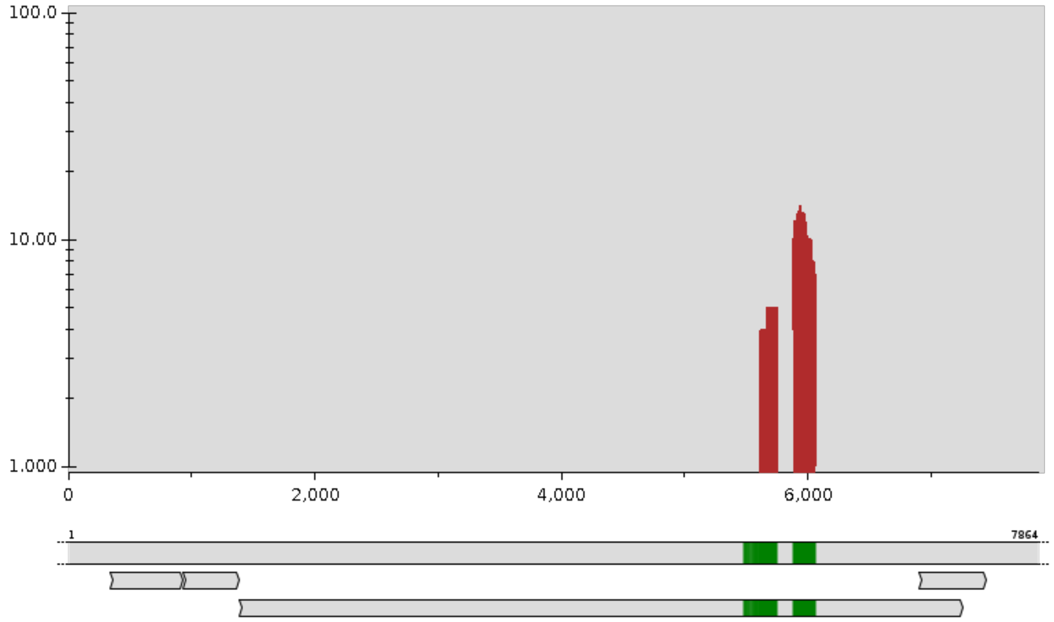

#### Assignment

| | |
|---|---|
| **Type** | Badnavirus volubetulae (Taxonomy ID: 3047683) |
| **Reference Genome** | NC_040635.1 |
| **NT Identity (%)** | 50.9485 |
| **AA Identity (%)** | 47.0588 |
| **Number Of Stop Codons** | 0 |
| **Number Of CDS** | 4 |

#### Alignment

| | |
|---|---|
| **Alignment Score** | -35.0 (NT) + 45.0 (AA) = 10.0 |
| **Concordance (%)** | 0.896057 |

| **Alignment Method** | Global, seeded, nucleotide + amino acids (AGA) |
|---|---|

#### Genome Region

Sequence starts at position 5482 and ends at position 6063 relative to NC_040635.1 reference sequence.

#### Alignment Detailed Statistics

| | Begin | End | Coverage | Score | Concordance | Matches | Identities | I/D/M/F* | Stop Codons |
|---|---|---|---|---|---|---|---|---|---|
| **NT** | **5482** | **6063** | **5.0%** | **-35** | **-5.1%** | **366 (92.9%)** | **188 (47.7%)** | **3/25** | |
| Mutations: | 5482T>A, 5484C>A, 5487T>A, 5490T>A, 5499A>C, 5500T>C, 5503C>A, 5504A>T, 5512T>G, 5513C>G, 5515C>G, 5516T>C, 5523G>A, 5526T>A, 5527C>A, 5528A>T, 5529A>G, 5538G>A, 5544C>A, 5547C>A, 5554C>G, 5556C>A, 5557C>T, 5558T>A, 5562T>A, 5566G>A, 5569T>A, 5570G>T, 5571T>A, 5572A>G, 5573G>C, 5575C>G, 5578C>A, 5580G>A, 5581A>G, 5583G>A, 5584C>G, 5585A>C, 5587G>A, 5588T>C, 5591C>A, 5592A>T, 5594C>A, 5597C>A, 5598A>C, 5601G>A, 5602A>G, 5604G>A, 5608_5610delCAG, 5613C>T, 5614A>C, 5618G>T, 5622C>A, 5624C>T, 5625T>A, 5626G>A, 5628C>A, 5629C>G, 5631G>A, 5632C>T, 5634G>A, 5635C>T, 5637C>A, 5638A>G, 5640G>T, 5641C>T, 5643C>A, 5646G>A, 5649T>A, 5652C>A, 5656C>A, 5657C>G, 5659A>T, 5660G>C, 5661C>A, 5662T>A, 5665A>T, 5666G>C, 5667C>T, 5670G>A, 5673T>C, 5677A>T, 5680A>G, 5681T>C, 5682G>A, 5685G>A, 5686A>T, 5691A>G, 5694T>A, 5695C>A, 5697G>A, 5698T>A, 5700A>C, 5701G>A, 5702G>T, 5706C>G, 5707_5728delAGTATAGATCCTAAAACAGGCA, 5731G>A, 5732A>T, 5736T>C, 5737C>A, 5747A>C, 5748A>C, 5757C>A, 5758C>A, 5877T>G, 5880C>T, 5882T>A, 5886T>C, 5890G>A, 5892C>T, 5893A>C, 5894C>T, 5895C>G, 5908T>G, 5909C>A, 5910A>T, 5911A>G, 5912T>C, 5914C>A, 5915C>A, 5917T>A, 5918G>A, 5923G>T, 5925A>T, 5929C>A, 5934A>C, 5938G>T, 5939A>G, 5940T>G, 5943G>A, 5944T>G, 5949C>T, 5950G>C, 5952A>C, 5954G>A, 5955G>T, 5957T>G, 5958A>G, 5961T>A, 5967A>G, 5979A>G, 5985C>T, 5986C>G, 5987C>G, 5988A>T, 5991A>T, 5992G>A, 5994A>G, 5996T>A, 5998C>A, 5999A>T, 6004A>G, 6005A>C, 6006A>T, 6010G>A, 6011A>C, 6012T>C, 6014A>C, 6015T>C, 6016T>A, 6017G>T, 6022A>C, 6024A>T, 6026G>A, 6029C>T, 6030_6031insATC, 6031G>C, 6033A>T, 6034G>A, 6036A>G, 6037T>G, 6038T>A, 6039C>G, 6042A>T, 6044C>A, 6045T>A, 6051C>T, 6052A>G, 6054C>G, 6063T>C | | | | | | | | |

##### CDS

| | | | | | | | | | |
|---|---|---|---|---|---|---|---|---|---|
| BLRaV_gp3 | 1380 | 1558 | 6.4% | 45 | 4.7% | 118 (92.9%) | 56 (44.1%) | 1/8/11/1 | 0 |
| Protein mutations: | Q1380M (5527C>A 5528A>T 5529A>G), C1394I (5569T>A 5570G>T 5571T>A), K1405E (5602A>G 5604G>A), Q1407del (5608_5610delCAG), K1409Q (5614A>C), R1410I (5618G>T), H1411Q (5622C>A), T1412I (5624C>T 5625T>A), D1413K (5626G>A 5628C>A), Q1414E (5629C>G 5631G>A), K1417D (5638A>G 5640G>T), P1423R (5656C>A 5657C>G), S1425T (5662T>A), T1430S (5677A>T), M1431A (5680A>G 5681T>C 5682G>A), I1433F (5686A>T), I1434M (5691A>G), Q1436K (5695C>A 5697G>A), S1437T (5698T>A 5700A>C), G1438I (5701G>A 5702G>T), S1440_G1446del (5707_5728delAGTATAGATCCTAAAACAGGCA), E1448I (5731G>A 5732A>T), E1453A (5747A>C 5748A>C), F1498Y (5882T>A), V1501I (5890G>A 5892C>T), T1502L (5893A>C 5894C>T 5895C>G), S1507D (5908T>G 5909C>A 5910A>T), I1508A (5911A>G 5912T>C), P1509K (5914C>A 5915C>A), W1510K (5917T>A 5918G>A), A1512S (5923G>T 5925A>T), L1514I (5929C>A), D1517W (5938G>T 5939A>G 5940T>G), L1519V (5944T>G), E1521H (5950G>C 5952A>C), W1522Y (5954G>A 5955G>T), L1523R (5957T>G 5958A>G), P1533G (5986C>G 5987C>G 5988A>T), V1535M (5992G>A 5994A>G), F1536Y (5996T>A), Q1537M (5998C>A 5999A>T), K1539A (6004A>G 6005A>C 6006A>T), D1541T (6010G>A 6011A>C 6012T>C), N1542T (6014A>C 6015T>C), C1543I (6016T>A 6017G>T), K1545H (6022A>C 6024A>T), G1546D (6026G>A), T1547M (6029C>T), T1547_E1548insI (6030_6031insATC), E1548H (6031G>C 6033A>T), E1549K (6034G>A 6036A>G), F1550E (6037T>G 6038T>A 6039C>G), A1552E (6044C>A 6045T>A), I1555V (6052A>G 6054C>G) | | | | | | | | |
| Codon mutations: | TTC1365A.A (5482T>A 5484C>A), ATT1366..A (5487T>A), GGT1367..A (5490T>A), CCA1370..C (5499A>C), TTA1371C.. (5500T>C), CAA1372AT. (5503C>A 5504A>T), TCA1375GG. (5512T>G 5513C>G), CTC1376GC. (5515C>G 5516T>C), AGG1378..A (5523G>A), ATT1379..A (5526T>A), CAA1380ATG (5527C>A 5528A>T 5529A>G), CTG1383..A (5538G>A), ATC1385..A (5544C>A), ATC1386..A (5547C>A), CAC1389G.A (5554C>G 5556C>A), CTT1390TA. (5557C>T 5558T>A), ACT1391..A (5562T>A), GAA1393A.. (5566G>A), TGT1394ATA (5569T>A 5570G>T 5571T>A), AGA1395GC. (5572A>G 5573G>C), CCA1396G.. (5575C>G), CTG1397A.A (5578C>A 5580G>A), AAG1398G.A (5581A>G 5583G>A), CAT1399GC. (5584C>G 5585A>C), GTA1400AC. (5587G>A 5588T>C), ACA1401.AT (5591C>A 5592A>T), CCA1402.A. (5594C>A), GCA1403.AC (5597C>A 5598A>C), ATG1404..A (5601G>A), AAG1405GAA (5602A>G 5604G>A), CAG1407del (5608_5610delCAG), TTC1408TTT (5613C>T), AAG1409CAG (5614A>C), AGA1410ATA (5618G>T), CAC1411CAA (5622C>A), ACT1412ATA (5624C>T 5625T>A), GAC1413AAA (5626G>A 5628C>A), CAG1414GAA (5629C>G 5631G>A), CTG1415TTA (5632C>T 5634G>A), CTC1416TTA (5635C>T 5637C>A), AAG1417GAT (5638A>G 5640G>T), CTC1418TTA (5641C>T 5643C>A), GGG1419GGA (5646G>A), GTT1420GTA (5649T>A), ATC1421ATA (5652C>A), CCA1423AGA (5656C>A 5657C>G), AGC1424TCA (5659A>T 5660G>C 5661C>A), TCA1425ACA (5662T>A), AGC1426TCT (5665A>T 5666G>C 5667C>T), AGG1427AGA (5670G>A), CAT1428CAC (5673T>C), ACA1430TCA (5677A>T), ATG1431GCA (5680A>G 5681T>C 5682G>A), GCG1432GCA (5685G>A), ATC1433TTC (5686A>T), ATA1434ATG (5691A>G), GTT1435GTA (5694T>A), CAG1436AAA (5695C>A 5697G>A), TCA1437ACC (5698T>A 5700A>C), GGA1438ATA (5701G>A 5702G>T), ACC1439ACG (5706C>G), AGT1440_GGC1446del (5707_5728delAGTATAGATCCTAAAACAGGCA), AAA1447-AA (5707_5728delAGTATAGATCCTAAAACAGGCA), GAA1448ATA (5731G>A 5732A>T), GTT1449GTC (5736T>C), CGA1450AGA (5737C>A), GAA1453GCC (5747A>C 5748A>C), GTC1456GTA (5757C>A), CTC1457AT. (5758C>A), AGT1496..G (5877T>G), GGC1497GGT (5880C>T), TTT1498TAT (5882T>A), CAT1499CAC (5886T>C), GTC1501ATT (5890G>A 5892C>T), ACC1502CTG (5893A>C 5894C>T 5895C>G), TCA1507GAT (5908T>G 5909C>A 5910A>T), ATA1508GCA (5911A>G 5912T>C), CCA1509AAA (5914C>A 5915C>A), TGG1510AAG (5917T>A 5918G>A), GCA1512TCT (5923G>T 5925A>T), CTC1514ATC (5929C>A), ACA1515ACC (5934A>C), GAT1517TGG (5938G>T 5939A>G 5940T>G), GGG1518GGA (5943G>A), TTG1519GTG (5944T>G), TAC1520TAT (5949C>T), GAA1521CAC (5950G>C 5952A>C), TGG1522TAT (5954G>A 5955G>T), CTA1523CGG (5957T>G 5958A>G), GTT1524GTA (5961T>A), CCA1526CCG (5967A>G), AAA1530AAG (5979A>G), GCC1532GCT (5985C>T), CCA1533GGT (5986C>G 5987C>G 5988A>T), GCA1534GCT (5991A>T), GTA1535ATG (5992G>A 5994A>G), TTC1536TAC (5996T>A), CAG1537ATG (5998C>A 5999A>T), AAA1539GCT (6004A>G 6005A>C 6006A>T), GAT1541ACC (6010G>A 6011A>C 6012T>C), AAT1542ACC (6014A>C 6015T>C), TGC1543ATC (6016T>A 6017G>T), AAA1545CAT (6022A>C 6024A>T), GGT1546GAT (6026G>A), ACG1547ATG (6029C>T), ACG1547_GAA1548insATC (6030_6031insATC), GAA1548CAT (6031G>C 6033A>T), GAA1549AAG (6034G>A 6036A>G), TTC1550GAG (6037T>G 6038T>A 6039C>G), ATA1551ATT (6042A>T), GCT1552GAA (6044C>A 6045T>A), TAC1554TAT (6051C>T), ATC1555GTG (6052A>G 6054C>G), ATT1558ATC (6063T>C) | | | | | | | | |

##### Proteins

| | | | | | | | | | |
|---|---|---|---|---|---|---|---|---|---|
| polyprotein (YP_009552737.1) | 1380 | 1558 | 6.4% | 45 | 4.7% | 118 (92.9%) | 56 (44.1%) | 1/8/11/1 | 0 |
| Protein mutations: | Q1380M (5527C>A 5528A>T 5529A>G), C1394I (5569T>A 5570G>T 5571T>A), K1405E (5602A>G 5604G>A), Q1407del (5608_5610delCAG), K1409Q (5614A>C), R1410I (5618G>T), H1411Q (5622C>A), T1412I (5624C>T 5625T>A), D1413K (5626G>A 5628C>A), Q1414E (5629C>G 5631G>A), K1417D (5638A>G 5640G>T), P1423R (5656C>A 5657C>G), S1425T (5662T>A), T1430S (5677A>T), M1431A (5680A>G 5681T>C 5682G>A), I1433F (5686A>T), I1434M (5691A>G), Q1436K (5695C>A 5697G>A), S1437T (5698T>A 5700A>C), G1438I (5701G>A 5702G>T), S1440_G1446del (5707_5728delAGTATAGATCCTAAAACAGGCA), E1448I (5731G>A 5732A>T), E1453A (5747A>C 5748A>C), F1498Y (5882T>A), V1501I (5890G>A 5892C>T), T1502L (5893A>C 5894C>T 5895C>G), S1507D (5908T>G 5909C>A 5910A>T), I1508A (5911A>G 5912T>C), P1509K (5914C>A 5915C>A), W1510K (5917T>A 5918G>A), A1512S (5923G>T 5925A>T), L1514I (5929C>A), D1517W (5938G>T 5939A>G 5940T>G), L1519V (5944T>G), E1521H (5950G>C 5952A>C), W1522Y (5954G>A 5955G>T), L1523R (5957T>G 5958A>G), P1533G (5986C>G 5987C>G 5988A>T), V1535M (5992G>A 5994A>G), F1536Y (5996T>A), Q1537M (5998C>A 5999A>T), K1539A (6004A>G 6005A>C 6006A>T), D1541T (6010G>A 6011A>C 6012T>C), N1542T (6014A>C 6015T>C), C1543I (6016T>A 6017G>T), K1545H (6022A>C 6024A>T), G1546D (6026G>A), T1547M (6029C>T), T1547_E1548insI (6030_6031insATC), E1548H (6031G>C 6033A>T), E1549K (6034G>A 6036A>G), F1550E (6037T>G 6038T>A 6039C>G), A1552E (6044C>A 6045T>A), I1555V (6052A>G 6054C>G) | | | | | | | | |
| Codon mutations: | TTC1365A.A (5482T>A 5484C>A), ATT1366..A (5487T>A), GGT1367..A (5490T>A), CCA1370..C (5499A>C), TTA1371C.. (5500T>C), CAA1372AT. (5503C>A 5504A>T), TCA1375GG. (5512T>G 5513C>G), CTC1376GC. (5515C>G 5516T>C), AGG1378..A (5523G>A), ATT1379..A (5526T>A), CAA1380ATG (5527C>A 5528A>T 5529A>G), CTG1383..A (5538G>A), ATC1385..A (5544C>A), ATC1386..A (5547C>A), CAC1389G.A (5554C>G 5556C>A), CTT1390TA. (5557C>T 5558T>A), ACT1391..A (5562T>A), GAA1393A.. (5566G>A), TGT1394ATA (5569T>A 5570G>T 5571T>A), AGA1395GC. (5572A>G 5573G>C), CCA1396G.. (5575C>G), CTG1397A.A (5578C>A 5580G>A), AAG1398G.A (5581A>G 5583G>A), CAT1399GC. (5584C>G 5585A>C), GTA1400AC. (5587G>A 5588T>C), ACA1401.AT (5591C>A 5592A>T), CCA1402.A. (5594C>A), GCA1403.AC (5597C>A 5598A>C), ATG1404..A (5601G>A), AAG1405GAA (5602A>G 5604G>A), CAG1407del (5608_5610delCAG), TTC1408TTT (5613C>T), AAG1409CAG (5614A>C), AGA1410ATA (5618G>T), CAC1411CAA (5622C>A), ACT1412ATA (5624C>T 5625T>A), GAC1413AAA (5626G>A 5628C>A), CAG1414GAA (5629C>G 5631G>A), CTG1415TTA (5632C>T 5634G>A), CTC1416TTA (5635C>T 5637C>A), AAG1417GAT (5638A>G 5640G>T), CTC1418TTA (5641C>T 5643C>A), GGG1419GGA (5646G>A), GTT1420GTA (5649T>A), ATC1421ATA (5652C>A), CCA1423AGA (5656C>A 5657C>G), AGC1424TCA (5659A>T 5660G>C 5661C>A), TCA1425ACA (5662T>A), AGC1426TCT (5665A>T 5666G>C 5667C>T), AGG1427AGA (5670G>A), CAT1428CAC (5673T>C), ACA1430TCA (5677A>T), ATG1431GCA (5680A>G 5681T>C 5682G>A), GCG1432GCA (5685G>A), ATC1433TTC (5686A>T), ATA1434ATG (5691A>G), GTT1435GTA (5694T>A), CAG1436AAA (5695C>A 5697G>A), TCA1437ACC (5698T>A 5700A>C), GGA1438ATA (5701G>A 5702G>T), ACC1439ACG (5706C>G), AGT1440_GGC1446del (5707_5728delAGTATAGATCCTAAAACAGGCA), AAA1447-AA (5707_5728delAGTATAGATCCTAAAACAGGCA), GAA1448ATA (5731G>A 5732A>T), GTT1449GTC (5736T>C), CGA1450AGA (5737C>A), GAA1453GCC (5747A>C 5748A>C), GTC1456GTA (5757C>A), CTC1457AT. (5758C>A), AGT1496..G (5877T>G), GGC1497GGT (5880C>T), TTT1498TAT (5882T>A), CAT1499CAC (5886T>C), GTC1501ATT (5890G>A 5892C>T), ACC1502CTG (5893A>C 5894C>T 5895C>G), TCA1507GAT (5908T>G 5909C>A 5910A>T), ATA1508GCA (5911A>G 5912T>C), CCA1509AAA (5914C>A 5915C>A), TGG1510AAG (5917T>A 5918G>A), GCA1512TCT (5923G>T 5925A>T), CTC1514ATC (5929C>A), ACA1515ACC (5934A>C), GAT1517TGG (5938G>T 5939A>G 5940T>G), GGG1518GGA (5943G>A), TTG1519GTG (5944T>G), TAC1520TAT (5949C>T), GAA1521CAC (5950G>C 5952A>C), TGG1522TAT (5954G>A 5955G>T), CTA1523CGG (5957T>G 5958A>G), GTT1524GTA (5961T>A), CCA1526CCG (5967A>G), AAA1530AAG (5979A>G), GCC1532GCT (5985C>T), CCA1533GGT (5986C>G 5987C>G 5988A>T), GCA1534GCT (5991A>T), GTA1535ATG (5992G>A 5994A>G), TTC1536TAC (5996T>A), CAG1537ATG (5998C>A 5999A>T), AAA1539GCT (6004A>G 6005A>C 6006A>T), GAT1541ACC (6010G>A 6011A>C 6012T>C), AAT1542ACC (6014A>C 6015T>C), TGC1543ATC (6016T>A 6017G>T), AAA1545CAT (6022A>C 6024A>T), GGT1546GAT (6026G>A), ACG1547ATG (6029C>T), ACG1547_GAA1548insATC (6030_6031insATC), GAA1548CAT (6031G>C 6033A>T), GAA1549AAG (6034G>A 6036A>G), TTC1550GAG (6037T>G 6038T>A 6039C>G), ATA1551ATT (6042A>T), GCT1552GAA (6044C>A 6045T>A), TAC1554TAT (6051C>T), ATC1555GTG (6052A>G 6054C>G), ATT1558ATC (6063T>C) | | | | | | | | |

*: Inserts / Deletes / Misaligned / Frameshifts

#### Analysis details

This analysis was performed with panviral2.64

### NGS Details (UN59): Petuvirus venapetuniae

#### Assembly

| | |
|---|---|
| **Coverage Length** | 610 (1 contig(s)) |
| **Depth Of Coverage** | 5.3 |
| **Number Of Reads** | 25 |
| **Reads Per Million** | 0.54 rpm (after QC) |
| **Ambiguities** | 0 |
| **Assembly Method** | de novo + reference guided assembly |
| **Consensus Caller** | Bcf Tools |

#### Coverage Map

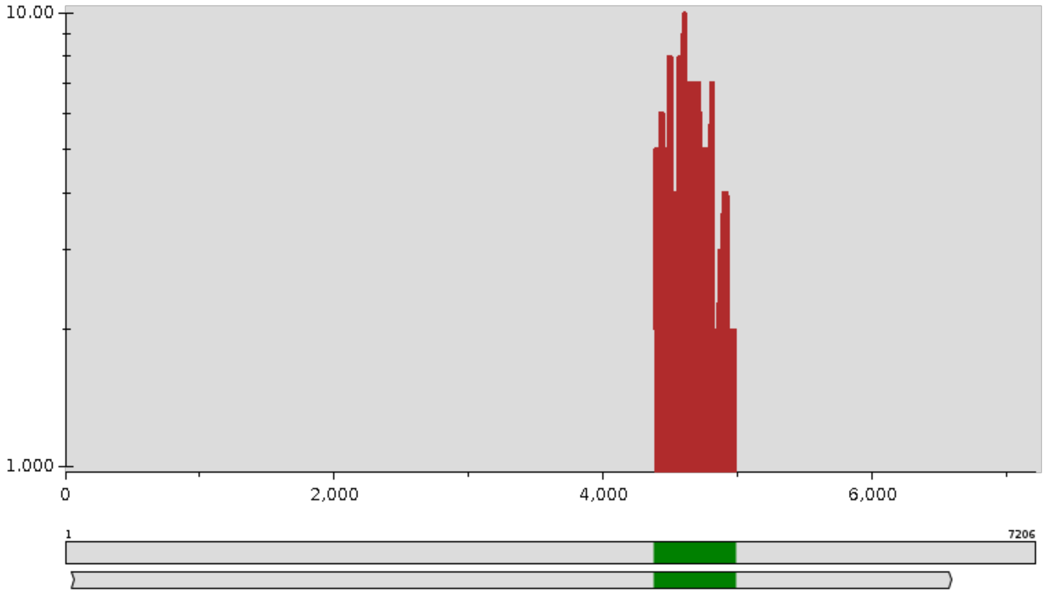

#### Assignment

| | |
|---|---|
| **Type** | Petuvirus venapetuniae (Taxonomy ID: 3048265) |
| **Reference Genome** | NC_001839.2 |
| **NT Identity (%)** | 51.8822 |
| **AA Identity (%)** | 42.9268 |
| **Number Of Stop Codons** | 1 |
| **Number Of CDS** | 1 |

#### Alignment

| | |
|---|---|
| **Alignment Score** | 38.0 (NT) + 491.0 (AA) = 529.0 |
| **Concordance (%)** | 20.983 |

| Alignment Method | Global, seeded, nucleotide + amino acids (AGA) |
|---|---|

#### Genome Region

Sequence starts at position 4380 and ends at position 4989 relative to NC_001839.2 reference sequence.

#### Alignment Detailed Statistics

| | Begin | End | Coverage | Score | Concordance | Matches | Identities | I/D/M/F* | Stop Codons |
|---|---|---|---|---|---|---|---|---|---|
| **NT** | **4380** | **4989** | **8.5%** | **38** | **3.1%** | **610 (99.8%)** | **317 (51.9%)** | **1/0** | |
| Mutations: | 4386C>A, 4391A>T, 4394A>G, 4397C>T, 4398C>A, 4399A>C, 4401T>G, 4402T>C, 4403C>T, 4404C>T, 4406C>A, 4407C>A, 4410G>T, 4411A>G, 4412T>G, 4413G>A, 4414A>T, 4415C>T, 4417A>G, 4418G>A, 4420T>A, 4421T>C, 4424C>T, 4430A>T, 4433T>C, 4436G>A, 4437C>A, 4438T>A, 4439C>G, 4440A>G, 4441C>A, 4442A>T, 4445C>T, 4446T>C, 4448C>A, 4449T>A, 4450C>A, 4451T>A, 4452C>A, 4453A>G, 4454C>A, 4455T>A, 4456T>C, 4457A>C, 4459C>T, 4460C>T, 4469G>C, 4470C>A, 4472C>T, 4474T>A, 4475T>C, 4476T>A, 4477C>G, 4478A>T, 4481G>A, 4488C>A, 4490T>G, 4493G>A, 4499A>G, 4509T>A, 4511A>T, 4512G>C, 4513G>A, 4514A>G, 4517C>T, 4518C>T, 4519A>C, 4521C>G, 4522C>A, 4523T>A, 4526T>A, 4529G>C, 4531G>A, 4533C>T, 4534C>A, 4535A>G, 4538A>G, 4539A>T, 4541A>C, 4543G>C, 4544G>T, 4548T>A, 4549G>A, 4550C>T, 4551A>G, 4556T>G, 4557G>T, 4558A>T, 4560A>G, 4562A>T, 4565T>A, 4566T>C, 4567T>A, 4569C>G, 4577G>C, 4580C>T, 4585C>T, 4586A>T, 4589C>T, 4592T>G, 4600C>A, 4601A>T, 4607T>C, 4610T>A, 4611T>G, 4612T>A, 4613G>A, 4616C>T, 4623G>A, 4624C>A, 4625C>T, 4625_4626insT, 4630T>A, 4631C>T, 4632A>G, 4634A>T, 4638T>A, 4640C>T, 4641C>A, 4643G>T, 4645C>A, 4646C>T, 4647A>T, 4649C>G, 4650C>A, 4651T>G, 4652T>C, 4653T>C, 4654T>A, 4656T>A, 4657C>T, 4658A>G, 4659G>T, 4661T>A, 4662T>A, 4664G>A, 4667C>T, 4670C>T, 4677G>A, 4680A>G, 4682C>T, 4683C>T, 4685T>A, 4686C>A, 4688T>C, 4691T>C, 4697A>G, 4698A>G, 4699C>A, 4700A>C, 4701C>A, 4703G>C, 4706A>T, 4707G>T, 4708A>C, 4712C>T, 4713A>T, 4715A>G, 4719T>C, 4720T>A, 4721G>C, 4724T>A, 4727C>T, 4728C>A, 4729A>T, 4730G>T, 4733C>A, 4734A>T, 4736A>T, 4738G>A, 4739T>C, 4740T>A, 4741T>C, 4742G>C, 4745T>C, 4750A>G, 4752T>A, 4753T>A, 4757A>T, 4758G>T, 4760C>G, 4761A>G, 4763G>A, 4764T>G, 4767T>A, 4768C>G, 4769G>C, 4770G>A, 4772A>T, 4781G>T, 4783T>A, 4784C>A, 4788G>T, 4789C>T, 4795A>C, 4796T>A, 4803C>A, 4804A>G, 4808C>T, 4811A>G, 4814A>T, 4815A>C, 4816T>A, 4817G>T, 4823T>A, 4824G>T, 4825C>A, 4826T>C, 4827G>C, 4829T>A, 4832A>T, 4833A>T, 4837T>G, 4838T>C, 4839T>A, 4840C>G, 4844T>A, 4845G>A, 4846C>T, 4848G>T, 4850T>C, 4851C>A, 4852A>G, 4853C>A, 4854A>G, 4855T>C, 4857T>A, 4858C>T, 4860T>G, 4861T>A, 4863G>T, 4864A>T, 4865A>T, 4866C>T, 4867T>C, 4868C>T, 4869C>A, 4871A>T, 4874G>A, 4880T>C, 4881G>A, 4884A>G, 4885C>T, 4886A>T, 4888A>T, 4890C>A, 4891T>C, 4892C>T, 4893T>G, 4894C>A, 4895G>C, 4896G>A, 4897T>A, 4900A>C, 4901G>C, 4904G>A, 4905A>C, 4907A>G, 4911C>A, 4912A>G, 4913A>G, 4917C>T, 4919T>G, 4922G>A, 4924T>G, 4926G>C, 4931T>C, 4935A>G, 4938A>G, 4939G>C, 4940A>T, 4943T>C, 4946T>C, 4952A>T, 4953G>A, 4960C>G, 4961T>A, 4962G>C, 4964G>A, 4965C>G, 4966A>T, 4968A>T, 4969T>G, 4971T>A, 4972C>A, 4973C>A, 4974C>T, 4977C>T, 4979T>G, 4980A>T, 4981G>T, 4985T>C, 4987T>G, 4989C>T | | | | | | | | |

##### CDS

| | Begin | End | Coverage | Score | Concordance | Matches | Identities | I/D/M/F* | Stop Codons |
|---|---|---|---|---|---|---|---|---|---|
| PVCVgp1 | 1444 | 1647 | 9.4% | 491 | 34.4% | 204 (99.5%) | 88 (42.9%) | 1/0/1/1 | 1 |
| Protein mutations: | Q1446K (4386C>A), H1450T (4398C>A 4399A>C), F1451A (4401T>G 4402T>C 4403C>T), Q1453K (4407C>A), D1454W (4410G>T 4411A>G 4412T>G), D1455I (4413G>A 4414A>T 4415C>T), K1456R (4417A>G 4418G>A), F1457Y (4420T>A 4421T>C), L1463K (4437C>A 4438T>A 4439C>G), T1464D (4440A>G 4441C>A 4442A>T), F1466L (4446T>C 4448C>A), S1467K (4449T>A 4450C>A 4451T>A), H1468R (4452C>A 4453A>G 4454C>A), L1469T (4455T>A 4456T>C 4457A>C), S1470F (4459C>T 4460C>T), K1473N (4469G>C), L1474I (4470C>A 4472C>T), F1475Y (4474T>A 4475T>C), L1480M (4488C>A 4490T>G), L1487I (4509T>A 4511A>T), G1488Q (4512G>C 4513G>A 4514A>G), H1490S (4518C>T 4519A>C), P1491E (4521C>G 4522C>A 4523T>A), N1492K (4526T>A), E1493D (4529G>C), R1494K (4531G>A), P1495* (4533C>T 4534C>A 4535A>G), T1497S (4539A>T 4541A>C), G1498A (4543G>C 4544G>T), C1500N (4548T>A 4549G>A 4550C>T), I1501V (4551A>G), D1503F (4557G>T 4558A>T), R1504G (4560A>G 4562A>T), H1505Q (4565T>A), F1506H (4566T>C 4567T>A), Q1507E (4569C>G), K1509N (4577G>C), P1512L (4585C>T 4586A>T), T1517N (4600C>A 4601A>T), L1521E (4611T>G 4612T>A 4613G>A), A1525N (4623G>A 4624C>A 4625C>T), A1525_M1526insX (4625_4626insT), I1527N (4630T>A 4631C>T), K1528D (4632A>G 4634A>T), F1530I (4638T>A 4640C>T), Q1531N (4641C>A 4643G>T), P1532H (4645C>A 4646C>T), I1533L (4647A>T 4649C>G), L1534S (4650C>A 4651T>G 4652T>C), F1535H (4653T>C 4654T>A), S1536M (4656T>A 4657C>T 4658A>G), A1537S (4659G>T 4661T>A), L1538I (4662T>A 4664G>A), D1543N (4677G>A), I1544V (4680A>G 4682C>T), L1546I (4686C>A 4688T>C), T1550D (4698A>G 4699C>A 4700A>C), L1551I (4701C>A 4703G>C), E1552D (4706A>T), D1553S (4707G>T 4708A>C), I1555L (4713A>T 4715A>G), L1557H (4719T>C 4720T>A 4721G>C), Q1560I (4728C>A 4729A>T 4730G>T), F1561L (4733C>A), I1562F (4734A>T 4736A>T), S1563N (4738G>A 4739T>C), L1564T (4740T>A 4741T>C 4742G>C), K1567R (4750A>G), F1568N (4752T>A 4753T>A), V1570L (4758G>T 4760C>G), M1571V (4761A>G 4763G>A), L1572V (4764T>G), A1574T (4770G>A 4772A>T), M1577I (4781G>T), I1578K (4783T>A 4784C>A), A1580F (4788G>T 4789C>T), N1582T (4795A>C 4796T>A), Q1585R (4803C>A 4804A>G), M1589H (4815A>C 4816T>A 4817G>T), F1591L (4823T>A), A1592Y (4824G>T 4825C>A 4826T>C), D1593Q (4827G>C 4829T>A), T1595S (4833A>T), F1596C (4837T>G 4838T>C), S1597R (4839T>A 4840C>G), A1599I (4845G>A 4846C>T), G1600C (4848G>T 4850T>C), H1601R (4851C>A 4852A>G 4853C>A), I1602A (4854A>G 4855T>C), S1603I (4857T>A 4858C>T), L1604E (4860T>G 4861T>A), E1605F (4863G>T 4864A>T 4865A>T), L1606S (4866C>T 4867T>C 4868C>T), Q1607N (4869C>A 4871A>T), D1611N (4881G>A), T1612V (4884A>G 4885C>T 4886A>T), N1613I (4888A>T), L1614T (4890C>A 4891T>C 4892C>T), S1615D (4893T>G 4894C>A 4895G>C), V1616K (4896G>A 4897T>A), K1617T (4900A>C 4901G>C), I1619L (4905A>C 4907A>G), Q1621R (4911C>A 4912A>G 4913A>G), I1625S (4924T>G), V1626L (4926G>C), I1629V (4935A>G), R1630A (4938A>G 4939G>C 4940A>T), E1635K (4953G>A), T1637R (4960C>G 4961T>A), E1638Q (4962G>C 4964G>A), H1639V (4965C>G 4966A>T), I1640C (4968A>T 4969T>G), S1641K (4971T>A 4972C>A 4973C>A), P1642S (4974C>T), S1644F (4980A>T 4981G>T), M1646R (4987T>G) | | | | | | | | |
| Codon mutations: | CAA1446AAA (4386C>A), CCA1447CCT (4391A>T), TTA1448TTG (4394A>G), AAC1449AAT (4397C>T), CAT1450ACT (4398C>A 4399A>C), TTC1451GCT (4401T>G 4402T>C 4403C>T), CTC1452TTA (4404C>T 4406C>A), CAA1453AAA (4407C>A), GAT1454TGG (4410G>T 4411A>G 4412T>G), GAC1455ATT (4413G>A 4414A>T 4415C>T), AAG1456AGA (4417A>G 4418G>A), TTT1457TAC (4420T>A 4421T>C), CCC1458CCT (4424C>T), CCA1460CCT (4430A>T), AAT1461AAC (4433T>C), AAG1462AAA (4436G>A), CTC1463AAG (4437C>A 4438T>A 4439C>G), ACA1464GAT (4440A>G 4441C>A 4442A>T), CTC1465CTT (4445C>T), TTC1466CTA (4446T>C 4448C>A), TCT1467AAA (4449T>A 4450C>A 4451T>A), CAC1468AGA (4452C>A 4453A>G 4454C>A), TTA1469ACC (4455T>A 4456T>C 4457A>C), TCC1470TTT (4459C>T 4460C>T), AAG1473AAC (4469G>C), CTC1474ATT (4470C>A 4472C>T), TTT1475TAC (4474T>A 4475T>C), TCA1476AGT (4476T>A 4477C>G 4478A>T), AAG1477AAA (4481G>A), CTT1480ATG (4488C>A 4490T>G), AAG1481AAA (4493G>A), GGA1483GGG (4499A>G), TTA1487ATT (4509T>A 4511A>T), GGA1488CAG (4512G>C 4513G>A 4514A>G), ATC1489ATT (4517C>T), CAT1490TCT (4518C>T 4519A>C), CCT1491GAA (4521C>G 4522C>A 4523T>A), AAT1492AAA (4526T>A), GAG1493GAC (4529G>C), AGA1494AAA (4531G>A), CCA1495TAG (4533C>T 4534C>A 4535A>G), AAA1496AAG (4538A>G), ACA1497TCC (4539A>T 4541A>C), GGG1498GCT (4543G>C 4544G>T), TGC1500AAT (4548T>A 4549G>A 4550C>T), ATT1501GTT (4551A>G), CCT1502CCG (4556T>G), GAT1503TTT (4557G>T 4558A>T), AGA1504GGT (4560A>G 4562A>T), CAT1505CAA (4565T>A), TTT1506CAT (4566T>C 4567T>A), CAA1507GAA (4569C>G), AAG1509AAC (4577G>C), GTC1510GTT (4580C>T), CCA1512CTT (4585C>T 4586A>T), TTC1513TTT (4589C>T), GGT1514GGG (4592T>G), ACA1517AAT (4600C>A 4601A>T), CCT1519CCC (4607T>C), TCT1520TCA (4610T>A), TTG1521GAA (4611T>G 4612T>A 4613G>A), TTC1522TTT (4616C>T), GCC1525AAT (4623G>A 4624C>A 4625C>T), GCC1525_ATG1526insT-- (4625_4626insT), ATC1527AAT (4630T>A 4631C>T), AAA1528GAT (4632A>G 4634A>T), TTC1530ATT (4638T>A 4640C>T), CAG1531AAT (4641C>A 4643G>T), CCC1532CAT (4645C>A 4646C>T), ATC1533TTG (4647A>T 4649C>G), CTT1534AGC (4650C>A 4651T>G 4652T>C), TTT1535CAT (4653T>C 4654T>A), TCA1536ATG (4656T>A 4657C>T 4658A>G), GCT1537TCA (4659G>T 4661T>A), TTG1538ATA (4662T>A 4664G>A), GTC1539GTT (4667C>T), TAC1540TAT (4670C>T), GAT1543AAT (4677G>A), ATC1544GTT (4680A>G 4682C>T), CTT1545TTA (4683C>T 4685T>A), CTT1546ATC (4686C>A 4688T>C), TTT1547TTC (4691T>C), GAA1549GAG (4697A>G), ACA1550GAC (4698A>G 4699C>A 4700A>C), CTG1551ATC (4701C>A 4703G>C), GAA1552GAT (4706A>T), GAT1553TCT (4707G>T 4708A>C), CAC1554CAT (4712C>T), ATA1555TTG (4713A>T 4715A>G), TTG1557CAC (4719T>C 4720T>A 4721G>C), CTT1558CTA (4724T>A), AAC1559AAT (4727C>T), CAG1560ATT (4728C>A 4729A>T 4730G>T), TTC1561TTA (4733C>A), ATA1562TTT (4734A>T 4736A>T), AGT1563AAC (4738G>A 4739T>C), TTG1564ACC (4740T>A 4741T>C 4742G>C), GTT1565GTC (4745T>C), AAA1567AGA (4750A>G), TTT1568AAT (4752T>A 4753T>A), GGA1569GGT (4757A>T), GTC1570TTG (4758G>T 4760C>G), ATG1571GTA (4761A>G 4763G>A), TTA1572GTA (4764T>G), TCG1573AGC (4767T>A 4768C>G 4769G>C), GCA1574ACT (4770G>A 4772A>T), ATG1577ATT (4781G>T), ATC1578AAA (4783T>A 4784C>A), GCT1580TTT (4788G>T 4789C>T), AAT1582ACA (4795A>C 4796T>A), CAA1585AGA (4803C>A 4804A>G), TTC1586TTT (4808C>T), TTA1587TTG (4811A>G), GGA1588GGT (4814A>T), ATG1589CAT (4815A>C 4816T>A 4817G>T), TTT1591TTA (4823T>A), GCT1592TAC (4824G>T 4825C>A 4826T>C), GAT1593CAA (4827G>C 4829T>A), GGA1594GGT (4832A>T), ACT1595TCT (4833A>T), TTT1596TGC (4837T>G 4838T>C), TCA1597AGA (4839T>A 4840C>G), CCT1598CCA (4844T>A), GCT1599ATT (4845G>A 4846C>T), GGT1600TGC (4848G>T 4850T>C), CAC1601AGA (4851C>A 4852A>G 4853C>A), ATC1602GCC (4854A>G 4855T>C), TCT1603ATT (4857T>A 4858C>T), TTG1604GAG (4860T>G 4861T>A), GAA1605TTT (4863G>T 4864A>T 4865A>T), CTC1606TCT (4866C>T 4867T>C 4868C>T), CAA1607AAT (4869C>A 4871A>T), AAG1608AAA (4874G>A), CCT1610CCC (4880T>C), GAT1611AAT (4881G>A), ACA1612GTT (4884A>G 4885C>T 4886A>T), AAT1613ATT (4888A>T), CTC1614ACT (4890C>A 4891T>C 4892C>T), TCG1615GAC (4893T>G 4894C>A 4895G>C), GTA1616AAA (4896G>A 4897T>A), AAG1617ACC (4900A>C 4901G>C), CAG1618CAA (4904G>A), ATA1619CTG (4905A>C 4907A>G), CAA1621AGG (4911C>A 4912A>G 4913A>G), CTT1623TTG (4917C>T 4919T>G), GGG1624GGA (4922G>A), ATT1625AGT (4924T>G), GTT1626CTT (4926G>C), AAT1627AAC (4931T>C), ATC1629GTC (4935A>G), AGA1630GCT (4938A>G 4939G>C 4940A>T), GAT1631GAC (4943T>C), TTT1632TTC (4946T>C), CCA1634CCT (4952A>T), GAG1635AAG (4953G>A), ACT1637AGA (4960C>G 4961T>A), GAG1638CAA (4962G>C 4964G>A), CAT1639GTT (4965C>G 4966A>T), ATT1640TGT (4968A>T 4969T>G), TCC1641AAA (4971T>A 4972C>A 4973C>A), CCT1642TCT (4974C>T), CTT1643TTG (4977C>T 4979T>G), AGT1644TTT (4980A>T 4981G>T), GAT1645GAC (4985T>C), ATG1646AGG (4987T>G), CTA1647T.. (4989C>T) | | | | | | | | |

##### Proteins

| | Begin | End | Coverage | Score | Concordance | Matches | Identities | I/D/M/F* | Stop Codons |
|---|---|---|---|---|---|---|---|---|---|
| ORF I polyprotein (NP_127504.1) | 1444 | 1647 | 9.4% | 491 | 34.4% | 204 (99.5%) | 88 (42.9%) | 1/0/1/1 | 1 |

| | Begin | End | Coverage | Score | Concordance | Matches | Identities | I/D/M/F* | Stop Codons |
|---|---|---|---|---|---|---|---|---|---|
| **NT** | **4380** | **4989** | **8.5%** | **38** | **3.1%** | **610 (99.8%)** | **317 (51.9%)** | **1/0** | |
| Protein mutations: | Q1446K (4386C>A), H1450T (4398C>A 4399A>C), F1451A (4401T>G 4402T>C 4403C>T), Q1453K (4407C>A), D1454W (4410G>T 4411A>G 4412T>G), D1455I (4413G>A 4414A>T 4415C>T), K1456R (4417A>G 4418G>A), F1457Y (4420T>A 4421T>C), L1463K (4437C>A 4438T>A 4439C>G), T1464D (4440A>G 4441C>A 4442A>T), F1466L (4446T>C 4448C>A), S1467K (4449T>A 4450C>A 4451T>A), H1468R (4452C>A 4453A>G 4454C>A), L1469T (4455T>A 4456T>C 4457A>C), S1470F (4459C>T 4460C>T), K1473N (4469G>C), L1474I (4470C>A 4472C>T), F1475Y (4474T>A 4475T>C), L1480M (4488C>A 4490T>G), L1487I (4509T>A 4511A>T), G1488Q (4512G>C 4513G>A 4514A>G), H1490S (4518C>T 4519A>C), P1491E (4521C>G 4522C>A 4523T>A), N1492K (4526T>A), E1493D (4529G>C), R1494K (4531G>A), P1495* (4533C>T 4534C>A 4535A>G), T1497S (4539A>T 4541A>C), G1498A (4543G>C 4544G>T), C1500N (4548T>A 4549G>A 4550C>T), I1501V (4551A>G), D1503F (4557G>T 4558A>T), R1504G (4560A>G 4562A>T), H1505Q (4565T>A), F1506H (4566T>C 4567T>A), Q1507E (4569C>G), K1509N (4577G>C), P1512L (4585C>T 4586A>T), T1517N (4600C>A 4601A>T), L1521E (4611T>G 4612T>A 4613G>A), A1525N (4623G>A 4624C>A 4625C>T), A1525_M1526insX (4625_4626insT), I1527N (4630T>A 4631C>T), K1528D (4632A>G 4634A>T), F1530I (4638T>A 4640C>T), Q1531N (4641C>A 4643G>T), P1532H (4645C>A 4646C>T), I1533L (4647A>T 4649C>G), L1534S (4650C>A 4651T>G 4652T>C), F1535H (4653T>C 4654T>A), S1536M (4656T>A 4657C>T 4658A>G), A1537S (4659G>T 4661T>A), L1538I (4662T>A 4664G>A), D1543N (4677G>A), I1544V (4680A>G 4682C>T), L1546I (4686C>A 4688T>C), T1550D (4698A>G 4699C>A 4700A>C), L1551I (4701C>A 4703G>C), E1552D (4706A>T), D1553S (4707G>T 4708A>C), I1555L (4713A>T 4715A>G), L1557H (4719T>C 4720T>A 4721G>C), Q1560I (4728C>A 4729A>T 4730G>T), F1561L (4733C>A), I1562F (4734A>T 4736A>T), S1563N (4738G>A 4739T>C), L1564T (4740T>A 4741T>C 4742G>C), K1567R (4750A>G), F1568N (4752T>A 4753T>A), V1570L (4758G>T 4760C>G), M1571V (4761A>G 4763G>A), L1572V (4764T>G), A1574T (4770G>A 4772A>T), M1577I (4781G>T), I1578K (4783T>A 4784C>A), A1580F (4788G>T 4789C>T), N1582T (4795A>C 4796T>A), Q1585R (4803C>A 4804A>G), M1589H (4815A>C 4816T>A 4817G>T), F1591L (4823T>A), A1592Y (4824G>T 4825C>A 4826T>C), D1593Q (4827G>C 4829T>A), T1595S (4833A>T), F1596C (4837T>G 4838T>C), S1597R (4839T>A 4840C>G), A1599I (4845G>A 4846C>T), G1600C (4848G>T 4850T>C), H1601R (4851C>A 4852A>G 4853C>A), I1602A (4854A>G 4855T>C), S1603I (4857T>A 4858C>T), L1604E (4860T>G 4861T>A), E1605F (4863G>T 4864A>T 4865A>T), L1606S (4866C>T 4867T>C 4868C>T), Q1607N (4869C>A 4871A>T), D1611N (4881G>A), T1612V (4884A>G 4885C>T 4886A>T), N1613I (4888A>T), L1614T (4890C>A 4891T>C 4892C>T), S1615D (4893T>G 4894C>A 4895G>C), V1616K (4896G>A 4897T>A), K1617T (4900A>C 4901G>C), I1619L (4905A>C 4907A>G), Q1621R (4911C>A 4912A>G 4913A>G), I1625S (4924T>G), V1626L (4926G>C), I1629V (4935A>G), R1630A (4938A>G 4939G>C 4940A>T), E1635K (4953G>A), T1637R (4960C>G 4961T>A), E1638Q (4962G>C 4964G>A), H1639V (4965C>G 4966A>T), I1640C (4968A>T 4969T>G), S1641K (4971T>A 4972C>A 4973C>A), P1642S (4974C>T), S1644F (4980A>T 4981G>T), M1646R (4987T>G) | | | | | | | | |
| Codon mutations: | CAA1446AAA (4386C>A), CCA1447CCT (4391A>T), TTA1448TTG (4394A>G), AAC1449AAT (4397C>T), CAT1450ACT (4398C>A 4399A>C), TTC1451GCT (4401T>G 4402T>C 4403C>T), CTC1452TTA (4404C>T 4406C>A), CAA1453AAA (4407C>A), GAT1454TGG (4410G>T 4411A>G 4412T>G), GAC1455ATT (4413G>A 4414A>T 4415C>T), AAG1456AGA (4417A>G 4418G>A), TTT1457TAC (4420T>A 4421T>C), CCC1458CCT (4424C>T), CCA1460CCT (4430A>T), AAT1461AAC (4433T>C), AAG1462AAA (4436G>A), CTC1463AAG (4437C>A 4438T>A 4439C>G), ACA1464GAT (4440A>G 4441C>A 4442A>T), CTC1465CTT (4445C>T), TTC1466CTA (4446T>C 4448C>A), TCT1467AAA (4449T>A 4450C>A 4451T>A), CAC1468AGA (4452C>A 4453A>G 4454C>A), TTA1469ACC (4455T>A 4456T>C 4457A>C), TCC1470TTT (4459C>T 4460C>T), AAG1473AAC (4469G>C), CTC1474ATT (4470C>A 4472C>T), TTT1475TAC (4474T>A 4475T>C), TCA1476AGT (4476T>A 4477C>G 4478A>T), AAG1477AAA (4481G>A), CTT1480ATG (4488C>A 4490T>G), AAG1481AAA (4493G>A), GGA1483GGG (4499A>G), TTA1487ATT (4509T>A 4511A>T), GGA1488CAG (4512G>C 4513G>A 4514A>G), ATC1489ATT (4517C>T), CAT1490TCT (4518C>T 4519A>C), CCT1491GAA (4521C>G 4522C>A 4523T>A), AAT1492AAA (4526T>A), GAG1493GAC (4529G>C), AGA1494AAA (4531G>A), CCA1495TAG (4533C>T 4534C>A 4535A>G), AAA1496AAG (4538A>G), ACA1497TCC (4539A>T 4541A>C), GGG1498GCT (4543G>C 4544G>T), TGC1500AAT (4548T>A 4549G>A 4550C>T), ATT1501GTT (4551A>G), CCT1502CCG (4556T>G), GAT1503TTT (4557G>T 4558A>T), AGA1504GGT (4560A>G 4562A>T), CAT1505CAA (4565T>A), TTT1506CAT (4566T>C 4567T>A), CAA1507GAA (4569C>G), AAG1509AAC (4577G>C), GTC1510GTT (4580C>T), CCA1512CTT (4585C>T 4586A>T), TTC1513TTT (4589C>T), GGT1514GGG (4592T>G), ACA1517AAT (4600C>A 4601A>T), CCT1519CCC (4607T>C), TCT1520TCA (4610T>A), TTG1521GAA (4611T>G 4612T>A 4613G>A), TTC1522TTT (4616C>T), GCC1525AAT (4623G>A 4624C>A 4625C>T), GCC1525_ATG1526insT-- (4625_4626insT), ATC1527AAT (4630T>A 4631C>T), AAA1528GAT (4632A>G 4634A>T), TTC1530ATT (4638T>A 4640C>T), CAG1531AAT (4641C>A 4643G>T), CCC1532CAT (4645C>A 4646C>T), ATC1533TTG (4647A>T 4649C>G), CTT1534AGC (4650C>A 4651T>G 4652T>C), TTT1535CAT (4653T>C 4654T>A), TCA1536ATG (4656T>A 4657C>T 4658A>G), GCT1537TCA (4659G>T 4661T>A), TTG1538ATA (4662T>A 4664G>A), GTC1539GTT (4667C>T), TAC1540TAT (4670C>T), GAT1543AAT (4677G>A), ATC1544GTT (4680A>G 4682C>T), CTT1545TTA (4683C>T 4685T>A), CTT1546ATC (4686C>A 4688T>C), TTT1547TTC (4691T>C), GAA1549GAG (4697A>G), ACA1550GAC (4698A>G 4699C>A 4700A>C), CTG1551ATC (4701C>A 4703G>C), GAA1552GAT (4706A>T), GAT1553TCT (4707G>T 4708A>C), CAC1554CAT (4712C>T), ATA1555TTG (4713A>T 4715A>G), TTG1557CAC (4719T>C 4720T>A 4721G>C), CTT1558CTA (4724T>A), AAC1559AAT (4727C>T), CAG1560ATT (4728C>A 4729A>T 4730G>T), TTC1561TTA (4733C>A), ATA1562TTT (4734A>T 4736A>T), AGT1563AAC (4738G>A 4739T>C), TTG1564ACC (4740T>A 4741T>C 4742G>C), GTT1565GTC (4745T>C), AAA1567AGA (4750A>G), TTT1568AAT (4752T>A 4753T>A), GGA1569GGT (4757A>T), GTC1570TTG (4758G>T 4760C>G), ATG1571GTA (4761A>G 4763G>A), TTA1572GTA (4764T>G), TCG1573AGC (4767T>A 4768C>G 4769G>C), GCA1574ACT (4770G>A 4772A>T), ATG1577ATT (4781G>T), ATC1578AAA (4783T>A 4784C>A), GCT1580TTT (4788G>T 4789C>T), AAT1582ACA (4795A>C 4796T>A), CAA1585AGA (4803C>A 4804A>G), TTC1586TTT (4808C>T), TTA1587TTG (4811A>G), GGA1588GGT (4814A>T), ATG1589CAT (4815A>C 4816T>A 4817G>T), TTT1591TTA (4823T>A), GCT1592TAC (4824G>T 4825C>A 4826T>C), GAT1593CAA (4827G>C 4829T>A), GGA1594GGT (4832A>T), ACT1595TCT (4833A>T), TTT1596TGC (4837T>G 4838T>C), TCA1597AGA (4839T>A 4840C>G), CCT1598CCA (4844T>A), GCT1599ATT (4845G>A 4846C>T), GGT1600TGC (4848G>T 4850T>C), CAC1601AGA (4851C>A 4852A>G 4853C>A), ATC1602GCC (4854A>G 4855T>C), TCT1603ATT (4857T>A 4858C>T), TTG1604GAG (4860T>G 4861T>A), GAA1605TTT (4863G>T 4864A>T 4865A>T), CTC1606TCT (4866C>T 4867T>C 4868C>T), CAA1607AAT (4869C>A 4871A>T), AAG1608AAA (4874G>A), CCT1610CCC (4880T>C), GAT1611AAT (4881G>A), ACA1612GTT (4884A>G 4885C>T 4886A>T), AAT1613ATT (4888A>T), CTC1614ACT (4890C>A 4891T>C 4892C>T), TCG1615GAC (4893T>G 4894C>A 4895G>C), GTA1616AAA (4896G>A 4897T>A), AAG1617ACC (4900A>C 4901G>C), CAG1618CAA (4904G>A), ATA1619CTG (4905A>C 4907A>G), CAA1621AGG (4911C>A 4912A>G 4913A>G), CTT1623TTG (4917C>T 4919T>G), GGG1624GGA (4922G>A), ATT1625AGT (4924T>G), GTT1626CTT (4926G>C), AAT1627AAC (4931T>C), ATC1629GTC (4935A>G), AGA1630GCT (4938A>G 4939G>C 4940A>T), GAT1631GAC (4943T>C), TTT1632TTC (4946T>C), CCA1634CCT (4952A>T), GAG1635AAG (4953G>A), ACT1637AGA (4960C>G 4961T>A), GAG1638CAA (4962G>C 4964G>A), CAT1639GTT (4965C>G 4966A>T), ATT1640TGT (4968A>T 4969T>G), TCC1641AAA (4971T>A 4972C>A 4973C>A), CCT1642TCT (4974C>T), CTT1643TTG (4977C>T 4979T>G), AGT1644TTT (4980A>T 4981G>T), GAT1645GAC (4985T>C), ATG1646AGG (4987T>G), CTA1647T.. (4989C>T) | | | | | | | | |

*: Inserts / Deletes / Misaligned / Frameshifts

#### Analysis details

This analysis was performed with panviral2.64

### NGS Details (UN59): Badnavirus maculasmallanthi

#### Assembly

| | |
|---|---|
| **Coverage Length** | 481 (1 contig(s)) |
| **Depth Of Coverage** | 5.8 |
| **Number Of Reads** | 23 |
| **Reads Per Million** | 0.50 rpm (after QC) |
| **Ambiguities** | 0 |
| **Assembly Method** | de novo + reference guided assembly |
| **Consensus Caller** | Bcf Tools |

#### Coverage Map

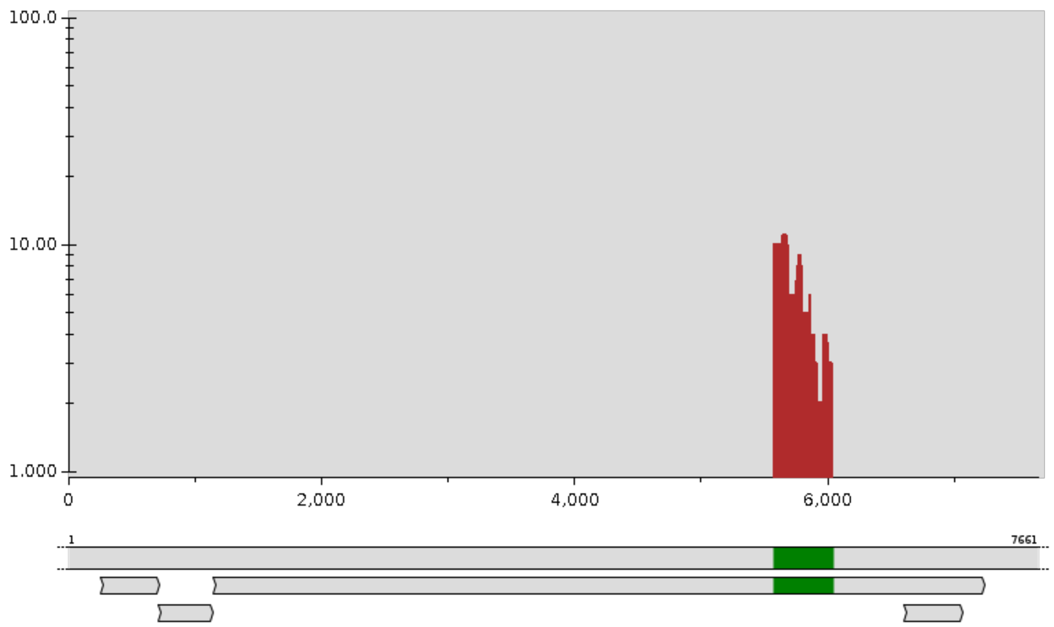

#### Assignment

| | |
|---|---|
| **Type** | Badnavirus maculasmallanthi (Taxonomy ID: 3048453) |
| **Reference Genome** | NC_026472.1 |
| **NT Identity (%)** | 58.7368 |
| **AA Identity (%)** | 60.3774 |
| **Number Of Stop Codons** | 0 |
| **Number Of CDS** | 4 |

#### Alignment

| | |
|---|---|
| **Alignment Score** | 142.0 (NT) + 654.0 (AA) = 796.0 |
| **Concordance (%)** | 39.602 |

| Alignment Method | Global, seeded, nucleotide + amino acids (AGA) |
|---|---|

#### Genome Region

Sequence starts at position 5569 and ends at position 6049 relative to NC_026472.1 reference sequence.

#### Alignment Detailed Statistics

| | Begin | End | Coverage | Score | Concordance | Matches | Identities | I/D/M/F* | Stop Codons |
|---|---|---|---|---|---|---|---|---|---|
| **NT** | **5569** | **6049** | **6.3%** | **142** | **15.3%** | **475 (98.8%)** | **279 (58.0%)** | **0/6** | |

Mutations: 5571C>T, 5577G>A, 5584C>T, 5585T>G, 5586G>C, 5589G>A, 5590T>A, 5591C>G, 5599C>T, 5600A>G, 5601T>G, 5605G>A, 5607T>A, 5608G>A, 5609C>A, 5610C>A, 5611A>C, 5613G>A, 5620G>A, 5622A>T, 5625C>T, 5629C>G, 5631C>T, 5633G>T, 5634G>A, 5637G>A, 5640C>A, 5643T>C, 5644T>A, 5645T>G, 5646A>T, 5648T>C, 5649C>A, 5653G>A, 5654G>A, 5655A>T, 5659T>C, 5660T>A, 5661A>C, 5671C>T, 5673T>A, 5676C>G, 5682A>T, 5685C>T, 5689C>T, 5691C>A, 5700A>C, 5703C>T, 5704G>C, 5705C>A, 5706T>A, 5709T>C, 5718A>G, 5731T>A, 5732G>T, 5733C>A, 5736C>T, 5737_5739delAAA, 5740G>T, 5741A>C, 5743A>C, 5744C>A, 5746G>A, 5748A>G, 5749G>A, 5751G>T, 5755A>C, 5757C>A, 5758G>A, 5759C>T, 5763A>C, 5769C>A, 5772C>T, 5775C>T, 5778C>A, 5779C>T, 5781G>A, 5782G>A, 5786A>G, 5787C>T, 5794A>T, 5795G>C, 5796T>A, 5797G>T, 5799A>T, 5800G>C, 5801C>A, 5802G>A, 5805T>A, 5809G>T, 5810A>T, 5813G>C, 5814G>T, 5818T>C, 5820A>T, 5823G>A, 5824A>C, 5825T>A, 5826T>A, 5827A>T, 5829G>T, 5830C>T, 5831T>C, 5832G>T, 5833G>C, 5834G>A, 5835T>A, 5837A>T, 5841C>T, 5842C>T, 5843A>G, 5844A>C, 5854T>A, 5856A>T, 5860C>T, 5863T>A, 5864C>G, 5865T>C, 5867C>A, 5868A>G, 5870C>A, 5874A>G, 5875A>G, 5876T>C, 5877G>A, 5878A>G, 5880G>C, 5884G>A, 5885C>A, 5886C>T, 5887G>A, 5888T>A, 5889C>A, 5890C>A, 5891C>A, 5892C>T, 5896G>A, 5898G>T, 5904C>T, 5907G>C, 5911G>A, 5912C>T, 5913A>G, 5919A>T, 5920G>T, 5921G>C, 5924G>A, 5925A>G, 5928C>T, 5929A>G, 5930A>G, 5931G>C, 5934A>T, 5935A>G, 5937G>A, 5938C>T, 5940G>A, 5943G>A, 5944C>A, 5946A>T, 5952T>A, 5953A>T, 5954T>C, 5955T>A, 5961A>G, 5964A>C, 5970C>A, 5974_5976delGAT, 5977G>C, 5978A>C, 5979G>A, 5982A>T, 5985G>A, 5991G>T, 5994T>A, 5998G>C, 5999C>A, 6000C>A, 6002G>A, 6003C>A, 6004C>A, 6006T>C, 6007A>C, 6008G>A, 6010T>A, 6011C>G, 6012C>G, 6015C>T, 6022A>T, 6023T>G, 6024C>T, 6030C>T, 6036C>A, 6039A>T, 6040A>G, 6042C>A, 6044A>T, 6048T>A

##### CDS

| | Begin | End | Coverage | Score | Concordance | Matches | Identities | I/D/M/F* | Stop Codons |
|---|---|---|---|---|---|---|---|---|---|
| UF61_gp3 | 1476 | 1636 | 7.9% | 654 | 59.7% | 159 (98.8%) | 96 (59.6%) | 0/2/0/0 | 0 |

Protein mutations: L1481C (5584C>T 5585T>G 5586G>C), H1486W (5599C>T 5600A>G 5601T>G), V1488I (5605G>A 5607T>A), A1489K (5608G>A 5609C>A 5610C>A), M1490L (5611A>C 5613G>A), E1493N (5620G>A 5622A>T), P1496A (5629C>G 5631C>T), W1497L (5633G>T 5634G>A), L1501S (5644T>A 5645T>G 5646A>T), I1502T (5648T>C 5649C>A), G1504N (5653G>A 5654G>A 5655A>T), L1506H (5659T>C 5660T>A 5661A>C), A1521Q (5704G>C 5705C>A 5706T>A), C1530I (5731T>A 5732G>T 5733C>A), K1532del (5737_5739delAAA), D1533S (5740G>T 5741A>C), T1534Q (5743A>C 5744C>A), E1535K (5746G>A 5748A>G), E1536N (5749G>A 5751G>T), I1538L (5755A>C 5757C>A), A1539I (5758G>A 5759C>T), V1547I (5782G>A), Y1548C (5786A>G 5787C>T), E1552Y (5797G>T 5799A>T), A1553Q (5800G>C 5801C>A 5802G>A), D1554E (5805T>A), E1556L (5809G>T 5810A>T), R1557T (5813G>C 5814G>T), I1561Q (5824A>C 5825T>A 5826T>A), M1562F (5827A>T 5829G>T), L1563S (5830C>T 5831T>C 5832G>T), G1564Q (5833G>C 5834G>A 5835T>A), K1565I (5837A>T), Q1567C (5842C>T 5843A>G 5844A>C), L1571I (5854T>A 5856A>T), P1575Q (5867C>A 5868A>G), T1576K (5870C>A), M1578A (5875A>G 5876T>C 5877G>A), K1579D (5878A>G 5880G>C), A1581N (5884G>A 5885C>A 5886C>T), V1582K (5887G>A 5888T>A 5889C>A), P1583N (5890C>A 5891C>A 5892C>T), V1585I (5896G>A 5898G>T), A1590M (5911G>A 5912C>T 5913A>G), G1593S (5920G>T 5921G>C), R1594K (5924G>A 5925A>G), K1596G (5929A>G 5930A>G 5931G>C), K1598E (5935A>G 5937G>A), P1601T (5944C>A 5946A>T), I1604S (5953A>T 5954T>C 5955T>A), D1609E (5970C>A), D1611del (5974_5976delGAT), E1612P (5977G>C 5978A>C 5979G>A), E1613D (5982A>T), K1616N (5991G>T), A1619Q (5998G>C 5999C>A 6000C>A), G1620E (6002G>A 6003C>A), L1621I (6004C>A 6006T>C), R1622Q (6007A>C 6008G>A), S1623R (6010T>A 6011C>G 6012C>G), I1627C (6022A>T 6023T>G 6024C>T), R1632S (6039A>T), N1633E (6040A>G 6042C>A), Y1634F (6044A>T)

Codon mutations: TTC1476TTT (5571C>T), AAG1478AAA (5577G>A), CTG1481TGC (5584C>T 5585T>G 5586G>C), AAG1482AAA (5589G>A), TCT1483AGT (5590T>A 5591C>G), CAT1486TGG (5599C>T 5600A>G 5601T>G), GTT1488ATA (5605G>A 5607T>A), GCC1489AAA (5608G>A 5609C>A 5610C>A), ATG1490CTA (5611A>C 5613G>A), GAA1493AAT (5620G>A 5622A>T), TCC1494TCT (5625C>T), CCC1496GCT (5629C>G 5631C>T), TGG1497TTA (5633G>T 5634G>A), ACG1498ACA (5637G>A), GCC1499GCA (5640C>A), TTT1500TTC (5643T>C), TTA1501AGT (5644T>A 5645T>G 5646A>T), ATC1502ACA (5648T>C 5649C>A), GGA1504AAT (5653G>A 5654G>A 5655A>T), TTA1506CAC (5659T>C 5660T>A 5661A>C), CTT1510TTA (5671C>T 5673T>A), GTC1511GTG (5676C>G), CCA1513CCT (5682A>T), TTC1514TTT (5685C>T), CTC1516TTA (5689C>T 5691C>A), GCA1519GCC (5700A>C), CCC1520CCT (5703C>T), GCT1521CAA (5704G>C 5705C>A 5706T>A), ATT1522ATC (5709T>C), AGA1525AGG (5718A>G), TGC1530ATA (5731T>A 5732G>T 5733C>A), TTC1531TTT (5736C>T), AAA1532del (5737_5739delAAA), GAC1533TCC (5740G>T 5741A>C), ACA1534CAA (5743A>C 5744C>A), GAA1535AAG (5746G>A 5748A>G), GAG1536AAT (5749G>A 5751G>T), ATC1538CTA (5755A>C 5757C>A), GCC1539ATC (5758G>A 5759C>T), GTA1540GTC (5763A>C), ATC1542ATA (5769C>A), GAC1543GAT (5772C>T), GAC1544GAT (5775C>T), ATC1545ATA (5778C>A), CTG1546TTA (5779C>T 5781G>A), GTA1547ATA (5782G>A), TAC1548TGT (5786A>G 5787C>T), AGT1551TCA (5794A>T 5795G>C 5796T>A), GAA1552TAT (5797G>T 5799A>T), GCG1553CAA (5800G>C 5801C>A 5802G>A), GAT1554GAA (5805T>A), GAA1556TTA (5809G>T 5810A>T), AGG1557ACT (5813G>C 5814G>T), TTA1559CTT (5818T>C 5820A>T), AAG1560AAA (5823G>A), ATT1561CAA (5824A>C 5825T>A 5826T>A), ATG1562TTT (5827A>T 5829G>T), CTG1563TCT (5830C>T 5831T>C 5832G>T), GGT1564CAA (5833G>C 5834G>A 5835T>A), AAA1565ATA (5837A>T), TGC1566TGT (5841C>T), CAA1567TGC (5842C>T 5843A>G 5844A>C), TTA1571ATT (5854T>A 5856A>T), CTA1573TTA (5860C>T), TCT1574AGC (5863T>A 5864C>G 5865T>C), CCA1575CAG (5867C>A 5868A>G), ACA1576AAA (5870C>A), AAA1577AAG (5874A>G), ATG1578GCA (5875A>G 5876T>C 5877G>A), AAG1579GAC (5878A>G 5880G>C), GCC1581AAT (5884G>A 5885C>A 5886C>T), GTC1582AAA (5887G>A 5888T>A 5889C>A), CCC1583AAT (5890C>A 5891C>A 5892C>T), GTG1585ATT (5896G>A 5898G>T), TTC1587TTT (5904C>T), CTG1588CTC (5907G>C), GCA1590ATG (5911G>A 5912C>T 5913A>G), ATA1592ATT (5919A>T), GGA1593TCA (5920G>T 5921G>C), AGA1594AAG (5924G>A 5925A>G), AAC1595AAT (5928C>T), AAG1596GGC (5929A>G 5930A>G 5931G>C), ATA1597ATT (5934A>T), AAG1598GAA (5935A>G 5937G>A), CTG1599TTA (5938C>T 5940G>A), CAG1600CAA (5943G>A), CCA1601ACT (5944C>A 5946A>T), ATT1603ATA (5952T>A), ATT1604TCA (5953A>T 5954T>C 5955T>A), AAA1606AAG (5961A>G), ATA1607ATC (5964A>C), GAC1609GAA (5970C>A), GAT1611del (5974_5976delGAT), GAG1612CCA (5977G>C 5978A>C 5979G>A), GAA1613GAT (5982A>T), AAG1614AAA (5985G>A), AAG1616AAT (5991G>T), ACT1617ACA (5994T>A), GCC1619CAA (5998G>C 5999C>A 6000C>A), GGC1620GAA (6002G>A 6003C>A), CTT1621ATC (6004C>A 6006T>C), AGA1622CAA (6007A>C 6008G>A), TCC1623AGG (6010T>A 6011C>G 6012C>G), TTC1624TTT (6015C>T), ATC1627TGT (6022A>T 6023T>G 6024C>T), AAC1629AAT (6030C>T), GCC1631GCA (6036C>A), AGA1632AGT (6039A>T), AAC1633GAA (6040A>G 6042C>A), TAC1634TTC (6044A>T), ATT1635ATA (6048T>A)

##### Proteins

| | Begin | End | Coverage | Score | Concordance | Matches | Identities | I/D/M/F* | Stop Codons |
|---|---|---|---|---|---|---|---|---|---|
| ORF3 (YP_009121747.1) | 1476 | 1636 | 7.9% | 654 | 59.7% | 159 (98.8%) | 96 (59.6%) | 0/2/0/0 | 0 |

Protein mutations: L1481C (5584C>T 5585T>G 5586G>C), H1486W (5599C>T 5600A>G 5601T>G), V1488I (5605G>A 5607T>A), A1489K (5608G>A 5609C>A 5610C>A), M1490L (5611A>C 5613G>A), E1493N (5620G>A 5622A>T), P1496A (5629C>G 5631C>T), W1497L (5633G>T 5634G>A), L1501S (5644T>A 5645T>G 5646A>T), I1502T (5648T>C 5649C>A), G1504N (5653G>A 5654G>A 5655A>T), L1506H (5659T>C 5660T>A 5661A>C), A1521Q (5704G>C 5705C>A 5706T>A), C1530I (5731T>A 5732G>T 5733C>A), K1532del (5737_5739delAAA), D1533S (5740G>T 5741A>C), T1534Q (5743A>C 5744C>A), E1535K (5746G>A 5748A>G), E1536N (5749G>A 5751G>T), I1538L (5755A>C 5757C>A), A1539I (5758G>A 5759C>T), V1547I (5782G>A), Y1548C (5786A>G 5787C>T), E1552Y (5797G>T 5799A>T), A1553Q (5800G>C 5801C>A 5802G>A), D1554E (5805T>A), E1556L (5809G>T 5810A>T), R1557T (5813G>C 5814G>T), I1561Q (5824A>C 5825T>A 5826T>A), M1562F (5827A>T 5829G>T), L1563S (5830C>T 5831T>C 5832G>T), G1564Q (5833G>C 5834G>A 5835T>A), K1565I (5837A>T), Q1567C (5842C>T 5843A>G 5844A>C), L1571I (5854T>A 5856A>T), P1575Q (5867C>A 5868A>G), T1576K (5870C>A), M1578A (5875A>G 5876T>C 5877G>A), K1579D (5878A>G 5880G>C), A1581N (5884G>A 5885C>A 5886C>T), V1582K (5887G>A 5888T>A 5889C>A), P1583N (5890C>A 5891C>A 5892C>T), V1585I (5896G>A 5898G>T), A1590M (5911G>A 5912C>T 5913A>G), G1593S (5920G>T 5921G>C), R1594K (5924G>A 5925A>G), K1596G (5929A>G 5930A>G 5931G>C), K1598E (5935A>G 5937G>A), P1601T (5944C>A 5946A>T), I1604S (5953A>T 5954T>C 5955T>A), D1609E (5970C>A), D1611del (5974_5976delGAT), E1612P (5977G>C 5978A>C 5979G>A), E1613D (5982A>T), K1616N (5991G>T), A1619Q (5998G>C 5999C>A 6000C>A), G1620E (6002G>A 6003C>A), L1621I (6004C>A 6006T>C), R1622Q (6007A>C 6008G>A), S1623R (6010T>A 6011C>G 6012C>G), I1627C (6022A>T 6023T>G 6024C>T), R1632S (6039A>T), N1633E (6040A>G 6042C>A), Y1634F (6044A>T)

Codon mutations: TTC1476TTT (5571C>T), AAG1478AAA (5577G>A), CTG1481TGC (5584C>T 5585T>G 5586G>C), AAG1482AAA (5589G>A), TCT1483AGT (5590T>A 5591C>G), CAT1486TGG (5599C>T 5600A>G 5601T>G), GTT1488ATA (5605G>A 5607T>A), GCC1489AAA (5608G>A 5609C>A 5610C>A), ATG1490CTA (5611A>C 5613G>A), GAA1493AAT (5620G>A 5622A>T), TCC1494TCT (5625C>T), CCC1496GCT (5629C>G 5631C>T), TGG1497TTA (5633G>T 5634G>A), ACG1498ACA (5637G>A), GCC1499GCA (5640C>A), TTT1500TTC (5643T>C), TTA1501AGT (5644T>A 5645T>G 5646A>T), ATC1502ACA (5648T>C 5649C>A), GGA1504AAT (5653G>A 5654G>A 5655A>T), TTA1506CAC (5659T>C 5660T>A 5661A>C), CTT1510TTA (5671C>T 5673T>A), GTC1511GTG (5676C>G), CCA1513CCT (5682A>T), TTC1514TTT (5685C>T), CTC1516TTA (5689C>T 5691C>A), GCA1519GCC (5700A>C), CCC1520CCT (5703C>T), GCT1521CAA (5704G>C 5705C>A 5706T>A), ATT1522ATC (5709T>C), AGA1525AGG (5718A>G), TGC1530ATA (5731T>A 5732G>T 5733C>A), TTC1531TTT (5736C>T), AAA1532del (5737_5739delAAA), GAC1533TCC (5740G>T 5741A>C), ACA1534CAA (5743A>C 5744C>A), GAA1535AAG (5746G>A 5748A>G), GAG1536AAT (5749G>A 5751G>T), ATC1538CTA (5755A>C 5757C>A), GCC1539ATC (5758G>A 5759C>T), GTA1540GTC (5763A>C), ATC1542ATA (5769C>A), GAC1543GAT (5772C>T), GAC1544GAT (5775C>T), ATC1545ATA (5778C>A), CTG1546TTA (5779C>T 5781G>A), GTA1547ATA (5782G>A), TAC1548TGT (5786A>G 5787C>T), AGT1551TCA (5794A>T 5795G>C 5796T>A), GAA1552TAT (5797G>T 5799A>T), GCG1553CAA (5800G>C 5801C>A 5802G>A), GAT1554GAA (5805T>A), GAA1556TTA (5809G>T 5810A>T), AGG1557ACT (5813G>C 5814G>T), TTA1559CTT (5818T>C 5820A>T), AAG1560AAA (5823G>A), ATT1561CAA (5824A>C 5825T>A 5826T>A), ATG1562TTT (5827A>T 5829G>T), CTG1563TCT (5830C>T 5831T>C 5832G>T), GGT1564CAA (5833G>C 5834G>A 5835T>A), AAA1565ATA (5837A>T), TGC1566TGT (5841C>T), CAA1567TGC (5842C>T 5843A>G 5844A>C), TTA1571ATT (5854T>A 5856A>T), CTA1573TTA (5860C>T), TCT1574AGC (5863T>A 5864C>G 5865T>C), CCA1575CAG (5867C>A 5868A>G), ACA1576AAA (5870C>A), AAA1577AAG (5874A>G), ATG1578GCA (5875A>G 5876T>C 5877G>A), AAG1579GAC (5878A>G 5880G>C), GCC1581AAT (5884G>A 5885C>A 5886C>T), GTC1582AAA (5887G>A 5888T>A 5889C>A), CCC1583AAT (5890C>A 5891C>A 5892C>T), GTG1585ATT (5896G>A 5898G>T), TTC1587TTT (5904C>T), CTG1588CTC (5907G>C), GCA1590ATG (5911G>A 5912C>T 5913A>G), ATA1592ATT (5919A>T), GGA1593TCA (5920G>T 5921G>C), AGA1594AAG (5924G>A 5925A>G), AAC1595AAT (5928C>T), AAG1596GGC (5929A>G 5930A>G 5931G>C), ATA1597ATT (5934A>T), AAG1598GAA (5935A>G 5937G>A), CTG1599TTA (5938C>T 5940G>A), CAG1600CAA (5943G>A), CCA1601ACT (5944C>A 5946A>T), ATT1603ATA (5952T>A), ATT1604TCA (5953A>T 5954T>C 5955T>A), AAA1606AAG (5961A>G), ATA1607ATC (5964A>C), GAC1609GAA (5970C>A), GAT1611del (5974_5976delGAT), GAG1612CCA (5977G>C 5978A>C 5979G>A), GAA1613GAT (5982A>T), AAG1614AAA (5985G>A), AAG1616AAT (5991G>T), ACT1617ACA (5994T>A), GCC1619CAA (5998G>C 5999C>A 6000C>A), GGC1620GAA (6002G>A 6003C>A), CTT1621ATC (6004C>A 6006T>C), AGA1622CAA (6007A>C 6008G>A), TCC1623AGG (6010T>A 6011C>G 6012C>G), TTC1624TTT (6015C>T), ATC1627TGT (6022A>T 6023T>G 6024C>T), AAC1629AAT (6030C>T), GCC1631GCA (6036C>A), AGA1632AGT (6039A>T), AAC1633GAA (6040A>G 6042C>A), TAC1634TTC (6044A>T), ATT1635ATA (6048T>A)

*: Inserts / Deletes / Misaligned / Frameshifts

#### Analysis details

This analysis was performed with panviral2.64

### NGS Details (UN59): Epiphyllum badnavirus 1

#### Assembly

| | |
|---|---|
| **Coverage Length** | 354 (1 contig(s)) |
| **Depth Of Coverage** | 7.4 |
| **Number Of Reads** | 22 |
| **Reads Per Million** | 0.48 rpm (after QC) |
| **Ambiguities** | 0 |
| **Assembly Method** | de novo + reference guided assembly |
| **Consensus Caller** | Bcf Tools |

#### Coverage Map

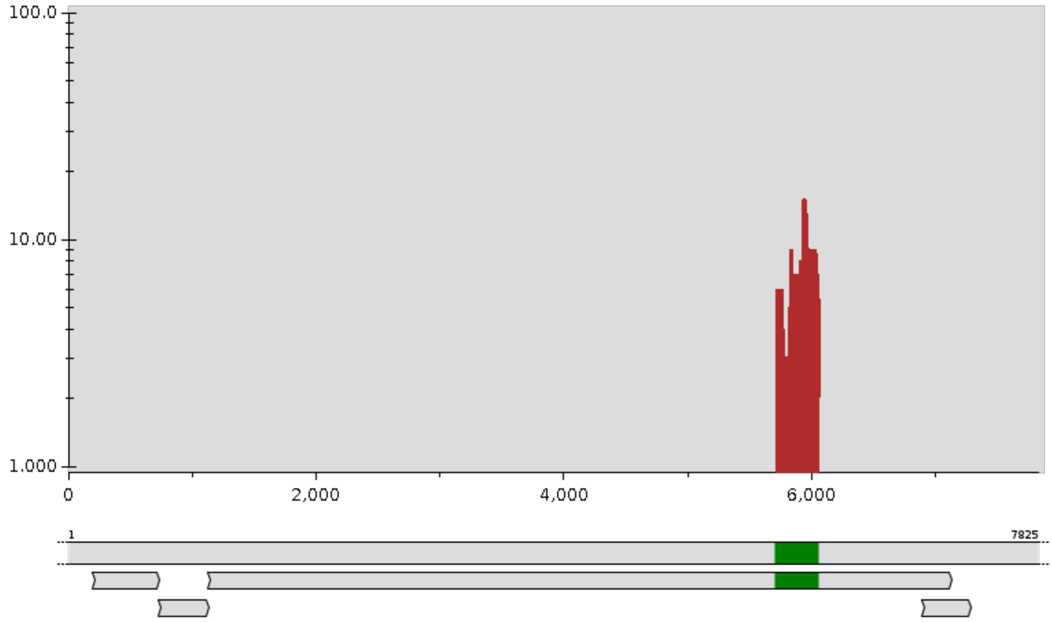

#### Assignment

| | |
|---|---|
| **Type** | Epiphyllum badnavirus 1 (Taxonomy ID: 2518008) |
| **Reference Genome** | NC_076247.1 |
| **NT Identity (%)** | 50.9804 |
| **AA Identity (%)** | 42.0168 |
| **Number Of Stop Codons** | 0 |
| **Number Of CDS** | 4 |

#### Alignment

| | |
|---|---|
| **Alignment Score** | 8.0 (NT) + 335.0 (AA) = 343.0 |
| **Concordance (%)** | 22.215 |

| | |
|---|---|
| **Alignment Method** | Global, seeded, nucleotide + amino acids (AGA) |

#### Genome Region

Sequence starts at position 5709 and ends at position 6062 relative to NC_076247.1 reference sequence.

#### Alignment Detailed Statistics

| | Begin | End | Coverage | Score | Concordance | Matches | Identities | I/D/M/F* | Stop Codons |
|---|---|---|---|---|---|---|---|---|---|
| **NT** | **5709** | **6062** | **4.5%** | **8** | **1.1%** | **354 (99.2%)** | **182 (51.0%)** | **3/0** | |
| Mutations: | 5709A>G, 5710A>T, 5711C>T, 5714G>T, 5715A>T, 5716G>A, 5718A>G, 5719G>A, 5721G>A, 5722A>C, 5724G>A, 5725A>G, 5727A>T, 5730A>T, 5732A>G, 5733G>T, 5734G>A, 5735A>T, 5738G>A, 5742G>A, 5743G>T, 5744T>G, 5745C>T, 5746T>A, 5748C>A, 5752T>A, 5755A>C, 5756A>G, 5758A>C, 5759C>A, 5760C>A, 5763G>A, 5766C>T, 5767G>A, 5768A>C, 5769C>G, 5770A>G, 5771A>T, 5775G>C, 5776T>A, 5778C>T, 5779A>C, 5780A>G, 5782G>A, 5784T>C, 5785C>A, 5793T>A, 5794C>T, 5796A>G, 5800G>C, 5802C>T, 5805T>A, 5806A>G, 5808C>T, 5809A>G, 5810C>A, 5811A>T, 5812A>C, 5815A>T, 5817A>T, 5818C>G, 5820G>C, 5821A>C, 5824G>T, 5826C>A, 5827T>C, 5828G>A, 5829T>G, 5835C>A, 5836A>T, 5837G>C, 5838T>A, 5841C>G, 5847C>A, 5850G>A, 5851T>A, 5853C>T, 5854G>A, 5859G>A, 5861A>G, 5862G>A, 5863A>T, 5864G>C, 5865C>T, 5868A>T, 5870T>A, 5874C>T, 5878A>T, 5882T>A, 5886G>T, 5887G>A, 5889C>G, 5890G>C, 5891A>C, 5892A>G, 5893G>A, 5895A>G, 5896T>G, 5897C>A, 5898C>T, 5899A>G, 5901C>G, 5905T>A, 5910C>T, 5913C>T, 5916C>T, 5917C>A, 5918T>G, 5919G>A, 5924C>G, 5925A>G, 5926G>T, 5931A>G, 5933T>A, 5934T>C, 5942G>T, 5943G>C, 5946A>G, 5947G>A, 5949T>G, 5953C>T, 5955A>G, 5956T>A, 5958C>T, 5961A>G, 5966A>C, 5967G>C, 5973C>G, 5976T>A, 5979T>A, 5981T>C, 5982G>C, 5986C>A, 5987A>T, 5988A>G, 5989C>A, 5991A>T, 5992A>T, 5993A>T, 5998G>A, 6000C>T, 6001A>G, 6002A>G, 6003C>G, 6004T>G, 6005G>T, 6006T>G, 6009C>T, 6011G>A, 6012A>G, 6013G>C, 6014A>C, 6015C>A, 6015_6016insTTT, 6016C>T, 6019T>G, 6020C>A, 6021A>T, 6022G>T, 6023G>C, 6024C>G, 6030A>G, 6031G>A, 6032C>T, 6033A>T, 6035T>G, 6036T>G, 6038A>T, 6039C>T, 6051C>T, 6054G>T, 6055G>A, 6057C>T | | | | | | | | |

##### CDS

| | Begin | End | Coverage | Score | Concordance | Matches | Identities | I/D/M/F* | Stop Codons |
|---|---|---|---|---|---|---|---|---|---|
| QKM20_gp3 | 1528 | 1645 | 5.9% | 335 | 39.7% | 118 (99.2%) | 50 (42.0%) | 1/0/0/0 | 0 |
| Protein mutations: | T1528F (5710A>T 5711C>T), G1529V (5714G>T 5715A>T), E1530K (5716G>A 5718A>G), E1531K (5719G>A 5721G>A), K1532Q (5722A>C 5724G>A), K1533D (5725A>G 5727A>T), K1535S (5732A>G 5733G>T), E1536M (5734G>A 5735A>T), R1537K (5738G>A), M1538I (5742G>A), V1539C (5743G>T 5744T>G 5745C>T), F1540I (5746T>A 5748C>A), Y1542N (5752T>A), K1543R (5755A>C 5756A>G), T1544Q (5758A>C 5759C>A 5760C>A), D1547T (5767G>A 5768A>C 5769C>G), N1548V (5770A>G 5771A>T), F1550I (5776T>A 5778C>T), K1551R (5779A>C 5780A>G), D1552N (5782G>A 5784T>C), Q1553K (5785C>A), G1558R (5800G>C 5802C>T), N1560D (5806A>G 5808C>T), T1561D (5809A>G 5810C>A 5811A>T), I1562L (5812A>C), I1563F (5815A>T 5817A>T), Q1564D (5818C>G 5820G>C), K1565Q (5821A>C), V1566L (5824G>T 5826C>A), C1567Q (5827T>C 5828G>A 5829T>G), F1575I (5851T>A 5853C>T), D1576N (5854G>A), K1578R (5861A>G 5862G>A), F1581Y (5870T>A), I1584F (5878A>T), M1585K (5882T>A), M1586I (5886G>T), D1587K (5887G>A 5889C>G), E1588P (5890G>C 5891A>C 5892A>G), E1589K (5893G>A 5895A>G), S1590D (5896T>G 5897C>A 5898C>T), I1591V (5899A>G 5901C>G), W1593R (5905T>A), L1597R (5917C>A 5918T>G 5919G>A), P1599R (5924C>G 5925A>G), D1600Y (5926G>T), L1602H (5933T>A 5934T>C), W1605F (5942G>T 5943G>C), V1607M (5947G>A 5949T>G), P1609S (5953C>T 5955A>G), F1610I (5956T>A 5958C>T), K1613T (5966A>C 5967G>C), V1618A (5981T>C 5982G>C), Q1620M (5986C>A 5987A>T 5988A>G), R1621S (5989C>A 5991A>T), K1622L (5992A>T 5993A>T), D1624N (5998G>A 6000C>T), N1625G (6001A>G 6002A>G 6003C>G), C1626V (6004T>G 6005G>T 6006T>G), R1628K (6011G>A 6012A>G), D1629P (6013G>C 6014A>C 6015C>A), D1629_L1630insF (6015_6016insTTT), S1631D (6019T>G 6020C>A 6021A>T), G1632S (6022G>T 6023G>C 6024C>G), A1635I (6031G>A 6032C>T 6033A>T), V1636G (6035T>G 6036T>G), Y1637F (6038A>T 6039C>T), V1643I (6055G>A 6057C>T) | | | | | | | | |
| Codon mutations: | GCA1527..G (5709A>G), ACT1528TTT (5710A>T 5711C>T), GGA1529GTT (5714G>T 5715A>T), GAA1530AAG (5716G>A 5718A>G), GAG1531AAA (5719G>A 5721G>A), AAG1532CAA (5722A>C 5724G>A), AAA1533GAT (5725A>G 5727A>T), GGA1534GGT (5730A>T), AAG1535AGT (5732A>G 5733G>T), GAG1536ATG (5734G>A 5735A>T), AGA1537AAA (5738G>A), ATG1538ATA (5742G>A), GTC1539TGT (5743G>T 5744T>G 5745C>T), TTC1540ATA (5746T>A 5748C>A), TAC1542AAC (5752T>A), AAA1543CGA (5755A>C 5756A>G), ACC1544CAA (5758A>C 5759C>A 5760C>A), TTG1545TTA (5763G>A), AAC1546AAT (5766C>T), GAC1547ACG (5767G>A 5768A>C 5769C>G), AAC1548GTC (5770A>G 5771A>T), ACG1549ACC (5775G>C), TTC1550ATT (5776T>A 5778C>T), AAA1551CGA (5779A>C 5780A>G), GAT1552AAC (5782G>A 5784T>C), CAG1553AAG (5785C>A), TCT1555TCA (5793T>A), CTA1556TTG (5794C>T 5796A>G), GGC1558CGT (5800G>C 5802C>T), ATT1559ATA (5805T>A), AAC1560GAT (5806A>G 5808C>T), ACA1561GAT (5809A>G 5810C>A 5811A>T), ATC1562CTC (5812A>C), ATA1563TTT (5815A>T 5817A>T), CAG1564GAC (5818C>G 5820G>C), AAG1565CAG (5821A>C), GTC1566TTA (5824G>T 5826C>A), TGT1567CAG (5827T>C 5828G>A 5829T>G), GCC1569GCA (5835C>A), AGT1570TCA (5836A>T 5837G>C 5838T>A), GTC1571GTG (5841C>G), TCC1573TCA (5847C>A), AAG1574AAA (5850G>A), TTC1575ATT (5851T>A 5853C>T), GAT1576AAT (5854G>A), CTG1577CTA (5859G>A), AAG1578AGA (5861A>G 5862G>A), AGC1579TCT (5863A>T 5864G>C 5865C>T), GGA1580GGT (5868A>T), TTC1581TAC (5870T>A), CAC1582CAT (5874C>T), ATT1584TTT (5878A>T), ATG1585AAG (5882T>A), ATG1586ATT (5886G>T), GAC1587AAG (5887G>A 5889C>G), GAA1588CCG (5890G>C 5891A>C 5892A>G), GAA1589AAG (5893G>A 5895A>G), TCC1590GAT (5896T>G 5897C>A 5898C>T), ATC1591GTG (5899A>G 5901C>G), TGG1593AGG (5905T>A), ACC1594ACT (5910C>T), GCC1595GCT (5913C>T), TTC1596TTT (5916C>T), CTG1597AGA (5917C>A 5918T>G 5919G>A), CCA1599CGG (5924C>G 5925A>G), GAT1600TAT (5926G>T), GGA1601GGG (5931A>G), CTT1602CAC (5933T>A 5934T>C), TGG1605TTC (5942G>T 5943G>C), TTA1606TTG (5946A>G), GTT1607ATG (5947G>A 5949T>G), CCA1609TCG (5953C>T 5955A>G), TTC1610ATT (5956T>A 5958C>T), GGA1611GGG (5961A>G), AAG1613ACC (5966A>C 5967G>C), GCC1615GCG (5973C>G), CCT1616CCA (5976T>A), GCT1617GCA (5979T>A), GTG1618GCC (5981T>C 5982G>C), CAA1620ATG (5986C>A 5987A>T 5988A>G), CGA1621AGT (5989C>A 5991A>T), AAG1622TTG (5992A>T 5993A>T), GAC1624AAT (5998G>A 6000C>T), AAC1625GGG (6001A>G 6002A>G 6003C>G), TGT1626GTG (6004T>G 6005G>T 6006T>G), TTC1627TTT (6009C>T), AGA1628AAG (6011G>A 6012A>G), GAC1629CCA (6013G>C 6014A>C 6015C>A), GAC1629_CTG1630insTTT (6015_6016insTTT), CTG1630TTG (6016C>T), TCA1631GAT (6019T>G 6020C>A 6021A>T), GGC1632TCG (6022G>T 6023G>C 6024C>G), GTA1634GTG (6030A>G), GCA1635ATT (6031G>A 6032C>T 6033A>T), GTT1636GGG (6035T>G 6036T>G), TAC1637TTT (6038A>T 6039C>T), ATC1641ATT (6051C>T), CTG1642CTT (6054G>T), GTC1643ATT (6055G>A 6057C>T) | | | | | | | | |

##### Proteins

| | Begin | End | Coverage | Score | Concordance | Matches | Identities | I/D/M/F* | Stop Codons |
|---|---|---|---|---|---|---|---|---|---|
| polyprotein (YP_010797894.1) | 1528 | 1645 | 5.9% | 335 | 39.7% | 118 (99.2%) | 50 (42.0%) | 1/0/0/0 | 0 |
| Protein mutations: | T1528F (5710A>T 5711C>T), G1529V (5714G>T 5715A>T), E1530K (5716G>A 5718A>G), E1531K (5719G>A 5721G>A), K1532Q (5722A>C 5724G>A), K1533D (5725A>G 5727A>T), K1535S (5732A>G 5733G>T), E1536M (5734G>A 5735A>T), R1537K (5738G>A), M1538I (5742G>A), V1539C (5743G>T 5744T>G 5745C>T), F1540I (5746T>A 5748C>A), Y1542N (5752T>A), K1543R (5755A>C 5756A>G), T1544Q (5758A>C 5759C>A 5760C>A), D1547T (5767G>A 5768A>C 5769C>G), N1548V (5770A>G 5771A>T), F1550I (5776T>A 5778C>T), K1551R (5779A>C 5780A>G), D1552N (5782G>A 5784T>C), Q1553K (5785C>A), G1558R (5800G>C 5802C>T), N1560D (5806A>G 5808C>T), T1561D (5809A>G 5810C>A 5811A>T), I1562L (5812A>C), I1563F (5815A>T 5817A>T), Q1564D (5818C>G 5820G>C), K1565Q (5821A>C), V1566L (5824G>T 5826C>A), C1567Q (5827T>C 5828G>A 5829T>G), F1575I (5851T>A 5853C>T), D1576N (5854G>A), K1578R (5861A>G 5862G>A), F1581Y (5870T>A), I1584F (5878A>T), M1585K (5882T>A), M1586I (5886G>T), D1587K (5887G>A 5889C>G), E1588P (5890G>C 5891A>C 5892A>G), E1589K (5893G>A 5895A>G), S1590D (5896T>G 5897C>A 5898C>T), I1591V (5899A>G 5901C>G), W1593R (5905T>A), L1597R (5917C>A 5918T>G 5919G>A), P1599R (5924C>G 5925A>G), D1600Y (5926G>T), L1602H (5933T>A 5934T>C), W1605F (5942G>T 5943G>C), V1607M (5947G>A 5949T>G), P1609S (5953C>T 5955A>G), F1610I (5956T>A 5958C>T), K1613T (5966A>C 5967G>C), V1618A (5981T>C 5982G>C), Q1620M (5986C>A 5987A>T 5988A>G), R1621S (5989C>A 5991A>T), K1622L (5992A>T 5993A>T), D1624N (5998G>A 6000C>T), N1625G (6001A>G 6002A>G 6003C>G), C1626V (6004T>G 6005G>T 6006T>G), R1628K (6011G>A 6012A>G), D1629P (6013G>C 6014A>C 6015C>A), D1629_L1630insF (6015_6016insTTT), S1631D (6019T>G 6020C>A 6021A>T), G1632S (6022G>T 6023G>C 6024C>G), A1635I (6031G>A 6032C>T 6033A>T), V1636G (6035T>G 6036T>G), Y1637F (6038A>T 6039C>T), V1643I (6055G>A 6057C>T) | | | | | | | | |
| Codon mutations: | GCA1527..G (5709A>G), ACT1528TTT (5710A>T 5711C>T), GGA1529GTT (5714G>T 5715A>T), GAA1530AAG (5716G>A 5718A>G), GAG1531AAA (5719G>A 5721G>A), AAG1532CAA (5722A>C 5724G>A), AAA1533GAT (5725A>G 5727A>T), GGA1534GGT (5730A>T), AAG1535AGT (5732A>G 5733G>T), GAG1536ATG (5734G>A 5735A>T), AGA1537AAA (5738G>A), ATG1538ATA (5742G>A), GTC1539TGT (5743G>T 5744T>G 5745C>T), TTC1540ATA (5746T>A 5748C>A), TAC1542AAC (5752T>A), AAA1543CGA (5755A>C 5756A>G), ACC1544CAA (5758A>C 5759C>A 5760C>A), TTG1545TTA (5763G>A), AAC1546AAT (5766C>T), GAC1547ACG (5767G>A 5768A>C 5769C>G), AAC1548GTC (5770A>G 5771A>T), ACG1549ACC (5775G>C), TTC1550ATT (5776T>A 5778C>T), AAA1551CGA (5779A>C 5780A>G), GAT1552AAC (5782G>A 5784T>C), CAG1553AAG (5785C>A), TCT1555TCA (5793T>A), CTA1556TTG (5794C>T 5796A>G), GGC1558CGT (5800G>C 5802C>T), ATT1559ATA (5805T>A), AAC1560GAT (5806A>G 5808C>T), ACA1561GAT (5809A>G 5810C>A 5811A>T), ATC1562CTC (5812A>C), ATA1563TTT (5815A>T 5817A>T), CAG1564GAC (5818C>G 5820G>C), AAG1565CAG (5821A>C), GTC1566TTA (5824G>T 5826C>A), TGT1567CAG (5827T>C 5828G>A 5829T>G), GCC1569GCA (5835C>A), AGT1570TCA (5836A>T 5837G>C 5838T>A), GTC1571GTG (5841C>G), TCC1573TCA (5847C>A), AAG1574AAA (5850G>A), TTC1575ATT (5851T>A 5853C>T), GAT1576AAT (5854G>A), CTG1577CTA (5859G>A), AAG1578AGA (5861A>G 5862G>A), AGC1579TCT (5863A>T 5864G>C 5865C>T), GGA1580GGT (5868A>T), TTC1581TAC (5870T>A), CAC1582CAT (5874C>T), ATT1584TTT (5878A>T), ATG1585AAG (5882T>A), ATG1586ATT (5886G>T), GAC1587AAG (5887G>A 5889C>G), GAA1588CCG (5890G>C 5891A>C 5892A>G), GAA1589AAG (5893G>A 5895A>G), TCC1590GAT (5896T>G 5897C>A 5898C>T), ATC1591GTG (5899A>G 5901C>G), TGG1593AGG (5905T>A), ACC1594ACT (5910C>T), GCC1595GCT (5913C>T), TTC1596TTT (5916C>T), CTG1597AGA (5917C>A 5918T>G 5919G>A), CCA1599CGG (5924C>G 5925A>G), GAT1600TAT (5926G>T), GGA1601GGG (5931A>G), CTT1602CAC (5933T>A 5934T>C), TGG1605TTC (5942G>T 5943G>C), TTA1606TTG (5946A>G), GTT1607ATG (5947G>A 5949T>G), CCA1609TCG (5953C>T 5955A>G), TTC1610ATT (5956T>A 5958C>T), GGA1611GGG (5961A>G), AAG1613ACC (5966A>C 5967G>C), GCC1615GCG (5973C>G), CCT1616CCA (5976T>A), GCT1617GCA (5979T>A), GTG1618GCC (5981T>C 5982G>C), CAA1620ATG (5986C>A 5987A>T 5988A>G), CGA1621AGT (5989C>A 5991A>T), AAG1622TTG (5992A>T 5993A>T), GAC1624AAT (5998G>A 6000C>T), AAC1625GGG (6001A>G 6002A>G 6003C>G), TGT1626GTG (6004T>G 6005G>T 6006T>G), TTC1627TTT (6009C>T), AGA1628AAG (6011G>A 6012A>G), GAC1629CCA (6013G>C 6014A>C 6015C>A), GAC1629_CTG1630insTTT (6015_6016insTTT), CTG1630TTG (6016C>T), TCA1631GAT (6019T>G 6020C>A 6021A>T), GGC1632TCG (6022G>T 6023G>C 6024C>G), GTA1634GTG (6030A>G), GCA1635ATT (6031G>A 6032C>T 6033A>T), GTT1636GGG (6035T>G 6036T>G), TAC1637TTT (6038A>T 6039C>T), ATC1641ATT (6051C>T), CTG1642CTT (6054G>T), GTC1643ATT (6055G>A 6057C>T) | | | | | | | | |

*: Inserts / Deletes / Misaligned / Frameshifts

#### Analysis details

This analysis was performed with panviral2.64

### NGS Details (UN59): Badnavirus rutilanscamelliae

#### Assembly

| | |
|---|---|
| **Coverage Length** | 283 (1 contig(s)) |
| **Depth Of Coverage** | 9.1 |
| **Number Of Reads** | 21 |
| **Reads Per Million** | 0.46 rpm (after QC) |
| **Ambiguities** | 0 |
| **Assembly Method** | de novo + reference guided assembly |
| **Consensus Caller** | Bcf Tools |

#### Coverage Map

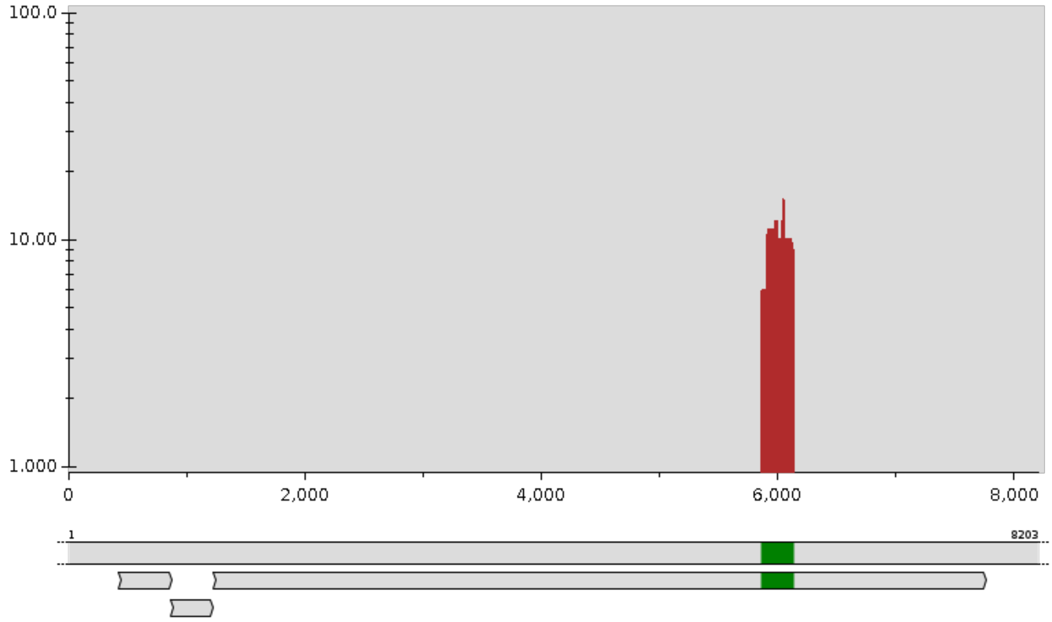

#### Assignment

| | |
|---|---|
| **Type** | Badnavirus rutilanscamelliae (Taxonomy ID: 3047719) |
| **Reference Genome** | NC_055598.1 |
| **NT Identity (%)** | 52.4476 |
| **AA Identity (%)** | 38.5417 |
| **Number Of Stop Codons** | 0 |
| **Number Of CDS** | 3 |

#### Alignment

| | |
|---|---|
| **Alignment Score** | 22.0 (NT) + 272.0 (AA) = 294.0 |
| **Concordance (%)** | 23.0588 |

| Alignment Method | Global, seeded, nucleotide + amino acids (AGA) |
|---|---|

#### Genome Region

Sequence starts at position 5861 and ends at position 6143 relative to NC_055598.1 reference sequence.

#### Alignment Detailed Statistics

| | Begin | End | Coverage | Score | Concordance | Matches | Identities | I/D/M/F* | Stop Codons |
|---|---|---|---|---|---|---|---|---|---|
| **NT** | **5861** | **6143** | **3.4%** | **22** | **3.9%** | **283 (99.0%)** | **150 (52.4%)** | **3/0** | |
| Mutations: | 5861A>C, 5862G>A, 5864C>T, 5866T>G, 5869C>G, 5870G>A, 5872C>G, 5873A>G, 5874A>T, 5875C>G, 5877C>T, 5878T>A, 5879C>G, 5880A>T, 5884A>G, 5885G>A, 5888C>A, 5890A>G, 5894A>C, 5895G>C, 5896C>T, 5902A>T, 5903G>A, 5911C>T, 5912A>G, 5913C>A, 5915A>C, 5917T>G, 5918G>T, 5920G>T, 5922C>A, 5923G>C, 5924A>C, 5925G>A, 5927A>C, 5929A>T, 5930G>C, 5933C>G, 5934A>G, 5936A>G, 5937G>C, 5938C>A, 5939A>G, 5940A>C, 5941A>T, 5942A>G, 5944C>G, 5946A>T, 5947T>C, 5950G>C, 5954T>G, 5957G>A, 5959C>T, 5960C>T, 5964A>G, 5968A>T, 5971G>A, 5973T>A, 5980A>G, 5981G>T, 5983T>G, 5984G>A, 5985C>G, 5986T>A, 5989G>C, 5990G>A, 5991A>G, 5992T>G, 5993C>G, 5996C>T, 5997A>C, 5998A>T, 5999C>G, 6005G>C, 6006A>C, 6007A>T, 6008T>A, 6009G>A, 6012C>T, 6013C>A, 6014G>T, 6026C>A, 6027C>G, 6028A>G, 6029G>T, 6030T>A, 6031G>T, 6034C>A, 6035T>C, 6036T>A, 6037A>C, 6043A>G, 6045G>T, 6046G>C, 6049G>T, 6050G>A, 6052T>G, 6056C>T, 6058A>C, 6064G>A, 6067T>A, 6069A>T, 6070G>T, 6082A>C, 6084T>C, 6085A>G, 6089C>A, 6090A>T, 6091A>G, 6092A>G, 6093G>A, 6095A>T, 6096A>T, 6101G>A, 6103C>T, 6104C>G, 6106A>T, 6107A>G, 6109C>A, 6110T>C, 6113A>C, 6114A>G, 6115T>A, 6121C>T, 6121_6122insTTG, 6123C>A, 6125G>T, 6126G>T, 6127A>C, 6134G>A, 6135C>T, 6139G>C, 6141A>T, 6142T>C | | | | | | | | |

##### CDS

| | Begin | End | Coverage | Score | Concordance | Matches | Identities | I/D/M/F* | Stop Codons |
|---|---|---|---|---|---|---|---|---|---|
| KM754_gp3 | 1547 | 1641 | 4.4% | 272 | 38.0% | 95 (99.0%) | 37 (38.5%) | 1/0/0/0 | 0 |
| Protein mutations: | R1547Q (5861A>C 5862G>A), N1549K (5869C>G), D1550K (5870G>A 5872C>G), N1551V (5873A>G 5874A>T 5875C>G), T1552I (5877C>T 5878T>A), H1553V (5879C>G 5880A>T), D1555N (5885G>A), Q1556K (5888C>A 5890A>G), S1558P (5894A>C 5895G>C 5896C>T), G1561R (5903G>A), T1564D (5912A>G 5913C>A), I1565L (5915A>C 5917T>G), V1566F (5918G>T 5920G>T), A1567D (5922C>A 5923G>C), R1568Q (5924A>C 5925G>A), I1569L (5927A>C 5929A>T), G1570R (5930G>C), H1571G (5933C>G 5934A>G), S1572A (5936A>G 5937G>C 5938C>A), K1573A (5939A>G 5940A>C 5941A>T), I1574V (5942A>G 5944C>G), Y1575F (5946A>T 5947T>C), F1578V (5954T>G), D1579N (5957G>A 5959C>T), K1581R (5964A>G), F1584Y (5973T>A), V1587L (5981G>T 5983T>G), A1588R (5984G>A 5985C>G 5986T>A), M1589I (5989G>C), D1590R (5990G>A 5991A>G 5992T>G), P1591A (5993C>G), Q1592S (5996C>T 5997A>C 5998A>T), H1593D (5999C>G), E1595P (6005G>C 6006A>C 6007A>T), W1596K (6008T>A 6009G>A), T1597I (6012C>T 6013C>A), A1598S (6014G>T), P1602R (6026C>A 6027C>G 6028A>G), V1603Y (6029G>T 6030T>A 6031G>T), L1605H (6035T>C 6036T>A 6037A>C), W1608F (6045G>T 6046G>C), V1610M (6050G>A 6052T>G), P1612S (6056C>T 6058A>C), K1616I (6069A>T 6070G>T), V1621A (6084T>C 6085A>G), Q1623M (6089C>A 6090A>T 6091A>G), R1624E (6092A>G 6093G>A), K1625L (6095A>T 6096A>T), D1627N (6101G>A 6103C>T), R1628G (6104C>G 6106A>T), I1629V (6107A>G 6109C>A), F1630L (6110T>C), N1631R (6113A>C 6114A>G 6115T>A), Y1633_A1634insL (6121_6122insTTG), A1634D (6123C>A), G1635F (6125G>T 6126G>T 6127A>C), A1638I (6134G>A 6135C>T), Y1640F (6141A>T 6142T>C) | | | | | | | | |
| Codon mutations: | AGG1547CAG (5861A>C 5862G>A), CTT1548TTG (5864C>T 5866T>G), AAC1549AAG (5869C>G), GAC1550AAG (5870G>A 5872C>G), AAC1551GTG (5873A>G 5874A>T 5875C>G), ACT1552ATA (5877C>T 5878T>A), CAC1553GTC (5879C>G 5880A>T), AAA1554AAG (5884A>G), GAC1555AAC (5885G>A), CAA1556AAG (5888C>A 5890A>G), AGC1558CCT (5894A>C 5895G>C 5896C>T), CCA1560CCT (5902A>T), GGG1561AGG (5903G>A), GAC1563GAT (5911C>T), ACT1564GAT (5912A>G 5913C>A), ATT1565CTG (5915A>C 5917T>G), GTG1566TTT (5918G>T 5920G>T), GCG1567GAC (5922C>A 5923G>C), AGG1568CAG (5924A>C 5925G>A), ATA1569CTT (5927A>C 5929A>T), GGG1570CGG (5930G>C), CAT1571GGT (5933C>G 5934A>G), AGC1572GCA (5936A>G 5937G>C 5938C>A), AAA1573GCT (5939A>G 5940A>C 5941A>T), ATC1574GTG (5942A>G 5944C>G), TAT1575TTC (5946A>T 5947T>C), TCG1576TCC (5950G>C), TTC1578GTC (5954T>G), GAC1579AAT (5957G>A 5959C>T), CTG1580TTG (5960C>T), AAG1581AGG (5964A>G), TCA1582TCT (5968A>T), GGG1583GGA (5971G>A), TTT1584TAT (5973T>A), CAA1586CAG (5980A>G), GTT1587TTG (5981G>T 5983T>G), GCT1588AGA (5984G>A 5985C>G 5986T>A), ATG1589ATC (5989G>C), GAT1590AGG (5990G>A 5991A>G 5992T>G), CCT1591GCT (5993C>G), CAA1592TCT (5996C>T 5997A>C 5998A>T), CAC1593GAC (5999C>G), GAA1595CCT (6005G>C 6006A>C 6007A>T), TGG1596AAG (6008T>A 6009G>A), ACC1597ATA (6012C>T 6013C>A), GCT1598TCT (6014G>T), CCA1602AGG (6026C>A 6027C>G 6028A>G), GTG1603TAT (6029G>T 6030T>A 6031G>T), GGC1604GGA (6034C>A), TTA1605CAC (6035T>C 6036T>A 6037A>C), GAA1607GAG (6043A>G), TGG1608TTC (6045G>T 6046G>C), CTG1609CTT (6049G>T), GTT1610ATG (6050G>A 6052T>G), CCA1612TCC (6056C>T 6058A>C), GGG1614GGA (6064G>A), CTT1615CTA (6067T>A), AAG1616ATT (6069A>T 6070G>T), GCA1620GCC (6082A>C), GTA1621GCG (6084T>C 6085A>G), CAA1623ATG (6089C>A 6090A>T 6091A>G), AGG1624GAG (6092A>G 6093G>A), AAG1625TTG (6095A>T 6096A>T), GAC1627AAT (6101G>A 6103C>T), CGA1628GGT (6104C>G 6106A>T), ATC1629GTA (6107A>G 6109C>A), TTT1630CTT (6110T>C), AAT1631CGA (6113A>C 6114A>G 6115T>A), TAC1633TAT (6121C>T), TAC1633_GCT1634insTTG (6121_6122insTTG), GCT1634GAT (6123C>A), GGA1635TTC (6125G>T 6126G>T 6127A>C), GCT1638ATT (6134G>A 6135C>T), GTG1639GTC (6139G>C), TAT1640TTC (6141A>T 6142T>C) | | | | | | | | |

##### Proteins

| | Begin | End | Coverage | Score | Concordance | Matches | Identities | I/D/M/F* | Stop Codons |
|---|---|---|---|---|---|---|---|---|---|
| polyprotein (YP_010087856.1) | 1547 | 1641 | 4.4% | 272 | 38.0% | 95 (99.0%) | 37 (38.5%) | 1/0/0/0 | 0 |
| Protein mutations: | R1547Q (5861A>C 5862G>A), N1549K (5869C>G), D1550K (5870G>A 5872C>G), N1551V (5873A>G 5874A>T 5875C>G), T1552I (5877C>T 5878T>A), H1553V (5879C>G 5880A>T), D1555N (5885G>A), Q1556K (5888C>A 5890A>G), S1558P (5894A>C 5895G>C 5896C>T), G1561R (5903G>A), T1564D (5912A>G 5913C>A), I1565L (5915A>C 5917T>G), V1566F (5918G>T 5920G>T), A1567D (5922C>A 5923G>C), R1568Q (5924A>C 5925G>A), I1569L (5927A>C 5929A>T), G1570R (5930G>C), H1571G (5933C>G 5934A>G), S1572A (5936A>G 5937G>C 5938C>A), K1573A (5939A>G 5940A>C 5941A>T), I1574V (5942A>G 5944C>G), Y1575F (5946A>T 5947T>C), F1578V (5954T>G), D1579N (5957G>A 5959C>T), K1581R (5964A>G), F1584Y (5973T>A), V1587L (5981G>T 5983T>G), A1588R (5984G>A 5985C>G 5986T>A), M1589I (5989G>C), D1590R (5990G>A 5991A>G 5992T>G), P1591A (5993C>G), Q1592S (5996C>T 5997A>C 5998A>T), H1593D (5999C>G), E1595P (6005G>C 6006A>C 6007A>T), W1596K (6008T>A 6009G>A), T1597I (6012C>T 6013C>A), A1598S (6014G>T), P1602R (6026C>A 6027C>G 6028A>G), V1603Y (6029G>T 6030T>A 6031G>T), L1605H (6035T>C 6036T>A 6037A>C), W1608F (6045G>T 6046G>C), V1610M (6050G>A 6052T>G), P1612S (6056C>T 6058A>C), K1616I (6069A>T 6070G>T), V1621A (6084T>C 6085A>G), Q1623M (6089C>A 6090A>T 6091A>G), R1624E (6092A>G 6093G>A), K1625L (6095A>T 6096A>T), D1627N (6101G>A 6103C>T), R1628G (6104C>G 6106A>T), I1629V (6107A>G 6109C>A), F1630L (6110T>C), N1631R (6113A>C 6114A>G 6115T>A), Y1633_A1634insL (6121_6122insTTG), A1634D (6123C>A), G1635F (6125G>T 6126G>T 6127A>C), A1638I (6134G>A 6135C>T), Y1640F (6141A>T 6142T>C) | | | | | | | | |
| Codon mutations: | AGG1547CAG (5861A>C 5862G>A), CTT1548TTG (5864C>T 5866T>G), AAC1549AAG (5869C>G), GAC1550AAG (5870G>A 5872C>G), AAC1551GTG (5873A>G 5874A>T 5875C>G), ACT1552ATA (5877C>T 5878T>A), CAC1553GTC (5879C>G 5880A>T), AAA1554AAG (5884A>G), GAC1555AAC (5885G>A), CAA1556AAG (5888C>A 5890A>G), AGC1558CCT (5894A>C 5895G>C 5896C>T), CCA1560CCT (5902A>T), GGG1561AGG (5903G>A), GAC1563GAT (5911C>T), ACT1564GAT (5912A>G 5913C>A), ATT1565CTG (5915A>C 5917T>G), GTG1566TTT (5918G>T 5920G>T), GCG1567GAC (5922C>A 5923G>C), AGG1568CAG (5924A>C 5925G>A), ATA1569CTT (5927A>C 5929A>T), GGG1570CGG (5930G>C), CAT1571GGT (5933C>G 5934A>G), AGC1572GCA (5936A>G 5937G>C 5938C>A), AAA1573GCT (5939A>G 5940A>C 5941A>T), ATC1574GTG (5942A>G 5944C>G), TAT1575TTC (5946A>T 5947T>C), TCG1576TCC (5950G>C), TTC1578GTC (5954T>G), GAC1579AAT (5957G>A 5959C>T), CTG1580TTG (5960C>T), AAG1581AGG (5964A>G), TCA1582TCT (5968A>T), GGG1583GGA (5971G>A), TTT1584TAT (5973T>A), CAA1586CAG (5980A>G), GTT1587TTG (5981G>T 5983T>G), GCT1588AGA (5984G>A 5985C>G 5986T>A), ATG1589ATC (5989G>C), GAT1590AGG (5990G>A 5991A>G 5992T>G), CCT1591GCT (5993C>G), CAA1592TCT (5996C>T 5997A>C 5998A>T), CAC1593GAC (5999C>G), GAA1595CCT (6005G>C 6006A>C 6007A>T), TGG1596AAG (6008T>A 6009G>A), ACC1597ATA (6012C>T 6013C>A), GCT1598TCT (6014G>T), CCA1602AGG (6026C>A 6027C>G 6028A>G), GTG1603TAT (6029G>T 6030T>A 6031G>T), GGC1604GGA (6034C>A), TTA1605CAC (6035T>C 6036T>A 6037A>C), GAA1607GAG (6043A>G), TGG1608TTC (6045G>T 6046G>C), CTG1609CTT (6049G>T), GTT1610ATG (6050G>A 6052T>G), CCA1612TCC (6056C>T 6058A>C), GGG1614GGA (6064G>A), CTT1615CTA (6067T>A), AAG1616ATT (6069A>T 6070G>T), GCA1620GCC (6082A>C), GTA1621GCG (6084T>C 6085A>G), CAA1623ATG (6089C>A 6090A>T 6091A>G), AGG1624GAG (6092A>G 6093G>A), AAG1625TTG (6095A>T 6096A>T), GAC1627AAT (6101G>A 6103C>T), CGA1628GGT (6104C>G 6106A>T), ATC1629GTA (6107A>G 6109C>A), TTT1630CTT (6110T>C), AAT1631CGA (6113A>C 6114A>G 6115T>A), TAC1633TAT (6121C>T), TAC1633_GCT1634insTTG (6121_6122insTTG), GCT1634GAT (6123C>A), GGA1635TTC (6125G>T 6126G>T 6127A>C), GCT1638ATT (6134G>A 6135C>T), GTG1639GTC (6139G>C), TAT1640TTC (6141A>T 6142T>C) | | | | | | | | |

*: Inserts / Deletes / Misaligned / Frameshifts

#### Analysis details

This analysis was performed with panviral2.64

### NGS Details (UN59): Unknown

#### Assembly

| | |
|---|---|
| **Coverage Length** | 420 (1 contig(s)) |
| **Depth Of Coverage** | 5.6 |
| **Number Of Reads** | 20 |
| **Reads Per Million** | 0.43 rpm (after QC) |
| **Ambiguities** | 0 |
| **Assembly Method** | de novo + reference guided assembly |
| **Consensus Caller** | Bcf Tools |

#### Coverage Map

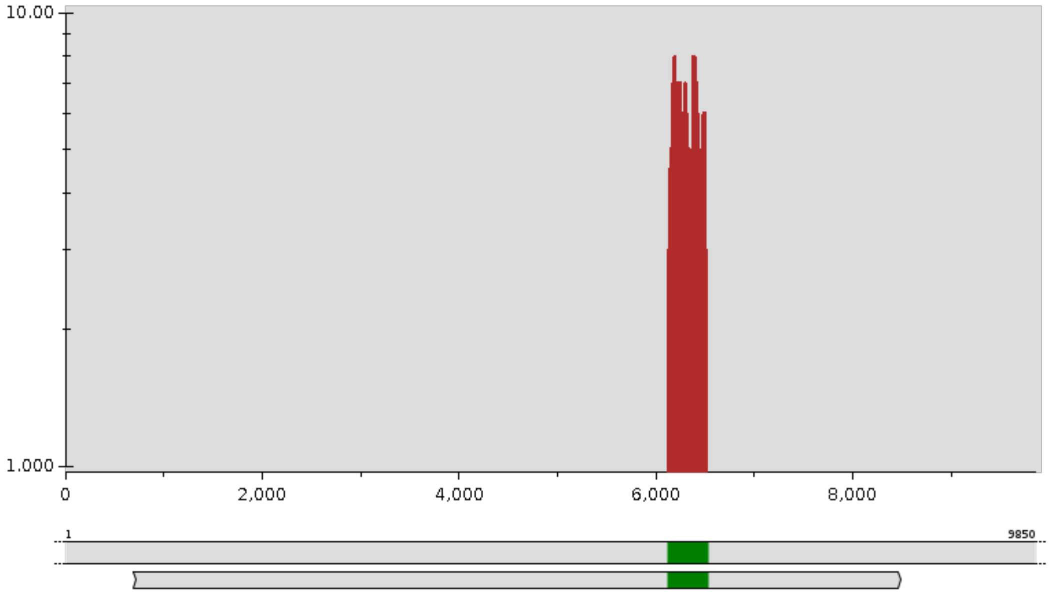

#### Assignment

| | |
|---|---|
| **Type** | Unknown (Taxonomy ID: -1) |
| **Reference Genome** | NC_028462.1 |
| **NT Identity (%)** | 56.6986 |
| **AA Identity (%)** | 53.5714 |
| **Number Of Stop Codons** | 0 |
| **Number Of CDS** | 1 |

#### Alignment

| | |
|---|---|
| **Alignment Score** | 101.0 (NT) + 432.0 (AA) = 533.0 |
| **Concordance (%)** | 31.585 |

| **Alignment Method** | Global, seeded, nucleotide + amino acids (AGA) |
|---|---|

#### Genome Region

Sequence starts at position 6114 and ends at position 6533 relative to NC_028462.1 reference sequence.

#### Alignment Detailed Statistics

| | Begin | End | Coverage | Score | Concordance | Matches | Identities | I/D/M/F* | Stop Codons |
|---|---|---|---|---|---|---|---|---|---|
| **NT** | **6114** | **6533** | **4.3%** | **101** | **12.2%** | **418 (99.5%)** | **237 (56.4%)** | **0/2** | |
| Mutations: | 6116C>T, 6119A>T, 6122C>A, 6125T>A, 6126G>A, 6131C>T, 6134C>T, 6137C>T, 6140C>T, 6143G>T, 6146T>C, 6149G>T, 6159A>T, 6160A>T, 6161G>A, 6164C>A, 6165C>A, 6167T>G, 6168C>T, 6182G>A, 6192A>C, 6194C>A, 6197A>T, 6200C>T, 6203G>A, 6204C>A, 6208G>A, 6209G>A, 6213G>T, 6215C>T, 6221C>A, 6224C>T, 6225A>T, 6226A>C, 6227A>T, 6230T>A, 6231_6232delTG, 6233T>A, 6234T>A, 6236T>G, 6237C>A, 6239G>A, 6242C>T, 6244G>T, 6245T>G, 6246G>A, 6247G>T, 6248T>A, 6255G>A, 6257C>A, 6260C>T, 6263T>C, 6266A>T, 6267C>T, 6269C>G, 6272C>A, 6274T>G, 6278C>A, 6280A>C, 6282G>A, 6283A>C, 6284C>A, 6285A>T, 6287A>T, 6288G>C, 6290A>G, 6296C>T, 6299A>T, 6300C>A, 6301T>C, 6302G>A, 6308G>T, 6309G>A, 6311T>A, 6312A>G, 6313T>A, 6314C>A, 6315A>T, 6317C>T, 6318A>T, 6319T>C, 6320C>A, 6323C>A, 6324A>C, 6325A>T, 6326A>C, 6327A>T, 6328T>G, 6333C>A, 6335G>T, 6340G>A, 6341A>C, 6344C>A, 6345A>G, 6347C>A, 6348A>T, 6350C>A, 6351G>A, 6353A>T, 6354A>C, 6355G>A, 6356A>G, 6358C>A, 6359C>A, 6362G>A, 6363T>G, 6364A>C, 6365C>A, 6369T>A, 6371G>A, 6372G>A, 6373T>A, 6374C>T, 6376G>A, 6378G>A, 6380A>T, 6382G>A, 6383C>A, 6386T>A, 6389C>A, 6392T>C, 6393T>C, 6395A>T, 6399G>A, 6401C>G, 6403C>T, 6404C>T, 6407A>T, 6409G>C, 6410C>A, 6414G>A, 6415G>A, 6416A>T, 6417A>G, 6418A>G, 6419G>A, 6422C>A, 6423A>G, 6425G>A, 6428G>A, 6431G>A, 6432C>A, 6440C>A, 6441C>T, 6442T>C, 6443T>C, 6445C>A, 6446T>A, 6447C>A, 6449C>G, 6450C>A, 6452C>A, 6453A>T, 6455G>A, 6470A>G, 6471C>T, 6473C>A, 6475A>T, 6476G>A, 6477G>A, 6478A>C, 6479T>C, 6480C>A, 6481G>A, 6484T>A, 6485T>A, 6486C>G, 6488G>A, 6489G>A, 6491C>A, 6494G>A, 6495C>A, 6496A>G, 6497G>A, 6503C>T, 6509T>C, 6510G>T, 6512C>A, 6518C>T, 6521C>A, 6525A>G, 6527G>A | | | | | | | | |
| **CDS** | | | | | | | | | |
| ATL80_gp1 | 1808 | 1947 | 5.4% | 432 | 42.8% | 140 (100%) | 75 (53.6%) | 0/0/1/1 | 0 |
| Protein mutations: | A1812T (6126G>A), K1817N (6143G>T), Q1819H (6149G>T), K1823L (6159A>T 6160A>T 6161G>A), L1825M (6165C>A 6167T>G), P1826S (6168C>T), N1834Q (6192A>C 6194C>A), R1839K (6208G>A 6209G>A), D1841Y (6213G>T 6215C>T), K1845S (6225A>T 6226A>C 6227A>T), D1846E (6230T>A), Y1848K (6234T>A 6236T>G), Q1849K (6237C>A 6239G>A), C1851L (6244G>T 6245T>G), G1852I (6246G>A 6247G>T 6248T>A), V1855I (6255G>A 6257C>A), F1861C (6274T>G), K1863T (6280A>C), D1864T (6282G>A 6283A>C 6284C>A), K1865Y (6285A>T 6287A>T), E1866Q (6288G>C 6290A>G), L1870T (6300C>A 6301T>C 6302G>A), D1873K (6309G>A 6311T>A), I1874E (6312A>G 6313T>A 6314C>A), I1875F (6315A>T 6317C>T), I1876S (6318A>T 6319T>C 6320C>A), N1877K (6323C>A), K1878L (6324A>C 6325A>T 6326A>C), I1879C (6327A>T 6328T>G), Q1881N (6333C>A 6335G>T), G1883D (6340G>A 6341A>C), I1885V (6345A>G 6347C>A), I1886L (6348A>T 6350C>A), G1887S (6351G>A 6353A>T), R1888Q (6354A>C 6355G>A 6356A>G), T1889K (6358C>A 6359C>A), Y1891A (6363T>G 6364A>C 6365C>A), L1893I (6369T>A 6371G>A), V1894N (6372G>A 6373T>A 6374C>T), R1895K (6376G>A), E1896N (6378G>A 6380A>T), S1897K (6382G>A 6383C>A), D1899E (6389C>A), V1903M (6399G>A 6401C>G), T1904I (6403C>T 6404C>T), C1906S (6409G>C 6410C>A), G1908N (6414G>A 6415G>A 6416A>T), K1909G (6417A>G 6418A>G 6419G>A), K1911E (6423A>G 6425G>A), P1914T (6432C>A), L1917S (6441C>T 6442T>C 6443T>C), T1918K (6445C>A 6446T>A), H1919K (6447C>A 6449C>G), L1920I (6450C>A 6452C>A), M1921L (6453A>T 6455G>A), K1928I (6475A>T 6476G>A), D1929T (6477G>A 6478A>C 6479T>C), R1930K (6480C>A 6481G>A), L1931Q (6484T>A 6485T>A), Q1932E (6486C>G 6488G>A), V1933I (6489G>A 6491C>A), Q1935R (6495C>A 6496A>G 6497G>A), V1940L (6510G>T 6512C>A), K1945E (6525A>G 6527G>A) | | | | | | | | |
| Codon mutations: | ATC1808ATT (6116C>T), CCA1809CCT (6119A>T), CTC1810CTA (6122C>A), ACT1811ACA (6125T>A), GCA1812ACA (6126G>A), TTC1813TTT (6131C>T), AGC1814AGT (6134C>T), ACC1815ACT (6137C>T), CCC1816CCT (6140C>T), AAG1817AAT (6143G>T), GGT1818GGC (6146T>C), CAG1819CAT (6149G>T), AAG1823TTA (6159A>T 6160A>T 6161G>A), GTC1824GTA (6164C>A), CTT1825ATG (6165C>A 6167T>G), CCA1826TCA (6168C>T), AAG1830AAA (6182G>A), AAC1834CAA (6192A>C 6194C>A), ATA1835ATT (6197A>T), TTC1836TTT (6200C>T), CAG1837CAA (6203G>A), CGA1838AGA (6204C>A), AGG1839AAA (6208G>A 6209G>A), GAC1841TAT (6213G>T 6215C>T), ATC1843ATA (6221C>A), TTC1844TTT (6224C>T), AAA1845TCT (6225A>T 6226A>C 6227A>T), GAT1846GAA (6230T>A), TGT1847--A (6231_6232delTG 6233T>A), TAT1848AAG (6234T>A 6236T>G), CAG1849AAA (6237C>A 6239G>A), TTC1850TTT (6242C>T), TGT1851TTG (6244G>T 6245T>G), GGT1852ATA (6246G>A 6247G>T 6248T>A), GTC1855ATA (6255G>A 6257C>A), GAC1856GAT (6260C>T), GAT1857GAC (6263T>C), ATA1858ATT (6266A>T), CTC1859TTG (6267C>T 6269C>G), GTC1860GTA (6272C>A), TTC1861TGC (6274T>G), TCC1862TCA (6278C>A), AAA1863ACA (6280A>C), GAC1864ACA (6282G>A 6283A>C 6284C>A), AAA1865TAT (6285A>T 6287A>T), GAA1866CAG (6288G>C 6290A>G), CAC1868CAT (6296C>T), CTA1869CTT (6299A>T), CTG1870ACA (6300C>A 6301T>C 6302G>A), CTG1872CTT (6308G>T), GAT1873AAA (6309G>A 6311T>A), ATC1874GAA (6312A>G 6313T>A 6314C>A), ATC1875TTT (6315A>T 6317C>T), ATC1876TCA (6318A>T 6319T>C 6320C>A), AAC1877AAA (6323C>A), AAA1878CTC (6324A>C 6325A>T 6326A>C), ATT1879TGT (6327A>T 6328T>G), CAG1881AAT (6333C>A 6335G>T), GGA1883GAC (6340G>A 6341A>C), ATC1884ATA (6344C>A), ATC1885GTA (6345A>G 6347C>A), ATC1886TTA (6348A>T 6350C>A), GGA1887AGT (6351G>A 6353A>T), AGA1888CAG (6354A>C 6355G>A 6356A>G), ACC1889AAA (6358C>A 6359C>A), AAG1890AAA (6362G>A), TAC1891GCA (6363T>G 6364A>C 6365C>A), TTG1893ATA (6369T>A 6371G>A), GTC1894AAT (6372G>A 6373T>A 6374C>T), AGA1895AAA (6376G>A), GAA1896AAT (6378G>A 6380A>T), AGC1897AAA (6382G>A 6383C>A), ATT1898ATA (6386T>A), GAC1899GAA (6389C>A), TTT1900TTC (6392T>C), TTA1901CTT (6393T>C 6395A>T), GTC1903ATG (6399G>A 6401C>G), ACC1904ATT (6403C>T 6404C>T), ATA1905ATT (6407A>T), TGC1906TCA (6409G>C 6410C>A), GGA1908AAT (6414G>A 6415G>A 6416A>T), AAG1909GGA (6417A>G 6418A>G 6419G>A), ATC1910ATA (6422C>A), AAG1911GAA (6423A>G 6425G>A), TTG1912TTA (6428G>A), CAG1913CAA (6431G>A), CCT1914ACT (6432C>A), ATC1916ATA (6440C>A), CTT1917TCC (6441C>T 6442T>C 6443T>C), ACT1918AAA (6445C>A 6446T>A), CAC1919AAG (6447C>A 6449C>G), CTC1920ATA (6450C>A 6452C>A), ATG1921TTA (6453A>T 6455G>A), AAA1926AAG (6470A>G), CTC1927TTA (6471C>T 6473C>A), AAG1928ATA (6475A>T 6476G>A), GAT1929ACC (6477G>A 6478A>C 6479T>C), CGA1930AAA (6480C>A 6481G>A), CTT1931CAA (6484T>A 6485T>A), CAG1932GAA (6486C>G 6488G>A), GTC1933ATA (6489G>A 6491C>A), CAG1934CAA (6494G>A), CAG1935AGA (6495C>A 6496A>G 6497G>A), CTC1937CTT (6503C>T), TGT1939TGC (6509T>C), GTC1940TTA (6510G>T 6512C>A), TAC1942TAT (6518C>T), GCC1943GCA (6521C>A), AAG1945GAA (6525A>G 6527G>A) | | | | | | | | |
| **Proteins** | | | | | | | | | |
| polyprotein (YP_009182100.1) | 1808 | 1947 | 5.4% | 432 | 42.8% | 140 (100%) | 75 (53.6%) | 0/0/1/1 | 0 |
| Protein mutations: | A1812T (6126G>A), K1817N (6143G>T), Q1819H (6149G>T), K1823L (6159A>T 6160A>T 6161G>A), L1825M (6165C>A 6167T>G), P1826S (6168C>T), N1834Q (6192A>C 6194C>A), R1839K (6208G>A 6209G>A), D1841Y (6213G>T 6215C>T), K1845S (6225A>T 6226A>C 6227A>T), D1846E (6230T>A), Y1848K (6234T>A 6236T>G), Q1849K (6237C>A 6239G>A), C1851L (6244G>T 6245T>G), G1852I (6246G>A 6247G>T 6248T>A), V1855I (6255G>A 6257C>A), F1861C (6274T>G), K1863T (6280A>C), D1864T (6282G>A 6283A>C 6284C>A), K1865Y (6285A>T 6287A>T), E1866Q (6288G>C 6290A>G), L1870T (6300C>A 6301T>C 6302G>A), D1873K (6309G>A 6311T>A), I1874E (6312A>G 6313T>A 6314C>A), I1875F (6315A>T 6317C>T), I1876S (6318A>T 6319T>C 6320C>A), N1877K (6323C>A), K1878L (6324A>C 6325A>T 6326A>C), I1879C (6327A>T 6328T>G), Q1881N (6333C>A 6335G>T), G1883D (6340G>A 6341A>C), I1885V (6345A>G 6347C>A), I1886L (6348A>T 6350C>A), G1887S (6351G>A 6353A>T), R1888Q (6354A>C 6355G>A 6356A>G), T1889K (6358C>A 6359C>A), Y1891A (6363T>G 6364A>C 6365C>A), L1893I (6369T>A 6371G>A), V1894N (6372G>A 6373T>A 6374C>T), R1895K (6376G>A), E1896N (6378G>A 6380A>T), S1897K (6382G>A 6383C>A), D1899E (6389C>A), V1903M (6399G>A 6401C>G), T1904I (6403C>T 6404C>T), C1906S (6409G>C 6410C>A), G1908N (6414G>A 6415G>A 6416A>T), K1909G (6417A>G 6418A>G 6419G>A), K1911E (6423A>G 6425G>A), P1914T (6432C>A), L1917S (6441C>T 6442T>C 6443T>C), T1918K (6445C>A 6446T>A), H1919K (6447C>A 6449C>G), L1920I (6450C>A 6452C>A), M1921L (6453A>T 6455G>A), K1928I (6475A>T 6476G>A), D1929T (6477G>A 6478A>C 6479T>C), R1930K (6480C>A 6481G>A), L1931Q (6484T>A 6485T>A), Q1932E (6486C>G 6488G>A), V1933I (6489G>A 6491C>A), Q1935R (6495C>A 6496A>G 6497G>A), V1940L (6510G>T 6512C>A), K1945E (6525A>G 6527G>A) | | | | | | | | |
| Codon mutations: | ATC1808ATT (6116C>T), CCA1809CCT (6119A>T), CTC1810CTA (6122C>A), ACT1811ACA (6125T>A), GCA1812ACA (6126G>A), TTC1813TTT (6131C>T), AGC1814AGT (6134C>T), ACC1815ACT (6137C>T), CCC1816CCT (6140C>T), AAG1817AAT (6143G>T), GGT1818GGC (6146T>C), CAG1819CAT (6149G>T), AAG1823TTA (6159A>T 6160A>T 6161G>A), GTC1824GTA (6164C>A), CTT1825ATG (6165C>A 6167T>G), CCA1826TCA (6168C>T), AAG1830AAA (6182G>A), AAC1834CAA (6192A>C 6194C>A), ATA1835ATT (6197A>T), TTC1836TTT (6200C>T), CAG1837CAA (6203G>A), CGA1838AGA (6204C>A), AGG1839AAA (6208G>A 6209G>A), GAC1841TAT (6213G>T 6215C>T), ATC1843ATA (6221C>A), TTC1844TTT (6224C>T), AAA1845TCT (6225A>T 6226A>C 6227A>T), GAT1846GAA (6230T>A), TGT1847--A (6231_6232delTG 6233T>A), TAT1848AAG (6234T>A 6236T>G), CAG1849AAA (6237C>A 6239G>A), TTC1850TTT (6242C>T), TGT1851TTG (6244G>T 6245T>G), GGT1852ATA (6246G>A 6247G>T 6248T>A), GTC1855ATA (6255G>A 6257C>A), GAC1856GAT (6260C>T), GAT1857GAC (6263T>C), ATA1858ATT (6266A>T), CTC1859TTG (6267C>T 6269C>G), GTC1860GTA (6272C>A), TTC1861TGC (6274T>G), TCC1862TCA (6278C>A), AAA1863ACA (6280A>C), GAC1864ACA (6282G>A 6283A>C 6284C>A), AAA1865TAT (6285A>T 6287A>T), GAA1866CAG (6288G>C 6290A>G), CAC1868CAT (6296C>T), CTA1869CTT (6299A>T), CTG1870ACA (6300C>A 6301T>C 6302G>A), CTG1872CTT (6308G>T), GAT1873AAA (6309G>A 6311T>A), ATC1874GAA (6312A>G 6313T>A 6314C>A), ATC1875TTT (6315A>T 6317C>T), ATC1876TCA (6318A>T 6319T>C 6320C>A), AAC1877AAA (6323C>A), AAA1878CTC (6324A>C 6325A>T 6326A>C), ATT1879TGT (6327A>T 6328T>G), CAG1881AAT (6333C>A 6335G>T), GGA1883GAC (6340G>A 6341A>C), ATC1884ATA (6344C>A), ATC1885GTA (6345A>G 6347C>A), ATC1886TTA (6348A>T 6350C>A), GGA1887AGT (6351G>A 6353A>T), AGA1888CAG (6354A>C 6355G>A 6356A>G), ACC1889AAA (6358C>A 6359C>A), AAG1890AAA (6362G>A), TAC1891GCA (6363T>G 6364A>C 6365C>A), TTG1893ATA (6369T>A 6371G>A), GTC1894AAT (6372G>A 6373T>A 6374C>T), AGA1895AAA (6376G>A), GAA1896AAT (6378G>A 6380A>T), AGC1897AAA (6382G>A 6383C>A), ATT1898ATA (6386T>A), GAC1899GAA (6389C>A), TTT1900TTC (6392T>C), TTA1901CTT (6393T>C 6395A>T), GTC1903ATG (6399G>A 6401C>G), ACC1904ATT (6403C>T 6404C>T), ATA1905ATT (6407A>T), TGC1906TCA (6409G>C 6410C>A), GGA1908AAT (6414G>A 6415G>A 6416A>T), AAG1909GGA (6417A>G 6418A>G 6419G>A), ATC1910ATA (6422C>A), AAG1911GAA (6423A>G 6425G>A), TTG1912TTA (6428G>A), CAG1913CAA (6431G>A), CCT1914ACT (6432C>A), ATC1916ATA (6440C>A), CTT1917TCC (6441C>T 6442T>C 6443T>C), ACT1918AAA (6445C>A 6446T>A), CAC1919AAG (6447C>A 6449C>G), CTC1920ATA (6450C>A 6452C>A), ATG1921TTA (6453A>T 6455G>A), AAA1926AAG (6470A>G), CTC1927TTA (6471C>T 6473C>A), AAG1928ATA (6475A>T 6476G>A), GAT1929ACC (6477G>A 6478A>C 6479T>C), CGA1930AAA (6480C>A 6481G>A), CTT1931CAA (6484T>A 6485T>A), CAG1932GAA (6486C>G 6488G>A), GTC1933ATA (6489G>A 6491C>A), CAG1934CAA (6494G>A), CAG1935AGA (6495C>A 6496A>G 6497G>A), CTC1937CTT (6503C>T), TGT1939TGC (6509T>C), GTC1940TTA (6510G>T 6512C>A), TAC1942TAT (6518C>T), GCC1943GCA (6521C>A), AAG1945GAA (6525A>G 6527G>A) | | | | | | | | |

*: Inserts / Deletes / Misaligned / Frameshifts

#### Analysis details

This analysis was performed with panviral2.64

### NGS Details (UN59): Badnavirus venabougainvilleae

#### Assembly

| | |
|---|---|
| **Coverage Length** | 303 (1 contig(s)) |
| **Depth Of Coverage** | 6.9 |
| **Number Of Reads** | 18 |
| **Reads Per Million** | 0.39 rpm (after QC) |
| **Ambiguities** | 0 |
| **Assembly Method** | de novo + reference guided assembly |
| **Consensus Caller** | Bcf Tools |

#### Coverage Map

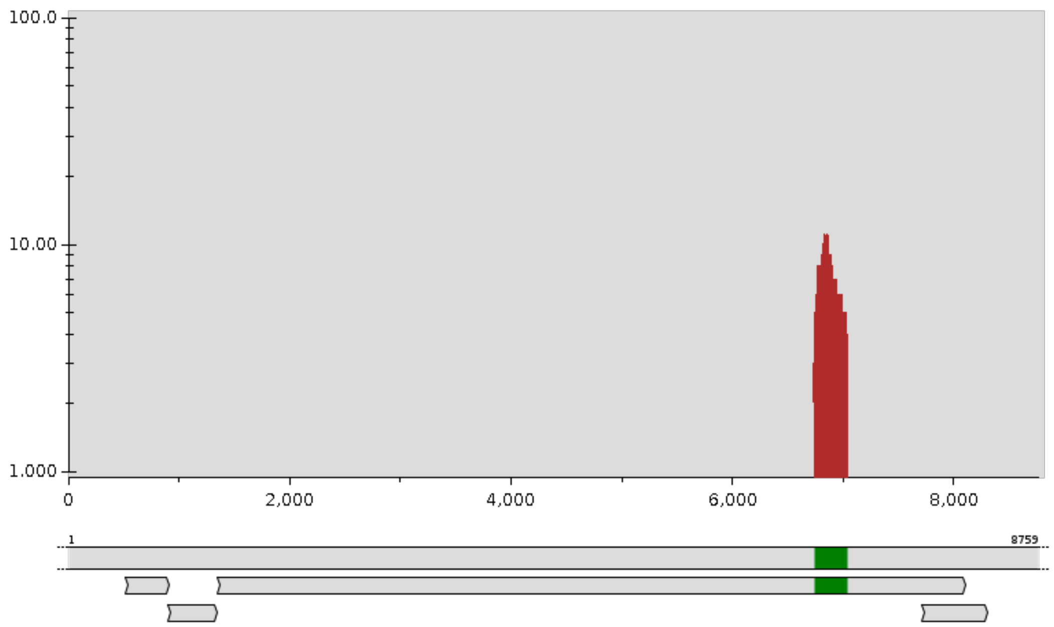

#### Assignment

| | |
|---|---|
| **Type** | Badnavirus venabougainvilleae (Taxonomy ID: 3047697) |
| **Reference Genome** | NC_011592.1 |
| **NT Identity (%)** | 53.268 |
| **AA Identity (%)** | 46.0784 |
| **Number Of Stop Codons** | 0 |
| **Number Of CDS** | 4 |

#### Alignment

| | |
|---|---|
| **Alignment Score** | 34.0 (NT) + 315.0 (AA) = 349.0 |
| **Concordance (%)** | 26.5198 |

| Alignment Method | Global, seeded, nucleotide + amino acids (AGA) |
|---|---|

#### Genome Region

Sequence starts at position 6741 and ends at position 7043 relative to NC_011592.1 reference sequence.

#### Alignment Detailed Statistics

| | Begin | End | Coverage | Score | Concordance | Matches | Identities | I/D/M/F* | Stop Codons |
|---|---|---|---|---|---|---|---|---|---|
| **NT** | **6741** | **7043** | **3.5%** | **34** | **5.6%** | **303 (99.0%)** | **163 (53.3%)** | **3/0** | |
| Mutations: | 6741C>T, 6747A>C, 6748T>A, 6749G>A, 6759C>T, 6761T>G, 6762T>A, 6765T>A, 6767C>G, 6768A>G, 6769C>T, 6771A>T, 6774A>T, 6775T>C, 6776T>A, 6777G>T, 6783A>G, 6785G>T, 6786G>T, 6789A>G, 6792C>T, 6796C>T, 6804T>G, 6809A>C, 6810G>A, 6819A>G, 6822T>A, 6823A>G, 6824T>C, 6828C>T, 6829C>A, 6830A>T, 6831A>G, 6832A>G, 6833G>A, 6834G>T, 6835A>T, 6836A>T, 6837G>A, 6841G>A, 6843C>T, 6845A>G, 6846C>G, 6847T>G, 6848G>T, 6849C>G, 6853T>A, 6854C>A, 6855A>G, 6857G>C, 6860A>T, 6861C>T, 6862A>T, 6863G>T, 6864C>G, 6867_6868insGTG, 6870C>T, 6871A>G, 6873T>G, 6874G>A, 6875C>T, 6876T>A, 6881A>T, 6885T>A, 6894A>T, 6902T>A, 6906T>A, 6907G>A, 6908A>G, 6909G>A, 6910A>T, 6911G>C, 6912T>C, 6916A>G, 6917G>A, 6918A>G, 6926A>T, 6927G>T, 6930G>T, 6933T>C, 6934C>T, 6936C>G, 6937A>C, 6940G>C, 6941T>A, 6942T>A, 6943A>G, 6946C>T, 6948T>A, 6952G>A, 6955T>C, 6956G>T, 6959A>G, 6960A>T, 6963A>T, 6964A>C, 6965A>G, 6967G>A, 6968G>A, 6969C>G, 6973G>T, 6974T>A, 6975T>C, 6976C>G, 6977T>C, 6979T>A, 6980C>A, 6981C>A, 6982C>T, 6983C>T, 6985A>T, 6991A>T, 6992T>G, 6993G>T, 6994A>G, 6996G>T, 6997G>T, 7000G>T, 7001C>G, 7003G>T, 7005C>A, 7007A>G, 7009A>T, 7011T>A, 7012A>G, 7014T>G, 7016A>C, 7017G>A, 7021C>T, 7023T>A, 7026A>T, 7027G>C, 7028C>A, 7029A>T, 7032A>T, 7033C>G, 7036G>T, 7037G>C, 7038G>T, 7039A>G, 7042C>G | | | | | | | | |

##### CDS

| | Begin | End | Coverage | Score | Concordance | Matches | Identities | I/D/M/F* | Stop Codons |
|---|---|---|---|---|---|---|---|---|---|
| BSCVV_gp3 | 1800 | 1900 | 4.5% | 315 | 43.6% | 101 (99.0%) | 47 (46.1%) | 1/0/0/0 | 0 |
| Protein mutations: | W1802K (6748T>A 6749G>A), L1806R (6761T>G 6762T>A), P1808R (6767C>G 6768A>G), Q1809Y (6769C>T 6771A>T), L1811H (6775T>C 6776T>A 6777G>T), W1814F (6785G>T 6786G>T), P1818S (6796C>T), K1822T (6809A>C 6810G>A), I1827A (6823A>G 6824T>C), Q1829M (6829C>A 6830A>T 6831A>G), R1830D (6832A>G 6833G>A 6834G>T), K1831L (6835A>T 6836A>T 6837G>A), D1833N (6841G>A 6843C>T), N1834R (6845A>G 6846C>G), C1835V (6847T>G 6848G>T 6849C>G), S1837K (6853T>A 6854C>A 6855A>G), R1838P (6857G>C), Y1839F (6860A>T 6861C>T), S1840L (6862A>T 6863G>T 6864C>G), D1841_F1842insV (6867_6868insGTG), I1843V (6871A>G 6873T>G), A1844I (6874G>A 6875C>T 6876T>A), Y1846F (6881A>T), F1853Y (6902T>A), E1855R (6907G>A 6908A>G 6909G>A), R1858E (6916A>G 6917G>A 6918A>G), E1861V (6926A>T 6927G>T), K1862N (6930G>T), V1866Q (6940G>C 6941T>A 6942T>A), M1867V (6943A>G), V1870I (6952G>A), C1871L (6955T>C 6956G>T), Q1872R (6959A>G 6960A>T), E1873D (6963A>T), N1874R (6964A>C 6965A>G), G1875K (6967G>A 6968G>A 6969C>G), V1877Y (6973G>T 6974T>A 6975T>C), L1878A (6976C>G 6977T>C), S1879K (6979T>A 6980C>A 6981C>A), P1880F (6982C>T 6983C>T), T1881S (6985A>T), M1883C (6991A>T 6992T>G 6993G>T), K1884D (6994A>G 6996G>T), V1885F (6997G>T), A1886W (7000G>T 7001C>G), V1887L (7003G>T 7005C>A), K1888R (7007A>G), T1889S (7009A>T 7011T>A), I1890V (7012A>G 7014T>G), E1891A (7016A>C 7017G>A), A1895H (7027G>C 7028C>A 7029A>T), L1897V (7033C>G), G1898S (7036G>T 7037G>C 7038G>T), N1899D (7039A>G) | | | | | | | | |
| Codon mutations: | TCC1799..T (6741C>T), CCA1801CCC (6747A>C), TGG1802AAG (6748T>A 6749G>A), TTC1805TTT (6759C>T), CTT1806CGA (6761T>G 6762T>A), ACT1807ACA (6765T>A), CCA1808CGG (6767C>G 6768A>G), CAA1809TAT (6769C>T 6771A>T), GGA1810GGT (6774A>T), TTG1811CAT (6775T>C 6776T>A 6777G>T), GAA1813GAG (6783A>G), TGG1814TTT (6785G>T 6786G>T), CTA1815CTG (6789A>G), GTC1816GTT (6792C>T), CCA1818TCA (6796C>T), GGT1820GGG (6804T>G), AAG1822ACA (6809A>C 6810G>A), CCA1825CCG (6819A>G), GCT1826GCA (6822T>A), ATC1827GCC (6823A>G 6824T>C), TTC1828TTT (6828C>T), CAA1829ATG (6829C>A 6830A>T 6831A>G), AGG1830GAT (6832A>G 6833G>A 6834G>T), AAG1831TTA (6835A>T 6836A>T 6837G>A), GAC1833AAT (6841G>A 6843C>T), AAC1834AGG (6845A>G 6846C>G), TGC1835GTG (6847T>G 6848G>T 6849C>G), TCA1837AAG (6853T>A 6854C>A 6855A>G), CGA1838CCA (6857G>C), TAC1839TTT (6860A>T 6861C>T), AGC1840TTG (6862A>T 6863G>T 6864C>G), GAT1841_TTC1842insGTG (6867_6868insGTG), TTC1842TTT (6870C>T), ATT1843GTG (6871A>G 6873T>G), GCT1844ATA (6874G>A 6875C>T 6876T>A), TAC1846TTC (6881A>T), ATT1847ATA (6885T>A), ATA1850ATT (6894A>T), TTT1853TAT (6902T>A), TCT1854TCA (6906T>A), GAG1855AGA (6907G>A 6908A>G 6909G>A), AGT1856TCC (6910A>T 6911G>C 6912T>C), AGA1858GAG (6916A>G 6917G>A 6918A>G), GAG1861GTT (6926A>T 6927G>T), AAG1862AAT (6930G>T), CAT1863CAC (6933T>C), CTC1864TTG (6934C>T 6936C>G), AGA1865CGA (6937A>C), GTT1866CAA (6940G>C 6941T>A 6942T>A), ATG1867GTG (6943A>G), CTT1868TTA (6946C>T 6948T>A), GTT1870ATT (6952G>A), TGT1871CTT (6955T>C 6956G>T), CAA1872CGT (6959A>G 6960A>T), GAA1873GAT (6963A>T), AAT1874CGT (6964A>C 6965A>G), GGC1875AAG (6967G>A 6968G>A 6969C>G), GTT1877TAC (6973G>T 6974T>A 6975T>C), CTA1878GCA (6976C>G 6977T>C), TCC1879AAA (6979T>A 6980C>A 6981C>A), CCC1880TTC (6982C>T 6983C>T), ACT1881TCT (6985A>T), ATG1883TGT (6991A>T 6992T>G 6993G>T), AAG1884GAT (6994A>G 6996G>T), GTT1885TTT (6997G>T), GCG1886TGG (7000G>T 7001C>G), GTC1887TTA (7003G>T 7005C>A), AAA1888AGA (7007A>G), ACT1889TCA (7009A>T 7011T>A), ATT1890GTG (7012A>G 7014T>G), GAG1891GCA (7016A>C 7017G>A), CTT1893TTA (7021C>T 7023T>A), GGA1894GGT (7026A>T), GCA1895CAT (7027G>C 7028C>A 7029A>T), ATA1896ATT (7032A>T), CTA1897GTA (7033C>G), GGG1898TCT (7036G>T 7037G>C 7038G>T), AAT1899GAT (7039A>G), CAA1900GA. (7042C>G) | | | | | | | | |

##### Proteins

| | Begin | End | Coverage | Score | Concordance | Matches | Identities | I/D/M/F* | Stop Codons |
|---|---|---|---|---|---|---|---|---|---|
| polyprotein (YP_002321513.1) | 1800 | 1900 | 4.5% | 315 | 43.6% | 101 (99.0%) | 47 (46.1%) | 1/0/0/0 | 0 |
| Protein mutations: | W1802K (6748T>A 6749G>A), L1806R (6761T>G 6762T>A), P1808R (6767C>G 6768A>G), Q1809Y (6769C>T 6771A>T), L1811H (6775T>C 6776T>A 6777G>T), W1814F (6785G>T 6786G>T), P1818S (6796C>T), K1822T (6809A>C 6810G>A), I1827A (6823A>G 6824T>C), Q1829M (6829C>A 6830A>T 6831A>G), R1830D (6832A>G 6833G>A 6834G>T), K1831L (6835A>T 6836A>T 6837G>A), D1833N (6841G>A 6843C>T), N1834R (6845A>G 6846C>G), C1835V (6847T>G 6848G>T 6849C>G), S1837K (6853T>A 6854C>A 6855A>G), R1838P (6857G>C), Y1839F (6860A>T 6861C>T), S1840L (6862A>T 6863G>T 6864C>G), D1841_F1842insV (6867_6868insGTG), I1843V (6871A>G 6873T>G), A1844I (6874G>A 6875C>T 6876T>A), Y1846F (6881A>T), F1853Y (6902T>A), E1855R (6907G>A 6908A>G 6909G>A), R1858E (6916A>G 6917G>A 6918A>G), E1861V (6926A>T 6927G>T), K1862N (6930G>T), V1866Q (6940G>C 6941T>A 6942T>A), M1867V (6943A>G), V1870I (6952G>A), C1871L (6955T>C 6956G>T), Q1872R (6959A>G 6960A>T), E1873D (6963A>T), N1874R (6964A>C 6965A>G), G1875K (6967G>A 6968G>A 6969C>G), V1877Y (6973G>T 6974T>A 6975T>C), L1878A (6976C>G 6977T>C), S1879K (6979T>A 6980C>A 6981C>A), P1880F (6982C>T 6983C>T), T1881S (6985A>T), M1883C (6991A>T 6992T>G 6993G>T), K1884D (6994A>G 6996G>T), V1885F (6997G>T), A1886W (7000G>T 7001C>G), V1887L (7003G>T 7005C>A), K1888R (7007A>G), T1889S (7009A>T 7011T>A), I1890V (7012A>G 7014T>G), E1891A (7016A>C 7017G>A), A1895H (7027G>C 7028C>A 7029A>T), L1897V (7033C>G), G1898S (7036G>T 7037G>C 7038G>T), N1899D (7039A>G) | | | | | | | | |
| Codon mutations: | TCC1799..T (6741C>T), CCA1801CCC (6747A>C), TGG1802AAG (6748T>A 6749G>A), TTC1805TTT (6759C>T), CTT1806CGA (6761T>G 6762T>A), ACT1807ACA (6765T>A), CCA1808CGG (6767C>G 6768A>G), CAA1809TAT (6769C>T 6771A>T), GGA1810GGT (6774A>T), TTG1811CAT (6775T>C 6776T>A 6777G>T), GAA1813GAG (6783A>G), TGG1814TTT (6785G>T 6786G>T), CTA1815CTG (6789A>G), GTC1816GTT (6792C>T), CCA1818TCA (6796C>T), GGT1820GGG (6804T>G), AAG1822ACA (6809A>C 6810G>A), CCA1825CCG (6819A>G), GCT1826GCA (6822T>A), ATC1827GCC (6823A>G 6824T>C), TTC1828TTT (6828C>T), CAA1829ATG (6829C>A 6830A>T 6831A>G), AGG1830GAT (6832A>G 6833G>A 6834G>T), AAG1831TTA (6835A>T 6836A>T 6837G>A), GAC1833AAT (6841G>A 6843C>T), AAC1834AGG (6845A>G 6846C>G), TGC1835GTG (6847T>G 6848G>T 6849C>G), TCA1837AAG (6853T>A 6854C>A 6855A>G), CGA1838CCA (6857G>C), TAC1839TTT (6860A>T 6861C>T), AGC1840TTG (6862A>T 6863G>T 6864C>G), GAT1841_TTC1842insGTG (6867_6868insGTG), TTC1842TTT (6870C>T), ATT1843GTG (6871A>G 6873T>G), GCT1844ATA (6874G>A 6875C>T 6876T>A), TAC1846TTC (6881A>T), ATT1847ATA (6885T>A), ATA1850ATT (6894A>T), TTT1853TAT (6902T>A), TCT1854TCA (6906T>A), GAG1855AGA (6907G>A 6908A>G 6909G>A), AGT1856TCC (6910A>T 6911G>C 6912T>C), AGA1858GAG (6916A>G 6917G>A 6918A>G), GAG1861GTT (6926A>T 6927G>T), AAG1862AAT (6930G>T), CAT1863CAC (6933T>C), CTC1864TTG (6934C>T 6936C>G), AGA1865CGA (6937A>C), GTT1866CAA (6940G>C 6941T>A 6942T>A), ATG1867GTG (6943A>G), CTT1868TTA (6946C>T 6948T>A), GTT1870ATT (6952G>A), TGT1871CTT (6955T>C 6956G>T), CAA1872CGT (6959A>G 6960A>T), GAA1873GAT (6963A>T), AAT1874CGT (6964A>C 6965A>G), GGC1875AAG (6967G>A 6968G>A 6969C>G), GTT1877TAC (6973G>T 6974T>A 6975T>C), CTA1878GCA (6976C>G 6977T>C), TCC1879AAA (6979T>A 6980C>A 6981C>A), CCC1880TTC (6982C>T 6983C>T), ACT1881TCT (6985A>T), ATG1883TGT (6991A>T 6992T>G 6993G>T), AAG1884GAT (6994A>G 6996G>T), GTT1885TTT (6997G>T), GCG1886TGG (7000G>T 7001C>G), GTC1887TTA (7003G>T 7005C>A), AAA1888AGA (7007A>G), ACT1889TCA (7009A>T 7011T>A), ATT1890GTG (7012A>G 7014T>G), GAG1891GCA (7016A>C 7017G>A), CTT1893TTA (7021C>T 7023T>A), GGA1894GGT (7026A>T), GCA1895CAT (7027G>C 7028C>A 7029A>T), ATA1896ATT (7032A>T), CTA1897GTA (7033C>G), GGG1898TCT (7036G>T 7037G>C 7038G>T), AAT1899GAT (7039A>G), CAA1900GA. (7042C>G) | | | | | | | | |

*: Inserts / Deletes / Misaligned / Frameshifts

#### Analysis details

This analysis was performed with panviral2.64

### NGS Details (UN59): Epiphyllum badnavirus 1

#### Assembly

| | |
|---|---|
| **Coverage Length** | 275 (1 contig(s)) |
| **Depth Of Coverage** | 6.7 |
| **Number Of Reads** | 16 |
| **Reads Per Million** | 0.35 rpm (after QC) |
| **Ambiguities** | 0 |
| **Assembly Method** | de novo + reference guided assembly |
| **Consensus Caller** | Bcf Tools |

#### Coverage Map

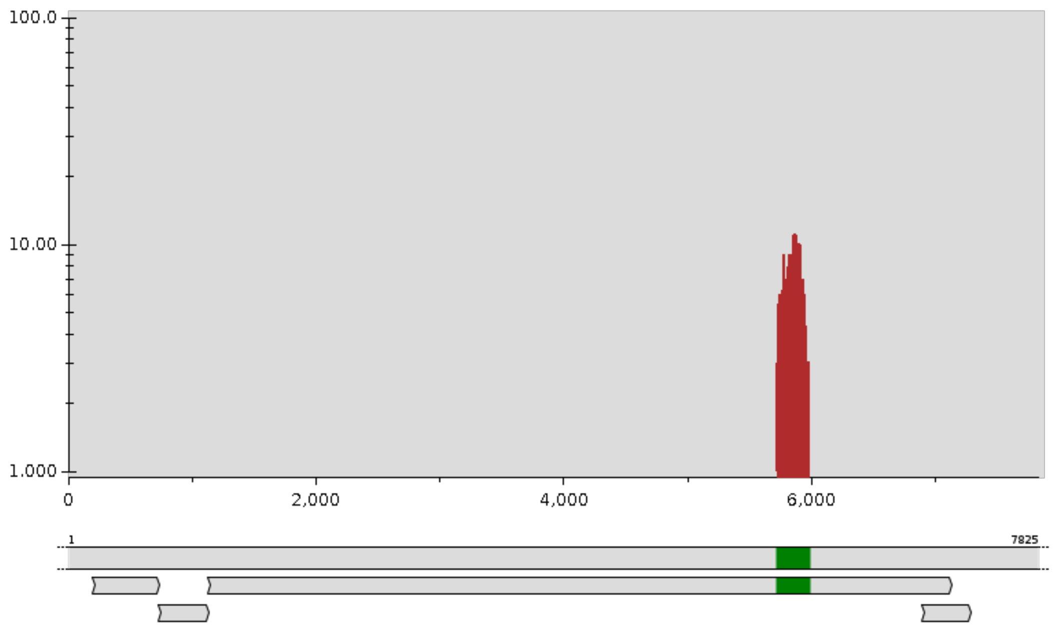

#### Assignment

| | |
|---|---|
| **Type** | Epiphyllum badnavirus 1 (Taxonomy ID: 2518008) |
| **Reference Genome** | NC_076247.1 |
| **NT Identity (%)** | 57.971 |
| **AA Identity (%)** | 50.0 |
| **Number Of Stop Codons** | 0 |
| **Number Of CDS** | 4 |

#### Alignment

| | |
|---|---|
| **Alignment Score** | 80.0 (NT) + 187.0 (AA) = 267.0 |
| **Concordance (%)** | 24.4983 |

| Alignment Method | Global, seeded, nucleotide + amino acids (AGA) |
|---|---|

#### Genome Region

Sequence starts at position 5714 and ends at position 5988 relative to NC_076247.1 reference sequence.

#### Alignment Detailed Statistics

| | Begin | End | Coverage | Score | Concordance | Matches | Identities | I/D/M/F* | Stop Codons |
|---|---|---|---|---|---|---|---|---|---|
| **NT** | **5714** | **5988** | **3.5%** | **80** | **14.5%** | **275 (99.6%)** | **160 (58.0%)** | **1/0** | |
| Mutations: | 5719G>A, 5725A>C, 5727A>T, 5730A>T, 5732A>G, 5733G>C, 5734G>A, 5735A>T, 5743G>T, 5744T>G, 5745C>T, 5746T>A, 5748C>T, 5749A>G, 5754C>T, 5755A>C, 5756A>G, 5757A>T, 5758A>C, 5759C>A, 5760C>G, 5767G>A, 5769C>A, 5770A>G, 5771A>T, 5772C>G, 5775G>C, 5776T>A, 5781A>G, 5782G>A, 5784T>C, 5785C>A, 5790C>T, 5791T>A, 5794C>T, 5796A>G, 5800G>C, 5802C>T, 5806A>G, 5808C>T, 5809A>G, 5810C>A, 5811A>T, 5812A>T, 5814C>A, 5815A>T, 5817A>T, 5818C>G, 5820G>T, 5821A>C, 5824G>C, 5826C>T, 5827T>C, 5828G>A, 5829T>G, 5830A>G, 5835C>T, 5836A>T, 5837G>C, 5838T>A, 5841C>G, 5847C>T, 5850G>A, 5851T>A, 5853C>T, 5857C>T, 5861A>G, 5863A>T, 5864G>C, 5865C>T, 5868A>G, 5870T>A, 5871C>T, 5878A>T, 5880T>G, 5882T>A, 5884A>G, 5886G>T, 5887G>C, 5888A>G, 5889C>G, 5892A>G, 5892_5893insT, 5893G>C, 5896T>A, 5897C>A, 5904T>G, 5905T>A, 5906G>A, 5910C>T, 5913C>T, 5917C>A, 5918T>A, 5922C>A, 5924C>G, 5926G>T, 5933T>A, 5934T>C, 5940A>G, 5942G>T, 5943G>T, 5944T>A, 5946A>G, 5953C>T, 5955A>T, 5958C>T, 5960G>C, 5961A>G, 5962T>C, 5964G>T, 5966A>C, 5967G>T, 5970T>C, 5979T>C, 5981T>C, 5982G>T | | | | | | | | |

##### CDS

| | Begin | End | Coverage | Score | Concordance | Matches | Identities | I/D/M/F* | Stop Codons |
|---|---|---|---|---|---|---|---|---|---|
| QKM20_gp3 | 1530 | 1620 | 4.5% | 187 | 28.6% | 91 (98.9%) | 46 (50.0%) | 1/0/1/1 | 0 |
| Protein mutations: | E1531K (5719G>A), K1533H (5725A>C 5727A>T), K1535S (5732A>G 5733G>C), E1536M (5734G>A 5735A>T), V1539C (5743G>T 5744T>G 5745C>T), F1540I (5746T>A 5748C>T), N1541D (5749A>G), K1543R (5755A>C 5756A>G 5757A>T), T1544Q (5758A>C 5759C>A 5760C>G), D1547K (5767G>A 5769C>A), N1548V (5770A>G 5771A>T 5772C>G), F1550I (5776T>A), D1552N (5782G>A 5784T>C), Q1553K (5785C>A), S1555T (5791T>A), G1558R (5800G>C 5802C>T), N1560D (5806A>G 5808C>T), T1561D (5809A>G 5810C>A 5811A>T), I1562L (5812A>T 5814C>A), I1563F (5815A>T 5817A>T), Q1564D (5818C>G 5820G>T), K1565Q (5821A>C), V1566L (5824G>C 5826C>T), C1567Q (5827T>C 5828G>A 5829T>G), N1568D (5830A>G), F1575I (5851T>A 5853C>T), K1578R (5861A>G), F1581Y (5870T>A 5871C>T), I1584L (5878A>T 5880T>G), M1585K (5882T>A), M1586V (5884A>G 5886G>T), D1587R (5887G>C 5888A>G 5889C>G), E1588_E1589insX (5892_5893insT), E1589Q (5893G>C), S1590N (5896T>A 5897C>A), W1593K (5905T>A 5906G>A), L1597K (5917C>A 5918T>A), P1599R (5924C>G), D1600Y (5926G>T), L1602H (5933T>A 5934T>C), W1605F (5942G>T 5943G>T), L1606M (5944T>A 5946A>G), P1609S (5953C>T 5955A>T), G1611A (5960G>C 5961A>G), K1613T (5966A>C 5967G>T), V1618A (5981T>C 5982G>T) | | | | | | | | |
| Codon mutations: | GAG1531AAG (5719G>A), AAA1533CAT (5725A>C 5727A>T), GGA1534GGT (5730A>T), AAG1535AGC (5732A>G 5733G>C), GAG1536ATG (5734G>A 5735A>T), GTC1539TGT (5743G>T 5744T>G 5745C>T), TTC1540ATT (5746T>A 5748C>T), AAC1541GAC (5749A>G), TAC1542TAT (5754C>T), AAA1543CGT (5755A>C 5756A>G 5757A>T), ACC1544CAG (5758A>C 5759C>A 5760C>G), GAC1547AAA (5767G>A 5769C>A), AAC1548GTG (5770A>G 5771A>T 5772C>G), ACG1549ACC (5775G>C), TTC1550ATC (5776T>A), AAA1551AAG (5781A>G), GAT1552AAC (5782G>A 5784T>C), CAG1553AAG (5785C>A), TAC1554TAT (5790C>T), TCT1555ACT (5791T>A), CTA1556TTG (5794C>T 5796A>G), GGC1558CGT (5800G>C 5802C>T), AAC1560GAT (5806A>G 5808C>T), ACA1561GAT (5809A>G 5810C>A 5811A>T), ATC1562TTA (5812A>T 5814C>A), ATA1563TTT (5815A>T 5817A>T), CAG1564GAT (5818C>G 5820G>T), AAG1565CAG (5821A>C), GTC1566CTT (5824G>C 5826C>T), TGT1567CAG (5827T>C 5828G>A 5829T>G), AAT1568GAT (5830A>G), GCC1569GCT (5835C>T), AGT1570TCA (5836A>T 5837G>C 5838T>A), GTC1571GTG (5841C>G), TCC1573TCT (5847C>T), AAG1574AAA (5850G>A), TTC1575ATT (5851T>A 5853C>T), CTG1577TTG (5857C>T), AAG1578AGG (5861A>G), AGC1579TCT (5863A>T 5864G>C 5865C>T), GGA1580GGG (5868A>G), TTC1581TAT (5870T>A 5871C>T), ATT1584TTG (5878A>T 5880T>G), ATG1585AAG (5882T>A), ATG1586GTT (5884A>G 5886G>T), GAC1587CGG (5887G>C 5888A>G 5889C>G), GAA1588GAG (5892A>G), GAA1588_GAA1589insT-- (5892_5893insT), GAA1589CAA (5893G>C), TCC1590AAC (5896T>A 5897C>A), CCT1592CCG (5904T>G), TGG1593AAG (5905T>A 5906G>A), ACC1594ACT (5910C>T), GCC1595GCT (5913C>T), CTG1597AAG (5917C>A 5918T>A), ACC1598ACA (5922C>A), CCA1599CGA (5924C>G), GAT1600TAT (5926G>T), CTT1602CAC (5933T>A 5934T>C), GAA1604GAG (5940A>G), TGG1605TTT (5942G>T 5943G>T), TTA1606ATG (5944T>A 5946A>G), CCA1609TCT (5953C>T 5955A>T), TTC1610TTT (5958C>T), GGA1611GCG (5960G>C 5961A>G), TTG1612CTT (5962T>C 5964G>T), AAG1613ACT (5966A>C 5967G>T), AAT1614AAC (5970T>C), GCT1617GCC (5979T>C), GTG1618GCT (5981T>C 5982G>T) | | | | | | | | |

##### Proteins

| | Begin | End | Coverage | Score | Concordance | Matches | Identities | I/D/M/F* | Stop Codons |
|---|---|---|---|---|---|---|---|---|---|
| polyprotein (YP_010797894.1) | 1530 | 1620 | 4.5% | 187 | 28.6% | 91 (98.9%) | 46 (50.0%) | 1/0/1/1 | 0 |
| Protein mutations: | E1531K (5719G>A), K1533H (5725A>C 5727A>T), K1535S (5732A>G 5733G>C), E1536M (5734G>A 5735A>T), V1539C (5743G>T 5744T>G 5745C>T), F1540I (5746T>A 5748C>T), N1541D (5749A>G), K1543R (5755A>C 5756A>G 5757A>T), T1544Q (5758A>C 5759C>A 5760C>G), D1547K (5767G>A 5769C>A), N1548V (5770A>G 5771A>T 5772C>G), F1550I (5776T>A), D1552N (5782G>A 5784T>C), Q1553K (5785C>A), S1555T (5791T>A), G1558R (5800G>C 5802C>T), N1560D (5806A>G 5808C>T), T1561D (5809A>G 5810C>A 5811A>T), I1562L (5812A>T 5814C>A), I1563F (5815A>T 5817A>T), Q1564D (5818C>G 5820G>T), K1565Q (5821A>C), V1566L (5824G>C 5826C>T), C1567Q (5827T>C 5828G>A 5829T>G), N1568D (5830A>G), F1575I (5851T>A 5853C>T), K1578R (5861A>G), F1581Y (5870T>A 5871C>T), I1584L (5878A>T 5880T>G), M1585K (5882T>A), M1586V (5884A>G 5886G>T), D1587R (5887G>C 5888A>G 5889C>G), E1588_E1589insX (5892_5893insT), E1589Q (5893G>C), S1590N (5896T>A 5897C>A), W1593K (5905T>A 5906G>A), L1597K (5917C>A 5918T>A), P1599R (5924C>G), D1600Y (5926G>T), L1602H (5933T>A 5934T>C), W1605F (5942G>T 5943G>T), L1606M (5944T>A 5946A>G), P1609S (5953C>T 5955A>T), G1611A (5960G>C 5961A>G), K1613T (5966A>C 5967G>T), V1618A (5981T>C 5982G>T) | | | | | | | | |
| Codon mutations: | GAG1531AAG (5719G>A), AAA1533CAT (5725A>C 5727A>T), GGA1534GGT (5730A>T), AAG1535AGC (5732A>G 5733G>C), GAG1536ATG (5734G>A 5735A>T), GTC1539TGT (5743G>T 5744T>G 5745C>T), TTC1540ATT (5746T>A 5748C>T), AAC1541GAC (5749A>G), TAC1542TAT (5754C>T), AAA1543CGT (5755A>C 5756A>G 5757A>T), ACC1544CAG (5758A>C 5759C>A 5760C>G), GAC1547AAA (5767G>A 5769C>A), AAC1548GTG (5770A>G 5771A>T 5772C>G), ACG1549ACC (5775G>C), TTC1550ATC (5776T>A), AAA1551AAG (5781A>G), GAT1552AAC (5782G>A 5784T>C), CAG1553AAG (5785C>A), TAC1554TAT (5790C>T), TCT1555ACT (5791T>A), CTA1556TTG (5794C>T 5796A>G), GGC1558CGT (5800G>C 5802C>T), AAC1560GAT (5806A>G 5808C>T), ACA1561GAT (5809A>G 5810C>A 5811A>T), ATC1562TTA (5812A>T 5814C>A), ATA1563TTT (5815A>T 5817A>T), CAG1564GAT (5818C>G 5820G>T), AAG1565CAG (5821A>C), GTC1566CTT (5824G>C 5826C>T), TGT1567CAG (5827T>C 5828G>A 5829T>G), AAT1568GAT (5830A>G), GCC1569GCT (5835C>T), AGT1570TCA (5836A>T 5837G>C 5838T>A), GTC1571GTG (5841C>G), TCC1573TCT (5847C>T), AAG1574AAA (5850G>A), TTC1575ATT (5851T>A 5853C>T), CTG1577TTG (5857C>T), AAG1578AGG (5861A>G), AGC1579TCT (5863A>T 5864G>C 5865C>T), GGA1580GGG (5868A>G), TTC1581TAT (5870T>A 5871C>T), ATT1584TTG (5878A>T 5880T>G), ATG1585AAG (5882T>A), ATG1586GTT (5884A>G 5886G>T), GAC1587CGG (5887G>C 5888A>G 5889C>G), GAA1588GAG (5892A>G), GAA1588_GAA1589insT-- (5892_5893insT), GAA1589CAA (5893G>C), TCC1590AAC (5896T>A 5897C>A), CCT1592CCG (5904T>G), TGG1593AAG (5905T>A 5906G>A), ACC1594ACT (5910C>T), GCC1595GCT (5913C>T), CTG1597AAG (5917C>A 5918T>A), ACC1598ACA (5922C>A), CCA1599CGA (5924C>G), GAT1600TAT (5926G>T), CTT1602CAC (5933T>A 5934T>C), GAA1604GAG (5940A>G), TGG1605TTT (5942G>T 5943G>T), TTA1606ATG (5944T>A 5946A>G), CCA1609TCT (5953C>T 5955A>T), TTC1610TTT (5958C>T), GGA1611GCG (5960G>C 5961A>G), TTG1612CTT (5962T>C 5964G>T), AAG1613ACT (5966A>C 5967G>T), AAT1614AAC (5970T>C), GCT1617GCC (5979T>C), GTG1618GCT (5981T>C 5982G>T) | | | | | | | | |

*: Inserts / Deletes / Misaligned / Frameshifts

#### Analysis details

This analysis was performed with panviral2.64

### NGS Details (UN59): Cavemovirus deltaepiphylli

#### Assembly

| | |
|---|---|
| **Coverage Length** | 413 (1 contig(s)) |
| **Depth Of Coverage** | 4.4 |
| **Number Of Reads** | 15 |
| **Reads Per Million** | 0.33 rpm (after QC) |
| **Ambiguities** | 0 |
| **Assembly Method** | de novo + reference guided assembly |
| **Consensus Caller** | Bcf Tools |

#### Coverage Map

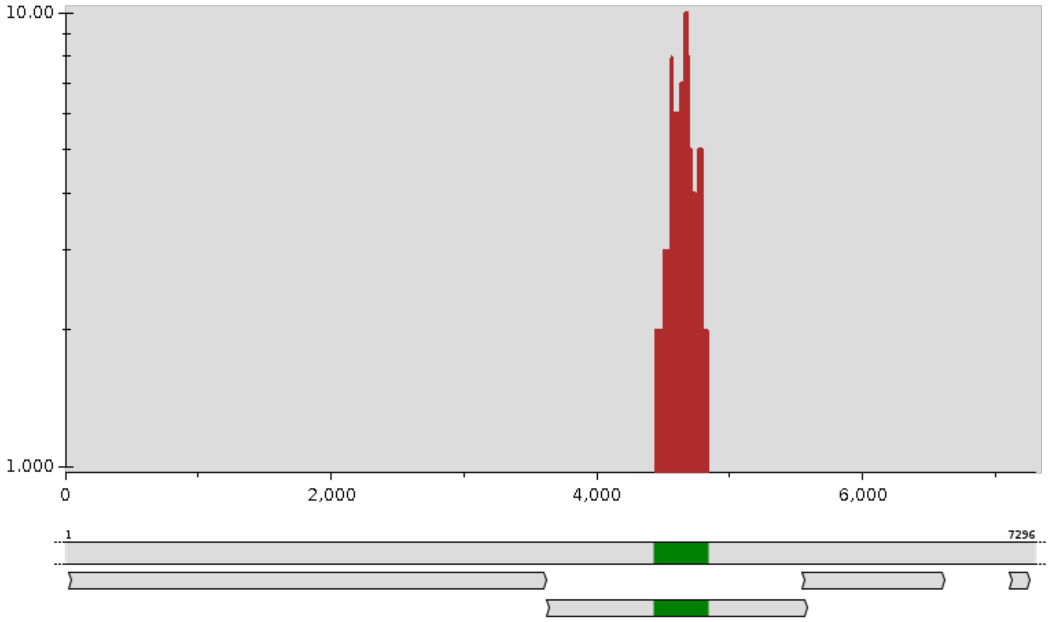

#### Assignment

| | |
|---|---|
| **Type** | Cavemovirus deltaepiphylli (Taxonomy ID: 3047827) |
| **Reference Genome** | NC_055588.1 |
| **NT Identity (%)** | 60.0484 |
| **AA Identity (%)** | 52.8986 |
| **Number Of Stop Codons** | 0 |
| **Number Of CDS** | 4 |

#### Alignment

| | |
|---|---|
| **Alignment Score** | 166.0 (NT) + 526.0 (AA) = 692.0 |
| **Concordance (%)** | 38.1898 |

| Alignment Method | Global, seeded, nucleotide + amino acids (AGA) |
|---|---|

#### Genome Region

Sequence starts at position 4433 and ends at position 4845 relative to NC_055588.1 reference sequence.

#### Alignment Detailed Statistics

| | Begin | End | Coverage | Score | Concordance | Matches | Identities | I/D/M/F* | Stop Codons |
|---|---|---|---|---|---|---|---|---|---|
| **NT** | **4433** | **4845** | **5.7%** | **166** | **20.1%** | **413 (100%)** | **248 (60.0%)** | **0/0** | |
| Mutations: | 4439C>G, 4443A>T, 4444T>G, 4445A>G, 4450A>G, 4451A>G, 4452C>T, 4454C>T, 4457A>T, 4460A>T, 4463A>C, 4469A>G, 4470G>A, 4472T>G, 4473A>G, 4474T>A, 4475A>T, 4479A>T, 4481C>A, 4482A>T, 4483T>C, 4484A>C, 4488G>T, 4489G>T, 4491A>G, 4492T>A, 4493A>T, 4496A>T, 4499C>T, 4503T>A, 4504A>T, 4505T>A, 4509A>C, 4510G>C, 4511C>A, 4517C>T, 4520C>T, 4522G>T, 4523T>A, 4534T>A, 4537A>G, 4538C>G, 4541C>A, 4544C>A, 4545A>C, 4548C>A, 4551G>T, 4552A>T, 4559A>G, 4560A>C, 4561G>A, 4564A>C, 4565A>T, 4566A>T, 4568A>T, 4569T>A, 4570A>G, 4571T>A, 4574A>T, 4577A>T, 4582C>A, 4583G>T, 4586A>C, 4590C>T, 4591A>T, 4592A>T, 4596T>C, 4598C>A, 4601C>T, 4602C>G, 4610A>T, 4613A>C, 4614T>A, 4616A>G, 4619C>A, 4622C>T, 4626T>C, 4627A>T, 4628T>A, 4629C>A, 4631T>A, 4635T>G, 4637A>T, 4640A>T, 4641A>T, 4642G>C, 4644A>G, 4645T>A, 4649C>T, 4652A>G, 4653C>A, 4656T>A, 4658T>C, 4662G>A, 4665A>G, 4667A>T, 4670A>T, 4675G>A, 4676G>T, 4683T>A, 4684A>T, 4685T>G, 4688A>T, 4691T>C, 4695T>A, 4697A>T, 4700A>C, 4712T>C, 4719A>G, 4721C>T, 4723T>A, 4724C>T, 4728C>A, 4730T>A, 4733A>T, 4734G>T, 4735A>T, 4736G>T, 4740G>A, 4741A>T, 4742T>G, 4746T>A, 4748A>T, 4751A>G, 4755A>T, 4758A>G, 4760A>G, 4763A>C, 4769A>G, 4770G>A, 4773T>A, 4774G>T, 4775C>T, 4776A>G, 4778A>T, 4779T>A, 4781G>A, 4790A>T, 4791A>T, 4793A>G, 4794A>G, 4795C>T, 4796C>A, 4797T>A, 4800A>T, 4801G>C, 4805T>A, 4806A>C, 4807A>C, 4811G>A, 4813G>T, 4814C>G, 4815G>A, 4816A>G, 4817A>T, 4821G>T, 4822G>T, 4824A>C, 4829A>T, 4830A>G, 4831C>A, 4837A>G, 4838C>G, 4841C>T, 4842C>T | | | | | | | | |

##### CDS

| | Begin | End | Coverage | Score | Concordance | Matches | Identities | I/D/M/F* | Stop Codons |
|---|---|---|---|---|---|---|---|---|---|
| KM744_gp2 | 271 | 408 | 21.1% | 526 | 52.7% | 138 (100%) | 73 (52.9%) | 0/0/0/0 | 0 |
| Protein mutations: | I274W (4443A>T 4444T>G 4445A>G), K276R (4450A>G 4451A>G), H277Y (4452C>T 4454C>T), D283K (4470G>A 4472T>G), I284D (4473A>G 4474T>A 4475A>T), I286L (4479A>T 4481C>A), I287S (4482A>T 4483T>C 4484A>C), G289L (4488G>T 4489G>T), I290D (4491A>G 4492T>A 4493A>T), K291N (4496A>T), Y294I (4503T>A 4504A>T 4505T>A), S296P (4509A>C 4510G>C 4511C>A), C300L (4522G>T 4523T>A), F304Y (4534T>A), Y305W (4537A>G 4538C>G), H306Q (4541C>A), K308Q (4545A>C), L309I (4548C>A), D310F (4551G>T 4552A>T), S313H (4560A>C 4561G>A), K314T (4564A>C 4565A>T), K315Y (4566A>T 4568A>T), Y316R (4569T>A 4570A>G 4571T>A), T320N (4582C>A 4583G>T), Q323F (4590C>T 4591A>T 4592A>T), Y325Q (4596T>C 4598C>A), Q327E (4602C>G), K329N (4610A>T), L331M (4614T>A 4616A>G), Y335L (4626T>C 4627A>T 4628T>A), H336K (4629C>A 4631T>A), S338A (4635T>G 4637A>T), I341E (4644A>G 4645T>A), Q344K (4653C>A), F345I (4656T>A 4658T>C), D347N (4662G>A), K348D (4665A>G 4667A>T), R351N (4675G>A 4676G>T), Y354M (4683T>A 4684A>T 4685T>G), E355D (4688A>T), L358I (4695T>A 4697A>T), I366V (4719A>G 4721C>T), F367Y (4723T>A 4724C>T), H369K (4728C>A 4730T>A), E371F (4734G>T 4735A>T 4736G>T), D373M (4740G>A 4741A>T 4742T>G), L375I (4746T>A 4748A>T), I378L (4755A>T), K379E (4758A>G 4760A>G), E383K (4770G>A), C384I (4773T>A 4774G>T 4775C>T), I385V (4776A>G 4778A>T), L386I (4779T>A 4781G>A), I390L (4791A>T 4793A>G), T391V (4794A>G 4795C>T 4796C>A), L392I (4797T>A), N394K (4805T>A), K395P (4806A>C 4807A>C), S397M (4813G>T 4814C>G), E398S (4815G>A 4816A>G 4817A>T), G400F (4821G>T 4822G>T), K401Q (4824A>C), T403D (4830A>G 4831C>A), H405R (4837A>G 4838C>G) | | | | | | | | |
| Codon mutations: | GCC272GCG (4439C>G), ATA274TGG (4443A>T 4444T>G 4445A>G), AAA276AGG (4450A>G 4451A>G), CAC277TAT (4452C>T 4454C>T), CCA278CCT (4457A>T), ATA279ATT (4460A>T), CCA280CCC (4463A>C), AAA282AAG (4469A>G), GAT283AAG (4470G>A 4472T>G), ATA284GAT (4473A>G 4474T>A 4475A>T), ATC286TTA (4479A>T 4481C>A), ATA287TCC (4482A>T 4483T>C 4484A>C), GGA289TTA (4488G>T 4489G>T), ATA290GAT (4491A>G 4492T>A 4493A>T), AAA291AAT (4496A>T), GCC292GCT (4499C>T), TAT294ATA (4503T>A 4504A>T 4505T>A), AGC296CCA (4509A>C 4510G>C 4511C>A), TTC298TTT (4517C>T), GAC299GAT (4520C>T), TGT300TTA (4522G>T 4523T>A), TTT304TAT (4534T>A), TAC305TGG (4537A>G 4538C>G), CAC306CAA (4541C>A), ATC307ATA (4544C>A), AAG308CAG (4545A>C), CTA309ATA (4548C>A), GAT310TTT (4551G>T 4552A>T), GAA312GAG (4559A>G), AGT313CAT (4560A>C 4561G>A), AAA314ACT (4564A>C 4565A>T), AAA315TAT (4566A>T 4568A>T), TAT316AGA (4569T>A 4570A>G 4571T>A), ACA317ACT (4574A>T), GCA318GCT (4577A>T), ACG320AAT (4582C>A 4583G>T), GTA321GTC (4586A>C), CAA323TTT (4590C>T 4591A>T 4592A>T), TAC325CAA (4596T>C 4598C>A), TAC326TAT (4601C>T), CAA327GAA (4602C>G), AAA329AAT (4610A>T), GTA330GTC (4613A>C), TTA331ATG (4614T>A 4616A>G), CCC332CCA (4619C>A), TTC333TTT (4622C>T), TAT335CTA (4626T>C 4627A>T 4628T>A), CAT336AAA (4629C>A 4631T>A), TCA338GCT (4635T>G 4637A>T), CCA339CCT (4640A>T), AGT340TCT (4641A>T 4642G>C), ATA341GAA (4644A>G 4645T>A), TTC342TTT (4649C>T), CAA343CAG (4652A>G), CAA344AAA (4653C>A), TTT345ATC (4656T>A 4658T>C), GAT347AAT (4662G>A), AAA348GAT (4665A>G 4667A>T), ATA349ATT (4670A>T), AGG351AAT (4675G>A 4676G>T), TAT354ATG (4683T>A 4684A>T 4685T>G), GAA355GAT (4688A>T), TTT356TTC (4691T>C), TTA358ATT (4695T>A 4697A>T), GTA359GTC (4700A>C), GAT363GAC (4712T>C), ATC366GTT (4719A>G 4721C>T), TTC367TAT (4723T>A 4724C>T), CAT369AAA (4728C>A 4730T>A), ACA370ACT (4733A>T), GAG371TTT (4734G>T 4735A>T 4736G>T), GAT373ATG (4740G>A 4741A>T 4742T>G), TTA375ATT (4746T>A 4748A>T), AAA376AAG (4751A>G), ATA378TTA (4755A>T), AAA379GAG (4758A>G 4760A>G), ATA380ATC (4763A>C), AAA382AAG (4769A>G), GAA383AAA (4770G>A), TGC384ATT (4773T>A 4774G>T 4775C>T), ATA385GTT (4776A>G 4778A>T), TTG386ATA (4779T>A 4781G>A), GGA389GGT (4790A>T), ATA390TTG (4791A>T 4793A>G), ACC391GTA (4794A>G 4795C>T 4796C>A), TTA392ATA (4797T>A), AGT393TCT (4800A>T 4801G>C), AAT394AAA (4805T>A), AAA395CCA (4806A>C 4807A>C), AAG396AAA (4811G>A), AGC397ATG (4813G>T 4814C>G), GAA398AGT (4815G>A 4816A>G 4817A>T), GGC400TTC (4821G>T 4822G>T), AAA401CAA (4824A>C), ACA402ACT (4829A>T), ACT403GAT (4830A>G 4831C>A), CAC405CGG (4837A>G 4838C>G), TTC406TTT (4841C>T), CTA407TTA (4842C>T) | | | | | | | | |

##### Proteins

| | Begin | End | Coverage | Score | Concordance | Matches | Identities | I/D/M/F* | Stop Codons |
|---|---|---|---|---|---|---|---|---|---|
| replicase (YP_010087807.1) | 271 | 408 | 21.1% | 526 | 52.7% | 138 (100%) | 73 (52.9%) | 0/0/0/0 | 0 |
| Protein mutations: | I274W (4443A>T 4444T>G 4445A>G), K276R (4450A>G 4451A>G), H277Y (4452C>T 4454C>T), D283K (4470G>A 4472T>G), I284D (4473A>G 4474T>A 4475A>T), I286L (4479A>T 4481C>A), I287S (4482A>T 4483T>C 4484A>C), G289L (4488G>T 4489G>T), I290D (4491A>G 4492T>A 4493A>T), K291N (4496A>T), Y294I (4503T>A 4504A>T 4505T>A), S296P (4509A>C 4510G>C 4511C>A), C300L (4522G>T 4523T>A), F304Y (4534T>A), Y305W (4537A>G 4538C>G), H306Q (4541C>A), K308Q (4545A>C), L309I (4548C>A), D310F (4551G>T 4552A>T), S313H (4560A>C 4561G>A), K314T (4564A>C 4565A>T), K315Y (4566A>T 4568A>T), Y316R (4569T>A 4570A>G 4571T>A), T320N (4582C>A 4583G>T), Q323F (4590C>T 4591A>T 4592A>T), Y325Q (4596T>C 4598C>A), Q327E (4602C>G), K329N (4610A>T), L331M (4614T>A 4616A>G), Y335L (4626T>C 4627A>T 4628T>A), H336K (4629C>A 4631T>A), S338A (4635T>G 4637A>T), I341E (4644A>G 4645T>A), Q344K (4653C>A), F345I (4656T>A 4658T>C), D347N (4662G>A), K348D (4665A>G 4667A>T), R351N (4675G>A 4676G>T), Y354M (4683T>A 4684A>T 4685T>G), E355D (4688A>T), L358I (4695T>A 4697A>T), I366V (4719A>G 4721C>T), F367Y (4723T>A 4724C>T), H369K (4728C>A 4730T>A), E371F (4734G>T 4735A>T 4736G>T), D373M (4740G>A 4741A>T 4742T>G), L375I (4746T>A 4748A>T), I378L (4755A>T), K379E (4758A>G 4760A>G), E383K (4770G>A), C384I (4773T>A 4774G>T 4775C>T), I385V (4776A>G 4778A>T), L386I (4779T>A 4781G>A), I390L (4791A>T 4793A>G), T391V (4794A>G 4795C>T 4796C>A), L392I (4797T>A), N394K (4805T>A), K395P (4806A>C 4807A>C), S397M (4813G>T 4814C>G), E398S (4815G>A 4816A>G 4817A>T), G400F (4821G>T 4822G>T), K401Q (4824A>C), T403D (4830A>G 4831C>A), H405R (4837A>G 4838C>G) | | | | | | | | |
| Codon mutations: | GCC272GCG (4439C>G), ATA274TGG (4443A>T 4444T>G 4445A>G), AAA276AGG (4450A>G 4451A>G), CAC277TAT (4452C>T 4454C>T), CCA278CCT (4457A>T), ATA279ATT (4460A>T), CCA280CCC (4463A>C), AAA282AAG (4469A>G), GAT283AAG (4470G>A 4472T>G), ATA284GAT (4473A>G 4474T>A 4475A>T), ATC286TTA (4479A>T 4481C>A), ATA287TCC (4482A>T 4483T>C 4484A>C), GGA289TTA (4488G>T 4489G>T), ATA290GAT (4491A>G 4492T>A 4493A>T), AAA291AAT (4496A>T), GCC292GCT (4499C>T), TAT294ATA (4503T>A 4504A>T 4505T>A), AGC296CCA (4509A>C 4510G>C 4511C>A), TTC298TTT (4517C>T), GAC299GAT (4520C>T), TGT300TTA (4522G>T 4523T>A), TTT304TAT (4534T>A), TAC305TGG (4537A>G 4538C>G), CAC306CAA (4541C>A), ATC307ATA (4544C>A), AAG308CAG (4545A>C), CTA309ATA (4548C>A), GAT310TTT (4551G>T 4552A>T), GAA312GAG (4559A>G), AGT313CAT (4560A>C 4561G>A), AAA314ACT (4564A>C 4565A>T), AAA315TAT (4566A>T 4568A>T), TAT316AGA (4569T>A 4570A>G 4571T>A), ACA317ACT (4574A>T), GCA318GCT (4577A>T), ACG320AAT (4582C>A 4583G>T), GTA321GTC (4586A>C), CAA323TTT (4590C>T 4591A>T 4592A>T), TAC325CAA (4596T>C 4598C>A), TAC326TAT (4601C>T), CAA327GAA (4602C>G), AAA329AAT (4610A>T), GTA330GTC (4613A>C), TTA331ATG (4614T>A 4616A>G), CCC332CCA (4619C>A), TTC333TTT (4622C>T), TAT335CTA (4626T>C 4627A>T 4628T>A), CAT336AAA (4629C>A 4631T>A), TCA338GCT (4635T>G 4637A>T), CCA339CCT (4640A>T), AGT340TCT (4641A>T 4642G>C), ATA341GAA (4644A>G 4645T>A), TTC342TTT (4649C>T), CAA343CAG (4652A>G), CAA344AAA (4653C>A), TTT345ATC (4656T>A 4658T>C), GAT347AAT (4662G>A), AAA348GAT (4665A>G 4667A>T), ATA349ATT (4670A>T), AGG351AAT (4675G>A 4676G>T), TAT354ATG (4683T>A 4684A>T 4685T>G), GAA355GAT (4688A>T), TTT356TTC (4691T>C), TTA358ATT (4695T>A 4697A>T), GTA359GTC (4700A>C), GAT363GAC (4712T>C), ATC366GTT (4719A>G 4721C>T), TTC367TAT (4723T>A 4724C>T), CAT369AAA (4728C>A 4730T>A), ACA370ACT (4733A>T), GAG371TTT (4734G>T 4735A>T 4736G>T), GAT373ATG (4740G>A 4741A>T 4742T>G), TTA375ATT (4746T>A 4748A>T), AAA376AAG (4751A>G), ATA378TTA (4755A>T), AAA379GAG (4758A>G 4760A>G), ATA380ATC (4763A>C), AAA382AAG (4769A>G), GAA383AAA (4770G>A), TGC384ATT (4773T>A 4774G>T 4775C>T), ATA385GTT (4776A>G 4778A>T), TTG386ATA (4779T>A 4781G>A), GGA389GGT (4790A>T), ATA390TTG (4791A>T 4793A>G), ACC391GTA (4794A>G 4795C>T 4796C>A), TTA392ATA (4797T>A), AGT393TCT (4800A>T 4801G>C), AAT394AAA (4805T>A), AAA395CCA (4806A>C 4807A>C), AAG396AAA (4811G>A), AGC397ATG (4813G>T 4814C>G), GAA398AGT (4815G>A 4816A>G 4817A>T), GGC400TTC (4821G>T 4822G>T), AAA401CAA (4824A>C), ACA402ACT (4829A>T), ACT403GAT (4830A>G 4831C>A), CAC405CGG (4837A>G 4838C>G), TTC406TTT (4841C>T), CTA407TTA (4842C>T) | | | | | | | | |

*: Inserts / Deletes / Misaligned / Frameshifts

#### Analysis details

This analysis was performed with panviral2.64

### NGS Details (UN59): Dioscorea bacilliform virus

#### Assembly

| | |
|---|---|
| **Coverage Length** | 421 (1 contig(s)) |
| **Depth Of Coverage** | 3.5 |
| **Number Of Reads** | 11 |
| **Reads Per Million** | 0.24 rpm (after QC) |
| **Ambiguities** | 0 |
| **Assembly Method** | de novo + reference guided assembly |
| **Consensus Caller** | Bcf Tools |

#### Coverage Map

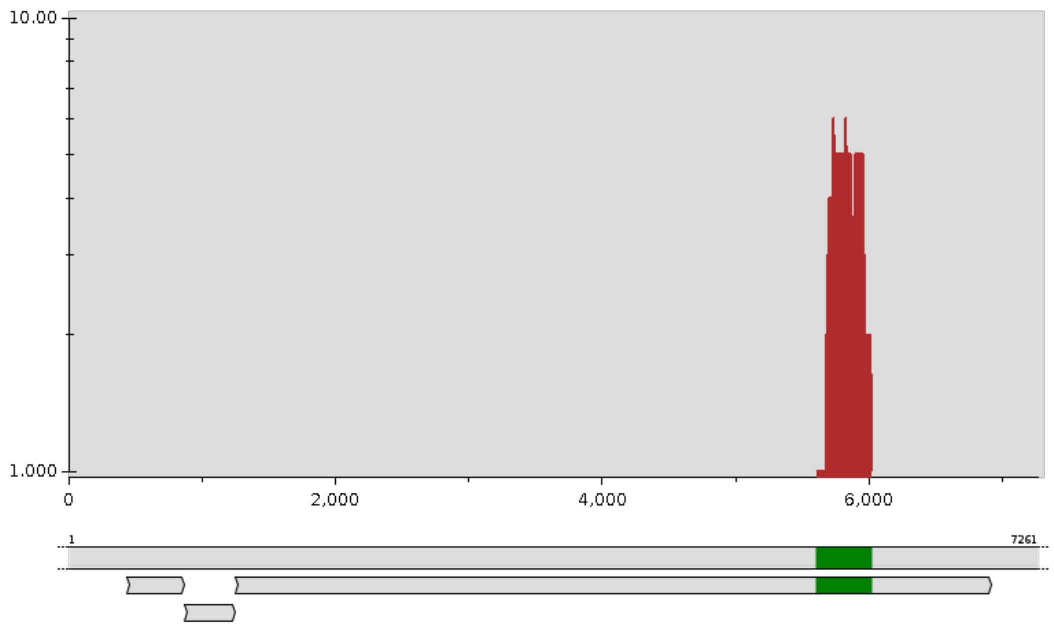

#### Assignment

| | |
|---|---|
| **Type** | Dioscorea bacilliform virus (Taxonomy ID: 52996) |
| **Reference Genome** | NC_009010.1 |
| **NT Identity (%)** | 53.066 |
| **AA Identity (%)** | 39.7163 |
| **Number Of Stop Codons** | 1 |
| **Number Of CDS** | 3 |

#### Alignment

| | |
|---|---|
| **Alignment Score** | 46.0 (NT) + 394.0 (AA) = 440.0 |
| **Concordance (%)** | 24.3363 |

| Alignment Method | Global, seeded, nucleotide + amino acids (AGA) |
|---|---|

#### Genome Region

Sequence starts at position 5600 and ends at position 6020 relative to NC_009010.1 reference sequence.

#### Alignment Detailed Statistics

| | Begin | End | Coverage | Score | Concordance | Matches | Identities | I/D/M/F* | Stop Codons |
|---|---|---|---|---|---|---|---|---|---|
| **NT** | **5600** | **6020** | **5.8%** | **46** | **5.5%** | **421 (99.3%)** | **225 (53.1%)** | **3/0** | |
| Mutations: | 5601G>A, 5602G>A, 5607C>G, 5608A>G, 5610A>T, 5615A>T, 5616G>C, 5617G>C, 5618A>T, 5619G>A, 5620A>T, 5626G>T, 5627T>G, 5628C>T, 5629T>A, 5631C>T, 5632A>G, 5639A>G, 5640A>G, 5641A>C, 5642G>A, 5643A>G, 5650G>A, 5652C>G, 5653A>G, 5654A>T, 5655C>G, 5660A>T, 5665G>A, 5669A>G, 5670A>T, 5673T>C, 5674A>C, 5675G>C, 5677C>A, 5679T>G, 5680C>A, 5682A>T, 5683G>T, 5684G>A, 5685G>T, 5688C>T, 5689A>G, 5691C>T, 5692A>G, 5693C>A, 5694A>T, 5695A>T, 5697T>A, 5698C>T, 5700G>C, 5701A>C, 5703G>C, 5704A>C, 5705G>A, 5706A>G, 5707G>C, 5710G>T, 5711G>A, 5712T>G, 5713C>G, 5715A>T, 5716A>G, 5717G>C, 5720A>C, 5722A>G, 5724A>G, 5726A>T, 5727T>C, 5730A>T, 5733A>G, 5734T>G, 5742A>G, 5744A>G, 5746A>T, 5747G>C, 5751C>T, 5753T>A, 5757T>C, 5761G>T, 5764G>A, 5765C>G, 5766A>G, 5769G>T, 5770G>A, 5771C>G, 5772G>A, 5773C>G, 5774C>T, 5776C>G, 5777A>C, 5778A>T, 5779T>G, 5780C>A, 5781A>C, 5782G>A, 5784T>A, 5785G>C, 5786A>C, 5787A>T, 5788T>A, 5789G>A, 5793G>T, 5794G>A, 5795C>T, 5796G>T, 5801T>G, 5802A>C, 5803G>A, 5805G>C, 5807C>G, 5808A>T, 5809G>T, 5810G>A, 5811A>T, 5814A>C, 5816T>A, 5820C>T, 5823A>G, 5825G>T, 5826G>C, 5827T>C, 5829G>A, 5836C>T, 5838A>T, 5844A>G, 5847G>T, 5849A>G, 5850G>T, 5856A>C, 5858C>T, 5861C>T, 5864T>C, 5865G>T, 5868C>T, 5869C>A, 5870A>T, 5872A>G, 5873G>A, 5874G>C, 5875A>C, 5876A>T, 5877G>T, 5881G>A, 5882A>C, 5883C>T, 5884A>C, 5885A>G, 5886C>A, 5889C>A, 5893C>A, 5895A>G, 5896G>C, 5897G>C, 5898_5899insTAT, 5899A>C, 5900C>T, 5902G>A, 5904A>C, 5905G>T, 5906A>C, 5907A>T, 5910T>C, 5913T>G, 5914G>A, 5915C>T, 5919A>G, 5921A>T, 5931T>C, 5937G>T, 5939T>C, 5942T>A, 5943T>C, 5946T>A, 5947G>C, 5951C>G, 5952A>T, 5953G>A, 5954A>G, 5965T>C, 5968A>C, 5970G>A, 5974C>T, 5979G>A, 5980A>G, 5982A>T, 5983C>G, 5991A>G, 5993T>C, 5996G>T, 5997T>G, 5998C>A, 5999A>G, 6001C>G, 6003A>G, 6006C>T, 6007G>C | | | | | | | | |

##### CDS

| | Begin | End | Coverage | Score | Concordance | Matches | Identities | I/D/M/F* | Stop Codons |
|---|---|---|---|---|---|---|---|---|---|
| DBV_gp3 | 1453 | 1592 | 7.4% | 394 | 40.5% | 140 (99.3%) | 56 (39.7%) | 1/0/0/0 | 1 |
| Protein mutations: | E1453K (5602G>A), N1454K (5607C>G), K1455D (5608A>G 5610A>T), K1457I (5615A>T 5616G>C), E1458L (5617G>C 5618A>T 5619G>A), R1459W (5620A>T), V1461C (5626G>T 5627T>G 5628C>T), F1462I (5629T>A 5631C>T), N1463D (5632A>G), K1465R (5639A>G 5640A>G), R1466Q (5641A>C 5642G>A 5643A>G), D1469K (5650G>A 5652C>G), N1470V (5653A>G 5654A>T 5655C>G), E1472V (5660A>T), D1474N (5665G>A), Q1475R (5669A>G 5670A>T), S1477P (5674A>C 5675G>C), L1478M (5677C>A 5679T>G), P1479T (5680C>A 5682A>T), G1480Y (5683G>T 5684G>A 5685G>T), N1482D (5689A>G 5691C>T), T1483D (5692A>G 5693C>A 5694A>T), I1484L (5695A>T 5697T>A), L1485F (5698C>T 5700G>C), K1486H (5701A>C 5703G>C), R1487Q (5704A>C 5705G>A 5706A>G), V1488L (5707G>C), G1489* (5710G>T 5711G>A 5712T>G), Q1490D (5713C>G 5715A>T), S1491A (5716A>G 5717G>C), K1492T (5720A>C), I1493V (5722A>G 5724A>G), Y1494F (5726A>T 5727T>C), F1497V (5734T>G), K1500R (5744A>G), F1503Y (5753T>A), V1506L (5761G>T), A1507R (5764G>A 5765C>G 5766A>G), M1508I (5769G>T), A1509R (5770G>A 5771C>G 5772G>A), P1510V (5773C>G 5774C>T), Q1511A (5776C>G 5777A>C 5778A>T), S1512D (5779T>G 5780C>A 5781A>C), V1513I (5782G>A 5784T>A), E1514P (5785G>C 5786A>C 5787A>T), W1515K (5788T>A 5789G>A), A1517I (5794G>A 5795C>T 5796G>T), L1519R (5801T>G 5802A>C), A1520T (5803G>A 5805G>C), P1521R (5807C>G 5808A>T), G1522Y (5809G>T 5810G>A 5811A>T), L1524H (5816T>A), W1527F (5825G>T 5826G>C), P1531S (5836C>T 5838A>T), K1535S (5849A>G 5850G>T), P1538L (5858C>T), A1539V (5861C>T), V1540A (5864T>C 5865G>T), Q1542M (5869C>A 5870A>T), R1543D (5872A>G 5873G>A 5874G>C), K1544L (5875A>C 5876A>T 5877G>T), D1546T (5881G>A 5882A>C 5883C>T), N1547R (5884A>C 5885A>G 5886C>A), G1551P (5896G>C 5897G>C), G1551_T1552insY (5898_5899insTAT), T1552L (5899A>C 5900C>T), E1553N (5902G>A 5904A>C), E1554S (5905G>T 5906A>C 5907A>T), I1556M (5913T>G), A1557I (5914G>A 5915C>T), Y1559F (5921A>T), V1565A (5939T>C), F1566Y (5942T>A 5943T>C), E1568Q (5947G>C), T1569S (5951C>G 5952A>T), E1570R (5953G>A 5954A>G), K1575Q (5968A>C 5970G>A), I1579V (5980A>G 5982A>T), L1580V (5983C>G), I1583T (5993T>C), C1584L (5996G>T 5997T>G), Q1585R (5998C>A 5999A>G), Q1586E (6001C>G 6003A>G), G1588R (6007G>C) | | | | | | | | |
| Codon mutations: | AAG1452.AA (5601G>A), GAG1453AAG (5602G>A), AAC1454AAG (5607C>G), AAA1455GAT (5608A>G 5610A>T), AAG1457ATC (5615A>T 5616G>C), GAG1458CTA (5617G>C 5618A>T 5619G>A), AGG1459TGG (5620A>T), GTC1461TGT (5626G>T 5627T>G 5628C>T), TTC1462ATT (5629T>A 5631C>T), AAC1463GAC (5632A>G), AAA1465AGG (5639A>G 5640A>G), AGA1466CAG (5641A>C 5642G>A 5643A>G), GAC1469AAG (5650G>A 5652C>G), AAC1470GTG (5653A>G 5654A>T 5655C>G), GAG1472GTG (5660A>T), GAC1474AAC (5665G>A), CAA1475CGT (5669A>G 5670A>T), TAT1476TAC (5673T>C), AGC1477CCC (5674A>C 5675G>C), CTT1478ATG (5677C>A 5679T>G), CCA1479ACT (5680C>A 5682A>T), GGG1480TAT (5683G>T 5684G>A 5685G>T), ATC1481ATT (5688C>T), AAC1482GAT (5689A>G 5691C>T), ACA1483GAT (5692A>G 5693C>A 5694A>T), ATT1484TTA (5695A>T 5697T>A), CTG1485TTC (5698C>T 5700G>C), AAG1486CAC (5701A>C 5703G>C), AGA1487CAG (5704A>C 5705G>A 5706A>G), GTC1488CTC (5707G>C), GGT1489TAG (5710G>T 5711G>A 5712T>G), CAA1490GAT (5713C>G 5715A>T), AGC1491GCC (5716A>G 5717G>C), AAA1492ACA (5720A>C), ATA1493GTG (5722A>G 5724A>G), TAT1494TTC (5726A>T 5727T>C), TCA1495TCT (5730A>T), AAA1496AAG (5733A>G), TTT1497GTT (5734T>G), TTA1499TTG (5742A>G), AAG1500AGG (5744A>G), AGC1501TCC (5746A>T 5747G>C), GGC1502GGT (5751C>T), TTT1503TAT (5753T>A), CAT1504CAC (5757T>C), GTG1506TTG (5761G>T), GCA1507AGG (5764G>A 5765C>G 5766A>G), ATG1508ATT (5769G>T), GCG1509AGA (5770G>A 5771C>G 5772G>A), CCA1510GTA (5773C>G 5774C>T), CAA1511GCT (5776C>G 5777A>C 5778A>T), TCA1512GAC (5779T>G 5780C>A 5781A>C), GTT1513ATA (5782G>A 5784T>A), GAA1514CCT (5785G>C 5786A>C 5787A>T), TGG1515AAG (5788T>A 5789G>A), ACG1516ACT (5793G>T), GCG1517ATT (5794G>A 5795C>T 5796G>T), CTA1519CGC (5801T>G 5802A>C), GCG1520ACC (5803G>A 5805G>C), CCA1521CGT (5807C>G 5808A>T), GGA1522TAT (5809G>T 5810G>A 5811A>T), GGA1523GGC (5814A>C), CTT1524CAT (5816T>A), TAC1525TAT (5820C>T), GAA1526GAG (5823A>G), TGG1527TTC (5825G>T 5826G>C), TTG1528CTA (5827T>C 5829G>A), CCA1531TCT (5836C>T 5838A>T), GGA1533GGG (5844A>G), CTG1534CTT (5847G>T), AAG1535AGT (5849A>G 5850G>T), GCA1537GCC (5856A>C), CCA1538CTA (5858C>T), GCA1539GTA (5861C>T), GTG1540GCT (5864T>C 5865G>T), TTC1541TTT (5868C>T), CAG1542ATG (5869C>A 5870A>T), AGG1543GAC (5872A>G 5873G>A 5874G>C), AAG1544CTT (5875A>C 5876A>T 5877G>T), GAC1546ACT (5881G>A 5882A>C 5883C>T), AAC1547CGA (5884A>C 5885A>G 5886C>A), GTC1548GTA (5889C>A), CGA1550AGG (5893C>A 5895A>G), GGT1551CCT (5896G>C 5897G>C), GGT1551_ACT1552insTAT (5898_5899insTAT), ACT1552CTT (5899A>C 5900C>T), GAA1553AAC (5902G>A 5904A>C), GAA1554TCT (5905G>T 5906A>C 5907A>T), TTT1555TTC (5910T>C), ATT1556ATG (5913T>G), GCT1557ATT (5914G>A 5915C>T), GTA1558GTG (5919A>G), TAC1559TTC (5921A>T), GAT1562GAC (5931T>C), CTG1564CTT (5937G>T), GTG1565GCG (5939T>C), TTT1566TAC (5942T>A 5943T>C), TCT1567TCA (5946T>A), GAA1568CAA (5947G>C), ACA1569AGT (5951C>G 5952A>T), GAA1570AGA (5953G>A 5954A>G), TTG1574CTG (5965T>C), AAG1575CAA (5968A>C 5970G>A), CTG1577TTG (5974C>T), AGG1578AGA (5979G>A), ATA1579GTT (5980A>G 5982A>T), CTG1580GTG (5983C>G), CAA1582CAG (5991A>G), ATC1583ACC (5993T>C), TGT1584TTG (5996G>T 5997T>G), CAA1585AGA (5998C>A 5999A>G), CAA1586GAG (6001C>G 6003A>G), CAC1587CAT (6006C>T), GGG1588CGG (6007G>C) | | | | | | | | |

##### Proteins

| | Begin | End | Coverage | Score | Concordance | Matches | Identities | I/D/M/F* | Stop Codons |
|---|---|---|---|---|---|---|---|---|---|
| ORF3 protein (YP_001036293.1) | 1453 | 1592 | 7.4% | 394 | 40.5% | 140 (99.3%) | 56 (39.7%) | 1/0/0/0 | 1 |
| Protein mutations: | E1453K (5602G>A), N1454K (5607C>G), K1455D (5608A>G 5610A>T), K1457I (5615A>T 5616G>C), E1458L (5617G>C 5618A>T 5619G>A), R1459W (5620A>T), V1461C (5626G>T 5627T>G 5628C>T), F1462I (5629T>A 5631C>T), N1463D (5632A>G), K1465R (5639A>G 5640A>G), R1466Q (5641A>C 5642G>A 5643A>G), D1469K (5650G>A 5652C>G), N1470V (5653A>G 5654A>T 5655C>G), E1472V (5660A>T), D1474N (5665G>A), Q1475R (5669A>G 5670A>T), S1477P (5674A>C 5675G>C), L1478M (5677C>A 5679T>G), P1479T (5680C>A 5682A>T), G1480Y (5683G>T 5684G>A 5685G>T), N1482D (5689A>G 5691C>T), T1483D (5692A>G 5693C>A 5694A>T), I1484L (5695A>T 5697T>A), L1485F (5698C>T 5700G>C), K1486H (5701A>C 5703G>C), R1487Q (5704A>C 5705G>A 5706A>G), V1488L (5707G>C), G1489* (5710G>T 5711G>A 5712T>G), Q1490D (5713C>G 5715A>T), S1491A (5716A>G 5717G>C), K1492T (5720A>C), I1493V (5722A>G 5724A>G), Y1494F (5726A>T 5727T>C), F1497V (5734T>G), K1500R (5744A>G), F1503Y (5753T>A), V1506L (5761G>T), A1507R (5764G>A 5765C>G 5766A>G), M1508I (5769G>T), A1509R (5770G>A 5771C>G 5772G>A), P1510V (5773C>G 5774C>T), Q1511A (5776C>G 5777A>C 5778A>T), S1512D (5779T>G 5780C>A 5781A>C), V1513I (5782G>A 5784T>A), E1514P (5785G>C 5786A>C 5787A>T), W1515K (5788T>A 5789G>A), A1517I (5794G>A 5795C>T 5796G>T), L1519R (5801T>G 5802A>C), A1520T (5803G>A 5805G>C), P1521R (5807C>G 5808A>T), G1522Y (5809G>T 5810G>A 5811A>T), L1524H (5816T>A), W1527F (5825G>T 5826G>C), P1531S (5836C>T 5838A>T), K1535S (5849A>G 5850G>T), P1538L (5858C>T), A1539V (5861C>T), V1540A (5864T>C 5865G>T), Q1542M (5869C>A 5870A>T), R1543D (5872A>G 5873G>A 5874G>C), K1544L (5875A>C 5876A>T 5877G>T), D1546T (5881G>A 5882A>C 5883C>T), N1547R (5884A>C 5885A>G 5886C>A), G1551P (5896G>C 5897G>C), G1551_T1552insY (5898_5899insTAT), T1552L (5899A>C 5900C>T), E1553N (5902G>A 5904A>C), E1554S (5905G>T 5906A>C 5907A>T), I1556M (5913T>G), A1557I (5914G>A 5915C>T), Y1559F (5921A>T), V1565A (5939T>C), F1566Y (5942T>A 5943T>C), E1568Q (5947G>C), T1569S (5951C>G 5952A>T), E1570R (5953G>A 5954A>G), K1575Q (5968A>C 5970G>A), I1579V (5980A>G 5982A>T), L1580V (5983C>G), I1583T (5993T>C), C1584L (5996G>T 5997T>G), Q1585R (5998C>A 5999A>G), Q1586E (6001C>G 6003A>G), G1588R (6007G>C) | | | | | | | | |

| | Begin | End | Coverage | Score | Concordance | Matches | Identities | I/D/M/F* | Stop Codons |
|---|---|---|---|---|---|---|---|---|---|
| **NT** | **5600** | **6020** | **5.8%** | **46** | **5.5%** | **421 (99.3%)** | **225 (53.1%)** | **3/0** | |
| Codon mutations: | AAG1452.AA (5601G>A), GAG1453AAG (5602G>A), AAC1454AAG (5607C>G), AAA1455GAT (5608A>G 5610A>T), AAG1457ATC (5615A>T 5616G>C), GAG1458CTA (5617G>C 5618A>T 5619G>A), AGG1459TGG (5620A>T), GTC1461TGT (5626G>T 5627T>G 5628C>T), TTC1462ATT (5629T>A 5631C>T), AAC1463GAC (5632A>G), AAA1465AGG (5639A>G 5640A>G), AGA1466CAG (5641A>C 5642G>A 5643A>G), GAC1469AAG (5650G>A 5652C>G), AAC1470GTG (5653A>G 5654A>T 5655C>G), GAG1472GTG (5660A>T), GAC1474AAC (5665G>A), CAA1475CGT (5669A>G 5670A>T), TAT1476TAC (5673T>C), AGC1477CCC (5674A>C 5675G>C), CTT1478ATG (5677C>A 5679T>G), CCA1479ACT (5680C>A 5682A>T), GGG1480TAT (5683G>T 5684G>A 5685G>T), ATC1481ATT (5688C>T), AAC1482GAT (5689A>G 5691C>T), ACA1483GAT (5692A>G 5693C>A 5694A>T), ATT1484TTA (5695A>T 5697T>A), CTG1485TTC (5698C>T 5700G>C), AAG1486CAC (5701A>C 5703G>C), AGA1487CAG (5704A>C 5705G>A 5706A>G), GTC1488CTC (5707G>C), GGT1489TAG (5710G>T 5711G>A 5712T>G), CAA1490GAT (5713C>G 5715A>T), AGC1491GCC (5716A>G 5717G>C), AAA1492ACA (5720A>C), ATA1493GTG (5722A>G 5724A>G), TAT1494TTC (5726A>T 5727T>C), TCA1495TCT (5730A>T), AAA1496AAG (5733A>G), TTT1497GTT (5734T>G), TTA1499TTG (5742A>G), AAG1500AGG (5744A>G), AGC1501TCC (5746A>T 5747G>C), GGC1502GGT (5751C>T), TTT1503TAT (5753T>A), CAT1504CAC (5757T>C), GTG1506TTG (5761G>T), GCA1507AGG (5764G>A 5765C>G 5766A>G), ATG1508ATT (5769G>T), GCG1509AGA (5770G>A 5771C>G 5772G>A), CCA1510GTA (5773C>G 5774C>T), CAA1511GCT (5776C>G 5777A>C 5778A>T), TCA1512GAC (5779T>G 5780C>A 5781A>C), GTT1513ATA (5782G>A 5784T>A), GAA1514CCT (5785G>C 5786A>C 5787A>T), TGG1515AAG (5788T>A 5789G>A), ACG1516ACT (5793G>T), GCG1517ATT (5794G>A 5795C>T 5796G>T), CTA1519CGC (5801T>G 5802A>C), GCG1520ACC (5803G>A 5805G>C), CCA1521CGT (5807C>G 5808A>T), GGA1522TAT (5809G>T 5810G>A 5811A>T), GGA1523GGC (5814A>C), CTT1524CAT (5816T>A), TAC1525TAT (5820C>T), GAA1526GAG (5823A>G), TGG1527TTC (5825G>T 5826G>C), TTG1528CTA (5827T>C 5829G>A), CCA1531TCT (5836C>T 5838A>T), GGA1533GGG (5844A>G), CTG1534CTT (5847G>T), AAG1535AGT (5849A>G 5850G>T), GCA1537GCC (5856A>C), CCA1538CTA (5858C>T), GCA1539GTA (5861C>T), GTG1540GCT (5864T>C 5865G>T), TTC1541TTT (5868C>T), CAG1542ATG (5869C>A 5870A>T), AGG1543GAC (5872A>G 5873G>A 5874G>C), AAG1544CTT (5875A>C 5876A>T 5877G>T), GAC1546ACT (5881G>A 5882A>C 5883C>T), AAC1547CGA (5884A>C 5885A>G 5886C>A), GTC1548GTA (5889C>A), CGA1550AGG (5893C>A 5895A>G), GGT1551CCT (5896G>C 5897G>C), GGT1551_ACT1552insTAT (5898_5899insTAT), ACT1552CTT (5899A>C 5900C>T), GAA1553AAC (5902G>A 5904A>C), GAA1554TCT (5905G>T 5906A>C 5907A>T), TTT1555TTC (5910T>C), ATT1556ATG (5913T>G), GCT1557ATT (5914G>A 5915C>T), GTA1558GTG (5919A>G), TAC1559TTC (5921A>T), GAT1562GAC (5931T>C), CTG1564CTT (5937G>T), GTG1565GCG (5939T>C), TTT1566TAC (5942T>A 5943T>C), TCT1567TCA (5946T>A), GAA1568CAA (5947G>C), ACA1569AGT (5951C>G 5952A>T), GAA1570AGA (5953G>A 5954A>G), TTG1574CTG (5965T>C), AAG1575CAA (5968A>C 5970G>A), CTG1577TTG (5974C>T), AGG1578AGA (5979G>A), ATA1579GTT (5980A>G 5982A>T), CTG1580GTG (5983C>G), CAA1582CAG (5991A>G), ATC1583ACC (5993T>C), TGT1584TTG (5996G>T 5997T>G), CAA1585AGA (5998C>A 5999A>G), CAA1586GAG (6001C>G 6003A>G), CAC1587CAT (6006C>T), GGG1588CGG (6007G>C) | | | | | | | | |

*: Inserts / Deletes / Misaligned / Frameshifts

#### Analysis details

This analysis was performed with panviral2.64

### NGS Details (UN59): Alphabaculovirus lydisparis

#### Assembly

| | |
|---|---|
| **Coverage Length** | 192 (1 contig(s)) |
| **Depth Of Coverage** | 4.3 |
| **Number Of Reads** | 8 |
| **Reads Per Million** | 0.17 rpm (after QC) |
| **Ambiguities** | 0 |
| **Assembly Method** | de novo + reference guided assembly |
| **Consensus Caller** | Bcf Tools |

#### Coverage Map

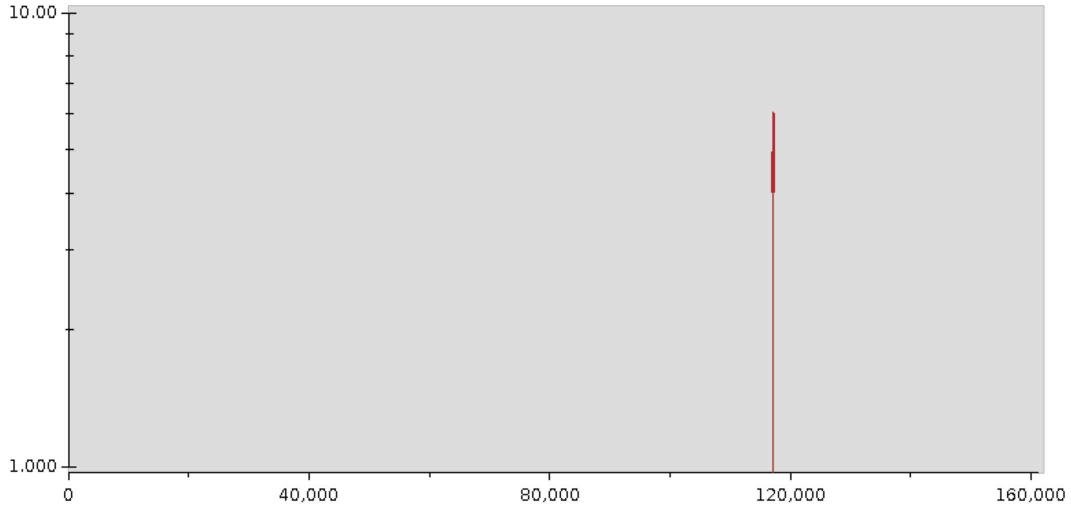

#### Assignment

| | |
|---|---|
| **Type** | Alphabaculovirus lydisparis (Taxonomy ID: 3048188) |
| **Reference Genome** | NC_001973.1 |
| **NT Identity (%)** | 78.6458 |
| **AA Identity (%)** | 89.0625 |
| **Number Of Stop Codons** | 0 |
| **Number Of CDS** | 164 |

#### Alignment

| | |
|---|---|
| **Alignment Score** | 220.0 (NT) + 423.0 (AA) = 643.0 |
| **Concordance (%)** | 75.915 |
| **Alignment Method** | Local, heuristic, nucleotide (BLASTN) |

#### Genome Region

Sequence starts at position 117029 and ends at position 117220 relative to NC_001973.1 reference sequence.

#### Alignment Detailed Statistics

| | Begin | End | Coverage | Score | Concordance | Matches | Identities | I/D/M/F* | Stop Codons |
|---|---|---|---|---|---|---|---|---|---|
| **NT** | **117029** | **117220** | **0.1%** | **220** | **57.3%** | **192 (100%)** | **151 (78.6%)** | **0/0** | |
| Mutations: | 117044G>A, 117046G>T, 117049C>G, 117055G>C, 117068G>A, 117070C>T, 117076T>C, 117083A>T, 117084G>C, 117085T>G, 117088C>T, 117091C>G, 117092G>T, 117097A>C, 117112A>G, 117130C>A, 117131C>T, 117136T>G, 117139C>T, 117140A>T, 117141G>C, 117142C>G, 117157C>G, 117158A>C, 117160G>C, 117169T>G, 117175T>C, 117176T>A, 117177G>C, 117187C>G, 117190T>C, 117191T>C, 117199C>T, 117201A>C, 117205T>C, 117206T>C, 117209G>C, 117210T>G, 117212C>A, 117213A>G, 117217A>G | | | | | | | | |

*: Inserts / Deletes / Misaligned / Frameshifts

#### Analysis details

This analysis was performed with panviral2.64

### NGS Details (UN59): Trialeurodes vaporariorum mononega-like virus 2

#### Assembly

| | |
|---|---|
| **Coverage Length** | 268 (2 contig(s)) |
| **Depth Of Coverage** | 4.0 |
| **Number Of Reads** | 8 |
| **Reads Per Million** | 0.17 rpm (after QC) |
| **Ambiguities** | 0 |
| **Assembly Method** | de novo + reference guided assembly |
| **Consensus Caller** | Bcf Tools |

#### Coverage Map

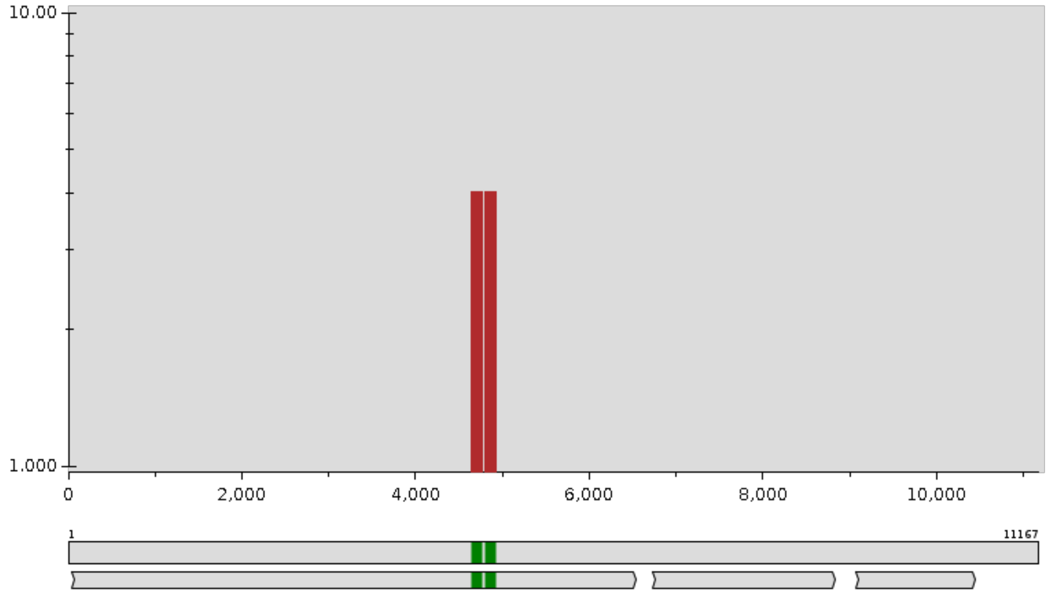

#### Assignment

| | |
|---|---|
| **Type** | Trialeurodes vaporariorum mononega-like virus 2 (Taxonomy ID: 2973797) |
| **Reference Genome** | NC_076992.1 |
| **NT Identity (%)** | 99.6269 |
| **AA Identity (%)** | 97.7528 |
| **Number Of Stop Codons** | 0 |
| **Number Of CDS** | 3 |

#### Alignment

| | |
|---|---|
| **Alignment Score** | 532.0 (NT) + 602.0 (AA) = 1134.0 |
| **Concordance (%)** | 98.7805 |

| Alignment Method | Global, seeded, nucleotide + amino acids (AGA) |
|---|---|

#### Genome Region

Sequence starts at position 4641 and ends at position 4933 relative to NC_076992.1 reference sequence.

#### Alignment Detailed Statistics

| | Begin | End | Coverage | Score | Concordance | Matches | Identities | I/D/M/F * | Stop Codons |
|---|---|---|---|---|---|---|---|---|---|
| **NT** | **4641** | **4933** | **2.4%** | **532** | **99.3%** | **268 (100%)** | **267 (99.6%)** | **0/0** | |
| Mutations: | 4763C>T | | | | | | | | |
| **CDS** | | | | | | | | | |
| QKT73_gp1 | 1537 | 1633 | 4.1% | 602 | 96.6% | 89 (100%) | 87 (97.8%) | 0/0/0/0 | 0 |
| Protein mutations: | R1577C (4763C>T) | | | | | | | | |
| Codon mutations: | CGT1577TGT (4763C>T) | | | | | | | | |
| **Proteins** | | | | | | | | | |
| RNA-dependent RNA polymerase (YP_010802337.1) | 1537 | 1633 | 4.1% | 602 | 96.6% | 89 (100%) | 87 (97.8%) | 0/0/0/0 | 0 |
| Protein mutations: | R1577C (4763C>T) | | | | | | | | |
| Codon mutations: | CGT1577TGT (4763C>T) | | | | | | | | |

*: Inserts / Deletes / Misaligned / Frameshifts

#### Analysis details

This analysis was performed with panviral2.64

### NGS Details (UN59): Caulimovirus venafragariae

#### Assembly

| | |
|---|---|
| **Coverage Length** | 396 (2 contig(s)) |
| **Depth Of Coverage** | 2.2 |
| **Number Of Reads** | 7 |
| **Reads Per Million** | 0.15 rpm (after QC) |
| **Ambiguities** | 0 |
| **Assembly Method** | de novo + reference guided assembly |
| **Consensus Caller** | Bcf Tools |

#### Coverage Map

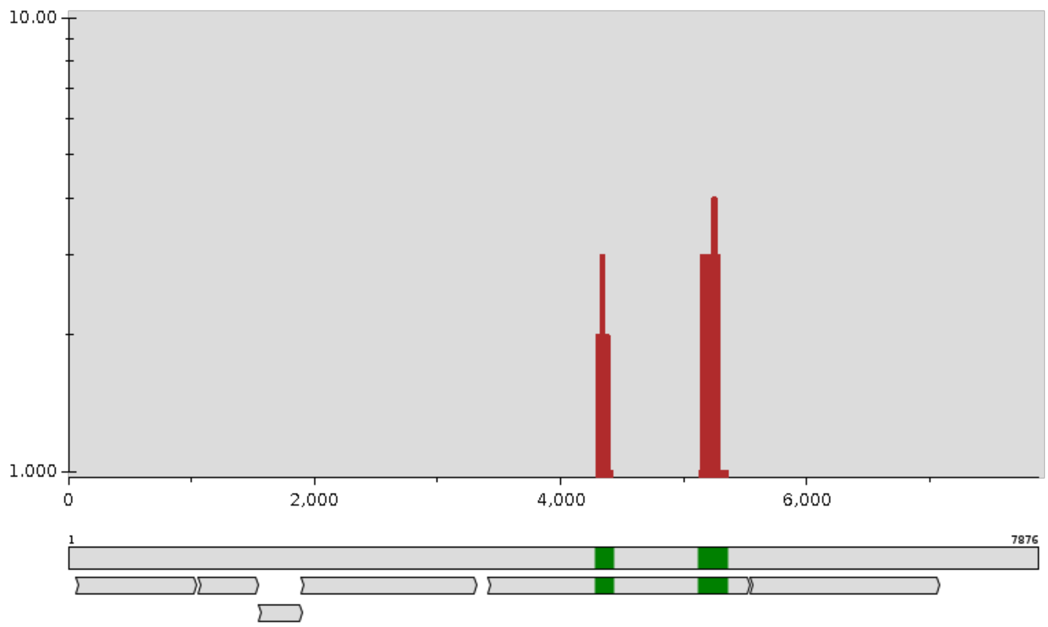

#### Assignment

| | |
|---|---|
| **Type** | Caulimovirus venafragariae (Taxonomy ID: 3048344) |
| **Reference Genome** | NC_001725.1 |
| **NT Identity (%)** | 65.1042 |
| **AA Identity (%)** | 60.9375 |
| **Number Of Stop Codons** | 1 |
| **Number Of CDS** | 6 |

#### Alignment

| | |
|---|---|
| **Alignment Score** | 211.0 (NT) + 577.0 (AA) = 788.0 |
| **Concordance (%)** | 49.4354 |

| Alignment Method | Global, seeded, nucleotide + amino acids (AGA) |
|---|---|

#### Genome Region

Sequence starts at position 4281 and ends at position 5363 relative to NC_001725.1 reference sequence.

#### Alignment Detailed Statistics

| | Begin | End | Coverage | Score | Concordance | Matches | Identities | I/D/M/F* | Stop Codons |
|---|---|---|---|---|---|---|---|---|---|
| **NT** | **4281** | **5363** | **5.0%** | **211** | **28.2%** | **384 (97.0%)** | **250 (63.1%)** | **0/12** | |
| Mutations: | 4281A>T, 4284G>A, 4287T>C, 4289A>G, 4292C>T, 4299C>T, 4300G>C, 4302A>G, 4308A>T, 4314G>A, 4315C>A, 4317T>G, 4318G>A, 4319G>A, 4320T>G, 4321A>T, 4323C>A, 4330C>G, 4332T>A, 4337A>G, 4338G>A, 4347T>C, 4348T>A, 4349C>A, 4351T>A, 4352C>G, 4353A>T, 4356A>C, 4359C>G, 4371G>A, 4374T>C, 4380C>G, 4383G>A, 4384A>C, 4385T>A, 4386C>A, 4395C>T, 4398A>G, 4401A>T, 4402A>C, 4410A>C, 4413T>A, 4416C>T, 4423A>G, 4426T>C, 4428A>T, 5126C>A, 5133G>A, 5139G>C, 5141C>T, 5142T>G, 5143A>G, 5145C>A, 5146A>T, 5148C>A, 5149G>C, 5152T>A, 5153G>C, 5154C>A, 5160A>C, 5163A>C, 5166T>C, 5167G>T, 5182A>T, 5184C>G, 5185C>T, 5187C>A, 5188A>C, 5191G>A, 5194A>G, 5196A>T, 5197C>T, 5199T>G, 5200C>A, 5201C>A, 5202A>T, 5206_5217delGGTAAAGAGGTA, 5220A>T, 5223C>T, 5226G>A, 5230G>A, 5233T>A, 5234C>G, 5235A>T, 5241C>T, 5247A>T, 5248C>G, 5249C>A, 5253T>A, 5256G>A, 5258A>C, 5259G>T, 5262T>A, 5265C>T, 5266C>T, 5267A>C, 5270G>C, 5271T>C, 5284A>T, 5286C>A, 5287C>T, 5289T>A, 5290T>G, 5292C>A, 5296A>G, 5299A>C, 5300A>G, 5301G>A, 5303C>G, 5309A>G, 5311G>A, 5312C>A, 5313T>A, 5317A>T, 5318G>C, 5320G>T, 5324A>T, 5325T>C, 5326A>C, 5328C>A, 5334T>A, 5335T>A, 5337T>A, 5338A>C, 5343T>C, 5344C>A, 5346T>C, 5347G>A, 5349A>C, 5350A>C, 5351G>A, 5352G>A, 5353A>T, 5355T>A, 5358T>C | | | | | | | | |

##### CDS

| | Begin | End | Coverage | Score | Concordance | Matches | Identities | I/D/M/F* | Stop Codons |
|---|---|---|---|---|---|---|---|---|---|
| ORF_V | 293 | 653 | 18.6% | 577 | 65.9% | 128 (97.0%) | 78 (59.1%) | 0/4/0/0 | 1 |
| Protein mutations: | K295R (4289A>G), T296I (4292C>T), E299Q (4300G>C 4302A>G), L304M (4315C>A 4317T>G), G305K (4318G>A 4319G>A 4320T>G), I306L (4321A>T 4323C>A), P309A (4330C>G 4332T>A), K311R (4337A>G 4338G>A), S315N (4348T>A 4349C>A), I327Q (4384A>C 4385T>A 4386C>A), K340E (4423A>G), S574* (5126C>A), K578N (5139G>C), P579L (5141C>T 5142T>G), I580V (5143A>G 5145C>A), I581L (5146A>T 5148C>A), E582Q (5149G>C), C583T (5152T>A 5153G>C 5154C>A), D588Y (5167G>T), I593L (5182A>T 5184C>G), K595Q (5188A>C), A596T (5191G>A), K597D (5194A>G 5196A>T), P599N (5200C>A 5201C>A 5202A>T), G601_V604del (5206_5217delGGTAAAGAGGTA), A609T (5230G>A), K614N (5247A>T), P615E (5248C>G 5249C>A), K618T (5258A>C 5259G>T), N619K (5262T>A), H621S (5266C>T 5267A>C), S622T (5270G>C 5271T>C), I627L (5284A>T 5286C>A), S629A (5290T>G 5292C>A), I631V (5296A>G), K632R (5299A>C 5300A>G 5301G>A), A633G (5303C>G), K635R (5309A>G), A636K (5311G>A 5312C>A 5313T>A), R638S (5317A>T 5318G>C), A639S (5320G>T), Y640F (5324A>T 5325T>C), I641L (5326A>C 5328C>A), Y644K (5335T>A 5337T>A), K645Q (5338A>C), L647I (5344C>A 5346T>C), V648I (5347G>A 5349A>C), R649Q (5350A>C 5351G>A 5352G>A), T650S (5353A>T 5355T>A) | | | | | | | | |
| Codon mutations: | GAA292..T (4281A>T), GAG293GAA (4284G>A), TTT294TTC (4287T>C), AAA295AGA (4289A>G), ACT296ATT (4292C>T), ATC298ATT (4299C>T), GAA299CAG (4300G>C 4302A>G), CTA301CTT (4308A>T), AAG303AAA (4314G>A), CTT304ATG (4315C>A 4317T>G), GGT305AAG (4318G>A 4319G>A 4320T>G), ATC306TTA (4321A>T 4323C>A), CCT309GCA (4330C>G 4332T>A), AAG311AGA (4337A>G 4338G>A), CAT314CAC (4347T>C), TCC315AAC (4348T>A 4349C>A), TCA316AGT (4351T>A 4352C>G 4353A>T), CCA317CCC (4356A>C), GCC318GCG (4359C>G), AGG322AGA (4371G>A), AAT323AAC (4374T>C), GCC325GCG (4380C>G), GAG326GAA (4383G>A), ATC327CAA (4384A>C 4385T>A 4386C>A), GGC330GGT (4395C>T), AAA331AAG (4398A>G), GCA332GCT (4401A>T), AGA333CGA (4402A>C), GTA335GTC (4410A>C), ATT336ATA (4413T>A), AAC337AAT (4416C>T), AAG340GAG (4423A>G), TTA341CTT (4426T>C 4428A>T), TCA574TAA (5126C>A), GAG576GAA (5133G>A), AAG578AAC (5139G>C), CCT579CTG (5141C>T 5142T>G), ATC580GTA (5143A>G 5145C>A), ATC581TTA (5146A>T 5148C>A), GAA582CAA (5149G>C), TGC583ACA (5152T>A 5153G>C 5154C>A), GCA585GCC (5160A>C), TCA586TCC (5163A>C), GAT587GAC (5166T>C), GAC588TAC (5167G>T), ATC593TTG (5182A>T 5184C>G), CTC594TTA (5185C>T 5187C>A), AAA595CAA (5188A>C), GCA596ACA (5191G>A), AAA597GAT (5194A>G 5196A>T), CTT598TTG (5197C>T 5199T>G), CCA599AAT (5200C>A 5201C>A 5202A>T), GGT601_GTA604del (5206_5217delGGTAAAGAGGTA), ATA605ATT (5220A>T), TGC606TGT (5223C>T), AGG607AGA (5226G>A), GCT609ACT (5230G>A), TCA610AGT (5233T>A 5234C>G 5235A>T), ACC612ACT (5241C>T), AAA614AAT (5247A>T), CCA615GAA (5248C>G 5249C>A), GCT616GCA (5253T>A), GAG617GAA (5256G>A), AAG618ACT (5258A>C 5259G>T), AAT619AAA (5262T>A), TAC620TAT (5265C>T), CAT621TCT (5266C>T 5267A>C), AGT622ACC (5270G>C 5271T>C), ATC627TTA (5284A>T 5286C>A), CTT628TTA (5287C>T 5289T>A), TCC629GCA (5290T>G 5292C>A), ATT631GTT (5296A>G), AAG632CGA (5299A>C 5300A>G 5301G>A), GCA633GGA (5303C>G), AAA635AGA (5309A>G), GCT636AAA (5311G>A 5312C>A 5313T>A), AGA638TCA (5317A>T 5318G>C), GCT639TCT (5320G>T), TAT640TTC (5324A>T 5325T>C), ATC641CTA (5326A>C 5328C>A), CCT643CCA (5334T>A), TAT644AAA (5335T>A 5337T>A), AAA645CAA (5338A>C), TTT646TTC (5343T>C), CTT647ATC (5344C>A 5346T>C), GTA648ATC (5347G>A 5349A>C), AGG649CAA (5350A>C 5351G>A 5352G>A), ACT650TCA (5353A>T 5355T>A), GAT651GAC (5358T>C) | | | | | | | | |

##### Proteins

| | Begin | End | Coverage | Score | Concordance | Matches | Identities | I/D/M/F* | Stop Codons |
|---|---|---|---|---|---|---|---|---|---|
| hypothetical protein (NP_043933.1) | 293 | 653 | 18.6% | 577 | 65.9% | 128 (97.0%) | 78 (59.1%) | 0/4/0/0 | 1 |
| Protein mutations: | K295R (4289A>G), T296I (4292C>T), E299Q (4300G>C 4302A>G), L304M (4315C>A 4317T>G), G305K (4318G>A 4319G>A 4320T>G), I306L (4321A>T 4323C>A), P309A (4330C>G 4332T>A), K311R (4337A>G 4338G>A), S315N (4348T>A 4349C>A), I327Q (4384A>C 4385T>A 4386C>A), K340E (4423A>G), S574* (5126C>A), K578N (5139G>C), P579L (5141C>T 5142T>G), I580V (5143A>G 5145C>A), I581L (5146A>T 5148C>A), E582Q (5149G>C), C583T (5152T>A 5153G>C 5154C>A), D588Y (5167G>T), I593L (5182A>T 5184C>G), K595Q (5188A>C), A596T (5191G>A), K597D (5194A>G 5196A>T), P599N (5200C>A 5201C>A 5202A>T), G601_V604del (5206_5217delGGTAAAGAGGTA), A609T (5230G>A), K614N (5247A>T), P615E (5248C>G 5249C>A), K618T (5258A>C 5259G>T), N619K (5262T>A), H621S (5266C>T 5267A>C), S622T (5270G>C 5271T>C), I627L (5284A>T 5286C>A), S629A (5290T>G 5292C>A), I631V (5296A>G), K632R (5299A>C 5300A>G 5301G>A), A633G (5303C>G), K635R (5309A>G), A636K (5311G>A 5312C>A 5313T>A), R638S (5317A>T 5318G>C), A639S (5320G>T), Y640F (5324A>T 5325T>C), I641L (5326A>C 5328C>A), Y644K (5335T>A 5337T>A), K645Q (5338A>C), L647I (5344C>A 5346T>C), V648I (5347G>A 5349A>C), R649Q (5350A>C 5351G>A 5352G>A), T650S (5353A>T 5355T>A) | | | | | | | | |
| Codon mutations: | GAA292..T (4281A>T), GAG293GAA (4284G>A), TTT294TTC (4287T>C), AAA295AGA (4289A>G), ACT296ATT (4292C>T), ATC298ATT (4299C>T), GAA299CAG (4300G>C 4302A>G), CTA301CTT (4308A>T), AAG303AAA (4314G>A), CTT304ATG (4315C>A 4317T>G), GGT305AAG (4318G>A 4319G>A 4320T>G), ATC306TTA (4321A>T 4323C>A), CCT309GCA (4330C>G 4332T>A), AAG311AGA (4337A>G 4338G>A), CAT314CAC (4347T>C), TCC315AAC (4348T>A 4349C>A), TCA316AGT (4351T>A 4352C>G 4353A>T), CCA317CCC (4356A>C), GCC318GCG (4359C>G), AGG322AGA (4371G>A), AAT323AAC (4374T>C), GCC325GCG (4380C>G), GAG326GAA (4383G>A), ATC327CAA (4384A>C 4385T>A 4386C>A), GGC330GGT (4395C>T), AAA331AAG (4398A>G), GCA332GCT (4401A>T), AGA333CGA (4402A>C), GTA335GTC (4410A>C), ATT336ATA (4413T>A), AAC337AAT (4416C>T), AAG340GAG (4423A>G), TTA341CTT (4426T>C 4428A>T), TCA574TAA (5126C>A), GAG576GAA (5133G>A), AAG578AAC (5139G>C), CCT579CTG (5141C>T 5142T>G), ATC580GTA (5143A>G 5145C>A), ATC581TTA (5146A>T 5148C>A), GAA582CAA (5149G>C), TGC583ACA (5152T>A 5153G>C 5154C>A), GCA585GCC (5160A>C), TCA586TCC (5163A>C), GAT587GAC (5166T>C), GAC588TAC (5167G>T), ATC593TTG (5182A>T 5184C>G), CTC594TTA (5185C>T 5187C>A), AAA595CAA (5188A>C), GCA596ACA (5191G>A), AAA597GAT (5194A>G 5196A>T), CTT598TTG (5197C>T 5199T>G), CCA599AAT (5200C>A 5201C>A 5202A>T), GGT601_GTA604del (5206_5217delGGTAAAGAGGTA), ATA605ATT (5220A>T), TGC606TGT (5223C>T), AGG607AGA (5226G>A), GCT609ACT (5230G>A), TCA610AGT (5233T>A 5234C>G 5235A>T), ACC612ACT (5241C>T), AAA614AAT (5247A>T), CCA615GAA (5248C>G 5249C>A), GCT616GCA (5253T>A), GAG617GAA (5256G>A), AAG618ACT (5258A>C 5259G>T), AAT619AAA (5262T>A), TAC620TAT (5265C>T), CAT621TCT (5266C>T 5267A>C), AGT622ACC (5270G>C 5271T>C), ATC627TTA (5284A>T 5286C>A), CTT628TTA (5287C>T 5289T>A), TCC629GCA (5290T>G 5292C>A), ATT631GTT (5296A>G), AAG632CGA (5299A>C 5300A>G 5301G>A), GCA633GGA (5303C>G), AAA635AGA (5309A>G), GCT636AAA (5311G>A 5312C>A 5313T>A), AGA638TCA (5317A>T 5318G>C), GCT639TCT (5320G>T), TAT640TTC (5324A>T 5325T>C), ATC641CTA (5326A>C 5328C>A), CCT643CCA (5334T>A), TAT644AAA (5335T>A 5337T>A), AAA645CAA (5338A>C), TTT646TTC (5343T>C), CTT647ATC (5344C>A 5346T>C), GTA648ATC (5347G>A 5349A>C), AGG649CAA (5350A>C 5351G>A 5352G>A), ACT650TCA (5353A>T 5355T>A), GAT651GAC (5358T>C) | | | | | | | | |

*: Inserts / Deletes / Misaligned / Frameshifts

#### Analysis details

This analysis was performed with panviral2.64

### NGS Details (UN59): Burkholderia phage Magia

#### Assembly

| | |
|---|---|
| **Coverage Length** | 197 (1 contig(s)) |
| **Depth Of Coverage** | 2.6 |
| **Number Of Reads** | 4 |
| **Reads Per Million** | 0.09 rpm (after QC) |
| **Ambiguities** | 0 |
| **Assembly Method** | de novo + reference guided assembly |
| **Consensus Caller** | Bcf Tools |

#### Coverage Map

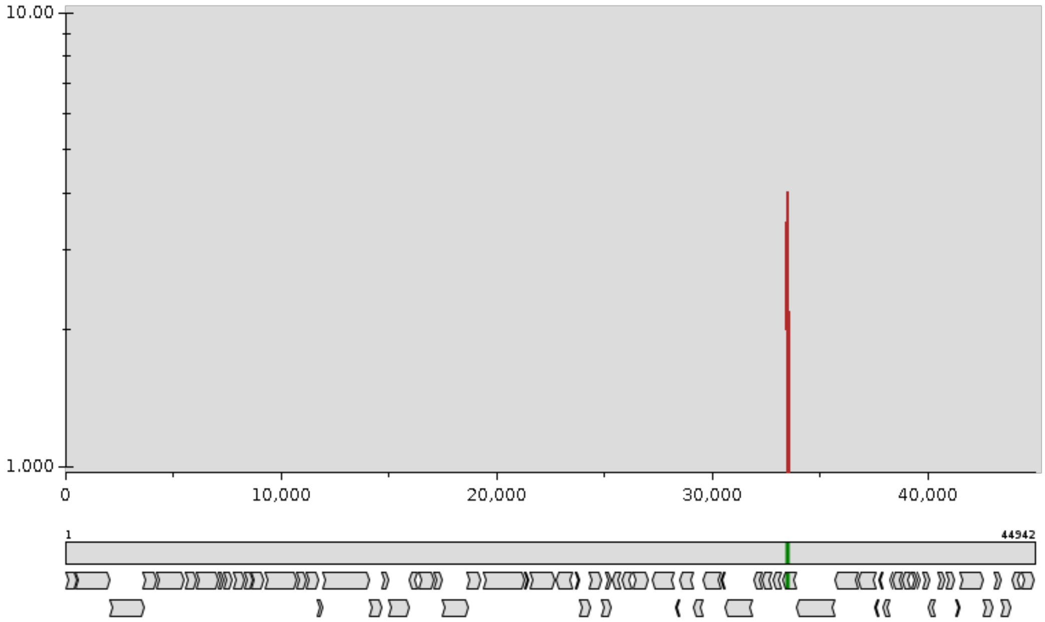

#### Assignment

| | |
|---|---|
| **Type** | Burkholderia phage Magia (Taxonomy ID: 2767577) |
| **Reference Genome** | NC_070954.1 |
| **NT Identity (%)** | 86.802 |
| **AA Identity (%)** | 84.8485 |
| **Number Of Stop Codons** | 0 |
| **Number Of CDS** | 70 |

#### Alignment

| | |
|---|---|
| **Alignment Score** | 290.0 (NT) + 381.0 (AA) = 671.0 |
| **Concordance (%)** | 82.2304 |

| Alignment Method | Local, heuristic, nucleotide (BLASTN) |
|---|---|

#### Genome Region

Sequence starts at position 33387 and ends at position 33583 relative to NC_070954.1 reference sequence.

#### Alignment Detailed Statistics

| | Begin | End | Coverage | Score | Concordance | Matches | Identities | I/D/M/F* | Stop Codons |
|---|---|---|---|---|---|---|---|---|---|
| **NT** | **33387** | **33583** | **0.4%** | **290** | **73.6%** | **197 (100%)** | **171 (86.8%)** | **0/0** | |
| Mutations: | 33394T>G, 33398T>C, 33404T>G, 33407G>T, 33415T>C, 33416C>G, 33422A>G, 33434A>G, 33437G>T, 33464A>G, 33473G>C, 33476T>C, 33482C>T, 33483G>A, 33509C>T, 33518T>C, 33527G>A, 33536T>G, 33538C>T, 33544T>G, 33545C>G, 33551A>G, 33567T>C, 33569T>G, 33570G>C, 33579C>T | | | | | | | | |
| **CDS** | | | | | | | | | |
| PQC04_gp48 | 102 | 167 | 31.7% | 381 | 87.2% | 66 (100%) | 56 (84.8%) | 0/0/0/0 | 0 |
| Protein mutations: | R103Q (33579C>T), A106G (33569T>G 33570G>C), D107G (33567T>C), I115L (33544T>G), A117T (33536T>G 33538C>T), T135I (33482C>T 33483G>A), I158V (33415T>C), K161N (33404T>G), K165Q (33394T>G) | | | | | | | | |
| Codon mutations: | CGG103CAG (33579C>T), GCA106GGC (33569T>G 33570G>C), GAC107GGC (33567T>C), GTT112GTC (33551A>G), CGG114CGC (33545C>G), ATC115CTC (33544T>G), GCA117ACC (33536T>G 33538C>T), GAC120GAT (33527G>A), CAA123CAG (33518T>C), GCG126GCA (33509C>T), ACG135ATA (33482C>T 33483G>A), GAA137GAG (33476T>C), GCC138GCG (33473G>C), GAT141GAC (33464A>G), GCC150GCA (33437G>T), CGT151CGC (33434A>G), TTT155TTC (33422A>G), ACG157ACC (33416C>G), ATC158GTC (33415T>C), CGC160CGA (33407G>T), AAA161AAC (33404T>G), AAA163AAG (33398T>C), AAG165CAG (33394T>G) | | | | | | | | |
| **Proteins** | | | | | | | | | |
| transcriptional regulator (YP_010668144.1) | 102 | 167 | 31.7% | 381 | 87.2% | 66 (100%) | 56 (84.8%) | 0/0/0/0 | 0 |
| Protein mutations: | R103Q (33579C>T), A106G (33569T>G 33570G>C), D107G (33567T>C), I115L (33544T>G), A117T (33536T>G 33538C>T), T135I (33482C>T 33483G>A), I158V (33415T>C), K161N (33404T>G), K165Q (33394T>G) | | | | | | | | |
| Codon mutations: | CGG103CAG (33579C>T), GCA106GGC (33569T>G 33570G>C), GAC107GGC (33567T>C), GTT112GTC (33551A>G), CGG114CGC (33545C>G), ATC115CTC (33544T>G), GCA117ACC (33536T>G 33538C>T), GAC120GAT (33527G>A), CAA123CAG (33518T>C), GCG126GCA (33509C>T), ACG135ATA (33482C>T 33483G>A), GAA137GAG (33476T>C), GCC138GCG (33473G>C), GAT141GAC (33464A>G), GCC150GCA (33437G>T), CGT151CGC (33434A>G), TTT155TTC (33422A>G), ACG157ACC (33416C>G), ATC158GTC (33415T>C), CGC160CGA (33407G>T), AAA161AAC (33404T>G), AAA163AAG (33398T>C), AAG165CAG (33394T>G) | | | | | | | | |

*: Inserts / Deletes / Misaligned / Frameshifts

#### Analysis details

This analysis was performed with panviral2.64

### NGS Details (UN59): Sugarcane chlorotic streak virus

#### Assembly

| | |
|---|---|
| **Coverage Length** | 200 (1 contig(s)) |
| **Depth Of Coverage** | 1.6 |
| **Number Of Reads** | 3 |
| **Reads Per Million** | 0.07 rpm (after QC) |
| **Ambiguities** | 0 |
| **Assembly Method** | de novo + reference guided assembly |
| **Consensus Caller** | Bcf Tools |

#### Coverage Map

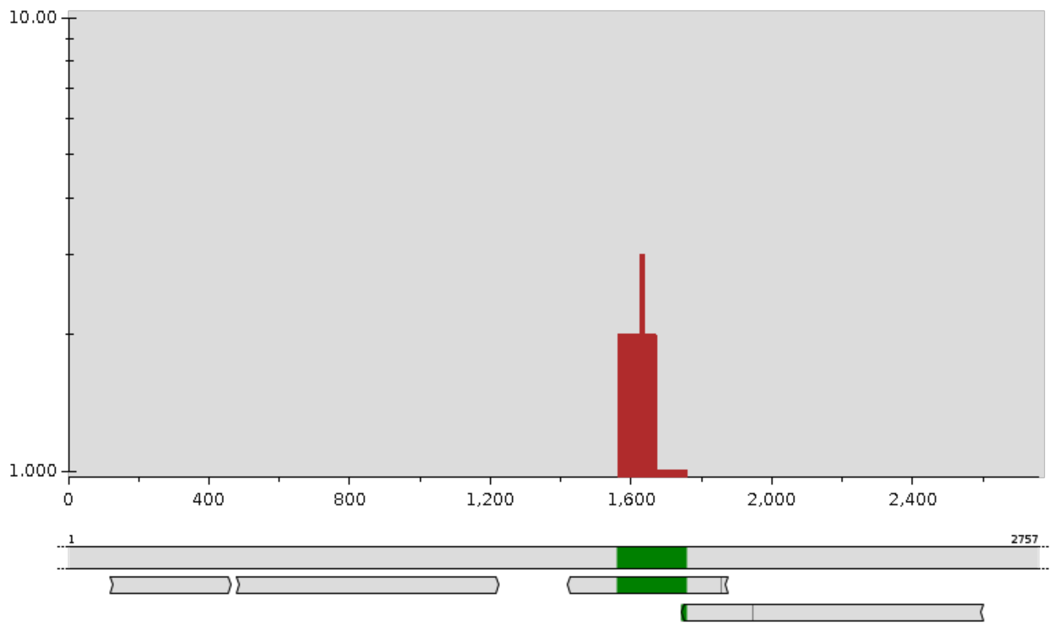

#### Assignment

| | |
|---|---|
| **Type** | Sugarcane chlorotic streak virus (Taxonomy ID: 1919062) |
| **Reference Genome** | NC_032004.1 |
| **NT Identity (%)** | 59.8985 |
| **AA Identity (%)** | 56.2044 |
| **Number Of Stop Codons** | 1 |
| **Number Of CDS** | 5 |

#### Alignment

| | |
|---|---|
| **Alignment Score** | 66.0 (NT) + 560.0 (AA) = 626.0 |
| **Concordance (%)** | 45.3623 |

| Alignment Method | Global, seeded, nucleotide + amino acids (AGA) |
|---|---|

#### Genome Region

Sequence starts at position 1562 and ends at position 1761 relative to NC_032004.1 reference sequence.

#### Alignment Detailed Statistics

| | Begin | End | Coverage | Score | Concordance | Matches | Identities | I/D/M/F* | Stop Codons |
|---|---|---|---|---|---|---|---|---|---|
| **NT** | **1562** | **1761** | **7.3%** | **66** | **17.3%** | **197 (98.5%)** | **118 (59.0%)** | **0/3** | |
| Mutations: | 1562C>T, 1566C>A, 1567T>G, 1573G>C, 1574C>A, 1578C>A, 1584A>C, 1585G>C, 1587T>C, 1593G>T, 1595T>C, 1596T>C, 1598T>C, 1599T>C, 1600G>T, 1601A>T, 1602_1604delTGC, 1605G>A, 1607C>T, 1609T>C, 1611C>T, 1614C>T, 1615C>T, 1617T>C, 1620G>T, 1622C>T, 1626C>T, 1629C>T, 1630G>T, 1631G>C, 1632G>A, 1638A>T, 1639A>G, 1641G>A, 1644A>G, 1645T>C, 1647C>G, 1648T>A, 1649T>A, 1650C>T, 1656G>A, 1659A>T, 1661T>A, 1662T>C, 1664A>G, 1665C>T, 1667G>C, 1672C>T, 1673A>T, 1675C>T, 1676A>T, 1677A>T, 1680A>T, 1681C>A, 1683G>A, 1684T>A, 1689G>A, 1692A>C, 1695T>G, 1701A>G, 1704A>T, 1710G>A, 1713G>A, 1714A>T, 1715C>A, 1716G>C, 1718C>A, 1721C>T, 1722C>G, 1723T>A, 1724C>T, 1725T>A, 1734A>G, 1735G>T, 1736A>T, 1737G>A, 1738G>C, 1740C>G, 1741C>A, 1744G>T | | | | | | | | |

##### CDS

| | Begin | End | Coverage | Score | Concordance | Matches | Identities | I/D/M/F* | Stop Codons |
|---|---|---|---|---|---|---|---|---|---|
| BS327_gp3 | 252 | 318 | 18.4% | 268 | 53.3% | 66 (98.5%) | 36 (53.7%) | 0/1/0/0 | 0 |
| Protein mutations: | A257D (1744G>T), W258F (1740C>G 1741C>A), T259S (1737G>A 1738G>C), S260N (1734A>G 1735G>T 1736A>T), E263D (1725T>A), E264I (1722C>G 1723T>A 1724C>T), A265T (1721C>T), V266L (1716G>C 1718C>A), V267Y (1713G>A 1714A>T 1715C>A), Y277F (1683G>A 1684T>A), C278L (1680A>T 1681C>A), C280N (1675C>T 1676A>T), W281K (1672C>T 1673A>T), Q283E (1665C>T 1667G>C), I285L (1659A>T 1661T>A), K289F (1647C>G 1648T>A 1649T>A), D290G (1644A>G 1645T>C), I292T (1638A>T 1639A>G), P295E (1629C>T 1630G>T 1631G>C), G298R (1620G>T 1622C>T), R300K (1614C>T 1615C>T), K302R (1609T>C), V303I (1605G>A 1607C>T), A304del (1602_1604delTGC), S305K (1599T>C 1600G>T 1601A>T), R306G (1596T>C 1598T>C), S307G (1593G>T 1595T>C), T310R (1584A>C 1585G>C), A314C (1573G>C 1574C>A), E316A (1566C>A 1567T>G) | | | | | | | | |
| Codon mutations: | GCC257GAC (1744G>T), TGG258TTC (1740C>G 1741C>A), ACC259AGT (1737G>A 1738G>C), TCT260AAC (1734A>G 1735G>T 1736A>T), GAA263GAT (1725T>A), GAG264ATC (1722C>G 1723T>A 1724C>T), GCA265ACA (1721C>T), GTC266TTG (1716G>C 1718C>A), GTC267TAT (1713G>A 1714A>T 1715C>A), AAC268AAT (1710G>A), ATT270ATA (1704A>T), GAT271GAC (1701A>G), ATA273ATC (1695T>G), CCT274CCG (1692A>C), TTC275TTT (1689G>A), TAC277TTT (1683G>A 1684T>A), TGT278TTA (1680A>T 1681C>A), CCT279CCA (1677A>T), TGC280AAC (1675C>T 1676A>T), TGG281AAG (1672C>T 1673A>T), CAG283GAA (1665C>T 1667G>C), TTA284CTG (1662T>C 1664A>G), ATT285TTA (1659A>T 1661T>A), GGC286GGT (1656G>A), CAG288CAA (1650C>T), AAG289TTC (1647C>G 1648T>A 1649T>A), GAT290GGC (1644A>G 1645T>C), TAC291TAT (1641G>A), ATT292ACA (1638A>T 1639A>G), AAC294AAT (1632G>A), CCG295GAA (1629C>T 1630G>T 1631G>C), AAG296AAA (1626C>T), GGC298AGA (1620G>T 1622C>T), AAA299AAG (1617T>C), AGG300AAA (1614C>T 1615C>T), AAG301AAA (1611C>T), AAG302AGG (1609T>C), GTC303ATT (1605G>A 1607C>T), GCA304del (1602_1604delTGC), TCA305AAG (1599T>C 1600G>T 1601A>T), AGA306GGG (1596T>C 1598T>C), AGC307GGA (1593G>T 1595T>C), TCA309TCG (1587T>C), ACT310AGG (1584A>C 1585G>C), GTG312GTT (1578C>A), GCC314TGC (1573G>C 1574C>A), GAG316GCT (1566C>A 1567T>G), GAG318A.. (1562C>T) | | | | | | | | |
| BS327_gp4 | 40 | 106 | 44.1% | 268 | 53.3% | 66 (98.5%) | 36 (53.7%) | 0/1/0/0 | 0 |
| Protein mutations: | A45D (1744G>T), W46F (1740C>G 1741C>A), T47S (1737G>A 1738G>C), S48N (1734A>G 1735G>T 1736A>T), E51D (1725T>A), E52I (1722C>G 1723T>A 1724C>T), A53T (1721C>T), V54L (1716G>C 1718C>A), V55Y (1713G>A 1714A>T 1715C>A), Y65F (1683G>A 1684T>A), C66L (1680A>T 1681C>A), C68N (1675C>T 1676A>T), W69K (1672C>T 1673A>T), Q71E (1665C>T 1667G>C), I73L (1659A>T 1661T>A), K77F (1647C>G 1648T>A 1649T>A), D78G (1644A>G 1645T>C), I80T (1638A>T 1639A>G), P83E (1629C>T 1630G>T 1631G>C), G86R (1620G>T 1622C>T), R88K (1614C>T 1615C>T), K90R (1609T>C), V91I (1605G>A 1607C>T), A92del (1602_1604delTGC), S93K (1599T>C 1600G>T 1601A>T), R94G (1596T>C 1598T>C), S95G (1593G>T 1595T>C), T98R (1584A>C 1585G>C), A102C (1573G>C 1574C>A), E104A (1566C>A 1567T>G) | | | | | | | | |
| Codon mutations: | GCC45GAC (1744G>T), TGG46TTC (1740C>G 1741C>A), ACC47AGT (1737G>A 1738G>C), TCT48AAC (1734A>G 1735G>T 1736A>T), GAA51GAT (1725T>A), GAG52ATC (1722C>G 1723T>A 1724C>T), GCA53ACA (1721C>T), GTC54TTG (1716G>C 1718C>A), GTC55TAT (1713G>A 1714A>T 1715C>A), AAC56AAT (1710G>A), ATT58ATA (1704A>T), GAT59GAC (1701A>G), ATA61ATC (1695T>G), CCT62CCG (1692A>C), TTC63TTT (1689G>A), TAC65TTT (1683G>A 1684T>A), TGT66TTA (1680A>T 1681C>A), CCT67CCA (1677A>T), TGC68AAC (1675C>T 1676A>T), TGG69AAG (1672C>T 1673A>T), CAG71GAA (1665C>T 1667G>C), TTA72CTG (1662T>C 1664A>G), ATT73TTA (1659A>T 1661T>A), GGC74GGT (1656G>A), CAG76CAA (1650C>T), AAG77TTC (1647C>G 1648T>A 1649T>A), GAT78GGC (1644A>G 1645T>C), TAC79TAT (1641G>A), ATT80ACA (1638A>T 1639A>G), AAC82AAT (1632G>A), CCG83GAA (1629C>T 1630G>T 1631G>C), AAG84AAA (1626C>T), GGC86AGA (1620G>T 1622C>T), AAA87AAG (1617T>C), AGG88AAA (1614C>T 1615C>T), AAG89AAA (1611C>T), AAG90AGG (1609T>C), GTC91ATT (1605G>A 1607C>T), GCA92del (1602_1604delTGC), TCA93AAG (1599T>C 1600G>T 1601A>T), AGA94GGG (1596T>C 1598T>C), AGC95GGA (1593G>T 1595T>C), TCA97TCG (1587T>C), ACT98AGG (1584A>C 1585G>C), GTG100GTT (1578C>A), GCC102TGC (1573G>C 1574C>A), GAG104GCT (1566C>A 1567T>G), GAG106A.. (1562C>T) | | | | | | | | |
| BS327_gp5 | 283 | 287 | 1.7% | 24 | 82.8% | 5 (100%) | 5 (100%) | 0/0/0/0 | 1 |

##### Proteins

| | Begin | End | Coverage | Score | Concordance | Matches | Identities | I/D/M/F* | Stop Codons |
|---|---|---|---|---|---|---|---|---|---|
| Rep-associated protein (YP_009325925.1) | 252 | 318 | 18.4% | 268 | 53.3% | 66 (98.5%) | 36 (53.7%) | 0/1/0/0 | 0 |
| Protein mutations: | A257D (1744G>T), W258F (1740C>G 1741C>A), T259S (1737G>A 1738G>C), S260N (1734A>G 1735G>T 1736A>T), E263D (1725T>A), E264I (1722C>G 1723T>A 1724C>T), A265T (1721C>T), V266L (1716G>C 1718C>A), V267Y (1713G>A 1714A>T 1715C>A), Y277F (1683G>A 1684T>A), C278L (1680A>T 1681C>A), C280N (1675C>T 1676A>T), W281K (1672C>T 1673A>T), Q283E (1665C>T 1667G>C), I285L (1659A>T 1661T>A), K289F (1647C>G 1648T>A 1649T>A), D290G (1644A>G 1645T>C), I292T (1638A>T 1639A>G), P295E (1629C>T 1630G>T 1631G>C), G298R (1620G>T 1622C>T), R300K (1614C>T 1615C>T), K302R (1609T>C), V303I (1605G>A 1607C>T), A304del (1602_1604delTGC), S305K (1599T>C 1600G>T 1601A>T), R306G (1596T>C 1598T>C), S307G (1593G>T 1595T>C), T310R (1584A>C 1585G>C), A314C (1573G>C 1574C>A), E316A (1566C>A 1567T>G) | | | | | | | | |
| Codon mutations: | GCC257GAC (1744G>T), TGG258TTC (1740C>G 1741C>A), ACC259AGT (1737G>A 1738G>C), TCT260AAC (1734A>G 1735G>T 1736A>T), GAA263GAT (1725T>A), GAG264ATC (1722C>G 1723T>A 1724C>T), GCA265ACA (1721C>T), GTC266TTG (1716G>C 1718C>A), GTC267TAT (1713G>A 1714A>T 1715C>A), AAC268AAT (1710G>A), ATT270ATA (1704A>T), GAT271GAC (1701A>G), ATA273ATC (1695T>G), CCT274CCG (1692A>C), TTC275TTT (1689G>A), TAC277TTT (1683G>A 1684T>A), TGT278TTA (1680A>T 1681C>A), CCT279CCA (1677A>T), TGC280AAC (1675C>T 1676A>T), TGG281AAG (1672C>T 1673A>T), CAG283GAA (1665C>T 1667G>C), TTA284CTG (1662T>C 1664A>G), ATT285TTA (1659A>T 1661T>A), GGC286GGT (1656G>A), CAG288CAA (1650C>T), AAG289TTC (1647C>G 1648T>A 1649T>A), GAT290GGC (1644A>G 1645T>C), TAC291TAT (1641G>A), ATT292ACA (1638A>T 1639A>G), AAC294AAT (1632G>A), CCG295GAA (1629C>T 1630G>T 1631G>C), AAG296AAA (1626C>T), GGC298AGA (1620G>T 1622C>T), AAA299AAG (1617T>C), AGG300AAA (1614C>T 1615C>T), AAG301AAA (1611C>T), AAG302AGG (1609T>C), GTC303ATT (1605G>A 1607C>T), GCA304del (1602_1604delTGC), TCA305AAG (1599T>C 1600G>T 1601A>T), AGA306GGG (1596T>C 1598T>C), AGC307GGA (1593G>T 1595T>C), TCA309TCG (1587T>C), ACT310AGG (1584A>C 1585G>C), GTG312GTT (1578C>A), GCC314TGC (1573G>C 1574C>A), GAG316GCT (1566C>A 1567T>G), GAG318A.. (1562C>T) | | | | | | | | |
| RepB (YP_009325926.1) | 40 | 106 | 44.1% | 268 | 53.3% | 66 (98.5%) | 36 (53.7%) | 0/1/0/0 | 0 |
| Protein mutations: | A45D (1744G>T), W46F (1740C>G 1741C>A), T47S (1737G>A 1738G>C), S48N (1734A>G 1735G>T 1736A>T), E51D (1725T>A), E52I (1722C>G 1723T>A 1724C>T), A53T (1721C>T), V54L (1716G>C 1718C>A), V55Y (1713G>A 1714A>T 1715C>A), Y65F (1683G>A 1684T>A), C66L (1680A>T 1681C>A), C68N (1675C>T 1676A>T), W69K (1672C>T 1673A>T), Q71E (1665C>T 1667G>C), I73L (1659A>T 1661T>A), K77F (1647C>G 1648T>A 1649T>A), D78G (1644A>G 1645T>C), I80T (1638A>T 1639A>G), P83E (1629C>T 1630G>T 1631G>C), G86R (1620G>T 1622C>T), R88K (1614C>T 1615C>T), K90R (1609T>C), V91I (1605G>A 1607C>T), A92del (1602_1604delTGC), S93K (1599T>C 1600G>T 1601A>T), R94G (1596T>C 1598T>C), S95G (1593G>T 1595T>C), T98R (1584A>C 1585G>C), A102C (1573G>C 1574C>A), E104A (1566C>A 1567T>G) | | | | | | | | |
| Codon mutations: | GCC45GAC (1744G>T), TGG46TTC (1740C>G 1741C>A), ACC47AGT (1737G>A 1738G>C), TCT48AAC (1734A>G 1735G>T 1736A>T), GAA51GAT (1725T>A), GAG52ATC (1722C>G 1723T>A 1724C>T), GCA53ACA (1721C>T), GTC54TTG (1716G>C 1718C>A), GTC55TAT (1713G>A 1714A>T 1715C>A), AAC56AAT (1710G>A), ATT58ATA (1704A>T), GAT59GAC (1701A>G), ATA61ATC (1695T>G), CCT62CCG (1692A>C), TTC63TTT (1689G>A), TAC65TTT (1683G>A 1684T>A), TGT66TTA (1680A>T 1681C>A), CCT67CCA (1677A>T), TGC68AAC (1675C>T 1676A>T), TGG69AAG (1672C>T 1673A>T), CAG71GAA (1665C>T 1667G>C), TTA72CTG (1662T>C 1664A>G), ATT73TTA (1659A>T 1661T>A), GGC74GGT (1656G>A), CAG76CAA (1650C>T), AAG77TTC (1647C>G 1648T>A 1649T>A), GAT78GGC (1644A>G 1645T>C), TAC79TAT (1641G>A), ATT80ACA (1638A>T 1639A>G), AAC82AAT (1632G>A), CCG83GAA (1629C>T 1630G>T 1631G>C), AAG84AAA (1626C>T), GGC86AGA (1620G>T 1622C>T), AAA87AAG (1617T>C), AGG88AAA (1614C>T 1615C>T), AAG89AAA (1611C>T), AAG90AGG (1609T>C), GTC91ATT (1605G>A 1607C>T), GCA92del (1602_1604delTGC), TCA93AAG (1599T>C 1600G>T 1601A>T), AGA94GGG (1596T>C 1598T>C), AGC95GGA (1593G>T 1595T>C), TCA97TCG (1587T>C), ACT98AGG (1584A>C 1585G>C), GTG100GTT (1578C>A), GCC102TGC (1573G>C 1574C>A), GAG104GCT (1566C>A 1567T>G), GAG106A.. (1562C>T) | | | | | | | | |
| RepA (YP_009325927.1) | 283 | 287 | 1.7% | 24 | 82.8% | 5 (100%) | 5 (100%) | 0/0/0/0 | 1 |
| Protein mutations: | *none* | | | | | | | | |
| Codon mutations: | *none* | | | | | | | | |

*: Inserts / Deletes / Misaligned / Frameshifts

#### Analysis details

This analysis was performed with panviral2.64

### NGS Details (UN59): Alphafusarivirus pleosporae

#### Assembly

| | |
|---|---|
| **Coverage Length** | 254 (1 contig(s)) |
| **Depth Of Coverage** | 1.1 |
| **Number Of Reads** | 2 |
| **Reads Per Million** | 0.04 rpm (after QC) |
| **Ambiguities** | 0 |
| **Assembly Method** | de novo + reference guided assembly |
| **Consensus Caller** | Bcf Tools |

#### Coverage Map

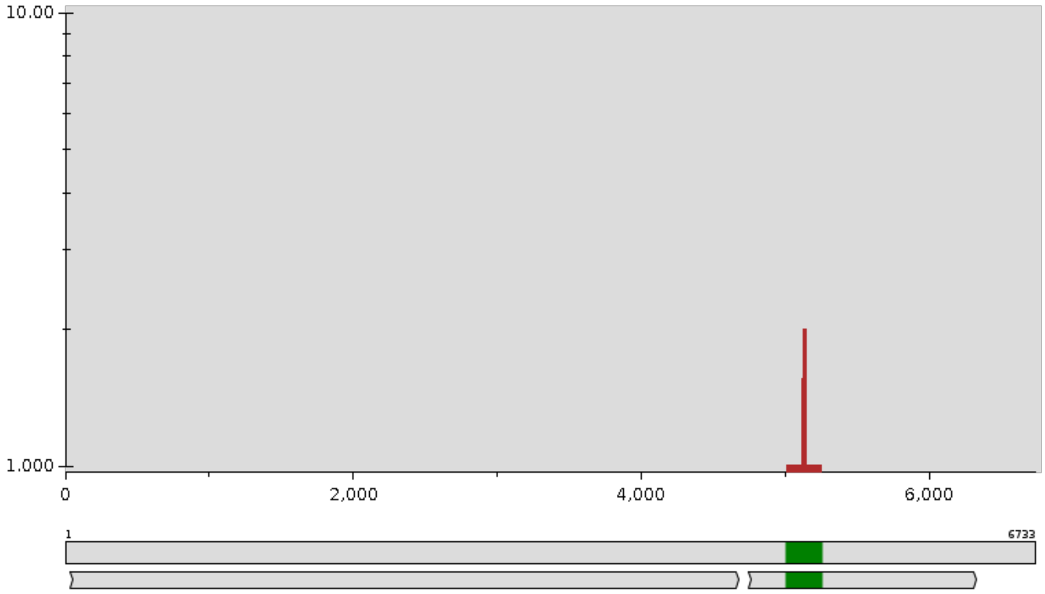

#### Assignment

| | |
|---|---|
| **Type** | Alphafusarivirus pleosporae (Taxonomy ID: 2955271) |
| **Reference Genome** | NC_028470.1 |
| **NT Identity (%)** | 75.1969 |
| **AA Identity (%)** | 85.7143 |
| **Number Of Stop Codons** | 0 |
| **Number Of CDS** | 2 |

#### Alignment

| | |
|---|---|
| **Alignment Score** | 256.0 (NT) + 417.0 (AA) = 673.0 |
| **Concordance (%)** | 68.4639 |

| Alignment Method | Global, seeded, nucleotide + amino acids (AGA) |
|---|---|

#### Genome Region

Sequence starts at position 5007 and ends at position 5260 relative to NC_028470.1 reference sequence.

#### Alignment Detailed Statistics

| | Begin | End | Coverage | Score | Concordance | Matches | Identities | I/D/M/F* | Stop Codons |
|---|---|---|---|---|---|---|---|---|---|
| **NT** | **5007** | **5260** | **3.8%** | **256** | **50.4%** | **254 (100%)** | **191 (75.2%)** | **0/0** | |
| Mutations: | 5011C>T, 5014T>A, 5027T>C, 5029G>T, 5035T>A, 5039T>C, 5041G>T, 5048C>T, 5050T>G, 5051C>A, 5053T>G, 5059A>G, 5062A>C, 5074A>G, 5075A>T, 5077C>T, 5095G>A, 5101T>C, 5104G>A, 5105G>C, 5106C>A, 5107C>A, 5110C>T, 5116A>T, 5119G>A, 5123T>C, 5125G>C, 5128T>C, 5134A>C, 5140A>C, 5146A>G, 5154C>T, 5155A>T, 5158A>T, 5159A>G, 5165T>G, 5168C>G, 5173T>A, 5174G>A, 5176T>C, 5177C>T, 5179G>A, 5182C>T, 5188T>C, 5189A>T, 5190G>C, 5191C>T, 5192A>G, 5193A>G, 5197G>T, 5200A>G, 5206G>A, 5207C>T, 5209C>G, 5210A>C, 5211A>G, 5224A>T, 5225A>T, 5226A>C, 5230T>C, 5236T>C, 5239T>C, 5244G>A | | | | | | | | |

##### CDS

| | Begin | End | Coverage | Score | Concordance | Matches | Identities | I/D/M/F* | Stop Codons |
|---|---|---|---|---|---|---|---|---|---|
| AT135_gp2 | 90 | 173 | 15.9% | 417 | 85.8% | 84 (100%) | 72 (85.7%) | 0/0/0/0 | 0 |
| Protein mutations: | T112S (5075A>T 5077C>T), A122Q (5105G>C 5106C>A 5107C>A), A138V (5154C>T 5155A>T), I140V (5159A>G), S142A (5165T>G), Q143E (5168C>G), D144E (5173T>A), G145S (5174G>A 5176T>C), N151G (5192A>G 5193A>G), K157R (5210A>C 5211A>G), K162S (5225A>T 5226A>C), R168K (5244G>A) | | | | | | | | |
| Codon mutations: | GTC90GTT (5011C>T), TCT91TCA (5014T>A), TTG96CTT (5027T>C 5029G>T), CGT98CGA (5035T>A), TTG100CTT (5039T>C 5041G>T), CTT103TTG (5048C>T 5050T>G), CGT104AGG (5051C>A 5053T>G), AAA106AAG (5059A>G), GCA107GCC (5062A>C), AAA111AAG (5074A>G), ACC112TCT (5075A>T 5077C>T), GAG118GAA (5095G>A), GCT120GCC (5101T>C), CAG121CAA (5104G>A), GCC122CAA (5105G>C 5106C>A 5107C>A), AAC123AAT (5110C>T), ACA125ACT (5116A>T), AAG126AAA (5119G>A), TTG128CTC (5123T>C 5125G>C), AAT129AAC (5128T>C), GCA131GCC (5134A>C), GCA133GCC (5140A>C), GAA135GAG (5146A>G), GCA138GTT (5154C>T 5155A>T), CGA139CGT (5158A>T), ATT140GTT (5159A>G), TCT142GCT (5165T>G), CAG143GAG (5168C>G), GAT144GAA (5173T>A), GGT145AGC (5174G>A 5176T>C), CTG146TTA (5177C>T 5179G>A), GCC147GCT (5182C>T), CTT149CTC (5188T>C), AGC150TCT (5189A>T 5190G>C 5191C>T), AAC151GGC (5192A>G 5193A>G), ACG152ACT (5197G>T), CAA153CAG (5200A>G), GAG155GAA (5206G>A), CTC156TTG (5207C>T 5209C>G), AAG157CGG (5210A>C 5211A>G), GCA161GCT (5224A>T), AAA162TCA (5225A>T 5226A>C), GTT163GTC (5230T>C), GCT165GCC (5236T>C), TCT166TCC (5239T>C), AGG168AAG (5244G>A) | | | | | | | | |

##### Proteins

| | Begin | End | Coverage | Score | Concordance | Matches | Identities | I/D/M/F* | Stop Codons |
|---|---|---|---|---|---|---|---|---|---|
| 59 kDa protein (YP_009182159.1) | 90 | 173 | 15.9% | 417 | 85.8% | 84 (100%) | 72 (85.7%) | 0/0/0/0 | 0 |
| Protein mutations: | T112S (5075A>T 5077C>T), A122Q (5105G>C 5106C>A 5107C>A), A138V (5154C>T 5155A>T), I140V (5159A>G), S142A (5165T>G), Q143E (5168C>G), D144E (5173T>A), G145S (5174G>A 5176T>C), N151G (5192A>G 5193A>G), K157R (5210A>C 5211A>G), K162S (5225A>T 5226A>C), R168K (5244G>A) | | | | | | | | |
| Codon mutations: | GTC90GTT (5011C>T), TCT91TCA (5014T>A), TTG96CTT (5027T>C 5029G>T), CGT98CGA (5035T>A), TTG100CTT (5039T>C 5041G>T), CTT103TTG (5048C>T 5050T>G), CGT104AGG (5051C>A 5053T>G), AAA106AAG (5059A>G), GCA107GCC (5062A>C), AAA111AAG (5074A>G), ACC112TCT (5075A>T 5077C>T), GAG118GAA (5095G>A), GCT120GCC (5101T>C), CAG121CAA (5104G>A), GCC122CAA (5105G>C 5106C>A 5107C>A), AAC123AAT (5110C>T), ACA125ACT (5116A>T), AAG126AAA (5119G>A), TTG128CTC (5123T>C 5125G>C), AAT129AAC (5128T>C), GCA131GCC (5134A>C), GCA133GCC (5140A>C), GAA135GAG (5146A>G), GCA138GTT (5154C>T 5155A>T), CGA139CGT (5158A>T), ATT140GTT (5159A>G), TCT142GCT (5165T>G), CAG143GAG (5168C>G), GAT144GAA (5173T>A), GGT145AGC (5174G>A 5176T>C), CTG146TTA (5177C>T 5179G>A), GCC147GCT (5182C>T), CTT149CTC (5188T>C), AGC150TCT (5189A>T 5190G>C 5191C>T), AAC151GGC (5192A>G 5193A>G), ACG152ACT (5197G>T), CAA153CAG (5200A>G), GAG155GAA (5206G>A), CTC156TTG (5207C>T 5209C>G), AAG157CGG (5210A>C 5211A>G), GCA161GCT (5224A>T), AAA162TCA (5225A>T 5226A>C), GTT163GTC (5230T>C), GCT165GCC (5236T>C), TCT166TCC (5239T>C), AGG168AAG (5244G>A) | | | | | | | | |

*: Inserts / Deletes / Misaligned / Frameshifts

#### Analysis details

This analysis was performed with panviral2.64

### NGS Details (UN59): Potato virus X

#### Assembly

| | |
|---|---|
| **Coverage Length** | 225 (1 contig(s)) |
| **Depth Of Coverage** | 1.2 |
| **Number Of Reads** | 2 |
| **Reads Per Million** | 0.04 rpm (after QC) |
| **Ambiguities** | 0 |
| **Assembly Method** | de novo + reference guided assembly |
| **Consensus Caller** | Bcf Tools |

#### Coverage Map

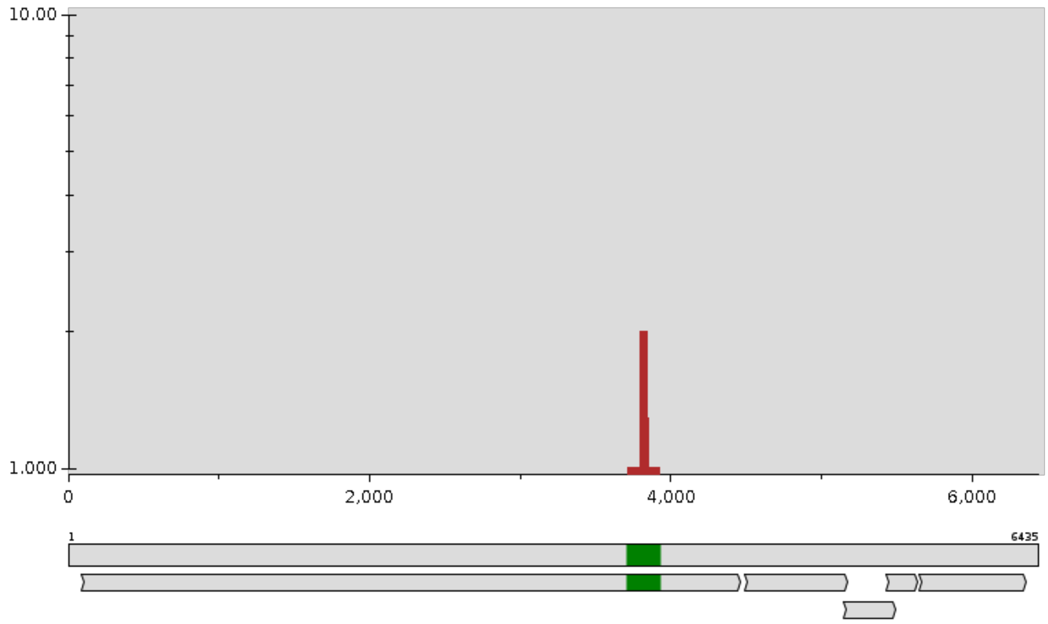

#### Assignment

| | |
|---|---|
| **Type** | Potato virus X (Taxonomy ID: 12183) |
| **Reference Genome** | NC_011620.1 |
| **NT Identity (%)** | 79.1111 |
| **AA Identity (%)** | 97.3333 |
| **Number Of Stop Codons** | 0 |
| **Number Of CDS** | 5 |

#### Alignment

| | |
|---|---|
| **Alignment Score** | 262.0 (NT) + 544.0 (AA) = 806.0 |
| **Concordance (%)** | 80.7615 |

| Alignment Method | Global, seeded, nucleotide + amino acids (AGA) |
|---|---|

#### Genome Region

Sequence starts at position 3708 and ends at position 3932 relative to NC_011620.1 reference sequence.

#### Alignment Detailed Statistics

| | Begin | End | Coverage | Score | Concordance | Matches | Identities | I/D/M/F* | Stop Codons |
|---|---|---|---|---|---|---|---|---|---|
| **NT** | **3708** | **3932** | **3.5%** | **262** | **58.2%** | **225 (100%)** | **178 (79.1%)** | **0/0** | |
| Mutations: | 3711G>A, 3714A>C, 3717A>G, 3723C>G, 3729A>C, 3735T>C, 3738G>A, 3741T>A, 3747A>T, 3750A>G, 3761T>C, 3768C>T, 3771G>A, 3783T>C, 3787A>T, 3788G>C, 3795T>C, 3799T>C, 3801A>G, 3804T>C, 3816A>T, 3819T>A, 3822C>T, 3825C>T, 3831T>A, 3834G>A, 3840A>T, 3843T>C, 3847C>T, 3849G>A, 3861G>A, 3864C>A, 3867A>G, 3873G>A, 3876C>T, 3879C>T, 3885A>C, 3894A>G, 3900C>A, 3903A>G, 3906A>G, 3912A>T, 3918T>C, 3921G>A, 3924C>T, 3927T>C, 3930A>C | | | | | | | | |

##### CDS

| | Begin | End | Coverage | Score | Concordance | Matches | Identities | I/D/M/F* | Stop Codons |
|---|---|---|---|---|---|---|---|---|---|
| PVX_gp1 | 1209 | 1283 | 5.1% | 544 | 96.1% | 75 (100%) | 73 (97.3%) | 0/0/0/0 | 0 |
| Protein mutations: | V1226A (3761T>C) | | | | | | | | |
| Codon mutations: | CAG1209CAA (3711G>A), CCA1210CCC (3714A>C), AAA1211AAG (3717A>G), GTC1213GTG (3723C>G), ATA1215ATC (3729A>C), TGT1217TGC (3735T>C), GAG1218GAA (3738G>A), ACT1219ACA (3741T>A), CCA1221CCT (3747A>T), GAA1222GAG (3750A>G), GTA1226GCA (3761T>C), GCC1228GCT (3768C>T), TTG1229TTA (3771G>A), AAT1233AAC (3783T>C), AGC1235TCC (3787A>T 3788G>C), CCT1237CCC (3795T>C), TTA1239CTG (3799T>C 3801A>G), GCT1240GCC (3804T>C), ACA1244ACT (3816A>T), GCT1245GCA (3819T>A), TTC1246TTT (3822C>T), GAC1247GAT (3825C>T), TCT1249TCA (3831T>A), CAG1250CAA (3834G>A), GGA1252GGT (3840A>T), GCT1253GCC (3843T>C), CTG1255TTA (3847C>T 3849G>A), GTG1259GTA (3861G>A), CTC1260CTA (3864C>A), AAA1261AAG (3867A>G), AAG1263AAA (3873G>A), CAC1264CAT (3876C>T), CAC1265CAT (3879C>T), ATA1267ATC (3885A>C), GAA1270GAG (3894A>G), ATC1272ATA (3900C>A), CAA1273CAG (3903A>G), GCA1274GCG (3906A>G), ATA1276ATT (3912A>T), ATT1278ATC (3918T>C), AAG1279AAA (3921G>A), ACC1280ACT (3924C>T), AAT1281AAC (3927T>C), GCA1282GCC (3930A>C) | | | | | | | | |

##### Proteins

| | Begin | End | Coverage | Score | Concordance | Matches | Identities | I/D/M/F* | Stop Codons |
|---|---|---|---|---|---|---|---|---|---|
| replicase (YP_002332929.1) | 1209 | 1283 | 5.1% | 544 | 96.1% | 75 (100%) | 73 (97.3%) | 0/0/0/0 | 0 |
| Protein mutations: | V1226A (3761T>C) | | | | | | | | |
| Codon mutations: | CAG1209CAA (3711G>A), CCA1210CCC (3714A>C), AAA1211AAG (3717A>G), GTC1213GTG (3723C>G), ATA1215ATC (3729A>C), TGT1217TGC (3735T>C), GAG1218GAA (3738G>A), ACT1219ACA (3741T>A), CCA1221CCT (3747A>T), GAA1222GAG (3750A>G), GTA1226GCA (3761T>C), GCC1228GCT (3768C>T), TTG1229TTA (3771G>A), AAT1233AAC (3783T>C), AGC1235TCC (3787A>T 3788G>C), CCT1237CCC (3795T>C), TTA1239CTG (3799T>C 3801A>G), GCT1240GCC (3804T>C), ACA1244ACT (3816A>T), GCT1245GCA (3819T>A), TTC1246TTT (3822C>T), GAC1247GAT (3825C>T), TCT1249TCA (3831T>A), CAG1250CAA (3834G>A), GGA1252GGT (3840A>T), GCT1253GCC (3843T>C), CTG1255TTA (3847C>T 3849G>A), GTG1259GTA (3861G>A), CTC1260CTA (3864C>A), AAA1261AAG (3867A>G), AAG1263AAA (3873G>A), CAC1264CAT (3876C>T), CAC1265CAT (3879C>T), ATA1267ATC (3885A>C), GAA1270GAG (3894A>G), ATC1272ATA (3900C>A), CAA1273CAG (3903A>G), GCA1274GCG (3906A>G), ATA1276ATT (3912A>T), ATT1278ATC (3918T>C), AAG1279AAA (3921G>A), ACC1280ACT (3924C>T), AAT1281AAC (3927T>C), GCA1282GCC (3930A>C) | | | | | | | | |

*: Inserts / Deletes / Misaligned / Frameshifts

#### Analysis details

This analysis was performed with panviral2.64

### NGS Details (UN59): Colombian datura virus

#### Assembly

| | |
|---|---|
| **Coverage Length** | 269 (2 contig(s)) |
| **Depth Of Coverage** | 1.0 |
| **Number Of Reads** | 2 |
| **Reads Per Million** | 0.04 rpm (after QC) |
| **Ambiguities** | 0 |
| **Assembly Method** | de novo + reference guided assembly |
| **Consensus Caller** | Bcf Tools |

#### Coverage Map

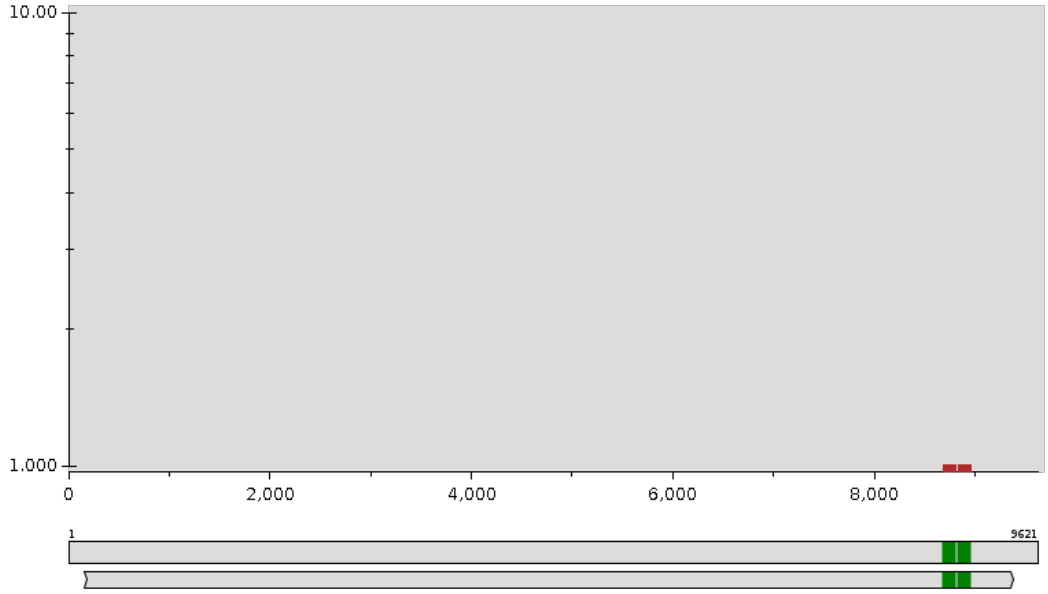

#### Assignment

| | |
|---|---|
| **Type** | Colombian datura virus (Taxonomy ID: 91613) |
| **Reference Genome** | NC_020072.1 |
| **NT Identity (%)** | 98.1413 |
| **AA Identity (%)** | 97.7778 |
| **Number Of Stop Codons** | 0 |
| **Number Of CDS** | 1 |

#### Alignment

| | |
|---|---|
| **Alignment Score** | 518.0 (NT) + 640.0 (AA) = 1158.0 |
| **Concordance (%)** | 98.3022 |

| Alignment Method | Global, seeded, nucleotide + amino acids (AGA) |
|---|---|

#### Genome Region

Sequence starts at position 8680 and ends at position 8961 relative to NC_020072.1 reference sequence.

#### Alignment Detailed Statistics

| | Begin | End | Coverage | Score | Concordance | Matches | Identities | I/D/M/F* | Stop Codons |
|---|---|---|---|---|---|---|---|---|---|
| **NT** | **8680** | **8961** | **2.8%** | **518** | **96.3%** | **269 (100%)** | **264 (98.1%)** | **0/0** | |
| Mutations: | 8761T>C, 8767A>G, 8788G>T, 8893T>C, 8914G>A | | | | | | | | |
| **CDS** | | | | | | | | | |
| G357_gp1 | 2843 | 2936 | 2.9% | 640 | 96.1% | 90 (100%) | 88 (97.8%) | 0/0/0/0 | 0 |
| **Proteins** | | | | | | | | | |
| polyprotein (YP_007346986.1) | 2843 | 2936 | 2.9% | 640 | 96.1% | 90 (100%) | 88 (97.8%) | 0/0/0/0 | 0 |
| Protein mutations: | *none* | | | | | | | | |
| Codon mutations: | AAT2869AAC (8761T>C), CAA2871CAG (8767A>G), CTG2878CTT (8788G>T), GAT2913GAC (8893T>C), TTG2920TTA (8914G>A) | | | | | | | | |

*: Inserts / Deletes / Misaligned / Frameshifts

#### Analysis details

This analysis was performed with panviral2.64

#### NGS Details (UN59): Rahariannevirus raharianne

##### Assembly

| | |
|---|---|
| **Coverage Length** | 251 (2 contig(s)) |
| **Depth Of Coverage** | 1.0 |
| **Number Of Reads** | 2 |
| **Reads Per Million** | 0.04 rpm (after QC) |
| **Ambiguities** | 0 |
| **Assembly Method** | de novo + reference guided assembly |
| **Consensus Caller** | Bcf Tools |

##### Coverage Map

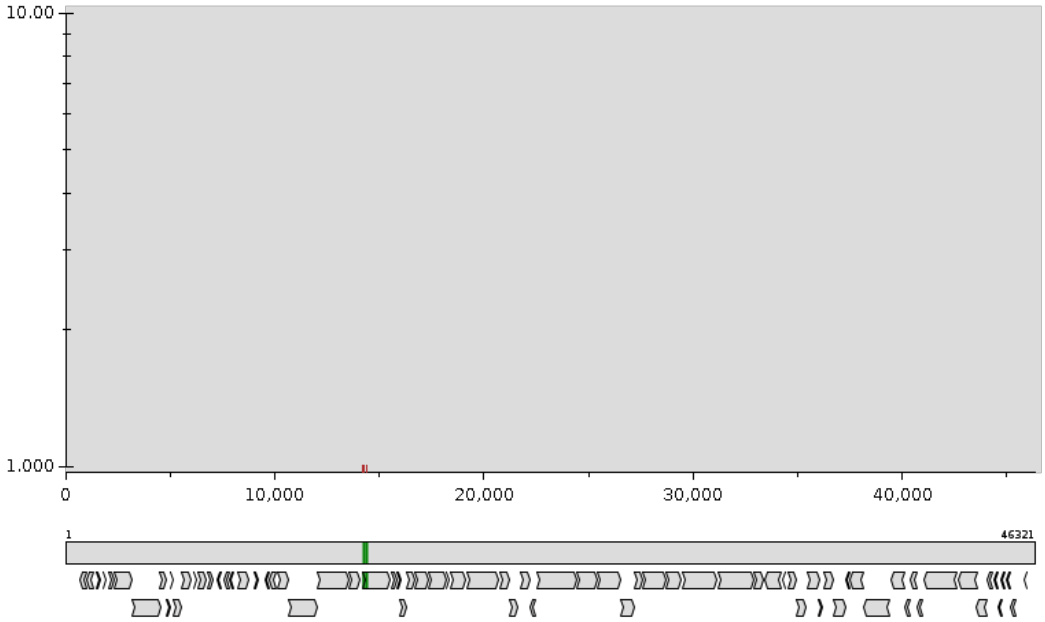

##### Assignment

| | |
|---|---|
| **Type** | Rahariannevirus raharianne (Taxonomy ID: 2846050) |
| **Reference Genome** | NC_054955.1 |
| **NT Identity (%)** | 82.0717 |
| **AA Identity (%)** | 82.1429 |
| **Number Of Stop Codons** | 0 |
| **Number Of CDS** | 75 |

##### Alignment

| | |
|---|---|
| **Alignment Score** | 322.0 (NT) + 482.0 (AA) = 804.0 |
| **Concordance (%)** | 76.1364 |

| Alignment Method | Local, heuristic, nucleotide (BLASTN) |
|---|---|

#### Genome Region

Sequence starts at position 14196 and ends at position 14467 relative to NC_054955.1 reference sequence.

#### Alignment Detailed Statistics

| | Begin | End | Coverage | Score | Concordance | Matches | Identities | I/D/M/F* | Stop Codons |
|---|---|---|---|---|---|---|---|---|---|
| **NT** | **14196** | **14467** | **0.5%** | **322** | **64.1%** | **251 (100%)** | **206 (82.1%)** | **0/0** | |
| Mutations: | 14204C>T, 14205A>C, 14206G>A, 14207C>A, 14210T>C, 14219C>T, 14222T>C, 14231G>C, 14264T>C, 14270T>A, 14272C>G, 14285C>G, 14288C>G, 14289G>A, 14293T>A, 14294C>T, 14298C>T, 14299A>C, 14306A>C, 14309T>C, 14336T>C, 14339A>G, 14342C>A, 14349A>C, 14357T>C, 14363T>C, 14366G>C, 14372G>T, 14375G>C, 14378C>G, 14381C>G, 14394C>G, 14397T>G, 14399G>C, 14400G>A, 14401C>G, 14402A>C, 14427T>A, 14432G>C, 14433T>A, 14434C>G, 14435G>C, 14447G>C, 14457T>A, 14462C>G | | | | | | | | |
| **CDS** | | | | | | | | | |
| KMC43_gp27 | 1 | 91 | 19.0% | 482 | 83.8% | 84 (100%) | 69 (82.1%) | 0/0/0/0 | 0 |
| Protein mutations: | S4Q (14205A>C 14206G>A 14207C>A), D25E (14270T>A), A26G (14272C>G), I30M (14285C>G), G32S (14289G>A), F33Y (14293T>A 14294C>T), Q35S (14298C>T 14299A>C), K52Q (14349A>C), P67A (14394C>G), S68A (14397T>G 14399G>C), A69S (14400G>A 14401C>G 14402A>C), S78T (14427T>A), S88T (14457T>A) | | | | | | | | |
| Codon mutations: | ATC3ATT (14204C>T), AGC4CAA (14205A>C 14206G>A 14207C>A), AAT5AAC (14210T>C), GCC8GCT (14219C>T), GCT9GCC (14222T>C), CCG12CCC (14231G>C), TTT23TTC (14264T>C), GAT25GAA (14270T>A), GCC26GGC (14272C>G), ATC30ATG (14285C>G), CTC31CTG (14288C>G), GGC32AGC (14289G>A), TTC33TAT (14293T>A 14294C>T), CAG35TCG (14298C>T 14299A>C), GCA37GCC (14306A>C), CGT38CGC (14309T>C), GGT47GGC (14336T>C), GAA48GAG (14339A>G), ACC49ACA (14342C>A), AAG52CAG (14349A>C), CGT54CGC (14357T>C), GGT56GGC (14363T>C), CTG57CTC (14366G>C), GCG59GCT (14372G>T), CCG60CCC (14375G>C), GTC61GTG (14378C>G), ACC62ACG (14381C>G), CCG67GCG (14394C>G), TCG68GCC (14397T>G 14399G>C), GCA69AGC (14400G>A 14401C>G 14402A>C), TCG78ACG (14427T>A), CCG79CCC (14432G>C), TCG80AGC (14433T>A 14434C>G 14435G>C), GTG84GTC (14447G>C), TCG88ACG (14457T>A), CTC89CTG (14462C>G) | | | | | | | | |
| **Proteins** | | | | | | | | | |
| DUF4043 domain-containing protein (YP_010078151.1) | 1 | 91 | 19.0% | 482 | 83.8% | 84 (100%) | 69 (82.1%) | 0/0/0/0 | 0 |
| Protein mutations: | S4Q (14205A>C 14206G>A 14207C>A), D25E (14270T>A), A26G (14272C>G), I30M (14285C>G), G32S (14289G>A), F33Y (14293T>A 14294C>T), Q35S (14298C>T 14299A>C), K52Q (14349A>C), P67A (14394C>G), S68A (14397T>G 14399G>C), A69S (14400G>A 14401C>G 14402A>C), S78T (14427T>A), S88T (14457T>A) | | | | | | | | |
| Codon mutations: | ATC3ATT (14204C>T), AGC4CAA (14205A>C 14206G>A 14207C>A), AAT5AAC (14210T>C), GCC8GCT (14219C>T), GCT9GCC (14222T>C), CCG12CCC (14231G>C), TTT23TTC (14264T>C), GAT25GAA (14270T>A), GCC26GGC (14272C>G), ATC30ATG (14285C>G), CTC31CTG (14288C>G), GGC32AGC (14289G>A), TTC33TAT (14293T>A 14294C>T), CAG35TCG (14298C>T 14299A>C), GCA37GCC (14306A>C), CGT38CGC (14309T>C), GGT47GGC (14336T>C), GAA48GAG (14339A>G), ACC49ACA (14342C>A), AAG52CAG (14349A>C), CGT54CGC (14357T>C), GGT56GGC (14363T>C), CTG57CTC (14366G>C), GCG59GCT (14372G>T), CCG60CCC (14375G>C), GTC61GTG (14378C>G), ACC62ACG (14381C>G), CCG67GCG (14394C>G), TCG68GCC (14397T>G 14399G>C), GCA69AGC (14400G>A 14401C>G 14402A>C), TCG78ACG (14427T>A), CCG79CCC (14432G>C), TCG80AGC (14433T>A 14434C>G 14435G>C), GTG84GTC (14447G>C), TCG88ACG (14457T>A), CTC89CTG (14462C>G) | | | | | | | | |

*: Inserts / Deletes / Misaligned / Frameshifts

#### Analysis details

This analysis was performed with panviral2.64

### NGS Details (UN59): Euphorbia caput-medusae latent virus

#### Assembly

| | |
|---|---|
| **Coverage Length** | 123 (1 contig(s)) |
| **Depth Of Coverage** | 1.0 |
| **Number Of Reads** | 1 |
| **Reads Per Million** | 0.02 rpm (after QC) |
| **Ambiguities** | 0 |
| **Assembly Method** | de novo + reference guided assembly |
| **Consensus Caller** | Bcf Tools |

#### Coverage Map

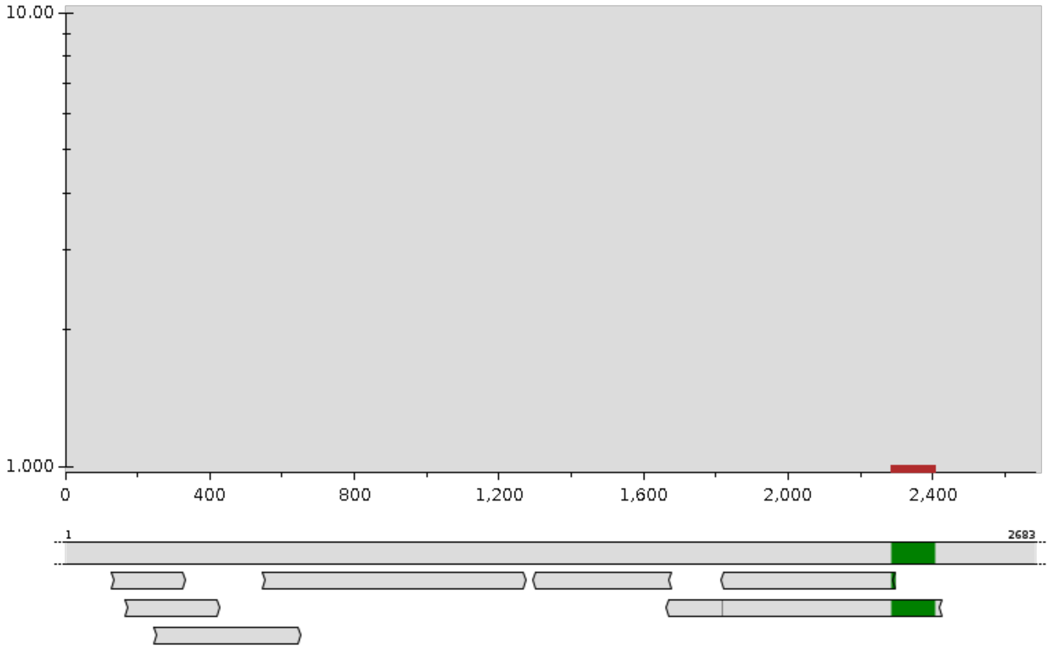

#### Assignment

| | |
|---|---|
| **Type** | Euphorbia caput-medusae latent virus (Taxonomy ID: 1853865) |
| **Reference Genome** | NC_030238.1 |
| **NT Identity (%)** | 64.2276 |
| **AA Identity (%)** | 66.2791 |
| **Number Of Stop Codons** | 0 |
| **Number Of CDS** | 7 |

#### Alignment

| | |
|---|---|
| **Alignment Score** | 70.0 (NT) + 438.0 (AA) = 508.0 |
| **Concordance (%)** | 59.4152 |

| **Alignment Method** | Global, seeded, nucleotide + amino acids (AGA) |
|---|---|

#### Genome Region

Sequence starts at position 2286 and ends at position 2408 relative to NC_030238.1 reference sequence.

#### Alignment Detailed Statistics

| | Begin | End | Coverage | Score | Concordance | Matches | Identities | I/D/M/F* | Stop Codons |
|---|---|---|---|---|---|---|---|---|---|
| **NT** | **2286** | **2408** | **4.6%** | **70** | **28.5%** | **123 (100%)** | **79 (64.2%)** | **0/0** | |
| Mutations: | 2286T>A, 2289C>T, 2293G>A, 2294C>T, 2298T>A, 2299G>T, 2307A>G, 2308A>T, 2312T>C, 2313A>T, 2315T>G, 2318A>G, 2319T>A, 2321G>T, 2324G>T, 2325A>T, 2327A>T, 2328T>G, 2330A>G, 2331A>G, 2332T>A, 2334A>G, 2337A>T, 2340T>G, 2342A>C, 2343A>G, 2346A>T, 2349A>T, 2351G>C, 2352A>G, 2354G>T, 2355A>T, 2364T>C, 2367T>G, 2376C>G, 2382A>T, 2384T>G, 2385A>T, 2388C>T, 2391A>T, 2397T>A, 2399A>G, 2400G>T, 2402G>T | | | | | | | | |

##### CDS

| | Begin | End | Coverage | Score | Concordance | Matches | Identities | I/D/M/F* | Stop Codons |
|---|---|---|---|---|---|---|---|---|---|
| A9K86_gp5 | 8 | 48 | 12.3% | 218 | 73.6% | 41 (100%) | 28 (68.3%) | 0/0/0/0 | 0 |
| Protein mutations: | I16L (2382A>T 2384T>G), L26I (2352A>G 2354G>T), P27A (2349A>T 2351G>C), L30V (2340T>G 2342A>C), Y33F (2331A>G 2332T>A), Y35K (2325A>T 2327A>T), Q36K (2324G>T), L37I (2319T>A 2321G>T), N39Q (2313A>T 2315T>G), N40D (2312T>C), F41Y (2307A>G 2308A>T), T44N (2298T>A 2299G>T), A46I (2293G>A 2294C>T) | | | | | | | | |
| Codon mutations: | CGC10AGA (2400G>T 2402G>T), TTA11CTT (2397T>A 2399A>G), GGT13GGA (2391A>T), AAG14AAA (2388C>T), TCT15TCA (2385A>T), ATT16CTA (2382A>T 2384T>G), CTG18CTC (2376C>G), CCA21CCC (2367T>G), AAA22AAG (2364T>C), CTT25CTA (2355A>T), CTT26ATC (2352A>G 2354G>T), CCT27GCA (2349A>T 2351G>C), ATT28ATA (2346A>T), TTT29TTC (2343A>G), TTA30GTC (2340T>G 2342A>C), ATT31ATA (2337A>T), GAT32GAC (2334A>G), TAT33TTC (2331A>G 2332T>A), TTA34CTC (2328T>G 2330A>G), TAT35AAA (2325A>T 2327A>T), CAA36AAA (2324G>T), CTA37ATT (2319T>A 2321G>T), TTA38CTA (2318A>G), AAT39CAA (2313A>T 2315T>G), AAT40GAT (2312T>C), TTT41TAC (2307A>G 2308A>T), ACA44AAT (2298T>A 2299G>T), GCT46ATT (2293G>A 2294C>T), AGG47AGA (2289C>T), GTA48GTT (2286T>A) | | | | | | | | |
| A9K86_gp6 | 8 | 48 | 16.1% | 218 | 73.6% | 41 (100%) | 28 (68.3%) | 0/0/0/0 | 0 |
| Protein mutations: | I16L (2382A>T 2384T>G), L26I (2352A>G 2354G>T), P27A (2349A>T 2351G>C), L30V (2340T>G 2342A>C), Y33F (2331A>G 2332T>A), Y35K (2325A>T 2327A>T), Q36K (2324G>T), L37I (2319T>A 2321G>T), N39Q (2313A>T 2315T>G), N40D (2312T>C), F41Y (2307A>G 2308A>T), T44N (2298T>A 2299G>T), A46I (2293G>A 2294C>T) | | | | | | | | |
| Codon mutations: | CGC10AGA (2400G>T 2402G>T), TTA11CTT (2397T>A 2399A>G), GGT13GGA (2391A>T), AAG14AAA (2388C>T), TCT15TCA (2385A>T), ATT16CTA (2382A>T 2384T>G), CTG18CTC (2376C>G), CCA21CCC (2367T>G), AAA22AAG (2364T>C), CTT25CTA (2355A>T), CTT26ATC (2352A>G 2354G>T), CCT27GCA (2349A>T 2351G>C), ATT28ATA (2346A>T), TTT29TTC (2343A>G), TTA30GTC (2340T>G 2342A>C), ATT31ATA (2337A>T), GAT32GAC (2334A>G), TAT33TTC (2331A>G 2332T>A), TTA34CTC (2328T>G 2330A>G), TAT35AAA (2325A>T 2327A>T), CAA36AAA (2324G>T), CTA37ATT (2319T>A 2321G>T), TTA38CTA (2318A>G), AAT39CAA (2313A>T 2315T>G), AAT40GAT (2312T>C), TTT41TAC (2307A>G 2308A>T), ACA44AAT (2298T>A 2299G>T), GCT46ATT (2293G>A 2294C>T), AGG47AGA (2289C>T), GTA48GTT (2286T>A) | | | | | | | | |
| A9K86_gp7 | 1 | 4 | 2.5% | 2 | 7.4% | 4 (100%) | 1 (25.0%) | 0/0/0/0 | 0 |
| Protein mutations: | M1I (2294C>T), G3E (2289C>T) | | | | | | | | |
| Codon mutations: | ATG1ATA (2294C>T), CTA2TTA (2293G>A), GGG3GAG (2289C>T), TAT4TT. (2286T>A) | | | | | | | | |

##### Proteins

| | Begin | End | Coverage | Score | Concordance | Matches | Identities | I/D/M/F* | Stop Codons |
|---|---|---|---|---|---|---|---|---|---|
| replication-associated protein (YP_009255247.1) | 8 | 48 | 12.3% | 218 | 73.6% | 41 (100%) | 28 (68.3%) | 0/0/0/0 | 0 |
| Protein mutations: | I16L (2382A>T 2384T>G), L26I (2352A>G 2354G>T), P27A (2349A>T 2351G>C), L30V (2340T>G 2342A>C), Y33F (2331A>G 2332T>A), Y35K (2325A>T 2327A>T), Q36K (2324G>T), L37I (2319T>A 2321G>T), N39Q (2313A>T 2315T>G), N40D (2312T>C), F41Y (2307A>G 2308A>T), T44N (2298T>A 2299G>T), A46I (2293G>A 2294C>T) | | | | | | | | |
| Codon mutations: | CGC10AGA (2400G>T 2402G>T), TTA11CTT (2397T>A 2399A>G), GGT13GGA (2391A>T), AAG14AAA (2388C>T), TCT15TCA (2385A>T), ATT16CTA (2382A>T 2384T>G), CTG18CTC (2376C>G), CCA21CCC (2367T>G), AAA22AAG (2364T>C), CTT25CTA (2355A>T), CTT26ATC (2352A>G 2354G>T), CCT27GCA (2349A>T 2351G>C), ATT28ATA (2346A>T), TTT29TTC (2343A>G), TTA30GTC (2340T>G 2342A>C), ATT31ATA (2337A>T), GAT32GAC (2334A>G), TAT33TTC (2331A>G 2332T>A), TTA34CTC (2328T>G 2330A>G), TAT35AAA (2325A>T 2327A>T), CAA36AAA (2324G>T), CTA37ATT (2319T>A 2321G>T), TTA38CTA (2318A>G), AAT39CAA (2313A>T 2315T>G), AAT40GAT (2312T>C), TTT41TAC (2307A>G 2308A>T), ACA44AAT (2298T>A 2299G>T), GCT46ATT (2293G>A 2294C>T), AGG47AGA (2289C>T), GTA48GTT (2286T>A) | | | | | | | | |
| RepA (YP_009255248.1) | 8 | 48 | 16.1% | 218 | 73.6% | 41 (100%) | 28 (68.3%) | 0/0/0/0 | 0 |
| Protein mutations: | I16L (2382A>T 2384T>G), L26I (2352A>G 2354G>T), P27A (2349A>T 2351G>C), L30V (2340T>G 2342A>C), Y33F (2331A>G 2332T>A), Y35K (2325A>T 2327A>T), Q36K (2324G>T), L37I (2319T>A 2321G>T), N39Q (2313A>T 2315T>G), N40D (2312T>C), F41Y (2307A>G 2308A>T), T44N (2298T>A 2299G>T), A46I (2293G>A 2294C>T) | | | | | | | | |
| Codon mutations: | CGC10AGA (2400G>T 2402G>T), TTA11CTT (2397T>A 2399A>G), GGT13GGA (2391A>T), AAG14AAA (2388C>T), TCT15TCA (2385A>T), ATT16CTA (2382A>T 2384T>G), CTG18CTC (2376C>G), CCA21CCC (2367T>G), AAA22AAG (2364T>C), CTT25CTA (2355A>T), CTT26ATC (2352A>G 2354G>T), CCT27GCA (2349A>T 2351G>C), ATT28ATA (2346A>T), TTT29TTC (2343A>G), TTA30GTC (2340T>G 2342A>C), ATT31ATA (2337A>T), GAT32GAC (2334A>G), TAT33TTC (2331A>G 2332T>A), TTA34CTC (2328T>G 2330A>G), TAT35AAA (2325A>T 2327A>T), CAA36AAA (2324G>T), CTA37ATT (2319T>A 2321G>T), TTA38CTA (2318A>G), AAT39CAA (2313A>T 2315T>G), AAT40GAT (2312T>C), TTT41TAC (2307A>G 2308A>T), ACA44AAT (2298T>A 2299G>T), GCT46ATT (2293G>A 2294C>T), AGG47AGA (2289C>T), GTA48GTT (2286T>A) | | | | | | | | |
| C3 (YP_009255249.1) | 1 | 4 | 2.5% | 2 | 7.4% | 4 (100%) | 1 (25.0%) | 0/0/0/0 | 0 |
| Protein mutations: | M1I (2294C>T), G3E (2289C>T) | | | | | | | | |
| Codon mutations: | ATG1ATA (2294C>T), CTA2TTA (2293G>A), GGG3GAG (2289C>T), TAT4TT. (2286T>A) | | | | | | | | |

*: Inserts / Deletes / Misaligned / Frameshifts

#### Analysis details

This analysis was performed with panviral2.64

### NGS Details (UN59): Escherichia virus DE3

#### Assembly

| | |
|---|---|
| **Coverage Length** | 135 (1 contig(s)) |
| **Depth Of Coverage** | 1.0 |
| **Number Of Reads** | 1 |
| **Reads Per Million** | 0.02 rpm (after QC) |
| **Ambiguities** | 0 |
| **Assembly Method** | de novo + reference guided assembly |
| **Consensus Caller** | Bcf Tools |

#### Coverage Map

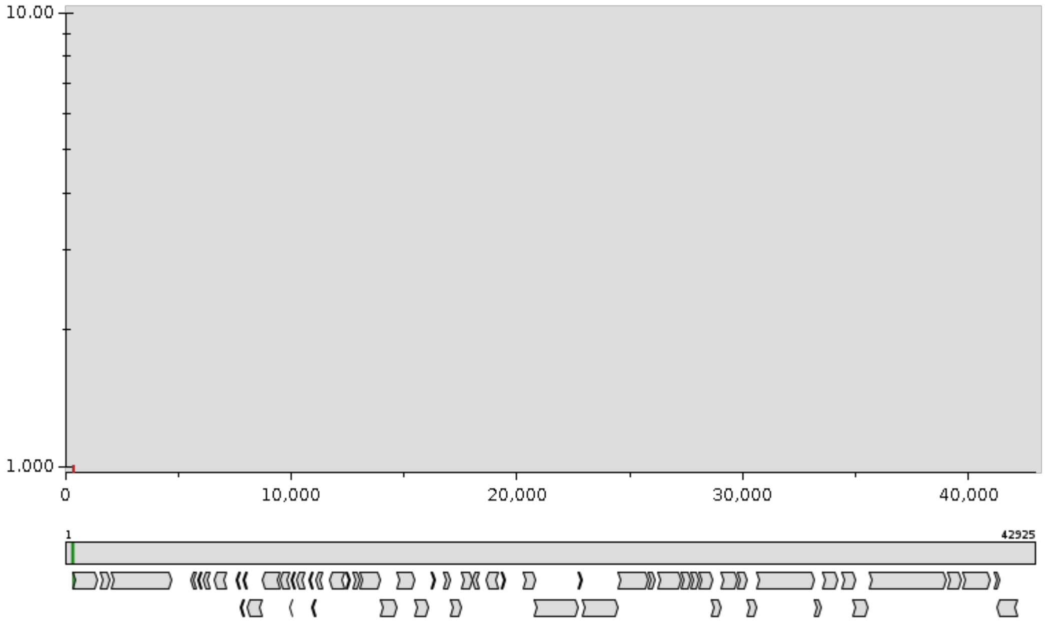

#### Assignment

| | |
|---|---|
| **Type** | Escherichia virus DE3 (Taxonomy ID: 2169967) |
| **Reference Genome** | NC_042057.1 |
| **NT Identity (%)** | 100.0 |
| **AA Identity (%)** | 96.2963 |
| **Number Of Stop Codons** | 0 |
| **Number Of CDS** | 57 |

#### Alignment

| | |
|---|---|
| **Alignment Score** | 270.0 (NT) + 162.0 (AA) = 432.0 |
| **Concordance (%)** | 100.0 |

| Alignment Method | Local, heuristic, nucleotide (BLASTN) |
|---|---|

#### Genome Region

Sequence starts at position 285 and ends at position 419 relative to NC_042057.1 reference sequence.

#### Alignment Detailed Statistics

| | Begin | End | Coverage | Score | Concordance | Matches | Identities | I/D/M/F * | Stop Codons |
|---|---|---|---|---|---|---|---|---|---|
| **NT** | **285** | **419** | **0.3%** | **270** | **100%** | **135 (100%)** | **135 (100%)** | **0/0** | |
| Mutations: | *none* | | | | | | | | |
| **CDS** | | | | | | | | | |
| lacI | 1 | 27 | 7.5% | 162 | 97.6% | 27 (100%) | 26 (96.3%) | 0/0/0/0 | 0 |
| **Proteins** | | | | | | | | | |
| DNA-binding transcriptional repressor LacI (YP_009617199.1) | 1 | 27 | 7.5% | 162 | 97.6% | 27 (100%) | 26 (96.3%) | 0/0/0/0 | 0 |
| Protein mutations: | *none* | | | | | | | | |
| Codon mutations: | *none* | | | | | | | | |

*: Inserts / Deletes / Misaligned / Frameshifts

#### Analysis details

This analysis was performed with panviral2.64

### Analysis of UN60

| | |
|---|---|
| **Technology** | Paired-end short reads |
| **Input Files** | UN60_R1.fq.gz (1.51 GB), UN60_R2.fq.gz (1.55 GB) |
| **Submitted On** | 2023-09-26 13:17:18 UTC |
| **Duration** | 3h 13m 35s |
| **Tool Version** | panviral2.64 |
| **Location** | https://www.genomedetective.com/db/ui/analysis/0a18e193-0210-4915-bede-4b932408d4fe |

#### Statistics

| | |
|---|---|
| **Original Read Length** | 20 - 150 |
| **Trimmed Read Length** | 50 - 135 |

| | # Reads | % of Reads |
|---|---|---|
| **Input file** | 47463978 | 100.0% |
| **After QC** | 47148532 | 99.3% |
| **After filtering** | 5923550 | 12.5% |
| **Mapped back to contigs** | 3180271 | 6.7% |

#### Assignments

| Assignment | No. of Reads | Depth of Coverage | Identity | | Genome Coverage | |
|---|---|---|---|---|---|---|
| | | | NT | AA | | |
| Potato virus Y | 1048341 | 14480.8 | 82.7% | 90.0% | 99.8% | 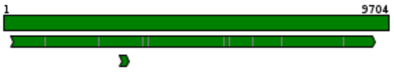 |
| Torradovirus lycopersici (2 segments out of 2) | 1041429 | 10584.1 | 88.8% | 94.7% | 99.3% | |
| Torradovirus lycopersici (segment RNA 2) | 685969 | 16080.2 | 85.3% | 89.6% | 99.6% | 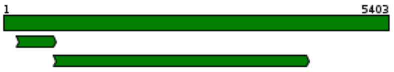 |
| Torradovirus lycopersici (segment RNA 1) | 355460 | 6761.0 | 91.2% | 98.0% | 99.1% | 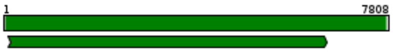 |
| Tomato chocolate spot virus (2 segments out of 2) | 367028 | 13358.9 | 67.5% | 69.9% | 28.2% | |
| Tomato chocolate spot virus (segment RNA2) | 239197 | 15922.8 | 67.3% | 70.6% | 33.1% | 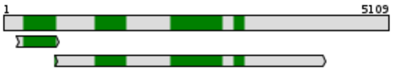 |
| Tomato chocolate spot virus (segment RNA 1) | 127831 | 11026.4 | 67.7% | 68.8% | 24.8% | 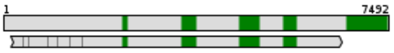 |
| Torradovirus marchitezum (2 segments out of 2) | 348393 | 5946.0 | 64.9% | 66.8% | 61.4% | |

| Assignment | No. of Reads | Depth of Coverage | Identity | | Genome Coverage | |
|---|---|---|---|---|---|---|
| | | | NT | AA | | |
| Torradovirus marchitezum (segment RNA 2) | 209020 | 18973.4 | 67.3% | 67.5% | 27.8% | 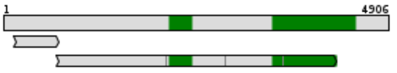 |
| Torradovirus marchitezum (segment RNA 1) | 139373 | 3023.2 | 64.3% | 66.7% | 84.2% | 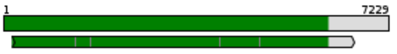 |
| Tomato necrotic dwarf virus (segment RNA1) | 244103 | 5251.7 | 64.2% | 65.6% | 85.3% | 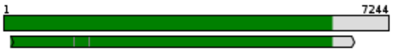 |
| Diachasmimorpha longicaudata entomopoxvirus (segment NC_043455.1) | 7189 | 803.2 | 60.5% | 55.2% | 83.9% | 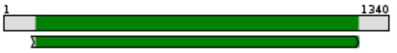 |
| Bracoviriform glomeratae (segment NC_043292.1) | 6143 | 2381.0 | 72.0% | 83.5% | 73.0% | 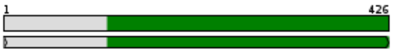 |
| Potato leafroll virus | 5542 | 126.6 | 97.5% | 96.5% | 97.3% | 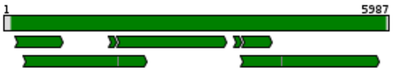 |
| Duamitovirus soch1 | 351 | 22.1 | 67.5% | 68.2% | 70.6% | 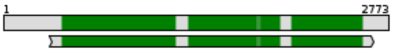 |
| Solendovirus venanicotianae | 130 | 11.4 | 77.9% | 75.6% | 17.7% | 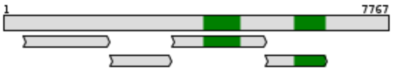 |
| Macrosiphum euphorbiae virus 1 | 106 | 2.6 | 92.8% | 96.5% | 24.3% | 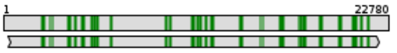 |
| Duamitovirus peex1 | 26 | 4.4 | 68.7% | 67.1% | 27.0% | 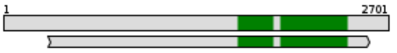 |

#### Discoveries

| Similar to | No. of Reads | Depth of Coverage | Identity | | Genome Coverage | |
|---|---|---|---|---|---|---|
| | | | NT | AA | | |
| Brazilian marseillevirus | 20377 | 8722.3 | 82.6% | 93.1% | 0.1% | 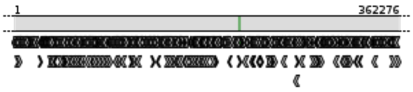 |
| Brazilian marseillevirus | 18320 | 8619.2 | 79.5% | 95.6% | 0.1% | 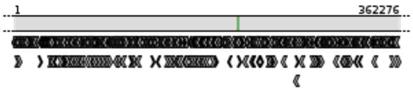 |
| Brazilian marseillevirus | 12747 | 5214.8 | 79.2% | 94.7% | 0.1% | 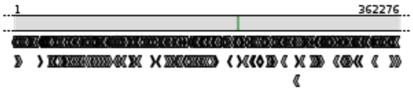 |
| Brazilian marseillevirus | 11487 | 6274.7 | 81.4% | 92.6% | 0.0% | 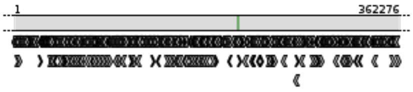 |
| Makelovirus prm1 | 10355 | 1317.0 | 76.8% | 87.5% | 0.7% | 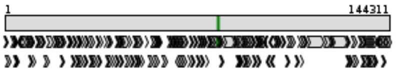 |

| Similar to | No. of Reads | Depth of Coverage | Identity | | Genome Coverage | |
|---|---|---|---|---|---|---|
| | | | NT | AA | | |
| Brazilian marseillevirus | 9741 | 4288.3 | 82.0% | 97.1% | 0.0% | 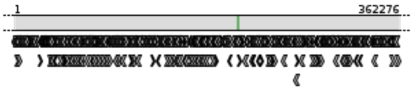 |
| Yellowstone lake phycodnavirus 1 | 5848 | 3350.0 | 82.1% | 93.1% | 0.1% | 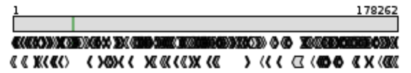 |
| Brazilian marseillevirus | 5218 | 2083.3 | 79.6% | 94.7% | 0.1% | 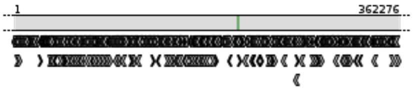 |
| Yellowstone lake phycodnavirus 1 | 5137 | 2919.0 | 80.0% | 90.4% | 0.1% | 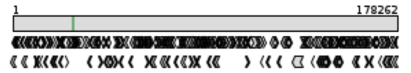 |
| Brazilian marseillevirus | 2553 | 1024.8 | 80.9% | 94.7% | 0.1% | 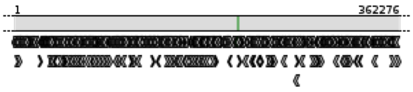 |
| Noumeavirus | 2495 | 1143.9 | 78.7% | 95.9% | 0.1% | 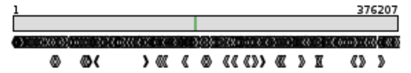 |
| Betabaculovirus disaccharalis | 2005 | 1183.5 | 82.0% | 93.0% | 0.1% | 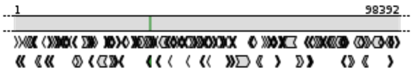 |
| Betabaculovirus disaccharalis | 1706 | 1065.4 | 84.7% | 92.0% | 0.2% | 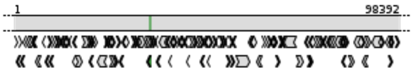 |
| Cladosporium fulvum T-1 virus | 605 | 46.5 | 52.2% | 47.5% | 20.9% | 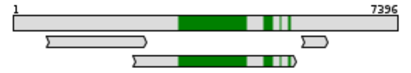 |
| Vellamovirus rhodeisland44 | 604 | 154.1 | 75.0% | 86.8% | 0.2% | 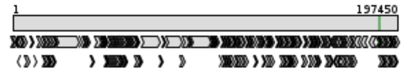 |
| Cassava brown streak virus | 356 | 82.0 | 60.3% | 57.2% | 5.8% | 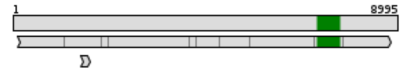 |
| Lowelvirus tuscon4d | 355 | 188.4 | 80.9% | 90.6% | 0.1% | 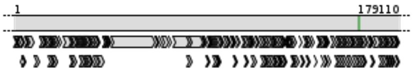 |
| Badnavirus occultipomeae | 353 | 112.5 | 51.7% | 45.7% | 4.3% | 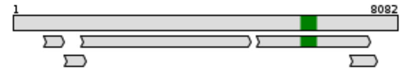 |
| Errantivirus | 227 | 29.2 | 55.2% | 43.5% | 8.5% | 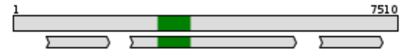 |
| Ichnoviriform fugitivi (4 segments out of 56) | 173 | 18.4 | 57.0% | 47.2% | 32.8% | |
| Ichnoviriform fugitivi (segment B17) | 173 | 18.4 | 57.0% | 47.2% | 32.8% | 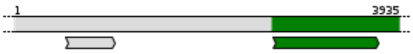 |
| Ichnoviriform fugitivi (segment B17) | 20 | 8.2 | 58.3% | 47.9% | 7.4% | 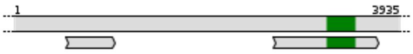 |

| Similar to | No. of Reads | Depth of Coverage | Identity | | Genome Coverage | |
|---|---|---|---|---|---|---|
| | | | NT | AA | | |
| Ichnoviriform fugitivi (segment B17) | 13 | 5.5 | 57.2% | 54.3% | 7.3% | 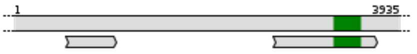 |
| Ichnoviriform fugitivi (segment B17) | 1 | 1.0 | 64.2% | 62.2% | 3.4% | 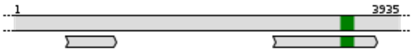 |
| Badnavirus maculasmallanthi | 110 | 43.1 | 56.5% | 55.2% | 3.4% | 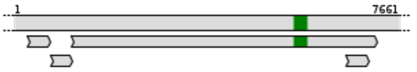 |
| Caulimovirus venafragariae | 77 | 9.9 | 56.3% | 52.3% | 13.2% | 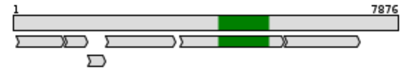 |
| Caulimovirus tesselloscrophulariae | 48 | 14.5 | 53.6% | 37.9% | 4.8% | 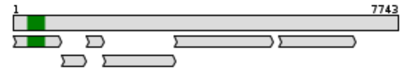 |
| Brazilian marseillevirus | 47 | 25.7 | 81.2% | 95.7% | 0.0% | 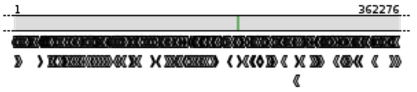 |
| Errantivirus | 46 | 16.8 | 53.0% | 41.0% | 4.0% | 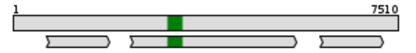 |
| Caulimovirus venafragariae | 45 | 6.3 | 58.5% | 54.0% | 10.8% | 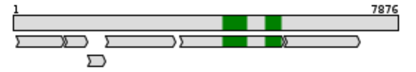 |
| Badnavirus rutilanscamelliae | 44 | 15.0 | 54.0% | 45.0% | 3.6% | 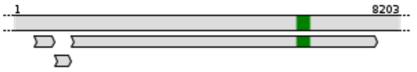 |
| Synechococcus phage S-SRM01 | 43 | 25.2 | 76.8% | 88.4% | 0.1% | 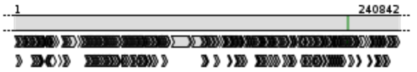 |
| Marseillevirus marseillevirus | 38 | 25.6 | 81.4% | 94.7% | 0.0% | 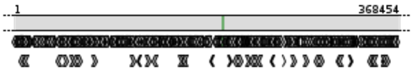 |
| Petuvirus venapetuniae | 36 | 5.1 | 58.6% | 53.7% | 13.0% | 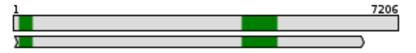 |
| Cavemovirus venamanihotis | 33 | 6.4 | 59.7% | 48.3% | 7.6% | 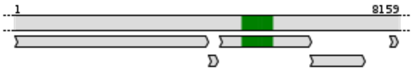 |
| Cavemovirus collusipomeae | 32 | 7.0 | 69.0% | 57.8% | 6.4% | 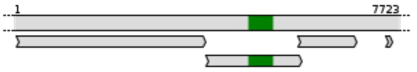 |
| Dioscovirus dioscoreae | 26 | 3.7 | 58.5% | 48.6% | 11.5% | 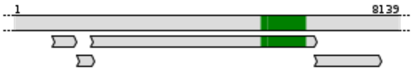 |
| Rahariannevirus raharianne | 26 | 4.0 | 79.5% | 81.2% | 1.9% | 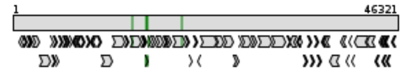 |
| Badnavirus maculasmallanthi | 21 | 9.4 | 56.5% | 51.6% | 3.5% | 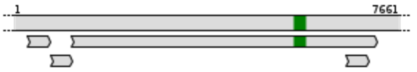 |

| Similar to | No. of Reads | Depth of Coverage | Identity | | Genome Coverage | |
|---|---|---|---|---|---|---|
| | | | NT | AA | | |
| Blunervirus vaccinii (segment RNA 2) | 14 | 6.4 | 61.7% | 61.7% | 4.6% | 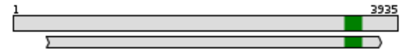 |
| Alfalfa leaf curl virus | 14 | 6.0 | 71.5% | 58.2% | 9.8% | 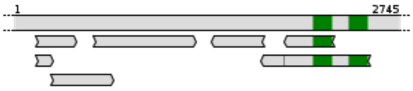 |
| Duamitovirus peex1 | 13 | 4.7 | 70.8% | 73.6% | 10.1% | 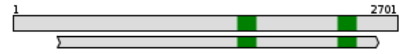 |
| Pseudomonas phage YMC11/07/P54_PAE_BP | 13 | 4.8 | 78.4% | 83.3% | 0.7% | 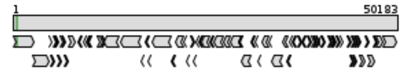 |
| Pinus nigra virus 1 | 11 | 3.1 | 56.8% | 45.9% | 6.7% | 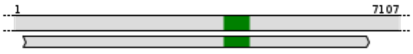 |
| Badnavirus rutilanscamelliae | 10 | 3.9 | 61.5% | 56.2% | 3.2% | 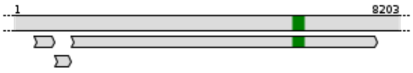 |
| Duamitovirus dapi1 | 8 | 6.5 | 68.9% | 66.7% | 4.8% | 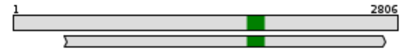 |
| Alphabaculovirus lyxylinae | 8 | 6.5 | 80.0% | 83.8% | 0.1% | 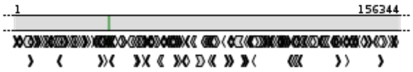 |
| Caulimovirus tessellobrassicae | 6 | 2.6 | 62.7% | 52.6% | 3.2% | 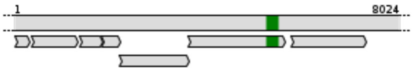 |
| Badnavirus occultipomeae | 6 | 3.2 | 54.3% | 43.5% | 3.1% | 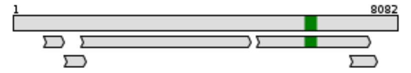 |
| Moloney murine leukemia virus | 6 | 3.0 | 75.6% | 100.0% | 3.2% | 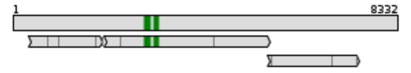 |
| Marseillevirus marseillevirus | 6 | 4.4 | 81.9% | 88.6% | 0.0% | 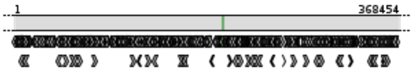 |
| Yaba monkey tumor virus | 4 | 3.3 | 81.3% | 84.1% | 0.1% | 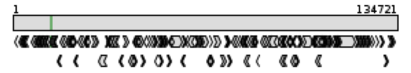 |
| Alphabaculovirus agipsilonis | 4 | 3.5 | 78.5% | 77.5% | 0.1% | 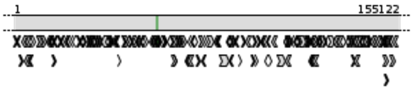 |
| Badnavirus tessellocastaneae | 4 | 2.1 | 59.4% | 53.7% | 3.4% | 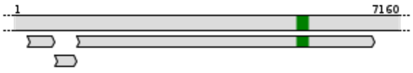 |
| Escherichia virus DE3 | 4 | 2.5 | 100.0% | 98.6% | 0.5% | 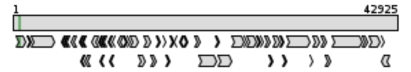 |
| Epiphyllum badnavirus 1 | 4 | 2.0 | 54.8% | 50.0% | 3.4% | 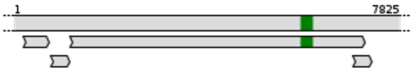 |

| Similar to | No. of Reads | Depth of Coverage | Identity | | Genome Coverage | |
|---|---|---|---|---|---|---|
| | | | NT | AA | | |
| Goose dicistrovirus | 4 | 2.2 | 85.0% | 95.0% | 2.6% | 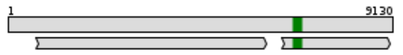 |
| Melanoplus sanguinipes entomopoxvirus | 2 | 1.4 | 81.7% | 76.6% | 0.1% | 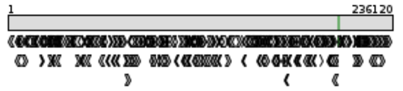 |
| Cotia virus | 2 | 1.8 | 80.0% | 93.3% | 0.1% | 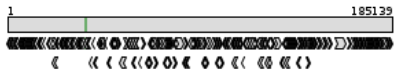 |
| Scottvirus scott | 2 | 1.1 | 82.0% | 92.6% | 0.6% | 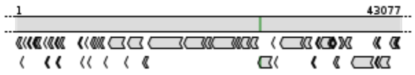 |
| Punavirus P1 | 2 | 1.2 | 100.0% | 98.3% | 0.2% | 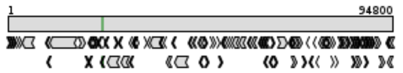 |
| Harvey murine sarcoma virus | 2 | 1.2 | 69.5% | 85.1% | 14.1% | 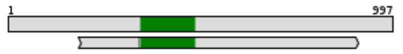 |
| Ochrobactrum phage POA1180 | 2 | 2.0 | 74.8% | 80.0% | 0.3% | 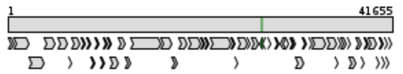 |
| Pseudomonas phage PA8 | 2 | 2.0 | 83.0% | 80.0% | 0.2% | 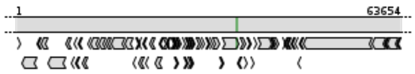 |
| Tomato associated geminivirus 1 | 2 | 1.2 | 73.0% | 75.3% | 8.9% | 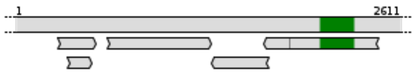 |
| Euphorbia caput-medusae latent virus | 1 | 1.0 | 47.6% | 45.4% | 5.5% | 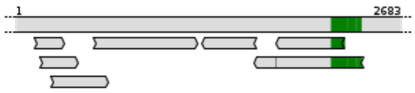 |

#### NGS Details (UN60): Potato virus Y

##### Assembly

| | |
|---|---|
| **Coverage Length** | 9683 (1 contig(s)) |
| **Depth Of Coverage** | 14480.8 |
| **Number Of Reads** | 1048341 |
| **Reads Per Million** | 22234.86 rpm (after QC) |
| **Ambiguities** | 0 |
| **Assembly Method** | de novo + reference guided assembly |
| **Consensus Caller** | Bcf Tools |

##### Coverage Map

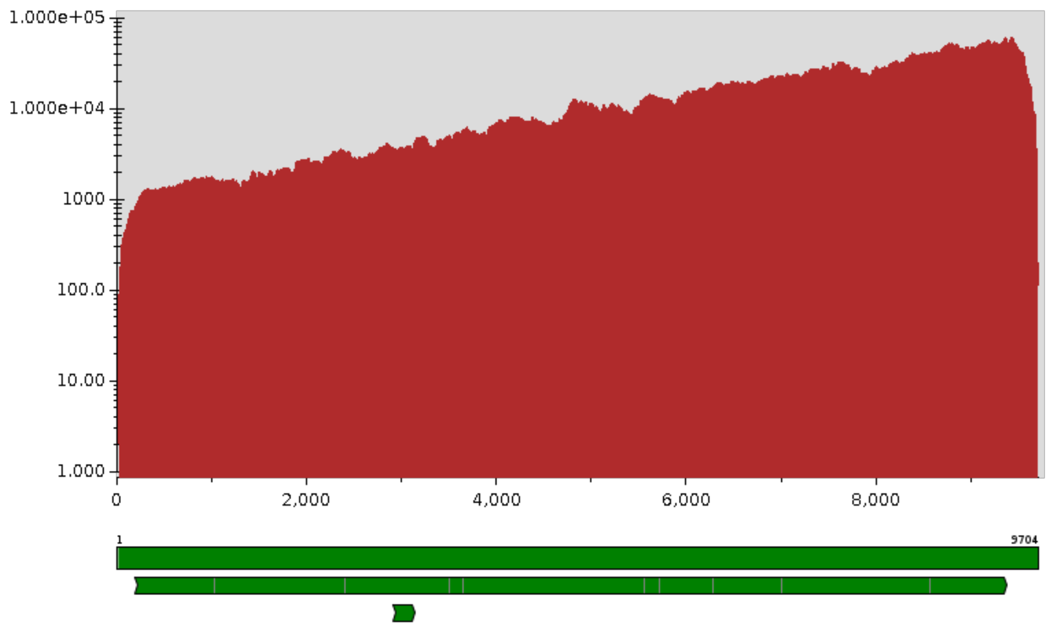

##### Assignment

| | |
|---|---|
| **Type** | Potato virus Y (Taxonomy ID: 12216) |
| **Reference Genome** | NC_001616.1 |
| **NT Identity (%)** | 82.6671 |
| **AA Identity (%)** | 89.9682 |
| **Number Of Stop Codons** | 3 |
| **Number Of CDS** | 2 |

##### Alignment

| | |
|---|---|
| **Alignment Score** | 12522.0 (NT) + 19018.0 (AA) = 31540.0 |
| **Concordance (%)** | 78.1255 |

| | |
|---|---|
| **Alignment Method** | Global, seeded, nucleotide + amino acids (AGA) |

#### Genome Region

Sequence starts at position 22 and ends at position 9704 relative to NC_001616.1 reference sequence.

#### Alignment Detailed Statistics

| | Begin | End | Coverage | Score | Concordance | Matches | Identities | I/D/M/F* | Stop Codons |
|---|---|---|---|---|---|---|---|---|---|
| **NT** | **22** | **9704** | **99.8%** | **12522** | **65.0%** | **9671 (99.8%)** | **8003 (82.6%)** | **10/12** | |

Mutations:

31A>C, 32A>G, 36C>A, 37G>A, 38C>G, 42A>T, 44C>T, 50T>C, 52A>G, 54C>A, 56C>T, 58C>T, 59A>T, 60_61insAAA, 75C>T, 79C>T, 83G>A, 85T>C, 87C>T, 89G>A, 90T>C, 93A>T, 94A>G, 95A>T, 100T>C, 101T>A, 102C>T, 104T>C, 107C>T, 110T>G, 112C>T, 114C>T, 115T>C, 116A>T, 117G>C, 119A>G, 120C>T, 121A>T, 125T>G, 126T>C, 127G>A, 128G>T, 131A>G, 132C>T, 139A>T, 139_140insG, 141C>T, 143C>T, 146C>T, 150C>T, 150_151insT, 154T>C, 156C>A, 158T>A, 160A>C, 164C>T, 168C>A, 169C>G, 171A>C, 172T>C, 173T>A, 174T>G, 177G>C, 178A>G, 179T>C, 180C>T, 181C>A, 182T>G, 183C>T, 193T>C, 194T>C, 196C>A, 202A>G, 208C>T, 209T>C, 210G>T, 211T>G, 220G>A, 221T>A, 223T>G, 226A>G, 235A>T, 241C>T, 247A>T, 251T>C, 255G>T, 256C>T, 258A>G, 262T>C, 263A>G, 265T>G, 267T>C, 273A>T, 277A>T, 280A>G, 283G>T, 286G>A, 287G>A, 289T>C, 293G>A, 294T>C, 301T>C, 304C>T, 305G>A, 306C>T, 309A>G, 311C>T, 316A>T, 317A>G, 318C>T, 319A>G, 322A>G, 331A>T, 339T>A, 340C>A, 342A>G, 347A>G, 348A>C, 349A>G, 350T>C, 355T>A, 356A>G, 357C>T, 364T>C, 365G>A, 366T>C, 367G>A, 368C>T, 369T>C, 373A>G, 376C>T, 384T>G, 388G>A, 391C>T, 394A>G, 400G>A, 406T>C, 408C>T, 409C>T, 412G>A, 415A>T, 416A>G, 417T>A, 424T>C, 427G>A, 433A>G, 436G>C, 437G>A, 440A>G, 441G>A, 443A>C, 444A>G, 445G>A, 448T>A, 454A>G, 456A>C, 460T>C, 461C>A, 469A>G, 475C>G, 481T>A, 502T>C, 505A>T, 511C>T, 517T>A, 520T>A, 523A>T, 531C>T, 535G>A, 536C>T, 539C>T, 540A>G, 541A>G, 549G>A, 550A>G, 556C>T, 559T>C, 562T>C, 568T>C, 571A>G, 576T>C, 580T>C, 585T>C, 586C>G, 593C>T, 594G>A, 595C>T, 597C>G, 599A>G, 601A>G, 602A>C, 603T>C, 604A>G, 613A>G, 616A>G, 619C>G, 622G>A, 625G>A, 628T>C, 631C>T, 635A>G, 637T>C, 644A>G, 646A>G, 655T>C, 661G>A, 663A>C, 664G>C, 667A>G, 679C>T, 682C>A, 683T>C, 685A>G, 691T>C, 699C>G, 703T>A, 706T>A, 719A>G, 721G>A, 722A>G, 724A>G, 725C>T, 727T>G, 730T>A, 731G>T, 734T>C, 735A>C, 736C>T, 737T>C, 738C>G, 739C>T, 744C>T, 745G>C, 749C>T, 751A>C, 754T>C, 765T>A, 770T>C, 773C>A, 774G>A, 775A>G, 776C>A, 790C>T, 801T>A, 813G>C, 814A>G, 816T>A, 818T>C, 823A>G, 825G>A, 826T>A, 833C>A, 835G>A, 841C>T, 851A>G, 853T>C, 859C>T, 860C>T, 862C>T, 863A>G, 865T>C, 867T>C, 869A>G, 871C>T, 872A>T, 874A>G, 876G>A, 879G>A, 886T>C, 898C>T, 903A>G, 906C>A, 907A>C, 909A>C, 910A>C, 912G>A, 913C>T, 922C>A, 931T>A, 938T>C, 939C>T, 940A>T, 943A>G, 952C>T, 967A>G, 973A>G, 976G>C, 980T>A, 982G>C, 991A>G, 994T>C, 999G>A, 1003T>C, 1007C>A, 1008A>G, 1009G>A, 1010A>G, 1012T>G, 1013A>G, 1018A>G, 1019A>C, 1021C>G, 1030C>T, 1039G>A, 1042T>C, 1045C>A, 1048C>A, 1051T>C, 1060G>A, 1063T>A, 1064C>T, 1069C>T, 1083G>A, 1093T>C, 1097T>G, 1099G>A, 1105C>T, 1114A>G, 1117T>G, 1123A>G, 1126T>A, 1129C>T, 1143G>A, 1147A>T, 1150T>A, 1154T>A, 1156G>A, 1165C>T, 1168T>C, 1172C>T, 1174T>A, 1201C>T, 1221G>A, 1229G>A, 1230T>C, 1231T>C, 1233G>A, 1234C>T, 1240G>A, 1246G>A, 1250T>C, 1252G>A, 1255T>C, 1257A>C, 1258A>G, 1267A>G, 1273T>C, 1276G>A, 1278A>G, 1283T>C, 1290C>T, 1291G>A, 1298G>A, 1299A>C, 1300C>T, 1306T>C, 1307A>G, 1309A>G, 1312T>C, 1316A>G, 1318T>A, 1327G>A, 1329T>C, 1333G>T, 1336A>G, 1343C>T, 1345A>G, 1354G>A, 1357G>T, 1360C>T, 1367C>T, 1369C>G, 1373C>A, 1378C>T, 1384G>A, 1385A>G, 1393A>G, 1396C>T, 1405G>T, 1408A>G, 1411G>A, 1415G>T, 1417A>T, 1420G>C, 1429T>C, 1432A>G, 1436G>A, 1438C>T, 1439T>C, 1441A>G, 1444T>C, 1453C>T, 1456G>A, 1468A>G, 1492A>G, 1499T>C, 1501G>A, 1504T>C, 1507G>A, 1510C>T, 1513A>G, 1516A>G, 1519A>T, 1522G>A, 1525C>T, 1528G>A, 1534C>T, 1540T>A, 1546C>T, 1555A>G, 1558T>C, 1561T>C, 1564A>C, 1567T>A, 1573C>T, 1576A>G, 1583T>C, 1588T>A, 1594G>A, 1600C>T, 1609G>A, 1618G>A, 1621C>T, 1624C>T, 1631C>T, 1639A>G, 1651C>T, 1654C>G, 1675T>C, 1693A>G, 1702T>C, 1707A>G, 1718G>A, 1726A>G, 1729C>T, 1732A>C, 1738T>C, 1742A>G, 1744C>T, 1747C>T, 1753T>C, 1758G>A, 1768G>A, 1786C>T, 1789A>G, 1790G>A, 1795C>T, 1805T>C, 1807A>G, 1813G>A, 1819G>A, 1825G>A, 1837C>T, 1840T>C, 1849A>G, 1858C>G, 1864A>G, 1870C>T, 1876T>C, 1882A>G, 1888T>A, 1891T>C, 1906T>C, 1915A>T, 1930T>C, 1936C>T, 1937A>G, 1939T>C, 1945A>G, 1951C>T, 1960A>T, 1978A>G, 1981T>A, 1984C>T, 1987T>C, 1996C>T, 2004T>A, 2020A>G, 2023G>A, 2026T>C, 2029G>T, 2032G>A, 2059T>C, 2068T>G, 2071C>T, 2072G>A, 2074G>A, 2087C>T, 2089G>A, 2101C>T, 2104G>A, 2107G>A, 2116G>A, 2122C>T, 2137C>T, 2158T>C, 2164C>T, 2170A>G, 2182T>C, 2188G>A, 2191C>A, 2208G>A, 2218T>C, 2224C>T, 2227A>T, 2242G>A, 2245C>T, 2254G>A, 2257T>C, 2260C>T, 2266C>G, 2272A>G, 2275G>A, 2293A>G, 2299C>T, 2302G>A, 2305G>A, 2314A>T, 2317T>C, 2326A>G, 2341G>A, 2347A>G, 2350T>C, 2356G>A, 2365T>C, 2371A>G, 2374A>G, 2383T>C, 2386A>T, 2389A>G, 2392T>C, 2395T>C, 2414A>G, 2415A>G, 2416T>G, 2419T>A, 2420A>T, 2440A>T, 2467T>C, 2476T>C, 2482G>A, 2491G>A, 2497G>T, 2503G>A, 2518A>G, 2521T>C, 2523A>G, 2533A>G, 2539G>A, 2545A>G, 2551G>A, 2554T>A, 2558C>T, 2561T>C, 2563G>C, 2572A>G, 2584T>C, 2587C>T, 2593G>C, 2599T>C, 2608T>C, 2620T>C, 2629G>A, 2632G>A, 2634G>A, 2638G>A, 2644T>A, 2653A>G, 2665T>C, 2678C>T, 2683A>G, 2689T>C, 2698A>C, 2707A>G, 2710G>A, 2716A>G, 2719A>G, 2722C>T, 2723G>A, 2725C>T, 2728A>G, 2734G>A, 2737T>A, 2746T>C, 2747G>A, 2749T>A, 2752A>T, 2755T>A, 2761C>T, 2764C>T, 2779T>C, 2785G>T, 2791C>T, 2795C>A, 2806C>T, 2816T>C, 2818G>C, 2822G>A, 2824G>A, 2827G>A, 2836T>C, 2839G>A, 2842T>C, 2863C>A, 2864C>T, 2882C>A, 2885A>C, 2886G>A, 2893C>T, 2895C>T, 2902C>T, 2905A>G, 2911T>C, 2923T>C, 2931A>G, 2940A>G, 2947T>C, 2962T>C, 2971_2972insA, 2989T>C, 2991C>T, 2994A>G, 2999G>A, 3000A>G, 3016delC, 3021G>A, 3024A>T, 3025C>T, 3036C>T, 3039A>G, 3040A>C, 3042A>C, 3043G>A, 3052G>A, 3061A>G, 3082G>A, 3091C>T, 3099C>T, 3100C>A, 3104G>A, 3105T>G, 3106G>A, 3109C>T, 3112A>G, 3118T>C, 3121C>T, 3127A>G, 3132G>A, 3137C>A, 3138G>A, 3139A>G, 3151T>C, 3152T>A, 3154C>T, 3165G>A, 3169A>G, 3199C>T, 3203A>C, 3208T>C, 3214A>G, 3217C>T, 3218T>C, 3223C>A, 3225C>A, 3230G>A, 3235C>T, 3257T>C, 3262C>T, 3265C>T, 3266G>A, 3268A>G, 3271G>A, 3275G>A, 3284G>A, 3301G>A, 3315G>A, 3316A>G, 3318A>C, 3322C>T, 3325G>A, 3328G>A, 3337A>T, 3349G>A, 3355C>T, 3358C>T, 3359A>G, 3367A>G, 3370T>C, 3377T>C, 3379A>G, 3394A>G, 3397C>T, 3400T>G, 3406A>G, 3418C>T, 3421T>A, 3427T>C, 3433G>A, 3436A>T, 3439A>G, 3442C>T, 3445T>C, 3457G>A, 3460T>C, 3463T>A, 3469G>T, 3472G>A, 3496C>T, 3499G>A, 3505C>T, 3513T>G, 3520A>G, 3529A>G, 3532A>G, 3545A>G, 3548G>A, 3551T>C, 3554G>A, 3556C>T, 3583T>C, 3586T>C, 3589C>T, 3595C>T, 3607T>C, 3613T>C, 3616G>A, 3619T>C, 3623C>G, 3628C>T, 3632C>A, 3637T>C, 3638T>C, 3643A>G, 3661A>G, 3670G>A, 3682T>C, 3688G>A, 3697G>A, 3698A>G, 3706T>C, 3709T>C, 3713T>C, 3715G>A, 3717G>A, 3721G>A, 3731C>A, 3732G>A, 3736T>A, 3746C>T, 3748A>G, 3754A>G, 3763T>C, 3778A>G, 3781A>G, 3784C>T, 3796T>C, 3805A>T, 3811C>T, 3814A>G, 3829C>T, 3838T>C, 3844A>G, 3847A>C, 3856T>A, 3862G>T, 3871C>T, 3877C>A, 3883C>T, 3886A>G, 3892A>G, 3895C>T, 3898T>C, 3904A>G, 3905C>A, 3907G>A, 3913T>C, 3928G>A, 3931A>T, 3937G>A, 3955T>C, 3958G>A, 3961C>T, 3964A>G, 3967T>C, 3976A>G, 3979T>C, 3985A>T, 3997A>T, 4000G>A, 4006C>T, 4009T>G, 4012C>T, 4015A>G, 4024T>C, 4030A>G, 4033A>T, 4042G>A, 4052C>T, 4054G>A, 4057T>C, 4063T>A, 4066A>G, 4073A>G, 4078T>C, 4081C>T, 4084T>C, 4096C>T, 4108C>T, 4111A>T, 4117G>A, 4118C>T, 4120A>G, 4126C>T, 4128T>A, 4129C>T, 4132C>T, 4135T>C, 4147A>G, 4148T>C, 4151G>T, 4162C>T, 4166G>A, 4171A>C, 4180G>A, 4190C>T, 4198T>C, 4201C>T, 4210G>T, 4213C>T, 4219T>C, 4220C>T, 4223C>A, 4225G>A, 4231T>G, 4237T>C, 4252A>G, 4253T>C, 4255A>G, 4258A>G, 4270T>C, 4282A>G, 4285G>A, 4291A>G, 4303G>A, 4312C>T, 4319A>G, 4321A>G, 4324G>T, 4327G>A, 4336G>T, 4339T>C, 4348A>T, 4360A>G, 4366T>C, 4369T>A, 4375T>C, 4387T>C, 4395T>A, 4406G>A, 4411T>C, 4438T>C, 4444C>A, 4450C>T, 4456C>T, 4465C>T, 4475A>G, 4477G>T, 4480C>T, 4481A>T, 4483A>G, 4495C>T, 4507G>A, 4510C>T, 4513T>G, 4517C>T, 4528C>T, 4534A>G, 4540C>T, 4544G>T, 4545C>T, 4546G>A, 4548G>A, 4558T>C, 4564A>T, 4570C>T, 4576A>T, 4597G>A, 4600C>T, 4606C>T, 4609G>A, 4615G>A, 4618C>T, 4625T>C, 4627G>T, 4648C>T, 4651T>C, 4660G>A, 4663C>T, 4669T>A, 4679G>A, 4681G>C, 4684T>C, 4687T>G, 4699G>A, 4702A>G, 4708A>G, 4711G>A, 4712C>T, 4738A>G, 4744A>T, 4747A>G, 4748T>C, 4765T>C, 4796G>A, 4798C>T, 4803C>G, 4804T>C, 4807G>A, 4810G>T, 4813T>G, 4816T>C, 4831T>C, 4834C>T, 4840A>T, 4852A>G, 4855C>T, 4867T>C, 4870G>C, 4873T>A, 4882T>C, 4903A>T, 4909G>A, 4915T>C, 4918A>G, 4919T>C, 4921G>A, 4924T>C, 4927C>G, 4933T>C, 4936C>A, 4939G>A, 4942T>C, 4945C>T, 4963A>G, 4972T>C, 4990T>C, 5002A>G, 5023T>C, 5024T>C, 5029C>T, 5032T>C, 5038T>C, 5044A>T, 5053A>T, 5056G>A, 5057A>G, 5060A>G, 5071G>T, 5090C>A, 5095A>G, 5098G>T, 5100C>T, 5104A>G, 5107A>T, 5110T>C, 5113A>G, 5116G>A, 5119A>G, 5120G>T, 5122C>T, 5126A>G, 5137T>C, 5140C>T, 5143A>G, 5146G>A, 5161C>A, 5167A>G, 5173T>G, 5182G>A, 5183G>A, 5188C>T, 5191G>A, 5197A>G, 5200C>T, 5206C>T, 5209A>G, 5218C>T, 5230G>A, 5236C>T, 5239C>T, 5245C>T, 5263A>T, 5269C>T, 5272C>T, 5275C>A, 5278C>T, 5281A>C, 5284A>G, 5287T>C, 5288C>T, 5293A>T, 5294T>C, 5296G>A, 5302G>A, 5311T>A, 5314A>G, 5323A>T, 5347T>C, 5353C>T, 5365C>T, 5386T>C, 5395C>A, 5422A>G, 5425T>C, 5431C>T, 5437G>A, 5452T>C, 5458G>A, 5467T>C, 5470A>G, 5471C>T, 5473A>G, 5479A>G, 5482C>T, 5485A>C, 5488T>C, 5489T>C, 5491G>A, 5497T>C, 5506T>C, 5509G>A, 5512G>A, 5516T>C, 5530T>C, 5533G>A, 5536G>T, 5539G>A, 5548T>G, 5554C>T, 5560T>A, 5561G>A, 5563G>A, 5566G>C, 5569A>G, 5584C>G, 5593G>A, 5596G>A, 5597A>G, 5605C>A, 5617G>T, 5619C>T, 5620T>G, 5629G>A, 5630A>C, 5641C>T, 5650A>T, 5662A>T, 5665A>G, 5689A>G, 5693G>A, 5698G>A, 5713A>G, 5716G>C, 5719A>G, 5743C>A, 5755C>A, 5758T>C, 5764T>C, 5770A>G, 5779C>T, 5782T>C, 5791C>T, 5803C>T, 5836G>A, 5839A>G, 5845A>G, 5848A>G, 5853A>G, 5857T>C, 5860C>T, 5863A>T, 5869T>A, 5875T>C, 5884C>T, 5893C>T, 5894A>G, 5896C>G, 5899C>T, 5908G>A, 5911T>C, 5923G>A, 5938A>G, 5941C>T, 5944T>C, 5947T>C, 5959G>A, 5960C>G, 5961G>C, 5962G>C, 5974A>G, 5986T>A, 5989C>T, 5990A>G, 5998T>C, 6008A>C, 6010A>T, 6019A>T, 6022G>T, 6025A>T, 6030A>C, 6049C>A, 6058G>A, 6061A>G, 6064C>A, 6070T>C, 6072G>A, 6073T>C, 6074A>C, 6079G>A, 6081C>A, 6085A>T, 6091A>T, 6094C>T, 6111G>C, 6112T>C, 6118A>G, 6124G>A, 6127G>A, 6128A>G, 6130T>C, 6133T>C, 6136A>G, 6148C>T, 6157A>G, 6158G>A, 6163T>C, 6169A>G, 6170A>T, 6172A>G, 6178C>T, 6187A>G, 6190T>C, 6196G>A, 6203C>T, 6205C>T, 6208A>G, 6209C>T, 6211A>G, 6217G>A, 6229T>A, 6230G>A, 6238C>T, 6247G>A, 6253A>T, 6257G>A, 6258C>A, 6274T>C, 6282A>G, 6286G>A, 6298C>T, 6301G>A, 6304A>G, 6307C>T, 6322C>T, 6328A>T, 6331T>G, 6334T>C, 6337G>A, 6338C>T, 6358T>C, 6361G>A, 6362G>A, 6367A>T, 6370G>A, 6376C>T, 6379T>G, 6385A>C, 6391A>T, 6394A>G, 6397C>T, 6401G>A, 6405C>T, 6412C>T, 6418A>G, 6421T>C, 6429A>T, 6436T>A, 6445G>A, 6450A>G, 6454C>G, 6460C>T, 6463T>A, 6469C>T, 6472G>A, 6475G>A, 6481T>C, 6485C>A, 6487C>T, 6490T>C, 6496C>T, 6502G>A, 6514T>C, 6517G>A, 6527C>A, 6532C>A, 6541G>A, 6544A>G, 6556C>T, 6559T>C, 6565G>A, 6569T>C, 6571G>A, 6586T>A, 6588C>T, 6598A>G, 6601A>G, 6604T>C, 6607T>C, 6616A>G, 6622C>T, 6631G>A, 6634G>A, 6635T>C, 6640T>A, 6646G>T, 6647A>G, 6652C>T, 6664C>T, 6670T>G, 6677A>G, 6679A>G, 6685C>T, 6691A>T, 6694C>T, 6715A>G, 6716G>A, 6719A>G, 6724A>G, 6734C>T, 6742G>A, 6745G>A, 6754C>T, 6760A>G, 6764A>T, 6766A>G, 6775T>A, 6782C>T, 6787A>T, 6790C>T, 6795C>T, 6796A>G, 6799C>A, 6802C>A, 6808C>T, 6811C>T, 6814C>T, 6823C>T, 6829A>G, 6847C>T, 6853A>C, 6857A>G, 6862G>A, 6871A>G, 6875G>A, 6876T>C, 6877C>A, 6880G>A, 6883T>A, 6889T>G, 6901C>T, 6916C>T, 6919G>A, 6925A>G, 6928T>C, 6934C>A, 6949G>C, 6955C>T, 6958A>G, 6976A>G, 6980C>T, 6985C>T, 6997A>T, 7000G>T, 7001G>A, 7003G>A, 7012T>C, 7015G>A, 7018T>C, 7032T>A, 7039C>T, 7040T>C, 7042G>A, 7048A>G, 7060T>A, 7063C>A, 7078C>T, 7082T>C, 7084A>G, 7087A>G, 7090C>A, 7096T>C, 7099A>G, 7102T>C, 7111G>A, 7117A>G, 7125C>A, 7129A>G, 7132T>C, 7133C>T, 7135G>A, 7141G>A, 7144T>C, 7145G>T, 7147A>G, 7150G>A, 7159A>T, 7160_7165delGAGGCA, 7171C>T, 7189G>T, 7198A>G, 7201C>T, 7205C>T, 7210T>C, 7216T>A, 7219G>A, 7225C>A, 7228G>A, 7240G>A, 7243T>C, 7249A>G, 7252T>G, 7258T>A, 7261T>C, 7264T>A, 7267C>G, 7270G>A, 7273_7274insT, 7274C>G, 7275G>T, 7281A>T, 7284T>C, 7296C>T, 7311C>T, 7314C>T, 7317G>A, 7321T>A, 7322delG, 7327C>T, 7330C>T, 7333C>T, 7334A>C, 7342T>C, 7343G>A, 7344C>A, 7345A>C, 7354T>C, 7363A>G, 7369T>C, 7372C>T, 7378G>T, 7381C>T, 7396A>T, 7399C>T, 7405C>T, 7407G>T, 7408T>G, 7414G>T, 7415T>G, 7420A>G, 7429C>T, 7432T>C, 7441T>C, 7459G>A, 7462A>G, 7468A>T, 7471C>A, 7477A>G, 7480C>T, 7486G>T, 7495T>C, 7498A>G, 7501T>C, 7503T>C, 7504G>A, 7513T>A, 7519C>T, 7522A>G, 7531G>A, 7537G>A, 7540C>T, 7546T>C, 7570G>A, 7573G>A, 7579G>A, 7588T>C, 7595C>T, 7604T>C, 7606G>A, 7615T>A, 7624T>C, 7633C>T, 7636C>T, 7639T>C, 7648T>C, 7666A>G, 7672T>C, 7678A>T, 7684G>A, 7690T>C, 7696T>C, 7702T>G, 7709G>A, 7713A>G, 7714A>G, 7715C>T, 7726T>C, 7735G>A, 7744A>G, 7747C>T, 7762C>T, 7768G>A, 7771T>C, 7780A>G, 7783A>C, 7786T>C, 7793C>T, 7795A>G, 7798C>T, 7804T>C, 7812C>T, 7834A>G, 7837C>T, 7844G>T, 7846G>A, 7855G>A, 7861G>A, 7870A>G, 7873C>T, 7876T>A, 7879G>A, 7882T>A, 7883G>A, 7894A>T, 7897T>C, 7921C>T, 7927G>A, 7936T>C, 7939T>C, 7942T>C, 7966G>A, 7969T>C, 7975T>C, 7987C>T, 7990T>C, 8005T>C, 8032A>G, 8038C>T, 8041C>T, 8043G>A, 8080G>A, 8089G>A, 8095G>A, 8098G>A, 8107C>T, 8110T>C, 8116T>C, 8128A>G, 8146T>C, 8151A>G, 8152C>T, 8164G>A, 8178G>A, 8182G>A, 8191G>A, 8197C>T, 8203C>T, 8208G>A, 8213C>T, 8215G>A, 8218A>G, 8221C>T, 8224G>A, 8239A>G, 8254G>A, 8260T>C, 8263A>G, 8281T>C, 8284A>G, 8294C>G, 8296A>G, 8299G>A, 8302C>T, 8317T>C, 8320C>T, 8326T>A, 8335G>A, 8338C>T, 8349C>T, 8353A>G, 8362C>T, 8365A>G, 8368C>T, 8380C>T, 8383A>G, 8389A>G, 8404T>C, 8408G>T, 8413A>G, 8419G>T, 8434T>C, 8440T>C, 8452G>A, 8458A>G, 8460G>A, 8464A>G, 8467G>A, 8479G>A, 8480G>A, 8482T>A, 8497G>A, 8498C>T, 8500A>G, 8502G>A, 8503A>G, 8506C>T, 8515A>T, 8524C>T, 8527A>C, 8528T>C, 8530A>G, 8539G>C, 8546C>T, 8547T>G, 8552T>A, 8563A>G, 8574C>G, 8585A>G, 8598G>A, 8602C>T, 8603A>T, 8604A>C, 8605C>T, 8620A>G, 8622C>A, 8625A>G, 8626G>A, 8629G>A, 8632C>T, 8638C>T, 8641G>A, 8647C>T, 8648C>T, 8649C>T, 8650G>C, 8653C>T, 8656A>G, 8658G>A, 8664A>C, 8665T>G, 8674T>G, 8679C>T, 8680A>T, 8683C>A, 8692G>A, 8695A>T, 8707G>A, 8708A>C, 8716G>A, 8722C>T, 8725G>A, 8731A>G, 8737A>G, 8743C>T, 8745C>A, 8746A>G, 8752G>A, 8755A>T, 8770C>T, 8771T>C, 8776A>G, 8779T>C, 8782G>A, 8791T>C, 8797A>G, 8800A>G, 8803A>G, 8809T>C, 8812T>C, 8818T>C, 8824G>A, 8827A>G, 8848G>A, 8854T>C, 8856A>G, 8857G>A, 8863G>A, 8865G>A, 8867A>C, 8869G>T, 8872A>G, 8883G>A, 8888A>T, 8929C>T, 8954G>A, 8959C>T, 8962A>T, 8983G>A, 8984A>G, 8986T>C, 8988A>T, 8998G>A, 9004G>A, 9007G>A, 9016C>T, 9037C>A, 9085A>T, 9097C>T, 9131A>G, 9136A>T, 9145G>T, 9149A>G, 9150T>G, 9151G>A, 9152G>A, 9157A>G, 9160G>T, 9163T>C, 9184G>A, 9196A>G, 9214G>A, 9217A>G, 9220G>A, 9244A>T, 9247G>A, 9256C>T, 9260C>T, 9262T>C, 9274G>A, 9280C>T, 9289C>T, 9298A>C, 9304G>A, 9334C>T, 9364C>G, 9377_9378insT, 9387C>T, 9404G>A, 9411C>T, 9412A>G, 9414A>T, 9437C>T, 9454A>G, 9458T>C, 9467G>A, 9470_9471insT, 9482G>A, 9483C>T, 9498T>C, 9516T>G, 9517delA, 9533G>A, 9534G>T, 9548A>T, 9551C>T, 9553_9554delTA, 9556A>T, 9557T>A, 9558A>T, 9574T>A, 9581G>T, 9582T>G, 9583G>C, 9584_9585insG, 9590C>T, 9591T>C, 9607C>T, 9610T>C, 9615delC, 9620A>G, 9648A>G, 9674T>C, 9690C>T

| | Begin | End | Coverage | Score | Concordance | Matches | Identities | I/D/M/F* | Stop Codons |
|---|---|---|---|---|---|---|---|---|---|
| **NT** | **22** | **9704** | **99.8%** | **12522** | **65.0%** | **9671 (99.8%)** | **8003 (82.6%)** | **10/12** | |
| **CDS** | | | | | | | | | |
| PVYgp1 | 1 | 3064 | 100% | 18923 | 89.7% | 3062 (99.9%) | 2771 (90.4%) | 2/2/4/4 | 1 |
| Protein mutations: | Y4Q (194T>C 196C>A), C9L (209T>C 210G>T 211T>G), F13M (221T>A 223T>G), S23P (251T>C), C24F (255G>T 256C>T), E25G (258A>G), I27V (263A>G 265T>G), V28A (267T>C), E30V (273A>T), A35T (287G>A 289T>C), V37T (293G>A 294T>C), A41I (305G>A 306C>T), D42G (309A>G), E44D (316A>T), T45V (317A>G 318C>T 319A>G), L52Q (339T>A 340C>A), K53R (342A>G), K55A (347A>G 348A>C 349A>G), Y56H (350T>C), T58V (356A>G 357C>T), V61T (365G>A 366T>C 367G>A), L62S (368C>T 369T>C), F67C (384T>G), A75V (408C>T 409C>T), M78E (416A>G 417T>A), E85K (437G>A), R86E (440A>G 441G>A), K87R (443A>C 444A>G 445G>A), D88E (448T>A), E91A (456A>C), H93N (461C>A), D111E (517T>A), S116F (531C>T), P118S (536C>T), Q119W (539C>T 540A>G 541A>G), R122K (549G>A 550A>G), M131T (576T>C), V134A (585T>C 586C>G), R137Y (593C>T 594G>A 595C>T), P138R (597C>G), I139V (599A>G 601A>G), I140P (602A>C 603T>C 604A>G), M147I (625G>A), I151V (635A>G 637T>C), I154V (644A>G 646A>G), E160A (663A>C 664G>C), H166Q (682C>A), T172S (699C>G), H174Q (706T>A), K179E (719A>G 721G>A), I180V (722A>G 724A>G), A183S (731G>T), Y184P (734T>C 735A>C 736C>T), S185R (737T>C 738C>G 739C>T), A187V (744C>T 745G>C), R189C (749C>T 751A>C), M194K (765T>A), R197K (773C>A 774G>A 775A>G), M206K (801T>A), G210A (813G>C 814A>G), L211H (816T>A), R214Q (825G>A 826T>A), N223D (851A>G 853T>C), R226C (860C>T 862C>T), T227A (863A>G 865T>C), I228T (867T>C), N229D (869A>G 871C>T), I230L (872A>T 874A>G), R231Q (876G>A), R232K (879G>A), N240S (903A>G), T241N (906C>A 907A>C), K242T (909A>C 910A>C), S243N (912G>A 913C>T), S252L (938T>C 939C>T 940A>T), L266I (980T>A 982G>C), R272K (999G>A), Q275R (1007C>A 1008A>G 1009G>A), S276G (1010A>G 1012T>G), I277V (1013A>G), N279Q (1019A>C 1021C>G), D288E (1048C>A), R300Q (1083G>A), S305A (1097T>G 1099G>A), R320K (1143G>A), L324I (1154T>A 1156G>A), S346N (1221G>A), V349T (1229G>A 1230T>C 1231T>C), S350N (1233G>A 1234C>T), K358T (1257A>C 1258A>G), N365S (1278A>G), A369V (1290C>T 1291G>A), D372T (1298G>A 1299A>C 1300C>T), I375V (1307A>G 1309A>G), N378E (1316A>G 1318T>A), I382T (1329T>C), L397I (1373C>A), I401V (1385A>G), E407D (1405G>T), A411S (1415G>T 1417A>T), V418I (1436G>A 1438C>T), E508G (1707A>G), A512T (1718G>A), I520V (1742A>G 1744C>T), S525N (1758G>A), V536I (1790G>A), I585V (1937A>G 1939T>C), F607Y (2004T>A), I628M (2068T>G), V630I (2072G>A 2074G>A), R675K (2208G>A), D694E (2266C>G), N744G (2414A>G 2415A>G 2416T>G), S746C (2420A>T), K780R (2523A>G), R817K (2634G>A), V847I (2723G>A 2725C>T), A855T (2747G>A 2749T>A), H871N (2795C>A), V880I (2822G>A 2824G>A), P900T (2882C>A), S901H (2885A>C 2886G>A), T904M (2895C>T), N916S (2931A>G), N919S (2940A>G), E929_N930insX (2971_2972insA), T936I (2991C>T), H937R (2994A>G), E939R (2999G>A 3000A>G), R946Q (3021G>A), Y947F (3024A>T 3025C>T), T951I (3036C>T), E952G (3039A>G 3040A>C), K953T (3042A>C 3043G>A), A972V (3099C>T 3100C>A), V974R (3104G>A 3105T>G 3106G>A), S983N (3132G>A), R985K (3137C>A 3138G>A 3139A>G), F990I (3152T>A 3154C>T), C994Y (3165G>A), F1012L (3218T>C), T1014N (3225C>A), V1016I (3230G>A), V1028M (3266G>A 3268A>G), V1031M (3275G>A), A1034T (3284G>A), R1044K (3315G>A 3316A>G), E1045A (3318A>C), M1059V (3359A>G), D1072E (3400T>G), V1110G (3513T>G), M1121V (3545A>G), A1122T (3548G>A), V1124I (3554G>A 3556C>T), L1147V (3623C>G), L1150M (3632C>A), Y1152H (3638T>C), I1172V (3698A>G), S1178N (3717G>A), R1183K (3731C>A 3732G>A), I1297V (4073A>G), F1315Y (4128T>A 4129C>T), A1323S (4151G>T), V1328I (4166G>A), L1347I (4223C>A 4225G>A), I1379V (4319A>G 4321A>G), F1404Y (4395T>A), V1408I (4406G>A), M1431V (4475A>G 4477G>T), T1433S (4481A>T 4483A>G), A1454L (4544G>T 4545C>T 4546G>A), R1455K (4548G>A), V1499I (4679G>A 4681G>C), V1538I (4796G>A 4798C>T), T1540S (4803C>G 4804T>C), S1625G (5057A>G), T1626A (5060A>G), L1636I (5090C>A), A1639V (5100C>T), E1641D (5107A>T), V1646F (5120G>T 5122C>T), I1648V (5126A>G), V1667I (5183G>A), A1793T (5561G>A 5563G>A), I1805V (5597A>G), N1807K (5605C>A), A1812V (5619C>T 5620T>G), I1816L (5630A>C), V1837I (5693G>A), K1890R (5853A>G), I1904V (5894A>G 5896C>G), R1926A (5960C>G 5961G>C 5962G>C), I1936V (5990A>G), E1945D (6019A>T), K1949T (6030A>C), D1955E (6049C>A), M1958I (6058G>A), S1963N (6072G>A 6073T>C), N1964H (6074A>C), T1966N (6081C>A), C1976S (6111G>C 6112T>C), I1982V (6128A>G 6130T>C), V1992I (6158G>A), T1996S (6170A>T 6172A>G), L2007F (6203C>T 6205C>T), V2016I (6230G>A), A2025K (6257G>A 6258C>A), K2033R (6282A>G), A2060T (6362G>A), V2073I (6401G>A), A2074V (6405C>T), Y2082F (6429A>T), Q2089R (6450A>G), H2101N (6485C>A 6487C>T), L2115I (6527C>A), T2135I (6588C>T), Y2151H (6635T>C), I2155V (6647A>G), I2165V (6677A>G 6679A>G), D2178N (6716G>A), N2179D (6719A>G), I2194L (6764A>T 6766A>G), A2204V (6795C>T 6796A>G), H2205Q (6799C>A), N2225D (6857A>G), V2231T (6875G>A 6876T>C 6877C>A), D2250E (6934C>A), V2273I (7001G>A 7003G>A), F2283Y (7032T>A), T2314K (7125C>A), A2321S (7145G>T 7147A>G), E2326_A2327del (7160_7165delGAGGCA), D2344E (7216T>A), D2358E (7258T>A), D2363_R2364insX (7273_7274insT), R2364V (7274C>G 7275G>T), H2366L (7281A>T), L2367S (7284T>C), S2371L (7296C>T), S2376F (7311C>T), T2377I (7314C>T), C2378Y (7317G>A), N2379K (7321T>A), K2384Q (7334A>C), A2387N (7343G>A 7344C>A 7345A>C), S2408M (7407G>T 7408T>G), C2411G (7415T>G), L2440S (7503T>C 7504G>A), D2509N (7709G>A), K2510R (7713A>G 7714A>G), T2543I (7812C>T), V2554L (7844G>T 7846G>A), V2567I (7883G>A), S2620N (8043G>A), N2656S (8151A>G 8152C>T), R2665K (8178G>A), R2675K (8208G>A), P2704A (8294C>G 8296A>G), S2722F (8349C>T), A2742S (8408G>T), M2756I (8452G>A), R2759K (8460G>A), A2766T (8480G>A 8482T>A), R2773K (8502G>A 8503A>G), E2777D (8515A>T), E2785D (8539G>C), L2788C (8546C>T 8547T>G), S2790T (8552T>A), A2797G (8574C>G), I2801V (8585A>G), G2805E (8598G>A), N2807S (8603A>T 8604A>C 8605C>T), P2813Q (8622C>A), E2814G (8625A>G 8626G>A), P2822F (8648C>T 8649C>T 8650G>C), G2825E (8658G>A), D2827A (8664A>C 8665T>G), A2832V (8679C>T 8680A>T), T2854K (8745C>A 8746A>G), E2891G (8856A>G 8857G>A), R2894Q (8865G>A), M2895L (8867A>C 8869G>T), G2900E (8883G>A), T2902S (8888A>T), V2924I (8954G>A), N2934D (8984A>G 8986T>C), E2935V (8988A>T), I2983V (9131A>G), M2989G (9149A>G 9150T>G 9151G>A), G2990S (9152G>A), P3026S (9260C>T 9262T>C), Q3038H (9298A>C) | | | | | | | | |

| | Begin | End | Coverage | Score | Concordance | Matches | Identities | I/D/M/F* | Stop Codons |
|---|---|---|---|---|---|---|---|---|---|
| NT | 22 | 9704 | 99.8% | 12522 | 65.0% | 9671 (99.8%) | 8003 (82.6%) | 10/12 | |

Codon mutations: ACT3ACC (193T>C), TAC4CAA (194T>C 196C>A), TCA6TCG (202A>G), ATC8ATT (208C>T), TGT9CTG (209T>C 210G>T 211T>G), TCG12TCA (220G>A), TTT13ATG (221T>A 223T>G), GAA14GAG (226A>G), CTA17CTT (235A>T), TAC19TAT (241C>T), CCA21CCT (247A>T), TCT23CCT (251T>C), TGC24TTT (255G>T 256C>T), GAG25GGG (258A>G), CAT26CAC (262T>C), ATT27GTG (263A>G 265T>G), GTG28GCG (267T>C), GAA30GTA (273A>T), CGA31CGT (277A>T), GAA32GAG (280A>G), GTG33GTT (283G>T), CCG34CCA (286G>A), GCT35ACC (287G>A 289T>C), GTT37ACT (293G>A 294T>C), CCT39CCC (301T>C), TTC40TTT (304C>T), GCA41ATA (305G>A 306C>T), GAT42GGT (309A>G), CTG43TTG (311C>T), GAA44GAT (316A>T), ACA45GTG (317A>G 318C>T 319A>G), CAA46CAG (322A>G), GCA49GCT (331A>T), CTC52CAA (339T>A 340C>A), AAG53AGG (342A>G), AAA55GCG (347A>G 348A>C 349A>G), TAT56CAT (350T>C), GCT57GCA (355T>A), ACT58GTT (356A>G 357C>T), CGT60CGC (364T>C), GTG61ACA (365G>A 366T>C 367G>A), CTC62TCC (368C>T 369T>C), AAA63AAG (373A>G), AAC64AAT (376C>T), TTT67TGT (384T>G), ACG68ACA (388G>A), TAC69TAT (391C>T), CGA70CGG (394A>G), AAG72AAA (400G>A), GAT74GAC (406T>C), GCC75GTT (408C>T 409C>T), CAG76CAA (412G>A), ATA77ATT (415A>T), ATG78GAG (416A>G 417T>A), ATT80ATC (424T>C), CAG81CAA (427G>A), AAA83AAG (433A>G), CTG84CTC (436G>C), GAG85AAG (437G>A), AGG86GAG (440A>G 441G>A), AAG87CGA (443A>C 444A>G 445G>A), GAT88GAA (448T>A), GAA90GAG (454A>G), GAA91GCA (456A>C), TAT92TAC (460T>C), CAC93AAC (461C>A), CAA95CAG (469A>G), GCC97GCG (475C>G), CCT99CCA (481T>A), ACT106ACC (502T>C), ATA107ATT (505A>T), GGC109GGT (511C>T), GAT111GAA (517T>A), CCT112CCA (520T>A), CCA113CCT (523A>T), TCT116TTT (531C>T), GAG117GAA (535G>A), CCA118TCA (536C>T), CAA119TGG (539C>T 540A>G 541A>G), AGA122AAG (549G>A 550A>G), ATC124ATT (556C>T), ATT125ATC (559T>C), CAT126CAC (562T>C), ACT128ACC (568T>C), CCA129CCG (571A>G), ATG131ACG (576T>C), CGT132CGC (580T>C), GTC134GCG (585T>C 586C>G), CGC137TAT (593C>T 594G>A 595C>T), CCC138CGC (597C>G), ATA139GTG (599A>G 601A>G), ATA140CCG (602A>C 603T>C 604A>G), ACA143ACG (613A>G), GAA144GAG (616A>G), GGC145GGG (619C>G), CAG146CAA (622G>A), ATG147ATA (625G>A), AAT148AAC (628T>C), CAC149CAT (631C>T), ATT151GTC (635A>G 637T>C), ATA154GTG (644A>G 646A>G), ATT157ATC (655T>C), TCG159TCA (661G>A), GAG160GCC (663A>C 664G>C), AAA161AAG (667A>G), GTC165GTT (679C>T), CAC166CAA (682C>A), TTA167CTG (683T>C 685A>G), AGT169AGC (691T>C), ACC172AGC (699C>G), ACT173ACA (703T>A), CAT174CAA (706T>A), AAG179GAA (719A>G 721G>A), ATA180GTG (722A>G 724A>G), CTT181TTG (725C>T 727T>G), GGT182GGA (730T>A), GCA183TCA (731G>T), TAC184CCT (734T>C 735A>C 736C>T), TCC185CGT (737T>C 738C>G 739C>T), GCG187GTC (744C>T 745G>C), CGA189TGC (749C>T 751A>C), ACT190ACC (754T>C), ATG194AAG (765T>A), TTG196CTG (770T>C), CGA197AAG (773C>A 774G>A 775A>G), CGG198AGG (776C>A), TTC202TTT (790C>T), ATG206AAG (801T>A), GGA210GCG (813G>C 814A>G), CTT211CAT (816T>A), TTG212CTG (818T>C), CAA213CAG (823A>G), CGT214CAA (825G>A 826T>A), CGG217AGA (833C>A 835G>A), GAC219GAT (841C>T), AAT223GAC (851A>G 853T>C), GTC225GTT (859C>T), CGC226TGT (860C>T 862C>T), ACT227GCC (863A>G 865T>C), ATC228ACC (867T>C), AAC229GAT (869A>G 871C>T), ATA230TTG (872A>T 874A>G), CGA231CAA (876G>A), AGG232AAG (879G>A), GAT234GAC (886T>C), ATC238ATT (898C>T), AAC240AGC (903A>G), ACA241AAC (906C>A 907A>C), AAA242ACC (909A>C 910A>C), AGC243AAT (912G>A 913C>T), GGC246GGA (922C>A), GGT249GGA (931T>A), TCA252CTT (938T>C 939C>T 940A>T), GGA253GGG (943A>G), TTC256TTT (952C>T), TCA261TCG (967A>G), GAA263GAG (973A>G), GGG264GGC (976G>C), TTG266ATC (980T>A 982G>C), GCA269GCG (991A>G), CGT270CGC (994T>C), AGA272AAA (999G>A), GTT273GTC (1003T>C), CAG275AGA (1007C>A 1008A>G 1009G>A), AGT276GGG (1010A>G 1012T>G), ATT277GTT (1013A>G), TTA278TTG (1018A>G), AAC279CAG (1019A>C 1021C>G), ATC282ATT (1030C>T), TCG285TCA (1039G>A), AAT286AAC (1042T>C), GCC287GCA (1045C>A), GAC288GAA (1048C>A), AAT289AAC (1051T>C), AAG292AAA (1060G>A), GGT293GGA (1063T>A), CTG294TTG (1064C>T), GAC295GAT (1069C>T), CGA300CAA (1083G>A), TAT303TAC (1093T>C), TCG305GCA (1097T>G 1099G>A), CAC307CAT (1105C>T), GTA310GTG (1114A>G), GCT311GCG (1117T>G), TTA313TTG (1123A>G), CCT314CCA (1126T>A), GTC315GTT (1129C>T), AGG320AAG (1143G>A), GTA321GTT (1147A>T), GCT322GCA (1150T>A), TTG324ATA (1154T>A 1156G>A), CAC327CAT (1165C>T), AGT328AGC (1168T>C), CTT330TTA (1172C>T 1174T>A), ACC339ACT (1201C>T), AGC346AAC (1221G>A), GTT349ACC (1229G>A 1230T>C 1231T>C), AGC350AAT (1233G>A 1234C>T), CTG352CTA (1240G>A), AAG354AAA (1246G>A), TTG356CTA (1250T>C 1252G>A), CAT357CAC (1255T>C), AAA358ACG (1257A>C 1258A>G), AGA361AGG (1267A>G), GGT363GGC (1273T>C), TTG364TTA (1276G>A), AAT365AGT (1278A>G), TTG367CTG (1283T>C), GCG369GTA (1290C>T 1291G>A), GAC372ACT (1298G>A 1299A>C 1300C>T), TTT374TTC (1306T>C), ATA375GTG (1307A>G 1309A>G), CAT376CAC (1312T>C), AAT378GAA (1316A>G 1318T>A), TTG381TTA (1327G>A), ATA382ACA (1329T>C), GCG383GCT (1333G>T), TTA384TTG (1336A>G), CTA387TTG (1343C>T 1345A>G), CCG390CCA (1354G>A), GTG391GTT (1357G>T), GAC392GAT (1360C>T), CTC395TTG (1367C>T 1369C>G), CTT397ATT (1373C>A), TTC398TTT (1378C>T), GAG400GAA (1384G>A), ATA401GTA (1385A>G), AAA403AAG (1393A>G), TCC404TCT (1396C>T), GAG407GAT (1405G>T), AAA408AAG (1408A>G), CAG409CAA (1411G>A), GCA411TCT (1415G>T 1417A>T), CCG412CCC (1420G>C), AAT415AAC (1429T>C), TTA416TTG (1432A>G), GTC418ATT (1436G>A 1438C>T), TTA419CTG (1439T>C 1441A>G), AAT420AAC (1444T>C), TTC423TTT (1453C>T), CTG424CTA (1456G>A), GAA428GAG (1468A>G), GTA436GTG (1492A>G), TTG439CTA (1499T>C 1501G>A), AGT440AGC (1504T>C), TTG441TTA (1507G>A), CTC442CTT (1510C>T), GAA443GAG (1513A>G), TTA444TTG (1516A>G), GCA445GCT (1519A>T), AGG446AGA (1522G>A), TTC447TTT (1525C>T), CAG448CAA (1528G>A), AAC450AAT (1534C>T), ACT452ACA (1540T>A), AAC454AAT (1546C>T), AAA457AAG (1555A>G), GGT458GGC (1558T>C), GAT459GAC (1561T>C), ATA460ATC (1564A>C), TCT461TCA (1567T>A), TTC463TTT (1573C>T), AGA464AGG (1576A>G), TTA467CTA (1583T>C), TCT468TCA (1588T>A), AAG470AAA (1594G>A), AAC472AAT (1600C>T), CTG475CTA (1609G>A), TCG478TCA (1618G>A), TGC479TGT (1621C>T), GAC480GAT (1624C>T), CTG483TTG (1631C>T), AAA485AAG (1639A>G), TTC489TTT (1651C>T), CTC490CTG (1654C>G), CAT497CAC (1675T>C), TCA503TCG (1693A>G), TTT506TTC (1702T>C), GAA508GGA (1707A>G), GCA512ACA (1718G>A), GGA514GGG (1726A>G), TAC515TAT (1729C>T), TCA516TCC (1732A>C), TAT518TAC (1738T>C), ATC520GTT (1742A>G 1744C>T), CGC521CGT (1747C>T), CAT523CAC (1753T>C), AGT525AAT (1758G>A), AGG528AGA (1768G>A), AAC534AAT (1786C>T), TTA535TTG (1789A>G), GTT536ATT (1790G>A), GTC537GTT (1795C>T), TTA541CTG (1805T>C 1807A>G), GAG543GAA (1813G>A), AGG545AGA (1819G>A), AAG547AAA (1825G>A), GAC551GAT (1837C>T), TAT552TAC (1840T>C), CAA555CAG (1849A>G), GTC558GTG (1858C>G), AAA560AAG (1864A>G), TGC562TGT (1870C>T), AGT564AGC (1876T>C), AAA566AAG (1882A>G), GGT568GGA (1888T>A), AAT569AAC (1891T>C), TGT574TGC (1906T>C), ACA577ACT (1915A>T), GGT582GGC (1930T>C), GCC584GCT (1936C>T), ATT585GTC (1937A>G 1939T>C), TCA587TCG (1945A>G), TTC589TTT (1951C>T), CCA592CCT (1960A>T), GTA598GTG (1978A>G), ATT599ATA (1981T>A), GGC600GGT (1984C>T), AAT601AAC (1987T>C), GAC604GAT (1996C>T), TTT607TAT (2004T>A), AAA612AAG (2020A>G), GGG613GGA (2023G>A), GAT614GAC (2026T>C), TCG615TCT (2029G>T), GAG616GAA (2032G>A), TAT625TAC (2059T>C), ATT628ATG (2068T>G), AAC629AAT (2071C>T), GTG630ATA (2072G>A 2074G>A), CTG635TTA (2087C>T 2089G>A), AGC639AGT (2101C>T), GAG640GAA (2104G>A), GAG641GAA (2107G>A), AAG644AAA (2116G>A), TTC646TTT (2122C>T), CGC651CGT (2137C>T), CTT658CTC (2158T>C), ACC660ACT (2164C>T), CCA662CCG (2170A>G), GAT666GAC (2182T>C), GCG668GCA (2188G>A), ACC669ACA (2191C>A), AGA675AAA (2208G>A), TAT678TAC (2218T>C), GAC680GAT (2224C>T), GTA681GTT (2227A>T), TTG686TTA (2242G>A), CCC687CCT (2245C>T), TTG690TTA (2254G>A), GTT691GTC (2257T>C), GAC692GAT (2260C>T), GAC694GAG (2266C>G), CAA696CAG (2272A>G), ACG697ACA (2275G>A), TCA703TCG (2293A>G), GGC705GGT (2299C>T), TCG706TCA (2302G>A), CAG707CAA (2305G>A), GGA710GGT (2314A>T), TAT711TAC (2317T>C), CTA714CTG (2326A>G), GTG719GTA (2341G>A), CAA721CAG (2347A>G), CTT722CTC (2350T>C), TTG724TTA (2356G>A), AAT727AAC (2365T>C), GAA729GAG (2371A>G), TTA730TTG (2374A>G), GAT733GAC (2383T>C), ATA734ATT (2386A>T), AAA735AAG (2389A>G), CAT736CAC (2392T>C), TAT737TAC (2395T>C), AAT744GGG (2414A>G 2415A>G 2416T>G), GCT745GCA (2419T>A), AGC746TGC (2420A>T), ACA752ACT (2440A>T), GTT761GTC (2467T>C), TCT764TCC (2476T>C), TCG766TCA (2482G>A), CTG769CTA (2491G>A), CTG771CTT (2497G>T), TTG773TTA (2503G>A), AGA778AGG (2518A>G), CCT779CCC (2521T>C), AAG780AGG (2523A>G), AGA783AGG (2533A>G), TTG785TTA (2539G>A), TTA787TTG (2545A>G), GAG789GAA (2551G>A), CCT790CCA (2554T>A), CTG792TTG (2558C>T), TTG793CTC (2561T>C 2563G>C), TCA796TCG (2572A>G), CCT800CCC (2584T>C), GGC801GGT (2587C>T), CTG803CTC (2593G>C), GCT805GCC (2599T>C), AAT808AAC (2608T>C), TTT812TTC (2620T>C), GCG815GCA (2629G>A), GTG816GTA (2632G>A), AGG817AAG (2634G>A), TTG818TTA (2638G>A), ATT820ATA (2644T>A), AAA823AAG (2653A>G), GCT827GCC (2665T>C), CTA832TTA (2678C>T), CTA833CTG (2683A>G), GCT835GCC (2689T>C), CTA838CTC (2698A>C), TCA841TCG (2707A>G), GCG842GCA (2710G>A), GAA844GAG (2716A>G), ACA845ACG (2719A>G), CTC846CTT (2722C>T), GTC847ATT (2723G>A 2725C>T), GCA848GCG (2728A>G), AGG850AGA (2734G>A), ATT851ATA (2737T>A), GAT854GAC (2746T>C), GCT855ACA (2747G>A 2749T>A), GCA856GCT (2752A>T), GCT857GCA (2755T>A), GAC859GAT (2761C>T), CTC860CTT (2764C>T), TGT865TGC (2779T>C), GGG867GGT (2785G>T), AAC869AAT (2791C>T), CAT871AAT (2795C>A), TAC874TAT (2806C>T), TTG878CTC (2816T>C 2818G>C), GTG880ATA (2822G>A 2824G>A), TTG881TTA (2827G>A), GTT884GTC (2836T>C), AAG885AAA (2839G>A), AAT886AAC (2842T>C), ACC893ACA (2863C>A), CTA894TTA (2864C>T), CCA900ACA (2882C>A), AGT901CAT (2885A>C 2886G>A), AAC903AAT (2893C>T), ACG904ATG (2895C>T), GTC906GTT (2902C>T), GTA907GTG (2905A>G), ATT909ATC (2911T>C), AAT913AAC (2923T>C), AAT916AGT (2931A>G), AAC919AGC (2940A>G), GCT921GCC (2947T>C), ACT926ACC (2962T>C), GAA929_AAT930insA-- (2971_2972insA), GGT935GGC (2989T>C), ACT936ATT (2991C>T), CAT937CGT (2994A>G), GAG939AGG (2999G>A 3000A>G), TCC944TC- (3016delC), CGG946CAG (3021G>A), TAC947TTT (3024A>T 3025C>T), ACA951ATA (3036C>T), GAA952GGC (3039A>G 3040A>C), AAG953ACA (3042A>C 3043G>A), TTG956TTA (3052G>A), TTA959TTG (3061A>G), GCG966GCA (3082G>A), GGC969GGT (3091C>T), GCC972GTA (3099C>T 3100C>A), GTG974AGA (3104G>A 3105T>G 3106G>A), GTC975GTT (3109C>T), AAA976AAG (3112A>G), ACT978ACC (3118T>C), GCC979GCT (3121C>T), GGA981GGG (3127A>G), AGC983AAC (3132G>A), CGA985AAG (3137C>A 3138G>A 3139A>G), TAT989TAC (3151T>C), TTC990ATT (3152T>A 3154C>T), TGT994TAT (3165G>A), GTA995GTG (3169A>G), ATC1005ATT (3199C>T), AGG1007CGG (3203A>C), CGT1008CGC (3208T>C), CCA1010CCG (3214A>G), ACC1011ACT (3217C>T), TTT1012CTT (3218T>C), GTC1013GTA (3223C>A), ACT1014AAT (3225C>A), GTT1016ATT (3230G>A), AAC1017AAT (3235C>T), TTA1025CTA (3257T>C), ACC1026ACT (3262C>T), AGC1027AGT (3265C>T), GTA1028ATG (3266G>A 3268A>G), GTG1029GTA (3271G>A), GTG1031ATG (3275G>A), GCA1034ACA (3284G>A), CAG1039CAA (3301G>A), AGA1044AAG (3315G>A 3316A>G), GAA1045GCA (3318A>C), ATC1046ATT (3322C>T), GAG1047GAA (3325G>A), TTG1048TTA (3328G>A), ATA1051ATT (3337A>T), GAG1055GAA (3349G>A), GTC1057GTT (3355C>T), TGC1058TGT (3358C>T), ATG1059GTG (3359A>G), CTA1061CTG (3367A>G), TAT1062TAC (3370T>C), TTA1065CTG (3377T>C 3379A>G), GAA1070GAG (3394A>G), CGC1071CGT (3397C>T), GAT1072GAG (3400T>G), ACA1074ACG (3406A>G), TAC1078TAT (3418C>T), ATT1079ATA (3421T>A), TAT1081TAC (3427T>C), AAG1083AAA (3433G>A), TCA1084TCT (3436A>T), GTA1085GTG (3439A>G), AAC1086AAT (3442C>T), CCT1087CCC (3445T>C), CAG1091CAA (3457G>A), TTT1092TTC (3460T>C), GCT1093GCA (3463T>A), GCG1095GCT (3469G>T), CAG1096CAA (3472G>A), CAC1104CAT (3496C>T), CAG1105CAA (3499G>A), TCC1107TCT (3505C>T), GTT1110GGT (3513T>G), AAA1112AAG (3520A>G), GAA1115GAG (3529A>G), CAA1116CAG (3532A>G), ATG1121GTG (3545A>G), GCT1122ACT (3548G>A), TTA1123CTA (3551T>C), GTC1124ATT (3554G>A 3556C>T), AGT1133AGC (3583T>C), GAT1134GAC (3586T>C), TGC1135TGT (3589C>T), TTC1137TTT (3595C>T), AAT1141AAC (3607T>C), TTT1143TTC (3613T>C), AAG1144AAA (3616G>A), GGT1145GGC (3619T>C), CTT1147GTT (3623C>G), TCC1148TCT (3628C>T), CTG1150ATG (3632C>A), GAT1151GAC (3637T>C), TAT1152CAT (3638T>C), GAA1153GAG (3643A>G), TTA1159TTG (3661A>G), GTG1162GTA (3670G>A), TTT1166TTC (3682T>C), GAG1168GAA (3688G>A), GAG1171GAA (3697G>A), ATT1172GTT (3698A>G), GAT1174GAC (3706T>C), TTT1175TTC (3709T>C), TTG1177CTA (3713T>C 3715G>A), AGT1178AAT (3717G>A), GAG1179GAA (3721G>A), CGA1183AAA (3731C>A 3732G>A), ACT1184ACA (3736T>A), CTA1188TTG (3746C>T 3748A>G), ACA1190ACG (3754A>G), AGT1193AGC (3763T>C), CGA1198CGG (3778A>G), CAA1199CAG (3781A>G), ATC1200ATT (3784C>T), CAT1204CAC (3796T>C), CCA1207CCT (3805A>T), TAC1209TAT (3811C>T), AGA1210AGG (3814A>G), TTC1215TTT (3829C>T), TTT1218TTC (3838T>C), AGA1220AGG (3844A>G), GCA1221GCC (3847A>C), GTT1224GTA (3856T>A), GTG1226GTT (3862G>T), GAC1229GAT (3871C>T), GCC1231GCA (3877C>A), AGC1233AGT (3883C>T), GAA1234GAG (3886A>G), CTA1236CTG (3892A>G), GAC1237GAT (3895C>T), TTT1238TTC (3898T>C), GTA1240GTG (3904A>G), CGG1241AGA (3905C>A 3907G>A), GCT1243GCC (3913T>C), AAG1248AAA (3928G>A), TCA1249TCT (3931A>T), GGG1251GGA (3937G>A), AGT1257AGC (3955T>C), GTG1258GTA (3958G>A), GCC1259GCT (3961C>T), GGA1260GGG (3964A>G), TCT1261TCC (3967T>C), TTA1264TTG (3976A>G), ATT1265ATC (3979T>C), CCA1267CCT (3985A>T), CTA1271CTT (3997A>T), GCG1272GCA (4000G>A), AAC1274AAT (4006C>T), GTT1275GTG (4009T>G), TTC1276TTT (4012C>T), AAA1277AAG (4015A>G), TCT1280TCC (4024T>C), GAA1282GAG (4030A>G), CCA1283CCT (4033A>T), AAG1286AAA (4042G>A), CTG1290TTA (4052C>T 4054G>A), CGT1291CGC (4057T>C), CGT1293CGA (4063T>A), GGA1294GGG (4066A>G), ATA1297GTA (4073A>G), TTT1298TTC (4078T>C), GGC1299GGT (4081C>T), TCT1300TCC (4084T>C), TCC1304TCT (4096C>T), AGC1308AGT (4108C>T), GGA1309GGT (4111A>T), GCG1311GCA (4117G>A), CTA1312TTG (4118C>T 4120A>G), TAC1314TAT (4126C>T), TTC1315TAT (4128T>A 4129C>T), GCC1316GCT (4132C>T), AAT1317AAC (4135T>C), CAA1321CAG (4147A>G), TTA1322CTA (4148T>C), GCT1323TCT (4151G>T), AAC1326AAT (4162C>T), GTA1328ATA (4166G>A), ATA1329ATC (4171A>C), GAG1332GAA (4180G>A), CTG1336TTG (4190C>T), CCT1338CCC (4198T>C), TCC1339TCT (4201C>T), GCG1342GCT (4210G>T), TTC1343TTT (4213C>T), AGT1345AGC (4219T>C), CTG1346TTG (4220C>T), CTG1347ATA (4223C>A 4225G>A), GTT1349GTG (4231T>G), CAT1351CAC (4237T>C), GTA1356GTG (4252A>G), TTA1357CTG (4253T>C 4255A>G), AAA1358AAG (4258A>G), ACT1362ACC (4270T>C), AGA1366AGG (4282A>G), GAG1367GAA (4285G>A), GAA1369GAG (4291A>G), CAG1373CAA (4303G>A), GTC1376GTT (4312C>T), ATA1379GTG (4319A>G 4321A>G), GTG1380GTT (4324G>T), GAG1381GAA (4327G>A), CTG1384CTT (4336G>T), TCT1385TCC (4339T>C), TCA1388TCT (4348A>T), GCA1392GCG (4360A>G), GGT1394GGC (4366T>C), TCT1395TCA (4369T>A), ACT1397ACC (4375T>C), GTT1401GTC (4387T>C), TTT1404TAT (4395T>A), GTA1408ATA (4406G>A), CTT1409CTC (4411T>C), GTT1418GTC (4438T>C), ACC1420ACA (4444C>A), GCC1422GCT (4450C>T), CTC1424CTT (4456C>T), GAC1427GAT (4465C>T), ATG1431GTT (4475A>G 4477G>T), GTC1432GTT (4480C>T), ACA1433TCG (4481A>T 4483A>G), GGC1437GGT (4495C>T), AAG1441AAA (4507G>A), CAC1442CAT (4510C>T), GGT1443GGG (4513T>G), CTA1445TTA (4517C>T), GTC1448GTT (4528C>T), AAA1450AAG (4534A>G), ACC1452ACT (4540C>T), GCG1454TTA (4544G>T 4545C>T 4546G>A), AGA1455AAA (4548G>A), TTT1458TTC (4558T>C), GTA1460GTT (4564A>T), ACC1462ACT (4570C>T), ATA1464ATT (4576A>T), TTG1471TTA (4597G>A), GAC1472GAT (4600C>T), GAC1474GAT (4606C>T), GTG1475GTA (4609G>A), GTG1477GTA (4615G>A), GAC1478GAT (4618C>T), TTG1481CTT (4625T>C 4627G>T), GAC1488GAT (4648C>T), ATT1489ATC (4651T>C), AGG1492AGA (4660G>A), AGC1493AGT (4663C>T), GCT1495GCA (4669T>A), GTG1499ATC (4679G>A 4681G>C), AGT1500AGC (4684T>C), GTT1501GTG (4687T>G), GAG1505GAA (4699G>A), AGA1506AGG (4702A>G), CAA1508CAG (4708A>G), AGG1509AGA (4711G>A), CTG1510TTG (4712C>T), AAA1518AAG (4738A>G), GTA1520GTT (4744A>T), GCA1521GCG (4747A>G), TTG1522CTG (4748T>C), ACT1527ACC (4765T>C), GTC1538ATT (4796G>A 4798C>T), ACT1540AGC (4803C>G 4804T>C), GAG1541GAA (4807G>A), GCG1542GCT (4810G>T), GCT1543GCG (4813T>G), CTT1544CTC (4816T>C), TAT1549TAC

| | | | | | | | | | |
|---|---|---|---|---|---|---|---|---|---|
| | (4831T>C), AAC1550AAT (4834C>T), CCA1552CCT (4840A>T), GGA1556GGG (4852A>G), GGC1557GGT (4855C>T), AGT1561AGC (4867T>C), CTG1562CTC (4870G>C), ATT1563ATA (4873T>A), TGT1566TGC (4882T>C), ACA1573ACT (4903A>T), CAG1575CAA (4909G>A), TTT1577TTC (4915T>C), GAA1578GAG (4918A>G), TTG1579CTA (4919T>C 4921G>A), AGT1580AGC (4924T>C), CCC1581CCG (4927C>G), TTT1583TTC (4933T>C), ATC1584ATA (4936C>A), CAG1585CAA (4939G>A), AAT1586AAC (4942T>C), TTC1587TTT (4945C>T), TCA1593TCG (4963A>G), CCT1596CCC (4972T>C), CTT1602CTC (4990T>C), AAA1606AAG (5002A>G), CCT1613CCC (5023T>C), TTG1614CTG (5024T>C), TGC1615TGT (5029C>T), GAT1616GAC (5032T>C), TCT1618TCC (5038T>C), CCA1620CCT (5044A>T), GCA1623GCT (5053A>T), TCG1624TCA (5056G>A), AGT1625GGT (5057A>G), ACT1626GCT (5060A>G), TCG1629TCT (5071G>T), CTT1636ATT (5090C>A), GGA1637GGG (5095A>G), GTG1638GTT (5098G>T), GCC1639GTC (5100C>T), TTA1640TTG (5104A>G), GAA1641GAT (5107A>T), ATT1642ATC (5110T>C), CCA1643CCG (5113A>G), AAG1644AAA (5116G>A), CAA1645CAG (5119A>G), GTC1646TTT (5120G>T 5122C>T), ATT1648GTT (5126A>G), CAT1651CAC (5137T>C), ATC1652ATT (5140C>T), AAA1653AAG (5143A>G), GAG1654GAA (5146G>A), CTC1659CTA (5161C>A), GAA1661GAG (5167A>G), CTT1663CTG (5173T>G), ACG1666ACA (5182G>A), GTT1667ATT (5183G>A), GTC1668GTT (5188C>T), AAG1669AAA (5191G>A), AAA1671AAG (5197A>G), GAC1672GAT (5200C>T), TGC1674TGT (5206C>T), TTA1675TTG (5209A>G), AGC1678AGT (5218C>T), TCG1682TCA (5230G>A), ATC1684ATT (5236C>T), AGC1685AGT (5239C>T), ATC1687ATT (5245C>T), ACA1693ACT (5263A>T), CTC1695CTT (5269C>T), TTC1696TTT (5272C>T), GCC1697GCA (5275C>A), ATC1698ATT (5278C>T), CCA1699CCC (5281A>C), AGA1700AGG (5284A>G), ACT1701ACC (5287T>C), CTA1702TTA (5288C>T), ATA1703ATT (5293A>T), TTG1704CTA (5294T>C 5296G>A), GAG1706GAA (5302G>A), CTT1709CTA (5311T>A), GAA1710GAG (5314A>G), CGA1713CGT (5323A>T), AGT1721AGC (5347T>C), ATC1723ATT (5353C>T), TGC1727TGT (5365C>T), GTT1734GTC (5386T>C), ACC1737ACA (5395C>A), AAA1746AAG (5422A>G), GAT1747GAC (5425T>C), ACC1749ACT (5431C>T), GAG1751GAA (5437G>A), CTT1756CTC (5452T>C), AAG1758AAA (5458G>A), AGT1761AGC (5467T>C), CAA1762CAG (5470A>G), CTA1763TTG (5471C>T 5473A>G), GAA1765GAG (5479A>G), TTC1766TTT (5482C>T), TCA1767TCC (5485A>C), AAT1768AAC (5488T>C), TTG1769CTA (5489T>C 5491G>A), GGT1771GGC (5497T>C), TGT1774TGC (5506T>C), GAG1775GAA (5509G>A), GAG1776GAA (5512G>A), TTA1778CTA (5516T>C), TAT1782TAC (5530T>C), GAG1783GAA (5533G>A), TCG1784TCT (5536G>T), TTG1785TTA (5539G>A), GTT1788GTG (5548T>G), CAC1790CAT (5554C>T), GCT1792GCA (5560T>A), GCG1793ACA (5561G>A 5563G>A), ACG1794ACC (5566G>C), TCA1795TCG (5569A>G), CTC1800CTG (5584C>G), AAG1803AAA (5593G>A), GGG1804GGA (5596G>A), ATT1805GTT (5597A>G), AAC1807AAA (5605C>A), GTG1811GTT (5617G>T), GCT1812GTG (5619C>T 5620T>G), TTG1815TTA (5629G>A), ATC1816CTC (5630A>C), GGC1819GGT (5641C>T), GCA1822GCT (5650A>T), ATA1826ATT (5662A>T), GGA1827GGG (5665A>G), CAA1835CAG (5689A>G), GTT1837ATT (5693G>A), GAG1838GAA (5698G>A), CAA1843CAG (5713A>G), GGG1844GGC (5716G>C), AAA1845AAG (5719A>G), GCC1853GCA (5743C>A), CGC1857CGA (5755C>A), CAT1858CAC (5758T>C), CGT1860CGC (5764T>C), AAA1862AAG (5770A>G), GGC1865GGT (5779C>T), TTT1866TTC (5782T>C), GAC1869GAT (5791C>T), GAC1873GAT (5803C>T), AGG1884AGA (5836G>A), AAA1885AAG (5839A>G), GGA1887GGG (5845A>G), AAA1888AAG (5848A>G), AAA1890AGA (5853A>G), GGT1891GGC (5857T>C), ACC1892ACT (5860C>T), ACA1893ACT (5863A>T), GGT1895GGA (5869T>A), GGT1897GGC (5875T>C), AGC1900AGT (5884C>T), TTC1903TTT (5893C>T), ATC1904GTG (5894A>G 5896C>G), AAC1905AAT (5899C>T), GGG1908GGA (5908G>A), TTT1909TTC (5911T>C), GAG1913GAA (5923G>A), CAA1918CAG (5938A>G), TTC1919TTT (5941C>T), GTT1920GTC (5944T>C), GAT1921GAC (5947T>C), GGG1925GGA (5959G>A), CGG1926GCC (5960C>G 5961G>C 5962G>C), GAA1930GAG (5974A>G), GCT1934GCA (5986T>A), GAC1935GAT (5989C>T), ATT1936GTT (5990A>G), GAT1938GAC (5998T>C), AGA1942CGT (6008A>C 6010A>T), GAA1945GAT (6019A>T), GTG1946GTT (6022G>T), CGA1947CGT (6025A>T), AAA1949ACA (6030A>C), GAC1955GAA (6049C>A), ATG1958ATA (6058G>A), CAA1959CAG (6061A>G), GCC1960GCA (6064C>A), GGT1962GGC (6070T>C), AGT1963AAC (6072G>A 6073T>C), AAC1964CAC (6074A>C), ACG1965ACA (6079G>A), ACC1966AAC (6081C>A), ATA1967ATT (6085A>T), GCA1969GCT (6091A>T), TAC1970TAT (6094C>T), TGT1976TCC (6111G>C 6112T>C), AAA1978AAG (6118A>G), TTG1980TTA (6124G>A), AAG1981AAA (6127G>A), ATT1982GTC (6128A>G 6130T>C), GAT1983GAC (6133T>C), TTA1984TTG (6136A>G), AAC1988AAT (6148C>T), AAA1991AAG (6157A>G), GTT1992ATT (6158G>A), TGT1993TGC (6163T>C), AAA1995AAG (6169A>G), ACA1996TCG (6170A>T 6172A>G), GGC1998GGT (6178C>T), AAA2001AAG (6187A>G), TTT2002TTC (6190T>C), GAG2004GAA (6196G>A), CTC2007TTT (6203C>T 6205C>T), GAA2008GAG (6208A>G), CTA2009TTG (6209C>T 6211A>G), CAG2011CAA (6217G>A), GCT2015GCA (6229T>A), GTA2016ATA (6230G>A), GTC2018GTT (6238C>T), AAG2021AAA (6247G>A), ATA2023ATT (6253A>T), GCA2025AAA (6257G>A 6258C>A), CAT2030CAC (6274T>C), AAA2033AGA (6282A>G), TCG2034TCA (6286G>A), GGC2038GGT (6298C>T), TTG2039TTA (6301G>A), AGA2040AGG (6304A>G), GAC2041GAT (6307C>T), GCC2046GCT (6322C>T), ACA2048ACT (6328A>T), GTT2049GTG (6331T>G), TGT2050TGC (6334T>C), AGG2051AGA (6337G>A), CTG2052TTG (6338C>T), TAT2058TAC (6358T>C), GGG2059GGA (6361G>A), GCA2060ACA (6362G>A), TCA2061TCT (6367A>T), GAG2062GAA (6370G>A), TAC2064TAT (6376C>T), GGT2065GGG (6379T>G), GGA2067GGC (6385A>C), GGA2069GGT (6391A>T), GCA2070GCG (6394A>G), TAC2071TAT (6397C>T), GTA2073ATA (6401G>A), GCG2074GTG (6405C>T), CAC2076CAT (6412C>T), TTA2078TTG (6418A>G), TTT2079TTC (6421T>C), TAC2082TTC (6429A>T), GGT2084GGA (6436T>A), GAG2087GAA (6445G>A), CAA2089CGA (6450A>G), TCC2090TCG (6454C>G), CAC2092CAT (6460C>T), GGT2093GGA (6463T>A), TTC2095TTT (6469C>T), AGG2096AGA (6472G>A), GTG2097GTA (6475G>A), AAT2099AAC (6481T>C), CAC2101AAT (6485C>A 6487C>T), AGT2102AGC (6490T>C), AGC2104AGT (6496C>T), CTG2106CTA (6502G>A), GGT2110GGC (6514T>C), AGG2111AGA (6517G>A), CTC2115ATC (6527C>A), ATC2116ATA (6532C>A), CCG2119CCA (6541G>A), AAA2120AAG (6544A>G), GTC2124GTT (6556C>T), TTT2125TTC (6559T>C), CAG2127CAA (6565G>A), TTG2129CTA (6569T>C 6571G>A), CCT2134CCA (6586T>A), ACA2135ATA (6588C>T), GAA2138GAG (6598A>G), AGA2139AGG (6601A>G), ATT2140ATC (6604T>C), TGT2141TGC (6607T>C), GGA2144GGG (6616A>G), AAC2146AAT (6622C>T), GAG2149GAA (6631G>A), AAG2150AAA (6634G>A), TAT2151CAT (6635T>C), GCT2152GCA (6640T>A), TCG2154TCT (6646G>T), ATC2155GTC (6647A>G), ATC2156ATT (6652C>T), AGC2160AGT (6664C>T), ACT2162ACG (6670T>G), ATA2165GTG (6677A>G 6679A>G), GGC2167GGT (6685C>T), ACA2169ACT (6691A>T), TTC2170TTT (6694C>T), ACA2177ACG (6715A>G), GAT2178AAT (6716G>A), AAT2179GAT (6719A>G), GGA2180GGG (6724A>G), CTA2184TTA (6734C>T), GTG2186GTA (6742G>A), GTG2187GTA (6745G>A), GCC2190GCT (6754C>T), GGA2192GGG (6760A>G), ATA2194TTG (6764A>T 6766A>G), ATT2197ATA (6775T>A), CTG2200TTG (6782C>T), GCA2201GCT (6787A>T), AAC2202AAT (6790C>T), GCA2204GTG (6795C>T 6796A>G), CAC2205CAA (6799C>A), ACC2206ACA (6802C>A), AAC2208AAT (6808C>T), TAC2209TAT (6811C>T), TAC2210TAT (6814C>T), TTC2213TTT (6823C>T), GAA2215GAG (6829A>G), TAC2221TAT (6847C>T), CGA2223CGC (6853A>C), AAT2225GAT (6857A>G), GAG2226GAA (6862G>A), GAA2229GAG (6871A>G), GTC2231ACA (6875G>A 6876T>C 6877C>A), AAG2232AAA (6880G>A), TCT2233TCA (6883T>A), GTT2235GTG (6889T>G), GAC2239GAT (6901C>T), GGC2244GGT (6916C>T), CCG2245CCA (6919G>A), AAA2247AAG (6925A>G), CTT2248CTC (6928T>C), GAC2250GAA (6934C>A), GGG2255GGC (6949G>C), TTC2257TTT (6955C>T), AAA2258AAG (6958A>G), CAA2264CAG (6976A>G), CTA2266TTA (6980C>T), ATC2267ATT (6985C>T), GTA2271GTT (6997A>T), GTG2272GTT (7000G>T), GTG2273ATA (7001G>A 7003G>A), GCT2276GCC (7012T>C), AAG2277AAA (7015G>A), CAT2278CAC (7018T>C), TTT2283TAT (7032T>A), GCC2285GCT (7039C>T), TTG2286CTA (7040T>C 7042G>A), GGA2288GGG (7048A>G), GCT2292GCA (7060T>A), GTC2293GTA (7063C>A), AGC2298AGT (7078C>T), TTA2300CTG (7082T>C 7084A>G), GTA2301GTG (7087A>G), ACC2302ACA (7090C>A), CAT2304CAC (7096T>C), GTA2305GTG (7099A>G), GTT2306GTC (7102T>C), GAG2309GAA (7111G>A), CGA2311CGG (7117A>G), ACA2314AAA (7125C>A), GAA2315GAG (7129A>G), TTT2316TTC (7132T>C), CTG2317TTA (7133C>T 7135G>A), GTG2319GTA (7141G>A), GAT2320GAC (7144T>C), GCA2321TCG (7145G>T 7147A>G), GAG2322GAA (7150G>A), GCA2325GCT (7159A>T), GAG2326_GCA2327del (7160_7165delGAGGCA), TTC2329TTT (7171C>T), GCG2335GCT (7189G>T), AAA2338AAG (7198A>G), AGC2339AGT (7201C>T), CTA2341TTA (7205C>T), AAT2342AAC (7210T>C), GAT2344GAA (7216T>A), GCG2345GCA (7219G>A), ATC2347ATA (7225C>A), AAG2348AAA (7228G>A), AAG2352AAA (7240G>A), TAT2353TAC (7243T>C), AAA2355AAG (7249A>G), CCT2356CCG (7252T>G), GAT2358GAA (7258T>A), GTT2359GTC (7261T>C), GGT2360GGA (7264T>A), GTC2361GTG (7267C>G), GTG2362GTA (7270G>A), GAT2363_CGG2364insT-- (7273_7274insT), CGG2364GTG (7274C>G 7275G>T), CAT2366CTT (7281A>T), TTG2367TCG (7284T>C), TCA2371TTA (7296C>T), TCT2376TTT (7311C>T), ACC2377ATC (7314C>T), TGC2378TAC (7317G>A), AAT2379AAA (7321T>A), GTG2380-TG (7322delG), CAC2381CAT (7327C>T), GGC2382GGT (7330C>T), TTC2383TTT (7333C>T), AAG2384CAG (7334A>C), TGT2386TGC (7342T>C), GCA2387AAC (7343G>A 7344C>A 7345A>C), ACT2390ACC (7354T>C), CAA2393CAG (7363A>G), ATT2395ATC (7369T>C), TTC2396TTT (7372C>T), GCG2398GCT (7378G>T), CTC2399CTT (7381C>T), GCA2404GCT (7396A>T), GTC2405GTT (7399C>T), GCC2407GCT (7405C>T), AGT2408ATG (7407G>T 7408T>G), GGG2410GGT (7414G>T), TGC2411GGC (7415T>G), AAA2412AAG (7420A>G), GAC2415GAT (7429C>T), TAT2416TAC (7432T>C), CAT2419CAC (7441T>C), AAG2425AAA (7459G>A), GAA2426GAG (7462A>G), ATA2428ATT (7468A>T), GTC2429GTA (7471C>A), CAA2431CAG (7477A>G), AGC2432AGT (7480C>T), CTG2434CTT (7486G>T), TAT2437TAC (7495T>C), AAA2438AAG (7498A>G), GGT2439GGC (7501T>C), TTG2440TCA (7503T>C 7504G>A), ATT2443ATA (7513T>A), AAC2445AAT (7519C>T), GGA2446GGG (7522A>G), AAG2449AAA (7531G>A), GAG2451GAA (7537G>A), CTC2452CTT (7540C>T), TGT2454TGC (7546T>C), AAG2462AAA (7570G>A), ACG2463ACA (7573G>A), ACG2465ACA (7579G>A), GCT2468GCC (7588T>C), CTA2471TTA (7595C>T), TTG2474CTA (7604T>C 7606G>A), GGT2477GGA (7615T>A), TGT2480TGC (7624T>C), GAC2483GAT (7633C>T), TTC2484TTT (7636C>T), AAT2485AAC (7639T>C), TTT2488TTC (7648T>C), GAA2494GAG (7666A>G), TGT2496TGC (7672T>C), ACA2498ACT (7678A>T), GGG2500GGA (7684G>A), ACT2502ACC (7690T>C), TTT2504TTC (7696T>C), GGT2506GGG (7702T>G), GAT2509AAT (7709G>A), AAA2510AGG (7713A>G 7714A>G), CTG2511TTG (7715C>T), CGT2514CGC (7726T>C), GAG2517GAA (7735G>A), GTA2520GTG (7744A>G), TAC2521TAT (7747C>T), GGC2526GGT (7762C>T), CAG2528CAA (7768G>A), TTT2529TTC (7771T>C), TCA2532TCG (7780A>G), CTA2533CTC (7783A>C), ACT2534ACC (7786T>C), CTA2537TTG (7793C>T 7795A>G), ATC2538ATT (7798C>T), GCT2540GCC (7804T>C), ACC2543ATC (7812C>T), GAA2550GAG (7834A>G), GAC2551GAT (7837C>T), GTG2554TTA (7844G>T 7846G>A), CAG2557CAA (7855G>A), CTG2559CTA (7861G>A), TTA2562TTG (7870A>G), TAC2563TAT (7873C>T), ACT2564ACA (7876T>A), GAG2565GAA (7879G>A), ATT2566ATA (7882T>A), GTT2567ATT (7883G>A), CCA2570CCT (7894A>T), ATT2571ATC (7897T>C), GTC2579GTT (7921C>T), AAG2581AAA (7927G>A), GGT2584GGC (7936T>C), AAT2585AAC (7939T>C), AAT2586AAC (7942T>C), GTG2594GTA (7966G>A), GAT2595GAC (7969T>C), TCT2597TCC (7975T>C), GTC2601GTT (7987C>T), CTT2602CTC (7990T>C), GCT2607GCC (8005T>C), GAA2616GAG (8032A>G), ATC2618ATT (8038C>T), GAC2619GAT (8041C>T), AGC2620AAC (8043G>A), TTG2632TTA (8080G>A), GTG2635GTA (8089G>A), CCG2637CCA (8095G>A), GAG2638GAA (8098G>A), AGC2641AGT (8107C>T), ATT2642ATC (8110T>C), GAT2644GAC (8116T>C), CAA2648CAG (8128A>G), GGT2654GGC (8146T>C), AAC2656AGT (8151A>G 8152C>T), TCG2660TCA (8164G>A), AGG2665AAG (8178G>A), AAG2666AAA (8182G>A), TTG2669TTA (8191G>A), TTC2671TTT (8197C>T), TCC2673TCT (8203C>T), AGA2675AAA (8208G>A), CTG2677TTA (8213C>T 8215G>A), CTA2678CTG (8218A>G), ATC2679ATT (8221C>T), GAG2680GAA (8224G>A), CCA2685CCG (8239A>G), GAG2690GAA (8254G>A), ATT2692ATC (8260T>C), GTA2693GTG (8263A>G), GAT2699GAC (8281T>C), AGA2700AGG (8284A>G), CCA2704GCG (8294C>G 8296A>G), GAG2705GAA (8299G>A), CAC2706CAT (8302C>T), ATT2711ATC (8317T>C), TGC2712TGT (8320C>T), GCT2714GCA (8326T>A), GAG2717GAA (8335G>A), TCC2718TCT (8338C>T), TCT2722TTT (8349C>T), GAA2723GAG (8353A>G), CAC2726CAT (8362C>T), CAA2727CAG (8365A>G), ATC2728ATT (8368C>T), TAC2732TAT (8380C>T), TCA2733TCG (8383A>G), TTA2735TTG (8389A>G), CCT2740CCC (8404T>C), GCA2742TCA (8408G>T), ACA2743ACG (8413A>G), GCG2745GCT (8419G>T), GCT2750GCC (8434T>C), TAT2752TAC (8440T>C), ATG2756ATA (8452G>A), CTA2758CTG (8458A>G), AGG2759AAG (8460G>A), AAA2760AAG (8464A>G), CTG2761CTA (8467G>A), AGG2765AGA (8479G>A), GCT2766ACA (8480G>A 8482T>A), GAG2771GAA (8497G>A), CTA2772TTG (8498C>T 8500A>G), AGA2773AAG (8502G>A 8503A>G), GCC2774GCT (8506C>T), GAA2777GAT (8515A>T), GTC2780GTT (8524C>T), GCA2781GCC (8527A>C), TTA2782CTG (8528T>C 8530A>G), GAG2785GAC (8539G>C), CTT2788TGT (8546C>T 8547T>G), TCT2790ACT (8552T>A), GTA2793GTG (8563A>G), GCA2797GGA (8574C>G), ATT2801GTT (8585A>G), GGA2805GAA (8598G>A), AGC2806AGT (8602C>T), AAC2807TCT (8603A>T 8604A>C 8605C>T), AAA2812AAG (8620A>G), CCA2813CAA (8622C>A), GAG2814GGA (8625A>G 8626G>A), CAG2815CAA (8629G>A), GGC2816GGT (8632C>T), ATC2818ATT (8638C>T), CAG2819CAA (8641G>A), AAC2821AAT (8647C>T), CCG2822TTC (8648C>T 8649C>T 8650G>C), AAC2823AAT (8653C>T), AAA2824AAG (8656A>G), GGA2825GAA (8658G>A), GAT2827GCG (8664A>C 8665T>G), GTT2830GTG (8674T>G), GCA2832GTT (8679C>T 8680A>T), GGC2833GGA (8683C>A), GGG2836GGA (8692G>A), ACA2837ACT (8695A>T), CCG2841CCA (8707G>A), AGA2842CGA (8708A>C), AAG2844AAA (8716G>A), ATC2846ATT (8722C>T), ACG2847ACA (8725G>A), AAA2849AAG (8731A>G), AGA2851AGG (8737A>G), CCC2853CCT (8743C>T), ACA2854AAG (8745C>A 8746A>G), AAG2856AAA (8752G>A), GGA2857GGT (8755A>T), AAC2862AAT (8770C>T), TTA2863CTA (8771T>C), GAA2864GAG (8776A>G), CAT2865CAC (8779T>C), TTG2866TTA (8782G>A), TAT2869TAC (8791T>C), CCA2871CCG (8797A>G), CAA2872CAG (8800A>G), CAA2873CAG (8803A>G), GAT2875GAC (8809T>C), ATT2876ATC (8812T>C), AAT2878AAC (8818T>C), CGG2880CGA (8824G>A), GCA2881GCG (8827A>G), ACG2888ACA (8848G>A), TAT2890TAC (8854T>C), GAG2891GGA (8856A>G 8857G>A), GTG2893GTA (8863G>A), CGG2894CAG (8865G>A), ATG2895CTT (8867A>C 8869G>T), GCA2896GCG (8872A>G), GGA2900GAA (8883G>A), ACT2902TCT (8888A>T), TGC2915TGT (8929C>T), GTC2924ATC (8954G>A), AAC2925AAT (8959C>T), GGA2926GGT (8962A>T), GGG2933GGA (8983G>A), AAT2934GAC (8984A>G 8986T>C), GAA2935GTA (8988A>T), GAG2938GAA (8998G>A), CCG2940CCA (9004G>A), TTG2941TTA (9007G>A), ATC2944ATT (9016C>T), ACC2951ACA (9037C>A), ATA2967ATT (9085A>T), AAC2971AAT (9097C>T), ATT2983GTT (9131A>G), CGA2984CGT (9136A>T), CGG2987CGT (9145G>T), ATG2989GGA (9149A>G 9150T>G 9151G>A), GGT2990AGT (9152G>A), TTA2991TTG (9157A>G), GCG2992GCT (9160G>T), CGT2993CGC (9163T>C), GAG3000GAA (9184G>A), CGA3004CGG (9196A>G), AGG3010AGA (9214G>A), GAA3011GAG (9217A>G), GCG3012GCA (9220G>A), GCA3020GCT (9244A>T), TTG3021TTA (9247G>A), GCC3024GCT (9256C>T), CCT3026TCC (9260C>T 9262T>C), GGG3030GGA (9274G>A), GAC3032GAT (9280C>T), ATC3035ATT (9289C>T), CAA3038CAC (9298A>C), GAG3040GAA (9304G>A), GTC3050GTT (9334C>T), GTC3060GTG (9364C>G) | | | | | | | | |
| PVYgp2 | 1 | 75 | 100% | 95 | 18.1% | 75 (98.7%) | 54 (71.1%) | 1/0/3/2 | 2 |
| Protein mutations: | R17_K17insX (2971_2972insA), Y23H (2989T>C 2991C>T), R26K (2999G>A 3000A>G), H35Y (3025C>T), K40H (3040A>C 3042A>C), G41S (3043G>A), E44K (3052G>A), I47V (3061A>G), V54I (3082G>A), P57S (3091C>T), P60T (3100C>A), G61E (3104G>A 3105T>G), G62S (3106G>A), Q63* (3109C>T), R64G (3112A>G), C66R (3118T>C), L67F (3121C>T), I69V (3127A>G), A72E (3137C>A 3138G>A), I73V (3139A>G) | | | | | | | | |
| Codon mutations: | TTA2CTA (2923T>C), AAA4AAG (2931A>G), GAA7GAG (2940A>G), TTG10CTG (2947T>C), TTG15CTG (2962T>C), AGA17_AAA17insAA- (2971_2972insA), TAC23CAT (2989T>C 2991C>T), TCA24TCG (2994A>G), AGA26AAG (2999G>A 3000A>G), CAC32-AC (3016delC), TCG33TCA (3021G>A), GTA34GTT (3024A>T), CAT35TAT (3025C>T), CAC38CAT (3036C>T), AGA39AGG (3039A>G), AAA40CAC (3040A>C 3042A>C), GGC41AGC (3043G>A), GAA44AAA (3052G>A), ATA47GTA (3061A>G), GTT54ATT (3082G>A), CCG57TCG (3091C>T), CGC59CGT (3099C>T), CCA60ACA (3100C>A), GGT61GAG (3104G>A 3105T>G), GGT62AGT (3106G>A), CAA63TAA (3109C>T), AGG64GGG (3112A>G), TGC66CGC (3118T>C), CTC67TTC (3121C>T), ATT69GTT (3127A>G), GAG70GAA (3132G>A), GCG72GAA (3137C>A 3138G>A), ATT73GTT (3139A>G) | | | | | | | | |

| | Begin | End | Coverage | Score | Concordance | Matches | Identities | I/D/M/F* | Stop Codons |
|---|---|---|---|---|---|---|---|---|---|
| **NT** | **22** | **9704** | **99.8%** | **12522** | **65.0%** | **9671 (99.8%)** | **8003 (82.6%)** | **10/12** | |
| **Proteins** | | | | | | | | | |
| polyprotein (NP_056759.1) | 1 | 3064 | 100% | 18923 | 89.7% | 3062 (99.9%) | 2771 (90.4%) | 2/2/4/4 | 1 |
| Protein mutations: | Y4Q (194T>C 196C>A), C9L (209T>C 210G>T 211T>G), F13M (221T>A 223T>G), S23P (251T>C), C24F (255G>T 256C>T), E25G (258A>G), I27V (263A>G 265T>G), V28A (267T>C), E30V (273A>T), A35T (287G>A 289T>C), V37T (293G>A 294T>C), A41I (305G>A 306C>T), D42G (309A>G), E44D (316A>T), T45V (317A>G 318C>T 319A>G), L52Q (339T>A 340C>A), K53R (342A>G), K55A (347A>G 348A>C 349A>G), Y56H (350T>C), T58V (356A>G 357C>T), V61T (365G>A 366T>C 367G>A), L62S (368C>T 369T>C), F67C (384T>G), A75V (408C>T 409C>T), M78E (416A>G 417T>A), E85K (437G>A), R86E (440A>G 441G>A), K87R (443A>C 444A>G 445G>A), D88E (448T>A), E91A (456A>C), H93N (461C>A), D111E (517T>A), S116F (531C>T), P118S (536C>T), Q119W (539C>T 540A>G 541A>G), R122K (549G>A 550A>G), M131T (576T>C), V134A (585T>C 586C>G), R137Y (593C>T 594G>A 595C>T), P138R (597C>G), I139V (599A>G 601A>G), I140P (602A>C 603T>C 604A>G), M147I (625G>A), I151V (635A>G 637T>C), I154V (644A>G 646A>G), E160A (663A>C 664G>C), H166Q (682C>A), T172S (699C>G), H174Q (706T>A), K179E (719A>G 721G>A), I180V (722A>G 724A>G), A183S (731G>T), Y184P (734T>C 735A>C 736C>T), S185R (737T>C 738C>G 739C>T), A187V (744C>T 745G>C), R189C (749C>T 751A>C), M194K (765T>A), R197K (773C>A 774G>A 775A>G), M206K (801T>A), G210A (813G>C 814A>G), L211H (816T>A), R214Q (825G>A 826T>A), N223D (851A>G 853T>C), R226C (860C>T 862C>T), T227A (863A>G 865T>C), I228T (867T>C), N229D (869A>G 871C>T), I230L (872A>T 874A>G), R231Q (876G>A), R232K (879G>A), N240S (903A>G), T241N (906C>A 907A>C), K242T (909A>C 910A>C), S243N (912G>A 913C>T), S252L (938T>C 939C>T 940A>T), L266I (980T>A 982G>C), R272K (999G>A), Q275R (1007C>A 1008A>G 1009G>A), S276G (1010A>G 1012T>G), I277V (1013A>G), N279Q (1019A>C 1021C>G), D288E (1048C>A), R300Q (1083G>A), S305A (1097T>G 1099G>A), R320K (1143G>A), L324I (1154T>A 1156G>A), S346N (1221G>A), V349T (1229G>A 1230T>C 1231T>C), S350N (1233G>A 1234C>T), K358T (1257A>C 1258A>G), N365S (1278A>G), A369V (1290C>T 1291G>A), D372T (1298G>A 1299A>C 1300C>T), I375V (1307A>G 1309A>G), N378E (1316A>G 1318T>A), I382T (1329T>C), L397I (1373C>A), I401V (1385A>G), E407D (1405G>T), A411S (1415G>T 1417A>T), V418I (1436G>A 1438C>T), E508G (1707A>G), A512T (1718G>A), I520V (1742A>G 1744C>T), S525N (1758G>A), V536I (1790G>A), I585V (1937A>G 1939T>C), F607Y (2004T>A), I628M (2068T>G), V630I (2072G>A 2074G>A), R675K (2208G>A), D694E (2266C>G), N744G (2414A>G 2415A>G 2416T>G), S746C (2420A>T), K780R (2523A>G), R817K (2634G>A), V847I (2723G>A 2725C>T), A855T (2747G>A 2749T>A), H871N (2795C>A), V880I (2822G>A 2824G>A), P900T (2882C>A), S901H (2885A>C 2886G>A), T904M (2895C>T), N916S (2931A>G), N919S (2940A>G), E929_N930insX (2971_2972insA), T936I (2991C>T), H937R (2994A>G), E939R (2999G>A 3000A>G), R946Q (3021G>A), Y947F (3024A>T 3025C>T), T951I (3036C>T), E952G (3039A>G 3040A>C), K953T (3042A>C 3043G>A), A972V (3099C>T 3100C>A), V974R (3104G>A 3105T>G 3106G>A), S983N (3132G>A), R985K (3137C>A 3138G>A 3139A>G), F990I (3152T>A 3154C>T), C994Y (3165G>A), F1012L (3218T>C), T1014N (3225C>A), V1016I (3230G>A), V1028M (3266G>A 3268A>G), V1031M (3275G>A), A1034T (3284G>A), R1044K (3315G>A 3316A>G), E1045A (3318A>C), M1059V (3359A>G), D1072E (3400T>G), V1110G (3513T>G), M1121V (3545A>G), A1122T (3548G>A), V1124I (3554G>A 3556C>T), L1147V (3623C>G), L1150M (3632C>A), Y1152H (3638T>C), I1172V (3698A>G), S1178N (3717G>A), R1183K (3731C>A 3732G>A), I1297V (4073A>G), F1315Y (4128T>A 4129C>T), A1323S (4151G>T), V1328I (4166G>A), L1347I (4223C>A 4225G>A), I1379V (4319A>G 4321A>G), F1404Y (4395T>A), V1408I (4406G>A), M1431V (4475A>G 4477G>T), T1433S (4481A>T 4483A>G), A1454L (4544G>T 4545C>T 4546G>A), R1455K (4548G>A), V1499I (4679G>A 4681G>C), V1538I (4796G>A 4798C>T), T1540S (4803C>G 4804T>C), S1625G (5057A>G), T1626A (5060A>G), L1636I (5090C>A), A1639V (5100C>T), E1641D (5107A>T), V1646F (5120G>T 5122C>T), I1648V (5126A>G), V1667I (5183G>A), A1793T (5561G>A 5563G>A), I1805V (5597A>G), N1807K (5605C>A), A1812V (5619C>T 5620T>G), I1816L (5630A>C), V1837I (5693G>A), K1890R (5853A>G), I1904V (5894A>G 5896C>G), R1926A (5960C>G 5961G>C 5962G>C), I1936V (5990A>G), E1945D (6019A>T), K1949T (6030A>C), D1955E (6049C>A), M1958I (6058G>A), S1963N (6072G>A 6073T>C), N1964H (6074A>C), T1966N (6081C>A), C1976S (6111G>C 6112T>C), I1982V (6128A>G 6130T>C), V1992I (6158G>A), T1996S (6170A>T 6172A>G), L2007F (6203C>T 6205C>T), V2016I (6230G>A), A2025K (6257G>A 6258C>A), K2033R (6282A>G), A2060T (6362G>A), V2073I (6401G>A), A2074V (6405C>T), Y2082F (6429A>T), Q2089R (6450A>G), H2101N (6485C>A 6487C>T), L2115I (6527C>A), T2135I (6588C>T), Y2151H (6635T>C), I2155V (6647A>G), I2165V (6677A>G 6679A>G), D2178N (6716G>A), N2179D (6719A>G), I2194L (6764A>T 6766A>G), A2204V (6795C>T 6796A>G), H2205Q (6799C>A), N2225D (6857A>G), V2231T (6875G>A 6876T>C 6877C>A), D2250E (6934C>A), V2273I (7001G>A 7003G>A), F2283Y (7032T>A), T2314K (7125C>A), A2321S (7145G>T 7147A>G), E2326_A2327del (7160_7165delGAGGCA), D2344E (7216T>A), D2358E (7258T>A), D2363_R2364insX (7273_7274insT), R2364V (7274C>G 7275G>T), H2366L (7281A>T), L2367S (7284T>C), S2371L (7296C>T), S2376F (7311C>T), T2377I (7314C>T), C2378Y (7317G>A), N2379K (7321T>A), K2384Q (7334A>C), A2387N (7343G>A 7344C>A 7345A>C), S2408M (7407G>T 7408T>G), C2411G (7415T>G), L2440S (7503T>C 7504G>A), D2509N (7709G>A), K2510R (7713A>G 7714A>G), T2543I (7812C>T), V2554L (7844G>T 7846G>A), V2567I (7883G>A), S2620N (8043G>A), N2656S (8151A>G 8152C>T), R2665K (8178G>A), R2675K (8208G>A), P2704A (8294C>G 8296A>G), S2722F (8349C>T), A2742S (8408G>T), M2756I (8452G>A), R2759K (8460G>A), A2766T (8480G>A 8482T>A), R2773K (8502G>A 8503A>G), E2777D (8515A>T), E2785D (8539G>C), L2788C (8546C>T 8547T>G), S2790T (8552T>A), A2797G (8574C>G), I2801V (8585A>G), G2805E (8598G>A), N2807S (8603A>T 8604A>C 8605C>T), P2813Q (8622C>A), E2814G (8625A>G 8626G>A), P2822F (8648C>T 8649C>T 8650G>C), G2825E (8658G>A), D2827A (8664A>C 8665T>G), A2832V (8679C>T 8680A>T), T2854K (8745C>A 8746A>G), E2891G (8856A>G 8857G>A), R2894Q (8865G>A), M2895L (8867A>C 8869G>T), G2900E (8883G>A), T2902S (8888A>T), V2924I (8954G>A), N2934D (8984A>G 8986T>C), E2935V (8988A>T), I2983V (9131A>G), M2989G (9149A>G 9150T>G 9151G>A), G2990S (9152G>A), P3026S (9260C>T 9262T>C), Q3038H (9298A>C) | | | | | | | | |

| | Begin | End | Coverage | Score | Concordance | Matches | Identities | I/D/M/F* | Stop Codons |
|---|---|---|---|---|---|---|---|---|---|
| **NT** | **22** | **9704** | **99.8%** | **12522** | **65.0%** | **9671 (99.8%)** | **8003 (82.6%)** | **10/12** | |

Codon mutations:

ACT3ACC (193T>C), TAC4CAA (194T>C 196C>A), TCA6TCG (202A>G), ATC8ATT (208C>T), TGT9CTG (209T>C 210G>T 211T>G), TCG12TCA (220G>A), TTT13ATG (221T>A 223T>G), GAA14GAG (226A>G), CTA17CTT (235A>T), TAC19TAT (241C>T), CCA21CCT (247A>T), TCT23CCT (251T>C), TGC24TTT (255G>T 256C>T), GAG25GGG (258A>G), CAT26CAC (262T>C), ATT27GTG (263A>G 265T>G), GTG28GCG (267T>C), GAA30GTA (273A>T), CGA31CGT (277A>T), GAA32GAG (280A>G), GTG33GTT (283G>T), CCG34CCA (286G>A), GCT35ACC (287G>A 289T>C), GTT37ACT (293G>A 294T>C), CCT39CCC (301T>C), TTC40TTT (304C>T), GCA41ATA (305G>A 306C>T), GAT42GGT (309A>G), CTG43TTG (311C>T), GAA44GAT (316A>T), ACA45GTG (317A>G 318C>T 319A>G), CAA46CAG (322A>G), GCA49GCT (331A>T), CTC52CAA (339T>A 340C>A), AAG53AGG (342A>G), AAA55GCG (347A>G 348A>C 349A>G), TAT56CAT (350T>C), GCT57GCA (355T>A), ACT58GTT (356A>G 357C>T), CGT60CGC (364T>C), GTG61ACA (365G>A 366T>C 367G>A), CTC62TCC (368C>T 369T>C), AAA63AAG (373A>G), AAC64AAT (376C>T), TTT67TGT (384T>G), ACG68ACA (388G>A), TAC69TAT (391C>T), CGA70CGG (394A>G), AAG72AAA (400G>A), GAT74GAC (406T>C), GCC75GTT (408C>T 409C>T), CAG76CAA (412G>A), ATA77ATT (415A>T), ATG78GAG (416A>G 417T>A), ATT80ATC (424T>C), CAG81CAA (427G>A), AAA83AAG (433A>G), CTG84CTC (436G>C), GAG85AAG (437G>A), AGG86GAG (440A>G 441G>A), AAG87CGA (443A>C 444A>G 445G>A), GAT88GAA (448T>A), GAA90GAG (454A>G), GAA91GCA (456A>C), TAT92TAC (460T>C), CAC93AAC (461C>A), CAA95CAG (469A>G), GCC97GCG (475C>G), CCT99CCA (481T>A), ACT106ACC (502T>C), ATA107ATT (505A>T), GGC109GGT (511C>T), GAT111GAA (517T>A), CCT112CCA (520T>A), CCA113CCT (523A>T), TCT116TTT (531C>T), GAG117GAA (535G>A), CCA118TCA (536C>T), CAA119TGG (539C>T 540A>G 541A>G), AGA122AAG (549G>A 550A>G), ATC124ATT (556C>T), ATT125ATC (559T>C), CAT126CAC (562T>C), ACT128ACC (568T>C), CCA129CCG (571A>G), ATG131ACG (576T>C), CGT132CGC (580T>C), GTC134GCG (585T>C 586C>G), CGC137TAT (593C>T 594G>A 595C>T), CCC138CGC (597C>G), ATA139GTG (599A>G 601A>G), ATA140CCG (602A>C 603T>C 604A>G), ACA143ACG (613A>G), GAA144GAG (616A>G), GGC145GGG (619C>G), CAG146CAA (622G>A), ATG147ATA (625G>A), AAT148AAC (628T>C), CAC149CAT (631C>T), ATT151GTC (635A>G 637T>C), ATA154GTG (644A>G 646A>G), ATT157ATC (655T>C), TCG159TCA (661G>A), GAG160GCC (663A>C 664G>C), AAA161AAG (667A>G), GTC165GTT (679C>T), CAC166CAA (682C>A), TTA167CTG (683T>C 685A>G), AGT169AGC (691T>C), ACC172AGC (699C>G), ACT173ACA (703T>A), CAT174CAA (706T>A), AAG179GAA (719A>G 721G>A), ATA180GTG (722A>G 724A>G), CTT181TTG (725C>T 727T>G), GGT182GGA (730T>A), GCA183TCA (731G>T), TAC184CCT (734T>C 735A>C 736C>T), TCC185CGT (737T>C 738C>G 739C>T), GCG187GTC (744C>T 745G>C), CGA189TGC (749C>T 751A>C), ACT190ACC (754T>C), ATG194AAG (765T>A), TTG196CTG (770T>C), CGA197AAG (773C>A 774G>A 775A>G), CGG198AGG (776C>A), TTC202TTT (790C>T), ATG206AAG (801T>A), GGA210GCG (813G>C 814A>G), CTT211CAT (816T>A), TTG212CTG (818T>C), CAA213CAG (823A>G), CGT214CAA (825G>A 826T>A), CGG217AGA (833C>A 835G>A), GAC219GAT (841C>T), AAT223GAC (851A>G 853T>C), GTC225GTT (859C>T), CGC226TGT (860C>T 862C>T), ACT227GCC (863A>G 865T>C), ATC228ACC (867T>C), AAC229GAT (869A>G 871C>T), ATA230TTG (872A>T 874A>G), CGA231CAA (876G>A), AGG232AAG (879G>A), GAT234GAC (886T>C), ATC238ATT (898C>T), AAC240AGC (903A>G), ACA241AAC (906C>A 907A>C), AAA242ACC (909A>C 910A>C), AGC243AAT (912G>A 913C>T), GGC246GGA (922C>A), GGT249GGA (931T>A), TCA252CTT (938T>C 939C>T 940A>T), GGA253GGG (943A>G), TTC256TTT (952C>T), TCA261TCG (967A>G), GAA263GAG (973A>G), GGG264GGC (976G>C), TTG266ATC (980T>A 982G>C), GCA269GCG (991A>G), CGT270CGC (994T>C), AGA272AAA (999G>A), GTT273GTC (1003T>C), CAG275AGA (1007C>A 1008A>G 1009G>A), AGT276GGG (1010A>G 1012T>G), ATT277GTT (1013A>G), TTA278TTG (1018A>G), AAC279CAG (1019A>C 1021C>G), ATC282ATT (1030C>T), TCG285TCA (1039G>A), AAT286AAC (1042T>C), GCC287GCA (1045C>A), GAC288GAA (1048C>A), AAT289AAC (1051T>C), AAG292AAA (1060G>A), GGT293GGA (1063T>A), CTG294TTG (1064C>T), GAC295GAT (1069C>T), CGA300CAA (1083G>A), TAT303TAC (1093T>C), TCG305GCA (1097T>G 1099G>A), CAC307CAT (1105C>T), GTA310GTG (1114A>G), GCT311GCG (1117T>G), TTA313TTG (1123A>G), CCT314CCA (1126T>A), GTC315GTT (1129C>T), AGG320AAG (1143G>A), GTA321GTT (1147A>T), GCT322GCA (1150T>A), TTG324ATA (1154T>A 1156G>A), CAC327CAT (1165C>T), AGT328AGC (1168T>C), CTT330TTA (1172C>T 1174T>A), ACC339ACT (1201C>T), AGC346AAC (1221G>A), GTT349ACC (1229G>A 1230T>C 1231T>C), AGC350AAT (1233G>A 1234C>T), CTG352CTA (1240G>A), AAG354AAA (1246G>A), TTG356CTA (1250T>C 1252G>A), CAT357CAC (1255T>C), AAA358ACG (1257A>C 1258A>G), AGA361AGG (1267A>G), GGT363GGC (1273T>C), TTG364TTA (1276G>A), AAT365AGT (1278A>G), TTG367CTG (1283T>C), GCG369GTA (1290C>T 1291G>A), GAC372ACT (1298G>A 1299A>C 1300C>T), TTT374TTC (1306T>C), ATA375GTG (1307A>G 1309A>G), CAT376CAC (1312T>C), AAT378GAA (1316A>G 1318T>A), TTG381TTA (1327G>A), ATA382ACA (1329T>C), GCG383GCT (1333G>T), TTA384TTG (1336A>G), CTA387TTG (1343C>T 1345A>G), CCG390CCA (1354G>A), GTG391GTT (1357G>T), GAC392GAT (1360C>T), CTC395TTG (1367C>T 1369C>G), CTT397ATT (1373C>A), TTC398TTT (1378C>T), GAG400GAA (1384G>A), ATA401GTA (1385A>G), AAA403AAG (1393A>G), TCC404TCT (1396C>T), GAG407GAT (1405G>T), AAA408AAG (1408A>G), CAG409CAA (1411G>A), GCA411TCT (1415G>T 1417A>T), CCG412CCC (1420G>C), AAT415AAC (1429T>C), TTA416TTG (1432A>G), GTC418ATT (1436G>A 1438C>T), TTA419CTG (1439T>C 1441A>G), AAT420AAC (1444T>C), TTC423TTT (1453C>T), CTG424CTA (1456G>A), GAA428GAG (1468A>G), GTA436GTG (1492A>G), TTG439CTA (1499T>C 1501G>A), AGT440AGC (1504T>C), TTG441TTA (1507G>A), CTC442CTT (1510C>T), GAA443GAG (1513A>G), TTA444TTG (1516A>G), GCA445GCT (1519A>T), AGG446AGA (1522G>A), TTC447TTT (1525C>T), CAG448CAA (1528G>A), AAC450AAT (1534C>T), ACT452ACA (1540T>A), AAC454AAT (1546C>T), AAA457AAG (1555A>G), GGT458GGC (1558T>C), GAT459GAC (1561T>C), ATA460ATC (1564A>C), TCT461TCA (1567T>A), TTC463TTT (1573C>T), AGA464AGG (1576A>G), TTA467CTA (1583T>C), TCT468TCA (1588T>A), AAG470AAA (1594G>A), AAC472AAT (1600C>T), CTG475CTA (1609G>A), TCG478TCA (1618G>A), TGC479TGT (1621C>T), GAC480GAT (1624C>T), CTG483TTG (1631C>T), AAA485AAG (1639A>G), TTC489TTT (1651C>T), CTC490CTG (1654C>G), CAT497CAC (1675T>C), TCA503TCG (1693A>G), TTT506TTC (1702T>C), GAA508GGA (1707A>G), GCA512ACA (1718G>A), GGA514GGG (1726A>G), TAC515TAT (1729C>T), TCA516TCC (1732A>C), TAT518TAC (1738T>C), ATC520GTT (1742A>G 1744C>T), CGC521CGT (1747C>T), CAT523CAC (1753T>C), AGT525AAT (1758G>A), AGG528AGA (1768G>A), AAC534AAT (1786C>T), TTA535TTG (1789A>G), GTT536ATT (1790G>A), GTC537GTT (1795C>T), TTA541CTG (1805T>C 1807A>G), GAG543GAA (1813G>A), AGG545AGA (1819G>A), AAG547AAA (1825G>A), GAC551GAT (1837C>T), TAT552TAC (1840T>C), CAA555CAG (1849A>G), GTC558GTG (1858C>G), AAA560AAG (1864A>G), TGC562TGT (1870C>T), AGT564AGC (1876T>C), AAA566AAG (1882A>G), GGT568GGA (1888T>A), AAT569AAC (1891T>C), TGT574TGC (1906T>C), ACA577ACT (1915A>T), GGT582GGC (1930T>C), GCC584GCT (1936C>T), ATT585GTC (1937A>G 1939T>C), TCA587TCG (1945A>G), TTC589TTT (1951C>T), CCA592CCT (1960A>T), GTA598GTG (1978A>G), ATT599ATA (1981T>A), GGC600GGT (1984C>T), AAT601AAC (1987T>C), GAC604GAT (1996C>T), TTT607TAT (2004T>A), AAA612AAG (2020A>G), GGG613GGA (2023G>A), GAT614GAC (2026T>C), TCG615TCT (2029G>T), GAG616GAA (2032G>A), TAT625TAC (2059T>C), ATT628ATG (2068T>G), AAC629AAT (2071C>T), GTG630ATA (2072G>A 2074G>A), CTG635TTA (2087C>T 2089G>A), AGC639AGT (2101C>T), GAG640GAA (2104G>A), GAG641GAA (2107G>A), AAG644AAA (2116G>A), TTC646TTT (2122C>T), CGC651CGT (2137C>T), CTT658CTC (2158T>C), ACC660ACT (2164C>T), CCA662CCG (2170A>G), GAT666GAC (2182T>C), GCG668GCA (2188G>A), ACC669ACA (2191C>A), AGA675AAA (2208G>A), TAT678TAC (2218T>C), GAC680GAT (2224C>T), GTA681GTT (2227A>T), TTG686TTA (2242G>A), CCC687CCT (2245C>T), TTG690TTA (2254G>A), GTT691GTC (2257T>C), GAC692GAT (2260C>T), GAC694GAG (2266C>G), CAA696CAG (2272A>G), ACG697ACA (2275G>A), TCA703TCG (2293A>G), GGC705GGT (2299C>T), TCG706TCA (2302G>A), CAG707CAA (2305G>A), GGA710GGT (2314A>T), TAT711TAC (2317T>C), CTA714CTG (2326A>G), GTG719GTA (2341G>A), CAA721CAG (2347A>G), CTT722CTC (2350T>C), TTG724TTA (2356G>A), AAT727AAC (2365T>C), GAA729GAG (2371A>G), TTA730TTG (2374A>G), GAT733GAC (2383T>C), ATA734ATT (2386A>T), AAA735AAG (2389A>G), CAT736CAC (2392T>C), TAT737TAC (2395T>C), AAT744GGG (2414A>G 2415A>G 2416T>G), GCT745GCA (2419T>A), AGC746TGC (2420A>T), ACA752ACT (2440A>T), GTT761GTC (2467T>C), TCT764TCC (2476T>C), TCG766TCA (2482G>A), CTG769CTA (2491G>A), CTG771CTT (2497G>T), TTG773TTA (2503G>A), AGA778AGG (2518A>G), CCT779CCC (2521T>C), AAG780AGG (2523A>G), AGA783AGG (2533A>G), TTG785TTA (2539G>A), TTA787TTG (2545A>G), GAG789GAA (2551G>A), CCT790CCA (2554T>A), CTG792TTG (2558C>T), TTG793CTC (2561T>C 2563G>C), TCA796TCG (2572A>G), CCT800CCC (2584T>C), GGC801GGT (2587C>T), CTG803CTC (2593G>C), GCT805GCC (2599T>C), AAT808AAC (2608T>C), TTT812TTC (2620T>C), GCG815GCA (2629G>A), GTG816GTA (2632G>A), AGG817AAG (2634G>A), TTG818TTA (2638G>A), ATT820ATA (2644T>A), AAA823AAG (2653A>G), GCT827GCC (2665T>C), CTA832TTA (2678C>T), CTA833CTG (2683A>G), GCT835GCC (2689T>C), CTA838CTC (2698A>C), TCA841TCG (2707A>G), GCG842GCA (2710G>A), GAA844GAG (2716A>G), ACA845ACG (2719A>G), CTC846CTT (2722C>T), GTC847ATT (2723G>A 2725C>T), GCA848GCG (2728A>G), AGG850AGA (2734G>A), ATT851ATA (2737T>A), GAT854GAC (2746T>C), GCT855ACA (2747G>A 2749T>A), GCA856GCT (2752A>T), GCT857GCA (2755T>A), GAC859GAT (2761C>T), CTC860CTT (2764C>T), TGT865TGC (2779T>C), GGG867GGT (2785G>T), AAC869AAT (2791C>T), CAT871AAT (2795C>A), TAC874TAT (2806C>T), TTG878CTC (2816T>C 2818G>C), GTG880ATA (2822G>A 2824G>A), TTG881TTA (2827G>A), GTT884GTC (2836T>C), AAG885AAA (2839G>A), AAT886AAC (2842T>C), ACC893ACA (2863C>A), CTA894TTA (2864C>T), CCA900ACA (2882C>A), AGT901CAT (2885A>C 2886G>A), AAC903AAT (2893C>T), ACG904ATG (2895C>T), GTC906GTT (2902C>T), GTA907GTG (2905A>G), ATT909ATC (2911T>C), AAT913AAC (2923T>C), AAT916AGT (2931A>G), AAC919AGC (2940A>G), GCT921GCC (2947T>C), ACT926ACC (2962T>C), GAA929_AAT930insA-- (2971_2972insA), GGT935GGC (2989T>C), ACT936ATT (2991C>T), CAT937CGT (2994A>G), GAG939AGG (2999G>A 3000A>G), TCC944TC- (3016delC), CGG946CAG (3021G>A), TAC947TTT (3024A>T 3025C>T), ACA951ATA (3036C>T), GAA952GGC (3039A>G 3040A>C), AAG953ACA (3042A>C 3043G>A), TTG956TTA (3052G>A), TTA959TTG (3061A>G), GCG966GCA (3082G>A), GGC969GGT (3091C>T), GCC972GTA (3099C>T 3100C>A), GTG974AGA (3104G>A 3105T>G 3106G>A), GTC975GTT (3109C>T), AAA976AAG (3112A>G), ACT978ACC (3118T>C), GCC979GCT (3121C>T), GGA981GGG (3127A>G), AGC983AAC (3132G>A), CGA985AAG (3137C>A 3138G>A 3139A>G), TAT989TAC (3151T>C), TTC990ATT (3152T>A 3154C>T), TGT994TAT (3165G>A), GTA995GTG (3169A>G), ATC1005ATT (3199C>T), AGG1007CGG (3203A>C), CGT1008CGC (3208T>C), CCA1010CCG (3214A>G), ACC1011ACT (3217C>T), TTT1012CTT (3218T>C), GTC1013GTA (3223C>A), ACT1014AAT (3225C>A), GTT1016ATT (3230G>A), AAC1017AAT (3235C>T), TTA1025CTA (3257T>C), ACC1026ACT (3262C>T), AGC1027AGT (3265C>T), GTA1028ATG (3266G>A 3268A>G), GTG1029GTA (3271G>A), GTG1031ATG (3275G>A), GCA1034ACA (3284G>A), CAG1039CAA (3301G>A), AGA1044AAG (3315G>A 3316A>G), GAA1045GCA (3318A>C), ATC1046ATT (3322C>T), GAG1047GAA (3325G>A), TTG1048TTA (3328G>A), ATA1051ATT (3337A>T), GAG1055GAA (3349G>A), GTC1057GTT (3355C>T), TGC1058TGT (3358C>T), ATG1059GTG (3359A>G), CTA1061CTG (3367A>G), TAT1062TAC (3370T>C), TTA1065CTG (3377T>C 3379A>G), GAA1070GAG (3394A>G), CGC1071CGT (3397C>T), GAT1072GAG (3400T>G), ACA1074ACG (3406A>G), TAC1078TAT (3418C>T), ATT1079ATA (3421T>A), TAT1081TAC (3427T>C), AAG1083AAA (3433G>A), TCA1084TCT (3436A>T), GTA1085GTG (3439A>G), AAC1086AAT (3442C>T), CCT1087CCC (3445T>C), CAG1091CAA (3457G>A), TTT1092TTC (3460T>C), GCT1093GCA (3463T>A), GCG1095GCT (3469G>T), CAG1096CAA (3472G>A), CAC1104CAT (3496C>T), CAG1105CAA (3499G>A), TCC1107TCT (3505C>T), GTT1110GGT (3513T>G), AAA1112AAG (3520A>G), GAA1115GAG (3529A>G), CAA1116CAG (3532A>G), ATG1121GTG (3545A>G), GCT1122ACT (3548G>A), TTA1123CTA (3551T>C), GTC1124ATT (3554G>A 3556C>T), AGT1133AGC (3583T>C), GAT1134GAC (3586T>C), TGC1135TGT (3589C>T), TTC1137TTT (3595C>T), AAT1141AAC (3607T>C), TTT1143TTC (3613T>C), AAG1144AAA (3616G>A), GGT1145GGC (3619T>C), CTT1147GTT (3623C>G), TCC1148TCT (3628C>T), CTG1150ATG (3632C>A), GAT1151GAC (3637T>C), TAT1152CAT (3638T>C), GAA1153GAG (3643A>G), TTA1159TTG (3661A>G), GTG1162GTA (3670G>A), TTT1166TTC (3682T>C), GAG1168GAA (3688G>A), GAG1171GAA (3697G>A), ATT1172GTT (3698A>G), GAT1174GAC (3706T>C), TTT1175TTC (3709T>C), TTG1177CTA (3713T>C 3715G>A), AGT1178AAT (3717G>A), GAG1179GAA (3721G>A), CGA1183AAA (3731C>A 3732G>A), ACT1184ACA (3736T>A), CTA1188TTG (3746C>T 3748A>G), ACA1190ACG (3754A>G), AGT1193AGC (3763T>C), CGA1198CGG (3778A>G), CAA1199CAG (3781A>G), ATC1200ATT (3784C>T), CAT1204CAC (3796T>C), CCA1207CCT (3805A>T), TAC1209TAT (3811C>T), AGA1210AGG (3814A>G), TTC1215TTT (3829C>T), TTT1218TTC (3838T>C), AGA1220AGG (3844A>G), GCA1221GCC (3847A>C), GTT1224GTA (3856T>A), GTG1226GTT (3862G>T), GAC1229GAT (3871C>T), GCC1231GCA (3877C>A), AGC1233AGT (3883C>T), GAA1234GAG (3886A>G), CTA1236CTG (3892A>G), GAC1237GAT (3895C>T), TTT1238TTC (3898T>C), GTA1240GTG (3904A>G), CGG1241AGA (3905C>A 3907G>A), GCT1243GCC (3913T>C), AAG1248AAA (3928G>A), TCA1249TCT (3931A>T), GGG1251GGA (3937G>A), AGT1257AGC (3955T>C), GTG1258GTA (3958G>A), GCC1259GCT (3961C>T), GGA1260GGG (3964A>G), TCT1261TCC (3967T>C), TTA1264TTG (3976A>G), ATT1265ATC (3979T>C), CCA1267CCT (3985A>T), CTA1271CTT (3997A>T), GCG1272GCA (4000G>A), AAC1274AAT (4006C>T), GTT1275GTG (4009T>G), TTC1276TTT (4012C>T), AAA1277AAG (4015A>G), TCT1280TCC (4024T>C), GAA1282GAG (4030A>G), CCA1283CCT (4033A>T), AAG1286AAA (4042G>A), CTG1290TTA (4052C>T 4054G>A), CGT1291CGC (4057T>C), CGT1293CGA (4063T>A), GGA1294GGG (4066A>G), ATA1297GTA (4073A>G), TTT1298TTC (4078T>C), GGC1299GGT (4081C>T), TCT1300TCC (4084T>C), TCC1304TCT (4096C>T), AGC1308AGT (4108C>T), GGA1309GGT (4111A>T), GCG1311GCA (4117G>A), CTA1312TTG (4118C>T 4120A>G), TAC1314TAT (4126C>T), TTC1315TAT (4128T>A 4129C>T), GCC1316GCT (4132C>T), AAT1317AAC (4135T>C), CAA1321CAG (4147A>G), TTA1322CTA (4148T>C), GCT1323TCT (4151G>T), AAC1326AAT (4162C>T), GTA1328ATA (4166G>A), ATA1329ATC (4171A>C), GAG1332GAA (4180G>A), CTG1336TTG (4190C>T), CCT1338CCC (4198T>C), TCC1339TCT (4201C>T), GCG1342GCT (4210G>T), TTC1343TTT (4213C>T), AGT1345AGC (4219T>C), CTG1346TTG (4220C>T), CTG1347ATA (4223C>A 4225G>A), GTT1349GTG (4231T>G), CAT1351CAC (4237T>C), GTA1356GTG (4252A>G), TTA1357CTG (4253T>C 4255A>G), AAA1358AAG (4258A>G), ACT1362ACC (4270T>C), AGA1366AGG (4282A>G), GAG1367GAA (4285G>A), GAA1369GAG (4291A>G), CAG1373CAA (4303G>A), GTC1376GTT (4312C>T), ATA1379GTG (4319A>G 4321A>G), GTG1380GTT (4324G>T), GAG1381GAA (4327G>A), CTG1384CTT (4336G>T), TCT1385TCC (4339T>C), TCA1388TCT (4348A>T), GCA1392GCG (4360A>G), GGT1394GGC (4366T>C), TCT1395TCA (4369T>A), ACT1397ACC (4375T>C), GTT1401GTC (4387T>C), TTT1404TAT (4395T>A), GTA1408ATA (4406G>A), CTT1409CTC (4411T>C), GTT1418GTC (4438T>C), ACC1420ACA (4444C>A), GCC1422GCT (4450C>T), CTC1424CTT (4456C>T), GAC1427GAT (4465C>T), ATG1431GTT (4475A>G 4477G>T), GTC1432GTT (4480C>T), ACA1433TCG (4481A>T 4483A>G), GGC1437GGT (4495C>T), AAG1441AAA (4507G>A), CAC1442CAT (4510C>T), GGT1443GGG (4513T>G), CTA1445TTA (4517C>T), GTC1448GTT (4528C>T), AAA1450AAG (4534A>G), ACC1452ACT (4540C>T), GCG1454TTA (4544G>T 4545C>T 4546G>A), AGA1455AAA (4548G>A), TTT1458TTC (4558T>C), GTA1460GTT (4564A>T), ACC1462ACT (4570C>T), ATA1464ATT (4576A>T), TTG1471TTA (4597G>A), GAC1472GAT (4600C>T), GAC1474GAT (4606C>T), GTG1475GTA (4609G>A), GTG1477GTA (4615G>A), GAC1478GAT (4618C>T), TTG1481CTT (4625T>C 4627G>T), GAC1488GAT (4648C>T), ATT1489ATC (4651T>C), AGG1492AGA (4660G>A), AGC1493AGT (4663C>T), GCT1495GCA (4669T>A), GTG1499ATC (4679G>A 4681G>C), AGT1500AGC (4684T>C), GTT1501GTG (4687T>G), GAG1505GAA (4699G>A), AGA1506AGG (4702A>G), CAA1508CAG (4708A>G), AGG1509AGA (4711G>A), CTG1510TTG (4712C>T), AAA1518AAG (4738A>G), GTA1520GTT (4744A>T), GCA1521GCG (4747A>G), TTG1522CTG (4748T>C), ACT1527ACC (4765T>C), GTC1538ATT (4796G>A 4798C>T), ACT1540AGC (4803C>G 4804T>C), GAG1541GAA (4807G>A), GCG1542GCT (4810G>T), GCT1543GCG (4813T>G), CTT1544CTC (4816T>C), TAT1549TAC

| | | | | | | | | | |
|---|---|---|---|---|---|---|---|---|---|
| | (4831T>C), AAC1550AAT (4834C>T), CCA1552CCT (4840A>T), GGA1556GGG (4852A>G), GGC1557GGT (4855C>T), AGT1561AGC (4867T>C), CTG1562CTC (4870G>C), ATT1563ATA (4873T>A), TGT1566TGC (4882T>C), ACA1573ACT (4903A>T), CAG1575CAA (4909G>A), TTT1577TTC (4915T>C), GAA1578GAG (4918A>G), TTG1579CTA (4919T>C 4921G>A), AGT1580AGC (4924T>C), CCC1581CCG (4927C>G), TTT1583TTC (4933T>C), ATC1584ATA (4936C>A), CAG1585CAA (4939G>A), AAT1586AAC (4942T>C), TTC1587TTT (4945C>T), TCA1593TCG (4963A>G), CCT1596CCC (4972T>C), CTT1602CTC (4990T>C), AAA1606AAG (5002A>G), CCT1613CCC (5023T>C), TTG1614CTG (5024T>C), TGC1615TGT (5029C>T), GAT1616GAC (5032T>C), TCT1618TCC (5038T>C), CCA1620CCT (5044A>T), GCA1623GCT (5053A>T), TCG1624TCA (5056G>A), AGT1625GGT (5057A>G), ACT1626GCT (5060A>G), TCG1629TCT (5071G>T), CTT1636ATT (5090C>A), GGA1637GGG (5095A>G), GTG1638GTT (5098G>T), GCC1639GTC (5100C>T), TTA1640TTG (5104A>G), GAA1641GAT (5107A>T), ATT1642ATC (5110T>C), CCA1643CCG (5113A>G), AAG1644AAA (5116G>A), CAA1645CAG (5119A>G), GTC1646TTT (5120G>T 5122C>T), ATT1648GTT (5126A>G), CAT1651CAC (5137T>C), ATC1652ATT (5140C>T), AAA1653AAG (5143A>G), GAG1654GAA (5146G>A), CTC1659CTA (5161C>A), GAA1661GAG (5167A>G), CTT1663CTG (5173T>G), ACG1666ACA (5182G>A), GTT1667ATT (5183G>A), GTC1668GTT (5188C>T), AAG1669AAA (5191G>A), AAA1671AAG (5197A>G), GAC1672GAT (5200C>T), TGC1674TGT (5206C>T), TTA1675TTG (5209A>G), AGC1678AGT (5218C>T), TCG1682TCA (5230G>A), ATC1684ATT (5236C>T), AGC1685AGT (5239C>T), ATC1687ATT (5245C>T), ACA1693ACT (5263A>T), CTC1695CTT (5269C>T), TTC1696TTT (5272C>T), GCC1697GCA (5275C>A), ATC1698ATT (5278C>T), CCA1699CCC (5281A>C), AGA1700AGG (5284A>G), ACT1701ACC (5287T>C), CTA1702TTA (5288C>T), ATA1703ATT (5293A>T), TTG1704CTA (5294T>C 5296G>A), GAG1706GAA (5302G>A), CTT1709CTA (5311T>A), GAA1710GAG (5314A>G), CGA1713CGT (5323A>T), AGT1721AGC (5347T>C), ATC1723ATT (5353C>T), TGC1727TGT (5365C>T), GTT1734GTC (5386T>C), ACC1737ACA (5395C>A), AAA1746AAG (5422A>G), GAT1747GAC (5425T>C), ACC1749ACT (5431C>T), GAG1751GAA (5437G>A), CTT1756CTC (5452T>C), AAG1758AAA (5458G>A), AGT1761AGC (5467T>C), CAA1762CAG (5470A>G), CTA1763TTG (5471C>T 5473A>G), GAA1765GAG (5479A>G), TTC1766TTT (5482C>T), TCA1767TCC (5485A>C), AAT1768AAC (5488T>C), TTG1769CTA (5489T>C 5491G>A), GGT1771GGC (5497T>C), TGT1774TGC (5506T>C), GAG1775GAA (5509G>A), GAG1776GAA (5512G>A), TTA1778CTA (5516T>C), TAT1782TAC (5530T>C), GAG1783GAA (5533G>A), TCG1784TCT (5536G>T), TTG1785TTA (5539G>A), GTT1788GTG (5548T>G), CAC1790CAT (5554C>T), GCT1792GCA (5560T>A), GCG1793ACA (5561G>A 5563G>A), ACG1794ACC (5566G>C), TCA1795TCG (5569A>G), CTC1800CTG (5584C>G), AAG1803AAA (5593G>A), GGG1804GGA (5596G>A), ATT1805GTT (5597A>G), AAC1807AAA (5605C>A), GTG1811GTT (5617G>T), GCT1812GTG (5619C>T 5620T>G), TTG1815TTA (5629G>A), ATC1816CTC (5630A>C), GGC1819GGT (5641C>T), GCA1822GCT (5650A>T), ATA1826ATT (5662A>T), GGA1827GGG (5665A>G), CAA1835CAG (5689A>G), GTT1837ATT (5693G>A), GAG1838GAA (5698G>A), CAA1843CAG (5713A>G), GGG1844GGC (5716G>C), AAA1845AAG (5719A>G), GCC1853GCA (5743C>A), CGC1857CGA (5755C>A), CAT1858CAC (5758T>C), CGT1860CGC (5764T>C), AAA1862AAG (5770A>G), GGC1865GGT (5779C>T), TTT1866TTC (5782T>C), GAC1869GAT (5791C>T), GAC1873GAT (5803C>T), AGG1884AGA (5836G>A), AAA1885AAG (5839A>G), GGA1887GGG (5845A>G), AAA1888AAG (5848A>G), AAA1890AGA (5853A>G), GGT1891GGC (5857T>C), ACC1892ACT (5860C>T), ACA1893ACT (5863A>T), GGT1895GGA (5869T>A), GGT1897GGC (5875T>C), AGC1900AGT (5884C>T), TTC1903TTT (5893C>T), ATC1904GTG (5894A>G 5896C>G), AAC1905AAT (5899C>T), GGG1908GGA (5908G>A), TTT1909TTC (5911T>C), GAG1913GAA (5923G>A), CAA1918CAG (5938A>G), TTC1919TTT (5941C>T), GTT1920GTC (5944T>C), GAT1921GAC (5947T>C), GGG1925GGA (5959G>A), CGG1926GCC (5960C>G 5961G>C 5962G>C), GAA1930GAG (5974A>G), GCT1934GCA (5986T>A), GAC1935GAT (5989C>T), ATT1936GTT (5990A>G), GAT1938GAC (5998T>C), AGA1942CGT (6008A>C 6010A>T), GAA1945GAT (6019A>T), GTG1946GTT (6022G>T), CGA1947CGT (6025A>T), AAA1949ACA (6030A>C), GAC1955GAA (6049C>A), ATG1958ATA (6058G>A), CAA1959CAG (6061A>G), GCC1960GCA (6064C>A), GGT1962GGC (6070T>C), AGT1963AAC (6072G>A 6073T>C), AAC1964CAC (6074A>C), ACG1965ACA (6079G>A), ACC1966AAC (6081C>A), ATA1967ATT (6085A>T), GCA1969GCT (6091A>T), TAC1970TAT (6094C>T), TGT1976TCC (6111G>C 6112T>C), AAA1978AAG (6118A>G), TTG1980TTA (6124G>A), AAG1981AAA (6127G>A), ATT1982GTC (6128A>G 6130T>C), GAT1983GAC (6133T>C), TTA1984TTG (6136A>G), AAC1988AAT (6148C>T), AAA1991AAG (6157A>G), GTT1992ATT (6158G>A), TGT1993TGC (6163T>C), AAA1995AAG (6169A>G), ACA1996TCG (6170A>T 6172A>G), GGC1998GGT (6178C>T), AAA2001AAG (6187A>G), TTT2002TTC (6190T>C), GAG2004GAA (6196G>A), CTC2007TTT (6203C>T 6205C>T), GAA2008GAG (6208A>G), CTA2009TTG (6209C>T 6211A>G), CAG2011CAA (6217G>A), GCT2015GCA (6229T>A), GTA2016ATA (6230G>A), GTC2018GTT (6238C>T), AAG2021AAA (6247G>A), ATA2023ATT (6253A>T), GCA2025AAA (6257G>A 6258C>A), CAT2030CAC (6274T>C), AAA2033AGA (6282A>G), TCG2034TCA (6286G>A), GGC2038GGT (6298C>T), TTG2039TTA (6301G>A), AGA2040AGG (6304A>G), GAC2041GAT (6307C>T), GCC2046GCT (6322C>T), ACA2048ACT (6328A>T), GTT2049GTG (6331T>G), TGT2050TGC (6334T>C), AGG2051AGA (6337G>A), CTG2052TTG (6338C>T), TAT2058TAC (6358T>C), GGG2059GGA (6361G>A), GCA2060ACA (6362G>A), TCA2061TCT (6367A>T), GAG2062GAA (6370G>A), TAC2064TAT (6376C>T), GGT2065GGG (6379T>G), GGA2067GGC (6385A>C), GGA2069GGT (6391A>T), GCA2070GCG (6394A>G), TAC2071TAT (6397C>T), GTA2073ATA (6401G>A), GCG2074GTG (6405C>T), CAC2076CAT (6412C>T), TTA2078TTG (6418A>G), TTT2079TTC (6421T>C), TAC2082TTC (6429A>T), GGT2084GGA (6436T>A), GAG2087GAA (6445G>A), CAA2089CGA (6450A>G), TCC2090TCG (6454C>G), CAC2092CAT (6460C>T), GGT2093GGA (6463T>A), TTC2095TTT (6469C>T), AGG2096AGA (6472G>A), GTG2097GTA (6475G>A), AAT2099AAC (6481T>C), CAC2101AAT (6485C>A 6487C>T), AGT2102AGC (6490T>C), AGC2104AGT (6496C>T), CTG2106CTA (6502G>A), GGT2110GGC (6514T>C), AGG2111AGA (6517G>A), CTC2115ATC (6527C>A), ATC2116ATA (6532C>A), CCG2119CCA (6541G>A), AAA2120AAG (6544A>G), GTC2124GTT (6556C>T), TTT2125TTC (6559T>C), CAG2127CAA (6565G>A), TTG2129CTA (6569T>C 6571G>A), CCT2134CCA (6586T>A), ACA2135ATA (6588C>T), GAA2138GAG (6598A>G), AGA2139AGG (6601A>G), ATT2140ATC (6604T>C), TGT2141TGC (6607T>C), GGA2144GGG (6616A>G), AAC2146AAT (6622C>T), GAG2149GAA (6631G>A), AAG2150AAA (6634G>A), TAT2151CAT (6635T>C), GCT2152GCA (6640T>A), TCG2154TCT (6646G>T), ATC2155GTC (6647A>G), ATC2156ATT (6652C>T), AGC2160AGT (6664C>T), ACT2162ACG (6670T>G), ATA2165GTG (6677A>G 6679A>G), GGC2167GGT (6685C>T), ACA2169ACT (6691A>T), TTC2170TTT (6694C>T), ACA2177ACG (6715A>G), GAT2178AAT (6716G>A), AAT2179GAT (6719A>G), GGA2180GGG (6724A>G), CTA2184TTA (6734C>T), GTG2186GTA (6742G>A), GTG2187GTA (6745G>A), GCC2190GCT (6754C>T), GGA2192GGG (6760A>G), ATA2194TTG (6764A>T 6766A>G), ATT2197ATA (6775T>A), CTG2200TTG (6782C>T), GCA2201GCT (6787A>T), AAC2202AAT (6790C>T), GCA2204GTG (6795C>T 6796A>G), CAC2205CAA (6799C>A), ACC2206ACA (6802C>A), AAC2208AAT (6808C>T), TAC2209TAT (6811C>T), TAC2210TAT (6814C>T), TTC2213TTT (6823C>T), GAA2215GAG (6829A>G), TAC2221TAT (6847C>T), CGA2223CGC (6853A>C), AAT2225GAT (6857A>G), GAG2226GAA (6862G>A), GAA2229GAG (6871A>G), GTC2231ACA (6875G>A 6876T>C 6877C>A), AAG2232AAA (6880G>A), TCT2233TCA (6883T>A), GTT2235GTG (6889T>G), GAC2239GAT (6901C>T), GGC2244GGT (6916C>T), CCG2245CCA (6919G>A), AAA2247AAG (6925A>G), CTT2248CTC (6928T>C), GAC2250GAA (6934C>A), GGG2255GGC (6949G>C), TTC2257TTT (6955C>T), AAA2258AAG (6958A>G), CAA2264CAG (6976A>G), CTA2266TTA (6980C>T), ATC2267ATT (6985C>T), GTA2271GTT (6997A>T), GTG2272GTT (7000G>T), GTG2273ATA (7001G>A 7003G>A), GCT2276GCC (7012T>C), AAG2277AAA (7015G>A), CAT2278CAC (7018T>C), TTT2283TAT (7032T>A), GCC2285GCT (7039C>T), TTG2286CTA (7040T>C 7042G>A), GGA2288GGG (7048A>G), GCT2292GCA (7060T>A), GTC2293GTA (7063C>A), AGC2298AGT (7078C>T), TTA2300CTG (7082T>C 7084A>G), GTA2301GTG (7087A>G), ACC2302ACA (7090C>A), CAT2304CAC (7096T>C), GTA2305GTG (7099A>G), GTT2306GTC (7102T>C), GAG2309GAA (7111G>A), CGA2311CGG (7117A>G), ACA2314AAA (7125C>A), GAA2315GAG (7129A>G), TTT2316TTC (7132T>C), CTG2317TTA (7133C>T 7135G>A), GTG2319GTA (7141G>A), GAT2320GAC (7144T>C), GCA2321TCG (7145G>T 7147A>G), GAG2322GAA (7150G>A), GCA2325GCT (7159A>T), GAG2326_GCA2327del (7160_7165delGAGGCA), TTC2329TTT (7171C>T), GCG2335GCT (7189G>T), AAA2338AAG (7198A>G), AGC2339AGT (7201C>T), CTA2341TTA (7205C>T), AAT2342AAC (7210T>C), GAT2344GAA (7216T>A), GCG2345GCA (7219G>A), ATC2347ATA (7225C>A), AAG2348AAA (7228G>A), AAG2352AAA (7240G>A), TAT2353TAC (7243T>C), AAA2355AAG (7249A>G), CCT2356CCG (7252T>G), GAT2358GAA (7258T>A), GTT2359GTC (7261T>C), GGT2360GGA (7264T>A), GTC2361GTG (7267C>G), GTG2362GTA (7270G>A), GAT2363_CGG2364insT-- (7273_7274insT), CGG2364GTG (7274C>G 7275G>T), CAT2366CTT (7281A>T), TTG2367TCG (7284T>C), TCA2371TTA (7296C>T), TCT2376TTT (7311C>T), ACC2377ATC (7314C>T), TGC2378TAC (7317G>A), AAT2379AAA (7321T>A), GTG2380-TG (7322delG), CAC2381CAT (7327C>T), GGC2382GGT (7330C>T), TTC2383TTT (7333C>T), AAG2384CAG (7334A>C), TGT2386TGC (7342T>C), GCA2387AAC (7343G>A 7344C>A 7345A>C), ACT2390ACC (7354T>C), CAA2393CAG (7363A>G), ATT2395ATC (7369T>C), TTC2396TTT (7372C>T), GCG2398GCT (7378G>T), CTC2399CTT (7381C>T), GCA2404GCT (7396A>T), GTC2405GTT (7399C>T), GCC2407GCT (7405C>T), AGT2408ATG (7407G>T 7408T>G), GGG2410GGT (7414G>T), TGC2411GGC (7415T>G), AAA2412AAG (7420A>G), GAC2415GAT (7429C>T), TAT2416TAC (7432T>C), CAT2419CAC (7441T>C), AAG2425AAA (7459G>A), GAA2426GAG (7462A>G), ATA2428ATT (7468A>T), GTC2429GTA (7471C>A), CAA2431CAG (7477A>G), AGC2432AGT (7480C>T), CTG2434CTT (7486G>T), TAT2437TAC (7495T>C), AAA2438AAG (7498A>G), GGT2439GGC (7501T>C), TTG2440TCA (7503T>C 7504G>A), ATT2443ATA (7513T>A), AAC2445AAT (7519C>T), GGA2446GGG (7522A>G), AAG2449AAA (7531G>A), GAG2451GAA (7537G>A), CTC2452CTT (7540C>T), TGT2454TGC (7546T>C), AAG2462AAA (7570G>A), ACG2463ACA (7573G>A), ACG2465ACA (7579G>A), GCT2468GCC (7588T>C), CTA2471TTA (7595C>T), TTG2474CTA (7604T>C 7606G>A), GGT2477GGA (7615T>A), TGT2480TGC (7624T>C), GAC2483GAT (7633C>T), TTC2484TTT (7636C>T), AAT2485AAC (7639T>C), TTT2488TTC (7648T>C), GAA2494GAG (7666A>G), TGT2496TGC (7672T>C), ACA2498ACT (7678A>T), GGG2500GGA (7684G>A), ACT2502ACC (7690T>C), TTT2504TTC (7696T>C), GGT2506GGG (7702T>G), GAT2509AAT (7709G>A), AAA2510AGG (7713A>G 7714A>G), CTG2511TTG (7715C>T), CGT2514CGC (7726T>C), GAG2517GAA (7735G>A), GTA2520GTG (7744A>G), TAC2521TAT (7747C>T), GGC2526GGT (7762C>T), CAG2528CAA (7768G>A), TTT2529TTC (7771T>C), TCA2532TCG (7780A>G), CTA2533CTC (7783A>C), ACT2534ACC (7786T>C), CTA2537TTG (7793C>T 7795A>G), ATC2538ATT (7798C>T), GCT2540GCC (7804T>C), ACC2543ATC (7812C>T), GAA2550GAG (7834A>G), GAC2551GAT (7837C>T), GTG2554TTA (7844G>T 7846G>A), CAG2557CAA (7855G>A), CTG2559CTA (7861G>A), TTA2562TTG (7870A>G), TAC2563TAT (7873C>T), ACT2564ACA (7876T>A), GAG2565GAA (7879G>A), ATT2566ATA (7882T>A), GTT2567ATT (7883G>A), CCA2570CCT (7894A>T), ATT2571ATC (7897T>C), GTC2579GTT (7921C>T), AAG2581AAA (7927G>A), GGT2584GGC (7936T>C), AAT2585AAC (7939T>C), AAT2586AAC (7942T>C), GTG2594GTA (7966G>A), GAT2595GAC (7969T>C), TCT2597TCC (7975T>C), GTC2601GTT (7987C>T), CTT2602CTC (7990T>C), GCT2607GCC (8005T>C), GAA2616GAG (8032A>G), ATC2618ATT (8038C>T), GAC2619GAT (8041C>T), AGC2620AAC (8043G>A), TTG2632TTA (8080G>A), GTG2635GTA (8089G>A), CCG2637CCA (8095G>A), GAG2638GAA (8098G>A), AGC2641AGT (8107C>T), ATT2642ATC (8110T>C), GAT2644GAC (8116T>C), CAA2648CAG (8128A>G), GGT2654GGC (8146T>C), AAC2656AGT (8151A>G 8152C>T), TCG2660TCA (8164G>A), AGG2665AAG (8178G>A), AAG2666AAA (8182G>A), TTG2669TTA (8191G>A), TTC2671TTT (8197C>T), TCC2673TCT (8203C>T), AGA2675AAA (8208G>A), CTG2677TTA (8213C>T 8215G>A), CTA2678CTG (8218A>G), ATC2679ATT (8221C>T), GAG2680GAA (8224G>A), CCA2685CCG (8239A>G), GAG2690GAA (8254G>A), ATT2692ATC (8260T>C), GTA2693GTG (8263A>G), GAT2699GAC (8281T>C), AGA2700AGG (8284A>G), CCA2704GCG (8294C>G 8296A>G), GAG2705GAA (8299G>A), CAC2706CAT (8302C>T), ATT2711ATC (8317T>C), TGC2712TGT (8320C>T), GCT2714GCA (8326T>A), GAG2717GAA (8335G>A), TCC2718TCT (8338C>T), TCT2722TTT (8349C>T), GAA2723GAG (8353A>G), CAC2726CAT (8362C>T), CAA2727CAG (8365A>G), ATC2728ATT (8368C>T), TAC2732TAT (8380C>T), TCA2733TCG (8383A>G), TTA2735TTG (8389A>G), CCT2740CCC (8404T>C), GCA2742TCA (8408G>T), ACA2743ACG (8413A>G), GCG2745GCT (8419G>T), GCT2750GCC (8434T>C), TAT2752TAC (8440T>C), ATG2756ATA (8452G>A), CTA2758CTG (8458A>G), AGG2759AAG (8460G>A), AAA2760AAG (8464A>G), CTG2761CTA (8467G>A), AGG2765AGA (8479G>A), GCT2766ACA (8480G>A 8482T>A), GAG2771GAA (8497G>A), CTA2772TTG (8498C>T 8500A>G), AGA2773AAG (8502G>A 8503A>G), GCC2774GCT (8506C>T), GAA2777GAT (8515A>T), GTC2780GTT (8524C>T), GCA2781GCC (8527A>C), TTA2782CTG (8528T>C 8530A>G), GAG2785GAC (8539G>C), CTT2788TGT (8546C>T 8547T>G), TCT2790ACT (8552T>A), GTA2793GTG (8563A>G), GCA2797GGA (8574C>G), ATT2801GTT (8585A>G), GGA2805GAA (8598G>A), AGC2806AGT (8602C>T), AAC2807TCT (8603A>T 8604A>C 8605C>T), AAA2812AAG (8620A>G), CCA2813CAA (8622C>A), GAG2814GGA (8625A>G 8626G>A), CAG2815CAA (8629G>A), GGC2816GGT (8632C>T), ATC2818ATT (8638C>T), CAG2819CAA (8641G>A), AAC2821AAT (8647C>T), CCG2822TTC (8648C>T 8649C>T 8650G>C), AAC2823AAT (8653C>T), AAA2824AAG (8656A>G), GGA2825GAA (8658G>A), GAT2827GCG (8664A>C 8665T>G), GTT2830GTG (8674T>G), GCA2832GTT (8679C>T 8680A>T), GGC2833GGA (8683C>A), GGG2836GGA (8692G>A), ACA2837ACT (8695A>T), CCG2841CCA (8707G>A), AGA2842CGA (8708A>C), AAG2844AAA (8716G>A), ATC2846ATT (8722C>T), ACG2847ACA (8725G>A), AAA2849AAG (8731A>G), AGA2851AGG (8737A>G), CCC2853CCT (8743C>T), ACA2854AAG (8745C>A 8746A>G), AAG2856AAA (8752G>A), GGA2857GGT (8755A>T), AAC2862AAT (8770C>T), TTA2863CTA (8771T>C), GAA2864GAG (8776A>G), CAT2865CAC (8779T>C), TTG2866TTA (8782G>A), TAT2869TAC (8791T>C), CCA2871CCG (8797A>G), CAA2872CAG (8800A>G), CAA2873CAG (8803A>G), GAT2875GAC (8809T>C), ATT2876ATC (8812T>C), AAT2878AAC (8818T>C), CGG2880CGA (8824G>A), GCA2881GCG (8827A>G), ACG2888ACA (8848G>A), TAT2890TAC (8854T>C), GAG2891GGA (8856A>G 8857G>A), GTG2893GTA (8863G>A), CGG2894CAG (8865G>A), ATG2895CTT (8867A>C 8869G>T), GCA2896GCG (8872A>G), GGA2900GAA (8883G>A), ACT2902TCT (8888A>T), TGC2915TGT (8929C>T), GTC2924ATC (8954G>A), AAC2925AAT (8959C>T), GGA2926GGT (8962A>T), GGG2933GGA (8983G>A), AAT2934GAC (8984A>G 8986T>C), GAA2935GTA (8988A>T), GAG2938GAA (8998G>A), CCG2940CCA (9004G>A), TTG2941TTA (9007G>A), ATC2944ATT (9016C>T), ACC2951ACA (9037C>A), ATA2967ATT (9085A>T), AAC2971AAT (9097C>T), ATT2983GTT (9131A>G), CGA2984CGT (9136A>T), CGG2987CGT (9145G>T), ATG2989GGA (9149A>G 9150T>G 9151G>A), GGT2990AGT (9152G>A), TTA2991TTG (9157A>G), GCG2992GCT (9160G>T), CGT2993CGC (9163T>C), GAG3000GAA (9184G>A), CGA3004CGG (9196A>G), AGG3010AGA (9214G>A), GAA3011GAG (9217A>G), GCG3012GCA (9220G>A), GCA3020GCT (9244A>T), TTG3021TTA (9247G>A), GCC3024GCT (9256C>T), CCT3026TCC (9260C>T 9262T>C), GGG3030GGA (9274G>A), GAC3032GAT (9280C>T), ATC3035ATT (9289C>T), CAA3038CAC (9298A>C), GAG3040GAA (9304G>A), GTC3050GTT (9334C>T), GTC3060GTG (9364C>G) | | | | | | | | |
| P1 protein (NP_734243.1) | 1 | 284 | 100% | 1394 | 72.7% | 284 (100%) | 203 (71.5%) | 0/0/0/0 | 0 |

| | Begin | End | Coverage | Score | Concordance | Matches | Identities | I/D/M/F* | Stop Codons |
|---|---|---|---|---|---|---|---|---|---|
| **NT** | **22** | **9704** | **99.8%** | **12522** | **65.0%** | **9671 (99.8%)** | **8003 (82.6%)** | **10/12** | |
| Protein mutations: | Y4Q (194T>C 196C>A), C9L (209T>C 210G>T 211T>G), F13M (221T>A 223T>G), S23P (251T>C), C24F (255G>T 256C>T), E25G (258A>G), I27V (263A>G 265T>G), V28A (267T>C), E30V (273A>T), A35T (287G>A 289T>C), V37T (293G>A 294T>C), A41I (305G>A 306C>T), D42G (309A>G), E44D (316A>T), T45V (317A>G 318C>T 319A>G), L52Q (339T>A 340C>A), K53R (342A>G), K55A (347A>G 348A>C 349A>G), Y56H (350T>C), T58V (356A>G 357C>T), V61T (365G>A 366T>C 367G>A), L62S (368C>T 369T>C), F67C (384T>G), A75V (408C>T 409C>T), M78E (416A>G 417T>A), E85K (437G>A), R86E (440A>G 441G>A), K87R (443A>C 444A>G 445G>A), D88E (448T>A), E91A (456A>C), H93N (461C>A), D111E (517T>A), S116F (531C>T), P118S (536C>T), Q119W (539C>T 540A>G 541A>G), R122K (549G>A 550A>G), M131T (576T>C), V134A (585T>C 586C>G), R137Y (593C>T 594G>A 595C>T), P138R (597C>G), I139V (599A>G 601A>G), I140P (602A>C 603T>C 604A>G), M147I (625G>A), I151V (635A>G 637T>C), I154V (644A>G 646A>G), E160A (663A>C 664G>C), H166Q (682C>A), T172S (699C>G), H174Q (706T>A), K179E (719A>G 721G>A), I180V (722A>G 724A>G), A183S (731G>T), Y184P (734T>C 735A>C 736C>T), S185R (737T>C 738C>G 739C>T), A187V (744C>T 745G>C), R189C (749C>T 751A>C), M194K (765T>A), R197K (773C>A 774G>A 775A>G), M206K (801T>A), G210A (813G>C 814A>G), L211H (816T>A), R214Q (825G>A 826T>A), N223D (851A>G 853T>C), R226C (860C>T 862C>T), T227A (863A>G 865T>C), I228T (867T>C), N229D (869A>G 871C>T), I230L (872A>T 874A>G), R231Q (876G>A), R232K (879G>A), N240S (903A>G), T241N (906C>A 907A>C), K242T (909A>C 910A>C), S243N (912G>A 913C>T), S252L (938T>C 939C>T 940A>T), L266I (980T>A 982G>C), R272K (999G>A), Q275R (1007C>A 1008A>G 1009G>A), S276G (1010A>G 1012T>G), I277V (1013A>G), N279Q (1019A>C 1021C>G) | | | | | | | | |
| Codon mutations: | ACT3ACC (193T>C), TAC4CAA (194T>C 196C>A), TCA6TCG (202A>G), ATC8ATT (208C>T), TGT9CTG (209T>C 210G>T 211T>G), TCG12TCA (220G>A), TTT13ATG (221T>A 223T>G), GAA14GAG (226A>G), CTA17CTT (235A>T), TAC19TAT (241C>T), CCA21CCT (247A>T), TCT23CCT (251T>C), TGC24TTT (255G>T 256C>T), GAG25GGG (258A>G), CAT26CAC (262T>C), ATT27GTG (263A>G 265T>G), GTG28GCG (267T>C), GAA30GTA (273A>T), CGA31CGT (277A>T), GAA32GAG (280A>G), GTG33GTT (283G>T), CCG34CCA (286G>A), GCT35ACC (287G>A 289T>C), GTT37ACT (293G>A 294T>C), CCT39CCC (301T>C), TTC40TTT (304C>T), GCA41ATA (305G>A 306C>T), GAT42GGT (309A>G), CTG43TTG (311C>T), GAA44GAT (316A>T), ACA45GTG (317A>G 318C>T 319A>G), CAA46CAG (322A>G), GCA49GCT (331A>T), CTC52CAA (339T>A 340C>A), AAG53AGG (342A>G), AAA55GCG (347A>G 348A>C 349A>G), TAT56CAT (350T>C), GCT57GCA (355T>A), ACT58GTT (356A>G 357C>T), CGT60CGC (364T>C), GTG61ACA (365G>A 366T>C 367G>A), CTC62TCC (368C>T 369T>C), AAA63AAG (373A>G), AAC64AAT (376C>T), TTT67TGT (384T>G), ACG68ACA (388G>A), TAC69TAT (391C>T), CGA70CGG (394A>G), AAG72AAA (400G>A), GAT74GAC (406T>C), GCC75GTT (408C>T 409C>T), CAG76CAA (412G>A), ATA77ATT (415A>T), ATG78GAG (416A>G 417T>A), ATT80ATC (424T>C), CAG81CAA (427G>A), AAA83AAG (433A>G), CTG84CTC (436G>C), GAG85AAG (437G>A), AGG86GAG (440A>G 441G>A), AAG87CGA (443A>C 444A>G 445G>A), GAT88GAA (448T>A), GAA90GAG (454A>G), GAA91GCA (456A>C), TAT92TAC (460T>C), CAC93AAC (461C>A), CAA95CAG (469A>G), GCC97GCG (475C>G), CCT99CCA (481T>A), ACT106ACC (502T>C), ATA107ATT (505A>T), GGC109GGT (511C>T), GAT111GAA (517T>A), CCT112CCA (520T>A), CCA113CCT (523A>T), TCT116TTT (531C>T), GAG117GAA (535G>A), CCA118TCA (536C>T), CAA119TGG (539C>T 540A>G 541A>G), AGA122AAG (549G>A 550A>G), ATC124ATT (556C>T), ATT125ATC (559T>C), CAT126CAC (562T>C), ACT128ACC (568T>C), CCA129CCG (571A>G), ATG131ACG (576T>C), CGT132CGC (580T>C), GTC134GCG (585T>C 586C>G), CGC137TAT (593C>T 594G>A 595C>T), CCC138CGC (597C>G), ATA139GTG (599A>G 601A>G), ATA140CCG (602A>C 603T>C 604A>G), ACA143ACG (613A>G), GAA144GAG (616A>G), GGC145GGG (619C>G), CAG146CAA (622G>A), ATG147ATA (625G>A), AAT148AAC (628T>C), CAC149CAT (631C>T), ATT151GTC (635A>G 637T>C), ATA154GTG (644A>G 646A>G), ATT157ATC (655T>C), TCG159TCA (661G>A), GAG160GCC (663A>C 664G>C), AAA161AAG (667A>G), GTC165GTT (679C>T), CAC166CAA (682C>A), TTA167CTG (683T>C 685A>G), AGT169AGC (691T>C), ACC172AGC (699C>G), ACT173ACA (703T>A), CAT174CAA (706T>A), AAG179GAA (719A>G 721G>A), ATA180GTG (722A>G 724A>G), CTT181TTG (725C>T 727T>G), GGT182GGA (730T>A), GCA183TCA (731G>T), TAC184CCT (734T>C 735A>C 736C>T), TCC185CGT (737T>C 738C>G 739C>T), GCG187GTC (744C>T 745G>C), CGA189TGC (749C>T 751A>C), ACT190ACC (754T>C), ATG194AAG (765T>A), TTG196CTG (770T>C), CGA197AAG (773C>A 774G>A 775A>G), CGG198AGG (776C>A), TTC202TTT (790C>T), ATG206AAG (801T>A), GGA210GCG (813G>C 814A>G), CTT211CAT (816T>A), TTG212CTG (818T>C), CAA213CAG (823A>G), CGT214CAA (825G>A 826T>A), CGG217AGA (833C>A 835G>A), GAC219GAT (841C>T), AAT223GAC (851A>G 853T>C), GTC225GTT (859C>T), CGC226TGT (860C>T 862C>T), ACT227GCC (863A>G 865T>C), ATC228ACC (867T>C), AAC229GAT (869A>G 871C>T), ATA230TTG (872A>T 874A>G), CGA231CAA (876G>A), AGG232AAG (879G>A), GAT234GAC (886T>C), ATC238ATT (898C>T), AAC240AGC (903A>G), ACA241AAC (906C>A 907A>C), AAA242ACC (909A>C 910A>C), AGC243AAT (912G>A 913C>T), GGC246GGA (922C>A), GGT249GGA (931T>A), TCA252CTT (938T>C 939C>T 940A>T), GGA253GGG (943A>G), TTC256TTT (952C>T), TCA261TCG (967A>G), GAA263GAG (973A>G), GGG264GGC (976G>C), TTG266ATC (980T>A 982G>C), GCA269GCG (991A>G), CGT270CGC (994T>C), AGA272AAA (999G>A), GTT273GTC (1003T>C), CAG275AGA (1007C>A 1008A>G 1009G>A), AGT276GGG (1010A>G 1012T>G), ATT277GTT (1013A>G), TTA278TTG (1018A>G), AAC279CAG (1019A>C 1021C>G), ATC282ATT (1030C>T) | | | | | | | | |
| HC-Pro protein (NP_734244.1) | 1 | 456 | 100% | 3144 | 95.7% | 456 (100%) | 425 (93.2%) | 0/0/0/0 | 0 |
| Protein mutations: | D4E (1048C>A), R16Q (1083G>A), S21A (1097T>G 1099G>A), R36K (1143G>A), L40I (1154T>A 1156G>A), S62N (1221G>A), V65T (1229G>A 1230T>C 1231T>C), S66N (1233G>A 1234C>T), K74T (1257A>C 1258A>G), N81S (1278A>G), A85V (1290C>T 1291G>A), D88T (1298G>A 1299A>C 1300C>T), I91V (1307A>G 1309A>G), N94E (1316A>G 1318T>A), I98T (1329T>C), L113I (1373C>A), I117V (1385A>G), E123D (1405G>T), A127S (1415G>T 1417A>T), V134I (1436G>A 1438C>T), E224G (1707A>G), A228T (1718G>A), I236V (1742A>G 1744C>T), S241N (1758G>A), V252I (1790G>A), I301V (1937A>G 1939T>C), F323Y (2004T>A), I344M (2068T>G), V346I (2072G>A 2074G>A), R391K (2208G>A), D410E (2266C>G) | | | | | | | | |
| Codon mutations: | TCG1TCA (1039G>A), AAT2AAC (1042T>C), GCC3GCA (1045C>A), GAC4GAA (1048C>A), AAT5AAC (1051T>C), AAG8AAA (1060G>A), GGT9GGA (1063T>A), CTG10TTG (1064C>T), GAC11GAT (1069C>T), CGA16CAA (1083G>A), TAT19TAC (1093T>C), TCG21GCA (1097T>G 1099G>A), CAC23CAT (1105C>T), GTA26GTG (1114A>G), GCT27GCG (1117T>G), TTA29TTG (1123A>G), CCT30CCA (1126T>A), GTC31GTT (1129C>T), AGG36AAG (1143G>A), GTA37GTT (1147A>T), GCT38GCA (1150T>A), TTG40ATA (1154T>A 1156G>A), CAC43CAT (1165C>T), AGT44AGC (1168T>C), CTT46TTA (1172C>T 1174T>A), ACC55ACT (1201C>T), AGC62AAC (1221G>A), GTT65ACC (1229G>A 1230T>C 1231T>C), AGC66AAT (1233G>A 1234C>T), CTG68CTA (1240G>A), AAG70AAA (1246G>A), TTG72CTA (1250T>C 1252G>A), CAT73CAC (1255T>C), AAA74ACG (1257A>C 1258A>G), AGA77AGG (1267A>G), GGT79GGC (1273T>C), TTG80TTA (1276G>A), AAT81AGT (1278A>G), TTG83CTG (1283T>C), GCG85GTA (1290C>T 1291G>A), GAC88ACT (1298G>A 1299A>C 1300C>T), TTT90TTC (1306T>C), ATA91GTG (1307A>G 1309A>G), CAT92CAC (1312T>C), AAT94GAA (1316A>G 1318T>A), TTG97TTA (1327G>A), ATA98ACA (1329T>C), GCG99GCT (1333G>T), TTA100TTG (1336A>G), CTA103TTG (1343C>T 1345A>G), CCG106CCA (1354G>A), GTG107GTT (1357G>T), GAC108GAT (1360C>T), CTC111TTG (1367C>T 1369C>G), CTT113ATT (1373C>A), TTC114TTT (1378C>T), GAG116GAA (1384G>A), ATA117GTA (1385A>G), AAA119AAG (1393A>G), TCC120TCT (1396C>T), GAG123GAT (1405G>T), AAA124AAG (1408A>G), CAG125CAA (1411G>A), GCA127TCT (1415G>T 1417A>T), CCG128CCC (1420G>C), AAT131AAC (1429T>C), TTA132TTG (1432A>G), GTC134ATT (1436G>A 1438C>T), TTA135CTG (1439T>C 1441A>G), AAT136AAC (1444T>C), TTC139TTT (1453C>T), CTG140CTA (1456G>A), GAA144GAG (1468A>G), GTA152GTG (1492A>G), TTG155CTA (1499T>C 1501G>A), AGT156AGC (1504T>C), TTG157TTA (1507G>A), CTC158CTT (1510C>T), GAA159GAG (1513A>G), TTA160TTG (1516A>G), GCA161GCT (1519A>T), AGG162AGA (1522G>A), TTC163TTT (1525C>T), CAG164CAA (1528G>A), AAC166AAT (1534C>T), ACT168ACA (1540T>A), AAC170AAT (1546C>T), AAA173AAG (1555A>G), GGT174GGC (1558T>C), GAT175GAC (1561T>C), ATA176ATC (1564A>C), TCT177TCA (1567T>A), TTC179TTT (1573C>T), AGA180AGG (1576A>G), TTA183CTA (1583T>C), TCT184TCA (1588T>A), AAG186AAA (1594G>A), AAC188AAT (1600C>T), CTG191CTA (1609G>A), TCG194TCA (1618G>A), TGC195TGT (1621C>T), GAC196GAT (1624C>T), CTG199TTG (1631C>T), AAA201AAG (1639A>G), TTC205TTT (1651C>T), CTC206CTG (1654C>G), CAT213CAC (1675T>C), TCA219TCG (1693A>G), TTT222TTC (1702T>C), GAA224GGA (1707A>G), GCA228ACA (1718G>A), GGA230GGG (1726A>G), TAC231TAT (1729C>T), TCA232TCC (1732A>C), TAT234TAC (1738T>C), ATC236GTT (1742A>G 1744C>T), CGC237CGT (1747C>T), CAT239CAC (1753T>C), AGT241AAT (1758G>A), AGG244AGA (1768G>A), AAC250AAT (1786C>T), TTA251TTG (1789A>G), GTT252ATT (1790G>A), GTC253GTT (1795C>T), TTA257CTG (1805T>C 1807A>G), GAG259GAA (1813G>A), AGG261AGA (1819G>A), AAG263AAA (1825G>A), GAC267GAT (1837C>T), TAT268TAC (1840T>C), CAA271CAG (1849A>G), GTC274GTG (1858C>G), AAA276AAG (1864A>G), TGC278TGT (1870C>T), AGT280AGC (1876T>C), AAA282AAG (1882A>G), GGT284GGA (1888T>A), AAT285AAC (1891T>C), TGT290TGC (1906T>C), ACA293ACT (1915A>T), GGT298GGC (1930T>C), GCC300GCT (1936C>T), ATT301GTC (1937A>G 1939T>C), TCA303TCG (1945A>G), TTC305TTT (1951C>T), CCA308CCT (1960A>T), GTA314GTG (1978A>G), ATT315ATA (1981T>A), GGC316GGT (1984C>T), AAT317AAC (1987T>C), GAC320GAT (1996C>T), TTT323TAT (2004T>A), AAA328AAG (2020A>G), GGG329GGA (2023G>A), GAT330GAC (2026T>C), TCG331TCT (2029G>T), GAG332GAA (2032G>A), TAT341TAC (2059T>C), ATT344ATG (2068T>G), AAC345AAT (2071C>T), GTG346ATA (2072G>A 2074G>A), CTG351TTA (2087C>T 2089G>A), AGC355AGT (2101C>T), GAG356GAA (2104G>A), GAG357GAA (2107G>A), AAG360AAA (2116G>A), TTC362TTT (2122C>T), CGC367CGT (2137C>T), CTT374CTC (2158T>C), ACC376ACT (2164C>T), CCA378CCG (2170A>G), GAT382GAC (2182T>C), GCG384GCA (2188G>A), ACC385ACA (2191C>A), AGA391AAA (2208G>A), TAT394TAC (2218T>C), GAC396GAT (2224C>T), GTA397GTT (2227A>T), TTG402TTA (2242G>A), CCC403CCT (2245C>T), TTG406TTA (2254G>A), GTT407GTC (2257T>C), GAC408GAT (2260C>T), GAC410GAG (2266C>G), CAA412CAG (2272A>G), ACG413ACA (2275G>A), TCA419TCG (2293A>G), GGC421GGT (2299C>T), TCG422TCA (2302G>A), CAG423CAA (2305G>A), GGA426GGT (2314A>T), TAT427TAC (2317T>C), CTA430CTG (2326A>G), GTG435GTA (2341G>A), CAA437CAG (2347A>G), CTT438CTC (2350T>C), TTG440TTA (2356G>A), AAT443AAC (2365T>C), GAA445GAG (2371A>G), TTA446TTG (2374A>G), GAT449GAC (2383T>C), ATA450ATT (2386A>T), AAA451AAG (2389A>G), CAT452CAC (2392T>C), TAT453TAC (2395T>C) | | | | | | | | |
| P3 protein (NP_734251.1) | 1 | 365 | 100% | 1924 | 79.6% | 365 (99.7%) | 327 (89.3%) | 1/0/2/2 | 0 |
| Protein mutations: | N4G (2414A>G 2415A>G 2416T>G), S6C (2420A>T), K40R (2523A>G), R77K (2634G>A), V107I (2723G>A 2725C>T), A115T (2747G>A 2749T>A), H131N (2795C>A), V140I (2822G>A 2824G>A), P160T (2882C>A), S161H (2885A>C 2886G>A), T164M (2895C>T), N176S (2931A>G), N179S (2940A>G), E189_N190insX (2971_2972insA), T196I (2991C>T), H197R (2994A>G), E199R (2999G>A 3000A>G), R206Q (3021G>A), Y207F (3024A>T 3025C>T), T211I (3036C>T), E212G (3039A>G 3040A>C), K213T (3042A>C 3043G>A), A232V (3099C>T 3100C>A), V234R (3104G>A 3105T>G 3106G>A), S243N (3132G>A), R245K (3137C>A 3138G>A 3139A>G), F250I (3152T>A 3154C>T), C254Y (3165G>A), F272L (3218T>C), T274N (3225C>A), V276I (3230G>A), V288M (3266G>A 3268A>G), V291M (3275G>A), A294T (3284G>A), R304K (3315G>A 3316A>G), E305A (3318A>C), M319V (3359A>G), D332E (3400T>G) | | | | | | | | |

| | Begin | End | Coverage | Score | Concordance | Matches | Identities | I/D/M/F* | Stop Codons |
|---|---|---|---|---|---|---|---|---|---|
| **NT** | **22** | **9704** | **99.8%** | **12522** | **65.0%** | **9671 (99.8%)** | **8003 (82.6%)** | **10/12** | |
| Codon mutations: | AAT4GGG (2414A>G 2415A>G 2416T>G), GCT5GCA (2419T>A), AGC6TGC (2420A>T), ACA12ACT (2440A>T), GTT21GTC (2467T>C), TCT24TCC (2476T>C), TCG26TCA (2482G>A), CTG29CTA (2491G>A), CTG31CTT (2497G>T), TTG33TTA (2503G>A), AGA38AGG (2518A>G), CCT39CCC (2521T>C), AAG40AGG (2523A>G), AGA43AGG (2533A>G), TTG45TTA (2539G>A), TTA47TTG (2545A>G), GAG49GAA (2551G>A), CCT50CCA (2554T>A), CTG52TTG (2558C>T), TTG53CTC (2561T>C 2563G>C), TCA56TCG (2572A>G), CCT60CCC (2584T>C), GGC61GGT (2587C>T), CTG63CTC (2593G>C), GCT65GCC (2599T>C), AAT68AAC (2608T>C), TTT72TTC (2620T>C), GCG75GCA (2629G>A), GTG76GTA (2632G>A), AGG77AAG (2634G>A), TTG78TTA (2638G>A), ATT80ATA (2644T>A), AAA83AAG (2653A>G), GCT87GCC (2665T>C), CTA92TTA (2678C>T), CTA93CTG (2683A>G), GCT95GCC (2689T>C), CTA98CTC (2698A>C), TCA101TCG (2707A>G), GCG102GCA (2710G>A), GAA104GAG (2716A>G), ACA105ACG (2719A>G), CTC106CTT (2722C>T), GTC107ATT (2723G>A 2725C>T), GCA108GCG (2728A>G), AGG110AGA (2734G>A), ATT111ATA (2737T>A), GAT114GAC (2746T>C), GCT115ACA (2747G>A 2749T>A), GCA116GCT (2752A>T), GCT117GCA (2755T>A), GAC119GAT (2761C>T), CTC120CTT (2764C>T), TGT125TGC (2779T>C), GGG127GGT (2785G>T), AAC129AAT (2791C>T), CAT131AAT (2795C>A), TAC134TAT (2806C>T), TTG138CTC (2816T>C 2818G>C), GTG140ATA (2822G>A 2824G>A), TTG141TTA (2827G>A), GTT144GTC (2836T>C), AAG145AAA (2839G>A), AAT146AAC (2842T>C), ACC153ACA (2863C>A), CTA154TTA (2864C>T), CCA160ACA (2882C>A), AGT161CAT (2885A>C 2886G>A), AAC163AAT (2893C>T), ACG164ATG (2895C>T), GTC166GTT (2902C>T), GTA167GTG (2905A>G), ATT169ATC (2911T>C), AAT173AAC (2923T>C), AAT176AGT (2931A>G), AAC179AGC (2940A>G), GCT181GCC (2947T>C), ACT186ACC (2962T>C), GAA189_AAT190insA-- (2971_2972insA), GGT195GGC (2989T>C), ACT196ATT (2991C>T), CAT197CGT (2994A>G), GAG199AGG (2999G>A 3000A>G), TCC204TC- (3016delC), CGG206CAG (3021G>A), TAC207TTT (3024A>T 3025C>T), ACA211ATA (3036C>T), GAA212GGC (3039A>G 3040A>C), AAG213ACA (3042A>C 3043G>A), TTG216TTA (3052G>A), TTA219TTG (3061A>G), GCG226GCA (3082G>A), GGC229GGT (3091C>T), GCC232GTA (3099C>T 3100C>A), GTG234AGA (3104G>A 3105T>G 3106G>A), GTC235GTT (3109C>T), AAA236AAG (3112A>G), ACT238ACC (3118T>C), GCC239GCT (3121C>T), GGA241GGG (3127A>G), AGC243AAC (3132G>A), CGA245AAG (3137C>A 3138G>A 3139A>G), TAT249TAC (3151T>C), TTC250ATT (3152T>A 3154C>T), TGT254TAT (3165G>A), GTA255GTG (3169A>G), ATC265ATT (3199C>T), AGG267CGG (3203A>C), CGT268CGC (3208T>C), CCA270CCG (3214A>G), ACC271ACT (3217C>T), TTT272CTT (3218T>C), GTC273GTA (3223C>A), ACT274AAT (3225C>A), GTT276ATT (3230G>A), AAC277AAT (3235C>T), TTA285CTA (3257T>C), ACC286ACT (3262C>T), AGC287AGT (3265C>T), GTA288ATG (3266G>A 3268A>G), GTG289GTA (3271G>A), GTG291ATG (3275G>A), GCA294ACA (3284G>A), CAG299CAA (3301G>A), AGA304AAG (3315G>A 3316A>G), GAA305GCA (3318A>C), ATC306ATT (3322C>T), GAG307GAA (3325G>A), TTG308TTA (3328G>A), ATA311ATT (3337A>T), GAG315GAA (3349G>A), GTC317GTT (3355C>T), TGC318TGT (3358C>T), ATG319GTG (3359A>G), CTA321CTG (3367A>G), TAT322TAC (3370T>C), TTA325CTG (3377T>C 3379A>G), GAA330GAG (3394A>G), CGC331CGT (3397C>T), GAT332GAG (3400T>G), ACA334ACG (3406A>G), TAC338TAT (3418C>T), ATT339ATA (3421T>A), TAT341TAC (3427T>C), AAG343AAA (3433G>A), TCA344TCT (3436A>T), GTA345GTG (3439A>G), AAC346AAT (3442C>T), CCT347CCC (3445T>C), CAG351CAA (3457G>A), TTT352TTC (3460T>C), GCT353GCA (3463T>A), GCG355GCT (3469G>T), CAG356CAA (3472G>A), CAC364CAT (3496C>T), CAG365CAA (3499G>A) | | | | | | | | |
| 6K1 protein (NP_734245.1) | 1 | 52 | 100% | 305 | 90.5% | 52 (100%) | 45 (86.5%) | 0/0/0/0 | 0 |
| Protein mutations: | V5G (3513T>G), M16V (3545A>G), A17T (3548G>A), V19I (3554G>A 3556C>T), L42V (3623C>G), L45M (3632C>A), Y47H (3638T>C) | | | | | | | | |
| Codon mutations: | TCC2TCT (3505C>T), GTT5GGT (3513T>G), AAA7AAG (3520A>G), GAA10GAG (3529A>G), CAA11CAG (3532A>G), ATG16GTG (3545A>G), GCT17ACT (3548G>A), TTA18CTA (3551T>C), GTC19ATT (3554G>A 3556C>T), AGT28AGC (3583T>C), GAT29GAC (3586T>C), TGC30TGT (3589C>T), TTC32TTT (3595C>T), AAT36AAC (3607T>C), TTT38TTC (3613T>C), AAG39AAA (3616G>A), GGT40GGC (3619T>C), CTT42GTT (3623C>G), TCC43TCT (3628C>T), CTG45ATG (3632C>A), GAT46GAC (3637T>C), TAT47CAT (3638T>C), GAA48GAG (3643A>G) | | | | | | | | |
| CI protein (NP_734246.1) | 1 | 634 | 100% | 4190 | 97.9% | 634 (100%) | 608 (95.9%) | 0/0/0/0 | 0 |
| Protein mutations: | I15V (3698A>G), S21N (3717G>A), R26K (3731C>A 3732G>A), I140V (4073A>G), F158Y (4128T>A 4129C>T), A166S (4151G>T), V171I (4166G>A), L190I (4223C>A 4225G>A), I222V (4319A>G 4321A>G), F247Y (4395T>A), V251I (4406G>A), M274V (4475A>G 4477G>T), T276S (4481A>T 4483A>G), A297L (4544G>T 4545C>T 4546G>A), R298K (4548G>A), V342I (4679G>A 4681G>C), V381I (4796G>A 4798C>T), T383S (4803C>G 4804T>C), S468G (5057A>G), T469A (5060A>G), L479I (5090C>A), A482V (5100C>T), E484D (5107A>T), V489F (5120G>T 5122C>T), I491V (5126A>G), V510I (5183G>A) | | | | | | | | |
| Codon mutations: | TTA2TTG (3661A>G), GTG5GTA (3670G>A), TTT9TTC (3682T>C), GAG11GAA (3688G>A), GAG14GAA (3697G>A), ATT15GTT (3698A>G), GAT17GAC (3706T>C), TTT18TTC (3709T>C), TTG20CTA (3713T>C 3715G>A), AGT21AAT (3717G>A), GAG22GAA (3721G>A), CGA26AAA (3731C>A 3732G>A), ACT27ACA (3736T>A), CTA31TTG (3746C>T 3748A>G), ACA33ACG (3754A>G), AGT36AGC (3763T>C), CGA41CGG (3778A>G), CAA42CAG (3781A>G), ATC43ATT (3784C>T), CAT47CAC (3796T>C), CCA50CCT (3805A>T), TAC52TAT (3811C>T), AGA53AGG (3814A>G), TTC58TTT (3829C>T), TTT61TTC (3838T>C), AGA63AGG (3844A>G), GCA64GCC (3847A>C), GTT67GTA (3856T>A), GTG69GTT (3862G>T), GAC72GAT (3871C>T), GCC74GCA (3877C>A), AGC76AGT (3883C>T), GAA77GAG (3886A>G), CTA79CTG (3892A>G), GAC80GAT (3895C>T), TTT81TTC (3898T>C), GTA83GTG (3904A>G), CGG84AGA (3905C>A 3907G>A), GCT86GCC (3913T>C), AAG91AAA (3928G>A), TCA92TCT (3931A>T), GGG94GGA (3937G>A), AGT100AGC (3955T>C), GTG101GTA (3958G>A), GCC102GCT (3961C>T), GGA103GGG (3964A>G), TCT104TCC (3967T>C), TTA107TTG (3976A>G), ATT108ATC (3979T>C), CCA110CCT (3985A>T), CTA114CTT (3997A>T), GCG115GCA (4000G>A), AAC117AAT (4006C>T), GTT118GTG (4009T>G), TTC119TTT (4012C>T), AAA120AAG (4015A>G), TCT123TCC (4024T>C), GAA125GAG (4030A>G), CCA126CCT (4033A>T), AAG129AAA (4042G>A), CTG133TTA (4052C>T 4054G>A), CGT134CGC (4057T>C), CGT136CGA (4063T>A), GGA137GGG (4066A>G), ATA140GTA (4073A>G), TTT141TTC (4078T>C), GGC142GGT (4081C>T), TCT143TCC (4084T>C), TCC147TCT (4096C>T), AGC151AGT (4108C>T), GGA152GGT (4111A>T), GCG154GCA (4117G>A), CTA155TTG (4118C>T 4120A>G), TAC157TAT (4126C>T), TTC158TAT (4128T>A 4129C>T), GCC159GCT (4132C>T), AAT160AAC (4135T>C), CAA164CAG (4147A>G), TTA165CTA (4148T>C), GCT166TCT (4151G>T), AAC169AAT (4162C>T), GTA171ATA (4166G>A), ATA172ATC (4171A>C), GAG175GAA (4180G>A), CTG179TTG (4190C>T), CCT181CCC (4198T>C), TCC182TCT (4201C>T), GCG185GCT (4210G>T), TTC186TTT (4213C>T), AGT188AGC (4219T>C), CTG189TTG (4220C>T), CTG190ATA (4223C>A 4225G>A), GTT192GTG (4231T>G), CAT194CAC (4237T>C), GTA199GTG (4252A>G), TTA200CTG (4253T>C 4255A>G), AAA201AAG (4258A>G), ACT205ACC (4270T>C), AGA209AGG (4282A>G), GAG210GAA (4285G>A), GAA212GAG (4291A>G), CAG216CAA (4303G>A), GTC219GTT (4312C>T), ATA222GTG (4319A>G 4321A>G), GTG223GTT (4324G>T), GAG224GAA (4327G>A), CTG227CTT (4336G>T), TCT228TCC (4339T>C), TCA231TCT (4348A>T), GCA235GCG (4360A>G), GGT237GGC (4366T>C), TCT238TCA (4369T>A), ACT240ACC (4375T>C), GTT244GTC (4387T>C), TTT247TAT (4395T>A), GTA251ATA (4406G>A), CTT252CTC (4411T>C), GTT261GTC (4438T>C), ACC263ACA (4444C>A), GCC265GCT (4450C>T), CTC267CTT (4456C>T), GAC270GAT (4465C>T), ATG274GTT (4475A>G 4477G>T), GTC275GTT (4480C>T), ACA276TCG (4481A>T 4483A>G), GGC280GGT (4495C>T), AAG284AAA (4507G>A), CAC285CAT (4510C>T), GGT286GGG (4513T>G), CTA288TTA (4517C>T), GTC291GTT (4528C>T), AAA293AAG (4534A>G), ACC295ACT (4540C>T), GCG297TTA (4544G>T 4545C>T 4546G>A), AGA298AAA (4548G>A), TTT301TTC (4558T>C), GTA303GTT (4564A>T), ACC305ACT (4570C>T), ATA307ATT (4576A>T), TTG314TTA (4597G>A), GAC315GAT (4600C>T), GAC317GAT (4606C>T), GTG318GTA (4609G>A), GTG320GTA (4615G>A), GAC321GAT (4618C>T), TTG324CTT (4625T>C 4627G>T), GAC331GAT (4648C>T), ATT332ATC (4651T>C), AGG335AGA (4660G>A), AGC336AGT (4663C>T), GCT338GCA (4669T>A), GTG342ATC (4679G>A 4681G>C), AGT343AGC (4684T>C), GTT344GTG (4687T>G), GAG348GAA (4699G>A), AGA349AGG (4702A>G), CAA351CAG (4708A>G), AGG352AGA (4711G>A), CTG353TTG (4712C>T), AAA361AAG (4738A>G), GTA363GTT (4744A>T), GCA364GCG (4747A>G), TTG365CTG (4748T>C), ACT370ACC (4765T>C), GTC381ATT (4796G>A 4798C>T), ACT383AGC (4803C>G 4804T>C), GAG384GAA (4807G>A), GCG385GCT (4810G>T), GCT386GCG (4813T>G), CTT387CTC (4816T>C), TAT392TAC (4831T>C), AAC393AAT (4834C>T), CCA395CCT (4840A>T), GGA399GGG (4852A>G), GGC400GGT (4855C>T), AGT404AGC (4867T>C), CTG405CTC (4870G>C), ATT406ATA (4873T>A), TGT409TGC (4882T>C), ACA416ACT (4903A>T), CAG418CAA (4909G>A), TTT420TTC (4915T>C), GAA421GAG (4918A>G), TTG422CTA (4919T>C 4921G>A), AGT423AGC (4924T>C), CCC424CCG (4927C>G), TTT426TTC (4933T>C), ATC427ATA (4936C>A), CAG428CAA (4939G>A), AAT429AAC (4942T>C), TTC430TTT (4945C>T), TCA436TCG (4963A>G), CCT439CCC (4972T>C), CTT445CTC (4990T>C), AAA449AAG (5002A>G), CCT456CCC (5023T>C), TTG457CTG (5024T>C), TGC458TGT (5029C>T), GAT459GAC (5032T>C), TCT461TCC (5038T>C), CCA463CCT (5044A>T), GCA466GCT (5053A>T), TCG467TCA (5056G>A), AGT468GGT (5057A>G), ACT469GCT (5060A>G), TCG472TCT (5071G>T), CTT479ATT (5090C>A), GGA480GGG (5095A>G), GTG481GTT (5098G>T), GCC482GTC (5100C>T), TTA483TTG (5104A>G), GAA484GAT (5107A>T), ATT485ATC (5110T>C), CCA486CCG (5113A>G), AAG487AAA (5116G>A), CAA488CAG (5119A>G), GTC489TTT (5120G>T 5122C>T), ATT491GTT (5126A>G), CAT494CAC (5137T>C), ATC495ATT (5140C>T), AAA496AAG (5143A>G), GAG497GAA (5146G>A), CTC502CTA (5161C>A), GAA504GAG (5167A>G), CTT506CTG (5173T>G), ACG509ACA (5182G>A), GTT510ATT (5183G>A), GTC511GTT (5188C>T), AAG512AAA (5191G>A), AAA514AAG (5197A>G), GAC515GAT (5200C>T), TGC517TGT (5206C>T), TTA518TTG (5209A>G), AGC521AGT (5218C>T), TCG525TCA (5230G>A), ATC527ATT (5236C>T), AGC528AGT (5239C>T), ATC530ATT (5245C>T), ACA536ACT (5263A>T), CTC538CTT (5269C>T), TTC539TTT (5272C>T), GCC540GCA (5275C>A), ATC541ATT (5278C>T), CCA542CCC (5281A>C), AGA543AGG (5284A>G), ACT544ACC (5287T>C), CTA545TTA (5288C>T), ATA546ATT (5293A>T), TTG547CTA (5294T>C 5296G>A), GAG549GAA (5302G>A), CTT552CTA (5311T>A), GAA553GAG (5314A>G), CGA556CGT (5323A>T), AGT564AGC (5347T>C), ATC566ATT (5353C>T), TGC570TGT (5365C>T), GTT577GTC (5386T>C), ACC580ACA (5395C>A), AAA589AAG (5422A>G), GAT590GAC (5425T>C), ACC592ACT (5431C>T), GAG594GAA (5437G>A), CTT599CTC (5452T>C), AAG601AAA (5458G>A), AGT604AGC (5467T>C), CAA605CAG (5470A>G), CTA606TTG (5471C>T 5473A>G), GAA608GAG (5479A>G), TTC609TTT (5482C>T), TCA610TCC (5485A>C), AAT611AAC (5488T>C), TTG612CTA (5489T>C 5491G>A), GGT614GGC (5497T>C), TGT617TGC (5506T>C), GAG618GAA (5509G>A), GAG619GAA (5512G>A), TTA621CTA (5516T>C), TAT625TAC (5530T>C), GAG626GAA (5533G>A), TCG627TCT (5536G>T), TTG628TTA (5539G>A), GTT631GTG (5548T>G), CAC633CAT (5554C>T) | | | | | | | | |
| 6K2 protein (NP_734247.1) | 1 | 52 | 100% | 303 | 93.5% | 52 (100%) | 46 (88.5%) | 0/0/0/0 | 0 |
| Protein mutations: | A2T (5561G>A 5563G>A), I14V (5597A>G), N16K (5605C>A), A21V (5619C>T 5620T>G), I25L (5630A>C), V46I (5693G>A) | | | | | | | | |
| Codon mutations: | GCT1GCA (5560T>A), GCG2ACA (5561G>A 5563G>A), ACG3ACC (5566G>C), TCA4TCG (5569A>G), CTC9CTG (5584C>G), AAG12AAA (5593G>A), GGG13GGA (5596G>A), ATT14GTT (5597A>G), AAC16AAA (5605C>A), GTG20GTT (5617G>T), GCT21GTG (5619C>T 5620T>G), TTG24TTA (5629G>A), ATC25CTC (5630A>C), GGC28GGT (5641C>T), GCA31GCT (5650A>T), ATA35ATT (5662A>T), GGA36GGG (5665A>G), CAA44CAG (5689A>G), GTT46ATT (5693G>A), GAG47GAA (5698G>A), CAA52CAG (5713A>G) | | | | | | | | |
| NIa-VPg protein (NP_734252.1) | 1 | 188 | 100% | 1227 | 93.3% | 188 (100%) | 170 (90.4%) | 0/0/0/0 | 0 |
| Protein mutations: | K47R (5853A>G), I61V (5894A>G 5896C>G), R83A (5960C>G 5961G>C 5962G>C), I93V (5990A>G), E102D (6019A>T), K106T (6030A>C), D112E (6049C>A), M115I (6058G>A), S120N (6072G>A 6073T>C), N121H (6074A>C), T123N (6081C>A), C133S (6111G>C 6112T>C), I139V (6128A>G 6130T>C), V149I (6158G>A), T153S (6170A>T 6172A>G), L164F (6203C>T 6205C>T), V173I (6230G>A), A182K (6257G>A 6258C>A) | | | | | | | | |

| | Begin | End | Coverage | Score | Concordance | Matches | Identities | I/D/M/F* | Stop Codons |
|---|---|---|---|---|---|---|---|---|---|
| **NT** | **22** | **9704** | **99.8%** | **12522** | **65.0%** | **9671 (99.8%)** | **8003 (82.6%)** | **10/12** | |
| Codon mutations: | GGG1GGC (5716G>C), AAA2AAG (5719A>G), GCC10GCA (5743C>A), CGC14CGA (5755C>A), CAT15CAC (5758T>C), CGT17CGC (5764T>C), AAA19AAG (5770A>G), GGC22GGT (5779C>T), TTT23TTC (5782T>C), GAC26GAT (5791C>T), GAC30GAT (5803C>T), AGG41AGA (5836G>A), AAA42AAG (5839A>G), GGA44GGG (5845A>G), AAA45AAG (5848A>G), AAA47AGA (5853A>G), GGT48GGC (5857T>C), ACC49ACT (5860C>T), ACA50ACT (5863A>T), GGT52GGA (5869T>A), GGT54GGC (5875T>C), AGC57AGT (5884C>T), TTC60TTT (5893C>T), ATC61GTG (5894A>G 5896C>G), AAC62AAT (5899C>T), GGG65GGA (5908G>A), TTT66TTC (5911T>C), GAG70GAA (5923G>A), CAA75CAG (5938A>G), TTC76TTT (5941C>T), GTT77GTC (5944T>C), GAT78GAC (5947T>C), GGG82GGA (5959G>A), CGG83GCC (5960C>G 5961G>C 5962G>C), GAA87GAG (5974A>G), GCT91GCA (5986T>A), GAC92GAT (5989C>T), ATT93GTT (5990A>G), GAT95GAC (5998T>C), AGA99CGT (6008A>C 6010A>T), GAA102GAT (6019A>T), GTG103GTT (6022G>T), CGA104CGT (6025A>T), AAA106ACA (6030A>C), GAC112GAA (6049C>A), ATG115ATA (6058G>A), CAA116CAG (6061A>G), GCC117GCA (6064C>A), GGT119GGC (6070T>C), AGT120AAC (6072G>A 6073T>C), AAC121CAC (6074A>C), ACG122ACA (6079G>A), ACC123AAC (6081C>A), ATA124ATT (6085A>T), GCA126GCT (6091A>T), TAC127TAT (6094C>T), TGT133TCC (6111G>C 6112T>C), AAA135AAG (6118A>G), TTG137TTA (6124G>A), AAG138AAA (6127G>A), ATT139GTC (6128A>G 6130T>C), GAT140GAC (6133T>C), TTA141TTG (6136A>G), AAC145AAT (6148C>T), AAA148AAG (6157A>G), GTT149ATT (6158G>A), TGT150TGC (6163T>C), AAA152AAG (6169A>G), ACA153TCG (6170A>T 6172A>G), GGC155GGT (6178C>T), AAA158AAG (6187A>G), TTT159TTC (6190T>C), GAG161GAA (6196G>A), CTC164TTT (6203C>T 6205C>T), GAA165GAG (6208A>G), CTA166TTG (6209C>T 6211A>G), CAG168CAA (6217G>A), GCT172GCA (6229T>A), GTA173ATA (6230G>A), GTC175GTT (6238C>T), AAG178AAA (6247G>A), ATA180ATT (6253A>T), GCA182AAA (6257G>A 6258C>A), CAT187CAC (6274T>C) | | | | | | | | |
| NIa-Pro protein (NP_734248.1) | 1 | 244 | 100% | 1654 | 93.7% | 244 (100%) | 223 (91.4%) | 0/0/0/0 | 0 |
| Protein mutations: | K2R (6282A>G), A29T (6362G>A), V42I (6401G>A), A43V (6405C>T), Y51F (6429A>T), Q58R (6450A>G), H70N (6485C>A 6487C>T), L84I (6527C>A), T104I (6588C>T), Y120H (6635T>C), I124V (6647A>G), I134V (6677A>G 6679A>G), D147N (6716G>A), N148D (6719A>G), I163L (6764A>T 6766A>G), A173V (6795C>T 6796A>G), H174Q (6799C>A), N194D (6857A>G), V200T (6875G>A 6876T>C 6877C>A), D219E (6934C>A), V242I (7001G>A 7003G>A) | | | | | | | | |
| Codon mutations: | AAA2AGA (6282A>G), TCG3TCA (6286G>A), GGC7GGT (6298C>T), TTG8TTA (6301G>A), AGA9AGG (6304A>G), GAC10GAT (6307C>T), GCC15GCT (6322C>T), ACA17ACT (6328A>T), GTT18GTG (6331T>G), TGT19TGC (6334T>C), AGG20AGA (6337G>A), CTG21TTG (6338C>T), TAT27TAC (6358T>C), GGG28GGA (6361G>A), GCA29ACA (6362G>A), TCA30TCT (6367A>T), GAG31GAA (6370G>A), TAC33TAT (6376C>T), GGT34GGG (6379T>G), GGA36GGC (6385A>C), GGA38GGT (6391A>T), GCA39GCG (6394A>G), TAC40TAT (6397C>T), GTA42ATA (6401G>A), GCG43GTG (6405C>T), CAC45CAT (6412C>T), TTA47TTG (6418A>G), TTT48TTC (6421T>C), TAC51TTC (6429A>T), GGT53GGA (6436T>A), GAG56GAA (6445G>A), CAA58CGA (6450A>G), TCC59TCG (6454C>G), CAC61CAT (6460C>T), GGT62GGA (6463T>A), TTC64TTT (6469C>T), AGG65AGA (6472G>A), GTG66GTA (6475G>A), AAT68AAC (6481T>C), CAC70AAT (6485C>A 6487C>T), AGT71AGC (6490T>C), AGC73AGT (6496C>T), CTG75CTA (6502G>A), GGT79GGC (6514T>C), AGG80AGA (6517G>A), CTC84ATC (6527C>A), ATC85ATA (6532C>A), CCG88CCA (6541G>A), AAA89AAG (6544A>G), GTC93GTT (6556C>T), TTT94TTC (6559T>C), CAG96CAA (6565G>A), TTG98CTA (6569T>C 6571G>A), CCT103CCA (6586T>A), ACA104ATA (6588C>T), GAA107GAG (6598A>G), AGA108AGG (6601A>G), ATT109ATC (6604T>C), TGT110TGC (6607T>C), GGA113GGG (6616A>G), AAC115AAT (6622C>T), GAG118GAA (6631G>A), AAG119AAA (6634G>A), TAT120CAT (6635T>C), GCT121GCA (6640T>A), TCG123TCT (6646G>T), ATC124GTC (6647A>G), ATC125ATT (6652C>T), AGC129AGT (6664C>T), ACT131ACG (6670T>G), ATA134GTG (6677A>G 6679A>G), GGC136GGT (6685C>T), ACA138ACT (6691A>T), TTC139TTT (6694C>T), ACA146ACG (6715A>G), GAT147AAT (6716G>A), AAT148GAT (6719A>G), GGA149GGG (6724A>G), CTA153TTA (6734C>T), GTG155GTA (6742G>A), GTG156GTA (6745G>A), GCC159GCT (6754C>T), GGA161GGG (6760A>G), ATA163TTG (6764A>T 6766A>G), ATT166ATA (6775T>A), CTG169TTG (6782C>T), GCA170GCT (6787A>T), AAC171AAT (6790C>T), GCA173GTG (6795C>T 6796A>G), CAC174CAA (6799C>A), ACC175ACA (6802C>A), AAC177AAT (6808C>T), TAC178TAT (6811C>T), TAC179TAT (6814C>T), TTC182TTT (6823C>T), GAA184GAG (6829A>G), TAC190TAT (6847C>T), CGA192CGC (6853A>C), AAT194GAT (6857A>G), GAG195GAA (6862G>A), GAA198GAG (6871A>G), GTC200ACA (6875G>A 6876T>C 6877C>A), AAG201AAA (6880G>A), TCT202TCA (6883T>A), GTT204GTG (6889T>G), GAC208GAT (6901C>T), GGC213GGT (6916C>T), CCG214CCA (6919G>A), AAA216AAG (6925A>G), CTT217CTC (6928T>C), GAC219GAA (6934C>A), GGG224GGC (6949G>C), TTC226TTT (6955C>T), AAA227AAG (6958A>G), CAA233CAG (6976A>G), CTA235TTA (6980C>T), ATC236ATT (6985C>T), GTA240GTT (6997A>T), GTG241GTT (7000G>T), GTG242ATA (7001G>A 7003G>A) | | | | | | | | |
| NIb protein (NP_734249.1) | 1 | 521 | 100% | 3129 | 86.1% | 519 (99.4%) | 480 (92.0%) | 1/2/2/2 | 0 |
| Protein mutations: | F8Y (7032T>A), T39K (7125C>A), A46S (7145G>T 7147A>G), E51_A52del (7160_7165delGAGGCA), D69E (7216T>A), D83E (7258T>A), D88_R89insX (7273_7274insT), R89V (7274C>G 7275G>T), H91L (7281A>T), L92S (7284T>C), S96L (7296C>T), S101F (7311C>T), T102I (7314C>T), C103Y (7317G>A), N104K (7321T>A), K109Q (7334A>C), A112N (7343G>A 7344C>A 7345A>C), S133M (7407G>T 7408T>G), C136G (7415T>G), L165S (7503T>C 7504G>A), D234N (7709G>A), K235R (7713A>G 7714A>G), T268I (7812C>T), V279L (7844G>T 7846G>A), V292I (7883G>A), S345N (8043G>A), N381S (8151A>G 8152C>T), R390K (8178G>A), R400K (8208G>A), P429A (8294C>G 8296A>G), S447F (8349C>T), A467S (8408G>T), M481I (8452G>A), R484K (8460G>A), A491T (8480G>A 8482T>A), R498K (8502G>A 8503A>G), E502D (8515A>T), E510D (8539G>C), L513C (8546C>T 8547T>G), S515T (8552T>A) | | | | | | | | |
| Codon mutations: | GCT1GCC (7012T>C), AAG2AAA (7015G>A), CAT3CAC (7018T>C), TTT8TAT (7032T>A), GCC10GCT (7039C>T), TTG11CTA (7040T>C 7042G>A), GGA13GGG (7048A>G), GCT17GCA (7060T>A), GTC18GTA (7063C>A), AGC23AGT (7078C>T), TTA25CTG (7082T>C 7084A>G), GTA26GTG (7087A>G), ACC27ACA (7090C>A), CAT29CAC (7096T>C), GTA30GTG (7099A>G), GTT31GTC (7102T>C), GAG34GAA (7111G>A), CGA36CGG (7117A>G), ACA39AAA (7125C>A), GAA40GAG (7129A>G), TTT41TTC (7132T>C), CTG42TTA (7133C>T 7135G>A), GTG44GTA (7141G>A), GAT45GAC (7144T>C), GCA46TCG (7145G>T 7147A>G), GAG47GAA (7150G>A), GCA50GCT (7159A>T), GAG51_GCA52del (7160_7165delGAGGCA), TTC54TTT (7171C>T), GCG60GCT (7189G>T), AAA63AAG (7198A>G), AGC64AGT (7201C>T), CTA66TTA (7205C>T), AAT67AAC (7210T>C), GAT69GAA (7216T>A), GCG70GCA (7219G>A), ATC72ATA (7225C>A), AAG73AAA (7228G>A), AAG77AAA (7240G>A), TAT78TAC (7243T>C), AAA80AAG (7249A>G), CCT81CCG (7252T>G), GAT83GAA (7258T>A), GTT84GTC (7261T>C), GGT85GGA (7264T>A), GTC86GTG (7267C>G), GTG87GTA (7270G>A), GAT88_CGG89insT-- (7273_7274insT), CGG89GTG (7274C>G 7275G>T), CAT91CTT (7281A>T), TTG92TCG (7284T>C), TCA96TTA (7296C>T), TCT101TTT (7311C>T), ACC102ATC (7314C>T), TGC103TAC (7317G>A), AAT104AAA (7321T>A), GTG105-TG (7322delG), CAC106CAT (7327C>T), GGC107GGT (7330C>T), TTC108TTT (7333C>T), AAG109CAG (7334A>C), TGT111TGC (7342T>C), GCA112AAC (7343G>A 7344C>A 7345A>C), ACT115ACC (7354T>C), CAA118CAG (7363A>G), ATT120ATC (7369T>C), TTC121TTT (7372C>T), GCG123GCT (7378G>T), CTC124CTT (7381C>T), GCA129GCT (7396A>T), GTC130GTT (7399C>T), GCC132GCT (7405C>T), AGT133ATG (7407G>T 7408T>G), GGG135GGT (7414G>T), TGC136GGC (7415T>G), AAA137AAG (7420A>G), GAC140GAT (7429C>T), TAT141TAC (7432T>C), CAT144CAC (7441T>C), AAG150AAA (7459G>A), GAA151GAG (7462A>G), ATA153ATT (7468A>T), GTC154GTA (7471C>A), CAA156CAG (7477A>G), AGC157AGT (7480C>T), CTG159CTT (7486G>T), TAT162TAC (7495T>C), AAA163AAG (7498A>G), GGT164GGC (7501T>C), TTG165TCA (7503T>C 7504G>A), ATT168ATA (7513T>A), AAC170AAT (7519C>T), GGA171GGG (7522A>G), AAG174AAA (7531G>A), GAG176GAA (7537G>A), CTC177CTT (7540C>T), TGT179TGC (7546T>C), AAG187AAA (7570G>A), ACG188ACA (7573G>A), ACG190ACA (7579G>A), GCT193GCC (7588T>C), CTA196TTA (7595C>T), TTG199CTA (7604T>C 7606G>A), GGT202GGA (7615T>A), TGT205TGC (7624T>C), GAC208GAT (7633C>T), TTC209TTT (7636C>T), AAT210AAC (7639T>C), TTT213TTC (7648T>C), GAA219GAG (7666A>G), TGT221TGC (7672T>C), ACA223ACT (7678A>T), GGG225GGA (7684G>A), ACT227ACC (7690T>C), TTT229TTC (7696T>C), GGT231GGG (7702T>G), GAT234AAT (7709G>A), AAA235AGG (7713A>G 7714A>G), CTG236TTG (7715C>T), CGT239CGC (7726T>C), GAG242GAA (7735G>A), GTA245GTG (7744A>G), TAC246TAT (7747C>T), GGC251GGT (7762C>T), CAG253CAA (7768G>A), TTT254TTC (7771T>C), TCA257TCG (7780A>G), CTA258CTC (7783A>C), ACT259ACC (7786T>C), CTA262TTG (7793C>T 7795A>G), ATC263ATT (7798C>T), GCT265GCC (7804T>C), ACC268ATC (7812C>T), GAA275GAG (7834A>G), GAC276GAT (7837C>T), GTG279TTA (7844G>T 7846G>A), CAG282CAA (7855G>A), CTG284CTA (7861G>A), TTA287TTG (7870A>G), TAC288TAT (7873C>T), ACT289ACA (7876T>A), GAG290GAA (7879G>A), ATT291ATA (7882T>A), GTT292ATT (7883G>A), CCA295CCT (7894A>T), ATT296ATC (7897T>C), GTC304GTT (7921C>T), AAG306AAA (7927G>A), GGT309GGC (7936T>C), AAT310AAC (7939T>C), AAT311AAC (7942T>C), GTG319GTA (7966G>A), GAT320GAC (7969T>C), TCT322TCC (7975T>C), GTC326GTT (7987C>T), CTT327CTC (7990T>C), GCT332GCC (8005T>C), GAA341GAG (8032A>G), ATC343ATT (8038C>T), GAC344GAT (8041C>T), AGC345AAC (8043G>A), TTG357TTA (8080G>A), GTG360GTA (8089G>A), CCG362CCA (8095G>A), GAG363GAA (8098G>A), AGC366AGT (8107C>T), ATT367ATC (8110T>C), GAT369GAC (8116T>C), CAA373CAG (8128A>G), GGT379GGC (8146T>C), AAC381AGT (8151A>G 8152C>T), TCG385TCA (8164G>A), AGG390AAG (8178G>A), AAG391AAA (8182G>A), TTG394TTA (8191G>A), TTC396TTT (8197C>T), TCC398TCT (8203C>T), AGA400AAA (8208G>A), CTG402TTA (8213C>T 8215G>A), CTA403CTG (8218A>G), ATC404ATT (8221C>T), GAG405GAA (8224G>A), CCA410CCG (8239A>G), GAG415GAA (8254G>A), ATT417ATC (8260T>C), GTA418GTG (8263A>G), GAT424GAC (8281T>C), AGA425AGG (8284A>G), CCA429GCG (8294C>G 8296A>G), GAG430GAA (8299G>A), CAC431CAT (8302C>T), ATT436ATC (8317T>C), TGC437TGT (8320C>T), GCT439GCA (8326T>A), GAG442GAA (8335G>A), TCC443TCT (8338C>T), TCT447TTT (8349C>T), GAA448GAG (8353A>G), CAC451CAT (8362C>T), CAA452CAG (8365A>G), ATC453ATT (8368C>T), TAC457TAT (8380C>T), TCA458TCG (8383A>G), TTA460TTG (8389A>G), CCT465CCC (8404T>C), GCA467TCA (8408G>T), ACA468ACG (8413A>G), GCG470GCT (8419G>T), GCT475GCC (8434T>C), TAT477TAC (8440T>C), ATG481ATA (8452G>A), CTA483CTG (8458A>G), AGG484AAG (8460G>A), AAA485AAG (8464A>G), CTG486CTA (8467G>A), AGG490AGA (8479G>A), GCT491ACA (8480G>A 8482T>A), GAG496GAA (8497G>A), CTA497TTG (8498C>T 8500A>G), AGA498AAG (8502G>A 8503A>G), GCC499GCT (8506C>T), GAA502GAT (8515A>T), GTC505GTT (8524C>T), GCA506GCC (8527A>C), TTA507CTG (8528T>C 8530A>G), GAG510GAC (8539G>C), CTT513TGT (8546C>T 8547T>G), TCT515ACT (8552T>A), GTA518GTG (8563A>G) | | | | | | | | |
| coat protein (NP_734250.1) | 1 | 267 | 100% | 1652 | 90.6% | 267 (100%) | 243 (91.0%) | 0/0/0/0 | 0 |
| Protein mutations: | A1G (8574C>G), I5V (8585A>G), G9E (8598G>A), N11S (8603A>T 8604A>C 8605C>T), P17Q (8622C>A), E18G (8625A>G 8626G>A), P26F (8648C>T 8649C>T 8650G>C), G29E (8658G>A), D31A (8664A>C 8665T>G), A36V (8679C>T 8680A>T), T58K (8745C>A 8746A>G), E95G (8856A>G 8857G>A), R98Q (8865G>A), M99L (8867A>C 8869G>T), G104E (8883G>A), T106S (8888A>T), V128I (8954G>A), N138D (8984A>G 8986T>C), E139V (8988A>T), I187V (9131A>G), M193G (9149A>G 9150T>G 9151G>A), G194S (9152G>A), P230S (9260C>T 9262T>C), Q242H (9298A>C) | | | | | | | | |
| Codon mutations: | GCA1GGA (8574C>G), ATT5GTT (8585A>G), GGA9GAA (8598G>A), AGC10AGT (8602C>T), AAC11TCT (8603A>T 8604A>C 8605C>T), AAA16AAG (8620A>G), CCA17CAA (8622C>A), GAG18GGA (8625A>G 8626G>A), CAG19CAA (8629G>A), GGC20GGT (8632C>T), ATC22ATT (8638C>T), CAG23CAA (8641G>A), AAC25AAT (8647C>T), CCG26TTC (8648C>T 8649C>T 8650G>C), AAC27AAT (8653C>T), AAA28AAG (8656A>G), GGA29GAA (8658G>A), GAT31GCG (8664A>C 8665T>G), GTT34GTG (8674T>G), GCA36GTT (8679C>T 8680A>T), GGC37GGA (8683C>A), GGG40GGA (8692G>A), ACA41ACT (8695A>T), CCG45CCA (8707G>A), AGA46CGA (8708A>C), AAG48AAA (8716G>A), ATC50ATT (8722C>T), ACG51ACA (8725G>A), AAA53AAG (8731A>G), AGA55AGG (8737A>G), CCC57CCT (8743C>T), ACA58AAG (8745C>A 8746A>G), AAG60AAA (8752G>A), GGA61GGT (8755A>T), AAC66AAT (8770C>T), TTA67CTA (8771T>C), GAA68GAG (8776A>G), CAT69CAC (8779T>C), TTG70TTA (8782G>A), TAT73TAC (8791T>C), CCA75CCG (8797A>G), CAA76CAG (8800A>G), CAA77CAG (8803A>G), GAT79GAC (8809T>C), ATT80ATC (8812T>C), AAT82AAC (8818T>C), CGG84CGA (8824G>A), GCA85GCG (8827A>G), ACG92ACA (8848G>A), TAT94TAC (8854T>C), GAG95GGA (8856A>G 8857G>A), GTG97GTA (8863G>A), CGG98CAG (8865G>A), ATG99CTT (8867A>C 8869G>T), GCA100GCG (8872A>G), GGA104GAA (8883G>A), ACT106TCT (8888A>T), TGC119TGT (8929C>T), GTC128ATC (8954G>A), AAC129AAT (8959C>T), GGA130GGT (8962A>T), GGG137GGA (8983G>A), AAT138GAC (8984A>G 8986T>C), GAA139GTA (8988A>T), GAG142GAA (8998G>A), CCG144CCA (9004G>A), TTG145TTA (9007G>A), ATC148ATT (9016C>T), ACC155ACA (9037C>A), ATA171ATT (9085A>T), AAC175AAT (9097C>T), ATT187GTT (9131A>G), CGA188CGT (9136A>T), CGG191CGT (9145G>T), ATG193GGA (9149A>G 9150T>G 9151G>A), GGT194AGT (9152G>A), TTA195TTG (9157A>G), GCG196GCT (9160G>T), CGT197CGC (9163T>C), GAG204GAA (9184G>A), CGA208CGG (9196A>G), AGG214AGA (9214G>A), GAA215GAG (9217A>G), GCG216GCA (9220G>A), GCA224GCT (9244A>T), TTG225TTA (9247G>A), GCC228GCT (9256C>T), CCT230TCC (9260C>T 9262T>C), GGG234GGA (9274G>A), GAC236GAT (9280C>T), ATC239ATT (9289C>T), CAA242CAC (9298A>C), GAG244GAA (9304G>A), GTC254GTT (9334C>T), GTC264GTG (9364C>G) | | | | | | | | |

| | Begin | End | Coverage | Score | Concordance | Matches | Identities | I/D/M/F* | Stop Codons |
|---|---|---|---|---|---|---|---|---|---|
| **NT** | **22** | **9704** | **99.8%** | **12522** | **65.0%** | **9671 (99.8%)** | **8003 (82.6%)** | **10/12** | |
| PIPO (YP_006393460.1) | 1 | 75 | 100% | 95 | 18.1% | 75 (98.7%) | 54 (71.1%) | 1/0/3/2 | 2 |
| Protein mutations: | R17_K17insX (2971_2972insA), Y23H (2989T>C 2991C>T), R26K (2999G>A 3000A>G), H35Y (3025C>T), K40H (3040A>C 3042A>C), G41S (3043G>A), E44K (3052G>A), I47V (3061A>G), V54I (3082G>A), P57S (3091C>T), P60T (3100C>A), G61E (3104G>A 3105T>G), G62S (3106G>A), Q63* (3109C>T), R64G (3112A>G), C66R (3118T>C), L67F (3121C>T), I69V (3127A>G), A72E (3137C>A 3138G>A), I73V (3139A>G) | | | | | | | | |
| Codon mutations: | TTA2CTA (2923T>C), AAA4AAG (2931A>G), GAA7GAG (2940A>G), TTG10CTG (2947T>C), TTG15CTG (2962T>C), AGA17_AAA17insAA- (2971_2972insA), TAC23CAT (2989T>C 2991C>T), TCA24TCG (2994A>G), AGA26AAG (2999G>A 3000A>G), CAC32-AC (3016delC), TCG33TCA (3021G>A), GTA34GTT (3024A>T), CAT35TAT (3025C>T), CAC38CAT (3036C>T), AGA39AGG (3039A>G), AAA40CAC (3040A>C 3042A>C), GGC41AGC (3043G>A), GAA44AAA (3052G>A), ATA47GTA (3061A>G), GTT54ATT (3082G>A), CCG57TCG (3091C>T), CGC59CGT (3099C>T), CCA60ACA (3100C>A), GGT61GAG (3104G>A 3105T>G), GGT62AGT (3106G>A), CAA63TAA (3109C>T), AGG64GGG (3112A>G), TGC66CGC (3118T>C), CTC67TTC (3121C>T), ATT69GTT (3127A>G), GAG70GAA (3132G>A), GCG72GAA (3137C>A 3138G>A), ATT73GTT (3139A>G) | | | | | | | | |

*: Inserts / Deletes / Misaligned / Frameshifts

#### Analysis details

This analysis was performed with panviral2.64

### NGS Details (UN60): Torradovirus lycopersici (segment RNA 1)

#### Assembly

| | |
|---|---|
| **Coverage Length** | 7740 (1 contig(s)) |
| **Depth Of Coverage** | 6761.0 |
| **Number Of Reads** | 355460 |
| **Reads Per Million** | 7539.15 rpm (after QC) |
| **Ambiguities** | 0 |
| **Assembly Method** | de novo + reference guided assembly |
| **Consensus Caller** | Bcf Tools |

#### Coverage Map

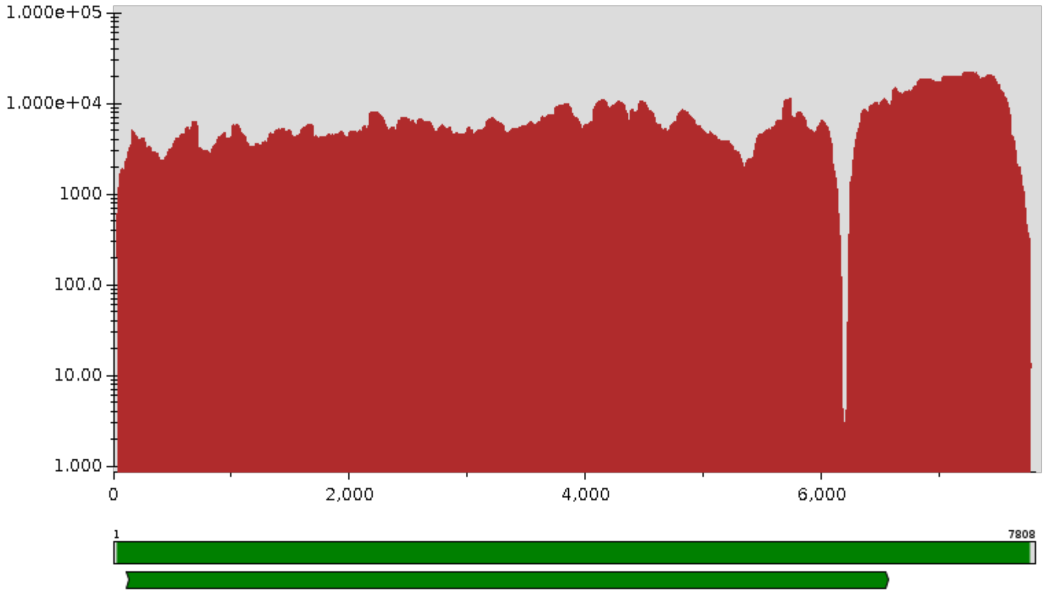

#### Assignment

| | |
|---|---|
| **Type** | Torradovirus lycopersici (Taxonomy ID: 3048378) |
| **Reference Genome** | NC_009013.1 |
| **NT Identity (%)** | 91.2359 |
| **AA Identity (%)** | 97.962 |
| **Number Of Stop Codons** | 1 |
| **Number Of CDS** | 1 |

#### Alignment

| | |
|---|---|
| **Alignment Score** | 11591.0 (NT) + 14866.0 (AA) = 26457.0 |
| **Concordance (%)** | 90.408 |

| Alignment Method | Global, seeded, nucleotide + amino acids (AGA) |
|---|---|

#### Genome Region

Sequence starts at position 30 and ends at position 7769 relative to NC_009013.1 reference sequence.

#### Alignment Detailed Statistics

| | Begin | End | Coverage | Score | Concordance | Matches | Identities | I/D/M/F* | Stop Codons |
|---|---|---|---|---|---|---|---|---|---|
| **NT** | **30** | **7769** | **99.1%** | **11591** | **81.7%** | **7347 (94.6%)** | **6725 (86.6%)** | **24/393** | |
| Mutations: | 31C>A, 69C>T, 76_77insT, 83C>T, 83_84insT, 85T>C, 87C>T, 101G>A, 102G>C, 115T>C, 127C>T, 128C>T, 130C>T, 135T>C, 148T>C, 151A>G, 158G>T, 161A>G, 178T>G, 187A>G, 199T>A, 208T>C, 220C>T, 224C>T, 265C>T, 328T>C, 334T>C, 340G>A, 349G>A, 373C>T, 376C>T, 433G>A, 487C>T, 502G>A, 541T>C, 559C>T, 578C>T, 628C>T, 655C>T, 676A>G, 694A>G, 736A>G, 781C>A, 790C>T, 799A>G, 805T>C, 818C>T, 820A>G, 832A>T, 835A>C, 841T>C, 842C>T, 847C>T, 883G>A, 886T>C, 901T>C, 925C>T, 952A>G, 964T>G, 970G>A, 997C>T, 1006T>C, 1033G>A, 1060T>C, 1078A>T, 1081C>T, 1093T>C, 1105A>G, 1108T>C, 1109C>T, 1165C>T, 1180G>A, 1196A>G, 1231G>A, 1240C>T, 1255G>A, 1267T>C, 1276A>G, 1300G>A, 1351A>G, 1372C>T, 1408T>C, 1411T>C, 1417T>C, 1423T>C, 1426T>C, 1429C>T, 1430C>T, 1441C>T, 1450T>C, 1451C>T, 1480C>T, 1481A>G, 1489T>C, 1522C>T, 1552C>T, 1555C>T, 1570A>G, 1573T>C, 1582C>T, 1585A>T, 1591C>T, 1594T>C, 1607G>T, 1612G>A, 1618T>G, 1619C>T, 1624C>A, 1645A>G, 1669C>T, 1681T>A, 1684G>A, 1699T>A, 1705T>C, 1738A>G, 1771C>T, 1781T>C, 1789A>G, 1897A>G, 1910T>C, 1954T>C, 1960A>G, 1984C>T, 2014A>G, 2042G>A, 2056T>C, 2082T>A, 2092A>G, 2107A>G, 2128T>C, 2138C>T, 2143T>C, 2149T>C, 2155T>C, 2174C>T, 2179A>C, 2182G>A, 2194T>A, 2200T>C, 2227C>T, 2241A>G, 2250G>A, 2251C>T, 2257C>T, 2263G>A, 2275T>C, 2287G>A, 2290C>T, 2314C>T, 2323A>T, 2347C>T, 2350A>G, 2356C>T, 2359G>A, 2362C>T, 2377G>A, 2380T>C, 2392A>G, 2407T>C, 2409G>A, 2443T>C, 2446G>A, 2479G>A, 2485C>T, 2497G>A, 2500T>C, 2503G>A, 2509C>T, 2512T>C, 2542G>A, 2548C>T, 2557C>T, 2620G>A, 2626T>C, 2650G>A, 2665T>C, 2675T>C, 2683A>G, 2692G>A, 2698T>C, 2702C>T, 2710A>G, 2725T>C, 2731C>T, 2734C>T, 2743A>G, 2744T>C, 2752A>G, 2765T>C, 2773A>G, 2785C>T, 2788C>T, 2809C>T, 2821G>T, 2827A>G, 2842T>C, 2860C>T, 2866T>A, 2869C>T, 2872A>T, 2926G>A, 2929C>T, 2930T>C, 2935G>A, 2941A>G, 2950C>T, 2951C>T, 2998A>G, 3031T>A, 3034G>A, 3037C>T, 3040T>C, 3049A>G, 3064A>G, 3073G>A, 3079T>C, 3109T>C, 3115G>A, 3145A>G, 3151C>T, 3163T>C, 3169C>T, 3172T>C, 3202T>C, 3241T>C, 3253C>T, 3256G>A, 3265A>G, 3418C>T, 3463A>G, 3466G>A, 3481C>T, 3484A>T, 3508G>A, 3559G>A, 3580T>C, 3583C>T, 3610A>G, 3613T>C, 3625C>T, 3628G>A, 3643A>G, 3646C>T, 3661A>G, 3664T>C, 3673G>A, 3676C>T, 3686C>T, 3706T>C, 3716T>C, 3721T>A, 3739A>T, 3754G>A, 3758G>A, 3766G>A, 3784C>G, 3808T>C, 3814C>T, 3817G>C, 3824T>C, 3835T>C, 3844T>C, 3857T>C, 3859G>A, 3868C>T, 3871T>C, 3883G>A, 3898T>C, 3907A>T, 3916C>T, 3925A>G, 3928G>A, 3949C>T, 3955G>A, 3961G>A, 3970A>G, 3973T>C, 4034T>C, 4066T>C, 4075A>G, 4084C>T, 4100T>C, 4118C>T, 4129C>T, 4147C>A, 4159G>A, 4162A>T, 4168A>G, 4195A>G, 4207G>A, 4210T>C, 4219C>T, 4228C>T, 4294A>G, 4348C>T, 4354T>C, 4396T>C, 4408C>T, 4414T>G, 4417T>C, 4420G>A, 4459T>C, 4462C>T, 4465C>T, 4468C>T, 4471T>C, 4474T>C, 4486T>C, 4510T>A, 4511C>T, 4514T>C, 4555A>G, 4570C>T, 4576A>T, 4588C>A, 4591T>C, 4625T>C, 4636T>C, 4648T>A, 4660A>G, 4684A>C, 4696C>T, 4702G>A, 4705C>T, 4714T>C, 4735C>T, 4747A>G, 4756A>T, 4759C>T, 4768T>C, 4774G>A, 4777C>T, 4783A>G, 4786A>G, 4789C>T, 4813A>G, 4816T>C, 4843G>A, 4898G>A, 4906G>A, 4969G>A, 4988A>G, 4993A>T, 5005T>C, 5012T>C, 5023C>T, 5029G>T, 5044A>C, 5059G>T, 5074T>C, 5077T>C, 5079T>C, 5102T>C, 5119T>C, 5120G>A, 5122T>A, 5123C>T, 5132T>A, 5146T>C, 5148C>A, 5200G>A, 5218T>G, 5244G>A, 5266C>T, 5281C>T, 5329C>T, 5341T>A, 5344A>G, 5347C>T, 5369G>A, 5371G>A, 5386C>T, 5392G>A, 5407A>T, 5419A>T, 5437T>C, 5440C>T, 5461T>C, 5471T>C, 5482A>G, 5533T>A, 5575A>G, 5578T>C, 5591G>T, 5603A>C, 5608A>G, 5611A>T, 5617A>T, 5641T>C, 5647T>C, 5674A>T, 5677G>T, 5686G>A, 5690T>C, 5710T>A, 5716G>T, 5719A>G, 5731C>T, 5734T>C, 5737T>C, 5739G>A, 5746T>C, 5750T>C, 5752G>A, 5758C>T, 5773G>A, 5782A>G, 5784T>C, 5786T>C, 5797A>G, 5806C>T, 5836A>T, 5849T>C, 5850T>A, 5851G>A, 5869G>A, 5875A>G, 5878G>T, 5884T>C, 5887C>T, 5890C>T, 5902G>A, 5914T>C, 5923T>C, 5927C>T, 5932C>T, 5955A>G, 5965G>A, 5998A>G, 6004C>T, 6016A>G, 6028C>T, 6031A>T, 6033C>T, 6035G>T, 6055T>A, 6067C>A, 6070T>C, 6092C>T, 6094T>A, 6100T>G, 6103G>A, 6106G>A, 6112A>G, 6121A>T, 6139T>C, 6145T>C, 6148T>C, 6154C>G, 6163C>G, 6184A>G, 6187T>G, 6193C>T, 6196A>G, 6202G>C, 6208A>G, 6217G>T, 6223A>T, 6226G>A, 6230A>G, 6232A>G, 6235G>A, 6244A>G, 6253A>C, 6256C>A, 6262A>T, 6265G>A, 6268G>T, 6273G>T, 6274C>G, 6296G>C, 6298A>G, 6304A>G, 6308T>C, 6313T>C, 6319A>G, 6322G>A, 6323A>G, 6344C>T, 6358T>C, 6382C>T, 6385C>T, 6406T>C, 6412C>T, 6414G>A, 6427A>G, 6442C>G, 6445C>T, 6448G>A, 6451T>C, 6457C>A, 6466T>C, 6484A>C, 6490A>G, 6493T>C, 6500A>T, 6517C>T, 6532C>T, 6545C>T, 6571C>T, 6587G>A, 6643A>G, 6670C>T, 6685A>G, 6715C>T, 6721A>T, 6742A>T, 6766delC, 6786T>C, 6798_6799insA, 6807A>G, 6808A>G, 6809C>T, 6810G>C, 6811G>T, 6814A>T, 6815G>A, 6817A>G, 6818C>A, 6821A>G, 6822C>T, 6823T>G, 6825T>A, 6826A>T, 6828C>T, 6831_6897delGTTGGGAGTCCGGCTCCATTTGGAGTACCAATGAATTTACATTGGTTAAAGAGATTTGCACGCCTCT, 6905T>C, 6907T>A, 6908G>T, 6911T>C, 6912_6916delCGTGT, 6921T>G, 6923C>G, 6924T>G, 6927_6941delGTTAGAAAGTTCCTT, 6947A>C, 6948_6971delGGTTTACAAAGTACGGGGAGCACT, 6976G>A, 6979T>A, 6983_7024delATAGTGCAGGTGCTATCCCATGATAGTCCTTTAACTCAAGGG, 7026T>C, 7031T>A, 7033G>A, 7035T>G, 7038G>A, 7040_7072delATCTTTTGCCGTGATGAAAGATGAGGTAGCTTC, 7074C>A, 7077T>C, 7083G>T, 7085_7196delAGGCTGAAACTACACATATGTAGTGGGTTTGACTGAGTCCTATAATCAGTCCGTTTGAAATTCGATAATTTTCCGTAGCTTGCGTCAAGCTGCTCACGTTAGGGGTGTGAGT, 7198A>T, 7204_7273delCGCCGTACCACGTCTTCCCCGGCAATGCCAGTGGTTCAGAGCGGGCCCTCAGAATAGAGGTTAAAACTAG, 7292G>T, 7293A>T, 7297_7298insC, 7299G>A, 7311T>A, 7318G>T, 7320G>T, 7342T>G, 7343C>A, 7344C>A, 7346A>T, 7346_7347insG, 7354G>A, 7354_7355insCAGTAGTGCCATTTA, 7358A>C, 7360G>A, 7361A>C, 7362A>C, 7364C>T, 7368G>A, 7372A>T, 7375T>G, 7376T>G, 7377T>A, 7379A>T, 7383_7384delAA, 7388C>T, 7397G>A, 7398C>A, 7399G>A, 7400A>T, 7404C>T, 7407A>T, 7408G>T, 7423A>T, 7425G>A, 7435T>C, 7436G>A, 7444C>T, 7445A>G, 7457A>G, 7458T>C, 7460G>A, 7498C>G, 7503C>T, 7505A>G, 7506T>C, 7511_7512insC, 7514A>C, 7525_7526insC, 7526_7527insG, 7569G>A, 7577C>T, 7591G>A, 7596C>T, 7600T>G, 7601T>A, 7602T>A, 7603_7604insC, 7610C>T, 7612G>A, 7615A>G, 7616A>T, 7617A>T, 7618C>T, 7624T>A, 7625A>T, 7628_7633delGCTTTA, 7637G>A, 7642T>G, 7644G>T, 7647T>G, 7648G>T, 7649T>C, 7654C>A, 7657_7672delGTTTTATTGTGTGTTT, 7686T>A, 7687T>C, 7689G>T, 7698G>A, 7701T>C, 7702C>T, 7705T>A, 7717A>T, 7730G>A, 7731A>G, 7739T>A, 7747C>T, 7750T>C, 7752G>A, 7757G>T, 7759T>A | | | | | | | | |
| **CDS** | | | | | | | | | |
| ToTV_sRNA1gp1 | 1 | 2159 | 100% | 14866 | 98.6% | 2159 (100%) | 2115 (98.0%) | 0/0/0/0 | 1 |
| Protein mutations: | P8S (128C>T 130C>T), F10S (135T>C), A18S (158G>T), T19A (161A>G), T364A (1196A>G), I459V (1481A>G), A501S (1607G>T), A646T (2042G>A), F659Y (2082T>A), N712S (2241A>G), S715N (2250G>A 2251C>T), R768K (2409G>A), V1218I (3758G>A), V1598I (4898G>A), T1628A (4988A>G), I1658T (5079T>C), V1672I (5120G>A 5122T>A), S1676T (5132T>A), T1681N (5148C>A), R1713K (5244G>A), V1755I (5369G>A 5371G>A), I1792M (5482A>G), A1829S (5591G>T), T1833P (5603A>C), Q1857H (5677G>T), R1878K (5739G>A), I1893T (5784T>C), L1915Q (5849T>C 5850T>A 5851G>A), N1950S (5955A>G), T1976I (6033C>T), A1977S (6035G>T), I2002M (6112A>G), M2032I (6202G>C), E2039D (6223A>T), I2042V (6230A>G 6232A>G), I2046M (6244A>G), L2049F (6253A>C), S2056M (6273G>T 6274C>G), V2064L (6296G>C 6298A>G), I2071M (6319A>G), I2073V (6323A>G), R2103Q (6414G>A), I2132F (6500A>T), L2147F (6545C>T) | | | | | | | | |

| | Begin | End | Coverage | Score | Concordance | Matches | Identities | I/D/M/F* | Stop Codons |
|---|---|---|---|---|---|---|---|---|---|
| **NT** | **30** | **7769** | **99.1%** | **11591** | **81.7%** | **7347 (94.6%)** | **6725 (86.6%)** | **24/393** | |

Codon mutations:

TTT3TTC (115T>C), TTC7TTT (127C>T), CCC8TCT (128C>T 130C>T), TTC10TCC (135T>C), ACT14ACC (148T>C), GAA15GAG (151A>G), GCT18TCT (158G>T), ACC19GCC (161A>G), TCT24TCG (178T>G), TCA27TCG (187A>G), GCT31GCA (199T>A), AGT34AGC (208T>C), CGC38CGT (220C>T), CTG40TTG (224C>T), CAC53CAT (265C>T), TCT74TCC (328T>C), ACT76ACC (334T>C), TTG78TTA (340G>A), AAG81AAA (349G>A), TTC89TTT (373C>T), ATC90ATT (376C>T), AAG109AAA (433G>A), GAC127GAT (487C>T), GTG132GTA (502G>A), TCT145TCC (541T>C), TTC151TTT (559C>T), CTA158TTA (578C>T), AAC174AAT (628C>T), GAC183GAT (655C>T), CAA190CAG (676A>G), GCA196GCG (694A>G), CCA210CCG (736A>G), ACC225ACA (781C>A), TTC228TTT (790C>T), GTA231GTG (799A>G), GGT233GGC (805T>C), CTA238TTG (818C>T 820A>G), CTA242CTT (832A>T), TCA243TCC (835A>C), ATT245ATC (841T>C), CTG246TTG (842C>T), TAC247TAT (847C>T), TTG259TTA (883G>A), TTT260TTC (886T>C), AAT265AAC (901T>C), GCC273GCT (925C>T), GTA282GTG (952A>G), GGT286GGG (964T>G), AAG288AAA (970G>A), GCC297GCT (997C>T), TCT300TCC (1006T>C), GAG309GAA (1033G>A), CAT318CAC (1060T>C), GCA324GCT (1078A>T), AAC325AAT (1081C>T), ACT329ACC (1093T>C), CAA333CAG (1105A>G), CAT334CAC (1108T>C), CTA335TTA (1109C>T), GCC353GCT (1165C>T), AGG358AGA (1180G>A), ACC364GCC (1196A>G), CAG375CAA (1231G>A), GGC378GGT (1240C>T), AGG383AGA (1255G>A), TTT387TTC (1267T>C), CAA390CAG (1276A>G), GGG398GGA (1300G>A), GAA415GAG (1351A>G), CTC422CTT (1372C>T), AAT434AAC (1408T>C), TAT435TAC (1411T>C), CAT437CAC (1417T>C), ACT439ACC (1423T>C), GCT440GCC (1426T>C), ATC441ATT (1429C>T), CTG442TTG (1430C>T), GAC445GAT (1441C>T), GCT448GCC (1450T>C), CTG449TTG (1451C>T), GAC458GAT (1480C>T), ATT459GTT (1481A>G), GAT461GAC (1489T>C), CCC472CCT (1522C>T), GGC482GGT (1552C>T), CGC483CGT (1555C>T), CGA488CGG (1570A>G), TAT489TAC (1573T>C), GCC492GCT (1582C>T), ACA493ACT (1585A>T), AAC495AAT (1591C>T), CTT496CTC (1594T>C), GCT501TCT (1607G>T), CAG502CAA (1612G>A), GGT504GGG (1618T>G), CTA505TTA (1619C>T), ACC506ACA (1624C>A), AGA513AGG (1645A>G), TGC521TGT (1669C>T), GGT525GGA (1681T>A), GAG526GAA (1684G>A), GCT531GCA (1699T>A), ACT533ACC (1705T>C), AAA544AAG (1738A>G), AGC555AGT (1771C>T), TTG559CTG (1781T>C), GAA561GAG (1789A>G), GCA597GCG (1897A>G), TTG602CTG (1910T>C), GTT616GTC (1954T>C), GTA618GTG (1960A>G), TAC626TAT (1984C>T), AAA636AAG (2014A>G), GCA646ACA (2042G>A), CCT650CCC (2056T>C), TTT659TAT (2082T>A), CTA662CTG (2092A>G), AAA667AAG (2107A>G), GTT674GTC (2128T>C), CTG678TTG (2138C>T), TAT679TAC (2143T>C), GGT681GGC (2149T>C), TGT683TGC (2155T>C), CTG690TTG (2174C>T), ATA691ATC (2179A>C), AAG692AAA (2182G>A), GCT696GCA (2194T>A), TAT698TAC (2200T>C), TAC707TAT (2227C>T), AAT712AGT (2241A>G), AGC715AAT (2250G>A 2251C>T), GAC717GAT (2257C>T), CAG719CAA (2263G>A), CCT723CCC (2275T>C), GCG727GCA (2287G>A), CAC728CAT (2290C>T), GAC736GAT (2314C>T), ATA739ATT (2323A>T), CAC747CAT (2347C>T), GTA748GTG (2350A>G), TTC750TTT (2356C>T), TTG751TTA (2359G>A), TAC752TAT (2362C>T), AGG757AGA (2377G>A), TAT758TAC (2380T>C), AGA762AGG (2392A>G), CCT767CCC (2407T>C), AGG768AAG (2409G>A), ATT779ATC (2443T>C), GTG780GTA (2446G>A), CCG791CCA (2479G>A), ATC793ATT (2485C>T), CTG797CTA (2497G>A), ATT798ATC (2500T>C), AAG799AAA (2503G>A), GGC801GGT (2509C>T), CTT802CTC (2512T>C), GGG812GGA (2542G>A), GTC814GTT (2548C>T), TTC817TTT (2557C>T), GAG838GAA (2620G>A), AAT840AAC (2626T>C), AAG848AAA (2650G>A), TTT853TTC (2665T>C), TTG857CTG (2675T>C), GCA859GCG (2683A>G), AAG862AAA (2692G>A), GAT864GAC (2698T>C), CTG866TTG (2702C>T), AAA868AAG (2710A>G), GAT873GAC (2725T>C), TTC875TTT (2731C>T), ATC876ATT (2734C>T), GGA879GGG (2743A>G), TTG880CTG (2744T>C), AAA882AAG (2752A>G), TTG887CTG (2765T>C), CTA889CTG (2773A>G), GTC893GTT (2785C>T), TAC894TAT (2788C>T), TGC901TGT (2809C>T), GTG905GTT (2821G>T), ACA907ACG (2827A>G), TTT912TTC (2842T>C), GGC918GGT (2860C>T), ACT920ACA (2866T>A), TTC921TTT (2869C>T), TCA922TCT (2872A>T), CGG940CGA (2926G>A), TTC941TTT (2929C>T), TTG942CTG (2930T>C), AAG943AAA (2935G>A), CAA945CAG (2941A>G), GTC948GTT (2950C>T), CTG949TTG (2951C>T), AAA964AAG (2998A>G), TCT975TCA (3031T>A), AAG976AAA (3034G>A), GGC977GGT (3037C>T), ACT978ACC (3040T>C), GTA981GTG (3049A>G), GTA986GTG (3064A>G), AAG989AAA (3073G>A), GCT991GCC (3079T>C), TTT1001TTC (3109T>C), GTG1003GTA (3115G>A), GTA1013GTG (3145A>G), TAC1015TAT (3151C>T), TCT1019TCC (3163T>C), TCC1021TCT (3169C>T), TTT1022TTC (3172T>C), TAT1032TAC (3202T>C), CTT1045CTC (3241T>C), AAC1049AAT (3253C>T), GTG1050GTA (3256G>A), AAA1053AAG (3265A>G), CTC1104CTT (3418C>T), AAA1119AAG (3463A>G), CAG1120CAA (3466G>A), GGC1125GGT (3481C>T), ACA1126ACT (3484A>T), ACG1134ACA (3508G>A), GGG1151GGA (3559G>A), TCT1158TCC (3580T>C), ATC1159ATT (3583C>T), GAA1168GAG (3610A>G), GGT1169GGC (3613T>C), TAC1173TAT (3625C>T), GTG1174GTA (3628G>A), AAA1179AAG (3643A>G), TAC1180TAT (3646C>T), CAA1185CAG (3661A>G), AAT1186AAC (3664T>C), GAG1189GAA (3673G>A), GAC1190GAT (3676C>T), CTA1194TTA (3686C>T), CGT1200CGC (3706T>C), TTG1204CTG (3716T>C), CCT1205CCA (3721T>A), CGA1211CGT (3739A>T), AAG1216AAA (3754G>A), GTC1218ATC (3758G>A), CAG1220CAA (3766G>A), CTC1226CTG (3784C>G), TAT1234TAC (3808T>C), GAC1236GAT (3814C>T), GGG1237GGC (3817G>C), TTG1240CTG (3824T>C), TCT1243TCC (3835T>C), TAT1246TAC (3844T>C), TTG1251CTA (3857T>C 3859G>A), GGC1254GGT (3868C>T), ACT1255ACC (3871T>C), GAG1259GAA (3883G>A), AGT1264AGC (3898T>C), GGA1267GGT (3907A>T), GTC1270GTT (3916C>T), AAA1273AAG (3925A>G), AGG1274AGA (3928G>A), GAC1281GAT (3949C>T), GAG1283GAA (3955G>A), TTG1285TTA (3961G>A), GAA1288GAG (3970A>G), ATT1289ATC (3973T>C), TTG1310CTG (4034T>C), CGT1320CGC (4066T>C), GAA1323GAG (4075A>G), CCC1326CCT (4084C>T), TTG1332CTG (4100T>C), CTG1338TTG (4118C>T), TCC1341TCT (4129C>T), TCC1347TCA (4147C>A), CTG1351CTA (4159G>A), CCA1352CCT (4162A>T), AAA1354AAG (4168A>G), GAA1363GAG (4195A>G), TTG1367TTA (4207G>A), GCT1368GCC (4210T>C), TTC1371TTT (4219C>T), GTC1374GTT (4228C>T), CAA1396CAG (4294A>G), GAC1414GAT (4348C>T), AAT1416AAC (4354T>C), ACT1430ACC (4396T>C), AAC1434AAT (4408C>T), GTT1436GTG (4414T>G), TTT1437TTC (4417T>C), AGG1438AGA (4420G>A), GTT1451GTC (4459T>C), ATC1452ATT (4462C>T), TTC1453TTT (4465C>T), AAC1454AAT (4468C>T), TCT1455TCC (4471T>C), CTT1456CTC (4474T>C), TTT1460TTC (4486T>C), TCT1468TCA (4510T>A), CTA1469TTA (4511C>T), TTG1470CTG (4514T>C), AAA1483AAG (4555A>G), AAC1488AAT (4570C>T), CTA1490CTT (4576A>T), GTC1494GTA (4588C>A), TAT1495TAC (4591T>C), TTG1507CTG (4625T>C), TAT1510TAC (4636T>C), GCT1514GCA (4648T>A), GAA1518GAG (4660A>G), ATA1526ATC (4684A>C), ATC1530ATT (4696C>T), AAG1532AAA (4702G>A), AAC1533AAT (4705C>T), GTT1536GTC (4714T>C), GAC1543GAT (4735C>T), CTA1547CTG (4747A>G), CTA1550CTT (4756A>T), AGC1551AGT (4759C>T), TTT1554TTC (4768T>C), AGG1556AGA (4774G>A), CAC1557CAT (4777C>T), CTA1559CTG (4783A>G), GGA1560GGG (4786A>G), TAC1561TAT (4789C>T), GTA1569GTG (4813A>G), TCT1570TCC (4816T>C), AGG1579AGA (4843G>A), GTT1598ATT (4898G>A), GAG1600GAA (4906G>A), AAG1621AAA (4969G>A), ACC1628GCC (4988A>G), ACA1629ACT (4993A>T), TAT1633TAC (5005T>C), TTG1636CTG (5012T>C), GAC1639GAT (5023C>T), CTG1641CTT (5029G>T), GGA1646GGC (5044A>C), GGG1651GGT (5059G>T), CTT1656CTC (5074T>C), CAT1657CAC (5077T>C), ATT1658ACT (5079T>C), TTG1666CTG (5102T>C), ACT1671ACC (5119T>C), GTT1672ATA (5120G>A 5122T>A), CTG1673TTG (5123C>T), TCT1676ACT (5132T>A), CAT1680CAC (5146T>C), ACT1681AAT (5148C>A), CAG1698CAA (5200G>A), GTT1704GTG (5218T>G), AGA1713AAA (5244G>A), CCC1720CCT (5266C>T), GAC1725GAT (5281C>T), CAC1741CAT (5329C>T), ATT1745ATA (5341T>A), GTA1746GTG (5344A>G), ACC1747ACT (5347C>T), GTG1755ATA (5369G>A 5371G>A), GGC1760GGT (5386C>T), GGG1762GGA (5392G>A), CCA1767CCT (5407A>T), GCA1771GCT (5419A>T), ACT1777ACC (5437T>C), GGC1778GGT (5440C>T), GTT1785GTC (5461T>C), TTG1789CTG (5471T>C), ATA1792ATG (5482A>G), CTT1809CTA (5533T>A), ACA1823ACG (5575A>G), TGT1824TGC (5578T>C), GCC1829TCC (5591G>T), ACA1833CCA (5603A>C), CCA1834CCG (5608A>G), GCA1835GCT (5611A>T), ATA1837ATT (5617A>T), GGT1845GGC (5641T>C), TTT1847TTC (5647T>C), CTA1856CTT (5674A>T), CAG1857CAT (5677G>T), CAG1860CAA (5686G>A), TTG1862CTG (5690T>C), CCT1868CCA (5710T>A), GGG1870GGT (5716G>T), GGA1871GGG (5719A>G), CAC1875CAT (5731C>T), ACT1876ACC (5734T>C), ACT1877ACC (5737T>C), AGA1878AAA (5739G>A), CTT1880CTC (5746T>C), TTG1882CTA (5750T>C 5752G>A), CTC1884CTT (5758C>T), AAG1889AAA (5773G>A), CAA1892CAG (5782A>G), ATT1893ACT (5784T>C), TTA1894CTA (5786T>C), AGA1897AGG (5797A>G), TTC1900TTT (5806C>T), TCA1910TCT (5836A>T), TTG1915CAA (5849T>C 5850T>A 5851G>A), AGG1921AGA (5869G>A), CAA1923CAG (5875A>G), GGG1924GGT (5878G>T), TTT1926TTC (5884T>C), CCC1927CCT (5887C>T), CGC1928CGT (5890C>T), GTG1932GTA (5902G>A), CTT1936CTC (5914T>C), TTT1939TTC (5923T>C), CTG1941TTG (5927C>T), AGC1942AGT (5932C>T), AAC1950AGC (5955A>G), ACG1953ACA (5965G>A), AAA1964AAG (5998A>G), ACC1966ACT (6004C>T), AAA1970AAG (6016A>G), TCC1974TCT (6028C>T), GCA1975GCT (6031A>T), ACA1976ATA (6033C>T), GCT1977TCT (6035G>T), GGT1983GGA (6055T>A), TCC1987TCA (6067C>A), GCT1988GCC (6070T>C), CTT1996TTA (6092C>T 6094T>A), CTT1998CTG (6100T>G), CAG1999CAA (6103G>A), AAG2000AAA (6106G>A), ATA2002ATG (6112A>G), TCA2005TCT (6121A>T), TTT2011TTC (6139T>C), ATT2013ATC (6145T>C), TAT2014TAC (6148T>C), GTC2016GTG (6154C>G), GGC2019GGG (6163C>G), GTA2026GTG (6184A>G), ACT2027ACG (6187T>G), TTC2029TTT (6193C>T), CAA2030CAG (6196A>G), ATG2032ATC (6202G>C), CAA2034CAG (6208A>G), CTG2037CTT (6217G>T), GAA2039GAT (6223A>T), GGG2040GGA (6226G>A), ATA2042GTG (6230A>G 6232A>G), GAG2043GAA (6235G>A), ATA2046ATG (6244A>G), TTA2049TTC (6253A>C), ACC2050ACA (6256C>A), ACA2052ACT (6262A>T), CTG2053CTA (6265G>A), TCG2054TCT (6268G>T), AGC2056ATG (6273G>T 6274C>G), GTA2064CTG (6296G>C 6298A>G), TTA2066TTG (6304A>G), TTG2068CTG (6308T>C), GAT2069GAC (6313T>C), ATA2071ATG (6319A>G), GAG2072GAA (6322G>A), ATC2073GTC (6323A>G), CTA2080TTA (6344C>T), AAT2084AAC (6358T>C), GCC2092GCT (6382C>T), ATC2093ATT (6385C>T), TGT2100TGC (6406T>C), AAC2102AAT (6412C>T), CGA2103CAA (6414G>A), AAA2107AAG (6427A>G), CGC2112CGG (6442C>G), TAC2113TAT (6445C>T), ACG2114ACA (6448G>A), ACT2115ACC (6451T>C), TCC2117TCA (6457C>A), GTT2120GTC (6466T>C), GCA2126GCC (6484A>C), AGA2128AGG (6490A>G), CAT2129CAC (6493T>C), ATT2132TTT (6500A>T), TCC2137TCT (6517C>T), CAC2142CAT (6532C>T), CTT2147TTT (6545C>T), TAC2155TAT (6571C>T)

#### Proteins

| | Begin | End | Coverage | Score | Concordance | Matches | Identities | I/D/M/F* | Stop Codons |
|---|---|---|---|---|---|---|---|---|---|
| polyprotein (YP_001039627.1) | 1 | 2159 | 100% | 14866 | 98.6% | 2159 (100%) | 2115 (98.0%) | 0/0/0/0 | 1 |

Protein mutations:

P8S (128C>T 130C>T), F10S (135T>C), A18S (158G>T), T19A (161A>G), T364A (1196A>G), I459V (1481A>G), A501S (1607G>T), A646T (2042G>A), F659Y (2082T>A), N712S (2241A>G), S715N (2250G>A 2251C>T), R768K (2409G>A), V1218I (3758G>A), V1598I (4898G>A), T1628A (4988A>G), I1658T (5079T>C), V1672I (5120G>A 5122T>A), S1676T (5132T>A), T1681N (5148C>A), R1713K (5244G>A), V1755I (5369G>A 5371G>A), I1792M (5482A>G), A1829S (5591G>T), T1833P (5603A>C), Q1857H (5677G>T), R1878K (5739G>A), I1893T (5784T>C), L1915Q (5849T>C 5850T>A 5851G>A), N1950S (5955A>G), T1976I (6033C>T), A1977S (6035G>T), I2002M (6112A>G), M2032I (6202G>C), E2039D (6223A>T), I2042V (6230A>G 6232A>G), I2046M (6244A>G), L2049F (6253A>C), S2056M (6273G>T 6274C>G), V2064L (6296G>C 6298A>G), I2071M (6319A>G), I2073V (6323A>G), R2103Q (6414G>A), I2132F (6500A>T), L2147F (6545C>T)

| | Begin | End | Coverage | Score | Concordance | Matches | Identities | I/D/M/F* | Stop Codons |
|---|---|---|---|---|---|---|---|---|---|
| **NT** | **30** | **7769** | **99.1%** | **11591** | **81.7%** | **7347 (94.6%)** | **6725 (86.6%)** | **24/393** | |
| Codon mutations: | TTT3TTC (115T>C), TTC7TTT (127C>T), CCC8TCT (128C>T 130C>T), TTC10TCC (135T>C), ACT14ACC (148T>C), GAA15GAG (151A>G), GCT18TCT (158G>T), ACC19GCC (161A>G), TCT24TCG (178T>G), TCA27TCG (187A>G), GCT31GCA (199T>A), AGT34AGC (208T>C), CGC38CGT (220C>T), CTG40TTG (224C>T), CAC53CAT (265C>T), TCT74TCC (328T>C), ACT76ACC (334T>C), TTG78TTA (340G>A), AAG81AAA (349G>A), TTC89TTT (373C>T), ATC90ATT (376C>T), AAG109AAA (433G>A), GAC127GAT (487C>T), GTG132GTA (502G>A), TCT145TCC (541T>C), TTC151TTT (559C>T), CTA158TTA (578C>T), AAC174AAT (628C>T), GAC183GAT (655C>T), CAA190CAG (676A>G), GCA196GCG (694A>G), CCA210CCG (736A>G), ACC225ACA (781C>A), TTC228TTT (790C>T), GTA231GTG (799A>G), GGT233GGC (805T>C), CTA238TTG (818C>T 820A>G), CTA242CTT (832A>T), TCA243TCC (835A>C), ATT245ATC (841T>C), CTG246TTG (842C>T), TAC247TAT (847C>T), TTG259TTA (883G>A), TTT260TTC (886T>C), AAT265AAC (901T>C), GCC273GCT (925C>T), GTA282GTG (952A>G), GGT286GGG (964T>G), AAG288AAA (970G>A), GCC297GCT (997C>T), TCT300TCC (1006T>C), GAG309GAA (1033G>A), CAT318CAC (1060T>C), GCA324GCT (1078A>T), AAC325AAT (1081C>T), ACT329ACC (1093T>C), CAA333CAG (1105A>G), CAT334CAC (1108T>C), CTA335TTA (1109C>T), GCC353GCT (1165C>T), AGG358AGA (1180G>A), ACC364GCC (1196A>G), CAG375CAA (1231G>A), GGC378GGT (1240C>T), AGG383AGA (1255G>A), TTT387TTC (1267T>C), CAA390CAG (1276A>G), GGG398GGA (1300G>A), GAA415GAG (1351A>G), CTC422CTT (1372C>T), AAT434AAC (1408T>C), TAT435TAC (1411T>C), CAT437CAC (1417T>C), ACT439ACC (1423T>C), GCT440GCC (1426T>C), ATC441ATT (1429C>T), CTG442TTG (1430C>T), GAC445GAT (1441C>T), GCT448GCC (1450T>C), CTG449TTG (1451C>T), GAC458GAT (1480C>T), ATT459GTT (1481A>G), GAT461GAC (1489T>C), CCC472CCT (1522C>T), GGC482GGT (1552C>T), CGC483CGT (1555C>T), CGA488CGG (1570A>G), TAT489TAC (1573T>C), GCC492GCT (1582C>T), ACA493ACT (1585A>T), AAC495AAT (1591C>T), CTT496CTC (1594T>C), GCT501TCT (1607G>T), CAG502CAA (1612G>A), GGT504GGG (1618T>G), CTA505TTA (1619C>T), ACC506ACA (1624C>A), AGA513AGG (1645A>G), TGC521TGT (1669C>T), GGT525GGA (1681T>A), GAG526GAA (1684G>A), GCT531GCA (1699T>A), ACT533ACC (1705T>C), AAA544AAG (1738A>G), AGC555AGT (1771C>T), TTG559CTG (1781T>C), GAA561GAG (1789A>G), GCA597GCG (1897A>G), TTG602CTG (1910T>C), GTT616GTC (1954T>C), GTA618GTG (1960A>G), TAC626TAT (1984C>T), AAA636AAG (2014A>G), GCA646ACA (2042G>A), CCT650CCC (2056T>C), TTT659TAT (2082T>A), CTA662CTG (2092A>G), AAA667AAG (2107A>G), GTT674GTC (2128T>C), CTG678TTG (2138C>T), TAT679TAC (2143T>C), GGT681GGC (2149T>C), TGT683TGC (2155T>C), CTG690TTG (2174C>T), ATA691ATC (2179A>C), AAG692AAA (2182G>A), GCT696GCA (2194T>A), TAT698TAC (2200T>C), TAC707TAT (2227C>T), AAT712AGT (2241A>G), AGC715AAT (2250G>A 2251C>T), GAC717GAT (2257C>T), CAG719CAA (2263G>A), CCT723CCC (2275T>C), GCG727GCA (2287G>A), CAC728CAT (2290C>T), GAC736GAT (2314C>T), ATA739ATT (2323A>T), CAC747CAT (2347C>T), GTA748GTG (2350A>G), TTC750TTT (2356C>T), TTG751TTA (2359G>A), TAC752TAT (2362C>T), AGG757AGA (2377G>A), TAT758TAC (2380T>C), AGA762AGG (2392A>G), CCT767CCC (2407T>C), AGG768AAG (2409G>A), ATT779ATC (2443T>C), GTG780GTA (2446G>A), CCG791CCA (2479G>A), ATC793ATT (2485C>T), CTG797CTA (2497G>A), ATT798ATC (2500T>C), AAG799AAA (2503G>A), GGC801GGT (2509C>T), CTT802CTC (2512T>C), GGG812GGA (2542G>A), GTC814GTT (2548C>T), TTC817TTT (2557C>T), GAG838GAA (2620G>A), AAT840AAC (2626T>C), AAG848AAA (2650G>A), TTT853TTC (2665T>C), TTG857CTG (2675T>C), GCA859GCG (2683A>G), AAG862AAA (2692G>A), GAT864GAC (2698T>C), CTG866TTG (2702C>T), AAA868AAG (2710A>G), GAT873GAC (2725T>C), TTC875TTT (2731C>T), ATC876ATT (2734C>T), GGA879GGG (2743A>G), TTG880CTG (2744T>C), AAA882AAG (2752A>G), TTG887CTG (2765T>C), CTA889CTG (2773A>G), GTC893GTT (2785C>T), TAC894TAT (2788C>T), TGC901TGT (2809C>T), GTG905GTT (2821G>T), ACA907ACG (2827A>G), TTT912TTC (2842T>C), GGC918GGT (2860C>T), ACT920ACA (2866T>A), TTC921TTT (2869C>T), TCA922TCT (2872A>T), CGG940CGA (2926G>A), TTC941TTT (2929C>T), TTG942CTG (2930T>C), AAG943AAA (2935G>A), CAA945CAG (2941A>G), GTC948GTT (2950C>T), CTG949TTG (2951C>T), AAA964AAG (2998A>G), TCT975TCA (3031T>A), AAG976AAA (3034G>A), GGC977GGT (3037C>T), ACT978ACC (3040T>C), GTA981GTG (3049A>G), GTA986GTG (3064A>G), AAG989AAA (3073G>A), GCT991GCC (3079T>C), TTT1001TTC (3109T>C), GTG1003GTA (3115G>A), GTA1013GTG (3145A>G), TAC1015TAT (3151C>T), TCT1019TCC (3163T>C), TCC1021TCT (3169C>T), TTT1022TTC (3172T>C), TAT1032TAC (3202T>C), CTT1045CTC (3241T>C), AAC1049AAT (3253C>T), GTG1050GTA (3256G>A), AAA1053AAG (3265A>G), CTC1104CTT (3418C>T), AAA1119AAG (3463A>G), CAG1120CAA (3466G>A), GGC1125GGT (3481C>T), ACA1126ACT (3484A>T), ACG1134ACA (3508G>A), GGG1151GGA (3559G>A), TCT1158TCC (3580T>C), ATC1159ATT (3583C>T), GAA1168GAG (3610A>G), GGT1169GGC (3613T>C), TAC1173TAT (3625C>T), GTG1174GTA (3628G>A), AAA1179AAG (3643A>G), TAC1180TAT (3646C>T), CAA1185CAG (3661A>G), AAT1186AAC (3664T>C), GAG1189GAA (3673G>A), GAC1190GAT (3676C>T), CTA1194TTA (3686C>T), CGT1200CGC (3706T>C), TTG1204CTG (3716T>C), CCT1205CCA (3721T>A), CGA1211CGT (3739A>T), AAG1216AAA (3754G>A), GTC1218ATC (3758G>A), CAG1220CAA (3766G>A), CTC1226CTG (3784C>G), TAT1234TAC (3808T>C), GAC1236GAT (3814C>T), GGG1237GGC (3817G>C), TTG1240CTG (3824T>C), TCT1243TCC (3835T>C), TAT1246TAC (3844T>C), TTG1251CTA (3857T>C 3859G>A), GGC1254GGT (3868C>T), ACT1255ACC (3871T>C), GAG1259GAA (3883G>A), AGT1264AGC (3898T>C), GGA1267GGT (3907A>T), GTC1270GTT (3916C>T), AAA1273AAG (3925A>G), AGG1274AGA (3928G>A), GAC1281GAT (3949C>T), GAG1283GAA (3955G>A), TTG1285TTA (3961G>A), GAA1288GAG (3970A>G), ATT1289ATC (3973T>C), TTG1310CTG (4034T>C), CGT1320CGC (4066T>C), GAA1323GAG (4075A>G), CCC1326CCT (4084C>T), TTG1332CTG (4100T>C), CTG1338TTG (4118C>T), TCC1341TCT (4129C>T), TCC1347TCA (4147C>A), CTG1351CTA (4159G>A), CCA1352CCT (4162A>T), AAA1354AAG (4168A>G), GAA1363GAG (4195A>G), TTG1367TTA (4207G>A), GCT1368GCC (4210T>C), TTC1371TTT (4219C>T), GTC1374GTT (4228C>T), CAA1396CAG (4294A>G), GAC1414GAT (4348C>T), AAT1416AAC (4354T>C), ACT1430ACC (4396T>C), AAC1434AAT (4408C>T), GTT1436GTG (4414T>G), TTT1437TTC (4417T>C), AGG1438AGA (4420G>A), GTT1451GTC (4459T>C), ATC1452ATT (4462C>T), TTC1453TTT (4465C>T), AAC1454AAT (4468C>T), TCT1455TCC (4471T>C), CTT1456CTC (4474T>C), TTT1460TTC (4486T>C), TCT1468TCA (4510T>A), CTA1469TTA (4511C>T), TTG1470CTG (4514T>C), AAA1483AAG (4555A>G), AAC1488AAT (4570C>T), CTA1490CTT (4576A>T), GTC1494GTA (4588C>A), TAT1495TAC (4591T>C), TTG1507CTG (4625T>C), TAT1510TAC (4636T>C), GCT1514GCA (4648T>A), GAA1518GAG (4660A>G), ATA1526ATC (4684A>C), ATC1530ATT (4696C>T), AAG1532AAA (4702G>A), AAC1533AAT (4705C>T), GTT1536GTC (4714T>C), GAC1543GAT (4735C>T), CTA1547CTG (4747A>G), CTA1550CTT (4756A>T), AGC1551AGT (4759C>T), TTT1554TTC (4768T>C), AGG1556AGA (4774G>A), CAC1557CAT (4777C>T), CTA1559CTG (4783A>G), GGA1560GGG (4786A>G), TAC1561TAT (4789C>T), GTA1569GTG (4813A>G), TCT1570TCC (4816T>C), AGG1579AGA (4843G>A), GTT1598ATT (4898G>A), GAG1600GAA (4906G>A), AAG1621AAA (4969G>A), ACC1628GCC (4988A>G), ACA1629ACT (4993A>T), TAT1633TAC (5005T>C), TTG1636CTG (5012T>C), GAC1639GAT (5023C>T), CTG1641CTT (5029G>T), GGA1646GGC (5044A>C), GGG1651GGT (5059G>T), CTT1656CTC (5074T>C), CAT1657CAC (5077T>C), ATT1658ACT (5079T>C), TTG1666CTG (5102T>C), ACT1671ACC (5119T>C), GTT1672ATA (5120G>A 5122T>A), CTG1673TTG (5123C>T), TCT1676ACT (5132T>A), CAT1680CAC (5146T>C), ACT1681AAT (5148C>A), CAG1698CAA (5200G>A), GTT1704GTG (5218T>G), AGA1713AAA (5244G>A), CCC1720CCT (5266C>T), GAC1725GAT (5281C>T), CAC1741CAT (5329C>T), ATT1745ATA (5341T>A), GTA1746GTG (5344A>G), ACC1747ACT (5347C>T), GTG1755ATA (5369G>A 5371G>A), GGC1760GGT (5386C>T), GGG1762GGA (5392G>A), CCA1767CCT (5407A>T), GCA1771GCT (5419A>T), ACT1777ACC (5437T>C), GGC1778GGT (5440C>T), GTT1785GTC (5461T>C), TTG1789CTG (5471T>C), ATA1792ATG (5482A>G), CTT1809CTA (5533T>A), ACA1823ACG (5575A>G), TGT1824TGC (5578T>C), GCC1829TCC (5591G>T), ACA1833CCA (5603A>C), CCA1834CCG (5608A>G), GCA1835GCT (5611A>T), ATA1837ATT (5617A>T), GGT1845GGC (5641T>C), TTT1847TTC (5647T>C), CTA1856CTT (5674A>T), CAG1857CAT (5677G>T), CAG1860CAA (5686G>A), TTG1862CTG (5690T>C), CCT1868CCA (5710T>A), GGG1870GGT (5716G>T), GGA1871GGG (5719A>G), CAC1875CAT (5731C>T), ACT1876ACC (5734T>C), ACT1877ACC (5737T>C), AGA1878AAA (5739G>A), CTT1880CTC (5746T>C), TTG1882CTA (5750T>C 5752G>A), CTC1884CTT (5758C>T), AAG1889AAA (5773G>A), CAA1892CAG (5782A>G), ATT1893ACT (5784T>C), TTA1894CTA (5786T>C), AGA1897AGG (5797A>G), TTC1900TTT (5806C>T), TCA1910TCT (5836A>T), TTG1915CAA (5849T>C 5850T>A 5851G>A), AGG1921AGA (5869G>A), CAA1923CAG (5875A>G), GGG1924GGT (5878G>T), TTT1926TTC (5884T>C), CCC1927CCT (5887C>T), CGC1928CGT (5890C>T), GTG1932GTA (5902G>A), CTT1936CTC (5914T>C), TTT1939TTC (5923T>C), CTG1941TTG (5927C>T), AGC1942AGT (5932C>T), AAC1950AGC (5955A>G), ACG1953ACA (5965G>A), AAA1964AAG (5998A>G), ACC1966ACT (6004C>T), AAA1970AAG (6016A>G), TCC1974TCT (6028C>T), GCA1975GCT (6031A>T), ACA1976ATA (6033C>T), GCT1977TCT (6035G>T), GGT1983GGA (6055T>A), TCC1987TCA (6067C>A), GCT1988GCC (6070T>C), CTT1996TTA (6092C>T 6094T>A), CTT1998CTG (6100T>G), CAG1999CAA (6103G>A), AAG2000AAA (6106G>A), ATA2002ATG (6112A>G), TCA2005TCT (6121A>T), TTT2011TTC (6139T>C), ATT2013ATC (6145T>C), TAT2014TAC (6148T>C), GTC2016GTG (6154C>G), GGC2019GGG (6163C>G), GTA2026GTG (6184A>G), ACT2027ACG (6187T>G), TTC2029TTT (6193C>T), CAA2030CAG (6196A>G), ATG2032ATC (6202G>C), CAA2034CAG (6208A>G), CTG2037CTT (6217G>T), GAA2039GAT (6223A>T), GGG2040GGA (6226G>A), ATA2042GTG (6230A>G 6232A>G), GAG2043GAA (6235G>A), ATA2046ATG (6244A>G), TTA2049TTC (6253A>C), ACC2050ACA (6256C>A), ACA2052ACT (6262A>T), CTG2053CTA (6265G>A), TCG2054TCT (6268G>T), AGC2056ATG (6273G>T 6274C>G), GTA2064CTG (6296G>C 6298A>G), TTA2066TTG (6304A>G), TTG2068CTG (6308T>C), GAT2069GAC (6313T>C), ATA2071ATG (6319A>G), GAG2072GAA (6322G>A), ATC2073GTC (6323A>G), CTA2080TTA (6344C>T), AAT2084AAC (6358T>C), GCC2092GCT (6382C>T), ATC2093ATT (6385C>T), TGT2100TGC (6406T>C), AAC2102AAT (6412C>T), CGA2103CAA (6414G>A), AAA2107AAG (6427A>G), CGC2112CGG (6442C>G), TAC2113TAT (6445C>T), ACG2114ACA (6448G>A), ACT2115ACC (6451T>C), TCC2117TCA (6457C>A), GTT2120GTC (6466T>C), GCA2126GCC (6484A>C), AGA2128AGG (6490A>G), CAT2129CAC (6493T>C), ATT2132TTT (6500A>T), TCC2137TCT (6517C>T), CAC2142CAT (6532C>T), CTT2147TTT (6545C>T), TAC2155TAT (6571C>T) | | | | | | | | |

*: Inserts / Deletes / Misaligned / Frameshifts

#### Analysis details

This analysis was performed with panviral2.64

### NGS Details (UN60): Torradovirus lycopersici (segment RNA 2)

#### Assembly

| | |
|---|---|
| **Coverage Length** | 5384 (1 contig(s)) |
| **Depth Of Coverage** | 16080.2 |
| **Number Of Reads** | 685969 |
| **Reads Per Million** | 14549.11 rpm (after QC) |
| **Ambiguities** | 0 |
| **Assembly Method** | de novo + reference guided assembly |
| **Consensus Caller** | Bcf Tools |

#### Coverage Map

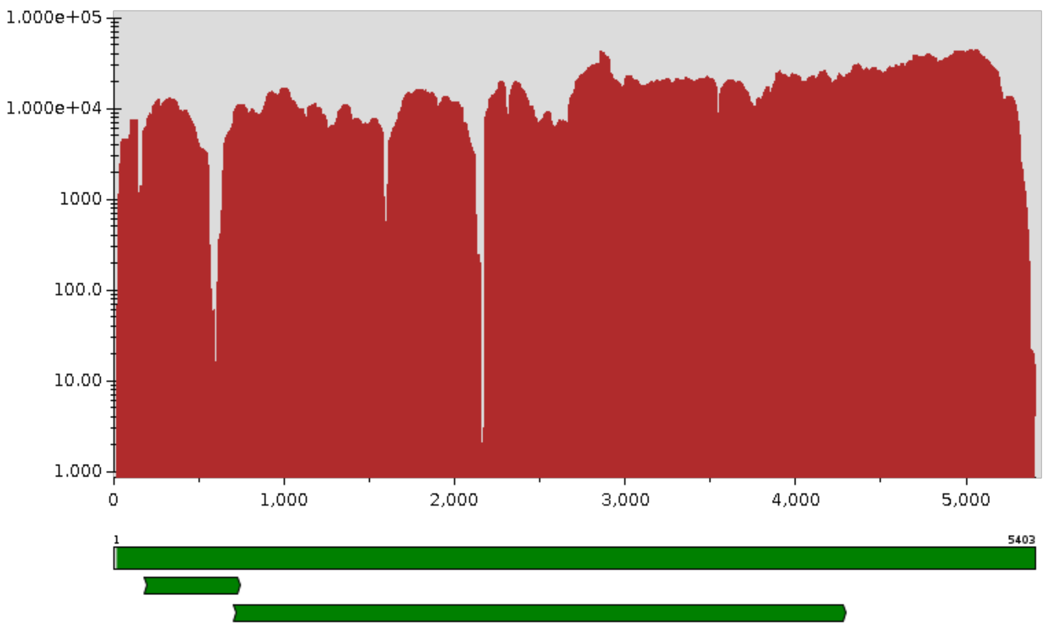

#### Assignment

| | |
|---|---|
| **Type** | Torradovirus lycopersici (Taxonomy ID: 3048378) |
| **Reference Genome** | NC_009032.1 |
| **NT Identity (%)** | 85.2794 |
| **AA Identity (%)** | 89.6179 |
| **Number Of Stop Codons** | 5 |
| **Number Of CDS** | 2 |

#### Alignment

| | |
|---|---|
| **Alignment Score** | 7578.0 (NT) + 8662.0 (AA) = 16240.0 |
| **Concordance (%)** | 80.1777 |

| Alignment Method | Global, seeded, nucleotide + amino acids (AGA) |
|---|---|

#### Genome Region

Sequence starts at position 20 and ends at position 5403 relative to NC_009032.1 reference sequence.

#### Alignment Detailed Statistics

| | Begin | End | Coverage | Score | Concordance | Matches | Identities | I/D/M/F* | Stop Codons |
|---|---|---|---|---|---|---|---|---|---|
| **NT** | **20** | **5403** | **99.6%** | **7578** | **70.4%** | **5384 (99.9%)** | **4594 (85.3%)** | **3/0** | |
| Mutations: | 52C>T, 58T>C, 60G>A, 69A>G, 79A>G, 91T>A, 96C>T, 99T>G, 103G>C, 104A>G, 105G>T, 107G>A, 108C>G, 111C>A, 112G>A, 117C>T, 120A>G, 121A>C, 132C>T, 137T>C, 149G>T, 161A>T, 164T>A, 170G>T, 181T>C, 187A>C, 212C>G, 223G>A, 232C>T, 235G>A, 238A>G, 244A>T, 247C>T, 265G>A, 268T>A, 277C>T, 280T>C, 286A>C, 289A>C, 292C>T, 301A>T, 304T>C, 305C>T, 307A>G, 313C>T, 316G>A, 325C>T, 329A>G, 334T>A, 340A>T, 352A>T, 356G>A, 370C>T, 372A>C, 376C>T, 383C>T, 385G>A, 388T>A, 400T>C, 401T>C, 403A>C, 418T>G, 433C>T, 439G>A, 445T>A, 448C>A, 451A>T, 461A>C, 481C>T, 484C>A, 487A>G, 496A>G, 502T>A, 505G>C, 507G>A, 508G>A, 509A>G, 511T>A, 515G>A, 520C>T, 541C>T, 547T>G, 552C>A, 553T>G, 554G>A, 555C>A, 564A>G, 565A>T, 568A>T, 570A>G, 576C>A, 578C>G, 586A>G, 587T>C, 589A>T, 595G>T, 599C>T, 601T>G, 603A>C, 604C>T, 607G>A, 610C>T, 613A>G, 618G>A, 619C>A, 622A>G, 631A>G, 632G>T, 637A>G, 644C>G, 645T>C, 654T>A, 658C>A, 665T>G, 670A>G, 673A>G, 679G>A, 683T>A, 685G>A, 689A>T, 690A>C, 696T>C, 706T>C, 731G>A, 733A>G, 750A>G, 755A>T, 758T>A, 761A>G, 788A>T, 818G>A, 821T>C, 824C>G, 833C>T, 842A>T, 844A>T, 851T>C, 854C>A, 860C>G, 863C>T, 866A>G, 878C>T, 884A>G, 887G>T, 893C>T, 911T>C, 929T>A, 932A>T, 968A>G, 1010T>G, 1061T>C, 1085G>A, 1095G>A, 1100G>A, 1109C>T, 1115G>A, 1118A>C, 1130A>G, 1139G>A, 1142C>T, 1151A>G, 1163A>G, 1169C>G, 1178C>T, 1187A>T, 1205T>A, 1220T>C, 1235C>A, 1247A>G, 1262T>C, 1313T>C, 1316C>T, 1317C>A, 1319A>G, 1325A>G, 1328T>C, 1331C>T, 1343G>A, 1355G>A, 1361A>T, 1367A>T, 1370A>T, 1376T>A, 1383A>G, 1385C>T, 1388C>T, 1390C>G, 1394A>C, 1395A>T, 1397T>A, 1400T>G, 1403A>T, 1406G>A, 1407C>T, 1409T>A, 1412G>T, 1418G>A, 1426A>C, 1427G>T, 1434G>A, 1439A>C, 1451G>T, 1454A>T, 1456G>C, 1457C>T, 1460A>G, 1461G>T, 1465G>A, 1472C>G, 1475G>C, 1478C>G, 1481C>T, 1487C>G, 1490G>A, 1493G>A, 1496C>T, 1502C>T, 1514T>C, 1524T>C, 1526C>T, 1529A>T, 1538A>T, 1543C>T, 1545A>G, 1547C>T, 1553T>C, 1559C>A, 1560C>T, 1562C>A, 1568C>G, 1574G>A, 1575C>A, 1577T>G, 1589A>G, 1593C>G, 1595G>T, 1603G>A, 1605A>G, 1608T>C, 1610C>G, 1613A>G, 1616G>A, 1621G>C, 1622T>A, 1626G>T, 1627A>C, 1628T>C, 1631G>A, 1634T>A, 1643C>T, 1646T>C, 1651A>G, 1655T>A, 1661T>C, 1662C>T, 1663T>A, 1664G>T, 1667C>T, 1674C>T, 1676T>C, 1677G>A, 1679G>T, 1685T>C, 1688T>C, 1695A>G, 1700G>A, 1721G>C, 1724C>T, 1727A>G, 1730C>T, 1740C>T, 1745G>A, 1748G>A, 1749A>C, 1754G>A, 1757A>G, 1760T>G, 1763T>A, 1766A>G, 1769A>T, 1778A>C, 1784A>T, 1790G>A, 1793T>A, 1794G>A, 1797T>G, 1799C>G, 1805C>T, 1811A>C, 1814T>C, 1817A>T, 1819T>A, 1820C>G, 1823C>T, 1832G>C, 1841C>T, 1843A>G, 1845T>C, 1847G>T, 1856C>T, 1859C>T, 1863C>T, 1865C>T, 1875G>A, 1878A>G, 1892C>T, 1898T>G, 1899G>A, 1900T>C, 1901G>A, 1906C>T, 1907C>G, 1913T>A, 1928A>C, 1931C>T, 1938A>G, 1941C>T, 1943T>G, 1946C>A, 1947T>C, 1952A>C, 1955G>A, 1958A>G, 1967G>A, 1970A>G, 1979A>G, 1986G>A, 1994T>C, 2002T>A, 2012C>T, 2024T>C, 2027T>A, 2049T>A, 2051A>T, 2054G>T, 2059A>G, 2060A>C, 2066C>G, 2074T>A, 2083T>C, 2084G>A, 2087A>C, 2094C>G, 2096G>C, 2098C>A, 2099G>A, 2100C>A, 2101G>T, 2102C>G, 2106A>G, 2107G>A, 2111T>G, 2114C>T, 2115A>C, 2116A>G, 2119A>C, 2120G>A, 2123G>T, 2125C>T, 2126C>T, 2127A>G, 2128G>T, 2130G>A, 2132T>C, 2135G>A, 2136G>C, 2138G>T, 2144C>G, 2159T>A, 2162T>C, 2164T>G, 2170A>T, 2171A>T, 2174A>T, 2177A>T, 2178T>G, 2183C>T, 2186A>C, 2189C>T, 2195A>C, 2198G>A, 2199G>A, 2200T>C, 2204C>T, 2205A>G, 2210G>A, 2216C>G, 2219T>C, 2225T>A, 2228C>T, 2229A>G, 2230T>A, 2231G>T, 2240T>A, 2243T>C, 2252A>C, 2256G>A, 2259A>C, 2261T>A, 2262G>C, 2270A>G, 2285T>C, 2294A>C, 2303A>G, 2312T>C, 2315T>A, 2320G>C, 2321T>C, 2324G>C, 2326C>T, 2339A>G, 2345A>G, 2375C>T, 2393G>A, 2414T>C, 2420T>G, 2429G>A, 2432G>A, 2436T>G, 2444G>C, 2447C>A, 2453G>A, 2459C>G, 2469T>A, 2471C>G, 2474G>A, 2489A>C, 2492G>A, 2495A>G, 2501A>G, 2503G>A, 2507T>C, 2516C>T, 2523A>C, 2546T>C, 2552C>T, 2561T>G, 2567T>A, 2570G>A, 2573T>A, 2588T>A, 2609C>A, 2627C>T, 2630C>G, 2645G>C, 2660T>C, 2672A>T, 2693G>A, 2702T>A, 2708A>G, 2717T>C, 2720C>A, 2729C>T, 2732T>G, 2747C>T, 2753G>C, 2792C>T, 2795C>T, 2798G>T, 2819G>A, 2835A>G, 2837C>T, 2840T>C, 2841C>T, 2846G>T, 2852A>G, 2861G>C, 2864C>T, 2870C>A, 2879T>A, 2882T>G, 2885T>G, 2886T>C, 2894A>G, 2900A>T, 2909T>C, 2915A>G, 2921C>T, 2927A>G, 2936A>C, 2939A>G, 2948A>G, 2954G>A, 2957A>T, 2960G>A, 2966C>T, 2972T>C, 2981G>T, 2982G>A, 2996T>G, 2999T>A, 3003T>C, 3011C>T, 3012A>G, 3014C>T, 3020T>C, 3023G>C, 3029C>T, 3032A>T, 3044C>T, 3047T>A, 3050G>C, 3051C>T, 3059A>T, 3062C>T, 3063T>C, 3065G>A, 3071A>G, 3077A>T, 3083C>A, 3101T>G, 3104C>T, 3107T>G, 3113G>C, 3122T>G, 3128T>C, 3131G>A, 3143C>T, 3146T>C, 3152T>A, 3158G>A, 3167A>T, 3168C>T, 3176A>C, 3179G>T, 3182C>T, 3187G>A, 3197C>T, 3206C>T, 3227T>G, 3230T>A, 3236C>T, 3239T>G, 3242G>A, 3248A>C, 3251A>T, 3253A>G, 3265G>A, 3269T>C, 3270A>T, 3272G>T, 3278C>T, 3293A>G, 3299T>A, 3303A>T, 3305C>T, 3311T>A, 3324C>T, 3329G>T, 3350C>T, 3354A>C, 3359G>A, 3368A>T, 3371C>G, 3372T>C, 3374G>C, 3386A>T, 3395A>C, 3404A>G, 3410G>A, 3413C>T, 3428G>A, 3429C>T, 3431T>G, 3434A>G, 3437C>T, 3440A>T, 3447T>C, 3449G>A, 3452C>A, 3455T>A, 3458G>A, 3461C>T, 3470T>C, 3473T>C, 3479C>T, 3482T>C, 3485C>T, 3488A>G, 3491C>T, 3503A>G, 3508A>G, 3512T>C, 3515G>T, 3522C>T, 3524A>G, 3527C>T, 3533G>A, 3543T>A, 3551C>T, 3560T>C, 3564A>C, 3567T>A, 3569A>G, 3575G>A, 3578A>G, 3581C>T, 3590A>C, 3605A>G, 3611G>A, 3617A>T, 3620G>A, 3623A>T, 3629G>C, 3631C>A, 3634T>C, 3635C>T, 3638A>G, 3641C>T, 3650A>T, 3656G>T, 3659T>C, 3662A>G, 3674G>C, 3689C>T, 3698A>C, 3702T>C, 3704G>A, 3707T>G, 3708T>C, 3713A>C, 3716C>T, 3719A>G, 3722C>A, 3728C>G, 3742A>C, 3743G>C, 3746G>A, 3749A>T, 3752G>A, 3755A>G, 3767C>A, 3770A>C, 3773C>T, 3776A>T, 3779C>T, 3800C>T, 3815A>T, 3827C>T, 3833T>A, 3839A>G, 3845A>G, 3848G>A, 3851G>A, 3852A>G, 3854T>C, 3857C>A, 3866A>C, 3869A>T, 3872A>T, 3878G>C, 3881C>T, 3882A>G, 3884C>A, 3885A>G, 3890A>G, 3893T>C, 3896A>T, 3914T>A, 3920A>T, 3923C>T, 3927A>C, 3928A>G, 3929G>A, 3932C>A, 3933A>G, 3935C>T, 3941T>G, 3944C>A, 3947T>C, 3950T>A, 3953T>A, 3960G>A, 3971T>A, 3983T>A, 3984A>G, 3986G>T, 3988G>A, 3989C>T, 3990G>A, 3991A>G, 3995C>T, 3998C>T, 4001T>C, 4016T>G, 4025G>A, 4028T>C, 4031T>C, 4034T>G, 4035A>C, 4037A>T, 4040C>T, 4052C>T, 4059A>G, 4061A>T, 4067A>T, 4068A>G, 4069A>G, 4070T>C, 4072C>T, 4073T>G, 4077A>C, 4088T>A, 4091G>A, 4100A>C, 4103A>T, 4106A>C, 4112C>T, 4133A>G, 4142A>C, 4145C>T, 4149C>A, 4151G>A, 4154A>G, 4157C>A, 4166G>A, 4181C>T, 4187G>A, 4188C>G, 4196T>A, 4208A>T, 4211A>T, 4223G>A, 4224C>T, 4227T>C, 4229G>T, 4250C>T, 4256T>G, 4262T>C, 4265T>A, 4274A>T, 4281A>C, 4284A>G, 4286T>A, 4289T>C, 4300T>C, 4302T>G, 4316T>C, 4354G>A, 4355T>G, 4382A>T, 4383C>T, 4388A>T, 4391A>C, 4393A>G, 4424C>T, 4425T>C, 4429T>C, 4457A>G, 4468T>C, 4496T>A, 4506A>T, 4525T>C, 4567_4568insT, 4605T>C, 4633T>A, 4634G>A, 4643G>A, 4648G>A, 4650T>C, 4651G>A, 4675A>T, 4812C>T, 4823A>G, 4974_4975insT, 5089T>C, 5192C>T, 5206C>T, 5213A>G, 5214C>T, 5222C>T, 5224G>A, 5225C>A, 5226T>G, 5233G>T, 5236G>T, 5239T>A, 5242T>G, 5244G>A, 5249A>G, 5258A>G, 5259T>A, 5262A>T, 5282T>C, 5313A>C, 5315T>A, 5369G>T, 5371G>A, 5374T>A, 5378A>T, 5379A>C, 5380_5381insT, 5400A>C | | | | | | | | |
| **CDS** | | | | | | | | | |
| ToTV_sRNA2gp1 | 1 | 188 | 100% | 1080 | 89.3% | 188 (100%) | 162 (86.2%) | 0/0/0/0 | 2 |
| Protein mutations: | L11V (212C>G), T50A (329A>G), V59I (356G>A), N64T (372A>C), L90F (451A>T), Q108H (505G>C), R109K (507G>A 508G>A), I110V (509A>G 511T>A), V112I (515G>A), A124E (552C>A 553T>G), A125K (554G>A 555C>A), Q128R (564A>G 565A>T), Q130R (570A>G), T132K (576C>A), L133V (578C>G), E138D (595G>T), D141A (603A>C 604C>T), R146Q (618G>A 619C>A), E151* (632G>T), L155A (644C>G 645T>C), L158Q (654T>A), S162A (665T>G), L168I (683T>A 685G>A), N170S (689A>T 690A>C), L172P (696T>C), E184K (731G>A 733A>G) | | | | | | | | |
| Codon mutations: | TCA2TCC (187A>C), CTT11GTT (212C>G), GAG14GAA (223G>A), CAC17CAT (232C>T), AAG18AAA (235G>A), CAA19CAG (238A>G), GCA21GCT (244A>T), GAC22GAT (247C>T), TCG28TCA (265G>A), GTT29GTA (268T>A), GGC32GGT (277C>T), TCT33TCC (280T>C), ATA35ATC (286A>C), ATA36ATC (289A>C), AAC37AAT (292C>T), CCA40CCT (301A>T), ACT41ACC (304T>C), CTA42TTG (305C>T 307A>G), TTC44TTT (313C>T), AAG45AAA (316G>A), CCC48CCT (325C>T), ACA50GCA (329A>G), GGT51GGA (334T>A), GCA53GCT (340A>T), CTA57CTT (352A>T), GTT59ATT (356G>A), AAC63AAT (370C>T), AAT64ACT (372A>C), AGC65AGT (376C>T), CTG68TTA (383C>T 385G>A), GTT69GTA (388T>A), TAT73TAC (400T>C), TTA74CTC (401T>C 403A>C), ACT79ACG (418T>G), GGC84GGT (433C>T), AAG86AAA (439G>A), TCT88TCA (445T>A), GGC89GGA (448C>A), TTA90TTT (451A>T), AGA94CGA (461A>C), ACC100ACT (481C>T), ACC101ACA (484C>A), AGA102AGG (487A>G), CTA105CTG (496A>G), GCT107GCA (502T>A), CAG108CAC (505G>C), AGG109AAA (507G>A 508G>A), ATT110GTA (509A>G 511T>A), GTA112ATA (515G>A), TCC113TCT (520C>T), TCC120TCT (541C>T), GCT122GCG (547T>G), GCT124GAG (552C>A 553T>G), GCA125AAA (554G>A 555C>A), CAA128CGT (564A>G 565A>T), CCA129CCT (568A>T), CAA130CGA (570A>G), ACG132AAG (576C>A), CTT133GTT (578C>G), CAA135CAG (586A>G), TTA136CTT (587T>C 589A>T), GAG138GAT (595G>T), CTT140TTG (599C>T 601T>G), GAC141GCT (603A>C 604C>T), GAG142GAA (607G>A), GCC143GCT (610C>T), AAA144AAG (613A>G), CGC146CAA (618G>A 619C>A), TTA147TTG (622A>G), AAA150AAG (631A>G), GAG151TAG (632G>T), AAA152AAG (637A>G), CTT155GCT (644C>G 645T>C), CTA158CAA (654T>A), TCC159TCA (658C>A), TCC162GCC (665T>G), AAA163AAG (670A>G), TTA164TTG (673A>G), TTG166TTA (679G>A), TTG168ATA (683T>A 685G>A), AAC170TCC (689A>T 690A>C), CTA172CCA (696T>C), GAT175GAC (706T>C), GAA184AAG (731G>A 733A>G) | | | | | | | | |
| ToTV_sRNA2gp2 | 1 | 1199 | 100% | 7582 | 91.6% | 1199 (100%) | 1081 (90.2%) | 0/0/0/0 | 3 |
| Protein mutations: | M2T (706T>C), K11R (733A>G), S17G (750A>G), Y48F (844A>T), V132I (1095G>A), T228A (1383A>G 1385C>T), S230* (1390C>G), T232S (1395A>T 1397T>A), D233E (1400T>G), Q242P (1426A>C 1427G>T), D245N (1434G>A), S252T (1456G>C 1457C>T), A254S (1461G>T), R255K (1465G>A), I257M (1472C>G), S275P (1524T>C 1526C>T), S281F (1543C>T), T282A (1545A>G 1547C>T), H286Q (1559C>A), Q298D (1593C>G 1595G>T), S301N (1603G>A), R302G (1605A>G), Y303Q (1608T>C 1610C>G), G307A (1621G>C 1622T>A), D309S (1626G>T 1627A>C 1628T>C), K317R (1651A>G), L321Y (1662C>T 1663T>A 1664G>T), H325Y (1674C>T 1676T>C), V326I (1677G>A 1679G>T), I332V (1695A>G), V365I (1794G>A), C366G (1797T>G 1799C>G), K370N (1811A>C), I373K (1819T>A 1820C>G), K377N (1832G>C), H381R (1843A>G), H388Y (1863C>T 1865C>T), V392M (1875G>A), I393V (1878A>G), I399M (1898T>G), V400T (1899G>A 1900T>C 1901G>A), A402V (1906C>T 1907C>G), E409D (1928A>C), N413D (1938A>G), D429N (1986G>A), F434Y (2002T>A), L450I (2049T>A 2051A>T), K453S (2059A>G 2060A>C), L458Q (2074T>A), V461A (2083T>C 2084G>A), Q465D (2094C>G 2096G>C), A466E (2098C>A 2099G>A), R467M (2100C>A 2101G>T 2102C>G), R469E (2106A>G 2107G>A), F470L (2111T>G), K472R (2115A>C 2116A>G), E473A (2119A>C 2120G>A), A475V (2125C>T 2126C>T), R476V (2127A>G 2128G>T), A477T (2130G>A 2132T>C), V479L (2136G>C 2138G>T), N481K (2144C>G), V488G (2164T>G), E490V (2170A>T 2171A>T), S493A (2178T>G), V500T (2199G>A 2200T>C), M502V (2205A>G), D508E (2225T>A), M510D (2229A>G 2230T>A 2231G>T), E517D (2252A>C), A519T (2256G>A), N520Q (2259A>C 2261T>A), E521Q (2262G>C), E531D (2294A>C), N538K (2315T>A), G540A (2320G>C 2321T>C), T542I (2326C>T), S579A (2436T>G), S590T (2469T>A 2471C>G), R601K (2503G>A), I608L (2523A>C), I643M (2630C>G), C667* (2702T>A), L699F (2798G>T), I712V (2835A>G 2837C>T), Q715H (2846G>T), A761T (2982G>A), I771V (3012A>G 3014C>T), I807M (3122T>G), S829N (3187G>A), K851R (3253A>G), R855K (3265G>A), T857S (3270A>T 3272G>T), T868S (3303A>T 3305C>T), N936S (3508A>G), S948T (3543T>A), K955Q (3564A>C), L956M (3567T>A 3569A>G), P977Q (3631C>A), L978P (3634T>C 3635C>T), Q1014P (3742A>C 3743G>C), T1051A (3852A>G 3854T>C), I1061V (3882A>G 3884C>A), T1062A (3885A>G), K1076R (3927A>C 3928A>G 3929G>A), T1078A (3933A>G 3935C>T), A1087T (3960G>A), M1095V (3984A>G 3986G>T), S1096N (3988G>A 3989C>T), D1097S (3990G>A 3991A>G), H1105Q (4016T>G), I1120V (4059A>G 4061A>T), N1123G (4068A>G 4069A>G 4070T>C), A1124V (4072C>T 4073T>G), L1150I (4149C>A 4151G>A), Q1163E (4188C>G), H1175Y (4224C>T), S1195G (4284A>G 4286T>A) | | | | | | | | |

| | Begin | End | Coverage | Score | Concordance | Matches | Identities | I/D/M/F* | Stop Codons |
|---|---|---|---|---|---|---|---|---|---|
| **NT** | **20** | **5403** | **99.6%** | **7578** | **70.4%** | **5384 (99.9%)** | **4594 (85.3%)** | **3/0** | |
| Codon mutations: | ATG2ACG (706T>C), AGG10AGA (731G>A), AAG11AGG (733A>G), AGT17GGT (750A>G), GCA18GCT (755A>T), GCT19GCA (758T>A), GTA20GTG (761A>G), ACA29ACT (788A>T), CTG39CTA (818G>A), CTT40CTC (821T>C), CGC41CGG (824C>G), GGC44GGT (833C>T), TCA47TCT (842A>T), TAT48TTT (844A>T), ATT50ATC (851T>C), CTC51CTA (854C>A), GGC53GGG (860C>G), GCC54GCT (863C>T), CAA55CAG (866A>G), GCC59GCT (878C>T), GAA61GAG (884A>G), CTG62CTT (887G>T), GAC64GAT (893C>T), AAT70AAC (911T>C), TCT76TCA (929T>A), TCA77TCT (932A>T), GGA89GGG (968A>G), GTT103GTG (1010T>G), AGT120AGC (1061T>C), CTG128CTA (1085G>A), GTT132ATT (1095G>A), TTG133TTA (1100G>A), TCC136TCT (1109C>T), GGG138GGA (1115G>A), GCA139GCC (1118A>C), TTA143TTG (1130A>G), CCG146CCA (1139G>A), ACC147ACT (1142C>T), CTA150CTG (1151A>G), AAA154AAG (1163A>G), GGC156GGG (1169C>G), TTC159TTT (1178C>T), ACA162ACT (1187A>T), ACT168ACA (1205T>A), CCT173CCC (1220T>C), GCC178GCA (1235C>A), CAA182CAG (1247A>G), CGT187CGC (1262T>C), CTT204CTC (1313T>C), CAC205CAT (1316C>T), CGA206AGG (1317C>A 1319A>G), CAA208CAG (1325A>G), TTT209TTC (1328T>C), TTC210TTT (1331C>T), CAG214CAA (1343G>A), CAG218CAA (1355G>A), ATA220ATT (1361A>T), TCA222TCT (1367A>T), GTA223GTT (1370A>T), TCT225TCA (1376T>A), ACC228GCT (1383A>G 1385C>T), TCC229TCT (1388C>T), TCA230TGA (1390C>G), GCA231GCC (1394A>C), ACT232TCA (1395A>T 1397T>A), GAT233GAG (1400T>G), TCA234TCT (1403A>T), GTG235GTA (1406G>A), CTT236TTA (1407C>T 1409T>A), CCG237CCT (1412G>T), GTG239GTA (1418G>A), CAG242CCT (1426A>C 1427G>T), GAC245AAC (1434G>A), ACA246ACC (1439A>C), GTG250GTT (1451G>T), CCA251CCT (1454A>T), AGC252ACT (1456G>C 1457C>T), GAA253GAG (1460A>G), GCA254TCA (1461G>T), AGA255AAA (1465G>A), ATC257ATG (1472C>G), CGG258CGC (1475G>C), TCC259TCG (1478C>G), TCC260TCT (1481C>T), CTC262CTG (1487C>G), CGG263CGA (1490G>A), GTG264GTA (1493G>A), CCC265CCT (1496C>T), CTC267CTT (1502C>T), AGT271AGC (1514T>C), TCC275CCT (1524T>C 1526C>T), TCA276TCT (1529A>T), ATA279ATT (1538A>T), TCT281TTT (1543C>T), ACC282GCT (1545A>G 1547C>T), ACT284ACC (1553T>C), CAC286CAA (1559C>A), CTC287TTA (1560C>T 1562C>A), GGC289GGG (1568C>G), ACG291ACA (1574G>A), CGT292AGG (1575C>A 1577T>G), GTA296GTG (1589A>G), CAG298GAT (1593C>G 1595G>T), AGT301AAT (1603G>A), AGG302GGG (1605A>G), TAC303CAG (1608T>C 1610C>G), GAA304GAG (1613A>G), GAG305GAA (1616G>A), GGT307GCA (1621G>C 1622T>A), GAT309TCC (1626G>T 1627A>C 1628T>C), GGG310GGA (1631G>A), GTT311GTA (1634T>A), CCC314CCT (1643C>T), GTT315GTC (1646T>C), AAG317AGG (1651A>G), GCT318GCA (1655T>A), GGT320GGC (1661T>C), CTG321TAT (1662C>T 1663T>A 1664G>T), AAC322AAT (1667C>T), CAT325TAC (1674C>T 1676T>C), GTG326ATT (1677G>A 1679G>T), GAT328GAC (1685T>C), AAT329AAC (1688T>C), ATT332GTT (1695A>G), GAG333GAA (1700G>A), CTG340CTC (1721G>C), CCC341CCT (1724C>T), GAA342GAG (1727A>G), AAC343AAT (1730C>T), CTG347TTG (1740C>T), TCG348TCA (1745G>A), GTG349GTA (1748G>A), AGA350CGA (1749A>C), CAG351CAA (1754G>A), CAA352CAG (1757A>G), GTT353GTG (1760T>G), GCT354GCA (1763T>A), GCA355GCG (1766A>G), GCA356GCT (1769A>T), GCA359GCC (1778A>C), GCA361GCT (1784A>T), AAG363AAA (1790G>A), GGT364GGA (1793T>A), GTT365ATT (1794G>A), TGC366GGG (1797T>G 1799C>G), GTC368GTT (1805C>T), AAA370AAC (1811A>C), CAT371CAC (1814T>C), ATA372ATT (1817A>T), ATC373AAG (1819T>A 1820C>G), AAC374AAT (1823C>T), AAG377AAC (1832G>C), ATC380ATT (1841C>T), CAC381CGC (1843A>G), TTG382CTT (1845T>C 1847G>T), ATC385ATT (1856C>T), ACC386ACT (1859C>T), CAC388TAT (1863C>T 1865C>T), GTG392ATG (1875G>A), ATA393GTA (1878A>G), TAC397TAT (1892C>T), ATT399ATG (1898T>G), GTG400ACA (1899G>A 1900T>C 1901G>A), GCC402GTG (1906C>T 1907C>G), TCT404TCA (1913T>A), GAA409GAC (1928A>C), AAC410AAT (1931C>T), AAT413GAT (1938A>G), CTT414TTG (1941C>T 1943T>G), ACC415ACA (1946C>A), TTG416CTG (1947T>C), GCA417GCC (1952A>C), CAG418CAA (1955G>A), AGA419AGG (1958A>G), GAG422GAA (1967G>A), CTA423CTG (1970A>G), CAA426CAG (1979A>G), GAT429AAT (1986G>A), TAT431TAC (1994T>C), TTT434TAT (2002T>A), AAC437AAT (2012C>T), TAT441TAC (2024T>C), CCT442CCA (2027T>A), TTA450ATT (2049T>A 2051A>T), ACG451ACT (2054G>T), AAA453AGC (2059A>G 2060A>C), CGC455CGG (2066C>G), CTA458CAA (2074T>A), GTG461GCA (2083T>C 2084G>A), GCA462GCC (2087A>C), CAG465GAC (2094C>G 2096G>C), GCG466GAA (2098C>A 2099G>A), CGC467ATG (2100C>A 2101G>T 2102C>G), AGA469GAA (2106A>G 2107G>A), TTT470TTG (2111T>G), TAC471TAT (2114C>T), AAG472CGG (2115A>C 2116A>G), GAG473GCA (2119A>C 2120G>A), GCG474GCT (2123G>T), GCC475GTT (2125C>T 2126C>T), AGG476GTG (2127A>G 2128G>T), GCT477ACC (2130G>A 2132T>C), CAG478CAA (2135G>A), GTG479CTT (2136G>C 2138G>T), AAC481AAG (2144C>G), ACT486ACA (2159T>A), AGT487AGC (2162T>C), GTG488GGG (2164T>G), GAA490GTT (2170A>T 2171A>T), ATA491ATT (2174A>T), CCA492CCT (2177A>T), TCC493GCC (2178T>G), ACC494ACT (2183C>T), TCA495TCC (2186A>C), TTC496TTT (2189C>T), ACA498ACC (2195A>C), AAG499AAA (2198G>A), GTT500ACT (2199G>A 2200T>C), GCC501GCT (2204C>T), ATG502GTG (2205A>G), GAG503GAA (2210G>A), GGC505GGG (2216C>G), AGT506AGC (2219T>C), GAT508GAA (2225T>A), AGC509AGT (2228C>T), ATG510GAT (2229A>G 2230T>A 2231G>T), GCT513GCA (2240T>A), ATT514ATC (2243T>C), GAA517GAC (2252A>C), GCC519ACC (2256G>A), AAT520CAA (2259A>C 2261T>A), GAA521CAA (2262G>C), GTA523GTG (2270A>G), GTT528GTC (2285T>C), GAA531GAC (2294A>C), CAA534CAG (2303A>G), AAT537AAC (2312T>C), AAT538AAA (2315T>A), GGT540GCC (2320G>C 2321T>C), GCG541GCC (2324G>C), ACA542ATA (2326C>T), TTA546TTG (2339A>G), CAA548CAG (2345A>G), TTC558TTT (2375C>T), GAG564GAA (2393G>A), AGT571AGC (2414T>C), GTT573GTG (2420T>G), GAG576GAA (2429G>A), GTG577GTA (2432G>A), TCA579GCA (2436T>G), GCG581GCC (2444G>C), CTC582CTA (2447C>A), GCG584GCA (2453G>A), CTC586CTG (2459C>G), TCC590ACG (2469T>A 2471C>G), AAG591AAA (2474G>A), GGA596GGC (2489A>C), TCG597TCA (2492G>A), CAA598CAG (2495A>G), CTA600CTG (2501A>G), AGG601AAG (2503G>A), TAT602TAC (2507T>C), GCC605GCT (2516C>T), ATT608CTT (2523A>C), TAT615TAC (2546T>C), TCC617TCT (2552C>T), GGT620GGG (2561T>G), ATT622ATA (2567T>A), TCG623TCA (2570G>A), GGT624GGA (2573T>A), GTT629GTA (2588T>A), CTC636CTA (2609C>A), TTC642TTT (2627C>T), ATC643ATG (2630C>G), CTG648CTC (2645G>C), CAT653CAC (2660T>C), TCA657TCT (2672A>T), GGG664GGA (2693G>A), TGT667TGA (2702T>A), ACA669ACG (2708A>G), GGT672GGC (2717T>C), ATC673ATA (2720C>A), TTC676TTT (2729C>T), GTT677GTG (2732T>G), GCC682GCT (2747C>T), GGG684GGC (2753G>C), CAC697CAT (2792C>T), CCC698CCT (2795C>T), TTG699TTT (2798G>T), GAG706GAA (2819G>A), ATC712GTT (2835A>G 2837C>T), CAT713CAC (2840T>C), CTG714TTG (2841C>T), CAG715CAT (2846G>T), AAA717AAG (2852A>G), TCG720TCC (2861G>C), ACC721ACT (2864C>T), ATC723ATA (2870C>A), CCT726CCA (2879T>A), CCT727CCG (2882T>G), CGT728CGG (2885T>G), TTG729CTG (2886T>C), GCA731GCG (2894A>G), GCA733GCT (2900A>T), GGT736GGC (2909T>C), AAA738AAG (2915A>G), GAC740GAT (2921C>T), ACA742ACG (2927A>G), CCA745CCC (2936A>C), CGA746CGG (2939A>G), ACA749ACG (2948A>G), CAG751CAA (2954G>A), GTA752GTT (2957A>T), CTG753CTA (2960G>A), CAC755CAT (2966C>T), AAT757AAC (2972T>C), GTG760GTT (2981G>T), GCT761ACT (2982G>A), GGT765GGG (2996T>G), ACT766ACA (2999T>A), TTA768CTA (3003T>C), AGC770AGT (3011C>T), ATC771GTT (3012A>G 3014C>T), TCT773TCC (3020T>C), CCG774CCC (3023G>C), GGC776GGT (3029C>T), ATA777ATT (3032A>T), GAC781GAT (3044C>T), GGT782GGA (3047T>A), ACG783ACC (3050G>C), CTG784TTG (3051C>T), CCA786CCT (3059A>T), TCC787TCT (3062C>T), TTG788CTA (3063T>C 3065G>A), GGA790GGG (3071A>G), ATA792ATT (3077A>T), CGC794CGA (3083C>A), ACT800ACG (3101T>G), GGC801GGT (3104C>T), ACT802ACG (3107T>G), GTG804GTC (3113G>C), ATT807ATG (3122T>G), ATT809ATC (3128T>C), GAG810GAA (3131G>A), TTC814TTT (3143C>T), CAT815CAC (3146T>C), GGT817GGA (3152T>A), TTG819TTA (3158G>A), GGA822GGT (3167A>T), CTG823TTG (3168C>T), ACA825ACC (3176A>C), CTG826CTT (3179G>T), AAC827AAT (3182C>T), AGC829AAC (3187G>A), ACC832ACT (3197C>T), GAC835GAT (3206C>T), GTT842GTG (3227T>G), ATT843ATA (3230T>A), AAC845AAT (3236C>T), CTT846CTG (3239T>G), GAG847GAA (3242G>A), GGA849GGC (3248A>C), CGA850CGT (3251A>T), AAG851AGG (3253A>G), AGG855AAG (3265G>A), TGT856TGC (3269T>C), ACG857TCT (3270A>T 3272G>T), ACC859ACT (3278C>T), AAA864AAG (3293A>G), CTT866CTA (3299T>A), ACC868TCT (3303A>T 3305C>T), GGT870GGA (3311T>A), CTA875TTA (3324C>T), CCG876CCT (3329G>T), CAC883CAT (3350C>T), AGG885CGG (3354A>C), TTG886TTA (3359G>A), ACA889ACT (3368A>T), GTC890GTG (3371C>G), TTG891CTC (3372T>C 3374G>C), GTA895GTT (3386A>T), TCA898TCC (3395A>C), CTA901CTG (3404A>G), ACG903ACA (3410G>A), GTC904GTT (3413C>T), CAG909CAA (3428G>A), CTT910TTG (3429C>T 3431T>G), AAA911AAG (3434A>G), TGC912TGT (3437C>T), ATA913ATT (3440A>T), TTG916CTA (3447T>C 3449G>A), GTC917GTA (3452C>A), CTT918CTA (3455T>A), GGG919GGA (3458G>A), GGC920GGT (3461C>T), TCT923TCC (3470T>C), GTT924GTC (3473T>C), CCC926CCT (3479C>T), ATT927ATC (3482T>C), TAC928TAT (3485C>T), GGA929GGG (3488A>G), CAC930CAT (3491C>T), GGA934GGG (3503A>G), AAT936AGT (3508A>G), GCT937GCC (3512T>C), GTG938GTT (3515G>T), CTA941TTG (3522C>T 3524A>G), TTC942TTT (3527C>T), GAG944GAA (3533G>A), TCT948ACT (3543T>A), CGC950CGT (3551C>T), ATT953ATC (3560T>C), AAA955CAA (3564A>C), TTA956ATG (3567T>A 3569A>G), AAG958AAA (3575G>A), GAA959GAG (3578A>G), AAC960AAT (3581C>T), ACA963ACC (3590A>C), GGA968GGG (3605A>G), AAG970AAA (3611G>A), CCA972CCT (3617A>T), CAG973CAA (3620G>A), GTA974GTT (3623A>T), GTG976GTC (3629G>C), CCG977CAG (3631C>A), CTC978CCT (3634T>C 3635C>T), AGA979AGG (3638A>G), GAC980GAT (3641C>T), TCA983TCT (3650A>T), GGG985GGT (3656G>T), GCT986GCC (3659T>C), GTA987GTG (3662A>G), GTG991GTC (3674G>C), GAC996GAT (3689C>T), CGA999CGC (3698A>C), TTG1001CTA (3702T>C 3704G>A), GTT1002GTG (3707T>G), TTA1003CTA (3708T>C), CCA1004CCC (3713A>C), TGC1005TGT (3716C>T), GCA1006GCG (3719A>G), CCC1007CCA (3722C>A), TCC1009TCG (3728C>G), CAG1014CCC (3742A>C 3743G>C), GGG1015GGA (3746G>A), GCA1016GCT (3749A>T), CTG1017CTA (3752G>A), GTA1018GTG (3755A>G), ATC1022ATA (3767C>A), ACA1023ACC (3770A>C), TGC1024TGT (3773C>T), CCA1025CCT (3776A>T), TTC1026TTT (3779C>T), TTC1033TTT (3800C>T), GGA1038GGT (3815A>T), TAC1042TAT (3827C>T), ATT1044ATA (3833T>A), GTA1046GTG (3839A>G), AGA1048AGG (3845A>G), GTG1049GTA (3848G>A), CAG1050CAA (3851G>A), ACT1051GCC (3852A>G 3854T>C), TCC1052TCA (3857C>A), ATA1055ATC (3866A>C), GGA1056GGT (3869A>T), GGA1057GGT (3872A>T), CTG1059CTC (3878G>C), AAC1060AAT (3881C>T), ATC1061GTA (3882A>G 3884C>A), ACC1062GCC (3885A>G), TTA1063TTG (3890A>G), GAT1064GAC (3893T>C), TCA1065TCT (3896A>T), CCT1071CCA (3914T>A), GGA1073GGT (3920A>T), ATC1074ATT (3923C>T), AAG1076CGA (3927A>C 3928A>G 3929G>A), GGC1077GGA (3932C>A), ACC1078GCT (3933A>G 3935C>T), GTT1080GTG (3941T>G), GTC1081GTA (3944C>A), TCT1082TCC (3947T>C), GCT1083GCA (3950T>A), GGT1084GGA (3953T>A), GCA1087ACA (3960G>A), GCT1090GCA (3971T>A), GGT1094GGA (3983T>A), ATG1095GTT (3984A>G 3986G>T), AGC1096AAT (3988G>A 3989C>T), GAC1097AGC (3990G>A 3991A>G), AAC1098AAT (3995C>T), ATC1099ATT (3998C>T), TTT1100TTC (4001T>C), CAT1105CAG (4016T>G), GAG1108GAA (4025G>A), TTT1109TTC (4028T>C), TTT1110TTC (4031T>C), CCT1111CCG (4034T>G), AGA1112CGT (4035A>C 4037A>T), CGC1113CGT (4040C>T), GCC1117GCT (4052C>T), ATA1120GTT (4059A>G 4061A>T), CCA1122CCT (4067A>T), AAT1123GGC (4068A>G 4069A>G 4070T>C), GCT1124GTG (4072C>T 4073T>G), AGA1126CGA (4077A>C), ACT1129ACA (4088T>A), CTG1130CTA (4091G>A), CGA1133CGC (4100A>C), CTA1134CTT (4103A>T), GGA1135GGC (4106A>C), CTC1137CTT (4112C>T), AAA1144AAG (4133A>G), ATA1147ATC (4142A>C), AGC1148AGT (4145C>T), CTG1150ATA (4149C>A 4151G>A), GAA1151GAG (4154A>G), ATC1152ATA (4157C>A), AAG1155AAA (4166G>A), TTC1160TTT (4181C>T), TTG1162TTA (4187G>A), CAA1163GAA (4188C>G), GCT1165GCA (4196T>A), TCA1169TCT (4208A>T), GCA1170GCT (4211A>T), AAG1174AAA (4223G>A), CAT1175TAT (4224C>T), TTG1176CTT (4227T>C 4229G>T), ACC1183ACT (4250C>T), CTT1185CTG (4256T>G), ACT1187ACC (4262T>C), CCT1188CCA (4265T>A), TCA1191TCT (4274A>T), AGG1194CGG (4281A>C), AGT1195GGA (4284A>G 4286T>A), TTT1196TTC (4289T>C) | | | | | | | | |

#### Proteins

| | Begin | End | Coverage | Score | Concordance | Matches | Identities | I/D/M/F* | Stop Codons |
|---|---|---|---|---|---|---|---|---|---|
| Orf1 (YP_001040017.1) | 1 | 188 | 100% | 1080 | 89.3% | 188 (100%) | 162 (86.2%) | 0/0/0/0 | 2 |
| Protein mutations: | L11V (212C>G), T50A (329A>G), V59I (356G>A), N64T (372A>C), L90F (451A>T), Q108H (505G>C), R109K (507G>A 508G>A), I110V (509A>G 511T>A), V112I (515G>A), A124E (552C>A 553T>G), A125K (554G>A 555C>A), Q128R (564A>G 565A>T), Q130R (570A>G), T132K (576C>A), L133V (578C>G), E138D (595G>T), D141A (603A>C 604C>T), R146Q (618G>A 619C>A), E151* (632G>T), L155A (644C>G 645T>C), L158Q (654T>A), S162A (665T>G), L168I (683T>A 685G>A), N170S (689A>T 690A>C), L172P (696T>C), E184K (731G>A 733A>G) | | | | | | | | |
| Codon mutations: | TCA2TCC (187A>C), CTT11GTT (212C>G), GAG14GAA (223G>A), CAC17CAT (232C>T), AAG18AAA (235G>A), CAA19CAG (238A>G), GCA21GCT (244A>T), GAC22GAT (247C>T), TCG28TCA (265G>A), GTT29GTA (268T>A), GGC32GGT (277C>T), TCT33TCC (280T>C), ATA35ATC (286A>C), ATA36ATC (289A>C), AAC37AAT (292C>T), CCA40CCT (301A>T), ACT41ACC (304T>C), CTA42TTG (305C>T 307A>G), TTC44TTT (313C>T), AAG45AAA (316G>A), CCC48CCT (325C>T), ACA50GCA (329A>G), GGT51GGA (334T>A), GCA53GCT (340A>T), CTA57CTT (352A>T), GTT59ATT (356G>A), AAC63AAT (370C>T), AAT64ACT (372A>C), AGC65AGT (376C>T), CTG68TTA (383C>T 385G>A), GTT69GTA (388T>A), TAT73TAC (400T>C), TTA74CTC (401T>C 403A>C), ACT79ACG (418T>G), GGC84GGT (433C>T), AAG86AAA (439G>A), TCT88TCA (445T>A), GGC89GGA (448C>A), TTA90TTT (451A>T), AGA94CGA (461A>C), ACC100ACT (481C>T), ACC101ACA (484C>A), AGA102AGG (487A>G), CTA105CTG (496A>G), GCT107GCA (502T>A), CAG108CAC (505G>C), AGG109AAA (507G>A 508G>A), ATT110GTA (509A>G 511T>A), GTA112ATA (515G>A), TCC113TCT (520C>T), TCC120TCT (541C>T), GCT122GCG (547T>G), GCT124GAG (552C>A 553T>G), GCA125AAA (554G>A 555C>A), CAA128CGT (564A>G 565A>T), CCA129CCT (568A>T), CAA130CGA (570A>G), ACG132AAG (576C>A), CTT133GTT (578C>G), CAA135CAG (586A>G), TTA136CTT (587T>C 589A>T), GAG138GAT (595G>T), CTT140TTG (599C>T 601T>G), GAC141GCT (603A>C 604C>T), GAG142GAA (607G>A), GCC143GCT (610C>T), AAA144AAG (613A>G), CGC146CAA (618G>A 619C>A), TTA147TTG (622A>G), AAA150AAG (631A>G), GAG151TAG (632G>T), AAA152AAG (637A>G), CTT155GCT (644C>G 645T>C), CTA158CAA (654T>A), TCC159TCA (658C>A), TCC162GCC (665T>G), AAA163AAG (670A>G), TTA164TTG (673A>G), TTG166TTA (679G>A), TTG168ATA (683T>A 685G>A), AAC170TCC (689A>T 690A>C), CTA172CCA (696T>C), GAT175GAC (706T>C), GAA184AAG (731G>A 733A>G) | | | | | | | | |
| polyprotein (YP_001040018.1) | 1 | 1199 | 100% | 7582 | 91.6% | 1199 (100%) | 1081 (90.2%) | 0/0/0/0 | 3 |

| | Begin | End | Coverage | Score | Concordance | Matches | Identities | I/D/M/F* | Stop Codons |
|---|---|---|---|---|---|---|---|---|---|
| **NT** | **20** | **5403** | **99.6%** | **7578** | **70.4%** | **5384 (99.9%)** | **4594 (85.3%)** | **3/0** | |

Protein mutations:

M2T (706T>C), K11R (733A>G), S17G (750A>G), Y48F (844A>T), V132I (1095G>A), T228A (1383A>G 1385C>T), S230* (1390C>G), T232S (1395A>T 1397T>A), D233E (1400T>G), Q242P (1426A>C 1427G>T), D245N (1434G>A), S252T (1456G>C 1457C>T), A254S (1461G>T), R255K (1465G>A), I257M (1472C>G), S275P (1524T>C 1526C>T), S281F (1543C>T), T282A (1545A>G 1547C>T), H286Q (1559C>A), Q298D (1593C>G 1595G>T), S301N (1603G>A), R302G (1605A>G), Y303Q (1608T>C 1610C>G), G307A (1621G>C 1622T>A), D309S (1626G>T 1627A>C 1628T>C), K317R (1651A>G), L321Y (1662C>T 1663T>A 1664G>T), H325Y (1674C>T 1676T>C), V326I (1677G>A 1679G>T), I332V (1695A>G), V365I (1794G>A), C366G (1797T>G 1799C>G), K370N (1811A>C), I373K (1819T>A 1820C>G), K377N (1832G>C), H381R (1843A>G), H388Y (1863C>T 1865C>T), V392M (1875G>A), I393V (1878A>G), I399M (1898T>G), V400T (1899G>A 1900T>C 1901G>A), A402V (1906C>T 1907C>G), E409D (1928A>C), N413D (1938A>G), D429N (1986G>A), F434Y (2002T>A), L450I (2049T>A 2051A>T), K453S (2059A>G 2060A>C), L458Q (2074T>A), V461A (2083T>C 2084G>A), Q465D (2094C>G 2096G>C), A466E (2098C>A 2099G>A), R467M (2100C>A 2101G>T 2102C>G), R469E (2106A>G 2107G>A), F470L (2111T>G), K472R (2115A>C 2116A>G), E473A (2119A>C 2120G>A), A475V (2125C>T 2126C>T), R476V (2127A>G 2128G>T), A477T (2130G>A 2132T>C), V479L (2136G>C 2138G>T), N481K (2144C>G), V488G (2164T>G), E490V (2170A>T 2171A>T), S493A (2178T>G), V500T (2199G>A 2200T>C), M502V (2205A>G), D508E (2225T>A), M510D (2229A>G 2230T>A 2231G>T), E517D (2252A>C), A519T (2256G>A), N520Q (2259A>C 2261T>A), E521Q (2262G>C), E531D (2294A>C), N538K (2315T>A), G540A (2320G>C 2321T>C), T542I (2326C>T), S579A (2436T>G), S590T (2469T>A 2471C>G), R601K (2503G>A), I608L (2523A>C), I643M (2630C>G), C667* (2702T>A), L699F (2798G>T), I712V (2835A>G 2837C>T), Q715H (2846G>T), A761T (2982G>A), I771V (3012A>G 3014C>T), I807M (3122T>G), S829N (3187G>A), K851R (3253A>G), R855K (3265G>A), T857S (3270A>T 3272G>T), T868S (3303A>T 3305C>T), N936S (3508A>G), S948T (3543T>A), K955Q (3564A>C), L956M (3567T>A 3569A>G), P977Q (3631C>A), L978P (3634T>C 3635C>T), Q1014P (3742A>C 3743G>C), T1051A (3852A>G 3854T>C), I1061V (3882A>G 3884C>A), T1062A (3885A>G), K1076R (3927A>C 3928A>G 3929G>A), T1078A (3933A>G 3935C>T), A1087T (3960G>A), M1095V (3984A>G 3986G>T), S1096N (3988G>A 3989C>T), D1097S (3990G>A 3991A>G), H1105Q (4016T>G), I1120V (4059A>G 4061A>T), N1123G (4068A>G 4069A>G 4070T>C), A1124V (4072C>T 4073T>G), L1150I (4149C>A 4151G>A), Q1163E (4188C>G), H1175Y (4224C>T), S1195G (4284A>G 4286T>A)

Codon mutations:

ATG2ACG (706T>C), AGG10AGA (731G>A), AAG11AGG (733A>G), AGT17GGT (750A>G), GCA18GCT (755A>T), GCT19GCA (758T>A), GTA20GTG (761A>G), ACA29ACT (788A>T), CTG39CTA (818G>A), CTT40CTC (821T>C), CGC41CGG (824C>G), GGC44GGT (833C>T), TCA47TCT (842A>T), TAT48TTT (844A>T), ATT50ATC (851T>C), CTC51CTA (854C>A), GGC53GGG (860C>G), GCC54GCT (863C>T), CAA55CAG (866A>G), GCC59GCT (878C>T), GAA61GAG (884A>G), CTG62CTT (887G>T), GAC64GAT (893C>T), AAT70AAC (911T>C), TCT76TCA (929T>A), TCA77TCT (932A>T), GGA89GGG (968A>G), GTT103GTG (1010T>G), AGT120AGC (1061T>C), CTG128CTA (1085G>A), GTT132ATT (1095G>A), TTG133TTA (1100G>A), TCC136TCT (1109C>T), GGG138GGA (1115G>A), GCA139GCC (1118A>C), TTA143TTG (1130A>G), CCG146CCA (1139G>A), ACC147ACT (1142C>T), CTA150CTG (1151A>G), AAA154AAG (1163A>G), GGC156GGG (1169C>G), TTC159TTT (1178C>T), ACA162ACT (1187A>T), ACT168ACA (1205T>A), CCT173CCC (1220T>C), GCC178GCA (1235C>A), CAA182CAG (1247A>G), CGT187CGC (1262T>C), CTT204CTC (1313T>C), CAC205CAT (1316C>T), CGA206AGG (1317C>A 1319A>G), CAA208CAG (1325A>G), TTT209TTC (1328T>C), TTC210TTT (1331C>T), CAG214CAA (1343G>A), CAG218CAA (1355G>A), ATA220ATT (1361A>T), TCA222TCT (1367A>T), GTA223GTT (1370A>T), TCT225TCA (1376T>A), ACC228GCT (1383A>G 1385C>T), TCC229TCT (1388C>T), TCA230TGA (1390C>G), GCA231GCC (1394A>C), ACT232TCA (1395A>T 1397T>A), GAT233GAG (1400T>G), TCA234TCT (1403A>T), GTG235GTA (1406G>A), CTT236TTA (1407C>T 1409T>A), CCG237CCT (1412G>T), GTG239GTA (1418G>A), CAG242CCT (1426A>C 1427G>T), GAC245AAC (1434G>A), ACA246ACC (1439A>C), GTG250GTT (1451G>T), CCA251CCT (1454A>T), AGC252ACT (1456G>C 1457C>T), GAA253GAG (1460A>G), GCA254TCA (1461G>T), AGA255AAA (1465G>A), ATC257ATG (1472C>G), CGG258CGC (1475G>C), TCC259TCG (1478C>G), TCC260TCT (1481C>T), CTC262CTG (1487C>G), CGG263CGA (1490G>A), GTG264GTA (1493G>A), CCC265CCT (1496C>T), CTC267CTT (1502C>T), AGT271AGC (1514T>C), TCC275CCT (1524T>C 1526C>T), TCA276TCT (1529A>T), ATA279ATT (1538A>T), TCT281TTT (1543C>T), ACC282GCT (1545A>G 1547C>T), ACT284ACC (1553T>C), CAC286CAA (1559C>A), CTC287TTA (1560C>T 1562C>A), GGC289GGG (1568C>G), ACG291ACA (1574G>A), CGT292AGG (1575C>A 1577T>G), GTA296GTG (1589A>G), CAG298GAT (1593C>G 1595G>T), AGT301AAT (1603G>A), AGG302GGG (1605A>G), TAC303CAG (1608T>C 1610C>G), GAA304GAG (1613A>G), GAG305GAA (1616G>A), GGT307GCA (1621G>C 1622T>A), GAT309TCC (1626G>T 1627A>C 1628T>C), GGG310GGA (1631G>A), GTT311GTA (1634T>A), CCC314CCT (1643C>T), GTT315GTC (1646T>C), AAG317AGG (1651A>G), GCT318GCA (1655T>A), GGT320GGC (1661T>C), CTG321TAT (1662C>T 1663T>A 1664G>T), AAC322AAT (1667C>T), CAT325TAC (1674C>T 1676T>C), GTG326ATT (1677G>A 1679G>T), GAT328GAC (1685T>C), AAT329AAC (1688T>C), ATT332GTT (1695A>G), GAG333GAA (1700G>A), CTG340CTC (1721G>C), CCC341CCT (1724C>T), GAA342GAG (1727A>G), AAC343AAT (1730C>T), CTG347TTG (1740C>T), TCG348TCA (1745G>A), GTG349GTA (1748G>A), AGA350CGA (1749A>C), CAG351CAA (1754G>A), CAA352CAG (1757A>G), GTT353GTG (1760T>G), GCT354GCA (1763T>A), GCA355GCG (1766A>G), GCA356GCT (1769A>T), GCA359GCC (1778A>C), GCA361GCT (1784A>T), AAG363AAA (1790G>A), GGT364GGA (1793T>A), GTT365ATT (1794G>A), TGC366GGG (1797T>G 1799C>G), GTC368GTT (1805C>T), AAA370AAC (1811A>C), CAT371CAC (1814T>C), ATA372ATT (1817A>T), ATC373AAG (1819T>A 1820C>G), AAC374AAT (1823C>T), AAG377AAC (1832G>C), ATC380ATT (1841C>T), CAC381CGC (1843A>G), TTG382CTT (1845T>C 1847G>T), ATC385ATT (1856C>T), ACC386ACT (1859C>T), CAC388TAT (1863C>T 1865C>T), GTG392ATG (1875G>A), ATA393GTA (1878A>G), TAC397TAT (1892C>T), ATT399ATG (1898T>G), GTG400ACA (1899G>A 1900T>C 1901G>A), GCC402GTG (1906C>T 1907C>G), TCT404TCA (1913T>A), GAA409GAC (1928A>C), AAC410AAT (1931C>T), AAT413GAT (1938A>G), CTT414TTG (1941C>T 1943T>G), ACC415ACA (1946C>A), TTG416CTG (1947T>C), GCA417GCC (1952A>C), CAG418CAA (1955G>A), AGA419AGG (1958A>G), GAG422GAA (1967G>A), CTA423CTG (1970A>G), CAA426CAG (1979A>G), GAT429AAT (1986G>A), TAT431TAC (1994T>C), TTT434TAT (2002T>A), AAC437AAT (2012C>T), TAT441TAC (2024T>C), CCT442CCA (2027T>A), TTA450ATT (2049T>A 2051A>T), ACG451ACT (2054G>T), AAA453AGC (2059A>G 2060A>C), CGC455CGG (2066C>G), CTA458CAA (2074T>A), GTG461GCA (2083T>C 2084G>A), GCA462GCC (2087A>C), CAG465GAC (2094C>G 2096G>C), GCG466GAA (2098C>A 2099G>A), CGC467ATG (2100C>A 2101G>T 2102C>G), AGA469GAA (2106A>G 2107G>A), TTT470TTG (2111T>G), TAC471TAT (2114C>T), AAG472CGG (2115A>C 2116A>G), GAG473GCA (2119A>C 2120G>A), GCG474GCT (2123G>T), GCC475GTT (2125C>T 2126C>T), AGG476GTG (2127A>G 2128G>T), GCT477ACC (2130G>A 2132T>C), CAG478CAA (2135G>A), GTG479CTT (2136G>C 2138G>T), AAC481AAG (2144C>G), ACT486ACA (2159T>A), AGT487AGC (2162T>C), GTG488GGG (2164T>G), GAA490GTT (2170A>T 2171A>T), ATA491ATT (2174A>T), CCA492CCT (2177A>T), TCC493GCC (2178T>G), ACC494ACT (2183C>T), TCA495TCC (2186A>C), TTC496TTT (2189C>T), ACA498ACC (2195A>C), AAG499AAA (2198G>A), GTT500ACT (2199G>A 2200T>C), GCC501GCT (2204C>T), ATG502GTG (2205A>G), GAG503GAA (2210G>A), GGC505GGG (2216C>G), AGT506AGC (2219T>C), GAT508GAA (2225T>A), AGC509AGT (2228C>T), ATG510GAT (2229A>G 2230T>A 2231G>T), GCT513GCA (2240T>A), ATT514ATC (2243T>C), GAA517GAC (2252A>C), GCC519ACC (2256G>A), AAT520CAA (2259A>C 2261T>A), GAA521CAA (2262G>C), GTA523GTG (2270A>G), GTT528GTC (2285T>C), GAA531GAC (2294A>C), CAA534CAG (2303A>G), AAT537AAC (2312T>C), AAT538AAA (2315T>A), GGT540GCC (2320G>C 2321T>C), GCG541GCC (2324G>C), ACA542ATA (2326C>T), TTA546TTG (2339A>G), CAA548CAG (2345A>G), TTC558TTT (2375C>T), GAG564GAA (2393G>A), AGT571AGC (2414T>C), GTT573GTG (2420T>G), GAG576GAA (2429G>A), GTG577GTA (2432G>A), TCA579GCA (2436T>G), GCG581GCC (2444G>C), CTC582CTA (2447C>A), GCG584GCA (2453G>A), CTC586CTG (2459C>G), TCC590ACG (2469T>A 2471C>G), AAG591AAA (2474G>A), GGA596GGC (2489A>C), TCG597TCA (2492G>A), CAA598CAG (2495A>G), CTA600CTG (2501A>G), AGG601AAG (2503G>A), TAT602TAC (2507T>C), GCC605GCT (2516C>T), ATT608CTT (2523A>C), TAT615TAC (2546T>C), TCC617TCT (2552C>T), GGT620GGG (2561T>G), ATT622ATA (2567T>A), TCG623TCA (2570G>A), GGT624GGA (2573T>A), GTT629GTA (2588T>A), CTC636CTA (2609C>A), TTC642TTT (2627C>T), ATC643ATG (2630C>G), CTG648CTC (2645G>C), CAT653CAC (2660T>C), TCA657TCT (2672A>T), GGG664GGA (2693G>A), TGT667TGA (2702T>A), ACA669ACG (2708A>G), GGT672GGC (2717T>C), ATC673ATA (2720C>A), TTC676TTT (2729C>T), GTT677GTG (2732T>G), GCC682GCT (2747C>T), GGG684GGC (2753G>C), CAC697CAT (2792C>T), CCC698CCT (2795C>T), TTG699TTT (2798G>T), GAG706GAA (2819G>A), ATC712GTT (2835A>G 2837C>T), CAT713CAC (2840T>C), CTG714TTG (2841C>T), CAG715CAT (2846G>T), AAA717AAG (2852A>G), TCG720TCC (2861G>C), ACC721ACT (2864C>T), ATC723ATA (2870C>A), CCT726CCA (2879T>A), CCT727CCG (2882T>G), CGT728CGG (2885T>G), TTG729CTG (2886T>C), GCA731GCG (2894A>G), GCA733GCT (2900A>T), GGT736GGC (2909T>C), AAA738AAG (2915A>G), GAC740GAT (2921C>T), ACA742ACG (2927A>G), CCA745CCC (2936A>C), CGA746CGG (2939A>G), ACA749ACG (2948A>G), CAG751CAA (2954G>A), GTA752GTT (2957A>T), CTG753CTA (2960G>A), CAC755CAT (2966C>T), AAT757AAC (2972T>C), GTG760GTT (2981G>T), GCT761ACT (2982G>A), GGT765GGG (2996T>G), ACT766ACA (2999T>A), TTA768CTA (3003T>C), AGC770AGT (3011C>T), ATC771GTT (3012A>G 3014C>T), TCT773TCC (3020T>C), CCG774CCC (3023G>C), GGC776GGT (3029C>T), ATA777ATT (3032A>T), GAC781GAT (3044C>T), GGT782GGA (3047T>A), ACG783ACC (3050G>C), CTG784TTG (3051C>T), CCA786CCT (3059A>T), TCC787TCT (3062C>T), TTG788CTA (3063T>C 3065G>A), GGA790GGG (3071A>G), ATA792ATT (3077A>T), CGC794CGA (3083C>A), ACT800ACG (3101T>G), GGC801GGT (3104C>T), ACT802ACG (3107T>G), GTG804GTC (3113G>C), ATT807ATG (3122T>G), ATT809ATC (3128T>C), GAG810GAA (3131G>A), TTC814TTT (3143C>T), CAT815CAC (3146T>C), GGT817GGA (3152T>A), TTG819TTA (3158G>A), GGA822GGT (3167A>T), CTG823TTG (3168C>T), ACA825ACC (3176A>C), CTG826CTT (3179G>T), AAC827AAT (3182C>T), AGC829AAC (3187G>A), ACC832ACT (3197C>T), GAC835GAT (3206C>T), GTT842GTG (3227T>G), ATT843ATA (3230T>A), AAC845AAT (3236C>T), CTT846CTG (3239T>G), GAG847GAA (3242G>A), GGA849GGC (3248A>C), CGA850CGT (3251A>T), AAG851AGG (3253A>G), AGG855AAG (3265G>A), TGT856TGC (3269T>C), ACG857TCT (3270A>T 3272G>T), ACC859ACT (3278C>T), AAA864AAG (3293A>G), CTT866CTA (3299T>A), ACC868TCT (3303A>T 3305C>T), GGT870GGA (3311T>A), CTA875TTA (3324C>T), CCG876CCT (3329G>T), CAC883CAT (3350C>T), AGG885CGG (3354A>C), TTG886TTA (3359G>A), ACA889ACT (3368A>T), GTC890GTG (3371C>G), TTG891CTC (3372T>C 3374G>C), GTA895GTT (3386A>T), TCA898TCC (3395A>C), CTA901CTG (3404A>G), ACG903ACA (3410G>A), GTC904GTT (3413C>T), CAG909CAA (3428G>A), CTT910TTG (3429C>T 3431T>G), AAA911AAG (3434A>G), TGC912TGT (3437C>T), ATA913ATT (3440A>T), TTG916CTA (3447T>C 3449G>A), GTC917GTA (3452C>A), CTT918CTA (3455T>A), GGG919GGA (3458G>A), GGC920GGT (3461C>T), TCT923TCC (3470T>C), GTT924GTC (3473T>C), CCC926CCT (3479C>T), ATT927ATC (3482T>C), TAC928TAT (3485C>T), GGA929GGG (3488A>G), CAC930CAT (3491C>T), GGA934GGG (3503A>G), AAT936AGT (3508A>G), GCT937GCC (3512T>C), GTG938GTT (3515G>T), CTA941TTG (3522C>T 3524A>G), TTC942TTT (3527C>T), GAG944GAA (3533G>A), TCT948ACT (3543T>A), CGC950CGT (3551C>T), ATT953ATC (3560T>C), AAA955CAA (3564A>C), TTA956ATG (3567T>A 3569A>G), AAG958AAA (3575G>A), GAA959GAG (3578A>G), AAC960AAT (3581C>T), ACA963ACC (3590A>C), GGA968GGG (3605A>G), AAG970AAA (3611G>A), CCA972CCT (3617A>T), CAG973CAA (3620G>A), GTA974GTT (3623A>T), GTG976GTC (3629G>C), CCG977CAG (3631C>A), CTC978CCT (3634T>C 3635C>T), AGA979AGG (3638A>G), GAC980GAT (3641C>T), TCA983TCT (3650A>T), GGG985GGT (3656G>T), GCT986GCC (3659T>C), GTA987GTG (3662A>G), GTG991GTC (3674G>C), GAC996GAT (3689C>T), CGA999CGC (3698A>C), TTG1001CTA (3702T>C 3704G>A), GTT1002GTG (3707T>G), TTA1003CTA (3708T>C), CCA1004CCC (3713A>C), TGC1005TGT (3716C>T), GCA1006GCG (3719A>G), CCC1007CCA (3722C>A), TCC1009TCG (3728C>G), CAG1014CCC (3742A>C 3743G>C), GGG1015GGA (3746G>A), GCA1016GCT (3749A>T), CTG1017CTA (3752G>A), GTA1018GTG (3755A>G), ATC1022ATA (3767C>A), ACA1023ACC (3770A>C), TGC1024TGT (3773C>T), CCA1025CCT (3776A>T), TTC1026TTT (3779C>T), TTC1033TTT (3800C>T), GGA1038GGT (3815A>T), TAC1042TAT (3827C>T), ATT1044ATA (3833T>A), GTA1046GTG (3839A>G), AGA1048AGG (3845A>G), GTG1049GTA (3848G>A), CAG1050CAA (3851G>A), ACT1051GCC (3852A>G 3854T>C), TCC1052TCA (3857C>A), ATA1055ATC (3866A>C), GGA1056GGT (3869A>T), GGA1057GGT (3872A>T), CTG1059CTC (3878G>C), AAC1060AAT (3881C>T), ATC1061GTA (3882A>G 3884C>A), ACC1062GCC (3885A>G), TTA1063TTG (3890A>G), GAT1064GAC (3893T>C), TCA1065TCT (3896A>T), CCT1071CCA (3914T>A), GGA1073GGT (3920A>T), ATC1074ATT (3923C>T), AAG1076CGA (3927A>C 3928A>G 3929G>A), GGC1077GGA (3932C>A), ACC1078GCT (3933A>G 3935C>T), GTT1080GTG (3941T>G), GTC1081GTA (3944C>A), TCT1082TCC (3947T>C), GCT1083GCA (3950T>A), GGT1084GGA (3953T>A), GCA1087ACA (3960G>A), GCT1090GCA (3971T>A), GGT1094GGA (3983T>A), ATG1095GTT (3984A>G 3986G>T), AGC1096AAT (3988G>A 3989C>T), GAC1097AGC (3990G>A 3991A>G), AAC1098AAT (3995C>T), ATC1099ATT (3998C>T), TTT1100TTC (4001T>C), CAT1105CAG (4016T>G), GAG1108GAA (4025G>A), TTT1109TTC (4028T>C), TTT1110TTC (4031T>C), CCT1111CCG (4034T>G), AGA1112CGT (4035A>C 4037A>T), CGC1113CGT (4040C>T), GCC1117GCT (4052C>T), ATA1120GTT (4059A>G 4061A>T), CCA1122CCT (4067A>T), AAT1123GGC (4068A>G 4069A>G 4070T>C), GCT1124GTG (4072C>T 4073T>G), AGA1126CGA (4077A>C), ACT1129ACA (4088T>A), CTG1130CTA (4091G>A), CGA1133CGC (4100A>C), CTA1134CTT (4103A>T), GGA1135GGC (4106A>C), CTC1137CTT (4112C>T), AAA1144AAG (4133A>G), ATA1147ATC (4142A>C), AGC1148AGT (4145C>T), CTG1150ATA (4149C>A 4151G>A), GAA1151GAG (4154A>G), ATC1152ATA (4157C>A), AAG1155AAA (4166G>A), TTC1160TTT (4181C>T), TTG1162TTA (4187G>A), CAA1163GAA (4188C>G), GCT1165GCA (4196T>A), TCA1169TCT (4208A>T), GCA1170GCT (4211A>T), AAG1174AAA (4223G>A), CAT1175TAT (4224C>T), TTG1176CTT (4227T>C 4229G>T), ACC1183ACT (4250C>T), CTT1185CTG (4256T>G), ACT1187ACC (4262T>C), CCT1188CCA (4265T>A), TCA1191TCT (4274A>T), AGG1194CGG (4281A>C), AGT1195GGA (4284A>G 4286T>A), TTT1196TTC (4289T>C)

*: Inserts / Deletes / Misaligned / Frameshifts

#### Analysis details

This analysis was performed with panviral2.64

### NGS Details (UN60): Tomato necrotic dwarf virus (segment RNA1)

#### Assembly

| | |
|---|---|
| **Coverage Length** | 6176 (1 contig(s)) |
| **Depth Of Coverage** | 5251.7 |
| **Number Of Reads** | 244103 |
| **Reads Per Million** | 5177.32 rpm (after QC) |
| **Ambiguities** | 0 |
| **Assembly Method** | de novo + reference guided assembly |
| **Consensus Caller** | Bcf Tools |

#### Coverage Map

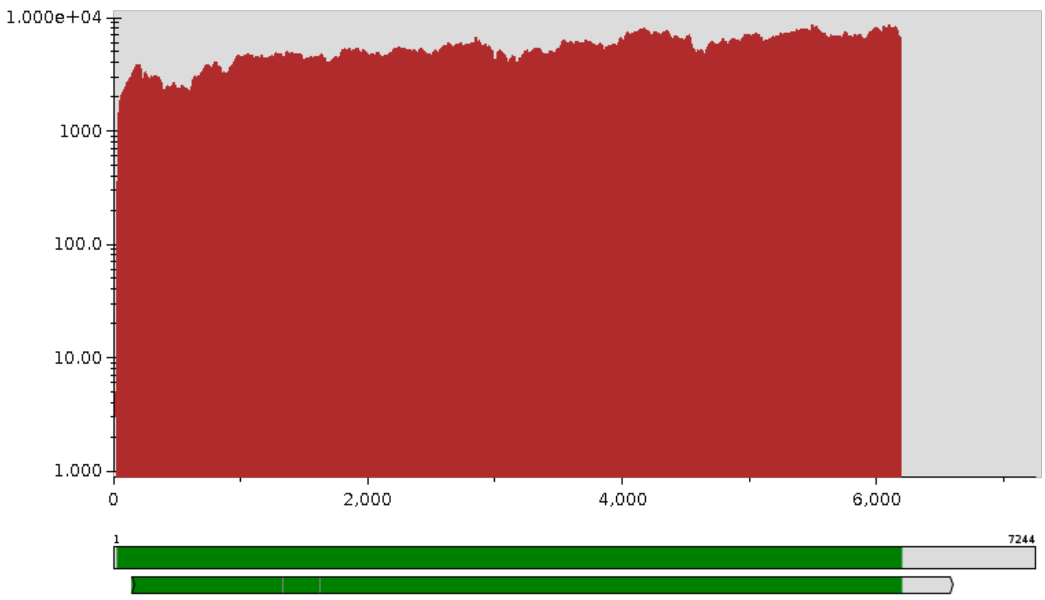

#### Assignment

| | |
|---|---|
| **Type** | Tomato necrotic dwarf virus (Taxonomy ID: 1481465) |
| **Reference Genome** | NC_027926.1 |
| **NT Identity (%)** | 64.2066 |
| **AA Identity (%)** | 65.5893 |
| **Number Of Stop Codons** | 0 |
| **Number Of CDS** | 1 |

#### Alignment

| | |
|---|---|
| **Alignment Score** | 3315.0 (NT) + 9642.0 (AA) = 12957.0 |
| **Concordance (%)** | 49.8691 |

| Alignment Method | Global, seeded, nucleotide + amino acids (AGA) |
|---|---|

#### Genome Region

Sequence starts at position 25 and ends at position 6200 relative to NC_027926.1 reference sequence.

#### Alignment Detailed Statistics

| | Begin | End | Coverage | Score | Concordance | Matches | Identities | I/D/M/F* | Stop Codons |
|---|---|---|---|---|---|---|---|---|---|
| NT | 25 | 6200 | 85.3% | 3315 | 27.4% | 6117 (98.7%) | 3941 (63.6%) | 21/59 | |

| | Begin | End | Coverage | Score | Concordance | Matches | Identities | I/D/M/F* | Stop Codons |
|---|---|---|---|---|---|---|---|---|---|
| **NT** | **25** | **6200** | **85.3%** | **3315** | **27.4%** | **6117 (98.7%)** | **3941 (63.6%)** | **21/59** | |

Mutations:

35T>C, 40G>A, 42C>T, 43C>T, 49T>C, 50C>T, 52A>C, 54T>C, 62A>C, 64T>C, 65T>A, 66G>T, 67G>A, 69A>C, 70T>A, 71T>A, 74A>C, 77G>T, 79T>C, 82A>T, 84G>T,
85_93delCTAAACTTC, 99C>A, 101T>C, 109_119delGAAACTTTTAA, 121A>C, 122C>T, 123A>C, 125T>A, 129T>C, 130T>C, 131C>T, 134C>G, 135T>A, 136T>A, 137A>T, 138T>C,
142G>A, 143T>C, 144G>A, 145A>T, 146T>A, 148A>C, 156T>A, 157C>T, 165A>G, 170T>G, 171T>G, 172G>A, 175T>C, 178T>C, 180A>T, 181T>G,
184_201delGAGAAGACTGGATCAGTT, 202C>T, 204C>T, 207T>A, 208A>C, 213T>A, 221G>C, 222T>A, 226C>G, 228G>T, 232G>T, 233G>C, 238A>T, 240C>T, 243A>T, 244A>T,
245A>C, 246A>T, 247T>C, 249C>T, 255A>G, 256G>A, 259A>C, 261T>C, 262A>T, 263G>C, 267C>T, 268A>G, 270C>T, 272C>A, 273A>T, 274T>A, 275C>A, 279A>T, 287A>C,
288A>C, 292T>G, 293C>T, 294T>A, 300T>C, 301T>A, 306T>G, 310T>G, 312G>C, 315A>G, 318A>G, 319T>C, 321T>A, 324G>T, 326C>T, 327T>G, 329A>T, 330T>A, 334T>G,
336T>A, 337T>A, 339T>A, 342T>A, 347T>C, 348T>C, 357T>C, 358T>A, 359C>A, 360C>T, 363A>T, 364T>G, 366G>A, 367A>G, 369T>A, 371A>G, 375G>T, 376C>T, 378T>A,
387A>T, 389G>C, 390T>A, 391T>A, 392C>A, 393C>A, 394T>A, 396G>A, 399T>G, 405T>C, 408T>C, 409C>A, 411G>C, 414A>G, 416C>G, 417A>T, 420C>T, 421T>G, 422C>A,
427T>G, 429G>A, 430G>A, 431C>A, 435C>T, 445T>A, 450C>A, 451A>G, 452A>T, 453G>A, 457A>C, 459G>T, 462T>A, 463T>C, 465A>T, 466C>A, 467G>A, 469T>A, 472T>G,
474C>T, 475G>T, 476A>T, 481T>G, 483G>C, 484T>A, 485C>A, 486C>G, 487A>G, 493G>T, 495T>C, 497G>T, 498A>C, 499A>C, 506T>A, 507T>C, 508C>G, 510G>A, 514G>T,
516T>A, 522G>A, 525G>A, 526A>G, 527G>T, 532G>T, 533A>C, 534T>A, 535G>T, 543T>A, 546A>G, 547G>T, 550T>A, 551A>T, 552T>A, 554G>C, 561G>A, 570A>T, 571A>T,
573A>G, 574A>G, 576T>C, 577C>A, 580C>T, 582T>G, 583G>C, 585T>A, 586T>C, 588G>A, 589T>C, 593G>A, 597T>A, 604T>G, 605C>T, 609C>T, 612A>T, 613A>G, 615T>A,
616G>A, 617A>C, 618A>G, 619G>C, 620G>C, 621A>T, 622G>T, 624A>T, 625T>A, 626C>A, 629G>A, 631A>G, 632C>T, 633C>A, 634A>G, 637A>G, 638A>T, 639A>T, 641A>C,
643A>T, 645T>G, 648C>T, 650T>C, 652A>G, 654A>T, 657A>T, 660G>A, 664A>T, 666G>T, 669T>C, 673A>T, 674G>C, 675T>A, 684C>T, 687G>A, 690C>T, 699C>A, 701A>C,
702G>A, 706G>T, 710A>G, 717C>T, 718A>T, 720A>C, 721A>C, 722T>A, 724G>C, 726A>G, 729A>G, 736A>G, 737G>C, 738T>A, 739A>G, 741G>A, 744A>T, 753G>A, 754T>A,
755C>G, 756T>C, 757G>T, 759C>A, 760T>A, 762A>G, 765C>T, 768T>A, 774A>T, 775T>A, 776C>G, 780G>A, 781T>C, 783G>T, 784A>T, 786G>A, 789A>C, 790A>T, 792T>A,
793A>T, 795T>G, 796A>T, 801A>T, 810T>C, 813C>A, 816G>T, 817C>T, 819C>A, 822G>A, 826A>G, 828A>T, 831A>T, 834G>T, 837A>T, 840A>G, 846C>A, 853T>C, 855G>T,
856A>T, 857G>C, 858C>A, 861T>C, 864C>A, 867C>G, 876T>C, 885A>T, 886A>G, 891G>A, 897G>T, 900T>G, 915A>G, 916T>C, 921T>C, 922A>C, 923C>A, 924A>G, 933A>G,
936T>C, 942A>G, 948C>T, 951T>A, 954A>T, 960A>C, 963T>C, 966T>A, 972G>T, 978A>G, 981T>A, 982A>G, 984A>G, 988G>A, 990T>C, 991G>A, 992C>T, 1002C>T, 1005G>A,
1014G>A, 1015C>T, 1017T>G, 1020T>A, 1023A>G, 1026T>A, 1028G>A, 1029C>T, 1032A>T, 1035A>C, 1038G>A, 1041T>C, 1044T>C, 1048T>C, 1050G>T, 1054G>C, 1056A>G,
1063C>A, 1064A>C, 1071G>T, 1072G>T, 1073A>C, 1074T>G, 1076C>A, 1077T>G, 1081T>G, 1082G>C, 1086T>A, 1090G>T, 1094C>T, 1095C>G, 1096A>C, 1098G>C, 1099T>C,
1101G>T, 1104G>A, 1107C>T, 1113C>A, 1116T>C, 1117A>C, 1118A>C, 1119A>T, 1122C>A, 1125C>T, 1128T>A, 1129G>A, 1130A>T, 1131C>T, 1134A>G, 1138A>C, 1140T>A,
1143C>T, 1144C>T, 1146T>A, 1148G>A, 1149G>A, 1152G>A, 1158A>C, 1159C>G, 1161G>C, 1164G>A, 1167T>A, 1173A>T, 1185A>C, 1188C>T, 1194T>C, 1197C>A, 1204A>T,
1212T>C, 1213C>A, 1215A>G, 1218G>T, 1219A>T, 1220G>C, 1224A>T, 1227T>C, 1230C>A, 1233T>G, 1239T>A, 1242A>T, 1245G>A, 1246G>C, 1248C>G, 1249T>G, 1250A>T,
1254G>T, 1258A>G, 1259G>C, 1260T>C, 1261T>A, 1264A>C, 1265G>A, 1270G>T, 1271T>C, 1278T>G, 1281T>C, 1284G>A, 1290A>G, 1296A>G, 1299C>T, 1308T>G, 1316A>T,
1320T>A, 1323C>T, 1329G>A, 1344A>G, 1345G>T, 1346C>G, 1347A>T, 1353A>C, 1354T>C, 1356G>T, 1372T>C, 1374G>C, 1377T>C, 1378C>G, 1379G>C, 1380C>A, 1386G>T,
1389G>A, 1392T>A, 1398T>G, 1402A>C, 1404A>C, 1405C>T, 1407T>G, 1413T>G, 1416T>A, 1419C>T, 1422G>A, 1423C>A, 1425C>A, 1431C>A, 1440T>C, 1443T>C, 1449C>A,
1452C>T, 1455G>A, 1461T>A, 1465T>C, 1467G>T, 1476C>T, 1482A>T, 1485A>T, 1486T>C, 1488G>T, 1489A>C, 1491A>G, 1503A>G, 1506A>G, 1512A>G, 1518T>A, 1536T>C,
1542C>T, 1545A>C, 1551A>T, 1557A>T, 1560G>T, 1563C>A, 1569A>T, 1572T>A, 1575G>C, 1578A>G, 1581C>T, 1584G>A, 1587G>A, 1593T>C, 1599T>A, 1608T>C, 1617A>C,
1627C>T, 1629T>G, 1630G>A, 1635A>T, 1638A>T, 1641T>A, 1644C>T, 1656T>A, 1659C>G, 1671T>A, 1677G>A, 1678C>A, 1680T>G, 1683A>G, 1686G>A, 1692C>T, 1704T>C,
1706T>G, 1707C>A, 1712A>T, 1713A>G, 1716T>C, 1719A>T, 1723T>G, 1724C>A, 1726A>C, 1729G>A, 1731G>T, 1732G>A, 1733A>C, 1734C>T, 1740C>A, 1742G>C, 1743A>C,
1749G>A, 1752G>T, 1756A>G, 1758C>T, 1759A>G, 1761T>G, 1771C>A, 1773G>A, 1776G>A, 1788A>C, 1798C>G, 1799A>C, 1800C>T, 1804A>T, 1805G>C, 1806T>C, 1809C>T,
1810T>G, 1811C>A, 1812G>T, 1821A>G, 1822G>C, 1824C>A, 1836C>A, 1842T>C, 1844A>G, 1848T>G, 1851T>C, 1852G>T, 1853C>A, 1858A>G, 1860A>G, 1865A>G, 1866G>C,
1869G>A, 1872G>T, 1874T>A, 1875T>C, 1876G>A, 1878A>T, 1879G>A, 1881A>T, 1882G>A, 1883A>C, 1885A>T, 1887T>G, 1890C>A, 1891A>T, 1893T>G, 1894T>G, 1896A>T,
1897A>G, 1899T>C, 1903A>G, 1904C>T, 1905A>T, 1908C>T, 1914T>C, 1915G>A, 1916C>A, 1920A>T, 1921T>G, 1924T>G, 1925C>T, 1929A>T, 1932G>A, 1935T>G, 1936G>C,
1940C>A, 1941A>T, 1945G>T, 1947T>C, 1948G>C, 1950T>A, 1952G>A, 1953G>T, 1956T>C, 1957_1959delCGC, 1962C>A, 1963A>C, 1966A>G, 1967C>T, 1968A>T, 1969A>T,
1970A>T, 1971A>G, 1973A>C, 1974A>T, 1975G>C, 1976A>C, 1979T>A, 1981C>T, 1984A>G, 1985T>C, 1988T>C, 1989G>T, 1990A>T, 1991C>A, 1993C>G, 1994C>A, 1995C>G,
1996A>C, 1997C>A, 1998A>G, 1999G>A, 2000T>C, 2001T>A, 2004A>T, 2010G>A, 2012A>C, 2013A>T, 2016T>A, 2018G>A, 2020G>A, 2022G>T, 2023A>G, 2031T>C, 2032A>C,
2034A>G, 2038T>C, 2040A>C, 2046G>T, 2049A>G, 2055T>C, 2060T>A, 2067T>C, 2070G>A, 2073C>G, 2077T>A, 2080G>A, 2082C>T, 2083C>A, 2084C>A, 2085G>A, 2086G>T,
2087A>T, 2088G>T, 2089A>G, 2090G>A, 2091T>G, 2094G>T, 2097T>C, 2100T>A, 2107T>C, 2108A>T, 2109T>C, 2110C>T, 2111A>C, 2112A>C, 2115A>G, 2118A>G, 2122C>A,
2126T>A, 2131T>G, 2132C>T, 2133C>G, 2137G>T, 2138A>C, 2140G>T, 2142A>T, 2145G>T, 2146C>G, 2147A>C, 2149C>A, 2151G>A, 2154A>G, 2155G>A, 2156T>A, 2157C>T,
2160C>T, 2161A>G, 2162T>A, 2163G>A, 2165G>T, 2166A>G, 2170A>G, 2172G>A, 2174C>A, 2178C>T, 2179A>T, 2182A>T, 2184C>G, 2185T>G, 2186A>T, 2193T>A, 2196C>A,
2198A>G, 2200G>C, 2203G>C, 2205A>G, 2208A>G, 2211C>T, 2212A>G, 2213G>C, 2214G>T, 2225A>G, 2229T>C, 2245C>A, 2256T>C, 2259A>G, 2262C>T, 2268C>T, 2272G>A,
2273C>G, 2274T>C, 2275T>A, 2277C>A, 2283C>T, 2285G>C, 2290A>G, 2293A>C, 2294A>T, 2300C>A, 2304A>C, 2308A>C, 2309C>G, 2310A>C, 2311G>A, 2313G>T, 2314A>C,
2316G>T, 2317C>G, 2318G>A, 2319C>G, 2320A>C, 2321C>A, 2325A>G, 2326T>A, 2327T>A, 2328T>A, 2331A>G, 2332C>T, 2337T>G, 2338T>G, 2339G>C, 2340T>G, 2341A>C,
2344G>C, 2346T>G, 2354T>C, 2355T>A, 2362T>C, 2364G>T, 2366T>C, 2367T>G, 2370C>T, 2373C>T, 2374C>G, 2376G>A, 2389T>A, 2390T>G, 2391T>A, 2394G>A, 2400T>C,
2401T>C, 2409A>T, 2412C>T, 2421T>C, 2424T>C, 2432C>G, 2433C>G, 2440T>G, 2442C>A, 2442_2443insGGT, 2444A>C, 2445A>T, 2448G>C, 2449G>A, 2450C>A, 2451C>G,
2454A>G, 2455G>A, 2456A>G, 2457C>A, 2458T>A, 2459C>A, 2464G>A, 2466C>T, 2468A>T, 2469G>C, 2470A>G, 2471T>A, 2475T>C, 2476T>A, 2477A>T, 2478C>G, 2479C>A,
2480A>T, 2481C>T, 2482A>G, 2483A>G, 2484A>T, 2487T>C, 2488G>A, 2489T>A, 2493T>C, 2494T>A, 2496C>A, 2501A>T, 2502A>C, 2503G>C, 2505A>G, 2507C>T, 2508T>G,
2512T>A, 2513G>A, 2514G>T, 2515A>G, 2520A>G, 2523A>C, 2525A>T, 2526T>C, 2527G>A, 2528C>T, 2529A>C, 2530C>T, 2534G>A, 2553C>A, 2556A>G, 2560A>T, 2562C>G,
2565G>A, 2568G>A, 2577A>C, 2578T>A, 2579T>C, 2580G>C, 2581T>A, 2583T>G, 2584G>A, 2585G>A, 2586G>T, 2588G>C, 2589A>C, 2590A>G, 2592A>T, 2595A>T, 2601T>C,
2602C>A, 2605G>A, 2610T>C, 2611T>C, 2612G>T, 2613G>C, 2614C>A, 2616A>G, 2617A>G, 2618G>C, 2619C>A, 2620A>G, 2621A>C, 2622T>A, 2623T>A, 2627T>C, 2628T>C,
2631A>T, 2632A>T, 2633G>T, 2634T>A, 2638A>G, 2639G>A, 2643A>C, 2649T>G, 2652T>C, 2655C>A, 2658A>C, 2673C>T, 2679T>C, 2685A>G, 2688G>A, 2691C>T, 2694G>A,
2697C>G, 2698C>A, 2699C>A, 2700C>G, 2701T>A, 2705G>A, 2707C>T, 2709G>C, 2710A>G, 2711A>C, 2712G>A, 2715T>A, 2717T>C, 2718G>C, 2722G>C, 2724G>A, 2730G>A,
2731A>G, 2732G>C, 2733T>G, 2734A>G, 2735G>C, 2736C>A, 2738C>A, 2744T>A, 2745T>C, 2746A>C, 2747C>A, 2748A>T, 2749C>G, 2750A>T, 2751A>G, 2754A>C,
2754_2755insCGG, 2762C>A, 2763T>A, 2764G>C, 2766A>G, 2772G>A, 2778C>T, 2781C>T, 2782C>G, 2785G>A, 2786C>G, 2787A>T, 2790C>G, 2791A>C, 2793A>C, 2796G>A,
2799T>C, 2800A>T, 2803A>G, 2808A>T, 2809C>T, 2812C>A, 2814G>A, 2815C>A, 2819G>C, 2820A>T, 2823G>A, 2829G>C, 2832C>T, 2844A>G, 2850G>T, 2856A>T, 2859G>T,
2865A>T, 2866A>C, 2868G>C, 2883A>C, 2886C>T, 2887C>A, 2888A>G, 2889C>A, 2892T>C, 2893G>A, 2898A>G, 2904A>C, 2907A>T, 2914A>T, 2919T>C, 2922C>T, 2926C>G,
2927C>A, 2928T>G, 2929A>C, 2931C>T, 2934A>C, 2936G>T, 2937G>T, 2939C>G, 2940A>T, 2943G>T, 2951A>T, 2955C>T, 2959C>A, 2960G>C, 2961G>C, 2962G>C, 2967C>A,
2969A>G, 2971G>T, 2973C>G, 2976T>A, 2982A>T, 2989A>G, 2991C>G, 2995C>T, 2998C>T, 3000A>G, 3001C>A, 3003T>G, 3006G>A, 3009T>G, 3012C>T, 3015T>A, 3028C>T,
3030T>A, 3036T>A, 3037C>A, 3039C>A, 3042G>A, 3051A>T, 3063G>T, 3066C>G, 3069C>T, 3071T>A, 3073A>T, 3075C>T, 3078A>G, 3081T>C, 3084T>A, 3085C>A, 3086A>T,
3093A>G, 3096T>C, 3097T>G, 3098G>T, 3099T>G, 3102A>G, 3105C>T, 3108G>A, 3111G>T, 3116C>A, 3117C>G, 3123A>C, 3125G>A, 3126A>G, 3127C>A, 3128T>A, 3129G>A,
3130G>T, 3131G>C, 3135C>T, 3136C>T, 3138A>G, 3141A>G, 3142T>G, 3145T>C, 3147G>T, 3152T>A, 3168C>A, 3171A>G, 3177T>C, 3180A>C, 3182T>A, 3183C>A, 3186G>A,
3189G>C, 3190G>A, 3191A>C, 3192G>A, 3197A>G, 3198G>A, 3204C>T, 3207T>G, 3208G>A, 3209C>A, 3210C>T, 3211A>G, 3212G>C, 3213T>A, 3217A>G, 3218A>C, 3219A>T,
3220T>C, 3222G>T, 3225T>C, 3228G>A, 3231G>A, 3232C>G, 3233A>T, 3234A>T, 3235T>G, 3243A>G, 3246T>C, 3249T>C, 3252G>T, 3256G>A, 3258A>T, 3267T>A, 3276A>T,
3283T>C, 3285A>C, 3286C>T, 3288T>A, 3294C>A, 3296G>C, 3300T>G, 3309A>G, 3312A>G, 3315T>G, 3316G>T, 3317T>G, 3318T>C, 3324G>T, 3327C>A, 3330T>A, 3339T>A,
3342A>T, 3350G>A, 3353T>C, 3354T>C, 3357A>T, 3363A>G, 3369A>T, 3371T>A, 3373A>G, 3375T>G, 3376T>G, 3381C>A, 3384T>C, 3387C>A, 3390G>A, 3393C>A, 3394A>G,
3396C>G, 3399C>T, 3400C>A, 3402G>A, 3405C>T, 3408G>A, 3409C>T, 3411T>G, 3415A>G, 3417C>T, 3429G>A, 3432T>C, 3433G>A, 3440C>T, 3444A>T, 3453A>C, 3459C>A,
3460A>C, 3462T>G, 3463C>T, 3465C>G, 3468A>G, 3469G>C, 3470C>A, 3472A>C, 3474T>G, 3477G>A, 3478A>T, 3479G>C, 3480T>C, 3482G>A, 3483A>G, 3486A>G, 3487A>G,
3488C>A, 3489C>A, 3491C>G, 3492T>A, 3493C>T, 3495T>C, 3501A>G, 3507C>A, 3510A>G, 3513C>G, 3514T>G, 3516C>T, 3519T>C, 3520C>T, 3522T>G, 3531T>A, 3534T>G,
3535C>T, 3536A>T, 3537A>G, 3541T>A, 3546C>T, 3549A>T, 3555A>G, 3558C>G, 3561T>C, 3567T>A, 3570A>C, 3573C>T, 3575G>A, 3577A>C, 3578A>G, 3579G>C, 3584A>C,
3585A>C, 3586T>C, 3597A>T, 3603T>G, 3604C>G, 3605C>A, 3606T>C, 3610G>C, 3612A>G, 3615C>T, 3618T>C, 3621C>A, 3624A>C, 3627C>T, 3630C>A, 3631A>T, 3633C>T,
3635T>C, 3639T>A, 3646C>A, 3654G>A, 3657A>G, 3660C>T, 3666A>C, 3675A>T, 3681G>C, 3682A>G, 3683T>A, 3687A>G, 3690C>T, 3699T>A, 3703C>T, 3704A>C, 3705T>C,
3706G>A, 3707G>A, 3708T>C, 3714T>G, 3717T>G, 3718G>C, 3719A>C, 3720T>A, 3726T>A, 3729A>G, 3730T>C, 3733T>G, 3735C>T, 3741A>G, 3742A>G, 3743C>G, 3747C>T,
3751A>G, 3752G>C, 3753T>A, 3756A>C, 3757C>A, 3759G>T, 3762G>T, 3765A>T, 3766A>G, 3767C>T, 3768A>G, 3769A>C, 3771G>T, 3774A>G, 3776A>T, 3777C>G, 3780A>G,
3781A>C, 3782A>G, 3795C>G, 3801A>T, 3802C>G, 3803A>C, 3804G>A, 3805G>A, 3806A>T, 3807G>C, 3811G>A, 3819T>C, 3822G>A, 3828G>C, 3837T>A, 3845G>T, 3854A>T,
3860G>C, 3861C>A, 3868C>T, 3870C>G, 3876A>G, 3879C>A, 3888C>T, 3894C>T, 3901A>T, 3906C>A, 3907A>C, 3909G>T, 3912A>T, 3913A>T, 3914T>C, 3915G>T, 3921G>A,
3922C>A, 3927G>A, 3933C>G, 3935T>A, 3937C>G, 3939A>C, 3940A>C, 3941C>A, 3943G>C, 3944G>C, 3946G>A, 3948A>C, 3949G>T, 3950G>C, 3951T>G, 3952G>C, 3954T>G,
3956A>G, 3957A>G, 3960A>T, 3961G>C, 3962A>T, 3963G>T, 3966T>C, 3968G>T, 3973G>A, 3981A>T, 3984G>C, 3990A>T, 3993T>A, 3999G>A, 4002A>G, 4003C>T, 4009A>T,
4010T>C, 4011T>G, 4014A>G, 4015A>G, 4017T>A, 4023G>A, 4028T>C, 4030C>T, 4031A>T, 4035G>T, 4038T>C, 4041A>C, 4042A>G, 4047C>A, 4050A>C, 4053A>G, 4059C>G,
4062G>A, 4068G>A, 4071C>G, 4072C>T, 4074T>G, 4077T>A, 4083A>G, 4104A>G, 4107A>C, 4108A>C, 4110A>C, 4111T>C, 4113A>G, 4116T>C, 4119G>A, 4122A>C, 4137C>T,
4149C>G, 4150A>C, 4161T>C, 4173C>T, 4176C>T, 4177A>T, 4178C>G, 4179C>T, 4188A>T, 4189T>G, 4190T>C, 4197T>C, 4200C>A, 4201A>C, 4206A>T, 4209T>C, 4215T>A,
4218A>T, 4221A>C, 4222G>A, 4230C>G, 4233T>C, 4234T>C, 4235C>A, 4243T>A, 4245T>A, 4246C>G, 4248T>G, 4251T>G, 4254C>T, 4256A>G, 4257C>T, 4259A>G, 4263C>T,
4267G>T, 4269T>A, 4279A>G, 4280C>A, 4281T>G, 4284G>A, 4287T>C, 4293T>C, 4302A>G, 4305T>C, 4311C>T, 4314G>A, 4315C>A, 4317C>A, 4323A>C, 4324G>A, 4326G>C,
4329C>T, 4332T>A, 4335T>C, 4338G>A, 4351T>C, 4356T>G, 4361A>C, 4362G>T, 4364A>T, 4370A>G, 4372A>G, 4373C>A, 4374C>T, 4375A>G, 4383A>G, 4384G>A, 4385A>G,
4386G>T, 4391C>A, 4392T>G, 4395C>A, 4401A>T, 4404A>G, 4407T>C, 4408G>A, 4410T>A, 4411C>T, 4413T>A, 4414C>T, 4416C>A, 4425A>C, 4428A>T, 4434C>T, 4437C>T,
4441T>C, 4443A>C, 4450G>A, 4452T>C, 4455A>G, 4456G>T, 4458T>G, 4461C>T, 4464T>C, 4465T>C, 4467G>A, 4470T>G, 4476C>A, 4482G>C, 4494A>T, 4495C>T, 4497T>G,
4500C>T, 4509C>T, 4512C>T, 4516C>T, 4519T>C, 4521G>A, 4537C>A, 4542T>C, 4545T>G, 4548C>T, 4551C>A, 4557T>G, 4560A>G, 4563A>G, 4566G>A, 4569T>A, 4572C>T,
4573A>G, 4575T>C, 4576G>A, 4577C>A, 4578C>A, 4581T>C, 4583G>A, 4585G>T, 4588A>G, 4590A>G, 4593T>C, 4594G>A, 4596A>T, 4602T>A, 4603G>A, 4604G>A, 4608T>C,
4614C>T, 4618C>T, 4620T>G, 4638A>T, 4647T>C, 4648C>T, 4653G>T, 4656T>G, 4659G>T, 4662T>A, 4663C>A, 4669C>T, 4680T>C, 4681T>G, 4682C>A, 4683A>T, 4686G>A,
4689T>A, 4690A>T, 4691A>C, 4692T>C, 4698C>A, 4699C>A, 4700A>G, 4701G>A, 4707T>G, 4710C>A, 4711A>G, 4712T>C, 4713G>T, 4716A>T, 4719T>C, 4728C>T, 4731A>G,
4737G>T, 4738C>T, 4743T>C, 4749T>A, 4759A>G, 4761C>G, 4762C>A, 4765G>A, 4766T>C, 4767G>A, 4768C>A, 4770G>A, 4773C>T, 4775A>T, 4779C>T, 4785A>T, 4788A>G,
4794T>C, 4803C>T, 4809A>T, 4812T>C, 4815A>G, 4818T>A, 4824C>A, 4825C>T, 4827T>G, 4830G>A, 4836C>A, 4857G>T, 4861A>G, 4863A>C, 4866T>C, 4872T>C, 4875G>T,
4878C>T, 4884C>T, 4887A>G, 4890T>A, 4894C>G, 4895C>T, 4899T>A, 4909A>G, 4910C>T, 4911T>A, 4912G>C, 4913T>A, 4914G>A, 4917A>T, 4921A>T, 4922T>A, 4923G>T,
4926G>A, 4930G>A, 4932G>T, 4938T>A, 4942C>A, 4944T>C, 4945C>A, 4947T>A, 4950G>A, 4956T>C, 4966A>C, 4967G>A, 4968A>G, 4970T>C, 4971T>C, 4972G>A, 4974A>T,
4980G>A, 4982A>G, 4983G>T, 4986A>T, 4987A>G, 4988C>T, 4989T>A, 4990T>A, 4999C>T, 5000A>T, 5001G>T, 5002T>G, 5004T>A, 5007C>T, 5012G>A, 5013A>G, 5014G>C,
5017C>A, 5019T>A, 5020G>A, 5021G>A, 5022A>T, 5026G>C, 5027G>C, 5029G>A, 5030C>T, 5031A>C, 5033C>T, 5034A>G, 5038T>A, 5040T>G, 5043T>C, 5044T>A, 5049T>C,
5050A>T, 5051T>A, 5052G>T, 5056G>C, 5058G>T, 5061G>A, 5062C>A, 5063A>G, 5064G>T, 5065G>C, 5071A>T, 5073C>G, 5074G>A, 5077A>G, 5093C>A, 5094A>G, 5095T>A,
5096C>G, 5097A>C, 5098G>A, 5099C>T, 5100G>T, 5101T>G, 5102T>G, 5104A>G, 5105C>T, 5106A>T, 5107A>T, 5109C>G, 5110T>A, 5111G>T, 5112T>G, 5115A>G, 5116A>C,
5118G>A, 5119G>T, 5121C>A, 5122A>G, 5124A>T, 5128T>A, 5129C>A, 5130T>A, 5132T>A, 5133C>T, 5136T>A, 5138T>A, 5142A>C, 5148G>A, 5151A>T, 5152C>G, 5158A>G,
5160T>A, 5164G>A, 5167_5169delGAA, 5178A>G, 5179A>G, 5180G>A, 5181A>G, 5183A>G, 5185C>G, 5188A>C, 5191T>G, 5192G>A, 5193G>A, 5198C>T, 5199C>G, 5202A>T,
5203A>G, 5205C>T, 5206A>G, 5208T>A, 5209G>T, 5210G>T, 5217C>T, 5218T>A, 5219C>G, 5220A>T, 5223C>T, 5226T>G, 5233G>T, 5235C>A, 5238G>C, 5239A>C, 5240C>A,
5241T>A, 5242G>C, 5248G>T, 5249A>C, 5250G>A, 5251C>T, 5253T>A, 5256G>T, 5257T>A, 5258G>A, 5262A>C, 5263G>C, 5265A>G, 5271C>T, 5274T>C, 5277A>T, 5280G>A,
5283G>C, 5284A>C, 5285A>G, 5286G>C, 5288A>G, 5289G>A, 5291G>C, 5292C>A, 5294A>C, 5301T>G, 5302C>A, 5304C>A, 5307A>G, 5313T>C, 5315G>A, 5325T>C, 5340G>A,
5346A>C, 5349T>C, 5352C>T, 5353A>T, 5356T>G, 5364T>C, 5366T>A, 5371G>C, 5372C>A, 5377A>C, 5379A>C, 5380G>A, 5381G>A, 5382C>G, 5384A>G, 5385A>T, 5386A>C,
5389T>G, 5390C>A, 5393A>T, 5394G>T, 5396A>G, 5397C>T, 5398A>G, 5399T>A, 5402C>G, 5403T>G, 5410G>T, 5411A>C, 5415T>C, 5416G>A, 5417T>C, 5418G>C, 5424G>T,
5430G>T, 5442T>A, 5443G>A, 5445G>A, 5446A>G, 5447G>C, 5448C>T, 5463T>A, 5466A>G, 5467A>T, 5469G>T, 5472T>A, 5473G>T, 5483G>A, 5487G>A, 5490T>C, 5491T>A,
5493C>G, 5494A>G, 5495T>C, 5496T>C, 5497G>A, 5498A>C, 5499G>A, 5502T>C, 5503A>C, 5505C>T, 5512A>C, 5514G>C, 5515T>C, 5517G>A, 5520C>A, 5523T>C, 5526T>G,
5528G>C, 5529C>T, 5530A>G, 5531A>G, 5532T>G, 5536T>C, 5550A>C, 5553T>C, 5556C>T, 5567T>C, 5568T>G, 5571G>A, 5577T>G, 5581C>A, 5589G>T, 5590C>G, 5591C>A,
5592T>A, 5593C>G, 5595G>C, 5595_5596insACC, 5596G>T, 5597A>C, 5599G>C, 5600G>A, 5601G>C, 5604A>G, 5605A>T, 5609C>G, 5614G>A, 5615T>G, 5620T>A, 5622G>A,
5623_5625delTCT, 5627T>A, 5628G>T, 5629C>G, 5630A>G, 5631A>C, 5633G>C, 5634T>C, 5636G>C, 5642T>A, 5643G>C, 5649A>T, 5649_5650insCACAGT, 5651G>C, 5652T>A,

| | |
|---|---|
| | 5656G>T, 5657A>C, 5658G>A, 5659C>G, 5664C>A, 5666A>T, 5667A>T, 5668A>C, 5670T>A, 5671C>A, 5673T>G, 5674G>A, 5676C>A, 5677A>G, 5679C>T, 5681A>C, 5682C>T, 5683G>T, 5684G>C, 5688T>C, 5689T>C, 5691A>C, 5694A>T, 5695A>G, 5696T>A, 5697G>A, 5698G>A, 5700A>G, 5707T>G, 5709G>T, 5712C>T, 5713A>T, 5721T>C, 5722G>A, 5723T>C, 5724T>A, 5725G>A, 5727G>T, 5730C>T, 5733A>T, 5734C>G, 5736T>A, 5739T>A, 5742A>T, 5747G>C, 5749A>C, 5751C>G, 5753T>G, 5754T>C, 5755A>T, 5757C>A, 5760A>G, 5763A>G, 5766T>C, 5769T>G, 5770G>A, 5772T>G, 5775A>T, 5776T>G, 5777T>C, 5778G>T, 5781T>G, 5783C>A, 5784A>G, 5786G>C, 5787T>A, 5788C>T, 5790T>G, 5791G>T, 5792A>C, 5793A>T, 5794A>G, 5795A>T, 5796G>C, 5797T>A, 5806G>A, 5807A>C, 5808A>T, 5809T>C, 5811G>T, 5812C>G, 5813T>C, 5814T>C, 5819A>G, 5820G>A, 5821A>G, 5822A>G, 5823C>A, 5824A>G, 5825C>T, 5826G>C, 5830C>G, 5835C>A, 5845G>A, 5847G>A, 5848A>G, 5851A>T, 5853G>C, 5854A>T, 5856T>C, 5860A>C, 5862A>C, 5866G>T, 5867T>G, 5868G>T, 5870C>A, 5872C>A, 5873A>T, 5874A>G, 5878G>C, 5880C>T, 5881A>G, 5882G>A, 5883T>G, 5887G>C, 5889A>G, 5890A>G, 5893C>A, 5894A>G, 5895G>T, 5902T>G, 5904C>A, 5905T>G, 5906A>T, 5907T>A, 5908C>T, 5909C>T, 5912A>T, 5915C>G, 5916C>G, 5918C>G, 5919A>C, 5920A>G, 5921C>T, 5922C>T, 5923G>A, 5924T>A, 5926G>A, 5931G>A, 5932C>A, 5934C>G, 5935A>T, 5936A>T, 5938G>T, 5939T>C, 5940A>C, 5943T>C, 5944G>A, 5945C>T, 5946T>G, 5947G>A, 5950G>C, 5951C>A, 5952T>A, 5955T>G, 5955_5956insATG, 5956A>T, 5958C>A, 5960T>A, 5961T>A, 5962G>A, 5963T>C, 5965A>C, 5968A>G, 5969C>T, 5973T>C, 5975A>G, 5976A>C, 5979T>A, 5981T>G, 5982C>T, 5983C>T, 5985G>T, 5986_5988delGGA, 5991A>C, 5992G>A, 5993T>C, 5994T>A, 5997C>T, 5998C>A, 6000G>T, 6004A>G, 6005A>C, 6006T>A, 6008T>G, 6009T>G, 6012C>A, 6014G>C, 6016C>A, 6017C>A, 6018T>G, 6021G>C, 6022T>A, 6023G>T, 6024G>T, 6024_6025insGTC, 6026A>T, 6027A>G, 6029T>C, 6030T>A, 6031G>A, 6032G>A, 6037G>A, 6042C>G, 6043A>C, 6046_6048delATG, 6050C>G, 6052A>T, 6054A>T, 6055C>A, 6057C>T, 6058T>C, 6060A>T, 6066G>A, 6068G>C, 6069T>A, 6070C>G, 6072T>G, 6073C>T, 6077C>T, 6078A>T, 6086T>C, 6087G>T, 6090G>A, 6094T>C, 6095C>T, 6098C>T, 6099C>T, 6100A>G, 6104A>G, 6105G>A, 6109T>A, 6112C>T, 6114C>G, 6115A>C, 6119G>A, 6120A>T, 6121T>A, 6122A>G, 6123C>T, 6124A>G, 6125A>G, 6128G>A, 6129A>G, 6131A>G, 6133A>T, 6134T>A, 6135G>C, 6138T>A, 6142C>A, 6143A>C, 6144T>A, 6145T>A, 6146G>C, 6147C>A, 6148A>C, 6150A>G, 6151A>G, 6154T>C, 6156G>T, 6159A>T, 6160C>T, 6162C>T, 6164A>G, 6165G>A, 6168A>G, 6171T>C, 6172G>A, 6174T>G, 6175_6180delCTCAAC, 6182T>G, 6183C>T, 6186G>A, 6187A>T, 6188T>G, 6190C>G, 6192G>A, 6193A>C, 6195T>C, 6199G>C, 6200A>C |

**CDS**

| | | | | | | | | | |
|---|---|---|---|---|---|---|---|---|---|
| APL33_sRA11gp1 | 1 | 2017 | 93.8% | 9642 | 69.4% | 2004 (99.0%) | 1319 (65.2%) | 7/13/0/0 | 0 |

| | Begin | End | Coverage | Score | Concordance | Matches | Identities | I/D/M/F* | Stop Codons |
|---|---|---|---|---|---|---|---|---|---|
| **NT** | **25** | **6200** | **85.3%** | **3315** | **27.4%** | **6117 (98.7%)** | **3941 (63.6%)** | **21/59** | |

Protein mutations:

L3F (157C>T), F7W (170T>G 171T>G), G8S (172G>A), S9P (175T>C), S10P (178T>C 180A>T), S11A (181T>G), E12_V17del (184_201delGAGAAGACTGGATCAGTT), P18S (202C>T 204C>T), T20P (208A>C), G24A (221G>C 222T>A), L26V (226C>G 228G>T), G28S (232G>T 233G>C), I30F (238A>T 240C>T), K32S (244A>T 245A>C 246A>T), F33L (247T>C 249C>T), A36T (256G>A), I37L (259A>C 261T>C), I40V (268A>G 270C>T), T41N (272C>A 273A>T), S42K (274T>A 275C>A), K46T (287A>C 288A>C), S48V (292T>G 293C>T 294T>A), L51I (301T>A), D52E (306T>G), L54V (310T>G 312G>C), S57P (319T>C 321T>A), T59M (326C>T 327T>G), D60V (329A>T 330T>A), S62A (334T>G 336T>A), S63T (337T>A 339T>A), V66A (347T>C 348T>C), S70N (358T>A 359C>A 360C>T), L72V (364T>G 366G>A), N73E (367A>G 369T>A), K74R (371A>G), G80A (389G>C 390T>A), S81K (391T>A 392C>A 393C>A), L82I (394T>A 396G>A), I83M (399T>G), L87I (409C>A 411G>C), A89G (416C>G 417A>T), S91D (421T>G 422C>A), L93V (427T>G 429G>A), A94K (430G>A 431C>A), S99T (445T>A), K101V (451A>G 452A>T 453G>A), M103L (457A>C 459G>T), R106K (466C>A 467G>A), L107M (469T>A), F108V (472T>G 474C>T), E109L (475G>T 476A>T), L111V (481T>G 483G>C), S112K (484T>A 485C>A 486C>G), N113D (487A>G), V115F (493G>T 495T>C), R116I (497G>T 498A>C), I117L (499A>C), F119Y (506T>A 507T>C), Q120E (508C>G 510G>A), V122L (514G>T 516T>A), S126V (526A>G 527G>T), D128S (532G>T 533A>C 534T>A), V129L (535G>T), V133F (547G>T), Y134I (550T>A 551A>T 552T>A), G135A (554G>C), I141L (571A>T 573A>G), T142A (574A>G 576T>C), L143I (577C>A), V145L (583G>C 585T>A), C148Y (593G>A), S152V (604T>G 605C>T), I155V (613A>G 615T>A), E156T (616G>A 617A>C 618A>G), G157P (619G>C 620G>C 621A>T), V158F (622G>T 624A>T), S159N (625T>A 626C>A), S160N (629G>A), T161V (631A>G 632C>T 633C>A), I162V (634A>G), K163V (637A>G 638A>T 639A>T), N164T (641A>C), I165L (643A>T 645T>G), L167S (650T>C), I168V (652A>G 654A>T), M172F (664A>T 666G>T), K184T (701A>C 702G>A), A186S (706G>T), E187G (710A>G), T190S (718A>T 720A>C), M191Q (721A>C 722T>A), V192L (724G>C 726A>G), S196A (736A>G 737G>C 738T>A), M197V (739A>G 741G>A), V203L (757G>T 759C>A), S204T (760T>A 762A>G), M212L (784A>T 786G>A), T214S (790A>T 792T>A), I215L (793A>T 795T>G), M216L (796A>T), I226V (826A>G 828A>T), T246A (886A>G), T258Q (922A>C 923C>A 924A>G), I278V (982A>G 984A>G), G280S (988G>A 990T>C), A281I (991G>A 992C>T), S293N (1028G>A 1029C>T), E302Q (1054G>C 1056A>G), Q305T (1063C>A 1064A>C), D308S (1072G>T 1073A>C 1074T>G), A309E (1076C>A 1077T>G), C311A (1081T>G 1082G>C), D312E (1086T>A), A314S (1090G>T), T315M (1094C>T 1095C>G), K323P (1117A>C 1118A>C 1119A>T), D327I (1129G>A 1130A>T 1131C>T), N330Q (1138A>C 1140T>A), R333K (1148G>A 1149G>A), L337V (1159C>G 1161G>C), M352L (1204A>T), E364D (1242A>T), V366L (1246G>C 1248C>G), Y367V (1249T>G 1250A>T), S370A (1258A>G 1259G>C 1260T>C), C371S (1261T>A), R372Q (1264A>C 1265G>A), V374S (1270G>T 1271T>C), Y389F (1316A>T), A399C (1345G>T 1346C>G 1347A>T), R410A (1378C>G 1379G>C 1380C>A), V494I (1630G>A), I519R (1706T>G 1707C>A), E521V (1712A>T 1713A>G), E523D (1719A>T), S525D (1723T>G 1724C>A), T526P (1726A>C), E527N (1729G>A 1731G>T), D528T (1732G>A 1733A>C 1734C>T), G531A (1742G>C 1743A>C), M534I (1752G>T), N536D (1756A>G 1758C>T), I537V (1759A>G 1761T>G), H550A (1798C>G 1799A>C 1800C>T), S554D (1810T>G 1811C>A 1812G>T), D558Q (1822G>C 1824C>A), Q565R (1844A>G), A568Y (1852G>T 1853C>A), I570V (1858A>G 1860A>G), K572S (1865A>G 1866G>C), F575Y (1874T>A 1875T>C), E576N (1876G>A 1878A>T), A577T (1879G>A 1881A>T), E578T (1882G>A 1883A>C), T579S (1885A>T 1887T>G), I581L (1891A>T 1893T>G), S582A (1894T>G 1896A>T), N583D (1897A>G 1899T>C), T585V (1903A>G 1904C>T 1905A>T), A589N (1915G>A 1916C>A), S591A (1921T>G), S592V (1924T>G 1925C>T), E593D (1929A>T), V596L (1936G>C), A597D (1940C>A 1941A>T), V599F (1945G>T 1947T>C), D600Q (1948G>C 1950T>A), G601D (1952G>A 1953G>T), R603del (1957_1959delCGC), F604L (1962C>A), K605Q (1963A>C), T606V (1966A>G 1967C>T 1968A>T), K607L (1969A>T 1970A>T 1971A>G), Q608P (1973A>C 1974A>T), D609P (1975G>C 1976A>C), V610E (1979T>A), P611S (1981C>T), I612A (1984A>G 1985T>C), V613A (1988T>C 1989G>T), T614Y (1990A>T 1991C>A), P615E (1993C>G 1994C>A 1995C>G), T616Q (1996A>C 1997C>A 1998A>G), V617T (1999G>A 2000T>C 2001T>A), E621A (2012A>C 2013A>T), D622E (2016T>A), R623K (2018G>A), V624I (2020G>A 2022G>T), I625V (2023A>G), K628Q (2032A>C 2034A>G), F637Y (2060T>A), L643M (2077T>A), D644N (2080G>A 2082C>T), P645K (2083C>A 2084C>A 2085G>A), E646F (2086G>T 2087A>T 2088G>T), S647E (2089A>G 2090G>A 2091T>G), L648F (2094G>T), Y653L (2107T>C 2108A>T 2109T>C), Q654S (2110C>T 2111A>C 2112A>C), L658I (2122C>A), F659Y (2126T>A), S661V (2131T>G 2132C>T 2133C>G), D663S (2137G>T 2138A>C), A664S (2140G>T 2142A>T), E665D (2145G>T), H666A (2146C>G 2147A>C), Q667K (2149C>A 2151G>A), V669N (2155G>A 2156T>A 2157C>T), M671E (2161A>G 2162T>A 2163G>A), R672M (2165G>T 2166A>G), M674V (2170A>G 2172G>A), T675N (2174C>A), M677L (2179A>T), I678L (2182A>T 2184C>G), Y679V (2185T>G 2186A>T), N683S (2198A>G), A684P (2200G>C), E685Q (2203G>C 2205A>G), R688A (2212A>G 2213G>C 2214G>T), Q692R (2225A>G), L699I (2245C>A), A708S (2272G>A 2273C>G 2274T>C), S709T (2275T>A 2277C>A), C712S (2285G>C), K714E (2290A>G), K715L (2293A>C 2294A>T), A717D (2300C>A), T720R (2308A>C 2309C>G 2310A>C), E721N (2311G>A 2313G>T), M722L (2314A>C 2316G>T), R723E (2317C>G 2318G>A 2319C>G), T724Q (2320A>C 2321C>A), F726K (2326T>A 2327T>A 2328T>A), H728Y (2332C>T), N729K (2337T>G), C730A (2338T>G 2339G>C 2340T>G), I731L (2341A>C), D732Q (2344G>C 2346T>G), V735A (2354T>C 2355T>A), I739T (2366T>C 2367T>G), Q742E (2374C>G 2376G>A), F747R (2389T>A 2390T>G 2391T>A), T761R (2432C>G 2433C>G), S764A (2440T>G 2442C>A), S764_K765insG (2442_2443insGGT), K765T (2444A>C 2445A>T), E766D (2448G>C), A767K (2449G>A 2450C>A 2451C>G), D769R (2455G>A 2456A>G 2457C>A), S770N (2458T>A 2459C>A), V772I (2464G>A 2466C>T), K773I (2468A>T 2469G>C), I774E (2470A>G 2471T>A), Y776M (2476T>A 2477A>T 2478C>G), H777I (2479C>A 2480A>T 2481C>T), K778G (2482A>G 2483A>G 2484A>T), V780K (2488G>A 2489T>A), Y782K (2494T>A 2496C>A), K784I (2501A>T 2502A>C), E785Q (2503G>C 2505A>G), T786M (2507C>T 2508T>G), W788N (2512T>A 2513G>A 2514G>T), N789D (2515A>G), Y792F (2525A>T 2526T>C), A793I (2527G>A 2528C>T 2529A>C), R795K (2534G>A), I804L (2560A>T 2562C>G), M805I (2565G>A), L810T (2578T>A 2579T>C 2580G>C), F811M (2581T>A 2583T>G), G812N (2584G>A 2585G>A 2586G>T), G813A (2588G>C 2589A>C), I814V (2590A>G 2592A>T), L818I (2602C>A), A819T (2605G>A), W821L (2611T>C 2612G>T 2613G>C), Q822K (2614C>A 2616A>G), S823A (2617A>G 2618G>C 2619C>A), N824A (2620A>G 2621A>C 2622T>A), S825T (2623T>A), F826S (2627T>C 2628T>C), S828L (2632A>T 2633G>T 2634T>A), S830D (2638A>G 2639G>A), P850K (2698C>A 2699C>A 2700C>G), S851T (2701T>A), R852H (2705G>A), L853F (2707C>T 2709G>C), K854A (2710A>G 2711A>C 2712G>A), N855K (2715T>A), L856S (2717T>C 2718G>C), V858L (2722G>C 2724G>A), S861A (2731A>G 2732G>C 2733T>G), S862A (2734A>G 2735G>C 2736C>A), A863D (2738C>A), I865N (2744T>A 2745T>C), T866H (2746A>C 2747C>A 2748A>T), Q867V (2749C>G 2750A>T 2751A>G), E868D (2754A>C), E868_W869insR (2754_2755insCGG), A871E (2762C>A 2763T>A), E872Q (2764G>C 2766A>G), Q878E (2782C>G), A879S (2785G>A 2786C>G 2787A>T), K881H (2791A>C 2793A>C), M884L (2800A>T), I885V (2803A>G), L889I (2815C>A), G890A (2819G>C 2820A>T), M906L (2866A>C 2868G>C), H913R (2887C>A 2888A>G 2889C>A), V915M (2893G>A), T922S (2914A>T), P926E (2926C>G 2927C>A 2928T>G), N927H (2929A>C 2931C>T), R929I (2936G>T 2937G>T), S930C (2939C>G 2940A>T), K931N (2943G>T), Y934F (2951A>T), R937T (2959C>A 2960G>C 2961G>C), E938Q (2962G>C), K940R (2969A>G), V941L (2971G>T 2973C>G), I947V (2989A>G 2991C>G), P960S (3028C>T 3030T>A), E971D (3063G>T), F974Y (3071T>A), T975S (3073A>T 3075C>T), Q979M (3085C>A 3086A>T), C983V (3097T>G 3098G>T 3099T>G), T989K (3116C>A 3117C>G), R992K (3125G>A 3126A>G), L993K (3127C>A 3128T>A 3129G>A), G994S (3130G>T 3131G>C), F998V (3142T>G), F1001Y (3152T>A), V1011E (3182T>A 3183C>A), E1014T (3190G>A 3191A>C 3192G>A), E1016G (3197A>G 3198G>A), A1020N (3208G>A 3209C>A 3210C>T), S1021A (3211A>G 3212G>C 3213T>A), K1023A (3217A>G 3218A>C 3219A>T), Q1028V (3232C>G 3233A>T 3234A>T), S1029A (3235T>G), M1034I (3252G>T), G1036S (3256G>A 3258A>T), S1049T (3296G>C), V1056C (3316G>T 3317T>G 3318T>C), G1067D (3350G>A), I1068T (3353T>C 3354T>C), F1074Y (3371T>A), I1075V (3373A>G 3375T>G), S1076A (3376T>G), I1082V (3394A>G 3396C>G), N1089D (3415A>G 3417C>T), V1095I (3433G>A), A1097V (3440C>T), D1103E (3459C>A), I1104L (3460A>C 3462T>G), A1107H (3469G>C 3470C>A), I1108L (3472A>C 3474T>G), R1111K (3482G>A 3483A>G), T1113E (3487A>G 3488C>A 3489C>A), T1114R (3491C>G 3492T>A), L1115F (3493C>T 3495T>C), N1119K (3507C>A), S1122A (3514T>G 3516C>T), Q1129L (3535C>T 3536A>T 3537A>G), L1131M (3541T>A), R1142K (3575G>A), K1143R (3577A>C 3578A>G 3579G>C), Q1145P (3584A>C 3585A>C), P1152D (3604C>G 3605C>A 3606T>C), E1154Q (3610G>C 3612A>G), T1161S (3631A>T 3633C>T), V1162A (3635T>C), N1163K (3639T>A), M1178E (3682A>G 3683T>A), H1185S (3703C>T 3704A>C 3705T>C), G1186N (3706G>A 3707G>A 3708T>C), D1188E (3714T>G), D1189E (3717T>G), D1190P (3718G>C 3719A>C 3720T>A), S1195A (3733T>G 3735C>T), T1198G (3742A>G 3743C>G), S1201A (3751A>G 3752G>C 3753T>A), L1203I (3757C>A 3759G>T), T1206V (3766A>G 3767C>T 3768A>G), M1207L (3769A>C 3771G>T), N1209M (3776A>T 3777C>G), K1211R (3781A>C 3782A>G), Q1218A (3802C>G 3803A>C 3804G>A), E1219I (3805G>A 3806A>T 3807G>C), D1221N (3811G>A), C1232F (3845G>T), Y1235F (3854A>T), G1237A (3860G>C 3861C>A), M1251L (3901A>T), M1253L (3907A>C 3909G>T), M1255S (3913A>T 3914T>C 3915G>T), V1262E (3935T>A), P1263A (3937C>G 3939A>C), T1264Q (3940A>C 3941C>A), G1265P (3943G>C 3944G>C), E1266N (3946G>A 3948A>C), G1267S (3949G>T 3950G>C 3951T>G), D1268Q (3952G>C 3954T>G), K1269R (3956A>G 3957A>G), E1271L (3961G>C 3962A>T 3963G>T), R1273I (3968G>T), D1275N (3973G>A), E1280D (3990A>T), I1287S (4009A>T 4010T>C 4011T>G), I1289V (4015A>G 4017T>A), I1293T (4028T>C), H1294F (4030C>T 4031A>T), I1298V (4042A>G), T1343C (4177A>T 4178C>G 4179C>T), L1347A (4189T>G 4190T>C), I1351L (4201A>C), D1358N (4222G>A), S1362H (4234T>C 4235C>A), S1365T (4243T>A 4245T>A), L1366V (4246C>G 4248T>G), N1369S (4256A>G 4257C>T), N1370S (4259A>G), A1373S (4267G>T 4269T>A), T1377E (4279A>G 4280C>A 4281T>G), V1392I (4324G>A 4326G>C), E1404A (4361A>C 4362G>T), Y1405F (4364A>T), D1407G (4370A>G), T1408D (4372A>G 4373C>A 4374C>T), K1409E (4375A>G), E1412S (4384G>A 4385A>G 4386G>T), A1414E (4391C>A 4392T>G), A1420T (4408G>A 4410T>A), D1434N (4450G>A 4452T>C), V1436L (4456G>T 4458T>G), I1475V (4573A>G 4575T>C), A1476K (4576G>A 4577C>A 4578C>A), R1478K (4583G>A), A1479S (4585G>T), I1480V (4588A>G 4590A>G), V1482I (4594G>A 4596A>T), G1485N (4603G>A 4604G>A), L1505M (4663C>A), S1511D (4681T>G 4682C>A 4683A>T), N1514S (4690A>T 4691A>C 4692T>C), Q1517R (4699C>A 4700A>G 4701G>A), D1520E (4710C>A), M1521A (4711A>G 4712T>C 4713G>T), N1537E (4759A>G 4761C>G), Q1538K (4762C>A), V1539T (4765G>A 4766T>C 4767G>A), Q1540K (4768C>A 4770G>A), Q1542L (4775A>T), H1562Q (4836C>A), I1571V (4861A>G 4863A>C), M1575I (4875G>T), P1582V (4894C>G 4895C>T), T1587V (4909A>G 4910C>T 4911T>A), V1588Q (4912G>C 4913T>A 4914G>A), M1591Y (4921A>T 4922T>A 4923G>T), V1594I (4930G>A 4932G>T), L1598I (4942C>A 4944T>C), R1606Q (4966A>C 4967G>A 4968A>G), V1607A (4970T>C 4971T>C), A1608T (4972G>A 4974A>T), K1611S (4982A>G 4983G>T), L1612F (4986A>T), T1613V (4987A>G 4988C>T 4989T>A), Y1614N (4990T>A), Q1617F (4999C>T 5000A>T 5001G>T), C1618G (5002T>G 5004T>A), R1621K (5012G>A 5013A>G), E1622Q (5014G>C), G1624N (5020G>A 5021G>A 5022A>T), G1626P (5026G>C 5027G>C), A1627I (5029G>A 5030C>T 5031A>C), T1628M (5033C>T 5034A>G), F1630M (5038T>A 5040T>G), S1632T (5044T>A), M1634Y (5050A>T 5051T>A 5052G>T), V1636L (5056G>C 5058G>T), Q1638S (5062C>A 5063A>G 5064G>T), E1639Q (5065G>C), I1641L (5071A>T 5073C>G), A1642T (5074G>A), K1643E (5077A>G), A1648E (5093C>A 5094A>G), A1650I (5098G>A 5099C>T 5100G>T), L1651G (5101T>G 5102T>G), T1652V (5104A>G 5105C>T 5106A>T), I1653L (5107A>T 5109C>G), C1654M (5110T>A 5111G>T 5112T>G), M1656L (5116A>C 5118G>A), A1657S (5119G>T 5121C>A), K1658D (5122A>G 5124A>T), S1660K (5128T>A 5129C>A 5130T>A), I1661N (5132T>A 5133C>T), F1663Y (5138T>A), P1668A (5152C>G), N1670E (5158A>G 5160T>A), V1672I (5164G>A), E1673del (5167_5169delGAA), R1677E (5179A>G 5180G>A 5181A>G), D1678G (5183A>G), L1679V (5185C>G), N1680H (5188A>C), W1681E (5191T>G 5192G>A 5193G>A), A1683V (5198C>T 5199C>G), N1685D (5203A>G 5205C>T), I1686V (5206A>G 5208T>A), G1687F (5209G>T 5210G>T), A1695S (5233G>T 5235C>A), K1696N (5238G>C), T1697Q (5239A>C 5240C>A 5241T>A), E1698Q (5242G>C), E1700S (5248G>T 5249A>C 5250G>A), P1701S (5251C>T 5253T>A), C1703N (5257T>A 5258G>A), V1705L (5263G>C 5265A>G), K1712R (5284A>C 5285A>G 5286G>C), K1713R (5288A>G 5289G>A), S1714T (5291G>C 5292C>A), K1715T (5294A>C), S1722N (5315G>A), M1730I (5340G>A), M1735L (5353A>T), S1736A (5356T>G), L1739Q (5366T>A), A1741H (5371G>C 5372C>A), G1744K (5380G>A 5381G>A 5382C>G), K1745S (5384A>G 5385A>T), M1746L (5386A>C), S1747E (5389T>G 5390C>A), E1748V (5393A>T 5394G>T), D1749G (5396A>G 5397C>T), I1750D (5398A>G 5399T>A), A1751G (5402C>G 5403T>G), D1754S (5410G>T 5411A>C), V1756T (5416G>A 5417T>C 5418G>C), V1765I (5443G>A 5445G>A), S1766A (5446A>G 5447G>C 5448C>T), M1773F (5467A>T 5469G>T), V1775L (5473G>T), R1778H (5483G>A), C1781R (5491T>A 5493C>G), I1782A (5494A>G 5495T>C 5496T>C), E1783T (5497G>A 5498A>C 5499G>A), I1785L (5503A>C 5505C>T), I1792M (5526T>G), G1793A (5528G>C 5529C>T), N1794G (5530A>G 5531A>G 5532T>G), V1806A (5567T>C 5568T>G), F1809L (5577T>G), Q1811K (5581C>A), P1814E (5590C>G 5591C>A 5592T>A), Q1815D (5593C>G 5595G>C), Q1815_E1816insT (5595_5596insACC), E1816S (5596G>T 5597A>C), G1817H (5599G>C 5600G>A 5601G>C), M1819L (5605A>T), A1820G (5609C>G), V1822S (5614G>A 5615T>G), S1824T (5620T>A 5622G>A), S1825del (5623_5625delTCT), L1826Y (5627T>A 5628G>T), Q1827G (5629C>G 5630A>G 5631A>C), S1828T (5633G>C 5634T>C), S1829T (5636G>C), M1831N (5642T>A 5643G>C), Q1833H (5649A>T), Q1833_G1834insHS (5649_5650insCACAGT), G1834A (5651G>C 5652T>A), E1836S (5656G>T 5657A>C 5658G>A), L1837V (5659C>G), Q1839L (5666A>T 5667A>T), I1840L (5668A>C 5670T>A), V1842I (5674G>A 5676C>A), I1843V (5677A>G 5679C>T), H1844P (5681A>C 5682C>T), G1845S (5683G>T 5684G>C), M1849E (5695A>G 5696T>A 5697G>A), E1850K (5698G>A 5700A>G), L1853V (5707T>G 5709G>T), M1855L (5713A>T), V1858T (5722G>A 5723T>C 5724T>A), V1859I (5725G>A 5727G>T), H1862E (5734C>G 5736T>A), G1866A (5747G>C), N1867Q (5749A>C 5751C>G), F1868C (5753T>G 5754T>C), T1869S (5755A>T 5757C>A), D1874K (5770G>A 5772T>G), Q1875H (5775A>T), L1876A (5776T>G 5777T>C 5778G>T), F1877L (5781T>G), A1878E (5783C>A 5784A>G), S1879T (5786G>C 5787T>A), E1881S (5791G>T 5792A>C 5793A>T), K1882V (5794A>G 5795A>T 5796G>C), C1883S (5797T>A), E1886T (5806G>A 5807A>C 5808A>T), L1888A (5812C>G 5813T>C 5814T>C), K1890R (5819A>G 5820G>A), N1891G (5821A>G 5822A>G 5823C>A), T1892V (5824A>G 5825C>T 5826G>C), L1894V (5830C>G), E1899K (5845G>A 5847G>A), K1900E (5848A>G), M1901F (5851A>T 5853G>C), T1902S (5854A>T 5856T>C), K1904H (5860A>C 5862A>C), V1906C (5866G>T 5867T>G 5868G>T), A1907E (5870C>A), Q1908M (5872C>A 5873A>T 5874A>G), V1910L (5878G>C 5880C>T), S1911E (5881A>G 5882G>A 5883T>G), V1913L (5887G>C 5889A>G), K1914E (5890A>G), Q1915S (5893C>A 5894A>G 5895G>T), F1918V (5902T>G 5904C>A), Y1919V (5905T>G 5906A>T 5907T>A), P1920F (5908C>T 5909C>T), K1921M (5912A>T), A1922G (5915C>G 5916C>G), T1923S (5918C>G 5919A>C), T1924V (5920A>G 5921C>T 5922C>T), V1925K (5923G>A 5924T>A), E1926K (5926G>A), L1928M (5932C>A 5934C>G), K1929L (5935A>T 5936A>T), V1930S (5938G>T 5939T>C 5940A>C), A1932M (5944G>A 5945C>T 5946T>G), D1933N (5947G>A), A1934Q (5950G>C 5951C>A 5952T>A), H1935Q (5955T>G), H1935_T1936insM (5955_5956insATG), T1936S (5956A>T 5958C>A), I1937K (5960T>A 5961T>A), V1938T (5962G>A 5963T>C), K1939Q (5965A>C), T1940V (5968A>G 5969C>T), K1942S (5975A>G 5976A>C), F1944C (5981T>G 5982C>T), R1945C (5983C>T 5985G>T), G1946del (5986_5988delGGA), V1948T (5992G>A 5993T>C 5994T>A), R1950S (5998C>A 6000G>T), N1952A (6004A>G 6005A>C 6006T>A), F1953W (6008T>G 6009T>G), S1955T (6014G>C), P1956K (6016C>A 6017C>A 6018T>G), E1957D (6021G>C), W1958I (6022T>A 6023G>T 6024G>T), W1958_K1959insV (6024_6025insGTC), K1959M (6026A>T 6027A>G), I1960T (6029T>C 6030T>A), G1961K (6031G>A 6032G>A), E1963K (6037G>A), N1964K (6042C>G), T1965P (6043A>C), M1966del (6046_6048delATG), A1967G (6050C>G), T1968S (6052A>T 6054A>T), L1969I (6055C>A 6057C>T), S1970P (6058T>C 6060A>T), S1973T (6068G>C 6069T>A), L1974V (6070C>G 6072T>G), L1975F (6073C>T), P1976L (6077C>T 6078A>T), L1979P (6086T>C 6087G>T), S1982L (6094T>C 6095C>T), A1983V (6098C>T 6099C>T), M1984V (6100A>G), K1985R (6104A>G 6105G>A), L1987M (6109T>A), S1989R (6115A>C), R1990N (6119G>A 6120A>T), Y1991S (6121T>A 6122A>G 6123C>T), K1992G (6124A>G

| | |
|---|---|
| | 6125A>G), R1993K (6128G>A 6129A>G), N1994S (6131A>G), M1995Y (6133A>T 6134T>A 6135G>C), H1998T (6142C>A 6143A>C 6144T>A), C1999T (6145T>A 6146G>C 6147C>A), I2000L (6148A>C 6150A>G), K2001E (6151A>G), L2004F (6160C>T 6162C>T), K2005R (6164A>G 6165G>A), I2006M (6168A>G), V2008M (6172G>A 6174T>G), L2009_N2010del (6175_6180delCTCAAC), F2011C (6182T>G 6183C>T), M2013W (6187A>T 6188T>G), L2014V (6190C>G 6192G>A), N2015H (6193A>C 6195T>C) |

| | Begin | End | Coverage | Score | Concordance | Matches | Identities | I/D/M/F* | Stop Codons |
|---|---|---|---|---|---|---|---|---|---|
| **NT** | **25** | **6200** | **85.3%** | **3315** | **27.4%** | **6117 (98.7%)** | **3941 (63.6%)** | **21/59** | |

GCT2GCA (156T>A), CTT3TTT (157C>T), AAA5AAG (165A>G), TTT7TGG (170T>G 171T>G), GGT8AGT (172G>A), TCT9CCT (175T>C), TCA10CCT (178T>C 180A>T), TCT11GCT (181T>G), GAG12_GTT17del (184_201delGAGAAGACTGGATCAGTT), CCC18TCT (202C>T 204C>T), TCT19TCA (207T>A), ACT20CCT (208A>C), TCT21TCA (213T>A), GGT24GCA (221G>C 222T>A), CTG26GTT (226C>G 228G>T), GGT28TCT (232G>T 233G>C), ATC30TTT (238A>T 240C>T), TCA31TCT (243A>T), AAA32TCT (244A>T 245A>C 246A>T), TTC33CTT (247T>C 249C>T), AAA35AAG (255A>G), GCT36ACT (256G>A), ATT37CTC (259A>C 261T>C), AGC38TCC (262A>T 263G>C), AAC39AAT (267C>T), ATC40GTT (268A>G 270C>T), ACA41AAT (272C>A 273A>T), TCA42AAA (274T>A 275C>A), GTA43GTT (279A>T), AAA46ACC (287A>C 288A>C), TCT48GTA (292T>G 293C>T 294T>A), CAT50CAC (300T>C), TTA51ATA (301T>A), GAT52GAG (306T>G), TTG54GTC (310T>G 312G>C), AAA55AAG (315A>G), CCA56CCG (318A>G), TCT57CCA (319T>C 321T>A), GTG58GTT (324G>T), ACT59ATG (326C>T 327T>G), GAT60GTA (329A>T 330T>A), TCT62GCA (334T>G 336T>A), TCT63ACA (337T>A 339T>A), TCT64TCA (342T>A), GTT66GCC (347T>C 348T>C), TGT69TGC (357T>C), TCC70AAT (358T>A 359C>A 360C>T), GCA71GCT (363A>T), TTG72GTA (364T>G 366G>A), AAT73GAA (367A>G 369T>A), AAA74AGA (371A>G), GTG75GTT (375G>T), CTT76TTA (376C>T 378T>A), ATA79ATT (387A>T), GGT80GCA (389G>C 390T>A), TCC81AAA (391T>A 392C>A 393C>A), TTG82ATA (394T>A 396G>A), ATT83ATG (399T>G), CCT85CCC (405T>C), TTT86TTC (408T>C), CTG87ATC (409C>A 411G>C), AAA88AAG (414A>G), GCA89GGT (416C>G 417A>T), TAC90TAT (420C>T), TCT91GAT (421T>G 422C>A), TTG93GTA (427T>G 429G>A), GCA94AAA (430G>A 431C>A), ACC95ACT (435C>T), TCT99ACT (445T>A), ATC100ATA (450C>A), AAG101GTA (451A>G 452A>T 453G>A), ATG103CTT (457A>C 459G>T), GTT104GTA (462T>A), TTA105CTT (463T>C 465A>T), CGG106AAG (466C>A 467G>A), TTG107ATG (469T>A), TTC108GTT (472T>G 474C>T), GAA109TTA (475G>T 476A>T), TTG111GTC (481T>G 483G>C), TCC112AAG (484T>A 485C>A 486C>G), AAT113GAT (487A>G), GTT115TTC (493G>T 495T>C), AGA116ATC (497G>T 498A>C), ATA117CTA (499A>C), TTT119TAC (506T>A 507T>C), CAG120GAA (508C>G 510G>A), GTT122TTA (514G>T 516T>A), GAG124GAA (522G>A), AAG125AAA (525G>A), AGT126GTT (526A>G 527G>T), GAT128TCA (532G>T 533A>C 534T>A), GTA129TTA (535G>T), ATT131ATA (543T>A), GTA132GTG (546A>G), GTT133TTT (547G>T), TAT134ATA (550T>A 551A>T 552T>A), GGT135GCT (554G>C), TTG137TTA (561G>A), GCA140GCT (570A>T), ATA141TTG (571A>T 573A>G), ACT142GCC (574A>G 576T>C), CTT143ATT (577C>A), CTT144TTG (580C>T 582T>G), GTT145CTA (583G>C 585T>A), TTG146CTA (586T>C 588G>A), TTG147CTG (589T>C), TGT148TAT (593G>A), GTT149GTA (597T>A), TCA152GTA (604T>G 605C>T), GAC153GAT (609C>T), ATA154ATT (612A>T), ATT155GTA (613A>G 615T>A), GAA156ACG (616G>A 617A>C 618A>G), GGA157CCT (619G>C 620G>C 621A>T), GTA158TTT (622G>T 624A>T), TCT159AAT (625T>A 626C>A), AGT160AAT (629G>A), ACC161GTA (631A>G 632C>T 633C>A), ATT162GTT (634A>G), AAA163GTT (637A>G 638A>T 639A>T), AAT164ACT (641A>C), ATT165TTG (643A>T 645T>G), TTC166TTT (648C>T), TTA167TCA (650T>C), ATA168GTT (652A>G 654A>T), GTA169GTT (657A>T), GGG170GGA (660G>A), ATG172TTT (664A>T 666G>T), TTT173TTC (669T>C), AGT175TCA (673A>T 674G>C 675T>A), AAC178AAT (684C>T), TTG179TTA (687G>A), GAC180GAT (690C>T), CCC183CCA (699C>A), AAG184ACA (701A>C 702G>A), GCT186TCT (706G>T), GAG187GGG (710A>G), TTC189TTT (717C>T), ACA190TCC (718A>T 720A>C), ATG191CAG (721A>C 722T>A), GTA192CTG (724G>C 726A>G), GCA193GCG (729A>G), AGT196GCA (736A>G 737G>C 738T>A), ATG197GTA (739A>G 741G>A), CTA198CTT (744A>T), GAG201GAA (753G>A), TCT202AGC (754T>A 755C>G 756T>C), GTC203TTA (757G>T 759C>A), TCA204ACG (760T>A 762A>G), CAC205CAT (765C>T), ACT206ACA (768T>A), ACA208ACT (774A>T), TCC209AGC (775T>A 776C>G), CAG210CAA (780G>A), TTG211CTT (781T>C 783G>T), ATG212TTA (784A>T 786G>A), TCA213TCC (789A>C), ACT214TCA (790A>T 792T>A), ATT215TTG (793A>T 795T>G), ATG216TTG (796A>T), GCA217GCT (801A>T), ATT220ATC (810T>C), TCC221TCA (813C>A), ACG222ACT (816G>T), CTC223TTA (817C>T 819C>A), GTG224GTA (822G>A), ATA226GTT (826A>G 828A>T), GCA227GCT (831A>T), GTG228GTT (834G>T), CCA229CCT (837A>T), GGA230GGG (840A>G), CCC232CCA (846C>A), TTG235CTT (853T>C 855G>T), AGC236TCA (856A>T 857G>C 858C>A), AAT237AAC (861T>C), CCC238CCA (864C>A), CTC239CTG (867C>G), ATT242ATC (876T>C), TCA245TCT (885A>T), ACT246GCT (886A>G), GGG247GGA (891G>A), GGG249GGT (897G>T), GCT250GCG (900T>G), CAA255CAG (915A>G), TTG256CTG (916T>C), TTT257TTC (921T>C), ACA258CAG (922A>C 923C>A 924A>G), AGA261AGG (933A>G), AAT262AAC (936T>C), AAA264AAG (942A>G), TGC266TGT (948C>T), ACT267ACA (951T>A), TCA268TCT (954A>T), GCA270GCC (960A>C), TTT271TTC (963T>C), TCT272TCA (966T>A), GTG274GTT (972G>T), GAA276GAG (978A>G), ATT277ATA (981T>A), ATA278GTG (982A>G 984A>G), GGT280AGC (988G>A 990T>C), GCT281ATT (991G>A 992C>T), TTC284TTT (1002C>T), AAG285AAA (1005G>A), GTG288GTA (1014G>A), CTT289TTG (1015C>T 1017T>G), TCT290TCA (1020T>A), GCA291GCG (1023A>G), ATT292ATA (1026T>A), AGC293AAT (1028G>A 1029C>T), GCA294GCT (1032A>T), ACA295ACC (1035A>C), TTG296TTA (1038G>A), TCT297TCC (1041T>C), ACT298ACC (1044T>C), TTG300CTT (1048T>C 1050G>T), GAA302CAG (1054G>C 1056A>G), CAG305ACG (1063C>A 1064A>C), GTG307GTT (1071G>T), GAT308TCG (1072G>T 1073A>C 1074T>G), GCT309GAG (1076C>A 1077T>G), TGT311GCT (1081T>G 1082G>C), GAT312GAA (1086T>A), GCT314TCT (1090G>T), ACC315ATG (1094C>T 1095C>G), AGG316CGC (1096A>C 1098G>C), TTG317CTT (1099T>C 1101G>T), GAG318GAA (1104G>A), AAC319AAT (1107C>T), GCC321GCA (1113C>A), AAT322AAC (1116T>C), AAA323CCT (1117A>C 1118A>C 1119A>T), GCC324GCA (1122C>A), TTC325TTT (1125C>T), CCT326CCA (1128T>A), GAC327ATT (1129G>A 1130A>T 1131C>T), AAA328AAG (1134A>G), AAT330CAA (1138A>C 1140T>A), CAC331CAT (1143C>T), CTT332TTA (1144C>T 1146T>A), AGG333AAA (1148G>A 1149G>A), GAG334GAA (1152G>A), GCA336GCC (1158A>C), CTG337GTC (1159C>G 1161G>C), AAG338AAA (1164G>A), ATT339ATA (1167T>A), GCA341GCT (1173A>T), ACA345ACC (1185A>C), CAC346CAT (1188C>T), GTT348GTC (1194T>C), GCC349GCA (1197C>A), ATG352TTG (1204A>T), CAT354CAC (1212T>C), CGA355AGG (1213C>A 1215A>G), GTG356GTT (1218G>T), AGT357TCT (1219A>T 1220G>C), GCA358GCT (1224A>T), GCT359GCC (1227T>C), ATC360ATA (1230C>A), ACT361ACG (1233T>G), CTT363CTA (1239T>A), GAA364GAT (1242A>T), AAG365AAA (1245G>A), GTC366CTG (1246G>C 1248C>G), TAT367GTT (1249T>G 1250A>T), GCG368GCT (1254G>T), AGT370GCC (1258A>G 1259G>C 1260T>C), TGC371AGC (1261T>A), AGA372CAA (1264A>C 1265G>A), GTT374TCT (1270G>T 1271T>C), GTT376GTG (1278T>G), GGT377GGC (1281T>C), CAG378CAA (1284G>A), AGA380AGG (1290A>G), GAA382GAG (1296A>G), CCC383CCT (1299C>T), GTT386GTG (1308T>G), TAT389TTT (1316A>T), GGT390GGA (1320T>A), GCC391GCT (1323C>T), GGG393GGA (1329G>A), ACA398ACG (1344A>G), GCA399TGT (1345G>T 1346C>G 1347A>T), CGA401CGC (1353A>C), TTG402CTT (1354T>C 1356G>T), TTG408CTC (1372T>C 1374G>C), GAT409GAC (1377T>C), CGC410GCA (1378C>G 1379G>C 1380C>A), GGG412GGT (1386G>T), GAG413GAA (1389G>A), CCT414CCA (1392T>A), CTT416CTG (1398T>G), AGA418CGC (1402A>C 1404A>C), CTT419TTG (1405C>T 1407T>G), GCT421GCG (1413T>G), GTT422GTA (1416T>A), AGC423AGT (1419C>T), AAG424AAA (1422G>A), CGC425AGA (1423C>A 1425C>A), GCC427GCA (1431C>A), TCT430TCC (1440T>C), AAT431AAC (1443T>C), GCC433GCA (1449C>A), CAC434CAT (1452C>T), CAG435CAA (1455G>A), GCT437GCA (1461T>A), TTG439CTT (1465T>C 1467G>T), GAC442GAT (1476C>T), GGA444GGT (1482A>T), GCA445GCT (1485A>T), TTG446CTT (1486T>C 1488G>T), AGA447CGG (1489A>C 1491A>G), GGA451GGG (1503A>G), CAA452CAG (1506A>G), CAA454CAG (1512A>G), ATT456ATA (1518T>A), ATT462ATC (1536T>C), TCC464TCT (1542C>T), ACA465ACC (1545A>C), CCA467CCT (1551A>T), CCA469CCT (1557A>T), CTG470CTT (1560G>T), CCC471CCA (1563C>A), GCA473GCT (1569A>T), GCT474GCA (1572T>A), GTG475GTC (1575G>C), GAA476GAG (1578A>G), GAC477GAT (1581C>T), AAG478AAA (1584G>A), GGG479GGA (1587G>A), CAT481CAC (1593T>C), ACT483ACA (1599T>A), TAT486TAC (1608T>C), GCA489GCC (1617A>C), CTT493TTG (1627C>T 1629T>G), GTA494ATA (1630G>A), TCA495TCT (1635A>T), GCA496GCT (1638A>T), CCT497CCA (1641T>A), GCC498GCT (1644C>T), CTT502CTA (1656T>A), ACC503ACG (1659C>G), GCT507GCA (1671T>A), GAG509GAA (1677G>A), CGT510AGG (1678C>A 1680T>G), AGA511AGG (1683A>G), AGG512AGA (1686G>A), ATC514ATT (1692C>T), TGT518TGC (1704T>C), ATC519AGA (1706T>G 1707C>A), GAA521GTG (1712A>T 1713A>G), GGT522GGC (1716T>C), GAA523GAT (1719A>T), TCT525GAT (1723T>G 1724C>A), ACA526CCA (1726A>C), GAG527AAT (1729G>A 1731G>T), GAC528ACT (1732G>A 1733A>C 1734C>T), ACC530ACA (1740C>A), GGA531GCC (1742G>C 1743A>C), CTG533CTA (1749G>A), ATG534ATT (1752G>T), AAC536GAT (1756A>G 1758C>T), ATT537GTG (1759A>G 1761T>G), CGG541AGA (1771C>A 1773G>A), AGG542AGA (1776G>A), ATA546ATC (1788A>C), CAC550GCT (1798C>G 1799A>C 1800C>T), AGT552TCC (1804A>T 1805G>C 1806T>C), TAC553TAT (1809C>T), TCG554GAT (1810T>G 1811C>A 1812G>T), TTA557TTG (1821A>G), GAC558CAA (1822G>C 1824C>A), GCC562GCA (1836C>A), TGT564TGC (1842T>C), CAG565CGG (1844A>G), GTT566GTG (1848T>G), CAT567CAC (1851T>C), GCT568TAT (1852G>T 1853C>A), ATA570GTG (1858A>G 1860A>G), AAG572AGC (1865A>G 1866G>C), CAG573CAA (1869G>A), CTG574CTT (1872G>T), TTT575TAC (1874T>A 1875T>C), GAA576AAT (1876G>A 1878A>T), GCA577ACT (1879G>A 1881A>T), GAA578ACA (1882G>A 1883A>C), ACT579TCG (1885A>T 1887T>G), GGC580GGA (1890C>A), ATT581TTG (1891A>T 1893T>G), TCA582GCT (1894T>G 1896A>T), AAT583GAC (1897A>G 1899T>C), ACA585GTT (1903A>G 1904C>T 1905A>T), GCC586GCT (1908C>T), GTT588GTC (1914T>C), GCT589AAT (1915G>A 1916C>A), GTA590GTT (1920A>T), TCT591GCT (1921T>G), TCT592GTT (1924T>G 1925C>T), GAA593GAT (1929A>T), GAG594GAA (1932G>A), GTT595GTG (1935T>G), GTG596CTG (1936G>C), GCA597GAT (1940C>A 1941A>T), GTT599TTC (1945G>T 1947T>C), GAT600CAA (1948G>C 1950T>A), GGG601GAT (1952G>A 1953G>T), GCT602GCC (1956T>C), CGC603del (1957_1959delCGC), TTC604TTA (1962C>A), AAA605CAA (1963A>C), ACA606GTT (1966A>G 1967C>T 1968A>T), AAA607TTG (1969A>T 1970A>T 1971A>G), CAA608CCT (1973A>C 1974A>T), GAT609CCT (1975G>C 1976A>C), GTG610GAG (1979T>A), CCA611TCA (1981C>T), ATA612GCA (1984A>G 1985T>C), GTG613GCT (1988T>C 1989G>T), ACT614TAT (1990A>T 1991C>A), CCC615GAG (1993C>G 1994C>A 1995C>G), ACA616CAG (1996A>C 1997C>A 1998A>G), GTT617ACA (1999G>A 2000T>C 2001T>A), ATA618ATT (2004A>T), GAG620GAA (2010G>A), GAA621GCT (2012A>C 2013A>T), GAT622GAA (2016T>A), AGA623AAA (2018G>A), GTG624ATT (2020G>A 2022G>T), ATT625GTT (2023A>G), TCT627TCC (2031T>C), AAA628CAG (2032A>C 2034A>G), TTA630CTC (2038T>C 2040A>C), GTG632GTT (2046G>T), GAA633GAG (2049A>G), CTT635CTC (2055T>C), TTT637TAT (2060T>A), TAT639TAC (2067T>C), CAG640CAA (2070G>A), GGC641GGG (2073C>G), TTG643ATG (2077T>A), GAC644AAT (2080G>A 2082C>T), CCG645AAA (2083C>A 2084C>A 2085G>A), GAG646TTT (2086G>T 2087A>T 2088G>T), AGT647GAG (2089A>G 2090G>A 2091T>G), TTG648TTT (2094G>T), TTT649TTC (2097T>C), CCT650CCA (2100T>A), TAT653CTC (2107T>C 2108A>T 2109T>C), CAA654TCC (2110C>T 2111A>C 2112A>C), AAA655AAG (2115A>G), CAA656CAG (2118A>G), CTC658ATC (2122C>A), TTT659TAT (2126T>A), TCC661GTG (2131T>G 2132C>T 2133C>G), GAT663TCT (2137G>T 2138A>C), GCA664TCT (2140G>T 2142A>T), GAG665GAT (2145G>T), CAC666GCC (2146C>G 2147A>C), CAG667AAA (2149C>A 2151G>A), CAA668CAG (2154A>G), GTC669AAT (2155G>A 2156T>A 2157C>T), TTC670TTT (2160C>T), ATG671GAA (2161A>G 2162T>A 2163G>A), AGA672ATG (2165G>T 2166A>G), ATG674GTA (2170A>G 2172G>A), ACT675AAT (2174C>A), GAC676GAT (2178C>T), ATG677TTG (2179A>T), ATC678TTG (2182A>T 2184C>G), TAC679GTC (2185T>G 2186A>T), GGT681GGA (2193T>A), GCC682GCA (2196C>A), AAT683AGT (2198A>G), GCT684CCT (2200G>C), GAA685CAG (2203G>C 2205A>G), CAA686CAG (2208A>G), TAC687TAT (2211C>T), AGG688GCT (2212A>G 2213G>C 2214G>T), CAA692CGA (2225A>G), AAT693AAC (2229T>C), CTC699ATC (2245C>A), TTT702TTC (2256T>C), AAA703AAG (2259A>G), AGC704AGT (2262C>T), ATC706ATT (2268C>T), GCT708AGC (2272G>A 2273C>G 2274T>C), TCC709ACA (2275T>A 2277C>A), ATC711ATT (2283C>T), TGT712TCT (2285G>C), AAG714GAG (2290A>G), AAG715CTG (2293A>C 2294A>T), GCT717GAT (2300C>A), GTA718GTC (2304A>C), ACA720CGC (2308A>C 2309C>G 2310A>C), GAG721AAT (2311G>A 2313G>T), ATG722CTT (2314A>C 2316G>T), CGC723GAG (2317C>G 2318G>A 2319C>G), ACA724CAA (2320A>C 2321C>A), GGA725GGG (2325A>G), TTT726AAA (2326T>A 2327T>A 2328T>A), GCA727GCG (2331A>G), CAT728TAT (2332C>T), AAT729AAG (2337T>G), TGT730GCG (2338T>G 2339G>C 2340T>G), ATT731CTT (2341A>C), GAT732CAG (2344G>C 2346T>G), GTT735GCA (2354T>C 2355T>A), TTG738CTT (2362T>C 2364G>T), ATT739ACG (2366T>C 2367T>G), TGC740TGT (2370C>T), ATC741ATT (2373C>T), CAG742GAA (2374C>G 2376G>A), TTT747AGA (2389T>A 2390T>G 2391T>A), GTG748GTA (2394G>A), TTT750TTC (2400T>C), TTG751CTG (2401T>C), ACA753ACT (2409A>T), GCC754GCT (2412C>T), CGT757CGC (2421T>C), TAT758TAC (2424T>C), ACC761AGG (2432C>G 2433C>G), TCC764GCA (2440T>G 2442C>A), TCC764_AAA765insGGT (2442_2443insGGT), AAA765ACT (2444A>C 2445A>T), GAG766GAC (2448G>C), GCC767AAG (2449G>A 2450C>A 2451C>G), AAA768AAG (2454A>G), GAC769AGA (2455G>A 2456A>G 2457C>A), TCT770AAT (2458T>A 2459C>A), GTC772ATT (2464G>A 2466C>T), AAG773ATC (2468A>T 2469G>C), ATA774GAA (2470A>G 2471T>A), TGT775TGC (2475T>C), TAC776ATG (2476T>A 2477A>T 2478C>G), CAC777ATT (2479C>A 2480A>T 2481C>T), AAA778GGT (2482A>G 2483A>G 2484A>T), ATT779ATC (2487T>C), GTG780AAG (2488G>A 2489T>A), GAT781GAC (2493T>C), TAC782AAA (2494T>A 2496C>A), AAA784ATC (2501A>T 2502A>C), GAA785CAG (2503G>C 2505A>G), ACT786ATG (2507C>T 2508T>G), TGG788AAT (2512T>A 2513G>A 2514G>T), AAC789GAC (2515A>G), TTA790TTG (2520A>G), CCA791CCC (2523A>C), TAT792TTC (2525A>T 2526T>C), GCA793ATC (2527G>A 2528C>T 2529A>C), CTG794TTG (2530C>T), AGG795AAG (2534G>A), GGC801GGA (2553C>A), CTA802CTG (2556A>G), ATC804TTG (2560A>T 2562C>G), ATG805ATA (2565G>A), GCG806GCA (2568G>A), GGA809GGC (2577A>C), TTG810ACC (2578T>A 2579T>C 2580G>C), TTT811ATG (2581T>A 2583T>G), GGG812AAT (2584G>A 2585G>A 2586G>T), GGA813GCC (2588G>C 2589A>C), ATA814GTT (2590A>G 2592A>T), ACA815ACT (2595A>T), TTT817TTC (2601T>C), CTT818ATT (2602C>A), GCT819ACT (2605G>A), TGT820TGC (2610T>C), TGG821CTC (2611T>C 2612G>T 2613G>C), CAA822AAG (2614C>A 2616A>G), AGC823GCA (2617A>G 2618G>C 2619C>A), AAT824GCA (2620A>G 2621A>C 2622T>A), TCT825ACT (2623T>A), TTT826TCC (2627T>C 2628T>C), CCA827CCT (2631A>T), AGT828TTA (2632A>T 2633G>T 2634T>A), AGT830GAT (2638A>G 2639G>A), GGA831GGC (2643A>C), GGT833GGG (2649T>G), GGT834GGC (2652T>C), GTC835GTA (2655C>A), ACA836ACC (2658A>C), AGC841AGT (2673C>T), TCT843TCC (2679T>C), AAA845AAG (2685A>G), AAG846AAA (2688G>A), AAC847AAT (2691C>T), AAG848AAA (2694G>A), GGC849GGG (2697C>G), CCC850AAG (2698C>A 2699C>A 2700C>G), TCT851ACT (2701T>A), CGT852CAT (2705G>A), CTG853TTC (2707C>T 2709G>C), AAG854GCA (2710A>G 2711A>C 2712G>A), AAT855AAA (2715T>A), TTG856TCC (2717T>C 2718G>C), GTG858CTA (2722G>C 2724G>A), CAG860CAA (2730G>A), AGT861GCG (2731A>G 2732G>C 2733T>G), AGC862GCA (2734A>G 2735G>C 2736C>A), GCT863GAT (2738C>A), ATT865AAC (2744T>A 2745T>C), ACA866CAT (2746A>C 2747C>A 2748A>T), CAA867GTG (2749C>G 2750A>T 2751A>G), GAA868GAC (2754A>C), GAA868_TGG869insCGG (2754_2755insCGG), GCT871GAA (2762C>A 2763T>A), GAA872CAG (2764G>C 2766A>G), GGG874GGA (2772G>A), GTC876GTT (2778C>T), AAC877AAT (2781C>T), CAA878GAA (2782C>G), GCA879AGT (2785G>A 2786C>G 2787A>T), CTC880CTG (2790C>G), AAA881CAC (2791A>C 2793A>C), AAG882AAA (2796G>A), AAT883AAC (2799T>C), ATG884TTG (2800A>T), ATT885GTT (2803A>G), GTA886GTT (2808A>T), CTG887TTG (2809C>T), CGG888AGA (2812C>A 2814G>A), CTT889ATT (2815C>A), GGA890GCT (2819G>C 2820A>T), GAG891GAA (2823G>A), GTG893GTC (2829G>C), TAC894TAT (2832C>T), ACA898ACG (2844A>G), GTG900GTT (2850G>T), TCA902TCT (2856A>T), GGG903GGT (2859G>T), ATA905ATT (2865A>T), ATG906CTC (2866A>C 2868G>C), GCA911GCC (2883A>C), TTC912TTT (2886C>T), CAC913AGA (2887C>A 2888A>G 2889C>A), AAT914AAC (2892T>C), GTG915ATG (2893G>A), AGA916AGG (2898A>G), GGA918GGC (2904A>C), ACA919ACT (2907A>T), ACC922TCC (2914A>T), ATT923ATC (2919T>C), ATC924ATT (2922C>T), CCT926GAG (2926C>G 2927C>A 2928T>G), AAC927CAT (2929A>C 2931C>T), TCA928TCC (2934A>C), AGG929ATT (2936G>T 2937G>T), TCA930TGT (2939C>G 2940A>T), AAG931AAT (2943G>T), TAC934TTC (2951A>T), AAC935AAT (2955C>T), CGG937ACC (2959C>A 2960G>C 2961G>C), GAA938CAA (2962G>C), TCC939TCA (2967C>A), AAG940AGG (2969A>G), GTC941TTG (2971G>T 2973C>G), ATT942ATA (2976T>A), GGA944GGT (2982A>T), ATC947GTG (2989A>G 2991C>G), CTG949TTG (2995C>T),

Codon mutations: CTA950TTG (2998C>T 3000A>G), CGT951AGG (3001C>A 3003T>G), GTG952GTA (3006G>A), GGT953GGG (3009T>G), GAC954GAT (3012C>T), CCT955CCA (3015T>A), CCT960TCA (3028C>T 3030T>A), ATT962ATA (3036T>A), CGC963AGA (3037C>A 3039C>A), AAG964AAA (3042G>A), CCA967CCT (3051A>T), GAG971GAT (3063G>T), GTC972GTG (3066C>G), TGC973TGT (3069C>T), TTC974TAC (3071T>A), ACC975TCT (3073A>T 3075C>T), AAA976AAG (3078A>G), GGT977GGC (3081T>C), TCT978TCA (3084T>A), CAG979ATG (3085C>A 3086A>T), TTA981TTG (3093A>G), TGT982TGC (3096T>C), TGT983GTG (3097T>G 3098G>T 3099T>G), AGA984AGG (3102A>G), GCC985GCT (3105C>T), GTG986GTA (3108G>A), GCG987GCT (3111G>T), ACC989AAG (3116C>A 3117C>G), CCA991CCC (3123A>C), AGA992AAG (3125G>A 3126A>G), CTG993AAA (3127C>A 3128T>A 3129G>A), GGC994TCC (3130G>T 3131G>C), AAC995AAT (3135C>T), CTA996TTG (3136C>T 3138A>G), GAA997GAG (3141A>G), TTT998GTT (3142T>G), TTG999CTT (3145T>C 3147G>T), TTT1001TAT (3152T>A), TCC1006TCA (3168C>A), AAA1007AAG (3171A>G), TAT1009TAC (3177T>C), ACA1010ACC (3180A>C), GTC1011GAA (3182T>A 3183C>A), AAG1012AAA (3186G>A), GTG1013GTC (3189G>C), GAG1014ACA (3190G>A 3191A>C 3192G>A), GAG1016GGA (3197A>G 3198G>A), GAC1018GAT (3204C>T), TCT1019TCG (3207T>G), GCC1020AAT (3208G>A 3209C>A 3210C>T), AGT1021GCA (3211A>G 3212G>C 3213T>A), AAA1023GCT (3217A>G 3218A>C 3219A>T), TTG1024CTT (3220T>C 3222G>T), AGT1025AGC (3225T>C), TCG1026TCA (3228G>A), CAG1027CAA (3231G>A), CAA1028GTT (3232C>G 3233A>T 3234A>T), TCA1029GCA (3235T>G), GAA1031GAG (3243A>G), TAT1032TAC (3246T>C), CAT1033CAC (3249T>C), ATG1034ATT (3252G>T), GGA1036AGT (3256G>A 3258A>T), GGT1039GGA (3267T>A), GGA1042GGT (3276A>T), TTA1045CTC (3283T>C 3285A>C), CTT1046TTA (3286C>T 3288T>A), CCC1048CCA (3294C>A), AGT1049ACT (3296G>C), GTT1050GTG (3300T>G), AAA1053AAG (3309A>G), CAA1054CAG (3312A>G), CCT1055CCG (3315T>G), GTT1056TGC (3316G>T 3317T>G 3318T>C), GTG1058GTT (3324G>T), GGC1059GGA (3327C>A), ATT1060ATA (3330T>A), GCT1063GCA (3339T>A), TCA1064TCT (3342A>T), GGT1067GAT (3350G>A), ATT1068ACC (3353T>C 3354T>C), GCA1069GCT (3357A>T), GAA1071GAG (3363A>G), GGA1073GGT (3369A>T), TTC1074TAC (3371T>A), ATT1075GTG (3373A>G 3375T>G), TCT1076GCT (3376T>G), TCC1077TCA (3381C>A), AAT1078AAC (3384T>C), GCC1079GCA (3387C>A), ACG1080ACA (3390G>A), GCC1081GCA (3393C>A), ATC1082GTG (3394A>G 3396C>G), TTC1083TTT (3399C>T), CGG1084AGA (3400C>A 3402G>A), GAC1085GAT (3405C>T), CAG1086CAA (3408G>A), CTT1087TTG (3409C>T 3411T>G), AAC1089GAT (3415A>G 3417C>T), GGG1093GGA (3429G>A), CCT1094CCC (3432T>C), GTT1095ATT (3433G>A), GCG1097GTG (3440C>T), GCA1098GCT (3444A>T), CGA1101CGC (3453A>C), GAC1103GAA (3459C>A), ATT1104CTG (3460A>C 3462T>G), CTC1105TTG (3463C>T 3465C>G), AAA1106AAG (3468A>G), GCC1107CAC (3469G>C 3470C>A), ATT1108CTG (3472A>C 3474T>G), AGG1109AGA (3477G>A), AGT1110TCC (3478A>T 3479G>C 3480T>C), AGA1111AAG (3482G>A 3483A>G), GAA1112GAG (3486A>G), ACC1113GAA (3487A>G 3488C>A 3489C>A), ACT1114AGA (3491C>G 3492T>A), CTT1115TTC (3493C>T 3495T>C), GAA1117GAG (3501A>G), AAC1119AAA (3507C>A), CAA1120CAG (3510A>G), GTC1121GTG (3513C>G), TCC1122GCT (3514T>G 3516C>T), TTT1123TTC (3519T>C), CTT1124TTG (3520C>T 3522T>G), GTT1127GTA (3531T>A), CCT1128CCG (3534T>G), CAA1129TTG (3535C>T 3536A>T 3537A>G), TTG1131ATG (3541T>A), GCC1132GCT (3546C>T), GCA1133GCT (3549A>T), GTA1135GTG (3555A>G), CCC1136CCG (3558C>G), CAT1137CAC (3561T>C), ACT1139ACA (3567T>A), ACA1140ACC (3570A>C), CTC1141CTT (3573C>T), AGA1142AAA (3575G>A), AAG1143CGC (3577A>C 3578A>G 3579G>C), CAA1145CCC (3584A>C 3585A>C), TTG1146CTG (3586T>C), GCA1149GCT (3597A>T), GGT1151GGG (3603T>G), CCT1152GAC (3604C>G 3605C>A 3606T>C), GAA1154CAG (3610G>C 3612A>G), ACC1155ACT (3615C>T), GCT1156GCC (3618T>C), CCC1157CCA (3621C>A), TCA1158TCC (3624A>C), ATC1159ATT (3627C>T), CTC1160CTA (3630C>A), ACC1161TCT (3631A>T 3633C>T), GTC1162GCC (3635T>C), AAT1163AAA (3639T>A), CGG1166AGG (3646C>A), GAG1168GAA (3654G>A), GGA1169GGG (3657A>G), TTC1170TTT (3660C>T), CCA1172CCC (3666A>C), GCA1175GCT (3675A>T), GTG1177GTC (3681G>C), ATG1178GAG (3682A>G 3683T>A), AAA1179AAG (3687A>G), TAC1180TAT (3690C>T), ACT1183ACA (3699T>A), CAT1185TCC (3703C>T 3704A>C 3705T>C), GGT1186AAC (3706G>A 3707G>A 3708T>C), GAT1188GAG (3714T>G), GAT1189GAG (3717T>G), GAT1190CCA (3718G>C 3719A>C 3720T>A), GCT1192GCA (3726T>A), AAA1193AAG (3729A>G), TTG1194CTG (3730T>C), TCC1195GCT (3733T>G 3735C>T), GAA1197GAG (3741A>G), ACT1198GGT (3742A>G 3743C>G), CTC1199CTT (3747C>T), AGT1201GCA (3751A>G 3752G>C 3753T>A), TCA1202TCC (3756A>C), CTG1203ATT (3757C>A 3759G>T), CTG1204CTT (3762G>T), CCA1205CCT (3765A>T), ACA1206GTG (3766A>G 3767C>T 3768A>G), ATG1207CTT (3769A>C 3771G>T), AAA1208AAG (3774A>G), AAC1209ATG (3776A>T 3777C>G), CAA1210CAG (3780A>G), AAA1211CGA (3781A>C 3782A>G), GGC1215GGG (3795C>G), CCA1217CCT (3801A>T), CAG1218GCA (3802C>G 3803A>C 3804G>A), GAG1219ATC (3805G>A 3806A>T 3807G>C), GAT1221AAT (3811G>A), GAT1223GAC (3819T>C), GTG1224GTA (3822G>A), CTG1226CTC (3828G>C), ATT1229ATA (3837T>A), TGT1232TTT (3845G>T), TAT1235TTT (3854A>T), GGC1237GCA (3860G>C 3861C>A), CTC1240TTG (3868C>T 3870C>G), ACA1242ACG (3876A>G), TCC1243TCA (3879C>A), TAC1246TAT (3888C>T), TTC1248TTT (3894C>T), ATG1251TTG (3901A>T), GGC1252GGA (3906C>A), ATG1253CTT (3907A>C 3909G>T), GGA1254GGT (3912A>T), ATG1255TCT (3913A>T 3914T>C 3915G>T), AAG1257AAA (3921G>A), CGG1258AGG (3922C>A), GAG1259GAA (3927G>A), GTC1261GTG (3933C>G), GTA1262GAA (3935T>A), CCA1263GCC (3937C>G 3939A>C), ACA1264CAA (3940A>C 3941C>A), GGA1265CCA (3943G>C 3944G>C), GAA1266AAC (3946G>A 3948A>C), GGT1267TCG (3949G>T 3950G>C 3951T>G), GAT1268CAG (3952G>C 3954T>G), AAA1269AGG (3956A>G 3957A>G), GTA1270GTT (3960A>T), GAG1271CTT (3961G>C 3962A>T 3963G>T), CTT1272CTC (3966T>C), AGA1273ATA (3968G>T), GAC1275AAC (3973G>A), CCA1277CCT (3981A>T), GTG1278GTC (3984G>C), GAA1280GAT (3990A>T), GCT1281GCA (3993T>A), GAG1283GAA (3999G>A), GAA1284GAG (4002A>G), CTG1285TTG (4003C>T), ATT1287TCG (4009A>T 4010T>C 4011T>G), CAA1288CAG (4014A>G), ATT1289GTA (4015A>G 4017T>A), CAG1291CAA (4023G>A), ATA1293ACA (4028T>C), CAT1294TTT (4030C>T 4031A>T), GTG1295GTT (4035G>T), GAT1296GAC (4038T>C), CTA1297CTC (4041A>C), ATC1298GTC (4042A>G), ACC1299ACA (4047C>A), ACA1300ACC (4050A>C), CAA1301CAG (4053A>G), GCC1303GCG (4059C>G), AAG1304AAA (4062G>A), GAG1306GAA (4068G>A), CGC1307CGG (4071C>G), CTT1308TTG (4072C>T 4074T>G), CCT1309CCA (4077T>A), GAA1311GAG (4083A>G), AAA1318AAG (4104A>G), ACA1319ACC (4107A>C), AGA1320CGC (4108A>C 4110A>C), TTA1321CTG (4111T>C 4113A>G), TTT1322TTC (4116T>C), GAG1323GAA (4119G>A), ATA1324ATC (4122A>C), TAC1329TAT (4137C>T), GTC1333GTG (4149C>G), AGG1334CGG (4150A>C), TTT1337TTC (4161T>C), TCC1341TCT (4173C>T), GCC1342GCT (4176C>T), ACC1343TGT (4177A>T 4178C>G 4179C>T), GCA1346GCT (4188A>T), TTA1347GCA (4189T>G 4190T>C), AAT1349AAC (4197T>C), GTC1350GTA (4200C>A), ATC1351CTC (4201A>C), CCA1352CCT (4206A>T), TGT1353TGC (4209T>C), GTT1355GTA (4215T>A), GGA1356GGT (4218A>T), ATA1357ATC (4221A>C), GAT1358AAT (4222G>A), ACC1360ACG (4230C>G), AGT1361AGC (4233T>C), TCT1362CAT (4234T>C 4235C>A), TCT1365ACA (4243T>A 4245T>A), CTT1366GTG (4246C>G 4248T>G), CTT1367CTG (4251T>G), GCC1368GCT (4254C>T), AAC1369AGT (4256A>G 4257C>T), AAC1370AGC (4259A>G), TTC1371TTT (4263C>T), GCT1373TCA (4267G>T 4269T>A), ACT1377GAG (4279A>G 4280C>A 4281T>G), GGG1378GGA (4284G>A), TTT1379TTC (4287T>C), GCT1381GCC (4293T>C), TCA1384TCG (4302A>G), AGT1385AGC (4305T>C), GAC1387GAT (4311C>T), GGG1388GGA (4314G>A), CGC1389AGA (4315C>A 4317C>A), CCA1391CCC (4323A>C), GTG1392ATC (4324G>A 4326G>C), TTC1393TTT (4329C>T), GCT1394GCA (4332T>A), TTT1395TTC (4335T>C), CAG1396CAA (4338G>A), TTG1401CTG (4351T>C), GTT1402GTG (4356T>G), GAG1404GCT (4361A>C 4362G>T), TAT1405TTT (4364A>T), GAT1407GGT (4370A>G), ACC1408GAT (4372A>G 4373C>A 4374C>T), AAG1409GAG (4375A>G), GGA1411GGG (4383A>G), GAG1412AGT (4384G>A 4385A>G 4386G>T), GCT1414GAG (4391C>A 4392T>G), TCC1415TCA (4395C>A), GCA1417GCT (4401A>T), AGA1418AGG (4404A>G), CAT1419CAC (4407T>C), GCT1420ACA (4408G>A 4410T>A), CTT1421TTA (4411C>T 4413T>A), CTC1422TTA (4414C>T 4416C>A), GCA1425GCC (4425A>C), TCA1426TCT (4428A>T), CAC1428CAT (4434C>T), TAC1429TAT (4437C>T), TTA1431CTC (4441T>C 4443A>C), GAT1434AAC (4450G>A 4452T>C), AAA1435AAG (4455A>G), GTT1436TTG (4456G>T 4458T>G), TTC1437TTT (4461C>T), CGT1438CGC (4464T>C), TTG1439CTA (4465T>C 4467G>A), GTT1440GTG (4470T>G), GGC1442GGA (4476C>A), CCG1444CCC (4482G>C), GCA1448GCT (4494A>T), CTT1449TTG (4495C>T 4497T>G), ACC1450ACT (4500C>T), TTC1453TTT (4509C>T), AAC1454AAT (4512C>T), CTG1456TTG (4516C>T), TTG1457CTA (4519T>C 4521G>A), CGG1463AGG (4537C>A), TAT1464TAC (4542T>C), GCT1465GCG (4545T>G), TTC1466TTT (4548C>T), ATC1467ATA (4551C>A), CTT1469CTG (4557T>G), TTA1470TTG (4560A>G), AGA1471AGG (4563A>G), AGG1472AGA (4566G>A), CCT1473CCA (4569T>A), CAC1474CAT (4572C>T), ATT1475GTC (4573A>G 4575T>C), GCC1476AAA (4576G>A 4577C>A 4578C>A), GCT1477GCC (4581T>C), AGA1478AAA (4583G>A), GCT1479TCT (4585G>T), ATA1480GTG (4588A>G 4590A>G), GGT1481GGC (4593T>C), GTA1482ATT (4594G>A 4596A>T), CCT1484CCA (4602T>A), GGT1485AAT (4603G>A 4604G>A), GAT1486GAC (4608T>C), AAC1488AAT (4614C>T), CTT1490TTG (4618C>T 4620T>G), GGA1496GGT (4638A>T), AAT1499AAC (4647T>C), CTG1500TTG (4648C>T), GTG1501GTT (4653G>T), GCT1502GCG (4656T>G), GTG1503GTT (4659G>T), CCT1504CCA (4662T>A), CTG1505ATG (4663C>A), CTG1507TTG (4669C>T), TAT1510TAC (4680T>C), TCA1511GAT (4681T>G 4682C>A 4683A>T), CTG1512CTA (4686G>A), CCT1513CCA (4689T>A), AAT1514TCC (4690A>T 4691A>C 4692T>C), GCC1516GCA (4698C>A), CAG1517AGA (4699C>A 4700A>G 4701G>A), CTT1519CTG (4707T>G), GAC1520GAA (4710C>A), ATG1521GCT (4711A>G 4712T>C 4713G>T), GTA1522GTT (4716A>T), AAT1523AAC (4719T>C), ATC1526ATT (4728C>T), AAA1527AAG (4731A>G), GGG1529GGT (4737G>T), CTG1530TTG (4738C>T), GAT1531GAC (4743T>C), TCT1533TCA (4749T>A), AAC1537GAG (4759A>G 4761C>G), CAG1538AAG (4762C>A), GTG1539ACA (4765G>A 4766T>C 4767G>A), CAG1540AAA (4768C>A 4770G>A), TTC1541TTT (4773C>T), CAG1542CTG (4775A>T), GAC1543GAT (4779C>T), TCA1545TCT (4785A>T), GAA1546GAG (4788A>G), ACT1548ACC (4794T>C), AGC1551AGT (4803C>T), GGA1553GGT (4809A>T), TTT1554TTC (4812T>C), AAA1555AAG (4815A>G), CGT1556CGA (4818T>A), GCC1558GCA (4824C>A), CTT1559TTG (4825C>T 4827T>G), GGG1560GGA (4830G>A), CAC1562CAA (4836C>A), GTG1569GTT (4857G>T), ATA1571GTC (4861A>G 4863A>C), ATT1572ATC (4866T>C), CCT1574CCC (4872T>C), ATG1575ATT (4875G>T), TAC1576TAT (4878C>T), ATC1578ATT (4884C>T), AGA1579AGG (4887A>G), CCT1580CCA (4890T>A), CCG1582GTG (4894C>G 4895C>T), GGT1583GGA (4899T>A), ACT1587GTA (4909A>G 4910C>T 4911T>A), GTG1588CAA (4912G>C 4913T>A 4914G>A), GCA1589GCT (4917A>T), ATG1591TAT (4921A>T 4922T>A 4923G>T), GAG1592GAA (4926G>A), GTG1594ATT (4930G>A 4932G>T), ACT1596ACA (4938T>A), CTT1598ATC (4942C>A 4944T>C), CGT1599AGA (4945C>A 4947T>A), GAG1600GAA (4950G>A), TTT1602TTC (4956T>C), AGA1606CAG (4966A>C 4967G>A 4968A>G), GTT1607GCC (4970T>C 4971T>C), GCA1608ACT (4972G>A 4974A>T), GAG1610GAA (4980G>A), AAG1611AGT (4982A>G 4983G>T), TTA1612TTT (4986A>T), ACT1613GTA (4987A>G 4988C>T 4989T>A), TAT1614AAT (4990T>A), CAG1617TTT (4999C>T 5000A>T 5001G>T), TGT1618GGA (5002T>G 5004T>A), GCC1619GCT (5007C>T), AGA1621AAG (5012G>A 5013A>G), GAG1622CAG (5014G>C), CGT1623AGA (5017C>A 5019T>A), GGA1624AAT (5020G>A 5021G>A 5022A>T), GGC1626CCC (5026G>C 5027G>C), GCA1627ATC (5029G>A 5030C>T 5031A>C), ACA1628ATG (5033C>T 5034A>G), TTT1630ATG (5038T>A 5040T>G), CCT1631CCC (5043T>C), TCT1632ACT (5044T>A), TAT1633TAC (5049T>C), ATG1634TAT (5050A>T 5051T>A 5052G>T), GTG1636CTT (5056G>C 5058G>T), GAG1637GAA (5061G>A), CAG1638AGT (5062C>A 5063A>G 5064G>T), GAA1639CAA (5065G>C), ATC1641TTG (5071A>T 5073C>G), GCA1642ACA (5074G>A), AAG1643GAG (5077A>G), GCA1648GAG (5093C>A 5094A>G), TCA1649AGC (5095T>A 5096C>G 5097A>C), GCG1650ATT (5098G>A 5099C>T 5100G>T), TTG1651GGG (5101T>G 5102T>G), ACA1652GTT (5104A>G 5105C>T 5106A>T), ATC1653TTG (5107A>T 5109C>G), TGT1654ATG (5110T>A 5111G>T 5112T>G), GAA1655GAG (5115A>G), ATG1656CTA (5116A>C 5118G>A), GCC1657TCA (5119G>T 5121C>A), AAA1658GAT (5122A>G 5124A>T), TCT1660AAA (5128T>A 5129C>A 5130T>A), ATC1661AAT (5132T>A 5133C>T), ACT1662ACA (5136T>A), TTC1663TAC (5138T>A), ACA1664ACC (5142A>C), TTG1666TTA (5148G>A), CCA1667CCT (5151A>T), CCT1668GCT (5152C>G), AAT1670GAA (5158A>G 5160T>A), GTT1672ATT (5164G>A), GAA1673del (5167_5169delGAA), GAA1676GAG (5178A>G), AGA1677GAG (5179A>G 5180G>A 5181A>G), GAT1678GGT (5183A>G), CTC1679GTC (5185C>G), AAT1680CAT (5188A>C), TGG1681GAA (5191T>G 5192G>A 5193G>A), GCC1683GTG (5198C>T 5199C>G), CCA1684CCT (5202A>T), AAC1685GAT (5203A>G 5205C>T), ATT1686GTA (5206A>G 5208T>A), GGT1687TTT (5209G>T 5210G>T), TGC1689TGT (5217C>T), TCA1690AGT (5218T>A 5219C>G 5220A>T), GCC1691GCT (5223C>T), CGT1692CGG (5226T>G), GCC1695TCA (5233G>T 5235C>A), AAG1696AAC (5238G>C), ACT1697CAA (5239A>C 5240C>A 5241T>A), GAA1698CAA (5242G>C), GAG1700TCA (5248G>T 5249A>C 5250G>A), CCT1701TCA (5251C>T 5253T>A), GGG1702GGT (5256G>T), TGC1703AAC (5257T>A 5258G>A), ATA1704ATC (5262A>C), GTA1705CTG (5263G>C 5265A>G), AAC1707AAT (5271C>T), TGT1708TGC (5274T>C), ACA1709ACT (5277A>T), GGG1710GGA (5280G>A), GCG1711GCC (5283G>C), AAG1712CGC (5284A>C 5285A>G 5286G>C), AAG1713AGA (5288A>G 5289G>A), AGC1714ACA (5291G>C 5292C>A), AAA1715ACA (5294A>C), GTT1717GTG (5301T>G), CGC1718AGA (5302C>A 5304C>A), GGA1719GGG (5307A>G), GCT1721GCC (5313T>C), AGT1722AAT (5315G>A), GAT1725GAC (5325T>C), ATG1730ATA (5340G>A), CCA1732CCC (5346A>C), TAT1733TAC (5349T>C), ACC1734ACT (5352C>T), ATG1735TTG (5353A>T), TCA1736GCA (5356T>G), ATT1738ATC (5364T>C), CTG1739CAG (5366T>A), GCT1741CAT (5371G>C 5372C>A), AGA1743CGC (5377A>C 5379A>C), GGC1744AAG (5380G>A 5381G>A 5382C>G), AAA1745AGT (5384A>G 5385A>T), ATG1746CTG (5386A>C), TCA1747GAA (5389T>G 5390C>A), GAG1748GTT (5393A>T 5394G>T), GAC1749GGT (5396A>G 5397C>T), ATT1750GAT (5398A>G 5399T>A), GCT1751GGG (5402C>G 5403T>G), GAT1754TCT (5410G>T 5411A>C), GTT1755GTC (5415T>C), GTG1756ACC (5416G>A 5417T>C 5418G>C), GTG1758GTT (5424G>T), GGG1760GGT (5430G>T), GCT1764GCA (5442T>A), GTG1765ATA (5443G>A 5445G>A), AGC1766GCT (5446A>G 5447G>C 5448C>T), GCT1771GCA (5463T>A), TTA1772TTG (5466A>G), ATG1773TTT (5467A>T 5469G>T), GCT1774GCA (5472T>A), GTG1775TTG (5473G>T), CGT1778CAT (5483G>A), CAG1779CAA (5487G>A), TAT1780TAC (5490T>C), TGC1781AGG (5491T>A 5493C>G), ATT1782GCC (5494A>G 5495T>C 5496T>C), GAG1783ACA (5497G>A 5498A>C 5499G>A), GAT1784GAC (5502T>C), ATC1785CTT (5503A>C 5505C>T), AGG1788CGC (5512A>C 5514G>C), TTG1789CTA (5515T>C 5517G>A), CGC1790CGA (5520C>A), ACT1791ACC (5523T>C), ATT1792ATG (5526T>G), GGC1793GCT (5528G>C 5529C>T), AAT1794GGG (5530A>G 5531A>G 5532T>G), TTG1796CTG (5536T>C), ACA1800ACC (5550A>C), TAT1801TAC (5553T>C), CCC1802CCT (5556C>T), GTT1806GCG (5567T>C 5568T>G), CAG1807CAA (5571G>A), TTT1809TTG (5577T>G), CAA1811AAA (5581C>A), GTG1813GTT (5589G>T), CCT1814GAA (5590C>G 5591C>A 5592T>A), CAG1815GAC (5593C>G 5595G>C), CAG1815_GAA1816insACC (5595_5596insACC), GAA1816TCA (5596G>T 5597A>C), GGG1817CAC (5599G>C 5600G>A 5601G>C), AAA1818AAG (5604A>G), ATG1819TTG (5605A>T), GCA1820GGA (5609C>G), GTT1822AGT (5614G>A 5615T>G), TCG1824ACA (5620T>A 5622G>A), TCT1825del (5623_5625delTCT), TTG1826TAT (5627T>A 5628G>T), CAA1827GGC (5629C>G 5630A>G 5631A>C), AGT1828ACC (5633G>C 5634T>C), AGT1829ACT (5636G>C), ATG1831AAC (5642T>A 5643G>C), CAA1833CAT (5649A>T), CAA1833_GGT1834insCACAGT (5649_5650insCACAGT), GGT1834GCA (5651G>C 5652T>A), GAG1836TCA (5656G>T 5657A>C 5658G>A), CTT1837GTT (5659C>G), GGC1838GGA (5664C>A), CAA1839CTT (5666A>T 5667A>T), ATT1840CTA (5668A>C 5670T>A), CGT1841AGG (5671C>A 5673T>G), GTC1842ATA (5674G>A 5676C>A), ATC1843GTT (5677A>G 5679C>T), CAC1844CCT (5681A>C 5682C>T), GGT1845TCT (5683G>T 5684G>C), GAT1846GAC (5688T>C), TTA1847CTC (5689T>C 5691A>C), TCA1848TCT (5694A>T), ATG1849GAA (5695A>G 5696T>A 5697G>A), GAA1850AAG (5698G>A 5700A>G), TTG1853GTT (5707T>G 5709G>T), CGC1854CGT (5712C>T), ATG1855TTG (5713A>T), TAT1857TAC (5721T>C), GTT1858ACA (5722G>A 5723T>C 5724T>A), GTG1859ATT (5725G>A 5727G>T), GGC1860GGT (5730C>T), CCA1861CCT (5733A>T), CAT1862GAA (5734C>G 5736T>A), GGT1863GGA (5739T>A), GGA1864GGT (5742A>T), GGC1866GCC (5747G>C), AAC1867CAG (5749A>C 5751C>G), TTT1868TGC (5753T>G 5754T>C), ACC1869TCA (5755A>T 5757C>A), ACA1870ACG (5760A>G), AAA1871AAG (5763A>G), AAT1872AAC (5766T>C), CTT1873CTG (5769T>G), GAT1874AAG (5770G>A 5772T>G), CAA1875CAT (5775A>T), TTG1876GCT (5776T>G 5777T>C 5778G>T), TTT1877TTG (5781T>G), GCA1878GAG (5783C>A 5784A>G), AGT1879ACA (5786G>C 5787T>A), CTT1880TTG (5788C>T 5790T>G), GAA1881TCT (5791G>T 5792A>C 5793A>T), AAG1882GTC (5794A>G 5795A>T 5796G>C), TGT1883AGT (5797T>A), GAA1886ACT (5806G>A 5807A>C 5808A>T), TTG1887CTT (5809T>C 5811G>T), CTT1888GCC (5812C>G 5813T>C 5814T>C), AAG1890AGA (5819A>G 5820G>A), AAC1891GGA (5821A>G 5822A>G 5823C>A), ACG1892GTC (5824A>G 5825C>T 5826G>C), CTG1894GTG (5830C>G), ACC1895ACA (5835C>A), GAG1899AAA (5845G>A 5847G>A), AAA1900GAA (5848A>G), ATG1901TTC (5851A>T 5853G>C), ACT1902TCC (5854A>T 5856T>C), AAA1904CAC (5860A>C 5862A>C), GTG1906TGT (5866G>T 5867T>G 5868G>T), GCA1907GAA (5870C>A), CAA1908ATG (5872C>A 5873A>T 5874A>G), GTC1910CTT (5878G>C 5880C>T), AGT1911GAG (5881A>G 5882G>A 5883T>G), GTA1913CTG (5887G>C 5889A>G), AAA1914GAA (5890A>G),

| | |
|---|---|
| | CAG1915AGT (5893C>A 5894A>G 5895G>T), TTC1918GTA (5902T>G 5904C>A), TAT1919GTA (5905T>G 5906A>T 5907T>A), CCC1920TTC (5908C>T 5909C>T), AAG1921ATG (5912A>T), GCC1922GGG (5915C>G 5916C>G), ACA1923AGC (5918C>G 5919A>C), ACC1924GTT (5920A>G 5921C>T 5922C>T), GTG1925AAG (5923G>A 5924T>A), GAA1926AAA (5926G>A), CAG1927CAA (5931G>A), CTC1928ATG (5932C>A 5934C>G), AAG1929TTG (5935A>T 5936A>T), GTA1930TCC (5938G>T 5939T>C 5940A>C), TTT1931TTC (5943T>C), GCT1932ATG (5944G>A 5945C>T 5946T>G), GAT1933AAT (5947G>A), GCT1934CAA (5950G>C 5951C>A 5952T>A), CAT1935CAG (5955T>G), CAT1935_ACC1936insATG (5955_5956insATG), ACC1936TCA (5956A>T 5958C>A), ATT1937AAA (5960T>A 5961T>A), GTC1938ACC (5962G>A 5963T>C), AAA1939CAA (5965A>C), ACT1940GTT (5968A>G 5969C>T), GCT1941GCC (5973T>C), AAA1942AGC (5975A>G 5976A>C), CCT1943CCA (5979T>A), TTC1944TGT (5981T>G 5982C>T), CGG1945TGT (5983C>T 5985G>T), GGA1946del (5986_5988delGGA), GTA1947GTC (5991A>C), GTT1948ACA (5992G>A 5993T>C 5994T>A), TTC1949TTT (5997C>T), CGG1950AGT (5998C>A 6000G>T), AAT1952GCA (6004A>G 6005A>C 6006T>A), TTT1953TGG (6008T>G 6009T>G), CTC1954CTA (6012C>A), AGT1955ACT (6014G>C), CCT1956AAG (6016C>A 6017C>A 6018T>G), GAG1957GAC (6021G>C), TGG1958ATT (6022T>A 6023G>T 6024G>T), TGG1958_AAA1959insGTC (6024_6025insGTC), AAA1959ATG (6026A>T 6027A>G), ATT1960ACA (6029T>C 6030T>A), GGA1961AAA (6031G>A 6032G>A), GAG1963AAG (6037G>A), AAC1964AAG (6042C>G), ACA1965CCA (6043A>C), ATG1966del (6046_6048delATG), GCA1967GGA (6050C>G), ACA1968TCT (6052A>T 6054A>T), CTC1969ATT (6055C>A 6057C>T), TCA1970CCT (6058T>C 6060A>T), GAG1972GAA (6066G>A), AGT1973ACA (6068G>C 6069T>A), CTT1974GTG (6070C>G 6072T>G), CTT1975TTT (6073C>T), CCA1976CTT (6077C>T 6078A>T), CTG1979CCT (6086T>C 6087G>T), TCG1980TCA (6090G>A), TCA1982CTA (6094T>C 6095C>T), GCC1983GTT (6098C>T 6099C>T), ATG1984GTG (6100A>G), AAG1985AGA (6104A>G 6105G>A), TTG1987ATG (6109T>A), CTC1988TTG (6112C>T 6114C>G), AGC1989CGC (6115A>C), AGA1990AAT (6119G>A 6120A>T), TAC1991AGT (6121T>A 6122A>G 6123C>T), AAA1992GGA (6124A>G 6125A>G), AGA1993AAG (6128G>A 6129A>G), AAC1994AGC (6131A>G), ATG1995TAC (6133A>T 6134T>A 6135G>C), TCT1996TCA (6138T>A), CAT1998ACA (6142C>A 6143A>C 6144T>A), TGC1999ACA (6145T>A 6146G>C 6147C>A), ATA2000CTG (6148A>C 6150A>G), AAA2001GAA (6151A>G), TTG2002CTT (6154T>C 6156G>T), GCA2003GCT (6159A>T), CTC2004TTT (6160C>T 6162C>T), AAG2005AGA (6164A>G 6165G>A), ATA2006ATG (6168A>G), TAT2007TAC (6171T>C), GTT2008ATG (6172G>A 6174T>G), CTC2009_AAC2010del (6175_6180delCTCAAC), TTC2011TGT (6182T>G 6183C>T), CAG2012CAA (6186G>A), ATG2013TGG (6187A>T 6188T>G), CTG2014GTA (6190C>G 6192G>A), AAT2015CAC (6193A>C 6195T>C), GAA2017CC. (6199G>C 6200A>C) |

**Proteins**

| | | | | | | | | | |
|---|---|---|---|---|---|---|---|---|---|
| polyprotein (YP_009165993.1) | 1 | 2017 | 93.8% | 9642 | 69.4% | 2004 (99.0%) | 1319 (65.2%) | 7/13/0/0 | 0 |

| | Begin | End | Coverage | Score | Concordance | Matches | Identities | I/D/M/F* | Stop Codons |
|---|---|---|---|---|---|---|---|---|---|
| **NT** | **25** | **6200** | **85.3%** | **3315** | **27.4%** | **6117 (98.7%)** | **3941 (63.6%)** | **21/59** | |

Protein mutations:

L3F (157C>T), F7W (170T>G 171T>G), G8S (172G>A), S9P (175T>C), S10P (178T>C 180A>T), S11A (181T>G), E12_V17del (184_201delGAGAAGACTGGATCAGTT), P18S (202C>T 204C>T), T20P (208A>C), G24A (221G>C 222T>A), L26V (226C>G 228G>T), G28S (232G>T 233G>C), I30F (238A>T 240C>T), K32S (244A>T 245A>C 246A>T), F33L (247T>C 249C>T), A36T (256G>A), I37L (259A>C 261T>C), I40V (268A>G 270C>T), T41N (272C>A 273A>T), S42K (274T>A 275C>A), K46T (287A>C 288A>C), S48V (292T>G 293C>T 294T>A), L51I (301T>A), D52E (306T>G), L54V (310T>G 312G>C), S57P (319T>C 321T>A), T59M (326C>T 327T>G), D60V (329A>T 330T>A), S62A (334T>G 336T>A), S63T (337T>A 339T>A), V66A (347T>C 348T>C), S70N (358T>A 359C>A 360C>T), L72V (364T>G 366G>A), N73E (367A>G 369T>A), K74R (371A>G), G80A (389G>C 390T>A), S81K (391T>A 392C>A 393C>A), L82I (394T>A 396G>A), I83M (399T>G), L87I (409C>A 411G>C), A89G (416C>G 417A>T), S91D (421T>G 422C>A), L93V (427T>G 429G>A), A94K (430G>A 431C>A), S99T (445T>A), K101V (451A>G 452A>T 453G>A), M103L (457A>C 459G>T), R106K (466C>A 467G>A), L107M (469T>A), F108V (472T>G 474C>T), E109L (475G>T 476A>T), L111V (481T>G 483G>C), S112K (484T>A 485C>A 486C>G), N113D (487A>G), V115F (493G>T 495T>C), R116I (497G>T 498A>C), I117L (499A>C), F119Y (506T>A 507T>C), Q120E (508C>G 510G>A), V122L (514G>T 516T>A), S126V (526A>G 527G>T), D128S (532G>T 533A>C 534T>A), V129L (535G>T), V133F (547G>T), Y134I (550T>A 551A>T 552T>A), G135A (554G>C), I141L (571A>T 573A>G), T142A (574A>G 576T>C), L143I (577C>A), V145L (583G>C 585T>A), C148Y (593G>A), S152V (604T>G 605C>T), I155V (613A>G 615T>A), E156T (616G>A 617A>C 618A>G), G157P (619G>C 620G>C 621A>T), V158F (622G>T 624A>T), S159N (625T>A 626C>A), S160N (629G>A), T161V (631A>G 632C>T 633C>A), I162V (634A>G), K163V (637A>G 638A>T 639A>T), N164T (641A>C), I165L (643A>T 645T>G), L167S (650T>C), I168V (652A>G 654A>T), M172F (664A>T 666G>T), K184T (701A>C 702G>A), A186S (706G>T), E187G (710A>G), T190S (718A>T 720A>C), M191Q (721A>C 722T>A), V192L (724G>C 726A>G), S196A (736A>G 737G>C 738T>A), M197V (739A>G 741G>A), V203L (757G>T 759C>A), S204T (760T>A 762A>G), M212L (784A>T 786G>A), T214S (790A>T 792T>A), I215L (793A>T 795T>G), M216L (796A>T), I226V (826A>G 828A>T), T246A (886A>G), T258Q (922A>C 923C>A 924A>G), I278V (982A>G 984A>G), G280S (988G>A 990T>C), A281I (991G>A 992C>T), S293N (1028G>A 1029C>T), E302Q (1054G>C 1056A>G), Q305T (1063C>A 1064A>C), D308S (1072G>T 1073A>C 1074T>G), A309E (1076C>A 1077T>G), C311A (1081T>G 1082G>C), D312E (1086T>A), A314S (1090G>T), T315M (1094C>T 1095C>G), K323P (1117A>C 1118A>C 1119A>T), D327I (1129G>A 1130A>T 1131C>T), N330Q (1138A>C 1140T>A), R333K (1148G>A 1149G>A), L337V (1159C>G 1161G>C), M352L (1204A>T), E364D (1242A>T), V366L (1246G>C 1248C>G), Y367V (1249T>G 1250A>T), S370A (1258A>G 1259G>C 1260T>C), C371S (1261T>A), R372Q (1264A>C 1265G>A), V374S (1270G>T 1271T>C), Y389F (1316A>T), A399C (1345G>T 1346C>G 1347A>T), R410A (1378C>G 1379G>C 1380C>A), V494I (1630G>A), I519R (1706T>G 1707C>A), E521V (1712A>T 1713A>G), E523D (1719A>T), S525D (1723T>G 1724C>A), T526P (1726A>C), E527N (1729G>A 1731G>T), D528T (1732G>A 1733A>C 1734C>T), G531A (1742G>C 1743A>C), M534I (1752G>T), N536D (1756A>G 1758C>T), I537V (1759A>G 1761T>G), H550A (1798C>G 1799A>C 1800C>T), S554D (1810T>G 1811C>A 1812G>T), D558Q (1822G>C 1824C>A), Q565R (1844A>G), A568Y (1852G>T 1853C>A), I570V (1858A>G 1860A>G), K572S (1865A>G 1866G>C), F575Y (1874T>A 1875T>C), E576N (1876G>A 1878A>T), A577T (1879G>A 1881A>T), E578T (1882G>A 1883A>C), T579S (1885A>T 1887T>G), I581L (1891A>T 1893T>G), S582A (1894T>G 1896A>T), N583D (1897A>G 1899T>C), T585V (1903A>G 1904C>T 1905A>T), A589N (1915G>A 1916C>A), S591A (1921T>G), S592V (1924T>G 1925C>T), E593D (1929A>T), V596L (1936G>C), A597D (1940C>A 1941A>T), V599F (1945G>T 1947T>C), D600Q (1948G>C 1950T>A), G601D (1952G>A 1953G>T), R603del (1957_1959delCGC), F604L (1962C>A), K605Q (1963A>C), T606V (1966A>G 1967C>T 1968A>T), K607L (1969A>T 1970A>T 1971A>G), Q608P (1973A>C 1974A>T), D609P (1975G>C 1976A>C), V610E (1979T>A), P611S (1981C>T), I612A (1984A>G 1985T>C), V613A (1988T>C 1989G>T), T614Y (1990A>T 1991C>A), P615E (1993C>G 1994C>A 1995C>G), T616Q (1996A>C 1997C>A 1998A>G), V617T (1999G>A 2000T>C 2001T>A), E621A (2012A>C 2013A>T), D622E (2016T>A), R623K (2018G>A), V624I (2020G>A 2022G>T), I625V (2023A>G), K628Q (2032A>C 2034A>G), F637Y (2060T>A), L643M (2077T>A), D644N (2080G>A 2082C>T), P645K (2083C>A 2084C>A 2085G>A), E646F (2086G>T 2087A>T 2088G>T), S647E (2089A>G 2090G>A 2091T>G), L648F (2094G>T), Y653L (2107T>C 2108A>T 2109T>C), Q654S (2110C>T 2111A>C 2112A>C), L658I (2122C>A), F659Y (2126T>A), S661V (2131T>G 2132C>T 2133C>G), D663S (2137G>T 2138A>C), A664S (2140G>T 2142A>T), E665D (2145G>T), H666A (2146C>G 2147A>C), Q667K (2149C>A 2151G>A), V669N (2155G>A 2156T>A 2157C>T), M671E (2161A>G 2162T>A 2163G>A), R672M (2165G>T 2166A>G), M674V (2170A>G 2172G>A), T675N (2174C>A), M677L (2179A>T), I678L (2182A>T 2184C>G), Y679V (2185T>G 2186A>T), N683S (2198A>G), A684P (2200G>C), E685Q (2203G>C 2205A>G), R688A (2212A>G 2213G>C 2214G>T), Q692R (2225A>G), L699I (2245C>A), A708S (2272G>A 2273C>G 2274T>C), S709T (2275T>A 2277C>A), C712S (2285G>C), K714E (2290A>G), K715L (2293A>C 2294A>T), A717D (2300C>A), T720R (2308A>C 2309C>G 2310A>C), E721N (2311G>A 2313G>T), M722L (2314A>C 2316G>T), R723E (2317C>G 2318G>A 2319C>G), T724Q (2320A>C 2321C>A), F726K (2326T>A 2327T>A 2328T>A), H728Y (2332C>T), N729K (2337T>G), C730A (2338T>G 2339G>C 2340T>G), I731L (2341A>C), D732Q (2344G>C 2346T>G), V735A (2354T>C 2355T>A), I739T (2366T>C 2367T>G), Q742E (2374C>G 2376G>A), F747R (2389T>A 2390T>G 2391T>A), T761R (2432C>G 2433C>G), S764A (2440T>G 2442C>A), S764_K765insG (2442_2443insGGT), K765T (2444A>C 2445A>T), E766D (2448G>C), A767K (2449G>A 2450C>A 2451C>G), D769R (2455G>A 2456A>G 2457C>A), S770N (2458T>A 2459C>A), V772I (2464G>A 2466C>T), K773I (2468A>T 2469G>C), I774E (2470A>G 2471T>A), Y776M (2476T>A 2477A>T 2478C>G), H777I (2479C>A 2480A>T 2481C>T), K778G (2482A>G 2483A>G 2484A>T), V780K (2488G>A 2489T>A), Y782K (2494T>A 2496C>A), K784I (2501A>T 2502A>C), E785Q (2503G>C 2505A>G), T786M (2507C>T 2508T>G), W788N (2512T>A 2513G>A 2514G>T), N789D (2515A>G), Y792F (2525A>T 2526T>C), A793I (2527G>A 2528C>T 2529A>C), R795K (2534G>A), I804L (2560A>T 2562C>G), M805I (2565G>A), L810T (2578T>A 2579T>C 2580G>C), F811M (2581T>A 2583T>G), G812N (2584G>A 2585G>A 2586G>T), G813A (2588G>C 2589A>C), I814V (2590A>G 2592A>T), L818I (2602C>A), A819T (2605G>A), W821L (2611T>C 2612G>T 2613G>C), Q822K (2614C>A 2616A>G), S823A (2617A>G 2618G>C 2619C>A), N824A (2620A>G 2621A>C 2622T>A), S825T (2623T>A), F826S (2627T>C 2628T>C), S828L (2632A>T 2633G>T 2634T>A), S830D (2638A>G 2639G>A), P850K (2698C>A 2699C>A 2700C>G), S851T (2701T>A), R852H (2705G>A), L853F (2707C>T 2709G>C), K854A (2710A>G 2711A>C 2712G>A), N855K (2715T>A), L856S (2717T>C 2718G>C), V858L (2722G>C 2724G>A), S861A (2731A>G 2732G>C 2733T>G), S862A (2734A>G 2735G>C 2736C>A), A863D (2738C>A), I865N (2744T>A 2745T>C), T866H (2746A>C 2747C>A 2748A>T), Q867V (2749C>G 2750A>T 2751A>G), E868D (2754A>C), E868_W869insR (2754_2755insCGG), A871E (2762C>A 2763T>A), E872Q (2764G>C 2766A>G), Q878E (2782C>G), A879S (2785G>A 2786C>G 2787A>T), K881H (2791A>C 2793A>C), M884L (2800A>T), I885V (2803A>G), L889I (2815C>A), G890A (2819G>C 2820A>T), M906L (2866A>C 2868G>C), H913R (2887C>A 2888A>G 2889C>A), V915M (2893G>A), T922S (2914A>T), P926E (2926C>G 2927C>A 2928T>G), N927H (2929A>C 2931C>T), R929I (2936G>T 2937G>T), S930C (2939C>G 2940A>T), K931N (2943G>T), Y934F (2951A>T), R937T (2959C>A 2960G>C 2961G>C), E938Q (2962G>C), K940R (2969A>G), V941L (2971G>T 2973C>G), I947V (2989A>G 2991C>G), P960S (3028C>T 3030T>A), E971D (3063G>T), F974Y (3071T>A), T975S (3073A>T 3075C>T), Q979M (3085C>A 3086A>T), C983V (3097T>G 3098G>T 3099T>G), T989K (3116C>A 3117C>G), R992K (3125G>A 3126A>G), L993K (3127C>A 3128T>A 3129G>A), G994S (3130G>T 3131G>C), F998V (3142T>G), F1001Y (3152T>A), V1011E (3182T>A 3183C>A), E1014T (3190G>A 3191A>C 3192G>A), E1016G (3197A>G 3198G>A), A1020N (3208G>A 3209C>A 3210C>T), S1021A (3211A>G 3212G>C 3213T>A), K1023A (3217A>G 3218A>C 3219A>T), Q1028V (3232C>G 3233A>T 3234A>T), S1029A (3235T>G), M1034I (3252G>T), G1036S (3256G>A 3258A>T), S1049T (3296G>C), V1056C (3316G>T 3317T>G 3318T>C), G1067D (3350G>A), I1068T (3353T>C 3354T>C), F1074Y (3371T>A), I1075V (3373A>G 3375T>G), S1076A (3376T>G), I1082V (3394A>G 3396C>G), N1089D (3415A>G 3417C>T), V1095I (3433G>A), A1097V (3440C>T), D1103E (3459C>A), I1104L (3460A>C 3462T>G), A1107H (3469G>C 3470C>A), I1108L (3472A>C 3474T>G), R1111K (3482G>A 3483A>G), T1113E (3487A>G 3488C>A 3489C>A), T1114R (3491C>G 3492T>A), L1115F (3493C>T 3495T>C), N1119K (3507C>A), S1122A (3514T>G 3516C>T), Q1129L (3535C>T 3536A>T 3537A>G), L1131M (3541T>A), R1142K (3575G>A), K1143R (3577A>C 3578A>G 3579G>C), Q1145P (3584A>C 3585A>C), P1152D (3604C>G 3605C>A 3606T>C), E1154Q (3610G>C 3612A>G), T1161S (3631A>T 3633C>T), V1162A (3635T>C), N1163K (3639T>A), M1178E (3682A>G 3683T>A), H1185S (3703C>T 3704A>C 3705T>C), G1186N (3706G>A 3707G>A 3708T>C), D1188E (3714T>G), D1189E (3717T>G), D1190P (3718G>C 3719A>C 3720T>A), S1195A (3733T>G 3735C>T), T1198G (3742A>G 3743C>G), S1201A (3751A>G 3752G>C 3753T>A), L1203I (3757C>A 3759G>T), T1206V (3766A>G 3767C>T 3768A>G), M1207L (3769A>C 3771G>T), N1209M (3776A>T 3777C>G), K1211R (3781A>C 3782A>G), Q1218A (3802C>G 3803A>C 3804G>A), E1219I (3805G>A 3806A>T 3807G>C), D1221N (3811G>A), C1232F (3845G>T), Y1235F (3854A>T), G1237A (3860G>C 3861C>A), M1251L (3901A>T), M1253L (3907A>C 3909G>T), M1255S (3913A>T 3914T>C 3915G>T), V1262E (3935T>A), P1263A (3937C>G 3939A>C), T1264Q (3940A>C 3941C>A), G1265P (3943G>C 3944G>C), E1266N (3946G>A 3948A>C), G1267S (3949G>T 3950G>C 3951T>G), D1268Q (3952G>C 3954T>G), K1269R (3956A>G 3957A>G), E1271L (3961G>C 3962A>T 3963G>T), R1273I (3968G>T), D1275N (3973G>A), E1280D (3990A>T), I1287S (4009A>T 4010T>C 4011T>G), I1289V (4015A>G 4017T>A), I1293T (4028T>C), H1294F (4030C>T 4031A>T), I1298V (4042A>G), T1343C (4177A>T 4178C>G 4179C>T), L1347A (4189T>G 4190T>C), I1351L (4201A>C), D1358N (4222G>A), S1362H (4234T>C 4235C>A), S1365T (4243T>A 4245T>A), L1366V (4246C>G 4248T>G), N1369S (4256A>G 4257C>T), N1370S (4259A>G), A1373S (4267G>T 4269T>A), T1377E (4279A>G 4280C>A 4281T>G), V1392I (4324G>A 4326G>C), E1404A (4361A>C 4362G>T), Y1405F (4364A>T), D1407G (4370A>G), T1408D (4372A>G 4373C>A 4374C>T), K1409E (4375A>G), E1412S (4384G>A 4385A>G 4386G>T), A1414E (4391C>A 4392T>G), A1420T (4408G>A 4410T>A), D1434N (4450G>A 4452T>C), V1436L (4456G>T 4458T>G), I1475V (4573A>G 4575T>C), A1476K (4576G>A 4577C>A 4578C>A), R1478K (4583G>A), A1479S (4585G>T), I1480V (4588A>G 4590A>G), V1482I (4594G>A 4596A>T), G1485N (4603G>A 4604G>A), L1505M (4663C>A), S1511D (4681T>G 4682C>A 4683A>T), N1514S (4690A>T 4691A>C 4692T>C), Q1517R (4699C>A 4700A>G 4701G>A), D1520E (4710C>A), M1521A (4711A>G 4712T>C 4713G>T), N1537E (4759A>G 4761C>G), Q1538K (4762C>A), V1539T (4765G>A 4766T>C 4767G>A), Q1540K (4768C>A 4770G>A), Q1542L (4775A>T), H1562Q (4836C>A), I1571V (4861A>G 4863A>C), M1575I (4875G>T), P1582V (4894C>G 4895C>T), T1587V (4909A>G 4910C>T 4911T>A), V1588Q (4912G>C 4913T>A 4914G>A), M1591Y (4921A>T 4922T>A 4923G>T), V1594I (4930G>A 4932G>T), L1598I (4942C>A 4944T>C), R1606Q (4966A>C 4967G>A 4968A>G), V1607A (4970T>C 4971T>C), A1608T (4972G>A 4974A>T), K1611S (4982A>G 4983G>T), L1612F (4986A>T), T1613V (4987A>G 4988C>T 4989T>A), Y1614N (4990T>A), Q1617F (4999C>T 5000A>T 5001G>T), C1618G (5002T>G 5004T>A), R1621K (5012G>A 5013A>G), E1622Q (5014G>C), G1624N (5020G>A 5021G>A 5022A>T), G1626P (5026G>C 5027G>C), A1627I (5029G>A 5030C>T 5031A>C), T1628M (5033C>T 5034A>G), F1630M (5038T>A 5040T>G), S1632T (5044T>A), M1634Y (5050A>T 5051T>A 5052G>T), V1636L (5056G>C 5058G>T), Q1638S (5062C>A 5063A>G 5064G>T), E1639Q (5065G>C), I1641L (5071A>T 5073C>G), A1642T (5074G>A), K1643E (5077A>G), A1648E (5093C>A 5094A>G), A1650I (5098G>A 5099C>T 5100G>T), L1651G (5101T>G 5102T>G), T1652V (5104A>G 5105C>T 5106A>T), I1653L (5107A>T 5109C>G), C1654M (5110T>A 5111G>T 5112T>G), M1656L (5116A>C 5118G>A), A1657S (5119G>T 5121C>A), K1658D (5122A>G 5124A>T), S1660K (5128T>A 5129C>A 5130T>A), I1661N (5132T>A 5133C>T), F1663Y (5138T>A), P1668A (5152C>G), N1670E (5158A>G 5160T>A), V1672I (5164G>A), E1673del (5167_5169delGAA), R1677E (5179A>G 5180G>A 5181A>G), D1678G (5183A>G), L1679V (5185C>G), N1680H (5188A>C), W1681E (5191T>G 5192G>A 5193G>A), A1683V (5198C>T 5199C>G), N1685D (5203A>G 5205C>T), I1686V (5206A>G 5208T>A), G1687F (5209G>T 5210G>T), A1695S (5233G>T 5235C>A), K1696N (5238G>C), T1697Q (5239A>C 5240C>A 5241T>A), E1698Q (5242G>C), E1700S (5248G>T 5249A>C 5250G>A), P1701S (5251C>T 5253T>A), C1703N (5257T>A 5258G>A), V1705L (5263G>C 5265A>G), K1712R (5284A>C 5285A>G 5286G>C), K1713R (5288A>G 5289G>A), S1714T (5291G>C 5292C>A), K1715T (5294A>C), S1722N (5315G>A), M1730I (5340G>A), M1735L (5353A>T), S1736A (5356T>G), L1739Q (5366T>A), A1741H (5371G>C 5372C>A), G1744K (5380G>A 5381G>A 5382C>G), K1745S (5384A>G 5385A>T), M1746L (5386A>C), S1747E (5389T>G 5390C>A), E1748V (5393A>T 5394G>T), D1749G (5396A>G 5397C>T), I1750D (5398A>G 5399T>A), A1751G (5402C>G 5403T>G), D1754S (5410G>T 5411A>C), V1756T (5416G>A 5417T>C 5418G>C), V1765I (5443G>A 5445G>A), S1766A (5446A>G 5447G>C 5448C>T), M1773F (5467A>T 5469G>T), V1775L (5473G>T), R1778H (5483G>A), C1781R (5491T>A 5493C>G), I1782A (5494A>G 5495T>C 5496T>C), E1783T (5497G>A 5498A>C 5499G>A), I1785L (5503A>C 5505C>T), I1792M (5526T>G), G1793A (5528G>C 5529C>T), N1794G (5530A>G 5531A>G 5532T>G), V1806A (5567T>C 5568T>G), F1809L (5577T>G), Q1811K (5581C>A), P1814E (5590C>G 5591C>A 5592T>A), Q1815D (5593C>G 5595G>C), Q1815_E1816insT (5595_5596insACC), E1816S (5596G>T 5597A>C), G1817H (5599G>C 5600G>A 5601G>C), M1819L (5605A>T), A1820G (5609C>G), V1822S (5614G>A 5615T>G), S1824T (5620T>A 5622G>A), S1825del (5623_5625delTCT), L1826Y (5627T>A 5628G>T), Q1827G (5629C>G 5630A>G 5631A>C), S1828T (5633G>C 5634T>C), S1829T (5636G>C), M1831N (5642T>A 5643G>C), Q1833H (5649A>T), Q1833_G1834insHS (5649_5650insCACAGT), G1834A (5651G>C 5652T>A), E1836S (5656G>T 5657A>C 5658G>A), L1837V (5659C>G), Q1839L (5666A>T 5667A>T), I1840L (5668A>C 5670T>A), V1842I (5674G>A 5676C>A), I1843V (5677A>G 5679C>T), H1844P (5681A>C 5682C>T), G1845S (5683G>T 5684G>C), M1849E (5695A>G 5696T>A 5697G>A), E1850K (5698G>A 5700A>G), L1853V (5707T>G 5709G>T), M1855L (5713A>T), V1858T (5722G>A 5723T>C 5724T>A), V1859I (5725G>A 5727G>T), H1862E (5734C>G 5736T>A), G1866A (5747G>C), N1867Q (5749A>C 5751C>G), F1868C (5753T>G 5754T>C), T1869S (5755A>T 5757C>A), D1874K (5770G>A 5772T>G), Q1875H (5775A>T), L1876A (5776T>G 5777T>C 5778G>T), F1877L (5781T>G), A1878E (5783C>A 5784A>G), S1879T (5786G>C 5787T>A), E1881S (5791G>T 5792A>C 5793A>T), K1882V (5794A>G 5795A>T 5796G>C), C1883S (5797T>A), E1886T (5806G>A 5807A>C 5808A>T), L1888A (5812C>G 5813T>C 5814T>C), K1890R (5819A>G 5820G>A), N1891G (5821A>G 5822A>G 5823C>A), T1892V (5824A>G 5825C>T 5826G>C), L1894V (5830C>G), E1899K (5845G>A 5847G>A), K1900E (5848A>G), M1901F (5851A>T 5853G>C), T1902S (5854A>T 5856T>C), K1904H (5860A>C 5862A>C), V1906C (5866G>T 5867T>G 5868G>T), A1907E (5870C>A), Q1908M (5872C>A 5873A>T 5874A>G), V1910L (5878G>C 5880C>T), S1911E (5881A>G 5882G>A 5883T>G), V1913L (5887G>C 5889A>G), K1914E (5890A>G), Q1915S (5893C>A 5894A>G 5895G>T), F1918V (5902T>G 5904C>A), Y1919V (5905T>G 5906A>T 5907T>A), P1920F (5908C>T 5909C>T), K1921M (5912A>T), A1922G (5915C>G 5916C>G), T1923S (5918C>G 5919A>C), T1924V (5920A>G 5921C>T 5922C>T), V1925K (5923G>A 5924T>A), E1926K (5926G>A), L1928M (5932C>A 5934C>G), K1929L (5935A>T 5936A>T), V1930S (5938G>T 5939T>C 5940A>C), A1932M (5944G>A 5945C>T 5946T>G), D1933N (5947G>A), A1934Q (5950G>C 5951C>A 5952T>A), H1935Q (5955T>G), H1935_T1936insM (5955_5956insATG), T1936S (5956A>T 5958C>A), I1937K (5960T>A 5961T>A), V1938T (5962G>A 5963T>C), K1939Q (5965A>C), T1940V (5968A>G 5969C>T), K1942S (5975A>G 5976A>C), F1944C (5981T>G 5982C>T), R1945C (5983C>T 5985G>T), G1946del (5986_5988delGGA), V1948T (5992G>A 5993T>C 5994T>A), R1950S (5998C>A 6000G>T), N1952A (6004A>G 6005A>C 6006T>A), F1953W (6008T>G 6009T>G), S1955T (6014G>C), P1956K (6016C>A 6017C>A 6018T>G), E1957D (6021G>C), W1958I (6022T>A 6023G>T 6024G>T), W1958_K1959insV (6024_6025insGTC), K1959M (6026A>T 6027A>G), I1960T (6029T>C 6030T>A), G1961K (6031G>A 6032G>A), E1963K (6037G>A), N1964K (6042C>G), T1965P (6043A>C), M1966del (6046_6048delATG), A1967G (6050C>G), T1968S (6052A>T 6054A>T), L1969I (6055C>A 6057C>T), S1970P (6058T>C 6060A>T), S1973T (6068G>C 6069T>A), L1974V (6070C>G 6072T>G), L1975F (6073C>T), P1976L (6077C>T 6078A>T), L1979P (6086T>C 6087G>T), S1982L (6094T>C 6095C>T), A1983V (6098C>T 6099C>T), M1984V (6100A>G), K1985R (6104A>G 6105G>A), L1987M (6109T>A), S1989R (6115A>C), R1990N (6119G>A 6120A>T), Y1991S (6121T>A 6122A>G 6123C>T), K1992G (6124A>G

| | |
|---|---|
| | 6125A>G), R1993K (6128G>A 6129A>G), N1994S (6131A>G), M1995Y (6133A>T 6134T>A 6135G>C), H1998T (6142C>A 6143A>C 6144T>A), C1999T (6145T>A 6146G>C 6147C>A), I2000L (6148A>C 6150A>G), K2001E (6151A>G), L2004F (6160C>T 6162C>T), K2005R (6164A>G 6165G>A), I2006M (6168A>G), V2008M (6172G>A 6174T>G), L2009_N2010del (6175_6180delCTCAAC), F2011C (6182T>G 6183C>T), M2013W (6187A>T 6188T>G), L2014V (6190C>G 6192G>A), N2015H (6193A>C 6195T>C) |

| | Begin | End | Coverage | Score | Concordance | Matches | Identities | I/D/M/F* | Stop Codons |
|---|---|---|---|---|---|---|---|---|---|
| **NT** | **25** | **6200** | **85.3%** | **3315** | **27.4%** | **6117 (98.7%)** | **3941 (63.6%)** | **21/59** | |

GCT2GCA (156T>A), CTT3TTT (157C>T), AAA5AAG (165A>G), TTT7TGG (170T>G 171T>G), GGT8AGT (172G>A), TCT9CCT (175T>C), TCA10CCT (178T>C 180A>T), TCT11GCT (181T>G), GAG12_GTT17del (184_201delGAGAAGACTGGATCAGTT), CCC18TCT (202C>T 204C>T), TCT19TCA (207T>A), ACT20CCT (208A>C), TCT21TCA (213T>A), GGT24GCA (221G>C 222T>A), CTG26GTT (226C>G 228G>T), GGT28TCT (232G>T 233G>C), ATC30TTT (238A>T 240C>T), TCA31TCT (243A>T), AAA32TCT (244A>T 245A>C 246A>T), TTC33CTT (247T>C 249C>T), AAA35AAG (255A>G), GCT36ACT (256G>A), ATT37CTC (259A>C 261T>C), AGC38TCC (262A>T 263G>C), AAC39AAT (267C>T), ATC40GTT (268A>G 270C>T), ACA41AAT (272C>A 273A>T), TCA42AAA (274T>A 275C>A), GTA43GTT (279A>T), AAA46ACC (287A>C 288A>C), TCT48GTA (292T>G 293C>T 294T>A), CAT50CAC (300T>C), TTA51ATA (301T>A), GAT52GAG (306T>G), TTG54GTC (310T>G 312G>C), AAA55AAG (315A>G), CCA56CCG (318A>G), TCT57CCA (319T>C 321T>A), GTG58GTT (324G>T), ACT59ATG (326C>T 327T>G), GAT60GTA (329A>T 330T>A), TCT62GCA (334T>G 336T>A), TCT63ACA (337T>A 339T>A), TCT64TCA (342T>A), GTT66GCC (347T>C 348T>C), TGT69TGC (357T>C), TCC70AAT (358T>A 359C>A 360C>T), GCA71GCT (363A>T), TTG72GTA (364T>G 366G>A), AAT73GAA (367A>G 369T>A), AAA74AGA (371A>G), GTG75GTT (375G>T), CTT76TTA (376C>T 378T>A), ATA79ATT (387A>T), GGT80GCA (389G>C 390T>A), TCC81AAA (391T>A 392C>A 393C>A), TTG82ATA (394T>A 396G>A), ATT83ATG (399T>G), CCT85CCC (405T>C), TTT86TTC (408T>C), CTG87ATC (409C>A 411G>C), AAA88AAG (414A>G), GCA89GGT (416C>G 417A>T), TAC90TAT (420C>T), TCT91GAT (421T>G 422C>A), TTG93GTA (427T>G 429G>A), GCA94AAA (430G>A 431C>A), ACC95ACT (435C>T), TCT99ACT (445T>A), ATC100ATA (450C>A), AAG101GTA (451A>G 452A>T 453G>A), ATG103CTT (457A>C 459G>T), GTT104GTA (462T>A), TTA105CTT (463T>C 465A>T), CGG106AAG (466C>A 467G>A), TTG107ATG (469T>A), TTC108GTT (472T>G 474C>T), GAA109TTA (475G>T 476A>T), TTG111GTC (481T>G 483G>C), TCC112AAG (484T>A 485C>A 486C>G), AAT113GAT (487A>G), GTT115TTC (493G>T 495T>C), AGA116ATC (497G>T 498A>C), ATA117CTA (499A>C), TTT119TAC (506T>A 507T>C), CAG120GAA (508C>G 510G>A), GTT122TTA (514G>T 516T>A), GAG124GAA (522G>A), AAG125AAA (525G>A), AGT126GTT (526A>G 527G>T), GAT128TCA (532G>T 533A>C 534T>A), GTA129TTA (535G>T), ATT131ATA (543T>A), GTA132GTG (546A>G), GTT133TTT (547G>T), TAT134ATA (550T>A 551A>T 552T>A), GGT135GCT (554G>C), TTG137TTA (561G>A), GCA140GCT (570A>T), ATA141TTG (571A>T 573A>G), ACT142GCC (574A>G 576T>C), CTT143ATT (577C>A), CTT144TTG (580C>T 582T>G), GTT145CTA (583G>C 585T>A), TTG146CTA (586T>C 588G>A), TTG147CTG (589T>C), TGT148TAT (593G>A), GTT149GTA (597T>A), TCA152GTA (604T>G 605C>T), GAC153GAT (609C>T), ATA154ATT (612A>T), ATT155GTA (613A>G 615T>A), GAA156ACG (616G>A 617A>C 618A>G), GGA157CCT (619G>C 620G>C 621A>T), GTA158TTT (622G>T 624A>T), TCT159AAT (625T>A 626C>A), AGT160AAT (629G>A), ACC161GTA (631A>G 632C>T 633C>A), ATT162GTT (634A>G), AAA163GTT (637A>G 638A>T 639A>T), AAT164ACT (641A>C), ATT165TTG (643A>T 645T>G), TTC166TTT (648C>T), TTA167TCA (650T>C), ATA168GTT (652A>G 654A>T), GTA169GTT (657A>T), GGG170GGA (660G>A), ATG172TTT (664A>T 666G>T), TTT173TTC (669T>C), AGT175TCA (673A>T 674G>C 675T>A), AAC178AAT (684C>T), TTG179TTA (687G>A), GAC180GAT (690C>T), CCC183CCA (699C>A), AAG184ACA (701A>C 702G>A), GCT186TCT (706G>T), GAG187GGG (710A>G), TTC189TTT (717C>T), ACA190TCC (718A>T 720A>C), ATG191CAG (721A>C 722T>A), GTA192CTG (724G>C 726A>G), GCA193GCG (729A>G), AGT196GCA (736A>G 737G>C 738T>A), ATG197GTA (739A>G 741G>A), CTA198CTT (744A>T), GAG201GAA (753G>A), TCT202AGC (754T>A 755C>G 756T>C), GTC203TTA (757G>T 759C>A), TCA204ACG (760T>A 762A>G), CAC205CAT (765C>T), ACT206ACA (768T>A), ACA208ACT (774A>T), TCC209AGC (775T>A 776C>G), CAG210CAA (780G>A), TTG211CTT (781T>C 783G>T), ATG212TTA (784A>T 786G>A), TCA213TCC (789A>C), ACT214TCA (790A>T 792T>A), ATT215TTG (793A>T 795T>G), ATG216TTG (796A>T), GCA217GCT (801A>T), ATT220ATC (810T>C), TCC221TCA (813C>A), ACG222ACT (816G>T), CTC223TTA (817C>T 819C>A), GTG224GTA (822G>A), ATA226GTT (826A>G 828A>T), GCA227GCT (831A>T), GTG228GTT (834G>T), CCA229CCT (837A>T), GGA230GGG (840A>G), CCC232CCA (846C>A), TTG235CTT (853T>C 855G>T), AGC236TCA (856A>T 857G>C 858C>A), AAT237AAC (861T>C), CCC238CCA (864C>A), CTC239CTG (867C>G), ATT242ATC (876T>C), TCA245TCT (885A>T), ACT246GCT (886A>G), GGG247GGA (891G>A), GGG249GGT (897G>T), GCT250GCG (900T>G), CAA255CAG (915A>G), TTG256CTG (916T>C), TTT257TTC (921T>C), ACA258CAG (922A>C 923C>A 924A>G), AGA261AGG (933A>G), AAT262AAC (936T>C), AAA264AAG (942A>G), TGC266TGT (948C>T), ACT267ACA (951T>A), TCA268TCT (954A>T), GCA270GCC (960A>C), TTT271TTC (963T>C), TCT272TCA (966T>A), GTG274GTT (972G>T), GAA276GAG (978A>G), ATT277ATA (981T>A), ATA278GTG (982A>G 984A>G), GGT280AGC (988G>A 990T>C), GCT281ATT (991G>A 992C>T), TTC284TTT (1002C>T), AAG285AAA (1005G>A), GTG288GTA (1014G>A), CTT289TTG (1015C>T 1017T>G), TCT290TCA (1020T>A), GCA291GCG (1023A>G), ATT292ATA (1026T>A), AGC293AAT (1028G>A 1029C>T), GCA294GCT (1032A>T), ACA295ACC (1035A>C), TTG296TTA (1038G>A), TCT297TCC (1041T>C), ACT298ACC (1044T>C), TTG300CTT (1048T>C 1050G>T), GAA302CAG (1054G>C 1056A>G), CAG305ACG (1063C>A 1064A>C), GTG307GTT (1071G>T), GAT308TCG (1072G>T 1073A>C 1074T>G), GCT309GAG (1076C>A 1077T>G), TGT311GCT (1081T>G 1082G>C), GAT312GAA (1086T>A), GCT314TCT (1090G>T), ACC315ATG (1094C>T 1095C>G), AGG316CGC (1096A>C 1098G>C), TTG317CTT (1099T>C 1101G>T), GAG318GAA (1104G>A), AAC319AAT (1107C>T), GCC321GCA (1113C>A), AAT322AAC (1116T>C), AAA323CCT (1117A>C 1118A>C 1119A>T), GCC324GCA (1122C>A), TTC325TTT (1125C>T), CCT326CCA (1128T>A), GAC327ATT (1129G>A 1130A>T 1131C>T), AAA328AAG (1134A>G), AAT330CAA (1138A>C 1140T>A), CAC331CAT (1143C>T), CTT332TTA (1144C>T 1146T>A), AGG333AAA (1148G>A 1149G>A), GAG334GAA (1152G>A), GCA336GCC (1158A>C), CTG337GTC (1159C>G 1161G>C), AAG338AAA (1164G>A), ATT339ATA (1167T>A), GCA341GCT (1173A>T), ACA345ACC (1185A>C), CAC346CAT (1188C>T), GTT348GTC (1194T>C), GCC349GCA (1197C>A), ATG352TTG (1204A>T), CAT354CAC (1212T>C), CGA355AGG (1213C>A 1215A>G), GTG356GTT (1218G>T), AGT357TCT (1219A>T 1220G>C), GCA358GCT (1224A>T), GCT359GCC (1227T>C), ATC360ATA (1230C>A), ACT361ACG (1233T>G), CTT363CTA (1239T>A), GAA364GAT (1242A>T), AAG365AAA (1245G>A), GTC366CTG (1246G>C 1248C>G), TAT367GTT (1249T>G 1250A>T), GCG368GCT (1254G>T), AGT370GCC (1258A>G 1259G>C 1260T>C), TGC371AGC (1261T>A), AGA372CAA (1264A>C 1265G>A), GTT374TCT (1270G>T 1271T>C), GTT376GTG (1278T>G), GGT377GGC (1281T>C), CAG378CAA (1284G>A), AGA380AGG (1290A>G), GAA382GAG (1296A>G), CCC383CCT (1299C>T), GTT386GTG (1308T>G), TAT389TTT (1316A>T), GGT390GGA (1320T>A), GCC391GCT (1323C>T), GGG393GGA (1329G>A), ACA398ACG (1344A>G), GCA399TGT (1345G>T 1346C>G 1347A>T), CGA401CGC (1353A>C), TTG402CTT (1354T>C 1356G>T), TTG408CTC (1372T>C 1374G>C), GAT409GAC (1377T>C), CGC410GCA (1378C>G 1379G>C 1380C>A), GGG412GGT (1386G>T), GAG413GAA (1389G>A), CCT414CCA (1392T>A), CTT416CTG (1398T>G), AGA418CGC (1402A>C 1404A>C), CTT419TTG (1405C>T 1407T>G), GCT421GCG (1413T>G), GTT422GTA (1416T>A), AGC423AGT (1419C>T), AAG424AAA (1422G>A), CGC425AGA (1423C>A 1425C>A), GCC427GCA (1431C>A), TCT430TCC (1440T>C), AAT431AAC (1443T>C), GCC433GCA (1449C>A), CAC434CAT (1452C>T), CAG435CAA (1455G>A), GCT437GCA (1461T>A), TTG439CTT (1465T>C 1467G>T), GAC442GAT (1476C>T), GGA444GGT (1482A>T), GCA445GCT (1485A>T), TTG446CTT (1486T>C 1488G>T), AGA447CGG (1489A>C 1491A>G), GGA451GGG (1503A>G), CAA452CAG (1506A>G), CAA454CAG (1512A>G), ATT456ATA (1518T>A), ATT462ATC (1536T>C), TCC464TCT (1542C>T), ACA465ACC (1545A>C), CCA467CCT (1551A>T), CCA469CCT (1557A>T), CTG470CTT (1560G>T), CCC471CCA (1563C>A), GCA473GCT (1569A>T), GCT474GCA (1572T>A), GTG475GTC (1575G>C), GAA476GAG (1578A>G), GAC477GAT (1581C>T), AAG478AAA (1584G>A), GGG479GGA (1587G>A), CAT481CAC (1593T>C), ACT483ACA (1599T>A), TAT486TAC (1608T>C), GCA489GCC (1617A>C), CTT493TTG (1627C>T 1629T>G), GTA494ATA (1630G>A), TCA495TCT (1635A>T), GCA496GCT (1638A>T), CCT497CCA (1641T>A), GCC498GCT (1644C>T), CTT502CTA (1656T>A), ACC503ACG (1659C>G), GCT507GCA (1671T>A), GAG509GAA (1677G>A), CGT510AGG (1678C>A 1680T>G), AGA511AGG (1683A>G), AGG512AGA (1686G>A), ATC514ATT (1692C>T), TGT518TGC (1704T>C), ATC519AGA (1706T>G 1707C>A), GAA521GTG (1712A>T 1713A>G), GGT522GGC (1716T>C), GAA523GAT (1719A>T), TCT525GAT (1723T>G 1724C>A), ACA526CCA (1726A>C), GAG527AAT (1729G>A 1731G>T), GAC528ACT (1732G>A 1733A>C 1734C>T), ACC530ACA (1740C>A), GGA531GCC (1742G>C 1743A>C), CTG533CTA (1749G>A), ATG534ATT (1752G>T), AAC536GAT (1756A>G 1758C>T), ATT537GTG (1759A>G 1761T>G), CGG541AGA (1771C>A 1773G>A), AGG542AGA (1776G>A), ATA546ATC (1788A>C), CAC550GCT (1798C>G 1799A>C 1800C>T), AGT552TCC (1804A>T 1805G>C 1806T>C), TAC553TAT (1809C>T), TCG554GAT (1810T>G 1811C>A 1812G>T), TTA557TTG (1821A>G), GAC558CAA (1822G>C 1824C>A), GCC562GCA (1836C>A), TGT564TGC (1842T>C), CAG565CGG (1844A>G), GTT566GTG (1848T>G), CAT567CAC (1851T>C), GCT568TAT (1852G>T 1853C>A), ATA570GTG (1858A>G 1860A>G), AAG572AGC (1865A>G 1866G>C), CAG573CAA (1869G>A), CTG574CTT (1872G>T), TTT575TAC (1874T>A 1875T>C), GAA576AAT (1876G>A 1878A>T), GCA577ACT (1879G>A 1881A>T), GAA578ACA (1882G>A 1883A>C), ACT579TCG (1885A>T 1887T>G), GGC580GGA (1890C>A), ATT581TTG (1891A>T 1893T>G), TCA582GCT (1894T>G 1896A>T), AAT583GAC (1897A>G 1899T>C), ACA585GTT (1903A>G 1904C>T 1905A>T), GCC586GCT (1908C>T), GTT588GTC (1914T>C), GCT589AAT (1915G>A 1916C>A), GTA590GTT (1920A>T), TCT591GCT (1921T>G), TCT592GTT (1924T>G 1925C>T), GAA593GAT (1929A>T), GAG594GAA (1932G>A), GTT595GTG (1935T>G), GTG596CTG (1936G>C), GCA597GAT (1940C>A 1941A>T), GTT599TTC (1945G>T 1947T>C), GAT600CAA (1948G>C 1950T>A), GGG601GAT (1952G>A 1953G>T), GCT602GCC (1956T>C), CGC603del (1957_1959delCGC), TTC604TTA (1962C>A), AAA605CAA (1963A>C), ACA606GTT (1966A>G 1967C>T 1968A>T), AAA607TTG (1969A>T 1970A>T 1971A>G), CAA608CCT (1973A>C 1974A>T), GAT609CCT (1975G>C 1976A>C), GTG610GAG (1979T>A), CCA611TCA (1981C>T), ATA612GCA (1984A>G 1985T>C), GTG613GCT (1988T>C 1989G>T), ACT614TAT (1990A>T 1991C>A), CCC615GAG (1993C>G 1994C>A 1995C>G), ACA616CAG (1996A>C 1997C>A 1998A>G), GTT617ACA (1999G>A 2000T>C 2001T>A), ATA618ATT (2004A>T), GAG620GAA (2010G>A), GAA621GCT (2012A>C 2013A>T), GAT622GAA (2016T>A), AGA623AAA (2018G>A), GTG624ATT (2020G>A 2022G>T), ATT625GTT (2023A>G), TCT627TCC (2031T>C), AAA628CAG (2032A>C 2034A>G), TTA630CTC (2038T>C 2040A>C), GTG632GTT (2046G>T), GAA633GAG (2049A>G), CTT635CTC (2055T>C), TTT637TAT (2060T>A), TAT639TAC (2067T>C), CAG640CAA (2070G>A), GGC641GGG (2073C>G), TTG643ATG (2077T>A), GAC644AAT (2080G>A 2082C>T), CCG645AAA (2083C>A 2084C>A 2085G>A), GAG646TTT (2086G>T 2087A>T 2088G>T), AGT647GAG (2089A>G 2090G>A 2091T>G), TTG648TTT (2094G>T), TTT649TTC (2097T>C), CCT650CCA (2100T>A), TAT653CTC (2107T>C 2108A>T 2109T>C), CAA654TCC (2110C>T 2111A>C 2112A>C), AAA655AAG (2115A>G), CAA656CAG (2118A>G), CTC658ATC (2122C>A), TTT659TAT (2126T>A), TCC661GTG (2131T>G 2132C>T 2133C>G), GAT663TCT (2137G>T 2138A>C), GCA664TCT (2140G>T 2142A>T), GAG665GAT (2145G>T), CAC666GCC (2146C>G 2147A>C), CAG667AAA (2149C>A 2151G>A), CAA668CAG (2154A>G), GTC669AAT (2155G>A 2156T>A 2157C>T), TTC670TTT (2160C>T), ATG671GAA (2161A>G 2162T>A 2163G>A), AGA672ATG (2165G>T 2166A>G), ATG674GTA (2170A>G 2172G>A), ACT675AAT (2174C>A), GAC676GAT (2178C>T), ATG677TTG (2179A>T), ATC678TTG (2182A>T 2184C>G), TAC679GTC (2185T>G 2186A>T), GGT681GGA (2193T>A), GCC682GCA (2196C>A), AAT683AGT (2198A>G), GCT684CCT (2200G>C), GAA685CAG (2203G>C 2205A>G), CAA686CAG (2208A>G), TAC687TAT (2211C>T), AGG688GCT (2212A>G 2213G>C 2214G>T), CAA692CGA (2225A>G), AAT693AAC (2229T>C), CTC699ATC (2245C>A), TTT702TTC (2256T>C), AAA703AAG (2259A>G), AGC704AGT (2262C>T), ATC706ATT (2268C>T), GCT708AGC (2272G>A 2273C>G 2274T>C), TCC709ACA (2275T>A 2277C>A), ATC711ATT (2283C>T), TGT712TCT (2285G>C), AAG714GAG (2290A>G), AAG715CTG (2293A>C 2294A>T), GCT717GAT (2300C>A), GTA718GTC (2304A>C), ACA720CGC (2308A>C 2309C>G 2310A>C), GAG721AAT (2311G>A 2313G>T), ATG722CTT (2314A>C 2316G>T), CGC723GAG (2317C>G 2318G>A 2319C>G), ACA724CAA (2320A>C 2321C>A), GGA725GGG (2325A>G), TTT726AAA (2326T>A 2327T>A 2328T>A), GCA727GCG (2331A>G), CAT728TAT (2332C>T), AAT729AAG (2337T>G), TGT730GCG (2338T>G 2339G>C 2340T>G), ATT731CTT (2341A>C), GAT732CAG (2344G>C 2346T>G), GTT735GCA (2354T>C 2355T>A), TTG738CTT (2362T>C 2364G>T), ATT739ACG (2366T>C 2367T>G), TGC740TGT (2370C>T), ATC741ATT (2373C>T), CAG742GAA (2374C>G 2376G>A), TTT747AGA (2389T>A 2390T>G 2391T>A), GTG748GTA (2394G>A), TTT750TTC (2400T>C), TTG751CTG (2401T>C), ACA753ACT (2409A>T), GCC754GCT (2412C>T), CGT757CGC (2421T>C), TAT758TAC (2424T>C), ACC761AGG (2432C>G 2433C>G), TCC764GCA (2440T>G 2442C>A), TCC764_AAA765insGGT (2442_2443insGGT), AAA765ACT (2444A>C 2445A>T), GAG766GAC (2448G>C), GCC767AAG (2449G>A 2450C>A 2451C>G), AAA768AAG (2454A>G), GAC769AGA (2455G>A 2456A>G 2457C>A), TCT770AAT (2458T>A 2459C>A), GTC772ATT (2464G>A 2466C>T), AAG773ATC (2468A>T 2469G>C), ATA774GAA (2470A>G 2471T>A), TGT775TGC (2475T>C), TAC776ATG (2476T>A 2477A>T 2478C>G), CAC777ATT (2479C>A 2480A>T 2481C>T), AAA778GGT (2482A>G 2483A>G 2484A>T), ATT779ATC (2487T>C), GTG780AAG (2488G>A 2489T>A), GAT781GAC (2493T>C), TAC782AAA (2494T>A 2496C>A), AAA784ATC (2501A>T 2502A>C), GAA785CAG (2503G>C 2505A>G), ACT786ATG (2507C>T 2508T>G), TGG788AAT (2512T>A 2513G>A 2514G>T), AAC789GAC (2515A>G), TTA790TTG (2520A>G), CCA791CCC (2523A>C), TAT792TTC (2525A>T 2526T>C), GCA793ATC (2527G>A 2528C>T 2529A>C), CTG794TTG (2530C>T), AGG795AAG (2534G>A), GGC801GGA (2553C>A), CTA802CTG (2556A>G), ATC804TTG (2560A>T 2562C>G), ATG805ATA (2565G>A), GCG806GCA (2568G>A), GGA809GGC (2577A>C), TTG810ACC (2578T>A 2579T>C 2580G>C), TTT811ATG (2581T>A 2583T>G), GGG812AAT (2584G>A 2585G>A 2586G>T), GGA813GCC (2588G>C 2589A>C), ATA814GTT (2590A>G 2592A>T), ACA815ACT (2595A>T), TTT817TTC (2601T>C), CTT818ATT (2602C>A), GCT819ACT (2605G>A), TGT820TGC (2610T>C), TGG821CTC (2611T>C 2612G>T 2613G>C), CAA822AAG (2614C>A 2616A>G), AGC823GCA (2617A>G 2618G>C 2619C>A), AAT824GCA (2620A>G 2621A>C 2622T>A), TCT825ACT (2623T>A), TTT826TCC (2627T>C 2628T>C), CCA827CCT (2631A>T), AGT828TTA (2632A>T 2633G>T 2634T>A), AGT830GAT (2638A>G 2639G>A), GGA831GGC (2643A>C), GGT833GGG (2649T>G), GGT834GGC (2652T>C), GTC835GTA (2655C>A), ACA836ACC (2658A>C), AGC841AGT (2673C>T), TCT843TCC (2679T>C), AAA845AAG (2685A>G), AAG846AAA (2688G>A), AAC847AAT (2691C>T), AAG848AAA (2694G>A), GGC849GGG (2697C>G), CCC850AAG (2698C>A 2699C>A 2700C>G), TCT851ACT (2701T>A), CGT852CAT (2705G>A), CTG853TTC (2707C>T 2709G>C), AAG854GCA (2710A>G 2711A>C 2712G>A), AAT855AAA (2715T>A), TTG856TCC (2717T>C 2718G>C), GTG858CTA (2722G>C 2724G>A), CAG860CAA (2730G>A), AGT861GCG (2731A>G 2732G>C 2733T>G), AGC862GCA (2734A>G 2735G>C 2736C>A), GCT863GAT (2738C>A), ATT865AAC (2744T>A 2745T>C), ACA866CAT (2746A>C 2747C>A 2748A>T), CAA867GTG (2749C>G 2750A>T 2751A>G), GAA868GAC (2754A>C), GAA868_TGG869insCGG (2754_2755insCGG), GCT871GAA (2762C>A 2763T>A), GAA872CAG (2764G>C 2766A>G), GGG874GGA (2772G>A), GTC876GTT (2778C>T), AAC877AAT (2781C>T), CAA878GAA (2782C>G), GCA879AGT (2785G>A 2786C>G 2787A>T), CTC880CTG (2790C>G), AAA881CAC (2791A>C 2793A>C), AAG882AAA (2796G>A), AAT883AAC (2799T>C), ATG884TTG (2800A>T), ATT885GTT (2803A>G), GTA886GTT (2808A>T), CTG887TTG (2809C>T), CGG888AGA (2812C>A 2814G>A), CTT889ATT (2815C>A), GGA890GCT (2819G>C 2820A>T), GAG891GAA (2823G>A), GTG893GTC (2829G>C), TAC894TAT (2832C>T), ACA898ACG (2844A>G), GTG900GTT (2850G>T), TCA902TCT (2856A>T), GGG903GGT (2859G>T), ATA905ATT (2865A>T), ATG906CTC (2866A>C 2868G>C), GCA911GCC (2883A>C), TTC912TTT (2886C>T), CAC913AGA (2887C>A 2888A>G 2889C>A), AAT914AAC (2892T>C), GTG915ATG (2893G>A), AGA916AGG (2898A>G), GGA918GGC (2904A>C), ACA919ACT (2907A>T), ACC922TCC (2914A>T), ATT923ATC (2919T>C), ATC924ATT (2922C>T), CCT926GAG (2926C>G 2927C>A 2928T>G), AAC927CAT (2929A>C 2931C>T), TCA928TCC (2934A>C), AGG929ATT (2936G>T 2937G>T), TCA930TGT (2939C>G 2940A>T), AAG931AAT (2943G>T), TAC934TTC (2951A>T), AAC935AAT (2955C>T), CGG937ACC (2959C>A 2960G>C 2961G>C), GAA938CAA (2962G>C), TCC939TCA (2967C>A), AAG940AGG (2969A>G), GTC941TTG (2971G>T 2973C>G), ATT942ATA (2976T>A), GGA944GGT (2982A>T), ATC947GTG (2989A>G 2991C>G), CTG949TTG (2995C>T),

Codon mutations:

CTA950TTG (2998C>T 3000A>G), CGT951AGG (3001C>A 3003T>G), GTG952GTA (3006G>A), GGT953GGG (3009T>G), GAC954GAT (3012C>T), CCT955CCA (3015T>A), CCT960TCA (3028C>T 3030T>A), ATT962ATA (3036T>A), CGC963AGA (3037C>A 3039C>A), AAG964AAA (3042G>A), CCA967CCT (3051A>T), GAG971GAT (3063G>T), GTC972GTG (3066C>G), TGC973TGT (3069C>T), TTC974TAC (3071T>A), ACC975TCT (3073A>T 3075C>T), AAA976AAG (3078A>G), GGT977GGC (3081T>C), TCT978TCA (3084T>A), CAG979ATG (3085C>A 3086A>T), TTA981TTG (3093A>G), TGT982TGC (3096T>C), TGT983GTG (3097T>G 3098G>T 3099T>G), AGA984AGG (3102A>G), GCC985GCT (3105C>T), GTG986GTA (3108G>A), GCG987GCT (3111G>T), ACC989AAG (3116C>A 3117C>G), CCA991CCC (3123A>C), AGA992AAG (3125G>A 3126A>G), CTG993AAA (3127C>A 3128T>A 3129G>A), GGC994TCC (3130G>T 3131G>C), AAC995AAT (3135C>T), CTA996TTG (3136C>T 3138A>G), GAA997GAG (3141A>G), TTT998GTT (3142T>G), TTG999CTT (3145T>C 3147G>T), TTT1001TAT (3152T>A), TCC1006TCA (3168C>A), AAA1007AAG (3171A>G), TAT1009TAC (3177T>C), ACA1010ACC (3180A>C), GTC1011GAA (3182T>A 3183C>A), AAG1012AAA (3186G>A), GTG1013GTC (3189G>C), GAG1014ACA (3190G>A 3191A>C 3192G>A), GAG1016GGA (3197A>G 3198G>A), GAC1018GAT (3204C>T), TCT1019TCG (3207T>G), GCC1020AAT (3208G>A 3209C>A 3210C>T), AGT1021GCA (3211A>G 3212G>C 3213T>A), AAA1023GCT (3217A>G 3218A>C 3219A>T), TTG1024CTT (3220T>C 3222G>T), AGT1025AGC (3225T>C), TCG1026TCA (3228G>A), CAG1027CAA (3231G>A), CAA1028GTT (3232C>G 3233A>T 3234A>T), TCA1029GCA (3235T>G), GAA1031GAG (3243A>G), TAT1032TAC (3246T>C), CAT1033CAC (3249T>C), ATG1034ATT (3252G>T), GGA1036AGT (3256G>A 3258A>T), GGT1039GGA (3267T>A), GGA1042GGT (3276A>T), TTA1045CTC (3283T>C 3285A>C), CTT1046TTA (3286C>T 3288T>A), CCC1048CCA (3294C>A), AGT1049ACT (3296G>C), GTT1050GTG (3300T>G), AAA1053AAG (3309A>G), CAA1054CAG (3312A>G), CCT1055CCG (3315T>G), GTT1056TGC (3316G>T 3317T>G 3318T>C), GTG1058GTT (3324G>T), GGC1059GGA (3327C>A), ATT1060ATA (3330T>A), GCT1063GCA (3339T>A), TCA1064TCT (3342A>T), GGT1067GAT (3350G>A), ATT1068ACC (3353T>C 3354T>C), GCA1069GCT (3357A>T), GAA1071GAG (3363A>G), GGA1073GGT (3369A>T), TTC1074TAC (3371T>A), ATT1075GTG (3373A>G 3375T>G), TCT1076GCT (3376T>G), TCC1077TCA (3381C>A), AAT1078AAC (3384T>C), GCC1079GCA (3387C>A), ACG1080ACA (3390G>A), GCC1081GCA (3393C>A), ATC1082GTG (3394A>G 3396C>G), TTC1083TTT (3399C>T), CGG1084AGA (3400C>A 3402G>A), GAC1085GAT (3405C>T), CAG1086CAA (3408G>A), CTT1087TTG (3409C>T 3411T>G), AAC1089GAT (3415A>G 3417C>T), GGG1093GGA (3429G>A), CCT1094CCC (3432T>C), GTT1095ATT (3433G>A), GCG1097GTG (3440C>T), GCA1098GCT (3444A>T), CGA1101CGC (3453A>C), GAC1103GAA (3459C>A), ATT1104CTG (3460A>C 3462T>G), CTC1105TTG (3463C>T 3465C>G), AAA1106AAG (3468A>G), GCC1107CAC (3469G>C 3470C>A), ATT1108CTG (3472A>C 3474T>G), AGG1109AGA (3477G>A), AGT1110TCC (3478A>T 3479G>C 3480T>C), AGA1111AAG (3482G>A 3483A>G), GAA1112GAG (3486A>G), ACC1113GAA (3487A>G 3488C>A 3489C>A), ACT1114AGA (3491C>G 3492T>A), CTT1115TTC (3493C>T 3495T>C), GAA1117GAG (3501A>G), AAC1119AAA (3507C>A), CAA1120CAG (3510A>G), GTC1121GTG (3513C>G), TCC1122GCT (3514T>G 3516C>T), TTT1123TTC (3519T>C), CTT1124TTG (3520C>T 3522T>G), GTT1127GTA (3531T>A), CCT1128CCG (3534T>G), CAA1129TTG (3535C>T 3536A>T 3537A>G), TTG1131ATG (3541T>A), GCC1132GCT (3546C>T), GCA1133GCT (3549A>T), GTA1135GTG (3555A>G), CCC1136CCG (3558C>G), CAT1137CAC (3561T>C), ACT1139ACA (3567T>A), ACA1140ACC (3570A>C), CTC1141CTT (3573C>T), AGA1142AAA (3575G>A), AAG1143CGC (3577A>C 3578A>G 3579G>C), CAA1145CCC (3584A>C 3585A>C), TTG1146CTG (3586T>C), GCA1149GCT (3597A>T), GGT1151GGG (3603T>G), CCT1152GAC (3604C>G 3605C>A 3606T>C), GAA1154CAG (3610G>C 3612A>G), ACC1155ACT (3615C>T), GCT1156GCC (3618T>C), CCC1157CCA (3621C>A), TCA1158TCC (3624A>C), ATC1159ATT (3627C>T), CTC1160CTA (3630C>A), ACC1161TCT (3631A>T 3633C>T), GTC1162GCC (3635T>C), AAT1163AAA (3639T>A), CGG1166AGG (3646C>A), GAG1168GAA (3654G>A), GGA1169GGG (3657A>G), TTC1170TTT (3660C>T), CCA1172CCC (3666A>C), GCA1175GCT (3675A>T), GTG1177GTC (3681G>C), ATG1178GAG (3682A>G 3683T>A), AAA1179AAG (3687A>G), TAC1180TAT (3690C>T), ACT1183ACA (3699T>A), CAT1185TCC (3703C>T 3704A>C 3705T>C), GGT1186AAC (3706G>A 3707G>A 3708T>C), GAT1188GAG (3714T>G), GAT1189GAG (3717T>G), GAT1190CCA (3718G>C 3719A>C 3720T>A), GCT1192GCA (3726T>A), AAA1193AAG (3729A>G), TTG1194CTG (3730T>C), TCC1195GCT (3733T>G 3735C>T), GAA1197GAG (3741A>G), ACT1198GGT (3742A>G 3743C>G), CTC1199CTT (3747C>T), AGT1201GCA (3751A>G 3752G>C 3753T>A), TCA1202TCC (3756A>C), CTG1203ATT (3757C>A 3759G>T), CTG1204CTT (3762G>T), CCA1205CCT (3765A>T), ACA1206GTG (3766A>G 3767C>T 3768A>G), ATG1207CTT (3769A>C 3771G>T), AAA1208AAG (3774A>G), AAC1209ATG (3776A>T 3777C>G), CAA1210CAG (3780A>G), AAA1211CGA (3781A>C 3782A>G), GGC1215GGG (3795C>G), CCA1217CCT (3801A>T), CAG1218GCA (3802C>G 3803A>C 3804G>A), GAG1219ATC (3805G>A 3806A>T 3807G>C), GAT1221AAT (3811G>A), GAT1223GAC (3819T>C), GTG1224GTA (3822G>A), CTG1226CTC (3828G>C), ATT1229ATA (3837T>A), TGT1232TTT (3845G>T), TAT1235TTT (3854A>T), GGC1237GCA (3860G>C 3861C>A), CTC1240TTG (3868C>T 3870C>G), ACA1242ACG (3876A>G), TCC1243TCA (3879C>A), TAC1246TAT (3888C>T), TTC1248TTT (3894C>T), ATG1251TTG (3901A>T), GGC1252GGA (3906C>A), ATG1253CTT (3907A>C 3909G>T), GGA1254GGT (3912A>T), ATG1255TCT (3913A>T 3914T>C 3915G>T), AAG1257AAA (3921G>A), CGG1258AGG (3922C>A), GAG1259GAA (3927G>A), GTC1261GTG (3933C>G), GTA1262GAA (3935T>A), CCA1263GCC (3937C>G 3939A>C), ACA1264CAA (3940A>C 3941C>A), GGA1265CCA (3943G>C 3944G>C), GAA1266AAC (3946G>A 3948A>C), GGT1267TCG (3949G>T 3950G>C 3951T>G), GAT1268CAG (3952G>C 3954T>G), AAA1269AGG (3956A>G 3957A>G), GTA1270GTT (3960A>T), GAG1271CTT (3961G>C 3962A>T 3963G>T), CTT1272CTC (3966T>C), AGA1273ATA (3968G>T), GAC1275AAC (3973G>A), CCA1277CCT (3981A>T), GTG1278GTC (3984G>C), GAA1280GAT (3990A>T), GCT1281GCA (3993T>A), GAG1283GAA (3999G>A), GAA1284GAG (4002A>G), CTG1285TTG (4003C>T), ATT1287TCG (4009A>T 4010T>C 4011T>G), CAA1288CAG (4014A>G), ATT1289GTA (4015A>G 4017T>A), CAG1291CAA (4023G>A), ATA1293ACA (4028T>C), CAT1294TTT (4030C>T 4031A>T), GTG1295GTT (4035G>T), GAT1296GAC (4038T>C), CTA1297CTC (4041A>C), ATC1298GTC (4042A>G), ACC1299ACA (4047C>A), ACA1300ACC (4050A>C), CAA1301CAG (4053A>G), GCC1303GCG (4059C>G), AAG1304AAA (4062G>A), GAG1306GAA (4068G>A), CGC1307CGG (4071C>G), CTT1308TTG (4072C>T 4074T>G), CCT1309CCA (4077T>A), GAA1311GAG (4083A>G), AAA1318AAG (4104A>G), ACA1319ACC (4107A>C), AGA1320CGC (4108A>C 4110A>C), TTA1321CTG (4111T>C 4113A>G), TTT1322TTC (4116T>C), GAG1323GAA (4119G>A), ATA1324ATC (4122A>C), TAC1329TAT (4137C>T), GTC1333GTG (4149C>G), AGG1334CGG (4150A>C), TTT1337TTC (4161T>C), TCC1341TCT (4173C>T), GCC1342GCT (4176C>T), ACC1343TGT (4177A>T 4178C>G 4179C>T), GCA1346GCT (4188A>T), TTA1347GCA (4189T>G 4190T>C), AAT1349AAC (4197T>C), GTC1350GTA (4200C>A), ATC1351CTC (4201A>C), CCA1352CCT (4206A>T), TGT1353TGC (4209T>C), GTT1355GTA (4215T>A), GGA1356GGT (4218A>T), ATA1357ATC (4221A>C), GAT1358AAT (4222G>A), ACC1360ACG (4230C>G), AGT1361AGC (4233T>C), TCT1362CAT (4234T>C 4235C>A), TCT1365ACA (4243T>A 4245T>A), CTT1366GTG (4246C>G 4248T>G), CTT1367CTG (4251T>G), GCC1368GCT (4254C>T), AAC1369AGT (4256A>G 4257C>T), AAC1370AGC (4259A>G), TTC1371TTT (4263C>T), GCT1373TCA (4267G>T 4269T>A), ACT1377GAG (4279A>G 4280C>A 4281T>G), GGG1378GGA (4284G>A), TTT1379TTC (4287T>C), GCT1381GCC (4293T>C), TCA1384TCG (4302A>G), AGT1385AGC (4305T>C), GAC1387GAT (4311C>T), GGG1388GGA (4314G>A), CGC1389AGA (4315C>A 4317C>A), CCA1391CCC (4323A>C), GTG1392ATC (4324G>A 4326G>C), TTC1393TTT (4329C>T), GCT1394GCA (4332T>A), TTT1395TTC (4335T>C), CAG1396CAA (4338G>A), TTG1401CTG (4351T>C), GTT1402GTG (4356T>G), GAG1404GCT (4361A>C 4362G>T), TAT1405TTT (4364A>T), GAT1407GGT (4370A>G), ACC1408GAT (4372A>G 4373C>A 4374C>T), AAG1409GAG (4375A>G), GGA1411GGG (4383A>G), GAG1412AGT (4384G>A 4385A>G 4386G>T), GCT1414GAG (4391C>A 4392T>G), TCC1415TCA (4395C>A), GCA1417GCT (4401A>T), AGA1418AGG (4404A>G), CAT1419CAC (4407T>C), GCT1420ACA (4408G>A 4410T>A), CTT1421TTA (4411C>T 4413T>A), CTC1422TTA (4414C>T 4416C>A), GCA1425GCC (4425A>C), TCA1426TCT (4428A>T), CAC1428CAT (4434C>T), TAC1429TAT (4437C>T), TTA1431CTC (4441T>C 4443A>C), GAT1434AAC (4450G>A 4452T>C), AAA1435AAG (4455A>G), GTT1436TTG (4456G>T 4458T>G), TTC1437TTT (4461C>T), CGT1438CGC (4464T>C), TTG1439CTA (4465T>C 4467G>A), GTT1440GTG (4470T>G), GGC1442GGA (4476C>A), CCG1444CCC (4482G>C), GCA1448GCT (4494A>T), CTT1449TTG (4495C>T 4497T>G), ACC1450ACT (4500C>T), TTC1453TTT (4509C>T), AAC1454AAT (4512C>T), CTG1456TTG (4516C>T), TTG1457CTA (4519T>C 4521G>A), CGG1463AGG (4537C>A), TAT1464TAC (4542T>C), GCT1465GCG (4545T>G), TTC1466TTT (4548C>T), ATC1467ATA (4551C>A), CTT1469CTG (4557T>G), TTA1470TTG (4560A>G), AGA1471AGG (4563A>G), AGG1472AGA (4566G>A), CCT1473CCA (4569T>A), CAC1474CAT (4572C>T), ATT1475GTC (4573A>G 4575T>C), GCC1476AAA (4576G>A 4577C>A 4578C>A), GCT1477GCC (4581T>C), AGA1478AAA (4583G>A), GCT1479TCT (4585G>T), ATA1480GTG (4588A>G 4590A>G), GGT1481GGC (4593T>C), GTA1482ATT (4594G>A 4596A>T), CCT1484CCA (4602T>A), GGT1485AAT (4603G>A 4604G>A), GAT1486GAC (4608T>C), AAC1488AAT (4614C>T), CTT1490TTG (4618C>T 4620T>G), GGA1496GGT (4638A>T), AAT1499AAC (4647T>C), CTG1500TTG (4648C>T), GTG1501GTT (4653G>T), GCT1502GCG (4656T>G), GTG1503GTT (4659G>T), CCT1504CCA (4662T>A), CTG1505ATG (4663C>A), CTG1507TTG (4669C>T), TAT1510TAC (4680T>C), TCA1511GAT (4681T>G 4682C>A 4683A>T), CTG1512CTA (4686G>A), CCT1513CCA (4689T>A), AAT1514TCC (4690A>T 4691A>C 4692T>C), GCC1516GCA (4698C>A), CAG1517AGA (4699C>A 4700A>G 4701G>A), CTT1519CTG (4707T>G), GAC1520GAA (4710C>A), ATG1521GCT (4711A>G 4712T>C 4713G>T), GTA1522GTT (4716A>T), AAT1523AAC (4719T>C), ATC1526ATT (4728C>T), AAA1527AAG (4731A>G), GGG1529GGT (4737G>T), CTG1530TTG (4738C>T), GAT1531GAC (4743T>C), TCT1533TCA (4749T>A), AAC1537GAG (4759A>G 4761C>G), CAG1538AAG (4762C>A), GTG1539ACA (4765G>A 4766T>C 4767G>A), CAG1540AAA (4768C>A 4770G>A), TTC1541TTT (4773C>T), CAG1542CTG (4775A>T), GAC1543GAT (4779C>T), TCA1545TCT (4785A>T), GAA1546GAG (4788A>G), ACT1548ACC (4794T>C), AGC1551AGT (4803C>T), GGA1553GGT (4809A>T), TTT1554TTC (4812T>C), AAA1555AAG (4815A>G), CGT1556CGA (4818T>A), GCC1558GCA (4824C>A), CTT1559TTG (4825C>T 4827T>G), GGG1560GGA (4830G>A), CAC1562CAA (4836C>A), GTG1569GTT (4857G>T), ATA1571GTC (4861A>G 4863A>C), ATT1572ATC (4866T>C), CCT1574CCC (4872T>C), ATG1575ATT (4875G>T), TAC1576TAT (4878C>T), ATC1578ATT (4884C>T), AGA1579AGG (4887A>G), CCT1580CCA (4890T>A), CCG1582GTG (4894C>G 4895C>T), GGT1583GGA (4899T>A), ACT1587GTA (4909A>G 4910C>T 4911T>A), GTG1588CAA (4912G>C 4913T>A 4914G>A), GCA1589GCT (4917A>T), ATG1591TAT (4921A>T 4922T>A 4923G>T), GAG1592GAA (4926G>A), GTG1594ATT (4930G>A 4932G>T), ACT1596ACA (4938T>A), CTT1598ATC (4942C>A 4944T>C), CGT1599AGA (4945C>A 4947T>A), GAG1600GAA (4950G>A), TTT1602TTC (4956T>C), AGA1606CAG (4966A>C 4967G>A 4968A>G), GTT1607GCC (4970T>C 4971T>C), GCA1608ACT (4972G>A 4974A>T), GAG1610GAA (4980G>A), AAG1611AGT (4982A>G 4983G>T), TTA1612TTT (4986A>T), ACT1613GTA (4987A>G 4988C>T 4989T>A), TAT1614AAT (4990T>A), CAG1617TTT (4999C>T 5000A>T 5001G>T), TGT1618GGA (5002T>G 5004T>A), GCC1619GCT (5007C>T), AGA1621AAG (5012G>A 5013A>G), GAG1622CAG (5014G>C), CGT1623AGA (5017C>A 5019T>A), GGA1624AAT (5020G>A 5021G>A 5022A>T), GGC1626CCC (5026G>C 5027G>C), GCA1627ATC (5029G>A 5030C>T 5031A>C), ACA1628ATG (5033C>T 5034A>G), TTT1630ATG (5038T>A 5040T>G), CCT1631CCC (5043T>C), TCT1632ACT (5044T>A), TAT1633TAC (5049T>C), ATG1634TAT (5050A>T 5051T>A 5052G>T), GTG1636CTT (5056G>C 5058G>T), GAG1637GAA (5061G>A), CAG1638AGT (5062C>A 5063A>G 5064G>T), GAA1639CAA (5065G>C), ATC1641TTG (5071A>T 5073C>G), GCA1642ACA (5074G>A), AAG1643GAG (5077A>G), GCA1648GAG (5093C>A 5094A>G), TCA1649AGC (5095T>A 5096C>G 5097A>C), GCG1650ATT (5098G>A 5099C>T 5100G>T), TTG1651GGG (5101T>G 5102T>G), ACA1652GTT (5104A>G 5105C>T 5106A>T), ATC1653TTG (5107A>T 5109C>G), TGT1654ATG (5110T>A 5111G>T 5112T>G), GAA1655GAG (5115A>G), ATG1656CTA (5116A>C 5118G>A), GCC1657TCA (5119G>T 5121C>A), AAA1658GAT (5122A>G 5124A>T), TCT1660AAA (5128T>A 5129C>A 5130T>A), ATC1661AAT (5132T>A 5133C>T), ACT1662ACA (5136T>A), TTC1663TAC (5138T>A), ACA1664ACC (5142A>C), TTG1666TTA (5148G>A), CCA1667CCT (5151A>T), CCT1668GCT (5152C>G), AAT1670GAA (5158A>G 5160T>A), GTT1672ATT (5164G>A), GAA1673del (5167_5169delGAA), GAA1676GAG (5178A>G), AGA1677GAG (5179A>G 5180G>A 5181A>G), GAT1678GGT (5183A>G), CTC1679GTC (5185C>G), AAT1680CAT (5188A>C), TGG1681GAA (5191T>G 5192G>A 5193G>A), GCC1683GTG (5198C>T 5199C>G), CCA1684CCT (5202A>T), AAC1685GAT (5203A>G 5205C>T), ATT1686GTA (5206A>G 5208T>A), GGT1687TTT (5209G>T 5210G>T), TGC1689TGT (5217C>T), TCA1690AGT (5218T>A 5219C>G 5220A>T), GCC1691GCT (5223C>T), CGT1692CGG (5226T>G), GCC1695TCA (5233G>T 5235C>A), AAG1696AAC (5238G>C), ACT1697CAA (5239A>C 5240C>A 5241T>A), GAA1698CAA (5242G>C), GAG1700TCA (5248G>T 5249A>C 5250G>A), CCT1701TCA (5251C>T 5253T>A), GGG1702GGT (5256G>T), TGC1703AAC (5257T>A 5258G>A), ATA1704ATC (5262A>C), GTA1705CTG (5263G>C 5265A>G), AAC1707AAT (5271C>T), TGT1708TGC (5274T>C), ACA1709ACT (5277A>T), GGG1710GGA (5280G>A), GCG1711GCC (5283G>C), AAG1712CGC (5284A>C 5285A>G 5286G>C), AAG1713AGA (5288A>G 5289G>A), AGC1714ACA (5291G>C 5292C>A), AAA1715ACA (5294A>C), GTT1717GTG (5301T>G), CGC1718AGA (5302C>A 5304C>A), GGA1719GGG (5307A>G), GCT1721GCC (5313T>C), AGT1722AAT (5315G>A), GAT1725GAC (5325T>C), ATG1730ATA (5340G>A), CCA1732CCC (5346A>C), TAT1733TAC (5349T>C), ACC1734ACT (5352C>T), ATG1735TTG (5353A>T), TCA1736GCA (5356T>G), ATT1738ATC (5364T>C), CTG1739CAG (5366T>A), GCT1741CAT (5371G>C 5372C>A), AGA1743CGC (5377A>C 5379A>C), GGC1744AAG (5380G>A 5381G>A 5382C>G), AAA1745AGT (5384A>G 5385A>T), ATG1746CTG (5386A>C), TCA1747GAA (5389T>G 5390C>A), GAG1748GTT (5393A>T 5394G>T), GAC1749GGT (5396A>G 5397C>T), ATT1750GAT (5398A>G 5399T>A), GCT1751GGG (5402C>G 5403T>G), GAT1754TCT (5410G>T 5411A>C), GTT1755GTC (5415T>C), GTG1756ACC (5416G>A 5417T>C 5418G>C), GTG1758GTT (5424G>T), GGG1760GGT (5430G>T), GCT1764GCA (5442T>A), GTG1765ATA (5443G>A 5445G>A), AGC1766GCT (5446A>G 5447G>C 5448C>T), GCT1771GCA (5463T>A), TTA1772TTG (5466A>G), ATG1773TTT (5467A>T 5469G>T), GCT1774GCA (5472T>A), GTG1775TTG (5473G>T), CGT1778CAT (5483G>A), CAG1779CAA (5487G>A), TAT1780TAC (5490T>C), TGC1781AGG (5491T>A 5493C>G), ATT1782GCC (5494A>G 5495T>C 5496T>C), GAG1783ACA (5497G>A 5498A>C 5499G>A), GAT1784GAC (5502T>C), ATC1785CTT (5503A>C 5505C>T), AGG1788CGC (5512A>C 5514G>C), TTG1789CTA (5515T>C 5517G>A), CGC1790CGA (5520C>A), ACT1791ACC (5523T>C), ATT1792ATG (5526T>G), GGC1793GCT (5528G>C 5529C>T), AAT1794GGG (5530A>G 5531A>G 5532T>G), TTG1796CTG (5536T>C), ACA1800ACC (5550A>C), TAT1801TAC (5553T>C), CCC1802CCT (5556C>T), GTT1806GCG (5567T>C 5568T>G), CAG1807CAA (5571G>A), TTT1809TTG (5577T>G), CAA1811AAA (5581C>A), GTG1813GTT (5589G>T), CCT1814GAA (5590C>G 5591C>A 5592T>A), CAG1815GAC (5593C>G 5595G>C), CAG1815_GAA1816insACC (5595_5596insACC), GAA1816TCA (5596G>T 5597A>C), GGG1817CAC (5599G>C 5600G>A 5601G>C), AAA1818AAG (5604A>G), ATG1819TTG (5605A>T), GCA1820GGA (5609C>G), GTT1822AGT (5614G>A 5615T>G), TCG1824ACA (5620T>A 5622G>A), TCT1825del (5623_5625delTCT), TTG1826TAT (5627T>A 5628G>T), CAA1827GGC (5629C>G 5630A>G 5631A>C), AGT1828ACC (5633G>C 5634T>C), AGT1829ACT (5636G>C), ATG1831AAC (5642T>A 5643G>C), CAA1833CAT (5649A>T), CAA1833_GGT1834insCACAGT (5649_5650insCACAGT), GGT1834GCA (5651G>C 5652T>A), GAG1836TCA (5656G>T 5657A>C 5658G>A), CTT1837GTT (5659C>G), GGC1838GGA (5664C>A), CAA1839CTT (5666A>T 5667A>T), ATT1840CTA (5668A>C 5670T>A), CGT1841AGG (5671C>A 5673T>G), GTC1842ATA (5674G>A 5676C>A), ATC1843GTT (5677A>G 5679C>T), CAC1844CCT (5681A>C 5682C>T), GGT1845TCT (5683G>T 5684G>C), GAT1846GAC (5688T>C), TTA1847CTC (5689T>C 5691A>C), TCA1848TCT (5694A>T), ATG1849GAA (5695A>G 5696T>A 5697G>A), GAA1850AAG (5698G>A 5700A>G), TTG1853GTT (5707T>G 5709G>T), CGC1854CGT (5712C>T), ATG1855TTG (5713A>T), TAT1857TAC (5721T>C), GTT1858ACA (5722G>A 5723T>C 5724T>A), GTG1859ATT (5725G>A 5727G>T), GGC1860GGT (5730C>T), CCA1861CCT (5733A>T), CAT1862GAA (5734C>G 5736T>A), GGT1863GGA (5739T>A), GGA1864GGT (5742A>T), GGC1866GCC (5747G>C), AAC1867CAG (5749A>C 5751C>G), TTT1868TGC (5753T>G 5754T>C), ACC1869TCA (5755A>T 5757C>A), ACA1870ACG (5760A>G), AAA1871AAG (5763A>G), AAT1872AAC (5766T>C), CTT1873CTG (5769T>G), GAT1874AAG (5770G>A 5772T>G), CAA1875CAT (5775A>T), TTG1876GCT (5776T>G 5777T>C 5778G>T), TTT1877TTG (5781T>G), GCA1878GAG (5783C>A 5784A>G), AGT1879ACA (5786G>C 5787T>A), CTT1880TTG (5788C>T 5790T>G), GAA1881TCT (5791G>T 5792A>C 5793A>T), AAG1882GTC (5794A>G 5795A>T 5796G>C), TGT1883AGT (5797T>A), GAA1886ACT (5806G>A 5807A>C 5808A>T), TTG1887CTT (5809T>C 5811G>T), CTT1888GCC (5812C>G 5813T>C 5814T>C), AAG1890AGA (5819A>G 5820G>A), AAC1891GGA (5821A>G 5822A>G 5823C>A), ACG1892GTC (5824A>G 5825C>T 5826G>C), CTG1894GTG (5830C>G), ACC1895ACA (5835C>A), GAG1899AAA (5845G>A 5847G>A), AAA1900GAA (5848A>G), ATG1901TTC (5851A>T 5853G>C), ACT1902TCC (5854A>T 5856T>C), AAA1904CAC (5860A>C 5862A>C), GTG1906TGT (5866G>T 5867T>G 5868G>T), GCA1907GAA (5870C>A), CAA1908ATG (5872C>A 5873A>T 5874A>G), GTC1910CTT (5878G>C 5880C>T), AGT1911GAG (5881A>G 5882G>A 5883T>G), GTA1913CTG (5887G>C 5889A>G), AAA1914GAA (5890A>G),

| | | | | | | | | | |
|---|---|---|---|---|---|---|---|---|---|
| | CAG1915AGT (5893C>A 5894A>G 5895G>T), TTC1918GTA (5902T>G 5904C>A), TAT1919GTA (5905T>G 5906A>T 5907T>A), CCC1920TTC (5908C>T 5909C>T), AAG1921ATG (5912A>T), GCC1922GGG (5915C>G 5916C>G), ACA1923AGC (5918C>G 5919A>C), ACC1924GTT (5920A>G 5921C>T 5922C>T), GTG1925AAG (5923G>A 5924T>A), GAA1926AAA (5926G>A), CAG1927CAA (5931G>A), CTC1928ATG (5932C>A 5934C>G), AAG1929TTG (5935A>T 5936A>T), GTA1930TCC (5938G>T 5939T>C 5940A>C), TTT1931TTC (5943T>C), GCT1932ATG (5944G>A 5945C>T 5946T>G), GAT1933AAT (5947G>A), GCT1934CAA (5950G>C 5951C>A 5952T>A), CAT1935CAG (5955T>G), CAT1935_ACC1936insATG (5955_5956insATG), ACC1936TCA (5956A>T 5958C>A), ATT1937AAA (5960T>A 5961T>A), GTC1938ACC (5962G>A 5963T>C), AAA1939CAA (5965A>C), ACT1940GTT (5968A>G 5969C>T), GCT1941GCC (5973T>C), AAA1942AGC (5975A>G 5976A>C), CCT1943CCA (5979T>A), TTC1944TGT (5981T>G 5982C>T), CGG1945TGT (5983C>T 5985G>T), GGA1946del (5986_5988delGGA), GTA1947GTC (5991A>C), GTT1948ACA (5992G>A 5993T>C 5994T>A), TTC1949TTT (5997C>T), CGG1950AGT (5998C>A 6000G>T), AAT1952GCA (6004A>G 6005A>C 6006T>A), TTT1953TGG (6008T>G 6009T>G), CTC1954CTA (6012C>A), AGT1955ACT (6014G>C), CCT1956AAG (6016C>A 6017C>A 6018T>G), GAG1957GAC (6021G>C), TGG1958ATT (6022T>A 6023G>T 6024G>T), TGG1958_AAA1959insGTC (6024_6025insGTC), AAA1959ATG (6026A>T 6027A>G), ATT1960ACA (6029T>C 6030T>A), GGA1961AAA (6031G>A 6032G>A), GAG1963AAG (6037G>A), AAC1964AAG (6042C>G), ACA1965CCA (6043A>C), ATG1966del (6046_6048delATG), GCA1967GGA (6050C>G), ACA1968TCT (6052A>T 6054A>T), CTC1969ATT (6055C>A 6057C>T), TCA1970CCT (6058T>C 6060A>T), GAG1972GAA (6066G>A), AGT1973ACA (6068G>C 6069T>A), CTT1974GTG (6070C>G 6072T>G), CTT1975TTT (6073C>T), CCA1976CTT (6077C>T 6078A>T), CTG1979CCT (6086T>C 6087G>T), TCG1980TCA (6090G>A), TCA1982CTA (6094T>C 6095C>T), GCC1983GTT (6098C>T 6099C>T), ATG1984GTG (6100A>G), AAG1985AGA (6104A>G 6105G>A), TTG1987ATG (6109T>A), CTC1988TTG (6112C>T 6114C>G), AGC1989CGC (6115A>C), AGA1990AAT (6119G>A 6120A>T), TAC1991AGT (6121T>A 6122A>G 6123C>T), AAA1992GGA (6124A>G 6125A>G), AGA1993AAG (6128G>A 6129A>G), AAC1994AGC (6131A>G), ATG1995TAC (6133A>T 6134T>A 6135G>C), TCT1996TCA (6138T>A), CAT1998ACA (6142C>A 6143A>C 6144T>A), TGC1999ACA (6145T>A 6146G>C 6147C>A), ATA2000CTG (6148A>C 6150A>G), AAA2001GAA (6151A>G), TTG2002CTT (6154T>C 6156G>T), GCA2003GCT (6159A>T), CTC2004TTT (6160C>T 6162C>T), AAG2005AGA (6164A>G 6165G>A), ATA2006ATG (6168A>G), TAT2007TAC (6171T>C), GTT2008ATG (6172G>A 6174T>G), CTC2009_AAC2010del (6175_6180delCTCAAC), TTC2011TGT (6182T>G 6183C>T), CAG2012CAA (6186G>A), ATG2013TGG (6187A>T 6188T>G), CTG2014GTA (6190C>G 6192G>A), AAT2015CAC (6193A>C 6195T>C), GAA2017CC. (6199G>C 6200A>C) | | | | | | | | |
| helicase motif A-C (YP_009167369.1) | 1 | 98 | 100% | 654 | 97.6% | 98 (100%) | 96 (98.0%) | 0/0/0/0 | 0 |
| Protein mutations: | A5C (1345G>T 1346C>G 1347A>T), R16A (1378C>G 1379G>C 1380C>A) | | | | | | | | |
| Codon mutations: | ACA4ACG (1344A>G), GCA5TGT (1345G>T 1346C>G 1347A>T), CGA7CGC (1353A>C), TTG8CTT (1354T>C 1356G>T), TTG14CTC (1372T>C 1374G>C), GAT15GAC (1377T>C), CGC16GCA (1378C>G 1379G>C 1380C>A), GGG18GGT (1386G>T), GAG19GAA (1389G>A), CCT20CCA (1392T>A), CTT22CTG (1398T>G), AGA24CGC (1402A>C 1404A>C), CTT25TTG (1405C>T 1407T>G), GCT27GCG (1413T>G), GTT28GTA (1416T>A), AGC29AGT (1419C>T), AAG30AAA (1422G>A), CGC31AGA (1423C>A 1425C>A), GCC33GCA (1431C>A), TCT36TCC (1440T>C), AAT37AAC (1443T>C), GCC39GCA (1449C>A), CAC40CAT (1452C>T), CAG41CAA (1455G>A), GCT43GCA (1461T>A), TTG45CTT (1465T>C 1467G>T), GAC48GAT (1476C>T), GGA50GGT (1482A>T), GCA51GCT (1485A>T), TTG52CTT (1486T>C 1488G>T), AGA53CGG (1489A>C 1491A>G), GGA57GGG (1503A>G), CAA58CAG (1506A>G), CAA60CAG (1512A>G), ATT62ATA (1518T>A), ATT68ATC (1536T>C), TCC70TCT (1542C>T), ACA71ACC (1545A>C), CCA73CCT (1551A>T), CCA75CCT (1557A>T), CTG76CTT (1560G>T), CCC77CCA (1563C>A), GCA79GCT (1569A>T), GCT80GCA (1572T>A), GTG81GTC (1575G>C), GAA82GAG (1578A>G), GAC83GAT (1581C>T), AAG84AAA (1584G>A), GGG85GGA (1587G>A), CAT87CAC (1593T>C), ACT89ACA (1599T>A), TAT92TAC (1608T>C), GCA95GCC (1617A>C) | | | | | | | | |

*: Inserts / Deletes / Misaligned / Frameshifts

#### Analysis details

This analysis was performed with panviral2.64

### NGS Details (UN60): Tomato chocolate spot virus (segment RNA 1)

#### Assembly

| | |
|---|---|
| **Coverage Length** | 1861 (5 contig(s)) |
| **Depth Of Coverage** | 11026.4 |
| **Number Of Reads** | 127831 |
| **Reads Per Million** | 2711.24 rpm (after QC) |
| **Ambiguities** | 0 |
| **Assembly Method** | de novo + reference guided assembly |
| **Consensus Caller** | Bcf Tools |

#### Coverage Map

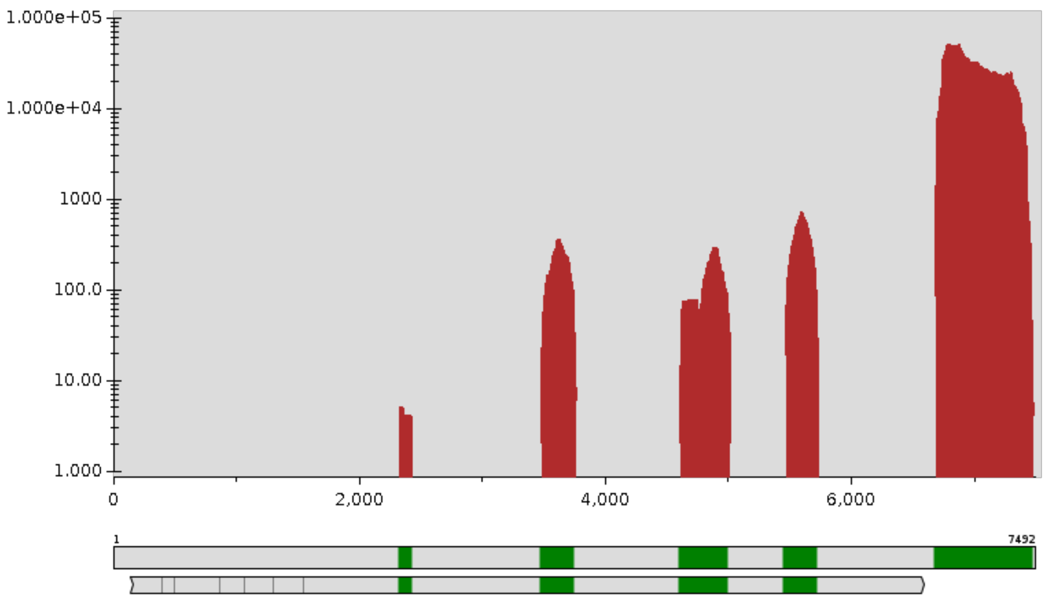

#### Assignment

| | |
|---|---|
| **Type** | Tomato chocolate spot virus (Taxonomy ID: 661101) |
| **Reference Genome** | NC_013075.1 |
| **NT Identity (%)** | 67.7475 |
| **AA Identity (%)** | 68.8022 |
| **Number Of Stop Codons** | 0 |
| **Number Of CDS** | 1 |

#### Alignment

| | |
|---|---|
| **Alignment Score** | 777.0 (NT) + 1773.0 (AA) = 2550.0 |
| **Concordance (%)** | 46.3047 |

| Alignment Method | Global, seeded, nucleotide + amino acids (AGA) |
|---|---|

#### Genome Region

Sequence starts at position 2324 and ends at position 7477 relative to NC_013075.1 reference sequence.

#### Alignment Detailed Statistics

| | Begin | End | Coverage | Score | Concordance | Matches | Identities | I/D/M/F* | Stop Codons |
|---|---|---|---|---|---|---|---|---|---|
| **NT** | **2324** | **7477** | **24.8%** | **777** | **26.0%** | **1698 (89.8%)** | **1170 (61.9%)** | **29/163** | |
| Mutations: | 2325T>A, 2329C>T, 2331C>T, 2335T>G, 2336G>C, 2337C>G, 2338A>C, 2341G>C, 2343T>G, 2351T>C, 2352T>A, 2355C>T, 2361C>T, 2363T>C, 2364A>G, 2367C>T, 2382C>A, 2385T>C, 2386T>A, 2387T>G, 2388T>A, 2391C>A, 2394A>G, 2397T>C, 2400T>G, 2403C>T, 2409C>T, 2415T>G, 2416A>C, 2418A>C, 2429A>G, 2430C>G, 2437T>G, 3484G>A, 3489C>T, 3490T>C, 3492G>C, 3498G>C, 3501G>A, 3504T>G, 3511T>G, 3513A>T, 3517C>T, 3519T>G, 3531A>T, 3537A>G, 3538T>A, 3549C>G, 3552C>G, 3562T>A, 3564A>C, 3567A>C, 3573A>G, 3575G>A, 3579C>T, 3581A>C, 3582G>A, 3594T>A, 3600A>T, 3601C>G, 3602C>A, 3603T>A, 3607G>C, 3612A>T, 3615A>T, 3618A>T, 3622A>C, 3624A>G, 3628A>T, 3630C>T, 3633C>G, 3636T>A, 3639C>T, 3642G>A, 3645G>A, 3648T>G, 3651G>T, 3654C>A, 3663A>T, 3675G>C, 3676G>A, 3678C>T, 3679A>G, 3680T>A, 3699A>C, 3700C>T, 3701A>C, 3702C>A, 3711T>G, 3714A>C, 3715G>A, 3717A>C, 3723C>A, 3724A>C, 3725A>G, 3727A>G, 3728T>C, 3729G>T, 3732C>A, 3738G>A, 3739A>G, 3740C>G, 3741C>T, 3747A>G, 3749G>C, 3753T>C, 3754T>A, 3756G>T, 3759G>A, 4615C>T, 4617T>A, 4620C>T, 4621G>A, 4626T>A, 4635A>G, 4644C>T, 4650T>G, 4656T>G, 4659C>A, 4660T>A, 4666T>C, 4668G>T, 4671G>A, 4677T>C, 4678T>G, 4679C>A, 4680A>G, 4683G>A, 4686C>T, 4687A>G, 4688A>C, 4689C>A, 4692T>A, 4701A>G, 4704G>A, 4707T>A, 4708A>C, 4709T>A, 4710G>A, 4713A>G, 4716T>C, 4720G>A, 4722A>T, 4725T>C, 4728A>G, 4731C>T, 4734T>C, 4735T>C, 4737G>T, 4740C>T, 4744T>A, 4745C>A, 4746A>C, 4750G>A, 4755G>C, 4761G>T, 4764G>C, 4765C>A, 4767G>A, 4768T>C, 4770T>C, 4785A>G, 4786C>T, 4791T>G, 4795C>T, 4806G>C, 4813A>C, 4815A>C, 4822C>T, 4827C>T, 4833T>A, 4834A>C, 4836G>A, 4839A>C, 4842G>C, 4845T>C, 4854A>T, 4860A>T, 4861A>G, 4863A>G, 4881A>C, 4882C>A, 4891C>G, 4892C>T, 4893C>T, 4896T>G, 4902T>A, 4906G>A, 4907C>T, 4908A>C, 4909C>A, 4910A>C, 4911G>T, 4914A>C, 4918A>T, 4919T>A, 4920G>T, 4929T>G, 4935T>G, 4938C>T, 4939A>G, 4950C>T, 4953T>C, 4956T>G, 4962T>A, 4965A>G, 4968T>C, 4970C>T, 4971A>T, 4977G>C, 4979A>C, 4984G>A, 4985T>C, 4986G>A, 4988C>A, 4989A>C, 4990G>A, 4995G>C, 4999G>A, 5005C>A, 5007G>A, 5009A>T, 5010T>G, 5011G>C, 5013A>G, 5016T>A, 5019A>C, 5469T>A, 5472G>A, 5473G>T, 5475G>A, 5478T>C, 5480G>A, 5481G>T, 5486T>A, 5488T>A, 5490T>A, 5497G>C, 5499C>A, 5500A>C, 5505T>A, 5508A>C, 5511G>A, 5515C>A, 5517C>A, 5521A>T, 5523T>G, 5526T>G, 5527A>G, 5528A>G, 5529T>C, 5530G>T, 5544C>T, 5545A>C, 5546C>A, 5547A>G, 5550C>T, 5563G>A, 5564C>T, 5565A>T, 5568G>A, 5571C>T, 5577A>C, 5578C>A, 5579A>C, 5580G>A, 5583C>T, 5586_5587insGTG, 5588C>A, 5589C>G, 5590C>A, 5592T>A, 5593G>A, 5596G>T, 5597A>G, 5598T>C, 5601A>G, 5602G>A, 5603T>C, 5604G>A, 5607A>T, 5614G>A, 5617_5619delATT, 5620T>G, 5621C>G, 5624T>G, 5625G>C, 5626C>G, 5628C>T, 5629A>G, 5631C>A, 5633G>T, 5634T>G, 5637T>G, 5638G>A, 5639C>A, 5640A>C, 5643T>C, 5646A>T, 5646_5647insCATGCT, 5648G>C, 5651T>A, 5654A>C, 5655A>C, 5656T>C, 5658G>T, 5661C>A, 5664T>C, 5665T>C, 5667C>T, 5668A>C, 5674A>G, 5676C>G, 5677C>G, 5679T>C, 5680G>T, 5683G>T, 5684A>C, 5685C>A, 5688C>T, 5693A>G, 5694G>A, 5696C>A, 5697T>G, 5699C>T, 5700T>G, 5703C>A, 5704T>G, 5710A>T, 5715A>C, 5718T>C, 5719G>A, 5720T>C, 5721G>T, 5727A>G, 5729C>A, 5730C>G, 5733T>A, 5735G>T, 5736A>T, 6706A>G, 6707T>A, 6721G>C, 6722G>C, 6729T>C, 6731G>A, 6735G>A, 6736T>G, 6740C>A, 6741G>A, 6753G>T, 6757T>G, 6758_6759insT, 6759_6760insG, 6772C>T, 6777T>C, 6795A>G, 6806C>T, 6814C>T, 6815_6834delAATTGTTTGGTGTTGTCAAC, 6837G>T, 6840G>A, 6842C>A, 6843T>C, 6845T>G, 6847_6850delGCCA, 6853A>G, 6855A>T, 6857C>A, 6859C>T, 6860C>G, 6861C>G, 6862C>T, 6863T>C, 6867_6868delGC, 6870T>G, 6872A>G, 6873T>A, 6878A>G, 6879A>T, 6881G>A, 6885A>C, 6886G>T, 6887A>T, 6889_6894delGTGGAG, 6897G>A, 6903C>A, 6906G>C, 6908A>C, 6909A>G, 6911C>G, 6912A>G, 6914A>G, 6915A>T, 6916G>A, 6918A>T, 6919_6921delGAG, 6923C>T, 6924T>C, 6926T>C, 6930_6937delGGTCGGGC, 6942G>C, 6944_6946delTGG, 6949G>A, 6950C>G, 6951C>T, 6953G>C, 6955A>G, 6956T>G, 6958A>T, 6960A>T, 6961G>T, 6962C>T, 6964T>C, 6965T>C, 6967G>A, 6969G>A, 6970T>G, 6972T>A, 6973G>A, 6976T>A, 6978T>A, 6982_6985delCATC, 6987G>T, 6989T>A, 6991T>C, 6996A>C, 6998C>T, 7001_7012delCTCGTTTGGAGT, 7014T>G, 7016A>G, 7018C>G, 7020A>T, 7022delT, 7026G>T, 7030_7035delTGGTGT, 7040A>C, 7041C>G, 7042G>A, 7047_7067delGCAATGACACCCGGGGTGTGC, 7073C>G, 7077T>C, 7079A>T, 7080A>G, 7084_7088delAAGTA, 7091G>A, 7093T>C, 7097delT, 7100C>G, 7102T>A, 7103A>T, 7108_7116delAACTAGAGC, 7118T>G, 7121_7129delGTCTCAACT, 7132A>G, 7134G>T, 7135A>G, 7136G>A, 7140_7145delCCCGTT, 7148A>T, 7151A>C, 7152T>A, 7154G>A, 7155A>C, 7156T>C, 7158A>C, 7159A>T, 7161T>C, 7161_7162insC, 7165G>C, 7169C>G, 7172delG, 7177A>C, 7180_7183delCTGC, 7186T>G, 7188G>T, 7190T>C, 7193G>A, 7195_7218delGCAACAGTGAAGAATCATCACACC, 7220C>G, 7223C>A, 7224T>C, 7225T>C, 7227C>T, 7228C>G, 7230A>C, 7231G>T, 7234T>G, 7235C>G, 7238C>T, 7241G>C, 7244G>A, 7245T>A, 7247_7248delTC, 7251A>T, 7259_7260delCT, 7262C>G, 7264T>A, 7266G>A, 7270A>C, 7274G>C, 7288G>C, 7289T>C, 7291T>G, 7292T>G, 7293G>A, 7294C>T, 7296_7297insGCTTCTCCTAGTTAA, 7298G>A, 7299A>C, 7300C>G, 7302A>G, 7303T>C, 7304G>A, 7308C>T, 7310T>A, 7311A>T, 7314G>C, 7315T>C, 7316T>A, 7317T>G, 7318G>T, 7319G>A, 7321T>A, 7325C>A, 7327T>A, 7328A>T, 7330A>T, 7335_7336insT, 7342T>G, 7346A>T, 7348A>T, 7350G>C, 7351G>T, 7352_7353insC, 7356A>G, 7358C>G, 7367C>T, 7369G>A, 7372G>T, 7373delA, 7375T>C, 7376G>C, 7378A>C, 7381C>G, 7385C>T, 7387T>G, 7388T>C, 7389C>A, 7390A>T, 7393_7396delTATT, 7407G>T, 7410delG, 7425A>C, 7428delC, 7430T>C, 7431C>A, 7447A>C, 7453A>G, 7454G>A, 7455A>T, 7459T>C, 7460C>G | | | | | | | | |

**CDS**

| | Begin | End | Coverage | Score | Concordance | Matches | Identities | I/D/M/F* | Stop Codons |
|---|---|---|---|---|---|---|---|---|---|
| ToChSV_s1_gp1 | 730 | 1866 | 16.6% | 1773 | 69.6% | 356 (98.9%) | 247 (68.6%) | 3/1/0/0 | 0 |
| Protein mutations: | H731Y (2329C>T 2331C>T), C733A (2335T>G 2336G>C 2337C>G), I734L (2338A>C), D735Q (2341G>C 2343T>G), V738A (2351T>C 2352T>A), I742T (2363T>C 2364A>G), F750R (2386T>A 2387T>G 2388T>A), N764R (2429A>G 2430C>G), A1116T (3484G>A), E1120D (3498G>C), N1122K (3504T>G), S1125A (3511T>G 3513A>T), L1134M (3538T>A), S1142T (3562T>A 3564A>C), R1146K (3575G>A), Q1148P (3581A>C 3582G>A), P1155E (3601C>G 3602C>A 3603T>A), E1157Q (3607G>C), I1162L (3622A>C 3624A>G), T1164S (3628A>T 3630C>T), N1166K (3636T>A), E1171D (3651G>T), V1180I (3676G>A 3678C>T), M1181E (3679A>G 3680T>A), H1188S (3700C>T 3701A>C 3702C>A), D1191E (3711T>G), E1192D (3714A>C), G1193S (3715G>A 3717A>C), K1196R (3724A>C 3725A>G), M1197A (3727A>G 3728T>C 3729G>T), T1201G (3739A>G 3740C>G 3741C>T), C1204S (3749G>C), L1206I (3754T>A 3756G>T), V1495I (4621G>A), L1508M (4660T>A), S1514E (4678T>G 4679C>A 4680A>G), N1517A (4687A>G 4688A>C 4689C>A), D1523E (4707T>A), M1524Q (4708A>C 4709T>A 4710G>A), V1528I (4720G>A 4722A>T), S1536N (4744T>A 4745C>A 4746A>C), D1538N (4750G>A), E1541D (4761G>T), Q1543K (4765C>A 4767G>A), F1544L (4768T>C 4770T>C), H1565Q (4833T>A), M1566L (4834A>C 4836G>A), I1575V (4861A>G 4863A>G), P1585V (4891C>G 4892C>T 4893C>T), A1590I (4906G>A 4907C>T 4908A>C), Q1591T (4909C>A 4910A>C 4911G>T), M1594Y (4918A>T 4919T>A 4920G>T), I1601V (4939A>G), H1606Q (4956T>G), A1611V (4970C>T 4971A>T), E1613D (4977G>C), K1614T (4979A>C), V1616T (4984G>A 4985T>C 4986G>A), A1617D (4988C>A 4989A>C), D1618N (4990G>A), E1621K (4999G>A), L1623I (5005C>A 5007G>A), N1624M (5009A>T 5010T>G), E1625Q (5011G>C 5013A>G), A1779S (5473G>T 5475G>A), R1781N (5480G>A 5481G>T), F1783Y (5486T>A), C1784R (5488T>A 5490T>A), D1787Q (5497G>C 5499C>A), I1788L (5500A>C), I1795L (5521A>T 5523T>G), N1797G (5527A>G 5528A>G 5529T>C), V1798L (5530G>T), T1803Q (5545A>C 5546C>A 5547A>G), A1809I (5563G>A 5564C>T 5565A>T), Q1814T (5578C>A 5579A>C 5580G>A), V1816_P1817insV (5586_5587insGTG), P1817Q (5588C>A 5589C>G), H1818K (5590C>A 5592T>A), G1819S (5593G>A), D1820C (5596G>T 5597A>G 5598T>C), V1822T (5602G>A 5603T>C 5604G>A), G1826S (5614G>A), I1827del (5617_5619delATT), S1828G (5620T>G 5621C>G), L1829R (5624T>G 5625G>C), H1830D (5626C>G 5628C>T), S1831G (5629A>G 5631C>A), S1832M (5633G>T 5634T>G), F1833L (5637T>G), A1834N (5638G>A 5639C>A 5640A>C), Q1836H (5646A>T), Q1836_G1837insHA (5646_5647insCATGCT), G1837A (5648G>C), F1838Y (5651T>A), E1839A (5654A>C 5655A>C), F1843L (5665T>C 5667C>T), I1846V (5674A>G 5676C>G), R1847G (5677C>G 5679T>C), G1848C (5680G>T), D1849S (5683G>T 5684A>C 5685C>A), K1852R (5693A>G 5694G>A), T1853K (5696C>A 5697T>G), T1854M (5699C>T 5700T>G), L1856V (5704T>G), M1858L (5710A>T), V1861T (5719G>A 5720T>C 5721G>T), P1864Q (5729C>A 5730C>G), N1865K (5733T>A), G1866V (5735G>T 5736A>T) | | | | | | | | |

| | Begin | End | Coverage | Score | Concordance | Matches | Identities | I/D/M/F* | Stop Codons |
|---|---|---|---|---|---|---|---|---|---|
| **NT** | **2324** | **7477** | **24.8%** | **777** | **26.0%** | **1698 (89.8%)** | **1170 (61.9%)** | **29/163** | |
| Codon mutations: | TTT729.TA (2325T>A), CAC731TAT (2329C>T 2331C>T), TGC733GCG (2335T>G 2336G>C 2337C>G), ATT734CTT (2338A>C), GAT735CAG (2341G>C 2343T>G), GTT738GCA (2351T>C 2352T>A), GAC739GAT (2355C>T), CTC741CTT (2361C>T), ATA742ACG (2363T>C 2364A>G), TGC743TGT (2367C>T), CCC748CCA (2382C>A), CCT749CCC (2385T>C), TTT750AGA (2386T>A 2387T>G 2388T>A), GTC751GTA (2391C>A), CAA752CAG (2394A>G), TTT753TTC (2397T>C), CTT754CTG (2400T>G), TAC755TAT (2403C>T), GCC757GCT (2409C>T), GTT759GTG (2415T>G), AGA760CGC (2416A>C 2418A>C), AAC764AGG (2429A>G 2430C>G), TCA767GC. (2437T>G), GCT1116ACT (3484G>A), GCC1117GCT (3489C>T), TTG1118CTC (3490T>C 3492G>C), GAG1120GAC (3498G>C), GAG1121GAA (3501G>A), AAT1122AAG (3504T>G), TCA1125GCT (3511T>G 3513A>T), CTT1127TTG (3517C>T 3519T>G), CCA1131CCT (3531A>T), GAA1133GAG (3537A>G), TTG1134ATG (3538T>A), ACC1137ACG (3549C>G), GTC1138GTG (3552C>G), TCA1142ACC (3562T>A 3564A>C), ACA1143ACC (3567A>C), AGA1145AGG (3573A>G), AGA1146AAA (3575G>A), AGC1147AGT (3579C>T), CAG1148CCA (3581A>C 3582G>A), GCT1152GCA (3594T>A), GGA1154GGT (3600A>T), CCT1155GAA (3601C>G 3602C>A 3603T>A), GAA1157CAA (3607G>C), ACA1158ACT (3612A>T), GCA1159GCT (3615A>T), CCA1160CCT (3618A>T), ATA1162CTG (3622A>C 3624A>G), ACC1164TCT (3628A>T 3630C>T), GTC1165GTG (3633C>G), AAT1166AAA (3636T>A), GAC1167GAT (3639C>T), AAG1168AAA (3642G>A), AGG1169AGA (3645G>A), GGT1170GGG (3648T>G), GAG1171GAT (3651G>T), GGC1172GGA (3654C>A), CCA1175CCT (3663A>T), GGG1179GGC (3675G>C), GTC1180ATT (3676G>A 3678C>T), ATG1181GAG (3679A>G 3680T>A), GCA1187GCC (3699A>C), CAC1188TCA (3700C>T 3701A>C 3702C>A), GAT1191GAG (3711T>G), GAA1192GAC (3714A>C), GGA1193AGC (3715G>A 3717A>C), GCC1195GCA (3723C>A), AAG1196CGG (3724A>C 3725A>G), ATG1197GCT (3727A>G 3728T>C 3729G>T), GCC1198GCA (3732C>A), GAG1200GAA (3738G>A), ACC1201GGT (3739A>G 3740C>G 3741C>T), AAA1203AAG (3747A>G), TGT1204TCT (3749G>C), GCT1205GCC (3753T>C), TTG1206ATT (3754T>A 3756G>T), TTG1207TTA (3759G>A), CTT1493TTA (4615C>T 4617T>A), TTC1494TTT (4620C>T), GTT1495ATT (4621G>A), GCT1496GCA (4626T>A), GGA1499GGG (4635A>G), AAC1502AAT (4644C>T), GTT1504GTG (4650T>G), GTT1506GTG (4656T>G), CCC1507CCA (4659C>A), TTG1508ATG (4660T>A), TTG1510CTT (4666T>C 4668G>T), CAG1511CAA (4671G>A), TAT1513TAC (4677T>C), TCA1514GAG (4678T>G 4679C>A 4680A>G), TTG1515TTA (4683G>A), CCC1516CCT (4686C>T), AAC1517GCA (4687A>G 4688A>C 4689C>A), ATT1518ATA (4692T>A), GAA1521GAG (4701A>G), CTG1522CTA (4704G>A), GAT1523GAA (4707T>A), ATG1524CAA (4708A>C 4709T>A 4710G>A), GTA1525GTG (4713A>G), AAT1526AAC (4716T>C), GTA1528ATT (4720G>A 4722A>T), ATT1529ATC (4725T>C), AAA1530AAG (4728A>G), AAC1531AAT (4731C>T), GGT1532GGC (4734T>C), TTG1533CTT (4735T>C 4737G>T), GAC1534GAT (4740C>T), TCA1536AAC (4744T>A 4745C>A 4746A>C), GAT1538AAT (4750G>A), GTG1539GTC (4755G>C), GAG1541GAT (4761G>T), GTG1542GTC (4764G>C), CAG1543AAA (4765C>A 4767G>A), TTT1544CTC (4768T>C 4770T>C), GAA1549GAG (4785A>G), CTG1550TTG (4786C>T), ACT1551ACG (4791T>G), CTG1553TTG (4795C>T), GGG1556GGC (4806G>C), AGA1559CGC (4813A>C 4815A>C), CTG1562TTG (4822C>T), GGC1563GGT (4827C>T), CAT1565CAA (4833T>A), ATG1566CTA (4834A>C 4836G>A), GCA1567GCC (4839A>C), CCG1568CCC (4842G>C), CTT1569CTC (4845T>C), GTA1572GTT (4854A>T), ATA1574ATT (4860A>T), ATA1575GTG (4861A>G 4863A>G), ATA1581ATC (4881A>C), CGA1582AGA (4882C>A), CCC1585GTT (4891C>G 4892C>T 4893C>T), GGT1586GGG (4896T>G), CCT1588CCA (4902T>A), GCA1590ATC (4906G>A 4907C>T 4908A>C), CAG1591ACT (4909C>A 4910A>C 4911G>T), GCA1592GCC (4914A>C), ATG1594TAT (4918A>T 4919T>A 4920G>T), GTT1597GTG (4929T>G), ACT1599ACG (4935T>G), GGC1600GGT (4938C>T), ATA1601GTA (4939A>G), GCC1604GCT (4950C>T), TTT1605TTC (4953T>C), CAT1606CAG (4956T>G), GGT1608GGA (4962T>A), CAA1609CAG (4965A>G), GTT1610GTC (4968T>C), GCA1611GTT (4970C>T 4971A>T), GAG1613GAC (4977G>C), AAA1614ACA (4979A>C), GTG1616ACA (4984G>A 4985T>C 4986G>A), GCA1617GAC (4988C>A 4989A>C), GAT1618AAT (4990G>A), GTG1619GTC (4995G>C), GAA1621AAA (4999G>A), CTG1623ATA (5005C>A 5007G>A), AAT1624ATG (5009A>T 5010T>G), GAA1625CAG (5011G>C 5013A>G), CGT1626CGA (5016T>A), GGA1627GGC (5019A>C), GCT1777..A (5469T>A), TTG1778TTA (5472G>A), GCG1779TCA (5473G>T 5475G>A), ACT1780ACC (5478T>C), AGG1781AAT (5480G>A 5481G>T), TTT1783TAT (5486T>A), TGT1784AGA (5488T>A 5490T>A), GAC1787CAA (5497G>C 5499C>A), ATT1788CTT (5500A>C), ATT1789ATA (5505T>A), GTA1790GTC (5508A>C), AGG1791AGA (5511G>A), CGC1793AGA (5515C>A 5517C>A), ATT1795TTG (5521A>T 5523T>G), GCT1796GCG (5526T>G), AAT1797GGC (5527A>G 5528A>G 5529T>C), GTG1798TTG (5530G>T), AAC1802AAT (5544C>T), ACA1803CAG (5545A>C 5546C>A 5547A>G), TAC1804TAT (5550C>T), GCA1809ATT (5563G>A 5564C>T 5565A>T), CAG1810CAA (5568G>A), TAC1811TAT (5571C>T), CTA1813CTC (5577A>C), CAG1814ACA (5578C>A 5579A>C 5580G>A), TGC1815TGT (5583C>T), GTC1816_CCC1817insGTG (5586_5587insGTG), CCC1817CAG (5588C>A 5589C>G), CAT1818AAA (5590C>A 5592T>A), GGT1819AGT (5593G>A), GAT1820TGC (5596G>T 5597A>G 5598T>C), AAA1821AAG (5601A>G), GTG1822ACA (5602G>A 5603T>C 5604G>A), GCA1823GCT (5607A>T), GGT1826AGT (5614G>A), ATT1827del (5617_5619delATT), TCG1828GGG (5620T>G 5621C>G), CTG1829CGC (5624T>G 5625G>C), CAC1830GAT (5626C>G 5628C>T), AGC1831GGA (5629A>G 5631C>A), AGT1832ATG (5633G>T 5634T>G), TTT1833TTG (5637T>G), GCA1834AAC (5638G>A 5639C>A 5640A>C), CAT1835CAC (5643T>C), CAA1836CAT (5646A>T), CAA1836_GGT1837insCATGCT (5646_5647insCATGCT), GGT1837GCT (5648G>C), TTT1838TAT (5651T>A), GAA1839GCC (5654A>C 5655A>C), TTG1840CTT (5656T>C 5658G>T), GGC1841GGA (5661C>A), AAT1842AAC (5664T>C), TTC1843CTT (5665T>C 5667C>T), AGA1844CGA (5668A>C), ATC1846GTG (5674A>G 5676C>G), CGT1847GGC (5677C>G 5679T>C), GGT1848TGT (5680G>T), GAC1849TCA (5683G>T 5684A>C 5685C>A), ATC1850ATT (5688C>T), AAG1852AGA (5693A>G 5694G>A), ACT1853AAG (5696C>A 5697T>G), ACT1854ATG (5699C>T 5700T>G), GCC1855GCA (5703C>A), TTG1856GTG (5704T>G), ATG1858TTG (5710A>T), CCA1859CCC (5715A>C), TAT1860TAC (5718T>C), GTG1861ACT (5719G>A 5720T>C 5721G>T), GGA1863GGG (5727A>G), CCC1864CAG (5729C>A 5730C>G), AAT1865AAA (5733T>A), GGA1866GTT (5735G>T 5736A>T) | | | | | | | | |

#### Proteins

| | | | | | | | | | |
|---|---|---|---|---|---|---|---|---|---|
| polyprotein (YP_003097229.1) | 730 | 1866 | 16.6% | 1773 | 69.6% | 356 (98.9%) | 247 (68.6%) | 3/1/0/0 | 0 |
| Protein mutations: | H731Y (2329C>T 2331C>T), C733A (2335T>G 2336G>C 2337C>G), I734L (2338A>C), D735Q (2341G>C 2343T>G), V738A (2351T>C 2352T>A), I742T (2363T>C 2364A>G), F750R (2386T>A 2387T>G 2388T>A), N764R (2429A>G 2430C>G), A1116T (3484G>A), E1120D (3498G>C), N1122K (3504T>G), S1125A (3511T>G 3513A>T), L1134M (3538T>A), S1142T (3562T>A 3564A>C), R1146K (3575G>A), Q1148P (3581A>C 3582G>A), P1155E (3601C>G 3602C>A 3603T>A), E1157Q (3607G>C), I1162L (3622A>C 3624A>G), T1164S (3628A>T 3630C>T), N1166K (3636T>A), E1171D (3651G>T), V1180I (3676G>A 3678C>T), M1181E (3679A>G 3680T>A), H1188S (3700C>T 3701A>C 3702C>A), D1191E (3711T>G), E1192D (3714A>C), G1193S (3715G>A 3717A>C), K1196R (3724A>C 3725A>G), M1197A (3727A>G 3728T>C 3729G>T), T1201G (3739A>G 3740C>G 3741C>T), C1204S (3749G>C), L1206I (3754T>A 3756G>T), V1495I (4621G>A), L1508M (4660T>A), S1514E (4678T>G 4679C>A 4680A>G), N1517A (4687A>G 4688A>C 4689C>A), D1523E (4707T>A), M1524Q (4708A>C 4709T>A 4710G>A), V1528I (4720G>A 4722A>T), S1536N (4744T>A 4745C>A 4746A>C), D1538N (4750G>A), E1541D (4761G>T), Q1543K (4765C>A 4767G>A), F1544L (4768T>C 4770T>C), H1565Q (4833T>A), M1566L (4834A>C 4836G>A), I1575V (4861A>G 4863A>G), P1585V (4891C>G 4892C>T 4893C>T), A1590I (4906G>A 4907C>T 4908A>C), Q1591T (4909C>A 4910A>C 4911G>T), M1594Y (4918A>T 4919T>A 4920G>T), I1601V (4939A>G), H1606Q (4956T>G), A1611V (4970C>T 4971A>T), E1613D (4977G>C), K1614T (4979A>C), V1616T (4984G>A 4985T>C 4986G>A), A1617D (4988C>A 4989A>C), D1618N (4990G>A), E1621K (4999G>A), L1623I (5005C>A 5007G>A), N1624M (5009A>T 5010T>G), E1625Q (5011G>C 5013A>G), A1779S (5473G>T 5475G>A), R1781N (5480G>A 5481G>T), F1783Y (5486T>A), C1784R (5488T>A 5490T>A), D1787Q (5497G>C 5499C>A), I1788L (5500A>C), I1795L (5521A>T 5523T>G), N1797G (5527A>G 5528A>G 5529T>C), V1798L (5530G>T), T1803Q (5545A>C 5546C>A 5547A>G), A1809I (5563G>A 5564C>T 5565A>T), Q1814T (5578C>A 5579A>C 5580G>A), V1816_P1817insV (5586_5587insGTG), P1817Q (5588C>A 5589C>G), H1818K (5590C>A 5592T>A), G1819S (5593G>A), D1820C (5596G>T 5597A>G 5598T>C), V1822T (5602G>A 5603T>C 5604G>A), G1826S (5614G>A), I1827del (5617_5619delATT), S1828G (5620T>G 5621C>G), L1829R (5624T>G 5625G>C), H1830D (5626C>G 5628C>T), S1831G (5629A>G 5631C>A), S1832M (5633G>T 5634T>G), F1833L (5637T>G), A1834N (5638G>A 5639C>A 5640A>C), Q1836H (5646A>T), Q1836_G1837insHA (5646_5647insCATGCT), G1837A (5648G>C), F1838Y (5651T>A), E1839A (5654A>C 5655A>C), F1843L (5665T>C 5667C>T), I1846V (5674A>G 5676C>G), R1847G (5677C>G 5679T>C), G1848C (5680G>T), D1849S (5683G>T 5684A>C 5685C>A), K1852R (5693A>G 5694G>A), T1853K (5696C>A 5697T>G), T1854M (5699C>T 5700T>G), L1856V (5704T>G), M1858L (5710A>T), V1861T (5719G>A 5720T>C 5721G>T), P1864Q (5729C>A 5730C>G), N1865K (5733T>A), G1866V (5735G>T 5736A>T) | | | | | | | | |

| | Begin | End | Coverage | Score | Concordance | Matches | Identities | I/D/M/F* | Stop Codons |
|---|---|---|---|---|---|---|---|---|---|
| **NT** | **2324** | **7477** | **24.8%** | **777** | **26.0%** | **1698 (89.8%)** | **1170 (61.9%)** | **29/163** | |

Codon mutations: TTT729.TA (2325T>A), CAC731TAT (2329C>T 2331C>T), TGC733GCG (2335T>G 2336G>C 2337C>G), ATT734CTT (2338A>C), GAT735CAG (2341G>C 2343T>G), GTT738GCA (2351T>C 2352T>A), GAC739GAT (2355C>T), CTC741CTT (2361C>T), ATA742ACG (2363T>C 2364A>G), TGC743TGT (2367C>T), CCC748CCA (2382C>A), CCT749CCC (2385T>C), TTT750AGA (2386T>A 2387T>G 2388T>A), GTC751GTA (2391C>A), CAA752CAG (2394A>G), TTT753TTC (2397T>C), CTT754CTG (2400T>G), TAC755TAT (2403C>T), GCC757GCT (2409C>T), GTT759GTG (2415T>G), AGA760CGC (2416A>C 2418A>C), AAC764AGG (2429A>G 2430C>G), TCA767GC. (2437T>G), GCT1116ACT (3484G>A), GCC1117GCT (3489C>T), TTG1118CTC (3490T>C 3492G>C), GAG1120GAC (3498G>C), GAG1121GAA (3501G>A), AAT1122AAG (3504T>G), TCA1125GCT (3511T>G 3513A>T), CTT1127TTG (3517C>T 3519T>G), CCA1131CCT (3531A>T), GAA1133GAG (3537A>G), TTG1134ATG (3538T>A), ACC1137ACG (3549C>G), GTC1138GTG (3552C>G), TCA1142ACC (3562T>A 3564A>C), ACA1143ACC (3567A>C), AGA1145AGG (3573A>G), AGA1146AAA (3575G>A), AGC1147AGT (3579C>T), CAG1148CCA (3581A>C 3582G>A), GCT1152GCA (3594T>A), GGA1154GGT (3600A>T), CCT1155GAA (3601C>G 3602C>A 3603T>A), GAA1157CAA (3607G>C), ACA1158ACT (3612A>T), GCA1159GCT (3615A>T), CCA1160CCT (3618A>T), ATA1162CTG (3622A>C 3624A>G), ACC1164TCT (3628A>T 3630C>T), GTC1165GTG (3633C>G), AAT1166AAA (3636T>A), GAC1167GAT (3639C>T), AAG1168AAA (3642G>A), AGG1169AGA (3645G>A), GGT1170GGG (3648T>G), GAG1171GAT (3651G>T), GGC1172GGA (3654C>A), CCA1175CCT (3663A>T), GGG1179GGC (3675G>C), GTC1180ATT (3676G>A 3678C>T), ATG1181GAG (3679A>G 3680T>A), GCA1187GCC (3699A>C), CAC1188TCA (3700C>T 3701A>C 3702C>A), GAT1191GAG (3711T>G), GAA1192GAC (3714A>C), GGA1193AGC (3715G>A 3717A>C), GCC1195GCA (3723C>A), AAG1196CGG (3724A>C 3725A>G), ATG1197GCT (3727A>G 3728T>C 3729G>T), GCC1198GCA (3732C>A), GAG1200GAA (3738G>A), ACC1201GGT (3739A>G 3740C>G 3741C>T), AAA1203AAG (3747A>G), TGT1204TCT (3749G>C), GCT1205GCC (3753T>C), TTG1206ATT (3754T>A 3756G>T), TTG1207TTA (3759G>A), CTT1493TTA (4615C>T 4617T>A), TTC1494TTT (4620C>T), GTT1495ATT (4621G>A), GCT1496GCA (4626T>A), GGA1499GGG (4635A>G), AAC1502AAT (4644C>T), GTT1504GTG (4650T>G), GTT1506GTG (4656T>G), CCC1507CCA (4659C>A), TTG1508ATG (4660T>A), TTG1510CTT (4666T>C 4668G>T), CAG1511CAA (4671G>A), TAT1513TAC (4677T>C), TCA1514GAG (4678T>G 4679C>A 4680A>G), TTG1515TTA (4683G>A), CCC1516CCT (4686C>T), AAC1517GCA (4687A>G 4688A>C 4689C>A), ATT1518ATA (4692T>A), GAA1521GAG (4701A>G), CTG1522CTA (4704G>A), GAT1523GAA (4707T>A), ATG1524CAA (4708A>C 4709T>A 4710G>A), GTA1525GTG (4713A>G), AAT1526AAC (4716T>C), GTA1528ATT (4720G>A 4722A>T), ATT1529ATC (4725T>C), AAA1530AAG (4728A>G), AAC1531AAT (4731C>T), GGT1532GGC (4734T>C), TTG1533CTT (4735T>C 4737G>T), GAC1534GAT (4740C>T), TCA1536AAC (4744T>A 4745C>A 4746A>C), GAT1538AAT (4750G>A), GTG1539GTC (4755G>C), GAG1541GAT (4761G>T), GTG1542GTC (4764G>C), CAG1543AAA (4765C>A 4767G>A), TTT1544CTC (4768T>C 4770T>C), GAA1549GAG (4785A>G), CTG1550TTG (4786C>T), ACT1551ACG (4791T>G), CTG1553TTG (4795C>T), GGG1556GGC (4806G>C), AGA1559CGC (4813A>C 4815A>C), CTG1562TTG (4822C>T), GGC1563GGT (4827C>T), CAT1565CAA (4833T>A), ATG1566CTA (4834A>C 4836G>A), GCA1567GCC (4839A>C), CCG1568CCC (4842G>C), CTT1569CTC (4845T>C), GTA1572GTT (4854A>T), ATA1574ATT (4860A>T), ATA1575GTG (4861A>G 4863A>G), ATA1581ATC (4881A>C), CGA1582AGA (4882C>A), CCC1585GTT (4891C>G 4892C>T 4893C>T), GGT1586GGG (4896T>G), CCT1588CCA (4902T>A), GCA1590ATC (4906G>A 4907C>T 4908A>C), CAG1591ACT (4909C>A 4910A>C 4911G>T), GCA1592GCC (4914A>C), ATG1594TAT (4918A>T 4919T>A 4920G>T), GTT1597GTG (4929T>G), ACT1599ACG (4935T>G), GGC1600GGT (4938C>T), ATA1601GTA (4939A>G), GCC1604GCT (4950C>T), TTT1605TTC (4953T>C), CAT1606CAG (4956T>G), GGT1608GGA (4962T>A), CAA1609CAG (4965A>G), GTT1610GTC (4968T>C), GCA1611GTT (4970C>T 4971A>T), GAG1613GAC (4977G>C), AAA1614ACA (4979A>C), GTG1616ACA (4984G>A 4985T>C 4986G>A), GCA1617GAC (4988C>A 4989A>C), GAT1618AAT (4990G>A), GTG1619GTC (4995G>C), GAA1621AAA (4999G>A), CTG1623ATA (5005C>A 5007G>A), AAT1624ATG (5009A>T 5010T>G), GAA1625CAG (5011G>C 5013A>G), CGT1626CGA (5016T>A), GGA1627GGC (5019A>C), GCT1777..A (5469T>A), TTG1778TTA (5472G>A), GCG1779TCA (5473G>T 5475G>A), ACT1780ACC (5478T>C), AGG1781AAT (5480G>A 5481G>T), TTT1783TAT (5486T>A), TGT1784AGA (5488T>A 5490T>A), GAC1787CAA (5497G>C 5499C>A), ATT1788CTT (5500A>C), ATT1789ATA (5505T>A), GTA1790GTC (5508A>C), AGG1791AGA (5511G>A), CGC1793AGA (5515C>A 5517C>A), ATT1795TTG (5521A>T 5523T>G), GCT1796GCG (5526T>G), AAT1797GGC (5527A>G 5528A>G 5529T>C), GTG1798TTG (5530G>T), AAC1802AAT (5544C>T), ACA1803CAG (5545A>C 5546C>A 5547A>G), TAC1804TAT (5550C>T), GCA1809ATT (5563G>A 5564C>T 5565A>T), CAG1810CAA (5568G>A), TAC1811TAT (5571C>T), CTA1813CTC (5577A>C), CAG1814ACA (5578C>A 5579A>C 5580G>A), TGC1815TGT (5583C>T), GTC1816_CCC1817insGTG (5586_5587insGTG), CCC1817CAG (5588C>A 5589C>G), CAT1818AAA (5590C>A 5592T>A), GGT1819AGT (5593G>A), GAT1820TGC (5596G>T 5597A>G 5598T>C), AAA1821AAG (5601A>G), GTG1822ACA (5602G>A 5603T>C 5604G>A), GCA1823GCT (5607A>T), GGT1826AGT (5614G>A), ATT1827del (5617_5619delATT), TCG1828GGG (5620T>G 5621C>G), CTG1829CGC (5624T>G 5625G>C), CAC1830GAT (5626C>G 5628C>T), AGC1831GGA (5629A>G 5631C>A), AGT1832ATG (5633G>T 5634T>G), TTT1833TTG (5637T>G), GCA1834AAC (5638G>A 5639C>A 5640A>C), CAT1835CAC (5643T>C), CAA1836CAT (5646A>T), CAA1836_GGT1837insCATGCT (5646_5647insCATGCT), GGT1837GCT (5648G>C), TTT1838TAT (5651T>A), GAA1839GCC (5654A>C 5655A>C), TTG1840CTT (5656T>C 5658G>T), GGC1841GGA (5661C>A), AAT1842AAC (5664T>C), TTC1843CTT (5665T>C 5667C>T), AGA1844CGA (5668A>C), ATC1846GTG (5674A>G 5676C>G), CGT1847GGC (5677C>G 5679T>C), GGT1848TGT (5680G>T), GAC1849TCA (5683G>T 5684A>C 5685C>A), ATC1850ATT (5688C>T), AAG1852AGA (5693A>G 5694G>A), ACT1853AAG (5696C>A 5697T>G), ACT1854ATG (5699C>T 5700T>G), GCC1855GCA (5703C>A), TTG1856GTG (5704T>G), ATG1858TTG (5710A>T), CCA1859CCC (5715A>C), TAT1860TAC (5718T>C), GTG1861ACT (5719G>A 5720T>C 5721G>T), GGA1863GGG (5727A>G), CCC1864CAG (5729C>A 5730C>G), AAT1865AAA (5733T>A), GGA1866GTT (5735G>T 5736A>T)

*: Inserts / Deletes / Misaligned / Frameshifts

#### Analysis details

This analysis was performed with panviral2.64

#### NGS Details (UN60): Tomato chocolate spot virus (segment RNA2)

##### Assembly

| | |
|---|---|
| **Coverage Length** | 1693 (4 contig(s)) |
| **Depth Of Coverage** | 15922.8 |
| **Number Of Reads** | 239197 |
| **Reads Per Million** | 5073.27 rpm (after QC) |
| **Ambiguities** | 0 |
| **Assembly Method** | de novo + reference guided assembly |
| **Consensus Caller** | Bcf Tools |

##### Coverage Map

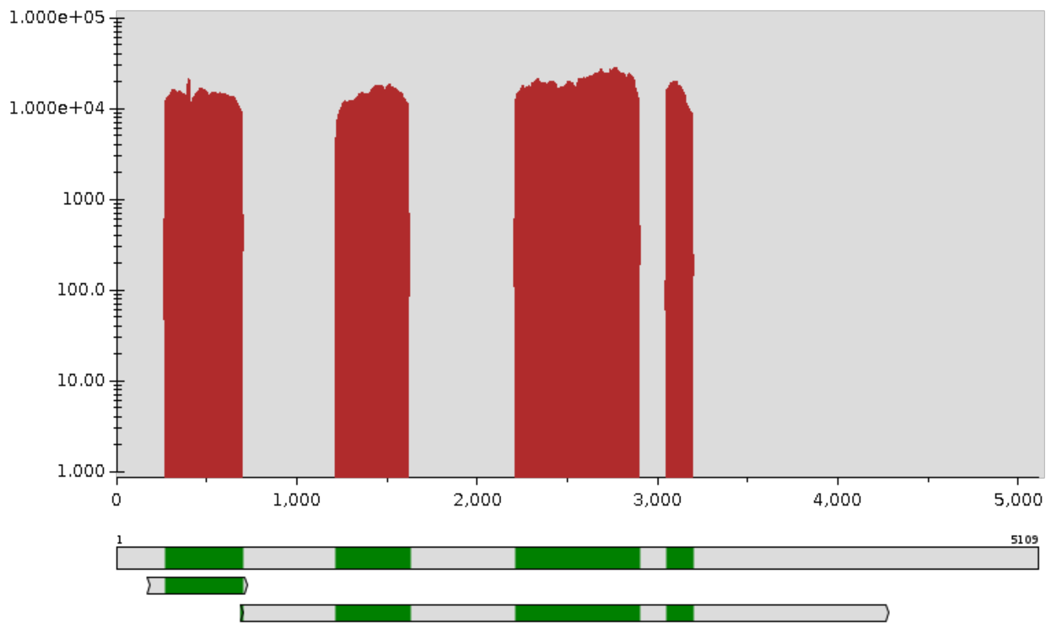

##### Assignment

| | |
|---|---|
| **Type** | Tomato chocolate spot virus (Taxonomy ID: 661101) |
| **Reference Genome** | NC_013076.1 |
| **NT Identity (%)** | 67.2803 |
| **AA Identity (%)** | 70.5986 |
| **Number Of Stop Codons** | 0 |
| **Number Of CDS** | 2 |

##### Alignment

| | |
|---|---|
| **Alignment Score** | 1110.0 (NT) + 2795.0 (AA) = 3905.0 |
| **Concordance (%)** | 55.524 |

| | |
|---|---|
| **Alignment Method** | Global, seeded, nucleotide + amino acids (AGA) |

#### Genome Region

Sequence starts at position 268 and ends at position 3198 relative to NC_013076.1 reference sequence.

#### Alignment Detailed Statistics

| | Begin | End | Coverage | Score | Concordance | Matches | Identities | I/D/M/F* | Stop Codons |
|---|---|---|---|---|---|---|---|---|---|
| **NT** | **268** | **3198** | **33.1%** | **1110** | **33.6%** | **1675 (98.4%)** | **1133 (66.6%)** | **9/18** | |
| Mutations: | 277A>G, 288C>T, 291C>G, 294T>G, 297A>T, 303T>C, 307G>A, 309G>C, 310A>T, 311T>G, 312T>C, 315C>G, 321A>T, 331G>A, 336A>T, 339G>C, 340C>A, 342A>C, 343A>T, 345T>C, 346G>A, 348T>C, 351G>C, 357G>A, 363A>T, 366A>T, 369A>G, 372G>A, 373T>C, 378T>A, 381G>T, 384C>T, 390T>C, 402T>G, 404G>A, 406_414delACTTGGCCT, 415G>A, 417T>C, 421A>G, 422A>C, 423C>A, 426G>A, 430T>C, 432A>T, 435T>A, 438A>C, 439C>T, 441C>G, 442T>A, 444G>T, 449A>G, 450T>A, 456T>C, 457G>A, 460C>A, 461G>A, 462T>A, 471C>A, 474C>A, 475A>C, 480A>G, 481G>T, 483T>G, 484C>T, 486A>G, 490C>G, 491A>C, 492G>C, 501T>G, 502T>C, 503C>A, 504A>T, 507A>T, 508G>T, 511_516delCAACAA, 517A>T, 520G>A, 521A>C, 522A>C, 523A>T, 524A>C, 529G>T, 530T>C, 534T>C, 535G>A, 538A>C, 539A>C, 540A>C, 541G>T, 542G>C, 543G>T, 544A>T, 545A>C, 546A>T, 549_550insTTA, 550A>T, 552C>A, 558A>T, 561T>A, 562T>A, 564G>T, 565A>T, 566A>T, 567G>A, 568T>A, 569T>G, 571A>G, 572A>G, 577G>A, 579A>C, 581T>A, 582T>G, 585T>A, 586_588delCAT, 591C>T, 594C>G, 596C>A, 597T>A, 599A>C, 600G>C, 606G>A, 607C>G, 608G>T, 609A>T, 612A>C, 618G>A, 622A>C, 624A>G, 625G>C, 626C>A, 631A>G, 632A>C, 635A>G, 636A>G, 639A>T, 645A>T, 646T>G, 647C>T, 651G>A, 652T>C, 654G>C, 655C>A, 656A>G, 657A>G, 658C>T, 660T>G, 663G>A, 664C>A, 666C>T, 667T>C, 668C>A, 669C>A, 670A>G, 671A>G, 675G>A, 678T>A, 681T>C, 685A>G, 687T>G, 691A>G, 1219G>T, 1222A>T, 1228A>G, 1231A>G, 1238G>A, 1239A>G, 1240T>C, 1241A>C, 1246G>A, 1252C>T, 1255A>T, 1258C>T, 1259T>A, 1261T>G, 1262C>G, 1263A>C, 1267G>A, 1273T>A, 1277T>A, 1278C>G, 1285C>A, 1291A>G, 1294A>T, 1295T>C, 1297G>C, 1300C>T, 1306T>A, 1309A>G, 1312T>C, 1315C>T, 1316C>T, 1317A>C, 1318A>T, 1327G>A, 1333T>A, 1336C>A, 1351A>T, 1354A>C, 1360T>G, 1366G>A, 1367C>G, 1369A>G, 1372T>G, 1373T>G, 1375C>G, 1376C>T, 1377A>C, 1378G>T, 1379A>C, 1380T>A, 1381G>A, 1385T>C, 1387G>A, 1394C>A, 1396T>A, 1397G>C, 1405T>A, 1408A>T, 1410A>C, 1411G>A, 1412A>C, 1413G>T, 1414G>T, 1423T>C, 1426A>T, 1428G>A, 1429A>C, 1433G>A, 1438A>T, 1440T>C, 1441G>C, 1442A>C, 1443G>A, 1444T>G, 1448A>T, 1449G>C, 1450C>T, 1453C>A, 1454A>T, 1456G>A, 1458C>T, 1459T>G, 1463A>G, 1464G>C, 1465C>G, 1468T>A, 1471C>G, 1474G>T, 1477A>C, 1483G>A, 1489C>A, 1493G>A, 1495G>C, 1501T>C, 1503G>A, 1505C>A, 1513A>T, 1516C>G, 1522A>G, 1525A>G, 1531C>T, 1532G>T, 1534G>T, 1535C>T, 1537T>A, 1539A>G, 1540C>G, 1541A>G, 1547C>T, 1548G>C, 1551T>C, 1552A>G, 1555_1556insGTAGGT, 1556A>G, 1559C>A, 1572C>G, 1573T>G, 1574C>G, 1575A>T, 1576C>A, 1582A>T, 1585C>T, 1586A>T, 1588T>C, 1589G>A, 1590G>A, 1592G>T, 1593G>T, 1595T>C, 1603A>T, 1606T>C, 1610A>G, 2218C>G, 2221A>T, 2224G>A, 2225C>G, 2228A>C, 2229T>C, 2230T>G, 2235A>G, 2236T>G, 2237G>A, 2239T>A, 2243A>G, 2244C>A, 2245T>G, 2249G>A, 2250G>A, 2254T>C, 2255G>A, 2258A>T, 2260T>C, 2266C>T, 2269C>T, 2272T>G, 2273T>G, 2274C>A, 2275A>G, 2278C>A, 2282G>A, 2283A>T, 2284C>G, 2285T>G, 2286C>G, 2287A>T, 2288A>G, 2290C>T, 2291C>G, 2293A>C, 2294G>C, 2296C>G, 2297A>T, 2300G>A, 2303T>C, 2305A>G, 2309A>T, 2317A>G, 2322T>C, 2332A>T, 2335C>T, 2341A>G, 2356T>G, 2357G>A, 2359G>T, 2365G>A, 2368C>T, 2369G>C, 2375G>A, 2386A>C, 2392G>T, 2395C>T, 2401A>G, 2404C>T, 2405A>C, 2407G>T, 2408T>G, 2411T>C, 2413G>A, 2414G>T, 2416A>C, 2419A>C, 2423G>A, 2424C>T, 2425T>G, 2435G>A, 2436C>G, 2437A>C, 2446A>G, 2449G>A, 2450A>G, 2452A>T, 2462G>T, 2464A>C, 2479T>C, 2485T>C, 2491T>A, 2492C>A, 2493T>A, 2495A>C, 2497C>T, 2502A>T, 2503C>T, 2506G>A, 2518T>C, 2519G>A, 2521G>T, 2524C>T, 2528G>A, 2530T>G, 2533T>A, 2536C>G, 2537A>G, 2539A>G, 2542A>G, 2548A>G, 2549T>C, 2551G>A, 2554T>C, 2557A>G, 2560C>G, 2577T>C, 2578T>A, 2581C>T, 2589A>C, 2590G>C, 2593A>G, 2599T>C, 2602A>C, 2605C>T, 2608T>A, 2614T>C, 2615C>A, 2626C>T, 2629G>A, 2632T>C, 2633A>C, 2634C>A, 2635C>T, 2638T>G, 2641T>G, 2644T>A, 2647A>T, 2650C>A, 2651C>G, 2653G>A, 2656A>G, 2657C>T, 2658A>C, 2659G>C, 2664A>G, 2666G>A, 2668A>G, 2672T>G, 2673G>T, 2674T>G, 2677T>C, 2678A>T, 2683A>T, 2684A>C, 2685C>A, 2689G>T, 2695G>T, 2697A>C, 2698G>C, 2700A>T, 2704T>A, 2708C>T, 2710T>G, 2715A>C, 2716G>T, 2717G>T, 2718G>C, 2719T>A, 2720A>C, 2721C>T, 2722T>G, 2727C>T, 2728C>T, 2729A>C, 2731G>T, 2734C>A, 2735T>C, 2737A>T, 2740A>C, 2743C>T, 2744A>G, 2746A>T, 2747C>A, 2749T>A, 2758A>G, 2761A>C, 2768T>C, 2770A>G, 2771T>G, 2782A>T, 2785G>A, 2789T>G, 2790C>A, 2791A>G, 2792A>T, 2793A>C, 2794T>A, 2795G>A, 2797C>A, 2800A>T, 2803C>T, 2806T>C, 2809C>A, 2812T>C, 2813T>C, 2815G>A, 2816C>T, 2817A>C, 2818G>C, 2827T>G, 2833C>A, 2842C>A, 2843C>A, 2848G>A, 2854T>A, 2855C>A, 2858C>T, 2860A>G, 2862T>G, 2863T>A, 2865C>G, 2866A>G, 2870T>G, 2872G>T, 2885A>C, 2886A>G, 2890G>A, 3055G>A, 3072C>G, 3073T>A, 3076G>C, 3079G>C, 3085C>T, 3091G>A, 3092A>G, 3094T>G, 3098A>G, 3109G>T, 3112C>G, 3115C>T, 3118C>T, 3121A>T, 3124A>G, 3128T>C, 3130G>T, 3133C>T, 3139A>G, 3140C>T, 3142C>G, 3145G>C, 3148A>G, 3152A>G, 3153A>C, 3154C>A, 3158A>G, 3160A>G, 3166A>C, 3168A>C, 3169T>A, 3172A>G, 3181T>A, 3184T>G | | | | | | | | |

##### CDS

| | Begin | End | Coverage | Score | Concordance | Matches | Identities | I/D/M/F* | Stop Codons |
|---|---|---|---|---|---|---|---|---|---|
| ToChV_s2_gp1 | 33 | 177 | 78.4% | 565 | 66.8% | 139 (95.2%) | 90 (61.6%) | 1/6/0/0 | 0 |
| Protein mutations: | I36V (277A>G), V46I (307G>A 309G>C), I47C (310A>T 311T>G 312T>C), V54I (331G>A), L57I (340C>A 342A>C), T58S (343A>T 345T>C), V59I (346G>A 348T>C), S77R (402T>G), G78D (404G>A), T79_P81del (406_414delACTTGGCCT), V82I (415G>A 417T>C), N84A (421A>G 422A>C 423C>A), L91I (442T>A 444G>T), H93R (449A>G 450T>A), V96I (457G>A), R97K (460C>A 461G>A 462T>A), V104L (481G>T 483T>G), Q107A (490C>G 491A>C 492G>C), S111H (502T>C 503C>A 504A>T), A113S (508G>T), Q114_Q115del (511_516delCAACAA), T116S (517A>T), E117T (520G>A 521A>C 522A>C), N118S (523A>T 524A>C), V120S (529G>T 530T>C), D122N (535G>A), K123P (538A>C 539A>C 540A>C), G124S (541G>T 542G>C 543G>T), K125S (544A>T 545A>C 546A>T), T126_T127insL (549_550insTTA), T127S (550A>T 552C>A), W131S (562T>A 564G>T), K132L (565A>T 566A>T 567G>A), F133S (568T>A 569T>G), K134G (571A>G 572A>G), E136N (577G>A 579A>C), I137K (581T>A 582T>G), H139del (586_588delCAT), N141K (594C>G), A142E (596C>A 597T>A), E143A (599A>C 600G>C), R146V (607C>G 608G>T 609A>T), I151L (622A>C 624A>G), A152Q (625G>C 626C>A), K154A (631A>G 632A>C), Q155R (635A>G 636A>G), S159V (646T>G 647C>T), Q162R (655C>A 656A>G 657A>G), L165I (664C>A 666C>T), S166Q (667T>C 668C>A 669C>A), N167G (670A>G 671A>G), N172E (685A>G 687T>G), I174V (691A>G) | | | | | | | | |
| Codon mutations: | ATT36GTT (277A>G), AAC39AAT (288C>T), CCC40CCG (291C>G), ACT41ACG (294T>G), CTA42CTT (297A>T), TTT44TTC (303T>C), GTG46ATC (307G>A 309G>C), ATT47TGC (310A>T 311T>G 312T>C), CCC48CCG (315C>G), ACA50ACT (321A>T), GTT54ATT (331G>A), TCA55TCT (336A>T), GTG56GTC (339G>C), CTA57ATC (340C>A 342A>C), ACT58TCC (343A>T 345T>C), GTT59ATC (346G>A 348T>C), TCG60TCC (351G>C), GAG62GAA (357G>A), TCA64TCT (363A>T), ACA65ACT (366A>T), CCA66CCG (369A>G), CAG67CAA (372G>A), TTG68CTG (373T>C), GTT69GTA (378T>A), CCG70CCT (381G>T), GGC71GGT (384C>T), TAT73TAC (390T>C), AGT77AGG (402T>G), GGT78GAT (404G>A), ACT79_CCT81del (406_414delACTTGGCCT), GTT82ATC (415G>A 417T>C), AAC84GCA (421A>G 422A>C 423C>A), GTG85GTA (426G>A), TTA87CTT (430T>C 432A>T), TCT88TCA (435T>A), GGA89GGC (438A>C), CTC90TTG (439C>T 441C>G), TTG91ATT (442T>A 444G>T), CAT93CGA (449A>G 450T>A), TCT95TCC (456T>C), GTT96ATT (457G>A), CGT97AAA (460C>A 461G>A 462T>A), ACC100ACA (471C>A), ACC101ACA (474C>A), AGG102CGG (475A>C), AAA103AAG (480A>G), GTT104TTG (481G>T 483T>G), CTA105TTG (484C>T 486A>G), CAG107GCC (490C>G 491A>C 492G>C), GTT110GTG (501T>G), TCA111CAT (502T>C 503C>A 504A>T), ATA112ATT (507A>T), GCT113TCT (508G>T), CAA114_CAA115del (511_516delCAACAA), ACT116TCT (517A>T), GAA117ACC (520G>A 521A>C 522A>C), AAT118TCT (523A>T 524A>C), GTA120TCA (529G>T 530T>C), TCT121TCC (534T>C), GAC122AAC (535G>A), AAA123CCC (538A>C 539A>C 540A>C), GGG124TCT (541G>T 542G>C 543G>T), AAA125TCT (544A>T 545A>C 546A>T), ACA126_ACC127insTTA (549_550insTTA), ACC127TCA (550A>T 552C>A), ACA129ACT (558A>T), GCT130GCA (561T>A), TGG131AGT (562T>A 564G>T), AAG132TTA (565A>T 566A>T 567G>A), TTC133AGC (568T>A 569T>G), AAA134GGA (571A>G 572A>G), GAA136AAC (577G>A 579A>C), ATT137AAG (581T>A 582T>G), GCT138GCA (585T>A), CAT139del (586_588delCAT), CTC140CTT (591C>T), AAC141AAG (594C>G), GCT142GAA (596C>A 597T>A), GAG143GCC (599A>C 600G>C), GAG145GAA (606G>A), CGA146GTT (607C>G 608G>T 609A>T), GCA147GCC (612A>C), AAG149AAA (618G>A), ATA151CTG (622A>C 624A>G), GCA152CAA (625G>C 626C>A), AAA154GCA (631A>G 632A>C), CAA155CGG (635A>G 636A>G), TCA156TCT (639A>T), ATA158ATT (645A>T), TCC159GTC (646T>G 647C>T), AAG160AAA (651G>A), TTG161CTC (652T>C 654G>C), CAA162AGG (655C>A 656A>G 657A>G), CTT163TTG (658C>T 660T>G), CAG164CAA (663G>A), CTC165ATT (664C>A 666C>T), TCC166CAA (667T>C 668C>A 669C>A), AAT167GGT (670A>G 671A>G), CAG168CAA (675G>A), CCT169CCA (678T>A), AGT170AGC (681T>C), AAT172GAG (685A>G 687T>G), ATC174GTC (691A>G) | | | | | | | | |
| ToChV_s2_gp2 | 1 | 838 | 35.6% | 2230 | 76.3% | 426 (99.5%) | 311 (72.7%) | 2/0/0/0 | 0 |
| Protein mutations: | M1R (687T>G), I2M (691A>G), D185S (1238G>A 1239A>G 1240T>C), T186P (1241A>C), S192T (1259T>A 1261T>G), Q193A (1262C>G 1263A>C), Q211S (1316C>T 1317A>C 1318A>T), Q228E (1367C>G 1369A>G), S229R (1372T>G), S230A (1373T>G 1375C>G), Q231S (1376C>T 1377A>C 1378G>T), M232Q (1379A>C 1380T>A 1381G>A), A238P (1397G>C), E242A (1410A>C 1411G>A), R243L (1412A>C 1413G>T 1414G>T), R248H (1428G>A 1429A>C), D250N (1433G>A), L252P (1440T>C 1441G>C), S253Q (1442A>C 1443G>A 1444T>G), M257L (1454A>T 1456G>A), T258M (1458C>T 1459T>G), S260A (1463A>G 1464G>C 1465C>G), V270I (1493G>A 1495G>C), R273K (1503G>A), V283F (1532G>T 1534G>T), P284S (1535C>T 1537T>A), N285R (1539A>G 1540C>G), T286A (1541A>G), R288S (1547C>T 1548G>C), L289S (1551T>C 1552A>G), T290_M291insVG (1555_1556insGTAGGT), M291V (1556A>G), A296G (1572C>G 1573T>G), H297V (1574C>G 1575A>T 1576C>A), E299D (1582A>T), T301S (1586A>T 1588T>C), G302N (1589G>A 1590G>A), G303L (1592G>T 1593G>T), K309E (1610A>G), H514D (2225C>G), I515P (2228A>C 2229T>C 2230T>G), D517G (2235A>G 2236T>G), D518K (2237G>A 2239T>A), T520E (2243A>G 2244C>A 2245T>G), G522N (2249G>A 2250G>A), V524I (2255G>A), I525F (2258A>T 2260T>C), S530E (2273T>G 2274C>A 2275A>G), D531E (2278C>A), D533M (2282G>A 2283A>T 2284C>G), S534G (2285T>G 2286C>G 2287A>T), N535D (2288A>G 2290C>T), P536A (2291C>G 2293A>C), A537P (2294G>C 2296C>G), I538L (2297A>T), D539N (2300G>A), M542L (2309A>T), V546A (2322T>C), V558I (2357G>A 2359G>T), V562L (2369G>C), E564K (2375G>A), K567N (2386A>C), M574L (2405A>C 2407G>T), S575A (2408T>G), A577S (2414G>T 2416A>C), A580M (2423G>A 2424C>T 2425T>G), A584S (2435G>A 2436C>G 2437A>C), T589A (2450A>G 2452A>T), A593S (2462G>T 2464A>C), L603N (2492C>A 2493T>A), I604L (2495A>C 2497C>T), Y606F (2502A>T 2503C>T), V612I (2519G>A 2521G>T), V615M (2528G>A 2530T>G), I618V (2537A>G 2539A>G), V631A (2577T>C 2578T>A), K635T (2589A>C 2590G>C), L644I (2615C>A), T650H (2633A>C 2634C>A 2635C>T), Q656E (2651C>G 2653G>A), Q658S (2657C>T 2658A>C 2659G>C), E660G (2664A>G), E661K (2666G>A 2668A>G), C663V (2672T>G 2673G>T 2674T>G), T665S (2678A>T), T667Q (2684A>C 2685C>A), K671T (2697A>C 2698G>C), Y672F (2700A>T), E677A (2715A>C 2716G>T), G678S (2717G>T 2718G>C 2719T>A), T679L (2720A>C 2721C>T 2722T>G), A681V (2727C>T 2728C>T), T682P (2729A>C 2731G>T), D683E (2734C>A), I687V (2744A>G 2746A>T), S696A (2771T>G), S702E (2789T>G 2790C>A 2791A>G), N703S (2792A>T 2793A>C 2794T>A), V704I (2795G>A 2797C>A), Q711S (2816C>T 2817A>C 2818G>C), L720M (2843C>A), I726R (2862T>G 2863T>A), A727G (2865C>G 2866A>G), S729A (2870T>G 2872G>T), K734R (2885A>C 2886A>G), T796R (3072C>G 3073T>A), I803V (3092A>G 3094T>G), I805V (3098A>G), N823A (3152A>G 3153A>C 3154C>A), K825E (3158A>G 3160A>G), N828T (3168A>C 3169T>A), F833L (3184T>G) | | | | | | | | |

| | Begin | End | Coverage | Score | Concordance | Matches | Identities | I/D/M/F* | Stop Codons |
|---|---|---|---|---|---|---|---|---|---|
| **NT** | **268** | **3198** | **33.1%** | **1110** | **33.6%** | **1675 (98.4%)** | **1133 (66.6%)** | **9/18** | |

Codon mutations: ATG1AGG (687T>G), ATA2ATG (691A>G), GCG178GCT (1219G>T), ATA179ATT (1222A>T), AAA181AAG (1228A>G), CAA182CAG (1231A>G), GAT185AGC (1238G>A 1239A>G 1240T>C), ACA186CCA (1241A>C), AGG187AGA (1246G>A), GCC189GCT (1252C>T), GTA190GTT (1255A>T), TAC191TAT (1258C>T), TCT192ACG (1259T>A 1261T>G), CAA193GCA (1262C>G 1263A>C), CAG194CAA (1267G>A), ACT196ACA (1273T>A), TCT198AGT (1277T>A 1278C>G), GCC200GCA (1285C>A), GAA202GAG (1291A>G), GCA203GCT (1294A>T), TTG204CTC (1295T>C 1297G>C), AAC205AAT (1300C>T), CCT207CCA (1306T>A), CAA208CAG (1309A>G), TTT209TTC (1312T>C), TTC210TTT (1315C>T), CAA211TCT (1316C>T 1317A>C 1318A>T), CAG214CAA (1327G>A), GCT216GCA (1333T>A), GCC217GCA (1336C>A), TCA222TCT (1351A>T), GTA223GTC (1354A>C), TCT225TCG (1360T>G), GGG227GGA (1366G>A), CAA228GAG (1367C>G 1369A>G), AGT229AGG (1372T>G), TCC230GCG (1373T>G 1375C>G), CAG231TCT (1376C>T 1377A>C 1378G>T), ATG232CAA (1379A>C 1380T>A 1381G>A), TTG234CTA (1385T>C 1387G>A), CGT237AGA (1394C>A 1396T>A), GCT238CCT (1397G>C), GTT240GTA (1405T>A), GGA241GGT (1408A>T), GAG242GCA (1410A>C 1411G>A), AGG243CTT (1412A>C 1413G>T 1414G>T), AAT246AAC (1423T>C), ACA247ACT (1426A>T), CGA248CAC (1428G>A 1429A>C), GAT250AAT (1433G>A), GTA251GTT (1438A>T), CTG252CCC (1440T>C 1441G>C), AGT253CAG (1442A>C 1443G>A 1444T>G), AGC255TCT (1448A>T 1449G>C 1450C>T), TCC256TCA (1453C>A), ATG257TTA (1454A>T 1456G>A), ACT258ATG (1458C>T 1459T>G), AGC260GCG (1463A>G 1464G>C 1465C>G), TCT261TCA (1468T>A), TCC262TCG (1471C>G), CTG263CTT (1474G>T), CGA264CGC (1477A>C), CCG266CCA (1483G>A), CTC268CTA (1489C>A), GTG270ATC (1493G>A 1495G>C), AGT272AGC (1501T>C), AGG273AAG (1503G>A), CGG274AGG (1505C>A), CCA276CCT (1513A>T), TCC277TCG (1516C>G), GAA279GAG (1522A>G), CTA280CTG (1525A>G), GAC282GAT (1531C>T), GTG283TTT (1532G>T 1534G>T), CCT284TCA (1535C>T 1537T>A), AAC285AGG (1539A>G 1540C>G), ACC286GCC (1541A>G), CGA288TCA (1547C>T 1548G>C), TTA289TCG (1551T>C 1552A>G), ACA290_ATG291insGTAGGT (1555_1556insGTAGGT), ATG291GTG (1556A>G), CGA292AGA (1559C>A), GCT296GGG (1572C>G 1573T>G), CAC297GTA (1574C>G 1575A>T 1576C>A), GAA299GAT (1582A>T), GAC300GAT (1585C>T), ACT301TCC (1586A>T 1588T>C), GGT302AAT (1589G>A 1590G>A), GGA303TTA (1592G>T 1593G>T), TTG304CTG (1595T>C), CCA306CCT (1603A>T), GTT307GTC (1606T>C), AAG309GAG (1610A>G), ACC511ACG (2218C>G), GCA512GCT (2221A>T), GAG513GAA (2224G>A), CAT514GAT (2225C>G), ATT515CCG (2228A>C 2229T>C 2230T>G), GAT517GGG (2235A>G 2236T>G), GAT518AAA (2237G>A 2239T>A), ACT520GAG (2243A>G 2244C>A 2245T>G), GGT522AAT (2249G>A 2250G>A), GAT523GAC (2254T>C), GTT524ATT (2255G>A), ATT525TTC (2258A>T 2260T>C), GAC527GAT (2266C>T), TTC528TTT (2269C>T), GGT529GGG (2272T>G), TCA530GAG (2273T>G 2274C>A 2275A>G), GAC531GAA (2278C>A), GAC533ATG (2282G>A 2283A>T 2284C>G), TCA534GGT (2285T>G 2286C>G 2287A>T), AAC535GAT (2288A>G 2290C>T), CCA536GCC (2291C>G 2293A>C), GCC537CCG (2294G>C 2296C>G), ATA538TTA (2297A>T), GAT539AAT (2300G>A), TTA540CTG (2303T>C 2305A>G), ATG542TTG (2309A>T), CAA544CAG (2317A>G), GTA546GCA (2322T>C), GCA549GCT (2332A>T), AGC550AGT (2335C>T), GAA552GAG (2341A>G), GGT557GGG (2356T>G), GTG558ATT (2357G>A 2359G>T), GAG560GAA (2365G>A), TTC561TTT (2368C>T), GTG562CTG (2369G>C), GAA564AAA (2375G>A), AAA567AAC (2386A>C), GTG569GTT (2392G>T), GCC570GCT (2395C>T), CAA572CAG (2401A>G), GTC573GTT (2404C>T), ATG574CTT (2405A>C 2407G>T), TCA575GCA (2408T>G), TTG576CTA (2411T>C 2413G>A), GCA577TCC (2414G>T 2416A>C), CTA578CTC (2419A>C), GCT580ATG (2423G>A 2424C>T 2425T>G), GCA584AGC (2435G>A 2436C>G 2437A>C), AAA587AAG (2446A>G), GAG588GAA (2449G>A), ACA589GCT (2450A>G 2452A>T), GCA593TCC (2462G>T 2464A>C), TAT598TAC (2479T>C), GAT600GAC (2485T>C), GCT602GCA (2491T>A), CTC603AAC (2492C>A 2493T>A), ATC604CTT (2495A>C 2497C>T), TAC606TTT (2502A>T 2503C>T), AAG607AAA (2506G>A), TAT611TAC (2518T>C), GTG612ATT (2519G>A 2521G>T), TCC613TCT (2524C>T), GTT615ATG (2528G>A 2530T>G), GGT616GGA (2533T>A), GCC617GCG (2536C>G), ATA618GTG (2537A>G 2539A>G), TCA619TCG (2542A>G), CAA621CAG (2548A>G), TTG622CTA (2549T>C 2551G>A), GCT623GCC (2554T>C), TTA624TTG (2557A>G), GTC625GTG (2560C>G), GTT631GCA (2577T>C 2578T>A), CTC632CTT (2581C>T), AAG635ACC (2589A>C 2590G>C), AAA636AAG (2593A>G), TTT638TTC (2599T>C), ATA639ATC (2602A>C), AAC640AAT (2605C>T), ATT641ATA (2608T>A), ACT643ACC (2614T>C), CTT644ATT (2615C>A), AGC647AGT (2626C>T), AAG648AAA (2629G>A), CAT649CAC (2632T>C), ACC650CAT (2633A>C 2634C>A 2635C>T), CTT651CTG (2638T>G), GTT652GTG (2641T>G), TCT653TCA (2644T>A), GCA654GCT (2647A>T), TCC655TCA (2650C>A), CAG656GAA (2651C>G 2653G>A), CAA657CAG (2656A>G), CAG658TCC (2657C>T 2658A>C 2659G>C), GAG660GGG (2664A>G), GAA661AAG (2666G>A 2668A>G), TGT663GTG (2672T>G 2673G>T 2674T>G), TTT664TTC (2677T>C), ACA665TCA (2678A>T), CCA666CCT (2683A>T), ACA667CAA (2684A>C 2685C>A), GGG668GGT (2689G>T), GGG670GGT (2695G>T), AAG671ACC (2697A>C 2698G>C), TAT672TTT (2700A>T), GTT673GTA (2704T>A), CTT675TTG (2708C>T 2710T>G), GAG677GCT (2715A>C 2716G>T), GGT678TCA (2717G>T 2718G>C 2719T>A), ACT679CTG (2720A>C 2721C>T 2722T>G), GCC681GTT (2727C>T 2728C>T), ACG682CCT (2729A>C 2731G>T), GAC683GAA (2734C>A), TTA684CTT (2735T>C 2737A>T), GGA685GGC (2740A>C), AGC686AGT (2743C>T), ATA687GTT (2744A>G 2746A>T), CGT688AGA (2747C>A 2749T>A), GTA691GTG (2758A>G), ACA692ACC (2761A>C), TTA695CTG (2768T>C 2770A>G), TCA696GCA (2771T>G), ACA699ACT (2782A>T), GAG700GAA (2785G>A), TCA702GAG (2789T>G 2790C>A 2791A>G), AAT703TCA (2792A>T 2793A>C 2794T>A), GTC704ATA (2795G>A 2797C>A), CCA705CCT (2800A>T), TGC706TGT (2803C>T), CAT707CAC (2806T>C), ATC708ATA (2809C>A), CAT709CAC (2812T>C), TTG710CTA (2813T>C 2815G>A), CAG711TCC (2816C>T 2817A>C 2818G>C), GTT714GTG (2827T>G), TCC716TCA (2833C>A), ATC719ATA (2842C>A), CTG720ATG (2843C>A), CAG721CAA (2848G>A), CCT723CCA (2854T>A), CGG724AGG (2855C>A), CTA725TTG (2858C>T 2860A>G), ATT726AGA (2862T>G 2863T>A), GCA727GGG (2865C>G 2866A>G), TCG729GCT (2870T>G 2872G>T), AAG734CGG (2885A>C 2886A>G), CCG735CCA (2890G>A), CGG790CGA (3055G>A), ACT796AGA (3072C>G 3073T>A), GGG797GGC (3076G>C), ACG798ACC (3079G>C), GTC800GTT (3085C>T), GAG802GAA (3091G>A), ATT803GTG (3092A>G 3094T>G), ATT805GTT (3098A>G), ACG808ACT (3109G>T), CTC809CTG (3112C>G), TTC810TTT (3115C>T), CAC811CAT (3118C>T), TCA812TCT (3121A>T), GGA813GGG (3124A>G), TTG815CTT (3128T>C 3130G>T), GCC816GCT (3133C>T), GGA818GGG (3139A>G), CTC819TTG (3140C>T 3142C>G), GGG820GGC (3145G>C), ACA821ACG (3148A>G), AAC823GCA (3152A>G 3153A>C 3154C>A), AAA825GAG (3158A>G 3160A>G), TCA827TCC (3166A>C), AAT828ACA (3168A>C 3169T>A), GCA829GCG (3172A>G), ATT832ATA (3181T>A), TTT833TTG (3184T>G)

#### Proteins

| | Begin | End | Coverage | Score | Concordance | Matches | Identities | I/D/M/F* | Stop Codons |
|---|---|---|---|---|---|---|---|---|---|
| hypothetical protein (YP_003097230.1) | 33 | 177 | 78.4% | 565 | 66.8% | 139 (95.2%) | 90 (61.6%) | 1/6/0/0 | 0 |

Protein mutations: I36V (277A>G), V46I (307G>A 309G>C), I47C (310A>T 311T>G 312T>C), V54I (331G>A), L57I (340C>A 342A>C), T58S (343A>T 345T>C), V59I (346G>A 348T>C), S77R (402T>G), G78D (404G>A), T79_P81del (406_414delACTTGGCCT), V82I (415G>A 417T>C), N84A (421A>G 422A>C 423C>A), L91I (442T>A 444G>T), H93R (449A>G 450T>A), V96I (457G>A), R97K (460C>A 461G>A 462T>A), V104L (481G>T 483T>G), Q107A (490C>G 491A>C 492G>C), S111H (502T>C 503C>A 504A>T), A113S (508G>T), Q114_Q115del (511_516delCAACAA), T116S (517A>T), E117T (520G>A 521A>C 522A>C), N118S (523A>T 524A>C), V120S (529G>T 530T>C), D122N (535G>A), K123P (538A>C 539A>C 540A>C), G124S (541G>T 542G>C 543G>T), K125S (544A>T 545A>C 546A>T), T126_T127insL (549_550insTTA), T127S (550A>T 552C>A), W131S (562T>A 564G>T), K132L (565A>T 566A>T 567G>A), F133S (568T>A 569T>G), K134G (571A>G 572A>G), E136N (577G>A 579A>C), I137K (581T>A 582T>G), H139del (586_588delCAT), N141K (594C>G), A142E (596C>A 597T>A), E143A (599A>C 600G>C), R146V (607C>G 608G>T 609A>T), I151L (622A>C 624A>G), A152Q (625G>C 626C>A), K154A (631A>G 632A>C), Q155R (635A>G 636A>G), S159V (646T>G 647C>T), Q162R (655C>A 656A>G 657A>G), L165I (664C>A 666C>T), S166Q (667T>C 668C>A 669C>A), N167G (670A>G 671A>G), N172E (685A>G 687T>G), I174V (691A>G)

Codon mutations: ATT36GTT (277A>G), AAC39AAT (288C>T), CCC40CCG (291C>G), ACT41ACG (294T>G), CTA42CTT (297A>T), TTT44TTC (303T>C), GTG46ATC (307G>A 309G>C), ATT47TGC (310A>T 311T>G 312T>C), CCC48CCG (315C>G), ACA50ACT (321A>T), GTT54ATT (331G>A), TCA55TCT (336A>T), GTG56GTC (339G>C), CTA57ATC (340C>A 342A>C), ACT58TCC (343A>T 345T>C), GTT59ATC (346G>A 348T>C), TCG60TCC (351G>C), GAG62GAA (357G>A), TCA64TCT (363A>T), ACA65ACT (366A>T), CCA66CCG (369A>G), CAG67CAA (372G>A), TTG68CTG (373T>C), GTT69GTA (378T>A), CCG70CCT (381G>T), GGC71GGT (384C>T), TAT73TAC (390T>C), AGT77AGG (402T>G), GGT78GAT (404G>A), ACT79_CCT81del (406_414delACTTGGCCT), GTT82ATC (415G>A 417T>C), AAC84GCA (421A>G 422A>C 423C>A), GTG85GTA (426G>A), TTA87CTT (430T>C 432A>T), TCT88TCA (435T>A), GGA89GGC (438A>C), CTC90TTG (439C>T 441C>G), TTG91ATT (442T>A 444G>T), CAT93CGA (449A>G 450T>A), TCT95TCC (456T>C), GTT96ATT (457G>A), CGT97AAA (460C>A 461G>A 462T>A), ACC100ACA (471C>A), ACC101ACA (474C>A), AGG102CGG (475A>C), AAA103AAG (480A>G), GTT104TTG (481G>T 483T>G), CTA105TTG (484C>T 486A>G), CAG107GCC (490C>G 491A>C 492G>C), GTT110GTG (501T>G), TCA111CAT (502T>C 503C>A 504A>T), ATA112ATT (507A>T), GCT113TCT (508G>T), CAA114_CAA115del (511_516delCAACAA), ACT116TCT (517A>T), GAA117ACC (520G>A 521A>C 522A>C), AAT118TCT (523A>T 524A>C), GTA120TCA (529G>T 530T>C), TCT121TCC (534T>C), GAC122AAC (535G>A), AAA123CCC (538A>C 539A>C 540A>C), GGG124TCT (541G>T 542G>C 543G>T), AAA125TCT (544A>T 545A>C 546A>T), ACA126_ACC127insTTA (549_550insTTA), ACC127TCA (550A>T 552C>A), ACA129ACT (558A>T), GCT130GCA (561T>A), TGG131AGT (562T>A 564G>T), AAG132TTA (565A>T 566A>T 567G>A), TTC133AGC (568T>A 569T>G), AAA134GGA (571A>G 572A>G), GAA136AAC (577G>A 579A>C), ATT137AAG (581T>A 582T>G), GCT138GCA (585T>A), CAT139del (586_588delCAT), CTC140CTT (591C>T), AAC141AAG (594C>G), GCT142GAA (596C>A 597T>A), GAG143GCC (599A>C 600G>C), GAG145GAA (606G>A), CGA146GTT (607C>G 608G>T 609A>T), GCA147GCC (612A>C), AAG149AAA (618G>A), ATA151CTG (622A>C 624A>G), GCA152CAA (625G>C 626C>A), AAA154GCA (631A>G 632A>C), CAA155CGG (635A>G 636A>G), TCA156TCT (639A>T), ATA158ATT (645A>T), TCC159GTC (646T>G 647C>T), AAG160AAA (651G>A), TTG161CTC (652T>C 654G>C), CAA162AGG (655C>A 656A>G 657A>G), CTT163TTG (658C>T 660T>G), CAG164CAA (663G>A), CTC165ATT (664C>A 666C>T), TCC166CAA (667T>C 668C>A 669C>A), AAT167GGT (670A>G 671A>G), CAG168CAA (675G>A), CCT169CCA (678T>A), AGT170AGC (681T>C), AAT172GAG (685A>G 687T>G), ATC174GTC (691A>G)

| | Begin | End | Coverage | Score | Concordance | Matches | Identities | I/D/M/F* | Stop Codons |
|---|---|---|---|---|---|---|---|---|---|
| polyprotein (YP_003097231.1) | 1 | 838 | 35.6% | 2230 | 76.3% | 426 (99.5%) | 311 (72.7%) | 2/0/0/0 | 0 |

Protein mutations: M1R (687T>G), I2M (691A>G), D185S (1238G>A 1239A>G 1240T>C), T186P (1241A>C), S192T (1259T>A 1261T>G), Q193A (1262C>G 1263A>C), Q211S (1316C>T 1317A>C 1318A>T), Q228E (1367C>G 1369A>G), S229R (1372T>G), S230A (1373T>G 1375C>G), Q231S (1376C>T 1377A>C 1378G>T), M232Q (1379A>C 1380T>A 1381G>A), A238P (1397G>C), E242A (1410A>C 1411G>A), R243L (1412A>C 1413G>T 1414G>T), R248H (1428G>A 1429A>C), D250N (1433G>A), L252P (1440T>C 1441G>C), S253Q (1442A>C 1443G>A 1444T>G), M257L (1454A>T 1456G>A), T258M (1458C>T 1459T>G), S260A (1463A>G 1464G>C 1465C>G), V270I (1493G>A 1495G>C), R273K (1503G>A), V283F (1532G>T 1534G>T), P284S (1535C>T 1537T>A), N285R (1539A>G 1540C>G), T286A (1541A>G), R288S (1547C>T 1548G>C), L289S (1551T>C 1552A>G), T290_M291insVG (1555_1556insGTAGGT), M291V (1556A>G), A296G (1572C>G 1573T>G), H297V (1574C>G 1575A>T 1576C>A), E299D (1582A>T), T301S (1586A>T 1588T>C), G302N (1589G>A 1590G>A), G303L (1592G>T 1593G>T), K309E (1610A>G), H514D (2225C>G), I515P (2228A>C 2229T>C 2230T>G), D517G (2235A>G 2236T>G), D518K (2237G>A 2239T>A), T520E (2243A>G 2244C>A 2245T>G), G522N (2249G>A 2250G>A), V524I (2255G>A), I525F (2258A>T 2260T>C), S530E (2273T>G 2274C>A 2275A>G), D531E (2278C>A), D533M (2282G>A 2283A>T 2284C>G), S534G (2285T>G 2286C>G 2287A>T), N535D (2288A>G 2290C>T), P536A (2291C>G 2293A>C), A537P (2294G>C 2296C>G), I538L (2297A>T), D539N (2300G>A), M542L (2309A>T), V546A (2322T>C), V558I (2357G>A 2359G>T), V562L (2369G>C), E564K (2375G>A), K567N (2386A>C), M574L (2405A>C 2407G>T), S575A (2408T>G), A577S (2414G>T 2416A>C), A580M (2423G>A 2424C>T 2425T>G), A584S (2435G>A 2436C>G 2437A>C), T589A (2450A>G 2452A>T), A593S (2462G>T 2464A>C), L603N (2492C>A 2493T>A), I604L (2495A>C 2497C>T), Y606F (2502A>T 2503C>T), V612I (2519G>A 2521G>T), V615M (2528G>A 2530T>G), I618V (2537A>G 2539A>G), V631A (2577T>C 2578T>A), K635T (2589A>C 2590G>C), L644I (2615C>A), T650H (2633A>C 2634C>A 2635C>T), Q656E (2651C>G 2653G>A), Q658S (2657C>T 2658A>C 2659G>C), E660G (2664A>G), E661K (2666G>A 2668A>G), C663V (2672T>G 2673G>T 2674T>G), T665S (2678A>T), T667Q (2684A>C 2685C>A), K671T (2697A>C 2698G>C), Y672F (2700A>T), E677A (2715A>C 2716G>T), G678S (2717G>T 2718G>C 2719T>A), T679L (2720A>C 2721C>T 2722T>G), A681V (2727C>T 2728C>T), T682P (2729A>C 2731G>T), D683E (2734C>A), I687V (2744A>G 2746A>T), S696A (2771T>G), S702E (2789T>G 2790C>A 2791A>G), N703S (2792A>T 2793A>C 2794T>A), V704I (2795G>A 2797C>A), Q711S (2816C>T 2817A>C 2818G>C), L720M (2843C>A), I726R (2862T>G 2863T>A), A727G (2865C>G 2866A>G), S729A (2870T>G 2872G>T), K734R (2885A>C 2886A>G), T796R (3072C>G 3073T>A), I803V (3092A>G 3094T>G), I805V (3098A>G), N823A (3152A>G 3153A>C 3154C>A), K825E (3158A>G 3160A>G), N828T (3168A>C 3169T>A), F833L (3184T>G)

| | Begin | End | Coverage | Score | Concordance | Matches | Identities | I/D/M/F* | Stop Codons |
|---|---|---|---|---|---|---|---|---|---|
| **NT** | **268** | **3198** | **33.1%** | **1110** | **33.6%** | **1675 (98.4%)** | **1133 (66.6%)** | **9/18** | |
| Codon mutations: | ATG1AGG (687T>G), ATA2ATG (691A>G), GCG178GCT (1219G>T), ATA179ATT (1222A>T), AAA181AAG (1228A>G), CAA182CAG (1231A>G), GAT185AGC (1238G>A 1239A>G 1240T>C), ACA186CCA (1241A>C), AGG187AGA (1246G>A), GCC189GCT (1252C>T), GTA190GTT (1255A>T), TAC191TAT (1258C>T), TCT192ACG (1259T>A 1261T>G), CAA193GCA (1262C>G 1263A>C), CAG194CAA (1267G>A), ACT196ACA (1273T>A), TCT198AGT (1277T>A 1278C>G), GCC200GCA (1285C>A), GAA202GAG (1291A>G), GCA203GCT (1294A>T), TTG204CTC (1295T>C 1297G>C), AAC205AAT (1300C>T), CCT207CCA (1306T>A), CAA208CAG (1309A>G), TTT209TTC (1312T>C), TTC210TTT (1315C>T), CAA211TCT (1316C>T 1317A>C 1318A>T), CAG214CAA (1327G>A), GCT216GCA (1333T>A), GCC217GCA (1336C>A), TCA222TCT (1351A>T), GTA223GTC (1354A>C), TCT225TCG (1360T>G), GGG227GGA (1366G>A), CAA228GAG (1367C>G 1369A>G), AGT229AGG (1372T>G), TCC230GCG (1373T>G 1375C>G), CAG231TCT (1376C>T 1377A>C 1378G>T), ATG232CAA (1379A>C 1380T>A 1381G>A), TTG234CTA (1385T>C 1387G>A), CGT237AGA (1394C>A 1396T>A), GCT238CCT (1397G>C), GTT240GTA (1405T>A), GGA241GGT (1408A>T), GAG242GCA (1410A>C 1411G>A), AGG243CTT (1412A>C 1413G>T 1414G>T), AAT246AAC (1423T>C), ACA247ACT (1426A>T), CGA248CAC (1428G>A 1429A>C), GAT250AAT (1433G>A), GTA251GTT (1438A>T), CTG252CCC (1440T>C 1441G>C), AGT253CAG (1442A>C 1443G>A 1444T>G), AGC255TCT (1448A>T 1449G>C 1450C>T), TCC256TCA (1453C>A), ATG257TTA (1454A>T 1456G>A), ACT258ATG (1458C>T 1459T>G), AGC260GCG (1463A>G 1464G>C 1465C>G), TCT261TCA (1468T>A), TCC262TCG (1471C>G), CTG263CTT (1474G>T), CGA264CGC (1477A>C), CCG266CCA (1483G>A), CTC268CTA (1489C>A), GTG270ATC (1493G>A 1495G>C), AGT272AGC (1501T>C), AGG273AAG (1503G>A), CGG274AGG (1505C>A), CCA276CCT (1513A>T), TCC277TCG (1516C>G), GAA279GAG (1522A>G), CTA280CTG (1525A>G), GAC282GAT (1531C>T), GTG283TTT (1532G>T 1534G>T), CCT284TCA (1535C>T 1537T>A), AAC285AGG (1539A>G 1540C>G), ACC286GCC (1541A>G), CGA288TCA (1547C>T 1548G>C), TTA289TCG (1551T>C 1552A>G), ACA290_ATG291insGTAGGT (1555_1556insGTAGGT), ATG291GTG (1556A>G), CGA292AGA (1559C>A), GCT296GGG (1572C>G 1573T>G), CAC297GTA (1574C>G 1575A>T 1576C>A), GAA299GAT (1582A>T), GAC300GAT (1585C>T), ACT301TCC (1586A>T 1588T>C), GGT302AAT (1589G>A 1590G>A), GGA303TTA (1592G>T 1593G>T), TTG304CTG (1595T>C), CCA306CCT (1603A>T), GTT307GTC (1606T>C), AAG309GAG (1610A>G), ACC511ACG (2218C>G), GCA512GCT (2221A>T), GAG513GAA (2224G>A), CAT514GAT (2225C>G), ATT515CCG (2228A>C 2229T>C 2230T>G), GAT517GGG (2235A>G 2236T>G), GAT518AAA (2237G>A 2239T>A), ACT520GAG (2243A>G 2244C>A 2245T>G), GGT522AAT (2249G>A 2250G>A), GAT523GAC (2254T>C), GTT524ATT (2255G>A), ATT525TTC (2258A>T 2260T>C), GAC527GAT (2266C>T), TTC528TTT (2269C>T), GGT529GGG (2272T>G), TCA530GAG (2273T>G 2274C>A 2275A>G), GAC531GAA (2278C>A), GAC533ATG (2282G>A 2283A>T 2284C>G), TCA534GGT (2285T>G 2286C>G 2287A>T), AAC535GAT (2288A>G 2290C>T), CCA536GCC (2291C>G 2293A>C), GCC537CCG (2294G>C 2296C>G), ATA538TTA (2297A>T), GAT539AAT (2300G>A), TTA540CTG (2303T>C 2305A>G), ATG542TTG (2309A>T), CAA544CAG (2317A>G), GTA546GCA (2322T>C), GCA549GCT (2332A>T), AGC550AGT (2335C>T), GAA552GAG (2341A>G), GGT557GGG (2356T>G), GTG558ATT (2357G>A 2359G>T), GAG560GAA (2365G>A), TTC561TTT (2368C>T), GTG562CTG (2369G>C), GAA564AAA (2375G>A), AAA567AAC (2386A>C), GTG569GTT (2392G>T), GCC570GCT (2395C>T), CAA572CAG (2401A>G), GTC573GTT (2404C>T), ATG574CTT (2405A>C 2407G>T), TCA575GCA (2408T>G), TTG576CTA (2411T>C 2413G>A), GCA577TCC (2414G>T 2416A>C), CTA578CTC (2419A>C), GCT580ATG (2423G>A 2424C>T 2425T>G), GCA584AGC (2435G>A 2436C>G 2437A>C), AAA587AAG (2446A>G), GAG588GAA (2449G>A), ACA589GCT (2450A>G 2452A>T), GCA593TCC (2462G>T 2464A>C), TAT598TAC (2479T>C), GAT600GAC (2485T>C), GCT602GCA (2491T>A), CTC603AAC (2492C>A 2493T>A), ATC604CTT (2495A>C 2497C>T), TAC606TTT (2502A>T 2503C>T), AAG607AAA (2506G>A), TAT611TAC (2518T>C), GTG612ATT (2519G>A 2521G>T), TCC613TCT (2524C>T), GTT615ATG (2528G>A 2530T>G), GGT616GGA (2533T>A), GCC617GCG (2536C>G), ATA618GTG (2537A>G 2539A>G), TCA619TCG (2542A>G), CAA621CAG (2548A>G), TTG622CTA (2549T>C 2551G>A), GCT623GCC (2554T>C), TTA624TTG (2557A>G), GTC625GTG (2560C>G), GTT631GCA (2577T>C 2578T>A), CTC632CTT (2581C>T), AAG635ACC (2589A>C 2590G>C), AAA636AAG (2593A>G), TTT638TTC (2599T>C), ATA639ATC (2602A>C), AAC640AAT (2605C>T), ATT641ATA (2608T>A), ACT643ACC (2614T>C), CTT644ATT (2615C>A), AGC647AGT (2626C>T), AAG648AAA (2629G>A), CAT649CAC (2632T>C), ACC650CAT (2633A>C 2634C>A 2635C>T), CTT651CTG (2638T>G), GTT652GTG (2641T>G), TCT653TCA (2644T>A), GCA654GCT (2647A>T), TCC655TCA (2650C>A), CAG656GAA (2651C>G 2653G>A), CAA657CAG (2656A>G), CAG658TCC (2657C>T 2658A>C 2659G>C), GAG660GGG (2664A>G), GAA661AAG (2666G>A 2668A>G), TGT663GTG (2672T>G 2673G>T 2674T>G), TTT664TTC (2677T>C), ACA665TCA (2678A>T), CCA666CCT (2683A>T), ACA667CAA (2684A>C 2685C>A), GGG668GGT (2689G>T), GGG670GGT (2695G>T), AAG671ACC (2697A>C 2698G>C), TAT672TTT (2700A>T), GTT673GTA (2704T>A), CTT675TTG (2708C>T 2710T>G), GAG677GCT (2715A>C 2716G>T), GGT678TCA (2717G>T 2718G>C 2719T>A), ACT679CTG (2720A>C 2721C>T 2722T>G), GCC681GTT (2727C>T 2728C>T), ACG682CCT (2729A>C 2731G>T), GAC683GAA (2734C>A), TTA684CTT (2735T>C 2737A>T), GGA685GGC (2740A>C), AGC686AGT (2743C>T), ATA687GTT (2744A>G 2746A>T), CGT688AGA (2747C>A 2749T>A), GTA691GTG (2758A>G), ACA692ACC (2761A>C), TTA695CTG (2768T>C 2770A>G), TCA696GCA (2771T>G), ACA699ACT (2782A>T), GAG700GAA (2785G>A), TCA702GAG (2789T>G 2790C>A 2791A>G), AAT703TCA (2792A>T 2793A>C 2794T>A), GTC704ATA (2795G>A 2797C>A), CCA705CCT (2800A>T), TGC706TGT (2803C>T), CAT707CAC (2806T>C), ATC708ATA (2809C>A), CAT709CAC (2812T>C), TTG710CTA (2813T>C 2815G>A), CAG711TCC (2816C>T 2817A>C 2818G>C), GTT714GTG (2827T>G), TCC716TCA (2833C>A), ATC719ATA (2842C>A), CTG720ATG (2843C>A), CAG721CAA (2848G>A), CCT723CCA (2854T>A), CGG724AGG (2855C>A), CTA725TTG (2858C>T 2860A>G), ATT726AGA (2862T>G 2863T>A), GCA727GGG (2865C>G 2866A>G), TCG729GCT (2870T>G 2872G>T), AAG734CGG (2885A>C 2886A>G), CCG735CCA (2890G>A), CGG790CGA (3055G>A), ACT796AGA (3072C>G 3073T>A), GGG797GGC (3076G>C), ACG798ACC (3079G>C), GTC800GTT (3085C>T), GAG802GAA (3091G>A), ATT803GTG (3092A>G 3094T>G), ATT805GTT (3098A>G), ACG808ACT (3109G>T), CTC809CTG (3112C>G), TTC810TTT (3115C>T), CAC811CAT (3118C>T), TCA812TCT (3121A>T), GGA813GGG (3124A>G), TTG815CTT (3128T>C 3130G>T), GCC816GCT (3133C>T), GGA818GGG (3139A>G), CTC819TTG (3140C>T 3142C>G), GGG820GGC (3145G>C), ACA821ACG (3148A>G), AAC823GCA (3152A>G 3153A>C 3154C>A), AAA825GAG (3158A>G 3160A>G), TCA827TCC (3166A>C), AAT828ACA (3168A>C 3169T>A), GCA829GCG (3172A>G), ATT832ATA (3181T>A), TTT833TTG (3184T>G) | | | | | | | | |

*: Inserts / Deletes / Misaligned / Frameshifts

#### Analysis details

This analysis was performed with panviral2.64

### NGS Details (UN60): Torradovirus marchitezum (segment RNA 1)

#### Assembly

| | |
|---|---|
| **Coverage Length** | 6084 (1 contig(s)) |
| **Depth Of Coverage** | 3023.2 |
| **Number Of Reads** | 139373 |
| **Reads Per Million** | 2956.04 rpm (after QC) |
| **Ambiguities** | 0 |
| **Assembly Method** | de novo + reference guided assembly |
| **Consensus Caller** | Bcf Tools |

#### Coverage Map

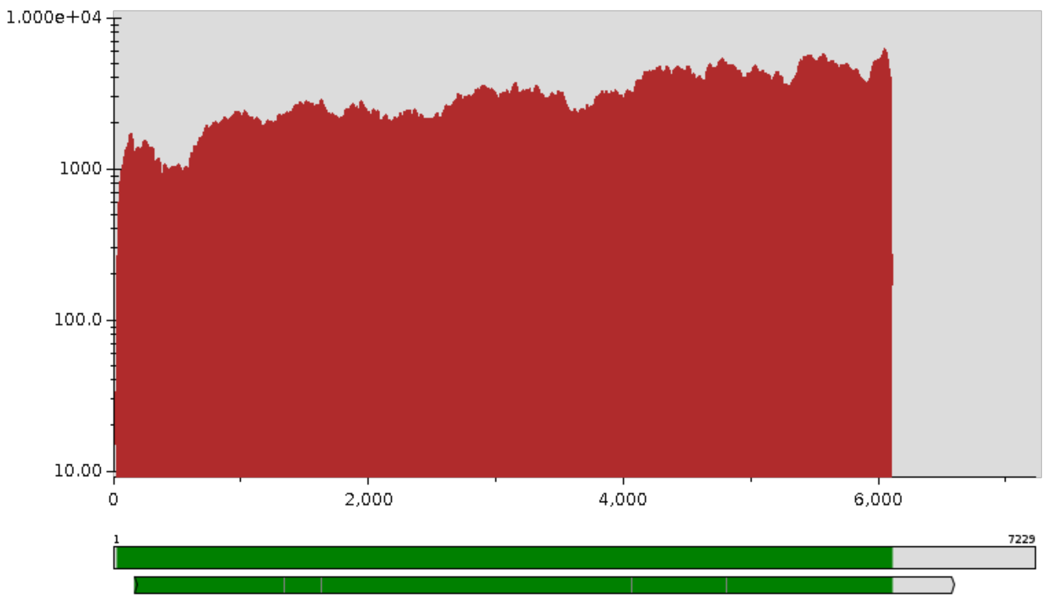

#### Assignment

| | |
|---|---|
| **Type** | Torradovirus marchitezum (Taxonomy ID: 3048376) |
| **Reference Genome** | NC_010987.1 |
| **NT Identity (%)** | 64.3453 |
| **AA Identity (%)** | 66.7002 |
| **Number Of Stop Codons** | 0 |
| **Number Of CDS** | 1 |

#### Alignment

| | |
|---|---|
| **Alignment Score** | 3417.0 (NT) + 9624.0 (AA) = 13041.0 |
| **Concordance (%)** | 50.3981 |

| Alignment Method | Global, seeded, nucleotide + amino acids (AGA) |
|---|---|

#### Genome Region

Sequence starts at position 24 and ends at position 6107 relative to NC_010987.1 reference sequence.

#### Alignment Detailed Statistics

| | Begin | End | Coverage | Score | Concordance | Matches | Identities | I/D/M/F* | Stop Codons |
|---|---|---|---|---|---|---|---|---|---|
| NT | 24 | 6107 | 84.2% | 3417 | 28.3% | 6063 (98.8%) | 3936 (64.1%) | 54/21 | |

| | Begin | End | Coverage | Score | Concordance | Matches | Identities | I/D/M/F* | Stop Codons |
|---|---|---|---|---|---|---|---|---|---|
| **NT** | **24** | **6107** | **84.2%** | **3417** | **28.3%** | **6063 (98.8%)** | **3936 (64.1%)** | **54/21** | |

Mutations: 37G>A, 40C>T, 51C>T, 59A>C, 60_61insCAAGCTACCTT, 63G>A, 65T>A, 66A>T, 69T>C, 72T>G, 74C>T, 76T>A, 77G>C, 79A>T, 80A>G, 81A>T, 84T>C, 86T>C, 88_89insCATTCTACCTTTCAGA, 91C>A, 92A>T, 93T>C, 95T>A, 97C>T, 98T>G, 100C>T, 102C>T, 104T>A, 105C>A, 108A>T, 109C>G, 111C>G, 112G>T, 113A>T, 119A>T, 121T>C, 123A>G, 126C>T, 128C>T, 130T>C, 133A>T, 135T>C, 137T>C, 138C>T, 139C>T, 140A>T, 141A>C, 145C>T, 149T>C, 150T>C, 151C>A, 152A>C, 155G>T, 157T>C, 158G>C, 159T>A, 160T>C, 162T>A, 163C>T, 164C>T, 165A>C, 167G>A, 168G>A, 170T>G, 173T>A, 174_176delGGT, 177G>T, 180G>T, 181A>C, 182T>A, 183G>A, 185C>A, 186G>A, 187C>T, 189A>G, 190G>T, 198T>G, 200C>T, 203C>T, 204A>C, 209A>T, 212C>T, 213T>C, 216G>T, 217G>C, 221A>G, 222C>T, 224T>G, 227A>T, 228A>C, 230T>G, 231T>A, 233T>A, 234A>G, 236T>G, 239C>A, 240A>T, 241A>C, 243T>C, 245T>G, 248A>C, 251A>G, 252G>A, 254G>A, 255G>A, 257T>G, 259G>A, 260T>G, 262A>G, 266T>G, 268C>G, 269C>T, 271C>A, 272T>A, 275G>A, 278A>T, 279C>G, 284G>A, 285A>T, 287T>C, 292A>G, 293T>C, 297C>A, 302G>A, 305T>A, 306C>T, 308G>A, 314A>T, 317T>A, 318G>A, 320G>T, 321A>G, 322C>T, 324G>A, 326T>A, 327G>A, 328T>C, 329G>T, 330T>G, 332C>A, 333A>G, 334C>A, 335A>T, 338T>C, 342G>A, 343T>A, 344G>T, 345G>A, 346C>G, 347C>T, 348A>G, 349C>T, 359A>T, 360C>G, 362T>A, 363A>G, 365T>A, 369G>T, 371T>A, 374G>A, 376A>G, 377C>G, 385G>C, 386T>C, 387G>A, 389A>G, 390C>G, 395A>T, 398T>A, 401A>T, 405C>G, 412C>G, 413G>T, 414T>A, 415A>T, 417T>A, 418C>A, 425G>C, 426T>A, 427C>A, 428C>A, 437C>T, 439A>G, 441T>A, 443T>C, 446C>A, 448A>T, 449G>T, 459T>C, 461A>G, 465T>G, 467G>T, 468T>G, 471G>T, 472A>T, 474A>G, 476T>C, 477T>A, 479G>C, 480T>A, 481C>A, 482C>A, 488A>T, 489G>T, 493A>T, 494A>T, 495T>A, 496T>G, 502T>A, 504G>T, 506T>A, 510G>C, 515A>G, 516A>G, 518T>A, 520A>C, 523G>C, 524T>A, 526A>G, 527A>T, 530T>A, 531G>T, 532T>A, 533A>T, 537A>G, 539A>T, 542T>A, 545A>T, 546T>A, 547A>T, 548T>A, 550G>C, 557T>A, 560A>C, 564G>T, 566T>C, 567A>C, 569A>T, 570A>T, 573T>C, 575G>A, 578A>G, 579G>T, 581G>A, 582C>T, 584T>A, 586T>A, 588T>A, 590T>A, 591G>A, 593T>C, 600A>G, 601G>T, 602T>A, 611T>A, 614G>A, 615G>A, 616G>T, 617T>G, 621T>A, 622C>G, 623T>A, 625A>G, 626C>T, 627A>G, 628C>T, 629T>A, 632C>T, 633A>T, 634A>G, 635A>T, 641C>A, 645T>C, 647G>A, 648A>G, 650A>T, 653C>A, 656A>T, 660A>T, 662G>C, 665T>C, 668A>T, 669A>T, 670G>C, 674A>G, 677C>T, 681T>A, 683A>T, 684G>A, 685A>G, 695A>C, 696A>C, 697A>T, 698A>C, 702G>T, 704C>T, 707A>G, 708C>A, 709G>A, 713C>T, 714A>T, 716C>T, 717A>C, 718T>A, 720G>A, 725C>A, 728A>G, 732A>T, 733G>C, 735A>G, 740T>A, 746G>T, 749A>G, 750T>A, 751C>G, 752A>C, 753G>T, 755T>G, 756T>A, 757C>A, 758A>T, 770G>C, 771T>A, 772C>G, 776G>A, 780A>C, 785C>T, 786A>T, 788A>T, 789A>T, 791A>G, 792T>A, 794G>A, 797T>A, 803G>C, 806T>C, 809G>A, 818G>A, 821C>T, 822A>G, 824T>G, 830G>A, 839A>G, 849T>C, 851G>T, 853G>C, 854C>A, 860A>G, 861T>C, 863G>A, 873C>T, 875C>G, 881G>A, 882A>G, 884A>G, 887A>C, 893T>A, 899G>A, 914A>C, 918A>C, 919C>A, 920T>A, 921C>T, 923T>G, 926C>T, 927C>A, 928G>A, 929T>G, 932T>C, 938A>G, 947C>A, 962A>T, 968T>G, 969C>T, 971T>G, 977A>T, 978A>G, 980A>T, 982T>C, 983C>T, 985G>A, 986C>T, 988C>T, 989C>A, 992T>C, 995A>G, 998C>T, 1001A>G, 1004T>C, 1010G>T, 1011T>C, 1016A>T, 1019T>A, 1022A>T, 1031G>A, 1032C>A, 1034G>T, 1046G>C, 1049C>T, 1059C>A, 1060A>C, 1061G>T, 1064G>A, 1068G>T, 1069A>C, 1077T>G, 1078G>C, 1091G>A, 1092C>A, 1094C>G, 1100G>A, 1109C>T, 1113A>C, 1114A>C, 1115G>T, 1121C>T, 1122C>T, 1124T>A, 1126C>T, 1142G>T, 1144G>A, 1145G>A, 1148A>G, 1149G>C, 1151A>G, 1154T>A, 1163C>A, 1169T>C, 1172C>T, 1178A>T, 1187A>T, 1190T>G, 1199C>T, 1200A>T, 1205C>T, 1211G>A, 1215A>T, 1216G>C, 1221G>A, 1223C>A, 1228C>T, 1232G>A, 1235A>G, 1238A>T, 1242G>C, 1244T>C, 1245T>A, 1246A>T, 1247T>A, 1250T>A, 1255G>A, 1256T>C, 1260A>C, 1261G>A, 1262G>A, 1265C>T, 1267T>C, 1268G>A, 1271C>A, 1277C>T, 1283T>C, 1284C>A, 1286T>A, 1295T>C, 1304A>T, 1313C>T, 1316G>T, 1317T>G, 1318C>A, 1319C>G, 1325G>A, 1328T>C, 1331T>C, 1337T>C, 1340A>G, 1341T>A, 1342C>G, 1349C>A, 1350T>C, 1352A>G, 1355T>C, 1358T>C, 1367G>A, 1373C>T, 1374A>G, 1375G>C, 1376A>C, 1380G>C, 1381G>A, 1382T>A, 1388A>C, 1391A>G, 1394C>G, 1397T>C, 1398C>A, 1401C>T, 1403A>G, 1412A>T, 1419A>C, 1421G>C, 1427A>T, 1430T>C, 1436C>A, 1439C>T, 1463G>C, 1472C>T, 1478A>T, 1482T>C, 1487A>G, 1493G>A, 1499G>C, 1502G>A, 1508A>G, 1511T>C, 1517A>G, 1520T>C, 1523G>C, 1529T>C, 1538A>C, 1553C>A, 1556A>G, 1559A>C, 1565C>T, 1571T>A, 1574G>A, 1583C>G, 1584A>C, 1595C>A, 1598C>A, 1604T>C, 1616A>C, 1622T>C, 1623C>T, 1626A>G, 1628C>T, 1634T>A, 1637T>A, 1640C>A, 1646T>C, 1649T>G, 1652T>G, 1661A>T, 1673G>A, 1674C>A, 1679A>G, 1680A>C, 1688C>T, 1689C>T, 1697G>A, 1700T>C, 1701A>C, 1702T>G, 1706G>A, 1708A>T, 1712C>G, 1715G>A, 1718T>C, 1721C>A, 1724T>A, 1727A>G, 1728G>A, 1730T>C, 1733T>A, 1736G>A, 1738G>C, 1748A>C, 1751T>C, 1757A>C, 1758G>A, 1760G>T, 1762A>G, 1763G>A, 1769A>G, 1772A>G, 1781T>C, 1784C>A, 1787T>A, 1793G>A, 1794A>G, 1796T>C, 1797T>C, 1799G>C, 1802C>T, 1806A>T, 1807G>C, 1811T>C, 1812C>T, 1814T>G, 1820G>A, 1829A>G, 1835A>G, 1838T>C, 1839C>A, 1844A>C, 1847T>C, 1848G>C, 1849C>A, 1850A>C, 1854G>A, 1856A>T, 1857T>G, 1861A>C, 1862G>C, 1866T>C, 1868G>C, 1870T>A, 1871T>C, 1872G>A, 1874A>C, 1875G>C, 1876C>A, 1877T>G, 1878G>T, 1879A>C, 1881T>A, 1883T>A, 1886C>T, 1887A>T, 1888T>G, 1889A>T, 1890A>G, 1891G>A, 1892T>A, 1893C>G, 1894C>A, 1895T>G, 1898G>A, 1899A>G, 1901A>T, 1904T>C, 1910T>G, 1911C>G, 1921C>T, 1922A>T, 1925T>A, 1928T>A, 1934A>G, 1935G>A, 1937G>A, 1940T>C, 1941G>T, 1943G>T, 1944G>C, 1948G>A, 1949G>T, 1953A>C, 1954G>A, 1956T>A, 1958G>A, 1959C>A, 1962A>T, 1963C>T, 1964C>T, 1965A>G, 1967G>C, 1968C>G, 1972A>T, 1974G>A, 1975A>G, 1976A>C, 1979A>C, 1981T>G, 1982A>T, 1983T>G, 1985G>T, 1986_1991delGTGCCC, 1994T>A, 1998G>C, 2000G>C, 2001A>T, 2002G>C, 2006A>G, 2007A>G, 2019A>G, 2020T>G, 2021C>A, 2025G>T, 2028A>C, 2029G>A, 2030A>G, 2033G>A, 2036C>A, 2039G>T, 2042A>G, 2045G>A, 2049T>C, 2054A>G, 2060A>T, 2063C>T, 2066G>A, 2069C>A, 2070A>G, 2071G>C, 2072T>A, 2075G>A, 2076G>A, 2081T>A, 2082G>T, 2083A>T, 2084G>T, 2088C>T, 2089T>A, 2096T>C, 2102C>T, 2103C>G, 2104A>C, 2106C>T, 2107A>C, 2108T>A, 2117C>A, 2120G>T, 2122A>T, 2127T>G, 2128C>T, 2129T>C, 2132T>G, 2133G>T, 2134A>C, 2135T>A, 2136G>A, 2137A>G, 2142C>G, 2143A>C, 2144T>C, 2152T>G, 2153T>C, 2156C>T, 2158A>C, 2159C>T, 2160A>C, 2161A>G, 2162A>C, 2166A>G, 2167G>T, 2169G>A, 2170T>A, 2171C>T, 2172A>G, 2174C>G, 2178T>A, 2180G>T, 2183T>C, 2184A>G, 2185G>A, 2186A>G, 2189A>G, 2192A>T, 2193G>T, 2194A>C, 2195T>G, 2196G>A, 2200A>C, 2202C>G, 2203A>G, 2204G>A, 2208C>G, 2209G>C, 2210A>C, 2217G>A, 2219A>C, 2220C>A, 2222A>G, 2223A>G, 2224A>C, 2228T>C, 2231T>A, 2241A>G, 2243A>C, 2255G>A, 2258T>C, 2268G>A, 2269C>G, 2270A>T, 2271T>A, 2277A>G, 2279C>G, 2282T>A, 2285G>A, 2286A>G, 2287A>C, 2288G>T, 2289A>C, 2291A>C, 2292C>A, 2294A>G, 2296C>A, 2297A>T, 2300G>A, 2303G>A, 2305C>G, 2306A>G, 2307G>A, 2308A>T, 2310A>T, 2313A>T, 2314G>A, 2315G>C, 2317C>A, 2321A>C, 2322T>A, 2323T>G, 2324T>A, 2331T>A, 2332C>G, 2333T>A, 2334T>G, 2335G>C, 2336C>A, 2337A>C, 2339T>G, 2342T>A, 2345A>T, 2350T>C, 2357C>T, 2358C>T, 2360C>A, 2363A>G, 2369C>T, 2372G>A, 2375G>A, 2381A>T, 2385T>C, 2386T>A, 2390T>C, 2393G>A, 2396T>C, 2397T>C, 2402T>C, 2415A>C, 2417A>G, 2426_2427insAGAAGG, 2429C>G, 2431A>T, 2432A>G, 2435A>C, 2436T>A, 2438A>T, 2441A>G, 2444G>A, 2445C>A, 2448_2453delAAGGAA, 2456A>T, 2460A>G, 2462C>A, 2463A>C, 2466A>T, 2467T>C, 2468T>G, 2469T>C, 2470G>T, 2474C>T, 2475C>G, 2477T>C, 2478A>T, 2479A>C, 2480A>C, 2482T>C, 2483C>A, 2484G>A, 2488A>G, 2489A>G, 2490T>A, 2491A>G, 2492T>A, 2495C>G, 2496A>C, 2498G>A, 2501T>A, 2503C>T, 2504A>G, 2508T>A, 2509G>A, 2510G>T, 2511A>G, 2512G>A, 2513C>T, 2516T>G, 2522T>C, 2523G>A, 2524C>T, 2525A>T, 2526C>T, 2530G>A, 2537C>T, 2549T>G, 2556A>C, 2561G>A, 2567T>C, 2573A>G, 2574G>A, 2575T>C, 2577T>A, 2579T>G, 2582A>T, 2584G>C, 2586A>G, 2588T>A, 2591A>T, 2597T>C, 2598T>A, 2600G>T, 2601G>T, 2603G>T, 2610C>A, 2612G>A, 2615T>C, 2617A>G, 2620C>T, 2621T>G, 2622T>A, 2624C>T, 2629A>C, 2631G>T, 2633A>C, 2634T>C, 2635C>A, 2641G>A, 2651G>T, 2663C>G, 2669T>C, 2675C>A, 2678T>C, 2693T>A, 2695A>G, 2696T>A, 2696_2697insTCC, 2697A>C, 2699G>T, 2700C>G, 2702C>A, 2703A>T, 2704G>C, 2708T>G, 2709C>A, 2711T>G, 2714C>G, 2715G>T, 2718G>A, 2720T>C, 2729T>A, 2733T>C, 2736T>A, 2737T>A, 2738G>T, 2740C>A, 2741A>T, 2744A>G, 2744_2745insGGA, 2745G>A, 2747T>A, 2754G>T, 2756T>C, 2757G>A, 2758A>G, 2759A>G, 2765A>T, 2771A>G, 2774T>C, 2775C>G, 2777G>A, 2778A>T, 2779G>C, 2780C>T, 2783C>T, 2784A>G, 2785A>C, 2786G>C, 2798G>A, 2802T>C, 2807A>G, 2808C>G, 2810T>G, 2812G>C, 2813T>A, 2822G>T, 2829A>C, 2831A>T, 2834C>A, 2837C>T, 2846T>C, 2847T>A, 2848C>G, 2849T>C, 2851G>A, 2852T>A, 2864C>T, 2870T>A, 2873T>C, 2876A>T, 2880C>A, 2881A>C, 2882C>G, 2883A>C, 2885T>G, 2886G>C, 2887C>T, 2888T>C, 2891A>T, 2897T>A, 2900T>C, 2907A>T, 2915C>T, 2920C>A, 2921C>A, 2922A>C, 2928C>G, 2929G>T, 2930A>C, 2932C>G, 2933T>C, 2934A>C, 2936A>G, 2939T>G, 2948C>T, 2953G>A, 2955G>C, 2957A>T, 2958T>G, 2960A>C, 2962G>A, 2964A>T, 2967A>T, 2969T>G, 2970G>C, 2971A>C, 2972G>T, 2974G>A, 2975C>A, 2981C>T, 2982A>G, 2984C>T, 2985A>G, 2987T>A, 2988C>T, 2990G>A, 2991T>C, 2994C>A, 2996C>G, 3002T>C, 3003G>T, 3004A>G, 3008A>T, 3011T>C, 3014T>G, 3017A>T, 3020G>A, 3021C>G, 3022C>T, 3023T>C, 3030C>A, 3032T>A, 3035A>G, 3041C>T, 3042C>G, 3044A>T, 3045A>G, 3046G>A, 3055A>C, 3056G>A, 3071G>A, 3074C>A, 3077A>T, 3078C>G, 3083A>C, 3089C>T, 3090T>G, 3091G>T, 3092T>C, 3098T>A, 3101T>G, 3104G>A, 3107T>A, 3109C>A, 3116A>T, 3117C>G, 3118G>A, 3119T>G, 3120C>A, 3121T>A, 3122T>A, 3123G>C, 3124G>A, 3125C>G, 3128T>C, 3131A>G, 3134G>C, 3144A>T, 3145T>A, 3146G>T, 3147C>T, 3161A>C, 3164G>A, 3170C>T, 3175T>A, 3176T>G, 3182G>C, 3183G>A, 3184A>C, 3188T>C, 3189G>A, 3190A>G, 3191A>G, 3192C>A, 3196A>G, 3198T>A, 3200C>A, 3201T>A, 3202C>A, 3203C>T, 3204A>T, 3205G>C, 3206T>A, 3210A>C, 3212G>C, 3212_3213insTCA, 3215T>C, 3216T>A, 3218C>T, 3219T>G, 3225C>G, 3228T>G, 3230T>C, 3242C>T, 3243A>G, 3245A>C, 3249G>A, 3251G>C, 3263C>T, 3269C>A, 3272G>T, 3279C>T, 3281G>A, 3284A>G, 3288A>T, 3289G>C, 3291G>A, 3293T>C, 3296G>A, 3299T>C, 3308T>A, 3309G>T, 3310T>G, 3311G>C, 3314C>T, 3320C>T, 3323C>T, 3326C>T, 3332A>T, 3335A>C, 3343G>A, 3344C>A, 3346T>C, 3347A>T, 3351G>A, 3353T>A, 3356A>G, 3357C>T, 3358G>T, 3359T>G, 3366A>G, 3368C>A, 3369T>G, 3371T>A, 3380T>C, 3383A>T, 3387A>G, 3389C>A, 3398A>C, 3401A>G, 3404A>G, 3405G>C, 3407G>A, 3414C>G, 3416A>T, 3417A>C, 3418C>A, 3419T>A, 3425G>A, 3428C>T, 3432G>A, 3433C>T, 3434A>G, 3437A>C, 3443A>G, 3444C>A, 3446C>A, 3452T>C, 3453A>T, 3455T>G, 3458G>C, 3461G>A, 3462T>C, 3463C>A, 3464A>T, 3465A>G, 3468A>C, 3471A>T, 3472G>C, 3473C>A, 3482A>T, 3483C>G, 3484A>C, 3485G>T, 3488T>C, 3494G>C, 3500C>G, 3506G>T, 3507T>G, 3508A>C, 3509C>T, 3511A>T, 3512C>T, 3513C>T, 3515T>G, 3518A>C, 3521A>T, 3527A>T, 3530G>A, 3534T>A, 3539C>A, 3542C>A, 3545A>G, 3548T>G, 3551C>A, 3563T>C, 3566G>C, 3567C>A, 3575C>T, 3577A>C, 3579T>C, 3587A>G, 3593C>T, 3596A>T, 3597C>G, 3598C>A, 3599T>A, 3603G>C, 3605G>A, 3608A>T, 3611A>T, 3614A>T, 3617C>A, 3618A>C, 3620T>G, 3623A>C, 3624A>T, 3626A>T, 3629T>G, 3630C>A, 3632T>A, 3635C>T, 3644T>G, 3650T>A, 3656C>T, 3659C>T, 3665G>T, 3671G>C, 3672G>A, 3674A>T, 3675A>G, 3676T>A, 3695T>C, 3697G>C, 3698T>A, 3699G>A, 3700G>A, 3701A>T, 3707T>G, 3710T>C, 3711G>A, 3712A>G, 3719C>A, 3720A>C, 3721A>G, 3722A>G, 3723C>G, 3724T>C, 3725A>T, 3731C>T, 3735A>G, 3736A>G, 3745G>C, 3746C>T, 3747T>G, 3749G>C, 3750C>A, 3752G>T, 3753C>T, 3758T>A, 3759A>G, 3761C>T, 3762A>C, 3764G>C, 3765C>A, 3766G>A, 3767T>G, 3770C>T, 3773G>A, 3775A>G, 3777A>G, 3779C>G, 3782T>A, 3785G>T, 3788A>T, 3789C>A, 3790G>T, 3791T>C, 3794A>C, 3795T>A, 3796G>C, 3797T>A, 3798G>A, 3799A>C, 3801A>C, 3803G>T, 3804G>T, 3805A>G, 3812T>C, 3813G>A, 3815A>C, 3818G>T, 3821C>G, 3827A>T, 3828A>G, 3830A>T, 3836A>T, 3838G>T, 3842T>C, 3844A>T, 3847A>T, 3852G>T, 3853G>C, 3860G>A, 3863G>C, 3872T>A, 3875C>T, 3881T>C, 3884C>T, 3890C>T, 3893G>A, 3894A>C, 3899G>T, 3900A>T, 3905G>T, 3907T>C, 3908G>C, 3911C>T, 3914G>A, 3917A>G, 3923T>C, 3926G>T, 3927C>G, 3930T>G, 3932C>A, 3933A>C, 3934C>A, 3935T>G, 3937G>A, 3938C>G, 3939G>A, 3940A>G, 3941A>T, 3942G>A, 3943G>C, 3944A>C, 3945G>C, 3947G>A, 3950A>G, 3953G>A, 3954G>C, 3955A>T, 3962A>G, 3966G>A, 3968C>T, 3969A>T, 3977A>T, 3980T>C, 3981G>C, 3992A>G, 3995G>A, 3996C>T, 4005C>G, 4007G>A, 4010T>A, 4016A>G, 4019C>T, 4021T>A, 4023C>T, 4031T>A, 4032C>A, 4037C>T, 4043C>T, 4049C>T, 4052C>T, 4055A>G, 4058T>C, 4062C>A, 4064C>G, 4067C>A, 4070A>T, 4073T>G, 4076G>C, 4079A>G, 4082C>A, 4085T>C, 4088C>G, 4091G>A, 4093G>A, 4094A>G, 4097G>A, 4101A>C, 4103G>T, 4106C>T, 4115A>T, 4116C>T, 4118T>G, 4121T>A, 4130C>T, 4137T>C, 4142C>G, 4143A>C, 4145G>C, 4148G>A, 4154T>C, 4163C>T, 4166A>T, 4169C>A, 4170A>T, 4171C>G, 4172A>T, 4175G>A, 4182T>G, 4183T>C, 4184G>A, 4187C>T, 4192C>T, 4194A>C, 4196A>G, 4199A>C, 4208T>G, 4214T>C, 4215G>A, 4220T>C, 4222C>T, 4223A>T, 4224A>T, 4225G>C, 4226T>A, 4227T>C, 4228C>A, 4238A>C, 4239T>A, 4240T>C, 4241G>C, 4242T>C, 4250T>C, 4251G>C, 4252G>A, 4256T>C, 4257A>C, 4259A>G, 4260G>A, 4261C>G, 4268A>T, 4271C>T, 4274G>T, 4277A>T, 4283A>T, 4286T>A, 4289T>C, 4292T>C, 4310A>G, 4313A>G, 4316T>A, 4317G>A, 4319T>A, 4325T>A, 4331G>A, 4354A>C, 4361C>T, 4364A>G, 4368A>G, 4369A>C, 4370G>T, 4373T>A, 4376C>A, 4379T>C, 4380C>G, 4381C>A, 4382T>G, 4388C>A, 4394T>A, 4395C>A, 4403A>T, 4406T>G, 4409A>G, 4418A>T, 4424T>C, 4433A>T, 4436G>A, 4449G>C, 4451G>C, 4454T>C, 4460G>A, 4466G>T, 4469C>A, 4475A>T, 4478A>T, 4487A>T, 4490A>C, 4493G>C, 4499C>A, 4502C>T, 4505T>C, 4508T>C, 4511C>T, 4512C>T, 4514C>A, 4517T>C, 4532A>G, 4538C>T, 4541T>C, 4547A>G, 4548T>C, 4550G>C, 4551T>C, 4553A>C, 4559A>G, 4565T>C, 4568T>C, 4569G>A, 4570C>A, 4571T>A, 4574C>A, 4576G>A, 4580T>A, 4581A>T, 4583A>G, 4586A>T, 4589T>A, 4592A>G, 4593C>T, 4604C>T, 4608C>G, 4610G>C, 4611C>T, 4616C>T, 4619A>T, 4622T>A, 4625T>G, 4631A>G, 4646T>G, 4652A>G, 4656T>A, 4658A>G, 4661T>C, 4664C>T, 4667G>A, 4673T>C, 4674T>G, 4675C>A, 4676T>G, 4677C>T, 4679G>A, 4682A>T, 4683A>G, 4684A>C, 4685T>A, 4694T>A, 4698T>C, 4705T>A, 4706G>A, 4709C>G, 4712T>C, 4715A>T, 4718C>T, 4721T>C, 4730T>C, 4736C>T, 4740T>A, 4741C>A, 4742G>C, 4746G>A, 4751T>C, 4757G>T, 4760A>C, 4761C>A, 4766T>C, 4769A>G, 4782C>T, 4784A>G, 4787T>G, 4791C>T, 4796T>C, 4802T>C, 4805T>C, 4808A>G, 4811A>C, 4818C>T, 4820T>G, 4823A>T, 4829C>A, 4830A>C, 4832G>A, 4835T>C, 4838T>C, 4853A>C, 4856C>T, 4857A>G, 4859T>G, 4862G>A, 4865C>A, 4877A>C, 4883T>A, 4886C>A, 4887C>G, 4888C>T, 4889A>T, 4892T>G, 4898T>A, 4901C>T, 4903C>T, 4904T>C, 4905C>A, 4906A>C, 4907G>T, 4914A>T, 4915T>A, 4916G>T, 4931A>G, 4934A>T, 4935A>G, 4938C>A, 4940T>A, 4943A>G, 4952T>G, 4955C>T, 4958T>A, 4960G>A, 4961T>G, 4964G>C, 4966C>T, 4975A>C, 4976G>A, 4977C>T, 4979T>G, 4980G>A, 4981T>C, 4982C>A, 4983T>G, 4984T>A, 4985A>C, 4986G>A, 4991T>C, 4994G>A, 4996C>A, 4997G>A, 5000G>A, 5001T>A, 5003G>A, 5004G>A, 5005A>T, 5006T>G, 5007G>C, 5009A>G, 5010A>C, 5012G>A, 5015T>C, 5018C>G, 5019A>C, 5020G>C, 5022G>A, 5023C>T, 5024T>C, 5028A>C, 5029T>A, 5030C>T, 5036T>A, 5039C>T, 5043T>G, 5044T>C, 5045G>T, 5049G>C, 5050T>A, 5051G>A, 5055C>A, 5056A>G, 5057G>T, 5060A>G, 5064A>C, 5066T>A, 5067G>C, 5068C>A, 5070A>G, 5071A>C, 5072G>C, 5075A>T, 5078A>T, 5081G>A, 5084T>C, 5085T>A, 5086C>G, 5087A>T, 5089G>A, 5090T>C, 5091G>A, 5092C>T, 5093C>A, 5094C>G, 5095T>G, 5097A>G, 5099A>T, 5100A>C, 5102T>G, 5103T>C, 5104G>T, 5105T>G, 5109A>C, 5112G>A, 5113C>G, 5114A>C, 5117A>G, 5119C>A, 5120A>T, 5127T>A, 5129C>T, 5132T>C, 5133A>G, 5134C>T, 5138A>T, 5139T>C, 5141G>T, 5142G>C, 5143A>C, 5144C>A, 5146C>A, 5151G>A, 5153G>A, 5156G>A, 5159C>T, 5160A>G, 5161C>A, 5163A>G, 5169G>T, 5170A>C, 5171G>A, 5173G>A, 5174T>A, 5176A>G, 5178C>G, 5180G>C, 5181A>C, 5184T>G, 5185G>A, 5186G>A, 5190G>T, 5191C>G, 5192A>T, 5193C>T, 5195A>T, 5196A>G, 5197A>G, 5198C>A, 5199A>G, 5202G>T, 5203G>T, 5204T>C, 5207T>C, 5214G>T, 5216A>G, 5219T>A, 5222T>A, 5224C>A, 5227C>T, 5228C>T, 5229C>A, 5231C>T, 5232T>C, 5234C>T, 5236C>A, 5237G>A, 5246A>G, 5248G>A, 5249G>C, 5250T>A, 5251A>C, 5252C>A, 5256A>C, 5264C>T, 5267T>C, 5279G>A, 5282G>A, 5283T>A, 5285C>A, 5287A>C, 5288T>A, 5294T>G, 5300T>C, 5339G>C, 5342C>T, 5345T>A, 5349T>G, 5354T>C, 5357A>C, 5359T>A, 5363T>C, 5364G>A, 5365C>A, 5366C>T, 5370A>C, 5372G>A, 5374A>G, 5376G>C, 5377T>G, 5378G>C, 5382G>T, 5385G>A, 5386G>T, 5387A>G, 5390A>G, 5391C>A, 5392A>T, 5395T>G, 5397G>A, 5399A>C, 5400A>C, 5401C>A, 5402C>A, 5405T>A, 5408C>T, 5411T>C, 5415G>A, 5417G>T, 5423A>G, 5429A>T, 5438C>G, 5439G>A, 5441C>T, 5444A>C, 5453A>T, 5456G>T, 5460A>T, 5462G>T, 5465C>A, 5469G>T, 5474A>C, 5476G>A, 5477G>T, 5480A>G, 5484T>A, 5486T>A, 5488T>C, 5489G>T, 5493G>C, 5495C>A, 5496A>C, 5498A>T, 5504T>C, 5511C>A, 5513C>A, 5517A>T, 5519T>G, 5521G>C, 5522A>G, 5523A>G, 5524A>G, 5525T>C, 5526G>T, 5531T>G, 5534C>T, 5535C>T, 5537C>G, 5541A>C, 5542C>A, 5543C>G, 5555C>T, 5559G>A, 5561G>T, 5564G>A, 5570T>C, 5573T>C, 5574C>A, 5575A>C, 5579_5580insGTC, 5584C>A, 5585T>G, 5586C>A, 5588T>A, 5589G>A, 5591A>T, 5592G>T, 5593A>G, 5597A>G, 5598G>A, 5599T>C, 5600G>A, 5607A>G, 5608G>C, 5610G>A, 5614C>G, 5615A>G, 5616_5618delTCG, 5620T>G, 5622C>G, 5624C>T, 5625A>G, 5627T>A, 5629G>T, 5630T>G, 5633T>G, 5635T>A, 5636G>C,

| | |
|---|---|
| | 5642A>T, 5642_5643insCATGCT, 5644G>C, 5645G>T, 5647T>A, 5650A>C, 5651A>C, 5652T>C, 5654G>T, 5657C>A, 5663C>T, 5666C>A, 5669T>A, 5670A>G, 5672A>G, 5673C>G, 5674A>G, 5675T>C, 5676G>T, 5679G>T, 5680A>C, 5681T>A, 5682T>A, 5684A>T, 5685G>T, 5689A>G, 5691C>A, 5695C>T, 5696A>G, 5700A>G, 5703C>A, 5705G>A, 5706A>T, 5711A>C, 5714T>C, 5715G>A, 5716T>C, 5717G>T, 5720A>T, 5723A>G, 5727C>A, 5728A>T, 5729T>G, 5732A>T, 5735A>T, 5740G>C, 5742A>G, 5746T>G, 5747T>C, 5750C>A, 5754C>A, 5755A>G, 5756G>A, 5757G>A, 5765G>T, 5766A>T, 5767G>C, 5769T>C, 5771G>C, 5772C>A, 5774A>C, 5775C>A, 5777C>A, 5778T>G, 5779A>C, 5780T>C, 5781T>C, 5783G>A, 5787C>T, 5788A>G, 5789G>C, 5790G>T, 5791G>C, 5792A>T, 5798A>T, 5800A>C, 5801G>A, 5803T>G, 5804A>T, 5805A>G, 5807T>G, 5808C>G, 5809A>C, 5810A>T, 5811A>C, 5812A>G, 5813G>T, 5814A>G, 5815A>G, 5816T>G, 5817A>T, 5818C>G, 5819A>T, 5823C>A, 5834T>C, 5837C>T, 5838A>G, 5840A>G, 5841G>A, 5843G>T, 5844C>T, 5846G>T, 5848G>A, 5850A>G, 5851T>C, 5852G>A, 5853G>C, 5857A>T, 5858T>A, 5860T>C, 5861G>T, 5862C>A, 5863A>C, 5864A>C, 5865C>A, 5866A>G, 5867A>G, 5868C>T, 5870G>A, 5871A>G, 5880G>C, 5885G>A, 5886C>A, 5887T>G, 5894G>T, 5895T>G, 5897T>G, 5900C>T, 5901C>A, 5902C>T, 5903A>C, 5905A>C, 5908A>C, 5909A>C, 5912T>A, 5913A>G, 5914C>G, 5916A>C, 5917T>A, 5918T>C, 5919C>A, 5921G>A, 5923A>T, 5925C>T, 5927C>G, 5928A>G, 5929A>T, 5930A>G, 5931A>C, 5933T>A, 5934T>A, 5935T>C, 5937G>C, 5939T>G, 5940G>A, 5941A>G, 5942T>C, 5943G>A, 5944C>A, 5946G>C, 5951T>G, 5953T>A, 5955A>T, 5957A>C, 5958T>C, 5959T>A, 5961A>G, 5964G>C, 5965C>A, 5969G>T, 5970A>T, 5972C>T, 5973T>G, 5975T>G, 5977G>A, 5979C>A, 5980A>G, 5981C>A, 5984A>C, 5985G>T, 5986T>A, 5993A>G, 5994A>C, 5995A>G, 5996G>A, 5999A>T, 6003T>C, 6005G>C, 6006A>T, 6007G>C, 6008T>A, 6009T>A, 6010C>A, 6012A>G, 6015T>A, 6016G>T, 6017G>A, 6019A>C, 6023_6024insTAC, 6024T>C, 6032A>T, 6033A>G, 6034G>C, 6035T>C, 6037T>A, 6038T>A, 6040T>A, 6045T>A, 6046C>T, 6048A>G, 6049G>T, 6050G>A, 6054_6056delGCA, 6063C>G, 6065A>G, 6069C>T, 6070C>T, 6071T>G, 6074C>T, 6078T>C, 6079T>C, 6080G>A, 6086T>C, 6087T>A, 6088C>T, 6089C>G, 6092A>T, 6097A>G, 6098A>G |

**CDS**

| | | | | | | | | | |
|---|---|---|---|---|---|---|---|---|---|
| ToMarV_RNA1gp1 | 1 | 1989 | 92.4% | 9624 | 69.7% | 1982 (99.2%) | 1328 (66.5%) | 9/7/0/0 | 0 |

| | Begin | End | Coverage | Score | Concordance | Matches | Identities | I/D/M/F* | Stop Codons |
|---|---|---|---|---|---|---|---|---|---|
| **NT** | **24** | **6107** | **84.2%** | **3417** | **28.3%** | **6063 (98.8%)** | **3936 (64.1%)** | **54/21** | |

Protein mutations:

M1L (141A>C), A2V (145C>T), S4H (150T>C 151C>A 152A>C), K5N (155G>T), M6T (157T>C 158G>C), F7T (159T>A 160T>C), S8I (162T>A 163C>T 164C>T), K9Q (165A>C 167G>A), V10M (168G>A 170T>G), G12del (174_176delGGT), V13F (177G>T), D14S (180G>T 181A>C 182T>A), D15K (183G>A 185C>A), A16I (186G>A 187C>T), S17V (189A>G 190G>T), S20A (198T>G 200C>T), T22P (204A>C), F25L (213T>C), G26S (216G>T 217G>C), N30Q (228A>C 230T>G), S31T (231T>A 233T>A), I32V (234A>G 236T>G), K34S (240A>T 241A>C), F35L (243T>C 245T>G), A38T (252G>A 254G>A), V39M (255G>A 257T>G), S40K (259G>A 260T>G), N41S (262A>G), I42M (266T>G), T43S (268C>G 269C>T), T44K (271C>A 272T>A), M45I (275G>A), Q47E (279C>G), I49F (285A>T 287T>C), D51G (292A>G 293T>C), L53I (297C>A), D55E (305T>A), V60I (318G>A 320G>T), T61V (321A>G 322C>T), D62K (324G>A 326T>A), V63T (327G>A 328T>C 329G>T), S64A (330T>G 332C>A), T65D (333A>G 334C>A 335A>T), V68N (342G>A 343T>A 344G>T), A69S (345G>A 346C>G 347C>T), T70V (348A>G 349C>T), L74V (360C>G 362T>A), N75E (363A>G 365T>A), V77L (369G>T 371T>A), D79G (376A>G 377C>G), G82A (385G>C 386T>C), A83T (387G>A 389A>G), L84V (390C>G), D86E (398T>A), L89V (405C>G), A91G (412C>G 413G>T), Y92I (414T>A 415A>T), S93N (417T>A 418C>A), L95F (425G>C), S96K (426T>A 427C>A 428C>A), K100R (439A>G), S101T (441T>A 443T>C), K103I (448A>T 449G>T), L109V (465T>G 467G>T), F110V (468T>G), E111L (471G>T 472A>T), S112G (474A>G 476T>C), L113I (477T>A 479C>G), S114K (480T>A 481C>A 482C>A), K116N (488A>T), V117F (489G>T), K118I (493A>T 494A>T), L119R (495T>A 496T>G), F121Y (502T>A), A122S (504G>T 506T>A), V124L (510G>C), N126E (516A>G 518T>A), K127T (520A>C), S128T (523G>C 524T>A), E129G (526A>G 527A>T), D130E (530T>A), V131Y (531G>T 532T>A 533A>T), I133V (537A>G 539A>T), Y136I (546T>A 547A>T 548T>A), G137A (550G>C), A142S (564G>T 566T>C), I143L (567A>C 569A>T), T144S (570A>T), V147L (579G>T 581G>A), L149Q (586T>A), F150I (588T>A 590T>A), V151I (591G>A 593T>C), S154V (600A>G 601G>T 602T>A), G159M (615G>A 616G>T 617T>G), S161R (621T>A 622C>G 623T>A), N162S (625A>G 626C>T), T163V (627A>G 628C>T 629T>A), K165C (633A>T 634A>G 635A>T), I170V (648A>G 650A>T), M174F (660A>T 662G>C), L181I (681T>A 683A>T), D182S (684G>A 685A>G), K186L (696A>C 697A>T 698A>C), A188S (702G>T 704C>T), R190K (708C>A 709G>A), T192S (714A>T 716C>T), M193Q (717A>C 718T>A), V194M (720G>A), M199V (735A>G), V205L (753G>T 755T>G), S206N (756T>A 757C>A 758A>T), M214L (780A>C), T216S (786A>T 788A>T), I217L (789A>T 791A>G), L218I (792T>A 794G>A), I228V (822A>G 824T>G), S238T (853G>C 854C>A), T248A (882A>G 884A>G), T260Q (918A>C 919C>A 920T>A), R263K (927C>A 928G>A 929T>G), I280V (978A>G 980A>T), V281A (982T>C 983C>T), G282D (985G>A 986C>T), T283I (988C>T 989C>A), L298I (1032C>A 1034G>T), Q307T (1059C>A 1060A>C 1061G>T), D310S (1068G>T 1069A>C), C313A (1077T>G 1078G>C), K325P (1113A>C 1114A>C 1115G>T), P328S (1122C>T 1124T>A), T329I (1126C>T), R335K (1144G>A 1145G>A), E337Q (1149G>C 1151A>G), M354L (1200A>T), A361T (1221G>A 1223C>A), A363V (1228C>T), E366D (1238A>T), V368L (1242G>C 1244T>C), Y369I (1245T>A 1246A>T 1247T>A), S372N (1255G>A 1256T>C), R374Q (1260A>C 1261G>A 1262G>A), M376T (1267T>C 1268G>A), S393E (1317T>G 1318C>A 1319C>G), R412A (1374A>G 1375G>C 1376A>C), G414Q (1380G>C 1381G>A 1382T>A), I496V (1626A>G 1628C>T), M521R (1701A>C 1702T>G), E523V (1708A>T), D530N (1728G>A 1730T>C), G533A (1738G>C), R536S (1748A>C), V540I (1758G>A 1760G>T), E541G (1762A>G 1763G>A), N552D (1794A>G 1796T>C), Q567K (1839C>A), A570H (1848G>C 1849C>A 1850A>C), V572I (1854G>A 1856A>T), S573A (1857T>G), K574T (1861A>C 1862G>C), F577Y (1870T>A 1871T>C), E578N (1872G>A 1874A>C), A579Q (1875G>C 1876C>A 1877T>G), E580S (1878G>T 1879A>C), S581T (1881T>A 1883T>A), I583C (1887A>T 1888T>G 1889A>T), S584E (1890A>G 1891G>A 1892T>A), P585E (1893C>G 1894C>A 1895T>G), I587V (1899A>G 1901A>T), Q591E (1911C>G), A594V (1921C>T 1922A>T), D595E (1925T>A), D596E (1928T>A), I598M (1934A>G), A599T (1935G>A 1937G>A), V601F (1941G>T 1943G>T), D602H (1944G>C), G603D (1948G>A 1949G>T), R605Q (1953A>C 1954G>A), L606I (1956T>A 1958G>A), R607S (1959C>A), T608F (1962A>T 1963C>T 1964C>T), K609D (1965A>G 1967G>C), Q610E (1968C>G), D611V (1972A>T), E612S (1974G>A 1975A>G 1976A>C), I614S (1981T>G 1982A>T), L615V (1983T>G 1985G>T), V616_P617del (1986_1991delGTGCCC), V620L (1998G>C 2000G>C), N623D (2007A>G), I627G (2019A>G 2020T>G 2021C>A), A629S (2025G>T), R630Q (2028A>C 2029G>A 2030A>G), S644A (2070A>G 2071G>C 2072T>A), D646N (2076G>A), E648F (2082G>T 2083A>T 2084G>T), L650Y (2088C>T 2089T>A), H655A (2103C>G 2104A>C), H656S (2106C>T 2107A>C 2108T>A), M660I (2120G>T), Y661F (2122A>T), S663V (2127T>G 2128C>T 2129T>C), D665S (2133G>T 2134A>C 2135T>A), D666S (2136G>A 2137A>G), H668A (2142C>G 2143A>C 2144T>C), I671S (2152T>G 2153T>C), N673T (2158A>C 2159C>T), K674R (2160A>C 2161A>G 2162A>C), R676V (2166A>G 2167G>T), V677N (2169G>A 2170T>A 2171C>T), N678E (2172A>G 2174C>G), L680I (2178T>A 2180G>T), R682E (2184A>G 2185G>A 2186A>G), D685S (2193G>T 2194A>C 2195T>G), A686T (2196G>A), E687A (2200A>C), Q688G (2202C>G 2203A>G 2204G>A), R690A (2208C>G 2209G>C 2210A>C), V693I (2217G>A 2219A>C), Q694K (2220C>A 2222A>G), N695A (2223A>G 2224A>C), I701V (2241A>G 2243A>C), A710S (2268G>A 2269C>G 2270A>T), S711T (2271T>A), I713V (2277A>G 2279C>G), K716A (2286A>G 2287A>C 2288G>T), K717H (2289A>C 2291A>C), L718M (2292C>A 2294A>G), A719D (2296C>A 2297A>T), T722R (2305C>G 2306A>G), E723M (2307G>A 2308A>T), M724L (2310A>T), R725Y (2313A>T 2314G>A 2315G>C), T726N (2317C>A), F728R (2322T>A 2323T>G 2324T>A), S731R (2331T>A 2332C>G 2333T>A), C732A (2334T>G 2335G>C 2336C>A), I733L (2337A>C 2339T>G), D734E (2342T>A), V737A (2350T>C), I741M (2363A>G), F749H (2385T>C 2386T>A), C762_N763insRR (2426_2427insAGAAGG), N763K (2429C>G), K764M (2431A>T 2432A>G), S766T (2436T>A 2438A>T), P769T (2445C>A), K770_E771del (2448_2453delAAGGAA), I774V (2460A>G 2462C>A), K775Q (2463A>C), I776S (2466A>T 2467T>C 2468T>G), C777L (2469T>C 2470G>T), H779D (2475C>G 2477T>C), K780S (2478A>T 2479A>C 2480A>C), I781T (2482T>C 2483C>A), V782I (2484G>A), E783G (2488A>G 2489A>G), Y784R (2490T>A 2491A>G 2492T>A), I785M (2495C>G), K786Q (2496A>C 2498G>A), D787E (2501T>A), T788M (2503C>T 2504A>G), W790N (2508T>A 2509G>A 2510G>T), S791D (2511A>G 2512G>A 2513C>T), A795I (2523G>A 2524C>T 2525A>T), R797K (2530G>A), I806L (2556A>C), M807I (2561G>A), V812T (2574G>A 2575T>C), F813M (2577T>A 2579T>G), G815A (2584G>C), I816V (2586A>G 2588T>A), L820I (2598T>A 2600G>T), A821S (2601G>T 2603G>T), Q824K (2610C>A 2612G>A), N826S (2617A>G), S827L (2620C>T 2621T>G), F828I (2622T>A 2624C>T), N830T (2629A>C), A831S (2631G>T 2633A>C), S832Q (2634T>C 2635C>A), R834K (2641G>A), N852R (2695A>G 2696T>A), N852_K853insS (2696_2697insTCC), K853H (2697A>C 2699G>T), L854V (2700C>G 2702C>A), R855S (2703A>T 2704G>C), N856K (2708T>G), L857M (2709C>A 2711T>G), V859F (2715G>T), G860S (2718G>A 2720T>C), S865P (2733T>C), L866N (2736T>A 2737T>A 2738G>T), A867D (2740C>A 2741A>T), Q868_D869insG (2744_2745insGGA), D869K (2745G>A 2747T>A), A872S (2754G>T 2756T>C), E873R (2757G>A 2758A>G 2759A>G), Q879E (2775C>G 2777G>A), K882A (2784A>G 2785A>C 2786G>C), L890V (2808C>G 2810T>G), G891A (2812G>C 2813T>A), G904E (2851G>A 2852T>A), H914T (2880C>A 2881A>C 2882C>G), N915Q (2883A>C 2885T>G), A916L (2886G>C 2887C>T 2888T>C), T923S (2907A>T), A927E (2920C>A 2921C>A), N928H (2922A>C), R930V (2928C>G 2929G>T 2930A>C), S931C (2932C>G 2933T>C), K932Q (2934A>C 2936A>G), R938K (2953G>A), E939H (2955G>C 2957A>T), S940A (2958T>G 2960A>C), R941K (2962G>A), I942F (2964A>T), I943L (2967A>T 2969T>G), E944P (2970G>C 2971A>C 2972G>T), G945E (2974G>A 2975C>A), I948V (2982A>G 2984C>T), I949V (2985A>G 2987T>A), D955C (3003G>T 3004A>G), P961V (3021C>G 3022C>T 3023T>C), P968A (3042C>G 3044A>T), R969E (3045A>G 3046G>A), E972A (3055A>C 3056G>A), Q980E (3078C>G), C984V (3090T>G 3091G>T 3092T>C), T990K (3109C>A), R993E (3117C>G 3118G>A 3119T>G), L994K (3120C>A 3121T>A 3122T>A), G995Q (3123G>C 3124G>A 3125C>G), E998D (3134G>C), M1002Y (3144A>T 3145T>A 3146G>T), P1003S (3147C>T), V1012E (3175T>A 3176T>G), E1015T (3183G>A 3184A>C), E1017R (3189G>A 3190A>G 3191A>G), L1018M (3192C>A), N1019S (3196A>G), S1020T (3198T>A 3200C>A), S1021N (3201T>A 3202C>A 3203C>T), K1024H (3210A>C 3212G>C), K1024_I1025insS (3212_3213insTCA), C1026S (3216T>A 3218C>T), S1027A (3219T>G), Q1029E (3225C>G), S1030A (3228T>G 3230T>C), I1035V (3243A>G 3245A>C), G1037S (3249G>A 3251G>C), V1051I (3291G>A 3293T>C), V1057C (3309G>T 3310T>G 3311G>C), G1068E (3343G>A 3344C>A), V1069A (3346T>C 3347A>T), A1071T (3351G>A 3353T>A), R1073L (3357C>T 3358G>T 3359T>G), I1076V (3366A>G 3368C>A), S1077A (3369T>G 3371T>A), I1083V (3387A>G 3389C>A), E1086D (3398A>C), E1089Q (3405G>C 3407G>A), P1092A (3414C>G 3416A>T), T1093Q (3417A>C 3418C>A 3419T>A), A1098M (3432G>A 3433C>T 3434A>G), I1105L (3453A>T 3455T>G), S1108H (3462T>C 3463C>A 3464A>T), I1109V (3465A>G), Q1115A (3483C>G 3484A>C 3485G>T), E1118D (3494G>C), N1120K (3500C>G), Y1123A (3507T>G 3508A>C 3509C>T), Y1124F (3511A>T 3512C>T), L1132M (3534T>A), Q1146P (3577A>C), P1153E (3597C>G 3598C>A 3599T>A), E1155Q (3603G>C 3605G>A), I1160L (3618A>C 3620T>G), T1162S (3624A>T 3626A>T), H1164K (3630C>A 3632T>A), V1178I (3672G>A 3674A>T), M1179E (3675A>G 3676T>A), C1186S (3697G>C 3698T>A), G1187N (3699G>A 3700G>A 3701A>T), D1189E (3707T>G), D1191S (3711G>A 3712A>G), K1194R (3720A>C 3721A>G 3722A>G), L1195A (3723C>G 3724T>C 3725A>T), N1199G (3735A>G 3736A>G), C1202S (3745G>C 3746C>T), S1203A (3747T>G 3749G>C), L1204I (3750C>A 3752G>T), I1207V (3759A>G 3761C>T), M1208L (3762A>C 3764G>C), R1209K (3765C>A 3766G>A 3767T>G), K1212R (3775A>G), I1213V (3777A>G 3779C>G), R1217I (3789C>A 3790G>T 3791T>C), C1219T (3795T>A 3796G>C 3797T>A), E1220T (3798G>A 3799A>C), D1222C (3804G>T 3805A>G), V1225I (3813G>A 3815A>C), I1230V (3828A>G 3830A>T), C1233F (3838G>T), Y1235F (3844A>T), Y1236F (3847A>T), G1238S (3852G>T 3853G>C), M1252L (3894A>C), M1254L (3900A>T), M1256T (3907T>C 3908G>C), Q1263E (3927C>G), S1264A (3930T>G 3932C>A), T1265Q (3933A>C 3934C>A 3935T>G), G1266E (3937G>A 3938C>G), E1267S (3939G>A 3940A>G 3941A>T), G1268T (3942G>A 3943G>C 3944A>C), E1269Q (3945G>C 3947G>A), E1272L (3954G>C 3955A>T), D1276N (3966G>A 3968C>T), T1277S (3969A>T), E1281Q (3981G>C), Q1289E (4005C>G 4007G>A), I1294N (4021T>A), H1295Y (4023C>T), D1297E (4031T>A), L1298M (4032C>A), E1312D (4076G>C), R1318K (4093G>A 4094A>G), T1344C (4170A>T 4171C>G 4172A>T), L1348A (4182T>G 4183T>C 4184G>A), A1351V (4192C>T), I1352L (4194A>C 4196A>G), D1359N (4215G>A), T1361I (4222C>T 4223A>T), S1363H (4227T>C 4228C>A), L1367T (4239T>A 4240T>C 4241G>C), G1371Q (4251G>C 4252G>A), A1374S (4260G>A 4261C>G), V1393I (4317G>A 4319T>A), E1405A (4354A>C), K1410A (4368A>G 4369A>C 4370G>T), P1414E (4380C>G 4381C>A 4382T>G), V1437L (4449G>C 4451G>C), A1477K (4569G>A 4570C>A 4571T>A), R1479K (4576G>A), I1481L (4581A>T 4583A>G), P1485S (4593C>T), Q1490D (4608C>G 4610G>C), L1506M (4656T>A 4658A>G), S1512E (4674T>G 4675C>A 4676T>G), N1515A (4683A>G 4684A>C 4685T>A), H1518Q (4694T>A), L1522Q (4705T>A 4706A>G), S1534N (4740T>A 4741C>A 4742G>C), D1536N (4746G>A), E1539D (4757G>T), Q1541K (4761C>A), H1563Q (4829C>A), M1564L (4830A>C 4832G>A), I1573V (4857A>G 4859T>G), P1583V (4887C>G 4888C>T 4889A>T), T1588I (4903C>T 4904T>C), Q1589T (4905C>A 4906A>C 4907G>T), M1592Y (4914A>T 4915T>A 4916G>T), I1599V (4935A>G), H1604Q (4952T>G), R1607Q (4960G>A 4961T>G), A1609V (4966C>T), K1612T (4975A>C 4976G>A), V1614T (4980G>A 4981T>C 4982C>A), L1615D (4983T>G 4984T>A 4985A>C), D1616N (4986G>A), T1619K (4996C>A 4997G>A), L1621I (5001T>A 5003G>A), D1622M (5004G>A 5005A>T 5006T>G), E1623Q (5007G>C 5009A>G), F1626L (5018C>G), R1627P (5019A>C 5020G>C), A1628I (5022G>A 5023C>T 5024T>C), I1630H (5028A>C 5029T>A 5030C>T), L1635A (5043T>G 5044T>C 5045G>T), V1637Q (5049G>C 5050T>A 5051G>A), Q1639S (5055C>A 5056A>G 5057G>T), I1642L (5064A>C 5066T>A), A1643Q (5067G>C 5068C>A), K1644A (5070A>G 5071A>C 5072G>C), S1650N (5089G>A 5090T>C), A1651I (5091G>A 5092C>T 5093C>A), L1652G (5094C>G 5095T>G), T1653A (5097A>G 5099A>T), I1654L (5100A>C 5102T>G), C1655L (5103T>C 5104G>T 5105T>G), M1657L (5109A>C), A1658S (5112G>A 5113C>G 5114A>C), A1660D (5119C>A 5120A>T), S1663T (5127T>A 5129C>T), T1665V (5133A>G 5134C>T), D1668P (5142G>C 5143A>C 5144C>A), A1669E (5146C>A), E1671K (5151G>A 5153G>A), T1674E (5160A>G 5161C>A), N1675D (5163A>G), E1677S (5169G>T 5170A>C 5171G>A), R1678Q (5173G>A 5174T>A), D1679G (5176A>G), L1680V (5178C>G 5180G>C), N1681H (5181A>C), W1682E (5184T>G 5185G>A 5186G>A), A1684C (5190G>T 5191C>G 5192A>T), P1685S (5193C>T 5195A>T), N1686G (5196A>G 5197A>G 5198C>A), I1687V (5199A>G), G1688F (5202G>T 5203G>T 5204T>C), A1692S (5214G>T 5216A>G), A1695E (5224C>A), A1696V (5227C>T 5228C>T), H1697N (5229C>A 5231C>T), Y1698H (5232T>C 5234C>T), T1699K (5236C>A 5237G>A), G1703D (5248G>A 5249G>C), Y1704T (5250T>A 5251A>C 5252C>A), I1706L (5256A>C), S1715T (5283T>A 5285C>A), N1716T (5287A>C 5288T>A), S1737A (5349T>G), M1740K (5359T>A), A1742N (5364G>A 5365C>A 5366C>T), K1744Q (5370A>C 5372G>A), N1745S (5374A>G), V1746R (5376G>C 5377T>G 5378G>C), A1748S (5382G>T), G1749M (5385G>A 5386G>T 5387A>G), H1751I (5391C>A 5392A>T), V1752G (5395T>G), A1753T (5397G>A 5399A>C), T1754Q (5400A>C 5401C>A 5402C>A), N1755K (5405T>A), V1759I (5415G>A 5417G>T), G1767S (5439G>A 5441C>T), M1774F (5460A>T 5462G>T), A1777S (5469G>T), R1779N (5476G>A 5477G>T), C1782R (5484T>A 5486T>A), V1783A (5488T>C 5489G>T), D1785Q (5493G>C 5495C>A), I1786L (5496A>C 5498A>T), I1793L (5517A>T 5519T>G), G1794A (5521G>C 5522A>G), N1795G (5523A>G 5524A>G 5525T>C), V1796L (5526G>T), T1801Q (5541A>C 5542C>A 5543C>G), V1807I (5559G>A 5561G>T), Q1812T (5574C>A 5575A>C), C1813_V1814insV (5579_5580insGTC), P1815Q (5584C>A 5585T>G), H1816K (5586C>A 5588T>A), G1817S (5589G>A 5591A>T), D1818C (5592G>T 5593A>G), V1820T (5598G>A 5599T>C 5600G>A), S1823A (5607A>G 5608G>C), G1824S (5610G>A), A1825G (5614C>G 5615A>G), S1826del (5616_5618delTCG), L1827R (5620T>G), H1828D (5622C>G 5624C>T), S1829G (5625A>G 5627T>A), S1830M (5629G>T 5630T>G), F1831L (5633T>G), M1832N (5635T>A 5636G>C), Q1834H (5642A>T), Q1834_G1835insHA (5642_5643insCATGCT), G1835A (5644G>C 5645G>T), F1836Y (5647T>A), E1837A (5650A>C 5651A>C), I1844V (5670A>G 5672A>G), H1845G (5673C>G 5674A>G 5675T>C), G1846C (5676G>T), D1847S (5679G>T 5680A>C 5681T>A), L1848I (5682T>A 5684A>T), A1849S (5685G>T), K1850R (5689A>G), Q1851K (5691C>A), T1852M (5695C>T 5696A>G), M1854V (5700A>G), M1856L (5706A>T), V1859T (5715G>A 5716T>C 5717G>T), H1863M (5727C>A 5728A>T 5729T>G), G1867A (5740G>C), N1868D (5742A>G), F1869C (5746T>G 5747T>C), Q1872R (5754C>A 5755A>G 5756G>A), D1873N (5757G>A), E1875D (5765G>T), L1878I (5772C>A 5774A>C), H1879K (5775C>A 5777C>A), Y1880A (5778T>G 5779A>C 5780T>C), Q1883C (5787C>T 5788A>G 5789G>C), G1884S (5790G>T 5791G>C 5792A>T), E1887A (5800A>C 5801G>A), L1888C (5803T>G 5804A>T), I1889V (5805A>G 5807T>G), Q1890A (5808C>G 5809A>C 5810A>T), K1891R (5811A>C 5812A>G 5813G>T), N1892G (5814A>G 5815A>G 5816T>G), T1893C (5817A>T 5818C>G 5819A>T), L1895I (5823C>A), K1900E (5838A>G 5840A>G), E1901N (5841G>A 5843G>T), L1902F (5844C>T 5846G>T), S1903N (5848G>A), M1904A (5850A>G 5851T>C 5852G>A), E1905Q (5853G>C), N1906I (5857A>T 5858T>A), V1907A (5860T>C 5861G>T), Q1908T (5862C>A 5863A>C 5864A>C), Q1909R (5865C>A 5866A>G 5867A>G), I1911V (5871A>G), V1914L (5880G>C), L1916S (5886C>A 5887T>G), F1919V (5895T>G 5897T>G), P1921I (5901C>A 5902C>T 5903A>C), K1922T (5905A>C), E1923A (5908A>C 5909A>C), T1925G (5913A>G 5914C>G), I1926H (5916A>C 5917T>A 5918T>C), Q1927K (5919C>A 5921G>A), K1928M (5923A>T), K1930V (5928A>G 5929A>T 5930A>G), N1931Q (5931A>C 5933T>A), F1932T (5934T>A 5935T>C), V1933L (5937G>C 5939T>G), D1934S (5940G>A 5941A>G 5942T>C), A1935N (5943G>A 5944C>A), E1936Q (5946G>C), C1937W (5951T>G), L1938Q (5953T>A), T1939S (5955A>T 5957A>C), F1940H (5958T>C 5959T>A), K1941E (5961A>G), A1942Q (5964G>C 5965C>A), R1943S (5969G>T), S1944C (5970A>T 5972C>T), F1945V (5973T>G 5975T>G), R1946K (5977G>A), H1947R (5979C>A 5980A>G 5981C>A), V1949Y (5985G>T 5986T>A), K1952R (5994A>C 5995A>G 5996G>A), K1953N (5999A>T), S1957N (6009T>A 6010C>A), T1958A (6012A>G), W1959I (6015T>A 6016G>T 6017G>A), K1960T (6019A>C), M1961_C1962insY (6023_6024insTAC), C1962R (6024T>C), E1964D (6032A>T), S1965A (6033A>G 6034G>C 6035T>C), I1966K (6037T>A 6038T>A), V1967D (6040T>A), S1969I (6045T>A 6046C>T), R1970V (6048A>G 6049G>T 6050G>A), A1972del (6054_6056delGCA), L1975V (6063C>G 6065A>G), P1977L (6069C>T 6070C>T 6071T>G), L1980P (6078T>C 6079T>C 6080G>A), S1983M (6087T>A 6088C>T 6089C>G), K1986R (6097A>G 6098A>G)

| | Begin | End | Coverage | Score | Concordance | Matches | Identities | I/D/M/F* | Stop Codons |
|---|---|---|---|---|---|---|---|---|---|
| **NT** | **24** | **6107** | **84.2%** | **3417** | **28.3%** | **6063 (98.8%)** | **3936 (64.1%)** | **54/21** | |

ATG1CTG (141A>C), GCT2GTT (145C>T), TTT3TTC (149T>C), TCA4CAC (150T>C 151C>A 152A>C), AAG5AAT (155G>T), ATG6ACC (157T>C 158G>C), TTC7ACC (159T>A 160T>C), TCC8ATT (162T>A 163C>T 164C>T), AAG9CAA (165A>C 167G>A), GTT10ATG (168G>A 170T>G), GCT11GCA (173T>A), GGT12del (174_176delGGT), GTT13TTT (177G>T), GAT14TCA (180G>T 181A>C 182T>A), GAC15AAA (183G>A 185C>A), GCC16ATC (186G>A 187C>T), AGT17GTT (189A>G 190G>T), TCC20GCT (198T>G 200C>T), ACC21ACT (203C>T), ACT22CCT (204A>C), TCA23TCT (209A>T), TTC24TTT (212C>T), TTT25CTT (213T>C), GGT26TCT (216G>T 217G>C), GAA27GAG (221A>G), CTT28TTG (222C>T 224T>G), TCA29TCT (227A>T), AAT30CAG (228A>C 230T>G), TCT31ACA (231T>A 233T>A), ATT32GTG (234A>G 236T>G), TCC33TCA (239C>A), AAA34TCA (240A>T 241A>C), TTT35CTG (243T>C 245T>G), TCA36TCC (248A>C), AAA37AAG (251A>G), GCG38ACA (252G>A 254G>A), GTT39ATG (255G>A 257T>G), AGT40AAG (259G>A 260T>G), AAT41AGT (262A>G), ATT42ATG (266T>G), ACC43AGT (268C>G 269C>T), ACT44AAA (271C>A 272T>A), ATG45ATA (275G>A), TCA46TCT (278A>T), CAG47GAG (279C>G), AAG48AAA (284G>A), ATT49TTC (285A>T 287T>C), GAT51GGC (292A>G 293T>C), CTT53ATT (297C>A), GAG54GAA (302G>A), GAT55GAA (305T>A), CTG56TTA (306C>T 308G>A), CCA58CCT (314A>T), TCT59TCA (317T>A), GTG60ATT (318G>A 320G>T), ACT61GTT (321A>G 322C>T), GAT62AAA (324G>A 326T>A), GTG63ACT (327G>A 328T>C 329G>T), TCC64GCA (330T>G 332C>A), ACA65GAT (333A>G 334C>A 335A>T), TCT66TCC (338T>C), GTG68AAT (342G>A 343T>A 344G>T), GCC69AGT (345G>A 346C>G 347C>T), ACT70GTT (348A>G 349C>T), TCA73TCT (359A>T), CTT74GTA (360C>G 362T>A), AAT75GAA (363A>G 365T>A), GTT77TTA (369G>T 371T>A), TTG78TTA (374G>A), GAC79GGG (376A>G 377C>G), GGT82GCC (385G>C 386T>C), GCA83ACG (387G>A 389A>G), CTT84GTT (390C>G), ATA85ATT (395A>T), GAT86GAA (398T>A), CCA87CCT (401A>T), CTT89GTT (405C>G), GCG91GGT (412C>G 413G>T), TAC92ATC (414T>A 415A>T), TCT93AAT (417T>A 418C>A), TTG95TTC (425G>C), TCC96AAA (426T>A 427C>A 428C>A), TAC99TAT (437C>T), AAA100AGA (439A>G), TCT101ACC (441T>A 443T>C), ATC102ATA (446C>A), AAG103ATT (448A>T 449G>T), TTA107CTG (459T>C 461A>G), TTG109GTT (465T>G 467G>T), TTT110GTT (468T>G), GAG111TTG (471G>T 472A>T), AGT112GGC (474A>G 476T>C), TTG113ATC (477T>A 479G>C), TCC114AAA (480T>A 481C>A 482C>A), AAA116AAT (488A>T), GTT117TTT (489G>T), AAA118ATT (493A>T 494A>T), TTA119AGA (495T>A 496T>G), TTT121TAT (502T>A), GCT122TCA (504G>T 506T>A), GTG124CTG (510G>C), TTA125TTG (515A>G), AAT126GAA (516A>G 518T>A), AAA127ACA (520A>C), AGT128ACA (523G>C 524T>A), GAA129GGT (526A>G 527A>T), GAT130GAA (530T>A), GTA131TAT (531G>T 532T>A 533A>T), ATA133GTT (537A>G 539A>T), GTT134GTA (542T>A), GTA135GTT (545A>T), TAT136ATA (546T>A 547A>T 548T>A), GGT137GCT (550G>C), CTT139CTA (557T>A), GTA140GTC (560A>C), GCT142TCC (564G>T 566T>C), ATA143CTT (567A>C 569A>T), ACA144TCA (570A>T), TTG145CTA (573T>C 575G>A), TTA146TTG (578A>G), GTG147TTA (579G>T 581G>A), CTT148TTA (582C>T 584T>A), CTA149CAA (586T>A), TTT150ATA (588T>A 590T>A), GTT151ATC (591G>A 593T>C), AGT154GTA (600A>G 601G>T 602T>A), GTT157GTA (611T>A), GAG158GAA (614G>A), GGT159ATG (615G>A 616G>T 617T>G), TCT161AGA (621T>A 622C>G 623T>A), AAC162AGT (625A>G 626C>T), ACT163GTA (627A>G 628C>T 629T>A), GTC164GTT (632C>T), AAA165TGT (633A>T 634A>G 635A>T), ATC167ATA (641C>A), TTG169CTA (645T>C 647G>A), ATA170GTT (648A>G 650A>T), GTC171GTA (653C>A), GGA172GGT (656A>T), ATG174TTC (660A>T 662G>C), TTT175TTC (665T>C), TCA176TCT (668A>T), AGT177TCT (669A>T 670G>C), CTA178CTG (674A>G), TAC179TAT (677C>T), TTA181ATT (681T>A 683A>T), GAT182AGT (684G>A 685A>G), CCA185CCC (695A>C), AAA186CTC (696A>C 697A>T 698A>C), GCC188TCT (702G>T 704C>T), GAA189GAG (707A>G), CGA190AAA (708C>A 709G>A), TTC191TTT (713C>T), ACC192TCT (714A>T 716C>T), ATG193CAG (717A>C 718T>A), GTG194ATG (720G>A), GCC195GCA (725C>A), CAA196CAG (728A>G), AGT198TCT (732A>T 733G>C), ATG199GTG (735A>G), CTT200CTA (740T>A), GGG202GGT (746G>T), GAA203GAG (749A>G), TCA204AGC (750T>A 751C>G 752A>C), GTT205TTG (753G>T 755T>G), TCA206AAT (756T>A 757C>A 758A>T), ACG210ACC (770G>C), TCT211AGT (771T>A 772C>G), CAG212CAA (776G>A), ATG214CTG (780A>C), TCC215TCT (785C>T), ACA216TCT (786A>T 788A>T), ATA217TTG (789A>T 791A>G), TTG218ATA (792T>A 794G>A), GCT219GCA (797T>A), GGG221GGC (803G>C), ATT222ATC (806T>C), TCG223TCA (809G>A), GTG226GTA (818G>A), TTC227TTT (821C>T), ATT228GTG (822A>G 824T>G), GTG230GTA (830G>A), AGA233AGG (839A>G), TTG237CTT (849T>C 851G>T), AGC238ACA (853G>C 854C>A), CCA240CCG (860A>G), TTG241CTA (861T>C 863G>A), CTC245TTG (873C>T 875C>G), TCG247TCA (881G>A), ACA248GCG (882A>G 884A>G), GGA249GGC (887A>C), GGT251GGA (893T>A), CAG253CAA (899G>A), CTA258CTC (914A>C), ACT260CAA (918A>C 919C>A 920T>A), CTT261TTG (921C>T 923T>G), TAC262TAT (926C>T), CGT263AAG (927C>A 928G>A 929T>G), AAT264AAC (932T>C), AAA266AAG (938A>G), ACC269ACA (947C>A), TCA274TCT (962A>T), GTT276GTG (968T>G), CTT277TTG (969C>T 971T>G), ATA279ATT (977A>T), ATA280GTT (978A>G 980A>T), GTC281GCT (982T>C 983C>T), GGC282GAT (985G>A 986C>T), ACC283ATA (988C>T 989C>A), TTT284TTC (992T>C), GGA285GGG (995A>G), TTC286TTT (998C>T), AAA287AAG (1001A>G), AAT288AAC (1004T>C), GTG290GTT (1010G>T), TTG291CTG (1011T>C), TCA292TCT (1016A>T), GCT293GCA (1019T>A), ATA294ATT (1022A>T), ACG297ACA (1031G>A), CTG298ATT (1032C>A 1034G>T), CTG302CTC (1046G>C), TTC303TTT (1049C>T), CAG307ACT (1059C>A 1060A>C 1061G>T), GAG308GAA (1064G>A), GAT310TCT (1068G>T 1069A>C), TGT313GCT (1077T>G 1078G>C), ACG317ACA (1091G>A), CGC318AGG (1092C>A 1094C>G), GAG320GAA (1100G>A), GCC323GCT (1109C>T), AAG325CCT (1113A>C 1114A>C 1115G>T), TTC327TTT (1121C>T), CCT328TCA (1122C>T 1124T>A), ACC329ATC (1126C>T), CTG334CTT (1142G>T), AGG335AAA (1144G>A 1145G>A), GAA336GAG (1148A>G), GAA337CAG (1149G>C 1151A>G), GCT338GCA (1154T>A), ATC341ATA (1163C>A), GCT343GCC (1169T>C), TAC344TAT (1172C>T), GCA346GCT (1178A>T), CCA349CCT (1187A>T), GTT350GTG (1190T>G), TTC353TTT (1199C>T), ATG354TTG (1200A>T), AGC355AGT (1205C>T), AGG357AGA (1211G>A), AGT359TCT (1215A>T 1216G>C), GCC361ACA (1221G>A 1223C>A), GCG363GTG (1228C>T), CAG364CAA (1232G>A), TTA365TTG (1235A>G), GAA366GAT (1238A>T), GTT368CTC (1242G>C 1244T>C), TAT369ATA (1245T>A 1246A>T 1247T>A), GCT370GCA (1250T>A), AGT372AAC (1255G>A 1256T>C), AGG374CAA (1260A>C 1261G>A 1262G>A), CAC375CAT (1265C>T), ATG376ACA (1267T>C 1268G>A), GGC377GGA (1271C>A), GGC379GGT (1277C>T), TAT381TAC (1283T>C), CGT382AGA (1284C>A 1286T>A), CCT385CCC (1295T>C), GTA388GTT (1304A>T), TTC391TTT (1313C>T), GGG392GGT (1316G>T), TCC393GAG (1317T>G 1318C>A 1319C>G), GGG395GGA (1325G>A), TGT396TGC (1328T>C), GGT397GGC (1331T>C), TCT399TCC (1337T>C), ACA400ACG (1340A>G), TCC401AGC (1341T>A 1342C>G), CGC403CGA (1349C>A), TTA404CTG (1350T>C 1352A>G), TTT405TTC (1355T>C), ATT406ATC (1358T>C), GTG409GTA (1367G>A), GAC411GAT (1373C>T), AGA412GCC (1374A>G 1375G>C 1376A>C), GGT414CAA (1380G>C 1381G>A 1382T>A), CCA416CCC (1388A>C), AAA417AAG (1391A>G), CTC418CTG (1394C>G), AAT419AAC (1397T>C), CGG420AGG (1398C>A), CTA421TTG (1401C>T 1403A>G), GTA424GTT (1412A>T), AGG427CGC (1419A>C 1421G>C), GCA429GCT (1427A>T), TAT430TAC (1430T>C), TCC432TCA (1436C>A), AAC433AAT (1439C>T), CTG441CTC (1463G>C), GAC444GAT (1472C>T), GGA446GGT (1478A>T), TTG448CTG (1482T>C), CGA449CGG (1487A>G), GGG451GGA (1493G>A), GGG453GGC (1499G>C), CAG454CAA (1502G>A), CAA456CAG (1508A>G), GAT457GAC (1511T>C), AAA459AAG (1517A>G), GAT460GAC (1520T>C), CTG461CTC (1523G>C), GAT463GAC (1529T>C), TCA466TCC (1538A>C), CCC471CCA (1553C>A), TTA472TTG (1556A>G), CCA473CCC (1559A>C), GCC475GCT (1565C>T), GTT477GTA (1571T>A), GAG478GAA (1574G>A), GGC481GGG (1583C>G), AGG482CGG (1584A>C), ACC485ACA (1595C>A), TCC486TCA (1598C>A), TAT488TAC (1604T>C), ACA492ACC (1616A>C), AAT494AAC (1622T>C), CTG495TTG (1623C>T), ATC496GTT (1626A>G 1628C>T), GCT498GCA (1634T>A), CCT499CCA (1637T>A), GCC500GCA (1640C>A), TGT502TGC (1646T>C), GGT503GGG (1649T>G), CTT504CTG (1652T>G), CCA507CCT (1661A>T), GAG511GAA (1673G>A), CGG512AGG (1674C>A), AGA513AGG (1679A>G), AGG514CGG (1680A>C), ATC516ATT (1688C>T), CTG517TTG (1689C>T), GAG519GAA (1697G>A), TGT520TGC (1700T>C), ATG521CGG (1701A>C 1702T>G), AAG522AAA (1706G>A), GAG523GTG (1708A>T), GGC524GGG (1712C>G), GAG525GAA (1715G>A), TTT526TTC (1718T>C), TCC527TCA (1721C>A), ACT528ACA (1724T>A), GAA529GAG (1727A>G), GAT530AAC (1728G>A 1730T>C), CCT531CCA (1733T>A), ACG532ACA (1736G>A), GGA533GCA (1738G>C), AGA536AGC (1748A>C), TTT537TTC (1751T>C), ATA539ATC (1757A>C), GTG540ATT (1758G>A 1760G>T), GAG541GGA (1762A>G 1763G>A), AGA543AGG (1769A>G), AGA544AGG (1772A>G), GCT547GCC (1781T>C), ATC548ATA (1784C>A), ACT549ACA (1787T>A), AGG551AGA (1793G>A), AAT552GAC (1794A>G 1796T>C), TTG553CTC (1797T>C 1799G>C), ACC554ACT (1802C>T), AGT556TCT (1806A>T 1807G>C), GAT557GAC (1811T>C), CTT558TTG (1812C>T 1814T>G), GAG560GAA (1820G>A), GTA563GTG (1829A>G), AAA565AAG (1835A>G), TGT566TGC (1838T>C), CAG567AAG (1839C>A), GTA568GTC (1844A>C), CAT569CAC (1847T>C), GCA570CAC (1848G>C 1849C>A 1850A>C), GTA572ATT (1854G>A 1856A>T), TCA573GCA (1857T>G), AAG574ACC (1861A>C 1862G>C), TTG576CTC (1866T>C 1868G>C), TTT577TAC (1870T>A 1871T>C), GAA578AAC (1872G>A 1874A>C), GCT579CAG (1875G>C 1876C>A 1877T>G), GAA580TCA (1878G>T 1879A>C), TCT581ACA (1881T>A 1883T>A), GGC582GGT (1886C>T), ATA583TGT (1887A>T 1888T>G 1889A>T), AGT584GAA (1890A>G 1891G>A 1892T>A), CCT585GAG (1893C>G 1894C>A 1895T>G), AAG586AAA (1898G>A), ATA587GTT (1899A>G 1901A>T), GCT588GCC (1904T>C), GTT590GTG (1910T>G), CAA591GAA (1911C>G), GCA594GTT (1921C>T 1922A>T), GAT595GAA (1925T>A), GAT596GAA (1928T>A), ATA598ATG (1934A>G), GCG599ACA (1935G>A 1937G>A), TCT600TCC (1940T>C), GTG601TTT (1941G>T 1943G>T), GAT602CAT (1944G>C), GGG603GAT (1948G>A 1949G>T), AGA605CAA (1953A>C 1954G>A), TTG606ATA (1956T>A 1958G>A), CGC607AGC (1959C>A), ACC608TTT (1962A>T 1963C>T 1964C>T), AAG609GAC (1965A>G 1967G>C), CAA610GAA (1968C>G), GAT611GTT (1972A>T), GAA612AGC (1974G>A 1975A>G 1976A>C), CCA613CCC (1979A>C), ATA614AGT (1981T>G 1982A>T), TTG615GTT (1983T>G 1985G>T), GTG616_CCC617del (1986_1991delGTGCCC), ACT618ACA (1994T>A), GTG620CTC (1998G>C 2000G>C), AGT621TCT (2001A>T 2002G>C), GAA622GAG (2006A>G), AAT623GAT (2007A>G), ATC627GGA (2019A>G 2020T>G 2021C>A), GCA629TCA (2025G>T), AGA630CAG (2028A>C 2029G>A 2030A>G), GAG631GAA (2033G>A), CTC632CTA (2036C>A), ACG633ACT (2039G>T), GTA634GTG (2042A>G), GAG635GAA (2045G>A), TTG637CTG (2049T>C), AAA638AAG (2054A>G), GCA640GCT (2060A>T), TAC641TAT (2063C>T), CAG642CAA (2066G>A), GGC643GGA (2069C>A), AGT644GCA (2070A>G 2071G>C 2072T>A), TTG645TTA (2075G>A), GAC646AAC (2076G>A), CCT647CCA (2081T>A), GAG648TTT (2082G>T 2083A>T 2084G>T), CTT650TAT (2088C>T 2089T>A), CCT652CCC (2096T>C), GAC654GAT (2102C>T), CAT655GCT (2103C>G 2104A>C), CAT656TCA (2106C>T 2107A>C 2108T>A), GCC659GCA (2117C>A), ATG660ATT (2120G>T), TAT661TTT (2122A>T), TCT663GTC (2127T>G 2128C>T 2129T>C), CTT664CTG (2132T>G), GAT665TCA (2133G>T 2134A>C 2135T>A), GAT666AGT (2136G>A 2137A>G), CAT668GCC (2142C>G 2143A>C 2144T>C), ATT671AGC (2152T>G 2153T>C), TTC672TTT (2156C>T), AAC673ACT (2158A>C 2159C>T), AAA674CGC (2160A>C 2161A>G 2162A>C), AGA676GTA (2166A>G 2167G>T), GTC677AAT (2169G>A 2170T>A 2171C>T), AAC678GAG (2172A>G 2174C>G), TTG680ATT (2178T>A 2180G>T), TAT681TAC (2183T>C), AGA682GAG (2184A>G 2185G>A 2186A>G), GGA683GGG (2189A>G), GCA684GCT (2192A>T), GAT685TCG (2193G>T 2194A>C 2195T>G), GCT686ACT (2196G>A), GAA687GCA (2200A>C), CAG688GGA (2202C>G 2203A>G 2204G>A), CGA690GCC (2208C>G 2209G>C 2210A>C), GTA693ATC (2217G>A 2219A>C), CAA694AAG (2220C>A 2222A>G), AAC695GCC (2223A>G 2224A>C), ATT696ATC (2228T>C), CCT697CCA (2231T>A), ATA701GTC (2241A>G 2243A>C), AAG705AAA (2255G>A), AGT706AGC (2258T>C), GCA710AGT (2268G>A 2269C>G 2270A>T), TCA711ACA (2271T>A), ATC713GTG (2277A>G 2279C>G), TCT714TCA (2282T>A), GAG715GAA (2285G>A), AAG716GCT (2286A>G 2287A>C 2288G>T), AAA717CAC (2289A>C 2291A>C), CTA718ATG (2292C>A 2294A>G), GCA719GAT (2296C>A 2297A>T), GTG720GTA (2300G>A), CAG721CAA (2303G>A), ACA722AGG (2305C>G 2306A>G), GAG723ATG (2307G>A 2308A>T), ATG724TTG (2310A>T), AGG725TAC (2313A>T 2314G>A 2315G>C), ACT726AAT (2317C>A), GGA727GGC (2321A>C), TTT728AGA (2322T>A 2323T>G 2324T>A), TCT731AGA (2331T>A 2332C>G 2333T>A), TGC732GCA (2334T>G 2335G>C 2336C>A), ATT733CTG (2337A>C 2339T>G), GAT734GAA (2342T>A), GCA735GCT (2345A>T), GTT737GCT (2350T>C), ACC739ACT (2357C>T), CTC740TTA (2358C>T 2360C>A), ATA741ATG (2363A>G), ATC743ATT (2369C>T), GAG744GAA (2372G>A), CAG745CAA (2375G>A), CCA747CCT (2381A>T), TTT749CAT (2385T>C 2386T>A), GTT750GTC (2390T>C), CAG751CAA (2393G>A), TTT752TTC (2396T>C), TTA753CTA (2397T>C), TAT754TAC (2402T>C), AGA759CGG (2415A>C 2417A>G), TGC762_AAC763insAGAAGG (2426_2427insAGAAGG), AAC763AAG (2429C>G), AAA764ATG (2431A>T 2432A>G), GTA765GTC (2435A>C), TCA766ACT (2436T>A 2438A>T), AAA767AAG (2441A>G), GAG768GAA (2444G>A), CCA769ACA (2445C>A), AAG770_GAA771del (2448_2453delAAGGAA), TCA772TCT (2456A>T), ATC774GTA (2460A>G 2462C>A), AAA775CAA (2463A>C), ATT776TCG (2466A>T 2467T>C 2468T>G), TGC777CTC (2469T>C 2470G>T), TAC778TAT (2474C>T), CAT779GAC (2475C>G 2477T>C), AAA780TCC (2478A>T 2479A>C 2480A>C), ATC781ACA (2482T>C 2483C>A), GTT782ATT (2484G>A), GAA783GGG (2488A>G 2489A>G), TAT784AGA (2490T>A 2491A>G 2492T>A), ATC785ATG (2495C>G), AAG786CAA (2496A>C 2498G>A), GAT787GAA (2501T>A), ACA788ATG (2503C>T 2504A>G), TGG790AAT (2508T>A 2509G>A 2510G>T), AGC791GAT (2511A>G 2512G>A 2513C>T), CTT792CTG (2516T>G), TAT794TAC (2522T>C), GCA795ATT (2523G>A 2524C>T 2525A>T), CTG796TTG (2526C>T), AGA797AAA (2530G>A), CTC799CTT (2537C>T), GGT803GGG (2549T>G), ATA806CTA (2556A>C), ATG807ATA (2561G>A), CTT809CTC (2567T>C), GGA811GGG (2573A>G), GTT812ACT (2574G>A 2575T>C), TTT813ATG (2577T>A 2579T>G), GGA814GGT (2582A>T), GGC815GCC (2584G>C), ATT816GTA (2586A>G 2588T>A), ACA817ACT (2591A>T), TTT819TTC (2597T>C), TTG820ATT (2598T>A 2600G>T), GCG821TCT (2601G>T 2603G>T), CAG824AAA (2610C>A 2612G>A), AGT825AGC (2615T>C), AAC826AGC (2617A>G), TCT827TTG (2620C>T 2621T>G), TTC828ATT (2622T>A 2624C>T), AAT830ACT (2629A>C), GCA831TCC (2631G>T 2633A>C), TCG832CAG (2634T>C 2635C>A), AGA834AAA (2641G>A), GTG837GTT (2651G>T), TCC841TCG (2663C>G), AGT843AGC (2669T>C), TCC845TCA (2675C>A), AGT846AGC (2678T>C), GGT851GGA (2693T>A), AAT852AGA (2695A>G 2696T>A), AAT852_AAG853insTCC (2696_2697insTCC), AAG853CAT (2697A>C 2699G>T), CTC854GTA (2700C>G 2702C>A), AGA855TCA (2703A>T 2704G>C), AAT856AAG (2708T>G), CTT857ATG (2709C>A 2711T>G), CTC858CTG (2714C>G), GTT859TTT (2715G>T), GGT860AGC (2718G>A 2720T>C), TCT863TCA (2729T>A), TCA865CCA (2733T>C), TTG866AAT (2736T>A 2737T>A 2738G>T), GCA867GAT (2740C>A 2741A>T), CAA868CAG (2744A>G), CAA868_GAT869insGGA (2744_2745insGGA), GAT869AAA (2745G>A 2747T>A), GCT872TCC (2754G>T 2756T>C), GAA873AGG (2757G>A 2758A>G 2759A>G), GGA875GGT (2765A>T), GTA877GTG (2771A>G), AAT878AAC (2774T>C), CAG879GAA (2775C>G 2777G>A), AGC880TCT (2778A>T 2779G>C 2780C>T), CTC881CTT (2783C>T), AAG882GCC (2784A>G 2785A>C 2786G>C), GTG886GTA (2798G>A), TTA888CTA (2802T>C), AGA889AGG (2807A>G), CTT890GTG (2808C>G 2810T>G), GGT891GCA (2812G>C 2813T>A), GTG894GTT (2822G>T), AGA897CGT (2829A>C 2831A>T), GGC898GGA (2834C>A), ACC899ACT (2837C>T), TGT902TGC (2846T>C), TCT903AGC (2847T>A 2848C>G 2849T>C), GGT904GAA (2851G>A 2852T>A), ACC908ACT (2864C>T), GCT910GCA (2870T>A), CAT911CAC (2873T>C), GCA912GCT (2876A>T), CAC914ACG (2880C>A 2881A>C 2882C>G), AAT915CAG (2883A>C 2885T>G), GCT916CTC (2886G>C 2887C>T 2888T>C), CGA917CGT (2891A>T), GGT919GGA (2897T>A), ACT920ACC (2900T>C), ACA923TCA (2907A>T), ATC925ATT (2915C>T), GCC927GAA (2920C>A 2921C>A), AAT928CAT (2922A>C), CGA930GTC (2928C>G 2929G>T 2930A>C), TCT931TGC (2932C>G 2933T>C), AAA932CAG (2934A>C 2936A>G), GTT933GTG (2939T>G), AAC936AAT (2948C>T), AGA938AAA (2953G>A), GAA939CAT (2955G>C 2957A>T), TCA940GCC (2958T>G 2960A>C), AGG941AAG (2962G>A), ATT942TTT (2964A>T), ATT943TTG (2967A>T 2969T>G), GAG944CCT (2970G>C 2971A>C 2972G>T), GGC945GAA (2974G>A 2975C>A), GAC947GAT (2981C>T), ATC948GTT (2982A>G 2984C>T), ATT949GTA (2985A>G 2987T>A), CTG950TTA (2988C>T 2990G>A), TTG951CTG (2991T>C), CGC952AGG (2994C>A 2996C>G), GGT954GGC (3002T>C), GAT955TGT (3003G>T 3004A>G), CCA956CCT (3008A>T), GAT957GAC (3011T>C),

| | |
|---|---|
| Codon mutations: | GGT958GGG (3014T>G), CCA959CCT (3017A>T), AAG960AAA (3020G>A), CCT961GTC (3021C>G 3022C>T 3023T>C), CGT964AGA (3030C>A 3032T>A), AAA965AAG (3035A>G), TTC967TTT (3041C>T), CCA968GCT (3042C>G 3044A>T), AGA969GAA (3045A>G 3046G>A), GAG972GCA (3055A>C 3056G>A), AAG977AAA (3071G>A), GGC978GGA (3074C>A), TCA979TCT (3077A>T), CAA980GAA (3078C>G), GGA981GGC (3083A>C), TGC983TGT (3089C>T), TGT984GTC (3090T>G 3091G>T 3092T>C), GCT986GCA (3098T>A), GTT987GTG (3101T>G), GCG988GCA (3104G>A), TCT989TCA (3107T>A), ACA990AAA (3109C>A), CCA992CCT (3116A>T), CGT993GAG (3117C>G 3118G>A 3119T>G), CTT994AAA (3120C>A 3121T>A 3122T>A), GGC995CAG (3123G>C 3124G>A 3125C>G), AAT996AAC (3128T>C), TTA997TTG (3131A>G), GAG998GAC (3134G>C), ATG1002TAT (3144A>T 3145T>A 3146G>T), CCA1003TCA (3147C>T), TCA1007TCC (3161A>C), AAG1008AAA (3164G>A), TAC1010TAT (3170C>T), GTT1012GAG (3175T>A 3176T>G), GTG1014GTC (3182G>C), GAA1015ACA (3183G>A 3184A>C), TAT1016TAC (3188T>C), GAA1017AGG (3189G>A 3190A>G 3191A>G), CTG1018ATG (3192C>A), AAC1019AGC (3196A>G), TCC1020ACA (3198T>A 3200C>A), TCC1021AAT (3201T>A 3202C>A 3203C>T), AGT1022TCA (3204A>T 3205G>C 3206T>A), AAG1024CAC (3210A>C 3212G>C), AAG1024_ATT1025insTCA (3212_3213insTCA), ATT1025ATC (3215T>C), TGC1026AGT (3216T>A 3218C>T), TCT1027GCT (3219T>G), CAA1029GAA (3225C>G), TCT1030GCC (3228T>G 3230T>C), CAC1034CAT (3242C>T), ATA1035GTC (3243A>G 3245A>C), GGG1037AGC (3249G>A 3251G>C), GAC1041GAT (3263C>T), GGC1043GGA (3269C>A), ACG1044ACT (3272G>T), CTG1047TTA (3279C>T 3281G>A), TTA1048TTG (3284A>G), AGT1050TCT (3288A>T 3289G>C), GTT1051ATC (3291G>A 3293T>C), CAG1052CAA (3296G>A), AAT1053AAC (3299T>C), CCT1056CCA (3308T>A), GTG1057TGC (3309G>T 3310T>G 3311G>C), ATC1058ATT (3314C>T), GGC1060GGT (3320C>T), ATC1061ATT (3323C>T), CAC1062CAT (3326C>T), GCA1064GCT (3332A>T), TCA1065TCC (3335A>C), GGC1068GAA (3343G>A 3344C>A), GTA1069GCT (3346T>C 3347A>T), GCT1071ACA (3351G>A 3353T>A), GAA1072GAG (3356A>G), CGT1073TTG (3357C>T 3358G>T 3359T>G), ATC1076GTA (3366A>G 3368C>A), TCT1077GCA (3369T>G 3371T>A), GCT1080GCC (3380T>C), ACA1081ACT (3383A>T), ATC1083GTA (3387A>G 3389C>A), GAA1086GAC (3398A>C), CAA1087CAG (3401A>G), CTA1088CTG (3404A>G), GAG1089CAA (3405G>C 3407G>A), CCA1092GCT (3414C>G 3416A>T), ACT1093CAA (3417A>C 3418C>A 3419T>A), CCG1095CCA (3425G>A), GTC1096GTT (3428C>T), GCA1098ATG (3432G>A 3433C>T 3434A>G), GCA1099GCC (3437A>C), GTA1101GTG (3443A>G), CGC1102AGA (3444C>A 3446C>A), GAT1104GAC (3452T>C), ATT1105TTG (3453A>T 3455T>G), CTG1106CTC (3458G>C), AAG1107AAA (3461G>A), TCA1108CAT (3462T>C 3463C>A 3464A>T), ATT1109GTT (3465A>G), AGA1110CGA (3468A>C), AGC1111TCA (3471A>T 3472G>C 3473C>A), ACA1114ACT (3482A>T), CAG1115GCT (3483C>G 3484A>C 3485G>T), CTT1116CTC (3488T>C), GAG1118GAC (3494G>C), AAC1120AAG (3500C>G), GTG1122GTT (3506G>T), TAC1123GCT (3507T>G 3508A>C 3509C>T), TAC1124TTT (3511A>T 3512C>T), CTT1125TTG (3513C>T 3515T>G), GGA1126GGC (3518A>C), ACA1127ACT (3521A>T), CCA1129CCT (3527A>T), CAG1130CAA (3530G>A), TTG1132ATG (3534T>A), GCC1133GCA (3539C>A), GCC1134GCA (3542C>A), ACA1135ACG (3545A>G), GTT1136GTG (3548T>G), CCC1137CCA (3551C>A), ACT1141ACC (3563T>C), CTG1142CTC (3566G>C), CGG1143AGG (3567C>A), AGC1145AGT (3575C>T), CAA1146CCA (3577A>C), TTG1147CTG (3579T>C), GAA1149GAG (3587A>G), TTC1151TTT (3593C>T), GGA1152GGT (3596A>T), CCT1153GAA (3597C>G 3598C>A 3599T>A), GAG1155CAA (3603G>C 3605G>A), ACA1156ACT (3608A>T), GCA1157GCT (3611A>T), CCA1158CCT (3614A>T), TCC1159TCA (3617C>A), ATT1160CTG (3618A>C 3620T>G), CTA1161CTC (3623A>C), ACA1162TCT (3624A>T 3626A>T), GTT1163GTG (3629T>G), CAT1164AAA (3630C>A 3632T>A), GAC1165GAT (3635C>T), GGT1168GGG (3644T>G), GGT1170GGA (3650T>A), GAC1172GAT (3656C>T), CCC1173CCT (3659C>T), GTG1175GTT (3665G>T), GGG1177GGC (3671G>C), GTA1178ATT (3672G>A 3674A>T), ATG1179GAG (3675A>G 3676T>A), GCT1185GCC (3695T>C), TGT1186TCA (3697G>C 3698T>A), GGA1187AAT (3699G>A 3700G>A 3701A>T), GAT1189GAG (3707T>G), GAT1190GAC (3710T>C), GAC1191AGC (3711G>A 3712A>G), GCC1193GCA (3719C>A), AAA1194CGG (3720A>C 3721A>G 3722A>G), CTA1195GCT (3723C>G 3724T>C 3725A>T), TTC1197TTT (3731C>T), AAT1199GGT (3735A>G 3736A>G), TGC1202TCT (3745G>C 3746C>T), TCG1203GCC (3747T>G 3749G>C), CTG1204ATT (3750C>A 3752G>T), CTA1205TTA (3753C>T), CCT1206CCA (3758T>A), ATC1207GTT (3759A>G 3761C>T), ATG1208CTC (3762A>C 3764G>C), CGT1209AAG (3765C>A 3766G>A 3767T>G), AGC1210AGT (3770C>T), CAG1211CAA (3773G>A), AAG1212AGG (3775A>G), ATC1213GTG (3777A>G 3779C>G), CCT1214CCA (3782T>A), GGG1215GGT (3785G>T), GGA1216GGT (3788A>T), CGT1217ATC (3789C>A 3790G>T 3791T>C), CCA1218CCC (3794A>C), TGT1219ACA (3795T>A 3796G>C 3797T>A), GAA1220ACA (3798G>A 3799A>C), AGG1221CGT (3801A>C 3803G>T), GAT1222TGT (3804G>T 3805A>G), GAT1224GAC (3812T>C), GTA1225ATC (3813G>A 3815A>C), GTG1226GTT (3818G>T), CTC1227CTG (3821C>G), GGA1229GGT (3827A>T), ATA1230GTT (3828A>G 3830A>T), GGA1232GGT (3836A>T), TGT1233TTT (3838G>T), GAT1234GAC (3842T>C), TAC1235TTC (3844A>T), TAT1236TTT (3847A>T), GGC1238TCC (3852G>T 3853G>C), GAG1240GAA (3860G>A), CTG1241CTC (3863G>C), TCT1244TCA (3872T>A), TGC1245TGT (3875C>T), TAT1247TAC (3881T>C), CCC1248CCT (3884C>T), AAC1250AAT (3890C>T), AAG1251AAA (3893G>A), ATG1252CTG (3894A>C), GGG1253GGT (3899G>T), ATG1254TTG (3900A>T), GGG1255GGT (3905G>T), ATG1256ACC (3907T>C 3908G>C), AAC1257AAT (3911C>T), AAG1258AAA (3914G>A), AGA1259AGG (3917A>G), TTT1261TTC (3923T>C), GTG1262GTT (3926G>T), CAA1263GAA (3927C>G), TCC1264GCA (3930T>G 3932C>A), ACT1265CAG (3933A>C 3934C>A 3935T>G), GGC1266GAG (3937G>A 3938C>G), GAA1267AGT (3939G>A 3940A>G 3941A>T), GGA1268ACC (3942G>A 3943G>C 3944A>C), GAG1269CAA (3945G>C 3947G>A), AGA1270AGG (3950A>G), GTG1271GTA (3953G>A), GAA1272CTA (3954G>C 3955A>T), AAA1274AAG (3962A>G), GAC1276AAT (3966G>A 3968C>T), ACT1277TCT (3969A>T), GTA1279GTT (3977A>T), TTT1280TTC (3980T>C), GAA1281CAA (3981G>C), GAA1284GAG (3992A>G), GAG1285GAA (3995G>A), CTA1286TTA (3996C>T), CAG1289GAA (4005C>G 4007G>A), ATT1290ATA (4010T>A), AAA1292AAG (4016A>G), GGC1293GGT (4019C>T), ATC1294AAC (4021T>A), CAT1295TAT (4023C>T), GAT1297GAA (4031T>A), CTG1298ATG (4032C>A), GTC1299GTT (4037C>T), ACC1301ACT (4043C>T), TGC1303TGT (4049C>T), GCC1304GCT (4052C>T), AAA1305AAG (4055A>G), GAT1306GAC (4058T>C), CGC1308AGG (4062C>A 4064C>G), CTC1309CTA (4067C>A), CCA1310CCT (4070A>T), CTT1311CTG (4073T>G), GAG1312GAC (4076G>C), AAA1313AAG (4079A>G), ATC1314ATA (4082C>A), TAT1315TAC (4085T>C), GGC1316GGG (4088C>G), AAG1317AAA (4091G>A), AGA1318AAG (4093G>A 4094A>G), AAG1319AAA (4097G>A), AGG1321CGT (4101A>C 4103G>T), CTC1322CTT (4106C>T), ATA1325ATT (4115A>T), CTT1326TTG (4116C>T 4118T>G), CCT1327CCA (4121T>A), TAC1330TAT (4130C>T), TTG1333CTG (4137T>C), GTC1334GTG (4142C>G), AGG1335CGC (4143A>C 4145G>C), AAG1336AAA (4148G>A), TTT1338TTC (4154T>C), TTC1341TTT (4163C>T), TCA1342TCT (4166A>T), GCC1343GCA (4169C>A), ACA1344TGT (4170A>T 4171C>G 4172A>T), TTG1345TTA (4175G>A), TTG1348GCA (4182T>G 4183T>C 4184G>A), CAC1349CAT (4187C>T), GCT1351GTT (4192C>T), ATA1352CTG (4194A>C 4196A>G), CCA1353CCC (4199A>C), GTT1356GTG (4208T>G), ATT1358ATC (4214T>C), GAT1359AAT (4215G>A), CCT1360CCC (4220T>C), ACA1361ATT (4222C>T 4223A>T), AGT1362TCA (4224A>T 4225G>C 4226T>A), TCT1363CAT (4227T>C 4228C>A), ACA1366ACC (4238A>C), TTG1367ACC (4239T>A 4240T>C 4241G>C), TTG1368CTG (4242T>C), AAT1370AAC (4250T>C), GGG1371CAG (4251G>C 4252G>A), TTT1372TTC (4256T>C), AGA1373CGG (4257A>C 4259A>G), GCT1374AGT (4260G>A 4261C>G), TCA1376TCT (4268A>T), GAC1377GAT (4271C>T), GTG1378GTT (4274G>T), GGA1379GGT (4277A>T), TCA1381TCT (4283A>T), GCT1382GCA (4286T>A), GAT1383GAC (4289T>C), TAT1384TAC (4292T>C), AGA1390AGG (4310A>G), GCA1391GCG (4313A>G), CCT1392CCA (4316T>A), GTT1393ATA (4317G>A 4319T>A), GCT1395GCA (4325T>A), CAG1397CAA (4331G>A), GAA1405GCA (4354A>C), TAC1407TAT (4361C>T), GGA1408GGG (4364A>G), AAG1410GCT (4368A>G 4369A>C 4370G>T), CCT1411CCA (4373T>A), GGC1412GGA (4376C>A), AGT1413AGC (4379T>C), CCT1414GAG (4380C>G 4381C>A 4382T>G), TCC1416TCA (4388C>A), GCT1418GCA (4394T>A), CGA1419AGA (4395C>A), GCA1421GCT (4403A>T), CTT1422CTG (4406T>G), TTA1423TTG (4409A>G), GCA1426GCT (4418A>T), TGT1428TGC (4424T>C), ACA1431ACT (4433A>T), CTG1432CTA (4436G>A), GTG1437CTC (4449G>C 4451G>C), TTT1438TTC (4454T>C), TTG1440TTA (4460G>A), GGG1442GGT (4466G>T), GGC1443GGA (4469C>A), CCA1445CCT (4475A>T), TCA1446TCT (4478A>T), GCA1449GCT (4487A>T), CTA1450CTC (4490A>C), ACG1451ACC (4493G>C), ATC1453ATA (4499C>A), TTC1454TTT (4502C>T), AAT1455AAC (4505T>C), TCT1456TCC (4508T>C), CTC1457CTT (4511C>T), CTC1458TTA (4512C>T 4514C>A), AAT1459AAC (4517T>C), CGA1464CGG (4532A>G), GCC1466GCT (4538C>T), TTT1467TTC (4541T>C), TCA1469TCG (4547A>G), TTG1470CTC (4548T>C 4550G>C), TTA1471CTC (4551T>C 4553A>C), AGA1473AGG (4559A>G), CAT1475CAC (4565T>C), ATT1476ATC (4568T>C), GCT1477AAA (4569G>A 4570C>A 4571T>A), GCC1478GCA (4574C>A), AGG1479AAG (4576G>A), GCT1480GCA (4580T>A), ATA1481TTG (4581A>T 4583A>G), GGA1482GGT (4586A>T), GTT1483GTA (4589T>A), AAA1484AAG (4592A>G), CCA1485TCA (4593C>T), TTC1488TTT (4604C>T), CAG1490GAC (4608C>G 4610G>C), CTA1491TTA (4611C>T), TTC1492TTT (4616C>T), ATA1493ATT (4619A>T), GCT1494GCA (4622T>A), GTT1495GTG (4625T>G), GGA1497GGG (4631A>G), GTT1502GTG (4646T>G), GTA1504GTG (4652A>G), TTA1506ATG (4656T>A 4658A>G), CAT1507CAC (4661T>C), CTC1508CTT (4664C>T), CAG1509CAA (4667G>A), TAT1511TAC (4673T>C), TCT1512GAG (4674T>G 4675C>A 4676T>G), CTG1513TTA (4677C>T 4679G>A), CCA1514CCT (4682A>T), AAT1515GCA (4683A>G 4684A>C 4685T>A), CAT1518CAA (4694T>A), TTA1520CTA (4698T>C), CTG1522CAA (4705T>A 4706G>A), GTC1523GTG (4709C>G), AAT1524AAC (4712T>C), GTA1525GTT (4715A>T), ATC1526ATT (4718C>T), ATT1527ATC (4721T>C), GGT1530GGC (4730T>C), GAC1532GAT (4736C>T), TCG1534AAC (4740T>A 4741C>A 4742G>C), GAT1536AAT (4746G>A), GTT1537GTC (4751T>C), GAG1539GAT (4757G>T), GTA1540GTC (4760A>C), CAA1541AAA (4761C>A), TTT1542TTC (4766T>C), CAA1543CAG (4769A>G), CTA1548TTG (4782C>T 4784A>G), ACT1549ACG (4787T>G), CTG1551TTG (4791C>T), AGT1552AGC (4796T>C), GGT1554GGC (4802T>C), TTT1555TTC (4805T>C), AAA1556AAG (4808A>G), CGA1557CGC (4811A>C), CTT1560TTG (4818C>T 4820T>G), GGA1561GGT (4823A>T), CAC1563CAA (4829C>A), ATG1564CTA (4830A>C 4832G>A), GCT1565GCC (4835T>C), CCT1566CCC (4838T>C), TCA1571TCC (4853A>C), ATC1572ATT (4856C>T), ATT1573GTG (4857A>G 4859T>G), GAG1574GAA (4862G>A), CCC1575CCA (4865C>A), ATA1579ATC (4877A>C), CCT1581CCA (4883T>A), GCC1582GCA (4886C>A), CCA1583GTT (4887C>G 4888C>T 4889A>T), GGT1584GGG (4892T>G), CCT1586CCA (4898T>A), GAC1587GAT (4901C>T), ACT1588ATC (4903C>T 4904T>C), CAG1589ACT (4905C>A 4906A>C 4907G>T), ATG1592TAT (4914A>T 4915T>A 4916G>T), ACA1597ACG (4931A>G), GGA1598GGT (4934A>T), ATA1599GTA (4935A>G), CGT1600AGA (4938C>A 4940T>A), GAA1601GAG (4943A>G), CAT1604CAG (4952T>G), CAC1605CAT (4955C>T), GGT1606GGA (4958T>A), CGT1607CAG (4960G>A 4961T>G), GTG1608GTC (4964G>C), GCT1609GTT (4966C>T), AAG1612ACA (4975A>C 4976G>A), CTT1613TTG (4977C>T 4979T>G), GTC1614ACA (4980G>A 4981T>C 4982C>A), TTA1615GAC (4983T>G 4984T>A 4985A>C), GAT1616AAT (4986G>A), GTT1617GTC (4991T>C), CAG1618CAA (4994G>A), ACG1619AAA (4996C>A 4997G>A), GCG1620GCA (5000G>A), TTG1621ATA (5001T>A 5003G>A), GAT1622ATG (5004G>A 5005A>T 5006T>G), GAA1623CAG (5007G>C 5009A>G), AGG1624CGA (5010A>C 5012G>A), GGT1625GGC (5015T>C), TTC1626TTG (5018C>G), AGA1627CCA (5019A>C 5020G>C), GCT1628ATC (5022G>A 5023C>T 5024T>C), ATC1630CAT (5028A>C 5029T>A 5030C>T), CCT1632CCA (5036T>A), TCC1633TCT (5039C>T), TTG1635GCT (5043T>G 5044T>C 5045G>T), GTG1637CAA (5049G>C 5050T>A 5051G>A), CAG1639AGT (5055C>A 5056A>G 5057G>T), GAA1640GAG (5060A>G), ATT1642CTA (5064A>C 5066T>A), GCA1643CAA (5067G>C 5068C>A), AAG1644GCC (5070A>G 5071A>C 5072G>C), GTA1645GTT (5075A>T), ACA1646ACT (5078A>T), GGG1647GGA (5081G>A), GAT1648GAC (5084T>C), TCA1649AGT (5085T>A 5086C>G 5087A>T), AGT1650AAC (5089G>A 5090T>C), GCC1651ATA (5091G>A 5092C>T 5093C>A), CTG1652GGG (5094C>G 5095T>G), ACA1653GCT (5097A>G 5099A>T), ATT1654CTG (5100A>C 5102T>G), TGT1655CTG (5103T>C 5104G>T 5105T>G), ATG1657CTG (5109A>C), GCA1658AGC (5112G>A 5113C>G 5114A>C), AAA1659AAG (5117A>G), GCA1660GAT (5119C>A 5120A>T), TCC1663ACT (5127T>A 5129C>T), TAT1664TAC (5132T>C), ACG1665GTG (5133A>G 5134C>T), CCA1666CCT (5138A>T), TTG1667CTT (5139T>C 5141G>T), GAC1668CCA (5142G>C 5143A>C 5144C>A), GCA1669GAA (5146C>A), GAG1671AAA (5151G>A 5153G>A), AAG1672AAA (5156G>A), ATC1673ATT (5159C>T), ACA1674GAA (5160A>G 5161C>A), AAT1675GAT (5163A>G), GAG1677TCA (5169G>T 5170A>C 5171G>A), CGT1678CAA (5173G>A 5174T>A), GAT1679GGT (5176A>G), CTG1680GTC (5178C>G 5180G>C), AAT1681CAT (5181A>C), TGG1682GAA (5184T>G 5185G>A 5186G>A), GCA1684TGT (5190G>T 5191C>G 5192A>T), CCA1685TCT (5193C>T 5195A>T), AAC1686GGA (5196A>G 5197A>G 5198C>A), ATT1687GTT (5199A>G), GGT1688TTC (5202G>T 5203G>T 5204T>C), TTT1689TTC (5207T>C), GCA1692TCG (5214G>T 5216A>G), CGT1693CGA (5219T>A), ACT1694ACA (5222T>A), GCA1695GAA (5224C>A), GCC1696GTT (5227C>T 5228C>T), CAC1697AAT (5229C>A 5231C>T), TAC1698CAT (5232T>C 5234C>T), ACG1699AAA (5236C>A 5237G>A), GAA1702GAG (5246A>G), GGG1703GAC (5248G>A 5249G>C), TAC1704ACA (5250T>A 5251A>C 5252C>A), ATT1706CTT (5256A>C), AAC1708AAT (5264C>T), TGT1709TGC (5267T>C), AAG1713AAA (5279G>A), AAG1714AAA (5282G>A), TCC1715ACA (5283T>A 5285C>A), AAT1716ACA (5287A>C 5288T>A), GTT1718GTG (5294T>G), GGT1720GGC (5300T>C), CCG1733CCC (5339G>C), TAC1734TAT (5342C>T), ACT1735ACA (5345T>A), TCA1737GCA (5349T>G), GCT1738GCC (5354T>C), ATA1739ATC (5357A>C), ATG1740AAG (5359T>A), GAT1741GAC (5363T>C), GCC1742AAT (5364G>A 5365C>A 5366C>T), AAG1744CAA (5370A>C 5372G>A), AAT1745AGT (5374A>G), GTG1746CGC (5376G>C 5377T>G 5378G>C), GCA1748TCA (5382G>T), GGA1749ATG (5385G>A 5386G>T 5387A>G), GGA1750GGG (5390A>G), CAT1751ATT (5391C>A 5392A>T), GTA1752GGA (5395T>G), GCA1753ACC (5397G>A 5399A>C), ACC1754CAA (5400A>C 5401C>A 5402C>A), AAT1755AAA (5405T>A), GTC1756GTT (5408C>T), GTT1757GTC (5411T>C), GTG1759ATT (5415G>A 5417G>T), GGA1761GGG (5423A>G), GGA1763GGT (5429A>T), GTC1766GTG (5438C>G), GGC1767AGT (5439G>A 5441C>T), CCA1768CCC (5444A>C), GCA1771GCT (5453A>T), GCG1772GCT (5456G>T), ATG1774TTT (5460A>T 5462G>T), GCC1775GCA (5465C>A), GCA1777TCA (5469G>T), ACA1778ACC (5474A>C), AGG1779AAT (5476G>A 5477G>T), CAA1780CAG (5480A>G), TGT1782AGA (5484T>A 5486T>A), GTG1783GCT (5488T>C 5489G>T), GAC1785CAA (5493G>C 5495C>A), ATA1786CTT (5496A>C 5498A>T), GTT1788GTC (5504T>C), CGC1791AGA (5511C>A 5513C>A), ATT1793TTG (5517A>T 5519T>G), GGA1794GCG (5521G>C 5522A>G), AAT1795GGC (5523A>G 5524A>G 5525T>C), GTG1796TTG (5526G>T), CTT1797CTG (5531T>G), GAC1798GAT (5534C>T), CTC1799TTG (5535C>T 5537C>G), ACC1801CAG (5541A>C 5542C>A 5543C>G), GGC1805GGT (5555C>T), GTG1807ATT (5559G>A 5561G>T), CAG1808CAA (5564G>A), TTT1810TTC (5570T>C), CTT1811CTC (5573T>C), CAA1812ACA (5574C>A 5575A>C), TGT1813_GTG1814insGTC (5579_5580insGTC), CCT1815CAG (5584C>A 5585T>G), CAT1816AAA (5586C>A 5588T>A), GGA1817AGT (5589G>A 5591A>T), GAC1818TGC (5592G>T 5593A>G), AAA1819AAG (5597A>G), GTG1820ACA (5598G>A 5599T>C 5600G>A), AGT1823GCT (5607A>G 5608G>C), GGT1824AGT (5610G>A), GCA1825GGG (5614C>G 5615A>G), TCG1826del (5616_5618delTCG), CTC1827CGC (5620T>G), CAC1828GAT (5622C>G 5624C>T), AGT1829GGA (5625A>G 5627T>A), AGT1830ATG (5629G>T 5630T>G), TTT1831TTG (5633T>G), ATG1832AAC (5635T>A 5636G>C), CAA1834CAT (5642A>T), CAA1834_GGG1835insCATGCT (5642_5643insCATGCT), GGG1835GCT (5644G>C 5645G>T), TTT1836TAT (5647T>A), GAA1837GCC (5650A>C 5651A>C), TTG1838CTT (5652T>C 5654G>T), GGC1839GGA (5657C>A), CTC1841CTT (5663C>T), CGC1842CGA (5666C>A), ATT1843ATA (5669T>A), ATA1844GTG (5670A>G 5672A>G), CAT1845GGC (5673C>G 5674A>G 5675T>C), GGT1846TGT (5676G>T), GAT1847TCA (5679G>T 5680A>C 5681T>A), TTA1848ATT (5682T>A 5684A>T), GCA1849TCA (5685G>T), AAA1850AGA (5689A>G), CAG1851AAG (5691C>A), ACA1852ATG (5695C>T 5696A>G), ATG1854GTG (5700A>G), CGG1855AGA (5703C>A 5705G>A), ATG1856TTG (5706A>T), CCA1857CCC (5711A>C), TAT1858TAC (5714T>C), GTG1859ACT (5715G>A 5716T>C 5717G>T), GTA1860GTT (5720A>T), GGA1861GGG (5723A>G), CAT1863ATG (5727C>A 5728A>T 5729T>G), GGA1864GGT (5732A>T), GGA1865GGT (5735A>T), GGA1867GCA (5740G>C), AAT1868GAT (5742A>G), TTT1869TGC (5746T>G 5747T>C), TCC1870TCA (5750C>A), CAG1872AGA (5754C>A 5755A>G 5756G>A), GAT1873AAT (5757G>A), GAG1875GAT (5765G>T), AGT1876TCT (5766A>T 5767G>C), TTG1877CTC (5769T>C 5771G>C), CTA1878ATC (5772C>A 5774A>C), CAC1879AAA (5775C>A 5777C>A), TAT1880GCC (5778T>G 5779A>C 5780T>C), TTG1881CTA (5781T>C 5783G>A), CAG1883TGC (5787C>T 5788A>G 5789G>C), GGA1884TCT (5790G>T 5791G>C 5792A>T), GCA1886GCT (5798A>T), GAG1887GCA (5800A>C 5801G>A), TTA1888TGT (5803T>G 5804A>T), ATT1889GTG (5805A>G 5807T>G), CAA1890GCT (5808C>G 5809A>C 5810A>T), AAG1891CGT (5811A>C 5812A>G 5813G>T), AAT1892GGG (5814A>G 5815A>G 5816T>G), ACA1893TGT (5817A>T 5818C>G 5819A>T), CTA1895ATA (5823C>A), TAT1898TAC (5834T>C), TTC1899TTT (5837C>T), AAA1900GAG (5838A>G 5840A>G), GAG1901AAT (5841G>A 5843G>T), CTG1902TTT (5844C>T 5846G>T), AGT1903AAT (5848G>A), ATG1904GCA (5850A>G 5851T>C 5852G>A), GAA1905CAA (5853G>C), AAT1906ATA (5857A>T 5858T>A), GTG1907GCT (5860T>C 5861G>T), CAA1908ACC (5862C>A 5863A>C 5864A>C), CAA1909AGG (5865C>A 5866A>G 5867A>G), CTG1910TTA (5868C>T 5870G>A), ATA1911GTA (5871A>G), GTT1914CTT (5880G>C), |

| | |
|---|---|
| | AAG1915AAA (5885G>A), CTT1916AGT (5886C>A 5887T>G), GGG1918GGT (5894G>T), TTT1919GTG (5895T>G 5897T>G), TTC1920TTT (5900C>T), CCA1921ATC (5901C>A 5902C>T 5903A>C), AAA1922ACA (5905A>C), GAA1923GCC (5908A>C 5909A>C), ACT1924ACA (5912T>A), ACC1925GGC (5913A>G 5914C>G), ATT1926CAC (5916A>C 5917T>A 5918T>C), CAG1927AAA (5919C>A 5921G>A), AAG1928ATG (5923A>T), CTC1929TTG (5925C>T 5927C>G), AAA1930GTG (5928A>G 5929A>T 5930A>G), AAT1931CAA (5931A>C 5933T>A), TTT1932ACT (5934T>A 5935T>C), GTT1933CTG (5937G>C 5939T>G), GAT1934AGC (5940G>A 5941A>G 5942T>C), GCT1935AAT (5943G>A 5944C>A), GAA1936CAA (5946G>C), TGT1937TGG (5951T>G), CTA1938CAA (5953T>A), ACA1939TCC (5955A>T 5957A>C), TTC1940CAC (5958T>C 5959T>A), AAA1941GAA (5961A>G), GCA1942CAA (5964G>C 5965C>A), AGG1943AGT (5969G>T), AGC1944TGT (5970A>T 5972C>T), TTT1945GTG (5973T>G 5975T>G), AGG1946AAG (5977G>A), CAC1947AGA (5979C>A 5980A>G 5981C>A), GTA1948GTC (5984A>C), GTT1949TAT (5985G>T 5986T>A), AAA1951AAG (5993A>G), AAG1952CGA (5994A>C 5995A>G 5996G>A), AAA1953AAT (5999A>T), TTG1955CTC (6003T>C 6005G>C), AGT1956TCA (6006A>T 6007G>C 6008T>A), TCC1957AAC (6009T>A 6010C>A), ACA1958GCA (6012A>G), TGG1959ATA (6015T>A 6016G>T 6017G>A), AAA1960ACA (6019A>C), ATG1961_TGT1962insTAC (6023_6024insTAC), TGT1962CGT (6024T>C), GAA1964GAT (6032A>T), AGT1965GCC (6033A>G 6034G>C 6035T>C), ATT1966AAA (6037T>A 6038T>A), GTT1967GAT (6040T>A), TCA1969ATA (6045T>A 6046C>T), AGG1970GTA (6048A>G 6049G>T 6050G>A), GCA1972del (6054_6056delGCA), CTA1975GTG (6063C>G 6065A>G), CCT1977TTG (6069C>T 6070C>T 6071T>G), GGC1978GGT (6074C>T), TTG1980CCA (6078T>C 6079T>C 6080G>A), GCT1982GCC (6086T>C), TCC1983ATG (6087T>A 6088C>T 6089C>G), GTA1984GTT (6092A>T), AAA1986AGG (6097A>G 6098A>G) |

**Proteins**

| | | | | | | | | | |
|---|---|---|---|---|---|---|---|---|---|
| polyprotein (YP_001976147.1) | 1 | 1989 | 92.4% | 9624 | 69.7% | 1982 (99.2%) | 1328 (66.5%) | 9/7/0/0 | 0 |

| | Begin | End | Coverage | Score | Concordance | Matches | Identities | I/D/M/F* | Stop Codons |
|---|---|---|---|---|---|---|---|---|---|
| **NT** | **24** | **6107** | **84.2%** | **3417** | **28.3%** | **6063 (98.8%)** | **3936 (64.1%)** | **54/21** | |
| Protein mutations: | M1L (141A>C), A2V (145C>T), S4H (150T>C 151C>A 152A>C), K5N (155G>T), M6T (157T>C 158G>C), F7T (159T>A 160T>C), S8I (162T>A 163C>T 164C>T), K9Q (165A>C 167G>A), V10M (168G>A 170T>G), G12del (174_176delGGT), V13F (177G>T), D14S (180G>T 181A>C 182T>A), D15K (183G>A 185C>A), A16I (186G>A 187C>T), S17V (189A>G 190G>T), S20A (198T>G 200C>T), T22P (204A>C), F25L (213T>C), G26S (216G>T 217G>C), N30Q (228A>C 230T>G), S31T (231T>A 233T>A), I32V (234A>G 236T>G), K34S (240A>T 241A>C), F35L (243T>C 245T>G), A38T (252G>A 254G>A), V39M (255G>A 257T>G), S40K (259G>A 260T>G), N41S (262A>G), I42M (266T>G), T43S (268C>G 269C>T), T44K (271C>A 272T>A), M45I (275G>A), Q47E (279C>G), I49F (285A>T 287T>C), D51G (292A>G 293T>C), L53I (297C>A), D55E (305T>A), V60I (318G>A 320G>T), T61V (321A>G 322C>T), D62K (324G>A 326T>A), V63T (327G>A 328T>C 329G>T), S64A (330T>G 332C>A), T65D (333A>G 334C>A 335A>T), V68N (342G>A 343T>A 344G>T), A69S (345G>A 346C>G 347C>T), T70V (348A>G 349C>T), L74V (360C>G 362T>A), N75E (363A>G 365T>A), V77L (369G>T 371T>A), D79G (376A>G 377C>G), G82A (385G>C 386T>C), A83T (387G>A 389A>G), L84V (390C>G), D86E (398T>A), L89V (405C>G), A91G (412C>G 413G>T), Y92I (414T>A 415A>T), S93N (417T>A 418C>A), L95F (425G>C), S96K (426T>A 427C>A 428C>A), K100R (439A>G), S101T (441T>A 443T>C), K103I (448A>T 449G>T), L109V (465T>G 467G>T), F110V (468T>G), E111L (471G>T 472A>T), S112G (474A>G 476T>C), L113I (477T>A 479G>C), S114K (480T>A 481C>A 482C>A), K116N (488A>T), V117F (489G>T), K118I (493A>T 494A>T), L119R (495T>A 496T>G), F121Y (502T>A), A122S (504G>T 506T>A), V124L (510G>C), N126E (516A>G 518T>A), K127T (520A>C), S128T (523G>C 524T>A), E129G (526A>G 527A>T), D130E (530T>A), V131Y (531G>T 532T>A 533A>T), I133V (537A>G 539A>T), Y136I (546T>A 547A>T 548T>A), G137A (550G>C), A142S (564G>T 566T>C), I143L (567A>C 569A>T), T144S (570A>T), V147L (579G>T 581G>A), L149Q (586T>A), F150I (588T>A 590T>A), V151I (591G>A 593T>C), S154V (600A>G 601G>T 602T>A), G159M (615G>A 616G>T 617T>G), S161R (621T>A 622C>G 623T>A), N162S (625A>G 626C>T), T163V (627A>G 628C>T 629T>A), K165C (633A>T 634A>G 635A>T), I170V (648A>G 650A>T), M174F (660A>T 662G>C), L181I (681T>A 683A>T), D182S (684G>A 685A>G), K186L (696A>C 697A>T 698A>C), A188S (702G>T 704C>T), R190K (708C>A 709G>A), T192S (714A>T 716C>T), M193Q (717A>C 718T>A), V194M (720G>A), M199V (735A>G), V205L (753G>T 755T>G), S206N (756T>A 757C>A 758A>T), M214L (780A>C), T216S (786A>T 788A>T), I217L (789A>T 791A>G), L218I (792T>A 794G>A), I228V (822A>G 824T>G), S238T (853G>C 854C>A), T248A (882A>G 884A>G), T260Q (918A>C 919C>A 920T>A), R263K (927C>A 928G>A 929T>G), I280V (978A>G 980A>T), V281A (982T>C 983C>T), G282D (985G>A 986C>T), T283I (988C>T 989C>A), L298I (1032C>A 1034G>T), Q307T (1059C>A 1060A>C 1061G>T), D310S (1068G>T 1069A>C), C313A (1077T>G 1078G>C), K325P (1113A>C 1114A>C 1115G>T), P328S (1122C>T 1124T>A), T329I (1126C>T), R335K (1144G>A 1145G>A), E337Q (1149G>C 1151A>G), M354L (1200A>T), A361T (1221G>A 1223C>A), A363V (1228C>T), E366D (1238A>T), V368L (1242G>C 1244T>C), Y369I (1245T>A 1246A>T 1247T>A), S372N (1255G>A 1256T>C), R374Q (1260A>C 1261G>A 1262G>A), M376T (1267T>C 1268G>A), S393E (1317T>G 1318C>A 1319C>G), R412A (1374A>G 1375G>C 1376A>C), G414Q (1380G>C 1381G>A 1382T>A), I496V (1626A>G 1628C>T), M521R (1701A>C 1702T>G), E523V (1708A>T), D530N (1728G>A 1730T>C), G533A (1738G>C), R536S (1748A>C), V540I (1758G>A 1760G>T), E541G (1762A>G 1763G>A), N552D (1794A>G 1796T>C), Q567K (1839C>A), A570H (1848G>C 1849C>A 1850A>C), V572I (1854G>A 1856A>T), S573A (1857T>G), K574T (1861A>C 1862G>C), F577Y (1870T>A 1871T>C), E578N (1872G>A 1874A>C), A579Q (1875G>C 1876C>A 1877T>G), E580S (1878G>T 1879A>C), S581T (1881T>A 1883T>A), I583C (1887A>T 1888T>G 1889A>T), S584E (1890A>G 1891G>A 1892T>A), P585E (1893C>G 1894C>A 1895T>G), I587V (1899A>G 1901A>T), Q591E (1911C>G), A594V (1921C>T 1922A>T), D595E (1925T>A), D596E (1928T>A), I598M (1934A>G), A599T (1935G>A 1937G>A), V601F (1941G>T 1943G>T), D602H (1944G>C), G603D (1948G>A 1949G>T), R605Q (1953A>C 1954G>A), L606I (1956T>A 1958G>A), R607S (1959C>A), T608F (1962A>T 1963C>T 1964C>T), K609D (1965A>G 1967G>C), Q610E (1968C>G), D611V (1972A>T), E612S (1974G>A 1975A>G 1976A>C), I614S (1981T>G 1982A>T), L615V (1983T>G 1985G>T), V616_P617del (1986_1991delGTGCCC), V620L (1998G>C 2000G>C), N623D (2007A>G), I627G (2019A>G 2020T>G 2021C>A), A629S (2025G>T), R630Q (2028A>C 2029G>A 2030A>G), S644A (2070A>G 2071G>C 2072T>A), D646N (2076G>A), E648F (2082G>T 2083A>T 2084G>T), L650Y (2088C>T 2089T>A), H655A (2103C>G 2104A>C), H656S (2106C>T 2107A>C 2108T>A), M660I (2120G>T), Y661F (2122A>T), S663V (2127T>G 2128C>T 2129T>C), D665S (2133G>T 2134A>C 2135T>A), D666S (2136G>A 2137A>G), H668A (2142C>G 2143A>C 2144T>C), I671S (2152T>G 2153T>C), N673T (2158A>C 2159C>T), K674R (2160A>C 2161A>G 2162A>C), R676V (2166A>G 2167G>T), V677N (2169G>A 2170T>A 2171C>T), N678E (2172A>G 2174C>G), L680I (2178T>A 2180G>T), R682E (2184A>G 2185G>A 2186A>G), D685S (2193G>T 2194A>C 2195T>G), A686T (2196G>A), E687A (2200A>C), Q688G (2202C>G 2203A>G 2204G>A), R690A (2208C>G 2209G>C 2210A>C), V693I (2217G>A 2219A>C), Q694K (2220C>A 2222A>G), N695A (2223A>G 2224A>C), I701V (2241A>G 2243A>C), A710S (2268G>A 2269C>G 2270A>T), S711T (2271T>A), I713V (2277A>G 2279C>G), K716A (2286A>G 2287A>C 2288G>T), K717H (2289A>C 2291A>C), L718M (2292C>A 2294A>G), A719D (2296C>A 2297A>T), T722R (2305C>G 2306A>G), E723M (2307G>A 2308A>T), M724L (2310A>T), R725Y (2313A>T 2314G>A 2315G>C), T726N (2317C>A), F728R (2322T>A 2323T>G 2324T>A), S731R (2331T>A 2332C>G 2333T>A), C732A (2334T>G 2335G>C 2336C>A), I733L (2337A>C 2339T>G), D734E (2342T>A), V737A (2350T>C), I741M (2363A>G), F749H (2385T>C 2386T>A), C762_N763insRR (2426_2427insAGAAGG), N763K (2429C>G), K764M (2431A>T 2432A>G), S766T (2436T>A 2438A>T), P769T (2445C>A), K770_E771del (2448_2453delAAGGAA), I774V (2460A>G 2462C>A), K775Q (2463A>C), I776S (2466A>T 2467T>C 2468T>G), C777L (2469T>C 2470G>T), H779D (2475C>G 2477T>C), K780S (2478A>T 2479A>C 2480A>C), I781T (2482T>C 2483C>A), V782I (2484G>A), E783G (2488A>G 2489A>G), Y784R (2490T>A 2491A>G 2492T>A), I785M (2495C>G), K786Q (2496A>C 2498G>A), D787E (2501T>A), T788M (2503C>T 2504A>G), W790N (2508T>A 2509G>A 2510G>T), S791D (2511A>G 2512G>A 2513C>T), A795I (2523G>A 2524C>T 2525A>T), R797K (2530G>A), I806L (2556A>C), M807I (2561G>A), V812T (2574G>A 2575T>C), F813M (2577T>A 2579T>G), G815A (2584G>C), I816V (2586A>G 2588T>A), L820I (2598T>A 2600G>T), A821S (2601G>T 2603G>T), Q824K (2610C>A 2612G>A), N826S (2617A>G), S827L (2620C>T 2621T>G), F828I (2622T>A 2624C>T), N830T (2629A>C), A831S (2631G>T 2633A>C), S832Q (2634T>C 2635C>A), R834K (2641G>A), N852R (2695A>G 2696T>A), N852_K853insS (2696_2697insTCC), K853H (2697A>C 2699G>T), L854V (2700C>G 2702C>A), R855S (2703A>T 2704G>C), N856K (2708T>G), L857M (2709C>A 2711T>G), V859F (2715G>T), G860S (2718G>A 2720T>C), S865P (2733T>C), L866N (2736T>A 2737T>A 2738G>T), A867D (2740C>A 2741A>T), Q868_D869insG (2744_2745insGGA), D869K (2745G>A 2747T>A), A872S (2754G>T 2756T>C), E873R (2757G>A 2758A>G 2759A>G), Q879E (2775C>G 2777G>A), K882A (2784A>G 2785A>C 2786G>C), L890V (2808C>G 2810T>G), G891A (2812G>C 2813T>A), G904E (2851G>A 2852T>A), H914T (2880C>A 2881A>C 2882C>G), N915Q (2883A>C 2885T>G), A916L (2886G>C 2887C>T 2888T>C), T923S (2907A>T), A927E (2920C>A 2921C>A), N928H (2922A>C), R930V (2928C>G 2929G>T 2930A>C), S931C (2932C>G 2933T>C), K932Q (2934A>C 2936A>G), R938K (2953G>A), E939H (2955G>C 2957A>T), S940A (2958T>G 2960A>C), R941K (2962G>A), I942F (2964A>T), I943L (2967A>T 2969T>G), E944P (2970G>C 2971A>C 2972G>T), G945E (2974G>A 2975C>A), I948V (2982A>G 2984C>T), I949V (2985A>G 2987T>A), D955C (3003G>T 3004A>G), P961V (3021C>G 3022C>T 3023T>C), P968A (3042C>G 3044A>T), R969E (3045A>G 3046G>A), E972A (3055A>C 3056G>A), Q980E (3078C>G), C984V (3090T>G 3091G>T 3092T>C), T990K (3109C>A), R993E (3117C>G 3118G>A 3119T>G), L994K (3120C>A 3121T>A 3122T>A), G995Q (3123G>C 3124G>A 3125C>G), E998D (3134G>C), M1002Y (3144A>T 3145T>A 3146G>T), P1003S (3147C>T), V1012E (3175T>A 3176T>G), E1015T (3183G>A 3184A>C), E1017R (3189G>A 3190A>G 3191A>G), L1018M (3192C>A), N1019S (3196A>G), S1020T (3198T>A 3200C>A), S1021N (3201T>A 3202C>A 3203C>T), K1024H (3210A>C 3212G>C), K1024_I1025insS (3212_3213insTCA), C1026S (3216T>A 3218C>T), S1027A (3219T>G), Q1029E (3225C>G), S1030A (3228T>G 3230T>C), I1035V (3243A>G 3245A>C), G1037S (3249G>A 3251G>C), V1051I (3291G>A 3293T>C), V1057C (3309G>T 3310T>G 3311G>C), G1068E (3343G>A 3344C>A), V1069A (3346T>C 3347A>T), A1071T (3351G>A 3353T>A), R1073L (3357C>T 3358G>T 3359T>G), I1076V (3366A>G 3368C>A), S1077A (3369T>G 3371T>A), I1083V (3387A>G 3389C>A), E1086D (3398A>C), E1089Q (3405G>C 3407G>A), P1092A (3414C>G 3416A>T), T1093Q (3417A>C 3418C>A 3419T>A), A1098M (3432G>A 3433C>T 3434A>G), I1105L (3453A>T 3455T>G), S1108H (3462T>C 3463C>A 3464A>T), I1109V (3465A>G), Q1115A (3483C>G 3484A>C 3485G>T), E1118D (3494G>C), N1120K (3500C>G), Y1123A (3507T>G 3508A>C 3509C>T), Y1124F (3511A>T 3512C>T), L1132M (3534T>A), Q1146P (3577A>C), P1153E (3597C>G 3598C>A 3599T>A), E1155Q (3603G>C 3605G>A), I1160L (3618A>C 3620T>G), T1162S (3624A>T 3626A>T), H1164K (3630C>A 3632T>A), V1178I (3672G>A 3674A>T), M1179E (3675A>G 3676T>A), C1186S (3697G>C 3698T>A), G1187N (3699G>A 3700G>A 3701A>T), D1189E (3707T>G), D1191S (3711G>A 3712A>G), K1194R (3720A>C 3721A>G 3722A>G), L1195A (3723C>G 3724T>C 3725A>T), N1199G (3735A>G 3736A>G), C1202S (3745G>C 3746C>T), S1203A (3747T>G 3749G>C), L1204I (3750C>A 3752G>T), I1207V (3759A>G 3761C>T), M1208L (3762A>C 3764G>C), R1209K (3765C>A 3766G>A 3767T>G), K1212R (3775A>G), I1213V (3777A>G 3779C>G), R1217I (3789C>A 3790G>T 3791T>C), C1219T (3795T>A 3796G>C 3797T>A), E1220T (3798G>A 3799A>C), D1222C (3804G>T 3805A>G), V1225I (3813G>A 3815A>C), I1230V (3828A>G 3830A>T), C1233F (3838G>T), Y1235F (3844A>T), Y1236F (3847A>T), G1238S (3852G>T 3853G>C), M1252L (3894A>C), M1254L (3900A>T), M1256T (3907T>C 3908G>C), Q1263E (3927C>G), S1264A (3930T>G 3932C>A), T1265Q (3933A>C 3934C>A 3935T>G), G1266E (3937G>A 3938C>G), E1267S (3939G>A 3940A>G 3941A>T), G1268T (3942G>A 3943G>C 3944A>C), E1269Q (3945G>C 3947G>A), E1272L (3954G>C 3955A>T), D1276N (3966G>A 3968C>T), T1277S (3969A>T), E1281Q (3981G>C), Q1289E (4005C>G 4007G>A), I1294N (4021T>A), H1295Y (4023C>T), D1297E (4031T>A), L1298M (4032C>A), E1312D (4076G>C), R1318K (4093G>A 4094A>G), T1344C (4170A>T 4171C>G 4172A>T), L1348A (4182T>G 4183T>C 4184G>A), A1351V (4192C>T), I1352L (4194A>C 4196A>G), D1359N (4215G>A), T1361I (4222C>T 4223A>T), S1363H (4227T>C 4228C>A), L1367T (4239T>A 4240T>C 4241G>C), G1371Q (4251G>C 4252G>A), A1374S (4260G>A 4261C>G), V1393I (4317G>A 4319T>A), E1405A (4354A>C), K1410A (4368A>G 4369A>C 4370G>T), P1414E (4380C>G 4381C>A 4382T>G), V1437L (4449G>C 4451G>C), A1477K (4569G>A 4570C>A 4571T>A), R1479K (4576G>A), I1481L (4581A>T 4583A>G), P1485S (4593C>T), Q1490D (4608C>G 4610G>C), L1506M (4656T>A 4658A>G), S1512E (4674T>G 4675C>A 4676T>G), N1515A (4683A>G 4684A>C 4685T>A), H1518Q (4694T>A), L1522Q (4705T>A 4706G>A), S1534N (4740T>A 4741C>A 4742G>C), D1536N (4746G>A), E1539D (4757G>T), Q1541K (4761C>A), H1563Q (4829C>A), M1564L (4830A>C 4832G>A), I1573V (4857A>G 4859T>G), P1583V (4887C>G 4888C>T 4889A>T), T1588I (4903C>T 4904T>C), Q1589T (4905C>A 4906A>C 4907G>T), M1592Y (4914A>T 4915T>A 4916G>T), I1599V (4935A>G), H1604Q (4952T>G), R1607Q (4960G>A 4961T>G), A1609V (4966C>T), K1612T (4975A>C 4976G>A), V1614T (4980G>A 4981T>C 4982C>A), L1615D (4983T>G 4984T>A 4985A>C), D1616N (4986G>A), T1619K (4996C>A 4997G>A), L1621I (5001T>A 5003G>A), D1622M (5004G>A 5005A>T 5006T>G), E1623Q (5007G>C 5009A>G), F1626L (5018C>G), R1627P (5019A>C 5020G>C), A1628I (5022G>A 5023C>T 5024T>C), I1630H (5028A>C 5029T>A 5030C>T), L1635A (5043T>G 5044T>C 5045G>T), V1637Q (5049G>C 5050T>A 5051G>A), Q1639S (5055C>A 5056A>G 5057G>T), I1642L (5064A>C 5066T>A), A1643Q (5067G>C 5068C>A), K1644A (5070A>G 5071A>C 5072G>C), S1650N (5089G>A 5090T>C), A1651I (5091G>A 5092C>T 5093C>A), L1652G (5094C>G 5095T>G), T1653A (5097A>G 5099A>T), I1654L (5100A>C 5102T>G), C1655L (5103T>C 5104G>T 5105T>G), M1657L (5109A>C), A1658S (5112G>A 5113C>G 5114A>C), A1660D (5119C>A 5120A>T), S1663T (5127T>A 5129C>T), T1665V (5133A>G 5134C>T), D1668P (5142G>C 5143A>C 5144C>A), A1669E (5146C>A), E1671K (5151G>A 5153G>A), T1674E (5160A>G 5161C>A), N1675D (5163A>G), E1677S (5169G>T 5170A>C 5171G>A), R1678Q (5173G>A 5174T>A), D1679G (5176A>G), L1680V (5178C>G 5180G>C), N1681H (5181A>C), W1682E (5184T>G 5185G>A 5186G>A), A1684C (5190G>T 5191C>G 5192A>T), P1685S (5193C>T 5195A>T), N1686G (5196A>G 5197A>G 5198C>A), I1687V (5199A>G), G1688F (5202G>T 5203G>T 5204T>C), A1692S (5214G>T 5216A>G), A1695E (5224C>A), A1696V (5227C>T 5228C>T), H1697N (5229C>A 5231C>T), Y1698H (5232T>C 5234C>T), T1699K (5236C>A 5237G>A), G1703D (5248G>A 5249G>C), Y1704T (5250T>A 5251A>C 5252C>A), I1706L (5256A>C), S1715T (5283T>A 5285C>A), N1716T (5287A>C 5288T>A), S1737A (5349T>G), M1740K (5359T>A), A1742N (5364G>A 5365C>A 5366C>T), K1744Q (5370A>C 5372G>A), N1745S (5374A>G), V1746R (5376G>C 5377T>G 5378G>C), A1748S (5382G>T), G1749M (5385G>A 5386G>T 5387A>G), H1751I (5391C>A 5392A>T), V1752G (5395T>G), A1753T (5397G>A 5399A>C), T1754Q (5400A>C 5401C>A 5402C>A), N1755K (5405T>A), V1759I (5415G>A 5417G>T), G1767S (5439G>A 5441C>T), M1774F (5460A>T 5462G>T), A1777S (5469G>T), R1779N (5476G>A 5477G>T), C1782R (5484T>A 5486T>A), V1783A (5488T>C 5489G>T), D1785Q (5493G>C 5495C>A), I1786L (5496A>C 5498A>T), I1793L (5517A>T 5519T>G), G1794A (5521G>C 5522A>G), N1795G (5523A>G 5524A>G 5525T>C), V1796L (5526G>T), T1801Q (5541A>C 5542C>A 5543C>G), V1807I (5559G>A 5561G>T), Q1812T (5574C>A 5575A>C), C1813_V1814insV (5579_5580insGTC), P1815Q (5584C>A 5585T>G), H1816K (5586C>A 5588T>A), G1817S (5589G>A 5591A>T), D1818C (5592G>T 5593A>G), V1820T (5598G>A 5599T>C 5600G>A), S1823A (5607A>G 5608G>C), G1824S (5610G>A), A1825G (5614C>G 5615A>G), S1826del (5616_5618delTCG), L1827R (5620T>G), H1828D (5622C>G 5624C>T), S1829G (5625A>G 5627T>A), S1830M (5629G>T 5630T>G), F1831L (5633T>G), M1832N (5635T>A 5636G>C), Q1834H (5642A>T), Q1834_G1835insHA (5642_5643insCATGCT), G1835A (5644G>C 5645G>T), F1836Y (5647T>A), E1837A (5650A>C 5651A>C), I1844V (5670A>G 5672A>G), H1845G (5673C>G 5674A>G 5675T>C), G1846C (5676G>T), D1847S (5679G>T 5680A>C 5681T>A), L1848I (5682T>A 5684A>T), A1849S (5685G>T), K1850R (5689A>G), Q1851K (5691C>A), T1852M (5695C>T 5696A>G), M1854V (5700A>G), M1856L (5706A>T), V1859T (5715G>A 5716T>C 5717G>T), H1863M (5727C>A 5728A>T 5729T>G), G1867A (5740G>C), N1868D (5742A>G), F1869C (5746T>G 5747T>C), Q1872R (5754C>A 5755A>G 5756G>A), D1873N (5757G>A), E1875D (5765G>T), L1878I (5772C>A 5774A>C), H1879K (5775C>A 5777C>A), Y1880A (5778T>G 5779A>C 5780T>C), Q1883C (5787C>T 5788A>G 5789G>C), G1884S (5790G>T 5791G>C 5792A>T), E1887A (5800A>C 5801G>A), L1888C (5803T>G 5804A>T), I1889V (5805A>G 5807T>G), Q1890A (5808C>G 5809A>C 5810A>T), K1891R (5811A>C 5812A>G 5813G>T), N1892G (5814A>G 5815A>G 5816T>G), T1893C (5817A>T 5818C>G 5819A>T), L1895I (5823C>A), K1900E (5838A>G 5840A>G), E1901N (5841G>A 5843G>T), L1902F (5844C>T 5846G>T), S1903N (5848G>A), M1904A (5850A>G 5851T>C 5852G>A), E1905Q (5853G>C), N1906I (5857A>T 5858T>A), V1907A (5860T>C 5861G>T), Q1908T (5862C>A 5863A>C 5864A>C), Q1909R (5865C>A 5866A>G 5867A>G), I1911V (5871A>G), V1914L (5880G>C), L1916S (5886C>A 5887T>G), F1919V (5895T>G 5897T>G), P1921I (5901C>A 5902C>T 5903A>C), K1922T (5905A>C), E1923A (5908A>C 5909A>C), T1925G (5913A>G 5914C>G), I1926H (5916A>C 5917T>A 5918T>C), Q1927K (5919C>A 5921G>A), K1928M (5923A>T), K1930V (5928A>G 5929A>T 5930A>G), N1931Q (5931A>C 5933T>A), F1932T (5934T>A 5935T>C), V1933L (5937G>C 5939T>G), D1934S (5940G>A 5941A>G 5942T>C), A1935N (5943G>A 5944C>A), E1936Q (5946G>C), C1937W (5951T>G), L1938Q (5953T>A), T1939S (5955A>T 5957A>C), F1940H (5958T>C 5959T>A), K1941E (5961A>G), A1942Q (5964G>C 5965C>A), R1943S (5969G>T), S1944C (5970A>T 5972C>T), F1945V (5973T>G 5975T>G), R1946K (5977G>A), H1947R (5979C>A 5980A>G 5981C>A), V1949Y (5985G>T 5986T>A), K1952R (5994A>C 5995A>G 5996G>A), K1953N (5999A>T), S1957N (6009T>A 6010C>A), T1958A (6012A>G), W1959I (6015T>A 6016G>T 6017G>A), K1960T (6019A>C), M1961_C1962insY (6023_6024insTAC), C1962R (6024T>C), E1964D (6032A>T), S1965A (6033A>G 6034G>C 6035T>C), I1966K (6037T>A 6038T>A), V1967D (6040T>A), S1969I (6045T>A 6046C>T), R1970V (6048A>G 6049G>T 6050G>A), A1972del (6054_6056delGCA), L1975V (6063C>G 6065A>G), P1977L (6069C>T 6070C>T 6071T>G), L1980P (6078T>C 6079T>C 6080G>A), S1983M (6087T>A 6088C>T 6089C>G), K1986R (6097A>G 6098A>G) | | | | | | | | |

| | Begin | End | Coverage | Score | Concordance | Matches | Identities | I/D/M/F* | Stop Codons |
|---|---|---|---|---|---|---|---|---|---|
| **NT** | **24** | **6107** | **84.2%** | **3417** | **28.3%** | **6063 (98.8%)** | **3936 (64.1%)** | **54/21** | |

ATG1CTG (141A>C), GCT2GTT (145C>T), TTT3TTC (149T>C), TCA4CAC (150T>C 151C>A 152A>C), AAG5AAT (155G>T), ATG6ACC (157T>C 158G>C), TTC7ACC (159T>A 160T>C), TCC8ATT (162T>A 163C>T 164C>T), AAG9CAA (165A>C 167G>A), GTT10ATG (168G>A 170T>G), GCT11GCA (173T>A), GGT12del (174_176delGGT), GTT13TTT (177G>T), GAT14TCA (180G>T 181A>C 182T>A), GAC15AAA (183G>A 185C>A), GCC16ATC (186G>A 187C>T), AGT17GTT (189A>G 190G>T), TCC20GCT (198T>G 200C>T), ACC21ACT (203C>T), ACT22CCT (204A>C), TCA23TCT (209A>T), TTC24TTT (212C>T), TTT25CTT (213T>C), GGT26TCT (216G>T 217G>C), GAA27GAG (221A>G), CTT28TTG (222C>T 224T>G), TCA29TCT (227A>T), AAT30CAG (228A>C 230T>G), TCT31ACA (231T>A 233T>A), ATT32GTG (234A>G 236T>G), TCC33TCA (239C>A), AAA34TCA (240A>T 241A>C), TTT35CTG (243T>C 245T>G), TCA36TCC (248A>C), AAA37AAG (251A>G), GCG38ACA (252G>A 254G>A), GTT39ATG (255G>A 257T>G), AGT40AAG (259G>A 260T>G), AAT41AGT (262A>G), ATT42ATG (266T>G), ACC43AGT (268C>G 269C>T), ACT44AAA (271C>A 272T>A), ATG45ATA (275G>A), TCA46TCT (278A>T), CAG47GAG (279C>G), AAG48AAA (284G>A), ATT49TTC (285A>T 287T>C), GAT51GGC (292A>G 293T>C), CTT53ATT (297C>A), GAG54GAA (302G>A), GAT55GAA (305T>A), CTG56TTA (306C>T 308G>A), CCA58CCT (314A>T), TCT59TCA (317T>A), GTG60ATT (318G>A 320G>T), ACT61GTT (321A>G 322C>T), GAT62AAA (324G>A 326T>A), GTG63ACT (327G>A 328T>C 329G>T), TCC64GCA (330T>G 332C>A), ACA65GAT (333A>G 334C>A 335A>T), TCT66TCC (338T>C), GTG68AAT (342G>A 343T>A 344G>T), GCC69AGT (345G>A 346C>G 347C>T), ACT70GTT (348A>G 349C>T), TCA73TCT (359A>T), CTT74GTA (360C>G 362T>A), AAT75GAA (363A>G 365T>A), GTT77TTA (369G>T 371T>A), TTG78TTA (374G>A), GAC79GGG (376A>G 377C>G), GGT82GCC (385G>C 386T>C), GCA83ACG (387G>A 389A>G), CTT84GTT (390C>G), ATA85ATT (395A>T), GAT86GAA (398T>A), CCA87CCT (401A>T), CTT89GTT (405C>G), GCG91GGT (412C>G 413G>T), TAC92ATC (414T>A 415A>T), TCT93AAT (417T>A 418C>A), TTG95TTC (425G>C), TCC96AAA (426T>A 427C>A 428C>A), TAC99TAT (437C>T), AAA100AGA (439A>G), TCT101ACC (441T>A 443T>C), ATC102ATA (446C>A), AAG103ATT (448A>T 449G>T), TTA107CTG (459T>C 461A>G), TTG109GTT (465T>G 467G>T), TTT110GTT (468T>G), GAG111TTG (471G>T 472A>T), AGT112GGC (474A>G 476T>C), TTG113ATC (477T>A 479G>C), TCC114AAA (480T>A 481C>A 482C>A), AAA116AAT (488A>T), GTT117TTT (489G>T), AAA118ATT (493A>T 494A>T), TTA119AGA (495T>A 496T>G), TTT121TAT (502T>A), GCT122TCA (504G>T 506T>A), GTG124CTG (510G>C), TTA125TTG (515A>G), AAT126GAA (516A>G 518T>A), AAA127ACA (520A>C), AGT128ACA (523G>C 524T>A), GAA129GGT (526A>G 527A>T), GAT130GAA (530T>A), GTA131TAT (531G>T 532T>A 533A>T), ATA133GTT (537A>G 539A>T), GTT134GTA (542T>A), GTA135GTT (545A>T), TAT136ATA (546T>A 547A>T 548T>A), GGT137GCT (550G>C), CTT139CTA (557T>A), GTA140GTC (560A>C), GCT142TCC (564G>T 566T>C), ATA143CTT (567A>C 569A>T), ACA144TCA (570A>T), TTG145CTA (573T>C 575G>A), TTA146TTG (578A>G), GTG147TTA (579G>T 581G>A), CTT148TTA (582C>T 584T>A), CTA149CAA (586T>A), TTT150ATA (588T>A 590T>A), GTT151ATC (591G>A 593T>C), AGT154GTA (600A>G 601G>T 602T>A), GTT157GTA (611T>A), GAG158GAA (614G>A), GGT159ATG (615G>A 616G>T 617T>G), TCT161AGA (621T>A 622C>G 623T>A), AAC162AGT (625A>G 626C>T), ACT163GTA (627A>G 628C>T 629T>A), GTC164GTT (632C>T), AAA165TGT (633A>T 634A>G 635A>T), ATC167ATA (641C>A), TTG169CTA (645T>C 647G>A), ATA170GTT (648A>G 650A>T), GTC171GTA (653C>A), GGA172GGT (656A>T), ATG174TTC (660A>T 662G>C), TTT175TTC (665T>C), TCA176TCT (668A>T), AGT177TCT (669A>T 670G>C), CTA178CTG (674A>G), TAC179TAT (677C>T), TTA181ATT (681T>A 683A>T), GAT182AGT (684G>A 685A>G), CCA185CCC (695A>C), AAA186CTC (696A>C 697A>T 698A>C), GCC188TCT (702G>T 704C>T), GAA189GAG (707A>G), CGA190AAA (708C>A 709G>A), TTC191TTT (713C>T), ACC192TCT (714A>T 716C>T), ATG193CAG (717A>C 718T>A), GTG194ATG (720G>A), GCC195GCA (725C>A), CAA196CAG (728A>G), AGT198TCT (732A>T 733G>C), ATG199GTG (735A>G), CTT200CTA (740T>A), GGG202GGT (746G>T), GAA203GAG (749A>G), TCA204AGC (750T>A 751C>G 752A>C), GTT205TTG (753G>T 755T>G), TCA206AAT (756T>A 757C>A 758A>T), ACG210ACC (770G>C), TCT211AGT (771T>A 772C>G), CAG212CAA (776G>A), ATG214CTG (780A>C), TCC215TCT (785C>T), ACA216TCT (786A>T 788A>T), ATA217TTG (789A>T 791A>G), TTG218ATA (792T>A 794G>A), GCT219GCA (797T>A), GGG221GGC (803G>C), ATT222ATC (806T>C), TCG223TCA (809G>A), GTG226GTA (818G>A), TTC227TTT (821C>T), ATT228GTG (822A>G 824T>G), GTG230GTA (830G>A), AGA233AGG (839A>G), TTG237CTT (849T>C 851G>T), AGC238ACA (853G>C 854C>A), CCA240CCG (860A>G), TTG241CTA (861T>C 863G>A), CTC245TTG (873C>T 875C>G), TCG247TCA (881G>A), ACA248GCG (882A>G 884A>G), GGA249GGC (887A>C), GGT251GGA (893T>A), CAG253CAA (899G>A), CTA258CTC (914A>C), ACT260CAA (918A>C 919C>A 920T>A), CTT261TTG (921C>T 923T>G), TAC262TAT (926C>T), CGT263AAG (927C>A 928G>A 929T>G), AAT264AAC (932T>C), AAA266AAG (938A>G), ACC269ACA (947C>A), TCA274TCT (962A>T), GTT276GTG (968T>G), CTT277TTG (969C>T 971T>G), ATA279ATT (977A>T), ATA280GTT (978A>G 980A>T), GTC281GCT (982T>C 983C>T), GGC282GAT (985G>A 986C>T), ACC283ATA (988C>T 989C>A), TTT284TTC (992T>C), GGA285GGG (995A>G), TTC286TTT (998C>T), AAA287AAG (1001A>G), AAT288AAC (1004T>C), GTG290GTT (1010G>T), TTG291CTG (1011T>C), TCA292TCT (1016A>T), GCT293GCA (1019T>A), ATA294ATT (1022A>T), ACG297ACA (1031G>A), CTG298ATT (1032C>A 1034G>T), CTG302CTC (1046G>C), TTC303TTT (1049C>T), CAG307ACT (1059C>A 1060A>C 1061G>T), GAG308GAA (1064G>A), GAT310TCT (1068G>T 1069A>C), TGT313GCT (1077T>G 1078G>C), ACG317ACA (1091G>A), CGC318AGG (1092C>A 1094C>G), GAG320GAA (1100G>A), GCC323GCT (1109C>T), AAG325CCT (1113A>C 1114A>C 1115G>T), TTC327TTT (1121C>T), CCT328TCA (1122C>T 1124T>A), ACC329ATC (1126C>T), CTG334CTT (1142G>T), AGG335AAA (1144G>A 1145G>A), GAA336GAG (1148A>G), GAA337CAG (1149G>C 1151A>G), GCT338GCA (1154T>A), ATC341ATA (1163C>A), GCT343GCC (1169T>C), TAC344TAT (1172C>T), GCA346GCT (1178A>T), CCA349CCT (1187A>T), GTT350GTG (1190T>G), TTC353TTT (1199C>T), ATG354TTG (1200A>T), AGC355AGT (1205C>T), AGG357AGA (1211G>A), AGT359TCT (1215A>T 1216G>C), GCC361ACA (1221G>A 1223C>A), GCG363GTG (1228C>T), CAG364CAA (1232G>A), TTA365TTG (1235A>G), GAA366GAT (1238A>T), GTT368CTC (1242G>C 1244T>C), TAT369ATA (1245T>A 1246A>T 1247T>A), GCT370GCA (1250T>A), AGT372AAC (1255G>A 1256T>C), AGG374CAA (1260A>C 1261G>A 1262G>A), CAC375CAT (1265C>T), ATG376ACA (1267T>C 1268G>A), GGC377GGA (1271C>A), GGC379GGT (1277C>T), TAT381TAC (1283T>C), CGT382AGA (1284C>A 1286T>A), CCT385CCC (1295T>C), GTA388GTT (1304A>T), TTC391TTT (1313C>T), GGG392GGT (1316G>T), TCC393GAG (1317T>G 1318C>A 1319C>G), GGG395GGA (1325G>A), TGT396TGC (1328T>C), GGT397GGC (1331T>C), TCT399TCC (1337T>C), ACA400ACG (1340A>G), TCC401AGC (1341T>A 1342C>G), CGC403CGA (1349C>A), TTA404CTG (1350T>C 1352A>G), TTT405TTC (1355T>C), ATT406ATC (1358T>C), GTG409GTA (1367G>A), GAC411GAT (1373C>T), AGA412GCC (1374A>G 1375G>C 1376A>C), GGT414CAA (1380G>C 1381G>A 1382T>A), CCA416CCC (1388A>C), AAA417AAG (1391A>G), CTC418CTG (1394C>G), AAT419AAC (1397T>C), CGG420AGG (1398C>A), CTA421TTG (1401C>T 1403A>G), GTA424GTT (1412A>T), AGG427CGC (1419A>C 1421G>C), GCA429GCT (1427A>T), TAT430TAC (1430T>C), TCC432TCA (1436C>A), AAC433AAT (1439C>T), CTG441CTC (1463G>C), GAC444GAT (1472C>T), GGA446GGT (1478A>T), TTG448CTG (1482T>C), CGA449CGG (1487A>G), GGG451GGA (1493G>A), GGG453GGC (1499G>C), CAG454CAA (1502G>A), CAA456CAG (1508A>G), GAT457GAC (1511T>C), AAA459AAG (1517A>G), GAT460GAC (1520T>C), CTG461CTC (1523G>C), GAT463GAC (1529T>C), TCA466TCC (1538A>C), CCC471CCA (1553C>A), TTA472TTG (1556A>G), CCA473CCC (1559A>C), GCC475GCT (1565C>T), GTT477GTA (1571T>A), GAG478GAA (1574G>A), GGC481GGG (1583C>G), AGG482CGG (1584A>C), ACC485ACA (1595C>A), TCC486TCA (1598C>A), TAT488TAC (1604T>C), ACA492ACC (1616A>C), AAT494AAC (1622T>C), CTG495TTG (1623C>T), ATC496GTT (1626A>G 1628C>T), GCT498GCA (1634T>A), CCT499CCA (1637T>A), GCC500GCA (1640C>A), TGT502TGC (1646T>C), GGT503GGG (1649T>G), CTT504CTG (1652T>G), CCA507CCT (1661A>T), GAG511GAA (1673G>A), CGG512AGG (1674C>A), AGA513AGG (1679A>G), AGG514CGG (1680A>C), ATC516ATT (1688C>T), CTG517TTG (1689C>T), GAG519GAA (1697G>A), TGT520TGC (1700T>C), ATG521CGG (1701A>C 1702T>G), AAG522AAA (1706G>A), GAG523GTG (1708A>T), GGC524GGG (1712C>G), GAG525GAA (1715G>A), TTT526TTC (1718T>C), TCC527TCA (1721C>A), ACT528ACA (1724T>A), GAA529GAG (1727A>G), GAT530AAC (1728G>A 1730T>C), CCT531CCA (1733T>A), ACG532ACA (1736G>A), GGA533GCA (1738G>C), AGA536AGC (1748A>C), TTT537TTC (1751T>C), ATA539ATC (1757A>C), GTG540ATT (1758G>A 1760G>T), GAG541GGA (1762A>G 1763G>A), AGA543AGG (1769A>G), AGA544AGG (1772A>G), GCT547GCC (1781T>C), ATC548ATA (1784C>A), ACT549ACA (1787T>A), AGG551AGA (1793G>A), AAT552GAC (1794A>G 1796T>C), TTG553CTC (1797T>C 1799G>C), ACC554ACT (1802C>T), AGT556TCT (1806A>T 1807G>C), GAT557GAC (1811T>C), CTT558TTG (1812C>T 1814T>G), GAG560GAA (1820G>A), GTA563GTG (1829A>G), AAA565AAG (1835A>G), TGT566TGC (1838T>C), CAG567AAG (1839C>A), GTA568GTC (1844A>C), CAT569CAC (1847T>C), GCA570CAC (1848G>C 1849C>A 1850A>C), GTA572ATT (1854G>A 1856A>T), TCA573GCA (1857T>G), AAG574ACC (1861A>C 1862G>C), TTG576CTC (1866T>C 1868G>C), TTT577TAC (1870T>A 1871T>C), GAA578AAC (1872G>A 1874A>C), GCT579CAG (1875G>C 1876C>A 1877T>G), GAA580TCA (1878G>T 1879A>C), TCT581ACA (1881T>A 1883T>A), GGC582GGT (1886C>T), ATA583TGT (1887A>T 1888T>G 1889A>T), AGT584GAA (1890A>G 1891G>A 1892T>A), CCT585GAG (1893C>G 1894C>A 1895T>G), AAG586AAA (1898G>A), ATA587GTT (1899A>G 1901A>T), GCT588GCC (1904T>C), GTT590GTG (1910T>G), CAA591GAA (1911C>G), GCA594GTT (1921C>T 1922A>T), GAT595GAA (1925T>A), GAT596GAA (1928T>A), ATA598ATG (1934A>G), GCG599ACA (1935G>A 1937G>A), TCT600TCC (1940T>C), GTG601TTT (1941G>T 1943G>T), GAT602CAT (1944G>C), GGG603GAT (1948G>A 1949G>T), AGA605CAA (1953A>C 1954G>A), TTG606ATA (1956T>A 1958G>A), CGC607AGC (1959C>A), ACC608TTT (1962A>T 1963C>T 1964C>T), AAG609GAC (1965A>G 1967G>C), CAA610GAA (1968C>G), GAT611GTT (1972A>T), GAA612AGC (1974G>A 1975A>G 1976A>C), CCA613CCC (1979A>C), ATA614AGT (1981T>G 1982A>T), TTG615GTT (1983T>G 1985G>T), GTG616_CCC617del (1986_1991delGTGCCC), ACT618ACA (1994T>A), GTG620CTC (1998G>C 2000G>C), AGT621TCT (2001A>T 2002G>C), GAA622GAG (2006A>G), AAT623GAT (2007A>G), ATC627GGA (2019A>G 2020T>G 2021C>A), GCA629TCA (2025G>T), AGA630CAG (2028A>C 2029G>A 2030A>G), GAG631GAA (2033G>A), CTC632CTA (2036C>A), ACG633ACT (2039G>T), GTA634GTG (2042A>G), GAG635GAA (2045G>A), TTG637CTG (2049T>C), AAA638AAG (2054A>G), GCA640GCT (2060A>T), TAC641TAT (2063C>T), CAG642CAA (2066G>A), GGC643GGA (2069C>A), AGT644GCA (2070A>G 2071G>C 2072T>A), TTG645TTA (2075G>A), GAC646AAC (2076G>A), CCT647CCA (2081T>A), GAG648TTT (2082G>T 2083A>T 2084G>T), CTT650TAT (2088C>T 2089T>A), CCT652CCC (2096T>C), GAC654GAT (2102C>T), CAT655GCT (2103C>G 2104A>C), CAT656TCA (2106C>T 2107A>C 2108T>A), GCC659GCA (2117C>A), ATG660ATT (2120G>T), TAT661TTT (2122A>T), TCT663GTC (2127T>G 2128C>T 2129T>C), CTT664CTG (2132T>G), GAT665TCA (2133G>T 2134A>C 2135T>A), GAT666AGT (2136G>A 2137A>G), CAT668GCC (2142C>G 2143A>C 2144T>C), ATT671AGC (2152T>G 2153T>C), TTC672TTT (2156C>T), AAC673ACT (2158A>C 2159C>T), AAA674CGC (2160A>C 2161A>G 2162A>C), AGA676GTA (2166A>G 2167G>T), GTC677AAT (2169G>A 2170T>A 2171C>T), AAC678GAG (2172A>G 2174C>G), TTG680ATT (2178T>A 2180G>T), TAT681TAC (2183T>C), AGA682GAG (2184A>G 2185G>A 2186A>G), GGA683GGG (2189A>G), GCA684GCT (2192A>T), GAT685TCG (2193G>T 2194A>C 2195T>G), GCT686ACT (2196G>A), GAA687GCA (2200A>C), CAG688GGA (2202C>G 2203A>G 2204G>A), CGA690GCC (2208C>G 2209G>C 2210A>C), GTA693ATC (2217G>A 2219A>C), CAA694AAG (2220C>A 2222A>G), AAC695GCC (2223A>G 2224A>C), ATT696ATC (2228T>C), CCT697CCA (2231T>A), ATA701GTC (2241A>G 2243A>C), AAG705AAA (2255G>A), AGT706AGC (2258T>C), GCA710AGT (2268G>A 2269C>G 2270A>T), TCA711ACA (2271T>A), ATC713GTG (2277A>G 2279C>G), TCT714TCA (2282T>A), GAG715GAA (2285G>A), AAG716GCT (2286A>G 2287A>C 2288G>T), AAA717CAC (2289A>C 2291A>C), CTA718ATG (2292C>A 2294A>G), GCA719GAT (2296C>A 2297A>T), GTG720GTA (2300G>A), CAG721CAA (2303G>A), ACA722AGG (2305C>G 2306A>G), GAG723ATG (2307G>A 2308A>T), ATG724TTG (2310A>T), AGG725TAC (2313A>T 2314G>A 2315G>C), ACT726AAT (2317C>A), GGA727GGC (2321A>C), TTT728AGA (2322T>A 2323T>G 2324T>A), TCT731AGA (2331T>A 2332C>G 2333T>A), TGC732GCA (2334T>G 2335G>C 2336C>A), ATT733CTG (2337A>C 2339T>G), GAT734GAA (2342T>A), GCA735GCT (2345A>T), GTT737GCT (2350T>C), ACC739ACT (2357C>T), CTC740TTA (2358C>T 2360C>A), ATA741ATG (2363A>G), ATC743ATT (2369C>T), GAG744GAA (2372G>A), CAG745CAA (2375G>A), CCA747CCT (2381A>T), TTT749CAT (2385T>C 2386T>A), GTT750GTC (2390T>C), CAG751CAA (2393G>A), TTT752TTC (2396T>C), TTA753CTA (2397T>C), TAT754TAC (2402T>C), AGA759CGG (2415A>C 2417A>G), TGC762_AAC763insAGAAGG (2426_2427insAGAAGG), AAC763AAG (2429C>G), AAA764ATG (2431A>T 2432A>G), GTA765GTC (2435A>C), TCA766ACT (2436T>A 2438A>T), AAA767AAG (2441A>G), GAG768GAA (2444G>A), CCA769ACA (2445C>A), AAG770_GAA771del (2448_2453delAAGGAA), TCA772TCT (2456A>T), ATC774GTA (2460A>G 2462C>A), AAA775CAA (2463A>C), ATT776TCG (2466A>T 2467T>C 2468T>G), TGC777CTC (2469T>C 2470G>T), TAC778TAT (2474C>T), CAT779GAC (2475C>G 2477T>C), AAA780TCC (2478A>T 2479A>C 2480A>C), ATC781ACA (2482T>C 2483C>A), GTT782ATT (2484G>A), GAA783GGG (2488A>G 2489A>G), TAT784AGA (2490T>A 2491A>G 2492T>A), ATC785ATG (2495C>G), AAG786CAA (2496A>C 2498G>A), GAT787GAA (2501T>A), ACA788ATG (2503C>T 2504A>G), TGG790AAT (2508T>A 2509G>A 2510G>T), AGC791GAT (2511A>G 2512G>A 2513C>T), CTT792CTG (2516T>G), TAT794TAC (2522T>C), GCA795ATT (2523G>A 2524C>T 2525A>T), CTG796TTG (2526C>T), AGA797AAA (2530G>A), CTC799CTT (2537C>T), GGT803GGG (2549T>G), ATA806CTA (2556A>C), ATG807ATA (2561G>A), CTT809CTC (2567T>C), GGA811GGG (2573A>G), GTT812ACT (2574G>A 2575T>C), TTT813ATG (2577T>A 2579T>G), GGA814GGT (2582A>T), GGC815GCC (2584G>C), ATT816GTA (2586A>G 2588T>A), ACA817ACT (2591A>T), TTT819TTC (2597T>C), TTG820ATT (2598T>A 2600G>T), GCG821TCT (2601G>T 2603G>T), CAG824AAA (2610C>A 2612G>A), AGT825AGC (2615T>C), AAC826AGC (2617A>G), TCT827TTG (2620C>T 2621T>G), TTC828ATT (2622T>A 2624C>T), AAT830ACT (2629A>C), GCA831TCC (2631G>T 2633A>C), TCG832CAG (2634T>C 2635C>A), AGA834AAA (2641G>A), GTG837GTT (2651G>T), TCC841TCG (2663C>G), AGT843AGC (2669T>C), TCC845TCA (2675C>A), AGT846AGC (2678T>C), GGT851GGA (2693T>A), AAT852AGA (2695A>G 2696T>A), AAT852_AAG853insTCC (2696_2697insTCC), AAG853CAT (2697A>C 2699G>T), CTC854GTA (2700C>G 2702C>A), AGA855TCA (2703A>T 2704G>C), AAT856AAG (2708T>G), CTT857ATG (2709C>A 2711T>G), CTC858CTG (2714C>G), GTT859TTT (2715G>T), GGT860AGC (2718G>A 2720T>C), TCT863TCA (2729T>A), TCA865CCA (2733T>C), TTG866AAT (2736T>A 2737T>A 2738G>T), GCA867GAT (2740C>A 2741A>T), CAA868CAG (2744A>G), CAA868_GAT869insGGA (2744_2745insGGA), GAT869AAA (2745G>A 2747T>A), GCT872TCC (2754G>T 2756T>C), GAA873AGG (2757G>A 2758A>G 2759A>G), GGA875GGT (2765A>T), GTA877GTG (2771A>G), AAT878AAC (2774T>C), CAG879GAA (2775C>G 2777G>A), AGC880TCT (2778A>T 2779G>C 2780C>T), CTC881CTT (2783C>T), AAG882GCC (2784A>G 2785A>C 2786G>C), GTG886GTA (2798G>A), TTA888CTA (2802T>C), AGA889AGG (2807A>G), CTT890GTG (2808C>G 2810T>G), GGT891GCA (2812G>C 2813T>A), GTG894GTT (2822G>T), AGA897CGT (2829A>C 2831A>T), GGC898GGA (2834C>A), ACC899ACT (2837C>T), TGT902TGC (2846T>C), TCT903AGC (2847T>A 2848C>G 2849T>C), GGT904GAA (2851G>A 2852T>A), ACC908ACT (2864C>T), GCT910GCA (2870T>A), CAT911CAC (2873T>C), GCA912GCT (2876A>T), CAC914ACG (2880C>A 2881A>C 2882C>G), AAT915CAG (2883A>C 2885T>G), GCT916CTC (2886G>C 2887C>T 2888T>C), CGA917CGT (2891A>T), GGT919GGA (2897T>A), ACT920ACC (2900T>C), ACA923TCA (2907A>T), ATC925ATT (2915C>T), GCC927GAA (2920C>A 2921C>A), AAT928CAT (2922A>C), CGA930GTC (2928C>G 2929G>T 2930A>C), TCT931TGC (2932C>G 2933T>C), AAA932CAG (2934A>C 2936A>G), GTT933GTG (2939T>G), AAC936AAT (2948C>T), AGA938AAA (2953G>A), GAA939CAT (2955G>C 2957A>T), TCA940GCC (2958T>G 2960A>C), AGG941AAG (2962G>A), ATT942TTT (2964A>T), ATT943TTG (2967A>T 2969T>G), GAG944CCT (2970G>C 2971A>C 2972G>T), GGC945GAA (2974G>A 2975C>A), GAC947GAT (2981C>T), ATC948GTT (2982A>G 2984C>T), ATT949GTA (2985A>G 2987T>A), CTG950TTA (2988C>T 2990G>A), TTG951CTG (2991T>C), CGC952AGG (2994C>A 2996C>G), GGT954GGC (3002T>C), GAT955TGT (3003G>T 3004A>G), CCA956CCT (3008A>T), GAT957GAC (3011T>C),

Codon mutations:

GGT958GGG (3014T>G), CCA959CCT (3017A>T), AAG960AAA (3020G>A), CCT961GTC (3021C>G 3022C>T 3023T>C), CGT964AGA (3030C>A 3032T>A), AAA965AAG (3035A>G), TTC967TTT (3041C>T), CCA968GCT (3042C>G 3044A>T), AGA969GAA (3045A>G 3046G>A), GAG972GCA (3055A>C 3056G>A), AAG977AAA (3071G>A), GGC978GGA (3074C>A), TCA979TCT (3077A>T), CAA980GAA (3078C>G), GGA981GGC (3083A>C), TGC983TGT (3089C>T), TGT984GTC (3090T>G 3091G>T 3092T>C), GCT986GCA (3098T>A), GTT987GTG (3101T>G), GCG988GCA (3104G>A), TCT989TCA (3107T>A), ACA990AAA (3109C>A), CCA992CCT (3116A>T), CGT993GAG (3117C>G 3118G>A 3119T>G), CTT994AAA (3120C>A 3121T>A 3122T>A), GGC995CAG (3123G>C 3124G>A 3125C>G), AAT996AAC (3128T>C), TTA997TTG (3131A>G), GAG998GAC (3134G>C), ATG1002TAT (3144A>T 3145T>A 3146G>T), CCA1003TCA (3147C>T), TCA1007TCC (3161A>C), AAG1008AAA (3164G>A), TAC1010TAT (3170C>T), GTT1012GAG (3175T>A 3176T>G), GTG1014GTC (3182G>C), GAA1015ACA (3183G>A 3184A>C), TAT1016TAC (3188T>C), GAA1017AGG (3189G>A 3190A>G 3191A>G), CTG1018ATG (3192C>A), AAC1019AGC (3196A>G), TCC1020ACA (3198T>A 3200C>A), TCC1021AAT (3201T>A 3202C>A 3203C>T), AGT1022TCA (3204A>T 3205G>C 3206T>A), AAG1024CAC (3210A>C 3212G>C), AAG1024_ATT1025insTCA (3212_3213insTCA), ATT1025ATC (3215T>C), TGC1026AGT (3216T>A 3218C>T), TCT1027GCT (3219T>G), CAA1029GAA (3225C>G), TCT1030GCC (3228T>G 3230T>C), CAC1034CAT (3242C>T), ATA1035GTC (3243A>G 3245A>C), GGG1037AGC (3249G>A 3251G>C), GAC1041GAT (3263C>T), GGC1043GGA (3269C>A), ACG1044ACT (3272G>T), CTG1047TTA (3279C>T 3281G>A), TTA1048TTG (3284A>G), AGT1050TCT (3288A>T 3289G>C), GTT1051ATC (3291G>A 3293T>C), CAG1052CAA (3296G>A), AAT1053AAC (3299T>C), CCT1056CCA (3308T>A), GTG1057TGC (3309G>T 3310T>G 3311G>C), ATC1058ATT (3314C>T), GGC1060GGT (3320C>T), ATC1061ATT (3323C>T), CAC1062CAT (3326C>T), GCA1064GCT (3332A>T), TCA1065TCC (3335A>C), GGC1068GAA (3343G>A 3344C>A), GTA1069GCT (3346T>C 3347A>T), GCT1071ACA (3351G>A 3353T>A), GAA1072GAG (3356A>G), CGT1073TTG (3357C>T 3358G>T 3359T>G), ATC1076GTA (3366A>G 3368C>A), TCT1077GCA (3369T>G 3371T>A), GCT1080GCC (3380T>C), ACA1081ACT (3383A>T), ATC1083GTA (3387A>G 3389C>A), GAA1086GAC (3398A>C), CAA1087CAG (3401A>G), CTA1088CTG (3404A>G), GAG1089CAA (3405G>C 3407G>A), CCA1092GCT (3414C>G 3416A>T), ACT1093CAA (3417A>C 3418C>A 3419T>A), CCG1095CCA (3425G>A), GTC1096GTT (3428C>T), GCA1098ATG (3432G>A 3433C>T 3434A>G), GCA1099GCC (3437A>C), GTA1101GTG (3443A>G), CGC1102AGA (3444C>A 3446C>A), GAT1104GAC (3452T>C), ATT1105TTG (3453A>T 3455T>G), CTG1106CTC (3458G>C), AAG1107AAA (3461G>A), TCA1108CAT (3462T>C 3463C>A 3464A>T), ATT1109GTT (3465A>G), AGA1110CGA (3468A>C), AGC1111TCA (3471A>T 3472G>C 3473C>A), ACA1114ACT (3482A>T), CAG1115GCT (3483C>G 3484A>C 3485G>T), CTT1116CTC (3488T>C), GAG1118GAC (3494G>C), AAC1120AAG (3500C>G), GTG1122GTT (3506G>T), TAC1123GCT (3507T>G 3508A>C 3509C>T), TAC1124TTT (3511A>T 3512C>T), CTT1125TTG (3513C>T 3515T>G), GGA1126GGC (3518A>C), ACA1127ACT (3521A>T), CCA1129CCT (3527A>T), CAG1130CAA (3530G>A), TTG1132ATG (3534T>A), GCC1133GCA (3539C>A), GCC1134GCA (3542C>A), ACA1135ACG (3545A>G), GTT1136GTG (3548T>G), CCC1137CCA (3551C>A), ACT1141ACC (3563T>C), CTG1142CTC (3566G>C), CGG1143AGG (3567C>A), AGC1145AGT (3575C>T), CAA1146CCA (3577A>C), TTG1147CTG (3579T>C), GAA1149GAG (3587A>G), TTC1151TTT (3593C>T), GGA1152GGT (3596A>T), CCT1153GAA (3597C>G 3598C>A 3599T>A), GAG1155CAA (3603G>C 3605G>A), ACA1156ACT (3608A>T), GCA1157GCT (3611A>T), CCA1158CCT (3614A>T), TCC1159TCA (3617C>A), ATT1160CTG (3618A>C 3620T>G), CTA1161CTC (3623A>C), ACA1162TCT (3624A>T 3626A>T), GTT1163GTG (3629T>G), CAT1164AAA (3630C>A 3632T>A), GAC1165GAT (3635C>T), GGT1168GGG (3644T>G), GGT1170GGA (3650T>A), GAC1172GAT (3656C>T), CCC1173CCT (3659C>T), GTG1175GTT (3665G>T), GGG1177GGC (3671G>C), GTA1178ATT (3672G>A 3674A>T), ATG1179GAG (3675A>G 3676T>A), GCT1185GCC (3695T>C), TGT1186TCA (3697G>C 3698T>A), GGA1187AAT (3699G>A 3700G>A 3701A>T), GAT1189GAG (3707T>G), GAT1190GAC (3710T>C), GAC1191AGC (3711G>A 3712A>G), GCC1193GCA (3719C>A), AAA1194CGG (3720A>C 3721A>G 3722A>G), CTA1195GCT (3723C>G 3724T>C 3725A>T), TTC1197TTT (3731C>T), AAT1199GGT (3735A>G 3736A>G), TGC1202TCT (3745G>C 3746C>T), TCG1203GCC (3747T>G 3749G>C), CTG1204ATT (3750C>A 3752G>T), CTA1205TTA (3753C>T), CCT1206CCA (3758T>A), ATC1207GTT (3759A>G 3761C>T), ATG1208CTC (3762A>C 3764G>C), CGT1209AAG (3765C>A 3766G>A 3767T>G), AGC1210AGT (3770C>T), CAG1211CAA (3773G>A), AAG1212AGG (3775A>G), ATC1213GTG (3777A>G 3779C>G), CCT1214CCA (3782T>A), GGG1215GGT (3785G>T), GGA1216GGT (3788A>T), CGT1217ATC (3789C>A 3790G>T 3791T>C), CCA1218CCC (3794A>C), TGT1219ACA (3795T>A 3796G>C 3797T>A), GAA1220ACA (3798G>A 3799A>C), AGG1221CGT (3801A>C 3803G>T), GAT1222TGT (3804G>T 3805A>G), GAT1224GAC (3812T>C), GTA1225ATC (3813G>A 3815A>C), GTG1226GTT (3818G>T), CTC1227CTG (3821C>G), GGA1229GGT (3827A>T), ATA1230GTT (3828A>G 3830A>T), GGA1232GGT (3836A>T), TGT1233TTT (3838G>T), GAT1234GAC (3842T>C), TAC1235TTC (3844A>T), TAT1236TTT (3847A>T), GGC1238TCC (3852G>T 3853G>C), GAG1240GAA (3860G>A), CTG1241CTC (3863G>C), TCT1244TCA (3872T>A), TGC1245TGT (3875C>T), TAT1247TAC (3881T>C), CCC1248CCT (3884C>T), AAC1250AAT (3890C>T), AAG1251AAA (3893G>A), ATG1252CTG (3894A>C), GGG1253GGT (3899G>T), ATG1254TTG (3900A>T), GGG1255GGT (3905G>T), ATG1256ACC (3907T>C 3908G>C), AAC1257AAT (3911C>T), AAG1258AAA (3914G>A), AGA1259AGG (3917A>G), TTT1261TTC (3923T>C), GTG1262GTT (3926G>T), CAA1263GAA (3927C>G), TCC1264GCA (3930T>G 3932C>A), ACT1265CAG (3933A>C 3934C>A 3935T>G), GGC1266GAG (3937G>A 3938C>G), GAA1267AGT (3939G>A 3940A>G 3941A>T), GGA1268ACC (3942G>A 3943G>C 3944A>C), GAG1269CAA (3945G>C 3947G>A), AGA1270AGG (3950A>G), GTG1271GTA (3953G>A), GAA1272CTA (3954G>C 3955A>T), AAA1274AAG (3962A>G), GAC1276AAT (3966G>A 3968C>T), ACT1277TCT (3969A>T), GTA1279GTT (3977A>T), TTT1280TTC (3980T>C), GAA1281CAA (3981G>C), GAA1284GAG (3992A>G), GAG1285GAA (3995G>A), CTA1286TTA (3996C>T), CAG1289GAA (4005C>G 4007G>A), ATT1290ATA (4010T>A), AAA1292AAG (4016A>G), GGC1293GGT (4019C>T), ATC1294AAC (4021T>A), CAT1295TAT (4023C>T), GAT1297GAA (4031T>A), CTG1298ATG (4032C>A), GTC1299GTT (4037C>T), ACC1301ACT (4043C>T), TGC1303TGT (4049C>T), GCC1304GCT (4052C>T), AAA1305AAG (4055A>G), GAT1306GAC (4058T>C), CGC1308AGG (4062C>A 4064C>G), CTC1309CTA (4067C>A), CCA1310CCT (4070A>T), CTT1311CTG (4073T>G), GAG1312GAC (4076G>C), AAA1313AAG (4079A>G), ATC1314ATA (4082C>A), TAT1315TAC (4085T>C), GGC1316GGG (4088C>G), AAG1317AAA (4091G>A), AGA1318AAG (4093G>A 4094A>G), AAG1319AAA (4097G>A), AGG1321CGT (4101A>C 4103G>T), CTC1322CTT (4106C>T), ATA1325ATT (4115A>T), CTT1326TTG (4116C>T 4118T>G), CCT1327CCA (4121T>A), TAC1330TAT (4130C>T), TTG1333CTG (4137T>C), GTC1334GTG (4142C>G), AGG1335CGC (4143A>C 4145G>C), AAG1336AAA (4148G>A), TTT1338TTC (4154T>C), TTC1341TTT (4163C>T), TCA1342TCT (4166A>T), GCC1343GCA (4169C>A), ACA1344TGT (4170A>T 4171C>G 4172A>T), TTG1345TTA (4175G>A), TTG1348GCA (4182T>G 4183T>C 4184G>A), CAC1349CAT (4187C>T), GCT1351GTT (4192C>T), ATA1352CTG (4194A>C 4196A>G), CCA1353CCC (4199A>C), GTT1356GTG (4208T>G), ATT1358ATC (4214T>C), GAT1359AAT (4215G>A), CCT1360CCC (4220T>C), ACA1361ATT (4222C>T 4223A>T), AGT1362TCA (4224A>T 4225G>C 4226T>A), TCT1363CAT (4227T>C 4228C>A), ACA1366ACC (4238A>C), TTG1367ACC (4239T>A 4240T>C 4241G>C), TTG1368CTG (4242T>C), AAT1370AAC (4250T>C), GGG1371CAG (4251G>C 4252G>A), TTT1372TTC (4256T>C), AGA1373CGG (4257A>C 4259A>G), GCT1374AGT (4260G>A 4261C>G), TCA1376TCT (4268A>T), GAC1377GAT (4271C>T), GTG1378GTT (4274G>T), GGA1379GGT (4277A>T), TCA1381TCT (4283A>T), GCT1382GCA (4286T>A), GAT1383GAC (4289T>C), TAT1384TAC (4292T>C), AGA1390AGG (4310A>G), GCA1391GCG (4313A>G), CCT1392CCA (4316T>A), GTT1393ATA (4317G>A 4319T>A), GCT1395GCA (4325T>A), CAG1397CAA (4331G>A), GAA1405GCA (4354A>C), TAC1407TAT (4361C>T), GGA1408GGG (4364A>G), AAG1410GCT (4368A>G 4369A>C 4370G>T), CCT1411CCA (4373T>A), GGC1412GGA (4376C>A), AGT1413AGC (4379T>C), CCT1414GAG (4380C>G 4381C>A 4382T>G), TCC1416TCA (4388C>A), GCT1418GCA (4394T>A), CGA1419AGA (4395C>A), GCA1421GCT (4403A>T), CTT1422CTG (4406T>G), TTA1423TTG (4409A>G), GCA1426GCT (4418A>T), TGT1428TGC (4424T>C), ACA1431ACT (4433A>T), CTG1432CTA (4436G>A), GTG1437CTC (4449G>C 4451G>C), TTT1438TTC (4454T>C), TTG1440TTA (4460G>A), GGG1442GGT (4466G>T), GGC1443GGA (4469C>A), CCA1445CCT (4475A>T), TCA1446TCT (4478A>T), GCA1449GCT (4487A>T), CTA1450CTC (4490A>C), ACG1451ACC (4493G>C), ATC1453ATA (4499C>A), TTC1454TTT (4502C>T), AAT1455AAC (4505T>C), TCT1456TCC (4508T>C), CTC1457CTT (4511C>T), CTC1458TTA (4512C>T 4514C>A), AAT1459AAC (4517T>C), CGA1464CGG (4532A>G), GCC1466GCT (4538C>T), TTT1467TTC (4541T>C), TCA1469TCG (4547A>G), TTG1470CTC (4548T>C 4550G>C), TTA1471CTC (4551T>C 4553A>C), AGA1473AGG (4559A>G), CAT1475CAC (4565T>C), ATT1476ATC (4568T>C), GCT1477AAA (4569G>A 4570C>A 4571T>A), GCC1478GCA (4574C>A), AGG1479AAG (4576G>A), GCT1480GCA (4580T>A), ATA1481TTG (4581A>T 4583A>G), GGA1482GGT (4586A>T), GTT1483GTA (4589T>A), AAA1484AAG (4592A>G), CCA1485TCA (4593C>T), TTC1488TTT (4604C>T), CAG1490GAC (4608C>G 4610G>C), CTA1491TTA (4611C>T), TTC1492TTT (4616C>T), ATA1493ATT (4619A>T), GCT1494GCA (4622T>A), GTT1495GTG (4625T>G), GGA1497GGG (4631A>G), GTT1502GTG (4646T>G), GTA1504GTG (4652A>G), TTA1506ATG (4656T>A 4658A>G), CAT1507CAC (4661T>C), CTC1508CTT (4664C>T), CAG1509CAA (4667G>A), TAT1511TAC (4673T>C), TCT1512GAG (4674T>G 4675C>A 4676T>G), CTG1513TTA (4677C>T 4679G>A), CCA1514CCT (4682A>T), AAT1515GCA (4683A>G 4684A>C 4685T>A), CAT1518CAA (4694T>A), TTA1520CTA (4698T>C), CTG1522CAA (4705T>A 4706G>A), GTC1523GTG (4709C>G), AAT1524AAC (4712T>C), GTA1525GTT (4715A>T), ATC1526ATT (4718C>T), ATT1527ATC (4721T>C), GGT1530GGC (4730T>C), GAC1532GAT (4736C>T), TCG1534AAC (4740T>A 4741C>A 4742G>C), GAT1536AAT (4746G>A), GTT1537GTC (4751T>C), GAG1539GAT (4757G>T), GTA1540GTC (4760A>C), CAA1541AAA (4761C>A), TTT1542TTC (4766T>C), CAA1543CAG (4769A>G), CTA1548TTG (4782C>T 4784A>G), ACT1549ACG (4787T>G), CTG1551TTG (4791C>T), AGT1552AGC (4796T>C), GGT1554GGC (4802T>C), TTT1555TTC (4805T>C), AAA1556AAG (4808A>G), CGA1557CGC (4811A>C), CTT1560TTG (4818C>T 4820T>G), GGA1561GGT (4823A>T), CAC1563CAA (4829C>A), ATG1564CTA (4830A>C 4832G>A), GCT1565GCC (4835T>C), CCT1566CCC (4838T>C), TCA1571TCC (4853A>C), ATC1572ATT (4856C>T), ATT1573GTG (4857A>G 4859T>G), GAG1574GAA (4862G>A), CCC1575CCA (4865C>A), ATA1579ATC (4877A>C), CCT1581CCA (4883T>A), GCC1582GCA (4886C>A), CCA1583GTT (4887C>G 4888C>T 4889A>T), GGT1584GGG (4892T>G), CCT1586CCA (4898T>A), GAC1587GAT (4901C>T), ACT1588ATC (4903C>T 4904T>C), CAG1589ACT (4905C>A 4906A>C 4907G>T), ATG1592TAT (4914A>T 4915T>A 4916G>T), ACA1597ACG (4931A>G), GGA1598GGT (4934A>T), ATA1599GTA (4935A>G), CGT1600AGA (4938C>A 4940T>A), GAA1601GAG (4943A>G), CAT1604CAG (4952T>G), CAC1605CAT (4955C>T), GGT1606GGA (4958T>A), CGT1607CAG (4960G>A 4961T>G), GTG1608GTC (4964G>C), GCT1609GTT (4966C>T), AAG1612ACA (4975A>C 4976G>A), CTT1613TTG (4977C>T 4979T>G), GTC1614ACA (4980G>A 4981T>C 4982C>A), TTA1615GAC (4983T>G 4984T>A 4985A>C), GAT1616AAT (4986G>A), GTT1617GTC (4991T>C), CAG1618CAA (4994G>A), ACG1619AAA (4996C>A 4997G>A), GCG1620GCA (5000G>A), TTG1621ATA (5001T>A 5003G>A), GAT1622ATG (5004G>A 5005A>T 5006T>G), GAA1623CAG (5007G>C 5009A>G), AGG1624CGA (5010A>C 5012G>A), GGT1625GGC (5015T>C), TTC1626TTG (5018C>G), AGA1627CCA (5019A>C 5020G>C), GCT1628ATC (5022G>A 5023C>T 5024T>C), ATC1630CAT (5028A>C 5029T>A 5030C>T), CCT1632CCA (5036T>A), TCC1633TCT (5039C>T), TTG1635GCT (5043T>G 5044T>C 5045G>T), GTG1637CAA (5049G>C 5050T>A 5051G>A), CAG1639AGT (5055C>A 5056A>G 5057G>T), GAA1640GAG (5060A>G), ATT1642CTA (5064A>C 5066T>A), GCA1643CAA (5067G>C 5068C>A), AAG1644GCC (5070A>G 5071A>C 5072G>C), GTA1645GTT (5075A>T), ACA1646ACT (5078A>T), GGG1647GGA (5081G>A), GAT1648GAC (5084T>C), TCA1649AGT (5085T>A 5086C>G 5087A>T), AGT1650AAC (5089G>A 5090T>C), GCC1651ATA (5091G>A 5092C>T 5093C>A), CTG1652GGG (5094C>G 5095T>G), ACA1653GCT (5097A>G 5099A>T), ATT1654CTG (5100A>C 5102T>G), TGT1655CTG (5103T>C 5104G>T 5105T>G), ATG1657CTG (5109A>C), GCA1658AGC (5112G>A 5113C>G 5114A>C), AAA1659AAG (5117A>G), GCA1660GAT (5119C>A 5120A>T), TCC1663ACT (5127T>A 5129C>T), TAT1664TAC (5132T>C), ACG1665GTG (5133A>G 5134C>T), CCA1666CCT (5138A>T), TTG1667CTT (5139T>C 5141G>T), GAC1668CCA (5142G>C 5143A>C 5144C>A), GCA1669GAA (5146C>A), GAG1671AAA (5151G>A 5153G>A), AAG1672AAA (5156G>A), ATC1673ATT (5159C>T), ACA1674GAA (5160A>G 5161C>A), AAT1675GAT (5163A>G), GAG1677TCA (5169G>T 5170A>C 5171G>A), CGT1678CAA (5173G>A 5174T>A), GAT1679GGT (5176A>G), CTG1680GTC (5178C>G 5180G>C), AAT1681CAT (5181A>C), TGG1682GAA (5184T>G 5185G>A 5186G>A), GCA1684TGT (5190G>T 5191C>G 5192A>T), CCA1685TCT (5193C>T 5195A>T), AAC1686GGA (5196A>G 5197A>G 5198C>A), ATT1687GTT (5199A>G), GGT1688TTC (5202G>T 5203G>T 5204T>C), TTT1689TTC (5207T>C), GCA1692TCG (5214G>T 5216A>G), CGT1693CGA (5219T>A), ACT1694ACA (5222T>A), GCA1695GAA (5224C>A), GCC1696GTT (5227C>T 5228C>T), CAC1697AAT (5229C>A 5231C>T), TAC1698CAT (5232T>C 5234C>T), ACG1699AAA (5236C>A 5237G>A), GAA1702GAG (5246A>G), GGG1703GAC (5248G>A 5249G>C), TAC1704ACA (5250T>A 5251A>C 5252C>A), ATT1706CTT (5256A>C), AAC1708AAT (5264C>T), TGT1709TGC (5267T>C), AAG1713AAA (5279G>A), AAG1714AAA (5282G>A), TCC1715ACA (5283T>A 5285C>A), AAT1716ACA (5287A>C 5288T>A), GTT1718GTG (5294T>G), GGT1720GGC (5300T>C), CCG1733CCC (5339G>C), TAC1734TAT (5342C>T), ACT1735ACA (5345T>A), TCA1737GCA (5349T>G), GCT1738GCC (5354T>C), ATA1739ATC (5357A>C), ATG1740AAG (5359T>A), GAT1741GAC (5363T>C), GCC1742AAT (5364G>A 5365C>A 5366C>T), AAG1744CAA (5370A>C 5372G>A), AAT1745AGT (5374A>G), GTG1746CGC (5376G>C 5377T>G 5378G>C), GCA1748TCA (5382G>T), GGA1749ATG (5385G>A 5386G>T 5387A>G), GGA1750GGG (5390A>G), CAT1751ATT (5391C>A 5392A>T), GTA1752GGA (5395T>G), GCA1753ACC (5397G>A 5399A>C), ACC1754CAA (5400A>C 5401C>A 5402C>A), AAT1755AAA (5405T>A), GTC1756GTT (5408C>T), GTT1757GTC (5411T>C), GTG1759ATT (5415G>A 5417G>T), GGA1761GGG (5423A>G), GGA1763GGT (5429A>T), GTC1766GTG (5438C>G), GGC1767AGT (5439G>A 5441C>T), CCA1768CCC (5444A>C), GCA1771GCT (5453A>T), GCG1772GCT (5456G>T), ATG1774TTT (5460A>T 5462G>T), GCC1775GCA (5465C>A), GCA1777TCA (5469G>T), ACA1778ACC (5474A>C), AGG1779AAT (5476G>A 5477G>T), CAA1780CAG (5480A>G), TGT1782AGA (5484T>A 5486T>A), GTG1783GCT (5488T>C 5489G>T), GAC1785CAA (5493G>C 5495C>A), ATA1786CTT (5496A>C 5498A>T), GTT1788GTC (5504T>C), CGC1791AGA (5511C>A 5513C>A), ATT1793TTG (5517A>T 5519T>G), GGA1794GCG (5521G>C 5522A>G), AAT1795GGC (5523A>G 5524A>G 5525T>C), GTG1796TTG (5526G>T), CTT1797CTG (5531T>G), GAC1798GAT (5534C>T), CTC1799TTG (5535C>T 5537C>G), ACC1801CAG (5541A>C 5542C>A 5543C>G), GGC1805GGT (5555C>T), GTG1807ATT (5559G>A 5561G>T), CAG1808CAA (5564G>A), TTT1810TTC (5570T>C), CTT1811CTC (5573T>C), CAA1812ACA (5574C>A 5575A>C), TGT1813_GTG1814insGTC (5579_5580insGTC), CCT1815CAG (5584C>A 5585T>G), CAT1816AAA (5586C>A 5588T>A), GGA1817AGT (5589G>A 5591A>T), GAC1818TGC (5592G>T 5593A>G), AAA1819AAG (5597A>G), GTG1820ACA (5598G>A 5599T>C 5600G>A), AGT1823GCT (5607A>G 5608G>C), GGT1824AGT (5610G>A), GCA1825GGG (5614C>G 5615A>G), TCG1826del (5616_5618delTCG), CTC1827CGC (5620T>G), CAC1828GAT (5622C>G 5624C>T), AGT1829GGA (5625A>G 5627T>A), AGT1830ATG (5629G>T 5630T>G), TTT1831TTG (5633T>G), ATG1832AAC (5635T>A 5636G>C), CAA1834CAT (5642A>T), CAA1834_GGG1835insCATGCT (5642_5643insCATGCT), GGG1835GCT (5644G>C 5645G>T), TTT1836TAT (5647T>A), GAA1837GCC (5650A>C 5651A>C), TTG1838CTT (5652T>C 5654G>T), GGC1839GGA (5657C>A), CTC1841CTT (5663C>T), CGC1842CGA (5666C>A), ATT1843ATA (5669T>A), ATA1844GTG (5670A>G 5672A>G), CAT1845GGC (5673C>G 5674A>G 5675T>C), GGT1846TGT (5676G>T), GAT1847TCA (5679G>T 5680A>C 5681T>A), TTA1848ATT (5682T>A 5684A>T), GCA1849TCA (5685G>T), AAA1850AGA (5689A>G), CAG1851AAG (5691C>A), ACA1852ATG (5695C>T 5696A>G), ATG1854GTG (5700A>G), CGG1855AGA (5703C>A 5705G>A), ATG1856TTG (5706A>T), CCA1857CCC (5711A>C), TAT1858TAC (5714T>C), GTG1859ACT (5715G>A 5716T>C 5717G>T), GTA1860GTT (5720A>T), GGA1861GGG (5723A>G), CAT1863ATG (5727C>A 5728A>T 5729T>G), GGA1864GGT (5732A>T), GGA1865GGT (5735A>T), GGA1867GCA (5740G>C), AAT1868GAT (5742A>G), TTT1869TGC (5746T>G 5747T>C), TCC1870TCA (5750C>A), CAG1872AGA (5754C>A 5755A>G 5756G>A), GAT1873AAT (5757G>A), GAG1875GAT (5765G>T), AGT1876TCT (5766A>T 5767G>C), TTG1877CTC (5769T>C 5771G>C), CTA1878ATC (5772C>A 5774A>C), CAC1879AAA (5775C>A 5777C>A), TAT1880GCC (5778T>G 5779A>C 5780T>C), TTG1881CTA (5781T>C 5783G>A), CAG1883TGC (5787C>T 5788A>G 5789G>C), GGA1884TCT (5790G>T 5791G>C 5792A>T), GCA1886GCT (5798A>T), GAG1887GCA (5800A>C 5801G>A), TTA1888TGT (5803T>G 5804A>T), ATT1889GTG (5805A>G 5807T>G), CAA1890GCT (5808C>G 5809A>C 5810A>T), AAG1891CGT (5811A>C 5812A>G 5813G>T), AAT1892GGG (5814A>G 5815A>G 5816T>G), ACA1893TGT (5817A>T 5818C>G 5819A>T), CTA1895ATA (5823C>A), TAT1898TAC (5834T>C), TTC1899TTT (5837C>T), AAA1900GAG (5838A>G 5840A>G), GAG1901AAT (5841G>A 5843G>T), CTG1902TTT (5844C>T 5846G>T), AGT1903AAT (5848G>A), ATG1904GCA (5850A>G 5851T>C 5852G>A), GAA1905CAA (5853G>C), AAT1906ATA (5857A>T 5858T>A), GTG1907GCT (5860T>C 5861G>T), CAA1908ACC (5862C>A 5863A>C 5864A>C), CAA1909AGG (5865C>A 5866A>G 5867A>G), CTG1910TTA (5868C>T 5870G>A), ATA1911GTA (5871A>G), GTT1914CTT (5880G>C),

| | | | | | | | | | |
|---|---|---|---|---|---|---|---|---|---|
| | AAG1915AAA (5885G>A), CTT1916AGT (5886C>A 5887T>G), GGG1918GGT (5894G>T), TTT1919GTG (5895T>G 5897T>G), TTC1920TTT (5900C>T), CCA1921ATC (5901C>A 5902C>T 5903A>C), AAA1922ACA (5905A>C), GAA1923GCC (5908A>C 5909A>C), ACT1924ACA (5912T>A), ACC1925GGC (5913A>G 5914C>G), ATT1926CAC (5916A>C 5917T>A 5918T>C), CAG1927AAA (5919C>A 5921G>A), AAG1928ATG (5923A>T), CTC1929TTG (5925C>T 5927C>G), AAA1930GTG (5928A>G 5929A>T 5930A>G), AAT1931CAA (5931A>C 5933T>A), TTT1932ACT (5934T>A 5935T>C), GTT1933CTG (5937G>C 5939T>G), GAT1934AGC (5940G>A 5941A>G 5942T>C), GCT1935AAT (5943G>A 5944C>A), GAA1936CAA (5946G>C), TGT1937TGG (5951T>G), CTA1938CAA (5953T>A), ACA1939TCC (5955A>T 5957A>C), TTC1940CAC (5958T>C 5959T>A), AAA1941GAA (5961A>G), GCA1942CAA (5964G>C 5965C>A), AGG1943AGT (5969G>T), AGC1944TGT (5970A>T 5972C>T), TTT1945GTG (5973T>G 5975T>G), AGG1946AAG (5977G>A), CAC1947AGA (5979C>A 5980A>G 5981C>A), GTA1948GTC (5984A>C), GTT1949TAT (5985G>T 5986T>A), AAA1951AAG (5993A>G), AAG1952CGA (5994A>C 5995A>G 5996G>A), AAA1953AAT (5999A>T), TTG1955CTC (6003T>C 6005G>C), AGT1956TCA (6006A>T 6007G>C 6008T>A), TCC1957AAC (6009T>A 6010C>A), ACA1958GCA (6012A>G), TGG1959ATA (6015T>A 6016G>T 6017G>A), AAA1960ACA (6019A>C), ATG1961_TGT1962insTAC (6023_6024insTAC), TGT1962CGT (6024T>C), GAA1964GAT (6032A>T), AGT1965GCC (6033A>G 6034G>C 6035T>C), ATT1966AAA (6037T>A 6038T>A), GTT1967GAT (6040T>A), TCA1969ATA (6045T>A 6046C>T), AGG1970GTA (6048A>G 6049G>T 6050G>A), GCA1972del (6054_6056delGCA), CTA1975GTG (6063C>G 6065A>G), CCT1977TTG (6069C>T 6070C>T 6071T>G), GGC1978GGT (6074C>T), TTG1980CCA (6078T>C 6079T>C 6080G>A), GCT1982GCC (6086T>C), TCC1983ATG (6087T>A 6088C>T 6089C>G), GTA1984GTT (6092A>T), AAA1986AGG (6097A>G 6098A>G) | | | | | | | | |
| helicase (YP_001976154.1) | 1 | 98 | 100% | 652 | 97.2% | 98 (100%) | 96 (98.0%) | 0/0/0/0 | 0 |
| Protein mutations: | R16A (1374A>G 1375G>C 1376A>C), G18Q (1380G>C 1381G>A 1382T>A) | | | | | | | | |
| Codon mutations: | GGT1GGC (1331T>C), TCT3TCC (1337T>C), ACA4ACG (1340A>G), TCC5AGC (1341T>A 1342C>G), CGC7CGA (1349C>A), TTA8CTG (1350T>C 1352A>G), TTT9TTC (1355T>C), ATT10ATC (1358T>C), GTG13GTA (1367G>A), GAC15GAT (1373C>T), AGA16GCC (1374A>G 1375G>C 1376A>C), GGT18CAA (1380G>C 1381G>A 1382T>A), CCA20CCC (1388A>C), AAA21AAG (1391A>G), CTC22CTG (1394C>G), AAT23AAC (1397T>C), CGG24AGG (1398C>A), CTA25TTG (1401C>T 1403A>G), GTA28GTT (1412A>T), AGG31CGC (1419A>C 1421G>C), GCA33GCT (1427A>T), TAT34TAC (1430T>C), TCC36TCA (1436C>A), AAC37AAT (1439C>T), CTG45CTC (1463G>C), GAC48GAT (1472C>T), GGA50GGT (1478A>T), TTG52CTG (1482T>C), CGA53CGG (1487A>G), GGG55GGA (1493G>A), GGG57GGC (1499G>C), CAG58CAA (1502G>A), CAA60CAG (1508A>G), GAT61GAC (1511T>C), AAA63AAG (1517A>G), GAT64GAC (1520T>C), CTG65CTC (1523G>C), GAT67GAC (1529T>C), TCA70TCC (1538A>C), CCC75CCA (1553C>A), TTA76TTG (1556A>G), CCA77CCC (1559A>C), GCC79GCT (1565C>T), GTT81GTA (1571T>A), GAG82GAA (1574G>A), GGC85GGG (1583C>G), AGG86CGG (1584A>C), ACC89ACA (1595C>A), TCC90TCA (1598C>A), TAT92TAC (1604T>C), ACA96ACC (1616A>C), AAT98AAC (1622T>C) | | | | | | | | |
| RNA-dependent RNA polymerase (YP_001976153.1) | 1 | 249 | 100% | 1569 | 89.4% | 249 (100%) | 218 (87.6%) | 0/0/0/0 | 0 |
| Protein mutations: | E8D (4076G>C), R14K (4093G>A 4094A>G), T40C (4170A>T 4171C>G 4172A>T), L44A (4182T>G 4183T>C 4184G>A), A47V (4192C>T), I48L (4194A>C 4196A>G), D55N (4215G>A), T57I (4222C>T 4223A>T), S59H (4227T>C 4228C>A), L63T (4239T>A 4240T>C 4241G>C), G67Q (4251G>C 4252G>A), A70S (4260G>A 4261C>G), V89I (4317G>A 4319T>A), E101A (4354A>C), K106A (4368A>G 4369A>C 4370G>T), P110E (4380C>G 4381C>A 4382T>G), V133L (4449G>C 4451G>C), A173K (4569G>A 4570C>A 4571T>A), R175K (4576G>A), I177L (4581A>T 4583A>G), P181S (4593C>T), Q186D (4608C>G 4610G>C), L202M (4656T>A 4658A>G), S208E (4674T>G 4675C>A 4676T>G), N211A (4683A>G 4684A>C 4685T>A), H214Q (4694T>A), L218Q (4705T>A 4706G>A), S230N (4740T>A 4741C>A 4742G>C), D232N (4746G>A), E235D (4757G>T), Q237K (4761C>A) | | | | | | | | |
| Codon mutations: | AAA1AAG (4055A>G), GAT2GAC (4058T>C), CGC4AGG (4062C>A 4064C>G), CTC5CTA (4067C>A), CCA6CCT (4070A>T), CTT7CTG (4073T>G), GAG8GAC (4076G>C), AAA9AAG (4079A>G), ATC10ATA (4082C>A), TAT11TAC (4085T>C), GGC12GGG (4088C>G), AAG13AAA (4091G>A), AGA14AAG (4093G>A 4094A>G), AAG15AAA (4097G>A), AGG17CGT (4101A>C 4103G>T), CTC18CTT (4106C>T), ATA21ATT (4115A>T), CTT22TTG (4116C>T 4118T>G), CCT23CCA (4121T>A), TAC26TAT (4130C>T), TTG29CTG (4137T>C), GTC30GTG (4142C>G), AGG31CGC (4143A>C 4145G>C), AAG32AAA (4148G>A), TTT34TTC (4154T>C), TTC37TTT (4163C>T), TCA38TCT (4166A>T), GCC39GCA (4169C>A), ACA40TGT (4170A>T 4171C>G 4172A>T), TTG41TTA (4175G>A), TTG44GCA (4182T>G 4183T>C 4184G>A), CAC45CAT (4187C>T), GCT47GTT (4192C>T), ATA48CTG (4194A>C 4196A>G), CCA49CCC (4199A>C), GTT52GTG (4208T>G), ATT54ATC (4214T>C), GAT55AAT (4215G>A), CCT56CCC (4220T>C), ACA57ATT (4222C>T 4223A>T), AGT58TCA (4224A>T 4225G>C 4226T>A), TCT59CAT (4227T>C 4228C>A), ACA62ACC (4238A>C), TTG63ACC (4239T>A 4240T>C 4241G>C), TTG64CTG (4242T>C), AAT66AAC (4250T>C), GGG67CAG (4251G>C 4252G>A), TTT68TTC (4256T>C), AGA69CGG (4257A>C 4259A>G), GCT70AGT (4260G>A 4261C>G), TCA72TCT (4268A>T), GAC73GAT (4271C>T), GTG74GTT (4274G>T), GGA75GGT (4277A>T), TCA77TCT (4283A>T), GCT78GCA (4286T>A), GAT79GAC (4289T>C), TAT80TAC (4292T>C), AGA86AGG (4310A>G), GCA87GCG (4313A>G), CCT88CCA (4316T>A), GTT89ATA (4317G>A 4319T>A), GCT91GCA (4325T>A), CAG93CAA (4331G>A), GAA101GCA (4354A>C), TAC103TAT (4361C>T), GGA104GGG (4364A>G), AAG106GCT (4368A>G 4369A>C 4370G>T), CCT107CCA (4373T>A), GGC108GGA (4376C>A), AGT109AGC (4379T>C), CCT110GAG (4380C>G 4381C>A 4382T>G), TCC112TCA (4388C>A), GCT114GCA (4394T>A), CGA115AGA (4395C>A), GCA117GCT (4403A>T), CTT118CTG (4406T>G), TTA119TTG (4409A>G), GCA122GCT (4418A>T), TGT124TGC (4424T>C), ACA127ACT (4433A>T), CTG128CTA (4436G>A), GTG133CTC (4449G>C 4451G>C), TTT134TTC (4454T>C), TTG136TTA (4460G>A), GGG138GGT (4466G>T), GGC139GGA (4469C>A), CCA141CCT (4475A>T), TCA142TCT (4478A>T), GCA145GCT (4487A>T), CTA146CTC (4490A>C), ACG147ACC (4493G>C), ATC149ATA (4499C>A), TTC150TTT (4502C>T), AAT151AAC (4505T>C), TCT152TCC (4508T>C), CTC153CTT (4511C>T), CTC154TTA (4512C>T 4514C>A), AAT155AAC (4517T>C), CGA160CGG (4532A>G), GCC162GCT (4538C>T), TTT163TTC (4541T>C), TCA165TCG (4547A>G), TTG166CTC (4548T>C 4550G>C), TTA167CTC (4551T>C 4553A>C), AGA169AGG (4559A>G), CAT171CAC (4565T>C), ATT172ATC (4568T>C), GCT173AAA (4569G>A 4570C>A 4571T>A), GCC174GCA (4574C>A), AGG175AAG (4576G>A), GCT176GCA (4580T>A), ATA177TTG (4581A>T 4583A>G), GGA178GGT (4586A>T), GTT179GTA (4589T>A), AAA180AAG (4592A>G), CCA181TCA (4593C>T), TTC184TTT (4604C>T), CAG186GAC (4608C>G 4610G>C), CTA187TTA (4611C>T), TTC188TTT (4616C>T), ATA189ATT (4619A>T), GCT190GCA (4622T>A), GTT191GTG (4625T>G), GGA193GGG (4631A>G), GTT198GTG (4646T>G), GTA200GTG (4652A>G), TTA202ATG (4656T>A 4658A>G), CAT203CAC (4661T>C), CTC204CTT (4664C>T), CAG205CAA (4667G>A), TAT207TAC (4673T>C), TCT208GAG (4674T>G 4675C>A 4676T>G), CTG209TTA (4677C>T 4679G>A), CCA210CCT (4682A>T), AAT211GCA (4683A>G 4684A>C 4685T>A), CAT214CAA (4694T>A), TTA216CTA (4698T>C), CTG218CAA (4705T>A 4706G>A), GTC219GTG (4709C>G), AAT220AAC (4712T>C), GTA221GTT (4715A>T), ATC222ATT (4718C>T), ATT223ATC (4721T>C), GGT226GGC (4730T>C), GAC228GAT (4736C>T), TCG230AAC (4740T>A 4741C>A 4742G>C), GAT232AAT (4746G>A), GTT233GTC (4751T>C), GAG235GAT (4757G>T), GTA236GTC (4760A>C), CAA237AAA (4761C>A), TTT238TTC (4766T>C), CAA239CAG (4769A>G), CTA244TTG (4782C>T 4784A>G), ACT245ACG (4787T>G), CTG247TTG (4791C>T), AGT248AGC (4796T>C) | | | | | | | | |

*: Inserts / Deletes / Misaligned / Frameshifts

#### Analysis details

This analysis was performed with panviral2.64

### NGS Details (UN60): Torradovirus marchitezum (segment RNA 2)

#### Assembly

| | |
|---|---|
| **Coverage Length** | 1365 (2 contig(s)) |
| **Depth Of Coverage** | 18973.4 |
| **Number Of Reads** | 209020 |
| **Reads Per Million** | 4433.22 rpm (after QC) |
| **Ambiguities** | 0 |
| **Assembly Method** | de novo + reference guided assembly |
| **Consensus Caller** | Bcf Tools |

#### Coverage Map

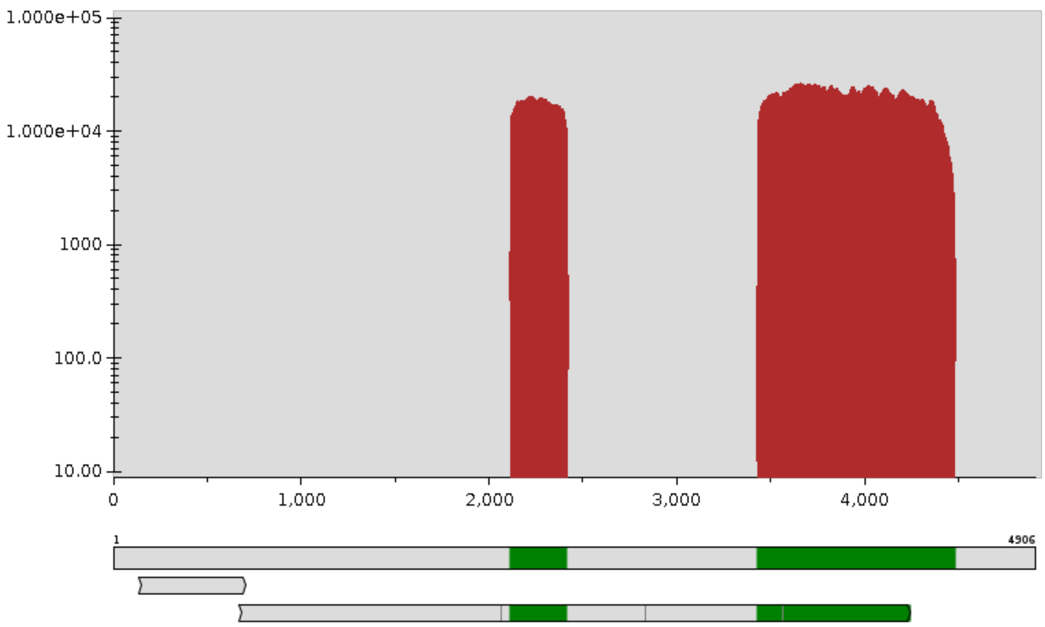

#### Assignment

| | |
|---|---|
| **Type** | Torradovirus marchitezum (Taxonomy ID: 3048376) |
| **Reference Genome** | NC_010988.1 |
| **NT Identity (%)** | 67.2951 |
| **AA Identity (%)** | 67.5462 |
| **Number Of Stop Codons** | 1 |
| **Number Of CDS** | 2 |

#### Alignment

| | |
|---|---|
| **Alignment Score** | 905.0 (NT) + 1799.0 (AA) = 2704.0 |
| **Concordance (%)** | 50.9423 |

| Alignment Method | Global, seeded, nucleotide + amino acids (AGA) |
|---|---|

#### Genome Region

Sequence starts at position 2113 and ends at position 4485 relative to NC_010988.1 reference sequence.

#### Alignment Detailed Statistics

| | Begin | End | Coverage | Score | Concordance | Matches | Identities | I/D/M/F* | Stop Codons |
|---|---|---|---|---|---|---|---|---|---|
| **NT** | **2113** | **4485** | **27.8%** | **905** | **33.2%** | **1365 (99.0%)** | **928 (67.3%)** | **14/0** | |
| Mutations: | 2122T>G, 2133T>C, 2134G>A, 2135T>A, 2137A>T, 2140T>A, 2143C>T, 2144A>G, 2146G>A, 2150G>C, 2152_2153insAGC, 2155T>G, 2156C>G, 2157C>G, 2158C>A, 2160C>T, 2164G>C, 2165C>A, 2166C>A, 2167A>G, 2170T>C, 2171A>C, 2173G>A, 2176A>T, 2178A>T, 2182T>A, 2184T>A, 2185T>G, 2187C>T, 2188A>T, 2190A>C, 2191G>T, 2193G>A, 2194A>C, 2195G>C, 2200A>G, 2201G>C, 2203A>G, 2207A>G, 2208C>T, 2209T>C, 2210C>T, 2211A>C, 2217G>A, 2218T>C, 2224T>C, 2225A>T, 2227T>C, 2239A>G, 2240C>G, 2241C>A, 2242A>G, 2249G>A, 2250A>T, 2251C>G, 2252A>G, 2253C>A, 2255T>G, 2256C>A, 2257C>T, 2258A>G, 2259T>C, 2260G>C, 2261G>C, 2263A>C, 2264G>A, 2267G>C, 2270C>T, 2275T>A, 2276A>T, 2281G>A, 2284A>G, 2287G>A, 2289T>C, 2290G>C, 2293T>C, 2296A>T, 2299T>C, 2305T>G, 2308T>G, 2314C>T, 2315A>C, 2320T>C, 2323A>C, 2326T>C, 2335C>T, 2336G>A, 2337T>A, 2338T>C, 2342G>A, 2347G>A, 2350C>T, 2353T>C, 2356T>C, 2359T>G, 2363G>C, 2365G>A, 2366C>G, 2368A>G, 2369G>C, 2372A>C, 2374G>A, 2375A>G, 2376G>C, 2377C>T, 2378T>C, 2383T>C, 2386G>A, 2389C>T, 2392T>A, 2395C>A, 2398A>G, 2402T>A, 2403C>G, 2404A>T, 2409G>A, 2410C>G, 3442C>G, 3445A>G, 3448C>T, 3449T>G, 3451A>C, 3457T>C, 3460C>T, 3461C>T, 3463T>G, 3466C>T, 3476G>A, 3478T>C, 3481A>G, 3482T>A, 3484A>T, 3493G>A, 3497A>C, 3506T>C, 3514G>A, 3517C>A, 3520C>T, 3521A>G, 3528A>C, 3529G>C, 3532C>T, 3533C>G, 3535G>T, 3538G>T, 3544C>G, 3544_3545insAAGAAG, 3550T>C, 3551C>A, 3552C>A, 3554C>G, 3555C>A, 3556C>G, 3559G>A, 3560A>G, 3561A>T, 3564C>G, 3568A>G, 3569A>G, 3570G>T, 3572A>G, 3573T>C, 3574T>A, 3577G>T, 3578T>G, 3579C>G, 3580C>A, 3581A>C, 3583A>T, 3584A>C, 3586A>G, 3589G>A, 3595T>C, 3598C>T, 3604T>A, 3610A>T, 3619C>T, 3622C>T, 3623A>T, 3624T>C, 3625G>T, 3627G>C, 3628T>C, 3638G>A, 3639A>C, 3641G>A, 3642A>G, 3643A>C, 3644A>C, 3646A>C, 3647T>C, 3650T>G, 3655A>T, 3656A>C, 3661T>A, 3667A>T, 3670A>G, 3676C>T, 3677A>G, 3680A>C, 3682G>A, 3685T>C, 3686G>T, 3687A>C, 3689G>C, 3690G>C, 3691G>C, 3692C>A, 3693A>C, 3700T>C, 3712A>C, 3715C>A, 3718T>G, 3724A>C, 3730A>C, 3739T>C, 3740A>T, 3742A>T, 3751T>C, 3760A>T, 3763T>C, 3767T>C, 3769G>T, 3770A>G, 3772T>G, 3775T>C, 3776T>A, 3778A>C, 3779A>C, 3781T>C, 3782G>A, 3784G>C, 3787A>C, 3790C>T, 3793A>G, 3796A>C, 3800T>G, 3801C>G, 3803A>T, 3804G>C, 3806C>A, 3807C>A, 3811T>C, 3812G>A, 3817A>T, 3826A>T, 3832T>G, 3838T>G, 3841T>C, 3842G>A, 3860C>T, 3862A>T, 3863G>T, 3864C>A, 3868G>T, 3869C>G, 3871T>G, 3872A>G, 3873A>C, 3874T>A, 3875A>G, 3877A>G, 3880A>G, 3881A>G, 3882A>C, 3883T>A, 3884T>C, 3890G>C, 3892A>T, 3901T>C, 3904A>T, 3907C>T, 3908A>T, 3910A>C, 3917G>A, 3922T>C, 3925A>C, 3931T>A, 3934G>T, 3935G>A, 3937A>C, 3940A>T, 3943T>C, 3945C>T, 3946G>T, 3949C>T, 3952A>C, 3955C>T, 3956A>G, 3957C>T, 3958T>G, 3967T>C, 3971G>A, 3973G>A, 3982A>T, 3983A>C, 3986C>A, 3989C>T, 3994A>T, 3996G>A, 3998A>G, 3999T>C, 4000G>A, 4001A>C, 4003G>A, 4009C>T, 4010T>G, 4011C>A, 4012A>C, 4014G>A, 4015C>T, 4019C>A, 4020A>G, 4021A>T, 4024C>T, 4027C>T, 4030C>T, 4031A>G, 4033G>T, 4036A>T, 4037C>T, 4039A>G, 4045T>C, 4047G>A, 4048G>A, 4049C>T, 4051T>G, 4054A>T, 4055A>C, 4056A>T, 4060G>T, 4061A>T, 4063C>G, 4066A>C, 4070T>C, 4072G>A, 4075T>A, 4078T>C, 4079T>G, 4080C>T, 4081C>A, 4083C>A, 4085A>T, 4088G>A, 4090G>T, 4093T>C, 4102G>A, 4105A>C, 4106C>A, 4108T>G, 4109A>G, 4111A>T, 4112T>A, 4113C>A, 4114T>G, 4117T>A, 4120A>G, 4122T>C, 4123T>A, 4129C>T, 4131A>G, 4135G>A, 4138G>A, 4139T>C, 4141G>T, 4154T>C, 4155C>T, 4156T>G, 4159C>T, 4162C>T, 4165T>A, 4165_4166insT, 4180C>T, 4181A>G, 4192T>G, 4195C>T, 4198C>T, 4201T>C, 4204G>A, 4213A>T, 4219T>C, 4222T>A, 4225G>A, 4228A>G, 4229C>A, 4231T>A, 4234T>C, 4237T>C, 4238G>T, 4242T>A, 4243T>C, 4248A>C, 4252T>A, 4279A>T, 4279_4280insT, 4280A>T, 4280_4281insC, 4283A>G, 4286C>T, 4287T>A, 4288T>C, 4291C>A, 4294C>T, 4306G>A, 4309G>A, 4312G>T, 4313C>A, 4313_4314insG, 4315A>G, 4316A>T, 4318T>C, 4319G>T, 4320G>A, 4323T>G, 4324T>A, 4328T>C, 4330G>A, 4333A>G, 4335G>T, 4336A>C, 4338A>G, 4339A>G, 4340G>T, 4341G>C, 4342G>T, 4343A>T, 4345A>T, 4346G>C, 4349A>C, 4364C>T, 4365T>C, 4369A>G, 4372A>G, 4373G>A, 4392T>C, 4393C>T, 4419_4420insG, 4422A>G, 4426G>A, 4430G>A, 4437C>T, 4441T>C, 4449A>T, 4451T>G | | | | | | | | |

##### CDS

| | | | | | | | | | |
|---|---|---|---|---|---|---|---|---|---|
| ToMarV_RNA2gp2 | 482 | 1192 | 31.5% | 1799 | 67.7% | 375 (98.9%) | 256 (67.5%) | 4/0/1/1 | 1 |
| Protein mutations: | V488A (2133T>C 2134G>A), S489T (2135T>A 2137A>T), T492A (2144A>G 2146G>A), E494Q (2150G>C), E494_D495insS (2152_2153insAGC), D495E (2155T>G), P496G (2156C>G 2157C>G 2158C>A), T497I (2160C>T), K498N (2164G>C), P499K (2165C>A 2166C>A 2167A>G), K501Q (2171A>C 2173G>A), E503V (2178A>T), V505E (2184T>A 2185T>G), A506V (2187C>T 2188A>T), E507A (2190A>C 2191G>T), G508D (2193G>A 2194A>C), A509P (2195G>C), E511Q (2201G>C 2203A>G), T513V (2207A>G 2208C>T 2209T>C), Q514S (2210C>T 2211A>C), G516D (2217G>A 2218T>C), I519F (2225A>T 2227T>C), P524E (2240C>G 2241C>A 2242A>G), D527M (2249G>A 2250A>T 2251C>G), T528E (2252A>G 2253C>A), S529D (2255T>G 2256C>A 2257C>T), M530A (2258A>G 2259T>C 2260G>C), A531P (2261G>C 2263A>C), V532I (2264G>A), E533Q (2267G>C), D535E (2275T>A), M536L (2276A>T), V540A (2289T>C 2290G>C), N545K (2305T>G), D546E (2308T>G), N549H (2315A>C), V556N (2336G>A 2337T>A 2338T>C), E558K (2342G>A), E565Q (2363G>C 2365G>A), Q566E (2366C>G 2368A>G), V567L (2369G>C), M568L (2372A>C 2374G>A), S569A (2375A>G 2376G>C 2377C>T), S580K (2409G>A 2410C>G), S927A (3449T>G 3451A>C), D936N (3476G>A 3478T>C), S938T (3482T>A 3484A>T), I943L (3497A>C), D949E (3517C>A), I951V (3521A>G), E953A (3528A>C 3529G>C), Q955D (3533C>G 3535G>T), E956D (3538G>T), G958_K959insKK (3544_3545insAAGAAG), P961K (3551C>A 3552C>A), P962E (3554C>G 3555C>A 3556C>G), K964V (3560A>G 3561A>T), A965G (3564C>G), S967V (3569A>G 3570G>T), I968A (3572A>G 3573T>C 3574T>A), S970G (3578T>G 3579C>G 3580C>A), I971L (3581A>C 3583A>T), M985S (3623A>T 3624T>C 3625G>T), G986A (3627G>C 3628T>C), D990T (3638G>A 3639A>C), E991S (3641G>A 3642A>G 3643A>C), L994V (3650T>G), I996L (3656A>C), I1003V (3677A>G), E1006S (3686G>T 3687A>C), G1007P (3689G>C 3690G>C 3691G>C), H1008T (3692C>A 3693A>C), T1024S (3740A>T 3742A>T), N1034E (3770A>G 3772T>G), S1036T (3776T>A 3778A>C), I1037L (3779A>C 3781T>C), V1038I (3782G>A 3784G>C), S1044G (3800T>G 3801C>G), P1046N (3806C>A 3807C>A), V1048I (3812G>A), F1056L (3838T>G), A1058T (3842G>A), P1064S (3860C>T 3862A>T), A1065Y (3863G>T 3864C>A), L1067V (3869C>G 3871T>G), N1068A (3872A>G 3873A>C 3874T>A), K1069E (3875A>G 3877A>G), N1071A (3881A>G 3882A>C 3883T>A), Y1072H (3884T>C), V1074L (3890G>C 3892A>T), T1080S (3908A>T 3910A>C), D1083N (3917G>A), A1089T (3935G>A 3937A>C), T1092I (3945C>T 3946G>T), T1096V (3956A>G 3957C>T 3958T>G), E1101K (3971G>A 3973G>A), H1107Y (3989C>T), R1109K (3996G>A), M1110A (3998A>G 3999T>C 4000G>A), S1114D (4010T>G 4011C>A 4012A>C), S1115N (4014G>A 4015C>T), Q1117S (4019C>A 4020A>G 4021A>T), M1121V (4031A>G 4033G>T), R1126K (4047G>A 4048G>A), N1129L (4055A>C 4056A>T), I1131L (4061A>T 4063C>G), S1137V (4079T>G 4080C>T 4081C>A), A1138D (4083C>A), I1139L (4085A>T), V1140I (4088G>A 4090G>T), L1146M (4106C>A 4108T>G), I1147V (4109A>G 4111A>T), S1148K (4112T>A 4113C>A 4114T>G), L1151P (4122T>C 4123T>A), K1154R (4131A>G), S1162L (4154T>C 4155C>T 4156T>G), H1165Q (4165T>A), H1165_E1166insX (4165_4166insT), N1171D (4181A>G), A1190S (4238G>T), I1191N (4242T>A 4243T>C) | | | | | | | | |

| | Begin | End | Coverage | Score | Concordance | Matches | Identities | I/D/M/F* | Stop Codons |
|---|---|---|---|---|---|---|---|---|---|
| **NT** | **2113** | **4485** | **27.8%** | **905** | **33.2%** | **1365 (99.0%)** | **928 (67.3%)** | **14/0** | |
| Codon mutations: | ACT484ACG (2122T>G), GTG488GCA (2133T>C 2134G>A), TCA489ACT (2135T>A 2137A>T), ACT490ACA (2140T>A), ACC491ACT (2143C>T), ACG492GCA (2144A>G 2146G>A), GAA494CAA (2150G>C), GAA494_GAT495insAGC (2152_2153insAGC), GAT495GAG (2155T>G), CCC496GGA (2156C>G 2157C>G 2158C>A), ACC497ATC (2160C>T), AAG498AAC (2164G>C), CCA499AAG (2165C>A 2166C>A 2167A>G), GAT500GAC (2170T>C), AAG501CAA (2171A>C 2173G>A), ATA502ATT (2176A>T), GAA503GTA (2178A>T), ATT504ATA (2182T>A), GTT505GAG (2184T>A 2185T>G), GCA506GTT (2187C>T 2188A>T), GAG507GCT (2190A>C 2191G>T), GGA508GAC (2193G>A 2194A>C), GCA509CCA (2195G>C), GAA510GAG (2200A>G), GAA511CAG (2201G>C 2203A>G), ACT513GTC (2207A>G 2208C>T 2209T>C), CAA514TCA (2210C>T 2211A>C), GGT516GAC (2217G>A 2218T>C), GTT518GTC (2224T>C), ATT519TTC (2225A>T 2227T>C), GGA523GGG (2239A>G), CCA524GAG (2240C>G 2241C>A 2242A>G), GAC527ATG (2249G>A 2250A>T 2251C>G), ACA528GAA (2252A>G 2253C>A), TCC529GAT (2255T>G 2256C>A 2257C>T), ATG530GCC (2258A>G 2259T>C 2260G>C), GCA531CCC (2261G>C 2263A>C), GTT532ATT (2264G>A), GAA533CAA (2267G>C), CTG534TTG (2270C>T), GAT535GAA (2275T>A), ATG536TTG (2276A>T), CAG537CAA (2281G>A), CAA538CAG (2284A>G), CCG539CCA (2287G>A), GTG540GCC (2289T>C 2290G>C), TGT541TGC (2293T>C), GTA542GTT (2296A>T), GCT543GCC (2299T>C), AAT545AAG (2305T>G), GAT546GAG (2308T>G), TTC548TTT (2314C>T), AAT549CAT (2315A>C), GTT550GTC (2320T>C), GGA551GGC (2323A>C), GTT552GTC (2326T>C), TTC555TTT (2335C>T), GTT556AAC (2336G>A 2337T>A 2338T>C), GAG558AAG (2342G>A), AAG559AAA (2347G>A), TCC560TCT (2350C>T), GCT561GCC (2353T>C), AAT562AAC (2356T>C), GTT563GTG (2359T>G), GAG565CAA (2363G>C 2365G>A), CAA566GAG (2366C>G 2368A>G), GTA567CTA (2369G>C), ATG568CTA (2372A>C 2374G>A), AGC569GCT (2375A>G 2376G>C 2377C>T), TTG570CTG (2378T>C), GCT571GCC (2383T>C), TTG572TTA (2386G>A), CCC573CCT (2389C>T), GCT574GCA (2392T>A), GCC575GCA (2395C>A), CTA576CTG (2398A>G), TCA578AGT (2402T>A 2403C>G 2404A>T), AGC580AAG (2409G>A 2410C>G), GGC924GGG (3442C>G), AAA925AAG (3445A>G), AGC926AGT (3448C>T), TCA927GCC (3449T>G 3451A>C), GAT929GAC (3457T>C), TTC930TTT (3460C>T), CTT931TTG (3461C>T 3463T>G), TTC932TTT (3466C>T), GAT936AAC (3476G>A 3478T>C), TTA937TTG (3481A>G), TCA938ACT (3482T>A 3484A>T), AAG941AAA (3493G>A), ATT943CTT (3497A>C), TTG946CTG (3506T>C), AAG948AAA (3514G>A), GAC949GAA (3517C>A), AAC950AAT (3520C>T), ATT951GTT (3521A>G), GAG953GCC (3528A>C 3529G>C), TAC954TAT (3532C>T), CAG955GAT (3533C>G 3535G>T), GAG956GAT (3538G>T), GGC958GGG (3544C>G), GGC958_AAA959insAAGAAG (3544_3545insAAGAAG), GAT960GAC (3550T>C), CCG961AAG (3551C>A 3552C>A), CCC962GAG (3554C>G 3555C>A 3556C>G), AAG963AAA (3559G>A), AAG964GTG (3560A>G 3561A>T), GCT965GGT (3564C>G), CAA966CAG (3568A>G), AGT967GTT (3569A>G 3570G>T), ATT968GCA (3572A>G 3573T>C 3574T>A), CTG969CTT (3577G>T), TCC970GGA (3578T>G 3579C>G 3580C>A), ATA971CTT (3581A>C 3583A>T), AGA972CGG (3584A>C 3586A>G), GAG973GAA (3589G>A), TTT975TTC (3595T>C), TCC976TCT (3598C>T), GGT978GGA (3604T>A), GTA980GTT (3610A>T), TTC983TTT (3619C>T), TGC984TGT (3622C>T), ATG985TCT (3623A>T 3624T>C 3625G>T), GGT986GCC (3627G>C 3628T>C), GAC990ACC (3638G>A 3639A>C), GAA991AGC (3641G>A 3642A>G 3643A>C), AGA992CGC (3644A>C 3646A>C), TTG993CTG (3647T>C), TTG994GTG (3650T>G), GTA995GTT (3655A>T), ATT996CTT (3656A>C), CCT997CCA (3661T>A), GCA999GCT (3667A>T), CCA1000CCG (3670A>G), TCC1002TCT (3676C>T), ATA1003GTA (3677A>G), AGG1004CGA (3680A>C 3682G>A), TTT1005TTC (3685T>C), GAA1006TCA (3686G>T 3687A>C), GGG1007CCC (3689G>C 3690G>C 3691G>C), CAC1008ACC (3692C>A 3693A>C), CCT1010CCC (3700T>C), GCA1014GCC (3712A>C), ATC1015ATA (3715C>A), ACT1016ACG (3718T>G), CCA1018CCC (3724A>C), ATA1020ATC (3730A>C), TGT1023TGC (3739T>C), ACA1024TCT (3740A>T 3742A>T), TGT1027TGC (3751T>C), TCA1030TCT (3760A>T), GGT1031GGC (3763T>C), TTG1033CTT (3767T>C 3769G>T), AAT1034GAG (3770A>G 3772T>G), TAT1035TAC (3775T>C), TCA1036ACC (3776T>A 3778A>C), ATT1037CTC (3779A>C 3781T>C), GTG1038ATC (3782G>A 3784G>C), ATA1039ATC (3787A>C), CAC1040CAT (3790C>T), AGA1041AGG (3793A>G), GTA1042GTC (3796A>C), TCC1044GGC (3800T>G 3801C>G), AGT1045TCT (3803A>T 3804G>C), CCT1046AAT (3806C>A 3807C>A), AAT1047AAC (3811T>C), GTT1048ATT (3812G>A), GGA1049GGT (3817A>T), CTA1052CTT (3826A>T), GTT1054GTG (3832T>G), TTT1056TTG (3838T>G), GAT1057GAC (3841T>C), GCC1058ACC (3842G>A), CCA1064TCT (3860C>T 3862A>T), GCT1065TAT (3863G>T 3864C>A), GGG1066GGT (3868G>T), CTT1067GTG (3869C>G 3871T>G), AAT1068GCA (3872A>G 3873A>C 3874T>A), AAA1069GAG (3875A>G 3877A>G), GGA1070GGG (3880A>G), AAT1071GCA (3881A>G 3882A>C 3883T>A), TAT1072CAT (3884T>C), GTA1074CTT (3890G>C 3892A>T), GGT1077GGC (3901T>C), GGA1078GGT (3904A>T), GGC1079GGT (3907C>T), ACA1080TCC (3908A>T 3910A>C), GAT1083AAT (3917G>A), TTT1084TTC (3922T>C), TCA1085TCC (3925A>C), GGT1087GGA (3931T>A), GTG1088GTT (3934G>T), GCA1089ACC (3935G>A 3937A>C), ACA1090ACT (3940A>T), AAT1091AAC (3943T>C), ACG1092ATT (3945C>T 3946G>T), TTC1093TTT (3949C>T), TCA1094TCC (3952A>C), TTC1095TTT (3955C>T), ACT1096GTG (3956A>G 3957C>T 3958T>G), GAT1099GAC (3967T>C), GAG1101AAA (3971G>A 3973G>A), CCA1104CCT (3982A>T), AGG1105CGG (3983A>C), CGG1106AGG (3986C>A), CAT1107TAT (3989C>T), ACA1108ACT (3994A>T), AGG1109AAG (3996G>A), ATG1110GCA (3998A>G 3999T>C 4000G>A), AGG1111CGA (4001A>C 4003G>A), TTC1113TTT (4009C>T), TCA1114GAC (4010T>G 4011C>A 4012A>C), AGC1115AAT (4014G>A 4015C>T), CAA1117AGT (4019C>A 4020A>G 4021A>T), TCC1118TCT (4024C>T), CGC1119CGT (4027C>T), ATC1120ATT (4030C>T), ATG1121GTT (4031A>G 4033G>T), TCA1122TCT (4036A>T), CTA1123TTG (4037C>T 4039A>G), GAT1125GAC (4045T>C), AGG1126AAA (4047G>A 4048G>A), CTT1127TTG (4049C>T 4051T>G), GGA1128GGT (4054A>T), AAT1129CTT (4055A>C 4056A>T), CTG1130CTT (4060G>T), ATC1131TTG (4061A>T 4063C>G), ATA1132ATC (4066A>C), TTG1134CTA (4070T>C 4072G>A), CCT1135CCA (4075T>A), CCT1136CCC (4078T>C), TCC1137GTA (4079T>G 4080C>T 4081C>A), GCC1138GAC (4083C>A), ATA1139TTA (4085A>T), GTG1140ATT (4088G>A 4090G>T), AGT1141AGC (4093T>C), GAG1144GAA (4102G>A), ATA1145ATC (4105A>C), CTT1146ATG (4106C>A 4108T>G), ATA1147GTT (4109A>G 4111A>T), TCT1148AAG (4112T>A 4113C>A 4114T>G), CCT1149CCA (4117T>A), GGA1150GGG (4120A>G), CTT1151CCA (4122T>C 4123T>A), TTC1153TTT (4129C>T), AAG1154AGG (4131A>G), TTG1155TTA (4135G>A), GAG1156GAA (4138G>A), TTG1157CTT (4139T>C 4141G>T), TCT1162CTG (4154T>C 4155C>T 4156T>G), GCC1163GCT (4159C>T), AAC1164AAT (4162C>T), CAT1165CAA (4165T>A), CAT1165_GAA1166insT-- (4165_4166insT), GGC1170GGT (4180C>T), AAT1171GAT (4181A>G), ACT1174ACG (4192T>G), CAC1175CAT (4195C>T), ACC1176ACT (4198C>T), TAT1177TAC (4201T>C), CAG1178CAA (4204G>A), TCA1181TCT (4213A>T), TTT1183TTC (4219T>C), TCT1184TCA (4222T>A), GAG1185GAA (4225G>A), CTA1186CTG (4228A>G), CGT1187AGA (4229C>A 4231T>A), GAT1188GAC (4234T>C), TTT1189TTC (4237T>C), GCG1190TCG (4238G>T), ATT1191AAC (4242T>A 4243T>C) | | | | | | | | |

#### Proteins

| | Begin | End | Coverage | Score | Concordance | Matches | Identities | I/D/M/F* | Stop Codons |
|---|---|---|---|---|---|---|---|---|---|
| polyprotein (YP_001976149.1) | 482 | 1192 | 31.5% | 1799 | 67.7% | 375 (98.9%) | 256 (67.5%) | 4/0/1/1 | 1 |
| Protein mutations: | V488A (2133T>C 2134G>A), S489T (2135T>A 2137A>T), T492A (2144A>G 2146G>A), E494Q (2150G>C), E494_D495insS (2152_2153insAGC), D495E (2155T>G), P496G (2156C>G 2157C>G 2158C>A), T497I (2160C>T), K498N (2164G>C), P499K (2165C>A 2166C>A 2167A>G), K501Q (2171A>C 2173G>A), E503V (2178A>T), V505E (2184T>A 2185T>G), A506V (2187C>T 2188A>T), E507A (2190A>C 2191G>T), G508D (2193G>A 2194A>C), A509P (2195G>C), E511Q (2201G>C 2203A>G), T513V (2207A>G 2208C>T 2209T>C), Q514S (2210C>T 2211A>C), G516D (2217G>A 2218T>C), I519F (2225A>T 2227T>C), P524E (2240C>G 2241C>A 2242A>G), D527M (2249G>A 2250A>T 2251C>G), T528E (2252A>G 2253C>A), S529D (2255T>G 2256C>A 2257C>T), M530A (2258A>G 2259T>C 2260G>C), A531P (2261G>C 2263A>C), V532I (2264G>A), E533Q (2267G>C), D535E (2275T>A), M536L (2276A>T), V540A (2289T>C 2290G>C), N545K (2305T>G), D546E (2308T>G), N549H (2315A>C), V556N (2336G>A 2337T>A 2338T>C), E558K (2342G>A), E565Q (2363G>C 2365G>A), Q566E (2366C>G 2368A>G), V567L (2369G>C), M568L (2372A>C 2374G>A), S569A (2375A>G 2376G>C 2377C>T), S580K (2409G>A 2410C>G), S927A (3449T>G 3451A>C), D936N (3476G>A 3478T>C), S938T (3482T>A 3484A>T), I943L (3497A>C), D949E (3517C>A), I951V (3521A>G), E953A (3528A>C 3529G>C), Q955D (3533C>G 3535G>T), E956D (3538G>T), G958_K959insKK (3544_3545insAAGAAG), P961K (3551C>A 3552C>A), P962E (3554C>G 3555C>A 3556C>G), K964V (3560A>G 3561A>T), A965G (3564C>G), S967V (3569A>G 3570G>T), I968A (3572A>G 3573T>C 3574T>A), S970G (3578T>G 3579C>G 3580C>A), I971L (3581A>C 3583A>T), M985S (3623A>T 3624T>C 3625G>T), G986A (3627G>C 3628T>C), D990T (3638G>A 3639A>C), E991S (3641G>A 3642A>G 3643A>C), L994V (3650T>G), I996L (3656A>C), I1003V (3677A>G), E1006S (3686G>T 3687A>C), G1007P (3689G>C 3690G>C 3691G>C), H1008T (3692C>A 3693A>C), T1024S (3740A>T 3742A>T), N1034E (3770A>G 3772T>G), S1036T (3776T>A 3778A>C), I1037L (3779A>C 3781T>C), V1038I (3782G>A 3784G>C), S1044G (3800T>G 3801C>G), P1046N (3806C>A 3807C>A), V1048I (3812G>A), F1056L (3838T>G), A1058T (3842G>A), P1064S (3860C>T 3862A>T), A1065Y (3863G>T 3864C>A), L1067V (3869C>G 3871T>G), N1068A (3872A>G 3873A>C 3874T>A), K1069E (3875A>G 3877A>G), N1071A (3881A>G 3882A>C 3883T>A), Y1072H (3884T>C), V1074L (3890G>C 3892A>T), T1080S (3908A>T 3910A>C), D1083N (3917G>A), A1089T (3935G>A 3937A>C), T1092I (3945C>T 3946G>T), T1096V (3956A>G 3957C>T 3958T>G), E1101K (3971G>A 3973G>A), H1107Y (3989C>T), R1109K (3996G>A), M1110A (3998A>G 3999T>C 4000G>A), S1114D (4010T>G 4011C>A 4012A>C), S1115N (4014G>A 4015C>T), Q1117S (4019C>A 4020A>G 4021A>T), M1121V (4031A>G 4033G>T), R1126K (4047G>A 4048G>A), N1129L (4055A>C 4056A>T), I1131L (4061A>T 4063C>G), S1137V (4079T>G 4080C>T 4081C>A), A1138D (4083C>A), I1139L (4085A>T), V1140I (4088G>A 4090G>T), L1146M (4106C>A 4108T>G), I1147V (4109A>G 4111A>T), S1148K (4112T>A 4113C>A 4114T>G), L1151P (4122T>C 4123T>A), K1154R (4131A>G), S1162L (4154T>C 4155C>T 4156T>G), H1165Q (4165T>A), H1165_E1166insX (4165_4166insT), N1171D (4181A>G), A1190S (4238G>T), I1191N (4242T>A 4243T>C) | | | | | | | | |

| | Begin | End | Coverage | Score | Concordance | Matches | Identities | I/D/M/F* | Stop Codons |
|---|---|---|---|---|---|---|---|---|---|
| **NT** | **2113** | **4485** | **27.8%** | **905** | **33.2%** | **1365 (99.0%)** | **928 (67.3%)** | **14/0** | |
| Codon mutations: | ACT484ACG (2122T>G), GTG488GCA (2133T>C 2134G>A), TCA489ACT (2135T>A 2137A>T), ACT490ACA (2140T>A), ACC491ACT (2143C>T), ACG492GCA (2144A>G 2146G>A), GAA494CAA (2150G>C), GAA494_GAT495insAGC (2152_2153insAGC), GAT495GAG (2155T>G), CCC496GGA (2156C>G 2157C>G 2158C>A), ACC497ATC (2160C>T), AAG498AAC (2164G>C), CCA499AAG (2165C>A 2166C>A 2167A>G), GAT500GAC (2170T>C), AAG501CAA (2171A>C 2173G>A), ATA502ATT (2176A>T), GAA503GTA (2178A>T), ATT504ATA (2182T>A), GTT505GAG (2184T>A 2185T>G), GCA506GTT (2187C>T 2188A>T), GAG507GCT (2190A>C 2191G>T), GGA508GAC (2193G>A 2194A>C), GCA509CCA (2195G>C), GAA510GAG (2200A>G), GAA511CAG (2201G>C 2203A>G), ACT513GTC (2207A>G 2208C>T 2209T>C), CAA514TCA (2210C>T 2211A>C), GGT516GAC (2217G>A 2218T>C), GTT518GTC (2224T>C), ATT519TTC (2225A>T 2227T>C), GGA523GGG (2239A>G), CCA524GAG (2240C>G 2241C>A 2242A>G), GAC527ATG (2249G>A 2250A>T 2251C>G), ACA528GAA (2252A>G 2253C>A), TCC529GAT (2255T>G 2256C>A 2257C>T), ATG530GCC (2258A>G 2259T>C 2260G>C), GCA531CCC (2261G>C 2263A>C), GTT532ATT (2264G>A), GAA533CAA (2267G>C), CTG534TTG (2270C>T), GAT535GAA (2275T>A), ATG536TTG (2276A>T), CAG537CAA (2281G>A), CAA538CAG (2284A>G), CCG539CCA (2287G>A), GTG540GCC (2289T>C 2290G>C), TGT541TGC (2293T>C), GTA542GTT (2296A>T), GCT543GCC (2299T>C), AAT545AAG (2305T>G), GAT546GAG (2308T>G), TTC548TTT (2314C>T), AAT549CAT (2315A>C), GTT550GTC (2320T>C), GGA551GGC (2323A>C), GTT552GTC (2326T>C), TTC555TTT (2335C>T), GTT556AAC (2336G>A 2337T>A 2338T>C), GAG558AAG (2342G>A), AAG559AAA (2347G>A), TCC560TCT (2350C>T), GCT561GCC (2353T>C), AAT562AAC (2356T>C), GTT563GTG (2359T>G), GAG565CAA (2363G>C 2365G>A), CAA566GAG (2366C>G 2368A>G), GTA567CTA (2369G>C), ATG568CTA (2372A>C 2374G>A), AGC569GCT (2375A>G 2376G>C 2377C>T), TTG570CTG (2378T>C), GCT571GCC (2383T>C), TTG572TTA (2386G>A), CCC573CCT (2389C>T), GCT574GCA (2392T>A), GCC575GCA (2395C>A), CTA576CTG (2398A>G), TCA578AGT (2402T>A 2403C>G 2404A>T), AGC580AAG (2409G>A 2410C>G), GGC924GGG (3442C>G), AAA925AAG (3445A>G), AGC926AGT (3448C>T), TCA927GCC (3449T>G 3451A>C), GAT929GAC (3457T>C), TTC930TTT (3460C>T), CTT931TTG (3461C>T 3463T>G), TTC932TTT (3466C>T), GAT936AAC (3476G>A 3478T>C), TTA937TTG (3481A>G), TCA938ACT (3482T>A 3484A>T), AAG941AAA (3493G>A), ATT943CTT (3497A>C), TTG946CTG (3506T>C), AAG948AAA (3514G>A), GAC949GAA (3517C>A), AAC950AAT (3520C>T), ATT951GTT (3521A>G), GAG953GCC (3528A>C 3529G>C), TAC954TAT (3532C>T), CAG955GAT (3533C>G 3535G>T), GAG956GAT (3538G>T), GGC958GGG (3544C>G), GGC958_AAA959insAAGAAG (3544_3545insAAGAAG), GAT960GAC (3550T>C), CCG961AAG (3551C>A 3552C>A), CCC962GAG (3554C>G 3555C>A 3556C>G), AAG963AAA (3559G>A), AAG964GTG (3560A>G 3561A>T), GCT965GGT (3564C>G), CAA966CAG (3568A>G), AGT967GTT (3569A>G 3570G>T), ATT968GCA (3572A>G 3573T>C 3574T>A), CTG969CTT (3577G>T), TCC970GGA (3578T>G 3579C>G 3580C>A), ATA971CTT (3581A>C 3583A>T), AGA972CGG (3584A>C 3586A>G), GAG973GAA (3589G>A), TTT975TTC (3595T>C), TCC976TCT (3598C>T), GGT978GGA (3604T>A), GTA980GTT (3610A>T), TTC983TTT (3619C>T), TGC984TGT (3622C>T), ATG985TCT (3623A>T 3624T>C 3625G>T), GGT986GCC (3627G>C 3628T>C), GAC990ACC (3638G>A 3639A>C), GAA991AGC (3641G>A 3642A>G 3643A>C), AGA992CGC (3644A>C 3646A>C), TTG993CTG (3647T>C), TTG994GTG (3650T>G), GTA995GTT (3655A>T), ATT996CTT (3656A>C), CCT997CCA (3661T>A), GCA999GCT (3667A>T), CCA1000CCG (3670A>G), TCC1002TCT (3676C>T), ATA1003GTA (3677A>G), AGG1004CGA (3680A>C 3682G>A), TTT1005TTC (3685T>C), GAA1006TCA (3686G>T 3687A>C), GGG1007CCC (3689G>C 3690G>C 3691G>C), CAC1008ACC (3692C>A 3693A>C), CCT1010CCC (3700T>C), GCA1014GCC (3712A>C), ATC1015ATA (3715C>A), ACT1016ACG (3718T>G), CCA1018CCC (3724A>C), ATA1020ATC (3730A>C), TGT1023TGC (3739T>C), ACA1024TCT (3740A>T 3742A>T), TGT1027TGC (3751T>C), TCA1030TCT (3760A>T), GGT1031GGC (3763T>C), TTG1033CTT (3767T>C 3769G>T), AAT1034GAG (3770A>G 3772T>G), TAT1035TAC (3775T>C), TCA1036ACC (3776T>A 3778A>C), ATT1037CTC (3779A>C 3781T>C), GTG1038ATC (3782G>A 3784G>C), ATA1039ATC (3787A>C), CAC1040CAT (3790C>T), AGA1041AGG (3793A>G), GTA1042GTC (3796A>C), TCC1044GGC (3800T>G 3801C>G), AGT1045TCT (3803A>T 3804G>C), CCT1046AAT (3806C>A 3807C>A), AAT1047AAC (3811T>C), GTT1048ATT (3812G>A), GGA1049GGT (3817A>T), CTA1052CTT (3826A>T), GTT1054GTG (3832T>G), TTT1056TTG (3838T>G), GAT1057GAC (3841T>C), GCC1058ACC (3842G>A), CCA1064TCT (3860C>T 3862A>T), GCT1065TAT (3863G>T 3864C>A), GGG1066GGT (3868G>T), CTT1067GTG (3869C>G 3871T>G), AAT1068GCA (3872A>G 3873A>C 3874T>A), AAA1069GAG (3875A>G 3877A>G), GGA1070GGG (3880A>G), AAT1071GCA (3881A>G 3882A>C 3883T>A), TAT1072CAT (3884T>C), GTA1074CTT (3890G>C 3892A>T), GGT1077GGC (3901T>C), GGA1078GGT (3904A>T), GGC1079GGT (3907C>T), ACA1080TCC (3908A>T 3910A>C), GAT1083AAT (3917G>A), TTT1084TTC (3922T>C), TCA1085TCC (3925A>C), GGT1087GGA (3931T>A), GTG1088GTT (3934G>T), GCA1089ACC (3935G>A 3937A>C), ACA1090ACT (3940A>T), AAT1091AAC (3943T>C), ACG1092ATT (3945C>T 3946G>T), TTC1093TTT (3949C>T), TCA1094TCC (3952A>C), TTC1095TTT (3955C>T), ACT1096GTG (3956A>G 3957C>T 3958T>G), GAT1099GAC (3967T>C), GAG1101AAA (3971G>A 3973G>A), CCA1104CCT (3982A>T), AGG1105CGG (3983A>C), CGG1106AGG (3986C>A), CAT1107TAT (3989C>T), ACA1108ACT (3994A>T), AGG1109AAG (3996G>A), ATG1110GCA (3998A>G 3999T>C 4000G>A), AGG1111CGA (4001A>C 4003G>A), TTC1113TTT (4009C>T), TCA1114GAC (4010T>G 4011C>A 4012A>C), AGC1115AAT (4014G>A 4015C>T), CAA1117AGT (4019C>A 4020A>G 4021A>T), TCC1118TCT (4024C>T), CGC1119CGT (4027C>T), ATC1120ATT (4030C>T), ATG1121GTT (4031A>G 4033G>T), TCA1122TCT (4036A>T), CTA1123TTG (4037C>T 4039A>G), GAT1125GAC (4045T>C), AGG1126AAA (4047G>A 4048G>A), CTT1127TTG (4049C>T 4051T>G), GGA1128GGT (4054A>T), AAT1129CTT (4055A>C 4056A>T), CTG1130CTT (4060G>T), ATC1131TTG (4061A>T 4063C>G), ATA1132ATC (4066A>C), TTG1134CTA (4070T>C 4072G>A), CCT1135CCA (4075T>A), CCT1136CCC (4078T>C), TCC1137GTA (4079T>G 4080C>T 4081C>A), GCC1138GAC (4083C>A), ATA1139TTA (4085A>T), GTG1140ATT (4088G>A 4090G>T), AGT1141AGC (4093T>C), GAG1144GAA (4102G>A), ATA1145ATC (4105A>C), CTT1146ATG (4106C>A 4108T>G), ATA1147GTT (4109A>G 4111A>T), TCT1148AAG (4112T>A 4113C>A 4114T>G), CCT1149CCA (4117T>A), GGA1150GGG (4120A>G), CTT1151CCA (4122T>C 4123T>A), TTC1153TTT (4129C>T), AAG1154AGG (4131A>G), TTG1155TTA (4135G>A), GAG1156GAA (4138G>A), TTG1157CTT (4139T>C 4141G>T), TCT1162CTG (4154T>C 4155C>T 4156T>G), GCC1163GCT (4159C>T), AAC1164AAT (4162C>T), CAT1165CAA (4165T>A), CAT1165_GAA1166insT-- (4165_4166insT), GGC1170GGT (4180C>T), AAT1171GAT (4181A>G), ACT1174ACG (4192T>G), CAC1175CAT (4195C>T), ACC1176ACT (4198C>T), TAT1177TAC (4201T>C), CAG1178CAA (4204G>A), TCA1181TCT (4213A>T), TTT1183TTC (4219T>C), TCT1184TCA (4222T>A), GAG1185GAA (4225G>A), CTA1186CTG (4228A>G), CGT1187AGA (4229C>A 4231T>A), GAT1188GAC (4234T>C), TTT1189TTC (4237T>C), GCG1190TCG (4238G>T), ATT1191AAC (4242T>A 4243T>C) | | | | | | | | |
| Coat protein A (YP_001976155.1) | 14 | 116 | 40.6% | 415 | 60.3% | 103 (99.0%) | 59 (56.7%) | 1/0/0/0 | 0 |
| Protein mutations: | V20A (2133T>C 2134G>A), S21T (2135T>A 2137A>T), T24A (2144A>G 2146G>A), E26Q (2150G>C), E26_D27insS (2152_2153insAGC), D27E (2155T>G), P28G (2156C>G 2157C>G 2158C>A), T29I (2160C>T), K30N (2164G>C), P31K (2165C>A 2166C>A 2167A>G), K33Q (2171A>C 2173G>A), E35V (2178A>T), V37E (2184T>A 2185T>G), A38V (2187C>T 2188A>T), E39A (2190A>C 2191G>T), G40D (2193G>A 2194A>C), A41P (2195G>C), E43Q (2201G>C 2203A>G), T45V (2207A>G 2208C>T 2209T>C), Q46S (2210C>T 2211A>C), G48D (2217G>A 2218T>C), I51F (2225A>T 2227T>C), P56E (2240C>G 2241C>A 2242A>G), D59M (2249G>A 2250A>T 2251C>G), T60E (2252A>G 2253C>A), S61D (2255T>G 2256C>A 2257C>T), M62A (2258A>G 2259T>C 2260G>C), A63P (2261G>C 2263A>C), V64I (2264G>A), E65Q (2267G>C), D67E (2275T>A), M68L (2276A>T), V72A (2289T>C 2290G>C), N77K (2305T>G), D78E (2308T>G), N81H (2315A>C), V88N (2336G>A 2337T>A 2338T>C), E90K (2342G>A), E97Q (2363G>C 2365G>A), Q98E (2366C>G 2368A>G), V99L (2369G>C), M100L (2372A>C 2374G>A), S101A (2375A>G 2376G>C 2377C>T), S112K (2409G>A 2410C>G) | | | | | | | | |
| Codon mutations: | ACT16ACG (2122T>G), GTG20GCA (2133T>C 2134G>A), TCA21ACT (2135T>A 2137A>T), ACT22ACA (2140T>A), ACC23ACT (2143C>T), ACG24GCA (2144A>G 2146G>A), GAA26CAA (2150G>C), GAA26_GAT27insAGC (2152_2153insAGC), GAT27GAG (2155T>G), CCC28GGA (2156C>G 2157C>G 2158C>A), ACC29ATC (2160C>T), AAG30AAC (2164G>C), CCA31AAG (2165C>A 2166C>A 2167A>G), GAT32GAC (2170T>C), AAG33CAA (2171A>C 2173G>A), ATA34ATT (2176A>T), GAA35GTA (2178A>T), ATT36ATA (2182T>A), GTT37GAG (2184T>A 2185T>G), GCA38GTT (2187C>T 2188A>T), GAG39GCT (2190A>C 2191G>T), GGA40GAC (2193G>A 2194A>C), GCA41CCA (2195G>C), GAA42GAG (2200A>G), GAA43CAG (2201G>C 2203A>G), ACT45GTC (2207A>G 2208C>T 2209T>C), CAA46TCA (2210C>T 2211A>C), GGT48GAC (2217G>A 2218T>C), GTT50GTC (2224T>C), ATT51TTC (2225A>T 2227T>C), GGA55GGG (2239A>G), CCA56GAG (2240C>G 2241C>A 2242A>G), GAC59ATG (2249G>A 2250A>T 2251C>G), ACA60GAA (2252A>G 2253C>A), TCC61GAT (2255T>G 2256C>A 2257C>T), ATG62GCC (2258A>G 2259T>C 2260G>C), GCA63CCC (2261G>C 2263A>C), GTT64ATT (2264G>A), GAA65CAA (2267G>C), CTG66TTG (2270C>T), GAT67GAA (2275T>A), ATG68TTG (2276A>T), CAG69CAA (2281G>A), CAA70CAG (2284A>G), CCG71CCA (2287G>A), GTG72GCC (2289T>C 2290G>C), TGT73TGC (2293T>C), GTA74GTT (2296A>T), GCT75GCC (2299T>C), AAT77AAG (2305T>G), GAT78GAG (2308T>G), TTC80TTT (2314C>T), AAT81CAT (2315A>C), GTT82GTC (2320T>C), GGA83GGC (2323A>C), GTT84GTC (2326T>C), TTC87TTT (2335C>T), GTT88AAC (2336G>A 2337T>A 2338T>C), GAG90AAG (2342G>A), AAG91AAA (2347G>A), TCC92TCT (2350C>T), GCT93GCC (2353T>C), AAT94AAC (2356T>C), GTT95GTG (2359T>G), GAG97CAA (2363G>C 2365G>A), CAA98GAG (2366C>G 2368A>G), GTA99CTA (2369G>C), ATG100CTA (2372A>C 2374G>A), AGC101GCT (2375A>G 2376G>C 2377C>T), TTG102CTG (2378T>C), GCT103GCC (2383T>C), TTG104TTA (2386G>A), CCC105CCT (2389C>T), GCT106GCA (2392T>A), GCC107GCA (2395C>A), CTA108CTG (2398A>G), TCA110AGT (2402T>A 2403C>G 2404A>T), AGC112AAG (2409G>A 2410C>G) | | | | | | | | |
| Coat protein B (YP_001976156.1) | 199 | 244 | 18.9% | 218 | 68.8% | 46 (95.8%) | 33 (68.8%) | 2/0/0/0 | 0 |
| Protein mutations: | S205A (3449T>G 3451A>C), D214N (3476G>A 3478T>C), S216T (3482T>A 3484A>T), I221L (3497A>C), D227E (3517C>A), I229V (3521A>G), E231A (3528A>C 3529G>C), Q233D (3533C>G 3535G>T), E234D (3538G>T), G236_K237insKK (3544_3545insAAGAAG), P239K (3551C>A 3552C>A), P240E (3554C>G 3555C>A 3556C>G), K242V (3560A>G 3561A>T), A243G (3564C>G) | | | | | | | | |
| Codon mutations: | GGC202GGG (3442C>G), AAA203AAG (3445A>G), AGC204AGT (3448C>T), TCA205GCC (3449T>G 3451A>C), GAT207GAC (3457T>C), TTC208TTT (3460C>T), CTT209TTG (3461C>T 3463T>G), TTC210TTT (3466C>T), GAT214AAC (3476G>A 3478T>C), TTA215TTG (3481A>G), TCA216ACT (3482T>A 3484A>T), AAG219AAA (3493G>A), ATT221CTT (3497A>C), TTG224CTG (3506T>C), AAG226AAA (3514G>A), GAC227GAA (3517C>A), AAC228AAT (3520C>T), ATT229GTT (3521A>G), GAG231GCC (3528A>C 3529G>C), TAC232TAT (3532C>T), CAG233GAT (3533C>G 3535G>T), GAG234GAT (3538G>T), GGC236GGG (3544C>G), GGC236_AAA237insAAGAAG (3544_3545insAAGAAG), GAT238GAC (3550T>C), CCG239AAG (3551C>A 3552C>A), CCC240GAG (3554C>G 3555C>A 3556C>G), AAG241AAA (3559G>A), AAG242GTG (3560A>G 3561A>T), GCT243GGT (3564C>G), CAA244CAG (3568A>G) | | | | | | | | |
| Coat protein C (YP_001976157.1) | 1 | 225 | 100% | 1165 | 70.6% | 225 (99.6%) | 163 (72.1%) | 1/0/1/1 | 0 |
| Protein mutations: | S1V (3569A>G 3570G>T), I2A (3572A>G 3573T>C 3574T>A), S4G (3578T>G 3579C>G 3580C>A), I5L (3581A>C 3583A>T), M19S (3623A>T 3624T>C 3625G>T), G20A (3627G>C 3628T>C), D24T (3638G>A 3639A>C), E25S (3641G>A 3642A>G 3643A>C), L28V (3650T>G), I30L (3656A>C), I37V (3677A>G), E40S (3686G>T 3687A>C), G41P (3689G>C 3690G>C 3691G>C), H42T (3692C>A 3693A>C), T58S (3740A>T 3742A>T), N68E (3770A>G 3772T>G), S70T (3776T>A 3778A>C), I71L (3779A>C 3781T>C), V72I (3782G>A 3784G>C), S78G (3800T>G 3801C>G), P80N (3806C>A 3807C>A), V82I (3812G>A), F90L (3838T>G), A92T (3842G>A), P98S (3860C>T 3862A>T), A99Y (3863G>T 3864C>A), L101V (3869C>G 3871T>G), N102A (3872A>G 3873A>C 3874T>A), K103E (3875A>G 3877A>G), N105A (3881A>G 3882A>C 3883T>A), Y106H (3884T>C), V108L (3890G>C 3892A>T), T114S (3908A>T 3910A>C), D117N (3917G>A), A123T (3935G>A 3937A>C), T126I (3945C>T 3946G>T), T130V (3956A>G 3957C>T 3958T>G), E135K (3971G>A 3973G>A), H141Y (3989C>T), R143K (3996G>A), M144A (3998A>G 3999T>C 4000G>A), S148D (4010T>G 4011C>A 4012A>C), S149N (4014G>A 4015C>T), Q151S (4019C>A 4020A>G 4021A>T), M155V (4031A>G 4033G>T), R160K (4047G>A 4048G>A), N163L (4055A>C 4056A>T), I165L (4061A>T 4063C>G), S171V (4079T>G 4080C>T 4081C>A), A172D (4083C>A), I173L (4085A>T), V174I (4088G>A 4090G>T), L180M (4106C>A 4108T>G), I181V (4109A>G 4111A>T), S182K (4112T>A 4113C>A 4114T>G), L185P (4122T>C 4123T>A), K188R (4131A>G), S196L (4154T>C 4155C>T 4156T>G), H199Q (4165T>A), H199_E200insX (4165_4166insT), N205D (4181A>G), A224S (4238G>T), I225N (4242T>A 4243T>C) | | | | | | | | |

| | Begin | End | Coverage | Score | Concordance | Matches | Identities | I/D/M/F* | Stop Codons |
|---|---|---|---|---|---|---|---|---|---|
| **NT** | **2113** | **4485** | **27.8%** | **905** | **33.2%** | **1365 (99.0%)** | **928 (67.3%)** | **14/0** | |
| Codon mutations: | AGT1GTT (3569A>G 3570G>T), ATT2GCA (3572A>G 3573T>C 3574T>A), CTG3CTT (3577G>T), TCC4GGA (3578T>G 3579C>G 3580C>A), ATA5CTT (3581A>C 3583A>T), AGA6CGG (3584A>C 3586A>G), GAG7GAA (3589G>A), TTT9TTC (3595T>C), TCC10TCT (3598C>T), GGT12GGA (3604T>A), GTA14GTT (3610A>T), TTC17TTT (3619C>T), TGC18TGT (3622C>T), ATG19TCT (3623A>T 3624T>C 3625G>T), GGT20GCC (3627G>C 3628T>C), GAC24ACC (3638G>A 3639A>C), GAA25AGC (3641G>A 3642A>G 3643A>C), AGA26CGC (3644A>C 3646A>C), TTG27CTG (3647T>C), TTG28GTG (3650T>G), GTA29GTT (3655A>T), ATT30CTT (3656A>C), CCT31CCA (3661T>A), GCA33GCT (3667A>T), CCA34CCG (3670A>G), TCC36TCT (3676C>T), ATA37GTA (3677A>G), AGG38CGA (3680A>C 3682G>A), TTT39TTC (3685T>C), GAA40TCA (3686G>T 3687A>C), GGG41CCC (3689G>C 3690G>C 3691G>C), CAC42ACC (3692C>A 3693A>C), CCT44CCC (3700T>C), GCA48GCC (3712A>C), ATC49ATA (3715C>A), ACT50ACG (3718T>G), CCA52CCC (3724A>C), ATA54ATC (3730A>C), TGT57TGC (3739T>C), ACA58TCT (3740A>T 3742A>T), TGT61TGC (3751T>C), TCA64TCT (3760A>T), GGT65GGC (3763T>C), TTG67CTT (3767T>C 3769G>T), AAT68GAG (3770A>G 3772T>G), TAT69TAC (3775T>C), TCA70ACC (3776T>A 3778A>C), ATT71CTC (3779A>C 3781T>C), GTG72ATC (3782G>A 3784G>C), ATA73ATC (3787A>C), CAC74CAT (3790C>T), AGA75AGG (3793A>G), GTA76GTC (3796A>C), TCC78GGC (3800T>G 3801C>G), AGT79TCT (3803A>T 3804G>C), CCT80AAT (3806C>A 3807C>A), AAT81AAC (3811T>C), GTT82ATT (3812G>A), GGA83GGT (3817A>T), CTA86CTT (3826A>T), GTT88GTG (3832T>G), TTT90TTG (3838T>G), GAT91GAC (3841T>C), GCC92ACC (3842G>A), CCA98TCT (3860C>T 3862A>T), GCT99TAT (3863G>T 3864C>A), GGG100GGT (3868G>T), CTT101GTG (3869C>G 3871T>G), AAT102GCA (3872A>G 3873A>C 3874T>A), AAA103GAG (3875A>G 3877A>G), GGA104GGG (3880A>G), AAT105GCA (3881A>G 3882A>C 3883T>A), TAT106CAT (3884T>C), GTA108CTT (3890G>C 3892A>T), GGT111GGC (3901T>C), GGA112GGT (3904A>T), GGC113GGT (3907C>T), ACA114TCC (3908A>T 3910A>C), GAT117AAT (3917G>A), TTT118TTC (3922T>C), TCA119TCC (3925A>C), GGT121GGA (3931T>A), GTG122GTT (3934G>T), GCA123ACC (3935G>A 3937A>C), ACA124ACT (3940A>T), AAT125AAC (3943T>C), ACG126ATT (3945C>T 3946G>T), TTC127TTT (3949C>T), TCA128TCC (3952A>C), TTC129TTT (3955C>T), ACT130GTG (3956A>G 3957C>T 3958T>G), GAT133GAC (3967T>C), GAG135AAA (3971G>A 3973G>A), CCA138CCT (3982A>T), AGG139CGG (3983A>C), CGG140AGG (3986C>A), CAT141TAT (3989C>T), ACA142ACT (3994A>T), AGG143AAG (3996G>A), ATG144GCA (3998A>G 3999T>C 4000G>A), AGG145CGA (4001A>C 4003G>A), TTC147TTT (4009C>T), TCA148GAC (4010T>G 4011C>A 4012A>C), AGC149AAT (4014G>A 4015C>T), CAA151AGT (4019C>A 4020A>G 4021A>T), TCC152TCT (4024C>T), CGC153CGT (4027C>T), ATC154ATT (4030C>T), ATG155GTT (4031A>G 4033G>T), TCA156TCT (4036A>T), CTA157TTG (4037C>T 4039A>G), GAT159GAC (4045T>C), AGG160AAA (4047G>A 4048G>A), CTT161TTG (4049C>T 4051T>G), GGA162GGT (4054A>T), AAT163CTT (4055A>C 4056A>T), CTG164CTT (4060G>T), ATC165TTG (4061A>T 4063C>G), ATA166ATC (4066A>C), TTG168CTA (4070T>C 4072G>A), CCT169CCA (4075T>A), CCT170CCC (4078T>C), TCC171GTA (4079T>G 4080C>T 4081C>A), GCC172GAC (4083C>A), ATA173TTA (4085A>T), GTG174ATT (4088G>A 4090G>T), AGT175AGC (4093T>C), GAG178GAA (4102G>A), ATA179ATC (4105A>C), CTT180ATG (4106C>A 4108T>G), ATA181GTT (4109A>G 4111A>T), TCT182AAG (4112T>A 4113C>A 4114T>G), CCT183CCA (4117T>A), GGA184GGG (4120A>G), CTT185CCA (4122T>C 4123T>A), TTC187TTT (4129C>T), AAG188AGG (4131A>G), TTG189TTA (4135G>A), GAG190GAA (4138G>A), TTG191CTT (4139T>C 4141G>T), TCT196CTG (4154T>C 4155C>T 4156T>G), GCC197GCT (4159C>T), AAC198AAT (4162C>T), CAT199CAA (4165T>A), CAT199_GAA200insT-- (4165_4166insT), GGC204GGT (4180C>T), AAT205GAT (4181A>G), ACT208ACG (4192T>G), CAC209CAT (4195C>T), ACC210ACT (4198C>T), TAT211TAC (4201T>C), CAG212CAA (4204G>A), TCA215TCT (4213A>T), TTT217TTC (4219T>C), TCT218TCA (4222T>A), GAG219GAA (4225G>A), CTA220CTG (4228A>G), CGT221AGA (4229C>A 4231T>A), GAT222GAC (4234T>C), TTT223TTC (4237T>C), GCG224TCG (4238G>T), ATT225AAC (4242T>A 4243T>C) | | | | | | | | |

*: Inserts / Deletes / Misaligned / Frameshifts

#### Analysis details

This analysis was performed with panviral2.64

### NGS Details (UN60): Diachasmimorpha longicaudata entomopoxvirus (segment NC_043455.1)

#### Assembly

| | |
|---|---|
| **Coverage Length** | 1124 (1 contig(s)) |
| **Depth Of Coverage** | 803.2 |
| **Number Of Reads** | 7189 |
| **Reads Per Million** | 152.48 rpm (after QC) |
| **Ambiguities** | 0 |
| **Assembly Method** | de novo + reference guided assembly |
| **Consensus Caller** | Bcf Tools |

#### Coverage Map

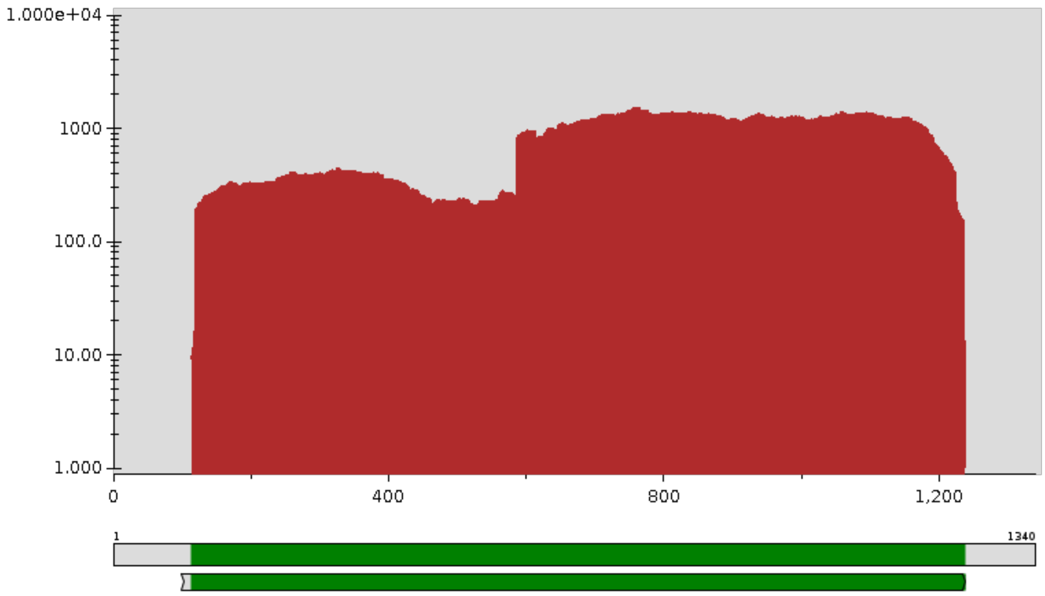

#### Assignment

| | |
|---|---|
| **Type** | Diachasmimorpha longicaudata entomopoxvirus (Taxonomy ID: 109981) |
| **Reference Genome** | NC_043455.1 |
| **NT Identity (%)** | 60.4982 |
| **AA Identity (%)** | 55.2 |
| **Number Of Stop Codons** | 0 |
| **Number Of CDS** | 1 |

#### Alignment

| | |
|---|---|
| **Alignment Score** | 454.0 (NT) + 1463.0 (AA) = 1917.0 |
| **Concordance (%)** | 40.8394 |

| Alignment Method | Global, seeded, nucleotide + amino acids (AGA) |
|---|---|

#### Genome Region

Sequence starts at position 115 and ends at position 1238 relative to NC_043455.1 reference sequence.

#### Alignment Detailed Statistics

| | Begin | End | Coverage | Score | Concordance | Matches | Identities | I/D/M/F* | Stop Codons |
|---|---|---|---|---|---|---|---|---|---|
| **NT** | **115** | **1238** | **83.9%** | **454** | **20.4%** | **1121 (99.5%)** | **680 (60.3%)** | **3/3** | |
| Mutations: | 121A>T, 126A>C, 127C>T, 128C>G, 130T>G, 136C>G, 137C>A, 139C>T, 142G>A, 143A>G, 145A>T, 146A>G, 148T>C, 149A>C, 151C>T, 154A>T, 156A>G, 157G>A, 160T>G, 163A>T, 166T>C, 167T>C, 168C>A, 169T>G, 171G>A, 172T>C, 175A>G, 178T>C, 187T>A, 190A>T, 191A>G, 193T>C, 203A>C, 204A>G, 208C>T, 209A>G, 211T>A, 214T>G, 217A>T, 218T>A, 219G>T, 223T>A, 226T>G, 230A>C, 231A>G, 232A>T, 238G>C, 241A>T, 243T>C, 244C>T, 247A>G, 256G>C, 259A>T, 262A>C, 271A>C, 272G>T, 274A>C, 276C>T, 277A>G, 278T>A, 279A>T, 283A>T, 284A>C, 286C>T, 288G>C, 289T>C, 294T>G, 295A>C, 298G>A, 299C>G, 300A>T, 301A>T, 302A>G, 304A>G, 310T>C, 311T>A, 312C>A, 313T>A, 320A>G, 322T>G, 323A>G, 325C>A, 331T>A, 337C>A, 341A>T, 343C>T, 346A>T, 352T>G, 355A>G, 356T>C, 358G>T, 361T>A, 362C>G, 363T>C, 364C>T, 368G>A, 370T>G, 371C>G, 374C>A, 375G>A, 376T>G, 379A>G, 380T>A, 382G>T, 383C>T, 384A>T, 385A>G, 386A>G, 394G>A, 395A>G, 400T>C, 401C>A, 403A>T, 404_406delTAT, 409T>C, 410T>A, 412T>A, 413A>C, 415A>G, 416T>G, 417G>C, 418T>A, 421A>T, 423T>C, 424T>G, 438C>A, 442T>C, 443A>G, 445C>G, 446A>G, 447A>G, 448G>T, 452A>G, 453G>A, 454C>T, 455C>A, 456A>T, 457G>C, 458G>A, 459A>G, 462C>A, 463T>G, 467A>G, 469_470insCAT, 470A>G, 471A>G, 474C>T, 475C>A, 478G>A, 482T>G, 484G>C, 485A>T, 486T>C, 500C>A, 503A>G, 505G>C, 506A>T, 507T>G, 511T>C, 512C>A, 514A>G, 515C>A, 517C>T, 519C>A, 520C>A, 521C>A, 523A>G, 525A>G, 528G>C, 530A>T, 532C>A, 533G>C, 534A>G, 535T>A, 540A>G, 541A>G, 544A>C, 547T>A, 551A>T, 554G>T, 556T>G, 559A>T, 560A>C, 562A>T, 569G>T, 585T>G, 586A>C, 587G>C, 588A>G, 589T>A, 590A>G, 591A>G, 595T>C, 596T>A, 597T>A, 602A>C, 604A>G, 607A>T, 608C>T, 610A>T, 616T>A, 620G>C, 622A>G, 623T>C, 628T>G, 629G>C, 630A>C, 632A>C, 633G>C, 634T>A, 635C>A, 637T>A, 638G>A, 646T>G, 647A>G, 648T>G, 650T>G, 652A>T, 655A>C, 658A>T, 661A>C, 664C>T, 665G>A, 667T>G, 671T>C, 674A>G, 675G>A, 678T>C, 680A>C, 682C>T, 683A>G, 685T>G, 687C>T, 688A>T, 689T>A, 692C>A, 693A>G, 694G>A, 695G>A, 696T>A, 697C>G, 700T>C, 705G>A, 706A>C, 707G>A, 709T>G, 715A>G, 717A>G, 718A>G, 724G>T, 727A>G, 730A>G, 731A>C, 732A>G, 735C>A, 739A>G, 740T>C, 742G>C, 745A>T, 748C>T, 754C>T, 759G>A, 765A>T, 770A>G, 772A>C, 778A>C, 779A>G, 781G>T, 784A>G, 785A>G, 787T>A, 790T>G, 792T>G, 793T>G, 796A>G, 797G>C, 798C>T, 799A>T, 810T>G, 820T>C, 823C>G, 824C>A, 825A>C, 830A>G, 831G>C, 833C>A, 838T>C, 841A>G, 843C>G, 844A>T, 845C>G, 847A>C, 850A>C, 854T>G, 855G>T, 856T>G, 859T>C, 862A>C, 863C>A, 865T>G, 866A>C, 868A>T, 869C>A, 878G>T, 879T>G, 880A>G, 885T>C, 886G>T, 889A>T, 890T>A, 891G>A, 892T>G, 893T>A, 895A>G, 896A>C, 897C>G, 898A>C, 900A>G, 901T>C, 902A>C, 904A>T, 905A>G, 908T>C, 909T>A, 915C>T, 919A>T, 920A>G, 921G>C, 922C>A, 924T>C, 925C>T, 934T>C, 938T>G, 939C>A, 943A>G, 944C>A, 946A>C, 947G>A, 948A>C, 949A>T, 950C>A, 952T>A, 956T>A, 958T>C, 965A>C, 966A>G, 970A>G, 973T>C, 974A>C, 977G>T, 978A>C, 982T>G, 983A>T, 984A>C, 986A>T, 989A>C, 991A>T, 992A>G, 994A>G, 997T>C, 998C>A, 1000T>C, 1001T>A, 1003A>T, 1007A>G, 1012A>T, 1015A>G, 1018A>T, 1019A>C, 1021A>T, 1024C>T, 1033T>A, 1036A>G, 1040A>G, 1042A>C, 1045T>C, 1046T>C, 1048A>T, 1051T>C, 1054C>T, 1057T>C, 1063T>C, 1064T>C, 1070C>A, 1071A>C, 1073A>C, 1075T>A, 1076A>C, 1077G>C, 1087T>C, 1088A>C, 1090A>G, 1103A>C, 1105A>T, 1111T>A, 1112A>C, 1120C>A, 1121C>A, 1123A>G, 1126A>G, 1129G>T, 1130A>G, 1141T>C, 1145A>G, 1147A>C, 1150T>C, 1151G>A, 1153A>G, 1156C>T, 1159A>T, 1160T>G, 1161T>A, 1163C>A, 1164C>G, 1165A>G, 1167A>T, 1172G>T, 1173C>T, 1177A>C, 1178T>A, 1181G>C, 1183A>G, 1185C>A, 1186G>A, 1192T>C, 1195T>C, 1196A>G, 1197C>T, 1198C>T, 1199A>G, 1200A>T, 1204A>C, 1207T>G, 1210A>G, 1211A>C, 1218A>C, 1219G>C, 1223A>G, 1227T>C, 1232T>C, 1234G>C, 1235A>C, 1237A>G | | | | | | | | |

##### CDS

| | Begin | End | Coverage | Score | Concordance | Matches | Identities | I/D/M/F* | Stop Codons |
|---|---|---|---|---|---|---|---|---|---|
| FLA14_p101 | 6 | 380 | 98.7% | 1463 | 59.0% | 374 (99.5%) | 207 (55.1%) | 1/1/0/0 | 0 |
| Protein mutations: | E7D (121A>T), D9A (126A>C 127C>T), H10E (128C>G 130T>G), L13I (137C>A 139C>T), K15D (143A>G 145A>T), N16D (146A>G 148T>C), I17L (149A>C 151C>T), K19R (156A>G 157G>A), S23Q (167T>C 168C>A 169T>G), C24Y (171G>A 172T>C), T31A (191A>G 193T>C), K35R (203A>C 204A>G), I37V (209A>G 211T>A), F38L (214T>G), C40I (218T>A 219G>T), K44R (230A>C 231A>G 232A>T), V48A (243T>C 244C>T), A58S (272G>T 274A>C), T59M (276C>T 277A>G), Y60I (278T>A 279A>T), I62L (284A>C 286C>T), S63T (288G>C 289T>C), L65C (294T>G 295A>C), Q67V (299C>G 300A>T 301A>T), I68V (302A>G 304A>G), S71K (311T>A 312C>A 313T>A), N74E (320A>G 322T>G), I75V (323A>G 325C>A), T81S (341A>T 343C>T), L88A (362C>G 363T>C 364C>T), A90T (368G>A 370T>G), Q91E (371C>G), R92K (374C>A 375G>A 376T>G), L94I (380T>A 382G>T), Q95L (383C>T 384A>T 385A>G), T96A (386A>G), N99D (395A>G), L101I (401C>A 403A>T), Y102del (404_406delTAT), F104I (410T>A 412T>A), K105Q (413A>C 415A>G), C106A (416T>G 417G>C 418T>A), Q107H (421A>T), V108A (423T>C 424T>G), T113K (438C>A), I115V (443A>G 445C>G), K116G (446A>G 447A>G 448G>T), S118D (452A>G 453G>A 454C>T), Q119I (455C>A 456A>T 457G>C), E120R (458G>A 459A>G), T121K (462C>A 463T>G), K123E (467A>G), K123_K124insH (469_470insCAT), K124G (470A>G 471A>G), A125V (474C>T 475C>A), L128V (482T>G 484G>C), I129S (485A>T 486T>C), M135V (503A>G 505G>C), I136C (506A>T 507T>G), L138M (512C>A 514A>G), L139I (515C>A 517C>T), T140K (519C>A 520C>A), K142R (525A>G), S143T (528G>C), I144L (530A>T 532C>A), D145R (533G>C 534A>G 535T>A), K147R (540A>G 541A>G), I151L (551A>T), V152L (554G>T 556T>G), I154L (560A>C 562A>T), A157S (569G>T), I162S (585T>G 586A>C), D163R (587G>C 588A>G 589T>A), N164G (590A>G 591A>G), L166K (596T>A 597T>A), K168Q (602A>C 604A>G), Q170Y (608C>T 610A>T), E174Q (620G>C 622A>G), F175L (623T>C), F176L (628T>G), E177P (629G>C 630A>C), S178P (632A>C 633G>C 634T>A), H179K (635C>A 637T>A), V180I (638G>A), I183G (647A>G 648T>G), L184V (650T>G 652A>T), L185F (655A>C), V189M (665G>A 667T>G), S191P (671T>C), R192E (674A>G 675G>A), V193A (678T>C), I194L (680A>C 682C>T), N195E (683A>G 685T>G), T196I (687C>T 688A>T), S197T (689T>A), Q198R (692C>A 693A>G 694G>A), V199K (695G>A 696T>A 697C>G), R202N (705G>A 706A>C), D203K (707G>A 709T>G), K206R (717A>G 718A>G), N211R (731A>C 732A>G), A212D (735C>A), R220K (759G>A), Y222F (765A>T), I224V (770A>G 772A>C), K227D (779A>G 781G>T), N229E (785A>G 787T>A), D230E (790T>G), F231W (792T>G 793T>G), A233L (797G>C 798C>T 799A>T), F237C (810T>G), D241E (823C>G), H242T (824C>A 825A>C), S244A (830A>G 831G>C), L245I (833C>A), T248S (843C>G 844A>T), L249V (845C>G 847A>C), C252V (854T>G 855G>T 856T>G), Q257K (869C>A), V260W (878G>T 879T>G 880A>G), M262T (885T>C 886G>T), E263D (889A>T), C264K (890T>A 891G>A 892T>G), L265M (893T>A 895A>G), T266R (896A>C 897C>G 898A>C), N267S (900A>G 901T>C), N269D (905A>G), F270H (908T>C 909T>A), A272V (915C>T), S274A (920A>G 921G>C 922C>A), I275T (924T>C 925C>T), S280D (938T>G 939C>A), Q282N (944C>A 946A>C), E283T (947G>A 948A>C 949A>T), F286I (956T>A 958T>C), K289R (965A>C 966A>G), D293S (977G>T 978A>C), K295S (983A>T 984A>C), T296S (986A>T), I298V (992A>G 994A>G), L300I (998C>A 1000T>C), S301T (1001T>A 1003A>T), N303D (1007A>G), I314V (1040A>G 1042A>C), H324T (1070C>A 1071A>C), N325Q (1073A>C 1075T>A), R326P (1076A>C 1077G>C), I330L (1088A>C 1090A>G), I344V (1130A>G), I349V (1145A>G 1147A>C), E351K (1151G>A 1153A>G), E353D (1159A>T), L354E (1160T>G 1161T>A), P355R (1163C>A 1164C>G 1165A>G), K356M (1167A>T), A358F (1172G>T 1173C>T), E359D (1177A>C), L360I (1178T>A), E361Q (1181G>C 1183A>G), T362K (1185C>A 1186G>A), T366V (1196A>G 1197C>T 1198C>T), K367V (1199A>G 1200A>T), D369E (1207T>G), M371L (1211A>C), E373A (1218A>C 1219G>C), I375V (1223A>G), V376A (1227T>C), I379L (1235A>C 1237A>G) | | | | | | | | |

| | Begin | End | Coverage | Score | Concordance | Matches | Identities | I/D/M/F* | Stop Codons |
|---|---|---|---|---|---|---|---|---|---|
| **NT** | **115** | **1238** | **83.9%** | **454** | **20.4%** | **1121 (99.5%)** | **680 (60.3%)** | **3/3** | |
| Codon mutations: | GAA7GAT (121A>T), GAC9GCT (126A>C 127C>T), CAT10GAG (128C>G 130T>G), GGC12GGG (136C>G), CTC13ATT (137C>A 139C>T), AAG14AAA (142G>A), AAA15GAT (143A>G 145A>T), AAT16GAC (146A>G 148T>C), ATC17CTT (149A>C 151C>T), CTA18CTT (154A>T), AAG19AGA (156A>G 157G>A), GGT20GGG (160T>G), ATA21ATT (163A>T), TAT22TAC (166T>C), TCT23CAG (167T>C 168C>A 169T>G), TGT24TAC (171G>A 172T>C), GGA25GGG (175A>G), TTT26TTC (178T>C), CCT29CCA (187T>A), TCA30TCT (190A>T), ACT31GCC (191A>G 193T>C), AAA35CGA (203A>C 204A>G), GCC36GCT (208C>T), ATT37GTA (209A>G 211T>A), TTT38TTG (214T>G), CCA39CCT (217A>T), TGT40ATT (218T>A 219G>T), ATT41ATA (223T>A), TCT42TCG (226T>G), AAA44CGT (230A>C 231A>G 232A>T), GTG46GTC (238G>C), ATA47ATT (241A>T), GTC48GCT (243T>C 244C>T), CAA49CAG (247A>G), TCG52TCC (256G>C), GGA53GGT (259A>T), ACA54ACC (262A>C), ACA57ACC (271A>C), GCA58TCC (272G>T 274A>C), ACA59ATG (276C>T 277A>G), TAT60ATT (278T>A 279A>T), GCA61GCT (283A>T), ATC62CTT (284A>C 286C>T), AGT63ACC (288G>C 289T>C), TTA65TGC (294T>G 295A>C), CAG66CAA (298G>A), CAA67GTT (299C>G 300A>T 301A>T), ATA68GTG (302A>G 304A>G), ACT70ACC (310T>C), TCT71AAA (311T>A 312C>A 313T>A), AAT74GAG (320A>G 322T>G), ATC75GTA (323A>G 325C>A), GCT77GCA (331T>A), ATC79ATA (337C>A), ACC81TCT (341A>T 343C>T), CCA82CCT (346A>T), CGT84CGG (352T>G), GAA85GAG (355A>G), TTG86CTT (356T>C 358G>T), GCT87GCA (361T>A), CTC88GCT (362C>G 363T>C 364C>T), GCT90ACG (368G>A 370T>G), CAA91GAA (371C>G), CGT92AAG (374C>A 375G>A 376T>G), GTA93GTG (379A>G), TTG94ATT (380T>A 382G>T), CAA95TTG (383C>T 384A>T 385A>G), ACA96GCA (386A>G), GGG98GGA (394G>A), AAT99GAT (395A>G), TAT100TAC (400T>C), CTA101ATT (401C>A 403A>T), TAT102del (404_406delTAT), AAT103AAC (409T>C), TTT104ATA (410T>A 412T>A), AAA105CAG (413A>C 415A>G), TGT106GCA (416T>G 417G>C 418T>A), CAA107CAT (421A>T), GTT108GCG (423T>C 424T>G), ACA113AAA (438C>A), AGT114AGC (442T>C), ATC115GTG (443A>G 445C>G), AAG116GGT (446A>G 447A>G 448G>T), AGC118GAT (452A>G 453G>A 454C>T), CAG119ATC (455C>A 456A>T 457G>C), GAA120AGA (458G>A 459A>G), ACT121AAG (462C>A 463T>G), AAG123GAG (467A>G), AAG123_AAA124insCAT (469_470insCAT), AAA124GGA (470A>G 471A>G), GCC125GTA (474C>T 475C>A), CAG126CAA (478G>A), TTG128GTC (482T>G 484G>C), ATT129TCT (485A>T 486T>C), CGA134AGA (500C>A), ATG135GTC (503A>G 505G>C), ATT136TGT (506A>T 507T>G), GAT137GAC (511T>C), CTA138ATG (512C>A 514A>G), CTC139ATT (515C>A 517C>T), ACC140AAA (519C>A 520C>A), CGA141AGG (521C>A 523A>G), AAA142AGA (525A>G), AGT143ACT (528G>C), ATC144TTA (530A>T 532C>A), GAT145CGA (533G>C 534A>G 535T>A), AAA147AGG (540A>G 541A>G), GCA148GCC (544A>C), ATT149ATA (547T>A), ATA151TTA (551A>T), GTT152TTG (554G>T 556T>G), GTA153GTT (559A>T), ATA154CTT (560A>C 562A>T), GCT157TCT (569G>T), ATA162AGC (585T>G 586A>C), GAT163CGA (587G>C 588A>G 589T>A), AAT164GGT (590A>G 591A>G), TTT165TTC (595T>C), TTG166AAG (596T>A 597T>A), AAA168CAG (602A>C 604A>G), ATA169ATT (607A>T), CAA170TAT (608C>T 610A>T), ATT172ATA (616T>A), GAA174CAG (620G>C 622A>G), TTT175CTT (623T>C), TTT176TTG (628T>G), GAA177CCA (629G>C 630A>C), AGT178CCA (632A>C 633G>C 634T>A), CAT179AAA (635C>A 637T>A), GTC180ATC (638G>A), GTT182GTG (646T>G), ATT183GGT (647A>G 648T>G), TTA184GTT (650T>G 652A>T), TTA185TTC (655A>C), TCA186TCT (658A>T), GCA187GCC (661A>C), ACC188ACT (664C>T), GTT189ATG (665G>A 667T>G), TCA191CCA (671T>C), AGG192GAG (674A>G 675G>A), GTT193GCT (678T>C), ATC194CTT (680A>C 682C>T), AAT195GAG (683A>G 685T>G), ACA196ATT (687C>T 688A>T), TCT197ACT (689T>A), CAG198AGA (692C>A 693A>G 694G>A), GTC199AAG (695G>A 696T>A 697C>G), TTT200TTC (700T>C), AGA202AAC (705G>A 706A>C), GAT203AAG (707G>A 709T>G), GTA205GTG (715A>G), AAA206AGG (717A>G 718A>G), CTG208CTT (724G>T), GTA209GTG (727A>G), AAA210AAG (730A>G), AAT211CGT (731A>C 732A>G), GCT212GAT (735C>A), GAA213GAG (739A>G), TTG214CTC (740T>C 742G>C), ACA215ACT (745A>T), CTC216CTT (748C>T), GGC218GGT (754C>T), AGG220AAG (759G>A), TAT222TTT (765A>T), ATA224GTC (770A>G 772A>C), GTA226GTC (778A>C), AAG227GAT (779A>G 781G>T), AAA228AAG (784A>G), AAT229GAA (785A>G 787T>A), GAT230GAG (790T>G), TTT231TGG (792T>G 793T>G), AAA232AAG (796A>G), GCA233CTT (797G>C 798C>T 799A>T), TTT237TGT (810T>G), TAT240TAC (820T>C), GAC241GAG (823C>G), CAC242ACC (824C>A 825A>C), AGC244GCC (830A>G 831G>C), CTC245ATC (833C>A), ACT246ACC (838T>C), CAA247CAG (841A>G), ACA248AGT (843C>G 844A>T), CTA249GTC (845C>G 847A>C), ATA250ATC (850A>C), TGT252GTG (854T>G 855G>T 856T>G), AAT253AAC (859T>C), ACA254ACC (862A>C), CGT255AGG (863C>A 865T>G), AGA256CGT (866A>C 868A>T), CAA257AAA (869C>A), GTA260TGG (878G>T 879T>G 880A>G), ATG262ACT (885T>C 886G>T), GAA263GAT (889A>T), TGT264AAG (890T>A 891G>A 892T>G), TTA265ATG (893T>A 895A>G), ACA266CGC (896A>C 897C>G 898A>C), AAT267AGC (900A>G 901T>C), AGA268CGT (902A>C 904A>T), AAT269GAT (905A>G), TTC270CAC (908T>C 909T>A), GCA272GTA (915C>T), TCA273TCT (919A>T), AGC274GCA (920A>G 921G>C 922C>A), ATC275ACT (924T>C 925C>T), GAT278GAC (934T>C), TCC280GAC (938T>G 939C>A), CAA281CAG (943A>G), CAA282AAC (944C>A 946A>C), GAA283ACT (947G>A 948A>C 949A>T), CGT284AGA (950C>A 952T>A), TTT286ATC (956T>A 958T>C), AAA289CGA (965A>C 966A>G), GAA290GAG (970A>G), TTT291TTC (973T>C), AGA292CGA (974A>C), GAT293TCT (977G>T 978A>C), GGT294GGG (982T>G), AAA295TCA (983A>T 984A>C), ACT296TCT (986A>T), AGA297CGT (989A>C 991A>T), ATA298GTG (992A>G 994A>G), CTT299CTC (997T>C), CTT300ATC (998C>A 1000T>C), TCA301ACT (1001T>A 1003A>T), AAT303GAT (1007A>G), CTA304CTT (1012A>T), TTA305TTG (1015A>G), GCA306GCT (1018A>T), AGA307CGT (1019A>C 1021A>T), GGC308GGT (1024C>T), GTT311GTA (1033T>A), CAA312CAG (1036A>G), ATA314GTC (1040A>G 1042A>C), TCT315TCC (1045T>C), TTA316CTT (1046T>C 1048A>T), GTT317GTC (1051T>C), ATC318ATT (1054C>T), AAT319AAC (1057T>C), GAT321GAC (1063T>C), TTG322CTG (1064T>C), CAT324ACT (1070C>A 1071A>C), AAT325CAA (1073A>C 1075T>A), AGA326CCA (1076A>C 1077G>C), TAT329TAC (1087T>C), ATA330CTG (1088A>C 1090A>G), AGA335CGT (1103A>C 1105A>T), GGT337GGA (1111T>A), AGA338CGA (1112A>C), GGC340GGA (1120C>A), CGA341AGG (1121C>A 1123A>G), AAA342AAG (1126A>G), GGG343GGT (1129G>T), ATT344GTT (1130A>G), AAT347AAC (1141T>C), ATA349GTC (1145A>G 1147A>C), ACT350ACC (1150T>C), GAA351AAG (1151G>A 1153A>G), GAC352GAT (1156C>T), GAA353GAT (1159A>T), TTA354GAA (1160T>G 1161T>A), CCA355AGG (1163C>A 1164C>G 1165A>G), AAG356ATG (1167A>T), GCT358TTT (1172G>T 1173C>T), GAA359GAC (1177A>C), TTA360ATA (1178T>A), GAA361CAG (1181G>C 1183A>G), ACG362AAA (1185C>A 1186G>A), TAT364TAC (1192T>C), AAT365AAC (1195T>C), ACC366GTT (1196A>G 1197C>T 1198C>T), AAA367GTA (1199A>G 1200A>T), ATA368ATC (1204A>C), GAT369GAG (1207T>G), GAA370GAG (1210A>G), ATG371CTG (1211A>C), GAG373GCC (1218A>C 1219G>C), ATT375GTT (1223A>G), GTT376GCT (1227T>C), TTG378CTC (1232T>C 1234G>C), ATA379CTG (1235A>C 1237A>G) | | | | | | | | |

#### Proteins

| | Begin | End | Coverage | Score | Concordance | Matches | Identities | I/D/M/F* | Stop Codons |
|---|---|---|---|---|---|---|---|---|---|
| hypothetical protein (YP_009666148.1) | 6 | 380 | 98.7% | 1463 | 59.0% | 374 (99.5%) | 207 (55.1%) | 1/1/0/0 | 0 |
| Protein mutations: | E7D (121A>T), D9A (126A>C 127C>T), H10E (128C>G 130T>G), L13I (137C>A 139C>T), K15D (143A>G 145A>T), N16D (146A>G 148T>C), I17L (149A>C 151C>T), K19R (156A>G 157G>A), S23Q (167T>C 168C>A 169T>G), C24Y (171G>A 172T>C), T31A (191A>G 193T>C), K35R (203A>C 204A>G), I37V (209A>G 211T>A), F38L (214T>G), C40I (218T>A 219G>T), K44R (230A>C 231A>G 232A>T), V48A (243T>C 244C>T), A58S (272G>T 274A>C), T59M (276C>T 277A>G), Y60I (278T>A 279A>T), I62L (284A>C 286C>T), S63T (288G>C 289T>C), L65C (294T>G 295A>C), Q67V (299C>G 300A>T 301A>T), I68V (302A>G 304A>G), S71K (311T>A 312C>A 313T>A), N74E (320A>G 322T>G), I75V (323A>G 325C>A), T81S (341A>T 343C>T), L88A (362C>G 363T>C 364C>T), A90T (368G>A 370T>G), Q91E (371C>G), R92K (374C>A 375G>A 376T>G), L94I (380T>A 382G>T), Q95L (383C>T 384A>T 385A>G), T96A (386A>G), N99D (395A>G), L101I (401C>A 403A>T), Y102del (404_406delTAT), F104I (410T>A 412T>A), K105Q (413A>C 415A>G), C106A (416T>G 417G>C 418T>A), Q107H (421A>T), V108A (423T>C 424T>G), T113K (438C>A), I115V (443A>G 445C>G), K116G (446A>G 447A>G 448G>T), S118D (452A>G 453G>A 454C>T), Q119I (455C>A 456A>T 457G>C), E120R (458G>A 459A>G), T121K (462C>A 463T>G), K123E (467A>G), K123_K124insH (469_470insCAT), K124G (470A>G 471A>G), A125V (474C>T 475C>A), L128V (482T>G 484G>C), I129S (485A>T 486T>C), M135V (503A>G 505G>C), I136C (506A>T 507T>G), L138M (512C>A 514A>G), L139I (515C>A 517C>T), T140K (519C>A 520C>A), K142R (525A>G), S143T (528G>C), I144L (530A>T 532C>A), D145R (533G>C 534A>G 535T>A), K147R (540A>G 541A>G), I151L (551A>T), V152L (554G>T 556T>G), I154L (560A>C 562A>T), A157S (569G>T), I162S (585T>G 586A>C), D163R (587G>C 588A>G 589T>A), N164G (590A>G 591A>G), L166K (596T>A 597T>A), K168Q (602A>C 604A>G), Q170Y (608C>T 610A>T), E174Q (620G>C 622A>G), F175L (623T>C), F176L (628T>G), E177P (629G>C 630A>C), S178P (632A>C 633G>C 634T>A), H179K (635C>A 637T>A), V180I (638G>A), I183G (647A>G 648T>G), L184V (650T>G 652A>T), L185F (655A>C), V189M (665G>A 667T>G), S191P (671T>C), R192E (674A>G 675G>A), V193A (678T>C), I194L (680A>C 682C>T), N195E (683A>G 685T>G), T196I (687C>T 688A>T), S197T (689T>A), Q198R (692C>A 693A>G 694G>A), V199K (695G>A 696T>A 697C>G), R202N (705G>A 706A>C), D203K (707G>A 709T>G), K206R (717A>G 718A>G), N211R (731A>C 732A>G), A212D (735C>A), R220K (759G>A), Y222F (765A>T), I224V (770A>G 772A>C), K227D (779A>G 781G>T), N229E (785A>G 787T>A), D230E (790T>G), F231W (792T>G 793T>G), A233L (797G>C 798C>T 799A>T), F237C (810T>G), D241E (823C>G), H242T (824C>A 825A>C), S244A (830A>G 831G>C), L245I (833C>A), T248S (843C>G 844A>T), L249V (845C>G 847A>C), C252V (854T>G 855G>T 856T>G), Q257K (869C>A), V260W (878G>T 879T>G 880A>G), M262T (885T>C 886G>T), E263D (889A>T), C264K (890T>A 891G>A 892T>G), L265M (893T>A 895A>G), T266R (896A>C 897C>G 898A>C), N267S (900A>G 901T>C), N269D (905A>G), F270H (908T>C 909T>A), A272V (915C>T), S274A (920A>G 921G>C 922C>A), I275T (924T>C 925C>T), S280D (938T>G 939C>A), Q282N (944C>A 946A>C), E283T (947G>A 948A>C 949A>T), F286I (956T>A 958T>C), K289R (965A>C 966A>G), D293S (977G>T 978A>C), K295S (983A>T 984A>C), T296S (986A>T), I298V (992A>G 994A>G), L300I (998C>A 1000T>C), S301T (1001T>A 1003A>T), N303D (1007A>G), I314V (1040A>G 1042A>C), H324T (1070C>A 1071A>C), N325Q (1073A>C 1075T>A), R326P (1076A>C 1077G>C), I330L (1088A>C 1090A>G), I344V (1130A>G), I349V (1145A>G 1147A>C), E351K (1151G>A 1153A>G), E353D (1159A>T), L354E (1160T>G 1161T>A), P355R (1163C>A 1164C>G 1165A>G), K356M (1167A>T), A358F (1172G>T 1173C>T), E359D (1177A>C), L360I (1178T>A), E361Q (1181G>C 1183A>G), T362K (1185C>A 1186G>A), T366V (1196A>G 1197C>T 1198C>T), K367V (1199A>G 1200A>T), D369E (1207T>G), M371L (1211A>C), E373A (1218A>C 1219G>C), I375V (1223A>G), V376A (1227T>C), I379L (1235A>C 1237A>G) | | | | | | | | |

| | Begin | End | Coverage | Score | Concordance | Matches | Identities | I/D/M/F* | Stop Codons |
|---|---|---|---|---|---|---|---|---|---|
| **NT** | **115** | **1238** | **83.9%** | **454** | **20.4%** | **1121 (99.5%)** | **680 (60.3%)** | **3/3** | |

Codon mutations: GAA7GAT (121A>T), GAC9GCT (126A>C 127C>T), CAT10GAG (128C>G 130T>G), GGC12GGG (136C>G), CTC13ATT (137C>A 139C>T), AAG14AAA (142G>A), AAA15GAT (143A>G 145A>T), AAT16GAC (146A>G 148T>C), ATC17CTT (149A>C 151C>T), CTA18CTT (154A>T), AAG19AGA (156A>G 157G>A), GGT20GGG (160T>G), ATA21ATT (163A>T), TAT22TAC (166T>C), TCT23CAG (167T>C 168C>A 169T>G), TGT24TAC (171G>A 172T>C), GGA25GGG (175A>G), TTT26TTC (178T>C), CCT29CCA (187T>A), TCA30TCT (190A>T), ACT31GCC (191A>G 193T>C), AAA35CGA (203A>C 204A>G), GCC36GCT (208C>T), ATT37GTA (209A>G 211T>A), TTT38TTG (214T>G), CCA39CCT (217A>T), TGT40ATT (218T>A 219G>T), ATT41ATA (223T>A), TCT42TCG (226T>G), AAA44CGT (230A>C 231A>G 232A>T), GTG46GTC (238G>C), ATA47ATT (241A>T), GTC48GCT (243T>C 244C>T), CAA49CAG (247A>G), TCG52TCC (256G>C), GGA53GGT (259A>T), ACA54ACC (262A>C), ACA57ACC (271A>C), GCA58TCC (272G>T 274A>C), ACA59ATG (276C>T 277A>G), TAT60ATT (278T>A 279A>T), GCA61GCT (283A>T), ATC62CTT (284A>C 286C>T), AGT63ACC (288G>C 289T>C), TTA65TGC (294T>G 295A>C), CAG66CAA (298G>A), CAA67GTT (299C>G 300A>T 301A>T), ATA68GTG (302A>G 304A>G), ACT70ACC (310T>C), TCT71AAA (311T>A 312C>A 313T>A), AAT74GAG (320A>G 322T>G), ATC75GTA (323A>G 325C>A), GCT77GCA (331T>A), ATC79ATA (337C>A), ACC81TCT (341A>T 343C>T), CCA82CCT (346A>T), CGT84CGG (352T>G), GAA85GAG (355A>G), TTG86CTT (356T>C 358G>T), GCT87GCA (361T>A), CTC88GCT (362C>G 363T>C 364C>T), GCT90ACG (368G>A 370T>G), CAA91GAA (371C>G), CGT92AAG (374C>A 375G>A 376T>G), GTA93GTG (379A>G), TTG94ATT (380T>A 382G>T), CAA95TTG (383C>T 384A>T 385A>G), ACA96GCA (386A>G), GGG98GGA (394G>A), AAT99GAT (395A>G), TAT100TAC (400T>C), CTA101ATT (401C>A 403A>T), TAT102del (404_406delTAT), AAT103AAC (409T>C), TTT104ATA (410T>A 412T>A), AAA105CAG (413A>C 415A>G), TGT106GCA (416T>G 417G>C 418T>A), CAA107CAT (421A>T), GTT108GCG (423T>C 424T>G), ACA113AAA (438C>A), AGT114AGC (442T>C), ATC115GTG (443A>G 445C>G), AAG116GGT (446A>G 447A>G 448G>T), AGC118GAT (452A>G 453G>A 454C>T), CAG119ATC (455C>A 456A>T 457G>C), GAA120AGA (458G>A 459A>G), ACT121AAG (462C>A 463T>G), AAG123GAG (467A>G), AAG123_AAA124insCAT (469_470insCAT), AAA124GGA (470A>G 471A>G), GCC125GTA (474C>T 475C>A), CAG126CAA (478G>A), TTG128GTC (482T>G 484G>C), ATT129TCT (485A>T 486T>C), CGA134AGA (500C>A), ATG135GTC (503A>G 505G>C), ATT136TGT (506A>T 507T>G), GAT137GAC (511T>C), CTA138ATG (512C>A 514A>G), CTC139ATT (515C>A 517C>T), ACC140AAA (519C>A 520C>A), CGA141AGG (521C>A 523A>G), AAA142AGA (525A>G), AGT143ACT (528G>C), ATC144TTA (530A>T 532C>A), GAT145CGA (533G>C 534A>G 535T>A), AAA147AGG (540A>G 541A>G), GCA148GCC (544A>C), ATT149ATA (547T>A), ATA151TTA (551A>T), GTT152TTG (554G>T 556T>G), GTA153GTT (559A>T), ATA154CTT (560A>C 562A>T), GCT157TCT (569G>T), ATA162AGC (585T>G 586A>C), GAT163CGA (587G>C 588A>G 589T>A), AAT164GGT (590A>G 591A>G), TTT165TTC (595T>C), TTG166AAG (596T>A 597T>A), AAA168CAG (602A>C 604A>G), ATA169ATT (607A>T), CAA170TAT (608C>T 610A>T), ATT172ATA (616T>A), GAA174CAG (620G>C 622A>G), TTT175CTT (623T>C), TTT176TTG (628T>G), GAA177CCA (629G>C 630A>C), AGT178CCA (632A>C 633G>C 634T>A), CAT179AAA (635C>A 637T>A), GTC180ATC (638G>A), GTT182GTG (646T>G), ATT183GGT (647A>G 648T>G), TTA184GTT (650T>G 652A>T), TTA185TTC (655A>C), TCA186TCT (658A>T), GCA187GCC (661A>C), ACC188ACT (664C>T), GTT189ATG (665G>A 667T>G), TCA191CCA (671T>C), AGG192GAG (674A>G 675G>A), GTT193GCT (678T>C), ATC194CTT (680A>C 682C>T), AAT195GAG (683A>G 685T>G), ACA196ATT (687C>T 688A>T), TCT197ACT (689T>A), CAG198AGA (692C>A 693A>G 694G>A), GTC199AAG (695G>A 696T>A 697C>G), TTT200TTC (700T>C), AGA202AAC (705G>A 706A>C), GAT203AAG (707G>A 709T>G), GTA205GTG (715A>G), AAA206AGG (717A>G 718A>G), CTG208CTT (724G>T), GTA209GTG (727A>G), AAA210AAG (730A>G), AAT211CGT (731A>C 732A>G), GCT212GAT (735C>A), GAA213GAG (739A>G), TTG214CTC (740T>C 742G>C), ACA215ACT (745A>T), CTC216CTT (748C>T), GGC218GGT (754C>T), AGG220AAG (759G>A), TAT222TTT (765A>T), ATA224GTC (770A>G 772A>C), GTA226GTC (778A>C), AAG227GAT (779A>G 781G>T), AAA228AAG (784A>G), AAT229GAA (785A>G 787T>A), GAT230GAG (790T>G), TTT231TGG (792T>G 793T>G), AAA232AAG (796A>G), GCA233CTT (797G>C 798C>T 799A>T), TTT237TGT (810T>G), TAT240TAC (820T>C), GAC241GAG (823C>G), CAC242ACC (824C>A 825A>C), AGC244GCC (830A>G 831G>C), CTC245ATC (833C>A), ACT246ACC (838T>C), CAA247CAG (841A>G), ACA248AGT (843C>G 844A>T), CTA249GTC (845C>G 847A>C), ATA250ATC (850A>C), TGT252GTG (854T>G 855G>T 856T>G), AAT253AAC (859T>C), ACA254ACC (862A>C), CGT255AGG (863C>A 865T>G), AGA256CGT (866A>C 868A>T), CAA257AAA (869C>A), GTA260TGG (878G>T 879T>G 880A>G), ATG262ACT (885T>C 886G>T), GAA263GAT (889A>T), TGT264AAG (890T>A 891G>A 892T>G), TTA265ATG (893T>A 895A>G), ACA266CGC (896A>C 897C>G 898A>C), AAT267AGC (900A>G 901T>C), AGA268CGT (902A>C 904A>T), AAT269GAT (905A>G), TTC270CAC (908T>C 909T>A), GCA272GTA (915C>T), TCA273TCT (919A>T), AGC274GCA (920A>G 921G>C 922C>A), ATC275ACT (924T>C 925C>T), GAT278GAC (934T>C), TCC280GAC (938T>G 939C>A), CAA281CAG (943A>G), CAA282AAC (944C>A 946A>C), GAA283ACT (947G>A 948A>C 949A>T), CGT284AGA (950C>A 952T>A), TTT286ATC (956T>A 958T>C), AAA289CGA (965A>C 966A>G), GAA290GAG (970A>G), TTT291TTC (973T>C), AGA292CGA (974A>C), GAT293TCT (977G>T 978A>C), GGT294GGG (982T>G), AAA295TCA (983A>T 984A>C), ACT296TCT (986A>T), AGA297CGT (989A>C 991A>T), ATA298GTG (992A>G 994A>G), CTT299CTC (997T>C), CTT300ATC (998C>A 1000T>C), TCA301ACT (1001T>A 1003A>T), AAT303GAT (1007A>G), CTA304CTT (1012A>T), TTA305TTG (1015A>G), GCA306GCT (1018A>T), AGA307CGT (1019A>C 1021A>T), GGC308GGT (1024C>T), GTT311GTA (1033T>A), CAA312CAG (1036A>G), ATA314GTC (1040A>G 1042A>C), TCT315TCC (1045T>C), TTA316CTT (1046T>C 1048A>T), GTT317GTC (1051T>C), ATC318ATT (1054C>T), AAT319AAC (1057T>C), GAT321GAC (1063T>C), TTG322CTG (1064T>C), CAT324ACT (1070C>A 1071A>C), AAT325CAA (1073A>C 1075T>A), AGA326CCA (1076A>C 1077G>C), TAT329TAC (1087T>C), ATA330CTG (1088A>C 1090A>G), AGA335CGT (1103A>C 1105A>T), GGT337GGA (1111T>A), AGA338CGA (1112A>C), GGC340GGA (1120C>A), CGA341AGG (1121C>A 1123A>G), AAA342AAG (1126A>G), GGG343GGT (1129G>T), ATT344GTT (1130A>G), AAT347AAC (1141T>C), ATA349GTC (1145A>G 1147A>C), ACT350ACC (1150T>C), GAA351AAG (1151G>A 1153A>G), GAC352GAT (1156C>T), GAA353GAT (1159A>T), TTA354GAA (1160T>G 1161T>A), CCA355AGG (1163C>A 1164C>G 1165A>G), AAG356ATG (1167A>T), GCT358TTT (1172G>T 1173C>T), GAA359GAC (1177A>C), TTA360ATA (1178T>A), GAA361CAG (1181G>C 1183A>G), ACG362AAA (1185C>A 1186G>A), TAT364TAC (1192T>C), AAT365AAC (1195T>C), ACC366GTT (1196A>G 1197C>T 1198C>T), AAA367GTA (1199A>G 1200A>T), ATA368ATC (1204A>C), GAT369GAG (1207T>G), GAA370GAG (1210A>G), ATG371CTG (1211A>C), GAG373GCC (1218A>C 1219G>C), ATT375GTT (1223A>G), GTT376GCT (1227T>C), TTG378CTC (1232T>C 1234G>C), ATA379CTG (1235A>C 1237A>G)

*: Inserts / Deletes / Misaligned / Frameshifts

#### Analysis details

This analysis was performed with panviral2.64

### NGS Details (UN60): Bracoviriform glomeratae (segment NC_043292.1)

#### Assembly

| | |
|---|---|
| **Coverage Length** | 311 (1 contig(s)) |
| **Depth Of Coverage** | 2381.0 |
| **Number Of Reads** | 6143 |
| **Reads Per Million** | 130.29 rpm (after QC) |
| **Ambiguities** | 0 |
| **Assembly Method** | de novo + reference guided assembly |
| **Consensus Caller** | Bcf Tools |

#### Coverage Map

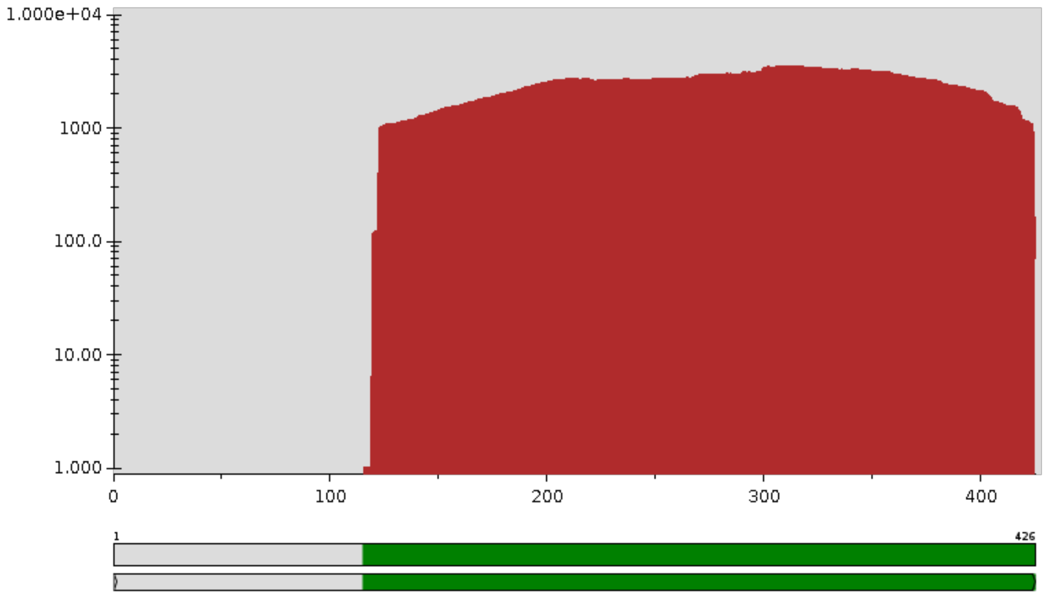

#### Assignment

| | |
|---|---|
| **Type** | Bracoviriform glomeratae (Taxonomy ID: 257816) |
| **Reference Genome** | NC_043292.1 |
| **NT Identity (%)** | 72.0257 |
| **AA Identity (%)** | 83.4951 |
| **Number Of Stop Codons** | 1 |
| **Number Of CDS** | 1 |

#### Alignment

| | |
|---|---|
| **Alignment Score** | 274.0 (NT) + 552.0 (AA) = 826.0 |
| **Concordance (%)** | 64.5817 |

| Alignment Method | Global, seeded, nucleotide + amino acids (AGA) |
|---|---|

#### Genome Region

Sequence starts at position 116 and ends at position 426 relative to NC_043292.1 reference sequence.

#### Alignment Detailed Statistics

| | Begin | End | Coverage | Score | Concordance | Matches | Identities | I/D/M/F* | Stop Codons |
|---|---|---|---|---|---|---|---|---|---|
| **NT** | **116** | **426** | **73.0%** | **274** | **44.1%** | **311 (100%)** | **224 (72.0%)** | **0/0** | |
| Mutations: | 119G>C, 124T>C, 125T>G, 126G>T, 129C>A, 132A>G, 136A>G, 137C>G, 138G>C, 150A>C, 153A>G, 155C>G, 156G>A, 159C>A, 166C>A, 168T>G, 169T>C, 172C>A, 174C>G, 177A>G, 183T>A, 184C>A, 189T>C, 198A>G, 201A>C, 207C>T, 214A>G, 219C>T, 222T>C, 223C>A, 225T>G, 228G>T, 235C>A, 237T>G, 243A>T, 246C>G, 255C>T, 261A>G, 262T>C, 264A>G, 265G>A, 273A>G, 278T>C, 279T>C, 284A>G, 285T>A, 288T>G, 289T>C, 300C>T, 301C>T, 303C>G, 309C>T, 312T>G, 315C>T, 322G>T, 329T>C, 333T>C, 336C>T, 339A>G, 345A>C, 347A>G, 348A>G, 349C>A, 351T>A, 360C>T, 363C>T, 366C>T, 375C>T, 378C>T, 384A>C, 385T>C, 387G>C, 390A>G, 391C>A, 392A>G, 393T>A, 394A>C, 396A>G, 399C>A, 400C>A, 402T>G, 404T>C, 405G>T, 406A>C, 414T>G, 416A>T, 420A>T | | | | | | | | |

##### CDS

| | Begin | End | Coverage | Score | Concordance | Matches | Identities | I/D/M/F* | Stop Codons |
|---|---|---|---|---|---|---|---|---|---|
| FK954_p501 | 40 | 142 | 72.5% | 552 | 83.0% | 103 (100%) | 86 (83.5%) | 0/0/0/0 | 1 |
| Protein mutations: | R40T (119G>C), L42R (124T>C 125T>G 126G>T), T46G (136A>G 137C>G 138G>C), A52G (155C>G 156G>A), Y57H (169T>C), T72A (214A>G), V89I (265G>A), I93T (278T>C 279T>C), D95G (284A>G 285T>A), A108S (322G>T), I110T (329T>C), K116R (347A>G 348A>G), H131R (391C>A 392A>G 393T>A), K132Q (394A>C 396A>G), M135T (404T>C 405G>T), I136L (406A>C), Y139F (416A>T) | | | | | | | | |
| Codon mutations: | AGA40ACA (119G>C), TTG42CGT (124T>C 125T>G 126G>T), GGC43GGA (129C>A), AAA44AAG (132A>G), ACG46GGC (136A>G 137C>G 138G>C), GGA50GGC (150A>C), AAA51AAG (153A>G), GCG52GGA (155C>G 156G>A), GGC53GGA (159C>A), CGT56AGG (166C>A 168T>G), TAT57CAT (169T>C), CGC58AGG (172C>A 174C>G), AAA59AAG (177A>G), CTT61CTA (183T>A), CGA62AGA (184C>A), GAT63GAC (189T>C), CAA66CAG (198A>G), GGA67GGC (201A>C), ACC69ACT (207C>T), ACT72GCT (214A>G), ATC73ATT (219C>T), CGT74CGC (222T>C), CGT75AGG (223C>A 225T>G), CTG76CTT (228G>T), CGT79AGG (235C>A 237T>G), GGA81GGT (243A>T), GTC82GTG (246C>G), ATC85ATT (255C>T), GGA87GGG (261A>G), TTA88CTG (262T>C 264A>G), GTC89ATC (265G>A), GAA91GAG (273A>G), ATT93ACC (278T>C 279T>C), GAT95GGA (284A>G 285T>A), GTT96GTG (288T>G), TTG97CTG (289T>C), TTC100TTT (300C>T), CTC101TTG (301C>T 303C>G), AAC103AAT (309C>T), GTT104GTG (312T>G), ATC105ATT (315C>T), GCT108TCT (322G>T), ATC110ACC (329T>C), TAT111TAC (333T>C), ACC112ACT (336C>T), GAA113GAG (339A>G), GCA115GCC (345A>C), AAA116AGG (347A>G 348A>G), CGT117AGA (349C>A 351T>A), GTC120GTT (360C>T), ACC121ACT (363C>T), GCC122GCT (366C>T), GTC125GTT (375C>T), GTC126GTT (378C>T), GCA128GCC (384A>C), TTG129CTC (385T>C 387G>C), AAA130AAG (390A>G), CAT131AGA (391C>A 392A>G 393T>A), AAA132CAG (394A>C 396A>G), GGC133GGA (399C>A), CGT134AGG (400C>A 402T>G), ATG135ACT (404T>C 405G>T), ATC136CTC (406A>C), GGT138GGG (414T>G), TAT139TTT (416A>T), GGA140GGT (420A>T) | | | | | | | | |

##### Proteins

| | Begin | End | Coverage | Score | Concordance | Matches | Identities | I/D/M/F* | Stop Codons |
|---|---|---|---|---|---|---|---|---|---|
| putative histone 4 (YP_009665791.1) | 40 | 142 | 72.5% | 552 | 83.0% | 103 (100%) | 86 (83.5%) | 0/0/0/0 | 1 |
| Protein mutations: | R40T (119G>C), L42R (124T>C 125T>G 126G>T), T46G (136A>G 137C>G 138G>C), A52G (155C>G 156G>A), Y57H (169T>C), T72A (214A>G), V89I (265G>A), I93T (278T>C 279T>C), D95G (284A>G 285T>A), A108S (322G>T), I110T (329T>C), K116R (347A>G 348A>G), H131R (391C>A 392A>G 393T>A), K132Q (394A>C 396A>G), M135T (404T>C 405G>T), I136L (406A>C), Y139F (416A>T) | | | | | | | | |
| Codon mutations: | AGA40ACA (119G>C), TTG42CGT (124T>C 125T>G 126G>T), GGC43GGA (129C>A), AAA44AAG (132A>G), ACG46GGC (136A>G 137C>G 138G>C), GGA50GGC (150A>C), AAA51AAG (153A>G), GCG52GGA (155C>G 156G>A), GGC53GGA (159C>A), CGT56AGG (166C>A 168T>G), TAT57CAT (169T>C), CGC58AGG (172C>A 174C>G), AAA59AAG (177A>G), CTT61CTA (183T>A), CGA62AGA (184C>A), GAT63GAC (189T>C), CAA66CAG (198A>G), GGA67GGC (201A>C), ACC69ACT (207C>T), ACT72GCT (214A>G), ATC73ATT (219C>T), CGT74CGC (222T>C), CGT75AGG (223C>A 225T>G), CTG76CTT (228G>T), CGT79AGG (235C>A 237T>G), GGA81GGT (243A>T), GTC82GTG (246C>G), ATC85ATT (255C>T), GGA87GGG (261A>G), TTA88CTG (262T>C 264A>G), GTC89ATC (265G>A), GAA91GAG (273A>G), ATT93ACC (278T>C 279T>C), GAT95GGA (284A>G 285T>A), GTT96GTG (288T>G), TTG97CTG (289T>C), TTC100TTT (300C>T), CTC101TTG (301C>T 303C>G), AAC103AAT (309C>T), GTT104GTG (312T>G), ATC105ATT (315C>T), GCT108TCT (322G>T), ATC110ACC (329T>C), TAT111TAC (333T>C), ACC112ACT (336C>T), GAA113GAG (339A>G), GCA115GCC (345A>C), AAA116AGG (347A>G 348A>G), CGT117AGA (349C>A 351T>A), GTC120GTT (360C>T), ACC121ACT (363C>T), GCC122GCT (366C>T), GTC125GTT (375C>T), GTC126GTT (378C>T), GCA128GCC (384A>C), TTG129CTC (385T>C 387G>C), AAA130AAG (390A>G), CAT131AGA (391C>A 392A>G 393T>A), AAA132CAG (394A>C 396A>G), GGC133GGA (399C>A), CGT134AGG (400C>A 402T>G), ATG135ACT (404T>C 405G>T), ATC136CTC (406A>C), GGT138GGG (414T>G), TAT139TTT (416A>T), GGA140GGT (420A>T) | | | | | | | | |

*: Inserts / Deletes / Misaligned / Frameshifts

#### Analysis details

This analysis was performed with panviral2.64

### NGS Details (UN60): Potato leafroll virus

#### Assembly

| | |
|---|---|
| **Coverage Length** | 5826 (1 contig(s)) |
| **Depth Of Coverage** | 126.6 |
| **Number Of Reads** | 5542 |
| **Reads Per Million** | 117.54 rpm (after QC) |
| **Ambiguities** | 0 |
| **Assembly Method** | read mapping against reference + variant calling |
| **Consensus Caller** | Bcf Tools |

#### Coverage Map

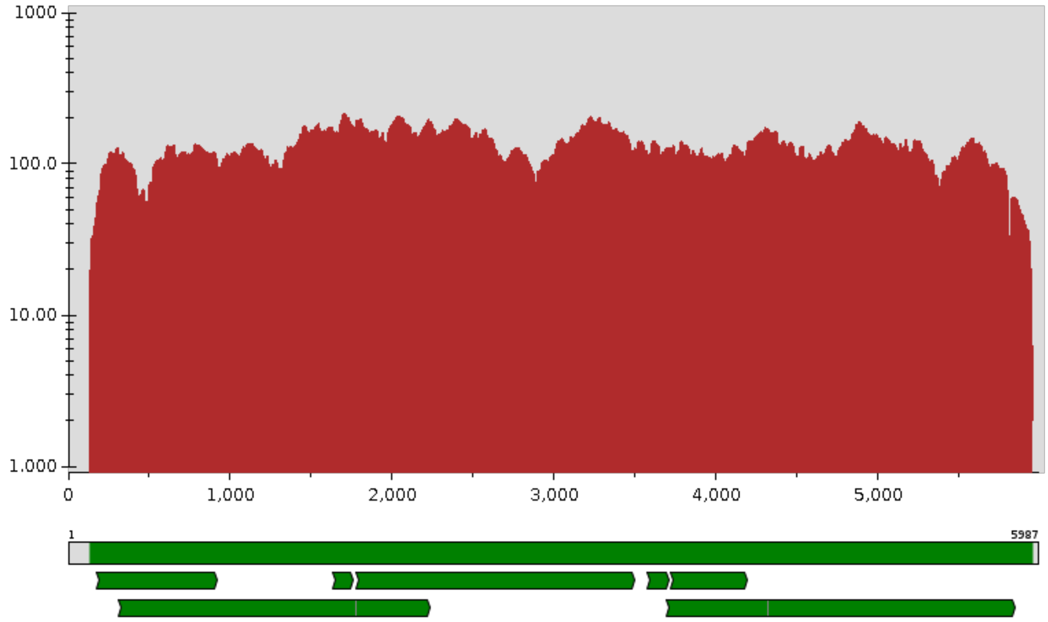

#### Assignment

| | |
|---|---|
| **Type** | Potato leafroll virus (Taxonomy ID: 12045) |
| **Reference Genome** | NC_001747.1 |
| **NT Identity (%)** | 97.5455 |
| **AA Identity (%)** | 96.4823 |
| **Number Of Stop Codons** | 8 |
| **Number Of CDS** | 8 |

#### Alignment

| | |
|---|---|
| **Alignment Score** | 11044.0 (NT) + 20020.0 (AA) = 31064.0 |
| **Concordance (%)** | 95.2114 |

| **Alignment Method** | Global, seeded, nucleotide + amino acids (AGA) |
|---|---|

#### Genome Region

Sequence starts at position 129 and ends at position 5954 relative to NC_001747.1 reference sequence.

#### Alignment Detailed Statistics

| | Begin | End | Coverage | Score | Concordance | Matches | Identities | I/D/M/F* | Stop Codons |
|---|---|---|---|---|---|---|---|---|---|
| **NT** | **129** | **5954** | **97.3%** | **11044** | **95.0%** | **5823 (99.9%)** | **5683 (97.5%)** | **3/3** | |
| Mutations: | 191A>C, 221T>C, 228C>T, 269T>C, 276G>A, 300C>T, 343C>T, 462T>A, 476_477insTT, 479_480delTT, 492T>C, 525C>T, 561C>T, 600T>C, 665A>G, 697C>A, 699A>C, 822C>T, 825T>G, 922G>A, 931A>T, 958G>A, 1054C>T, 1132T>G, 1147C>T, 1171C>T, 1183T>C, 1204A>G, 1319G>A, 1356G>A, 1381T>C, 1411T>G, 1434A>G, 1465C>T, 1474G>A, 1534G>A, 1577G>T, 1593G>A, 1701A>G, 1716A>G, 1755T>C, 1757A>G, 1774T>C, 1814G>T, 1830C>T, 1894C>A, 1924G>A, 2007T>C, 2021A>G, 2047G>A, 2064A>G, 2079C>T, 2154C>T, 2187C>T, 2265A>G, 2283G>A, 2316T>G, 2400C>T, 2433T>C, 2470G>A, 2550G>A, 2574C>T, 2580A>G, 2586A>G, 2673C>T, 2704C>T, 2729A>G, 2850A>G, 2856G>A, 2869G>A, 2877C>T, 2898T>C, 2931T>C, 3000T>C, 3033C>T, 3036C>T, 3054T>C, 3069C>T, 3096C>T, 3105C>T, 3117C>T, 3195C>T, 3321C>T, 3379A>G, 3420A>G, 3437A>G, 3441C>T, 3572T>A, 3584_3585insT, 3591delT, 3619C>T, 3652T>A, 3712A>G, 3713A>G, 3721T>G, 3748T>G, 3750C>A, 3777G>T, 3788T>A, 3939A>T, 4013C>T, 4031A>G, 4034C>T, 4132A>T, 4151T>C, 4214T>G, 4265T>C, 4280C>T, 4295G>A, 4341A>G, 4379G>A, 4385C>T, 4421A>G, 4430G>A, 4433C>T, 4511T>C, 4520G>T, 4521T>G, 4661T>C, 4700C>T, 4733C>G, 4736A>G, 4817T>C, 4835C>T, 4883C>T, 4886T>C, 4949G>A, 4964A>G, 4979C>T, 5062C>T, 5072C>T, 5095G>A, 5129G>A, 5162C>T, 5199C>T, 5210G>A, 5215G>A, 5375T>G, 5385G>A, 5432T>C, 5493G>A, 5517A>G, 5530T>C, 5925T>C | | | | | | | | |
| **CDS** | | | | | | | | | |
| PLRVgp1 | 1 | 248 | 100% | 1428 | 82.4% | 248 (99.6%) | 241 (96.8%) | 1/0/2/2 | 1 |
| Protein mutations: | Q6P (191A>C), F16S (221T>C), L32P (269T>C), H57Y (343C>T), G101_L102insX (476_477insTT), Q164R (665A>G), Q175N (697C>A 699A>C) | | | | | | | | |
| Codon mutations: | CAG6CCG (191A>C), TTT16TCT (221T>C), CTC18CTT (228C>T), CTT32CCT (269T>C), CTG34CTA (276G>A), GGC42GGT (300C>T), CAT57TAT (343C>T), GCT96GCA (462T>A), GGC101GGT (476_477insTT), GGC101_CTT102insT-C (476_477insTT), CTT102C-- (479_480delTT), ATT106ATC (492T>C), TAC117TAT (525C>T), TAC129TAT (561C>T), CAT142CAC (600T>C), CAA164CGA (665A>G), CAA175AAC (697C>A 699A>C), CGC216CGT (822C>T), GCT217GCG (825T>G) | | | | | | | | |
| PLRVgp2 | 1 | 1063 | 100% | 7021 | 94.1% | 1063 (99.9%) | 1031 (96.9%) | 1/0/2/2 | 1 |
| Protein mutations: | L52Q (462T>A), W56_A56insV (476_477insTT), L61S (492T>C), T72I (525C>T), T84I (561C>T), M97T (600T>C), K119E (665A>G), N130T (699A>C), A171V (822C>T), L172R (825T>G), E207D (931A>T), A337T (1319G>A), G349E (1356G>A), E375G (1434A>G), V423F (1577G>T), R428K (1593G>A), Q464R (1701A>G), E469G (1716A>G), L482P (1755T>C), T483A (1757A>G), W489R (1774T>C), R502L (1814G>T), Q529K (1894C>A), E539K (1924G>A), N571S (2021A>G), E580K (2047G>A), A721T (2470G>A), N807S (2729A>G), A854T (2869G>A), I1024V (3379A>G), E1043G (3437A>G) | | | | | | | | |
| Codon mutations: | TTC12TTT (343C>T), CTA52CAA (462T>A), TGG56_GCC56insGTT (476_477insTT), TTA57--A (479_480delTT), TTA61TCA (492T>C), ACA72ATA (525C>T), ACA84ATA (561C>T), ATG97ACG (600T>C), AAA119GAA (665A>G), ATC129ATA (697C>A), AAC130ACC (699A>C), GCG171GTG (822C>T), CTA172CGA (825T>G), AGG204AGA (922G>A), GAA207GAT (931A>T), CAG216CAA (958G>A), GCC248GCT (1054C>T), ACT274ACG (1132T>G), TTC279TTT (1147C>T), TCC287TCT (1171C>T), GGT291GGC (1183T>C), CTA298CTG (1204A>G), GCC337ACC (1319G>A), GGA349GAA (1356G>A), TTT357TTC (1381T>C), CTT367CTG (1411T>G), GAG375GGG (1434A>G), CCC385CCT (1465C>T), CCG388CCA (1474G>A), TCG408TCA (1534G>A), GTC423TTC (1577G>T), AGA428AAA (1593G>A), CAA464CGA (1701A>G), GAG469GGG (1716A>G), CTA482CCA (1755T>C), ACA483GCA (1757A>G), AAT488AAC (1774T>C), TGG489CGG (1774T>C), CGA502CTA (1814G>T), CGC507CGT (1830C>T), CAA529AAA (1894C>A), GAG539AAG (1924G>A), TTT566TTC (2007T>C), AAC571AGC (2021A>G), GAA580AAA (2047G>A), CCA585CCG (2064A>G), TGC590TGT (2079C>T), CAC615CAT (2154C>T), GGC626GGT (2187C>T), TCA652TCG (2265A>G), GCG658GCA (2283G>A), GTT669GTG (2316T>G), ATC697ATT (2400C>T), GCT708GCC (2433T>C), GCA721ACA (2470G>A), TCG747TCA (2550G>A), AGC755AGT (2574C>T), GAA757GAG (2580A>G), CTA759CTG (2586A>G), CGC788CGT (2673C>T), CTG799TTG (2704C>T), AAT807AGT (2729A>G), CCA847CCG (2850A>G), GTG849GTA (2856G>A), GCA854ACA (2869G>A), CAC856CAT (2877C>T), ACT863ACC (2898T>C), TAT874TAC (2931T>C), CGT897CGC (3000T>C), AAC908AAT (3033C>T), TCC909TCT (3036C>T), GAT915GAC (3054T>C), GCC920GCT (3069C>T), AGC929AGT (3096C>T), TAC932TAT (3105C>T), TCC936TCT (3117C>T), CTC962CTT (3195C>T), AAC1004AAT (3321C>T), ATT1024GTT (3379A>G), GAA1037GAG (3420A>G), GAG1043GGG (3437A>G), CTC1044CTT (3441C>T) | | | | | | | | |
| PLRVgp3 | 1 | 640 | 100% | 3892 | 90.7% | 640 (99.8%) | 611 (95.3%) | 1/0/2/2 | 1 |
| Protein mutations: | L52Q (462T>A), W56_A56insV (476_477insTT), L61S (492T>C), T72I (525C>T), T84I (561C>T), M97T (600T>C), K119E (665A>G), N130T (699A>C), A171V (822C>T), L172R (825T>G), E207D (931A>T), A337T (1319G>A), G349E (1356G>A), E375G (1434A>G), V423F (1577G>T), R428K (1593G>A), Q464R (1701A>G), E469G (1716A>G), L482P (1755T>C), T483A (1757A>G), D502Y (1814G>T), A507V (1830C>T), L566S (2007T>C), T571A (2021A>G), Q585R (2064A>G), A590V (2079C>T), T615I (2154C>T), A626V (2187C>T) | | | | | | | | |
| Codon mutations: | TTC12TTT (343C>T), CTA52CAA (462T>A), TGG56_GCC56insGTT (476_477insTT), TTA57--A (479_480delTT), TTA61TCA (492T>C), ACA72ATA (525C>T), ACA84ATA (561C>T), ATG97ACG (600T>C), AAA119GAA (665A>G), ATC129ATA (697C>A), AAC130ACC (699A>C), GCG171GTG (822C>T), CTA172CGA (825T>G), AGG204AGA (922G>A), GAA207GAT (931A>T), CAG216CAA (958G>A), GCC248GCT (1054C>T), ACT274ACG (1132T>G), TTC279TTT (1147C>T), TCC287TCT (1171C>T), GGT291GGC (1183T>C), CTA298CTG (1204A>G), GCC337ACC (1319G>A), GGA349GAA (1356G>A), TTT357TTC (1381T>C), CTT367CTG (1411T>G), GAG375GGG (1434A>G), CCC385CCT (1465C>T), CCG388CCA (1474G>A), TCG408TCA (1534G>A), GTC423TTC (1577G>T), AGA428AAA (1593G>A), CAA464CGA (1701A>G), GAG469GGG (1716A>G), CTA482CCA (1755T>C), ACA483GCA (1757A>G), AAT488AAC (1774T>C), GAC502TAC (1814G>T), GCT507GTT (1830C>T), ATC528ATA (1894C>A), GTG538GTA (1924G>A), TTA566TCA (2007T>C), ACG571GCG (2021A>G), AAG579AAA (2047G>A), CAG585CGG (2064A>G), GCG590GTG (2079C>T), ACC615ATC (2154C>T), GCT626GTT (2187C>T) | | | | | | | | |
| Rap1 | 1 | 42 | 100% | 267 | 93.0% | 42 (100%) | 39 (92.9%) | 0/0/0/0 | 0 |
| Protein mutations: | K24E (1701A>G), R29G (1716A>G), *42Q (1755T>C 1757A>G) | | | | | | | | |
| Codon mutations: | AAA24GAA (1701A>G), AGA29GGA (1716A>G), TAA42CAG (1755T>C 1757A>G) | | | | | | | | |
| ORF3a | 1 | 46 | 100% | 17 | 5.7% | 46 (97.9%) | 43 (91.5%) | 1/0/3/2 | 1 |
| Protein mutations: | Y3_K3insX (3584_3585insT), F25L (3652T>A) | | | | | | | | |
| Codon mutations: | TAT3_AAA3insAT- (3584_3585insT), TTA5T-A (3591delT), TCC14TCT (3619C>T), TTT25TTA (3652T>A), TAA45TAG (3712A>G) | | | | | | | | |
| PLRVgp4 | 1 | 718 | 100% | 4940 | 98.1% | 718 (100%) | 700 (97.5%) | 0/0/0/0 | 2 |
| Protein mutations: | K7R (3712A>G 3713A>G), V10G (3721T>G), M19R (3748T>G), A29S (3777G>T), T83S (3939A>T), Y147F (4132A>T), S217G (4341A>G), K276N (4520G>T), F277V (4521T>G), A457V (5062C>T), G468E (5095G>A), P503S (5199C>T), R508Q (5215G>A), N561K (5375T>G), E565K (5385G>A), V601I (5493G>A), K609E (5517A>G), L613P (5530T>C) | | | | | | | | |
| Codon mutations: | AAA7AGG (3712A>G 3713A>G), GTC10GGC (3721T>G), ATG19AGG (3748T>G), CGA20AGA (3750C>A), GCT29TCT (3777G>T), GTT32GTA (3788T>A), ACC83TCC (3939A>T), TAC107TAT (4013C>T), ACA113ACG (4031A>G), AGC114AGT (4034C>T), TAC147TTC (4132A>T), ATT153ATC (4151T>C), TCT174TCG (4214T>G), GAT191GAC (4265T>C), TTC196TTT (4280C>T), AAG201AAA (4295G>A), AGT217GGT (4341A>G), CAG229CAA (4379G>A), CAC231CAT (4385C>T), CTA243CTG (4421A>G), CAG246CAA (4430G>A), GCC247GCT (4433C>T), AAT273AAC (4511T>C), AAG276AAT (4520G>T), TTT277GTT (4521T>G), GGT323GGC (4661T>C), TCC336TCT (4700C>T), GTC347GTG (4733C>G), AAA348AAG (4736A>G), GGT375GGC (4817T>C), GAC381GAT (4835C>T), TTC397TTT (4883C>T), CTT398CTC (4886T>C), ACG419ACA (4949G>A), GAA424GAG (4964A>G), ACC429ACT (4979C>T), GCC457GTC (5062C>T), CAC460CAT (5072C>T), GGA468GAA (5095G>A), ACG479ACA (5129G>A), GAC490GAT (5162C>T), CCA503TCA (5199C>T), TCG506TCA (5210G>A), CGA508CAA (5215G>A), AAT561AAG (5375T>G), GAA565AAA (5385G>A), ACT580ACC (5432T>C), GTC601ATC (5493G>A), AAG609GAG (5517A>G), CTT613CCT (5530T>C) | | | | | | | | |
| PLRVgp5 | 1 | 209 | 100% | 1406 | 98.0% | 209 (100%) | 203 (97.1%) | 0/0/0/0 | 1 |
| Protein mutations: | K7R (3712A>G 3713A>G), V10G (3721T>G), M19R (3748T>G), A29S (3777G>T), T83S (3939A>T), Y147F (4132A>T) | | | | | | | | |
| Codon mutations: | AAA7AGG (3712A>G 3713A>G), GTC10GGC (3721T>G), ATG19AGG (3748T>G), CGA20AGA (3750C>A), GCT29TCT (3777G>T), GTT32GTA (3788T>A), ACC83TCC (3939A>T), TAC107TAT (4013C>T), ACA113ACG (4031A>G), AGC114AGT (4034C>T), TAC147TTC (4132A>T), ATT153ATC (4151T>C), TCT174TCG (4214T>G), GAT191GAC (4265T>C), TTC196TTT (4280C>T), AAG201AAA (4295G>A) | | | | | | | | |
| PLRVgp6 | 1 | 157 | 100% | 1049 | 95.2% | 157 (100%) | 149 (94.9%) | 0/0/0/0 | 1 |
| Protein mutations: | S2A (3721T>G), C11G (3748T>G 3750C>A), F24Y (3788T>A), T99I (4013C>T), Q105R (4031A>G), A106V (4034C>T), T139S (4132A>T), L145S (4151T>C) | | | | | | | | |
| Codon mutations: | TCA2GCA (3721T>G), TGC11GGA (3748T>G 3750C>A), GCG20GCT (3777G>T), TTC24TAC (3788T>A), ACA74ACT (3939A>T), ACC99ATC (4013C>T), CAA105CGA (4031A>G), GCA106GTA (4034C>T), ACG139TCG (4132A>T), TTA145TCA (4151T>C) | | | | | | | | |

| | Begin | End | Coverage | Score | Concordance | Matches | Identities | I/D/M/F* | Stop Codons |
|---|---|---|---|---|---|---|---|---|---|
| **NT** | **129** | **5954** | **97.3%** | **11044** | **95.0%** | **5823 (99.9%)** | **5683 (97.5%)** | **3/3** | |
| **Proteins** | | | | | | | | | |
| P0 protein (NP_056746.1) | 1 | 248 | 100% | 1428 | 82.4% | 248 (99.6%) | 241 (96.8%) | 1/0/2/2 | 1 |
| Protein mutations: | Q6P (191A>C), F16S (221T>C), L32P (269T>C), H57Y (343C>T), G101_L102insX (476_477insTT), Q164R (665A>G), Q175N (697C>A 699A>C) | | | | | | | | |
| Codon mutations: | CAG6CCG (191A>C), TTT16TCT (221T>C), CTC18CTT (228C>T), CTT32CCT (269T>C), CTG34CTA (276G>A), GGC42GGT (300C>T), CAT57TAT (343C>T), GCT96GCA (462T>A), GGC101GGT (476_477insTT), GGC101_CTT102insT-C (476_477insTT), CTT102C-- (479_480delTT), ATT106ATC (492T>C), TAC117TAT (525C>T), TAC129TAT (561C>T), CAT142CAC (600T>C), CAA164CGA (665A>G), CAA175AAC (697C>A 699A>C), CGC216CGT (822C>T), GCT217GCG (825T>G) | | | | | | | | |
| RNA-dependent RNA polymerase (NP_056748.3) | 1 | 1063 | 100% | 7021 | 94.1% | 1063 (99.9%) | 1031 (96.9%) | 1/0/2/2 | 1 |
| Protein mutations: | L52Q (462T>A), W56_A56insV (476_477insTT), L61S (492T>C), T72I (525C>T), T84I (561C>T), M97T (600T>C), K119E (665A>G), N130T (699A>C), A171V (822C>T), L172R (825T>G), E207D (931A>T), A337T (1319G>A), G349E (1356G>A), E375G (1434A>G), V423F (1577G>T), R428K (1593G>A), Q464R (1701A>G), E469G (1716A>G), L482P (1755T>C), T483A (1757A>G), W489R (1774T>C), R502L (1814G>T), Q529K (1894C>A), E539K (1924G>A), N571S (2021A>G), E580K (2047G>A), A721T (2470G>A), N807S (2729A>G), A854T (2869G>A), I1024V (3379A>G), E1043G (3437A>G) | | | | | | | | |
| Codon mutations: | TTC12TTT (343C>T), CTA52CAA (462T>A), TGG56_GCC56insGTT (476_477insTT), TTA57--A (479_480delTT), TTA61TCA (492T>C), ACA72ATA (525C>T), ACA84ATA (561C>T), ATG97ACG (600T>C), AAA119GAA (665A>G), ATC129ATA (697C>A), AAC130ACC (699A>C), GCG171GTG (822C>T), CTA172CGA (825T>G), AGG204AGA (922G>A), GAA207GAT (931A>T), CAG216CAA (958G>A), GCC248GCT (1054C>T), ACT274ACG (1132T>G), TTC279TTT (1147C>T), TCC287TCT (1171C>T), GGT291GGC (1183T>C), CTA298CTG (1204A>G), GCC337ACC (1319G>A), GGA349GAA (1356G>A), TTT357TTC (1381T>C), CTT367CTG (1411T>G), GAG375GGG (1434A>G), CCC385CCT (1465C>T), CCG388CCA (1474G>A), TCG408TCA (1534G>A), GTC423TTC (1577G>T), AGA428AAA (1593G>A), CAA464CGA (1701A>G), GAG469GGG (1716A>G), CTA482CCA (1755T>C), ACA483GCA (1757A>G), AAT488AAC (1774T>C), TGG489CGG (1774T>C), CGA502CTA (1814G>T), CGC507CGT (1830C>T), CAA529AAA (1894C>A), GAG539AAG (1924G>A), TTT566TTC (2007T>C), AAC571AGC (2021A>G), GAA580AAA (2047G>A), CCA585CCG (2064A>G), TGC590TGT (2079C>T), CAC615CAT (2154C>T), GGC626GGT (2187C>T), TCA652TCG (2265A>G), GCG658GCA (2283G>A), GTT669GTG (2316T>G), ATC697ATT (2400C>T), GCT708GCC (2433T>C), GCA721ACA (2470G>A), TCG747TCA (2550G>A), AGC755AGT (2574C>T), GAA757GAG (2580A>G), CTA759CTG (2586A>G), CGC788CGT (2673C>T), CTG799TTG (2704C>T), AAT807AGT (2729A>G), CCA847CCG (2850A>G), GTG849GTA (2856G>A), GCA854ACA (2869G>A), CAC856CAT (2877C>T), ACT863ACC (2898T>C), TAT874TAC (2931T>C), CGT897CGC (3000T>C), AAC908AAT (3033C>T), TCC909TCT (3036C>T), GAT915GAC (3054T>C), GCC920GCT (3069C>T), AGC929AGT (3096C>T), TAC932TAT (3105C>T), TCC936TCT (3117C>T), CTC962CTT (3195C>T), AAC1004AAT (3321C>T), ATT1024GTT (3379A>G), GAA1037GAG (3420A>G), GAG1043GGG (3437A>G), CTC1044CTT (3441C>T) | | | | | | | | |
| P1 protein (NP_056747.1) | 1 | 640 | 100% | 3892 | 90.7% | 640 (99.8%) | 611 (95.3%) | 1/0/2/2 | 1 |
| Protein mutations: | L52Q (462T>A), W56_A56insV (476_477insTT), L61S (492T>C), T72I (525C>T), T84I (561C>T), M97T (600T>C), K119E (665A>G), N130T (699A>C), A171V (822C>T), L172R (825T>G), E207D (931A>T), A337T (1319G>A), G349E (1356G>A), E375G (1434A>G), V423F (1577G>T), R428K (1593G>A), Q464R (1701A>G), E469G (1716A>G), L482P (1755T>C), T483A (1757A>G), D502Y (1814G>T), A507V (1830C>T), L566S (2007T>C), T571A (2021A>G), Q585R (2064A>G), A590V (2079C>T), T615I (2154C>T), A626V (2187C>T) | | | | | | | | |
| Codon mutations: | TTC12TTT (343C>T), CTA52CAA (462T>A), TGG56_GCC56insGTT (476_477insTT), TTA57--A (479_480delTT), TTA61TCA (492T>C), ACA72ATA (525C>T), ACA84ATA (561C>T), ATG97ACG (600T>C), AAA119GAA (665A>G), ATC129ATA (697C>A), AAC130ACC (699A>C), GCG171GTG (822C>T), CTA172CGA (825T>G), AGG204AGA (922G>A), GAA207GAT (931A>T), CAG216CAA (958G>A), GCC248GCT (1054C>T), ACT274ACG (1132T>G), TTC279TTT (1147C>T), TCC287TCT (1171C>T), GGT291GGC (1183T>C), CTA298CTG (1204A>G), GCC337ACC (1319G>A), GGA349GAA (1356G>A), TTT357TTC (1381T>C), CTT367CTG (1411T>G), GAG375GGG (1434A>G), CCC385CCT (1465C>T), CCG388CCA (1474G>A), TCG408TCA (1534G>A), GTC423TTC (1577G>T), AGA428AAA (1593G>A), CAA464CGA (1701A>G), GAG469GGG (1716A>G), CTA482CCA (1755T>C), ACA483GCA (1757A>G), AAT488AAC (1774T>C), GAC502TAC (1814G>T), GCT507GTT (1830C>T), ATC528ATA (1894C>A), GTG538GTA (1924G>A), TTA566TCA (2007T>C), ACG571GCG (2021A>G), AAG579AAA (2047G>A), CAG585CGG (2064A>G), GCG590GTG (2079C>T), ACC615ATC (2154C>T), GCT626GTT (2187C>T) | | | | | | | | |
| Replication-associated protein (YP_006355442.1) | 1 | 42 | 100% | 267 | 93.0% | 42 (100%) | 39 (92.9%) | 0/0/0/0 | 0 |
| Protein mutations: | K24E (1701A>G), R29G (1716A>G), *42Q (1755T>C 1757A>G) | | | | | | | | |
| Codon mutations: | AAA24GAA (1701A>G), AGA29GGA (1716A>G), TAA42CAG (1755T>C 1757A>G) | | | | | | | | |
| protein 3a (YP_009179365.2) | 1 | 46 | 100% | 17 | 5.7% | 46 (97.9%) | 43 (91.5%) | 1/0/3/2 | 1 |
| Protein mutations: | Y3_K3insX (3584_3585insT), F25L (3652T>A) | | | | | | | | |
| Codon mutations: | TAT3_AAA3insAT- (3584_3585insT), TTA5T-A (3591delT), TCC14TCT (3619C>T), TTT25TTA (3652T>A), TAA45TAG (3712A>G) | | | | | | | | |
| CP read-through protein (NP_056751.2) | 1 | 718 | 100% | 4940 | 98.1% | 718 (100%) | 700 (97.5%) | 0/0/0/0 | 2 |
| Protein mutations: | K7R (3712A>G 3713A>G), V10G (3721T>G), M19R (3748T>G), A29S (3777G>T), T83S (3939A>T), Y147F (4132A>T), S217G (4341A>G), K276N (4520G>T), F277V (4521T>G), A457V (5062C>T), G468E (5095G>A), P503S (5199C>T), R508Q (5215G>A), N561K (5375T>G), E565K (5385G>A), V601I (5493G>A), K609E (5517A>G), L613P (5530T>C) | | | | | | | | |
| Codon mutations: | AAA7AGG (3712A>G 3713A>G), GTC10GGC (3721T>G), ATG19AGG (3748T>G), CGA20AGA (3750C>A), GCT29TCT (3777G>T), GTT32GTA (3788T>A), ACC83TCC (3939A>T), TAC107TAT (4013C>T), ACA113ACG (4031A>G), AGC114AGT (4034C>T), TAC147TTC (4132A>T), ATT153ATC (4151T>C), TCT174TCG (4214T>G), GAT191GAC (4265T>C), TTC196TTT (4280C>T), AAG201AAA (4295G>A), AGT217GGT (4341A>G), CAG229CAA (4379G>A), CAC231CAT (4385C>T), CTA243CTG (4421A>G), CAG246CAA (4430G>A), GCC247GCT (4433C>T), AAT273AAC (4511T>C), AAG276AAT (4520G>T), TTT277GTT (4521T>G), GGT323GGC (4661T>C), TCC336TCT (4700C>T), GTC347GTG (4733C>G), AAA348AAG (4736A>G), GGT375GGC (4817T>C), GAC381GAT (4835C>T), TTC397TTT (4883C>T), CTT398CTC (4886T>C), ACG419ACA (4949G>A), GAA424GAG (4964A>G), ACC429ACT (4979C>T), GCC457GTC (5062C>T), CAC460CAT (5072C>T), GGA468GAA (5095G>A), ACG479ACA (5129G>A), GAC490GAT (5162C>T), CCA503TCA (5199C>T), TCG506TCA (5210G>A), CGA508CAA (5215G>A), AAT561AAG (5375T>G), GAA565AAA (5385G>A), ACT580ACC (5432T>C), GTC601ATC (5493G>A), AAG609GAG (5517A>G), CTT613CCT (5530T>C) | | | | | | | | |
| coat protein (NP_056749.1) | 1 | 209 | 100% | 1406 | 98.0% | 209 (100%) | 203 (97.1%) | 0/0/0/0 | 1 |
| Protein mutations: | K7R (3712A>G 3713A>G), V10G (3721T>G), M19R (3748T>G), A29S (3777G>T), T83S (3939A>T), Y147F (4132A>T) | | | | | | | | |
| Codon mutations: | AAA7AGG (3712A>G 3713A>G), GTC10GGC (3721T>G), ATG19AGG (3748T>G), CGA20AGA (3750C>A), GCT29TCT (3777G>T), GTT32GTA (3788T>A), ACC83TCC (3939A>T), TAC107TAT (4013C>T), ACA113ACG (4031A>G), AGC114AGT (4034C>T), TAC147TTC (4132A>T), ATT153ATC (4151T>C), TCT174TCG (4214T>G), GAT191GAC (4265T>C), TTC196TTT (4280C>T), AAG201AAA (4295G>A) | | | | | | | | |
| movement protein (NP_056750.1) | 1 | 157 | 100% | 1049 | 95.2% | 157 (100%) | 149 (94.9%) | 0/0/0/0 | 1 |
| Protein mutations: | S2A (3721T>G), C11G (3748T>G 3750C>A), F24Y (3788T>A), T99I (4013C>T), Q105R (4031A>G), A106V (4034C>T), T139S (4132A>T), L145S (4151T>C) | | | | | | | | |
| Codon mutations: | TCA2GCA (3721T>G), TGC11GGA (3748T>G 3750C>A), GCG20GCT (3777G>T), TTC24TAC (3788T>A), ACA74ACT (3939A>T), ACC99ATC (4013C>T), CAA105CGA (4031A>G), GCA106GTA (4034C>T), ACG139TCG (4132A>T), TTA145TCA (4151T>C) | | | | | | | | |

*: Inserts / Deletes / Misaligned / Frameshifts

#### Analysis details

This analysis was performed with panviral2.64

### NGS Details (UN60): Duamitovirus soch1

#### Assembly

| | |
|---|---|
| **Coverage Length** | 1957 (6 contig(s)) |
| **Depth Of Coverage** | 22.1 |
| **Number Of Reads** | 351 |
| **Reads Per Million** | 7.44 rpm (after QC) |
| **Ambiguities** | 0 |
| **Assembly Method** | de novo + reference guided assembly |
| **Consensus Caller** | Bcf Tools |

#### Coverage Map

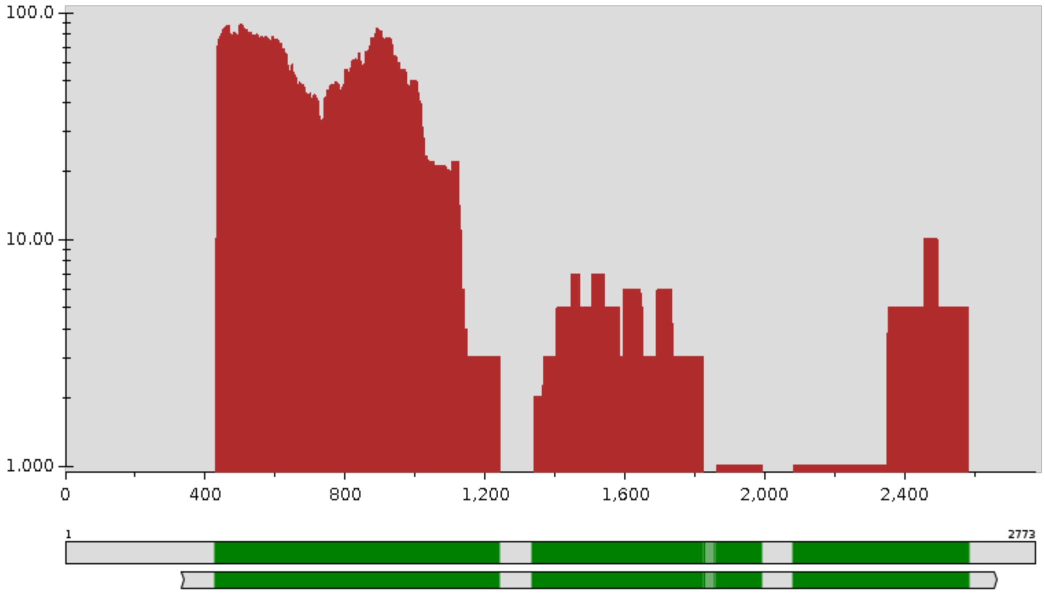

#### Assignment

| | |
|---|---|
| **Type** | Duamitovirus soch1 (Taxonomy ID: 2955838) |
| **Reference Genome** | NC_076524.1 |
| **NT Identity (%)** | 67.4872 |
| **AA Identity (%)** | 68.2099 |
| **Number Of Stop Codons** | 4 |
| **Number Of CDS** | 1 |

#### Alignment

| | |
|---|---|
| **Alignment Score** | 1294.0 (NT) + 2702.0 (AA) = 3996.0 |
| **Concordance (%)** | 50.319 |

| | |
|---|---|
| **Alignment Method** | Global, seeded, nucleotide + amino acids (AGA) |

#### Genome Region

Sequence starts at position 428 and ends at position 2585 relative to NC_076524.1 reference sequence.

#### Alignment Detailed Statistics

| | Begin | End | Coverage | Score | Concordance | Matches | Identities | I/D/M/F* | Stop Codons |
|---|---|---|---|---|---|---|---|---|---|
| **NT** | **428** | **2585** | **70.6%** | **1294** | **33.9%** | **1940 (98.6%)** | **1316 (66.9%)** | **10/17** | |
| Mutations: | 437A>G, 441A>C, 443C>T, 447C>T, 459C>T, 466C>T, 470C>T, 471G>A, 472C>T, 473T>G, 477T>A, 480A>C, 481C>A, 482A>T, 485A>G, 489A>G, 494T>C, 497T>A, 500T>G, 503A>C, 506T>C, 509C>T, 512T>C, 513G>A, 516T>A, 517C>A, 518T>A, 520A>C, 524G>T, 525A>G, 526C>T, 527T>G, 529C>A, 530C>A, 531T>C, 533A>G, 534C>A, 535G>A, 536C>A, 539G>A, 540A>C, 541A>G, 542G>T, 545G>A, 548C>T, 550A>T, 556T>A, 557C>T, 560A>T, 569C>T, 575G>A, 581C>T, 582G>A, 587A>T, 589G>C, 591T>C, 593A>T, 599A>T, 602C>T, 612T>A, 613C>G, 614C>T, 616T>A, 618C>T, 619G>C, 620A>T, 624G>A, 626A>T, 629A>C, 630C>A, 632A>C, 633C>A, 635T>G, 638G>T, 641G>T, 644T>A, 647G>T, 651C>T, 653C>G, 656A>C, 657A>C, 661C>G, 662G>T, 665T>A, 668C>T, 672C>A, 673G>A, 674T>G, 680C>A, 683C>T, 684A>G, 685A>C, 686A>C, 692G>A, 695T>A, 699G>C, 700C>A, 701A>C, 702A>G, 705C>A, 707T>G, 708A>G, 709G>C, 710G>A, 713A>G, 714G>T, 715A>C, 719A>T, 722C>T, 737T>C, 746C>T, 752A>T, 761G>A, 765G>T, 767T>A, 770G>A, 771G>T, 772T>A, 773T>G, 774T>A, 776G>T, 777A>G, 778A>C, 779A>G, 782T>G, 796G>A, 797T>G, 803A>C, 809T>C, 810G>C, 815C>A, 818A>T, 821T>G, 824C>T, 827A>T, 831C>T, 833T>A, 841G>A, 842G>T, 850T>G, 851A>T, 857G>T, 860G>T, 862A>G, 869T>G, 875G>T, 880T>G, 881C>A, 885C>A, 887T>G, 892A>G, 905A>T, 908C>A, 909T>C, 910T>A, 916G>A, 918G>T, 919G>C, 923T>A, 926A>T, 927A>C, 929G>T, 934T>A, 935T>G, 938G>T, 942T>A, 943C>A, 948C>A, 971C>A, 972C>T, 975A>C, 976A>T, 977T>C, 983T>C, 990A>C, 992G>C, 993T>C, 994C>G, 995A>G, 997A>T, 998C>T, 999G>C, 1000G>A, 1001T>A, 1003T>A, 1004A>T, 1005G>T, 1007C>G, 1010A>G, 1016C>T, 1017A>G, 1018A>T, 1019A>T, 1022T>A, 1025T>C, 1026C>G, 1027A>G, 1030A>C, 1031G>A, 1034T>G, 1035A>G, 1036G>C, 1044T>G, 1046C>T, 1049G>A, 1050T>G, 1051T>A, 1052T>A, 1054C>A, 1055A>C, 1061T>C, 1067T>G, 1073G>A, 1076A>G, 1083C>A, 1088G>T, 1091C>G, 1099G>T, 1104A>G, 1105C>T, 1106A>G, 1115T>G, 1121A>C, 1122T>A, 1124C>T, 1127G>A, 1130C>T, 1133C>G, 1142A>C, 1145C>T, 1148G>T, 1151G>T, 1152T>C, 1154G>T, 1160C>T, 1161A>C, 1164T>A, 1165C>T, 1166T>G, 1169C>T, 1172C>A, 1174T>A, 1175T>C, 1178T>A, 1181A>T, 1184C>T, 1185A>T, 1186G>C, 1187G>T, 1188G>A, 1189C>A, 1191G>A, 1192C>A, 1193A>T, 1196T>C, 1197_1199delGAA, 1205A>T, 1211C>T, 1212T>G, 1213T>A, 1217T>C, 1220C>T, 1223A>T, 1224C>T, 1225A>T, 1226C>A, 1232G>A, 1236A>C, 1237G>A, 1238T>A, 1241C>A, 1340A>C, 1345T>C, 1349G>A, 1355C>T, 1359A>G, 1364_1365insAAAA, 1369G>A, 1370A>G, 1371C>T, 1372T>G, 1373T>A, 1376C>G, 1377C>T, 1378A>T, 1379C>T, 1385T>C, 1389G>C, 1391T>A, 1397T>A, 1402C>A, 1409G>A, 1413A>C, 1414A>G, 1415A>C, 1416A>C, 1418A>T, 1419A>C, 1420A>C, 1422A>G, 1424A>G, 1427C>T, 1430G>A, 1431A>G, 1433T>A, 1445G>A, 1446A>C, 1447A>G, 1448A>T, 1451T>G, 1455T>G, 1457G>A, 1458G>C, 1459C>G, 1460T>A, 1464A>C, 1465A>G, 1466G>A, 1467C>T, 1468A>C, 1469A>T, 1470A>C, 1472G>A, 1475C>G, 1476T>A, 1477T>A, 1478T>A, 1479A>G, 1480A>G, 1481A>T, 1486G>C, 1489A>T, 1490T>C, 1502A>C, 1505G>A, 1508A>G, 1526G>A, 1538G>A, 1541T>C, 1545C>T, 1547T>C, 1553T>C, 1554A>T, 1555T>G, 1556G>C, 1559A>G, 1566C>A, 1567C>G, 1571C>A, 1572T>C, 1580C>A, 1589T>C, 1593_1597delAGCTC, 1598T>A, 1607T>C, 1613G>T, 1614A>T, 1617C>T, 1619T>G, 1620G>A, 1622T>C, 1623A>G, 1624A>T, 1626A>C, 1627A>C, 1628A>T, 1632T>A, 1634A>G, 1635A>G, 1636C>T, 1637A>C, 1643A>G, 1647T>A, 1648A>G, 1664G>A, 1670G>A, 1676T>A, 1679G>A, 1682C>T, 1686C>T, 1691T>A, 1698G>T, 1700A>T, 1703T>C, 1707T>G, 1709G>T, 1710T>C, 1712G>A, 1724T>C, 1736A>G, 1738C>G, 1742G>A, 1744G>A, 1745C>G, 1759G>A, 1760A>C, 1761T>C, 1766C>A, 1767T>A, 1769T>C, 1772A>C, 1775T>C, 1776G>A, 1777G>A, 1778G>A, 1781T>C, 1784C>T, 1787T>C, 1790A>G, 1808A>G, 1814C>T, 1818A>G, 1819G>C, 1820T>A, 1823A>G, 1829C>G, 1830A>C, 1831A>G, 1832T>G, 1833G>C, 1840A>G, 1847A>G, 1848T>C, 1856A>G, 1859C>T, 1860A>G, 1861A>G, 1865C>T, 1867A>C, 1871T>C, 1874G>T, 1875delA, 1878C>G, 1880A>T, 1886T>G, 1887T>A, 1888T>G, 1889G>A, 1895T>C, 1898T>G, 1901T>C, 1910T>C, 1919G>T, 1925G>A, 1934T>C, 1937A>G, 1938T>G, 1939C>A, 1940A>C, 1950G>T, 1952T>C, 1953T>C, 1955A>T, 1964T>C, 1967C>T, 1976T>A, 1980T>A, 1981T>C, 1982A>T, 1985T>C, 1989C>A, 1991T>G, 1994T>A, 2084G>T, 2088T>C, 2090A>G, 2091T>A, 2093C>G, 2094A>T, 2099C>T, 2102A>G, 2103A>T, 2104G>C, 2105C>A, 2106A>C, 2107A>C, 2108G>A, 2111T>A, 2117G>A, 2118C>A, 2120T>A, 2126A>G, 2129T>G, 2133C>G, 2134A>C, 2136C>T, 2137T>G, 2141G>A, 2145G>A, 2146T>A, 2147T>C, 2148C>A, 2152C>T, 2156C>T, 2157T>C, 2160C>T, 2166T>C, 2167A>T, 2168C>T, 2173G>T, 2174G>T, 2187G>T, 2188G>T, 2190A>G, 2191T>G, 2192G>A, 2193C>A, 2204C>T, 2213A>G, 2215G>C, 2217_2219delAAG, 2223G>A, 2226A>G, 2227G>A, 2231C>T, 2234C>G, 2236T>G, 2238A>G, 2240A>G, 2242A>C, 2243A>T, 2244T>G, 2249A>G, 2250G>C, 2251T>C, 2256G>C, 2259T>A, 2264G>A, 2276G>A, 2280C>T, 2285G>A, 2288C>T, 2294T>A, 2298G>C, 2299T>G, 2300T>G, 2306A>C, 2309A>T, 2312A>G, 2315T>G, 2322C>A, 2324A>C, 2326G>T, 2327C>T, 2328C>A, 2330A>T, 2334G>A, 2336A>G, 2337G>A, 2348A>G, 2350A>C, 2351G>T, 2355_2359delCTCAA, 2360C>T, 2366C>T, 2372G>T, 2373A>G, 2375C>A, 2376T>G, 2377G>C, 2378T>A, 2379A>T, 2381G>A, 2382A>T, 2383A>C, 2384C>A, 2393A>G, 2394T>A, 2400A>G, 2402G>T, 2403A>C, 2404G>A, 2405T>A, 2408C>A, 2409A>T, 2411C>T, 2414C>T, 2415T>G, 2416C>T, 2417C>T, 2423A>G, 2426C>T, 2429A>C, 2432A>T, 2435C>T, 2442A>C, 2444G>A, 2445C>T, 2446T>C, 2447C>A, 2448G>A, 2450A>C, 2451A>G, 2452C>A, 2453T>A, 2458T>A, 2462T>A, 2474C>T, 2477C>G, 2478C>T, 2479T>A, 2480A>C, 2481A>G, 2483C>T, 2489A>G, 2504T>A, 2516T>C, 2518C>T, 2519C>T, 2525C>T, 2528T>A, 2531A>T, 2532C>A, 2533A>G, 2545A>G, 2550A>C, 2551A>C, 2552C>A, 2555_2556insCTCAGT, 2556G>T, 2561T>A, 2562C>T, 2563A>T, 2564A>G, 2565T>G, 2568A>G, 2571A>C, 2575G>A, 2577A>G, 2582A>T, 2585G>T | | | | | | | | |

##### CDS

| | Begin | End | Coverage | Score | Concordance | Matches | Identities | I/D/M/F* | Stop Codons |
|---|---|---|---|---|---|---|---|---|---|
| RdRp | 32 | 750 | 83.4% | 2702 | 58.7% | 644 (98.8%) | 442 (67.8%) | 4/4/5/4 | 4 |
| Protein mutations: | I36L (441A>C 443C>T), L38F (447C>T), A44V (466C>T), A46M (471G>A 472C>T 473T>G), L48M (477T>A), T49H (480A>C 481C>A 482A>T), I52V (489A>G), V60I (513G>A), S61K (516T>A 517C>A 518T>A), K62T (520A>C), T64V (525A>G 526C>T 527T>G), T65K (529C>A 530C>A), R67K (534C>A 535G>A 536C>A), K69R (540A>C 541A>G 542G>T), Y72F (550A>T), F74Y (556T>A 557C>T), A83T (582G>A), C85S (589G>C), F94Y (616T>A), R95S (618C>T 619G>C 620A>T), E97N (624G>A 626A>T), E98D (629A>C), P99T (630C>A 632A>C), L100M (633C>A 635T>G), S109C (661C>G 662G>T), R113K (672C>A 673G>A 674T>G), K117A (684A>G 685A>C 686A>C), A122H (699G>C 700C>A 701A>C), I123V (702A>G), R125A (708A>G 709G>C 710G>A), D127S (714G>T 715A>C), E128D (719A>T), A144S (765G>T 767T>A), V146* (771G>T 772T>A 773T>G), L147I (774T>A 776G>T), K148A (777A>G 778A>C 779A>G), S154K (796G>A 797T>G), E159Q (810G>C), P166S (831C>T 833T>A), G169D (841G>A 842G>T), L172R (850T>G 851A>T), E176G (862A>G), I178M (869T>G), F182* (880T>G 881C>A), H184K (885C>A 887T>G), Q186R (892A>G), F192H (909T>C 910T>A), R194Q (916G>A), G195S (918G>T 919G>C), M198L (927A>C 929G>T), I200K (934T>A 935T>G), S203N (942T>A 943C>A), Q205K (948C>A), N214L (975A>C 976A>T 977T>C), S220R (993T>C 994C>G 995A>G), Y221F (997A>T 998C>T), G222Q (999G>C 1000G>A 1001T>A), L223Y (1003T>A 1004A>T), A224S (1005G>T 1007C>G), K228V (1017A>G 1018A>T 1019A>T), Q231G (1026C>G 1027A>G), E232A (1030A>C 1031G>A), S234A (1035A>G 1036G>C), F237V (1044T>G 1046C>T), F239E (1050T>G 1051T>A 1052T>A), P240H (1054C>A 1055A>C), L250I (1083C>A), W255L (1099G>T), T257V (1104A>G 1105C>T 1106A>G), I260M (1115T>G), F263I (1122T>A 1124C>T), M272I (1151G>T), K276Q (1161A>C), S277M (1164T>A 1165C>T 1166T>G), F279L (1172C>A), F280Y (1174T>A 1175T>C), L282F (1181A>T), R284S (1185A>T 1186G>C 1187G>T), A285N (1188G>A 1189C>A), A286N (1191G>A 1192C>A 1193A>T), E288del (1197_1199delGAA), L290F (1205A>T), L293E (1212T>G 1213T>A), E296D (1223A>T), H297L (1224C>T 1225A>T 1226C>A), S301Q (1236A>C 1237G>A 1238T>A), V337A (1345T>C), M338I (1349G>A), I342V (1359A>G), K343_Q344insKX (1364_1365insAAAA), R345K (1369G>A 1370A>G), L346* (1371C>T 1372T>G 1373T>A), H348F (1377C>T 1378A>T 1379C>T), D352Q (1389G>C 1391T>A), T356K (1402C>A), K360R (1413A>C 1414A>G 1415A>C), I361L (1416A>C 1418A>T), K362P (1419A>C 1420A>C), T363A (1422A>G 1424A>G), T366A (1431A>G 1433T>A), K371R (1446A>C 1447A>G 1448A>T), L374V (1455T>G 1457G>A), A375R (1458G>C 1459C>G 1460T>A), K377R (1464A>C 1465A>G 1466G>A), Q378S (1467C>T 1468A>C 1469A>T), K379Q (1470A>C 1472G>A), N380K (1475C>G), F381K (1476T>A 1477T>A 1478T>A), K382G (1479A>G 1480A>G 1481A>T), C384S (1486G>C), Y385F (1489A>T 1490T>C), L389F (1502A>C), M401I (1538G>A), L404F (1545C>T 1547T>C), M407C (1554A>T 1555T>G 1556G>C), I408M (1559A>G), P411S (1566C>A 1567C>G), S420del (1593_1597delAGCTC), I427F (1614A>T), V429I (1620G>A 1622T>C), N430V (1623A>G 1624A>T), K431P (1626A>C 1627A>C 1628A>T), L433M (1632T>A 1634A>G), T434V (1635A>G 1636C>T 1637A>C), Y438S (1647T>A 1648A>G), H451Y (1686C>T), A455S (1698G>T 1700A>T), S458A (1707T>G 1709G>T), A468G (1738C>G), S470K (1744G>A 1745C>G), G475D (1759G>A 1760A>C), C476R (1761T>C), N477K (1766C>A), S478T (1767T>A 1769T>C), G481K (1776G>A 1777G>A 1778G>A), S495A (1818A>G 1819G>C 1820T>A), N499R (1830A>C 1831A>G 1832T>G), K511T (1867A>C), P515A (1878C>G 1880A>T), L518R (1887T>A 1888T>G 1889G>A), D521E (1898T>G), S535D (1938T>G 1939C>A 1940A>C), D539Y (1950G>T 1952T>C), L549T (1980T>A 1981T>C 1982A>T), F586M (2091T>A 2093C>G), T587S (2094A>T), K591P (2106A>C 2107A>C 2108G>A), H600A (2133C>G 2134A>C), L601C (2136C>T 2137T>G), V604N (2145G>A 2146T>A 2147T>C), S606L (2152C>T), Y611L (2166T>C 2167A>T 2168C>T), W613F (2173G>T 2174G>T), G618F (2187G>T 2188G>T), M619G (2190A>G 2191T>G 2192G>A), P620T (2193C>A), G627A (2215G>C), K628del (2217_2219delAAG), V630I (2223G>A), S631D (2226A>G 2227G>A), L634R (2236T>G), K635E (2238A>G 2240A>G), E636A (2242A>C 2243A>T), L637V (2244T>G), V639P (2250G>C 2251T>C), E641Q (2256G>C), L642I (2259T>A), V655R (2298G>C 2299T>G 2300T>G), L663I (2322C>A 2324A>C), R664L (2326G>T 2327C>T), Q665N (2328C>A 2330A>T), V667M (2334G>A 2336A>G), E668K (2337G>A), K672T (2350A>C 2351G>T), L674del (2355_2359delCTCAA), I680V (2373A>G 2375C>A), C681A (2376T>G 2377G>C 2378T>A), M682L (2379A>T 2381G>A), N683S (2382A>T 2383A>C 2384C>A), S687T (2394T>A), K689D (2400A>G 2402G>T), S690Q (2403A>C 2404G>A 2405T>A), I692F (2409A>T 2411C>T), S694V (2415T>G 2416C>T 2417C>T), I696M (2423A>G), L704S (2445C>T 2446T>C 2447C>A), E705N (2448G>A 2450A>C), T706E (2451A>G 2452C>A 2453T>A), F708Y (2458T>A), D709E (2462T>A), L715Y (2478C>T 2479T>A 2480A>C), I716V (2481A>G 2483C>T), P728L (2518C>T 2519C>T), Q733R (2532C>A 2533A>G), D737G (2545A>G), N739P (2550A>C 2551A>C 2552C>A), T740_V741insLS (2555_2556insCTCAGT), V741F (2556G>T), Q743L (2562C>T 2563A>T 2564A>G), F744V (2565T>G), N745D (2568A>G), K746Q (2571A>C), W747* (2575G>A), I748V (2577A>G), L749F (2582A>T) | | | | | | | | |

| | Begin | End | Coverage | Score | Concordance | Matches | Identities | I/D/M/F* | Stop Codons |
|---|---|---|---|---|---|---|---|---|---|
| **NT** | **428** | **2585** | **70.6%** | **1294** | **33.9%** | **1940 (98.6%)** | **1316 (66.9%)** | **10/17** | |
| Codon mutations: | AAA34AAG (437A>G), ATC36CTT (441A>C 443C>T), CTT38TTT (447C>T), CTA42TTA (459C>T), GCC44GTC (466C>T), TCC45TCT (470C>T), GCT46ATG (471G>A 472C>T 473T>G), TTG48ATG (477T>A), ACA49CAT (480A>C 481C>A 482A>T), AAA50AAG (485A>G), ATC52GTC (489A>G), GCT53GCC (494T>C), CTT54CTA (497T>A), GCT55GCG (500T>G), GGA56GGC (503A>C), TGT57TGC (506T>C), GAC58GAT (509C>T), TTT59TTC (512T>C), GTA60ATA (513G>A), TCT61AAA (516T>A 517C>A 518T>A), AAA62ACA (520A>C), GTG63GTT (524G>T), ACT64GTG (525A>G 526C>T 527T>G), ACC65AAA (529C>A 530C>A), TTA66CTG (531T>C 533A>G), CGC67AAA (534C>A 535G>A 536C>A), CGG68CGA (539G>A), AAG69CGT (540A>C 541A>G 542G>T), TCG70TCA (545G>A), GGC71GGT (548C>T), TAT72TTT (550A>T), TTC74TAT (556T>A 557C>T), ACA75ACT (560A>T), TAC78TAT (569C>T), AAG80AAA (575G>A), TGC82TGT (581C>T), GCA83ACA (582G>A), GTA84GTT (587A>T), TGC85TCC (589G>C), TTA86CTT (591T>C 593A>T), CGA88CGT (599A>T), TAC89TAT (602C>T), TCC93AGT (612T>A 613C>G 614C>T), TTT94TAT (616T>A), CGA95TCT (618C>T 619G>C 620A>T), GAA97AAT (624G>A 626A>T), GAA98GAC (629A>C), CCA99ACC (630C>A 632A>C), CTT100ATG (633C>A 635T>G), TCG101TCT (638G>T), GTG102GTT (641G>T), CCT103CCA (644T>A), GTG104GTT (647G>T), CTC106TTG (651C>T 653C>G), ACA107ACC (656A>C), AGA108CGA (657A>C), TCG109TGT (661C>G 662G>T), GGT110GGA (665T>A), ATC111ATT (668C>T), CGT113AAG (672C>A 673G>A 674T>G), ATC115ATA (680C>A), CCC116CCT (683C>T), AAA117GCC (684A>G 685A>C 686A>C), TTG119TTA (692G>A), CGT120CGA (695T>A), GCA122CAC (699G>C 700C>A 701A>C), ATA123GTA (702A>G), CGT124AGG (705C>A 707T>G), AGG125GCA (708A>G 709G>C 710G>A), AAA126AAG (713A>G), GAT127TCT (714G>T 715A>C), GAA128GAT (719A>T), CAC129CAT (722C>T), GTT134GTC (737T>C), TAC137TAT (746C>T), TCA139TCT (752A>T), GGG142GGA (761G>A), GCT144TCA (765G>T 767T>A), AAG145AAA (770G>A), GTT146TAG (771G>T 772T>A 773T>G), TTG147ATT (774T>A 776G>T), AAA148GCG (777A>G 778A>C 779A>G), GTT149GTG (782T>G), AGT154AAG (796G>A 797T>G), GCA156GCC (803A>C), TTT158TTC (809T>C), GAA159CAA (810G>C), TCC160TCA (815C>A), ATA161ATT (818A>T), ACT162ACG (821T>G), ACC163ACT (824C>T), CCA164CCT (827A>T), CCT166TCA (831C>T 833T>A), GGG169GAT (841G>A 842G>T), CTA172CGT (850T>G 851A>T), GTG174GTT (857G>T), CTG175CTT (860G>T), GAA176GGA (862A>G), ATT178ATG (869T>G), ACG180ACT (875G>T), TTC182TGA (880T>G 881C>A), CAT184AAG (885C>A 887T>G), CAA186CGA (892A>G), CTA190CTT (905A>T), CCC191CCA (908C>A), TTT192CAT (909T>C 910T>A), CGA194CAA (916G>A), GGC195TCC (918G>T 919G>C), ATT196ATA (923T>A), CCA197CCT (926A>T), ATG198CTT (927A>C 929G>T), ATT200AAG (934T>A 935T>G), GGG201GGT (938G>T), TCT203AAT (942T>A 943C>A), CAA205AAA (948C>A), CCC212CCA (971C>A), CTA213TTA (972C>T), AAT214CTC (975A>C 976A>T 977T>C), AAT216AAC (983T>C), AGG219CGC (990A>C 992G>C), TCA220CGG (993T>C 994C>G 995A>G), TAC221TTT (997A>T 998C>T), GGT222CAA (999G>C 1000G>A 1001T>A), TTA223TAT (1003T>A 1004A>T), GCC224TCG (1005G>T 1007C>G), GTA225GTG (1010A>G), GAC227GAT (1016C>T), AAA228GTT (1017A>G 1018A>T 1019A>T), GTT229GTA (1022T>A), GAT230GAC (1025T>C), CAA231GGA (1026C>G 1027A>G), GAG232GCA (1030A>C 1031G>A), GCT233GCG (1034T>G), AGT234GCT (1035A>G 1036G>C), TTC237GTT (1044T>G 1046C>T), AAG238AAA (1049G>A), TTT239GAA (1050T>G 1051T>A 1052T>A), CCA240CAC (1054C>A 1055A>C), ATT242ATC (1061T>C), GTT244GTG (1067T>G), TTG246TTA (1073G>A), AAA247AAG (1076A>G), CTA250ATA (1083C>A), GCG251GCT (1088G>T), GCC252GCG (1091C>G), TGG255TTG (1099G>T), ACA257GTG (1104A>G 1105C>T 1106A>G), ATT260ATG (1115T>G), TCA262TCC (1121A>C), TTC263ATT (1122T>A 1124C>T), CAG264CAA (1127G>A), GAC265GAT (1130C>T), GGC266GGG (1133C>G), TCA269TCC (1142A>C), CCC270CCT (1145C>T), GGG271GGT (1148G>T), ATG272ATT (1151G>T), TTG273CTT (1152T>C 1154G>T), TCC275TCT (1160C>T), AAA276CAA (1161A>C), TCT277ATG (1164T>A 1165C>T 1166T>G), GTC278GTT (1169C>T), TTC279TTA (1172C>A), TTT280TAC (1174T>A 1175T>C), CCT281CCA (1178T>A), TTA282TTT (1181A>T), GAC283GAT (1184C>T), AGG284TCT (1185A>T 1186G>C 1187G>T), GCT285AAT (1188G>A 1189C>A), GCA286AAT (1191G>A 1192C>A 1193A>T), ACT287ACC (1196T>C), GAA288del (1197_1199delGAA), TTA290TTT (1205A>T), AAC292AAT (1211C>T), TTA293GAA (1212T>G 1213T>A), TCT294TCC (1217T>C), CTC295CTT (1220C>T), GAA296GAT (1223A>T), CAC297TTA (1224C>T 1225A>T 1226C>A), GAG299GAA (1232G>A), AGT301CAA (1236A>C 1237G>A 1238T>A), GTC302GTA (1241C>A), GTA335..C (1340A>C), GTA337GCA (1345T>C), ATG338ATA (1349G>A), AAC340AAT (1355C>T), ATA342GTA (1359A>G), AAA343_CAA344insAAAA-- (1364_1365insAAAA), AGA345AAG (1369G>A 1370A>G), CTT346TGA (1371C>T 1372T>G 1373T>A), CTC347CTG (1376C>G), CAC348TTT (1377C>T 1378A>T 1379C>T), GTT350GTC (1385T>C), GAT352CAA (1389G>C 1391T>A), GCT354GCA (1397T>A), ACA356AAA (1402C>A), CTG358CTA (1409G>A), AAA360CGC (1413A>C 1414A>G 1415A>C), ATA361CTT (1416A>C 1418A>T), AAG362CCG (1419A>C 1420A>C), ACA363GCG (1422A>G 1424A>G), GAC364GAT (1427C>T), GGG365GGA (1430G>A), ACT366GCA (1431A>G 1433T>A), GAG370GAA (1445G>A), AAA371CGT (1446A>C 1447A>G 1448A>T), CCT372CCG (1451T>G), TTG374GTA (1455T>G 1457G>A), GCT375CGA (1458G>C 1459C>G 1460T>A), AAG377CGA (1464A>C 1465A>G 1466G>A), CAA378TCT (1467C>T 1468A>C 1469A>T), AAG379CAA (1470A>C 1472G>A), AAC380AAG (1475C>G), TTT381AAA (1476T>A 1477T>A 1478T>A), AAA382GGT (1479A>G 1480A>G 1481A>T), TGT384TCT (1486G>C), TAT385TTC (1489A>T 1490T>C), TTA389TTC (1502A>C), AAG390AAA (1505G>A), TCA391TCG (1508A>G), CCG397CCA (1526G>A), ATG401ATA (1538G>A), TAT402TAC (1541T>C), CTT404TTC (1545C>T 1547T>C), TCT406TCC (1553T>C), ATG407TGC (1554A>T 1555T>G 1556G>C), ATA408ATG (1559A>G), CCT411AGT (1566C>A 1567C>G), ACC412ACA (1571C>A), TTG413CTG (1572T>C), TCC415TCA (1580C>A), GTT418GTC (1589T>C), AGC420del (1593_1597delAGCTC), TCT421--A (1593_1597delAGCTC 1598T>A), CTT424CTC (1607T>C), ACG426ACT (1613G>T), ATC427TTC (1614A>T), CTT428TTG (1617C>T 1619T>G), GTT429ATC (1620G>A 1622T>C), AAC430GTC (1623A>G 1624A>T), AAA431CCT (1626A>C 1627A>C 1628A>T), TTA433ATG (1632T>A 1634A>G), ACA434GTC (1635A>G 1636C>T 1637A>C), AGA436AGG (1643A>G), TAT438AGT (1647T>A 1648A>G), CTG443CTA (1664G>A), GGG445GGA (1670G>A), CCT447CCA (1676T>A), TTG448TTA (1679G>A), GGC449GGT (1682C>T), CAC451TAC (1686C>T), GGT452GGA (1691T>A), GCA455TCT (1698G>T 1700A>T), CTT456CTC (1703T>C), TCG458GCT (1707T>G 1709G>T), TTG459CTA (1710T>C 1712G>A), TAT463TAC (1724T>C), TTA467TTG (1736A>G), GCA468GGA (1738C>G), GCG469GCA (1742G>A), AGC470AAG (1744G>A 1745C>G), GGA475GAC (1759G>A 1760A>C), TGT476CGT (1761T>C), AAC477AAA (1766C>A), TCT478ACC (1767T>A 1769T>C), CCA479CCC (1772A>C), TTT480TTC (1775T>C), GGG481AAA (1776G>A 1777G>A 1778G>A), GAT482GAC (1781T>C), TAC483TAT (1784C>T), GCT484GCC (1787T>C), TTA485TTG (1790A>G), CTA491CTG (1808A>G), ACC493ACT (1814C>T), AGT495GCA (1818A>G 1819G>C 1820T>A), AAA496AAG (1823A>G), GCC498..G (1829C>G), AAT499CGG (1830A>C 1831A>G 1832T>G), GAG500C.. (1833G>C), AAG502.G. (1840A>G), TTA504..G (1847A>G), TTG505C.. (1848T>C), AGA507..G (1856A>G), CTC508..T (1859C>T), AAT509GG. (1860A>G 1861A>G), GTC510GTT (1865C>T), AAA511ACA (1867A>C), ATT512ATC (1871T>C), TCG513TCT (1874G>T), ATA514-TA (1875delA), CCA515GCT (1878C>G 1880A>T), TCT517TCG (1886T>G), TTG518AGA (1887T>A 1888T>G 1889G>A), TCT520TCC (1895T>C), GAT521GAG (1898T>G), AAT522AAC (1901T>C), ATT525ATC (1910T>C), GCG528GCT (1919G>T), AGG530AGA (1925G>A), ACT533ACC (1934T>C), AAA534AAG (1937A>G), TCA535GAC (1938T>G 1939C>A 1940A>C), GAT539TAC (1950G>T 1952T>C), TTA540CTT (1953T>C 1955A>T), ATT543ATC (1964T>C), TCC544TCT (1967C>T), GCT547GCA (1976T>A), TTA549ACT (1980T>A 1981T>C 1982A>T), TCT550TCC (1985T>C), CGT552AGG (1989C>A 1991T>G), ACT553ACA (1994T>A), GCG583.CT (2084G>T), TTA585CTG (2088T>C 2090A>G), TTC586ATG (2091T>A 2093C>G), ACT587TCT (2094A>T), ACC588ACT (2099C>T), CAA589CAG (2102A>G), AGC590TCA (2103A>T 2104G>C 2105C>A), AAG591CCA (2106A>C 2107A>C 2108G>A), CGT592CGA (2111T>A), GAG594GAA (2117G>A), CGT595AGA (2118C>A 2120T>A), AAA597AAG (2126A>G), GCT598GCG (2129T>G), CAT600GCT (2133C>G 2134A>C), CTT601TGT (2136C>T 2137T>G), AAG602AAA (2141G>A), GTT604AAC (2145G>A 2146T>A 2147T>C), CGG605AGG (2148C>A), TCA606TTA (2152C>T), CAC607CAT (2156C>T), TTA608CTA (2157T>C), CTA609TTA (2160C>T), TAC611CTT (2166T>C 2167A>T 2168C>T), TGG613TTT (2173G>T 2174G>T), GGT618TTT (2187G>T 2188G>T), ATG619GGA (2190A>G 2191T>G 2192G>A), CCC620ACC (2193C>A), CCC623CCT (2204C>T), AAA626AAG (2213A>G), GGG627GCG (2215G>C), AAG628del (2217_2219delAAG), GTA630ATA (2223G>A), AGT631GAT (2226A>G 2227G>A), TAC632TAT (2231C>T), CTC633CTG (2234C>G), CTT634CGT (2236T>G), AAA635GAG (2238A>G 2240A>G), GAA636GCT (2242A>C 2243A>T), TTA637GTA (2244T>G), AAA638AAG (2249A>G), GTT639CCT (2250G>C 2251T>C), GAG641CAG (2256G>C), TTA642ATA (2259T>A), CAG643CAA (2264G>A), AAG647AAA (2276G>A), CTG649TTG (2280C>T), GTG650GTA (2285G>A), TTC651TTT (2288C>T), GGT653GGA (2294T>A), GTT655CGG (2298G>C 2299T>G 2300T>G), ATA657ATC (2306A>C), CTA658CTT (2309A>T), GAA659GAG (2312A>G), CGT660CGG (2315T>G), CTA663ATC (2322C>A 2324A>C), CGC664CTT (2326G>T 2327C>T), CAA665AAT (2328C>A 2330A>T), GTA667ATG (2334G>A 2336A>G), GAA668AAA (2337G>A), TTA671TTG (2348A>G), AAG672ACT (2350A>C 2351G>T), CTC674del (2355_2359delCTCAA), AAC675--T (2355_2359delCTCAA 2360C>T), TAC677TAT (2366C>T), ACG679ACT (2372G>T), ATC680GTA (2373A>G 2375C>A), TGT681GCA (2376T>G 2377G>C 2378T>A), ATG682TTA (2379A>T 2381G>A), AAC683TCA (2382A>T 2383A>C 2384C>A), GTA686GTG (2393A>G), TCC687ACC (2394T>A), AAG689GAT (2400A>G 2402G>T), AGT690CAA (2403A>C 2404G>A 2405T>A), CTC691CTA (2408C>A), ATC692TTT (2409A>T 2411C>T), GAC693GAT (2414C>T), TCC694GTT (2415T>G 2416C>T 2417C>T), ATA696ATG (2423A>G), TGC697TGT (2426C>T), GCA698GCC (2429A>C), ACA699ACT (2432A>T), TCC700TCT (2435C>T), AGG703CGA (2442A>C 2444G>A), CTC704TCA (2445C>T 2446T>C 2447C>A), GAA705AAC (2448G>A 2450A>C), ACT706GAA (2451A>G 2452C>A 2453T>A), TTT708TAT (2458T>A), GAT709GAA (2462T>A), TTC713TTT (2474C>T), GGC714GGG (2477C>G), CTA715TAC (2478C>T 2479T>A 2480A>C), ATC716GTT (2481A>G 2483C>T), AAA718AAG (2489A>G), GGT723GGA (2504T>A), AAT727AAC (2516T>C), CCC728CTT (2518C>T 2519C>T), ACC730ACT (2525C>T), ACT731ACA (2528T>A), CCA732CCT (2531A>T), CAG733AGG (2532C>A 2533A>G), GAT737GGT (2545A>G), AAC739CCA (2550A>C 2551A>C 2552C>A), ACT740_GTT741insCTCAGT (2555_2556insCTCAGT), GTT741TTT (2556G>T), ATT742ATA (2561T>A), CAA743TTG (2562C>T 2563A>T 2564A>G), TTT744GTT (2565T>G), AAT745GAT (2568A>G), AAA746CAA (2571A>C), TGG747TAG (2575G>A), ATA748GTA (2577A>G), TTA749TTT (2582A>T), GGG750GGT (2585G>T) | | | | | | | | |

#### Proteins

| | Begin | End | Coverage | Score | Concordance | Matches | Identities | I/D/M/F* | Stop Codons |
|---|---|---|---|---|---|---|---|---|---|
| RNA-dependent RNA polymerase (YP_010798874.1) | 32 | 750 | 83.4% | 2702 | 58.7% | 644 (98.8%) | 442 (67.8%) | 4/4/5/4 | 4 |

| | Begin | End | Coverage | Score | Concordance | Matches | Identities | I/D/M/F* | Stop Codons |
|---|---|---|---|---|---|---|---|---|---|
| **NT** | **428** | **2585** | **70.6%** | **1294** | **33.9%** | **1940 (98.6%)** | **1316 (66.9%)** | **10/17** | |

Protein mutations:

I36L (441A>C 443C>T), L38F (447C>T), A44V (466C>T), A46M (471G>A 472C>T 473T>G), L48M (477T>A), T49H (480A>C 481C>A 482A>T), I52V (489A>G), V60I (513G>A), S61K (516T>A 517C>A 518T>A), K62T (520A>C), T64V (525A>G 526C>T 527T>G), T65K (529C>A 530C>A), R67K (534C>A 535G>A 536C>A), K69R (540A>C 541A>G 542G>T), Y72F (550A>T), F74Y (556T>A 557C>T), A83T (582G>A), C85S (589G>C), F94Y (616T>A), R95S (618C>T 619G>C 620A>T), E97N (624G>A 626A>T), E98D (629A>C), P99T (630C>A 632A>C), L100M (633C>A 635T>G), S109C (661C>G 662G>T), R113K (672C>A 673G>A 674T>G), K117A (684A>G 685A>C 686A>C), A122H (699G>C 700C>A 701A>C), I123V (702A>G), R125A (708A>G 709G>C 710G>A), D127S (714G>T 715A>C), E128D (719A>T), A144S (765G>T 767T>A), V146* (771G>T 772T>A 773T>G), L147I (774T>A 776G>T), K148A (777A>G 778A>C 779A>G), S154K (796G>A 797T>G), E159Q (810G>C), P166S (831C>T 833T>A), G169D (841G>A 842G>T), L172R (850T>G 851A>T), E176G (862A>G), I178M (869T>G), F182* (880T>G 881C>A), H184K (885C>A 887T>G), Q186R (892A>G), F192H (909T>C 910T>A), R194Q (916G>A), G195S (918G>T 919G>C), M198L (927A>C 929G>T), I200K (934T>A 935T>G), S203N (942T>A 943C>A), Q205K (948C>A), N214L (975A>C 976A>T 977T>C), S220R (993T>C 994C>G 995A>G), Y221F (997A>T 998C>T), G222Q (999G>C 1000G>A 1001T>A), L223Y (1003T>A 1004A>T), A224S (1005G>T 1007C>G), K228V (1017A>G 1018A>T 1019A>T), Q231G (1026C>G 1027A>G), E232A (1030A>C 1031G>A), S234A (1035A>G 1036G>C), F237V (1044T>G 1046C>T), F239E (1050T>G 1051T>A 1052T>A), P240H (1054C>A 1055A>C), L250I (1083C>A), W255L (1099G>T), T257V (1104A>G 1105C>T 1106A>G), I260M (1115T>G), F263I (1122T>A 1124C>T), M272I (1151G>T), K276Q (1161A>C), S277M (1164T>A 1165C>T 1166T>G), F279L (1172C>A), F280Y (1174T>A 1175T>C), L282F (1181A>T), R284S (1185A>T 1186G>C 1187G>T), A285N (1188G>A 1189C>A), A286N (1191G>A 1192C>A 1193A>T), E288del (1197_1199delGAA), L290F (1205A>T), L293E (1212T>G 1213T>A), E296D (1223A>T), H297L (1224C>T 1225A>T 1226C>A), S301Q (1236A>C 1237G>A 1238T>A), V337A (1345T>C), M338I (1349G>A), I342V (1359A>G), K343_Q344insKX (1364_1365insAAAA), R345K (1369G>A 1370A>G), L346* (1371C>T 1372T>G 1373T>A), H348F (1377C>T 1378A>T 1379C>T), D352Q (1389G>C 1391T>A), T356K (1402C>A), K360R (1413A>C 1414A>G 1415A>C), I361L (1416A>C 1418A>T), K362P (1419A>C 1420A>C), T363A (1422A>G 1424A>G), T366A (1431A>G 1433T>A), K371R (1446A>C 1447A>G 1448A>T), L374V (1455T>G 1457G>A), A375R (1458G>C 1459C>G 1460T>A), K377R (1464A>C 1465A>G 1466G>A), Q378S (1467C>T 1468A>C 1469A>T), K379Q (1470A>C 1472G>A), N380K (1475C>G), F381K (1476T>A 1477T>A 1478T>A), K382G (1479A>G 1480A>G 1481A>T), C384S (1486G>C), Y385F (1489A>T 1490T>C), L389F (1502A>C), M401I (1538G>A), L404F (1545C>T 1547T>C), M407C (1554A>T 1555T>G 1556G>C), I408M (1559A>G), P411S (1566C>A 1567C>G), S420del (1593_1597delAGCTC), I427F (1614A>T), V429I (1620G>A 1622T>C), N430V (1623A>G 1624A>T), K431P (1626A>C 1627A>C 1628A>T), L433M (1632T>A 1634A>G), T434V (1635A>G 1636C>T 1637A>C), Y438S (1647T>A 1648A>G), H451Y (1686C>T), A455S (1698G>T 1700A>T), S458A (1707T>G 1709G>T), A468G (1738C>G), S470K (1744G>A 1745C>G), G475D (1759G>A 1760A>C), C476R (1761T>C), N477K (1766C>A), S478T (1767T>A 1769T>C), G481K (1776G>A 1777G>A 1778G>A), S495A (1818A>G 1819G>C 1820T>A), N499R (1830A>C 1831A>G 1832T>G), K511T (1867A>C), P515A (1878C>G 1880A>T), L518R (1887T>A 1888T>G 1889G>A), D521E (1898T>G), S535D (1938T>G 1939C>A 1940A>C), D539Y (1950G>T 1952T>C), L549T (1980T>A 1981T>C 1982A>T), F586M (2091T>A 2093C>G), T587S (2094A>T), K591P (2106A>C 2107A>C 2108G>A), H600A (2133C>G 2134A>C), L601C (2136C>T 2137T>G), V604N (2145G>A 2146T>A 2147T>C), S606L (2152C>T), Y611L (2166T>C 2167A>T 2168C>T), W613F (2173G>T 2174G>T), G618F (2187G>T 2188G>T), M619G (2190A>G 2191T>G 2192G>A), P620T (2193C>A), G627A (2215G>C), K628del (2217_2219delAAG), V630I (2223G>A), S631D (2226A>G 2227G>A), L634R (2236T>G), K635E (2238A>G 2240A>G), E636A (2242A>C 2243A>T), L637V (2244T>G), V639P (2250G>C 2251T>C), E641Q (2256G>C), L642I (2259T>A), V655R (2298G>C 2299T>G 2300T>G), L663I (2322C>A 2324A>C), R664L (2326G>T 2327C>T), Q665N (2328C>A 2330A>T), V667M (2334G>A 2336A>G), E668K (2337G>A), K672T (2350A>C 2351G>T), L674del (2355_2359delCTCAA), I680V (2373A>G 2375C>A), C681A (2376T>G 2377G>C 2378T>A), M682L (2379A>T 2381G>A), N683S (2382A>T 2383A>C 2384C>A), S687T (2394T>A), K689D (2400A>G 2402G>T), S690Q (2403A>C 2404G>A 2405T>A), I692F (2409A>T 2411C>T), S694V (2415T>G 2416C>T 2417C>T), I696M (2423A>G), L704S (2445C>T 2446T>C 2447C>A), E705N (2448G>A 2450A>C), T706E (2451A>G 2452C>A 2453T>A), F708Y (2458T>A), D709E (2462T>A), L715Y (2478C>T 2479T>A 2480A>C), I716V (2481A>G 2483C>T), P728L (2518C>T 2519C>T), Q733R (2532C>A 2533A>G), D737G (2545A>G), N739P (2550A>C 2551A>C 2552C>A), T740_V741insLS (2555_2556insCTCAGT), V741F (2556G>T), Q743L (2562C>T 2563A>T 2564A>G), F744V (2565T>G), N745D (2568A>G), K746Q (2571A>C), W747* (2575G>A), I748V (2577A>G), L749F (2582A>T)

Codon mutations:

AAA34AAG (437A>G), ATC36CTT (441A>C 443C>T), CTT38TTT (447C>T), CTA42TTA (459C>T), GCC44GTC (466C>T), TCC45TCT (470C>T), GCT46ATG (471G>A 472C>T 473T>G), TTG48ATG (477T>A), ACA49CAT (480A>C 481C>A 482A>T), AAA50AAG (485A>G), ATC52GTC (489A>G), GCT53GCC (494T>C), CTT54CTA (497T>A), GCT55GCG (500T>G), GGA56GGC (503A>C), TGT57TGC (506T>C), GAC58GAT (509C>T), TTT59TTC (512T>C), GTA60ATA (513G>A), TCT61AAA (516T>A 517C>A 518T>A), AAA62ACA (520A>C), GTG63GTT (524G>T), ACT64GTG (525A>G 526C>T 527T>G), ACC65AAA (529C>A 530C>A), TTA66CTG (531T>C 533A>G), CGC67AAA (534C>A 535G>A 536C>A), CGG68CGA (539G>A), AAG69CGT (540A>C 541A>G 542G>T), TCG70TCA (545G>A), GGC71GGT (548C>T), TAT72TTT (550A>T), TTC74TAT (556T>A 557C>T), ACA75ACT (560A>T), TAC78TAT (569C>T), AAG80AAA (575G>A), TGC82TGT (581C>T), GCA83ACA (582G>A), GTA84GTT (587A>T), TGC85TCC (589G>C), TTA86CTT (591T>C 593A>T), CGA88CGT (599A>T), TAC89TAT (602C>T), TCC93AGT (612T>A 613C>G 614C>T), TTT94TAT (616T>A), CGA95TCT (618C>T 619G>C 620A>T), GAA97AAT (624G>A 626A>T), GAA98GAC (629A>C), CCA99ACC (630C>A 632A>C), CTT100ATG (633C>A 635T>G), TCG101TCT (638G>T), GTG102GTT (641G>T), CCT103CCA (644T>A), GTG104GTT (647G>T), CTC106TTG (651C>T 653C>G), ACA107ACC (656A>C), AGA108CGA (657A>C), TCG109TGT (661C>G 662G>T), GGT110GGA (665T>A), ATC111ATT (668C>T), CGT113AAG (672C>A 673G>A 674T>G), ATC115ATA (680C>A), CCC116CCT (683C>T), AAA117GCC (684A>G 685A>C 686A>C), TTG119TTA (692G>A), CGT120CGA (695T>A), GCA122CAC (699G>C 700C>A 701A>C), ATA123GTA (702A>G), CGT124AGG (705C>A 707T>G), AGG125GCA (708A>G 709G>C 710G>A), AAA126AAG (713A>G), GAT127TCT (714G>T 715A>C), GAA128GAT (719A>T), CAC129CAT (722C>T), GTT134GTC (737T>C), TAC137TAT (746C>T), TCA139TCT (752A>T), GGG142GGA (761G>A), GCT144TCA (765G>T 767T>A), AAG145AAA (770G>A), GTT146TAG (771G>T 772T>A 773T>G), TTG147ATT (774T>A 776G>T), AAA148GCG (777A>G 778A>C 779A>G), GTT149GTG (782T>G), AGT154AAG (796G>A 797T>G), GCA156GCC (803A>C), TTT158TTC (809T>C), GAA159CAA (810G>C), TCC160TCA (815C>A), ATA161ATT (818A>T), ACT162ACG (821T>G), ACC163ACT (824C>T), CCA164CCT (827A>T), CCT166TCA (831C>T 833T>A), GGG169GAT (841G>A 842G>T), CTA172CGT (850T>G 851A>T), GTG174GTT (857G>T), CTG175CTT (860G>T), GAA176GGA (862A>G), ATT178ATG (869T>G), ACG180ACT (875G>T), TTC182TGA (880T>G 881C>A), CAT184AAG (885C>A 887T>G), CAA186CGA (892A>G), CTA190CTT (905A>T), CCC191CCA (908C>A), TTT192CAT (909T>C 910T>A), CGA194CAA (916G>A), GGC195TCC (918G>T 919G>C), ATT196ATA (923T>A), CCA197CCT (926A>T), ATG198CTT (927A>C 929G>T), ATT200AAG (934T>A 935T>G), GGG201GGT (938G>T), TCT203AAT (942T>A 943C>A), CAA205AAA (948C>A), CCC212CCA (971C>A), CTA213TTA (972C>T), AAT214CTC (975A>C 976A>T 977T>C), AAT216AAC (983T>C), AGG219CGC (990A>C 992G>C), TCA220CGG (993T>C 994C>G 995A>G), TAC221TTT (997A>T 998C>T), GGT222CAA (999G>C 1000G>A 1001T>A), TTA223TAT (1003T>A 1004A>T), GCC224TCG (1005G>T 1007C>G), GTA225GTG (1010A>G), GAC227GAT (1016C>T), AAA228GTT (1017A>G 1018A>T 1019A>T), GTT229GTA (1022T>A), GAT230GAC (1025T>C), CAA231GGA (1026C>G 1027A>G), GAG232GCA (1030A>C 1031G>A), GCT233GCG (1034T>G), AGT234GCT (1035A>G 1036G>C), TTC237GTT (1044T>G 1046C>T), AAG238AAA (1049G>A), TTT239GAA (1050T>G 1051T>A 1052T>A), CCA240CAC (1054C>A 1055A>C), ATT242ATC (1061T>C), GTT244GTG (1067T>G), TTG246TTA (1073G>A), AAA247AAG (1076A>G), CTA250ATA (1083C>A), GCG251GCT (1088G>T), GCC252GCG (1091C>G), TGG255TTG (1099G>T), ACA257GTG (1104A>G 1105C>T 1106A>G), ATT260ATG (1115T>G), TCA262TCC (1121A>C), TTC263ATT (1122T>A 1124C>T), CAG264CAA (1127G>A), GAC265GAT (1130C>T), GGC266GGG (1133C>G), TCA269TCC (1142A>C), CCC270CCT (1145C>T), GGG271GGT (1148G>T), ATG272ATT (1151G>T), TTG273CTT (1152T>C 1154G>T), TCC275TCT (1160C>T), AAA276CAA (1161A>C), TCT277ATG (1164T>A 1165C>T 1166T>G), GTC278GTT (1169C>T), TTC279TTA (1172C>A), TTT280TAC (1174T>A 1175T>C), CCT281CCA (1178T>A), TTA282TTT (1181A>T), GAC283GAT (1184C>T), AGG284TCT (1185A>T 1186G>C 1187G>T), GCT285AAT (1188G>A 1189C>A), GCA286AAT (1191G>A 1192C>A 1193A>T), ACT287ACC (1196T>C), GAA288del (1197_1199delGAA), TTA290TTT (1205A>T), AAC292AAT (1211C>T), TTA293GAA (1212T>G 1213T>A), TCT294TCC (1217T>C), CTC295CTT (1220C>T), GAA296GAT (1223A>T), CAC297TTA (1224C>T 1225A>T 1226C>A), GAG299GAA (1232G>A), AGT301CAA (1236A>C 1237G>A 1238T>A), GTC302GTA (1241C>A), GTA335..C (1340A>C), GTA337GCA (1345T>C), ATG338ATA (1349G>A), AAC340AAT (1355C>T), ATA342GTA (1359A>G), AAA343_CAA344insAAAA-- (1364_1365insAAAA), AGA345AAG (1369G>A 1370A>G), CTT346TGA (1371C>T 1372T>G 1373T>A), CTC347CTG (1376C>G), CAC348TTT (1377C>T 1378A>T 1379C>T), GTT350GTC (1385T>C), GAT352CAA (1389G>C 1391T>A), GCT354GCA (1397T>A), ACA356AAA (1402C>A), CTG358CTA (1409G>A), AAA360CGC (1413A>C 1414A>G 1415A>C), ATA361CTT (1416A>C 1418A>T), AAG362CCG (1419A>C 1420A>C), ACA363GCG (1422A>G 1424A>G), GAC364GAT (1427C>T), GGG365GGA (1430G>A), ACT366GCA (1431A>G 1433T>A), GAG370GAA (1445G>A), AAA371CGT (1446A>C 1447A>G 1448A>T), CCT372CCG (1451T>G), TTG374GTA (1455T>G 1457G>A), GCT375CGA (1458G>C 1459C>G 1460T>A), AAG377CGA (1464A>C 1465A>G 1466G>A), CAA378TCT (1467C>T 1468A>C 1469A>T), AAG379CAA (1470A>C 1472G>A), AAC380AAG (1475C>G), TTT381AAA (1476T>A 1477T>A 1478T>A), AAA382GGT (1479A>G 1480A>G 1481A>T), TGT384TCT (1486G>C), TAT385TTC (1489A>T 1490T>C), TTA389TTC (1502A>C), AAG390AAA (1505G>A), TCA391TCG (1508A>G), CCG397CCA (1526G>A), ATG401ATA (1538G>A), TAT402TAC (1541T>C), CTT404TTC (1545C>T 1547T>C), TCT406TCC (1553T>C), ATG407TGC (1554A>T 1555T>G 1556G>C), ATA408ATG (1559A>G), CCT411AGT (1566C>A 1567C>G), ACC412ACA (1571C>A), TTG413CTG (1572T>C), TCC415TCA (1580C>A), GTT418GTC (1589T>C), AGC420del (1593_1597delAGCTC), TCT421--A (1593_1597delAGCTC 1598T>A), CTT424CTC (1607T>C), ACG426ACT (1613G>T), ATC427TTC (1614A>T), CTT428TTG (1617C>T 1619T>G), GTT429ATC (1620G>A 1622T>C), AAC430GTC (1623A>G 1624A>T), AAA431CCT (1626A>C 1627A>C 1628A>T), TTA433ATG (1632T>A 1634A>G), ACA434GTC (1635A>G 1636C>T 1637A>C), AGA436AGG (1643A>G), TAT438AGT (1647T>A 1648A>G), CTG443CTA (1664G>A), GGG445GGA (1670G>A), CCT447CCA (1676T>A), TTG448TTA (1679G>A), GGC449GGT (1682C>T), CAC451TAC (1686C>T), GGT452GGA (1691T>A), GCA455TCT (1698G>T 1700A>T), CTT456CTC (1703T>C), TCG458GCT (1707T>G 1709G>T), TTG459CTA (1710T>C 1712G>A), TAT463TAC (1724T>C), TTA467TTG (1736A>G), GCA468GGA (1738C>G), GCG469GCA (1742G>A), AGC470AAG (1744G>A 1745C>G), GGA475GAC (1759G>A 1760A>C), TGT476CGT (1761T>C), AAC477AAA (1766C>A), TCT478ACC (1767T>A 1769T>C), CCA479CCC (1772A>C), TTT480TTC (1775T>C), GGG481AAA (1776G>A 1777G>A 1778G>A), GAT482GAC (1781T>C), TAC483TAT (1784C>T), GCT484GCC (1787T>C), TTA485TTG (1790A>G), CTA491CTG (1808A>G), ACC493ACT (1814C>T), AGT495GCA (1818A>G 1819G>C 1820T>A), AAA496AAG (1823A>G), GCC498..G (1829C>G), AAT499CGG (1830A>C 1831A>G 1832T>G), GAG500C.. (1833G>C), AAG502.G. (1840A>G), TTA504..G (1847A>G), TTG505C.. (1848T>C), AGA507..G (1856A>G), CTC508..T (1859C>T), AAT509GG. (1860A>G 1861A>G), GTC510GTT (1865C>T), AAA511ACA (1867A>C), ATT512ATC (1871T>C), TCG513TCT (1874G>T), ATA514-TA (1875delA), CCA515GCT (1878C>G 1880A>T), TCT517TCG (1886T>G), TTG518AGA (1887T>A 1888T>G 1889G>A), TCT520TCC (1895T>C), GAT521GAG (1898T>G), AAT522AAC (1901T>C), ATT525ATC (1910T>C), GCG528GCT (1919G>T), AGG530AGA (1925G>A), ACT533ACC (1934T>C), AAA534AAG (1937A>G), TCA535GAC (1938T>G 1939C>A 1940A>C), GAT539TAC (1950G>T 1952T>C), TTA540CTT (1953T>C 1955A>T), ATT543ATC (1964T>C), TCC544TCT (1967C>T), GCT547GCA (1976T>A), TTA549ACT (1980T>A 1981T>C 1982A>T), TCT550TCC (1985T>C), CGT552AGG (1989C>A 1991T>G), ACT553ACA (1994T>A), GCG583.CT (2084G>T), TTA585CTG (2088T>C 2090A>G), TTC586ATG (2091T>A 2093C>G), ACT587TCT (2094A>T), ACC588ACT (2099C>T), CAA589CAG (2102A>G), AGC590TCA (2103A>T 2104G>C 2105C>A), AAG591CCA (2106A>C 2107A>C 2108G>A), CGT592CGA (2111T>A), GAG594GAA (2117G>A), CGT595AGA (2118C>A 2120T>A), AAA597AAG (2126A>G), GCT598GCG (2129T>G), CAT600GCT (2133C>G 2134A>C), CTT601TGT (2136C>T 2137T>G), AAG602AAA (2141G>A), GTT604AAC (2145G>A 2146T>A 2147T>C), CGG605AGG (2148C>A), TCA606TTA (2152C>T), CAC607CAT (2156C>T), TTA608CTA (2157T>C), CTA609TTA (2160C>T), TAC611CTT (2166T>C 2167A>T 2168C>T), TGG613TTT (2173G>T 2174G>T), GGT618TTT (2187G>T 2188G>T), ATG619GGA (2190A>G 2191T>G 2192G>A), CCC620ACC (2193C>A), CCC623CCT (2204C>T), AAA626AAG (2213A>G), GGG627GCG (2215G>C), AAG628del (2217_2219delAAG), GTA630ATA (2223G>A), AGT631GAT (2226A>G 2227G>A), TAC632TAT (2231C>T), CTC633CTG (2234C>G), CTT634CGT (2236T>G), AAA635GAG (2238A>G 2240A>G), GAA636GCT (2242A>C 2243A>T), TTA637GTA (2244T>G), AAA638AAG (2249A>G), GTT639CCT (2250G>C 2251T>C), GAG641CAG (2256G>C), TTA642ATA (2259T>A), CAG643CAA (2264G>A), AAG647AAA (2276G>A), CTG649TTG (2280C>T), GTG650GTA (2285G>A), TTC651TTT (2288C>T), GGT653GGA (2294T>A), GTT655CGG (2298G>C 2299T>G 2300T>G), ATA657ATC (2306A>C), CTA658CTT (2309A>T), GAA659GAG (2312A>G), CGT660CGG (2315T>G), CTA663ATC (2322C>A 2324A>C), CGC664CTT (2326G>T 2327C>T), CAA665AAT (2328C>A 2330A>T), GTA667ATG (2334G>A 2336A>G), GAA668AAA (2337G>A), TTA671TTG (2348A>G), AAG672ACT (2350A>C 2351G>T), CTC674del (2355_2359delCTCAA), AAC675--T (2355_2359delCTCAA 2360C>T), TAC677TAT (2366C>T), ACG679ACT (2372G>T), ATC680GTA (2373A>G 2375C>A), TGT681GCA (2376T>G 2377G>C 2378T>A), ATG682TTA (2379A>T 2381G>A), AAC683TCA (2382A>T 2383A>C 2384C>A), GTA686GTG (2393A>G), TCC687ACC (2394T>A), AAG689GAT (2400A>G 2402G>T), AGT690CAA (2403A>C 2404G>A 2405T>A), CTC691CTA (2408C>A), ATC692TTT (2409A>T 2411C>T), GAC693GAT (2414C>T), TCC694GTT (2415T>G 2416C>T 2417C>T), ATA696ATG (2423A>G), TGC697TGT (2426C>T), GCA698GCC (2429A>C), ACA699ACT (2432A>T), TCC700TCT (2435C>T), AGG703CGA (2442A>C 2444G>A), CTC704TCA (2445C>T 2446T>C 2447C>A), GAA705AAC (2448G>A 2450A>C), ACT706GAA (2451A>G 2452C>A 2453T>A), TTT708TAT (2458T>A), GAT709GAA (2462T>A), TTC713TTT (2474C>T), GGC714GGG (2477C>G), CTA715TAC (2478C>T 2479T>A 2480A>C), ATC716GTT (2481A>G 2483C>T), AAA718AAG (2489A>G), GGT723GGA (2504T>A), AAT727AAC (2516T>C), CCC728CTT (2518C>T 2519C>T), ACC730ACT (2525C>T), ACT731ACA (2528T>A), CCA732CCT (2531A>T), CAG733AGG (2532C>A 2533A>G), GAT737GGT (2545A>G), AAC739CCA (2550A>C 2551A>C 2552C>A), ACT740_GTT741insCTCAGT (2555_2556insCTCAGT), GTT741TTT (2556G>T), ATT742ATA (2561T>A), CAA743TTG (2562C>T 2563A>T 2564A>G), TTT744GTT (2565T>G), AAT745GAT (2568A>G), AAA746CAA (2571A>C), TGG747TAG (2575G>A), ATA748GTA (2577A>G), TTA749TTT (2582A>T), GGG750GGT (2585G>T)

*: Inserts / Deletes / Misaligned / Frameshifts

#### Analysis details

This analysis was performed with panviral2.64

### NGS Details (UN60): Solendovirus venanicotianae

#### Assembly

| | |
|---|---|
| **Coverage Length** | 1373 (2 contig(s)) |
| **Depth Of Coverage** | 11.4 |
| **Number Of Reads** | 130 |
| **Reads Per Million** | 2.76 rpm (after QC) |
| **Ambiguities** | 0 |
| **Assembly Method** | de novo + reference guided assembly |
| **Consensus Caller** | Bcf Tools |

#### Coverage Map

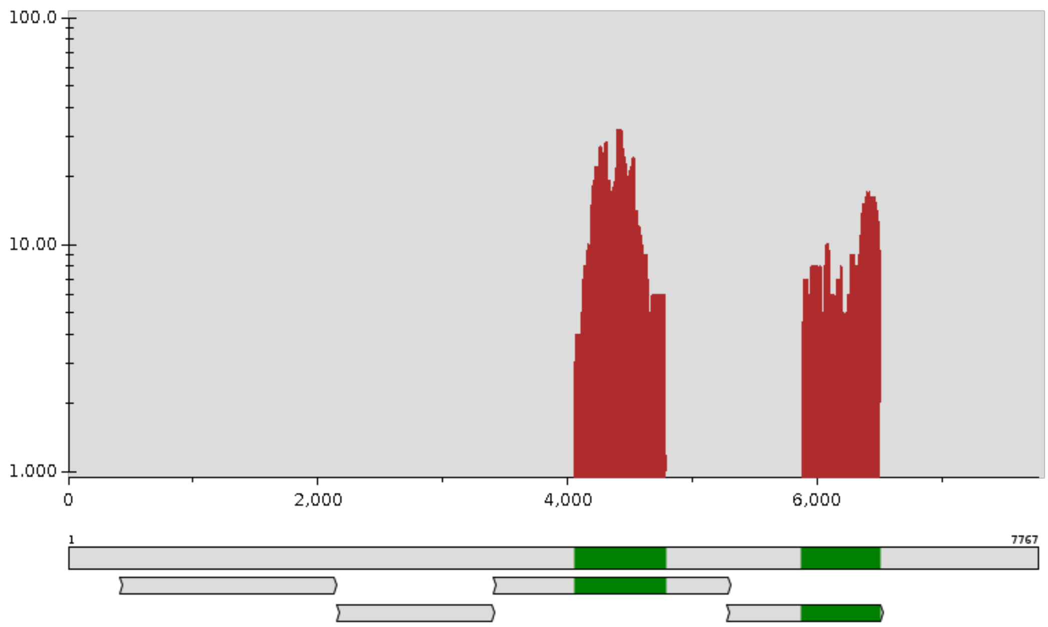

#### Assignment

| | |
|---|---|
| **Type** | Solendovirus venanicotianae (Taxonomy ID: 3048371) |
| **Reference Genome** | NC_003378.1 |
| **NT Identity (%)** | 77.8592 |
| **AA Identity (%)** | 75.6044 |
| **Number Of Stop Codons** | 1 |
| **Number Of CDS** | 4 |

#### Alignment

| | |
|---|---|
| **Alignment Score** | 1493.0 (NT) + 2180.0 (AA) = 3673.0 |
| **Concordance (%)** | 66.1114 |

| **Alignment Method** | Global, seeded, nucleotide + amino acids (AGA) |
|---|---|

#### Genome Region

Sequence starts at position 4050 and ends at position 6507 relative to NC_003378.1 reference sequence.

#### Alignment Detailed Statistics

| | Begin | End | Coverage | Score | Concordance | Matches | Identities | I/D/M/F* | Stop Codons |
|---|---|---|---|---|---|---|---|---|---|
| **NT** | **4050** | **6507** | **17.7%** | **1493** | **55.3%** | **1364 (99.3%)** | **1062 (77.3%)** | **0/9** | |
| Mutations: | 4052C>T, 4056A>G, 4061T>C, 4065A>G, 4066C>A, 4073T>C, 4079A>G, 4088A>C, 4091T>A, 4095T>C, 4097G>A, 4098G>A, 4100T>A, 4103T>G, 4104A>G, 4105A>G, 4115A>G, 4116G>A, 4121T>C, 4122A>G, 4124T>A, 4125A>G, 4127C>T, 4133T>C, 4161C>T, 4165G>A, 4181A>T, 4190G>A, 4196C>T, 4199A>T, 4202C>T, 4212T>C, 4235T>C, 4247A>C, 4250C>A, 4253T>C, 4260T>C, 4265G>A, 4292C>T, 4298T>C, 4310T>C, 4313G>A, 4325T>C, 4328T>C, 4329T>C, 4331G>A, 4340G>A, 4343T>A, 4355A>G, 4364T>A, 4376A>C, 4385T>C, 4400T>A, 4401T>C, 4406A>T, 4436C>T, 4440C>T, 4451C>T, 4453A>G, 4467C>T, 4470G>C, 4471A>C, 4475T>C, 4478T>C, 4482G>A, 4484A>T, 4497A>G, 4500C>T, 4503C>T, 4513G>A, 4515A>C, 4517A>T, 4518C>A, 4523T>A, 4530A>T, 4545A>C, 4550C>T, 4551G>A, 4552C>T, 4554C>A, 4559T>A, 4560A>G, 4568C>A, 4571T>A, 4574T>A, 4581T>C, 4591C>A, 4592C>G, 4598A>T, 4601A>G, 4604T>A, 4605A>T, 4607G>T, 4626T>C, 4643T>C, 4653A>G, 4667T>A, 4688C>T, 4691T>C, 4692C>T, 4693T>C, 4694T>A, 4703C>T, 4706T>A, 4730A>C, 4734C>T, 4740A>C, 4745A>T, 4748C>T, 4755A>C, 4757G>A, 4772A>G, 4784C>T, 4785T>C, 5872T>C, 5874G>A, 5880T>A, 5881T>C, 5882T>A, 5889T>A, 5892A>T, 5894G>A, 5898A>C, 5901A>T, 5907A>T, 5908G>A, 5909C>G, 5910C>A, 5913C>A, 5916C>T, 5919C>T, 5921A>G, 5932C>T, 5935G>A, 5940T>C, 5949A>C, 5950C>T, 5952G>T, 5953A>G, 5954C>A, 5955A>C, 5959C>G, 5960A>G, 5964G>A, 5968G>T, 5970C>A, 5971A>G, 5972A>C, 5979T>A, 5982G>A, 5983T>C, 5988C>T, 5991G>A, 5994A>C, 5997C>T, 6001C>A, 6002A>T, 6006T>C, 6009G>A, 6011G>A, 6012G>A, 6015A>T, 6018T>A, 6020A>G, 6023G>A, 6024G>A, 6026C>T, 6027A>T, 6033C>T, 6035A>T, 6036C>T, 6037G>A, 6040C>A, 6041G>A, 6044T>G, 6045C>T, 6049T>A, 6051G>A, 6054T>C, 6055A>C, 6067C>T, 6075A>T, 6078G>A, 6084G>A, 6087T>A, 6088A>G, 6090C>T, 6096A>G, 6099T>G, 6105A>G, 6108T>A, 6109G>T, 6112C>T, 6114C>G, 6120G>A, 6123A>T, 6129C>T, 6133C>G, 6139A>G, 6141G>T, 6144A>T, 6146A>G, 6148G>C, 6154G>C, 6155A>C, 6163G>C, 6164A>C, 6169G>A, 6171A>T, 6172A>G, 6174T>G, 6177T>A, 6181G>A, 6182A>G, 6183A>T, 6186C>T, 6189T>C, 6190G>A, 6192T>C, 6198A>C, 6201G>A, 6204A>T, 6211A>C, 6214T>G, 6219T>A, 6223C>A, 6225A>C, 6232C>T, 6236C>G, 6240C>T, 6246T>C, 6247C>A, 6248T>C, 6253C>G, 6254A>G, 6255G>A, 6258T>C, 6259G>C, 6260C>T, 6264T>G, 6269G>T, 6270C>T, 6277T>A, 6279A>C, 6282G>A, 6285A>T, 6288G>A, 6290C>T, 6308G>C, 6309C>A, 6315T>A, 6318A>G, 6319A>C, 6322G>A, 6325G>C, 6326C>A, 6334G>A, 6340C>G, 6341T>C, 6345A>G, 6352G>A, 6354C>A, 6355T>A, 6357A>G, 6359G>C, 6360T>A, 6363T>G, 6369A>T, 6371C>T, 6376C>A, 6377A>G, 6380C>A, 6381T>A, 6382A>C, 6384A>T, 6401G>A, 6402G>A, 6404G>T, 6406delA, 6407A>G, 6408T>A, 6411C>T, 6414C>T, 6417C>A, 6418C>G, 6419C>A, 6422A>G, 6423T>A, 6424C>T, 6426G>A, 6429A>T, 6444G>A, 6445C>G, 6446T>A, 6447A>T, 6451A>G, 6453T>A, 6456C>G, 6462T>C, 6464C>A, 6466_6473delGACCACCT, 6477C>T, 6480A>T, 6489A>C | | | | | | | | |

##### CDS

| | Begin | End | Coverage | Score | Concordance | Matches | Identities | I/D/M/F* | Stop Codons |
|---|---|---|---|---|---|---|---|---|---|
| Tvvgp3 | 214 | 459 | 38.7% | 1475 | 87.0% | 246 (100%) | 211 (85.8%) | 0/0/0/0 | 0 |
| Protein mutations: | N216D (4056A>G), T219E (4065A>G 4066C>A), I223M (4079A>G), K226N (4088A>C), D227E (4091T>A), D230K (4098G>A 4100T>A), N231K (4103T>G), K232G (4104A>G 4105A>G), E236K (4116G>A), N238E (4122A>G 4124T>A), S239G (4125A>G 4127C>T), H251Y (4161C>T), S252N (4165G>A), D311E (4343T>A), H344Y (4440C>T), N348S (4453A>G), E354P (4470G>C 4471A>C), V358I (4482G>A 4484A>T), I363V (4497A>G), R368K (4513G>A), T369P (4515A>C 4517A>T), Q370K (4518C>A), D371E (4523T>A), I374L (4530A>T), K379Q (4545A>C), A381I (4551G>A 4552C>T), H382N (4554C>A), I384V (4560A>G), N386K (4568C>A), T394K (4591C>A 4592C>G), M399F (4605A>T 4607G>T), I415V (4653A>G), L428S (4692C>T 4693T>C 4694T>A), I444L (4740A>C) | | | | | | | | |
| Codon mutations: | AAC214AAT (4052C>T), AAT216GAT (4056A>G), GAT217GAC (4061T>C), ACA219GAA (4065A>G 4066C>A), TTT221TTC (4073T>C), ATA223ATG (4079A>G), AAA226AAC (4088A>C), GAT227GAA (4091T>A), TTG229CTA (4095T>C 4097G>A), GAT230AAA (4098G>A 4100T>A), AAT231AAG (4103T>G), AAA232GGA (4104A>G 4105A>G), CAA235CAG (4115A>G), GAA236AAA (4116G>A), AGT237AGC (4121T>C), AAT238GAA (4122A>G 4124T>A), AGC239GGT (4125A>G 4127C>T), CAT241CAC (4133T>C), CAT251TAT (4161C>T), AGT252AAT (4165G>A), GGA257GGT (4181A>T), AGG260AGA (4190G>A), GTC262GTT (4196C>T), ATA263ATT (4199A>T), GAC264GAT (4202C>T), TTA268CTA (4212T>C), TAT275TAC (4235T>C), ATA279ATC (4247A>C), CCC280CCA (4250C>A), AAT281AAC (4253T>C), TTA284CTA (4260T>C), AAG285AAA (4265G>A), TAC294TAT (4292C>T), AGT296AGC (4298T>C), TGT300TGC (4310T>C), AAG301AAA (4313G>A), TAT305TAC (4325T>C), CAT306CAC (4328T>C), TTG307CTA (4329T>C 4331G>A), GAG310GAA (4340G>A), GAT311GAA (4343T>A), AAA315AAG (4355A>G), GCT318GCA (4364T>A), CCA322CCC (4376A>C), TTT325TTC (4385T>C), GTT330GTA (4400T>A), TTA331CTA (4401T>C), CCA332CCT (4406A>T), TAC342TAT (4436C>T), CAT344TAT (4440C>T), GAC347GAT (4451C>T), AAC348AGC (4453A>G), CTA353TTA (4467C>T), GAA354CCA (4470G>C 4471A>C), AAT355AAC (4475T>C), TGT356TGC (4478T>C), GTA358ATT (4482G>A 4484A>T), ATA363GTA (4497A>G), CTA364TTA (4500C>T), CTA365TTA (4503C>T), AGA368AAA (4513G>A), ACA369CCT (4515A>C 4517A>T), CAA370AAA (4518C>A), GAT371GAA (4523T>A), ATA374TTA (4530A>T), AAA379CAA (4545A>C), TTC380TTT (4550C>T), GCA381ATA (4551G>A 4552C>T), CAT382AAT (4554C>A), ATT383ATA (4559T>A), ATA384GTA (4560A>G), AAC386AAA (4568C>A), TCT387TCA (4571T>A), GGT388GGA (4574T>A), TTA391CTA (4581T>C), ACC394AAG (4591C>A 4592C>G), GCA396GCT (4598A>T), GAA397GAG (4601A>G), ATT398ATA (4604T>A), ATG399TTT (4605A>T 4607G>T), TTA406CTA (4626T>C), GAT411GAC (4643T>C), ATA415GTA (4653A>G), ACT419ACA (4667T>A), ATC426ATT (4688C>T), AAT427AAC (4691T>C), CTT428TCA (4692C>T 4693T>C 4694T>A), AAC431AAT (4703C>T), ATT432ATA (4706T>A), TCA440TCC (4730A>C), CTA442TTA (4734C>T), ATA444CTA (4740A>C), GTA445GTT (4745A>T), AAC446AAT (4748C>T), AGG449CGA (4755A>C 4757G>A), AAA454AAG (4772A>G), AAC458AAT (4784C>T), TTA459CT. (4785T>C) | | | | | | | | |
| Tvvgp4 | 200 | 411 | 50.5% | 705 | 49.9% | 209 (99.1%) | 133 (63.0%) | 0/2/2/2 | 1 |
| Protein mutations: | L203Q (5881T>C 5882T>A), N205K (5889T>A), S207N (5894G>A), K209N (5901A>T), A212R (5908G>A 5909C>G 5910C>A), N216S (5921A>G), D221N (5935G>A), Q226Y (5950C>T 5952G>T), T227D (5953A>G 5954C>A 5955A>C), Q229G (5959C>G 5960A>G), A232S (5968G>T 5970C>A), N233A (5971A>G 5972A>C), Q243I (6001C>A 6002A>T), R246K (6011G>A 6012G>A), K249R (6020A>G), G250E (6023G>A 6024G>A), T251I (6026C>T 6027A>T), Y254F (6035A>T 6036C>T), V255I (6037G>A), R256K (6040C>A 6041G>A), F257C (6044T>G 6045C>T), S259T (6049T>A 6051G>A), T261P (6055A>C), E267D (6075A>T), I272V (6088A>G 6090C>T), G279* (6109G>T), E283D (6123A>T), P287A (6133C>G), K289D (6139A>G 6141G>T), E291G (6146A>G), E292Q (6148G>C), E294P (6154G>C 6155A>C), E297P (6163G>C 6164A>C), V299I (6169G>A 6171A>T), N300E (6172A>G 6174T>G), E303S (6181G>A 6182A>G 6183A>T), A306T (6190G>A 6192T>C), K308N (6198A>C), I313L (6211A>C), S314A (6214T>G), L317I (6223C>A 6225A>C), A321G (6236C>G), L325T (6247C>A 6248T>C), Q327G (6253C>G 6254A>G 6255G>A), A329L (6259G>C 6260C>T), I330M (6264T>G), S332I (6269G>T 6270C>T), S335T (6277T>A 6279A>C), E337D (6285A>T), T339I (6290C>T), C345S (6308G>C 6309C>A), D347E (6315T>A), I348M (6318A>G), E350K (6322G>A), A351Q (6325G>C 6326C>A), E354K (6334G>A), L356A (6340C>G 6341T>C), V360I (6352G>A 6354C>A), L361M (6355T>A 6357A>G), S362T (6359G>C 6360T>A), K365N (6369A>T), P366L (6371C>T), Q368R (6376C>A 6377A>G), P369Q (6380C>A 6381T>A), T370P (6382A>C 6384A>T), R376K (6401G>A 6402G>A), R377M (6404G>T), P382E (6418C>G 6419C>A), D383G (6422A>G 6423T>A), L385F (6429A>T), L391D (6445C>G 6446T>A 6447A>T), S393G (6451A>G 6453T>A), H394Q (6456C>G), P397Q (6464C>A), D398_H399del (6466_6473delGACCACCT), K405N (6489A>C) | | | | | | | | |
| Codon mutations: | TTG200CTA (5872T>C 5874G>A), GCT202GCA (5880T>A), TTA203CAA (5881T>C 5882T>A), AAT205AAA (5889T>A), ACA206ACT (5892A>T), AGT207AAT (5894G>A), GCA208GCC (5898A>C), AAA209AAT (5901A>T), GTA211GTT (5907A>T), GCC212AGA (5908G>A 5909C>G 5910C>A), ACC213ACA (5913C>A), TGC214TGT (5916C>T), TAC215TAT (5919C>T), AAC216AGC (5921A>G), CTA220TTA (5932C>T), GAT221AAT (5935G>A), ACT222ACC (5940T>C), ACA225ACC (5949A>C), CAG226TAT (5950C>T 5952G>T), ACA227GAC (5953A>G 5954C>A 5955A>C), CAA229GGA (5959C>G 5960A>G), GAG230GAA (5964G>A), GCC232TCA (5968G>T 5970C>A), AAT233GCT (5971A>G 5972A>C), CCT235CCA (5979T>A), GAG236GAA (5982G>A), TTA237CTA (5983T>C), TAC238TAT (5988C>T), AAG239AAA (5991G>A), GCA240GCC (5994A>C), TTC241TTT (5997C>T), CAA243ATA (6001C>A 6002A>T), TAT244TAC (6006T>C), AAG245AAA (6009G>A), AGG246AAA (6011G>A 6012G>A), ATA247ATT (6015A>T), ACT248ACA (6018T>A), AAA249AGA (6020A>G), GGG250GAA (6023G>A 6024G>A), ACA251ATT (6026C>T 6027A>T), TTC253TTT (6033C>T), TAC254TTT (6035A>T 6036C>T), GTA255ATA (6037G>A), CGA256AAA (6040C>A 6041G>A), TTC257TGT (6044T>G 6045C>T), TCG259ACA (6049T>A 6051G>A), GCT260GCC (6054T>C), ACA261CCA (6055A>C), CTA265TTA (6067C>T), GAA267GAT (6075A>T), GAG268GAA (6078G>A), AAG270AAA (6084G>A), CCT271CCA (6087T>A), ATC272GTT (6088A>G 6090C>T), CAA274CAG (6096A>G), GTT275GTG (6099T>G), AAA277AAG (6105A>G), ATT278ATA (6108T>A), GGA279TGA (6109G>T), CTC280TTG (6112C>T 6114C>G), AGG282AGA (6120G>A), GAA283GAT (6123A>T), ATC285ATT (6129C>T), CCA287GCA (6133C>G), AAG289GAT (6139A>G 6141G>T), ATA290ATT (6144A>T), GAA291GGA (6146A>G), GAA292CAA (6148G>C), GAA294CCA (6154G>C 6155A>C), GAA297CCA (6163G>C 6164A>C), GTA299ATT (6169G>A 6171A>T), AAT300GAG (6172A>G 6174T>G), ATT301ATA (6177T>A), GAA303AGT (6181G>A 6182A>G 6183A>T), TTC304TTT (6186C>T), TAT305TAC (6189T>C), GCT306ACC (6190G>A 6192T>C), AAA308AAC (6198A>C), AGG309AGA (6201G>A), ATA310ATT (6204A>T), ATA313CTA (6211A>C), TCT314GCT (6214T>G), ACT315ACA (6219T>A), CTA317ATC (6223C>A 6225A>C), CTA320TTA (6232C>T), GCA321GGA (6236C>G), AAC322AAT (6240C>T), TAT324TAC (6246T>C), CTA325ACA (6247C>A 6248T>C), CAG327GGA (6253C>G 6254A>G 6255G>A), AAT328AAC (6258T>C), GCA329CTA (6259G>C 6260C>T), ATT330ATG (6264T>G), AGC332ATT (6269G>T 6270C>T), TCA335ACC (6277T>A 6279A>C), AGG336AGA (6282G>A), GAA337GAT (6285A>T), CAG338CAA (6288G>A), ACA339ATA (6290C>T), TGC345TCA (6308G>C 6309C>A), GAT347GAA (6315T>A), ATA348ATG (6318A>G), AGA349CGA (6319A>C), GAA350AAA (6322G>A), GCA351CAA (6325G>C 6326C>A), GAA354AAA (6334G>A), CTA356GCA (6340C>G 6341T>C), AGA357AGG (6345A>G), GTC360ATA (6352G>A 6354C>A), TTA361ATG (6355T>A 6357A>G), AGT362ACA (6359G>C 6360T>A), CTT363CTG (6363T>G), AAA365AAT (6369A>T), CCA366CTA (6371C>T), CAA368AGA (6376C>A 6377A>G), CCT369CAA (6380C>A 6381T>A), ACA370CCT (6382A>C 6384A>T), AGG376AAA (6401G>A 6402G>A), AGG377ATG (6404G>T), AAT378-GA (6406delA 6407A>G 6408T>A), TTC379TTT (6411C>T), ATC380ATT (6414C>T), TCC381TCA (6417C>A), CCA382GAA (6418C>G 6419C>A), GAT383GGA (6422A>G 6423T>A), CTG384TTA (6424C>T 6426G>A), TTA385TTT (6429A>T), AAG390AAA (6444G>A), CTA391GAT (6445C>G 6446T>A 6447A>T), AGT393GGA (6451A>G 6453T>A), CAC394CAG (6456C>G), TAT396TAC (6462T>C), CCA397CAA (6464C>A), GAC398_CAC399del (6466_6473delGACCACCT), CTA400--A (6466_6473delGACCACCT), TGC401TGT (6477C>T), TCA402TCT (6480A>T), AAA405AAC (6489A>C) | | | | | | | | |

##### Proteins

| | Begin | End | Coverage | Score | Concordance | Matches | Identities | I/D/M/F* | Stop Codons |
|---|---|---|---|---|---|---|---|---|---|
| polyprotein (NP_569141.1) | 214 | 459 | 38.7% | 1475 | 87.0% | 246 (100%) | 211 (85.8%) | 0/0/0/0 | 0 |
| Protein mutations: | N216D (4056A>G), T219E (4065A>G 4066C>A), I223M (4079A>G), K226N (4088A>C), D227E (4091T>A), D230K (4098G>A 4100T>A), N231K (4103T>G), K232G (4104A>G 4105A>G), E236K (4116G>A), N238E (4122A>G 4124T>A), S239G (4125A>G 4127C>T), H251Y (4161C>T), S252N (4165G>A), D311E (4343T>A), H344Y (4440C>T), N348S (4453A>G), E354P (4470G>C 4471A>C), V358I (4482G>A 4484A>T), I363V (4497A>G), R368K (4513G>A), T369P (4515A>C 4517A>T), Q370K (4518C>A), D371E (4523T>A), I374L (4530A>T), K379Q (4545A>C), A381I (4551G>A 4552C>T), H382N (4554C>A), I384V (4560A>G), N386K (4568C>A), T394K (4591C>A 4592C>G), M399F (4605A>T 4607G>T), I415V (4653A>G), L428S (4692C>T 4693T>C 4694T>A), I444L (4740A>C) | | | | | | | | |

| | Begin | End | Coverage | Score | Concordance | Matches | Identities | I/D/M/F* | Stop Codons |
|---|---|---|---|---|---|---|---|---|---|
| **NT** | **4050** | **6507** | **17.7%** | **1493** | **55.3%** | **1364 (99.3%)** | **1062 (77.3%)** | **0/9** | |
| Codon mutations: | AAC214AAT (4052C>T), AAT216GAT (4056A>G), GAT217GAC (4061T>C), ACA219GAA (4065A>G 4066C>A), TTT221TTC (4073T>C), ATA223ATG (4079A>G), AAA226AAC (4088A>C), GAT227GAA (4091T>A), TTG229CTA (4095T>C 4097G>A), GAT230AAA (4098G>A 4100T>A), AAT231AAG (4103T>G), AAA232GGA (4104A>G 4105A>G), CAA235CAG (4115A>G), GAA236AAA (4116G>A), AGT237AGC (4121T>C), AAT238GAA (4122A>G 4124T>A), AGC239GGT (4125A>G 4127C>T), CAT241CAC (4133T>C), CAT251TAT (4161C>T), AGT252AAT (4165G>A), GGA257GGT (4181A>T), AGG260AGA (4190G>A), GTC262GTT (4196C>T), ATA263ATT (4199A>T), GAC264GAT (4202C>T), TTA268CTA (4212T>C), TAT275TAC (4235T>C), ATA279ATC (4247A>C), CCC280CCA (4250C>A), AAT281AAC (4253T>C), TTA284CTA (4260T>C), AAG285AAA (4265G>A), TAC294TAT (4292C>T), AGT296AGC (4298T>C), TGT300TGC (4310T>C), AAG301AAA (4313G>A), TAT305TAC (4325T>C), CAT306CAC (4328T>C), TTG307CTA (4329T>C 4331G>A), GAG310GAA (4340G>A), GAT311GAA (4343T>A), AAA315AAG (4355A>G), GCT318GCA (4364T>A), CCA322CCC (4376A>C), TTT325TTC (4385T>C), GTT330GTA (4400T>A), TTA331CTA (4401T>C), CCA332CCT (4406A>T), TAC342TAT (4436C>T), CAT344TAT (4440C>T), GAC347GAT (4451C>T), AAC348AGC (4453A>G), CTA353TTA (4467C>T), GAA354CCA (4470G>C 4471A>C), AAT355AAC (4475T>C), TGT356TGC (4478T>C), GTA358ATT (4482G>A 4484A>T), ATA363GTA (4497A>G), CTA364TTA (4500C>T), CTA365TTA (4503C>T), AGA368AAA (4513G>A), ACA369CCT (4515A>C 4517A>T), CAA370AAA (4518C>A), GAT371GAA (4523T>A), ATA374TTA (4530A>T), AAA379CAA (4545A>C), TTC380TTT (4550C>T), GCA381ATA (4551G>A 4552C>T), CAT382AAT (4554C>A), ATT383ATA (4559T>A), ATA384GTA (4560A>G), AAC386AAA (4568C>A), TCT387TCA (4571T>A), GGT388GGA (4574T>A), TTA391CTA (4581T>C), ACC394AAG (4591C>A 4592C>G), GCA396GCT (4598A>T), GAA397GAG (4601A>G), ATT398ATA (4604T>A), ATG399TTT (4605A>T 4607G>T), TTA406CTA (4626T>C), GAT411GAC (4643T>C), ATA415GTA (4653A>G), ACT419ACA (4667T>A), ATC426ATT (4688C>T), AAT427AAC (4691T>C), CTT428TCA (4692C>T 4693T>C 4694T>A), AAC431AAT (4703C>T), ATT432ATA (4706T>A), TCA440TCC (4730A>C), CTA442TTA (4734C>T), ATA444CTA (4740A>C), GTA445GTT (4745A>T), AAC446AAT (4748C>T), AGG449CGA (4755A>C 4757G>A), AAA454AAG (4772A>G), AAC458AAT (4784C>T), TTA459CT. (4785T>C) | | | | | | | | |
| putative transactivator factor (NP_569142.1) | 200 | 411 | 50.5% | 705 | 49.9% | 209 (99.1%) | 133 (63.0%) | 0/2/2/2 | 1 |
| Protein mutations: | L203Q (5881T>C 5882T>A), N205K (5889T>A), S207N (5894G>A), K209N (5901A>T), A212R (5908G>A 5909C>G 5910C>A), N216S (5921A>G), D221N (5935G>A), Q226Y (5950C>T 5952G>T), T227D (5953A>G 5954C>A 5955A>C), Q229G (5959C>G 5960A>G), A232S (5968G>T 5970C>A), N233A (5971A>G 5972A>C), Q243I (6001C>A 6002A>T), R246K (6011G>A 6012G>A), K249R (6020A>G), G250E (6023G>A 6024G>A), T251I (6026C>T 6027A>T), Y254F (6035A>T 6036C>T), V255I (6037G>A), R256K (6040C>A 6041G>A), F257C (6044T>G 6045C>T), S259T (6049T>A 6051G>A), T261P (6055A>C), E267D (6075A>T), I272V (6088A>G 6090C>T), G279* (6109G>T), E283D (6123A>T), P287A (6133C>G), K289D (6139A>G 6141G>T), E291G (6146A>G), E292Q (6148G>C), E294P (6154G>C 6155A>C), E297P (6163G>C 6164A>C), V299I (6169G>A 6171A>T), N300E (6172A>G 6174T>G), E303S (6181G>A 6182A>G 6183A>T), A306T (6190G>A 6192T>C), K308N (6198A>C), I313L (6211A>C), S314A (6214T>G), L317I (6223C>A 6225A>C), A321G (6236C>G), L325T (6247C>A 6248T>C), Q327G (6253C>G 6254A>G 6255G>A), A329L (6259G>C 6260C>T), I330M (6264T>G), S332I (6269G>T 6270C>T), S335T (6277T>A 6279A>C), E337D (6285A>T), T339I (6290C>T), C345S (6308G>C 6309C>A), D347E (6315T>A), I348M (6318A>G), E350K (6322G>A), A351Q (6325G>C 6326C>A), E354K (6334G>A), L356A (6340C>G 6341T>C), V360I (6352G>A 6354C>A), L361M (6355T>A 6357A>G), S362T (6359G>C 6360T>A), K365N (6369A>T), P366L (6371C>T), Q368R (6376C>A 6377A>G), P369Q (6380C>A 6381T>A), T370P (6382A>C 6384A>T), R376K (6401G>A 6402G>A), R377M (6404G>T), P382E (6418C>G 6419C>A), D383G (6422A>G 6423T>A), L385F (6429A>T), L391D (6445C>G 6446T>A 6447A>T), S393G (6451A>G 6453T>A), H394Q (6456C>G), P397Q (6464C>A), D398_H399del (6466_6473delGACCACCT), K405N (6489A>C) | | | | | | | | |
| Codon mutations: | TTG200CTA (5872T>C 5874G>A), GCT202GCA (5880T>A), TTA203CAA (5881T>C 5882T>A), AAT205AAA (5889T>A), ACA206ACT (5892A>T), AGT207AAT (5894G>A), GCA208GCC (5898A>C), AAA209AAT (5901A>T), GTA211GTT (5907A>T), GCC212AGA (5908G>A 5909C>G 5910C>A), ACC213ACA (5913C>A), TGC214TGT (5916C>T), TAC215TAT (5919C>T), AAC216AGC (5921A>G), CTA220TTA (5932C>T), GAT221AAT (5935G>A), ACT222ACC (5940T>C), ACA225ACC (5949A>C), CAG226TAT (5950C>T 5952G>T), ACA227GAC (5953A>G 5954C>A 5955A>C), CAA229GGA (5959C>G 5960A>G), GAG230GAA (5964G>A), GCC232TCA (5968G>T 5970C>A), AAT233GCT (5971A>G 5972A>C), CCT235CCA (5979T>A), GAG236GAA (5982G>A), TTA237CTA (5983T>C), TAC238TAT (5988C>T), AAG239AAA (5991G>A), GCA240GCC (5994A>C), TTC241TTT (5997C>T), CAA243ATA (6001C>A 6002A>T), TAT244TAC (6006T>C), AAG245AAA (6009G>A), AGG246AAA (6011G>A 6012G>A), ATA247ATT (6015A>T), ACT248ACA (6018T>A), AAA249AGA (6020A>G), GGG250GAA (6023G>A 6024G>A), ACA251ATT (6026C>T 6027A>T), TTC253TTT (6033C>T), TAC254TTT (6035A>T 6036C>T), GTA255ATA (6037G>A), CGA256AAA (6040C>A 6041G>A), TTC257TGT (6044T>G 6045C>T), TCG259ACA (6049T>A 6051G>A), GCT260GCC (6054T>C), ACA261CCA (6055A>C), CTA265TTA (6067C>T), GAA267GAT (6075A>T), GAG268GAA (6078G>A), AAG270AAA (6084G>A), CCT271CCA (6087T>A), ATC272GTT (6088A>G 6090C>T), CAA274CAG (6096A>G), GTT275GTG (6099T>G), AAA277AAG (6105A>G), ATT278ATA (6108T>A), GGA279TGA (6109G>T), CTC280TTG (6112C>T 6114C>G), AGG282AGA (6120G>A), GAA283GAT (6123A>T), ATC285ATT (6129C>T), CCA287GCA (6133C>G), AAG289GAT (6139A>G 6141G>T), ATA290ATT (6144A>T), GAA291GGA (6146A>G), GAA292CAA (6148G>C), GAA294CCA (6154G>C 6155A>C), GAA297CCA (6163G>C 6164A>C), GTA299ATT (6169G>A 6171A>T), AAT300GAG (6172A>G 6174T>G), ATT301ATA (6177T>A), GAA303AGT (6181G>A 6182A>G 6183A>T), TTC304TTT (6186C>T), TAT305TAC (6189T>C), GCT306ACC (6190G>A 6192T>C), AAA308AAC (6198A>C), AGG309AGA (6201G>A), ATA310ATT (6204A>T), ATA313CTA (6211A>C), TCT314GCT (6214T>G), ACT315ACA (6219T>A), CTA317ATC (6223C>A 6225A>C), CTA320TTA (6232C>T), GCA321GGA (6236C>G), AAC322AAT (6240C>T), TAT324TAC (6246T>C), CTA325ACA (6247C>A 6248T>C), CAG327GGA (6253C>G 6254A>G 6255G>A), AAT328AAC (6258T>C), GCA329CTA (6259G>C 6260C>T), ATT330ATG (6264T>G), AGC332ATT (6269G>T 6270C>T), TCA335ACC (6277T>A 6279A>C), AGG336AGA (6282G>A), GAA337GAT (6285A>T), CAG338CAA (6288G>A), ACA339ATA (6290C>T), TGC345TCA (6308G>C 6309C>A), GAT347GAA (6315T>A), ATA348ATG (6318A>G), AGA349CGA (6319A>C), GAA350AAA (6322G>A), GCA351CAA (6325G>C 6326C>A), GAA354AAA (6334G>A), CTA356GCA (6340C>G 6341T>C), AGA357AGG (6345A>G), GTC360ATA (6352G>A 6354C>A), TTA361ATG (6355T>A 6357A>G), AGT362ACA (6359G>C 6360T>A), CTT363CTG (6363T>G), AAA365AAT (6369A>T), CCA366CTA (6371C>T), CAA368AGA (6376C>A 6377A>G), CCT369CAA (6380C>A 6381T>A), ACA370CCT (6382A>C 6384A>T), AGG376AAA (6401G>A 6402G>A), AGG377ATG (6404G>T), AAT378-GA (6406delA 6407A>G 6408T>A), TTC379TTT (6411C>T), ATC380ATT (6414C>T), TCC381TCA (6417C>A), CCA382GAA (6418C>G 6419C>A), GAT383GGA (6422A>G 6423T>A), CTG384TTA (6424C>T 6426G>A), TTA385TTT (6429A>T), AAG390AAA (6444G>A), CTA391GAT (6445C>G 6446T>A 6447A>T), AGT393GGA (6451A>G 6453T>A), CAC394CAG (6456C>G), TAT396TAC (6462T>C), CCA397CAA (6464C>A), GAC398_CAC399del (6466_6473delGACCACCT), CTA400--A (6466_6473delGACCACCT), TGC401TGT (6477C>T), TCA402TCT (6480A>T), AAA405AAC (6489A>C) | | | | | | | | |

*: Inserts / Deletes / Misaligned / Frameshifts

#### Analysis details

This analysis was performed with panviral2.64

### NGS Details (UN60): Macrosiphum euphorbiae virus 1

#### Assembly

| | |
|---|---|
| **Coverage Length** | 5542 (29 contig(s)) |
| **Depth Of Coverage** | 2.6 |
| **Number Of Reads** | 106 |
| **Reads Per Million** | 2.25 rpm (after QC) |
| **Ambiguities** | 3 |
| **Assembly Method** | de novo + reference guided assembly |
| **Consensus Caller** | Bcf Tools |

#### Coverage Map

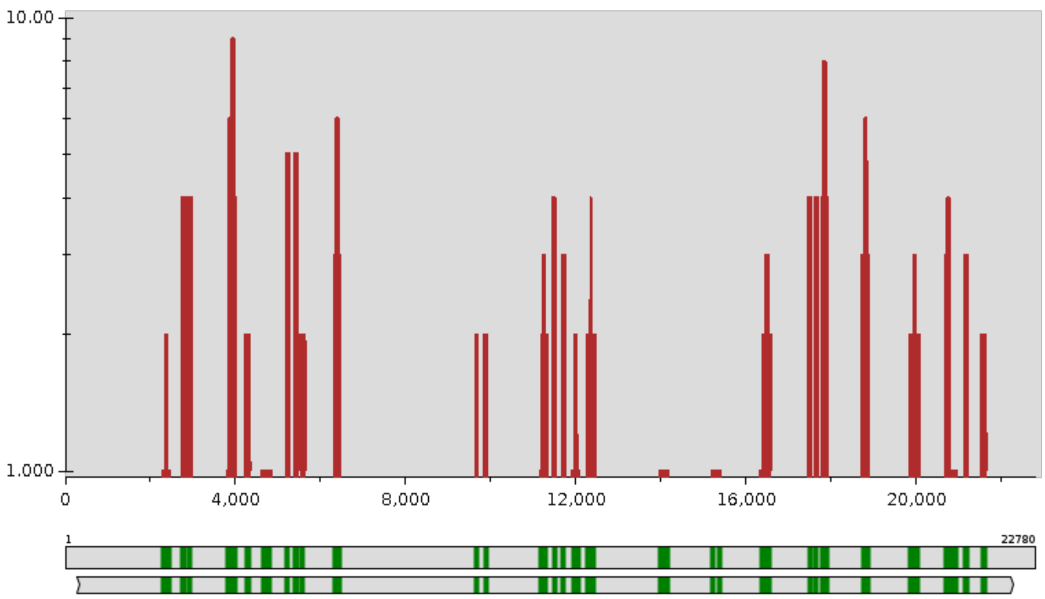

#### Assignment

| | |
|---|---|
| **Type** | Macrosiphum euphorbiae virus 1 (Taxonomy ID: 1685953) |
| **Reference Genome** | NC_028137.1 |
| **NT Identity (%)** | 92.8378 |
| **AA Identity (%)** | 96.4655 |
| **Number Of Stop Codons** | 0 |
| **Number Of CDS** | 1 |

#### Alignment

| | |
|---|---|
| **Alignment Score** | 9449.0 (NT) + 12248.0 (AA) = 21697.0 |
| **Concordance (%)** | 91.9092 |

| Alignment Method | Global, seeded, nucleotide + amino acids (AGA) |
|---|---|

#### Genome Region

Sequence starts at position 2251 and ends at position 21653 relative to NC_028137.1 reference sequence.

#### Alignment Detailed Statistics

| | Begin | End | Coverage | Score | Concordance | Matches | Identities | I/D/M/F* | Stop Codons |
|---|---|---|---|---|---|---|---|---|---|
| **NT** | **2251** | **21653** | **24.3%** | **9449** | **85.7%** | **5529 (99.8%)** | **5133 (92.6%)** | **0/13** | |
| Mutations: | 2262C>T, 2277G>A, 2279G>T, 2282C>T, 2291T>C, 2303T>C, 2312C>A, 2318T>C, 2321A>G, 2324T>C, 2327C>T, 2336T>G, 2343A>G, 2344G>A, 2354T>C, 2363T>C, 2369T>C, 2381A>G, 2399T>C, 2405T>C, 2411C>T, 2420A>G, 2429C>G, 2432A>G, 2438A>G, 2456T>C, 2465A>G, 2466C>T, 2471C>T, 2474C>T, 2475C>T, 2480T>C, 2711A>G, 2717T>C, 2720G>T, 2735C>T, 2738T>A, 2747G>A, 2748G>A, 2759C>T, 2762G>A, 2765C>T, 2769T>G, 2771T>C, 2774G>A, 2777T>C, 2780A>T, 2783A>G, 2789A>G, 2798A>G, 2802C>T, 2804A>G, 2808A>G, 2810T>G, 2825G>A, 2831G>T, 2836G>A, 2837C>T, 2855C>A, 2867A>C, 2870A>T, 2871G>A, 2872T>C, 2876A>G, 2879T>C, 2881G>K, 2882G>C, 2883C>A, 2885C>G, 2888G>A, 2895C>G, 2900A>G, 2903C>A, 2904A>G, 2906C>T, 2912A>C, 2918A>G, 2930G>A, 2939A>G, 2942T>C, 2945A>G, 2948C>T, 2950T>C, 2951T>A, 2954G>T, 2957C>T, 2965A>T, 2972C>T, 3770G>A, 3776G>C, 3779A>T, 3782C>T, 3785T>C, 3788A>G, 3791A>G, 3794C>T, 3797T>C, 3806C>T, 3815T>C, 3821C>T, 3824C>T, 3827C>T, 3836C>T, 3842C>T, 3845G>A, 3860T>C, 3866C>T, 3872T>C, 3884C>T, 3885A>G, 3886C>T, 3887A>G, 3888C>T, 3890T>A, 3893C>T, 3896A>G, 3905A>G, 3911T>C, 3914G>A, 3916A>T, 3917G>A, 3920A>G, 3925A>G, 3929A>G, 3931T>G, 3932T>C, 3938C>T, 3947T>C, 3953T>C, 3956C>T, 3959T>C, 3962A>G, 3963C>T, 3971C>T, 3974G>T, 3983C>T, 3992G>C, 4004A>G, 4007A>G, 4008T>C, 4016T>C, 4022T>C, 4031T>C, 4034T>C, 4208T>C, 4212C>T, 4217T>C, 4223G>A, 4232T>C, 4237T>C, 4238C>T, 4247C>T, 4250A>G, 4259C>A, 4265C>T, 4268T>C, 4271C>T, 4289G>A, 4304T>C, 4316T>C, 4334T>C, 4337T>A, 4340C>T, 4343A>G, 4346G>A, 4355G>A, 4358G>A, 4598T>A, 4601A>G, 4607T>C, 4610A>T, 4620C>T, 4624A>G, 4625C>T, 4628T>C, 4631C>T, 4634G>A, 4638A>C, 4643A>G, 4661T>C, 4667A>G, 4673T>C, 4674C>T, 4676A>G, 4677A>G, 4685T>C, 4691C>T, 4694T>C, 4700T>C, 4705C>T, 4751T>C, 4784T>C, 4786C>T, 4807G>A, 4828A>G, 4838C>T, 4854T>C, 4855T>C, 5156C>T, 5165T>C, 5166G>A, 5174C>T, 5183T>C, 5203T>G, 5210A>G, 5217T>C, 5228C>T, 5246C>T, 5264C>T, 5285C>T, 5358G>A, 5363G>A, 5366G>A, 5390A>C, 5397T>C, 5411T>C, 5414T>C, 5417C>T, 5438A>G, 5439C>T, 5450C>T, 5454G>A, 5459T>C, 5462A>G, 5474T>C, 5489A>G, 5507C>T, 5528C>T, 5537C>T, 5540C>T, 5543G>A, 5546T>C, 5561C>T, 5579G>A, 5582A>G, 5591T>C, 5612A>T, 5621T>C, 5636T>C, 5640G>A, 5642C>T, 5645A>G, 6296C>T, 6302A>C, 6303A>G, 6309A>G, 6326C>T, 6344C>G, 6347T>C, 6356C>T, 6365T>C, 6371C>T, 6377T>C, 6401G>A, 6402C>T, 6449T>C, 6458G>A, 6488T>A, 9674T>C, 9701T>C, 9720G>K, 9823A>W, 9878G>A, 9910T>C, 11189T>C, 11231T>C, 11296T>G, 11513C>T, 11549T>C, 11555C>T, 11646T>C, 11681T>C, 11756G>C, 11915C>T, 11922A>G, 12257T>C, 12350C>T, 12383G>A, 12443T>C, 13928T>C, 13940T>C, 13946A>G, 13955T>A, 14058_14064delGAAGATA, 14084C>T, 15161T>C, 15183C>T, 15317T>C, 16307T>C, 16349C>T, 16369A>G, 16376C>T, 16430T>C, 16481C>T, 16523G>T, 16553T>C, 16577T>C, 17471T>C, 17516C>T, 17635G>A, 17684T>C, 17838T>G, 17912T>C, 17942C>T, 18701T>C, 18722C>T, 18743A>C, 18752A>G, 18755T>G, 18758C>T, 18761T>C, 18764G>A, 18794C>T, 18797G>A, 18833C>T, 18864T>C, 18878T>C, 18887T>C, 18890A>G, 18896G>T, 18905G>A, 18908C>T, 18920G>A, 19808A>G, 19818G>T, 19829C>T, 19835T>C, 19859C>T, 19904T>C, 19907G>T, 19926C>A, 19961C>T, 19982C>T, 20000T>C, 20036C>T, 20078T>C, 20082G>A, 20083T>C, 20084T>G, 20648T>C, 20654C>T, 20663G>A, 20666T>C, 20672G>A, 20675C>T, 20711T>C, 20716A>G, 20720A>G, 20732T>C, 20735T>C, 20756T>C, 20759T>C, 20771T>A, 20780C>T, 20783T>C, 20786C>T, 20807G>A, 20810T>C, 20816T>C, 20822C>T, 20828G>A, 20840T>C, 20844_20849delTCCATT, 20855G>A, 20867A>C, 20873C>T, 20876G>A, 20882A>G, 20891A>G, 20909C>T, 20912T>C, 20924T>C, 20939T>G, 21122T>C, 21137T>C, 21143T>C, 21156T>C, 21158G>T, 21164C>T, 21170C>T, 21179C>T, 21182C>T, 21185A>G, 21188T>C, 21191A>G, 21194C>T, 21200C>T, 21210A>G, 21218G>A, 21503T>C, 21518C>T, 21525A>G, 21526G>A, 21599C>T, 21617T>C, 21618G>A, 21635C>T, 21647G>A | | | | | | | | |
| **CDS** | | | | | | | | | |
| ASK90_gp1 | 655 | 7121 | 25.1% | 12248 | 94.7% | 1839 (99.8%) | 1774 (96.3%) | 0/4/4/1 | 0 |
| Protein mutations: | V663I (2277G>A 2279G>T), S685D (2343A>G 2344G>A), A820T (2748G>A), S827A (2769T>G 2771T>C), I840V (2808A>G 2810T>G), R849H (2836G>A 2837C>T), V861T (2871G>A 2872T>C), Q869E (2895C>G), I872V (2904A>G 2906C>T), V887A (2950T>C 2951T>A), N892I (2965A>T), T1199V (3885A>G 3886C>T 3887A>G), Q1209L (3916A>T 3917G>A), N1212S (3925A>G), V1214G (3931T>G 3932T>C), L1308F (4212C>T), V1316A (4237T>C 4238C>T), E1440D (4610A>T), H1444Y (4620C>T), D1445G (4624A>G 4625C>T), M1450L (4638A>C), K1463E (4677A>G), A1472V (4705C>T), T1499I (4786C>T), R1506Q (4807G>A), D1513G (4828A>G), L1522P (4854T>C 4855T>C), D1626N (5166G>A), V1638G (5203T>G), A1690T (5358G>A), V1722I (5454G>A), V1784I (5640G>A 5642C>T), I2005V (6303A>G), I2007V (6309A>G), V3207A (9910T>C), I3669R (11296T>G), C3786R (11646T>C), I3878V (11922A>G), E4590_D4591del (14058_14064delGAAGATA), D5360G (16369A>G), R5782K (17635G>A), S5850A (17838T>G), V6510L (19818G>T), K6539N (19907G>T), V6598T (20082G>A 20083T>C 20084T>G), K6809R (20716A>G), S6852_I6853del (20844_20849delTCCATT), I6974V (21210A>G), S7079D (21525A>G 21526G>A), D7110N (21618G>A) | | | | | | | | |

| | Begin | End | Coverage | Score | Concordance | Matches | Identities | I/D/M/F* | Stop Codons |
|---|---|---|---|---|---|---|---|---|---|
| **NT** | **2251** | **21653** | **24.3%** | **9449** | **85.7%** | **5529 (99.8%)** | **5133 (92.6%)** | **0/13** | |
| Codon mutations: | CTG658TTG (2262C>T), GTG663ATT (2277G>A 2279G>T), TTC664TTT (2282C>T), GCT667GCC (2291T>C), GTT671GTC (2303T>C), TCC674TCA (2312C>A), GCT676GCC (2318T>C), AAA677AAG (2321A>G), TAT678TAC (2324T>C), CTC679CTT (2327C>T), ACT682ACG (2336T>G), AGC685GAC (2343A>G 2344G>A), CCT688CCC (2354T>C), ATT691ATC (2363T>C), GAT693GAC (2369T>C), GAA697GAG (2381A>G), CGT703CGC (2399T>C), CTT705CTC (2405T>C), TAC707TAT (2411C>T), GCA710GCG (2420A>G), TCC713TCG (2429C>G), AAA714AAG (2432A>G), AAA716AAG (2438A>G), ACT722ACC (2456T>C), CAA725CAG (2465A>G), CTG726TTG (2466C>T), ATC727ATT (2471C>T), GGC728GGT (2474C>T), CTA729TTA (2475C>T), GCT730GCC (2480T>C), GGA807GGG (2711A>G), TCT809TCC (2717T>C), TCG810TCT (2720G>T), TAC815TAT (2735C>T), GGT816GGA (2738T>A), CCG819CCA (2747G>A), GCC820ACC (2748G>A), CAC823CAT (2759C>T), GGG824GGA (2762G>A), TTC825TTT (2765C>T), TCT827GCC (2769T>G 2771T>C), TCG828TCA (2774G>A), CGT829CGC (2777T>C), GGA830GGT (2780A>T), GGA831GGG (2783A>G), AGA833AGG (2789A>G), GGA836GGG (2798A>G), CTA838TTG (2802C>T 2804A>G), ATT840GTG (2808A>G 2810T>G), GAG845GAA (2825G>A), GTG847GTT (2831G>T), CGC849CAT (2836G>A 2837C>T), GTC855..A (2855C>A), CCA859CCC (2867A>C), TCA860TCT (2870A>T), GTG861ACG (2871G>A 2872T>C), CCA862CCG (2876A>G), ATT863ATC (2879T>C), CGG864CKC (2881G>K 2882G>C), CGC865AGG (2883C>A 2885C>G), CTG866CTA (2888G>A), CAG869GAG (2895C>G), AAA870AAG (2900A>G), CCC871CCA (2903C>A), ATC872GTT (2904A>G 2906C>T), TCA874TCC (2912A>C), GAA876GAG (2918A>G), GAG880GAA (2930G>A), AAA883AAG (2939A>G), ACT884ACC (2942T>C), TTA885TTG (2945A>G), AAC886AAT (2948C>T), GTT887GCA (2950T>C 2951T>A), GGG888GGT (2954G>T), GTC889GTT (2957C>T), AAC892ATC (2965A>T), TCC894TCT (2972C>T), TTG1160TTA (3770G>A), GGG1162GGC (3776G>C), GCA1163GCT (3779A>T), GCC1164GCT (3782C>T), TTT1165TTC (3785T>C), GGA1166GGG (3788A>G), CAA1167CAG (3791A>G), TGC1168TGT (3794C>T), TTT1169TTC (3797T>C), TTC1172TTT (3806C>T), TCT1175TCC (3815T>C), TTC1177TTT (3821C>T), ATC1178ATT (3824C>T), AGC1179AGT (3827C>T), GTC1182GTT (3836C>T), TGC1184TGT (3842C>T), GGG1185GGA (3845G>A), CTT1190CTC (3860T>C), GAC1192GAT (3866C>T), AAT1194AAC (3872T>C), ACC1198ACT (3884C>T), ACA1199GTG (3885A>G 3886C>T 3887A>G), CTT1200TTA (3888C>T 3890T>A), AGC1201AGT (3893C>T), GAA1202GAG (3896A>G), AGA1205AGG (3905A>G), TGT1207TGC (3911T>C), TTG1208TTA (3914G>A), CAG1209CTA (3916A>T 3917G>A), ACA1210ACG (3920A>G), AAC1212AGC (3925A>G), ACA1213ACG (3929A>G), GTT1214GGC (3931T>G 3932T>C), CGC1216CGT (3938C>T), TAT1219TAC (3947T>C), GTT1221GTC (3953T>C), AGC1222AGT (3956C>T), AAT1223AAC (3959T>C), AAA1224AAG (3962A>G), CTG1225TTG (3963C>T), GAC1227GAT (3971C>T), GTG1228GTT (3974G>T), CCC1231CCT (3983C>T), GTG1234GTC (3992G>C), AAA1238AAG (4004A>G), GAA1239GAG (4007A>G), TTG1240CTG (4008T>C), ACT1242ACC (4016T>C), GCT1244GCC (4022T>C), GAT1247GAC (4031T>C), TAT1248TAC (4034T>C), GTT1306GTC (4208T>C), CTT1308TTT (4212C>T), TGT1309TGC (4217T>C), GTG1311GTA (4223G>A), AAT1314AAC (4232T>C), GTC1316GCT (4237T>C 4238C>T), GTC1319GTT (4247C>T), GAA1320GAG (4250A>G), CCC1323CCA (4259C>A), GAC1325GAT (4265C>T), ATT1326ATC (4268T>C), GTC1327GTT (4271C>T), GAG1333GAA (4289G>A), GCT1338GCC (4304T>C), GAT1342GAC (4316T>C), GCT1348GCC (4334T>C), GCT1349GCA (4337T>A), GCC1350GCT (4340C>T), TTA1351TTG (4343A>G), AAG1352AAA (4346G>A), GCG1355GCA (4355G>A), GTG1356GTA (4358G>A), TCT1436TCA (4598T>A), AGA1437AGG (4601A>G), GCT1439GCC (4607T>C), GAA1440GAT (4610A>T), CAT1444TAT (4620C>T), GAC1445GGT (4624A>G 4625C>T), GAT1446GAC (4628T>C), CCC1447CCT (4631C>T), GAG1448GAA (4634G>A), ATG1450CTG (4638A>C), TCA1451TCG (4643A>G), GCT1457GCC (4661T>C), CAA1459CAG (4667A>G), AAT1461AAC (4673T>C), CTA1462TTG (4674C>T 4676A>G), AAA1463GAA (4677A>G), CGT1465CGC (4685T>C), GAC1467GAT (4691C>T), GAT1468GAC (4694T>C), ATT1470ATC (4700T>C), GCT1472GTT (4705C>T), GCT1487GCC (4751T>C), GAT1498GAC (4784T>C), ACT1499ATT (4786C>T), CGA1506CAA (4807G>A), GAC1513GGC (4828A>G), ACC1516ACT (4838C>T), TTG1522CCG (4854T>C 4855T>C), TTC1622TTT (5156C>T), AAT1625AAC (5165T>C), GAC1626AAC (5166G>A), TAC1628TAT (5174C>T), GCT1631GCC (5183T>C), GTC1638GGC (5203T>G), TTA1640TTG (5210A>G), TTG1643CTG (5217T>C), GCC1646GCT (5228C>T), AAC1652AAT (5246C>T), ATC1658ATT (5264C>T), GGC1665GGT (5285C>T), GCC1690ACC (5358G>A), GAG1691GAA (5363G>A), AAG1692AAA (5366G>A), TCA1700TCC (5390A>C), TTA1703CTA (5397T>C), AAT1707AAC (5411T>C), AGT1708AGC (5414T>C), GAC1709GAT (5417C>T), CTA1716CTG (5438A>G), CTG1717TTG (5439C>T), GAC1720GAT (5450C>T), GTT1722ATT (5454G>A), TAT1723TAC (5459T>C), GAA1724GAG (5462A>G), GGT1728GGC (5474T>C), ACA1733ACG (5489A>G), TCC1739.CT (5507C>T), CTC1746CTT (5528C>T), GCC1749GCT (5537C>T), AAC1750AAT (5540C>T), TCG1751TCA (5543G>A), GCT1752GCC (5546T>C), CGC1757CGT (5561C>T), GTG1763GTA (5579G>A), GAA1764GAG (5582A>G), TCT1767TCC (5591T>C), GGA1774GGT (5612A>T), AGT1777AGC (5621T>C), GGT1782GGC (5636T>C), GTC1784ATT (5640G>A 5642C>T), GCA1785GCG (5645A>G), CGC2002CGT (6296C>T), CGA2004CGC (6302A>C), ATT2005GTT (6303A>G), ATT2007GTT (6309A>G), GCC2012GCT (6326C>T), TCC2018TCG (6344C>G), TAT2019TAC (6347T>C), AAC2022AAT (6356C>T), AAT2025AAC (6365T>C), TTC2027TTT (6371C>T), CAT2029CAC (6377T>C), AAG2037AAA (6401G>A), CTG2038TTG (6402C>T), GAT2053GAC (6449T>C), AAG2056AAA (6458G>A), ATT2066ATA (6488T>A), TCT3128TCC (9674T>C), CGT3137CGC (9701T>C), GCC3144KCC (9720G>K), AAA3178AWA (9823A>W), CAG3196CAA (9878G>A), GTT3207GCT (9910T>C), TTT3633TTC (11189T>C), AAT3647AAC (11231T>C), ATA3669AGA (11296T>G), GAC3741GAT (11513C>T), TTT3753TTC (11549T>C), AAC3755AAT (11555C>T), TGC3786CGC (11646T>C), AGT3797AGC (11681T>C), CCG3822CCC (11756G>C), TCC3875TCT (11915C>T), ATA3878GTA (11922A>G), GCT3989GCC (12257T>C), CAC4020CAT (12350C>T), AGG4031AGA (12383G>A), TTT4051TTC (12443T>C), AGT4546AGC (13928T>C), TTT4550TTC (13940T>C), GTA4552GTG (13946A>G), GTT4555GTA (13955T>A), GAA4590_GAT4591del (14058_14064delGAAGATA), ATG4592-TG (14058_14064delGAAGATA), TAC4598TAT (14084C>T), GTT4957GTC (15161T>C), CTG4965TTG (15183C>T), GAT5009GAC (15317T>C), GTT5339GTC (16307T>C), GAC5353GAT (16349C>T), GAT5360GGT (16369A>G), TAC5362TAT (16376C>T), AAT5380AAC (16430T>C), GTC5397GTT (16481C>T), GCG5411GCT (16523G>T), CTT5421CTC (16553T>C), GAT5429GAC (16577T>C), GAT5727GAC (17471T>C), AAC5742AAT (17516C>T), AGG5782AAG (17635G>A), CAT5798CAC (17684T>C), TCT5850GCT (17838T>G), GCT5874GCC (17912T>C), CAC5884CAT (17942C>T), AGT6137AGC (18701T>C), GAC6144GAT (18722C>T), ACA6151ACC (18743A>C), GAA6154GAG (18752A>G), GTT6155GTG (18755T>G), AGC6156AGT (18758C>T), GAT6157GAC (18761T>C), GAG6158GAA (18764G>A), AAC6168AAT (18794C>T), AAG6169AAA (18797G>A), TCC6181TCT (18833C>T), TTG6192CTG (18864T>C), GAT6196GAC (18878T>C), AAT6199AAC (18887T>C), GTA6200GTG (18890A>G), GCG6202GCT (18896G>T), GCG6205GCA (18905G>A), GCC6206GCT (18908C>T), CAG6210CAA (18920G>A), CCA6506CCG (19808A>G), GTG6510TTG (19818G>T), GAC6513GAT (19829C>T), GAT6515GAC (19835T>C), CGC6523CGT (19859C>T), TCT6538TCC (19904T>C), AAG6539AAT (19907G>T), CGA6546AGA (19926C>A), AGC6557AGT (19961C>T), GGC6564GGT (19982C>T), GTT6570GTC (20000T>C), AAC6582AAT (20036C>T), AAT6596AAC (20078T>C), GTT6598ACG (20082G>A 20083T>C 20084T>G), ATT6786.TC (20648T>C), GGC6788GGT (20654C>T), CCG6791CCA (20663G>A), CAT6792CAC (20666T>C), CAG6794CAA (20672G>A), GGC6795GGT (20675C>T), CCT6807CCC (20711T>C), AAG6809AGG (20716A>G), AAA6810AAG (20720A>G), CAT6814CAC (20732T>C), GTT6815GTC (20735T>C), GAT6822GAC (20756T>C), ATT6823ATC (20759T>C), CCT6827CCA (20771T>A), GGC6830GGT (20780C>T), TAT6831TAC (20783T>C), CTC6832CTT (20786C>T), AAG6839AAA (20807G>A), ATT6840ATC (20810T>C), TCT6842TCC (20816T>C), GAC6844GAT (20822C>T), GAG6846GAA (20828G>A), GGT6850GGC (20840T>C), TCC6852_ATT6853del (20844_20849delTCCATT), GAG6855GAA (20855G>A), ACA6859ACC (20867A>C), AAC6861AAT (20873C>T), GAG6862GAA (20876G>A), CAA6864CAG (20882A>G), TTA6867TTG (20891A>G), ATC6873ATT (20909C>T), GAT6874GAC (20912T>C), CTT6878CTC (20924T>C), ACT6883ACG (20939T>G), AAT6944AAC (21122T>C), ACT6949ACC (21137T>C), AAT6951AAC (21143T>C), TTG6956CTT (21156T>C 21158G>T), CAC6958CAT (21164C>T), CTC6960CTT (21170C>T), AGC6963AGT (21179C>T), ATC6964ATT (21182C>T), GAA6965GAG (21185A>G), GAT6966GAC (21188T>C), CTA6967CTG (21191A>G), ACC6968ACT (21194C>T), ACC6970ACT (21200C>T), ATC6974GTC (21210A>G), GTG6976GTA (21218G>A), TAT7071TAC (21503T>C), ATC7076ATT (21518C>T), AGC7079GAC (21525A>G 21526G>A), CTC7103CTT (21599C>T), AAT7109AAC (21617T>C), GAC7110AAC (21618G>A), TTC7115TTT (21635C>T), AGG7119AGA (21647G>A) | | | | | | | | |

#### Proteins

| | Begin | End | Coverage | Score | Concordance | Matches | Identities | I/D/M/F* | Stop Codons |
|---|---|---|---|---|---|---|---|---|---|
| polyprotein (YP_009175071.1) | 655 | 7121 | 25.1% | 12248 | 94.7% | 1839 (99.8%) | 1774 (96.3%) | 0/4/4/1 | 0 |
| Protein mutations: | V663I (2277G>A 2279G>T), S685D (2343A>G 2344G>A), A820T (2748G>A), S827A (2769T>G 2771T>C), I840V (2808A>G 2810T>G), R849H (2836G>A 2837C>T), V861T (2871G>A 2872T>C), Q869E (2895C>G), I872V (2904A>G 2906C>T), V887A (2950T>C 2951T>A), N892I (2965A>T), T1199V (3885A>G 3886C>T 3887A>G), Q1209L (3916A>T 3917G>A), N1212S (3925A>G), V1214G (3931T>G 3932T>C), L1308F (4212C>T), V1316A (4237T>C 4238C>T), E1440D (4610A>T), H1444Y (4620C>T), D1445G (4624A>G 4625C>T), M1450L (4638A>C), K1463E (4677A>G), A1472V (4705C>T), T1499I (4786C>T), R1506Q (4807G>A), D1513G (4828A>G), L1522P (4854T>C 4855T>C), D1626N (5166G>A), V1638G (5203T>G), A1690T (5358G>A), V1722I (5454G>A), V1784I (5640G>A 5642C>T), I2005V (6303A>G), I2007V (6309A>G), V3207A (9910T>C), I3669R (11296T>G), C3786R (11646T>C), I3878V (11922A>G), E4590_D4591del (14058_14064delGAAGATA), D5360G (16369A>G), R5782K (17635G>A), S5850A (17838T>G), V6510L (19818G>T), K6539N (19907G>T), V6598T (20082G>A 20083T>C 20084T>G), K6809R (20716A>G), S6852_I6853del (20844_20849delTCCATT), I6974V (21210A>G), S7079D (21525A>G 21526G>A), D7110N (21618G>A) | | | | | | | | |

| | Begin | End | Coverage | Score | Concordance | Matches | Identities | I/D/M/F* | Stop Codons |
|---|---|---|---|---|---|---|---|---|---|
| **NT** | **2251** | **21653** | **24.3%** | **9449** | **85.7%** | **5529 (99.8%)** | **5133 (92.6%)** | **0/13** | |
| Codon mutations: | CTG658TTG (2262C>T), GTG663ATT (2277G>A 2279G>T), TTC664TTT (2282C>T), GCT667GCC (2291T>C), GTT671GTC (2303T>C), TCC674TCA (2312C>A), GCT676GCC (2318T>C), AAA677AAG (2321A>G), TAT678TAC (2324T>C), CTC679CTT (2327C>T), ACT682ACG (2336T>G), AGC685GAC (2343A>G 2344G>A), CCT688CCC (2354T>C), ATT691ATC (2363T>C), GAT693GAC (2369T>C), GAA697GAG (2381A>G), CGT703CGC (2399T>C), CTT705CTC (2405T>C), TAC707TAT (2411C>T), GCA710GCG (2420A>G), TCC713TCG (2429C>G), AAA714AAG (2432A>G), AAA716AAG (2438A>G), ACT722ACC (2456T>C), CAA725CAG (2465A>G), CTG726TTG (2466C>T), ATC727ATT (2471C>T), GGC728GGT (2474C>T), CTA729TTA (2475C>T), GCT730GCC (2480T>C), GGA807GGG (2711A>G), TCT809TCC (2717T>C), TCG810TCT (2720G>T), TAC815TAT (2735C>T), GGT816GGA (2738T>A), CCG819CCA (2747G>A), GCC820ACC (2748G>A), CAC823CAT (2759C>T), GGG824GGA (2762G>A), TTC825TTT (2765C>T), TCT827GCC (2769T>G 2771T>C), TCG828TCA (2774G>A), CGT829CGC (2777T>C), GGA830GGT (2780A>T), GGA831GGG (2783A>G), AGA833AGG (2789A>G), GGA836GGG (2798A>G), CTA838TTG (2802C>T 2804A>G), ATT840GTG (2808A>G 2810T>G), GAG845GAA (2825G>A), GTG847GTT (2831G>T), CGC849CAT (2836G>A 2837C>T), GTC855..A (2855C>A), CCA859CCC (2867A>C), TCA860TCT (2870A>T), GTG861ACG (2871G>A 2872T>C), CCA862CCG (2876A>G), ATT863ATC (2879T>C), CGG864CKC (2881G>K 2882G>C), CGC865AGG (2883C>A 2885C>G), CTG866CTA (2888G>A), CAG869GAG (2895C>G), AAA870AAG (2900A>G), CCC871CCA (2903C>A), ATC872GTT (2904A>G 2906C>T), TCA874TCC (2912A>C), GAA876GAG (2918A>G), GAG880GAA (2930G>A), AAA883AAG (2939A>G), ACT884ACC (2942T>C), TTA885TTG (2945A>G), AAC886AAT (2948C>T), GTT887GCA (2950T>C 2951T>A), GGG888GGT (2954G>T), GTC889GTT (2957C>T), AAC892ATC (2965A>T), TCC894TCT (2972C>T), TTG1160TTA (3770G>A), GGG1162GGC (3776G>C), GCA1163GCT (3779A>T), GCC1164GCT (3782C>T), TTT1165TTC (3785T>C), GGA1166GGG (3788A>G), CAA1167CAG (3791A>G), TGC1168TGT (3794C>T), TTT1169TTC (3797T>C), TTC1172TTT (3806C>T), TCT1175TCC (3815T>C), TTC1177TTT (3821C>T), ATC1178ATT (3824C>T), AGC1179AGT (3827C>T), GTC1182GTT (3836C>T), TGC1184TGT (3842C>T), GGG1185GGA (3845G>A), CTT1190CTC (3860T>C), GAC1192GAT (3866C>T), AAT1194AAC (3872T>C), ACC1198ACT (3884C>T), ACA1199GTG (3885A>G 3886C>T 3887A>G), CTT1200TTA (3888C>T 3890T>A), AGC1201AGT (3893C>T), GAA1202GAG (3896A>G), AGA1205AGG (3905A>G), TGT1207TGC (3911T>C), TTG1208TTA (3914G>A), CAG1209CTA (3916A>T 3917G>A), ACA1210ACG (3920A>G), AAC1212AGC (3925A>G), ACA1213ACG (3929A>G), GTT1214GGC (3931T>G 3932T>C), CGC1216CGT (3938C>T), TAT1219TAC (3947T>C), GTT1221GTC (3953T>C), AGC1222AGT (3956C>T), AAT1223AAC (3959T>C), AAA1224AAG (3962A>G), CTG1225TTG (3963C>T), GAC1227GAT (3971C>T), GTG1228GTT (3974G>T), CCC1231CCT (3983C>T), GTG1234GTC (3992G>C), AAA1238AAG (4004A>G), GAA1239GAG (4007A>G), TTG1240CTG (4008T>C), ACT1242ACC (4016T>C), GCT1244GCC (4022T>C), GAT1247GAC (4031T>C), TAT1248TAC (4034T>C), GTT1306GTC (4208T>C), CTT1308TTT (4212C>T), TGT1309TGC (4217T>C), GTG1311GTA (4223G>A), AAT1314AAC (4232T>C), GTC1316GCT (4237T>C 4238C>T), GTC1319GTT (4247C>T), GAA1320GAG (4250A>G), CCC1323CCA (4259C>A), GAC1325GAT (4265C>T), ATT1326ATC (4268T>C), GTC1327GTT (4271C>T), GAG1333GAA (4289G>A), GCT1338GCC (4304T>C), GAT1342GAC (4316T>C), GCT1348GCC (4334T>C), GCT1349GCA (4337T>A), GCC1350GCT (4340C>T), TTA1351TTG (4343A>G), AAG1352AAA (4346G>A), GCG1355GCA (4355G>A), GTG1356GTA (4358G>A), TCT1436TCA (4598T>A), AGA1437AGG (4601A>G), GCT1439GCC (4607T>C), GAA1440GAT (4610A>T), CAT1444TAT (4620C>T), GAC1445GGT (4624A>G 4625C>T), GAT1446GAC (4628T>C), CCC1447CCT (4631C>T), GAG1448GAA (4634G>A), ATG1450CTG (4638A>C), TCA1451TCG (4643A>G), GCT1457GCC (4661T>C), CAA1459CAG (4667A>G), AAT1461AAC (4673T>C), CTA1462TTG (4674C>T 4676A>G), AAA1463GAA (4677A>G), CGT1465CGC (4685T>C), GAC1467GAT (4691C>T), GAT1468GAC (4694T>C), ATT1470ATC (4700T>C), GCT1472GTT (4705C>T), GCT1487GCC (4751T>C), GAT1498GAC (4784T>C), ACT1499ATT (4786C>T), CGA1506CAA (4807G>A), GAC1513GGC (4828A>G), ACC1516ACT (4838C>T), TTG1522CCG (4854T>C 4855T>C), TTC1622TTT (5156C>T), AAT1625AAC (5165T>C), GAC1626AAC (5166G>A), TAC1628TAT (5174C>T), GCT1631GCC (5183T>C), GTC1638GGC (5203T>G), TTA1640TTG (5210A>G), TTG1643CTG (5217T>C), GCC1646GCT (5228C>T), AAC1652AAT (5246C>T), ATC1658ATT (5264C>T), GGC1665GGT (5285C>T), GCC1690ACC (5358G>A), GAG1691GAA (5363G>A), AAG1692AAA (5366G>A), TCA1700TCC (5390A>C), TTA1703CTA (5397T>C), AAT1707AAC (5411T>C), AGT1708AGC (5414T>C), GAC1709GAT (5417C>T), CTA1716CTG (5438A>G), CTG1717TTG (5439C>T), GAC1720GAT (5450C>T), GTT1722ATT (5454G>A), TAT1723TAC (5459T>C), GAA1724GAG (5462A>G), GGT1728GGC (5474T>C), ACA1733ACG (5489A>G), TCC1739.CT (5507C>T), CTC1746CTT (5528C>T), GCC1749GCT (5537C>T), AAC1750AAT (5540C>T), TCG1751TCA (5543G>A), GCT1752GCC (5546T>C), CGC1757CGT (5561C>T), GTG1763GTA (5579G>A), GAA1764GAG (5582A>G), TCT1767TCC (5591T>C), GGA1774GGT (5612A>T), AGT1777AGC (5621T>C), GGT1782GGC (5636T>C), GTC1784ATT (5640G>A 5642C>T), GCA1785GCG (5645A>G), CGC2002CGT (6296C>T), CGA2004CGC (6302A>C), ATT2005GTT (6303A>G), ATT2007GTT (6309A>G), GCC2012GCT (6326C>T), TCC2018TCG (6344C>G), TAT2019TAC (6347T>C), AAC2022AAT (6356C>T), AAT2025AAC (6365T>C), TTC2027TTT (6371C>T), CAT2029CAC (6377T>C), AAG2037AAA (6401G>A), CTG2038TTG (6402C>T), GAT2053GAC (6449T>C), AAG2056AAA (6458G>A), ATT2066ATA (6488T>A), TCT3128TCC (9674T>C), CGT3137CGC (9701T>C), GCC3144KCC (9720G>K), AAA3178AWA (9823A>W), CAG3196CAA (9878G>A), GTT3207GCT (9910T>C), TTT3633TTC (11189T>C), AAT3647AAC (11231T>C), ATA3669AGA (11296T>G), GAC3741GAT (11513C>T), TTT3753TTC (11549T>C), AAC3755AAT (11555C>T), TGC3786CGC (11646T>C), AGT3797AGC (11681T>C), CCG3822CCC (11756G>C), TCC3875TCT (11915C>T), ATA3878GTA (11922A>G), GCT3989GCC (12257T>C), CAC4020CAT (12350C>T), AGG4031AGA (12383G>A), TTT4051TTC (12443T>C), AGT4546AGC (13928T>C), TTT4550TTC (13940T>C), GTA4552GTG (13946A>G), GTT4555GTA (13955T>A), GAA4590_GAT4591del (14058_14064delGAAGATA), ATG4592-TG (14058_14064delGAAGATA), TAC4598TAT (14084C>T), GTT4957GTC (15161T>C), CTG4965TTG (15183C>T), GAT5009GAC (15317T>C), GTT5339GTC (16307T>C), GAC5353GAT (16349C>T), GAT5360GGT (16369A>G), TAC5362TAT (16376C>T), AAT5380AAC (16430T>C), GTC5397GTT (16481C>T), GCG5411GCT (16523G>T), CTT5421CTC (16553T>C), GAT5429GAC (16577T>C), GAT5727GAC (17471T>C), AAC5742AAT (17516C>T), AGG5782AAG (17635G>A), CAT5798CAC (17684T>C), TCT5850GCT (17838T>G), GCT5874GCC (17912T>C), CAC5884CAT (17942C>T), AGT6137AGC (18701T>C), GAC6144GAT (18722C>T), ACA6151ACC (18743A>C), GAA6154GAG (18752A>G), GTT6155GTG (18755T>G), AGC6156AGT (18758C>T), GAT6157GAC (18761T>C), GAG6158GAA (18764G>A), AAC6168AAT (18794C>T), AAG6169AAA (18797G>A), TCC6181TCT (18833C>T), TTG6192CTG (18864T>C), GAT6196GAC (18878T>C), AAT6199AAC (18887T>C), GTA6200GTG (18890A>G), GCG6202GCT (18896G>T), GCG6205GCA (18905G>A), GCC6206GCT (18908C>T), CAG6210CAA (18920G>A), CCA6506CCG (19808A>G), GTG6510TTG (19818G>T), GAC6513GAT (19829C>T), GAT6515GAC (19835T>C), CGC6523CGT (19859C>T), TCT6538TCC (19904T>C), AAG6539AAT (19907G>T), CGA6546AGA (19926C>A), AGC6557AGT (19961C>T), GGC6564GGT (19982C>T), GTT6570GTC (20000T>C), AAC6582AAT (20036C>T), AAT6596AAC (20078T>C), GTT6598ACG (20082G>A 20083T>C 20084T>G), ATT6786.TC (20648T>C), GGC6788GGT (20654C>T), CCG6791CCA (20663G>A), CAT6792CAC (20666T>C), CAG6794CAA (20672G>A), GGC6795GGT (20675C>T), CCT6807CCC (20711T>C), AAG6809AGG (20716A>G), AAA6810AAG (20720A>G), CAT6814CAC (20732T>C), GTT6815GTC (20735T>C), GAT6822GAC (20756T>C), ATT6823ATC (20759T>C), CCT6827CCA (20771T>A), GGC6830GGT (20780C>T), TAT6831TAC (20783T>C), CTC6832CTT (20786C>T), AAG6839AAA (20807G>A), ATT6840ATC (20810T>C), TCT6842TCC (20816T>C), GAC6844GAT (20822C>T), GAG6846GAA (20828G>A), GGT6850GGC (20840T>C), TCC6852_ATT6853del (20844_20849delTCCATT), GAG6855GAA (20855G>A), ACA6859ACC (20867A>C), AAC6861AAT (20873C>T), GAG6862GAA (20876G>A), CAA6864CAG (20882A>G), TTA6867TTG (20891A>G), ATC6873ATT (20909C>T), GAT6874GAC (20912T>C), CTT6878CTC (20924T>C), ACT6883ACG (20939T>G), AAT6944AAC (21122T>C), ACT6949ACC (21137T>C), AAT6951AAC (21143T>C), TTG6956CTT (21156T>C 21158G>T), CAC6958CAT (21164C>T), CTC6960CTT (21170C>T), AGC6963AGT (21179C>T), ATC6964ATT (21182C>T), GAA6965GAG (21185A>G), GAT6966GAC (21188T>C), CTA6967CTG (21191A>G), ACC6968ACT (21194C>T), ACC6970ACT (21200C>T), ATC6974GTC (21210A>G), GTG6976GTA (21218G>A), TAT7071TAC (21503T>C), ATC7076ATT (21518C>T), AGC7079GAC (21525A>G 21526G>A), CTC7103CTT (21599C>T), AAT7109AAC (21617T>C), GAC7110AAC (21618G>A), TTC7115TTT (21635C>T), AGG7119AGA (21647G>A) | | | | | | | | |

*: Inserts / Deletes / Misaligned / Frameshifts

#### Analysis details

This analysis was performed with panviral2.64

#### NGS Details (UN60): Duamitovirus peex1

##### Assembly

| | |
|---|---|
| **Coverage Length** | 729 (2 contig(s)) |
| **Depth Of Coverage** | 4.4 |
| **Number Of Reads** | 26 |
| **Reads Per Million** | 0.55 rpm (after QC) |
| **Ambiguities** | 0 |
| **Assembly Method** | de novo + reference guided assembly |
| **Consensus Caller** | Bcf Tools |

##### Coverage Map

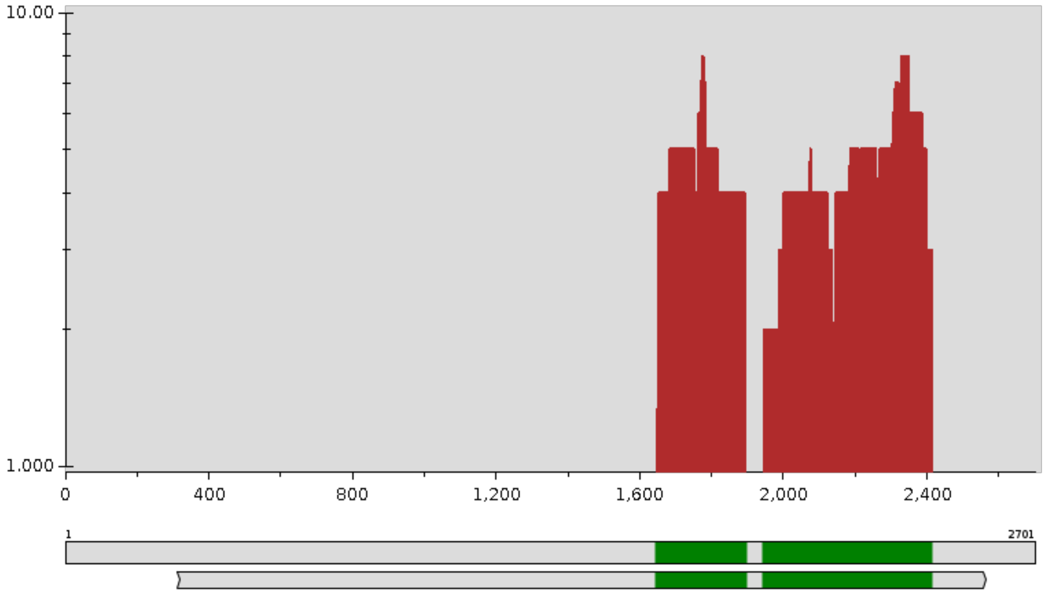

##### Assignment

| | |
|---|---|
| **Type** | Duamitovirus peex1 (Taxonomy ID: 2955799) |
| **Reference Genome** | NC_076525.1 |
| **NT Identity (%)** | 68.6813 |
| **AA Identity (%)** | 67.0782 |
| **Number Of Stop Codons** | 1 |
| **Number Of CDS** | 1 |

##### Alignment

| | |
|---|---|
| **Alignment Score** | 534.0 (NT) + 1035.0 (AA) = 1569.0 |
| **Concordance (%)** | 51.9883 |

| **Alignment Method** | Global, seeded, nucleotide + amino acids (AGA) |
|---|---|

#### Genome Region

Sequence starts at position 1645 and ends at position 2417 relative to NC_076525.1 reference sequence.

#### Alignment Detailed Statistics

| | Begin | End | Coverage | Score | Concordance | Matches | Identities | I/D/M/F* | Stop Codons |
|---|---|---|---|---|---|---|---|---|---|
| **NT** | **1645** | **2417** | **27.0%** | **534** | **36.9%** | **728 (99.9%)** | **500 (68.6%)** | **0/1** | |
| Mutations: | 1655C>G, 1658G>T, 1661A>C, 1671T>A, 1676A>G, 1680T>C, 1682G>A, 1685G>T, 1689G>A, 1691G>A, 1692A>T, 1694G>T, 1696T>C, 1697C>A, 1703G>A, 1704G>A, 1705G>C, 1706C>G, 1707C>A, 1711G>C, 1713C>A, 1714G>C, 1715G>C, 1721C>T, 1722T>G, 1723G>C, 1724G>T, 1727A>T, 1730T>C, 1736T>G, 1737C>T, 1739C>A, 1751T>C, 1754C>A, 1757T>C, 1758G>A, 1764G>A, 1765A>C, 1766G>A, 1772C>G, 1774G>C, 1775C>G, 1776A>C, 1777A>G, 1781A>G, 1784T>C, 1785G>A, 1786C>G, 1788G>A, 1789A>T, 1790A>T, 1792A>T, 1793C>G, 1799G>T, 1800C>A, 1808A>T, 1814C>A, 1817T>C, 1820G>T, 1821delA, 1824C>G, 1825A>C, 1834T>G, 1841G>C, 1842A>G, 1844T>G, 1847T>C, 1850T>A, 1851T>A, 1853T>C, 1854C>A, 1856T>C, 1862C>T, 1865A>T, 1871G>A, 1880T>C, 1884T>G, 1885T>A, 1886A>C, 1893G>A, 1944A>G, 1946T>G, 1958G>A, 1959A>C, 1961T>C, 1963G>C, 1964G>A, 1966T>C, 1967G>C, 1974G>T, 1975G>C, 1976A>G, 1977C>A, 1980C>T, 1981A>C, 1984T>C, 1985G>A, 1988C>G, 1989G>A, 1990C>T, 1996A>C, 1997A>C, 1998C>A, 1999G>T, 2001C>T, 2003A>T, 2004G>T, 2005C>G, 2006T>G, 2009C>G, 2012A>C, 2015C>A, 2017A>T, 2018T>C, 2021G>A, 2027A>T, 2033C>T, 2034C>T, 2036T>G, 2040A>T, 2042G>T, 2048G>A, 2057C>T, 2066G>A, 2067A>C, 2069T>G, 2072G>A, 2075T>G, 2077T>C, 2081T>A, 2082T>A, 2084T>C, 2088T>C, 2090T>A, 2091T>C, 2095T>G, 2096A>G, 2102G>T, 2104G>T, 2105G>A, 2109T>C, 2114T>C, 2119G>A, 2120G>T, 2129A>G, 2132G>A, 2133G>T, 2134G>T, 2135A>T, 2138G>A, 2141C>T, 2145G>A, 2152A>T, 2156A>G, 2158G>A, 2159A>G, 2162T>A, 2171C>A, 2173C>A, 2180T>C, 2183T>G, 2189G>A, 2194A>C, 2209A>G, 2210G>A, 2211T>C, 2213G>C, 2214A>T, 2216A>T, 2222A>T, 2223C>G, 2225A>G, 2226C>T, 2234A>T, 2243A>G, 2244C>A, 2245A>G, 2246T>A, 2249G>A, 2255A>T, 2268G>T, 2272G>T, 2273T>C, 2274C>A, 2275A>G, 2276T>A, 2280A>G, 2283G>A, 2294T>G, 2295C>A, 2296G>A, 2301C>G, 2303T>C, 2306A>T, 2312T>C, 2315C>T, 2321G>T, 2325A>G, 2326A>C, 2327T>A, 2330C>A, 2331C>G, 2333G>A, 2339G>A, 2345A>T, 2348C>T, 2351A>G, 2352C>T, 2354T>G, 2363G>T, 2373G>A, 2374C>G, 2375G>T, 2377C>A, 2378T>A, 2381T>C, 2384G>A, 2388C>A, 2390T>A, 2393T>A, 2394C>A, 2396G>C, 2397A>G, 2398C>A, 2399T>A, 2406C>G, 2411A>G, 2412G>A, 2414G>C | | | | | | | | |

##### CDS

| | Begin | End | Coverage | Score | Concordance | Matches | Identities | I/D/M/F* | Stop Codons |
|---|---|---|---|---|---|---|---|---|---|
| RdRp | 446 | 702 | 32.4% | 1035 | 59.8% | 243 (100%) | 163 (67.1%) | 0/0/1/1 | 1 |
| Protein mutations: | L454M (1671T>A), E460K (1689G>A 1691G>A), K461Y (1692A>T 1694G>T), V462A (1696T>C 1697C>A), G465T (1704G>A 1705G>C 1706C>G), R466S (1707C>A), S467T (1711G>C), R468T (1713C>A 1714G>C 1715G>C), W471A (1722T>G 1723G>C 1724G>T), E472D (1727A>T), A483T (1758G>A), E485T (1764G>A 1765A>C 1766G>A), G488A (1774G>C 1775C>G), K489R (1776A>C 1777A>G), A492R (1785G>A 1786C>G), E493I (1788G>A 1789A>T 1790A>T), Y494L (1792A>T 1793C>G), E496D (1799G>T), H505A (1824C>G 1825A>C), I508R (1834T>G), N511E (1842A>G 1844T>G), S514T (1851T>A 1853T>C), L515I (1854C>A 1856T>C), L525D (1884T>G 1885T>A 1886A>C), V528I (1893G>A), I545V (1944A>G 1946T>G), I550L (1959A>C 1961T>C), G551A (1963G>C 1964G>A), M552T (1966T>C 1967G>C), G555S (1974G>T 1975G>C 1976A>G), L556I (1977C>A), Q557S (1980C>T 1981A>C), M558T (1984T>C 1985G>A), A560I (1989G>A 1990C>T), Q562P (1996A>C 1997A>C), R563I (1998C>A 1999G>T), L564F (2001C>T 2003A>T), A565W (2004G>T 2005C>G 2006T>G), Y569F (2017A>T 2018T>C), T577S (2040A>T 2042G>T), I586L (2067A>C 2069T>G), V589A (2077T>C), C591S (2082T>A 2084T>C), S593P (2088T>C 2090T>A), Y594H (2091T>C), I595R (2095T>G 2096A>G), Q597H (2102G>T), R598L (2104G>T 2105G>A), S600P (2109T>C), W603Y (2119G>A 2120G>T), G608F (2133G>T 2134G>T 2135A>T), D612N (2145G>A), Y614F (2152A>T), R616K (2158G>A 2159A>G), A621D (2173C>A), K628T (2194A>C), Q633R (2209A>G 2210G>A), I635F (2214A>T 2216A>T), K637N (2222A>T), Q638E (2223C>G 2225A>G), L641F (2234A>T), H645R (2244C>A 2245A>G 2246T>A), V653L (2268G>T), R654L (2272G>T 2273T>C), H655R (2274C>A 2275A>G 2276T>A), M657V (2280A>G), E658K (2283G>A), R662K (2295C>A 2296G>A), L664V (2301C>G 2303T>C), N672A (2325A>G 2326A>C 2327T>A), P674A (2331C>G 2333G>A), A688S (2373G>A 2374C>G 2375G>T), T689K (2377C>A 2378T>A), W691* (2384G>A), Q695N (2394C>A 2396G>C), T696E (2397A>G 2398C>A 2399T>A), H699D (2406C>G), V701I (2412G>A 2414G>C) | | | | | | | | |
| Codon mutations: | GCC448GCG (1655C>G), CTG449CTT (1658G>T), TCA450TCC (1661A>C), TTG454ATG (1671T>A), GTA455GTG (1676A>G), TTG457CTA (1680T>C 1682G>A), GCG458GCT (1685G>T), GAG460AAA (1689G>A 1691G>A), AAG461TAT (1692A>T 1694G>T), GTC462GCA (1696T>C 1697C>A), CCG464CCA (1703G>A), GGC465ACG (1704G>A 1705G>C 1706C>G), CGT466AGT (1707C>A), AGT467ACT (1711G>C), CGG468ACC (1713C>A 1714G>C 1715G>C), TTC470TTT (1721C>T), TGG471GCT (1722T>G 1723G>C 1724G>T), GAA472GAT (1727A>T), TAT473TAC (1730T>C), CTT475CTG (1736T>G), CTC476TTA (1737C>T 1739C>A), ATT480ATC (1751T>C), CTC481CTA (1754C>A), ATT482ATC (1757T>C), GCC483ACC (1758G>A), GAG485ACA (1764G>A 1765A>C 1766G>A), GTC487GTG (1772C>G), GGC488GCG (1774G>C 1775C>G), AAG489CGG (1776A>C 1777A>G), CAA490CAG (1781A>G), TAT491TAC (1784T>C), GCG492AGG (1785G>A 1786C>G), GAA493ATT (1788G>A 1789A>T 1790A>T), TAC494TTG (1792A>T 1793C>G), GAG496GAT (1799G>T), CGG497AGG (1800C>A), GGA499GGT (1808A>T), ACC501ACA (1814C>A), ATT502ATC (1817T>C), TCG503TCT (1820G>T), ATA504-TA (1821delA), CAT505GCT (1824C>G 1825A>C), ATA508AGA (1834T>G), TCG510TCC (1841G>C), AAT511GAG (1842A>G 1844T>G), AAT512AAC (1847T>C), GGT513GGA (1850T>A), TCT514ACC (1851T>A 1853T>C), CTT515ATC (1854C>A 1856T>C), TTC517TTT (1862C>T), GCA518GCT (1865A>T), AGG520AGA (1871G>A), ACT523ACC (1880T>C), TTA525GAC (1884T>G 1885T>A 1886A>C), GTT528ATT (1893G>A), ATT545GTG (1944A>G 1946T>G), CAG549CAA (1958G>A), ATT550CTC (1959A>C 1961T>C), GGG551GCA (1963G>C 1964G>A), ATG552ACC (1966T>C 1967G>C), GGA555TCG (1974G>T 1975G>C 1976A>G), CTT556ATT (1977C>A), CAA557TCA (1980C>T 1981A>C), ATG558ACA (1984T>C 1985G>A), TCC559TCG (1988C>G), GCT560ATT (1989G>A 1990C>T), CAA562CCC (1996A>C 1997A>C), CGT563ATT (1998C>A 1999G>T), CTA564TTT (2001C>T 2003A>T), GCT565TGG (2004G>T 2005C>G 2006T>G), GGC566GGG (2009C>G), GCA567GCC (2012A>C), GGC568GGA (2015C>A), TAT569TTC (2017A>T 2018T>C), AGG570AGA (2021G>A), CGA572CGT (2027A>T), CGC574CGT (2033C>T), CTT575TTG (2034C>T 2036T>G), ACG577TCT (2040A>T 2042G>T), CAG579CAA (2048G>A), CGC582CGT (2057C>T), AGG585AGA (2066G>A), ATT586CTG (2067A>C 2069T>G), AAG587AAA (2072G>A), GCT588GCG (2075T>G), GTT589GCT (2077T>C), GCT590GCA (2081T>A), TGT591AGC (2082T>A 2084T>C), TCT593CCA (2088T>C 2090T>A), TAC594CAC (2091T>C), ATA595AGG (2095T>G 2096A>G), CAG597CAT (2102G>T), CGG598CTA (2104G>T 2105G>A), TCT600CCT (2109T>C), CTT601CTC (2114T>C), TGG603TAT (2119G>A 2120G>T), GGA606GGG (2129A>G), AGG607AGA (2132G>A), GGA608TTT (2133G>T 2134G>T 2135A>T), AAG609AAA (2138G>A), CCC610CCT (2141C>T), GAT612AAT (2145G>A), TAT614TTT (2152A>T), TTA615TTG (2156A>G), AGA616AAG (2158G>A 2159A>G), GGT617GGA (2162T>A), ATC620ATA (2171C>A), GCT621GAT (2173C>A), CTT623CTC (2180T>C), CGT624CGG (2183T>G), GAG626GAA (2189G>A), AAA628ACA (2194A>C), CAG633CGA (2209A>G 2210G>A), TTG634CTC (2211T>C 2213G>C), ATA635TTT (2214A>T 2216A>T), AAA637AAT (2222A>T), CAA638GAG (2223C>G 2225A>G), CTG639TTG (2226C>T), TTA641TTT (2234A>T), GAA644GAG (2243A>G), CAT645AGA (2244C>A 2245A>G 2246T>A), GAG646GAA (2249G>A), CTA648CTT (2255A>T), GTA653TTA (2268G>T), CGT654CTC (2272G>T 2273T>C), CAT655AGA (2274C>A 2275A>G 2276T>A), ATG657GTG (2280A>G), GAA658AAA (2283G>A), CTT661CTG (2294T>G), CGA662AAA (2295C>A 2296G>A), CTT664GTC (2301C>G 2303T>C), TCA665TCT (2306A>T), TAT667TAC (2312T>C), CAC668CAT (2315C>T), GTG670GTT (2321G>T), AAT672GCA (2325A>G 2326A>C 2327T>A), TCC673TCA (2330C>A), CCG674GCA (2331C>G 2333G>A), GTG676GTA (2339G>A), ATA678ATT (2345A>T), GAC679GAT (2348C>T), CAA680CAG (2351A>G), CTT681TTG (2352C>T 2354T>G), GTG684GTT (2363G>T), GCG688AGT (2373G>A 2374C>G 2375G>T), ACT689AAA (2377C>A 2378T>A), TCT690TCC (2381T>C), TGG691TGA (2384G>A), CGT693AGA (2388C>A 2390T>A), ACT694ACA (2393T>A), CAG695AAC (2394C>A 2396G>C), ACT696GAA (2397A>G 2398C>A 2399T>A), CAT699GAT (2406C>G), TTA700TTG (2411A>G), GTG701ATC (2412G>A 2414G>C) | | | | | | | | |

##### Proteins

| | Begin | End | Coverage | Score | Concordance | Matches | Identities | I/D/M/F* | Stop Codons |
|---|---|---|---|---|---|---|---|---|---|
| RNA-dependent RNA polymerase (YP_010798875.1) | 446 | 702 | 32.4% | 1035 | 59.8% | 243 (100%) | 163 (67.1%) | 0/0/1/1 | 1 |
| Protein mutations: | L454M (1671T>A), E460K (1689G>A 1691G>A), K461Y (1692A>T 1694G>T), V462A (1696T>C 1697C>A), G465T (1704G>A 1705G>C 1706C>G), R466S (1707C>A), S467T (1711G>C), R468T (1713C>A 1714G>C 1715G>C), W471A (1722T>G 1723G>C 1724G>T), E472D (1727A>T), A483T (1758G>A), E485T (1764G>A 1765A>C 1766G>A), G488A (1774G>C 1775C>G), K489R (1776A>C 1777A>G), A492R (1785G>A 1786C>G), E493I (1788G>A 1789A>T 1790A>T), Y494L (1792A>T 1793C>G), E496D (1799G>T), H505A (1824C>G 1825A>C), I508R (1834T>G), N511E (1842A>G 1844T>G), S514T (1851T>A 1853T>C), L515I (1854C>A 1856T>C), L525D (1884T>G 1885T>A 1886A>C), V528I (1893G>A), I545V (1944A>G 1946T>G), I550L (1959A>C 1961T>C), G551A (1963G>C 1964G>A), M552T (1966T>C 1967G>C), G555S (1974G>T 1975G>C 1976A>G), L556I (1977C>A), Q557S (1980C>T 1981A>C), M558T (1984T>C 1985G>A), A560I (1989G>A 1990C>T), Q562P (1996A>C 1997A>C), R563I (1998C>A 1999G>T), L564F (2001C>T 2003A>T), A565W (2004G>T 2005C>G 2006T>G), Y569F (2017A>T 2018T>C), T577S (2040A>T 2042G>T), I586L (2067A>C 2069T>G), V589A (2077T>C), C591S (2082T>A 2084T>C), S593P (2088T>C 2090T>A), Y594H (2091T>C), I595R (2095T>G 2096A>G), Q597H (2102G>T), R598L (2104G>T 2105G>A), S600P (2109T>C), W603Y (2119G>A 2120G>T), G608F (2133G>T 2134G>T 2135A>T), D612N (2145G>A), Y614F (2152A>T), R616K (2158G>A 2159A>G), A621D (2173C>A), K628T (2194A>C), Q633R (2209A>G 2210G>A), I635F (2214A>T 2216A>T), K637N (2222A>T), Q638E (2223C>G 2225A>G), L641F (2234A>T), H645R (2244C>A 2245A>G 2246T>A), V653L (2268G>T), R654L (2272G>T 2273T>C), H655R (2274C>A 2275A>G 2276T>A), M657V (2280A>G), E658K (2283G>A), R662K (2295C>A 2296G>A), L664V (2301C>G 2303T>C), N672A (2325A>G 2326A>C 2327T>A), P674A (2331C>G 2333G>A), A688S (2373G>A 2374C>G 2375G>T), T689K (2377C>A 2378T>A), W691* (2384G>A), Q695N (2394C>A 2396G>C), T696E (2397A>G 2398C>A 2399T>A), H699D (2406C>G), V701I (2412G>A 2414G>C) | | | | | | | | |

| | Begin | End | Coverage | Score | Concordance | Matches | Identities | I/D/M/F* | Stop Codons |
|---|---|---|---|---|---|---|---|---|---|
| **NT** | **1645** | **2417** | **27.0%** | **534** | **36.9%** | **728 (99.9%)** | **500 (68.6%)** | **0/1** | |
| Codon mutations: | GCC448GCG (1655C>G), CTG449CTT (1658G>T), TCA450TCC (1661A>C), TTG454ATG (1671T>A), GTA455GTG (1676A>G), TTG457CTA (1680T>C 1682G>A), GCG458GCT (1685G>T), GAG460AAA (1689G>A 1691G>A), AAG461TAT (1692A>T 1694G>T), GTC462GCA (1696T>C 1697C>A), CCG464CCA (1703G>A), GGC465ACG (1704G>A 1705G>C 1706C>G), CGT466AGT (1707C>A), AGT467ACT (1711G>C), CGG468ACC (1713C>A 1714G>C 1715G>C), TTC470TTT (1721C>T), TGG471GCT (1722T>G 1723G>C 1724G>T), GAA472GAT (1727A>T), TAT473TAC (1730T>C), CTT475CTG (1736T>G), CTC476TTA (1737C>T 1739C>A), ATT480ATC (1751T>C), CTC481CTA (1754C>A), ATT482ATC (1757T>C), GCC483ACC (1758G>A), GAG485ACA (1764G>A 1765A>C 1766G>A), GTC487GTG (1772C>G), GGC488GCG (1774G>C 1775C>G), AAG489CGG (1776A>C 1777A>G), CAA490CAG (1781A>G), TAT491TAC (1784T>C), GCG492AGG (1785G>A 1786C>G), GAA493ATT (1788G>A 1789A>T 1790A>T), TAC494TTG (1792A>T 1793C>G), GAG496GAT (1799G>T), CGG497AGG (1800C>A), GGA499GGT (1808A>T), ACC501ACA (1814C>A), ATT502ATC (1817T>C), TCG503TCT (1820G>T), ATA504-TA (1821delA), CAT505GCT (1824C>G 1825A>C), ATA508AGA (1834T>G), TCG510TCC (1841G>C), AAT511GAG (1842A>G 1844T>G), AAT512AAC (1847T>C), GGT513GGA (1850T>A), TCT514ACC (1851T>A 1853T>C), CTT515ATC (1854C>A 1856T>C), TTC517TTT (1862C>T), GCA518GCT (1865A>T), AGG520AGA (1871G>A), ACT523ACC (1880T>C), TTA525GAC (1884T>G 1885T>A 1886A>C), GTT528ATT (1893G>A), ATT545GTG (1944A>G 1946T>G), CAG549CAA (1958G>A), ATT550CTC (1959A>C 1961T>C), GGG551GCA (1963G>C 1964G>A), ATG552ACC (1966T>C 1967G>C), GGA555TCG (1974G>T 1975G>C 1976A>G), CTT556ATT (1977C>A), CAA557TCA (1980C>T 1981A>C), ATG558ACA (1984T>C 1985G>A), TCC559TCG (1988C>G), GCT560ATT (1989G>A 1990C>T), CAA562CCC (1996A>C 1997A>C), CGT563ATT (1998C>A 1999G>T), CTA564TTT (2001C>T 2003A>T), GCT565TGG (2004G>T 2005C>G 2006T>G), GGC566GGG (2009C>G), GCA567GCC (2012A>C), GGC568GGA (2015C>A), TAT569TTC (2017A>T 2018T>C), AGG570AGA (2021G>A), CGA572CGT (2027A>T), CGC574CGT (2033C>T), CTT575TTG (2034C>T 2036T>G), ACG577TCT (2040A>T 2042G>T), CAG579CAA (2048G>A), CGC582CGT (2057C>T), AGG585AGA (2066G>A), ATT586CTG (2067A>C 2069T>G), AAG587AAA (2072G>A), GCT588GCG (2075T>G), GTT589GCT (2077T>C), GCT590GCA (2081T>A), TGT591AGC (2082T>A 2084T>C), TCT593CCA (2088T>C 2090T>A), TAC594CAC (2091T>C), ATA595AGG (2095T>G 2096A>G), CAG597CAT (2102G>T), CGG598CTA (2104G>T 2105G>A), TCT600CCT (2109T>C), CTT601CTC (2114T>C), TGG603TAT (2119G>A 2120G>T), GGA606GGG (2129A>G), AGG607AGA (2132G>A), GGA608TTT (2133G>T 2134G>T 2135A>T), AAG609AAA (2138G>A), CCC610CCT (2141C>T), GAT612AAT (2145G>A), TAT614TTT (2152A>T), TTA615TTG (2156A>G), AGA616AAG (2158G>A 2159A>G), GGT617GGA (2162T>A), ATC620ATA (2171C>A), GCT621GAT (2173C>A), CTT623CTC (2180T>C), CGT624CGG (2183T>G), GAG626GAA (2189G>A), AAA628ACA (2194A>C), CAG633CGA (2209A>G 2210G>A), TTG634CTC (2211T>C 2213G>C), ATA635TTT (2214A>T 2216A>T), AAA637AAT (2222A>T), CAA638GAG (2223C>G 2225A>G), CTG639TTG (2226C>T), TTA641TTT (2234A>T), GAA644GAG (2243A>G), CAT645AGA (2244C>A 2245A>G 2246T>A), GAG646GAA (2249G>A), CTA648CTT (2255A>T), GTA653TTA (2268G>T), CGT654CTC (2272G>T 2273T>C), CAT655AGA (2274C>A 2275A>G 2276T>A), ATG657GTG (2280A>G), GAA658AAA (2283G>A), CTT661CTG (2294T>G), CGA662AAA (2295C>A 2296G>A), CTT664GTC (2301C>G 2303T>C), TCA665TCT (2306A>T), TAT667TAC (2312T>C), CAC668CAT (2315C>T), GTG670GTT (2321G>T), AAT672GCA (2325A>G 2326A>C 2327T>A), TCC673TCA (2330C>A), CCG674GCA (2331C>G 2333G>A), GTG676GTA (2339G>A), ATA678ATT (2345A>T), GAC679GAT (2348C>T), CAA680CAG (2351A>G), CTT681TTG (2352C>T 2354T>G), GTG684GTT (2363G>T), GCG688AGT (2373G>A 2374C>G 2375G>T), ACT689AAA (2377C>A 2378T>A), TCT690TCC (2381T>C), TGG691TGA (2384G>A), CGT693AGA (2388C>A 2390T>A), ACT694ACA (2393T>A), CAG695AAC (2394C>A 2396G>C), ACT696GAA (2397A>G 2398C>A 2399T>A), CAT699GAT (2406C>G), TTA700TTG (2411A>G), GTG701ATC (2412G>A 2414G>C) | | | | | | | | |

*: Inserts / Deletes / Misaligned / Frameshifts

#### Analysis details

This analysis was performed with panviral2.64

### NGS Details (UN60): Brazilian marseillevirus

#### Assembly

| | |
|---|---|
| **Coverage Length** | 224 (1 contig(s)) |
| **Depth Of Coverage** | 8722.3 |
| **Number Of Reads** | 20377 |
| **Reads Per Million** | 432.19 rpm (after QC) |
| **Ambiguities** | 0 |
| **Assembly Method** | de novo + reference guided assembly |
| **Consensus Caller** | Bcf Tools |

#### Coverage Map

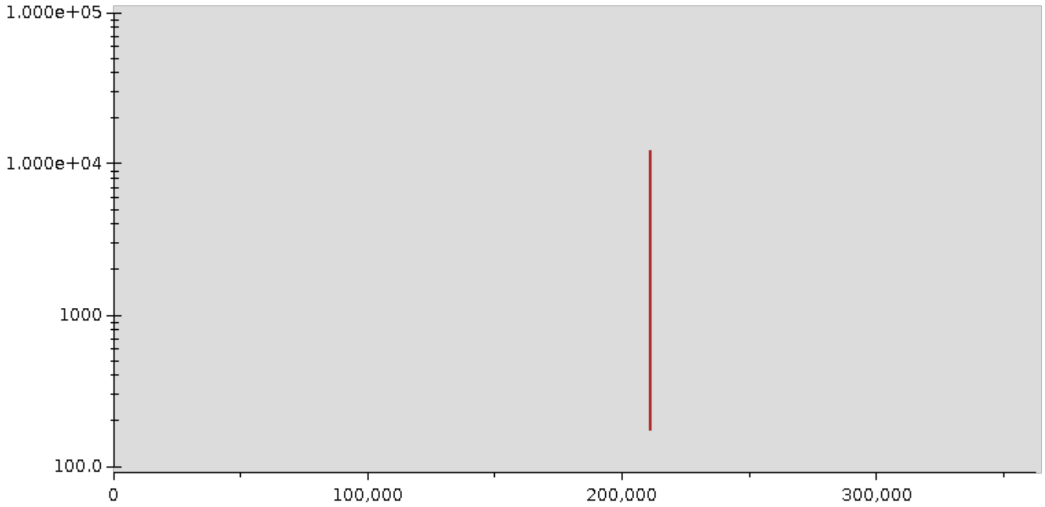

#### Assignment

| | |
|---|---|
| **Type** | Brazilian marseillevirus (Taxonomy ID: 1813599) |
| **Reference Genome** | NC_029692.1 |
| **NT Identity (%)** | 82.5893 |
| **AA Identity (%)** | 93.0556 |
| **Number Of Stop Codons** | 0 |
| **Number Of CDS** | 491 |

#### Alignment

| | |
|---|---|
| **Alignment Score** | 292.0 (NT) + 432.0 (AA) = 724.0 |
| **Concordance (%)** | 80.9843 |
| **Alignment Method** | Local, heuristic, nucleotide (BLASTN) |

#### Genome Region

Sequence starts at position 210900 and ends at position 211123 relative to NC_029692.1 reference sequence.

#### Alignment Detailed Statistics

| | Begin | End | Coverage | Score | Concordance | Matches | Identities | I/D/M/F* | Stop Codons |
|---|---|---|---|---|---|---|---|---|---|
| **NT** | **210900** | **211123** | **0.1%** | **292** | **65.2%** | **224 (100%)** | **185 (82.6%)** | **0/0** | |
| Mutations: | 210914A>C, 210920G>T, 210926T>C, 210929T>C, 210932G>A, 210938A>G, 210953C>G, 210955T>G, 210956T>G, 210962T>C, 210965C>A, 210974G>A, 210977T>A, 210986A>C, 210988G>A, 210989G>C, 210995T>C, 211001A>T, 211004A>T, 211007A>G, 211009C>T, 211013T>C, 211031T>G, 211032T>G, 211033G>C, 211034T>C, 211037C>A, 211040G>A, 211043T>G, 211046G>A, 211061T>C, 211070G>C, 211079C>G, 211088C>G, 211091G>A, 211100T>A, 211103A>G, 211106C>G, 211108C>T | | | | | | | | |

*: Inserts / Deletes / Misaligned / Frameshifts

#### Analysis details

This analysis was performed with panviral2.64

### NGS Details (UN60): Brazilian marseillevirus

#### Assembly

| | |
|---|---|
| **Coverage Length** | 210 (1 contig(s)) |
| **Depth Of Coverage** | 8619.2 |
| **Number Of Reads** | 18320 |
| **Reads Per Million** | 388.56 rpm (after QC) |
| **Ambiguities** | 0 |
| **Assembly Method** | de novo + reference guided assembly |
| **Consensus Caller** | Bcf Tools |

#### Coverage Map

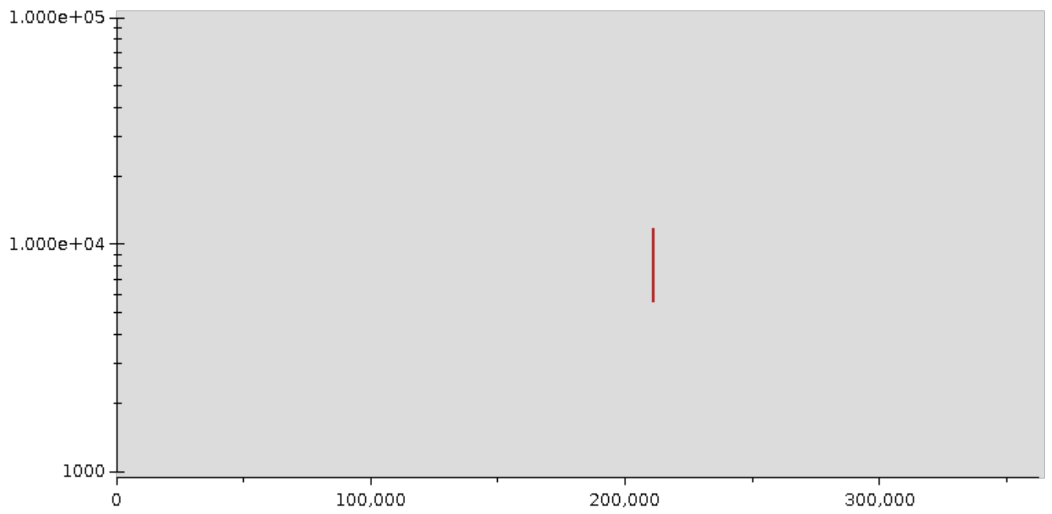

#### Assignment

| | |
|---|---|
| **Type** | Brazilian marseillevirus (Taxonomy ID: 1813599) |
| **Reference Genome** | NC_029692.1 |
| **NT Identity (%)** | 79.5238 |
| **AA Identity (%)** | 95.5882 |
| **Number Of Stop Codons** | 1 |
| **Number Of CDS** | 491 |

#### Alignment

| | |
|---|---|
| **Alignment Score** | 248.0 (NT) + 412.0 (AA) = 660.0 |
| **Concordance (%)** | 78.1065 |
| **Alignment Method** | Local, heuristic, nucleotide (BLASTN) |

#### Genome Region

Sequence starts at position 210884 and ends at position 211093 relative to NC_029692.1 reference sequence.

#### Alignment Detailed Statistics

| | Begin | End | Coverage | Score | Concordance | Matches | Identities | I/D/M/F* | Stop Codons |
|---|---|---|---|---|---|---|---|---|---|
| **NT** | **210884** | **211093** | **0.1%** | **248** | **59.0%** | **210 (100%)** | **167 (79.5%)** | **0/0** | |

Mutations: 210896A>G, 210899T>A, 210901T>G, 210902T>G, 210914A>C, 210920G>T, 210926T>C, 210929T>C, 210935G>A, 210938A>G, 210941A>G, 210953C>G, 210955T>G, 210956T>G, 210965C>G, 210974G>A, 210983G>T, 210986A>C, 210989G>C, 210991G>T, 210992T>C, 210995T>C, 211001A>T, 211004A>T, 211007A>G, 211009C>T, 211010T>C, 211013T>C, 211025G>A, 211031T>A, 211032T>G, 211033G>C, 211040G>A, 211043T>G, 211046G>A, 211052G>A, 211058G>A, 211061T>C, 211064A>C, 211070G>C, 211072G>A, 211079C>G, 211082C>T

*: Inserts / Deletes / Misaligned / Frameshifts

#### Analysis details

This analysis was performed with panviral2.64

### NGS Details (UN60): Brazilian marseillevirus

#### Assembly

| | |
|---|---|
| **Coverage Length** | 236 (1 contig(s)) |
| **Depth Of Coverage** | 5214.8 |
| **Number Of Reads** | 12747 |
| **Reads Per Million** | 270.36 rpm (after QC) |
| **Ambiguities** | 0 |
| **Assembly Method** | de novo + reference guided assembly |
| **Consensus Caller** | Bcf Tools |

#### Coverage Map

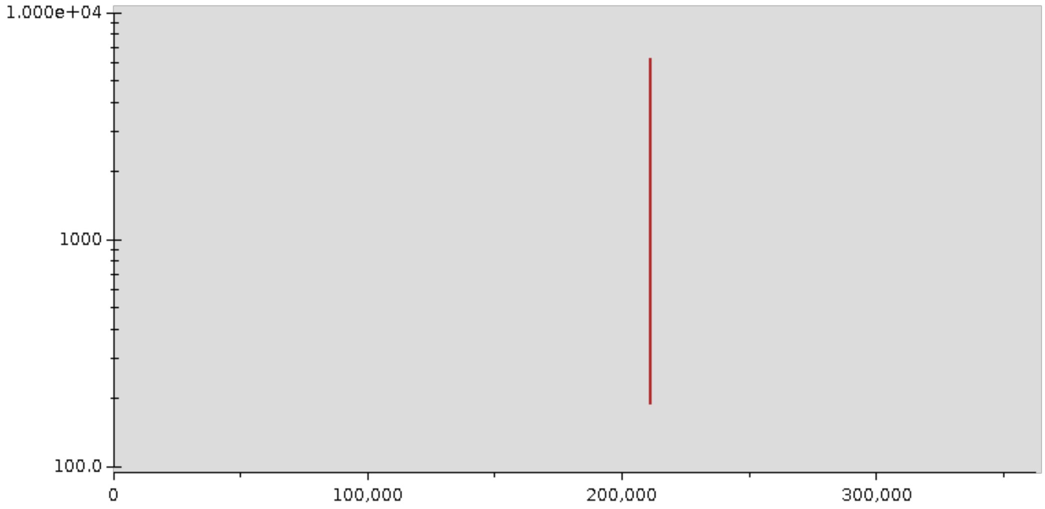

#### Assignment

| | |
|---|---|
| **Type** | Brazilian marseillevirus (Taxonomy ID: 1813599) |
| **Reference Genome** | NC_029692.1 |
| **NT Identity (%)** | 79.2373 |
| **AA Identity (%)** | 94.6667 |
| **Number Of Stop Codons** | 1 |
| **Number Of CDS** | 491 |

#### Alignment

| | |
|---|---|
| **Alignment Score** | 276.0 (NT) + 454.0 (AA) = 730.0 |
| **Concordance (%)** | 77.6596 |
| **Alignment Method** | Local, heuristic, nucleotide (BLASTN) |

#### Genome Region

Sequence starts at position 210888 and ends at position 211123 relative to NC_029692.1 reference sequence.

#### Alignment Detailed Statistics

| | Begin | End | Coverage | Score | Concordance | Matches | Identities | I/D/M/F* | Stop Codons |
|---|---|---|---|---|---|---|---|---|---|
| **NT** | **210888** | **211123** | **0.1%** | **276** | **58.5%** | **236 (100%)** | **187 (79.2%)** | **0/0** | |

Mutations: 210899T>C, 210902T>C, 210905G>A, 210908A>C, 210910G>A, 210914A>G, 210917A>C, 210920G>C, 210926T>C, 210929T>C, 210932G>A, 210938A>G, 210947A>G, 210953C>G, 210955T>G, 210956T>G, 210962T>C, 210965C>T, 210974G>A, 210977T>A, 210980G>A, 210986A>C, 210988G>A, 210989G>C, 210991G>T, 210992T>C, 210995T>C, 211001A>T, 211004A>T, 211007A>G, 211009C>T, 211010T>C, 211013T>C, 211031T>G, 211032T>G, 211033G>C, 211034T>C, 211037C>G, 211040G>A, 211043T>G, 211046G>A, 211061T>C, 211070G>C, 211079C>G, 211088C>G, 211094A>G, 211100T>A, 211106C>A, 211108C>T

*: Inserts / Deletes / Misaligned / Frameshifts

#### Analysis details

This analysis was performed with panviral2.64

### NGS Details (UN60): Brazilian marseillevirus

#### Assembly

| | |
|---|---|
| **Coverage Length** | 161 (1 contig(s)) |
| **Depth Of Coverage** | 6274.7 |
| **Number Of Reads** | 11487 |
| **Reads Per Million** | 243.63 rpm (after QC) |
| **Ambiguities** | 0 |
| **Assembly Method** | de novo + reference guided assembly |
| **Consensus Caller** | Bcf Tools |

#### Coverage Map

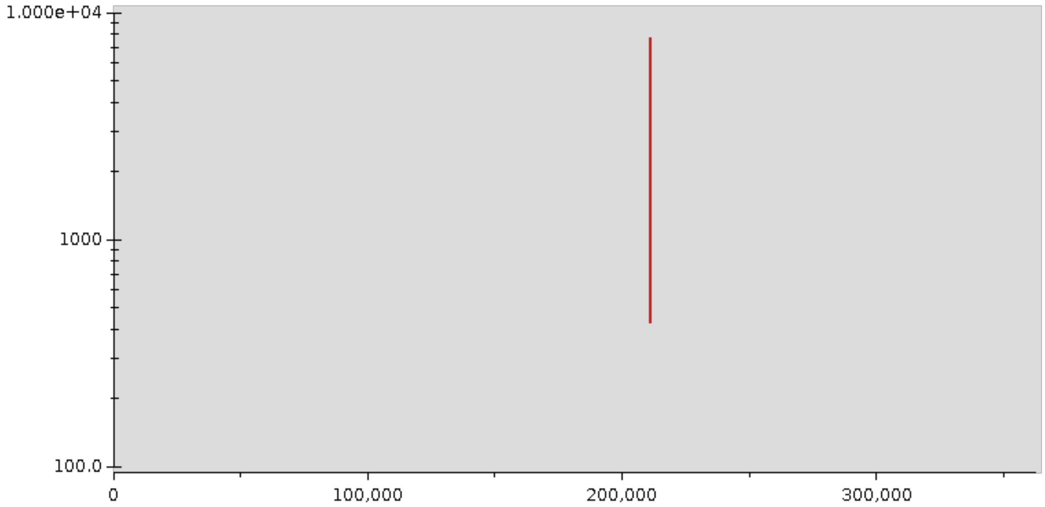

#### Assignment

| | |
|---|---|
| **Type** | Brazilian marseillevirus (Taxonomy ID: 1813599) |
| **Reference Genome** | NC_029692.1 |
| **NT Identity (%)** | 81.3665 |
| **AA Identity (%)** | 92.5926 |
| **Number Of Stop Codons** | 0 |
| **Number Of CDS** | 491 |

#### Alignment

| | |
|---|---|
| **Alignment Score** | 202.0 (NT) + 335.0 (AA) = 537.0 |
| **Concordance (%)** | 80.0298 |
| **Alignment Method** | Local, heuristic, nucleotide (BLASTN) |

#### Genome Region

Sequence starts at position 210894 and ends at position 211054 relative to NC_029692.1 reference sequence.

#### Alignment Detailed Statistics

| | Begin | End | Coverage | Score | Concordance | Matches | Identities | I/D/M/F* | Stop Codons |
|---|---|---|---|---|---|---|---|---|---|
| **NT** | **210894** | **211054** | **0.1%** | **202** | **62.7%** | **161 (100%)** | **131 (81.4%)** | **0/0** | |

Mutations: 210908A>C, 210910G>A, 210917A>G, 210920G>A, 210926T>C, 210929T>C, 210938A>G, 210944C>A, 210956T>A, 210965C>G, 210971C>T, 210974G>C, 210977T>A, 210980G>A, 210986A>C, 210989G>T, 210991G>T, 210992T>C, 210995T>C, 210998G>A, 211001A>T, 211004A>G, 211009C>T, 211010T>C, 211031T>G, 211032T>G, 211033G>T, 211034T>C, 211037C>G, 211043T>G

*: Inserts / Deletes / Misaligned / Frameshifts

#### Analysis details

This analysis was performed with panviral2.64

### NGS Details (UN60): Makelovirus prm1

#### Assembly

| | |
|---|---|
| **Coverage Length** | 1013 (1 contig(s)) |
| **Depth Of Coverage** | 1317.0 |
| **Number Of Reads** | 10355 |
| **Reads Per Million** | 219.63 rpm (after QC) |
| **Ambiguities** | 0 |
| **Assembly Method** | de novo + reference guided assembly |
| **Consensus Caller** | Bcf Tools |

#### Coverage Map

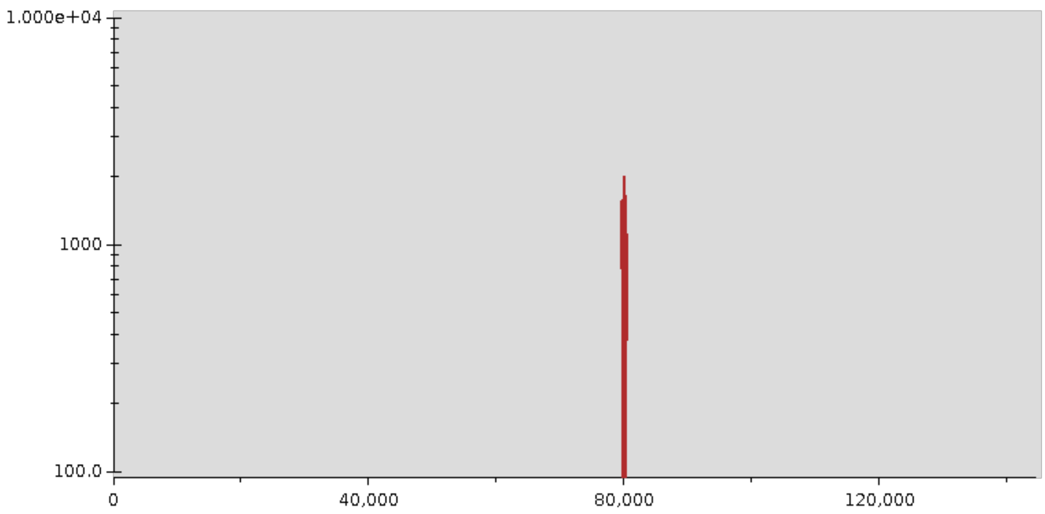

#### Assignment

| | |
|---|---|
| **Type** | Makelovirus prm1 (Taxonomy ID: 2956181) |
| **Reference Genome** | NC_055761.1 |
| **NT Identity (%)** | 76.8016 |
| **AA Identity (%)** | 87.5371 |
| **Number Of Stop Codons** | 0 |
| **Number Of CDS** | 190 |

#### Alignment

| | |
|---|---|
| **Alignment Score** | 1086.0 (NT) + 2250.0 (AA) = 3336.0 |
| **Concordance (%)** | 74.3979 |
| **Alignment Method** | Local, heuristic, nucleotide (BLASTN) |

#### Genome Region

Sequence starts at position 79507 and ends at position 80519 relative to NC_055761.1 reference sequence.

#### Alignment Detailed Statistics

| | Begin | End | Coverage | Score | Concordance | Matches | Identities | I/D/M/F* | Stop Codons |
|---|---|---|---|---|---|---|---|---|---|
| **NT** | **79507** | **80519** | **0.7%** | **1086** | **53.6%** | **1013 (100%)** | **778 (76.8%)** | **0/0** | |
| Mutations: | 79519A>G, 79520A>T, 79522A>G, 79523A>C, 79529C>T, 79530G>A, 79532G>C, 79536G>A, 79541G>T, 79542T>A, 79543C>G, 79544T>C, 79547A>T, 79548A>G, 79550C>A, 79563G>A, 79565C>T, 79568T>A, 79574C>T, 79580G>T, 79581C>T, 79592A>T, 79595T>C, 79596C>T, 79598G>A, 79602G>A, 79604A>G, 79610C>T, 79611G>T, 79612T>C, 79613C>T, 79614T>G, 79615G>T, 79616C>A, 79619C>T, 79624C>T, 79628G>C, 79632G>A, 79643C>T, 79646T>A, 79649G>A, 79655C>T, 79658C>T, 79664C>T, 79673A>T, 79676C>T, 79677G>T, 79679T>A, 79682T>G, 79688G>A, 79689A>C, 79691G>T, 79694T>C, 79703C>T, 79712T>C, 79718T>C, 79719G>A, 79722G>A, 79724A>T, 79728T>A, 79733C>T, 79734A>G, 79735A>C, 79736C>A, 79739A>T, 79742T>A, 79746C>T, 79748T>A, 79751C>T, 79754C>T, 79760C>A, 79763T>C, 79772T>G, 79779C>G, 79791C>T, 79793C>A, 79799T>C, 79811C>T, 79812C>G, 79817G>A, 79818G>A, 79820C>T, 79823A>T, 79824T>C, 79826T>A, 79833C>T, 79835G>A, 79836A>C, 79838C>T, 79842A>G, 79844C>A, 79845T>G, 79846T>C, 79847C>T, 79850C>T, 79871A>G, 79874C>T, 79875T>A, 79876C>G, 79879A>T, 79880T>C, 79895T>A, 79898C>T, 79904C>T, 79905T>G, 79906G>C, 79907C>T, 79916C>T, 79919T>A, 79922A>T, 79931T>A, 79937A>T, 79938T>A, 79940T>C, 79946C>T, 79950C>T, 79953G>A, 79958T>C, 79961T>A, 79962T>A, 79973T>A, 79974T>A, 79975C>G, 79976G>T, 79979C>T, 79985C>T, 79994C>T, 79997C>A, 80000C>A, 80018C>T, 80025C>A, 80030C>A, 80039A>T, 80042A>G, 80054G>T, 80063C>A, 80066C>T, 80069T>C, 80073C>T, 80075T>A, 80078T>C, 80090C>A, 80096T>C, 80099T>C, 80105G>A, 80114A>T, 80120C>T, 80126T>C, 80127C>T, 80132T>A, 80139T>A, 80140C>G, 80141G>T, 80142C>T, 80145G>A, 80147T>C, 80148C>A, 80150T>G, 80159C>A, 80162G>A, 80165C>T, 80168G>A, 80171C>T, 80172C>G, 80173A>C, 80174G>T, 80177C>T, 80178T>G, 80180T>A, 80188A>G, 80189G>A, 80201A>G, 80207G>A, 80210C>T, 80213C>T, 80216C>T, 80222G>A, 80225T>C, 80237C>T, 80240C>T, 80243T>C, 80246C>A, 80247C>T, 80261C>T, 80265T>A, 80266C>G, 80267C>T, 80279C>T, 80286C>T, 80288G>A, 80291T>C, 80300T>A, 80306A>T, 80315T>A, 80318C>A, 80333T>C, 80336A>T, 80337C>T, 80339T>A, 80343G>A, 80346T>A, 80347C>G, 80351C>T, 80357A>T, 80366G>A, 80369C>T, 80378C>T, 80387G>A, 80390C>T, 80391A>G, 80393C>A, 80394A>G, 80396G>T, 80399T>C, 80400G>A, 80409A>C, 80410A>G, 80411A>T, 80414C>A, 80415C>A, 80417C>T, 80433G>A, 80435T>C, 80436C>A, 80438A>C, 80447C>T, 80453G>T, 80456G>T, 80462G>A, 80465A>T, 80471C>T, 80474G>A, 80486C>T, 80498G>A, 80504G>A | | | | | | | | |

*: Inserts / Deletes / Misaligned / Frameshifts

#### Analysis details

This analysis was performed with panviral2.64

### NGS Details (UN60): Brazilian marseillevirus

#### Assembly

| | |
|---|---|
| **Coverage Length** | 111 (1 contig(s)) |
| **Depth Of Coverage** | 4288.3 |
| **Number Of Reads** | 9741 |
| **Reads Per Million** | 206.60 rpm (after QC) |
| **Ambiguities** | 0 |
| **Assembly Method** | de novo + reference guided assembly |
| **Consensus Caller** | Bcf Tools |

#### Coverage Map

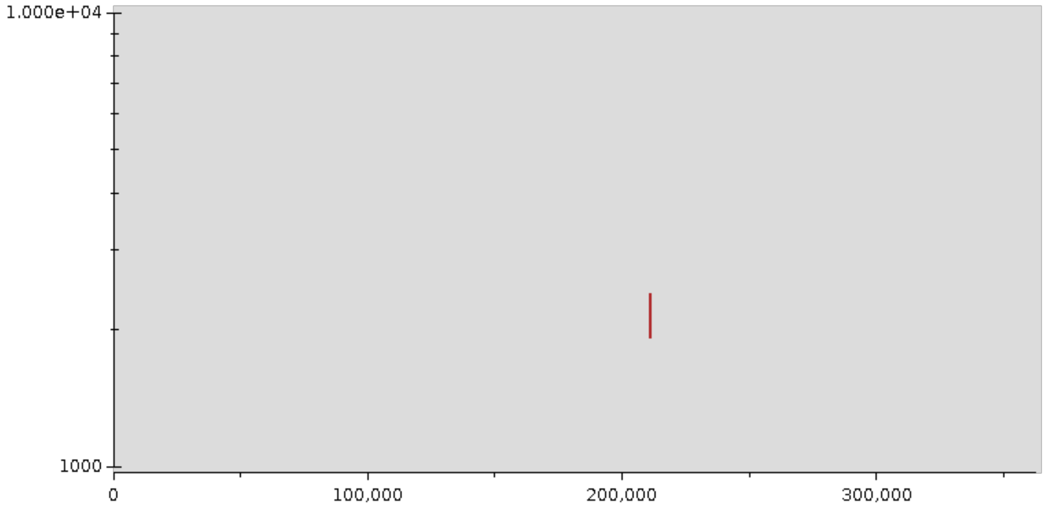

#### Assignment

| | |
|---|---|
| **Type** | Brazilian marseillevirus (Taxonomy ID: 1813599) |
| **Reference Genome** | NC_029692.1 |
| **NT Identity (%)** | 81.982 |
| **AA Identity (%)** | 97.1429 |
| **Number Of Stop Codons** | 1 |
| **Number Of CDS** | 491 |

#### Alignment

| | |
|---|---|
| **Alignment Score** | 142.0 (NT) + 215.0 (AA) = 357.0 |
| **Concordance (%)** | 80.9524 |
| **Alignment Method** | Local, heuristic, nucleotide (BLASTN) |

#### Genome Region

Sequence starts at position 210884 and ends at position 210994 relative to NC_029692.1 reference sequence.

#### Alignment Detailed Statistics

| | Begin | End | Coverage | Score | Concordance | Matches | Identities | I/D/M/F* | Stop Codons |
|---|---|---|---|---|---|---|---|---|---|
| NT | 210884 | 210994 | 0.1% | 142 | 64.0% | 111 (100%) | 91 (82.0%) | 0/0 | |

Mutations: 210896A>G, 210899T>A, 210901T>G, 210902T>A, 210907C>A, 210914A>G, 210917A>G, 210920G>A, 210926T>C, 210929T>C, 210938A>G, 210941A>G, 210944C>A, 210947A>T, 210950G>T, 210953C>T, 210955T>G, 210962T>C, 210965C>A, 210983G>A

*: Inserts / Deletes / Misaligned / Frameshifts

#### Analysis details

This analysis was performed with panviral2.64

### NGS Details (UN60): Yellowstone lake phycodnavirus 1

#### Assembly

| | |
|---|---|
| **Coverage Length** | 173 (1 contig(s)) |
| **Depth Of Coverage** | 3350.0 |
| **Number Of Reads** | 5848 |
| **Reads Per Million** | 124.03 rpm (after QC) |
| **Ambiguities** | 0 |
| **Assembly Method** | de novo + reference guided assembly |
| **Consensus Caller** | Bcf Tools |

#### Coverage Map

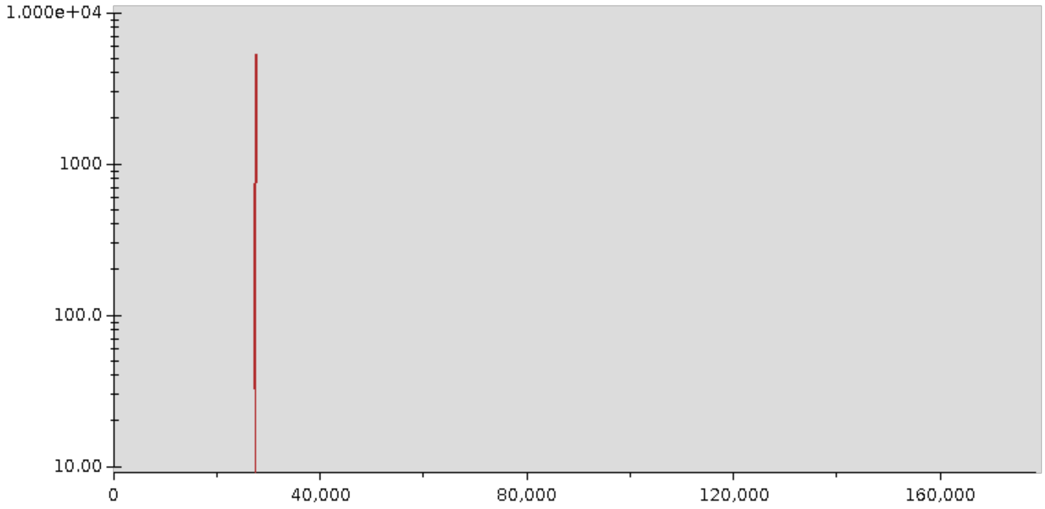

#### Assignment

| | |
|---|---|
| **Type** | Yellowstone lake phycodnavirus 1 (Taxonomy ID: 1586713) |
| **Reference Genome** | NC_028112.1 |
| **NT Identity (%)** | 82.0809 |
| **AA Identity (%)** | 93.1034 |
| **Number Of Stop Codons** | 0 |
| **Number Of CDS** | 248 |

#### Alignment

| | |
|---|---|
| **Alignment Score** | 222.0 (NT) + 350.0 (AA) = 572.0 |
| **Concordance (%)** | 80.5634 |
| **Alignment Method** | Local, heuristic, nucleotide (BLASTN) |

#### Genome Region

Sequence starts at position 27469 and ends at position 27641 relative to NC_028112.1 reference sequence.

#### Alignment Detailed Statistics

| | Begin | End | Coverage | Score | Concordance | Matches | Identities | I/D/M/F* | Stop Codons |
|---|---|---|---|---|---|---|---|---|---|
| **NT** | **27469** | **27641** | **0.1%** | **222** | **64.2%** | **173 (100%)** | **142 (82.1%)** | **0/0** | |

Mutations: 27481T>G, 27486G>A, 27492A>G, 27510A>G, 27513T>C, 27516T>A, 27529C>A, 27531T>G, 27534A>G, 27537T>A, 27540T>C, 27543T>A, 27552A>G, 27558G>A, 27563A>G, 27564T>C, 27567T>C, 27573A>T, 27576A>G, 27582T>C, 27586G>A, 27591A>G, 27597A>G, 27600C>A, 27606T>G, 27609T>C, 27612A>T, 27616T>C, 27618G>C, 27624T>C, 27627C>T

*: Inserts / Deletes / Misaligned / Frameshifts

#### Analysis details

This analysis was performed with panviral2.64

### NGS Details (UN60): Brazilian marseillevirus

#### Assembly

| | |
|---|---|
| **Coverage Length** | 240 (1 contig(s)) |
| **Depth Of Coverage** | 2083.3 |
| **Number Of Reads** | 5218 |
| **Reads Per Million** | 110.67 rpm (after QC) |
| **Ambiguities** | 0 |
| **Assembly Method** | de novo + reference guided assembly |
| **Consensus Caller** | Bcf Tools |

#### Coverage Map

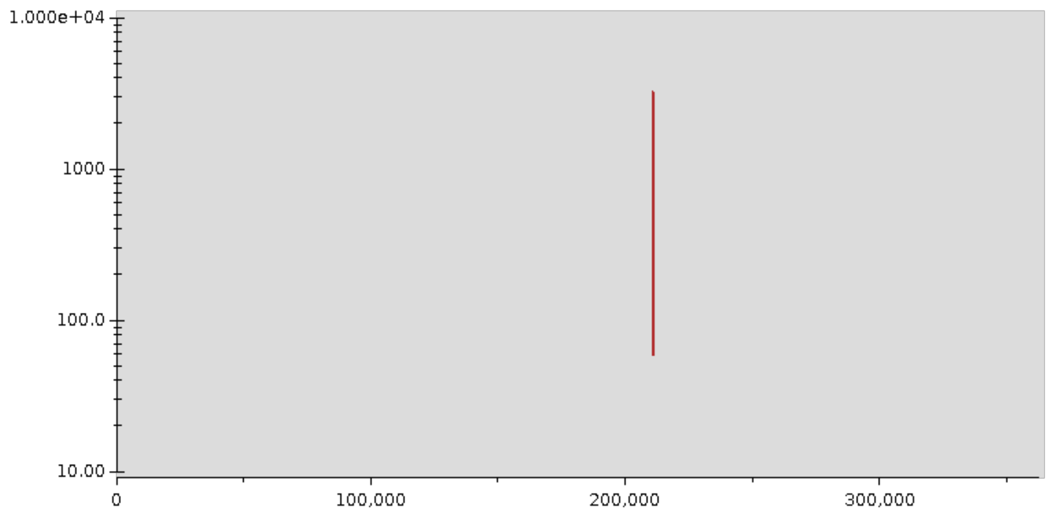

#### Assignment

| | |
|---|---|
| **Type** | Brazilian marseillevirus (Taxonomy ID: 1813599) |
| **Reference Genome** | NC_029692.1 |
| **NT Identity (%)** | 79.5833 |
| **AA Identity (%)** | 94.6667 |
| **Number Of Stop Codons** | 1 |
| **Number Of CDS** | 491 |

#### Alignment

| | |
|---|---|
| **Alignment Score** | 284.0 (NT) + 454.0 (AA) = 738.0 |
| **Concordance (%)** | 77.8481 |
| **Alignment Method** | Local, heuristic, nucleotide (BLASTN) |

#### Genome Region

Sequence starts at position 210884 and ends at position 211123 relative to NC_029692.1 reference sequence.

#### Alignment Detailed Statistics

| | Begin | End | Coverage | Score | Concordance | Matches | Identities | I/D/M/F* | Stop Codons |
|---|---|---|---|---|---|---|---|---|---|
| **NT** | **210884** | **211123** | **0.1%** | **284** | **59.2%** | **240 (100%)** | **191 (79.6%)** | **0/0** | |

Mutations: 210896A>G, 210899T>A, 210901T>G, 210902T>G, 210914A>C, 210916G>A, 210920G>T, 210926T>C, 210938A>G, 210941A>G, 210944C>T, 210947A>C, 210950G>A, 210974G>C, 210980G>A, 210986A>C, 210989G>C, 210991G>T, 210992T>C, 210995T>C, 211001A>T, 211004A>G, 211009C>T, 211010T>G, 211025G>A, 211031T>A, 211032T>G, 211033G>C, 211034T>C, 211040G>A, 211043T>G, 211046G>A, 211049G>A, 211052G>A, 211058G>A, 211064A>C, 211070G>C, 211072G>A, 211076G>A, 211079C>G, 211082C>T, 211085T>C, 211088C>A, 211091G>C, 211093G>A, 211094A>T, 211100T>G, 211106C>G, 211108C>T

*: Inserts / Deletes / Misaligned / Frameshifts

#### Analysis details

This analysis was performed with panviral2.64

### NGS Details (UN60): Yellowstone lake phycodnavirus 1

#### Assembly

| | |
|---|---|
| **Coverage Length** | 155 (1 contig(s)) |
| **Depth Of Coverage** | 2919.0 |
| **Number Of Reads** | 5137 |
| **Reads Per Million** | 108.95 rpm (after QC) |
| **Ambiguities** | 0 |
| **Assembly Method** | de novo + reference guided assembly |
| **Consensus Caller** | Bcf Tools |

#### Coverage Map

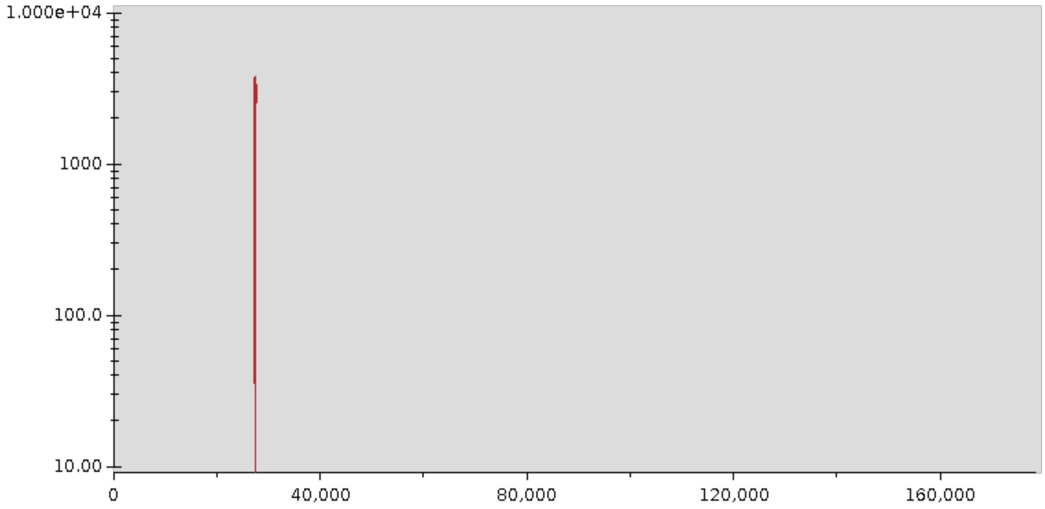

#### Assignment

| | |
|---|---|
| **Type** | Yellowstone lake phycodnavirus 1 (Taxonomy ID: 1586713) |
| **Reference Genome** | NC_028112.1 |
| **NT Identity (%)** | 80.0 |
| **AA Identity (%)** | 90.3846 |
| **Number Of Stop Codons** | 0 |
| **Number Of CDS** | 248 |

#### Alignment

| | |
|---|---|
| **Alignment Score** | 186.0 (NT) + 299.0 (AA) = 485.0 |
| **Concordance (%)** | 77.476 |
| **Alignment Method** | Local, heuristic, nucleotide (BLASTN) |

#### Genome Region

Sequence starts at position 27397 and ends at position 27551 relative to NC_028112.1 reference sequence.

#### Alignment Detailed Statistics

| | Begin | End | Coverage | Score | Concordance | Matches | Identities | I/D/M/F* | Stop Codons |
|---|---|---|---|---|---|---|---|---|---|
| NT | 27397 | 27551 | 0.1% | 186 | 60.0% | 155 (100%) | 124 (80.0%) | 0/0 | |

Mutations: 27417T>C, 27420C>G, 27426G>C, 27427C>T, 27429T>G, 27432C>T, 27447T>C, 27456T>G, 27459G>A, 27460T>A, 27461C>G, 27462A>T, 27463G>T, 27464G>C, 27468T>C, 27470G>C, 27476C>A, 27477A>C, 27481T>G, 27483G>T, 27489T>C, 27492A>G, 27504G>A, 27510A>T, 27513T>C, 27516T>G, 27529C>A, 27531T>G, 27532C>T, 27534A>G, 27537T>A

*: Inserts / Deletes / Misaligned / Frameshifts

#### Analysis details

This analysis was performed with panviral2.64

### NGS Details (UN60): Brazilian marseillevirus

#### Assembly

| | |
|---|---|
| **Coverage Length** | 241 (1 contig(s)) |
| **Depth Of Coverage** | 1024.8 |
| **Number Of Reads** | 2553 |
| **Reads Per Million** | 54.15 rpm (after QC) |
| **Ambiguities** | 0 |
| **Assembly Method** | de novo + reference guided assembly |
| **Consensus Caller** | Bcf Tools |

#### Coverage Map

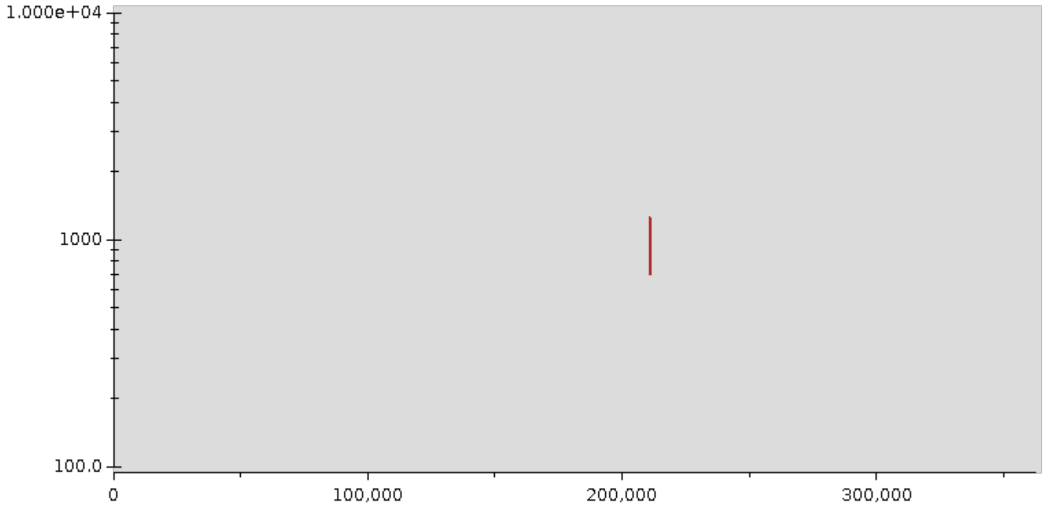

#### Assignment

| | |
|---|---|
| **Type** | Brazilian marseillevirus (Taxonomy ID: 1813599) |
| **Reference Genome** | NC_029692.1 |
| **NT Identity (%)** | 80.9129 |
| **AA Identity (%)** | 94.6667 |
| **Number Of Stop Codons** | 1 |
| **Number Of CDS** | 491 |

#### Alignment

| | |
|---|---|
| **Alignment Score** | 298.0 (NT) + 454.0 (AA) = 752.0 |
| **Concordance (%)** | 79.1579 |
| **Alignment Method** | Local, heuristic, nucleotide (BLASTN) |

#### Genome Region

Sequence starts at position 210884 and ends at position 211124 relative to NC_029692.1 reference sequence.

#### Alignment Detailed Statistics

| | Begin | End | Coverage | Score | Concordance | Matches | Identities | I/D/M/F* | Stop Codons |
|---|---|---|---|---|---|---|---|---|---|
| NT | 210884 | 211124 | 0.1% | 298 | 61.8% | 241 (100%) | 195 (80.9%) | 0/0 | |

Mutations: 210899T>G, 210901T>G, 210902T>G, 210911G>A, 210917A>G, 210926T>C, 210938A>G, 210941A>G, 210944C>G, 210947A>T, 210950G>T, 210953C>T, 210955T>G, 210956T>G, 210965C>A, 210971C>T, 210974G>T, 210977T>G, 210980G>A, 210986A>C, 210989G>C, 210991G>T, 210992T>C, 210995T>C, 211004A>G, 211009C>T, 211010T>G, 211022C>T, 211025G>A, 211031T>A, 211032T>G, 211033G>C, 211037C>G, 211040G>A, 211043T>A, 211058G>A, 211061T>C, 211064A>G, 211070G>A, 211079C>G, 211088C>G, 211094A>C, 211100T>A, 211106C>G, 211108C>T, 211115T>C

*: Inserts / Deletes / Misaligned / Frameshifts

#### Analysis details

This analysis was performed with panviral2.64

### NGS Details (UN60): Noumeavirus

#### Assembly

| | |
|---|---|
| **Coverage Length** | 225 (1 contig(s)) |
| **Depth Of Coverage** | 1143.9 |
| **Number Of Reads** | 2495 |
| **Reads Per Million** | 52.92 rpm (after QC) |
| **Ambiguities** | 0 |
| **Assembly Method** | de novo + reference guided assembly |
| **Consensus Caller** | Bcf Tools |

#### Coverage Map

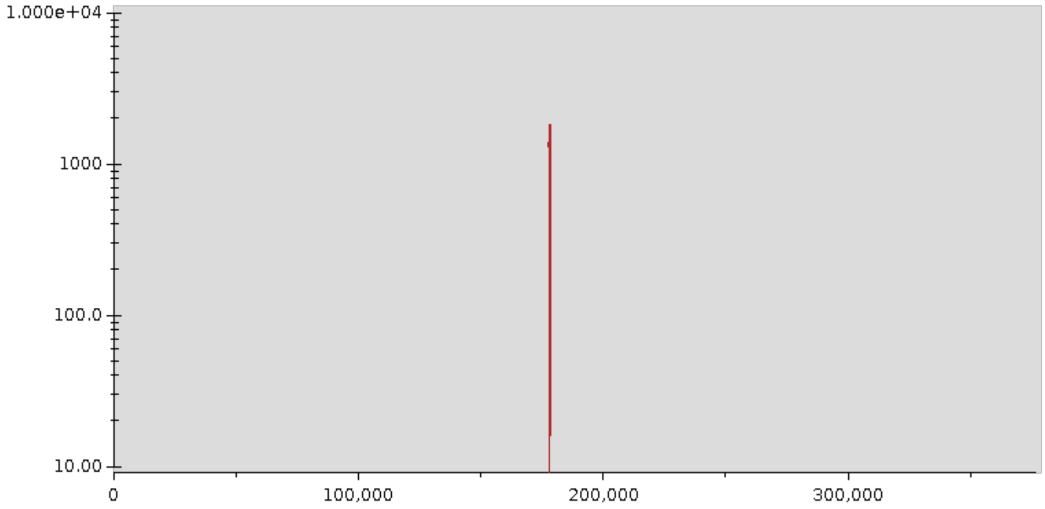

#### Assignment

| | |
|---|---|
| **Type** | Noumeavirus (Taxonomy ID: 1955558) |
| **Reference Genome** | NC_033775.1 |
| **NT Identity (%)** | 78.6667 |
| **AA Identity (%)** | 95.9459 |
| **Number Of Stop Codons** | 1 |
| **Number Of CDS** | 452 |

#### Alignment

| | |
|---|---|
| **Alignment Score** | 258.0 (NT) + 450.0 (AA) = 708.0 |
| **Concordance (%)** | 77.5465 |
| **Alignment Method** | Local, heuristic, nucleotide (BLASTN) |

#### Genome Region

Sequence starts at position 177923 and ends at position 178147 relative to NC_033775.1 reference sequence.

#### Alignment Detailed Statistics

| | Begin | End | Coverage | Score | Concordance | Matches | Identities | I/D/M/F* | Stop Codons |
|---|---|---|---|---|---|---|---|---|---|
| **NT** | **177923** | **178147** | **0.1%** | **258** | **57.3%** | **225 (100%)** | **177 (78.7%)** | **0/0** | |

Mutations: 177947A>C, 177950A>T, 177956G>A, 177959C>T, 177962C>T, 177965C>G, 177968G>A, 177971G>T, 177975A>T, 177976G>C, 177978T>A, 177979C>G, 177980A>C, 177983C>T, 177986C>A, 177992G>C, 177995C>T, 178001A>G, 178002C>G, 178003A>C, 178004A>C, 178010T>A, 178019A>G, 178028C>T, 178031T>A, 178044C>A, 178049T>G, 178052C>T, 178055C>T, 178058C>T, 178061A>T, 178064A>G, 178079A>C, 178080A>C, 178082G>C, 178085T>C, 178086C>T, 178088T>G, 178091C>G, 178097T>C, 178101G>A, 178106A>G, 178118A>T, 178121G>A, 178127T>G, 178130T>G, 178131T>C, 178136A>G

*: Inserts / Deletes / Misaligned / Frameshifts

#### Analysis details

This analysis was performed with panviral2.64

### NGS Details (UN60): Betabaculovirus disaccharalis

#### Assembly

| | |
|---|---|
| **Coverage Length** | 128 (1 contig(s)) |
| **Depth Of Coverage** | 1183.5 |
| **Number Of Reads** | 2005 |
| **Reads Per Million** | 42.53 rpm (after QC) |
| **Ambiguities** | 0 |
| **Assembly Method** | de novo + reference guided assembly |
| **Consensus Caller** | Bcf Tools |

#### Coverage Map

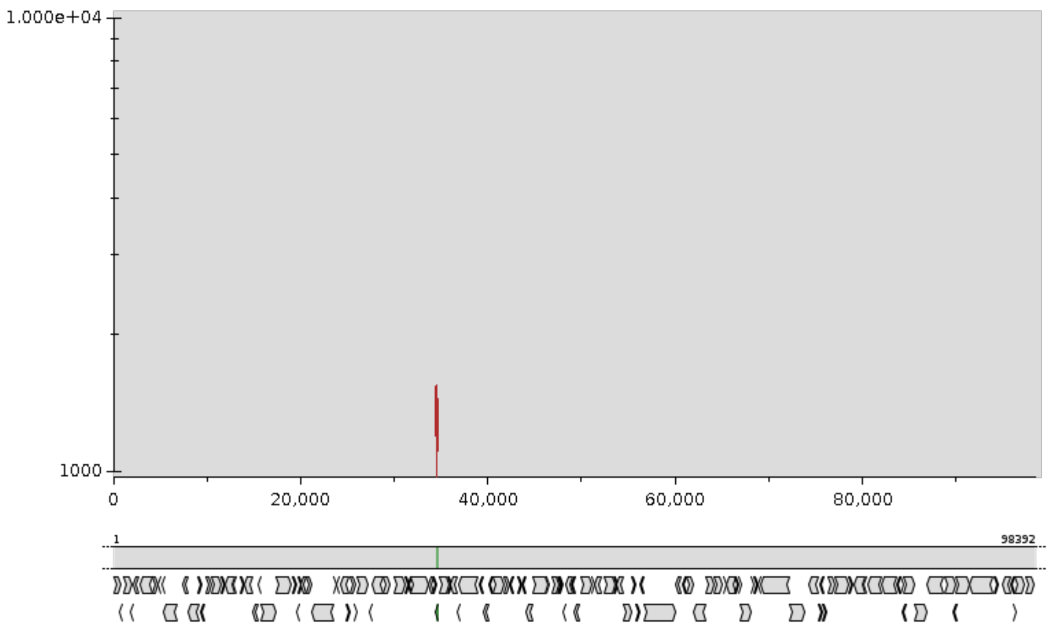

#### Assignment

| | |
|---|---|
| **Type** | Betabaculovirus disaccharalis (Taxonomy ID: 3047795) |
| **Reference Genome** | NC_028491.1 |
| **NT Identity (%)** | 82.0312 |
| **AA Identity (%)** | 93.0233 |
| **Number Of Stop Codons** | 0 |
| **Number Of CDS** | 125 |

#### Alignment

| | |
|---|---|
| **Alignment Score** | 164.0 (NT) + 276.0 (AA) = 440.0 |
| **Concordance (%)** | 81.0313 |

| **Alignment Method** | Local, heuristic, nucleotide (BLASTN) |
|---|---|

#### Genome Region

Sequence starts at position 34471 and ends at position 34598 relative to NC_028491.1 reference sequence.

#### Alignment Detailed Statistics

| | Begin | End | Coverage | Score | Concordance | Matches | Identities | I/D/M/F* | Stop Codons |
|---|---|---|---|---|---|---|---|---|---|
| **NT** | **34471** | **34598** | **0.1%** | **164** | **64.1%** | **128 (100%)** | **105 (82.0%)** | **0/0** | |
| Mutations: | 34482G>T, 34485T>C, 34488A>C, 34490G>A, 34491A>C, 34512G>T, 34513C>G, 34514T>A, 34521T>C, 34524A>G, 34533A>G, 34538A>C, 34545T>C, 34547G>T, 34548C>G, 34549T>C, 34551G>A, 34554T>C, 34569G>A, 34575T>A, 34578T>C, 34580A>G, 34587T>C | | | | | | | | |
| **CDS** | | | | | | | | | |
| v-ubq | 60 | 102 | 36.1% | 276 | 89.3% | 43 (100%) | 40 (93.0%) | 0/0/0/0 | 0 |
| Protein mutations: | E76G (34548C>G 34549T>C), S80A (34538A>C) | | | | | | | | |
| Codon mutations: | CAA63CAG (34587T>C), TTA66CTG (34578T>C 34580A>G), ATA67ATT (34575T>A), GCC69GCT (34569G>A), GAA74GAG (34554T>C), GAC75GAT (34551G>A), GAG76GGC (34548C>G 34549T>C), CGA77AGG (34545T>C 34547G>T), TCA80GCA (34538A>C), GAT81GAC (34533A>G), ATT84ATC (34524A>G), CAA85CAG (34521T>C), AGC88TCA (34512G>T 34513C>G 34514T>A), CGT95CGG (34491A>C), CTT96TTG (34488A>C 34490G>A), AGA97AGG (34485T>C), GGC98GGA (34482G>T) | | | | | | | | |
| **Proteins** | | | | | | | | | |
| ubiquitin-like protein (YP_009182246.1) | 60 | 102 | 36.1% | 276 | 89.3% | 43 (100%) | 40 (93.0%) | 0/0/0/0 | 0 |
| Protein mutations: | E76G (34548C>G 34549T>C), S80A (34538A>C) | | | | | | | | |
| Codon mutations: | CAA63CAG (34587T>C), TTA66CTG (34578T>C 34580A>G), ATA67ATT (34575T>A), GCC69GCT (34569G>A), GAA74GAG (34554T>C), GAC75GAT (34551G>A), GAG76GGC (34548C>G 34549T>C), CGA77AGG (34545T>C 34547G>T), TCA80GCA (34538A>C), GAT81GAC (34533A>G), ATT84ATC (34524A>G), CAA85CAG (34521T>C), AGC88TCA (34512G>T 34513C>G 34514T>A), CGT95CGG (34491A>C), CTT96TTG (34488A>C 34490G>A), AGA97AGG (34485T>C), GGC98GGA (34482G>T) | | | | | | | | |

*: Inserts / Deletes / Misaligned / Frameshifts

#### Analysis details

This analysis was performed with panviral2.64

### NGS Details (UN60): Betabaculovirus disaccharalis

#### Assembly

| | |
|---|---|
| **Coverage Length** | 150 (1 contig(s)) |
| **Depth Of Coverage** | 1065.4 |
| **Number Of Reads** | 1706 |
| **Reads Per Million** | 36.18 rpm (after QC) |
| **Ambiguities** | 0 |
| **Assembly Method** | de novo + reference guided assembly |
| **Consensus Caller** | Bcf Tools |

#### Coverage Map

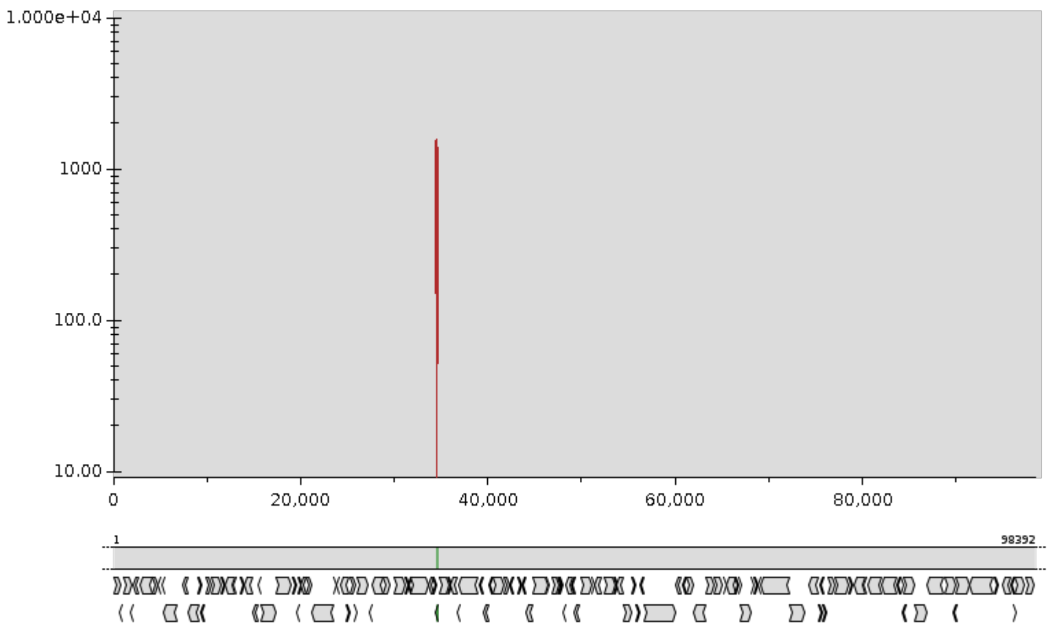

#### Assignment

| | |
|---|---|
| **Type** | Betabaculovirus disaccharalis (Taxonomy ID: 3047795) |
| **Reference Genome** | NC_028491.1 |
| **NT Identity (%)** | 84.6667 |
| **AA Identity (%)** | 92.0 |
| **Number Of Stop Codons** | 0 |
| **Number Of CDS** | 125 |

#### Alignment

| | |
|---|---|
| **Alignment Score** | 208.0 (NT) + 314.0 (AA) = 522.0 |
| **Concordance (%)** | 83.3866 |

| Alignment Method | Local, heuristic, nucleotide (BLASTN) |
| --- | --- |

#### Genome Region

Sequence starts at position 34474 and ends at position 34623 relative to NC_028491.1 reference sequence.

#### Alignment Detailed Statistics

| | Begin | End | Coverage | Score | Concordance | Matches | Identities | I/D/M/F* | Stop Codons |
| --- | --- | --- | --- | --- | --- | --- | --- | --- | --- |
| **NT** | **34474** | **34623** | **0.2%** | **208** | **69.3%** | **150 (100%)** | **127 (84.7%)** | **0/0** | |
| Mutations: | 34485T>C, 34488A>C, 34490G>A, 34513C>G, 34514T>A, 34521T>C, 34524A>G, 34533A>G, 34538A>C, 34542T>A, 34545T>C, 34547G>T, 34549T>C, 34554T>C, 34569G>A, 34575T>G, 34578T>C, 34587T>C, 34593C>T, 34596C>G, 34599C>A, 34601C>T, 34602T>A | | | | | | | | |
| **CDS** | | | | | | | | | |
| v-ubq | 52 | 101 | 42.0% | 314 | 91.3% | 50 (100%) | 46 (92.0%) | 0/0/0/0 | 0 |
| Protein mutations: | V59I (34599C>A 34601C>T), E76G (34549T>C), S80A (34538A>C) | | | | | | | | |
| Codon mutations: | GGA58GGT (34602T>A), GTG59ATT (34599C>A 34601C>T), CCG60CCC (34596C>G), CCG61CCA (34593C>T), CAA63CAG (34587T>C), TTA66TTG (34578T>C), ATA67ATC (34575T>G), GCC69GCT (34569G>A), GAA74GAG (34554T>C), GAG76GGG (34549T>C), CGA77AGG (34545T>C 34547G>T), ACA78ACT (34542T>A), TCA80GCA (34538A>C), GAT81GAC (34533A>G), ATT84ATC (34524A>G), CAA85CAG (34521T>C), AGC88TCC (34513C>G 34514T>A), CTT96TTG (34488A>C 34490G>A), AGA97AGG (34485T>C) | | | | | | | | |
| **Proteins** | | | | | | | | | |
| ubiquitin-like protein (YP_009182246.1) | 52 | 101 | 42.0% | 314 | 91.3% | 50 (100%) | 46 (92.0%) | 0/0/0/0 | 0 |
| Protein mutations: | V59I (34599C>A 34601C>T), E76G (34549T>C), S80A (34538A>C) | | | | | | | | |
| Codon mutations: | GGA58GGT (34602T>A), GTG59ATT (34599C>A 34601C>T), CCG60CCC (34596C>G), CCG61CCA (34593C>T), CAA63CAG (34587T>C), TTA66TTG (34578T>C), ATA67ATC (34575T>G), GCC69GCT (34569G>A), GAA74GAG (34554T>C), GAG76GGG (34549T>C), CGA77AGG (34545T>C 34547G>T), ACA78ACT (34542T>A), TCA80GCA (34538A>C), GAT81GAC (34533A>G), ATT84ATC (34524A>G), CAA85CAG (34521T>C), AGC88TCC (34513C>G 34514T>A), CTT96TTG (34488A>C 34490G>A), AGA97AGG (34485T>C) | | | | | | | | |

*: Inserts / Deletes / Misaligned / Frameshifts

#### Analysis details

This analysis was performed with panviral2.64

### NGS Details (UN60): Cladosporium fulvum T-1 virus

#### Assembly

| | |
|---|---|
| **Coverage Length** | 1543 (4 contig(s)) |
| **Depth Of Coverage** | 46.5 |
| **Number Of Reads** | 605 |
| **Reads Per Million** | 12.83 rpm (after QC) |
| **Ambiguities** | 0 |
| **Assembly Method** | de novo + reference guided assembly |
| **Consensus Caller** | Bcf Tools |

#### Coverage Map

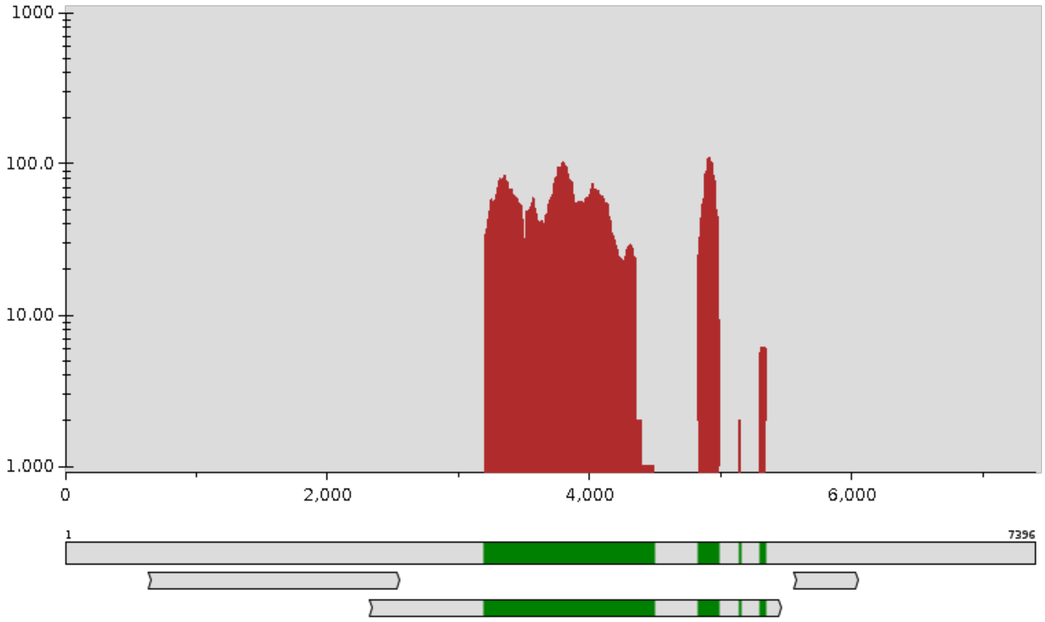

#### Assignment

| | |
|---|---|
| **Type** | Cladosporium fulvum T-1 virus (Taxonomy ID: 2052899) |
| **Reference Genome** | NC_043491.1 |
| **NT Identity (%)** | 52.2078 |
| **AA Identity (%)** | 47.4708 |
| **Number Of Stop Codons** | 0 |
| **Number Of CDS** | 3 |

#### Alignment

| | |
|---|---|
| **Alignment Score** | 88.0 (NT) + 1723.0 (AA) = 1811.0 |
| **Concordance (%)** | 27.5563 |

| Alignment Method | Global, seeded, nucleotide + amino acids (AGA) |
|---|---|

#### Genome Region

Sequence starts at position 3196 and ends at position 5342 relative to NC_043491.1 reference sequence.

#### Alignment Detailed Statistics

| | Begin | End | Coverage | Score | Concordance | Matches | Identities | I/D/M/F* | Stop Codons |
|---|---|---|---|---|---|---|---|---|---|
| **NT** | **3196** | **5342** | **20.9%** | **88** | **2.9%** | **1528 (98.3%)** | **804 (51.7%)** | **12/15** | |
| Mutations: | 3204T>A, 3205C>G, 3207T>A, 3208T>C, 3210A>C, 3211A>T, 3214G>T, 3216A>T, 3217A>G, 3219G>T, 3220A>G, 3221G>C, 3222G>C, 3223C>A, 3224A>C, 3225A>C, 3226G>T, 3229C>G, 3230G>T, 3231C>T, 3232C>G, 3233T>C, 3234T>C, 3236C>A, 3237C>A, 3238T>G, 3243A>T, 3244C>G, 3246A>C, 3251T>A, 3253_3254insAGGGATATT, 3261A>G, 3262G>A, 3265A>T, 3266A>G, 3268C>G, 3269A>T, 3271T>G, 3272C>A, 3273A>T, 3280G>T, 3281A>T, 3282A>C, 3283A>T, 3284G>A, 3285A>C, 3289T>G, 3292A>T, 3293T>G, 3294G>C, 3295G>A, 3296G>C, 3297G>A, 3299C>G, 3301C>T, 3303T>C, 3304A>T, 3307T>C, 3308C>A, 3309A>G, 3314T>G, 3317G>C, 3318A>C, 3319G>T, 3326C>T, 3328A>G, 3329C>G, 3330A>C, 3331G>T, 3332A>G, 3333C>A, 3334C>A, 3337A>G, 3338C>A, 3340A>G, 3341G>A, 3343A>G, 3344T>C, 3345G>A, 3352G>C, 3355G>A, 3356A>T, 3357A>T, 3361A>T, 3363C>A, 3364C>T, 3365A>T, 3366A>C, 3371T>C, 3372G>T, 3376A>T, 3378G>A, 3379A>G, 3381G>C, 3382A>T, 3385C>T, 3387C>A, 3388C>G, 3389T>G, 3391A>T, 3392A>C, 3393G>C, 3394T>C, 3395G>T, 3396C>A, 3397A>T, 3400A>T, 3401A>G, 3403T>A, 3406A>T, 3407T>G, 3408G>T, 3409C>T, 3410A>T, 3412G>A, 3415C>T, 3416G>C, 3417T>A, 3418T>G, 3419C>A, 3420C>A, 3425G>C, 3426C>A, 3427A>G, 3428A>G, 3430C>T, 3434A>T, 3435A>C, 3437C>T, 3440C>A, 3442A>G, 3443C>A, 3445C>G, 3446G>T, 3447T>G, 3448A>C, 3449C>A, 3450A>T, 3451A>G, 3457C>T, 3460A>G, 3461A>G, 3462A>C, 3464T>C, 3470G>A, 3473A>G, 3478G>T, 3479A>G, 3481C>G, 3488C>A, 3489G>A, 3490A>G, 3496G>A, 3499A>T, 3502C>T, 3504A>G, 3505C>G, 3508C>T, 3511A>T, 3514A>T, 3515G>T, 3516C>T, 3517A>G, 3518C>A, 3519A>T, 3520A>G, 3526A>G, 3529A>G, 3531C>G, 3533G>A, 3534G>A, 3536T>G, 3538A>G, 3539G>T, 3540A>G, 3546A>T, 3550G>T, 3554A>C, 3556T>A, 3559C>T, 3562A>T, 3563C>A, 3565A>G, 3566G>T, 3567A>C, 3568C>A, 3570C>G, 3573T>A, 3574C>T, 3576A>G, 3577T>G, 3578G>C, 3579C>A, 3580T>G, 3581A>G, 3583C>T, 3584C>A, 3586A>G, 3589G>A, 3598A>G, 3601A>T, 3604A>G, 3605T>C, 3606G>C, 3607G>T, 3610A>G, 3613C>T, 3614G>A, 3618T>G, 3619C>T, 3620A>G, 3621G>T, 3622G>A, 3625A>T, 3628A>G, 3631C>T, 3634A>G, 3635C>T, 3636T>C, 3637C>A, 3640C>T, 3647T>C, 3649G>T, 3652C>A, 3658A>G, 3659A>T, 3661G>T, 3664A>G, 3670C>T, 3673C>T, 3676A>C, 3679C>G, 3682A>T, 3683T>A, 3687G>T, 3688C>T, 3689C>T, 3690A>G, 3691G>C, 3692G>A, 3694C>T, 3695C>T, 3697T>A, 3698G>A, 3700C>G, 3703C>T, 3706A>C, 3707A>G, 3708C>T, 3712T>G, 3713A>T, 3714G>T, 3715A>T, 3719C>T, 3720T>A, 3721A>T, 3724C>T, 3727C>T, 3729T>A, 3730G>T, 3732G>T, 3733C>T, 3736C>A, 3739T>A, 3741C>T, 3742T>C, 3745C>T, 3746A>C, 3748G>A, 3751C>T, 3757A>T, 3758C>G, 3760G>C, 3761G>A, 3763C>A, 3766C>T, 3768C>G, 3769A>T, 3770A>C, 3773_3775delGGA, 3776T>A, 3778C>G, 3779C>T, 3781C>G, 3782C>G, 3784G>T, 3791A>G, 3792C>T, 3793C>T, 3795A>G, 3796G>C, 3799A>T, 3800G>T, 3802T>G, 3803C>A, 3804A>G, 3805A>T, 3806G>T, 3807A>T, 3808T>G, 3811G>A, 3814C>A, 3815G>T, 3816A>C, 3817A>T, 3820A>T, 3821C>T, 3823C>A, 3825C>G, 3827A>C, 3829G>A, 3831C>A, 3833G>A, 3834G>C, 3835A>T, 3836T>C, 3839A>T, 3841G>T, 3842A>G, 3843C>T, 3844A>T, 3845G>A, 3846C>A, 3848C>A, 3849C>T, 3850C>G, 3856A>G, 3859C>T, 3862A>G, 3865C>T, 3866C>G, 3867A>C, 3869A>C, 3871G>A, 3877A>G, 3878G>A, 3880C>A, 3886T>C, 3887T>C, 3892C>G, 3893T>C, 3894T>A, 3896A>C, 3898C>A, 3899A>G, 3901C>T, 3905A>G, 3906C>A, 3909C>A, 3910A>C, 3911G>C, 3912G>A, 3913G>A, 3914A>G, 3918C>G, 3922C>G, 3925C>T, 3929G>A, 3930C>A, 3931A>G, 3934G>A, 3935A>G, 3936C>T, 3937A>G, 3940G>A, 3941T>G, 3943A>C, 3946C>T, 3947A>G, 3948G>T, 3949A>T, 3952A>T, 3957C>A, 3960A>C, 3964G>C, 3967G>A, 3970A>T, 3973C>G, 3980G>C, 3982A>G, 3983C>A, 3984A>G, 3988A>T, 3992C>T, 3994T>G, 3997A>G, 4000C>A, 4003C>T, 4010A>T, 4012C>T, 4025A>G, 4026A>C, 4027G>T, 4029A>G, 4030C>A, 4033T>C, 4041C>A, 4048A>T, 4049C>G, 4052A>C, 4057G>T, 4058A>G, 4059T>A, 4060G>T, 4061C>T, 4063T>G, 4064A>C, 4065C>T, 4066A>G, 4068G>A, 4069A>G, 4072A>G, 4075C>A, 4081C>G, 4085A>G, 4086A>T, 4091G>T, 4092G>C, 4093A>G, 4094A>G, 4095A>T, 4097G>A, 4098A>G, 4100C>T, 4101A>G, 4102G>C, 4103A>G, 4104C>A, 4105C>A, 4111G>T, 4114C>T, 4115A>C, 4117A>G, 4120A>T, 4121C>T, 4123C>G, 4130C>G, 4131A>C, 4132G>T, 4133T>A, 4134G>T, 4135C>T, 4138T>A, 4143C>A, 4144C>A, 4149C>T, 4150G>A, 4153T>G, 4154C>A, 4155G>A, 4156A>G, 4159A>G, 4160T>C, 4161T>C, 4162C>A, 4165T>C, 4166G>T, 4167G>T, 4169A>G, 4170G>A, 4171C>G, 4172A>C, 4173A>T, 4174G>A, 4175G>C, 4176A>C, 4177A>T, 4178G>T, 4180C>T, 4181C>G, 4183C>G, 4184A>G, 4186C>A, 4187G>C, 4189G>C, 4192C>T, 4195C>T, 4201T>G, 4204T>C, 4206T>A, 4207G>A, 4210A>T, 4211A>G, 4213A>G, 4216C>A, 4218C>G, 4219A>C, 4220T>G, 4221G>T, 4222T>A, 4223C>T, 4225A>G, 4226A>G, 4227C>T, 4228A>G, 4231G>A, 4232_4237delACACAC, 4240T>A, 4243G>T, 4244_4249delAAAAGA, 4252C>T, 4255A>G, 4263A>T, 4265T>G, 4267T>A, 4268T>A, 4269C>G, 4271C>A, 4277A>T, 4279G>A, 4281C>A, 4282C>T, 4283A>G, 4284C>A, 4285A>T, 4288G>T, 4291A>G, 4296A>G, 4297C>G, 4301G>T, 4302A>C, 4303C>T, 4305T>C, 4306C>T, 4312C>A, 4319C>A, 4321T>G, 4322C>G, 4324A>T, 4327C>A, 4328A>G, 4330T>G, 4333T>G, 4334G>C, 4335C>A, 4337G>T, 4338C>G, 4339C>T, 4340A>C, 4345A>G, 4348T>A, 4352A>C, 4354A>T, 4356T>C, 4357G>C, 4358T>G, 4360C>G, 4361G>C, 4363C>T, 4364G>C, 4366G>A, 4369C>A, 4369_4370insTCC, 4372A>T, 4374C>A, 4375G>C, 4378G>A, 4379T>A, 4381A>T, 4382A>G, 4383C>A, 4384G>A, 4390T>C, 4391T>A, 4396C>T, 4399C>T, 4402G>A, 4403A>G, 4404A>C, 4406C>T, 4408C>A, 4409A>G, 4410C>A, 4411G>A, 4417C>T, 4419C>T, 4421A>T, 4423G>A, 4426G>T, 4429G>A, 4430G>A, 4431A>C, 4432A>T, 4433C>T, 4435T>A, 4437C>A, 4439C>T, 4440G>C, 4441A>T, 4442A>C, 4444A>T, 4447A>G, 4450C>A, 4453C>T, 4457T>G, 4459G>A, 4462G>C, 4463C>T, 4465G>T, 4466C>T, 4468T>A, 4469G>T, 4470G>C, 4471G>C, 4472C>G, 4474G>A, 4477C>T, 4480G>C, 4483C>T, 4484G>A, 4485A>T, 4486A>T, 4832C>G, 4833G>A, 4834C>G, 4836A>T, 4837C>T, 4838A>G, 4840C>G, 4841A>G, 4842A>C, 4843G>C, 4846A>G, 4849C>T, 4850G>C, 4851T>C, 4853C>A, 4855C>T, 4865A>G, 4866A>T, 4867C>T, 4870A>G, 4872C>T, 4873T>A, 4875C>A, 4878G>A, 4879G>T, 4882C>A, 4883G>A, 4884C>G, 4886A>C, 4887A>C, 4889T>G, 4890A>G, 4894T>C, 4896A>T, 4897C>A, 4900A>T, 4904T>G, 4905T>A, 4906C>T, 4908G>T, 4909G>T, 4910A>G, 4911C>A, 4912A>G, 4913C>A, 4914C>T, 4915A>T, 4918A>T, 4921G>T, 4922A>T, 4923A>G, 4924A>G, 4925C>A, 4926C>A, 4927A>G, 4933C>A, 4939T>G, 4941C>A, 4942G>C, 4951C>T, 4952A>G, 4954T>C, 4955A>G, 4956C>T, 4958A>G, 4959A>G, 4961C>T, 4963C>A, 4966G>T, 4968G>A, 4970T>A, 4972A>T, 4974A>G, 4976G>A, 4977A>G, 4978T>G, 4980G>A, 5132G>T, 5133T>C, 5134A>C, 5135T>A, 5137C>T, 5141A>T, 5143A>T, 5146C>T, 5151A>G, 5152C>T, 5154A>C, 5155G>T, 5157T>A, 5294A>G, 5296C>G, 5298A>C, 5299C>G, 5300T>G, 5301A>C, 5302C>G, 5303G>A, 5304C>G, 5305A>C, 5308A>G, 5309G>A, 5310A>G, 5311C>A, 5314C>T, 5320T>G, 5321T>G, 5322C>A, 5323A>T, 5329G>T, 5330C>G, 5331C>A, 5332A>T, 5334T>C, 5335G>T, 5341G>A, 5342A>T | | | | | | | | |
| **CDS** | | | | | | | | | |
| homologue_of_retroviral_POL_genes | 291 | 1006 | 49.2% | 1723 | 47.7% | 510 (98.3%) | 244 (47.0%) | 4/5/0/0 | 0 |

| | Begin | End | Coverage | Score | Concordance | Matches | Identities | I/D/M/F* | Stop Codons |
|---|---|---|---|---|---|---|---|---|---|
| **NT** | **3196** | **5342** | **20.9%** | **88** | **2.9%** | **1528 (98.3%)** | **804 (51.7%)** | **12/15** | |
| Protein mutations: | L293Q (3204T>A 3205C>G), F294Y (3207T>A 3208T>C), E295A (3210A>C 3211A>T), E296D (3214G>T), E297V (3216A>T 3217A>G), R298M (3219G>T 3220A>G), G299P (3221G>C 3222G>C 3223C>A), K300P (3224A>C 3225A>C 3226G>T), D301E (3229C>G), A302L (3230G>T 3231C>T 3232C>G), L303P (3233T>C 3234T>C), P304K (3236C>A 3237C>A 3238T>G), H306L (3243A>T 3244C>G), Q307P (3246A>C), W309R (3251T>A), W309_D310insRDI (3253_3254insAGGGATATT), K312R (3261A>G 3262G>A), N314E (3266A>G 3268C>G), I315L (3269A>T 3271T>G), Q316M (3272C>A 3273A>T), K319S (3281A>T 3282A>C 3283A>T), E320T (3284G>A 3285A>C), W323A (3293T>G 3294G>C 3295G>A), G324Q (3296G>C 3297G>A), P325A (3299C>G 3301C>T), L326P (3303T>C 3304A>T), Q328R (3308C>A 3309A>G), S330A (3314T>G), E331P (3317G>C 3318A>C 3319G>T), Q335A (3329C>G 3330A>C 3331G>T), T336E (3332A>G 3333C>A 3334C>A), E339K (3341G>A 3343A>G), W340Q (3344T>C 3345G>A), K342N (3352G>C), K344L (3356A>T 3357A>T), A346D (3363C>A 3364C>T), K347S (3365A>T 3366A>C), W349L (3371T>C 3372G>T), R351Q (3378G>A 3379A>G), R352P (3381G>C 3382A>T), T354K (3387C>A 3388C>G), S355A (3389T>G 3391A>T), S356P (3392A>C 3393G>C 3394T>C), A357Y (3395G>T 3396C>A 3397A>T), T359A (3401A>G 3403T>A), C361V (3407T>G 3408G>T 3409C>T), M362L (3410A>T 3412G>A), V364Q (3416G>C 3417T>A 3418T>G), P365K (3419C>A 3420C>A), A367Q (3425G>C 3426C>A 3427A>G), N368D (3428A>G 3430C>T), K370S (3434A>T 3435A>C), L373M (3443C>A 3445C>G), V374C (3446G>T 3447T>G 3448A>C), Q375M (3449C>A 3450A>T 3451A>G), K379A (3461A>G 3462A>C), E382K (3470G>A), I383V (3473A>G), I385V (3479A>G 3481C>G), R388K (3488C>A 3489G>A 3490A>G), N393R (3504A>G 3505C>G), E395D (3511A>T), E396D (3514A>T), A397L (3515G>T 3516C>T 3517A>G), Q398M (3518C>A 3519A>T 3520A>G), T402S (3531C>G), G403K (3533G>A 3534G>A), S404A (3536T>G 3538A>G), D405C (3539G>T 3540A>G), Y407F (3546A>T), I410L (3554A>C 3556T>A), D414S (3566G>T 3567A>C 3568C>A), A415G (3570C>G), F416Y (3573T>A 3574C>T), Y417W (3576A>G 3577T>G), A418Q (3578G>C 3579C>A 3580T>G), I419V (3581A>G 3583C>T), M421I (3589G>A), E425D (3601A>T), W427P (3605T>C 3606G>C 3607G>T), A430T (3614G>A), F431C (3618T>G 3619C>T), R432V (3620A>G 3621G>T 3622G>A), L437S (3635C>T 3636T>C 3637C>A), M445F (3659A>T 3661G>T), S453T (3683T>A), C454F (3687G>T 3688C>T), Q455C (3689C>T 3690A>G 3691G>C), D456N (3692G>A 3694C>T), V458M (3698G>A 3700C>G), E460D (3706A>C), T461V (3707A>G 3708C>T), R463F (3713A>T 3714G>T 3715A>T), L465Y (3719C>T 3720T>A 3721A>T), V468D (3729T>A 3730G>T), C469F (3732G>T 3733C>T), A472V (3741C>T 3742T>C), M474L (3746A>C 3748G>A), L478V (3758C>G 3760G>C), V479I (3761G>A 3763C>A), T481S (3768C>G 3769A>T), K482Q (3770A>C), G483del (3773_3775delGGA), S484T (3776T>A 3778C>G), Q486D (3782C>G 3784G>T), T489V (3791A>G 3792C>T 3793C>T), K490S (3795A>G 3796G>C), Q491H (3799A>T), V492L (3800G>T 3802T>G), Q493S (3803C>A 3804A>G 3805A>T), D494L (3806G>T 3807A>T 3808T>G), F496L (3814C>A), E497S (3815G>T 3816A>C 3817A>T), T500R (3825C>G), K501Q (3827A>C 3829G>A), S502Y (3831C>A), G503T (3833G>A 3834G>C 3835A>T), F504L (3836T>C), K505Y (3839A>T 3841G>T), T506V (3842A>G 3843C>T 3844A>T), A507K (3845G>A 3846C>A), P508M (3848C>A 3849C>T 3850C>G), H514A (3866C>G 3867A>C), K515Q (3869A>C 3871G>A), V518I (3878G>A 3880C>A), F523H (3893T>C 3894T>A), I524L (3896A>C 3898C>A), I525V (3899A>G 3901C>T), T527E (3905A>G 3906C>A), T528N (3909C>A 3910A>C), G529Q (3911G>C 3912G>A 3913G>A), I530V (3914A>G), T531R (3918C>G), I532M (3922C>G), A535K (3929G>A 3930C>A 3931A>G), T537V (3935A>G 3936C>T 3937A>G), S539A (3941T>G 3943A>C), R541V (3947A>G 3948G>T 3949A>T), E542D (3952A>T), P544Q (3957C>A), E545A (3960A>C), V552L (3980G>C 3982A>G), Q553R (3983C>A 3984A>G), N562Y (4010A>T 4012C>T), K567A (4025A>G 4026A>C 4027G>T), D568G (4029A>G 4030C>A), T572K (4041C>A), P575A (4049C>G), M576L (4052A>C), M578D (4058A>G 4059T>A 4060G>T), T580L (4064A>C 4065C>T 4066A>G), R581K (4068G>A 4069A>G), D583E (4075C>A), N585K (4081C>G), K587V (4085A>G 4086A>T), G589S (4091G>T 4092G>C 4093A>G), K590V (4094A>G 4095A>T), E591R (4097G>A 4098A>G), Q592C (4100C>T 4101A>G 4102G>C), T593E (4103A>G 4104C>A 4105C>A), K597Q (4115A>C 4117A>G), R598S (4120A>T), Q602A (4130C>G 4131A>C 4132G>T), C603I (4133T>A 4134G>T 4135C>T), A606E (4143C>A 4144C>A), T608I (4149C>T 4150G>A), R610K (4154C>A 4155G>A 4156A>G), F612P (4160T>C 4161T>C 4162C>A), G614F (4166G>T 4167G>T), S615E (4169A>G 4170G>A 4171C>G), K616L (4172A>C 4173A>T 4174G>A), E617P (4175G>C 4176A>C 4177A>T), V618F (4178G>T 4180C>T), H619E (4181C>G 4183C>G), I620V (4184A>G 4186C>A), E621H (4187G>C 4189G>C), M627K (4206T>A 4207G>A), I629V (4211A>G 4213A>G), A631G (4218C>G 4219A>C), C632V (4220T>G 4221G>T 4222T>A), T634V (4226A>G 4227C>T 4228A>G), T636_H637del (4232_4237delACACAC), D638E (4240T>A), K640_R641del (4244_4249delAAAAGA), Y646F (4263A>T), Y647E (4265T>G 4267T>A), M651L (4277A>T 4279G>A), T652N (4281C>A 4282C>T), T653D (4283A>G 4284C>A 4285A>T), N657R (4296A>G 4297C>G), D659S (4301G>T 4302A>C 4303C>T), I660T (4305T>C 4306C>T), D662E (4312C>A), L665M (4319C>A 4321T>G), L666V (4322C>G 4324A>T), I668V (4328A>G 4330T>G), A670H (4334G>C 4335C>A), A671C (4337G>T 4338C>G 4339C>T), M672L (4340A>C), H674Q (4348T>A), V677A (4356T>C 4357G>C), Y678E (4358T>G 4360C>G), V679L (4361G>C 4363C>T), E680Q (4364G>C 4366G>A), G681_P682insS (4369_4370insTCC), P683H (4374C>A 4375G>C), L685I (4379T>A 4381A>T), T686E (4382A>G 4383C>A 4384G>A), S689T (4391T>A), N693A (4403A>G 4404A>C), T695E (4409A>G 4410C>A 4411G>A), T698M (4419C>T), T699S (4421A>T 4423G>A), E702T (4430G>A 4431A>C 4432A>T), T704N (4437C>A), R705S (4439C>T 4440G>C 4441A>T), S711A (4457T>G 4459G>A), E712D (4462G>C), L713F (4463C>T 4465G>T), G715S (4469G>T 4470G>C 4471G>C), Q716E (4472C>G 4474G>A), K718N (4480G>C), E720I (4484G>A 4485A>T 4486A>T), R836E (4832C>G 4833G>A 4834C>G), Y837F (4836A>T 4837C>T), I838V (4838A>G 4840C>G), K839A (4841A>G 4842A>C 4843G>C), V842P (4850G>C 4851T>C), H843N (4853C>A 4855C>T), N847V (4865A>G 4866A>T 4867C>T), A849V (4872C>T 4873T>A), A850E (4875C>A), R851H (4878G>A 4879G>T), H852Q (4882C>A), A853R (4883G>A 4884C>G), K854P (4886A>C 4887A>C), Y855G (4889T>G 4890A>G), H857L (4896A>T 4897C>A), F860D (4904T>G 4905T>A 4906C>T), R861I (4908G>T 4909G>T), T862E (4910A>G 4911C>A 4912A>G), P863I (4913C>A 4914C>T 4915A>T), K866W (4922A>T 4923A>G 4924A>G), P867K (4925C>A 4926C>A 4927A>G), D869E (4933C>A), T872N (4941C>A 4942G>C), I876V (4952A>G 4954T>C), T877V (4955A>G 4956C>T), K878G (4958A>G 4959A>G), R881K (4968G>A), S882T (4970T>A 4972A>T), K883R (4974A>G), D884R (4976G>A 4977A>G 4978T>G), R885Q (4980G>A), V936S (5132G>T 5133T>C 5134A>C), F937I (5135T>A 5137C>T), T939S (5141A>T 5143A>T), D942G (5151A>G 5152C>T), K943T (5154A>C 5155G>T), I990V (5294A>G 5296C>G), N991T (5298A>C 5299C>G), Y992A (5300T>G 5301A>C 5302C>G), A993S (5303G>A 5304C>G 5305A>C), D995R (5309G>A 5310A>G 5311C>A), S999D (5321T>G 5322C>A 5323A>T), P1002D (5330C>G 5331C>A 5332A>T), M1003T (5334T>C 5335G>T) | | | | | | | | |

| | Begin | End | Coverage | Score | Concordance | Matches | Identities | I/D/M/F* | Stop Codons |
|---|---|---|---|---|---|---|---|---|---|
| **NT** | **3196** | **5342** | **20.9%** | **88** | **2.9%** | **1528 (98.3%)** | **804 (51.7%)** | **12/15** | |

Codon mutations:

CTC293CAG (3204T>A 3205C>G), TTT294TAC (3207T>A 3208T>C), GAA295GCT (3210A>C 3211A>T), GAG296GAT (3214G>T), GAA297GTG (3216A>T 3217A>G), AGA298ATG (3219G>T 3220A>G), GGC299CCA (3221G>C 3222G>C 3223C>A), AAG300CCT (3224A>C 3225A>C 3226G>T), GAC301GAG (3229C>G), GCC302TTG (3230G>T 3231C>T 3232C>G), TTA303CCA (3233T>C 3234T>C), CCT304AAG (3236C>A 3237C>A 3238T>G), CAC306CTG (3243A>T 3244C>G), CAA307CCA (3246A>C), TGG309AGG (3251T>A), TGG309_GAT310insAGGGATATT (3253_3254insAGGGATATT), AAG312AGA (3261A>G 3262G>A), ATA313ATT (3265A>T), AAC314GAG (3266A>G 3268C>G), ATT315TTG (3269A>T 3271T>G), CAG316ATG (3272C>A 3273A>T), GGG318GGT (3280G>T), AAA319TCT (3281A>T 3282A>C 3283A>T), GAG320ACG (3284G>A 3285A>C), CCT321CCG (3289T>G), CCA322CCT (3292A>T), TGG323GCA (3293T>G 3294G>C 3295G>A), GGA324CAA (3296G>C 3297G>A), CCC325GCT (3299C>G 3301C>T), CTA326CCT (3303T>C 3304A>T), TAT327TAC (3307T>C), CAA328AGA (3308C>A 3309A>G), TCT330GCT (3314T>G), GAG331CCT (3317G>C 3318A>C 3319G>T), CTA334TTG (3326C>T 3328A>G), CAG335GCT (3329C>G 3330A>C 3331G>T), ACC336GAA (3332A>G 3333C>A 3334C>A), CTA337CTG (3337A>G), CGA338AGG (3338C>A 3340A>G), GAA339AAG (3341G>A 3343A>G), TGG340CAG (3344T>C 3345G>A), AAG342AAC (3352G>C), GAG343GAA (3355G>A), AAG344TTG (3356A>T 3357A>T), CTA345CTT (3361A>T), GCC346GAT (3363C>A 3364C>T), AAA347TCA (3365A>T 3366A>C), TGG349CTG (3371T>C 3372G>T), ATA350ATT (3376A>T), CGA351CAG (3378G>A 3379A>G), CGA352CCT (3381G>C 3382A>T), TCC353TCT (3385C>T), ACC354AAG (3387C>A 3388C>G), TCA355GCT (3389T>G 3391A>T), AGT356CCC (3392A>C 3393G>C 3394T>C), GCA357TAT (3395G>T 3396C>A 3397A>T), GGA358GGT (3400A>T), ACT359GCA (3401A>G 3403T>A), CCA360CCT (3406A>T), TGC361GTT (3407T>G 3408G>T 3409C>T), ATG362TTA (3410A>T 3412G>A), TTC363TTT (3415C>T), GTT364CAG (3416G>C 3417T>A 3418T>G), CCA365AAA (3419C>A 3420C>A), GCA367CAG (3425G>C 3426C>A 3427A>G), AAC368GAT (3428A>G 3430C>T), AAA370TCA (3434A>T 3435A>C), CTA371TTA (3437C>T), CGA372AGG (3440C>A 3442A>G), CTC373ATG (3443C>A 3445C>G), GTA374TGC (3446G>T 3447T>G 3448A>C), CAA375ATG (3449C>A 3450A>T 3451A>G), TAC377TAT (3457C>T), CGA378CGG (3460A>G), AAG379GCG (3461A>G 3462A>C), TTG380CTG (3464T>C), GAG382AAG (3470G>A), ATC383GTC (3473A>G), ACG384ACT (3478G>T), ATC385GTG (3479A>G 3481C>G), CGA388AAG (3488C>A 3489G>A 3490A>G), CCG390CCA (3496G>A), CTA391CTT (3499A>T), CCC392CCT (3502C>T), AAC393AGG (3504A>G 3505C>G), ATC394ATT (3508C>T), GAA395GAT (3511A>T), GAA396GAT (3514A>T), GCA397TTG (3515G>T 3516C>T 3517A>G), CAA398ATG (3518C>A 3519A>T 3520A>G), AGA400AGG (3526A>G), TTA401TTG (3529A>G), ACC402AGC (3531C>G), GGA403AAA (3533G>A 3534G>A), TCA404GCG (3536T>G 3538A>G), GAC405TGC (3539G>T 3540A>G), TAC407TTC (3546A>T), ACG408ACT (3550G>T), ATT410CTA (3554A>C 3556T>A), GAC411GAT (3559C>T), CTA412CTT (3562A>T), CGA413AGG (3563C>A 3565A>G), GAC414TCA (3566G>T 3567A>C 3568C>A), GCC415GGC (3570C>G), TTC416TAT (3573T>A 3574C>T), TAT417TGG (3576A>G 3577T>G), GCT418CAG (3578G>C 3579C>A 3580T>G), ATC419GTT (3581A>G 3583C>T), CGA420AGG (3584C>A 3586A>G), ATG421ATA (3589G>A), GGA424GGG (3598A>G), GAA425GAT (3601A>T), GAA426GAG (3604A>G), TGG427CCT (3605T>C 3606G>C 3607G>T), AAA428AAG (3610A>G), ACC429ACT (3613C>T), GCT430ACT (3614G>A), TTC431TGT (3618T>G 3619C>T), AGG432GTA (3620A>G 3621G>T 3622G>A), ACA433ACT (3625A>T), AGA434AGG (3628A>G), TAC435TAT (3631C>T), GGA436GGG (3634A>G), CTC437TCA (3635C>T 3636T>C 3637C>A), TAC438TAT (3640C>T), TTG441CTT (3647T>C 3649G>T), GTC442GTA (3652C>A), CCA444CCG (3658A>G), ATG445TTT (3659A>T 3661G>T), GGA446GGG (3664A>G), ACC448ACT (3670C>T), AAC449AAT (3673C>T), GCA450GCC (3676A>C), CCC451CCG (3679C>G), GCA452GCT (3682A>T), TCC453ACC (3683T>A), TGC454TTT (3687G>T 3688C>T), CAG455TGC (3689C>T 3690A>G 3691G>C), GAC456AAT (3692G>A 3694C>T), CTT457TTA (3695C>T 3697T>A), GTC458ATG (3698G>A 3700C>G), AAC459AAT (3703C>T), GAA460GAC (3706A>C), ACA461GTA (3707A>G 3708C>T), CTT462CTG (3712T>G), AGA463TTT (3713A>T 3714G>T 3715A>T), CTA465TAT (3719C>T 3720T>A 3721A>T), CTC466CTT (3724C>T), GAC467GAT (3727C>T), GTG468GAT (3729T>A 3730G>T), TGC469TTT (3732G>T 3733C>T), GTC470GTA (3736C>A), GTT471GTA (3739T>A), GCT472GTC (3741C>T 3742T>C), TAC473TAT (3745C>T), ATG474CTA (3746A>C 3748G>A), GAC475GAT (3751C>T), ATA477ATT (3757A>T), CTG478GTC (3758C>G 3760G>C), GTC479ATA (3761G>A 3763C>A), TAC480TAT (3766C>T), ACA481AGT (3768C>G 3769A>T), AAA482CAA (3770A>C), GGA483del (3773_3775delGGA), TCC484ACG (3776T>A 3778C>G), CTC485TTG (3779C>T 3781C>G), CAG486GAT (3782C>G 3784G>T), ACC489GTT (3791A>G 3792C>T 3793C>T), AAG490AGC (3795A>G 3796G>C), CAA491CAT (3799A>T), GTT492TTG (3800G>T 3802T>G), CAA493AGT (3803C>A 3804A>G 3805A>T), GAT494TTG (3806G>T 3807A>T 3808T>G), GTG495GTA (3811G>A), TTC496TTA (3814C>A), GAA497TCT (3815G>T 3816A>C 3817A>T), CGA498CGT (3820A>T), CTC499TTA (3821C>T 3823C>A), ACG500AGG (3825C>G), AAG501CAA (3827A>C 3829G>A), TCC502TAC (3831C>A), GGA503ACT (3833G>A 3834G>C 3835A>T), TTC504CTC (3836T>C), AAG505TAT (3839A>T 3841G>T), ACA506GTT (3842A>G 3843C>T 3844A>T), GCA507AAA (3845G>A 3846C>A), CCC508ATG (3848C>A 3849C>T 3850C>G), AAA510AAG (3856A>G), TGC511TGT (3859C>T), GAA512GAG (3862A>G), TTC513TTT (3865C>T), CAC514GCC (3866C>G 3867A>C), AAG515CAA (3869A>C 3871G>A), GAA517GAG (3877A>G), GTC518ATA (3878G>A 3880C>A), TTT520TTC (3886T>C), TTA521CTA (3887T>C), GGC522GGG (3892C>G), TTT523CAT (3893T>C 3894T>A), ATC524CTA (3896A>C 3898C>A), ATC525GTT (3899A>G 3901C>T), ACA527GAA (3905A>G 3906C>A), ACA528AAC (3909C>A 3910A>C), GGG529CAA (3911G>C 3912G>A 3913G>A), ATA530GTA (3914A>G), ACG531AGG (3918C>G), ATC532ATG (3922C>G), GAC533GAT (3925C>T), GCA535AAG (3929G>A 3930C>A 3931A>G), AAG536AAA (3934G>A), ACA537GTG (3935A>G 3936C>T 3937A>G), CAG538CAA (3940G>A), TCA539GCC (3941T>G 3943A>C), ATC540ATT (3946C>T), AGA541GTT (3947A>G 3948G>T 3949A>T), GAA542GAT (3952A>T), CCA544CAA (3957C>A), GAA545GCA (3960A>C), CCG546CCC (3964G>C), AAG547AAA (3967G>A), ACA548ACT (3970A>T), GTC549GTG (3973C>G), GTA552CTG (3980G>C 3982A>G), CAG553AGG (3983C>A 3984A>G), TCA554TCT (3988A>T), CTT556TTG (3992C>T 3994T>G), GGA557GGG (3997A>G), CTC558CTA (4000C>A), GCC559GCT (4003C>T), AAC562TAT (4010A>T 4012C>T), AAG567GCT (4025A>G 4026A>C 4027G>T), GAC568GGA (4029A>G 4030C>A), TAT569TAC (4033T>C), ACA572AAA (4041C>A), GCA574GCT (4048A>T), CCA575GCA (4049C>G), ATG576CTG (4052A>C), ACG577ACT (4057G>T), ATG578GAT (4058A>G 4059T>A 4060G>T), CTT579TTG (4061C>T 4063T>G), ACA580CTG (4064A>C 4065C>T 4066A>G), AGA581AAG (4068G>A 4069A>G), AAA582AAG (4072A>G), GAC583GAA (4075C>A), AAC585AAG (4081C>G), AAA587GTA (4085A>G 4086A>T), GGA589TCG (4091G>T 4092G>C 4093A>G), AAA590GTA (4094A>G 4095A>T), GAA591AGA (4097G>A 4098A>G), CAG592TGC (4100C>T 4101A>G 4102G>C), ACC593GAA (4103A>G 4104C>A 4105C>A), GCG595GCT (4111G>T), TTC596TTT (4114C>T), AAA597CAG (4115A>C 4117A>G), AGA598AGT (4120A>T), CTC599TTG (4121C>T 4123C>G), CAG602GCT (4130C>G 4131A>C 4132G>T), TGC603ATT (4133T>A 4134G>T 4135C>T), GCT604GCA (4138T>A), GCC606GAA (4143C>A 4144C>A), ACG608ATA (4149C>T 4150G>A), CTT609CTG (4153T>G), CGA610AAG (4154C>A 4155G>A 4156A>G), CTA611CTG (4159A>G), TTC612CCA (4160T>C 4161T>C 4162C>A), GAT613GAC (4165T>C), GGT614TTT (4166G>T 4167G>T), AGC615GAG (4169A>G 4170G>A 4171C>G), AAG616CTA (4172A>C 4173A>T 4174G>A), GAA617CCT (4175G>C 4176A>C 4177A>T), GTC618TTT (4178G>T 4180C>T), CAC619GAG (4181C>G 4183C>G), ATC620GTA (4184A>G 4186C>A), GAG621CAC (4187G>C 4189G>C), ACC622ACT (4192C>T), GAC623GAT (4195C>T), TCT625TCG (4201T>G), GAT626GAC (4204T>C), ATG627AAA (4206T>A 4207G>A), GCA628GCT (4210A>T), ATA629GTG (4211A>G 4213A>G), GGC630GGA (4216C>A), GCA631GGC (4218C>G 4219A>C), TGT632GTA (4220T>G 4221G>T 4222T>A), CTA633TTG (4223C>T 4225A>G), ACA634GTG (4226A>G 4227C>T 4228A>G), CAG635CAA (4231G>A), ACA636_CAC637del (4232_4237delACACAC), GAT638GAA (4240T>A), GGG639GGT (4243G>T), AAA640_AGA641del (4244_4249delAAAAGA), CAC642CAT (4252C>T), CCA643CCG (4255A>G), TAT646TTT (4263A>T), TAT647GAA (4265T>G 4267T>A), TCC648AGC (4268T>A 4269C>G), CGG649AGG (4271C>A), ATG651TTA (4277A>T 4279G>A), ACC652AAT (4281C>A 4282C>T), ACA653GAT (4283A>G 4284C>A 4285A>T), GCG654GCT (4288G>T), GAA655GAG (4291A>G), AAC657AGG (4296A>G 4297C>G), GAC659TCT (4301G>T 4302A>C 4303C>T), ATC660ACT (4305T>C 4306C>T), GAC662GAA (4312C>A), CTT665ATG (4319C>A 4321T>G), CTA666GTT (4322C>G 4324A>T), GCC667GCA (4327C>A), ATT668GTG (4328A>G 4330T>G), GTT669GTG (4333T>G), GCC670CAC (4334G>C 4335C>A), GCC671TGT (4337G>T 4338C>G 4339C>T), ATG672CTG (4340A>C), CAA673CAG (4345A>G), CAT674CAA (4348T>A), AGA676CGT (4352A>C 4354A>T), GTG677GCC (4356T>C 4357G>C), TAC678GAG (4358T>G 4360C>G), GTC679CTT (4361G>C 4363C>T), GAG680CAA (4364G>C 4366G>A), GGC681GGA (4369C>A), GGC681_CCA682insTCC (4369_4370insTCC), CCA682CCT (4372A>T), CCG683CAC (4374C>A 4375G>C), AAG684AAA (4378G>A), TTA685ATT (4379T>A 4381A>T), ACG686GAA (4382A>G 4383C>A 4384G>A), CTT688CTC (4390T>C), TCA689ACA (4391T>A), GAC690GAT (4396C>T), CAC691CAT (4399C>T), AAG692AAA (4402G>A), AAT693GCT (4403A>G 4404A>C), CTC694TTA (4406C>T 4408C>A), ACG695GAA (4409A>G 4410C>A 4411G>A), TTC697TTT (4417C>T), ACG698ATG (4419C>T), ACG699TCA (4421A>T 4423G>A), ACG700ACT (4426G>T), AAG701AAA (4429G>A), GAA702ACT (4430G>A 4431A>C 4432A>T), CTT703TTA (4433C>T 4435T>A), ACC704AAC (4437C>A), CGA705TCT (4439C>T 4440G>C 4441A>T), AGA706CGT (4442A>C 4444A>T), CAA707CAG (4447A>G), GCC708GCA (4450C>A), CGC709CGT (4453C>T), TCG711GCA (4457T>G 4459G>A), GAG712GAC (4462G>C), CTG713TTT (4463C>T 4465G>T), CTT714TTA (4466C>T 4468T>A), GGG715TCC (4469G>T 4470G>C 4471G>C), CAG716GAA (4472C>G 4474G>A), TAC717TAT (4477C>T), AAG718AAC (4480G>C), TTC719TTT (4483C>T), GAA720ATT (4484G>A 4485A>T 4486A>T), CGC836GAG (4832C>G 4833G>A 4834C>G), TAC837TTT (4836A>T 4837C>T), ATC838GTG (4838A>G 4840C>G), AAG839GCC (4841A>G 4842A>C 4843G>C), AAA840AAG (4846A>G), TGC841TGT (4849C>T), GTA842CCA (4850G>C 4851T>C), CAC843AAT (4853C>A 4855C>T), AAC847GTT (4865A>G 4866A>T 4867C>T), AAA848AAG (4870A>G), GCT849GTA (4872C>T 4873T>A), GCA850GAA (4875C>A), CGG851CAT (4878G>A 4879G>T), CAC852CAA (4882C>A), GCG853AGG (4883G>A 4884C>G), AAA854CCA (4886A>C 4887A>C), TAC855GGC (4889T>G 4890A>G), GGT856GGC (4894T>C), CAC857CTA (4896A>T 4897C>A), CTA858CTT (4900A>T), TTC860GAT (4904T>G 4905T>A 4906C>T), AGG861ATT (4908G>T 4909G>T), ACA862GAG (4910A>G 4911C>A 4912A>G), CCA863ATT (4913C>A 4914C>T 4915A>T), CCA864CCT (4918A>T), ACG865ACT (4921G>T), AAA866TGG (4922A>T 4923A>G 4924A>G), CCA867AAG (4925C>A 4926C>A 4927A>G), GAC869GAA (4933C>A), GTT871GTG (4939T>G), ACG872AAC (4941C>A 4942G>C), TTC875TTT (4951C>T), ATT876GTC (4952A>G 4954T>C), ACG877GTG (4955A>G 4956C>T), AAA878GGA (4958A>G 4959A>G), CTC879TTA (4961C>T 4963C>A), CCG880CCT (4966G>T), AGG881AAG (4968G>A), TCA882ACT (4970T>A 4972A>T), AAG883AGG (4974A>G), GAT884AGG (4976G>A 4977A>G 4978T>G), CGA885CAA (4980G>A), GTA936TCC (5132G>T 5133T>C 5134A>C), TTC937ATT (5135T>A 5137C>T), ACA939TCT (5141A>T 5143A>T), GAC940GAT (5146C>T), GAC942GGT (5151A>G 5152C>T), AAG943ACT (5154A>C 5155G>T), CTC944CA. (5157T>A), ATC990GTG (5294A>G 5296C>G), AAC991ACG (5298A>C 5299C>G), TAC992GCG (5300T>G 5301A>C 5302C>G), GCA993AGC (5303G>A 5304C>G 5305A>C), CAA994CAG (5308A>G), GAC995AGA (5309G>A 5310A>G 5311C>A), AAC996AAT (5314C>T), GTT998GTG (5320T>G), TCA999GAT (5321T>G 5322C>A 5323A>T), CTG1001CTT (5329G>T), CCA1002GAT (5330C>G 5331C>A 5332A>T), ATG1003ACT (5334T>C 5335G>T), CAG1005CAA (5341G>A), ATC1006T.. (5342A>T)

#### Proteins

| | Begin | End | Coverage | Score | Concordance | Matches | Identities | I/D/M/F* | Stop Codons |
|---|---|---|---|---|---|---|---|---|---|
| Reverse Transcriptase (YP_009666308.1) | 291 | 1006 | 49.2% | 1723 | 47.7% | 510 (98.3%) | 244 (47.0%) | 4/5/0/0 | 0 |

| | Begin | End | Coverage | Score | Concordance | Matches | Identities | I/D/M/F* | Stop Codons |
|---|---|---|---|---|---|---|---|---|---|
| **NT** | **3196** | **5342** | **20.9%** | **88** | **2.9%** | **1528 (98.3%)** | **804 (51.7%)** | **12/15** | |
| Protein mutations: | L293Q (3204T>A 3205C>G), F294Y (3207T>A 3208T>C), E295A (3210A>C 3211A>T), E296D (3214G>T), E297V (3216A>T 3217A>G), R298M (3219G>T 3220A>G), G299P (3221G>C 3222G>C 3223C>A), K300P (3224A>C 3225A>C 3226G>T), D301E (3229C>G), A302L (3230G>T 3231C>T 3232C>G), L303P (3233T>C 3234T>C), P304K (3236C>A 3237C>A 3238T>G), H306L (3243A>T 3244C>G), Q307P (3246A>C), W309R (3251T>A), W309_D310insRDI (3253_3254insAGGGATATT), K312R (3261A>G 3262G>A), N314E (3266A>G 3268C>G), I315L (3269A>T 3271T>G), Q316M (3272C>A 3273A>T), K319S (3281A>T 3282A>C 3283A>T), E320T (3284G>A 3285A>C), W323A (3293T>G 3294G>C 3295G>A), G324Q (3296G>C 3297G>A), P325A (3299C>G 3301C>T), L326P (3303T>C 3304A>T), Q328R (3308C>A 3309A>G), S330A (3314T>G), E331P (3317G>C 3318A>C 3319G>T), Q335A (3329C>G 3330A>C 3331G>T), T336E (3332A>G 3333C>A 3334C>A), E339K (3341G>A 3343A>G), W340Q (3344T>C 3345G>A), K342N (3352G>C), K344L (3356A>T 3357A>T), A346D (3363C>A 3364C>T), K347S (3365A>T 3366A>C), W349L (3371T>C 3372G>T), R351Q (3378G>A 3379A>G), R352P (3381G>C 3382A>T), T354K (3387C>A 3388C>G), S355A (3389T>G 3391A>T), S356P (3392A>C 3393G>C 3394T>C), A357Y (3395G>T 3396C>A 3397A>T), T359A (3401A>G 3403T>A), C361V (3407T>G 3408G>T 3409C>T), M362L (3410A>T 3412G>A), V364Q (3416G>C 3417T>A 3418T>G), P365K (3419C>A 3420C>A), A367Q (3425G>C 3426C>A 3427A>G), N368D (3428A>G 3430C>T), K370S (3434A>T 3435A>C), L373M (3443C>A 3445C>G), V374C (3446G>T 3447T>G 3448A>C), Q375M (3449C>A 3450A>T 3451A>G), K379A (3461A>G 3462A>C), E382K (3470G>A), I383V (3473A>G), I385V (3479A>G 3481C>G), R388K (3488C>A 3489G>A 3490A>G), N393R (3504A>G 3505C>G), E395D (3511A>T), E396D (3514A>T), A397L (3515G>T 3516C>T 3517A>G), Q398M (3518C>A 3519A>T 3520A>G), T402S (3531C>G), G403K (3533G>A 3534G>A), S404A (3536T>G 3538A>G), D405C (3539G>T 3540A>G), Y407F (3546A>T), I410L (3554A>C 3556T>A), D414S (3566G>T 3567A>C 3568C>A), A415G (3570C>G), F416Y (3573T>A 3574C>T), Y417W (3576A>G 3577T>G), A418Q (3578G>C 3579C>A 3580T>G), I419V (3581A>G 3583C>T), M421I (3589G>A), E425D (3601A>T), W427P (3605T>C 3606G>C 3607G>T), A430T (3614G>A), F431C (3618T>G 3619C>T), R432V (3620A>G 3621G>T 3622G>A), L437S (3635C>T 3636T>C 3637C>A), M445F (3659A>T 3661G>T), S453T (3683T>A), C454F (3687G>T 3688C>T), Q455C (3689C>T 3690A>G 3691G>C), D456N (3692G>A 3694C>T), V458M (3698G>A 3700C>G), E460D (3706A>C), T461V (3707A>G 3708C>T), R463F (3713A>T 3714G>T 3715A>T), L465Y (3719C>T 3720T>A 3721A>T), V468D (3729T>A 3730G>T), C469F (3732G>T 3733C>T), A472V (3741C>T 3742T>C), M474L (3746A>C 3748G>A), L478V (3758C>G 3760G>C), V479I (3761G>A 3763C>A), T481S (3768C>G 3769A>T), K482Q (3770A>C), G483del (3773_3775delGGA), S484T (3776T>A 3778C>G), Q486D (3782C>G 3784G>T), T489V (3791A>G 3792C>T 3793C>T), K490S (3795A>G 3796G>C), Q491H (3799A>T), V492L (3800G>T 3802T>G), Q493S (3803C>A 3804A>G 3805A>T), D494L (3806G>T 3807A>T 3808T>G), F496L (3814C>A), E497S (3815G>T 3816A>C 3817A>T), T500R (3825C>G), K501Q (3827A>C 3829G>A), S502Y (3831C>A), G503T (3833G>A 3834G>C 3835A>T), F504L (3836T>C), K505Y (3839A>T 3841G>T), T506V (3842A>G 3843C>T 3844A>T), A507K (3845G>A 3846C>A), P508M (3848C>A 3849C>T 3850C>G), H514A (3866C>G 3867A>C), K515Q (3869A>C 3871G>A), V518I (3878G>A 3880C>A), F523H (3893T>C 3894T>A), I524L (3896A>C 3898C>A), I525V (3899A>G 3901C>T), T527E (3905A>G 3906C>A), T528N (3909C>A 3910A>C), G529Q (3911G>C 3912G>A 3913G>A), I530V (3914A>G), T531R (3918C>G), I532M (3922C>G), A535K (3929G>A 3930C>A 3931A>G), T537V (3935A>G 3936C>T 3937A>G), S539A (3941T>G 3943A>C), R541V (3947A>G 3948G>T 3949A>T), E542D (3952A>T), P544Q (3957C>A), E545A (3960A>C), V552L (3980G>C 3982A>G), Q553R (3983C>A 3984A>G), N562Y (4010A>T 4012C>T), K567A (4025A>G 4026A>C 4027G>T), D568G (4029A>G 4030C>A), T572K (4041C>A), P575A (4049C>G), M576L (4052A>C), M578D (4058A>G 4059T>A 4060G>T), T580L (4064A>C 4065C>T 4066A>G), R581K (4068G>A 4069A>G), D583E (4075C>A), N585K (4081C>G), K587V (4085A>G 4086A>T), G589S (4091G>T 4092G>C 4093A>G), K590V (4094A>G 4095A>T), E591R (4097G>A 4098A>G), Q592C (4100C>T 4101A>G 4102G>C), T593E (4103A>G 4104C>A 4105C>A), K597Q (4115A>C 4117A>G), R598S (4120A>T), Q602A (4130C>G 4131A>C 4132G>T), C603I (4133T>A 4134G>T 4135C>T), A606E (4143C>A 4144C>A), T608I (4149C>T 4150G>A), R610K (4154C>A 4155G>A 4156A>G), F612P (4160T>C 4161T>C 4162C>A), G614F (4166G>T 4167G>T), S615E (4169A>G 4170G>A 4171C>G), K616L (4172A>C 4173A>T 4174G>A), E617P (4175G>C 4176A>C 4177A>T), V618F (4178G>T 4180C>T), H619E (4181C>G 4183C>G), I620V (4184A>G 4186C>A), E621H (4187G>C 4189G>C), M627K (4206T>A 4207G>A), I629V (4211A>G 4213A>G), A631G (4218C>G 4219A>C), C632V (4220T>G 4221G>T 4222T>A), T634V (4226A>G 4227C>T 4228A>G), T636_H637del (4232_4237delACACAC), D638E (4240T>A), K640_R641del (4244_4249delAAAAGA), Y646F (4263A>T), Y647E (4265T>G 4267T>A), M651L (4277A>T 4279G>A), T652N (4281C>A 4282C>T), T653D (4283A>G 4284C>A 4285A>T), N657R (4296A>G 4297C>G), D659S (4301G>T 4302A>C 4303C>T), I660T (4305T>C 4306C>T), D662E (4312C>A), L665M (4319C>A 4321T>G), L666V (4322C>G 4324A>T), I668V (4328A>G 4330T>G), A670H (4334G>C 4335C>A), A671C (4337G>T 4338C>G 4339C>T), M672L (4340A>C), H674Q (4348T>A), V677A (4356T>C 4357G>C), Y678E (4358T>G 4360C>G), V679L (4361G>C 4363C>T), E680Q (4364G>C 4366G>A), G681_P682insS (4369_4370insTCC), P683H (4374C>A 4375G>C), L685I (4379T>A 4381A>T), T686E (4382A>G 4383C>A 4384G>A), S689T (4391T>A), N693A (4403A>G 4404A>C), T695E (4409A>G 4410C>A 4411G>A), T698M (4419C>T), T699S (4421A>T 4423G>A), E702T (4430G>A 4431A>C 4432A>T), T704N (4437C>A), R705S (4439C>T 4440G>C 4441A>T), S711A (4457T>G 4459G>A), E712D (4462G>C), L713F (4463C>T 4465G>T), G715S (4469G>T 4470G>C 4471G>C), Q716E (4472C>G 4474G>A), K718N (4480G>C), E720I (4484G>A 4485A>T 4486A>T), R836E (4832C>G 4833G>A 4834C>G), Y837F (4836A>T 4837C>T), I838V (4838A>G 4840C>G), K839A (4841A>G 4842A>C 4843G>C), V842P (4850G>C 4851T>C), H843N (4853C>A 4855C>T), N847V (4865A>G 4866A>T 4867C>T), A849V (4872C>T 4873T>A), A850E (4875C>A), R851H (4878G>A 4879G>T), H852Q (4882C>A), A853R (4883G>A 4884C>G), K854P (4886A>C 4887A>C), Y855G (4889T>G 4890A>G), H857L (4896A>T 4897C>A), F860D (4904T>G 4905T>A 4906C>T), R861I (4908G>T 4909G>T), T862E (4910A>G 4911C>A 4912A>G), P863I (4913C>A 4914C>T 4915A>T), K866W (4922A>T 4923A>G 4924A>G), P867K (4925C>A 4926C>A 4927A>G), D869E (4933C>A), T872N (4941C>A 4942G>C), I876V (4952A>G 4954T>C), T877V (4955A>G 4956C>T), K878G (4958A>G 4959A>G), R881K (4968G>A), S882T (4970T>A 4972A>T), K883R (4974A>G), D884R (4976G>A 4977A>G 4978T>G), R885Q (4980G>A), V936S (5132G>T 5133T>C 5134A>C), F937I (5135T>A 5137C>T), T939S (5141A>T 5143A>T), D942G (5151A>G 5152C>T), K943T (5154A>C 5155G>T), I990V (5294A>G 5296C>G), N991T (5298A>C 5299C>G), Y992A (5300T>G 5301A>C 5302C>G), A993S (5303G>A 5304C>G 5305A>C), D995R (5309G>A 5310A>G 5311C>A), S999D (5321T>G 5322C>A 5323A>T), P1002D (5330C>G 5331C>A 5332A>T), M1003T (5334T>C 5335G>T) | | | | | | | | |

| | Begin | End | Coverage | Score | Concordance | Matches | Identities | I/D/M/F* | Stop Codons |
|---|---|---|---|---|---|---|---|---|---|
| **NT** | **3196** | **5342** | **20.9%** | **88** | **2.9%** | **1528 (98.3%)** | **804 (51.7%)** | **12/15** | |
| Codon mutations: | CTC293CAG (3204T>A 3205C>G), TTT294TAC (3207T>A 3208T>C), GAA295GCT (3210A>C 3211A>T), GAG296GAT (3214G>T), GAA297GTG (3216A>T 3217A>G), AGA298ATG (3219G>T 3220A>G), GGC299CCA (3221G>C 3222G>C 3223C>A), AAG300CCT (3224A>C 3225A>C 3226G>T), GAC301GAG (3229C>G), GCC302TTG (3230G>T 3231C>T 3232C>G), TTA303CCA (3233T>C 3234T>C), CCT304AAG (3236C>A 3237C>A 3238T>G), CAC306CTG (3243A>T 3244C>G), CAA307CCA (3246A>C), TGG309AGG (3251T>A), TGG309_GAT310insAGGGATATT (3253_3254insAGGGATATT), AAG312AGA (3261A>G 3262G>A), ATA313ATT (3265A>T), AAC314GAG (3266A>G 3268C>G), ATT315TTG (3269A>T 3271T>G), CAG316ATG (3272C>A 3273A>T), GGG318GGT (3280G>T), AAA319TCT (3281A>T 3282A>C 3283A>T), GAG320ACG (3284G>A 3285A>C), CCT321CCG (3289T>G), CCA322CCT (3292A>T), TGG323GCA (3293T>G 3294G>C 3295G>A), GGA324CAA (3296G>C 3297G>A), CCC325GCT (3299C>G 3301C>T), CTA326CCT (3303T>C 3304A>T), TAT327TAC (3307T>C), CAA328AGA (3308C>A 3309A>G), TCT330GCT (3314T>G), GAG331CCT (3317G>C 3318A>C 3319G>T), CTA334TTG (3326C>T 3328A>G), CAG335GCT (3329C>G 3330A>C 3331G>T), ACC336GAA (3332A>G 3333C>A 3334C>A), CTA337CTG (3337A>G), CGA338AGG (3338C>A 3340A>G), GAA339AAG (3341G>A 3343A>G), TGG340CAG (3344T>C 3345G>A), AAG342AAC (3352G>C), GAG343GAA (3355G>A), AAG344TTG (3356A>T 3357A>T), CTA345CTT (3361A>T), GCC346GAT (3363C>A 3364C>T), AAA347TCA (3365A>T 3366A>C), TGG349CTG (3371T>C 3372G>T), ATA350ATT (3376A>T), CGA351CAG (3378G>A 3379A>G), CGA352CCT (3381G>C 3382A>T), TCC353TCT (3385C>T), ACC354AAG (3387C>A 3388C>G), TCA355GCT (3389T>G 3391A>T), AGT356CCC (3392A>C 3393G>C 3394T>C), GCA357TAT (3395G>T 3396C>A 3397A>T), GGA358GGT (3400A>T), ACT359GCA (3401A>G 3403T>A), CCA360CCT (3406A>T), TGC361GTT (3407T>G 3408G>T 3409C>T), ATG362TTA (3410A>T 3412G>A), TTC363TTT (3415C>T), GTT364CAG (3416G>C 3417T>A 3418T>G), CCA365AAA (3419C>A 3420C>A), GCA367CAG (3425G>C 3426C>A 3427A>G), AAC368GAT (3428A>G 3430C>T), AAA370TCA (3434A>T 3435A>C), CTA371TTA (3437C>T), CGA372AGG (3440C>A 3442A>G), CTC373ATG (3443C>A 3445C>G), GTA374TGC (3446G>T 3447T>G 3448A>C), CAA375ATG (3449C>A 3450A>T 3451A>G), TAC377TAT (3457C>T), CGA378CGG (3460A>G), AAG379GCG (3461A>G 3462A>C), TTG380CTG (3464T>C), GAG382AAG (3470G>A), ATC383GTC (3473A>G), ACG384ACT (3478G>T), ATC385GTG (3479A>G 3481C>G), CGA388AAG (3488C>A 3489G>A 3490A>G), CCG390CCA (3496G>A), CTA391CTT (3499A>T), CCC392CCT (3502C>T), AAC393AGG (3504A>G 3505C>G), ATC394ATT (3508C>T), GAA395GAT (3511A>T), GAA396GAT (3514A>T), GCA397TTG (3515G>T 3516C>T 3517A>G), CAA398ATG (3518C>A 3519A>T 3520A>G), AGA400AGG (3526A>G), TTA401TTG (3529A>G), ACC402AGC (3531C>G), GGA403AAA (3533G>A 3534G>A), TCA404GCG (3536T>G 3538A>G), GAC405TGC (3539G>T 3540A>G), TAC407TTC (3546A>T), ACG408ACT (3550G>T), ATT410CTA (3554A>C 3556T>A), GAC411GAT (3559C>T), CTA412CTT (3562A>T), CGA413AGG (3563C>A 3565A>G), GAC414TCA (3566G>T 3567A>C 3568C>A), GCC415GGC (3570C>G), TTC416TAT (3573T>A 3574C>T), TAT417TGG (3576A>G 3577T>G), GCT418CAG (3578G>C 3579C>A 3580T>G), ATC419GTT (3581A>G 3583C>T), CGA420AGG (3584C>A 3586A>G), ATG421ATA (3589G>A), GGA424GGG (3598A>G), GAA425GAT (3601A>T), GAA426GAG (3604A>G), TGG427CCT (3605T>C 3606G>C 3607G>T), AAA428AAG (3610A>G), ACC429ACT (3613C>T), GCT430ACT (3614G>A), TTC431TGT (3618T>G 3619C>T), AGG432GTA (3620A>G 3621G>T 3622G>A), ACA433ACT (3625A>T), AGA434AGG (3628A>G), TAC435TAT (3631C>T), GGA436GGG (3634A>G), CTC437TCA (3635C>T 3636T>C 3637C>A), TAC438TAT (3640C>T), TTG441CTT (3647T>C 3649G>T), GTC442GTA (3652C>A), CCA444CCG (3658A>G), ATG445TTT (3659A>T 3661G>T), GGA446GGG (3664A>G), ACC448ACT (3670C>T), AAC449AAT (3673C>T), GCA450GCC (3676A>C), CCC451CCG (3679C>G), GCA452GCT (3682A>T), TCC453ACC (3683T>A), TGC454TTT (3687G>T 3688C>T), CAG455TGC (3689C>T 3690A>G 3691G>C), GAC456AAT (3692G>A 3694C>T), CTT457TTA (3695C>T 3697T>A), GTC458ATG (3698G>A 3700C>G), AAC459AAT (3703C>T), GAA460GAC (3706A>C), ACA461GTA (3707A>G 3708C>T), CTT462CTG (3712T>G), AGA463TTT (3713A>T 3714G>T 3715A>T), CTA465TAT (3719C>T 3720T>A 3721A>T), CTC466CTT (3724C>T), GAC467GAT (3727C>T), GTG468GAT (3729T>A 3730G>T), TGC469TTT (3732G>T 3733C>T), GTC470GTA (3736C>A), GTT471GTA (3739T>A), GCT472GTC (3741C>T 3742T>C), TAC473TAT (3745C>T), ATG474CTA (3746A>C 3748G>A), GAC475GAT (3751C>T), ATA477ATT (3757A>T), CTG478GTC (3758C>G 3760G>C), GTC479ATA (3761G>A 3763C>A), TAC480TAT (3766C>T), ACA481AGT (3768C>G 3769A>T), AAA482CAA (3770A>C), GGA483del (3773_3775delGGA), TCC484ACG (3776T>A 3778C>G), CTC485TTG (3779C>T 3781C>G), CAG486GAT (3782C>G 3784G>T), ACC489GTT (3791A>G 3792C>T 3793C>T), AAG490AGC (3795A>G 3796G>C), CAA491CAT (3799A>T), GTT492TTG (3800G>T 3802T>G), CAA493AGT (3803C>A 3804A>G 3805A>T), GAT494TTG (3806G>T 3807A>T 3808T>G), GTG495GTA (3811G>A), TTC496TTA (3814C>A), GAA497TCT (3815G>T 3816A>C 3817A>T), CGA498CGT (3820A>T), CTC499TTA (3821C>T 3823C>A), ACG500AGG (3825C>G), AAG501CAA (3827A>C 3829G>A), TCC502TAC (3831C>A), GGA503ACT (3833G>A 3834G>C 3835A>T), TTC504CTC (3836T>C), AAG505TAT (3839A>T 3841G>T), ACA506GTT (3842A>G 3843C>T 3844A>T), GCA507AAA (3845G>A 3846C>A), CCC508ATG (3848C>A 3849C>T 3850C>G), AAA510AAG (3856A>G), TGC511TGT (3859C>T), GAA512GAG (3862A>G), TTC513TTT (3865C>T), CAC514GCC (3866C>G 3867A>C), AAG515CAA (3869A>C 3871G>A), GAA517GAG (3877A>G), GTC518ATA (3878G>A 3880C>A), TTT520TTC (3886T>C), TTA521CTA (3887T>C), GGC522GGG (3892C>G), TTT523CAT (3893T>C 3894T>A), ATC524CTA (3896A>C 3898C>A), ATC525GTT (3899A>G 3901C>T), ACA527GAA (3905A>G 3906C>A), ACA528AAC (3909C>A 3910A>C), GGG529CAA (3911G>C 3912G>A 3913G>A), ATA530GTA (3914A>G), ACG531AGG (3918C>G), ATC532ATG (3922C>G), GAC533GAT (3925C>T), GCA535AAG (3929G>A 3930C>A 3931A>G), AAG536AAA (3934G>A), ACA537GTG (3935A>G 3936C>T 3937A>G), CAG538CAA (3940G>A), TCA539GCC (3941T>G 3943A>C), ATC540ATT (3946C>T), AGA541GTT (3947A>G 3948G>T 3949A>T), GAA542GAT (3952A>T), CCA544CAA (3957C>A), GAA545GCA (3960A>C), CCG546CCC (3964G>C), AAG547AAA (3967G>A), ACA548ACT (3970A>T), GTC549GTG (3973C>G), GTA552CTG (3980G>C 3982A>G), CAG553AGG (3983C>A 3984A>G), TCA554TCT (3988A>T), CTT556TTG (3992C>T 3994T>G), GGA557GGG (3997A>G), CTC558CTA (4000C>A), GCC559GCT (4003C>T), AAC562TAT (4010A>T 4012C>T), AAG567GCT (4025A>G 4026A>C 4027G>T), GAC568GGA (4029A>G 4030C>A), TAT569TAC (4033T>C), ACA572AAA (4041C>A), GCA574GCT (4048A>T), CCA575GCA (4049C>G), ATG576CTG (4052A>C), ACG577ACT (4057G>T), ATG578GAT (4058A>G 4059T>A 4060G>T), CTT579TTG (4061C>T 4063T>G), ACA580CTG (4064A>C 4065C>T 4066A>G), AGA581AAG (4068G>A 4069A>G), AAA582AAG (4072A>G), GAC583GAA (4075C>A), AAC585AAG (4081C>G), AAA587GTA (4085A>G 4086A>T), GGA589TCG (4091G>T 4092G>C 4093A>G), AAA590GTA (4094A>G 4095A>T), GAA591AGA (4097G>A 4098A>G), CAG592TGC (4100C>T 4101A>G 4102G>C), ACC593GAA (4103A>G 4104C>A 4105C>A), GCG595GCT (4111G>T), TTC596TTT (4114C>T), AAA597CAG (4115A>C 4117A>G), AGA598AGT (4120A>T), CTC599TTG (4121C>T 4123C>G), CAG602GCT (4130C>G 4131A>C 4132G>T), TGC603ATT (4133T>A 4134G>T 4135C>T), GCT604GCA (4138T>A), GCC606GAA (4143C>A 4144C>A), ACG608ATA (4149C>T 4150G>A), CTT609CTG (4153T>G), CGA610AAG (4154C>A 4155G>A 4156A>G), CTA611CTG (4159A>G), TTC612CCA (4160T>C 4161T>C 4162C>A), GAT613GAC (4165T>C), GGT614TTT (4166G>T 4167G>T), AGC615GAG (4169A>G 4170G>A 4171C>G), AAG616CTA (4172A>C 4173A>T 4174G>A), GAA617CCT (4175G>C 4176A>C 4177A>T), GTC618TTT (4178G>T 4180C>T), CAC619GAG (4181C>G 4183C>G), ATC620GTA (4184A>G 4186C>A), GAG621CAC (4187G>C 4189G>C), ACC622ACT (4192C>T), GAC623GAT (4195C>T), TCT625TCG (4201T>G), GAT626GAC (4204T>C), ATG627AAA (4206T>A 4207G>A), GCA628GCT (4210A>T), ATA629GTG (4211A>G 4213A>G), GGC630GGA (4216C>A), GCA631GGC (4218C>G 4219A>C), TGT632GTA (4220T>G 4221G>T 4222T>A), CTA633TTG (4223C>T 4225A>G), ACA634GTG (4226A>G 4227C>T 4228A>G), CAG635CAA (4231G>A), ACA636_CAC637del (4232_4237delACACAC), GAT638GAA (4240T>A), GGG639GGT (4243G>T), AAA640_AGA641del (4244_4249delAAAAGA), CAC642CAT (4252C>T), CCA643CCG (4255A>G), TAT646TTT (4263A>T), TAT647GAA (4265T>G 4267T>A), TCC648AGC (4268T>A 4269C>G), CGG649AGG (4271C>A), ATG651TTA (4277A>T 4279G>A), ACC652AAT (4281C>A 4282C>T), ACA653GAT (4283A>G 4284C>A 4285A>T), GCG654GCT (4288G>T), GAA655GAG (4291A>G), AAC657AGG (4296A>G 4297C>G), GAC659TCT (4301G>T 4302A>C 4303C>T), ATC660ACT (4305T>C 4306C>T), GAC662GAA (4312C>A), CTT665ATG (4319C>A 4321T>G), CTA666GTT (4322C>G 4324A>T), GCC667GCA (4327C>A), ATT668GTG (4328A>G 4330T>G), GTT669GTG (4333T>G), GCC670CAC (4334G>C 4335C>A), GCC671TGT (4337G>T 4338C>G 4339C>T), ATG672CTG (4340A>C), CAA673CAG (4345A>G), CAT674CAA (4348T>A), AGA676CGT (4352A>C 4354A>T), GTG677GCC (4356T>C 4357G>C), TAC678GAG (4358T>G 4360C>G), GTC679CTT (4361G>C 4363C>T), GAG680CAA (4364G>C 4366G>A), GGC681GGA (4369C>A), GGC681_CCA682insTCC (4369_4370insTCC), CCA682CCT (4372A>T), CCG683CAC (4374C>A 4375G>C), AAG684AAA (4378G>A), TTA685ATT (4379T>A 4381A>T), ACG686GAA (4382A>G 4383C>A 4384G>A), CTT688CTC (4390T>C), TCA689ACA (4391T>A), GAC690GAT (4396C>T), CAC691CAT (4399C>T), AAG692AAA (4402G>A), AAT693GCT (4403A>G 4404A>C), CTC694TTA (4406C>T 4408C>A), ACG695GAA (4409A>G 4410C>A 4411G>A), TTC697TTT (4417C>T), ACG698ATG (4419C>T), ACG699TCA (4421A>T 4423G>A), ACG700ACT (4426G>T), AAG701AAA (4429G>A), GAA702ACT (4430G>A 4431A>C 4432A>T), CTT703TTA (4433C>T 4435T>A), ACC704AAC (4437C>A), CGA705TCT (4439C>T 4440G>C 4441A>T), AGA706CGT (4442A>C 4444A>T), CAA707CAG (4447A>G), GCC708GCA (4450C>A), CGC709CGT (4453C>T), TCG711GCA (4457T>G 4459G>A), GAG712GAC (4462G>C), CTG713TTT (4463C>T 4465G>T), CTT714TTA (4466C>T 4468T>A), GGG715TCC (4469G>T 4470G>C 4471G>C), CAG716GAA (4472C>G 4474G>A), TAC717TAT (4477C>T), AAG718AAC (4480G>C), TTC719TTT (4483C>T), GAA720ATT (4484G>A 4485A>T 4486A>T), CGC836GAG (4832C>G 4833G>A 4834C>G), TAC837TTT (4836A>T 4837C>T), ATC838GTG (4838A>G 4840C>G), AAG839GCC (4841A>G 4842A>C 4843G>C), AAA840AAG (4846A>G), TGC841TGT (4849C>T), GTA842CCA (4850G>C 4851T>C), CAC843AAT (4853C>A 4855C>T), AAC847GTT (4865A>G 4866A>T 4867C>T), AAA848AAG (4870A>G), GCT849GTA (4872C>T 4873T>A), GCA850GAA (4875C>A), CGG851CAT (4878G>A 4879G>T), CAC852CAA (4882C>A), GCG853AGG (4883G>A 4884C>G), AAA854CCA (4886A>C 4887A>C), TAC855GGC (4889T>G 4890A>G), GGT856GGC (4894T>C), CAC857CTA (4896A>T 4897C>A), CTA858CTT (4900A>T), TTC860GAT (4904T>G 4905T>A 4906C>T), AGG861ATT (4908G>T 4909G>T), ACA862GAG (4910A>G 4911C>A 4912A>G), CCA863ATT (4913C>A 4914C>T 4915A>T), CCA864CCT (4918A>T), ACG865ACT (4921G>T), AAA866TGG (4922A>T 4923A>G 4924A>G), CCA867AAG (4925C>A 4926C>A 4927A>G), GAC869GAA (4933C>A), GTT871GTG (4939T>G), ACG872AAC (4941C>A 4942G>C), TTC875TTT (4951C>T), ATT876GTC (4952A>G 4954T>C), ACG877GTG (4955A>G 4956C>T), AAA878GGA (4958A>G 4959A>G), CTC879TTA (4961C>T 4963C>A), CCG880CCT (4966G>T), AGG881AAG (4968G>A), TCA882ACT (4970T>A 4972A>T), AAG883AGG (4974A>G), GAT884AGG (4976G>A 4977A>G 4978T>G), CGA885CAA (4980G>A), GTA936TCC (5132G>T 5133T>C 5134A>C), TTC937ATT (5135T>A 5137C>T), ACA939TCT (5141A>T 5143A>T), GAC940GAT (5146C>T), GAC942GGT (5151A>G 5152C>T), AAG943ACT (5154A>C 5155G>T), CTC944CA. (5157T>A), ATC990GTG (5294A>G 5296C>G), AAC991ACG (5298A>C 5299C>G), TAC992GCG (5300T>G 5301A>C 5302C>G), GCA993AGC (5303G>A 5304C>G 5305A>C), CAA994CAG (5308A>G), GAC995AGA (5309G>A 5310A>G 5311C>A), AAC996AAT (5314C>T), GTT998GTG (5320T>G), TCA999GAT (5321T>G 5322C>A 5323A>T), CTG1001CTT (5329G>T), CCA1002GAT (5330C>G 5331C>A 5332A>T), ATG1003ACT (5334T>C 5335G>T), CAG1005CAA (5341G>A), ATC1006T.. (5342A>T) | | | | | | | | |

*: Inserts / Deletes / Misaligned / Frameshifts

#### Analysis details

This analysis was performed with panviral2.64

### NGS Details (UN60): Vellamovirus rhodeisland44

#### Assembly

| | |
|---|---|
| **Coverage Length** | 432 (1 contig(s)) |
| **Depth Of Coverage** | 154.1 |
| **Number Of Reads** | 604 |
| **Reads Per Million** | 12.81 rpm (after QC) |
| **Ambiguities** | 0 |
| **Assembly Method** | de novo + reference guided assembly |
| **Consensus Caller** | Bcf Tools |

#### Coverage Map

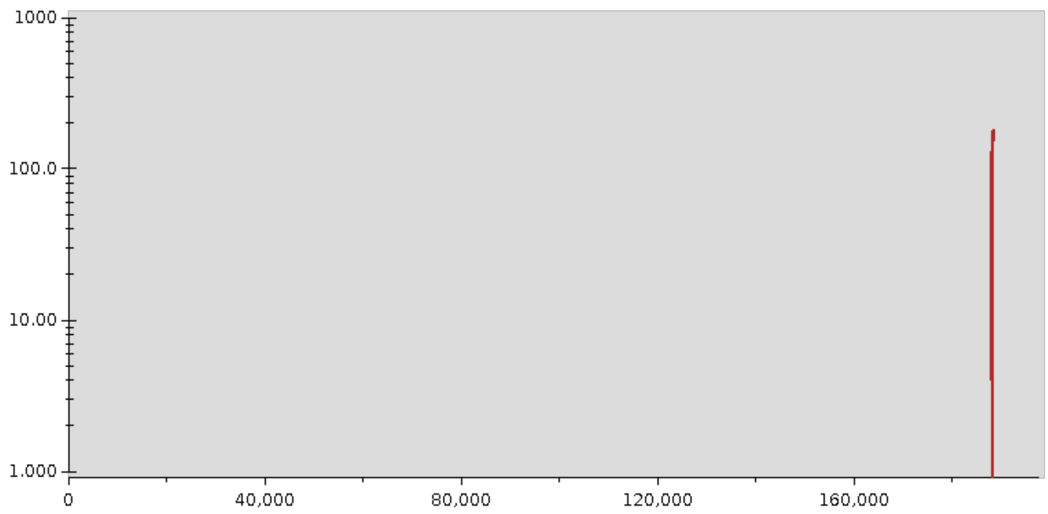

#### Assignment

| | |
|---|---|
| **Type** | Vellamovirus rhodeisland44 (Taxonomy ID: 2734152) |
| **Reference Genome** | NC_047734.1 |
| **NT Identity (%)** | 75.0 |
| **AA Identity (%)** | 86.8056 |
| **Number Of Stop Codons** | 0 |
| **Number Of CDS** | 235 |

#### Alignment

| | |
|---|---|
| **Alignment Score** | 432.0 (NT) + 958.0 (AA) = 1390.0 |
| **Concordance (%)** | 72.6607 |
| **Alignment Method** | Local, heuristic, nucleotide (BLASTN) |

#### Genome Region

Sequence starts at position 188039 and ends at position 188470 relative to NC_047734.1 reference sequence.

#### Alignment Detailed Statistics

| | Begin | End | Coverage | Score | Concordance | Matches | Identities | I/D/M/F* | Stop Codons |
|---|---|---|---|---|---|---|---|---|---|
| **NT** | **188039** | **188470** | **0.2%** | **432** | **50.0%** | **432 (100%)** | **324 (75.0%)** | **0/0** | |
| Mutations: | 188048A>C, 188051T>G, 188054C>T, 188057G>T, 188060G>T, 188067G>A, 188069C>T, 188070G>A, 188071C>G, 188073A>T, 188075G>A, 188077G>C, 188078T>A, 188084T>G, 188085C>T, 188087T>G, 188088C>T, 188096T>A, 188108G>A, 188109T>G, 188111T>A, 188117G>A, 188120C>T, 188124C>A, 188125A>C, 188126A>T, 188129C>T, 188135C>T, 188139C>T, 188141T>G, 188147A>T, 188150C>G, 188155A>C, 188162G>T, 188165A>T, 188171C>T, 188174T>A, 188177C>T, 188182C>G, 188183T>C, 188186C>A, 188189T>A, 188199C>T, 188201T>A, 188204A>T, 188207G>A, 188210T>C, 188213A>G, 188222C>A, 188224T>C, 188225G>T, 188226A>G, 188229G>C, 188230G>A, 188231T>A, 188232A>T, 188234C>G, 188235C>A, 188237T>A, 188243C>T, 188249T>A, 188252T>C, 188255G>A, 188267T>A, 188282C>T, 188283A>T, 188284G>C, 188291C>T, 188294T>G, 188297C>T, 188306G>A, 188316T>G, 188317C>G, 188318A>T, 188339C>T, 188342G>T, 188351A>T, 188354C>A, 188358A>C, 188364C>A, 188366A>C, 188367T>C, 188369G>A, 188372C>T, 188373C>T, 188378G>A, 188381T>G, 188384C>T, 188387C>T, 188390C>T, 188397C>T, 188399C>G, 188408C>T, 188426T>C, 188430A>G, 188433C>T, 188435A>G, 188438A>C, 188440G>C, 188441A>T, 188444A>T, 188445C>T, 188447A>G, 188450C>A, 188451A>T, 188453T>C, 188456T>C, 188459C>T | | | | | | | | |

*: Inserts / Deletes / Misaligned / Frameshifts

#### Analysis details

This analysis was performed with panviral2.64

### NGS Details (UN60): Cassava brown streak virus

#### Assembly

| | |
|---|---|
| **Coverage Length** | 519 (1 contig(s)) |
| **Depth Of Coverage** | 82.0 |
| **Number Of Reads** | 356 |
| **Reads Per Million** | 7.55 rpm (after QC) |
| **Ambiguities** | 0 |
| **Assembly Method** | de novo + reference guided assembly |
| **Consensus Caller** | Bcf Tools |

#### Coverage Map

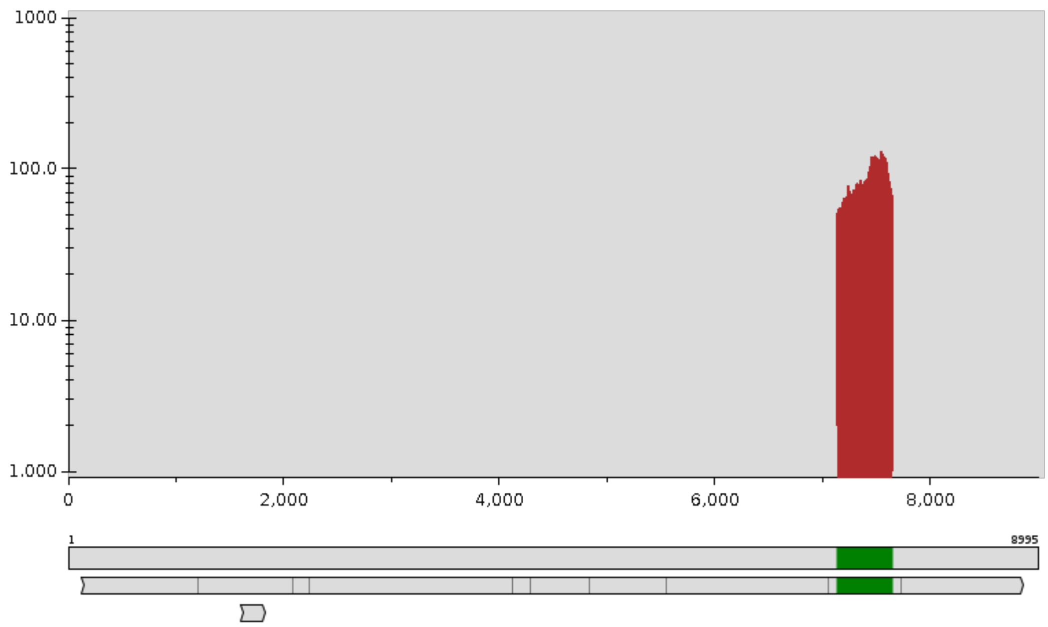

#### Assignment

| | |
|---|---|
| **Type** | Cassava brown streak virus (Taxonomy ID: 137758) |
| **Reference Genome** | NC_012698.2 |
| **NT Identity (%)** | 60.3083 |
| **AA Identity (%)** | 57.2254 |
| **Number Of Stop Codons** | 0 |
| **Number Of CDS** | 2 |

#### Alignment

| | |
|---|---|
| **Alignment Score** | 214.0 (NT) + 728.0 (AA) = 942.0 |
| **Concordance (%)** | 41.9599 |

| Alignment Method | Global, seeded, nucleotide + amino acids (AGA) |
|---|---|

#### Genome Region

Sequence starts at position 7129 and ends at position 7647 relative to NC_012698.2 reference sequence.

#### Alignment Detailed Statistics

| | Begin | End | Coverage | Score | Concordance | Matches | Identities | I/D/M/F* | Stop Codons |
|---|---|---|---|---|---|---|---|---|---|
| **NT** | **7129** | **7647** | **5.8%** | **214** | **20.6%** | **519 (100%)** | **313 (60.3%)** | **0/0** | |
| Mutations: | 7130A>G, 7133A>G, 7134A>G, 7136T>G, 7142C>T, 7145A>T, 7148A>C, 7151G>A, 7157A>T, 7158C>A, 7163A>G, 7167A>G, 7175A>T, 7177A>G, 7178A>G, 7179C>G, 7180A>C, 7184A>T, 7185T>C, 7190T>C, 7192C>A, 7193C>G, 7194A>T, 7202A>C, 7203A>T, 7205C>T, 7206A>C, 7207T>A, 7208C>G, 7211T>C, 7213G>T, 7214G>T, 7217A>G, 7218G>C, 7229A>T, 7234C>T, 7235A>T, 7241A>G, 7242A>G, 7243C>A, 7244A>G, 7245G>C, 7246T>C, 7250G>A, 7253A>T, 7256T>A, 7257A>T, 7258T>C, 7270T>C, 7271A>T, 7272C>A, 7273G>A, 7274T>A, 7275G>A, 7277A>T, 7280G>T, 7281T>G, 7283T>C, 7284G>A, 7286G>A, 7287T>G, 7288T>A, 7290A>G, 7292T>G, 7293G>A, 7294G>A, 7295A>T, 7298G>A, 7304T>G, 7307C>A, 7310A>T, 7313A>G, 7319A>T, 7320A>T, 7323T>C, 7325G>C, 7331T>C, 7332G>A, 7334T>C, 7337T>C, 7340C>T, 7343T>G, 7355A>G, 7358A>T, 7361C>T, 7364T>C, 7374A>T, 7377G>C, 7380G>A, 7381G>A, 7385T>C, 7388A>T, 7390T>A, 7391A>T, 7397A>T, 7398T>C, 7400A>T, 7401T>A, 7403T>C, 7406G>C, 7409G>A, 7410G>T, 7413G>A, 7414A>T, 7416C>G, 7418G>T, 7422C>G, 7424A>G, 7425A>G, 7432T>C, 7433G>A, 7436T>A, 7437A>T, 7438G>A, 7442T>C, 7443C>A, 7445C>G, 7448C>T, 7449G>A, 7451G>C, 7455G>T, 7457T>A, 7458T>C, 7462T>C, 7463A>T, 7464A>C, 7465A>T, 7467A>G, 7468A>G, 7469A>G, 7470G>C, 7471T>C, 7472T>A, 7473G>A, 7474G>A, 7476G>A, 7477A>C, 7481T>G, 7484T>G, 7487A>G, 7489T>C, 7490C>T, 7494A>G, 7495A>T, 7496G>T, 7499T>A, 7500G>A, 7501T>A, 7502G>A, 7503C>A, 7504T>C, 7505A>G, 7506A>C, 7507G>T, 7508A>G, 7511T>A, 7512G>A, 7517T>A, 7520A>G, 7521A>C, 7522T>C, 7523G>A, 7524C>G, 7526A>T, 7527C>A, 7529C>G, 7535A>C, 7539T>A, 7540C>A, 7541A>T, 7544T>C, 7547G>A, 7553C>T, 7556A>G, 7559T>A, 7562C>T, 7568A>T, 7570T>A, 7572A>G, 7573A>G, 7576G>A, 7577G>T, 7578A>G, 7579G>A, 7580A>C, 7581A>C, 7586A>T, 7588T>A, 7589T>C, 7599A>C, 7600T>C, 7601G>C, 7602G>A, 7603C>A, 7604C>G, 7607G>A, 7610G>A, 7616T>C, 7618T>A, 7622A>T, 7628T>C, 7631A>C, 7632T>G, 7633T>G, 7635C>A, 7636G>A | | | | | | | | |

##### CDS

| | Begin | End | Coverage | Score | Concordance | Matches | Identities | I/D/M/F* | Stop Codons |
|---|---|---|---|---|---|---|---|---|---|
| CBSV_gp1 | 2336 | 2508 | 5.9% | 728 | 59.9% | 173 (100%) | 99 (57.2%) | 0/0/0/0 | 0 |
| Protein mutations: | I2337V (7134A>G 7136T>G), Q2345K (7158C>A), K2348E (7167A>G), K2351R (7177A>G 7178A>G), Q2352A (7179C>G 7180A>C), F2354L (7185T>C), P2356Q (7192C>A 7193C>G), T2357S (7194A>T), I2360F (7203A>T 7205C>T), I2361Q (7206A>C 7207T>A 7208C>G), R2363L (7213G>T 7214G>T), V2365L (7218G>C), P2370L (7234C>T 7235A>T), T2373E (7242A>G 7243C>A 7244A>G), V2374P (7245G>C 7246T>C), E2376D (7253A>T), I2378S (7257A>T 7258T>C), V2382A (7270T>C 7271A>T), R2383K (7272C>A 7273G>A 7274T>A), V2384I (7275G>A 7277A>T), S2386A (7281T>G 7283T>C), E2387K (7284G>A 7286G>A), L2388E (7287T>G 7288T>A), I2389V (7290A>G 7292T>G), G2390N (7293G>A 7294G>A 7295A>T), S2399C (7320A>T), D2403N (7332G>A 7334T>C), N2406K (7343T>G), M2417L (7374A>T), E2418Q (7377G>C), G2419K (7380G>A 7381G>A), L2422H (7390T>A 7391A>T), Y2426N (7401T>A 7403T>C), K2427N (7406G>C), V2429L (7410G>T), E2430M (7413G>A 7414A>T), P2431A (7416C>G 7418G>T), Q2433E (7422C>G 7424A>G), N2434D (7425A>G), M2436T (7432T>C 7433G>A), S2438Y (7437A>T 7438G>A), L2440M (7443C>A 7445C>G), V2442I (7449G>A 7451G>C), A2444S (7455G>T 7457T>A), F2445L (7458T>C), V2446A (7462T>C 7463A>T), N2447L (7464A>C 7465A>T), K2448G (7467A>G 7468A>G 7469A>G), V2449P (7470G>C 7471T>C 7472T>A), G2450N (7473G>A 7474G>A), D2451T (7476G>A 7477A>C), D2452E (7481T>G), I2454M (7487A>G), I2455T (7489T>C 7490C>T), K2457V (7494A>G 7495A>T 7496G>T), V2459K (7500G>A 7501T>A 7502G>A), L2460T (7503C>A 7504T>C 7505A>G), R2461L (7506A>C 7507G>T 7508A>G), E2463K (7512G>A), M2466P (7521A>C 7522T>C 7523G>A), P2467A (7524C>G 7526A>T), S2472N (7539T>A 7540C>A 7541A>T), L2482H (7570T>A), N2483G (7572A>G 7573A>G), W2484Y (7576G>A 7577G>T), R2485D (7578A>G 7579G>A 7580A>C), K2486Q (7581A>C), F2488Y (7588T>A 7589T>C), M2492P (7599A>C 7600T>C 7601G>C), A2493K (7602G>A 7603C>A 7604C>G), M2498K (7618T>A), F2503G (7632T>G 7633T>G), R2504K (7635C>A 7636G>A) | | | | | | | | |
| Codon mutations: | GCA2335.CG (7130A>G), CCA2336CCG (7133A>G), ATT2337GTG (7134A>G 7136T>G), TTC2339TTT (7142C>T), GTA2340GTT (7145A>T), ACA2341ACC (7148A>C), GGG2342GGA (7151G>A), GCA2344GCT (7157A>T), CAA2345AAA (7158C>A), AAA2346AAG (7163A>G), AAG2348GAG (7167A>G), GTA2350GTT (7175A>T), AAA2351AGG (7177A>G 7178A>G), CAA2352GCA (7179C>G 7180A>C), ATA2353ATT (7184A>T), TTT2354CTT (7185T>C), GGT2355GGC (7190T>C), CCC2356CAG (7192C>A 7193C>G), ACT2357TCT (7194A>T), CCA2359CCC (7202A>C), ATC2360TTT (7203A>T 7205C>T), ATC2361CAG (7206A>C 7207T>A 7208C>G), TCT2362TCC (7211T>C), CGG2363CTT (7213G>T 7214G>T), AAA2364AAG (7217A>G), GTT2365CTT (7218G>C), CCA2368CCT (7229A>T), CCA2370CTT (7234C>T 7235A>T), GGA2372GGG (7241A>G), ACA2373GAG (7242A>G 7243C>A 7244A>G), GTT2374CCT (7245G>C 7246T>C), GAG2375GAA (7250G>A), GAA2376GAT (7253A>T), ATT2377ATA (7256T>A), ATC2378TCC (7257A>T 7258T>C), GTA2382GCT (7270T>C 7271A>T), CGT2383AAA (7272C>A 7273G>A 7274T>A), GTA2384ATT (7275G>A 7277A>T), GCG2385GCT (7280G>T), TCT2386GCC (7281T>G 7283T>C), GAG2387AAA (7284G>A 7286G>A), TTG2388GAG (7287T>G 7288T>A), ATT2389GTG (7290A>G 7292T>G), GGA2390AAT (7293G>A 7294G>A 7295A>T), GGG2391GGA (7298G>A), GTT2393GTG (7304T>G), CTC2394CTA (7307C>A), GTA2395GTT (7310A>T), GAA2396GAG (7313A>G), ACA2398ACT (7319A>T), AGT2399TGT (7320A>T), TTG2400CTC (7323T>C 7325G>C), TTT2402TTC (7331T>C), GAT2403AAC (7332G>A 7334T>C), GCT2404GCC (7337T>C), CTC2405CTT (7340C>T), AAT2406AAG (7343T>G), GGA2410GGG (7355A>G), CCA2411CCT (7358A>T), TAC2412TAT (7361C>T), ATT2413ATC (7364T>C), ATG2417TTG (7374A>T), GAA2418CAA (7377G>C), GGG2419AAG (7380G>A 7381G>A), ATT2420ATC (7385T>C), GGA2421GGT (7388A>T), CTA2422CAT (7390T>A 7391A>T), GGA2424GGT (7397A>T), TTA2425CTT (7398T>C 7400A>T), TAT2426AAC (7401T>A 7403T>C), AAG2427AAC (7406G>C), TTG2428TTA (7409G>A), GTG2429TTG (7410G>T), GAG2430ATG (7413G>A 7414A>T), CCG2431GCT (7416C>G 7418G>T), CAA2433GAG (7422C>G 7424A>G), AAT2434GAT (7425A>G), ATG2436ACA (7432T>C 7433G>A), GCT2437GCA (7436T>A), AGT2438TAT (7437A>T 7438G>A), GCT2439GCC (7442T>C), CTC2440ATG (7443C>A 7445C>G), TGC2441TGT (7448C>T), GTG2442ATC (7449G>A 7451G>C), GCT2444TCA (7455G>T 7457T>A), TTT2445CTT (7458T>C), GTA2446GCT (7462T>C 7463A>T), AAT2447CTT (7464A>C 7465A>T), AAA2448GGG (7467A>G 7468A>G 7469A>G), GTT2449CCA (7470G>C 7471T>C 7472T>A), GGT2450AAT (7473G>A 7474G>A), GAT2451ACT (7476G>A 7477A>C), GAT2452GAG (7481T>G), CCT2453CCG (7484T>G), ATA2454ATG (7487A>G), ATC2455ACT (7489T>C 7490C>T), AAG2457GTT (7494A>G 7495A>T 7496G>T), GGT2458GGA (7499T>A), GTG2459AAA (7500G>A 7501T>A 7502G>A), CTA2460ACG (7503C>A 7504T>C 7505A>G), AGA2461CTG (7506A>C 7507G>T 7508A>G), GGT2462GGA (7511T>A), GAG2463AAG (7512G>A), ATT2464ATA (7517T>A), GTA2465GTG (7520A>G), ATG2466CCA (7521A>C 7522T>C 7523G>A), CCA2467GCT (7524C>G 7526A>T), CGC2468AGG (7527C>A 7529C>G), CCA2470CCC (7535A>C), TCA2472AAT (7539T>A 7540C>A 7541A>T), TTT2473TTC (7544T>C), GGG2474GGA (7547G>A), GAC2476GAT (7553C>T), CCA2477CCG (7556A>G), ATT2478ATA (7559T>A), TTC2479TTT (7562C>T), CCA2481CCT (7568A>T), CTT2482CAT (7570T>A), AAC2483GGC (7572A>G 7573A>G), TGG2484TAT (7576G>A 7577G>T), AGA2485GAC (7578A>G 7579G>A 7580A>C), AAG2486CAG (7581A>C), ACA2487ACT (7586A>T), TTT2488TAC (7588T>A 7589T>C), ATG2492CCC (7599A>C 7600T>C 7601G>C), GCC2493AAG (7602G>A 7603C>A 7604C>G), GAG2494GAA (7607G>A), GAG2495GAA (7610G>A), AAT2497AAC (7616T>C), ATG2498AAG (7618T>A), ATA2499ATT (7622A>T), CAT2501CAC (7628T>C), CGA2502CGC (7631A>C), TTT2503GGT (7632T>G 7633T>G), CGA2504AAA (7635C>A 7636G>A) | | | | | | | | |

##### Proteins

| | Begin | End | Coverage | Score | Concordance | Matches | Identities | I/D/M/F* | Stop Codons |
|---|---|---|---|---|---|---|---|---|---|
| polyprotein (YP_007027011.1) | 2336 | 2508 | 5.9% | 728 | 59.9% | 173 (100%) | 99 (57.2%) | 0/0/0/0 | 0 |
| Protein mutations: | I2337V (7134A>G 7136T>G), Q2345K (7158C>A), K2348E (7167A>G), K2351R (7177A>G 7178A>G), Q2352A (7179C>G 7180A>C), F2354L (7185T>C), P2356Q (7192C>A 7193C>G), T2357S (7194A>T), I2360F (7203A>T 7205C>T), I2361Q (7206A>C 7207T>A 7208C>G), R2363L (7213G>T 7214G>T), V2365L (7218G>C), P2370L (7234C>T 7235A>T), T2373E (7242A>G 7243C>A 7244A>G), V2374P (7245G>C 7246T>C), E2376D (7253A>T), I2378S (7257A>T 7258T>C), V2382A (7270T>C 7271A>T), R2383K (7272C>A 7273G>A 7274T>A), V2384I (7275G>A 7277A>T), S2386A (7281T>G 7283T>C), E2387K (7284G>A 7286G>A), L2388E (7287T>G 7288T>A), I2389V (7290A>G 7292T>G), G2390N (7293G>A 7294G>A 7295A>T), S2399C (7320A>T), D2403N (7332G>A 7334T>C), N2406K (7343T>G), M2417L (7374A>T), E2418Q (7377G>C), G2419K (7380G>A 7381G>A), L2422H (7390T>A 7391A>T), Y2426N (7401T>A 7403T>C), K2427N (7406G>C), V2429L (7410G>T), E2430M (7413G>A 7414A>T), P2431A (7416C>G 7418G>T), Q2433E (7422C>G 7424A>G), N2434D (7425A>G), M2436T (7432T>C 7433G>A), S2438Y (7437A>T 7438G>A), L2440M (7443C>A 7445C>G), V2442I (7449G>A 7451G>C), A2444S (7455G>T 7457T>A), F2445L (7458T>C), V2446A (7462T>C 7463A>T), N2447L (7464A>C 7465A>T), K2448G (7467A>G 7468A>G 7469A>G), V2449P (7470G>C 7471T>C 7472T>A), G2450N (7473G>A 7474G>A), D2451T (7476G>A 7477A>C), D2452E (7481T>G), I2454M (7487A>G), I2455T (7489T>C 7490C>T), K2457V (7494A>G 7495A>T 7496G>T), V2459K (7500G>A 7501T>A 7502G>A), L2460T (7503C>A 7504T>C 7505A>G), R2461L (7506A>C 7507G>T 7508A>G), E2463K (7512G>A), M2466P (7521A>C 7522T>C 7523G>A), P2467A (7524C>G 7526A>T), S2472N (7539T>A 7540C>A 7541A>T), L2482H (7570T>A), N2483G (7572A>G 7573A>G), W2484Y (7576G>A 7577G>T), R2485D (7578A>G 7579G>A 7580A>C), K2486Q (7581A>C), F2488Y (7588T>A 7589T>C), M2492P (7599A>C 7600T>C 7601G>C), A2493K (7602G>A 7603C>A 7604C>G), M2498K (7618T>A), F2503G (7632T>G 7633T>G), R2504K (7635C>A 7636G>A) | | | | | | | | |

| | Begin | End | Coverage | Score | Concordance | Matches | Identities | I/D/M/F* | Stop Codons |
|---|---|---|---|---|---|---|---|---|---|
| **NT** | **7129** | **7647** | **5.8%** | **214** | **20.6%** | **519 (100%)** | **313 (60.3%)** | **0/0** | |
| Codon mutations: | GCA2335.CG (7130A>G), CCA2336CCG (7133A>G), ATT2337GTG (7134A>G 7136T>G), TTC2339TTT (7142C>T), GTA2340GTT (7145A>T), ACA2341ACC (7148A>C), GGG2342GGA (7151G>A), GCA2344GCT (7157A>T), CAA2345AAA (7158C>A), AAA2346AAG (7163A>G), AAG2348GAG (7167A>G), GTA2350GTT (7175A>T), AAA2351AGG (7177A>G 7178A>G), CAA2352GCA (7179C>G 7180A>C), ATA2353ATT (7184A>T), TTT2354CTT (7185T>C), GGT2355GGC (7190T>C), CCC2356CAG (7192C>A 7193C>G), ACT2357TCT (7194A>T), CCA2359CCC (7202A>C), ATC2360TTT (7203A>T 7205C>T), ATC2361CAG (7206A>C 7207T>A 7208C>G), TCT2362TCC (7211T>C), CGG2363CTT (7213G>T 7214G>T), AAA2364AAG (7217A>G), GTT2365CTT (7218G>C), CCA2368CCT (7229A>T), CCA2370CTT (7234C>T 7235A>T), GGA2372GGG (7241A>G), ACA2373GAG (7242A>G 7243C>A 7244A>G), GTT2374CCT (7245G>C 7246T>C), GAG2375GAA (7250G>A), GAA2376GAT (7253A>T), ATT2377ATA (7256T>A), ATC2378TCC (7257A>T 7258T>C), GTA2382GCT (7270T>C 7271A>T), CGT2383AAA (7272C>A 7273G>A 7274T>A), GTA2384ATT (7275G>A 7277A>T), GCG2385GCT (7280G>T), TCT2386GCC (7281T>G 7283T>C), GAG2387AAA (7284G>A 7286G>A), TTG2388GAG (7287T>G 7288T>A), ATT2389GTG (7290A>G 7292T>G), GGA2390AAT (7293G>A 7294G>A 7295A>T), GGG2391GGA (7298G>A), GTT2393GTG (7304T>G), CTC2394CTA (7307C>A), GTA2395GTT (7310A>T), GAA2396GAG (7313A>G), ACA2398ACT (7319A>T), AGT2399TGT (7320A>T), TTG2400CTC (7323T>C 7325G>C), TTT2402TTC (7331T>C), GAT2403AAC (7332G>A 7334T>C), GCT2404GCC (7337T>C), CTC2405CTT (7340C>T), AAT2406AAG (7343T>G), GGA2410GGG (7355A>G), CCA2411CCT (7358A>T), TAC2412TAT (7361C>T), ATT2413ATC (7364T>C), ATG2417TTG (7374A>T), GAA2418CAA (7377G>C), GGG2419AAG (7380G>A 7381G>A), ATT2420ATC (7385T>C), GGA2421GGT (7388A>T), CTA2422CAT (7390T>A 7391A>T), GGA2424GGT (7397A>T), TTA2425CTT (7398T>C 7400A>T), TAT2426AAC (7401T>A 7403T>C), AAG2427AAC (7406G>C), TTG2428TTA (7409G>A), GTG2429TTG (7410G>T), GAG2430ATG (7413G>A 7414A>T), CCG2431GCT (7416C>G 7418G>T), CAA2433GAG (7422C>G 7424A>G), AAT2434GAT (7425A>G), ATG2436ACA (7432T>C 7433G>A), GCT2437GCA (7436T>A), AGT2438TAT (7437A>T 7438G>A), GCT2439GCC (7442T>C), CTC2440ATG (7443C>A 7445C>G), TGC2441TGT (7448C>T), GTG2442ATC (7449G>A 7451G>C), GCT2444TCA (7455G>T 7457T>A), TTT2445CTT (7458T>C), GTA2446GCT (7462T>C 7463A>T), AAT2447CTT (7464A>C 7465A>T), AAA2448GGG (7467A>G 7468A>G 7469A>G), GTT2449CCA (7470G>C 7471T>C 7472T>A), GGT2450AAT (7473G>A 7474G>A), GAT2451ACT (7476G>A 7477A>C), GAT2452GAG (7481T>G), CCT2453CCG (7484T>G), ATA2454ATG (7487A>G), ATC2455ACT (7489T>C 7490C>T), AAG2457GTT (7494A>G 7495A>T 7496G>T), GGT2458GGA (7499T>A), GTG2459AAA (7500G>A 7501T>A 7502G>A), CTA2460ACG (7503C>A 7504T>C 7505A>G), AGA2461CTG (7506A>C 7507G>T 7508A>G), GGT2462GGA (7511T>A), GAG2463AAG (7512G>A), ATT2464ATA (7517T>A), GTA2465GTG (7520A>G), ATG2466CCA (7521A>C 7522T>C 7523G>A), CCA2467GCT (7524C>G 7526A>T), CGC2468AGG (7527C>A 7529C>G), CCA2470CCC (7535A>C), TCA2472AAT (7539T>A 7540C>A 7541A>T), TTT2473TTC (7544T>C), GGG2474GGA (7547G>A), GAC2476GAT (7553C>T), CCA2477CCG (7556A>G), ATT2478ATA (7559T>A), TTC2479TTT (7562C>T), CCA2481CCT (7568A>T), CTT2482CAT (7570T>A), AAC2483GGC (7572A>G 7573A>G), TGG2484TAT (7576G>A 7577G>T), AGA2485GAC (7578A>G 7579G>A 7580A>C), AAG2486CAG (7581A>C), ACA2487ACT (7586A>T), TTT2488TAC (7588T>A 7589T>C), ATG2492CCC (7599A>C 7600T>C 7601G>C), GCC2493AAG (7602G>A 7603C>A 7604C>G), GAG2494GAA (7607G>A), GAG2495GAA (7610G>A), AAT2497AAC (7616T>C), ATG2498AAG (7618T>A), ATA2499ATT (7622A>T), CAT2501CAC (7628T>C), CGA2502CGC (7631A>C), TTT2503GGT (7632T>G 7633T>G), CGA2504AAA (7635C>A 7636G>A) | | | | | | | | |
| HAM1-like protein (YP_007032446.1) | 28 | 200 | 76.5% | 728 | 59.9% | 173 (100%) | 99 (57.2%) | 0/0/0/0 | 0 |
| Protein mutations: | I29V (7134A>G 7136T>G), Q37K (7158C>A), K40E (7167A>G), K43R (7177A>G 7178A>G), Q44A (7179C>G 7180A>C), F46L (7185T>C), P48Q (7192C>A 7193C>G), T49S (7194A>T), I52F (7203A>T 7205C>T), I53Q (7206A>C 7207T>A 7208C>G), R55L (7213G>T 7214G>T), V57L (7218G>C), P62L (7234C>T 7235A>T), T65E (7242A>G 7243C>A 7244A>G), V66P (7245G>C 7246T>C), E68D (7253A>T), I70S (7257A>T 7258T>C), V74A (7270T>C 7271A>T), R75K (7272C>A 7273G>A 7274T>A), V76I (7275G>A 7277A>T), S78A (7281T>G 7283T>C), E79K (7284G>A 7286G>A), L80E (7287T>G 7288T>A), I81V (7290A>G 7292T>G), G82N (7293G>A 7294G>A 7295A>T), S91C (7320A>T), D95N (7332G>A 7334T>C), N98K (7343T>G), M109L (7374A>T), E110Q (7377G>C), G111K (7380G>A 7381G>A), L114H (7390T>A 7391A>T), Y118N (7401T>A 7403T>C), K119N (7406G>C), V121L (7410G>T), E122M (7413G>A 7414A>T), P123A (7416C>G 7418G>T), Q125E (7422C>G 7424A>G), N126D (7425A>G), M128T (7432T>C 7433G>A), S130Y (7437A>T 7438G>A), L132M (7443C>A 7445C>G), V134I (7449G>A 7451G>C), A136S (7455G>T 7457T>A), F137L (7458T>C), V138A (7462T>C 7463A>T), N139L (7464A>C 7465A>T), K140G (7467A>G 7468A>G 7469A>G), V141P (7470G>C 7471T>C 7472T>A), G142N (7473G>A 7474G>A), D143T (7476G>A 7477A>C), D144E (7481T>G), I146M (7487A>G), I147T (7489T>C 7490C>T), K149V (7494A>G 7495A>T 7496G>T), V151K (7500G>A 7501T>A 7502G>A), L152T (7503C>A 7504T>C 7505A>G), R153L (7506A>C 7507G>T 7508A>G), E155K (7512G>A), M158P (7521A>C 7522T>C 7523G>A), P159A (7524C>G 7526A>T), S164N (7539T>A 7540C>A 7541A>T), L174H (7570T>A), N175G (7572A>G 7573A>G), W176Y (7576G>A 7577G>T), R177D (7578A>G 7579G>A 7580A>C), K178Q (7581A>C), F180Y (7588T>A 7589T>C), M184P (7599A>C 7600T>C 7601G>C), A185K (7602G>A 7603C>A 7604C>G), M190K (7618T>A), F195G (7632T>G 7633T>G), R196K (7635C>A 7636G>A) | | | | | | | | |
| Codon mutations: | GCA27.CG (7130A>G), CCA28CCG (7133A>G), ATT29GTG (7134A>G 7136T>G), TTC31TTT (7142C>T), GTA32GTT (7145A>T), ACA33ACC (7148A>C), GGG34GGA (7151G>A), GCA36GCT (7157A>T), CAA37AAA (7158C>A), AAA38AAG (7163A>G), AAG40GAG (7167A>G), GTA42GTT (7175A>T), AAA43AGG (7177A>G 7178A>G), CAA44GCA (7179C>G 7180A>C), ATA45ATT (7184A>T), TTT46CTT (7185T>C), GGT47GGC (7190T>C), CCC48CAG (7192C>A 7193C>G), ACT49TCT (7194A>T), CCA51CCC (7202A>C), ATC52TTT (7203A>T 7205C>T), ATC53CAG (7206A>C 7207T>A 7208C>G), TCT54TCC (7211T>C), CGG55CTT (7213G>T 7214G>T), AAA56AAG (7217A>G), GTT57CTT (7218G>C), CCA60CCT (7229A>T), CCA62CTT (7234C>T 7235A>T), GGA64GGG (7241A>G), ACA65GAG (7242A>G 7243C>A 7244A>G), GTT66CCT (7245G>C 7246T>C), GAG67GAA (7250G>A), GAA68GAT (7253A>T), ATT69ATA (7256T>A), ATC70TCC (7257A>T 7258T>C), GTA74GCT (7270T>C 7271A>T), CGT75AAA (7272C>A 7273G>A 7274T>A), GTA76ATT (7275G>A 7277A>T), GCG77GCT (7280G>T), TCT78GCC (7281T>G 7283T>C), GAG79AAA (7284G>A 7286G>A), TTG80GAG (7287T>G 7288T>A), ATT81GTG (7290A>G 7292T>G), GGA82AAT (7293G>A 7294G>A 7295A>T), GGG83GGA (7298G>A), GTT85GTG (7304T>G), CTC86CTA (7307C>A), GTA87GTT (7310A>T), GAA88GAG (7313A>G), ACA90ACT (7319A>T), AGT91TGT (7320A>T), TTG92CTC (7323T>C 7325G>C), TTT94TTC (7331T>C), GAT95AAC (7332G>A 7334T>C), GCT96GCC (7337T>C), CTC97CTT (7340C>T), AAT98AAG (7343T>G), GGA102GGG (7355A>G), CCA103CCT (7358A>T), TAC104TAT (7361C>T), ATT105ATC (7364T>C), ATG109TTG (7374A>T), GAA110CAA (7377G>C), GGG111AAG (7380G>A 7381G>A), ATT112ATC (7385T>C), GGA113GGT (7388A>T), CTA114CAT (7390T>A 7391A>T), GGA116GGT (7397A>T), TTA117CTT (7398T>C 7400A>T), TAT118AAC (7401T>A 7403T>C), AAG119AAC (7406G>C), TTG120TTA (7409G>A), GTG121TTG (7410G>T), GAG122ATG (7413G>A 7414A>T), CCG123GCT (7416C>G 7418G>T), CAA125GAG (7422C>G 7424A>G), AAT126GAT (7425A>G), ATG128ACA (7432T>C 7433G>A), GCT129GCA (7436T>A), AGT130TAT (7437A>T 7438G>A), GCT131GCC (7442T>C), CTC132ATG (7443C>A 7445C>G), TGC133TGT (7448C>T), GTG134ATC (7449G>A 7451G>C), GCT136TCA (7455G>T 7457T>A), TTT137CTT (7458T>C), GTA138GCT (7462T>C 7463A>T), AAT139CTT (7464A>C 7465A>T), AAA140GGG (7467A>G 7468A>G 7469A>G), GTT141CCA (7470G>C 7471T>C 7472T>A), GGT142AAT (7473G>A 7474G>A), GAT143ACT (7476G>A 7477A>C), GAT144GAG (7481T>G), CCT145CCG (7484T>G), ATA146ATG (7487A>G), ATC147ACT (7489T>C 7490C>T), AAG149GTT (7494A>G 7495A>T 7496G>T), GGT150GGA (7499T>A), GTG151AAA (7500G>A 7501T>A 7502G>A), CTA152ACG (7503C>A 7504T>C 7505A>G), AGA153CTG (7506A>C 7507G>T 7508A>G), GGT154GGA (7511T>A), GAG155AAG (7512G>A), ATT156ATA (7517T>A), GTA157GTG (7520A>G), ATG158CCA (7521A>C 7522T>C 7523G>A), CCA159GCT (7524C>G 7526A>T), CGC160AGG (7527C>A 7529C>G), CCA162CCC (7535A>C), TCA164AAT (7539T>A 7540C>A 7541A>T), TTT165TTC (7544T>C), GGG166GGA (7547G>A), GAC168GAT (7553C>T), CCA169CCG (7556A>G), ATT170ATA (7559T>A), TTC171TTT (7562C>T), CCA173CCT (7568A>T), CTT174CAT (7570T>A), AAC175GGC (7572A>G 7573A>G), TGG176TAT (7576G>A 7577G>T), AGA177GAC (7578A>G 7579G>A 7580A>C), AAG178CAG (7581A>C), ACA179ACT (7586A>T), TTT180TAC (7588T>A 7589T>C), ATG184CCC (7599A>C 7600T>C 7601G>C), GCC185AAG (7602G>A 7603C>A 7604C>G), GAG186GAA (7607G>A), GAG187GAA (7610G>A), AAT189AAC (7616T>C), ATG190AAG (7618T>A), ATA191ATT (7622A>T), CAT193CAC (7628T>C), CGA194CGC (7631A>C), TTT195GGT (7632T>G 7633T>G), CGA196AAA (7635C>A 7636G>A) | | | | | | | | |

*: Inserts / Deletes / Misaligned / Frameshifts

#### Analysis details

This analysis was performed with panviral2.64

### NGS Details (UN60): Lowelvirus tuscon4d

#### Assembly

| | |
|---|---|
| **Coverage Length** | 164 (1 contig(s)) |
| **Depth Of Coverage** | 188.4 |
| **Number Of Reads** | 355 |
| **Reads Per Million** | 7.53 rpm (after QC) |
| **Ambiguities** | 0 |
| **Assembly Method** | de novo + reference guided assembly |
| **Consensus Caller** | Bcf Tools |

#### Coverage Map

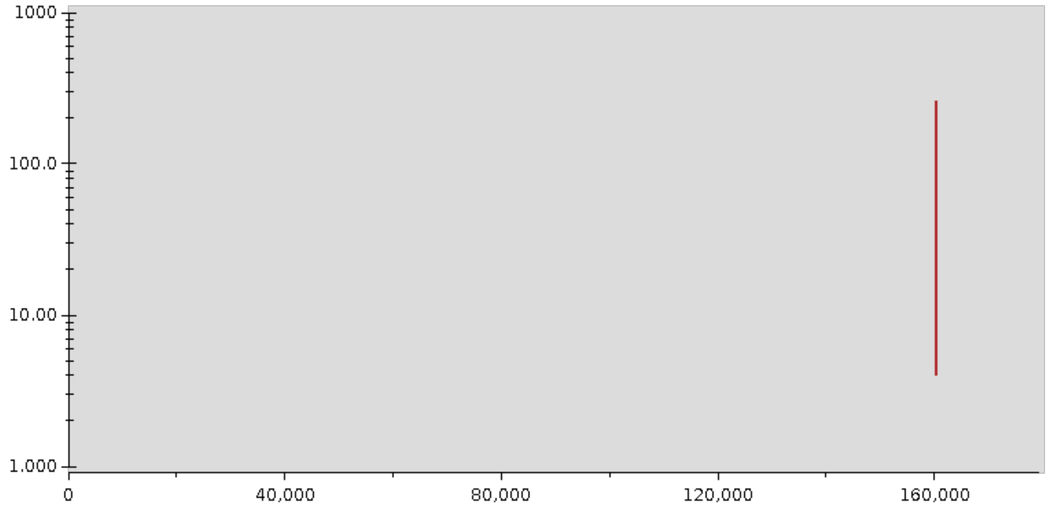

#### Assignment

| | |
|---|---|
| **Type** | Lowelvirus tuscon4d (Taxonomy ID: 2956131) |
| **Reference Genome** | NC_026923.1 |
| **NT Identity (%)** | 80.8642 |
| **AA Identity (%)** | 90.566 |
| **Number Of Stop Codons** | 1 |
| **Number Of CDS** | 218 |

#### Alignment

| | |
|---|---|
| **Alignment Score** | 189.0 (NT) + 233.0 (AA) = 422.0 |
| **Concordance (%)** | 75.5481 |
| **Alignment Method** | Local, heuristic, nucleotide (BLASTN) |

#### Genome Region

Sequence starts at position 160235 and ends at position 160398 relative to NC_026923.1 reference sequence.

#### Alignment Detailed Statistics

| | Begin | End | Coverage | Score | Concordance | Matches | Identities | I/D/M/F* | Stop Codons |
|---|---|---|---|---|---|---|---|---|---|
| **NT** | **160235** | **160398** | **0.1%** | **189** | **60.4%** | **162 (98.8%)** | **131 (79.9%)** | **0/2** | |
| Mutations: | 160246A>C, 160248A>C, 160254A>G, 160270G>A, 160275C>T, 160287G>A, 160290C>T, 160291_160292delAT, 160293C>A, 160296A>C, 160299G>A, 160302T>C, 160308A>T, 160309C>A, 160326A>G, 160327C>G, 160330G>C, 160331C>A, 160332T>G, 160338A>G, 160341A>T, 160350C>T, 160356C>A, 160359C>T, 160368A>G, 160371A>T, 160377T>A, 160378A>C, 160380A>T, 160383T>A, 160389A>T, 160392G>T | | | | | | | | |

*: Inserts / Deletes / Misaligned / Frameshifts

#### Analysis details

This analysis was performed with panviral2.64

### NGS Details (UN60): Badnavirus occultipomeae

#### Assembly

| | |
|---|---|
| **Coverage Length** | 345 (1 contig(s)) |
| **Depth Of Coverage** | 112.5 |
| **Number Of Reads** | 353 |
| **Reads Per Million** | 7.49 rpm (after QC) |
| **Ambiguities** | 0 |
| **Assembly Method** | de novo + reference guided assembly |
| **Consensus Caller** | Bcf Tools |

#### Coverage Map

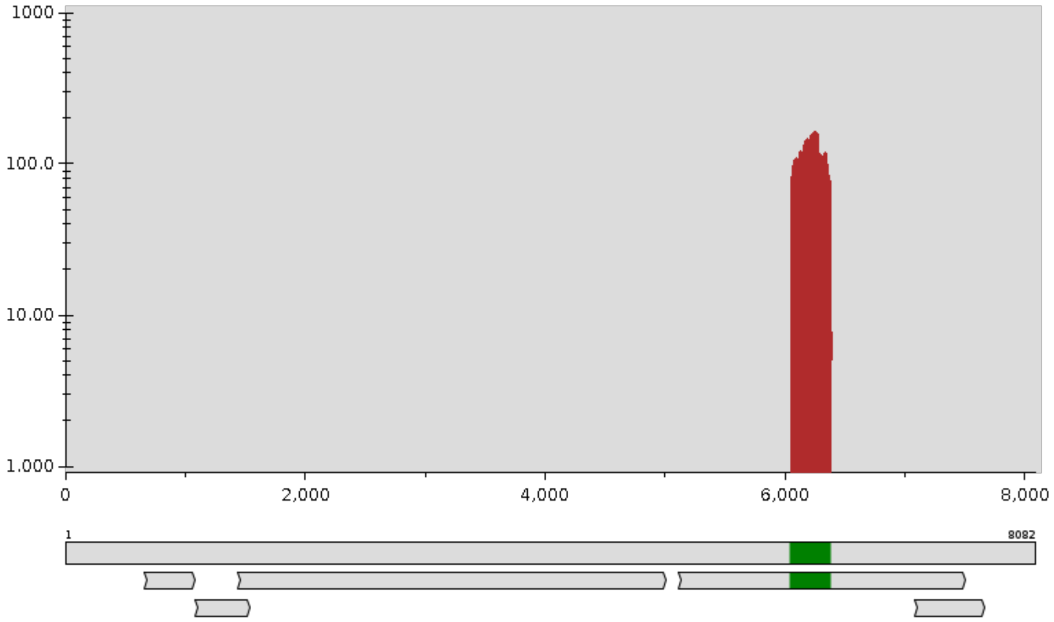

#### Assignment

| | |
|---|---|
| **Type** | Badnavirus occultipomeae (Taxonomy ID: 3048353) |
| **Reference Genome** | NC_015655.1 |
| **NT Identity (%)** | 51.7241 |
| **AA Identity (%)** | 45.6897 |
| **Number Of Stop Codons** | 0 |
| **Number Of CDS** | 5 |

#### Alignment

| | |
|---|---|
| **Alignment Score** | 18.0 (NT) + 370.0 (AA) = 388.0 |
| **Concordance (%)** | 25.9532 |

| **Alignment Method** | Global, seeded, nucleotide + amino acids (AGA) |
|---|---|

#### Genome Region

Sequence starts at position 6040 and ends at position 6384 relative to NC_015655.1 reference sequence.

#### Alignment Detailed Statistics

| | Begin | End | Coverage | Score | Concordance | Matches | Identities | I/D/M/F* | Stop Codons |
|---|---|---|---|---|---|---|---|---|---|
| **NT** | **6040** | **6384** | **4.3%** | **18** | **2.6%** | **345 (99.1%)** | **180 (51.7%)** | **3/0** | |
| Mutations: | 6042A>G, 6043T>A, 6051C>T, 6052A>C, 6053A>G, 6055G>T, 6057A>G, 6060C>T, 6062T>A, 6063T>C, 6070A>T, 6072A>G, 6076A>G, 6078G>T, 6079G>C, 6080A>G, 6084A>G, 6085A>T, 6086A>C, 6087G>A, 6088T>G, 6089C>A, 6090C>T, 6092A>T, 6093A>T, 6096A>G, 6097T>A, 6098G>A, 6105T>G, 6108T>C, 6109T>A, 6114T>C, 6116C>G, 6117A>C, 6118G>T, 6120A>T, 6123G>T, 6125T>A, 6126C>T, 6132A>G, 6137A>T, 6138A>G, 6141C>T, 6145C>T, 6147C>T, 6153A>G, 6154C>T, 6158T>C, 6159G>T, 6165A>C, 6168A>T, 6171A>T, 6173A>C, 6177T>C, 6178C>A, 6179A>T, 6181A>G, 6182G>A, 6183A>G, 6184A>T, 6185A>T, 6190G>A, 6192T>C, 6193A>C, 6194A>G, 6195T>A, 6197C>T, 6198A>G, 6201T>C, 6203G>A, 6204G>A, 6205G>T, 6206G>C, 6207A>T, 6207_6208insTAT, 6208A>C, 6209C>T, 6210A>T, 6214G>C, 6215C>A, 6216A>G, 6220A>G, 6222C>G, 6223G>A, 6224C>T, 6225A>T, 6228A>G, 6230A>T, 6231T>C, 6234T>A, 6240C>T, 6243A>C, 6249A>T, 6251T>A, 6255T>C, 6256G>A, 6258A>G, 6260A>G, 6264A>G, 6265G>A, 6266A>G, 6267A>T, 6270G>T, 6275A>T, 6277G>A, 6279T>A, 6282T>C, 6285C>G, 6286T>A, 6287T>G, 6289A>T, 6290A>T, 6292T>G, 6294G>A, 6295G>T, 6296C>T, 6297C>A, 6300A>G, 6302T>G, 6303T>G, 6304G>T, 6309A>G, 6310A>G, 6311G>A, 6312A>T, 6315A>G, 6316G>A, 6317G>A, 6319C>T, 6321T>G, 6322A>T, 6323T>A, 6324T>C, 6325C>G, 6326T>C, 6327A>C, 6329G>A, 6330C>G, 6331C>T, 6332C>T, 6333A>C, 6334A>T, 6339A>G, 6340A>T, 6341T>G, 6342G>T, 6343A>G, 6345A>G, 6346A>T, 6349G>T, 6351A>G, 6352G>C, 6354C>A, 6355A>C, 6357A>G, 6358G>T, 6359A>C, 6363G>A, 6364G>T, 6365A>C, 6369C>T, 6375A>T, 6376A>C, 6377T>A, 6378A>T, 6379A>G | | | | | | | | |
| **CDS** | | | | | | | | | |
| SPBVa_gp4 | 309 | 423 | 14.5% | 370 | 46.0% | 115 (99.1%) | 53 (45.7%) | 1/0/0/0 | 0 |
| Protein mutations: | F310I (6043T>A), K313R (6052A>C 6053A>G), A314S (6055G>T 6057A>G), F316Y (6062T>A 6063T>C), I319L (6070A>T 6072A>G), M321V (6076A>G 6078G>T), E322R (6079G>C 6080A>G), K324S (6085A>T 6086A>C 6087G>A), S325D (6088T>G 6089C>A 6090C>T), K326I (6092A>T 6093A>T), W328K (6097T>A 6098G>A), W332R (6109T>A), P334R (6116C>G 6117A>C), E335Y (6118G>T 6120A>T), L337H (6125T>A 6126C>T), E341V (6137A>T 6138A>G), P344S (6145C>T 6147C>T), M348T (6158T>C 6159G>T), D353A (6173A>C), Q355M (6178C>A 6179A>T), R356E (6181A>G 6182G>A 6183A>G), K357L (6184A>T 6185A>T), D359N (6190G>A 6192T>C), N360R (6193A>C 6194A>G 6195T>A), A361V (6197C>T 6198A>G), R363K (6203G>A 6204G>A), G364S (6205G>T 6206G>C 6207A>T), G364_T365insY (6207_6208insTAT), T365L (6208A>C 6209C>T 6210A>T), A367Q (6214G>C 6215C>A 6216A>G), I369V (6220A>G 6222C>G), A370I (6223G>A 6224C>T 6225A>T), Y372F (6230A>T 6231T>C), F379Y (6251T>A), E381K (6256G>A 6258A>G), N382S (6260A>G), E384S (6265G>A 6266A>G 6267A>T), E385D (6270G>T), E387V (6275A>T), D388K (6277G>A 6279T>A), L391R (6286T>A 6287T>G), N392F (6289A>T 6290A>T), L393V (6292T>G 6294G>A), A394L (6295G>T 6296C>T 6297C>A), I396R (6302T>G 6303T>G), V397L (6304G>T), R399D (6310A>G 6311G>A 6312A>T), G401K (6316G>A 6317G>A), I403Y (6322A>T 6323T>A 6324T>C), L404A (6325C>G 6326T>C 6327A>C), S405K (6329G>A 6330C>G), P406F (6331C>T 6332C>T 6333A>C), T407S (6334A>T), M409C (6340A>T 6341T>G 6342G>T), K410E (6343A>G 6345A>G), I411F (6346A>T), G412W (6349G>T 6351A>G), V413L (6352G>C 6354C>A), K414Q (6355A>C 6357A>G), E415S (6358G>T 6359A>C), D417S (6364G>T 6365A>C), I421H (6376A>C 6377T>A 6378A>T), K422E (6379A>G) | | | | | | | | |
| Codon mutations: | AAA309AAG (6042A>G), TTT310ATT (6043T>A), CTC312CTT (6051C>T), AAA313CGA (6052A>C 6053A>G), GCA314TCG (6055G>T 6057A>G), GGC315GGT (6060C>T), TTT316TAC (6062T>A 6063T>C), ATA319TTG (6070A>T 6072A>G), ATG321GTT (6076A>G 6078G>T), GAG322CGG (6079G>C 6080A>G), GAA323GAG (6084A>G), AAG324TCA (6085A>T 6086A>C 6087G>A), TCC325GAT (6088T>G 6089C>A 6090C>T), AAA326ATT (6092A>T 6093A>T), CCA327CCG (6096A>G), TGG328AAG (6097T>A 6098G>A), GCT330GCG (6105T>G), TTT331TTC (6108T>C), TGG332AGG (6109T>A), ACT333ACC (6114T>C), CCA334CGC (6116C>G 6117A>C), GAA335TAT (6118G>T 6120A>T), GGG336GGT (6123G>T), CTC337CAT (6125T>A 6126C>T), GAA339GAG (6132A>G), GAA341GTG (6137A>T 6138A>G), GTC342GTT (6141C>T), CCC344TCT (6145C>T 6147C>T), GGA346GGG (6153A>G), CTG347TTG (6154C>T), ATG348ACT (6158T>C 6159G>T), GCA350GCC (6165A>C), CCA351CCT (6168A>T), GCA352GCT (6171A>T), GAT353GCT (6173A>C), TTT354TTC (6177T>C), CAG355ATG (6178C>A 6179A>T), AGA356GAG (6181A>G 6182G>A 6183A>G), AAG357TTG (6184A>T 6185A>T), GAT359AAC (6190G>A 6192T>C), AAT360CGA (6193A>C 6194A>G 6195T>A), GCA361GTG (6197C>T 6198A>G), TTT362TTC (6201T>C), AGG363AAA (6203G>A 6204G>A), GGA364TCT (6205G>T 6206G>C 6207A>T), GGA364_ACA365insTAT (6207_6208insTAT), ACA365CTT (6208A>C 6209C>T 6210A>T), GCA367CAG (6214G>C 6215C>A 6216A>G), ATC369GTG (6220A>G 6222C>G), GCA370ATT (6223G>A 6224C>T 6225A>T), GTA371GTG (6228A>G), TAT372TTC (6230A>T 6231T>C), ATT373ATA (6234T>A), GAC375GAT (6240C>T), ATA376ATC (6243A>C), GTA378GTT (6249A>T), TTC379TAC (6251T>A), TCT380TCC (6255T>C), GAA381AAG (6256G>A 6258A>G), AAT382AGT (6260A>G), GAA383GAG (6264A>G), GAA384AGT (6265G>A 6266A>G 6267A>T), GAG385GAT (6270G>T), GAG387GTG (6275A>T), GAT388AAA (6277G>A 6279T>A), CAT389CAC (6282T>C), CTC390CTG (6285C>G), TTA391AGA (6286T>A 6287T>G), AAT392TTT (6289A>T 6290A>T), TTG393GTA (6292T>G 6294G>A), GCC394TTA (6295G>T 6296C>T 6297C>A), CAA395CAG (6300A>G), ATT396AGG (6302T>G 6303T>G), GTG397TTG (6304G>T), AGA398AGG (6309A>G), AGA399GAT (6310A>G 6311G>A 6312A>T), GAA400GAG (6315A>G), GGG401AAG (6316G>A 6317G>A), CTT402TTG (6319C>T 6321T>G), ATT403TAC (6322A>T 6323T>A 6324T>C), CTA404GCC (6325C>G 6326T>C 6327A>C), AGC405AAG (6329G>A 6330C>G), CCA406TTC (6331C>T 6332C>T 6333A>C), ACA407TCA (6334A>T), AAA408AAG (6339A>G), ATG409TGT (6340A>T 6341T>G 6342G>T), AAA410GAG (6343A>G 6345A>G), ATT411TTT (6346A>T), GGA412TGG (6349G>T 6351A>G), GTC413CTA (6352G>C 6354C>A), AAA414CAG (6355A>C 6357A>G), GAA415TCA (6358G>T 6359A>C), GTG416GTA (6363G>A), GAT417TCT (6364G>T 6365A>C), TTC418TTT (6369C>T), GGA420GGT (6375A>T), ATA421CAT (6376A>C 6377T>A 6378A>T), AAG422GAG (6379A>G) | | | | | | | | |
| **Proteins** | | | | | | | | | |
| RNaseH/reverse transcriptase (YP_004581513.1) | 309 | 423 | 14.5% | 370 | 46.0% | 115 (99.1%) | 53 (45.7%) | 1/0/0/0 | 0 |
| Protein mutations: | F310I (6043T>A), K313R (6052A>C 6053A>G), A314S (6055G>T 6057A>G), F316Y (6062T>A 6063T>C), I319L (6070A>T 6072A>G), M321V (6076A>G 6078G>T), E322R (6079G>C 6080A>G), K324S (6085A>T 6086A>C 6087G>A), S325D (6088T>G 6089C>A 6090C>T), K326I (6092A>T 6093A>T), W328K (6097T>A 6098G>A), W332R (6109T>A), P334R (6116C>G 6117A>C), E335Y (6118G>T 6120A>T), L337H (6125T>A 6126C>T), E341V (6137A>T 6138A>G), P344S (6145C>T 6147C>T), M348T (6158T>C 6159G>T), D353A (6173A>C), Q355M (6178C>A 6179A>T), R356E (6181A>G 6182G>A 6183A>G), K357L (6184A>T 6185A>T), D359N (6190G>A 6192T>C), N360R (6193A>C 6194A>G 6195T>A), A361V (6197C>T 6198A>G), R363K (6203G>A 6204G>A), G364S (6205G>T 6206G>C 6207A>T), G364_T365insY (6207_6208insTAT), T365L (6208A>C 6209C>T 6210A>T), A367Q (6214G>C 6215C>A 6216A>G), I369V (6220A>G 6222C>G), A370I (6223G>A 6224C>T 6225A>T), Y372F (6230A>T 6231T>C), F379Y (6251T>A), E381K (6256G>A 6258A>G), N382S (6260A>G), E384S (6265G>A 6266A>G 6267A>T), E385D (6270G>T), E387V (6275A>T), D388K (6277G>A 6279T>A), L391R (6286T>A 6287T>G), N392F (6289A>T 6290A>T), L393V (6292T>G 6294G>A), A394L (6295G>T 6296C>T 6297C>A), I396R (6302T>G 6303T>G), V397L (6304G>T), R399D (6310A>G 6311G>A 6312A>T), G401K (6316G>A 6317G>A), I403Y (6322A>T 6323T>A 6324T>C), L404A (6325C>G 6326T>C 6327A>C), S405K (6329G>A 6330C>G), P406F (6331C>T 6332C>T 6333A>C), T407S (6334A>T), M409C (6340A>T 6341T>G 6342G>T), K410E (6343A>G 6345A>G), I411F (6346A>T), G412W (6349G>T 6351A>G), V413L (6352G>C 6354C>A), K414Q (6355A>C 6357A>G), E415S (6358G>T 6359A>C), D417S (6364G>T 6365A>C), I421H (6376A>C 6377T>A 6378A>T), K422E (6379A>G) | | | | | | | | |
| Codon mutations: | AAA309AAG (6042A>G), TTT310ATT (6043T>A), CTC312CTT (6051C>T), AAA313CGA (6052A>C 6053A>G), GCA314TCG (6055G>T 6057A>G), GGC315GGT (6060C>T), TTT316TAC (6062T>A 6063T>C), ATA319TTG (6070A>T 6072A>G), ATG321GTT (6076A>G 6078G>T), GAG322CGG (6079G>C 6080A>G), GAA323GAG (6084A>G), AAG324TCA (6085A>T 6086A>C 6087G>A), TCC325GAT (6088T>G 6089C>A 6090C>T), AAA326ATT (6092A>T 6093A>T), CCA327CCG (6096A>G), TGG328AAG (6097T>A 6098G>A), GCT330GCG (6105T>G), TTT331TTC (6108T>C), TGG332AGG (6109T>A), ACT333ACC (6114T>C), CCA334CGC (6116C>G 6117A>C), GAA335TAT (6118G>T 6120A>T), GGG336GGT (6123G>T), CTC337CAT (6125T>A 6126C>T), GAA339GAG (6132A>G), GAA341GTG (6137A>T 6138A>G), GTC342GTT (6141C>T), CCC344TCT (6145C>T 6147C>T), GGA346GGG (6153A>G), CTG347TTG (6154C>T), ATG348ACT (6158T>C 6159G>T), GCA350GCC (6165A>C), CCA351CCT (6168A>T), GCA352GCT (6171A>T), GAT353GCT (6173A>C), TTT354TTC (6177T>C), CAG355ATG (6178C>A 6179A>T), AGA356GAG (6181A>G 6182G>A 6183A>G), AAG357TTG (6184A>T 6185A>T), GAT359AAC (6190G>A 6192T>C), AAT360CGA (6193A>C 6194A>G 6195T>A), GCA361GTG (6197C>T 6198A>G), TTT362TTC (6201T>C), AGG363AAA (6203G>A 6204G>A), GGA364TCT (6205G>T 6206G>C 6207A>T), GGA364_ACA365insTAT (6207_6208insTAT), ACA365CTT (6208A>C 6209C>T 6210A>T), GCA367CAG (6214G>C 6215C>A 6216A>G), ATC369GTG (6220A>G 6222C>G), GCA370ATT (6223G>A 6224C>T 6225A>T), GTA371GTG (6228A>G), TAT372TTC (6230A>T 6231T>C), ATT373ATA (6234T>A), GAC375GAT (6240C>T), ATA376ATC (6243A>C), GTA378GTT (6249A>T), TTC379TAC (6251T>A), TCT380TCC (6255T>C), GAA381AAG (6256G>A 6258A>G), AAT382AGT (6260A>G), GAA383GAG (6264A>G), GAA384AGT (6265G>A 6266A>G 6267A>T), GAG385GAT (6270G>T), GAG387GTG (6275A>T), GAT388AAA (6277G>A 6279T>A), CAT389CAC (6282T>C), CTC390CTG (6285C>G), TTA391AGA (6286T>A 6287T>G), AAT392TTT (6289A>T 6290A>T), TTG393GTA (6292T>G 6294G>A), GCC394TTA (6295G>T 6296C>T 6297C>A), CAA395CAG (6300A>G), ATT396AGG (6302T>G 6303T>G), GTG397TTG (6304G>T), AGA398AGG (6309A>G), AGA399GAT (6310A>G 6311G>A 6312A>T), GAA400GAG (6315A>G), GGG401AAG (6316G>A 6317G>A), CTT402TTG (6319C>T 6321T>G), ATT403TAC (6322A>T 6323T>A 6324T>C), CTA404GCC (6325C>G 6326T>C 6327A>C), AGC405AAG (6329G>A 6330C>G), CCA406TTC (6331C>T 6332C>T 6333A>C), ACA407TCA (6334A>T), AAA408AAG (6339A>G), ATG409TGT (6340A>T 6341T>G 6342G>T), AAA410GAG (6343A>G 6345A>G), ATT411TTT (6346A>T), GGA412TGG (6349G>T 6351A>G), GTC413CTA (6352G>C 6354C>A), AAA414CAG (6355A>C 6357A>G), GAA415TCA (6358G>T 6359A>C), GTG416GTA (6363G>A), GAT417TCT (6364G>T 6365A>C), TTC418TTT (6369C>T), GGA420GGT (6375A>T), ATA421CAT (6376A>C 6377T>A 6378A>T), AAG422GAG (6379A>G) | | | | | | | | |

*: Inserts / Deletes / Misaligned / Frameshifts

#### Analysis details

This analysis was performed with panviral2.64

#### NGS Details (UN60): Errantivirus

##### Assembly

| | |
|---|---|
| **Coverage Length** | 638 (1 contig(s)) |
| **Depth Of Coverage** | 29.2 |
| **Number Of Reads** | 227 |
| **Reads Per Million** | 4.81 rpm (after QC) |
| **Ambiguities** | 0 |
| **Assembly Method** | de novo + reference guided assembly |
| **Consensus Caller** | Bcf Tools |

##### Coverage Map

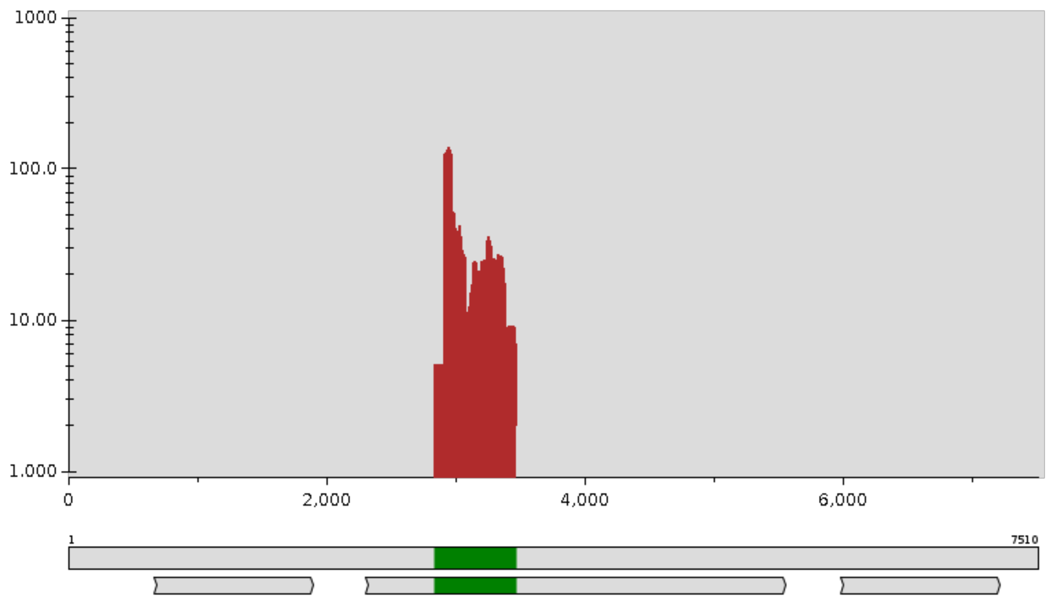

##### Assignment

| | |
|---|---|
| **Type** | Errantivirus (Taxonomy ID: 186666) |
| **Reference Genome** | NC_038512.1 |
| **NT Identity (%)** | 55.1834 |
| **AA Identity (%)** | 43.5407 |
| **Number Of Stop Codons** | 0 |
| **Number Of CDS** | 3 |

##### Alignment

| | |
|---|---|
| **Alignment Score** | 101.0 (NT) + 551.0 (AA) = 652.0 |
| **Concordance (%)** | 25.7402 |

| Alignment Method | Global, seeded, nucleotide + amino acids (AGA) |
|---|---|

#### Genome Region

Sequence starts at position 2834 and ends at position 3471 relative to NC_038512.1 reference sequence.

#### Alignment Detailed Statistics

| | Begin | End | Coverage | Score | Concordance | Matches | Identities | I/D/M/F* | Stop Codons |
|---|---|---|---|---|---|---|---|---|---|
| **NT** | **2834** | **3471** | **8.5%** | **101** | **8.2%** | **627 (98.3%)** | **346 (54.2%)** | **0/11** | |
| Mutations: | 2836C>A, 2837A>G, 2839A>G, 2841G>T, 2843G>T, 2845T>A, 2849_2851delCCC, 2859T>A, 2860C>A, 2863C>T, 2865C>G, 2866T>G, 2867T>A, 2870_2877delGGGAAACA, 2880A>C, 2881G>A, 2885C>A, 2887T>A, 2888C>T, 2891G>T, 2892T>A, 2893A>T, 2894G>A, 2901T>A, 2903C>A, 2905T>A, 2906A>G, 2909T>C, 2914C>T, 2915G>A, 2917G>C, 2919A>T, 2920G>C, 2924A>G, 2926C>G, 2927G>A, 2928A>G, 2929T>G, 2932C>T, 2934A>G, 2935A>G, 2937A>T, 2941G>A, 2944A>C, 2949A>C, 2950C>T, 2953A>T, 2954A>G, 2955G>A, 2959C>G, 2960G>C, 2963C>T, 2968C>T, 2969A>G, 2972T>G, 2975G>A, 2976G>A, 2978A>G, 2979A>G, 2980G>C, 2981T>G, 2982G>C, 2983C>T, 2984C>G, 2985A>T, 2986A>T, 2989C>T, 2993A>T, 2995C>A, 2997C>A, 2998C>G, 2999T>C, 3001A>T, 3008G>T, 3009C>T, 3011A>T, 3012G>C, 3016G>A, 3018T>A, 3019T>C, 3020T>G, 3025G>A, 3026G>A, 3028G>T, 3029G>C, 3030A>G, 3032A>G, 3034G>T, 3035G>A, 3037C>G, 3038C>G, 3039C>T, 3040T>G, 3041C>G, 3042A>C, 3050T>C, 3052G>C, 3058C>T, 3061G>T, 3066A>G, 3067C>G, 3068G>A, 3069T>C, 3071G>C, 3072A>G, 3074C>T, 3076C>T, 3080C>A, 3081A>G, 3084T>A, 3095C>G, 3096G>T, 3103T>G, 3104A>T, 3106G>T, 3109A>G, 3110T>C, 3114A>C, 3115A>T, 3118C>T, 3119T>G, 3121A>C, 3124A>G, 3125T>G, 3133T>C, 3134C>T, 3135A>G, 3136A>T, 3138G>A, 3139A>C, 3140G>T, 3142T>A, 3146G>A, 3157A>T, 3158A>T, 3159G>A, 3160A>T, 3162G>A, 3164C>T, 3165T>A, 3166C>T, 3168A>T, 3169A>T, 3170A>G, 3173A>G, 3176A>T, 3179T>G, 3180G>T, 3181T>A, 3182C>G, 3184C>A, 3191C>T, 3193T>A, 3196C>T, 3199T>C, 3202T>A, 3203A>G, 3205T>C, 3206G>A, 3208C>A, 3216C>G, 3217T>A, 3218T>A, 3220C>T, 3221C>T, 3223A>G, 3224C>G, 3226G>A, 3229A>C, 3232C>T, 3233C>T, 3236G>C, 3237A>T, 3239A>C, 3241C>T, 3242C>T, 3245G>A, 3246A>G, 3247A>G, 3248C>A, 3249G>A, 3250A>G, 3253T>G, 3256C>G, 3272A>T, 3273G>A, 3274T>C, 3275A>C, 3277C>A, 3278T>C, 3281A>T, 3283A>T, 3284A>G, 3286T>C, 3287C>A, 3289A>G, 3295C>G, 3300C>G, 3301C>T, 3307C>T, 3309T>C, 3310G>T, 3312A>G, 3313G>T, 3315T>G, 3316C>T, 3319A>G, 3321C>T, 3322T>C, 3323G>A, 3324C>A, 3325T>G, 3327A>T, 3329C>T, 3331T>A, 3334T>C, 3337C>T, 3338A>T, 3340C>G, 3341A>G, 3343A>T, 3346C>T, 3347A>G, 3348G>A, 3349G>A, 3350G>A, 3353G>C, 3354G>A, 3355T>A, 3356A>G, 3358C>G, 3359A>C, 3360A>G, 3361G>C, 3362C>A, 3363C>T, 3364T>G, 3365A>G, 3371G>A, 3373T>G, 3376G>A, 3377A>G, 3379T>G, 3380T>C, 3381C>A, 3382C>G, 3385T>A, 3386A>G, 3388T>A, 3389C>G, 3390A>T, 3392A>G, 3394A>T, 3396A>G, 3397T>G, 3399T>A, 3400G>A, 3401A>G, 3402T>C, 3403T>A, 3406A>C, 3408A>C, 3409G>T, 3410A>G, 3411C>G, 3412C>G, 3413C>G, 3414C>T, 3415T>C, 3422A>C, 3426A>G, 3428C>T, 3429A>C, 3430A>C, 3431T>A, 3439C>G, 3442T>G, 3443C>G, 3445C>A, 3446G>A, 3447G>A, 3448T>C, 3454C>T | | | | | | | | |

##### CDS

| | Begin | End | Coverage | Score | Concordance | Matches | Identities | I/D/M/F* | Stop Codons |
|---|---|---|---|---|---|---|---|---|---|
| D1R33_gp2 | 177 | 389 | 19.5% | 551 | 38.1% | 209 (98.6%) | 91 (42.9%) | 0/3/1/1 | 0 |
| Protein mutations: | I178V (2837A>G 2839A>G), W179L (2841G>T), V180L (2843G>T 2845T>A), P182del (2849_2851delCCC), I185K (2859T>A 2860C>A), A187G (2865C>G 2866T>G), S188T (2867T>A), G189_K190del (2870_2877delGGGAAACA), K192T (2880A>C 2881G>A), L195F (2888C>T), V196Y (2891G>T 2892T>A 2893A>T), V197I (2894G>A), F199Y (2901T>A), K201E (2906A>G), E204N (2915G>A 2917G>C), K205I (2919A>T 2920G>C), I207V (2924A>G 2926C>G), D208R (2927G>A 2928A>G 2929T>G), K210R (2934A>G 2935A>G), Y211F (2937A>T), N215T (2949A>C 2950C>T), S217D (2954A>G 2955G>A), D218E (2959C>G), V219L (2960G>C), L220F (2963C>T), K222E (2969A>G), L223V (2972T>G), G224N (2975G>A 2976G>A), K225G (2978A>G 2979A>G 2980G>C), C226A (2981T>G 2982G>C 2983C>T), Q227V (2984C>G 2985A>T 2986A>T), T230S (2993A>T 2995C>A), T231K (2997C>A 2998C>G), A235L (3008G>T 3009C>T), F238Y (3018T>A 3019T>C), Y239D (3020T>G), V241I (3026G>A 3028G>T), E242R (3029G>C 3030A>G), M243V (3032A>G 3034G>T), D244K (3035G>A 3037C>G), P245V (3038C>G 3039C>T 3040T>G), Q246A (3041C>G 3042A>C), S249P (3050T>C 3052G>C), N254R (3066A>G 3067C>G), V255T (3068G>A 3069T>C), E256R (3071G>C 3072A>G), H257Y (3074C>T 3076C>T), H259S (3080C>A 3081A>G), F260Y (3084T>A), R264V (3095C>G 3096G>T), M267F (3104A>T 3106G>T), K270T (3114A>C 3115A>T), S272A (3119T>G 3121A>C), S274A (3125T>G), Q277C (3134C>T 3135A>G 3136A>T), R278N (3138G>A 3139A>C), V279L (3140G>T 3142T>A), D281N (3146G>A), R285Y (3158A>T 3159G>A 3160A>T), G286D (3162G>A), L287Y (3164C>T 3165T>A 3166C>T), Q288L (3168A>T 3169A>T), N289D (3170A>G), N290D (3173A>G), I291F (3176A>T), C292V (3179T>G 3180G>T 3181T>A), L293V (3182C>G 3184C>A), I300V (3203A>G 3205T>C), V301I (3206G>A 3208C>A), T304R (3216C>G 3217T>A), S305T (3218T>A 3220C>T), Q307E (3224C>G 3226G>A), E308D (3229A>C), E311L (3236G>C 3237A>T), N312H (3239A>C 3241C>T), E314R (3245G>A 3246A>G 3247A>G), R315K (3248C>A 3249G>A 3250A>G), F317L (3256C>G), S323Y (3272A>T 3273G>A 3274T>C), N324Q (3275A>C 3277C>A), F325L (3278T>C), K326Y (3281A>T 3283A>T), I327V (3284A>G 3286T>C), Q328K (3287C>A 3289A>G), D330E (3295C>G), S332C (3300C>G 3301C>T), L335S (3309T>C 3310G>T), K336S (3312A>G 3313G>T), L337R (3315T>G 3316C>T), T339I (3321C>T 3322T>C), A340K (3323G>A 3324C>A 3325T>G), Y341F (3327A>T), I345L (3338A>T 3340C>G), I346V (3341A>G 3343A>T), R348E (3347A>G 3348G>A 3349G>A), D349N (3350G>A), G350Q (3353G>C 3354G>A 3355T>A), I351V (3356A>G 3358C>G), K352R (3359A>C 3360A>G 3361G>C), P353M (3362C>A 3363C>T 3364T>G), N354D (3365A>G), D356K (3371G>A 3373T>G), I358V (3377A>G 3379T>G), S359Q (3380T>C 3381C>A 3382C>G), I361V (3386A>G 3388T>A), Q362V (3389C>G 3390A>T), K363D (3392A>G 3394A>T), Y364W (3396A>G 3397T>G), L365Q (3399T>A 3400G>A), I366A (3401A>G 3402T>C 3403T>A), K368T (3408A>C 3409G>T), T369G (3410A>G 3411C>G 3412C>G), P370V (3413C>G 3414C>T 3415T>C), I373L (3422A>C), K374R (3426A>G), Q375S (3428C>T 3429A>C 3430A>C), F376I (3431T>A), L380V (3443C>G 3445C>A), G381N (3446G>A 3447G>A 3448T>C) | | | | | | | | |
| Codon mutations: | CCC177CCA (2836C>A), ATA178GTG (2837A>G 2839A>G), TGG179TTG (2841G>T), GTT180TTA (2843G>T 2845T>A), CCC182del (2849_2851delCCC), ATC185AAA (2859T>A 2860C>A), GAC186GAT (2863C>T), GCT187GGG (2865C>G 2866T>G), TCT188ACT (2867T>A), GGG189_AAA190del (2870_2877delGGGAAACA), CAA191--A (2870_2877delGGGAAACA), AAG192ACA (2880A>C 2881G>A), CGT194AGA (2885C>A 2887T>A), CTC195TTC (2888C>T), GTA196TAT (2891G>T 2892T>A 2893A>T), GTT197ATT (2894G>A), TTC199TAC (2901T>A), CGT200AGA (2903C>A 2905T>A), AAG201GAG (2906A>G), TTG202CTG (2909T>C), AAC203AAT (2914C>T), GAG204AAC (2915G>A 2917G>C), AAG205ATC (2919A>T 2920G>C), ATC207GTG (2924A>G 2926C>G), GAT208AGG (2927G>A 2928A>G 2929T>G), GAC209GAT (2932C>T), AAA210AGG (2934A>G 2935A>G), TAC211TTC (2937A>T), CCG212CCA (2941G>A), ATA213ATC (2944A>C), AAC215ACT (2949A>C 2950C>T), ATA216ATT (2953A>T), AGT217GAT (2954A>G 2955G>A), GAC218GAG (2959C>G), GTA219CTA (2960G>C), CTT220TTT (2963C>T), GAC221GAT (2968C>T), AAG222GAG (2969A>G), TTA223GTA (2972T>G), GGT224AAT (2975G>A 2976G>A), AAG225GGC (2978A>G 2979A>G 2980G>C), TGC226GCT (2981T>G 2982G>C 2983C>T), CAA227GTT (2984C>G 2985A>T 2986A>T), TAC228TAT (2989C>T), ACC230TCA (2993A>T 2995C>A), ACC231AAG (2997C>A 2998C>G), TTA232CTT (2999T>C 3001A>T), GCA235TTA (3008G>T 3009C>T), AGT236TCT (3011A>T 3012G>C), GGG237GGA (3016G>A), TTT238TAC (3018T>A 3019T>C), TAT239GAT (3020T>G), CAG240CAA (3025G>A), GTG241ATT (3026G>A 3028G>T), GAG242CGG (3029G>C 3030A>G), ATG243GTT (3032A>G 3034G>T), GAC244AAG (3035G>A 3037C>G), CCT245GTG (3038C>G 3039C>T 3040T>G), CAA246GCA (3041C>G 3042A>C), TCG249CCC (3050T>C 3052G>C), ACC251ACT (3058C>T), GCG252GCT (3061G>T), AAC254AGG (3066A>G 3067C>G), GTA255ACA (3068G>A 3069T>C), GAA256CGA (3071G>C 3072A>G), CAC257TAT (3074C>T 3076C>T), CAT259AGT (3080C>A 3081A>G), TTT260TAT (3084T>A), CGA264GTA (3095C>G 3096G>T), CCT266CCG (3103T>G), ATG267TTT (3104A>T 3106G>T), GGA268GGG (3109A>G), TTA269CTA (3110T>C), AAA270ACT (3114A>C 3115A>T), AAC271AAT (3118C>T), TCA272GCC (3119T>G 3121A>C), CCA273CCG (3124A>G), TCT274GCT (3125T>G), TTT276TTC (3133T>C), CAA277TGT (3134C>T 3135A>G 3136A>T), AGA278AAC (3138G>A 3139A>C), GTT279TTA (3140G>T 3142T>A), GAC281AAC (3146G>A), CTA284CTT (3157A>T), AGA285TAT (3158A>T 3159G>A 3160A>T), GGT286GAT (3162G>A), CTC287TAT (3164C>T 3165T>A 3166C>T), CAA288CTT (3168A>T 3169A>T), AAT289GAT (3170A>G), AAC290GAC (3173A>G), ATC291TTC (3176A>T), TGT292GTA (3179T>G 3180G>T 3181T>A), CTC293GTA (3182C>G 3184C>A), CTT296TTA (3191C>T 3193T>A), GAC297GAT (3196C>T), GAT298GAC (3199T>C), ATT299ATA (3202T>A), ATT300GTC (3203A>G 3205T>C), GTC301ATA (3206G>A 3208C>A), ACT304AGA (3216C>G 3217T>A), TCC305ACT (3218T>A 3220C>T), CTA306TTG (3221C>T 3223A>G), CAG307GAA (3224C>G 3226G>A), GAA308GAC (3229A>C), CAC309CAT (3232C>T), CTG310TTG (3233C>T), GAG311CTG (3236G>C 3237A>T), AAC312CAT (3239A>C 3241C>T), CTG313TTG (3242C>T), GAA314AGG (3245G>A 3246A>G 3247A>G), CGA315AAG (3248C>A 3249G>A 3250A>G), GTT316GTG (3253T>G), TTC317TTG (3256C>G), AGT323TAC (3272A>T 3273G>A 3274T>C), AAC324CAA (3275A>C 3277C>A), TTC325CTC (3278T>C), AAA326TAT (3281A>T 3283A>T), ATT327GTC (3284A>G 3286T>C), CAA328AAG (3287C>A 3289A>G), GAC330GAG (3295C>G), TCC332TGT (3300C>G 3301C>T), TTC334TTT (3307C>T), TTG335TCT (3309T>C 3310G>T), AAG336AGT (3312A>G 3313G>T), CTC337CGT (3315T>G 3316C>T), GAA338GAG (3319A>G), ACT339ATC (3321C>T 3322T>C), GCT340AAG (3323G>A 3324C>A 3325T>G), TAT341TTT (3327A>T), CTT342TTA (3329C>T 3331T>A), GGT343GGC (3334T>C), CAC344CAT (3337C>T), ATC345TTG (3338A>T 3340C>G), ATA346GTT (3341A>G 3343A>T), AGC347AGT (3346C>T), AGG348GAA (3347A>G 3348G>A 3349G>A), GAC349AAC (3350G>A), GGT350CAA (3353G>C 3354G>A 3355T>A), ATC351GTG (3356A>G 3358C>G), AAG352CGC (3359A>C 3360A>G 3361G>C), CCT353ATG (3362C>A 3363C>T 3364T>G), AAC354GAC (3365A>G), GAT356AAG (3371G>A 3373T>G), AAG357AAA (3376G>A), ATT358GTG (3377A>G 3379T>G), TCC359CAG (3380T>C 3381C>A 3382C>G), GCT360GCA (3385T>A), ATT361GTA (3386A>G 3388T>A), CAA362GTA (3389C>G 3390A>T), AAA363GAT (3392A>G 3394A>T), TAT364TGG (3396A>G 3397T>G), CTG365CAA (3399T>A 3400G>A), ATT366GCA (3401A>G 3402T>C 3403T>A), CCA367CCC (3406A>C), AAG368ACT (3408A>C 3409G>T), ACC369GGG (3410A>G 3411C>G 3412C>G), CCT370GTC (3413C>G 3414C>T 3415T>C), ATA373CTA (3422A>C), AAA374AGA (3426A>G), CAA375TCC (3428C>T 3429A>C 3430A>C), TTT376ATT (3431T>A), GGC378GGG (3439C>G), CTT379CTG (3442T>G), CTC380GTA (3443C>G 3445C>A), GGT381AAC (3446G>A 3447G>A 3448T>C), TAC383TAT (3454C>T) | | | | | | | | |

##### Proteins

| | Begin | End | Coverage | Score | Concordance | Matches | Identities | I/D/M/F* | Stop Codons |
|---|---|---|---|---|---|---|---|---|---|
| ORF B (YP_009507248.1) | 177 | 389 | 19.5% | 551 | 38.1% | 209 (98.6%) | 91 (42.9%) | 0/3/1/1 | 0 |

| | Begin | End | Coverage | Score | Concordance | Matches | Identities | I/D/M/F* | Stop Codons |
|---|---|---|---|---|---|---|---|---|---|
| **NT** | **2834** | **3471** | **8.5%** | **101** | **8.2%** | **627 (98.3%)** | **346 (54.2%)** | **0/11** | |
| Protein mutations: | I178V (2837A>G 2839A>G), W179L (2841G>T), V180L (2843G>T 2845T>A), P182del (2849_2851delCCC), I185K (2859T>A 2860C>A), A187G (2865C>G 2866T>G), S188T (2867T>A), G189_K190del (2870_2877delGGGAAACA), K192T (2880A>C 2881G>A), L195F (2888C>T), V196Y (2891G>T 2892T>A 2893A>T), V197I (2894G>A), F199Y (2901T>A), K201E (2906A>G), E204N (2915G>A 2917G>C), K205I (2919A>T 2920G>C), I207V (2924A>G 2926C>G), D208R (2927G>A 2928A>G 2929T>G), K210R (2934A>G 2935A>G), Y211F (2937A>T), N215T (2949A>C 2950C>T), S217D (2954A>G 2955G>A), D218E (2959C>G), V219L (2960G>C), L220F (2963C>T), K222E (2969A>G), L223V (2972T>G), G224N (2975G>A 2976G>A), K225G (2978A>G 2979A>G 2980G>C), C226A (2981T>G 2982G>C 2983C>T), Q227V (2984C>G 2985A>T 2986A>T), T230S (2993A>T 2995C>A), T231K (2997C>A 2998C>G), A235L (3008G>T 3009C>T), F238Y (3018T>A 3019T>C), Y239D (3020T>G), V241I (3026G>A 3028G>T), E242R (3029G>C 3030A>G), M243V (3032A>G 3034G>T), D244K (3035G>A 3037C>G), P245V (3038C>G 3039C>T 3040T>G), Q246A (3041C>G 3042A>C), S249P (3050T>C 3052G>C), N254R (3066A>G 3067C>G), V255T (3068G>A 3069T>C), E256R (3071G>C 3072A>G), H257Y (3074C>T 3076C>T), H259S (3080C>A 3081A>G), F260Y (3084T>A), R264V (3095C>G 3096G>T), M267F (3104A>T 3106G>T), K270T (3114A>C 3115A>T), S272A (3119T>G 3121A>C), S274A (3125T>G), Q277C (3134C>T 3135A>G 3136A>T), R278N (3138G>A 3139A>C), V279L (3140G>T 3142T>A), D281N (3146G>A), R285Y (3158A>T 3159G>A 3160A>T), G286D (3162G>A), L287Y (3164C>T 3165T>A 3166C>T), Q288L (3168A>T 3169A>T), N289D (3170A>G), N290D (3173A>G), I291F (3176A>T), C292V (3179T>G 3180G>T 3181T>A), L293V (3182C>G 3184C>A), I300V (3203A>G 3205T>C), V301I (3206G>A 3208C>A), T304R (3216C>G 3217T>A), S305T (3218T>A 3220C>T), Q307E (3224C>G 3226G>A), E308D (3229A>C), E311L (3236G>C 3237A>T), N312H (3239A>C 3241C>T), E314R (3245G>A 3246A>G 3247A>G), R315K (3248C>A 3249G>A 3250A>G), F317L (3256C>G), S323Y (3272A>T 3273G>A 3274T>C), N324Q (3275A>C 3277C>A), F325L (3278T>C), K326Y (3281A>T 3283A>T), I327V (3284A>G 3286T>C), Q328K (3287C>A 3289A>G), D330E (3295C>G), S332C (3300C>G 3301C>T), L335S (3309T>C 3310G>T), K336S (3312A>G 3313G>T), L337R (3315T>G 3316C>T), T339I (3321C>T 3322T>C), A340K (3323G>A 3324C>A 3325T>G), Y341F (3327A>T), I345L (3338A>T 3340C>G), I346V (3341A>G 3343A>T), R348E (3347A>G 3348G>A 3349G>A), D349N (3350G>A), G350Q (3353G>C 3354G>A 3355T>A), I351V (3356A>G 3358C>G), K352R (3359A>C 3360A>G 3361G>C), P353M (3362C>A 3363C>T 3364T>G), N354D (3365A>G), D356K (3371G>A 3373T>G), I358V (3377A>G 3379T>G), S359Q (3380T>C 3381C>A 3382C>G), I361V (3386A>G 3388T>A), Q362V (3389C>G 3390A>T), K363D (3392A>G 3394A>T), Y364W (3396A>G 3397T>G), L365Q (3399T>A 3400G>A), I366A (3401A>G 3402T>C 3403T>A), K368T (3408A>C 3409G>T), T369G (3410A>G 3411C>G 3412C>G), P370V (3413C>G 3414C>T 3415T>C), I373L (3422A>C), K374R (3426A>G), Q375S (3428C>T 3429A>C 3430A>C), F376I (3431T>A), L380V (3443C>G 3445C>A), G381N (3446G>A 3447G>A 3448T>C) | | | | | | | | |
| Codon mutations: | CCC177CCA (2836C>A), ATA178GTG (2837A>G 2839A>G), TGG179TTG (2841G>T), GTT180TTA (2843G>T 2845T>A), CCC182del (2849_2851delCCC), ATC185AAA (2859T>A 2860C>A), GAC186GAT (2863C>T), GCT187GGG (2865C>G 2866T>G), TCT188ACT (2867T>A), GGG189_AAA190del (2870_2877delGGGAAACA), CAA191--A (2870_2877delGGGAAACA), AAG192ACA (2880A>C 2881G>A), CGT194AGA (2885C>A 2887T>A), CTC195TTC (2888C>T), GTA196TAT (2891G>T 2892T>A 2893A>T), GTT197ATT (2894G>A), TTC199TAC (2901T>A), CGT200AGA (2903C>A 2905T>A), AAG201GAG (2906A>G), TTG202CTG (2909T>C), AAC203AAT (2914C>T), GAG204AAC (2915G>A 2917G>C), AAG205ATC (2919A>T 2920G>C), ATC207GTG (2924A>G 2926C>G), GAT208AGG (2927G>A 2928A>G 2929T>G), GAC209GAT (2932C>T), AAA210AGG (2934A>G 2935A>G), TAC211TTC (2937A>T), CCG212CCA (2941G>A), ATA213ATC (2944A>C), AAC215ACT (2949A>C 2950C>T), ATA216ATT (2953A>T), AGT217GAT (2954A>G 2955G>A), GAC218GAG (2959C>G), GTA219CTA (2960G>C), CTT220TTT (2963C>T), GAC221GAT (2968C>T), AAG222GAG (2969A>G), TTA223GTA (2972T>G), GGT224AAT (2975G>A 2976G>A), AAG225GGC (2978A>G 2979A>G 2980G>C), TGC226GCT (2981T>G 2982G>C 2983C>T), CAA227GTT (2984C>G 2985A>T 2986A>T), TAC228TAT (2989C>T), ACC230TCA (2993A>T 2995C>A), ACC231AAG (2997C>A 2998C>G), TTA232CTT (2999T>C 3001A>T), GCA235TTA (3008G>T 3009C>T), AGT236TCT (3011A>T 3012G>C), GGG237GGA (3016G>A), TTT238TAC (3018T>A 3019T>C), TAT239GAT (3020T>G), CAG240CAA (3025G>A), GTG241ATT (3026G>A 3028G>T), GAG242CGG (3029G>C 3030A>G), ATG243GTT (3032A>G 3034G>T), GAC244AAG (3035G>A 3037C>G), CCT245GTG (3038C>G 3039C>T 3040T>G), CAA246GCA (3041C>G 3042A>C), TCG249CCC (3050T>C 3052G>C), ACC251ACT (3058C>T), GCG252GCT (3061G>T), AAC254AGG (3066A>G 3067C>G), GTA255ACA (3068G>A 3069T>C), GAA256CGA (3071G>C 3072A>G), CAC257TAT (3074C>T 3076C>T), CAT259AGT (3080C>A 3081A>G), TTT260TAT (3084T>A), CGA264GTA (3095C>G 3096G>T), CCT266CCG (3103T>G), ATG267TTT (3104A>T 3106G>T), GGA268GGG (3109A>G), TTA269CTA (3110T>C), AAA270ACT (3114A>C 3115A>T), AAC271AAT (3118C>T), TCA272GCC (3119T>G 3121A>C), CCA273CCG (3124A>G), TCT274GCT (3125T>G), TTT276TTC (3133T>C), CAA277TGT (3134C>T 3135A>G 3136A>T), AGA278AAC (3138G>A 3139A>C), GTT279TTA (3140G>T 3142T>A), GAC281AAC (3146G>A), CTA284CTT (3157A>T), AGA285TAT (3158A>T 3159G>A 3160A>T), GGT286GAT (3162G>A), CTC287TAT (3164C>T 3165T>A 3166C>T), CAA288CTT (3168A>T 3169A>T), AAT289GAT (3170A>G), AAC290GAC (3173A>G), ATC291TTC (3176A>T), TGT292GTA (3179T>G 3180G>T 3181T>A), CTC293GTA (3182C>G 3184C>A), CTT296TTA (3191C>T 3193T>A), GAC297GAT (3196C>T), GAT298GAC (3199T>C), ATT299ATA (3202T>A), ATT300GTC (3203A>G 3205T>C), GTC301ATA (3206G>A 3208C>A), ACT304AGA (3216C>G 3217T>A), TCC305ACT (3218T>A 3220C>T), CTA306TTG (3221C>T 3223A>G), CAG307GAA (3224C>G 3226G>A), GAA308GAC (3229A>C), CAC309CAT (3232C>T), CTG310TTG (3233C>T), GAG311CTG (3236G>C 3237A>T), AAC312CAT (3239A>C 3241C>T), CTG313TTG (3242C>T), GAA314AGG (3245G>A 3246A>G 3247A>G), CGA315AAG (3248C>A 3249G>A 3250A>G), GTT316GTG (3253T>G), TTC317TTG (3256C>G), AGT323TAC (3272A>T 3273G>A 3274T>C), AAC324CAA (3275A>C 3277C>A), TTC325CTC (3278T>C), AAA326TAT (3281A>T 3283A>T), ATT327GTC (3284A>G 3286T>C), CAA328AAG (3287C>A 3289A>G), GAC330GAG (3295C>G), TCC332TGT (3300C>G 3301C>T), TTC334TTT (3307C>T), TTG335TCT (3309T>C 3310G>T), AAG336AGT (3312A>G 3313G>T), CTC337CGT (3315T>G 3316C>T), GAA338GAG (3319A>G), ACT339ATC (3321C>T 3322T>C), GCT340AAG (3323G>A 3324C>A 3325T>G), TAT341TTT (3327A>T), CTT342TTA (3329C>T 3331T>A), GGT343GGC (3334T>C), CAC344CAT (3337C>T), ATC345TTG (3338A>T 3340C>G), ATA346GTT (3341A>G 3343A>T), AGC347AGT (3346C>T), AGG348GAA (3347A>G 3348G>A 3349G>A), GAC349AAC (3350G>A), GGT350CAA (3353G>C 3354G>A 3355T>A), ATC351GTG (3356A>G 3358C>G), AAG352CGC (3359A>C 3360A>G 3361G>C), CCT353ATG (3362C>A 3363C>T 3364T>G), AAC354GAC (3365A>G), GAT356AAG (3371G>A 3373T>G), AAG357AAA (3376G>A), ATT358GTG (3377A>G 3379T>G), TCC359CAG (3380T>C 3381C>A 3382C>G), GCT360GCA (3385T>A), ATT361GTA (3386A>G 3388T>A), CAA362GTA (3389C>G 3390A>T), AAA363GAT (3392A>G 3394A>T), TAT364TGG (3396A>G 3397T>G), CTG365CAA (3399T>A 3400G>A), ATT366GCA (3401A>G 3402T>C 3403T>A), CCA367CCC (3406A>C), AAG368ACT (3408A>C 3409G>T), ACC369GGG (3410A>G 3411C>G 3412C>G), CCT370GTC (3413C>G 3414C>T 3415T>C), ATA373CTA (3422A>C), AAA374AGA (3426A>G), CAA375TCC (3428C>T 3429A>C 3430A>C), TTT376ATT (3431T>A), GGC378GGG (3439C>G), CTT379CTG (3442T>G), CTC380GTA (3443C>G 3445C>A), GGT381AAC (3446G>A 3447G>A 3448T>C), TAC383TAT (3454C>T) | | | | | | | | |

*: Inserts / Deletes / Misaligned / Frameshifts

#### Analysis details

This analysis was performed with panviral2.64

### NGS Details (UN60): Ichnoviriform fugitivi (segment B17)

#### Assembly

| | |
|---|---|
| **Coverage Length** | 134 (1 contig(s)) |
| **Depth Of Coverage** | 1.0 |
| **Number Of Reads** | 1 |
| **Reads Per Million** | 0.02 rpm (after QC) |
| **Ambiguities** | 0 |
| **Assembly Method** | de novo + reference guided assembly |
| **Consensus Caller** | Bcf Tools |

#### Coverage Map

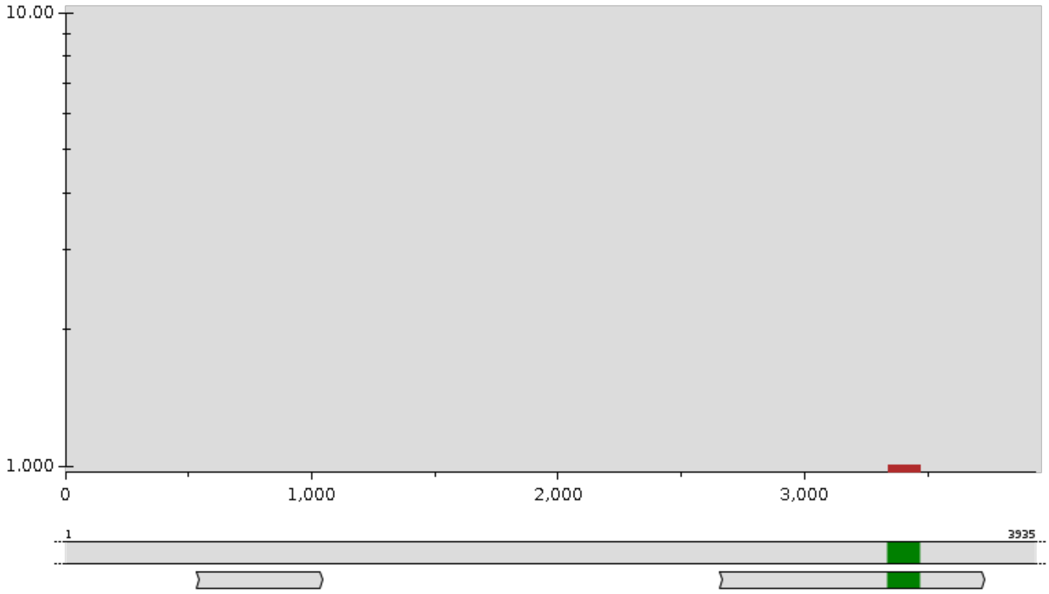

#### Assignment

| | |
|---|---|
| **Type** | Ichnoviriform fugitivi (Taxonomy ID: 265522) |
| **Reference Genome** | NC_008953.1 |
| **NT Identity (%)** | 64.1791 |
| **AA Identity (%)** | 62.2222 |
| **Number Of Stop Codons** | 0 |
| **Number Of CDS** | 2 |

#### Alignment

| | |
|---|---|
| **Alignment Score** | 76.0 (NT) + 243.0 (AA) = 319.0 |
| **Concordance (%)** | 52.9022 |

| Alignment Method | Global, seeded, nucleotide + amino acids (AGA) |
|---|---|

#### Genome Region

Sequence starts at position 3339 and ends at position 3472 relative to NC_008953.1 reference sequence.

#### Alignment Detailed Statistics

| | Begin | End | Coverage | Score | Concordance | Matches | Identities | I/D/M/F* | Stop Codons |
|---|---|---|---|---|---|---|---|---|---|
| **NT** | **3339** | **3472** | **3.4%** | **76** | **28.4%** | **134 (100%)** | **86 (64.2%)** | **0/0** | |
| Mutations: | 3344G>A, 3348C>A, 3354A>T, 3355A>C, 3356C>G, 3357A>T, 3360C>G, 3363T>A, 3364C>A, 3365G>A, 3375T>C, 3378C>T, 3379A>C, 3380A>G, 3384C>T, 3388C>T, 3390T>A, 3393G>A, 3396C>T, 3397T>G, 3398C>A, 3400C>G, 3402G>A, 3403G>C, 3405A>G, 3407A>G, 3408C>A, 3409T>C, 3411C>T, 3414G>T, 3416G>C, 3420C>G, 3423C>T, 3429C>T, 3432G>A, 3433G>T, 3434A>T, 3439G>A, 3441G>T, 3442G>A, 3447C>T, 3450G>A, 3453G>A, 3456C>T, 3459C>T, 3462T>C, 3466C>T, 3468C>G | | | | | | | | |
| **CDS** | | | | | | | | | |
| HfIV_sB17gp2 | 229 | 273 | 12.6% | 243 | 69.0% | 45 (100%) | 28 (62.2%) | 0/0/0/0 | 0 |
| Protein mutations: | R230Q (3344G>A), T234R (3355A>C 3356C>G 3357A>T), I235M (3360C>G), R237K (3364C>A 3365G>A), K242R (3379A>C 3380A>G), P245S (3388C>T 3390T>A), S248D (3397T>G 3398C>A), L249V (3400C>G 3402G>A), E250Q (3403G>C 3405A>G), N251R (3407A>G 3408C>A), Y252H (3409T>C 3411C>T), E253D (3414G>T), G254A (3416G>C), E260L (3433G>T 3434A>T), V262I (3439G>A 3441G>T), V263I (3442G>A) | | | | | | | | |
| Codon mutations: | CGA230CAA (3344G>A), GTC231GTA (3348C>A), CCA233CCT (3354A>T), ACA234CGT (3355A>C 3356C>G 3357A>T), ATC235ATG (3360C>G), ACT236ACA (3363T>A), CGA237AAA (3364C>A 3365G>A), TTT240TTC (3375T>C), CAC241CAT (3378C>T), AAA242CGA (3379A>C 3380A>G), TAC243TAT (3384C>T), CCT245TCA (3388C>T 3390T>A), TCG246TCA (3393G>A), GGC247GGT (3396C>T), TCC248GAC (3397T>G 3398C>A), CTG249GTA (3400C>G 3402G>A), GAA250CAG (3403G>C 3405A>G), AAC251AGA (3407A>G 3408C>A), TAC252CAT (3409T>C 3411C>T), GAG253GAT (3414G>T), GGA254GCA (3416G>C), CTC255CTG (3420C>G), TGC256TGT (3423C>T), CTC258CTT (3429C>T), CCG259CCA (3432G>A), GAG260TTG (3433G>T 3434A>T), GTG262ATT (3439G>A 3441G>T), GTC263ATC (3442G>A), AAC264AAT (3447C>T), GAG265GAA (3450G>A), AAG266AAA (3453G>A), ATC267ATT (3456C>T), TAC268TAT (3459C>T), ATT269ATC (3462T>C), CTC271TTG (3466C>T 3468C>G) | | | | | | | | |
| **Proteins** | | | | | | | | | |
| viral inexin-like protein (YP_001031233.1) | 229 | 273 | 12.6% | 243 | 69.0% | 45 (100%) | 28 (62.2%) | 0/0/0/0 | 0 |
| Protein mutations: | R230Q (3344G>A), T234R (3355A>C 3356C>G 3357A>T), I235M (3360C>G), R237K (3364C>A 3365G>A), K242R (3379A>C 3380A>G), P245S (3388C>T 3390T>A), S248D (3397T>G 3398C>A), L249V (3400C>G 3402G>A), E250Q (3403G>C 3405A>G), N251R (3407A>G 3408C>A), Y252H (3409T>C 3411C>T), E253D (3414G>T), G254A (3416G>C), E260L (3433G>T 3434A>T), V262I (3439G>A 3441G>T), V263I (3442G>A) | | | | | | | | |
| Codon mutations: | CGA230CAA (3344G>A), GTC231GTA (3348C>A), CCA233CCT (3354A>T), ACA234CGT (3355A>C 3356C>G 3357A>T), ATC235ATG (3360C>G), ACT236ACA (3363T>A), CGA237AAA (3364C>A 3365G>A), TTT240TTC (3375T>C), CAC241CAT (3378C>T), AAA242CGA (3379A>C 3380A>G), TAC243TAT (3384C>T), CCT245TCA (3388C>T 3390T>A), TCG246TCA (3393G>A), GGC247GGT (3396C>T), TCC248GAC (3397T>G 3398C>A), CTG249GTA (3400C>G 3402G>A), GAA250CAG (3403G>C 3405A>G), AAC251AGA (3407A>G 3408C>A), TAC252CAT (3409T>C 3411C>T), GAG253GAT (3414G>T), GGA254GCA (3416G>C), CTC255CTG (3420C>G), TGC256TGT (3423C>T), CTC258CTT (3429C>T), CCG259CCA (3432G>A), GAG260TTG (3433G>T 3434A>T), GTG262ATT (3439G>A 3441G>T), GTC263ATC (3442G>A), AAC264AAT (3447C>T), GAG265GAA (3450G>A), AAG266AAA (3453G>A), ATC267ATT (3456C>T), TAC268TAT (3459C>T), ATT269ATC (3462T>C), CTC271TTG (3466C>T 3468C>G) | | | | | | | | |

*: Inserts / Deletes / Misaligned / Frameshifts

#### Analysis details

This analysis was performed with panviral2.64

### NGS Details (UN60): Ichnoviriform fugitivi (segment B17)

#### Assembly

| | |
|---|---|
| **Coverage Length** | 286 (1 contig(s)) |
| **Depth Of Coverage** | 5.5 |
| **Number Of Reads** | 13 |
| **Reads Per Million** | 0.28 rpm (after QC) |
| **Ambiguities** | 0 |
| **Assembly Method** | de novo + reference guided assembly |
| **Consensus Caller** | Bcf Tools |

#### Coverage Map

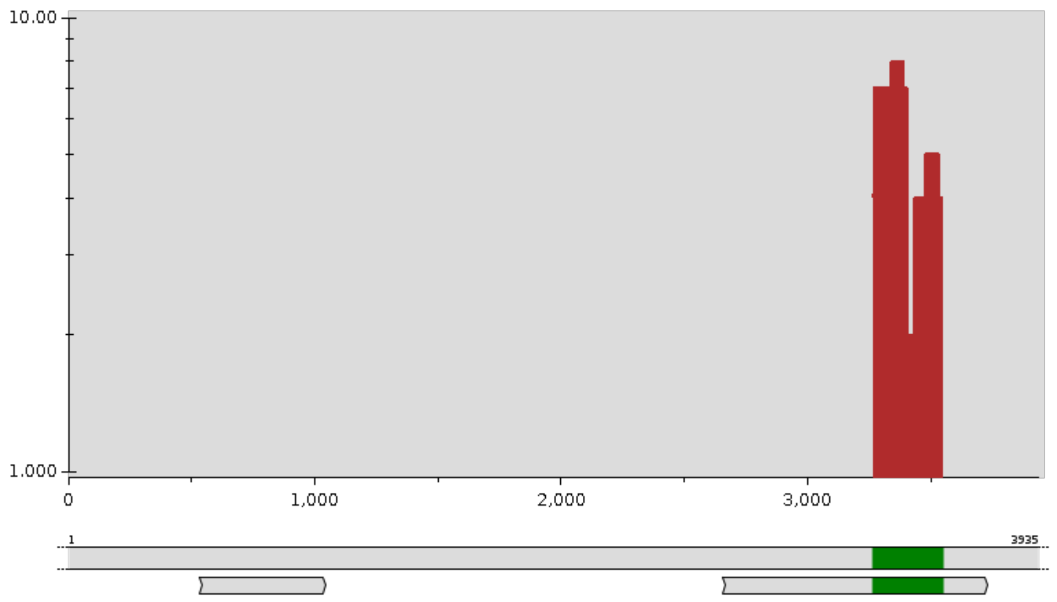

#### Assignment

| | |
|---|---|
| **Type** | Ichnoviriform fugitivi (Taxonomy ID: 265522) |
| **Reference Genome** | NC_008953.1 |
| **NT Identity (%)** | 57.2438 |
| **AA Identity (%)** | 54.2553 |
| **Number Of Stop Codons** | 0 |
| **Number Of CDS** | 2 |

#### Alignment

| | |
|---|---|
| **Alignment Score** | 70.0 (NT) + 421.0 (AA) = 491.0 |
| **Concordance (%)** | 40.0489 |

| | |
|---|---|
| **Alignment Method** | Global, seeded, nucleotide + amino acids (AGA) |

#### Genome Region

Sequence starts at position 3266 and ends at position 3551 relative to NC_008953.1 reference sequence.

#### Alignment Detailed Statistics

| | Begin | End | Coverage | Score | Concordance | Matches | Identities | I/D/M/F* | Stop Codons |
|---|---|---|---|---|---|---|---|---|---|
| **NT** | **3266** | **3551** | **7.3%** | **70** | **12.6%** | **283 (99.0%)** | **162 (56.6%)** | **0/3** | |
| Mutations: | 3267C>A, 3273G>C, 3274T>A, 3275T>C, 3276G>A, 3282A>T, 3285C>T, 3286T>A, 3288T>A, 3291T>G, 3292A>T, 3293C>T, 3294C>A, 3295T>G, 3296T>A, 3297C>A, 3298T>C, 3302A>G, 3303C>T, 3306G>A, 3309A>G, 3311C>A, 3312C>T, 3313_3315delGCC, 3318T>A, 3319C>G, 3320C>A, 3321G>A, 3326T>G, 3327G>A, 3329C>T, 3330G>T, 3331A>G, 3333C>T, 3340A>G, 3341A>T, 3342G>A, 3343C>T, 3344G>C, 3345A>T, 3351T>C, 3356C>G, 3358A>G, 3360C>A, 3363T>A, 3364C>A, 3365G>A, 3366A>G, 3369T>C, 3372T>A, 3383A>T, 3390T>A, 3393G>T, 3396C>A, 3399C>A, 3400C>T, 3403G>C, 3405A>G, 3407A>C, 3408C>A, 3409T>C, 3411C>T, 3414G>T, 3416G>C, 3420C>T, 3423C>T, 3424A>G, 3426C>A, 3427C>T, 3429C>G, 3430C>G, 3432G>A, 3433G>T, 3434A>T, 3435G>A, 3438C>T, 3439G>A, 3441G>C, 3442G>T, 3444C>G, 3447C>T, 3450G>A, 3453G>A, 3462T>A, 3465C>T, 3466C>T, 3468C>G, 3474C>T, 3481T>A, 3482A>T, 3483C>T, 3484G>A, 3486G>A, 3487C>T, 3489A>G, 3492C>A, 3498C>A, 3499A>T, 3500G>C, 3501C>T, 3504T>A, 3507C>G, 3509T>C, 3510T>G, 3511G>T, 3513T>G, 3514C>G, 3515T>C, 3516G>C, 3520C>A, 3522C>T, 3529C>G, 3531G>T, 3532C>A, 3535G>A, 3537G>A, 3539C>T, 3540C>G, 3544G>T, 3546G>A, 3547C>A, 3549C>T | | | | | | | | |
| **CDS** | | | | | | | | | |
| HfIV_sB17gp2 | 205 | 299 | 26.5% | 421 | 61.0% | 94 (98.9%) | 51 (53.7%) | 0/1/0/0 | 0 |
| Protein mutations: | L206F (3273G>C), L207T (3274T>A 3275T>C 3276G>A), Y211K (3286T>A 3288T>A), T213L (3292A>T 3293C>T 3294C>A), F214E (3295T>G 3296T>A 3297C>A), F215L (3298T>C), N216S (3302A>G 3303C>T), A219D (3311C>A 3312C>T), A220del (3313_3315delGCC), H221Q (3318T>A), P222E (3319C>G 3320C>A 3321G>A), M224R (3326T>G 3327G>A), T225I (3329C>T 3330G>T), N226D (3331A>G 3333C>T), K229V (3340A>G 3341A>T 3342G>A), R230S (3343C>T 3344G>C 3345A>T), T234R (3356C>G), I235V (3358A>G 3360C>A), R237K (3364C>A 3365G>A 3366A>G), Y243F (3383A>T), E250Q (3403G>C 3405A>G), N251T (3407A>C 3408C>A), Y252H (3409T>C 3411C>T), E253D (3414G>T), G254A (3416G>C), I257V (3424A>G 3426C>A), P259A (3430C>G 3432G>A), E260L (3433G>T 3434A>T 3435G>A), V262I (3439G>A 3441G>C), V263L (3442G>T 3444C>G), Y276I (3481T>A 3482A>T 3483C>T), V277I (3484G>A 3486G>A), I284M (3507C>G), V285A (3509T>C 3510T>G), V286L (3511G>T 3513T>G), L287A (3514C>G 3515T>C 3516G>C), R289S (3520C>A 3522C>T), L292V (3529C>G 3531G>T), L293I (3532C>A), A294T (3535G>A 3537G>A), S295L (3539C>T 3540C>G), A297S (3544G>T 3546G>A), L298I (3547C>A 3549C>T) | | | | | | | | |
| Codon mutations: | GAC204.AA (3267C>A), TTG206TTC (3273G>C), TTG207ACA (3274T>A 3275T>C 3276G>A), GGA209GGT (3282A>T), ATC210ATT (3285C>T), TAT211AAA (3286T>A 3288T>A), GTT212GTG (3291T>G), ACC213TTA (3292A>T 3293C>T 3294C>A), TTC214GAA (3295T>G 3296T>A 3297C>A), TTC215CTC (3298T>C), AAC216AGT (3302A>G 3303C>T), CAG217CAA (3306G>A), GAA218GAG (3309A>G), GCC219GAT (3311C>A 3312C>T), GCC220del (3313_3315delGCC), CAT221CAA (3318T>A), CCG222GAA (3319C>G 3320C>A 3321G>A), ATG224AGA (3326T>G 3327G>A), ACG225ATT (3329C>T 3330G>T), AAC226GAT (3331A>G 3333C>T), AAG229GTA (3340A>G 3341A>T 3342G>A), CGA230TCT (3343C>T 3344G>C 3345A>T), TTT232TTC (3351T>C), ACA234AGA (3356C>G), ATC235GTA (3358A>G 3360C>A), ACT236ACA (3363T>A), CGA237AAG (3364C>A 3365G>A 3366A>G), TGT238TGC (3369T>C), ACT239ACA (3372T>A), TAC243TTC (3383A>T), CCT245CCA (3390T>A), TCG246TCT (3393G>T), GGC247GGA (3396C>A), TCC248TCA (3399C>A), CTG249TTG (3400C>T), GAA250CAG (3403G>C 3405A>G), AAC251ACA (3407A>C 3408C>A), TAC252CAT (3409T>C 3411C>T), GAG253GAT (3414G>T), GGA254GCA (3416G>C), CTC255CTT (3420C>T), TGC256TGT (3423C>T), ATC257GTA (3424A>G 3426C>A), CTC258TTG (3427C>T 3429C>G), CCG259GCA (3430C>G 3432G>A), GAG260TTA (3433G>T 3434A>T 3435G>A), AAC261AAT (3438C>T), GTG262ATC (3439G>A 3441G>C), GTC263TTG (3442G>T 3444C>G), AAC264AAT (3447C>T), GAG265GAA (3450G>A), AAG266AAA (3453G>A), ATT269ATA (3462T>A), TTC270TTT (3465C>T), CTC271TTG (3466C>T 3468C>G), TTC273TTT (3474C>T), TAC276ATT (3481T>A 3482A>T 3483C>T), GTG277ATA (3484G>A 3486G>A), CTA278TTG (3487C>T 3489A>G), GCC279GCA (3492C>A), ATC281ATA (3498C>A), AGC282TCT (3499A>T 3500G>C 3501C>T), GGT283GGA (3504T>A), ATC284ATG (3507C>G), GTT285GCG (3509T>C 3510T>G), GTT286TTG (3511G>T 3513T>G), CTG287GCC (3514C>G 3515T>C 3516G>C), CGC289AGT (3520C>A 3522C>T), CTG292GTT (3529C>G 3531G>T), CTA293ATA (3532C>A), GCG294ACA (3535G>A 3537G>A), TCC295TTG (3539C>T 3540C>G), GCG297TCA (3544G>T 3546G>A), CTC298ATT (3547C>A 3549C>T) | | | | | | | | |
| **Proteins** | | | | | | | | | |
| viral inexin-like protein (YP_001031233.1) | 205 | 299 | 26.5% | 421 | 61.0% | 94 (98.9%) | 51 (53.7%) | 0/1/0/0 | 0 |
| Protein mutations: | L206F (3273G>C), L207T (3274T>A 3275T>C 3276G>A), Y211K (3286T>A 3288T>A), T213L (3292A>T 3293C>T 3294C>A), F214E (3295T>G 3296T>A 3297C>A), F215L (3298T>C), N216S (3302A>G 3303C>T), A219D (3311C>A 3312C>T), A220del (3313_3315delGCC), H221Q (3318T>A), P222E (3319C>G 3320C>A 3321G>A), M224R (3326T>G 3327G>A), T225I (3329C>T 3330G>T), N226D (3331A>G 3333C>T), K229V (3340A>G 3341A>T 3342G>A), R230S (3343C>T 3344G>C 3345A>T), T234R (3356C>G), I235V (3358A>G 3360C>A), R237K (3364C>A 3365G>A 3366A>G), Y243F (3383A>T), E250Q (3403G>C 3405A>G), N251T (3407A>C 3408C>A), Y252H (3409T>C 3411C>T), E253D (3414G>T), G254A (3416G>C), I257V (3424A>G 3426C>A), P259A (3430C>G 3432G>A), E260L (3433G>T 3434A>T 3435G>A), V262I (3439G>A 3441G>C), V263L (3442G>T 3444C>G), Y276I (3481T>A 3482A>T 3483C>T), V277I (3484G>A 3486G>A), I284M (3507C>G), V285A (3509T>C 3510T>G), V286L (3511G>T 3513T>G), L287A (3514C>G 3515T>C 3516G>C), R289S (3520C>A 3522C>T), L292V (3529C>G 3531G>T), L293I (3532C>A), A294T (3535G>A 3537G>A), S295L (3539C>T 3540C>G), A297S (3544G>T 3546G>A), L298I (3547C>A 3549C>T) | | | | | | | | |
| Codon mutations: | GAC204.AA (3267C>A), TTG206TTC (3273G>C), TTG207ACA (3274T>A 3275T>C 3276G>A), GGA209GGT (3282A>T), ATC210ATT (3285C>T), TAT211AAA (3286T>A 3288T>A), GTT212GTG (3291T>G), ACC213TTA (3292A>T 3293C>T 3294C>A), TTC214GAA (3295T>G 3296T>A 3297C>A), TTC215CTC (3298T>C), AAC216AGT (3302A>G 3303C>T), CAG217CAA (3306G>A), GAA218GAG (3309A>G), GCC219GAT (3311C>A 3312C>T), GCC220del (3313_3315delGCC), CAT221CAA (3318T>A), CCG222GAA (3319C>G 3320C>A 3321G>A), ATG224AGA (3326T>G 3327G>A), ACG225ATT (3329C>T 3330G>T), AAC226GAT (3331A>G 3333C>T), AAG229GTA (3340A>G 3341A>T 3342G>A), CGA230TCT (3343C>T 3344G>C 3345A>T), TTT232TTC (3351T>C), ACA234AGA (3356C>G), ATC235GTA (3358A>G 3360C>A), ACT236ACA (3363T>A), CGA237AAG (3364C>A 3365G>A 3366A>G), TGT238TGC (3369T>C), ACT239ACA (3372T>A), TAC243TTC (3383A>T), CCT245CCA (3390T>A), TCG246TCT (3393G>T), GGC247GGA (3396C>A), TCC248TCA (3399C>A), CTG249TTG (3400C>T), GAA250CAG (3403G>C 3405A>G), AAC251ACA (3407A>C 3408C>A), TAC252CAT (3409T>C 3411C>T), GAG253GAT (3414G>T), GGA254GCA (3416G>C), CTC255CTT (3420C>T), TGC256TGT (3423C>T), ATC257GTA (3424A>G 3426C>A), CTC258TTG (3427C>T 3429C>G), CCG259GCA (3430C>G 3432G>A), GAG260TTA (3433G>T 3434A>T 3435G>A), AAC261AAT (3438C>T), GTG262ATC (3439G>A 3441G>C), GTC263TTG (3442G>T 3444C>G), AAC264AAT (3447C>T), GAG265GAA (3450G>A), AAG266AAA (3453G>A), ATT269ATA (3462T>A), TTC270TTT (3465C>T), CTC271TTG (3466C>T 3468C>G), TTC273TTT (3474C>T), TAC276ATT (3481T>A 3482A>T 3483C>T), GTG277ATA (3484G>A 3486G>A), CTA278TTG (3487C>T 3489A>G), GCC279GCA (3492C>A), ATC281ATA (3498C>A), AGC282TCT (3499A>T 3500G>C 3501C>T), GGT283GGA (3504T>A), ATC284ATG (3507C>G), GTT285GCG (3509T>C 3510T>G), GTT286TTG (3511G>T 3513T>G), CTG287GCC (3514C>G 3515T>C 3516G>C), CGC289AGT (3520C>A 3522C>T), CTG292GTT (3529C>G 3531G>T), CTA293ATA (3532C>A), GCG294ACA (3535G>A 3537G>A), TCC295TTG (3539C>T 3540C>G), GCG297TCA (3544G>T 3546G>A), CTC298ATT (3547C>A 3549C>T) | | | | | | | | |

*: Inserts / Deletes / Misaligned / Frameshifts

#### Analysis details

This analysis was performed with panviral2.64

### NGS Details (UN60): Ichnoviriform fugitivi (segment B17)

#### Assembly

| | |
|---|---|
| **Coverage Length** | 291 (1 contig(s)) |
| **Depth Of Coverage** | 8.2 |
| **Number Of Reads** | 20 |
| **Reads Per Million** | 0.42 rpm (after QC) |
| **Ambiguities** | 0 |
| **Assembly Method** | de novo + reference guided assembly |
| **Consensus Caller** | Bcf Tools |

#### Coverage Map

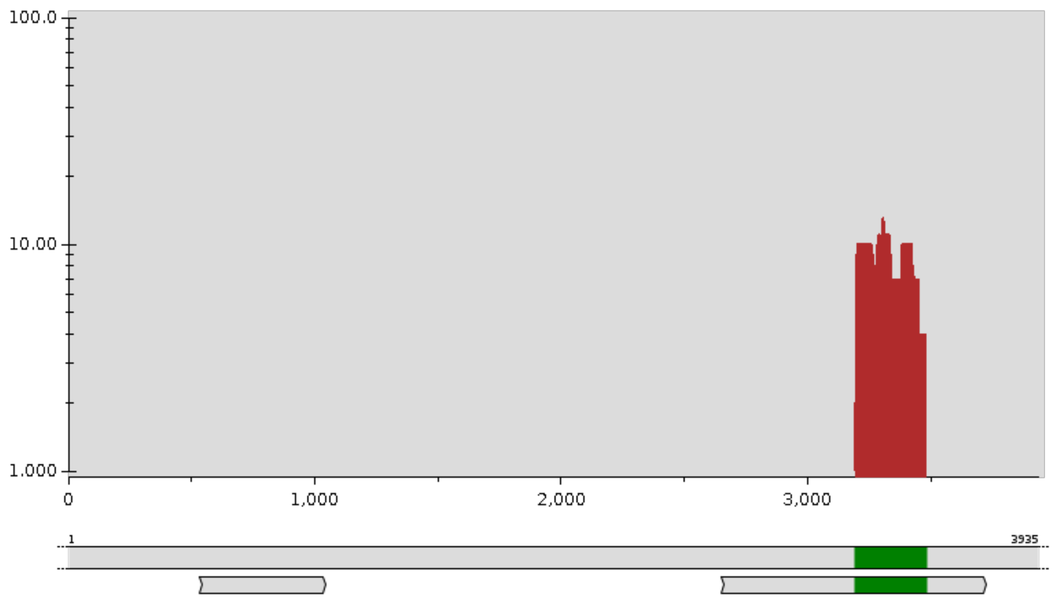

#### Assignment

| | |
|---|---|
| **Type** | Ichnoviriform fugitivi (Taxonomy ID: 265522) |
| **Reference Genome** | NC_008953.1 |
| **NT Identity (%)** | 58.3333 |
| **AA Identity (%)** | 47.9167 |
| **Number Of Stop Codons** | 0 |
| **Number Of CDS** | 2 |

#### Alignment

| | |
|---|---|
| **Alignment Score** | 75.0 (NT) + 415.0 (AA) = 490.0 |
| **Concordance (%)** | 38.6435 |

| Alignment Method | Global, seeded, nucleotide + amino acids (AGA) |
|---|---|

#### Genome Region

Sequence starts at position 3195 and ends at position 3485 relative to NC_008953.1 reference sequence.

#### Alignment Detailed Statistics

| | Begin | End | Coverage | Score | Concordance | Matches | Identities | I/D/M/F* | Stop Codons |
|---|---|---|---|---|---|---|---|---|---|
| **NT** | **3195** | **3485** | **7.4%** | **75** | **13.5%** | **285 (96.9%)** | **168 (57.1%)** | **3/6** | |
| Mutations: | 3201C>T, 3204G>A, 3205C>A, 3207G>T, 3208C>A, 3210A>C, 3211A>T, 3213C>T, 3217G>A, 3219C>T, 3220A>C, 3222C>T, 3223G>C, 3226G>A, 3228C>T, 3229G>T, 3230C>G, 3231A>C, 3234G>A, 3240C>G, 3245T>C, 3246G>C, 3250G>C, 3251C>T, 3252C>G, 3255T>C, 3256A>C, 3258A>T, 3261C>T, 3263A>G, 3265G>T, 3270T>C, 3273G>T, 3275T>C, 3276G>A, 3278A>T, 3279C>G, 3282A>C, 3285_3286insGAG, 3287A>G, 3288T>G, 3289G>A, 3291T>G, 3292A>T, 3296T>A, 3297C>T, 3299T>C, 3300C>A, 3301A>C, 3303C>T, 3304C>A, 3305A>T, 3311C>A, 3312C>G, 3314C>A, 3315C>T, 3316_3321delCATCCG, 3324C>T, 3325A>C, 3326T>G, 3327G>T, 3328A>C, 3329C>A, 3330G>T, 3331A>G, 3333C>T, 3336C>T, 3337A>T, 3341A>T, 3342G>C, 3346G>A, 3347T>C, 3348C>T, 3354A>T, 3355A>C, 3356C>G, 3357A>T, 3358A>T, 3360C>A, 3363T>C, 3364C>A, 3365G>A, 3369T>C, 3370A>T, 3372T>A, 3375T>C, 3378C>T, 3384C>T, 3388C>T, 3389C>A, 3396C>T, 3397T>A, 3399C>A, 3400C>A, 3402G>T, 3405A>G, 3406A>G, 3407A>C, 3408C>T, 3409T>C, 3414G>T, 3416G>T, 3417A>T, 3423C>T, 3424A>T, 3426C>T, 3429C>T, 3432G>T, 3433G>C, 3434A>T, 3435G>C, 3439G>A, 3441G>T, 3450G>A, 3459C>T, 3460A>G, 3462T>C, 3463T>A, 3468C>T | | | | | | | | |

##### CDS

| | Begin | End | Coverage | Score | Concordance | Matches | Identities | I/D/M/F* | Stop Codons |
|---|---|---|---|---|---|---|---|---|---|
| HfIV_sB17gp2 | 181 | 277 | 27.1% | 415 | 57.0% | 95 (96.9%) | 46 (46.9%) | 1/2/0/0 | 0 |
| Protein mutations: | L184I (3205C>A 3207G>T), L185I (3208C>A 3210A>C), N186Y (3211A>T 3213C>T), V188I (3217G>A 3219C>T), N189H (3220A>C 3222C>T), V190L (3223G>C), V191I (3226G>A 3228C>T), A192C (3229G>T 3230C>G 3231A>C), C195W (3240C>G), M197T (3245T>C 3246G>C), A199L (3250G>C 3251C>T 3252C>G), I201L (3256A>C 3258A>T), E203G (3263A>G), D204Y (3265G>T), L206F (3273G>T), L207S (3275T>C 3276G>A), Y208L (3278A>T 3279C>G), I210_Y211insE (3285_3286insGAG), Y211W (3287A>G 3288T>G), V212M (3289G>A 3291T>G), T213S (3292A>T), F214Y (3296T>A 3297C>T), F215S (3299T>C 3300C>A), N216H (3301A>C 3303C>T), Q217M (3304C>A 3305A>T), A219E (3311C>A 3312C>G), A220D (3314C>A 3315C>T), H221_P222del (3316_3321delCATCCG), M224R (3325A>C 3326T>G 3327G>T), T225H (3328A>C 3329C>A 3330G>T), N226D (3331A>G 3333C>T), M228L (3337A>T), K229I (3341A>T 3342G>C), V231T (3346G>A 3347T>C 3348C>T), T234R (3355A>C 3356C>G 3357A>T), I235L (3358A>T 3360C>A), R237K (3364C>A 3365G>A), T239S (3370A>T 3372T>A), P245Y (3388C>T 3389C>A), S248T (3397T>A 3399C>A), L249I (3400C>A 3402G>T), N251A (3406A>G 3407A>C 3408C>T), Y252H (3409T>C), E253D (3414G>T), G254V (3416G>T 3417A>T), I257F (3424A>T 3426C>T), E260L (3433G>C 3434A>T 3435G>C), V262I (3439G>A 3441G>T), I269V (3460A>G 3462T>C), F270I (3463T>A) | | | | | | | | |
| Codon mutations: | TGC182TGT (3201C>T), GAG183GAA (3204G>A), CTG184ATT (3205C>A 3207G>T), CTA185ATC (3208C>A 3210A>C), AAC186TAT (3211A>T 3213C>T), GTC188ATT (3217G>A 3219C>T), AAC189CAT (3220A>C 3222C>T), GTT190CTT (3223G>C), GTC191ATT (3226G>A 3228C>T), GCA192TGC (3229G>T 3230C>G 3231A>C), CAG193CAA (3234G>A), TGC195TGG (3240C>G), ATG197ACC (3245T>C 3246G>C), GCC199CTG (3250G>C 3251C>T 3252C>G), TTT200TTC (3255T>C), ATA201CTT (3256A>C 3258A>T), GGC202GGT (3261C>T), GAA203GGA (3263A>G), GAC204TAC (3265G>T), TTT205TTC (3270T>C), TTG206TTT (3273G>T), TTG207TCA (3275T>C 3276G>A), TAC208TTG (3278A>T 3279C>G), GGA209GGC (3282A>C), ATC210_TAT211insGAG (3285_3286insGAG), TAT211TGG (3287A>G 3288T>G), GTT212ATG (3289G>A 3291T>G), ACC213TCC (3292A>T), TTC214TAT (3296T>A 3297C>T), TTC215TCA (3299T>C 3300C>A), AAC216CAT (3301A>C 3303C>T), CAG217ATG (3304C>A 3305A>T), GCC219GAG (3311C>A 3312C>G), GCC220GAT (3314C>A 3315C>T), CAT221_CCG222del (3316_3321delCATCCG), AAC223AAT (3324C>T), ATG224CGT (3325A>C 3326T>G 3327G>T), ACG225CAT (3328A>C 3329C>A 3330G>T), AAC226GAT (3331A>G 3333C>T), CCC227CCT (3336C>T), ATG228TTG (3337A>T), AAG229ATC (3341A>T 3342G>C), GTC231ACT (3346G>A 3347T>C 3348C>T), CCA233CCT (3354A>T), ACA234CGT (3355A>C 3356C>G 3357A>T), ATC235TTA (3358A>T 3360C>A), ACT236ACC (3363T>C), CGA237AAA (3364C>A 3365G>A), TGT238TGC (3369T>C), ACT239TCA (3370A>T 3372T>A), TTT240TTC (3375T>C), CAC241CAT (3378C>T), TAC243TAT (3384C>T), CCT245TAT (3388C>T 3389C>A), GGC247GGT (3396C>T), TCC248ACA (3397T>A 3399C>A), CTG249ATT (3400C>A 3402G>T), GAA250GAG (3405A>G), AAC251GCT (3406A>G 3407A>C 3408C>T), TAC252CAC (3409T>C), GAG253GAT (3414G>T), GGA254GTT (3416G>T 3417A>T), TGC256TGT (3423C>T), ATC257TTT (3424A>T 3426C>T), CTC258CTT (3429C>T), CCG259CCT (3432G>T), GAG260CTC (3433G>C 3434A>T 3435G>C), GTG262ATT (3439G>A 3441G>T), GAG265GAA (3450G>A), TAC268TAT (3459C>T), ATT269GTC (3460A>G 3462T>C), TTC270ATC (3463T>A), CTC271CTT (3468C>T) | | | | | | | | |

##### Proteins

| | Begin | End | Coverage | Score | Concordance | Matches | Identities | I/D/M/F* | Stop Codons |
|---|---|---|---|---|---|---|---|---|---|
| viral inexin-like protein (YP_001031233.1) | 181 | 277 | 27.1% | 415 | 57.0% | 95 (96.9%) | 46 (46.9%) | 1/2/0/0 | 0 |
| Protein mutations: | L184I (3205C>A 3207G>T), L185I (3208C>A 3210A>C), N186Y (3211A>T 3213C>T), V188I (3217G>A 3219C>T), N189H (3220A>C 3222C>T), V190L (3223G>C), V191I (3226G>A 3228C>T), A192C (3229G>T 3230C>G 3231A>C), C195W (3240C>G), M197T (3245T>C 3246G>C), A199L (3250G>C 3251C>T 3252C>G), I201L (3256A>C 3258A>T), E203G (3263A>G), D204Y (3265G>T), L206F (3273G>T), L207S (3275T>C 3276G>A), Y208L (3278A>T 3279C>G), I210_Y211insE (3285_3286insGAG), Y211W (3287A>G 3288T>G), V212M (3289G>A 3291T>G), T213S (3292A>T), F214Y (3296T>A 3297C>T), F215S (3299T>C 3300C>A), N216H (3301A>C 3303C>T), Q217M (3304C>A 3305A>T), A219E (3311C>A 3312C>G), A220D (3314C>A 3315C>T), H221_P222del (3316_3321delCATCCG), M224R (3325A>C 3326T>G 3327G>T), T225H (3328A>C 3329C>A 3330G>T), N226D (3331A>G 3333C>T), M228L (3337A>T), K229I (3341A>T 3342G>C), V231T (3346G>A 3347T>C 3348C>T), T234R (3355A>C 3356C>G 3357A>T), I235L (3358A>T 3360C>A), R237K (3364C>A 3365G>A), T239S (3370A>T 3372T>A), P245Y (3388C>T 3389C>A), S248T (3397T>A 3399C>A), L249I (3400C>A 3402G>T), N251A (3406A>G 3407A>C 3408C>T), Y252H (3409T>C), E253D (3414G>T), G254V (3416G>T 3417A>T), I257F (3424A>T 3426C>T), E260L (3433G>C 3434A>T 3435G>C), V262I (3439G>A 3441G>T), I269V (3460A>G 3462T>C), F270I (3463T>A) | | | | | | | | |
| Codon mutations: | TGC182TGT (3201C>T), GAG183GAA (3204G>A), CTG184ATT (3205C>A 3207G>T), CTA185ATC (3208C>A 3210A>C), AAC186TAT (3211A>T 3213C>T), GTC188ATT (3217G>A 3219C>T), AAC189CAT (3220A>C 3222C>T), GTT190CTT (3223G>C), GTC191ATT (3226G>A 3228C>T), GCA192TGC (3229G>T 3230C>G 3231A>C), CAG193CAA (3234G>A), TGC195TGG (3240C>G), ATG197ACC (3245T>C 3246G>C), GCC199CTG (3250G>C 3251C>T 3252C>G), TTT200TTC (3255T>C), ATA201CTT (3256A>C 3258A>T), GGC202GGT (3261C>T), GAA203GGA (3263A>G), GAC204TAC (3265G>T), TTT205TTC (3270T>C), TTG206TTT (3273G>T), TTG207TCA (3275T>C 3276G>A), TAC208TTG (3278A>T 3279C>G), GGA209GGC (3282A>C), ATC210_TAT211insGAG (3285_3286insGAG), TAT211TGG (3287A>G 3288T>G), GTT212ATG (3289G>A 3291T>G), ACC213TCC (3292A>T), TTC214TAT (3296T>A 3297C>T), TTC215TCA (3299T>C 3300C>A), AAC216CAT (3301A>C 3303C>T), CAG217ATG (3304C>A 3305A>T), GCC219GAG (3311C>A 3312C>G), GCC220GAT (3314C>A 3315C>T), CAT221_CCG222del (3316_3321delCATCCG), AAC223AAT (3324C>T), ATG224CGT (3325A>C 3326T>G 3327G>T), ACG225CAT (3328A>C 3329C>A 3330G>T), AAC226GAT (3331A>G 3333C>T), CCC227CCT (3336C>T), ATG228TTG (3337A>T), AAG229ATC (3341A>T 3342G>C), GTC231ACT (3346G>A 3347T>C 3348C>T), CCA233CCT (3354A>T), ACA234CGT (3355A>C 3356C>G 3357A>T), ATC235TTA (3358A>T 3360C>A), ACT236ACC (3363T>C), CGA237AAA (3364C>A 3365G>A), TGT238TGC (3369T>C), ACT239TCA (3370A>T 3372T>A), TTT240TTC (3375T>C), CAC241CAT (3378C>T), TAC243TAT (3384C>T), CCT245TAT (3388C>T 3389C>A), GGC247GGT (3396C>T), TCC248ACA (3397T>A 3399C>A), CTG249ATT (3400C>A 3402G>T), GAA250GAG (3405A>G), AAC251GCT (3406A>G 3407A>C 3408C>T), TAC252CAC (3409T>C), GAG253GAT (3414G>T), GGA254GTT (3416G>T 3417A>T), TGC256TGT (3423C>T), ATC257TTT (3424A>T 3426C>T), CTC258CTT (3429C>T), CCG259CCT (3432G>T), GAG260CTC (3433G>C 3434A>T 3435G>C), GTG262ATT (3439G>A 3441G>T), GAG265GAA (3450G>A), TAC268TAT (3459C>T), ATT269GTC (3460A>G 3462T>C), TTC270ATC (3463T>A), CTC271CTT (3468C>T) | | | | | | | | |

*: Inserts / Deletes / Misaligned / Frameshifts

#### Analysis details

This analysis was performed with panviral2.64

### NGS Details (UN60): Ichnoviriform fugitivi (segment B17)

#### Assembly

| | |
|---|---|
| **Coverage Length** | 1290 (1 contig(s)) |
| **Depth Of Coverage** | 18.4 |
| **Number Of Reads** | 173 |
| **Reads Per Million** | 3.67 rpm (after QC) |
| **Ambiguities** | 0 |
| **Assembly Method** | de novo + reference guided assembly |
| **Consensus Caller** | Bcf Tools |

#### Coverage Map

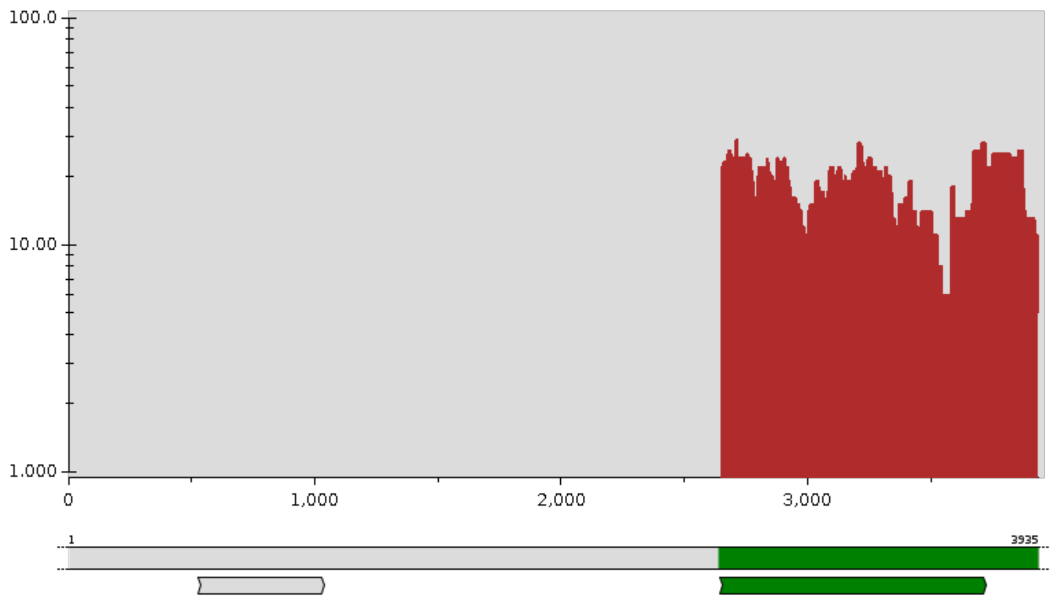

#### Assignment

| | |
|---|---|
| **Type** | Ichnoviriform fugitivi (Taxonomy ID: 265522) |
| **Reference Genome** | NC_008953.1 |
| **NT Identity (%)** | 56.9547 |
| **AA Identity (%)** | 47.191 |
| **Number Of Stop Codons** | 0 |
| **Number Of CDS** | 2 |

#### Alignment

| | |
|---|---|
| **Alignment Score** | 164.0 (NT) + 1334.0 (AA) = 1498.0 |
| **Concordance (%)** | 31.124 |

| Alignment Method | Global, seeded, nucleotide + amino acids (AGA) |
|---|---|

#### Genome Region

Sequence starts at position 2646 and ends at position 3935 relative to NC_008953.1 reference sequence.

#### Alignment Detailed Statistics

| | Begin | End | Coverage | Score | Concordance | Matches | Identities | I/D/M/F* | Stop Codons |
|---|---|---|---|---|---|---|---|---|---|
| **NT** | **2646** | **3935** | **32.8%** | **164** | **7.3%** | **1203 (92.4%)** | **692 (53.1%)** | **12/87** | |
| Mutations: | 2659C>T, 2660G>T, 2661G>C, 2662A>G, 2665C>G, 2667A>G, 2668A>T, 2671A>G, 2672A>G, 2674G>T, 2679G>C, 2685C>G, 2686T>C, 2688A>G, 2689A>C, 2691A>G, 2694G>A, 2695C>A, 2697T>C, 2698C>G, 2699C>A, 2700A>C, 2702C>G, 2703A>C, 2706C>G, 2708C>G, 2718T>C, 2719G>A, 2720T>A, 2721A>C, 2722T>A, 2724C>A, 2736T>C, 2740C>A, 2743T>G, 2745C>G, 2751G>C, 2760C>G, 2761A>G, 2763C>G, 2764G>T, 2769T>C, 2772A>G, 2778G>C, 2781G>C, 2784T>G, 2787G>C, 2790G>C, 2797T>A, 2802A>C, 2803A>G, 2805G>C, 2807T>C, 2808C>G, 2811A>C, 2818C>A, 2819A>T, 2820T>C, 2821T>G, 2823T>C, 2824C>G, 2825C>A, 2826G>C, 2829C>T, 2830T>G, 2831A>T, 2832T>G, 2836T>C, 2837A>T, 2839G>A, 2840G>A, 2842T>A, 2843C>T, 2845C>A, 2847C>G, 2848A>G, 2850T>C, 2851G>A, 2852A>C, 2853C>G, 2855T>A, 2861C>G, 2862C>G, 2863G>A, 2865G>A, 2866C>T, 2868A>C, 2869C>T, 2874C>G, 2876A>T, 2877_2878insACCATA, 2879T>C, 2880A>C, 2881G>A, 2883A>C, 2884G>C, 2885T>G, 2886G>T, 2887A>C, 2892_2893insGGCCGC, 2894C>T, 2896A>G, 2897C>G, 2898A>C, 2899C>A, 2901T>A, 2904T>C, 2907T>G, 2908A>G, 2910A>C, 2911A>C, 2912G>A, 2913C>G, 2916C>G, 2917A>G, 2918T>G, 2920T>G, 2921C>T, 2922G>C, 2923C>G, 2925T>G, 2926C>T, 2927A>C, 2928T>C, 2931A>C, 2934T>G, 2935_2943delCGTACATCG, 2944A>G, 2946T>C, 2947C>G, 2948A>G, 2949G>C, 2950C>G, 2952A>G, 2953C>G, 2954G>A, 2955A>C, 2958A>G, 2959A>G, 2967T>C, 2968T>C, 2970T>C, 2971G>A, 2972G>A, 2973C>G, 2976T>C, 2982A>G, 2988T>G, 2990T>G, 2992A>T, 2997A>G, 2998T>C, 3000G>C, 3004A>T, 3012G>A, 3013G>A, 3023C>A, 3025A>C, 3027T>G, 3030G>C, 3031C>A, 3032A>G, 3039C>G, 3045G>A, 3046G>A, 3047T>C, 3048A>G, 3051T>G, 3054A>G, 3057A>T, 3060G>C, 3061A>C, 3062A>G, 3063A>C, 3066G>C, 3071C>T, 3072A>G, 3073T>C, 3075G>C, 3077C>T, 3078T>G, 3080A>T, 3081T>C, 3084T>C, 3087G>C, 3089C>A, 3090T>C, 3092C>G, 3093A>C, 3097T>A, 3100C>A, 3102G>C, 3106A>G, 3108A>G, 3116T>A, 3117C>G, 3119C>A, 3120A>C, 3123G>C, 3124A>C, 3125A>G, 3126G>C, 3127G>A, 3128T>A, 3130G>A, 3132T>G, 3134A>T, 3135T>G, 3136T>C, 3141G>C, 3144T>C, 3147T>C, 3151T>G, 3152T>C, 3155T>C, 3156G>C, 3157C>A, 3159G>T, 3166G>A, 3171A>G, 3176C>T, 3177T>C, 3184T>A, 3185A>T, 3186C>A, 3187A>C, 3188A>G, 3189A>C, 3191A>T, 3195T>C, 3196G>C, 3197G>T, 3198C>G, 3205C>T, 3207G>C, 3210A>C, 3216T>C, 3225T>C, 3226G>A, 3231A>G, 3237A>T, 3239G>T, 3247A>G, 3249T>C, 3250G>T, 3251C>A, 3255T>C, 3256A>C, 3258A>C, 3260G>A, 3261C>A, 3263A>G, 3264A>T, 3267C>G, 3270T>C, 3271T>A, 3272T>G, 3273G>T, 3274T>A, 3275T>C, 3282A>C, 3283A>T, 3284T>C, 3285C>G, 3286T>G, 3291T>G, 3292A>C, 3293C>T, 3294C>G, 3295T>C, 3296T>G, 3297C>G, 3302A>C, 3304_3306delCAG, 3310G>A, 3311C>T, 3312C>G, 3314C>A, 3315C>G, 3317A>C, 3318T>A, 3319C>G, 3320C>A, 3321G>A, 3322A>G, 3324C>A, 3325A>C, 3326T>G, 3327G>T, 3328A>G, 3329C>A, 3330G>A, 3331A>G, 3333C>T, 3340A>G, 3341A>C, 3342G>C, 3345A>G, 3348C>G, 3351T>C, 3354A>G, 3356C>A, 3358A>G, 3360C>T, 3363T>A, 3364C>A, 3365G>A, 3366A>G, 3372T>A, 3375T>C, 3384C>T, 3390T>A, 3393G>T, 3396C>A, 3400C>A, 3402G>C, 3403G>C, 3405A>G, 3408C>G, 3410A>T, 3411C>A, 3414G>C, 3417A>T, 3420C>T, 3423C>T, 3424A>G, 3426C>G, 3427C>T, 3429C>G, 3432G>A, 3433G>C, 3434A>T, 3435G>C, 3439G>A, 3441G>T, 3450G>A, 3453G>A, 3459C>T, 3460A>G, 3462T>G, 3468C>G, 3482A>T, 3483C>G, 3484G>T, 3486G>C, 3487C>A, 3489A>T, 3493A>G, 3495C>A, 3496A>C, 3501C>T, 3505A>C, 3508G>A, 3509T>A, 3510T>C, 3511G>T, 3513T>G, 3514C>G, 3524T>C, 3525T>A, 3528T>C, 3529C>G, 3531G>T, 3532C>G, 3536C>T, 3538T>A, 3539C>T, 3540C>G, 3543T>C, 3544G>A, 3545C>A, 3546G>A, 3547C>T, 3550A>C, 3552A>T, 3556T>C, 3557A>T, 3558C>G, 3559A>T, 3564C>G, 3565C>A, 3568A>G, 3569A>C, 3572C>G, 3573G>A, 3575G>C, 3576C>T, 3578T>G, 3579C>T, 3580A>T, 3583A>G, 3584A>C, 3585C>T, 3586T>C, 3587T>C, 3588C>T, 3590C>A, 3591C>G, 3594T>C, 3597C>A, 3600C>T, 3601C>G, 3604C>A, 3605T>C, 3606G>C, 3607G>A, 3609A>T, 3610C>A, 3611A>C, 3613G>A, 3615A>G, 3616C>A, 3619C>T, 3620T>G, 3621A>C, 3624A>G, 3627A>C, 3630A>T, 3637C>T, 3640C>G, 3642G>A, 3643G>T, 3645A>G, 3646C>T, 3648T>C, 3649G>C, 3650G>A, 3651T>G, 3652C>T, 3654T>G, 3655T>G, 3657G>A, 3660A>G, 3665C>T, 3666G>T, 3667A>G, 3672A>C, 3673A>T, 3675G>A, 3680A>T, 3684A>G, 3687A>G, 3690C>T, 3691A>G, 3694A>T, 3696G>A, 3697C>G, 3698G>A, 3700A>T, 3702T>A, 3705A>C, 3706C>A, 3708T>G, 3709_3714delCGTATC, 3720C>G, 3723T>C, 3725T>G, 3727T>A, 3730T>C, 3731delT, 3733C>G, 3734T>A, 3735T>G, 3736T>C, 3741T>A, 3742G>A, 3743T>A, 3745delC, 3748T>G, 3749T>A, 3753T>G, 3754T>A, 3756C>T, 3757T>C, 3758T>A, 3760T>C, 3762G>T, 3763T>C, 3767C>T, 3768G>T, 3769A>T, 3774_3809delACTCCCGTACCAAGAGTGCGGGTGTGTGTGTGTGTG, 3813T>A, 3814T>A, 3816G>A, 3817A>C, 3820T>A, 3822C>T, 3824C>A, 3826C>A, 3830T>A, 3833G>A, 3836T>G, 3838_3853delCCGTCATGTACCCTAG, 3860T>A, 3862T>G, 3863_3869delAATTAAT, 3875G>C, 3877G>A, 3878T>A, 3880G>A, 3883T>G, 3885T>A, 3886A>T, 3889C>G, 3890G>C, 3891_3898delTGTTCATT, 3901A>T, 3905A>T, 3906C>T, 3909A>G, 3910C>A, 3911A>C, 3913T>A, 3914A>T, 3915A>T, 3916A>T, 3917T>G, 3922T>A, 3930G>A, 3932C>T, 3935C>A | | | | | | | | |

**CDS**

| | Begin | End | Coverage | Score | Concordance | Matches | Identities | I/D/M/F* | Stop Codons |
|---|---|---|---|---|---|---|---|---|---|
| HfIV_sB17gp2 | 1 | 358 | 100% | 1334 | 51.8% | 352 (97.2%) | 168 (46.4%) | 4/6/0/0 | 0 |
| Protein mutations: | R2F (2659C>T 2660G>T 2661G>C), N3D (2662A>G), L4V (2665C>G 2667A>G), I5F (2668A>T), N6G (2671A>G 2672A>G), A7S (2674G>T), I12L (2689A>C 2691A>G), L14I (2695C>A 2697T>C), P15D (2698C>G 2699C>A 2700A>C), T16S (2702C>G 2703A>C), S18C (2708C>G), V22N (2719G>A 2720T>A 2721A>C), F23I (2722T>A 2724C>A), Q29K (2740C>A), F30V (2743T>G 2745C>G), I36V (2761A>G 2763C>G), A37S (2764G>T), F48I (2797T>A), K50D (2803A>G 2805G>C), L51P (2807T>C 2808C>G), H55I (2818C>A 2819A>T 2820T>C), F56V (2821T>G 2823T>C), P57D (2824C>G 2825C>A 2826G>C), Y59V (2830T>G 2831A>T 2832T>G), Y61L (2836T>C 2837A>T), G62N (2839G>A 2840G>A), S63I (2842T>A 2843C>T), L64M (2845C>A 2847C>G), N65D (2848A>G 2850T>C), D66T (2851G>A 2852A>C 2853C>G), F67Y (2855T>A), S69W (2861C>G 2862C>G), V70I (2863G>A 2865G>A), Q71Y (2866C>T 2868A>C), P72S (2869C>T), Y74F (2876A>T), Y74_L75insTI (2877_2878insACCATA), L75P (2879T>C 2880A>C), E76N (2881G>A 2883A>C), V77R (2884G>C 2885T>G 2886G>T), I78L (2887A>C), G79_T80insGR (2892_2893insGGCCGC), T80I (2894C>T), T81G (2896A>G 2897C>G 2898A>C), H82K (2899C>A 2901T>A), I85V (2908A>G 2910A>C), S86Q (2911A>C 2912G>A 2913C>G), I88G (2917A>G 2918T>G), S89V (2920T>G 2921C>T 2922G>C), P90A (2923C>G 2925T>G), H91S (2926C>T 2927A>C 2928T>C), Q92H (2931A>C), R94_S96del (2935_2943delCGTACATCG), N97D (2944A>G 2946T>C), Q98G (2947C>G 2948A>G 2949G>C), Q99E (2950C>G 2952A>G), R100D (2953C>G 2954G>A 2955A>C), I102V (2959A>G), Y105H (2968T>C 2970T>C), G106K (2971G>A 2972G>A 2973C>G), F112C (2990T>G), I113F (2992A>T), I117F (3004A>T), V120M (3013G>A), S123Y (3023C>A), I124L (3025A>C 3027T>G), Q126R (3031C>A 3032A>G), I128M (3039C>G), V131T (3046G>A 3047T>C 3048A>G), C132W (3051T>G), K136R (3061A>C 3062A>G 3063A>C), M137I (3066G>C), T139M (3071C>T 3072A>G), A141V (3077C>T 3078T>G), H142L (3080A>T 3081T>C), T145N (3089C>A 3090T>C), S146C (3092C>G 3093A>C), F148I (3097T>A), L149I (3100C>A 3102G>C), K151E (3106A>G 3108A>G), I154K (3116T>A 3117C>G), T155N (3119C>A 3120A>C), E156D (3123G>C), K157R (3124A>C 3125A>G 3126G>C), V158K (3127G>A 3128T>A), D159K (3130G>A 3132T>G), H160L (3134A>T 3135T>G), M162I (3141G>C), F166A (3151T>G 3152T>C), M167T (3155T>C 3156G>C), Q168N (3157C>A 3159G>T), A171T (3166G>A), S174F (3176C>T 3177T>C), Y177I (3184T>A 3185A>T 3186C>A), K178R (3187A>C 3188A>G 3189A>C), Y179F (3191A>T), G181L (3196G>C 3197G>T 3198C>G), L184F (3205C>T 3207G>C), V191I (3226G>A), C195F (3239G>T), N198D (3247A>G 3249T>C), A199Y (3250G>T 3251C>A), I201L (3256A>C 3258A>C), G202E (3260G>A 3261C>A), E203G (3263A>G 3264A>T), D204E (3267C>G), L206S (3271T>A 3272T>G 3273G>T), L207T (3274T>A 3275T>C), I210S (3283A>T 3284T>C 3285C>G), Y211D (3286T>G), T213L (3292A>C 3293C>T 3294C>G), F214R (3295T>C 3296T>G 3297C>G), N216T (3302A>C), Q217del (3304_3306delCAG), A219M (3310G>A 3311C>T 3312C>G), A220E (3314C>A 3315C>G), H221P (3317A>C 3318T>A), P222E (3319C>G 3320C>A 3321G>A), N223E (3322A>G 3324C>A), M224R (3325A>C 3326T>G 3327G>T), T225E (3328A>G 3329C>A 3330G>A), N226D (3331A>G 3333C>T), K229A (3340A>G 3341A>C 3342G>C), T234K (3356C>A), I235V (3358A>G 3360C>T), R237K (3364C>A 3365G>A 3366A>G), L249I (3400C>A 3402G>C), E250Q (3403G>C 3405A>G), N251K (3408C>G), Y252L (3410A>T 3411C>A), E253D (3414G>C), I257V (3424A>G 3426C>G), E260L (3433G>C 3434A>T 3435G>C), V262I (3439G>A 3441G>T), I269V (3460A>G 3462T>G), Y276L (3482A>T 3483C>G), V277F (3484G>T 3486G>C), L278I (3487C>A 3489A>T), I280V (3493A>G 3495C>A), I281L (3496A>C), I284L (3505A>C), V285N (3508G>A 3509T>A 3510T>C), V286L (3511G>T 3513T>G), L287V (3514C>G), I290T (3524T>C 3525T>A), L292V (3529C>G 3531G>T), L293V (3532C>G), A294V (3536C>T), S295M (3538T>A 3539C>T 3540C>G), A297K (3544G>A 3545C>A 3546G>A), L298F (3547C>T), Y301L (3556T>C 3557A>T 3558C>G), M302L (3559A>T), F303L (3564C>G), K305A (3568A>G 3569A>C), T306R (3572C>G 3573G>A), C307S (3575G>C 3576C>T), L308R (3578T>G 3579C>T), M309L (3580A>T), N310A (3583A>G 3584A>C 3585C>T), F311P (3586T>C 3587T>C 3588C>T), P312Q (3590C>A 3591C>G), D314E (3597C>A), Q316E (3601C>G), L317T (3604C>A 3605T>C 3606G>C), V318I (3607G>A 3609A>T), H319T (3610C>A 3611A>C), E320K (3613G>A 3615A>G), Q321K (3616C>A), L322C (3619C>T 3620T>G 3621A>C), L328F (3637C>T), L329V (3640C>G 3642G>A), V330L (3643G>T 3645A>G), H331Y (3646C>T 3648T>C), G332Q (3649G>C 3650G>A 3651T>G), W334G (3655T>G 3657G>A), T337I (3665C>T 3666G>T), N338D (3667A>G), M340L (3673A>T 3675G>A), Y342F (3680A>T), I346V (3691A>G), T347S (3694A>T 3696G>A), R348D (3697C>G 3698G>A), I349L (3700A>T 3702T>A), H351K (3706C>A 3708T>G), R352_I353del (3709_3714delCGTATC), F355L (3720C>G), V357G (3725T>G), *358K (3727T>A) | | | | | | | | |

| | Begin | End | Coverage | Score | Concordance | Matches | Identities | I/D/M/F* | Stop Codons |
|---|---|---|---|---|---|---|---|---|---|
| **NT** | **2646** | **3935** | **32.8%** | **164** | **7.3%** | **1203 (92.4%)** | **692 (53.1%)** | **12/87** | |

| | |
|---|---|
| Codon mutations: | CGG2TTC (2659C>T 2660G>T 2661G>C), AAC3GAC (2662A>G), CTA4GTG (2665C>G 2667A>G), ATC5TTC (2668A>T), AAC6GGC (2671A>G 2672A>G), GCC7TCC (2674G>T), GTG8GTC (2679G>C), GGC10GGG (2685C>G), TTA11CTG (2686T>C 2688A>G), ATA12CTG (2689A>C 2691A>G), AAG13AAA (2694G>A), CTT14ATC (2695C>A 2697T>C), CCA15GAC (2698C>G 2699C>A 2700A>C), ACA16AGC (2702C>G 2703A>C), GTC17GTG (2706C>G), TCC18TGC (2708C>G), AAT21AAC (2718T>C), GTA22AAC (2719G>A 2720T>A 2721A>C), TTC23ATA (2722T>A 2724C>A), CAT27CAC (2736T>C), CAG29AAG (2740C>A), TTC30GTG (2743T>G 2745C>G), GTG32GTC (2751G>C), CTC35CTG (2760C>G), ATC36GTG (2761A>G 2763C>G), GCG37TCG (2764G>T), TTT38TTC (2769T>C), TCA39TCG (2772A>G), CTG41CTC (2778G>C), GTG42GTC (2781G>C), ACT43ACG (2784T>G), TCG44TCC (2787G>C), CGG45CGC (2790G>C), TTC48ATC (2797T>A), GGA49GGC (2802A>C), AAG50GAC (2803A>G 2805G>C), CTC51CCG (2807T>C 2808C>G), ATA52ATC (2811A>C), CAT55ATC (2818C>A 2819A>T 2820T>C), TTT56GTC (2821T>G 2823T>C), CCG57GAC (2824C>G 2825C>A 2826G>C), GAC58GAT (2829C>T), TAT59GTG (2830T>G 2831A>T 2832T>G), TAC61CTC (2836T>C 2837A>T), GGC62AAC (2839G>A 2840G>A), TCC63ATC (2842T>A 2843C>T), CTC64ATG (2845C>A 2847C>G), AAT65GAC (2848A>G 2850T>C), GAC66ACG (2851G>A 2852A>C 2853C>G), TTC67TAC (2855T>A), TCC69TGG (2861C>G 2862C>G), GTG70ATA (2863G>A 2865G>A), CAA71TAC (2866C>T 2868A>C), CCG72TCG (2869C>T), ACC73ACG (2874C>G), TAC74TTC (2876A>T), TAC74_CTA75insACCATA (2877_2878insACCATA), CTA75CCC (2879T>C 2880A>C), GAA76AAC (2881G>A 2883A>C), GTG77CGT (2884G>C 2885T>G 2886G>T), ATC78CTC (2887A>C), GGT79_ACC80insGGCCGC (2892_2893insGGCCGC), ACC80ATC (2894C>T), ACA81GGC (2896A>G 2897C>G 2898A>C), CAT82AAA (2899C>A 2901T>A), GAT83GAC (2904T>C), GTT84GTG (2907T>G), ATA85GTC (2908A>G 2910A>C), AGC86CAG (2911A>C 2912G>A 2913C>G), CCC87CCG (2916C>G), ATC88GGC (2917A>G 2918T>G), TCG89GTC (2920T>G 2921C>T 2922G>C), CCT90GCG (2923C>G 2925T>G), CAT91TCC (2926C>T 2927A>C 2928T>C), CAA92CAC (2931A>C), GTT93GTG (2934T>G), CGT94_TCG96del (2935_2943delCGTACATCG), AAT97GAC (2944A>G 2946T>C), CAG98GGC (2947C>G 2948A>G 2949G>C), CAA99GAG (2950C>G 2952A>G), CGA100GAC (2953C>G 2954G>A 2955A>C), GAA101GAG (2958A>G), ATC102GTC (2959A>G), TAT104TAC (2967T>C), TAT105CAC (2968T>C 2970T>C), GGC106AAG (2971G>A 2972G>A 2973C>G), TAT107TAC (2976T>C), CAA109CAG (2982A>G), GTT111GTG (2988T>G), TTC112TGC (2990T>G), ATC113TTC (2992A>T), GTA114GTG (2997A>G), TTG115CTC (2998T>C 3000G>C), ATC117TTC (3004A>T), GCG119GCA (3012G>A), GTG120ATG (3013G>A), TCC123TAC (3023C>A), ATT124CTG (3025A>C 3027T>G), CCG125CCC (3030G>C), CAG126AGG (3031C>A 3032A>G), ATC128ATG (3039C>G), AAG130AAA (3045G>A), GTA131ACG (3046G>A 3047T>C 3048A>G), TGT132TGG (3051T>G), GAA133GAG (3054A>G), GGA134GGT (3057A>T), GGG135GGC (3060G>C), AAA136CGC (3061A>C 3062A>G 3063A>C), ATG137ATC (3066G>C), ACA139ATG (3071C>T 3072A>G), TTG140CTC (3073T>C 3075G>C), GCT141GTG (3077C>T 3078T>G), CAT142CTC (3080A>T 3081T>C), GAT143GAC (3084T>C), CTG144CTC (3087G>C), ACT145AAC (3089C>A 3090T>C), TCA146TGC (3092C>G 3093A>C), TTC148ATC (3097T>A), CTG149ATC (3100C>A 3102G>C), AAA151GAG (3106A>G 3108A>G), ATC154AAG (3116T>A 3117C>G), ACA155AAC (3119C>A 3120A>C), GAG156GAC (3123G>C), AAG157CGC (3124A>C 3125A>G 3126G>C), GTG158AAG (3127G>A 3128T>A), GAT159AAG (3130G>A 3132T>G), CAT160CTG (3134A>T 3135T>G), TTG161CTG (3136T>C), ATG162ATC (3141G>C), GAT163GAC (3144T>C), TAT164TAC (3147T>C), TTC166GCC (3151T>G 3152T>C), ATG167ACC (3155T>C 3156G>C), CAG168AAT (3157C>A 3159G>T), GCG171ACG (3166G>A), CAA172CAG (3171A>G), TCT174TTC (3176C>T 3177T>C), TAC177ATA (3184T>A 3185A>T 3186C>A), AAA178CGC (3187A>C 3188A>G 3189A>C), TAC179TTC (3191A>T), TTT180TTC (3195T>C), GGC181CTG (3196G>C 3197G>T 3198C>G), CTG184TTC (3205C>T 3207G>C), CTA185CTC (3210A>C), TTT187TTC (3216T>C), GTT190GTC (3225T>C), GTC191ATC (3226G>A), GCA192GCG (3231A>G), ATA194ATT (3237A>T), TGC195TTC (3239G>T), AAT198GAC (3247A>G 3249T>C), GCC199TAC (3250G>T 3251C>A), TTT200TTC (3255T>C), ATA201CTC (3256A>C 3258A>C), GGC202GAA (3260G>A 3261C>A), GAA203GGT (3263A>G 3264A>T), GAC204GAG (3267C>G), TTT205TTC (3270T>C), TTG206AGT (3271T>A 3272T>G 3273G>T), TTG207ACG (3274T>A 3275T>C), GGA209GGC (3282A>C), ATC210TCG (3283A>T 3284T>C 3285C>G), TAT211GAT (3286T>G), GTT212GTG (3291T>G), ACC213CTG (3292A>C 3293C>T 3294C>G), TTC214CGG (3295T>C 3296T>G 3297C>G), AAC216ACC (3302A>C), CAG217del (3304_3306delCAG), GCC219ATG (3310G>A 3311C>T 3312C>G), GCC220GAG (3314C>A 3315C>G), CAT221CCA (3317A>C 3318T>A), CCG222GAA (3319C>G 3320C>A 3321G>A), AAC223GAA (3322A>G 3324C>A), ATG224CGT (3325A>C 3326T>G 3327G>T), ACG225GAA (3328A>G 3329C>A 3330G>A), AAC226GAT (3331A>G 3333C>T), AAG229GCC (3340A>G 3341A>C 3342G>C), CGA230CGG (3345A>G), GTC231GTG (3348C>G), TTT232TTC (3351T>C), CCA233CCG (3354A>G), ACA234AAA (3356C>A), ATC235GTT (3358A>G 3360C>T), ACT236ACA (3363T>A), CGA237AAG (3364C>A 3365G>A 3366A>G), ACT239ACA (3372T>A), TTT240TTC (3375T>C), TAC243TAT (3384C>T), CCT245CCA (3390T>A), TCG246TCT (3393G>T), GGC247GGA (3396C>A), CTG249ATC (3400C>A 3402G>C), GAA250CAG (3403G>C 3405A>G), AAC251AAG (3408C>G), TAC252TTA (3410A>T 3411C>A), GAG253GAC (3414G>C), GGA254GGT (3417A>T), CTC255CTT (3420C>T), TGC256TGT (3423C>T), ATC257GTG (3424A>G 3426C>G), CTC258TTG (3427C>T 3429C>G), CCG259CCA (3432G>A), GAG260CTC (3433G>C 3434A>T 3435G>C), GTG262ATT (3439G>A 3441G>T), GAG265GAA (3450G>A), AAG266AAA (3453G>A), TAC268TAT (3459C>T), ATT269GTG (3460A>G 3462T>G), CTC271CTG (3468C>G), TAC276TTG (3482A>T 3483C>G), GTG277TTC (3484G>T 3486G>C), CTA278ATT (3487C>A 3489A>T), ATC280GTA (3493A>G 3495C>A), ATC281CTC (3496A>C), AGC282AGT (3501C>T), ATC284CTC (3505A>C), GTT285AAC (3508G>A 3509T>A 3510T>C), GTT286TTG (3511G>T 3513T>G), CTG287GTG (3514C>G), ATT290ACA (3524T>C 3525T>A), GCT291GCC (3528T>C), CTG292GTT (3529C>G 3531G>T), CTA293GTA (3532C>G), GCG294GTG (3536C>T), TCC295ATG (3538T>A 3539C>T 3540C>G), CCT296CCC (3543T>C), GCG297AAA (3544G>A 3545C>A 3546G>A), CTC298TTC (3547C>T), AGA299CGT (3550A>C 3552A>T), TAC301CTG (3556T>C 3557A>T 3558C>G), ATG302TTG (3559A>T), TTC303TTG (3564C>G), CGA304AGA (3565C>A), AAA305GCA (3568A>G 3569A>C), ACG306AGA (3572C>G 3573G>A), TGC307TCT (3575G>C 3576C>T), CTC308CGT (3578T>G 3579C>T), ATG309TTG (3580A>T), AAC310GCT (3583A>G 3584A>C 3585C>T), TTC311CCT (3586T>C 3587T>C 3588C>T), CCC312CAG (3590C>A 3591C>G), GAT313GAC (3594T>C), GAC314GAA (3597C>A), GTC315GTT (3600C>T), CAA316GAA (3601C>G), CTG317ACC (3604C>A 3605T>C 3606G>C), GTA318ATT (3607G>A 3609A>T), CAC319ACC (3610C>A 3611A>C), GAA320AAG (3613G>A 3615A>G), CAG321AAG (3616C>A), CTA322TGC (3619C>T 3620T>G 3621A>C), CAA323CAG (3624A>G), ATA324ATC (3627A>C), GGA325GGT (3630A>T), CTT328TTT (3637C>T), CTG329GTA (3640C>G 3642G>A), GTA330TTG (3643G>T 3645A>G), CAT331TAC (3646C>T 3648T>C), GGT332CAG (3649G>C 3650G>A 3651T>G), CTT333TTG (3652C>T 3654T>G), TGG334GGA (3655T>G 3657G>A), AAA335AAG (3660A>G), ACG337ATT (3665C>T 3666G>T), AAC338GAC (3667A>G), CCA339CCC (3672A>C), ATG340TTA (3673A>T 3675G>A), TAC342TTC (3680A>T), AAA343AAG (3684A>G), GAA344GAG (3687A>G), CTC345CTT (3690C>T), ATA346GTA (3691A>G), ACG347TCA (3694A>T 3696G>A), CGC348GAC (3697C>G 3698G>A), ATT349TTA (3700A>T 3702T>A), GCA350GCC (3705A>C), CAT351AAG (3706C>A 3708T>G), CGT352_ATC353del (3709_3714delCGTATC), TTC355TTG (3720C>G), GAT356GAC (3723T>C), GTT357GGT (3725T>G), TAG358AAG (3727T>A) |

#### Proteins

| | Begin | End | Coverage | Score | Concordance | Matches | Identities | I/D/M/F* | Stop Codons |
|---|---|---|---|---|---|---|---|---|---|
| viral inexin-like protein (YP_001031233.1) | 1 | 358 | 100% | 1334 | 51.8% | 352 (97.2%) | 168 (46.4%) | 4/6/0/0 | 0 |

| | |
|---|---|
| Protein mutations: | R2F (2659C>T 2660G>T 2661G>C), N3D (2662A>G), L4V (2665C>G 2667A>G), I5F (2668A>T), N6G (2671A>G 2672A>G), A7S (2674G>T), I12L (2689A>C 2691A>G), L14I (2695C>A 2697T>C), P15D (2698C>G 2699C>A 2700A>C), T16S (2702C>G 2703A>C), S18C (2708C>G), V22N (2719G>A 2720T>A 2721A>C), F23I (2722T>A 2724C>A), Q29K (2740C>A), F30V (2743T>G 2745C>G), I36V (2761A>G 2763C>G), A37S (2764G>T), F48I (2797T>A), K50D (2803A>G 2805G>C), L51P (2807T>C 2808C>G), H55I (2818C>A 2819A>T 2820T>C), F56V (2821T>G 2823T>C), P57D (2824C>G 2825C>A 2826G>C), Y59V (2830T>G 2831A>T 2832T>G), Y61L (2836T>C 2837A>T), G62N (2839G>A 2840G>A), S63I (2842T>A 2843C>T), L64M (2845C>A 2847C>G), N65D (2848A>G 2850T>C), D66T (2851G>A 2852A>C 2853C>G), F67Y (2855T>A), S69W (2861C>G 2862C>G), V70I (2863G>A 2865G>A), Q71Y (2866C>T 2868A>C), P72S (2869C>T), Y74F (2876A>T), Y74_L75insTI (2877_2878insACCATA), L75P (2879T>C 2880A>C), E76N (2881G>A 2883A>C), V77R (2884G>C 2885T>G 2886G>T), I78L (2887A>C), G79_T80insGR (2892_2893insGGCCGC), T80I (2894C>T), T81G (2896A>G 2897C>G 2898A>C), H82K (2899C>A 2901T>A), I85V (2908A>G 2910A>C), S86Q (2911A>C 2912G>A 2913C>G), I88G (2917A>G 2918T>G), S89V (2920T>G 2921C>T 2922G>C), P90A (2923C>G 2925T>G), H91S (2926C>T 2927A>C 2928T>C), Q92H (2931A>C), R94_S96del (2935_2943delCGTACATCG), N97D (2944A>G 2946T>C), Q98G (2947C>G 2948A>G 2949G>C), Q99E (2950C>G 2952A>G), R100D (2953C>G 2954G>A 2955A>C), I102V (2959A>G), Y105H (2968T>C 2970T>C), G106K (2971G>A 2972G>A 2973C>G), F112C (2990T>G), I113F (2992A>T), I117F (3004A>T), V120M (3013G>A), S123Y (3023C>A), I124L (3025A>C 3027T>G), Q126R (3031C>A 3032A>G), I128M (3039C>G), V131T (3046G>A 3047T>C 3048A>G), C132W (3051T>G), K136R (3061A>C 3062A>G 3063A>C), M137I (3066G>C), T139M (3071C>T 3072A>G), A141V (3077C>T 3078T>G), H142L (3080A>T 3081T>C), T145N (3089C>A 3090T>C), S146C (3092C>G 3093A>C), F148I (3097T>A), L149I (3100C>A 3102G>C), K151E (3106A>G 3108A>G), I154K (3116T>A 3117C>G), T155N (3119C>A 3120A>C), E156D (3123G>C), K157R (3124A>C 3125A>G 3126G>C), V158K (3127G>A 3128T>A), D159K (3130G>A 3132T>G), H160L (3134A>T 3135T>G), M162I (3141G>C), F166A (3151T>G 3152T>C), M167T (3155T>C 3156G>C), Q168N (3157C>A 3159G>T), A171T (3166G>A), S174F (3176C>T 3177T>C), Y177I (3184T>A 3185A>T 3186C>A), K178R (3187A>C 3188A>G 3189A>C), Y179F (3191A>T), G181L (3196G>C 3197G>T 3198C>G), L184F (3205C>T 3207G>C), V191I (3226G>A), C195F (3239G>T), N198D (3247A>G 3249T>C), A199Y (3250G>T 3251C>A), I201L (3256A>C 3258A>C), G202E (3260G>A 3261C>A), E203G (3263A>G 3264A>T), D204E (3267C>G), L206S (3271T>A 3272T>G 3273G>T), L207T (3274T>A 3275T>C), I210S (3283A>T 3284T>C 3285C>G), Y211D (3286T>G), T213L (3292A>C 3293C>T 3294C>G), F214R (3295T>C 3296T>G 3297C>G), N216T (3302A>C), Q217del (3304_3306delCAG), A219M (3310G>A 3311C>T 3312C>G), A220E (3314C>A 3315C>G), H221P (3317A>C 3318T>A), P222E (3319C>G 3320C>A 3321G>A), N223E (3322A>G 3324C>A), M224R (3325A>C 3326T>G 3327G>T), T225E (3328A>G 3329C>A 3330G>A), N226D (3331A>G 3333C>T), K229A (3340A>G 3341A>C 3342G>C), T234K (3356C>A), I235V (3358A>G 3360C>T), R237K (3364C>A 3365G>A 3366A>G), L249I (3400C>A 3402G>C), E250Q (3403G>C 3405A>G), N251K (3408C>G), Y252L (3410A>T 3411C>A), E253D (3414G>C), I257V (3424A>G 3426C>G), E260L (3433G>C 3434A>T 3435G>C), V262I (3439G>A 3441G>T), I269V (3460A>G 3462T>G), Y276L (3482A>T 3483C>G), V277F (3484G>T 3486G>C), L278I (3487C>A 3489A>T), I280V (3493A>G 3495C>A), I281L (3496A>C), I284L (3505A>C), V285N (3508G>A 3509T>A 3510T>C), V286L (3511G>T 3513T>G), L287V (3514C>G), I290T (3524T>C 3525T>A), L292V (3529C>G 3531G>T), L293V (3532C>G), A294V (3536C>T), S295M (3538T>A 3539C>T 3540C>G), A297K (3544G>A 3545C>A 3546G>A), L298F (3547C>T), Y301L (3556T>C 3557A>T 3558C>G), M302L (3559A>T), F303L (3564C>G), K305A (3568A>G 3569A>C), T306R (3572C>G 3573G>A), C307S (3575G>C 3576C>T), L308R (3578T>G 3579C>T), M309L (3580A>T), N310A (3583A>G 3584A>C 3585C>T), F311P (3586T>C 3587T>C 3588C>T), P312Q (3590C>A 3591C>G), D314E (3597C>A), Q316E (3601C>G), L317T (3604C>A 3605T>C 3606G>C), V318I (3607G>A 3609A>T), H319T (3610C>A 3611A>C), E320K (3613G>A 3615A>G), Q321K (3616C>A), L322C (3619C>T 3620T>G 3621A>C), L328F (3637C>T), L329V (3640C>G 3642G>A), V330L (3643G>T 3645A>G), H331Y (3646C>T 3648T>C), G332Q (3649G>C 3650G>A 3651T>G), W334G (3655T>G 3657G>A), T337I (3665C>T 3666G>T), N338D (3667A>G), M340L (3673A>T 3675G>A), Y342F (3680A>T), I346V (3691A>G), T347S (3694A>T 3696G>A), R348D (3697C>G 3698G>A), I349L (3700A>T 3702T>A), H351K (3706C>A 3708T>G), R352_I353del (3709_3714delCGTATC), F355L (3720C>G), V357G (3725T>G), *358K (3727T>A) |

| | Begin | End | Coverage | Score | Concordance | Matches | Identities | I/D/M/F* | Stop Codons |
|---|---|---|---|---|---|---|---|---|---|
| **NT** | **2646** | **3935** | **32.8%** | **164** | **7.3%** | **1203 (92.4%)** | **692 (53.1%)** | **12/87** | |
| Codon mutations: | CGG2TTC (2659C>T 2660G>T 2661G>C), AAC3GAC (2662A>G), CTA4GTG (2665C>G 2667A>G), ATC5TTC (2668A>T), AAC6GGC (2671A>G 2672A>G), GCC7TCC (2674G>T), GTG8GTC (2679G>C), GGC10GGG (2685C>G), TTA11CTG (2686T>C 2688A>G), ATA12CTG (2689A>C 2691A>G), AAG13AAA (2694G>A), CTT14ATC (2695C>A 2697T>C), CCA15GAC (2698C>G 2699C>A 2700A>C), ACA16AGC (2702C>G 2703A>C), GTC17GTG (2706C>G), TCC18TGC (2708C>G), AAT21AAC (2718T>C), GTA22AAC (2719G>A 2720T>A 2721A>C), TTC23ATA (2722T>A 2724C>A), CAT27CAC (2736T>C), CAG29AAG (2740C>A), TTC30GTG (2743T>G 2745C>G), GTG32GTC (2751G>C), CTC35CTG (2760C>G), ATC36GTG (2761A>G 2763C>G), GCG37TCG (2764G>T), TTT38TTC (2769T>C), TCA39TCG (2772A>G), CTG41CTC (2778G>C), GTG42GTC (2781G>C), ACT43ACG (2784T>G), TCG44TCC (2787G>C), CGG45CGC (2790G>C), TTC48ATC (2797T>A), GGA49GGC (2802A>C), AAG50GAC (2803A>G 2805G>C), CTC51CCG (2807T>C 2808C>G), ATA52ATC (2811A>C), CAT55ATC (2818C>A 2819A>T 2820T>C), TTT56GTC (2821T>G 2823T>C), CCG57GAC (2824C>G 2825C>A 2826G>C), GAC58GAT (2829C>T), TAT59GTG (2830T>G 2831A>T 2832T>G), TAC61CTC (2836T>C 2837A>T), GGC62AAC (2839G>A 2840G>A), TCC63ATC (2842T>A 2843C>T), CTC64ATG (2845C>A 2847C>G), AAT65GAC (2848A>G 2850T>C), GAC66ACG (2851G>A 2852A>C 2853C>G), TTC67TAC (2855T>A), TCC69TGG (2861C>G 2862C>G), GTG70ATA (2863G>A 2865G>A), CAA71TAC (2866C>T 2868A>C), CCG72TCG (2869C>T), ACC73ACG (2874C>G), TAC74TTC (2876A>T), TAC74_CTA75insACCATA (2877_2878insACCATA), CTA75CCC (2879T>C 2880A>C), GAA76AAC (2881G>A 2883A>C), GTG77CGT (2884G>C 2885T>G 2886G>T), ATC78CTC (2887A>C), GGT79_ACC80insGGCCGC (2892_2893insGGCCGC), ACC80ATC (2894C>T), ACA81GGC (2896A>G 2897C>G 2898A>C), CAT82AAA (2899C>A 2901T>A), GAT83GAC (2904T>C), GTT84GTG (2907T>G), ATA85GTC (2908A>G 2910A>C), AGC86CAG (2911A>C 2912G>A 2913C>G), CCC87CCG (2916C>G), ATC88GGC (2917A>G 2918T>G), TCG89GTC (2920T>G 2921C>T 2922G>C), CCT90GCG (2923C>G 2925T>G), CAT91TCC (2926C>T 2927A>C 2928T>C), CAA92CAC (2931A>C), GTT93GTG (2934T>G), CGT94_TCG96del (2935_2943delCGTACATCG), AAT97GAC (2944A>G 2946T>C), CAG98GGC (2947C>G 2948A>G 2949G>C), CAA99GAG (2950C>G 2952A>G), CGA100GAC (2953C>G 2954G>A 2955A>C), GAA101GAG (2958A>G), ATC102GTC (2959A>G), TAT104TAC (2967T>C), TAT105CAC (2968T>C 2970T>C), GGC106AAG (2971G>A 2972G>A 2973C>G), TAT107TAC (2976T>C), CAA109CAG (2982A>G), GTT111GTG (2988T>G), TTC112TGC (2990T>G), ATC113TTC (2992A>T), GTA114GTG (2997A>G), TTG115CTC (2998T>C 3000G>C), ATC117TTC (3004A>T), GCG119GCA (3012G>A), GTG120ATG (3013G>A), TCC123TAC (3023C>A), ATT124CTG (3025A>C 3027T>G), CCG125CCC (3030G>C), CAG126AGG (3031C>A 3032A>G), ATC128ATG (3039C>G), AAG130AAA (3045G>A), GTA131ACG (3046G>A 3047T>C 3048A>G), TGT132TGG (3051T>G), GAA133GAG (3054A>G), GGA134GGT (3057A>T), GGG135GGC (3060G>C), AAA136CGC (3061A>C 3062A>G 3063A>C), ATG137ATC (3066G>C), ACA139ATG (3071C>T 3072A>G), TTG140CTC (3073T>C 3075G>C), GCT141GTG (3077C>T 3078T>G), CAT142CTC (3080A>T 3081T>C), GAT143GAC (3084T>C), CTG144CTC (3087G>C), ACT145AAC (3089C>A 3090T>C), TCA146TGC (3092C>G 3093A>C), TTC148ATC (3097T>A), CTG149ATC (3100C>A 3102G>C), AAA151GAG (3106A>G 3108A>G), ATC154AAG (3116T>A 3117C>G), ACA155AAC (3119C>A 3120A>C), GAG156GAC (3123G>C), AAG157CGC (3124A>C 3125A>G 3126G>C), GTG158AAG (3127G>A 3128T>A), GAT159AAG (3130G>A 3132T>G), CAT160CTG (3134A>T 3135T>G), TTG161CTG (3136T>C), ATG162ATC (3141G>C), GAT163GAC (3144T>C), TAT164TAC (3147T>C), TTC166GCC (3151T>G 3152T>C), ATG167ACC (3155T>C 3156G>C), CAG168AAT (3157C>A 3159G>T), GCG171ACG (3166G>A), CAA172CAG (3171A>G), TCT174TTC (3176C>T 3177T>C), TAC177ATA (3184T>A 3185A>T 3186C>A), AAA178CGC (3187A>C 3188A>G 3189A>C), TAC179TTC (3191A>T), TTT180TTC (3195T>C), GGC181CTG (3196G>C 3197G>T 3198C>G), CTG184TTC (3205C>T 3207G>C), CTA185CTC (3210A>C), TTT187TTC (3216T>C), GTT190GTC (3225T>C), GTC191ATC (3226G>A), GCA192GCG (3231A>G), ATA194ATT (3237A>T), TGC195TTC (3239G>T), AAT198GAC (3247A>G 3249T>C), GCC199TAC (3250G>T 3251C>A), TTT200TTC (3255T>C), ATA201CTC (3256A>C 3258A>C), GGC202GAA (3260G>A 3261C>A), GAA203GGT (3263A>G 3264A>T), GAC204GAG (3267C>G), TTT205TTC (3270T>C), TTG206AGT (3271T>A 3272T>G 3273G>T), TTG207ACG (3274T>A 3275T>C), GGA209GGC (3282A>C), ATC210TCG (3283A>T 3284T>C 3285C>G), TAT211GAT (3286T>G), GTT212GTG (3291T>G), ACC213CTG (3292A>C 3293C>T 3294C>G), TTC214CGG (3295T>C 3296T>G 3297C>G), AAC216ACC (3302A>C), CAG217del (3304_3306delCAG), GCC219ATG (3310G>A 3311C>T 3312C>G), GCC220GAG (3314C>A 3315C>G), CAT221CCA (3317A>C 3318T>A), CCG222GAA (3319C>G 3320C>A 3321G>A), AAC223GAA (3322A>G 3324C>A), ATG224CGT (3325A>C 3326T>G 3327G>T), ACG225GAA (3328A>G 3329C>A 3330G>A), AAC226GAT (3331A>G 3333C>T), AAG229GCC (3340A>G 3341A>C 3342G>C), CGA230CGG (3345A>G), GTC231GTG (3348C>G), TTT232TTC (3351T>C), CCA233CCG (3354A>G), ACA234AAA (3356C>A), ATC235GTT (3358A>G 3360C>T), ACT236ACA (3363T>A), CGA237AAG (3364C>A 3365G>A 3366A>G), ACT239ACA (3372T>A), TTT240TTC (3375T>C), TAC243TAT (3384C>T), CCT245CCA (3390T>A), TCG246TCT (3393G>T), GGC247GGA (3396C>A), CTG249ATC (3400C>A 3402G>C), GAA250CAG (3403G>C 3405A>G), AAC251AAG (3408C>G), TAC252TTA (3410A>T 3411C>A), GAG253GAC (3414G>C), GGA254GGT (3417A>T), CTC255CTT (3420C>T), TGC256TGT (3423C>T), ATC257GTG (3424A>G 3426C>G), CTC258TTG (3427C>T 3429C>G), CCG259CCA (3432G>A), GAG260CTC (3433G>C 3434A>T 3435G>C), GTG262ATT (3439G>A 3441G>T), GAG265GAA (3450G>A), AAG266AAA (3453G>A), TAC268TAT (3459C>T), ATT269GTG (3460A>G 3462T>G), CTC271CTG (3468C>G), TAC276TTG (3482A>T 3483C>G), GTG277TTC (3484G>T 3486G>C), CTA278ATT (3487C>A 3489A>T), ATC280GTA (3493A>G 3495C>A), ATC281CTC (3496A>C), AGC282AGT (3501C>T), ATC284CTC (3505A>C), GTT285AAC (3508G>A 3509T>A 3510T>C), GTT286TTG (3511G>T 3513T>G), CTG287GTG (3514C>G), ATT290ACA (3524T>C 3525T>A), GCT291GCC (3528T>C), CTG292GTT (3529C>G 3531G>T), CTA293GTA (3532C>G), GCG294GTG (3536C>T), TCC295ATG (3538T>A 3539C>T 3540C>G), CCT296CCC (3543T>C), GCG297AAA (3544G>A 3545C>A 3546G>A), CTC298TTC (3547C>T), AGA299CGT (3550A>C 3552A>T), TAC301CTG (3556T>C 3557A>T 3558C>G), ATG302TTG (3559A>T), TTC303TTG (3564C>G), CGA304AGA (3565C>A), AAA305GCA (3568A>G 3569A>C), ACG306AGA (3572C>G 3573G>A), TGC307TCT (3575G>C 3576C>T), CTC308CGT (3578T>G 3579C>T), ATG309TTG (3580A>T), AAC310GCT (3583A>G 3584A>C 3585C>T), TTC311CCT (3586T>C 3587T>C 3588C>T), CCC312CAG (3590C>A 3591C>G), GAT313GAC (3594T>C), GAC314GAA (3597C>A), GTC315GTT (3600C>T), CAA316GAA (3601C>G), CTG317ACC (3604C>A 3605T>C 3606G>C), GTA318ATT (3607G>A 3609A>T), CAC319ACC (3610C>A 3611A>C), GAA320AAG (3613G>A 3615A>G), CAG321AAG (3616C>A), CTA322TGC (3619C>T 3620T>G 3621A>C), CAA323CAG (3624A>G), ATA324ATC (3627A>C), GGA325GGT (3630A>T), CTT328TTT (3637C>T), CTG329GTA (3640C>G 3642G>A), GTA330TTG (3643G>T 3645A>G), CAT331TAC (3646C>T 3648T>C), GGT332CAG (3649G>C 3650G>A 3651T>G), CTT333TTG (3652C>T 3654T>G), TGG334GGA (3655T>G 3657G>A), AAA335AAG (3660A>G), ACG337ATT (3665C>T 3666G>T), AAC338GAC (3667A>G), CCA339CCC (3672A>C), ATG340TTA (3673A>T 3675G>A), TAC342TTC (3680A>T), AAA343AAG (3684A>G), GAA344GAG (3687A>G), CTC345CTT (3690C>T), ATA346GTA (3691A>G), ACG347TCA (3694A>T 3696G>A), CGC348GAC (3697C>G 3698G>A), ATT349TTA (3700A>T 3702T>A), GCA350GCC (3705A>C), CAT351AAG (3706C>A 3708T>G), CGT352_ATC353del (3709_3714delCGTATC), TTC355TTG (3720C>G), GAT356GAC (3723T>C), GTT357GGT (3725T>G), TAG358AAG (3727T>A) | | | | | | | | |

*: Inserts / Deletes / Misaligned / Frameshifts

#### Analysis details

This analysis was performed with panviral2.64

### NGS Details (UN60): Badnavirus maculasmallanthi

#### Assembly

| | |
|---|---|
| **Coverage Length** | 257 (1 contig(s)) |
| **Depth Of Coverage** | 43.1 |
| **Number Of Reads** | 110 |
| **Reads Per Million** | 2.33 rpm (after QC) |
| **Ambiguities** | 0 |
| **Assembly Method** | de novo + reference guided assembly |
| **Consensus Caller** | Bcf Tools |

#### Coverage Map

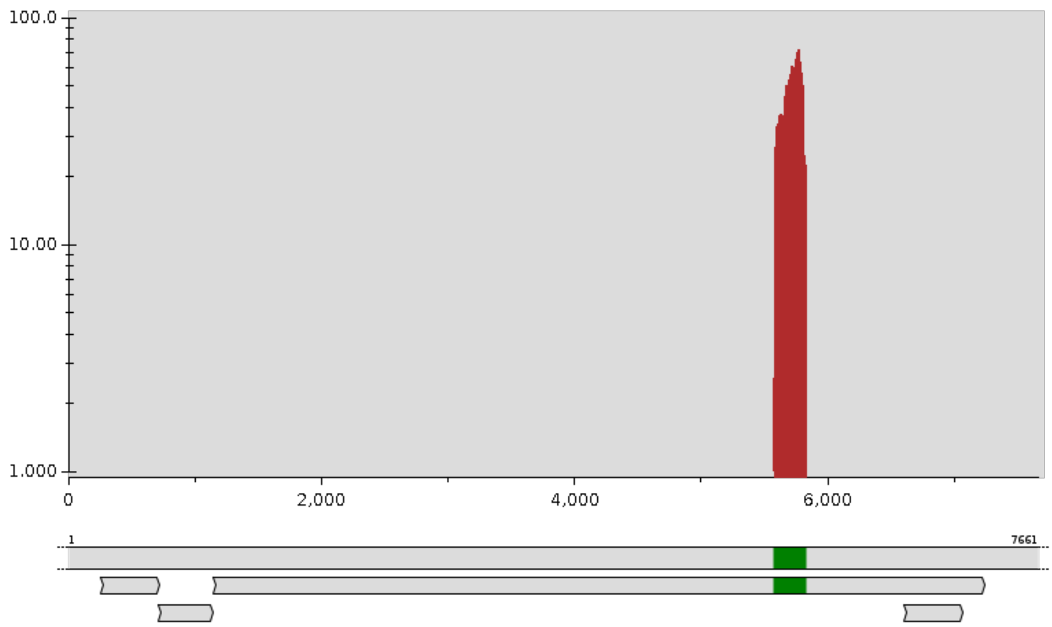

#### Assignment

| | |
|---|---|
| **Type** | Badnavirus maculasmallanthi (Taxonomy ID: 3048453) |
| **Reference Genome** | NC_026472.1 |
| **NT Identity (%)** | 56.5385 |
| **AA Identity (%)** | 55.1724 |
| **Number Of Stop Codons** | 0 |
| **Number Of CDS** | 4 |

#### Alignment

| | |
|---|---|
| **Alignment Score** | 62.0 (NT) + 334.0 (AA) = 396.0 |
| **Concordance (%)** | 35.1375 |

| Alignment Method | Global, seeded, nucleotide + amino acids (AGA) |
|---|---|

#### Genome Region

Sequence starts at position 5577 and ends at position 5833 relative to NC_026472.1 reference sequence.

#### Alignment Detailed Statistics

| | Begin | End | Coverage | Score | Concordance | Matches | Identities | I/D/M/F* | Stop Codons |
|---|---|---|---|---|---|---|---|---|---|
| **NT** | **5577** | **5833** | **3.4%** | **62** | **12.1%** | **257 (98.8%)** | **147 (56.5%)** | **3/0** | |
| Mutations: | 5588A>G, 5589G>A, 5592T>G, 5595A>T, 5597T>A, 5604A>G, 5605G>T, 5607T>G, 5608G>A, 5609C>G, 5610C>A, 5611A>G, 5613G>T, 5614G>C, 5615A>G, 5616A>G, 5619A>G, 5620G>T, 5621A>C, 5623T>G, 5624C>A, 5625C>T, 5628A>C, 5631C>G, 5632T>A, 5633G>A, 5637G>T, 5640C>T, 5644T>C, 5645T>G, 5646A>G, 5648T>C, 5649C>A, 5650C>T, 5651C>G, 5652A>C, 5653G>T, 5654G>A, 5655A>T, 5658A>T, 5659T>C, 5660T>A, 5661A>T, 5664C>T, 5667A>G, 5669G>T, 5670G>T, 5671C>T, 5673T>G, 5676C>T, 5680C>T, 5682A>T, 5685C>T, 5688A>G, 5691C>T, 5693A>C, 5700A>C, 5703C>T, 5707A>G, 5708T>C, 5712T>C, 5713C>A, 5714A>T, 5716A>G, 5717G>A, 5718A>G, 5719A>T, 5720A>T, 5721A>G, 5725G>A, 5728A>C, 5729A>G, 5730A>G, 5731T>G, 5732G>T, 5733C>G, 5736C>T, 5739A>G, 5740G>T, 5741A>C, 5742_5743insTAT, 5743A>T, 5744C>T, 5745A>G, 5748A>T, 5749G>C, 5755A>G, 5757C>G, 5758G>A, 5759C>T, 5765A>T, 5766C>T, 5769C>A, 5772C>T, 5775C>T, 5779C>T, 5784A>T, 5787C>T, 5793C>G, 5799A>G, 5800G>A, 5801C>G, 5802G>T, 5810A>T, 5813G>A, 5814G>A, 5817T>C, 5820A>G, 5822A>G, 5823G>A, 5827A>G | | | | | | | | |
| **CDS** | | | | | | | | | |
| UF61_gp3 | 1479 | 1564 | 4.2% | 334 | 53.4% | 86 (98.9%) | 48 (55.2%) | 1/0/0/0 | 0 |
| Protein mutations: | K1482R (5588A>G 5589G>A), F1485Y (5597T>A), V1488L (5605G>T 5607T>G), A1489R (5608G>A 5609C>G 5610C>A), M1490V (5611A>G 5613G>T), E1491R (5614G>C 5615A>G 5616A>G), E1493S (5620G>T 5621A>C), S1494D (5623T>G 5624C>A 5625C>T), W1497K (5632T>A 5633G>A), L1501R (5644T>C 5645T>G 5646A>G), I1502T (5648T>C 5649C>A), P1503C (5650C>T 5651C>G 5652A>C), G1504Y (5653G>T 5654G>A 5655A>T), L1506H (5659T>C 5660T>A 5661A>T), W1509F (5669G>T 5670G>T), P1513S (5680C>T 5682A>T), K1517T (5693A>C), I1522A (5707A>G 5708T>C), Q1524M (5713C>A 5714A>T), R1525E (5716A>G 5717G>A 5718A>G), K1526L (5719A>T 5720A>T 5721A>G), D1528N (5725G>A), K1529R (5728A>C 5729A>G 5730A>G), C1530V (5731T>G 5732G>T 5733C>G), D1533S (5740G>T 5741A>C), D1533_T1534insY (5742_5743insTAT), T1534L (5743A>T 5744C>T 5745A>G), E1535D (5748A>T), E1536Q (5749G>C), I1538V (5755A>G 5757C>G), A1539I (5758G>A 5759C>T), Y1541F (5765A>T 5766C>T), N1550K (5793C>G), A1553S (5800G>A 5801C>G 5802G>T), E1556V (5810A>T), R1557K (5813G>A 5814G>A), K1560R (5822A>G 5823G>A), M1562V (5827A>G) | | | | | | | | |
| Codon mutations: | AAG1482AGA (5588A>G 5589G>A), TCT1483TCG (5592T>G), GGA1484GGT (5595A>T), TTT1485TAT (5597T>A), CAA1487CAG (5604A>G), GTT1488TTG (5605G>T 5607T>G), GCC1489AGA (5608G>A 5609C>G 5610C>A), ATG1490GTT (5611A>G 5613G>T), GAA1491CGG (5614G>C 5615A>G 5616A>G), GAA1492GAG (5619A>G), GAA1493TCA (5620G>T 5621A>C), TCC1494GAT (5623T>G 5624C>A 5625C>T), ATA1495ATC (5628A>C), CCC1496CCG (5631C>G), TGG1497AAG (5632T>A 5633G>A), ACG1498ACT (5637G>T), GCC1499GCT (5640C>T), TTA1501CGG (5644T>C 5645T>G 5646A>G), ATC1502ACA (5648T>C 5649C>A), CCA1503TGC (5650C>T 5651C>G 5652A>C), GGA1504TAT (5653G>T 5654G>A 5655A>T), GGA1505GGT (5658A>T), TTA1506CAT (5659T>C 5660T>A 5661A>T), TAC1507TAT (5664C>T), GAA1508GAG (5667A>G), TGG1509TTT (5669G>T 5670G>T), CTT1510TTG (5671C>T 5673T>G), GTC1511GTT (5676C>T), CCA1513TCT (5680C>T 5682A>T), TTC1514TTT (5685C>T), GGA1515GGG (5688A>G), CTC1516CTT (5691C>T), AAA1517ACA (5693A>C), GCA1519GCC (5700A>C), CCC1520CCT (5703C>T), ATT1522GCT (5707A>G 5708T>C), TTT1523TTC (5712T>C), CAG1524ATG (5713C>A 5714A>T), AGA1525GAG (5716A>G 5717G>A 5718A>G), AAA1526TTG (5719A>T 5720A>T 5721A>G), GAT1528AAT (5725G>A), AAA1529CGG (5728A>C 5729A>G 5730A>G), TGC1530GTG (5731T>G 5732G>T 5733C>G), TTC1531TTT (5736C>T), AAA1532AAG (5739A>G), GAC1533TCC (5740G>T 5741A>C), GAC1533_ACA1534insTAT (5742_5743insTAT), ACA1534TTG (5743A>T 5744C>T 5745A>G), GAA1535GAT (5748A>T), GAG1536CAG (5749G>C), ATC1538GTG (5755A>G 5757C>G), GCC1539ATC (5758G>A 5759C>T), TAC1541TTT (5765A>T 5766C>T), ATC1542ATA (5769C>A), GAC1543GAT (5772C>T), GAC1544GAT (5775C>T), CTG1546TTG (5779C>T), GTA1547GTT (5784A>T), TAC1548TAT (5787C>T), AAC1550AAG (5793C>G), GAA1552GAG (5799A>G), GCG1553AGT (5800G>A 5801C>G 5802G>T), GAA1556GTA (5810A>T), AGG1557AAA (5813G>A 5814G>A), CAT1558CAC (5817T>C), TTA1559TTG (5820A>G), AAG1560AGA (5822A>G 5823G>A), ATG1562GTG (5827A>G) | | | | | | | | |
| **Proteins** | | | | | | | | | |
| ORF3 (YP_009121747.1) | 1479 | 1564 | 4.2% | 334 | 53.4% | 86 (98.9%) | 48 (55.2%) | 1/0/0/0 | 0 |
| Protein mutations: | K1482R (5588A>G 5589G>A), F1485Y (5597T>A), V1488L (5605G>T 5607T>G), A1489R (5608G>A 5609C>G 5610C>A), M1490V (5611A>G 5613G>T), E1491R (5614G>C 5615A>G 5616A>G), E1493S (5620G>T 5621A>C), S1494D (5623T>G 5624C>A 5625C>T), W1497K (5632T>A 5633G>A), L1501R (5644T>C 5645T>G 5646A>G), I1502T (5648T>C 5649C>A), P1503C (5650C>T 5651C>G 5652A>C), G1504Y (5653G>T 5654G>A 5655A>T), L1506H (5659T>C 5660T>A 5661A>T), W1509F (5669G>T 5670G>T), P1513S (5680C>T 5682A>T), K1517T (5693A>C), I1522A (5707A>G 5708T>C), Q1524M (5713C>A 5714A>T), R1525E (5716A>G 5717G>A 5718A>G), K1526L (5719A>T 5720A>T 5721A>G), D1528N (5725G>A), K1529R (5728A>C 5729A>G 5730A>G), C1530V (5731T>G 5732G>T 5733C>G), D1533S (5740G>T 5741A>C), D1533_T1534insY (5742_5743insTAT), T1534L (5743A>T 5744C>T 5745A>G), E1535D (5748A>T), E1536Q (5749G>C), I1538V (5755A>G 5757C>G), A1539I (5758G>A 5759C>T), Y1541F (5765A>T 5766C>T), N1550K (5793C>G), A1553S (5800G>A 5801C>G 5802G>T), E1556V (5810A>T), R1557K (5813G>A 5814G>A), K1560R (5822A>G 5823G>A), M1562V (5827A>G) | | | | | | | | |
| Codon mutations: | AAG1482AGA (5588A>G 5589G>A), TCT1483TCG (5592T>G), GGA1484GGT (5595A>T), TTT1485TAT (5597T>A), CAA1487CAG (5604A>G), GTT1488TTG (5605G>T 5607T>G), GCC1489AGA (5608G>A 5609C>G 5610C>A), ATG1490GTT (5611A>G 5613G>T), GAA1491CGG (5614G>C 5615A>G 5616A>G), GAA1492GAG (5619A>G), GAA1493TCA (5620G>T 5621A>C), TCC1494GAT (5623T>G 5624C>A 5625C>T), ATA1495ATC (5628A>C), CCC1496CCG (5631C>G), TGG1497AAG (5632T>A 5633G>A), ACG1498ACT (5637G>T), GCC1499GCT (5640C>T), TTA1501CGG (5644T>C 5645T>G 5646A>G), ATC1502ACA (5648T>C 5649C>A), CCA1503TGC (5650C>T 5651C>G 5652A>C), GGA1504TAT (5653G>T 5654G>A 5655A>T), GGA1505GGT (5658A>T), TTA1506CAT (5659T>C 5660T>A 5661A>T), TAC1507TAT (5664C>T), GAA1508GAG (5667A>G), TGG1509TTT (5669G>T 5670G>T), CTT1510TTG (5671C>T 5673T>G), GTC1511GTT (5676C>T), CCA1513TCT (5680C>T 5682A>T), TTC1514TTT (5685C>T), GGA1515GGG (5688A>G), CTC1516CTT (5691C>T), AAA1517ACA (5693A>C), GCA1519GCC (5700A>C), CCC1520CCT (5703C>T), ATT1522GCT (5707A>G 5708T>C), TTT1523TTC (5712T>C), CAG1524ATG (5713C>A 5714A>T), AGA1525GAG (5716A>G 5717G>A 5718A>G), AAA1526TTG (5719A>T 5720A>T 5721A>G), GAT1528AAT (5725G>A), AAA1529CGG (5728A>C 5729A>G 5730A>G), TGC1530GTG (5731T>G 5732G>T 5733C>G), TTC1531TTT (5736C>T), AAA1532AAG (5739A>G), GAC1533TCC (5740G>T 5741A>C), GAC1533_ACA1534insTAT (5742_5743insTAT), ACA1534TTG (5743A>T 5744C>T 5745A>G), GAA1535GAT (5748A>T), GAG1536CAG (5749G>C), ATC1538GTG (5755A>G 5757C>G), GCC1539ATC (5758G>A 5759C>T), TAC1541TTT (5765A>T 5766C>T), ATC1542ATA (5769C>A), GAC1543GAT (5772C>T), GAC1544GAT (5775C>T), CTG1546TTG (5779C>T), GTA1547GTT (5784A>T), TAC1548TAT (5787C>T), AAC1550AAG (5793C>G), GAA1552GAG (5799A>G), GCG1553AGT (5800G>A 5801C>G 5802G>T), GAA1556GTA (5810A>T), AGG1557AAA (5813G>A 5814G>A), CAT1558CAC (5817T>C), TTA1559TTG (5820A>G), AAG1560AGA (5822A>G 5823G>A), ATG1562GTG (5827A>G) | | | | | | | | |

*: Inserts / Deletes / Misaligned / Frameshifts

#### Analysis details

This analysis was performed with panviral2.64

### NGS Details (UN60): Caulimovirus venafragariae

#### Assembly

| | |
|---|---|
| **Coverage Length** | 1041 (1 contig(s)) |
| **Depth Of Coverage** | 9.9 |
| **Number Of Reads** | 77 |
| **Reads Per Million** | 1.63 rpm (after QC) |
| **Ambiguities** | 0 |
| **Assembly Method** | de novo + reference guided assembly |
| **Consensus Caller** | Bcf Tools |

#### Coverage Map

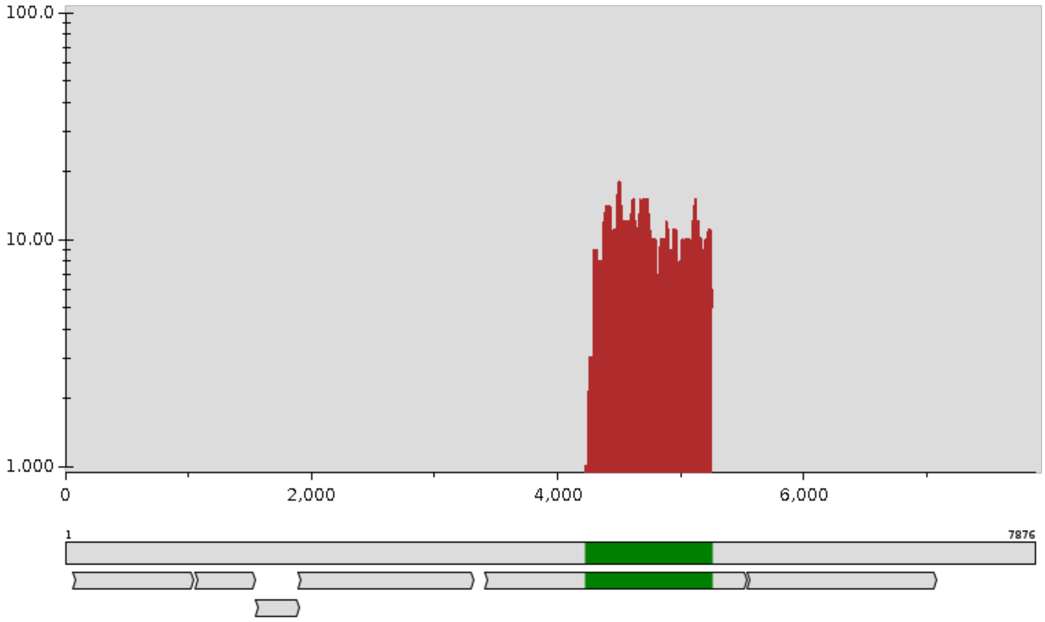

#### Assignment

| | |
|---|---|
| **Type** | Caulimovirus venafragariae (Taxonomy ID: 3048344) |
| **Reference Genome** | NC_001725.1 |
| **NT Identity (%)** | 56.3477 |
| **AA Identity (%)** | 52.3392 |
| **Number Of Stop Codons** | 0 |
| **Number Of CDS** | 6 |

#### Alignment

| | |
|---|---|
| **Alignment Score** | 216.0 (NT) + 1173.0 (AA) = 1389.0 |
| **Concordance (%)** | 32.3873 |

| Alignment Method | Global, seeded, nucleotide + amino acids (AGA) |
|---|---|

#### Genome Region

Sequence starts at position 4219 and ends at position 5259 relative to NC_001725.1 reference sequence.

#### Alignment Detailed Statistics

| | Begin | End | Coverage | Score | Concordance | Matches | Identities | I/D/M/F* | Stop Codons |
|---|---|---|---|---|---|---|---|---|---|
| **NT** | **4219** | **5259** | **13.2%** | **216** | **10.7%** | **1023 (98.2%)** | **577 (55.4%)** | **1/18** | |
| Mutations: | 4219T>C, 4222T>A, 4223T>A, 4227C>T, 4228A>C, 4230T>C, 4232A>G, 4235A>T, 4236A>C, 4239T>A, 4242T>A, 4244A>G, 4245G>A, 4246T>G, 4247G>T, 4254A>T, 4259A>T, 4260G>A, 4263T>C, 4264T>A, 4265C>A, 4266A>T, 4267C>G, 4268C>A, 4269T>A, 4275C>T, 4276C>A, 4277G>A, 4279G>A, 4280A>C, 4281A>T, 4284G>A, 4292C>T, 4299C>T, 4300G>C, 4306C>T, 4309C>T, 4311C>A, 4314G>T, 4315C>A, 4317T>G, 4318G>A, 4319G>A, 4320T>A, 4321A>T, 4323C>A, 4326C>A, 4331C>A, 4335C>T, 4338G>A, 4348T>A, 4349C>G, 4350C>T, 4351T>A, 4352C>G, 4353A>C, 4356A>C, 4359C>A, 4362C>T, 4366G>A, 4371G>A, 4374T>C, 4379C>G, 4383G>A, 4384A>C, 4385T>A, 4386C>A, 4392A>T, 4395C>A, 4401A>T, 4402A>C, 4413T>A, 4423A>G, 4426T>C, 4428A>T, 4432G>A, 4434C>A, 4435C>A, 4438A>T, 4439C>G, 4440A>T, 4441A>G, 4442A>T, 4443G>C, 4444G>T, 4445G>T, 4446A>C, 4449T>C, 4452C>A, 4455C>T, 4456C>T, 4458A>C, 4459C>A, 4461T>C, 4464T>A, 4467C>T, 4470G>A, 4471G>A, 4473G>T, 4474C>A, 4476A>C, 4479G>T, 4480C>A, 4482T>A, 4483C>A, 4485A>T, 4486A>C, 4487G>T, 4488A>T, 4489A>G, 4490T>C, 4492G>A, 4493G>A, 4497T>A, 4500G>A, 4505T>A, 4506T>C, 4508A>T, 4509C>T, 4512T>A, 4513T>A, 4514C>A, 4515C>A, 4521C>T, 4524T>C, 4528T>A, 4529C>G, 4542G>A, 4543G>A, 4546C>A, 4547G>A, 4548C>A, 4551T>A, 4553C>A, 4554T>A, 4555C>G, 4556C>A, 4558G>A, 4560A>T, 4561A>T, 4563A>T, 4566T>A, 4567C>G, 4568A>C, 4569G>T, 4570C>T, 4575C>A, 4578T>G, 4584C>T, 4585T>A, 4586G>C, 4587T>G, 4590C>A, 4591C>A, 4593A>T, 4599C>T, 4614C>G, 4633C>A, 4635A>T, 4642G>C, 4643C>A, 4644T>A, 4647T>C, 4650C>T, 4653A>G, 4657C>A, 4659C>A, 4665C>T, 4666G>A, 4670G>T, 4671T>A, 4672C>T, 4674C>T, 4675A>T, 4676G>C, 4678A>C, 4680C>A, 4682T>A, 4684T>A, 4687_4692delCCACAG, 4696T>C, 4697G>T, 4698T>A, 4699G>A, 4700C>T, 4701T>C, 4704A>C, 4707T>C, 4708G>A, 4710T>A, 4716C>T, 4719C>A, 4720A>T, 4722C>A, 4723G>A, 4725C>A, 4727T>G, 4728C>T, 4729A>T, 4730G>C, 4734A>C, 4735A>T, 4737T>A, 4738G>T, 4740A>T, 4741G>C, 4753G>A, 4754G>C, 4755A>T, 4759G>C, 4761C>T, 4765A>C, 4766T>A, 4767T>A, 4768G>T, 4771C>T, 4772T>C, 4773A>T, 4774A>C, 4776C>A, 4778G>T, 4782C>T, 4783A>T, 4784A>G, 4785A>C, 4787C>A, 4788C>A, 4789C>A, 4790T>A, 4791A>T, 4797A>T, 4801C>T, 4803T>A, 4807A>C, 4809A>G, 4812G>A, 4815A>G, 4818C>A, 4819C>G, 4821G>C, 4822T>A, 4824G>A, 4825T>A, 4826G>A, 4827C>T, 4832C>A, 4833A>T, 4834A>G, 4835C>A, 4836T>A, 4839C>T, 4840A>G, 4842T>A, 4845C>T, 4848T>C, 4851T>A, 4852C>A, 4854A>G, 4855G>A, 4860A>T, 4861G>T, 4862A>C, 4865G>A, 4867G>A, 4868G>A, 4869A>T, 4870A>G, 4871A>G, 4872T>C, 4873C>A, 4875C>T, 4876A>G, 4879G>T, 4881C>A, 4884G>A, 4886G>C, 4890T>C, 4893C>A, 4894G>T, 4895G>C, 4897C>A, 4898T>A, 4900C>A, 4902C>A, 4902_4903insG, 4903T>A, 4905A>C, 4906G>T, 4907T>G, 4910C>A, 4911T>A, 4915C>T, 4921C>A, 4926T>A, 4928G>A, 4929T>C, 4930G>A, 4931A>C, 4932T>A, 4934G>A, 4935G>A, 4936A>C, 4938T>A, 4940C>A, 4941C>A, 4942T>A, 4944A>C, 4950A>G, 4954T>C, 4956A>T, 4959C>A, 4960C>T, 4961T>G, 4962C>T, 4963C>T, 4971T>C, 4972A>G, 4973T>C, 4974C>A, 4975T>A, 4976C>G, 4977C>T, 4979C>A, 4980T>A, 4982A>T, 4984T>A, 4986T>A, 4990A>G, 4992A>C, 4993A>C, 5001C>A, 5002C>A, 5003T>A, 5004T>G, 5007G>A, 5008T>A, 5009C>A, 5010A>T, 5012C>T, 5019G>A, 5020G>A, 5021T>G, 5022T>G, 5023A>C, 5024A>T, 5025A>T, 5026C>A, 5029A>C, 5030A>G, 5031G>A, 5035G>T, 5036A>C, 5037G>C, 5039T>A, 5040A>C, 5041A>C, 5042C>G, 5043T>A, 5044T>A, 5045G>T, 5047T>G, 5048C>G, 5049T>A, 5055T>A, 5058G>T, 5059A>G, 5062G>C, 5064C>T, 5067G>T, 5068G>A, 5071A>T, 5072C>G, 5076C>T, 5077A>C, 5078G>A, 5081A>G, 5082A>C, 5083A>C, 5088G>A, 5089A>C, 5090G>A, 5091T>A, 5092C>G, 5093T>A, 5095G>T, 5096T>G, 5097A>T, 5098A>T, 5099A>C, 5100A>C, 5102C>A, 5103T>A, 5110G>A, 5112C>G, 5115T>A, 5116T>A, 5117A>G, 5118C>A, 5119A>C, 5120A>T, 5124T>G, 5125T>G, 5126C>A, 5133G>A, 5136T>C, 5139G>C, 5140C>T, 5141C>T, 5142T>A, 5143A>G, 5145C>T, 5146A>T, 5148C>A, 5149G>C, 5152T>A, 5153G>C, 5154C>A, 5160A>T, 5167G>T, 5182A>T, 5184C>A, 5185C>T, 5187C>A, 5188A>C, 5191G>A, 5193A>T, 5194A>G, 5196A>C, 5200C>A, 5201C>A, 5202A>T, 5206_5217delGGTAAAGAGGTA, 5220A>T, 5223C>T, 5230G>A, 5232T>A, 5233T>A, 5234C>G, 5235A>T, 5238A>T, 5241C>A, 5247A>T, 5248C>G, 5249C>A, 5251G>A, 5253T>C, 5256G>A, 5258A>C, 5259G>T | | | | | | | | |

##### CDS

| | Begin | End | Coverage | Score | Concordance | Matches | Identities | I/D/M/F* | Stop Codons |
|---|---|---|---|---|---|---|---|---|---|
| ORF_V | 272 | 618 | 48.9% | 1173 | 49.7% | 341 (98.0%) | 179 (51.4%) | 1/6/1/1 | 0 |
| Protein mutations: | L273K (4222T>A 4223T>A), T275P (4228A>C 4230T>C), N276S (4232A>G), K277I (4235A>T 4236A>C), K280R (4244A>G 4245G>A), C281V (4246T>G 4247G>T), Q285L (4259A>T 4260G>A), S287N (4264T>A 4265C>A 4266A>T), P288E (4267C>G 4268C>A 4269T>A), R291K (4276C>A 4277G>A), E292T (4279G>A 4280A>C 4281A>T), T296I (4292C>T), E299Q (4300G>C), K303N (4314G>T), L304M (4315C>A 4317T>G), G305K (4318G>A 4319G>A 4320T>A), I306L (4321A>T 4323C>A), P309H (4331C>A), V321I (4366G>A), A325G (4379C>G), I327Q (4384A>C 4385T>A 4386C>A), K340E (4423A>G), D343K (4432G>A 4434C>A), H344N (4435C>A), T345C (4438A>T 4439C>G 4440A>T), K346V (4441A>G 4442A>T 4443G>C), G347F (4444G>T 4445G>T 4446A>C), L351F (4456C>T 4458A>C), L352I (4459C>A 4461T>C), E356N (4471G>A 4473G>T), Q357N (4474C>A 4476A>C), L359I (4480C>A 4482T>A), Q360N (4483C>A 4485A>T), R361L (4486A>C 4487G>T 4488A>T), I362A (4489A>G 4490T>C), G363K (4492G>A 4493G>A), F367Y (4505T>A 4506T>C), Y368F (4508A>T 4509C>T), S370K (4513T>A 4514C>A 4515C>A), V380I (4543G>A), R381K (4546C>A 4547G>A 4548C>A), A383E (4553C>A 4554T>A), P384E (4555C>G 4556C>A), E385N (4558G>A 4560A>T), T386S (4561A>T 4563A>T), Q388A (4567C>G 4568A>C 4569G>T), C394T (4585T>A 4586G>C 4587T>G), Q396N (4591C>A 4593A>T), Q410N (4633C>A 4635A>T), A413Q (4642G>C 4643C>A 4644T>A), H418K (4657C>A 4659C>A), E421K (4666G>A), S422I (4670G>T 4671T>A), L423F (4672C>T 4674C>T), N425Q (4678A>C 4680C>A), M426K (4682T>A), Y427N (4684T>A), P428_Q429del (4687_4692delCCACAG), C431L (4696T>C 4697G>T 4698T>A), A432I (4699G>A 4700C>T 4701T>C), V435I (4708G>A 4710T>A), I439L (4720A>T 4722C>A), V440I (4723G>A 4725C>A), F441C (4727T>G 4728C>T), K443N (4734A>C), T444S (4735A>T 4737T>A), E445Y (4738G>T 4740A>T), E446Q (4741G>C), G450T (4753G>A 4754G>C 4755A>T), V452L (4759G>C 4761C>T), I454Q (4765A>C 4766T>A 4767T>A), V455F (4768G>T), L456S (4771C>T 4772T>C 4773A>T), N457Q (4774A>C 4776C>A), R458I (4778G>T), K460C (4783A>T 4784A>G 4785A>C), A461E (4787C>A 4788C>A), L462N (4789C>A 4790T>A 4791A>T), K468Q (4807A>C 4809A>G), Q472D (4819C>G 4821G>C), L473I (4822T>A 4824G>A), C474N (4825T>A 4826G>A 4827C>T), T476N (4832C>A 4833A>T), T477E (4834A>G 4835C>A 4836T>A), N479E (4840A>G 4842T>A), L483M (4852C>A 4854A>G), V484I (4855G>A), E486S (4861G>T 4862A>C), R487K (4865G>A), G488N (4867G>A 4868G>A 4869A>T), N489G (4870A>G 4871A>G 4872T>C), L490I (4873C>A 4875C>T), K491E (4876A>G), V492L (4879G>T 4881C>A), S494T (4886G>C), G497S (4894G>T 4895G>C), L498K (4897C>A 4898T>A), H499K (4900C>A 4902C>A), H499_L500insX (4902_4903insG), L500I (4903T>A 4905A>C), V501C (4906G>T 4907T>G), A502E (4910C>A 4911T>A), P504S (4915C>T), Q506K (4921C>A), S508N (4928G>A 4929T>C), D509T (4930G>A 4931A>C 4932T>A), R510K (4934G>A 4935G>A), N511Q (4936A>C 4938T>A), A512E (4940C>A 4941C>A), L513I (4942T>A 4944A>C), L519C (4960C>T 4961T>G 4962C>T), I523A (4972A>G 4973T>C 4974C>A), A525E (4979C>A 4980T>A), Y526F (4982A>T), F527I (4984T>A 4986T>A), K529D (4990A>G 4992A>C), I530L (4993A>C), N532K (5001C>A), L533K (5002C>A 5003T>A 5004T>G), S535N (5008T>A 5009C>A 5010A>T), P536L (5012C>T), V539R (5020G>A 5021T>G 5022T>G), K540L (5023A>C 5024A>T 5025A>T), L541I (5026C>A), K542R (5029A>C 5030A>G 5031G>A), E544S (5035G>T 5036A>C 5037G>C), I545N (5039T>A 5040A>C), T546R (5041A>C 5042C>G 5043T>A), W547M (5044T>A 5045G>T), S548G (5047T>G 5048C>G 5049T>A), E551D (5058G>T), K552E (5059A>G), D553H (5062G>C 5064C>T), E555K (5068G>A), T556C (5071A>T 5072C>G), R558Q (5077A>C 5078G>A), K559S (5081A>G 5082A>C), I560L (5083A>C), S562Q (5089A>C 5090G>A 5091T>A), L563E (5092C>G 5093T>A), V564C (5095G>T 5096T>G 5097A>T), K565S (5098A>T 5099A>C 5100A>C), T566K (5102C>A 5103T>A), D569K (5110G>A 5112C>G), Y571R (5116T>A 5117A>G 5118C>A), N572L (5119A>C 5120A>T), S574E (5125T>G 5126C>A), K578N (5139G>C), P579L (5140C>T 5141C>T 5142T>A), I580V (5143A>G 5145C>T), I581L (5146A>T 5148C>A), E582Q (5149G>C), C583T (5152T>A 5153G>C 5154C>A), D588Y (5167G>T), I593L (5182A>T 5184C>A), K595Q (5188A>C), A596T (5191G>A 5193A>T), K597D (5194A>G 5196A>C), P599N (5200C>A 5201C>A 5202A>T), G601_V604del (5206_5217delGGTAAAGAGGTA), A609T (5230G>A 5232T>A), K614N (5247A>T), P615E (5248C>G 5249C>A), A616T (5251G>A 5253T>C), K618T (5258A>C 5259G>T) | | | | | | | | |

| | Begin | End | Coverage | Score | Concordance | Matches | Identities | I/D/M/F* | Stop Codons |
|---|---|---|---|---|---|---|---|---|---|
| **NT** | **4219** | **5259** | **13.2%** | **216** | **10.7%** | **1023 (98.2%)** | **577 (55.4%)** | **1/18** | |
| Codon mutations: | TTA272CTA (4219T>C), TTA273AAA (4222T>A 4223T>A), GAC274GAT (4227C>T), ACT275CCC (4228A>C 4230T>C), AAC276AGC (4232A>G), AAA277ATC (4235A>T 4236A>C), ATT278ATA (4239T>A), ATT279ATA (4242T>A), AAG280AGA (4244A>G 4245G>A), TGT281GTT (4246T>G 4247G>T), CCA283CCT (4254A>T), CAG285CTA (4259A>T 4260G>A), TAT286TAC (4263T>C), TCA287AAT (4264T>A 4265C>A 4266A>T), CCT288GAA (4267C>G 4268C>A 4269T>A), GAC290GAT (4275C>T), CGA291AAA (4276C>A 4277G>A), GAA292ACT (4279G>A 4280A>C 4281A>T), GAG293GAA (4284G>A), ACT296ATT (4292C>T), ATC298ATT (4299C>T), GAA299CAA (4300G>C), CTA301TTA (4306C>T), CTC302TTA (4309C>T 4311C>A), AAG303AAT (4314G>T), CTT304ATG (4315C>A 4317T>G), GGT305AAA (4318G>A 4319G>A 4320T>A), ATC306TTA (4321A>T 4323C>A), ATC307ATA (4326C>A), CCT309CAT (4331C>A), AGC310AGT (4335C>T), AAG311AAA (4338G>A), TCC315AGT (4348T>A 4349C>G 4350C>T), TCA316AGC (4351T>A 4352C>G 4353A>C), CCA317CCC (4356A>C), GCC318GCA (4359C>A), TTC319TTT (4362C>T), GTA321ATA (4366G>A), AGG322AGA (4371G>A), AAT323AAC (4374T>C), GCC325GGC (4379C>G), GAG326GAA (4383G>A), ATC327CAA (4384A>C 4385T>A 4386C>A), CGA329CGT (4392A>T), GGC330GGA (4395C>A), GCA332GCT (4401A>T), AGA333CGA (4402A>C), ATT336ATA (4413T>A), AAG340GAG (4423A>G), TTA341CTT (4426T>C 4428A>T), GAC343AAA (4432G>A 4434C>A), CAT344AAT (4435C>A), ACA345TGT (4438A>T 4439C>G 4440A>T), AAG346GTC (4441A>G 4442A>T 4443G>C), GGA347TTC (4444G>T 4445G>T 4446A>C), GAT348GAC (4449T>C), GGC349GGA (4452C>A), TAC350TAT (4455C>T), CTA351TTC (4456C>T 4458A>C), CTT352ATC (4459C>A 4461T>C), CCT353CCA (4464T>A), AAC354AAT (4467C>T), AAG355AAA (4470G>A), GAG356AAT (4471G>A 4473G>T), CAA357AAC (4474C>A 4476A>C), CTG358CTT (4479G>T), CTT359ATA (4480C>A 4482T>A), CAA360AAT (4483C>A 4485A>T), AGA361CTT (4486A>C 4487G>T 4488A>T), ATC362GCC (4489A>G 4490T>C), GGA363AAA (4492G>A 4493G>A), GGT364GGA (4497T>A), AAG365AAA (4500G>A), TTT367TAC (4505T>A 4506T>C), TAC368TTT (4508A>T 4509C>T), TCT369TCA (4512T>A), TCC370AAA (4513T>A 4514C>A 4515C>A), GAC372GAT (4521C>T), TGT373TGC (4524T>C), TCT375AGT (4528T>A 4529C>G), CAG379CAA (4542G>A), GTA380ATA (4543G>A), CGC381AAA (4546C>A 4547G>A 4548C>A), CTT382CTA (4551T>A), GCT383GAA (4553C>A 4554T>A), CCA384GAA (4555C>G 4556C>A), GAA385AAT (4558G>A 4560A>T), ACA386TCT (4561A>T 4563A>T), ATT387ATA (4566T>A), CAG388GCT (4567C>G 4568A>C 4569G>T), CTA389TTA (4570C>T), ACC390ACA (4575C>A), GCT391GCG (4578T>G), AGC393AGT (4584C>T), TGT394ACG (4585T>A 4586G>C 4587T>G), CCC395CCA (4590C>A), CAA396AAT (4591C>A 4593A>T), CAC398CAT (4599C>T), GTC403GTG (4614C>G), CAA410AAT (4633C>A 4635A>T), GCT413CAA (4642G>C 4643C>A 4644T>A), ATT414ATC (4647T>C), TTC415TTT (4650C>T), CAA416CAG (4653A>G), CAC418AAA (4657C>A 4659C>A), GAC420GAT (4665C>T), GAA421AAA (4666G>A), AGT422ATA (4670G>T 4671T>A), CTC423TTT (4672C>T 4674C>T), AGC424TCC (4675A>T 4676G>C), AAC425CAA (4678A>C 4680C>A), ATG426AAG (4682T>A), TAT427AAT (4684T>A), CCA428_CAG429del (4687_4692delCCACAG), TGT431CTA (4696T>C 4697G>T 4698T>A), GCT432ATC (4699G>A 4700C>T 4701T>C), GTA433GTC (4704A>C), TAT434TAC (4707T>C), GTT435ATA (4708G>A 4710T>A), GAC437GAT (4716C>T), ATC438ATA (4719C>A), ATC439TTA (4720A>T 4722C>A), GTC440ATA (4723G>A 4725C>A), TTC441TGT (4727T>G 4728C>T), AGC442TCC (4729A>T 4730G>C), AAA443AAC (4734A>C), ACT444TCA (4735A>T 4737T>A), GAA445TAT (4738G>T 4740A>T), GAA446CAA (4741G>C), GGA450ACT (4753G>A 4754G>C 4755A>T), GTC452CTT (4759G>C 4761C>T), ATT454CAA (4765A>C 4766T>A 4767T>A), GTT455TTT (4768G>T), CTA456TCT (4771C>T 4772T>C 4773A>T), AAC457CAA (4774A>C 4776C>A), AGA458ATA (4778G>T), TGC459TGT (4782C>T), AAA460TGC (4783A>T 4784A>G 4785A>C), GCC461GAA (4787C>A 4788C>A), CTA462AAT (4789C>A 4790T>A 4791A>T), ATA464ATT (4797A>T), CTT466TTA (4801C>T 4803T>A), AAA468CAG (4807A>C 4809A>G), AAG469AAA (4812G>A), AAA470AAG (4815A>G), GCC471GCA (4818C>A), CAG472GAC (4819C>G 4821G>C), TTG473ATA (4822T>A 4824G>A), TGC474AAT (4825T>A 4826G>A 4827C>T), ACA476AAT (4832C>A 4833A>T), ACT477GAA (4834A>G 4835C>A 4836T>A), ATC478ATT (4839C>T), AAT479GAA (4840A>G 4842T>A), TTC480TTT (4845C>T), CTT481CTC (4848T>C), GGT482GGA (4851T>A), CTA483ATG (4852C>A 4854A>G), GTA484ATA (4855G>A), ATA485ATT (4860A>T), GAA486TCA (4861G>T 4862A>C), AGA487AAA (4865G>A), GGA488AAT (4867G>A 4868G>A 4869A>T), AAT489GGC (4870A>G 4871A>G 4872T>C), CTC490ATT (4873C>A 4875C>T), AAA491GAA (4876A>G), GTC492TTA (4879G>T 4881C>A), CAG493CAA (4884G>A), AGT494ACT (4886G>C), CAT495CAC (4890T>C), ATC496ATA (4893C>A), GGA497TCA (4894G>T 4895G>C), CTA498AAA (4897C>A 4898T>A), CAC499AAA (4900C>A 4902C>A), CAC499_TTA500insG-- (4902_4903insG), TTA500ATC (4903T>A 4905A>C), GTT501TGT (4906G>T 4907T>G), GCT502GAA (4910C>A 4911T>A), CCA504TCA (4915C>T), CAA506AAA (4921C>A), CTT507CTA (4926T>A), AGT508AAC (4928G>A 4929T>C), GAT509ACA (4930G>A 4931A>C 4932T>A), AGG510AAA (4934G>A 4935G>A), AAT511CAA (4936A>C 4938T>A), GCC512GAA (4940C>A 4941C>A), TTA513ATC (4942T>A 4944A>C), AGA515AGG (4950A>G), TTA517CTT (4954T>C 4956A>T), GGC518GGA (4959C>A), CTC519TGT (4960C>T 4961T>G 4962C>T), CTA520TTA (4963C>T), TAT522TAC (4971T>C), ATC523GCA (4972A>G 4973T>C 4974C>A), TCC524AGT (4975T>A 4976C>G 4977C>T), GCT525GAA (4979C>A 4980T>A), TAT526TTT (4982A>T), TTT527ATA (4984T>A 4986T>A), AAA529GAC (4990A>G 4992A>C), ATA530CTA (4993A>C), AAC532AAA (5001C>A), CTT533AAG (5002C>A 5003T>A 5004T>G), AGG534AGA (5007G>A), TCA535AAT (5008T>A 5009C>A 5010A>T), CCT536CTT (5012C>T), CAG538CAA (5019G>A), GTT539AGG (5020G>A 5021T>G 5022T>G), AAA540CTT (5023A>C 5024A>T 5025A>T), CTT541ATT (5026C>A), AAG542CGA (5029A>C 5030A>G 5031G>A), GAG544TCC (5035G>T 5036A>C 5037G>C), ATA545AAC (5039T>A 5040A>C), ACT546CGA (5041A>C 5042C>G 5043T>A), TGG547ATG (5044T>A 5045G>T), TCT548GGA (5047T>G 5048C>G 5049T>A), ACT550ACA (5055T>A), GAG551GAT (5058G>T), AAA552GAA (5059A>G), GAC553CAT (5062G>C 5064C>T), ACG554ACT (5067G>T), GAA555AAA (5068G>A), ACT556TGT (5071A>T 5072C>G), GTC557GTT (5076C>T), AGA558CAA (5077A>C 5078G>A), AAA559AGC (5081A>G 5082A>C), ATA560CTA (5083A>C), AAG561AAA (5088G>A), AGT562CAA (5089A>C 5090G>A 5091T>A), CTA563GAA (5092C>G 5093T>A), GTA564TGT (5095G>T 5096T>G 5097A>T), AAA565TCC (5098A>T 5099A>C 5100A>C), ACT566AAA (5102C>A 5103T>A), GAC569AAG (5110G>A 5112C>G), CTT570CTA (5115T>A), TAC571AGA (5116T>A 5117A>G 5118C>A), AAT572CTT (5119A>C 5120A>T), CCT573CCG (5124T>G), TCA574GAA (5125T>G 5126C>A), GAG576GAA (5133G>A), GAT577GAC (5136T>C), AAG578AAC (5139G>C), CCT579TTA (5140C>T 5141C>T 5142T>A), ATC580GTT (5143A>G 5145C>T), ATC581TTA (5146A>T 5148C>A), GAA582CAA (5149G>C), TGC583ACA (5152T>A 5153G>C 5154C>A), GCA585GCT (5160A>T), GAC588TAC (5167G>T), ATC593TTA (5182A>T 5184C>A), CTC594TTA (5185C>T 5187C>A), AAA595CAA (5188A>C), GCA596ACT (5191G>A 5193A>T), AAA597GAC (5194A>G 5196A>C), CCA599AAT (5200C>A 5201C>A 5202A>T), GGT601_GTA604del (5206_5217delGGTAAAGAGGTA), ATA605ATT (5220A>T), TGC606TGT (5223C>T), GCT609ACA (5230G>A 5232T>A), TCA610AGT (5233T>A 5234C>G 5235A>T), GGA611GGT (5238A>T), ACC612ACA (5241C>A), AAA614AAT (5247A>T), CCA615GAA (5248C>G 5249C>A), GCT616ACC (5251G>A 5253T>C), GAG617GAA (5256G>A), AAG618ACT (5258A>C 5259G>T) | | | | | | | | |

#### Proteins

| | Begin | End | Coverage | Score | Concordance | Matches | Identities | I/D/M/F* | Stop Codons |
|---|---|---|---|---|---|---|---|---|---|
| hypothetical protein (NP_043933.1) | 272 | 618 | 48.9% | 1173 | 49.7% | 341 (98.0%) | 179 (51.4%) | 1/6/1/1 | 0 |
| Protein mutations: | L273K (4222T>A 4223T>A), T275P (4228A>C 4230T>C), N276S (4232A>G), K277I (4235A>T 4236A>C), K280R (4244A>G 4245G>A), C281V (4246T>G 4247G>T), Q285L (4259A>T 4260G>A), S287N (4264T>A 4265C>A 4266A>T), P288E (4267C>G 4268C>A 4269T>A), R291K (4276C>A 4277G>A), E292T (4279G>A 4280A>C 4281A>T), T296I (4292C>T), E299Q (4300G>C), K303N (4314G>T), L304M (4315C>A 4317T>G), G305K (4318G>A 4319G>A 4320T>A), I306L (4321A>T 4323C>A), P309H (4331C>A), V321I (4366G>A), A325G (4379C>G), I327Q (4384A>C 4385T>A 4386C>A), K340E (4423A>G), D343K (4432G>A 4434C>A), H344N (4435C>A), T345C (4438A>T 4439C>G 4440A>T), K346V (4441A>G 4442A>T 4443G>C), G347F (4444G>T 4445G>T 4446A>C), L351F (4456C>T 4458A>C), L352I (4459C>A 4461T>C), E356N (4471G>A 4473G>T), Q357N (4474C>A 4476A>C), L359I (4480C>A 4482T>A), Q360N (4483C>A 4485A>T), R361L (4486A>C 4487G>T 4488A>T), I362A (4489A>G 4490T>C), G363K (4492G>A 4493G>A), F367Y (4505T>A 4506T>C), Y368F (4508A>T 4509C>T), S370K (4513T>A 4514C>A 4515C>A), V380I (4543G>A), R381K (4546C>A 4547G>A 4548C>A), A383E (4553C>A 4554T>A), P384E (4555C>G 4556C>A), E385N (4558G>A 4560A>T), T386S (4561A>T 4563A>T), Q388A (4567C>G 4568A>C 4569G>T), C394T (4585T>A 4586G>C 4587T>G), Q396N (4591C>A 4593A>T), Q410N (4633C>A 4635A>T), A413Q (4642G>C 4643C>A 4644T>A), H418K (4657C>A 4659C>A), E421K (4666G>A), S422I (4670G>T 4671T>A), L423F (4672C>T 4674C>T), N425Q (4678A>C 4680C>A), M426K (4682T>A), Y427N (4684T>A), P428_Q429del (4687_4692delCCACAG), C431L (4696T>C 4697G>T 4698T>A), A432I (4699G>A 4700C>T 4701T>C), V435I (4708G>A 4710T>A), I439L (4720A>T 4722C>A), V440I (4723G>A 4725C>A), F441C (4727T>G 4728C>T), K443N (4734A>C), T444S (4735A>T 4737T>A), E445Y (4738G>T 4740A>T), E446Q (4741G>C), G450T (4753G>A 4754G>C 4755A>T), V452L (4759G>C 4761C>T), I454Q (4765A>C 4766T>A 4767T>A), V455F (4768G>T), L456S (4771C>T 4772T>C 4773A>T), N457Q (4774A>C 4776C>A), R458I (4778G>T), K460C (4783A>T 4784A>G 4785A>C), A461E (4787C>A 4788C>A), L462N (4789C>A 4790T>A 4791A>T), K468Q (4807A>C 4809A>G), Q472D (4819C>G 4821G>C), L473I (4822T>A 4824G>A), C474N (4825T>A 4826G>A 4827C>T), T476N (4832C>A 4833A>T), T477E (4834A>G 4835C>A 4836T>A), N479E (4840A>G 4842T>A), L483M (4852C>A 4854A>G), V484I (4855G>A), E486S (4861G>T 4862A>C), R487K (4865G>A), G488N (4867G>A 4868G>A 4869A>T), N489G (4870A>G 4871A>G 4872T>C), L490I (4873C>A 4875C>T), K491E (4876A>G), V492L (4879G>T 4881C>A), S494T (4886G>C), G497S (4894G>T 4895G>C), L498K (4897C>A 4898T>A), H499K (4900C>A 4902C>A), H499_L500insX (4902_4903insG), L500I (4903T>A 4905A>C), V501C (4906G>T 4907T>G), A502E (4910C>A 4911T>A), P504S (4915C>T), Q506K (4921C>A), S508N (4928G>A 4929T>C), D509T (4930G>A 4931A>C 4932T>A), R510K (4934G>A 4935G>A), N511Q (4936A>C 4938T>A), A512E (4940C>A 4941C>A), L513I (4942T>A 4944A>C), L519C (4960C>T 4961T>G 4962C>T), I523A (4972A>G 4973T>C 4974C>A), A525E (4979C>A 4980T>A), Y526F (4982A>T), F527I (4984T>A 4986T>A), K529D (4990A>G 4992A>C), I530L (4993A>C), N532K (5001C>A), L533K (5002C>A 5003T>A 5004T>G), S535N (5008T>A 5009C>A 5010A>T), P536L (5012C>T), V539R (5020G>A 5021T>G 5022T>G), K540L (5023A>C 5024A>T 5025A>T), L541I (5026C>A), K542R (5029A>C 5030A>G 5031G>A), E544S (5035G>T 5036A>C 5037G>C), I545N (5039T>A 5040A>C), T546R (5041A>C 5042C>G 5043T>A), W547M (5044T>A 5045G>T), S548G (5047T>G 5048C>G 5049T>A), E551D (5058G>T), K552E (5059A>G), D553H (5062G>C 5064C>T), E555K (5068G>A), T556C (5071A>T 5072C>G), R558Q (5077A>C 5078G>A), K559S (5081A>G 5082A>C), I560L (5083A>C), S562Q (5089A>C 5090G>A 5091T>A), L563E (5092C>G 5093T>A), V564C (5095G>T 5096T>G 5097A>T), K565S (5098A>T 5099A>C 5100A>C), T566K (5102C>A 5103T>A), D569K (5110G>A 5112C>G), Y571R (5116T>A 5117A>G 5118C>A), N572L (5119A>C 5120A>T), S574E (5125T>G 5126C>A), K578N (5139G>C), P579L (5140C>T 5141C>T 5142T>A), I580V (5143A>G 5145C>T), I581L (5146A>T 5148C>A), E582Q (5149G>C), C583T (5152T>A 5153G>C 5154C>A), D588Y (5167G>T), I593L (5182A>T 5184C>A), K595Q (5188A>C), A596T (5191G>A 5193A>T), K597D (5194A>G 5196A>C), P599N (5200C>A 5201C>A 5202A>T), G601_V604del (5206_5217delGGTAAAGAGGTA), A609T (5230G>A 5232T>A), K614N (5247A>T), P615E (5248C>G 5249C>A), A616T (5251G>A 5253T>C), K618T (5258A>C 5259G>T) | | | | | | | | |

| | Begin | End | Coverage | Score | Concordance | Matches | Identities | I/D/M/F* | Stop Codons |
|---|---|---|---|---|---|---|---|---|---|
| **NT** | **4219** | **5259** | **13.2%** | **216** | **10.7%** | **1023 (98.2%)** | **577 (55.4%)** | **1/18** | |

Codon mutations: TTA272CTA (4219T>C), TTA273AAA (4222T>A 4223T>A), GAC274GAT (4227C>T), ACT275CCC (4228A>C 4230T>C), AAC276AGC (4232A>G), AAA277ATC (4235A>T 4236A>C), ATT278ATA (4239T>A), ATT279ATA (4242T>A), AAG280AGA (4244A>G 4245G>A), TGT281GTT (4246T>G 4247G>T), CCA283CCT (4254A>T), CAG285CTA (4259A>T 4260G>A), TAT286TAC (4263T>C), TCA287AAT (4264T>A 4265C>A 4266A>T), CCT288GAA (4267C>G 4268C>A 4269T>A), GAC290GAT (4275C>T), CGA291AAA (4276C>A 4277G>A), GAA292ACT (4279G>A 4280A>C 4281A>T), GAG293GAA (4284G>A), ACT296ATT (4292C>T), ATC298ATT (4299C>T), GAA299CAA (4300G>C), CTA301TTA (4306C>T), CTC302TTA (4309C>T 4311C>A), AAG303AAT (4314G>T), CTT304ATG (4315C>A 4317T>G), GGT305AAA (4318G>A 4319G>A 4320T>A), ATC306TTA (4321A>T 4323C>A), ATC307ATA (4326C>A), CCT309CAT (4331C>A), AGC310AGT (4335C>T), AAG311AAA (4338G>A), TCC315AGT (4348T>A 4349C>G 4350C>T), TCA316AGC (4351T>A 4352C>G 4353A>C), CCA317CCC (4356A>C), GCC318GCA (4359C>A), TTC319TTT (4362C>T), GTA321ATA (4366G>A), AGG322AGA (4371G>A), AAT323AAC (4374T>C), GCC325GGC (4379C>G), GAG326GAA (4383G>A), ATC327CAA (4384A>C 4385T>A 4386C>A), CGA329CGT (4392A>T), GGC330GGA (4395C>A), GCA332GCT (4401A>T), AGA333CGA (4402A>C), ATT336ATA (4413T>A), AAG340GAG (4423A>G), TTA341CTT (4426T>C 4428A>T), GAC343AAA (4432G>A 4434C>A), CAT344AAT (4435C>A), ACA345TGT (4438A>T 4439C>G 4440A>T), AAG346GTC (4441A>G 4442A>T 4443G>C), GGA347TTC (4444G>T 4445G>T 4446A>C), GAT348GAC (4449T>C), GGC349GGA (4452C>A), TAC350TAT (4455C>T), CTA351TTC (4456C>T 4458A>C), CTT352ATC (4459C>A 4461T>C), CCT353CCA (4464T>A), AAC354AAT (4467C>T), AAG355AAA (4470G>A), GAG356AAT (4471G>A 4473G>T), CAA357AAC (4474C>A 4476A>C), CTG358CTT (4479G>T), CTT359ATA (4480C>A 4482T>A), CAA360AAT (4483C>A 4485A>T), AGA361CTT (4486A>C 4487G>T 4488A>T), ATC362GCC (4489A>G 4490T>C), GGA363AAA (4492G>A 4493G>A), GGT364GGA (4497T>A), AAG365AAA (4500G>A), TTT367TAC (4505T>A 4506T>C), TAC368TTT (4508A>T 4509C>T), TCT369TCA (4512T>A), TCC370AAA (4513T>A 4514C>A 4515C>A), GAC372GAT (4521C>T), TGT373TGC (4524T>C), TCT375AGT (4528T>A 4529C>G), CAG379CAA (4542G>A), GTA380ATA (4543G>A), CGC381AAA (4546C>A 4547G>A 4548C>A), CTT382CTA (4551T>A), GCT383GAA (4553C>A 4554T>A), CCA384GAA (4555C>G 4556C>A), GAA385AAT (4558G>A 4560A>T), ACA386TCT (4561A>T 4563A>T), ATT387ATA (4566T>A), CAG388GCT (4567C>G 4568A>C 4569G>T), CTA389TTA (4570C>T), ACC390ACA (4575C>A), GCT391GCG (4578T>G), AGC393AGT (4584C>T), TGT394ACG (4585T>A 4586G>C 4587T>G), CCC395CCA (4590C>A), CAA396AAT (4591C>A 4593A>T), CAC398CAT (4599C>T), GTC403GTG (4614C>G), CAA410AAT (4633C>A 4635A>T), GCT413CAA (4642G>C 4643C>A 4644T>A), ATT414ATC (4647T>C), TTC415TTT (4650C>T), CAA416CAG (4653A>G), CAC418AAA (4657C>A 4659C>A), GAC420GAT (4665C>T), GAA421AAA (4666G>A), AGT422ATA (4670G>T 4671T>A), CTC423TTT (4672C>T 4674C>T), AGC424TCC (4675A>T 4676G>C), AAC425CAA (4678A>C 4680C>A), ATG426AAG (4682T>A), TAT427AAT (4684T>A), CCA428_CAG429del (4687_4692delCCACAG), TGT431CTA (4696T>C 4697G>T 4698T>A), GCT432ATC (4699G>A 4700C>T 4701T>C), GTA433GTC (4704A>C), TAT434TAC (4707T>C), GTT435ATA (4708G>A 4710T>A), GAC437GAT (4716C>T), ATC438ATA (4719C>A), ATC439TTA (4720A>T 4722C>A), GTC440ATA (4723G>A 4725C>A), TTC441TGT (4727T>G 4728C>T), AGC442TCC (4729A>T 4730G>C), AAA443AAC (4734A>C), ACT444TCA (4735A>T 4737T>A), GAA445TAT (4738G>T 4740A>T), GAA446CAA (4741G>C), GGA450ACT (4753G>A 4754G>C 4755A>T), GTC452CTT (4759G>C 4761C>T), ATT454CAA (4765A>C 4766T>A 4767T>A), GTT455TTT (4768G>T), CTA456TCT (4771C>T 4772T>C 4773A>T), AAC457CAA (4774A>C 4776C>A), AGA458ATA (4778G>T), TGC459TGT (4782C>T), AAA460TGC (4783A>T 4784A>G 4785A>C), GCC461GAA (4787C>A 4788C>A), CTA462AAT (4789C>A 4790T>A 4791A>T), ATA464ATT (4797A>T), CTT466TTA (4801C>T 4803T>A), AAA468CAG (4807A>C 4809A>G), AAG469AAA (4812G>A), AAA470AAG (4815A>G), GCC471GCA (4818C>A), CAG472GAC (4819C>G 4821G>C), TTG473ATA (4822T>A 4824G>A), TGC474AAT (4825T>A 4826G>A 4827C>T), ACA476AAT (4832C>A 4833A>T), ACT477GAA (4834A>G 4835C>A 4836T>A), ATC478ATT (4839C>T), AAT479GAA (4840A>G 4842T>A), TTC480TTT (4845C>T), CTT481CTC (4848T>C), GGT482GGA (4851T>A), CTA483ATG (4852C>A 4854A>G), GTA484ATA (4855G>A), ATA485ATT (4860A>T), GAA486TCA (4861G>T 4862A>C), AGA487AAA (4865G>A), GGA488AAT (4867G>A 4868G>A 4869A>T), AAT489GGC (4870A>G 4871A>G 4872T>C), CTC490ATT (4873C>A 4875C>T), AAA491GAA (4876A>G), GTC492TTA (4879G>T 4881C>A), CAG493CAA (4884G>A), AGT494ACT (4886G>C), CAT495CAC (4890T>C), ATC496ATA (4893C>A), GGA497TCA (4894G>T 4895G>C), CTA498AAA (4897C>A 4898T>A), CAC499AAA (4900C>A 4902C>A), CAC499_TTA500insG-- (4902_4903insG), TTA500ATC (4903T>A 4905A>C), GTT501TGT (4906G>T 4907T>G), GCT502GAA (4910C>A 4911T>A), CCA504TCA (4915C>T), CAA506AAA (4921C>A), CTT507CTA (4926T>A), AGT508AAC (4928G>A 4929T>C), GAT509ACA (4930G>A 4931A>C 4932T>A), AGG510AAA (4934G>A 4935G>A), AAT511CAA (4936A>C 4938T>A), GCC512GAA (4940C>A 4941C>A), TTA513ATC (4942T>A 4944A>C), AGA515AGG (4950A>G), TTA517CTT (4954T>C 4956A>T), GGC518GGA (4959C>A), CTC519TGT (4960C>T 4961T>G 4962C>T), CTA520TTA (4963C>T), TAT522TAC (4971T>C), ATC523GCA (4972A>G 4973T>C 4974C>A), TCC524AGT (4975T>A 4976C>G 4977C>T), GCT525GAA (4979C>A 4980T>A), TAT526TTT (4982A>T), TTT527ATA (4984T>A 4986T>A), AAA529GAC (4990A>G 4992A>C), ATA530CTA (4993A>C), AAC532AAA (5001C>A), CTT533AAG (5002C>A 5003T>A 5004T>G), AGG534AGA (5007G>A), TCA535AAT (5008T>A 5009C>A 5010A>T), CCT536CTT (5012C>T), CAG538CAA (5019G>A), GTT539AGG (5020G>A 5021T>G 5022T>G), AAA540CTT (5023A>C 5024A>T 5025A>T), CTT541ATT (5026C>A), AAG542CGA (5029A>C 5030A>G 5031G>A), GAG544TCC (5035G>T 5036A>C 5037G>C), ATA545AAC (5039T>A 5040A>C), ACT546CGA (5041A>C 5042C>G 5043T>A), TGG547ATG (5044T>A 5045G>T), TCT548GGA (5047T>G 5048C>G 5049T>A), ACT550ACA (5055T>A), GAG551GAT (5058G>T), AAA552GAA (5059A>G), GAC553CAT (5062G>C 5064C>T), ACG554ACT (5067G>T), GAA555AAA (5068G>A), ACT556TGT (5071A>T 5072C>G), GTC557GTT (5076C>T), AGA558CAA (5077A>C 5078G>A), AAA559AGC (5081A>G 5082A>C), ATA560CTA (5083A>C), AAG561AAA (5088G>A), AGT562CAA (5089A>C 5090G>A 5091T>A), CTA563GAA (5092C>G 5093T>A), GTA564TGT (5095G>T 5096T>G 5097A>T), AAA565TCC (5098A>T 5099A>C 5100A>C), ACT566AAA (5102C>A 5103T>A), GAC569AAG (5110G>A 5112C>G), CTT570CTA (5115T>A), TAC571AGA (5116T>A 5117A>G 5118C>A), AAT572CTT (5119A>C 5120A>T), CCT573CCG (5124T>G), TCA574GAA (5125T>G 5126C>A), GAG576GAA (5133G>A), GAT577GAC (5136T>C), AAG578AAC (5139G>C), CCT579TTA (5140C>T 5141C>T 5142T>A), ATC580GTT (5143A>G 5145C>T), ATC581TTA (5146A>T 5148C>A), GAA582CAA (5149G>C), TGC583ACA (5152T>A 5153G>C 5154C>A), GCA585GCT (5160A>T), GAC588TAC (5167G>T), ATC593TTA (5182A>T 5184C>A), CTC594TTA (5185C>T 5187C>A), AAA595CAA (5188A>C), GCA596ACT (5191G>A 5193A>T), AAA597GAC (5194A>G 5196A>C), CCA599AAT (5200C>A 5201C>A 5202A>T), GGT601_GTA604del (5206_5217delGGTAAAGAGGTA), ATA605ATT (5220A>T), TGC606TGT (5223C>T), GCT609ACA (5230G>A 5232T>A), TCA610AGT (5233T>A 5234C>G 5235A>T), GGA611GGT (5238A>T), ACC612ACA (5241C>A), AAA614AAT (5247A>T), CCA615GAA (5248C>G 5249C>A), GCT616ACC (5251G>A 5253T>C), GAG617GAA (5256G>A), AAG618ACT (5258A>C 5259G>T)

*: Inserts / Deletes / Misaligned / Frameshifts

#### Analysis details

This analysis was performed with panviral2.64

### NGS Details (UN60): Caulimovirus tesselloscrophulariae

#### Assembly

| | |
|---|---|
| **Coverage Length** | 373 (1 contig(s)) |
| **Depth Of Coverage** | 14.5 |
| **Number Of Reads** | 48 |
| **Reads Per Million** | 1.02 rpm (after QC) |
| **Ambiguities** | 0 |
| **Assembly Method** | de novo + reference guided assembly |
| **Consensus Caller** | Bcf Tools |

#### Coverage Map

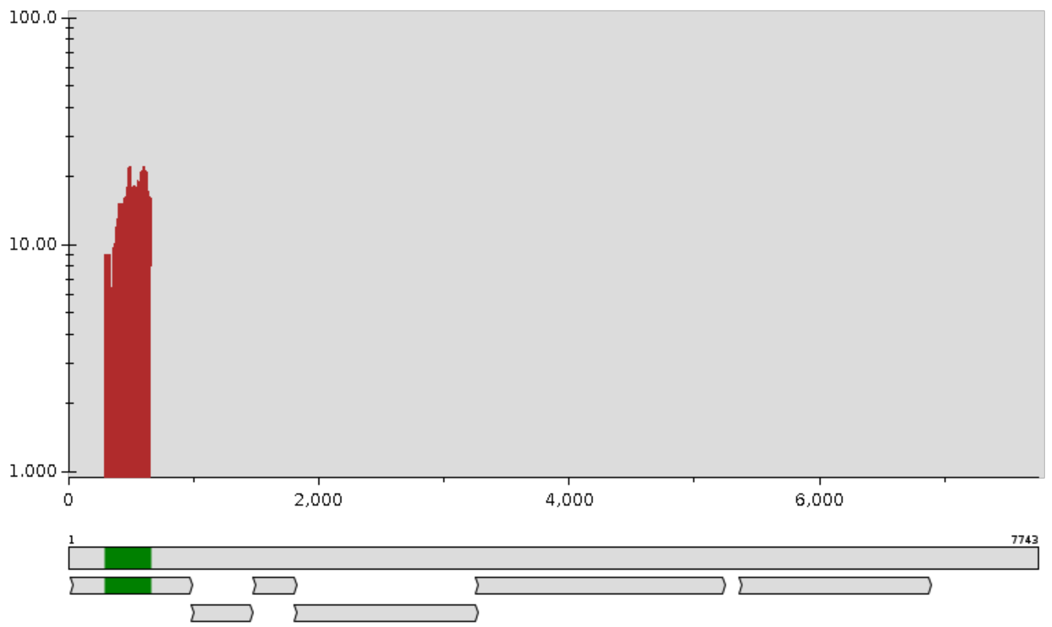

#### Assignment

| | |
|---|---|
| **Type** | Caulimovirus tesselloscrophulariae (Taxonomy ID: 3047834) |
| **Reference Genome** | NC_003554.1 |
| **NT Identity (%)** | 53.6193 |
| **AA Identity (%)** | 37.9032 |
| **Number Of Stop Codons** | 0 |
| **Number Of CDS** | 6 |

#### Alignment

| | |
|---|---|
| **Alignment Score** | 54.0 (NT) + 317.0 (AA) = 371.0 |
| **Concordance (%)** | 24.4079 |

| Alignment Method | Global, seeded, nucleotide + amino acids (AGA) |
|---|---|

#### Genome Region

Sequence starts at position 288 and ends at position 660 relative to NC_003554.1 reference sequence.

#### Alignment Detailed Statistics

| | Begin | End | Coverage | Score | Concordance | Matches | Identities | I/D/M/F* | Stop Codons |
|---|---|---|---|---|---|---|---|---|---|
| **NT** | **288** | **660** | **4.8%** | **54** | **7.2%** | **373 (100%)** | **200 (53.6%)** | **0/0** | |
| Mutations: | 288T>C, 289C>T, 290T>A, 296C>A, 299G>C, 300T>A, 302T>A, 304T>A, 305C>T, 307T>A, 308C>A, 309C>A, 311C>T, 313T>A, 314A>T, 322G>A, 324A>C, 325G>A, 326G>A, 327A>T, 328A>G, 331A>T, 332C>A, 334A>T, 339A>T, 340G>C, 343G>A, 344A>C, 345T>A, 346G>A, 348A>G, 353G>C, 354A>C, 355T>A, 356C>G, 358A>C, 359T>A, 360C>G, 361A>C, 362G>A, 363T>A, 368A>C, 369G>A, 373A>G, 376C>G, 377T>A, 378C>A, 379A>T, 380A>T, 381T>A, 382G>T, 383A>G, 385C>T, 388C>T, 389C>T, 391T>G, 396C>G, 398G>A, 401A>C, 403G>A, 406T>C, 407C>A, 409A>G, 410C>G, 412T>G, 414C>A, 416G>T, 417C>T, 419C>A, 420A>C, 421A>T, 425C>A, 426G>A, 427G>A, 428C>G, 433A>T, 436T>A, 437G>A, 440A>T, 441C>G, 442G>T, 443T>C, 445A>T, 446G>A, 448T>A, 450A>T, 454G>A, 455G>A, 456C>A, 457A>C, 461A>T, 462T>C, 463C>T, 469T>A, 473A>T, 475C>T, 476G>A, 478C>T, 482C>A, 483G>C, 484A>T, 486A>G, 490T>A, 491A>G, 493T>A, 494C>A, 495T>A, 496G>T, 499C>T, 503G>A, 504C>T, 507C>T, 508A>T, 509A>G, 510G>A, 518C>T, 520C>A, 526T>C, 527G>A, 528G>C, 535T>A, 536A>C, 538G>T, 544C>A, 545G>T, 546T>A, 550C>T, 553T>C, 555A>G, 558T>A, 560G>T, 561C>G, 562A>T, 563T>A, 565G>A, 572C>A, 575A>G, 576G>A, 577C>T, 578A>G, 579A>C, 580G>T, 583T>C, 586A>T, 587G>A, 590A>G, 593A>G, 598A>G, 601C>T, 604T>G, 605A>C, 606T>A, 608C>T, 609A>T, 613A>G, 614T>G, 616T>C, 617G>A, 628C>T, 632A>T, 634G>C, 637A>G, 638A>C, 643C>T, 644G>A, 647A>C, 649A>T, 650G>A, 653T>A, 655T>G, 657C>G, 658G>T, 659G>A | | | | | | | | |
| **CDS** | | | | | | | | | |
| FMVgp1 | 93 | 216 | 38.3% | 317 | 40.4% | 124 (100%) | 47 (37.9%) | 0/0/0/0 | 0 |
| Protein mutations: | S93T (290T>A), Q95K (296C>A), V96H (299G>C 300T>A), Y97K (302T>A 304T>A), P99N (308C>A 309C>A), I101L (314A>T), Q104P (324A>C 325G>A), E105M (326G>A 327A>T 328A>G), Q107N (332C>A 334A>T), K109I (339A>T 340G>C), M111Q (344A>C 345T>A 346G>A), K112R (348A>G), D114P (353G>C 354A>C 355T>A), P115A (356C>G 358A>C), V117K (362G>A 363T>A), S119H (368A>C 369G>A), I121M (376C>G), S122N (377T>A 378C>A 379A>T), M123Y (380A>T 381T>A 382G>T), I124V (383A>G 385C>T), A128G (396C>G), V129I (398G>A), K130Q (401A>C 403G>A), L132M (407C>A 409A>G), L133V (410C>G 412T>G), T134K (414C>A), A135F (416G>T 417C>T), Q136T (419C>A 420A>C 421A>T), R138K (425C>A 426G>A 427G>A), Q139E (428C>G), D142N (437G>A), T143C (440A>T 441C>G 442G>T), S144P (443T>C 445A>T), V145I (446G>A 448T>A), K146I (450A>T), M147I (454G>A), A148N (455G>A 456C>A 457A>C), I150S (461A>T 462T>C 463C>T), D152E (469T>A), I154F (473A>T 475C>T), V155I (476G>A 478C>T), R157T (482C>A 483G>C 484A>T), K158R (486A>G), D159E (490T>A), S160G (491A>G 493T>A), L161N (494C>A 495T>A 496G>T), A164I (503G>A 504C>T), A165V (507C>T 508A>T), R166E (509A>G 510G>A), G172T (527G>A 528G>C), F174L (535T>A), M175L (536A>C 538G>T), V178Y (545G>T 546T>A), K181R (555A>G), F182Y (558T>A), A183C (560G>T 561C>G 562A>T), L184I (563T>A 565G>A), Q187K (572C>A), S188D (575A>G 576G>A 577C>T), K189A (578A>G 579A>C 580G>T), L191F (586A>T), D192N (587G>A), K193E (590A>G), T194A (593A>G), F197L (604T>G), I198H (605A>C 606T>A), H199F (608C>T 609A>T), F201V (614T>G 616T>C), E202K (617G>A), M207F (632A>T 634G>C), T209P (638A>C), D211N (644G>A), K212H (647A>C 649A>T), V213I (650G>A), F214M (653T>A 655T>G), T215S (657C>G 658G>T) | | | | | | | | |
| Codon mutations: | GTC92.CT (288T>C 289C>T), TCT93ACT (290T>A), CAA95AAA (296C>A), GTT96CAT (299G>C 300T>A), TAT97AAA (302T>A 304T>A), CTT98TTA (305C>T 307T>A), CCT99AAT (308C>A 309C>A), CTT100TTA (311C>T 313T>A), ATA101TTA (314A>T), AAG103AAA (322G>A), CAG104CCA (324A>C 325G>A), GAA105ATG (326G>A 327A>T 328A>G), ATA106ATT (331A>T), CAA107AAT (332C>A 334A>T), AAG109ATC (339A>T 340G>C), CTG110CTA (343G>A), ATG111CAA (344A>C 345T>A 346G>A), AAA112AGA (348A>G), GAT114CCA (353G>C 354A>C 355T>A), CCA115GCC (356C>G 358A>C), TCA116AGC (359T>A 360C>G 361A>C), GTA117AAA (362G>A 363T>A), AGT119CAT (368A>C 369G>A), AAA120AAG (373A>G), ATC121ATG (376C>G), TCA122AAT (377T>A 378C>A 379A>T), ATG123TAT (380A>T 381T>A 382G>T), ATC124GTT (383A>G 385C>T), CAC125CAT (388C>T), CTT126TTG (389C>T 391T>G), GCA128GGA (396C>G), GTA129ATA (398G>A), AAG130CAA (401A>C 403G>A), ATT131ATC (406T>C), CTA132ATG (407C>A 409A>G), CTT133GTG (410C>G 412T>G), ACA134AAA (414C>A), GCC135TTC (416G>T 417C>T), CAA136ACT (419C>A 420A>C 421A>T), CGG138AAA (425C>A 426G>A 427G>A), CAA139GAA (428C>G), GGA140GGT (433A>T), ATT141ATA (436T>A), GAT142AAT (437G>A), ACG143TGT (440A>T 441C>G 442G>T), TCA144CCT (443T>C 445A>T), GTT145ATA (446G>A 448T>A), AAA146ATA (450A>T), ATG147ATA (454G>A), GCA148AAC (455G>A 456C>A 457A>C), ATC150TCT (461A>T 462T>C 463C>T), GAT152GAA (469T>A), ATC154TTT (473A>T 475C>T), GTC155ATT (476G>A 478C>T), CGA157ACT (482C>A 483G>C 484A>T), AAA158AGA (486A>G), GAT159GAA (490T>A), AGT160GGA (491A>G 493T>A), CTG161AAT (494C>A 495T>A 496G>T), CTC162CTT (499C>T), GCA164ATA (503G>A 504C>T), GCA165GTT (507C>T 508A>T), AGA166GAA (509A>G 510G>A), CTC169TTA (518C>T 520C>A), TAT171TAC (526T>C), GGT172ACT (527G>A 528G>C), TTT174TTA (535T>A), ATG175CTT (536A>C 538G>T), ACC177ACA (544C>A), GTT178TAT (545G>T 546T>A), TAC179TAT (550C>T), CCT180CCC (553T>C), AAA181AGA (555A>G), TTT182TAT (558T>A), GCA183TGT (560G>T 561C>G 562A>T), TTG184ATA (563T>A 565G>A), CAA187AAA (572C>A), AGC188GAT (575A>G 576G>A 577C>T), AAG189GCT (578A>G 579A>C 580G>T), AAT190AAC (583T>C), TTA191TTT (586A>T), GAT192AAT (587G>A), AAA193GAA (590A>G), ACA194GCA (593A>G), TTA195TTG (598A>G), AGC196AGT (601C>T), TTT197TTG (604T>G), ATT198CAT (605A>C 606T>A), CAT199TTT (608C>T 609A>T), CAA200CAG (613A>G), TTT201GTC (614T>G 616T>C), GAA202AAA (617G>A), GAC205GAT (628C>T), ATG207TTC (632A>T 634G>C), AAA208AAG (637A>G), ACA209CCA (638A>C), GGC210GGT (643C>T), GAT211AAT (644G>A), AAA212CAT (647A>C 649A>T), GTT213ATT (650G>A), TTT214ATG (653T>A 655T>G), ACG215AGT (657C>G 658G>T), GTC216AT. (659G>A) | | | | | | | | |
| **Proteins** | | | | | | | | | |
| hypothetical protein (NP_619544.1) | 93 | 216 | 38.3% | 317 | 40.4% | 124 (100%) | 47 (37.9%) | 0/0/0/0 | 0 |
| Protein mutations: | S93T (290T>A), Q95K (296C>A), V96H (299G>C 300T>A), Y97K (302T>A 304T>A), P99N (308C>A 309C>A), I101L (314A>T), Q104P (324A>C 325G>A), E105M (326G>A 327A>T 328A>G), Q107N (332C>A 334A>T), K109I (339A>T 340G>C), M111Q (344A>C 345T>A 346G>A), K112R (348A>G), D114P (353G>C 354A>C 355T>A), P115A (356C>G 358A>C), V117K (362G>A 363T>A), S119H (368A>C 369G>A), I121M (376C>G), S122N (377T>A 378C>A 379A>T), M123Y (380A>T 381T>A 382G>T), I124V (383A>G 385C>T), A128G (396C>G), V129I (398G>A), K130Q (401A>C 403G>A), L132M (407C>A 409A>G), L133V (410C>G 412T>G), T134K (414C>A), A135F (416G>T 417C>T), Q136T (419C>A 420A>C 421A>T), R138K (425C>A 426G>A 427G>A), Q139E (428C>G), D142N (437G>A), T143C (440A>T 441C>G 442G>T), S144P (443T>C 445A>T), V145I (446G>A 448T>A), K146I (450A>T), M147I (454G>A), A148N (455G>A 456C>A 457A>C), I150S (461A>T 462T>C 463C>T), D152E (469T>A), I154F (473A>T 475C>T), V155I (476G>A 478C>T), R157T (482C>A 483G>C 484A>T), K158R (486A>G), D159E (490T>A), S160G (491A>G 493T>A), L161N (494C>A 495T>A 496G>T), A164I (503G>A 504C>T), A165V (507C>T 508A>T), R166E (509A>G 510G>A), G172T (527G>A 528G>C), F174L (535T>A), M175L (536A>C 538G>T), V178Y (545G>T 546T>A), K181R (555A>G), F182Y (558T>A), A183C (560G>T 561C>G 562A>T), L184I (563T>A 565G>A), Q187K (572C>A), S188D (575A>G 576G>A 577C>T), K189A (578A>G 579A>C 580G>T), L191F (586A>T), D192N (587G>A), K193E (590A>G), T194A (593A>G), F197L (604T>G), I198H (605A>C 606T>A), H199F (608C>T 609A>T), F201V (614T>G 616T>C), E202K (617G>A), M207F (632A>T 634G>C), T209P (638A>C), D211N (644G>A), K212H (647A>C 649A>T), V213I (650G>A), F214M (653T>A 655T>G), T215S (657C>G 658G>T) | | | | | | | | |
| Codon mutations: | GTC92.CT (288T>C 289C>T), TCT93ACT (290T>A), CAA95AAA (296C>A), GTT96CAT (299G>C 300T>A), TAT97AAA (302T>A 304T>A), CTT98TTA (305C>T 307T>A), CCT99AAT (308C>A 309C>A), CTT100TTA (311C>T 313T>A), ATA101TTA (314A>T), AAG103AAA (322G>A), CAG104CCA (324A>C 325G>A), GAA105ATG (326G>A 327A>T 328A>G), ATA106ATT (331A>T), CAA107AAT (332C>A 334A>T), AAG109ATC (339A>T 340G>C), CTG110CTA (343G>A), ATG111CAA (344A>C 345T>A 346G>A), AAA112AGA (348A>G), GAT114CCA (353G>C 354A>C 355T>A), CCA115GCC (356C>G 358A>C), TCA116AGC (359T>A 360C>G 361A>C), GTA117AAA (362G>A 363T>A), AGT119CAT (368A>C 369G>A), AAA120AAG (373A>G), ATC121ATG (376C>G), TCA122AAT (377T>A 378C>A 379A>T), ATG123TAT (380A>T 381T>A 382G>T), ATC124GTT (383A>G 385C>T), CAC125CAT (388C>T), CTT126TTG (389C>T 391T>G), GCA128GGA (396C>G), GTA129ATA (398G>A), AAG130CAA (401A>C 403G>A), ATT131ATC (406T>C), CTA132ATG (407C>A 409A>G), CTT133GTG (410C>G 412T>G), ACA134AAA (414C>A), GCC135TTC (416G>T 417C>T), CAA136ACT (419C>A 420A>C 421A>T), CGG138AAA (425C>A 426G>A 427G>A), CAA139GAA (428C>G), GGA140GGT (433A>T), ATT141ATA (436T>A), GAT142AAT (437G>A), ACG143TGT (440A>T 441C>G 442G>T), TCA144CCT (443T>C 445A>T), GTT145ATA (446G>A 448T>A), AAA146ATA (450A>T), ATG147ATA (454G>A), GCA148AAC (455G>A 456C>A 457A>C), ATC150TCT (461A>T 462T>C 463C>T), GAT152GAA (469T>A), ATC154TTT (473A>T 475C>T), GTC155ATT (476G>A 478C>T), CGA157ACT (482C>A 483G>C 484A>T), AAA158AGA (486A>G), GAT159GAA (490T>A), AGT160GGA (491A>G 493T>A), CTG161AAT (494C>A 495T>A 496G>T), CTC162CTT (499C>T), GCA164ATA (503G>A 504C>T), GCA165GTT (507C>T 508A>T), AGA166GAA (509A>G 510G>A), CTC169TTA (518C>T 520C>A), TAT171TAC (526T>C), GGT172ACT (527G>A 528G>C), TTT174TTA (535T>A), ATG175CTT (536A>C 538G>T), ACC177ACA (544C>A), GTT178TAT (545G>T 546T>A), TAC179TAT (550C>T), CCT180CCC (553T>C), AAA181AGA (555A>G), TTT182TAT (558T>A), GCA183TGT (560G>T 561C>G 562A>T), TTG184ATA (563T>A 565G>A), CAA187AAA (572C>A), AGC188GAT (575A>G 576G>A 577C>T), AAG189GCT (578A>G 579A>C 580G>T), AAT190AAC (583T>C), TTA191TTT (586A>T), GAT192AAT (587G>A), AAA193GAA (590A>G), ACA194GCA (593A>G), TTA195TTG (598A>G), AGC196AGT (601C>T), TTT197TTG (604T>G), ATT198CAT (605A>C 606T>A), CAT199TTT (608C>T 609A>T), CAA200CAG (613A>G), TTT201GTC (614T>G 616T>C), GAA202AAA (617G>A), GAC205GAT (628C>T), ATG207TTC (632A>T 634G>C), AAA208AAG (637A>G), ACA209CCA (638A>C), GGC210GGT (643C>T), GAT211AAT (644G>A), AAA212CAT (647A>C 649A>T), GTT213ATT (650G>A), TTT214ATG (653T>A 655T>G), ACG215AGT (657C>G 658G>T), GTC216AT. (659G>A) | | | | | | | | |

*: Inserts / Deletes / Misaligned / Frameshifts

#### Analysis details

This analysis was performed with panviral2.64

### NGS Details (UN60): Brazilian marseillevirus

#### Assembly

| | |
|---|---|
| **Coverage Length** | 138 (1 contig(s)) |
| **Depth Of Coverage** | 25.7 |
| **Number Of Reads** | 47 |
| **Reads Per Million** | 1.00 rpm (after QC) |
| **Ambiguities** | 0 |
| **Assembly Method** | de novo + reference guided assembly |
| **Consensus Caller** | Bcf Tools |

#### Coverage Map

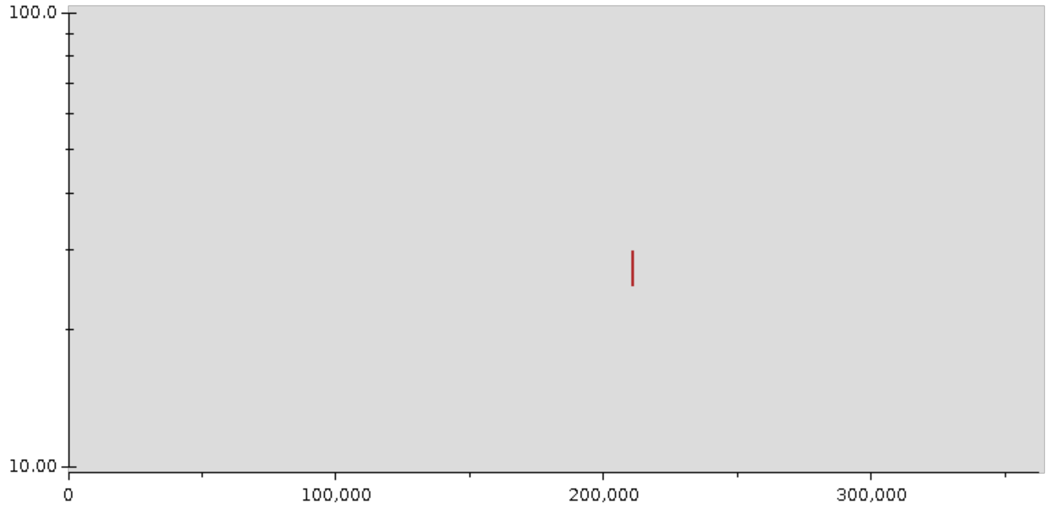

#### Assignment

| | |
|---|---|
| **Type** | Brazilian marseillevirus (Taxonomy ID: 1813599) |
| **Reference Genome** | NC_029692.1 |
| **NT Identity (%)** | 81.1594 |
| **AA Identity (%)** | 95.6522 |
| **Number Of Stop Codons** | 0 |
| **Number Of CDS** | 491 |

#### Alignment

| | |
|---|---|
| **Alignment Score** | 172.0 (NT) + 303.0 (AA) = 475.0 |
| **Concordance (%)** | 81.4751 |
| **Alignment Method** | Local, heuristic, nucleotide (BLASTN) |

#### Genome Region

Sequence starts at position 210893 and ends at position 211030 relative to NC_029692.1 reference sequence.

#### Alignment Detailed Statistics

| | Begin | End | Coverage | Score | Concordance | Matches | Identities | I/D/M/F* | Stop Codons |
|---|---|---|---|---|---|---|---|---|---|
| **NT** | **210893** | **211030** | **0.1%** | **172** | **62.3%** | **138 (100%)** | **112 (81.2%)** | **0/0** | |

Mutations: 210901T>G, 210904G>A, 210908A>C, 210910G>A, 210920G>A, 210932G>A, 210938A>G, 210944C>G, 210945G>C, 210946C>T, 210947A>C, 210949G>A, 210950G>A, 210953C>A, 210955T>G, 210965C>A, 210971C>T, 210974G>A, 210977T>A, 210989G>A, 210992T>C, 211001A>G, 211004A>T, 211007A>G, 211009C>T, 211010T>A

*: Inserts / Deletes / Misaligned / Frameshifts

#### Analysis details

This analysis was performed with panviral2.64

### NGS Details (UN60): Errantivirus

#### Assembly

| | |
|---|---|
| **Coverage Length** | 300 (1 contig(s)) |
| **Depth Of Coverage** | 16.8 |
| **Number Of Reads** | 46 |
| **Reads Per Million** | 0.98 rpm (after QC) |
| **Ambiguities** | 0 |
| **Assembly Method** | de novo + reference guided assembly |
| **Consensus Caller** | Bcf Tools |

#### Coverage Map

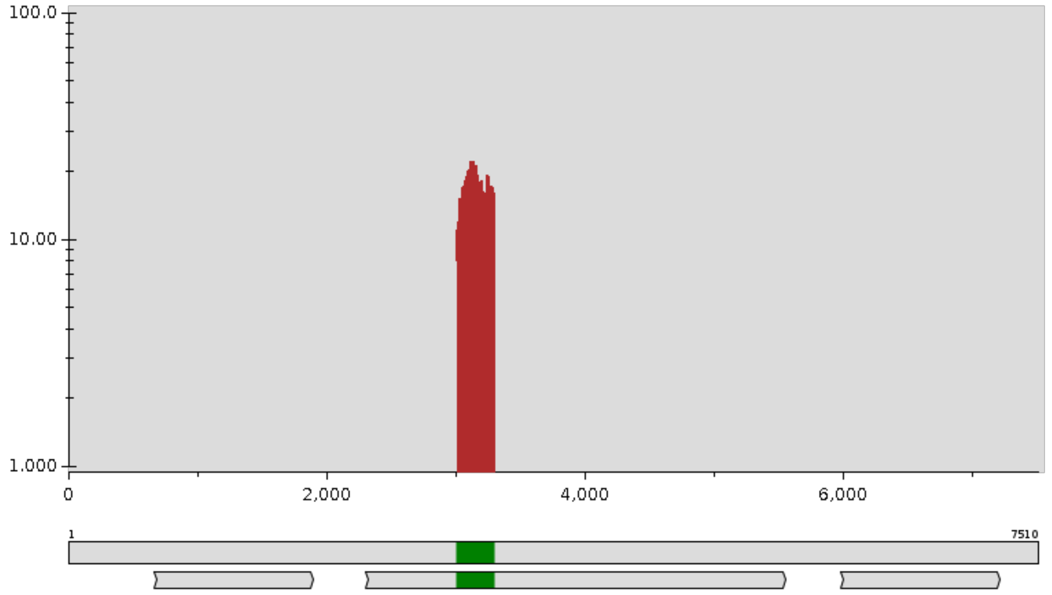

#### Assignment

| | |
|---|---|
| **Type** | Errantivirus (Taxonomy ID: 186666) |
| **Reference Genome** | NC_038512.1 |
| **NT Identity (%)** | 53.0 |
| **AA Identity (%)** | 41.0 |
| **Number Of Stop Codons** | 0 |
| **Number Of CDS** | 3 |

#### Alignment

| | |
|---|---|
| **Alignment Score** | 36.0 (NT) + 301.0 (AA) = 337.0 |
| **Concordance (%)** | 26.1646 |

| | |
|---|---|
| **Alignment Method** | Global, seeded, nucleotide + amino acids (AGA) |

#### Genome Region

Sequence starts at position 3007 and ends at position 3306 relative to NC_038512.1 reference sequence.

#### Alignment Detailed Statistics

| | Begin | End | Coverage | Score | Concordance | Matches | Identities | I/D/M/F* | Stop Codons |
|---|---|---|---|---|---|---|---|---|---|
| **NT** | **3007** | **3306** | **4.0%** | **36** | **6.0%** | **300 (100%)** | **159 (53.0%)** | **0/0** | |
| Mutations: | 3007G>T, 3008G>A, 3009C>G, 3011A>G, 3012G>C, 3016G>A, 3018T>A, 3020T>C, 3022T>C, 3026G>T, 3029G>A, 3030A>G, 3031G>A, 3035G>T, 3037C>T, 3038C>G, 3039C>A, 3040T>A, 3041C>A, 3043A>G, 3047A>G, 3048T>C, 3049A>T, 3050T>C, 3051C>A, 3052G>C, 3055A>G, 3058C>T, 3061G>C, 3067C>A, 3068G>A, 3069T>C, 3070A>T, 3071G>C, 3073A>T, 3074C>G, 3076C>A, 3082T>C, 3084T>A, 3088A>G, 3091C>T, 3094T>G, 3095C>G, 3096G>T, 3103T>C, 3104A>T, 3106G>T, 3109A>G, 3114A>C, 3115A>C, 3118C>T, 3119T>G, 3121A>C, 3124A>T, 3128A>T, 3133T>C, 3138G>A, 3139A>C, 3140G>C, 3142T>C, 3146G>A, 3149A>T, 3150A>C, 3151T>A, 3154C>A, 3155C>T, 3157A>T, 3161G>C, 3162G>C, 3163T>A, 3164C>T, 3166C>A, 3167C>T, 3168A>T, 3171A>G, 3172T>G, 3175C>A, 3178C>T, 3179T>G, 3180G>T, 3181T>C, 3182C>T, 3184C>G, 3187C>T, 3189A>T, 3191C>T, 3196C>T, 3199T>C, 3202T>C, 3203A>C, 3206G>A, 3211T>C, 3214T>C, 3216C>A, 3217T>G, 3221C>A, 3223A>G, 3224C>A, 3225A>C, 3226G>T, 3229A>T, 3233C>A, 3235G>T, 3238G>A, 3239A>C, 3241C>T, 3242C>G, 3244G>T, 3245G>A, 3248C>G, 3249G>C, 3253T>G, 3256C>T, 3257C>G, 3259A>T, 3260A>C, 3261G>T, 3262A>G, 3263C>A, 3265T>G, 3266A>T, 3267G>T, 3269G>A, 3271A>G, 3272A>T, 3273G>T, 3275A>C, 3277C>G, 3280C>A, 3281A>T, 3283A>T, 3284A>G, 3285T>C, 3287C>A, 3293G>T, 3294A>C, 3295C>T, 3298G>A, 3300C>G, 3301C>T, 3303A>C | | | | | | | | |

##### CDS

| | Begin | End | Coverage | Score | Concordance | Matches | Identities | I/D/M/F* | Stop Codons |
|---|---|---|---|---|---|---|---|---|---|
| D1R33_gp2 | 235 | 334 | 9.2% | 301 | 42.8% | 100 (100%) | 41 (41.0%) | 0/0/0/0 | 0 |
| Protein mutations: | A235R (3008G>A 3009C>G), S236A (3011A>G 3012G>C), F238Y (3018T>A), Y239H (3020T>C 3022T>C), V241L (3026G>T), E242R (3029G>A 3030A>G 3031G>A), D244Y (3035G>T 3037C>T), P245E (3038C>G 3039C>A 3040T>A), Q246K (3041C>A 3043A>G), I248A (3047A>G 3048T>C 3049A>T), S249H (3050T>C 3051C>A 3052G>C), N254K (3067C>A), V255T (3068G>A 3069T>C 3070A>T), E256H (3071G>C 3073A>T), H257E (3074C>G 3076C>A), F260Y (3084T>A), R264V (3095C>G 3096G>T), M267F (3104A>T 3106G>T), K270T (3114A>C 3115A>C), S272A (3119T>G 3121A>C), T275S (3128A>T), R278N (3138G>A 3139A>C), V279L (3140G>C 3142T>C), D281N (3146G>A), N282S (3149A>T 3150A>C 3151T>A), L284F (3155C>T 3157A>T), G286P (3161G>C 3162G>C 3163T>A), Q288L (3167C>T 3168A>T), N289R (3171A>G 3172T>G), N290K (3175C>A), C292V (3179T>G 3180G>T 3181T>C), Y295F (3189A>T), L296F (3191C>T), I300L (3203A>C), V301I (3206G>A), T304K (3216C>A 3217T>G), L306M (3221C>A 3223A>G), Q307T (3224C>A 3225A>C 3226G>T), E308D (3229A>T), L310I (3233C>A 3235G>T), N312H (3239A>C 3241C>T), L313V (3242C>G 3244G>T), E314K (3245G>A), R315A (3248C>G 3249G>C), Q318D (3257C>G 3259A>T), R319L (3260A>C 3261G>T 3262A>G), L320M (3263C>A 3265T>G), R321L (3266A>T 3267G>T), E322K (3269G>A 3271A>G), S323F (3272A>T 3273G>T), N324Q (3275A>C 3277C>G), F325L (3280C>A), K326Y (3281A>T 3283A>T), I327A (3284A>G 3285T>C), Q328K (3287C>A), D330S (3293G>T 3294A>C 3295C>T), S332C (3300C>G 3301C>T), E333A (3303A>C) | | | | | | | | |
| Codon mutations: | TTG234..T (3007G>T), GCA235AGA (3008G>A 3009C>G), AGT236GCT (3011A>G 3012G>C), GGG237GGA (3016G>A), TTT238TAT (3018T>A), TAT239CAC (3020T>C 3022T>C), GTG241TTG (3026G>T), GAG242AGA (3029G>A 3030A>G 3031G>A), GAC244TAT (3035G>T 3037C>T), CCT245GAA (3038C>G 3039C>A 3040T>A), CAA246AAG (3041C>A 3043A>G), ATA248GCT (3047A>G 3048T>C 3049A>T), TCG249CAC (3050T>C 3051C>A 3052G>C), AAA250AAG (3055A>G), ACC251ACT (3058C>T), GCG252GCC (3061G>C), AAC254AAA (3067C>A), GTA255ACT (3068G>A 3069T>C 3070A>T), GAA256CAT (3071G>C 3073A>T), CAC257GAA (3074C>G 3076C>A), CAT259CAC (3082T>C), TTT260TAT (3084T>A), GAA261GAG (3088A>G), TTC262TTT (3091C>T), CTT263CTG (3094T>G), CGA264GTA (3095C>G 3096G>T), CCT266CCC (3103T>C), ATG267TTT (3104A>T 3106G>T), GGA268GGG (3109A>G), AAA270ACC (3114A>C 3115A>C), AAC271AAT (3118C>T), TCA272GCC (3119T>G 3121A>C), CCA273CCT (3124A>T), ACT275TCT (3128A>T), TTT276TTC (3133T>C), AGA278AAC (3138G>A 3139A>C), GTT279CTC (3140G>C 3142T>C), GAC281AAC (3146G>A), AAT282TCA (3149A>T 3150A>C 3151T>A), GTC283GTA (3154C>A), CTA284TTT (3155C>T 3157A>T), GGT286CCA (3161G>C 3162G>C 3163T>A), CTC287TTA (3164C>T 3166C>A), CAA288TTA (3167C>T 3168A>T), AAT289AGG (3171A>G 3172T>G), AAC290AAA (3175C>A), ATC291ATT (3178C>T), TGT292GTC (3179T>G 3180G>T 3181T>C), CTC293TTG (3182C>T 3184C>G), GTC294GTT (3187C>T), TAC295TTC (3189A>T), CTT296TTT (3191C>T), GAC297GAT (3196C>T), GAT298GAC (3199T>C), ATT299ATC (3202T>C), ATT300CTT (3203A>C), GTC301ATC (3206G>A), TAT302TAC (3211T>C), AGT303AGC (3214T>C), ACT304AAG (3216C>A 3217T>G), CTA306ATG (3221C>A 3223A>G), CAG307ACT (3224C>A 3225A>C 3226G>T), GAA308GAT (3229A>T), CTG310ATT (3233C>A 3235G>T), GAG311GAA (3238G>A), AAC312CAT (3239A>C 3241C>T), CTG313GTT (3242C>G 3244G>T), GAA314AAA (3245G>A), CGA315GCA (3248C>G 3249G>C), GTT316GTG (3253T>G), TTC317TTT (3256C>T), CAA318GAT (3257C>G 3259A>T), AGA319CTG (3260A>C 3261G>T 3262A>G), CTT320ATG (3263C>A 3265T>G), AGA321TTA (3266A>T 3267G>T), GAA322AAG (3269G>A 3271A>G), AGT323TTT (3272A>T 3273G>T), AAC324CAG (3275A>C 3277C>G), TTC325TTA (3280C>A), AAA326TAT (3281A>T 3283A>T), ATT327GCT (3284A>G 3285T>C), CAA328AAA (3287C>A), GAC330TCT (3293G>T 3294A>C 3295C>T), AAG331AAA (3298G>A), TCC332TGT (3300C>G 3301C>T), GAA333GCA (3303A>C) | | | | | | | | |

##### Proteins

| | Begin | End | Coverage | Score | Concordance | Matches | Identities | I/D/M/F* | Stop Codons |
|---|---|---|---|---|---|---|---|---|---|
| ORF B (YP_009507248.1) | 235 | 334 | 9.2% | 301 | 42.8% | 100 (100%) | 41 (41.0%) | 0/0/0/0 | 0 |
| Protein mutations: | A235R (3008G>A 3009C>G), S236A (3011A>G 3012G>C), F238Y (3018T>A), Y239H (3020T>C 3022T>C), V241L (3026G>T), E242R (3029G>A 3030A>G 3031G>A), D244Y (3035G>T 3037C>T), P245E (3038C>G 3039C>A 3040T>A), Q246K (3041C>A 3043A>G), I248A (3047A>G 3048T>C 3049A>T), S249H (3050T>C 3051C>A 3052G>C), N254K (3067C>A), V255T (3068G>A 3069T>C 3070A>T), E256H (3071G>C 3073A>T), H257E (3074C>G 3076C>A), F260Y (3084T>A), R264V (3095C>G 3096G>T), M267F (3104A>T 3106G>T), K270T (3114A>C 3115A>C), S272A (3119T>G 3121A>C), T275S (3128A>T), R278N (3138G>A 3139A>C), V279L (3140G>C 3142T>C), D281N (3146G>A), N282S (3149A>T 3150A>C 3151T>A), L284F (3155C>T 3157A>T), G286P (3161G>C 3162G>C 3163T>A), Q288L (3167C>T 3168A>T), N289R (3171A>G 3172T>G), N290K (3175C>A), C292V (3179T>G 3180G>T 3181T>C), Y295F (3189A>T), L296F (3191C>T), I300L (3203A>C), V301I (3206G>A), T304K (3216C>A 3217T>G), L306M (3221C>A 3223A>G), Q307T (3224C>A 3225A>C 3226G>T), E308D (3229A>T), L310I (3233C>A 3235G>T), N312H (3239A>C 3241C>T), L313V (3242C>G 3244G>T), E314K (3245G>A), R315A (3248C>G 3249G>C), Q318D (3257C>G 3259A>T), R319L (3260A>C 3261G>T 3262A>G), L320M (3263C>A 3265T>G), R321L (3266A>T 3267G>T), E322K (3269G>A 3271A>G), S323F (3272A>T 3273G>T), N324Q (3275A>C 3277C>G), F325L (3280C>A), K326Y (3281A>T 3283A>T), I327A (3284A>G 3285T>C), Q328K (3287C>A), D330S (3293G>T 3294A>C 3295C>T), S332C (3300C>G 3301C>T), E333A (3303A>C) | | | | | | | | |
| Codon mutations: | TTG234..T (3007G>T), GCA235AGA (3008G>A 3009C>G), AGT236GCT (3011A>G 3012G>C), GGG237GGA (3016G>A), TTT238TAT (3018T>A), TAT239CAC (3020T>C 3022T>C), GTG241TTG (3026G>T), GAG242AGA (3029G>A 3030A>G 3031G>A), GAC244TAT (3035G>T 3037C>T), CCT245GAA (3038C>G 3039C>A 3040T>A), CAA246AAG (3041C>A 3043A>G), ATA248GCT (3047A>G 3048T>C 3049A>T), TCG249CAC (3050T>C 3051C>A 3052G>C), AAA250AAG (3055A>G), ACC251ACT (3058C>T), GCG252GCC (3061G>C), AAC254AAA (3067C>A), GTA255ACT (3068G>A 3069T>C 3070A>T), GAA256CAT (3071G>C 3073A>T), CAC257GAA (3074C>G 3076C>A), CAT259CAC (3082T>C), TTT260TAT (3084T>A), GAA261GAG (3088A>G), TTC262TTT (3091C>T), CTT263CTG (3094T>G), CGA264GTA (3095C>G 3096G>T), CCT266CCC (3103T>C), ATG267TTT (3104A>T 3106G>T), GGA268GGG (3109A>G), AAA270ACC (3114A>C 3115A>C), AAC271AAT (3118C>T), TCA272GCC (3119T>G 3121A>C), CCA273CCT (3124A>T), ACT275TCT (3128A>T), TTT276TTC (3133T>C), AGA278AAC (3138G>A 3139A>C), GTT279CTC (3140G>C 3142T>C), GAC281AAC (3146G>A), AAT282TCA (3149A>T 3150A>C 3151T>A), GTC283GTA (3154C>A), CTA284TTT (3155C>T 3157A>T), GGT286CCA (3161G>C 3162G>C 3163T>A), CTC287TTA (3164C>T 3166C>A), CAA288TTA (3167C>T 3168A>T), AAT289AGG (3171A>G 3172T>G), AAC290AAA (3175C>A), ATC291ATT (3178C>T), TGT292GTC (3179T>G 3180G>T 3181T>C), CTC293TTG (3182C>T 3184C>G), GTC294GTT (3187C>T), TAC295TTC (3189A>T), CTT296TTT (3191C>T), GAC297GAT (3196C>T), GAT298GAC (3199T>C), ATT299ATC (3202T>C), ATT300CTT (3203A>C), GTC301ATC (3206G>A), TAT302TAC (3211T>C), AGT303AGC (3214T>C), ACT304AAG (3216C>A 3217T>G), CTA306ATG (3221C>A 3223A>G), CAG307ACT (3224C>A 3225A>C 3226G>T), GAA308GAT (3229A>T), CTG310ATT (3233C>A 3235G>T), GAG311GAA (3238G>A), AAC312CAT (3239A>C 3241C>T), CTG313GTT (3242C>G 3244G>T), GAA314AAA (3245G>A), CGA315GCA (3248C>G 3249G>C), GTT316GTG (3253T>G), TTC317TTT (3256C>T), CAA318GAT (3257C>G 3259A>T), AGA319CTG (3260A>C 3261G>T 3262A>G), CTT320ATG (3263C>A 3265T>G), AGA321TTA (3266A>T 3267G>T), GAA322AAG (3269G>A 3271A>G), AGT323TTT (3272A>T 3273G>T), AAC324CAG (3275A>C 3277C>G), TTC325TTA (3280C>A), AAA326TAT (3281A>T 3283A>T), ATT327GCT (3284A>G 3285T>C), CAA328AAA (3287C>A), GAC330TCT (3293G>T 3294A>C 3295C>T), AAG331AAA (3298G>A), TCC332TGT (3300C>G 3301C>T), GAA333GCA (3303A>C) | | | | | | | | |

*: Inserts / Deletes / Misaligned / Frameshifts

#### Analysis details

This analysis was performed with panviral2.64

### NGS Details (UN60): Caulimovirus venafragariae

#### Assembly

| | |
|---|---|
| **Coverage Length** | 847 (2 contig(s)) |
| **Depth Of Coverage** | 6.3 |
| **Number Of Reads** | 45 |
| **Reads Per Million** | 0.95 rpm (after QC) |
| **Ambiguities** | 0 |
| **Assembly Method** | de novo + reference guided assembly |
| **Consensus Caller** | Bcf Tools |

#### Coverage Map

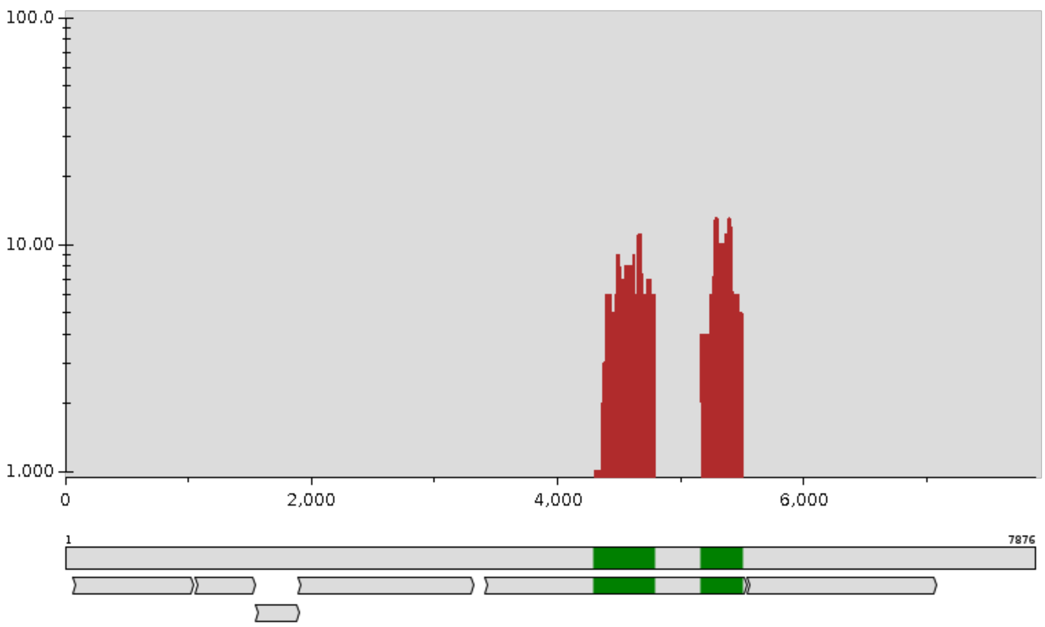

#### Assignment

| | |
|---|---|
| **Type** | Caulimovirus venafragariae (Taxonomy ID: 3048344) |
| **Reference Genome** | NC_001725.1 |
| **NT Identity (%)** | 58.5042 |
| **AA Identity (%)** | 53.9855 |
| **Number Of Stop Codons** | 0 |
| **Number Of CDS** | 6 |

#### Alignment

| | |
|---|---|
| **Alignment Score** | 246.0 (NT) + 1127.0 (AA) = 1373.0 |
| **Concordance (%)** | 38.8182 |

| Alignment Method | Global, seeded, nucleotide + amino acids (AGA) |
|---|---|

#### Genome Region

Sequence starts at position 4295 and ends at position 5508 relative to NC_001725.1 reference sequence.

#### Alignment Detailed Statistics

| | Begin | End | Coverage | Score | Concordance | Matches | Identities | I/D/M/F* | Stop Codons |
|---|---|---|---|---|---|---|---|---|---|
| **NT** | **4295** | **5508** | **10.8%** | **246** | **15.2%** | **829 (97.9%)** | **485 (57.3%)** | **0/18** | |

Mutations: 4299C>T, 4300G>C, 4302A>G, 4308A>T, 4314G>A, 4315C>A, 4317T>G, 4318G>A, 4319G>A, 4320T>G, 4321A>T, 4323C>A, 4330C>G, 4332T>A, 4337A>G, 4338G>A, 4347T>C, 4348T>A, 4349C>A, 4351T>A, 4352C>G, 4353A>T, 4356A>C, 4359C>G, 4371G>A, 4374T>C, 4380C>G, 4381G>A, 4383G>A, 4384A>C, 4385T>A, 4386C>A, 4395C>A, 4401A>T, 4402A>C, 4413T>A, 4423A>G, 4426T>C, 4428A>T, 4432G>A, 4434C>A, 4435C>A, 4438A>T, 4439C>G, 4440A>T, 4441A>G, 4442A>T, 4443G>C, 4444G>T, 4445G>T, 4446A>T, 4452C>A, 4455C>T, 4456C>T, 4458A>T, 4459C>A, 4464T>A, 4470G>A, 4471G>A, 4473G>T, 4474C>A, 4476A>T, 4479G>T, 4480C>A, 4482T>A, 4483C>A, 4485A>T, 4486A>C, 4487G>T, 4489A>G, 4490T>C, 4492G>A, 4493G>A, 4497T>A, 4500G>A, 4505T>A, 4506T>C, 4508A>T, 4509C>T, 4512T>A, 4513T>A, 4514C>A, 4515C>A, 4528T>A, 4529C>G, 4542G>A, 4543G>A, 4546C>A, 4547G>A, 4548C>A, 4551T>A, 4552G>A, 4553C>A, 4554T>A, 4555C>G, 4556C>A, 4558G>A, 4560A>T, 4561A>T, 4563A>T, 4566T>A, 4567C>G, 4568A>C, 4569G>T, 4570C>T, 4575C>A, 4578T>A, 4584C>T, 4585T>A, 4586G>C, 4587T>A, 4590C>A, 4591C>A, 4593A>T, 4599C>T, 4602C>T, 4614C>A, 4633C>A, 4635A>T, 4642G>C, 4643C>A, 4644T>A, 4647T>C, 4650C>T, 4656G>A, 4657C>A, 4659C>A, 4665C>T, 4670G>T, 4671T>A, 4672C>T, 4674C>T, 4675A>T, 4676G>C, 4677C>T, 4678A>G, 4680C>A, 4682T>A, 4684T>A, 4687_4692delCCACAG, 4696T>C, 4697G>T, 4698T>A, 4699G>A, 4700C>T, 4701T>C, 4704A>C, 4708G>A, 4710T>A, 4716C>T, 4719C>A, 4720A>C, 4722C>A, 4723G>A, 4725C>A, 4727T>G, 4728C>T, 4729A>T, 4730G>C, 4731C>T, 4734A>T, 4735A>T, 4737T>A, 4738G>T, 4740A>T, 4741G>C, 4746A>G, 4753G>A, 4754G>C, 4758T>C, 4759G>C, 4761C>T, 4765A>C, 4766T>A, 4767T>A, 4768G>T, 4771C>T, 4772T>C, 4773A>T, 4774A>C, 4776C>A, 4777A>T, 4778G>T, 4782C>T, 4783A>G, 4784A>T, 4785A>T, 4787C>A, 4788C>A, 4789C>A, 4790T>A, 4791A>T, 5163A>C, 5167G>T, 5169C>T, 5182A>T, 5184C>A, 5185C>T, 5187C>A, 5188A>C, 5191G>A, 5193A>T, 5194A>G, 5196A>T, 5199T>A, 5200C>A, 5201C>A, 5202A>T, 5206_5217delGGTAAAGAGGTA, 5220A>T, 5223C>T, 5226G>A, 5230G>A, 5233T>A, 5234C>G, 5235A>T, 5247A>T, 5248C>G, 5249C>A, 5251G>A, 5253T>A, 5256G>A, 5258A>C, 5259G>T, 5261A>G, 5262T>G, 5266C>T, 5267A>T, 5268T>A, 5270G>C, 5271T>A, 5280G>A, 5284A>T, 5286C>A, 5289T>A, 5290T>G, 5292C>T, 5295T>A, 5296A>G, 5298T>A, 5300A>G, 5301G>A, 5303C>G, 5304A>C, 5307C>A, 5308A>C, 5309A>G, 5311G>A, 5312C>A, 5313T>G, 5317A>T, 5318G>C, 5319A>T, 5322T>A, 5324A>T, 5325T>C, 5326A>T, 5328C>A, 5331T>A, 5334T>A, 5335T>A, 5337T>A, 5338A>C, 5343T>C, 5344C>A, 5346T>C, 5347G>A, 5349A>C, 5352G>A, 5353A>T, 5355T>G, 5358T>C, 5364T>G, 5365A>C, 5367T>G, 5369C>T, 5370T>A, 5371G>T, 5373T>A, 5374T>G, 5375A>G, 5377T>C, 5379T>A, 5380G>A, 5382G>A, 5383A>T, 5384G>T, 5385A>C, 5387C>A, 5388A>T, 5391T>A, 5392A>C, 5395G>C, 5398G>T, 5399G>C, 5403T>A, 5404T>C, 5405A>C, 5407A>C, 5409G>A, 5410C>T, 5412A>T, 5415C>A, 5416C>A, 5418G>A, 5419C>T, 5421T>A, 5422G>C, 5423T>A, 5424G>T, 5434A>G, 5436G>A, 5437G>T, 5438C>T, 5439T>G, 5443A>T, 5444G>C, 5445A>T, 5446G>T, 5447A>T, 5448A>T, 5452T>A, 5454T>C, 5457T>C, 5458G>A, 5460T>A, 5463A>T, 5467C>A, 5470G>A, 5473T>A, 5474C>A, 5475T>A, 5478T>A, 5479C>T, 5480A>C, 5481A>T, 5482A>G, 5484A>T, 5491C>T, 5493A>T, 5496T>A, 5500A>T, 5503A>T, 5505G>A, 5506A>T, 5508T>A

##### CDS

| | Begin | End | Coverage | Score | Concordance | Matches | Identities | I/D/M/F* | Stop Codons |
|---|---|---|---|---|---|---|---|---|---|
| ORF_V | 298 | 701 | 39.8% | 1127 | 58.3% | 276 (97.9%) | 149 (52.8%) | 0/6/0/0 | 0 |

Protein mutations: E299Q (4300G>C 4302A>G), L304M (4315C>A 4317T>G), G305K (4318G>A 4319G>A 4320T>G), I306L (4321A>T 4323C>A), P309A (4330C>G 4332T>A), K311R (4337A>G 4338G>A), S315N (4348T>A 4349C>A), E326K (4381G>A 4383G>A), I327Q (4384A>C 4385T>A 4386C>A), K340E (4423A>G), D343K (4432G>A 4434C>A), H344N (4435C>A), T345C (4438A>T 4439C>G 4440A>T), K346V (4441A>G 4442A>T 4443G>C), G347F (4444G>T 4445G>T 4446A>T), L351F (4456C>T 4458A>T), L352I (4459C>A), E356N (4471G>A 4473G>T), Q357N (4474C>A 4476A>T), L359I (4480C>A 4482T>A), Q360N (4483C>A 4485A>T), R361L (4486A>C 4487G>T), I362A (4489A>G 4490T>C), G363K (4492G>A 4493G>A), F367Y (4505T>A 4506T>C), Y368F (4508A>T 4509C>T), S370K (4513T>A 4514C>A 4515C>A), V380I (4543G>A), R381K (4546C>A 4547G>A 4548C>A), A383K (4552G>A 4553C>A 4554T>A), P384E (4555C>G 4556C>A), E385N (4558G>A 4560A>T), T386S (4561A>T 4563A>T), Q388A (4567C>G 4568A>C 4569G>T), C394T (4585T>A 4586G>C 4587T>A), Q396N (4591C>A 4593A>T), Q410N (4633C>A 4635A>T), A413Q (4642G>C 4643C>A 4644T>A), H418K (4657C>A 4659C>A), S422I (4670G>T 4671T>A), L423F (4672C>T 4674C>T), N425E (4678A>G 4680C>A), M426K (4682T>A), Y427N (4684T>A), P428_Q429del (4687_4692delCCACAG), C431L (4696T>C 4697G>T 4698T>A), A432I (4699G>A 4700C>T 4701T>C), V435I (4708G>A 4710T>A), I439L (4720A>C 4722C>A), V440I (4723G>A 4725C>A), F441C (4727T>G 4728C>T), K443N (4734A>T), T444S (4735A>T 4737T>A), E445Y (4738G>T 4740A>T), E446Q (4741G>C), G450T (4753G>A 4754G>C), V452L (4759G>C 4761C>T), I454Q (4765A>C 4766T>A 4767T>A), V455F (4768G>T), L456S (4771C>T 4772T>C 4773A>T), N457Q (4774A>C 4776C>A), R458L (4777A>T 4778G>T), K460V (4783A>G 4784A>T 4785A>T), A461E (4787C>A 4788C>A), L462N (4789C>A 4790T>A 4791A>T), D588Y (5167G>T 5169C>T), I593L (5182A>T 5184C>A), K595Q (5188A>C), A596T (5191G>A 5193A>T), K597D (5194A>G 5196A>T), P599N (5200C>A 5201C>A 5202A>T), G601_V604del (5206_5217delGGTAAAGAGGTA), A609T (5230G>A), K614N (5247A>T), P615E (5248C>G 5249C>A), A616T (5251G>A 5253T>A), K618T (5258A>C 5259G>T), N619R (5261A>G 5262T>G), H621L (5266C>T 5267A>T 5268T>A), S622T (5270G>C 5271T>A), I627L (5284A>T 5286C>A), S629A (5290T>G 5292C>T), I631V (5296A>G 5298T>A), K632R (5300A>G 5301G>A), A633G (5303C>G 5304A>C), K635R (5308A>C 5309A>G), A636K (5311G>A 5312C>A 5313T>G), R638S (5317A>T 5318G>C 5319A>T), Y640F (5324A>T 5325T>C), I641L (5326A>T 5328C>A), Y644K (5335T>A 5337T>A), K645Q (5338A>C), L647I (5344C>A 5346T>C), V648I (5347G>A 5349A>C), T650S (5353A>T 5355T>G), N654Q (5365A>C 5367T>G), A655V (5369C>T 5370T>A), A656S (5371G>T 5373T>A), Y657G (5374T>G 5375A>G), F658L (5377T>C 5379T>A), V659I (5380G>A 5382G>A), R660F (5383A>T 5384G>T 5385A>C), T661N (5387C>A 5388A>T), N662K (5391T>A), I663L (5392A>C), A664P (5395G>C), G665S (5398G>T 5399G>C), D666E (5403T>A), Y667P (5404T>C 5405A>C), K668Q (5407A>C 5409G>A), Q669Y (5410C>T 5412A>T), S670R (5415C>A), V673H (5422G>C 5423T>A 5424G>T), M677V (5434A>G 5436G>A), A678L (5437G>T 5438C>T 5439T>G), R680S (5443A>T 5444G>C 5445A>T), E681F (5446G>T 5447A>T 5448A>T), S683T (5452T>A 5454T>C), D685K (5458G>A 5460T>A), H688N (5467C>A), V689I (5470G>A), S690K (5473T>A 5474C>A 5475T>A), Q692S (5479C>T 5480A>C 5481A>T), K693D (5482A>G 5484A>T), L696F (5491C>T 5493A>T), I699F (5500A>T), M700L (5503A>T 5505G>A), T701S (5506A>T 5508T>A)

Codon mutations: ATC298ATT (4299C>T), GAA299CAG (4300G>C 4302A>G), CTA301CTT (4308A>T), AAG303AAA (4314G>A), CTT304ATG (4315C>A 4317T>G), GGT305AAG (4318G>A 4319G>A 4320T>G), ATC306TTA (4321A>T 4323C>A), CCT309GCA (4330C>G 4332T>A), AAG311AGA (4337A>G 4338G>A), CAT314CAC (4347T>C), TCC315AAC (4348T>A 4349C>A), TCA316AGT (4351T>A 4352C>G 4353A>T), CCA317CCC (4356A>C), GCC318GCG (4359C>G), AGG322AGA (4371G>A), AAT323AAC (4374T>C), GCC325GCG (4380C>G), GAG326AAA (4381G>A 4383G>A), ATC327CAA (4384A>C 4385T>A 4386C>A), GGC330GGA (4395C>A), GCA332GCT (4401A>T), AGA333CGA (4402A>C), ATT336ATA (4413T>A), AAG340GAG (4423A>G), TTA341CTT (4426T>C 4428A>T), GAC343AAA (4432G>A 4434C>A), CAT344AAT (4435C>A), ACA345TGT (4438A>T 4439C>G 4440A>T), AAG346GTC (4441A>G 4442A>T 4443G>C), GGA347TTT (4444G>T 4445G>T 4446A>T), GGC349GGA (4452C>A), TAC350TAT (4455C>T), CTA351TTT (4456C>T 4458A>T), CTT352ATT (4459C>A), CCT353CCA (4464T>A), AAG355AAA (4470G>A), GAG356AAT (4471G>A 4473G>T), CAA357AAT (4474C>A 4476A>T), CTG358CTT (4479G>T), CTT359ATA (4480C>A 4482T>A), CAA360AAT (4483C>A 4485A>T), AGA361CTA (4486A>C 4487G>T), ATC362GCC (4489A>G 4490T>C), GGA363AAA (4492G>A 4493G>A), GGT364GGA (4497T>A), AAG365AAA (4500G>A), TTT367TAC (4505T>A 4506T>C), TAC368TTT (4508A>T 4509C>T), TCT369TCA (4512T>A), TCC370AAA (4513T>A 4514C>A 4515C>A), TCT375AGT (4528T>A 4529C>G), CAG379CAA (4542G>A), GTA380ATA (4543G>A), CGC381AAA (4546C>A 4547G>A 4548C>A), CTT382CTA (4551T>A), GCT383AAA (4552G>A 4553C>A 4554T>A), CCA384GAA (4555C>G 4556C>A), GAA385AAT (4558G>A 4560A>T), ACA386TCT (4561A>T 4563A>T), ATT387ATA (4566T>A), CAG388GCT (4567C>G 4568A>C 4569G>T), CTA389TTA (4570C>T), ACC390ACA (4575C>A), GCT391GCA (4578T>A), AGC393AGT (4584C>T), TGT394ACA (4585T>A 4586G>C 4587T>A), CCC395CCA (4590C>A), CAA396AAT (4591C>A 4593A>T), CAC398CAT (4599C>T), TAC399TAT (4602C>T), GTC403GTA (4614C>A), CAA410AAT (4633C>A 4635A>T), GCT413CAA (4642G>C 4643C>A 4644T>A), ATT414ATC (4647T>C), TTC415TTT (4650C>T), AGG417AGA (4656G>A), CAC418AAA (4657C>A 4659C>A), GAC420GAT (4665C>T), AGT422ATA (4670G>T 4671T>A), CTC423TTT (4672C>T 4674C>T), AGC424TCT (4675A>T 4676G>C 4677C>T), AAC425GAA (4678A>G 4680C>A), ATG426AAG (4682T>A), TAT427AAT (4684T>A), CCA428_CAG429del (4687_4692delCCACAG), TGT431CTA (4696T>C 4697G>T 4698T>A), GCT432ATC (4699G>A 4700C>T 4701T>C), GTA433GTC (4704A>C), GTT435ATA (4708G>A 4710T>A), GAC437GAT (4716C>T), ATC438ATA (4719C>A), ATC439CTA (4720A>C 4722C>A), GTC440ATA (4723G>A 4725C>A), TTC441TGT (4727T>G 4728C>T), AGC442TCT (4729A>T 4730G>C 4731C>T), AAA443AAT (4734A>T), ACT444TCA (4735A>T 4737T>A), GAA445TAT (4738G>T 4740A>T), GAA446CAA (4741G>C), GAA447GAG (4746A>G), GGA450ACA (4753G>A 4754G>C), CAT451CAC (4758T>C), GTC452CTT (4759G>C 4761C>T), ATT454CAA (4765A>C 4766T>A 4767T>A), GTT455TTT (4768G>T), CTA456TCT (4771C>T 4772T>C 4773A>T), AAC457CAA (4774A>C 4776C>A), AGA458TTA (4777A>T 4778G>T), TGC459TGT (4782C>T), AAA460GTT (4783A>G 4784A>T 4785A>T), GCC461GAA (4787C>A 4788C>A), CTA462AAT (4789C>A 4790T>A 4791A>T), TCA586..C (5163A>C), GAC588TAT (5167G>T 5169C>T), ATC593TTA (5182A>T 5184C>A), CTC594TTA (5185C>T 5187C>A), AAA595CAA (5188A>C), GCA596ACT (5191G>A 5193A>T), AAA597GAT (5194A>G 5196A>T), CTT598CTA (5199T>A), CCA599AAT (5200C>A 5201C>A 5202A>T), GGT601_GTA604del (5206_5217delGGTAAAGAGGTA), ATA605ATT (5220A>T), TGC606TGT (5223C>T), AGG607AGA (5226G>A), GCT609ACT (5230G>A), TCA610AGT (5233T>A 5234C>G 5235A>T), AAA614AAT (5247A>T), CCA615GAA (5248C>G 5249C>A), GCT616ACA (5251G>A 5253T>A), GAG617GAA (5256G>A), AAG618ACT (5258A>C 5259G>T), AAT619AGG (5261A>G 5262T>G), CAT621TTA (5266C>T 5267A>T 5268T>A), AGT622ACA (5270G>C 5271T>A), AAG625AAA (5280G>A), ATC627TTA (5284A>T 5286C>A), CTT628CTA (5289T>A), TCC629GCT (5290T>G 5292C>T), ATT630ATA (5295T>A), ATT631GTA (5296A>G 5298T>A), AAG632AGA (5300A>G 5301G>A), GCA633GGC (5303C>G 5304A>C), ATC634ATA (5307C>A), AAA635CGA (5308A>C 5309A>G), GCT636AAG (5311G>A 5312C>A 5313T>G), AGA638TCT (5317A>T 5318G>C 5319A>T), GCT639GCA (5322T>A), TAT640TTC (5324A>T 5325T>C), ATC641TTA (5326A>T 5328C>A), CTT642CTA (5331T>A), CCT643CCA (5334T>A), TAT644AAA (5335T>A 5337T>A), AAA645CAA (5338A>C), TTT646TTC (5343T>C), CTT647ATC (5344C>A 5346T>C), GTA648ATC (5347G>A 5349A>C), AGG649AGA (5352G>A), ACT650TCG (5353A>T 5355T>G), GAT651GAC (5358T>C), ACT653ACG (5364T>G), AAT654CAG (5365A>C 5367T>G), GCT655GTA (5369C>T 5370T>A), GCT656TCA (5371G>T 5373T>A), TAT657GGT (5374T>G 5375A>G), TTT658CTA (5377T>C 5379T>A), GTG659ATA (5380G>A 5382G>A), AGA660TTC (5383A>T 5384G>T 5385A>C), ACA661AAT (5387C>A 5388A>T), AAT662AAA (5391T>A), ATT663CTT (5392A>C), GCA664CCA (5395G>C), GGT665TCT (5398G>T 5399G>C), GAT666GAA (5403T>A), TAT667CCT (5404T>C 5405A>C), AAG668CAA (5407A>C 5409G>A), CAA669TAT (5410C>T 5412A>T), AGC670AGA (5415C>A), CGG671AGA (5416C>A 5418G>A), CTT672TTA (5419C>T 5421T>A), GTG673CAT (5422G>C 5423T>A 5424G>T), ATG677GTA (5434A>G 5436G>A), GCT678TTG (5437G>T 5438C>T 5439T>G), AGA680TCT (5443A>T 5444G>C 5445A>T), GAA681TTT (5446G>T 5447A>T 5448A>T), TCT683ACC (5452T>A 5454T>C), TTT684TTC (5457T>C), GAT685AAA (5458G>A 5460T>A), ATA686ATT (5463A>T), CAT688AAT (5467C>A), GTT689ATT (5470G>A), TCT690AAA (5473T>A 5474C>A 5475T>A), GGT691GGA (5478T>A), CAA692TCT (5479C>T 5480A>C 5481A>T), AAA693GAT (5482A>G 5484A>T), CTA696TTT (5491C>T 5493A>T), GCT697GCA (5496T>A), ATC699TTC (5500A>T), ATG700TTA (5503A>T 5505G>A), ACT701TCA (5506A>T 5508T>A)

##### Proteins

| | Begin | End | Coverage | Score | Concordance | Matches | Identities | I/D/M/F* | Stop Codons |
|---|---|---|---|---|---|---|---|---|---|
| hypothetical protein (NP_043933.1) | 298 | 701 | 39.8% | 1127 | 58.3% | 276 (97.9%) | 149 (52.8%) | 0/6/0/0 | 0 |

| | Begin | End | Coverage | Score | Concordance | Matches | Identities | I/D/M/F* | Stop Codons |
|---|---|---|---|---|---|---|---|---|---|
| **NT** | **4295** | **5508** | **10.8%** | **246** | **15.2%** | **829 (97.9%)** | **485 (57.3%)** | **0/18** | |
| Protein mutations: | E299Q (4300G>C 4302A>G), L304M (4315C>A 4317T>G), G305K (4318G>A 4319G>A 4320T>G), I306L (4321A>T 4323C>A), P309A (4330C>G 4332T>A), K311R (4337A>G 4338G>A), S315N (4348T>A 4349C>A), E326K (4381G>A 4383G>A), I327Q (4384A>C 4385T>A 4386C>A), K340E (4423A>G), D343K (4432G>A 4434C>A), H344N (4435C>A), T345C (4438A>T 4439C>G 4440A>T), K346V (4441A>G 4442A>T 4443G>C), G347F (4444G>T 4445G>T 4446A>T), L351F (4456C>T 4458A>T), L352I (4459C>A), E356N (4471G>A 4473G>T), Q357N (4474C>A 4476A>T), L359I (4480C>A 4482T>A), Q360N (4483C>A 4485A>T), R361L (4486A>C 4487G>T), I362A (4489A>G 4490T>C), G363K (4492G>A 4493G>A), F367Y (4505T>A 4506T>C), Y368F (4508A>T 4509C>T), S370K (4513T>A 4514C>A 4515C>A), V380I (4543G>A), R381K (4546C>A 4547G>A 4548C>A), A383K (4552G>A 4553C>A 4554T>A), P384E (4555C>G 4556C>A), E385N (4558G>A 4560A>T), T386S (4561A>T 4563A>T), Q388A (4567C>G 4568A>C 4569G>T), C394T (4585T>A 4586G>C 4587T>A), Q396N (4591C>A 4593A>T), Q410N (4633C>A 4635A>T), A413Q (4642G>C 4643C>A 4644T>A), H418K (4657C>A 4659C>A), S422I (4670G>T 4671T>A), L423F (4672C>T 4674C>T), N425E (4678A>G 4680C>A), M426K (4682T>A), Y427N (4684T>A), P428_Q429del (4687_4692delCCACAG), C431L (4696T>C 4697G>T 4698T>A), A432I (4699G>A 4700C>T 4701T>C), V435I (4708G>A 4710T>A), I439L (4720A>C 4722C>A), V440I (4723G>A 4725C>A), F441C (4727T>G 4728C>T), K443N (4734A>T), T444S (4735A>T 4737T>A), E445Y (4738G>T 4740A>T), E446Q (4741G>C), G450T (4753G>A 4754G>C), V452L (4759G>C 4761C>T), I454Q (4765A>C 4766T>A 4767T>A), V455F (4768G>T), L456S (4771C>T 4772T>C 4773A>T), N457Q (4774A>C 4776C>A), R458L (4777A>T 4778G>T), K460V (4783A>G 4784A>T 4785A>T), A461E (4787C>A 4788C>A), L462N (4789C>A 4790T>A 4791A>T), D588Y (5167G>T 5169C>T), I593L (5182A>T 5184C>A), K595Q (5188A>C), A596T (5191G>A 5193A>T), K597D (5194A>G 5196A>T), P599N (5200C>A 5201C>A 5202A>T), G601_V604del (5206_5217delGGTAAAGAGGTA), A609T (5230G>A), K614N (5247A>T), P615E (5248C>G 5249C>A), A616T (5251G>A 5253T>A), K618T (5258A>C 5259G>T), N619R (5261A>G 5262T>G), H621L (5266C>T 5267A>T 5268T>A), S622T (5270G>C 5271T>A), I627L (5284A>T 5286C>A), S629A (5290T>G 5292C>T), I631V (5296A>G 5298T>A), K632R (5300A>G 5301G>A), A633G (5303C>G 5304A>C), K635R (5308A>C 5309A>G), A636K (5311G>A 5312C>A 5313T>G), R638S (5317A>T 5318G>C 5319A>T), Y640F (5324A>T 5325T>C), I641L (5326A>T 5328C>A), Y644K (5335T>A 5337T>A), K645Q (5338A>C), L647I (5344C>A 5346T>C), V648I (5347G>A 5349A>C), T650S (5353A>T 5355T>G), N654Q (5365A>C 5367T>G), A655V (5369C>T 5370T>A), A656S (5371G>T 5373T>A), Y657G (5374T>G 5375A>G), F658L (5377T>C 5379T>A), V659I (5380G>A 5382G>A), R660F (5383A>T 5384G>T 5385A>C), T661N (5387C>A 5388A>T), N662K (5391T>A), I663L (5392A>C), A664P (5395G>C), G665S (5398G>T 5399G>C), D666E (5403T>A), Y667P (5404T>C 5405A>C), K668Q (5407A>C 5409G>A), Q669Y (5410C>T 5412A>T), S670R (5415C>A), V673H (5422G>C 5423T>A 5424G>T), M677V (5434A>G 5436G>A), A678L (5437G>T 5438C>T 5439T>G), R680S (5443A>T 5444G>C 5445A>T), E681F (5446G>T 5447A>T 5448A>T), S683T (5452T>A 5454T>C), D685K (5458G>A 5460T>A), H688N (5467C>A), V689I (5470G>A), S690K (5473T>A 5474C>A 5475T>A), Q692S (5479C>T 5480A>C 5481A>T), K693D (5482A>G 5484A>T), L696F (5491C>T 5493A>T), I699F (5500A>T), M700L (5503A>T 5505G>A), T701S (5506A>T 5508T>A) | | | | | | | | |
| Codon mutations: | ATC298ATT (4299C>T), GAA299CAG (4300G>C 4302A>G), CTA301CTT (4308A>T), AAG303AAA (4314G>A), CTT304ATG (4315C>A 4317T>G), GGT305AAG (4318G>A 4319G>A 4320T>G), ATC306TTA (4321A>T 4323C>A), CCT309GCA (4330C>G 4332T>A), AAG311AGA (4337A>G 4338G>A), CAT314CAC (4347T>C), TCC315AAC (4348T>A 4349C>A), TCA316AGT (4351T>A 4352C>G 4353A>T), CCA317CCC (4356A>C), GCC318GCG (4359C>G), AGG322AGA (4371G>A), AAT323AAC (4374T>C), GCC325GCG (4380C>G), GAG326AAA (4381G>A 4383G>A), ATC327CAA (4384A>C 4385T>A 4386C>A), GGC330GGA (4395C>A), GCA332GCT (4401A>T), AGA333CGA (4402A>C), ATT336ATA (4413T>A), AAG340GAG (4423A>G), TTA341CTT (4426T>C 4428A>T), GAC343AAA (4432G>A 4434C>A), CAT344AAT (4435C>A), ACA345TGT (4438A>T 4439C>G 4440A>T), AAG346GTC (4441A>G 4442A>T 4443G>C), GGA347TTT (4444G>T 4445G>T 4446A>T), GGC349GGA (4452C>A), TAC350TAT (4455C>T), CTA351TTT (4456C>T 4458A>T), CTT352ATT (4459C>A), CCT353CCA (4464T>A), AAG355AAA (4470G>A), GAG356AAT (4471G>A 4473G>T), CAA357AAT (4474C>A 4476A>T), CTG358CTT (4479G>T), CTT359ATA (4480C>A 4482T>A), CAA360AAT (4483C>A 4485A>T), AGA361CTA (4486A>C 4487G>T), ATC362GCC (4489A>G 4490T>C), GGA363AAA (4492G>A 4493G>A), GGT364GGA (4497T>A), AAG365AAA (4500G>A), TTT367TAC (4505T>A 4506T>C), TAC368TTT (4508A>T 4509C>T), TCT369TCA (4512T>A), TCC370AAA (4513T>A 4514C>A 4515C>A), TCT375AGT (4528T>A 4529C>G), CAG379CAA (4542G>A), GTA380ATA (4543G>A), CGC381AAA (4546C>A 4547G>A 4548C>A), CTT382CTA (4551T>A), GCT383AAA (4552G>A 4553C>A 4554T>A), CCA384GAA (4555C>G 4556C>A), GAA385AAT (4558G>A 4560A>T), ACA386TCT (4561A>T 4563A>T), ATT387ATA (4566T>A), CAG388GCT (4567C>G 4568A>C 4569G>T), CTA389TTA (4570C>T), ACC390ACA (4575C>A), GCT391GCA (4578T>A), AGC393AGT (4584C>T), TGT394ACA (4585T>A 4586G>C 4587T>A), CCC395CCA (4590C>A), CAA396AAT (4591C>A 4593A>T), CAC398CAT (4599C>T), TAC399TAT (4602C>T), GTC403GTA (4614C>A), CAA410AAT (4633C>A 4635A>T), GCT413CAA (4642G>C 4643C>A 4644T>A), ATT414ATC (4647T>C), TTC415TTT (4650C>T), AGG417AGA (4656G>A), CAC418AAA (4657C>A 4659C>A), GAC420GAT (4665C>T), AGT422ATA (4670G>T 4671T>A), CTC423TTT (4672C>T 4674C>T), AGC424TCT (4675A>T 4676G>C 4677C>T), AAC425GAA (4678A>G 4680C>A), ATG426AAG (4682T>A), TAT427AAT (4684T>A), CCA428_CAG429del (4687_4692delCCACAG), TGT431CTA (4696T>C 4697G>T 4698T>A), GCT432ATC (4699G>A 4700C>T 4701T>C), GTA433GTC (4704A>C), GTT435ATA (4708G>A 4710T>A), GAC437GAT (4716C>T), ATC438ATA (4719C>A), ATC439CTA (4720A>C 4722C>A), GTC440ATA (4723G>A 4725C>A), TTC441TGT (4727T>G 4728C>T), AGC442TCT (4729A>T 4730G>C 4731C>T), AAA443AAT (4734A>T), ACT444TCA (4735A>T 4737T>A), GAA445TAT (4738G>T 4740A>T), GAA446CAA (4741G>C), GAA447GAG (4746A>G), GGA450ACA (4753G>A 4754G>C), CAT451CAC (4758T>C), GTC452CTT (4759G>C 4761C>T), ATT454CAA (4765A>C 4766T>A 4767T>A), GTT455TTT (4768G>T), CTA456TCT (4771C>T 4772T>C 4773A>T), AAC457CAA (4774A>C 4776C>A), AGA458TTA (4777A>T 4778G>T), TGC459TGT (4782C>T), AAA460GTT (4783A>G 4784A>T 4785A>T), GCC461GAA (4787C>A 4788C>A), CTA462AAT (4789C>A 4790T>A 4791A>T), TCA586..C (5163A>C), GAC588TAT (5167G>T 5169C>T), ATC593TTA (5182A>T 5184C>A), CTC594TTA (5185C>T 5187C>A), AAA595CAA (5188A>C), GCA596ACT (5191G>A 5193A>T), AAA597GAT (5194A>G 5196A>T), CTT598CTA (5199T>A), CCA599AAT (5200C>A 5201C>A 5202A>T), GGT601_GTA604del (5206_5217delGGTAAAGAGGTA), ATA605ATT (5220A>T), TGC606TGT (5223C>T), AGG607AGA (5226G>A), GCT609ACT (5230G>A), TCA610AGT (5233T>A 5234C>G 5235A>T), AAA614AAT (5247A>T), CCA615GAA (5248C>G 5249C>A), GCT616ACA (5251G>A 5253T>A), GAG617GAA (5256G>A), AAG618ACT (5258A>C 5259G>T), AAT619AGG (5261A>G 5262T>G), CAT621TTA (5266C>T 5267A>T 5268T>A), AGT622ACA (5270G>C 5271T>A), AAG625AAA (5280G>A), ATC627TTA (5284A>T 5286C>A), CTT628CTA (5289T>A), TCC629GCT (5290T>G 5292C>T), ATT630ATA (5295T>A), ATT631GTA (5296A>G 5298T>A), AAG632AGA (5300A>G 5301G>A), GCA633GGC (5303C>G 5304A>C), ATC634ATA (5307C>A), AAA635CGA (5308A>C 5309A>G), GCT636AAG (5311G>A 5312C>A 5313T>G), AGA638TCT (5317A>T 5318G>C 5319A>T), GCT639GCA (5322T>A), TAT640TTC (5324A>T 5325T>C), ATC641TTA (5326A>T 5328C>A), CTT642CTA (5331T>A), CCT643CCA (5334T>A), TAT644AAA (5335T>A 5337T>A), AAA645CAA (5338A>C), TTT646TTC (5343T>C), CTT647ATC (5344C>A 5346T>C), GTA648ATC (5347G>A 5349A>C), AGG649AGA (5352G>A), ACT650TCG (5353A>T 5355T>G), GAT651GAC (5358T>C), ACT653ACG (5364T>G), AAT654CAG (5365A>C 5367T>G), GCT655GTA (5369C>T 5370T>A), GCT656TCA (5371G>T 5373T>A), TAT657GGT (5374T>G 5375A>G), TTT658CTA (5377T>C 5379T>A), GTG659ATA (5380G>A 5382G>A), AGA660TTC (5383A>T 5384G>T 5385A>C), ACA661AAT (5387C>A 5388A>T), AAT662AAA (5391T>A), ATT663CTT (5392A>C), GCA664CCA (5395G>C), GGT665TCT (5398G>T 5399G>C), GAT666GAA (5403T>A), TAT667CCT (5404T>C 5405A>C), AAG668CAA (5407A>C 5409G>A), CAA669TAT (5410C>T 5412A>T), AGC670AGA (5415C>A), CGG671AGA (5416C>A 5418G>A), CTT672TTA (5419C>T 5421T>A), GTG673CAT (5422G>C 5423T>A 5424G>T), ATG677GTA (5434A>G 5436G>A), GCT678TTG (5437G>T 5438C>T 5439T>G), AGA680TCT (5443A>T 5444G>C 5445A>T), GAA681TTT (5446G>T 5447A>T 5448A>T), TCT683ACC (5452T>A 5454T>C), TTT684TTC (5457T>C), GAT685AAA (5458G>A 5460T>A), ATA686ATT (5463A>T), CAT688AAT (5467C>A), GTT689ATT (5470G>A), TCT690AAA (5473T>A 5474C>A 5475T>A), GGT691GGA (5478T>A), CAA692TCT (5479C>T 5480A>C 5481A>T), AAA693GAT (5482A>G 5484A>T), CTA696TTT (5491C>T 5493A>T), GCT697GCA (5496T>A), ATC699TTC (5500A>T), ATG700TTA (5503A>T 5505G>A), ACT701TCA (5506A>T 5508T>A) | | | | | | | | |

*: Inserts / Deletes / Misaligned / Frameshifts

#### Analysis details

This analysis was performed with panviral2.64

### NGS Details (UN60): Badnavirus rutilanscamelliae

#### Assembly

| | |
|---|---|
| **Coverage Length** | 297 (1 contig(s)) |
| **Depth Of Coverage** | 15.0 |
| **Number Of Reads** | 44 |
| **Reads Per Million** | 0.93 rpm (after QC) |
| **Ambiguities** | 0 |
| **Assembly Method** | de novo + reference guided assembly |
| **Consensus Caller** | Bcf Tools |

#### Coverage Map

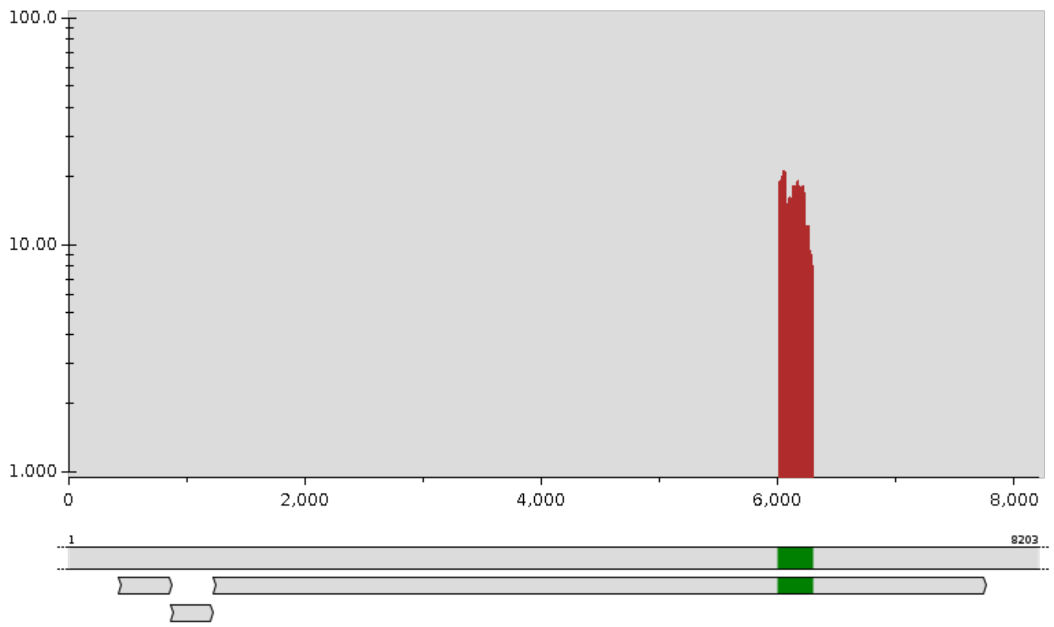

#### Assignment

| | |
|---|---|
| **Type** | Badnavirus rutilanscamelliae (Taxonomy ID: 3047719) |
| **Reference Genome** | NC_055598.1 |
| **NT Identity (%)** | 54.0 |
| **AA Identity (%)** | 45.0 |
| **Number Of Stop Codons** | 0 |
| **Number Of CDS** | 3 |

#### Alignment

| | |
|---|---|
| **Alignment Score** | 42.0 (NT) + 327.0 (AA) = 369.0 |
| **Concordance (%)** | 28.4942 |

| Alignment Method | Global, seeded, nucleotide + amino acids (AGA) |
|---|---|

#### Genome Region

Sequence starts at position 6010 and ends at position 6306 relative to NC_055598.1 reference sequence.

#### Alignment Detailed Statistics

| | Begin | End | Coverage | Score | Concordance | Matches | Identities | I/D/M/F* | Stop Codons |
|---|---|---|---|---|---|---|---|---|---|
| **NT** | **6010** | **6306** | **3.6%** | **42** | **7.1%** | **297 (99.0%)** | **162 (54.0%)** | **3/0** | |

Mutations: 6013C>T, 6019T>C, 6020T>A, 6022G>A, 6027C>A, 6028A>T, 6031G>A, 6034C>G, 6035T>C, 6036T>A, 6037A>T, 6040T>C, 6043A>G, 6045G>T, 6046G>C, 6047C>A, 6048T>A, 6052T>G, 6058A>G, 6064G>C, 6069A>C, 6070G>A, 6082A>T, 6083G>A, 6084T>C, 6085A>T, 6088T>C, 6092A>G, 6093G>C, 6094G>C, 6095A>T, 6096A>T, 6097G>C, 6101G>A, 6103C>T, 6105G>A, 6106A>G, 6107A>G, 6109C>T, 6112T>C, 6113A>G, 6115T>A, 6118A>T, 6120A>T, 6121C>T, 6122G>C, 6123C>T, 6124T>G, 6125G>A, 6127_6128insAAG, 6131A>G, 6134G>T, 6135C>T, 6136T>G, 6139G>A, 6141A>T, 6143A>T, 6154C>A, 6157C>T, 6158A>G, 6160A>T, 6162T>A, 6166C>T, 6167C>T, 6168C>T, 6169A>G, 6173C>G, 6174C>T, 6175T>A, 6176G>A, 6177A>G, 6178A>T, 6180C>A, 6181C>T, 6185A>G, 6186A>T, 6187A>T, 6188A>C, 6196G>A, 6197G>A, 6198C>G, 6199C>G, 6200C>G, 6201A>T, 6202A>T, 6203T>G, 6206G>T, 6208G>T, 6213T>C, 6214A>T, 6215T>C, 6216G>T, 6221G>T, 6222A>G, 6223A>T, 6227G>T, 6228G>C, 6229G>A, 6232T>C, 6233G>T, 6234T>A, 6235C>T, 6236T>G, 6237T>C, 6238G>T, 6240G>A, 6241T>G, 6242C>A, 6243C>G, 6244A>G, 6245A>T, 6251A>T, 6252T>G, 6253G>T, 6254A>T, 6255A>C, 6256G>C, 6257G>T, 6259G>T, 6261C>G, 6266G>C, 6268C>A, 6272A>G, 6277C>G, 6279T>A, 6280T>C, 6283G>A, 6286C>T, 6287T>C, 6288C>A, 6289A>C, 6291C>T, 6298A>C, 6301T>A, 6303G>A, 6304G>A

##### CDS

| | Begin | End | Coverage | Score | Concordance | Matches | Identities | I/D/M/F* | Stop Codons |
|---|---|---|---|---|---|---|---|---|---|
| KM754_gp3 | 1597 | 1695 | 4.5% | 327 | 44.8% | 99 (99.0%) | 45 (45.0%) | 1/0/0/0 | 0 |

Protein mutations: W1600R (6020T>A 6022G>A), P1602H (6027C>A 6028A>T), L1605H (6035T>C 6036T>A 6037A>T), W1608F (6045G>T 6046G>C), L1609K (6047C>A 6048T>A), K1616T (6069A>C 6070G>A), V1621T (6083G>A 6084T>C 6085A>T), R1624A (6092A>G 6093G>C 6094G>C), K1625F (6095A>T 6096A>T 6097G>C), D1627N (6101G>A 6103C>T), R1628Q (6105G>A 6106A>G), I1629V (6107A>G 6109C>T), N1631E (6113A>G 6115T>A), Y1633F (6120A>T 6121C>T), A1634L (6122G>C 6123C>T 6124T>G), G1635R (6125G>A), G1635_F1636insK (6127_6128insAAG), I1637V (6131A>G), A1638L (6134G>T 6135C>T 6136T>G), Y1640F (6141A>T), I1641F (6143A>T), I1646V (6158A>G 6160A>T), F1647Y (6162T>A), P1649L (6167C>T 6168C>T 6169A>G), P1651V (6173C>G 6174C>T 6175T>A), E1652S (6176G>A 6177A>G 6178A>T), A1653D (6180C>A 6181C>T), K1655V (6185A>G 6186A>T 6187A>T), K1656Q (6188A>C), A1659R (6197G>A 6198C>G 6199C>G), Q1660V (6200C>G 6201A>T 6202A>T), F1661V (6203T>G), V1662F (6206G>T 6208G>T), I1664T (6213T>C 6214A>T), C1665L (6215T>C 6216G>T), E1667C (6221G>T 6222A>G 6223A>T), G1669S (6227G>T 6228G>C 6229G>A), V1671Y (6233G>T 6234T>A 6235C>T), L1672A (6236T>G 6237T>C 6238G>T), S1673K (6240G>A 6241T>G), P1674R (6242C>A 6243C>G 6244A>G), T1675S (6245A>T), M1677C (6251A>T 6252T>G 6253G>T), K1678S (6254A>T 6255A>C 6256G>C), V1679F (6257G>T 6259G>T), A1680G (6261C>G), A1682P (6266G>C 6268C>A), I1684V (6272A>G), D1685E (6277C>G), F1686Y (6279T>A 6280T>C), S1689H (6287T>C 6288C>A 6289A>C), T1690I (6291C>T), N1693K (6301T>A), G1694E (6303G>A 6304G>A)

Codon mutations: ACC1597ACT (6013C>T), TTT1599TTC (6019T>C), TGG1600AGA (6020T>A 6022G>A), CCA1602CAT (6027C>A 6028A>T), GTG1603GTA (6031G>A), GGC1604GGG (6034C>G), TTA1605CAT (6035T>C 6036T>A 6037A>T), TAT1606TAC (6040T>C), GAA1607GAG (6043A>G), TGG1608TTC (6045G>T 6046G>C), CTG1609AAG (6047C>A 6048T>A), GTT1610GTG (6052T>G), CCA1612CCG (6058A>G), GGG1614GGC (6064G>C), AAG1616ACA (6069A>C 6070G>A), GCA1620GCT (6082A>T), GTA1621ACT (6083G>A 6084T>C 6085A>T), TTT1622TTC (6088T>C), AGG1624GCC (6092A>G 6093G>C 6094G>C), AAG1625TTC (6095A>T 6096A>T 6097G>C), GAC1627AAT (6101G>A 6103C>T), CGA1628CAG (6105G>A 6106A>G), ATC1629GTT (6107A>G 6109C>T), TTT1630TTC (6112T>C), AAT1631GAA (6113A>G 6115T>A), CCA1632CCT (6118A>T), TAC1633TTT (6120A>T 6121C>T), GCT1634CTG (6122G>C 6123C>T 6124T>G), GGA1635AGA (6125G>A), GGA1635_TTT1636insAAG (6127_6128insAAG), ATT1637GTT (6131A>G), GCT1638TTG (6134G>T 6135C>T 6136T>G), GTG1639GTA (6139G>A), TAT1640TTT (6141A>T), ATT1641TTT (6143A>T), ATC1644ATA (6154C>A), CTC1645CTT (6157C>T), ATA1646GTT (6158A>G 6160A>T), TTC1647TAC (6162T>A), AGC1648AGT (6166C>T), CCA1649TTG (6167C>T 6168C>T 6169A>G), CCT1651GTA (6173C>G 6174C>T 6175T>A), GAA1652AGT (6176G>A 6177A>G 6178A>T), GCC1653GAT (6180C>A 6181C>T), AAA1655GTT (6185A>G 6186A>T 6187A>T), AAG1656CAG (6188A>C), TTG1658TTA (6196G>A), GCC1659AGG (6197G>A 6198C>G 6199C>G), CAA1660GTT (6200C>G 6201A>T 6202A>T), TTC1661GTC (6203T>G), GTG1662TTT (6206G>T 6208G>T), ATA1664ACT (6213T>C 6214A>T), TGC1665CTC (6215T>C 6216G>T), GAA1667TGT (6221G>T 6222A>G 6223A>T), GGG1669TCA (6227G>T 6228G>C 6229G>A), CTT1670CTC (6232T>C), GTC1671TAT (6233G>T 6234T>A 6235C>T), TTG1672GCT (6236T>G 6237T>C 6238G>T), AGT1673AAG (6240G>A 6241T>G), CCA1674AGG (6242C>A 6243C>G 6244A>G), ACA1675TCA (6245A>T), ATG1677TGT (6251A>T 6252T>G 6253G>T), AAG1678TCC (6254A>T 6255A>C 6256G>C), GTG1679TTT (6257G>T 6259G>T), GCC1680GGC (6261C>G), GCC1682CCA (6266G>C 6268C>A), ATT1684GTT (6272A>G), GAC1685GAG (6277C>G), TTT1686TAC (6279T>A 6280T>C), TTG1687TTA (6283G>A), GGC1688GGT (6286C>T), TCA1689CAC (6287T>C 6288C>A 6289A>C), ACA1690ATA (6291C>T), ACA1692ACC (6298A>C), AAT1693AAA (6301T>A), GGG1694GAA (6303G>A 6304G>A)

##### Proteins

| | Begin | End | Coverage | Score | Concordance | Matches | Identities | I/D/M/F* | Stop Codons |
|---|---|---|---|---|---|---|---|---|---|
| polyprotein (YP_010087856.1) | 1597 | 1695 | 4.5% | 327 | 44.8% | 99 (99.0%) | 45 (45.0%) | 1/0/0/0 | 0 |

Protein mutations: W1600R (6020T>A 6022G>A), P1602H (6027C>A 6028A>T), L1605H (6035T>C 6036T>A 6037A>T), W1608F (6045G>T 6046G>C), L1609K (6047C>A 6048T>A), K1616T (6069A>C 6070G>A), V1621T (6083G>A 6084T>C 6085A>T), R1624A (6092A>G 6093G>C 6094G>C), K1625F (6095A>T 6096A>T 6097G>C), D1627N (6101G>A 6103C>T), R1628Q (6105G>A 6106A>G), I1629V (6107A>G 6109C>T), N1631E (6113A>G 6115T>A), Y1633F (6120A>T 6121C>T), A1634L (6122G>C 6123C>T 6124T>G), G1635R (6125G>A), G1635_F1636insK (6127_6128insAAG), I1637V (6131A>G), A1638L (6134G>T 6135C>T 6136T>G), Y1640F (6141A>T), I1641F (6143A>T), I1646V (6158A>G 6160A>T), F1647Y (6162T>A), P1649L (6167C>T 6168C>T 6169A>G), P1651V (6173C>G 6174C>T 6175T>A), E1652S (6176G>A 6177A>G 6178A>T), A1653D (6180C>A 6181C>T), K1655V (6185A>G 6186A>T 6187A>T), K1656Q (6188A>C), A1659R (6197G>A 6198C>G 6199C>G), Q1660V (6200C>G 6201A>T 6202A>T), F1661V (6203T>G), V1662F (6206G>T 6208G>T), I1664T (6213T>C 6214A>T), C1665L (6215T>C 6216G>T), E1667C (6221G>T 6222A>G 6223A>T), G1669S (6227G>T 6228G>C 6229G>A), V1671Y (6233G>T 6234T>A 6235C>T), L1672A (6236T>G 6237T>C 6238G>T), S1673K (6240G>A 6241T>G), P1674R (6242C>A 6243C>G 6244A>G), T1675S (6245A>T), M1677C (6251A>T 6252T>G 6253G>T), K1678S (6254A>T 6255A>C 6256G>C), V1679F (6257G>T 6259G>T), A1680G (6261C>G), A1682P (6266G>C 6268C>A), I1684V (6272A>G), D1685E (6277C>G), F1686Y (6279T>A 6280T>C), S1689H (6287T>C 6288C>A 6289A>C), T1690I (6291C>T), N1693K (6301T>A), G1694E (6303G>A 6304G>A)

Codon mutations: ACC1597ACT (6013C>T), TTT1599TTC (6019T>C), TGG1600AGA (6020T>A 6022G>A), CCA1602CAT (6027C>A 6028A>T), GTG1603GTA (6031G>A), GGC1604GGG (6034C>G), TTA1605CAT (6035T>C 6036T>A 6037A>T), TAT1606TAC (6040T>C), GAA1607GAG (6043A>G), TGG1608TTC (6045G>T 6046G>C), CTG1609AAG (6047C>A 6048T>A), GTT1610GTG (6052T>G), CCA1612CCG (6058A>G), GGG1614GGC (6064G>C), AAG1616ACA (6069A>C 6070G>A), GCA1620GCT (6082A>T), GTA1621ACT (6083G>A 6084T>C 6085A>T), TTT1622TTC (6088T>C), AGG1624GCC (6092A>G 6093G>C 6094G>C), AAG1625TTC (6095A>T 6096A>T 6097G>C), GAC1627AAT (6101G>A 6103C>T), CGA1628CAG (6105G>A 6106A>G), ATC1629GTT (6107A>G 6109C>T), TTT1630TTC (6112T>C), AAT1631GAA (6113A>G 6115T>A), CCA1632CCT (6118A>T), TAC1633TTT (6120A>T 6121C>T), GCT1634CTG (6122G>C 6123C>T 6124T>G), GGA1635AGA (6125G>A), GGA1635_TTT1636insAAG (6127_6128insAAG), ATT1637GTT (6131A>G), GCT1638TTG (6134G>T 6135C>T 6136T>G), GTG1639GTA (6139G>A), TAT1640TTT (6141A>T), ATT1641TTT (6143A>T), ATC1644ATA (6154C>A), CTC1645CTT (6157C>T), ATA1646GTT (6158A>G 6160A>T), TTC1647TAC (6162T>A), AGC1648AGT (6166C>T), CCA1649TTG (6167C>T 6168C>T 6169A>G), CCT1651GTA (6173C>G 6174C>T 6175T>A), GAA1652AGT (6176G>A 6177A>G 6178A>T), GCC1653GAT (6180C>A 6181C>T), AAA1655GTT (6185A>G 6186A>T 6187A>T), AAG1656CAG (6188A>C), TTG1658TTA (6196G>A), GCC1659AGG (6197G>A 6198C>G 6199C>G), CAA1660GTT (6200C>G 6201A>T 6202A>T), TTC1661GTC (6203T>G), GTG1662TTT (6206G>T 6208G>T), ATA1664ACT (6213T>C 6214A>T), TGC1665CTC (6215T>C 6216G>T), GAA1667TGT (6221G>T 6222A>G 6223A>T), GGG1669TCA (6227G>T 6228G>C 6229G>A), CTT1670CTC (6232T>C), GTC1671TAT (6233G>T 6234T>A 6235C>T), TTG1672GCT (6236T>G 6237T>C 6238G>T), AGT1673AAG (6240G>A 6241T>G), CCA1674AGG (6242C>A 6243C>G 6244A>G), ACA1675TCA (6245A>T), ATG1677TGT (6251A>T 6252T>G 6253G>T), AAG1678TCC (6254A>T 6255A>C 6256G>C), GTG1679TTT (6257G>T 6259G>T), GCC1680GGC (6261C>G), GCC1682CCA (6266G>C 6268C>A), ATT1684GTT (6272A>G), GAC1685GAG (6277C>G), TTT1686TAC (6279T>A 6280T>C), TTG1687TTA (6283G>A), GGC1688GGT (6286C>T), TCA1689CAC (6287T>C 6288C>A 6289A>C), ACA1690ATA (6291C>T), ACA1692ACC (6298A>C), AAT1693AAA (6301T>A), GGG1694GAA (6303G>A 6304G>A)

*: Inserts / Deletes / Misaligned / Frameshifts

#### Analysis details

This analysis was performed with panviral2.64

### NGS Details (UN60): Synechococcus phage S-SRM01

#### Assembly

| | |
|---|---|
| **Coverage Length** | 207 (1 contig(s)) |
| **Depth Of Coverage** | 25.2 |
| **Number Of Reads** | 43 |
| **Reads Per Million** | 0.91 rpm (after QC) |
| **Ambiguities** | 0 |
| **Assembly Method** | de novo + reference guided assembly |
| **Consensus Caller** | Bcf Tools |

#### Coverage Map

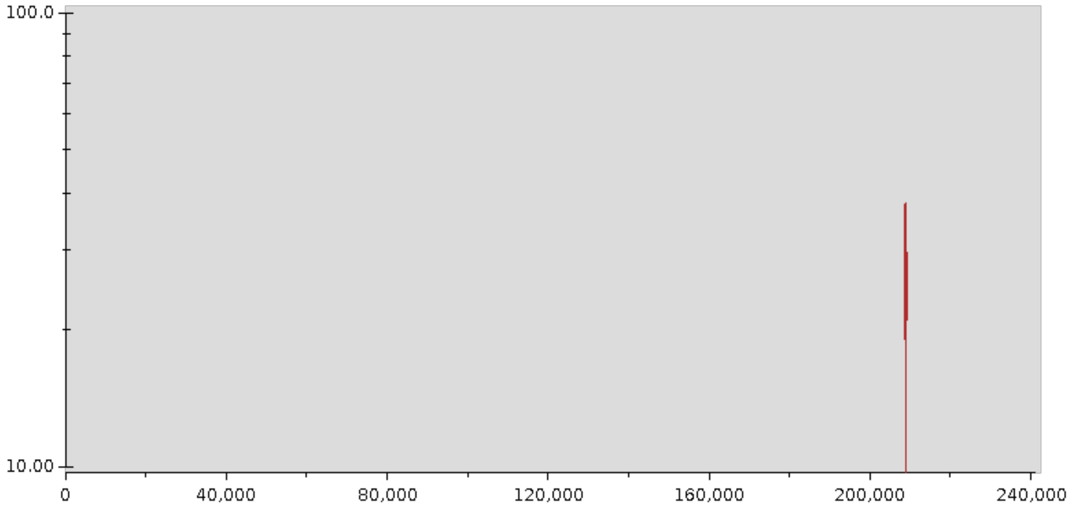

#### Assignment

| | |
|---|---|
| **Type** | Synechococcus phage S-SRM01 (Taxonomy ID: 2781608) |
| **Reference Genome** | NC_070963.1 |
| **NT Identity (%)** | 76.8116 |
| **AA Identity (%)** | 88.4058 |
| **Number Of Stop Codons** | 0 |
| **Number Of CDS** | 353 |

#### Alignment

| | |
|---|---|
| **Alignment Score** | 222.0 (NT) + 433.0 (AA) = 655.0 |
| **Concordance (%)** | 74.3473 |
| **Alignment Method** | Local, heuristic, nucleotide (BLASTN) |

#### Genome Region

Sequence starts at position 208802 and ends at position 209008 relative to NC_070963.1 reference sequence.

#### Alignment Detailed Statistics

| | Begin | End | Coverage | Score | Concordance | Matches | Identities | I/D/M/F* | Stop Codons |
|---|---|---|---|---|---|---|---|---|---|
| NT | 208802 | 209008 | 0.1% | 222 | 53.6% | 207 (100%) | 159 (76.8%) | 0/0 | |
| Mutations: | 208807T>C, 208809G>A, 208812T>A, 208816A>T, 208817G>C, 208821A>T, 208830C>T, 208832A>T, 208837C>A, 208842G>A, 208851A>T, 208866G>T, 208875C>A, 208878C>T, 208881T>C, 208885C>T, 208887T>A, 208890A>C, 208893T>G, 208902C>A, 208908T>C, 208911T>C, 208914T>C, 208917G>A, 208932C>T, 208938T>C, 208939C>T, 208944T>A, 208950G>T, 208951T>A, 208952C>G, 208953A>T, 208954C>T, 208957G>A, 208959T>C, 208960C>A, 208962T>G, 208971T>A, 208976C>A, 208983A>T, 208984C>G, 208985A>C, 208986G>T, 208990T>G, 208992T>A, 209000A>G, 209001G>A, 209004T>C | | | | | | | | |

*: Inserts / Deletes / Misaligned / Frameshifts

#### Analysis details

This analysis was performed with panviral2.64

### NGS Details (UN60): Marseillevirus marseillevirus

#### Assembly

| | |
|---|---|
| **Coverage Length** | 113 (1 contig(s)) |
| **Depth Of Coverage** | 25.6 |
| **Number Of Reads** | 38 |
| **Reads Per Million** | 0.81 rpm (after QC) |
| **Ambiguities** | 0 |
| **Assembly Method** | de novo + reference guided assembly |
| **Consensus Caller** | Bcf Tools |

#### Coverage Map

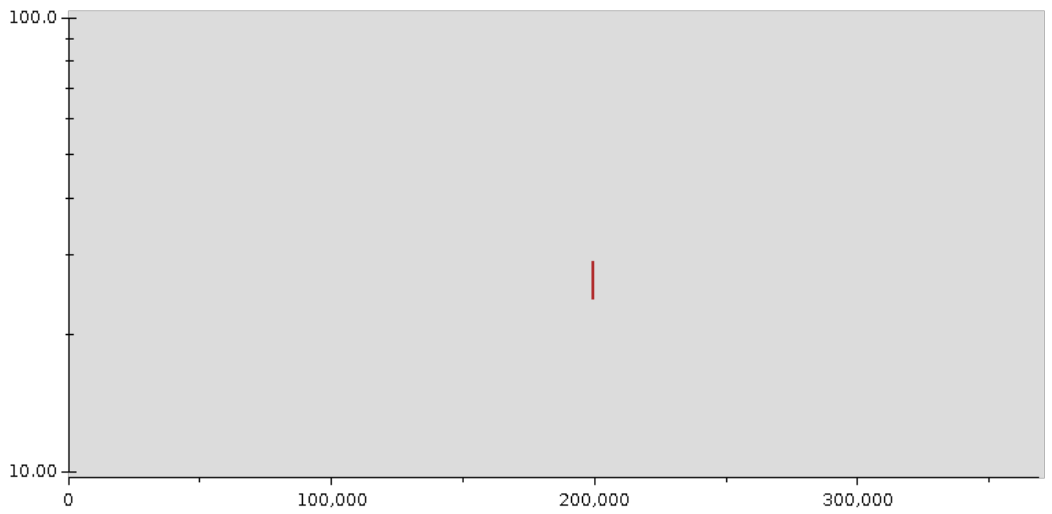

#### Assignment

| | |
|---|---|
| **Type** | Marseillevirus marseillevirus (Taxonomy ID: 694581) |
| **Reference Genome** | NC_013756.1 |
| **NT Identity (%)** | 81.4159 |
| **AA Identity (%)** | 94.7368 |
| **Number Of Stop Codons** | 0 |
| **Number Of CDS** | 428 |

#### Alignment

| | |
|---|---|
| **Alignment Score** | 142.0 (NT) + 245.0 (AA) = 387.0 |
| **Concordance (%)** | 81.9915 |
| **Alignment Method** | Local, heuristic, nucleotide (BLASTN) |

#### Genome Region

Sequence starts at position 199409 and ends at position 199521 relative to NC_013756.1 reference sequence.

#### Alignment Detailed Statistics

| | Begin | End | Coverage | Score | Concordance | Matches | Identities | I/D/M/F* | Stop Codons |
|---|---|---|---|---|---|---|---|---|---|
| **NT** | **199409** | **199521** | **0.1%** | **142** | **62.8%** | **113 (100%)** | **92 (81.4%)** | **0/0** | |

Mutations: 199411A>C, 199414T>A, 199416T>G, 199423A>G, 199432A>G, 199444T>C, 199449C>T, 199459A>C, 199462C>A, 199464A>G, 199465A>G, 199470T>G, 199471T>G, 199474A>G, 199477T>C, 199480C>G, 199489T>G, 199495G>A, 199501G>C, 199503G>A, 199516C>T

*: Inserts / Deletes / Misaligned / Frameshifts

#### Analysis details

This analysis was performed with panviral2.64

### NGS Details (UN60): Petuvirus venapetuniae

#### Assembly

| | |
|---|---|
| **Coverage Length** | 939 (2 contig(s)) |
| **Depth Of Coverage** | 5.1 |
| **Number Of Reads** | 36 |
| **Reads Per Million** | 0.76 rpm (after QC) |
| **Ambiguities** | 0 |
| **Assembly Method** | de novo + reference guided assembly |
| **Consensus Caller** | Bcf Tools |

#### Coverage Map

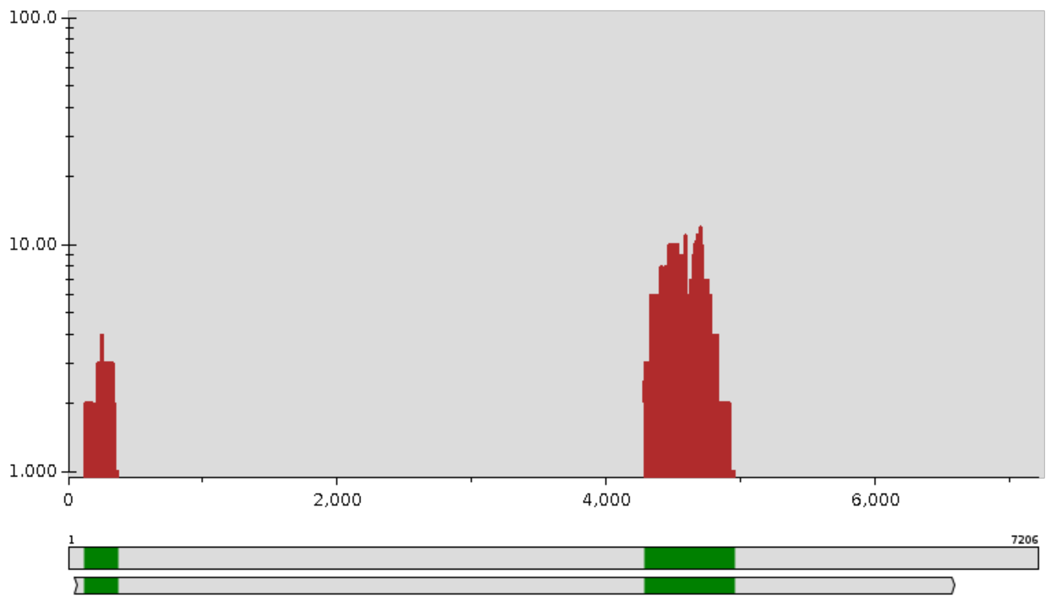

#### Assignment

| | |
|---|---|
| **Type** | Petuvirus venapetuniae (Taxonomy ID: 3048265) |
| **Reference Genome** | NC_001839.2 |
| **NT Identity (%)** | 58.5729 |
| **AA Identity (%)** | 53.6741 |
| **Number Of Stop Codons** | 3 |
| **Number Of CDS** | 1 |

#### Alignment

| | |
|---|---|
| **Alignment Score** | 304.0 (NT) + 1191.0 (AA) = 1495.0 |
| **Concordance (%)** | 37.4218 |

| Alignment Method | Global, seeded, nucleotide + amino acids (AGA) |
|---|---|

#### Genome Region

Sequence starts at position 116 and ends at position 4957 relative to NC_001839.2 reference sequence.

#### Alignment Detailed Statistics

| | Begin | End | Coverage | Score | Concordance | Matches | Identities | I/D/M/F* | Stop Codons |
|---|---|---|---|---|---|---|---|---|---|
| **NT** | **116** | **4957** | **13.0%** | **304** | **16.3%** | **936 (99.4%)** | **550 (58.4%)** | **3/3** | |

Mutations: 116A>T, 119C>T, 121C>G, 122T>A, 126G>A, 128T>C, 129T>C, 131G>T, 134T>A, 136A>G, 137T>A, 149C>A, 150C>T, 152T>G, 159G>A, 161A>T, 165A>G, 170T>A, 171A>C, 172A>C, 173A>C, 177A>G, 179C>G, 186A>G, 187T>C, 192G>T, 193A>C, 194T>C, 197C>G, 199A>C, 200T>C, 201C>T, 203T>A, 207T>C, 209G>T, 210T>A, 221C>T, 233A>C, 238G>A, 239A>T, 241C>A, 242C>A, 243G>T, 245T>A, 246A>T, 248C>G, 251T>A, 255T>A, 256C>T, 257T>C, 260G>T, 263C>T, 265G>A, 269A>C, 278G>A, 279T>A, 280C>A, 281T>C, 286C>T, 287T>C, 288C>A, 289G>T, 290C>A, 291C>G, 292A>G, 293T>A, 296G>A, 298A>G, 310C>T, 317C>A, 321T>C, 323A>C, 324G>A, 329G>A, 333C>T, 336G>A, 338C>A, 339T>C, 344T>A, 347A>T, 348G>C, 349G>C, 359G>A, 362T>C, 363G>T, 366A>T, 4287C>A, 4289A>T, 4293G>A, 4294A>G, 4298C>T, 4302G>A, 4304T>A, 4307A>T, 4309A>C, 4310A>T, 4314G>A, 4315C>G, 4316A>C, 4320G>T, 4321A>C, 4325T>C, 4328T>C, 4334C>T, 4337C>T, 4340G>A, 4342G>C, 4343G>T, 4344T>G, 4346A>T, 4349G>A, 4350C>A, 4354T>A, 4355G>A, 4361A>C, 4362A>T, 4363A>C, 4364A>C, 4365T>C, 4366T>C, 4367A>T, 4376C>T, 4379A>T, 4386C>A, 4388A>G, 4391A>T, 4397C>T, 4398C>T, 4399A>C, 4401T>G, 4403C>T, 4404C>T, 4406C>A, 4407C>A, 4410G>T, 4411A>G, 4412T>G, 4413G>A, 4414A>T, 4415C>T, 4417A>G, 4418G>A, 4420T>A, 4421T>C, 4424C>G, 4430A>T, 4436G>A, 4437C>A, 4439C>G, 4440A>G, 4441C>A, 4442A>T, 4445C>T, 4448C>G, 4449T>A, 4450C>A, 4451T>A, 4452C>A, 4453A>G, 4454C>A, 4455T>A, 4456T>C, 4457A>T, 4459C>T, 4460C>T, 4464G>A, 4466A>C, 4469G>T, 4470C>A, 4472C>T, 4474T>A, 4475T>C, 4476T>A, 4477C>G, 4478A>T, 4481G>A, 4488C>A, 4490T>G, 4493G>A, 4499A>T, 4502T>C, 4509T>A, 4511A>T, 4512G>C, 4513G>A, 4517C>T, 4518C>T, 4519A>C, 4520T>G, 4521C>A, 4522C>A, 4523T>A, 4526T>A, 4529G>T, 4531G>A, 4533C>T, 4534C>A, 4535A>T, 4541A>T, 4543G>C, 4544G>T, 4547T>C, 4548T>A, 4549G>A, 4551A>G, 4553T>C, 4556T>C, 4557G>T, 4558A>T, 4559T>C, 4560A>G, 4565T>A, 4567T>A, 4569C>G, 4577G>T, 4580C>T, 4586A>C, 4589C>T, 4592T>A, 4593C>T, 4595T>A, 4600C>A, 4601A>T, 4611T>G, 4612T>A, 4613G>A, 4616C>T, 4622A>C, 4624C>T, 4625C>T, 4630T>A, 4631C>T, 4632A>T, 4633A>C, 4634A>T, 4637C>T, 4640C>T, 4641C>A, 4643G>T, 4645C>A, 4650C>A, 4651T>G, 4652T>C, 4653T>A, 4654T>A, 4655T>C, 4656T>A, 4657C>T, 4658A>C, 4659G>T, 4662T>A, 4664G>T, 4667C>G, 4670C>T, 4680A>G, 4682C>T, 4683C>T, 4685T>G, 4686C>A, 4688T>A, 4697A>G, 4698A>G, 4699C>A, 4700A>T, 4701C>A, 4703G>T, 4706A>T, 4707G>T, 4708A>C, 4712C>T, 4713A>T, 4715A>T, 4719T>C, 4720T>A, 4721G>T, 4722C>T, 4724T>A, 4727C>T, 4728C>A, 4729A>C, 4730G>C, 4734A>T, 4736A>C, 4738G>A, 4739T>A, 4740T>A, 4742G>T, 4745T>C, 4750A>G, 4752T>A, 4753T>A, 4757A>T, 4758G>C, 4760C>T, 4761A>G, 4763G>C, 4764T>A, 4766A>C, 4767T>A, 4768C>G, 4769G>T, 4772A>T, 4775A>G, 4778G>A, 4783T>A, 4784C>A, 4785C>T, 4787T>G, 4788G>T, 4789C>T, 4795A>C, 4796T>A, 4802T>C, 4803C>A, 4804A>G, 4808C>T, 4809T>C, 4811A>T, 4814A>G, 4815A>C, 4816T>A, 4817G>T, 4823T>A, 4824G>T, 4825C>A, 4827G>C, 4829T>A, 4835T>A, 4837T>A, 4839T>A, 4840C>A, 4844T>C, 4845G>A, 4846C>T, 4847T>C, 4848G>T, 4850T>C, 4851_4853delCAC, 4855T>G, 4856C>A, 4857T>G, 4859T>C, 4860T>A, 4862G>C, 4866C>T, 4868C>T, 4869C>T, 4870A>C, 4871A>T, 4871_4872insGGT, 4874G>A, 4877T>C, 4881G>A, 4884A>G, 4885C>A, 4888A>T, 4889T>C, 4890C>A, 4891T>C, 4892C>T, 4893T>G, 4894C>A, 4895G>C, 4896G>A, 4897T>A, 4900A>C, 4901G>T, 4905A>T, 4910A>G, 4911C>A, 4912A>G, 4916T>C, 4917C>T, 4919T>G, 4922G>A, 4924T>G, 4925T>C, 4926G>C, 4928T>C, 4935A>G, 4937C>T, 4938A>G, 4939G>C, 4943T>C, 4946T>C, 4952A>T, 4953G>A, 4955G>A

##### CDS

| | Begin | End | Coverage | Score | Concordance | Matches | Identities | I/D/M/F* | Stop Codons |
|---|---|---|---|---|---|---|---|---|---|
| PVCVgp1 | 23 | 1636 | 14.4% | 1191 | 55.2% | 312 (99.4%) | 168 (53.5%) | 1/1/0/0 | 3 |

Protein mutations: S24* (121C>G 122T>A), A26T (126G>A 128T>C), N29R (136A>G 137T>A), Y33* (149C>A), V37I (159G>A 161A>T), I39V (165A>G), K41P (171A>C 172A>C 173A>C), N43E (177A>G 179C>G), I46A (186A>G 187T>C), D48S (192G>T 193A>C 194T>C), Y50S (199A>C 200T>C), L54I (210T>A), R63N (238G>A 239A>T), S64* (241C>A 242C>A), V65L (243G>T 245T>A), T66S (246A>T 248C>G), S69I (255T>A 256C>T 257T>C), Q70H (260G>T), R72H (265G>A), S77N (279T>A 280C>A 281T>C), P79L (286C>T 287T>C), R80I (288C>A 289G>T 290C>A), H81G (291C>G 292A>G 293T>A), K83R (298A>G), A87V (310C>T), D92N (324G>A), P95S (333C>T), V96I (336G>A 338C>A), F97L (339T>C), G100P (348G>C 349G>C), V105F (363G>T), T106S (366A>T), L1413I (4287C>A 4289A>T), E1415R (4293G>A 4294A>G), D1418K (4302G>A 4304T>A), Q1420P (4309A>C 4310A>T), A1422S (4314G>A 4315C>G 4316A>C), E1424S (4320G>T 4321A>C), R1431T (4342G>C 4343G>T), S1432A (4344T>G 4346A>T), Q1434K (4350C>A), V1435E (4354T>A 4355G>A), K1438S (4362A>T 4363A>C 4364A>C), L1439P (4365T>C 4366T>C 4367A>T), Q1446K (4386C>A 4388A>G), H1450S (4398C>T 4399A>C), F1451V (4401T>G 4403C>T), Q1453K (4407C>A), D1454W (4410G>T 4411A>G 4412T>G), D1455I (4413G>A 4414A>T 4415C>T), K1456R (4417A>G 4418G>A), F1457Y (4420T>A 4421T>C), L1463M (4437C>A 4439C>G), T1464D (4440A>G 4441C>A 4442A>T), F1466L (4448C>G), S1467K (4449T>A 4450C>A 4451T>A), H1468R (4452C>A 4453A>G 4454C>A), L1469T (4455T>A 4456T>C 4457A>T), S1470F (4459C>T 4460C>T), A1472T (4464G>A 4466A>C), K1473N (4469G>T), L1474I (4470C>A 4472C>T), F1475Y (4474T>A 4475T>C), L1480M (4488C>A 4490T>G), L1487I (4509T>A 4511A>T), G1488Q (4512G>C 4513G>A), H1490S (4518C>T 4519A>C 4520T>G), P1491K (4521C>A 4522C>A 4523T>A), N1492K (4526T>A), E1493D (4529G>T), R1494K (4531G>A), P1495Y (4533C>T 4534C>A 4535A>T), G1498A (4543G>C 4544G>T), C1500N (4548T>A 4549G>A), I1501V (4551A>G 4553T>C), D1503F (4557G>T 4558A>T 4559T>C), R1504G (4560A>G), H1505Q (4565T>A), F1506Y (4567T>A), Q1507E (4569C>G), K1509N (4577G>T), T1517N (4600C>A 4601A>T), L1521E (4611T>G 4612T>A 4613G>A), K1524N (4622A>C), A1525V (4624C>T 4625C>T), I1527N (4630T>A 4631C>T), K1528S (4632A>T 4633A>C 4634A>T), Q1531N (4641C>A 4643G>T), P1532H (4645C>A), L1534S (4650C>A 4651T>G 4652T>C), F1535N (4653T>A 4654T>A 4655T>C), S1536I (4656T>A 4657C>T 4658A>C), A1537S (4659G>T), L1538I (4662T>A 4664G>T), I1544V (4680A>G 4682C>T), L1546I (4686C>A 4688T>A), T1550D (4698A>G 4699C>A 4700A>T), L1551I (4701C>A 4703G>T), E1552D (4706A>T), D1553S (4707G>T 4708A>C), I1555F (4713A>T 4715A>T), L1557H (4719T>C 4720T>A 4721G>T), Q1560T (4728C>A 4729A>C 4730G>C), I1562F (4734A>T 4736A>C), S1563K (4738G>A 4739T>A), L1564I (4740T>A 4742G>T), K1567R (4750A>G), F1568N (4752T>A 4753T>A), V1570L (4758G>C 4760C>T), M1571V (4761A>G 4763G>C), L1572I (4764T>A 4766A>C), I1578K (4783T>A 4784C>A), A1580F (4788G>T 4789C>T), N1582T (4795A>C 4796T>A), Q1585R (4803C>A 4804A>G), M1589H (4815A>C 4816T>A 4817G>T), F1591L (4823T>A), A1592Y (4824G>T 4825C>A), D1593Q (4827G>C 4829T>A), F1596Y (4837T>A), S1597K (4839T>A 4840C>A), A1599I (4845G>A 4846C>T 4847T>C), G1600C (4848G>T 4850T>C), H1601del (4851_4853delCAC), I1602R (4855T>G 4856C>A), S1603A (4857T>G 4859T>C), L1604I (4860T>A 4862G>C), L1606F (4866C>T 4868C>T), Q1607S (4869C>T 4870A>C 4871A>T), Q1607_K1608insG (4871_4872insGGT), D1611N (4881G>A), T1612E (4884A>G 4885C>A), N1613I (4888A>T 4889T>C), L1614T (4890C>A 4891T>C 4892C>T), S1615D (4893T>G 4894C>A 4895G>C), V1616K (4896G>A 4897T>A), K1617T (4900A>C 4901G>T), I1619L (4905A>T), Q1621R (4911C>A 4912A>G), I1625S (4924T>G 4925T>C), V1626L (4926G>C 4928T>C), I1629V (4935A>G 4937C>T), R1630A (4938A>G 4939G>C), E1635K (4953G>A 4955G>A)

Codon mutations: AAA22..T (116A>T), CTC23CTT (119C>T), TCT24TGA (121C>G 122T>A), GCT26ACC (126G>A 128T>C), TTG27CTT (129T>C 131G>T), TCT28TCA (134T>A), AAT29AGA (136A>G 137T>A), TAC33TAA (149C>A), CTT34TTG (150C>T 152T>G), GTA37ATT (159G>A 161A>T), ATC39GTC (165A>G), CTT40CTA (170T>A), AAA41CCC (171A>C 172A>C 173A>C), AAC43GAG (177A>G 179C>G), ATA46GCA (186A>G 187T>C), GAT48TCC (192G>T 193A>C 194T>C), ACC49ACG (197C>G), TAT50TCC (199A>C 200T>C), CTT51TTA (201C>T 203T>A), TTG53CTT (207T>C 209G>T), TTA54ATA (210T>A), TAC57TAT (221C>T), GCA61GCC (233A>C), AGA63AAT (238G>A 239A>T), TCC64TAA (241C>A 242C>A), GTT65TTA (243G>T 245T>A), ACC66TCG (246A>T 248C>G), CCT67CCA (251T>A), TCT69ATC (255T>A 256C>T 257T>C), CAG70CAT (260G>T), ATC71ATT (263C>T), CGT72CAT (265G>A), TCA73TCC (269A>C), CAG76CAA (278G>A), TCT77AAC (279T>A 280C>A 281T>C), CCT79CTC (286C>T 287T>C), CGC80ATA (288C>A 289G>T 290C>A), CAT81GGA (291C>G 292A>G 293T>A), GTG82GTA (296G>A), AAA83AGA (298A>G), GCT87GTT (310C>T), TCC89TCA (317C>A), TTA91CTC (321T>C 323A>C), GAT92AAT (324G>A), CAG93CAA (329G>A), CCT95TCT (333C>T), GTC96ATA (336G>A 338C>A), TTT97CTT (339T>C), GCT98GCA (344T>A), ACA99ACT (347A>T), GGA100CCA (348G>C 349G>C), CAG103CAA (359G>A), TTT104TTC (362T>C), GTT105TTT (363G>T), ACT106TCT (366A>T), CTA1413ATT (4287C>A 4289A>T), GAA1415AGA (4293G>A 4294A>G), CCC1416CCT (4298C>T), GAT1418AAA (4302G>A 4304T>A), TCA1419TCT (4307A>T), CAA1420CCT (4309A>C 4310A>T), GCA1422AGC (4314G>A 4315C>G 4316A>C), GAA1424TCA (4320G>T 4321A>C), GCT1425GCC (4325T>C), TTT1426TTC (4328T>C), GTC1428GTT (4334C>T), AAC1429AAT (4337C>T), AAG1430AAA (4340G>A), AGG1431ACT (4342G>C 4343G>T), TCA1432GCT (4344T>G 4346A>T), GAG1433GAA (4349G>A), CAA1434AAA (4350C>A), GTG1435GAA (4354T>A 4355G>A), GGA1437GGC (4361A>C), AAA1438TCC (4362A>T 4363A>C 4364A>C), TTA1439CCT (4365T>C 4366T>C 4367A>T), GTC1442GTT (4376C>T), ATA1443ATT (4379A>T), CAA1446AAG (4386C>A 4388A>G), CCA1447CCT (4391A>T), AAC1449AAT (4397C>T), CAT1450TCT (4398C>T 4399A>C), TTC1451GTT (4401T>G 4403C>T), CTC1452TTA (4404C>T 4406C>A), CAA1453AAA (4407C>A), GAT1454TGG (4410G>T 4411A>G 4412T>G), GAC1455ATT (4413G>A 4414A>T 4415C>T), AAG1456AGA (4417A>G 4418G>A), TTT1457TAC (4420T>A 4421T>C), CCC1458CCG (4424C>G), CCA1460CCT (4430A>T), AAG1462AAA (4436G>A), CTC1463ATG (4437C>A 4439C>G), ACA1464GAT (4440A>G 4441C>A 4442A>T), CTC1465CTT (4445C>T), TTC1466TTG (4448C>G), TCT1467AAA (4449T>A 4450C>A 4451T>A), CAC1468AGA (4452C>A 4453A>G 4454C>A), TTA1469ACT (4455T>A 4456T>C 4457A>T), TCC1470TTT (4459C>T 4460C>T), GCA1472ACC (4464G>A 4466A>C), AAG1473AAT (4469G>T), CTC1474ATT (4470C>A 4472C>T), TTT1475TAC (4474T>A 4475T>C), TCA1476AGT (4476T>A 4477C>G 4478A>T), AAG1477AAA (4481G>A), CTT1480ATG (4488C>A 4490T>G), AAG1481AAA (4493G>A), GGA1483GGT (4499A>T), TTT1484TTC (4502T>C), TTA1487ATT (4509T>A 4511A>T), GGA1488CAA (4512G>C 4513G>A), ATC1489ATT (4517C>T), CAT1490TCG (4518C>T 4519A>C 4520T>G), CCT1491AAA (4521C>A 4522C>A 4523T>A), AAT1492AAA (4526T>A), GAG1493GAT (4529G>T), AGA1494AAA (4531G>A), CCA1495TAT (4533C>T 4534C>A 4535A>T), ACA1497ACT (4541A>T), GGG1498GCT (4543G>C 4544G>T), TTT1499TTC (4547T>C), TGC1500AAC (4548T>A 4549G>A), ATT1501GTC (4551A>G 4553T>C), CCT1502CCC (4556T>C), GAT1503TTC (4557G>T 4558A>T 4559T>C), AGA1504GGA (4560A>G), CAT1505CAA (4565T>A), TTT1506TAT (4567T>A), CAA1507GAA (4569C>G), AAG1509AAT (4577G>T), GTC1510GTT (4580C>T), CCA1512CCC (4586A>C), TTC1513TTT (4589C>T), GGT1514GGA (4592T>A), CTT1515TTA (4593C>T 4595T>A), ACA1517AAT (4600C>A 4601A>T), TTG1521GAA (4611T>G 4612T>A 4613G>A), TTC1522TTT (4616C>T), AAA1524AAC (4622A>C), GCC1525GTT (4624C>T 4625C>T), ATC1527AAT (4630T>A 4631C>T), AAA1528TCT (4632A>T 4633A>C 4634A>T), ATC1529ATT (4637C>T), TTC1530TTT (4640C>T), CAG1531AAT (4641C>A 4643G>T), CCC1532CAC (4645C>A), CTT1534AGC (4650C>A 4651T>G 4652T>C), TTT1535AAC (4653T>A 4654T>A 4655T>C), TCA1536ATC (4656T>A 4657C>T 4658A>C), GCT1537TCT (4659G>T), TTG1538ATT (4662T>A 4664G>T), GTC1539GTG (4667C>G), TAC1540TAT (4670C>T), ATC1544GTT (4680A>G 4682C>T), CTT1545TTG (4683C>T 4685T>G), CTT1546ATA (4686C>A 4688T>A), GAA1549GAG (4697A>G), ACA1550GAT (4698A>G 4699C>A 4700A>T), CTG1551ATT (4701C>A 4703G>T), GAA1552GAT (4706A>T), GAT1553TCT (4707G>T 4708A>C), CAC1554CAT (4712C>T), ATA1555TTT (4713A>T 4715A>T), TTG1557CAT (4719T>C 4720T>A 4721G>T), CTT1558TTA (4722C>T 4724T>A), AAC1559AAT (4727C>T), CAG1560ACC (4728C>A 4729A>C 4730G>C), ATA1562TTC (4734A>T 4736A>C), AGT1563AAA (4738G>A 4739T>A), TTG1564ATT (4740T>A 4742G>T), GTT1565GTC (4745T>C), AAA1567AGA (4750A>G), TTT1568AAT (4752T>A 4753T>A), GGA1569GGT (4757A>T), GTC1570CTT (4758G>C 4760C>T), ATG1571GTC (4761A>G 4763G>C), TTA1572ATC (4764T>A 4766A>C), TCG1573AGT (4767T>A 4768C>G 4769G>T), GCA1574GCT (4772A>T), AAA1575AAG (4775A>G), AAG1576AAA (4778G>A), ATC1578AAA (4783T>A 4784C>A), CTT1579TTG (4785C>T 4787T>G), GCT1580TTT (4788G>T 4789C>T), AAT1582ACA (4795A>C 4796T>A), ATT1584ATC (4802T>C), CAA1585AGA (4803C>A 4804A>G), TTC1586TTT (4808C>T), TTA1587CTT (4809T>C 4811A>T), GGA1588GGG (4814A>G), ATG1589CAT (4815A>C 4816T>A 4817G>T), TTT1591TTA (4823T>A), GCT1592TAT (4824G>T 4825C>A), GAT1593CAA (4827G>C 4829T>A), ACT1595ACA (4835T>A), TTT1596TAT (4837T>A), TCA1597AAA (4839T>A 4840C>A), CCT1598CCC (4844T>C), GCT1599ATC (4845G>A 4846C>T 4847T>C), GGT1600TGC (4848G>T 4850T>C), CAC1601del (4851_4853delCAC), ATC1602AGA (4855T>G 4856C>A), TCT1603GCC (4857T>G 4859T>C), TTG1604ATC (4860T>A 4862G>C), CTC1606TTT (4866C>T 4868C>T), CAA1607TCT (4869C>T 4870A>C 4871A>T), CAA1607_AAG1608insGGT (4871_4872insGGT), AAG1608AAA (4874G>A), TTT1609TTC (4877T>C), GAT1611AAT (4881G>A), ACA1612GAA (4884A>G 4885C>A), AAT1613ATC (4888A>T 4889T>C), CTC1614ACT (4890C>A 4891T>C 4892C>T), TCG1615GAC (4893T>G 4894C>A 4895G>C), GTA1616AAA (4896G>A 4897T>A), AAG1617ACT (4900A>C 4901G>T), ATA1619TTA (4905A>T), CAA1620CAG (4910A>G), CAA1621AGA (4911C>A 4912A>G), TTT1622TTC (4916T>C), CTT1623TTG (4917C>T 4919T>G), GGG1624GGA (4922G>A), ATT1625AGC (4924T>G 4925T>C), GTT1626CTC (4926G>C 4928T>C), ATC1629GTT (4935A>G 4937C>T), AGA1630GCA (4938A>G 4939G>C), GAT1631GAC (4943T>C), TTT1632TTC (4946T>C), CCA1634CCT (4952A>T), GAG1635AAA (4953G>A 4955G>A)

| | Begin | End | Coverage | Score | Concordance | Matches | Identities | I/D/M/F* | Stop Codons |
|---|---|---|---|---|---|---|---|---|---|
| **NT** | **116** | **4957** | **13.0%** | **304** | **16.3%** | **936 (99.4%)** | **550 (58.4%)** | **3/3** | |

**Proteins**

| | Begin | End | Coverage | Score | Concordance | Matches | Identities | I/D/M/F* | Stop Codons |
|---|---|---|---|---|---|---|---|---|---|
| ORF I polyprotein (NP_127504.1) | 23 | 1636 | 14.4% | 1191 | 55.2% | 312 (99.4%) | 168 (53.5%) | 1/1/0/0 | 3 |

| | |
|---|---|
| Protein mutations: | S24* (121C>G 122T>A), A26T (126G>A 128T>C), N29R (136A>G 137T>A), Y33* (149C>A), V37I (159G>A 161A>T), I39V (165A>G), K41P (171A>C 172A>C 173A>C), N43E (177A>G 179C>G), I46A (186A>G 187T>C), D48S (192G>T 193A>C 194T>C), Y50S (199A>C 200T>C), L54I (210T>A), R63N (238G>A 239A>T), S64* (241C>A 242C>A), V65L (243G>T 245T>A), T66S (246A>T 248C>G), S69I (255T>A 256C>T 257T>C), Q70H (260G>T), R72H (265G>A), S77N (279T>A 280C>A 281T>C), P79L (286C>T 287T>C), R80I (288C>A 289G>T 290C>A), H81G (291C>G 292A>G 293T>A), K83R (298A>G), A87V (310C>T), D92N (324G>A), P95S (333C>T), V96I (336G>A 338C>A), F97L (339T>C), G100P (348G>C 349G>C), V105F (363G>T), T106S (366A>T), L1413I (4287C>A 4289A>T), E1415R (4293G>A 4294A>G), D1418K (4302G>A 4304T>A), Q1420P (4309A>C 4310A>T), A1422S (4314G>A 4315C>G 4316A>C), E1424S (4320G>T 4321A>C), R1431T (4342G>C 4343G>T), S1432A (4344T>G 4346A>T), Q1434K (4350C>A), V1435E (4354T>A 4355G>A), K1438S (4362A>T 4363A>C 4364A>C), L1439P (4365T>C 4366T>C 4367A>T), Q1446K (4386C>A 4388A>G), H1450S (4398C>T 4399A>C), F1451V (4401T>G 4403C>T), Q1453K (4407C>A), D1454W (4410G>T 4411A>G 4412T>G), D1455I (4413G>A 4414A>T 4415C>T), K1456R (4417A>G 4418G>A), F1457Y (4420T>A 4421T>C), L1463M (4437C>A 4439C>G), T1464D (4440A>G 4441C>A 4442A>T), F1466L (4448C>G), S1467K (4449T>A 4450C>A 4451T>A), H1468R (4452C>A 4453A>G 4454C>A), L1469T (4455T>A 4456T>C 4457A>T), S1470F (4459C>T 4460C>T), A1472T (4464G>A 4466A>C), K1473N (4469G>T), L1474I (4470C>A 4472C>T), F1475Y (4474T>A 4475T>C), L1480M (4488C>A 4490T>G), L1487I (4509T>A 4511A>T), G1488Q (4512G>C 4513G>A), H1490S (4518C>T 4519A>C 4520T>G), P1491K (4521C>A 4522C>A 4523T>A), N1492K (4526T>A), E1493D (4529G>T), R1494K (4531G>A), P1495Y (4533C>T 4534C>A 4535A>T), G1498A (4543G>C 4544G>T), C1500N (4548T>A 4549G>A), I1501V (4551A>G 4553T>C), D1503F (4557G>T 4558A>T 4559T>C), R1504G (4560A>G), H1505Q (4565T>A), F1506Y (4567T>A), Q1507E (4569C>G), K1509N (4577G>T), T1517N (4600C>A 4601A>T), L1521E (4611T>G 4612T>A 4613G>A), K1524N (4622A>C), A1525V (4624C>T 4625C>T), I1527N (4630T>A 4631C>T), K1528S (4632A>T 4633A>C 4634A>T), Q1531N (4641C>A 4643G>T), P1532H (4645C>A), L1534S (4650C>A 4651T>G 4652T>C), F1535N (4653T>A 4654T>A 4655T>C), S1536I (4656T>A 4657C>T 4658A>C), A1537S (4659G>T), L1538I (4662T>A 4664G>T), I1544V (4680A>G 4682C>T), L1546I (4686C>A 4688T>A), T1550D (4698A>G 4699C>A 4700A>T), L1551I (4701C>A 4703G>T), E1552D (4706A>T), D1553S (4707G>T 4708A>C), I1555F (4713A>T 4715A>T), L1557H (4719T>C 4720T>A 4721G>T), Q1560T (4728C>A 4729A>C 4730G>C), I1562F (4734A>T 4736A>C), S1563K (4738G>A 4739T>A), L1564I (4740T>A 4742G>T), K1567R (4750A>G), F1568N (4752T>A 4753T>A), V1570L (4758G>C 4760C>T), M1571V (4761A>G 4763G>C), L1572I (4764T>A 4766A>C), I1578K (4783T>A 4784C>A), A1580F (4788G>T 4789C>T), N1582T (4795A>C 4796T>A), Q1585R (4803C>A 4804A>G), M1589H (4815A>C 4816T>A 4817G>T), F1591L (4823T>A), A1592Y (4824G>T 4825C>A), D1593Q (4827G>C 4829T>A), F1596Y (4837T>A), S1597K (4839T>A 4840C>A), A1599I (4845G>A 4846C>T 4847T>C), G1600C (4848G>T 4850T>C), H1601del (4851_4853delCAC), I1602R (4855T>G 4856C>A), S1603A (4857T>G 4859T>C), L1604I (4860T>A 4862G>C), L1606F (4866C>T 4868C>T), Q1607S (4869C>T 4870A>C 4871A>T), Q1607_K1608insG (4871_4872insGGT), D1611N (4881G>A), T1612E (4884A>G 4885C>A), N1613I (4888A>T 4889T>C), L1614T (4890C>A 4891T>C 4892C>T), S1615D (4893T>G 4894C>A 4895G>C), V1616K (4896G>A 4897T>A), K1617T (4900A>C 4901G>T), I1619L (4905A>T), Q1621R (4911C>A 4912A>G), I1625S (4924T>G 4925T>C), V1626L (4926G>C 4928T>C), I1629V (4935A>G 4937C>T), R1630A (4938A>G 4939G>C), E1635K (4953G>A 4955G>A) |
| Codon mutations: | AAA22..T (116A>T), CTC23CTT (119C>T), TCT24TGA (121C>G 122T>A), GCT26ACC (126G>A 128T>C), TTG27CTT (129T>C 131G>T), TCT28TCA (134T>A), AAT29AGA (136A>G 137T>A), TAC33TAA (149C>A), CTT34TTG (150C>T 152T>G), GTA37ATT (159G>A 161A>T), ATC39GTC (165A>G), CTT40CTA (170T>A), AAA41CCC (171A>C 172A>C 173A>C), AAC43GAG (177A>G 179C>G), ATA46GCA (186A>G 187T>C), GAT48TCC (192G>T 193A>C 194T>C), ACC49ACG (197C>G), TAT50TCC (199A>C 200T>C), CTT51TTA (201C>T 203T>A), TTG53CTT (207T>C 209G>T), TTA54ATA (210T>A), TAC57TAT (221C>T), GCA61GCC (233A>C), AGA63AAT (238G>A 239A>T), TCC64TAA (241C>A 242C>A), GTT65TTA (243G>T 245T>A), ACC66TCG (246A>T 248C>G), CCT67CCA (251T>A), TCT69ATC (255T>A 256C>T 257T>C), CAG70CAT (260G>T), ATC71ATT (263C>T), CGT72CAT (265G>A), TCA73TCC (269A>C), CAG76CAA (278G>A), TCT77AAC (279T>A 280C>A 281T>C), CCT79CTC (286C>T 287T>C), CGC80ATA (288C>A 289G>T 290C>A), CAT81GGA (291C>G 292A>G 293T>A), GTG82GTA (296G>A), AAA83AGA (298A>G), GCT87GTT (310C>T), TCC89TCA (317C>A), TTA91CTC (321T>C 323A>C), GAT92AAT (324G>A), CAG93CAA (329G>A), CCT95TCT (333C>T), GTC96ATA (336G>A 338C>A), TTT97CTT (339T>C), GCT98GCA (344T>A), ACA99ACT (347A>T), GGA100CCA (348G>C 349G>C), CAG103CAA (359G>A), TTT104TTC (362T>C), GTT105TTT (363G>T), ACT106TCT (366A>T), CTA1413ATT (4287C>A 4289A>T), GAA1415AGA (4293G>A 4294A>G), CCC1416CCT (4298C>T), GAT1418AAA (4302G>A 4304T>A), TCA1419TCT (4307A>T), CAA1420CCT (4309A>C 4310A>T), GCA1422AGC (4314G>A 4315C>G 4316A>C), GAA1424TCA (4320G>T 4321A>C), GCT1425GCC (4325T>C), TTT1426TTC (4328T>C), GTC1428GTT (4334C>T), AAC1429AAT (4337C>T), AAG1430AAA (4340G>A), AGG1431ACT (4342G>C 4343G>T), TCA1432GCT (4344T>G 4346A>T), GAG1433GAA (4349G>A), CAA1434AAA (4350C>A), GTG1435GAA (4354T>A 4355G>A), GGA1437GGC (4361A>C), AAA1438TCC (4362A>T 4363A>C 4364A>C), TTA1439CCT (4365T>C 4366T>C 4367A>T), GTC1442GTT (4376C>T), ATA1443ATT (4379A>T), CAA1446AAG (4386C>A 4388A>G), CCA1447CCT (4391A>T), AAC1449AAT (4397C>T), CAT1450TCT (4398C>T 4399A>C), TTC1451GTT (4401T>G 4403C>T), CTC1452TTA (4404C>T 4406C>A), CAA1453AAA (4407C>A), GAT1454TGG (4410G>T 4411A>G 4412T>G), GAC1455ATT (4413G>A 4414A>T 4415C>T), AAG1456AGA (4417A>G 4418G>A), TTT1457TAC (4420T>A 4421T>C), CCC1458CCG (4424C>G), CCA1460CCT (4430A>T), AAG1462AAA (4436G>A), CTC1463ATG (4437C>A 4439C>G), ACA1464GAT (4440A>G 4441C>A 4442A>T), CTC1465CTT (4445C>T), TTC1466TTG (4448C>G), TCT1467AAA (4449T>A 4450C>A 4451T>A), CAC1468AGA (4452C>A 4453A>G 4454C>A), TTA1469ACT (4455T>A 4456T>C 4457A>T), TCC1470TTT (4459C>T 4460C>T), GCA1472ACC (4464G>A 4466A>C), AAG1473AAT (4469G>T), CTC1474ATT (4470C>A 4472C>T), TTT1475TAC (4474T>A 4475T>C), TCA1476AGT (4476T>A 4477C>G 4478A>T), AAG1477AAA (4481G>A), CTT1480ATG (4488C>A 4490T>G), AAG1481AAA (4493G>A), GGA1483GGT (4499A>T), TTT1484TTC (4502T>C), TTA1487ATT (4509T>A 4511A>T), GGA1488CAA (4512G>C 4513G>A), ATC1489ATT (4517C>T), CAT1490TCG (4518C>T 4519A>C 4520T>G), CCT1491AAA (4521C>A 4522C>A 4523T>A), AAT1492AAA (4526T>A), GAG1493GAT (4529G>T), AGA1494AAA (4531G>A), CCA1495TAT (4533C>T 4534C>A 4535A>T), ACA1497ACT (4541A>T), GGG1498GCT (4543G>C 4544G>T), TTT1499TTC (4547T>C), TGC1500AAC (4548T>A 4549G>A), ATT1501GTC (4551A>G 4553T>C), CCT1502CCC (4556T>C), GAT1503TTC (4557G>T 4558A>T 4559T>C), AGA1504GGA (4560A>G), CAT1505CAA (4565T>A), TTT1506TAT (4567T>A), CAA1507GAA (4569C>G), AAG1509AAT (4577G>T), GTC1510GTT (4580C>T), CCA1512CCC (4586A>C), TTC1513TTT (4589C>T), GGT1514GGA (4592T>A), CTT1515TTA (4593C>T 4595T>A), ACA1517AAT (4600C>A 4601A>T), TTG1521GAA (4611T>G 4612T>A 4613G>A), TTC1522TTT (4616C>T), AAA1524AAC (4622A>C), GCC1525GTT (4624C>T 4625C>T), ATC1527AAT (4630T>A 4631C>T), AAA1528TCT (4632A>T 4633A>C 4634A>T), ATC1529ATT (4637C>T), TTC1530TTT (4640C>T), CAG1531AAT (4641C>A 4643G>T), CCC1532CAC (4645C>A), CTT1534AGC (4650C>A 4651T>G 4652T>C), TTT1535AAC (4653T>A 4654T>A 4655T>C), TCA1536ATC (4656T>A 4657C>T 4658A>C), GCT1537TCT (4659G>T), TTG1538ATT (4662T>A 4664G>T), GTC1539GTG (4667C>G), TAC1540TAT (4670C>T), ATC1544GTT (4680A>G 4682C>T), CTT1545TTG (4683C>T 4685T>G), CTT1546ATA (4686C>A 4688T>A), GAA1549GAG (4697A>G), ACA1550GAT (4698A>G 4699C>A 4700A>T), CTG1551ATT (4701C>A 4703G>T), GAA1552GAT (4706A>T), GAT1553TCT (4707G>T 4708A>C), CAC1554CAT (4712C>T), ATA1555TTT (4713A>T 4715A>T), TTG1557CAT (4719T>C 4720T>A 4721G>T), CTT1558TTA (4722C>T 4724T>A), AAC1559AAT (4727C>T), CAG1560ACC (4728C>A 4729A>C 4730G>C), ATA1562TTC (4734A>T 4736A>C), AGT1563AAA (4738G>A 4739T>A), TTG1564ATT (4740T>A 4742G>T), GTT1565GTC (4745T>C), AAA1567AGA (4750A>G), TTT1568AAT (4752T>A 4753T>A), GGA1569GGT (4757A>T), GTC1570CTT (4758G>C 4760C>T), ATG1571GTC (4761A>G 4763G>C), TTA1572ATC (4764T>A 4766A>C), TCG1573AGT (4767T>A 4768C>G 4769G>T), GCA1574GCT (4772A>T), AAA1575AAG (4775A>G), AAG1576AAA (4778G>A), ATC1578AAA (4783T>A 4784C>A), CTT1579TTG (4785C>T 4787T>G), GCT1580TTT (4788G>T 4789C>T), AAT1582ACA (4795A>C 4796T>A), ATT1584ATC (4802T>C), CAA1585AGA (4803C>A 4804A>G), TTC1586TTT (4808C>T), TTA1587CTT (4809T>C 4811A>T), GGA1588GGG (4814A>G), ATG1589CAT (4815A>C 4816T>A 4817G>T), TTT1591TTA (4823T>A), GCT1592TAT (4824G>T 4825C>A), GAT1593CAA (4827G>C 4829T>A), ACT1595ACA (4835T>A), TTT1596TAT (4837T>A), TCA1597AAA (4839T>A 4840C>A), CCT1598CCC (4844T>C), GCT1599ATC (4845G>A 4846C>T 4847T>C), GGT1600TGC (4848G>T 4850T>C), CAC1601del (4851_4853delCAC), ATC1602AGA (4855T>G 4856C>A), TCT1603GCC (4857T>G 4859T>C), TTG1604ATC (4860T>A 4862G>C), CTC1606TTT (4866C>T 4868C>T), CAA1607TCT (4869C>T 4870A>C 4871A>T), CAA1607_AAG1608insGGT (4871_4872insGGT), AAG1608AAA (4874G>A), TTT1609TTC (4877T>C), GAT1611AAT (4881G>A), ACA1612GAA (4884A>G 4885C>A), AAT1613ATC (4888A>T 4889T>C), CTC1614ACT (4890C>A 4891T>C 4892C>T), TCG1615GAC (4893T>G 4894C>A 4895G>C), GTA1616AAA (4896G>A 4897T>A), AAG1617ACT (4900A>C 4901G>T), ATA1619TTA (4905A>T), CAA1620CAG (4910A>G), CAA1621AGA (4911C>A 4912A>G), TTT1622TTC (4916T>C), CTT1623TTG (4917C>T 4919T>G), GGG1624GGA (4922G>A), ATT1625AGC (4924T>G 4925T>C), GTT1626CTC (4926G>C 4928T>C), ATC1629GTT (4935A>G 4937C>T), AGA1630GCA (4938A>G 4939G>C), GAT1631GAC (4943T>C), TTT1632TTC (4946T>C), CCA1634CCT (4952A>T), GAG1635AAA (4953G>A 4955G>A) |

*: Inserts / Deletes / Misaligned / Frameshifts

#### Analysis details

This analysis was performed with panviral2.64

### NGS Details (UN60): Cavemovirus venamanihotis

#### Assembly

| | |
|---|---|
| **Coverage Length** | 620 (2 contig(s)) |
| **Depth Of Coverage** | 6.4 |
| **Number Of Reads** | 33 |
| **Reads Per Million** | 0.70 rpm (after QC) |
| **Ambiguities** | 0 |
| **Assembly Method** | de novo + reference guided assembly |
| **Consensus Caller** | Bcf Tools |

#### Coverage Map

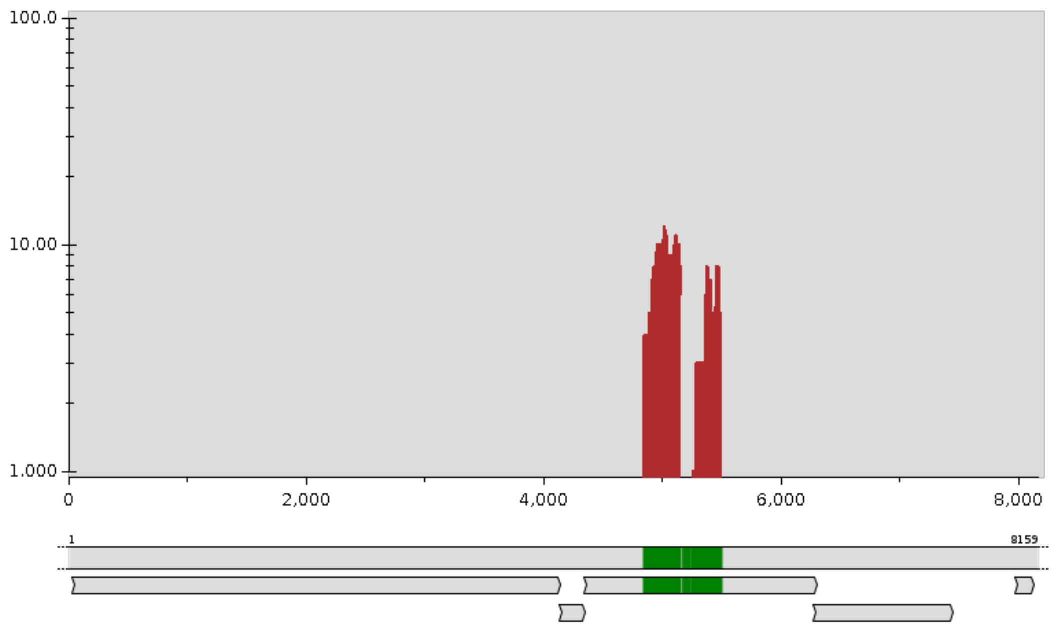

#### Assignment

| | |
|---|---|
| **Type** | Cavemovirus venamanihotis (Taxonomy ID: 3047736) |
| **Reference Genome** | NC_001648.1 |
| **NT Identity (%)** | 59.6694 |
| **AA Identity (%)** | 48.2587 |
| **Number Of Stop Codons** | 1 |
| **Number Of CDS** | 5 |

#### Alignment

| | |
|---|---|
| **Alignment Score** | 201.0 (NT) + 585.0 (AA) = 786.0 |
| **Concordance (%)** | 32.0685 |

| **Alignment Method** | Global, seeded, nucleotide + amino acids (AGA) |
|---|---|

#### Genome Region

Sequence starts at position 4833 and ends at position 5501 relative to NC_001648.1 reference sequence.

#### Alignment Detailed Statistics

| | Begin | End | Coverage | Score | Concordance | Matches | Identities | I/D/M/F* | Stop Codons |
|---|---|---|---|---|---|---|---|---|---|
| **NT** | **4833** | **5501** | **7.6%** | **201** | **17.1%** | **605 (97.6%)** | **361 (58.2%)** | **0/15** | |
| Mutations: | 4842G>A, 4845G>A, 4853T>A, 4857A>G, 4858T>A, 4859A>T, 4863G>C, 4864G>A, 4867C>A, 4868A>G, 4869G>T, 4870A>C, 4872_4874delATA, 4880C>A, 4889A>C, 4890T>C, 4904A>T, 4908C>A, 4909A>G, 4911A>C, 4913A>C, 4914A>G, 4915C>A, 4917T>A, 4918T>A, 4921C>G, 4924A>T, 4925A>T, 4926A>T, 4929G>A, 4930A>T, 4931A>C, 4932T>C, 4934A>T, 4938A>G, 4941G>C, 4942A>T, 4943A>T, 4946A>G, 4947G>A, 4948A>T, 4951A>T, 4952T>A, 4955T>A, 4956T>A, 4957A>G, 4958T>G, 4959A>G, 4960A>T, 4962C>A, 4963C>G, 4977C>A, 4979A>T, 4984C>A, 4989T>A, 4990T>A, 4992C>T, 4994A>T, 4995G>A, 5000T>C, 5002A>G, 5006G>T, 5009T>G, 5012A>T, 5013G>C, 5019A>C, 5021G>T, 5022A>C, 5024T>C, 5028G>T, 5029A>T, 5031G>A, 5032G>A, 5033T>A, 5035T>A, 5036T>A, 5040G>A, 5041A>G, 5047A>G, 5048A>T, 5050C>A, 5051T>G, 5053A>G, 5055_5066delTTTGAAGATAAA, 5067A>C, 5068A>C, 5070T>C, 5073T>A, 5074C>G, 5075T>C, 5078C>T, 5081A>T, 5084T>A, 5090A>G, 5091G>A, 5095A>G, 5096T>A, 5099A>T, 5103T>G, 5105A>G, 5123A>G, 5124A>G, 5127A>C, 5135A>T, 5138T>A, 5139G>A, 5141T>C, 5144T>C, 5150T>G, 5151T>C, 5153A>T, 5160A>G, 5161A>C, 5162A>T, 5163G>T, 5164C>T, 5165T>A, 5169G>T, 5170T>G, 5171A>G, 5172G>A, 5174A>T, 5176A>G, 5180T>C, 5186C>T, 5189A>T, 5192T>C, 5196G>A, 5198T>G, 5199A>G, 5200C>A, 5201A>T, 5202T>C, 5204A>T, 5205A>C, 5208C>A, 5210T>A, 5215G>C, 5216T>C, 5217A>T, 5219A>T, 5220C>A, 5222G>A, 5227G>A, 5228A>C, 5229T>A, 5230A>T, 5231C>T, 5234T>C, 5247T>A, 5248G>T, 5249T>G, 5258A>T, 5260T>A, 5263A>G, 5264T>G, 5267T>A, 5271A>C, 5274T>A, 5276A>C, 5277G>T, 5278A>T, 5279A>T, 5280G>A, 5285T>G, 5286A>C, 5287G>A, 5290A>C, 5291A>T, 5292A>T, 5294A>T, 5295T>A, 5296A>G, 5297T>A, 5300A>T, 5308C>A, 5309A>T, 5315G>T, 5316C>T, 5317A>T, 5318G>T, 5322T>C, 5324T>A, 5328C>G, 5330G>A, 5336A>C, 5339A>T, 5340T>A, 5342A>G, 5345A>T, 5349G>T, 5351A>T, 5353A>T, 5354T>G, 5355C>A, 5357T>A, 5360C>T, 5361T>G, 5362C>T, 5363A>C, 5366A>T, 5367A>T, 5368G>C, 5370A>G, 5371T>A, 5372T>A, 5375C>T, 5378A>G, 5379C>A, 5382T>A, 5384T>C, 5388G>A, 5391A>G, 5392G>A, 5393A>T, 5396A>T, 5401G>A, 5402A>T, 5410A>T, 5411T>A, 5420T>C, 5426G>T, 5438T>C, 5441A>T, 5444A>G, 5445G>A, 5447C>T, 5449T>A, 5456G>A, 5459T>G, 5460A>T, 5466G>A, 5467A>T, 5473A>T, 5474A>C, 5476T>A, 5481A>C, 5485C>A, 5486A>T, 5488A>T, 5489A>C, 5492T>C, 5494G>A, 5496G>A, 5498T>A, 5501A>T | | | | | | | | |

**CDS**

| | Begin | End | Coverage | Score | Concordance | Matches | Identities | I/D/M/F* | Stop Codons |
|---|---|---|---|---|---|---|---|---|---|
| CsVMVgp3 | 164 | 386 | 31.5% | 585 | 39.9% | 201 (97.6%) | 97 (47.1%) | 0/5/4/0 | 1 |
| Protein mutations: | E167K (4842G>A), E168K (4845G>A), N170K (4853T>A), I172D (4857A>G 4858T>A 4859A>T), G174Q (4863G>C 4864G>A), T175K (4867C>A 4868A>G), D176S (4869G>T 4870A>C), I177del (4872_4874delATA), E187D (4904A>T), H189S (4908C>A 4909A>G), K190H (4911A>C 4913A>C), T191E (4914A>G 4915C>A), L192K (4917T>A 4918T>A), A193G (4921C>G), K194I (4924A>T 4925A>T), I195L (4926A>T), E196I (4929G>A 4930A>T 4931A>C), N199D (4938A>G), E200L (4941G>C 4942A>T 4943A>T), D202I (4947G>A 4948A>T), N203I (4951A>T 4952T>A), Y205R (4956T>A 4957A>G 4958T>G), K206V (4959A>G 4960A>T), P207R (4962C>A 4963C>G), Q212N (4977C>A 4979A>T), T214K (4984C>A), L216K (4989T>A 4990T>A), P217S (4992C>T 4994A>T), E218K (4995G>A), K220R (5002A>G), M221I (5006G>T), H222Q (5009T>G), E224Q (5013G>C), M226L (5019A>C 5021G>T), I227L (5022A>C 5024T>C), E229L (5028G>T 5029A>T), G230K (5031G>A 5032G>A 5033T>A), F231* (5035T>A 5036T>A), E233R (5040G>A 5041A>G), K235S (5047A>G 5048A>T), T236K (5050C>A 5051T>G), N237S (5053A>G), F238_K241del (5055_5066delTTTGAAGATAAA), K242P (5067A>C 5068A>C), Y243H (5070T>C), I249M (5090A>G), V250I (5091G>A), N251R (5095A>G 5096T>A), K252N (5099A>T), S254A (5103T>G 5105A>G), T261A (5124A>G), D266N (5139G>A 5141T>C), D269E (5150T>G), K273A (5160A>G 5161A>C 5162A>T), A274L (5163G>T 5164C>T 5165T>A), V276W (5169G>T 5170T>G 5171A>G), T286D (5199A>G 5200C>A 5201A>T), Y296I (5229T>A 5230A>T 5231C>T), C302M (5247T>A 5248G>T 5249T>G), F306Y (5260T>A), Y307W (5263A>G 5264T>G), H308Q (5267T>A), K310Q (5271A>C), L311I (5274T>A 5276A>C), E312F (5277G>T 5278A>T 5279A>T), E313K (5280G>A), D314E (5285T>G), S315H (5286A>C 5287G>A), K316T (5290A>C 5291A>T), K317Y (5292A>T 5294A>T), Y318R (5295T>A 5296A>G 5297T>A), T322N (5308C>A 5309A>T), Q325F (5316C>T 5317A>T 5318G>T), Y327Q (5322T>C 5324T>A), Q329E (5328C>G 5330G>A), K331N (5336A>C), L333M (5340T>A 5342A>G), G336C (5349G>T 5351A>T), Y337L (5353A>T 5354T>G), H338K (5355C>A 5357T>A), S340V (5361T>G 5362C>T 5363A>C), I343E (5370A>G 5371T>A 5372T>A), Q346K (5379C>A), F347I (5382T>A 5384T>C), D349N (5388G>A), R350D (5391A>G 5392G>A 5393A>T), R353N (5401G>A 5402A>T), Y356L (5410A>T 5411T>A), V368I (5445G>A 5447C>T), F369Y (5449T>A), I373F (5460A>T), E375M (5466G>A 5467A>T), K377I (5473A>T 5474A>C), I378K (5476T>A), I380L (5481A>C), A381D (5485C>A 5486A>T), K382I (5488A>T 5489A>C), R384K (5494G>A), D385K (5496G>A 5498T>A) | | | | | | | | |
| Codon mutations: | GAA167AAA (4842G>A), GAA168AAA (4845G>A), AAT170AAA (4853T>A), ATA172GAT (4857A>G 4858T>A 4859A>T), GGA174CAA (4863G>C 4864G>A), ACA175AAG (4867C>A 4868A>G), GAT176TCT (4869G>T 4870A>C), ATA177del (4872_4874delATA), GGC179GGA (4880C>A), CCA182CCC (4889A>C), TTA183CTA (4890T>C), GAA187GAT (4904A>T), CAT189AGT (4908C>A 4909A>G), AAA190CAC (4911A>C 4913A>C), ACA191GAA (4914A>G 4915C>A), TTA192AAA (4917T>A 4918T>A), GCA193GGA (4921C>G), AAA194ATT (4924A>T 4925A>T), ATA195TTA (4926A>T), GAA196ATC (4929G>A 4930A>T 4931A>C), TTA197CTT (4932T>C 4934A>T), AAT199GAT (4938A>G), GAA200CTT (4941G>C 4942A>T 4943A>T), ACA201ACG (4946A>G), GAT202ATT (4947G>A 4948A>T), AAT203ATA (4951A>T 4952T>A), ATT204ATA (4955T>A), TAT205AGG (4956T>A 4957A>G 4958T>G), AAA206GTA (4959A>G 4960A>T), CCA207AGA (4962C>A 4963C>G), CAA212AAT (4977C>A 4979A>T), ACA214AAA (4984C>A), TTA216AAA (4989T>A 4990T>A), CCA217TCT (4992C>T 4994A>T), GAA218AAA (4995G>A), TTT219TTC (5000T>C), AAA220AGA (5002A>G), ATG221ATT (5006G>T), CAT222CAG (5009T>G), ATA223ATT (5012A>T), GAA224CAA (5013G>C), ATG226CTT (5019A>C 5021G>T), ATT227CTC (5022A>C 5024T>C), GAA229TTA (5028G>T 5029A>T), GGT230AAA (5031G>A 5032G>A 5033T>A), TTT231TAA (5035T>A 5036T>A), GAA233AGA (5040G>A 5041A>G), AAA235AGT (5047A>G 5048A>T), ACT236AAG (5050C>A 5051T>G), AAT237AGT (5053A>G), TTT238_AAA241del (5055_5066delTTTGAAGATAAA), AAA242CCA (5067A>C 5068A>C), TAT243CAT (5070T>C), TCT244AGC (5073T>A 5074C>G 5075T>C), AGC245AGT (5078C>T), CCA246CCT (5081A>T), GCT247GCA (5084T>A), ATA249ATG (5090A>G), GTA250ATA (5091G>A), AAT251AGA (5095A>G 5096T>A), AAA252AAT (5099A>T), TCA254GCG (5103T>G 5105A>G), AAA260AAG (5123A>G), ACT261GCT (5124A>G), AGA262CGA (5127A>C), GTA264GTT (5135A>T), ATT265ATA (5138T>A), GAT266AAC (5139G>A 5141T>C), TAT267TAC (5144T>C), GAT269GAG (5150T>G), TTA270CTT (5151T>C 5153A>T), AAA273GCT (5160A>G 5161A>C 5162A>T), GCT274TTA (5163G>T 5164C>T 5165T>A), GTA276TGG (5169G>T 5170T>G 5171A>G), GTA277A.T (5172G>A 5174A>T), AAA278.G. (5176A>G), TAT279..C (5180T>C), ATC281..T (5186C>T), CCA282..T (5189A>T), AAT283..C (5192T>C), GAT285A.G (5196G>A 5198T>G), ACA286GAT (5199A>G 5200C>A 5201A>T), TTA287C.T (5202T>C 5204A>T), ATA288C.. (5205A>C), CAT289A.A (5208C>A 5210T>A), AGT291.CC (5215G>C 5216T>C), ATA292T.T (5217A>T 5219A>T), CAG293A.A (5220C>A 5222G>A), AGA295.AC (5227G>A 5228A>C), TAC296ATT (5229T>A 5230A>T 5231C>T), TAT297..C (5234T>C), TGT302ATG (5247T>A 5248G>T 5249T>G), GGA305GGT (5258A>T), TTT306TAT (5260T>A), TAT307TGG (5263A>G 5264T>G), CAT308CAA (5267T>A), AAG310CAG (5271A>C), TTA311ATC (5274T>A 5276A>C), GAA312TTT (5277G>T 5278A>T 5279A>T), GAA313AAA (5280G>A), GAT314GAG (5285T>G), AGT315CAT (5286A>C 5287G>A), AAA316ACT (5290A>C 5291A>T), AAA317TAT (5292A>T 5294A>T), TAT318AGA (5295T>A 5296A>G 5297T>A), ACA319ACT (5300A>T), ACA322AAT (5308C>A 5309A>T), CCG324CCT (5315G>T), CAG325TTT (5316C>T 5317A>T 5318G>T), TAT327CAA (5322T>C 5324T>A), CAG329GAA (5328C>G 5330G>A), AAA331AAC (5336A>C), GTA332GTT (5339A>T), TTA333ATG (5340T>A 5342A>G), CCA334CCT (5345A>T), GGA336TGT (5349G>T 5351A>T), TAT337TTG (5353A>T 5354T>G), CAT338AAA (5355C>A 5357T>A), AAC339AAT (5360C>T), TCA340GTC (5361T>G 5362C>T 5363A>C), CCA341CCT (5366A>T), AGT342TCT (5367A>T 5368G>C), ATT343GAA (5370A>G 5371T>A 5372T>A), TTC344TTT (5375C>T), CAA345CAG (5378A>G), CAA346AAA (5379C>A), TTT347ATC (5382T>A 5384T>C), GAT349AAT (5388G>A), AGA350GAT (5391A>G 5392G>A 5393A>T), ATA351ATT (5396A>T), AGA353AAT (5401G>A 5402A>T), TAT356TTA (5410A>T 5411T>A), ATT359ATC (5420T>C), GTG361GTT (5426G>T), GAT365GAC (5438T>C), ATA366ATT (5441A>T), TTA367TTG (5444A>G), GTC368ATT (5445G>A 5447C>T), TTT369TAT (5449T>A), AAG371AAA (5456G>A), ACT372ACG (5459T>G), ATT373TTT (5460A>T), GAG375ATG (5466G>A 5467A>T), AAA377ATC (5473A>T 5474A>C), ATA378AAA (5476T>A), ATT380CTT (5481A>C), GCA381GAT (5485C>A 5486A>T), AAA382ATC (5488A>T 5489A>C), TTT383TTC (5492T>C), AGA384AAA (5494G>A), GAT385AAA (5496G>A 5498T>A), ATA386ATT (5501A>T) | | | | | | | | |

**Proteins**

| | Begin | End | Coverage | Score | Concordance | Matches | Identities | I/D/M/F* | Stop Codons |
|---|---|---|---|---|---|---|---|---|---|
| aspartic protease/reverse transcriptase (NP_056848.1) | 164 | 386 | 31.5% | 585 | 39.9% | 201 (97.6%) | 97 (47.1%) | 0/5/4/0 | 1 |
| Protein mutations: | E167K (4842G>A), E168K (4845G>A), N170K (4853T>A), I172D (4857A>G 4858T>A 4859A>T), G174Q (4863G>C 4864G>A), T175K (4867C>A 4868A>G), D176S (4869G>T 4870A>C), I177del (4872_4874delATA), E187D (4904A>T), H189S (4908C>A 4909A>G), K190H (4911A>C 4913A>C), T191E (4914A>G 4915C>A), L192K (4917T>A 4918T>A), A193G (4921C>G), K194I (4924A>T 4925A>T), I195L (4926A>T), E196I (4929G>A 4930A>T 4931A>C), N199D (4938A>G), E200L (4941G>C 4942A>T 4943A>T), D202I (4947G>A 4948A>T), N203I (4951A>T 4952T>A), Y205R (4956T>A 4957A>G 4958T>G), K206V (4959A>G 4960A>T), P207R (4962C>A 4963C>G), Q212N (4977C>A 4979A>T), T214K (4984C>A), L216K (4989T>A 4990T>A), P217S (4992C>T 4994A>T), E218K (4995G>A), K220R (5002A>G), M221I (5006G>T), H222Q (5009T>G), E224Q (5013G>C), M226L (5019A>C 5021G>T), I227L (5022A>C 5024T>C), E229L (5028G>T 5029A>T), G230K (5031G>A 5032G>A 5033T>A), F231* (5035T>A 5036T>A), E233R (5040G>A 5041A>G), K235S (5047A>G 5048A>T), T236K (5050C>A 5051T>G), N237S (5053A>G), F238_K241del (5055_5066delTTTGAAGATAAA), K242P (5067A>C 5068A>C), Y243H (5070T>C), I249M (5090A>G), V250I (5091G>A), N251R (5095A>G 5096T>A), K252N (5099A>T), S254A (5103T>G 5105A>G), T261A (5124A>G), D266N (5139G>A 5141T>C), D269E (5150T>G), K273A (5160A>G 5161A>C 5162A>T), A274L (5163G>T 5164C>T 5165T>A), V276W (5169G>T 5170T>G 5171A>G), T286D (5199A>G 5200C>A 5201A>T), Y296I (5229T>A 5230A>T 5231C>T), C302M (5247T>A 5248G>T 5249T>G), F306Y (5260T>A), Y307W (5263A>G 5264T>G), H308Q (5267T>A), K310Q (5271A>C), L311I (5274T>A 5276A>C), E312F (5277G>T 5278A>T 5279A>T), E313K (5280G>A), D314E (5285T>G), S315H (5286A>C 5287G>A), K316T (5290A>C 5291A>T), K317Y (5292A>T 5294A>T), Y318R (5295T>A 5296A>G 5297T>A), T322N (5308C>A 5309A>T), Q325F (5316C>T 5317A>T 5318G>T), Y327Q (5322T>C 5324T>A), Q329E (5328C>G 5330G>A), K331N (5336A>C), L333M (5340T>A 5342A>G), G336C (5349G>T 5351A>T), Y337L (5353A>T 5354T>G), H338K (5355C>A 5357T>A), S340V (5361T>G 5362C>T 5363A>C), I343E (5370A>G 5371T>A 5372T>A), Q346K (5379C>A), F347I (5382T>A 5384T>C), D349N (5388G>A), R350D (5391A>G 5392G>A 5393A>T), R353N (5401G>A 5402A>T), Y356L (5410A>T 5411T>A), V368I (5445G>A 5447C>T), F369Y (5449T>A), I373F (5460A>T), E375M (5466G>A 5467A>T), K377I (5473A>T 5474A>C), I378K (5476T>A), I380L (5481A>C), A381D (5485C>A 5486A>T), K382I (5488A>T 5489A>C), R384K (5494G>A), D385K (5496G>A 5498T>A) | | | | | | | | |

| | Begin | End | Coverage | Score | Concordance | Matches | Identities | I/D/M/F* | Stop Codons |
|---|---|---|---|---|---|---|---|---|---|
| **NT** | **4833** | **5501** | **7.6%** | **201** | **17.1%** | **605 (97.6%)** | **361 (58.2%)** | **0/15** | |
| Codon mutations: | GAA167AAA (4842G>A), GAA168AAA (4845G>A), AAT170AAA (4853T>A), ATA172GAT (4857A>G 4858T>A 4859A>T), GGA174CAA (4863G>C 4864G>A), ACA175AAG (4867C>A 4868A>G), GAT176TCT (4869G>T 4870A>C), ATA177del (4872_4874delATA), GGC179GGA (4880C>A), CCA182CCC (4889A>C), TTA183CTA (4890T>C), GAA187GAT (4904A>T), CAT189AGT (4908C>A 4909A>G), AAA190CAC (4911A>C 4913A>C), ACA191GAA (4914A>G 4915C>A), TTA192AAA (4917T>A 4918T>A), GCA193GGA (4921C>G), AAA194ATT (4924A>T 4925A>T), ATA195TTA (4926A>T), GAA196ATC (4929G>A 4930A>T 4931A>C), TTA197CTT (4932T>C 4934A>T), AAT199GAT (4938A>G), GAA200CTT (4941G>C 4942A>T 4943A>T), ACA201ACG (4946A>G), GAT202ATT (4947G>A 4948A>T), AAT203ATA (4951A>T 4952T>A), ATT204ATA (4955T>A), TAT205AGG (4956T>A 4957A>G 4958T>G), AAA206GTA (4959A>G 4960A>T), CCA207AGA (4962C>A 4963C>G), CAA212AAT (4977C>A 4979A>T), ACA214AAA (4984C>A), TTA216AAA (4989T>A 4990T>A), CCA217TCT (4992C>T 4994A>T), GAA218AAA (4995G>A), TTT219TTC (5000T>C), AAA220AGA (5002A>G), ATG221ATT (5006G>T), CAT222CAG (5009T>G), ATA223ATT (5012A>T), GAA224CAA (5013G>C), ATG226CTT (5019A>C 5021G>T), ATT227CTC (5022A>C 5024T>C), GAA229TTA (5028G>T 5029A>T), GGT230AAA (5031G>A 5032G>A 5033T>A), TTT231TAA (5035T>A 5036T>A), GAA233AGA (5040G>A 5041A>G), AAA235AGT (5047A>G 5048A>T), ACT236AAG (5050C>A 5051T>G), AAT237AGT (5053A>G), TTT238_AAA241del (5055_5066delTTTGAAGATAAA), AAA242CCA (5067A>C 5068A>C), TAT243CAT (5070T>C), TCT244AGC (5073T>A 5074C>G 5075T>C), AGC245AGT (5078C>T), CCA246CCT (5081A>T), GCT247GCA (5084T>A), ATA249ATG (5090A>G), GTA250ATA (5091G>A), AAT251AGA (5095A>G 5096T>A), AAA252AAT (5099A>T), TCA254GCG (5103T>G 5105A>G), AAA260AAG (5123A>G), ACT261GCT (5124A>G), AGA262CGA (5127A>C), GTA264GTT (5135A>T), ATT265ATA (5138T>A), GAT266AAC (5139G>A 5141T>C), TAT267TAC (5144T>C), GAT269GAG (5150T>G), TTA270CTT (5151T>C 5153A>T), AAA273GCT (5160A>G 5161A>C 5162A>T), GCT274TTA (5163G>T 5164C>T 5165T>A), GTA276TGG (5169G>T 5170T>G 5171A>G), GTA277A.T (5172G>A 5174A>T), AAA278.G. (5176A>G), TAT279..C (5180T>C), ATC281..T (5186C>T), CCA282..T (5189A>T), AAT283..C (5192T>C), GAT285A.G (5196G>A 5198T>G), ACA286GAT (5199A>G 5200C>A 5201A>T), TTA287C.T (5202T>C 5204A>T), ATA288C.. (5205A>C), CAT289A.A (5208C>A 5210T>A), AGT291.CC (5215G>C 5216T>C), ATA292T.T (5217A>T 5219A>T), CAG293A.A (5220C>A 5222G>A), AGA295.AC (5227G>A 5228A>C), TAC296ATT (5229T>A 5230A>T 5231C>T), TAT297..C (5234T>C), TGT302ATG (5247T>A 5248G>T 5249T>G), GGA305GGT (5258A>T), TTT306TAT (5260T>A), TAT307TGG (5263A>G 5264T>G), CAT308CAA (5267T>A), AAG310CAG (5271A>C), TTA311ATC (5274T>A 5276A>C), GAA312TTT (5277G>T 5278A>T 5279A>T), GAA313AAA (5280G>A), GAT314GAG (5285T>G), AGT315CAT (5286A>C 5287G>A), AAA316ACT (5290A>C 5291A>T), AAA317TAT (5292A>T 5294A>T), TAT318AGA (5295T>A 5296A>G 5297T>A), ACA319ACT (5300A>T), ACA322AAT (5308C>A 5309A>T), CCG324CCT (5315G>T), CAG325TTT (5316C>T 5317A>T 5318G>T), TAT327CAA (5322T>C 5324T>A), CAG329GAA (5328C>G 5330G>A), AAA331AAC (5336A>C), GTA332GTT (5339A>T), TTA333ATG (5340T>A 5342A>G), CCA334CCT (5345A>T), GGA336TGT (5349G>T 5351A>T), TAT337TTG (5353A>T 5354T>G), CAT338AAA (5355C>A 5357T>A), AAC339AAT (5360C>T), TCA340GTC (5361T>G 5362C>T 5363A>C), CCA341CCT (5366A>T), AGT342TCT (5367A>T 5368G>C), ATT343GAA (5370A>G 5371T>A 5372T>A), TTC344TTT (5375C>T), CAA345CAG (5378A>G), CAA346AAA (5379C>A), TTT347ATC (5382T>A 5384T>C), GAT349AAT (5388G>A), AGA350GAT (5391A>G 5392G>A 5393A>T), ATA351ATT (5396A>T), AGA353AAT (5401G>A 5402A>T), TAT356TTA (5410A>T 5411T>A), ATT359ATC (5420T>C), GTG361GTT (5426G>T), GAT365GAC (5438T>C), ATA366ATT (5441A>T), TTA367TTG (5444A>G), GTC368ATT (5445G>A 5447C>T), TTT369TAT (5449T>A), AAG371AAA (5456G>A), ACT372ACG (5459T>G), ATT373TTT (5460A>T), GAG375ATG (5466G>A 5467A>T), AAA377ATC (5473A>T 5474A>C), ATA378AAA (5476T>A), ATT380CTT (5481A>C), GCA381GAT (5485C>A 5486A>T), AAA382ATC (5488A>T 5489A>C), TTT383TTC (5492T>C), AGA384AAA (5494G>A), GAT385AAA (5496G>A 5498T>A), ATA386ATT (5501A>T) | | | | | | | | |

*: Inserts / Deletes / Misaligned / Frameshifts

#### Analysis details

This analysis was performed with panviral2.64

### NGS Details (UN60): Cavemovirus collusipomeae

#### Assembly

| | |
|---|---|
| **Coverage Length** | 497 (1 contig(s)) |
| **Depth Of Coverage** | 7.0 |
| **Number Of Reads** | 32 |
| **Reads Per Million** | 0.68 rpm (after QC) |
| **Ambiguities** | 0 |
| **Assembly Method** | de novo + reference guided assembly |
| **Consensus Caller** | Bcf Tools |

#### Coverage Map

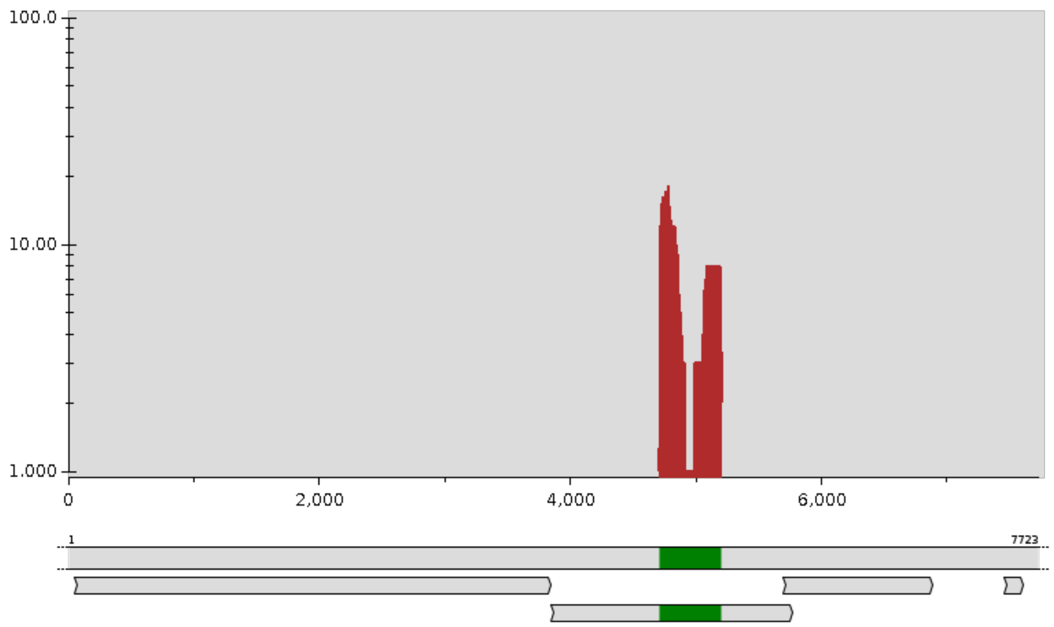

#### Assignment

| | |
|---|---|
| **Type** | Cavemovirus collusipomeae (Taxonomy ID: 3048352) |
| **Reference Genome** | NC_015328.1 |
| **NT Identity (%)** | 69.0141 |
| **AA Identity (%)** | 57.8313 |
| **Number Of Stop Codons** | 1 |
| **Number Of CDS** | 4 |

#### Alignment

| | |
|---|---|
| **Alignment Score** | 378.0 (NT) + 640.0 (AA) = 1018.0 |
| **Concordance (%)** | 47.3709 |

| **Alignment Method** | Global, seeded, nucleotide + amino acids (AGA) |
|---|---|

#### Genome Region

Sequence starts at position 4709 and ends at position 5205 relative to NC_015328.1 reference sequence.

#### Alignment Detailed Statistics

| | Begin | End | Coverage | Score | Concordance | Matches | Identities | I/D/M/F* | Stop Codons |
|---|---|---|---|---|---|---|---|---|---|
| **NT** | **4709** | **5205** | **6.4%** | **378** | **38.0%** | **497 (100%)** | **343 (69.0%)** | **0/0** | |
| Mutations: | 4723T>C, 4729C>T, 4732C>T, 4734G>A, 4750T>C, 4753T>C, 4754A>C, 4756T>A, 4759A>G, 4760C>T, 4763G>A, 4764A>C, 4771T>A, 4772A>T, 4773G>C, 4778A>C, 4780A>G, 4782A>T, 4783T>A, 4809A>T, 4813T>C, 4815T>A, 4816A>G, 4818G>A, 4821T>A, 4828A>G, 4831T>A, 4841C>G, 4843T>A, 4846T>C, 4847T>A, 4853A>G, 4855T>A, 4857T>G, 4865C>T, 4867A>T, 4873G>T, 4877G>A, 4879A>C, 4880A>T, 4881T>G, 4882A>T, 4887G>A, 4888A>T, 4890C>A, 4893A>T, 4894T>A, 4895T>C, 4896A>C, 4897T>A, 4898G>A, 4900T>C, 4902T>G, 4903T>C, 4907C>A, 4909T>A, 4912A>G, 4924T>C, 4927A>T, 4930A>G, 4931A>G, 4935T>G, 4936T>C, 4939G>C, 4941A>C, 4948C>T, 4949G>C, 4951A>G, 4958A>T, 4959A>T, 4960A>T, 4967T>C, 4969G>T, 4970G>A, 4973A>G, 4974T>A, 4979A>T, 4980G>T, 4984T>A, 4985A>T, 4987A>T, 4988A>T, 4989T>G, 4990A>T, 4991A>G, 4993A>C, 4996A>T, 4997C>A, 5001G>A, 5002A>C, 5015A>C, 5029G>A, 5033G>A, 5034G>A, 5035A>T, 5039C>A, 5041A>T, 5053T>C, 5056A>T, 5060G>A, 5062T>G, 5064A>T, 5065G>T, 5068A>T, 5069G>T, 5070A>C, 5072C>A, 5075G>A, 5076G>A, 5077A>T, 5092G>A, 5093C>A, 5098T>C, 5102A>T, 5103T>C, 5104A>C, 5105G>A, 5107T>A, 5110A>G, 5119A>G, 5120A>T, 5121A>T, 5122A>T, 5124A>C, 5126A>G, 5127T>A, 5128A>C, 5131A>G, 5132A>T, 5136A>T, 5139G>C, 5140T>C, 5144A>C, 5150T>A, 5156T>A, 5157C>G, 5159A>T, 5161A>T, 5162T>C, 5164A>T, 5169T>G, 5170A>C, 5171G>T, 5177C>T, 5179A>T, 5180A>G, 5181T>C, 5182T>A, 5183A>G, 5186A>G, 5188T>A, 5192T>A, 5198A>G, 5200T>A | | | | | | | | |
| **CDS** | | | | | | | | | |
| SPCV_gp2 | 290 | 455 | 25.9% | 640 | 55.2% | 166 (100%) | 96 (57.8%) | 0/0/0/0 | 1 |
| Protein mutations: | C298Y (4734G>A), I305L (4754A>C 4756T>A), E308T (4763G>A 4764A>C), D310E (4771T>A), K313Q (4778A>C 4780A>G), Y314L (4782A>T 4783T>A), Y323F (4809A>T), V325E (4815T>A 4816A>G), W326* (4818G>A), I327N (4821T>A), H334E (4841C>G 4843T>A), S336T (4847T>A), S338G (4853A>G 4855T>A), I339R (4857T>G), Q342Y (4865C>T 4867A>T), M344I (4873G>T), G346S (4877G>A 4879A>C), I347C (4880A>T 4881T>G 4882A>T), R349N (4887G>A 4888A>T), P350Q (4890C>A), Y351L (4893A>T 4894T>A), Y352P (4895T>C 4896A>C 4897T>A), D353N (4898G>A 4900T>C), F354C (4902T>G 4903T>C), L356I (4907C>A 4909T>A), I364V (4931A>G), F365C (4935T>G 4936T>C), K367T (4941A>C), E370Q (4949G>C 4951A>G), K373F (4958A>T 4959A>T 4960A>T), E377K (4970G>A), I378E (4973A>G 4974T>A), R380L (4979A>T 4980G>T), N381K (4984T>A), I382F (4985A>T 4987A>T), I383C (4988A>T 4989T>G 4990A>T), I384V (4991A>G 4993A>C), K385N (4996A>T), H386N (4997C>A), G387D (5001G>A 5002A>C), K392Q (5015A>C), G398N (5033G>A 5034G>A 5035A>T), Q400N (5039C>A 5041A>T), V407M (5060G>A 5062T>G), K408I (5064A>T 5065G>T), E410S (5069G>T 5070A>C), Q411K (5072C>A), G412N (5075G>A 5076G>A 5077A>T), P418T (5093C>A), I421S (5102A>T 5103T>C 5104A>C), D422K (5105G>A 5107T>A), K427F (5120A>T 5121A>T 5122A>T), H428P (5124A>C), I429D (5126A>G 5127T>A 5128A>C), I431L (5132A>T), K432I (5136A>T), S433T (5139G>C 5140T>C), K435Q (5144A>C), L437I (5150T>A), S439R (5156T>A 5157C>G), I440F (5159A>T 5161A>T), L443C (5169T>G 5170A>C), V444L (5171G>T), Q446Y (5177C>T 5179A>T), I447A (5180A>G 5181T>C 5182T>A), R448G (5183A>G), N449E (5186A>G 5188T>A), L451I (5192T>A), N453E (5198A>G 5200T>A) | | | | | | | | |
| Codon mutations: | AGT294AGC (4723T>C), TTC296TTT (4729C>T), GAC297GAT (4732C>T), TGT298TAT (4734G>A), TAT303TAC (4750T>C), CAT304CAC (4753T>C), ATT305CTA (4754A>C 4756T>A), AAA306AAG (4759A>G), CTA307TTA (4760C>T), GAA308ACA (4763G>A 4764A>C), GAT310GAA (4771T>A), AGT311TCT (4772A>T 4773G>C), AAA313CAG (4778A>C 4780A>G), TAT314TTA (4782A>T 4783T>A), TAT323TTT (4809A>T), TAT324TAC (4813T>C), GTA325GAG (4815T>A 4816A>G), TGG326TAG (4818G>A), ATT327AAT (4821T>A), TTA329TTG (4828A>G), CCT330CCA (4831T>A), CAT334GAA (4841C>G 4843T>A), AAT335AAC (4846T>C), TCA336ACA (4847T>A), AGT338GGA (4853A>G 4855T>A), ATA339AGA (4857T>G), CAA342TAT (4865C>T 4867A>T), ATG344ATT (4873G>T), GGA346AGC (4877G>A 4879A>C), ATA347TGT (4880A>T 4881T>G 4882A>T), AGA349AAT (4887G>A 4888A>T), CCA350CAA (4890C>A), TAT351TTA (4893A>T 4894T>A), TAT352CCA (4895T>C 4896A>C 4897T>A), GAT353AAC (4898G>A 4900T>C), TTT354TGC (4902T>G 4903T>C), CTT356ATA (4907C>A 4909T>A), GTA357GTG (4912A>G), GAT361GAC (4924T>C), ATA362ATT (4927A>T), TTA363TTG (4930A>G), ATA364GTA (4931A>G), TTT365TGC (4935T>G 4936T>C), TCG366TCC (4939G>C), AAA367ACA (4941A>C), TAC369TAT (4948C>T), GAA370CAG (4949G>C 4951A>G), AAA373TTT (4958A>T 4959A>T 4960A>T), TTG376CTT (4967T>C 4969G>T), GAA377AAA (4970G>A), ATA378GAA (4973A>G 4974T>A), AGA380TTA (4979A>T 4980G>T), AAT381AAA (4984T>A), ATA382TTT (4985A>T 4987A>T), ATA383TGT (4988A>T 4989T>G 4990A>T), ATA384GTC (4991A>G 4993A>C), AAA385AAT (4996A>T), CAT386AAT (4997C>A), GGA387GAC (5001G>A 5002A>C), AAG392CAG (5015A>C), GAG396GAA (5029G>A), GGA398AAT (5033G>A 5034G>A 5035A>T), CAA400AAT (5039C>A 5041A>T), TTT404TTC (5053T>C), CTA405CTT (5056A>T), GTT407ATG (5060G>A 5062T>G), AAG408ATT (5064A>T 5065G>T), ATA409ATT (5068A>T), GAA410TCA (5069G>T 5070A>C), CAA411AAA (5072C>A), GGA412AAT (5075G>A 5076G>A 5077A>T), CAG417CAA (5092G>A), CCT418ACT (5093C>A), CAT419CAC (5098T>C), ATA421TCC (5102A>T 5103T>C 5104A>C), GAT422AAA (5105G>A 5107T>A), AAA423AAG (5110A>G), GAA426GAG (5119A>G), AAA427TTT (5120A>T 5121A>T 5122A>T), CAT428CCT (5124A>C), ATA429GAC (5126A>G 5127T>A 5128A>C), AAA430AAG (5131A>G), ATA431TTA (5132A>T), AAA432ATA (5136A>T), AGT433ACC (5139G>C 5140T>C), AAA435CAA (5144A>C), TTA437ATA (5150T>A), TCA439AGA (5156T>A 5157C>G), ATA440TTT (5159A>T 5161A>T), TTA441CTT (5162T>C 5164A>T), TTA443TGC (5169T>G 5170A>C), GTA444TTA (5171G>T), CAA446TAT (5177C>T 5179A>T), ATT447GCA (5180A>G 5181T>C 5182T>A), AGA448GGA (5183A>G), AAT449GAA (5186A>G 5188T>A), TTA451ATA (5192T>A), AAT453GAA (5198A>G 5200T>A) | | | | | | | | |
| **Proteins** | | | | | | | | | |
| replicase (YP_004347415.1) | 290 | 455 | 25.9% | 640 | 55.2% | 166 (100%) | 96 (57.8%) | 0/0/0/0 | 1 |
| Protein mutations: | C298Y (4734G>A), I305L (4754A>C 4756T>A), E308T (4763G>A 4764A>C), D310E (4771T>A), K313Q (4778A>C 4780A>G), Y314L (4782A>T 4783T>A), Y323F (4809A>T), V325E (4815T>A 4816A>G), W326* (4818G>A), I327N (4821T>A), H334E (4841C>G 4843T>A), S336T (4847T>A), S338G (4853A>G 4855T>A), I339R (4857T>G), Q342Y (4865C>T 4867A>T), M344I (4873G>T), G346S (4877G>A 4879A>C), I347C (4880A>T 4881T>G 4882A>T), R349N (4887G>A 4888A>T), P350Q (4890C>A), Y351L (4893A>T 4894T>A), Y352P (4895T>C 4896A>C 4897T>A), D353N (4898G>A 4900T>C), F354C (4902T>G 4903T>C), L356I (4907C>A 4909T>A), I364V (4931A>G), F365C (4935T>G 4936T>C), K367T (4941A>C), E370Q (4949G>C 4951A>G), K373F (4958A>T 4959A>T 4960A>T), E377K (4970G>A), I378E (4973A>G 4974T>A), R380L (4979A>T 4980G>T), N381K (4984T>A), I382F (4985A>T 4987A>T), I383C (4988A>T 4989T>G 4990A>T), I384V (4991A>G 4993A>C), K385N (4996A>T), H386N (4997C>A), G387D (5001G>A 5002A>C), K392Q (5015A>C), G398N (5033G>A 5034G>A 5035A>T), Q400N (5039C>A 5041A>T), V407M (5060G>A 5062T>G), K408I (5064A>T 5065G>T), E410S (5069G>T 5070A>C), Q411K (5072C>A), G412N (5075G>A 5076G>A 5077A>T), P418T (5093C>A), I421S (5102A>T 5103T>C 5104A>C), D422K (5105G>A 5107T>A), K427F (5120A>T 5121A>T 5122A>T), H428P (5124A>C), I429D (5126A>G 5127T>A 5128A>C), I431L (5132A>T), K432I (5136A>T), S433T (5139G>C 5140T>C), K435Q (5144A>C), L437I (5150T>A), S439R (5156T>A 5157C>G), I440F (5159A>T 5161A>T), L443C (5169T>G 5170A>C), V444L (5171G>T), Q446Y (5177C>T 5179A>T), I447A (5180A>G 5181T>C 5182T>A), R448G (5183A>G), N449E (5186A>G 5188T>A), L451I (5192T>A), N453E (5198A>G 5200T>A) | | | | | | | | |
| Codon mutations: | AGT294AGC (4723T>C), TTC296TTT (4729C>T), GAC297GAT (4732C>T), TGT298TAT (4734G>A), TAT303TAC (4750T>C), CAT304CAC (4753T>C), ATT305CTA (4754A>C 4756T>A), AAA306AAG (4759A>G), CTA307TTA (4760C>T), GAA308ACA (4763G>A 4764A>C), GAT310GAA (4771T>A), AGT311TCT (4772A>T 4773G>C), AAA313CAG (4778A>C 4780A>G), TAT314TTA (4782A>T 4783T>A), TAT323TTT (4809A>T), TAT324TAC (4813T>C), GTA325GAG (4815T>A 4816A>G), TGG326TAG (4818G>A), ATT327AAT (4821T>A), TTA329TTG (4828A>G), CCT330CCA (4831T>A), CAT334GAA (4841C>G 4843T>A), AAT335AAC (4846T>C), TCA336ACA (4847T>A), AGT338GGA (4853A>G 4855T>A), ATA339AGA (4857T>G), CAA342TAT (4865C>T 4867A>T), ATG344ATT (4873G>T), GGA346AGC (4877G>A 4879A>C), ATA347TGT (4880A>T 4881T>G 4882A>T), AGA349AAT (4887G>A 4888A>T), CCA350CAA (4890C>A), TAT351TTA (4893A>T 4894T>A), TAT352CCA (4895T>C 4896A>C 4897T>A), GAT353AAC (4898G>A 4900T>C), TTT354TGC (4902T>G 4903T>C), CTT356ATA (4907C>A 4909T>A), GTA357GTG (4912A>G), GAT361GAC (4924T>C), ATA362ATT (4927A>T), TTA363TTG (4930A>G), ATA364GTA (4931A>G), TTT365TGC (4935T>G 4936T>C), TCG366TCC (4939G>C), AAA367ACA (4941A>C), TAC369TAT (4948C>T), GAA370CAG (4949G>C 4951A>G), AAA373TTT (4958A>T 4959A>T 4960A>T), TTG376CTT (4967T>C 4969G>T), GAA377AAA (4970G>A), ATA378GAA (4973A>G 4974T>A), AGA380TTA (4979A>T 4980G>T), AAT381AAA (4984T>A), ATA382TTT (4985A>T 4987A>T), ATA383TGT (4988A>T 4989T>G 4990A>T), ATA384GTC (4991A>G 4993A>C), AAA385AAT (4996A>T), CAT386AAT (4997C>A), GGA387GAC (5001G>A 5002A>C), AAG392CAG (5015A>C), GAG396GAA (5029G>A), GGA398AAT (5033G>A 5034G>A 5035A>T), CAA400AAT (5039C>A 5041A>T), TTT404TTC (5053T>C), CTA405CTT (5056A>T), GTT407ATG (5060G>A 5062T>G), AAG408ATT (5064A>T 5065G>T), ATA409ATT (5068A>T), GAA410TCA (5069G>T 5070A>C), CAA411AAA (5072C>A), GGA412AAT (5075G>A 5076G>A 5077A>T), CAG417CAA (5092G>A), CCT418ACT (5093C>A), CAT419CAC (5098T>C), ATA421TCC (5102A>T 5103T>C 5104A>C), GAT422AAA (5105G>A 5107T>A), AAA423AAG (5110A>G), GAA426GAG (5119A>G), AAA427TTT (5120A>T 5121A>T 5122A>T), CAT428CCT (5124A>C), ATA429GAC (5126A>G 5127T>A 5128A>C), AAA430AAG (5131A>G), ATA431TTA (5132A>T), AAA432ATA (5136A>T), AGT433ACC (5139G>C 5140T>C), AAA435CAA (5144A>C), TTA437ATA (5150T>A), TCA439AGA (5156T>A 5157C>G), ATA440TTT (5159A>T 5161A>T), TTA441CTT (5162T>C 5164A>T), TTA443TGC (5169T>G 5170A>C), GTA444TTA (5171G>T), CAA446TAT (5177C>T 5179A>T), ATT447GCA (5180A>G 5181T>C 5182T>A), AGA448GGA (5183A>G), AAT449GAA (5186A>G 5188T>A), TTA451ATA (5192T>A), AAT453GAA (5198A>G 5200T>A) | | | | | | | | |

*: Inserts / Deletes / Misaligned / Frameshifts

#### Analysis details

This analysis was performed with panviral2.64

### NGS Details (UN60): Dioscovirus dioscoreae

#### Assembly

| | |
|---|---|
| **Coverage Length** | 932 (1 contig(s)) |
| **Depth Of Coverage** | 3.7 |
| **Number Of Reads** | 26 |
| **Reads Per Million** | 0.55 rpm (after QC) |
| **Ambiguities** | 0 |
| **Assembly Method** | de novo + reference guided assembly |
| **Consensus Caller** | Bcf Tools |

#### Coverage Map

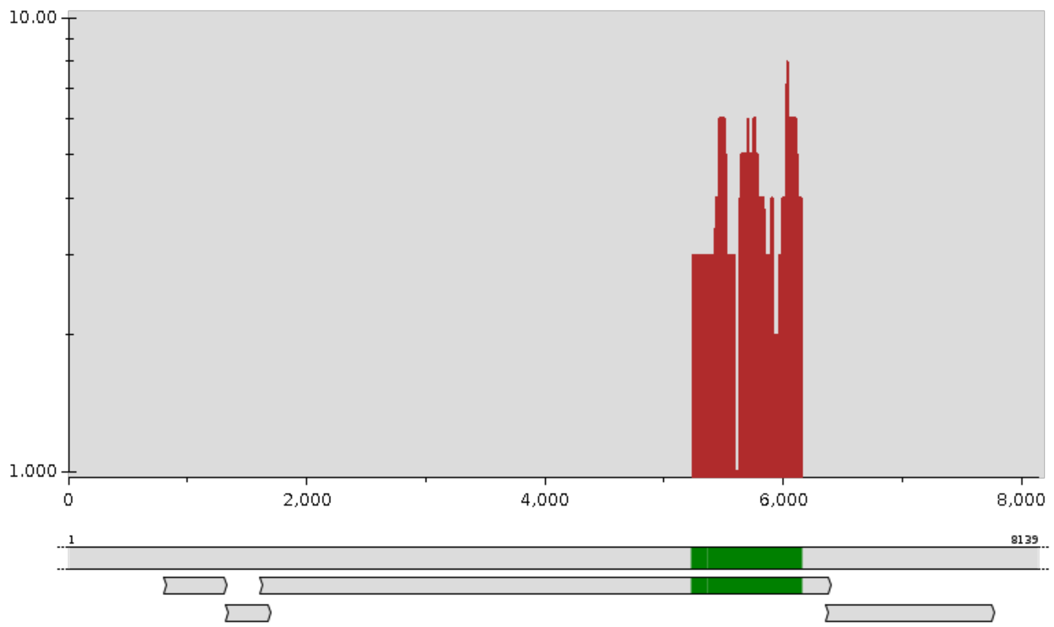

#### Assignment

| | |
|---|---|
| **Type** | Dioscovirus dioscoreae (Taxonomy ID: 3052184) |
| **Reference Genome** | NC_040712.1 |
| **NT Identity (%)** | 58.5288 |
| **AA Identity (%)** | 48.5623 |
| **Number Of Stop Codons** | 0 |
| **Number Of CDS** | 4 |

#### Alignment

| | |
|---|---|
| **Alignment Score** | 299.0 (NT) + 1031.0 (AA) = 1330.0 |
| **Concordance (%)** | 33.4087 |

| Alignment Method | Global, seeded, nucleotide + amino acids (AGA) |
|---|---|

#### Genome Region

Sequence starts at position 5232 and ends at position 6164 relative to NC_040712.1 reference sequence.

#### Alignment Detailed Statistics

| | Begin | End | Coverage | Score | Concordance | Matches | Identities | I/D/M/F* | Stop Codons |
|---|---|---|---|---|---|---|---|---|---|
| **NT** | **5232** | **6164** | **11.5%** | **299** | **16.2%** | **929 (98.7%)** | **549 (58.3%)** | **9/3** | |
| Mutations: | 5233A>T, 5246A>G, 5247A>G, 5248G>C, 5249T>A, 5252T>A, 5259T>A, 5260A>T, 5261C>A, 5264T>C, 5267T>C, 5270G>A, 5271C>A, 5276A>T, 5285T>C, 5288A>T, 5289T>A, 5290A>G, 5291C>A, 5293A>C, 5294A>G, 5295G>A, 5297C>T, 5298C>G, 5299A>C, 5300A>C, 5303C>T, 5305C>A, 5306A>G, 5309C>A, 5310C>T, 5313A>G, 5314G>A, 5317T>A, 5318T>A, 5319G>A, 5320A>G, 5321C>T, 5322T>G, 5324T>A, 5325C>T, 5327C>A, 5328C>A, 5331C>A, 5332T>A, 5335A>G, 5339T>A, 5340A>C, 5344A>G, 5345C>T, 5350T>A, 5351T>A, 5352G>A, 5354C>A, 5356A>T, 5360C>T, 5370C>T, 5371T>G, 5372A>T, 5374A>G, 5378A>T, 5383T>A, 5384C>T, 5387C>T, 5390G>A, 5391A>C, 5395T>G, 5396G>A, 5401A>C, 5402T>C, 5403C>G, 5406C>G, 5409A>G, 5410G>A, 5412A>G, 5414A>G, 5415G>C, 5416A>C, 5418T>A, 5419G>A, 5420G>A, 5423A>T, 5426A>C, 5430G>A, 5431T>G, 5432C>A, 5433T>A, 5434G>C, 5435C>T, 5437C>A, 5438A>C, 5439C>T, 5440A>C, 5441G>T, 5444T>G, 5453A>G, 5455G>A, 5456G>C, 5457A>T, 5462A>C, 5468A>C, 5471C>T, 5477C>G, 5478A>T, 5479A>C, 5485C>T, 5486A>C, 5489T>A, 5490T>G, 5492A>C, 5493G>A, 5494T>C, 5498C>T, 5502C>G, 5505A>T, 5506A>T, 5511G>A, 5514A>T, 5515A>C, 5517A>G, 5519A>G, 5523A>C, 5525A>G, 5530A>T, 5532T>C, 5533C>T, 5534_5535insAGG, 5535G>A, 5537A>G, 5538T>C, 5539T>A, 5543C>G, 5545G>T, 5546T>G, 5549T>A, 5551A>T, 5552C>T, 5553A>T, 5555A>T, 5558C>T, 5564T>A, 5565T>C, 5567G>T, 5572T>A, 5574T>A, 5575C>G, 5576T>C, 5577G>A, 5578A>G, 5581G>C, 5582T>C, 5586C>G, 5589C>G, 5591A>T, 5595G>C, 5598C>G, 5599A>G, 5600A>T, 5603T>C, 5604T>C, 5607T>C, 5608T>A, 5609A>G, 5610C>A, 5611A>C, 5612G>A, 5613T>A, 5615C>A, 5618C>T, 5619C>A, 5620A>G, 5622G>C, 5626G>T, 5628A>G, 5629A>T, 5630G>A, 5631G>A, 5632A>G, 5633A>T, 5634G>A, 5636A>C, 5639G>C, 5640C>A, 5642T>A, 5645T>A, 5646T>G, 5648G>A, 5649T>A, 5650C>G, 5651A>C, 5656C>A, 5657A>G, 5661T>A, 5667A>T, 5670G>T, 5672A>T, 5673G>A, 5674T>A, 5676G>A, 5678A>C, 5679A>C, 5685G>A, 5687A>T, 5691C>T, 5693T>A, 5696C>G, 5697C>A, 5699A>T, 5700G>A, 5701A>T, 5703A>T, 5705A>G, 5711A>T, 5714A>G, 5717A>G, 5718G>A, 5720T>A, 5721C>A, 5726C>A, 5739T>G, 5740T>C, 5741A>T, 5748A>G, 5756A>T, 5757T>A, 5759T>G, 5762T>A, 5763_5765delGAA, 5768C>T, 5769C>A, 5777A>T, 5779C>A, 5781T>A, 5782T>C, 5786G>A, 5788G>A, 5790C>T, 5795G>A, 5798G>A, 5807G>A, 5808A>T, 5811T>G, 5813A>G, 5816C>T, 5822C>T, 5828T>C, 5830A>T, 5834C>A, 5836C>A, 5843G>A, 5844A>G, 5846T>A, 5850T>A, 5851A>T, 5853A>G, 5855T>A, 5856A>G, 5859A>C, 5860T>C, 5864T>A, 5868A>G, 5869A>C, 5870T>A, 5873G>A, 5874T>A, 5875G>T, 5876T>A, 5877T>G, 5878C>G, 5879C>A, 5882C>T, 5889G>C, 5893G>A, 5895A>C, 5897A>C, 5903T>C, 5904T>A, 5906T>A, 5907C>G, 5910G>A, 5912C>T, 5913T>A, 5914G>T, 5915G>T, 5918G>A, 5919A>C, 5924T>A, 5926G>A, 5928A>C, 5933C>T, 5940G>A, 5941T>A, 5945T>A, 5947C>A, 5948C>A, 5951A>C, 5952T>A, 5957C>A, 5958C>G, 5959C>A, 5960T>C, 5963G>A, 5965T>G, 5966T>C, 5967T>A, 5973A>C, 5974A>T, 5976C>G, 5977C>A, 5979C>T, 5980A>C, 5982A>G, 5983G>A, 5988A>C, 5990T>A, 5999G>A, 6005T>C, 6009T>A, 6010C>G, 6011T>C, 6014A>C, 6015G>C, 6026G>A, 6028G>C, 6029A>T, 6030A>G, 6032T>A, 6039T>G, 6040G>C, 6041G>A, 6047C>A, 6048T>A, 6049C>A, 6050T>C, 6051C>A, 6054C>T, 6056A>C, 6057G>A, 6058A>G, 6060T>C, 6062A>T, 6063C>A, 6064C>A, 6065T>A, 6066T>A, 6067C>A, 6069A>G, 6070C>A, 6082C>G, 6083A>T, 6084C>G, 6085G>C, 6086G>T, 6089T>C, 6090T>C, 6091G>A, 6092T>A, 6100G>A, 6101C>A, 6104T>C, 6107_6108insGAAAAG, 6108T>G, 6109C>G, 6111G>A, 6112C>A, 6116C>G, 6117T>A, 6118C>G, 6120G>A, 6121C>G, 6122A>T, 6130C>T, 6131T>C, 6135A>G, 6138A>T, 6140G>A, 6144T>A, 6145G>T, 6146T>A, 6154T>G, 6155A>G, 6156T>C, 6159G>A | | | | | | | | |
| **CDS** | | | | | | | | | |
| EXK67_gp3 | 1208 | 1518 | 19.5% | 1031 | 47.8% | 310 (98.7%) | 152 (48.4%) | 3/1/1/0 | 0 |
| Protein mutations: | Q1208L (5233A>T), S1213A (5247A>G 5248G>C 5249T>A), Y1217I (5259T>A 5260A>T 5261C>A), Y1227R (5289T>A 5290A>G 5291C>A), K1228T (5293A>C 5294A>G), D1229N (5295G>A 5297C>T), Q1230A (5298C>G 5299A>C 5300A>C), T1232K (5305C>A 5306A>G), P1234S (5310C>T), S1235D (5313A>G 5314G>A), I1236K (5317T>A 5318T>A), D1237S (5319G>A 5320A>G 5321C>T), Y1238E (5322T>G 5324T>A), L1240I (5328C>A), L1241N (5331C>A 5332T>A), K1242R (5335A>G), K1244Q (5340A>C), D1245G (5344A>G 5345C>T), I1247K (5350T>A 5351T>A), V1248I (5352G>A 5354C>A), Y1249F (5356A>T), L1254C (5370C>T 5371T>G 5372A>T), F1258Y (5383T>A 5384C>T), I1261L (5391A>C), M1262R (5395T>G 5396G>A), D1264A (5401A>C 5402T>C), P1265A (5403C>G), Q1266E (5406C>G), S1267D (5409A>G 5410G>A), I1268V (5412A>G 5414A>G), E1269P (5415G>C 5416A>C), W1270K (5418T>A 5419G>A 5420G>A), V1274R (5430G>A 5431T>G 5432C>A), C1275T (5433T>A 5434G>C 5435C>T), P1276H (5437C>A 5438A>C), Q1277S (5439C>T 5440A>C 5441G>T), W1282Y (5455G>A 5456G>C), I1283L (5457A>T), K1290S (5478A>T 5479A>C), A1292V (5485C>T 5486A>C), S1294A (5490T>G 5492A>C), V1295T (5493G>A 5494T>C), R1298G (5502C>G), K1299L (5505A>T 5506A>T), D1301N (5511G>A), N1302S (5514A>T 5515A>C), I1303V (5517A>G 5519A>G), K1305Q (5523A>C 5525A>G), Y1307F (5530A>T), S1308L (5532T>C 5533C>T), S1308_E1309insR (5534_5535insAGG), E1309K (5535G>A 5537A>G), F1310H (5538T>C 5539T>A), C1312L (5545G>T 5546T>G), Y1314F (5551A>T 5552C>T), I1315F (5553A>T 5555A>T), F1321Y (5572T>A), E1323R (5577G>A 5578A>G), S1324T (5581G>C 5582T>C), Q1326E (5586C>G), Q1327D (5589C>G 5591A>T), V1329L (5595G>C), Q1330G (5598C>G 5599A>G 5600A>T), L1333Q (5607T>C 5608T>A 5609A>G), Q1334T (5610C>A 5611A>C 5612G>A), F1335I (5613T>A 5615C>A), Q1337R (5619C>A 5620A>G), V1338L (5622G>C), C1339F (5626G>T), K1340V (5628A>G 5629A>T 5630G>A), E1341S (5631G>A 5632A>G 5633A>T), E1342N (5634G>A 5636A>C), L1344I (5640C>A 5642T>A), L1346V (5646T>G 5648G>A), T1349K (5656C>A 5657A>G), L1351M (5661T>A), I1353L (5667A>T), G1354C (5670G>T 5672A>T), V1355K (5673G>A 5674T>A), A1356T (5676G>A 5678A>C), N1357H (5679A>C), E1359N (5685G>A 5687A>T), L1363I (5697C>A 5699A>T), E1364I (5700G>A 5701A>T), I1365L (5703A>T 5705A>G), E1367D (5711A>T), V1370I (5718G>A 5720T>A), Q1371K (5721C>A), L1377A (5739T>G 5740T>C 5741A>T), I1380V (5748A>G), E1382D (5756A>T), F1383M (5757T>A 5759T>G), E1385del (5763_5765delGAA), Q1387K (5769C>A), E1389D (5777A>T), T1390K (5779C>A), L1391T (5781T>A 5782T>C), G1393D (5788G>A), I1400L (5808A>T), L1401V (5811T>G 5813A>G), Y1407F (5830A>T), P1409Q (5836C>A), S1412G (5844A>G 5846T>A), Y1414I (5850T>A 5851A>T), T1415A (5853A>G 5855T>A), R1416G (5856A>G), I1417P (5859A>C 5860T>C), F1418L (5864T>A), N1420A (5868A>G 5869A>C 5870T>A), C1422I (5874T>A 5875G>T 5876T>A), S1423G (5877T>G 5878C>G 5879C>A), E1427Q (5889G>C), R1428K (5893G>A), K1429H (5895A>C 5897A>C), S1432T (5904T>A 5906T>A), Q1433E (5907C>G), D1434N (5910G>A 5912C>T), W1435I (5913T>A 5914G>T 5915G>T), M1437L (5919A>C), R1439K (5926G>A), K1440Q (5928A>C), V1444K (5940G>A 5941T>A), T1446K (5947C>A 5948C>A), K1447N (5951A>C), L1448I (5952T>A), P1450D (5958C>G 5959C>A 5960T>C), I1452S (5965T>G 5966T>C), L1453I (5967T>A), K1455L (5973A>C 5974A>T), P1456E (5976C>G 5977C>A), Q1457S (5979C>T 5980A>C), S1458D (5982A>G 5983G>A), I1460L (5988A>C 5990T>A), E1469Q (6015G>C), G1473A (6028G>C 6029A>T), I1474V (6030A>G 6032T>A), W1477A (6039T>G 6040G>C 6041G>A), S1480N (6048T>A 6049C>A 6050T>C), Q1481K (6051C>A), Q1482Y (6054C>T 6056A>C), D1483S (6057G>A 6058A>G), S1484P (6060T>C 6062A>T), P1485K (6063C>A 6064C>A 6065T>A), S1486K (6066T>A 6067C>A), T1487E (6069A>G 6070C>A), S1491C (6082C>G 6083A>T), R1492A (6084C>G 6085G>C 6086G>T), C1494Q (6090T>C 6091G>A 6092T>A), S1497K (6100G>A 6101C>A), K1499_S1500insEK (6107_6108insGAAAAG), S1500G (6108T>G 6109C>G), A1501N (6111G>A 6112C>A), I1502M (6116C>G), A1504S (6120G>A 6121C>G 6122A>T), A1507V (6130C>T 6131T>C), I1509V (6135A>G), M1510L (6138A>T 6140G>A), C1512I (6144T>A 6145G>T 6146T>A), V1515G (6154T>G 6155A>G), D1517N (6159G>A) | | | | | | | | |

| | Begin | End | Coverage | Score | Concordance | Matches | Identities | I/D/M/F* | Stop Codons |
|---|---|---|---|---|---|---|---|---|---|
| **NT** | **5232** | **6164** | **11.5%** | **299** | **16.2%** | **929 (98.7%)** | **549 (58.3%)** | **9/3** | |
| Codon mutations: | CAA1208CTA (5233A>T), AAA1212AAG (5246A>G), AGT1213GCA (5247A>G 5248G>C 5249T>A), CGT1214CGA (5252T>A), TAC1217ATA (5259T>A 5260A>T 5261C>A), AAT1218AAC (5264T>C), TAT1219TAC (5267T>C), AAG1220AAA (5270G>A), CGA1221AGA (5271C>A), CTA1222CTT (5276A>T), AAT1225AAC (5285T>C), ACA1226ACT (5288A>T), TAC1227AGA (5289T>A 5290A>G 5291C>A), AAA1228ACG (5293A>C 5294A>G), GAC1229AAT (5295G>A 5297C>T), CAA1230GCC (5298C>G 5299A>C 5300A>C), TAC1231TAT (5303C>T), ACA1232AAG (5305C>A 5306A>G), CTC1233CTA (5309C>A), CCA1234TCA (5310C>T), AGT1235GAT (5313A>G 5314G>A), ATT1236AAA (5317T>A 5318T>A), GAC1237AGT (5319G>A 5320A>G 5321C>T), TAT1238GAA (5322T>G 5324T>A), CTC1239TTA (5325C>T 5327C>A), CTA1240ATA (5328C>A), CTT1241AAT (5331C>A 5332T>A), AAA1242AGA (5335A>G), ATT1243ATA (5339T>A), AAA1244CAA (5340A>C), GAC1245GGT (5344A>G 5345C>T), ATT1247AAA (5350T>A 5351T>A), GTC1248ATA (5352G>A 5354C>A), TAC1249TTC (5356A>T), AGC1250AGT (5360C>T), CTA1254TGT (5370C>T 5371T>G 5372A>T), AAG1255.GG (5374A>G), TCA1256TCT (5378A>T), TTC1258TAT (5383T>A 5384C>T), CAC1259CAT (5387C>T), CAG1260CAA (5390G>A), ATT1261CTT (5391A>C), ATG1262AGA (5395T>G 5396G>A), GAT1264GCC (5401A>C 5402T>C), CCA1265GCA (5403C>G), CAG1266GAG (5406C>G), AGT1267GAT (5409A>G 5410G>A), ATA1268GTG (5412A>G 5414A>G), GAA1269CCA (5415G>C 5416A>C), TGG1270AAA (5418T>A 5419G>A 5420G>A), ACA1271ACT (5423A>T), GCA1272GCC (5426A>C), GTC1274AGA (5430G>A 5431T>G 5432C>A), TGC1275ACT (5433T>A 5434G>C 5435C>T), CCA1276CAC (5437C>A 5438A>C), CAG1277TCT (5439C>T 5440A>C 5441G>T), GGT1278GGG (5444T>G), GAA1281GAG (5453A>G), TGG1282TAC (5455G>A 5456G>C), ATA1283TTA (5457A>T), GTA1284GTC (5462A>C), CCA1286CCC (5468A>C), TTC1287TTT (5471C>T), CTC1289CTG (5477C>G), AAA1290TCA (5478A>T 5479A>C), GCA1292GTC (5485C>T 5486A>C), CCT1293CCA (5489T>A), TCA1294GCC (5490T>G 5492A>C), GTA1295ACA (5493G>A 5494T>C), TTC1296TTT (5498C>T), CGA1298GGA (5502C>G), AAA1299TTA (5505A>T 5506A>T), GAT1301AAT (5511G>A), AAT1302TCT (5514A>T 5515A>C), ATA1303GTG (5517A>G 5519A>G), AAA1305CAG (5523A>C 5525A>G), TAT1307TTT (5530A>T), TCA1308CTA (5532T>C 5533C>T), TCA1308_GAA1309insAGG (5534_5535insAGG), GAA1309AAG (5535G>A 5537A>G), TTT1310CAT (5538T>C 5539T>A), GTC1311GTG (5543C>G), TGT1312TTG (5545G>T 5546T>G), GTT1313GTA (5549T>A), TAC1314TTT (5551A>T 5552C>T), ATA1315TTT (5553A>T 5555A>T), GAC1316GAT (5558C>T), ATT1318ATA (5564T>A), TTG1319CTT (5565T>C 5567G>T), TTT1321TAT (5572T>A), TCT1322AGC (5574T>A 5575C>G 5576T>C), GAA1323AGA (5577G>A 5578A>G), AGT1324ACC (5581G>C 5582T>C), CAA1326GAA (5586C>G), CAA1327GAT (5589C>G 5591A>T), GTA1329CTA (5595G>C), CAA1330GGT (5598C>G 5599A>G 5600A>T), CAT1331CAC (5603T>C), TTA1332CTA (5604T>C), TTA1333CAG (5607T>C 5608T>A 5609A>G), CAG1334ACA (5610C>A 5611A>C 5612G>A), TTC1335ATA (5613T>A 5615C>A), TTC1336TTT (5618C>T), CAA1337AGA (5619C>A 5620A>G), GTA1338CTA (5622G>C), TGT1339TTT (5626G>T), AAG1340GTA (5628A>G 5629A>T 5630G>A), GAA1341AGT (5631G>A 5632A>G 5633A>T), GAA1342AAC (5634G>A 5636A>C), GGG1343GGC (5639G>C), CTT1344ATA (5640C>A 5642T>A), ATT1345ATA (5645T>A), TTG1346GTA (5646T>G 5648G>A), TCA1347AGC (5649T>A 5650C>G 5651A>C), ACA1349AAG (5656C>A 5657A>G), TTG1351ATG (5661T>A), ATA1353TTA (5667A>T), GGA1354TGT (5670G>T 5672A>T), GTA1355AAA (5673G>A 5674T>A), GCA1356ACC (5676G>A 5678A>C), AAT1357CAT (5679A>C), GAA1359AAT (5685G>A 5687A>T), CTT1361TTA (5691C>T 5693T>A), GGC1362GGG (5696C>G), CTA1363ATT (5697C>A 5699A>T), GAA1364ATA (5700G>A 5701A>T), ATA1365TTG (5703A>T 5705A>G), GAA1367GAT (5711A>T), GGA1368GGG (5714A>G), AAA1369AAG (5717A>G), GTT1370ATA (5718G>A 5720T>A), CAA1371AAA (5721C>A), CTC1372CTA (5726C>A), TTA1377GCT (5739T>G 5740T>C 5741A>T), ATA1380GTA (5748A>G), GAA1382GAT (5756A>T), TTT1383ATG (5757T>A 5759T>G), CCT1384CCA (5762T>A), GAA1385del (5763_5765delGAA), GAC1386GAT (5768C>T), CAA1387AAA (5769C>A), GAA1389GAT (5777A>T), ACA1390AAA (5779C>A), TTA1391ACA (5781T>A 5782T>C), AAG1392AAA (5786G>A), GGT1393GAT (5788G>A), CTA1394TTA (5790C>T), CAG1395CAA (5795G>A), AAG1396AAA (5798G>A), GGG1399GGA (5807G>A), ATA1400TTA (5808A>T), TTA1401GTG (5811T>G 5813A>G), AAC1402AAT (5816C>T), GCC1404GCT (5822C>T), AAT1406AAC (5828T>C), TAT1407TTT (5830A>T), ATC1408ATA (5834C>A), CCA1409CAA (5836C>A), TTG1411TTA (5843G>A), AGT1412GGA (5844A>G 5846T>A), TAT1414ATT (5850T>A 5851A>T), ACT1415GCA (5853A>G 5855T>A), AGA1416GGA (5856A>G), ATA1417CCA (5859A>C 5860T>C), TTT1418TTA (5864T>A), AAT1420GCA (5868A>G 5869A>C 5870T>A), AAG1421AAA (5873G>A), TGT1422ATA (5874T>A 5875G>T 5876T>A), TCC1423GGA (5877T>G 5878C>G 5879C>A), AGC1424AGT (5882C>T), GAA1427CAA (5889G>C), AGA1428AAA (5893G>A), AAA1429CAC (5895A>C 5897A>C), AAT1431AAC (5903T>C), TCT1432ACA (5904T>A 5906T>A), CAA1433GAA (5907C>G), GAC1434AAT (5910G>A 5912C>T), TGG1435ATT (5913T>A 5914G>T 5915G>T), AAG1436AAA (5918G>A), ATG1437CTG (5919A>C), GTT1438GTA (5924T>A), AGA1439AAA (5926G>A), AAA1440CAA (5928A>C), ATC1441ATT (5933C>T), GTA1444AAA (5940G>A 5941T>A), ATT1445ATA (5945T>A), ACC1446AAA (5947C>A 5948C>A), AAA1447AAC (5951A>C), TTA1448ATA (5952T>A), CCC1449CCA (5957C>A), CCT1450GAC (5958C>G 5959C>A 5960T>C), TTG1451TTA (5963G>A), ATT1452AGC (5965T>G 5966T>C), TTA1453ATA (5967T>A), AAA1455CTA (5973A>C 5974A>T), CCA1456GAA (5976C>G 5977C>A), CAA1457TCA (5979C>T 5980A>C), AGT1458GAT (5982A>G 5983G>A), ATT1460CTA (5988A>C 5990T>A), GAG1463GAA (5999G>A), GAT1465GAC (6005T>C), TCT1467AGC (6009T>A 6010C>G 6011T>C), CTA1468CTC (6014A>C), GAA1469CAA (6015G>C), GGG1472GGA (6026G>A), GGA1473GCT (6028G>C 6029A>T), ATT1474GTA (6030A>G 6032T>A), TGG1477GCA (6039T>G 6040G>C 6041G>A), CCC1479CCA (6047C>A), TCT1480AAC (6048T>A 6049C>A 6050T>C), CAA1481AAA (6051C>A), CAA1482TAC (6054C>T 6056A>C), GAT1483AGT (6057G>A 6058A>G), TCA1484CCT (6060T>C 6062A>T), CCT1485AAA (6063C>A 6064C>A 6065T>A), TCA1486AAA (6066T>A 6067C>A), ACA1487GAA (6069A>G 6070C>A), TCA1491TGT (6082C>G 6083A>T), CGG1492GCT (6084C>G 6085G>C 6086G>T), TAT1493TAC (6089T>C), TGT1494CAA (6090T>C 6091G>A 6092T>A), AGC1497AAA (6100G>A 6101C>A), TAT1498TAC (6104T>C), AAA1499_TCT1500insGAAAAG (6107_6108insGAAAAG), TCT1500GGT (6108T>G 6109C>G), GCC1501AAC (6111G>A 6112C>A), ATC1502ATG (6116C>G), TCT1503AGT (6117T>A 6118C>G), GCA1504AGT (6120G>A 6121C>G 6122A>T), GCT1507GTC (6130C>T 6131T>C), ATA1509GTA (6135A>G), ATG1510TTA (6138A>T 6140G>A), TGT1512ATA (6144T>A 6145G>T 6146T>A), GTA1515GGG (6154T>G 6155A>G), TTA1516CTA (6156T>C), GAT1517AAT (6159G>A) | | | | | | | | |

#### Proteins

| | Begin | End | Coverage | Score | Concordance | Matches | Identities | I/D/M/F* | Stop Codons |
|---|---|---|---|---|---|---|---|---|---|
| ORF3 (YP_009553219.1) | 1208 | 1518 | 19.5% | 1031 | 47.8% | 310 (98.7%) | 152 (48.4%) | 3/1/1/0 | 0 |
| Protein mutations: | Q1208L (5233A>T), S1213A (5247A>G 5248G>C 5249T>A), Y1217I (5259T>A 5260A>T 5261C>A), Y1227R (5289T>A 5290A>G 5291C>A), K1228T (5293A>C 5294A>G), D1229N (5295G>A 5297C>T), Q1230A (5298C>G 5299A>C 5300A>C), T1232K (5305C>A 5306A>G), P1234S (5310C>T), S1235D (5313A>G 5314G>A), I1236K (5317T>A 5318T>A), D1237S (5319G>A 5320A>G 5321C>T), Y1238E (5322T>G 5324T>A), L1240I (5328C>A), L1241N (5331C>A 5332T>A), K1242R (5335A>G), K1244Q (5340A>C), D1245G (5344A>G 5345C>T), I1247K (5350T>A 5351T>A), V1248I (5352G>A 5354C>A), Y1249F (5356A>T), L1254C (5370C>T 5371T>G 5372A>T), F1258Y (5383T>A 5384C>T), I1261L (5391A>C), M1262R (5395T>G 5396G>A), D1264A (5401A>C 5402T>C), P1265A (5403C>G), Q1266E (5406C>G), S1267D (5409A>G 5410G>A), I1268V (5412A>G 5414A>G), E1269P (5415G>C 5416A>C), W1270K (5418T>A 5419G>A 5420G>A), V1274R (5430G>A 5431T>G 5432C>A), C1275T (5433T>A 5434G>C 5435C>T), P1276H (5437C>A 5438A>C), Q1277S (5439C>T 5440A>C 5441G>T), W1282Y (5455G>A 5456G>C), I1283L (5457A>T), K1290S (5478A>T 5479A>C), A1292V (5485C>T 5486A>C), S1294A (5490T>G 5492A>C), V1295T (5493G>A 5494T>C), R1298G (5502C>G), K1299L (5505A>T 5506A>T), D1301N (5511G>A), N1302S (5514A>T 5515A>C), I1303V (5517A>G 5519A>G), K1305Q (5523A>C 5525A>G), Y1307F (5530A>T), S1308L (5532T>C 5533C>T), S1308_E1309insR (5534_5535insAGG), E1309K (5535G>A 5537A>G), F1310H (5538T>C 5539T>A), C1312L (5545G>T 5546T>G), Y1314F (5551A>T 5552C>T), I1315F (5553A>T 5555A>T), F1321Y (5572T>A), E1323R (5577G>A 5578A>G), S1324T (5581G>C 5582T>C), Q1326E (5586C>G), Q1327D (5589C>G 5591A>T), V1329L (5595G>C), Q1330G (5598C>G 5599A>G 5600A>T), L1333Q (5607T>C 5608T>A 5609A>G), Q1334T (5610C>A 5611A>C 5612G>A), F1335I (5613T>A 5615C>A), Q1337R (5619C>A 5620A>G), V1338L (5622G>C), C1339F (5626G>T), K1340V (5628A>G 5629A>T 5630G>A), E1341S (5631G>A 5632A>G 5633A>T), E1342N (5634G>A 5636A>C), L1344I (5640C>A 5642T>A), L1346V (5646T>G 5648G>A), T1349K (5656C>A 5657A>G), L1351M (5661T>A), I1353L (5667A>T), G1354C (5670G>T 5672A>T), V1355K (5673G>A 5674T>A), A1356T (5676G>A 5678A>C), N1357H (5679A>C), E1359N (5685G>A 5687A>T), L1363I (5697C>A 5699A>T), E1364I (5700G>A 5701A>T), I1365L (5703A>T 5705A>G), E1367D (5711A>T), V1370I (5718G>A 5720T>A), Q1371K (5721C>A), L1377A (5739T>G 5740T>C 5741A>T), I1380V (5748A>G), E1382D (5756A>T), F1383M (5757T>A 5759T>G), E1385del (5763_5765delGAA), Q1387K (5769C>A), E1389D (5777A>T), T1390K (5779C>A), L1391T (5781T>A 5782T>C), G1393D (5788G>A), I1400L (5808A>T), L1401V (5811T>G 5813A>G), Y1407F (5830A>T), P1409Q (5836C>A), S1412G (5844A>G 5846T>A), Y1414I (5850T>A 5851A>T), T1415A (5853A>G 5855T>A), R1416G (5856A>G), I1417P (5859A>C 5860T>C), F1418L (5864T>A), N1420A (5868A>G 5869A>C 5870T>A), C1422I (5874T>A 5875G>T 5876T>A), S1423G (5877T>G 5878C>G 5879C>A), E1427Q (5889G>C), R1428K (5893G>A), K1429H (5895A>C 5897A>C), S1432T (5904T>A 5906T>A), Q1433E (5907C>G), D1434N (5910G>A 5912C>T), W1435I (5913T>A 5914G>T 5915G>T), M1437L (5919A>C), R1439K (5926G>A), K1440Q (5928A>C), V1444K (5940G>A 5941T>A), T1446K (5947C>A 5948C>A), K1447N (5951A>C), L1448I (5952T>A), P1450D (5958C>G 5959C>A 5960T>C), I1452S (5965T>G 5966T>C), L1453I (5967T>A), K1455L (5973A>C 5974A>T), P1456E (5976C>G 5977C>A), Q1457S (5979C>T 5980A>C), S1458D (5982A>G 5983G>A), I1460L (5988A>C 5990T>A), E1469Q (6015G>C), G1473A (6028G>C 6029A>T), I1474V (6030A>G 6032T>A), W1477A (6039T>G 6040G>C 6041G>A), S1480N (6048T>A 6049C>A 6050T>C), Q1481K (6051C>A), Q1482Y (6054C>T 6056A>C), D1483S (6057G>A 6058A>G), S1484P (6060T>C 6062A>T), P1485K (6063C>A 6064C>A 6065T>A), S1486K (6066T>A 6067C>A), T1487E (6069A>G 6070C>A), S1491C (6082C>G 6083A>T), R1492A (6084C>G 6085G>C 6086G>T), C1494Q (6090T>C 6091G>A 6092T>A), S1497K (6100G>A 6101C>A), K1499_S1500insEK (6107_6108insGAAAAG), S1500G (6108T>G 6109C>G), A1501N (6111G>A 6112C>A), I1502M (6116C>G), A1504S (6120G>A 6121C>G 6122A>T), A1507V (6130C>T 6131T>C), I1509V (6135A>G), M1510L (6138A>T 6140G>A), C1512I (6144T>A 6145G>T 6146T>A), V1515G (6154T>G 6155A>G), D1517N (6159G>A) | | | | | | | | |

| | Begin | End | Coverage | Score | Concordance | Matches | Identities | I/D/M/F* | Stop Codons |
|---|---|---|---|---|---|---|---|---|---|
| **NT** | **5232** | **6164** | **11.5%** | **299** | **16.2%** | **929 (98.7%)** | **549 (58.3%)** | **9/3** | |
| Codon mutations: | CAA1208CTA (5233A>T), AAA1212AAG (5246A>G), AGT1213GCA (5247A>G 5248G>C 5249T>A), CGT1214CGA (5252T>A), TAC1217ATA (5259T>A 5260A>T 5261C>A), AAT1218AAC (5264T>C), TAT1219TAC (5267T>C), AAG1220AAA (5270G>A), CGA1221AGA (5271C>A), CTA1222CTT (5276A>T), AAT1225AAC (5285T>C), ACA1226ACT (5288A>T), TAC1227AGA (5289T>A 5290A>G 5291C>A), AAA1228ACG (5293A>C 5294A>G), GAC1229AAT (5295G>A 5297C>T), CAA1230GCC (5298C>G 5299A>C 5300A>C), TAC1231TAT (5303C>T), ACA1232AAG (5305C>A 5306A>G), CTC1233CTA (5309C>A), CCA1234TCA (5310C>T), AGT1235GAT (5313A>G 5314G>A), ATT1236AAA (5317T>A 5318T>A), GAC1237AGT (5319G>A 5320A>G 5321C>T), TAT1238GAA (5322T>G 5324T>A), CTC1239TTA (5325C>T 5327C>A), CTA1240ATA (5328C>A), CTT1241AAT (5331C>A 5332T>A), AAA1242AGA (5335A>G), ATT1243ATA (5339T>A), AAA1244CAA (5340A>C), GAC1245GGT (5344A>G 5345C>T), ATT1247AAA (5350T>A 5351T>A), GTC1248ATA (5352G>A 5354C>A), TAC1249TTC (5356A>T), AGC1250AGT (5360C>T), CTA1254TGT (5370C>T 5371T>G 5372A>T), AAG1255.GG (5374A>G), TCA1256TCT (5378A>T), TTC1258TAT (5383T>A 5384C>T), CAC1259CAT (5387C>T), CAG1260CAA (5390G>A), ATT1261CTT (5391A>C), ATG1262AGA (5395T>G 5396G>A), GAT1264GCC (5401A>C 5402T>C), CCA1265GCA (5403C>G), CAG1266GAG (5406C>G), AGT1267GAT (5409A>G 5410G>A), ATA1268GTG (5412A>G 5414A>G), GAA1269CCA (5415G>C 5416A>C), TGG1270AAA (5418T>A 5419G>A 5420G>A), ACA1271ACT (5423A>T), GCA1272GCC (5426A>C), GTC1274AGA (5430G>A 5431T>G 5432C>A), TGC1275ACT (5433T>A 5434G>C 5435C>T), CCA1276CAC (5437C>A 5438A>C), CAG1277TCT (5439C>T 5440A>C 5441G>T), GGT1278GGG (5444T>G), GAA1281GAG (5453A>G), TGG1282TAC (5455G>A 5456G>C), ATA1283TTA (5457A>T), GTA1284GTC (5462A>C), CCA1286CCC (5468A>C), TTC1287TTT (5471C>T), CTC1289CTG (5477C>G), AAA1290TCA (5478A>T 5479A>C), GCA1292GTC (5485C>T 5486A>C), CCT1293CCA (5489T>A), TCA1294GCC (5490T>G 5492A>C), GTA1295ACA (5493G>A 5494T>C), TTC1296TTT (5498C>T), CGA1298GGA (5502C>G), AAA1299TTA (5505A>T 5506A>T), GAT1301AAT (5511G>A), AAT1302TCT (5514A>T 5515A>C), ATA1303GTG (5517A>G 5519A>G), AAA1305CAG (5523A>C 5525A>G), TAT1307TTT (5530A>T), TCA1308CTA (5532T>C 5533C>T), TCA1308_GAA1309insAGG (5534_5535insAGG), GAA1309AAG (5535G>A 5537A>G), TTT1310CAT (5538T>C 5539T>A), GTC1311GTG (5543C>G), TGT1312TTG (5545G>T 5546T>G), GTT1313GTA (5549T>A), TAC1314TTT (5551A>T 5552C>T), ATA1315TTT (5553A>T 5555A>T), GAC1316GAT (5558C>T), ATT1318ATA (5564T>A), TTG1319CTT (5565T>C 5567G>T), TTT1321TAT (5572T>A), TCT1322AGC (5574T>A 5575C>G 5576T>C), GAA1323AGA (5577G>A 5578A>G), AGT1324ACC (5581G>C 5582T>C), CAA1326GAA (5586C>G), CAA1327GAT (5589C>G 5591A>T), GTA1329CTA (5595G>C), CAA1330GGT (5598C>G 5599A>G 5600A>T), CAT1331CAC (5603T>C), TTA1332CTA (5604T>C), TTA1333CAG (5607T>C 5608T>A 5609A>G), CAG1334ACA (5610C>A 5611A>C 5612G>A), TTC1335ATA (5613T>A 5615C>A), TTC1336TTT (5618C>T), CAA1337AGA (5619C>A 5620A>G), GTA1338CTA (5622G>C), TGT1339TTT (5626G>T), AAG1340GTA (5628A>G 5629A>T 5630G>A), GAA1341AGT (5631G>A 5632A>G 5633A>T), GAA1342AAC (5634G>A 5636A>C), GGG1343GGC (5639G>C), CTT1344ATA (5640C>A 5642T>A), ATT1345ATA (5645T>A), TTG1346GTA (5646T>G 5648G>A), TCA1347AGC (5649T>A 5650C>G 5651A>C), ACA1349AAG (5656C>A 5657A>G), TTG1351ATG (5661T>A), ATA1353TTA (5667A>T), GGA1354TGT (5670G>T 5672A>T), GTA1355AAA (5673G>A 5674T>A), GCA1356ACC (5676G>A 5678A>C), AAT1357CAT (5679A>C), GAA1359AAT (5685G>A 5687A>T), CTT1361TTA (5691C>T 5693T>A), GGC1362GGG (5696C>G), CTA1363ATT (5697C>A 5699A>T), GAA1364ATA (5700G>A 5701A>T), ATA1365TTG (5703A>T 5705A>G), GAA1367GAT (5711A>T), GGA1368GGG (5714A>G), AAA1369AAG (5717A>G), GTT1370ATA (5718G>A 5720T>A), CAA1371AAA (5721C>A), CTC1372CTA (5726C>A), TTA1377GCT (5739T>G 5740T>C 5741A>T), ATA1380GTA (5748A>G), GAA1382GAT (5756A>T), TTT1383ATG (5757T>A 5759T>G), CCT1384CCA (5762T>A), GAA1385del (5763_5765delGAA), GAC1386GAT (5768C>T), CAA1387AAA (5769C>A), GAA1389GAT (5777A>T), ACA1390AAA (5779C>A), TTA1391ACA (5781T>A 5782T>C), AAG1392AAA (5786G>A), GGT1393GAT (5788G>A), CTA1394TTA (5790C>T), CAG1395CAA (5795G>A), AAG1396AAA (5798G>A), GGG1399GGA (5807G>A), ATA1400TTA (5808A>T), TTA1401GTG (5811T>G 5813A>G), AAC1402AAT (5816C>T), GCC1404GCT (5822C>T), AAT1406AAC (5828T>C), TAT1407TTT (5830A>T), ATC1408ATA (5834C>A), CCA1409CAA (5836C>A), TTG1411TTA (5843G>A), AGT1412GGA (5844A>G 5846T>A), TAT1414ATT (5850T>A 5851A>T), ACT1415GCA (5853A>G 5855T>A), AGA1416GGA (5856A>G), ATA1417CCA (5859A>C 5860T>C), TTT1418TTA (5864T>A), AAT1420GCA (5868A>G 5869A>C 5870T>A), AAG1421AAA (5873G>A), TGT1422ATA (5874T>A 5875G>T 5876T>A), TCC1423GGA (5877T>G 5878C>G 5879C>A), AGC1424AGT (5882C>T), GAA1427CAA (5889G>C), AGA1428AAA (5893G>A), AAA1429CAC (5895A>C 5897A>C), AAT1431AAC (5903T>C), TCT1432ACA (5904T>A 5906T>A), CAA1433GAA (5907C>G), GAC1434AAT (5910G>A 5912C>T), TGG1435ATT (5913T>A 5914G>T 5915G>T), AAG1436AAA (5918G>A), ATG1437CTG (5919A>C), GTT1438GTA (5924T>A), AGA1439AAA (5926G>A), AAA1440CAA (5928A>C), ATC1441ATT (5933C>T), GTA1444AAA (5940G>A 5941T>A), ATT1445ATA (5945T>A), ACC1446AAA (5947C>A 5948C>A), AAA1447AAC (5951A>C), TTA1448ATA (5952T>A), CCC1449CCA (5957C>A), CCT1450GAC (5958C>G 5959C>A 5960T>C), TTG1451TTA (5963G>A), ATT1452AGC (5965T>G 5966T>C), TTA1453ATA (5967T>A), AAA1455CTA (5973A>C 5974A>T), CCA1456GAA (5976C>G 5977C>A), CAA1457TCA (5979C>T 5980A>C), AGT1458GAT (5982A>G 5983G>A), ATT1460CTA (5988A>C 5990T>A), GAG1463GAA (5999G>A), GAT1465GAC (6005T>C), TCT1467AGC (6009T>A 6010C>G 6011T>C), CTA1468CTC (6014A>C), GAA1469CAA (6015G>C), GGG1472GGA (6026G>A), GGA1473GCT (6028G>C 6029A>T), ATT1474GTA (6030A>G 6032T>A), TGG1477GCA (6039T>G 6040G>C 6041G>A), CCC1479CCA (6047C>A), TCT1480AAC (6048T>A 6049C>A 6050T>C), CAA1481AAA (6051C>A), CAA1482TAC (6054C>T 6056A>C), GAT1483AGT (6057G>A 6058A>G), TCA1484CCT (6060T>C 6062A>T), CCT1485AAA (6063C>A 6064C>A 6065T>A), TCA1486AAA (6066T>A 6067C>A), ACA1487GAA (6069A>G 6070C>A), TCA1491TGT (6082C>G 6083A>T), CGG1492GCT (6084C>G 6085G>C 6086G>T), TAT1493TAC (6089T>C), TGT1494CAA (6090T>C 6091G>A 6092T>A), AGC1497AAA (6100G>A 6101C>A), TAT1498TAC (6104T>C), AAA1499_TCT1500insGAAAAG (6107_6108insGAAAAG), TCT1500GGT (6108T>G 6109C>G), GCC1501AAC (6111G>A 6112C>A), ATC1502ATG (6116C>G), TCT1503AGT (6117T>A 6118C>G), GCA1504AGT (6120G>A 6121C>G 6122A>T), GCT1507GTC (6130C>T 6131T>C), ATA1509GTA (6135A>G), ATG1510TTA (6138A>T 6140G>A), TGT1512ATA (6144T>A 6145G>T 6146T>A), GTA1515GGG (6154T>G 6155A>G), TTA1516CTA (6156T>C), GAT1517AAT (6159G>A) | | | | | | | | |

*: Inserts / Deletes / Misaligned / Frameshifts

#### Analysis details

This analysis was performed with panviral2.64

#### NGS Details (UN60): Rahariannevirus raharianne

##### Assembly

| | |
|---|---|
| **Coverage Length** | 885 (3 contig(s)) |
| **Depth Of Coverage** | 4.0 |
| **Number Of Reads** | 26 |
| **Reads Per Million** | 0.55 rpm (after QC) |
| **Ambiguities** | 0 |
| **Assembly Method** | de novo + reference guided assembly |
| **Consensus Caller** | Bcf Tools |

##### Coverage Map

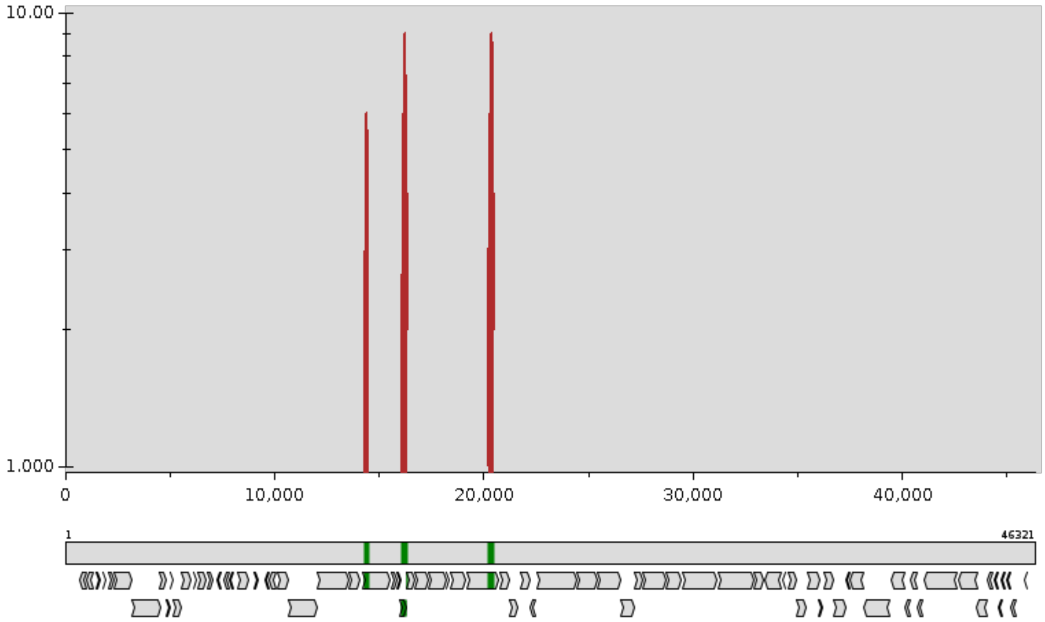

##### Assignment

| | |
|---|---|
| **Type** | Rahariannevirus raharianne (Taxonomy ID: 2846050) |
| **Reference Genome** | NC_054955.1 |
| **NT Identity (%)** | 79.5249 |
| **AA Identity (%)** | 81.1644 |
| **Number Of Stop Codons** | 1 |
| **Number Of CDS** | 75 |

##### Alignment

| | |
|---|---|
| **Alignment Score** | 1034.0 (NT) + 1598.0 (AA) = 2632.0 |
| **Concordance (%)** | 72.1887 |

| **Alignment Method** | Local, heuristic, nucleotide (BLASTN) |
|---|---|

#### Genome Region

Sequence starts at position 14270 and ends at position 20492 relative to NC_054955.1 reference sequence.

#### Alignment Detailed Statistics

| | Begin | End | Coverage | Score | Concordance | Matches | Identities | I/D/M/F* | Stop Codons |
|---|---|---|---|---|---|---|---|---|---|
| **NT** | **14270** | **20492** | **1.9%** | **1034** | **58.8%** | **884 (99.9%)** | **703 (79.4%)** | **0/1** | |
| Mutations: | 14285C>G, 14288C>G, 14289G>A, 14293T>A, 14294C>T, 14298C>T, 14299A>C, 14306A>C, 14309T>C, 14317G>A, 14318T>G, 14336T>C, 14339A>G, 14342C>A, 14357T>C, 14363T>C, 14366G>C, 14372G>T, 14375G>C, 14378C>G, 14381C>G, 14394C>G, 14397T>G, 14399G>C, 14400G>A, 14401C>G, 14402A>C, 14427T>A, 14432G>C, 14433T>A, 14434C>G, 14435G>C, 14447G>C, 14457T>A, 14462C>G, 14486G>C, 14500C>T, 14504G>C, 14516C>G, 16035C>G, 16036A>C, 16037A>G, 16038A>C, 16040G>A, 16045A>G, 16047C>G, 16053T>C, 16057A>G, 16059G>T, 16061C>G, 16062C>G, 16065G>C, 16066C>A, 16067A>C, 16071G>C, 16074T>C, 16078T>A, 16086C>A, 16087A>G, 16088A>C, 16089G>C, 16091A>G, 16095T>C, 16099A>C, 16101C>G, 16113A>C, 16126A>C, 16128T>G, 16131T>G, 16134C>G, 16137T>G, 16140G>C, 16143G>C, 16146T>C, 16147G>A, 16149G>T, 16150G>C, 16151G>C, 16158C>G, 16161G>C, 16164C>T, 16170G>A, 16176G>C, 16177A>G, 16178C>T, 16179G>C, 16182C>T, 16183A>G, 16184C>G, 16190C>T, 16191T>C, 16198A>G, 16199A>T, 16204A>G, 16208G>C, 16209C>G, 16212G>C, 16218G>C, 16227G>C, 16231T>A, 16233A>C, 16239C>G, 16245T>C, 16251G>C, 16255C>T, 16258A>G, 16259A>C, 16260C>A, 16263C>G, 16270A>G, 16278T>C, 16282G>A, 16284T>C, 16293C>G, 16296T>C, 16297T>G, 16302T>C, 16305T>C, 16317C>T, 16319G>C, 16321G>C, 16323delG, 16335A>C, 16338C>G, 16339G>C, 16340C>A, 16341C>G, 16345A>C, 16346A>G, 20211A>T, 20229T>C, 20231T>G, 20235G>A, 20237G>C, 20240G>C, 20253T>A, 20254C>G, 20255G>C, 20268A>C, 20269C>A, 20279C>G, 20282C>T, 20286T>C, 20292T>C, 20293C>A, 20298A>G, 20299T>G, 20301C>A, 20302A>T, 20303G>C, 20306G>C, 20309T>C, 20336C>G, 20343T>A, 20344C>G, 20345G>C, 20355T>G, 20356G>C, 20357C>G, 20360G>C, 20362C>T, 20363G>C, 20369T>C, 20385G>C, 20388C>G, 20394C>A, 20396G>C, 20411C>G, 20433A>T, 20434G>A, 20439G>C, 20444T>C, 20461A>T, 20463G>A, 20468G>T, 20477A>G, 20478T>A, 20479C>G, 20480G>C, 20481G>T, 20483C>G, 20489G>C | | | | | | | | |

##### CDS

| | Begin | End | Coverage | Score | Concordance | Matches | Identities | I/D/M/F* | Stop Codons |
|---|---|---|---|---|---|---|---|---|---|
| KMC43_gp27 | 26 | 109 | 19.0% | 489 | 86.5% | 84 (100%) | 72 (85.7%) | 0/0/0/0 | 0 |
| Protein mutations: | I30M (14285C>G), G32S (14289G>A), F33Y (14293T>A 14294C>T), Q35S (14298C>T 14299A>C), R41Q (14317G>A 14318T>G), P67A (14394C>G), S68A (14397T>G 14399G>C), A69S (14400G>A 14401C>G 14402A>C), S78T (14427T>A), S88T (14457T>A), T102M (14500C>T) | | | | | | | | |
| Codon mutations: | ATC30ATG (14285C>G), CTC31CTG (14288C>G), GGC32AGC (14289G>A), TTC33TAT (14293T>A 14294C>T), CAG35TCG (14298C>T 14299A>C), GCA37GCC (14306A>C), CGT38CGC (14309T>C), CGT41CAG (14317G>A 14318T>G), GGT47GGC (14336T>C), GAA48GAG (14339A>G), ACC49ACA (14342C>A), CGT54CGC (14357T>C), GGT56GGC (14363T>C), CTG57CTC (14366G>C), GCG59GCT (14372G>T), CCG60CCC (14375G>C), GTC61GTG (14378C>G), ACC62ACG (14381C>G), CCG67GCG (14394C>G), TCG68GCC (14397T>G 14399G>C), GCA69AGC (14400G>A 14401C>G 14402A>C), TCG78ACG (14427T>A), CCG79CCC (14432G>C), TCG80AGC (14433T>A 14434C>G 14435G>C), GTG84GTC (14447G>C), TCG88ACG (14457T>A), CTC89CTG (14462C>G), ACG97ACC (14486G>C), ACG102ATG (14500C>T), GTG103GTC (14504G>C), GTC107GTG (14516C>G) | | | | | | | | |
| KMC43_gp30 | 7 | 103 | 94.2% | 504 | 79.6% | 97 (100%) | 73 (75.3%) | 0/0/0/0 | 1 |
| Protein mutations: | K11R (16036A>C 16037A>G 16038A>C), S12N (16040G>A), T14A (16045A>G 16047C>G), T18A (16057A>G 16059G>T), A19G (16061C>G 16062C>G), Q21T (16066C>A 16067A>C), S25T (16078T>A), K28A (16087A>G 16088A>C 16089G>C), D29G (16091A>G), I32L (16099A>C 16101C>G), N41Q (16126A>C 16128T>G), A48T (16147G>A 16149G>T), G49P (16150G>C 16151G>C), T58V (16177A>G 16178C>T 16179G>C), T60G (16183A>G 16184C>G), A62V (16190C>T 16191T>C), N65V (16198A>G 16199A>T), I67V (16204A>G), G68A (16208G>C 16209C>G), S76T (16231T>A 16233A>C), N85A (16258A>G 16259A>C 16260C>A), I89V (16270A>G), A93T (16282G>A 16284T>C), S98A (16297T>G) | | | | | | | | |
| Codon mutations: | GGC10GGG (16035C>G), AAA11CGC (16036A>C 16037A>G 16038A>C), AGC12AAC (16040G>A), ACC14GCG (16045A>G 16047C>G), AAT16AAC (16053T>C), ACG18GCT (16057A>G 16059G>T), GCC19GGG (16061C>G 16062C>G), GCG20GCC (16065G>C), CAG21ACG (16066C>A 16067A>C), GTG22GTC (16071G>C), GTT23GTC (16074T>C), TCG25ACG (16078T>A), CCC27CCA (16086C>A), AAG28GCC (16087A>G 16088A>C 16089G>C), GAC29GGC (16091A>G), TTT30TTC (16095T>C), ATC32CTG (16099A>C 16101C>G), CGA36CGC (16113A>C), AAT41CAG (16126A>C 16128T>G), GTT42GTG (16131T>G), CTC43CTG (16134C>G), GTT44GTG (16137T>G), GCG45GCC (16140G>C), GGG46GGC (16143G>C), ACT47ACC (16146T>C), GCG48ACT (16147G>A 16149G>T), GGG49CCG (16150G>C 16151G>C), GCC51GCG (16158C>G), GCG52GCC (16161G>C), TAC53TAT (16164C>T), GCG55GCA (16170G>A), ACG57ACC (16176G>C), ACG58GTC (16177A>G 16178C>T 16179G>C), GCC59GCT (16182C>T), ACC60GGC (16183A>G 16184C>G), GCT62GTC (16190C>T 16191T>C), AAC65GTC (16198A>G 16199A>T), ATC67GTC (16204A>G), GGC68GCG (16208G>C 16209C>G), GCG69GCC (16212G>C), CCG71CCC (16218G>C), GTG74GTC (16227G>C), TCA76ACC (16231T>A 16233A>C), CTC78CTG (16239C>G), GAT80GAC (16245T>C), CCG82CCC (16251G>C), CTG84TTG (16255C>T), AAC85GCA (16258A>G 16259A>C 16260C>A), GGC86GGG (16263C>G), ATC89GTC (16270A>G), CCT91CCC (16278T>C), GCT93ACC (16282G>A 16284T>C), ACC96ACG (16293C>G), GTT97GTC (16296T>C), TCT98GCT (16297T>G), GTT99GTC (16302T>C), TCT100TCC (16305T>C) | | | | | | | | |
| KMC43_gp31 | 1 | 11 | 8.6% | 58 | 90.6% | 11 (100%) | 9 (81.8%) | 0/0/0/0 | 0 |
| Protein mutations: | A6Q (16339G>C 16340C>A 16341C>G), K8R (16345A>C 16346A>G) | | | | | | | | |
| Codon mutations: | ACA4ACC (16335A>C), ACC5ACG (16338C>G), GCC6CAG (16339G>C 16340C>A 16341C>G), AAG8CGG (16345A>C 16346A>G) | | | | | | | | |
| KMC43_gp36 | 329 | 428 | 19.6% | 547 | 84.2% | 100 (100%) | 83 (83.0%) | 0/0/0/0 | 0 |
| Protein mutations: | T335S (20211A>T), Y341Q (20229T>C 20231T>G), V343I (20235G>A 20237G>C), T354Q (20268A>C 20269C>A), S362Q (20292T>C 20293C>A), I364G (20298A>G 20299T>G), Q365I (20301C>A 20302A>T 20303G>C), C383A (20355T>G 20356G>C 20357C>G), S385F (20362C>T 20363G>C), A393P (20385G>C), L394V (20388C>G), Q396N (20394C>A 20396G>C), S409Y (20433A>T 20434G>A), V411L (20439G>C), Y418F (20461A>T), V419I (20463G>A), A425S (20481G>T 20483C>G) | | | | | | | | |
| Codon mutations: | ACG335TCG (20211A>T), TAT341CAG (20229T>C 20231T>G), GTG343ATC (20235G>A 20237G>C), GTG344GTC (20240G>C), TCG349AGC (20253T>A 20254C>G 20255G>C), ACG354CAG (20268A>C 20269C>A), TCC357TCG (20279C>G), GCC358GCT (20282C>T), TTG360CTG (20286T>C), TCG362CAG (20292T>C 20293C>A), ATC364GGC (20298A>G 20299T>G), CAG365ATC (20301C>A 20302A>T 20303G>C), GCG366GCC (20306G>C), GGT367GGC (20309T>C), CCC376CCG (20336C>G), TCG379AGC (20343T>A 20344C>G 20345G>C), TGC383GCG (20355T>G 20356G>C 20357C>G), CGG384CGC (20360G>C), TCG385TTC (20362C>T 20363G>C), CAT387CAC (20369T>C), GCG393CCG (20385G>C), CTG394GTG (20388C>G), CAG396AAC (20394C>A 20396G>C), ACC401ACG (20411C>G), AGC409TAC (20433A>T 20434G>A), GTC411CTC (20439G>C), AAT412AAC (20444T>C), TAC418TTC (20461A>T), GTC419ATC (20463G>A), GGG420GGT (20468G>T), CAA423CAG (20477A>G), TCG424AGC (20478T>A 20479C>G 20480G>C), GCC425TCG (20481G>T 20483C>G), GTG427GTC (20489G>C) | | | | | | | | |

##### Proteins

| | Begin | End | Coverage | Score | Concordance | Matches | Identities | I/D/M/F* | Stop Codons |
|---|---|---|---|---|---|---|---|---|---|
| DUF4043 domain-containing protein (YP_010078151.1) | 26 | 109 | 19.0% | 489 | 86.5% | 84 (100%) | 72 (85.7%) | 0/0/0/0 | 0 |
| Protein mutations: | I30M (14285C>G), G32S (14289G>A), F33Y (14293T>A 14294C>T), Q35S (14298C>T 14299A>C), R41Q (14317G>A 14318T>G), P67A (14394C>G), S68A (14397T>G 14399G>C), A69S (14400G>A 14401C>G 14402A>C), S78T (14427T>A), S88T (14457T>A), T102M (14500C>T) | | | | | | | | |
| Codon mutations: | ATC30ATG (14285C>G), CTC31CTG (14288C>G), GGC32AGC (14289G>A), TTC33TAT (14293T>A 14294C>T), CAG35TCG (14298C>T 14299A>C), GCA37GCC (14306A>C), CGT38CGC (14309T>C), CGT41CAG (14317G>A 14318T>G), GGT47GGC (14336T>C), GAA48GAG (14339A>G), ACC49ACA (14342C>A), CGT54CGC (14357T>C), GGT56GGC (14363T>C), CTG57CTC (14366G>C), GCG59GCT (14372G>T), CCG60CCC (14375G>C), GTC61GTG (14378C>G), ACC62ACG (14381C>G), CCG67GCG (14394C>G), TCG68GCC (14397T>G 14399G>C), GCA69AGC (14400G>A 14401C>G 14402A>C), TCG78ACG (14427T>A), CCG79CCC (14432G>C), TCG80AGC (14433T>A 14434C>G 14435G>C), GTG84GTC (14447G>C), TCG88ACG (14457T>A), CTC89CTG (14462C>G), ACG97ACC (14486G>C), ACG102ATG (14500C>T), GTG103GTC (14504G>C), GTC107GTG (14516C>G) | | | | | | | | |
| hypothetical protein (YP_010078154.1) | 7 | 103 | 94.2% | 504 | 79.6% | 97 (100%) | 73 (75.3%) | 0/0/0/0 | 1 |
| Protein mutations: | K11R (16036A>C 16037A>G 16038A>C), S12N (16040G>A), T14A (16045A>G 16047C>G), T18A (16057A>G 16059G>T), A19G (16061C>G 16062C>G), Q21T (16066C>A 16067A>C), S25T (16078T>A), K28A (16087A>G 16088A>C 16089G>C), D29G (16091A>G), I32L (16099A>C 16101C>G), N41Q (16126A>C 16128T>G), A48T (16147G>A 16149G>T), G49P (16150G>C 16151G>C), T58V (16177A>G 16178C>T 16179G>C), T60G (16183A>G 16184C>G), A62V (16190C>T 16191T>C), N65V (16198A>G 16199A>T), I67V (16204A>G), G68A (16208G>C 16209C>G), S76T (16231T>A 16233A>C), N85A (16258A>G 16259A>C 16260C>A), I89V (16270A>G), A93T (16282G>A 16284T>C), S98A (16297T>G) | | | | | | | | |

| | Begin | End | Coverage | Score | Concordance | Matches | Identities | I/D/M/F* | Stop Codons |
|---|---|---|---|---|---|---|---|---|---|
| **NT** | **14270** | **20492** | **1.9%** | **1034** | **58.8%** | **884 (99.9%)** | **703 (79.4%)** | **0/1** | |
| Codon mutations: | GGC10GGG (16035C>G), AAA11CGC (16036A>C 16037A>G 16038A>C), AGC12AAC (16040G>A), ACC14GCG (16045A>G 16047C>G), AAT16AAC (16053T>C), ACG18GCT (16057A>G 16059G>T), GCC19GGG (16061C>G 16062C>G), GCG20GCC (16065G>C), CAG21ACG (16066C>A 16067A>C), GTG22GTC (16071G>C), GTT23GTC (16074T>C), TCG25ACG (16078T>A), CCC27CCA (16086C>A), AAG28GCC (16087A>G 16088A>C 16089G>C), GAC29GGC (16091A>G), TTT30TTC (16095T>C), ATC32CTG (16099A>C 16101C>G), CGA36CGC (16113A>C), AAT41CAG (16126A>C 16128T>G), GTT42GTG (16131T>G), CTC43CTG (16134C>G), GTT44GTG (16137T>G), GCG45GCC (16140G>C), GGG46GGC (16143G>C), ACT47ACC (16146T>C), GCG48ACT (16147G>A 16149G>T), GGG49CCG (16150G>C 16151G>C), GCC51GCG (16158C>G), GCG52GCC (16161G>C), TAC53TAT (16164C>T), GCG55GCA (16170G>A), ACG57ACC (16176G>C), ACG58GTC (16177A>G 16178C>T 16179G>C), GCC59GCT (16182C>T), ACC60GGC (16183A>G 16184C>G), GCT62GTC (16190C>T 16191T>C), AAC65GTC (16198A>G 16199A>T), ATC67GTC (16204A>G), GGC68GCG (16208G>C 16209C>G), GCG69GCC (16212G>C), CCG71CCC (16218G>C), GTG74GTC (16227G>C), TCA76ACC (16231T>A 16233A>C), CTC78CTG (16239C>G), GAT80GAC (16245T>C), CCG82CCC (16251G>C), CTG84TTG (16255C>T), AAC85GCA (16258A>G 16259A>C 16260C>A), GGC86GGG (16263C>G), ATC89GTC (16270A>G), CCT91CCC (16278T>C), GCT93ACC (16282G>A 16284T>C), ACC96ACG (16293C>G), GTT97GTC (16296T>C), TCT98GCT (16297T>G), GTT99GTC (16302T>C), TCT100TCC (16305T>C) | | | | | | | | |
| hypothetical protein (YP_010078155.1) | 1 | 11 | 8.6% | 58 | 90.6% | 11 (100%) | 9 (81.8%) | 0/0/0/0 | 0 |
| Protein mutations: | A6Q (16339G>C 16340C>A 16341C>G), K8R (16345A>C 16346A>G) | | | | | | | | |
| Codon mutations: | ACA4ACC (16335A>C), ACC5ACG (16338C>G), GCC6CAG (16339G>C 16340C>A 16341C>G), AAG8CGG (16345A>C 16346A>G) | | | | | | | | |
| tail protein (YP_010078160.1) | 329 | 428 | 19.6% | 547 | 84.2% | 100 (100%) | 83 (83.0%) | 0/0/0/0 | 0 |
| Protein mutations: | T335S (20211A>T), Y341Q (20229T>C 20231T>G), V343I (20235G>A 20237G>C), T354Q (20268A>C 20269C>A), S362Q (20292T>C 20293C>A), I364G (20298A>G 20299T>G), Q365I (20301C>A 20302A>T 20303G>C), C383A (20355T>G 20356G>C 20357C>G), S385F (20362C>T 20363G>C), A393P (20385G>C), L394V (20388C>G), Q396N (20394C>A 20396G>C), S409Y (20433A>T 20434G>A), V411L (20439G>C), Y418F (20461A>T), V419I (20463G>A), A425S (20481G>T 20483C>G) | | | | | | | | |
| Codon mutations: | ACG335TCG (20211A>T), TAT341CAG (20229T>C 20231T>G), GTG343ATC (20235G>A 20237G>C), GTG344GTC (20240G>C), TCG349AGC (20253T>A 20254C>G 20255G>C), ACG354CAG (20268A>C 20269C>A), TCC357TCG (20279C>G), GCC358GCT (20282C>T), TTG360CTG (20286T>C), TCG362CAG (20292T>C 20293C>A), ATC364GGC (20298A>G 20299T>G), CAG365ATC (20301C>A 20302A>T 20303G>C), GCG366GCC (20306G>C), GGT367GGC (20309T>C), CCC376CCG (20336C>G), TCG379AGC (20343T>A 20344C>G 20345G>C), TGC383GCG (20355T>G 20356G>C 20357C>G), CGG384CGC (20360G>C), TCG385TTC (20362C>T 20363G>C), CAT387CAC (20369T>C), GCG393CCG (20385G>C), CTG394GTG (20388C>G), CAG396AAC (20394C>A 20396G>C), ACC401ACG (20411C>G), AGC409TAC (20433A>T 20434G>A), GTC411CTC (20439G>C), AAT412AAC (20444T>C), TAC418TTC (20461A>T), GTC419ATC (20463G>A), GGG420GGT (20468G>T), CAA423CAG (20477A>G), TCG424AGC (20478T>A 20479C>G 20480G>C), GCC425TCG (20481G>T 20483C>G), GTG427GTC (20489G>C) | | | | | | | | |

*: Inserts / Deletes / Misaligned / Frameshifts

#### Analysis details

This analysis was performed with panviral2.64

### NGS Details (UN60): Badnavirus maculasmallanthi

#### Assembly

| | |
|---|---|
| **Coverage Length** | 268 (1 contig(s)) |
| **Depth Of Coverage** | 9.4 |
| **Number Of Reads** | 21 |
| **Reads Per Million** | 0.45 rpm (after QC) |
| **Ambiguities** | 0 |
| **Assembly Method** | de novo + reference guided assembly |
| **Consensus Caller** | Bcf Tools |

#### Coverage Map

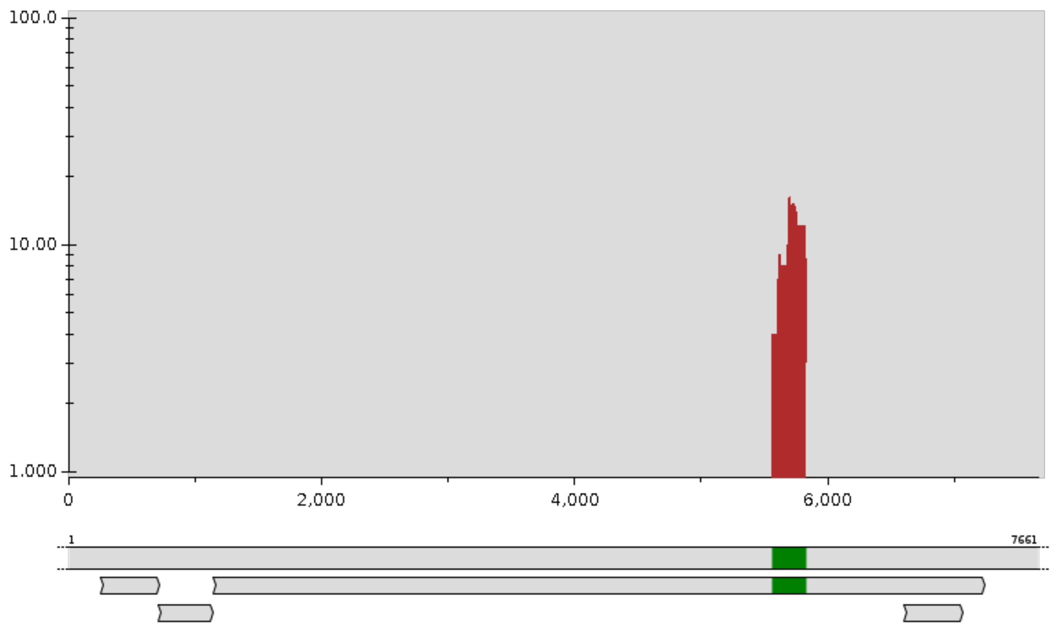

#### Assignment

| | |
|---|---|
| **Type** | Badnavirus maculasmallanthi (Taxonomy ID: 3048453) |
| **Reference Genome** | NC_026472.1 |
| **NT Identity (%)** | 56.4576 |
| **AA Identity (%)** | 51.6484 |
| **Number Of Stop Codons** | 0 |
| **Number Of CDS** | 4 |

#### Alignment

| | |
|---|---|
| **Alignment Score** | 64.0 (NT) + 333.0 (AA) = 397.0 |
| **Concordance (%)** | 33.8737 |

| Alignment Method | Global, seeded, nucleotide + amino acids (AGA) |
|---|---|

#### Genome Region

Sequence starts at position 5560 and ends at position 5827 relative to NC_026472.1 reference sequence.

#### Alignment Detailed Statistics

| | Begin | End | Coverage | Score | Concordance | Matches | Identities | I/D/M/F* | Stop Codons |
|---|---|---|---|---|---|---|---|---|---|
| **NT** | **5560** | **5827** | **3.5%** | **64** | **11.9%** | **268 (98.9%)** | **153 (56.5%)** | **3/0** | |
| Mutations: | 5560G>T, 5562A>C, 5574A>T, 5577G>A, 5578T>A, 5580T>A, 5583T>C, 5584C>T, 5586G>A, 5588A>G, 5589G>A, 5597T>A, 5598T>C, 5605G>C, 5608G>A, 5609C>G, 5610C>A, 5615A>C, 5616A>C, 5617G>A, 5618A>C, 5622A>G, 5623T>G, 5624C>A, 5625C>T, 5626A>G, 5628A>G, 5631C>A, 5632T>A, 5633G>A, 5634G>A, 5637G>T, 5644T>A, 5645T>G, 5648T>C, 5649C>T, 5651C>A, 5652A>C, 5653G>T, 5654G>C, 5655A>T, 5658A>G, 5659T>C, 5660T>A, 5661A>T, 5663A>T, 5664C>T, 5667A>G, 5669G>A, 5670G>C, 5671C>T, 5673T>A, 5682A>C, 5685C>T, 5691C>G, 5692A>T, 5693A>C, 5699C>T, 5700A>C, 5703C>A, 5706T>C, 5708T>C, 5709T>A, 5715G>A, 5716A>G, 5719A>T, 5720A>T, 5725G>A, 5728A>T, 5729A>C, 5730A>T, 5731T>G, 5732G>T, 5733C>G, 5736_5737insCAG, 5740G>T, 5741A>T, 5742C>T, 5743A>C, 5744C>T, 5746G>A, 5747A>G, 5748A>G, 5749G>A, 5752T>C, 5753T>A, 5755A>G, 5757C>G, 5758G>T, 5759C>T, 5760C>G, 5765A>T, 5766C>T, 5767A>T, 5769C>T, 5772C>T, 5775C>T, 5778C>A, 5781G>T, 5782G>A, 5787C>T, 5788T>A, 5789C>G, 5792A>G, 5793C>A, 5795G>C, 5796T>C, 5797G>A, 5798A>T, 5801C>A, 5802G>A, 5809G>A, 5810A>T, 5812A>T, 5813G>T, 5820A>G | | | | | | | | |

##### CDS

| | Begin | End | Coverage | Score | Concordance | Matches | Identities | I/D/M/F* | Stop Codons |
|---|---|---|---|---|---|---|---|---|---|
| UF61_gp3 | 1473 | 1562 | 4.4% | 333 | 51.9% | 90 (98.9%) | 47 (51.6%) | 1/0/0/0 | 0 |
| Protein mutations: | A1473S (5560G>T 5562A>C), F1479I (5578T>A 5580T>A), K1482R (5588A>G 5589G>A), F1485Y (5597T>A 5598T>C), V1488L (5605G>C), A1489R (5608G>A 5609C>G 5610C>A), E1491A (5615A>C 5616A>C), E1492T (5617G>A 5618A>C), S1494D (5623T>G 5624C>A 5625C>T), I1495V (5626A>G 5628A>G), W1497K (5632T>A 5633G>A 5634G>A), L1501R (5644T>A 5645T>G), I1502T (5648T>C 5649C>T), P1503H (5651C>A 5652A>C), G1504S (5653G>T 5654G>C 5655A>T), L1506H (5659T>C 5660T>A 5661A>T), Y1507F (5663A>T 5664C>T), W1509Y (5669G>A 5670G>C), K1517S (5692A>T 5693A>C), A1519V (5699C>T 5700A>C), I1522T (5708T>C 5709T>A), R1525G (5716A>G), K1526L (5719A>T 5720A>T), D1528N (5725G>A), K1529S (5728A>T 5729A>C 5730A>T), C1530V (5731T>G 5732G>T 5733C>G), F1531_K1532insQ (5736_5737insCAG), D1533F (5740G>T 5741A>T 5742C>T), T1534L (5743A>C 5744C>T), E1535R (5746G>A 5747A>G 5748A>G), E1536K (5749G>A), F1537H (5752T>C 5753T>A), I1538V (5755A>G 5757C>G), A1539L (5758G>T 5759C>T 5760C>G), Y1541F (5765A>T 5766C>T), I1542F (5767A>T 5769C>T), V1547I (5782G>A), N1550R (5792A>G 5793C>A), S1551T (5795G>C 5796T>C), E1552I (5797G>A 5798A>T), A1553E (5801C>A 5802G>A), E1556I (5809G>A 5810A>T), R1557L (5812A>T 5813G>T) | | | | | | | | |
| Codon mutations: | GCA1473TCC (5560G>T 5562A>C), TCA1477TCT (5574A>T), AAG1478AAA (5577G>A), TTT1479ATA (5578T>A 5580T>A), GAT1480GAC (5583T>C), CTG1481TTA (5584C>T 5586G>A), AAG1482AGA (5588A>G 5589G>A), TTT1485TAC (5597T>A 5598T>C), GTT1488CTT (5605G>C), GCC1489AGA (5608G>A 5609C>G 5610C>A), GAA1491GCC (5615A>C 5616A>C), GAA1492ACA (5617G>A 5618A>C), GAA1493GAG (5622A>G), TCC1494GAT (5623T>G 5624C>A 5625C>T), ATA1495GTG (5626A>G 5628A>G), CCC1496CCA (5631C>A), TGG1497AAA (5632T>A 5633G>A 5634G>A), ACG1498ACT (5637G>T), TTA1501AGA (5644T>A 5645T>G), ATC1502ACT (5648T>C 5649C>T), CCA1503CAC (5651C>A 5652A>C), GGA1504TCT (5653G>T 5654G>C 5655A>T), GGA1505GGG (5658A>G), TTA1506CAT (5659T>C 5660T>A 5661A>T), TAC1507TTT (5663A>T 5664C>T), GAA1508GAG (5667A>G), TGG1509TAC (5669G>A 5670G>C), CTT1510TTA (5671C>T 5673T>A), CCA1513CCC (5682A>C), TTC1514TTT (5685C>T), CTC1516CTG (5691C>G), AAA1517TCA (5692A>T 5693A>C), GCA1519GTC (5699C>T 5700A>C), CCC1520CCA (5703C>A), GCT1521GCC (5706T>C), ATT1522ACA (5708T>C 5709T>A), CAG1524CAA (5715G>A), AGA1525GGA (5716A>G), AAA1526TTA (5719A>T 5720A>T), GAT1528AAT (5725G>A), AAA1529TCT (5728A>T 5729A>C 5730A>T), TGC1530GTG (5731T>G 5732G>T 5733C>G), TTC1531_AAA1532insCAG (5736_5737insCAG), GAC1533TTT (5740G>T 5741A>T 5742C>T), ACA1534CTA (5743A>C 5744C>T), GAA1535AGG (5746G>A 5747A>G 5748A>G), GAG1536AAG (5749G>A), TTT1537CAT (5752T>C 5753T>A), ATC1538GTG (5755A>G 5757C>G), GCC1539TTG (5758G>T 5759C>T 5760C>G), TAC1541TTT (5765A>T 5766C>T), ATC1542TTT (5767A>T 5769C>T), GAC1543GAT (5772C>T), GAC1544GAT (5775C>T), ATC1545ATA (5778C>A), CTG1546CTT (5781G>T), GTA1547ATA (5782G>A), TAC1548TAT (5787C>T), TCC1549AGC (5788T>A 5789C>G), AAC1550AGA (5792A>G 5793C>A), AGT1551ACC (5795G>C 5796T>C), GAA1552ATA (5797G>A 5798A>T), GCG1553GAA (5801C>A 5802G>A), GAA1556ATA (5809G>A 5810A>T), AGG1557TTG (5812A>T 5813G>T), TTA1559TTG (5820A>G) | | | | | | | | |

##### Proteins

| | Begin | End | Coverage | Score | Concordance | Matches | Identities | I/D/M/F* | Stop Codons |
|---|---|---|---|---|---|---|---|---|---|
| ORF3 (YP_009121747.1) | 1473 | 1562 | 4.4% | 333 | 51.9% | 90 (98.9%) | 47 (51.6%) | 1/0/0/0 | 0 |
| Protein mutations: | A1473S (5560G>T 5562A>C), F1479I (5578T>A 5580T>A), K1482R (5588A>G 5589G>A), F1485Y (5597T>A 5598T>C), V1488L (5605G>C), A1489R (5608G>A 5609C>G 5610C>A), E1491A (5615A>C 5616A>C), E1492T (5617G>A 5618A>C), S1494D (5623T>G 5624C>A 5625C>T), I1495V (5626A>G 5628A>G), W1497K (5632T>A 5633G>A 5634G>A), L1501R (5644T>A 5645T>G), I1502T (5648T>C 5649C>T), P1503H (5651C>A 5652A>C), G1504S (5653G>T 5654G>C 5655A>T), L1506H (5659T>C 5660T>A 5661A>T), Y1507F (5663A>T 5664C>T), W1509Y (5669G>A 5670G>C), K1517S (5692A>T 5693A>C), A1519V (5699C>T 5700A>C), I1522T (5708T>C 5709T>A), R1525G (5716A>G), K1526L (5719A>T 5720A>T), D1528N (5725G>A), K1529S (5728A>T 5729A>C 5730A>T), C1530V (5731T>G 5732G>T 5733C>G), F1531_K1532insQ (5736_5737insCAG), D1533F (5740G>T 5741A>T 5742C>T), T1534L (5743A>C 5744C>T), E1535R (5746G>A 5747A>G 5748A>G), E1536K (5749G>A), F1537H (5752T>C 5753T>A), I1538V (5755A>G 5757C>G), A1539L (5758G>T 5759C>T 5760C>G), Y1541F (5765A>T 5766C>T), I1542F (5767A>T 5769C>T), V1547I (5782G>A), N1550R (5792A>G 5793C>A), S1551T (5795G>C 5796T>C), E1552I (5797G>A 5798A>T), A1553E (5801C>A 5802G>A), E1556I (5809G>A 5810A>T), R1557L (5812A>T 5813G>T) | | | | | | | | |
| Codon mutations: | GCA1473TCC (5560G>T 5562A>C), TCA1477TCT (5574A>T), AAG1478AAA (5577G>A), TTT1479ATA (5578T>A 5580T>A), GAT1480GAC (5583T>C), CTG1481TTA (5584C>T 5586G>A), AAG1482AGA (5588A>G 5589G>A), TTT1485TAC (5597T>A 5598T>C), GTT1488CTT (5605G>C), GCC1489AGA (5608G>A 5609C>G 5610C>A), GAA1491GCC (5615A>C 5616A>C), GAA1492ACA (5617G>A 5618A>C), GAA1493GAG (5622A>G), TCC1494GAT (5623T>G 5624C>A 5625C>T), ATA1495GTG (5626A>G 5628A>G), CCC1496CCA (5631C>A), TGG1497AAA (5632T>A 5633G>A 5634G>A), ACG1498ACT (5637G>T), TTA1501AGA (5644T>A 5645T>G), ATC1502ACT (5648T>C 5649C>T), CCA1503CAC (5651C>A 5652A>C), GGA1504TCT (5653G>T 5654G>C 5655A>T), GGA1505GGG (5658A>G), TTA1506CAT (5659T>C 5660T>A 5661A>T), TAC1507TTT (5663A>T 5664C>T), GAA1508GAG (5667A>G), TGG1509TAC (5669G>A 5670G>C), CTT1510TTA (5671C>T 5673T>A), CCA1513CCC (5682A>C), TTC1514TTT (5685C>T), CTC1516CTG (5691C>G), AAA1517TCA (5692A>T 5693A>C), GCA1519GTC (5699C>T 5700A>C), CCC1520CCA (5703C>A), GCT1521GCC (5706T>C), ATT1522ACA (5708T>C 5709T>A), CAG1524CAA (5715G>A), AGA1525GGA (5716A>G), AAA1526TTA (5719A>T 5720A>T), GAT1528AAT (5725G>A), AAA1529TCT (5728A>T 5729A>C 5730A>T), TGC1530GTG (5731T>G 5732G>T 5733C>G), TTC1531_AAA1532insCAG (5736_5737insCAG), GAC1533TTT (5740G>T 5741A>T 5742C>T), ACA1534CTA (5743A>C 5744C>T), GAA1535AGG (5746G>A 5747A>G 5748A>G), GAG1536AAG (5749G>A), TTT1537CAT (5752T>C 5753T>A), ATC1538GTG (5755A>G 5757C>G), GCC1539TTG (5758G>T 5759C>T 5760C>G), TAC1541TTT (5765A>T 5766C>T), ATC1542TTT (5767A>T 5769C>T), GAC1543GAT (5772C>T), GAC1544GAT (5775C>T), ATC1545ATA (5778C>A), CTG1546CTT (5781G>T), GTA1547ATA (5782G>A), TAC1548TAT (5787C>T), TCC1549AGC (5788T>A 5789C>G), AAC1550AGA (5792A>G 5793C>A), AGT1551ACC (5795G>C 5796T>C), GAA1552ATA (5797G>A 5798A>T), GCG1553GAA (5801C>A 5802G>A), GAA1556ATA (5809G>A 5810A>T), AGG1557TTG (5812A>T 5813G>T), TTA1559TTG (5820A>G) | | | | | | | | |

*: Inserts / Deletes / Misaligned / Frameshifts

#### Analysis details

This analysis was performed with panviral2.64

### NGS Details (UN60): Blunervirus vaccinii (segment RNA 2)

#### Assembly

| | |
|---|---|
| **Coverage Length** | 180 (1 contig(s)) |
| **Depth Of Coverage** | 6.4 |
| **Number Of Reads** | 14 |
| **Reads Per Million** | 0.30 rpm (after QC) |
| **Ambiguities** | 0 |
| **Assembly Method** | de novo + reference guided assembly |
| **Consensus Caller** | Bcf Tools |

#### Coverage Map

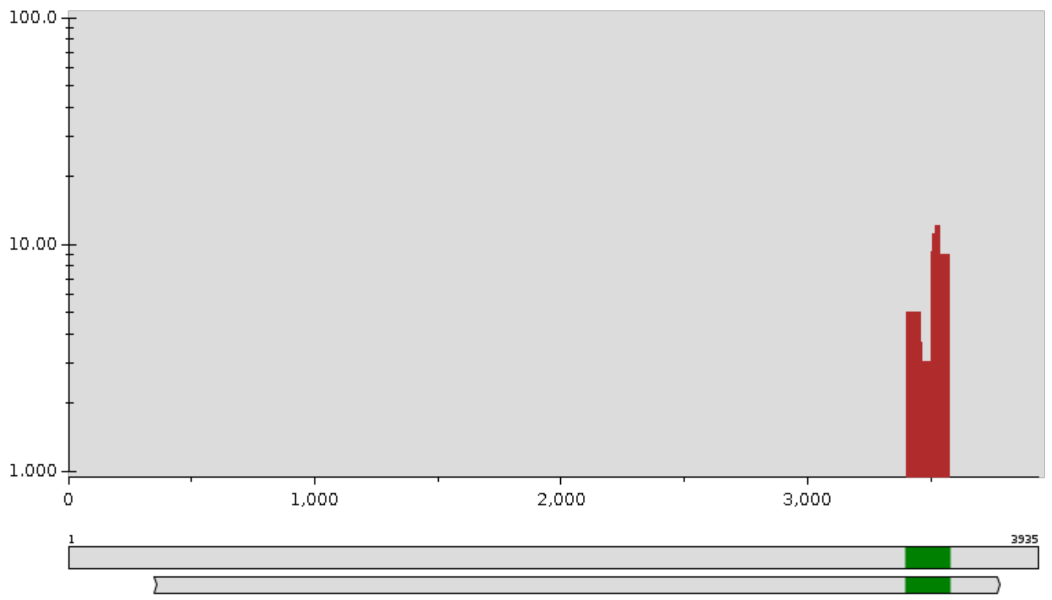

#### Assignment

| | |
|---|---|
| **Type** | Blunervirus vaccinii (Taxonomy ID: 3047693) |
| **Reference Genome** | NC_016085.1 |
| **NT Identity (%)** | 61.6667 |
| **AA Identity (%)** | 61.6667 |
| **Number Of Stop Codons** | 0 |
| **Number Of CDS** | 1 |

#### Alignment

| | |
|---|---|
| **Alignment Score** | 84.0 (NT) + 266.0 (AA) = 350.0 |
| **Concordance (%)** | 43.4783 |

| Alignment Method | Global, seeded, nucleotide + amino acids (AGA) |
|---|---|

#### Genome Region

Sequence starts at position 3400 and ends at position 3579 relative to NC_016085.1 reference sequence.

#### Alignment Detailed Statistics

| | Begin | End | Coverage | Score | Concordance | Matches | Identities | I/D/M/F* | Stop Codons |
|---|---|---|---|---|---|---|---|---|---|
| **NT** | **3400** | **3579** | **4.6%** | **84** | **23.3%** | **180 (100%)** | **111 (61.7%)** | **0/0** | |
| Mutations: | 3405A>T, 3406T>G, 3409C>T, 3412A>T, 3418G>A, 3421G>A, 3423T>G, 3424T>C, 3425G>A, 3426A>T, 3427C>G, 3430C>T, 3431C>A, 3433G>C, 3435A>C, 3437C>A, 3439C>G, 3445C>T, 3448C>T, 3451G>A, 3457T>C, 3458G>T, 3460C>A, 3461C>T, 3463T>G, 3469G>T, 3470A>G, 3472T>A, 3476C>A, 3477G>T, 3478T>C, 3480G>T, 3481G>A, 3482G>A, 3483T>G, 3484T>A, 3485G>T, 3487C>A, 3488A>G, 3489C>T, 3490A>G, 3493C>T, 3499G>T, 3500C>G, 3505G>A, 3506C>G, 3507G>T, 3509G>T, 3510C>T, 3512G>A, 3514C>T, 3517A>G, 3518C>T, 3520C>G, 3523A>C, 3524C>A, 3526T>G, 3527G>C, 3528C>A, 3533C>T, 3535A>G, 3542T>G, 3553G>T, 3556G>A, 3565G>A, 3568G>A, 3574C>G, 3575A>T, 3577T>G | | | | | | | | |
| **CDS** | | | | | | | | | |
| BNRBV_s2gp1 | 1019 | 1078 | 5.2% | 266 | 57.5% | 60 (100%) | 37 (61.7%) | 0/0/0/0 | 0 |
| Protein mutations: | N1020M (3405A>T 3406T>G), E1022D (3412A>T), F1026C (3423T>G 3424T>C), D1027M (3425G>A 3426A>T 3427C>G), Q1029N (3431C>A 3433G>C), Y1030S (3435A>C), H1031K (3437C>A 3439C>G), V1038L (3458G>T 3460C>A), N1042E (3470A>G 3472T>A), R1044I (3476C>A 3477G>T 3478T>C), W1045L (3480G>T 3481G>A), V1046R (3482G>A 3483T>G 3484T>A), V1047L (3485G>T 3487C>A), T1048V (3488A>G 3489C>T 3490A>G), L1052V (3500C>G), R1054V (3506C>G 3507G>T), A1055L (3509G>T 3510C>T), V1056I (3512G>A 3514C>T), A1061H (3527G>C 3528C>A), F1066V (3542T>G), F1076L (3574C>G), I1077L (3575A>T 3577T>G) | | | | | | | | |
| Codon mutations: | AAT1020ATG (3405A>T 3406T>G), TTC1021TTT (3409C>T), GAA1022GAT (3412A>T), AAG1024AAA (3418G>A), TTG1025TTA (3421G>A), TTT1026TGC (3423T>G 3424T>C), GAC1027ATG (3425G>A 3426A>T 3427C>G), TTC1028TTT (3430C>T), CAG1029AAC (3431C>A 3433G>C), TAT1030TCT (3435A>C), CAC1031AAG (3437C>A 3439C>G), TTC1033TTT (3445C>T), TGC1034TGT (3448C>T), TCG1035TCA (3451G>A), TTT1037TTC (3457T>C), GTC1038TTA (3458G>T 3460C>A), CTT1039TTG (3461C>T 3463T>G), GTG1041GTT (3469G>T), AAT1042GAA (3470A>G 3472T>A), CGT1044ATC (3476C>A 3477G>T 3478T>C), TGG1045TTA (3480G>T 3481G>A), GTT1046AGA (3482G>A 3483T>G 3484T>A), GTC1047TTA (3485G>T 3487C>A), ACA1048GTG (3488A>G 3489C>T 3490A>G), CCC1049CCT (3493C>T), CCG1051CCT (3499G>T), CTG1052GTG (3500C>G), AAG1053AAA (3505G>A), CGT1054GTT (3506C>G 3507G>T), GCG1055TTG (3509G>T 3510C>T), GTC1056ATT (3512G>A 3514C>T), AAA1057AAG (3517A>G), CTC1058TTG (3518C>T 3520C>G), GGA1059GGC (3523A>C), CGT1060AGG (3524C>A 3526T>G), GCT1061CAT (3527G>C 3528C>A), CTA1063TTG (3533C>T 3535A>G), TTT1066GTT (3542T>G), GTG1069GTT (3553G>T), GAG1070GAA (3556G>A), AGG1073AGA (3565G>A), GTG1074GTA (3568G>A), TTC1076TTG (3574C>G), ATT1077TTG (3575A>T 3577T>G) | | | | | | | | |
| **Proteins** | | | | | | | | | |
| polymerase (YP_004901701.1) | 1019 | 1078 | 5.2% | 266 | 57.5% | 60 (100%) | 37 (61.7%) | 0/0/0/0 | 0 |
| Protein mutations: | N1020M (3405A>T 3406T>G), E1022D (3412A>T), F1026C (3423T>G 3424T>C), D1027M (3425G>A 3426A>T 3427C>G), Q1029N (3431C>A 3433G>C), Y1030S (3435A>C), H1031K (3437C>A 3439C>G), V1038L (3458G>T 3460C>A), N1042E (3470A>G 3472T>A), R1044I (3476C>A 3477G>T 3478T>C), W1045L (3480G>T 3481G>A), V1046R (3482G>A 3483T>G 3484T>A), V1047L (3485G>T 3487C>A), T1048V (3488A>G 3489C>T 3490A>G), L1052V (3500C>G), R1054V (3506C>G 3507G>T), A1055L (3509G>T 3510C>T), V1056I (3512G>A 3514C>T), A1061H (3527G>C 3528C>A), F1066V (3542T>G), F1076L (3574C>G), I1077L (3575A>T 3577T>G) | | | | | | | | |
| Codon mutations: | AAT1020ATG (3405A>T 3406T>G), TTC1021TTT (3409C>T), GAA1022GAT (3412A>T), AAG1024AAA (3418G>A), TTG1025TTA (3421G>A), TTT1026TGC (3423T>G 3424T>C), GAC1027ATG (3425G>A 3426A>T 3427C>G), TTC1028TTT (3430C>T), CAG1029AAC (3431C>A 3433G>C), TAT1030TCT (3435A>C), CAC1031AAG (3437C>A 3439C>G), TTC1033TTT (3445C>T), TGC1034TGT (3448C>T), TCG1035TCA (3451G>A), TTT1037TTC (3457T>C), GTC1038TTA (3458G>T 3460C>A), CTT1039TTG (3461C>T 3463T>G), GTG1041GTT (3469G>T), AAT1042GAA (3470A>G 3472T>A), CGT1044ATC (3476C>A 3477G>T 3478T>C), TGG1045TTA (3480G>T 3481G>A), GTT1046AGA (3482G>A 3483T>G 3484T>A), GTC1047TTA (3485G>T 3487C>A), ACA1048GTG (3488A>G 3489C>T 3490A>G), CCC1049CCT (3493C>T), CCG1051CCT (3499G>T), CTG1052GTG (3500C>G), AAG1053AAA (3505G>A), CGT1054GTT (3506C>G 3507G>T), GCG1055TTG (3509G>T 3510C>T), GTC1056ATT (3512G>A 3514C>T), AAA1057AAG (3517A>G), CTC1058TTG (3518C>T 3520C>G), GGA1059GGC (3523A>C), CGT1060AGG (3524C>A 3526T>G), GCT1061CAT (3527G>C 3528C>A), CTA1063TTG (3533C>T 3535A>G), TTT1066GTT (3542T>G), GTG1069GTT (3553G>T), GAG1070GAA (3556G>A), AGG1073AGA (3565G>A), GTG1074GTA (3568G>A), TTC1076TTG (3574C>G), ATT1077TTG (3575A>T 3577T>G) | | | | | | | | |

*: Inserts / Deletes / Misaligned / Frameshifts

#### Analysis details

This analysis was performed with panviral2.64

### NGS Details (UN60): Alfalfa leaf curl virus

#### Assembly

| | |
|---|---|
| **Coverage Length** | 270 (2 contig(s)) |
| **Depth Of Coverage** | 6.0 |
| **Number Of Reads** | 14 |
| **Reads Per Million** | 0.30 rpm (after QC) |
| **Ambiguities** | 0 |
| **Assembly Method** | de novo + reference guided assembly |
| **Consensus Caller** | Bcf Tools |

#### Coverage Map

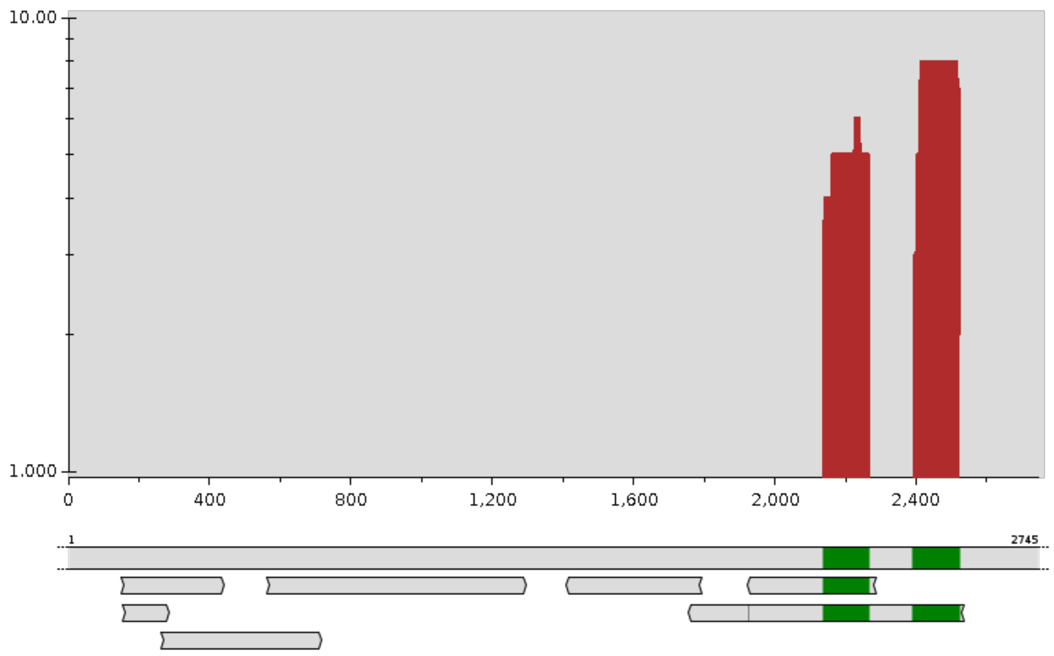

#### Assignment

| | |
|---|---|
| **Type** | Alfalfa leaf curl virus (Taxonomy ID: 1306546) |
| **Reference Genome** | NC_027714.1 |
| **NT Identity (%)** | 71.4815 |
| **AA Identity (%)** | 58.2222 |
| **Number Of Stop Codons** | 0 |
| **Number Of CDS** | 7 |

#### Alignment

| | |
|---|---|
| **Alignment Score** | 232.0 (NT) + 989.0 (AA) = 1221.0 |
| **Concordance (%)** | 57.297 |

| Alignment Method | Global, seeded, nucleotide + amino acids (AGA) |
|---|---|

#### Genome Region

Sequence starts at position 2136 and ends at position 2525 relative to NC_027714.1 reference sequence.

#### Alignment Detailed Statistics

| | Begin | End | Coverage | Score | Concordance | Matches | Identities | I/D/M/F* | Stop Codons |
|---|---|---|---|---|---|---|---|---|---|
| **NT** | **2136** | **2525** | **9.8%** | **232** | **43.0%** | **270 (100%)** | **193 (71.5%)** | **0/0** | |
| Mutations: | 2141C>G, 2144G>T, 2147G>A, 2152C>T, 2158A>T, 2161T>G, 2162C>T, 2170T>C, 2173T>G, 2175C>T, 2179A>T, 2181G>T, 2182C>T, 2183T>C, 2184C>T, 2188G>A, 2191A>C, 2195T>A, 2197G>T, 2209C>T, 2210C>T, 2219G>T, 2225G>A, 2228G>A, 2233C>A, 2237A>G, 2239C>G, 2241G>T, 2242C>T, 2243A>G, 2247C>G, 2248T>A, 2251G>T, 2252G>C, 2255A>T, 2261G>A, 2264A>G, 2267T>G, 2396G>A, 2402T>A, 2403G>A, 2404C>T, 2405G>A, 2409G>T, 2414A>G, 2416C>T, 2417C>G, 2418C>T, 2420G>A, 2422T>C, 2423C>T, 2425T>G, 2428A>G, 2429T>A, 2431A>T, 2434G>T, 2435A>T, 2437A>T, 2442T>A, 2444A>G, 2447G>T, 2450T>G, 2452A>C, 2453A>G, 2456C>T, 2461G>C, 2467A>G, 2476G>T, 2477C>G, 2486T>G, 2488A>G, 2492A>T, 2494T>G, 2495A>T, 2498C>T, 2501A>T, 2504C>T | | | | | | | | |

##### CDS

| | Begin | End | Coverage | Score | Concordance | Matches | Identities | I/D/M/F* | Stop Codons |
|---|---|---|---|---|---|---|---|---|---|
| AL349_gp5 | 5 | 134 | 27.2% | 440 | 64.9% | 90 (100%) | 56 (62.2%) | 0/0/0/0 | 0 |
| Protein mutations: | I15L (2492A>T 2494T>G), Q21K (2476G>T), P26A (2461G>C), M27I (2456C>T), L29V (2450T>G 2452A>C), Y32F (2442T>A), Y34K (2435A>T 2437A>T), Q35K (2434G>T), L36I (2429T>A 2431A>T), K38Q (2423C>T 2425T>G), N39D (2420G>A 2422T>C), W40Y (2417C>G 2418C>T), D41N (2414A>G 2416C>T), T43N (2409G>T), A45I (2402T>A 2403G>A 2404C>T), R97S (2247C>G 2248T>A), A99K (2241G>T 2242C>T), D100H (2237A>G 2239C>G), A102S (2233C>A), V110I (2209C>T), R114S (2195T>A 2197G>T), L116V (2191A>C), L117F (2188G>A), R118K (2183T>C 2184C>T), A119N (2181G>T 2182C>T), S120T (2179A>T), R121K (2175C>T), S123G (2170T>C), K126Q (2161T>G), S127T (2158A>T), D129N (2152C>T), W132C (2141C>G) | | | | | | | | |
| Codon mutations: | CAG11CAA (2504C>T), GGT12GGA (2501A>T), AAG13AAA (2498C>T), TCT14TCA (2495A>T), ATT15CTA (2492A>T 2494T>G), TTA17CTC (2486T>G 2488A>G), CCG20CCC (2477C>G), CAG21AAG (2476G>T), TTA24CTA (2467A>G), CCA26GCA (2461G>C), ATG27ATA (2456C>T), TTT28TTC (2453A>G), TTA29GTC (2450T>G 2452A>C), ATC30ATA (2447G>T), GAT31GAC (2444A>G), TAC32TTC (2442T>A), TAT34AAA (2435A>T 2437A>T), CAA35AAA (2434G>T), TTA36ATT (2429T>A 2431A>T), TTA37CTA (2428A>G), AAG38CAA (2423C>T 2425T>G), AAC39GAT (2420G>A 2422T>C), TGG40TAC (2417C>G 2418C>T), GAT41AAC (2414A>G 2416C>T), ACT43AAT (2409G>T), TAC44TAT (2405G>A), GCA45ATT (2402T>A 2403G>A 2404C>T), GTC47GTT (2396G>A), CCA90CCC (2267T>G), AAT91AAC (2264A>G), ACC92ACT (2261G>A), ATT94ATA (2255A>T), CCC95CCG (2252G>C), CGA96AGA (2251G>T), AGG97TCG (2247C>G 2248T>A), GAT98GAC (2243A>G), GCA99AAA (2241G>T 2242C>T), GAT100CAC (2237A>G 2239C>G), GCC102TCC (2233C>A), GAC103GAT (2228G>A), TAC104TAT (2225G>A), TCC106TCA (2219G>T), GGG109GGA (2210C>T), GTA110ATA (2209C>T), CGA114AGT (2195T>A 2197G>T), TTA116GTA (2191A>C), CTT117TTT (2188G>A), AGA118AAG (2183T>C 2184C>T), GCT119AAT (2181G>T 2182C>T), TCA120ACA (2179A>T), AGA121AAA (2175C>T), AGA122CGA (2173T>G), AGT123GGT (2170T>C), AAG125AAA (2162C>T), AAG126CAG (2161T>G), TCT127ACT (2158A>T), GAT129AAT (2152C>T), TCC130TCT (2147G>A), ATC131ATA (2144G>T), TGG132TGC (2141C>G) | | | | | | | | |
| repA | 5 | 134 | 34.6% | 440 | 64.9% | 90 (100%) | 56 (62.2%) | 0/0/0/0 | 0 |
| Protein mutations: | I15L (2492A>T 2494T>G), Q21K (2476G>T), P26A (2461G>C), M27I (2456C>T), L29V (2450T>G 2452A>C), Y32F (2442T>A), Y34K (2435A>T 2437A>T), Q35K (2434G>T), L36I (2429T>A 2431A>T), K38Q (2423C>T 2425T>G), N39D (2420G>A 2422T>C), W40Y (2417C>G 2418C>T), D41N (2414A>G 2416C>T), T43N (2409G>T), A45I (2402T>A 2403G>A 2404C>T), R97S (2247C>G 2248T>A), A99K (2241G>T 2242C>T), D100H (2237A>G 2239C>G), A102S (2233C>A), V110I (2209C>T), R114S (2195T>A 2197G>T), L116V (2191A>C), L117F (2188G>A), R118K (2183T>C 2184C>T), A119N (2181G>T 2182C>T), S120T (2179A>T), R121K (2175C>T), S123G (2170T>C), K126Q (2161T>G), S127T (2158A>T), D129N (2152C>T), W132C (2141C>G) | | | | | | | | |
| Codon mutations: | CAG11CAA (2504C>T), GGT12GGA (2501A>T), AAG13AAA (2498C>T), TCT14TCA (2495A>T), ATT15CTA (2492A>T 2494T>G), TTA17CTC (2486T>G 2488A>G), CCG20CCC (2477C>G), CAG21AAG (2476G>T), TTA24CTA (2467A>G), CCA26GCA (2461G>C), ATG27ATA (2456C>T), TTT28TTC (2453A>G), TTA29GTC (2450T>G 2452A>C), ATC30ATA (2447G>T), GAT31GAC (2444A>G), TAC32TTC (2442T>A), TAT34AAA (2435A>T 2437A>T), CAA35AAA (2434G>T), TTA36ATT (2429T>A 2431A>T), TTA37CTA (2428A>G), AAG38CAA (2423C>T 2425T>G), AAC39GAT (2420G>A 2422T>C), TGG40TAC (2417C>G 2418C>T), GAT41AAC (2414A>G 2416C>T), ACT43AAT (2409G>T), TAC44TAT (2405G>A), GCA45ATT (2402T>A 2403G>A 2404C>T), GTC47GTT (2396G>A), CCA90CCC (2267T>G), AAT91AAC (2264A>G), ACC92ACT (2261G>A), ATT94ATA (2255A>T), CCC95CCG (2252G>C), CGA96AGA (2251G>T), AGG97TCG (2247C>G 2248T>A), GAT98GAC (2243A>G), GCA99AAA (2241G>T 2242C>T), GAT100CAC (2237A>G 2239C>G), GCC102TCC (2233C>A), GAC103GAT (2228G>A), TAC104TAT (2225G>A), TCC106TCA (2219G>T), GGG109GGA (2210C>T), GTA110ATA (2209C>T), CGA114AGT (2195T>A 2197G>T), TTA116GTA (2191A>C), CTT117TTT (2188G>A), AGA118AAG (2183T>C 2184C>T), GCT119AAT (2181G>T 2182C>T), TCA120ACA (2179A>T), AGA121AAA (2175C>T), AGA122CGA (2173T>G), AGT123GGT (2170T>C), AAG125AAA (2162C>T), AAG126CAG (2161T>G), TCT127ACT (2158A>T), GAT129AAT (2152C>T), TCC130TCT (2147G>A), ATC131ATA (2144G>T), TGG132TGC (2141C>G) | | | | | | | | |
| C3 | 8 | 52 | 36.9% | 109 | 34.1% | 45 (100%) | 19 (42.2%) | 0/0/0/0 | 0 |
| Protein mutations: | Q8P (2267T>G), I9T (2264A>G), P10L (2261G>A), F12Y (2255A>T), P13R (2251G>T 2252G>C), E14D (2248T>A), G15R (2247C>G), M16T (2242C>T 2243A>G), Q17N (2239C>G 2241G>T), M18T (2237A>G), T21I (2228G>A), T22I (2225G>A), P24Q (2219G>T), G27E (2209C>T 2210C>T), N31K (2197G>T), E32V (2195T>A), D33E (2191A>C), E36R (2182C>T 2183T>C 2184C>T), L37I (2179A>T 2181G>T), E39N (2173T>G 2175C>T), R43N (2161T>G 2162C>T), S44R (2158A>T), P48L (2147G>A), S49Y (2144G>T), G50A (2141C>G) | | | | | | | | |
| Codon mutations: | CAA8CCA (2267T>G), ATA9ACA (2264A>G), CCC10CTC (2261G>A), TTC12TAC (2255A>T), CCC13CGA (2251G>T 2252G>C), GAA14GAT (2248T>A), GGG15CGG (2247C>G), ATG16ACA (2242C>T 2243A>G), CAG17AAC (2239C>G 2241G>T), ATG18ACG (2237A>G), TCG19TCT (2233C>A), ACT21ATT (2228G>A), ACA22ATA (2225G>A), CCA24CAA (2219G>T), GGG27GAA (2209C>T 2210C>T), AAC31AAA (2197G>T), GAG32GTG (2195T>A), GAT33GAG (2191A>C), TAC34TAT (2188G>A), GAG36AGA (2182C>T 2183T>C 2184C>T), CTT37ATA (2179A>T 2181G>T), GAA39AAC (2173T>G 2175C>T), GAA40GAG (2170T>C), AGA43AAC (2161T>G 2162C>T), AGT44AGA (2158A>T), GGG46GGA (2152C>T), CCA48CTA (2147G>A), TCT49TAT (2144G>T), GGA50GCA (2141C>G) | | | | | | | | |

##### Proteins

| | Begin | End | Coverage | Score | Concordance | Matches | Identities | I/D/M/F* | Stop Codons |
|---|---|---|---|---|---|---|---|---|---|
| replication associated protein (YP_009162635.2) | 5 | 134 | 27.2% | 440 | 64.9% | 90 (100%) | 56 (62.2%) | 0/0/0/0 | 0 |
| Protein mutations: | I15L (2492A>T 2494T>G), Q21K (2476G>T), P26A (2461G>C), M27I (2456C>T), L29V (2450T>G 2452A>C), Y32F (2442T>A), Y34K (2435A>T 2437A>T), Q35K (2434G>T), L36I (2429T>A 2431A>T), K38Q (2423C>T 2425T>G), N39D (2420G>A 2422T>C), W40Y (2417C>G 2418C>T), D41N (2414A>G 2416C>T), T43N (2409G>T), A45I (2402T>A 2403G>A 2404C>T), R97S (2247C>G 2248T>A), A99K (2241G>T 2242C>T), D100H (2237A>G 2239C>G), A102S (2233C>A), V110I (2209C>T), R114S (2195T>A 2197G>T), L116V (2191A>C), L117F (2188G>A), R118K (2183T>C 2184C>T), A119N (2181G>T 2182C>T), S120T (2179A>T), R121K (2175C>T), S123G (2170T>C), K126Q (2161T>G), S127T (2158A>T), D129N (2152C>T), W132C (2141C>G) | | | | | | | | |
| Codon mutations: | CAG11CAA (2504C>T), GGT12GGA (2501A>T), AAG13AAA (2498C>T), TCT14TCA (2495A>T), ATT15CTA (2492A>T 2494T>G), TTA17CTC (2486T>G 2488A>G), CCG20CCC (2477C>G), CAG21AAG (2476G>T), TTA24CTA (2467A>G), CCA26GCA (2461G>C), ATG27ATA (2456C>T), TTT28TTC (2453A>G), TTA29GTC (2450T>G 2452A>C), ATC30ATA (2447G>T), GAT31GAC (2444A>G), TAC32TTC (2442T>A), TAT34AAA (2435A>T 2437A>T), CAA35AAA (2434G>T), TTA36ATT (2429T>A 2431A>T), TTA37CTA (2428A>G), AAG38CAA (2423C>T 2425T>G), AAC39GAT (2420G>A 2422T>C), TGG40TAC (2417C>G 2418C>T), GAT41AAC (2414A>G 2416C>T), ACT43AAT (2409G>T), TAC44TAT (2405G>A), GCA45ATT (2402T>A 2403G>A 2404C>T), GTC47GTT (2396G>A), CCA90CCC (2267T>G), AAT91AAC (2264A>G), ACC92ACT (2261G>A), ATT94ATA (2255A>T), CCC95CCG (2252G>C), CGA96AGA (2251G>T), AGG97TCG (2247C>G 2248T>A), GAT98GAC (2243A>G), GCA99AAA (2241G>T 2242C>T), GAT100CAC (2237A>G 2239C>G), GCC102TCC (2233C>A), GAC103GAT (2228G>A), TAC104TAT (2225G>A), TCC106TCA (2219G>T), GGG109GGA (2210C>T), GTA110ATA (2209C>T), CGA114AGT (2195T>A 2197G>T), TTA116GTA (2191A>C), CTT117TTT (2188G>A), AGA118AAG (2183T>C 2184C>T), GCT119AAT (2181G>T 2182C>T), TCA120ACA (2179A>T), AGA121AAA (2175C>T), AGA122CGA (2173T>G), AGT123GGT (2170T>C), AAG125AAA (2162C>T), AAG126CAG (2161T>G), TCT127ACT (2158A>T), GAT129AAT (2152C>T), TCC130TCT (2147G>A), ATC131ATA (2144G>T), TGG132TGC (2141C>G) | | | | | | | | |
| RepA (YP_009162636.1) | 5 | 134 | 34.6% | 440 | 64.9% | 90 (100%) | 56 (62.2%) | 0/0/0/0 | 0 |
| Protein mutations: | I15L (2492A>T 2494T>G), Q21K (2476G>T), P26A (2461G>C), M27I (2456C>T), L29V (2450T>G 2452A>C), Y32F (2442T>A), Y34K (2435A>T 2437A>T), Q35K (2434G>T), L36I (2429T>A 2431A>T), K38Q (2423C>T 2425T>G), N39D (2420G>A 2422T>C), W40Y (2417C>G 2418C>T), D41N (2414A>G 2416C>T), T43N (2409G>T), A45I (2402T>A 2403G>A 2404C>T), R97S (2247C>G 2248T>A), A99K (2241G>T 2242C>T), D100H (2237A>G 2239C>G), A102S (2233C>A), V110I (2209C>T), R114S (2195T>A 2197G>T), L116V (2191A>C), L117F (2188G>A), R118K (2183T>C 2184C>T), A119N (2181G>T 2182C>T), S120T (2179A>T), R121K (2175C>T), S123G (2170T>C), K126Q (2161T>G), S127T (2158A>T), D129N (2152C>T), W132C (2141C>G) | | | | | | | | |

| | Begin | End | Coverage | Score | Concordance | Matches | Identities | I/D/M/F* | Stop Codons |
|---|---|---|---|---|---|---|---|---|---|
| **NT** | **2136** | **2525** | **9.8%** | **232** | **43.0%** | **270 (100%)** | **193 (71.5%)** | **0/0** | |
| Codon mutations: | CAG11CAA (2504C>T), GGT12GGA (2501A>T), AAG13AAA (2498C>T), TCT14TCA (2495A>T), ATT15CTA (2492A>T 2494T>G), TTA17CTC (2486T>G 2488A>G), CCG20CCC (2477C>G), CAG21AAG (2476G>T), TTA24CTA (2467A>G), CCA26GCA (2461G>C), ATG27ATA (2456C>T), TTT28TTC (2453A>G), TTA29GTC (2450T>G 2452A>C), ATC30ATA (2447G>T), GAT31GAC (2444A>G), TAC32TTC (2442T>A), TAT34AAA (2435A>T 2437A>T), CAA35AAA (2434G>T), TTA36ATT (2429T>A 2431A>T), TTA37CTA (2428A>G), AAG38CAA (2423C>T 2425T>G), AAC39GAT (2420G>A 2422T>C), TGG40TAC (2417C>G 2418C>T), GAT41AAC (2414A>G 2416C>T), ACT43AAT (2409G>T), TAC44TAT (2405G>A), GCA45ATT (2402T>A 2403G>A 2404C>T), GTC47GTT (2396G>A), CCA90CCC (2267T>G), AAT91AAC (2264A>G), ACC92ACT (2261G>A), ATT94ATA (2255A>T), CCC95CCG (2252G>C), CGA96AGA (2251G>T), AGG97TCG (2247C>G 2248T>A), GAT98GAC (2243A>G), GCA99AAA (2241G>T 2242C>T), GAT100CAC (2237A>G 2239C>G), GCC102TCC (2233C>A), GAC103GAT (2228G>A), TAC104TAT (2225G>A), TCC106TCA (2219G>T), GGG109GGA (2210C>T), GTA110ATA (2209C>T), CGA114AGT (2195T>A 2197G>T), TTA116GTA (2191A>C), CTT117TTT (2188G>A), AGA118AAG (2183T>C 2184C>T), GCT119AAT (2181G>T 2182C>T), TCA120ACA (2179A>T), AGA121AAA (2175C>T), AGA122CGA (2173T>G), AGT123GGT (2170T>C), AAG125AAA (2162C>T), AAG126CAG (2161T>G), TCT127ACT (2158A>T), GAT129AAT (2152C>T), TCC130TCT (2147G>A), ATC131ATA (2144G>T), TGG132TGC (2141C>G) | | | | | | | | |
| C3 (YP_009162637.1) | 8 | 52 | 36.9% | 109 | 34.1% | 45 (100%) | 19 (42.2%) | 0/0/0/0 | 0 |
| Protein mutations: | Q8P (2267T>G), I9T (2264A>G), P10L (2261G>A), F12Y (2255A>T), P13R (2251G>T 2252G>C), E14D (2248T>A), G15R (2247C>G), M16T (2242C>T 2243A>G), Q17N (2239C>G 2241G>T), M18T (2237A>G), T21I (2228G>A), T22I (2225G>A), P24Q (2219G>T), G27E (2209C>T 2210C>T), N31K (2197G>T), E32V (2195T>A), D33E (2191A>C), E36R (2182C>T 2183T>C 2184C>T), L37I (2179A>T 2181G>T), E39N (2173T>G 2175C>T), R43N (2161T>G 2162C>T), S44R (2158A>T), P48L (2147G>A), S49Y (2144G>T), G50A (2141C>G) | | | | | | | | |
| Codon mutations: | CAA8CCA (2267T>G), ATA9ACA (2264A>G), CCC10CTC (2261G>A), TTC12TAC (2255A>T), CCC13CGA (2251G>T 2252G>C), GAA14GAT (2248T>A), GGG15CGG (2247C>G), ATG16ACA (2242C>T 2243A>G), CAG17AAC (2239C>G 2241G>T), ATG18ACG (2237A>G), TCG19TCT (2233C>A), ACT21ATT (2228G>A), ACA22ATA (2225G>A), CCA24CAA (2219G>T), GGG27GAA (2209C>T 2210C>T), AAC31AAA (2197G>T), GAG32GTG (2195T>A), GAT33GAG (2191A>C), TAC34TAT (2188G>A), GAG36AGA (2182C>T 2183T>C 2184C>T), CTT37ATA (2179A>T 2181G>T), GAA39AAC (2173T>G 2175C>T), GAA40GAG (2170T>C), AGA43AAC (2161T>G 2162C>T), AGT44AGA (2158A>T), GGG46GGA (2152C>T), CCA48CTA (2147G>A), TCT49TAT (2144G>T), GGA50GCA (2141C>G) | | | | | | | | |

*: Inserts / Deletes / Misaligned / Frameshifts

#### Analysis details

This analysis was performed with panviral2.64

### NGS Details (UN60): Duamitovirus peex1

#### Assembly

| | |
|---|---|
| **Coverage Length** | 274 (2 contig(s)) |
| **Depth Of Coverage** | 4.7 |
| **Number Of Reads** | 13 |
| **Reads Per Million** | 0.28 rpm (after QC) |
| **Ambiguities** | 0 |
| **Assembly Method** | de novo + reference guided assembly |
| **Consensus Caller** | Bcf Tools |

#### Coverage Map

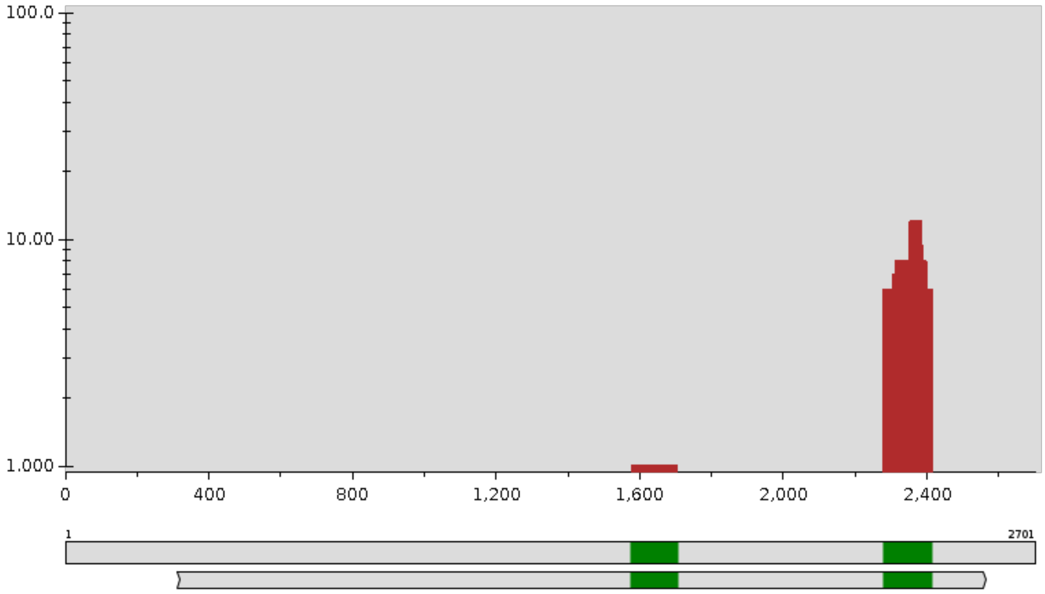

#### Assignment

| | |
|---|---|
| **Type** | Duamitovirus peex1 (Taxonomy ID: 2955799) |
| **Reference Genome** | NC_076525.1 |
| **NT Identity (%)** | 70.8029 |
| **AA Identity (%)** | 73.6264 |
| **Number Of Stop Codons** | 0 |
| **Number Of CDS** | 1 |

#### Alignment

| | |
|---|---|
| **Alignment Score** | 228.0 (NT) + 541.0 (AA) = 769.0 |
| **Concordance (%)** | 63.5537 |

| Alignment Method | Global, seeded, nucleotide + amino acids (AGA) |
|---|---|

#### Genome Region

Sequence starts at position 1575 and ends at position 2417 relative to NC_076525.1 reference sequence.

#### Alignment Detailed Statistics

| | Begin | End | Coverage | Score | Concordance | Matches | Identities | I/D/M/F* | Stop Codons |
|---|---|---|---|---|---|---|---|---|---|
| **NT** | **1575** | **2417** | **10.1%** | **228** | **41.6%** | **274 (100%)** | **194 (70.8%)** | **0/0** | |
| Mutations: | 1583C>G, 1590G>A, 1592G>T, 1593T>A, 1594C>G, 1598G>A, 1599A>G, 1601C>A, 1604T>C, 1610G>T, 1611G>A, 1616G>A, 1619G>A, 1622T>G, 1623C>T, 1628C>T, 1632C>T, 1633T>A, 1634G>C, 1637A>T, 1646C>A, 1649A>G, 1652T>C, 1653G>T, 1655C>G, 1661A>C, 1664C>T, 1667T>C, 1671T>A, 1673G>A, 1676A>G, 1688G>T, 1689G>T, 1690A>T, 1696T>C, 1697C>T, 1700C>T, 2278G>A, 2281T>A, 2283G>A, 2294T>G, 2295C>A, 2296G>A, 2301C>G, 2303T>C, 2306A>T, 2311A>G, 2312T>C, 2315C>T, 2321G>T, 2325A>G, 2326A>C, 2327T>A, 2330C>A, 2331C>G, 2333G>T, 2339G>A, 2345A>T, 2348C>T, 2351A>G, 2352C>T, 2354T>C, 2363G>T, 2373G>A, 2374C>G, 2375G>T, 2377C>A, 2378T>A, 2379T>A, 2388C>A, 2390T>A, 2393T>A, 2394C>A, 2396G>C, 2398C>A, 2399T>A, 2406C>G, 2411A>G, 2412G>A, 2414G>C | | | | | | | | |

##### CDS

| | Begin | End | Coverage | Score | Concordance | Matches | Identities | I/D/M/F* | Stop Codons |
|---|---|---|---|---|---|---|---|---|---|
| RdRp | 422 | 702 | 12.1% | 541 | 78.3% | 91 (100%) | 67 (73.6%) | 0/0/0/0 | 0 |
| Protein mutations: | V427I (1590G>A 1592G>T), I430V (1599A>G 1601C>A), A434T (1611G>A), L441Y (1632C>T 1633T>A 1634G>C), A448S (1653G>T 1655C>G), L454I (1671T>A 1673G>A), E460L (1689G>T 1690A>T), V462A (1696T>C 1697C>T), M657K (2281T>A), E658K (2283G>A), R662K (2295C>A 2296G>A), L664V (2301C>G 2303T>C), Y667C (2311A>G 2312T>C), N672A (2325A>G 2326A>C 2327T>A), P674A (2331C>G 2333G>T), L681F (2352C>T 2354T>C), A688S (2373G>A 2374C>G 2375G>T), T689K (2377C>A 2378T>A), S690T (2379T>A), Q695N (2394C>A 2396G>C), T696K (2398C>A 2399T>A), H699D (2406C>G), V701I (2412G>A 2414G>C) | | | | | | | | |
| Codon mutations: | ACC424ACG (1583C>G), GTG427ATT (1590G>A 1592G>T), TCT428AGT (1593T>A 1594C>G), GAG429GAA (1598G>A), ATC430GTA (1599A>G 1601C>A), GCT431GCC (1604T>C), CTG433CTT (1610G>T), GCT434ACT (1611G>A), GGG435GGA (1616G>A), CAG436CAA (1619G>A), CCT437CCG (1622T>G), CTA438TTA (1623C>T), GGC439GGT (1628C>T), CTG441TAC (1632C>T 1633T>A 1634G>C), GGA442GGT (1637A>T), TCC445TCA (1646C>A), CTA446CTG (1649A>G), TTT447TTC (1652T>C), GCC448TCG (1653G>T 1655C>G), TCA450TCC (1661A>C), CAC451CAT (1664C>T), CAT452CAC (1667T>C), TTG454ATA (1671T>A 1673G>A), GTA455GTG (1676A>G), GCG459GCT (1688G>T), GAG460TTG (1689G>T 1690A>T), GTC462GCT (1696T>C 1697C>T), TAC463TAT (1700C>T), TGG656.AG (2278G>A), ATG657AAG (2281T>A), GAA658AAA (2283G>A), CTT661CTG (2294T>G), CGA662AAA (2295C>A 2296G>A), CTT664GTC (2301C>G 2303T>C), TCA665TCT (2306A>T), TAT667TGC (2311A>G 2312T>C), CAC668CAT (2315C>T), GTG670GTT (2321G>T), AAT672GCA (2325A>G 2326A>C 2327T>A), TCC673TCA (2330C>A), CCG674GCT (2331C>G 2333G>T), GTG676GTA (2339G>A), ATA678ATT (2345A>T), GAC679GAT (2348C>T), CAA680CAG (2351A>G), CTT681TTC (2352C>T 2354T>C), GTG684GTT (2363G>T), GCG688AGT (2373G>A 2374C>G 2375G>T), ACT689AAA (2377C>A 2378T>A), TCT690ACT (2379T>A), CGT693AGA (2388C>A 2390T>A), ACT694ACA (2393T>A), CAG695AAC (2394C>A 2396G>C), ACT696AAA (2398C>A 2399T>A), CAT699GAT (2406C>G), TTA700TTG (2411A>G), GTG701ATC (2412G>A 2414G>C) | | | | | | | | |

##### Proteins

| | Begin | End | Coverage | Score | Concordance | Matches | Identities | I/D/M/F* | Stop Codons |
|---|---|---|---|---|---|---|---|---|---|
| RNA-dependent RNA polymerase (YP_010798875.1) | 422 | 702 | 12.1% | 541 | 78.3% | 91 (100%) | 67 (73.6%) | 0/0/0/0 | 0 |
| Protein mutations: | V427I (1590G>A 1592G>T), I430V (1599A>G 1601C>A), A434T (1611G>A), L441Y (1632C>T 1633T>A 1634G>C), A448S (1653G>T 1655C>G), L454I (1671T>A 1673G>A), E460L (1689G>T 1690A>T), V462A (1696T>C 1697C>T), M657K (2281T>A), E658K (2283G>A), R662K (2295C>A 2296G>A), L664V (2301C>G 2303T>C), Y667C (2311A>G 2312T>C), N672A (2325A>G 2326A>C 2327T>A), P674A (2331C>G 2333G>T), L681F (2352C>T 2354T>C), A688S (2373G>A 2374C>G 2375G>T), T689K (2377C>A 2378T>A), S690T (2379T>A), Q695N (2394C>A 2396G>C), T696K (2398C>A 2399T>A), H699D (2406C>G), V701I (2412G>A 2414G>C) | | | | | | | | |
| Codon mutations: | ACC424ACG (1583C>G), GTG427ATT (1590G>A 1592G>T), TCT428AGT (1593T>A 1594C>G), GAG429GAA (1598G>A), ATC430GTA (1599A>G 1601C>A), GCT431GCC (1604T>C), CTG433CTT (1610G>T), GCT434ACT (1611G>A), GGG435GGA (1616G>A), CAG436CAA (1619G>A), CCT437CCG (1622T>G), CTA438TTA (1623C>T), GGC439GGT (1628C>T), CTG441TAC (1632C>T 1633T>A 1634G>C), GGA442GGT (1637A>T), TCC445TCA (1646C>A), CTA446CTG (1649A>G), TTT447TTC (1652T>C), GCC448TCG (1653G>T 1655C>G), TCA450TCC (1661A>C), CAC451CAT (1664C>T), CAT452CAC (1667T>C), TTG454ATA (1671T>A 1673G>A), GTA455GTG (1676A>G), GCG459GCT (1688G>T), GAG460TTG (1689G>T 1690A>T), GTC462GCT (1696T>C 1697C>T), TAC463TAT (1700C>T), TGG656.AG (2278G>A), ATG657AAG (2281T>A), GAA658AAA (2283G>A), CTT661CTG (2294T>G), CGA662AAA (2295C>A 2296G>A), CTT664GTC (2301C>G 2303T>C), TCA665TCT (2306A>T), TAT667TGC (2311A>G 2312T>C), CAC668CAT (2315C>T), GTG670GTT (2321G>T), AAT672GCA (2325A>G 2326A>C 2327T>A), TCC673TCA (2330C>A), CCG674GCT (2331C>G 2333G>T), GTG676GTA (2339G>A), ATA678ATT (2345A>T), GAC679GAT (2348C>T), CAA680CAG (2351A>G), CTT681TTC (2352C>T 2354T>C), GTG684GTT (2363G>T), GCG688AGT (2373G>A 2374C>G 2375G>T), ACT689AAA (2377C>A 2378T>A), TCT690ACT (2379T>A), CGT693AGA (2388C>A 2390T>A), ACT694ACA (2393T>A), CAG695AAC (2394C>A 2396G>C), ACT696AAA (2398C>A 2399T>A), CAT699GAT (2406C>G), TTA700TTG (2411A>G), GTG701ATC (2412G>A 2414G>C) | | | | | | | | |

*: Inserts / Deletes / Misaligned / Frameshifts

#### Analysis details

This analysis was performed with panviral2.64

### NGS Details (UN60): Pseudomonas phage YMC11/07/P54_PAE_BP

#### Assembly

| | |
|---|---|
| **Coverage Length** | 334 (1 contig(s)) |
| **Depth Of Coverage** | 4.8 |
| **Number Of Reads** | 13 |
| **Reads Per Million** | 0.28 rpm (after QC) |
| **Ambiguities** | 0 |
| **Assembly Method** | de novo + reference guided assembly |
| **Consensus Caller** | Bcf Tools |

#### Coverage Map

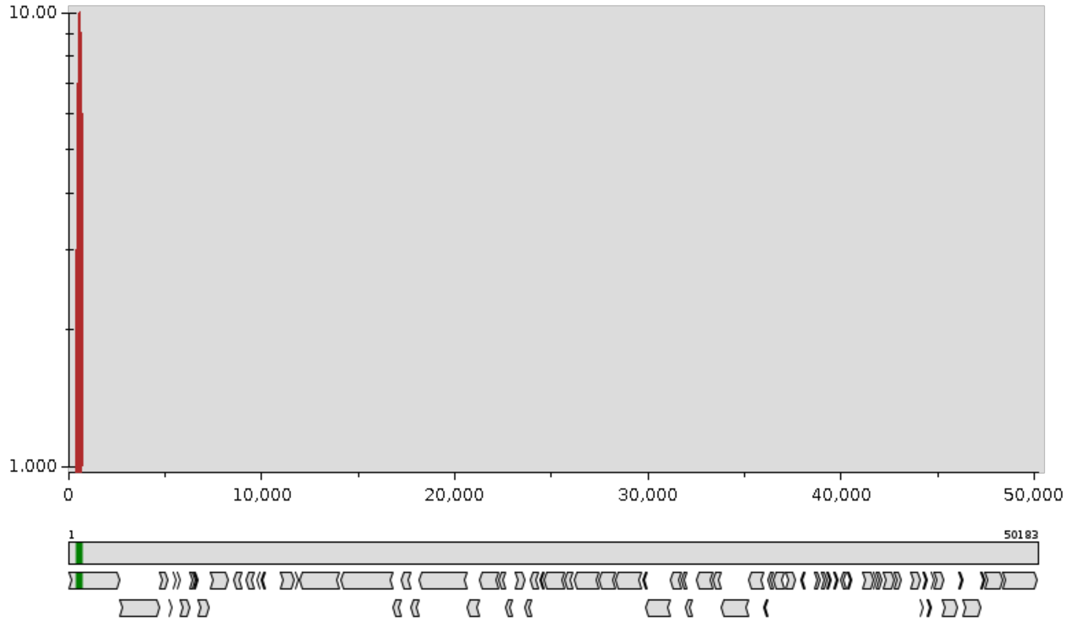

#### Assignment

| | |
|---|---|
| **Type** | Pseudomonas phage YMC11/07/P54_PAE_BP (Taxonomy ID: 1777064) |
| **Reference Genome** | NC_030909.1 |
| **NT Identity (%)** | 78.4257 |
| **AA Identity (%)** | 83.3333 |
| **Number Of Stop Codons** | 0 |
| **Number Of CDS** | 73 |

#### Alignment

| | |
|---|---|
| **Alignment Score** | 390.0 (NT) + 622.0 (AA) = 1012.0 |
| **Concordance (%)** | 71.9772 |

| **Alignment Method** | Local, heuristic, nucleotide (BLASTN) |
|---|---|

#### Genome Region

Sequence starts at position 373 and ends at position 706 relative to NC_030909.1 reference sequence.

#### Alignment Detailed Statistics

| | Begin | End | Coverage | Score | Concordance | Matches | Identities | I/D/M/F* | Stop Codons |
|---|---|---|---|---|---|---|---|---|---|
| **NT** | **373** | **706** | **0.7%** | **390** | **58.4%** | **334 (97.4%)** | **269 (78.4%)** | **9/0** | |
| Mutations: | 385G>C, 391T>C, 397G>A, 400C>G, 401G>A, 410G>C, 418C>G, 424T>G, 426G>T, 427C>G, 429G>T, 442A>C, 454A>G, 457C>G, 458A>G, 459G>C, 460T>A, 466T>G, 490G>C, 508T>G, 517T>C, 520T>C, 526T>C, 529C>G, 529_530insCGCGACATG, 533G>A, 534C>A, 536G>C, 539A>G, 541C>G, 542C>A, 543G>A, 544C>G, 545G>A, 549A>C, 556C>G, 558G>C, 562C>T, 565G>C, 567A>T, 577T>G, 586A>C, 589C>T, 592G>A, 598A>G, 600A>T, 601C>T, 602A>G, 610A>G, 616T>C, 620T>A, 621C>G, 622G>C, 631C>G, 643T>C, 646C>T, 661C>G, 662A>C, 664G>T, 673A>G, 680T>A, 681C>G, 682T>C, 695C>A, 697A>C, 700C>G | | | | | | | | |
| **CDS** | | | | | | | | | |
| BH775_gp01 | 114 | 224 | 12.7% | 622 | 83.7% | 111 (97.4%) | 95 (83.3%) | 3/0/0/0 | 0 |
| Protein mutations: | V123I (401G>A), G126R (410G>C), R131L (426G>T 427C>G), R132L (429G>T), S142A (458A>G 459G>C 460T>A), R165_A166insRDM (529_530insCGCGACATG), A167N (533G>A 534C>A), G168R (536G>C), I169V (539A>G 541C>G), R170K (542C>A 543G>A 544C>G), G171S (545G>A), D172A (549A>C), R175P (558G>C), Y178F (567A>T), Y189F (600A>T 601C>T), I190V (602A>G), Q221N (695C>A 697A>C) | | | | | | | | |
| Codon mutations: | CTG117CTC (385G>C), AAT119AAC (391T>C), GAG121GAA (397G>A), ACC122ACG (400C>G), GTC123ATC (401G>A), GGC126CGC (410G>C), CTC128CTG (418C>G), CCT130CCG (424T>G), CGC131CTG (426G>T 427C>G), CGG132CTG (429G>T), GCA136GCC (442A>C), GAA140GAG (454A>G), GCC141GCG (457C>G), AGT142GCA (458A>G 459G>C 460T>A), CTT144CTG (466T>G), ACG152ACC (490G>C), GCT158GCG (508T>G), GCT161GCC (517T>C), GGT162GGC (520T>C), GAT164GAC (526T>C), CGC165CGG (529C>G), CGC165_GCC166insCGCGACATG (529_530insCGCGACATG), GCC167AAC (533G>A 534C>A), GGC168CGC (536G>C), ATC169GTG (539A>G 541C>G), CGC170AAG (542C>A 543G>A 544C>G), GGC171AGC (545G>A), GAT172GCT (549A>C), GTC174GTG (556C>G), CGG175CCG (558G>C), TAC176TAT (562C>T), CGG177CGC (565G>C), TAC178TTC (567A>T), ACT181ACG (577T>G), CCA184CCC (586A>C), AGC185AGT (589C>T), CCG186CCA (592G>A), GAA188GAG (598A>G), TAC189TTT (600A>T 601C>T), ATC190GTC (602A>G), CTA192CTG (610A>G), GCT194GCC (616T>C), TCG196AGC (620T>A 621C>G 622G>C), CTC199CTG (631C>G), GAT203GAC (643T>C), GTC204GTT (646C>T), ACC209ACG (661C>G), AGG210CGT (662A>C 664G>T), AAA213AAG (673A>G), TCT216AGC (680T>A 681C>G 682T>C), CAA221AAC (695C>A 697A>C), CTC222CTG (700C>G) | | | | | | | | |
| **Proteins** | | | | | | | | | |
| DNA methylase N-4 (YP_009273683.1) | 114 | 224 | 12.7% | 622 | 83.7% | 111 (97.4%) | 95 (83.3%) | 3/0/0/0 | 0 |
| Protein mutations: | V123I (401G>A), G126R (410G>C), R131L (426G>T 427C>G), R132L (429G>T), S142A (458A>G 459G>C 460T>A), R165_A166insRDM (529_530insCGCGACATG), A167N (533G>A 534C>A), G168R (536G>C), I169V (539A>G 541C>G), R170K (542C>A 543G>A 544C>G), G171S (545G>A), D172A (549A>C), R175P (558G>C), Y178F (567A>T), Y189F (600A>T 601C>T), I190V (602A>G), Q221N (695C>A 697A>C) | | | | | | | | |
| Codon mutations: | CTG117CTC (385G>C), AAT119AAC (391T>C), GAG121GAA (397G>A), ACC122ACG (400C>G), GTC123ATC (401G>A), GGC126CGC (410G>C), CTC128CTG (418C>G), CCT130CCG (424T>G), CGC131CTG (426G>T 427C>G), CGG132CTG (429G>T), GCA136GCC (442A>C), GAA140GAG (454A>G), GCC141GCG (457C>G), AGT142GCA (458A>G 459G>C 460T>A), CTT144CTG (466T>G), ACG152ACC (490G>C), GCT158GCG (508T>G), GCT161GCC (517T>C), GGT162GGC (520T>C), GAT164GAC (526T>C), CGC165CGG (529C>G), CGC165_GCC166insCGCGACATG (529_530insCGCGACATG), GCC167AAC (533G>A 534C>A), GGC168CGC (536G>C), ATC169GTG (539A>G 541C>G), CGC170AAG (542C>A 543G>A 544C>G), GGC171AGC (545G>A), GAT172GCT (549A>C), GTC174GTG (556C>G), CGG175CCG (558G>C), TAC176TAT (562C>T), CGG177CGC (565G>C), TAC178TTC (567A>T), ACT181ACG (577T>G), CCA184CCC (586A>C), AGC185AGT (589C>T), CCG186CCA (592G>A), GAA188GAG (598A>G), TAC189TTT (600A>T 601C>T), ATC190GTC (602A>G), CTA192CTG (610A>G), GCT194GCC (616T>C), TCG196AGC (620T>A 621C>G 622G>C), CTC199CTG (631C>G), GAT203GAC (643T>C), GTC204GTT (646C>T), ACC209ACG (661C>G), AGG210CGT (662A>C 664G>T), AAA213AAG (673A>G), TCT216AGC (680T>A 681C>G 682T>C), CAA221AAC (695C>A 697A>C), CTC222CTG (700C>G) | | | | | | | | |

*: Inserts / Deletes / Misaligned / Frameshifts

#### Analysis details

This analysis was performed with panviral2.64

#### NGS Details (UN60): Pinus nigra virus 1

##### Assembly

| | |
|---|---|
| **Coverage Length** | 474 (1 contig(s)) |
| **Depth Of Coverage** | 3.1 |
| **Number Of Reads** | 11 |
| **Reads Per Million** | 0.23 rpm (after QC) |
| **Ambiguities** | 0 |
| **Assembly Method** | de novo + reference guided assembly |
| **Consensus Caller** | Bcf Tools |

##### Coverage Map

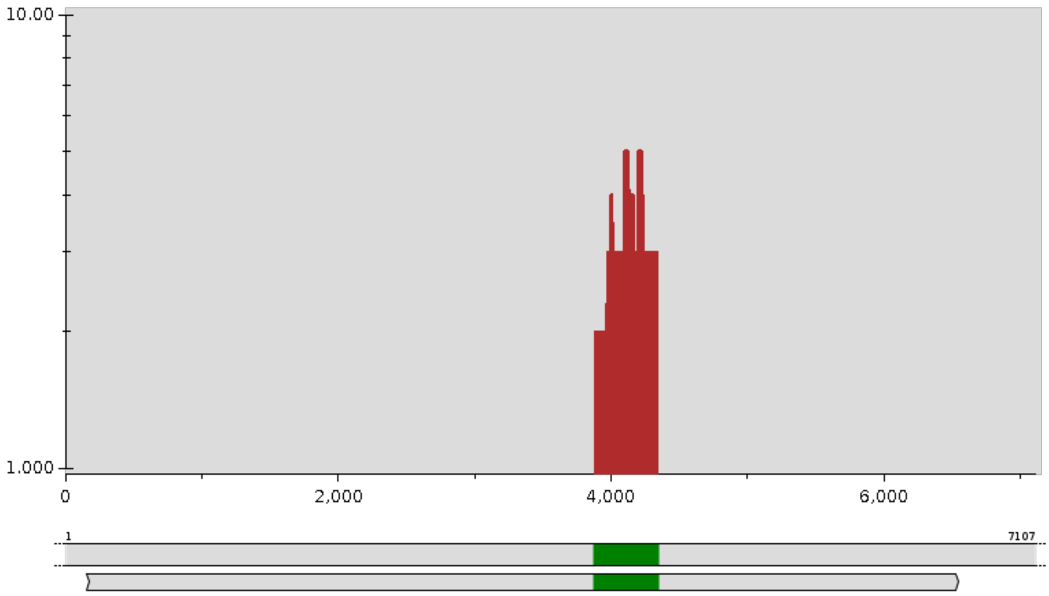

##### Assignment

| | |
|---|---|
| **Type** | Pinus nigra virus 1 (Taxonomy ID: 2267679) |
| **Reference Genome** | NC_040841.1 |
| **NT Identity (%)** | 56.8134 |
| **AA Identity (%)** | 45.9119 |
| **Number Of Stop Codons** | 1 |
| **Number Of CDS** | 1 |

##### Alignment

| | |
|---|---|
| **Alignment Score** | 115.0 (NT) + 496.0 (AA) = 611.0 |
| **Concordance (%)** | 29.5597 |

| Alignment Method | Global, seeded, nucleotide + amino acids (AGA) |
|---|---|

#### Genome Region

Sequence starts at position 3879 and ends at position 4352 relative to NC_040841.1 reference sequence.

#### Alignment Detailed Statistics

| | Begin | End | Coverage | Score | Concordance | Matches | Identities | I/D/M/F* | Stop Codons |
|---|---|---|---|---|---|---|---|---|---|
| **NT** | **3879** | **4352** | **6.7%** | **115** | **12.4%** | **471 (98.1%)** | **271 (56.5%)** | **6/3** | |
| Mutations: | 3880G>T, 3884C>G, 3885A>T, 3887T>G, 3892G>A, 3894T>A, 3897T>A, 3899A>T, 3901A>C, 3903G>A, 3904T>G, 3905A>T, 3906A>G, 3907A>T, 3914A>C, 3916G>T, 3919A>T, 3922A>C, 3935C>T, 3937T>G, 3941G>A, 3943C>T, 3944A>G, 3945T>C, 3946C>T, 3947A>T, 3948C>T, 3949T>A, 3950C>A, 3952G>A, 3953A>T, 3954A>G, 3955T>G, 3956T>A, 3960A>G, 3961T>A, 3967A>T, 3968T>A, 3970G>T, 3973A>T, 3975G>A, 3976A>C, 3977C>A, 3978C>A, 3980G>A, 3982A>G, 3984T>A, 3985G>T, 3986A>C, 3989A>C, 3991G>A, 3992C>A, 3996A>G, 3999T>C, 4000T>C, 4001C>T, 4002A>T, 4003G>T, 4004C>A, 4006T>A, 4007A>G, 4008G>C, 4009T>A, 4012G>C, 4013G>A, 4015G>T, 4017T>A, 4021C>T, 4024G>A, 4030C>T, 4036G>A, 4044A>T, 4047A>G, 4048T>G, 4051G>A, 4054A>T, 4057A>G, 4061C>T, 4062A>C, 4063G>T, 4064C>G, 4065C>A, 4067G>A, 4072T>C, 4074G>A, 4076C>T, 4078C>G, 4081A>G, 4082A>T, 4084A>C, 4090C>T, 4092T>A, 4093A>T, 4094T>G, 4095G>T, 4099A>G, 4100G>T, 4101C>T, 4102A>T, 4105A>T, 4106T>C, 4107T>A, 4108C>A, 4109T>C, 4112C>G, 4114G>A, 4120A>C, 4124G>A, 4126T>G, 4128C>T, 4129A>T, 4135T>G, 4136T>C, 4138G>T, 4141G>A, 4147G>C, 4150A>C, 4151G>T, 4153C>A, 4154T>G, 4155T>A, 4156C>A, 4164G>A, 4167G>T, 4168G>T, 4172G>A, 4175T>G, 4181T>A, 4184G>A, 4185C>A, 4187A>C, 4189A>T, 4191A>T, 4192T>G, 4193G>A, 4194A>G, 4195T>C, 4195_4196insCATATG, 4197T>C, 4198T>A, 4205_4207delACA, 4210C>T, 4213A>T, 4217G>A, 4220A>G, 4228T>C, 4229C>T, 4230A>T, 4231T>C, 4232A>T, 4233G>C, 4235C>G, 4236C>A, 4237A>G, 4240T>C, 4241G>A, 4243A>C, 4244C>G, 4246G>T, 4247A>T, 4248A>C, 4258G>A, 4262T>C, 4265G>A, 4267A>T, 4273T>A, 4274C>T, 4277G>A, 4279A>C, 4280G>A, 4281A>C, 4282A>C, 4288G>A, 4290A>G, 4292C>A, 4297A>T, 4298A>T, 4300T>G, 4303T>A, 4304T>G, 4306G>A, 4307T>A, 4308C>G, 4309A>C, 4310G>A, 4311A>C, 4312A>T, 4314G>A, 4321G>T, 4322T>A, 4323C>A, 4324C>A, 4325T>C, 4327A>T, 4333G>A, 4334G>A, 4335A>C, 4336T>A, 4339T>A, 4343G>A, 4344A>G, 4345T>A | | | | | | | | |

##### CDS

| | Begin | End | Coverage | Score | Concordance | Matches | Identities | I/D/M/F* | Stop Codons |
|---|---|---|---|---|---|---|---|---|---|
| EXL67_gp1 | 1243 | 1400 | 7.4% | 496 | 43.0% | 157 (98.1%) | 73 (45.6%) | 2/1/0/0 | 1 |
| Protein mutations: | H1244V (3884C>G 3885A>T), S1245A (3887T>G), I1247K (3894T>A), V1248E (3897T>A), R1249C (3899A>T 3901A>C), G1250E (3903G>A 3904T>G), K1251C (3905A>T 3906A>G 3907A>T), M1254L (3914A>C 3916G>T), A1263T (3941G>A 3943C>T), I1264A (3944A>G 3945T>C 3946C>T), T1265L (3947A>T 3948C>T 3949T>A), Q1266K (3950C>A 3952G>A), N1267W (3953A>T 3954A>G 3955T>G), F1268I (3956T>A), N1269R (3960A>G 3961T>A), L1272I (3968T>A 3970G>T), R1274N (3975G>A 3976A>C), P1275K (3977C>A 3978C>A), E1276K (3980G>A 3982A>G), V1277D (3984T>A 3985G>T), I1278L (3986A>C), M1279L (3989A>C 3991G>A), Q1280K (3992C>A), K1281R (3996A>G), I1282T (3999T>C 4000T>C), Q1283F (4001C>T 4002A>T 4003G>T), H1284K (4004C>A 4006T>A), S1285A (4007A>G 4008G>C 4009T>A), K1286N (4012G>C), V1287I (4013G>A 4015G>T), F1288Y (4017T>A), Y1297F (4044A>T), Y1298W (4047A>G 4048T>G), Q1303S (4061C>T 4062A>C 4063G>T), P1304E (4064C>G 4065C>A), E1305K (4067G>A), R1307K (4074G>A), H1308* (4076C>T 4078C>G), T1310S (4082A>T 4084A>C), I1313N (4092T>A 4093A>T), C1314V (4094T>G 4095G>T), A1316F (4100G>T 4101C>T 4102A>T), F1318Q (4106T>C 4107T>A 4108C>A), Y1319H (4109T>C), Q1320E (4112C>G 4114G>A), K1322N (4120A>C), V1324M (4124G>A 4126T>G), P1325L (4128C>T 4129A>T), A1333S (4151G>T 4153C>A), F1334E (4154T>G 4155T>A 4156C>A), R1337K (4164G>A), R1338I (4167G>T 4168G>T), D1340N (4172G>A), Y1341D (4175T>G), F1343I (4181T>A), A1344N (4184G>A 4185C>A), K1345H (4187A>C 4189A>T), Y1346L (4191A>T 4192T>G), D1347S (4193G>A 4194A>G 4195T>C), D1347_F1348insHM (4195_4196insCATATG), F1348S (4197T>C 4198T>A), T1351del (4205_4207delACA), D1355N (4217G>A), I1356V (4220A>G), H1359F (4229C>T 4230A>T 4231T>C), P1361E (4235C>G 4236C>A 4237A>G), V1363I (4241G>A 4243A>C), Q1364D (4244C>G 4246G>T), N1365S (4247A>T 4248A>C), E1371N (4265G>A 4267A>T), F1373L (4273T>A), L1374F (4274C>T), E1375N (4277G>A 4279A>C), E1376T (4280G>A 4281A>C 4282A>C), K1379R (4290A>G), H1380N (4292C>A), I1382L (4298A>T 4300T>G), L1384V (4304T>G 4306G>A), E1386T (4310G>A 4311A>C 4312A>T), R1387K (4314G>A), M1389I (4321G>T), S1390K (4322T>A 4323C>A 4324C>A), D1394T (4334G>A 4335A>C 4336T>A), N1395K (4339T>A), D1397R (4343G>A 4344A>G 4345T>A) | | | | | | | | |
| Codon mutations: | AGG1242.GT (3880G>T), CAT1244GTT (3884C>G 3885A>T), TCT1245GCT (3887T>G), GAG1246GAA (3892G>A), ATA1247AAA (3894T>A), GTA1248GAA (3897T>A), AGA1249TGC (3899A>T 3901A>C), GGT1250GAG (3903G>A 3904T>G), AAA1251TGT (3905A>T 3906A>G 3907A>T), ATG1254CTT (3914A>C 3916G>T), GTA1255GTT (3919A>T), ATA1256ATC (3922A>C), CTT1261TTG (3935C>T 3937T>G), GCC1263ACT (3941G>A 3943C>T), ATC1264GCT (3944A>G 3945T>C 3946C>T), ACT1265TTA (3947A>T 3948C>T 3949T>A), CAG1266AAA (3950C>A 3952G>A), AAT1267TGG (3953A>T 3954A>G 3955T>G), TTT1268ATT (3956T>A), AAT1269AGA (3960A>G 3961T>A), CCA1271CCT (3967A>T), TTG1272ATT (3968T>A 3970G>T), CCA1273CCT (3973A>T), AGA1274AAC (3975G>A 3976A>C), CCA1275AAA (3977C>A 3978C>A), GAA1276AAG (3980G>A 3982A>G), GTG1277GAT (3984T>A 3985G>T), ATT1278CTT (3986A>C), ATG1279CTA (3989A>C 3991G>A), CAA1280AAA (3992C>A), AAA1281AGA (3996A>G), ATT1282ACC (3999T>C 4000T>C), CAG1283TTT (4001C>T 4002A>T 4003G>T), CAT1284AAA (4004C>A 4006T>A), AGT1285GCA (4007A>G 4008G>C 4009T>A), AAG1286AAC (4012G>C), GTG1287ATT (4013G>A 4015G>T), TTC1288TAC (4017T>A), AGC1289AGT (4021C>T), AAG1290AAA (4024G>A), GAC1292GAT (4030C>T), AAG1294AAA (4036G>A), TAT1297TTT (4044A>T), TAT1298TGG (4047A>G 4048T>G), CAG1299CAA (4051G>A), ATA1300ATT (4054A>T), CAA1301CAG (4057A>G), CAG1303TCT (4061C>T 4062A>C 4063G>T), CCA1304GAA (4064C>G 4065C>A), GAA1305AAA (4067G>A), GAT1306GAC (4072T>C), AGA1307AAA (4074G>A), CAC1308TAG (4076C>T 4078C>G), AAA1309AAG (4081A>G), ACA1310TCC (4082A>T 4084A>C), TTC1312TTT (4090C>T), ATA1313AAT (4092T>A 4093A>T), TGT1314GTT (4094T>G 4095G>T), CCA1315CCG (4099A>G), GCA1316TTT (4100G>T 4101C>T 4102A>T), GGA1317GGT (4105A>T), TTC1318CAA (4106T>C 4107T>A 4108C>A), TAT1319CAT (4109T>C), CAG1320GAA (4112C>G 4114G>A), AAA1322AAC (4120A>C), GTT1324ATG (4124G>A 4126T>G), CCA1325CTT (4128C>T 4129A>T), GGT1327GGG (4135T>G), TTG1328CTT (4136T>C 4138G>T), AAG1329AAA (4141G>A), GCG1331GCC (4147G>C), CCA1332CCC (4150A>C), GCC1333TCA (4151G>T 4153C>A), TTC1334GAA (4154T>G 4155T>A 4156C>A), AGA1337AAA (4164G>A), AGG1338ATT (4167G>T 4168G>T), GAT1340AAT (4172G>A), TAT1341GAT (4175T>G), TTT1343ATT (4181T>A), GCT1344AAT (4184G>A 4185C>A), AAA1345CAT (4187A>C 4189A>T), TAT1346TTG (4191A>T 4192T>G), GAT1347AGC (4193G>A 4194A>G 4195T>C), GAT1347_TTT1348insCATATG (4195_4196insCATATG), TTT1348TCA (4197T>C 4198T>A), ACA1351del (4205_4207delACA), TAC1352TAT (4210C>T), ATA1353ATT (4213A>T), GAT1355AAT (4217G>A), ATT1356GTT (4220A>G), ATT1358ATC (4228T>C), CAT1359TTC (4229C>T 4230A>T 4231T>C), AGT1360TCT (4232A>T 4233G>C), CCA1361GAG (4235C>G 4236C>A 4237A>G), GAT1362GAC (4240T>C), GTA1363ATC (4241G>A 4243A>C), CAG1364GAT (4244C>G 4246G>T), AAT1365TCT (4247A>T 4248A>C), AAG1368AAA (4258G>A), TTA1370CTA (4262T>C), GAA1371AAT (4265G>A 4267A>T), TTT1373TTA (4273T>A), CTT1374TTT (4274C>T), GAA1375AAC (4277G>A 4279A>C), GAA1376ACC (4280G>A 4281A>C 4282A>C), AAG1378AAA (4288G>A), AAA1379AGA (4290A>G), CAT1380AAT (4292C>A), GGA1381GGT (4297A>T), ATT1382TTG (4298A>T 4300T>G), GTT1383GTA (4303T>A), TTG1384GTA (4304T>G 4306G>A), TCA1385AGC (4307T>A 4308C>G 4309A>C), GAA1386ACT (4310G>A 4311A>C 4312A>T), AGA1387AAA (4314G>A), ATG1389ATT (4321G>T), TCC1390AAA (4322T>A 4323C>A 4324C>A), TTA1391CTT (4325T>C 4327A>T), CAG1393CAA (4333G>A), GAT1394ACA (4334G>A 4335A>C 4336T>A), AAT1395AAA (4339T>A), GAT1397AGA (4343G>A 4344A>G 4345T>A) | | | | | | | | |

##### Proteins

| | Begin | End | Coverage | Score | Concordance | Matches | Identities | I/D/M/F* | Stop Codons |
|---|---|---|---|---|---|---|---|---|---|
| polyprotein (YP_009553669.1) | 1243 | 1400 | 7.4% | 496 | 43.0% | 157 (98.1%) | 73 (45.6%) | 2/1/0/0 | 1 |
| Protein mutations: | H1244V (3884C>G 3885A>T), S1245A (3887T>G), I1247K (3894T>A), V1248E (3897T>A), R1249C (3899A>T 3901A>C), G1250E (3903G>A 3904T>G), K1251C (3905A>T 3906A>G 3907A>T), M1254L (3914A>C 3916G>T), A1263T (3941G>A 3943C>T), I1264A (3944A>G 3945T>C 3946C>T), T1265L (3947A>T 3948C>T 3949T>A), Q1266K (3950C>A 3952G>A), N1267W (3953A>T 3954A>G 3955T>G), F1268I (3956T>A), N1269R (3960A>G 3961T>A), L1272I (3968T>A 3970G>T), R1274N (3975G>A 3976A>C), P1275K (3977C>A 3978C>A), E1276K (3980G>A 3982A>G), V1277D (3984T>A 3985G>T), I1278L (3986A>C), M1279L (3989A>C 3991G>A), Q1280K (3992C>A), K1281R (3996A>G), I1282T (3999T>C 4000T>C), Q1283F (4001C>T 4002A>T 4003G>T), H1284K (4004C>A 4006T>A), S1285A (4007A>G 4008G>C 4009T>A), K1286N (4012G>C), V1287I (4013G>A 4015G>T), F1288Y (4017T>A), Y1297F (4044A>T), Y1298W (4047A>G 4048T>G), Q1303S (4061C>T 4062A>C 4063G>T), P1304E (4064C>G 4065C>A), E1305K (4067G>A), R1307K (4074G>A), H1308* (4076C>T 4078C>G), T1310S (4082A>T 4084A>C), I1313N (4092T>A 4093A>T), C1314V (4094T>G 4095G>T), A1316F (4100G>T 4101C>T 4102A>T), F1318Q (4106T>C 4107T>A 4108C>A), Y1319H (4109T>C), Q1320E (4112C>G 4114G>A), K1322N (4120A>C), V1324M (4124G>A 4126T>G), P1325L (4128C>T 4129A>T), A1333S (4151G>T 4153C>A), F1334E (4154T>G 4155T>A 4156C>A), R1337K (4164G>A), R1338I (4167G>T 4168G>T), D1340N (4172G>A), Y1341D (4175T>G), F1343I (4181T>A), A1344N (4184G>A 4185C>A), K1345H (4187A>C 4189A>T), Y1346L (4191A>T 4192T>G), D1347S (4193G>A 4194A>G 4195T>C), D1347_F1348insHM (4195_4196insCATATG), F1348S (4197T>C 4198T>A), T1351del (4205_4207delACA), D1355N (4217G>A), I1356V (4220A>G), H1359F (4229C>T 4230A>T 4231T>C), P1361E (4235C>G 4236C>A 4237A>G), V1363I (4241G>A 4243A>C), Q1364D (4244C>G 4246G>T), N1365S (4247A>T 4248A>C), E1371N (4265G>A 4267A>T), F1373L (4273T>A), L1374F (4274C>T), E1375N (4277G>A 4279A>C), E1376T (4280G>A 4281A>C 4282A>C), K1379R (4290A>G), H1380N (4292C>A), I1382L (4298A>T 4300T>G), L1384V (4304T>G 4306G>A), E1386T (4310G>A 4311A>C 4312A>T), R1387K (4314G>A), M1389I (4321G>T), S1390K (4322T>A 4323C>A 4324C>A), D1394T (4334G>A 4335A>C 4336T>A), N1395K (4339T>A), D1397R (4343G>A 4344A>G 4345T>A) | | | | | | | | |

| | Begin | End | Coverage | Score | Concordance | Matches | Identities | I/D/M/F* | Stop Codons |
|---|---|---|---|---|---|---|---|---|---|
| **NT** | **3879** | **4352** | **6.7%** | **115** | **12.4%** | **471 (98.1%)** | **271 (56.5%)** | **6/3** | |
| Codon mutations: | AGG1242.GT (3880G>T), CAT1244GTT (3884C>G 3885A>T), TCT1245GCT (3887T>G), GAG1246GAA (3892G>A), ATA1247AAA (3894T>A), GTA1248GAA (3897T>A), AGA1249TGC (3899A>T 3901A>C), GGT1250GAG (3903G>A 3904T>G), AAA1251TGT (3905A>T 3906A>G 3907A>T), ATG1254CTT (3914A>C 3916G>T), GTA1255GTT (3919A>T), ATA1256ATC (3922A>C), CTT1261TTG (3935C>T 3937T>G), GCC1263ACT (3941G>A 3943C>T), ATC1264GCT (3944A>G 3945T>C 3946C>T), ACT1265TTA (3947A>T 3948C>T 3949T>A), CAG1266AAA (3950C>A 3952G>A), AAT1267TGG (3953A>T 3954A>G 3955T>G), TTT1268ATT (3956T>A), AAT1269AGA (3960A>G 3961T>A), CCA1271CCT (3967A>T), TTG1272ATT (3968T>A 3970G>T), CCA1273CCT (3973A>T), AGA1274AAC (3975G>A 3976A>C), CCA1275AAA (3977C>A 3978C>A), GAA1276AAG (3980G>A 3982A>G), GTG1277GAT (3984T>A 3985G>T), ATT1278CTT (3986A>C), ATG1279CTA (3989A>C 3991G>A), CAA1280AAA (3992C>A), AAA1281AGA (3996A>G), ATT1282ACC (3999T>C 4000T>C), CAG1283TTT (4001C>T 4002A>T 4003G>T), CAT1284AAA (4004C>A 4006T>A), AGT1285GCA (4007A>G 4008G>C 4009T>A), AAG1286AAC (4012G>C), GTG1287ATT (4013G>A 4015G>T), TTC1288TAC (4017T>A), AGC1289AGT (4021C>T), AAG1290AAA (4024G>A), GAC1292GAT (4030C>T), AAG1294AAA (4036G>A), TAT1297TTT (4044A>T), TAT1298TGG (4047A>G 4048T>G), CAG1299CAA (4051G>A), ATA1300ATT (4054A>T), CAA1301CAG (4057A>G), CAG1303TCT (4061C>T 4062A>C 4063G>T), CCA1304GAA (4064C>G 4065C>A), GAA1305AAA (4067G>A), GAT1306GAC (4072T>C), AGA1307AAA (4074G>A), CAC1308TAG (4076C>T 4078C>G), AAA1309AAG (4081A>G), ACA1310TCC (4082A>T 4084A>C), TTC1312TTT (4090C>T), ATA1313AAT (4092T>A 4093A>T), TGT1314GTT (4094T>G 4095G>T), CCA1315CCG (4099A>G), GCA1316TTT (4100G>T 4101C>T 4102A>T), GGA1317GGT (4105A>T), TTC1318CAA (4106T>C 4107T>A 4108C>A), TAT1319CAT (4109T>C), CAG1320GAA (4112C>G 4114G>A), AAA1322AAC (4120A>C), GTT1324ATG (4124G>A 4126T>G), CCA1325CTT (4128C>T 4129A>T), GGT1327GGG (4135T>G), TTG1328CTT (4136T>C 4138G>T), AAG1329AAA (4141G>A), GCG1331GCC (4147G>C), CCA1332CCC (4150A>C), GCC1333TCA (4151G>T 4153C>A), TTC1334GAA (4154T>G 4155T>A 4156C>A), AGA1337AAA (4164G>A), AGG1338ATT (4167G>T 4168G>T), GAT1340AAT (4172G>A), TAT1341GAT (4175T>G), TTT1343ATT (4181T>A), GCT1344AAT (4184G>A 4185C>A), AAA1345CAT (4187A>C 4189A>T), TAT1346TTG (4191A>T 4192T>G), GAT1347AGC (4193G>A 4194A>G 4195T>C), GAT1347_TTT1348insCATATG (4195_4196insCATATG), TTT1348TCA (4197T>C 4198T>A), ACA1351del (4205_4207delACA), TAC1352TAT (4210C>T), ATA1353ATT (4213A>T), GAT1355AAT (4217G>A), ATT1356GTT (4220A>G), ATT1358ATC (4228T>C), CAT1359TTC (4229C>T 4230A>T 4231T>C), AGT1360TCT (4232A>T 4233G>C), CCA1361GAG (4235C>G 4236C>A 4237A>G), GAT1362GAC (4240T>C), GTA1363ATC (4241G>A 4243A>C), CAG1364GAT (4244C>G 4246G>T), AAT1365TCT (4247A>T 4248A>C), AAG1368AAA (4258G>A), TTA1370CTA (4262T>C), GAA1371AAT (4265G>A 4267A>T), TTT1373TTA (4273T>A), CTT1374TTT (4274C>T), GAA1375AAC (4277G>A 4279A>C), GAA1376ACC (4280G>A 4281A>C 4282A>C), AAG1378AAA (4288G>A), AAA1379AGA (4290A>G), CAT1380AAT (4292C>A), GGA1381GGT (4297A>T), ATT1382TTG (4298A>T 4300T>G), GTT1383GTA (4303T>A), TTG1384GTA (4304T>G 4306G>A), TCA1385AGC (4307T>A 4308C>G 4309A>C), GAA1386ACT (4310G>A 4311A>C 4312A>T), AGA1387AAA (4314G>A), ATG1389ATT (4321G>T), TCC1390AAA (4322T>A 4323C>A 4324C>A), TTA1391CTT (4325T>C 4327A>T), CAG1393CAA (4333G>A), GAT1394ACA (4334G>A 4335A>C 4336T>A), AAT1395AAA (4339T>A), GAT1397AGA (4343G>A 4344A>G 4345T>A) | | | | | | | | |

*: Inserts / Deletes / Misaligned / Frameshifts

#### Analysis details

This analysis was performed with panviral2.64

### NGS Details (UN60): Badnavirus rutilanscamelliae

#### Assembly

| | |
|---|---|
| **Coverage Length** | 262 (1 contig(s)) |
| **Depth Of Coverage** | 3.9 |
| **Number Of Reads** | 10 |
| **Reads Per Million** | 0.21 rpm (after QC) |
| **Ambiguities** | 0 |
| **Assembly Method** | de novo + reference guided assembly |
| **Consensus Caller** | Bcf Tools |

#### Coverage Map

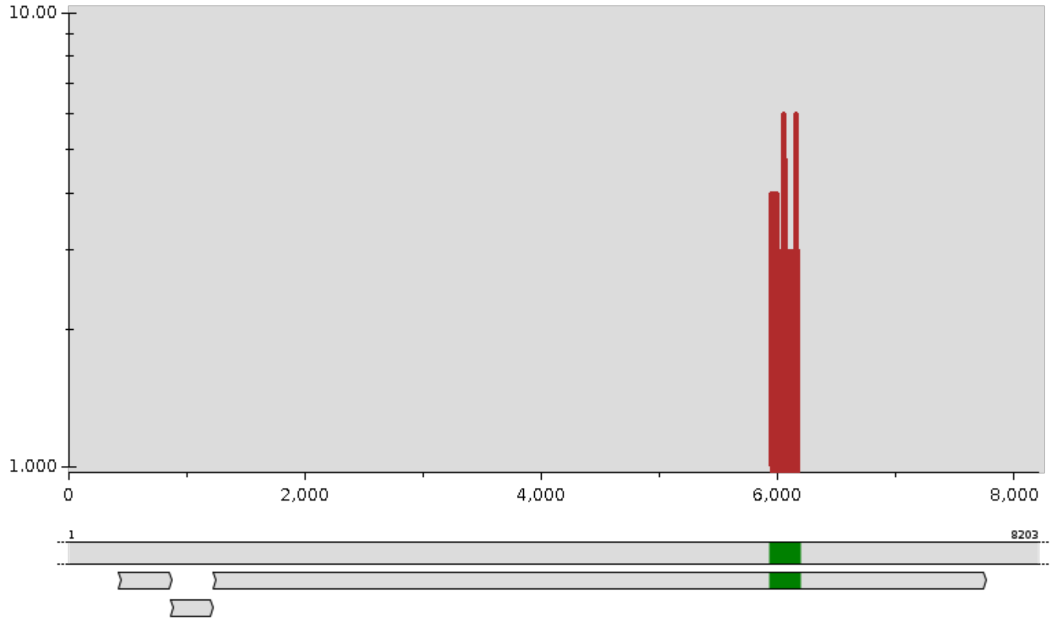

#### Assignment

| | |
|---|---|
| **Type** | Badnavirus rutilanscamelliae (Taxonomy ID: 3047719) |
| **Reference Genome** | NC_055598.1 |
| **NT Identity (%)** | 61.5094 |
| **AA Identity (%)** | 56.1798 |
| **Number Of Stop Codons** | 0 |
| **Number Of CDS** | 3 |

#### Alignment

| | |
|---|---|
| **Alignment Score** | 98.0 (NT) + 398.0 (AA) = 496.0 |
| **Concordance (%)** | 42.7586 |

| | |
|---|---|
| **Alignment Method** | Global, seeded, nucleotide + amino acids (AGA) |

#### Genome Region

Sequence starts at position 5933 and ends at position 6194 relative to NC_055598.1 reference sequence.

#### Alignment Detailed Statistics

| | Begin | End | Coverage | Score | Concordance | Matches | Identities | I/D/M/F* | Stop Codons |
|---|---|---|---|---|---|---|---|---|---|
| **NT** | **5933** | **6194** | **3.2%** | **98** | **19.4%** | **259 (96.6%)** | **163 (60.8%)** | **6/3** | |
| Mutations: | 5937G>C, 5946A>T, 5950G>T, 5953G>A, 5954T>A, 5956C>T, 5959C>T, 5965G>A, 5966T>A, 5967C>G, 5968A>T, 5971G>T, 5973T>A, 5974T>C, 5978C>G, 5980A>G, 5981G>A, 5984G>C, 5985C>G, 5986T>A, 5990G>A, 5996C>G, 5997A>G, 5999C>G, 6001C>T, 6002_6004delATT, 6010_6011insAAA, 6013C>T, 6020T>A, 6021G>A, 6025T>C, 6026C>A, 6027C>A, 6029G>T, 6030T>A, 6031G>T, 6034C>A, 6035T>C, 6037A>T, 6043A>G, 6047C>A, 6058A>C, 6064G>C, 6065C>T, 6067T>A, 6069A>C, 6070G>T, 6076T>A, 6080G>A, 6081C>G, 6082A>C, 6083G>A, 6084T>C, 6085A>T, 6089C>A, 6090A>T, 6091A>G, 6094G>A, 6095A>C, 6096A>T, 6097G>A, 6101G>A, 6103C>T, 6105G>A, 6106A>T, 6107A>G, 6109C>T, 6115T>A, 6116C>G, 6117C>A, 6118A>T, 6120A>T, 6122G>C, 6123C>A, 6127_6128insAAA, 6131A>G, 6135C>T, 6136T>G, 6143A>T, 6151T>C, 6155C>T, 6157C>G, 6160A>C, 6163C>T, 6164A>T, 6165G>C, 6166C>T, 6168C>A, 6171C>A, 6172A>T, 6174C>T, 6175T>A, 6178A>T, 6180C>A, 6181C>A, 6184T>C, 6185A>C, 6186A>T, 6188A>G | | | | | | | | |

##### CDS

| | Begin | End | Coverage | Score | Concordance | Matches | Identities | I/D/M/F* | Stop Codons |
|---|---|---|---|---|---|---|---|---|---|
| KM754_gp3 | 1571 | 1658 | 4.0% | 398 | 60.5% | 87 (96.7%) | 50 (55.6%) | 2/1/0/0 | 0 |
| Protein mutations: | S1572T (5937G>C), Y1575F (5946A>T), F1578I (5954T>A 5956C>T), F1584Y (5973T>A 5974T>C), Q1586E (5978C>G 5980A>G), V1587I (5981G>A), A1588R (5984G>C 5985C>G 5986T>A), D1590N (5990G>A), Q1592G (5996C>G 5997A>G), H1593D (5999C>G 6001C>T), I1594del (6002_6004delATT), W1596_T1597insK (6010_6011insAAA), W1600K (6020T>A 6021G>A), P1602K (6026C>A 6027C>A), V1603Y (6029G>T 6030T>A 6031G>T), L1609M (6047C>A), K1616T (6069A>C 6070G>T), A1620S (6080G>A 6081C>G 6082A>C), V1621T (6083G>A 6084T>C 6085A>T), Q1623M (6089C>A 6090A>T 6091A>G), K1625L (6095A>C 6096A>T 6097G>A), D1627N (6101G>A 6103C>T), R1628H (6105G>A 6106A>T), I1629V (6107A>G 6109C>T), N1631K (6115T>A), P1632D (6116C>G 6117C>A 6118A>T), Y1633F (6120A>T), A1634H (6122G>C 6123C>A), G1635_F1636insK (6127_6128insAAA), I1637V (6131A>G), A1638V (6135C>T 6136T>G), I1641F (6143A>T), P1649Q (6168C>A), T1650N (6171C>A 6172A>T), P1651L (6174C>T 6175T>A), E1652D (6178A>T), A1653E (6180C>A 6181C>A), K1655L (6185A>C 6186A>T), K1656E (6188A>G) | | | | | | | | |
| Codon mutations: | AGC1572ACC (5937G>C), TAT1575TTT (5946A>T), TCG1576TCT (5950G>T), AAG1577AAA (5953G>A), TTC1578ATT (5954T>A 5956C>T), GAC1579GAT (5959C>T), AAG1581AAA (5965G>A), TCA1582AGT (5966T>A 5967C>G 5968A>T), GGG1583GGT (5971G>T), TTT1584TAC (5973T>A 5974T>C), CAA1586GAG (5978C>G 5980A>G), GTT1587ATT (5981G>A), GCT1588CGA (5984G>C 5985C>G 5986T>A), GAT1590AAT (5990G>A), CAA1592GGA (5996C>G 5997A>G), CAC1593GAT (5999C>G 6001C>T), ATT1594del (6002_6004delATT), TGG1596_ACC1597insAAA (6010_6011insAAA), ACC1597ACT (6013C>T), TGG1600AAG (6020T>A 6021G>A), ACT1601ACC (6025T>C), CCA1602AAA (6026C>A 6027C>A), GTG1603TAT (6029G>T 6030T>A 6031G>T), GGC1604GGA (6034C>A), TTA1605CTT (6035T>C 6037A>T), GAA1607GAG (6043A>G), CTG1609ATG (6047C>A), CCA1612CCC (6058A>C), GGG1614GGC (6064G>C), CTT1615TTA (6065C>T 6067T>A), AAG1616ACT (6069A>C 6070G>T), GCT1618GCA (6076T>A), GCA1620AGC (6080G>A 6081C>G 6082A>C), GTA1621ACT (6083G>A 6084T>C 6085A>T), CAA1623ATG (6089C>A 6090A>T 6091A>G), AGG1624AGA (6094G>A), AAG1625CTA (6095A>C 6096A>T 6097G>A), GAC1627AAT (6101G>A 6103C>T), CGA1628CAT (6105G>A 6106A>T), ATC1629GTT (6107A>G 6109C>T), AAT1631AAA (6115T>A), CCA1632GAT (6116C>G 6117C>A 6118A>T), TAC1633TTC (6120A>T), GCT1634CAT (6122G>C 6123C>A), GGA1635_TTT1636insAAA (6127_6128insAAA), ATT1637GTT (6131A>G), GCT1638GTG (6135C>T 6136T>G), ATT1641TTT (6143A>T), GAT1643GAC (6151T>C), CTC1645TTG (6155C>T 6157C>G), ATA1646ATC (6160A>C), TTC1647TTT (6163C>T), AGC1648TCT (6164A>T 6165G>C 6166C>T), CCA1649CAA (6168C>A), ACA1650AAT (6171C>A 6172A>T), CCT1651CTA (6174C>T 6175T>A), GAA1652GAT (6178A>T), GCC1653GAA (6180C>A 6181C>A), CAT1654CAC (6184T>C), AAA1655CTA (6185A>C 6186A>T), AAG1656GAG (6188A>G) | | | | | | | | |

##### Proteins

| | Begin | End | Coverage | Score | Concordance | Matches | Identities | I/D/M/F* | Stop Codons |
|---|---|---|---|---|---|---|---|---|---|
| polyprotein (YP_010087856.1) | 1571 | 1658 | 4.0% | 398 | 60.5% | 87 (96.7%) | 50 (55.6%) | 2/1/0/0 | 0 |
| Protein mutations: | S1572T (5937G>C), Y1575F (5946A>T), F1578I (5954T>A 5956C>T), F1584Y (5973T>A 5974T>C), Q1586E (5978C>G 5980A>G), V1587I (5981G>A), A1588R (5984G>C 5985C>G 5986T>A), D1590N (5990G>A), Q1592G (5996C>G 5997A>G), H1593D (5999C>G 6001C>T), I1594del (6002_6004delATT), W1596_T1597insK (6010_6011insAAA), W1600K (6020T>A 6021G>A), P1602K (6026C>A 6027C>A), V1603Y (6029G>T 6030T>A 6031G>T), L1609M (6047C>A), K1616T (6069A>C 6070G>T), A1620S (6080G>A 6081C>G 6082A>C), V1621T (6083G>A 6084T>C 6085A>T), Q1623M (6089C>A 6090A>T 6091A>G), K1625L (6095A>C 6096A>T 6097G>A), D1627N (6101G>A 6103C>T), R1628H (6105G>A 6106A>T), I1629V (6107A>G 6109C>T), N1631K (6115T>A), P1632D (6116C>G 6117C>A 6118A>T), Y1633F (6120A>T), A1634H (6122G>C 6123C>A), G1635_F1636insK (6127_6128insAAA), I1637V (6131A>G), A1638V (6135C>T 6136T>G), I1641F (6143A>T), P1649Q (6168C>A), T1650N (6171C>A 6172A>T), P1651L (6174C>T 6175T>A), E1652D (6178A>T), A1653E (6180C>A 6181C>A), K1655L (6185A>C 6186A>T), K1656E (6188A>G) | | | | | | | | |
| Codon mutations: | AGC1572ACC (5937G>C), TAT1575TTT (5946A>T), TCG1576TCT (5950G>T), AAG1577AAA (5953G>A), TTC1578ATT (5954T>A 5956C>T), GAC1579GAT (5959C>T), AAG1581AAA (5965G>A), TCA1582AGT (5966T>A 5967C>G 5968A>T), GGG1583GGT (5971G>T), TTT1584TAC (5973T>A 5974T>C), CAA1586GAG (5978C>G 5980A>G), GTT1587ATT (5981G>A), GCT1588CGA (5984G>C 5985C>G 5986T>A), GAT1590AAT (5990G>A), CAA1592GGA (5996C>G 5997A>G), CAC1593GAT (5999C>G 6001C>T), ATT1594del (6002_6004delATT), TGG1596_ACC1597insAAA (6010_6011insAAA), ACC1597ACT (6013C>T), TGG1600AAG (6020T>A 6021G>A), ACT1601ACC (6025T>C), CCA1602AAA (6026C>A 6027C>A), GTG1603TAT (6029G>T 6030T>A 6031G>T), GGC1604GGA (6034C>A), TTA1605CTT (6035T>C 6037A>T), GAA1607GAG (6043A>G), CTG1609ATG (6047C>A), CCA1612CCC (6058A>C), GGG1614GGC (6064G>C), CTT1615TTA (6065C>T 6067T>A), AAG1616ACT (6069A>C 6070G>T), GCT1618GCA (6076T>A), GCA1620AGC (6080G>A 6081C>G 6082A>C), GTA1621ACT (6083G>A 6084T>C 6085A>T), CAA1623ATG (6089C>A 6090A>T 6091A>G), AGG1624AGA (6094G>A), AAG1625CTA (6095A>C 6096A>T 6097G>A), GAC1627AAT (6101G>A 6103C>T), CGA1628CAT (6105G>A 6106A>T), ATC1629GTT (6107A>G 6109C>T), AAT1631AAA (6115T>A), CCA1632GAT (6116C>G 6117C>A 6118A>T), TAC1633TTC (6120A>T), GCT1634CAT (6122G>C 6123C>A), GGA1635_TTT1636insAAA (6127_6128insAAA), ATT1637GTT (6131A>G), GCT1638GTG (6135C>T 6136T>G), ATT1641TTT (6143A>T), GAT1643GAC (6151T>C), CTC1645TTG (6155C>T 6157C>G), ATA1646ATC (6160A>C), TTC1647TTT (6163C>T), AGC1648TCT (6164A>T 6165G>C 6166C>T), CCA1649CAA (6168C>A), ACA1650AAT (6171C>A 6172A>T), CCT1651CTA (6174C>T 6175T>A), GAA1652GAT (6178A>T), GCC1653GAA (6180C>A 6181C>A), CAT1654CAC (6184T>C), AAA1655CTA (6185A>C 6186A>T), AAG1656GAG (6188A>G) | | | | | | | | |

*: Inserts / Deletes / Misaligned / Frameshifts

#### Analysis details

This analysis was performed with panviral2.64

### NGS Details (UN60): Duamitovirus dapi1

#### Assembly

| | |
|---|---|
| **Coverage Length** | 135 (1 contig(s)) |
| **Depth Of Coverage** | 6.5 |
| **Number Of Reads** | 8 |
| **Reads Per Million** | 0.17 rpm (after QC) |
| **Ambiguities** | 0 |
| **Assembly Method** | de novo + reference guided assembly |
| **Consensus Caller** | Bcf Tools |

#### Coverage Map

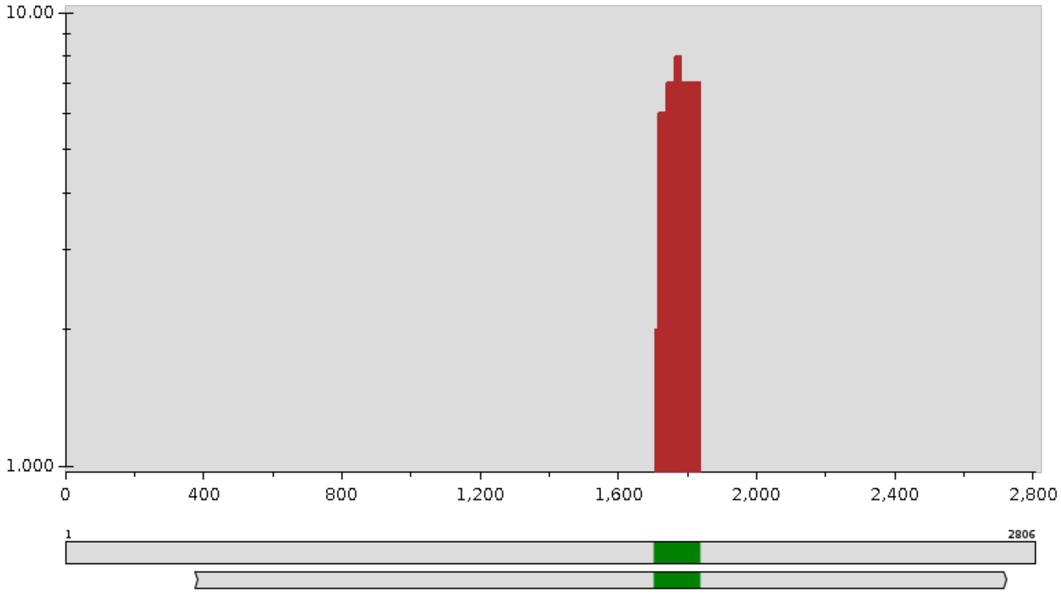

#### Assignment

| | |
|---|---|
| **Type** | Duamitovirus dapi1 (Taxonomy ID: 2955778) |
| **Reference Genome** | NC_076528.1 |
| **NT Identity (%)** | 68.8889 |
| **AA Identity (%)** | 66.6667 |
| **Number Of Stop Codons** | 0 |
| **Number Of CDS** | 1 |

#### Alignment

| | |
|---|---|
| **Alignment Score** | 102.0 (NT) + 249.0 (AA) = 351.0 |
| **Concordance (%)** | 57.1661 |

| Alignment Method | Global, seeded, nucleotide + amino acids (AGA) |
|---|---|

#### Genome Region

Sequence starts at position 1706 and ends at position 1840 relative to NC_076528.1 reference sequence.

#### Alignment Detailed Statistics

| | Begin | End | Coverage | Score | Concordance | Matches | Identities | I/D/M/F* | Stop Codons |
|---|---|---|---|---|---|---|---|---|---|
| **NT** | **1706** | **1840** | **4.8%** | **102** | **37.8%** | **135 (100%)** | **93 (68.9%)** | **0/0** | |
| Mutations: | 1706G>C, 1710C>T, 1713A>G, 1715G>A, 1725T>C, 1728G>C, 1734C>T, 1736A>T, 1737C>G, 1738T>A, 1739C>G, 1743C>T, 1749G>T, 1750T>C, 1751C>T, 1756G>A, 1763C>A, 1774G>C, 1776A>T, 1783C>A, 1785G>A, 1788G>T, 1789G>A, 1791C>T, 1792T>G, 1793G>A, 1794G>A, 1800A>G, 1804C>G, 1805C>G, 1806T>C, 1809T>G, 1810C>A, 1811A>C, 1816T>C, 1819A>T, 1820A>G, 1821G>C, 1827C>T, 1830T>C, 1831C>A, 1839T>C | | | | | | | | |
| **CDS** | | | | | | | | | |
| RdRp | 445 | 489 | 5.7% | 249 | 68.8% | 45 (100%) | 30 (66.7%) | 0/0/0/0 | 0 |
| Protein mutations: | R447K (1715G>A), Y454L (1736A>T 1737C>G), S459L (1750T>C 1751C>T), A461T (1756G>A), T463K (1763C>A), V467L (1774G>C 1776A>T), L470I (1783C>A 1785G>A), A472T (1789G>A 1791C>T), W473E (1792T>G 1793G>A 1794G>A), P477G (1804C>G 1805C>G 1806T>C), Q479T (1810C>A 1811A>C), F481L (1816T>C), K482C (1819A>T 1820A>G 1821G>C), L486I (1831C>A) | | | | | | | | |
| Codon mutations: | CGG444.CG (1706G>C), TTC445TTT (1710C>T), ACA446ACG (1713A>G), AGG447AAG (1715G>A), CCT450CCC (1725T>C), CTG451CTC (1728G>C), TTC453TTT (1734C>T), TAC454TTG (1736A>T 1737C>G), TCT455AGT (1738T>A 1739C>G), TCC456TCT (1743C>T), CCG458CCT (1749G>T), TCA459CTA (1750T>C 1751C>T), GCA461ACA (1756G>A), ACA463AAA (1763C>A), GTA467CTT (1774G>C 1776A>T), CTG470ATA (1783C>A 1785G>A), GCG471GCT (1788G>T), GCC472ACT (1789G>A 1791C>T), TGG473GAA (1792T>G 1793G>A 1794G>A), GTA475GTG (1800A>G), CCT477GGC (1804C>G 1805C>G 1806T>C), ACT478ACG (1809T>G), CAG479ACG (1810C>A 1811A>C), TTT481CTT (1816T>C), AAG482TGC (1819A>T 1820A>G 1821G>C), TAC484TAT (1827C>T), GCT485GCC (1830T>C), CTT486ATT (1831C>A), GGT488GGC (1839T>C) | | | | | | | | |
| **Proteins** | | | | | | | | | |
| RNA-dependent RNA polymerase (YP_010798878.1) | 445 | 489 | 5.7% | 249 | 68.8% | 45 (100%) | 30 (66.7%) | 0/0/0/0 | 0 |
| Protein mutations: | R447K (1715G>A), Y454L (1736A>T 1737C>G), S459L (1750T>C 1751C>T), A461T (1756G>A), T463K (1763C>A), V467L (1774G>C 1776A>T), L470I (1783C>A 1785G>A), A472T (1789G>A 1791C>T), W473E (1792T>G 1793G>A 1794G>A), P477G (1804C>G 1805C>G 1806T>C), Q479T (1810C>A 1811A>C), F481L (1816T>C), K482C (1819A>T 1820A>G 1821G>C), L486I (1831C>A) | | | | | | | | |
| Codon mutations: | CGG444.CG (1706G>C), TTC445TTT (1710C>T), ACA446ACG (1713A>G), AGG447AAG (1715G>A), CCT450CCC (1725T>C), CTG451CTC (1728G>C), TTC453TTT (1734C>T), TAC454TTG (1736A>T 1737C>G), TCT455AGT (1738T>A 1739C>G), TCC456TCT (1743C>T), CCG458CCT (1749G>T), TCA459CTA (1750T>C 1751C>T), GCA461ACA (1756G>A), ACA463AAA (1763C>A), GTA467CTT (1774G>C 1776A>T), CTG470ATA (1783C>A 1785G>A), GCG471GCT (1788G>T), GCC472ACT (1789G>A 1791C>T), TGG473GAA (1792T>G 1793G>A 1794G>A), GTA475GTG (1800A>G), CCT477GGC (1804C>G 1805C>G 1806T>C), ACT478ACG (1809T>G), CAG479ACG (1810C>A 1811A>C), TTT481CTT (1816T>C), AAG482TGC (1819A>T 1820A>G 1821G>C), TAC484TAT (1827C>T), GCT485GCC (1830T>C), CTT486ATT (1831C>A), GGT488GGC (1839T>C) | | | | | | | | |

*: Inserts / Deletes / Misaligned / Frameshifts

#### Analysis details

This analysis was performed with panviral2.64

### NGS Details (UN60): Alphabaculovirus lyxylinae

#### Assembly

| | |
|---|---|
| **Coverage Length** | 110 (1 contig(s)) |
| **Depth Of Coverage** | 6.5 |
| **Number Of Reads** | 8 |
| **Reads Per Million** | 0.17 rpm (after QC) |
| **Ambiguities** | 0 |
| **Assembly Method** | de novo + reference guided assembly |
| **Consensus Caller** | Bcf Tools |

#### Coverage Map

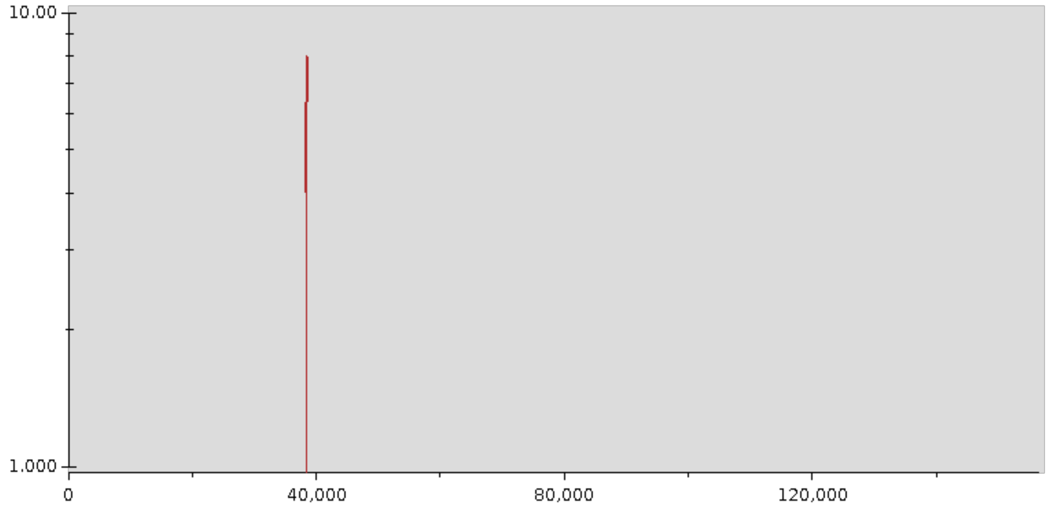

#### Assignment

| | |
|---|---|
| **Type** | Alphabaculovirus lyxylinae (Taxonomy ID: 3051419) |
| **Reference Genome** | NC_013953.1 |
| **NT Identity (%)** | 80.0 |
| **AA Identity (%)** | 83.7838 |
| **Number Of Stop Codons** | 0 |
| **Number Of CDS** | 157 |

#### Alignment

| | |
|---|---|
| **Alignment Score** | 132.0 (NT) + 204.0 (AA) = 336.0 |
| **Concordance (%)** | 73.523 |
| **Alignment Method** | Local, heuristic, nucleotide (BLASTN) |

#### Genome Region

Sequence starts at position 38361 and ends at position 38470 relative to NC_013953.1 reference sequence.

#### Alignment Detailed Statistics

| | Begin | End | Coverage | Score | Concordance | Matches | Identities | I/D/M/F* | Stop Codons |
|---|---|---|---|---|---|---|---|---|---|
| **NT** | **38361** | **38470** | **0.1%** | **132** | **60.0%** | **110 (100%)** | **88 (80.0%)** | **0/0** | |

Mutations: 38363A>G, 38369G>C, 38377A>T, 38387A>G, 38393G>C, 38396C>A, 38401A>G, 38402G>C, 38408C>G, 38412A>G, 38413A>C, 38426C>T, 38432G>A, 38435A>G, 38436G>T, 38447T>C, 38448T>C, 38450G>C, 38453G>C, 38463A>C, 38465A>C, 38468C>T

*: Inserts / Deletes / Misaligned / Frameshifts

#### Analysis details

This analysis was performed with panviral2.64

### NGS Details (UN60): Caulimovirus tessellobrassicae

#### Assembly

| | |
|---|---|
| **Coverage Length** | 254 (1 contig(s)) |
| **Depth Of Coverage** | 2.6 |
| **Number Of Reads** | 6 |
| **Reads Per Million** | 0.13 rpm (after QC) |
| **Ambiguities** | 0 |
| **Assembly Method** | de novo + reference guided assembly |
| **Consensus Caller** | Bcf Tools |

#### Coverage Map

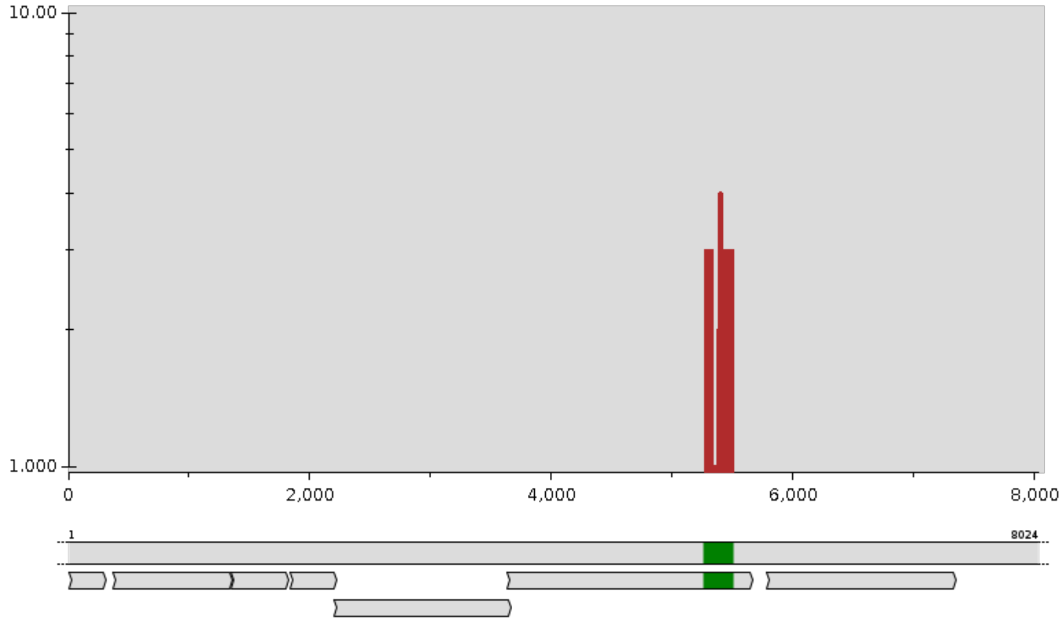

#### Assignment

| | |
|---|---|
| **Type** | Caulimovirus tessellobrassicae (Taxonomy ID: 3047738) |
| **Reference Genome** | NC_001497.2 |
| **NT Identity (%)** | 62.6609 |
| **AA Identity (%)** | 52.5641 |
| **Number Of Stop Codons** | 0 |
| **Number Of CDS** | 7 |

#### Alignment

| | |
|---|---|
| **Alignment Score** | 79.0 (NT) + 296.0 (AA) = 375.0 |
| **Concordance (%)** | 40.1929 |

| Alignment Method | Global, seeded, nucleotide + amino acids (AGA) |
|---|---|

#### Genome Region

Sequence starts at position 5256 and ends at position 5509 relative to NC_001497.2 reference sequence.

#### Alignment Detailed Statistics

| | Begin | End | Coverage | Score | Concordance | Matches | Identities | I/D/M/F* | Stop Codons |
|---|---|---|---|---|---|---|---|---|---|
| **NT** | **5256** | **5509** | **3.2%** | **79** | **18.5%** | **233 (91.7%)** | **146 (57.5%)** | **0/21** | |
| Mutations: | 5256A>T, 5262G>A, 5265G>C, 5268G>T, 5272A>G, 5274C>A, 5275A>T, 5277C>A, 5278G>C, 5280G>A, 5283C>A, 5284G>A, 5288C>T, 5289A>C, 5292A>C, 5295C>T, 5296G>T, 5298C>T, 5299T>C, 5301C>T, 5309G>C, 5310T>A, 5311A>T, 5314T>C, 5317A>C, 5320G>A, 5322T>A, 5323_5325delATC, 5326A>G, 5328A>C, 5329A>C, 5331T>A, 5338_5355delGGTACTAATACTGAGTTA, 5361C>T, 5364A>G, 5367C>T, 5368G>A, 5371T>A, 5372C>G, 5378G>C, 5379C>T, 5385A>C, 5387C>A, 5388T>A, 5392G>A, 5395A>G, 5396A>T, 5397G>T, 5403C>T, 5404C>T, 5405A>C, 5408G>C, 5409C>A, 5415C>A, 5421G>A, 5422A>T, 5423C>T, 5424A>G, 5427G>A, 5434A>G, 5438A>G, 5439T>A, 5440A>G, 5441C>G, 5442T>A, 5446A>C, 5448G>A, 5455A>T, 5456G>C, 5457T>G, 5458A>G, 5459T>C, 5462A>G, 5463T>C, 5464C>T, 5467A>T, 5468C>T, 5469T>A, 5472T>A, 5473G>A, 5474T>A, 5475T>A, 5476C>T, 5477A>C, 5478T>A, 5482C>G, 5484G>T, 5489G>A, 5490G>A | | | | | | | | |

##### CDS

| | Begin | End | Coverage | Score | Concordance | Matches | Identities | I/D/M/F* | Stop Codons |
|---|---|---|---|---|---|---|---|---|---|
| CaMVgp6 | 542 | 626 | 12.5% | 296 | 57.6% | 78 (91.8%) | 41 (48.2%) | 0/7/0/0 | 0 |
| Protein mutations: | E544D (5265G>C), K545N (5268G>T), I547V (5272A>G 5274C>A), I548L (5275A>T 5277C>A), E549Q (5278G>C 5280G>A), D551N (5284G>A), A552V (5288C>T 5289A>C), D555Y (5296G>T 5298C>T), Y556H (5299T>C 5301C>T), G559A (5309G>C 5310T>A), M560L (5311A>T), K562Q (5317A>C), A563T (5320G>A 5322T>A), I564del (5323_5325delATC), K565D (5326A>G 5328A>C), I566L (5329A>C 5331T>A), G569_L574del (5338_5355delGGTACTAATACTGAGTTA), A579T (5368G>A), S582T (5378G>C 5379C>T), K584N (5385A>C), A585E (5387C>A 5388T>A), E587K (5392G>A), K588V (5395A>G 5396A>T 5397G>T), H591S (5404C>T 5405A>C), S592T (5408G>C 5409C>A), D594E (5415C>A), T597L (5422A>T 5423C>T 5424A>G), I601V (5434A>G), N602R (5438A>G 5439T>A), T603G (5440A>G 5441C>G 5442T>A), K605Q (5446A>C 5448G>A), I609A (5458A>G 5459T>C), Y610C (5462A>G 5463T>C), T612L (5467A>T 5468C>T 5469T>A), V614K (5473G>A 5474T>A 5475T>A), H615S (5476C>T 5477A>C 5478T>A), L617V (5482C>G 5484G>T), R619K (5489G>A 5490G>A) | | | | | | | | |
| Codon mutations: | TTA541..T (5256A>T), GAG543GAA (5262G>A), GAG544GAC (5265G>C), AAG545AAT (5268G>T), ATC547GTA (5272A>G 5274C>A), ATC548TTA (5275A>T 5277C>A), GAG549CAA (5278G>C 5280G>A), ACC550ACA (5283C>A), GAT551AAT (5284G>A), GCA552GTC (5288C>T 5289A>C), TCA553TCC (5292A>C), GAC554GAT (5295C>T), GAC555TAT (5296G>T 5298C>T), TAC556CAT (5299T>C 5301C>T), GGT559GCA (5309G>C 5310T>A), ATG560TTG (5311A>T), TTA561CTA (5314T>C), AAA562CAA (5317A>C), GCT563ACA (5320G>A 5322T>A), ATC564del (5323_5325delATC), AAA565GAC (5326A>G 5328A>C), ATT566CTA (5329A>C 5331T>A), GGT569_TTA574del (5338_5355delGGTACTAATACTGAGTTA), TGC576TGT (5361C>T), AGA577AGG (5364A>G), TAC578TAT (5367C>T), GCA579ACA (5368G>A), TCT580AGT (5371T>A 5372C>G), AGC582ACT (5378G>C 5379C>T), AAA584AAC (5385A>C), GCT585GAA (5387C>A 5388T>A), GAA587AAA (5392G>A), AAG588GTT (5395A>G 5396A>T 5397G>T), TAC590TAT (5403C>T), CAC591TCC (5404C>T 5405A>C), AGC592ACA (5408G>C 5409C>A), GAC594GAA (5415C>A), GAG596GAA (5421G>A), ACA597TTG (5422A>T 5423C>T 5424A>G), TTG598TTA (5427G>A), ATA601GTA (5434A>G), AAT602AGA (5438A>G 5439T>A), ACT603GGA (5440A>G 5441C>G 5442T>A), AAG605CAA (5446A>C 5448G>A), AGT608TCG (5455A>T 5456G>C 5457T>G), ATT609GCT (5458A>G 5459T>C), TAT610TGC (5462A>G 5463T>C), CTA611TTA (5464C>T), ACT612TTA (5467A>T 5468C>T 5469T>A), CCT613CCA (5472T>A), GTT614AAA (5473G>A 5474T>A 5475T>A), CAT615TCA (5476C>T 5477A>C 5478T>A), CTG617GTT (5482C>G 5484G>T), AGG619AAA (5489G>A 5490G>A) | | | | | | | | |

##### Proteins

| | Begin | End | Coverage | Score | Concordance | Matches | Identities | I/D/M/F* | Stop Codons |
|---|---|---|---|---|---|---|---|---|---|
| reverse transcriptase (NP_056728.1) | 542 | 626 | 12.5% | 296 | 57.6% | 78 (91.8%) | 41 (48.2%) | 0/7/0/0 | 0 |
| Protein mutations: | E544D (5265G>C), K545N (5268G>T), I547V (5272A>G 5274C>A), I548L (5275A>T 5277C>A), E549Q (5278G>C 5280G>A), D551N (5284G>A), A552V (5288C>T 5289A>C), D555Y (5296G>T 5298C>T), Y556H (5299T>C 5301C>T), G559A (5309G>C 5310T>A), M560L (5311A>T), K562Q (5317A>C), A563T (5320G>A 5322T>A), I564del (5323_5325delATC), K565D (5326A>G 5328A>C), I566L (5329A>C 5331T>A), G569_L574del (5338_5355delGGTACTAATACTGAGTTA), A579T (5368G>A), S582T (5378G>C 5379C>T), K584N (5385A>C), A585E (5387C>A 5388T>A), E587K (5392G>A), K588V (5395A>G 5396A>T 5397G>T), H591S (5404C>T 5405A>C), S592T (5408G>C 5409C>A), D594E (5415C>A), T597L (5422A>T 5423C>T 5424A>G), I601V (5434A>G), N602R (5438A>G 5439T>A), T603G (5440A>G 5441C>G 5442T>A), K605Q (5446A>C 5448G>A), I609A (5458A>G 5459T>C), Y610C (5462A>G 5463T>C), T612L (5467A>T 5468C>T 5469T>A), V614K (5473G>A 5474T>A 5475T>A), H615S (5476C>T 5477A>C 5478T>A), L617V (5482C>G 5484G>T), R619K (5489G>A 5490G>A) | | | | | | | | |
| Codon mutations: | TTA541..T (5256A>T), GAG543GAA (5262G>A), GAG544GAC (5265G>C), AAG545AAT (5268G>T), ATC547GTA (5272A>G 5274C>A), ATC548TTA (5275A>T 5277C>A), GAG549CAA (5278G>C 5280G>A), ACC550ACA (5283C>A), GAT551AAT (5284G>A), GCA552GTC (5288C>T 5289A>C), TCA553TCC (5292A>C), GAC554GAT (5295C>T), GAC555TAT (5296G>T 5298C>T), TAC556CAT (5299T>C 5301C>T), GGT559GCA (5309G>C 5310T>A), ATG560TTG (5311A>T), TTA561CTA (5314T>C), AAA562CAA (5317A>C), GCT563ACA (5320G>A 5322T>A), ATC564del (5323_5325delATC), AAA565GAC (5326A>G 5328A>C), ATT566CTA (5329A>C 5331T>A), GGT569_TTA574del (5338_5355delGGTACTAATACTGAGTTA), TGC576TGT (5361C>T), AGA577AGG (5364A>G), TAC578TAT (5367C>T), GCA579ACA (5368G>A), TCT580AGT (5371T>A 5372C>G), AGC582ACT (5378G>C 5379C>T), AAA584AAC (5385A>C), GCT585GAA (5387C>A 5388T>A), GAA587AAA (5392G>A), AAG588GTT (5395A>G 5396A>T 5397G>T), TAC590TAT (5403C>T), CAC591TCC (5404C>T 5405A>C), AGC592ACA (5408G>C 5409C>A), GAC594GAA (5415C>A), GAG596GAA (5421G>A), ACA597TTG (5422A>T 5423C>T 5424A>G), TTG598TTA (5427G>A), ATA601GTA (5434A>G), AAT602AGA (5438A>G 5439T>A), ACT603GGA (5440A>G 5441C>G 5442T>A), AAG605CAA (5446A>C 5448G>A), AGT608TCG (5455A>T 5456G>C 5457T>G), ATT609GCT (5458A>G 5459T>C), TAT610TGC (5462A>G 5463T>C), CTA611TTA (5464C>T), ACT612TTA (5467A>T 5468C>T 5469T>A), CCT613CCA (5472T>A), GTT614AAA (5473G>A 5474T>A 5475T>A), CAT615TCA (5476C>T 5477A>C 5478T>A), CTG617GTT (5482C>G 5484G>T), AGG619AAA (5489G>A 5490G>A) | | | | | | | | |

*: Inserts / Deletes / Misaligned / Frameshifts

#### Analysis details

This analysis was performed with panviral2.64

### NGS Details (UN60): Badnavirus occultipomeae

#### Assembly

| | |
|---|---|
| **Coverage Length** | 251 (1 contig(s)) |
| **Depth Of Coverage** | 3.2 |
| **Number Of Reads** | 6 |
| **Reads Per Million** | 0.13 rpm (after QC) |
| **Ambiguities** | 0 |
| **Assembly Method** | de novo + reference guided assembly |
| **Consensus Caller** | Bcf Tools |

#### Coverage Map

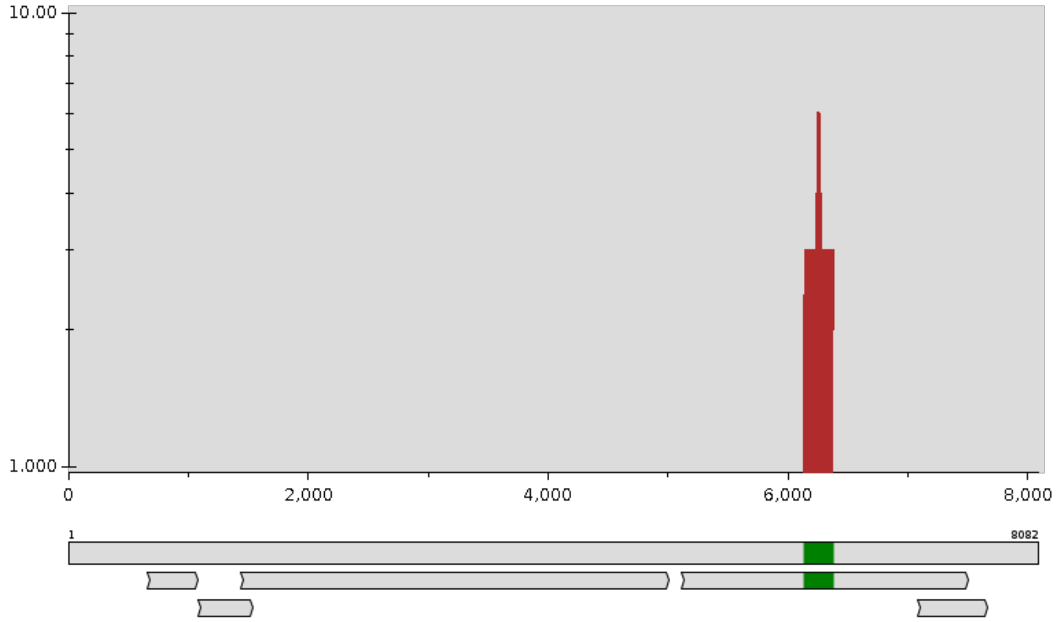

#### Assignment

| | |
|---|---|
| **Type** | Badnavirus occultipomeae (Taxonomy ID: 3048353) |
| **Reference Genome** | NC_015655.1 |
| **NT Identity (%)** | 54.3307 |
| **AA Identity (%)** | 43.5294 |
| **Number Of Stop Codons** | 0 |
| **Number Of CDS** | 5 |

#### Alignment

| | |
|---|---|
| **Alignment Score** | 38.0 (NT) + 264.0 (AA) = 302.0 |
| **Concordance (%)** | 28.3568 |

| **Alignment Method** | Global, seeded, nucleotide + amino acids (AGA) |
|---|---|

#### Genome Region

Sequence starts at position 6129 and ends at position 6379 relative to NC_015655.1 reference sequence.

#### Alignment Detailed Statistics

| | Begin | End | Coverage | Score | Concordance | Matches | Identities | I/D/M/F* | Stop Codons |
|---|---|---|---|---|---|---|---|---|---|
| **NT** | **6129** | **6379** | **3.1%** | **38** | **7.6%** | **251 (98.8%)** | **138 (54.3%)** | **3/0** | |
| Mutations: | 6129T>C, 6132A>G, 6137A>T, 6138A>G, 6141C>A, 6145C>T, 6147C>T, 6153A>G, 6154C>T, 6165A>G, 6169G>A, 6171A>T, 6172G>A, 6173A>T, 6174T>A, 6178C>A, 6179A>T, 6181A>G, 6182G>A, 6183A>T, 6184A>C, 6185A>T, 6186G>T, 6190G>A, 6194A>G, 6195T>G, 6197C>T, 6198A>T, 6201T>C, 6203G>A, 6204G>A, 6205G>C, 6206G>A, 6207A>C, 6207_6208insTAC, 6208A>C, 6209C>T, 6210A>T, 6214G>C, 6215C>T, 6216A>T, 6220A>G, 6222C>T, 6224C>T, 6225A>G, 6226G>T, 6230A>T, 6241A>G, 6244T>C, 6246G>A, 6247G>A, 6249A>C, 6251T>A, 6255T>C, 6256G>C, 6260A>G, 6264A>G, 6265G>A, 6267A>G, 6270G>A, 6276G>A, 6279T>A, 6283C>T, 6285C>G, 6286T>A, 6287T>G, 6290A>T, 6291T>A, 6292T>G, 6294G>C, 6295G>C, 6296C>T, 6297C>T, 6302T>C, 6304G>T, 6310A>G, 6311G>A, 6312A>T, 6313G>A, 6315A>G, 6316G>A, 6317G>A, 6319C>T, 6321T>G, 6322A>T, 6323T>A, 6325C>G, 6326T>C, 6327A>C, 6329G>A, 6330C>A, 6331C>T, 6332C>T, 6333A>C, 6335C>A, 6336A>C, 6340A>T, 6341T>G, 6342G>T, 6343A>G, 6346A>T, 6349G>T, 6351A>G, 6352G>A, 6354C>G, 6363G>T, 6364G>T, 6365A>C, 6372A>T, 6375A>C, 6376A>C, 6377T>A, 6378A>T, 6379A>G | | | | | | | | |

##### CDS

| | Begin | End | Coverage | Score | Concordance | Matches | Identities | I/D/M/F* | Stop Codons |
|---|---|---|---|---|---|---|---|---|---|
| SPBVa_gp4 | 339 | 422 | 10.6% | 264 | 45.8% | 84 (98.8%) | 37 (43.5%) | 1/0/0/0 | 0 |
| Protein mutations: | E341V (6137A>T 6138A>G), P344S (6145C>T 6147C>T), A352T (6169G>A 6171A>T), D353I (6172G>A 6173A>T 6174T>A), Q355M (6178C>A 6179A>T), R356D (6181A>G 6182G>A 6183A>T), K357L (6184A>C 6185A>T 6186G>T), D359N (6190G>A), N360R (6194A>G 6195T>G), A361V (6197C>T 6198A>T), R363K (6203G>A 6204G>A), G364H (6205G>C 6206G>A 6207A>C), G364_T365insY (6207_6208insTAC), T365L (6208A>C 6209C>T 6210A>T), A367L (6214G>C 6215C>T 6216A>T), I369V (6220A>G 6222C>T), A370V (6224C>T 6225A>G), V371L (6226G>T), Y372F (6230A>T), I376V (6241A>G), V378I (6247G>A 6249A>C), F379Y (6251T>A), E381Q (6256G>C), N382S (6260A>G), E384K (6265G>A 6267A>G), D388E (6279T>A), L391R (6286T>A 6287T>G), N392I (6290A>T 6291T>A), L393V (6292T>G 6294G>C), A394L (6295G>C 6296C>T 6297C>T), I396T (6302T>C), V397L (6304G>T), R399D (6310A>G 6311G>A 6312A>T), E400K (6313G>A 6315A>G), G401K (6316G>A 6317G>A), I403Y (6322A>T 6323T>A), L404A (6325C>G 6326T>C 6327A>C), S405K (6329G>A 6330C>A), P406F (6331C>T 6332C>T 6333A>C), T407N (6335C>A 6336A>C), M409C (6340A>T 6341T>G 6342G>T), K410E (6343A>G), I411F (6346A>T), G412W (6349G>T 6351A>G), V413M (6352G>A 6354C>G), D417S (6364G>T 6365A>C), I421H (6376A>C 6377T>A 6378A>T) | | | | | | | | |
| Codon mutations: | TAT338..C (6129T>C), GAA339GAG (6132A>G), GAA341GTG (6137A>T 6138A>G), GTC342GTA (6141C>A), CCC344TCT (6145C>T 6147C>T), GGA346GGG (6153A>G), CTG347TTG (6154C>T), GCA350GCG (6165A>G), GCA352ACT (6169G>A 6171A>T), GAT353ATA (6172G>A 6173A>T 6174T>A), CAG355ATG (6178C>A 6179A>T), AGA356GAT (6181A>G 6182G>A 6183A>T), AAG357CTT (6184A>C 6185A>T 6186G>T), GAT359AAT (6190G>A), AAT360AGG (6194A>G 6195T>G), GCA361GTT (6197C>T 6198A>T), TTT362TTC (6201T>C), AGG363AAA (6203G>A 6204G>A), GGA364CAC (6205G>C 6206G>A 6207A>C), GGA364_ACA365insTAC (6207_6208insTAC), ACA365CTT (6208A>C 6209C>T 6210A>T), GCA367CTT (6214G>C 6215C>T 6216A>T), ATC369GTT (6220A>G 6222C>T), GCA370GTG (6224C>T 6225A>G), GTA371TTA (6226G>T), TAT372TTT (6230A>T), ATA376GTA (6241A>G), TTG377CTA (6244T>C 6246G>A), GTA378ATC (6247G>A 6249A>C), TTC379TAC (6251T>A), TCT380TCC (6255T>C), GAA381CAA (6256G>C), AAT382AGT (6260A>G), GAA383GAG (6264A>G), GAA384AAG (6265G>A 6267A>G), GAG385GAA (6270G>A), GAG387GAA (6276G>A), GAT388GAA (6279T>A), CTC390TTG (6283C>T 6285C>G), TTA391AGA (6286T>A 6287T>G), AAT392ATA (6290A>T 6291T>A), TTG393GTC (6292T>G 6294G>C), GCC394CTT (6295G>C 6296C>T 6297C>T), ATT396ACT (6302T>C), GTG397TTG (6304G>T), AGA399GAT (6310A>G 6311G>A 6312A>T), GAA400AAG (6313G>A 6315A>G), GGG401AAG (6316G>A 6317G>A), CTT402TTG (6319C>T 6321T>G), ATT403TAT (6322A>T 6323T>A), CTA404GCC (6325C>G 6326T>C 6327A>C), AGC405AAA (6329G>A 6330C>A), CCA406TTC (6331C>T 6332C>T 6333A>C), ACA407AAC (6335C>A 6336A>C), ATG409TGT (6340A>T 6341T>G 6342G>T), AAA410GAA (6343A>G), ATT411TTT (6346A>T), GGA412TGG (6349G>T 6351A>G), GTC413ATG (6352G>A 6354C>G), GTG416GTT (6363G>T), GAT417TCT (6364G>T 6365A>C), CTA419CTT (6372A>T), GGA420GGC (6375A>C), ATA421CAT (6376A>C 6377T>A 6378A>T), AAG422G.. (6379A>G) | | | | | | | | |

##### Proteins

| | Begin | End | Coverage | Score | Concordance | Matches | Identities | I/D/M/F* | Stop Codons |
|---|---|---|---|---|---|---|---|---|---|
| RNaseH/reverse transcriptase (YP_004581513.1) | 339 | 422 | 10.6% | 264 | 45.8% | 84 (98.8%) | 37 (43.5%) | 1/0/0/0 | 0 |
| Protein mutations: | E341V (6137A>T 6138A>G), P344S (6145C>T 6147C>T), A352T (6169G>A 6171A>T), D353I (6172G>A 6173A>T 6174T>A), Q355M (6178C>A 6179A>T), R356D (6181A>G 6182G>A 6183A>T), K357L (6184A>C 6185A>T 6186G>T), D359N (6190G>A), N360R (6194A>G 6195T>G), A361V (6197C>T 6198A>T), R363K (6203G>A 6204G>A), G364H (6205G>C 6206G>A 6207A>C), G364_T365insY (6207_6208insTAC), T365L (6208A>C 6209C>T 6210A>T), A367L (6214G>C 6215C>T 6216A>T), I369V (6220A>G 6222C>T), A370V (6224C>T 6225A>G), V371L (6226G>T), Y372F (6230A>T), I376V (6241A>G), V378I (6247G>A 6249A>C), F379Y (6251T>A), E381Q (6256G>C), N382S (6260A>G), E384K (6265G>A 6267A>G), D388E (6279T>A), L391R (6286T>A 6287T>G), N392I (6290A>T 6291T>A), L393V (6292T>G 6294G>C), A394L (6295G>C 6296C>T 6297C>T), I396T (6302T>C), V397L (6304G>T), R399D (6310A>G 6311G>A 6312A>T), E400K (6313G>A 6315A>G), G401K (6316G>A 6317G>A), I403Y (6322A>T 6323T>A), L404A (6325C>G 6326T>C 6327A>C), S405K (6329G>A 6330C>A), P406F (6331C>T 6332C>T 6333A>C), T407N (6335C>A 6336A>C), M409C (6340A>T 6341T>G 6342G>T), K410E (6343A>G), I411F (6346A>T), G412W (6349G>T 6351A>G), V413M (6352G>A 6354C>G), D417S (6364G>T 6365A>C), I421H (6376A>C 6377T>A 6378A>T) | | | | | | | | |
| Codon mutations: | TAT338..C (6129T>C), GAA339GAG (6132A>G), GAA341GTG (6137A>T 6138A>G), GTC342GTA (6141C>A), CCC344TCT (6145C>T 6147C>T), GGA346GGG (6153A>G), CTG347TTG (6154C>T), GCA350GCG (6165A>G), GCA352ACT (6169G>A 6171A>T), GAT353ATA (6172G>A 6173A>T 6174T>A), CAG355ATG (6178C>A 6179A>T), AGA356GAT (6181A>G 6182G>A 6183A>T), AAG357CTT (6184A>C 6185A>T 6186G>T), GAT359AAT (6190G>A), AAT360AGG (6194A>G 6195T>G), GCA361GTT (6197C>T 6198A>T), TTT362TTC (6201T>C), AGG363AAA (6203G>A 6204G>A), GGA364CAC (6205G>C 6206G>A 6207A>C), GGA364_ACA365insTAC (6207_6208insTAC), ACA365CTT (6208A>C 6209C>T 6210A>T), GCA367CTT (6214G>C 6215C>T 6216A>T), ATC369GTT (6220A>G 6222C>T), GCA370GTG (6224C>T 6225A>G), GTA371TTA (6226G>T), TAT372TTT (6230A>T), ATA376GTA (6241A>G), TTG377CTA (6244T>C 6246G>A), GTA378ATC (6247G>A 6249A>C), TTC379TAC (6251T>A), TCT380TCC (6255T>C), GAA381CAA (6256G>C), AAT382AGT (6260A>G), GAA383GAG (6264A>G), GAA384AAG (6265G>A 6267A>G), GAG385GAA (6270G>A), GAG387GAA (6276G>A), GAT388GAA (6279T>A), CTC390TTG (6283C>T 6285C>G), TTA391AGA (6286T>A 6287T>G), AAT392ATA (6290A>T 6291T>A), TTG393GTC (6292T>G 6294G>C), GCC394CTT (6295G>C 6296C>T 6297C>T), ATT396ACT (6302T>C), GTG397TTG (6304G>T), AGA399GAT (6310A>G 6311G>A 6312A>T), GAA400AAG (6313G>A 6315A>G), GGG401AAG (6316G>A 6317G>A), CTT402TTG (6319C>T 6321T>G), ATT403TAT (6322A>T 6323T>A), CTA404GCC (6325C>G 6326T>C 6327A>C), AGC405AAA (6329G>A 6330C>A), CCA406TTC (6331C>T 6332C>T 6333A>C), ACA407AAC (6335C>A 6336A>C), ATG409TGT (6340A>T 6341T>G 6342G>T), AAA410GAA (6343A>G), ATT411TTT (6346A>T), GGA412TGG (6349G>T 6351A>G), GTC413ATG (6352G>A 6354C>G), GTG416GTT (6363G>T), GAT417TCT (6364G>T 6365A>C), CTA419CTT (6372A>T), GGA420GGC (6375A>C), ATA421CAT (6376A>C 6377T>A 6378A>T), AAG422G.. (6379A>G) | | | | | | | | |

*: Inserts / Deletes / Misaligned / Frameshifts

#### Analysis details

This analysis was performed with panviral2.64

### NGS Details (UN60): Moloney murine leukemia virus

#### Assembly

| | |
|---|---|
| **Coverage Length** | 270 (2 contig(s)) |
| **Depth Of Coverage** | 3.0 |
| **Number Of Reads** | 6 |
| **Reads Per Million** | 0.13 rpm (after QC) |
| **Ambiguities** | 0 |
| **Assembly Method** | de novo + reference guided assembly |
| **Consensus Caller** | Bcf Tools |

#### Coverage Map

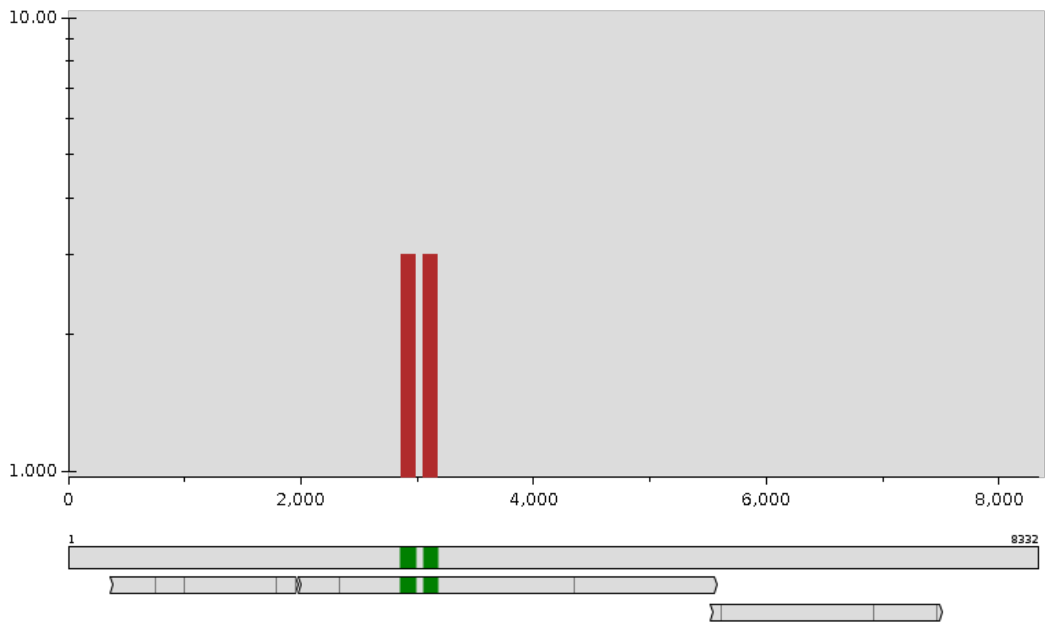

#### Assignment

| | |
|---|---|
| **Type** | Moloney murine leukemia virus (Taxonomy ID: 11801) |
| **Reference Genome** | NC_001501.1 |
| **NT Identity (%)** | 75.5556 |
| **AA Identity (%)** | 100.0 |
| **Number Of Stop Codons** | 0 |
| **Number Of CDS** | 3 |

#### Alignment

| | |
|---|---|
| **Alignment Score** | 276.0 (NT) + 627.0 (AA) = 903.0 |
| **Concordance (%)** | 77.3779 |

| Alignment Method | Global, seeded, nucleotide + amino acids (AGA) |
|---|---|

#### Genome Region

Sequence starts at position 2856 and ends at position 3182 relative to NC_001501.1 reference sequence.

#### Alignment Detailed Statistics

| | Begin | End | Coverage | Score | Concordance | Matches | Identities | I/D/M/F* | Stop Codons |
|---|---|---|---|---|---|---|---|---|---|
| **NT** | **2856** | **3182** | **3.2%** | **276** | **51.1%** | **270 (100%)** | **204 (75.6%)** | **0/0** | |
| Mutations: | 2858A>T, 2861G>A, 2867A>C, 2870C>T, 2871T>A, 2872C>G, 2873A>C, 2876A>C, 2879A>G, 2880T>C, 2892A>C, 2894A>C, 2897C>G, 2900A>G, 2906T>C, 2909C>T, 2918T>C, 2921C>G, 2936G>A, 2939A>G, 2945C>T, 2946A>C, 2948A>C, 2951C>T, 2954A>G, 2957A>G, 2960C>T, 2963C>T, 2966G>C, 2969C>T, 2975C>T, 2978A>G, 2981C>T, 2982T>C, 2987C>T, 3050T>C, 3053T>C, 3056G>C, 3059C>G, 3063T>C, 3065A>G, 3068A>G, 3074A>G, 3077G>C, 3083C>G, 3086G>C, 3092G>C, 3095C>G, 3096T>A, 3097C>G, 3098G>C, 3101C>G, 3104G>A, 3110C>G, 3113A>G, 3131C>G, 3134G>A, 3143G>C, 3149T>G, 3152A>G, 3158G>A, 3161T>C, 3165A>C, 3167A>C, 3176T>C, 3179G>A | | | | | | | | |
| **CDS** | | | | | | | | | |
| gag-pol | 833 | 941 | 5.2% | 627 | 100% | 90 (100%) | 90 (100%) | 0/0/0/0 | 0 |
| **Proteins** | | | | | | | | | |
| Pr180 (NP_057933.2) | 833 | 941 | 5.2% | 627 | 100% | 90 (100%) | 90 (100%) | 0/0/0/0 | 0 |
| Protein mutations: | *none* | | | | | | | | |
| Codon mutations: | CCA833CCT (2858A>T), GAG834GAA (2861G>A), GGA836GGC (2867A>C), ATC837ATT (2870C>T), TCA838AGC (2871T>A 2872C>G 2873A>C), GGA839GGC (2876A>C), CAA840CAG (2879A>G), TTG841CTG (2880T>C), AGA845CGC (2892A>C 2894A>C), CTC846CTG (2897C>G), CCA847CCG (2900A>G), GGT849GGC (2906T>C), TTC850TTT (2909C>T), AGT853AGC (2918T>C), CCC854CCG (2921C>G), GAG859GAA (2936G>A), GCA860GCG (2939A>G), CAC862CAT (2945C>T), AGA863CGC (2946A>C 2948A>C), GAC864GAT (2951C>T), CTA865CTG (2954A>G), GCA866GCG (2957A>G), GAC867GAT (2960C>T), TTC868TTT (2963C>T), CGG869CGC (2966G>C), ATC870ATT (2969C>T), CAC872CAT (2975C>T), CCA873CCG (2978A>G), GAC874GAT (2981C>T), TTG875CTG (2982T>C), ATC876ATT (2987C>T), GGT897GGC (3050T>C), ACT898ACC (3053T>C), CGG899CGC (3056G>C), GCC900GCG (3059C>G), TTA902CTG (3063T>C 3065A>G), CAA903CAG (3068A>G), CTA905CTG (3074A>G), GGG906GGC (3077G>C), CTC908CTG (3083C>G), GGG909GGC (3086G>C), CGG911CGC (3092G>C), GCC912GCG (3095C>G), TCG913AGC (3096T>A 3097C>G 3098G>C), GCC914GCG (3101C>G), AAG915AAA (3104G>A), GCC917GCG (3110C>G), CAA918CAG (3113A>G), GTC924GTG (3131C>G), AAG925AAA (3134G>A), GGG928GGC (3143G>C), CTT930CTG (3149T>G), CTA931CTG (3152A>G), GAG933GAA (3158G>A), GGT934GGC (3161T>C), AGA936CGC (3165A>C 3167A>C), ACT939ACC (3176T>C), GAG940GAA (3179G>A) | | | | | | | | |
| p80 RT (NP_955591.1) | 174 | 282 | 13.4% | 627 | 100% | 90 (100%) | 90 (100%) | 0/0/0/0 | 0 |
| Protein mutations: | *none* | | | | | | | | |
| Codon mutations: | CCA174CCT (2858A>T), GAG175GAA (2861G>A), GGA177GGC (2867A>C), ATC178ATT (2870C>T), TCA179AGC (2871T>A 2872C>G 2873A>C), GGA180GGC (2876A>C), CAA181CAG (2879A>G), TTG182CTG (2880T>C), AGA186CGC (2892A>C 2894A>C), CTC187CTG (2897C>G), CCA188CCG (2900A>G), GGT190GGC (2906T>C), TTC191TTT (2909C>T), AGT194AGC (2918T>C), CCC195CCG (2921C>G), GAG200GAA (2936G>A), GCA201GCG (2939A>G), CAC203CAT (2945C>T), AGA204CGC (2946A>C 2948A>C), GAC205GAT (2951C>T), CTA206CTG (2954A>G), GCA207GCG (2957A>G), GAC208GAT (2960C>T), TTC209TTT (2963C>T), CGG210CGC (2966G>C), ATC211ATT (2969C>T), CAC213CAT (2975C>T), CCA214CCG (2978A>G), GAC215GAT (2981C>T), TTG216CTG (2982T>C), ATC217ATT (2987C>T), GGT238GGC (3050T>C), ACT239ACC (3053T>C), CGG240CGC (3056G>C), GCC241GCG (3059C>G), TTA243CTG (3063T>C 3065A>G), CAA244CAG (3068A>G), CTA246CTG (3074A>G), GGG247GGC (3077G>C), CTC249CTG (3083C>G), GGG250GGC (3086G>C), CGG252CGC (3092G>C), GCC253GCG (3095C>G), TCG254AGC (3096T>A 3097C>G 3098G>C), GCC255GCG (3101C>G), AAG256AAA (3104G>A), GCC258GCG (3110C>G), CAA259CAG (3113A>G), GTC265GTG (3131C>G), AAG266AAA (3134G>A), GGG269GGC (3143G>C), CTT271CTG (3149T>G), CTA272CTG (3152A>G), GAG274GAA (3158G>A), GGT275GGC (3161T>C), AGA277CGC (3165A>C 3167A>C), ACT280ACC (3176T>C), GAG281GAA (3179G>A) | | | | | | | | |

*: Inserts / Deletes / Misaligned / Frameshifts

#### Analysis details

This analysis was performed with panviral2.64

### NGS Details (UN60): Marseillevirus marseillevirus

#### Assembly

| | |
|---|---|
| **Coverage Length** | 105 (1 contig(s)) |
| **Depth Of Coverage** | 4.4 |
| **Number Of Reads** | 6 |
| **Reads Per Million** | 0.13 rpm (after QC) |
| **Ambiguities** | 1 |
| **Assembly Method** | de novo + reference guided assembly |
| **Consensus Caller** | Bcf Tools |

#### Coverage Map

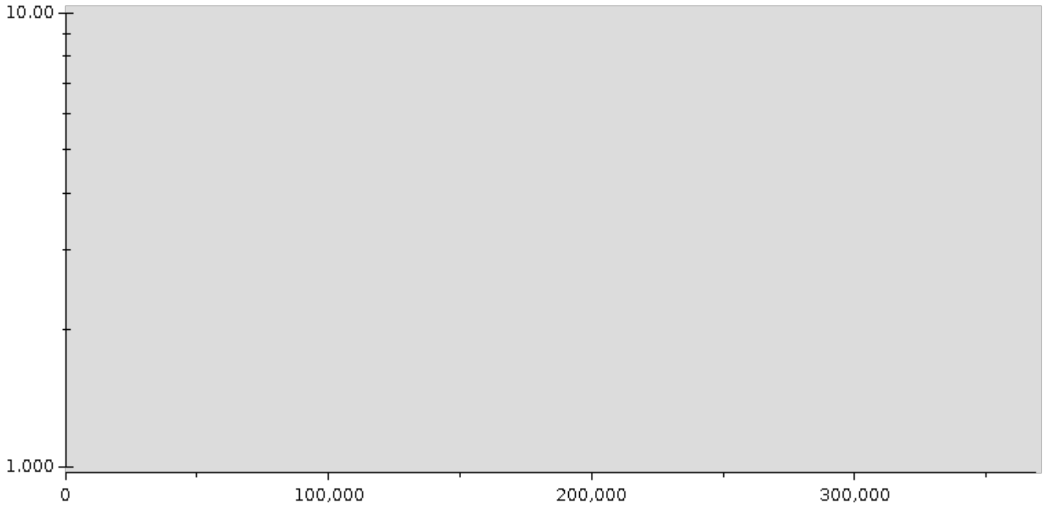

#### Assignment

| | |
|---|---|
| **Type** | Marseillevirus marseillevirus (Taxonomy ID: 694581) |
| **Reference Genome** | NC_013756.1 |
| **NT Identity (%)** | 81.9048 |
| **AA Identity (%)** | 88.5714 |
| **Number Of Stop Codons** | 0 |
| **Number Of CDS** | 428 |

#### Alignment

| | |
|---|---|
| **Alignment Score** | 136.0 (NT) + 183.0 (AA) = 319.0 |
| **Concordance (%)** | 73.8426 |
| **Alignment Method** | Local, heuristic, nucleotide (BLASTN) |

#### Genome Region

Sequence starts at position 199412 and ends at position 199516 relative to NC_013756.1 reference sequence.

#### Alignment Detailed Statistics

| | Begin | End | Coverage | Score | Concordance | Matches | Identities | I/D/M/F* | Stop Codons |
|---|---|---|---|---|---|---|---|---|---|
| **NT** | **199412** | **199516** | **0.1%** | **136** | **64.8%** | **105 (100%)** | **86 (81.9%)** | **0/0** | |

Mutations: 199414T>C, 199420G>A, 199423A>C, 199425G>A, 199426G>M, 199429G>C, 199432A>G, 199444T>C, 199449C>T, 199450G>A, 199461C>A, 199462C>G, 199464A>G, 199465A>G, 199477T>C, 199480C>G, 199492G>A, 199495G>A, 199513G>A

*: Inserts / Deletes / Misaligned / Frameshifts

#### Analysis details

This analysis was performed with panviral2.64

### NGS Details (UN60): Yaba monkey tumor virus

#### Assembly

| | |
|---|---|
| **Coverage Length** | 134 (1 contig(s)) |
| **Depth Of Coverage** | 3.3 |
| **Number Of Reads** | 4 |
| **Reads Per Million** | 0.08 rpm (after QC) |
| **Ambiguities** | 0 |
| **Assembly Method** | de novo + reference guided assembly |
| **Consensus Caller** | Bcf Tools |

#### Coverage Map

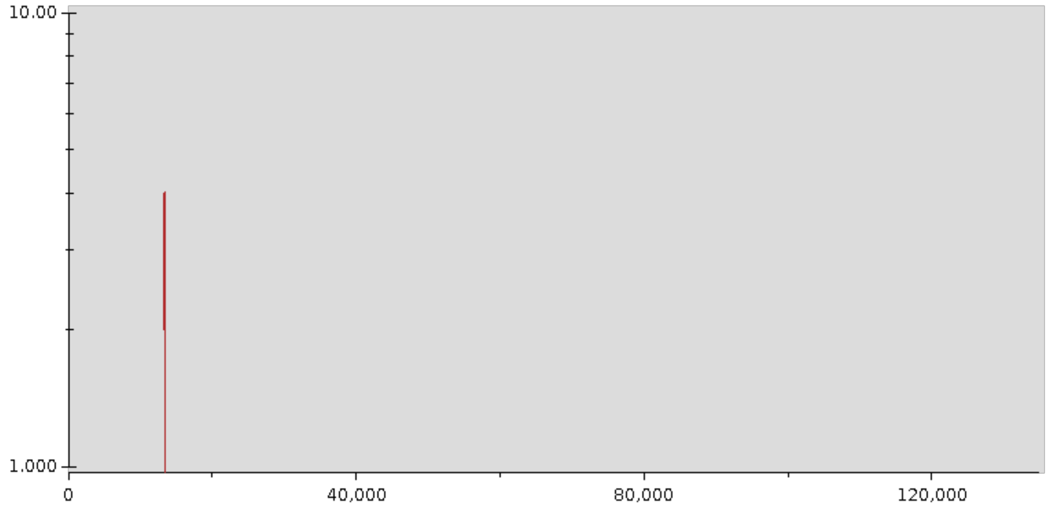

#### Assignment

| | |
|---|---|
| **Type** | Yaba monkey tumor virus (Taxonomy ID: 38804) |
| **Reference Genome** | NC_005179.1 |
| **NT Identity (%)** | 81.3433 |
| **AA Identity (%)** | 84.0909 |
| **Number Of Stop Codons** | 0 |
| **Number Of CDS** | 140 |

#### Alignment

| | |
|---|---|
| **Alignment Score** | 168.0 (NT) + 273.0 (AA) = 441.0 |
| **Concordance (%)** | 75.7732 |
| **Alignment Method** | Local, heuristic, nucleotide (BLASTN) |

#### Genome Region

Sequence starts at position 13363 and ends at position 13496 relative to NC_005179.1 reference sequence.

#### Alignment Detailed Statistics

| | Begin | End | Coverage | Score | Concordance | Matches | Identities | I/D/M/F* | Stop Codons |
|---|---|---|---|---|---|---|---|---|---|
| NT | 13363 | 13496 | 0.1% | 168 | 62.7% | 134 (100%) | 109 (81.3%) | 0/0 | |

Mutations: 13366G>A, 13369G>C, 13370T>G, 13376T>G, 13381T>A, 13384T>A, 13387G>A, 13397T>A, 13399A>G, 13401T>C, 13408C>T, 13411A>T, 13412C>G, 13413T>G, 13418A>T, 13419A>G, 13426A>T, 13429A>G, 13432T>A, 13435C>T, 13453T>C, 13459A>G, 13472G>A, 13473C>T, 13483A>C

*: Inserts / Deletes / Misaligned / Frameshifts

#### Analysis details

This analysis was performed with panviral2.64

### NGS Details (UN60): Alphabaculovirus agipsilonis

#### Assembly

| | |
|---|---|
| **Coverage Length** | 121 (1 contig(s)) |
| **Depth Of Coverage** | 3.5 |
| **Number Of Reads** | 4 |
| **Reads Per Million** | 0.08 rpm (after QC) |
| **Ambiguities** | 0 |
| **Assembly Method** | de novo + reference guided assembly |
| **Consensus Caller** | Bcf Tools |

#### Coverage Map

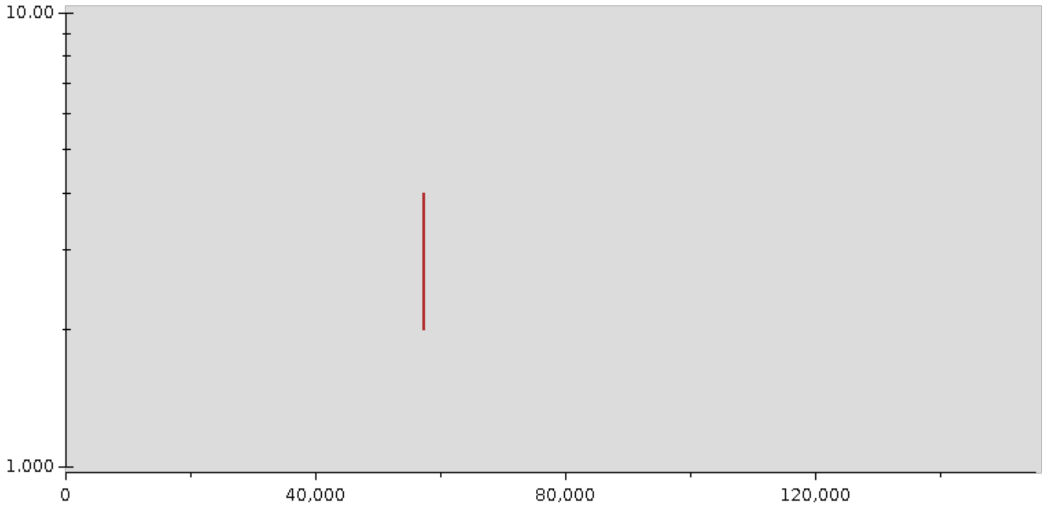

#### Assignment

| | |
|---|---|
| **Type** | Alphabaculovirus agipsilonis (Taxonomy ID: 3047352) |
| **Reference Genome** | NC_011345.1 |
| **NT Identity (%)** | 78.5124 |
| **AA Identity (%)** | 77.5 |
| **Number Of Stop Codons** | 0 |
| **Number Of CDS** | 163 |

#### Alignment

| | |
|---|---|
| **Alignment Score** | 138.0 (NT) + 272.0 (AA) = 410.0 |
| **Concordance (%)** | 72.1831 |
| **Alignment Method** | Local, heuristic, nucleotide (BLASTN) |

#### Genome Region

Sequence starts at position 57321 and ends at position 57441 relative to NC_011345.1 reference sequence.

#### Alignment Detailed Statistics

| | Begin | End | Coverage | Score | Concordance | Matches | Identities | I/D/M/F* | Stop Codons |
|---|---|---|---|---|---|---|---|---|---|
| **NT** | **57321** | **57441** | **0.1%** | **138** | **57.0%** | **121 (100%)** | **95 (78.5%)** | **0/0** | |

Mutations: 57330C>T, 57334A>T, 57336G>A, 57339C>G, 57344T>C, 57357T>C, 57359C>T, 57361C>G, 57362T>C, 57363G>T, 57368T>C, 57375A>G, 57381C>G, 57382T>G, 57383C>G, 57384G>T, 57387C>A, 57390C>A, 57393C>T, 57399G>C, 57409G>T, 57414G>A, 57422C>G, 57432A>G, 57435T>G, 57439T>C

*: Inserts / Deletes / Misaligned / Frameshifts

#### Analysis details

This analysis was performed with panviral2.64

#### NGS Details (UN60): Badnavirus tessellocastaneae

##### Assembly

| | |
|---|---|
| **Coverage Length** | 245 (1 contig(s)) |
| **Depth Of Coverage** | 2.1 |
| **Number Of Reads** | 4 |
| **Reads Per Million** | 0.08 rpm (after QC) |
| **Ambiguities** | 0 |
| **Assembly Method** | de novo + reference guided assembly |
| **Consensus Caller** | Bcf Tools |

##### Coverage Map

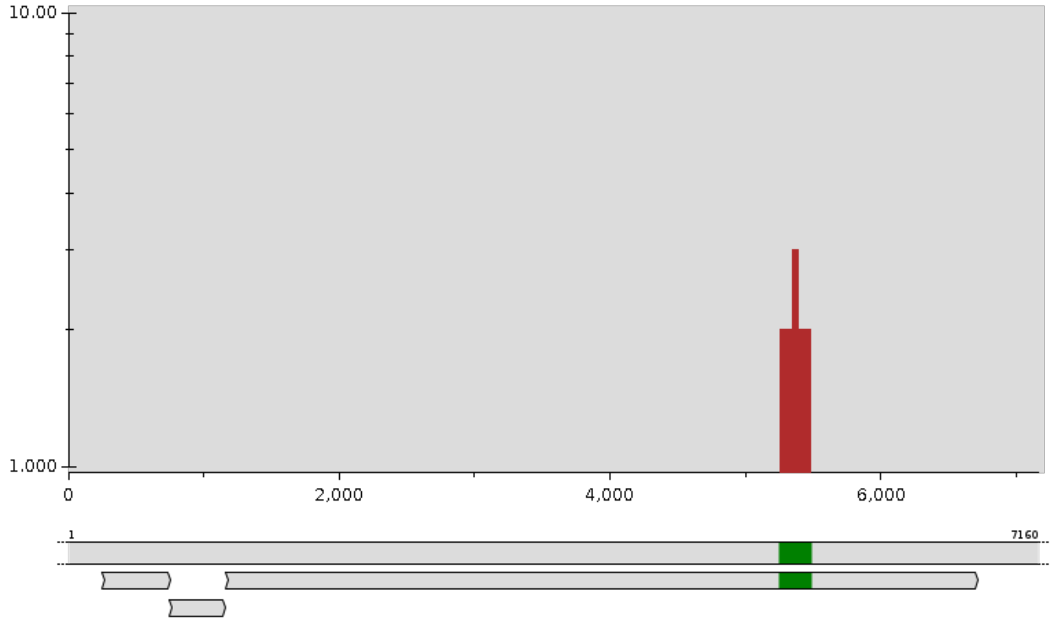

##### Assignment

| | |
|---|---|
| **Type** | Badnavirus tessellocastaneae (Taxonomy ID: 3051987) |
| **Reference Genome** | NC_076885.1 |
| **NT Identity (%)** | 59.4262 |
| **AA Identity (%)** | 53.6585 |
| **Number Of Stop Codons** | 0 |
| **Number Of CDS** | 3 |

##### Alignment

| | |
|---|---|
| **Alignment Score** | 82.0 (NT) + 195.0 (AA) = 277.0 |
| **Concordance (%)** | 30.9948 |

| **Alignment Method** | Global, seeded, nucleotide + amino acids (AGA) |
|---|---|

#### Genome Region

Sequence starts at position 5248 and ends at position 5492 relative to NC_076885.1 reference sequence.

#### Alignment Detailed Statistics

| | Begin | End | Coverage | Score | Concordance | Matches | Identities | I/D/M/F* | Stop Codons |
|---|---|---|---|---|---|---|---|---|---|
| **NT** | **5248** | **5492** | **3.4%** | **82** | **17.2%** | **244 (99.6%)** | **145 (59.2%)** | **0/1** | |
| Mutations: | 5248G>A, 5254G>T, 5258G>A, 5259T>A, 5260G>T, 5263A>C, 5266A>C, 5267G>T, 5268G>T, 5269A>T, 5273T>C, 5274T>A, 5281G>A, 5285C>A, 5286T>A, 5287G>T, 5296A>T, 5305A>T, 5307G>A, 5314A>C, 5321A>G, 5322C>A, 5323C>A, 5326C>T, 5332G>T, 5334A>T, 5335G>T, 5339G>A, 5341C>T, 5342C>T, 5343A>C, 5345delT, 5346G>T, 5352G>A, 5353G>T, 5354G>C, 5355G>A, 5356G>T, 5357A>C, 5358C>T, 5359A>T, 5360G>A, 5361A>G, 5362A>C, 5363G>C, 5365A>T, 5369A>T, 5370T>C, 5371A>T, 5372T>A, 5373C>T, 5374A>T, 5386C>T, 5389T>C, 5390A>G, 5392A>T, 5393C>T, 5395C>A, 5396G>A, 5398C>A, 5404C>A, 5405C>G, 5406C>A, 5411C>A, 5416A>T, 5417A>T, 5423G>T, 5424C>T, 5425A>C, 5432T>C, 5434G>C, 5435G>A, 5437G>T, 5438A>G, 5439A>C, 5440A>T, 5441A>T, 5443G>C, 5444C>T, 5446G>C, 5447G>A, 5448C>A, 5451T>C, 5452C>T, 5453T>A, 5454G>T, 5455C>T, 5459G>A, 5460A>T, 5464T>C, 5470A>T, 5474C>A, 5476G>T, 5477T>A, 5478C>G, 5479A>T, 5480C>G, 5484C>A, 5485A>G, 5491G>A | | | | | | | | |

##### CDS

| | Begin | End | Coverage | Score | Concordance | Matches | Identities | I/D/M/F* | Stop Codons |
|---|---|---|---|---|---|---|---|---|---|
| ORF3 | 1363 | 1444 | 4.4% | 195 | 32.4% | 82 (100%) | 44 (53.7%) | 0/0/1/1 | 0 |
| Protein mutations: | V1366N (5258G>A 5259T>A 5260G>T), G1369F (5267G>T 5268G>T 5269A>T), L1371Q (5273T>C 5274T>A), L1375N (5285C>A 5286T>A 5287G>T), R1382K (5307G>A), T1387E (5321A>G 5322C>A 5323C>A), R1390S (5332G>T), K1391I (5334A>T 5335G>T), D1393N (5339G>A 5341C>T), Q1394S (5342C>T 5343A>C), R1397N (5352G>A 5353G>T), G1398H (5354G>C 5355G>A 5356G>T), T1399L (5357A>C 5358C>T 5359A>T), E1400S (5360G>A 5361A>G 5362A>C), E1401H (5363G>C 5365A>T), I1403S (5369A>T 5370T>C 5371A>T), S1404I (5372T>A 5373C>T 5374A>T), I1410V (5390A>G 5392A>T), V1412I (5396G>A 5398C>A), P1415E (5405C>G 5406C>A), L1417I (5411C>A), E1418D (5416A>T), T1419S (5417A>T), A1421F (5423G>T 5424C>T 5425A>C), E1425N (5435G>A 5437G>T), K1426A (5438A>G 5439A>C 5440A>T), M1427F (5441A>T 5443G>C), L1428F (5444C>T 5446G>C), A1429K (5447G>A 5448C>A), I1430T (5451T>C 5452C>T), C1431I (5453T>A 5454G>T 5455C>T), D1433I (5459G>A 5460A>T), L1438I (5474C>A 5476G>T), P1440A (5480C>G), T1441K (5484C>A 5485A>G), M1443I (5491G>A) | | | | | | | | |
| Codon mutations: | TGG1362..A (5248G>A), GCG1364GCT (5254G>T), GTG1366AAT (5258G>A 5259T>A 5260G>T), GTA1367GTC (5263A>C), CCA1368CCC (5266A>C), GGA1369TTT (5267G>T 5268G>T 5269A>T), TTA1371CAA (5273T>C 5274T>A), GAG1373GAA (5281G>A), CTG1375AAT (5285C>A 5286T>A 5287G>T), CCA1378CCT (5296A>T), CTA1381CTT (5305A>T), AGA1382AAA (5307G>A), GCA1384GCC (5314A>C), ACC1387GAA (5321A>G 5322C>A 5323C>A), TTC1388TTT (5326C>T), AGG1390AGT (5332G>T), AAG1391ATT (5334A>T 5335G>T), GAC1393AAT (5339G>A 5341C>T), CAA1394TCA (5342C>T 5343A>C), TGT1395-TT (5345delT 5346G>T), AGG1397AAT (5352G>A 5353G>T), GGG1398CAT (5354G>C 5355G>A 5356G>T), ACA1399CTT (5357A>C 5358C>T 5359A>T), GAA1400AGC (5360G>A 5361A>G 5362A>C), GAA1401CAT (5363G>C 5365A>T), ATA1403TCT (5369A>T 5370T>C 5371A>T), TCA1404ATT (5372T>A 5373C>T 5374A>T), GAC1408GAT (5386C>T), GAT1409GAC (5389T>C), ATA1410GTT (5390A>G 5392A>T), CTC1411TTA (5393C>T 5395C>A), GTC1412ATA (5396G>A 5398C>A), TCC1414TCA (5404C>A), CCA1415GAA (5405C>G 5406C>A), CTT1417ATT (5411C>A), GAA1418GAT (5416A>T), ACT1419TCT (5417A>T), GCA1421TTC (5423G>T 5424C>T 5425A>C), TTG1424CTC (5432T>C 5434G>C), GAG1425AAT (5435G>A 5437G>T), AAA1426GCT (5438A>G 5439A>C 5440A>T), ATG1427TTC (5441A>T 5443G>C), CTG1428TTC (5444C>T 5446G>C), GCA1429AAA (5447G>A 5448C>A), ATC1430ACT (5451T>C 5452C>T), TGC1431ATT (5453T>A 5454G>T 5455C>T), GAC1433ATC (5459G>A 5460A>T), AAT1434AAC (5464T>C), CTA1436CTT (5470A>T), CTG1438ATT (5474C>A 5476G>T), TCA1439AGT (5477T>A 5478C>G 5479A>T), CCA1440GCA (5480C>G), ACA1441AAG (5484C>A 5485A>G), ATG1443ATA (5491G>A) | | | | | | | | |

##### Proteins

| | Begin | End | Coverage | Score | Concordance | Matches | Identities | I/D/M/F* | Stop Codons |
|---|---|---|---|---|---|---|---|---|---|
| polyprotein (YP_010800602.1) | 1363 | 1444 | 4.4% | 195 | 32.4% | 82 (100%) | 44 (53.7%) | 0/0/1/1 | 0 |
| Protein mutations: | V1366N (5258G>A 5259T>A 5260G>T), G1369F (5267G>T 5268G>T 5269A>T), L1371Q (5273T>C 5274T>A), L1375N (5285C>A 5286T>A 5287G>T), R1382K (5307G>A), T1387E (5321A>G 5322C>A 5323C>A), R1390S (5332G>T), K1391I (5334A>T 5335G>T), D1393N (5339G>A 5341C>T), Q1394S (5342C>T 5343A>C), R1397N (5352G>A 5353G>T), G1398H (5354G>C 5355G>A 5356G>T), T1399L (5357A>C 5358C>T 5359A>T), E1400S (5360G>A 5361A>G 5362A>C), E1401H (5363G>C 5365A>T), I1403S (5369A>T 5370T>C 5371A>T), S1404I (5372T>A 5373C>T 5374A>T), I1410V (5390A>G 5392A>T), V1412I (5396G>A 5398C>A), P1415E (5405C>G 5406C>A), L1417I (5411C>A), E1418D (5416A>T), T1419S (5417A>T), A1421F (5423G>T 5424C>T 5425A>C), E1425N (5435G>A 5437G>T), K1426A (5438A>G 5439A>C 5440A>T), M1427F (5441A>T 5443G>C), L1428F (5444C>T 5446G>C), A1429K (5447G>A 5448C>A), I1430T (5451T>C 5452C>T), C1431I (5453T>A 5454G>T 5455C>T), D1433I (5459G>A 5460A>T), L1438I (5474C>A 5476G>T), P1440A (5480C>G), T1441K (5484C>A 5485A>G), M1443I (5491G>A) | | | | | | | | |
| Codon mutations: | TGG1362..A (5248G>A), GCG1364GCT (5254G>T), GTG1366AAT (5258G>A 5259T>A 5260G>T), GTA1367GTC (5263A>C), CCA1368CCC (5266A>C), GGA1369TTT (5267G>T 5268G>T 5269A>T), TTA1371CAA (5273T>C 5274T>A), GAG1373GAA (5281G>A), CTG1375AAT (5285C>A 5286T>A 5287G>T), CCA1378CCT (5296A>T), CTA1381CTT (5305A>T), AGA1382AAA (5307G>A), GCA1384GCC (5314A>C), ACC1387GAA (5321A>G 5322C>A 5323C>A), TTC1388TTT (5326C>T), AGG1390AGT (5332G>T), AAG1391ATT (5334A>T 5335G>T), GAC1393AAT (5339G>A 5341C>T), CAA1394TCA (5342C>T 5343A>C), TGT1395-TT (5345delT 5346G>T), AGG1397AAT (5352G>A 5353G>T), GGG1398CAT (5354G>C 5355G>A 5356G>T), ACA1399CTT (5357A>C 5358C>T 5359A>T), GAA1400AGC (5360G>A 5361A>G 5362A>C), GAA1401CAT (5363G>C 5365A>T), ATA1403TCT (5369A>T 5370T>C 5371A>T), TCA1404ATT (5372T>A 5373C>T 5374A>T), GAC1408GAT (5386C>T), GAT1409GAC (5389T>C), ATA1410GTT (5390A>G 5392A>T), CTC1411TTA (5393C>T 5395C>A), GTC1412ATA (5396G>A 5398C>A), TCC1414TCA (5404C>A), CCA1415GAA (5405C>G 5406C>A), CTT1417ATT (5411C>A), GAA1418GAT (5416A>T), ACT1419TCT (5417A>T), GCA1421TTC (5423G>T 5424C>T 5425A>C), TTG1424CTC (5432T>C 5434G>C), GAG1425AAT (5435G>A 5437G>T), AAA1426GCT (5438A>G 5439A>C 5440A>T), ATG1427TTC (5441A>T 5443G>C), CTG1428TTC (5444C>T 5446G>C), GCA1429AAA (5447G>A 5448C>A), ATC1430ACT (5451T>C 5452C>T), TGC1431ATT (5453T>A 5454G>T 5455C>T), GAC1433ATC (5459G>A 5460A>T), AAT1434AAC (5464T>C), CTA1436CTT (5470A>T), CTG1438ATT (5474C>A 5476G>T), TCA1439AGT (5477T>A 5478C>G 5479A>T), CCA1440GCA (5480C>G), ACA1441AAG (5484C>A 5485A>G), ATG1443ATA (5491G>A) | | | | | | | | |

*: Inserts / Deletes / Misaligned / Frameshifts

#### Analysis details

This analysis was performed with panviral2.64

### NGS Details (UN60): Escherichia virus DE3

#### Assembly

| | |
|---|---|
| **Coverage Length** | 219 (1 contig(s)) |
| **Depth Of Coverage** | 2.5 |
| **Number Of Reads** | 4 |
| **Reads Per Million** | 0.08 rpm (after QC) |
| **Ambiguities** | 0 |
| **Assembly Method** | de novo + reference guided assembly |
| **Consensus Caller** | Bcf Tools |

#### Coverage Map

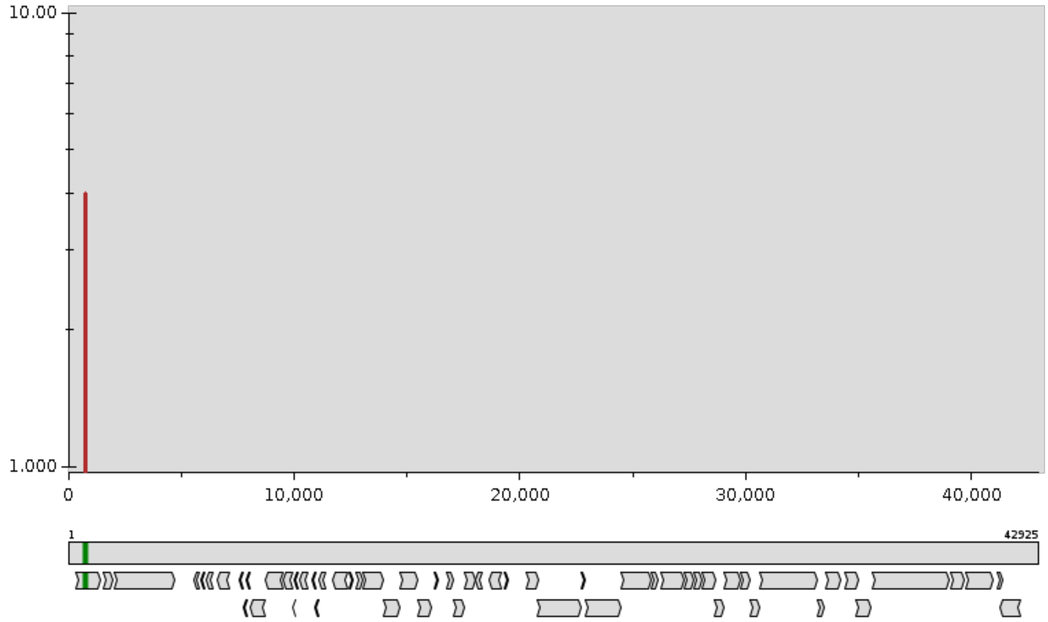

#### Assignment

| | |
|---|---|
| **Type** | Escherichia virus DE3 (Taxonomy ID: 2169967) |
| **Reference Genome** | NC_042057.1 |
| **NT Identity (%)** | 100.0 |
| **AA Identity (%)** | 98.6301 |
| **Number Of Stop Codons** | 0 |
| **Number Of CDS** | 57 |

#### Alignment

| | |
|---|---|
| **Alignment Score** | 438.0 (NT) + 491.0 (AA) = 929.0 |
| **Concordance (%)** | 100.0 |

| Alignment Method | Local, heuristic, nucleotide (BLASTN) |
|---|---|

#### Genome Region

Sequence starts at position 657 and ends at position 875 relative to NC_042057.1 reference sequence.

#### Alignment Detailed Statistics

| | Begin | End | Coverage | Score | Concordance | Matches | Identities | I/D/M/F * | Stop Codons |
|---|---|---|---|---|---|---|---|---|---|
| **NT** | **657** | **875** | **0.5%** | **438** | **100%** | **219 (100%)** | **219 (100%)** | **0/0** | |
| Mutations: | *none* | | | | | | | | |
| **CDS** | | | | | | | | | |
| lacI | 107 | 179 | 20.2% | 491 | 96.3% | 73 (100%) | 72 (98.6%) | 0/0/0/0 | 0 |
| **Proteins** | | | | | | | | | |
| DNA-binding transcriptional repressor LacI (YP_009617199.1) | 107 | 179 | 20.2% | 491 | 96.3% | 73 (100%) | 72 (98.6%) | 0/0/0/0 | 0 |
| Protein mutations: | *none* | | | | | | | | |
| Codon mutations: | *none* | | | | | | | | |

*: Inserts / Deletes / Misaligned / Frameshifts

#### Analysis details

This analysis was performed with panviral2.64

### NGS Details (UN60): Epiphyllum badnavirus 1

#### Assembly

| | |
|---|---|
| **Coverage Length** | 267 (1 contig(s)) |
| **Depth Of Coverage** | 2.0 |
| **Number Of Reads** | 4 |
| **Reads Per Million** | 0.08 rpm (after QC) |
| **Ambiguities** | 0 |
| **Assembly Method** | de novo + reference guided assembly |
| **Consensus Caller** | Bcf Tools |

#### Coverage Map

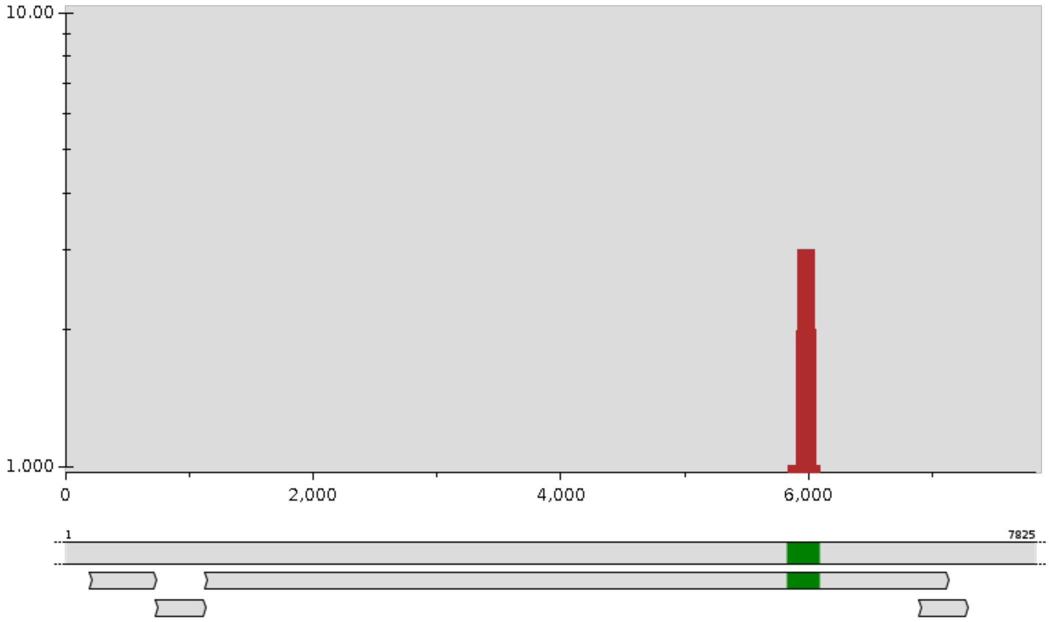

#### Assignment

| | |
|---|---|
| **Type** | Epiphyllum badnavirus 1 (Taxonomy ID: 2518008) |
| **Reference Genome** | NC_076247.1 |
| **NT Identity (%)** | 54.8148 |
| **AA Identity (%)** | 50.0 |
| **Number Of Stop Codons** | 0 |
| **Number Of CDS** | 4 |

#### Alignment

| | |
|---|---|
| **Alignment Score** | 46.0 (NT) + 315.0 (AA) = 361.0 |
| **Concordance (%)** | 30.1839 |

| Alignment Method | Global, seeded, nucleotide + amino acids (AGA) |
|---|---|

#### Genome Region

Sequence starts at position 5825 and ends at position 6091 relative to NC_076247.1 reference sequence.

#### Alignment Detailed Statistics

| | Begin | End | Coverage | Score | Concordance | Matches | Identities | I/D/M/F* | Stop Codons |
|---|---|---|---|---|---|---|---|---|---|
| **NT** | **5825** | **6091** | **3.4%** | **46** | **8.6%** | **267 (98.9%)** | **148 (54.8%)** | **3/0** | |
| Mutations: | 5826C>G, 5830A>G, 5831A>G, 5832T>A, 5835C>T, 5836A>T, 5837G>C, 5841C>G, 5851T>A, 5853C>A, 5859G>T, 5861A>G, 5862G>A, 5863A>G, 5864G>C, 5865C>T, 5868A>C, 5870T>A, 5871C>T, 5877G>A, 5878A>C, 5880T>A, 5882T>G, 5883G>A, 5887G>A, 5888A>T, 5892A>G, 5893G>T, 5894A>G, 5895A>T, 5896T>G, 5897C>A, 5900T>C, 5901C>A, 5903C>A, 5905T>A, 5906G>A, 5907G>A, 5910C>G, 5913C>T, 5917C>A, 5918T>A, 5919G>A, 5922C>T, 5924C>A, 5925A>T, 5928T>G, 5931A>T, 5933T>A, 5937C>T, 5942G>T, 5943G>T, 5955A>T, 5958C>T, 5961A>C, 5966A>C, 5967G>A, 5976T>A, 5977G>T, 5980G>T, 5981T>C, 5982G>A, 5985C>T, 5988A>G, 5989C>A, 5991A>C, 5992A>C, 5993A>T, 5994G>C, 5998G>A, 6000C>T, 6001A>G, 6002A>T, 6003C>T, 6004T>G, 6005G>T, 6011G>A, 6013G>C, 6014A>C, 6015C>A, 6016C>T, 6018G>A, 6020C>T, 6021A>G, 6022G>A, 6024C>G, 6024_6025insAAA, 6026T>C, 6027T>A, 6030A>G, 6031G>C, 6032C>T, 6033A>G, 6038A>T, 6039C>T, 6040A>T, 6043G>T, 6052C>T, 6055G>A, 6057C>T, 6059T>A, 6060C>T, 6063T>C, 6066C>G, 6069T>C, 6070G>A, 6071A>T, 6072A>G, 6073G>T, 6074A>C, 6076G>A, 6077A>C, 6078C>T, 6082G>A, 6083C>T, 6084A>T, 6085C>G, 6087G>T, 6090C>T, 6091C>G | | | | | | | | |
| **CDS** | | | | | | | | | |
| QKM20_gp3 | 1567 | 1655 | 4.4% | 315 | 46.9% | 89 (98.9%) | 45 (50.0%) | 1/0/0/0 | 0 |
| Protein mutations: | N1568G (5830A>G 5831A>G 5832T>A), F1575I (5851T>A 5853C>A), K1578R (5861A>G 5862G>A), S1579A (5863A>G 5864G>C 5865C>T), F1581Y (5870T>A 5871C>T), I1584L (5878A>C 5880T>A), M1585R (5882T>G 5883G>A), D1587I (5887G>A 5888A>T), E1589C (5893G>T 5894A>G 5895A>T), S1590D (5896T>G 5897C>A), I1591T (5900T>C 5901C>A), P1592H (5903C>A), W1593K (5905T>A 5906G>A 5907G>A), L1597K (5917C>A 5918T>A 5919G>A), P1599H (5924C>A 5925A>T), D1600E (5928T>G), L1602H (5933T>A), W1605F (5942G>T 5943G>T), K1613T (5966A>C 5967G>A), A1617S (5977G>T), V1618S (5980G>T 5981T>C 5982G>A), R1621S (5989C>A 5991A>C), K1622L (5992A>C 5993A>T 5994G>C), D1624N (5998G>A 6000C>T), N1625V (6001A>G 6002A>T 6003C>T), C1626V (6004T>G 6005G>T), R1628K (6011G>A), D1629P (6013G>C 6014A>C 6015C>A), S1631L (6020C>T 6021A>G), G1632R (6022G>A 6024C>G), G1632_F1633insK (6024_6025insAAA), F1633S (6026T>C 6027T>A), A1635L (6031G>C 6032C>T 6033A>G), Y1637F (6038A>T 6039C>T), I1638F (6040A>T), D1639Y (6043G>T), V1643I (6055G>A 6057C>T), F1644Y (6059T>A 6060C>T), N1646K (6066C>G), E1648M (6070G>A 6071A>T 6072A>G), D1649S (6073G>T 6074A>C), D1650T (6076G>A 6077A>C 6078C>T), A1652I (6082G>A 6083C>T 6084A>T), Q1653D (6085C>G 6087G>T) | | | | | | | | |
| Codon mutations: | GTC1566.TG (5826C>G), AAT1568GGA (5830A>G 5831A>G 5832T>A), GCC1569GCT (5835C>T), AGT1570TCT (5836A>T 5837G>C), GTC1571GTG (5841C>G), TTC1575ATA (5851T>A 5853C>A), CTG1577CTT (5859G>T), AAG1578AGA (5861A>G 5862G>A), AGC1579GCT (5863A>G 5864G>C 5865C>T), GGA1580GGC (5868A>C), TTC1581TAT (5870T>A 5871C>T), CAG1583CAA (5877G>A), ATT1584CTA (5878A>C 5880T>A), ATG1585AGA (5882T>G 5883G>A), GAC1587ATC (5887G>A 5888A>T), GAA1588GAG (5892A>G), GAA1589TGT (5893G>T 5894A>G 5895A>T), TCC1590GAC (5896T>G 5897C>A), ATC1591ACA (5900T>C 5901C>A), CCT1592CAT (5903C>A), TGG1593AAA (5905T>A 5906G>A 5907G>A), ACC1594ACG (5910C>G), GCC1595GCT (5913C>T), CTG1597AAA (5917C>A 5918T>A 5919G>A), ACC1598ACT (5922C>T), CCA1599CAT (5924C>A 5925A>T), GAT1600GAG (5928T>G), GGA1601GGT (5931A>T), CTT1602CAT (5933T>A), TAC1603TAT (5937C>T), TGG1605TTT (5942G>T 5943G>T), CCA1609CCT (5955A>T), TTC1610TTT (5958C>T), GGA1611GGC (5961A>C), AAG1613ACA (5966A>C 5967G>A), CCT1616CCA (5976T>A), GCT1617TCT (5977G>T), GTG1618TCA (5980G>T 5981T>C 5982G>A), TTC1619TTT (5985C>T), CAA1620CAG (5988A>G), CGA1621AGC (5989C>A 5991A>C), AAG1622CTC (5992A>C 5993A>T 5994G>C), GAC1624AAT (5998G>A 6000C>T), AAC1625GTT (6001A>G 6002A>T 6003C>T), TGT1626GTT (6004T>G 6005G>T), AGA1628AAA (6011G>A), GAC1629CCA (6013G>C 6014A>C 6015C>A), CTG1630TTA (6016C>T 6018G>A), TCA1631TTG (6020C>T 6021A>G), GGC1632AGG (6022G>A 6024C>G), GGC1632_TTT1633insAAA (6024_6025insAAA), TTT1633TCA (6026T>C 6027T>A), GTA1634GTG (6030A>G), GCA1635CTG (6031G>C 6032C>T 6033A>G), TAC1637TTT (6038A>T 6039C>T), ATT1638TTT (6040A>T), GAT1639TAT (6043G>T), CTG1642TTG (6052C>T), GTC1643ATT (6055G>A 6057C>T), TTC1644TAT (6059T>A 6060C>T), AGT1645AGC (6063T>C), AAC1646AAG (6066C>G), ACT1647ACC (6069T>C), GAA1648ATG (6070G>A 6071A>T 6072A>G), GAT1649TCT (6073G>T 6074A>C), GAC1650ACT (6076G>A 6077A>C 6078C>T), GCA1652ATT (6082G>A 6083C>T 6084A>T), CAG1653GAT (6085C>G 6087G>T), CAC1654CAT (6090C>T), CTG1655G.. (6091C>G) | | | | | | | | |
| **Proteins** | | | | | | | | | |
| polyprotein (YP_010797894.1) | 1567 | 1655 | 4.4% | 315 | 46.9% | 89 (98.9%) | 45 (50.0%) | 1/0/0/0 | 0 |
| Protein mutations: | N1568G (5830A>G 5831A>G 5832T>A), F1575I (5851T>A 5853C>A), K1578R (5861A>G 5862G>A), S1579A (5863A>G 5864G>C 5865C>T), F1581Y (5870T>A 5871C>T), I1584L (5878A>C 5880T>A), M1585R (5882T>G 5883G>A), D1587I (5887G>A 5888A>T), E1589C (5893G>T 5894A>G 5895A>T), S1590D (5896T>G 5897C>A), I1591T (5900T>C 5901C>A), P1592H (5903C>A), W1593K (5905T>A 5906G>A 5907G>A), L1597K (5917C>A 5918T>A 5919G>A), P1599H (5924C>A 5925A>T), D1600E (5928T>G), L1602H (5933T>A), W1605F (5942G>T 5943G>T), K1613T (5966A>C 5967G>A), A1617S (5977G>T), V1618S (5980G>T 5981T>C 5982G>A), R1621S (5989C>A 5991A>C), K1622L (5992A>C 5993A>T 5994G>C), D1624N (5998G>A 6000C>T), N1625V (6001A>G 6002A>T 6003C>T), C1626V (6004T>G 6005G>T), R1628K (6011G>A), D1629P (6013G>C 6014A>C 6015C>A), S1631L (6020C>T 6021A>G), G1632R (6022G>A 6024C>G), G1632_F1633insK (6024_6025insAAA), F1633S (6026T>C 6027T>A), A1635L (6031G>C 6032C>T 6033A>G), Y1637F (6038A>T 6039C>T), I1638F (6040A>T), D1639Y (6043G>T), V1643I (6055G>A 6057C>T), F1644Y (6059T>A 6060C>T), N1646K (6066C>G), E1648M (6070G>A 6071A>T 6072A>G), D1649S (6073G>T 6074A>C), D1650T (6076G>A 6077A>C 6078C>T), A1652I (6082G>A 6083C>T 6084A>T), Q1653D (6085C>G 6087G>T) | | | | | | | | |
| Codon mutations: | GTC1566.TG (5826C>G), AAT1568GGA (5830A>G 5831A>G 5832T>A), GCC1569GCT (5835C>T), AGT1570TCT (5836A>T 5837G>C), GTC1571GTG (5841C>G), TTC1575ATA (5851T>A 5853C>A), CTG1577CTT (5859G>T), AAG1578AGA (5861A>G 5862G>A), AGC1579GCT (5863A>G 5864G>C 5865C>T), GGA1580GGC (5868A>C), TTC1581TAT (5870T>A 5871C>T), CAG1583CAA (5877G>A), ATT1584CTA (5878A>C 5880T>A), ATG1585AGA (5882T>G 5883G>A), GAC1587ATC (5887G>A 5888A>T), GAA1588GAG (5892A>G), GAA1589TGT (5893G>T 5894A>G 5895A>T), TCC1590GAC (5896T>G 5897C>A), ATC1591ACA (5900T>C 5901C>A), CCT1592CAT (5903C>A), TGG1593AAA (5905T>A 5906G>A 5907G>A), ACC1594ACG (5910C>G), GCC1595GCT (5913C>T), CTG1597AAA (5917C>A 5918T>A 5919G>A), ACC1598ACT (5922C>T), CCA1599CAT (5924C>A 5925A>T), GAT1600GAG (5928T>G), GGA1601GGT (5931A>T), CTT1602CAT (5933T>A), TAC1603TAT (5937C>T), TGG1605TTT (5942G>T 5943G>T), CCA1609CCT (5955A>T), TTC1610TTT (5958C>T), GGA1611GGC (5961A>C), AAG1613ACA (5966A>C 5967G>A), CCT1616CCA (5976T>A), GCT1617TCT (5977G>T), GTG1618TCA (5980G>T 5981T>C 5982G>A), TTC1619TTT (5985C>T), CAA1620CAG (5988A>G), CGA1621AGC (5989C>A 5991A>C), AAG1622CTC (5992A>C 5993A>T 5994G>C), GAC1624AAT (5998G>A 6000C>T), AAC1625GTT (6001A>G 6002A>T 6003C>T), TGT1626GTT (6004T>G 6005G>T), AGA1628AAA (6011G>A), GAC1629CCA (6013G>C 6014A>C 6015C>A), CTG1630TTA (6016C>T 6018G>A), TCA1631TTG (6020C>T 6021A>G), GGC1632AGG (6022G>A 6024C>G), GGC1632_TTT1633insAAA (6024_6025insAAA), TTT1633TCA (6026T>C 6027T>A), GTA1634GTG (6030A>G), GCA1635CTG (6031G>C 6032C>T 6033A>G), TAC1637TTT (6038A>T 6039C>T), ATT1638TTT (6040A>T), GAT1639TAT (6043G>T), CTG1642TTG (6052C>T), GTC1643ATT (6055G>A 6057C>T), TTC1644TAT (6059T>A 6060C>T), AGT1645AGC (6063T>C), AAC1646AAG (6066C>G), ACT1647ACC (6069T>C), GAA1648ATG (6070G>A 6071A>T 6072A>G), GAT1649TCT (6073G>T 6074A>C), GAC1650ACT (6076G>A 6077A>C 6078C>T), GCA1652ATT (6082G>A 6083C>T 6084A>T), CAG1653GAT (6085C>G 6087G>T), CAC1654CAT (6090C>T), CTG1655G.. (6091C>G) | | | | | | | | |

*: Inserts / Deletes / Misaligned / Frameshifts

#### Analysis details

This analysis was performed with panviral2.64

### NGS Details (UN60): Goose dicistrovirus

#### Assembly

| | |
|---|---|
| **Coverage Length** | 240 (1 contig(s)) |
| **Depth Of Coverage** | 2.2 |
| **Number Of Reads** | 4 |
| **Reads Per Million** | 0.08 rpm (after QC) |
| **Ambiguities** | 0 |
| **Assembly Method** | de novo + reference guided assembly |
| **Consensus Caller** | Bcf Tools |

#### Coverage Map

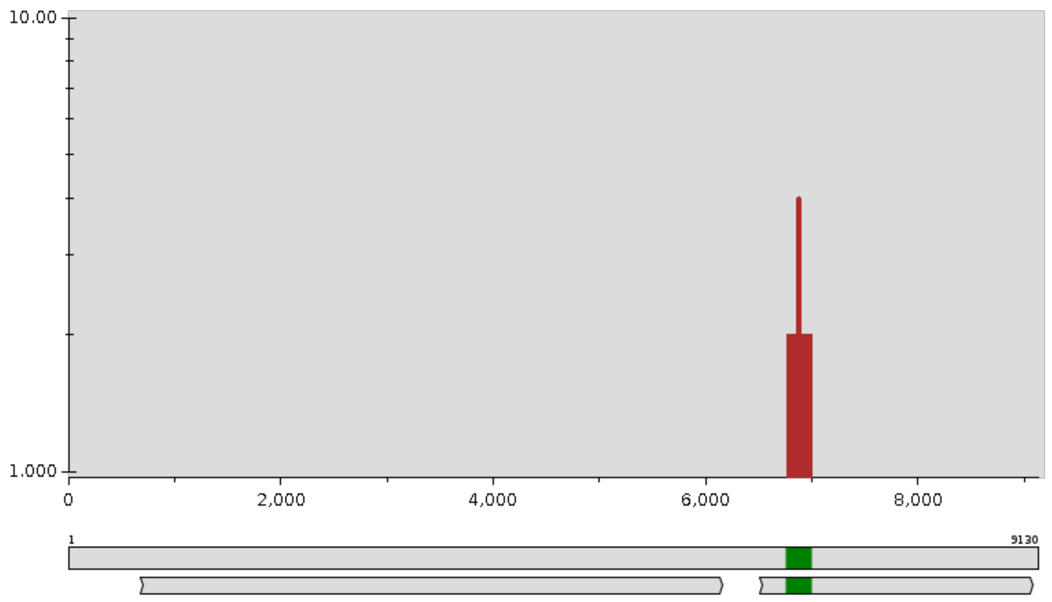

#### Assignment

| | |
|---|---|
| **Type** | Goose dicistrovirus (Taxonomy ID: 1776109) |
| **Reference Genome** | NC_029052.1 |
| **NT Identity (%)** | 85.0 |
| **AA Identity (%)** | 95.0 |
| **Number Of Stop Codons** | 0 |
| **Number Of CDS** | 2 |

#### Alignment

| | |
|---|---|
| **Alignment Score** | 336.0 (NT) + 540.0 (AA) = 876.0 |
| **Concordance (%)** | 84.8015 |

| Alignment Method | Global, seeded, nucleotide + amino acids (AGA) |
| --- | --- |

#### Genome Region

Sequence starts at position 6764 and ends at position 7003 relative to NC_029052.1 reference sequence.

#### Alignment Detailed Statistics

| | Begin | End | Coverage | Score | Concordance | Matches | Identities | I/D/M/F* | Stop Codons |
| --- | --- | --- | --- | --- | --- | --- | --- | --- | --- |
| **NT** | **6764** | **7003** | **2.6%** | **336** | **70.0%** | **240 (100%)** | **204 (85.0%)** | **0/0** | |
| Mutations: | 6774C>T, 6780G>A, 6792T>C, 6795T>C, 6801A>G, 6802G>A, 6805C>T, 6807T>A, 6819T>A, 6828G>A, 6837G>T, 6846A>C, 6852T>C, 6855A>T, 6864T>C, 6870T>G, 6873A>G, 6879G>A, 6882G>A, 6885A>T, 6886A>C, 6900C>G, 6903G>C, 6907G>A, 6912T>A, 6924G>A, 6927T>A, 6943T>C, 6945G>A, 6955T>C, 6960T>G, 6972A>G, 6978T>C, 6993C>T, 7000C>T, 7002T>A | | | | | | | | |

##### CDS

| | Begin | End | Coverage | Score | Concordance | Matches | Identities | I/D/M/F* | Stop Codons |
| --- | --- | --- | --- | --- | --- | --- | --- | --- | --- |
| AXF40_gp2 | 87 | 166 | 9.3% | 540 | 95.6% | 80 (100%) | 76 (95.0%) | 0/0/0/0 | 0 |
| Protein mutations: | A99T (6802G>A), V134I (6907G>A), Y150H (6955T>C) | | | | | | | | |
| Codon mutations: | CCC89CCT (6774C>T), CAG91CAA (6780G>A), GGT95GGC (6792T>C), AAT96AAC (6795T>C), ACA98ACG (6801A>G), GCA99ACA (6802G>A), CTT100TTA (6805C>T 6807T>A), ACT104ACA (6819T>A), GGG107GGA (6828G>A), TCG110TCT (6837G>T), GGA113GGC (6846A>C), GAT115GAC (6852T>C), ATA116ATT (6855A>T), TGT119TGC (6864T>C), GGT121GGG (6870T>G), TCA122TCG (6873A>G), CCG124CCA (6879G>A), CAG125CAA (6882G>A), TCA126TCT (6885A>T), AGA127CGA (6886A>C), GCC131GCG (6900C>G), ACG132ACC (6903G>C), GTT134ATT (6907G>A), ATT135ATA (6912T>A), TCG139TCA (6924G>A), CCT140CCA (6927T>A), TTG146CTA (6943T>C 6945G>A), TAT150CAT (6955T>C), GGT151GGG (6960T>G), CAA155CAG (6972A>G), AAT157AAC (6978T>C), AGC162AGT (6993C>T), CTT165TTA (7000C>T 7002T>A) | | | | | | | | |

##### Proteins

| | Begin | End | Coverage | Score | Concordance | Matches | Identities | I/D/M/F* | Stop Codons |
| --- | --- | --- | --- | --- | --- | --- | --- | --- | --- |
| capsid protein precursor (YP_009221982.1) | 87 | 166 | 9.3% | 540 | 95.6% | 80 (100%) | 76 (95.0%) | 0/0/0/0 | 0 |
| Protein mutations: | A99T (6802G>A), V134I (6907G>A), Y150H (6955T>C) | | | | | | | | |
| Codon mutations: | CCC89CCT (6774C>T), CAG91CAA (6780G>A), GGT95GGC (6792T>C), AAT96AAC (6795T>C), ACA98ACG (6801A>G), GCA99ACA (6802G>A), CTT100TTA (6805C>T 6807T>A), ACT104ACA (6819T>A), GGG107GGA (6828G>A), TCG110TCT (6837G>T), GGA113GGC (6846A>C), GAT115GAC (6852T>C), ATA116ATT (6855A>T), TGT119TGC (6864T>C), GGT121GGG (6870T>G), TCA122TCG (6873A>G), CCG124CCA (6879G>A), CAG125CAA (6882G>A), TCA126TCT (6885A>T), AGA127CGA (6886A>C), GCC131GCG (6900C>G), ACG132ACC (6903G>C), GTT134ATT (6907G>A), ATT135ATA (6912T>A), TCG139TCA (6924G>A), CCT140CCA (6927T>A), TTG146CTA (6943T>C 6945G>A), TAT150CAT (6955T>C), GGT151GGG (6960T>G), CAA155CAG (6972A>G), AAT157AAC (6978T>C), AGC162AGT (6993C>T), CTT165TTA (7000C>T 7002T>A) | | | | | | | | |

*: Inserts / Deletes / Misaligned / Frameshifts

#### Analysis details

This analysis was performed with panviral2.64

### NGS Details (UN60): Melanoplus sanguinipes entomopoxvirus

#### Assembly

| | |
|---|---|
| **Coverage Length** | 142 (1 contig(s)) |
| **Depth Of Coverage** | 1.4 |
| **Number Of Reads** | 2 |
| **Reads Per Million** | 0.04 rpm (after QC) |
| **Ambiguities** | 0 |
| **Assembly Method** | de novo + reference guided assembly |
| **Consensus Caller** | Bcf Tools |

#### Coverage Map

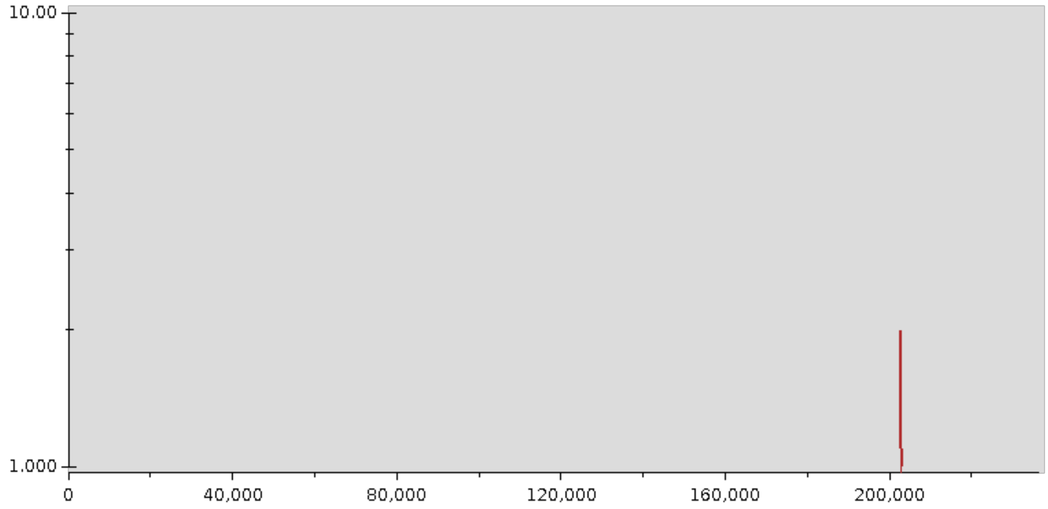

#### Assignment

| | |
|---|---|
| **Type** | Melanoplus sanguinipes entomopoxvirus (Taxonomy ID: 83191) |
| **Reference Genome** | NC_001993.1 |
| **NT Identity (%)** | 81.6901 |
| **AA Identity (%)** | 76.5957 |
| **Number Of Stop Codons** | 0 |
| **Number Of CDS** | 267 |

#### Alignment

| | |
|---|---|
| **Alignment Score** | 180.0 (NT) + 336.0 (AA) = 516.0 |
| **Concordance (%)** | 75.7709 |
| **Alignment Method** | Local, heuristic, nucleotide (BLASTN) |

#### Genome Region

Sequence starts at position 202763 and ends at position 202904 relative to NC_001993.1 reference sequence.

#### Alignment Detailed Statistics

| | Begin | End | Coverage | Score | Concordance | Matches | Identities | I/D/M/F* | Stop Codons |
|---|---|---|---|---|---|---|---|---|---|
| **NT** | **202763** | **202904** | **0.1%** | **180** | **63.4%** | **142 (100%)** | **116 (81.7%)** | **0/0** | |

Mutations: 202768T>C, 202771A>T, 202773G>T, 202774T>C, 202775C>T, 202777A>G, 202784T>G, 202789A>G, 202790C>T, 202807A>T, 202813T>A, 202817G>T, 202818G>C, 202831T>C, 202832G>C, 202834T>C, 202835C>T, 202836T>G, 202852A>T, 202862A>G, 202874T>A, 202876T>A, 202880T>C, 202885A>G, 202891T>A, 202897T>C

*: Inserts / Deletes / Misaligned / Frameshifts

#### Analysis details

This analysis was performed with panviral2.64

### NGS Details (UN60): Cotia virus

#### Assembly

| | |
|---|---|
| **Coverage Length** | 135 (1 contig(s)) |
| **Depth Of Coverage** | 1.8 |
| **Number Of Reads** | 2 |
| **Reads Per Million** | 0.04 rpm (after QC) |
| **Ambiguities** | 0 |
| **Assembly Method** | de novo + reference guided assembly |
| **Consensus Caller** | Bcf Tools |

#### Coverage Map

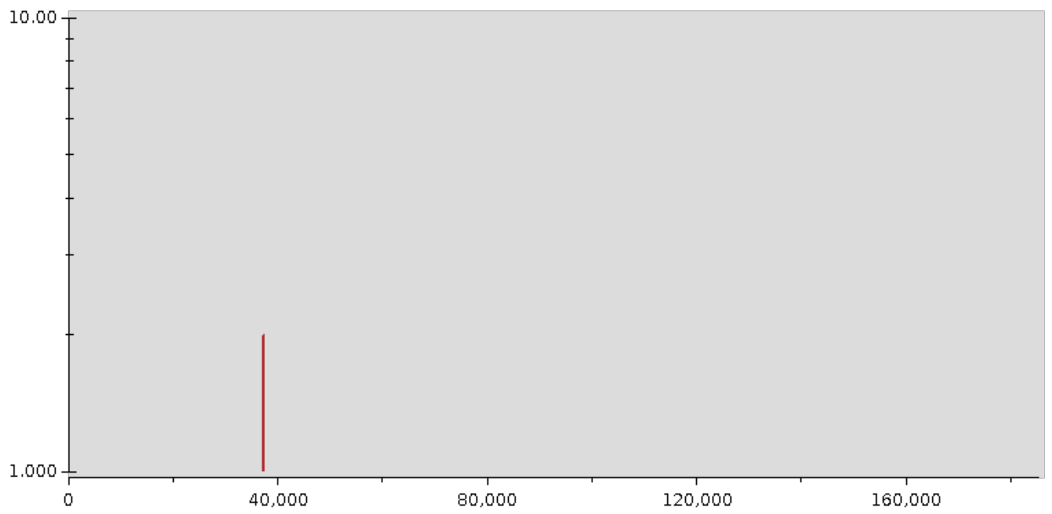

#### Assignment

| | |
|---|---|
| **Type** | Cotia virus (Taxonomy ID: 39444) |
| **Reference Genome** | NC_016924.1 |
| **NT Identity (%)** | 80.0 |
| **AA Identity (%)** | 93.3333 |
| **Number Of Stop Codons** | 0 |
| **Number Of CDS** | 185 |

#### Alignment

| | |
|---|---|
| **Alignment Score** | 162.0 (NT) + 285.0 (AA) = 447.0 |
| **Concordance (%)** | 78.4211 |
| **Alignment Method** | Local, heuristic, nucleotide (BLASTN) |

#### Genome Region

Sequence starts at position 37199 and ends at position 37333 relative to NC_016924.1 reference sequence.

#### Alignment Detailed Statistics

| | Begin | End | Coverage | Score | Concordance | Matches | Identities | I/D/M/F* | Stop Codons |
|---|---|---|---|---|---|---|---|---|---|
| NT | 37199 | 37333 | 0.1% | 162 | 60.0% | 135 (100%) | 108 (80.0%) | 0/0 | |

Mutations: 37209C>G, 37210T>A, 37211T>A, 37216A>G, 37223A>G, 37224G>C, 37225A>T, 37229G>A, 37235T>A, 37247A>T, 37256T>C, 37259A>C, 37261T>A, 37270T>A, 37274T>A, 37281G>C, 37282A>T, 37289A>G, 37292G>A, 37295A>T, 37307T>A, 37310T>A, 37319T>A, 37322A>T, 37325A>C, 37329G>T, 37330T>C

*: Inserts / Deletes / Misaligned / Frameshifts

#### Analysis details

This analysis was performed with panviral2.64

### NGS Details (UN60): Scottvirus scott

#### Assembly

| | |
|---|---|
| **Coverage Length** | 244 (1 contig(s)) |
| **Depth Of Coverage** | 1.1 |
| **Number Of Reads** | 2 |
| **Reads Per Million** | 0.04 rpm (after QC) |
| **Ambiguities** | 0 |
| **Assembly Method** | de novo + reference guided assembly |
| **Consensus Caller** | Bcf Tools |

#### Coverage Map

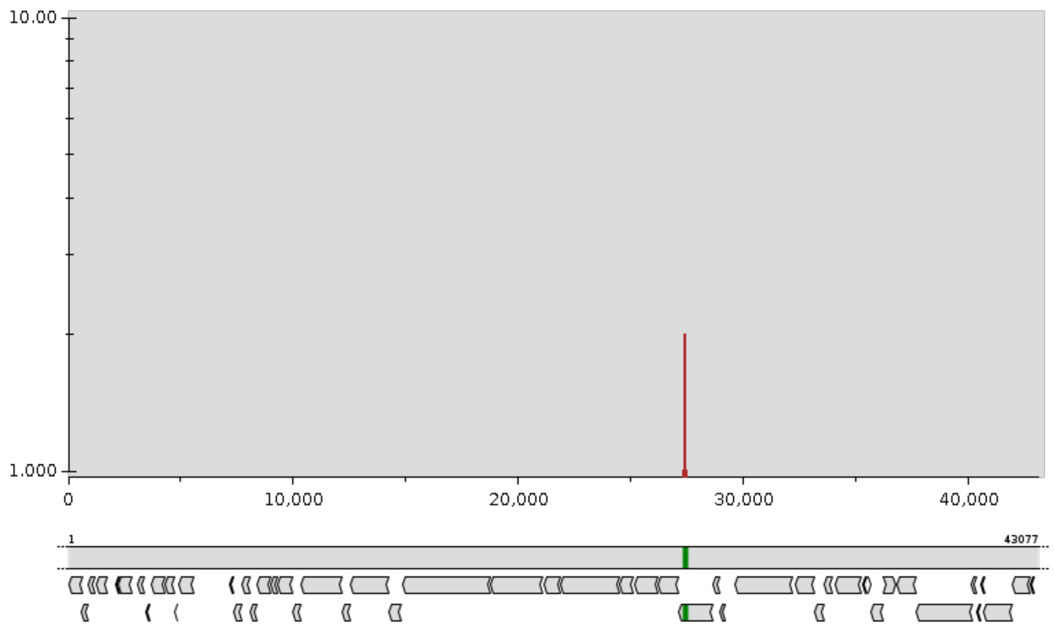

#### Assignment

| | |
|---|---|
| **Type** | Scottvirus scott (Taxonomy ID: 2733954) |
| **Reference Genome** | NC_048029.1 |
| **NT Identity (%)** | 81.9672 |
| **AA Identity (%)** | 92.5926 |
| **Number Of Stop Codons** | 0 |
| **Number Of CDS** | 51 |

#### Alignment

| | |
|---|---|
| **Alignment Score** | 312.0 (NT) + 506.0 (AA) = 818.0 |
| **Concordance (%)** | 80.1961 |

| Alignment Method | Local, heuristic, nucleotide (BLASTN) |
|---|---|

#### Genome Region

Sequence starts at position 27285 and ends at position 27528 relative to NC_048029.1 reference sequence.

#### Alignment Detailed Statistics

| | Begin | End | Coverage | Score | Concordance | Matches | Identities | I/D/M/F* | Stop Codons |
|---|---|---|---|---|---|---|---|---|---|
| **NT** | **27285** | **27528** | **0.6%** | **312** | **63.9%** | **244 (100%)** | **200 (82.0%)** | **0/0** | |
| Mutations: | 27287T>C, 27291T>C, 27296G>C, 27299T>G, 27317T>A, 27318T>G, 27319G>C, 27323A>T, 27326C>T, 27329T>C, 27344A>G, 27347C>T, 27350A>G, 27353C>T, 27362G>C, 27368C>G, 27380G>A, 27392T>A, 27398G>C, 27401G>T, 27404A>T, 27410C>G, 27412A>G, 27416G>C, 27419A>T, 27428A>T, 27434G>A, 27440C>T, 27443A>G, 27445T>C, 27446C>G, 27461C>G, 27462C>T, 27463G>T, 27464C>A, 27476A>T, 27482A>C, 27485A>C, 27487G>T, 27491G>A, 27509A>G, 27515T>C, 27521G>A, 27524A>G | | | | | | | | |

##### CDS

| | Begin | End | Coverage | Score | Concordance | Matches | Identities | I/D/M/F* | Stop Codons |
|---|---|---|---|---|---|---|---|---|---|
| HOT83_gp32 | 373 | 453 | 15.9% | 506 | 93.7% | 81 (100%) | 75 (92.6%) | 0/0/0/0 | 0 |
| Protein mutations: | L386M (27485A>C 27487G>T), R394N (27461C>G 27462C>T 27463G>T), T400A (27443A>G 27445T>C), Q442A (27317T>A 27318T>G 27319G>C), K451R (27291T>C) | | | | | | | | |
| Codon mutations: | CTT373CTC (27524A>G), GGC374GGT (27521G>A), GCA376GCG (27515T>C), TCT378TCC (27509A>G), TTC384TTT (27491G>A), CTT386ATG (27485A>C 27487G>T), CCT387CCG (27482A>C), GCT389GCA (27476A>T), CTG393CTT (27464C>A), CGG394AAC (27461C>G 27462C>T 27463G>T), CCG399CCC (27446C>G), ACT400GCC (27443A>G 27445T>C), AAG401AAA (27440C>T), TTC403TTT (27434G>A), CCT405CCA (27428A>T), GTT408GTA (27419A>T), ACC409ACG (27416G>C), TTG411CTC (27410C>G 27412A>G), GCT413GCA (27404A>T), CTC414CTA (27401G>T), TCC415TCG (27398G>C), GGA417GGT (27392T>A), GAC421GAT (27380G>A), CTG425CTC (27368C>G), CTC427CTG (27362G>C), GTG430GTA (27353C>T), GCT431GCC (27350A>G), GCG432GCA (27347C>T), CTT433CTC (27344A>G), GAA438GAG (27329T>C), CCG439CCA (27326C>T), GCT440GCA (27323A>T), CAA442GCT (27317T>A 27318T>G 27319G>C), GCA448GCC (27299T>G), CTC449CTG (27296G>C), AAG451AGG (27291T>C), GCA452GCG (27287T>C) | | | | | | | | |

##### Proteins

| | Begin | End | Coverage | Score | Concordance | Matches | Identities | I/D/M/F* | Stop Codons |
|---|---|---|---|---|---|---|---|---|---|
| head-to-tail joining protein (YP_009807875.1) | 373 | 453 | 15.9% | 506 | 93.7% | 81 (100%) | 75 (92.6%) | 0/0/0/0 | 0 |
| Protein mutations: | L386M (27485A>C 27487G>T), R394N (27461C>G 27462C>T 27463G>T), T400A (27443A>G 27445T>C), Q442A (27317T>A 27318T>G 27319G>C), K451R (27291T>C) | | | | | | | | |
| Codon mutations: | CTT373CTC (27524A>G), GGC374GGT (27521G>A), GCA376GCG (27515T>C), TCT378TCC (27509A>G), TTC384TTT (27491G>A), CTT386ATG (27485A>C 27487G>T), CCT387CCG (27482A>C), GCT389GCA (27476A>T), CTG393CTT (27464C>A), CGG394AAC (27461C>G 27462C>T 27463G>T), CCG399CCC (27446C>G), ACT400GCC (27443A>G 27445T>C), AAG401AAA (27440C>T), TTC403TTT (27434G>A), CCT405CCA (27428A>T), GTT408GTA (27419A>T), ACC409ACG (27416G>C), TTG411CTC (27410C>G 27412A>G), GCT413GCA (27404A>T), CTC414CTA (27401G>T), TCC415TCG (27398G>C), GGA417GGT (27392T>A), GAC421GAT (27380G>A), CTG425CTC (27368C>G), CTC427CTG (27362G>C), GTG430GTA (27353C>T), GCT431GCC (27350A>G), GCG432GCA (27347C>T), CTT433CTC (27344A>G), GAA438GAG (27329T>C), CCG439CCA (27326C>T), GCT440GCA (27323A>T), CAA442GCT (27317T>A 27318T>G 27319G>C), GCA448GCC (27299T>G), CTC449CTG (27296G>C), AAG451AGG (27291T>C), GCA452GCG (27287T>C) | | | | | | | | |

*: Inserts / Deletes / Misaligned / Frameshifts

#### Analysis details

This analysis was performed with panviral2.64

### NGS Details (UN60): Punavirus P1

#### Assembly

| | |
|---|---|
| **Coverage Length** | 224 (1 contig(s)) |
| **Depth Of Coverage** | 1.2 |
| **Number Of Reads** | 2 |
| **Reads Per Million** | 0.04 rpm (after QC) |
| **Ambiguities** | 0 |
| **Assembly Method** | de novo + reference guided assembly |
| **Consensus Caller** | Bcf Tools |

#### Coverage Map

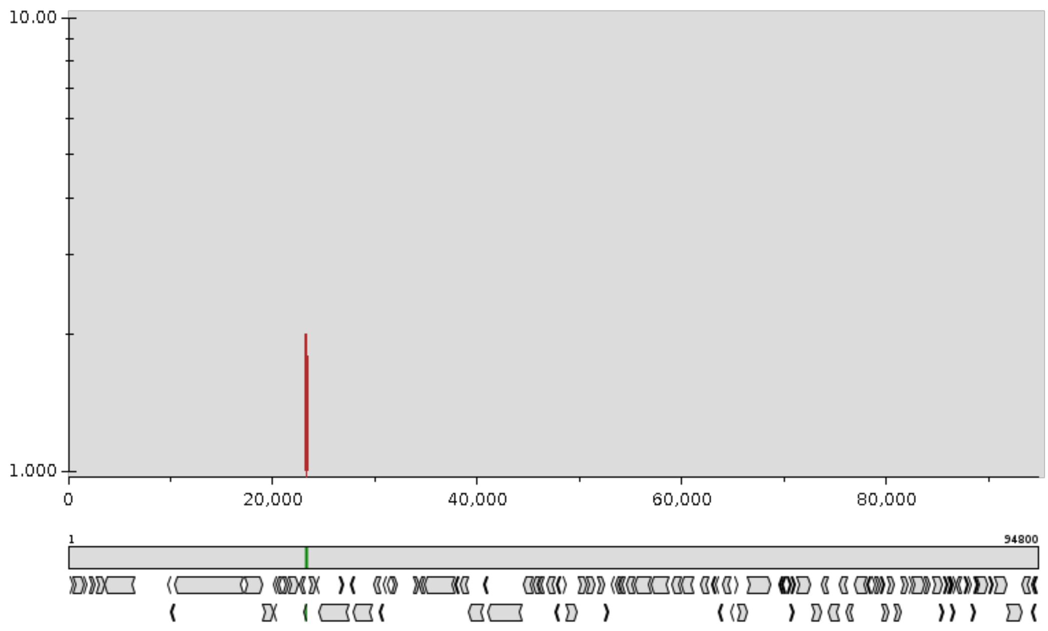

#### Assignment

| | |
|---|---|
| **Type** | Punavirus P1 (Taxonomy ID: 10678) |
| **Reference Genome** | NC_005856.1 |
| **NT Identity (%)** | 100.0 |
| **AA Identity (%)** | 98.3051 |
| **Number Of Stop Codons** | 0 |
| **Number Of CDS** | 110 |

#### Alignment

| | |
|---|---|
| **Alignment Score** | 448.0 (NT) + 438.0 (AA) = 886.0 |
| **Concordance (%)** | 100.0 |

| Alignment Method | Local, heuristic, nucleotide (BLASTN) |
|---|---|

#### Genome Region

Sequence starts at position 23164 and ends at position 23387 relative to NC_005856.1 reference sequence.

#### Alignment Detailed Statistics

| | Begin | End | Coverage | Score | Concordance | Matches | Identities | I/D/M/F* | Stop Codons |
|---|---|---|---|---|---|---|---|---|---|
| **NT** | **23164** | **23387** | **0.2%** | **448** | **100%** | **224 (100%)** | **224 (100%)** | **0/0** | |
| Mutations: | *none* | | | | | | | | |
| **CDS** | | | | | | | | | |
| insA | 1 | 59 | 64.1% | 438 | 98.6% | 59 (100%) | 58 (98.3%) | 0/0/0/0 | 0 |
| **Proteins** | | | | | | | | | |
| IS1 family transposase (YP_006488.1) | 1 | 59 | 64.1% | 438 | 98.6% | 59 (100%) | 58 (98.3%) | 0/0/0/0 | 0 |
| Protein mutations: | *none* | | | | | | | | |
| Codon mutations: | *none* | | | | | | | | |

*: Inserts / Deletes / Misaligned / Frameshifts

#### Analysis details

This analysis was performed with panviral2.64

### NGS Details (UN60): Harvey murine sarcoma virus

#### Assembly

| | |
|---|---|
| **Coverage Length** | 141 (1 contig(s)) |
| **Depth Of Coverage** | 1.2 |
| **Number Of Reads** | 2 |
| **Reads Per Million** | 0.04 rpm (after QC) |
| **Ambiguities** | 0 |
| **Assembly Method** | de novo + reference guided assembly |
| **Consensus Caller** | Bcf Tools |

#### Coverage Map

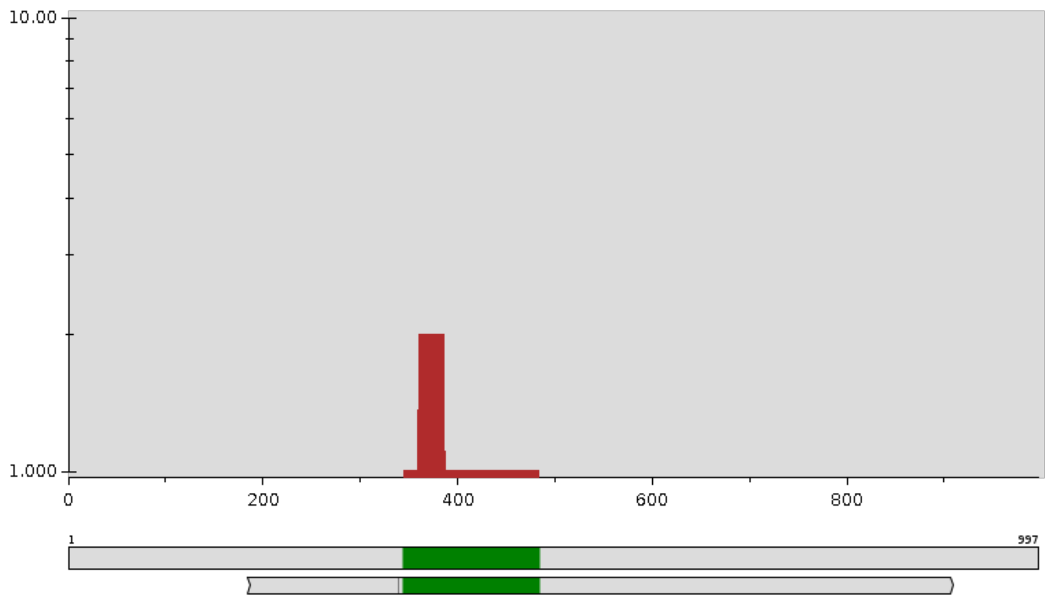

#### Assignment

| | |
|---|---|
| **Type** | Harvey murine sarcoma virus (Taxonomy ID: 11807) |
| **Reference Genome** | NC_038668.1 |
| **NT Identity (%)** | 69.5035 |
| **AA Identity (%)** | 85.1064 |
| **Number Of Stop Codons** | 0 |
| **Number Of CDS** | 2 |

#### Alignment

| | |
|---|---|
| **Alignment Score** | 110.0 (NT) + 516.0 (AA) = 626.0 |
| **Concordance (%)** | 70.3371 |

| Alignment Method | Global, seeded, nucleotide + amino acids (AGA) |
|---|---|

#### Genome Region

Sequence starts at position 345 and ends at position 485 relative to NC_038668.1 reference sequence.

#### Alignment Detailed Statistics

| | Begin | End | Coverage | Score | Concordance | Matches | Identities | I/D/M/F* | Stop Codons |
|---|---|---|---|---|---|---|---|---|---|
| **NT** | **345** | **485** | **14.1%** | **110** | **39.0%** | **141 (100%)** | **98 (69.5%)** | **0/0** | |
| Mutations: | 345C>G, 349A>G, 358T>C, 361G>T, 364G>C, 367G>C, 370C>A, 372C>G, 373T>C, 374A>G, 379C>G, 389A>T, 390G>C, 391T>A, 392G>T, 393C>G, 394C>T, 397G>T, 400C>G, 406G>A, 407C>T, 409G>A, 412C>T, 415G>A, 417A>G, 418C>T, 424T>C, 427G>A, 430C>T, 433G>A, 442C>G, 445T>A, 448A>C, 451G>A, 457C>A, 463G>T, 469G>A, 470G>T, 471T>G, 472A>C, 475C>G, 481T>C, 483G>A | | | | | | | | |
| **CDS** | | | | | | | | | |
| D1R95_gp1 | 55 | 101 | 19.4% | 258 | 81.6% | 47 (100%) | 40 (85.1%) | 0/0/0/0 | 0 |
| Protein mutations: | A63G (372C>G 373T>C), R64G (374A>G), A70C (392G>T 393C>G 394C>T), N78S (417A>G 418C>T), V96C (470G>T 471T>G 472A>C), G100E (483G>A) | | | | | | | | |
| Codon mutations: | ACA54.GA (345C>G), GAA55GAG (349A>G), CTT58CTC (358T>C), GTG59GTT (361G>T), GTG60GTC (364G>C), GTG61GTC (367G>C), GGC62GGA (370C>A), GCT63GGC (372C>G 373T>C), AGA64GGA (374A>G), GGC65GGG (379C>G), AGT69TCA (389A>T 390G>C 391T>A), GCC70TGT (392G>T 393C>G 394C>T), CTG71CTT (397G>T), ACC72ACG (400C>G), CAG74CAA (406G>A), CTG75TTA (407C>T 409G>A), ATC76ATT (412C>T), CAG77CAA (415G>A), AAC78AGT (417A>G 418C>T), TTT80TTC (424T>C), GTG81GTA (427G>A), GAC82GAT (430C>T), GAG83GAA (433G>A), CCC86CCG (442C>G), ACT87ACA (445T>A), ATA88ATC (448A>C), GAG89GAA (451G>A), TCC91TCA (457C>A), CGG93CGT (463G>T), CAG95CAA (469G>A), GTA96TGC (470G>T 471T>G 472A>C), GTC97GTG (475C>G), GAT99GAC (481T>C), GGG100GAG (483G>A) | | | | | | | | |
| D1R95_gp2 | 3 | 49 | 24.7% | 258 | 81.6% | 47 (100%) | 40 (85.1%) | 0/0/0/0 | 0 |
| Protein mutations: | A11G (372C>G 373T>C), R12G (374A>G), A18C (392G>T 393C>G 394C>T), N26S (417A>G 418C>T), V44C (470G>T 471T>G 472A>C), G48E (483G>A) | | | | | | | | |
| Codon mutations: | ACA2.GA (345C>G), GAA3GAG (349A>G), CTT6CTC (358T>C), GTG7GTT (361G>T), GTG8GTC (364G>C), GTG9GTC (367G>C), GGC10GGA (370C>A), GCT11GGC (372C>G 373T>C), AGA12GGA (374A>G), GGC13GGG (379C>G), AGT17TCA (389A>T 390G>C 391T>A), GCC18TGT (392G>T 393C>G 394C>T), CTG19CTT (397G>T), ACC20ACG (400C>G), CAG22CAA (406G>A), CTG23TTA (407C>T 409G>A), ATC24ATT (412C>T), CAG25CAA (415G>A), AAC26AGT (417A>G 418C>T), TTT28TTC (424T>C), GTG29GTA (427G>A), GAC30GAT (430C>T), GAG31GAA (433G>A), CCC34CCG (442C>G), ACT35ACA (445T>A), ATA36ATC (448A>C), GAG37GAA (451G>A), TCC39TCA (457C>A), CGG41CGT (463G>T), CAG43CAA (469G>A), GTA44TGC (470G>T 471T>G 472A>C), GTC45GTG (475C>G), GAT47GAC (481T>C), GGG48GAG (483G>A) | | | | | | | | |
| **Proteins** | | | | | | | | | |
| hypothetical protein (YP_009507788.1) | 55 | 101 | 19.4% | 258 | 81.6% | 47 (100%) | 40 (85.1%) | 0/0/0/0 | 0 |
| Protein mutations: | A63G (372C>G 373T>C), R64G (374A>G), A70C (392G>T 393C>G 394C>T), N78S (417A>G 418C>T), V96C (470G>T 471T>G 472A>C), G100E (483G>A) | | | | | | | | |
| Codon mutations: | ACA54.GA (345C>G), GAA55GAG (349A>G), CTT58CTC (358T>C), GTG59GTT (361G>T), GTG60GTC (364G>C), GTG61GTC (367G>C), GGC62GGA (370C>A), GCT63GGC (372C>G 373T>C), AGA64GGA (374A>G), GGC65GGG (379C>G), AGT69TCA (389A>T 390G>C 391T>A), GCC70TGT (392G>T 393C>G 394C>T), CTG71CTT (397G>T), ACC72ACG (400C>G), CAG74CAA (406G>A), CTG75TTA (407C>T 409G>A), ATC76ATT (412C>T), CAG77CAA (415G>A), AAC78AGT (417A>G 418C>T), TTT80TTC (424T>C), GTG81GTA (427G>A), GAC82GAT (430C>T), GAG83GAA (433G>A), CCC86CCG (442C>G), ACT87ACA (445T>A), ATA88ATC (448A>C), GAG89GAA (451G>A), TCC91TCA (457C>A), CGG93CGT (463G>T), CAG95CAA (469G>A), GTA96TGC (470G>T 471T>G 472A>C), GTC97GTG (475C>G), GAT99GAC (481T>C), GGG100GAG (483G>A) | | | | | | | | |
| hypothetical protein (YP_009507789.1) | 3 | 49 | 24.7% | 258 | 81.6% | 47 (100%) | 40 (85.1%) | 0/0/0/0 | 0 |
| Protein mutations: | A11G (372C>G 373T>C), R12G (374A>G), A18C (392G>T 393C>G 394C>T), N26S (417A>G 418C>T), V44C (470G>T 471T>G 472A>C), G48E (483G>A) | | | | | | | | |
| Codon mutations: | ACA2.GA (345C>G), GAA3GAG (349A>G), CTT6CTC (358T>C), GTG7GTT (361G>T), GTG8GTC (364G>C), GTG9GTC (367G>C), GGC10GGA (370C>A), GCT11GGC (372C>G 373T>C), AGA12GGA (374A>G), GGC13GGG (379C>G), AGT17TCA (389A>T 390G>C 391T>A), GCC18TGT (392G>T 393C>G 394C>T), CTG19CTT (397G>T), ACC20ACG (400C>G), CAG22CAA (406G>A), CTG23TTA (407C>T 409G>A), ATC24ATT (412C>T), CAG25CAA (415G>A), AAC26AGT (417A>G 418C>T), TTT28TTC (424T>C), GTG29GTA (427G>A), GAC30GAT (430C>T), GAG31GAA (433G>A), CCC34CCG (442C>G), ACT35ACA (445T>A), ATA36ATC (448A>C), GAG37GAA (451G>A), TCC39TCA (457C>A), CGG41CGT (463G>T), CAG43CAA (469G>A), GTA44TGC (470G>T 471T>G 472A>C), GTC45GTG (475C>G), GAT47GAC (481T>C), GGG48GAG (483G>A) | | | | | | | | |

*: Inserts / Deletes / Misaligned / Frameshifts

#### Analysis details

This analysis was performed with panviral2.64

### NGS Details (UN60): Ochrobactrum phage POA1180

#### Assembly

| | |
|---|---|
| Coverage Length | 119 (1 contig(s)) |
| Depth Of Coverage | 2.0 |
| Number Of Reads | 2 |
| Reads Per Million | 0.04 rpm (after QC) |
| Ambiguities | 0 |
| Assembly Method | de novo + reference guided assembly |
| Consensus Caller | Bcf Tools |

#### Coverage Map

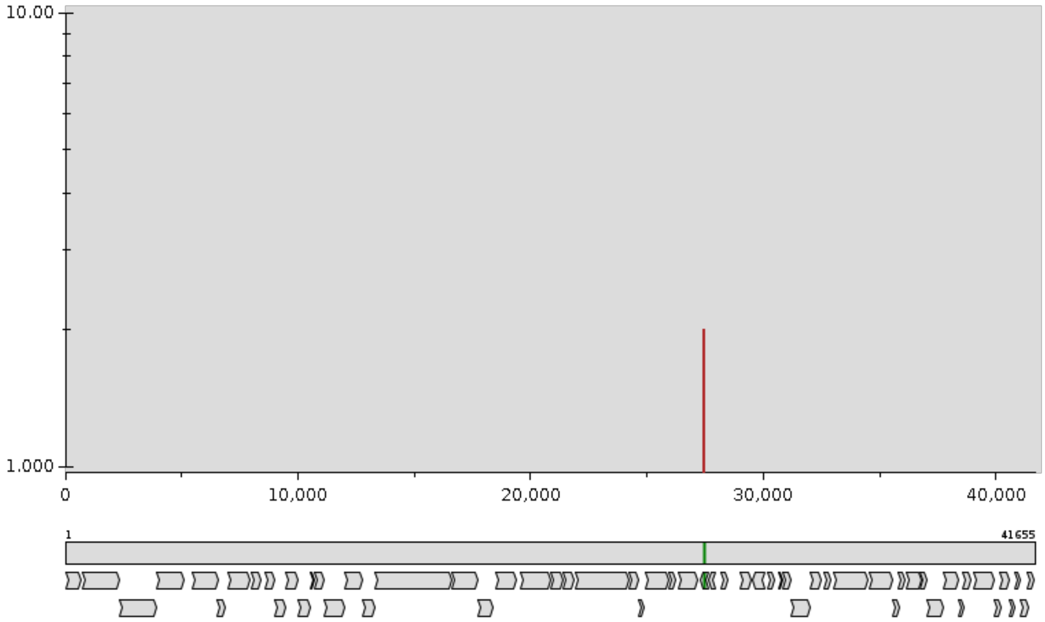

#### Assignment

| | |
|---|---|
| Type | Ochrobactrum phage POA1180 (Taxonomy ID: 1897640) |
| Reference Genome | NC_070931.1 |
| NT Identity (%) | 74.7899 |
| AA Identity (%) | 80.0 |
| Number Of Stop Codons | 0 |
| Number Of CDS | 58 |

#### Alignment

| | |
|---|---|
| Alignment Score | 118.0 (NT) + 250.0 (AA) = 368.0 |
| Concordance (%) | 70.0952 |

| Alignment Method | Local, heuristic, nucleotide (BLASTN) |
|---|---|

#### Genome Region

Sequence starts at position 27400 and ends at position 27518 relative to NC_070931.1 reference sequence.

#### Alignment Detailed Statistics

| | Begin | End | Coverage | Score | Concordance | Matches | Identities | I/D/M/F* | Stop Codons |
|---|---|---|---|---|---|---|---|---|---|
| **NT** | **27400** | **27518** | **0.3%** | **118** | **49.6%** | **119 (100%)** | **89 (74.8%)** | **0/0** | |
| Mutations: | 27417C>G, 27418G>C, 27419T>C, 27420G>C, 27423G>C, 27424C>G, 27425C>A, 27426G>C, 27428T>C, 27435T>C, 27436T>C, 27437T>G, 27447G>C, 27449T>G, 27453C>G, 27465G>C, 27468A>G, 27474G>C, 27477G>C, 27480G>C, 27492G>C, 27493C>G, 27494T>A, 27495C>G, 27498G>C, 27501C>G, 27507C>G, 27510C>A, 27511G>A, 27512C>G | | | | | | | | |
| **CDS** | | | | | | | | | |
| PQB33_gp31 | 30 | 69 | 36.7% | 250 | 85.9% | 40 (100%) | 32 (80.0%) | 0/0/0/0 | 0 |
| Protein mutations: | A32L (27510C>A 27511G>A 27512C>G), I53L (27447G>C 27449T>G), K57R (27435T>C 27436T>C 27437T>G), I60V (27426G>C 27428T>C), G61S (27423G>C 27424C>G 27425C>A), D62E (27420G>C), T63G (27417C>G 27418G>C 27419T>C) | | | | | | | | |
| Codon mutations: | GCG32CTT (27510C>A 27511G>A 27512C>G), CGG33CGC (27507C>G), GCG35GCC (27501C>G), GCC36GCG (27498G>C), CTG37CTC (27495C>G), AGC38TCG (27492G>C 27493C>G 27494T>A), CCC42CCG (27480G>C), GCC43GCG (27477G>C), GGC44GGG (27474G>C), ATT46ATC (27468A>G), GCC47GCG (27465G>C), CGG51CGC (27453C>G), ATC53CTG (27447G>C 27449T>G), AAA57CGG (27435T>C 27436T>C 27437T>G), ATC60GTG (27426G>C 27428T>C), GGC61TCG (27423G>C 27424C>G 27425C>A), GAC62GAG (27420G>C), ACG63GGC (27417C>G 27418G>C 27419T>C) | | | | | | | | |
| **Proteins** | | | | | | | | | |
| chromate transport protein (YP_010665114.1) | 30 | 69 | 36.7% | 250 | 85.9% | 40 (100%) | 32 (80.0%) | 0/0/0/0 | 0 |
| Protein mutations: | A32L (27510C>A 27511G>A 27512C>G), I53L (27447G>C 27449T>G), K57R (27435T>C 27436T>C 27437T>G), I60V (27426G>C 27428T>C), G61S (27423G>C 27424C>G 27425C>A), D62E (27420G>C), T63G (27417C>G 27418G>C 27419T>C) | | | | | | | | |
| Codon mutations: | GCG32CTT (27510C>A 27511G>A 27512C>G), CGG33CGC (27507C>G), GCG35GCC (27501C>G), GCC36GCG (27498G>C), CTG37CTC (27495C>G), AGC38TCG (27492G>C 27493C>G 27494T>A), CCC42CCG (27480G>C), GCC43GCG (27477G>C), GGC44GGG (27474G>C), ATT46ATC (27468A>G), GCC47GCG (27465G>C), CGG51CGC (27453C>G), ATC53CTG (27447G>C 27449T>G), AAA57CGG (27435T>C 27436T>C 27437T>G), ATC60GTG (27426G>C 27428T>C), GGC61TCG (27423G>C 27424C>G 27425C>A), GAC62GAG (27420G>C), ACG63GGC (27417C>G 27418G>C 27419T>C) | | | | | | | | |

*: Inserts / Deletes / Misaligned / Frameshifts

#### Analysis details

This analysis was performed with panviral2.64

### NGS Details (UN60): Pseudomonas phage PA8

#### Assembly

| | |
|---|---|
| **Coverage Length** | 135 (1 contig(s)) |
| **Depth Of Coverage** | 2.0 |
| **Number Of Reads** | 2 |
| **Reads Per Million** | 0.04 rpm (after QC) |
| **Ambiguities** | 0 |
| **Assembly Method** | de novo + reference guided assembly |
| **Consensus Caller** | Bcf Tools |

#### Coverage Map

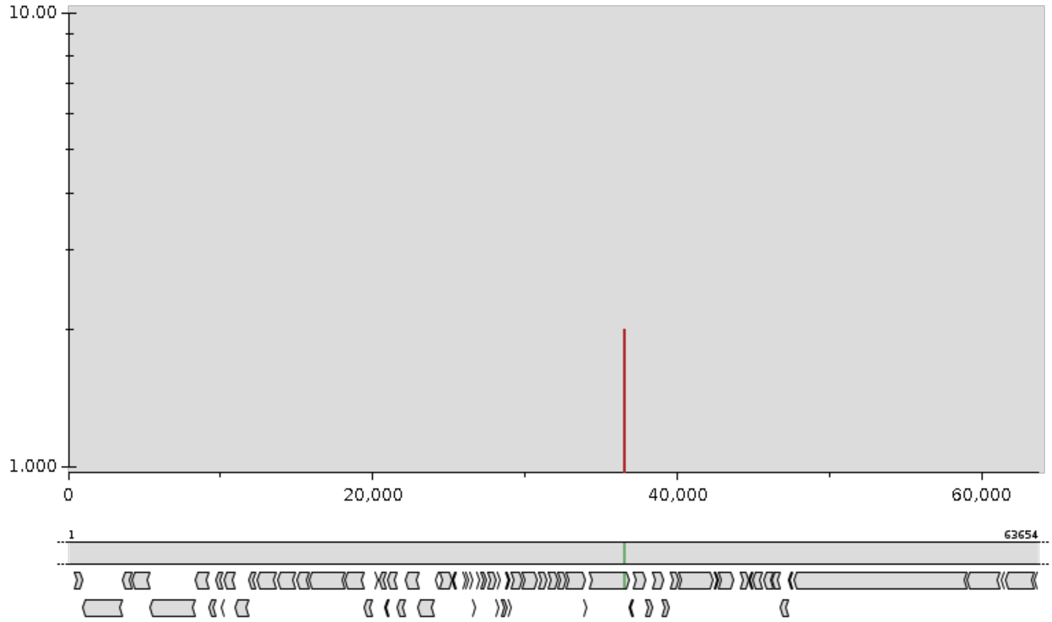

#### Assignment

| | |
|---|---|
| **Type** | Pseudomonas phage PA8 (Taxonomy ID: 2860997) |
| **Reference Genome** | NC_073672.1 |
| **NT Identity (%)** | 82.963 |
| **AA Identity (%)** | 80.0 |
| **Number Of Stop Codons** | 0 |
| **Number Of CDS** | 69 |

#### Alignment

| | |
|---|---|
| **Alignment Score** | 178.0 (NT) + 275.0 (AA) = 453.0 |
| **Concordance (%)** | 74.0196 |

| Alignment Method | Local, heuristic, nucleotide (BLASTN) |
|---|---|

#### Genome Region

Sequence starts at position 36438 and ends at position 36572 relative to NC_073672.1 reference sequence.

#### Alignment Detailed Statistics

| | Begin | End | Coverage | Score | Concordance | Matches | Identities | I/D/M/F* | Stop Codons |
|---|---|---|---|---|---|---|---|---|---|
| **NT** | **36438** | **36572** | **0.2%** | **178** | **65.9%** | **135 (100%)** | **112 (83.0%)** | **0/0** | |
| Mutations: | 36444C>G, 36449C>G, 36461A>G, 36462T>G, 36469A>C, 36470G>C, 36472T>A, 36473C>G, 36474A>C, 36475C>G, 36479G>C, 36481A>T, 36492G>A, 36493G>C, 36508C>A, 36516T>A, 36517C>G, 36524C>T, 36533A>C, 36545C>G, 36546G>A, 36548C>G, 36557G>C | | | | | | | | |
| **CDS** | | | | | | | | | |
| QGM55_gp48 | 733 | 777 | 5.1% | 275 | 80.4% | 45 (100%) | 36 (80.0%) | 0/0/0/0 | 0 |
| Protein mutations: | H735D (36444C>G), S741A (36462T>G), E743A (36469A>C 36470G>C), V744E (36472T>A 36473C>G), T745R (36474A>C 36475C>G), K747M (36481A>T), G751T (36492G>A 36493G>C), P756Q (36508C>A), A769T (36546G>A 36548C>G) | | | | | | | | |
| Codon mutations: | CAC735GAC (36444C>G), GTC736GTG (36449C>G), GAA740GAG (36461A>G), TCG741GCG (36462T>G), GAG743GCC (36469A>C 36470G>C), GTC744GAG (36472T>A 36473C>G), ACC745CGC (36474A>C 36475C>G), GGG746GGC (36479G>C), AAG747ATG (36481A>T), GGC751ACC (36492G>A 36493G>C), CCG756CAG (36508C>A), TCC759AGC (36516T>A 36517C>G), CAC761CAT (36524C>T), GTA764GTC (36533A>C), GTC768GTG (36545C>G), GCC769ACG (36546G>A 36548C>G), CGG772CGC (36557G>C) | | | | | | | | |
| **Proteins** | | | | | | | | | |
| hypothetical protein (YP_010765738.1) | 733 | 777 | 5.1% | 275 | 80.4% | 45 (100%) | 36 (80.0%) | 0/0/0/0 | 0 |
| Protein mutations: | H735D (36444C>G), S741A (36462T>G), E743A (36469A>C 36470G>C), V744E (36472T>A 36473C>G), T745R (36474A>C 36475C>G), K747M (36481A>T), G751T (36492G>A 36493G>C), P756Q (36508C>A), A769T (36546G>A 36548C>G) | | | | | | | | |
| Codon mutations: | CAC735GAC (36444C>G), GTC736GTG (36449C>G), GAA740GAG (36461A>G), TCG741GCG (36462T>G), GAG743GCC (36469A>C 36470G>C), GTC744GAG (36472T>A 36473C>G), ACC745CGC (36474A>C 36475C>G), GGG746GGC (36479G>C), AAG747ATG (36481A>T), GGC751ACC (36492G>A 36493G>C), CCG756CAG (36508C>A), TCC759AGC (36516T>A 36517C>G), CAC761CAT (36524C>T), GTA764GTC (36533A>C), GTC768GTG (36545C>G), GCC769ACG (36546G>A 36548C>G), CGG772CGC (36557G>C) | | | | | | | | |

*: Inserts / Deletes / Misaligned / Frameshifts

#### Analysis details

This analysis was performed with panviral2.64

#### NGS Details (UN60): Tomato associated geminivirus 1

##### Assembly

| | |
|---|---|
| **Coverage Length** | 233 (1 contig(s)) |
| **Depth Of Coverage** | 1.2 |
| **Number Of Reads** | 2 |
| **Reads Per Million** | 0.04 rpm (after QC) |
| **Ambiguities** | 0 |
| **Assembly Method** | de novo + reference guided assembly |
| **Consensus Caller** | Bcf Tools |

##### Coverage Map

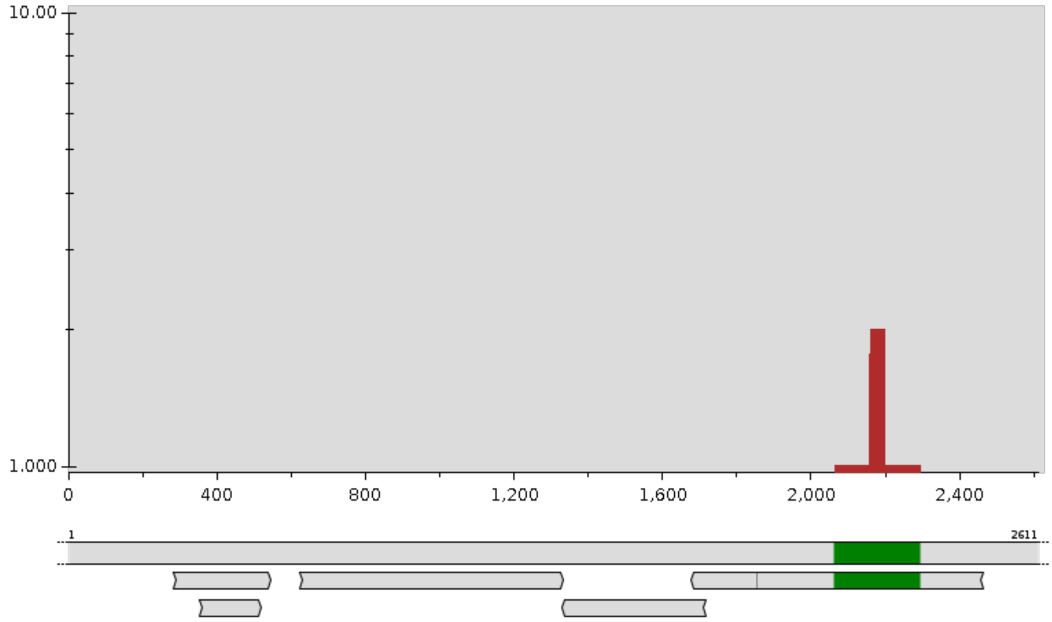

##### Assignment

| | |
|---|---|
| **Type** | Tomato associated geminivirus 1 (Taxonomy ID: 2027645) |
| **Reference Genome** | NC_035676.1 |
| **NT Identity (%)** | 72.9614 |
| **AA Identity (%)** | 75.3247 |
| **Number Of Stop Codons** | 4 |
| **Number Of CDS** | 5 |

##### Alignment

| | |
|---|---|
| **Alignment Score** | 214.0 (NT) + 844.0 (AA) = 1058.0 |
| **Concordance (%)** | 64.7491 |

| Alignment Method | Global, seeded, nucleotide + amino acids (AGA) |
|---|---|

#### Genome Region

Sequence starts at position 2064 and ends at position 2296 relative to NC_035676.1 reference sequence.

#### Alignment Detailed Statistics

| | Begin | End | Coverage | Score | Concordance | Matches | Identities | I/D/M/F* | Stop Codons |
|---|---|---|---|---|---|---|---|---|---|
| **NT** | **2064** | **2296** | **8.9%** | **214** | **45.9%** | **233 (100%)** | **170 (73.0%)** | **0/0** | |
| Mutations: | 2065C>G, 2067T>A, 2070C>T, 2073G>T, 2076A>T, 2078C>A, 2079G>A, 2082T>C, 2084T>G, 2086T>G, 2087T>A, 2088C>T, 2094C>A, 2102T>A, 2105G>T, 2106G>T, 2118A>T, 2120T>A, 2121A>T, 2126T>G, 2136G>C, 2137G>T, 2138T>G, 2150A>C, 2151A>G, 2157G>A, 2160C>G, 2163A>G, 2175C>T, 2177T>G, 2190G>A, 2191A>C, 2192T>A, 2196T>G, 2201G>C, 2205A>G, 2209C>G, 2210C>T, 2216C>T, 2220A>T, 2223G>A, 2227G>A, 2228T>C, 2229G>A, 2241C>A, 2243T>A, 2244C>T, 2247G>A, 2251T>C, 2252G>T, 2253C>G, 2256C>G, 2258G>T, 2263C>T, 2264T>G, 2268C>T, 2271A>C, 2272G>A, 2277T>G, 2280C>A, 2289G>A, 2291T>G, 2295T>C | | | | | | | | |
| **CDS** | | | | | | | | | |
| CK833_gp4 | 58 | 134 | 23.1% | 422 | 70.9% | 77 (100%) | 58 (75.3%) | 0/0/0/0 | 2 |
| Protein mutations: | I59L (2289G>A 2291T>G), T65M (2271A>C 2272G>A), R68Q (2263C>T 2264T>G), Q70N (2256C>G 2258G>T), Q72R (2251T>C 2252G>T), R75C (2241C>A 2243T>A), T80V (2227G>A 2228T>C), G84R (2216C>T), G86T (2209C>G 2210C>T), H89D (2201G>C), I92C (2190G>A 2191A>C 2192T>A), S106A (2150A>C), T110Q (2136G>C 2137G>T 2138T>G), I116L (2118A>T 2120T>A), R122* (2102T>A), N127S (2086T>G 2087T>A), A130S (2076A>T 2078C>A), W132* (2070C>T), R134T (2065C>G) | | | | | | | | |
| Codon mutations: | CCA57.CG (2295T>C), ATC59CTT (2289G>A 2291T>G), CTG62CTT (2280C>A), GTA63GTC (2277T>G), ACT65ATG (2271A>C 2272G>A), GAG66GAA (2268C>T), AGG68CAG (2263C>T 2264T>G), CAG70AAC (2256C>G 2258G>T), ACG71ACC (2253C>G), CAG72AGG (2251T>C 2252G>T), AAC73AAT (2247G>A), CAG74CAA (2244C>T), AGG75TGT (2241C>A 2243T>A), ATC79ATT (2229G>A), ACC80GTC (2227G>A 2228T>C), GAC81GAT (2223G>A), CCT82CCA (2220A>T), GGA84AGA (2216C>T), GGA86ACA (2209C>G 2210C>T), ATT87ATC (2205A>G), CAT89GAT (2201G>C), CCA90CCC (2196T>G), ATC92TGT (2190G>A 2191A>C 2192T>A), AGG97CGA (2175C>T 2177T>G), GTT101GTC (2163A>G), GCG102GCC (2160C>G), GAC103GAT (2157G>A), ATT105ATC (2151A>G), TCC106GCC (2150A>C), ACC110CAG (2136G>C 2137G>T 2138T>G), AGA114CGA (2126T>G), GGT115GGA (2121A>T), ATT116TTA (2118A>T 2120T>A), TCC120TCA (2106G>T), CGA121AGA (2105G>T), AGA122TGA (2102T>A), CCG124CCT (2094C>A), AAG126AAA (2088C>T), AAT127TCT (2086T>G 2087T>A), AGA128CGG (2082T>C 2084T>G), GAC129GAT (2079G>A), GCT130TCA (2076A>T 2078C>A), ATC131ATA (2073G>T), TGG132TGA (2070C>T), ACA133ACT (2067T>A), AGA134ACA (2065C>G) | | | | | | | | |
| CK833_gp5 | 58 | 134 | 29.4% | 422 | 70.9% | 77 (100%) | 58 (75.3%) | 0/0/0/0 | 2 |
| Protein mutations: | I59L (2289G>A 2291T>G), T65M (2271A>C 2272G>A), R68Q (2263C>T 2264T>G), Q70N (2256C>G 2258G>T), Q72R (2251T>C 2252G>T), R75C (2241C>A 2243T>A), T80V (2227G>A 2228T>C), G84R (2216C>T), G86T (2209C>G 2210C>T), H89D (2201G>C), I92C (2190G>A 2191A>C 2192T>A), S106A (2150A>C), T110Q (2136G>C 2137G>T 2138T>G), I116L (2118A>T 2120T>A), R122* (2102T>A), N127S (2086T>G 2087T>A), A130S (2076A>T 2078C>A), W132* (2070C>T), R134T (2065C>G) | | | | | | | | |
| Codon mutations: | CCA57.CG (2295T>C), ATC59CTT (2289G>A 2291T>G), CTG62CTT (2280C>A), GTA63GTC (2277T>G), ACT65ATG (2271A>C 2272G>A), GAG66GAA (2268C>T), AGG68CAG (2263C>T 2264T>G), CAG70AAC (2256C>G 2258G>T), ACG71ACC (2253C>G), CAG72AGG (2251T>C 2252G>T), AAC73AAT (2247G>A), CAG74CAA (2244C>T), AGG75TGT (2241C>A 2243T>A), ATC79ATT (2229G>A), ACC80GTC (2227G>A 2228T>C), GAC81GAT (2223G>A), CCT82CCA (2220A>T), GGA84AGA (2216C>T), GGA86ACA (2209C>G 2210C>T), ATT87ATC (2205A>G), CAT89GAT (2201G>C), CCA90CCC (2196T>G), ATC92TGT (2190G>A 2191A>C 2192T>A), AGG97CGA (2175C>T 2177T>G), GTT101GTC (2163A>G), GCG102GCC (2160C>G), GAC103GAT (2157G>A), ATT105ATC (2151A>G), TCC106GCC (2150A>C), ACC110CAG (2136G>C 2137G>T 2138T>G), AGA114CGA (2126T>G), GGT115GGA (2121A>T), ATT116TTA (2118A>T 2120T>A), TCC120TCA (2106G>T), CGA121AGA (2105G>T), AGA122TGA (2102T>A), CCG124CCT (2094C>A), AAG126AAA (2088C>T), AAT127TCT (2086T>G 2087T>A), AGA128CGG (2082T>C 2084T>G), GAC129GAT (2079G>A), GCT130TCA (2076A>T 2078C>A), ATC131ATA (2073G>T), TGG132TGA (2070C>T), ACA133ACT (2067T>A), AGA134ACA (2065C>G) | | | | | | | | |
| **Proteins** | | | | | | | | | |
| replication-associated protein (YP_009417307.1) | 58 | 134 | 23.1% | 422 | 70.9% | 77 (100%) | 58 (75.3%) | 0/0/0/0 | 2 |
| Protein mutations: | I59L (2289G>A 2291T>G), T65M (2271A>C 2272G>A), R68Q (2263C>T 2264T>G), Q70N (2256C>G 2258G>T), Q72R (2251T>C 2252G>T), R75C (2241C>A 2243T>A), T80V (2227G>A 2228T>C), G84R (2216C>T), G86T (2209C>G 2210C>T), H89D (2201G>C), I92C (2190G>A 2191A>C 2192T>A), S106A (2150A>C), T110Q (2136G>C 2137G>T 2138T>G), I116L (2118A>T 2120T>A), R122* (2102T>A), N127S (2086T>G 2087T>A), A130S (2076A>T 2078C>A), W132* (2070C>T), R134T (2065C>G) | | | | | | | | |
| Codon mutations: | CCA57.CG (2295T>C), ATC59CTT (2289G>A 2291T>G), CTG62CTT (2280C>A), GTA63GTC (2277T>G), ACT65ATG (2271A>C 2272G>A), GAG66GAA (2268C>T), AGG68CAG (2263C>T 2264T>G), CAG70AAC (2256C>G 2258G>T), ACG71ACC (2253C>G), CAG72AGG (2251T>C 2252G>T), AAC73AAT (2247G>A), CAG74CAA (2244C>T), AGG75TGT (2241C>A 2243T>A), ATC79ATT (2229G>A), ACC80GTC (2227G>A 2228T>C), GAC81GAT (2223G>A), CCT82CCA (2220A>T), GGA84AGA (2216C>T), GGA86ACA (2209C>G 2210C>T), ATT87ATC (2205A>G), CAT89GAT (2201G>C), CCA90CCC (2196T>G), ATC92TGT (2190G>A 2191A>C 2192T>A), AGG97CGA (2175C>T 2177T>G), GTT101GTC (2163A>G), GCG102GCC (2160C>G), GAC103GAT (2157G>A), ATT105ATC (2151A>G), TCC106GCC (2150A>C), ACC110CAG (2136G>C 2137G>T 2138T>G), AGA114CGA (2126T>G), GGT115GGA (2121A>T), ATT116TTA (2118A>T 2120T>A), TCC120TCA (2106G>T), CGA121AGA (2105G>T), AGA122TGA (2102T>A), CCG124CCT (2094C>A), AAG126AAA (2088C>T), AAT127TCT (2086T>G 2087T>A), AGA128CGG (2082T>C 2084T>G), GAC129GAT (2079G>A), GCT130TCA (2076A>T 2078C>A), ATC131ATA (2073G>T), TGG132TGA (2070C>T), ACA133ACT (2067T>A), AGA134ACA (2065C>G) | | | | | | | | |
| RepA (YP_009417308.1) | 58 | 134 | 29.4% | 422 | 70.9% | 77 (100%) | 58 (75.3%) | 0/0/0/0 | 2 |
| Protein mutations: | I59L (2289G>A 2291T>G), T65M (2271A>C 2272G>A), R68Q (2263C>T 2264T>G), Q70N (2256C>G 2258G>T), Q72R (2251T>C 2252G>T), R75C (2241C>A 2243T>A), T80V (2227G>A 2228T>C), G84R (2216C>T), G86T (2209C>G 2210C>T), H89D (2201G>C), I92C (2190G>A 2191A>C 2192T>A), S106A (2150A>C), T110Q (2136G>C 2137G>T 2138T>G), I116L (2118A>T 2120T>A), R122* (2102T>A), N127S (2086T>G 2087T>A), A130S (2076A>T 2078C>A), W132* (2070C>T), R134T (2065C>G) | | | | | | | | |
| Codon mutations: | CCA57.CG (2295T>C), ATC59CTT (2289G>A 2291T>G), CTG62CTT (2280C>A), GTA63GTC (2277T>G), ACT65ATG (2271A>C 2272G>A), GAG66GAA (2268C>T), AGG68CAG (2263C>T 2264T>G), CAG70AAC (2256C>G 2258G>T), ACG71ACC (2253C>G), CAG72AGG (2251T>C 2252G>T), AAC73AAT (2247G>A), CAG74CAA (2244C>T), AGG75TGT (2241C>A 2243T>A), ATC79ATT (2229G>A), ACC80GTC (2227G>A 2228T>C), GAC81GAT (2223G>A), CCT82CCA (2220A>T), GGA84AGA (2216C>T), GGA86ACA (2209C>G 2210C>T), ATT87ATC (2205A>G), CAT89GAT (2201G>C), CCA90CCC (2196T>G), ATC92TGT (2190G>A 2191A>C 2192T>A), AGG97CGA (2175C>T 2177T>G), GTT101GTC (2163A>G), GCG102GCC (2160C>G), GAC103GAT (2157G>A), ATT105ATC (2151A>G), TCC106GCC (2150A>C), ACC110CAG (2136G>C 2137G>T 2138T>G), AGA114CGA (2126T>G), GGT115GGA (2121A>T), ATT116TTA (2118A>T 2120T>A), TCC120TCA (2106G>T), CGA121AGA (2105G>T), AGA122TGA (2102T>A), CCG124CCT (2094C>A), AAG126AAA (2088C>T), AAT127TCT (2086T>G 2087T>A), AGA128CGG (2082T>C 2084T>G), GAC129GAT (2079G>A), GCT130TCA (2076A>T 2078C>A), ATC131ATA (2073G>T), TGG132TGA (2070C>T), ACA133ACT (2067T>A), AGA134ACA (2065C>G) | | | | | | | | |

*: Inserts / Deletes / Misaligned / Frameshifts

#### Analysis details

This analysis was performed with panviral2.64

### NGS Details (UN60): Euphorbia caput-medusae latent virus

#### Assembly

| | |
|---|---|
| **Coverage Length** | 147 (3 contig(s)) |
| **Depth Of Coverage** | 1.0 |
| **Number Of Reads** | 1 |
| **Reads Per Million** | 0.02 rpm (after QC) |
| **Ambiguities** | 0 |
| **Assembly Method** | de novo + reference guided assembly |
| **Consensus Caller** | Bcf Tools |

#### Coverage Map

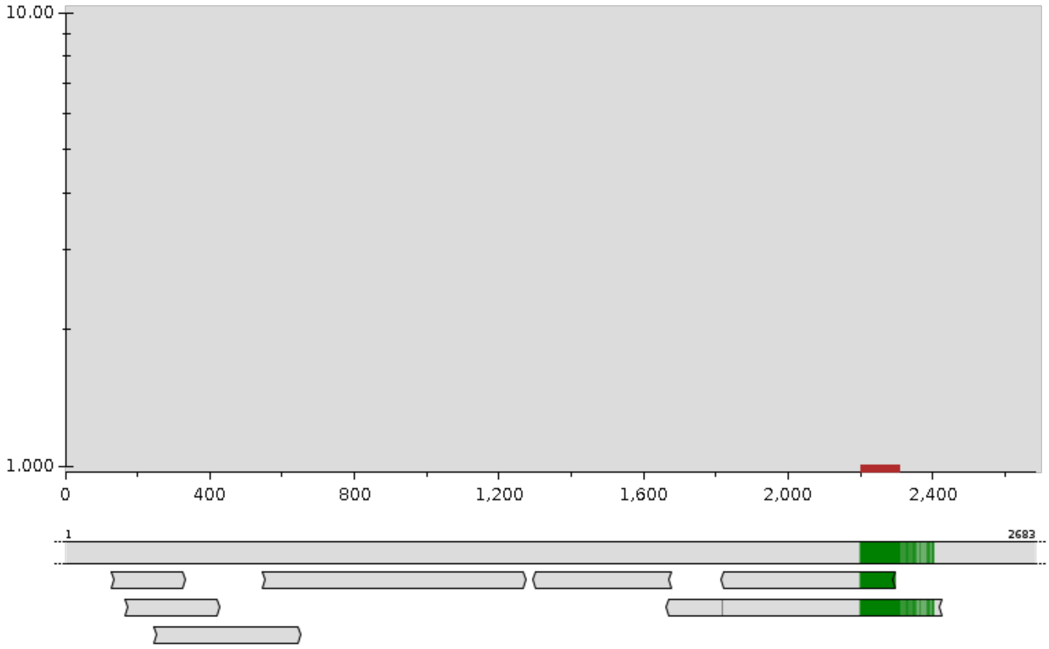

#### Assignment

| | |
|---|---|
| **Type** | Euphorbia caput-medusae latent virus (Taxonomy ID: 1853865) |
| **Reference Genome** | NC_030238.1 |
| **NT Identity (%)** | 47.619 |
| **AA Identity (%)** | 45.3704 |
| **Number Of Stop Codons** | 6 |
| **Number Of CDS** | 7 |

#### Alignment

| | |
|---|---|
| **Alignment Score** | -14.0 (NT) + 366.0 (AA) = 352.0 |
| **Concordance (%)** | 31.6832 |

| **Alignment Method** | Global, seeded, nucleotide + amino acids (AGA) |
|---|---|

#### Genome Region

Sequence starts at position 2201 and ends at position 2402 relative to NC_030238.1 reference sequence.

#### Alignment Detailed Statistics

| | Begin | End | Coverage | Score | Concordance | Matches | Identities | I/D/M/F* | Stop Codons |
|---|---|---|---|---|---|---|---|---|---|
| **NT** | **2201** | **2402** | **5.5%** | **-14** | **-4.8%** | **147 (100%)** | **70 (47.6%)** | **0/0** | |
| Mutations: | 2201A>G, 2204G>T, 2206T>A, 2211G>T, 2212G>A, 2214T>A, 2215G>A, 2217G>A, 2219T>G, 2222G>A, 2223C>T, 2224C>T, 2229G>C, 2231C>A, 2232C>G, 2233A>G, 2238G>A, 2239A>C, 2240C>A, 2241A>T, 2243G>T, 2253G>A, 2259C>T, 2262A>C, 2265G>T, 2271G>A, 2274A>T, 2275T>G, 2277C>T, 2279C>A, 2280C>T, 2283A>G, 2284C>G, 2286T>A, 2289C>T, 2293G>A, 2294C>T, 2298T>A, 2299G>T, 2307A>G, 2308A>T, 2312T>C, 2313A>T, 2315T>G, 2318A>G, 2319T>A, 2321G>T, 2324G>T, 2325A>T, 2327A>T, 2328T>G, 2330A>G, 2331A>G, 2332T>A, 2334A>G, 2337A>T, 2340T>G, 2342A>C, 2343A>G, 2346A>T, 2349A>T, 2351G>C, 2352A>G, 2354G>T, 2355A>T, 2364T>C, 2367T>G, 2376C>G, 2382A>T, 2384T>G, 2385A>T, 2388C>T, 2391A>T, 2397T>A, 2399A>G, 2400G>T, 2402G>T | | | | | | | | |

##### CDS

| | Begin | End | Coverage | Score | Concordance | Matches | Identities | I/D/M/F* | Stop Codons |
|---|---|---|---|---|---|---|---|---|---|
| A9K86_gp5 | 40 | 77 | 11.4% | 160 | 34.6% | 38 (100%) | 19 (50.0%) | 0/0/0/0 | 2 |
| Protein mutations: | N40D (2312T>C), F41Y (2307A>G 2308A>T), T44N (2298T>A 2299G>T), A46I (2293G>A 2294C>T), C49S (2283A>G 2284C>G), E51* (2277C>T 2279C>A), N52T (2274A>T 2275T>G), D55E (2265G>T), L63I (2241A>T 2243G>T), V64C (2238G>A 2239A>C 2240C>A), M66T (2232C>G 2233A>G), D67* (2229G>C 2231C>A), R69K (2223C>T 2224C>T), L70F (2222G>A), N71H (2217G>A 2219T>G), T72I (2214T>A 2215G>A), T73I (2211G>T 2212G>A), Q75L (2206T>A) | | | | | | | | |
| Codon mutations: | CGC10A.A (2400G>T 2402G>T), TTA11C.T (2397T>A 2399A>G), GGT13..A (2391A>T), AAG14..A (2388C>T), TCT15..A (2385A>T), ATT16C.A (2382A>T 2384T>G), CTG18..C (2376C>G), CCA21..C (2367T>G), AAA22..G (2364T>C), CTT25..A (2355A>T), CTT26A.C (2352A>G 2354G>T), CCT27G.A (2349A>T 2351G>C), ATT28..A (2346A>T), TTT29..C (2343A>G), TTA30G.C (2340T>G 2342A>C), ATT31..A (2337A>T), GAT32..C (2334A>G), TAT33.TC (2331A>G 2332T>A), TTA34C.C (2328T>G 2330A>G), TAT35A.A (2325A>T 2327A>T), CAA36A.. (2324G>T), CTA37A.T (2319T>A 2321G>T), TTA38C.. (2318A>G), AAT39C.A (2313A>T 2315T>G), AAT40GAT (2312T>C), TTT41TAC (2307A>G 2308A>T), ACA44AAT (2298T>A 2299G>T), GCT46ATT (2293G>A 2294C>T), AGG47AGA (2289C>T), GTA48GTT (2286T>A), TGT49TCC (2283A>G 2284C>G), ACG50ACA (2280C>T), GAG51TAA (2277C>T 2279C>A), AAT52ACA (2274A>T 2275T>G), CAC53CAT (2271G>A), GAC55GAA (2265G>T), GGT56GGG (2262A>C), GAG57GAA (2259C>T), CAC59CAT (2253G>A), CTT63ATA (2241A>T 2243G>T), GTC64TGT (2238G>A 2239A>C 2240C>A), ATG66ACC (2232C>G 2233A>G), GAC67TAG (2229G>C 2231C>A), AGG69AAA (2223C>T 2224C>T), CTC70TTC (2222G>A), AAC71CAT (2217G>A 2219T>G), ACA72ATT (2214T>A 2215G>A), ACC73ATA (2211G>T 2212G>A), CAA75CTA (2206T>A), CGG76AGG (2204G>T), TTC77C.. (2201A>G) | | | | | | | | |
| A9K86_gp6 | 40 | 77 | 14.9% | 160 | 34.6% | 38 (100%) | 19 (50.0%) | 0/0/0/0 | 2 |
| Protein mutations: | N40D (2312T>C), F41Y (2307A>G 2308A>T), T44N (2298T>A 2299G>T), A46I (2293G>A 2294C>T), C49S (2283A>G 2284C>G), E51* (2277C>T 2279C>A), N52T (2274A>T 2275T>G), D55E (2265G>T), L63I (2241A>T 2243G>T), V64C (2238G>A 2239A>C 2240C>A), M66T (2232C>G 2233A>G), D67* (2229G>C 2231C>A), R69K (2223C>T 2224C>T), L70F (2222G>A), N71H (2217G>A 2219T>G), T72I (2214T>A 2215G>A), T73I (2211G>T 2212G>A), Q75L (2206T>A) | | | | | | | | |
| Codon mutations: | CGC10A.A (2400G>T 2402G>T), TTA11C.T (2397T>A 2399A>G), GGT13..A (2391A>T), AAG14..A (2388C>T), TCT15..A (2385A>T), ATT16C.A (2382A>T 2384T>G), CTG18..C (2376C>G), CCA21..C (2367T>G), AAA22..G (2364T>C), CTT25..A (2355A>T), CTT26A.C (2352A>G 2354G>T), CCT27G.A (2349A>T 2351G>C), ATT28..A (2346A>T), TTT29..C (2343A>G), TTA30G.C (2340T>G 2342A>C), ATT31..A (2337A>T), GAT32..C (2334A>G), TAT33.TC (2331A>G 2332T>A), TTA34C.C (2328T>G 2330A>G), TAT35A.A (2325A>T 2327A>T), CAA36A.. (2324G>T), CTA37A.T (2319T>A 2321G>T), TTA38C.. (2318A>G), AAT39C.A (2313A>T 2315T>G), AAT40GAT (2312T>C), TTT41TAC (2307A>G 2308A>T), ACA44AAT (2298T>A 2299G>T), GCT46ATT (2293G>A 2294C>T), AGG47AGA (2289C>T), GTA48GTT (2286T>A), TGT49TCC (2283A>G 2284C>G), ACG50ACA (2280C>T), GAG51TAA (2277C>T 2279C>A), AAT52ACA (2274A>T 2275T>G), CAC53CAT (2271G>A), GAC55GAA (2265G>T), GGT56GGG (2262A>C), GAG57GAA (2259C>T), CAC59CAT (2253G>A), CTT63ATA (2241A>T 2243G>T), GTC64TGT (2238G>A 2239A>C 2240C>A), ATG66ACC (2232C>G 2233A>G), GAC67TAG (2229G>C 2231C>A), AGG69AAA (2223C>T 2224C>T), CTC70TTC (2222G>A), AAC71CAT (2217G>A 2219T>G), ACA72ATT (2214T>A 2215G>A), ACC73ATA (2211G>T 2212G>A), CAA75CTA (2206T>A), CGG76AGG (2204G>T), TTC77C.. (2201A>G) | | | | | | | | |
| A9K86_gp7 | 1 | 32 | 19.9% | 46 | 20.8% | 32 (100%) | 11 (34.4%) | 0/0/0/0 | 2 |
| Protein mutations: | M1I (2294C>T), G3E (2289C>T), Y4F (2286T>A), V5P (2283A>G 2284C>G), R6H (2279C>A 2280C>T), R7K (2277C>T), I8H (2274A>T 2275T>G), T9I (2271G>A), T11K (2265G>T), V12G (2262A>C), S13N (2259C>T), T15I (2253G>A), L19Y (2240C>A 2241A>T), S20V (2238G>A 2239A>C), W22P (2231C>A 2232C>G 2233A>G), T23R (2229G>C), G25N (2222G>A 2223C>T 2224C>T), T27I (2217G>A), Q28L (2214T>A 2215G>A), P29* (2211G>T 2212G>A), N31* (2204G>T 2206T>A) | | | | | | | | |
| Codon mutations: | ATG1ATA (2294C>T), CTA2TTA (2293G>A), GGG3GAG (2289C>T), TAT4TTT (2286T>A), GTA5CCA (2283A>G 2284C>G), CGG6CAT (2279C>A 2280C>T), AGA7AAA (2277C>T), ATC8CAC (2274A>T 2275T>G), ACC9ATC (2271G>A), ACG11AAG (2265G>T), GTG12GGG (2262A>C), AGC13AAC (2259C>T), ACC15ATC (2253G>A), GCC18GCA (2243G>T), TTG19TAT (2240C>A 2241A>T), TCC20GTC (2238G>A 2239A>C), TGG22CCT (2231C>A 2232C>G 2233A>G), ACA23AGA (2229G>C), GGC25AAT (2222G>A 2223C>T 2224C>T), TCA26TCC (2219T>G), ACA27ATA (2217G>A), CAA28TTA (2214T>A 2215G>A), CCA29TAA (2211G>T 2212G>A), AAC31TAA (2204G>T 2206T>A), GGT32GGC (2201A>G) | | | | | | | | |

##### Proteins

| | Begin | End | Coverage | Score | Concordance | Matches | Identities | I/D/M/F* | Stop Codons |
|---|---|---|---|---|---|---|---|---|---|
| replication-associated protein (YP_009255247.1) | 40 | 77 | 11.4% | 160 | 34.6% | 38 (100%) | 19 (50.0%) | 0/0/0/0 | 2 |
| Protein mutations: | N40D (2312T>C), F41Y (2307A>G 2308A>T), T44N (2298T>A 2299G>T), A46I (2293G>A 2294C>T), C49S (2283A>G 2284C>G), E51* (2277C>T 2279C>A), N52T (2274A>T 2275T>G), D55E (2265G>T), L63I (2241A>T 2243G>T), V64C (2238G>A 2239A>C 2240C>A), M66T (2232C>G 2233A>G), D67* (2229G>C 2231C>A), R69K (2223C>T 2224C>T), L70F (2222G>A), N71H (2217G>A 2219T>G), T72I (2214T>A 2215G>A), T73I (2211G>T 2212G>A), Q75L (2206T>A) | | | | | | | | |
| Codon mutations: | CGC10A.A (2400G>T 2402G>T), TTA11C.T (2397T>A 2399A>G), GGT13..A (2391A>T), AAG14..A (2388C>T), TCT15..A (2385A>T), ATT16C.A (2382A>T 2384T>G), CTG18..C (2376C>G), CCA21..C (2367T>G), AAA22..G (2364T>C), CTT25..A (2355A>T), CTT26A.C (2352A>G 2354G>T), CCT27G.A (2349A>T 2351G>C), ATT28..A (2346A>T), TTT29..C (2343A>G), TTA30G.C (2340T>G 2342A>C), ATT31..A (2337A>T), GAT32..C (2334A>G), TAT33.TC (2331A>G 2332T>A), TTA34C.C (2328T>G 2330A>G), TAT35A.A (2325A>T 2327A>T), CAA36A.. (2324G>T), CTA37A.T (2319T>A 2321G>T), TTA38C.. (2318A>G), AAT39C.A (2313A>T 2315T>G), AAT40GAT (2312T>C), TTT41TAC (2307A>G 2308A>T), ACA44AAT (2298T>A 2299G>T), GCT46ATT (2293G>A 2294C>T), AGG47AGA (2289C>T), GTA48GTT (2286T>A), TGT49TCC (2283A>G 2284C>G), ACG50ACA (2280C>T), GAG51TAA (2277C>T 2279C>A), AAT52ACA (2274A>T 2275T>G), CAC53CAT (2271G>A), GAC55GAA (2265G>T), GGT56GGG (2262A>C), GAG57GAA (2259C>T), CAC59CAT (2253G>A), CTT63ATA (2241A>T 2243G>T), GTC64TGT (2238G>A 2239A>C 2240C>A), ATG66ACC (2232C>G 2233A>G), GAC67TAG (2229G>C 2231C>A), AGG69AAA (2223C>T 2224C>T), CTC70TTC (2222G>A), AAC71CAT (2217G>A 2219T>G), ACA72ATT (2214T>A 2215G>A), ACC73ATA (2211G>T 2212G>A), CAA75CTA (2206T>A), CGG76AGG (2204G>T), TTC77C.. (2201A>G) | | | | | | | | |
| RepA (YP_009255248.1) | 40 | 77 | 14.9% | 160 | 34.6% | 38 (100%) | 19 (50.0%) | 0/0/0/0 | 2 |
| Protein mutations: | N40D (2312T>C), F41Y (2307A>G 2308A>T), T44N (2298T>A 2299G>T), A46I (2293G>A 2294C>T), C49S (2283A>G 2284C>G), E51* (2277C>T 2279C>A), N52T (2274A>T 2275T>G), D55E (2265G>T), L63I (2241A>T 2243G>T), V64C (2238G>A 2239A>C 2240C>A), M66T (2232C>G 2233A>G), D67* (2229G>C 2231C>A), R69K (2223C>T 2224C>T), L70F (2222G>A), N71H (2217G>A 2219T>G), T72I (2214T>A 2215G>A), T73I (2211G>T 2212G>A), Q75L (2206T>A) | | | | | | | | |
| Codon mutations: | CGC10A.A (2400G>T 2402G>T), TTA11C.T (2397T>A 2399A>G), GGT13..A (2391A>T), AAG14..A (2388C>T), TCT15..A (2385A>T), ATT16C.A (2382A>T 2384T>G), CTG18..C (2376C>G), CCA21..C (2367T>G), AAA22..G (2364T>C), CTT25..A (2355A>T), CTT26A.C (2352A>G 2354G>T), CCT27G.A (2349A>T 2351G>C), ATT28..A (2346A>T), TTT29..C (2343A>G), TTA30G.C (2340T>G 2342A>C), ATT31..A (2337A>T), GAT32..C (2334A>G), TAT33.TC (2331A>G 2332T>A), TTA34C.C (2328T>G 2330A>G), TAT35A.A (2325A>T 2327A>T), CAA36A.. (2324G>T), CTA37A.T (2319T>A 2321G>T), TTA38C.. (2318A>G), AAT39C.A (2313A>T 2315T>G), AAT40GAT (2312T>C), TTT41TAC (2307A>G 2308A>T), ACA44AAT (2298T>A 2299G>T), GCT46ATT (2293G>A 2294C>T), AGG47AGA (2289C>T), GTA48GTT (2286T>A), TGT49TCC (2283A>G 2284C>G), ACG50ACA (2280C>T), GAG51TAA (2277C>T 2279C>A), AAT52ACA (2274A>T 2275T>G), CAC53CAT (2271G>A), GAC55GAA (2265G>T), GGT56GGG (2262A>C), GAG57GAA (2259C>T), CAC59CAT (2253G>A), CTT63ATA (2241A>T 2243G>T), GTC64TGT (2238G>A 2239A>C 2240C>A), ATG66ACC (2232C>G 2233A>G), GAC67TAG (2229G>C 2231C>A), AGG69AAA (2223C>T 2224C>T), CTC70TTC (2222G>A), AAC71CAT (2217G>A 2219T>G), ACA72ATT (2214T>A 2215G>A), ACC73ATA (2211G>T 2212G>A), CAA75CTA (2206T>A), CGG76AGG (2204G>T), TTC77C.. (2201A>G) | | | | | | | | |
| C3 (YP_009255249.1) | 1 | 32 | 19.9% | 46 | 20.8% | 32 (100%) | 11 (34.4%) | 0/0/0/0 | 2 |
| Protein mutations: | M1I (2294C>T), G3E (2289C>T), Y4F (2286T>A), V5P (2283A>G 2284C>G), R6H (2279C>A 2280C>T), R7K (2277C>T), I8H (2274A>T 2275T>G), T9I (2271G>A), T11K (2265G>T), V12G (2262A>C), S13N (2259C>T), T15I (2253G>A), L19Y (2240C>A 2241A>T), S20V (2238G>A 2239A>C), W22P (2231C>A 2232C>G 2233A>G), T23R (2229G>C), G25N (2222G>A 2223C>T 2224C>T), T27I (2217G>A), Q28L (2214T>A 2215G>A), P29* (2211G>T 2212G>A), N31* (2204G>T 2206T>A) | | | | | | | | |

| | Begin | End | Coverage | Score | Concordance | Matches | Identities | I/D/M/F* | Stop Codons |
|---|---|---|---|---|---|---|---|---|---|
| **NT** | **2201** | **2402** | **5.5%** | **-14** | **-4.8%** | **147 (100%)** | **70 (47.6%)** | **0/0** | |
| Codon mutations: | ATG1ATA (2294C>T), CTA2TTA (2293G>A), GGG3GAG (2289C>T), TAT4TTT (2286T>A), GTA5CCA (2283A>G 2284C>G), CGG6CAT (2279C>A 2280C>T), AGA7AAA (2277C>T), ATC8CAC (2274A>T 2275T>G), ACC9ATC (2271G>A), ACG11AAG (2265G>T), GTG12GGG (2262A>C), AGC13AAC (2259C>T), ACC15ATC (2253G>A), GCC18GCA (2243G>T), TTG19TAT (2240C>A 2241A>T), TCC20GTC (2238G>A 2239A>C), TGG22CCT (2231C>A 2232C>G 2233A>G), ACA23AGA (2229G>C), GGC25AAT (2222G>A 2223C>T 2224C>T), TCA26TCC (2219T>G), ACA27ATA (2217G>A), CAA28TTA (2214T>A 2215G>A), CCA29TAA (2211G>T 2212G>A), AAC31TAA (2204G>T 2206T>A), GGT32GGC (2201A>G) | | | | | | | | |

*: Inserts / Deletes / Misaligned / Frameshifts

#### Analysis details

This analysis was performed with panviral2.64

### Analysis of UN62

| | |
|---|---|
| **Technology** | Paired-end short reads |
| **Input Files** | UN62_R1.fq.gz (1.38 GB), UN62_R2.fq.gz (1.41 GB) |
| **Submitted On** | 2023-09-26 14:21:19 UTC |
| **Duration** | 2h 33m 10s |
| **Tool Version** | panviral2.64 |
| **Location** | https://www.genomedetective.com/db/ui/analysis/1abb2807-2d0c-4c65-b599-46af9f9dd8ba |

#### Statistics

| | |
|---|---|
| **Original Read Length** | 20 - 150 |
| **Trimmed Read Length** | 50 - 135 |

| | # Reads | % of Reads |
|---|---|---|
| **Input file** | 43419312 | 100.0% |
| **After QC** | 43139000 | 99.4% |
| **After filtering** | 3469850 | 8.0% |
| **Mapped back to contigs** | 1265033 | 2.9% |

#### Assignments

| Assignment | No. of Reads | Depth of Coverage | Identity | | Genome Coverage | |
|---|---|---|---|---|---|---|
| | | | NT | AA | | |
| Torradovirus lycopersici (2 segments out of 2) | 464494 | 4623.0 | 88.8% | 93.7% | 99.8% | |
| Torradovirus lycopersici (segment RNA 2) | 333725 | 7488.0 | 91.5% | 94.3% | 99.6% | 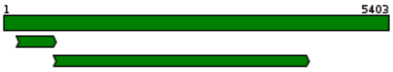  |
| Torradovirus lycopersici (segment RNA 1) | 130769 | 2646.1 | 87.0% | 93.2% | 99.9% | 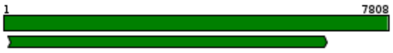  |
| Potato virus Y | 431062 | 5971.0 | 82.8% | 90.0% | 99.5% | 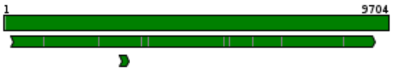  |
| Torradovirus marchitezum (2 segments out of 2) | 240631 | 4344.5 | 67.6% | 71.0% | 58.2% | |
| Torradovirus marchitezum (segment RNA 2) | 143680 | 14475.3 | 69.5% | 76.0% | 25.4% | 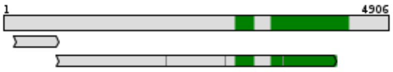  |
| Torradovirus marchitezum (segment RNA 1) | 96951 | 2179.8 | 67.2% | 70.0% | 80.5% | 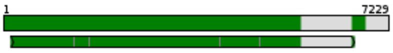  |
| Diachasmimorpha longicaudata entomopoxvirus (segment NC_043455.1) | 8833 | 993.3 | 60.0% | 55.5% | 84.0% | 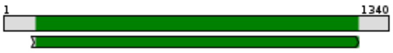  |

| Assignment | No. of Reads | Depth of Coverage | Identity | | Genome Coverage | |
|---|---|---|---|---|---|---|
| | | | NT | AA | | |
| Bracoviriform glomeratae (segment NC_043292.1) | 8084 | 3029.1 | 72.6% | 84.5% | 72.8% | 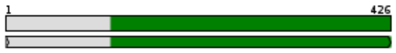 |
| Potato leafroll virus | 3283 | 75.3 | 98.1% | 97.2% | 97.2% | 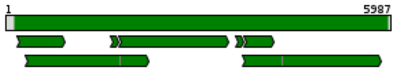 |
| Duamitovirus soch1 | 543 | 32.4 | 68.3% | 69.2% | 75.7% | 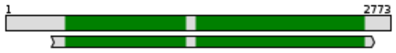 |
| Solendovirus venanicotianae | 124 | 8.5 | 77.0% | 73.5% | 20.1% | 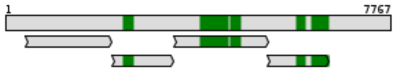 |

#### Discoveries

| Similar to | No. of Reads | Depth of Coverage | Identity | | Genome Coverage | |
|---|---|---|---|---|---|---|
| | | | NT | AA | | |
| Tomato chocolate spot virus (2 segments out of 2) | 79686 | 12924.2 | 73.1% | 70.5% | 5.5% | |
| Tomato chocolate spot virus (segment RNA 1) | 61095 | 24274.9 | 82.1% | 0.0% | 3.7% | 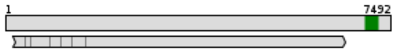 |
| Tomato chocolate spot virus (segment RNA2) | 18591 | 5209.1 | 67.0% | 70.5% | 8.0% | 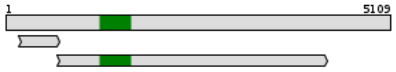 |
| Noumeavirus | 24245 | 10898.8 | 82.0% | 94.4% | 0.1% | 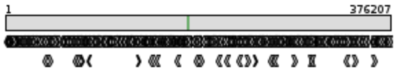 |
| Tomato necrotic dwarf virus (segment RNA1) | 2020 | 356.8 | 64.8% | 63.2% | 10.3% | 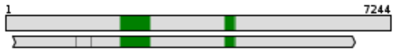 |
| Cladosporium fulvum T-1 virus | 511 | 39.2 | 53.0% | 47.2% | 18.5% | 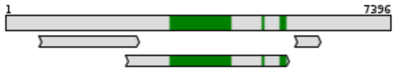 |
| Bracoviriform congregatae (3 segments out of 30) | 408 | 206.7 | 78.9% | 90.0% | 0.7% | |
| Bracoviriform congregatae (segment Circle 7) | 408 | 206.7 | 78.9% | 90.0% | 0.7% | 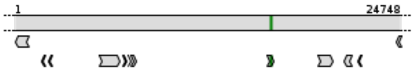 |
| Bracoviriform congregatae (segment Circle 7) | 277 | 198.1 | 78.6% | 86.8% | 0.5% | 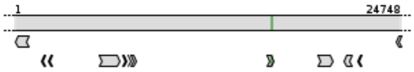 |
| Bracoviriform congregatae (segment Circle 7) | 5 | 3.1 | 75.5% | 85.4% | 0.6% | 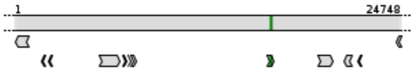 |
| Cassava brown streak virus | 391 | 85.7 | 60.8% | 57.7% | 5.8% | 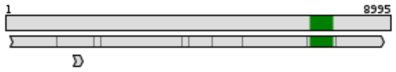 |
| Human gammaherpesvirus 8 (subtype: Could not assign) | 322 | 200.8 | 83.7% | 84.4% | 0.1% | 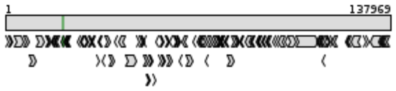 |

| Similar to | No. of Reads | Depth of Coverage | Identity | | Genome Coverage | |
|---|---|---|---|---|---|---|
| | | | NT | AA | | |
| Hibiscus bacilliform virus GD1 | 53 | 11.3 | 51.6% | 42.9% | 4.5% | 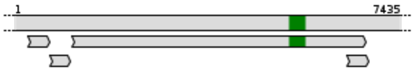 |
| Caulimovirus venafragariae | 51 | 6.8 | 57.6% | 51.7% | 11.1% | 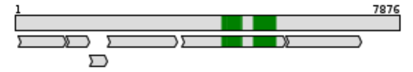 |
| Badnavirus occultipomeae | 46 | 8.1 | 49.5% | 38.4% | 5.3% | 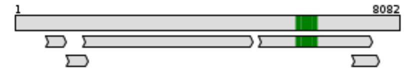 |
| Errantivirus | 38 | 8.9 | 37.9% | 33.3% | 7.7% | 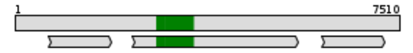 |
| Badnavirus volubetulae | 30 | 10.1 | 52.6% | 42.1% | 4.3% | 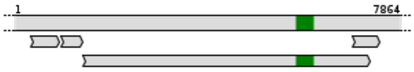 |
| Badnavirus zetavirgamusae | 29 | 8.4 | 53.9% | 40.4% | 6.3% | 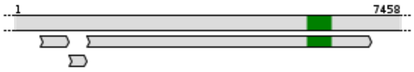 |
| Badnavirus maculasmallanthi | 15 | 6.2 | 57.1% | 48.8% | 3.2% | 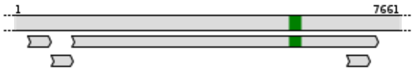 |
| Chinaberry tree badnavirus 1 | 13 | 6.5 | 55.4% | 45.8% | 3.5% | 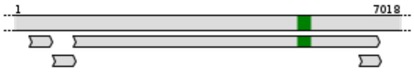 |
| Pinus nigra virus 1 | 12 | 4.5 | 55.3% | 47.4% | 4.1% | 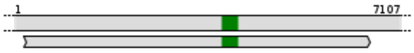 |
| Dioscovirus dioscoreae | 11 | 2.7 | 61.1% | 54.9% | 5.3% | 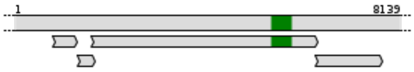 |
| Duamitovirus peex1 | 10 | 5.1 | 73.4% | 64.4% | 9.7% | 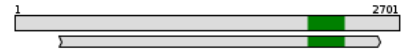 |
| Cacao swollen shoot Ghana J virus | 10 | 4.1 | 58.0% | 49.5% | 4.3% | 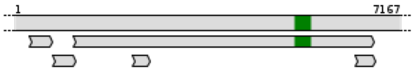 |
| Hibiscus bacilliform virus GD1 | 10 | 5.5 | 56.4% | 46.3% | 3.2% | 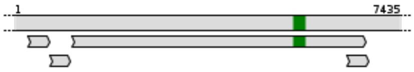 |
| Badnavirus maculasmallanthi | 9 | 2.8 | 51.8% | 45.0% | 5.5% | 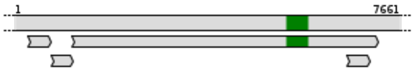 |
| Badnavirus venazanthoxyli | 8 | 3.8 | 58.4% | 53.2% | 3.5% | 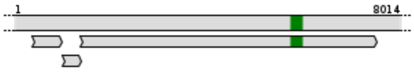 |
| Dioscovirus dioscoreae | 8 | 3.1 | 63.4% | 48.7% | 4.2% | 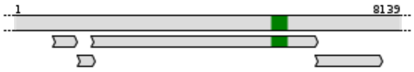 |
| Duamitovirus soch1 | 7 | 2.5 | 69.2% | 73.9% | 12.9% | 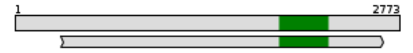 |

| Similar to | No. of Reads | Depth of Coverage | Identity | | Genome Coverage | |
|---|---|---|---|---|---|---|
| | | | NT | AA | | |
| Escherichia virus DE3 | 6 | 3.6 | 100.0% | 100.0% | 0.5% | 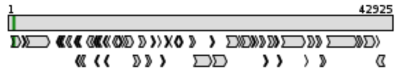 |
| Potato virus X | 4 | 2.4 | 73.7% | 88.0% | 3.5% | 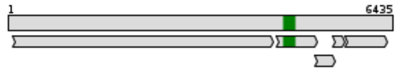 |
| Pandoravirus salinus | 4 | 2.7 | 75.4% | 80.5% | 0.0% | 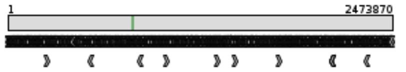 |
| Dioscorea bacilliform virus | 4 | 1.7 | 52.9% | 47.1% | 4.3% | 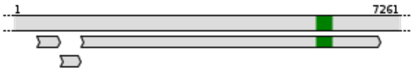 |
| Rhizobium phage RHEph10 | 3 | 2.9 | 80.0% | 82.2% | 0.1% | 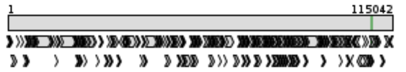 |
| Melanoplus sanguinipes entomopoxvirus | 2 | 2.0 | 80.0% | 80.0% | 0.1% | 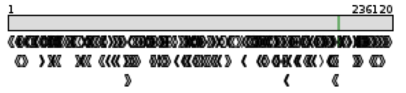 |
| Wadgaonvirus wv5004651 | 2 | 1.4 | 100.0% | 100.0% | 0.5% | 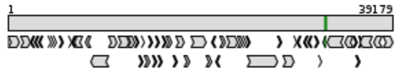 |
| Enterobacteria phage P7 | 2 | 1.6 | 100.0% | 100.0% | 0.2% | 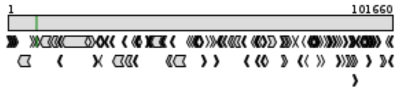 |
| Orthomarburgvirus marburgense | 2 | 1.0 | 70.4% | 98.9% | 1.4% | 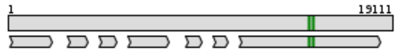 |
| Sclerotinia sclerotiorum deltaflexivirus 2 | 2 | 1.2 | 60.7% | 65.8% | 3.2% | 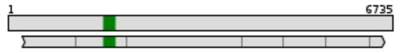 |
| Friend murine leukemia virus | 1 | 1.0 | 77.0% | 97.8% | 1.6% | 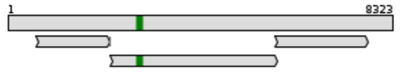 |
| Moloney murine leukemia virus | 1 | 1.0 | 76.3% | 100.0% | 1.6% | 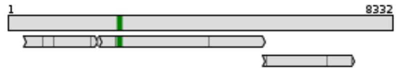 |
| Ochrobactrum phage POA1180 | 1 | 1.0 | 79.3% | 71.1% | 0.3% | 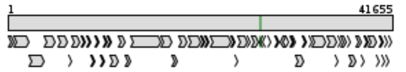 |

### NGS Details (UN62): Potato virus Y

#### Assembly

| | |
|---|---|
| **Coverage Length** | 9651 (1 contig(s)) |
| **Depth Of Coverage** | 5971.0 |
| **Number Of Reads** | 431062 |
| **Reads Per Million** | 9992.40 rpm (after QC) |
| **Ambiguities** | 0 |
| **Assembly Method** | de novo + reference guided assembly |
| **Consensus Caller** | Bcf Tools |

#### Coverage Map

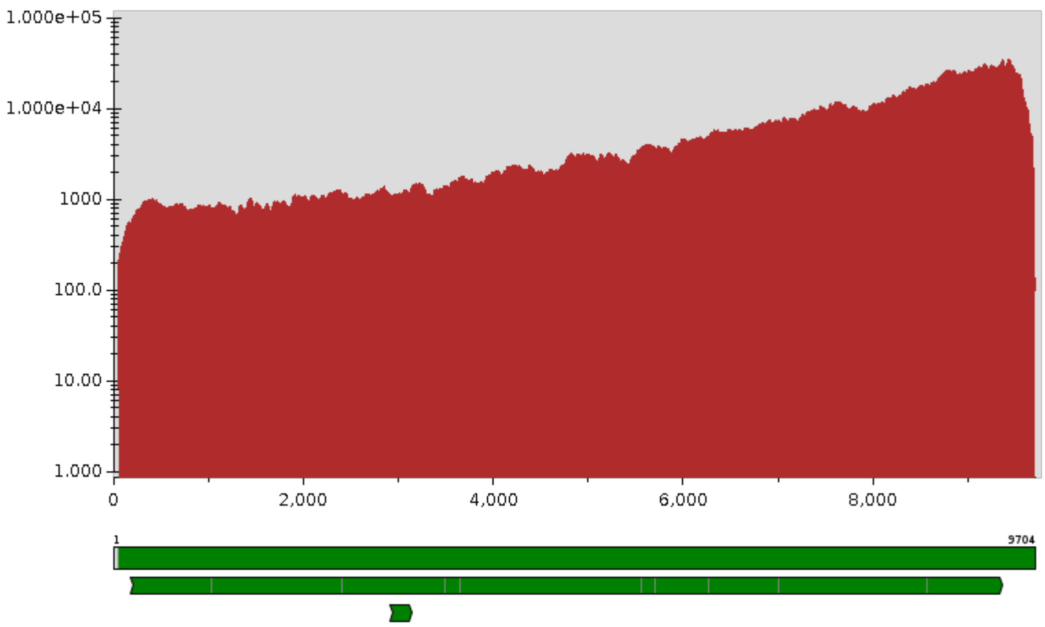

#### Assignment

| | |
|---|---|
| **Type** | Potato virus Y (Taxonomy ID: 12216) |
| **Reference Genome** | NC_001616.1 |
| **NT Identity (%)** | 82.7701 |
| **AA Identity (%)** | 89.9682 |
| **Number Of Stop Codons** | 3 |
| **Number Of CDS** | 2 |

#### Alignment

| | |
|---|---|
| **Alignment Score** | 12522.0 (NT) + 19019.0 (AA) = 31541.0 |
| **Concordance (%)** | 78.2514 |

| | |
|---|---|
| **Alignment Method** | Global, seeded, nucleotide + amino acids (AGA) |

#### Genome Region

Sequence starts at position 54 and ends at position 9704 relative to NC_001616.1 reference sequence.

#### Alignment Detailed Statistics

| | Begin | End | Coverage | Score | Concordance | Matches | Identities | I/D/M/F* | Stop Codons |
|---|---|---|---|---|---|---|---|---|---|
| **NT** | **54** | **9704** | **99.5%** | **12522** | **65.2%** | **9639 (99.8%)** | **7984 (82.7%)** | **7/12** | |

Mutations:

75C>T, 79C>T, 83G>A, 85T>C, 87C>T, 89G>A, 90T>C, 93A>T, 94A>G, 95A>T, 100T>C, 101T>A, 102C>T, 104T>C, 107C>T, 110T>G, 112C>T, 114C>T, 115T>C, 116A>T, 117G>C, 119A>G, 120C>T, 121A>T, 125T>G, 126T>C, 127G>A, 128G>T, 131A>G, 132C>T, 139A>T, 139_140insG, 141C>T, 143C>T, 146C>T, 150C>T, 150_151insT, 154T>C, 156C>A, 158T>A, 160A>C, 164C>T, 168C>A, 169C>G, 171A>C, 172T>C, 173T>A, 174T>G, 177G>C, 178A>G, 179T>C, 180C>T, 181C>A, 182T>G, 183C>T, 193T>C, 194T>C, 196C>A, 202A>G, 208C>T, 209T>C, 210G>T, 211T>G, 220G>A, 221T>A, 223T>G, 226A>G, 235A>T, 241C>T, 247A>T, 251T>C, 255G>T, 256C>T, 258A>G, 262T>C, 263A>G, 265T>G, 267T>C, 273A>T, 277A>T, 280A>G, 283G>T, 286G>A, 287G>A, 289T>C, 293G>A, 294T>C, 301T>C, 304C>T, 305G>A, 306C>T, 309A>G, 311C>T, 316A>T, 317A>G, 318C>T, 319A>G, 322A>G, 331A>T, 339T>A, 340C>A, 342A>G, 347A>G, 348A>C, 349A>G, 350T>C, 355T>A, 356A>G, 357C>T, 364T>C, 365G>A, 366T>C, 367G>A, 368C>T, 369T>C, 373A>G, 376C>T, 384T>G, 388G>A, 391C>T, 394A>G, 400G>A, 406T>C, 408C>T, 409C>T, 412G>A, 415A>T, 416A>G, 417T>A, 424T>C, 427G>A, 433A>G, 436G>C, 437G>A, 440A>G, 441G>A, 443A>C, 444A>G, 445G>A, 448T>A, 454A>G, 456A>C, 460T>C, 461C>A, 469A>G, 475C>G, 481T>A, 502T>C, 505A>T, 511C>T, 517T>A, 520T>A, 523A>T, 531C>T, 535G>A, 536C>T, 539C>T, 540A>G, 541A>G, 549G>A, 550A>G, 556C>T, 559T>C, 562T>C, 568T>C, 571A>G, 576T>C, 580T>C, 585T>C, 586C>G, 593C>T, 594G>A, 595C>T, 597C>G, 599A>G, 601A>G, 602A>C, 603T>C, 604A>G, 613A>G, 616A>G, 619C>G, 622G>A, 625G>A, 628T>C, 631C>T, 635A>G, 637T>C, 644A>G, 646A>G, 655T>C, 661G>A, 663A>C, 664G>C, 667A>G, 679C>T, 682C>A, 683T>C, 685A>G, 691T>C, 699C>G, 703T>A, 706T>A, 719A>G, 721G>A, 722A>G, 724A>G, 725C>T, 727T>G, 730T>A, 731G>T, 734T>C, 735A>C, 736C>T, 737T>C, 738C>G, 739C>T, 744C>T, 745G>C, 749C>T, 751A>C, 754T>C, 765T>A, 770T>C, 773C>A, 774G>A, 775A>G, 776C>A, 790C>T, 801T>A, 813G>C, 814A>G, 816T>A, 818T>C, 823A>G, 825G>A, 826T>A, 833C>A, 835G>A, 841C>T, 851A>G, 853T>C, 859C>T, 860C>T, 862C>T, 863A>G, 865T>C, 867T>C, 869A>G, 871C>T, 872A>T, 874A>G, 876G>A, 879G>A, 886T>C, 898C>T, 903A>G, 906C>A, 907A>C, 909A>C, 910A>C, 912G>A, 913C>T, 922C>A, 931T>A, 938T>C, 939C>T, 940A>T, 943A>G, 952C>T, 967A>G, 973A>G, 976G>C, 980T>A, 982G>C, 991A>G, 994T>C, 999G>A, 1003T>C, 1007C>A, 1008A>G, 1009G>A, 1010A>G, 1012T>G, 1013A>G, 1018A>G, 1019A>C, 1021C>G, 1030C>T, 1039G>A, 1042T>C, 1045C>A, 1048C>A, 1051T>C, 1060G>A, 1063T>A, 1064C>T, 1069C>T, 1083G>A, 1093T>C, 1097T>G, 1099G>A, 1105C>T, 1114A>G, 1117T>G, 1123A>G, 1126T>A, 1129C>T, 1143G>A, 1147A>T, 1150T>A, 1154T>A, 1156G>A, 1165C>T, 1168T>C, 1172C>T, 1174T>A, 1201C>T, 1221G>A, 1229G>A, 1230T>C, 1231T>C, 1233G>A, 1234C>T, 1240G>A, 1246G>A, 1250T>C, 1252G>A, 1255T>C, 1257A>C, 1258A>G, 1267A>G, 1273T>C, 1276G>A, 1278A>G, 1283T>C, 1290C>T, 1291G>A, 1298G>A, 1299A>C, 1300C>T, 1306T>C, 1307A>G, 1309A>G, 1312T>C, 1316A>G, 1318T>A, 1327G>A, 1329T>C, 1333G>T, 1336A>G, 1343C>T, 1345A>G, 1354G>A, 1357G>T, 1360C>T, 1367C>T, 1369C>G, 1373C>A, 1378C>T, 1384G>A, 1385A>G, 1393A>G, 1396C>T, 1405G>T, 1408A>G, 1411G>A, 1415G>T, 1417A>T, 1420G>C, 1429T>C, 1432A>G, 1436G>A, 1438C>T, 1439T>C, 1441A>G, 1444T>C, 1453C>T, 1456G>A, 1468A>G, 1492A>G, 1499T>C, 1501G>A, 1504T>C, 1507G>A, 1510C>T, 1513A>G, 1516A>G, 1519A>T, 1522G>A, 1525C>T, 1528G>A, 1534C>T, 1540T>A, 1546C>T, 1555A>G, 1558T>C, 1561T>C, 1564A>C, 1567T>A, 1573C>T, 1576A>G, 1583T>C, 1588T>A, 1594G>A, 1600C>T, 1609G>A, 1618G>A, 1621C>T, 1624C>T, 1631C>T, 1639A>G, 1651C>T, 1654C>G, 1675T>C, 1693A>G, 1702T>C, 1706G>A, 1707A>G, 1718G>A, 1726A>G, 1729C>T, 1732A>C, 1738T>C, 1742A>G, 1744C>T, 1747C>T, 1753T>C, 1758G>A, 1768G>A, 1786C>T, 1789A>G, 1790G>A, 1795C>T, 1805T>C, 1807A>G, 1813G>A, 1819G>A, 1825G>A, 1837C>T, 1840T>C, 1849A>G, 1858C>G, 1864A>G, 1870C>T, 1876T>C, 1882A>G, 1888T>A, 1891T>C, 1906T>C, 1915A>T, 1930T>C, 1936C>T, 1937A>G, 1939T>C, 1945A>G, 1951C>T, 1960A>T, 1978A>G, 1981T>A, 1984C>T, 1987T>C, 1996C>T, 2004T>A, 2020A>G, 2023G>A, 2026T>C, 2029G>T, 2032G>A, 2059T>C, 2068T>G, 2071C>T, 2072G>A, 2074G>A, 2087C>T, 2089G>A, 2101C>T, 2104G>A, 2107G>A, 2116G>A, 2122C>T, 2137C>T, 2158T>C, 2164C>T, 2170A>G, 2182T>C, 2188G>A, 2191C>A, 2208G>A, 2218T>C, 2224C>T, 2227A>T, 2242G>A, 2245C>T, 2254G>A, 2257T>C, 2260C>T, 2266C>G, 2272A>G, 2275G>A, 2293A>G, 2299C>T, 2302G>A, 2305G>A, 2314A>T, 2317T>C, 2326A>G, 2341G>A, 2347A>G, 2350T>C, 2356G>A, 2365T>C, 2371A>G, 2374A>G, 2383T>C, 2386A>T, 2389A>G, 2392T>C, 2395T>C, 2414A>G, 2415A>G, 2416T>G, 2419T>A, 2420A>T, 2440A>T, 2467T>C, 2476T>C, 2482G>A, 2491G>A, 2497G>T, 2503G>A, 2518A>G, 2521T>C, 2523A>G, 2533A>G, 2539G>A, 2545A>G, 2551G>A, 2554T>A, 2558C>T, 2561T>C, 2563G>C, 2572A>G, 2584T>C, 2587C>T, 2593G>C, 2599T>C, 2608T>C, 2620T>C, 2629G>A, 2632G>A, 2634G>A, 2638G>A, 2644T>A, 2653A>G, 2665T>C, 2678C>T, 2683A>G, 2689T>C, 2698A>C, 2707A>G, 2710G>A, 2716A>G, 2719A>G, 2722C>T, 2723G>A, 2725C>T, 2728A>G, 2734G>A, 2737T>A, 2746T>C, 2747G>A, 2749T>A, 2752A>T, 2755T>A, 2761C>T, 2764C>T, 2779T>C, 2785G>T, 2791C>T, 2795C>A, 2806C>T, 2816T>C, 2818G>C, 2822G>A, 2824G>A, 2827G>A, 2836T>C, 2839G>A, 2842T>C, 2863C>A, 2864C>T, 2882C>A, 2885A>C, 2886G>A, 2893C>T, 2895C>T, 2902C>T, 2905A>G, 2911T>C, 2923T>C, 2931A>G, 2940A>G, 2947T>C, 2962T>C, 2971_2972insA, 2989T>C, 2991C>T, 2994A>G, 2999G>A, 3000A>G, 3016delC, 3021G>A, 3024A>T, 3025C>T, 3036C>T, 3039A>G, 3040A>C, 3042A>C, 3043G>A, 3052G>A, 3061A>G, 3082G>A, 3091C>T, 3099C>T, 3100C>A, 3104G>A, 3105T>G, 3106G>A, 3109C>T, 3112A>G, 3118T>C, 3121C>T, 3127A>G, 3132G>A, 3137C>A, 3138G>A, 3139A>G, 3151T>C, 3152T>A, 3154C>T, 3165G>A, 3169A>G, 3199C>T, 3203A>C, 3208T>C, 3214A>G, 3217C>T, 3218T>C, 3223C>A, 3225C>A, 3230G>A, 3235C>T, 3257T>C, 3262C>T, 3265C>T, 3266G>A, 3268A>G, 3271G>A, 3275G>A, 3284G>A, 3301G>A, 3315G>A, 3316A>G, 3318A>C, 3322C>T, 3325G>A, 3328G>A, 3337A>T, 3349G>A, 3355C>T, 3358C>T, 3359A>G, 3367A>G, 3370T>C, 3377T>C, 3379A>G, 3394A>G, 3397C>T, 3400T>G, 3406A>G, 3418C>T, 3421T>A, 3427T>C, 3433G>A, 3436A>T, 3439A>G, 3442C>T, 3445T>C, 3457G>A, 3460T>C, 3463T>A, 3469G>T, 3472G>A, 3496C>T, 3499G>A, 3505C>T, 3513T>G, 3520A>G, 3529A>G, 3532A>G, 3545A>G, 3548G>A, 3551T>C, 3554G>A, 3556C>T, 3583T>C, 3586T>C, 3589C>T, 3595C>T, 3607T>C, 3613T>C, 3616G>A, 3619T>C, 3623C>G, 3628C>T, 3632C>A, 3637T>C, 3638T>C, 3643A>G, 3661A>G, 3670G>A, 3682T>C, 3688G>A, 3697G>A, 3698A>G, 3706T>C, 3709T>C, 3713T>C, 3715G>A, 3717G>A, 3721G>A, 3731C>A, 3732G>A, 3736T>A, 3746C>T, 3748A>G, 3754A>G, 3763T>C, 3778A>G, 3781A>G, 3784C>T, 3796T>C, 3805A>T, 3811C>T, 3814A>G, 3829C>T, 3838T>C, 3844A>G, 3847A>C, 3856T>A, 3862G>T, 3871C>T, 3877C>A, 3883C>T, 3886A>G, 3892A>G, 3895C>T, 3898T>C, 3904A>G, 3905C>A, 3907G>A, 3913T>C, 3928G>A, 3931A>T, 3937G>A, 3955T>C, 3958G>A, 3961C>T, 3964A>G, 3967T>C, 3976A>G, 3979T>C, 3985A>T, 3997A>T, 4000G>A, 4006C>T, 4009T>G, 4012C>T, 4015A>G, 4024T>C, 4030A>G, 4033A>T, 4042G>A, 4052C>T, 4054G>A, 4057T>C, 4063T>A, 4066A>G, 4073A>G, 4078T>C, 4081C>T, 4084T>C, 4096C>T, 4108C>T, 4111A>T, 4117G>A, 4118C>T, 4120A>G, 4126C>T, 4128T>A, 4129C>T, 4132C>T, 4135T>C, 4147A>G, 4148T>C, 4151G>T, 4162C>T, 4166G>A, 4171A>C, 4180G>A, 4190C>T, 4198T>C, 4201C>T, 4210G>T, 4213C>T, 4219T>C, 4220C>T, 4223C>A, 4225G>A, 4231T>G, 4237T>C, 4252A>G, 4253T>C, 4255A>G, 4258A>G, 4270T>C, 4282A>G, 4285G>A, 4291A>G, 4303G>A, 4312C>T, 4319A>G, 4321A>G, 4324G>T, 4327G>A, 4336G>T, 4339T>C, 4348A>T, 4360A>G, 4366T>C, 4369T>A, 4375T>C, 4387T>C, 4395T>A, 4406G>A, 4411T>C, 4438T>C, 4444C>A, 4450C>T, 4456C>T, 4465C>T, 4475A>G, 4477G>T, 4480C>T, 4481A>T, 4483A>G, 4495C>T, 4507G>A, 4510C>T, 4513T>G, 4517C>T, 4528C>T, 4534A>G, 4540C>T, 4544G>T, 4545C>T, 4546G>A, 4548G>A, 4558T>C, 4564A>T, 4570C>T, 4576A>T, 4597G>A, 4600C>T, 4606C>T, 4609G>A, 4615G>A, 4618C>T, 4625T>C, 4627G>T, 4648C>T, 4651T>C, 4660G>A, 4663C>T, 4669T>A, 4679G>A, 4681G>C, 4684T>C, 4687T>G, 4699G>A, 4702A>G, 4708A>G, 4711G>A, 4712C>T, 4738A>G, 4744A>T, 4747A>G, 4748T>C, 4765T>C, 4796G>A, 4798C>T, 4803C>G, 4804T>C, 4807G>A, 4810G>T, 4813T>G, 4816T>C, 4831T>C, 4834C>T, 4840A>T, 4852A>G, 4855C>T, 4867T>C, 4870G>C, 4873T>A, 4882T>C, 4903A>T, 4909G>A, 4915T>C, 4918A>G, 4919T>C, 4921G>A, 4924T>C, 4927C>G, 4933T>C, 4936C>A, 4939G>A, 4942T>C, 4945C>T, 4963A>G, 4972T>C, 4990T>C, 5002A>G, 5023T>C, 5024T>C, 5029C>T, 5032T>C, 5038T>C, 5044A>T, 5053A>T, 5056G>A, 5057A>G, 5060A>G, 5071G>T, 5090C>A, 5095A>G, 5098G>T, 5100C>T, 5104A>G, 5107A>T, 5110T>C, 5113A>G, 5116G>A, 5119A>G, 5120G>T, 5122C>T, 5126A>G, 5137T>C, 5140C>T, 5143A>G, 5146G>A, 5161C>A, 5167A>G, 5173T>G, 5182G>A, 5183G>A, 5188C>T, 5191G>A, 5197A>G, 5200C>T, 5206C>T, 5209A>G, 5218C>T, 5230G>A, 5236C>T, 5239C>T, 5245C>T, 5263A>T, 5269C>T, 5272C>T, 5275C>A, 5278C>T, 5281A>C, 5284A>G, 5287T>C, 5288C>T, 5293A>T, 5294T>C, 5296G>A, 5302G>A, 5311T>A, 5314A>G, 5323A>T, 5347T>C, 5353C>T, 5365C>T, 5386T>C, 5395C>A, 5422A>G, 5425T>C, 5431C>T, 5437G>A, 5452T>C, 5458G>A, 5467T>C, 5470A>G, 5471C>T, 5473A>G, 5479A>G, 5482C>T, 5485A>C, 5488T>C, 5489T>C, 5491G>A, 5497T>C, 5506T>C, 5509G>A, 5512G>A, 5516T>C, 5530T>C, 5533G>A, 5536G>T, 5539G>A, 5548T>G, 5554C>T, 5560T>A, 5561G>A, 5563G>A, 5566G>C, 5569A>G, 5584C>G, 5593G>A, 5596G>A, 5597A>G, 5605C>A, 5617G>T, 5619C>T, 5620T>G, 5629G>A, 5630A>C, 5641C>T, 5650A>T, 5662A>T, 5665A>G, 5689A>G, 5693G>A, 5698G>A, 5713A>G, 5716G>C, 5719A>G, 5743C>A, 5755C>A, 5758T>C, 5764T>C, 5770A>G, 5779C>T, 5782T>C, 5791C>T, 5803C>T, 5836G>A, 5839A>G, 5845A>G, 5848A>G, 5853A>G, 5857T>C, 5860C>T, 5863A>T, 5869T>A, 5875T>C, 5884C>T, 5893C>T, 5894A>G, 5896C>G, 5899C>T, 5908G>A, 5911T>C, 5923G>A, 5938A>G, 5941C>T, 5944T>C, 5947T>C, 5959G>A, 5960C>G, 5961G>C, 5962G>C, 5974A>G, 5986T>A, 5989C>T, 5990A>G, 5998T>C, 6008A>C, 6010A>T, 6019A>T, 6022G>T, 6025A>T, 6030A>C, 6049C>A, 6058G>A, 6061A>G, 6064C>A, 6070T>C, 6072G>A, 6073T>C, 6074A>C, 6079G>A, 6081C>A, 6085A>T, 6091A>T, 6094C>T, 6111G>C, 6112T>C, 6118A>G, 6124G>A, 6127G>A, 6128A>G, 6130T>C, 6133T>C, 6136A>G, 6148C>T, 6157A>G, 6158G>A, 6163T>C, 6169A>G, 6170A>T, 6172A>G, 6178C>T, 6187A>G, 6190T>C, 6196G>A, 6203C>T, 6205C>T, 6208A>G, 6209C>T, 6211A>G, 6217G>A, 6229T>A, 6230G>A, 6238C>T, 6247G>A, 6253A>T, 6257G>A, 6258C>A, 6274T>C, 6282A>G, 6286G>A, 6298C>T, 6301G>A, 6304A>G, 6307C>T, 6322C>T, 6328A>T, 6331T>G, 6334T>C, 6337G>A, 6338C>T, 6358T>C, 6361G>A, 6362G>A, 6367A>T, 6370G>A, 6376C>T, 6379T>G, 6385A>C, 6391A>T, 6394A>G, 6397C>T, 6401G>A, 6405C>T, 6412C>T, 6418A>G, 6421T>C, 6429A>T, 6436T>A, 6445G>A, 6450A>G, 6454C>G, 6460C>T, 6463T>A, 6469C>T, 6472G>A, 6475G>A, 6481T>C, 6485C>A, 6487C>T, 6490T>C, 6496C>T, 6502G>A, 6514T>C, 6517G>A, 6527C>A, 6532C>A, 6541G>A, 6544A>G, 6556C>T, 6559T>C, 6565G>A, 6569T>C, 6571G>A, 6586T>A, 6588C>T, 6598A>G, 6601A>G, 6604T>C, 6607T>C, 6616A>G, 6622C>T, 6631G>A, 6634G>A, 6635T>C, 6640T>A, 6646G>T, 6647A>G, 6652C>T, 6664C>T, 6670T>G, 6677A>G, 6679A>G, 6685C>T, 6691A>T, 6694C>T, 6715A>G, 6716G>A, 6719A>G, 6724A>G, 6734C>T, 6742G>A, 6745G>A, 6754C>T, 6760A>G, 6764A>T, 6766A>G, 6775T>A, 6782C>T, 6787A>T, 6790C>T, 6795C>T, 6796A>G, 6799C>A, 6802C>A, 6808C>T, 6811C>T, 6814C>T, 6823C>T, 6829A>G, 6847C>T, 6853A>C, 6857A>G, 6862G>A, 6871A>G, 6875G>A, 6876T>C, 6877C>A, 6880G>A, 6883T>A, 6889T>G, 6901C>T, 6916C>T, 6919G>A, 6925A>G, 6928T>C, 6934C>A, 6949G>C, 6955C>T, 6958A>G, 6976A>G, 6980C>T, 6985C>T, 6997A>T, 7000G>T, 7001G>A, 7003G>A, 7012T>C, 7015G>A, 7018T>C, 7032T>A, 7039C>T, 7040T>C, 7042G>A, 7048A>G, 7060T>A, 7063C>A, 7078C>T, 7082T>C, 7084A>G, 7087A>G, 7090C>A, 7096T>C, 7099A>G, 7102T>C, 7111G>A, 7117A>G, 7125C>A, 7129A>G, 7132T>C, 7133C>T, 7135G>A, 7141G>A, 7144T>C, 7145G>T, 7147A>G, 7150G>A, 7159A>T, 7160_7165delGAGGCA, 7171C>T, 7189G>T, 7198A>G, 7201C>T, 7205C>T, 7210T>C, 7216T>A, 7219G>A, 7225C>A, 7228G>A, 7240G>A, 7243T>C, 7249A>G, 7252T>G, 7258T>A, 7261T>C, 7264T>A, 7267C>G, 7270G>A, 7273_7274insT, 7274C>G, 7275G>T, 7281A>T, 7284T>C, 7296C>T, 7311C>T, 7314C>T, 7317G>A, 7321T>A, 7322delG, 7327C>T, 7330C>T, 7333C>T, 7334A>C, 7342T>C, 7343G>A, 7344C>A, 7345A>C, 7354T>C, 7363A>G, 7369T>C, 7372C>T, 7378G>T, 7381C>T, 7396A>T, 7399C>T, 7405C>T, 7407G>T, 7408T>G, 7414G>T, 7415T>G, 7420A>G, 7429C>T, 7432T>C, 7441T>C, 7459G>A, 7462A>G, 7468A>T, 7471C>A, 7477A>G, 7480C>T, 7486G>T, 7495T>C, 7498A>G, 7501T>C, 7503T>C, 7504G>A, 7513T>A, 7519C>T, 7522A>G, 7531G>A, 7537G>A, 7540C>T, 7546T>C, 7570G>A, 7573G>A, 7579G>A, 7588T>C, 7595C>T, 7604T>C, 7606G>A, 7615T>A, 7624T>C, 7633C>T, 7636C>T, 7639T>C, 7648T>C, 7666A>G, 7672T>C, 7678A>T, 7684G>A, 7690T>C, 7696T>C, 7702T>G, 7709G>A, 7713A>G, 7714A>G, 7715C>T, 7726T>C, 7735G>A, 7744A>G, 7747C>T, 7762C>T, 7768G>A, 7771T>C, 7780A>G, 7783A>C, 7786T>C, 7793C>T, 7795A>G, 7798C>T, 7804T>C, 7812C>T, 7834A>G, 7837C>T, 7844G>T, 7846G>A, 7855G>A, 7861G>A, 7870A>G, 7873C>T, 7876T>A, 7879G>A, 7882T>A, 7883G>A, 7894A>T, 7897T>C, 7921C>T, 7927G>A, 7936T>C, 7939T>C, 7942T>C, 7966G>A, 7969T>C, 7975T>C, 7987C>T, 7990T>C, 8005T>C, 8032A>G, 8038C>T, 8041C>T, 8043G>A, 8080G>A, 8089G>A, 8095G>A, 8098G>A, 8107C>T, 8110T>C, 8116T>C, 8128A>G, 8146T>C, 8151A>G, 8152C>T, 8164G>A, 8178G>A, 8182G>A, 8191G>A, 8197C>T, 8203C>T, 8208G>A, 8213C>T, 8215G>A, 8218A>G, 8221C>T, 8224G>A, 8239A>G, 8254G>A, 8260T>C, 8263A>G, 8281T>C, 8284A>G, 8294C>G, 8296A>G, 8299G>A, 8302C>T, 8317T>C, 8320C>T, 8326T>A, 8335G>A, 8338C>T, 8349C>T, 8353A>G, 8362C>T, 8365A>G, 8368C>T, 8380C>T, 8383A>G, 8389A>G, 8404T>C, 8408G>T, 8413A>G, 8419G>T, 8434T>C, 8440T>C, 8452G>A, 8458A>G, 8460G>A, 8464A>G, 8467G>A, 8479G>A, 8480G>A, 8482T>A, 8497G>A, 8498C>T, 8500A>G, 8502G>A, 8503A>G, 8506C>T, 8515A>T, 8524C>T, 8527A>C, 8528T>C, 8530A>G, 8539G>C, 8546C>T, 8547T>G, 8552T>A, 8563A>G, 8574C>G, 8585A>G, 8598G>A, 8602C>T, 8603A>T, 8604A>C, 8605C>T, 8620A>G, 8622C>A, 8625A>G, 8626G>A, 8629G>A, 8632C>T, 8638C>T, 8641G>A, 8647C>T, 8648C>T, 8649C>T, 8650G>C, 8653C>T, 8656A>G, 8658G>A, 8664A>C, 8665T>G, 8674T>G, 8679C>T, 8680A>T, 8683C>A, 8692G>A, 8695A>T, 8707G>A, 8708A>C, 8716G>A, 8722C>T, 8725G>A, 8731A>G, 8737A>G, 8743C>T, 8745C>A, 8752G>A, 8755A>T, 8770C>T, 8771T>C, 8776A>G, 8779T>C, 8782G>A, 8791T>C, 8797A>G, 8800A>G, 8803A>G, 8809T>C, 8812T>C, 8818T>C, 8824G>A, 8827A>G, 8848G>A, 8854T>C, 8856A>G, 8857G>A, 8863G>A, 8865G>A, 8867A>C, 8869G>T, 8872A>G, 8883G>A, 8888A>T, 8929C>T, 8954G>A, 8959C>T, 8962A>T, 8983G>A, 8984A>G, 8986T>C, 8988A>T, 8998G>A, 9004G>A, 9007G>A, 9016C>T, 9037C>A, 9085A>T, 9097C>T, 9131A>G, 9136A>T, 9145G>T, 9149A>G, 9150T>G, 9151G>A, 9152G>A, 9157A>G, 9160G>T, 9163T>C, 9184G>A, 9196A>G, 9214G>A, 9217A>G, 9220G>A, 9244A>T, 9247G>A, 9256C>T, 9260C>T, 9262T>C, 9274G>A, 9280C>T, 9289C>T, 9298A>C, 9304G>A, 9334C>T, 9364C>G, 9377_9378insT, 9387C>T, 9404G>A, 9411C>T, 9412A>G, 9414A>T, 9437C>T, 9454A>G, 9458T>C, 9467G>A, 9470_9471insT, 9482G>A, 9483C>T, 9498T>C, 9516T>G, 9517delA, 9533G>A, 9534G>T, 9548A>T, 9551C>T, 9553_9554delTA, 9556A>T, 9557T>A, 9558A>T, 9574T>A, 9581G>T, 9582T>G, 9583G>C, 9584_9585insG, 9590C>T, 9591T>C, 9607C>T, 9610T>C, 9615delC, 9620A>G, 9648A>G, 9674T>C, 9690C>T

| | Begin | End | Coverage | Score | Concordance | Matches | Identities | I/D/M/F* | Stop Codons |
|---|---|---|---|---|---|---|---|---|---|
| **NT** | **54** | **9704** | **99.5%** | **12522** | **65.2%** | **9639 (99.8%)** | **7984 (82.7%)** | **7/12** | |
| **CDS** | | | | | | | | | |
| PVYgp1 | 1 | 3064 | 100% | 18924 | 89.7% | 3062 (99.9%) | 2771 (90.4%) | 2/2/4/4 | 1 |
| Protein mutations: | Y4Q (194T>C 196C>A), C9L (209T>C 210G>T 211T>G), F13M (221T>A 223T>G), S23P (251T>C), C24F (255G>T 256C>T), E25G (258A>G), I27V (263A>G 265T>G), V28A (267T>C), E30V (273A>T), A35T (287G>A 289T>C), V37T (293G>A 294T>C), A41I (305G>A 306C>T), D42G (309A>G), E44D (316A>T), T45V (317A>G 318C>T 319A>G), L52Q (339T>A 340C>A), K53R (342A>G), K55A (347A>G 348A>C 349A>G), Y56H (350T>C), T58V (356A>G 357C>T), V61T (365G>A 366T>C 367G>A), L62S (368C>T 369T>C), F67C (384T>G), A75V (408C>T 409C>T), M78E (416A>G 417T>A), E85K (437G>A), R86E (440A>G 441G>A), K87R (443A>C 444A>G 445G>A), D88E (448T>A), E91A (456A>C), H93N (461C>A), D111E (517T>A), S116F (531C>T), P118S (536C>T), Q119W (539C>T 540A>G 541A>G), R122K (549G>A 550A>G), M131T (576T>C), V134A (585T>C 586C>G), R137Y (593C>T 594G>A 595C>T), P138R (597C>G), I139V (599A>G 601A>G), I140P (602A>C 603T>C 604A>G), M147I (625G>A), I151V (635A>G 637T>C), I154V (644A>G 646A>G), E160A (663A>C 664G>C), H166Q (682C>A), T172S (699C>G), H174Q (706T>A), K179E (719A>G 721G>A), I180V (722A>G 724A>G), A183S (731G>T), Y184P (734T>C 735A>C 736C>T), S185R (737T>C 738C>G 739C>T), A187V (744C>T 745G>C), R189C (749C>T 751A>C), M194K (765T>A), R197K (773C>A 774G>A 775A>G), M206K (801T>A), G210A (813G>C 814A>G), L211H (816T>A), R214Q (825G>A 826T>A), N223D (851A>G 853T>C), R226C (860C>T 862C>T), T227A (863A>G 865T>C), I228T (867T>C), N229D (869A>G 871C>T), I230L (872A>T 874A>G), R231Q (876G>A), R232K (879G>A), N240S (903A>G), T241N (906C>A 907A>C), K242T (909A>C 910A>C), S243N (912G>A 913C>T), S252L (938T>C 939C>T 940A>T), L266I (980T>A 982G>C), R272K (999G>A), Q275R (1007C>A 1008A>G 1009G>A), S276G (1010A>G 1012T>G), I277V (1013A>G), N279Q (1019A>C 1021C>G), D288E (1048C>A), R300Q (1083G>A), S305A (1097T>G 1099G>A), R320K (1143G>A), L324I (1154T>A 1156G>A), S346N (1221G>A), V349T (1229G>A 1230T>C 1231T>C), S350N (1233G>A 1234C>T), K358T (1257A>C 1258A>G), N365S (1278A>G), A369V (1290C>T 1291G>A), D372T (1298G>A 1299A>C 1300C>T), I375V (1307A>G 1309A>G), N378E (1316A>G 1318T>A), I382T (1329T>C), L397I (1373C>A), I401V (1385A>G), E407D (1405G>T), A411S (1415G>T 1417A>T), V418I (1436G>A 1438C>T), E508R (1706G>A 1707A>G), A512T (1718G>A), I520V (1742A>G 1744C>T), S525N (1758G>A), V536I (1790G>A), I585V (1937A>G 1939T>C), F607Y (2004T>A), I628M (2068T>G), V630I (2072G>A 2074G>A), R675K (2208G>A), D694E (2266C>G), N744G (2414A>G 2415A>G 2416T>G), S746C (2420A>T), K780R (2523A>G), R817K (2634G>A), V847I (2723G>A 2725C>T), A855T (2747G>A 2749T>A), H871N (2795C>A), V880I (2822G>A 2824G>A), P900T (2882C>A), S901H (2885A>C 2886G>A), T904M (2895C>T), N916S (2931A>G), N919S (2940A>G), E929_N930insX (2971_2972insA), T936I (2991C>T), H937R (2994A>G), E939R (2999G>A 3000A>G), R946Q (3021G>A), Y947F (3024A>T 3025C>T), T951I (3036C>T), E952G (3039A>G 3040A>C), K953T (3042A>C 3043G>A), A972V (3099C>T 3100C>A), V974R (3104G>A 3105T>G 3106G>A), S983N (3132G>A), R985K (3137C>A 3138G>A 3139A>G), F990I (3152T>A 3154C>T), C994Y (3165G>A), F1012L (3218T>C), T1014N (3225C>A), V1016I (3230G>A), V1028M (3266G>A 3268A>G), V1031M (3275G>A), A1034T (3284G>A), R1044K (3315G>A 3316A>G), E1045A (3318A>C), M1059V (3359A>G), D1072E (3400T>G), V1110G (3513T>G), M1121V (3545A>G), A1122T (3548G>A), V1124I (3554G>A 3556C>T), L1147V (3623C>G), L1150M (3632C>A), Y1152H (3638T>C), I1172V (3698A>G), S1178N (3717G>A), R1183K (3731C>A 3732G>A), I1297V (4073A>G), F1315Y (4128T>A 4129C>T), A1323S (4151G>T), V1328I (4166G>A), L1347I (4223C>A 4225G>A), I1379V (4319A>G 4321A>G), F1404Y (4395T>A), V1408I (4406G>A), M1431V (4475A>G 4477G>T), T1433S (4481A>T 4483A>G), A1454L (4544G>T 4545C>T 4546G>A), R1455K (4548G>A), V1499I (4679G>A 4681G>C), V1538I (4796G>A 4798C>T), T1540S (4803C>G 4804T>C), S1625G (5057A>G), T1626A (5060A>G), L1636I (5090C>A), A1639V (5100C>T), E1641D (5107A>T), V1646F (5120G>T 5122C>T), I1648V (5126A>G), V1667I (5183G>A), A1793T (5561G>A 5563G>A), I1805V (5597A>G), N1807K (5605C>A), A1812V (5619C>T 5620T>G), I1816L (5630A>C), V1837I (5693G>A), K1890R (5853A>G), I1904V (5894A>G 5896C>G), R1926A (5960C>G 5961G>C 5962G>C), I1936V (5990A>G), E1945D (6019A>T), K1949T (6030A>C), D1955E (6049C>A), M1958I (6058G>A), S1963N (6072G>A 6073T>C), N1964H (6074A>C), T1966N (6081C>A), C1976S (6111G>C 6112T>C), I1982V (6128A>G 6130T>C), V1992I (6158G>A), T1996S (6170A>T 6172A>G), L2007F (6203C>T 6205C>T), V2016I (6230G>A), A2025K (6257G>A 6258C>A), K2033R (6282A>G), A2060T (6362G>A), V2073I (6401G>A), A2074V (6405C>T), Y2082F (6429A>T), Q2089R (6450A>G), H2101N (6485C>A 6487C>T), L2115I (6527C>A), T2135I (6588C>T), Y2151H (6635T>C), I2155V (6647A>G), I2165V (6677A>G 6679A>G), D2178N (6716G>A), N2179D (6719A>G), I2194L (6764A>T 6766A>G), A2204V (6795C>T 6796A>G), H2205Q (6799C>A), N2225D (6857A>G), V2231T (6875G>A 6876T>C 6877C>A), D2250E (6934C>A), V2273I (7001G>A 7003G>A), F2283Y (7032T>A), T2314K (7125C>A), A2321S (7145G>T 7147A>G), E2326_A2327del (7160_7165delGAGGCA), D2344E (7216T>A), D2358E (7258T>A), D2363_R2364insX (7273_7274insT), R2364V (7274C>G 7275G>T), H2366L (7281A>T), L2367S (7284T>C), S2371L (7296C>T), S2376F (7311C>T), T2377I (7314C>T), C2378Y (7317G>A), N2379K (7321T>A), K2384Q (7334A>C), A2387N (7343G>A 7344C>A 7345A>C), S2408M (7407G>T 7408T>G), C2411G (7415T>G), L2440S (7503T>C 7504G>A), D2509N (7709G>A), K2510R (7713A>G 7714A>G), T2543I (7812C>T), V2554L (7844G>T 7846G>A), V2567I (7883G>A), S2620N (8043G>A), N2656S (8151A>G 8152C>T), R2665K (8178G>A), R2675K (8208G>A), P2704A (8294C>G 8296A>G), S2722F (8349C>T), A2742S (8408G>T), M2756I (8452G>A), R2759K (8460G>A), A2766T (8480G>A 8482T>A), R2773K (8502G>A 8503A>G), E2777D (8515A>T), E2785D (8539G>C), L2788C (8546C>T 8547T>G), S2790T (8552T>A), A2797G (8574C>G), I2801V (8585A>G), G2805E (8598G>A), N2807S (8603A>T 8604A>C 8605C>T), P2813Q (8622C>A), E2814G (8625A>G 8626G>A), P2822F (8648C>T 8649C>T 8650G>C), G2825E (8658G>A), D2827A (8664A>C 8665T>G), A2832V (8679C>T 8680A>T), T2854K (8745C>A), E2891G (8856A>G 8857G>A), R2894Q (8865G>A), M2895L (8867A>C 8869G>T), G2900E (8883G>A), T2902S (8888A>T), V2924I (8954G>A), N2934D (8984A>G 8986T>C), E2935V (8988A>T), I2983V (9131A>G), M2989G (9149A>G 9150T>G 9151G>A), G2990S (9152G>A), P3026S (9260C>T 9262T>C), Q3038H (9298A>C) | | | | | | | | |

| | Begin | End | Coverage | Score | Concordance | Matches | Identities | I/D/M/F* | Stop Codons |
|---|---|---|---|---|---|---|---|---|---|
| **NT** | **54** | **9704** | **99.5%** | **12522** | **65.2%** | **9639 (99.8%)** | **7984 (82.7%)** | **7/12** | |

Codon mutations:

ACT3ACC (193T>C), TAC4CAA (194T>C 196C>A), TCA6TCG (202A>G), ATC8ATT (208C>T), TGT9CTG (209T>C 210G>T 211T>G), TCG12TCA (220G>A), TTT13ATG (221T>A 223T>G), GAA14GAG (226A>G), CTA17CTT (235A>T), TAC19TAT (241C>T), CCA21CCT (247A>T), TCT23CCT (251T>C), TGC24TTT (255G>T 256C>T), GAG25GGG (258A>G), CAT26CAC (262T>C), ATT27GTG (263A>G 265T>G), GTG28GCG (267T>C), GAA30GTA (273A>T), CGA31CGT (277A>T), GAA32GAG (280A>G), GTG33GTT (283G>T), CCG34CCA (286G>A), GCT35ACC (287G>A 289T>C), GTT37ACT (293G>A 294T>C), CCT39CCC (301T>C), TTC40TTT (304C>T), GCA41ATA (305G>A 306C>T), GAT42GGT (309A>G), CTG43TTG (311C>T), GAA44GAT (316A>T), ACA45GTG (317A>G 318C>T 319A>G), CAA46CAG (322A>G), GCA49GCT (331A>T), CTC52CAA (339T>A 340C>A), AAG53AGG (342A>G), AAA55GCG (347A>G 348A>C 349A>G), TAT56CAT (350T>C), GCT57GCA (355T>A), ACT58GTT (356A>G 357C>T), CGT60CGC (364T>C), GTG61ACA (365G>A 366T>C 367G>A), CTC62TCC (368C>T 369T>C), AAA63AAG (373A>G), AAC64AAT (376C>T), TTT67TGT (384T>G), ACG68ACA (388G>A), TAC69TAT (391C>T), CGA70CGG (394A>G), AAG72AAA (400G>A), GAT74GAC (406T>C), GCC75GTT (408C>T 409C>T), CAG76CAA (412G>A), ATA77ATT (415A>T), ATG78GAG (416A>G 417T>A), ATT80ATC (424T>C), CAG81CAA (427G>A), AAA83AAG (433A>G), CTG84CTC (436G>C), GAG85AAG (437G>A), AGG86GAG (440A>G 441G>A), AAG87CGA (443A>C 444A>G 445G>A), GAT88GAA (448T>A), GAA90GAG (454A>G), GAA91GCA (456A>C), TAT92TAC (460T>C), CAC93AAC (461C>A), CAA95CAG (469A>G), GCC97GCG (475C>G), CCT99CCA (481T>A), ACT106ACC (502T>C), ATA107ATT (505A>T), GGC109GGT (511C>T), GAT111GAA (517T>A), CCT112CCA (520T>A), CCA113CCT (523A>T), TCT116TTT (531C>T), GAG117GAA (535G>A), CCA118TCA (536C>T), CAA119TGG (539C>T 540A>G 541A>G), AGA122AAG (549G>A 550A>G), ATC124ATT (556C>T), ATT125ATC (559T>C), CAT126CAC (562T>C), ACT128ACC (568T>C), CCA129CCG (571A>G), ATG131ACG (576T>C), CGT132CGC (580T>C), GTC134GCG (585T>C 586C>G), CGC137TAT (593C>T 594G>A 595C>T), CCC138CGC (597C>G), ATA139GTG (599A>G 601A>G), ATA140CCG (602A>C 603T>C 604A>G), ACA143ACG (613A>G), GAA144GAG (616A>G), GGC145GGG (619C>G), CAG146CAA (622G>A), ATG147ATA (625G>A), AAT148AAC (628T>C), CAC149CAT (631C>T), ATT151GTC (635A>G 637T>C), ATA154GTG (644A>G 646A>G), ATT157ATC (655T>C), TCG159TCA (661G>A), GAG160GCC (663A>C 664G>C), AAA161AAG (667A>G), GTC165GTT (679C>T), CAC166CAA (682C>A), TTA167CTG (683T>C 685A>G), AGT169AGC (691T>C), ACC172AGC (699C>G), ACT173ACA (703T>A), CAT174CAA (706T>A), AAG179GAA (719A>G 721G>A), ATA180GTG (722A>G 724A>G), CTT181TTG (725C>T 727T>G), GGT182GGA (730T>A), GCA183TCA (731G>T), TAC184CCT (734T>C 735A>C 736C>T), TCC185CGT (737T>C 738C>G 739C>T), GCG187GTC (744C>T 745G>C), CGA189TGC (749C>T 751A>C), ACT190ACC (754T>C), ATG194AAG (765T>A), TTG196CTG (770T>C), CGA197AAG (773C>A 774G>A 775A>G), CGG198AGG (776C>A), TTC202TTT (790C>T), ATG206AAG (801T>A), GGA210GCG (813G>C 814A>G), CTT211CAT (816T>A), TTG212CTG (818T>C), CAA213CAG (823A>G), CGT214CAA (825G>A 826T>A), CGG217AGA (833C>A 835G>A), GAC219GAT (841C>T), AAT223GAC (851A>G 853T>C), GTC225GTT (859C>T), CGC226TGT (860C>T 862C>T), ACT227GCC (863A>G 865T>C), ATC228ACC (867T>C), AAC229GAT (869A>G 871C>T), ATA230TTG (872A>T 874A>G), CGA231CAA (876G>A), AGG232AAG (879G>A), GAT234GAC (886T>C), ATC238ATT (898C>T), AAC240AGC (903A>G), ACA241AAC (906C>A 907A>C), AAA242ACC (909A>C 910A>C), AGC243AAT (912G>A 913C>T), GGC246GGA (922C>A), GGT249GGA (931T>A), TCA252CTT (938T>C 939C>T 940A>T), GGA253GGG (943A>G), TTC256TTT (952C>T), TCA261TCG (967A>G), GAA263GAG (973A>G), GGG264GGC (976G>C), TTG266ATC (980T>A 982G>C), GCA269GCG (991A>G), CGT270CGC (994T>C), AGA272AAA (999G>A), GTT273GTC (1003T>C), CAG275AGA (1007C>A 1008A>G 1009G>A), AGT276GGG (1010A>G 1012T>G), ATT277GTT (1013A>G), TTA278TTG (1018A>G), AAC279CAG (1019A>C 1021C>G), ATC282ATT (1030C>T), TCG285TCA (1039G>A), AAT286AAC (1042T>C), GCC287GCA (1045C>A), GAC288GAA (1048C>A), AAT289AAC (1051T>C), AAG292AAA (1060G>A), GGT293GGA (1063T>A), CTG294TTG (1064C>T), GAC295GAT (1069C>T), CGA300CAA (1083G>A), TAT303TAC (1093T>C), TCG305GCA (1097T>G 1099G>A), CAC307CAT (1105C>T), GTA310GTG (1114A>G), GCT311GCG (1117T>G), TTA313TTG (1123A>G), CCT314CCA (1126T>A), GTC315GTT (1129C>T), AGG320AAG (1143G>A), GTA321GTT (1147A>T), GCT322GCA (1150T>A), TTG324ATA (1154T>A 1156G>A), CAC327CAT (1165C>T), AGT328AGC (1168T>C), CTT330TTA (1172C>T 1174T>A), ACC339ACT (1201C>T), AGC346AAC (1221G>A), GTT349ACC (1229G>A 1230T>C 1231T>C), AGC350AAT (1233G>A 1234C>T), CTG352CTA (1240G>A), AAG354AAA (1246G>A), TTG356CTA (1250T>C 1252G>A), CAT357CAC (1255T>C), AAA358ACG (1257A>C 1258A>G), AGA361AGG (1267A>G), GGT363GGC (1273T>C), TTG364TTA (1276G>A), AAT365AGT (1278A>G), TTG367CTG (1283T>C), GCG369GTA (1290C>T 1291G>A), GAC372ACT (1298G>A 1299A>C 1300C>T), TTT374TTC (1306T>C), ATA375GTG (1307A>G 1309A>G), CAT376CAC (1312T>C), AAT378GAA (1316A>G 1318T>A), TTG381TTA (1327G>A), ATA382ACA (1329T>C), GCG383GCT (1333G>T), TTA384TTG (1336A>G), CTA387TTG (1343C>T 1345A>G), CCG390CCA (1354G>A), GTG391GTT (1357G>T), GAC392GAT (1360C>T), CTC395TTG (1367C>T 1369C>G), CTT397ATT (1373C>A), TTC398TTT (1378C>T), GAG400GAA (1384G>A), ATA401GTA (1385A>G), AAA403AAG (1393A>G), TCC404TCT (1396C>T), GAG407GAT (1405G>T), AAA408AAG (1408A>G), CAG409CAA (1411G>A), GCA411TCT (1415G>T 1417A>T), CCG412CCC (1420G>C), AAT415AAC (1429T>C), TTA416TTG (1432A>G), GTC418ATT (1436G>A 1438C>T), TTA419CTG (1439T>C 1441A>G), AAT420AAC (1444T>C), TTC423TTT (1453C>T), CTG424CTA (1456G>A), GAA428GAG (1468A>G), GTA436GTG (1492A>G), TTG439CTA (1499T>C 1501G>A), AGT440AGC (1504T>C), TTG441TTA (1507G>A), CTC442CTT (1510C>T), GAA443GAG (1513A>G), TTA444TTG (1516A>G), GCA445GCT (1519A>T), AGG446AGA (1522G>A), TTC447TTT (1525C>T), CAG448CAA (1528G>A), AAC450AAT (1534C>T), ACT452ACA (1540T>A), AAC454AAT (1546C>T), AAA457AAG (1555A>G), GGT458GGC (1558T>C), GAT459GAC (1561T>C), ATA460ATC (1564A>C), TCT461TCA (1567T>A), TTC463TTT (1573C>T), AGA464AGG (1576A>G), TTA467CTA (1583T>C), TCT468TCA (1588T>A), AAG470AAA (1594G>A), AAC472AAT (1600C>T), CTG475CTA (1609G>A), TCG478TCA (1618G>A), TGC479TGT (1621C>T), GAC480GAT (1624C>T), CTG483TTG (1631C>T), AAA485AAG (1639A>G), TTC489TTT (1651C>T), CTC490CTG (1654C>G), CAT497CAC (1675T>C), TCA503TCG (1693A>G), TTT506TTC (1702T>C), GAA508AGA (1706G>A 1707A>G), GCA512ACA (1718G>A), GGA514GGG (1726A>G), TAC515TAT (1729C>T), TCA516TCC (1732A>C), TAT518TAC (1738T>C), ATC520GTT (1742A>G 1744C>T), CGC521CGT (1747C>T), CAT523CAC (1753T>C), AGT525AAT (1758G>A), AGG528AGA (1768G>A), AAC534AAT (1786C>T), TTA535TTG (1789A>G), GTT536ATT (1790G>A), GTC537GTT (1795C>T), TTA541CTG (1805T>C 1807A>G), GAG543GAA (1813G>A), AGG545AGA (1819G>A), AAG547AAA (1825G>A), GAC551GAT (1837C>T), TAT552TAC (1840T>C), CAA555CAG (1849A>G), GTC558GTG (1858C>G), AAA560AAG (1864A>G), TGC562TGT (1870C>T), AGT564AGC (1876T>C), AAA566AAG (1882A>G), GGT568GGA (1888T>A), AAT569AAC (1891T>C), TGT574TGC (1906T>C), ACA577ACT (1915A>T), GGT582GGC (1930T>C), GCC584GCT (1936C>T), ATT585GTC (1937A>G 1939T>C), TCA587TCG (1945A>G), TTC589TTT (1951C>T), CCA592CCT (1960A>T), GTA598GTG (1978A>G), ATT599ATA (1981T>A), GGC600GGT (1984C>T), AAT601AAC (1987T>C), GAC604GAT (1996C>T), TTT607TAT (2004T>A), AAA612AAG (2020A>G), GGG613GGA (2023G>A), GAT614GAC (2026T>C), TCG615TCT (2029G>T), GAG616GAA (2032G>A), TAT625TAC (2059T>C), ATT628ATG (2068T>G), AAC629AAT (2071C>T), GTG630ATA (2072G>A 2074G>A), CTG635TTA (2087C>T 2089G>A), AGC639AGT (2101C>T), GAG640GAA (2104G>A), GAG641GAA (2107G>A), AAG644AAA (2116G>A), TTC646TTT (2122C>T), CGC651CGT (2137C>T), CTT658CTC (2158T>C), ACC660ACT (2164C>T), CCA662CCG (2170A>G), GAT666GAC (2182T>C), GCG668GCA (2188G>A), ACC669ACA (2191C>A), AGA675AAA (2208G>A), TAT678TAC (2218T>C), GAC680GAT (2224C>T), GTA681GTT (2227A>T), TTG686TTA (2242G>A), CCC687CCT (2245C>T), TTG690TTA (2254G>A), GTT691GTC (2257T>C), GAC692GAT (2260C>T), GAC694GAG (2266C>G), CAA696CAG (2272A>G), ACG697ACA (2275G>A), TCA703TCG (2293A>G), GGC705GGT (2299C>T), TCG706TCA (2302G>A), CAG707CAA (2305G>A), GGA710GGT (2314A>T), TAT711TAC (2317T>C), CTA714CTG (2326A>G), GTG719GTA (2341G>A), CAA721CAG (2347A>G), CTT722CTC (2350T>C), TTG724TTA (2356G>A), AAT727AAC (2365T>C), GAA729GAG (2371A>G), TTA730TTG (2374A>G), GAT733GAC (2383T>C), ATA734ATT (2386A>T), AAA735AAG (2389A>G), CAT736CAC (2392T>C), TAT737TAC (2395T>C), AAT744GGG (2414A>G 2415A>G 2416T>G), GCT745GCA (2419T>A), AGC746TGC (2420A>T), ACA752ACT (2440A>T), GTT761GTC (2467T>C), TCT764TCC (2476T>C), TCG766TCA (2482G>A), CTG769CTA (2491G>A), CTG771CTT (2497G>T), TTG773TTA (2503G>A), AGA778AGG (2518A>G), CCT779CCC (2521T>C), AAG780AGG (2523A>G), AGA783AGG (2533A>G), TTG785TTA (2539G>A), TTA787TTG (2545A>G), GAG789GAA (2551G>A), CCT790CCA (2554T>A), CTG792TTG (2558C>T), TTG793CTC (2561T>C 2563G>C), TCA796TCG (2572A>G), CCT800CCC (2584T>C), GGC801GGT (2587C>T), CTG803CTC (2593G>C), GCT805GCC (2599T>C), AAT808AAC (2608T>C), TTT812TTC (2620T>C), GCG815GCA (2629G>A), GTG816GTA (2632G>A), AGG817AAG (2634G>A), TTG818TTA (2638G>A), ATT820ATA (2644T>A), AAA823AAG (2653A>G), GCT827GCC (2665T>C), CTA832TTA (2678C>T), CTA833CTG (2683A>G), GCT835GCC (2689T>C), CTA838CTC (2698A>C), TCA841TCG (2707A>G), GCG842GCA (2710G>A), GAA844GAG (2716A>G), ACA845ACG (2719A>G), CTC846CTT (2722C>T), GTC847ATT (2723G>A 2725C>T), GCA848GCG (2728A>G), AGG850AGA (2734G>A), ATT851ATA (2737T>A), GAT854GAC (2746T>C), GCT855ACA (2747G>A 2749T>A), GCA856GCT (2752A>T), GCT857GCA (2755T>A), GAC859GAT (2761C>T), CTC860CTT (2764C>T), TGT865TGC (2779T>C), GGG867GGT (2785G>T), AAC869AAT (2791C>T), CAT871AAT (2795C>A), TAC874TAT (2806C>T), TTG878CTC (2816T>C 2818G>C), GTG880ATA (2822G>A 2824G>A), TTG881TTA (2827G>A), GTT884GTC (2836T>C), AAG885AAA (2839G>A), AAT886AAC (2842T>C), ACC893ACA (2863C>A), CTA894TTA (2864C>T), CCA900ACA (2882C>A), AGT901CAT (2885A>C 2886G>A), AAC903AAT (2893C>T), ACG904ATG (2895C>T), GTC906GTT (2902C>T), GTA907GTG (2905A>G), ATT909ATC (2911T>C), AAT913AAC (2923T>C), AAT916AGT (2931A>G), AAC919AGC (2940A>G), GCT921GCC (2947T>C), ACT926ACC (2962T>C), GAA929_AAT930insA-- (2971_2972insA), GGT935GGC (2989T>C), ACT936ATT (2991C>T), CAT937CGT (2994A>G), GAG939AGG (2999G>A 3000A>G), TCC944TC- (3016delC), CGG946CAG (3021G>A), TAC947TTT (3024A>T 3025C>T), ACA951ATA (3036C>T), GAA952GGC (3039A>G 3040A>C), AAG953ACA (3042A>C 3043G>A), TTG956TTA (3052G>A), TTA959TTG (3061A>G), GCG966GCA (3082G>A), GGC969GGT (3091C>T), GCC972GTA (3099C>T 3100C>A), GTG974AGA (3104G>A 3105T>G 3106G>A), GTC975GTT (3109C>T), AAA976AAG (3112A>G), ACT978ACC (3118T>C), GCC979GCT (3121C>T), GGA981GGG (3127A>G), AGC983AAC (3132G>A), CGA985AAG (3137C>A 3138G>A 3139A>G), TAT989TAC (3151T>C), TTC990ATT (3152T>A 3154C>T), TGT994TAT (3165G>A), GTA995GTG (3169A>G), ATC1005ATT (3199C>T), AGG1007CGG (3203A>C), CGT1008CGC (3208T>C), CCA1010CCG (3214A>G), ACC1011ACT (3217C>T), TTT1012CTT (3218T>C), GTC1013GTA (3223C>A), ACT1014AAT (3225C>A), GTT1016ATT (3230G>A), AAC1017AAT (3235C>T), TTA1025CTA (3257T>C), ACC1026ACT (3262C>T), AGC1027AGT (3265C>T), GTA1028ATG (3266G>A 3268A>G), GTG1029GTA (3271G>A), GTG1031ATG (3275G>A), GCA1034ACA (3284G>A), CAG1039CAA (3301G>A), AGA1044AAG (3315G>A 3316A>G), GAA1045GCA (3318A>C), ATC1046ATT (3322C>T), GAG1047GAA (3325G>A), TTG1048TTA (3328G>A), ATA1051ATT (3337A>T), GAG1055GAA (3349G>A), GTC1057GTT (3355C>T), TGC1058TGT (3358C>T), ATG1059GTG (3359A>G), CTA1061CTG (3367A>G), TAT1062TAC (3370T>C), TTA1065CTG (3377T>C 3379A>G), GAA1070GAG (3394A>G), CGC1071CGT (3397C>T), GAT1072GAG (3400T>G), ACA1074ACG (3406A>G), TAC1078TAT (3418C>T), ATT1079ATA (3421T>A), TAT1081TAC (3427T>C), AAG1083AAA (3433G>A), TCA1084TCT (3436A>T), GTA1085GTG (3439A>G), AAC1086AAT (3442C>T), CCT1087CCC (3445T>C), CAG1091CAA (3457G>A), TTT1092TTC (3460T>C), GCT1093GCA (3463T>A), GCG1095GCT (3469G>T), CAG1096CAA (3472G>A), CAC1104CAT (3496C>T), CAG1105CAA (3499G>A), TCC1107TCT (3505C>T), GTT1110GGT (3513T>G), AAA1112AAG (3520A>G), GAA1115GAG (3529A>G), CAA1116CAG (3532A>G), ATG1121GTG (3545A>G), GCT1122ACT (3548G>A), TTA1123CTA (3551T>C), GTC1124ATT (3554G>A 3556C>T), AGT1133AGC (3583T>C), GAT1134GAC (3586T>C), TGC1135TGT (3589C>T), TTC1137TTT (3595C>T), AAT1141AAC (3607T>C), TTT1143TTC (3613T>C), AAG1144AAA (3616G>A), GGT1145GGC (3619T>C), CTT1147GTT (3623C>G), TCC1148TCT (3628C>T), CTG1150ATG (3632C>A), GAT1151GAC (3637T>C), TAT1152CAT (3638T>C), GAA1153GAG (3643A>G), TTA1159TTG (3661A>G), GTG1162GTA (3670G>A), TTT1166TTC (3682T>C), GAG1168GAA (3688G>A), GAG1171GAA (3697G>A), ATT1172GTT (3698A>G), GAT1174GAC (3706T>C), TTT1175TTC (3709T>C), TTG1177CTA (3713T>C 3715G>A), AGT1178AAT (3717G>A), GAG1179GAA (3721G>A), CGA1183AAA (3731C>A 3732G>A), ACT1184ACA (3736T>A), CTA1188TTG (3746C>T 3748A>G), ACA1190ACG (3754A>G), AGT1193AGC (3763T>C), CGA1198CGG (3778A>G), CAA1199CAG (3781A>G), ATC1200ATT (3784C>T), CAT1204CAC (3796T>C), CCA1207CCT (3805A>T), TAC1209TAT (3811C>T), AGA1210AGG (3814A>G), TTC1215TTT (3829C>T), TTT1218TTC (3838T>C), AGA1220AGG (3844A>G), GCA1221GCC (3847A>C), GTT1224GTA (3856T>A), GTG1226GTT (3862G>T), GAC1229GAT (3871C>T), GCC1231GCA (3877C>A), AGC1233AGT (3883C>T), GAA1234GAG (3886A>G), CTA1236CTG (3892A>G), GAC1237GAT (3895C>T), TTT1238TTC (3898T>C), GTA1240GTG (3904A>G), CGG1241AGA (3905C>A 3907G>A), GCT1243GCC (3913T>C), AAG1248AAA (3928G>A), TCA1249TCT (3931A>T), GGG1251GGA (3937G>A), AGT1257AGC (3955T>C), GTG1258GTA (3958G>A), GCC1259GCT (3961C>T), GGA1260GGG (3964A>G), TCT1261TCC (3967T>C), TTA1264TTG (3976A>G), ATT1265ATC (3979T>C), CCA1267CCT (3985A>T), CTA1271CTT (3997A>T), GCG1272GCA (4000G>A), AAC1274AAT (4006C>T), GTT1275GTG (4009T>G), TTC1276TTT (4012C>T), AAA1277AAG (4015A>G), TCT1280TCC (4024T>C), GAA1282GAG (4030A>G), CCA1283CCT (4033A>T), AAG1286AAA (4042G>A), CTG1290TTA (4052C>T 4054G>A), CGT1291CGC (4057T>C), CGT1293CGA (4063T>A), GGA1294GGG (4066A>G), ATA1297GTA (4073A>G), TTT1298TTC (4078T>C), GGC1299GGT (4081C>T), TCT1300TCC (4084T>C), TCC1304TCT (4096C>T), AGC1308AGT (4108C>T), GGA1309GGT (4111A>T), GCG1311GCA (4117G>A), CTA1312TTG (4118C>T 4120A>G), TAC1314TAT (4126C>T), TTC1315TAT (4128T>A 4129C>T), GCC1316GCT (4132C>T), AAT1317AAC (4135T>C), CAA1321CAG (4147A>G), TTA1322CTA (4148T>C), GCT1323TCT (4151G>T), AAC1326AAT (4162C>T), GTA1328ATA (4166G>A), ATA1329ATC (4171A>C), GAG1332GAA (4180G>A), CTG1336TTG (4190C>T), CCT1338CCC (4198T>C), TCC1339TCT (4201C>T), GCG1342GCT (4210G>T), TTC1343TTT (4213C>T), AGT1345AGC (4219T>C), CTG1346TTG (4220C>T), CTG1347ATA (4223C>A 4225G>A), GTT1349GTG (4231T>G), CAT1351CAC (4237T>C), GTA1356GTG (4252A>G), TTA1357CTG (4253T>C 4255A>G), AAA1358AAG (4258A>G), ACT1362ACC (4270T>C), AGA1366AGG (4282A>G), GAG1367GAA (4285G>A), GAA1369GAG (4291A>G), CAG1373CAA (4303G>A), GTC1376GTT (4312C>T), ATA1379GTG (4319A>G 4321A>G), GTG1380GTT (4324G>T), GAG1381GAA (4327G>A), CTG1384CTT (4336G>T), TCT1385TCC (4339T>C), TCA1388TCT (4348A>T), GCA1392GCG (4360A>G), GGT1394GGC (4366T>C), TCT1395TCA (4369T>A), ACT1397ACC (4375T>C), GTT1401GTC (4387T>C), TTT1404TAT (4395T>A), GTA1408ATA (4406G>A), CTT1409CTC (4411T>C), GTT1418GTC (4438T>C), ACC1420ACA (4444C>A), GCC1422GCT (4450C>T), CTC1424CTT (4456C>T), GAC1427GAT (4465C>T), ATG1431GTT (4475A>G 4477G>T), GTC1432GTT (4480C>T), ACA1433TCG (4481A>T 4483A>G), GGC1437GGT (4495C>T), AAG1441AAA (4507G>A), CAC1442CAT (4510C>T), GGT1443GGG (4513T>G), CTA1445TTA (4517C>T), GTC1448GTT (4528C>T), AAA1450AAG (4534A>G), ACC1452ACT (4540C>T), GCG1454TTA (4544G>T 4545C>T 4546G>A), AGA1455AAA (4548G>A), TTT1458TTC (4558T>C), GTA1460GTT (4564A>T), ACC1462ACT (4570C>T), ATA1464ATT (4576A>T), TTG1471TTA (4597G>A), GAC1472GAT (4600C>T), GAC1474GAT (4606C>T), GTG1475GTA (4609G>A), GTG1477GTA (4615G>A), GAC1478GAT (4618C>T), TTG1481CTT (4625T>C 4627G>T), GAC1488GAT (4648C>T), ATT1489ATC (4651T>C), AGG1492AGA (4660G>A), AGC1493AGT (4663C>T), GCT1495GCA (4669T>A), GTG1499ATC (4679G>A 4681G>C), AGT1500AGC (4684T>C), GTT1501GTG (4687T>G), GAG1505GAA (4699G>A), AGA1506AGG (4702A>G), CAA1508CAG (4708A>G), AGG1509AGA (4711G>A), CTG1510TTG (4712C>T), AAA1518AAG (4738A>G), GTA1520GTT (4744A>T), GCA1521GCG (4747A>G), TTG1522CTG (4748T>C), ACT1527ACC (4765T>C), GTC1538ATT (4796G>A 4798C>T), ACT1540AGC (4803C>G 4804T>C), GAG1541GAA (4807G>A), GCG1542GCT (4810G>T), GCT1543GCG (4813T>G), CTT1544CTC

| | | | | | | | | | |
|---|---|---|---|---|---|---|---|---|---|
| | (4816T>C), TAT1549TAC (4831T>C), AAC1550AAT (4834C>T), CCA1552CCT (4840A>T), GGA1556GGG (4852A>G), GGC1557GGT (4855C>T), AGT1561AGC (4867T>C), CTG1562CTC (4870G>C), ATT1563ATA (4873T>A), TGT1566TGC (4882T>C), ACA1573ACT (4903A>T), CAG1575CAA (4909G>A), TTT1577TTC (4915T>C), GAA1578GAG (4918A>G), TTG1579CTA (4919T>C 4921G>A), AGT1580AGC (4924T>C), CCC1581CCG (4927C>G), TTT1583TTC (4933T>C), ATC1584ATA (4936C>A), CAG1585CAA (4939G>A), AAT1586AAC (4942T>C), TTC1587TTT (4945C>T), TCA1593TCG (4963A>G), CCT1596CCC (4972T>C), CTT1602CTC (4990T>C), AAA1606AAG (5002A>G), CCT1613CCC (5023T>C), TTG1614CTG (5024T>C), TGC1615TGT (5029C>T), GAT1616GAC (5032T>C), TCT1618TCC (5038T>C), CCA1620CCT (5044A>T), GCA1623GCT (5053A>T), TCG1624TCA (5056G>A), AGT1625GGT (5057A>G), ACT1626GCT (5060A>G), TCG1629TCT (5071G>T), CTT1636ATT (5090C>A), GGA1637GGG (5095A>G), GTG1638GTT (5098G>T), GCC1639GTC (5100C>T), TTA1640TTG (5104A>G), GAA1641GAT (5107A>T), ATT1642ATC (5110T>C), CCA1643CCG (5113A>G), AAG1644AAA (5116G>A), CAA1645CAG (5119A>G), GTC1646TTT (5120G>T 5122C>T), ATT1648GTT (5126A>G), CAT1651CAC (5137T>C), ATC1652ATT (5140C>T), AAA1653AAG (5143A>G), GAG1654GAA (5146G>A), CTC1659CTA (5161C>A), GAA1661GAG (5167A>G), CTT1663CTG (5173T>G), ACG1666ACA (5182G>A), GTT1667ATT (5183G>A), GTC1668GTT (5188C>T), AAG1669AAA (5191G>A), AAA1671AAG (5197A>G), GAC1672GAT (5200C>T), TGC1674TGT (5206C>T), TTA1675TTG (5209A>G), AGC1678AGT (5218C>T), TCG1682TCA (5230G>A), ATC1684ATT (5236C>T), AGC1685AGT (5239C>T), ATC1687ATT (5245C>T), ACA1693ACT (5263A>T), CTC1695CTT (5269C>T), TTC1696TTT (5272C>T), GCC1697GCA (5275C>A), ATC1698ATT (5278C>T), CCA1699CCC (5281A>C), AGA1700AGG (5284A>G), ACT1701ACC (5287T>C), CTA1702TTA (5288C>T), ATA1703ATT (5293A>T), TTG1704CTA (5294T>C 5296G>A), GAG1706GAA (5302G>A), CTT1709CTA (5311T>A), GAA1710GAG (5314A>G), CGA1713CGT (5323A>T), AGT1721AGC (5347T>C), ATC1723ATT (5353C>T), TGC1727TGT (5365C>T), GTT1734GTC (5386T>C), ACC1737ACA (5395C>A), AAA1746AAG (5422A>G), GAT1747GAC (5425T>C), ACC1749ACT (5431C>T), GAG1751GAA (5437G>A), CTT1756CTC (5452T>C), AAG1758AAA (5458G>A), AGT1761AGC (5467T>C), CAA1762CAG (5470A>G), CTA1763TTG (5471C>T 5473A>G), GAA1765GAG (5479A>G), TTC1766TTT (5482C>T), TCA1767TCC (5485A>C), AAT1768AAC (5488T>C), TTG1769CTA (5489T>C 5491G>A), GGT1771GGC (5497T>C), TGT1774TGC (5506T>C), GAG1775GAA (5509G>A), GAG1776GAA (5512G>A), TTA1778CTA (5516T>C), TAT1782TAC (5530T>C), GAG1783GAA (5533G>A), TCG1784TCT (5536G>T), TTG1785TTA (5539G>A), GTT1788GTG (5548T>G), CAC1790CAT (5554C>T), GCT1792GCA (5560T>A), GCG1793ACA (5561G>A 5563G>A), ACG1794ACC (5566G>C), TCA1795TCG (5569A>G), CTC1800CTG (5584C>G), AAG1803AAA (5593G>A), GGG1804GGA (5596G>A), ATT1805GTT (5597A>G), AAC1807AAA (5605C>A), GTG1811GTT (5617G>T), GCT1812GTG (5619C>T 5620T>G), TTG1815TTA (5629G>A), ATC1816CTC (5630A>C), GGC1819GGT (5641C>T), GCA1822GCT (5650A>T), ATA1826ATT (5662A>T), GGA1827GGG (5665A>G), CAA1835CAG (5689A>G), GTT1837ATT (5693G>A), GAG1838GAA (5698G>A), CAA1843CAG (5713A>G), GGG1844GGC (5716G>C), AAA1845AAG (5719A>G), GCC1853GCA (5743C>A), CGC1857CGA (5755C>A), CAT1858CAC (5758T>C), CGT1860CGC (5764T>C), AAA1862AAG (5770A>G), GGC1865GGT (5779C>T), TTT1866TTC (5782T>C), GAC1869GAT (5791C>T), GAC1873GAT (5803C>T), AGG1884AGA (5836G>A), AAA1885AAG (5839A>G), GGA1887GGG (5845A>G), AAA1888AAG (5848A>G), AAA1890AGA (5853A>G), GGT1891GGC (5857T>C), ACC1892ACT (5860C>T), ACA1893ACT (5863A>T), GGT1895GGA (5869T>A), GGT1897GGC (5875T>C), AGC1900AGT (5884C>T), TTC1903TTT (5893C>T), ATC1904GTG (5894A>G 5896C>G), AAC1905AAT (5899C>T), GGG1908GGA (5908G>A), TTT1909TTC (5911T>C), GAG1913GAA (5923G>A), CAA1918CAG (5938A>G), TTC1919TTT (5941C>T), GTT1920GTC (5944T>C), GAT1921GAC (5947T>C), GGG1925GGA (5959G>A), CGG1926GCC (5960C>G 5961G>C 5962G>C), GAA1930GAG (5974A>G), GCT1934GCA (5986T>A), GAC1935GAT (5989C>T), ATT1936GTT (5990A>G), GAT1938GAC (5998T>C), AGA1942CGT (6008A>C 6010A>T), GAA1945GAT (6019A>T), GTG1946GTT (6022G>T), CGA1947CGT (6025A>T), AAA1949ACA (6030A>C), GAC1955GAA (6049C>A), ATG1958ATA (6058G>A), CAA1959CAG (6061A>G), GCC1960GCA (6064C>A), GGT1962GGC (6070T>C), AGT1963AAC (6072G>A 6073T>C), AAC1964CAC (6074A>C), ACG1965ACA (6079G>A), ACC1966AAC (6081C>A), ATA1967ATT (6085A>T), GCA1969GCT (6091A>T), TAC1970TAT (6094C>T), TGT1976TCC (6111G>C 6112T>C), AAA1978AAG (6118A>G), TTG1980TTA (6124G>A), AAG1981AAA (6127G>A), ATT1982GTC (6128A>G 6130T>C), GAT1983GAC (6133T>C), TTA1984TTG (6136A>G), AAC1988AAT (6148C>T), AAA1991AAG (6157A>G), GTT1992ATT (6158G>A), TGT1993TGC (6163T>C), AAA1995AAG (6169A>G), ACA1996TCG (6170A>T 6172A>G), GGC1998GGT (6178C>T), AAA2001AAG (6187A>G), TTT2002TTC (6190T>C), GAG2004GAA (6196G>A), CTC2007TTT (6203C>T 6205C>T), GAA2008GAG (6208A>G), CTA2009TTG (6209C>T 6211A>G), CAG2011CAA (6217G>A), GCT2015GCA (6229T>A), GTA2016ATA (6230G>A), GTC2018GTT (6238C>T), AAG2021AAA (6247G>A), ATA2023ATT (6253A>T), GCA2025AAA (6257G>A 6258C>A), CAT2030CAC (6274T>C), AAA2033AGA (6282A>G), TCG2034TCA (6286G>A), GGC2038GGT (6298C>T), TTG2039TTA (6301G>A), AGA2040AGG (6304A>G), GAC2041GAT (6307C>T), GCC2046GCT (6322C>T), ACA2048ACT (6328A>T), GTT2049GTG (6331T>G), TGT2050TGC (6334T>C), AGG2051AGA (6337G>A), CTG2052TTG (6338C>T), TAT2058TAC (6358T>C), GGG2059GGA (6361G>A), GCA2060ACA (6362G>A), TCA2061TCT (6367A>T), GAG2062GAA (6370G>A), TAC2064TAT (6376C>T), GGT2065GGG (6379T>G), GGA2067GGC (6385A>C), GGA2069GGT (6391A>T), GCA2070GCG (6394A>G), TAC2071TAT (6397C>T), GTA2073ATA (6401G>A), GCG2074GTG (6405C>T), CAC2076CAT (6412C>T), TTA2078TTG (6418A>G), TTT2079TTC (6421T>C), TAC2082TTC (6429A>T), GGT2084GGA (6436T>A), GAG2087GAA (6445G>A), CAA2089CGA (6450A>G), TCC2090TCG (6454C>G), CAC2092CAT (6460C>T), GGT2093GGA (6463T>A), TTC2095TTT (6469C>T), AGG2096AGA (6472G>A), GTG2097GTA (6475G>A), AAT2099AAC (6481T>C), CAC2101AAT (6485C>A 6487C>T), AGT2102AGC (6490T>C), AGC2104AGT (6496C>T), CTG2106CTA (6502G>A), GGT2110GGC (6514T>C), AGG2111AGA (6517G>A), CTC2115ATC (6527C>A), ATC2116ATA (6532C>A), CCG2119CCA (6541G>A), AAA2120AAG (6544A>G), GTC2124GTT (6556C>T), TTT2125TTC (6559T>C), CAG2127CAA (6565G>A), TTG2129CTA (6569T>C 6571G>A), CCT2134CCA (6586T>A), ACA2135ATA (6588C>T), GAA2138GAG (6598A>G), AGA2139AGG (6601A>G), ATT2140ATC (6604T>C), TGT2141TGC (6607T>C), GGA2144GGG (6616A>G), AAC2146AAT (6622C>T), GAG2149GAA (6631G>A), AAG2150AAA (6634G>A), TAT2151CAT (6635T>C), GCT2152GCA (6640T>A), TCG2154TCT (6646G>T), ATC2155GTC (6647A>G), ATC2156ATT (6652C>T), AGC2160AGT (6664C>T), ACT2162ACG (6670T>G), ATA2165GTG (6677A>G 6679A>G), GGC2167GGT (6685C>T), ACA2169ACT (6691A>T), TTC2170TTT (6694C>T), ACA2177ACG (6715A>G), GAT2178AAT (6716G>A), AAT2179GAT (6719A>G), GGA2180GGG (6724A>G), CTA2184TTA (6734C>T), GTG2186GTA (6742G>A), GTG2187GTA (6745G>A), GCC2190GCT (6754C>T), GGA2192GGG (6760A>G), ATA2194TTG (6764A>T 6766A>G), ATT2197ATA (6775T>A), CTG2200TTG (6782C>T), GCA2201GCT (6787A>T), AAC2202AAT (6790C>T), GCA2204GTG (6795C>T 6796A>G), CAC2205CAA (6799C>A), ACC2206ACA (6802C>A), AAC2208AAT (6808C>T), TAC2209TAT (6811C>T), TAC2210TAT (6814C>T), TTC2213TTT (6823C>T), GAA2215GAG (6829A>G), TAC2221TAT (6847C>T), CGA2223CGC (6853A>C), AAT2225GAT (6857A>G), GAG2226GAA (6862G>A), GAA2229GAG (6871A>G), GTC2231ACA (6875G>A 6876T>C 6877C>A), AAG2232AAA (6880G>A), TCT2233TCA (6883T>A), GTT2235GTG (6889T>G), GAC2239GAT (6901C>T), GGC2244GGT (6916C>T), CCG2245CCA (6919G>A), AAA2247AAG (6925A>G), CTT2248CTC (6928T>C), GAC2250GAA (6934C>A), GGG2255GGC (6949G>C), TTC2257TTT (6955C>T), AAA2258AAG (6958A>G), CAA2264CAG (6976A>G), CTA2266TTA (6980C>T), ATC2267ATT (6985C>T), GTA2271GTT (6997A>T), GTG2272GTT (7000G>T), GTG2273ATA (7001G>A 7003G>A), GCT2276GCC (7012T>C), AAG2277AAA (7015G>A), CAT2278CAC (7018T>C), TTT2283TAT (7032T>A), GCC2285GCT (7039C>T), TTG2286CTA (7040T>C 7042G>A), GGA2288GGG (7048A>G), GCT2292GCA (7060T>A), GTC2293GTA (7063C>A), AGC2298AGT (7078C>T), TTA2300CTG (7082T>C 7084A>G), GTA2301GTG (7087A>G), ACC2302ACA (7090C>A), CAT2304CAC (7096T>C), GTA2305GTG (7099A>G), GTT2306GTC (7102T>C), GAG2309GAA (7111G>A), CGA2311CGG (7117A>G), ACA2314AAA (7125C>A), GAA2315GAG (7129A>G), TTT2316TTC (7132T>C), CTG2317TTA (7133C>T 7135G>A), GTG2319GTA (7141G>A), GAT2320GAC (7144T>C), GCA2321TCG (7145G>T 7147A>G), GAG2322GAA (7150G>A), GCA2325GCT (7159A>T), GAG2326_GCA2327del (7160_7165delGAGGCA), TTC2329TTT (7171C>T), GCG2335GCT (7189G>T), AAA2338AAG (7198A>G), AGC2339AGT (7201C>T), CTA2341TTA (7205C>T), AAT2342AAC (7210T>C), GAT2344GAA (7216T>A), GCG2345GCA (7219G>A), ATC2347ATA (7225C>A), AAG2348AAA (7228G>A), AAG2352AAA (7240G>A), TAT2353TAC (7243T>C), AAA2355AAG (7249A>G), CCT2356CCG (7252T>G), GAT2358GAA (7258T>A), GTT2359GTC (7261T>C), GGT2360GGA (7264T>A), GTC2361GTG (7267C>G), GTG2362GTA (7270G>A), GAT2363_CGG2364insT-- (7273_7274insT), CGG2364GTG (7274C>G 7275G>T), CAT2366CTT (7281A>T), TTG2367TCG (7284T>C), TCA2371TTA (7296C>T), TCT2376TTT (7311C>T), ACC2377ATC (7314C>T), TGC2378TAC (7317G>A), AAT2379AAA (7321T>A), GTG2380-TG (7322delG), CAC2381CAT (7327C>T), GGC2382GGT (7330C>T), TTC2383TTT (7333C>T), AAG2384CAG (7334A>C), TGT2386TGC (7342T>C), GCA2387AAC (7343G>A 7344C>A 7345A>C), ACT2390ACC (7354T>C), CAA2393CAG (7363A>G), ATT2395ATC (7369T>C), TTC2396TTT (7372C>T), GCG2398GCT (7378G>T), CTC2399CTT (7381C>T), GCA2404GCT (7396A>T), GTC2405GTT (7399C>T), GCC2407GCT (7405C>T), AGT2408ATG (7407G>T 7408T>G), GGG2410GGT (7414G>T), TGC2411GGC (7415T>G), AAA2412AAG (7420A>G), GAC2415GAT (7429C>T), TAT2416TAC (7432T>C), CAT2419CAC (7441T>C), AAG2425AAA (7459G>A), GAA2426GAG (7462A>G), ATA2428ATT (7468A>T), GTC2429GTA (7471C>A), CAA2431CAG (7477A>G), AGC2432AGT (7480C>T), CTG2434CTT (7486G>T), TAT2437TAC (7495T>C), AAA2438AAG (7498A>G), GGT2439GGC (7501T>C), TTG2440TCA (7503T>C 7504G>A), ATT2443ATA (7513T>A), AAC2445AAT (7519C>T), GGA2446GGG (7522A>G), AAG2449AAA (7531G>A), GAG2451GAA (7537G>A), CTC2452CTT (7540C>T), TGT2454TGC (7546T>C), AAG2462AAA (7570G>A), ACG2463ACA (7573G>A), ACG2465ACA (7579G>A), GCT2468GCC (7588T>C), CTA2471TTA (7595C>T), TTG2474CTA (7604T>C 7606G>A), GGT2477GGA (7615T>A), TGT2480TGC (7624T>C), GAC2483GAT (7633C>T), TTC2484TTT (7636C>T), AAT2485AAC (7639T>C), TTT2488TTC (7648T>C), GAA2494GAG (7666A>G), TGT2496TGC (7672T>C), ACA2498ACT (7678A>T), GGG2500GGA (7684G>A), ACT2502ACC (7690T>C), TTT2504TTC (7696T>C), GGT2506GGG (7702T>G), GAT2509AAT (7709G>A), AAA2510AGG (7713A>G 7714A>G), CTG2511TTG (7715C>T), CGT2514CGC (7726T>C), GAG2517GAA (7735G>A), GTA2520GTG (7744A>G), TAC2521TAT (7747C>T), GGC2526GGT (7762C>T), CAG2528CAA (7768G>A), TTT2529TTC (7771T>C), TCA2532TCG (7780A>G), CTA2533CTC (7783A>C), ACT2534ACC (7786T>C), CTA2537TTG (7793C>T 7795A>G), ATC2538ATT (7798C>T), GCT2540GCC (7804T>C), ACC2543ATC (7812C>T), GAA2550GAG (7834A>G), GAC2551GAT (7837C>T), GTG2554TTA (7844G>T 7846G>A), CAG2557CAA (7855G>A), CTG2559CTA (7861G>A), TTA2562TTG (7870A>G), TAC2563TAT (7873C>T), ACT2564ACA (7876T>A), GAG2565GAA (7879G>A), ATT2566ATA (7882T>A), GTT2567ATT (7883G>A), CCA2570CCT (7894A>T), ATT2571ATC (7897T>C), GTC2579GTT (7921C>T), AAG2581AAA (7927G>A), GGT2584GGC (7936T>C), AAT2585AAC (7939T>C), AAT2586AAC (7942T>C), GTG2594GTA (7966G>A), GAT2595GAC (7969T>C), TCT2597TCC (7975T>C), GTC2601GTT (7987C>T), CTT2602CTC (7990T>C), GCT2607GCC (8005T>C), GAA2616GAG (8032A>G), ATC2618ATT (8038C>T), GAC2619GAT (8041C>T), AGC2620AAC (8043G>A), TTG2632TTA (8080G>A), GTG2635GTA (8089G>A), CCG2637CCA (8095G>A), GAG2638GAA (8098G>A), AGC2641AGT (8107C>T), ATT2642ATC (8110T>C), GAT2644GAC (8116T>C), CAA2648CAG (8128A>G), GGT2654GGC (8146T>C), AAC2656AGT (8151A>G 8152C>T), TCG2660TCA (8164G>A), AGG2665AAG (8178G>A), AAG2666AAA (8182G>A), TTG2669TTA (8191G>A), TTC2671TTT (8197C>T), TCC2673TCT (8203C>T), AGA2675AAA (8208G>A), CTG2677TTA (8213C>T 8215G>A), CTA2678CTG (8218A>G), ATC2679ATT (8221C>T), GAG2680GAA (8224G>A), CCA2685CCG (8239A>G), GAG2690GAA (8254G>A), ATT2692ATC (8260T>C), GTA2693GTG (8263A>G), GAT2699GAC (8281T>C), AGA2700AGG (8284A>G), CCA2704GCG (8294C>G 8296A>G), GAG2705GAA (8299G>A), CAC2706CAT (8302C>T), ATT2711ATC (8317T>C), TGC2712TGT (8320C>T), GCT2714GCA (8326T>A), GAG2717GAA (8335G>A), TCC2718TCT (8338C>T), TCT2722TTT (8349C>T), GAA2723GAG (8353A>G), CAC2726CAT (8362C>T), CAA2727CAG (8365A>G), ATC2728ATT (8368C>T), TAC2732TAT (8380C>T), TCA2733TCG (8383A>G), TTA2735TTG (8389A>G), CCT2740CCC (8404T>C), GCA2742TCA (8408G>T), ACA2743ACG (8413A>G), GCG2745GCT (8419G>T), GCT2750GCC (8434T>C), TAT2752TAC (8440T>C), ATG2756ATA (8452G>A), CTA2758CTG (8458A>G), AGG2759AAG (8460G>A), AAA2760AAG (8464A>G), CTG2761CTA (8467G>A), AGG2765AGA (8479G>A), GCT2766ACA (8480G>A 8482T>A), GAG2771GAA (8497G>A), CTA2772TTG (8498C>T 8500A>G), AGA2773AAG (8502G>A 8503A>G), GCC2774GCT (8506C>T), GAA2777GAT (8515A>T), GTC2780GTT (8524C>T), GCA2781GCC (8527A>C), TTA2782CTG (8528T>C 8530A>G), GAG2785GAC (8539G>C), CTT2788TGT (8546C>T 8547T>G), TCT2790ACT (8552T>A), GTA2793GTG (8563A>G), GCA2797GGA (8574C>G), ATT2801GTT (8585A>G), GGA2805GAA (8598G>A), AGC2806AGT (8602C>T), AAC2807TCT (8603A>T 8604A>C 8605C>T), AAA2812AAG (8620A>G), CCA2813CAA (8622C>A), GAG2814GGA (8625A>G 8626G>A), CAG2815CAA (8629G>A), GGC2816GGT (8632C>T), ATC2818ATT (8638C>T), CAG2819CAA (8641G>A), AAC2821AAT (8647C>T), CCG2822TTC (8648C>T 8649C>T 8650G>C), AAC2823AAT (8653C>T), AAA2824AAG (8656A>G), GGA2825GAA (8658G>A), GAT2827GCG (8664A>C 8665T>G), GTT2830GTG (8674T>G), GCA2832GTT (8679C>T 8680A>T), GGC2833GGA (8683C>A), GGG2836GGA (8692G>A), ACA2837ACT (8695A>T), CCG2841CCA (8707G>A), AGA2842CGA (8708A>C), AAG2844AAA (8716G>A), ATC2846ATT (8722C>T), ACG2847ACA (8725G>A), AAA2849AAG (8731A>G), AGA2851AGG (8737A>G), CCC2853CCT (8743C>T), ACA2854AAA (8745C>A), AAG2856AAA (8752G>A), GGA2857GGT (8755A>T), AAC2862AAT (8770C>T), TTA2863CTA (8771T>C), GAA2864GAG (8776A>G), CAT2865CAC (8779T>C), TTG2866TTA (8782G>A), TAT2869TAC (8791T>C), CCA2871CCG (8797A>G), CAA2872CAG (8800A>G), CAA2873CAG (8803A>G), GAT2875GAC (8809T>C), ATT2876ATC (8812T>C), AAT2878AAC (8818T>C), CGG2880CGA (8824G>A), GCA2881GCG (8827A>G), ACG2888ACA (8848G>A), TAT2890TAC (8854T>C), GAG2891GGA (8856A>G 8857G>A), GTG2893GTA (8863G>A), CGG2894CAG (8865G>A), ATG2895CTT (8867A>C 8869G>T), GCA2896GCG (8872A>G), GGA2900GAA (8883G>A), ACT2902TCT (8888A>T), TGC2915TGT (8929C>T), GTC2924ATC (8954G>A), AAC2925AAT (8959C>T), GGA2926GGT (8962A>T), GGG2933GGA (8983G>A), AAT2934GAC (8984A>G 8986T>C), GAA2935GTA (8988A>T), GAG2938GAA (8998G>A), CCG2940CCA (9004G>A), TTG2941TTA (9007G>A), ATC2944ATT (9016C>T), ACC2951ACA (9037C>A), ATA2967ATT (9085A>T), AAC2971AAT (9097C>T), ATT2983GTT (9131A>G), CGA2984CGT (9136A>T), CGG2987CGT (9145G>T), ATG2989GGA (9149A>G 9150T>G 9151G>A), GGT2990AGT (9152G>A), TTA2991TTG (9157A>G), GCG2992GCT (9160G>T), CGT2993CGC (9163T>C), GAG3000GAA (9184G>A), CGA3004CGG (9196A>G), AGG3010AGA (9214G>A), GAA3011GAG (9217A>G), GCG3012GCA (9220G>A), GCA3020GCT (9244A>T), TTG3021TTA (9247G>A), GCC3024GCT (9256C>T), CCT3026TCC (9260C>T 9262T>C), GGG3030GGA (9274G>A), GAC3032GAT (9280C>T), ATC3035ATT (9289C>T), CAA3038CAC (9298A>C), GAG3040GAA (9304G>A), GTC3050GTT (9334C>T), GTC3060GTG (9364C>G) | | | | | | | | |
| PVYgp2 | 1 | 75 | 100% | 95 | 18.1% | 75 (98.7%) | 54 (71.1%) | 1/0/3/2 | 2 |
| Protein mutations: | R17_K17insX (2971_2972insA), Y23H (2989T>C 2991C>T), R26K (2999G>A 3000A>G), H35Y (3025C>T), K40H (3040A>C 3042A>C), G41S (3043G>A), E44K (3052G>A), I47V (3061A>G), V54I (3082G>A), P57S (3091C>T), P60T (3100C>A), G61E (3104G>A 3105T>G), G62S (3106G>A), Q63* (3109C>T), R64G (3112A>G), C66R (3118T>C), L67F (3121C>T), I69V (3127A>G), A72E (3137C>A 3138G>A), I73V (3139A>G) | | | | | | | | |
| Codon mutations: | TTA2CTA (2923T>C), AAA4AAG (2931A>G), GAA7GAG (2940A>G), TTG10CTG (2947T>C), TTG15CTG (2962T>C), AGA17_AAA17insAA- (2971_2972insA), TAC23CAT (2989T>C 2991C>T), TCA24TCG (2994A>G), AGA26AAG (2999G>A 3000A>G), CAC32-AC (3016delC), TCG33TCA (3021G>A), GTA34GTT (3024A>T), CAT35TAT (3025C>T), CAC38CAT (3036C>T), AGA39AGG (3039A>G), AAA40CAC (3040A>C 3042A>C), GGC41AGC (3043G>A), GAA44AAA (3052G>A), ATA47GTA (3061A>G), GTT54ATT (3082G>A), CCG57TCG (3091C>T), CGC59CGT (3099C>T), CCA60ACA (3100C>A), GGT61GAG (3104G>A 3105T>G), GGT62AGT (3106G>A), CAA63TAA (3109C>T), AGG64GGG (3112A>G), TGC66CGC (3118T>C), CTC67TTC (3121C>T), ATT69GTT (3127A>G), GAG70GAA (3132G>A), GCG72GAA (3137C>A 3138G>A), ATT73GTT (3139A>G) | | | | | | | | |

| | Begin | End | Coverage | Score | Concordance | Matches | Identities | I/D/M/F* | Stop Codons |
|---|---|---|---|---|---|---|---|---|---|
| **NT** | **54** | **9704** | **99.5%** | **12522** | **65.2%** | **9639 (99.8%)** | **7984 (82.7%)** | **7/12** | |
| **Proteins** | | | | | | | | | |
| polyprotein (NP_056759.1) | 1 | 3064 | 100% | 18924 | 89.7% | 3062 (99.9%) | 2771 (90.4%) | 2/2/4/4 | 1 |
| Protein mutations: | Y4Q (194T>C 196C>A), C9L (209T>C 210G>T 211T>G), F13M (221T>A 223T>G), S23P (251T>C), C24F (255G>T 256C>T), E25G (258A>G), I27V (263A>G 265T>G), V28A (267T>C), E30V (273A>T), A35T (287G>A 289T>C), V37T (293G>A 294T>C), A41I (305G>A 306C>T), D42G (309A>G), E44D (316A>T), T45V (317A>G 318C>T 319A>G), L52Q (339T>A 340C>A), K53R (342A>G), K55A (347A>G 348A>C 349A>G), Y56H (350T>C), T58V (356A>G 357C>T), V61T (365G>A 366T>C 367G>A), L62S (368C>T 369T>C), F67C (384T>G), A75V (408C>T 409C>T), M78E (416A>G 417T>A), E85K (437G>A), R86E (440A>G 441G>A), K87R (443A>C 444A>G 445G>A), D88E (448T>A), E91A (456A>C), H93N (461C>A), D111E (517T>A), S116F (531C>T), P118S (536C>T), Q119W (539C>T 540A>G 541A>G), R122K (549G>A 550A>G), M131T (576T>C), V134A (585T>C 586C>G), R137Y (593C>T 594G>A 595C>T), P138R (597C>G), I139V (599A>G 601A>G), I140P (602A>C 603T>C 604A>G), M147I (625G>A), I151V (635A>G 637T>C), I154V (644A>G 646A>G), E160A (663A>C 664G>C), H166Q (682C>A), T172S (699C>G), H174Q (706T>A), K179E (719A>G 721G>A), I180V (722A>G 724A>G), A183S (731G>T), Y184P (734T>C 735A>C 736C>T), S185R (737T>C 738C>G 739C>T), A187V (744C>T 745G>C), R189C (749C>T 751A>C), M194K (765T>A), R197K (773C>A 774G>A 775A>G), M206K (801T>A), G210A (813G>C 814A>G), L211H (816T>A), R214Q (825G>A 826T>A), N223D (851A>G 853T>C), R226C (860C>T 862C>T), T227A (863A>G 865T>C), I228T (867T>C), N229D (869A>G 871C>T), I230L (872A>T 874A>G), R231Q (876G>A), R232K (879G>A), N240S (903A>G), T241N (906C>A 907A>C), K242T (909A>C 910A>C), S243N (912G>A 913C>T), S252L (938T>C 939C>T 940A>T), L266I (980T>A 982G>C), R272K (999G>A), Q275R (1007C>A 1008A>G 1009G>A), S276G (1010A>G 1012T>G), I277V (1013A>G), N279Q (1019A>C 1021C>G), D288E (1048C>A), R300Q (1083G>A), S305A (1097T>G 1099G>A), R320K (1143G>A), L324I (1154T>A 1156G>A), S346N (1221G>A), V349T (1229G>A 1230T>C 1231T>C), S350N (1233G>A 1234C>T), K358T (1257A>C 1258A>G), N365S (1278A>G), A369V (1290C>T 1291G>A), D372T (1298G>A 1299A>C 1300C>T), I375V (1307A>G 1309A>G), N378E (1316A>G 1318T>A), I382T (1329T>C), L397I (1373C>A), I401V (1385A>G), E407D (1405G>T), A411S (1415G>T 1417A>T), V418I (1436G>A 1438C>T), E508R (1706G>A 1707A>G), A512T (1718G>A), I520V (1742A>G 1744C>T), S525N (1758G>A), V536I (1790G>A), I585V (1937A>G 1939T>C), F607Y (2004T>A), I628M (2068T>G), V630I (2072G>A 2074G>A), R675K (2208G>A), D694E (2266C>G), N744G (2414A>G 2415A>G 2416T>G), S746C (2420A>T), K780R (2523A>G), R817K (2634G>A), V847I (2723G>A 2725C>T), A855T (2747G>A 2749T>A), H871N (2795C>A), V880I (2822G>A 2824G>A), P900T (2882C>A), S901H (2885A>C 2886G>A), T904M (2895C>T), N916S (2931A>G), N919S (2940A>G), E929_N930insX (2971_2972insA), T936I (2991C>T), H937R (2994A>G), E939R (2999G>A 3000A>G), R946Q (3021G>A), Y947F (3024A>T 3025C>T), T951I (3036C>T), E952G (3039A>G 3040A>C), K953T (3042A>C 3043G>A), A972V (3099C>T 3100C>A), V974R (3104G>A 3105T>G 3106G>A), S983N (3132G>A), R985K (3137C>A 3138G>A 3139A>G), F990I (3152T>A 3154C>T), C994Y (3165G>A), F1012L (3218T>C), T1014N (3225C>A), V1016I (3230G>A), V1028M (3266G>A 3268A>G), V1031M (3275G>A), A1034T (3284G>A), R1044K (3315G>A 3316A>G), E1045A (3318A>C), M1059V (3359A>G), D1072E (3400T>G), V1110G (3513T>G), M1121V (3545A>G), A1122T (3548G>A), V1124I (3554G>A 3556C>T), L1147V (3623C>G), L1150M (3632C>A), Y1152H (3638T>C), I1172V (3698A>G), S1178N (3717G>A), R1183K (3731C>A 3732G>A), I1297V (4073A>G), F1315Y (4128T>A 4129C>T), A1323S (4151G>T), V1328I (4166G>A), L1347I (4223C>A 4225G>A), I1379V (4319A>G 4321A>G), F1404Y (4395T>A), V1408I (4406G>A), M1431V (4475A>G 4477G>T), T1433S (4481A>T 4483A>G), A1454L (4544G>T 4545C>T 4546G>A), R1455K (4548G>A), V1499I (4679G>A 4681G>C), V1538I (4796G>A 4798C>T), T1540S (4803C>G 4804T>C), S1625G (5057A>G), T1626A (5060A>G), L1636I (5090C>A), A1639V (5100C>T), E1641D (5107A>T), V1646F (5120G>T 5122C>T), I1648V (5126A>G), V1667I (5183G>A), A1793T (5561G>A 5563G>A), I1805V (5597A>G), N1807K (5605C>A), A1812V (5619C>T 5620T>G), I1816L (5630A>C), V1837I (5693G>A), K1890R (5853A>G), I1904V (5894A>G 5896C>G), R1926A (5960C>G 5961G>C 5962G>C), I1936V (5990A>G), E1945D (6019A>T), K1949T (6030A>C), D1955E (6049C>A), M1958I (6058G>A), S1963N (6072G>A 6073T>C), N1964H (6074A>C), T1966N (6081C>A), C1976S (6111G>C 6112T>C), I1982V (6128A>G 6130T>C), V1992I (6158G>A), T1996S (6170A>T 6172A>G), L2007F (6203C>T 6205C>T), V2016I (6230G>A), A2025K (6257G>A 6258C>A), K2033R (6282A>G), A2060T (6362G>A), V2073I (6401G>A), A2074V (6405C>T), Y2082F (6429A>T), Q2089R (6450A>G), H2101N (6485C>A 6487C>T), L2115I (6527C>A), T2135I (6588C>T), Y2151H (6635T>C), I2155V (6647A>G), I2165V (6677A>G 6679A>G), D2178N (6716G>A), N2179D (6719A>G), I2194L (6764A>T 6766A>G), A2204V (6795C>T 6796A>G), H2205Q (6799C>A), N2225D (6857A>G), V2231T (6875G>A 6876T>C 6877C>A), D2250E (6934C>A), V2273I (7001G>A 7003G>A), F2283Y (7032T>A), T2314K (7125C>A), A2321S (7145G>T 7147A>G), E2326_A2327del (7160_7165delGAGGCA), D2344E (7216T>A), D2358E (7258T>A), D2363_R2364insX (7273_7274insT), R2364V (7274C>G 7275G>T), H2366L (7281A>T), L2367S (7284T>C), S2371L (7296C>T), S2376F (7311C>T), T2377I (7314C>T), C2378Y (7317G>A), N2379K (7321T>A), K2384Q (7334A>C), A2387N (7343G>A 7344C>A 7345A>C), S2408M (7407G>T 7408T>G), C2411G (7415T>G), L2440S (7503T>C 7504G>A), D2509N (7709G>A), K2510R (7713A>G 7714A>G), T2543I (7812C>T), V2554L (7844G>T 7846G>A), V2567I (7883G>A), S2620N (8043G>A), N2656S (8151A>G 8152C>T), R2665K (8178G>A), R2675K (8208G>A), P2704A (8294C>G 8296A>G), S2722F (8349C>T), A2742S (8408G>T), M2756I (8452G>A), R2759K (8460G>A), A2766T (8480G>A 8482T>A), R2773K (8502G>A 8503A>G), E2777D (8515A>T), E2785D (8539G>C), L2788C (8546C>T 8547T>G), S2790T (8552T>A), A2797G (8574C>G), I2801V (8585A>G), G2805E (8598G>A), N2807S (8603A>T 8604A>C 8605C>T), P2813Q (8622C>A), E2814G (8625A>G 8626G>A), P2822F (8648C>T 8649C>T 8650G>C), G2825E (8658G>A), D2827A (8664A>C 8665T>G), A2832V (8679C>T 8680A>T), T2854K (8745C>A), E2891G (8856A>G 8857G>A), R2894Q (8865G>A), M2895L (8867A>C 8869G>T), G2900E (8883G>A), T2902S (8888A>T), V2924I (8954G>A), N2934D (8984A>G 8986T>C), E2935V (8988A>T), I2983V (9131A>G), M2989G (9149A>G 9150T>G 9151G>A), G2990S (9152G>A), P3026S (9260C>T 9262T>C), Q3038H (9298A>C) | | | | | | | | |

| | Begin | End | Coverage | Score | Concordance | Matches | Identities | I/D/M/F* | Stop Codons |
|---|---|---|---|---|---|---|---|---|---|
| **NT** | **54** | **9704** | **99.5%** | **12522** | **65.2%** | **9639 (99.8%)** | **7984 (82.7%)** | **7/12** | |

Codon mutations:

ACT3ACC (193T>C), TAC4CAA (194T>C 196C>A), TCA6TCG (202A>G), ATC8ATT (208C>T), TGT9CTG (209T>C 210G>T 211T>G), TCG12TCA (220G>A), TTT13ATG (221T>A 223T>G), GAA14GAG (226A>G), CTA17CTT (235A>T), TAC19TAT (241C>T), CCA21CCT (247A>T), TCT23CCT (251T>C), TGC24TTT (255G>T 256C>T), GAG25GGG (258A>G), CAT26CAC (262T>C), ATT27GTG (263A>G 265T>G), GTG28GCG (267T>C), GAA30GTA (273A>T), CGA31CGT (277A>T), GAA32GAG (280A>G), GTG33GTT (283G>T), CCG34CCA (286G>A), GCT35ACC (287G>A 289T>C), GTT37ACT (293G>A 294T>C), CCT39CCC (301T>C), TTC40TTT (304C>T), GCA41ATA (305G>A 306C>T), GAT42GGT (309A>G), CTG43TTG (311C>T), GAA44GAT (316A>T), ACA45GTG (317A>G 318C>T 319A>G), CAA46CAG (322A>G), GCA49GCT (331A>T), CTC52CAA (339T>A 340C>A), AAG53AGG (342A>G), AAA55GCG (347A>G 348A>C 349A>G), TAT56CAT (350T>C), GCT57GCA (355T>A), ACT58GTT (356A>G 357C>T), CGT60CGC (364T>C), GTG61ACA (365G>A 366T>C 367G>A), CTC62TCC (368C>T 369T>C), AAA63AAG (373A>G), AAC64AAT (376C>T), TTT67TGT (384T>G), ACG68ACA (388G>A), TAC69TAT (391C>T), CGA70CGG (394A>G), AAG72AAA (400G>A), GAT74GAC (406T>C), GCC75GTT (408C>T 409C>T), CAG76CAA (412G>A), ATA77ATT (415A>T), ATG78GAG (416A>G 417T>A), ATT80ATC (424T>C), CAG81CAA (427G>A), AAA83AAG (433A>G), CTG84CTC (436G>C), GAG85AAG (437G>A), AGG86GAG (440A>G 441G>A), AAG87CGA (443A>C 444A>G 445G>A), GAT88GAA (448T>A), GAA90GAG (454A>G), GAA91GCA (456A>C), TAT92TAC (460T>C), CAC93AAC (461C>A), CAA95CAG (469A>G), GCC97GCG (475C>G), CCT99CCA (481T>A), ACT106ACC (502T>C), ATA107ATT (505A>T), GGC109GGT (511C>T), GAT111GAA (517T>A), CCT112CCA (520T>A), CCA113CCT (523A>T), TCT116TTT (531C>T), GAG117GAA (535G>A), CCA118TCA (536C>T), CAA119TGG (539C>T 540A>G 541A>G), AGA122AAG (549G>A 550A>G), ATC124ATT (556C>T), ATT125ATC (559T>C), CAT126CAC (562T>C), ACT128ACC (568T>C), CCA129CCG (571A>G), ATG131ACG (576T>C), CGT132CGC (580T>C), GTC134GCG (585T>C 586C>G), CGC137TAT (593C>T 594G>A 595C>T), CCC138CGC (597C>G), ATA139GTG (599A>G 601A>G), ATA140CCG (602A>C 603T>C 604A>G), ACA143ACG (613A>G), GAA144GAG (616A>G), GGC145GGG (619C>G), CAG146CAA (622G>A), ATG147ATA (625G>A), AAT148AAC (628T>C), CAC149CAT (631C>T), ATT151GTC (635A>G 637T>C), ATA154GTG (644A>G 646A>G), ATT157ATC (655T>C), TCG159TCA (661G>A), GAG160GCC (663A>C 664G>C), AAA161AAG (667A>G), GTC165GTT (679C>T), CAC166CAA (682C>A), TTA167CTG (683T>C 685A>G), AGT169AGC (691T>C), ACC172AGC (699C>G), ACT173ACA (703T>A), CAT174CAA (706T>A), AAG179GAA (719A>G 721G>A), ATA180GTG (722A>G 724A>G), CTT181TTG (725C>T 727T>G), GGT182GGA (730T>A), GCA183TCA (731G>T), TAC184CCT (734T>C 735A>C 736C>T), TCC185CGT (737T>C 738C>G 739C>T), GCG187GTC (744C>T 745G>C), CGA189TGC (749C>T 751A>C), ACT190ACC (754T>C), ATG194AAG (765T>A), TTG196CTG (770T>C), CGA197AAG (773C>A 774G>A 775A>G), CGG198AGG (776C>A), TTC202TTT (790C>T), ATG206AAG (801T>A), GGA210GCG (813G>C 814A>G), CTT211CAT (816T>A), TTG212CTG (818T>C), CAA213CAG (823A>G), CGT214CAA (825G>A 826T>A), CGG217AGA (833C>A 835G>A), GAC219GAT (841C>T), AAT223GAC (851A>G 853T>C), GTC225GTT (859C>T), CGC226TGT (860C>T 862C>T), ACT227GCC (863A>G 865T>C), ATC228ACC (867T>C), AAC229GAT (869A>G 871C>T), ATA230TTG (872A>T 874A>G), CGA231CAA (876G>A), AGG232AAG (879G>A), GAT234GAC (886T>C), ATC238ATT (898C>T), AAC240AGC (903A>G), ACA241AAC (906C>A 907A>C), AAA242ACC (909A>C 910A>C), AGC243AAT (912G>A 913C>T), GGC246GGA (922C>A), GGT249GGA (931T>A), TCA252CTT (938T>C 939C>T 940A>T), GGA253GGG (943A>G), TTC256TTT (952C>T), TCA261TCG (967A>G), GAA263GAG (973A>G), GGG264GGC (976G>C), TTG266ATC (980T>A 982G>C), GCA269GCG (991A>G), CGT270CGC (994T>C), AGA272AAA (999G>A), GTT273GTC (1003T>C), CAG275AGA (1007C>A 1008A>G 1009G>A), AGT276GGG (1010A>G 1012T>G), ATT277GTT (1013A>G), TTA278TTG (1018A>G), AAC279CAG (1019A>C 1021C>G), ATC282ATT (1030C>T), TCG285TCA (1039G>A), AAT286AAC (1042T>C), GCC287GCA (1045C>A), GAC288GAA (1048C>A), AAT289AAC (1051T>C), AAG292AAA (1060G>A), GGT293GGA (1063T>A), CTG294TTG (1064C>T), GAC295GAT (1069C>T), CGA300CAA (1083G>A), TAT303TAC (1093T>C), TCG305GCA (1097T>G 1099G>A), CAC307CAT (1105C>T), GTA310GTG (1114A>G), GCT311GCG (1117T>G), TTA313TTG (1123A>G), CCT314CCA (1126T>A), GTC315GTT (1129C>T), AGG320AAG (1143G>A), GTA321GTT (1147A>T), GCT322GCA (1150T>A), TTG324ATA (1154T>A 1156G>A), CAC327CAT (1165C>T), AGT328AGC (1168T>C), CTT330TTA (1172C>T 1174T>A), ACC339ACT (1201C>T), AGC346AAC (1221G>A), GTT349ACC (1229G>A 1230T>C 1231T>C), AGC350AAT (1233G>A 1234C>T), CTG352CTA (1240G>A), AAG354AAA (1246G>A), TTG356CTA (1250T>C 1252G>A), CAT357CAC (1255T>C), AAA358ACG (1257A>C 1258A>G), AGA361AGG (1267A>G), GGT363GGC (1273T>C), TTG364TTA (1276G>A), AAT365AGT (1278A>G), TTG367CTG (1283T>C), GCG369GTA (1290C>T 1291G>A), GAC372ACT (1298G>A 1299A>C 1300C>T), TTT374TTC (1306T>C), ATA375GTG (1307A>G 1309A>G), CAT376CAC (1312T>C), AAT378GAA (1316A>G 1318T>A), TTG381TTA (1327G>A), ATA382ACA (1329T>C), GCG383GCT (1333G>T), TTA384TTG (1336A>G), CTA387TTG (1343C>T 1345A>G), CCG390CCA (1354G>A), GTG391GTT (1357G>T), GAC392GAT (1360C>T), CTC395TTG (1367C>T 1369C>G), CTT397ATT (1373C>A), TTC398TTT (1378C>T), GAG400GAA (1384G>A), ATA401GTA (1385A>G), AAA403AAG (1393A>G), TCC404TCT (1396C>T), GAG407GAT (1405G>T), AAA408AAG (1408A>G), CAG409CAA (1411G>A), GCA411TCT (1415G>T 1417A>T), CCG412CCC (1420G>C), AAT415AAC (1429T>C), TTA416TTG (1432A>G), GTC418ATT (1436G>A 1438C>T), TTA419CTG (1439T>C 1441A>G), AAT420AAC (1444T>C), TTC423TTT (1453C>T), CTG424CTA (1456G>A), GAA428GAG (1468A>G), GTA436GTG (1492A>G), TTG439CTA (1499T>C 1501G>A), AGT440AGC (1504T>C), TTG441TTA (1507G>A), CTC442CTT (1510C>T), GAA443GAG (1513A>G), TTA444TTG (1516A>G), GCA445GCT (1519A>T), AGG446AGA (1522G>A), TTC447TTT (1525C>T), CAG448CAA (1528G>A), AAC450AAT (1534C>T), ACT452ACA (1540T>A), AAC454AAT (1546C>T), AAA457AAG (1555A>G), GGT458GGC (1558T>C), GAT459GAC (1561T>C), ATA460ATC (1564A>C), TCT461TCA (1567T>A), TTC463TTT (1573C>T), AGA464AGG (1576A>G), TTA467CTA (1583T>C), TCT468TCA (1588T>A), AAG470AAA (1594G>A), AAC472AAT (1600C>T), CTG475CTA (1609G>A), TCG478TCA (1618G>A), TGC479TGT (1621C>T), GAC480GAT (1624C>T), CTG483TTG (1631C>T), AAA485AAG (1639A>G), TTC489TTT (1651C>T), CTC490CTG (1654C>G), CAT497CAC (1675T>C), TCA503TCG (1693A>G), TTT506TTC (1702T>C), GAA508AGA (1706G>A 1707A>G), GCA512ACA (1718G>A), GGA514GGG (1726A>G), TAC515TAT (1729C>T), TCA516TCC (1732A>C), TAT518TAC (1738T>C), ATC520GTT (1742A>G 1744C>T), CGC521CGT (1747C>T), CAT523CAC (1753T>C), AGT525AAT (1758G>A), AGG528AGA (1768G>A), AAC534AAT (1786C>T), TTA535TTG (1789A>G), GTT536ATT (1790G>A), GTC537GTT (1795C>T), TTA541CTG (1805T>C 1807A>G), GAG543GAA (1813G>A), AGG545AGA (1819G>A), AAG547AAA (1825G>A), GAC551GAT (1837C>T), TAT552TAC (1840T>C), CAA555CAG (1849A>G), GTC558GTG (1858C>G), AAA560AAG (1864A>G), TGC562TGT (1870C>T), AGT564AGC (1876T>C), AAA566AAG (1882A>G), GGT568GGA (1888T>A), AAT569AAC (1891T>C), TGT574TGC (1906T>C), ACA577ACT (1915A>T), GGT582GGC (1930T>C), GCC584GCT (1936C>T), ATT585GTC (1937A>G 1939T>C), TCA587TCG (1945A>G), TTC589TTT (1951C>T), CCA592CCT (1960A>T), GTA598GTG (1978A>G), ATT599ATA (1981T>A), GGC600GGT (1984C>T), AAT601AAC (1987T>C), GAC604GAT (1996C>T), TTT607TAT (2004T>A), AAA612AAG (2020A>G), GGG613GGA (2023G>A), GAT614GAC (2026T>C), TCG615TCT (2029G>T), GAG616GAA (2032G>A), TAT625TAC (2059T>C), ATT628ATG (2068T>G), AAC629AAT (2071C>T), GTG630ATA (2072G>A 2074G>A), CTG635TTA (2087C>T 2089G>A), AGC639AGT (2101C>T), GAG640GAA (2104G>A), GAG641GAA (2107G>A), AAG644AAA (2116G>A), TTC646TTT (2122C>T), CGC651CGT (2137C>T), CTT658CTC (2158T>C), ACC660ACT (2164C>T), CCA662CCG (2170A>G), GAT666GAC (2182T>C), GCG668GCA (2188G>A), ACC669ACA (2191C>A), AGA675AAA (2208G>A), TAT678TAC (2218T>C), GAC680GAT (2224C>T), GTA681GTT (2227A>T), TTG686TTA (2242G>A), CCC687CCT (2245C>T), TTG690TTA (2254G>A), GTT691GTC (2257T>C), GAC692GAT (2260C>T), GAC694GAG (2266C>G), CAA696CAG (2272A>G), ACG697ACA (2275G>A), TCA703TCG (2293A>G), GGC705GGT (2299C>T), TCG706TCA (2302G>A), CAG707CAA (2305G>A), GGA710GGT (2314A>T), TAT711TAC (2317T>C), CTA714CTG (2326A>G), GTG719GTA (2341G>A), CAA721CAG (2347A>G), CTT722CTC (2350T>C), TTG724TTA (2356G>A), AAT727AAC (2365T>C), GAA729GAG (2371A>G), TTA730TTG (2374A>G), GAT733GAC (2383T>C), ATA734ATT (2386A>T), AAA735AAG (2389A>G), CAT736CAC (2392T>C), TAT737TAC (2395T>C), AAT744GGG (2414A>G 2415A>G 2416T>G), GCT745GCA (2419T>A), AGC746TGC (2420A>T), ACA752ACT (2440A>T), GTT761GTC (2467T>C), TCT764TCC (2476T>C), TCG766TCA (2482G>A), CTG769CTA (2491G>A), CTG771CTT (2497G>T), TTG773TTA (2503G>A), AGA778AGG (2518A>G), CCT779CCC (2521T>C), AAG780AGG (2523A>G), AGA783AGG (2533A>G), TTG785TTA (2539G>A), TTA787TTG (2545A>G), GAG789GAA (2551G>A), CCT790CCA (2554T>A), CTG792TTG (2558C>T), TTG793CTC (2561T>C 2563G>C), TCA796TCG (2572A>G), CCT800CCC (2584T>C), GGC801GGT (2587C>T), CTG803CTC (2593G>C), GCT805GCC (2599T>C), AAT808AAC (2608T>C), TTT812TTC (2620T>C), GCG815GCA (2629G>A), GTG816GTA (2632G>A), AGG817AAG (2634G>A), TTG818TTA (2638G>A), ATT820ATA (2644T>A), AAA823AAG (2653A>G), GCT827GCC (2665T>C), CTA832TTA (2678C>T), CTA833CTG (2683A>G), GCT835GCC (2689T>C), CTA838CTC (2698A>C), TCA841TCG (2707A>G), GCG842GCA (2710G>A), GAA844GAG (2716A>G), ACA845ACG (2719A>G), CTC846CTT (2722C>T), GTC847ATT (2723G>A 2725C>T), GCA848GCG (2728A>G), AGG850AGA (2734G>A), ATT851ATA (2737T>A), GAT854GAC (2746T>C), GCT855ACA (2747G>A 2749T>A), GCA856GCT (2752A>T), GCT857GCA (2755T>A), GAC859GAT (2761C>T), CTC860CTT (2764C>T), TGT865TGC (2779T>C), GGG867GGT (2785G>T), AAC869AAT (2791C>T), CAT871AAT (2795C>A), TAC874TAT (2806C>T), TTG878CTC (2816T>C 2818G>C), GTG880ATA (2822G>A 2824G>A), TTG881TTA (2827G>A), GTT884GTC (2836T>C), AAG885AAA (2839G>A), AAT886AAC (2842T>C), ACC893ACA (2863C>A), CTA894TTA (2864C>T), CCA900ACA (2882C>A), AGT901CAT (2885A>C 2886G>A), AAC903AAT (2893C>T), ACG904ATG (2895C>T), GTC906GTT (2902C>T), GTA907GTG (2905A>G), ATT909ATC (2911T>C), AAT913AAC (2923T>C), AAT916AGT (2931A>G), AAC919AGC (2940A>G), GCT921GCC (2947T>C), ACT926ACC (2962T>C), GAA929_AAT930insA-- (2971_2972insA), GGT935GGC (2989T>C), ACT936ATT (2991C>T), CAT937CGT (2994A>G), GAG939AGG (2999G>A 3000A>G), TCC944TC- (3016delC), CGG946CAG (3021G>A), TAC947TTT (3024A>T 3025C>T), ACA951ATA (3036C>T), GAA952GGC (3039A>G 3040A>C), AAG953ACA (3042A>C 3043G>A), TTG956TTA (3052G>A), TTA959TTG (3061A>G), GCG966GCA (3082G>A), GGC969GGT (3091C>T), GCC972GTA (3099C>T 3100C>A), GTG974AGA (3104G>A 3105T>G 3106G>A), GTC975GTT (3109C>T), AAA976AAG (3112A>G), ACT978ACC (3118T>C), GCC979GCT (3121C>T), GGA981GGG (3127A>G), AGC983AAC (3132G>A), CGA985AAG (3137C>A 3138G>A 3139A>G), TAT989TAC (3151T>C), TTC990ATT (3152T>A 3154C>T), TGT994TAT (3165G>A), GTA995GTG (3169A>G), ATC1005ATT (3199C>T), AGG1007CGG (3203A>C), CGT1008CGC (3208T>C), CCA1010CCG (3214A>G), ACC1011ACT (3217C>T), TTT1012CTT (3218T>C), GTC1013GTA (3223C>A), ACT1014AAT (3225C>A), GTT1016ATT (3230G>A), AAC1017AAT (3235C>T), TTA1025CTA (3257T>C), ACC1026ACT (3262C>T), AGC1027AGT (3265C>T), GTA1028ATG (3266G>A 3268A>G), GTG1029GTA (3271G>A), GTG1031ATG (3275G>A), GCA1034ACA (3284G>A), CAG1039CAA (3301G>A), AGA1044AAG (3315G>A 3316A>G), GAA1045GCA (3318A>C), ATC1046ATT (3322C>T), GAG1047GAA (3325G>A), TTG1048TTA (3328G>A), ATA1051ATT (3337A>T), GAG1055GAA (3349G>A), GTC1057GTT (3355C>T), TGC1058TGT (3358C>T), ATG1059GTG (3359A>G), CTA1061CTG (3367A>G), TAT1062TAC (3370T>C), TTA1065CTG (3377T>C 3379A>G), GAA1070GAG (3394A>G), CGC1071CGT (3397C>T), GAT1072GAG (3400T>G), ACA1074ACG (3406A>G), TAC1078TAT (3418C>T), ATT1079ATA (3421T>A), TAT1081TAC (3427T>C), AAG1083AAA (3433G>A), TCA1084TCT (3436A>T), GTA1085GTG (3439A>G), AAC1086AAT (3442C>T), CCT1087CCC (3445T>C), CAG1091CAA (3457G>A), TTT1092TTC (3460T>C), GCT1093GCA (3463T>A), GCG1095GCT (3469G>T), CAG1096CAA (3472G>A), CAC1104CAT (3496C>T), CAG1105CAA (3499G>A), TCC1107TCT (3505C>T), GTT1110GGT (3513T>G), AAA1112AAG (3520A>G), GAA1115GAG (3529A>G), CAA1116CAG (3532A>G), ATG1121GTG (3545A>G), GCT1122ACT (3548G>A), TTA1123CTA (3551T>C), GTC1124ATT (3554G>A 3556C>T), AGT1133AGC (3583T>C), GAT1134GAC (3586T>C), TGC1135TGT (3589C>T), TTC1137TTT (3595C>T), AAT1141AAC (3607T>C), TTT1143TTC (3613T>C), AAG1144AAA (3616G>A), GGT1145GGC (3619T>C), CTT1147GTT (3623C>G), TCC1148TCT (3628C>T), CTG1150ATG (3632C>A), GAT1151GAC (3637T>C), TAT1152CAT (3638T>C), GAA1153GAG (3643A>G), TTA1159TTG (3661A>G), GTG1162GTA (3670G>A), TTT1166TTC (3682T>C), GAG1168GAA (3688G>A), GAG1171GAA (3697G>A), ATT1172GTT (3698A>G), GAT1174GAC (3706T>C), TTT1175TTC (3709T>C), TTG1177CTA (3713T>C 3715G>A), AGT1178AAT (3717G>A), GAG1179GAA (3721G>A), CGA1183AAA (3731C>A 3732G>A), ACT1184ACA (3736T>A), CTA1188TTG (3746C>T 3748A>G), ACA1190ACG (3754A>G), AGT1193AGC (3763T>C), CGA1198CGG (3778A>G), CAA1199CAG (3781A>G), ATC1200ATT (3784C>T), CAT1204CAC (3796T>C), CCA1207CCT (3805A>T), TAC1209TAT (3811C>T), AGA1210AGG (3814A>G), TTC1215TTT (3829C>T), TTT1218TTC (3838T>C), AGA1220AGG (3844A>G), GCA1221GCC (3847A>C), GTT1224GTA (3856T>A), GTG1226GTT (3862G>T), GAC1229GAT (3871C>T), GCC1231GCA (3877C>A), AGC1233AGT (3883C>T), GAA1234GAG (3886A>G), CTA1236CTG (3892A>G), GAC1237GAT (3895C>T), TTT1238TTC (3898T>C), GTA1240GTG (3904A>G), CGG1241AGA (3905C>A 3907G>A), GCT1243GCC (3913T>C), AAG1248AAA (3928G>A), TCA1249TCT (3931A>T), GGG1251GGA (3937G>A), AGT1257AGC (3955T>C), GTG1258GTA (3958G>A), GCC1259GCT (3961C>T), GGA1260GGG (3964A>G), TCT1261TCC (3967T>C), TTA1264TTG (3976A>G), ATT1265ATC (3979T>C), CCA1267CCT (3985A>T), CTA1271CTT (3997A>T), GCG1272GCA (4000G>A), AAC1274AAT (4006C>T), GTT1275GTG (4009T>G), TTC1276TTT (4012C>T), AAA1277AAG (4015A>G), TCT1280TCC (4024T>C), GAA1282GAG (4030A>G), CCA1283CCT (4033A>T), AAG1286AAA (4042G>A), CTG1290TTA (4052C>T 4054G>A), CGT1291CGC (4057T>C), CGT1293CGA (4063T>A), GGA1294GGG (4066A>G), ATA1297GTA (4073A>G), TTT1298TTC (4078T>C), GGC1299GGT (4081C>T), TCT1300TCC (4084T>C), TCC1304TCT (4096C>T), AGC1308AGT (4108C>T), GGA1309GGT (4111A>T), GCG1311GCA (4117G>A), CTA1312TTG (4118C>T 4120A>G), TAC1314TAT (4126C>T), TTC1315TAT (4128T>A 4129C>T), GCC1316GCT (4132C>T), AAT1317AAC (4135T>C), CAA1321CAG (4147A>G), TTA1322CTA (4148T>C), GCT1323TCT (4151G>T), AAC1326AAT (4162C>T), GTA1328ATA (4166G>A), ATA1329ATC (4171A>C), GAG1332GAA (4180G>A), CTG1336TTG (4190C>T), CCT1338CCC (4198T>C), TCC1339TCT (4201C>T), GCG1342GCT (4210G>T), TTC1343TTT (4213C>T), AGT1345AGC (4219T>C), CTG1346TTG (4220C>T), CTG1347ATA (4223C>A 4225G>A), GTT1349GTG (4231T>G), CAT1351CAC (4237T>C), GTA1356GTG (4252A>G), TTA1357CTG (4253T>C 4255A>G), AAA1358AAG (4258A>G), ACT1362ACC (4270T>C), AGA1366AGG (4282A>G), GAG1367GAA (4285G>A), GAA1369GAG (4291A>G), CAG1373CAA (4303G>A), GTC1376GTT (4312C>T), ATA1379GTG (4319A>G 4321A>G), GTG1380GTT (4324G>T), GAG1381GAA (4327G>A), CTG1384CTT (4336G>T), TCT1385TCC (4339T>C), TCA1388TCT (4348A>T), GCA1392GCG (4360A>G), GGT1394GGC (4366T>C), TCT1395TCA (4369T>A), ACT1397ACC (4375T>C), GTT1401GTC (4387T>C), TTT1404TAT (4395T>A), GTA1408ATA (4406G>A), CTT1409CTC (4411T>C), GTT1418GTC (4438T>C), ACC1420ACA (4444C>A), GCC1422GCT (4450C>T), CTC1424CTT (4456C>T), GAC1427GAT (4465C>T), ATG1431GTT (4475A>G 4477G>T), GTC1432GTT (4480C>T), ACA1433TCG (4481A>T 4483A>G), GGC1437GGT (4495C>T), AAG1441AAA (4507G>A), CAC1442CAT (4510C>T), GGT1443GGG (4513T>G), CTA1445TTA (4517C>T), GTC1448GTT (4528C>T), AAA1450AAG (4534A>G), ACC1452ACT (4540C>T), GCG1454TTA (4544G>T 4545C>T 4546G>A), AGA1455AAA (4548G>A), TTT1458TTC (4558T>C), GTA1460GTT (4564A>T), ACC1462ACT (4570C>T), ATA1464ATT (4576A>T), TTG1471TTA (4597G>A), GAC1472GAT (4600C>T), GAC1474GAT (4606C>T), GTG1475GTA (4609G>A), GTG1477GTA (4615G>A), GAC1478GAT (4618C>T), TTG1481CTT (4625T>C 4627G>T), GAC1488GAT (4648C>T), ATT1489ATC (4651T>C), AGG1492AGA (4660G>A), AGC1493AGT (4663C>T), GCT1495GCA (4669T>A), GTG1499ATC (4679G>A 4681G>C), AGT1500AGC (4684T>C), GTT1501GTG (4687T>G), GAG1505GAA (4699G>A), AGA1506AGG (4702A>G), CAA1508CAG (4708A>G), AGG1509AGA (4711G>A), CTG1510TTG (4712C>T), AAA1518AAG (4738A>G), GTA1520GTT (4744A>T), GCA1521GCG (4747A>G), TTG1522CTG (4748T>C), ACT1527ACC (4765T>C), GTC1538ATT (4796G>A 4798C>T), ACT1540AGC (4803C>G 4804T>C), GAG1541GAA (4807G>A), GCG1542GCT (4810G>T), GCT1543GCG (4813T>G), CTT1544CTC

| | | | | | | | | | |
|---|---|---|---|---|---|---|---|---|---|
| | (4816T>C), TAT1549TAC (4831T>C), AAC1550AAT (4834C>T), CCA1552CCT (4840A>T), GGA1556GGG (4852A>G), GGC1557GGT (4855C>T), AGT1561AGC (4867T>C), CTG1562CTC (4870G>C), ATT1563ATA (4873T>A), TGT1566TGC (4882T>C), ACA1573ACT (4903A>T), CAG1575CAA (4909G>A), TTT1577TTC (4915T>C), GAA1578GAG (4918A>G), TTG1579CTA (4919T>C 4921G>A), AGT1580AGC (4924T>C), CCC1581CCG (4927C>G), TTT1583TTC (4933T>C), ATC1584ATA (4936C>A), CAG1585CAA (4939G>A), AAT1586AAC (4942T>C), TTC1587TTT (4945C>T), TCA1593TCG (4963A>G), CCT1596CCC (4972T>C), CTT1602CTC (4990T>C), AAA1606AAG (5002A>G), CCT1613CCC (5023T>C), TTG1614CTG (5024T>C), TGC1615TGT (5029C>T), GAT1616GAC (5032T>C), TCT1618TCC (5038T>C), CCA1620CCT (5044A>T), GCA1623GCT (5053A>T), TCG1624TCA (5056G>A), AGT1625GGT (5057A>G), ACT1626GCT (5060A>G), TCG1629TCT (5071G>T), CTT1636ATT (5090C>A), GGA1637GGG (5095A>G), GTG1638GTT (5098G>T), GCC1639GTC (5100C>T), TTA1640TTG (5104A>G), GAA1641GAT (5107A>T), ATT1642ATC (5110T>C), CCA1643CCG (5113A>G), AAG1644AAA (5116G>A), CAA1645CAG (5119A>G), GTC1646TTT (5120G>T 5122C>T), ATT1648GTT (5126A>G), CAT1651CAC (5137T>C), ATC1652ATT (5140C>T), AAA1653AAG (5143A>G), GAG1654GAA (5146G>A), CTC1659CTA (5161C>A), GAA1661GAG (5167A>G), CTT1663CTG (5173T>G), ACG1666ACA (5182G>A), GTT1667ATT (5183G>A), GTC1668GTT (5188C>T), AAG1669AAA (5191G>A), AAA1671AAG (5197A>G), GAC1672GAT (5200C>T), TGC1674TGT (5206C>T), TTA1675TTG (5209A>G), AGC1678AGT (5218C>T), TCG1682TCA (5230G>A), ATC1684ATT (5236C>T), AGC1685AGT (5239C>T), ATC1687ATT (5245C>T), ACA1693ACT (5263A>T), CTC1695CTT (5269C>T), TTC1696TTT (5272C>T), GCC1697GCA (5275C>A), ATC1698ATT (5278C>T), CCA1699CCC (5281A>C), AGA1700AGG (5284A>G), ACT1701ACC (5287T>C), CTA1702TTA (5288C>T), ATA1703ATT (5293A>T), TTG1704CTA (5294T>C 5296G>A), GAG1706GAA (5302G>A), CTT1709CTA (5311T>A), GAA1710GAG (5314A>G), CGA1713CGT (5323A>T), AGT1721AGC (5347T>C), ATC1723ATT (5353C>T), TGC1727TGT (5365C>T), GTT1734GTC (5386T>C), ACC1737ACA (5395C>A), AAA1746AAG (5422A>G), GAT1747GAC (5425T>C), ACC1749ACT (5431C>T), GAG1751GAA (5437G>A), CTT1756CTC (5452T>C), AAG1758AAA (5458G>A), AGT1761AGC (5467T>C), CAA1762CAG (5470A>G), CTA1763TTG (5471C>T 5473A>G), GAA1765GAG (5479A>G), TTC1766TTT (5482C>T), TCA1767TCC (5485A>C), AAT1768AAC (5488T>C), TTG1769CTA (5489T>C 5491G>A), GGT1771GGC (5497T>C), TGT1774TGC (5506T>C), GAG1775GAA (5509G>A), GAG1776GAA (5512G>A), TTA1778CTA (5516T>C), TAT1782TAC (5530T>C), GAG1783GAA (5533G>A), TCG1784TCT (5536G>T), TTG1785TTA (5539G>A), GTT1788GTG (5548T>G), CAC1790CAT (5554C>T), GCT1792GCA (5560T>A), GCG1793ACA (5561G>A 5563G>A), ACG1794ACC (5566G>C), TCA1795TCG (5569A>G), CTC1800CTG (5584C>G), AAG1803AAA (5593G>A), GGG1804GGA (5596G>A), ATT1805GTT (5597A>G), AAC1807AAA (5605C>A), GTG1811GTT (5617G>T), GCT1812GTG (5619C>T 5620T>G), TTG1815TTA (5629G>A), ATC1816CTC (5630A>C), GGC1819GGT (5641C>T), GCA1822GCT (5650A>T), ATA1826ATT (5662A>T), GGA1827GGG (5665A>G), CAA1835CAG (5689A>G), GTT1837ATT (5693G>A), GAG1838GAA (5698G>A), CAA1843CAG (5713A>G), GGG1844GGC (5716G>C), AAA1845AAG (5719A>G), GCC1853GCA (5743C>A), CGC1857CGA (5755C>A), CAT1858CAC (5758T>C), CGT1860CGC (5764T>C), AAA1862AAG (5770A>G), GGC1865GGT (5779C>T), TTT1866TTC (5782T>C), GAC1869GAT (5791C>T), GAC1873GAT (5803C>T), AGG1884AGA (5836G>A), AAA1885AAG (5839A>G), GGA1887GGG (5845A>G), AAA1888AAG (5848A>G), AAA1890AGA (5853A>G), GGT1891GGC (5857T>C), ACC1892ACT (5860C>T), ACA1893ACT (5863A>T), GGT1895GGA (5869T>A), GGT1897GGC (5875T>C), AGC1900AGT (5884C>T), TTC1903TTT (5893C>T), ATC1904GTG (5894A>G 5896C>G), AAC1905AAT (5899C>T), GGG1908GGA (5908G>A), TTT1909TTC (5911T>C), GAG1913GAA (5923G>A), CAA1918CAG (5938A>G), TTC1919TTT (5941C>T), GTT1920GTC (5944T>C), GAT1921GAC (5947T>C), GGG1925GGA (5959G>A), CGG1926GCC (5960C>G 5961G>C 5962G>C), GAA1930GAG (5974A>G), GCT1934GCA (5986T>A), GAC1935GAT (5989C>T), ATT1936GTT (5990A>G), GAT1938GAC (5998T>C), AGA1942CGT (6008A>C 6010A>T), GAA1945GAT (6019A>T), GTG1946GTT (6022G>T), CGA1947CGT (6025A>T), AAA1949ACA (6030A>C), GAC1955GAA (6049C>A), ATG1958ATA (6058G>A), CAA1959CAG (6061A>G), GCC1960GCA (6064C>A), GGT1962GGC (6070T>C), AGT1963AAC (6072G>A 6073T>C), AAC1964CAC (6074A>C), ACG1965ACA (6079G>A), ACC1966AAC (6081C>A), ATA1967ATT (6085A>T), GCA1969GCT (6091A>T), TAC1970TAT (6094C>T), TGT1976TCC (6111G>C 6112T>C), AAA1978AAG (6118A>G), TTG1980TTA (6124G>A), AAG1981AAA (6127G>A), ATT1982GTC (6128A>G 6130T>C), GAT1983GAC (6133T>C), TTA1984TTG (6136A>G), AAC1988AAT (6148C>T), AAA1991AAG (6157A>G), GTT1992ATT (6158G>A), TGT1993TGC (6163T>C), AAA1995AAG (6169A>G), ACA1996TCG (6170A>T 6172A>G), GGC1998GGT (6178C>T), AAA2001AAG (6187A>G), TTT2002TTC (6190T>C), GAG2004GAA (6196G>A), CTC2007TTT (6203C>T 6205C>T), GAA2008GAG (6208A>G), CTA2009TTG (6209C>T 6211A>G), CAG2011CAA (6217G>A), GCT2015GCA (6229T>A), GTA2016ATA (6230G>A), GTC2018GTT (6238C>T), AAG2021AAA (6247G>A), ATA2023ATT (6253A>T), GCA2025AAA (6257G>A 6258C>A), CAT2030CAC (6274T>C), AAA2033AGA (6282A>G), TCG2034TCA (6286G>A), GGC2038GGT (6298C>T), TTG2039TTA (6301G>A), AGA2040AGG (6304A>G), GAC2041GAT (6307C>T), GCC2046GCT (6322C>T), ACA2048ACT (6328A>T), GTT2049GTG (6331T>G), TGT2050TGC (6334T>C), AGG2051AGA (6337G>A), CTG2052TTG (6338C>T), TAT2058TAC (6358T>C), GGG2059GGA (6361G>A), GCA2060ACA (6362G>A), TCA2061TCT (6367A>T), GAG2062GAA (6370G>A), TAC2064TAT (6376C>T), GGT2065GGG (6379T>G), GGA2067GGC (6385A>C), GGA2069GGT (6391A>T), GCA2070GCG (6394A>G), TAC2071TAT (6397C>T), GTA2073ATA (6401G>A), GCG2074GTG (6405C>T), CAC2076CAT (6412C>T), TTA2078TTG (6418A>G), TTT2079TTC (6421T>C), TAC2082TTC (6429A>T), GGT2084GGA (6436T>A), GAG2087GAA (6445G>A), CAA2089CGA (6450A>G), TCC2090TCG (6454C>G), CAC2092CAT (6460C>T), GGT2093GGA (6463T>A), TTC2095TTT (6469C>T), AGG2096AGA (6472G>A), GTG2097GTA (6475G>A), AAT2099AAC (6481T>C), CAC2101AAT (6485C>A 6487C>T), AGT2102AGC (6490T>C), AGC2104AGT (6496C>T), CTG2106CTA (6502G>A), GGT2110GGC (6514T>C), AGG2111AGA (6517G>A), CTC2115ATC (6527C>A), ATC2116ATA (6532C>A), CCG2119CCA (6541G>A), AAA2120AAG (6544A>G), GTC2124GTT (6556C>T), TTT2125TTC (6559T>C), CAG2127CAA (6565G>A), TTG2129CTA (6569T>C 6571G>A), CCT2134CCA (6586T>A), ACA2135ATA (6588C>T), GAA2138GAG (6598A>G), AGA2139AGG (6601A>G), ATT2140ATC (6604T>C), TGT2141TGC (6607T>C), GGA2144GGG (6616A>G), AAC2146AAT (6622C>T), GAG2149GAA (6631G>A), AAG2150AAA (6634G>A), TAT2151CAT (6635T>C), GCT2152GCA (6640T>A), TCG2154TCT (6646G>T), ATC2155GTC (6647A>G), ATC2156ATT (6652C>T), AGC2160AGT (6664C>T), ACT2162ACG (6670T>G), ATA2165GTG (6677A>G 6679A>G), GGC2167GGT (6685C>T), ACA2169ACT (6691A>T), TTC2170TTT (6694C>T), ACA2177ACG (6715A>G), GAT2178AAT (6716G>A), AAT2179GAT (6719A>G), GGA2180GGG (6724A>G), CTA2184TTA (6734C>T), GTG2186GTA (6742G>A), GTG2187GTA (6745G>A), GCC2190GCT (6754C>T), GGA2192GGG (6760A>G), ATA2194TTG (6764A>T 6766A>G), ATT2197ATA (6775T>A), CTG2200TTG (6782C>T), GCA2201GCT (6787A>T), AAC2202AAT (6790C>T), GCA2204GTG (6795C>T 6796A>G), CAC2205CAA (6799C>A), ACC2206ACA (6802C>A), AAC2208AAT (6808C>T), TAC2209TAT (6811C>T), TAC2210TAT (6814C>T), TTC2213TTT (6823C>T), GAA2215GAG (6829A>G), TAC2221TAT (6847C>T), CGA2223CGC (6853A>C), AAT2225GAT (6857A>G), GAG2226GAA (6862G>A), GAA2229GAG (6871A>G), GTC2231ACA (6875G>A 6876T>C 6877C>A), AAG2232AAA (6880G>A), TCT2233TCA (6883T>A), GTT2235GTG (6889T>G), GAC2239GAT (6901C>T), GGC2244GGT (6916C>T), CCG2245CCA (6919G>A), AAA2247AAG (6925A>G), CTT2248CTC (6928T>C), GAC2250GAA (6934C>A), GGG2255GGC (6949G>C), TTC2257TTT (6955C>T), AAA2258AAG (6958A>G), CAA2264CAG (6976A>G), CTA2266TTA (6980C>T), ATC2267ATT (6985C>T), GTA2271GTT (6997A>T), GTG2272GTT (7000G>T), GTG2273ATA (7001G>A 7003G>A), GCT2276GCC (7012T>C), AAG2277AAA (7015G>A), CAT2278CAC (7018T>C), TTT2283TAT (7032T>A), GCC2285GCT (7039C>T), TTG2286CTA (7040T>C 7042G>A), GGA2288GGG (7048A>G), GCT2292GCA (7060T>A), GTC2293GTA (7063C>A), AGC2298AGT (7078C>T), TTA2300CTG (7082T>C 7084A>G), GTA2301GTG (7087A>G), ACC2302ACA (7090C>A), CAT2304CAC (7096T>C), GTA2305GTG (7099A>G), GTT2306GTC (7102T>C), GAG2309GAA (7111G>A), CGA2311CGG (7117A>G), ACA2314AAA (7125C>A), GAA2315GAG (7129A>G), TTT2316TTC (7132T>C), CTG2317TTA (7133C>T 7135G>A), GTG2319GTA (7141G>A), GAT2320GAC (7144T>C), GCA2321TCG (7145G>T 7147A>G), GAG2322GAA (7150G>A), GCA2325GCT (7159A>T), GAG2326_GCA2327del (7160_7165delGAGGCA), TTC2329TTT (7171C>T), GCG2335GCT (7189G>T), AAA2338AAG (7198A>G), AGC2339AGT (7201C>T), CTA2341TTA (7205C>T), AAT2342AAC (7210T>C), GAT2344GAA (7216T>A), GCG2345GCA (7219G>A), ATC2347ATA (7225C>A), AAG2348AAA (7228G>A), AAG2352AAA (7240G>A), TAT2353TAC (7243T>C), AAA2355AAG (7249A>G), CCT2356CCG (7252T>G), GAT2358GAA (7258T>A), GTT2359GTC (7261T>C), GGT2360GGA (7264T>A), GTC2361GTG (7267C>G), GTG2362GTA (7270G>A), GAT2363_CGG2364insT-- (7273_7274insT), CGG2364GTG (7274C>G 7275G>T), CAT2366CTT (7281A>T), TTG2367TCG (7284T>C), TCA2371TTA (7296C>T), TCT2376TTT (7311C>T), ACC2377ATC (7314C>T), TGC2378TAC (7317G>A), AAT2379AAA (7321T>A), GTG2380-TG (7322delG), CAC2381CAT (7327C>T), GGC2382GGT (7330C>T), TTC2383TTT (7333C>T), AAG2384CAG (7334A>C), TGT2386TGC (7342T>C), GCA2387AAC (7343G>A 7344C>A 7345A>C), ACT2390ACC (7354T>C), CAA2393CAG (7363A>G), ATT2395ATC (7369T>C), TTC2396TTT (7372C>T), GCG2398GCT (7378G>T), CTC2399CTT (7381C>T), GCA2404GCT (7396A>T), GTC2405GTT (7399C>T), GCC2407GCT (7405C>T), AGT2408ATG (7407G>T 7408T>G), GGG2410GGT (7414G>T), TGC2411GGC (7415T>G), AAA2412AAG (7420A>G), GAC2415GAT (7429C>T), TAT2416TAC (7432T>C), CAT2419CAC (7441T>C), AAG2425AAA (7459G>A), GAA2426GAG (7462A>G), ATA2428ATT (7468A>T), GTC2429GTA (7471C>A), CAA2431CAG (7477A>G), AGC2432AGT (7480C>T), CTG2434CTT (7486G>T), TAT2437TAC (7495T>C), AAA2438AAG (7498A>G), GGT2439GGC (7501T>C), TTG2440TCA (7503T>C 7504G>A), ATT2443ATA (7513T>A), AAC2445AAT (7519C>T), GGA2446GGG (7522A>G), AAG2449AAA (7531G>A), GAG2451GAA (7537G>A), CTC2452CTT (7540C>T), TGT2454TGC (7546T>C), AAG2462AAA (7570G>A), ACG2463ACA (7573G>A), ACG2465ACA (7579G>A), GCT2468GCC (7588T>C), CTA2471TTA (7595C>T), TTG2474CTA (7604T>C 7606G>A), GGT2477GGA (7615T>A), TGT2480TGC (7624T>C), GAC2483GAT (7633C>T), TTC2484TTT (7636C>T), AAT2485AAC (7639T>C), TTT2488TTC (7648T>C), GAA2494GAG (7666A>G), TGT2496TGC (7672T>C), ACA2498ACT (7678A>T), GGG2500GGA (7684G>A), ACT2502ACC (7690T>C), TTT2504TTC (7696T>C), GGT2506GGG (7702T>G), GAT2509AAT (7709G>A), AAA2510AGG (7713A>G 7714A>G), CTG2511TTG (7715C>T), CGT2514CGC (7726T>C), GAG2517GAA (7735G>A), GTA2520GTG (7744A>G), TAC2521TAT (7747C>T), GGC2526GGT (7762C>T), CAG2528CAA (7768G>A), TTT2529TTC (7771T>C), TCA2532TCG (7780A>G), CTA2533CTC (7783A>C), ACT2534ACC (7786T>C), CTA2537TTG (7793C>T 7795A>G), ATC2538ATT (7798C>T), GCT2540GCC (7804T>C), ACC2543ATC (7812C>T), GAA2550GAG (7834A>G), GAC2551GAT (7837C>T), GTG2554TTA (7844G>T 7846G>A), CAG2557CAA (7855G>A), CTG2559CTA (7861G>A), TTA2562TTG (7870A>G), TAC2563TAT (7873C>T), ACT2564ACA (7876T>A), GAG2565GAA (7879G>A), ATT2566ATA (7882T>A), GTT2567ATT (7883G>A), CCA2570CCT (7894A>T), ATT2571ATC (7897T>C), GTC2579GTT (7921C>T), AAG2581AAA (7927G>A), GGT2584GGC (7936T>C), AAT2585AAC (7939T>C), AAT2586AAC (7942T>C), GTG2594GTA (7966G>A), GAT2595GAC (7969T>C), TCT2597TCC (7975T>C), GTC2601GTT (7987C>T), CTT2602CTC (7990T>C), GCT2607GCC (8005T>C), GAA2616GAG (8032A>G), ATC2618ATT (8038C>T), GAC2619GAT (8041C>T), AGC2620AAC (8043G>A), TTG2632TTA (8080G>A), GTG2635GTA (8089G>A), CCG2637CCA (8095G>A), GAG2638GAA (8098G>A), AGC2641AGT (8107C>T), ATT2642ATC (8110T>C), GAT2644GAC (8116T>C), CAA2648CAG (8128A>G), GGT2654GGC (8146T>C), AAC2656AGT (8151A>G 8152C>T), TCG2660TCA (8164G>A), AGG2665AAG (8178G>A), AAG2666AAA (8182G>A), TTG2669TTA (8191G>A), TTC2671TTT (8197C>T), TCC2673TCT (8203C>T), AGA2675AAA (8208G>A), CTG2677TTA (8213C>T 8215G>A), CTA2678CTG (8218A>G), ATC2679ATT (8221C>T), GAG2680GAA (8224G>A), CCA2685CCG (8239A>G), GAG2690GAA (8254G>A), ATT2692ATC (8260T>C), GTA2693GTG (8263A>G), GAT2699GAC (8281T>C), AGA2700AGG (8284A>G), CCA2704GCG (8294C>G 8296A>G), GAG2705GAA (8299G>A), CAC2706CAT (8302C>T), ATT2711ATC (8317T>C), TGC2712TGT (8320C>T), GCT2714GCA (8326T>A), GAG2717GAA (8335G>A), TCC2718TCT (8338C>T), TCT2722TTT (8349C>T), GAA2723GAG (8353A>G), CAC2726CAT (8362C>T), CAA2727CAG (8365A>G), ATC2728ATT (8368C>T), TAC2732TAT (8380C>T), TCA2733TCG (8383A>G), TTA2735TTG (8389A>G), CCT2740CCC (8404T>C), GCA2742TCA (8408G>T), ACA2743ACG (8413A>G), GCG2745GCT (8419G>T), GCT2750GCC (8434T>C), TAT2752TAC (8440T>C), ATG2756ATA (8452G>A), CTA2758CTG (8458A>G), AGG2759AAG (8460G>A), AAA2760AAG (8464A>G), CTG2761CTA (8467G>A), AGG2765AGA (8479G>A), GCT2766ACA (8480G>A 8482T>A), GAG2771GAA (8497G>A), CTA2772TTG (8498C>T 8500A>G), AGA2773AAG (8502G>A 8503A>G), GCC2774GCT (8506C>T), GAA2777GAT (8515A>T), GTC2780GTT (8524C>T), GCA2781GCC (8527A>C), TTA2782CTG (8528T>C 8530A>G), GAG2785GAC (8539G>C), CTT2788TGT (8546C>T 8547T>G), TCT2790ACT (8552T>A), GTA2793GTG (8563A>G), GCA2797GGA (8574C>G), ATT2801GTT (8585A>G), GGA2805GAA (8598G>A), AGC2806AGT (8602C>T), AAC2807TCT (8603A>T 8604A>C 8605C>T), AAA2812AAG (8620A>G), CCA2813CAA (8622C>A), GAG2814GGA (8625A>G 8626G>A), CAG2815CAA (8629G>A), GGC2816GGT (8632C>T), ATC2818ATT (8638C>T), CAG2819CAA (8641G>A), AAC2821AAT (8647C>T), CCG2822TTC (8648C>T 8649C>T 8650G>C), AAC2823AAT (8653C>T), AAA2824AAG (8656A>G), GGA2825GAA (8658G>A), GAT2827GCG (8664A>C 8665T>G), GTT2830GTG (8674T>G), GCA2832GTT (8679C>T 8680A>T), GGC2833GGA (8683C>A), GGG2836GGA (8692G>A), ACA2837ACT (8695A>T), CCG2841CCA (8707G>A), AGA2842CGA (8708A>C), AAG2844AAA (8716G>A), ATC2846ATT (8722C>T), ACG2847ACA (8725G>A), AAA2849AAG (8731A>G), AGA2851AGG (8737A>G), CCC2853CCT (8743C>T), ACA2854AAA (8745C>A), AAG2856AAA (8752G>A), GGA2857GGT (8755A>T), AAC2862AAT (8770C>T), TTA2863CTA (8771T>C), GAA2864GAG (8776A>G), CAT2865CAC (8779T>C), TTG2866TTA (8782G>A), TAT2869TAC (8791T>C), CCA2871CCG (8797A>G), CAA2872CAG (8800A>G), CAA2873CAG (8803A>G), GAT2875GAC (8809T>C), ATT2876ATC (8812T>C), AAT2878AAC (8818T>C), CGG2880CGA (8824G>A), GCA2881GCG (8827A>G), ACG2888ACA (8848G>A), TAT2890TAC (8854T>C), GAG2891GGA (8856A>G 8857G>A), GTG2893GTA (8863G>A), CGG2894CAG (8865G>A), ATG2895CTT (8867A>C 8869G>T), GCA2896GCG (8872A>G), GGA2900GAA (8883G>A), ACT2902TCT (8888A>T), TGC2915TGT (8929C>T), GTC2924ATC (8954G>A), AAC2925AAT (8959C>T), GGA2926GGT (8962A>T), GGG2933GGA (8983G>A), AAT2934GAC (8984A>G 8986T>C), GAA2935GTA (8988A>T), GAG2938GAA (8998G>A), CCG2940CCA (9004G>A), TTG2941TTA (9007G>A), ATC2944ATT (9016C>T), ACC2951ACA (9037C>A), ATA2967ATT (9085A>T), AAC2971AAT (9097C>T), ATT2983GTT (9131A>G), CGA2984CGT (9136A>T), CGG2987CGT (9145G>T), ATG2989GGA (9149A>G 9150T>G 9151G>A), GGT2990AGT (9152G>A), TTA2991TTG (9157A>G), GCG2992GCT (9160G>T), CGT2993CGC (9163T>C), GAG3000GAA (9184G>A), CGA3004CGG (9196A>G), AGG3010AGA (9214G>A), GAA3011GAG (9217A>G), GCG3012GCA (9220G>A), GCA3020GCT (9244A>T), TTG3021TTA (9247G>A), GCC3024GCT (9256C>T), CCT3026TCC (9260C>T 9262T>C), GGG3030GGA (9274G>A), GAC3032GAT (9280C>T), ATC3035ATT (9289C>T), CAA3038CAC (9298A>C), GAG3040GAA (9304G>A), GTC3050GTT (9334C>T), GTC3060GTG (9364C>G) | | | | | | | | |
| P1 protein (NP_734243.1) | 1 | 284 | 100% | 1394 | 72.7% | 284 (100%) | 203 (71.5%) | 0/0/0/0 | 0 |

| | Begin | End | Coverage | Score | Concordance | Matches | Identities | I/D/M/F* | Stop Codons |
|---|---|---|---|---|---|---|---|---|---|
| **NT** | **54** | **9704** | **99.5%** | **12522** | **65.2%** | **9639 (99.8%)** | **7984 (82.7%)** | **7/12** | |
| Protein mutations: | Y4Q (194T>C 196C>A), C9L (209T>C 210G>T 211T>G), F13M (221T>A 223T>G), S23P (251T>C), C24F (255G>T 256C>T), E25G (258A>G), I27V (263A>G 265T>G), V28A (267T>C), E30V (273A>T), A35T (287G>A 289T>C), V37T (293G>A 294T>C), A41I (305G>A 306C>T), D42G (309A>G), E44D (316A>T), T45V (317A>G 318C>T 319A>G), L52Q (339T>A 340C>A), K53R (342A>G), K55A (347A>G 348A>C 349A>G), Y56H (350T>C), T58V (356A>G 357C>T), V61T (365G>A 366T>C 367G>A), L62S (368C>T 369T>C), F67C (384T>G), A75V (408C>T 409C>T), M78E (416A>G 417T>A), E85K (437G>A), R86E (440A>G 441G>A), K87R (443A>C 444A>G 445G>A), D88E (448T>A), E91A (456A>C), H93N (461C>A), D111E (517T>A), S116F (531C>T), P118S (536C>T), Q119W (539C>T 540A>G 541A>G), R122K (549G>A 550A>G), M131T (576T>C), V134A (585T>C 586C>G), R137Y (593C>T 594G>A 595C>T), P138R (597C>G), I139V (599A>G 601A>G), I140P (602A>C 603T>C 604A>G), M147I (625G>A), I151V (635A>G 637T>C), I154V (644A>G 646A>G), E160A (663A>C 664G>C), H166Q (682C>A), T172S (699C>G), H174Q (706T>A), K179E (719A>G 721G>A), I180V (722A>G 724A>G), A183S (731G>T), Y184P (734T>C 735A>C 736C>T), S185R (737T>C 738C>G 739C>T), A187V (744C>T 745G>C), R189C (749C>T 751A>C), M194K (765T>A), R197K (773C>A 774G>A 775A>G), M206K (801T>A), G210A (813G>C 814A>G), L211H (816T>A), R214Q (825G>A 826T>A), N223D (851A>G 853T>C), R226C (860C>T 862C>T), T227A (863A>G 865T>C), I228T (867T>C), N229D (869A>G 871C>T), I230L (872A>T 874A>G), R231Q (876G>A), R232K (879G>A), N240S (903A>G), T241N (906C>A 907A>C), K242T (909A>C 910A>C), S243N (912G>A 913C>T), S252L (938T>C 939C>T 940A>T), L266I (980T>A 982G>C), R272K (999G>A), Q275R (1007C>A 1008A>G 1009G>A), S276G (1010A>G 1012T>G), I277V (1013A>G), N279Q (1019A>C 1021C>G) | | | | | | | | |
| Codon mutations: | ACT3ACC (193T>C), TAC4CAA (194T>C 196C>A), TCA6TCG (202A>G), ATC8ATT (208C>T), TGT9CTG (209T>C 210G>T 211T>G), TCG12TCA (220G>A), TTT13ATG (221T>A 223T>G), GAA14GAG (226A>G), CTA17CTT (235A>T), TAC19TAT (241C>T), CCA21CCT (247A>T), TCT23CCT (251T>C), TGC24TTT (255G>T 256C>T), GAG25GGG (258A>G), CAT26CAC (262T>C), ATT27GTG (263A>G 265T>G), GTG28GCG (267T>C), GAA30GTA (273A>T), CGA31CGT (277A>T), GAA32GAG (280A>G), GTG33GTT (283G>T), CCG34CCA (286G>A), GCT35ACC (287G>A 289T>C), GTT37ACT (293G>A 294T>C), CCT39CCC (301T>C), TTC40TTT (304C>T), GCA41ATA (305G>A 306C>T), GAT42GGT (309A>G), CTG43TTG (311C>T), GAA44GAT (316A>T), ACA45GTG (317A>G 318C>T 319A>G), CAA46CAG (322A>G), GCA49GCT (331A>T), CTC52CAA (339T>A 340C>A), AAG53AGG (342A>G), AAA55GCG (347A>G 348A>C 349A>G), TAT56CAT (350T>C), GCT57GCA (355T>A), ACT58GTT (356A>G 357C>T), CGT60CGC (364T>C), GTG61ACA (365G>A 366T>C 367G>A), CTC62TCC (368C>T 369T>C), AAA63AAG (373A>G), AAC64AAT (376C>T), TTT67TGT (384T>G), ACG68ACA (388G>A), TAC69TAT (391C>T), CGA70CGG (394A>G), AAG72AAA (400G>A), GAT74GAC (406T>C), GCC75GTT (408C>T 409C>T), CAG76CAA (412G>A), ATA77ATT (415A>T), ATG78GAG (416A>G 417T>A), ATT80ATC (424T>C), CAG81CAA (427G>A), AAA83AAG (433A>G), CTG84CTC (436G>C), GAG85AAG (437G>A), AGG86GAG (440A>G 441G>A), AAG87CGA (443A>C 444A>G 445G>A), GAT88GAA (448T>A), GAA90GAG (454A>G), GAA91GCA (456A>C), TAT92TAC (460T>C), CAC93AAC (461C>A), CAA95CAG (469A>G), GCC97GCG (475C>G), CCT99CCA (481T>A), ACT106ACC (502T>C), ATA107ATT (505A>T), GGC109GGT (511C>T), GAT111GAA (517T>A), CCT112CCA (520T>A), CCA113CCT (523A>T), TCT116TTT (531C>T), GAG117GAA (535G>A), CCA118TCA (536C>T), CAA119TGG (539C>T 540A>G 541A>G), AGA122AAG (549G>A 550A>G), ATC124ATT (556C>T), ATT125ATC (559T>C), CAT126CAC (562T>C), ACT128ACC (568T>C), CCA129CCG (571A>G), ATG131ACG (576T>C), CGT132CGC (580T>C), GTC134GCG (585T>C 586C>G), CGC137TAT (593C>T 594G>A 595C>T), CCC138CGC (597C>G), ATA139GTG (599A>G 601A>G), ATA140CCG (602A>C 603T>C 604A>G), ACA143ACG (613A>G), GAA144GAG (616A>G), GGC145GGG (619C>G), CAG146CAA (622G>A), ATG147ATA (625G>A), AAT148AAC (628T>C), CAC149CAT (631C>T), ATT151GTC (635A>G 637T>C), ATA154GTG (644A>G 646A>G), ATT157ATC (655T>C), TCG159TCA (661G>A), GAG160GCC (663A>C 664G>C), AAA161AAG (667A>G), GTC165GTT (679C>T), CAC166CAA (682C>A), TTA167CTG (683T>C 685A>G), AGT169AGC (691T>C), ACC172AGC (699C>G), ACT173ACA (703T>A), CAT174CAA (706T>A), AAG179GAA (719A>G 721G>A), ATA180GTG (722A>G 724A>G), CTT181TTG (725C>T 727T>G), GGT182GGA (730T>A), GCA183TCA (731G>T), TAC184CCT (734T>C 735A>C 736C>T), TCC185CGT (737T>C 738C>G 739C>T), GCG187GTC (744C>T 745G>C), CGA189TGC (749C>T 751A>C), ACT190ACC (754T>C), ATG194AAG (765T>A), TTG196CTG (770T>C), CGA197AAG (773C>A 774G>A 775A>G), CGG198AGG (776C>A), TTC202TTT (790C>T), ATG206AAG (801T>A), GGA210GCG (813G>C 814A>G), CTT211CAT (816T>A), TTG212CTG (818T>C), CAA213CAG (823A>G), CGT214CAA (825G>A 826T>A), CGG217AGA (833C>A 835G>A), GAC219GAT (841C>T), AAT223GAC (851A>G 853T>C), GTC225GTT (859C>T), CGC226TGT (860C>T 862C>T), ACT227GCC (863A>G 865T>C), ATC228ACC (867T>C), AAC229GAT (869A>G 871C>T), ATA230TTG (872A>T 874A>G), CGA231CAA (876G>A), AGG232AAG (879G>A), GAT234GAC (886T>C), ATC238ATT (898C>T), AAC240AGC (903A>G), ACA241AAC (906C>A 907A>C), AAA242ACC (909A>C 910A>C), AGC243AAT (912G>A 913C>T), GGC246GGA (922C>A), GGT249GGA (931T>A), TCA252CTT (938T>C 939C>T 940A>T), GGA253GGG (943A>G), TTC256TTT (952C>T), TCA261TCG (967A>G), GAA263GAG (973A>G), GGG264GGC (976G>C), TTG266ATC (980T>A 982G>C), GCA269GCG (991A>G), CGT270CGC (994T>C), AGA272AAA (999G>A), GTT273GTC (1003T>C), CAG275AGA (1007C>A 1008A>G 1009G>A), AGT276GGG (1010A>G 1012T>G), ATT277GTT (1013A>G), TTA278TTG (1018A>G), AAC279CAG (1019A>C 1021C>G), ATC282ATT (1030C>T) | | | | | | | | |
| HC-Pro protein (NP_734244.1) | 1 | 456 | 100% | 3145 | 95.7% | 456 (100%) | 425 (93.2%) | 0/0/0/0 | 0 |
| Protein mutations: | D4E (1048C>A), R16Q (1083G>A), S21A (1097T>G 1099G>A), R36K (1143G>A), L40I (1154T>A 1156G>A), S62N (1221G>A), V65T (1229G>A 1230T>C 1231T>C), S66N (1233G>A 1234C>T), K74T (1257A>C 1258A>G), N81S (1278A>G), A85V (1290C>T 1291G>A), D88T (1298G>A 1299A>C 1300C>T), I91V (1307A>G 1309A>G), N94E (1316A>G 1318T>A), I98T (1329T>C), L113I (1373C>A), I117V (1385A>G), E123D (1405G>T), A127S (1415G>T 1417A>T), V134I (1436G>A 1438C>T), E224R (1706G>A 1707A>G), A228T (1718G>A), I236V (1742A>G 1744C>T), S241N (1758G>A), V252I (1790G>A), I301V (1937A>G 1939T>C), F323Y (2004T>A), I344M (2068T>G), V346I (2072G>A 2074G>A), R391K (2208G>A), D410E (2266C>G) | | | | | | | | |
| Codon mutations: | TCG1TCA (1039G>A), AAT2AAC (1042T>C), GCC3GCA (1045C>A), GAC4GAA (1048C>A), AAT5AAC (1051T>C), AAG8AAA (1060G>A), GGT9GGA (1063T>A), CTG10TTG (1064C>T), GAC11GAT (1069C>T), CGA16CAA (1083G>A), TAT19TAC (1093T>C), TCG21GCA (1097T>G 1099G>A), CAC23CAT (1105C>T), GTA26GTG (1114A>G), GCT27GCG (1117T>G), TTA29TTG (1123A>G), CCT30CCA (1126T>A), GTC31GTT (1129C>T), AGG36AAG (1143G>A), GTA37GTT (1147A>T), GCT38GCA (1150T>A), TTG40ATA (1154T>A 1156G>A), CAC43CAT (1165C>T), AGT44AGC (1168T>C), CTT46TTA (1172C>T 1174T>A), ACC55ACT (1201C>T), AGC62AAC (1221G>A), GTT65ACC (1229G>A 1230T>C 1231T>C), AGC66AAT (1233G>A 1234C>T), CTG68CTA (1240G>A), AAG70AAA (1246G>A), TTG72CTA (1250T>C 1252G>A), CAT73CAC (1255T>C), AAA74ACG (1257A>C 1258A>G), AGA77AGG (1267A>G), GGT79GGC (1273T>C), TTG80TTA (1276G>A), AAT81AGT (1278A>G), TTG83CTG (1283T>C), GCG85GTA (1290C>T 1291G>A), GAC88ACT (1298G>A 1299A>C 1300C>T), TTT90TTC (1306T>C), ATA91GTG (1307A>G 1309A>G), CAT92CAC (1312T>C), AAT94GAA (1316A>G 1318T>A), TTG97TTA (1327G>A), ATA98ACA (1329T>C), GCG99GCT (1333G>T), TTA100TTG (1336A>G), CTA103TTG (1343C>T 1345A>G), CCG106CCA (1354G>A), GTG107GTT (1357G>T), GAC108GAT (1360C>T), CTC111TTG (1367C>T 1369C>G), CTT113ATT (1373C>A), TTC114TTT (1378C>T), GAG116GAA (1384G>A), ATA117GTA (1385A>G), AAA119AAG (1393A>G), TCC120TCT (1396C>T), GAG123GAT (1405G>T), AAA124AAG (1408A>G), CAG125CAA (1411G>A), GCA127TCT (1415G>T 1417A>T), CCG128CCC (1420G>C), AAT131AAC (1429T>C), TTA132TTG (1432A>G), GTC134ATT (1436G>A 1438C>T), TTA135CTG (1439T>C 1441A>G), AAT136AAC (1444T>C), TTC139TTT (1453C>T), CTG140CTA (1456G>A), GAA144GAG (1468A>G), GTA152GTG (1492A>G), TTG155CTA (1499T>C 1501G>A), AGT156AGC (1504T>C), TTG157TTA (1507G>A), CTC158CTT (1510C>T), GAA159GAG (1513A>G), TTA160TTG (1516A>G), GCA161GCT (1519A>T), AGG162AGA (1522G>A), TTC163TTT (1525C>T), CAG164CAA (1528G>A), AAC166AAT (1534C>T), ACT168ACA (1540T>A), AAC170AAT (1546C>T), AAA173AAG (1555A>G), GGT174GGC (1558T>C), GAT175GAC (1561T>C), ATA176ATC (1564A>C), TCT177TCA (1567T>A), TTC179TTT (1573C>T), AGA180AGG (1576A>G), TTA183CTA (1583T>C), TCT184TCA (1588T>A), AAG186AAA (1594G>A), AAC188AAT (1600C>T), CTG191CTA (1609G>A), TCG194TCA (1618G>A), TGC195TGT (1621C>T), GAC196GAT (1624C>T), CTG199TTG (1631C>T), AAA201AAG (1639A>G), TTC205TTT (1651C>T), CTC206CTG (1654C>G), CAT213CAC (1675T>C), TCA219TCG (1693A>G), TTT222TTC (1702T>C), GAA224AGA (1706G>A 1707A>G), GCA228ACA (1718G>A), GGA230GGG (1726A>G), TAC231TAT (1729C>T), TCA232TCC (1732A>C), TAT234TAC (1738T>C), ATC236GTT (1742A>G 1744C>T), CGC237CGT (1747C>T), CAT239CAC (1753T>C), AGT241AAT (1758G>A), AGG244AGA (1768G>A), AAC250AAT (1786C>T), TTA251TTG (1789A>G), GTT252ATT (1790G>A), GTC253GTT (1795C>T), TTA257CTG (1805T>C 1807A>G), GAG259GAA (1813G>A), AGG261AGA (1819G>A), AAG263AAA (1825G>A), GAC267GAT (1837C>T), TAT268TAC (1840T>C), CAA271CAG (1849A>G), GTC274GTG (1858C>G), AAA276AAG (1864A>G), TGC278TGT (1870C>T), AGT280AGC (1876T>C), AAA282AAG (1882A>G), GGT284GGA (1888T>A), AAT285AAC (1891T>C), TGT290TGC (1906T>C), ACA293ACT (1915A>T), GGT298GGC (1930T>C), GCC300GCT (1936C>T), ATT301GTC (1937A>G 1939T>C), TCA303TCG (1945A>G), TTC305TTT (1951C>T), CCA308CCT (1960A>T), GTA314GTG (1978A>G), ATT315ATA (1981T>A), GGC316GGT (1984C>T), AAT317AAC (1987T>C), GAC320GAT (1996C>T), TTT323TAT (2004T>A), AAA328AAG (2020A>G), GGG329GGA (2023G>A), GAT330GAC (2026T>C), TCG331TCT (2029G>T), GAG332GAA (2032G>A), TAT341TAC (2059T>C), ATT344ATG (2068T>G), AAC345AAT (2071C>T), GTG346ATA (2072G>A 2074G>A), CTG351TTA (2087C>T 2089G>A), AGC355AGT (2101C>T), GAG356GAA (2104G>A), GAG357GAA (2107G>A), AAG360AAA (2116G>A), TTC362TTT (2122C>T), CGC367CGT (2137C>T), CTT374CTC (2158T>C), ACC376ACT (2164C>T), CCA378CCG (2170A>G), GAT382GAC (2182T>C), GCG384GCA (2188G>A), ACC385ACA (2191C>A), AGA391AAA (2208G>A), TAT394TAC (2218T>C), GAC396GAT (2224C>T), GTA397GTT (2227A>T), TTG402TTA (2242G>A), CCC403CCT (2245C>T), TTG406TTA (2254G>A), GTT407GTC (2257T>C), GAC408GAT (2260C>T), GAC410GAG (2266C>G), CAA412CAG (2272A>G), ACG413ACA (2275G>A), TCA419TCG (2293A>G), GGC421GGT (2299C>T), TCG422TCA (2302G>A), CAG423CAA (2305G>A), GGA426GGT (2314A>T), TAT427TAC (2317T>C), CTA430CTG (2326A>G), GTG435GTA (2341G>A), CAA437CAG (2347A>G), CTT438CTC (2350T>C), TTG440TTA (2356G>A), AAT443AAC (2365T>C), GAA445GAG (2371A>G), TTA446TTG (2374A>G), GAT449GAC (2383T>C), ATA450ATT (2386A>T), AAA451AAG (2389A>G), CAT452CAC (2392T>C), TAT453TAC (2395T>C) | | | | | | | | |
| P3 protein (NP_734251.1) | 1 | 365 | 100% | 1924 | 79.6% | 365 (99.7%) | 327 (89.3%) | 1/0/2/2 | 0 |
| Protein mutations: | N4G (2414A>G 2415A>G 2416T>G), S6C (2420A>T), K40R (2523A>G), R77K (2634G>A), V107I (2723G>A 2725C>T), A115T (2747G>A 2749T>A), H131N (2795C>A), V140I (2822G>A 2824G>A), P160T (2882C>A), S161H (2885A>C 2886G>A), T164M (2895C>T), N176S (2931A>G), N179S (2940A>G), E189_N190insX (2971_2972insA), T196I (2991C>T), H197R (2994A>G), E199R (2999G>A 3000A>G), R206Q (3021G>A), Y207F (3024A>T 3025C>T), T211I (3036C>T), E212G (3039A>G 3040A>C), K213T (3042A>C 3043G>A), A232V (3099C>T 3100C>A), V234R (3104G>A 3105T>G 3106G>A), S243N (3132G>A), R245K (3137C>A 3138G>A 3139A>G), F250I (3152T>A 3154C>T), C254Y (3165G>A), F272L (3218T>C), T274N (3225C>A), V276I (3230G>A), V288M (3266G>A 3268A>G), V291M (3275G>A), A294T (3284G>A), R304K (3315G>A 3316A>G), E305A (3318A>C), M319V (3359A>G), D332E (3400T>G) | | | | | | | | |

| | Begin | End | Coverage | Score | Concordance | Matches | Identities | I/D/M/F* | Stop Codons |
|---|---|---|---|---|---|---|---|---|---|
| **NT** | **54** | **9704** | **99.5%** | **12522** | **65.2%** | **9639 (99.8%)** | **7984 (82.7%)** | **7/12** | |
| Codon mutations: | AAT4GGG (2414A>G 2415A>G 2416T>G), GCT5GCA (2419T>A), AGC6TGC (2420A>T), ACA12ACT (2440A>T), GTT21GTC (2467T>C), TCT24TCC (2476T>C), TCG26TCA (2482G>A), CTG29CTA (2491G>A), CTG31CTT (2497G>T), TTG33TTA (2503G>A), AGA38AGG (2518A>G), CCT39CCC (2521T>C), AAG40AGG (2523A>G), AGA43AGG (2533A>G), TTG45TTA (2539G>A), TTA47TTG (2545A>G), GAG49GAA (2551G>A), CCT50CCA (2554T>A), CTG52TTG (2558C>T), TTG53CTC (2561T>C 2563G>C), TCA56TCG (2572A>G), CCT60CCC (2584T>C), GGC61GGT (2587C>T), CTG63CTC (2593G>C), GCT65GCC (2599T>C), AAT68AAC (2608T>C), TTT72TTC (2620T>C), GCG75GCA (2629G>A), GTG76GTA (2632G>A), AGG77AAG (2634G>A), TTG78TTA (2638G>A), ATT80ATA (2644T>A), AAA83AAG (2653A>G), GCT87GCC (2665T>C), CTA92TTA (2678C>T), CTA93CTG (2683A>G), GCT95GCC (2689T>C), CTA98CTC (2698A>C), TCA101TCG (2707A>G), GCG102GCA (2710G>A), GAA104GAG (2716A>G), ACA105ACG (2719A>G), CTC106CTT (2722C>T), GTC107ATT (2723G>A 2725C>T), GCA108GCG (2728A>G), AGG110AGA (2734G>A), ATT111ATA (2737T>A), GAT114GAC (2746T>C), GCT115ACA (2747G>A 2749T>A), GCA116GCT (2752A>T), GCT117GCA (2755T>A), GAC119GAT (2761C>T), CTC120CTT (2764C>T), TGT125TGC (2779T>C), GGG127GGT (2785G>T), AAC129AAT (2791C>T), CAT131AAT (2795C>A), TAC134TAT (2806C>T), TTG138CTC (2816T>C 2818G>C), GTG140ATA (2822G>A 2824G>A), TTG141TTA (2827G>A), GTT144GTC (2836T>C), AAG145AAA (2839G>A), AAT146AAC (2842T>C), ACC153ACA (2863C>A), CTA154TTA (2864C>T), CCA160ACA (2882C>A), AGT161CAT (2885A>C 2886G>A), AAC163AAT (2893C>T), ACG164ATG (2895C>T), GTC166GTT (2902C>T), GTA167GTG (2905A>G), ATT169ATC (2911T>C), AAT173AAC (2923T>C), AAT176AGT (2931A>G), AAC179AGC (2940A>G), GCT181GCC (2947T>C), ACT186ACC (2962T>C), GAA189_AAT190insA-- (2971_2972insA), GGT195GGC (2989T>C), ACT196ATT (2991C>T), CAT197CGT (2994A>G), GAG199AGG (2999G>A 3000A>G), TCC204TC- (3016delC), CGG206CAG (3021G>A), TAC207TTT (3024A>T 3025C>T), ACA211ATA (3036C>T), GAA212GGC (3039A>G 3040A>C), AAG213ACA (3042A>C 3043G>A), TTG216TTA (3052G>A), TTA219TTG (3061A>G), GCG226GCA (3082G>A), GGC229GGT (3091C>T), GCC232GTA (3099C>T 3100C>A), GTG234AGA (3104G>A 3105T>G 3106G>A), GTC235GTT (3109C>T), AAA236AAG (3112A>G), ACT238ACC (3118T>C), GCC239GCT (3121C>T), GGA241GGG (3127A>G), AGC243AAC (3132G>A), CGA245AAG (3137C>A 3138G>A 3139A>G), TAT249TAC (3151T>C), TTC250ATT (3152T>A 3154C>T), TGT254TAT (3165G>A), GTA255GTG (3169A>G), ATC265ATT (3199C>T), AGG267CGG (3203A>C), CGT268CGC (3208T>C), CCA270CCG (3214A>G), ACC271ACT (3217C>T), TTT272CTT (3218T>C), GTC273GTA (3223C>A), ACT274AAT (3225C>A), GTT276ATT (3230G>A), AAC277AAT (3235C>T), TTA285CTA (3257T>C), ACC286ACT (3262C>T), AGC287AGT (3265C>T), GTA288ATG (3266G>A 3268A>G), GTG289GTA (3271G>A), GTG291ATG (3275G>A), GCA294ACA (3284G>A), CAG299CAA (3301G>A), AGA304AAG (3315G>A 3316A>G), GAA305GCA (3318A>C), ATC306ATT (3322C>T), GAG307GAA (3325G>A), TTG308TTA (3328G>A), ATA311ATT (3337A>T), GAG315GAA (3349G>A), GTC317GTT (3355C>T), TGC318TGT (3358C>T), ATG319GTG (3359A>G), CTA321CTG (3367A>G), TAT322TAC (3370T>C), TTA325CTG (3377T>C 3379A>G), GAA330GAG (3394A>G), CGC331CGT (3397C>T), GAT332GAG (3400T>G), ACA334ACG (3406A>G), TAC338TAT (3418C>T), ATT339ATA (3421T>A), TAT341TAC (3427T>C), AAG343AAA (3433G>A), TCA344TCT (3436A>T), GTA345GTG (3439A>G), AAC346AAT (3442C>T), CCT347CCC (3445T>C), CAG351CAA (3457G>A), TTT352TTC (3460T>C), GCT353GCA (3463T>A), GCG355GCT (3469G>T), CAG356CAA (3472G>A), CAC364CAT (3496C>T), CAG365CAA (3499G>A) | | | | | | | | |
| 6K1 protein (NP_734245.1) | 1 | 52 | 100% | 305 | 90.5% | 52 (100%) | 45 (86.5%) | 0/0/0/0 | 0 |
| Protein mutations: | V5G (3513T>G), M16V (3545A>G), A17T (3548G>A), V19I (3554G>A 3556C>T), L42V (3623C>G), L45M (3632C>A), Y47H (3638T>C) | | | | | | | | |
| Codon mutations: | TCC2TCT (3505C>T), GTT5GGT (3513T>G), AAA7AAG (3520A>G), GAA10GAG (3529A>G), CAA11CAG (3532A>G), ATG16GTG (3545A>G), GCT17ACT (3548G>A), TTA18CTA (3551T>C), GTC19ATT (3554G>A 3556C>T), AGT28AGC (3583T>C), GAT29GAC (3586T>C), TGC30TGT (3589C>T), TTC32TTT (3595C>T), AAT36AAC (3607T>C), TTT38TTC (3613T>C), AAG39AAA (3616G>A), GGT40GGC (3619T>C), CTT42GTT (3623C>G), TCC43TCT (3628C>T), CTG45ATG (3632C>A), GAT46GAC (3637T>C), TAT47CAT (3638T>C), GAA48GAG (3643A>G) | | | | | | | | |
| CI protein (NP_734246.1) | 1 | 634 | 100% | 4190 | 97.9% | 634 (100%) | 608 (95.9%) | 0/0/0/0 | 0 |
| Protein mutations: | I15V (3698A>G), S21N (3717G>A), R26K (3731C>A 3732G>A), I140V (4073A>G), F158Y (4128T>A 4129C>T), A166S (4151G>T), V171I (4166G>A), L190I (4223C>A 4225G>A), I222V (4319A>G 4321A>G), F247Y (4395T>A), V251I (4406G>A), M274V (4475A>G 4477G>T), T276S (4481A>T 4483A>G), A297L (4544G>T 4545C>T 4546G>A), R298K (4548G>A), V342I (4679G>A 4681G>C), V381I (4796G>A 4798C>T), T383S (4803C>G 4804T>C), S468G (5057A>G), T469A (5060A>G), L479I (5090C>A), A482V (5100C>T), E484D (5107A>T), V489F (5120G>T 5122C>T), I491V (5126A>G), V510I (5183G>A) | | | | | | | | |
| Codon mutations: | TTA2TTG (3661A>G), GTG5GTA (3670G>A), TTT9TTC (3682T>C), GAG11GAA (3688G>A), GAG14GAA (3697G>A), ATT15GTT (3698A>G), GAT17GAC (3706T>C), TTT18TTC (3709T>C), TTG20CTA (3713T>C 3715G>A), AGT21AAT (3717G>A), GAG22GAA (3721G>A), CGA26AAA (3731C>A 3732G>A), ACT27ACA (3736T>A), CTA31TTG (3746C>T 3748A>G), ACA33ACG (3754A>G), AGT36AGC (3763T>C), CGA41CGG (3778A>G), CAA42CAG (3781A>G), ATC43ATT (3784C>T), CAT47CAC (3796T>C), CCA50CCT (3805A>T), TAC52TAT (3811C>T), AGA53AGG (3814A>G), TTC58TTT (3829C>T), TTT61TTC (3838T>C), AGA63AGG (3844A>G), GCA64GCC (3847A>C), GTT67GTA (3856T>A), GTG69GTT (3862G>T), GAC72GAT (3871C>T), GCC74GCA (3877C>A), AGC76AGT (3883C>T), GAA77GAG (3886A>G), CTA79CTG (3892A>G), GAC80GAT (3895C>T), TTT81TTC (3898T>C), GTA83GTG (3904A>G), CGG84AGA (3905C>A 3907G>A), GCT86GCC (3913T>C), AAG91AAA (3928G>A), TCA92TCT (3931A>T), GGG94GGA (3937G>A), AGT100AGC (3955T>C), GTG101GTA (3958G>A), GCC102GCT (3961C>T), GGA103GGG (3964A>G), TCT104TCC (3967T>C), TTA107TTG (3976A>G), ATT108ATC (3979T>C), CCA110CCT (3985A>T), CTA114CTT (3997A>T), GCG115GCA (4000G>A), AAC117AAT (4006C>T), GTT118GTG (4009T>G), TTC119TTT (4012C>T), AAA120AAG (4015A>G), TCT123TCC (4024T>C), GAA125GAG (4030A>G), CCA126CCT (4033A>T), AAG129AAA (4042G>A), CTG133TTA (4052C>T 4054G>A), CGT134CGC (4057T>C), CGT136CGA (4063T>A), GGA137GGG (4066A>G), ATA140GTA (4073A>G), TTT141TTC (4078T>C), GGC142GGT (4081C>T), TCT143TCC (4084T>C), TCC147TCT (4096C>T), AGC151AGT (4108C>T), GGA152GGT (4111A>T), GCG154GCA (4117G>A), CTA155TTG (4118C>T 4120A>G), TAC157TAT (4126C>T), TTC158TAT (4128T>A 4129C>T), GCC159GCT (4132C>T), AAT160AAC (4135T>C), CAA164CAG (4147A>G), TTA165CTA (4148T>C), GCT166TCT (4151G>T), AAC169AAT (4162C>T), GTA171ATA (4166G>A), ATA172ATC (4171A>C), GAG175GAA (4180G>A), CTG179TTG (4190C>T), CCT181CCC (4198T>C), TCC182TCT (4201C>T), GCG185GCT (4210G>T), TTC186TTT (4213C>T), AGT188AGC (4219T>C), CTG189TTG (4220C>T), CTG190ATA (4223C>A 4225G>A), GTT192GTG (4231T>G), CAT194CAC (4237T>C), GTA199GTG (4252A>G), TTA200CTG (4253T>C 4255A>G), AAA201AAG (4258A>G), ACT205ACC (4270T>C), AGA209AGG (4282A>G), GAG210GAA (4285G>A), GAA212GAG (4291A>G), CAG216CAA (4303G>A), GTC219GTT (4312C>T), ATA222GTG (4319A>G 4321A>G), GTG223GTT (4324G>T), GAG224GAA (4327G>A), CTG227CTT (4336G>T), TCT228TCC (4339T>C), TCA231TCT (4348A>T), GCA235GCG (4360A>G), GGT237GGC (4366T>C), TCT238TCA (4369T>A), ACT240ACC (4375T>C), GTT244GTC (4387T>C), TTT247TAT (4395T>A), GTA251ATA (4406G>A), CTT252CTC (4411T>C), GTT261GTC (4438T>C), ACC263ACA (4444C>A), GCC265GCT (4450C>T), CTC267CTT (4456C>T), GAC270GAT (4465C>T), ATG274GTT (4475A>G 4477G>T), GTC275GTT (4480C>T), ACA276TCG (4481A>T 4483A>G), GGC280GGT (4495C>T), AAG284AAA (4507G>A), CAC285CAT (4510C>T), GGT286GGG (4513T>G), CTA288TTA (4517C>T), GTC291GTT (4528C>T), AAA293AAG (4534A>G), ACC295ACT (4540C>T), GCG297TTA (4544G>T 4545C>T 4546G>A), AGA298AAA (4548G>A), TTT301TTC (4558T>C), GTA303GTT (4564A>T), ACC305ACT (4570C>T), ATA307ATT (4576A>T), TTG314TTA (4597G>A), GAC315GAT (4600C>T), GAC317GAT (4606C>T), GTG318GTA (4609G>A), GTG320GTA (4615G>A), GAC321GAT (4618C>T), TTG324CTT (4625T>C 4627G>T), GAC331GAT (4648C>T), ATT332ATC (4651T>C), AGG335AGA (4660G>A), AGC336AGT (4663C>T), GCT338GCA (4669T>A), GTG342ATC (4679G>A 4681G>C), AGT343AGC (4684T>C), GTT344GTG (4687T>G), GAG348GAA (4699G>A), AGA349AGG (4702A>G), CAA351CAG (4708A>G), AGG352AGA (4711G>A), CTG353TTG (4712C>T), AAA361AAG (4738A>G), GTA363GTT (4744A>T), GCA364GCG (4747A>G), TTG365CTG (4748T>C), ACT370ACC (4765T>C), GTC381ATT (4796G>A 4798C>T), ACT383AGC (4803C>G 4804T>C), GAG384GAA (4807G>A), GCG385GCT (4810G>T), GCT386GCG (4813T>G), CTT387CTC (4816T>C), TAT392TAC (4831T>C), AAC393AAT (4834C>T), CCA395CCT (4840A>T), GGA399GGG (4852A>G), GGC400GGT (4855C>T), AGT404AGC (4867T>C), CTG405CTC (4870G>C), ATT406ATA (4873T>A), TGT409TGC (4882T>C), ACA416ACT (4903A>T), CAG418CAA (4909G>A), TTT420TTC (4915T>C), GAA421GAG (4918A>G), TTG422CTA (4919T>C 4921G>A), AGT423AGC (4924T>C), CCC424CCG (4927C>G), TTT426TTC (4933T>C), ATC427ATA (4936C>A), CAG428CAA (4939G>A), AAT429AAC (4942T>C), TTC430TTT (4945C>T), TCA436TCG (4963A>G), CCT439CCC (4972T>C), CTT445CTC (4990T>C), AAA449AAG (5002A>G), CCT456CCC (5023T>C), TTG457CTG (5024T>C), TGC458TGT (5029C>T), GAT459GAC (5032T>C), TCT461TCC (5038T>C), CCA463CCT (5044A>T), GCA466GCT (5053A>T), TCG467TCA (5056G>A), AGT468GGT (5057A>G), ACT469GCT (5060A>G), TCG472TCT (5071G>T), CTT479ATT (5090C>A), GGA480GGG (5095A>G), GTG481GTT (5098G>T), GCC482GTC (5100C>T), TTA483TTG (5104A>G), GAA484GAT (5107A>T), ATT485ATC (5110T>C), CCA486CCG (5113A>G), AAG487AAA (5116G>A), CAA488CAG (5119A>G), GTC489TTT (5120G>T 5122C>T), ATT491GTT (5126A>G), CAT494CAC (5137T>C), ATC495ATT (5140C>T), AAA496AAG (5143A>G), GAG497GAA (5146G>A), CTC502CTA (5161C>A), GAA504GAG (5167A>G), CTT506CTG (5173T>G), ACG509ACA (5182G>A), GTT510ATT (5183G>A), GTC511GTT (5188C>T), AAG512AAA (5191G>A), AAA514AAG (5197A>G), GAC515GAT (5200C>T), TGC517TGT (5206C>T), TTA518TTG (5209A>G), AGC521AGT (5218C>T), TCG525TCA (5230G>A), ATC527ATT (5236C>T), AGC528AGT (5239C>T), ATC530ATT (5245C>T), ACA536ACT (5263A>T), CTC538CTT (5269C>T), TTC539TTT (5272C>T), GCC540GCA (5275C>A), ATC541ATT (5278C>T), CCA542CCC (5281A>C), AGA543AGG (5284A>G), ACT544ACC (5287T>C), CTA545TTA (5288C>T), ATA546ATT (5293A>T), TTG547CTA (5294T>C 5296G>A), GAG549GAA (5302G>A), CTT552CTA (5311T>A), GAA553GAG (5314A>G), CGA556CGT (5323A>T), AGT564AGC (5347T>C), ATC566ATT (5353C>T), TGC570TGT (5365C>T), GTT577GTC (5386T>C), ACC580ACA (5395C>A), AAA589AAG (5422A>G), GAT590GAC (5425T>C), ACC592ACT (5431C>T), GAG594GAA (5437G>A), CTT599CTC (5452T>C), AAG601AAA (5458G>A), AGT604AGC (5467T>C), CAA605CAG (5470A>G), CTA606TTG (5471C>T 5473A>G), GAA608GAG (5479A>G), TTC609TTT (5482C>T), TCA610TCC (5485A>C), AAT611AAC (5488T>C), TTG612CTA (5489T>C 5491G>A), GGT614GGC (5497T>C), TGT617TGC (5506T>C), GAG618GAA (5509G>A), GAG619GAA (5512G>A), TTA621CTA (5516T>C), TAT625TAC (5530T>C), GAG626GAA (5533G>A), TCG627TCT (5536G>T), TTG628TTA (5539G>A), GTT631GTG (5548T>G), CAC633CAT (5554C>T) | | | | | | | | |
| 6K2 protein (NP_734247.1) | 1 | 52 | 100% | 303 | 93.5% | 52 (100%) | 46 (88.5%) | 0/0/0/0 | 0 |
| Protein mutations: | A2T (5561G>A 5563G>A), I14V (5597A>G), N16K (5605C>A), A21V (5619C>T 5620T>G), I25L (5630A>C), V46I (5693G>A) | | | | | | | | |
| Codon mutations: | GCT1GCA (5560T>A), GCG2ACA (5561G>A 5563G>A), ACG3ACC (5566G>C), TCA4TCG (5569A>G), CTC9CTG (5584C>G), AAG12AAA (5593G>A), GGG13GGA (5596G>A), ATT14GTT (5597A>G), AAC16AAA (5605C>A), GTG20GTT (5617G>T), GCT21GTG (5619C>T 5620T>G), TTG24TTA (5629G>A), ATC25CTC (5630A>C), GGC28GGT (5641C>T), GCA31GCT (5650A>T), ATA35ATT (5662A>T), GGA36GGG (5665A>G), CAA44CAG (5689A>G), GTT46ATT (5693G>A), GAG47GAA (5698G>A), CAA52CAG (5713A>G) | | | | | | | | |
| NIa-VPg protein (NP_734252.1) | 1 | 188 | 100% | 1227 | 93.3% | 188 (100%) | 170 (90.4%) | 0/0/0/0 | 0 |
| Protein mutations: | K47R (5853A>G), I61V (5894A>G 5896C>G), R83A (5960C>G 5961G>C 5962G>C), I93V (5990A>G), E102D (6019A>T), K106T (6030A>C), D112E (6049C>A), M115I (6058G>A), S120N (6072G>A 6073T>C), N121H (6074A>C), T123N (6081C>A), C133S (6111G>C 6112T>C), I139V (6128A>G 6130T>C), V149I (6158G>A), T153S (6170A>T 6172A>G), L164F (6203C>T 6205C>T), V173I (6230G>A), A182K (6257G>A 6258C>A) | | | | | | | | |

| | Begin | End | Coverage | Score | Concordance | Matches | Identities | I/D/M/F* | Stop Codons |
|---|---|---|---|---|---|---|---|---|---|
| **NT** | **54** | **9704** | **99.5%** | **12522** | **65.2%** | **9639 (99.8%)** | **7984 (82.7%)** | **7/12** | |
| Codon mutations: | GGG1GGC (5716G>C), AAA2AAG (5719A>G), GCC10GCA (5743C>A), CGC14CGA (5755C>A), CAT15CAC (5758T>C), CGT17CGC (5764T>C), AAA19AAG (5770A>G), GGC22GGT (5779C>T), TTT23TTC (5782T>C), GAC26GAT (5791C>T), GAC30GAT (5803C>T), AGG41AGA (5836G>A), AAA42AAG (5839A>G), GGA44GGG (5845A>G), AAA45AAG (5848A>G), AAA47AGA (5853A>G), GGT48GGC (5857T>C), ACC49ACT (5860C>T), ACA50ACT (5863A>T), GGT52GGA (5869T>A), GGT54GGC (5875T>C), AGC57AGT (5884C>T), TTC60TTT (5893C>T), ATC61GTG (5894A>G 5896C>G), AAC62AAT (5899C>T), GGG65GGA (5908G>A), TTT66TTC (5911T>C), GAG70GAA (5923G>A), CAA75CAG (5938A>G), TTC76TTT (5941C>T), GTT77GTC (5944T>C), GAT78GAC (5947T>C), GGG82GGA (5959G>A), CGG83GCC (5960C>G 5961G>C 5962G>C), GAA87GAG (5974A>G), GCT91GCA (5986T>A), GAC92GAT (5989C>T), ATT93GTT (5990A>G), GAT95GAC (5998T>C), AGA99CGT (6008A>C 6010A>T), GAA102GAT (6019A>T), GTG103GTT (6022G>T), CGA104CGT (6025A>T), AAA106ACA (6030A>C), GAC112GAA (6049C>A), ATG115ATA (6058G>A), CAA116CAG (6061A>G), GCC117GCA (6064C>A), GGT119GGC (6070T>C), AGT120AAC (6072G>A 6073T>C), AAC121CAC (6074A>C), ACG122ACA (6079G>A), ACC123AAC (6081C>A), ATA124ATT (6085A>T), GCA126GCT (6091A>T), TAC127TAT (6094C>T), TGT133TCC (6111G>C 6112T>C), AAA135AAG (6118A>G), TTG137TTA (6124G>A), AAG138AAA (6127G>A), ATT139GTC (6128A>G 6130T>C), GAT140GAC (6133T>C), TTA141TTG (6136A>G), AAC145AAT (6148C>T), AAA148AAG (6157A>G), GTT149ATT (6158G>A), TGT150TGC (6163T>C), AAA152AAG (6169A>G), ACA153TCG (6170A>T 6172A>G), GGC155GGT (6178C>T), AAA158AAG (6187A>G), TTT159TTC (6190T>C), GAG161GAA (6196G>A), CTC164TTT (6203C>T 6205C>T), GAA165GAG (6208A>G), CTA166TTG (6209C>T 6211A>G), CAG168CAA (6217G>A), GCT172GCA (6229T>A), GTA173ATA (6230G>A), GTC175GTT (6238C>T), AAG178AAA (6247G>A), ATA180ATT (6253A>T), GCA182AAA (6257G>A 6258C>A), CAT187CAC (6274T>C) | | | | | | | | |
| NIa-Pro protein (NP_734248.1) | 1 | 244 | 100% | 1654 | 93.7% | 244 (100%) | 223 (91.4%) | 0/0/0/0 | 0 |
| Protein mutations: | K2R (6282A>G), A29T (6362G>A), V42I (6401G>A), A43V (6405C>T), Y51F (6429A>T), Q58R (6450A>G), H70N (6485C>A 6487C>T), L84I (6527C>A), T104I (6588C>T), Y120H (6635T>C), I124V (6647A>G), I134V (6677A>G 6679A>G), D147N (6716G>A), N148D (6719A>G), I163L (6764A>T 6766A>G), A173V (6795C>T 6796A>G), H174Q (6799C>A), N194D (6857A>G), V200T (6875G>A 6876T>C 6877C>A), D219E (6934C>A), V242I (7001G>A 7003G>A) | | | | | | | | |
| Codon mutations: | AAA2AGA (6282A>G), TCG3TCA (6286G>A), GGC7GGT (6298C>T), TTG8TTA (6301G>A), AGA9AGG (6304A>G), GAC10GAT (6307C>T), GCC15GCT (6322C>T), ACA17ACT (6328A>T), GTT18GTG (6331T>G), TGT19TGC (6334T>C), AGG20AGA (6337G>A), CTG21TTG (6338C>T), TAT27TAC (6358T>C), GGG28GGA (6361G>A), GCA29ACA (6362G>A), TCA30TCT (6367A>T), GAG31GAA (6370G>A), TAC33TAT (6376C>T), GGT34GGG (6379T>G), GGA36GGC (6385A>C), GGA38GGT (6391A>T), GCA39GCG (6394A>G), TAC40TAT (6397C>T), GTA42ATA (6401G>A), GCG43GTG (6405C>T), CAC45CAT (6412C>T), TTA47TTG (6418A>G), TTT48TTC (6421T>C), TAC51TTC (6429A>T), GGT53GGA (6436T>A), GAG56GAA (6445G>A), CAA58CGA (6450A>G), TCC59TCG (6454C>G), CAC61CAT (6460C>T), GGT62GGA (6463T>A), TTC64TTT (6469C>T), AGG65AGA (6472G>A), GTG66GTA (6475G>A), AAT68AAC (6481T>C), CAC70AAT (6485C>A 6487C>T), AGT71AGC (6490T>C), AGC73AGT (6496C>T), CTG75CTA (6502G>A), GGT79GGC (6514T>C), AGG80AGA (6517G>A), CTC84ATC (6527C>A), ATC85ATA (6532C>A), CCG88CCA (6541G>A), AAA89AAG (6544A>G), GTC93GTT (6556C>T), TTT94TTC (6559T>C), CAG96CAA (6565G>A), TTG98CTA (6569T>C 6571G>A), CCT103CCA (6586T>A), ACA104ATA (6588C>T), GAA107GAG (6598A>G), AGA108AGG (6601A>G), ATT109ATC (6604T>C), TGT110TGC (6607T>C), GGA113GGG (6616A>G), AAC115AAT (6622C>T), GAG118GAA (6631G>A), AAG119AAA (6634G>A), TAT120CAT (6635T>C), GCT121GCA (6640T>A), TCG123TCT (6646G>T), ATC124GTC (6647A>G), ATC125ATT (6652C>T), AGC129AGT (6664C>T), ACT131ACG (6670T>G), ATA134GTG (6677A>G 6679A>G), GGC136GGT (6685C>T), ACA138ACT (6691A>T), TTC139TTT (6694C>T), ACA146ACG (6715A>G), GAT147AAT (6716G>A), AAT148GAT (6719A>G), GGA149GGG (6724A>G), CTA153TTA (6734C>T), GTG155GTA (6742G>A), GTG156GTA (6745G>A), GCC159GCT (6754C>T), GGA161GGG (6760A>G), ATA163TTG (6764A>T 6766A>G), ATT166ATA (6775T>A), CTG169TTG (6782C>T), GCA170GCT (6787A>T), AAC171AAT (6790C>T), GCA173GTG (6795C>T 6796A>G), CAC174CAA (6799C>A), ACC175ACA (6802C>A), AAC177AAT (6808C>T), TAC178TAT (6811C>T), TAC179TAT (6814C>T), TTC182TTT (6823C>T), GAA184GAG (6829A>G), TAC190TAT (6847C>T), CGA192CGC (6853A>C), AAT194GAT (6857A>G), GAG195GAA (6862G>A), GAA198GAG (6871A>G), GTC200ACA (6875G>A 6876T>C 6877C>A), AAG201AAA (6880G>A), TCT202TCA (6883T>A), GTT204GTG (6889T>G), GAC208GAT (6901C>T), GGC213GGT (6916C>T), CCG214CCA (6919G>A), AAA216AAG (6925A>G), CTT217CTC (6928T>C), GAC219GAA (6934C>A), GGG224GGC (6949G>C), TTC226TTT (6955C>T), AAA227AAG (6958A>G), CAA233CAG (6976A>G), CTA235TTA (6980C>T), ATC236ATT (6985C>T), GTA240GTT (6997A>T), GTG241GTT (7000G>T), GTG242ATA (7001G>A 7003G>A) | | | | | | | | |
| NIb protein (NP_734249.1) | 1 | 521 | 100% | 3129 | 86.1% | 519 (99.4%) | 480 (92.0%) | 1/2/2/2 | 0 |
| Protein mutations: | F8Y (7032T>A), T39K (7125C>A), A46S (7145G>T 7147A>G), E51_A52del (7160_7165delGAGGCA), D69E (7216T>A), D83E (7258T>A), D88_R89insX (7273_7274insT), R89V (7274C>G 7275G>T), H91L (7281A>T), L92S (7284T>C), S96L (7296C>T), S101F (7311C>T), T102I (7314C>T), C103Y (7317G>A), N104K (7321T>A), K109Q (7334A>C), A112N (7343G>A 7344C>A 7345A>C), S133M (7407G>T 7408T>G), C136G (7415T>G), L165S (7503T>C 7504G>A), D234N (7709G>A), K235R (7713A>G 7714A>G), T268I (7812C>T), V279L (7844G>T 7846G>A), V292I (7883G>A), S345N (8043G>A), N381S (8151A>G 8152C>T), R390K (8178G>A), R400K (8208G>A), P429A (8294C>G 8296A>G), S447F (8349C>T), A467S (8408G>T), M481I (8452G>A), R484K (8460G>A), A491T (8480G>A 8482T>A), R498K (8502G>A 8503A>G), E502D (8515A>T), E510D (8539G>C), L513C (8546C>T 8547T>G), S515T (8552T>A) | | | | | | | | |
| Codon mutations: | GCT1GCC (7012T>C), AAG2AAA (7015G>A), CAT3CAC (7018T>C), TTT8TAT (7032T>A), GCC10GCT (7039C>T), TTG11CTA (7040T>C 7042G>A), GGA13GGG (7048A>G), GCT17GCA (7060T>A), GTC18GTA (7063C>A), AGC23AGT (7078C>T), TTA25CTG (7082T>C 7084A>G), GTA26GTG (7087A>G), ACC27ACA (7090C>A), CAT29CAC (7096T>C), GTA30GTG (7099A>G), GTT31GTC (7102T>C), GAG34GAA (7111G>A), CGA36CGG (7117A>G), ACA39AAA (7125C>A), GAA40GAG (7129A>G), TTT41TTC (7132T>C), CTG42TTA (7133C>T 7135G>A), GTG44GTA (7141G>A), GAT45GAC (7144T>C), GCA46TCG (7145G>T 7147A>G), GAG47GAA (7150G>A), GCA50GCT (7159A>T), GAG51_GCA52del (7160_7165delGAGGCA), TTC54TTT (7171C>T), GCG60GCT (7189G>T), AAA63AAG (7198A>G), AGC64AGT (7201C>T), CTA66TTA (7205C>T), AAT67AAC (7210T>C), GAT69GAA (7216T>A), GCG70GCA (7219G>A), ATC72ATA (7225C>A), AAG73AAA (7228G>A), AAG77AAA (7240G>A), TAT78TAC (7243T>C), AAA80AAG (7249A>G), CCT81CCG (7252T>G), GAT83GAA (7258T>A), GTT84GTC (7261T>C), GGT85GGA (7264T>A), GTC86GTG (7267C>G), GTG87GTA (7270G>A), GAT88_CGG89insT-- (7273_7274insT), CGG89GTG (7274C>G 7275G>T), CAT91CTT (7281A>T), TTG92TCG (7284T>C), TCA96TTA (7296C>T), TCT101TTT (7311C>T), ACC102ATC (7314C>T), TGC103TAC (7317G>A), AAT104AAA (7321T>A), GTG105-TG (7322delG), CAC106CAT (7327C>T), GGC107GGT (7330C>T), TTC108TTT (7333C>T), AAG109CAG (7334A>C), TGT111TGC (7342T>C), GCA112AAC (7343G>A 7344C>A 7345A>C), ACT115ACC (7354T>C), CAA118CAG (7363A>G), ATT120ATC (7369T>C), TTC121TTT (7372C>T), GCG123GCT (7378G>T), CTC124CTT (7381C>T), GCA129GCT (7396A>T), GTC130GTT (7399C>T), GCC132GCT (7405C>T), AGT133ATG (7407G>T 7408T>G), GGG135GGT (7414G>T), TGC136GGC (7415T>G), AAA137AAG (7420A>G), GAC140GAT (7429C>T), TAT141TAC (7432T>C), CAT144CAC (7441T>C), AAG150AAA (7459G>A), GAA151GAG (7462A>G), ATA153ATT (7468A>T), GTC154GTA (7471C>A), CAA156CAG (7477A>G), AGC157AGT (7480C>T), CTG159CTT (7486G>T), TAT162TAC (7495T>C), AAA163AAG (7498A>G), GGT164GGC (7501T>C), TTG165TCA (7503T>C 7504G>A), ATT168ATA (7513T>A), AAC170AAT (7519C>T), GGA171GGG (7522A>G), AAG174AAA (7531G>A), GAG176GAA (7537G>A), CTC177CTT (7540C>T), TGT179TGC (7546T>C), AAG187AAA (7570G>A), ACG188ACA (7573G>A), ACG190ACA (7579G>A), GCT193GCC (7588T>C), CTA196TTA (7595C>T), TTG199CTA (7604T>C 7606G>A), GGT202GGA (7615T>A), TGT205TGC (7624T>C), GAC208GAT (7633C>T), TTC209TTT (7636C>T), AAT210AAC (7639T>C), TTT213TTC (7648T>C), GAA219GAG (7666A>G), TGT221TGC (7672T>C), ACA223ACT (7678A>T), GGG225GGA (7684G>A), ACT227ACC (7690T>C), TTT229TTC (7696T>C), GGT231GGG (7702T>G), GAT234AAT (7709G>A), AAA235AGG (7713A>G 7714A>G), CTG236TTG (7715C>T), CGT239CGC (7726T>C), GAG242GAA (7735G>A), GTA245GTG (7744A>G), TAC246TAT (7747C>T), GGC251GGT (7762C>T), CAG253CAA (7768G>A), TTT254TTC (7771T>C), TCA257TCG (7780A>G), CTA258CTC (7783A>C), ACT259ACC (7786T>C), CTA262TTG (7793C>T 7795A>G), ATC263ATT (7798C>T), GCT265GCC (7804T>C), ACC268ATC (7812C>T), GAA275GAG (7834A>G), GAC276GAT (7837C>T), GTG279TTA (7844G>T 7846G>A), CAG282CAA (7855G>A), CTG284CTA (7861G>A), TTA287TTG (7870A>G), TAC288TAT (7873C>T), ACT289ACA (7876T>A), GAG290GAA (7879G>A), ATT291ATA (7882T>A), GTT292ATT (7883G>A), CCA295CCT (7894A>T), ATT296ATC (7897T>C), GTC304GTT (7921C>T), AAG306AAA (7927G>A), GGT309GGC (7936T>C), AAT310AAC (7939T>C), AAT311AAC (7942T>C), GTG319GTA (7966G>A), GAT320GAC (7969T>C), TCT322TCC (7975T>C), GTC326GTT (7987C>T), CTT327CTC (7990T>C), GCT332GCC (8005T>C), GAA341GAG (8032A>G), ATC343ATT (8038C>T), GAC344GAT (8041C>T), AGC345AAC (8043G>A), TTG357TTA (8080G>A), GTG360GTA (8089G>A), CCG362CCA (8095G>A), GAG363GAA (8098G>A), AGC366AGT (8107C>T), ATT367ATC (8110T>C), GAT369GAC (8116T>C), CAA373CAG (8128A>G), GGT379GGC (8146T>C), AAC381AGT (8151A>G 8152C>T), TCG385TCA (8164G>A), AGG390AAG (8178G>A), AAG391AAA (8182G>A), TTG394TTA (8191G>A), TTC396TTT (8197C>T), TCC398TCT (8203C>T), AGA400AAA (8208G>A), CTG402TTA (8213C>T 8215G>A), CTA403CTG (8218A>G), ATC404ATT (8221C>T), GAG405GAA (8224G>A), CCA410CCG (8239A>G), GAG415GAA (8254G>A), ATT417ATC (8260T>C), GTA418GTG (8263A>G), GAT424GAC (8281T>C), AGA425AGG (8284A>G), CCA429GCG (8294C>G 8296A>G), GAG430GAA (8299G>A), CAC431CAT (8302C>T), ATT436ATC (8317T>C), TGC437TGT (8320C>T), GCT439GCA (8326T>A), GAG442GAA (8335G>A), TCC443TCT (8338C>T), TCT447TTT (8349C>T), GAA448GAG (8353A>G), CAC451CAT (8362C>T), CAA452CAG (8365A>G), ATC453ATT (8368C>T), TAC457TAT (8380C>T), TCA458TCG (8383A>G), TTA460TTG (8389A>G), CCT465CCC (8404T>C), GCA467TCA (8408G>T), ACA468ACG (8413A>G), GCG470GCT (8419G>T), GCT475GCC (8434T>C), TAT477TAC (8440T>C), ATG481ATA (8452G>A), CTA483CTG (8458A>G), AGG484AAG (8460G>A), AAA485AAG (8464A>G), CTG486CTA (8467G>A), AGG490AGA (8479G>A), GCT491ACA (8480G>A 8482T>A), GAG496GAA (8497G>A), CTA497TTG (8498C>T 8500A>G), AGA498AAG (8502G>A 8503A>G), GCC499GCT (8506C>T), GAA502GAT (8515A>T), GTC505GTT (8524C>T), GCA506GCC (8527A>C), TTA507CTG (8528T>C 8530A>G), GAG510GAC (8539G>C), CTT513TGT (8546C>T 8547T>G), TCT515ACT (8552T>A), GTA518GTG (8563A>G) | | | | | | | | |
| coat protein (NP_734250.1) | 1 | 267 | 100% | 1652 | 90.6% | 267 (100%) | 243 (91.0%) | 0/0/0/0 | 0 |
| Protein mutations: | A1G (8574C>G), I5V (8585A>G), G9E (8598G>A), N11S (8603A>T 8604A>C 8605C>T), P17Q (8622C>A), E18G (8625A>G 8626G>A), P26F (8648C>T 8649C>T 8650G>C), G29E (8658G>A), D31A (8664A>C 8665T>G), A36V (8679C>T 8680A>T), T58K (8745C>A), E95G (8856A>G 8857G>A), R98Q (8865G>A), M99L (8867A>C 8869G>T), G104E (8883G>A), T106S (8888A>T), V128I (8954G>A), N138D (8984A>G 8986T>C), E139V (8988A>T), I187V (9131A>G), M193G (9149A>G 9150T>G 9151G>A), G194S (9152G>A), P230S (9260C>T 9262T>C), Q242H (9298A>C) | | | | | | | | |
| Codon mutations: | GCA1GGA (8574C>G), ATT5GTT (8585A>G), GGA9GAA (8598G>A), AGC10AGT (8602C>T), AAC11TCT (8603A>T 8604A>C 8605C>T), AAA16AAG (8620A>G), CCA17CAA (8622C>A), GAG18GGA (8625A>G 8626G>A), CAG19CAA (8629G>A), GGC20GGT (8632C>T), ATC22ATT (8638C>T), CAG23CAA (8641G>A), AAC25AAT (8647C>T), CCG26TTC (8648C>T 8649C>T 8650G>C), AAC27AAT (8653C>T), AAA28AAG (8656A>G), GGA29GAA (8658G>A), GAT31GCG (8664A>C 8665T>G), GTT34GTG (8674T>G), GCA36GTT (8679C>T 8680A>T), GGC37GGA (8683C>A), GGG40GGA (8692G>A), ACA41ACT (8695A>T), CCG45CCA (8707G>A), AGA46CGA (8708A>C), AAG48AAA (8716G>A), ATC50ATT (8722C>T), ACG51ACA (8725G>A), AAA53AAG (8731A>G), AGA55AGG (8737A>G), CCC57CCT (8743C>T), ACA58AAA (8745C>A), AAG60AAA (8752G>A), GGA61GGT (8755A>T), AAC66AAT (8770C>T), TTA67CTA (8771T>C), GAA68GAG (8776A>G), CAT69CAC (8779T>C), TTG70TTA (8782G>A), TAT73TAC (8791T>C), CCA75CCG (8797A>G), CAA76CAG (8800A>G), CAA77CAG (8803A>G), GAT79GAC (8809T>C), ATT80ATC (8812T>C), AAT82AAC (8818T>C), CGG84CGA (8824G>A), GCA85GCG (8827A>G), ACG92ACA (8848G>A), TAT94TAC (8854T>C), GAG95GGA (8856A>G 8857G>A), GTG97GTA (8863G>A), CGG98CAG (8865G>A), ATG99CTT (8867A>C 8869G>T), GCA100GCG (8872A>G), GGA104GAA (8883G>A), ACT106TCT (8888A>T), TGC119TGT (8929C>T), GTC128ATC (8954G>A), AAC129AAT (8959C>T), GGA130GGT (8962A>T), GGG137GGA (8983G>A), AAT138GAC (8984A>G 8986T>C), GAA139GTA (8988A>T), GAG142GAA (8998G>A), CCG144CCA (9004G>A), TTG145TTA (9007G>A), ATC148ATT (9016C>T), ACC155ACA (9037C>A), ATA171ATT (9085A>T), AAC175AAT (9097C>T), ATT187GTT (9131A>G), CGA188CGT (9136A>T), CGG191CGT (9145G>T), ATG193GGA (9149A>G 9150T>G 9151G>A), GGT194AGT (9152G>A), TTA195TTG (9157A>G), GCG196GCT (9160G>T), CGT197CGC (9163T>C), GAG204GAA (9184G>A), CGA208CGG (9196A>G), AGG214AGA (9214G>A), GAA215GAG (9217A>G), GCG216GCA (9220G>A), GCA224GCT (9244A>T), TTG225TTA (9247G>A), GCC228GCT (9256C>T), CCT230TCC (9260C>T 9262T>C), GGG234GGA (9274G>A), GAC236GAT (9280C>T), ATC239ATT (9289C>T), CAA242CAC (9298A>C), GAG244GAA (9304G>A), GTC254GTT (9334C>T), GTC264GTG (9364C>G) | | | | | | | | |

| | Begin | End | Coverage | Score | Concordance | Matches | Identities | I/D/M/F* | Stop Codons |
|---|---|---|---|---|---|---|---|---|---|
| **NT** | **54** | **9704** | **99.5%** | **12522** | **65.2%** | **9639 (99.8%)** | **7984 (82.7%)** | **7/12** | |
| PIPO (YP_006393460.1) | 1 | 75 | 100% | 95 | 18.1% | 75 (98.7%) | 54 (71.1%) | 1/0/3/2 | 2 |
| Protein mutations: | R17_K17insX (2971_2972insA), Y23H (2989T>C 2991C>T), R26K (2999G>A 3000A>G), H35Y (3025C>T), K40H (3040A>C 3042A>C), G41S (3043G>A), E44K (3052G>A), I47V (3061A>G), V54I (3082G>A), P57S (3091C>T), P60T (3100C>A), G61E (3104G>A 3105T>G), G62S (3106G>A), Q63* (3109C>T), R64G (3112A>G), C66R (3118T>C), L67F (3121C>T), I69V (3127A>G), A72E (3137C>A 3138G>A), I73V (3139A>G) | | | | | | | | |
| Codon mutations: | TTA2CTA (2923T>C), AAA4AAG (2931A>G), GAA7GAG (2940A>G), TTG10CTG (2947T>C), TTG15CTG (2962T>C), AGA17_AAA17insAA- (2971_2972insA), TAC23CAT (2989T>C 2991C>T), TCA24TCG (2994A>G), AGA26AAG (2999G>A 3000A>G), CAC32-AC (3016delC), TCG33TCA (3021G>A), GTA34GTT (3024A>T), CAT35TAT (3025C>T), CAC38CAT (3036C>T), AGA39AGG (3039A>G), AAA40CAC (3040A>C 3042A>C), GGC41AGC (3043G>A), GAA44AAA (3052G>A), ATA47GTA (3061A>G), GTT54ATT (3082G>A), CCG57TCG (3091C>T), CGC59CGT (3099C>T), CCA60ACA (3100C>A), GGT61GAG (3104G>A 3105T>G), GGT62AGT (3106G>A), CAA63TAA (3109C>T), AGG64GGG (3112A>G), TGC66CGC (3118T>C), CTC67TTC (3121C>T), ATT69GTT (3127A>G), GAG70GAA (3132G>A), GCG72GAA (3137C>A 3138G>A), ATT73GTT (3139A>G) | | | | | | | | |

*: Inserts / Deletes / Misaligned / Frameshifts

#### Analysis details

This analysis was performed with panviral2.64

### NGS Details (UN62): Torradovirus lycopersici (segment RNA 1)

#### Assembly

| | |
|---|---|
| **Coverage Length** | 7803 (1 contig(s)) |
| **Depth Of Coverage** | 2646.1 |
| **Number Of Reads** | 130769 |
| **Reads Per Million** | 3031.34 rpm (after QC) |
| **Ambiguities** | 0 |
| **Assembly Method** | de novo + reference guided assembly |
| **Consensus Caller** | Bcf Tools |

#### Coverage Map

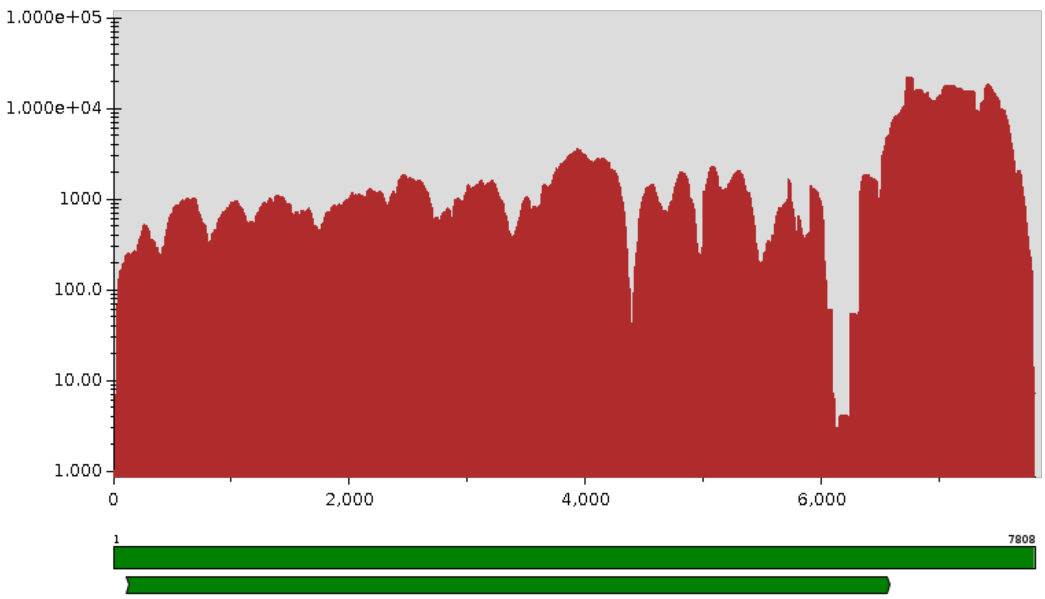

#### Assignment

| | |
|---|---|
| **Type** | Torradovirus lycopersici (Taxonomy ID: 3048378) |
| **Reference Genome** | NC_009013.1 |
| **NT Identity (%)** | 87.0179 |
| **AA Identity (%)** | 93.2376 |
| **Number Of Stop Codons** | 3 |
| **Number Of CDS** | 1 |

#### Alignment

| | |
|---|---|
| **Alignment Score** | 10368.0 (NT) + 14226.0 (AA) = 24594.0 |
| **Concordance (%)** | 83.7585 |

| Alignment Method | Global, seeded, nucleotide + amino acids (AGA) |
|---|---|

#### Genome Region

Sequence starts at position 1 and ends at position 7803 relative to NC_009013.1 reference sequence.

#### Alignment Detailed Statistics

| | Begin | End | Coverage | Score | Concordance | Matches | Identities | I/D/M/F* | Stop Codons |
|---|---|---|---|---|---|---|---|---|---|
| **NT** | **1** | **7803** | **99.9%** | **10368** | **72.5%** | **7431 (95.1%)** | **6475 (82.9%)** | **10/372** | |

Mutations: 1T>G, 69C>T, 72C>T, 77A>T, 87C>T, 101G>T, 115T>C, 127C>T, 128C>T, 130C>T, 135T>C, 138A>G, 144T>C, 147C>T, 155T>A, 157C>T, 159C>T, 161A>G, 178T>A, 187A>T, 197G>C, 198C>A, 199T>A, 220C>T, 262T>A, 265C>T, 289T>G, 292A>T, 301T>A, 302T>G, 307C>T, 310T>C, 311A>T, 313C>T, 322T>C, 324A>G, 325C>T, 328T>A, 340G>A, 349G>A, 358T>A, 359T>C, 361A>T, 367A>G, 370C>T, 373C>T, 376C>T, 382T>A, 388T>C, 394C>T, 409A>G, 412A>C, 427T>G, 433G>A, 443A>G, 451A>G, 487C>T, 502G>A, 541T>C, 559C>T, 578C>T, 599A>T, 606T>C, 615A>G, 618C>T, 619A>T, 655C>T, 661A>G, 676A>G, 694A>G, 707C>T, 715G>A, 718A>G, 719T>G, 727A>T, 748G>T, 751A>T, 760C>T, 766C>T, 778T>A, 781C>T, 790C>T, 799A>T, 802T>A, 806A>C, 821T>A, 829G>A, 832A>T, 835A>C, 842C>T, 847C>T, 883G>A, 886T>C, 889C>A, 901T>C, 913T>C, 925C>T, 938C>T, 952A>G, 964T>A, 970G>A, 977G>A, 991T>C, 997C>T, 1001T>C, 1006T>A, 1018C>T, 1019A>G, 1020C>T, 1033G>A, 1060T>C, 1063C>A, 1093T>C, 1109C>T, 1129A>G, 1135T>A, 1165C>A, 1180G>A, 1186G>A, 1192A>C, 1195A>T, 1196A>T, 1198C>T, 1204T>G, 1207T>G, 1219G>C, 1240C>T, 1249A>G, 1255G>A, 1276A>G, 1300G>A, 1309C>A, 1318G>A, 1319C>T, 1324C>T, 1336T>G, 1337T>C, 1339G>T, 1357A>C, 1361T>C, 1411T>C, 1429C>T, 1489T>C, 1522C>A, 1534A>T, 1552C>T, 1553C>A, 1555C>G, 1568C>A, 1570A>G, 1576A>C, 1582C>T, 1585A>C, 1591C>T, 1607G>T, 1612G>A, 1618T>G, 1619C>T, 1624C>A, 1630C>T, 1643A>C, 1669C>T, 1681T>A, 1684G>A, 1692C>T, 1699T>G, 1706T>G, 1708T>C, 1714T>G, 1737A>G, 1738A>G, 1739A>C, 1755C>T, 1756A>C, 1770G>A, 1771C>T, 1781T>C, 1789A>G, 1813T>G, 1825C>A, 1828A>T, 1834A>G, 1839A>T, 1844G>A, 1849A>C, 1850T>A, 1858G>A, 1867G>A, 1876G>C, 1877T>A, 1879A>T, 1885G>A, 1889A>G, 1891A>G, 1894A>T, 1906T>A, 1907A>C, 1908T>C, 1909G>A, 1911T>G, 1927C>A, 1938C>T, 1942G>A, 1957T>G, 1960A>C, 1966A>G, 1978G>A, 1984C>T, 2014A>G, 2092A>G, 2120A>C, 2128T>C, 2130A>C, 2134C>T, 2138C>T, 2141T>C, 2143T>A, 2146A>G, 2149T>A, 2155T>C, 2174C>T, 2179A>C, 2200T>C, 2224C>T, 2227C>T, 2230T>G, 2240A>T, 2241A>G, 2250G>A, 2251C>A, 2257C>T, 2263G>A, 2270C>T, 2281A>T, 2282A>T, 2287G>A, 2290C>T, 2299A>T, 2318C>T, 2323A>T, 2344C>A, 2347C>T, 2350A>G, 2356C>T, 2359G>A, 2365A>T, 2368A>T, 2375A>C, 2380T>C, 2387C>A, 2389C>A, 2392A>G, 2407T>C, 2409G>A, 2446G>A, 2479G>A, 2485C>T, 2488G>A, 2491A>G, 2494T>G, 2497G>C, 2500T>C, 2503G>A, 2506T>A, 2509C>T, 2512T>C, 2516T>C, 2524A>C, 2542G>T, 2605A>T, 2614C>A, 2620G>A, 2626T>C, 2639C>A, 2650G>A, 2662C>T, 2692G>A, 2716A>C, 2726G>A, 2731C>T, 2734C>T, 2743A>C, 2744T>C, 2765T>C, 2767G>C, 2768A>C, 2773A>G, 2776T>A, 2806C>T, 2809C>T, 2812G>A, 2821G>A, 2827A>T, 2848C>T, 2852C>A, 2854T>A, 2860C>G, 2866T>A, 2869C>T, 2875A>T, 2878A>C, 2881T>C, 2882G>T, 2890A>C, 2914C>T, 2926G>A, 2929C>T, 2932G>A, 2947T>A, 2950C>T, 2992T>C, 2998A>G, 3031T>A, 3049A>C, 3064A>T, 3079T>C, 3083C>A, 3085A>G, 3101T>C, 3103G>T, 3109T>C, 3124C>A, 3136T>G, 3146A>T, 3151C>T, 3166T>C, 3169C>T, 3172T>C, 3175A>G, 3176C>T, 3184T>A, 3202T>C, 3238T>G, 3241T>A, 3242C>T, 3253C>T, 3265A>G, 3277T>A, 3289T>C, 3298T>A, 3331A>T, 3349T>A, 3365T>C, 3376T>G, 3379G>A, 3382T>A, 3388G>T, 3400A>C, 3406A>G, 3413C>G, 3418C>A, 3421T>C, 3430A>G, 3431C>A, 3445G>T, 3451T>C, 3466G>A, 3476G>C, 3481C>T, 3484A>T, 3508G>T, 3520A>G, 3529G>C, 3530C>A, 3532A>G, 3534G>A, 3541C>T, 3542T>C, 3544G>T, 3553T>A, 3559G>A, 3580T>C, 3583C>A, 3589A>G, 3591C>T, 3592T>C, 3593T>C, 3594C>A, 3610A>G, 3613T>C, 3622G>A, 3625C>T, 3628G>A, 3629G>T, 3646C>T, 3658A>C, 3661A>G, 3676C>T, 3679T>A, 3682G>C, 3688A>G, 3691C>T, 3693A>T, 3701C>T, 3710G>T, 3713A>G, 3716T>C, 3739A>T, 3742T>G, 3754G>A, 3758G>A, 3766G>A, 3776G>A, 3778G>T, 3781G>T, 3784C>G, 3790T>A, 3814C>T, 3824T>C, 3847G>A, 3856G>A, 3868C>T, 3871T>C, 3883G>A, 3895A>C, 3907A>T, 3910T>C, 3911C>A, 3949C>T, 3955G>A, 3958T>A, 3959T>C, 3961G>A, 3973T>C, 3982A>C, 4018A>G, 4028T>C, 4033G>T, 4034T>C, 4055C>A, 4067T>C, 4078T>A, 4084C>T, 4093C>T, 4118C>T, 4129C>T, 4147C>A, 4157C>T, 4159G>A, 4160C>G, 4162A>T, 4189T>A, 4192T>G, 4207G>A, 4210T>C, 4225A>T, 4228C>T, 4282C>A, 4288C>A, 4291T>C, 4294A>G, 4309T>G, 4318C>T, 4327A>T, 4339T>C, 4348C>T, 4358A>C, 4360A>C, 4381T>C, 4384A>C, 4393T>C, 4414T>G, 4417T>C, 4421T>C, 4426G>T, 4429A>G, 4441A>T, 4459T>C, 4468C>T, 4471T>C, 4501A>T, 4510T>A, 4537T>A, 4549G>C, 4555A>G, 4588C>T, 4606A>T, 4618C>A, 4625T>C, 4636T>C, 4648T>A, 4651T>A, 4654C>T, 4661T>C, 4663G>A, 4678T>C, 4684A>T, 4696C>T, 4699C>T, 4714T>C, 4716G>A, 4719G>A, 4735C>T, 4736T>C, 4747A>G, 4753T>C, 4756A>T, 4759C>T, 4765T>C, 4768T>C, 4774G>A, 4780T>C, 4783A>G, 4786A>G, 4789C>T, 4801T>C, 4802C>T, 4813A>G, 4816T>C, 4828A>C, 4834C>T, 4843G>A, 4861C>T, 4870C>G, 4885C>T, 4894G>T, 4897A>C, 4898G>A, 4906G>A, 4909A>C, 4921G>T, 4947C>A, 4954T>G, 4958A>C, 4960C>T, 4969G>A, 4975G>A, 4981C>T, 4987C>T, 4988A>G, 4990C>A, 4999T>C, 5002A>T, 5005T>C, 5012T>C, 5017G>A, 5020G>A, 5023C>T, 5030G>A, 5035G>A, 5041A>T, 5044A>C, 5059G>T, 5065C>T, 5074T>C, 5077T>C, 5079T>C, 5083A>T, 5090G>T, 5102T>C, 5105C>G, 5113C>T, 5116T>C, 5119T>C, 5120G>A, 5123C>T, 5131T>C, 5132T>A, 5143G>T, 5146T>C, 5158C>T, 5167T>C, 5173C>T, 5176C>T, 5182A>G, 5194A>C, 5200G>A, 5206C>T, 5209C>A, 5218T>G, 5227C>T, 5239A>C, 5266C>A, 5269C>A, 5278G>A, 5290G>T, 5293T>C, 5302T>A, 5329C>T, 5344A>G, 5347C>A, 5357A>T, 5360A>C, 5366C>A, 5369G>A, 5371G>T, 5374T>A, 5386C>G, 5392G>A, 5399A>G, 5407A>T, 5410G>T, 5413C>T, 5419A>T, 5429C>T, 5437T>C, 5440C>G, 5459G>A, 5461T>C, 5462G>A, 5471T>C, 5482A>T, 5485A>T, 5486G>A, 5491T>C, 5492A>C, 5494A>T, 5498C>T, 5500C>G, 5512A>T, 5515A>T, 5531C>T, 5533T>G, 5534C>G, 5537A>C, 5543G>T, 5558G>C, 5566T>G, 5569A>T, 5572T>C, 5578T>C, 5582A>G, 5591G>T, 5602G>A, 5604C>T, 5611A>T, 5613A>T, 5617A>T, 5620T>A, 5626A>T, 5638T>G, 5641T>C, 5647T>C, 5664G>A, 5674A>T, 5677G>T, 5686G>A, 5689C>A, 5707A>T, 5710T>A, 5711T>C, 5716G>T, 5719A>T, 5731C>T, 5734T>C, 5737T>C, 5739G>A, 5746T>C, 5751T>A, 5752G>A, 5767G>T, 5773G>A, 5774A>G, 5776A>G, 5779T>C, 5786T>C, 5791C>A, 5797A>G, 5804T>A, 5805T>C, 5806C>T, 5815C>G, 5818C>T, 5826A>T, 5845G>A, 5849T>C, 5850T>A, 5857T>G, 5860C>A, 5866T>G, 5869G>A, 5872C>G, 5878G>T, 5881T>C, 5887C>T, 5890C>T, 5894A>C, 5907A>T, 5914T>C, 5923T>C, 5927C>T, 5942C>T, 5944C>T, 5955A>G, 5965G>A, 5968T>G, 5974T>C, 5975A>C, 5980G>A, 5983T>C, 5986T>C, 5998A>G, 6002A>G, 6003C>T, 6004C>T, 6009G>A, 6016A>G, 6028C>T, 6031A>T, 6033C>T, 6035G>T, 6040T>A, 6055T>G, 6070T>C, 6079C>G, 6085T>C, 6089C>T, 6091A>G, 6092C>T, 6097T>A, 6100T>G, 6103G>A, 6106G>A, 6112A>G, 6118T>C, 6121A>T, 6130G>T, 6139T>C, 6154C>A, 6163C>G, 6166C>T, 6184A>G, 6185A>G, 6186C>A, 6187T>C, 6189A>G, 6193C>T, 6196A>G, 6199C>T, 6202G>C, 6214T>G, 6215C>G, 6217G>T, 6218A>C, 6221G>C, 6222A>C, 6225G>A, 6227A>G, 6230A>G, 6232A>G, 6235G>A, 6239G>A, 6240T>C, 6251T>C, 6253A>T, 6256C>A, 6262A>T, 6263C>T, 6265G>A, 6266T>A, 6268G>T, 6273G>T, 6274C>G, 6291T>A, 6292T>A, 6296G>C, 6298A>T, 6301T>A, 6302T>A, 6304A>G, 6308T>C, 6311G>T, 6313T>A, 6317A>G, 6319A>T, 6322G>A, 6337T>C, 6344C>T, 6371G>A, 6382C>T, 6397C>G, 6404T>C, 6405G>A, 6414G>T, 6415A>G, 6416T>C, 6417C>A, 6418A>C, 6433G>T, 6436C>T, 6437T>A, 6445C>T, 6448G>A, 6457C>A, 6458T>A, 6461C>T, 6463T>A, 6481T>A, 6485C>A, 6490A>C, 6491C>A, 6493T>C, 6496G>A, 6497A>C, 6499A>T, 6500A>G, 6501T>A, 6502T>G, 6514G>T, 6515T>C, 6517C>A, 6520C>T, 6532C>T, 6545C>T, 6556A>C, 6562A>T, 6566G>A, 6567G>A, 6568C>T, 6571C>T, 6574A>G, 6587G>A, 6643A>G, 6670C>T, 6685A>G, 6721A>T, 6742A>T, 6766delC, 6786T>C, 6798_6799insA, 6806T>G, 6807_6832delAACGGAGAGTACTGACTTTAACTAGT, 6835G>T, 6838_6850delGTCCGGCTCCATT, 6854A>T, 6858_6867delCCAATGAATT, 6871_6895delATTGGTTAAAGAGATTTGCACGCCT, 6897T>A, 6905T>G, 6907T>A, 6908G>T, 6911T>C, 6912_6916delCGTGT, 6921T>G, 6923C>G, 6924T>A, 6927_6941delGTTAGAAAGTTCCTT, 6947_6970delAGGTTTACAAAGTACGGGGAGCAC, 6972C>T, 6976G>A, 6984T>C, 6984_6985insG, 6988G>A, 6994G>T, 6995C>A, 6997A>C, 6998T>A, 7002A>T, 7003_7023delTGATAGTCCTTTAACTCAAGG, 7028A>T, 7030T>G, 7032C>A, 7034G>A, 7036T>G, 7039_7050delCATCTTTTGCCG, 7056_7106delAAAGATGAGGTAGCTTCCCCCTTATTGGGAGGCTGAAACTACACATATGTA, 7110G>T, 7111G>A, 7113T>A, 7115G>C, 7119_7121delGAG, 7123C>T, 7124C>A, 7125T>A, 7131C>G, 7133G>C, 7134T>A, 7137G>C, 7138_7185delTTTGAAATTCGATAATTTTCCGTAGCTTGCGTCAAGCTGCTCACGTTA, 7189G>A, 7194_7216delAGTGAAGATGCGCCGTACCACGT, 7221C>T, 7224_7231delGGCAATGC, 7236_7285delGGTTCAGAGCGGGCCCTCAGAATAGAGGTTAAAACTAGTGTGATGGTGTA, 7288T>A, 7294T>G, 7295A>C, 7297A>C, 7300T>G, 7303C>A, 7304_7315delACCCGGGTTGTG, 7320G>C, 7322C>T, 7323_7328delTAGTTA, 7332C>A, 7333G>C, 7336C>T, 7338C>A, 7339A>T, 7341_7347delGTCCCAC, 7350T>A, 7351A>C, 7353A>T, 7354G>A, 7358A>C, 7360G>A, 7361A>C, 7362A>C, 7363A>G, 7364C>T, 7368G>A, 7372A>T, 7374T>C, 7375T>G, 7376T>G, 7377T>A, 7379A>T, 7383_7384delAA, 7388C>T, 7397G>A, 7398C>A, 7399G>A, 7400A>T, 7404C>T, 7407A>T, 7408G>T, 7423A>T, 7425G>A, 7435T>C, 7436G>A, 7444C>T, 7445A>G, 7457A>G, 7458T>C, 7460G>A, 7503C>T, 7505A>G, 7506T>C, 7511_7512insT, 7513A>C, 7514A>T, 7515A>C, 7525_7526insG, 7526T>A, 7526_7527insG, 7545T>A, 7554A>G, 7569G>A, 7585T>C, 7589T>A, 7600T>G, 7601T>A, 7602T>G, 7604A>C, 7604_7605insG, 7610C>T, 7612G>A, 7615A>G, 7616A>C, 7617A>T, 7618C>T, 7622C>T, 7625delA, 7627T>A, 7629C>A, 7634A>G, 7637G>T, 7644_7652delGTTTGTTTT, 7657G>C, 7660T>A, 7665G>T, 7666T>C, 7667G>T, 7678C>T, 7686T>G, 7687T>A, 7689G>T, 7698G>A, 7717A>C, 7719T>A, 7723T>C, 7739T>A, 7742_7743insC, 7749T>C, 7751T>C, 7752G>A, 7754_7755insG, 7756_7757insT, 7762C>T, 7763A>C, 7764A>T, 7767C>T, 7773G>T, 7775G>A, 7778T>A, 7782A>T, 7783A>C, 7784_7785insT

**CDS**

| | Begin | End | Coverage | Score | Concordance | Matches | Identities | I/D/M/F* | Stop Codons |
|---|---|---|---|---|---|---|---|---|---|
| ToTV_sRNA1gp1 | 1 | 2159 | 100% | 14226 | 94.4% | 2159 (100%) | 2013 (93.2%) | 0/0/0/0 | 3 |

Protein mutations: P8S (128C>T 130C>T), F10S (135T>C), N11S (138A>G), V13A (144T>C), T14I (147C>T), C17S (155T>A 157C>T), A18V (159C>T), T19A (161A>G), A31Q (197G>C 198C>A 199T>A), S66A (302T>G), T69S (311A>T 313C>T), N73S (324A>G 325C>T), S113G (443A>G), T165S (599A>T), I167T (606T>C), N170S (615A>G), T171I (618C>T 619A>T), S205A (719T>G), S239T (821T>A), V291I (977G>A), T305V (1019A>G 1020C>T), T364S (1196A>T 1198C>T), D367E (1207T>G), A501S (1607G>T), T529I (1692C>T), S534A (1706T>G 1708T>C), K544R (1737A>G 1738A>G), T550I (1755C>T 1756A>C), S555N (1770G>A 1771C>T), Y578F (1839A>T), A580T (1844G>A), S582T (1850T>A), S591T (1877T>A 1879A>T), I595V (1889A>G 1891A>G), D600E (1906T>A), M601P (1907A>C 1908T>C 1909G>A), L602W (1911T>G), A611V (1938C>T), N675T (2130A>C), Y679Q (2141T>C 2143T>A), N712C (2240A>T 2241A>G), S715K (2250G>A 2251C>A), M726L (2282A>T), R768K (2409G>A), G874S (2726G>A), A926S (2882G>T), Q993K (3083C>A 3085A>G), T1014S (3146A>T), L1046F (3242C>T), Q1103E (3413C>G), V1124L (3476G>C), R1143K (3534G>A), A1162V (3591C>T 3592T>C), S1163Q (3593T>C 3594C>A), A1175S (3629G>T), Y1196F (3693A>T), A1202S (3710G>T), I1203V (3713A>G), V1218I (3758G>A), V1224I (3776G>A 3778G>T), P1352A (4160C>G 4162A>T), I1362M (4192T>G), S1537N (4716G>A), S1538N (4719G>A), V1598I (4898G>A), T1614K (4947C>A), N1618H (4958A>C 4960C>T), T1628A (4988A>G 4990C>A), V1642M (5030G>A), I1658T (5079T>C), A1662S (5090G>T), P1667A (5105C>G), V1672I (5120G>A), S1676T (5132T>A), T1751S (5357A>T), K1752Q (5360A>C), H1754N (5366C>A), V1755I (5369G>A 5371G>T), I1765V (5399A>G), V1785I (5459G>A 5461T>C), V1786I (5462G>A), D1794N (5486G>A), T1796P (5492A>C 5494A>T), L1810V (5534C>G), K1811Q (5537A>C), A1813S (5543G>T), E1818Q (5558G>C), I1826V (5582A>G), A1829S (5591G>T), T1833I (5604C>T), Y1836F (5613A>T), R1853K (5664G>A), Q1857H (5677G>T), Y1869H (5711T>C), R1878K (5739G>A), L1882* (5751T>A 5752G>A), M1887I (5767G>T), I1890V (5774A>G 5776A>G), F1900T (5804T>A 5805T>C 5806C>T), Y1907F (5826A>T), L1915Q (5849T>C 5850T>A), D1918E (5860C>A), N1930H (5894A>C), Y1934F (5907A>T), H1946Y (5942C>T 5944C>T), N1950S (5955A>G), T1966V (6002A>G 6003C>T 6004C>T), G1968E (6009G>A), T1976I (6033C>T), A1977S (6035G>T), L1996F (6092C>T), H1997Q (6097T>A), I2002M (6112A>G), E2008D (6130G>T), T2027D (6185A>G 6186C>A 6187T>C), H2028R (6189A>G), M2032I (6202G>C), D2036E (6214T>G), L2037V (6215C>G 6217G>T), T2038P (6218A>C), E2039P (6221G>C 6222A>C), G2040E (6225G>A), I2041V (6227A>G), I2042V (6230A>G 6232A>G), V2045T (6239G>A 6240T>C), S2054T (6266T>A 6268G>T), S2056M (6273G>T 6274C>G), V2062E (6291T>A 6292T>A), V2064L (6296G>C 6298A>T), L2066M (6302T>A 6304A>G), D2069* (6311G>T 6313T>A), I2071V (6317A>G 6319A>T), A2089T (6371G>A), C2100H (6404T>C 6405G>A), R2103L (6414G>T 6415A>G), S2104H (6416T>C 6417C>A 6418A>C), C2111S (6437T>A), S2118T (6458T>A), H2127N (6485C>A), R2128S (6490A>C), H2129N (6491C>A 6493T>C), T2131P (6497A>C 6499A>T), I2132E (6500A>G 6501T>A 6502T>G), S2137P (6515T>C 6517C>A), L2147F (6545C>T), G2154N (6566G>A 6567G>A 6568C>T)

| | Begin | End | Coverage | Score | Concordance | Matches | Identities | I/D/M/F* | Stop Codons |
|---|---|---|---|---|---|---|---|---|---|
| **NT** | **1** | **7803** | **99.9%** | **10368** | **72.5%** | **7431 (95.1%)** | **6475 (82.9%)** | **10/372** | |

Codon mutations:

TTT3TTC (115T>C), TTC7TTT (127C>T), CCC8TCT (128C>T 130C>T), TTC10TCC (135T>C), AAC11AGC (138A>G), GTT13GCT (144T>C), ACT14ATT (147C>T), TGC17AGT (155T>A 157C>T), GCT18GTT (159C>T), ACC19GCC (161A>G), TCT24TCA (178T>A), TCA27TCT (187A>T), GCT31CAA (197G>C 198C>A 199T>A), CGC38CGT (220C>T), TCT52TCA (262T>A), CAC53CAT (265C>T), GTT61GTG (289T>G), ACA62ACT (292A>T), GCT65GCA (301T>A), TCT66GCT (302T>G), TCC67TCT (307C>T), TTT68TTC (310T>C), ACC69TCT (311A>T 313C>T), TGT72TGC (322T>C), AAC73AGT (324A>G 325C>T), TCT74TCA (328T>A), TTG78TTA (340G>A), AAG81AAA (349G>A), ACT84ACA (358T>A), TTA85CTT (359T>C 361A>T), GAA87GAG (367A>G), CCC88CCT (370C>T), TTC89TTT (373C>T), ATC90ATT (376C>T), GCT92GCA (382T>A), TCT94TCC (388T>C), GTC96GTT (394C>T), AAA101AAG (409A>G), TCA102TCC (412A>C), GTT107GTG (427T>G), AAG109AAA (433G>A), AGC113GGC (443A>G), AAA115AAG (451A>G), GAC127GAT (487C>T), GTG132GTA (502G>A), TCT145TCC (541T>C), TTC151TTT (559C>T), CTA158TTA (578C>T), ACT165TCT (599A>T), ATA167ACA (606T>C), AAT170AGT (615A>G), ACA171ATT (618C>T 619A>T), GAC183GAT (655C>T), TTA185TTG (661A>G), CAA190CAG (676A>G), GCA196GCG (694A>G), CTA201TTA (707C>T), GGG203GGA (715G>A), GAA204GAG (718A>G), TCC205GCC (719T>G), TCA207TCT (727A>T), GTG214GTT (748G>T), GTA215GTT (751A>T), CTC218CTT (760C>T), GCC220GCT (766C>T), TCT224TCA (778T>A), ACC225ACT (781C>T), TTC228TTT (790C>T), GTA231GTT (799A>T), CCT232CCA (802T>A), AGA234CGA (806A>C), TCC239ACC (821T>A), CCG241CCA (829G>A), CTA242CTT (832A>T), TCA243TCC (835A>C), CTG246TTG (842C>T), TAC247TAT (847C>T), TTG259TTA (883G>A), TTT260TTC (886T>C), ACC261ACA (889C>A), AAT265AAC (901T>C), TGT269TGC (913T>C), GCC273GCT (925C>T), CTG278TTG (938C>T), GTA282GTG (952A>G), GGT286GGA (964T>A), AAG288AAA (970G>A), GTT291ATT (977G>A), ATT295ATC (991T>C), GCC297GCT (997C>T), TTG299CTG (1001T>C), TCT300TCA (1006T>A), TTC304TTT (1018C>T), ACG305GTG (1019A>G 1020C>T), GAG309GAA (1033G>A), CAT318CAC (1060T>C), CGC319CGA (1063C>A), ACT329ACC (1093T>C), CTA335TTA (1109C>T), AAA341AAG (1129A>G), TCT343TCA (1135T>A), GCC353GCA (1165C>A), AGG358AGA (1180G>A), ACG360ACA (1186G>A), GCA362GCC (1192A>C), ATA363ATT (1195A>T), ACC364TCT (1196A>T 1198C>T), CTT366CTG (1204T>G), GAT367GAG (1207T>G), GGG371GGC (1219G>C), GGC378GGT (1240C>T), CAA381CAG (1249A>G), AGG383AGA (1255G>A), CAA390CAG (1276A>G), GGG398GGA (1300G>A), ACC401ACA (1309C>A), CGG404CGA (1318G>A), CTG405TTG (1319C>T), TTC406TTT (1324C>T), GTT410GTG (1336T>G), TTG411CTT (1337T>C 1339G>T), CCA417CCC (1357A>C), TTG419CTG (1361T>C), TAT435TAC (1411T>C), ATC441ATT (1429C>T), GAT461GAC (1489T>C), CCC472CCA (1522C>A), GCA476GCT (1534A>T), GGC482GGT (1552C>T), CGC483AGG (1553C>A 1555C>G), CGA488AGG (1568C>A 1570A>G), ATA490ATC (1576A>C), GCC492GCT (1582C>T), ACA493ACC (1585A>C), AAC495AAT (1591C>T), GCT501TCT (1607G>T), CAG502CAA (1612G>A), GGT504GGG (1618T>G), CTA505TTA (1619C>T), ACC506ACA (1624C>A), CCC508CCT (1630C>T), AGA513CGA (1643A>C), TGC521TGT (1669C>T), GGT525GGA (1681T>A), GAG526GAA (1684G>A), ACT529ATT (1692C>T), GCT531GCG (1699T>G), TCT534GCC (1706T>G 1708T>C), CTT536CTG (1714T>G), AAA544AGG (1737A>G 1738A>G), AGA545CGA (1739A>C), ACA550ATC (1755C>T 1756A>C), AGC555AAT (1770G>A 1771C>T), TTG559CTG (1781T>C), GAA561GAG (1789A>G), GTT569GTG (1813T>G), ATC573ATA (1825C>A), TCA574TCT (1828A>T), AAA576AAG (1834A>G), TAT578TTT (1839A>T), GCC580ACC (1844G>A), ACA581ACC (1849A>C), TCA582ACA (1850T>A), AAG584AAA (1858G>A), CAG587CAA (1867G>A), GTG590GTC (1876G>C), TCA591ACT (1877T>A 1879A>T), GAG593GAA (1885G>A), ATA595GTG (1889A>G 1891A>G), ATA596ATT (1894A>T), GAT600GAA (1906T>A), ATG601CCA (1907A>C 1908T>C 1909G>A), TTG602TGG (1911T>G), ACC607ACA (1927C>A), GCT611GTT (1938C>T), AAG612AAA (1942G>A), GTT617GTG (1957T>G), GTA618GTC (1960A>C), GAA620GAG (1966A>G), GTG624GTA (1978G>A), TAC626TAT (1984C>T), AAA636AAG (2014A>G), CTA662CTG (2092A>G), AGG672CGG (2120A>C), GTT674GTC (2128T>C), AAT675ACT (2130A>C), GAC676GAT (2134C>T), CTG678TTG (2138C>T), TAT679CAA (2141T>C 2143T>A), CAA680CAG (2146A>G), GGT681GGA (2149T>A), TGT683TGC (2155T>C), CTG690TTG (2174C>T), ATA691ATC (2179A>C), TAT698TAC (2200T>C), ATC706ATT (2224C>T), TAC707TAT (2227C>T), GCT708GCG (2230T>G), AAT712TGT (2240A>T 2241A>G), AGC715AAA (2250G>A 2251C>A), GAC717GAT (2257C>T), CAG719CAA (2263G>A), CTG722TTG (2270C>T), GGA725GGT (2281A>T), ATG726TTG (2282A>T), GCG727GCA (2287G>A), CAC728CAT (2290C>T), ATA731ATT (2299A>T), CTG738TTG (2318C>T), ATA739ATT (2323A>T), GCC746GCA (2344C>A), CAC747CAT (2347C>T), GTA748GTG (2350A>G), TTC750TTT (2356C>T), TTG751TTA (2359G>A), ACA753ACT (2365A>T), GCA754GCT (2368A>T), AGG757CGG (2375A>C), TAT758TAC (2380T>C), CGC761AGA (2387C>A 2389C>A), AGA762AGG (2392A>G), CCT767CCC (2407T>C), AGG768AAG (2409G>A), GTG780GTA (2446G>A), CCG791CCA (2479G>A), ATC793ATT (2485C>T), TTG794TTA (2488G>A), AGA795AGG (2491A>G), GTT796GTG (2494T>G), CTG797CTC (2497G>C), ATT798ATC (2500T>C), AAG799AAA (2503G>A), GCT800GCA (2506T>A), GGC801GGT (2509C>T), CTT802CTC (2512T>C), TTA804CTA (2516T>C), GCA806GCC (2524A>C), GGG812GGT (2542G>T), GGA833GGT (2605A>T), ACC836ACA (2614C>A), GAG838GAA (2620G>A), AAT840AAC (2626T>C), CGG845AGG (2639C>A), AAG848AAA (2650G>A), ATC852ATT (2662C>T), AAG862AAA (2692G>A), GCA870GCC (2716A>C), GGC874AGC (2726G>A), TTC875TTT (2731C>T), ATC876ATT (2734C>T), GGA879GGC (2743A>C), TTG880CTG (2744T>C), TTG887CTC (2765T>C 2767G>C), AGA888CGA (2768A>C), CTA889CTG (2773A>G), GGT890GGA (2776T>A), GTC900GTT (2806C>T), TGC901TGT (2809C>T), TCG902TCA (2812G>A), GTG905GTA (2821G>A), ACA907ACT (2827A>T), AGC914AGT (2848C>T), CGT916AGA (2852C>A 2854T>A), GGC918GGG (2860C>G), ACT920ACA (2866T>A), TTC921TTT (2869C>T), ATA923ATT (2875A>T), ATA924ATC (2878A>C), CAT925CAC (2881T>C), GCC926TCC (2882G>T), TCA928TCC (2890A>C), GCC936GCT (2914C>T), CGG940CGA (2926G>A), TTC941TTT (2929C>T), TTG942TTA (2932G>A), ATT947ATA (2947T>A), GTC948GTT (2950C>T), ATT962ATC (2992T>C), AAA964AAG (2998A>G), TCT975TCA (3031T>A), GTA981GTC (3049A>C), GTA986GTT (3064A>T), GCT991GCC (3079T>C), CAA993AAG (3083C>A 3085A>G), TTG999CTT (3101T>C 3103G>T), TTT1001TTC (3109T>C), TCC1006TCA (3124C>A), CTT1010CTG (3136T>G), ACG1014TCG (3146A>T), TAC1015TAT (3151C>T), AGT1020AGC (3166T>C), TCC1021TCT (3169C>T), TTT1022TTC (3172T>C), AAA1023AAG (3175A>G), CTG1024TTG (3176C>T), TCT1026TCA (3184T>A), TAT1032TAC (3202T>C), CTT1044CTG (3238T>G), CTT1045CTA (3241T>A), CTT1046TTT (3242C>T), AAC1049AAT (3253C>T), AAA1053AAG (3265A>G), ATT1057ATA (3277T>A), CAT1061CAC (3289T>C), TCT1064TCA (3298T>A), GTA1075GTT (3331A>T), GCT1081GCA (3349T>A), TTG1087CTG (3365T>C), CTT1090CTG (3376T>G), CCG1091CCA (3379G>A), ACT1092ACA (3382T>A), CCG1094CCT (3388G>T), GCA1098GCC (3400A>C), GTA1100GTG (3406A>G), CAG1103GAG (3413C>G), CTC1104CTA (3418C>A), CTT1105CTC (3421T>C), CTA1108CTG (3430A>G), CGA1109AGA (3431C>A), GCG1113GCT (3445G>T), CTT1115CTC (3451T>C), CAG1120CAA (3466G>A), GTT1124CTT (3476G>C), GGC1125GGT (3481C>T), ACA1126ACT (3484A>T), ACG1134ACT (3508G>T), AAA1138AAG (3520A>G), CTG1141CTC (3529G>C), CGA1142AGG (3530C>A 3532A>G), AGG1143AAG (3534G>A), GGC1145GGT (3541C>T), TTG1146CTT (3542T>C 3544G>T), GCT1149GCA (3553T>A), GGG1151GGA (3559G>A), TCT1158TCC (3580T>C), ATC1159ATA (3583C>A), TCA1161TCG (3589A>G), GCT1162GTC (3591C>T 3592T>C), TCA1163CAA (3593T>C 3594C>A), GAA1168GAG (3610A>G), GGT1169GGC (3613T>C), CCG1172CCA (3622G>A), TAC1173TAT (3625C>T), GTG1174GTA (3628G>A), GCT1175TCT (3629G>T), TAC1180TAT (3646C>T), GCA1184GCC (3658A>C), CAA1185CAG (3661A>G), GAC1190GAT (3676C>T), ATT1191ATA (3679T>A), GCG1192GCC (3682G>C), CTA1194CTG (3688A>G), GCC1195GCT (3691C>T), TAT1196TTT (3693A>T), CTA1199TTA (3701C>T), GCA1202TCA (3710G>T), ATT1203GTT (3713A>G), TTG1204CTG (3716T>C), CGA1211CGT (3739A>T), GTT1212GTG (3742T>G), AAG1216AAA (3754G>A), GTC1218ATC (3758G>A), CAG1220CAA (3766G>A), GTG1224ATT (3776G>A 3778G>T), GTG1225GTT (3781G>T), CTC1226CTG (3784C>G), GGT1228GGA (3790T>A), GAC1236GAT (3814C>T), TTG1240CTG (3824T>C), CCG1247CCA (3847G>A), AAG1250AAA (3856G>A), GGC1254GGT (3868C>T), ACT1255ACC (3871T>C), GAG1259GAA (3883G>A), CCA1263CCC (3895A>C), GGA1267GGT (3907A>T), GAT1268GAC (3910T>C), CGA1269AGA (3911C>A), GAC1281GAT (3949C>T), GAG1283GAA (3955G>A), GCT1284GCA (3958T>A), TTG1285CTA (3959T>C 3961G>A), ATT1289ATC (3973T>C), GGA1292GGC (3982A>C), AAA1304AAG (4018A>G), TTG1308CTG (4028T>C), CCG1309CCT (4033G>T), TTG1310CTG (4034T>C), CGG1317AGG (4055C>A), TTG1321CTG (4067T>C), ATT1324ATA (4078T>A), CCC1326CCT (4084C>T), TAC1329TAT (4093C>T), CTG1338TTG (4118C>T), TCC1341TCT (4129C>T), TCC1347TCA (4147C>A), CTG1351TTA (4157C>T 4159G>A), CCA1352GCT (4160C>G 4162A>T), GGT1361GGA (4189T>A), ATT1362ATG (4192T>G), TTG1367TTA (4207G>A), GCT1368GCC (4210T>C), GCA1373GCT (4225A>T), GTC1374GTT (4228C>T), ATC1392ATA (4282C>A), GCC1394GCA (4288C>A), TTT1395TTC (4291T>C), CAA1396CAG (4294A>G), CTT1401CTG (4309T>G), GAC1404GAT (4318C>T), GGA1407GGT (4327A>T), GGT1411GGC (4339T>C), GAC1414GAT (4348C>T), AGA1418CGC (4358A>C 4360A>C), GCT1425GCC (4381T>C), TCA1426TCC (4384A>C), TAT1429TAC (4393T>C), GTT1436GTG (4414T>G), TTT1437TTC (4417T>C), TTG1439CTG (4421T>C), GTG1440GTT (4426G>T), GGA1441GGG (4429A>G), TCA1445TCT (4441A>T), GTT1451GTC (4459T>C), AAC1454AAT (4468C>T), TCT1455TCC (4471T>C), GCA1465GCT (4501A>T), TCT1468TCA (4510T>A), GCT1477GCA (4537T>A), GGG1481GGC (4549G>C), AAA1483AAG (4555A>G), GTC1494GTT (4588C>T), CTA1500CTT (4606A>T), CCC1504CCA (4618C>A), TTG1507CTG (4625T>C), TAT1510TAC (4636T>C), GCT1514GCA (4648T>A), ATT1515ATA (4651T>A), GCC1516GCT (4654C>T), TTG1519CTA (4661T>C 4663G>A), GTT1524GTC (4678T>C), ATA1526ATT (4684A>T), ATC1530ATT (4696C>T), GAC1531GAT (4699C>T), GTT1536GTC (4714T>C), AGC1537AAC (4716G>A), AGT1538AAT (4719G>A), GAC1543GAT (4735C>T), TTG1544CTG (4736T>C), CTA1547CTG (4747A>G), TTT1549TTC (4753T>C), CTA1550CTT (4756A>T), AGC1551AGT (4759C>T), GGT1553GGC (4765T>C), TTT1554TTC (4768T>C), AGG1556AGA (4774G>A), CGT1558CGC (4780T>C), CTA1559CTG (4783A>G), GGA1560GGG (4786A>G), TAC1561TAT (4789C>T), CCT1565CCC (4801T>C), CTG1566TTG (4802C>T), GTA1569GTG (4813A>G), TCT1570TCC (4816T>C), CCA1574CCC (4828A>C), TAC1576TAT (4834C>T), AGG1579AGA (4843G>A), CCC1585CCT (4861C>T), CTC1588CTG (4870C>G), AAC1593AAT (4885C>T), ACG1596ACT (4894G>T), GGA1597GGC (4897A>C), GTT1598ATT (4898G>A), GAG1600GAA (4906G>A), GCA1601GCC (4909A>C), GGG1605GGT (4921G>T), ACA1614AAA (4947C>A), GTT1616GTG (4954T>G), AAC1618CAT (4958A>C 4960C>T), AAG1621AAA (4969G>A), CGG1623CGA (4975G>A), TTC1625TTT (4981C>T), GCC1627GCT (4987C>T), ACC1628GCA (4988A>G 4990C>A), CCT1631CCC (4999T>C), ACA1632ACT (5002A>T), TAT1633TAC (5005T>C), TTG1636CTG (5012T>C), GAG1637GAA (5017G>A), CAG1638CAA (5020G>A), GAC1639GAT (5023C>T), GTG1642ATG (5030G>A), GAG1643GAA (5035G>A), ACA1645ACT (5041A>T), GGA1646GGC (5044A>C), GGG1651GGT (5059G>T), ATC1653ATT (5065C>T), CTT1656CTC (5074T>C), CAT1657CAC (5077T>C), ATT1658ACT (5079T>C), GCA1659GCT (5083A>T), GCT1662TCT (5090G>T), TTG1666CTG (5102T>C), CCC1667GCC (5105C>G), GGC1669GGT (5113C>T), AAT1670AAC (5116T>C), ACT1671ACC (5119T>C), GTT1672ATT (5120G>A), CTG1673TTG (5123C>T), TTT1675TTC (5131T>C), TCT1676ACT (5132T>A), GTG1679GTT (5143G>T), CAT1680CAC (5146T>C), GAC1684GAT (5158C>T), AGT1687AGC (5167T>C), TGC1689TGT (5173C>T), TCC1690TCT (5176C>T), CGA1692CGG (5182A>G), GCA1696GCC (5194A>C), CAG1698CAA (5200G>A), GAC1700GAT (5206C>T), ACC1701ACA (5209C>A), GTT1704GTG (5218T>G), AAC1707AAT (5227C>T), GCA1711GCC (5239A>C), CCC1720CCA (5266C>A), ACC1721ACA (5269C>A), AGG1724AGA (5278G>A), GGG1728GGT (5290G>T), CTT1729CTC (5293T>C), CCT1732CCA (5302T>A), CAC1741CAT (5329C>T), GTA1746GTG (5344A>G), ACC1747ACA (5347C>A), ACC1751TCC (5357A>T), AAA1752CAA (5360A>C), CAT1754AAT (5366C>A), GTG1755ATT (5369G>A 5371G>T), GTT1756GTA (5374T>A), GGC1760GGG (5386C>G), GGG1762GGA (5392G>A), ATT1765GTT (5399A>G), CCA1767CCT (5407A>T), GTG1768GTT (5410G>T), TGC1769TGT (5413C>T), GCA1771GCT (5419A>T), CTG1775TTG (5429C>T), ACT1777ACC (5437T>C), GGC1778GGG (5440C>G), GTT1785ATC (5459G>A 5461T>C), GTT1786ATT (5462G>A), TTG1789CTG (5471T>C), ATA1792ATT (5482A>T), GCA1793GCT (5485A>T), GAT1794AAT (5486G>A), GTT1795GTC (5491T>C), ACA1796CCT (5492A>C 5494A>T), CTC1798TTG (5498C>T 5500C>G), CCA1802CCT (5512A>T), GGA1803GGT (5515A>T), CTT1809TTG (5531C>T 5533T>G), CTG1810GTG (5534C>G), AAA1811CAA (5537A>C), GCT1813TCT (5543G>T), GAA1818CAA (5558G>C), CTT1820CTG (5566T>G), GCA1821GCT (5569A>T), GAT1822GAC (5572T>C), TGT1824TGC (5578T>C), ATT1826GTT (5582A>G), GCC1829TCC (5591G>T), GAG1832GAA (5602G>A), ACA1833ATA (5604C>T), GCA1835GCT (5611A>T), TAC1836TTC (5613A>T), ATA1837ATT (5617A>T), CCT1838CCA (5620T>A), GGA1840GGT (5626A>T), CTT1844CTG (5638T>G), GGT1845GGC (5641T>C), TTT1847TTC (5647T>C), AGA1853AAA (5664G>A), CTA1856CTT (5674A>T), CAG1857CAT (5677G>T), CAG1860CAA (5686G>A), CGC1861CGA (5689C>A), GGA1867GGT (5707A>T), CCT1868CCA (5710T>A), TAT1869CAT (5711T>C), GGG1870GGT (5716G>T), GGA1871GGT (5719A>T), CAC1875CAT (5731C>T), ACT1876ACC (5734T>C), ACT1877ACC (5737T>C), AGA1878AAA (5739G>A), CTT1880CTC (5746T>C), TTG1882TAA (5751T>A 5752G>A), ATG1887ATT (5767G>T), AAG1889AAA (5773G>A), ATA1890GTG (5774A>G 5776A>G), TAT1891TAC (5779T>C), TTA1894CTA (5786T>C), GTC1895GTA (5791C>A), AGA1897AGG (5797A>G), TTC1900ACT (5804T>A 5805T>C 5806C>T), CTC1903CTG (5815C>G), TAC1904TAT (5818C>T), TAT1907TTT (5826A>T), GTG1913GTA (5845G>A), TTG1915CAG (5849T>C 5850T>A), GTT1917GTG (5857T>G), GAC1918GAA (5860C>A), CTT1920CTG (5866T>G), AGG1921AGA (5869G>A), CTC1922CTG (5872C>G), GGG1924GGT (5878G>T), TTT1925TTC (5881T>C), CCC1927CCT (5887C>T), CGC1928CGT (5890C>T), AAT1930CAT (5894A>C), TAC1934TTC (5907A>T), CTT1936CTC (5914T>C), TTT1939TTC (5923T>C), CTG1941TTG (5927C>T), CAC1946TAT (5942C>T 5944C>T), AAC1950AGC (5955A>G), ACG1953ACA (5965G>A), GTT1954GTG (5968T>G), TTT1956TTC (5974T>C), AGA1957CGA (5975A>C), AAG1958AAA (5980G>A), CCT1959CCC (5983T>C), TTT1960TTC (5986T>C), AAA1964AAG (5998A>G), ACC1966GTT (6002A>G 6003C>T 6004C>T), GGG1968GAG (6009G>A), AAA1970AAG (6016A>G), TCC1974TCT (6028C>T), GCA1975GCT (6031A>T), ACA1976ATA (6033C>T), GCT1977TCT (6035G>T), GCT1978GCA (6040T>A), GGT1983GGG (6055T>G), GCT1988GCC (6070T>C), CTC1991CTG (6079C>G), AGT1993AGC (6085T>C), CTA1995TTG (6089C>T 6091A>G), CTT1996TTT (6092C>T), CAT1997CAA (6097T>A), CTT1998CTG (6100T>G), CAG1999CAA (6103G>A), AAG2000AAA (6106G>A), ATA2002ATG (6112A>G), TGT2004TGC (6118T>C), TCA2005TCT (6121A>T), GAG2008GAT (6130G>T), TTT2011TTC (6139T>C), GTC2016GTA (6154C>A), GGC2019GGG (6163C>G), CAC2020CAT (6166C>T), GTA2026GTG (6184A>G), ACT2027GAC (6185A>G 6186C>A 6187T>C), CAC2028CGC (6189A>G), TTC2029TTT (6193C>T), CAA2030CAG (6196A>G), CGC2031CGT (6199C>T), ATG2032ATC (6202G>C), GAT2036GAG (6214T>G), CTG2037GTT (6215C>G 6217G>T), ACA2038CCA (6218A>C), GAA2039CCA (6221G>C 6222A>C), GGG2040GAG (6225G>A), ATC2041GTC (6227A>G), ATA2042GTG (6230A>G 6232A>G), GAG2043GAA (6235G>A), GTG2045ACG (6239G>A 6240T>C), TTA2049CTT (6251T>C 6253A>T), ACC2050ACA (6256C>A), ACA2052ACT (6262A>T), CTG2053TTA (6263C>T 6265G>A), TCG2054ACT (6266T>A 6268G>T), AGC2056ATG (6273G>T 6274C>G), GTT2062GAA (6291T>A 6292T>A), GTA2064CTT (6296G>C 6298A>T), CCT2065CCA (6301T>A), TTA2066ATG (6302T>A 6304A>G), TTG2068CTG (6308T>C), GAT2069TAA (6311G>T 6313T>A), ATA2071GTT (6317A>G 6319A>T), GAG2072GAA (6322G>A), GGT2077GGC (6337T>C), CTA2080TTA (6344C>T), GCA2089ACA (6371G>A), GCC2092GCT (6382C>T), CTC2097CTG (6397C>G), TGT2100CAT (6404T>C 6405G>A), CGA2103CTG (6414G>T 6415A>G), TCA2104CAC (6416T>C 6417C>A 6418A>C), GTG2109GTT (6433G>T), TTC2110TTT (6436C>T), TGT2111AGT (6437T>A), TAC2113TAT (6445C>T), ACG2114ACA (6448G>A), TCC2117TCA (6457C>A), TCT2118ACT (6458T>A), CTT2119TTA (6461C>T 6463T>A), CTT2125CTA (6481T>A), CAT2127AAT (6485C>A), AGA2128AGC (6490A>C), CAT2129AAC (6491C>A 6493T>C), CAG2130CAA (6496G>A), ACA2131CCT (6497A>C 6499A>T), ATT2132GAG (6500A>G 6501T>A 6502T>G), CCG2136CCT (6514G>T), TCC2137CCA (6515T>C 6517C>A), TTC2138TTT (6520C>T), CAC2142CAT (6532C>T), CTT2147TTT (6545C>T), CCA2150CCC (6556A>C), CTA2152CTT (6562A>T), GGC2154AAT (6566G>A 6567G>A 6568C>T), TAC2155TAT (6571C>T), AAA2156AAG (6574A>G)

| | Begin | End | Coverage | Score | Concordance | Matches | Identities | I/D/M/F* | Stop Codons |
|---|---|---|---|---|---|---|---|---|---|
| **NT** | **1** | **7803** | **99.9%** | **10368** | **72.5%** | **7431 (95.1%)** | **6475 (82.9%)** | **10/372** | |

**Proteins**

| | Begin | End | Coverage | Score | Concordance | Matches | Identities | I/D/M/F* | Stop Codons |
|---|---|---|---|---|---|---|---|---|---|
| polyprotein (YP_001039627.1) | 1 | 2159 | 100% | 14226 | 94.4% | 2159 (100%) | 2013 (93.2%) | 0/0/0/0 | 3 |

Protein mutations: P8S (128C>T 130C>T), F10S (135T>C), N11S (138A>G), V13A (144T>C), T14I (147C>T), C17S (155T>A 157C>T), A18V (159C>T), T19A (161A>G), A31Q (197G>C 198C>A 199T>A), S66A (302T>G), T69S (311A>T 313C>T), N73S (324A>G 325C>T), S113G (443A>G), T165S (599A>T), I167T (606T>C), N170S (615A>G), T171I (618C>T 619A>T), S205A (719T>G), S239T (821T>A), V291I (977G>A), T305V (1019A>G 1020C>T), T364S (1196A>T 1198C>T), D367E (1207T>G), A501S (1607G>T), T529I (1692C>T), S534A (1706T>G 1708T>C), K544R (1737A>G 1738A>G), T550I (1755C>T 1756A>C), S555N (1770G>A 1771C>T), Y578F (1839A>T), A580T (1844G>A), S582T (1850T>A), S591T (1877T>A 1879A>T), I595V (1889A>G 1891A>G), D600E (1906T>A), M601P (1907A>C 1908T>C 1909G>A), L602W (1911T>G), A611V (1938C>T), N675T (2130A>C), Y679Q (2141T>C 2143T>A), N712C (2240A>T 2241A>G), S715K (2250G>A 2251C>A), M726L (2282A>T), R768K (2409G>A), G874S (2726G>A), A926S (2882G>T), Q993K (3083C>A 3085A>G), T1014S (3146A>T), L1046F (3242C>T), Q1103E (3413C>G), V1124L (3476G>C), R1143K (3534G>A), A1162V (3591C>T 3592T>C), S1163Q (3593T>C 3594C>A), A1175S (3629G>T), Y1196F (3693A>T), A1202S (3710G>T), I1203V (3713A>G), V1218I (3758G>A), V1224I (3776G>A 3778G>T), P1352A (4160C>G 4162A>T), I1362M (4192T>G), S1537N (4716G>A), S1538N (4719G>A), V1598I (4898G>A), T1614K (4947C>A), N1618H (4958A>C 4960C>T), T1628A (4988A>G 4990C>A), V1642M (5030G>A), I1658T (5079T>C), A1662S (5090G>T), P1667A (5105C>G), V1672I (5120G>A), S1676T (5132T>A), T1751S (5357A>T), K1752Q (5360A>C), H1754N (5366C>A), V1755I (5369G>A 5371G>T), I1765V (5399A>G), V1785I (5459G>A 5461T>C), V1786I (5462G>A), D1794N (5486G>A), T1796P (5492A>C 5494A>T), L1810V (5534C>G), K1811Q (5537A>C), A1813S (5543G>T), E1818Q (5558G>C), I1826V (5582A>G), A1829S (5591G>T), T1833I (5604C>T), Y1836F (5613A>T), R1853K (5664G>A), Q1857H (5677G>T), Y1869H (5711T>C), R1878K (5739G>A), L1882* (5751T>A 5752G>A), M1887I (5767G>T), I1890V (5774A>G 5776A>G), F1900T (5804T>A 5805T>C 5806C>T), Y1907F (5826A>T), L1915Q (5849T>C 5850T>A), D1918E (5860C>A), N1930H (5894A>C), Y1934F (5907A>T), H1946Y (5942C>T 5944C>T), N1950S (5955A>G), T1966V (6002A>G 6003C>T 6004C>T), G1968E (6009G>A), T1976I (6033C>T), A1977S (6035G>T), L1996F (6092C>T), H1997Q (6097T>A), I2002M (6112A>G), E2008D (6130G>T), T2027D (6185A>G 6186C>A 6187T>C), H2028R (6189A>G), M2032I (6202G>C), D2036E (6214T>G), L2037V (6215C>G 6217G>T), T2038P (6218A>C), E2039P (6221G>C 6222A>C), G2040E (6225G>A), I2041V (6227A>G), I2042V (6230A>G 6232A>G), V2045T (6239G>A 6240T>C), S2054T (6266T>A 6268G>T), S2056M (6273G>T 6274C>G), V2062E (6291T>A 6292T>A), V2064L (6296G>C 6298A>T), L2066M (6302T>A 6304A>G), D2069* (6311G>T 6313T>A), I2071V (6317A>G 6319A>T), A2089T (6371G>A), C2100H (6404T>C 6405G>A), R2103L (6414G>T 6415A>G), S2104H (6416T>C 6417C>A 6418A>C), C2111S (6437T>A), S2118T (6458T>A), H2127N (6485C>A), R2128S (6490A>C), H2129N (6491C>A 6493T>C), T2131P (6497A>C 6499A>T), I2132E (6500A>G 6501T>A 6502T>G), S2137P (6515T>C 6517C>A), L2147F (6545C>T), G2154N (6566G>A 6567G>A 6568C>T)

| | Begin | End | Coverage | Score | Concordance | Matches | Identities | I/D/M/F* | Stop Codons |
|---|---|---|---|---|---|---|---|---|---|
| **NT** | **1** | **7803** | **99.9%** | **10368** | **72.5%** | **7431 (95.1%)** | **6475 (82.9%)** | **10/372** | |

Codon mutations:

TTT3TTC (115T>C), TTC7TTT (127C>T), CCC8TCT (128C>T 130C>T), TTC10TCC (135T>C), AAC11AGC (138A>G), GTT13GCT (144T>C), ACT14ATT (147C>T), TGC17AGT (155T>A 157C>T), GCT18GTT (159C>T), ACC19GCC (161A>G), TCT24TCA (178T>A), TCA27TCT (187A>T), GCT31CAA (197G>C 198C>A 199T>A), CGC38CGT (220C>T), TCT52TCA (262T>A), CAC53CAT (265C>T), GTT61GTG (289T>G), ACA62ACT (292A>T), GCT65GCA (301T>A), TCT66GCT (302T>G), TCC67TCT (307C>T), TTT68TTC (310T>C), ACC69TCT (311A>T 313C>T), TGT72TGC (322T>C), AAC73AGT (324A>G 325C>T), TCT74TCA (328T>A), TTG78TTA (340G>A), AAG81AAA (349G>A), ACT84ACA (358T>A), TTA85CTT (359T>C 361A>T), GAA87GAG (367A>G), CCC88CCT (370C>T), TTC89TTT (373C>T), ATC90ATT (376C>T), GCT92GCA (382T>A), TCT94TCC (388T>C), GTC96GTT (394C>T), AAA101AAG (409A>G), TCA102TCC (412A>C), GTT107GTG (427T>G), AAG109AAA (433G>A), AGC113GGC (443A>G), AAA115AAG (451A>G), GAC127GAT (487C>T), GTG132GTA (502G>A), TCT145TCC (541T>C), TTC151TTT (559C>T), CTA158TTA (578C>T), ACT165TCT (599A>T), ATA167ACA (606T>C), AAT170AGT (615A>G), ACA171ATT (618C>T 619A>T), GAC183GAT (655C>T), TTA185TTG (661A>G), CAA190CAG (676A>G), GCA196GCG (694A>G), CTA201TTA (707C>T), GGG203GGA (715G>A), GAA204GAG (718A>G), TCC205GCC (719T>G), TCA207TCT (727A>T), GTG214GTT (748G>T), GTA215GTT (751A>T), CTC218CTT (760C>T), GCC220GCT (766C>T), TCT224TCA (778T>A), ACC225ACT (781C>T), TTC228TTT (790C>T), GTA231GTT (799A>T), CCT232CCA (802T>A), AGA234CGA (806A>C), TCC239ACC (821T>A), CCG241CCA (829G>A), CTA242CTT (832A>T), TCA243TCC (835A>C), CTG246TTG (842C>T), TAC247TAT (847C>T), TTG259TTA (883G>A), TTT260TTC (886T>C), ACC261ACA (889C>A), AAT265AAC (901T>C), TGT269TGC (913T>C), GCC273GCT (925C>T), CTG278TTG (938C>T), GTA282GTG (952A>G), GGT286GGA (964T>A), AAG288AAA (970G>A), GTT291ATT (977G>A), ATT295ATC (991T>C), GCC297GCT (997C>T), TTG299CTG (1001T>C), TCT300TCA (1006T>A), TTC304TTT (1018C>T), ACG305GTG (1019A>G 1020C>T), GAG309GAA (1033G>A), CAT318CAC (1060T>C), CGC319CGA (1063C>A), ACT329ACC (1093T>C), CTA335TTA (1109C>T), AAA341AAG (1129A>G), TCT343TCA (1135T>A), GCC353GCA (1165C>A), AGG358AGA (1180G>A), ACG360ACA (1186G>A), GCA362GCC (1192A>C), ATA363ATT (1195A>T), ACC364TCT (1196A>T 1198C>T), CTT366CTG (1204T>G), GAT367GAG (1207T>G), GGG371GGC (1219G>C), GGC378GGT (1240C>T), CAA381CAG (1249A>G), AGG383AGA (1255G>A), CAA390CAG (1276A>G), GGG398GGA (1300G>A), ACC401ACA (1309C>A), CGG404CGA (1318G>A), CTG405TTG (1319C>T), TTC406TTT (1324C>T), GTT410GTG (1336T>G), TTG411CTT (1337T>C 1339G>T), CCA417CCC (1357A>C), TTG419CTG (1361T>C), TAT435TAC (1411T>C), ATC441ATT (1429C>T), GAT461GAC (1489T>C), CCC472CCA (1522C>A), GCA476GCT (1534A>T), GGC482GGT (1552C>T), CGC483AGG (1553C>A 1555C>G), CGA488AGG (1568C>A 1570A>G), ATA490ATC (1576A>C), GCC492GCT (1582C>T), ACA493ACC (1585A>C), AAC495AAT (1591C>T), GCT501TCT (1607G>T), CAG502CAA (1612G>A), GGT504GGG (1618T>G), CTA505TTA (1619C>T), ACC506ACA (1624C>A), CCC508CCT (1630C>T), AGA513CGA (1643A>C), TGC521TGT (1669C>T), GGT525GGA (1681T>A), GAG526GAA (1684G>A), ACT529ATT (1692C>T), GCT531GCG (1699T>G), TCT534GCC (1706T>G 1708T>C), CTT536CTG (1714T>G), AAA544AGG (1737A>G 1738A>G), AGA545CGA (1739A>C), ACA550ATC (1755C>T 1756A>C), AGC555AAT (1770G>A 1771C>T), TTG559CTG (1781T>C), GAA561GAG (1789A>G), GTT569GTG (1813T>G), ATC573ATA (1825C>A), TCA574TCT (1828A>T), AAA576AAG (1834A>G), TAT578TTT (1839A>T), GCC580ACC (1844G>A), ACA581ACC (1849A>C), TCA582ACA (1850T>A), AAG584AAA (1858G>A), CAG587CAA (1867G>A), GTG590GTC (1876G>C), TCA591ACT (1877T>A 1879A>T), GAG593GAA (1885G>A), ATA595GTG (1889A>G 1891A>G), ATA596ATT (1894A>T), GAT600GAA (1906T>A), ATG601CCA (1907A>C 1908T>C 1909G>A), TTG602TGG (1911T>G), ACC607ACA (1927C>A), GCT611GTT (1938C>T), AAG612AAA (1942G>A), GTT617GTG (1957T>G), GTA618GTC (1960A>C), GAA620GAG (1966A>G), GTG624GTA (1978G>A), TAC626TAT (1984C>T), AAA636AAG (2014A>G), CTA662CTG (2092A>G), AGG672CGG (2120A>C), GTT674GTC (2128T>C), AAT675ACT (2130A>C), GAC676GAT (2134C>T), CTG678TTG (2138C>T), TAT679CAA (2141T>C 2143T>A), CAA680CAG (2146A>G), GGT681GGA (2149T>A), TGT683TGC (2155T>C), CTG690TTG (2174C>T), ATA691ATC (2179A>C), TAT698TAC (2200T>C), ATC706ATT (2224C>T), TAC707TAT (2227C>T), GCT708GCG (2230T>G), AAT712TGT (2240A>T 2241A>G), AGC715AAA (2250G>A 2251C>A), GAC717GAT (2257C>T), CAG719CAA (2263G>A), CTG722TTG (2270C>T), GGA725GGT (2281A>T), ATG726TTG (2282A>T), GCG727GCA (2287G>A), CAC728CAT (2290C>T), ATA731ATT (2299A>T), CTG738TTG (2318C>T), ATA739ATT (2323A>T), GCC746GCA (2344C>A), CAC747CAT (2347C>T), GTA748GTG (2350A>G), TTC750TTT (2356C>T), TTG751TTA (2359G>A), ACA753ACT (2365A>T), GCA754GCT (2368A>T), AGG757CGG (2375A>C), TAT758TAC (2380T>C), CGC761AGA (2387C>A 2389C>A), AGA762AGG (2392A>G), CCT767CCC (2407T>C), AGG768AAG (2409G>A), GTG780GTA (2446G>A), CCG791CCA (2479G>A), ATC793ATT (2485C>T), TTG794TTA (2488G>A), AGA795AGG (2491A>G), GTT796GTG (2494T>G), CTG797CTC (2497G>C), ATT798ATC (2500T>C), AAG799AAA (2503G>A), GCT800GCA (2506T>A), GGC801GGT (2509C>T), CTT802CTC (2512T>C), TTA804CTA (2516T>C), GCA806GCC (2524A>C), GGG812GGT (2542G>T), GGA833GGT (2605A>T), ACC836ACA (2614C>A), GAG838GAA (2620G>A), AAT840AAC (2626T>C), CGG845AGG (2639C>A), AAG848AAA (2650G>A), ATC852ATT (2662C>T), AAG862AAA (2692G>A), GCA870GCC (2716A>C), GGC874AGC (2726G>A), TTC875TTT (2731C>T), ATC876ATT (2734C>T), GGA879GGC (2743A>C), TTG880CTG (2744T>C), TTG887CTC (2765T>C 2767G>C), AGA888CGA (2768A>C), CTA889CTG (2773A>G), GGT890GGA (2776T>A), GTC900GTT (2806C>T), TGC901TGT (2809C>T), TCG902TCA (2812G>A), GTG905GTA (2821G>A), ACA907ACT (2827A>T), AGC914AGT (2848C>T), CGT916AGA (2852C>A 2854T>A), GGC918GGG (2860C>G), ACT920ACA (2866T>A), TTC921TTT (2869C>T), ATA923ATT (2875A>T), ATA924ATC (2878A>C), CAT925CAC (2881T>C), GCC926TCC (2882G>T), TCA928TCC (2890A>C), GCC936GCT (2914C>T), CGG940CGA (2926G>A), TTC941TTT (2929C>T), TTG942TTA (2932G>A), ATT947ATA (2947T>A), GTC948GTT (2950C>T), ATT962ATC (2992T>C), AAA964AAG (2998A>G), TCT975TCA (3031T>A), GTA981GTC (3049A>C), GTA986GTT (3064A>T), GCT991GCC (3079T>C), CAA993AAG (3083C>A 3085A>G), TTG999CTT (3101T>C 3103G>T), TTT1001TTC (3109T>C), TCC1006TCA (3124C>A), CTT1010CTG (3136T>G), ACG1014TCG (3146A>T), TAC1015TAT (3151C>T), AGT1020AGC (3166T>C), TCC1021TCT (3169C>T), TTT1022TTC (3172T>C), AAA1023AAG (3175A>G), CTG1024TTG (3176C>T), TCT1026TCA (3184T>A), TAT1032TAC (3202T>C), CTT1044CTG (3238T>G), CTT1045CTA (3241T>A), CTT1046TTT (3242C>T), AAC1049AAT (3253C>T), AAA1053AAG (3265A>G), ATT1057ATA (3277T>A), CAT1061CAC (3289T>C), TCT1064TCA (3298T>A), GTA1075GTT (3331A>T), GCT1081GCA (3349T>A), TTG1087CTG (3365T>C), CTT1090CTG (3376T>G), CCG1091CCA (3379G>A), ACT1092ACA (3382T>A), CCG1094CCT (3388G>T), GCA1098GCC (3400A>C), GTA1100GTG (3406A>G), CAG1103GAG (3413C>G), CTC1104CTA (3418C>A), CTT1105CTC (3421T>C), CTA1108CTG (3430A>G), CGA1109AGA (3431C>A), GCG1113GCT (3445G>T), CTT1115CTC (3451T>C), CAG1120CAA (3466G>A), GTT1124CTT (3476G>C), GGC1125GGT (3481C>T), ACA1126ACT (3484A>T), ACG1134ACT (3508G>T), AAA1138AAG (3520A>G), CTG1141CTC (3529G>C), CGA1142AGG (3530C>A 3532A>G), AGG1143AAG (3534G>A), GGC1145GGT (3541C>T), TTG1146CTT (3542T>C 3544G>T), GCT1149GCA (3553T>A), GGG1151GGA (3559G>A), TCT1158TCC (3580T>C), ATC1159ATA (3583C>A), TCA1161TCG (3589A>G), GCT1162GTC (3591C>T 3592T>C), TCA1163CAA (3593T>C 3594C>A), GAA1168GAG (3610A>G), GGT1169GGC (3613T>C), CCG1172CCA (3622G>A), TAC1173TAT (3625C>T), GTG1174GTA (3628G>A), GCT1175TCT (3629G>T), TAC1180TAT (3646C>T), GCA1184GCC (3658A>C), CAA1185CAG (3661A>G), GAC1190GAT (3676C>T), ATT1191ATA (3679T>A), GCG1192GCC (3682G>C), CTA1194CTG (3688A>G), GCC1195GCT (3691C>T), TAT1196TTT (3693A>T), CTA1199TTA (3701C>T), GCA1202TCA (3710G>T), ATT1203GTT (3713A>G), TTG1204CTG (3716T>C), CGA1211CGT (3739A>T), GTT1212GTG (3742T>G), AAG1216AAA (3754G>A), GTC1218ATC (3758G>A), CAG1220CAA (3766G>A), GTG1224ATT (3776G>A 3778G>T), GTG1225GTT (3781G>T), CTC1226CTG (3784C>G), GGT1228GGA (3790T>A), GAC1236GAT (3814C>T), TTG1240CTG (3824T>C), CCG1247CCA (3847G>A), AAG1250AAA (3856G>A), GGC1254GGT (3868C>T), ACT1255ACC (3871T>C), GAG1259GAA (3883G>A), CCA1263CCC (3895A>C), GGA1267GGT (3907A>T), GAT1268GAC (3910T>C), CGA1269AGA (3911C>A), GAC1281GAT (3949C>T), GAG1283GAA (3955G>A), GCT1284GCA (3958T>A), TTG1285CTA (3959T>C 3961G>A), ATT1289ATC (3973T>C), GGA1292GGC (3982A>C), AAA1304AAG (4018A>G), TTG1308CTG (4028T>C), CCG1309CCT (4033G>T), TTG1310CTG (4034T>C), CGG1317AGG (4055C>A), TTG1321CTG (4067T>C), ATT1324ATA (4078T>A), CCC1326CCT (4084C>T), TAC1329TAT (4093C>T), CTG1338TTG (4118C>T), TCC1341TCT (4129C>T), TCC1347TCA (4147C>A), CTG1351TTA (4157C>T 4159G>A), CCA1352GCT (4160C>G 4162A>T), GGT1361GGA (4189T>A), ATT1362ATG (4192T>G), TTG1367TTA (4207G>A), GCT1368GCC (4210T>C), GCA1373GCT (4225A>T), GTC1374GTT (4228C>T), ATC1392ATA (4282C>A), GCC1394GCA (4288C>A), TTT1395TTC (4291T>C), CAA1396CAG (4294A>G), CTT1401CTG (4309T>G), GAC1404GAT (4318C>T), GGA1407GGT (4327A>T), GGT1411GGC (4339T>C), GAC1414GAT (4348C>T), AGA1418CGC (4358A>C 4360A>C), GCT1425GCC (4381T>C), TCA1426TCC (4384A>C), TAT1429TAC (4393T>C), GTT1436GTG (4414T>G), TTT1437TTC (4417T>C), TTG1439CTG (4421T>C), GTG1440GTT (4426G>T), GGA1441GGG (4429A>G), TCA1445TCT (4441A>T), GTT1451GTC (4459T>C), AAC1454AAT (4468C>T), TCT1455TCC (4471T>C), GCA1465GCT (4501A>T), TCT1468TCA (4510T>A), GCT1477GCA (4537T>A), GGG1481GGC (4549G>C), AAA1483AAG (4555A>G), GTC1494GTT (4588C>T), CTA1500CTT (4606A>T), CCC1504CCA (4618C>A), TTG1507CTG (4625T>C), TAT1510TAC (4636T>C), GCT1514GCA (4648T>A), ATT1515ATA (4651T>A), GCC1516GCT (4654C>T), TTG1519CTA (4661T>C 4663G>A), GTT1524GTC (4678T>C), ATA1526ATT (4684A>T), ATC1530ATT (4696C>T), GAC1531GAT (4699C>T), GTT1536GTC (4714T>C), AGC1537AAC (4716G>A), AGT1538AAT (4719G>A), GAC1543GAT (4735C>T), TTG1544CTG (4736T>C), CTA1547CTG (4747A>G), TTT1549TTC (4753T>C), CTA1550CTT (4756A>T), AGC1551AGT (4759C>T), GGT1553GGC (4765T>C), TTT1554TTC (4768T>C), AGG1556AGA (4774G>A), CGT1558CGC (4780T>C), CTA1559CTG (4783A>G), GGA1560GGG (4786A>G), TAC1561TAT (4789C>T), CCT1565CCC (4801T>C), CTG1566TTG (4802C>T), GTA1569GTG (4813A>G), TCT1570TCC (4816T>C), CCA1574CCC (4828A>C), TAC1576TAT (4834C>T), AGG1579AGA (4843G>A), CCC1585CCT (4861C>T), CTC1588CTG (4870C>G), AAC1593AAT (4885C>T), ACG1596ACT (4894G>T), GGA1597GGC (4897A>C), GTT1598ATT (4898G>A), GAG1600GAA (4906G>A), GCA1601GCC (4909A>C), GGG1605GGT (4921G>T), ACA1614AAA (4947C>A), GTT1616GTG (4954T>G), AAC1618CAT (4958A>C 4960C>T), AAG1621AAA (4969G>A), CGG1623CGA (4975G>A), TTC1625TTT (4981C>T), GCC1627GCT (4987C>T), ACC1628GCA (4988A>G 4990C>A), CCT1631CCC (4999T>C), ACA1632ACT (5002A>T), TAT1633TAC (5005T>C), TTG1636CTG (5012T>C), GAG1637GAA (5017G>A), CAG1638CAA (5020G>A), GAC1639GAT (5023C>T), GTG1642ATG (5030G>A), GAG1643GAA (5035G>A), ACA1645ACT (5041A>T), GGA1646GGC (5044A>C), GGG1651GGT (5059G>T), ATC1653ATT (5065C>T), CTT1656CTC (5074T>C), CAT1657CAC (5077T>C), ATT1658ACT (5079T>C), GCA1659GCT (5083A>T), GCT1662TCT (5090G>T), TTG1666CTG (5102T>C), CCC1667GCC (5105C>G), GGC1669GGT (5113C>T), AAT1670AAC (5116T>C), ACT1671ACC (5119T>C), GTT1672ATT (5120G>A), CTG1673TTG (5123C>T), TTT1675TTC (5131T>C), TCT1676ACT (5132T>A), GTG1679GTT (5143G>T), CAT1680CAC (5146T>C), GAC1684GAT (5158C>T), AGT1687AGC (5167T>C), TGC1689TGT (5173C>T), TCC1690TCT (5176C>T), CGA1692CGG (5182A>G), GCA1696GCC (5194A>C), CAG1698CAA (5200G>A), GAC1700GAT (5206C>T), ACC1701ACA (5209C>A), GTT1704GTG (5218T>G), AAC1707AAT (5227C>T), GCA1711GCC (5239A>C), CCC1720CCA (5266C>A), ACC1721ACA (5269C>A), AGG1724AGA (5278G>A), GGG1728GGT (5290G>T), CTT1729CTC (5293T>C), CCT1732CCA (5302T>A), CAC1741CAT (5329C>T), GTA1746GTG (5344A>G), ACC1747ACA (5347C>A), ACC1751TCC (5357A>T), AAA1752CAA (5360A>C), CAT1754AAT (5366C>A), GTG1755ATT (5369G>A 5371G>T), GTT1756GTA (5374T>A), GGC1760GGG (5386C>G), GGG1762GGA (5392G>A), ATT1765GTT (5399A>G), CCA1767CCT (5407A>T), GTG1768GTT (5410G>T), TGC1769TGT (5413C>T), GCA1771GCT (5419A>T), CTG1775TTG (5429C>T), ACT1777ACC (5437T>C), GGC1778GGG (5440C>G), GTT1785ATC (5459G>A 5461T>C), GTT1786ATT (5462G>A), TTG1789CTG (5471T>C), ATA1792ATT (5482A>T), GCA1793GCT (5485A>T), GAT1794AAT (5486G>A), GTT1795GTC (5491T>C), ACA1796CCT (5492A>C 5494A>T), CTC1798TTG (5498C>T 5500C>G), CCA1802CCT (5512A>T), GGA1803GGT (5515A>T), CTT1809TTG (5531C>T 5533T>G), CTG1810GTG (5534C>G), AAA1811CAA (5537A>C), GCT1813TCT (5543G>T), GAA1818CAA (5558G>C), CTT1820CTG (5566T>G), GCA1821GCT (5569A>T), GAT1822GAC (5572T>C), TGT1824TGC (5578T>C), ATT1826GTT (5582A>G), GCC1829TCC (5591G>T), GAG1832GAA (5602G>A), ACA1833ATA (5604C>T), GCA1835GCT (5611A>T), TAC1836TTC (5613A>T), ATA1837ATT (5617A>T), CCT1838CCA (5620T>A), GGA1840GGT (5626A>T), CTT1844CTG (5638T>G), GGT1845GGC (5641T>C), TTT1847TTC (5647T>C), AGA1853AAA (5664G>A), CTA1856CTT (5674A>T), CAG1857CAT (5677G>T), CAG1860CAA (5686G>A), CGC1861CGA (5689C>A), GGA1867GGT (5707A>T), CCT1868CCA (5710T>A), TAT1869CAT (5711T>C), GGG1870GGT (5716G>T), GGA1871GGT (5719A>T), CAC1875CAT (5731C>T), ACT1876ACC (5734T>C), ACT1877ACC (5737T>C), AGA1878AAA (5739G>A), CTT1880CTC (5746T>C), TTG1882TAA (5751T>A 5752G>A), ATG1887ATT (5767G>T), AAG1889AAA (5773G>A), ATA1890GTG (5774A>G 5776A>G), TAT1891TAC (5779T>C), TTA1894CTA (5786T>C), GTC1895GTA (5791C>A), AGA1897AGG (5797A>G), TTC1900ACT (5804T>A 5805T>C 5806C>T), CTC1903CTG (5815C>G), TAC1904TAT (5818C>T), TAT1907TTT (5826A>T), GTG1913GTA (5845G>A), TTG1915CAG (5849T>C 5850T>A), GTT1917GTG (5857T>G), GAC1918GAA (5860C>A), CTT1920CTG (5866T>G), AGG1921AGA (5869G>A), CTC1922CTG (5872C>G), GGG1924GGT (5878G>T), TTT1925TTC (5881T>C), CCC1927CCT (5887C>T), CGC1928CGT (5890C>T), AAT1930CAT (5894A>C), TAC1934TTC (5907A>T), CTT1936CTC (5914T>C), TTT1939TTC (5923T>C), CTG1941TTG (5927C>T), CAC1946TAT (5942C>T 5944C>T), AAC1950AGC (5955A>G), ACG1953ACA (5965G>A), GTT1954GTG (5968T>G), TTT1956TTC (5974T>C), AGA1957CGA (5975A>C), AAG1958AAA (5980G>A), CCT1959CCC (5983T>C), TTT1960TTC (5986T>C), AAA1964AAG (5998A>G), ACC1966GTT (6002A>G 6003C>T 6004C>T), GGG1968GAG (6009G>A), AAA1970AAG (6016A>G), TCC1974TCT (6028C>T), GCA1975GCT (6031A>T), ACA1976ATA (6033C>T), GCT1977TCT (6035G>T), GCT1978GCA (6040T>A), GGT1983GGG (6055T>G), GCT1988GCC (6070T>C), CTC1991CTG (6079C>G), AGT1993AGC (6085T>C), CTA1995TTG (6089C>T 6091A>G), CTT1996TTT (6092C>T), CAT1997CAA (6097T>A), CTT1998CTG (6100T>G), CAG1999CAA (6103G>A), AAG2000AAA (6106G>A), ATA2002ATG (6112A>G), TGT2004TGC (6118T>C), TCA2005TCT (6121A>T), GAG2008GAT (6130G>T), TTT2011TTC (6139T>C), GTC2016GTA (6154C>A), GGC2019GGG (6163C>G), CAC2020CAT (6166C>T), GTA2026GTG (6184A>G), ACT2027GAC (6185A>G 6186C>A 6187T>C), CAC2028CGC (6189A>G), TTC2029TTT (6193C>T), CAA2030CAG (6196A>G), CGC2031CGT (6199C>T), ATG2032ATC (6202G>C), GAT2036GAG (6214T>G), CTG2037GTT (6215C>G 6217G>T), ACA2038CCA (6218A>C), GAA2039CCA (6221G>C 6222A>C), GGG2040GAG (6225G>A), ATC2041GTC (6227A>G), ATA2042GTG (6230A>G 6232A>G), GAG2043GAA (6235G>A), GTG2045ACG (6239G>A 6240T>C), TTA2049CTT (6251T>C 6253A>T), ACC2050ACA (6256C>A), ACA2052ACT (6262A>T), CTG2053TTA (6263C>T 6265G>A), TCG2054ACT (6266T>A 6268G>T), AGC2056ATG (6273G>T 6274C>G), GTT2062GAA (6291T>A 6292T>A), GTA2064CTT (6296G>C 6298A>T), CCT2065CCA (6301T>A), TTA2066ATG (6302T>A 6304A>G), TTG2068CTG (6308T>C), GAT2069TAA (6311G>T 6313T>A), ATA2071GTT (6317A>G 6319A>T), GAG2072GAA (6322G>A), GGT2077GGC (6337T>C), CTA2080TTA (6344C>T), GCA2089ACA (6371G>A), GCC2092GCT (6382C>T), CTC2097CTG (6397C>G), TGT2100CAT (6404T>C 6405G>A), CGA2103CTG (6414G>T 6415A>G), TCA2104CAC (6416T>C 6417C>A 6418A>C), GTG2109GTT (6433G>T), TTC2110TTT (6436C>T), TGT2111AGT (6437T>A), TAC2113TAT (6445C>T), ACG2114ACA (6448G>A), TCC2117TCA (6457C>A), TCT2118ACT (6458T>A), CTT2119TTA (6461C>T 6463T>A), CTT2125CTA (6481T>A), CAT2127AAT (6485C>A), AGA2128AGC (6490A>C), CAT2129AAC (6491C>A 6493T>C), CAG2130CAA (6496G>A), ACA2131CCT (6497A>C 6499A>T), ATT2132GAG (6500A>G 6501T>A 6502T>G), CCG2136CCT (6514G>T), TCC2137CCA (6515T>C 6517C>A), TTC2138TTT (6520C>T), CAC2142CAT (6532C>T), CTT2147TTT (6545C>T), CCA2150CCC (6556A>C), CTA2152CTT (6562A>T), GGC2154AAT (6566G>A 6567G>A 6568C>T), TAC2155TAT (6571C>T), AAA2156AAG (6574A>G)

*: Inserts / Deletes / Misaligned / Frameshifts

#### Analysis details

This analysis was performed with panviral2.64

### NGS Details (UN62): Torradovirus lycopersici (segment RNA 2)

#### Assembly

| | |
|---|---|
| **Coverage Length** | 5384 (1 contig(s)) |
| **Depth Of Coverage** | 7488.0 |
| **Number Of Reads** | 333725 |
| **Reads Per Million** | 7736.04 rpm (after QC) |
| **Ambiguities** | 0 |
| **Assembly Method** | de novo + reference guided assembly |
| **Consensus Caller** | Bcf Tools |

#### Coverage Map

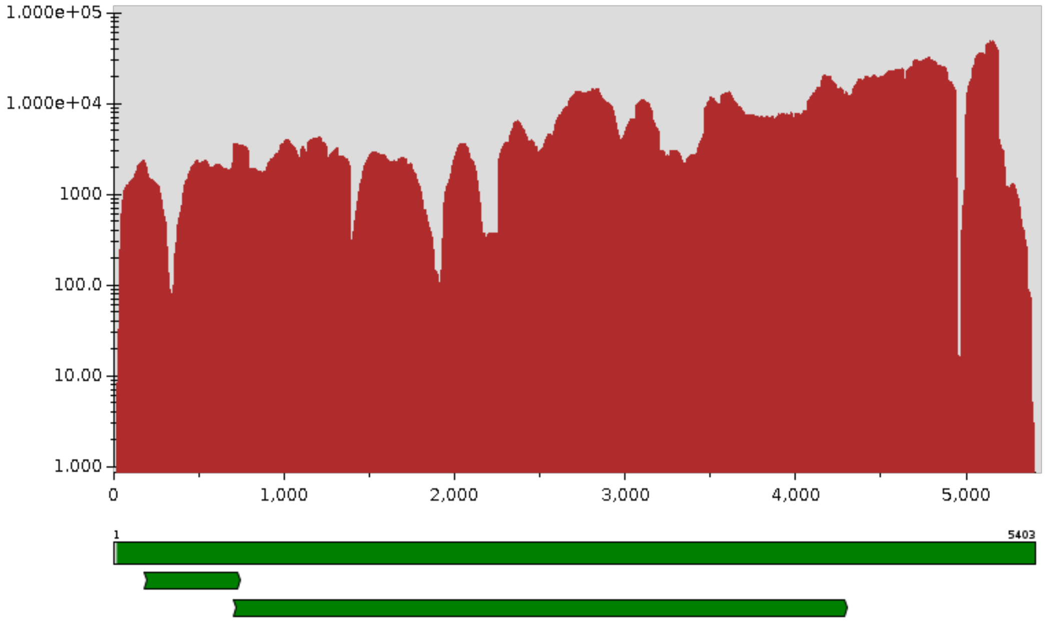

#### Assignment

| | |
|---|---|
| **Type** | Torradovirus lycopersici (Taxonomy ID: 3048378) |
| **Reference Genome** | NC_009032.1 |
| **NT Identity (%)** | 91.4763 |
| **AA Identity (%)** | 94.3043 |
| **Number Of Stop Codons** | 2 |
| **Number Of CDS** | 2 |

#### Alignment

| | |
|---|---|
| **Alignment Score** | 8926.0 (NT) + 9075.0 (AA) = 18001.0 |
| **Concordance (%)** | 88.8719 |

| Alignment Method | Global, seeded, nucleotide + amino acids (AGA) |
|---|---|

#### Genome Region

Sequence starts at position 20 and ends at position 5403 relative to NC_009032.1 reference sequence.

#### Alignment Detailed Statistics

| | Begin | End | Coverage | Score | Concordance | Matches | Identities | I/D/M/F* | Stop Codons |
|---|---|---|---|---|---|---|---|---|---|
| **NT** | **20** | **5403** | **99.6%** | **8926** | **82.9%** | **5384 (99.9%)** | **4926 (91.5%)** | **1/0** | |
| Mutations: | 89G>A, 90C>T, 97C>T, 107G>A, 111C>A, 116A>G, 137T>C, 144A>G, 161A>G, 179T>C, 180C>T, 271T>C, 277C>T, 280T>C, 283C>T, 295T>C, 301A>T, 310C>T, 313C>T, 316G>A, 340A>T, 356G>A, 370C>T, 372A>C, 376C>T, 388T>A, 400T>C, 401T>C, 418T>G, 433C>T, 439G>A, 445T>A, 448C>A, 460T>C, 461A>C, 481C>A, 487A>G, 502T>A, 507G>A, 508G>A, 509A>G, 511T>A, 515G>A, 552C>T, 565A>T, 568A>T, 576C>A, 583A>T, 586A>G, 587T>C, 589A>T, 603A>C, 607G>A, 613A>G, 622A>G, 634G>A, 637A>G, 644C>G, 645T>C, 654T>A, 670A>G, 679G>A, 683T>A, 690A>C, 696T>C, 704G>A, 721C>T, 733A>G, 748A>G, 760T>C, 797T>C, 806C>T, 827A>G, 844A>T, 968A>G, 1043A>G, 1061T>C, 1083C>T, 1100G>A, 1106T>C, 1109C>T, 1139G>A, 1151A>G, 1163A>G, 1169C>G, 1178C>T, 1187A>T, 1205T>A, 1226A>G, 1235C>A, 1247A>G, 1277A>G, 1313T>C, 1317C>A, 1319A>G, 1325A>G, 1328T>C, 1331C>T, 1343G>A, 1355G>A, 1361A>T, 1367A>T, 1383A>G, 1388C>T, 1397T>A, 1403A>T, 1406G>A, 1412G>T, 1418G>A, 1421A>G, 1427G>T, 1434G>A, 1451G>T, 1457C>T, 1460A>G, 1461G>T, 1472C>G, 1490G>A, 1502C>T, 1514T>C, 1524T>C, 1547C>T, 1550A>C, 1589A>G, 1608T>C, 1619G>A, 1634T>A, 1646T>C, 1651A>G, 1655T>A, 1661T>C, 1662C>T, 1664G>T, 1667C>T, 1677G>A, 1685T>C, 1721G>C, 1724C>T, 1740C>T, 1748G>A, 1757A>G, 1778A>C, 1794G>A, 1796T>A, 1814T>C, 1817A>T, 1819T>A, 1820C>G, 1832G>C, 1845T>C, 1856C>T, 1859C>T, 1863C>T, 1865C>T, 1892C>T, 1899G>A, 1900T>C, 1901G>A, 1906C>T, 1907C>G, 1928A>C, 1931C>T, 1937C>T, 1941C>T, 1947T>C, 1955G>A, 1958A>G, 1967G>A, 1970A>G, 1979A>G, 1985C>T, 1994T>C, 2072G>A, 2084G>A, 2094C>G, 2099G>A, 2106A>G, 2108A>G, 2126C>T, 2128G>T, 2135G>A, 2136G>C, 2140A>C, 2144C>G, 2159T>A, 2164T>G, 2183C>T, 2186A>C, 2189C>T, 2198G>A, 2199G>A, 2200T>C, 2205A>G, 2210G>A, 2216C>G, 2229A>G, 2230T>A, 2232A>G, 2240T>A, 2243T>A, 2252A>T, 2256G>C, 2259A>G, 2261T>G, 2262G>C, 2270A>G, 2285T>C, 2294A>C, 2303A>G, 2312T>C, 2315T>A, 2320G>C, 2321T>C, 2326C>T, 2339A>G, 2345A>G, 2375C>T, 2393G>A, 2414T>C, 2420T>G, 2432G>A, 2436T>G, 2444G>C, 2447C>A, 2453G>A, 2459C>G, 2469T>A, 2471C>G, 2474G>A, 2489A>C, 2492G>A, 2495A>G, 2501A>G, 2503G>A, 2507T>C, 2516C>T, 2523A>C, 2546T>C, 2552C>T, 2561T>G, 2567T>A, 2570G>A, 2573T>A, 2630C>T, 2645G>A, 2660T>C, 2693G>A, 2708A>G, 2720C>A, 2732T>C, 2765G>A, 2792C>T, 2819G>A, 2840T>C, 2841C>T, 2852A>G, 2870C>A, 2879T>A, 2885T>C, 2886T>C, 2888G>A, 2900A>T, 2915A>G, 2939A>G, 2957A>T, 2958C>T, 2966C>T, 2972T>C, 2982G>A, 2999T>A, 3003T>C, 3011C>T, 3012A>G, 3020T>C, 3032A>T, 3050G>A, 3063T>C, 3122T>C, 3131G>A, 3146T>C, 3152T>A, 3167A>T, 3187G>A, 3206C>T, 3227T>G, 3230T>A, 3236C>T, 3242G>A, 3248A>C, 3253A>G, 3265G>A, 3270A>T, 3303A>T, 3305C>T, 3324C>T, 3335C>T, 3350C>T, 3354A>C, 3359G>A, 3368A>G, 3371C>G, 3386A>T, 3404A>G, 3413C>T, 3422A>G, 3428G>A, 3431T>C, 3437C>T, 3446T>C, 3449G>A, 3452C>T, 3455T>A, 3470T>C, 3479C>T, 3482T>C, 3485C>T, 3491C>T, 3503A>G, 3508A>G, 3512T>C, 3515G>T, 3522C>T, 3524A>G, 3527C>T, 3533G>A, 3543T>A, 3551C>T, 3560T>A, 3569A>G, 3575G>A, 3581C>T, 3590A>C, 3626A>G, 3635C>T, 3638A>G, 3650A>T, 3662A>T, 3674G>A, 3722C>T, 3743G>A, 3755A>G, 3767C>T, 3773C>T, 3800C>T, 3827C>T, 3831A>G, 3881C>T, 3885A>G, 3890A>G, 3893T>C, 3933A>G, 3941T>G, 3950T>A, 3983T>A, 3984A>G, 3986G>T, 3990G>A, 3995C>T, 3998C>T, 4013G>C, 4031T>C, 4040C>T, 4059A>G, 4061A>G, 4069A>G, 4100A>T, 4112C>T, 4130C>T, 4223G>A, 4224C>T, 4250C>T, 4336T>A, 4457A>G, 4590G>A, 4635C>T, 4812C>T, 4814T>C, 4823A>G, 4940C>A, 4941C>A, 4943A>T, 4949T>C, 4955A>C, 4959A>C, 4960A>G, 4961C>T, 4965G>A, 4969A>T, 4971T>C, 4972T>G, 4974T>A, 4980A>G, 4984C>T, 4993G>A, 4994C>A, 4995G>A, 4996A>T, 5000C>T, 5003A>T, 5004G>T, 5019A>T, 5031T>C, 5032G>A, 5040C>T, 5041A>G, 5053A>G, 5054T>C, 5056G>A, 5099C>T, 5101A>G, 5107A>T, 5110A>C, 5112A>G, 5118T>A, 5153A>G, 5173C>T, 5184C>T, 5185T>G, 5187G>A, 5196T>G, 5197T>A, 5198T>A, 5199_5200insC, 5206C>T, 5208G>A, 5211A>G, 5212A>T, 5213A>T, 5214C>T, 5220T>A, 5221A>T, 5236G>A, 5239T>A, 5242T>A, 5250C>T, 5262A>T, 5265G>A, 5266T>A, 5268T>C, 5282T>A, 5285G>T, 5294G>A, 5297T>C, 5298C>T, 5301T>A, 5313A>C, 5326G>A, 5327A>G, 5335T>A, 5346T>C, 5348G>A, 5360A>T, 5363C>A, 5364A>T, 5369G>T, 5371G>T, 5373G>A, 5374T>G, 5377G>T | | | | | | | | |
| **CDS** | | | | | | | | | |
| ToTV_sRNA2gp1 | 1 | 188 | 100% | 1130 | 93.5% | 188 (100%) | 173 (92.0%) | 0/0/0/0 | 1 |
| Protein mutations: | V59I (356G>A), N64T (372A>C), R109K (507G>A 508G>A), I110V (509A>G 511T>A), V112I (515G>A), A124V (552C>T), Q128H (565A>T), T132K (576C>A), D141A (603A>C), L155A (644C>G 645T>C), L158Q (654T>A), L168M (683T>A), N170T (690A>C), L172P (696T>C), D175N (704G>A) | | | | | | | | |
| Codon mutations: | GAT30GAC (271T>C), GGC32GGT (277C>T), TCT33TCC (280T>C), GGC34GGT (283C>T), AGT38AGC (295T>C), CCA40CCT (301A>T), GAC43GAT (310C>T), TTC44TTT (313C>T), AAG45AAA (316G>A), GCA53GCT (340A>T), GTT59ATT (356G>A), AAC63AAT (370C>T), AAT64ACT (372A>C), AGC65AGT (376C>T), GTT69GTA (388T>A), TAT73TAC (400T>C), TTA74CTA (401T>C), ACT79ACG (418T>G), GGC84GGT (433C>T), AAG86AAA (439G>A), TCT88TCA (445T>A), GGC89GGA (448C>A), CAT93CAC (460T>C), AGA94CGA (461A>C), ACC100ACA (481C>A), AGA102AGG (487A>G), GCT107GCA (502T>A), AGG109AAA (507G>A 508G>A), ATT110GTA (509A>G 511T>A), GTA112ATA (515G>A), GCT124GTT (552C>T), CAA128CAT (565A>T), CCA129CCT (568A>T), ACG132AAG (576C>A), ACA134ACT (583A>T), CAA135CAG (586A>G), TTA136CTT (587T>C 589A>T), GAC141GCC (603A>C), GAG142GAA (607G>A), AAA144AAG (613A>G), TTA147TTG (622A>G), GAG151GAA (634G>A), AAA152AAG (637A>G), CTT155GCT (644C>G 645T>C), CTA158CAA (654T>A), AAA163AAG (670A>G), TTG166TTA (679G>A), TTG168ATG (683T>A), AAC170ACC (690A>C), CTA172CCA (696T>C), GAT175AAT (704G>A), GGC180GGT (721C>T), GAA184GAG (733A>G) | | | | | | | | |
| ToTV_sRNA2gp2 | 1 | 1199 | 100% | 7945 | 96.0% | 1199 (100%) | 1135 (94.7%) | 0/0/0/0 | 1 |
| Protein mutations: | M1I (704G>A), A7V (721C>T), K11R (733A>G), K16R (748A>G), V20A (760T>C), Y48F (844A>T), T228A (1383A>G), Q242H (1427G>T), D245N (1434G>A), A254S (1461G>T), I257M (1472C>G), S275P (1524T>C), Y303H (1608T>C), K317R (1651A>G), L321F (1662C>T 1664G>T), V326M (1677G>A), V365I (1794G>A 1796T>A), I373K (1819T>A 1820C>G), K377N (1832G>C), H388Y (1863C>T 1865C>T), V400T (1899G>A 1900T>C 1901G>A), A402V (1906C>T 1907C>G), E409D (1928A>C), L414F (1941C>T), Q465E (2094C>G), R469G (2106A>G 2108A>G), R476M (2128G>T), V479L (2136G>C), K480T (2140A>C), N481K (2144C>G), V488G (2164T>G), V500T (2199G>A 2200T>C), M502V (2205A>G), M510E (2229A>G 2230T>A), K511E (2232A>G), E517D (2252A>T), A519P (2256G>C), N520E (2259A>G 2261T>G), E521Q (2262G>C), E531D (2294A>C), N538K (2315T>A), G540A (2320G>C 2321T>C), T542I (2326C>T), S579A (2436T>G), S590T (2469T>A 2471C>G), R601K (2503G>A), I608L (2523A>C), A761T (2982G>A), I771V (3012A>G), S829N (3187G>A), K851R (3253A>G), R855K (3265G>A), T857S (3270A>T), T868S (3303A>T 3305C>T), N936S (3508A>G), S948T (3543T>A), I1044V (3831A>G), T1062A (3885A>G), T1078A (3933A>G), M1095V (3984A>G 3986G>T), D1097N (3990G>A), I1120V (4059A>G 4061A>G), N1123S (4069A>G), H1175Y (4224C>T) | | | | | | | | |

| | Begin | End | Coverage | Score | Concordance | Matches | Identities | I/D/M/F* | Stop Codons |
|---|---|---|---|---|---|---|---|---|---|
| **NT** | **20** | **5403** | **99.6%** | **8926** | **82.9%** | **5384 (99.9%)** | **4926 (91.5%)** | **1/0** | |
| Codon mutations: | ATG1ATA (704G>A), GCT7GTT (721C>T), AAG11AGG (733A>G), AAA16AGA (748A>G), GTA20GCA (760T>C), GTT32GTC (797T>C), GGC35GGT (806C>T), AAA42AAG (827A>G), TAT48TTT (844A>T), GGA89GGG (968A>G), AAA114AAG (1043A>G), AGT120AGC (1061T>C), CTG128TTG (1083C>T), TTG133TTA (1100G>A), TTT135TTC (1106T>C), TCC136TCT (1109C>T), CCG146CCA (1139G>A), CTA150CTG (1151A>G), AAA154AAG (1163A>G), GGC156GGG (1169C>G), TTC159TTT (1178C>T), ACA162ACT (1187A>T), ACT168ACA (1205T>A), TCA175TCG (1226A>G), GCC178GCA (1235C>A), CAA182CAG (1247A>G), TCA192TCG (1277A>G), CTT204CTC (1313T>C), CGA206AGG (1317C>A 1319A>G), CAA208CAG (1325A>G), TTT209TTC (1328T>C), TTC210TTT (1331C>T), CAG214CAA (1343G>A), CAG218CAA (1355G>A), ATA220ATT (1361A>T), TCA222TCT (1367A>T), ACC228GCC (1383A>G), TCC229TCT (1388C>T), ACT232ACA (1397T>A), TCA234TCT (1403A>T), GTG235GTA (1406G>A), CCG237CCT (1412G>T), GTG239GTA (1418G>A), GGA240GGG (1421A>G), CAG242CAT (1427G>T), GAC245AAC (1434G>A), GTG250GTT (1451G>T), AGC252AGT (1457C>T), GAA253GAG (1460A>G), GCA254TCA (1461G>T), ATC257ATG (1472C>G), CGG263CGA (1490G>A), CTC267CTT (1502C>T), AGT271AGC (1514T>C), TCC275CCC (1524T>C), ACC282ACT (1547C>T), TCA283TCC (1550A>C), GTA296GTG (1589A>G), TAC303CAC (1608T>C), GAG306GAA (1619G>A), GTT311GTA (1634T>A), GTT315GTC (1646T>C), AAG317AGG (1651A>G), GCT318GCA (1655T>A), GGT320GGC (1661T>C), CTG321TTT (1662C>T 1664G>T), AAC322AAT (1667C>T), GTG326ATG (1677G>A), GAT328GAC (1685T>C), CTG340CTC (1721G>C), CCC341CCT (1724C>T), CTG347TTG (1740C>T), GTG349GTA (1748G>A), CAA352CAG (1757A>G), GCA359GCC (1778A>C), GTT365ATA (1794G>A 1796T>A), CAT371CAC (1814T>C), ATA372ATT (1817A>T), ATC373AAG (1819T>A 1820C>G), AAG377AAC (1832G>C), TTG382CTG (1845T>C), ATC385ATT (1856C>T), ACC386ACT (1859C>T), CAC388TAT (1863C>T 1865C>T), TAC397TAT (1892C>T), GTG400ACA (1899G>A 1900T>C 1901G>A), GCC402GTG (1906C>T 1907C>G), GAA409GAC (1928A>C), AAC410AAT (1931C>T), TAC412TAT (1937C>T), CTT414TTT (1941C>T), TTG416CTG (1947T>C), CAG418CAA (1955G>A), AGA419AGG (1958A>G), GAG422GAA (1967G>A), CTA423CTG (1970A>G), CAA426CAG (1979A>G), GAC428GAT (1985C>T), TAT431TAC (1994T>C), GAG457GAA (2072G>A), GTG461GTA (2084G>A), CAG465GAG (2094C>G), GCG466GCA (2099G>A), AGA469GGG (2106A>G 2108A>G), GCC475GCT (2126C>T), AGG476ATG (2128G>T), CAG478CAA (2135G>A), GTG479CTG (2136G>C), AAA480ACA (2140A>C), AAC481AAG (2144C>G), ACT486ACA (2159T>A), GTG488GGG (2164T>G), ACC494ACT (2183C>T), TCA495TCC (2186A>C), TTC496TTT (2189C>T), AAG499AAA (2198G>A), GTT500ACT (2199G>A 2200T>C), ATG502GTG (2205A>G), GAG503GAA (2210G>A), GGC505GGG (2216C>G), ATG510GAG (2229A>G 2230T>A), AAA511GAA (2232A>G), GCT513GCA (2240T>A), ATT514ATA (2243T>A), GAA517GAT (2252A>T), GCC519CCC (2256G>C), AAT520GAG (2259A>G 2261T>G), GAA521CAA (2262G>C), GTA523GTG (2270A>G), GTT528GTC (2285T>C), GAA531GAC (2294A>C), CAA534CAG (2303A>G), AAT537AAC (2312T>C), AAT538AAA (2315T>A), GGT540GCC (2320G>C 2321T>C), ACA542ATA (2326C>T), TTA546TTG (2339A>G), CAA548CAG (2345A>G), TTC558TTT (2375C>T), GAG564GAA (2393G>A), AGT571AGC (2414T>C), GTT573GTG (2420T>G), GTG577GTA (2432G>A), TCA579GCA (2436T>G), GCG581GCC (2444G>C), CTC582CTA (2447C>A), GCG584GCA (2453G>A), CTC586CTG (2459C>G), TCC590ACG (2469T>A 2471C>G), AAG591AAA (2474G>A), GGA596GGC (2489A>C), TCG597TCA (2492G>A), CAA598CAG (2495A>G), CTA600CTG (2501A>G), AGG601AAG (2503G>A), TAT602TAC (2507T>C), GCC605GCT (2516C>T), ATT608CTT (2523A>C), TAT615TAC (2546T>C), TCC617TCT (2552C>T), GGT620GGG (2561T>G), ATT622ATA (2567T>A), TCG623TCA (2570G>A), GGT624GGA (2573T>A), ATC643ATT (2630C>T), CTG648CTA (2645G>A), CAT653CAC (2660T>C), GGG664GGA (2693G>A), ACA669ACG (2708A>G), ATC673ATA (2720C>A), GTT677GTC (2732T>C), CTG688CTA (2765G>A), CAC697CAT (2792C>T), GAG706GAA (2819G>A), CAT713CAC (2840T>C), CTG714TTG (2841C>T), AAA717AAG (2852A>G), ATC723ATA (2870C>A), CCT726CCA (2879T>A), CGT728CGC (2885T>C), TTG729CTA (2886T>C 2888G>A), GCA733GCT (2900A>T), AAA738AAG (2915A>G), CGA746CGG (2939A>G), GTA752GTT (2957A>T), CTG753TTG (2958C>T), CAC755CAT (2966C>T), AAT757AAC (2972T>C), GCT761ACT (2982G>A), ACT766ACA (2999T>A), TTA768CTA (3003T>C), AGC770AGT (3011C>T), ATC771GTC (3012A>G), TCT773TCC (3020T>C), ATA777ATT (3032A>T), ACG783ACA (3050G>A), TTG788CTG (3063T>C), ATT807ATC (3122T>C), GAG810GAA (3131G>A), CAT815CAC (3146T>C), GGT817GGA (3152T>A), GGA822GGT (3167A>T), AGC829AAC (3187G>A), GAC835GAT (3206C>T), GTT842GTG (3227T>G), ATT843ATA (3230T>A), AAC845AAT (3236C>T), GAG847GAA (3242G>A), GGA849GGC (3248A>C), AAG851AGG (3253A>G), AGG855AAG (3265G>A), ACG857TCG (3270A>T), ACC868TCT (3303A>T 3305C>T), CTA875TTA (3324C>T), CCC878CCT (3335C>T), CAC883CAT (3350C>T), AGG885CGG (3354A>C), TTG886TTA (3359G>A), ACA889ACG (3368A>G), GTC890GTG (3371C>G), GTA895GTT (3386A>T), CTA901CTG (3404A>G), GTC904GTT (3413C>T), ACA907ACG (3422A>G), CAG909CAA (3428G>A), CTT910CTC (3431T>C), TGC912TGT (3437C>T), GAT915GAC (3446T>C), TTG916TTA (3449G>A), GTC917GTT (3452C>T), CTT918CTA (3455T>A), TCT923TCC (3470T>C), CCC926CCT (3479C>T), ATT927ATC (3482T>C), TAC928TAT (3485C>T), CAC930CAT (3491C>T), GGA934GGG (3503A>G), AAT936AGT (3508A>G), GCT937GCC (3512T>C), GTG938GTT (3515G>T), CTA941TTG (3522C>T 3524A>G), TTC942TTT (3527C>T), GAG944GAA (3533G>A), TCT948ACT (3543T>A), CGC950CGT (3551C>T), ATT953ATA (3560T>A), TTA956TTG (3569A>G), AAG958AAA (3575G>A), AAC960AAT (3581C>T), ACA963ACC (3590A>C), CAA975CAG (3626A>G), CTC978CTT (3635C>T), AGA979AGG (3638A>G), TCA983TCT (3650A>T), GTA987GTT (3662A>T), GTG991GTA (3674G>A), CCC1007CCT (3722C>T), CAG1014CAA (3743G>A), GTA1018GTG (3755A>G), ATC1022ATT (3767C>T), TGC1024TGT (3773C>T), TTC1033TTT (3800C>T), TAC1042TAT (3827C>T), ATT1044GTT (3831A>G), AAC1060AAT (3881C>T), ACC1062GCC (3885A>G), TTA1063TTG (3890A>G), GAT1064GAC (3893T>C), ACC1078GCC (3933A>G), GTT1080GTG (3941T>G), GCT1083GCA (3950T>A), GGT1094GGA (3983T>A), ATG1095GTT (3984A>G 3986G>T), GAC1097AAC (3990G>A), AAC1098AAT (3995C>T), ATC1099ATT (3998C>T), GTG1104GTC (4013G>C), TTT1110TTC (4031T>C), CGC1113CGT (4040C>T), ATA1120GTG (4059A>G 4061A>G), AAT1123AGT (4069A>G), CGA1133CGT (4100A>T), CTC1137CTT (4112C>T), GCC1143GCT (4130C>T), AAG1174AAA (4223G>A), CAT1175TAT (4224C>T), ACC1183ACT (4250C>T) | | | | | | | | |

#### Proteins

| | Begin | End | Coverage | Score | Concordance | Matches | Identities | I/D/M/F* | Stop Codons |
|---|---|---|---|---|---|---|---|---|---|
| Orf1 (YP_001040017.1) | 1 | 188 | 100% | 1130 | 93.5% | 188 (100%) | 173 (92.0%) | 0/0/0/0 | 1 |
| Protein mutations: | V59I (356G>A), N64T (372A>C), R109K (507G>A 508G>A), I110V (509A>G 511T>A), V112I (515G>A), A124V (552C>T), Q128H (565A>T), T132K (576C>A), D141A (603A>C), L155A (644C>G 645T>C), L158Q (654T>A), L168M (683T>A), N170T (690A>C), L172P (696T>C), D175N (704G>A) | | | | | | | | |
| Codon mutations: | GAT30GAC (271T>C), GGC32GGT (277C>T), TCT33TCC (280T>C), GGC34GGT (283C>T), AGT38AGC (295T>C), CCA40CCT (301A>T), GAC43GAT (310C>T), TTC44TTT (313C>T), AAG45AAA (316G>A), GCA53GCT (340A>T), GTT59ATT (356G>A), AAC63AAT (370C>T), AAT64ACT (372A>C), AGC65AGT (376C>T), GTT69GTA (388T>A), TAT73TAC (400T>C), TTA74CTA (401T>C), ACT79ACG (418T>G), GGC84GGT (433C>T), AAG86AAA (439G>A), TCT88TCA (445T>A), GGC89GGA (448C>A), CAT93CAC (460T>C), AGA94CGA (461A>C), ACC100ACA (481C>A), AGA102AGG (487A>G), GCT107GCA (502T>A), AGG109AAA (507G>A 508G>A), ATT110GTA (509A>G 511T>A), GTA112ATA (515G>A), GCT124GTT (552C>T), CAA128CAT (565A>T), CCA129CCT (568A>T), ACG132AAG (576C>A), ACA134ACT (583A>T), CAA135CAG (586A>G), TTA136CTT (587T>C 589A>T), GAC141GCC (603A>C), GAG142GAA (607G>A), AAA144AAG (613A>G), TTA147TTG (622A>G), GAG151GAA (634G>A), AAA152AAG (637A>G), CTT155GCT (644C>G 645T>C), CTA158CAA (654T>A), AAA163AAG (670A>G), TTG166TTA (679G>A), TTG168ATG (683T>A), AAC170ACC (690A>C), CTA172CCA (696T>C), GAT175AAT (704G>A), GGC180GGT (721C>T), GAA184GAG (733A>G) | | | | | | | | |
| polyprotein (YP_001040018.1) | 1 | 1199 | 100% | 7945 | 96.0% | 1199 (100%) | 1135 (94.7%) | 0/0/0/0 | 1 |
| Protein mutations: | M1I (704G>A), A7V (721C>T), K11R (733A>G), K16R (748A>G), V20A (760T>C), Y48F (844A>T), T228A (1383A>G), Q242H (1427G>T), D245N (1434G>A), A254S (1461G>T), I257M (1472C>G), S275P (1524T>C), Y303H (1608T>C), K317R (1651A>G), L321F (1662C>T 1664G>T), V326M (1677G>A), V365I (1794G>A 1796T>A), I373K (1819T>A 1820C>G), K377N (1832G>C), H388Y (1863C>T 1865C>T), V400T (1899G>A 1900T>C 1901G>A), A402V (1906C>T 1907C>G), E409D (1928A>C), L414F (1941C>T), Q465E (2094C>G), R469G (2106A>G 2108A>G), R476M (2128G>T), V479L (2136G>C), K480T (2140A>C), N481K (2144C>G), V488G (2164T>G), V500T (2199G>A 2200T>C), M502V (2205A>G), M510E (2229A>G 2230T>A), K511E (2232A>G), E517D (2252A>T), A519P (2256G>C), N520E (2259A>G 2261T>G), E521Q (2262G>C), E531D (2294A>C), N538K (2315T>A), G540A (2320G>C 2321T>C), T542I (2326C>T), S579A (2436T>G), S590T (2469T>A 2471C>G), R601K (2503G>A), I608L (2523A>C), A761T (2982G>A), I771V (3012A>G), S829N (3187G>A), K851R (3253A>G), R855K (3265G>A), T857S (3270A>T), T868S (3303A>T 3305C>T), N936S (3508A>G), S948T (3543T>A), I1044V (3831A>G), T1062A (3885A>G), T1078A (3933A>G), M1095V (3984A>G 3986G>T), D1097N (3990G>A), I1120V (4059A>G 4061A>G), N1123S (4069A>G), H1175Y (4224C>T) | | | | | | | | |

| | Begin | End | Coverage | Score | Concordance | Matches | Identities | I/D/M/F* | Stop Codons |
|---|---|---|---|---|---|---|---|---|---|
| **NT** | **20** | **5403** | **99.6%** | **8926** | **82.9%** | **5384 (99.9%)** | **4926 (91.5%)** | **1/0** | |
| Codon mutations: | ATG1ATA (704G>A), GCT7GTT (721C>T), AAG11AGG (733A>G), AAA16AGA (748A>G), GTA20GCA (760T>C), GTT32GTC (797T>C), GGC35GGT (806C>T), AAA42AAG (827A>G), TAT48TTT (844A>T), GGA89GGG (968A>G), AAA114AAG (1043A>G), AGT120AGC (1061T>C), CTG128TTG (1083C>T), TTG133TTA (1100G>A), TTT135TTC (1106T>C), TCC136TCT (1109C>T), CCG146CCA (1139G>A), CTA150CTG (1151A>G), AAA154AAG (1163A>G), GGC156GGG (1169C>G), TTC159TTT (1178C>T), ACA162ACT (1187A>T), ACT168ACA (1205T>A), TCA175TCG (1226A>G), GCC178GCA (1235C>A), CAA182CAG (1247A>G), TCA192TCG (1277A>G), CTT204CTC (1313T>C), CGA206AGG (1317C>A 1319A>G), CAA208CAG (1325A>G), TTT209TTC (1328T>C), TTC210TTT (1331C>T), CAG214CAA (1343G>A), CAG218CAA (1355G>A), ATA220ATT (1361A>T), TCA222TCT (1367A>T), ACC228GCC (1383A>G), TCC229TCT (1388C>T), ACT232ACA (1397T>A), TCA234TCT (1403A>T), GTG235GTA (1406G>A), CCG237CCT (1412G>T), GTG239GTA (1418G>A), GGA240GGG (1421A>G), CAG242CAT (1427G>T), GAC245AAC (1434G>A), GTG250GTT (1451G>T), AGC252AGT (1457C>T), GAA253GAG (1460A>G), GCA254TCA (1461G>T), ATC257ATG (1472C>G), CGG263CGA (1490G>A), CTC267CTT (1502C>T), AGT271AGC (1514T>C), TCC275CCC (1524T>C), ACC282ACT (1547C>T), TCA283TCC (1550A>C), GTA296GTG (1589A>G), TAC303CAC (1608T>C), GAG306GAA (1619G>A), GTT311GTA (1634T>A), GTT315GTC (1646T>C), AAG317AGG (1651A>G), GCT318GCA (1655T>A), GGT320GGC (1661T>C), CTG321TTT (1662C>T 1664G>T), AAC322AAT (1667C>T), GTG326ATG (1677G>A), GAT328GAC (1685T>C), CTG340CTC (1721G>C), CCC341CCT (1724C>T), CTG347TTG (1740C>T), GTG349GTA (1748G>A), CAA352CAG (1757A>G), GCA359GCC (1778A>C), GTT365ATA (1794G>A 1796T>A), CAT371CAC (1814T>C), ATA372ATT (1817A>T), ATC373AAG (1819T>A 1820C>G), AAG377AAC (1832G>C), TTG382CTG (1845T>C), ATC385ATT (1856C>T), ACC386ACT (1859C>T), CAC388TAT (1863C>T 1865C>T), TAC397TAT (1892C>T), GTG400ACA (1899G>A 1900T>C 1901G>A), GCC402GTG (1906C>T 1907C>G), GAA409GAC (1928A>C), AAC410AAT (1931C>T), TAC412TAT (1937C>T), CTT414TTT (1941C>T), TTG416CTG (1947T>C), CAG418CAA (1955G>A), AGA419AGG (1958A>G), GAG422GAA (1967G>A), CTA423CTG (1970A>G), CAA426CAG (1979A>G), GAC428GAT (1985C>T), TAT431TAC (1994T>C), GAG457GAA (2072G>A), GTG461GTA (2084G>A), CAG465GAG (2094C>G), GCG466GCA (2099G>A), AGA469GGG (2106A>G 2108A>G), GCC475GCT (2126C>T), AGG476ATG (2128G>T), CAG478CAA (2135G>A), GTG479CTG (2136G>C), AAA480ACA (2140A>C), AAC481AAG (2144C>G), ACT486ACA (2159T>A), GTG488GGG (2164T>G), ACC494ACT (2183C>T), TCA495TCC (2186A>C), TTC496TTT (2189C>T), AAG499AAA (2198G>A), GTT500ACT (2199G>A 2200T>C), ATG502GTG (2205A>G), GAG503GAA (2210G>A), GGC505GGG (2216C>G), ATG510GAG (2229A>G 2230T>A), AAA511GAA (2232A>G), GCT513GCA (2240T>A), ATT514ATA (2243T>A), GAA517GAT (2252A>T), GCC519CCC (2256G>C), AAT520GAG (2259A>G 2261T>G), GAA521CAA (2262G>C), GTA523GTG (2270A>G), GTT528GTC (2285T>C), GAA531GAC (2294A>C), CAA534CAG (2303A>G), AAT537AAC (2312T>C), AAT538AAA (2315T>A), GGT540GCC (2320G>C 2321T>C), ACA542ATA (2326C>T), TTA546TTG (2339A>G), CAA548CAG (2345A>G), TTC558TTT (2375C>T), GAG564GAA (2393G>A), AGT571AGC (2414T>C), GTT573GTG (2420T>G), GTG577GTA (2432G>A), TCA579GCA (2436T>G), GCG581GCC (2444G>C), CTC582CTA (2447C>A), GCG584GCA (2453G>A), CTC586CTG (2459C>G), TCC590ACG (2469T>A 2471C>G), AAG591AAA (2474G>A), GGA596GGC (2489A>C), TCG597TCA (2492G>A), CAA598CAG (2495A>G), CTA600CTG (2501A>G), AGG601AAG (2503G>A), TAT602TAC (2507T>C), GCC605GCT (2516C>T), ATT608CTT (2523A>C), TAT615TAC (2546T>C), TCC617TCT (2552C>T), GGT620GGG (2561T>G), ATT622ATA (2567T>A), TCG623TCA (2570G>A), GGT624GGA (2573T>A), ATC643ATT (2630C>T), CTG648CTA (2645G>A), CAT653CAC (2660T>C), GGG664GGA (2693G>A), ACA669ACG (2708A>G), ATC673ATA (2720C>A), GTT677GTC (2732T>C), CTG688CTA (2765G>A), CAC697CAT (2792C>T), GAG706GAA (2819G>A), CAT713CAC (2840T>C), CTG714TTG (2841C>T), AAA717AAG (2852A>G), ATC723ATA (2870C>A), CCT726CCA (2879T>A), CGT728CGC (2885T>C), TTG729CTA (2886T>C 2888G>A), GCA733GCT (2900A>T), AAA738AAG (2915A>G), CGA746CGG (2939A>G), GTA752GTT (2957A>T), CTG753TTG (2958C>T), CAC755CAT (2966C>T), AAT757AAC (2972T>C), GCT761ACT (2982G>A), ACT766ACA (2999T>A), TTA768CTA (3003T>C), AGC770AGT (3011C>T), ATC771GTC (3012A>G), TCT773TCC (3020T>C), ATA777ATT (3032A>T), ACG783ACA (3050G>A), TTG788CTG (3063T>C), ATT807ATC (3122T>C), GAG810GAA (3131G>A), CAT815CAC (3146T>C), GGT817GGA (3152T>A), GGA822GGT (3167A>T), AGC829AAC (3187G>A), GAC835GAT (3206C>T), GTT842GTG (3227T>G), ATT843ATA (3230T>A), AAC845AAT (3236C>T), GAG847GAA (3242G>A), GGA849GGC (3248A>C), AAG851AGG (3253A>G), AGG855AAG (3265G>A), ACG857TCG (3270A>T), ACC868TCT (3303A>T 3305C>T), CTA875TTA (3324C>T), CCC878CCT (3335C>T), CAC883CAT (3350C>T), AGG885CGG (3354A>C), TTG886TTA (3359G>A), ACA889ACG (3368A>G), GTC890GTG (3371C>G), GTA895GTT (3386A>T), CTA901CTG (3404A>G), GTC904GTT (3413C>T), ACA907ACG (3422A>G), CAG909CAA (3428G>A), CTT910CTC (3431T>C), TGC912TGT (3437C>T), GAT915GAC (3446T>C), TTG916TTA (3449G>A), GTC917GTT (3452C>T), CTT918CTA (3455T>A), TCT923TCC (3470T>C), CCC926CCT (3479C>T), ATT927ATC (3482T>C), TAC928TAT (3485C>T), CAC930CAT (3491C>T), GGA934GGG (3503A>G), AAT936AGT (3508A>G), GCT937GCC (3512T>C), GTG938GTT (3515G>T), CTA941TTG (3522C>T 3524A>G), TTC942TTT (3527C>T), GAG944GAA (3533G>A), TCT948ACT (3543T>A), CGC950CGT (3551C>T), ATT953ATA (3560T>A), TTA956TTG (3569A>G), AAG958AAA (3575G>A), AAC960AAT (3581C>T), ACA963ACC (3590A>C), CAA975CAG (3626A>G), CTC978CTT (3635C>T), AGA979AGG (3638A>G), TCA983TCT (3650A>T), GTA987GTT (3662A>T), GTG991GTA (3674G>A), CCC1007CCT (3722C>T), CAG1014CAA (3743G>A), GTA1018GTG (3755A>G), ATC1022ATT (3767C>T), TGC1024TGT (3773C>T), TTC1033TTT (3800C>T), TAC1042TAT (3827C>T), ATT1044GTT (3831A>G), AAC1060AAT (3881C>T), ACC1062GCC (3885A>G), TTA1063TTG (3890A>G), GAT1064GAC (3893T>C), ACC1078GCC (3933A>G), GTT1080GTG (3941T>G), GCT1083GCA (3950T>A), GGT1094GGA (3983T>A), ATG1095GTT (3984A>G 3986G>T), GAC1097AAC (3990G>A), AAC1098AAT (3995C>T), ATC1099ATT (3998C>T), GTG1104GTC (4013G>C), TTT1110TTC (4031T>C), CGC1113CGT (4040C>T), ATA1120GTG (4059A>G 4061A>G), AAT1123AGT (4069A>G), CGA1133CGT (4100A>T), CTC1137CTT (4112C>T), GCC1143GCT (4130C>T), AAG1174AAA (4223G>A), CAT1175TAT (4224C>T), ACC1183ACT (4250C>T) | | | | | | | | |

*: Inserts / Deletes / Misaligned / Frameshifts

#### Analysis details

This analysis was performed with panviral2.64

### NGS Details (UN62): Torradovirus marchitezum (segment RNA 1)

#### Assembly

| | |
|---|---|
| **Coverage Length** | 5822 (2 contig(s)) |
| **Depth Of Coverage** | 2179.8 |
| **Number Of Reads** | 96951 |
| **Reads Per Million** | 2247.41 rpm (after QC) |
| **Ambiguities** | 0 |
| **Assembly Method** | de novo + reference guided assembly |
| **Consensus Caller** | Bcf Tools |

#### Coverage Map

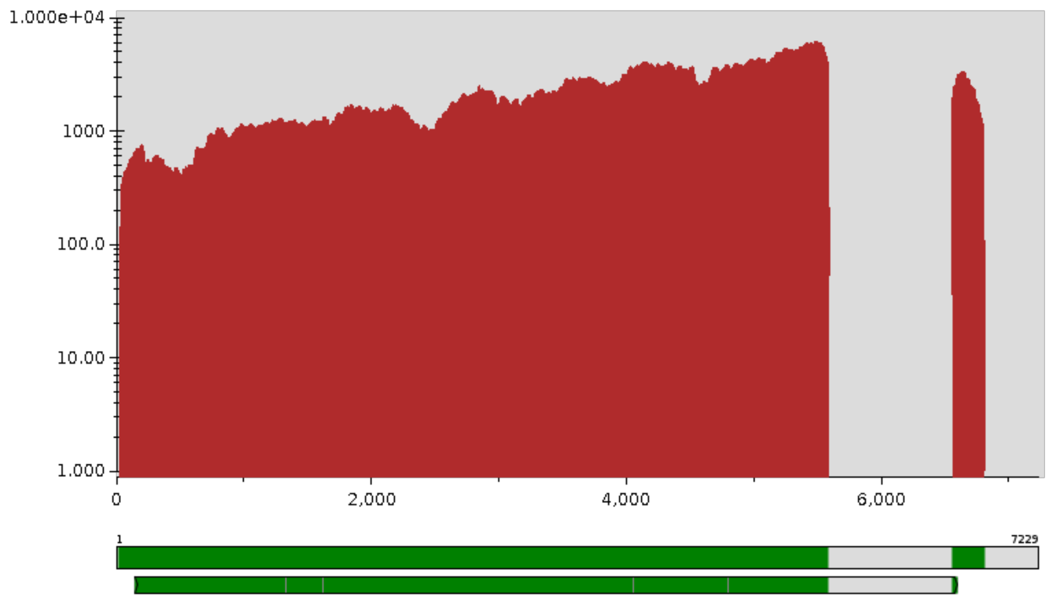

#### Assignment

| | |
|---|---|
| **Type** | Torradovirus marchitezum (Taxonomy ID: 3048376) |
| **Reference Genome** | NC_010987.1 |
| **NT Identity (%)** | 67.1567 |
| **AA Identity (%)** | 69.9945 |
| **Number Of Stop Codons** | 1 |
| **Number Of CDS** | 1 |

#### Alignment

| | |
|---|---|
| **Alignment Score** | 3829.0 (NT) + 9356.0 (AA) = 13185.0 |
| **Concordance (%)** | 54.5579 |

| | |
|---|---|
| **Alignment Method** | Global, seeded, nucleotide + amino acids (AGA) |

#### Genome Region

Sequence starts at position 22 and ends at position 6809 relative to NC_010987.1 reference sequence.

#### Alignment Detailed Statistics

| | Begin | End | Coverage | Score | Concordance | Matches | Identities | I/D/M/F* | Stop Codons |
|---|---|---|---|---|---|---|---|---|---|
| **NT** | **22** | **6809** | **80.5%** | **3829** | **33.5%** | **5772 (99.0%)** | **3883 (66.6%)** | **10/50** | |

| | Begin | End | Coverage | Score | Concordance | Matches | Identities | I/D/M/F* | Stop Codons |
|---|---|---|---|---|---|---|---|---|---|
| **NT** | **22** | **6809** | **80.5%** | **3829** | **33.5%** | **5772 (99.0%)** | **3883 (66.6%)** | **10/50** | |

Mutations:

32T>C, 37G>A, 40C>T, 59A>C, 61T>C, 62T>A, 63G>T, 66A>C, 67_78delTTTGATTCTTGT, 81A>T, 82C>A, 83T>C, 87C>G, 89A>T, 90A>T, 91C>T, 94C>T, 99T>A, 100_101insC, 108A>T, 112G>C, 114C>T, 115C>T, 116A>C, 117T>C, 118T>C, 119A>T, 122T>G, 125A>T, 128C>A, 129C>A, 130T>A, 133_134delAT, 136T>A, 146T>A, 160T>G, 161C>G, 162T>A, 163C>G, 164C>T, 165_188delAAGGTTGCTGGTGTTGATGACGCC, 189A>C, 190G>C, 192T>C, 195T>G, 200C>T, 201A>T, 203C>A, 204A>C, 217G>C, 218T>A, 222C>G, 227A>T, 228A>T, 229A>C, 234A>T, 239C>T, 240A>T, 241A>C, 242A>T, 243T>C, 251A>G, 252G>A, 254G>T, 255G>C, 257T>C, 258A>T, 259G>C, 260T>C, 264A>G, 268C>A, 269C>T, 271C>A, 272T>A, 273A>G, 275G>T, 283A>C, 284G>C, 288T>G, 289C>T, 296T>C, 297C>A, 299T>A, 306C>G, 308G>C, 311A>G, 314A>G, 315T>C, 317T>A, 320G>T, 322C>T, 323T>G, 325A>C, 326T>A, 328T>C, 329G>A, 330T>G, 332C>A, 338T>A, 343T>C, 344G>C, 345G>T, 347C>T, 350T>A, 353T>C, 355G>A, 357T>G, 359A>T, 360C>G, 362T>A, 363A>G, 365T>A, 367A>G, 374G>A, 377C>T, 380A>G, 385G>C, 386T>A, 387G>A, 388C>A, 390C>A, 392T>A, 395A>G, 398T>A, 401A>C, 404T>C, 405C>A, 407T>C, 410A>G, 412C>G, 413G>T, 416C>T, 417T>G, 418C>A, 422C>T, 423T>G, 425G>A, 426T>A, 427C>A, 428C>A, 431A>T, 437C>T, 441T>A, 446C>A, 447A>G, 448A>T, 449G>A, 453A>C, 455G>T, 458T>A, 459T>C, 461A>T, 464A>G, 465T>A, 468T>G, 471G>T, 472A>T, 473G>A, 477T>G, 479G>C, 480T>A, 481C>A, 482C>G, 483A>G, 489G>T, 491T>C, 493A>T, 494A>C, 495T>C, 502T>A, 503T>C, 505C>A, 506T>A, 510G>T, 512G>A, 516A>G, 518T>A, 522A>G, 523G>T, 527A>G, 528G>T, 529A>C, 530T>A, 531G>T, 542T>G, 543G>T, 545A>T, 546T>A, 547A>T, 548T>A, 550G>C, 555C>T, 557T>A, 560A>T, 567A>T, 569A>G, 570A>G, 572A>C, 573T>A, 575G>T, 578A>G, 579G>C, 581G>A, 584T>A, 587A>G, 589T>A, 593T>A, 596T>C, 600A>G, 601G>T, 602T>A, 611T>A, 612G>A, 613A>C, 615G>C, 616G>C, 618G>T, 621T>A, 622C>A, 626C>T, 627A>G, 628C>T, 629T>A, 632C>T, 633A>G, 634A>T, 635A>T, 637A>C, 639A>T, 641C>G, 646T>C, 647G>A, 648A>G, 650A>T, 653C>T, 659C>T, 660A>T, 662G>T, 665T>C, 668A>T, 669A>T, 670G>C, 671T>A, 672C>T, 674A>G, 677C>T, 697A>C, 702G>T, 704C>T, 706A>G, 707A>G, 708C>A, 709G>A, 710A>G, 713C>T, 714A>T, 717A>C, 718T>A, 720G>C, 725C>G, 731T>A, 732A>G, 733G>C, 734T>A, 735A>G, 737G>A, 746G>T, 750T>A, 751C>G, 752A>C, 753G>T, 755T>A, 756T>A, 758A>G, 770G>T, 771T>A, 772C>G, 773T>C, 776G>A, 777T>C, 779G>T, 780A>T, 782G>A, 786A>T, 789A>T, 791A>G, 803G>T, 806T>C, 809G>A, 812G>T, 815G>A, 818G>A, 821C>T, 822A>G, 827A>T, 830G>T, 836T>G, 848C>T, 849T>C, 851G>T, 852A>T, 853G>C, 854C>A, 861T>C, 866C>A, 872T>C, 875C>G, 878T>C, 881G>T, 882A>G, 884A>T, 896T>G, 899G>A, 911A>G, 914A>G, 918A>C, 919C>A, 920T>G, 921C>T, 923T>G, 927C>A, 929T>G, 932T>C, 938A>G, 944C>T, 947C>A, 950C>T, 953G>A, 956T>C, 974A>G, 978A>G, 980A>G, 983C>T, 984G>A, 988C>T, 989C>T, 995A>T, 998C>T, 1010G>A, 1019T>G, 1024G>A, 1028A>T, 1031G>C, 1032C>T, 1034G>A, 1040T>C, 1046G>T, 1050G>C, 1059C>A, 1060A>C, 1064G>A, 1068G>T, 1069A>C, 1070T>G, 1072C>A, 1073A>G, 1076G>T, 1077T>G, 1078G>C, 1082T>A, 1085T>A, 1086G>T, 1088A>T, 1090C>T, 1100G>A, 1103C>T, 1109C>A, 1112T>C, 1113A>C, 1114A>C, 1115G>T, 1118T>A, 1121C>T, 1124T>A, 1126C>T, 1127C>T, 1130A>G, 1134A>C, 1136C>A, 1140C>T, 1142G>A, 1144G>A, 1145G>A, 1148A>G, 1149G>C, 1151A>G, 1154T>C, 1155C>G, 1160G>A, 1163C>A, 1178A>C, 1190T>C, 1193G>A, 1199C>T, 1200A>T, 1215A>T, 1216G>C, 1226T>A, 1227G>A, 1233T>C, 1238A>T, 1242G>C, 1244T>G, 1245T>G, 1246A>T, 1253A>G, 1254A>G, 1255G>C, 1256T>C, 1257T>A, 1259T>C, 1260A>C, 1261G>A, 1262G>A, 1265C>T, 1266A>T, 1267T>C, 1268G>T, 1271C>T, 1280G>A, 1284C>A, 1286T>G, 1292A>G, 1298T>C, 1304A>G, 1313C>T, 1316G>A, 1317T>G, 1319C>T, 1325G>A, 1337T>A, 1340A>G, 1342C>G, 1343C>T, 1350T>C, 1352A>T, 1355T>C, 1358T>C, 1367G>T, 1368T>C, 1370G>C, 1374A>G, 1375G>C, 1385G>A, 1391A>G, 1394C>G, 1397T>C, 1400G>C, 1401C>T, 1403A>G, 1409A>G, 1418G>A, 1421G>A, 1445T>A, 1448C>T, 1457T>A, 1458A>G, 1460C>G, 1463G>T, 1472C>T, 1478A>T, 1481A>T, 1482T>C, 1484G>T, 1487A>G, 1493G>A, 1496T>A, 1505C>T, 1508A>G, 1511T>C, 1514T>A, 1517A>G, 1529T>C, 1538A>T, 1541A>C, 1544A>G, 1547A>T, 1553C>T, 1554T>C, 1556A>T, 1565C>T, 1571T>C, 1577C>T, 1583C>A, 1589T>C, 1592C>T, 1595C>A, 1598C>A, 1604T>C, 1607A>C, 1616A>C, 1622T>C, 1623C>T, 1628C>A, 1631A>T, 1637T>A, 1640C>T, 1643A>G, 1649T>G, 1652T>A, 1655A>G, 1661A>T, 1667T>A, 1673G>A, 1674C>A, 1679A>G, 1682G>A, 1685C>T, 1688C>T, 1689C>T, 1691G>A, 1694G>T, 1700T>C, 1702T>G, 1703G>A, 1708A>T, 1715G>T, 1719T>G, 1720C>A, 1721C>T, 1722A>C, 1724T>A, 1725G>A, 1727A>T, 1728G>A, 1729A>C, 1733T>A, 1736G>A, 1738G>C, 1739A>C, 1745C>A, 1747G>T, 1748A>T, 1752A>G, 1754C>T, 1755A>G, 1757A>G, 1763G>A, 1775T>C, 1781T>C, 1787T>A, 1790C>T, 1793G>A, 1794A>G, 1795A>C, 1799G>A, 1800A>T, 1806A>G, 1807G>A, 1811T>C, 1814T>C, 1818G>C, 1820G>A, 1829A>G, 1832C>A, 1835A>G, 1838T>C, 1840A>G, 1844A>G, 1847T>C, 1848G>T, 1849C>A, 1850A>T, 1853A>G, 1856A>G, 1861A>G, 1862G>C, 1866T>C, 1868G>T, 1870T>A, 1871T>C, 1872G>A, 1874A>T, 1875G>A, 1878G>A, 1879A>C, 1883T>G, 1886C>A, 1887A>T, 1889A>G, 1890A>G, 1891G>C, 1893C>G, 1894C>A, 1895T>C, 1899A>G, 1901A>T, 1910T>C, 1911C>A, 1913A>T, 1917T>G, 1921C>T, 1922A>T, 1928T>A, 1932A>C, 1934A>G, 1936C>A, 1937G>T, 1941G>T, 1943G>C, 1944G>C, 1946T>A, 1948G>A, 1949G>T, 1952T>C, 1953_1955delAGA, 1958G>A, 1960G>A, 1961C>A, 1962A>G, 1963C>T, 1964C>T, 1965A>T, 1966A>T, 1969A>C, 1970A>T, 1971G>C, 1972A>C, 1976A>G, 1980A>G, 1981T>C, 1983T>G, 1984T>C, 1985G>T, 1986G>T, 1987T>A, 1988G>T, 1989C>G, 1990C>A, 1991C>G, 1992A>C, 1993C>A, 1994T>G, 1995G>A, 1996T>C, 1997T>A, 1998G>A, 2000G>T, 2002G>C, 2003T>A, 2007A>G, 2008A>C, 2012T>A, 2014G>A, 2016G>A, 2018G>T, 2019A>G, 2021C>T, 2024T>C, 2025G>T, 2027A>C, 2028A>C, 2029G>A, 2030A>G, 2033G>A, 2039G>A, 2042A>T, 2048A>C, 2049T>C, 2051G>C, 2060A>T, 2066G>A, 2069C>G, 2073T>A, 2076G>A, 2079C>A, 2080C>A, 2081T>A, 2082G>T, 2083A>T, 2084G>T, 2087A>G, 2088C>T, 2093T>C, 2096T>A, 2104A>T, 2105T>C, 2106C>T, 2107A>C, 2108T>C, 2117C>T, 2120G>C, 2127T>G, 2128C>T, 2129T>G, 2132T>A, 2133G>T, 2134A>C, 2136G>T, 2137A>C, 2141A>T, 2142C>G, 2143A>C, 2144T>C, 2148G>C, 2152T>A, 2157A>G, 2159C>A, 2161A>T, 2162A>G, 2166A>G, 2167G>T, 2169G>A, 2170T>A, 2171C>T, 2172A>G, 2174C>T, 2175A>T, 2181T>G, 2182A>T, 2183T>C, 2185G>A, 2193G>A, 2194A>G, 2196G>C, 2199G>C, 2201A>G, 2208C>G, 2209G>C, 2210A>T, 2219A>G, 2221A>G, 2237C>T, 2243A>C, 2249C>T, 2252T>C, 2268G>A, 2269C>G, 2270A>C, 2271T>A, 2279C>T, 2285G>A, 2286A>G, 2289A>C, 2290A>T, 2291A>G, 2294A>G, 2296C>A, 2297A>T, 2300G>C, 2303G>A, 2304A>C, 2305C>G, 2306A>C, 2307G>A, 2309G>T, 2310A>C, 2312G>T, 2313A>G, 2314G>A, 2316A>C, 2317C>A, 2318T>A, 2321A>G, 2322T>A, 2323T>A, 2324T>A, 2327A>G, 2328C>T, 2331T>A, 2332C>A, 2333T>G, 2334T>G, 2335G>C, 2336C>G, 2337A>C, 2340G>C, 2342T>G, 2345A>T, 2350T>C, 2351T>A, 2354C>T, 2357C>A, 2360C>T, 2362T>C, 2363A>G, 2369C>T, 2372G>A, 2385T>A, 2386T>G, 2387T>A, 2390T>A, 2396T>C, 2397T>C, 2399A>G, 2405A>T, 2408A>T, 2415A>C, 2417A>C, 2426C>T, 2428A>G, 2429C>G, 2435A>T, 2436T>G, 2438_2439insGGT, 2440A>C, 2441A>T, 2444G>C, 2445C>A, 2446C>A, 2447A>G, 2451G>A, 2452A>G, 2454T>A, 2455C>A, 2456A>T, 2462C>T, 2464A>T, 2465A>C, 2466A>G, 2467T>A, 2468T>A, 2472T>A, 2473A>T, 2474C>G, 2475C>A, 2476A>T, 2478A>G, 2479A>G, 2480A>T, 2484G>A, 2485T>A, 2486T>G, 2489A>C, 2490T>A, 2492T>A, 2495C>G, 2497A>T, 2498G>C, 2499G>C, 2501T>G, 2503C>T, 2504A>G, 2508T>A, 2509G>A, 2510G>T, 2511A>G, 2512G>A, 2514C>T, 2516T>G, 2521A>T, 2522T>C, 2523G>A, 2524C>T, 2525A>C, 2526C>T, 2530G>A, 2531A>G, 2537C>T, 2546A>C, 2549T>A, 2556A>T, 2558A>G, 2561G>A, 2564T>A, 2573A>C, 2574G>A, 2575T>C, 2576T>C, 2577T>A, 2579T>G, 2580G>A, 2581G>A, 2582A>T, 2584G>C, 2586A>G, 2591A>T, 2594A>T, 2597T>C, 2598T>A, 2600G>T, 2601G>A, 2603G>T, 2607T>C, 2608G>T, 2609G>C, 2610C>A, 2613A>G, 2614G>C, 2615T>A, 2616A>G, 2617A>C, 2618C>A, 2619T>A, 2623T>C, 2627C>T, 2628A>T, 2629A>T, 2630T>A, 2633A>T, 2634T>G, 2635C>A, 2636G>T, 2645A>G, 2648T>C, 2651G>A, 2660A>G, 2672A>C, 2678T>C, 2684G>A, 2687C>T, 2690G>A, 2693T>G, 2696T>G, 2698A>C, 2699G>T, 2699_2700insCAT, 2700C>T, 2703A>G, 2704G>C, 2708T>A, 2709C>T, 2710T>C, 2711T>C, 2712C>T, 2714C>G, 2715G>C, 2717T>A, 2720T>G, 2724A>G, 2725G>C, 2726T>G, 2727T>G, 2729T>A, 2730C>G, 2732A>T, 2733T>A, 2734C>A, 2735A>C, 2736T>A, 2737T>A, 2738G>C, 2739G>C, 2740C>A, 2741A>T, 2742C>G, 2743A>T, 2744A>G, 2747T>C, 2747_2748insCGG, 2753T>A, 2755C>A, 2756T>A, 2757G>C, 2759A>G, 2771A>T, 2775C>G, 2777G>A, 2780C>T, 2783C>G, 2784A>C, 2786G>C, 2792T>C, 2798G>T, 2801G>T, 2804A>G, 2808C>A, 2812G>C, 2822G>C, 2825C>T, 2828T>C, 2834C>T, 2837C>G, 2840C>T, 2843G>T, 2858A>T, 2859A>C, 2861G>C, 2864C>T, 2867A>G, 2870T>A, 2876A>C, 2879C>T, 2880C>A, 2881A>G, 2882C>A, 2885T>C, 2886G>A, 2887C>T, 2888T>G, 2889C>A, 2891A>G, 2897T>C, 2903A>T, 2907A>T, 2909A>C, 2915C>T, 2920C>A, 2921C>G, 2922A>C, 2927T>C, 2928C>A, 2929G>T, 2930A>T, 2932C>G, 2936A>T, 2939T>G, 2942A>G, 2944A>T, 2948C>T, 2953G>C, 2954A>C, 2955G>C, 2964A>T, 2966T>G, 2969T>A, 2975C>T, 2981C>T, 2982A>G, 2984C>G, 2988C>T, 2994C>A, 2996C>G, 2999T>A, 3002T>G, 3017A>C, 3020G>A, 3021C>T, 3023T>A, 3026C>T, 3029C>A, 3030C>A, 3032T>A, 3038C>T, 3044A>T, 3045A>C, 3047A>C, 3050G>A, 3056G>T, 3062C>T, 3064T>A, 3066A>T, 3068A>T, 3078C>A, 3079A>T, 3080A>G, 3083A>C, 3090T>G, 3091G>T, 3092T>G, 3095A>G, 3101T>A, 3104G>T, 3107T>A, 3109C>A, 3110A>G, 3116A>C, 3117C>A, 3118G>A, 3119T>G, 3120C>A, 3121T>A, 3122T>A, 3123G>T, 3124G>C, 3131A>G, 3135T>G, 3140C>T, 3143G>A, 3144A>T, 3145T>A, 3146G>T, 3167A>C, 3173A>C, 3175T>A, 3176T>A, 3182G>C, 3183G>A, 3184A>C, 3190A>G, 3192C>T, 3194G>A, 3195A>G, 3197C>T, 3200C>G, 3201T>A, 3202C>A, 3203C>T, 3204A>G, 3205G>C, 3206T>A, 3210A>G, 3211A>C, 3212G>T, 3213A>C, 3216T>A, 3221T>A, 3225C>G, 3226A>T, 3227A>T, 3228T>G, 3230T>A, 3236A>G, 3245A>T, 3249G>A, 3251G>T, 3254A>G, 3260T>A, 3263C>T, 3269C>T, 3272G>T, 3273T>C, 3275G>T, 3278A>C, 3279C>T, 3281G>A, 3284A>G, 3289G>C, 3293T>G, 3299T>C, 3305A>G, 3308T>G, 3309G>T, 3310T>G, 3311G>C, 3320C>A, 3323C>A, 3326C>T, 3335A>T, 3343G>A, 3344C>T, 3345G>A, 3346T>C, 3347A>C, 3350A>T, 3353T>A, 3356A>G, 3357C>A, 3358G>A, 3359T>G, 3362A>T, 3364T>A, 3365T>C, 3366A>G, 3368C>G, 3369T>G, 3377T>C, 3380T>A, 3386T>A, 3387A>G, 3389C>G, 3391A>T, 3392C>T, 3395G>A, 3398A>T, 3402C>T, 3404A>G, 3407G>A, 3416A>C, 3419T>G, 3422G>A, 3425G>C, 3426G>A, 3428C>T, 3433C>T, 3434A>G, 3437A>T, 3443A>G, 3452T>A, 3453A>C, 3455T>G, 3456C>T, 3462T>C, 3463C>A, 3464A>C, 3465A>C, 3467T>G, 3471A>T, 3472G>C, 3475G>A, 3476A>G, 3479A>G, 3480A>G, 3481C>A, 3483C>A, 3484A>G, 3485G>A, 3486C>T, 3488T>C, 3497A>G, 3500C>A, 3503A>G, 3507T>G, 3508A>C, 3509C>T, 3511A>T, 3513C>T, 3515T>G, 3518A>C, 3521A>T, 3524T>A, 3527A>G, 3528C>T, 3529A>T, 3533G>A, 3534T>A, 3539C>T, 3542C>T, 3548T>G, 3551C>G, 3557G>A, 3560C>A, 3563T>C, 3566G>T, 3567C>A, 3568G>A, 3569G>A, 3570A>C, 3571A>G, 3572A>C, 3575C>T, 3577A>C, 3578A>C, 3579T>C, 3587A>G, 3590A>T, 3593C>T, 3596A>G, 3597C>G, 3598C>A, 3599T>C, 3602A>T, 3603G>C, 3608A>T, 3611A>C, 3624A>T, 3626A>T, 3628T>C, 3629T>C, 3630C>A, 3632T>A, 3641A>G, 3644T>A, 3647T>A, 3650T>G, 3656C>T, 3671G>T, 3674A>C, 3675A>G, 3676T>A, 3680A>G, 3683C>T, 3697G>C, 3698T>C, 3699G>A, 3700G>A, 3701A>C, 3707T>G, 3710T>G, 3711G>C, 3712A>C, 3713C>A, 3719C>A, 3722A>G, 3725A>G, 3728A>T, 3731C>T, 3734A>G, 3735A>G, 3736A>G, 3740C>T, 3743G>A, 3744T>G, 3745G>C, 3746C>A, 3749G>C, 3750C>A, 3752G>T, 3755A>T, 3759A>G, 3761C>G, 3762A>C, 3764G>T, 3765C>A, 3766G>A, 3767T>G, 3769G>T, 3770C>G, 3774A>C, 3775A>G, 3776G>A, 3777A>G, 3779C>G, 3782T>C, 3788A>G, 3789C>A, 3790G>A, 3791T>A, 3794A>T, 3795T>G, 3796G>C, 3797T>A, 3798G>A, 3799A>T, 3800A>C, 3803G>A, 3804G>A, 3827A>G, 3838G>T, 3845C>T, 3847A>T, 3853G>C, 3854C>A, 3860G>A, 3861C>T, 3869A>G, 3872T>A, 3875C>T, 3878A>T, 3884C>T, 3887C>T, 3894A>T, 3899G>A, 3900A>C, 3902G>T, 3905G>T, 3906A>T, 3907T>C, 3908G>T, 3914G>A, 3917A>G, 3927C>G, 3930T>G, 3933A>C, 3934C>A, 3935T>A, 3936G>C, 3937G>C, 3938C>A, 3939G>A, 3941A>C, 3942G>T, 3943G>C, 3944A>G, 3945G>C, 3950A>G, 3953G>T, 3954G>C, 3955A>T, 3956A>T, 3961A>T, 3965A>G, 3966G>A, 3971T>A, 3977A>C, 3983A>T, 3996C>T, 3998A>G, 4001T>C, 4002G>T, 4003T>C, 4008A>G, 4010T>A, 4014A>C, 4021T>C, 4022C>A, 4023C>T, 4024A>T, 4028G>T, 4031T>C, 4034G>C, 4040C>A, 4046A>G, 4049C>T, 4052C>G, 4064C>G, 4065C>T, 4067C>G, 4079A>G, 4082C>T, 4088C>G, 4094A>G, 4101A>C, 4103G>C, 4106C>G, 4109T>C, 4112G>A, 4115A>C, 4116C>T, 4118T>G, 4130C>T, 4137T>C, 4142C>G, 4143A>C, 4148G>A, 4151T>C, 4154T>C, 4160T>C, 4163C>T, 4166A>T, 4169C>T, 4170A>T, 4171C>G, 4172A>T, 4182T>G, 4183T>C, 4184G>A, 4190T>C, 4192C>T, 4193T>A, 4194A>C, 4196A>C, 4199A>T, 4205A>G, 4208T>A, 4214T>C, 4215G>A, 4220T>A, 4223A>G, 4226T>C, 4227T>C, 4228C>A, 4239T>G, 4242T>C, 4247A>T, 4249A>G, 4251G>A, 4253G>C, 4259A>G, 4260G>T, 4262T>A, 4265G>A, 4268A>T, 4271C>T, 4273T>A, 4280T>C, 4286T>C, 4292T>C, 4295C>G, 4316T>C, 4317G>A, 4319T>C, 4325T>A, 4328T>C, 4331G>A, 4344T>C, 4354A>C, 4355A>T, 4357A>T, 4358C>T, 4361C>T, 4364A>T, 4365T>G, 4366C>A, 4367A>T, 4368A>G, 4376C>G, 4385T>G, 4388C>A, 4395C>A, 4397A>G, 4400T>C, 4401G>A, 4404C>T, 4406T>A, 4418A>C, 4424T>C, 4430C>T, 4433A>C, 4436G>C, 4439C>T, 4443G>A, 4445T>C, 4448A>G, 4449G>T, 4455A>C, 4457G>C, 4458T>C, 4460G>A, 4463T>G, 4466G>T, 4469C>A, 4475A>C, 4478A>T, 4481A>T, 4487A>T, 4488C>T, 4490A>G, 4493G>T, 4499C>A, 4502C>T, 4508T>A, 4509C>T, 4511C>G, 4514C>A, 4530C>A, 4532A>G, 4535T>C, 4538C>G, 4548T>C, 4553A>G, 4556A>G, 4562C>A, 4566A>G, 4568T>C, 4569G>A, 4570C>A, 4571T>A, 4576G>A, 4577G>A, 4578G>T, 4581A>G, 4583A>G, 4586A>C, 4587G>A, 4592A>G, 4597G>A, 4601T>C, 4611C>T, 4613A>G, 4622T>A, 4625T>A, 4631A>T, 4637C>T, 4640T>C, 4641C>T, 4643T>G, 4649T>G, 4652A>T, 4656T>A, 4658A>G, 4662C>T, 4664C>G, 4665C>A, 4666A>G, 4673T>C, 4674T>G, 4675C>A, 4679G>A, 4683A>T, 4684A>C, 4685T>C, 4688A>T, 4692C>A, 4693A>G, 4694T>A, 4697G>A, 4698T>C, 4700A>G, 4704C>G, 4705T>C, 4706G>T, 4709C>T, 4712T>C, 4715A>C, 4716A>G, 4731C>T, 4733T>G, 4739A>G, 4742G>A, 4751T>A, 4752A>G, 4754T>G, 4755G>A, 4758G>A, 4759T>C, 4761C>A, 4768A>T, 4769A>G, 4784A>C, 4787T>C, 4793G>C, 4799A>G, 4805T>C, 4808A>G, 4817T>A, 4818C>T, 4820T>G, 4826C>T, 4829C>A, 4835T>A, 4838T>A, 4844G>A, 4854A>G, 4859T>C, 4868G>T, 4877A>T, 4880A>G, 4883T>A, 4886C>A, 4887C>G, 4888C>T, 4889A>G, 4892T>A, 4901C>T, 4902A>G, 4903C>T, 4904T>A, 4907G>A, 4910C>T, 4914A>T, 4915T>A, 4916G>T, 4923G>A, 4925G>T, 4934A>C, 4937A>C, 4938C>A, 4940T>A, 4946T>C, 4955C>T, 4958T>G, 4960G>A, 4961T>G, 4963T>C, 4964G>C, 4965G>A, 4973C>A, 4975A>G, 4976G>T, 4977C>T, 4982C>A, 4983T>A, 4984T>A, 4985A>T, 4992C>T, 4993A>T, 4994G>T, 4995A>G, 4996C>G, 4997G>A, 5000G>T, 5001T>C, 5003G>C, 5004G>A, 5006T>G, 5007G>C, 5009A>G, 5012G>A, 5013G>A, 5014G>A, 5016T>C, 5019A>C, 5020G>C, 5021A>C, 5022G>A, 5023C>T, 5024T>C, 5025G>A, 5031T>A, 5033T>G, 5036T>C, 5037T>A, 5039C>T, 5042T>C, 5044T>A, 5045G>T, 5049G>C, 5051G>T, 5055C>A, 5056A>G, 5057G>T, 5058G>C, 5064A>T, 5066T>G, 5067G>A, 5070A>G, 5075A>C, 5081G>A, 5085T>G, 5086C>A, 5087A>G, 5090T>C, 5091G>A, 5092C>T, 5093C>T, 5094C>G, 5095T>G, 5097A>G, 5098C>T, 5099A>T, 5100A>T, 5102T>G, 5103T>A, 5104G>T, 5105T>G, 5108A>G, 5109A>C, 5111G>A, 5112G>T, 5115A>G, 5117A>T, 5120A>T, 5121G>A, 5122C>A, 5123T>A, 5125T>A, 5127T>A, 5129C>A, 5132T>C, 5135G>C, 5138A>T, 5141G>A, 5142G>C, 5143A>C, 5144C>T, 5147A>T, 5150T>G, 5153G>A, 5159C>T, 5160_5162delACA, 5172C>G, 5173G>A, 5174T>G, 5176A>G, 5178C>G, 5180G>C, 5181A>C, 5184T>G, 5185G>A, 5186G>A, 5191C>T, 5192A>G, 5195A>T, 5196A>G, 5198C>T, 5199A>G, 5201T>A, 5202G>T, 5203G>T, 5207T>C, 5211T>A, 5212C>G, 5213A>T, 5216A>T, 5219T>G, 5226G>T, 5228C>A, 5229C>A, 5232T>C, 5234C>A, 5235A>C, 5236C>A, 5237G>A, 5241G>T, 5242A>C, 5243T>A, 5244G>T, 5245A>C, 5249G>T, 5250T>A, 5255T>C, 5256A>C, 5258T>G, 5264C>T, 5267T>C, 5270A>T, 5273T>A, 5276A>C, 5277A>C, 5278A>G, 5279G>C, 5281A>G, 5282G>A, 5283T>A, 5285C>A, 5287A>C, 5288T>A, 5294T>G, 5300T>G, 5303A>T, 5309C>T, 5333G>A, 5339G>C, 5346A>T, 5349T>G, 5354T>A, 5357A>C, 5358A>C, 5359T>A, 5364G>C, 5365C>A, 5366C>T, 5370A>C, 5371A>G, 5372G>C, 5375T>G, 5376G>A, 5377T>G, 5378G>T, 5379T>C, 5383C>A, 5386G>T, 5387A>T, 5390A>T, 5391C>G, 5395T>G, 5396A>G, 5399A>T, 5403A>T, 5404A>C, 5409G>A, 5410T>C, 5411T>C, 5417G>T, 5423A>T, 5429A>T, 5435T>A, 5436G>A, 5438C>A, 5440G>C, 5441C>T, 5447T>C, 5456G>A, 5457C>T, 5460A>T, 5462G>T, 5465C>A, 5468A>G, 5471A>C, 5474A>T, 5475A>C, 5476G>A, 5477G>T, 5483T>C, 5484T>A, 5486T>G, 5488T>C, 5489G>C, 5490G>A, 5491A>C, 5492G>A, 5496A>C, 5498A>T, 5501A>T, 5504T>G, 5505A>C, 5507A>C, 5508T>C, 5510G>A, 5513C>A, 5516A>C, 5519T>G, 5521G>C, 5522A>T, 5523A>G, 5524A>G, 5525T>G, 5531T>G, 5534C>T, 5537C>G, 5540T>C, 5546T>C, 5555C>T, 5560T>C, 5564G>A, 5567T>C, 5570T>G, 5573T>C, 5574C>A, 6565A>C, 6567A>C, 6572T>C, 6584C>T, 6589A>C, 6621G>A, 6623G>A, 6624A>T, 6625T>G, 6626A>C, 6627T>A, 6628G>A, 6641T>G, 6643G>A, 6650A>T, 6651T>G, 6652_6653delAT, 6655G>A, 6657T>G, 6664G>T, 6668_6671delTGTT, 6674G>C, 6686T>G, 6693A>T, 6697A>T, 6698T>C, 6707C>G, 6708C>G, 6709T>C, 6713C>T, 6716T>C, 6721A>G, 6722G>T, 6726A>T, 6727A>G, 6733C>T, 6737A>G, 6738T>C, 6739T>A, 6741G>T, 6743A>T, 6745A>G, 6749G>A, 6750T>C, 6759T>C, 6760C>T, 6764T>C, 6772A>T, 6774T>G, 6775T>A

| | Begin | End | Coverage | Score | Concordance | Matches | Identities | I/D/M/F* | Stop Codons |
|---|---|---|---|---|---|---|---|---|---|
| **NT** | **22** | **6809** | **80.5%** | **3829** | **33.5%** | **5772 (99.0%)** | **3883 (66.6%)** | **10/50** | |
| **CDS** | | | | | | | | | |
| ToMarV_RNA1gp1 | 1 | 2152 | 85.0% | 9356 | 73.4% | 1820 (99.3%) | 1276 (69.6%) | 3/10/0/0 | 1 |

Protein mutations:

F7W (160T>G 161C>G), K9_A16del (165_188delAAGGTTGCTGGTGTTGATGACGCC), S17P (189A>C 190G>C), S18P (192T>C), S19A (195T>G), T21S (201A>T 203C>A), T22P (204A>C), G26A (217G>C 218T>A), L28V (222C>G), N30S (228A>T 229A>C), I32F (234A>T), K34S (240A>T 241A>C 242A>T), F35L (243T>C), A38T (252G>A 254G>T), V39L (255G>C 257T>C), I42V (264A>G), T43N (268C>A 269C>T), T44K (271C>A 272T>A), M45V (273A>G 275G>T), K48T (283A>C 284G>C), S50V (288T>G 289C>T), L53I (297C>A 299T>A), L56V (306C>G 308G>C), S59P (315T>C 317T>A), T61M (322C>T 323T>G), D62A (325A>C 326T>A), V63A (328T>C 329G>A), S64A (330T>G 332C>A), V68A (343T>C 344G>C), A69S (345G>T 347C>T), S72N (355G>A), S73A (357T>G 359A>T), L74V (360C>G 362T>A), N75E (363A>G 365T>A), K76R (367A>G), G82A (385G>C 386T>A), A83K (387G>A 388C>A), L84I (390C>A 392T>A), I85M (395A>G), D86E (398T>A), L89I (405C>A 407T>C), A91G (412C>G 413G>T), S93D (417T>G 418C>A), L95V (423T>G 425G>A), S96K (426T>A 427C>A 428C>A), S101T (441T>A), K103V (447A>G 448A>T 449G>A), M105L (453A>C 455G>T), L109M (465T>A), F110V (468T>G), E111L (471G>T 472A>T 473G>A), L113V (477T>G 479G>C), S114K (480T>A 481C>A 482C>G), N115D (483A>G), V117F (489G>T 491T>C), K118I (493A>T 494A>C), F121Y (502T>A 503T>C), A122E (505C>A 506T>A), V124L (510G>T 512G>A), N126E (516A>G 518T>A), S128V (522A>G 523G>T), D130S (528G>T 529A>C 530T>A), V131L (531G>T), V135F (543G>T 545A>T), Y136I (546T>A 547A>T 548T>A), G137A (550G>C), I143L (567A>T 569A>G), T144A (570A>G 572A>C), L145I (573T>A 575G>T), V147L (579G>C 581G>A), F150Y (589T>A), S154V (600A>G 601G>T 602T>A), E158T (612G>A 613A>C), G159P (615G>C 616G>C), V160F (618G>T), S161N (621T>A 622C>A), T163V (627A>G 628C>T 629T>A), K165V (633A>G 634A>T 635A>T), N166T (637A>C), I167L (639A>T 641C>G), L169S (646T>C 647G>A), I170V (648A>G 650A>T), M174F (660A>T 662G>T), K186T (697A>C), A188S (702G>T 704C>T), E189G (706A>G 707A>G), R190K (708C>A 709G>A 710A>G), T192S (714A>T), M193Q (717A>C 718T>A), V194L (720G>C), S198A (732A>G 733G>C 734T>A), M199V (735A>G 737G>A), V205L (753G>T 755T>A), S206T (756T>A 758A>G), M214L (780A>T 782G>A), T216S (786A>T), I217L (789A>T 791A>G), I228V (822A>G), T248A (882A>G 884A>T), T260Q (918A>C 919C>A 920T>G), I280V (978A>G 980A>G), G282S (984G>A), T283I (988C>T 989C>T), S295N (1024G>A), E304Q (1050G>C), Q307T (1059C>A 1060A>C), D310S (1068G>T 1069A>C 1070T>G), A311E (1072C>A 1073A>G), C313A (1077T>G 1078G>C), D314E (1082T>A), A316S (1086G>T 1088A>T), T317M (1090C>T), K325P (1113A>C 1114A>C 1115G>T), T329I (1126C>T 1127C>T), N332Q (1134A>C 1136C>A), R335K (1144G>A 1145G>A), E337Q (1149G>C 1151A>G), L339V (1155C>G), M354L (1200A>T), A363T (1227G>A), E366D (1238A>T), V368L (1242G>C 1244T>G), Y369V (1245T>G 1246A>T), S372A (1254A>G 1255G>C 1256T>C), C373S (1257T>A 1259T>C), R374Q (1260A>C 1261G>A 1262G>A), M376S (1266A>T 1267T>C 1268G>T), S393A (1317T>G 1319C>T), S401C (1342C>G 1343C>T), R412A (1374A>G 1375G>C), I440V (1458A>G 1460C>G), M521R (1702T>G 1703G>A), E523V (1708A>T), E525D (1715G>T), S527D (1719T>G 1720C>A 1721C>T), T528P (1722A>C 1724T>A), E529N (1725G>A 1727A>T), D530T (1728G>A 1729A>C), G533A (1738G>C 1739A>C), R536I (1747G>T 1748A>T), N538D (1752A>G 1754C>T), I539V (1755A>G 1757A>G), N552A (1794A>G 1795A>C), T554S (1800A>T), S556D (1806A>G 1807G>A), E560Q (1818G>C 1820G>A), Q567R (1840A>G), A570Y (1848G>T 1849C>A 1850A>T), K574S (1861A>G 1862G>C), F577Y (1870T>A 1871T>C), E578N (1872G>A 1874A>T), A579T (1875G>A), E580T (1878G>A 1879A>C), I583L (1887A>T 1889A>G), S584A (1890A>G 1891G>C), P585D (1893C>G 1894C>A 1895T>C), I587V (1899A>G 1901A>T), Q591N (1911C>A 1913A>T), S593A (1917T>G), A594V (1921C>T 1922A>T), D596E (1928T>A), I598L (1932A>C 1934A>G), A599D (1936C>A 1937G>T), V601F (1941G>T 1943G>C), D602Q (1944G>C 1946T>A), G603D (1948G>A 1949G>T), R605del (1953_1955delAGA), R607Q (1960G>A 1961C>A), T608V (1962A>G 1963C>T 1964C>T), K609L (1965A>T 1966A>T), Q610P (1969A>C 1970A>T), D611P (1971G>C 1972A>C), I614A (1980A>G 1981T>C), L615A (1983T>G 1984T>C 1985G>T), V616Y (1986G>T 1987T>A 1988G>T), P617E (1989C>G 1990C>A 1991C>G), T618Q (1992A>C 1993C>A 1994T>G), V619T (1995G>A 1996T>C 1997T>A), V620I (1998G>A 2000G>T), S621T (2002G>C 2003T>A), N623A (2007A>G 2008A>C), D624E (2012T>A), R625K (2014G>A), V626I (2016G>A 2018G>T), I627V (2019A>G 2021C>T), A629S (2025G>T 2027A>C), R630Q (2028A>C 2029G>A 2030A>G), L645M (2073T>A), D646N (2076G>A), P647K (2079C>A 2080C>A 2081T>A), E648F (2082G>T 2083A>T 2084G>T), L650F (2088C>T), H655L (2104A>T 2105T>C), H656S (2106C>T 2107A>C 2108T>C), M660I (2120G>C), S663V (2127T>G 2128C>T 2129T>G), D665S (2133G>T 2134A>C), D666S (2136G>T 2137A>C), E667D (2141A>T), H668A (2142C>G 2143A>C 2144T>C), E670Q (2148G>C), I671N (2152T>A), N673E (2157A>G 2159C>A), K674M (2161A>T 2162A>G), R676V (2166A>G 2167G>T), V677N (2169G>A 2170T>A 2171C>T), N678D (2172A>G 2174C>T), M679L (2175A>T), Y681V (2181T>G 2182A>T 2183T>C), R682K (2185G>A), D685S (2193G>A 2194A>G), A686P (2196G>C), E687Q (2199G>C 2201A>G), R690A (2208C>G 2209G>C 2210A>T), Q694R (2221A>G), A710S (2268G>A 2269C>G 2270A>C), S711T (2271T>A), K716E (2286A>G), K717L (2289A>C 2290A>T 2291A>G), A719D (2296C>A 2297A>T), T722R (2304A>C 2305C>G 2306A>C), E723N (2307G>A 2309G>T), M724L (2310A>C 2312G>T), R725E (2313A>G 2314G>A), T726Q (2316A>C 2317C>A 2318T>A), F728K (2322T>A 2323T>A 2324T>A), H730Y (2328C>T), S731K (2331T>A 2332C>A 2333T>G), C732A (2334T>G 2335G>C 2336C>G), I733L (2337A>C), D734Q (2340G>C 2342T>G), V737A (2350T>C 2351T>A), I741T (2362T>C 2363A>G), F749R (2385T>A 2386T>G 2387T>A), N763R (2428A>G 2429C>G), S766A (2436T>G), S766_K767insG (2438_2439insGGT), K767T (2440A>C 2441A>T), E768D (2444G>C), P769K (2445C>A 2446C>A 2447A>G), E771R (2451G>A 2452A>G), S772N (2454T>A 2455C>A 2456A>T), K775I (2464A>T 2465A>C), I776E (2466A>G 2467T>A 2468T>A), Y778M (2472T>A 2473A>T 2474C>G), H779I (2475C>A 2476A>T), K780G (2478A>G 2479A>G 2480A>T), V782K (2484G>A 2485T>A 2486T>G), E783D (2489A>C), Y784K (2490T>A 2492T>A), I785M (2495C>G), K786I (2497A>T 2498G>C), D787Q (2499G>C 2501T>G), T788M (2503C>T 2504A>G), W790N (2508T>A 2509G>A 2510G>T), S791D (2511A>G 2512G>A), Y794F (2521A>T 2522T>C), A795I (2523G>A 2524C>T 2525A>C), R797K (2530G>A 2531A>G), I806L (2556A>T 2558A>G), M807I (2561G>A), V812T (2574G>A 2575T>C 2576T>C), F813M (2577T>A 2579T>G), G814N (2580G>A 2581G>A 2582A>T), G815A (2584G>C), I816V (2586A>G), L820I (2598T>A 2600G>T), A821T (2601G>A 2603G>T), W823L (2607T>C 2608G>T 2609G>C), Q824K (2610C>A), S825A (2613A>G 2614G>C 2615T>A), N826A (2616A>G 2617A>C 2618C>A), S827T (2619T>A), F828S (2623T>C), N830L (2628A>T 2629A>T 2630T>A), S832D (2634T>G 2635C>A 2636G>T), N852K (2696T>G), K853T (2698A>C 2699G>T), K853_L854insH (2699_2700insCAT), L854F (2700C>T), R855A (2703A>G 2704G>C), N856K (2708T>A), L857S (2709C>T 2710T>C 2711T>C), V859L (2715G>C 2717T>A), S862A (2724A>G 2725G>C 2726T>G), S863A (2727T>G 2729T>A), Q864D (2730C>G 2732A>T), S865N (2733T>A 2734C>A 2735A>C), L866N (2736T>A 2737T>A 2738G>C), A867H (2739G>C 2740C>A 2741A>T), Q868V (2742C>G 2743A>T 2744A>G), D869_W870insR (2747_2748insCGG), A872E (2755C>A 2756T>A), E873Q (2757G>C 2759A>G), Q879E (2775C>G 2777G>A), K882H (2784A>C 2786G>C), L890I (2808C>A), G891A (2812G>C), M907L (2859A>C 2861G>C), H914R (2880C>A 2881A>G 2882C>A), A916M (2886G>A 2887C>T 2888T>G), T923S (2907A>T 2909A>C), A927E (2920C>A 2921C>G), N928H (2922A>C), R930I (2928C>A 2929G>T 2930A>T), S931C (2932C>G), K932N (2936A>T), Y935F (2944A>T), R938T (2953G>C 2954A>C), E939Q (2955G>C), I942L (2964A>T 2966T>G), I948V (2982A>G 2984C>G), P961S (3021C>T 3023T>A), E972D (3056G>T), F975Y (3064T>A), T976S (3066A>T 3068A>T), Q980M (3078C>A 3079A>T 3080A>G), C984V (3090T>G 3091G>T 3092T>G), T990K (3109C>A 3110A>G), R993K (3117C>A 3118G>A 3119T>G), L994K (3120C>A 3121T>A 3122T>A), G995S (3123G>T 3124G>C), F999V (3135T>G), M1002Y (3144A>T 3145T>A 3146G>T), V1012E (3175T>A 3176T>A), E1015T (3183G>A 3184A>C), E1017G (3190A>G), N1019D (3195A>G 3197C>T), S1021N (3201T>A 3202C>A 3203C>T), S1022A (3204A>G 3205G>C 3206T>A), K1024A (3210A>G 3211A>C 3212G>T), I1025L (3213A>C), C1026S (3216T>A), Q1029V (3225C>G 3226A>T 3227A>T), S1030A (3228T>G 3230T>A), G1037S (3249G>A 3251G>T), S1050T (3289G>C), V1057C (3309G>T 3310T>G 3311G>C), G1068D (3343G>A 3344C>T), V1069T (3345G>A 3346T>C 3347A>C), R1073K (3357C>A 3358G>A 3359T>G), F1075Y (3364T>A 3365T>C), I1076V (3366A>G 3368C>G), S1077A (3369T>G), I1083V (3387A>G 3389C>G), Y1084F (3391A>T 3392C>T), E1086D (3398A>T), V1096I (3426G>A 3428C>T), A1098V (3433C>T 3434A>G), D1104E (3452T>A), I1105L (3453A>C 3455T>G), S1108H (3462T>C 3463C>A 3464A>C), I1109L (3465A>C 3467T>G), R1112K (3475G>A 3476A>G), T1114E (3480A>G 3481C>A), Q1115R (3483C>A 3484A>G 3485G>A), L1116F (3486C>T 3488T>C), N1120K (3500C>A), Y1123A (3507T>G 3508A>C 3509C>T), Y1124F (3511A>T), Q1130L (3528C>T 3529A>T), L1132M (3534T>A), R1143K (3567C>A 3568G>A 3569G>A), K1144R (3570A>C 3571A>G 3572A>C), Q1146P (3577A>C 3578A>C), P1153D (3597C>G 3598C>A 3599T>C), E1155Q (3603G>C), T1162S (3624A>T 3626A>T), V1163A (3628T>C 3629T>C), H1164K (3630C>A 3632T>A), D1169E (3647T>A), M1179E (3675A>G 3676T>A), C1186S (3697G>C 3698T>C), G1187N (3699G>A 3700G>A 3701A>C), D1189E (3707T>G), D1190E (3710T>G), D1191P (3711G>C 3712A>C 3713C>A), N1199G (3735A>G 3736A>G), C1202A (3744T>G 3745G>C 3746C>A), L1204I (3750C>A 3752G>T), I1207V (3759A>G 3761C>G), M1208L (3762A>C 3764G>T), R1209K (3765C>A 3766G>A 3767T>G), S1210M (3769G>T 3770C>G), K1212R (3774A>C 3775A>G 3776G>A), I1213V (3777A>G 3779C>G), R1217K (3789C>A 3790G>A 3791T>A), C1219A (3795T>G 3796G>C 3797T>A), E1220I (3798G>A 3799A>T 3800A>C), D1222N (3804G>A), C1233F (3838G>T), Y1236F (3847A>T), G1238A (3853G>C 3854C>A), M1252L (3894A>T), M1254L (3900A>C 3902G>T), M1256S (3906A>T 3907T>C 3908G>T), Q1263E (3927C>G), S1264A (3930T>G), T1265Q (3933A>C 3934C>A 3935T>A), G1266P (3936G>C 3937G>C 3938C>A), E1267N (3939G>A 3941A>C), G1268S (3942G>T 3943G>C 3944A>G), E1269Q (3945G>C), E1272L (3954G>C 3955A>T 3956A>T), K1274I (3961A>T), D1276N (3966G>A), E1281D (3983A>T), V1288S (4002G>T 4003T>C), I1290V (4008A>G 4010T>A), K1292Q (4014A>C), I1294T (4021T>C 4022C>A), H1295F (4023C>T 4024A>T), T1344C (4170A>T 4171C>G 4172A>T), L1348A (4182T>G 4183T>C 4184G>A), A1351V (4192C>T 4193T>A), I1352L (4194A>C 4196A>C), D1359N (4215G>A), S1363H (4227T>C 4228C>A), L1367V (4239T>G), N1370S (4249A>G), G1371S (4251G>A 4253G>C), A1374S (4260G>T 4262T>A), V1378E (4273T>A), V1393I (4317G>A 4319T>C), E1405A (4354A>C 4355A>T), Y1406F (4357A>T 4358C>T), S1409D (4365T>G 4366C>A 4367A>T), K1410E (4368A>G), D1415E (4385T>G), A1421T (4401G>A), D1435N (4443G>A 4445T>C), V1437L (4449G>T), I1476V (4566A>G 4568T>C), A1477K (4569G>A 4570C>A 4571T>A), R1479K (4576G>A 4577G>A), A1480S (4578G>T), I1481V (4581A>G 4583A>G), V1483I (4587G>A), S1486N (4597G>A), L1506M (4656T>A 4658A>G), Q1509R (4665C>A 4666A>G), S1512D (4674T>G 4675C>A), N1515S (4683A>T 4684A>C 4685T>C), H1518R (4692C>A 4693A>G 4694T>A), L1522A (4704C>G 4705T>C 4706G>T), I1526V (4716A>G), N1538E (4752A>G 4754T>G), E1539K (4755G>A), V1540T (4758G>A 4759T>C), Q1541K (4761C>A), Q1543L (4768A>T 4769A>G), H1563Q (4829C>A), I1572V (4854A>G), M1576I (4868G>T), P1583V (4887C>G 4888C>T 4889A>G), T1588V (4902A>G 4903C>T 4904T>A), M1592Y (4914A>T 4915T>A 4916G>T), V1595I (4923G>A 4925G>T), R1607Q (4960G>A 4961T>G), V1608A (4963T>C 4964G>C), A1609T (4965G>A), D1611E (4973C>A), K1612S (4975A>G 4976G>T), L1613F (4977C>T), L1615N (4983T>A 4984T>A 4985A>T), Q1618F (4992C>T 4993A>T 4994G>T), T1619G (4995A>G 4996C>G 4997G>A), D1622K (5004G>A 5006T>G), E1623Q (5007G>C 5009A>G), G1625N (5013G>A 5014G>A), F1626L (5016T>C), R1627P (5019A>C 5020G>C 5021A>C), A1628I (5022G>A 5023C>T 5024T>C), V1629M (5025G>A), F1631M (5031T>A 5033T>G), S1633T (5037T>A 5039C>T), L1635Y (5044T>A 5045G>T), V1637L (5049G>C 5051G>T), Q1639S (5055C>A 5056A>G 5057G>T), E1640Q (5058G>C), I1642L (5064A>T 5066T>G), A1643T (5067G>A), K1644E (5070A>G), S1649E (5085T>G 5086C>A 5087A>G), A1651I (5091G>A 5092C>T 5093C>T), L1652G (5094C>G 5095T>G), T1653V (5097A>G 5098C>T 5099A>T), I1654L (5100A>T 5102T>G), C1655M (5103T>A 5104G>T 5105T>G), M1657L (5109A>C 5111G>A), A1658S (5112G>T), K1659D (5115A>G 5117A>T), A1661K (5121G>A 5122C>A 5123T>A), I1662N (5125T>A), S1663T (5127T>A 5129C>A), D1668P (5142G>C 5143A>C 5144C>T), T1674del (5160_5162delACA), R1678E (5172C>G 5173G>A 5174T>G), D1679G (5176A>G), L1680V (5178C>G 5180G>C), N1681H (5181A>C), W1682E (5184T>G 5185G>A 5186G>A), A1684V (5191C>T 5192A>G), N1686D (5196A>G 5198C>T), I1687V (5199A>G 5201T>A), G1688F (5202G>T 5203G>T), A1696S (5226G>T 5228C>A), H1697N (5229C>A), Y1698Q (5232T>C 5234C>A), T1699Q (5235A>C 5236C>A 5237G>A), D1701S (5241G>T 5242A>C 5243T>A), E1702S (5244G>T 5245A>C), Y1704N (5250T>A), I1706L (5256A>C 5258T>G), K1713R (5277A>C 5278A>G 5279G>C), K1714R (5281A>G 5282G>A), S1715T (5283T>A 5285C>A), N1716T (5287A>C 5288T>A), M1731I (5333G>A), M1736L (5346A>T), S1737A (5349T>G), M1740Q (5358A>C 5359T>A), A1742H (5364G>C 5365C>A 5366C>T), K1744R (5370A>C 5371A>G 5372G>C), N1745K (5375T>G), V1746S (5376G>A 5377T>G 5378G>T), A1748E (5383C>A), G1749V (5386G>T 5387A>T), H1751D (5391C>G), V1752G (5395T>G 5396A>G), N1755S (5403A>T 5404A>C), V1757T (5409G>A 5410T>C 5411T>C), V1766I (5436G>A 5438C>A), G1767A (5440G>C 5441C>T), M1774F (5460A>T 5462G>T), R1779H (5475A>C 5476G>A 5477G>T), C1782R (5484T>A 5486T>G), V1783A (5488T>C 5489G>C), E1784T (5490G>A 5491A>C 5492G>A), I1786L (5496A>C 5498A>T), I1793M (5519T>G), G1794A (5521G>C 5522A>T), N1795G (5523A>G 5524A>G 5525T>G), V1807A (5560T>C), F1810L (5570T>G), Q1812K (5574C>A), E2142A (6565A>C), T2143P (6567A>C), E2150A (6589A>C)

| | Begin | End | Coverage | Score | Concordance | Matches | Identities | I/D/M/F* | Stop Codons |
|---|---|---|---|---|---|---|---|---|---|
| **NT** | **22** | **6809** | **80.5%** | **3829** | **33.5%** | **5772 (99.0%)** | **3883 (66.6%)** | **10/50** | |

Codon mutations:

GCT2GCA (146T>A), TTC7TGG (160T>G 161C>G), TCC8AGT (162T>A 163C>G 164C>T), AAG9_GCC16del (165_188delAAGGTTGCTGGTGTTGATGACGCC), AGT17CCT (189A>C 190G>C), TCT18CCT (192T>C), TCT19GCT (195T>G), TCC20TCT (200C>T), ACC21TCA (201A>T 203C>A), ACT22CCT (204A>C), GGT26GCA (217G>C 218T>A), CTT28GTT (222C>G), TCA29TCT (227A>T), AAT30TCT (228A>T 229A>C), ATT32TTT (234A>T), TCC33TCT (239C>T), AAA34TCT (240A>T 241A>C 242A>T), TTT35CTT (243T>C), AAA37AAG (251A>G), GCG38ACT (252G>A 254G>T), GTT39CTC (255G>C 257T>C), AGT40TCC (258A>T 259G>C 260T>C), ATT42GTT (264A>G), ACC43AAT (268C>A 269C>T), ACT44AAA (271C>A 272T>A), ATG45GTT (273A>G 275G>T), AAG48ACC (283A>C 284G>C), TCA50GTA (288T>G 289C>T), CAT52CAC (296T>C), CTT53ATA (297C>A 299T>A), CTG56GTC (306C>G 308G>C), AAA57AAG (311A>G), CCA58CCG (314A>G), TCT59CCA (315T>C 317T>A), GTG60GTT (320G>T), ACT61ATG (322C>T 323T>G), GAT62GCA (325A>C 326T>A), GTG63GCA (328T>C 329G>A), TCC64GCA (330T>G 332C>A), TCT66TCA (338T>A), GTG68GCC (343T>C 344G>C), GCC69TCT (345G>T 347C>T), ACT70ACA (350T>A), TGT71TGC (353T>C), AGT72AAT (355G>A), TCA73GCT (357T>G 359A>T), CTT74GTA (360C>G 362T>A), AAT75GAA (363A>G 365T>A), AAA76AGA (367A>G), TTG78TTA (374G>A), GAC79GAT (377C>T), AAA80AAG (380A>G), GGT82GCA (385G>C 386T>A), GCA83AAA (387G>A 388C>A), CTT84ATA (390C>A 392T>A), ATA85ATG (395A>G), GAT86GAA (398T>A), CCA87CCC (401A>C), TTT88TTC (404T>C), CTT89ATC (405C>A 407T>C), AAA90AAG (410A>G), GCG91GGT (412C>G 413G>T), TAC92TAT (416C>T), TCT93GAT (417T>G 418C>A), TTC94TTT (422C>T), TTG95GTA (423T>G 425G>A), TCC96AAA (426T>A 427C>A 428C>A), ACA97ACT (431A>T), TAC99TAT (437C>T), TCT101ACT (441T>A), ATC102ATA (446C>A), AAG103GTA (447A>G 448A>T 449G>A), ATG105CTT (453A>C 455G>T), GTT106GTA (458T>A), TTA107CTT (459T>C 461A>T), AAA108AAG (464A>G), TTG109ATG (465T>A), TTT110GTT (468T>G), GAG111TTA (471G>T 472A>T 473G>A), TTG113GTC (477T>G 479G>C), TCC114AAG (480T>A 481C>A 482C>G), AAT115GAT (483A>G), GTT117TTC (489G>T 491T>C), AAA118ATC (493A>T 494A>C), TTA119CTA (495T>C), TTT121TAC (502T>A 503T>C), GCT122GAA (505C>A 506T>A), GTG124TTA (510G>T 512G>A), AAT126GAA (516A>G 518T>A), AGT128GTT (522A>G 523G>T), GAA129GAG (527A>G), GAT130TCA (528G>T 529A>C 530T>A), GTA131TTA (531G>T), GTT134GTG (542T>G), GTA135TTT (543G>T 545A>T), TAT136ATA (546T>A 547A>T 548T>A), GGT137GCT (550G>C), CTT139TTA (555C>T 557T>A), GTA140GTT (560A>T), ATA143TTG (567A>T 569A>G), ACA144GCC (570A>G 572A>C), TTG145ATT (573T>A 575G>T), TTA146TTG (578A>G), GTG147CTA (579G>C 581G>A), CTT148CTA (584T>A), CTA149CTG (587A>G), TTT150TAT (589T>A), GTT151GTA (593T>A), TGT152TGC (596T>C), AGT154GTA (600A>G 601G>T 602T>A), GTT157GTA (611T>A), GAG158ACG (612G>A 613A>C), GGT159CCT (615G>C 616G>C), GTT160TTT (618G>T), TCT161AAT (621T>A 622C>A), AAC162AAT (626C>T), ACT163GTA (627A>G 628C>T 629T>A), GTC164GTT (632C>T), AAA165GTT (633A>G 634A>T 635A>T), AAT166ACT (637A>C), ATC167TTG (639A>T 641C>G), TTG169TCA (646T>C 647G>A), ATA170GTT (648A>G 650A>T), GTC171GTT (653C>T), AAC173AAT (659C>T), ATG174TTT (660A>T 662G>T), TTT175TTC (665T>C), TCA176TCT (668A>T), AGT177TCA (669A>T 670G>C 671T>A), CTA178TTG (672C>T 674A>G), TAC179TAT (677C>T), AAA186ACA (697A>C), GCC188TCT (702G>T 704C>T), GAA189GGG (706A>G 707A>G), CGA190AAG (708C>A 709G>A 710A>G), TTC191TTT (713C>T), ACC192TCC (714A>T), ATG193CAG (717A>C 718T>A), GTG194CTG (720G>C), GCC195GCG (725C>G), GCT197GCA (731T>A), AGT198GCA (732A>G 733G>C 734T>A), ATG199GTA (735A>G 737G>A), GGG202GGT (746G>T), TCA204AGC (750T>A 751C>G 752A>C), GTT205TTA (753G>T 755T>A), TCA206ACG (756T>A 758A>G), ACG210ACT (770G>T), TCT211AGC (771T>A 772C>G 773T>C), CAG212CAA (776G>A), TTG213CTT (777T>C 779G>T), ATG214TTA (780A>T 782G>A), ACA216TCA (786A>T), ATA217TTG (789A>T 791A>G), GGG221GGT (803G>T), ATT222ATC (806T>C), TCG223TCA (809G>A), ACG224ACT (812G>T), TTG225TTA (815G>A), GTG226GTA (818G>A), TTC227TTT (821C>T), ATT228GTT (822A>G), GCA229GCT (827A>T), GTG230GTT (830G>T), GGT232GGG (836T>G), GGC236GGT (848C>T), TTG237CTT (849T>C 851G>T), AGC238TCA (852A>T 853G>C 854C>A), TTG241CTG (861T>C), TCC242TCA (866C>A), ATT244ATC (872T>C), CTC245CTG (875C>G), TAT246TAC (878T>C), TCG247TCT (881G>T), ACA248GCT (882A>G 884A>T), GCT252GCG (896T>G), CAG253CAA (899G>A), CAA257CAG (911A>G), CTA258CTG (914A>G), ACT260CAG (918A>C 919C>A 920T>G), CTT261TTG (921C>T 923T>G), CGT263AGG (927C>A 929T>G), AAT264AAC (932T>C), AAA266AAG (938A>G), TGC268TGT (944C>T), ACC269ACA (947C>A), TCC270TCT (950C>T), CAG271CAA (953G>A), GCT272GCC (956T>C), GAA278GAG (974A>G), ATA280GTG (978A>G 980A>G), GTC281GTT (983C>T), GGC282AGC (984G>A), ACC283ATT (988C>T 989C>T), GGA285GGT (995A>T), TTC286TTT (998C>T), GTG290GTA (1010G>A), GCT293GCG (1019T>G), AGT295AAT (1024G>A), GCA296GCT (1028A>T), ACG297ACC (1031G>C), CTG298TTA (1032C>T 1034G>A), ACT300ACC (1040T>C), CTG302CTT (1046G>T), GAG304CAG (1050G>C), CAG307ACG (1059C>A 1060A>C), GAG308GAA (1064G>A), GAT310TCG (1068G>T 1069A>C 1070T>G), GCA311GAG (1072C>A 1073A>G), GTG312GTT (1076G>T), TGT313GCT (1077T>G 1078G>C), GAT314GAA (1082T>A), CCT315CCA (1085T>A), GCA316TCT (1086G>T 1088A>T), ACG317ATG (1090C>T), GAG320GAA (1100G>A), AAC321AAT (1103C>T), GCC323GCA (1109C>A), AAT324AAC (1112T>C), AAG325CCT (1113A>C 1114A>C 1115G>T), GCT326GCA (1118T>A), TTC327TTT (1121C>T), CCT328CCA (1124T>A), ACC329ATT (1126C>T 1127C>T), AAA330AAG (1130A>G), AAC332CAA (1134A>C 1136C>A), CTG334TTA (1140C>T 1142G>A), AGG335AAA (1144G>A 1145G>A), GAA336GAG (1148A>G), GAA337CAG (1149G>C 1151A>G), GCT338GCC (1154T>C), CTC339GTC (1155C>G), AAG340AAA (1160G>A), ATC341ATA (1163C>A), GCA346GCC (1178A>C), GTT350GTC (1190T>C), GCG351GCA (1193G>A), TTC353TTT (1199C>T), ATG354TTG (1200A>T), AGT359TCT (1215A>T 1216G>C), ATT362ATA (1226T>A), GCG363ACG (1227G>A), TTA365CTA (1233T>C), GAA366GAT (1238A>T), GTT368CTG (1242G>C 1244T>G), TAT369GTT (1245T>G 1246A>T), GAA371GAG (1253A>G), AGT372GCC (1254A>G 1255G>C 1256T>C), TGT373AGC (1257T>A 1259T>C), AGG374CAA (1260A>C 1261G>A 1262G>A), CAC375CAT (1265C>T), ATG376TCT (1266A>T 1267T>C 1268G>T), GGC377GGT (1271C>T), CAG380CAA (1280G>A), CGT382AGG (1284C>A 1286T>G), GAA384GAG (1292A>G), TTT386TTC (1298T>C), GTA388GTG (1304A>G), TTC391TTT (1313C>T), GGG392GGA (1316G>A), TCC393GCT (1317T>G 1319C>T), GGG395GGA (1325G>A), TCT399TCA (1337T>A), ACA400ACG (1340A>G), TCC401TGT (1342C>G 1343C>T), TTA404CTT (1350T>C 1352A>T), TTT405TTC (1355T>C), ATT406ATC (1358T>C), GTG409GTT (1367G>T), TTG410CTC (1368T>C 1370G>C), AGA412GCA (1374A>G 1375G>C), GAG415GAA (1385G>A), AAA417AAG (1391A>G), CTC418CTG (1394C>G), AAT419AAC (1397T>C), CGG420CGC (1400G>C), CTA421TTG (1401C>T 1403A>G), GCA423GCG (1409A>G), AAG426AAA (1418G>A), AGG427AGA (1421G>A), GCT435GCA (1445T>A), CAC436CAT (1448C>T), GCT439GCA (1457T>A), ATC440GTG (1458A>G 1460C>G), CTG441CTT (1463G>T), GAC444GAT (1472C>T), GGA446GGT (1478A>T), GCA447GCT (1481A>T), TTG448CTT (1482T>C 1484G>T), CGA449CGG (1487A>G), GGG451GGA (1493G>A), GCT452GCA (1496T>A), TGC455TGT (1505C>T), CAA456CAG (1508A>G), GAT457GAC (1511T>C), ATT458ATA (1514T>A), AAA459AAG (1517A>G), GAT463GAC (1529T>C), TCA466TCT (1538A>T), ACA467ACC (1541A>C), CAA468CAG (1544A>G), CCA469CCT (1547A>T), CCC471CCT (1553C>T), TTA472CTT (1554T>C 1556A>T), GCC475GCT (1565C>T), GTT477GTC (1571T>C), GAC479GAT (1577C>T), GGC481GGA (1583C>A), CAT483CAC (1589T>C), TTC484TTT (1592C>T), ACC485ACA (1595C>A), TCC486TCA (1598C>A), TAT488TAC (1604T>C), ATA489ATC (1607A>C), ACA492ACC (1616A>C), AAT494AAC (1622T>C), CTG495TTG (1623C>T), ATC496ATA (1628C>A), TCA497TCT (1631A>T), CCT499CCA (1637T>A), GCC500GCT (1640C>T), CAA501CAG (1643A>G), GGT503GGG (1649T>G), CTT504CTA (1652T>A), ACA505ACG (1655A>G), CCA507CCT (1661A>T), GCT509GCA (1667T>A), GAG511GAA (1673G>A), CGG512AGG (1674C>A), AGA513AGG (1679A>G), AGG514AGA (1682G>A), GAC515GAT (1685C>T), ATC516ATT (1688C>T), CTG517TTA (1689C>T 1691G>A), GTG518GTT (1694G>T), TGT520TGC (1700T>C), ATG521AGA (1702T>G 1703G>A), GAG523GTG (1708A>T), GAG525GAT (1715G>T), TCC527GAT (1719T>G 1720C>A 1721C>T), ACT528CCA (1722A>C 1724T>A), GAA529AAT (1725G>A 1727A>T), GAT530ACT (1728G>A 1729A>C), CCT531CCA (1733T>A), ACG532ACA (1736G>A), GGA533GCC (1738G>C 1739A>C), CTC535CTA (1745C>A), AGA536ATT (1747G>T 1748A>T), AAC538GAT (1752A>G 1754C>T), ATA539GTG (1755A>G 1757A>G), GAG541GAA (1763G>A), CCT545CCC (1775T>C), GCT547GCC (1781T>C), ACT549ACA (1787T>A), CAC550CAT (1790C>T), AGG551AGA (1793G>A), AAT552GCT (1794A>G 1795A>C), TTG553TTA (1799G>A), ACC554TCC (1800A>T), AGT556GAT (1806A>G 1807G>A), GAT557GAC (1811T>C), CTT558CTC (1814T>C), GAG560CAA (1818G>C 1820G>A), GTA563GTG (1829A>G), GCC564GCA (1832C>A), AAA565AAG (1835A>G), TGT566TGC (1838T>C), CAG567CGG (1840A>G), GTA568GTG (1844A>G), CAT569CAC (1847T>C), GCA570TAT (1848G>T 1849C>A 1850A>T), GAA571GAG (1853A>G), GTA572GTG (1856A>G), AAG574AGC (1861A>G 1862G>C), TTG576CTT (1866T>C 1868G>T), TTT577TAC (1870T>A 1871T>C), GAA578AAT (1872G>A 1874A>T), GCT579ACT (1875G>A), GAA580ACA (1878G>A 1879A>C), TCT581TCG (1883T>G), GGC582GGA (1886C>A), ATA583TTG (1887A>T 1889A>G), AGT584GCT (1890A>G 1891G>C), CCT585GAC (1893C>G 1894C>A 1895T>C), ATA587GTT (1899A>G 1901A>T), GTT590GTC (1910T>C), CAA591AAT (1911C>A 1913A>T), TCT593GCT (1917T>G), GCA594GTT (1921C>T 1922A>T), GAT596GAA (1928T>A), ATA598CTG (1932A>C 1934A>G), GCG599GAT (1936C>A 1937G>T), GTG601TTC (1941G>T 1943G>C), GAT602CAA (1944G>C 1946T>A), GGG603GAT (1948G>A 1949G>T), GCT604GCC (1952T>C), AGA605del (1953_1955delAGA), TTG606TTA (1958G>A), CGC607CAA (1960G>A 1961C>A), ACC608GTT (1962A>G 1963C>T 1964C>T), AAG609TTG (1965A>T 1966A>T), CAA610CCT (1969A>C 1970A>T), GAT611CCT (1971G>C 1972A>C), GAA612GAG (1976A>G), ATA614GCA (1980A>G 1981T>C), TTG615GCT (1983T>G 1984T>C 1985G>T), GTG616TAT (1986G>T 1987T>A 1988G>T), CCC617GAG (1989C>G 1990C>A 1991C>G), ACT618CAG (1992A>C 1993C>A 1994T>G), GTT619ACA (1995G>A 1996T>C 1997T>A), GTG620ATT (1998G>A 2000G>T), AGT621ACA (2002G>C 2003T>A), AAT623GCT (2007A>G 2008A>C), GAT624GAA (2012T>A), AGA625AAA (2014G>A), GTG626ATT (2016G>A 2018G>T), ATC627GTT (2019A>G 2021C>T), TAT628TAC (2024T>C), GCA629TCC (2025G>T 2027A>C), AGA630CAG (2028A>C 2029G>A 2030A>G), GAG631GAA (2033G>A), ACG633ACA (2039G>A), GTA634GTT (2042A>T), GCA636GCC (2048A>C), TTG637CTC (2049T>C 2051G>C), GCA640GCT (2060A>T), CAG642CAA (2066G>A), GGC643GGG (2069C>G), TTG645ATG (2073T>A), GAC646AAC (2076G>A), CCT647AAA (2079C>A 2080C>A 2081T>A), GAG648TTT (2082G>T 2083A>T 2084G>T), GAA649GAG (2087A>G), CTT650TTT (2088C>T), TTT651TTC (2093T>C), CCT652CCA (2096T>A), CAT655CTC (2104A>T 2105T>C), CAT656TCC (2106C>T 2107A>C 2108T>C), GCC659GCT (2117C>T), ATG660ATC (2120G>C), TCT663GTG (2127T>G 2128C>T 2129T>G), CTT664CTA (2132T>A), GAT665TCT (2133G>T 2134A>C), GAT666TCT (2136G>T 2137A>C), GAA667GAT (2141A>T), CAT668GCC (2142C>G 2143A>C 2144T>C), GAG670CAG (2148G>C), ATT671AAT (2152T>A), AAC673GAA (2157A>G 2159C>A), AAA674ATG (2161A>T 2162A>G), AGA676GTA (2166A>G 2167G>T), GTC677AAT (2169G>A 2170T>A 2171C>T), AAC678GAT (2172A>G 2174C>T), ATG679TTG (2175A>T), TAT681GTC (2181T>G 2182A>T 2183T>C), AGA682AAA (2185G>A), GAT685AGT (2193G>A 2194A>G), GCT686CCT (2196G>C), GAA687CAG (2199G>C 2201A>G), CGA690GCT (2208C>G 2209G>C 2210A>T), GTA693GTG (2219A>G), CAA694CGA (2221A>G), GAC699GAT (2237C>T), ATA701ATC (2243A>C), CAC703CAT (2249C>T), TTT704TTC (2252T>C), GCA710AGC (2268G>A 2269C>G 2270A>C), TCA711ACA (2271T>A), ATC713ATT (2279C>T), GAG715GAA (2285G>A), AAG716GAG (2286A>G), AAA717CTG (2289A>C 2290A>T 2291A>G), CTA718CTG (2294A>G), GCA719GAT (2296C>A 2297A>T), GTG720GTC (2300G>C), CAG721CAA (2303G>A), ACA722CGC (2304A>C 2305C>G 2306A>C), GAG723AAT (2307G>A 2309G>T), ATG724CTT (2310A>C 2312G>T), AGG725GAG (2313A>G 2314G>A), ACT726CAA (2316A>C 2317C>A 2318T>A), GGA727GGG (2321A>G), TTT728AAA (2322T>A 2323T>A 2324T>A), GCA729GCG (2327A>G), CAT730TAT (2328C>T), TCT731AAG (2331T>A 2332C>A 2333T>G), TGC732GCG (2334T>G 2335G>C 2336C>G), ATT733CTT (2337A>C), GAT734CAG (2340G>C 2342T>G), GCA735GCT (2345A>T), GTT737GCA (2350T>C 2351T>A), GAC738GAT (2354C>T), ACC739ACA (2357C>A), CTC740CTT (2360C>T), ATA741ACG (2362T>C 2363A>G), ATC743ATT (2369C>T), GAG744GAA (2372G>A), TTT749AGA (2385T>A 2386T>G 2387T>A), GTT750GTA (2390T>A), TTT752TTC (2396T>C), TTA753CTG (2397T>C 2399A>G), ACA755ACT (2405A>T), GCA756GCT (2408A>T), AGA759CGC (2415A>C 2417A>C), TGC762TGT (2426C>T), AAC763AGG (2428A>G 2429C>G), GTA765GTT (2435A>T), TCA766GCA (2436T>G), TCA766_AAA767insGGT (2438_2439insGGT), AAA767ACT (2440A>C 2441A>T), GAG768GAC (2444G>C), CCA769AAG (2445C>A 2446C>A 2447A>G), GAA771AGA (2451G>A 2452A>G), TCA772AAT (2454T>A 2455C>A 2456A>T), ATC774ATT (2462C>T), AAA775ATC (2464A>T 2465A>C), ATT776GAA (2466A>G 2467T>A 2468T>A), TAC778ATG (2472T>A 2473A>T 2474C>G), CAT779ATT (2475C>A 2476A>T), AAA780GGT (2478A>G 2479A>G 2480A>T), GTT782AAG (2484G>A 2485T>A 2486T>G), GAA783GAC (2489A>C), TAT784AAA (2490T>A 2492T>A), ATC785ATG (2495C>G), AAG786ATC (2497A>T 2498G>C), GAT787CAG (2499G>C 2501T>G), ACA788ATG (2503C>T 2504A>G), TGG790AAT (2508T>A 2509G>A 2510G>T), AGC791GAC (2511A>G 2512G>A), CTT792TTG (2514C>T 2516T>G), TAT794TTC (2521A>T 2522T>C), GCA795ATC (2523G>A 2524C>T 2525A>C), CTG796TTG (2526C>T), AGA797AAG (2530G>A 2531A>G), CTC799CTT (2537C>T), GCA802GCC (2546A>C), GGT803GGA (2549T>A), ATA806TTG (2556A>T 2558A>G), ATG807ATA (2561G>A), GCT808GCA (2564T>A), GGA811GGC (2573A>C), GTT812ACC (2574G>A 2575T>C 2576T>C), TTT813ATG (2577T>A 2579T>G), GGA814AAT (2580G>A 2581G>A 2582A>T), GGC815GCC (2584G>C), ATT816GTT (2586A>G), ACA817ACT (2591A>T), GCA818GCT (2594A>T), TTT819TTC (2597T>C), TTG820ATT (2598T>A 2600G>T), GCG821ACT (2601G>A 2603G>T), TGG823CTC (2607T>C 2608G>T 2609G>C), CAG824AAG (2610C>A), AGT825GCA (2613A>G 2614G>C 2615T>A), AAC826GCA (2616A>G 2617A>C 2618C>A), TCT827ACT (2619T>A), TTC828TCC (2623T>C), CCC829CCT (2627C>T), AAT830TTA (2628A>T 2629A>T 2630T>A), GCA831GCT (2633A>T), TCG832GAT (2634T>G 2635C>A 2636G>T), GGA835GGG (2645A>G), GGT836GGC (2648T>C), GTG837GTA (2651G>A), GAA840GAG (2660A>G), ATA844ATC (2672A>C), AGT846AGC (2678T>C), AAG848AAA (2684G>A), AAC849AAT (2687C>T), AAG850AAA (2690G>A), GGT851GGG (2693T>G), AAT852AAG (2696T>G), AAG853ACT (2698A>C 2699G>T), AAG853_CTC854insCAT (2699_2700insCAT), CTC854TTC (2700C>T), AGA855GCA (2703A>G 2704G>C), AAT856AAA (2708T>A), CTT857TCC (2709C>T 2710T>C 2711T>C), CTC858TTG (2712C>T 2714C>G), GTT859CTA (2715G>C 2717T>A), GGT860GGG (2720T>G), AGT862GCG (2724A>G 2725G>C 2726T>G), TCT863GCA (2727T>G 2729T>A), CAA864GAT (2730C>G 2732A>T), TCA865AAC (2733T>A 2734C>A 2735A>C), TTG866AAC (2736T>A 2737T>A 2738G>C), GCA867CAT (2739G>C 2740C>A 2741A>T), CAA868GTG (2742C>G 2743A>T 2744A>G), GAT869GAC (2747T>C), GAT869_TGG870insCGG (2747_2748insCGG), GCT871GCA (2753T>A), GCT872GAA (2755C>A 2756T>A), GAA873CAG (2757G>C 2759A>G), GTA877GTT (2771A>T), CAG879GAA (2775C>G 2777G>A), AGC880AGT (2780C>T), CTC881CTG (2783C>G), AAG882CAC (2784A>C 2786G>C), AAT884AAC (2792T>C), GTG886GTT (2798G>T), GTG887GTT (2801G>T), TTA888TTG (2804A>G), CTT890ATT (2808C>A), GGT891GCT (2812G>C), GTG894GTC (2822G>C), TAC895TAT (2825C>T), TTT896TTC (2828T>C), GGC898GGT (2834C>T), ACC899ACG (2837C>G), TAC900TAT (2840C>T), GTG901GTT (2843G>T), ATA906ATT (2858A>T), ATG907CTC (2859A>C 2861G>C), ACC908ACT (2864C>T), GTA909GTG (2867A>G), GCT910GCA (2870T>A), GCA912GCC (2876A>C), TTC913TTT (2879C>T), CAC914AGA (2880C>A 2881A>G 2882C>A), AAT915AAC (2885T>C), GCT916ATG (2886G>A 2887C>T 2888T>G), CGA917AGG (2889C>A 2891A>G), GGT919GGC (2897T>C), CCA921CCT (2903A>T), ACA923TCC (2907A>T 2909A>C), ATC925ATT (2915C>T), GCC927GAG (2920C>A 2921C>G), AAT928CAT (2922A>C), TCT929TCC (2927T>C), CGA930ATT (2928C>A 2929G>T 2930A>T), TCT931TGT (2932C>G), AAA932AAT (2936A>T), GTT933GTG (2939T>G), CAA934CAG (2942A>G), TAC935TTC (2944A>T), AAC936AAT (2948C>T), AGA938ACC (2953G>C 2954A>C), GAA939CAA (2955G>C), ATT942TTG (2964A>T 2966T>G), ATT943ATA (2969T>A), GGC945GGT (2975C>T), GAC947GAT (2981C>T), ATC948GTG (2982A>G 2984C>G), CTG950TTG (2988C>T), CGC952AGG (2994C>A 2996C>G), GTT953GTA (2999T>A), GGT954GGG (3002T>G), CCA959CCC (3017A>C), AAG960AAA (3020G>A), CCT961TCA (3021C>T 3023T>A), GAC962GAT (3026C>T), ATC963ATA (3029C>A), CGT964AGA (3030C>A 3032T>A), CAC966CAT (3038C>T), CCA968CCT (3044A>T), AGA969CGC (3045A>C 3047A>C), AGG970AGA (3050G>A),

| | |
|---|---|
| | GAG972GAT (3056G>T), TGC974TGT (3062C>T), TTC975TAC (3064T>A), ACA976TCT (3066A>T 3068A>T), CAA980ATG (3078C>A 3079A>T 3080A>G), GGA981GGC (3083A>C), TGT984GTG (3090T>G 3091G>T 3092T>G), AGA985AGG (3095A>G), GTT987GTA (3101T>A), GCG988GCT (3104G>T), TCT989TCA (3107T>A), ACA990AAG (3109C>A 3110A>G), CCA992CCC (3116A>C), CGT993AAG (3117C>A 3118G>A 3119T>G), CTT994AAA (3120C>A 3121T>A 3122T>A), GGC995TCC (3123G>T 3124G>C), TTA997TTG (3131A>G), TTT999GTT (3135T>G), CTC1000CTT (3140C>T), AAG1001AAA (3143G>A), ATG1002TAT (3144A>T 3145T>A 3146G>T), GGA1009GGC (3167A>C), ACA1011ACC (3173A>C), GTT1012GAA (3175T>A 3176T>A), GTG1014GTC (3182G>C), GAA1015ACA (3183G>A 3184A>C), GAA1017GGA (3190A>G), CTG1018TTA (3192C>T 3194G>A), AAC1019GAT (3195A>G 3197C>T), TCC1020TCG (3200C>G), TCC1021AAT (3201T>A 3202C>A 3203C>T), AGT1022GCA (3204A>G 3205G>C 3206T>A), AAG1024GCT (3210A>G 3211A>C 3212G>T), ATT1025CTT (3213A>C), TGC1026AGC (3216T>A), TCT1027TCA (3221T>A), CAA1029GTT (3225C>G 3226A>T 3227A>T), TCT1030GCA (3228T>G 3230T>A), GAA1032GAG (3236A>G), ATA1035ATT (3245A>T), GGG1037AGT (3249G>A 3251G>T), GAA1038GAG (3254A>G), GGT1040GGA (3260T>A), GAC1041GAT (3263C>T), GGC1043GGT (3269C>T), ACG1044ACT (3272G>T), TTG1045CTT (3273T>C 3275G>T), CTA1046CTC (3278A>C), CTG1047TTA (3279C>T 3281G>A), TTA1048TTG (3284A>G), AGT1050ACT (3289G>C), GTT1051GTG (3293T>G), AAT1053AAC (3299T>C), CAA1055CAG (3305A>G), CCT1056CCG (3308T>G), GTG1057TGC (3309G>T 3310T>G 3311G>C), GGC1060GGA (3320C>A), ATC1061ATA (3323C>A), CAC1062CAT (3326C>T), TCA1065TCT (3335A>T), GGC1068GAT (3343G>A 3344C>T), GTA1069ACC (3345G>A 3346T>C 3347A>C), GCA1070GCT (3350A>T), GCT1071GCA (3353T>A), GAA1072GAG (3356A>G), CGT1073AAG (3357C>A 3358G>A 3359T>G), GGA1074GGT (3362A>T), TTT1075TAC (3364T>A 3365T>C), ATC1076GTG (3366A>G 3368C>G), TCT1077GCT (3369T>G), AAT1079AAC (3377T>C), GCT1080GCA (3380T>A), GCT1082GCA (3386T>A), ATC1083GTG (3387A>G 3389C>G), TAC1084TTT (3391A>T 3392C>T), AGG1085AGA (3395G>A), GAA1086GAT (3398A>T), CTA1088TTG (3402C>T 3404A>G), GAG1089GAA (3407G>A), CCA1092CCC (3416A>C), ACT1093ACG (3419T>G), GGG1094GGA (3422G>A), CCG1095CCC (3425G>C), GTC1096ATT (3426G>A 3428C>T), GCA1098GTG (3433C>T 3434A>G), GCA1099GCT (3437A>T), GTA1101GTG (3443A>G), GAT1104GAA (3452T>A), ATT1105CTG (3453A>C 3455T>G), CTG1106TTG (3456C>T), TCA1108CAC (3462T>C 3463C>A 3464A>C), ATT1109CTG (3465A>C 3467T>G), AGC1111TCC (3471A>T 3472G>C), AGA1112AAG (3475G>A 3476A>G), GAA1113GAG (3479A>G), ACA1114GAA (3480A>G 3481C>A), CAG1115AGA (3483C>A 3484A>G 3485G>A), CTT1116TTC (3486C>T 3488T>C), GAA1119GAG (3497A>G), AAC1120AAA (3500C>A), CAA1121CAG (3503A>G), TAC1123GCT (3507T>G 3508A>C 3509C>T), TAC1124TTC (3511A>T), CTT1125TTG (3513C>T 3515T>G), GGA1126GGC (3518A>C), ACA1127ACT (3521A>T), GTT1128GTA (3524T>A), CCA1129CCG (3527A>G), CAG1130TTG (3528C>T 3529A>T), GAG1131GAA (3533G>A), TTG1132ATG (3534T>A), GCC1133GCT (3539C>T), GCC1134GCT (3542C>T), GTT1136GTG (3548T>G), CCC1137CCG (3551C>G), AAG1139AAA (3557G>A), ACC1140ACA (3560C>A), ACT1141ACC (3563T>C), CTG1142CTT (3566G>T), CGG1143AAA (3567C>A 3568G>A 3569G>A), AAA1144CGC (3570A>C 3571A>G 3572A>C), AGC1145AGT (3575C>T), CAA1146CCC (3577A>C 3578A>C), TTG1147CTG (3579T>C), GAA1149GAG (3587A>G), GCA1150GCT (3590A>T), TTC1151TTT (3593C>T), GGA1152GGG (3596A>G), CCT1153GAC (3597C>G 3598C>A 3599T>C), GCA1154GCT (3602A>T), GAG1155CAG (3603G>C), ACA1156ACT (3608A>T), GCA1157GCC (3611A>C), ACA1162TCT (3624A>T 3626A>T), GTT1163GCC (3628T>C 3629T>C), CAT1164AAA (3630C>A 3632T>A), AGA1167AGG (3641A>G), GGT1168GGA (3644T>A), GAT1169GAA (3647T>A), GGT1170GGG (3650T>G), GAC1172GAT (3656C>T), GGG1177GGT (3671G>T), GTA1178GTC (3674A>C), ATG1179GAG (3675A>G 3676T>A), AAA1180AAG (3680A>G), TAC1181TAT (3683C>T), TGT1186TCC (3697G>C 3698T>C), GGA1187AAC (3699G>A 3700G>A 3701A>C), GAT1189GAG (3707T>G), GAT1190GAG (3710T>G), GAC1191CCA (3711G>C 3712A>C 3713C>A), GCC1193GCA (3719C>A), AAA1194AAG (3722A>G), CTA1195CTG (3725A>G), GCA1196GCT (3728A>T), TTC1197TTT (3731C>T), GAA1198GAG (3734A>G), AAT1199GGT (3735A>G 3736A>G), CTC1200CTT (3740C>T), AAG1201AAA (3743G>A), TGC1202GCA (3744T>G 3745G>C 3746C>A), TCG1203TCC (3749G>C), CTG1204ATT (3750C>A 3752G>T), CTA1205CTT (3755A>T), ATC1207GTG (3759A>G 3761C>G), ATG1208CTT (3762A>C 3764G>T), CGT1209AAG (3765C>A 3766G>A 3767T>G), AGC1210ATG (3769G>T 3770C>G), AAG1212CGA (3774A>C 3775A>G 3776G>A), ATC1213GTG (3777A>G 3779C>G), CCT1214CCC (3782T>C), GGA1216GGG (3788A>G), CGT1217AAA (3789C>A 3790G>A 3791T>A), CCA1218CCT (3794A>T), TGT1219GCA (3795T>G 3796G>C 3797T>A), GAA1220ATC (3798G>A 3799A>T 3800A>C), AGG1221AGA (3803G>A), GAT1222AAT (3804G>A), GGA1229GGG (3827A>G), TGT1233TTT (3838G>T), TAC1235TAT (3845C>T), TAT1236TTT (3847A>T), GGC1238GCA (3853G>C 3854C>A), GAG1240GAA (3860G>A), CTG1241TTG (3861C>T), ACA1243ACG (3869A>G), TCT1244TCA (3872T>A), TGC1245TGT (3875C>T), GGA1246GGT (3878A>T), CCC1248CCT (3884C>T), TTC1249TTT (3887C>T), ATG1252TTG (3894A>T), GGG1253GGA (3899G>A), ATG1254CTT (3900A>C 3902G>T), GGG1255GGT (3905G>T), ATG1256TCT (3906A>T 3907T>C 3908G>T), AAG1258AAA (3914G>A), AGA1259AGG (3917A>G), CAA1263GAA (3927C>G), TCC1264GCC (3930T>G), ACT1265CAA (3933A>C 3934C>A 3935T>A), GGC1266CCA (3936G>C 3937G>C 3938C>A), GAA1267AAC (3939G>A 3941A>C), GGA1268TCG (3942G>T 3943G>C 3944A>G), GAG1269CAG (3945G>C), AGA1270AGG (3950A>G), GTG1271GTT (3953G>T), GAA1272CTT (3954G>C 3955A>T 3956A>T), AAA1274ATA (3961A>T), AGA1275AGG (3965A>G), GAC1276AAC (3966G>A), ACT1277ACA (3971T>A), GTA1279GTC (3977A>C), GAA1281GAT (3983A>T), CTA1286TTG (3996C>T 3998A>G), GAT1287GAC (4001T>C), GTG1288TCG (4002G>T 4003T>C), ATT1290GTA (4008A>G 4010T>A), AAA1292CAA (4014A>C), ATC1294ACA (4021T>C 4022C>A), CAT1295TTT (4023C>T 4024A>T), GTG1296GTT (4028G>T), GAT1297GAC (4031T>C), CTG1298CTC (4034G>C), ACC1300ACA (4040C>A), CAA1302CAG (4046A>G), TGC1303TGT (4049C>T), GCC1304GCG (4052C>G), CGC1308CGG (4064C>G), CTC1309TTG (4065C>T 4067C>G), AAA1313AAG (4079A>G), ATC1314ATT (4082C>T), GGC1316GGG (4088C>G), AGA1318AGG (4094A>G), AGG1321CGC (4101A>C 4103G>C), CTC1322CTG (4106C>G), TTT1323TTC (4109T>C), GAG1324GAA (4112G>A), ATA1325ATC (4115A>C), CTT1326TTG (4116C>T 4118T>G), TAC1330TAT (4130C>T), TTG1333CTG (4137T>C), GTC1334GTG (4142C>G), AGG1335CGG (4143A>C), AAG1336AAA (4148G>A), TAT1337TAC (4151T>C), TTT1338TTC (4154T>C), GAT1340GAC (4160T>C), TTC1341TTT (4163C>T), TCA1342TCT (4166A>T), GCC1343GCT (4169C>T), ACA1344TGT (4170A>T 4171C>G 4172A>T), TTG1348GCA (4182T>G 4183T>C 4184G>A), AAT1350AAC (4190T>C), GCT1351GTA (4192C>T 4193T>A), ATA1352CTC (4194A>C 4196A>C), CCA1353CCT (4199A>T), AAA1355AAG (4205A>G), GTT1356GTA (4208T>A), ATT1358ATC (4214T>C), GAT1359AAT (4215G>A), CCT1360CCA (4220T>A), ACA1361ACG (4223A>G), AGT1362AGC (4226T>C), TCT1363CAT (4227T>C 4228C>A), TTG1367GTG (4239T>G), TTG1368CTG (4242T>C), GCA1369GCT (4247A>T), AAT1370AGT (4249A>G), GGG1371AGC (4251G>A 4253G>C), AGA1373AGG (4259A>G), GCT1374TCA (4260G>T 4262T>A), GTG1375GTA (4265G>A), TCA1376TCT (4268A>T), GAC1377GAT (4271C>T), GTG1378GAG (4273T>A), TTT1380TTC (4280T>C), GCT1382GCC (4286T>C), TAT1384TAC (4292T>C), TCC1385TCG (4295C>G), CCT1392CCC (4316T>C), GTT1393ATC (4317G>A 4319T>C), GCT1395GCA (4325T>A), TTT1396TTC (4328T>C), CAG1397CAA (4331G>A), TTG1402CTG (4344T>C), GAA1405GCT (4354A>C 4355A>T), TAC1406TTT (4357A>T 4358C>T), TAC1407TAT (4361C>T), GGA1408GGT (4364A>T), TCA1409GAT (4365T>G 4366C>A 4367A>T), AAG1410GAG (4368A>G), GGC1412GGG (4376C>G), GAT1415GAG (4385T>G), TCC1416TCA (4388C>A), CGA1419AGG (4395C>A 4397A>G), CAT1420CAC (4400T>C), GCA1421ACA (4401G>A), CTT1422TTA (4404C>T 4406T>A), GCA1426GCC (4418A>C), TGT1428TGC (4424T>C), TAC1430TAT (4430C>T), ACA1431ACC (4433A>C), CTG1432CTC (4436G>C), TGC1433TGT (4439C>T), GAT1435AAC (4443G>A 4445T>C), AAA1436AAG (4448A>G), GTG1437TTG (4449G>T), AGG1439CGC (4455A>C 4457G>C), TTG1440CTA (4458T>C 4460G>A), GTT1441GTG (4463T>G), GGG1442GGT (4466G>T), GGC1443GGA (4469C>A), CCA1445CCC (4475A>C), TCA1446TCT (4478A>T), GGA1447GGT (4481A>T), GCA1449GCT (4487A>T), CTA1450TTG (4488C>T 4490A>G), ACG1451ACT (4493G>T), ATC1453ATA (4499C>A), TTC1454TTT (4502C>T), TCT1456TCA (4508T>A), CTC1457TTG (4509C>T 4511C>G), CTC1458CTA (4514C>A), CGA1464AGG (4530C>A 4532A>G), TAT1465TAC (4535T>C), GCC1466GCG (4538C>G), TTG1470CTG (4548T>C), TTA1471TTG (4553A>G), AGA1472AGG (4556A>G), CCC1474CCA (4562C>A), ATT1476GTC (4566A>G 4568T>C), GCT1477AAA (4569G>A 4570C>A 4571T>A), AGG1479AAA (4576G>A 4577G>A), GCT1480TCT (4578G>T), ATA1481GTG (4581A>G 4583A>G), GGA1482GGC (4586A>C), GTT1483ATT (4587G>A), AAA1484AAG (4592A>G), AGT1486AAT (4597G>A), GAT1487GAC (4601T>C), CTA1491TTG (4611C>T 4613A>G), GCT1494GCA (4622T>A), GTT1495GTA (4625T>A), GGA1497GGT (4631A>T), GAC1499GAT (4637C>T), AAT1500AAC (4640T>C), CTT1501TTG (4641C>T 4643T>G), GCT1503GCG (4649T>G), GTA1504GTT (4652A>T), TTA1506ATG (4656T>A 4658A>G), CTC1508TTG (4662C>T 4664C>G), CAG1509AGG (4665C>A 4666A>G), TAT1511TAC (4673T>C), TCT1512GAT (4674T>G 4675C>A), CTG1513CTA (4679G>A), AAT1515TCC (4683A>T 4684A>C 4685T>C), ATA1516ATT (4688A>T), CAT1518AGA (4692C>A 4693A>G 4694T>A), GAG1519GAA (4697G>A), TTA1520CTG (4698T>C 4700A>G), CTG1522GCT (4704C>G 4705T>C 4706G>T), GTC1523GTT (4709C>T), AAT1524AAC (4712T>C), GTA1525GTC (4715A>C), ATC1526GTC (4716A>G), CTT1531TTG (4731C>T 4733T>G), AAA1533AAG (4739A>G), TCG1534TCA (4742G>A), GTT1537GTA (4751T>A), AAT1538GAG (4752A>G 4754T>G), GAG1539AAG (4755G>A), GTA1540ACA (4758G>A 4759T>C), CAA1541AAA (4761C>A), CAA1543CTG (4768A>T 4769A>G), CTA1548CTC (4784A>C), ACT1549ACC (4787T>C), CTG1551CTC (4793G>C), AGA1553AGG (4799A>G), TTT1555TTC (4805T>C), AAA1556AAG (4808A>G), GCT1559GCA (4817T>A), CTT1560TTG (4818C>T 4820T>G), TAC1562TAT (4826C>T), CAC1563CAA (4829C>A), GCT1565GCA (4835T>A), CCT1566CCA (4838T>A), AAG1568AAA (4844G>A), ATC1572GTC (4854A>G), ATT1573ATC (4859T>C), ATG1576ATT (4868G>T), ATA1579ATT (4877A>T), AGA1580AGG (4880A>G), CCT1581CCA (4883T>A), GCC1582GCA (4886C>A), CCA1583GTG (4887C>G 4888C>T 4889A>G), GGT1584GGA (4892T>A), GAC1587GAT (4901C>T), ACT1588GTA (4902A>G 4903C>T 4904T>A), CAG1589CAA (4907G>A), GCC1590GCT (4910C>T), ATG1592TAT (4914A>T 4915T>A 4916G>T), GTG1595ATT (4923G>A 4925G>T), GGA1598GGC (4934A>C), ATA1599ATC (4937A>C), CGT1600AGA (4938C>A 4940T>A), GCT1602GCC (4946T>C), CAC1605CAT (4955C>T), GGT1606GGG (4958T>G), CGT1607CAG (4960G>A 4961T>G), GTG1608GCC (4963T>C 4964G>C), GCT1609ACT (4965G>A), GAC1611GAA (4973C>A), AAG1612AGT (4975A>G 4976G>T), CTT1613TTT (4977C>T), GTC1614GTA (4982C>A), TTA1615AAT (4983T>A 4984T>A 4985A>T), CAG1618TTT (4992C>T 4993A>T 4994G>T), ACG1619GGA (4995A>G 4996C>G 4997G>A), GCG1620GCT (5000G>T), TTG1621CTC (5001T>C 5003G>C), GAT1622AAG (5004G>A 5006T>G), GAA1623CAG (5007G>C 5009A>G), AGG1624AGA (5012G>A), GGT1625AAT (5013G>A 5014G>A), TTC1626CTC (5016T>C), AGA1627CCC (5019A>C 5020G>C 5021A>C), GCT1628ATC (5022G>A 5023C>T 5024T>C), GTG1629ATG (5025G>A), TTT1631ATG (5031T>A 5033T>G), CCT1632CCC (5036T>C), TCC1633ACT (5037T>A 5039C>T), TAT1634TAC (5042T>C), TTG1635TAT (5044T>A 5045G>T), GTG1637CTT (5049G>C 5051G>T), CAG1639AGT (5055C>A 5056A>G 5057G>T), GAA1640CAA (5058G>C), ATT1642TTG (5064A>T 5066T>G), GCA1643ACA (5067G>A), AAG1644GAG (5070A>G), GTA1645GTC (5075A>C), GGG1647GGA (5081G>A), TCA1649GAG (5085T>G 5086C>A 5087A>G), AGT1650AGC (5090T>C), GCC1651ATT (5091G>A 5092C>T 5093C>T), CTG1652GGG (5094C>G 5095T>G), ACA1653GTT (5097A>G 5098C>T 5099A>T), ATT1654TTG (5100A>T 5102T>G), TGT1655ATG (5103T>A 5104G>T 5105T>G), GAA1656GAG (5108A>G), ATG1657CTA (5109A>C 5111G>A), GCA1658TCA (5112G>T), AAA1659GAT (5115A>G 5117A>T), GCA1660GCT (5120A>T), GCT1661AAA (5121G>A 5122C>A 5123T>A), ATT1662AAT (5125T>A), TCC1663ACA (5127T>A 5129C>A), TAT1664TAC (5132T>C), ACG1665ACC (5135G>C), CCA1666CCT (5138A>T), TTG1667TTA (5141G>A), GAC1668CCT (5142G>C 5143A>C 5144C>T), GCA1669GCT (5147A>T), GGT1670GGG (5150T>G), GAG1671GAA (5153G>A), ATC1673ATT (5159C>T), ACA1674del (5160_5162delACA), CGT1678GAG (5172C>G 5173G>A 5174T>G), GAT1679GGT (5176A>G), CTG1680GTC (5178C>G 5180G>C), AAT1681CAT (5181A>C), TGG1682GAA (5184T>G 5185G>A 5186G>A), GCA1684GTG (5191C>T 5192A>G), CCA1685CCT (5195A>T), AAC1686GAT (5196A>G 5198C>T), ATT1687GTA (5199A>G 5201T>A), GGT1688TTT (5202G>T 5203G>T), TTT1689TTC (5207T>C), TCA1691AGT (5211T>A 5212C>G 5213A>T), GCA1692GCT (5216A>T), CGT1693CGG (5219T>G), GCC1696TCA (5226G>T 5228C>A), CAC1697AAC (5229C>A), TAC1698CAA (5232T>C 5234C>A), ACG1699CAA (5235A>C 5236C>A 5237G>A), GAT1701TCA (5241G>T 5242A>C 5243T>A), GAA1702TCA (5244G>T 5245A>C), GGG1703GGT (5249G>T), TAC1704AAC (5250T>A), ATT1705ATC (5255T>C), ATT1706CTG (5256A>C 5258T>G), AAC1708AAT (5264C>T), TGT1709TGC (5267T>C), ACA1710ACT (5270A>T), GGT1711GGA (5273T>A), GCA1712GCC (5276A>C), AAG1713CGC (5277A>C 5278A>G 5279G>C), AAG1714AGA (5281A>G 5282G>A), TCC1715ACA (5283T>A 5285C>A), AAT1716ACA (5287A>C 5288T>A), GTT1718GTG (5294T>G), GGT1720GGG (5300T>G), CCA1721CCT (5303A>T), AAC1723AAT (5309C>T), ATG1731ATA (5333G>A), CCG1733CCC (5339G>C), ATG1736TTG (5346A>T), TCA1737GCA (5349T>G), GCT1738GCA (5354T>A), ATA1739ATC (5357A>C), ATG1740CAG (5358A>C 5359T>A), GCC1742CAT (5364G>C 5365C>A 5366C>T), AAG1744CGC (5370A>C 5371A>G 5372G>C), AAT1745AAG (5375T>G), GTG1746AGT (5376G>A 5377T>G 5378G>T), TTG1747CTG (5379T>C), GCA1748GAA (5383C>A), GGA1749GTT (5386G>T 5387A>T), GGA1750GGT (5390A>T), CAT1751GAT (5391C>G), GTA1752GGG (5395T>G 5396A>G), GCA1753GCT (5399A>T), AAT1755TCT (5403A>T 5404A>C), GTT1757ACC (5409G>A 5410T>C 5411T>C), GTG1759GTT (5417G>T), GGA1761GGT (5423A>T), GGA1763GGT (5429A>T), GCT1765GCA (5435T>A), GTC1766ATA (5436G>A 5438C>A), GGC1767GCT (5440G>C 5441C>T), ATT1769ATC (5447T>C), GCG1772GCA (5456G>A), CTG1773TTG (5457C>T), ATG1774TTT (5460A>T 5462G>T), GCC1775GCA (5465C>A), TTA1776TTG (5468A>G), GCA1777GCC (5471A>C), ACA1778ACT (5474A>T), AGG1779CAT (5475A>C 5476G>A 5477G>T), TAT1781TAC (5483T>C), TGT1782AGG (5484T>A 5486T>G), GTG1783GCC (5488T>C 5489G>C), GAG1784ACA (5490G>A 5491A>C 5492G>A), ATA1786CTT (5496A>C 5498A>T), ATA1787ATT (5501A>T), GTT1788GTG (5504T>G), AGA1789CGC (5505A>C 5507A>C), TTG1790CTA (5508T>C 5510G>A), CGC1791CGA (5513C>A), ACA1792ACC (5516A>C), ATT1793ATG (5519T>G), GGA1794GCT (5521G>C 5522A>T), AAT1795GGG (5523A>G 5524A>G 5525T>G), CTT1797CTG (5531T>G), GAC1798GAT (5534C>T), CTC1799CTG (5537C>G), AAT1800AAC (5540T>C), TAT1802TAC (5546T>C), GGC1805GGT (5555C>T), GTG1807GCG (5560T>C), CAG1808CAA (5564G>A), TAT1809TAC (5567T>C), TTT1810TTG (5570T>G), CTT1811CTC (5573T>C), CAA1812AAA (5574C>A), GAA2142GCA (6565A>C), ACC2143CCC (6567A>C), ATT2144ATC (6572T>C), TAC2148TAT (6584C>T), GAG2150GCG (6589A>C) |

#### Proteins

| | | | | | | | | | |
|---|---|---|---|---|---|---|---|---|---|
| polyprotein (YP_001976147.1) | 1 | 2152 | 85.0% | 9356 | 73.4% | 1820 (99.3%) | 1276 (69.6%) | 3/10/0/0 | 1 |

| | Begin | End | Coverage | Score | Concordance | Matches | Identities | I/D/M/F* | Stop Codons |
|---|---|---|---|---|---|---|---|---|---|
| **NT** | **22** | **6809** | **80.5%** | **3829** | **33.5%** | **5772 (99.0%)** | **3883 (66.6%)** | **10/50** | |

Protein mutations:

F7W (160T>G 161C>G), K9_A16del (165_188delAAGGTTGCTGGTGTTGATGACGCC), S17P (189A>C 190G>C), S18P (192T>C), S19A (195T>G), T21S (201A>T 203C>A), T22P (204A>C), G26A (217G>C 218T>A), L28V (222C>G), N30S (228A>T 229A>C), I32F (234A>T), K34S (240A>T 241A>C 242A>T), F35L (243T>C), A38T (252G>A 254G>T), V39L (255G>C 257T>C), I42V (264A>G), T43N (268C>A 269C>T), T44K (271C>A 272T>A), M45V (273A>G 275G>T), K48T (283A>C 284G>C), S50V (288T>G 289C>T), L53I (297C>A 299T>A), L56V (306C>G 308G>C), S59P (315T>C 317T>A), T61M (322C>T 323T>G), D62A (325A>C 326T>A), V63A (328T>C 329G>A), S64A (330T>G 332C>A), V68A (343T>C 344G>C), A69S (345G>T 347C>T), S72N (355G>A), S73A (357T>G 359A>T), L74V (360C>G 362T>A), N75E (363A>G 365T>A), K76R (367A>G), G82A (385G>C 386T>A), A83K (387G>A 388C>A), L84I (390C>A 392T>A), I85M (395A>G), D86E (398T>A), L89I (405C>A 407T>C), A91G (412C>G 413G>T), S93D (417T>G 418C>A), L95V (423T>G 425G>A), S96K (426T>A 427C>A 428C>A), S101T (441T>A), K103V (447A>G 448A>T 449G>A), M105L (453A>C 455G>T), L109M (465T>A), F110V (468T>G), E111L (471G>T 472A>T 473G>A), L113V (477T>G 479G>C), S114K (480T>A 481C>A 482C>G), N115D (483A>G), V117F (489G>T 491T>C), K118I (493A>T 494A>C), F121Y (502T>A 503T>C), A122E (505C>A 506T>A), V124L (510G>T 512G>A), N126E (516A>G 518T>A), S128V (522A>G 523G>T), D130S (528G>T 529A>C 530T>A), V131L (531G>T), V135F (543G>T 545A>T), Y136I (546T>A 547A>T 548T>A), G137A (550G>C), I143L (567A>T 569A>G), T144A (570A>G 572A>C), L145I (573T>A 575G>T), V147L (579G>C 581G>A), F150Y (589T>A), S154V (600A>G 601G>T 602T>A), E158T (612G>A 613A>C), G159P (615G>C 616G>C), V160F (618G>T), S161N (621T>A 622C>A), T163V (627A>G 628C>T 629T>A), K165V (633A>G 634A>T 635A>T), N166T (637A>C), I167L (639A>T 641C>G), L169S (646T>C 647G>A), I170V (648A>G 650A>T), M174F (660A>T 662G>T), K186T (697A>C), A188S (702G>T 704C>T), E189G (706A>G 707A>G), R190K (708C>A 709G>A 710A>G), T192S (714A>T), M193Q (717A>C 718T>A), V194L (720G>C), S198A (732A>G 733G>C 734T>A), M199V (735A>G 737G>A), V205L (753G>T 755T>A), S206T (756T>A 758A>G), M214L (780A>T 782G>A), T216S (786A>T), I217L (789A>T 791A>G), I228V (822A>G), T248A (882A>G 884A>T), T260Q (918A>C 919C>A 920T>G), I280V (978A>G 980A>G), G282S (984G>A), T283I (988C>T 989C>T), S295N (1024G>A), E304Q (1050G>C), Q307T (1059C>A 1060A>C), D310S (1068G>T 1069A>C 1070T>G), A311E (1072C>A 1073A>G), C313A (1077T>G 1078G>C), D314E (1082T>A), A316S (1086G>T 1088A>T), T317M (1090C>T), K325P (1113A>C 1114A>C 1115G>T), T329I (1126C>T 1127C>T), N332Q (1134A>C 1136C>A), R335K (1144G>A 1145G>A), E337Q (1149G>C 1151A>G), L339V (1155C>G), M354L (1200A>T), A363T (1227G>A), E366D (1238A>T), V368L (1242G>C 1244T>G), Y369V (1245T>G 1246A>T), S372A (1254A>G 1255G>C 1256T>C), C373S (1257T>A 1259T>C), R374Q (1260A>C 1261G>A 1262G>A), M376S (1266A>T 1267T>C 1268G>T), S393A (1317T>G 1319C>T), S401C (1342C>G 1343C>T), R412A (1374A>G 1375G>C), I440V (1458A>G 1460C>G), M521R (1702T>G 1703G>A), E523V (1708A>T), E525D (1715G>T), S527D (1719T>G 1720C>A 1721C>T), T528P (1722A>C 1724T>A), E529N (1725G>A 1727A>T), D530T (1728G>A 1729A>C), G533A (1738G>C 1739A>C), R536I (1747G>T 1748A>T), N538D (1752A>G 1754C>T), I539V (1755A>G 1757A>G), N552A (1794A>G 1795A>C), T554S (1800A>T), S556D (1806A>G 1807G>A), E560Q (1818G>C 1820G>A), Q567R (1840A>G), A570Y (1848G>T 1849C>A 1850A>T), K574S (1861A>G 1862G>C), F577Y (1870T>A 1871T>C), E578N (1872G>A 1874A>T), A579T (1875G>A), E580T (1878G>A 1879A>C), I583L (1887A>T 1889A>G), S584A (1890A>G 1891G>C), P585D (1893C>G 1894C>A 1895T>C), I587V (1899A>G 1901A>T), Q591N (1911C>A 1913A>T), S593A (1917T>G), A594V (1921C>T 1922A>T), D596E (1928T>A), I598L (1932A>C 1934A>G), A599D (1936C>A 1937G>T), V601F (1941G>T 1943G>C), D602Q (1944G>C 1946T>A), G603D (1948G>A 1949G>T), R605del (1953_1955delAGA), R607Q (1960G>A 1961C>A), T608V (1962A>G 1963C>T 1964C>T), K609L (1965A>T 1966A>T), Q610P (1969A>C 1970A>T), D611P (1971G>C 1972A>C), I614A (1980A>G 1981T>C), L615A (1983T>G 1984T>C 1985G>T), V616Y (1986G>T 1987T>A 1988G>T), P617E (1989C>G 1990C>A 1991C>G), T618Q (1992A>C 1993C>A 1994T>G), V619T (1995G>A 1996T>C 1997T>A), V620I (1998G>A 2000G>T), S621T (2002G>C 2003T>A), N623A (2007A>G 2008A>C), D624E (2012T>A), R625K (2014G>A), V626I (2016G>A 2018G>T), I627V (2019A>G 2021C>T), A629S (2025G>T 2027A>C), R630Q (2028A>C 2029G>A 2030A>G), L645M (2073T>A), D646N (2076G>A), P647K (2079C>A 2080C>A 2081T>A), E648F (2082G>T 2083A>T 2084G>T), L650F (2088C>T), H655L (2104A>T 2105T>C), H656S (2106C>T 2107A>C 2108T>C), M660I (2120G>C), S663V (2127T>G 2128C>T 2129T>G), D665S (2133G>T 2134A>C), D666S (2136G>T 2137A>C), E667D (2141A>T), H668A (2142C>G 2143A>C 2144T>C), E670Q (2148G>C), I671N (2152T>A), N673E (2157A>G 2159C>A), K674M (2161A>T 2162A>G), R676V (2166A>G 2167G>T), V677N (2169G>A 2170T>A 2171C>T), N678D (2172A>G 2174C>T), M679L (2175A>T), Y681V (2181T>G 2182A>T 2183T>C), R682K (2185G>A), D685S (2193G>A 2194A>G), A686P (2196G>C), E687Q (2199G>C 2201A>G), R690A (2208C>G 2209G>C 2210A>T), Q694R (2221A>G), A710S (2268G>A 2269C>G 2270A>C), S711T (2271T>A), K716E (2286A>G), K717L (2289A>C 2290A>T 2291A>G), A719D (2296C>A 2297A>T), T722R (2304A>C 2305C>G 2306A>C), E723N (2307G>A 2309G>T), M724L (2310A>C 2312G>T), R725E (2313A>G 2314G>A), T726Q (2316A>C 2317C>A 2318T>A), F728K (2322T>A 2323T>A 2324T>A), H730Y (2328C>T), S731K (2331T>A 2332C>A 2333T>G), C732A (2334T>G 2335G>C 2336C>G), I733L (2337A>C), D734Q (2340G>C 2342T>G), V737A (2350T>C 2351T>A), I741T (2362T>C 2363A>G), F749R (2385T>A 2386T>G 2387T>A), N763R (2428A>G 2429C>G), S766A (2436T>G), S766_K767insG (2438_2439insGGT), K767T (2440A>C 2441A>T), E768D (2444G>C), P769K (2445C>A 2446C>A 2447A>G), E771R (2451G>A 2452A>G), S772N (2454T>A 2455C>A 2456A>T), K775I (2464A>T 2465A>C), I776E (2466A>G 2467T>A 2468T>A), Y778M (2472T>A 2473A>T 2474C>G), H779I (2475C>A 2476A>T), K780G (2478A>G 2479A>G 2480A>T), V782K (2484G>A 2485T>A 2486T>G), E783D (2489A>C), Y784K (2490T>A 2492T>A), I785M (2495C>G), K786I (2497A>T 2498G>C), D787Q (2499G>C 2501T>G), T788M (2503C>T 2504A>G), W790N (2508T>A 2509G>A 2510G>T), S791D (2511A>G 2512G>A), Y794F (2521A>T 2522T>C), A795I (2523G>A 2524C>T 2525A>C), R797K (2530G>A 2531A>G), I806L (2556A>T 2558A>G), M807I (2561G>A), V812T (2574G>A 2575T>C 2576T>C), F813M (2577T>A 2579T>G), G814N (2580G>A 2581G>A 2582A>T), G815A (2584G>C), I816V (2586A>G), L820I (2598T>A 2600G>T), A821T (2601G>A 2603G>T), W823L (2607T>C 2608G>T 2609G>C), Q824K (2610C>A), S825A (2613A>G 2614G>C 2615T>A), N826A (2616A>G 2617A>C 2618C>A), S827T (2619T>A), F828S (2623T>C), N830L (2628A>T 2629A>T 2630T>A), S832D (2634T>G 2635C>A 2636G>T), N852K (2696T>G), K853T (2698A>C 2699G>T), K853_L854insH (2699_2700insCAT), L854F (2700C>T), R855A (2703A>G 2704G>C), N856K (2708T>A), L857S (2709C>T 2710T>C 2711T>C), V859L (2715G>C 2717T>A), S862A (2724A>G 2725G>C 2726T>G), S863A (2727T>G 2729T>A), Q864D (2730C>G 2732A>T), S865N (2733T>A 2734C>A 2735A>C), L866N (2736T>A 2737T>A 2738G>C), A867H (2739G>C 2740C>A 2741A>T), Q868V (2742C>G 2743A>T 2744A>G), D869_W870insR (2747_2748insCGG), A872E (2755C>A 2756T>A), E873Q (2757G>C 2759A>G), Q879E (2775C>G 2777G>A), K882H (2784A>C 2786G>C), L890I (2808C>A), G891A (2812G>C), M907L (2859A>C 2861G>C), H914R (2880C>A 2881A>G 2882C>A), A916M (2886G>A 2887C>T 2888T>G), T923S (2907A>T 2909A>C), A927E (2920C>A 2921C>G), N928H (2922A>C), R930I (2928C>A 2929G>T 2930A>T), S931C (2932C>G), K932N (2936A>T), Y935F (2944A>T), R938T (2953G>C 2954A>C), E939Q (2955G>C), I942L (2964A>T 2966T>G), I948V (2982A>G 2984C>G), P961S (3021C>T 3023T>A), E972D (3056G>T), F975Y (3064T>A), T976S (3066A>T 3068A>T), Q980M (3078C>A 3079A>T 3080A>G), C984V (3090T>G 3091G>T 3092T>G), T990K (3109C>A 3110A>G), R993K (3117C>A 3118G>A 3119T>G), L994K (3120C>A 3121T>A 3122T>A), G995S (3123G>T 3124G>C), F999V (3135T>G), M1002Y (3144A>T 3145T>A 3146G>T), V1012E (3175T>A 3176T>A), E1015T (3183G>A 3184A>C), E1017G (3190A>G), N1019D (3195A>G 3197C>T), S1021N (3201T>A 3202C>A 3203C>T), S1022A (3204A>G 3205G>C 3206T>A), K1024A (3210A>G 3211A>C 3212G>T), I1025L (3213A>C), C1026S (3216T>A), Q1029V (3225C>G 3226A>T 3227A>T), S1030A (3228T>G 3230T>A), G1037S (3249G>A 3251G>T), S1050T (3289G>C), V1057C (3309G>T 3310T>G 3311G>C), G1068D (3343G>A 3344C>T), V1069T (3345G>A 3346T>C 3347A>C), R1073K (3357C>A 3358G>A 3359T>G), F1075Y (3364T>A 3365T>C), I1076V (3366A>G 3368C>G), S1077A (3369T>G), I1083V (3387A>G 3389C>G), Y1084F (3391A>T 3392C>T), E1086D (3398A>T), V1096I (3426G>A 3428C>T), A1098V (3433C>T 3434A>G), D1104E (3452T>A), I1105L (3453A>C 3455T>G), S1108H (3462T>C 3463C>A 3464A>C), I1109L (3465A>C 3467T>G), R1112K (3475G>A 3476A>G), T1114E (3480A>G 3481C>A), Q1115R (3483C>A 3484A>G 3485G>A), L1116F (3486C>T 3488T>C), N1120K (3500C>A), Y1123A (3507T>G 3508A>C 3509C>T), Y1124F (3511A>T), Q1130L (3528C>T 3529A>T), L1132M (3534T>A), R1143K (3567C>A 3568G>A 3569G>A), K1144R (3570A>C 3571A>G 3572A>C), Q1146P (3577A>C 3578A>C), P1153D (3597C>G 3598C>A 3599T>C), E1155Q (3603G>C), T1162S (3624A>T 3626A>T), V1163A (3628T>C 3629T>C), H1164K (3630C>A 3632T>A), D1169E (3647T>A), M1179E (3675A>G 3676T>A), C1186S (3697G>C 3698T>C), G1187N (3699G>A 3700G>A 3701A>C), D1189E (3707T>G), D1190E (3710T>G), D1191P (3711G>C 3712A>C 3713C>A), N1199G (3735A>G 3736A>G), C1202A (3744T>G 3745G>C 3746C>A), L1204I (3750C>A 3752G>T), I1207V (3759A>G 3761C>G), M1208L (3762A>C 3764G>T), R1209K (3765C>A 3766G>A 3767T>G), S1210M (3769G>T 3770C>G), K1212R (3774A>C 3775A>G 3776G>A), I1213V (3777A>G 3779C>G), R1217K (3789C>A 3790G>A 3791T>A), C1219A (3795T>G 3796G>C 3797T>A), E1220I (3798G>A 3799A>T 3800A>C), D1222N (3804G>A), C1233F (3838G>T), Y1236F (3847A>T), G1238A (3853G>C 3854C>A), M1252L (3894A>T), M1254L (3900A>C 3902G>T), M1256S (3906A>T 3907T>C 3908G>T), Q1263E (3927C>G), S1264A (3930T>G), T1265Q (3933A>C 3934C>A 3935T>A), G1266P (3936G>C 3937G>C 3938C>A), E1267N (3939G>A 3941A>C), G1268S (3942G>T 3943G>C 3944A>G), E1269Q (3945G>C), E1272L (3954G>C 3955A>T 3956A>T), K1274I (3961A>T), D1276N (3966G>A), E1281D (3983A>T), V1288S (4002G>T 4003T>C), I1290V (4008A>G 4010T>A), K1292Q (4014A>C), I1294T (4021T>C 4022C>A), H1295F (4023C>T 4024A>T), T1344C (4170A>T 4171C>G 4172A>T), L1348A (4182T>G 4183T>C 4184G>A), A1351V (4192C>T 4193T>A), I1352L (4194A>C 4196A>C), D1359N (4215G>A), S1363H (4227T>C 4228C>A), L1367V (4239T>G), N1370S (4249A>G), G1371S (4251G>A 4253G>C), A1374S (4260G>T 4262T>A), V1378E (4273T>A), V1393I (4317G>A 4319T>C), E1405A (4354A>C 4355A>T), Y1406F (4357A>T 4358C>T), S1409D (4365T>G 4366C>A 4367A>T), K1410E (4368A>G), D1415E (4385T>G), A1421T (4401G>A), D1435N (4443G>A 4445T>C), V1437L (4449G>T), I1476V (4566A>G 4568T>C), A1477K (4569G>A 4570C>A 4571T>A), R1479K (4576G>A 4577G>A), A1480S (4578G>T), I1481V (4581A>G 4583A>G), V1483I (4587G>A), S1486N (4597G>A), L1506M (4656T>A 4658A>G), Q1509R (4665C>A 4666A>G), S1512D (4674T>G 4675C>A), N1515S (4683A>T 4684A>C 4685T>C), H1518R (4692C>A 4693A>G 4694T>A), L1522A (4704C>G 4705T>C 4706G>T), I1526V (4716A>G), N1538E (4752A>G 4754T>G), E1539K (4755G>A), V1540T (4758G>A 4759T>C), Q1541K (4761C>A), Q1543L (4768A>T 4769A>G), H1563Q (4829C>A), I1572V (4854A>G), M1576I (4868G>T), P1583V (4887C>G 4888C>T 4889A>G), T1588V (4902A>G 4903C>T 4904T>A), M1592Y (4914A>T 4915T>A 4916G>T), V1595I (4923G>A 4925G>T), R1607Q (4960G>A 4961T>G), V1608A (4963T>C 4964G>C), A1609T (4965G>A), D1611E (4973C>A), K1612S (4975A>G 4976G>T), L1613F (4977C>T), L1615N (4983T>A 4984T>A 4985A>T), Q1618F (4992C>T 4993A>T 4994G>T), T1619G (4995A>G 4996C>G 4997G>A), D1622K (5004G>A 5006T>G), E1623Q (5007G>C 5009A>G), G1625N (5013G>A 5014G>A), F1626L (5016T>C), R1627P (5019A>C 5020G>C 5021A>C), A1628I (5022G>A 5023C>T 5024T>C), V1629M (5025G>A), F1631M (5031T>A 5033T>G), S1633T (5037T>A 5039C>T), L1635Y (5044T>A 5045G>T), V1637L (5049G>C 5051G>T), Q1639S (5055C>A 5056A>G 5057G>T), E1640Q (5058G>C), I1642L (5064A>T 5066T>G), A1643T (5067G>A), K1644E (5070A>G), S1649E (5085T>G 5086C>A 5087A>G), A1651I (5091G>A 5092C>T 5093C>T), L1652G (5094C>G 5095T>G), T1653V (5097A>G 5098C>T 5099A>T), I1654L (5100A>T 5102T>G), C1655M (5103T>A 5104G>T 5105T>G), M1657L (5109A>C 5111G>A), A1658S (5112G>T), K1659D (5115A>G 5117A>T), A1661K (5121G>A 5122C>A 5123T>A), I1662N (5125T>A), S1663T (5127T>A 5129C>A), D1668P (5142G>C 5143A>C 5144C>T), T1674del (5160_5162delACA), R1678E (5172C>G 5173G>A 5174T>G), D1679G (5176A>G), L1680V (5178C>G 5180G>C), N1681H (5181A>C), W1682E (5184T>G 5185G>A 5186G>A), A1684V (5191C>T 5192A>G), N1686D (5196A>G 5198C>T), I1687V (5199A>G 5201T>A), G1688F (5202G>T 5203G>T), A1696S (5226G>T 5228C>A), H1697N (5229C>A), Y1698Q (5232T>C 5234C>A), T1699Q (5235A>C 5236C>A 5237G>A), D1701S (5241G>T 5242A>C 5243T>A), E1702S (5244G>T 5245A>C), Y1704N (5250T>A), I1706L (5256A>C 5258T>G), K1713R (5277A>C 5278A>G 5279G>C), K1714R (5281A>G 5282G>A), S1715T (5283T>A 5285C>A), N1716T (5287A>C 5288T>A), M1731I (5333G>A), M1736L (5346A>T), S1737A (5349T>G), M1740Q (5358A>C 5359T>A), A1742H (5364G>C 5365C>A 5366C>T), K1744R (5370A>C 5371A>G 5372G>C), N1745K (5375T>G), V1746S (5376G>A 5377T>G 5378G>T), A1748E (5383C>A), G1749V (5386G>T 5387A>T), H1751D (5391C>G), V1752G (5395T>G 5396A>G), N1755S (5403A>T 5404A>C), V1757T (5409G>A 5410T>C 5411T>C), V1766I (5436G>A 5438C>A), G1767A (5440G>C 5441C>T), M1774F (5460A>T 5462G>T), R1779H (5475A>C 5476G>A 5477G>T), C1782R (5484T>A 5486T>G), V1783A (5488T>C 5489G>C), E1784T (5490G>A 5491A>C 5492G>A), I1786L (5496A>C 5498A>T), I1793M (5519T>G), G1794A (5521G>C 5522A>T), N1795G (5523A>G 5524A>G 5525T>G), V1807A (5560T>C), F1810L (5570T>G), Q1812K (5574C>A), E2142A (6565A>C), T2143P (6567A>C), E2150A (6589A>C)

| | Begin | End | Coverage | Score | Concordance | Matches | Identities | I/D/M/F* | Stop Codons |
|---|---|---|---|---|---|---|---|---|---|
| **NT** | **22** | **6809** | **80.5%** | **3829** | **33.5%** | **5772 (99.0%)** | **3883 (66.6%)** | **10/50** | |

Codon mutations:

GCT2GCA (146T>A), TTC7TGG (160T>G 161C>G), TCC8AGT (162T>A 163C>G 164C>T), AAG9_GCC16del (165_188delAAGGTTGCTGGTGTTGATGACGCC), AGT17CCT (189A>C 190G>C), TCT18CCT (192T>C), TCT19GCT (195T>G), TCC20TCT (200C>T), ACC21TCA (201A>T 203C>A), ACT22CCT (204A>C), GGT26GCA (217G>C 218T>A), CTT28GTT (222C>G), TCA29TCT (227A>T), AAT30TCT (228A>T 229A>C), ATT32TTT (234A>T), TCC33TCT (239C>T), AAA34TCT (240A>T 241A>C 242A>T), TTT35CTT (243T>C), AAA37AAG (251A>G), GCG38ACT (252G>A 254G>T), GTT39CTC (255G>C 257T>C), AGT40TCC (258A>T 259G>C 260T>C), ATT42GTT (264A>G), ACC43AAT (268C>A 269C>T), ACT44AAA (271C>A 272T>A), ATG45GTT (273A>G 275G>T), AAG48ACC (283A>C 284G>C), TCA50GTA (288T>G 289C>T), CAT52CAC (296T>C), CTT53ATA (297C>A 299T>A), CTG56GTC (306C>G 308G>C), AAA57AAG (311A>G), CCA58CCG (314A>G), TCT59CCA (315T>C 317T>A), GTG60GTT (320G>T), ACT61ATG (322C>T 323T>G), GAT62GCA (325A>C 326T>A), GTG63GCA (328T>C 329G>A), TCC64GCA (330T>G 332C>A), TCT66TCA (338T>A), GTG68GCC (343T>C 344G>C), GCC69TCT (345G>T 347C>T), ACT70ACA (350T>A), TGT71TGC (353T>C), AGT72AAT (355G>A), TCA73GCT (357T>G 359A>T), CTT74GTA (360C>G 362T>A), AAT75GAA (363A>G 365T>A), AAA76AGA (367A>G), TTG78TTA (374G>A), GAC79GAT (377C>T), AAA80AAG (380A>G), GGT82GCA (385G>C 386T>A), GCA83AAA (387G>A 388C>A), CTT84ATA (390C>A 392T>A), ATA85ATG (395A>G), GAT86GAA (398T>A), CCA87CCC (401A>C), TTT88TTC (404T>C), CTT89ATC (405C>A 407T>C), AAA90AAG (410A>G), GCG91GGT (412C>G 413G>T), TAC92TAT (416C>T), TTT93GAT (417T>G 418C>A), TTC94TTT (422C>T), TTG95GTA (423T>G 425G>A), TCC96AAA (426T>A 427C>A 428C>A), ACA97ACT (431A>T), TAC99TAT (437C>T), TCT101ACT (441T>A), ATC102ATA (446C>A), AAG103GTA (447A>G 448A>T 449G>A), ATG105CTT (453A>C 455G>T), GTT106GTA (458T>A), TTA107CTT (459T>C 461A>T), AAA108AAG (464A>G), TTG109ATG (465T>A), TTT110GTT (468T>G), GAG111TTA (471G>T 472A>T 473G>A), TTG113GTC (477T>G 479G>C), TCC114AAG (480T>A 481C>A 482C>G), AAT115GAT (483A>G), GTT117TTC (489G>T 491T>C), AAA118ATC (493A>T 494A>C), TTA119CTA (495T>C), TTT121TAC (502T>A 503T>C), GCT122GAA (505C>A 506T>A), GTG124TTA (510G>T 512G>A), AAT126GAA (516A>G 518T>A), AGT128GTT (522A>G 523G>T), GAA129GAG (527A>G), GAT130TCA (528G>T 529A>C 530T>A), GTA131TTA (531G>T), GTT134GTG (542T>G), GTA135TTT (543G>T 545A>T), TAT136ATA (546T>A 547A>T 548T>A), GGT137GCT (550G>C), CTT139TTA (555C>T 557T>A), GTA140GTT (560A>T), ATA143TTG (567A>T 569A>G), ACA144GCC (570A>G 572A>C), TTG145ATT (573T>A 575G>T), TTA146TTG (578A>G), GTG147CTA (579G>C 581G>A), CTT148CTA (584T>A), CTA149CTG (587A>G), TTT150TAT (589T>A), GTT151GTA (593T>A), TGT152TGC (596T>C), AGT154GTA (600A>G 601G>T 602T>A), GTT157GTA (611T>A), GAG158ACG (612G>A 613A>C), GGT159CCT (615G>C 616G>C), GTT160TTT (618G>T), TCT161AAT (621T>A 622C>A), AAC162AAT (626C>T), ACT163GTA (627A>G 628C>T 629T>A), GTC164GTT (632C>T), AAA165GTT (633A>G 634A>T 635A>T), AAT166ACT (637A>C), ATC167TTG (639A>T 641C>G), TTG169TCA (646T>C 647G>A), ATA170GTT (648A>G 650A>T), GTC171GTT (653C>T), AAC173AAT (659C>T), ATG174TTT (660A>T 662G>T), TTT175TTC (665T>C), TCA176TCT (668A>T), AGT177TCA (669A>T 670G>C 671T>A), CTA178TTG (672C>T 674A>G), TAC179TAT (677C>T), AAA186ACA (697A>C), GCC188TCT (702G>T 704C>T), GAA189GGG (706A>G 707A>G), CGA190AAG (708C>A 709G>A 710A>G), TTC191TTT (713C>T), ACC192TCC (714A>T), ATG193CAG (717A>C 718T>A), GTG194CTG (720G>C), GCC195GCG (725C>G), GCT197GCA (731T>A), AGT198GCA (732A>G 733G>C 734T>A), ATG199GTA (735A>G 737G>A), GGG202GGT (746G>T), TCA204AGC (750T>A 751C>G 752A>C), GTT205TTA (753G>T 755T>A), TCA206ACG (756T>A 758A>G), ACG210ACT (770G>T), TCT211AGC (771T>A 772C>G 773T>C), CAG212CAA (776G>A), TTG213CTT (777T>C 779G>T), ATG214TTA (780A>T 782G>A), ACA216TCA (786A>T), ATA217TTG (789A>T 791A>G), GGG221GGT (803G>T), ATT222ATC (806T>C), TCG223TCA (809G>A), ACG224ACT (812G>T), TTG225TTA (815G>A), GTG226GTA (818G>A), TTC227TTT (821C>T), ATT228GTT (822A>G), GCA229GCT (827A>T), GTG230GTT (830G>T), GGT232GGG (836T>G), GGC236GGT (848C>T), TTG237CTT (849T>C 851G>T), AGC238TCA (852A>T 853G>C 854C>A), TTG241CTG (861T>C), TCC242TCA (866C>A), ATT244ATC (872T>C), CTC245CTG (875C>G), TAT246TAC (878T>C), TCG247TCT (881G>T), ACA248GCT (882A>G 884A>T), GCT252GCG (896T>G), CAG253CAA (899G>A), CAA257CAG (911A>G), CTA258CTG (914A>G), ACT260CAG (918A>C 919C>A 920T>G), CTT261TTG (921C>T 923T>G), CGT263AGG (927C>A 929T>G), AAT264AAC (932T>C), AAA266AAG (938A>G), TGC268TGT (944C>T), ACC269ACA (947C>A), TCC270TCT (950C>T), CAG271CAA (953G>A), GCT272GCC (956T>C), GAA278GAG (974A>G), ATA280GTG (978A>G 980A>G), GTC281GTT (983C>T), GGC282AGC (984G>A), ACC283ATT (988C>T 989C>T), GGA285GGT (995A>T), TTC286TTT (998C>T), GTG290GTA (1010G>A), GCT293GCG (1019T>G), AGT295AAT (1024G>A), GCA296GCT (1028A>T), ACG297ACC (1031G>C), CTG298TTA (1032C>T 1034G>A), ACT300ACC (1040T>C), CTG302CTT (1046G>T), GAG304CAG (1050G>C), CAG307ACG (1059C>A 1060A>C), GAG308GAA (1064G>A), GAT310TCG (1068G>T 1069A>C 1070T>G), GCA311GAG (1072C>A 1073A>G), GTG312GTT (1076G>T), TGT313GCT (1077T>G 1078G>C), GAT314GAA (1082T>A), CCT315CCA (1085T>A), GCA316TCT (1086G>T 1088A>T), ACG317ATG (1090C>T), GAG320GAA (1100G>A), AAC321AAT (1103C>T), GCC323GCA (1109C>A), AAT324AAC (1112T>C), AAG325CCT (1113A>C 1114A>C 1115G>T), GCT326GCA (1118T>A), TTC327TTT (1121C>T), CCT328CCA (1124T>A), ACC329ATT (1126C>T 1127C>T), AAA330AAG (1130A>G), AAC332CAA (1134A>C 1136C>A), CTG334TTA (1140C>T 1142G>A), AGG335AAA (1144G>A 1145G>A), GAA336GAG (1148A>G), GAA337CAG (1149G>C 1151A>G), GCT338GCC (1154T>C), CTC339GTC (1155C>G), AAG340AAA (1160G>A), ATC341ATA (1163C>A), GCA346GCC (1178A>C), GTT350GTC (1190T>C), GCG351GCA (1193G>A), TTC353TTT (1199C>T), ATG354TTG (1200A>T), AGT359TCT (1215A>T 1216G>C), ATT362ATA (1226T>A), GCG363ACG (1227G>A), TTA365CTA (1233T>C), GAA366GAT (1238A>T), GTT368CTG (1242G>C 1244T>G), TAT369GTT (1245T>G 1246A>T), GAA371GAG (1253A>G), AGT372GCC (1254A>G 1255G>C 1256T>C), TGT373AGC (1257T>A 1259T>C), AGG374CAA (1260A>C 1261G>A 1262G>A), CAC375CAT (1265C>T), ATG376TCT (1266A>T 1267T>C 1268G>T), GGC377GGT (1271C>T), CAG380CAA (1280G>A), CGT382AGG (1284C>A 1286T>G), GAA384GAG (1292A>G), TTT386TTC (1298T>C), GTA388GTG (1304A>G), TTC391TTT (1313C>T), GGG392GGA (1316G>A), TCC393GCT (1317T>G 1319C>T), GGG395GGA (1325G>A), TCT399TCA (1337T>A), ACA400ACG (1340A>G), TCC401TGT (1342C>G 1343C>T), TTA404CTT (1350T>C 1352A>T), TTT405TTC (1355T>C), ATT406ATC (1358T>C), GTG409GTT (1367G>T), TTG410CTC (1368T>C 1370G>C), AGA412GCA (1374A>G 1375G>C), GAG415GAA (1385G>A), AAA417AAG (1391A>G), CTC418CTG (1394C>G), AAT419AAC (1397T>C), CGG420CGC (1400G>C), CTA421TTG (1401C>T 1403A>G), GCA423GCG (1409A>G), AAG426AAA (1418G>A), AGG427AGA (1421G>A), GCT435GCA (1445T>A), CAC436CAT (1448C>T), GCT439GCA (1457T>A), ATC440GTG (1458A>G 1460C>G), CTG441CTT (1463G>T), GAC444GAT (1472C>T), GGA446GGT (1478A>T), GCA447GCT (1481A>T), TTG448CTT (1482T>C 1484G>T), CGA449CGG (1487A>G), GGG451GGA (1493G>A), GCT452GCA (1496T>A), TGC455TGT (1505C>T), CAA456CAG (1508A>G), GAT457GAC (1511T>C), ATT458ATA (1514T>A), AAA459AAG (1517A>G), GAT463GAC (1529T>C), TCA466TCT (1538A>T), ACA467ACC (1541A>C), CAA468CAG (1544A>G), CCA469CCT (1547A>T), CCC471CCT (1553C>T), TTA472CTT (1554T>C 1556A>T), GCC475GCT (1565C>T), GTT477GTC (1571T>C), GAC479GAT (1577C>T), GGC481GGA (1583C>A), CAT483CAC (1589T>C), TTC484TTT (1592C>T), ACC485ACA (1595C>A), TCC486TCA (1598C>A), TAT488TAC (1604T>C), ATA489ATC (1607A>C), ACA492ACC (1616A>C), AAT494AAC (1622T>C), CTG495TTG (1623C>T), ATC496ATA (1628C>A), TCA497TCT (1631A>T), CCT499CCA (1637T>A), GCC500GCT (1640C>T), CAA501CAG (1643A>G), GGT503GGG (1649T>G), CTT504CTA (1652T>A), ACA505ACG (1655A>G), CCA507CCT (1661A>T), GCT509GCA (1667T>A), GGA511GAA (1673G>A), CGG512AGG (1674C>A), AGA513AGG (1679A>G), AGG514AGA (1682G>A), GAC515GAT (1685C>T), ATC516ATT (1688C>T), CTG517TTA (1689C>T 1691G>A), GTG518GTT (1694G>T), TGT520TGC (1700T>C), ATG521AGA (1702T>G 1703G>A), GAG523GTG (1708A>T), GAG525GAT (1715G>T), TCC527GAT (1719T>G 1720C>A 1721C>T), ACT528CCA (1722A>C 1724T>A), GAA529AAT (1725G>A 1727A>T), GAT530ACT (1728G>A 1729A>C), CCT531CCA (1733T>A), ACG532ACA (1736G>A), GGA533GCC (1738G>C 1739A>C), CTC535CTA (1745C>A), AGA536ATT (1747G>T 1748A>T), AAC538GAT (1752A>G 1754C>T), ATA539GTG (1755A>G 1757A>G), GAG541GAA (1763G>A), CCT545CCC (1775T>C), GCT547GCC (1781T>C), ACT549ACA (1787T>A), CAC550CAT (1790C>T), AGG551AGA (1793G>A), AAT552GCT (1794A>G 1795A>C), TTG553TTA (1799G>A), ACC554TCC (1800A>T), AGT556GAT (1806A>G 1807G>A), GAT557GAC (1811T>C), CTT558CTC (1814T>C), GAG560CAA (1818G>C 1820G>A), GTA563GTG (1829A>G), GCC564GCA (1832C>A), AAA565AAG (1835A>G), TGT566TGC (1838T>C), CAG567CGG (1840A>G), GTA568GTG (1844A>G), CAT569CAC (1847T>C), GCA570TAT (1848G>T 1849C>A 1850A>T), GAA571GAG (1853A>G), GTA572GTG (1856A>G), AAG574AGC (1861A>G 1862G>C), TTG576CTT (1866T>C 1868G>T), TTT577TAC (1870T>A 1871T>C), GAA578AAT (1872G>A 1874A>T), GCT579ACT (1875G>A), GAA580ACA (1878G>A 1879A>C), TCT581TCG (1883T>G), GGC582GGA (1886C>A), ATA583TTG (1887A>T 1889A>G), AGT584GCT (1890A>G 1891G>C), CCT585GAC (1893C>G 1894C>A 1895T>C), ATA587GTT (1899A>G 1901A>T), GTT590GTC (1910T>C), CAA591AAT (1911C>A 1913A>T), TCT593GCT (1917T>G), GCA594GTT (1921C>T 1922A>T), GAT596GAA (1928T>A), ATA598CTG (1932A>C 1934A>G), GCG599GAT (1936C>A 1937G>T), GTG601TTC (1941G>T 1943G>C), GAT602CAA (1944G>C 1946T>A), GGG603GAT (1948G>A 1949G>T), GCT604GCC (1952T>C), AGA605del (1953_1955delAGA), TTG606TTA (1958G>A), CGC607CAA (1960G>A 1961C>A), ACC608GTT (1962A>G 1963C>T 1964C>T), AAG609TTG (1965A>T 1966A>T), CAA610CCT (1969A>C 1970A>T), GAT611CCT (1971G>C 1972A>C), GAA612GAG (1976A>G), ATA614GCA (1980A>G 1981T>C), TTG615GCT (1983T>G 1984T>C 1985G>T), GTG616TAT (1986G>T 1987T>A 1988G>T), CCC617GAG (1989C>G 1990C>A 1991C>G), ACT618CAG (1992A>C 1993C>A 1994T>G), GTT619ACA (1995G>A 1996T>C 1997T>A), GTG620ATT (1998G>A 2000G>T), AGT621ACA (2002G>C 2003T>A), AAT623GCT (2007A>G 2008A>C), GAT624GAA (2012T>A), AGA625AAA (2014G>A), GTG626ATT (2016G>A 2018G>T), ATC627GTT (2019A>G 2021C>T), TAT628TAC (2024T>C), GCA629TCC (2025G>T 2027A>C), AGA630CAG (2028A>C 2029G>A 2030A>G), GAG631GAA (2033G>A), ACG633ACA (2039G>A), GTA634GTT (2042A>T), GCA636GCC (2048A>C), TTG637CTC (2049T>C 2051G>C), GCA640GCT (2060A>T), CAG642CAA (2066G>A), GGC643GGG (2069C>G), TTG645ATG (2073T>A), GAC646AAC (2076G>A), CCT647AAA (2079C>A 2080C>A 2081T>A), GAG648TTT (2082G>T 2083A>T 2084G>T), GAA649GAG (2087A>G), CTT650TTT (2088C>T), TTT651TTC (2093T>C), CCT652CCA (2096T>A), CAT655CTC (2104A>T 2105T>C), CAT656TCC (2106C>T 2107A>C 2108T>C), GCC659GCT (2117C>T), ATG660ATC (2120G>C), TCT663GTG (2127T>G 2128C>T 2129T>G), CTT664CTA (2132T>A), GAT665TCT (2133G>T 2134A>C), GAT666TCT (2136G>T 2137A>C), GAA667GAT (2141A>T), CAT668GCC (2142C>G 2143A>C 2144T>C), GAG670CAG (2148G>C), ATT671AAT (2152T>A), AAC673GAA (2157A>G 2159C>A), AAA674ATG (2161A>T 2162A>G), AGA676GTA (2166A>G 2167G>T), GTC677AAT (2169G>A 2170T>A 2171C>T), AAC678GAT (2172A>G 2174C>T), ATG679TTG (2175A>T), TAT681GTC (2181T>G 2182A>T 2183T>C), AGA682AAA (2185G>A), GAT685AGT (2193G>A 2194A>G), GCT686CCT (2196G>C), GAA687CAG (2199G>C 2201A>G), CGA690GCT (2208C>G 2209G>C 2210A>T), ATG693GTG (2219A>G), CAA694CGA (2221A>G), GAC699GAT (2237C>T), ATA701ATC (2243A>C), CAC703CAT (2249C>T), TTT704TTC (2252T>C), GCA710AGC (2268G>A 2269C>G 2270A>C), TCA711ACA (2271T>A), ATC713ATT (2279C>T), GAG715GAA (2285G>A), AAG716GAG (2286A>G), AAA717CTG (2289A>C 2290A>T 2291A>G), CTA718CTG (2294A>G), GCA719GAT (2296C>A 2297A>T), GTG720GTC (2300G>C), CAG721CAA (2303G>A), ACA722CGC (2304A>C 2305C>G 2306A>C), GAG723AAT (2307G>A 2309G>T), ATG724CTT (2310A>C 2312G>T), AGG725GAG (2313A>G 2314G>A), ACT726CAA (2316A>C 2317C>A 2318T>A), GGA727GGG (2321A>G), TTT728AAA (2322T>A 2323T>A 2324T>A), GCA729GCG (2327A>G), CAT730TAT (2328C>T), TCT731AAG (2331T>A 2332C>A 2333T>G), TGC732GCG (2334T>G 2335G>C 2336C>G), ATT733CTT (2337A>C), GAT734CAG (2340G>C 2342T>G), GCA735GCT (2345A>T), GTT737GCA (2350T>C 2351T>A), GAC738GAT (2354C>T), ACC739ACA (2357C>A), CTC740CTT (2360C>T), ATA741ACG (2362T>C 2363A>G), ATC743ATT (2369C>T), GAG744GAA (2372G>A), TTT749AGA (2385T>A 2386T>G 2387T>A), GTT750GTA (2390T>A), TTT752TTC (2396T>C), TTA753CTG (2397T>C 2399A>G), ACA755ACT (2405A>T), GCA756GCT (2408A>T), AGA759CGC (2415A>C 2417A>C), TGC762TGT (2426C>T), AAC763AGG (2428A>G 2429C>G), GTA765GTT (2435A>T), TCA766GCA (2436T>G), TCA766_AAA767insGGT (2438_2439insGGT), AAA767ACT (2440A>C 2441A>T), GAG768GAC (2444G>C), CCA769AAG (2445C>A 2446C>A 2447A>G), GAA771AGA (2451G>A 2452A>G), TCA772AAT (2454T>A 2455C>A 2456A>T), ATC774ATT (2462C>T), AAA775ACE (2464A>T 2465A>C), ATT776GAA (2466A>G 2467T>A 2468T>A), TAC778ATG (2472T>A 2473A>T 2474C>G), CAT779ATT (2475C>A 2476A>T), AAA780GGT (2478A>G 2479A>G 2480A>T), GTT782AAG (2484G>A 2485T>A 2486T>G), GAA783GAC (2489A>C), TAT784AAA (2490T>A 2492T>A), ATC785ATG (2495C>G), AAG786ATC (2497A>T 2498G>C), GAT787CAG (2499G>C 2501T>G), ACA788ATG (2503C>T 2504A>G), TGG790AAT (2508T>A 2509G>A 2510G>T), AGC791GAC (2511A>G 2512G>A), CTT792TTG (2514C>T 2516T>G), TAT794TTC (2521A>T 2522T>C), GCA795ATC (2523G>A 2524C>T 2525A>C), CTG796TTG (2526C>T), AGA797AAG (2530G>A 2531A>G), CTC799CTT (2537C>T), GCA802GCC (2546A>C), GGT803GGA (2549T>A), ATA806TTG (2556A>T 2558A>G), ATG807ATA (2561G>A), GCT808GCA (2564T>A), GGA811GGC (2573A>C), GTT812ACC (2574G>A 2575T>C 2576T>C), TTT813ATG (2577T>A 2579T>G), GGA814AAT (2580G>A 2581G>A 2582A>T), GGC815GCC (2584G>C), ATT816GTT (2586A>G), ACA817ACT (2591A>T), GCA818GCT (2594A>T), TTT819TTC (2597T>C), TTG820ATT (2598T>A 2600G>T), GCG821ACT (2601G>A 2603G>T), TGG823CTC (2607T>C 2608G>T 2609G>C), CAG824AAG (2610C>A), AGT825GCA (2613A>G 2614G>C 2615T>A), AAC826GCA (2616A>G 2617A>C 2618C>A), TCT827ACT (2619T>A), TTC828TCC (2623T>C), CCC829CCT (2627C>T), AAT830TTA (2628A>T 2629A>T 2630T>A), GCA831GCT (2633A>T), TCG832GAT (2634T>G 2635C>A 2636G>T), GGA835GGG (2645A>G), GGT836GGC (2648T>C), GTG837GTA (2651G>A), GAA840GAG (2660A>G), ATA844ATC (2672A>C), AGT846AGC (2678T>C), AAG848AAA (2684G>A), AAC849AAT (2687C>T), AAG850AAA (2690G>A), GGT851GGG (2693T>G), AAT852AAG (2696T>G), AAG853ACT (2698A>C 2699G>T), AAG853_CTC854insCAT (2699_2700insCAT), CTC854TTC (2700C>T), AGA855GCA (2703A>G 2704G>C), AAT856AAA (2708T>A), CTT857TCC (2709C>T 2710T>C 2711T>C), CTC858TTG (2712C>T 2714C>G), GTT859CTA (2715G>C 2717T>A), GGT860GGG (2720T>G), AGT862GCG (2724A>G 2725G>C 2726T>G), TCT863GCA (2727T>G 2729T>A), CAA864GAT (2730C>G 2732A>T), TCA865AAC (2733T>A 2734C>A 2735A>C), TTG866AAC (2736T>A 2737T>A 2738G>C), GCA867CAT (2739G>C 2740C>A 2741A>T), CAA868GTG (2742C>G 2743A>T 2744A>G), GAT869GAC (2747T>C), CAT869_TGG870insCGG (2747_2748insCGG), GCT871GCA (2753T>A), GCT872GAA (2755C>A 2756T>A), GAA873CAG (2757G>C 2759A>G), GTA877GTT (2771A>T), CAG879GAA (2775C>G 2777G>A), AGC880AGT (2780C>T), CTC881CTG (2783C>G), AAG882CAC (2784A>C 2786G>C), AAT884AAC (2792T>C), GTG886GTT (2798G>T), GTG887GTT (2801G>T), TTA888TTG (2804A>G), CTT890ATT (2808C>A), GGT891GCT (2812G>C), GTG894GTC (2822G>C), TAC895TAT (2825C>T), TTT896TTC (2828T>C), GGC898GGT (2834C>T), ACC899ACG (2837C>G), TAC900TAT (2840C>T), GTG901GTT (2843G>T), ATA906ATT (2858A>T), ATG907CTC (2859A>C 2861G>C), ACC908ACT (2864C>T), GTA909GTG (2867A>G), GCT910GCA (2870T>A), GCA912GCC (2876A>C), TTC913TTT (2879C>T), CAC914AGA (2880C>A 2881A>G 2882C>A), AAT915AAC (2885T>C), GCT916ATG (2886G>A 2887C>T 2888T>G), CGA917AGG (2889C>A 2891A>G), GGT919GGC (2897T>C), CCA921CCT (2903A>T), ACA923TCC (2907A>T 2909A>C), ATC925ATT (2915C>T), GCC927GAG (2920C>A 2921C>G), AAT928CAT (2922A>C), TCT929TCC (2927T>C), CGA930ATT (2928C>A 2929G>T 2930A>T), TCT931TGT (2932C>G), AAA932AAT (2936A>T), GTT933GTG (2939T>G), CAA934CAG (2942A>G), TAC935TTC (2944A>T), AAC936AAT (2948C>T), AGA938ACC (2953G>C 2954A>C), GAA939CAA (2955G>C), ATT942TTG (2964A>T 2966T>G), ATT943ATA (2969T>A), GGC945GGT (2975C>T), GAC947GAT (2981C>T), ATC948GTG (2982A>G 2984C>G), CTG950TTG (2988C>T), CGC952AGG (2994C>A 2996C>G), GTT953GTA (2999T>A), GGT954GGG (3002T>G), CCA959CCC (3017A>C), AAG960AAA (3020G>A), CCT961TCA (3021C>T 3023T>A), GAC962GAT (3026C>T), ATC963ATA (3029C>A), CGT964AGA (3030C>A 3032T>A), CAC966CAT (3038C>T), CCA968CCT (3044A>T), AGA969CGC (3045A>C 3047A>C), AGG970AGA (3050G>A),

| | | | | | | | | | |
|---|---|---|---|---|---|---|---|---|---|
| | GAG972GAT (3056G>T), TGC974TGT (3062C>T), TTC975TAC (3064T>A), ACA976TCT (3066A>T 3068A>T), CAA980ATG (3078C>A 3079A>T 3080A>G), GGA981GGC (3083A>C), TGT984GTG (3090T>G 3091G>T 3092T>G), AGA985AGG (3095A>G), GTT987GTA (3101T>A), GCG988GCT (3104G>T), TCT989TCA (3107T>A), ACA990AAG (3109C>A 3110A>G), CCA992CCC (3116A>C), CGT993AAG (3117C>A 3118G>A 3119T>G), CTT994AAA (3120C>A 3121T>A 3122T>A), GGC995TCC (3123G>T 3124G>C), TTA997TTG (3131A>G), TTT999GTT (3135T>G), CTC1000CTT (3140C>T), AAG1001AAA (3143G>A), ATG1002TAT (3144A>T 3145T>A 3146G>T), GGA1009GGC (3167A>C), ACA1011ACC (3173A>C), GTT1012GAA (3175T>A 3176T>A), GTG1014GTC (3182G>C), GAA1015ACA (3183G>A 3184A>C), GAA1017GGA (3190A>G), CTG1018TTA (3192C>T 3194G>A), AAC1019GAT (3195A>G 3197C>T), TCC1020TCG (3200C>G), TCC1021AAT (3201T>A 3202C>A 3203C>T), AGT1022GCA (3204A>G 3205G>C 3206T>A), AAG1024GCT (3210A>G 3211A>C 3212G>T), ATT1025CTT (3213A>C), TGC1026AGC (3216T>A), TCT1027TCA (3221T>A), CAA1029GTT (3225C>G 3226A>T 3227A>T), TCT1030GCA (3228T>G 3230T>A), GAA1032GAG (3236A>G), ATA1035ATT (3245A>T), GGG1037AGT (3249G>A 3251G>T), GAA1038GAG (3254A>G), GGT1040GGA (3260T>A), GAC1041GAT (3263C>T), GGC1043GGT (3269C>T), ACG1044ACT (3272G>T), TTG1045CTT (3273T>C 3275G>T), CTA1046CTC (3278A>C), CTG1047TTA (3279C>T 3281G>A), TTA1048TTG (3284A>G), AGT1050ACT (3289G>C), GTT1051GTG (3293T>G), AAT1053AAC (3299T>C), CAA1055CAG (3305A>G), CCT1056CCG (3308T>G), GTG1057TGC (3309G>T 3310T>G 3311G>C), GGC1060GGA (3320C>A), ATC1061ATA (3323C>A), CAC1062CAT (3326C>T), TCA1065TCT (3335A>T), GGC1068GAT (3343G>A 3344C>T), GTA1069ACC (3345G>A 3346T>C 3347A>C), GCA1070GCT (3350A>T), GCT1071GCA (3353T>A), GAA1072GAG (3356A>G), CGT1073AAG (3357C>A 3358G>A 3359T>G), GGA1074GGT (3362A>T), TTT1075TAC (3364T>A 3365T>C), ATC1076GTG (3366A>G 3368C>G), TCT1077GCT (3369T>G), AAT1079AAC (3377T>C), GCT1080GCA (3380T>A), GCT1082GCA (3386T>A), ATC1083GTG (3387A>G 3389C>G), TAC1084TTT (3391A>T 3392C>T), AGG1085AGA (3395G>A), GAA1086GAT (3398A>T), CTA1088TTG (3402C>T 3404A>G), GAG1089GAA (3407G>A), CCA1092CCC (3416A>C), ACT1093ACG (3419T>G), GGG1094GGA (3422G>A), CCG1095CCC (3425G>C), GTC1096ATT (3426G>A 3428C>T), GCA1098GTG (3433C>T 3434A>G), GCA1099GCT (3437A>T), GTA1101GTG (3443A>G), GAT1104GAA (3452T>A), ATT1105CTG (3453A>C 3455T>G), CTG1106TTG (3456C>T), TCA1108CAC (3462T>C 3463C>A 3464A>C), ATT1109CTG (3465A>C 3467T>G), AGC1111TCC (3471A>T 3472G>C), AGA1112AAG (3475G>A 3476A>G), GAA1113GAG (3479A>G), ACA1114GAA (3480A>G 3481C>A), CAG1115AGA (3483C>A 3484A>G 3485G>A), CTT1116TTC (3486C>T 3488T>C), GAA1119GAG (3497A>G), AAC1120AAA (3500C>A), CAA1121CAG (3503A>G), TAC1123GCT (3507T>G 3508A>C 3509C>T), TAC1124TTC (3511A>T), CTT1125TTG (3513C>T 3515T>G), GGA1126GGC (3518A>C), ACA1127ACT (3521A>T), GTT1128GTA (3524T>A), CCA1129CCG (3527A>G), CAG1130TTG (3528C>T 3529A>T), GAG1131GAA (3533G>A), TTG1132ATG (3534T>A), GCC1133GCT (3539C>T), GCC1134GCT (3542C>T), GTT1136GTG (3548T>G), CCC1137CCG (3551C>G), AAG1139AAA (3557G>A), ACC1140ACA (3560C>A), ACT1141ACC (3563T>C), CTG1142CTT (3566G>T), CGG1143AAA (3567C>A 3568G>A 3569G>A), AAA1144CGC (3570A>C 3571A>G 3572A>C), AGC1145AGT (3575C>T), CAA1146CCC (3577A>C 3578A>C), TTG1147CTG (3579T>C), GAA1149GAG (3587A>G), GCA1150GCT (3590A>T), TTC1151TTT (3593C>T), GGA1152GGG (3596A>G), CCT1153GAC (3597C>G 3598C>A 3599T>C), GCA1154GCT (3602A>T), GAG1155CAG (3603G>C), ACA1156ACT (3608A>T), GCA1157GCC (3611A>C), ACA1162TCT (3624A>T 3626A>T), GTT1163GCC (3628T>C 3629T>C), CAT1164AAA (3630C>A 3632T>A), AGA1167AGG (3641A>G), GGT1168GGA (3644T>A), GAT1169GAA (3647T>A), GGT1170GGG (3650T>G), GAC1172GAT (3656C>T), GGG1177GGT (3671G>T), GTA1178GTC (3674A>C), ATG1179GAG (3675A>G 3676T>A), AAA1180AAG (3680A>G), TAC1181TAT (3683C>T), TGT1186TCC (3697G>C 3698T>C), GGA1187AAC (3699G>A 3700G>A 3701A>C), GAT1189GAG (3707T>G), GAT1190GAG (3710T>G), GAC1191CCA (3711G>C 3712A>C 3713C>A), GCC1193GCA (3719C>A), AAA1194AAG (3722A>G), CTA1195CTG (3725A>G), GCA1196GCT (3728A>T), TTC1197TTT (3731C>T), GAA1198GAG (3734A>G), AAT1199GGT (3735A>G 3736A>G), CTC1200CTT (3740C>T), AAG1201AAA (3743G>A), TGC1202GCA (3744T>G 3745G>C 3746C>A), TCG1203TCC (3749G>C), CTG1204ATT (3750C>A 3752G>T), CTA1205CTT (3755A>T), ATC1207GTG (3759A>G 3761C>G), ATG1208CTT (3762A>C 3764G>T), CGT1209AAG (3765C>A 3766G>A 3767T>G), AGC1210ATG (3769G>T 3770C>G), AAG1212CGA (3774A>C 3775A>G 3776G>A), ATC1213GTG (3777A>G 3779C>G), CCT1214CCC (3782T>C), GGA1216GGG (3788A>G), CGT1217AAA (3789C>A 3790G>A 3791T>A), CCA1218CCT (3794A>T), TGT1219GCA (3795T>G 3796G>C 3797T>A), GAA1220ATC (3798G>A 3799A>T 3800A>C), AGG1221AGA (3803G>A), GAT1222AAT (3804G>A), GGA1229GGG (3827A>G), TGT1233TTT (3838G>T), TAC1235TAT (3845C>T), TAT1236TTT (3847A>T), GGC1238GCA (3853G>C 3854C>A), GAG1240GAA (3860G>A), CTG1241TTG (3861C>T), ACA1243ACG (3869A>G), TCT1244TCA (3872T>A), TGC1245TGT (3875C>T), GGA1246GGT (3878A>T), CCC1248CCT (3884C>T), TTC1249TTT (3887C>T), ATG1252TTG (3894A>T), GGG1253GGA (3899G>A), ATG1254CTT (3900A>C 3902G>T), GGG1255GGT (3905G>T), ATG1256TCT (3906A>T 3907T>C 3908G>T), AAG1258AAA (3914G>A), AGA1259AGG (3917A>G), CAA1263GAA (3927C>G), TCC1264GCC (3930T>G), ACT1265CAA (3933A>C 3934C>A 3935T>A), GGC1266CCA (3936G>C 3937G>C 3938C>A), GAA1267AAC (3939G>A 3941A>C), GGA1268TCG (3942G>T 3943G>C 3944A>G), GAG1269CAG (3945G>C), AGA1270AGG (3950A>G), GTG1271GTT (3953G>T), GAA1272CTT (3954G>C 3955A>T 3956A>T), AAA1274ATA (3961A>T), AGA1275AGG (3965A>G), GAC1276AAC (3966G>A), ACT1277ACA (3971T>A), GTA1279GTC (3977A>C), GAA1281GAT (3983A>T), CTA1286TTG (3996C>T 3998A>G), GAT1287GAC (4001T>C), GTG1288TCG (4002G>T 4003T>C), ATT1290GTA (4008A>G 4010T>A), AAA1292CAA (4014A>C), ATC1294ACA (4021T>C 4022C>A), CAT1295TTT (4023C>T 4024A>T), GTG1296GTT (4028G>T), GAT1297GAC (4031T>C), CTG1298CTC (4034G>C), ACC1300ACA (4040C>A), CAA1302CAG (4046A>G), TGC1303TGT (4049C>T), GCC1304GCG (4052C>G), CGC1308CGG (4064C>G), CTC1309TTG (4065C>T 4067C>G), AAA1313AAG (4079A>G), ATC1314ATT (4082C>T), GGC1316GGG (4088C>G), AGA1318AGG (4094A>G), AGG1321CGC (4101A>C 4103G>C), CTC1322CTG (4106C>G), TTT1323TTC (4109T>C), GAG1324GAA (4112G>A), ATA1325ATC (4115A>C), CTT1326TTG (4116C>T 4118T>G), TAC1330TAT (4130C>T), TTG1333CTG (4137T>C), GTC1334GTG (4142C>G), AGG1335CGG (4143A>C), AAG1336AAA (4148G>A), TAT1337TAC (4151T>C), TTT1338TTC (4154T>C), GAT1340GAC (4160T>C), TTC1341TTT (4163C>T), TCA1342TCT (4166A>T), GCC1343GCT (4169C>T), ACA1344TGT (4170A>T 4171C>G 4172A>T), TTG1348GCA (4182T>G 4183T>C 4184G>A), AAT1350AAC (4190T>C), GCT1351GTA (4192C>T 4193T>A), ATA1352CTC (4194A>C 4196A>C), CCA1353CCT (4199A>T), AAA1355AAG (4205A>G), GTT1356GTA (4208T>A), ATT1358ATC (4214T>C), GAT1359AAT (4215G>A), CCT1360CCA (4220T>A), ACA1361ACG (4223A>G), AGT1362AGC (4226T>C), TCT1363CAT (4227T>C 4228C>A), TTG1367GTG (4239T>G), TTG1368CTG (4242T>C), GCA1369GCT (4247A>T), AAT1370AGT (4249A>G), GGG1371AGC (4251G>A 4253G>C), AGA1373AGG (4259A>G), GCT1374TCA (4260G>T 4262T>A), GTG1375GTA (4265G>A), TCA1376TCT (4268A>T), GAC1377GAT (4271C>T), GTG1378GAG (4273T>A), TTT1380TTC (4280T>C), GCT1382GCC (4286T>C), TAT1384TAC (4292T>C), TCC1385TCG (4295C>G), CCT1392CCC (4316T>C), GTT1393ATC (4317G>A 4319T>C), GCT1395GCA (4325T>A), TTT1396TTC (4328T>C), CAG1397CAA (4331G>A), TTG1402CTG (4344T>C), GAA1405GCT (4354A>C 4355A>T), TAC1406TTT (4357A>T 4358C>T), TAC1407TAT (4361C>T), GGA1408GGT (4364A>T), TCA1409GAT (4365T>G 4366C>A 4367A>T), AAG1410GAG (4368A>G), GGC1412GGG (4376C>G), GAT1415GAG (4385T>G), TCC1416TCA (4388C>A), CGA1419AGG (4395C>A 4397A>G), CAT1420CAC (4400T>C), GCA1421ACA (4401G>A), CTT1422TTA (4404C>T 4406T>A), GCA1426GCC (4418A>C), TGT1428TGC (4424T>C), TAC1430TAT (4430C>T), ACA1431ACC (4433A>C), CTG1432CTC (4436G>C), TGC1433TGT (4439C>T), GAT1435AAC (4443G>A 4445T>C), AAA1436AAG (4448A>G), GTG1437TTG (4449G>T), AGG1439CGC (4455A>C 4457G>C), TTG1440CTA (4458T>C 4460G>A), GTT1441GTG (4463T>G), GGG1442GGT (4466G>T), GGC1443GGA (4469C>A), CCA1445CCC (4475A>C), TCA1446TCT (4478A>T), GGA1447GGT (4481A>T), GCA1449GCT (4487A>T), CTA1450TTG (4488C>T 4490A>G), ACG1451ACT (4493G>T), ATC1453ATA (4499C>A), TTC1454TTT (4502C>T), TCT1456TCA (4508T>A), CTC1457TTG (4509C>T 4511C>G), CTC1458CTA (4514C>A), CGA1464AGG (4530C>A 4532A>G), TAT1465TAC (4535T>C), GCC1466GCG (4538C>G), TTG1470CTG (4548T>C), TTA1471TTG (4553A>G), AGA1472AGG (4556A>G), CCC1474CCA (4562C>A), ATT1476GTC (4566A>G 4568T>C), GCT1477AAA (4569G>A 4570C>A 4571T>A), AGG1479AAA (4576G>A 4577G>A), GCT1480TCT (4578G>T), ATA1481GTG (4581A>G 4583A>G), GGA1482GGC (4586A>C), GTT1483ATT (4587G>A), AAA1484AAG (4592A>G), AGT1486AAT (4597G>A), GAT1487GAC (4601T>C), CTA1491TTG (4611C>T 4613A>G), GCT1494GCA (4622T>A), GTT1495GTA (4625T>A), GGA1497GGT (4631A>T), GAC1499GAT (4637C>T), AAT1500AAC (4640T>C), CTT1501TTG (4641C>T 4643T>G), GCT1503GCG (4649T>G), GTA1504GTT (4652A>T), TTA1506ATG (4656T>A 4658A>G), CTC1508TTG (4662C>T 4664C>G), CAG1509AGG (4665C>A 4666A>G), TAT1511TAC (4673T>C), TCT1512GAT (4674T>G 4675C>A), CTG1513CTA (4679G>A), AAT1515TCC (4683A>T 4684A>C 4685T>C), ATA1516ATT (4688A>T), CAT1518AGA (4692C>A 4693A>G 4694T>A), GAG1519GAA (4697G>A), TTA1520CTG (4698T>C 4700A>G), CTG1522GCT (4704C>G 4705T>C 4706G>T), GTC1523GTT (4709C>T), AAT1524AAC (4712T>C), GTA1525GTC (4715A>C), ATC1526GTC (4716A>G), CTT1531TTG (4731C>T 4733T>G), AAA1533AAG (4739A>G), TCG1534TCA (4742G>A), GTT1537GTA (4751T>A), AAT1538GAG (4752A>G 4754T>G), GAG1539AAG (4755G>A), GTA1540ACA (4758G>A 4759T>C), CAA1541AAA (4761C>A), CAA1543CTG (4768A>T 4769A>G), CTA1548CTC (4784A>C), ACT1549ACC (4787T>C), CTG1551CTC (4793G>C), AGA1553AGG (4799A>G), TTT1555TTC (4805T>C), AAA1556AAG (4808A>G), GCT1559GCA (4817T>A), CTT1560TTG (4818C>T 4820T>G), TAC1562TAT (4826C>T), CAC1563CAA (4829C>A), GCT1565GCA (4835T>A), CCT1566CCA (4838T>A), AAG1568AAA (4844G>A), ATC1572GTC (4854A>G), ATT1573ATC (4859T>C), ATG1576ATT (4868G>T), ATA1579ATT (4877A>T), AGA1580AGG (4880A>G), CCT1581CCA (4883T>A), GCC1582GCA (4886C>A), CCA1583GTG (4887C>G 4888C>T 4889A>G), GGT1584GGA (4892T>A), GAC1587GAT (4901C>T), ACT1588GTA (4902A>G 4903C>T 4904T>A), CAG1589CAA (4907G>A), GCC1590GCT (4910C>T), ATG1592TAT (4914A>T 4915T>A 4916G>T), GTG1595ATT (4923G>A 4925G>T), GGA1598GGC (4934A>C), ATA1599ATC (4937A>C), CGT1600AGA (4938C>A 4940T>A), GCT1602GCC (4946T>C), CAC1605CAT (4955C>T), GGT1606GGG (4958T>G), CGT1607CAG (4960G>A 4961T>G), GTG1608GCC (4963T>C 4964G>C), GCT1609ACT (4965G>A), GAC1611GAA (4973C>A), AAG1612AGT (4975A>G 4976G>T), CTT1613TTT (4977C>T), GTC1614GTA (4982C>A), TTA1615AAT (4983T>A 4984T>A 4985A>T), CAG1618TTT (4992C>T 4993A>T 4994G>T), ACG1619GGA (4995A>G 4996C>G 4997G>A), GCG1620GCT (5000G>T), TTG1621CTC (5001T>C 5003G>C), GAT1622AAG (5004G>A 5006T>G), GAA1623CAG (5007G>C 5009A>G), AGG1624AGA (5012G>A), GGT1625AAT (5013G>A 5014G>A), TTC1626CTC (5016T>C), AGA1627CCC (5019A>C 5020G>C 5021A>C), GCT1628ATC (5022G>A 5023C>T 5024T>C), GTG1629ATG (5025G>A), TTT1631ATG (5031T>A 5033T>G), CCT1632CCC (5036T>C), TCC1633ACT (5037T>A 5039C>T), TAT1634TAC (5042T>C), TTG1635TAT (5044T>A 5045G>T), GTG1637CTT (5049G>C 5051G>T), CAG1639AGT (5055C>A 5056A>G 5057G>T), GAA1640CAA (5058G>C), ATT1642TTG (5064A>T 5066T>G), GCA1643ACA (5067G>A), AAG1644GAG (5070A>G), GTA1645GTC (5075A>C), GGG1647GGA (5081G>A), TCA1649GAG (5085T>G 5086C>A 5087A>G), AGT1650AGC (5090T>C), GCC1651ATT (5091G>A 5092C>T 5093C>T), CTG1652GGG (5094C>G 5095T>G), ACA1653GTT (5097A>G 5098C>T 5099A>T), ATT1654TTG (5100A>T 5102T>G), TGT1655ATG (5103T>A 5104G>T 5105T>G), GAA1656GAG (5108A>G), ATG1657CTA (5109A>C 5111G>A), GCA1658TCA (5112G>T), AAA1659GAT (5115A>G 5117A>T), GCA1660GCT (5120A>T), GCT1661AAA (5121G>A 5122C>A 5123T>A), ATT1662AAT (5125T>A), TCC1663ACA (5127T>A 5129C>A), TAT1664TAC (5132T>C), ACG1665ACC (5135G>C), CCA1666CCT (5138A>T), TTG1667TTA (5141G>A), GAC1668CCT (5142G>C 5143A>C 5144C>T), GCA1669GCT (5147A>T), GGT1670GGG (5150T>G), GAG1671GAA (5153G>A), ATC1673ATT (5159C>T), ACA1674del (5160_5162delACA), CGT1678GAG (5172C>G 5173G>A 5174T>G), GAT1679GGT (5176A>G), CTG1680GTC (5178C>G 5180G>C), AAT1681CAT (5181A>C), TGG1682GAA (5184T>G 5185G>A 5186G>A), GCA1684GTG (5191C>T 5192A>G), CCA1685CCT (5195A>T), AAC1686GAT (5196A>G 5198C>T), ATT1687GTA (5199A>G 5201T>A), GGT1688TTT (5202G>T 5203G>T), TTT1689TTC (5207T>C), TCA1691AGT (5211T>A 5212C>G 5213A>T), GCA1692GCT (5216A>T), CGT1693CGG (5219T>G), GCC1696TCA (5226G>T 5228C>A), CAC1697AAC (5229C>A), TAC1698CAA (5232T>C 5234C>A), ACG1699CAA (5235A>C 5236C>A 5237G>A), GAT1701TCA (5241G>T 5242A>C 5243T>A), GAA1702TCA (5244G>T 5245A>C), GGG1703GGT (5249G>T), TAC1704AAC (5250T>A), ATT1705ATC (5255T>C), ATT1706CTG (5256A>C 5258T>G), AAC1708AAT (5264C>T), TGT1709TGC (5267T>C), ACA1710ACT (5270A>T), GGT1711GGA (5273T>A), GCA1712GCC (5276A>C), AAG1713CGC (5277A>C 5278A>G 5279G>C), AAG1714AGA (5281A>G 5282G>A), TCC1715ACA (5283T>A 5285C>A), AAT1716ACA (5287A>C 5288T>A), GTT1718GTG (5294T>G), GGT1720GGG (5300T>G), CCA1721CCT (5303A>T), AAC1723AAT (5309C>T), ATG1731ATA (5333G>A), CCG1733CCC (5339G>C), ATG1736TTG (5346A>T), TCA1737GCA (5349T>G), GCT1738GCA (5354T>A), ATA1739ATC (5357A>C), ATG1740CAG (5358A>C 5359T>A), GCC1742CAT (5364G>C 5365C>A 5366C>T), AAG1744CGC (5370A>C 5371A>G 5372G>C), AAT1745AAG (5375T>G), GTG1746AGT (5376G>A 5377T>G 5378G>T), TTG1747CTG (5379T>C), GCA1748GAA (5383C>A), GGA1749GTT (5386G>T 5387A>T), GGA1750GGT (5390A>T), CAT1751GAT (5391C>G), GTA1752GGG (5395T>G 5396A>G), GCA1753GCT (5399A>T), AAT1755TCT (5403A>T 5404A>C), GTT1757ACC (5409G>A 5410T>C 5411T>C), GTG1759GTT (5417G>T), GGA1761GGT (5423A>T), GGA1763GGT (5429A>T), GCT1765GCA (5435T>A), GTC1766ATA (5436G>A 5438C>A), GGC1767GCT (5440G>C 5441C>T), ATT1769ATC (5447T>C), GCG1772GCA (5456G>A), CTG1773TTG (5457C>T), ATG1774TTT (5460A>T 5462G>T), GCC1775GCA (5465C>A), TTA1776TTG (5468A>G), GCA1777GCC (5471A>C), ACA1778ACT (5474A>T), AGG1779CAT (5475A>C 5476G>A 5477G>T), TAT1781TAC (5483T>C), TGT1782AGG (5484T>A 5486T>G), GTG1783GCC (5488T>C 5489G>C), GAG1784ACA (5490G>A 5491A>C 5492G>A), ATA1786CTT (5496A>C 5498A>T), ATA1787ATT (5501A>T), GTT1788GTG (5504T>G), AGA1789CGC (5505A>C 5507A>C), TTG1790CTA (5508T>C 5510G>A), CGC1791CGA (5513C>A), ACA1792ACC (5516A>C), ATT1793ATG (5519T>G), GGA1794GCT (5521G>C 5522A>T), AAT1795GGG (5523A>G 5524A>G 5525T>G), CTT1797CTG (5531T>G), GAC1798GAT (5534C>T), CTC1799CTG (5537C>G), AAT1800AAC (5540T>C), TAT1802TAC (5546T>C), GGC1805GGT (5555C>T), GTG1807GCG (5560T>C), CAG1808CAA (5564G>A), TAT1809TAC (5567T>C), TTT1810TTG (5570T>G), CTT1811CTC (5573T>C), CAA1812AAA (5574C>A), GAA2142GCA (6565A>C), ACC2143CCC (6567A>C), ATT2144ATC (6572T>C), TAC2148TAT (6584C>T), GAG2150GCG (6589A>C) | | | | | | | | |
| helicase (YP_001976154.1) | 1 | 98 | 100% | 654 | 97.5% | 98 (100%) | 95 (96.9%) | 0/0/0/0 | 0 |
| Protein mutations: | S5C (1342C>G 1343C>T), R16A (1374A>G 1375G>C), I44V (1458A>G 1460C>G) | | | | | | | | |
| Codon mutations: | TCT3TCA (1337T>A), ACA4ACG (1340A>G), TCC5TGT (1342C>G 1343C>T), TTA8CTT (1350T>C 1352A>T), TTT9TTC (1355T>C), ATT10ATC (1358T>C), GTG13GTT (1367G>T), TTG14CTC (1368T>C 1370G>C), AGA16GCA (1374A>G 1375G>C), GAG19GAA (1385G>A), AAA21AAG (1391A>G), CTC22CTG (1394C>G), AAT23AAC (1397T>C), CGG24CGC (1400G>C), CTA25TTG (1401C>T 1403A>G), GCA27GCG (1409A>G), AAG30AAA (1418G>A), AGG31AGA (1421G>A), GCT39GCA (1445T>A), CAC40CAT (1448C>T), GCT43GCA (1457T>A), ATC44GTG (1458A>G 1460C>G), CTG45CTT (1463G>T), GAC48GAT (1472C>T), GGA50GGT (1478A>T), GCA51GCT (1481A>T), TTG52CTT (1482T>C 1484G>T), CGA53CGG (1487A>G), GGG55GGA (1493G>A), GCT56GCA (1496T>A), TGC59TGT (1505C>T), CAA60CAG (1508A>G), GAT61GAC (1511T>C), ATT62ATA (1514T>A), AAA63AAG (1517A>G), GAT67GAC (1529T>C), TCA70TCT (1538A>T), ACA71ACC (1541A>C), CAA72CAG (1544A>G), CCA73CCT (1547A>T), CCC75CCT (1553C>T), TTA76CTT (1554T>C 1556A>T), GCC79GCT (1565C>T), GTT81GTC (1571T>C), GAC83GAT (1577C>T), GGC85GGA (1583C>A), CAT87CAC (1589T>C), TTC88TTT (1592C>T), ACC89ACA (1595C>A), TCC90TCA (1598C>A), TAT92TAC (1604T>C), ATA93ATC (1607A>C), ACA96ACC (1616A>C), AAT98AAC (1622T>C) | | | | | | | | |

| | Begin | End | Coverage | Score | Concordance | Matches | Identities | I/D/M/F* | Stop Codons |
|---|---|---|---|---|---|---|---|---|---|
| **NT** | **22** | **6809** | **80.5%** | **3829** | **33.5%** | **5772 (99.0%)** | **3883 (66.6%)** | **10/50** | |
| RNA-dependent RNA polymerase (YP_001976153.1) | 1 | 249 | 100% | 1552 | 88.4% | 249 (100%) | 210 (84.3%) | 0/0/0/0 | 0 |
| Protein mutations: | T40C (4170A>T 4171C>G 4172A>T), L44A (4182T>G 4183T>C 4184G>A), A47V (4192C>T 4193T>A), I48L (4194A>C 4196A>C), D55N (4215G>A), S59H (4227T>C 4228C>A), L63V (4239T>G), N66S (4249A>G), G67S (4251G>A 4253G>C), A70S (4260G>T 4262T>A), V74E (4273T>A), V89I (4317G>A 4319T>C), E101A (4354A>C 4355A>T), Y102F (4357A>T 4358C>T), S105D (4365T>G 4366C>A 4367A>T), K106E (4368A>G), D111E (4385T>G), A117T (4401G>A), D131N (4443G>A 4445T>C), V133L (4449G>T), I172V (4566A>G 4568T>C), A173K (4569G>A 4570C>A 4571T>A), R175K (4576G>A 4577G>A), A176S (4578G>T), I177V (4581A>G 4583A>G), V179I (4587G>A), S182N (4597G>A), L202M (4656T>A 4658A>G), Q205R (4665C>A 4666A>G), S208D (4674T>G 4675C>A), N211S (4683A>T 4684A>C 4685T>C), H214R (4692C>A 4693A>G 4694T>A), L218A (4704C>G 4705T>C 4706G>T), I222V (4716A>G), N234E (4752A>G 4754T>G), E235K (4755G>A), V236T (4758G>A 4759T>C), Q237K (4761C>A), Q239L (4768A>T 4769A>G) | | | | | | | | |
| Codon mutations: | CGC4CGG (4064C>G), CTC5TTG (4065C>T 4067C>G), AAA9AAG (4079A>G), ATC10ATT (4082C>T), GGC12GGG (4088C>G), AGA14AGG (4094A>G), AGG17CGC (4101A>C 4103G>C), CTC18CTG (4106C>G), TTT19TTC (4109T>C), GAG20GAA (4112G>A), ATA21ATC (4115A>C), CTT22TTG (4116C>T 4118T>G), TAC26TAT (4130C>T), TTG29CTG (4137T>C), GTC30GTG (4142C>G), AGG31CGG (4143A>C), AAG32AAA (4148G>A), TAT33TAC (4151T>C), TTT34TTC (4154T>C), GAT36GAC (4160T>C), TTC37TTT (4163C>T), TCA38TCT (4166A>T), GCC39GCT (4169C>T), ACA40TGT (4170A>T 4171C>G 4172A>T), TTG44GCA (4182T>G 4183T>C 4184G>A), AAT46AAC (4190T>C), GCT47GTA (4192C>T 4193T>A), ATA48CTC (4194A>C 4196A>C), CCA49CCT (4199A>T), AAA51AAG (4205A>G), GTT52GTA (4208T>A), ATT54ATC (4214T>C), GAT55AAT (4215G>A), CCT56CCA (4220T>A), ACA57ACG (4223A>G), AGT58AGC (4226T>C), TCT59CAT (4227T>C 4228C>A), TTG63GTG (4239T>G), TTG64CTG (4242T>C), GCA65GCT (4247A>T), AAT66AGT (4249A>G), GGG67AGC (4251G>A 4253G>C), AGA69AGG (4259A>G), GCT70TCA (4260G>T 4262T>A), GTG71GTA (4265G>A), TCA72TCT (4268A>T), GAC73GAT (4271C>T), GTG74GAG (4273T>A), TTT76TTC (4280T>C), GCT78GCC (4286T>C), TAT80TAC (4292T>C), TCC81TCG (4295C>G), CCT88CCC (4316T>C), GTT89ATC (4317G>A 4319T>C), GCT91GCA (4325T>A), TTT92TTC (4328T>C), CAG93CAA (4331G>A), TTG98CTG (4344T>C), GAA101GCT (4354A>C 4355A>T), TAC102TTT (4357A>T 4358C>T), TAC103TAT (4361C>T), GGA104GGT (4364A>T), TCA105GAT (4365T>G 4366C>A 4367A>T), AAG106GAG (4368A>G), GGC108GGG (4376C>G), GAT111GAG (4385T>G), TCC112TCA (4388C>A), CGA115AGG (4395C>A 4397A>G), CAT116CAC (4400T>C), GCA117ACA (4401G>A), CTT118TTA (4404C>T 4406T>A), GCA122GCC (4418A>C), TGT124TGC (4424T>C), TAC126TAT (4430C>T), ACA127ACC (4433A>C), CTG128CTC (4436G>C), TGC129TGT (4439C>T), GAT131AAC (4443G>A 4445T>C), AAA132AAG (4448A>G), GTG133TTG (4449G>T), AGG135CGC (4455A>C 4457G>C), TTG136CTA (4458T>C 4460G>A), GTT137GTG (4463T>G), GGG138GGT (4466G>T), GGC139GGA (4469C>A), CCA141CCC (4475A>C), TCA142TCT (4478A>T), GGA143GGT (4481A>T), GCA145GCT (4487A>T), CTA146TTG (4488C>T 4490A>G), ACG147ACT (4493G>T), ATC149ATA (4499C>A), TTC150TTT (4502C>T), TCT152TCA (4508T>A), CTC153TTG (4509C>T 4511C>G), CTC154CTA (4514C>A), CGA160AGG (4530C>A 4532A>G), TAT161TAC (4535T>C), GCC162GCG (4538C>G), TTG166CTG (4548T>C), TTA167TTG (4553A>G), AGA168AGG (4556A>G), CCC170CCA (4562C>A), ATT172GTC (4566A>G 4568T>C), GCT173AAA (4569G>A 4570C>A 4571T>A), AGG175AAA (4576G>A 4577G>A), GCT176TCT (4578G>T), ATA177GTG (4581A>G 4583A>G), GGA178GGC (4586A>C), GTT179ATT (4587G>A), AAA180AAG (4592A>G), AGT182AAT (4597G>A), GAT183GAC (4601T>C), CTA187TTG (4611C>T 4613A>G), GCT190GCA (4622T>A), GTT191GTA (4625T>A), GGA193GGT (4631A>T), GAC195GAT (4637C>T), AAT196AAC (4640T>C), CTT197TTG (4641C>T 4643T>G), GCT199GCG (4649T>G), GTA200GTT (4652A>T), TTA202ATG (4656T>A 4658A>G), CTC204TTG (4662C>T 4664C>G), CAG205AGG (4665C>A 4666A>G), TAT207TAC (4673T>C), TCT208GAT (4674T>G 4675C>A), CTG209CTA (4679G>A), AAT211TCC (4683A>T 4684A>C 4685T>C), ATA212ATT (4688A>T), CAT214AGA (4692C>A 4693A>G 4694T>A), GAG215GAA (4697G>A), TTA216CTG (4698T>C 4700A>G), CTG218GCT (4704C>G 4705T>C 4706G>T), GTC219GTT (4709C>T), AAT220AAC (4712T>C), GTA221GTC (4715A>C), ATC222GTC (4716A>G), CTT227TTG (4731C>T 4733T>G), AAA229AAG (4739A>G), TCG230TCA (4742G>A), GTT233GTA (4751T>A), AAT234GAG (4752A>G 4754T>G), GAG235AAG (4755G>A), GTA236ACA (4758G>A 4759T>C), CAA237AAA (4761C>A), CAA239CTG (4768A>T 4769A>G), CTA244CTC (4784A>C), ACT245ACC (4787T>C), CTG247CTC (4793G>C), AGA249AGG (4799A>G) | | | | | | | | |

*: Inserts / Deletes / Misaligned / Frameshifts

#### Analysis details

This analysis was performed with panviral2.64

### NGS Details (UN62): Torradovirus marchitezum (segment RNA 2)

#### Assembly

| | |
|---|---|
| **Coverage Length** | 1244 (2 contig(s)) |
| **Depth Of Coverage** | 14475.3 |
| **Number Of Reads** | 143680 |
| **Reads Per Million** | 3330.63 rpm (after QC) |
| **Ambiguities** | 0 |
| **Assembly Method** | de novo + reference guided assembly |
| **Consensus Caller** | Bcf Tools |

#### Coverage Map

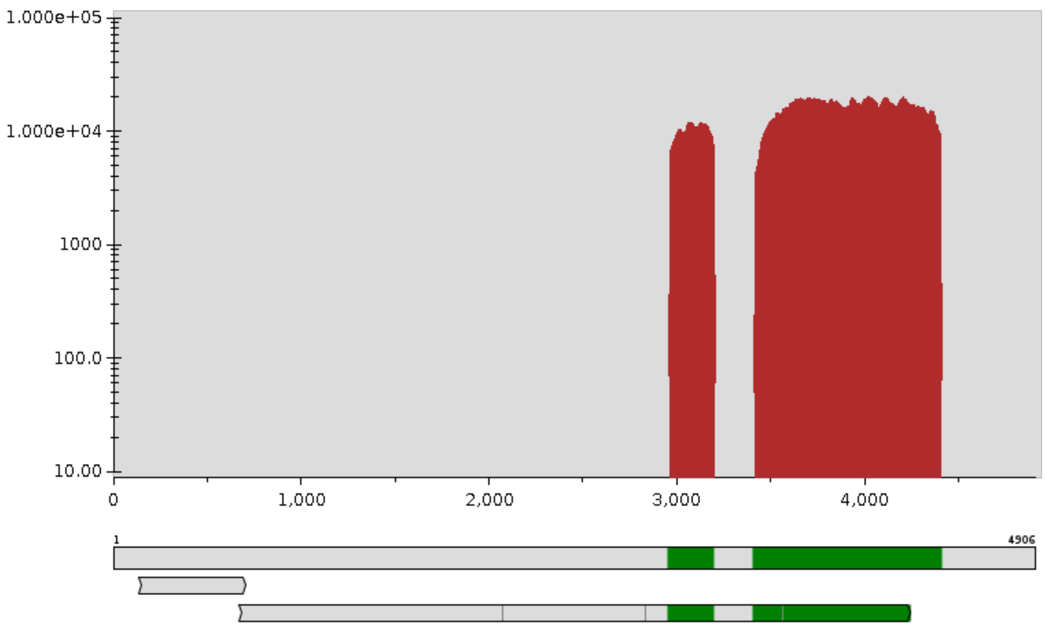

#### Assignment

| | |
|---|---|
| **Type** | Torradovirus marchitezum (Taxonomy ID: 3048376) |
| **Reference Genome** | NC_010988.1 |
| **NT Identity (%)** | 69.4712 |
| **AA Identity (%)** | 75.9669 |
| **Number Of Stop Codons** | 1 |
| **Number Of CDS** | 2 |

#### Alignment

| | |
|---|---|
| **Alignment Score** | 931.0 (NT) + 2123.0 (AA) = 3054.0 |
| **Concordance (%)** | 60.0708 |

| **Alignment Method** | Global, seeded, nucleotide + amino acids (AGA) |
|---|---|

#### Genome Region

Sequence starts at position 2960 and ends at position 4410 relative to NC_010988.1 reference sequence.

#### Alignment Detailed Statistics

| | Begin | End | Coverage | Score | Concordance | Matches | Identities | I/D/M/F* | Stop Codons |
|---|---|---|---|---|---|---|---|---|---|
| **NT** | **2960** | **4410** | **25.4%** | **931** | **37.8%** | **1239 (98.9%)** | **867 (69.2%)** | **9/5** | |
| Mutations: | 2971A>T, 2977G>A, 2989C>A, 2990T>C, 2998G>A, 3001T>A, 3002T>C, 3005C>T, 3007T>G, 3010T>A, 3013C>T, 3016A>T, 3019C>T, 3022G>A, 3028C>T, 3038A>C, 3039C>G, 3040T>G, 3043A>C, 3046T>A, 3049T>C, 3058A>G, 3061T>A, 3064C>T, 3065A>G, 3067T>C, 3070G>A, 3073A>G, 3079G>A, 3085T>C, 3088A>T, 3091T>A, 3097G>T, 3100A>C, 3106A>T, 3107C>T, 3109T>G, 3112A>T, 3118G>A, 3119A>G, 3120A>C, 3121C>A, 3124C>A, 3125A>G, 3131A>G, 3134A>G, 3135A>C, 3136T>A, 3148A>T, 3149T>C, 3151T>A, 3154C>T, 3160A>C, 3163T>C, 3178T>A, 3182A>G, 3187T>C, 3190A>C, 3191A>C, 3192A>G, 3193A>T, 3427G>T, 3429A>T, 3436A>T, 3442C>G, 3445A>G, 3448C>T, 3449T>G, 3451A>C, 3457T>C, 3460C>T, 3461C>T, 3463T>G, 3466C>T, 3476G>A, 3478T>C, 3481A>G, 3482T>A, 3484A>T, 3493G>A, 3497A>C, 3506T>C, 3514G>A, 3517C>A, 3520C>T, 3521A>G, 3528A>T, 3529G>C, 3532C>T, 3533C>G, 3535G>T, 3538G>T, 3544C>G, 3544_3545insAAGAAG, 3550T>C, 3551C>A, 3552C>A, 3554C>G, 3555C>A, 3556C>G, 3559G>A, 3560A>G, 3561A>T, 3564C>G, 3568A>G, 3569A>G, 3570G>T, 3572A>G, 3573T>C, 3574T>A, 3577G>T, 3578T>G, 3579C>G, 3580C>A, 3581A>C, 3583A>T, 3584A>C, 3586A>G, 3589G>A, 3595T>C, 3598C>T, 3604T>A, 3610A>T, 3619C>T, 3622C>T, 3623A>T, 3624T>C, 3625G>T, 3627G>C, 3628T>C, 3638G>A, 3639A>C, 3641G>A, 3642A>G, 3643A>C, 3644A>C, 3646A>C, 3647T>C, 3650T>G, 3655A>T, 3656A>C, 3661T>A, 3667A>T, 3670A>G, 3676C>T, 3677A>G, 3680A>C, 3682G>A, 3685T>C, 3686G>T, 3687A>C, 3689G>C, 3690G>C, 3691G>C, 3692C>A, 3693A>C, 3700T>C, 3712A>C, 3715C>A, 3718T>G, 3724A>C, 3730A>C, 3739T>C, 3740A>T, 3742A>T, 3751T>C, 3760A>T, 3763T>C, 3767T>C, 3769G>T, 3770A>G, 3772T>G, 3775T>C, 3776T>A, 3778A>T, 3779A>C, 3781T>C, 3782G>A, 3784G>C, 3787A>C, 3790C>T, 3793A>G, 3796A>C, 3800T>G, 3801C>G, 3803A>T, 3804G>C, 3806C>A, 3807C>A, 3811T>C, 3812G>A, 3817A>T, 3826A>T, 3832T>G, 3838T>G, 3841T>C, 3842G>A, 3860C>T, 3862A>T, 3863G>T, 3864C>A, 3868G>T, 3869C>G, 3871T>G, 3872A>G, 3873A>C, 3874T>A, 3875A>G, 3877A>G, 3880A>G, 3881A>G, 3882A>C, 3883T>A, 3884T>C, 3890G>C, 3892A>T, 3901T>C, 3904A>T, 3907C>T, 3908A>T, 3910A>C, 3917G>A, 3922T>C, 3925A>C, 3931T>A, 3934G>T, 3935G>A, 3937A>C, 3940A>T, 3943T>C, 3945C>T, 3946G>T, 3949C>T, 3952A>C, 3955C>T, 3956A>G, 3957C>T, 3958T>G, 3967T>C, 3971G>A, 3973G>A, 3982A>T, 3983A>C, 3986C>A, 3989C>T, 3994A>T, 3996G>A, 3998A>G, 3999T>C, 4000G>A, 4001A>C, 4003G>A, 4009C>T, 4010T>G, 4011C>A, 4012A>C, 4014G>A, 4015C>T, 4019C>A, 4020A>G, 4021A>T, 4024C>T, 4027C>T, 4030C>T, 4031A>G, 4033G>T, 4036A>T, 4037C>T, 4039A>G, 4045T>C, 4047G>A, 4048G>A, 4049C>T, 4051T>G, 4054A>T, 4055A>C, 4056A>T, 4060G>T, 4061A>T, 4063C>G, 4066A>C, 4070T>C, 4072G>A, 4075T>A, 4078T>C, 4079T>G, 4080C>T, 4081C>A, 4083C>A, 4085A>T, 4088G>A, 4090G>T, 4093T>C, 4102G>A, 4105A>C, 4106C>A, 4108T>G, 4109A>G, 4111A>T, 4112T>A, 4113C>A, 4114T>G, 4117T>A, 4120A>G, 4122T>C, 4123T>A, 4129C>T, 4131A>G, 4135G>A, 4138G>A, 4139T>C, 4141G>T, 4154T>C, 4155C>T, 4156T>G, 4159C>T, 4162C>T, 4180C>T, 4181A>G, 4192T>G, 4195C>T, 4198C>T, 4201T>C, 4204G>A, 4213A>T, 4219T>C, 4222T>A, 4225G>A, 4228A>G, 4229C>A, 4231T>A, 4234T>C, 4237T>C, 4238G>T, 4242T>A, 4243T>C, 4248A>C, 4252T>A, 4279A>T, 4279_4280insT, 4280A>T, 4280_4281insC, 4283A>G, 4286C>T, 4287T>A, 4288T>C, 4291C>A, 4294C>T, 4306G>A, 4309G>A, 4312G>T, 4313C>A, 4313_4314insG, 4315A>G, 4316A>T, 4318T>C, 4319G>T, 4320G>A, 4323T>G, 4324T>A, 4328T>C, 4330G>A, 4333A>G, 4335G>T, 4336A>C, 4338A>G, 4339_4343delAGGGA, 4345A>T, 4346G>C, 4349A>C, 4364C>T, 4365T>C, 4369A>G, 4372A>G, 4373G>A, 4392T>C, 4393C>T | | | | | | | | |
| **CDS** | | | | | | | | | |
| ToMarV_RNA2gp2 | 764 | 1192 | 30.2% | 2123 | 80.3% | 360 (99.4%) | 275 (76.0%) | 2/0/0/0 | 1 |
| Protein mutations: | T790R (3038A>C 3039C>G 3040T>G), I799V (3065A>G 3067T>C), N817A (3119A>G 3120A>C 3121C>A), K819E (3125A>G), T821A (3131A>G), N822A (3134A>G 3135A>C 3136T>A), F827L (3149T>C 3151T>A), M838V (3182A>G), K841R (3191A>C 3192A>G 3193A>T), H920L (3429A>T), S927A (3449T>G 3451A>C), D936N (3476G>A 3478T>C), S938T (3482T>A 3484A>T), I943L (3497A>C), D949E (3517C>A), I951V (3521A>G), E953V (3528A>T 3529G>C), Q955D (3533C>G 3535G>T), E956D (3538G>T), G958_K959insKK (3544_3545insAAGAAG), P961K (3551C>A 3552C>A), P962E (3554C>G 3555C>A 3556C>G), K964V (3560A>G 3561A>T), A965G (3564C>G), S967V (3569A>G 3570G>T), I968A (3572A>G 3573T>C 3574T>A), S970G (3578T>G 3579C>G 3580C>A), I971L (3581A>C 3583A>T), M985S (3623A>T 3624T>C 3625G>T), G986A (3627G>C 3628T>C), D990T (3638G>A 3639A>C), E991S (3641G>A 3642A>G 3643A>C), L994V (3650T>G), I996L (3656A>C), I1003V (3677A>G), E1006S (3686G>T 3687A>C), G1007P (3689G>C 3690G>C 3691G>C), H1008T (3692C>A 3693A>C), T1024S (3740A>T 3742A>T), N1034E (3770A>G 3772T>G), S1036T (3776T>A 3778A>T), I1037L (3779A>C 3781T>C), V1038I (3782G>A 3784G>C), S1044G (3800T>G 3801C>G), P1046N (3806C>A 3807C>A), V1048I (3812G>A), F1056L (3838T>G), A1058T (3842G>A), P1064S (3860C>T 3862A>T), A1065Y (3863G>T 3864C>A), L1067V (3869C>G 3871T>G), N1068A (3872A>G 3873A>C 3874T>A), K1069E (3875A>G 3877A>G), N1071A (3881A>G 3882A>C 3883T>A), Y1072H (3884T>C), V1074L (3890G>C 3892A>T), T1080S (3908A>T 3910A>C), D1083N (3917G>A), A1089T (3935G>A 3937A>C), T1092I (3945C>T 3946G>T), T1096V (3956A>G 3957C>T 3958T>G), E1101K (3971G>A 3973G>A), H1107Y (3989C>T), R1109K (3996G>A), M1110A (3998A>G 3999T>C 4000G>A), S1114D (4010T>G 4011C>A 4012A>C), S1115N (4014G>A 4015C>T), Q1117S (4019C>A 4020A>G 4021A>T), M1121V (4031A>G 4033G>T), R1126K (4047G>A 4048G>A), N1129L (4055A>C 4056A>T), I1131L (4061A>T 4063C>G), S1137V (4079T>G 4080C>T 4081C>A), A1138D (4083C>A), I1139L (4085A>T), V1140I (4088G>A 4090G>T), L1146M (4106C>A 4108T>G), I1147V (4109A>G 4111A>T), S1148K (4112T>A 4113C>A 4114T>G), L1151P (4122T>C 4123T>A), K1154R (4131A>G), S1162L (4154T>C 4155C>T 4156T>G), N1171D (4181A>G), A1190S (4238G>T), I1191N (4242T>A 4243T>C) | | | | | | | | |
| Codon mutations: | ATA767ATT (2971A>T), GAG769GAA (2977G>A), ACC773ACA (2989C>A), TTG774CTG (2990T>C), CCG776CCA (2998G>A), TCT777TCA (3001T>A), TTG778CTG (3002T>C), CTT779TTG (3005C>T 3007T>G), GGT780GGA (3010T>A), AAC781AAT (3013C>T), ATA782ATT (3016A>T), GCC783GCT (3019C>T), AGG784AGA (3022G>A), TGC786TGT (3028C>T), ACT790CGG (3038A>C 3039C>G 3040T>G), GGA791GGC (3043A>C), ACT792ACA (3046T>A), TGT793TGC (3049T>C), GAA796GAG (3058A>G), ATT797ATA (3061T>A), TGC798TGT (3064C>T), ATT799GTC (3065A>G 3067T>C), GAG800GAA (3070G>A), AAA801AAG (3073A>G), TTG803TTA (3079G>A), CAT805CAC (3085T>C), TCA806TCT (3088A>T), GGT807GGA (3091T>A), CTG809CTT (3097G>T), GCA810GCC (3100A>C), GGA812GGT (3106A>T), CTT813TTG (3107C>T 3109T>G), GGA814GGT (3112A>T), CTG816CTA (3118G>A), AAC817GCA (3119A>G 3120A>C 3121C>A), ACC818ACA (3124C>A), AAA819GAA (3125A>G), ACC821GCC (3131A>G), AAT822GCA (3134A>G 3135A>C 3136T>A), ATA826ATT (3148A>T), TTT827CTA (3149T>C 3151T>A), AAC828AAT (3154C>T), CCA830CCC (3160A>C), CAT831CAC (3163T>C), CTT836CTA (3178T>A), ATG838GTG (3182A>G), GGT839GGC (3187T>C), CGA840CGC (3190A>C), AAA841CGT (3191A>C 3192A>G 3193A>T), GGG919GGT (3427G>T), CAT920CTT (3429A>T), ACA922ACT (3436A>T), GGC924GGG (3442C>G), AAA925AAG (3445A>G), AGC926AGT (3448C>T), TCA927GCC (3449T>G 3451A>C), GAT929GAC (3457T>C), TTC930TTT (3460C>T), CTT931TTG (3461C>T 3463T>G), TTC932TTT (3466C>T), GAT936AAC (3476G>A 3478T>C), TTA937TTG (3481A>G), TCA938ACT (3482T>A 3484A>T), AAG941AAA (3493G>A), ATT943CTT (3497A>C), TTG946CTG (3506T>C), AAG948AAA (3514G>A), GAC949GAA (3517C>A), AAC950AAT (3520C>T), ATT951GTT (3521A>G), GAG953GTC (3528A>T 3529G>C), TAC954TAT (3532C>T), CAG955GAT (3533C>G 3535G>T), GAG956GAT (3538G>T), GGC958GGG (3544C>G), GGC958_AAA959insAAGAAG (3544_3545insAAGAAG), GAT960GAC (3550T>C), CCG961AAG (3551C>A 3552C>A), CCC962GAG (3554C>G 3555C>A 3556C>G), AAG963AAA (3559G>A), AAG964GTG (3560A>G 3561A>T), GCT965GGT (3564C>G), CAA966CAG (3568A>G), AGT967GTT (3569A>G 3570G>T), ATT968GCA (3572A>G 3573T>C 3574T>A), CTG969CTT (3577G>T), TCC970GGA (3578T>G 3579C>G 3580C>A), ATA971CTT (3581A>C 3583A>T), AGA972CGG (3584A>C 3586A>G), GAG973GAA (3589G>A), TTT975TTC (3595T>C), TCC976TCT (3598C>T), GGT978GGA (3604T>A), GTA980GTT (3610A>T), TTC983TTT (3619C>T), TGC984TGT (3622C>T), ATG985TCT (3623A>T 3624T>C 3625G>T), GGT986GCC (3627G>C 3628T>C), GAC990ACC (3638G>A 3639A>C), GAA991AGC (3641G>A 3642A>G 3643A>C), AGA992CGC (3644A>C 3646A>C), TTG993CTG (3647T>C), TTG994GTG (3650T>G), GTA995GTT (3655A>T), ATT996CTT (3656A>C), CCT997CCA (3661T>A), GCA999GCT (3667A>T), CCA1000CCG (3670A>G), TCC1002TCT (3676C>T), ATA1003GTA (3677A>G), AGG1004CGA (3680A>C 3682G>A), TTT1005TTC (3685T>C), GAA1006TCA (3686G>T 3687A>C), GGG1007CCC (3689G>C 3690G>C 3691G>C), CAC1008ACC (3692C>A 3693A>C), CCT1010CCC (3700T>C), GCA1014GCC (3712A>C), ATC1015ATA (3715C>A), ACT1016ACG (3718T>G), CCA1018CCC (3724A>C), ATA1020ATC (3730A>C), TGT1023TGC (3739T>C), ACA1024TCT (3740A>T 3742A>T), TGT1027TGC (3751T>C), TCA1030TCT (3760A>T), GGT1031GGC (3763T>C), TTG1033CTT (3767T>C 3769G>T), AAT1034GAG (3770A>G 3772T>G), TAT1035TAC (3775T>C), TCA1036ACT (3776T>A 3778A>T), ATT1037CTC (3779A>C 3781T>C), GTG1038ATC (3782G>A 3784G>C), ATA1039ATC (3787A>C), CAC1040CAT (3790C>T), AGA1041AGG (3793A>G), GTA1042GTC (3796A>C), TCC1044GGC (3800T>G 3801C>G), AGT1045TCT (3803A>T 3804G>C), CCT1046AAT (3806C>A 3807C>A), AAT1047AAC (3811T>C), GTT1048ATT (3812G>A), GGA1049GGT (3817A>T), CTA1052CTT (3826A>T), GTT1054GTG (3832T>G), TTT1056TTG (3838T>G), GAT1057GAC (3841T>C), GCC1058ACC (3842G>A), CCA1064TCT (3860C>T 3862A>T), GCT1065TAT (3863G>T 3864C>A), GGG1066GGT (3868G>T), CTT1067GTG (3869C>G 3871T>G), AAT1068GCA (3872A>G 3873A>C 3874T>A), AAA1069GAG (3875A>G 3877A>G), GGA1070GGG (3880A>G), AAT1071GCA (3881A>G 3882A>C 3883T>A), TAT1072CAT (3884T>C), GTA1074CTT (3890G>C 3892A>T), GGT1077GGC (3901T>C), GGA1078GGT (3904A>T), GGC1079GGT (3907C>T), ACA1080TCC (3908A>T 3910A>C), GAT1083AAT (3917G>A), TTT1084TTC (3922T>C), TCA1085TCC (3925A>C), GGT1087GGA (3931T>A), GTG1088GTT (3934G>T), GCA1089ACC (3935G>A 3937A>C), ACA1090ACT (3940A>T), AAT1091AAC (3943T>C), ACG1092ATT (3945C>T 3946G>T), TTC1093TTT (3949C>T), TCA1094TCC (3952A>C), TTC1095TTT (3955C>T), ACT1096GTG (3956A>G 3957C>T 3958T>G), GAT1099GAC (3967T>C), GAG1101AAA (3971G>A 3973G>A), CCA1104CCT (3982A>T), AGG1105CGG (3983A>C), CGG1106AGG (3986C>A), CAT1107TAT (3989C>T), ACA1108ACT (3994A>T), AGG1109AAG (3996G>A), ATG1110GCA (3998A>G 3999T>C 4000G>A), AGG1111CGA (4001A>C 4003G>A), TTC1113TTT (4009C>T), TCA1114GAC (4010T>G 4011C>A 4012A>C), AGC1115AAT (4014G>A 4015C>T), CAA1117AGT (4019C>A 4020A>G 4021A>T), TCC1118TCT (4024C>T), CGC1119CGT (4027C>T), ATC1120ATT (4030C>T), ATG1121GTT (4031A>G 4033G>T), TCA1122TCT (4036A>T), CTA1123TTG (4037C>T 4039A>G), GAT1125GAC (4045T>C), AGG1126AAA (4047G>A 4048G>A), CTT1127TTG (4049C>T 4051T>G), GGA1128GGT (4054A>T), AAT1129CTT (4055A>C 4056A>T), CTG1130CTT (4060G>T), ATC1131TTG (4061A>T 4063C>G), ATA1132ATC (4066A>C), TTG1134CTA (4070T>C 4072G>A), CCT1135CCA (4075T>A), CCT1136CCC (4078T>C), TCC1137GTA (4079T>G 4080C>T 4081C>A), GCC1138GAC (4083C>A), ATA1139TTA (4085A>T), GTG1140ATT (4088G>A 4090G>T), AGT1141AGC (4093T>C), GAG1144GAA (4102G>A), ATA1145ATC (4105A>C), CTT1146ATG (4106C>A 4108T>G), ATA1147GTT (4109A>G 4111A>T), TCT1148AAG (4112T>A 4113C>A 4114T>G), CCT1149CCA (4117T>A), GGA1150GGG (4120A>G), CTT1151CCA (4122T>C 4123T>A), TTC1153TTT (4129C>T), AAG1154AGG (4131A>G), TTG1155TTA (4135G>A), GAG1156GAA (4138G>A), TTG1157CTT (4139T>C 4141G>T), TCT1162CTG (4154T>C 4155C>T 4156T>G), GCC1163GCT (4159C>T), AAC1164AAT (4162C>T), GGC1170GGT (4180C>T), AAT1171GAT (4181A>G), ACT1174ACG (4192T>G), CAC1175CAT (4195C>T), ACC1176ACT (4198C>T), TAT1177TAC (4201T>C), CAG1178CAA (4204G>A), TCA1181TCT (4213A>T), TTT1183TTC (4219T>C), TCT1184TCA (4222T>A), GAG1185GAA (4225G>A), CTA1186CTG (4228A>G), CGT1187AGA (4229C>A 4231T>A), GAT1188GAC (4234T>C), TTT1189TTC (4237T>C), GCG1190TCG (4238G>T), ATT1191AAC (4242T>A 4243T>C) | | | | | | | | |
| **Proteins** | | | | | | | | | |
| polyprotein (YP_001976149.1) | 764 | 1192 | 30.2% | 2123 | 80.3% | 360 (99.4%) | 275 (76.0%) | 2/0/0/0 | 1 |

| | Begin | End | Coverage | Score | Concordance | Matches | Identities | I/D/M/F* | Stop Codons |
|---|---|---|---|---|---|---|---|---|---|
| **NT** | **2960** | **4410** | **25.4%** | **931** | **37.8%** | **1239 (98.9%)** | **867 (69.2%)** | **9/5** | |
| Protein mutations: | T790R (3038A>C 3039C>G 3040T>G), I799V (3065A>G 3067T>C), N817A (3119A>G 3120A>C 3121C>A), K819E (3125A>G), T821A (3131A>G), N822A (3134A>G 3135A>C 3136T>A), F827L (3149T>C 3151T>A), M838V (3182A>G), K841R (3191A>C 3192A>G 3193A>T), H920L (3429A>T), S927A (3449T>G 3451A>C), D936N (3476G>A 3478T>C), S938T (3482T>A 3484A>T), I943L (3497A>C), D949E (3517C>A), I951V (3521A>G), E953V (3528A>T 3529G>C), Q955D (3533C>G 3535G>T), E956D (3538G>T), G958_K959insKK (3544_3545insAAGAAG), P961K (3551C>A 3552C>A), P962E (3554C>G 3555C>A 3556C>G), K964V (3560A>G 3561A>T), A965G (3564C>G), S967V (3569A>G 3570G>T), I968A (3572A>G 3573T>C 3574T>A), S970G (3578T>G 3579C>G 3580C>A), I971L (3581A>C 3583A>T), M985S (3623A>T 3624T>C 3625G>T), G986A (3627G>C 3628T>C), D990T (3638G>A 3639A>C), E991S (3641G>A 3642A>G 3643A>C), L994V (3650T>G), I996L (3656A>C), I1003V (3677A>G), E1006S (3686G>T 3687A>C), G1007P (3689G>C 3690G>C 3691G>C), H1008T (3692C>A 3693A>C), T1024S (3740A>T 3742A>T), N1034E (3770A>G 3772T>G), S1036T (3776T>A 3778A>T), I1037L (3779A>C 3781T>C), V1038I (3782G>A 3784G>C), S1044G (3800T>G 3801C>G), P1046N (3806C>A 3807C>A), V1048I (3812G>A), F1056L (3838T>G), A1058T (3842G>A), P1064S (3860C>T 3862A>T), A1065Y (3863G>T 3864C>A), L1067V (3869C>G 3871T>G), N1068A (3872A>G 3873A>C 3874T>A), K1069E (3875A>G 3877A>G), N1071A (3881A>G 3882A>C 3883T>A), Y1072H (3884T>C), V1074L (3890G>C 3892A>T), T1080S (3908A>T 3910A>C), D1083N (3917G>A), A1089T (3935G>A 3937A>C), T1092I (3945C>T 3946G>T), T1096V (3956A>G 3957C>T 3958T>G), E1101K (3971G>A 3973G>A), H1107Y (3989C>T), R1109K (3996G>A), M1110A (3998A>G 3999T>C 4000G>A), S1114D (4010T>G 4011C>A 4012A>C), S1115N (4014G>A 4015C>T), Q1117S (4019C>A 4020A>G 4021A>T), M1121V (4031A>G 4033G>T), R1126K (4047G>A 4048G>A), N1129L (4055A>C 4056A>T), I1131L (4061A>T 4063C>G), S1137V (4079T>G 4080C>T 4081C>A), A1138D (4083C>A), I1139L (4085A>T), V1140I (4088G>A 4090G>T), L1146M (4106C>A 4108T>G), I1147V (4109A>G 4111A>T), S1148K (4112T>A 4113C>A 4114T>G), L1151P (4122T>C 4123T>A), K1154R (4131A>G), S1162L (4154T>C 4155C>T 4156T>G), N1171D (4181A>G), A1190S (4238G>T), I1191N (4242T>A 4243T>C) | | | | | | | | |
| Codon mutations: | ATA767ATT (2971A>T), GAG769GAA (2977G>A), ACC773ACA (2989C>A), TTG774CTG (2990T>C), CCG776CCA (2998G>A), TCT777TCA (3001T>A), TTG778CTG (3002T>C), CTT779TTG (3005C>T 3007T>G), GGT780GGA (3010T>A), AAC781AAT (3013C>T), ATA782ATT (3016A>T), GCC783GCT (3019C>T), AGG784AGA (3022G>A), TGC786TGT (3028C>T), ACT790CGG (3038A>C 3039C>G 3040T>G), GGA791GGC (3043A>C), ACT792ACA (3046T>A), TGT793TGC (3049T>C), GAA796GAG (3058A>G), ATT797ATA (3061T>A), TGC798TGT (3064C>T), ATT799GTC (3065A>G 3067T>C), GAG800GAA (3070G>A), AAA801AAG (3073A>G), TTG803TTA (3079G>A), CAT805CAC (3085T>C), TCA806TCT (3088A>T), GGT807GGA (3091T>A), CTG809CTT (3097G>T), GCA810GCC (3100A>C), GGA812GGT (3106A>T), CTT813TTG (3107C>T 3109T>G), GGA814GGT (3112A>T), CTG816CTA (3118G>A), AAC817GCA (3119A>G 3120A>C 3121C>A), ACC818ACA (3124C>A), AAA819GAA (3125A>G), ACC821GCC (3131A>G), AAT822GCA (3134A>G 3135A>C 3136T>A), ATA826ATT (3148A>T), TTT827CTA (3149T>C 3151T>A), AAC828AAT (3154C>T), CCA830CCC (3160A>C), CAT831CAC (3163T>C), CTT836CTA (3178T>A), ATG838GTG (3182A>G), GGT839GGC (3187T>C), CGA840CGC (3190A>C), AAA841CGT (3191A>C 3192A>G 3193A>T), GGG919GGT (3427G>T), CAT920CTT (3429A>T), ACA922ACT (3436A>T), GGC924GGG (3442C>G), AAA925AAG (3445A>G), AGC926AGT (3448C>T), TCA927GCC (3449T>G 3451A>C), GAT929GAC (3457T>C), TTC930TTT (3460C>T), CTT931TTG (3461C>T 3463T>G), TTC932TTT (3466C>T), GAT936AAC (3476G>A 3478T>C), TTA937TTG (3481A>G), TCA938ACT (3482T>A 3484A>T), AAG941AAA (3493G>A), ATT943CTT (3497A>C), TTG946CTG (3506T>C), AAG948AAA (3514G>A), GAC949GAA (3517C>A), AAC950AAT (3520C>T), ATT951GTT (3521A>G), GAG953GTC (3528A>T 3529G>C), TAC954TAT (3532C>T), CAG955GAT (3533C>G 3535G>T), GAG956GAT (3538G>T), GGC958GGG (3544C>G), GGC958_AAA959insAAGAAG (3544_3545insAAGAAG), GAT960GAC (3550T>C), CCG961AAG (3551C>A 3552C>A), CCC962GAG (3554C>G 3555C>A 3556C>G), AAG963AAA (3559G>A), AAG964GTG (3560A>G 3561A>T), GCT965GGT (3564C>G), CAA966CAG (3568A>G), AGT967GTT (3569A>G 3570G>T), ATT968GCA (3572A>G 3573T>C 3574T>A), CTG969CTT (3577G>T), TCC970GGA (3578T>G 3579C>G 3580C>A), ATA971CTT (3581A>C 3583A>T), AGA972CGG (3584A>C 3586A>G), GAG973GAA (3589G>A), TTT975TTC (3595T>C), TCC976TCT (3598C>T), GGT978GGA (3604T>A), GTA980GTT (3610A>T), TTC983TTT (3619C>T), TGC984TGT (3622C>T), ATG985TCT (3623A>T 3624T>C 3625G>T), GGT986GCC (3627G>C 3628T>C), GAC990ACC (3638G>A 3639A>C), GAA991AGC (3641G>A 3642A>G 3643A>C), AGA992CGC (3644A>C 3646A>C), TTG993CTG (3647T>C), TTG994GTG (3650T>G), GTA995GTT (3655A>T), ATT996CTT (3656A>C), CCT997CCA (3661T>A), GCA999GCT (3667A>T), CCA1000CCG (3670A>G), TCC1002TCT (3676C>T), ATA1003GTA (3677A>G), AGG1004CGA (3680A>C 3682G>A), TTT1005TTC (3685T>C), GAA1006TCA (3686G>T 3687A>C), GGG1007CCC (3689G>C 3690G>C 3691G>C), CAC1008ACC (3692C>A 3693A>C), CCT1010CCC (3700T>C), GCA1014GCC (3712A>C), ATC1015ATA (3715C>A), ACT1016ACG (3718T>G), CCA1018CCC (3724A>C), ATA1020ATC (3730A>C), TGT1023TGC (3739T>C), ACA1024TCT (3740A>T 3742A>T), TGT1027TGC (3751T>C), TCA1030TCT (3760A>T), GGT1031GGC (3763T>C), TTG1033CTT (3767T>C 3769G>T), AAT1034GAG (3770A>G 3772T>G), TAT1035TAC (3775T>C), TCA1036ACT (3776T>A 3778A>T), ATT1037CTC (3779A>C 3781T>C), GTG1038ATC (3782G>A 3784G>C), ATA1039ATC (3787A>C), CAC1040CAT (3790C>T), AGA1041AGG (3793A>G), GTA1042GTC (3796A>C), TCC1044GGC (3800T>G 3801C>G), AGT1045TCT (3803A>T 3804G>C), CCT1046AAT (3806C>A 3807C>A), AAT1047AAC (3811T>C), GTT1048ATT (3812G>A), GGA1049GGT (3817A>T), CTA1052CTT (3826A>T), GTT1054GTG (3832T>G), TTT1056TTG (3838T>G), GAT1057GAC (3841T>C), GCC1058ACC (3842G>A), CCA1064TCT (3860C>T 3862A>T), GCT1065TAT (3863G>T 3864C>A), GGG1066GGT (3868G>T), CTT1067GTG (3869C>G 3871T>G), AAT1068GCA (3872A>G 3873A>C 3874T>A), AAA1069GAG (3875A>G 3877A>G), GGA1070GGG (3880A>G), AAT1071GCA (3881A>G 3882A>C 3883T>A), TAT1072CAT (3884T>C), GTA1074CTT (3890G>C 3892A>T), GGT1077GGC (3901T>C), GGA1078GGT (3904A>T), GGC1079GGT (3907C>T), ACA1080TCC (3908A>T 3910A>C), GAT1083AAT (3917G>A), TTT1084TTC (3922T>C), TCA1085TCC (3925A>C), GGT1087GGA (3931T>A), GTG1088GTT (3934G>T), GCA1089ACC (3935G>A 3937A>C), ACA1090ACT (3940A>T), AAT1091AAC (3943T>C), ACG1092ATT (3945C>T 3946G>T), TTC1093TTT (3949C>T), TCA1094TCC (3952A>C), TTC1095TTT (3955C>T), ACT1096GTG (3956A>G 3957C>T 3958T>G), GAT1099GAC (3967T>C), GAG1101AAA (3971G>A 3973G>A), CCA1104CCT (3982A>T), AGG1105CGG (3983A>C), CGG1106AGG (3986C>A), CAT1107TAT (3989C>T), ACA1108ACT (3994A>T), AGG1109AAG (3996G>A), ATG1110GCA (3998A>G 3999T>C 4000G>A), AGG1111CGA (4001A>C 4003G>A), TTC1113TTT (4009C>T), TCA1114GAC (4010T>G 4011C>A 4012A>C), AGC1115AAT (4014G>A 4015C>T), CAA1117AGT (4019C>A 4020A>G 4021A>T), TCC1118TCT (4024C>T), CGC1119CGT (4027C>T), ATC1120ATT (4030C>T), ATG1121GTT (4031A>G 4033G>T), TCA1122TCT (4036A>T), CTA1123TTG (4037C>T 4039A>G), GAT1125GAC (4045T>C), AGG1126AAA (4047G>A 4048G>A), CTT1127TTG (4049C>T 4051T>G), GGA1128GGT (4054A>T), AAT1129CTT (4055A>C 4056A>T), CTG1130CTT (4060G>T), ATC1131TTG (4061A>T 4063C>G), ATA1132ATC (4066A>C), TTG1134CTA (4070T>C 4072G>A), CCT1135CCA (4075T>A), CCT1136CCC (4078T>C), TCC1137GTA (4079T>G 4080C>T 4081C>A), GCC1138GAC (4083C>A), ATA1139TTA (4085A>T), GTG1140ATT (4088G>A 4090G>T), AGT1141AGC (4093T>C), GAG1144GAA (4102G>A), ATA1145ATC (4105A>C), CTT1146ATG (4106C>A 4108T>G), ATA1147GTT (4109A>G 4111A>T), TCT1148AAG (4112T>A 4113C>A 4114T>G), CCT1149CCA (4117T>A), GGA1150GGG (4120A>G), CTT1151CCA (4122T>C 4123T>A), TTC1153TTT (4129C>T), AAG1154AGG (4131A>G), TTG1155TTA (4135G>A), GAG1156GAA (4138G>A), TTG1157CTT (4139T>C 4141G>T), TCT1162CTG (4154T>C 4155C>T 4156T>G), GCC1163GCT (4159C>T), AAC1164AAT (4162C>T), GGC1170GGT (4180C>T), AAT1171GAT (4181A>G), ACT1174ACG (4192T>G), CAC1175CAT (4195C>T), ACC1176ACT (4198C>T), TAT1177TAC (4201T>C), CAG1178CAA (4204G>A), TCA1181TCT (4213A>T), TTT1183TTC (4219T>C), TCT1184TCA (4222T>A), GAG1185GAA (4225G>A), CTA1186CTG (4228A>G), CGT1187AGA (4229C>A 4231T>A), GAT1188GAC (4234T>C), TTT1189TTC (4237T>C), GCG1190TCG (4238G>T), ATT1191AAC (4242T>A 4243T>C) | | | | | | | | |
| Coat protein B (YP_001976156.1) | 42 | 244 | 54.9% | 817 | 82.4% | 134 (98.5%) | 110 (80.9%) | 2/0/0/0 | 0 |
| Protein mutations: | T68R (3038A>C 3039C>G 3040T>G), I77V (3065A>G 3067T>C), N95A (3119A>G 3120A>C 3121C>A), K97E (3125A>G), T99A (3131A>G), N100A (3134A>G 3135A>C 3136T>A), F105L (3149T>C 3151T>A), M116V (3182A>G), K119R (3191A>C 3192A>G 3193A>T), H198L (3429A>T), S205A (3449T>G 3451A>C), D214N (3476G>A 3478T>C), S216T (3482T>A 3484A>T), I221L (3497A>C), D227E (3517C>A), I229V (3521A>G), E231V (3528A>T 3529G>C), Q233D (3533C>G 3535G>T), E234D (3538G>T), G236_K237insKK (3544_3545insAAGAAG), P239K (3551C>A 3552C>A), P240E (3554C>G 3555C>A 3556C>G), K242V (3560A>G 3561A>T), A243G (3564C>G) | | | | | | | | |
| Codon mutations: | ATA45ATT (2971A>T), GAG47GAA (2977G>A), ACC51ACA (2989C>A), TTG52CTG (2990T>C), CCG54CCA (2998G>A), TCT55TCA (3001T>A), TTG56CTG (3002T>C), CTT57TTG (3005C>T 3007T>G), GGT58GGA (3010T>A), AAC59AAT (3013C>T), ATA60ATT (3016A>T), GCC61GCT (3019C>T), AGG62AGA (3022G>A), TGC64TGT (3028C>T), ACT68CGG (3038A>C 3039C>G 3040T>G), GGA69GGC (3043A>C), ACT70ACA (3046T>A), TGT71TGC (3049T>C), GAA74GAG (3058A>G), ATT75ATA (3061T>A), TGC76TGT (3064C>T), ATT77GTC (3065A>G 3067T>C), GAG78GAA (3070G>A), AAA79AAG (3073A>G), TTG81TTA (3079G>A), CAT83CAC (3085T>C), TCA84TCT (3088A>T), GGT85GGA (3091T>A), CTG87CTT (3097G>T), GCA88GCC (3100A>C), GGA90GGT (3106A>T), CTT91TTG (3107C>T 3109T>G), GGA92GGT (3112A>T), CTG94CTA (3118G>A), AAC95GCA (3119A>G 3120A>C 3121C>A), ACC96ACA (3124C>A), AAA97GAA (3125A>G), ACC99GCC (3131A>G), AAT100GCA (3134A>G 3135A>C 3136T>A), ATA104ATT (3148A>T), TTT105CTA (3149T>C 3151T>A), AAC106AAT (3154C>T), CCA108CCC (3160A>C), CAT109CAC (3163T>C), CTT114CTA (3178T>A), ATG116GTG (3182A>G), GGT117GGC (3187T>C), CGA118CGC (3190A>C), AAA119CGT (3191A>C 3192A>G 3193A>T), GGG197GGT (3427G>T), CAT198CTT (3429A>T), ACA200ACT (3436A>T), GGC202GGG (3442C>G), AAA203AAG (3445A>G), AGC204AGT (3448C>T), TCA205GCC (3449T>G 3451A>C), GAT207GAC (3457T>C), TTC208TTT (3460C>T), CTT209TTG (3461C>T 3463T>G), TTC210TTT (3466C>T), GAT214AAC (3476G>A 3478T>C), TTA215TTG (3481A>G), TCA216ACT (3482T>A 3484A>T), AAG219AAA (3493G>A), ATT221CTT (3497A>C), TTG224CTG (3506T>C), AAG226AAA (3514G>A), GAC227GAA (3517C>A), AAC228AAT (3520C>T), ATT229GTT (3521A>G), GAG231GTC (3528A>T 3529G>C), TAC232TAT (3532C>T), CAG233GAT (3533C>G 3535G>T), GAG234GAT (3538G>T), GGC236GGG (3544C>G), GGC236_AAA237insAAGAAG (3544_3545insAAGAAG), GAT238GAC (3550T>C), CCG239AAG (3551C>A 3552C>A), CCC240GAG (3554C>G 3555C>A 3556C>G), AAG241AAA (3559G>A), AAG242GTG (3560A>G 3561A>T), GCT243GGT (3564C>G), CAA244CAG (3568A>G) | | | | | | | | |
| Coat protein C (YP_001976157.1) | 1 | 225 | 100% | 1305 | 79.1% | 225 (100%) | 164 (72.9%) | 0/0/0/0 | 0 |
| Protein mutations: | S1V (3569A>G 3570G>T), I2A (3572A>G 3573T>C 3574T>A), S4G (3578T>G 3579C>G 3580C>A), I5L (3581A>C 3583A>T), M19S (3623A>T 3624T>C 3625G>T), G20A (3627G>C 3628T>C), D24T (3638G>A 3639A>C), E25S (3641G>A 3642A>G 3643A>C), L28V (3650T>G), I30L (3656A>C), I37V (3677A>G), E40S (3686G>T 3687A>C), G41P (3689G>C 3690G>C 3691G>C), H42T (3692C>A 3693A>C), T58S (3740A>T 3742A>T), N68E (3770A>G 3772T>G), S70T (3776T>A 3778A>T), I71L (3779A>C 3781T>C), V72I (3782G>A 3784G>C), S78G (3800T>G 3801C>G), P80N (3806C>A 3807C>A), V82I (3812G>A), F90L (3838T>G), A92T (3842G>A), P98S (3860C>T 3862A>T), A99Y (3863G>T 3864C>A), L101V (3869C>G 3871T>G), N102A (3872A>G 3873A>C 3874T>A), K103E (3875A>G 3877A>G), N105A (3881A>G 3882A>C 3883T>A), Y106H (3884T>C), V108L (3890G>C 3892A>T), T114S (3908A>T 3910A>C), D117N (3917G>A), A123T (3935G>A 3937A>C), T126I (3945C>T 3946G>T), T130V (3956A>G 3957C>T 3958T>G), E135K (3971G>A 3973G>A), H141Y (3989C>T), R143K (3996G>A), M144A (3998A>G 3999T>C 4000G>A), S148D (4010T>G 4011C>A 4012A>C), S149N (4014G>A 4015C>T), Q151S (4019C>A 4020A>G 4021A>T), M155V (4031A>G 4033G>T), R160K (4047G>A 4048G>A), N163L (4055A>C 4056A>T), I165L (4061A>T 4063C>G), S171V (4079T>G 4080C>T 4081C>A), A172D (4083C>A), I173L (4085A>T), V174I (4088G>A 4090G>T), L180M (4106C>A 4108T>G), I181V (4109A>G 4111A>T), S182K (4112T>A 4113C>A 4114T>G), L185P (4122T>C 4123T>A), K188R (4131A>G), S196L (4154T>C 4155C>T 4156T>G), N205D (4181A>G), A224S (4238G>T), I225N (4242T>A 4243T>C) | | | | | | | | |

| | Begin | End | Coverage | Score | Concordance | Matches | Identities | I/D/M/F* | Stop Codons |
|---|---|---|---|---|---|---|---|---|---|
| **NT** | **2960** | **4410** | **25.4%** | **931** | **37.8%** | **1239 (98.9%)** | **867 (69.2%)** | **9/5** | |
| Codon mutations: | AGT1GTT (3569A>G 3570G>T), ATT2GCA (3572A>G 3573T>C 3574T>A), CTG3CTT (3577G>T), TCC4GGA (3578T>G 3579C>G 3580C>A), ATA5CTT (3581A>C 3583A>T), AGA6CGG (3584A>C 3586A>G), GAG7GAA (3589G>A), TTT9TTC (3595T>C), TCC10TCT (3598C>T), GGT12GGA (3604T>A), GTA14GTT (3610A>T), TTC17TTT (3619C>T), TGC18TGT (3622C>T), ATG19TCT (3623A>T 3624T>C 3625G>T), GGT20GCC (3627G>C 3628T>C), GAC24ACC (3638G>A 3639A>C), GAA25AGC (3641G>A 3642A>G 3643A>C), AGA26CGC (3644A>C 3646A>C), TTG27CTG (3647T>C), TTG28GTG (3650T>G), GTA29GTT (3655A>T), ATT30CTT (3656A>C), CCT31CCA (3661T>A), GCA33GCT (3667A>T), CCA34CCG (3670A>G), TCC36TCT (3676C>T), ATA37GTA (3677A>G), AGG38CGA (3680A>C 3682G>A), TTT39TTC (3685T>C), GAA40TCA (3686G>T 3687A>C), GGG41CCC (3689G>C 3690G>C 3691G>C), CAC42ACC (3692C>A 3693A>C), CCT44CCC (3700T>C), GCA48GCC (3712A>C), ATC49ATA (3715C>A), ACT50ACG (3718T>G), CCA52CCC (3724A>C), ATA54ATC (3730A>C), TGT57TGC (3739T>C), ACA58TCT (3740A>T 3742A>T), TGT61TGC (3751T>C), TCA64TCT (3760A>T), GGT65GGC (3763T>C), TTG67CTT (3767T>C 3769G>T), AAT68GAG (3770A>G 3772T>G), TAT69TAC (3775T>C), TCA70ACT (3776T>A 3778A>T), ATT71CTC (3779A>C 3781T>C), GTG72ATC (3782G>A 3784G>C), ATA73ATC (3787A>C), CAC74CAT (3790C>T), AGA75AGG (3793A>G), GTA76GTC (3796A>C), TCC78GGC (3800T>G 3801C>G), AGT79TCT (3803A>T 3804G>C), CCT80AAT (3806C>A 3807C>A), AAT81AAC (3811T>C), GTT82ATT (3812G>A), GGA83GGT (3817A>T), CTA86CTT (3826A>T), GTT88GTG (3832T>G), TTT90TTG (3838T>G), GAT91GAC (3841T>C), GCC92ACC (3842G>A), CCA98TCT (3860C>T 3862A>T), GCT99TAT (3863G>T 3864C>A), GGG100GGT (3868G>T), CTT101GTG (3869C>G 3871T>G), AAT102GCA (3872A>G 3873A>C 3874T>A), AAA103GAG (3875A>G 3877A>G), GGA104GGG (3880A>G), AAT105GCA (3881A>G 3882A>C 3883T>A), TAT106CAT (3884T>C), GTA108CTT (3890G>C 3892A>T), GGT111GGC (3901T>C), GGA112GGT (3904A>T), GGC113GGT (3907C>T), ACA114TCC (3908A>T 3910A>C), GAT117AAT (3917G>A), TTT118TTC (3922T>C), TCA119TCC (3925A>C), GGT121GGA (3931T>A), GTG122GTT (3934G>T), GCA123ACC (3935G>A 3937A>C), ACA124ACT (3940A>T), AAT125AAC (3943T>C), ACG126ATT (3945C>T 3946G>T), TTC127TTT (3949C>T), TCA128TCC (3952A>C), TTC129TTT (3955C>T), ACT130GTG (3956A>G 3957C>T 3958T>G), GAT133GAC (3967T>C), GAG135AAA (3971G>A 3973G>A), CCA138CCT (3982A>T), AGG139CGG (3983A>C), CGG140AGG (3986C>A), CAT141TAT (3989C>T), ACA142ACT (3994A>T), AGG143AAG (3996G>A), ATG144GCA (3998A>G 3999T>C 4000G>A), AGG145CGA (4001A>C 4003G>A), TTC147TTT (4009C>T), TCA148GAC (4010T>G 4011C>A 4012A>C), AGC149AAT (4014G>A 4015C>T), CAA151AGT (4019C>A 4020A>G 4021A>T), TCC152TCT (4024C>T), CGC153CGT (4027C>T), ATC154ATT (4030C>T), ATG155GTT (4031A>G 4033G>T), TCA156TCT (4036A>T), CTA157TTG (4037C>T 4039A>G), GAT159GAC (4045T>C), AGG160AAA (4047G>A 4048G>A), CTT161TTG (4049C>T 4051T>G), GGA162GGT (4054A>T), AAT163CTT (4055A>C 4056A>T), CTG164CTT (4060G>T), ATC165TTG (4061A>T 4063C>G), ATA166ATC (4066A>C), TTG168CTA (4070T>C 4072G>A), CCT169CCA (4075T>A), CCT170CCC (4078T>C), TCC171GTA (4079T>G 4080C>T 4081C>A), GCC172GAC (4083C>A), ATA173TTA (4085A>T), GTG174ATT (4088G>A 4090G>T), AGT175AGC (4093T>C), GAG178GAA (4102G>A), ATA179ATC (4105A>C), CTT180ATG (4106C>A 4108T>G), ATA181GTT (4109A>G 4111A>T), TCT182AAG (4112T>A 4113C>A 4114T>G), CCT183CCA (4117T>A), GGA184GGG (4120A>G), CTT185CCA (4122T>C 4123T>A), TTC187TTT (4129C>T), AAG188AGG (4131A>G), TTG189TTA (4135G>A), GAG190GAA (4138G>A), TTG191CTT (4139T>C 4141G>T), TCT196CTG (4154T>C 4155C>T 4156T>G), GCC197GCT (4159C>T), AAC198AAT (4162C>T), GGC204GGT (4180C>T), AAT205GAT (4181A>G), ACT208ACG (4192T>G), CAC209CAT (4195C>T), ACC210ACT (4198C>T), TAT211TAC (4201T>C), CAG212CAA (4204G>A), TCA215TCT (4213A>T), TTT217TTC (4219T>C), TCT218TCA (4222T>A), GAG219GAA (4225G>A), CTA220CTG (4228A>G), CGT221AGA (4229C>A 4231T>A), GAT222GAC (4234T>C), TTT223TTC (4237T>C), GCG224TCG (4238G>T), ATT225AAC (4242T>A 4243T>C) | | | | | | | | |

*: Inserts / Deletes / Misaligned / Frameshifts

#### Analysis details

This analysis was performed with panviral2.64

### NGS Details (UN62): Diachasmimorpha longicaudata entomopoxvirus (segment NC_043455.1)

#### Assembly

| | |
|---|---|
| **Coverage Length** | 1126 (1 contig(s)) |
| **Depth Of Coverage** | 993.3 |
| **Number Of Reads** | 8833 |
| **Reads Per Million** | 204.76 rpm (after QC) |
| **Ambiguities** | 0 |
| **Assembly Method** | de novo + reference guided assembly |
| **Consensus Caller** | Bcf Tools |

#### Coverage Map

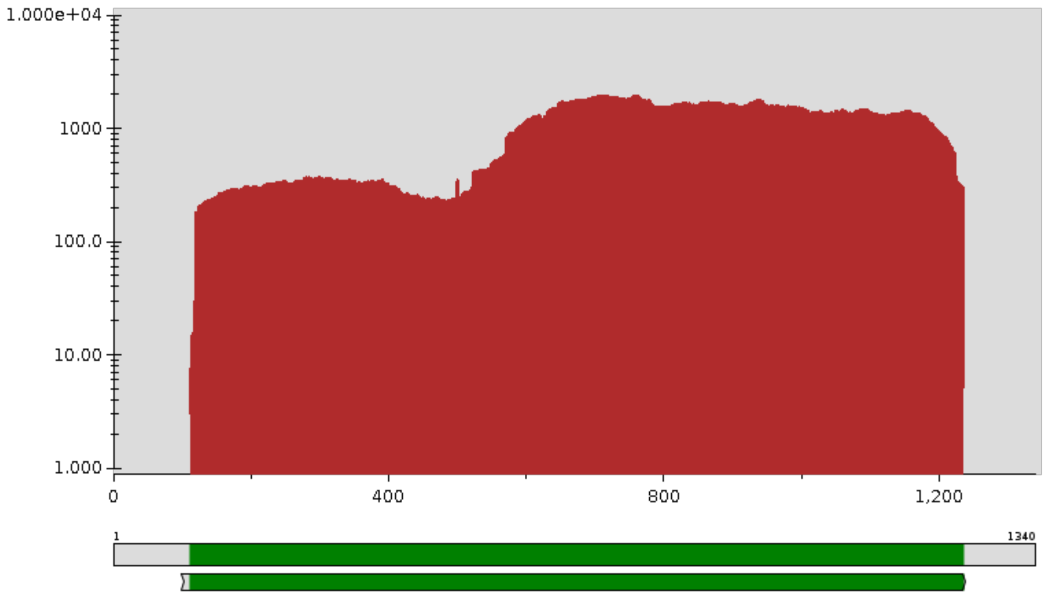

#### Assignment

| | |
|---|---|
| **Type** | Diachasmimorpha longicaudata entomopoxvirus (Taxonomy ID: 109981) |
| **Reference Genome** | NC_043455.1 |
| **NT Identity (%)** | 60.0355 |
| **AA Identity (%)** | 55.4667 |
| **Number Of Stop Codons** | 0 |
| **Number Of CDS** | 1 |

#### Alignment

| | |
|---|---|
| **Alignment Score** | 434.0 (NT) + 1463.0 (AA) = 1897.0 |
| **Concordance (%)** | 40.2418 |

| Alignment Method | Global, seeded, nucleotide + amino acids (AGA) |
|---|---|

#### Genome Region

Sequence starts at position 112 and ends at position 1237 relative to NC_043455.1 reference sequence.

#### Alignment Detailed Statistics

| | Begin | End | Coverage | Score | Concordance | Matches | Identities | I/D/M/F* | Stop Codons |
|---|---|---|---|---|---|---|---|---|---|
| **NT** | **112** | **1237** | **84.0%** | **434** | **19.4%** | **1123 (99.5%)** | **676 (59.9%)** | **3/3** | |
| Mutations: | 121A>T, 126A>C, 127C>T, 128C>G, 130T>G, 136C>G, 137C>A, 139C>T, 142G>A, 143A>G, 145A>T, 146A>G, 148T>C, 149A>C, 151C>T, 154A>T, 156A>G, 157G>A, 160T>G, 163A>T, 166T>C, 167T>C, 168C>A, 169T>G, 171G>A, 172T>C, 175A>G, 178T>C, 187T>A, 190A>T, 191A>G, 193T>C, 203A>C, 204A>G, 208C>T, 209A>G, 211T>A, 214T>G, 217A>T, 218T>A, 219G>T, 223T>A, 226T>G, 230A>C, 231A>G, 232A>T, 238G>C, 241A>T, 243T>C, 244C>T, 247A>G, 256G>C, 259A>T, 262A>C, 271A>C, 272G>T, 274A>C, 276C>T, 277A>G, 278T>A, 279A>T, 283A>T, 284A>C, 286C>T, 288G>C, 289T>C, 294T>G, 295A>C, 298G>A, 299C>G, 300A>T, 301A>T, 302A>G, 304A>G, 310T>C, 311T>A, 312C>A, 313T>A, 320A>G, 322T>G, 323A>G, 325C>A, 331T>A, 337C>A, 341A>T, 343C>T, 346A>T, 352T>G, 355A>G, 356T>C, 358G>T, 361T>A, 362C>G, 363T>C, 364C>T, 368G>A, 370T>G, 371C>G, 374C>A, 375G>A, 376T>G, 379A>G, 380T>A, 382G>T, 383C>T, 384A>T, 385A>G, 386A>G, 394G>A, 395A>G, 400T>C, 401C>A, 403A>T, 404_406delTAT, 409T>C, 410T>A, 412T>A, 413A>C, 415A>G, 416T>G, 417G>C, 418T>A, 421A>T, 423T>C, 424T>G, 438C>A, 442T>C, 443A>G, 445C>G, 446A>G, 447A>G, 448G>T, 452A>G, 453G>A, 454C>T, 455C>A, 456A>T, 457G>C, 458G>A, 459A>G, 462C>A, 463T>G, 467A>G, 469_470insCAT, 470A>G, 471A>G, 474C>T, 475C>A, 478G>A, 482T>G, 484G>C, 485A>T, 486T>C, 500C>A, 503A>G, 505G>C, 506A>T, 507T>G, 511T>C, 512C>A, 514A>G, 515C>T, 517C>G, 519C>G, 520C>A, 521C>A, 524A>C, 526A>G, 527A>T, 528G>C, 530A>C, 533G>C, 534A>G, 535T>C, 536A>C, 538C>T, 539A>G, 541A>T, 542G>T, 543C>A, 544A>C, 549A>C, 550A>G, 553A>G, 554G>T, 560A>T, 562A>G, 565T>C, 568A>G, 577G>A, 581T>C, 583G>C, 584A>T, 585T>C, 586A>C, 587G>A, 588A>G, 589T>A, 590A>G, 591A>G, 595T>C, 596T>A, 597T>A, 602A>C, 604A>G, 607A>T, 608C>T, 610A>T, 620G>C, 622A>G, 623T>C, 628T>G, 629G>C, 630A>C, 632A>C, 633G>C, 634T>A, 635C>A, 637T>G, 638G>A, 646T>G, 647A>G, 648T>G, 650T>G, 652A>T, 655A>C, 658A>T, 661A>C, 664C>T, 665G>A, 667T>G, 671T>C, 674A>G, 675G>A, 678T>C, 680A>C, 682C>T, 683A>G, 685T>G, 687C>T, 688A>T, 689T>A, 692C>A, 693A>G, 694G>A, 695G>A, 696T>A, 697C>G, 700T>C, 705G>A, 706A>C, 707G>A, 709T>G, 715A>G, 717A>G, 718A>G, 724G>T, 727A>G, 730A>G, 731A>C, 732A>G, 735C>A, 739A>G, 740T>C, 742G>C, 745A>T, 748C>T, 754C>T, 759G>A, 765A>T, 770A>G, 772A>C, 778A>C, 779A>G, 781G>T, 784A>G, 785A>G, 787T>A, 790T>A, 792T>G, 793T>G, 796A>G, 797G>C, 798C>T, 799A>T, 810T>G, 823C>G, 824C>A, 825A>C, 826C>T, 830A>G, 831G>C, 833C>A, 838T>C, 841A>G, 843C>G, 844A>T, 845C>G, 847A>C, 850A>C, 854T>G, 855G>T, 856T>G, 859T>C, 862A>C, 863C>A, 865T>G, 866A>C, 868A>T, 869C>A, 878G>T, 879T>G, 880A>G, 885T>C, 886G>T, 889A>T, 890T>A, 891G>A, 892T>G, 893T>A, 895A>G, 896A>C, 897C>G, 898A>C, 900A>G, 901T>C, 902A>C, 904A>T, 905A>G, 908T>C, 909T>A, 915C>T, 919A>T, 920A>G, 921G>C, 922C>A, 924T>C, 925C>T, 934T>C, 938T>G, 939C>A, 943A>G, 944C>A, 946A>C, 947G>A, 948A>C, 949A>T, 950C>A, 952T>A, 956T>A, 958T>C, 965A>C, 966A>G, 970A>G, 973T>C, 974A>C, 977G>T, 978A>C, 982T>G, 983A>T, 984A>C, 986A>T, 989A>C, 991A>T, 992A>G, 994A>G, 997T>C, 998C>A, 1000T>C, 1001T>A, 1003A>T, 1007A>G, 1012A>T, 1015A>G, 1018A>T, 1019A>C, 1021A>T, 1024C>T, 1033T>A, 1036A>G, 1040A>G, 1042A>C, 1045T>C, 1046T>C, 1048A>T, 1051T>C, 1054C>T, 1057T>C, 1063T>C, 1064T>C, 1070C>A, 1071A>C, 1073A>C, 1075T>A, 1076A>C, 1077G>C, 1087T>C, 1088A>C, 1090A>G, 1103A>C, 1105A>T, 1111T>A, 1112A>C, 1120C>A, 1121C>A, 1123A>G, 1126A>G, 1129G>T, 1130A>G, 1141T>C, 1145A>G, 1147A>C, 1150T>C, 1151G>A, 1153A>G, 1156C>T, 1159A>T, 1160T>G, 1161T>A, 1163C>A, 1164C>G, 1165A>G, 1167A>T, 1172G>T, 1173C>T, 1177A>C, 1178T>A, 1181G>C, 1183A>G, 1185C>A, 1186G>A, 1192T>C, 1195T>C, 1196A>G, 1197C>T, 1199A>G, 1200A>T, 1204A>C, 1207T>G, 1210A>G, 1211A>C, 1218A>C, 1219G>C, 1223A>G, 1227T>C, 1232T>C, 1234G>C, 1235A>C, 1237A>G | | | | | | | | |
| **CDS** | | | | | | | | | |
| FLA14_p101 | 5 | 379 | 98.7% | 1463 | 58.8% | 374 (99.5%) | 208 (55.3%) | 1/1/0/0 | 0 |
| Protein mutations: | E7D (121A>T), D9A (126A>C 127C>T), H10E (128C>G 130T>G), L13I (137C>A 139C>T), K15D (143A>G 145A>T), N16D (146A>G 148T>C), I17L (149A>C 151C>T), K19R (156A>G 157G>A), S23Q (167T>C 168C>A 169T>G), C24Y (171G>A 172T>C), T31A (191A>G 193T>C), K35R (203A>C 204A>G), I37V (209A>G 211T>A), F38L (214T>G), C40I (218T>A 219G>T), K44R (230A>C 231A>G 232A>T), V48A (243T>C 244C>T), A58S (272G>T 274A>C), T59M (276C>T 277A>G), Y60I (278T>A 279A>T), I62L (284A>C 286C>T), S63T (288G>C 289T>C), L65C (294T>G 295A>C), Q67V (299C>G 300A>T 301A>T), I68V (302A>G 304A>G), S71K (311T>A 312C>A 313T>A), N74E (320A>G 322T>G), I75V (323A>G 325C>A), T81S (341A>T 343C>T), L88A (362C>G 363T>C 364C>T), A90T (368G>A 370T>G), Q91E (371C>G), R92K (374C>A 375G>A 376T>G), L94I (380T>A 382G>T), Q95L (383C>T 384A>T 385A>G), T96A (386A>G), N99D (395A>G), L101I (401C>A 403A>T), Y102del (404_406delTAT), F104I (410T>A 412T>A), K105Q (413A>C 415A>G), C106A (416T>G 417G>C 418T>A), Q107H (421A>T), V108A (423T>C 424T>G), T113K (438C>A), I115V (443A>G 445C>G), K116G (446A>G 447A>G 448G>T), S118D (452A>G 453G>A 454C>T), Q119I (455C>A 456A>T 457G>C), E120R (458G>A 459A>G), T121K (462C>A 463T>G), K123E (467A>G), K123_K124insH (469_470insCAT), K124G (470A>G 471A>G), A125V (474C>T 475C>A), L128V (482T>G 484G>C), I129S (485A>T 486T>C), M135V (503A>G 505G>C), I136C (506A>T 507T>G), L138M (512C>A 514A>G), T140R (519C>G 520C>A), K142Q (524A>C 526A>G), I144L (530A>C), D145R (533G>C 534A>G 535T>C), T146P (536A>C 538C>T), K147D (539A>G 541A>T), A148Y (542G>T 543C>A 544A>C), K150T (549A>C 550A>G), I151M (553A>G), V152F (554G>T), I154L (560A>T 562A>G), I162S (584A>T 585T>C 586A>C), D163R (587G>A 588A>G 589T>A), N164G (590A>G 591A>G), L166K (596T>A 597T>A), K168Q (602A>C 604A>G), Q170Y (608C>T 610A>T), E174Q (620G>C 622A>G), F175L (623T>C), F176L (628T>G), E177P (629G>C 630A>C), S178P (632A>C 633G>C 634T>A), H179K (635C>A 637T>G), V180I (638G>A), I183G (647A>G 648T>G), L184V (650T>G 652A>T), L185F (655A>C), V189M (665G>A 667T>G), S191P (671T>C), R192E (674A>G 675G>A), V193A (678T>C), I194L (680A>C 682C>T), N195E (683A>G 685T>G), T196I (687C>T 688A>T), S197T (689T>A), Q198R (692C>A 693A>G 694G>A), V199K (695G>A 696T>A 697C>G), R202N (705G>A 706A>C), D203K (707G>A 709T>G), K206R (717A>G 718A>G), N211R (731A>C 732A>G), A212D (735C>A), R220K (759G>A), Y222F (765A>T), I224V (770A>G 772A>C), K227D (779A>G 781G>T), N229E (785A>G 787T>A), D230E (790T>A), F231W (792T>G 793T>G), A233L (797G>C 798C>T 799A>T), F237C (810T>G), D241E (823C>G), H242T (824C>A 825A>C 826C>T), S244A (830A>G 831G>C), L245I (833C>A), T248S (843C>G 844A>T), L249V (845C>G 847A>C), C252V (854T>G 855G>T 856T>G), Q257K (869C>A), V260W (878G>T 879T>G 880A>G), M262T (885T>C 886G>T), E263D (889A>T), C264K (890T>A 891G>A 892T>G), L265M (893T>A 895A>G), T266R (896A>C 897C>G 898A>C), N267S (900A>G 901T>C), N269D (905A>G), F270H (908T>C 909T>A), A272V (915C>T), S274A (920A>G 921G>C 922C>A), I275T (924T>C 925C>T), S280D (938T>G 939C>A), Q282N (944C>A 946A>C), E283T (947G>A 948A>C 949A>T), F286I (956T>A 958T>C), K289R (965A>C 966A>G), D293S (977G>T 978A>C), K295S (983A>T 984A>C), T296S (986A>T), I298V (992A>G 994A>G), L300I (998C>A 1000T>C), S301T (1001T>A 1003A>T), N303D (1007A>G), I314V (1040A>G 1042A>C), H324T (1070C>A 1071A>C), N325Q (1073A>C 1075T>A), R326P (1076A>C 1077G>C), I330L (1088A>C 1090A>G), I344V (1130A>G), I349V (1145A>G 1147A>C), E351K (1151G>A 1153A>G), E353D (1159A>T), L354E (1160T>G 1161T>A), P355R (1163C>A 1164C>G 1165A>G), K356M (1167A>T), A358F (1172G>T 1173C>T), E359D (1177A>C), L360I (1178T>A), E361Q (1181G>C 1183A>G), T362K (1185C>A 1186G>A), T366V (1196A>G 1197C>T), K367V (1199A>G 1200A>T), D369E (1207T>G), M371L (1211A>C), E373A (1218A>C 1219G>C), I375V (1223A>G), V376A (1227T>C), I379L (1235A>C 1237A>G) | | | | | | | | |

| | Begin | End | Coverage | Score | Concordance | Matches | Identities | I/D/M/F* | Stop Codons |
|---|---|---|---|---|---|---|---|---|---|
| **NT** | **112** | **1237** | **84.0%** | **434** | **19.4%** | **1123 (99.5%)** | **676 (59.9%)** | **3/3** | |
| Codon mutations: | GAA7GAT (121A>T), GAC9GCT (126A>C 127C>T), CAT10GAG (128C>G 130T>G), GGC12GGG (136C>G), CTC13ATT (137C>A 139C>T), AAG14AAA (142G>A), AAA15GAT (143A>G 145A>T), AAT16GAC (146A>G 148T>C), ATC17CTT (149A>C 151C>T), CTA18CTT (154A>T), AAG19AGA (156A>G 157G>A), GGT20GGG (160T>G), ATA21ATT (163A>T), TAT22TAC (166T>C), TCT23CAG (167T>C 168C>A 169T>G), TGT24TAC (171G>A 172T>C), GGA25GGG (175A>G), TTT26TTC (178T>C), CCT29CCA (187T>A), TCA30TCT (190A>T), ACT31GCC (191A>G 193T>C), AAA35CGA (203A>C 204A>G), GCC36GCT (208C>T), ATT37GTA (209A>G 211T>A), TTT38TTG (214T>G), CCA39CCT (217A>T), TGT40ATT (218T>A 219G>T), ATT41ATA (223T>A), TCT42TCG (226T>G), AAA44CGT (230A>C 231A>G 232A>T), GTG46GTC (238G>C), ATA47ATT (241A>T), GTC48GCT (243T>C 244C>T), CAA49CAG (247A>G), TCG52TCC (256G>C), GGA53GGT (259A>T), ACA54ACC (262A>C), ACA57ACC (271A>C), GCA58TCC (272G>T 274A>C), ACA59ATG (276C>T 277A>G), TAT60ATT (278T>A 279A>T), GCA61GCT (283A>T), ATC62CTT (284A>C 286C>T), AGT63ACC (288G>C 289T>C), TTA65TGC (294T>G 295A>C), CAG66CAA (298G>A), CAA67GTT (299C>G 300A>T 301A>T), ATA68GTG (302A>G 304A>G), ACT70ACC (310T>C), TCT71AAA (311T>A 312C>A 313T>A), AAT74GAG (320A>G 322T>G), ATC75GTA (323A>G 325C>A), GCT77GCA (331T>A), ATC79ATA (337C>A), ACC81TCT (341A>T 343C>T), CCA82CCT (346A>T), CGT84CGG (352T>G), GAA85GAG (355A>G), TTG86CTT (356T>C 358G>T), GCT87GCA (361T>A), CTC88GCT (362C>G 363T>C 364C>T), GCT90ACG (368G>A 370T>G), CAA91GAA (371C>G), CGT92AAG (374C>A 375G>A 376T>G), GTA93GTG (379A>G), TTG94ATT (380T>A 382G>T), CAA95TTG (383C>T 384A>T 385A>G), ACA96GCA (386A>G), GGG98GGA (394G>A), AAT99GAT (395A>G), TAT100TAC (400T>C), CTA101ATT (401C>A 403A>T), TAT102del (404_406delTAT), AAT103AAC (409T>C), TTT104ATA (410T>A 412T>A), AAA105CAG (413A>C 415A>G), TGT106GCA (416T>G 417G>C 418T>A), CAA107CAT (421A>T), GTT108GCG (423T>C 424T>G), ACA113AAA (438C>A), AGT114AGC (442T>C), ATC115GTG (443A>G 445C>G), AAG116GGT (446A>G 447A>G 448G>T), AGC118GAT (452A>G 453G>A 454C>T), CAG119ATC (455C>A 456A>T 457G>C), GAA120AGA (458G>A 459A>G), ACT121AAG (462C>A 463T>G), AAG123GAG (467A>G), AAG123_AAA124insCAT (469_470insCAT), AAA124GGA (470A>G 471A>G), GCC125GTA (474C>T 475C>A), CAG126CAA (478G>A), TTG128GTC (482T>G 484G>C), ATT129TCT (485A>T 486T>C), CGA134AGA (500C>A), ATG135GTC (503A>G 505G>C), ATT136TGT (506A>T 507T>G), GAT137GAC (511T>C), CTA138ATG (512C>A 514A>G), CTC139TTG (515C>T 517C>G), ACC140AGA (519C>G 520C>A), CGA141AGA (521C>A), AAA142CAG (524A>C 526A>G), AGT143TCT (527A>T 528G>C), ATC144CTC (530A>C), GAT145CGC (533G>C 534A>G 535T>C), ACC146CCT (536A>C 538C>T), AAA147GAT (539A>G 541A>T), GCA148TAC (542G>T 543C>A 544A>C), AAA150ACG (549A>C 550A>G), ATA151ATG (553A>G), GTT152TTT (554G>T), ATA154TTG (560A>T 562A>G), GAT155GAC (565T>C), GAA156GAG (568A>G), GAG159GAA (577G>A), TTG161CTC (581T>C 583G>C), ATA162TCC (584A>T 585T>C 586A>C), GAT163AGA (587G>A 588A>G 589T>A), AAT164GGT (590A>G 591A>G), TTT165TTC (595T>C), TTG166AAG (596T>A 597T>A), AAA168CAG (602A>C 604A>G), ATA169ATT (607A>T), CAA170TAT (608C>T 610A>T), GAA174CAG (620G>C 622A>G), TTT175CTT (623T>C), TTT176TTG (628T>G), GAA177CCA (629G>C 630A>C), AGT178CCA (632A>C 633G>C 634T>A), CAT179AAG (635C>A 637T>G), GTC180ATC (638G>A), GTT182GTG (646T>G), ATT183GGT (647A>G 648T>G), TTA184GTT (650T>G 652A>T), TTA185TTC (655A>C), TCA186TCT (658A>T), GCA187GCC (661A>C), ACC188ACT (664C>T), GTT189ATG (665G>A 667T>G), TCA191CCA (671T>C), AGG192GAG (674A>G 675G>A), GTT193GCT (678T>C), ATC194CTT (680A>C 682C>T), AAT195GAG (683A>G 685T>G), ACA196ATT (687C>T 688A>T), TCT197ACT (689T>A), CAG198AGA (692C>A 693A>G 694G>A), GTC199AAG (695G>A 696T>A 697C>G), TTT200TTC (700T>C), AGA202AAC (705G>A 706A>C), GAT203AAG (707G>A 709T>G), GTA205GTG (715A>G), AAA206AGG (717A>G 718A>G), CTG208CTT (724G>T), GTA209GTG (727A>G), AAA210AAG (730A>G), AAT211CGT (731A>C 732A>G), GCT212GAT (735C>A), GAA213GAG (739A>G), TTG214CTC (740T>C 742G>C), ACA215ACT (745A>T), CTC216CTT (748C>T), GGC218GGT (754C>T), AGG220AAG (759G>A), TAT222TTT (765A>T), ATA224GTC (770A>G 772A>C), GTA226GTC (778A>C), AAG227GAT (779A>G 781G>T), AAA228AAG (784A>G), AAT229GAA (785A>G 787T>A), GAT230GAA (790T>A), TTT231TGG (792T>G 793T>G), AAA232AAG (796A>G), GCA233CTT (797G>C 798C>T 799A>T), TTT237TGT (810T>G), GAC241GAG (823C>G), CAC242ACT (824C>A 825A>C 826C>T), AGC244GCC (830A>G 831G>C), CTC245ATC (833C>A), ACT246ACC (838T>C), CAA247CAG (841A>G), ACA248AGT (843C>G 844A>T), CTA249GTC (845C>G 847A>C), ATA250ATC (850A>C), TGT252GTG (854T>G 855G>T 856T>G), AAT253AAC (859T>C), ACA254ACC (862A>C), CGT255AGG (863C>A 865T>G), AGA256CGT (866A>C 868A>T), CAA257AAA (869C>A), GTA260TGG (878G>T 879T>G 880A>G), ATG262ACT (885T>C 886G>T), GAA263GAT (889A>T), TGT264AAG (890T>A 891G>A 892T>G), TTA265ATG (893T>A 895A>G), ACA266CGC (896A>C 897C>G 898A>C), AAT267AGC (900A>G 901T>C), AGA268CGT (902A>C 904A>T), AAT269GAT (905A>G), TTC270CAC (908T>C 909T>A), GCA272GTA (915C>T), TCA273TCT (919A>T), AGC274GCA (920A>G 921G>C 922C>A), ATC275ACT (924T>C 925C>T), GAT278GAC (934T>C), TCC280GAC (938T>G 939C>A), CAA281CAG (943A>G), CAA282AAC (944C>A 946A>C), GAA283ACT (947G>A 948A>C 949A>T), CGT284AGA (950C>A 952T>A), TTT286ATC (956T>A 958T>C), AAA289CGA (965A>C 966A>G), GAA290GAG (970A>G), TTT291TTC (973T>C), AGA292CGA (974A>C), GAT293TCT (977G>T 978A>C), GGT294GGG (982T>G), AAA295TCA (983A>T 984A>C), ACT296TCT (986A>T), AGA297CGT (989A>C 991A>T), ATA298GTG (992A>G 994A>G), CTT299CTC (997T>C), CTT300ATC (998C>A 1000T>C), TCA301ACT (1001T>A 1003A>T), AAT303GAT (1007A>G), CTA304CTT (1012A>T), TTA305TTG (1015A>G), GCA306GCT (1018A>T), AGA307CGT (1019A>C 1021A>T), GGC308GGT (1024C>T), GTT311GTA (1033T>A), CAA312CAG (1036A>G), ATA314GTC (1040A>G 1042A>C), TCT315TCC (1045T>C), TTA316CTT (1046T>C 1048A>T), GTT317GTC (1051T>C), ATC318ATT (1054C>T), AAT319AAC (1057T>C), GAT321GAC (1063T>C), TTG322CTG (1064T>C), CAT324ACT (1070C>A 1071A>C), AAT325CAA (1073A>C 1075T>A), AGA326CCA (1076A>C 1077G>C), TAT329TAC (1087T>C), ATA330CTG (1088A>C 1090A>G), AGA335CGT (1103A>C 1105A>T), GGT337GGA (1111T>A), AGA338CGA (1112A>C), GGC340GGA (1120C>A), CGA341AGG (1121C>A 1123A>G), AAA342AAG (1126A>G), GGG343GGT (1129G>T), ATT344GTT (1130A>G), AAT347AAC (1141T>C), ATA349GTC (1145A>G 1147A>C), ACT350ACC (1150T>C), GAA351AAG (1151G>A 1153A>G), GAC352GAT (1156C>T), GAA353GAT (1159A>T), TTA354GAA (1160T>G 1161T>A), CCA355AGG (1163C>A 1164C>G 1165A>G), AAG356ATG (1167A>T), GCT358TTT (1172G>T 1173C>T), GAA359GAC (1177A>C), TTA360ATA (1178T>A), GAA361CAG (1181G>C 1183A>G), ACG362AAA (1185C>A 1186G>A), TAT364TAC (1192T>C), AAT365AAC (1195T>C), ACC366GTC (1196A>G 1197C>T), AAA367GTA (1199A>G 1200A>T), ATA368ATC (1204A>C), GAT369GAG (1207T>G), GAA370GAG (1210A>G), ATG371CTG (1211A>C), GAG373GCC (1218A>C 1219G>C), ATT375GTT (1223A>G), GTT376GCT (1227T>C), TTG378CTC (1232T>C 1234G>C), ATA379CTG (1235A>C 1237A>G) | | | | | | | | |

#### Proteins

| | Begin | End | Coverage | Score | Concordance | Matches | Identities | I/D/M/F* | Stop Codons |
|---|---|---|---|---|---|---|---|---|---|
| hypothetical protein (YP_009666148.1) | 5 | 379 | 98.7% | 1463 | 58.8% | 374 (99.5%) | 208 (55.3%) | 1/1/0/0 | 0 |
| Protein mutations: | E7D (121A>T), D9A (126A>C 127C>T), H10E (128C>G 130T>G), L13I (137C>A 139C>T), K15D (143A>G 145A>T), N16D (146A>G 148T>C), I17L (149A>C 151C>T), K19R (156A>G 157G>A), S23Q (167T>C 168C>A 169T>G), C24Y (171G>A 172T>C), T31A (191A>G 193T>C), K35R (203A>C 204A>G), I37V (209A>G 211T>A), F38L (214T>G), C40I (218T>A 219G>T), K44R (230A>C 231A>G 232A>T), V48A (243T>C 244C>T), A58S (272G>T 274A>C), T59M (276C>T 277A>G), Y60I (278T>A 279A>T), I62L (284A>C 286C>T), S63T (288G>C 289T>C), L65C (294T>G 295A>C), Q67V (299C>G 300A>T 301A>T), I68V (302A>G 304A>G), S71K (311T>A 312C>A 313T>A), N74E (320A>G 322T>G), I75V (323A>G 325C>A), T81S (341A>T 343C>T), L88A (362C>G 363T>C 364C>T), A90T (368G>A 370T>G), Q91E (371C>G), R92K (374C>A 375G>A 376T>G), L94I (380T>A 382G>T), Q95L (383C>T 384A>T 385A>G), T96A (386A>G), N99D (395A>G), L101I (401C>A 403A>T), Y102del (404_406delTAT), F104I (410T>A 412T>A), K105Q (413A>C 415A>G), C106A (416T>G 417G>C 418T>A), Q107H (421A>T), V108A (423T>C 424T>G), T113K (438C>A), I115V (443A>G 445C>G), K116G (446A>G 447A>G 448G>T), S118D (452A>G 453G>A 454C>T), Q119I (455C>A 456A>T 457G>C), E120R (458G>A 459A>G), T121K (462C>A 463T>G), K123E (467A>G), K123_K124insH (469_470insCAT), K124G (470A>G 471A>G), A125V (474C>T 475C>A), L128V (482T>G 484G>C), I129S (485A>T 486T>C), M135V (503A>G 505G>C), I136C (506A>T 507T>G), L138M (512C>A 514A>G), T140R (519C>G 520C>A), K142Q (524A>C 526A>G), I144L (530A>C), D145R (533G>C 534A>G 535T>C), T146P (536A>C 538C>T), K147D (539A>G 541A>T), A148Y (542G>T 543C>A 544A>C), K150T (549A>C 550A>G), I151M (553A>G), V152F (554G>T), I154L (560A>T 562A>G), I162S (584A>T 585T>C 586A>C), D163R (587G>A 588A>G 589T>A), N164G (590A>G 591A>G), L166K (596T>A 597T>A), K168Q (602A>C 604A>G), Q170Y (608C>T 610A>T), E174Q (620G>C 622A>G), F175L (623T>C), F176L (628T>G), E177P (629G>C 630A>C), S178P (632A>C 633G>C 634T>A), H179K (635C>A 637T>G), V180I (638G>A), I183G (647A>G 648T>G), L184V (650T>G 652A>T), L185F (655A>C), V189M (665G>A 667T>G), S191P (671T>C), R192E (674A>G 675G>A), V193A (678T>C), I194L (680A>C 682C>T), N195E (683A>G 685T>G), T196I (687C>T 688A>T), S197T (689T>A), Q198R (692C>A 693A>G 694G>A), V199K (695G>A 696T>A 697C>G), R202N (705G>A 706A>C), D203K (707G>A 709T>G), K206R (717A>G 718A>G), N211R (731A>C 732A>G), A212D (735C>A), R220K (759G>A), Y222F (765A>T), I224V (770A>G 772A>C), K227D (779A>G 781G>T), N229E (785A>G 787T>A), D230E (790T>A), F231W (792T>G 793T>G), A233L (797G>C 798C>T 799A>T), F237C (810T>G), D241E (823C>G), H242T (824C>A 825A>C 826C>T), S244A (830A>G 831G>C), L245I (833C>A), T248S (843C>G 844A>T), L249V (845C>G 847A>C), C252V (854T>G 855G>T 856T>G), Q257K (869C>A), V260W (878G>T 879T>G 880A>G), M262T (885T>C 886G>T), E263D (889A>T), C264K (890T>A 891G>A 892T>G), L265M (893T>A 895A>G), T266R (896A>C 897C>G 898A>C), N267S (900A>G 901T>C), N269D (905A>G), F270H (908T>C 909T>A), A272V (915C>T), S274A (920A>G 921G>C 922C>A), I275T (924T>C 925C>T), S280D (938T>G 939C>A), Q282N (944C>A 946A>C), E283T (947G>A 948A>C 949A>T), F286I (956T>A 958T>C), K289R (965A>C 966A>G), D293S (977G>T 978A>C), K295S (983A>T 984A>C), T296S (986A>T), I298V (992A>G 994A>G), L300I (998C>A 1000T>C), S301T (1001T>A 1003A>T), N303D (1007A>G), I314V (1040A>G 1042A>C), H324T (1070C>A 1071A>C), N325Q (1073A>C 1075T>A), R326P (1076A>C 1077G>C), I330L (1088A>C 1090A>G), I344V (1130A>G), I349V (1145A>G 1147A>C), E351K (1151G>A 1153A>G), E353D (1159A>T), L354E (1160T>G 1161T>A), P355R (1163C>A 1164C>G 1165A>G), K356M (1167A>T), A358F (1172G>T 1173C>T), E359D (1177A>C), L360I (1178T>A), E361Q (1181G>C 1183A>G), T362K (1185C>A 1186G>A), T366V (1196A>G 1197C>T), K367V (1199A>G 1200A>T), D369E (1207T>G), M371L (1211A>C), E373A (1218A>C 1219G>C), I375V (1223A>G), V376A (1227T>C), I379L (1235A>C 1237A>G) | | | | | | | | |

| | Begin | End | Coverage | Score | Concordance | Matches | Identities | I/D/M/F* | Stop Codons |
|---|---|---|---|---|---|---|---|---|---|
| **NT** | **112** | **1237** | **84.0%** | **434** | **19.4%** | **1123 (99.5%)** | **676 (59.9%)** | **3/3** | |
| Codon mutations: | GAA7GAT (121A>T), GAC9GCT (126A>C 127C>T), CAT10GAG (128C>G 130T>G), GGC12GGG (136C>G), CTC13ATT (137C>A 139C>T), AAG14AAA (142G>A), AAA15GAT (143A>G 145A>T), AAT16GAC (146A>G 148T>C), ATC17CTT (149A>C 151C>T), CTA18CTT (154A>T), AAG19AGA (156A>G 157G>A), GGT20GGG (160T>G), ATA21ATT (163A>T), TAT22TAC (166T>C), TCT23CAG (167T>C 168C>A 169T>G), TGT24TAC (171G>A 172T>C), GGA25GGG (175A>G), TTT26TTC (178T>C), CCT29CCA (187T>A), TCA30TCT (190A>T), ACT31GCC (191A>G 193T>C), AAA35CGA (203A>C 204A>G), GCC36GCT (208C>T), ATT37GTA (209A>G 211T>A), TTT38TTG (214T>G), CCA39CCT (217A>T), TGT40ATT (218T>A 219G>T), ATT41ATA (223T>A), TCT42TCG (226T>G), AAA44CGT (230A>C 231A>G 232A>T), GTG46GTC (238G>C), ATA47ATT (241A>T), GTC48GCT (243T>C 244C>T), CAA49CAG (247A>G), TCG52TCC (256G>C), GGA53GGT (259A>T), ACA54ACC (262A>C), ACA57ACC (271A>C), GCA58TCC (272G>T 274A>C), ACA59ATG (276C>T 277A>G), TAT60ATT (278T>A 279A>T), GCA61GCT (283A>T), ATC62CTT (284A>C 286C>T), AGT63ACC (288G>C 289T>C), TTA65TGC (294T>G 295A>C), CAG66CAA (298G>A), CAA67GTT (299C>G 300A>T 301A>T), ATA68GTG (302A>G 304A>G), ACT70ACC (310T>C), TCT71AAA (311T>A 312C>A 313T>A), AAT74GAG (320A>G 322T>G), ATC75GTA (323A>G 325C>A), GCT77GCA (331T>A), ATC79ATA (337C>A), ACC81TCT (341A>T 343C>T), CCA82CCT (346A>T), CGT84CGG (352T>G), GAA85GAG (355A>G), TTG86CTT (356T>C 358G>T), GCT87GCA (361T>A), CTC88GCT (362C>G 363T>C 364C>T), GCT90ACG (368G>A 370T>G), CAA91GAA (371C>G), CGT92AAG (374C>A 375G>A 376T>G), GTA93GTG (379A>G), TTG94ATT (380T>A 382G>T), CAA95TTG (383C>T 384A>T 385A>G), ACA96GCA (386A>G), GGG98GGA (394G>A), AAT99GAT (395A>G), TAT100TAC (400T>C), CTA101ATT (401C>A 403A>T), TAT102del (404_406delTAT), AAT103AAC (409T>C), TTT104ATA (410T>A 412T>A), AAA105CAG (413A>C 415A>G), TGT106GCA (416T>G 417G>C 418T>A), CAA107CAT (421A>T), GTT108GCG (423T>C 424T>G), ACA113AAA (438C>A), AGT114AGC (442T>C), ATC115GTG (443A>G 445C>G), AAG116GGT (446A>G 447A>G 448G>T), AGC118GAT (452A>G 453G>A 454C>T), CAG119ATC (455C>A 456A>T 457G>C), GAA120AGA (458G>A 459A>G), ACT121AAG (462C>A 463T>G), AAG123GAG (467A>G), AAG123_AAA124insCAT (469_470insCAT), AAA124GGA (470A>G 471A>G), GCC125GTA (474C>T 475C>A), CAG126CAA (478G>A), TTG128GTC (482T>G 484G>C), ATT129TCT (485A>T 486T>C), CGA134AGA (500C>A), ATG135GTC (503A>G 505G>C), ATT136TGT (506A>T 507T>G), GAT137GAC (511T>C), CTA138ATG (512C>A 514A>G), CTC139TTG (515C>T 517C>G), ACC140AGA (519C>G 520C>A), CGA141AGA (521C>A), AAA142CAG (524A>C 526A>G), AGT143TCT (527A>T 528G>C), ATC144CTC (530A>C), GAT145CGC (533G>C 534A>G 535T>C), ACC146CCT (536A>C 538C>T), AAA147GAT (539A>G 541A>T), GCA148TAC (542G>T 543C>A 544A>C), AAA150ACG (549A>C 550A>G), ATA151ATG (553A>G), GTT152TTT (554G>T), ATA154TTG (560A>T 562A>G), GAT155GAC (565T>C), GAA156GAG (568A>G), GAG159GAA (577G>A), TTG161CTC (581T>C 583G>C), ATA162TCC (584A>T 585T>C 586A>C), GAT163AGA (587G>A 588A>G 589T>A), AAT164GGT (590A>G 591A>G), TTT165TTC (595T>C), TTG166AAG (596T>A 597T>A), AAA168CAG (602A>C 604A>G), ATA169ATT (607A>T), CAA170TAT (608C>T 610A>T), GAA174CAG (620G>C 622A>G), TTT175CTT (623T>C), TTT176TTG (628T>G), GAA177CCA (629G>C 630A>C), AGT178CCA (632A>C 633G>C 634T>A), CAT179AAG (635C>A 637T>G), GTC180ATC (638G>A), GTT182GTG (646T>G), ATT183GGT (647A>G 648T>G), TTA184GTT (650T>G 652A>T), TTA185TTC (655A>C), TCA186TCT (658A>T), GCA187GCC (661A>C), ACC188ACT (664C>T), GTT189ATG (665G>A 667T>G), TCA191CCA (671T>C), AGG192GAG (674A>G 675G>A), GTT193GCT (678T>C), ATC194CTT (680A>C 682C>T), AAT195GAG (683A>G 685T>G), ACA196ATT (687C>T 688A>T), TCT197ACT (689T>A), CAG198AGA (692C>A 693A>G 694G>A), GTC199AAG (695G>A 696T>A 697C>G), TTT200TTC (700T>C), AGA202AAC (705G>A 706A>C), GAT203AAG (707G>A 709T>G), GTA205GTG (715A>G), AAA206AGG (717A>G 718A>G), CTG208CTT (724G>T), GTA209GTG (727A>G), AAA210AAG (730A>G), AAT211CGT (731A>C 732A>G), GCT212GAT (735C>A), GAA213GAG (739A>G), TTG214CTC (740T>C 742G>C), ACA215ACT (745A>T), CTC216CTT (748C>T), GGC218GGT (754C>T), AGG220AAG (759G>A), TAT222TTT (765A>T), ATA224GTC (770A>G 772A>C), GTA226GTC (778A>C), AAG227GAT (779A>G 781G>T), AAA228AAG (784A>G), AAT229GAA (785A>G 787T>A), GAT230GAA (790T>A), TTT231TGG (792T>G 793T>G), AAA232AAG (796A>G), GCA233CTT (797G>C 798C>T 799A>T), TTT237TGT (810T>G), GAC241GAG (823C>G), CAC242ACT (824C>A 825A>C 826C>T), AGC244GCC (830A>G 831G>C), CTC245ATC (833C>A), ACT246ACC (838T>C), CAA247CAG (841A>G), ACA248AGT (843C>G 844A>T), CTA249GTC (845C>G 847A>C), ATA250ATC (850A>C), TGT252GTG (854T>G 855G>T 856T>G), AAT253AAC (859T>C), ACA254ACC (862A>C), CGT255AGG (863C>A 865T>G), AGA256CGT (866A>C 868A>T), CAA257AAA (869C>A), GTA260TGG (878G>T 879T>G 880A>G), ATG262ACT (885T>C 886G>T), GAA263GAT (889A>T), TGT264AAG (890T>A 891G>A 892T>G), TTA265ATG (893T>A 895A>G), ACA266CGC (896A>C 897C>G 898A>C), AAT267AGC (900A>G 901T>C), AGA268CGT (902A>C 904A>T), AAT269GAT (905A>G), TTC270CAC (908T>C 909T>A), GCA272GTA (915C>T), TCA273TCT (919A>T), AGC274GCA (920A>G 921G>C 922C>A), ATC275ACT (924T>C 925C>T), GAT278GAC (934T>C), TCC280GAC (938T>G 939C>A), CAA281CAG (943A>G), CAA282AAC (944C>A 946A>C), GAA283ACT (947G>A 948A>C 949A>T), CGT284AGA (950C>A 952T>A), TTT286ATC (956T>A 958T>C), AAA289CGA (965A>C 966A>G), GAA290GAG (970A>G), TTT291TTC (973T>C), AGA292CGA (974A>C), GAT293TCT (977G>T 978A>C), GGT294GGG (982T>G), AAA295TCA (983A>T 984A>C), ACT296TCT (986A>T), AGA297CGT (989A>C 991A>T), ATA298GTG (992A>G 994A>G), CTT299CTC (997T>C), CTT300ATC (998C>A 1000T>C), TCA301ACT (1001T>A 1003A>T), AAT303GAT (1007A>G), CTA304CTT (1012A>T), TTA305TTG (1015A>G), GCA306GCT (1018A>T), AGA307CGT (1019A>C 1021A>T), GGC308GGT (1024C>T), GTT311GTA (1033T>A), CAA312CAG (1036A>G), ATA314GTC (1040A>G 1042A>C), TCT315TCC (1045T>C), TTA316CTT (1046T>C 1048A>T), GTT317GTC (1051T>C), ATC318ATT (1054C>T), AAT319AAC (1057T>C), GAT321GAC (1063T>C), TTG322CTG (1064T>C), CAT324ACT (1070C>A 1071A>C), AAT325CAA (1073A>C 1075T>A), AGA326CCA (1076A>C 1077G>C), TAT329TAC (1087T>C), ATA330CTG (1088A>C 1090A>G), AGA335CGT (1103A>C 1105A>T), GGT337GGA (1111T>A), AGA338CGA (1112A>C), GGC340GGA (1120C>A), CGA341AGG (1121C>A 1123A>G), AAA342AAG (1126A>G), GGG343GGT (1129G>T), ATT344GTT (1130A>G), AAT347AAC (1141T>C), ATA349GTC (1145A>G 1147A>C), ACT350ACC (1150T>C), GAA351AAG (1151G>A 1153A>G), GAC352GAT (1156C>T), GAA353GAT (1159A>T), TTA354GAA (1160T>G 1161T>A), CCA355AGG (1163C>A 1164C>G 1165A>G), AAG356ATG (1167A>T), GCT358TTT (1172G>T 1173C>T), GAA359GAC (1177A>C), TTA360ATA (1178T>A), GAA361CAG (1181G>C 1183A>G), ACG362AAA (1185C>A 1186G>A), TAT364TAC (1192T>C), AAT365AAC (1195T>C), ACC366GTC (1196A>G 1197C>T), AAA367GTA (1199A>G 1200A>T), ATA368ATC (1204A>C), GAT369GAG (1207T>G), GAA370GAG (1210A>G), ATG371CTG (1211A>C), GAG373GCC (1218A>C 1219G>C), ATT375GTT (1223A>G), GTT376GCT (1227T>C), TTG378CTC (1232T>C 1234G>C), ATA379CTG (1235A>C 1237A>G) | | | | | | | | |

*: Inserts / Deletes / Misaligned / Frameshifts

#### Analysis details

This analysis was performed with panviral2.64

### NGS Details (UN62): Bracoviriform glomeratae (segment NC_043292.1)

#### Assembly

| | |
|---|---|
| **Coverage Length** | 310 (1 contig(s)) |
| **Depth Of Coverage** | 3029.1 |
| **Number Of Reads** | 8084 |
| **Reads Per Million** | 187.39 rpm (after QC) |
| **Ambiguities** | 0 |
| **Assembly Method** | de novo + reference guided assembly |
| **Consensus Caller** | Bcf Tools |

#### Coverage Map

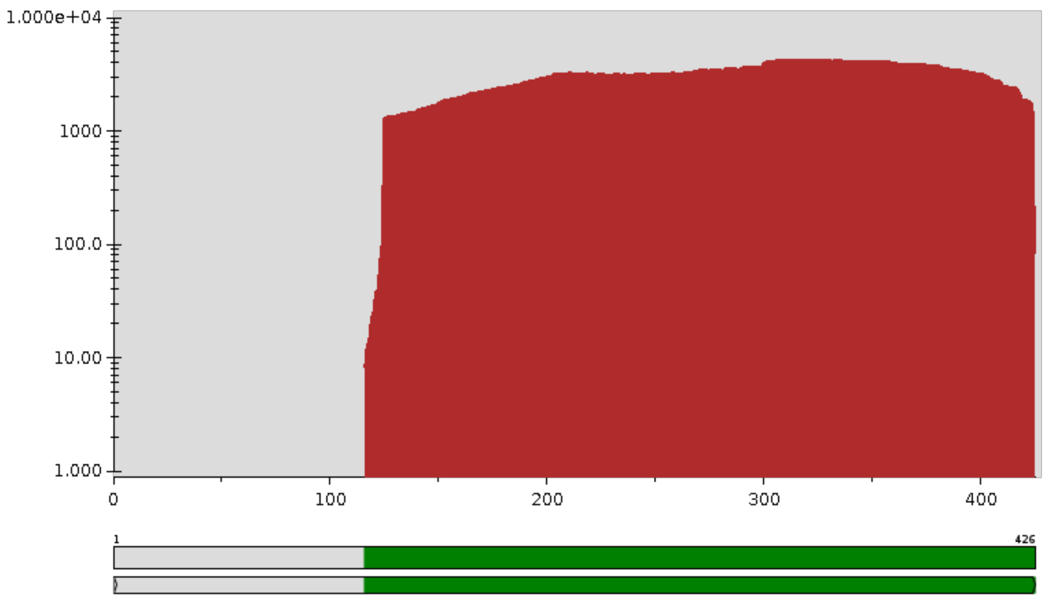

#### Assignment

| | |
|---|---|
| **Type** | Bracoviriform glomeratae (Taxonomy ID: 257816) |
| **Reference Genome** | NC_043292.1 |
| **NT Identity (%)** | 72.5806 |
| **AA Identity (%)** | 84.466 |
| **Number Of Stop Codons** | 1 |
| **Number Of CDS** | 1 |

#### Alignment

| | |
|---|---|
| **Alignment Score** | 280.0 (NT) + 567.0 (AA) = 847.0 |
| **Concordance (%)** | 66.3273 |

| Alignment Method | Global, seeded, nucleotide + amino acids (AGA) |
|---|---|

#### Genome Region

Sequence starts at position 117 and ends at position 426 relative to NC_043292.1 reference sequence.

#### Alignment Detailed Statistics

| | Begin | End | Coverage | Score | Concordance | Matches | Identities | I/D/M/F* | Stop Codons |
|---|---|---|---|---|---|---|---|---|---|
| **NT** | **117** | **426** | **72.8%** | **280** | **45.2%** | **310 (100%)** | **225 (72.6%)** | **0/0** | |
| Mutations: | 126G>T, 129C>A, 132A>G, 136A>G, 137C>G, 138G>C, 150A>C, 153A>G, 155C>G, 156G>A, 159C>A, 166C>A, 168T>G, 169T>C, 172C>A, 174C>G, 177A>G, 183T>A, 184C>A, 189T>C, 198A>G, 201A>C, 207C>T, 214A>G, 219C>T, 222T>C, 223C>A, 225T>G, 228G>T, 235C>A, 237T>G, 243A>T, 246C>G, 255C>T, 261A>G, 262T>C, 264A>G, 265G>A, 273A>G, 278T>C, 279T>C, 284A>G, 285T>A, 288T>G, 289T>C, 300C>T, 301C>T, 303C>G, 309C>T, 312T>G, 315C>T, 322G>T, 324T>C, 329T>C, 333T>C, 336C>T, 339A>G, 345A>C, 347A>G, 348A>G, 349C>A, 351T>A, 360C>T, 363C>T, 366C>T, 375C>T, 378C>T, 384A>C, 385T>C, 387G>C, 390A>G, 391C>A, 392A>G, 393T>A, 394A>C, 396A>G, 399C>A, 400C>A, 402T>G, 404T>C, 405G>T, 406A>C, 414T>G, 416A>T, 420A>T | | | | | | | | |
| **CDS** | | | | | | | | | |
| FK954_p501 | 40 | 142 | 72.5% | 567 | 85.3% | 103 (100%) | 87 (84.5%) | 0/0/0/0 | 1 |
| Protein mutations: | L42F (126G>T), T46G (136A>G 137C>G 138G>C), A52G (155C>G 156G>A), Y57H (169T>C), T72A (214A>G), V89I (265G>A), I93T (278T>C 279T>C), D95G (284A>G 285T>A), A108S (322G>T 324T>C), I110T (329T>C), K116R (347A>G 348A>G), H131R (391C>A 392A>G 393T>A), K132Q (394A>C 396A>G), M135T (404T>C 405G>T), I136L (406A>C), Y139F (416A>T) | | | | | | | | |
| Codon mutations: | TTG42TTT (126G>T), GGC43GGA (129C>A), AAA44AAG (132A>G), ACG46GGC (136A>G 137C>G 138G>C), GGA50GGC (150A>C), AAA51AAG (153A>G), GCG52GGA (155C>G 156G>A), GGC53GGA (159C>A), CGT56AGG (166C>A 168T>G), TAT57CAT (169T>C), CGC58AGG (172C>A 174C>G), AAA59AAG (177A>G), CTT61CTA (183T>A), CGA62AGA (184C>A), GAT63GAC (189T>C), CAA66CAG (198A>G), GGA67GGC (201A>C), ACC69ACT (207C>T), ACT72GCT (214A>G), ATC73ATT (219C>T), CGT74CGC (222T>C), CGT75AGG (223C>A 225T>G), CTG76CTT (228G>T), CGT79AGG (235C>A 237T>G), GGA81GGT (243A>T), GTC82GTG (246C>G), ATC85ATT (255C>T), GGA87GGG (261A>G), TTA88CTG (262T>C 264A>G), GTC89ATC (265G>A), GAA91GAG (273A>G), ATT93ACC (278T>C 279T>C), GAT95GGA (284A>G 285T>A), GTT96GTG (288T>G), TTG97CTG (289T>C), TTC100TTT (300C>T), CTC101TTG (301C>T 303C>G), AAC103AAT (309C>T), GTT104GTG (312T>G), ATC105ATT (315C>T), GCT108TCC (322G>T 324T>C), ATC110ACC (329T>C), TAT111TAC (333T>C), ACC112ACT (336C>T), GAA113GAG (339A>G), GCA115GCC (345A>C), AAA116AGG (347A>G 348A>G), CGT117AGA (349C>A 351T>A), GTC120GTT (360C>T), ACC121ACT (363C>T), GCC122GCT (366C>T), GTC125GTT (375C>T), GTC126GTT (378C>T), GCA128GCC (384A>C), TTG129CTC (385T>C 387G>C), AAA130AAG (390A>G), CAT131AGA (391C>A 392A>G 393T>A), AAA132CAG (394A>C 396A>G), GGC133GGA (399C>A), CGT134AGG (400C>A 402T>G), ATG135ACT (404T>C 405G>T), ATC136CTC (406A>C), GGT138GGG (414T>G), TAT139TTT (416A>T), GGA140GGT (420A>T) | | | | | | | | |
| **Proteins** | | | | | | | | | |
| putative histone 4 (YP_009665791.1) | 40 | 142 | 72.5% | 567 | 85.3% | 103 (100%) | 87 (84.5%) | 0/0/0/0 | 1 |
| Protein mutations: | L42F (126G>T), T46G (136A>G 137C>G 138G>C), A52G (155C>G 156G>A), Y57H (169T>C), T72A (214A>G), V89I (265G>A), I93T (278T>C 279T>C), D95G (284A>G 285T>A), A108S (322G>T 324T>C), I110T (329T>C), K116R (347A>G 348A>G), H131R (391C>A 392A>G 393T>A), K132Q (394A>C 396A>G), M135T (404T>C 405G>T), I136L (406A>C), Y139F (416A>T) | | | | | | | | |
| Codon mutations: | TTG42TTT (126G>T), GGC43GGA (129C>A), AAA44AAG (132A>G), ACG46GGC (136A>G 137C>G 138G>C), GGA50GGC (150A>C), AAA51AAG (153A>G), GCG52GGA (155C>G 156G>A), GGC53GGA (159C>A), CGT56AGG (166C>A 168T>G), TAT57CAT (169T>C), CGC58AGG (172C>A 174C>G), AAA59AAG (177A>G), CTT61CTA (183T>A), CGA62AGA (184C>A), GAT63GAC (189T>C), CAA66CAG (198A>G), GGA67GGC (201A>C), ACC69ACT (207C>T), ACT72GCT (214A>G), ATC73ATT (219C>T), CGT74CGC (222T>C), CGT75AGG (223C>A 225T>G), CTG76CTT (228G>T), CGT79AGG (235C>A 237T>G), GGA81GGT (243A>T), GTC82GTG (246C>G), ATC85ATT (255C>T), GGA87GGG (261A>G), TTA88CTG (262T>C 264A>G), GTC89ATC (265G>A), GAA91GAG (273A>G), ATT93ACC (278T>C 279T>C), GAT95GGA (284A>G 285T>A), GTT96GTG (288T>G), TTG97CTG (289T>C), TTC100TTT (300C>T), CTC101TTG (301C>T 303C>G), AAC103AAT (309C>T), GTT104GTG (312T>G), ATC105ATT (315C>T), GCT108TCC (322G>T 324T>C), ATC110ACC (329T>C), TAT111TAC (333T>C), ACC112ACT (336C>T), GAA113GAG (339A>G), GCA115GCC (345A>C), AAA116AGG (347A>G 348A>G), CGT117AGA (349C>A 351T>A), GTC120GTT (360C>T), ACC121ACT (363C>T), GCC122GCT (366C>T), GTC125GTT (375C>T), GTC126GTT (378C>T), GCA128GCC (384A>C), TTG129CTC (385T>C 387G>C), AAA130AAG (390A>G), CAT131AGA (391C>A 392A>G 393T>A), AAA132CAG (394A>C 396A>G), GGC133GGA (399C>A), CGT134AGG (400C>A 402T>G), ATG135ACT (404T>C 405G>T), ATC136CTC (406A>C), GGT138GGG (414T>G), TAT139TTT (416A>T), GGA140GGT (420A>T) | | | | | | | | |

*: Inserts / Deletes / Misaligned / Frameshifts

#### Analysis details

This analysis was performed with panviral2.64

### NGS Details (UN62): Potato leafroll virus

#### Assembly

| | |
|---|---|
| **Coverage Length** | 5822 (1 contig(s)) |
| **Depth Of Coverage** | 75.3 |
| **Number Of Reads** | 3283 |
| **Reads Per Million** | 76.10 rpm (after QC) |
| **Ambiguities** | 0 |
| **Assembly Method** | read mapping against reference + variant calling |
| **Consensus Caller** | Bcf Tools |

#### Coverage Map

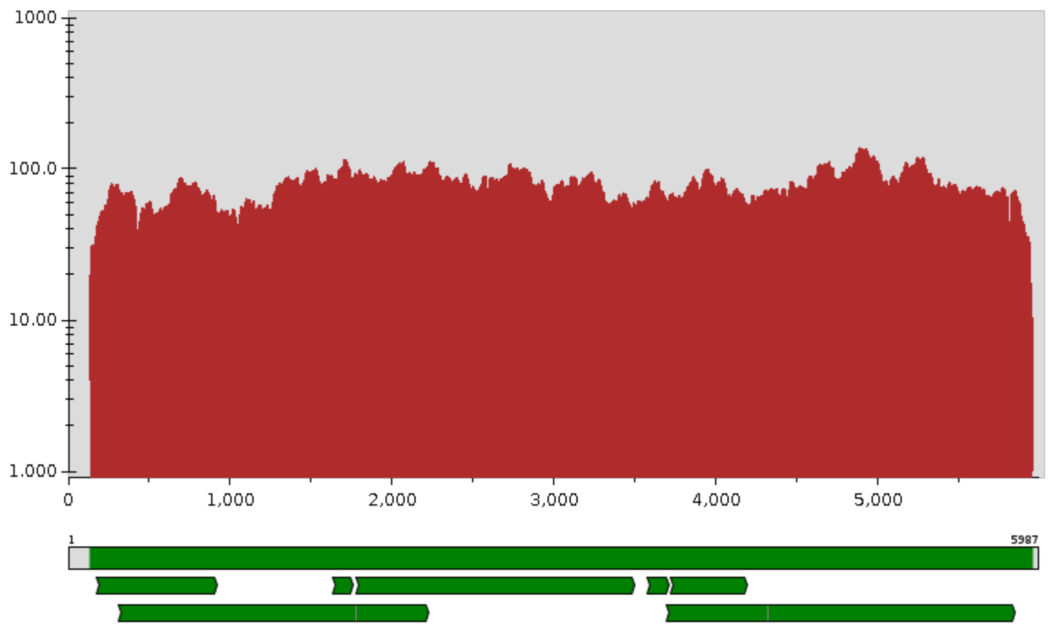

#### Assignment

| | |
|---|---|
| **Type** | Potato leafroll virus (Taxonomy ID: 12045) |
| **Reference Genome** | NC_001747.1 |
| **NT Identity (%)** | 98.1106 |
| **AA Identity (%)** | 97.2178 |
| **Number Of Stop Codons** | 9 |
| **Number Of CDS** | 8 |

#### Alignment

| | |
|---|---|
| **Alignment Score** | 11168.0 (NT) + 20158.0 (AA) = 31326.0 |
| **Concordance (%)** | 95.9828 |

| Alignment Method | Global, seeded, nucleotide + amino acids (AGA) |
|---|---|

#### Genome Region

Sequence starts at position 135 and ends at position 5956 relative to NC_001747.1 reference sequence.

#### Alignment Detailed Statistics

| | Begin | End | Coverage | Score | Concordance | Matches | Identities | I/D/M/F* | Stop Codons |
|---|---|---|---|---|---|---|---|---|---|
| **NT** | **135** | **5956** | **97.2%** | **11168** | **96.2%** | **5818 (99.9%)** | **5712 (98.0%)** | **4/4** | |
| Mutations: | 191A>C, 221T>C, 228C>T, 269T>C, 276G>A, 300C>T, 343C>T, 462T>A, 476_477insTT, 479_480delTT, 492T>C, 525C>T, 561C>T, 600T>C, 665A>G, 699A>C, 822C>T, 825T>G, 931A>G, 958G>A, 1054C>T, 1132T>G, 1147C>T, 1171C>T, 1204A>G, 1231T>C, 1319G>A, 1356G>A, 1465C>T, 1474G>A, 1534G>A, 1577G>T, 1593G>A, 1611A>G, 1716A>G, 1754C>T, 1779A>G, 1894C>A, 2007T>C, 2051G>A, 2154C>T, 2265A>G, 2283G>A, 2316T>G, 2470G>A, 2555C>T, 2586A>G, 2640C>T, 2649C>T, 2729A>G, 2856G>A, 2877C>T, 2898T>C, 2931T>C, 3000T>C, 3033C>T, 3036C>T, 3069C>T, 3117C>T, 3195C>T, 3321C>T, 3379A>G, 3420A>G, 3437A>G, 3441C>T, 3619C>T, 3652T>A, 3721T>G, 3748T>G, 3750C>A, 3777G>T, 3788T>A, 3939A>T, 4031A>G, 4132A>T, 4214T>G, 4433C>T, 4511T>C, 4520G>T, 4521T>G, 4661T>C, 4700C>T, 4733C>G, 4817T>C, 4883C>T, 4886T>C, 4895T>C, 4898C>T, 4949G>A, 4964A>G, 4979C>T, 5062C>T, 5072C>T, 5095G>A, 5129G>A, 5162C>T, 5199C>T, 5215G>A, 5375T>G, 5385G>A, 5432T>C, 5474G>A, 5493G>A, 5517A>G, 5689_5690insGA, 5692_5693delGT, 5741T>G, 5748A>G, 5781G>A, 5925T>C | | | | | | | | |
| **CDS** | | | | | | | | | |
| PLRVgp1 | 1 | 248 | 100% | 1429 | 82.5% | 248 (99.6%) | 241 (96.8%) | 1/0/2/2 | 1 |
| Protein mutations: | Q6P (191A>C), F16S (221T>C), L32P (269T>C), H57Y (343C>T), G101_L102insX (476_477insTT), Q164R (665A>G), Q175H (699A>C) | | | | | | | | |
| Codon mutations: | CAG6CCG (191A>C), TTT16TCT (221T>C), CTC18CTT (228C>T), CTT32CCT (269T>C), CTG34CTA (276G>A), GGC42GGT (300C>T), CAT57TAT (343C>T), GCT96GCA (462T>A), GGC101GGT (476_477insTT), GGC101_CTT102insT-C (476_477insTT), CTT102C-- (479_480delTT), ATT106ATC (492T>C), TAC117TAT (525C>T), TAC129TAT (561C>T), CAT142CAC (600T>C), CAA164CGA (665A>G), CAA175CAC (699A>C), CGC216CGT (822C>T), GCT217GCG (825T>G) | | | | | | | | |
| PLRVgp2 | 1 | 1063 | 100% | 7076 | 94.9% | 1063 (99.9%) | 1039 (97.7%) | 1/0/2/2 | 1 |
| Protein mutations: | L52Q (462T>A), W56_A56insV (476_477insTT), L61S (492T>C), T72I (525C>T), T84I (561C>T), M97T (600T>C), K119E (665A>G), N130T (699A>C), A171V (822C>T), L172R (825T>G), A337T (1319G>A), G349E (1356G>A), V423F (1577G>T), R428K (1593G>A), D434G (1611A>G), E469G (1716A>G), Q529K (1894C>A), G581E (2051G>A), A721T (2470G>A), A749V (2555C>T), N807S (2729A>G), I1024V (3379A>G), E1043G (3437A>G) | | | | | | | | |
| Codon mutations: | TTC12TTT (343C>T), CTA52CAA (462T>A), TGG56_GCC56insGTT (476_477insTT), TTA57--A (479_480delTT), TTA61TCA (492T>C), ACA72ATA (525C>T), ACA84ATA (561C>T), ATG97ACG (600T>C), AAA119GAA (665A>G), AAC130ACC (699A>C), GCG171GTG (822C>T), CTA172CGA (825T>G), GAA207GAG (931A>G), CAG216CAA (958G>A), GCC248GCT (1054C>T), ACT274ACG (1132T>G), TTC279TTT (1147C>T), TCC287TCT (1171C>T), CTA298CTG (1204A>G), ATT307ATC (1231T>C), GCC337ACC (1319G>A), GGA349GAA (1356G>A), CCC385CCT (1465C>T), CCG388CCA (1474G>A), TCG408TCA (1534G>A), GTC423TTC (1577G>T), AGA428AAA (1593G>A), GAC434GGC (1611A>G), GAG469GGG (1716A>G), CTA482TTA (1754C>T), GCA490GCG (1779A>G), CAA529AAA (1894C>A), TTT566TTC (2007T>C), GGA581GAA (2051G>A), CAC615CAT (2154C>T), TCA652TCG (2265A>G), GCG658GCA (2283G>A), GTT669GTG (2316T>G), GCA721ACA (2470G>A), GCC749GTC (2555C>T), CTA759CTG (2586A>G), CAC777CAT (2640C>T), AGC780AGT (2649C>T), AAT807AGT (2729A>G), GTG849GTA (2856G>A), CAC856CAT (2877C>T), ACT863ACC (2898T>C), TAT874TAC (2931T>C), CGT897CGC (3000T>C), AAC908AAT (3033C>T), TCC909TCT (3036C>T), GCC920GCT (3069C>T), TCC936TCT (3117C>T), CTC962CTT (3195C>T), AAC1004AAT (3321C>T), ATT1024GTT (3379A>G), GAA1037GAG (3420A>G), GAG1043GGG (3437A>G), CTC1044CTT (3441C>T) | | | | | | | | |
| PLRVgp3 | 1 | 640 | 100% | 3930 | 91.6% | 640 (99.8%) | 619 (96.6%) | 1/0/2/2 | 1 |
| Protein mutations: | L52Q (462T>A), W56_A56insV (476_477insTT), L61S (492T>C), T72I (525C>T), T84I (561C>T), M97T (600T>C), K119E (665A>G), N130T (699A>C), A171V (822C>T), L172R (825T>G), A337T (1319G>A), G349E (1356G>A), V423F (1577G>T), R428K (1593G>A), D434G (1611A>G), E469G (1716A>G), Q490R (1779A>G), L566S (2007T>C), E581K (2051G>A), T615I (2154C>T) | | | | | | | | |
| Codon mutations: | TTC12TTT (343C>T), CTA52CAA (462T>A), TGG56_GCC56insGTT (476_477insTT), TTA57--A (479_480delTT), TTA61TCA (492T>C), ACA72ATA (525C>T), ACA84ATA (561C>T), ATG97ACG (600T>C), AAA119GAA (665A>G), AAC130ACC (699A>C), GCG171GTG (822C>T), CTA172CGA (825T>G), GAA207GAG (931A>G), CAG216CAA (958G>A), GCC248GCT (1054C>T), ACT274ACG (1132T>G), TTC279TTT (1147C>T), TCC287TCT (1171C>T), CTA298CTG (1204A>G), ATT307ATC (1231T>C), GCC337ACC (1319G>A), GGA349GAA (1356G>A), CCC385CCT (1465C>T), CCG388CCA (1474G>A), TCG408TCA (1534G>A), GTC423TTC (1577G>T), AGA428AAA (1593G>A), GAC434GGC (1611A>G), GAG469GGG (1716A>G), CTA482TTA (1754C>T), CAA490CGA (1779A>G), ATC528ATA (1894C>A), TTA566TCA (2007T>C), GAG581AAG (2051G>A), ACC615ATC (2154C>T) | | | | | | | | |
| Rap1 | 1 | 42 | 100% | 277 | 96.5% | 42 (100%) | 41 (97.6%) | 0/0/0/0 | 1 |
| Protein mutations: | R29G (1716A>G) | | | | | | | | |
| Codon mutations: | AGA29GGA (1716A>G), CTC41CTT (1754C>T) | | | | | | | | |
| ORF3a | 1 | 46 | 100% | 291 | 97.3% | 46 (100%) | 45 (97.8%) | 0/0/0/0 | 1 |
| Protein mutations: | F26L (3652T>A) | | | | | | | | |
| Codon mutations: | TCC15TCT (3619C>T), TTT26TTA (3652T>A) | | | | | | | | |
| PLRVgp4 | 1 | 718 | 100% | 4683 | 93.0% | 718 (99.9%) | 699 (97.2%) | 1/0/2/2 | 2 |
| Protein mutations: | V10G (3721T>G), M19R (3748T>G), A29S (3777G>T), T83S (3939A>T), Y147F (4132A>T), K276N (4520G>T), F277V (4521T>G), A457V (5062C>T), G468E (5095G>A), P503S (5199C>T), R508Q (5215G>A), N561K (5375T>G), E565K (5385G>A), V601I (5493G>A), K609E (5517A>G), E666_R667insX (5689_5690insGA), F683L (5741T>G), T686A (5748A>G), E697K (5781G>A) | | | | | | | | |
| Codon mutations: | GTC10GGC (3721T>G), ATG19AGG (3748T>G), CGA20AGA (3750C>A), GCT29TCT (3777G>T), GTT32GTA (3788T>A), ACC83TCC (3939A>T), ACA113ACG (4031A>G), TAC147TTC (4132A>T), TCT174TCG (4214T>G), GCC247GCT (4433C>T), AAT273AAC (4511T>C), AAG276AAT (4520G>T), TTT277GTT (4521T>G), GGT323GGC (4661T>C), TCC336TCT (4700C>T), GTC347GTG (4733C>G), GGT375GGC (4817T>C), TTC397TTT (4883C>T), CTT398CTC (4886T>C), CCT401CCC (4895T>C), GCC402GCT (4898C>T), ACG419ACA (4949G>A), GAA424GAG (4964A>G), ACC429ACT (4979C>T), GCC457GTC (5062C>T), CAC460CAT (5072C>T), GGA468GAA (5095G>A), ACG479ACA (5129G>A), GAC490GAT (5162C>T), CCA503TCA (5199C>T), CGA508CAA (5215G>A), AAT561AAG (5375T>G), GAA565AAA (5385G>A), ACT580ACC (5432T>C), TTG594TTA (5474G>A), GTC601ATC (5493G>A), AAG609GAG (5517A>G), GAA666GAG (5689_5690insGA), GAA666_CGT667insA-A (5689_5690insGA), CGT667C-- (5692_5693delGT), TTT683TTG (5741T>G), ACT686GCT (5748A>G), GAA697AAA (5781G>A) | | | | | | | | |
| PLRVgp5 | 1 | 209 | 100% | 1409 | 98.2% | 209 (100%) | 204 (97.6%) | 0/0/0/0 | 1 |
| Protein mutations: | V10G (3721T>G), M19R (3748T>G), A29S (3777G>T), T83S (3939A>T), Y147F (4132A>T) | | | | | | | | |
| Codon mutations: | GTC10GGC (3721T>G), ATG19AGG (3748T>G), CGA20AGA (3750C>A), GCT29TCT (3777G>T), GTT32GTA (3788T>A), ACC83TCC (3939A>T), ACA113ACG (4031A>G), TAC147TTC (4132A>T), TCT174TCG (4214T>G) | | | | | | | | |
| PLRVgp6 | 1 | 157 | 100% | 1063 | 96.5% | 157 (100%) | 152 (96.8%) | 0/0/0/0 | 1 |
| Protein mutations: | S2A (3721T>G), C11G (3748T>G 3750C>A), F24Y (3788T>A), Q105R (4031A>G), T139S (4132A>T) | | | | | | | | |
| Codon mutations: | TCA2GCA (3721T>G), TGC11GGA (3748T>G 3750C>A), GCG20GCT (3777G>T), TTC24TAC (3788T>A), ACA74ACT (3939A>T), CAA105CGA (4031A>G), ACG139TCG (4132A>T) | | | | | | | | |
| **Proteins** | | | | | | | | | |
| P0 protein (NP_056746.1) | 1 | 248 | 100% | 1429 | 82.5% | 248 (99.6%) | 241 (96.8%) | 1/0/2/2 | 1 |
| Protein mutations: | Q6P (191A>C), F16S (221T>C), L32P (269T>C), H57Y (343C>T), G101_L102insX (476_477insTT), Q164R (665A>G), Q175H (699A>C) | | | | | | | | |

| | Begin | End | Coverage | Score | Concordance | Matches | Identities | I/D/M/F* | Stop Codons |
|---|---|---|---|---|---|---|---|---|---|
| **NT** | **135** | **5956** | **97.2%** | **11168** | **96.2%** | **5818 (99.9%)** | **5712 (98.0%)** | **4/4** | |
| Codon mutations: | CAG6CCG (191A>C), TTT16TCT (221T>C), CTC18CTT (228C>T), CTT32CCT (269T>C), CTG34CTA (276G>A), GGC42GGT (300C>T), CAT57TAT (343C>T), GCT96GCA (462T>A), GGC101GGT (476_477insTT), GGC101_CTT102insT-C (476_477insTT), CTT102C-- (479_480delTT), ATT106ATC (492T>C), TAC117TAT (525C>T), TAC129TAT (561C>T), CAT142CAC (600T>C), CAA164CGA (665A>G), CAA175CAC (699A>C), CGC216CGT (822C>T), GCT217GCG (825T>G) | | | | | | | | |
| RNA-dependent RNA polymerase (NP_056748.3) | 1 | 1063 | 100% | 7076 | 94.9% | 1063 (99.9%) | 1039 (97.7%) | 1/0/2/2 | 1 |
| Protein mutations: | L52Q (462T>A), W56_A56insV (476_477insTT), L61S (492T>C), T72I (525C>T), T84I (561C>T), M97T (600T>C), K119E (665A>G), N130T (699A>C), A171V (822C>T), L172R (825T>G), A337T (1319G>A), G349E (1356G>A), V423F (1577G>T), R428K (1593G>A), D434G (1611A>G), E469G (1716A>G), Q529K (1894C>A), G581E (2051G>A), A721T (2470G>A), A749V (2555C>T), N807S (2729A>G), I1024V (3379A>G), E1043G (3437A>G) | | | | | | | | |
| Codon mutations: | TTC12TTT (343C>T), CTA52CAA (462T>A), TGG56_GCC56insGTT (476_477insTT), TTA57--A (479_480delTT), TTA61TCA (492T>C), ACA72ATA (525C>T), ACA84ATA (561C>T), ATG97ACG (600T>C), AAA119GAA (665A>G), AAC130ACC (699A>C), GCG171GTG (822C>T), CTA172CGA (825T>G), GAA207GAG (931A>G), CAG216CAA (958G>A), GCC248GCT (1054C>T), ACT274ACG (1132T>G), TTC279TTT (1147C>T), TCC287TCT (1171C>T), CTA298CTG (1204A>G), ATT307ATC (1231T>C), GCC337ACC (1319G>A), GGA349GAA (1356G>A), CCC385CCT (1465C>T), CCG388CCA (1474G>A), TCG408TCA (1534G>A), GTC423TTC (1577G>T), AGA428AAA (1593G>A), GAC434GGC (1611A>G), GAG469GGG (1716A>G), CTA482TTA (1754C>T), GCA490GCG (1779A>G), CAA529AAA (1894C>A), TTT566TTC (2007T>C), GGA581GAA (2051G>A), CAC615CAT (2154C>T), TCA652TCG (2265A>G), GCG658GCA (2283G>A), GTT669GTG (2316T>G), GCA721ACA (2470G>A), GCC749GTC (2555C>T), CTA759CTG (2586A>G), CAC777CAT (2640C>T), AGC780AGT (2649C>T), AAT807AGT (2729A>G), GTG849GTA (2856G>A), CAC856CAT (2877C>T), ACT863ACC (2898T>C), TAT874TAC (2931T>C), CGT897CGC (3000T>C), AAC908AAT (3033C>T), TCC909TCT (3036C>T), GCC920GCT (3069C>T), TCC936TCT (3117C>T), CTC962CTT (3195C>T), AAC1004AAT (3321C>T), ATT1024GTT (3379A>G), GAA1037GAG (3420A>G), GAG1043GGG (3437A>G), CTC1044CTT (3441C>T) | | | | | | | | |
| P1 protein (NP_056747.1) | 1 | 640 | 100% | 3930 | 91.6% | 640 (99.8%) | 619 (96.6%) | 1/0/2/2 | 1 |
| Protein mutations: | L52Q (462T>A), W56_A56insV (476_477insTT), L61S (492T>C), T72I (525C>T), T84I (561C>T), M97T (600T>C), K119E (665A>G), N130T (699A>C), A171V (822C>T), L172R (825T>G), A337T (1319G>A), G349E (1356G>A), V423F (1577G>T), R428K (1593G>A), D434G (1611A>G), E469G (1716A>G), Q490R (1779A>G), L566S (2007T>C), E581K (2051G>A), T615I (2154C>T) | | | | | | | | |
| Codon mutations: | TTC12TTT (343C>T), CTA52CAA (462T>A), TGG56_GCC56insGTT (476_477insTT), TTA57--A (479_480delTT), TTA61TCA (492T>C), ACA72ATA (525C>T), ACA84ATA (561C>T), ATG97ACG (600T>C), AAA119GAA (665A>G), AAC130ACC (699A>C), GCG171GTG (822C>T), CTA172CGA (825T>G), GAA207GAG (931A>G), CAG216CAA (958G>A), GCC248GCT (1054C>T), ACT274ACG (1132T>G), TTC279TTT (1147C>T), TCC287TCT (1171C>T), CTA298CTG (1204A>G), ATT307ATC (1231T>C), GCC337ACC (1319G>A), GGA349GAA (1356G>A), CCC385CCT (1465C>T), CCG388CCA (1474G>A), TCG408TCA (1534G>A), GTC423TTC (1577G>T), AGA428AAA (1593G>A), GAC434GGC (1611A>G), GAG469GGG (1716A>G), CTA482TTA (1754C>T), CAA490CGA (1779A>G), ATC528ATA (1894C>A), TTA566TCA (2007T>C), GAG581AAG (2051G>A), ACC615ATC (2154C>T) | | | | | | | | |
| Replication-associated protein (YP_006355442.1) | 1 | 42 | 100% | 277 | 96.5% | 42 (100%) | 41 (97.6%) | 0/0/0/0 | 1 |
| Protein mutations: | R29G (1716A>G) | | | | | | | | |
| Codon mutations: | AGA29GGA (1716A>G), CTC41CTT (1754C>T) | | | | | | | | |
| protein 3a (YP_009179365.2) | 1 | 46 | 100% | 291 | 97.3% | 46 (100%) | 45 (97.8%) | 0/0/0/0 | 1 |
| Protein mutations: | F26L (3652T>A) | | | | | | | | |
| Codon mutations: | TCC15TCT (3619C>T), TTT26TTA (3652T>A) | | | | | | | | |
| CP read-through protein (NP_056751.2) | 1 | 718 | 100% | 4683 | 93.0% | 718 (99.9%) | 699 (97.2%) | 1/0/2/2 | 2 |
| Protein mutations: | V10G (3721T>G), M19R (3748T>G), A29S (3777G>T), T83S (3939A>T), Y147F (4132A>T), K276N (4520G>T), F277V (4521T>G), A457V (5062C>T), G468E (5095G>A), P503S (5199C>T), R508Q (5215G>A), N561K (5375T>G), E565K (5385G>A), V601I (5493G>A), K609E (5517A>G), E666_R667insX (5689_5690insGA), F683L (5741T>G), T686A (5748A>G), E697K (5781G>A) | | | | | | | | |
| Codon mutations: | GTC10GGC (3721T>G), ATG19AGG (3748T>G), CGA20AGA (3750C>A), GCT29TCT (3777G>T), GTT32GTA (3788T>A), ACC83TCC (3939A>T), ACA113ACG (4031A>G), TAC147TTC (4132A>T), TCT174TCG (4214T>G), GCC247GCT (4433C>T), AAT273AAC (4511T>C), AAG276AAT (4520G>T), TTT277GTT (4521T>G), GGT323GGC (4661T>C), TCC336TCT (4700C>T), GTC347GTG (4733C>G), GGT375GGC (4817T>C), TTC397TTT (4883C>T), CTT398CTC (4886T>C), CCT401CCC (4895T>C), GCC402GCT (4898C>T), ACG419ACA (4949G>A), GAA424GAG (4964A>G), ACC429ACT (4979C>T), GCC457GTC (5062C>T), CAC460CAT (5072C>T), GGA468GAA (5095G>A), ACG479ACA (5129G>A), GAC490GAT (5162C>T), CCA503TCA (5199C>T), CGA508CAA (5215G>A), AAT561AAG (5375T>G), GAA565AAA (5385G>A), ACT580ACC (5432T>C), TTG594TTA (5474G>A), GTC601ATC (5493G>A), AAG609GAG (5517A>G), GAA666GAG (5689_5690insGA), GAA666_CGT667insA-A (5689_5690insGA), CGT667C-- (5692_5693delGT), TTT683TTG (5741T>G), ACT686GCT (5748A>G), GAA697AAA (5781G>A) | | | | | | | | |
| coat protein (NP_056749.1) | 1 | 209 | 100% | 1409 | 98.2% | 209 (100%) | 204 (97.6%) | 0/0/0/0 | 1 |
| Protein mutations: | V10G (3721T>G), M19R (3748T>G), A29S (3777G>T), T83S (3939A>T), Y147F (4132A>T) | | | | | | | | |
| Codon mutations: | GTC10GGC (3721T>G), ATG19AGG (3748T>G), CGA20AGA (3750C>A), GCT29TCT (3777G>T), GTT32GTA (3788T>A), ACC83TCC (3939A>T), ACA113ACG (4031A>G), TAC147TTC (4132A>T), TCT174TCG (4214T>G) | | | | | | | | |
| movement protein (NP_056750.1) | 1 | 157 | 100% | 1063 | 96.5% | 157 (100%) | 152 (96.8%) | 0/0/0/0 | 1 |
| Protein mutations: | S2A (3721T>G), C11G (3748T>G 3750C>A), F24Y (3788T>A), Q105R (4031A>G), T139S (4132A>T) | | | | | | | | |
| Codon mutations: | TCA2GCA (3721T>G), TGC11GGA (3748T>G 3750C>A), GCG20GCT (3777G>T), TTC24TAC (3788T>A), ACA74ACT (3939A>T), CAA105CGA (4031A>G), ACG139TCG (4132A>T) | | | | | | | | |

*: Inserts / Deletes / Misaligned / Frameshifts

#### Analysis details

This analysis was performed with panviral2.64

### NGS Details (UN62): Duamitovirus soch1

#### Assembly

| | |
|---|---|
| **Coverage Length** | 2098 (2 contig(s)) |
| **Depth Of Coverage** | 32.4 |
| **Number Of Reads** | 543 |
| **Reads Per Million** | 12.59 rpm (after QC) |
| **Ambiguities** | 0 |
| **Assembly Method** | de novo + reference guided assembly |
| **Consensus Caller** | Bcf Tools |

#### Coverage Map

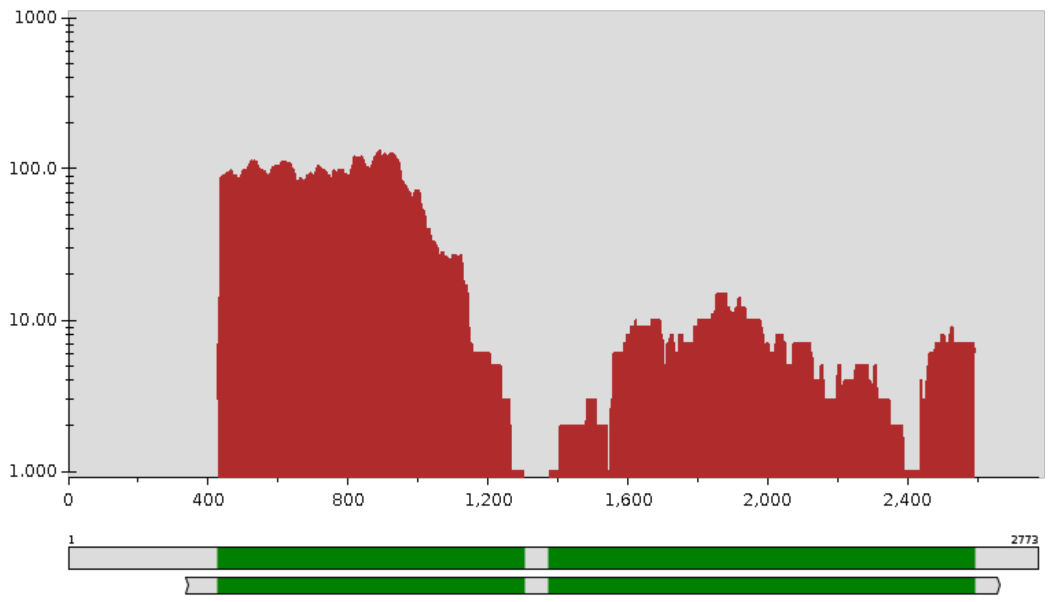

#### Assignment

| | |
|---|---|
| **Type** | Duamitovirus soch1 (Taxonomy ID: 2955838) |
| **Reference Genome** | NC_076524.1 |
| **NT Identity (%)** | 68.3102 |
| **AA Identity (%)** | 69.1535 |
| **Number Of Stop Codons** | 3 |
| **Number Of CDS** | 1 |

#### Alignment

| | |
|---|---|
| **Alignment Score** | 1494.0 (NT) + 3210.0 (AA) = 4704.0 |
| **Concordance (%)** | 53.7195 |

| Alignment Method | Global, seeded, nucleotide + amino acids (AGA) |
|---|---|

#### Genome Region

Sequence starts at position 428 and ends at position 2593 relative to NC_076524.1 reference sequence.

#### Alignment Detailed Statistics

| | Begin | End | Coverage | Score | Concordance | Matches | Identities | I/D/M/F* | Stop Codons |
|---|---|---|---|---|---|---|---|---|---|
| **NT** | **428** | **2593** | **75.7%** | **1494** | **36.1%** | **2089 (99.6%)** | **1427 (68.0%)** | **0/9** | |
| Mutations: | 437A>G, 441A>C, 443C>T, 447C>T, 459C>T, 466C>T, 470C>T, 471G>A, 472C>T, 473T>G, 477T>A, 480A>C, 481C>A, 482A>T, 485A>G, 489A>G, 494T>C, 497T>A, 500T>G, 503A>C, 506T>C, 509C>T, 512T>C, 513G>A, 516T>A, 517C>A, 518T>A, 520A>C, 524G>T, 525A>G, 526C>T, 527T>G, 529C>A, 530C>A, 531T>C, 533A>G, 534C>A, 535G>A, 536C>A, 539G>A, 540A>C, 541A>G, 542G>T, 545G>A, 548C>T, 550A>T, 556T>A, 557C>T, 560A>T, 569C>T, 575G>A, 581C>T, 582G>A, 587A>T, 589G>C, 591T>C, 593A>T, 599A>T, 602C>T, 612T>A, 613C>G, 614C>T, 616T>A, 618C>T, 619G>C, 620A>T, 624G>A, 626A>T, 629A>C, 630C>A, 632A>C, 633C>A, 635T>G, 638G>T, 641G>T, 644T>A, 647G>T, 651C>T, 653C>G, 656A>C, 657A>C, 661C>G, 662G>T, 665T>A, 668C>T, 672C>A, 673G>A, 674T>G, 680C>A, 683C>T, 684A>G, 685A>C, 686A>C, 692G>A, 695T>A, 699G>C, 700C>A, 701A>C, 702A>G, 705C>A, 707T>G, 708A>G, 709G>C, 710G>A, 713A>G, 714G>T, 715A>C, 719A>T, 722C>T, 737T>C, 746C>T, 752A>T, 761G>A, 765G>T, 767T>A, 770G>A, 771G>T, 772T>A, 773T>G, 774T>A, 776G>T, 777A>G, 778A>C, 779A>G, 782T>G, 796G>A, 797T>G, 803A>C, 809T>C, 810G>C, 815C>A, 818A>T, 821T>G, 824C>T, 827A>T, 831C>T, 833T>A, 841G>A, 842G>T, 850T>G, 851A>T, 857G>T, 860G>T, 862A>G, 869T>G, 875G>T, 880T>G, 881C>A, 885C>A, 887T>G, 892A>G, 905A>T, 908C>A, 909T>C, 910T>A, 916G>A, 918G>T, 919G>C, 923T>A, 926A>T, 927A>C, 929G>T, 934T>A, 935T>G, 938G>T, 942T>A, 943C>A, 948C>A, 971C>A, 972C>T, 975A>C, 976A>T, 977T>C, 983T>C, 990A>C, 992G>C, 993T>C, 994C>G, 995A>G, 997A>T, 998C>T, 999G>C, 1000G>A, 1001T>A, 1003T>A, 1004A>T, 1005G>T, 1007C>G, 1010A>G, 1016C>T, 1017A>G, 1018A>T, 1019A>T, 1022T>A, 1025T>C, 1026C>G, 1027A>G, 1030A>C, 1031G>A, 1034T>G, 1035A>G, 1036G>C, 1044T>G, 1046C>T, 1049G>A, 1050T>G, 1051T>A, 1052T>A, 1054C>A, 1055A>C, 1061T>C, 1067T>G, 1073G>A, 1076A>G, 1083C>A, 1088G>T, 1091C>G, 1099G>T, 1104A>G, 1105C>T, 1106A>G, 1115T>G, 1121A>C, 1122T>A, 1124C>T, 1127G>A, 1130C>T, 1133C>G, 1142A>C, 1145C>T, 1148G>T, 1151G>T, 1152T>C, 1154G>T, 1160C>T, 1161A>C, 1164T>A, 1165C>T, 1166T>G, 1169C>T, 1172C>A, 1174T>A, 1175T>C, 1178T>A, 1181A>T, 1184C>T, 1185A>T, 1186G>C, 1187G>T, 1188G>A, 1189C>A, 1191G>A, 1192C>A, 1193A>T, 1196T>C, 1197_1199delGAA, 1205A>T, 1211C>T, 1212T>G, 1213T>A, 1217T>C, 1220C>T, 1223A>T, 1224C>T, 1225A>T, 1226C>A, 1232G>A, 1236A>C, 1237G>A, 1238T>A, 1241C>A, 1247A>C, 1248T>C, 1249G>T, 1250G>T, 1253T>C, 1256A>T, 1258C>G, 1265G>A, 1266A>G, 1268T>G, 1269C>G, 1271T>C, 1274T>C, 1276G>A, 1280G>A, 1283G>T, 1292T>G, 1376C>G, 1379C>T, 1382T>C, 1385T>C, 1388C>T, 1389G>A, 1390A>T, 1403A>T, 1406C>A, 1409G>A, 1412T>A, 1414A>G, 1415A>T, 1416A>C, 1418A>T, 1424A>T, 1427C>T, 1430G>T, 1433T>A, 1437G>A, 1445G>A, 1446A>C, 1447A>G, 1448A>T, 1451T>G, 1455T>C, 1456T>A, 1458G>T, 1459C>A, 1461T>C, 1465A>G, 1466G>A, 1468A>T, 1469A>T, 1472G>A, 1474A>G, 1475C>A, 1477T>C, 1478T>G, 1481A>G, 1482A>T, 1483A>C, 1486G>C, 1489A>T, 1490T>C, 1502A>C, 1505G>A, 1508A>G, 1526G>A, 1538G>A, 1541T>C, 1545C>T, 1547T>C, 1553T>C, 1554A>T, 1555T>G, 1556G>C, 1559A>G, 1566C>A, 1567C>G, 1571C>A, 1572T>C, 1580C>A, 1589T>C, 1593_1597delAGCTC, 1598T>A, 1607T>C, 1613G>T, 1614A>T, 1617C>T, 1619T>G, 1620G>A, 1622T>C, 1623A>G, 1624A>T, 1626A>C, 1627A>C, 1628A>T, 1632T>A, 1634A>G, 1635A>G, 1636C>T, 1637A>C, 1643A>G, 1647T>A, 1648A>G, 1655A>T, 1661T>C, 1662C>T, 1664G>A, 1667T>A, 1670G>A, 1676T>G, 1679G>A, 1682C>T, 1684T>A, 1686C>T, 1691T>A, 1694T>A, 1698G>T, 1700A>C, 1703T>A, 1706C>T, 1707T>G, 1710T>C, 1712G>T, 1715T>C, 1721C>T, 1727A>G, 1734T>C, 1739A>T, 1744G>A, 1745C>A, 1746A>T, 1748A>T, 1754T>C, 1758G>A, 1759G>C, 1760A>G, 1761T>A, 1765A>C, 1766C>T, 1767T>A, 1769T>C, 1772A>T, 1777G>C, 1778G>T, 1787T>C, 1788T>C, 1790A>G, 1791C>T, 1819G>C, 1820T>A, 1821A>G, 1823A>G, 1829C>G, 1830A>C, 1831A>G, 1832T>G, 1833G>C, 1840A>G, 1847A>G, 1848T>C, 1856A>G, 1859C>T, 1860A>G, 1861A>G, 1865C>T, 1867A>C, 1871T>C, 1874G>T, 1875delA, 1878C>G, 1880A>T, 1886T>G, 1887T>A, 1888T>G, 1889G>A, 1895T>C, 1898T>G, 1901T>C, 1910T>C, 1919G>T, 1925G>A, 1934T>C, 1937A>G, 1938T>G, 1939C>A, 1940A>C, 1950G>T, 1952T>C, 1953T>C, 1955A>T, 1964T>C, 1967C>T, 1976T>A, 1980T>A, 1981T>C, 1982A>T, 1985T>C, 1989C>A, 1991T>G, 1994T>A, 2000T>G, 2003A>T, 2006T>G, 2009C>T, 2010G>C, 2011C>A, 2012C>A, 2020T>C, 2028G>T, 2029A>C, 2030T>G, 2034A>T, 2035A>C, 2036C>A, 2038T>C, 2042A>G, 2043G>A, 2050A>C, 2051A>C, 2053G>T, 2054A>T, 2057G>T, 2058G>T, 2063T>G, 2069G>A, 2071A>T, 2074A>G, 2075G>A, 2078A>T, 2082G>T, 2084G>T, 2087C>T, 2090A>G, 2091T>A, 2093C>G, 2094A>T, 2099C>T, 2103A>T, 2104G>C, 2105C>T, 2108G>A, 2117G>A, 2118C>A, 2120T>A, 2123A>G, 2129T>G, 2133C>G, 2134A>C, 2135T>A, 2136C>A, 2137T>G, 2138T>C, 2141G>A, 2145G>C, 2146T>A, 2147T>C, 2148C>A, 2156C>T, 2157T>C, 2160C>T, 2166T>C, 2167A>T, 2173G>A, 2174G>T, 2183A>G, 2186G>A, 2187G>T, 2188G>T, 2191T>A, 2192G>A, 2195C>T, 2204C>T, 2206A>T, 2213A>G, 2216G>A, 2223G>A, 2226A>G, 2227G>A, 2231C>T, 2236T>G, 2237T>G, 2240A>G, 2248A>C, 2250G>C, 2251T>C, 2259T>A, 2263A>G, 2264G>A, 2265A>C, 2276G>T, 2279T>G, 2280C>T, 2288C>T, 2297A>G, 2298G>A, 2299T>G, 2300T>A, 2306A>C, 2309A>T, 2315T>C, 2322C>T, 2326G>T, 2328C>A, 2329A>G, 2336A>G, 2337G>A, 2346T>C, 2348A>G, 2351G>A, 2355C>G, 2358A>T, 2359A>C, 2360C>T, 2372G>T, 2373A>G, 2375C>T, 2376T>G, 2377G>C, 2378T>A, 2379A>G, 2380T>C, 2381G>A, 2382A>T, 2383A>C, 2384C>A, 2385C>G, 2390T>A, 2394T>A, 2400A>G, 2402G>T, 2403A>C, 2404G>A, 2405T>G, 2406C>T, 2408C>G, 2411C>G, 2414C>A, 2415T>G, 2416C>T, 2417C>T, 2423A>G, 2426C>T, 2427G>A, 2428C>G, 2429A>T, 2431C>A, 2438G>A, 2441A>G, 2442A>C, 2444G>A, 2445C>T, 2446T>C, 2447C>G, 2448G>A, 2450A>T, 2451A>G, 2452C>A, 2453T>A, 2456T>C, 2457T>C, 2459T>G, 2460G>A, 2462T>A, 2465A>G, 2466A>G, 2477C>T, 2478C>T, 2489A>G, 2492T>C, 2495C>T, 2504T>A, 2510T>C, 2514A>G, 2518C>T, 2519C>G, 2525C>T, 2528T>A, 2532C>A, 2533A>G, 2536G>A, 2537G>T, 2541T>C, 2543T>G, 2551A>G, 2552C>T, 2555T>C, 2556G>T, 2557T>C, 2561T>C, 2568A>G, 2571A>C, 2572A>G, 2582A>G, 2585G>A | | | | | | | | |
| **CDS** | | | | | | | | | |
| RdRp | 32 | 753 | 90.0% | 3210 | 65.4% | 697 (99.7%) | 482 (69.0%) | 0/2/2/2 | 3 |
| Protein mutations: | I36L (441A>C 443C>T), L38F (447C>T), A44V (466C>T), A46M (471G>A 472C>T 473T>G), L48M (477T>A), T49H (480A>C 481C>A 482A>T), I52V (489A>G), V60I (513G>A), S61K (516T>A 517C>A 518T>A), K62T (520A>C), T64V (525A>G 526C>T 527T>G), T65K (529C>A 530C>A), R67K (534C>A 535G>A 536C>A), K69R (540A>C 541A>G 542G>T), Y72F (550A>T), F74Y (556T>A 557C>T), A83T (582G>A), C85S (589G>C), F94Y (616T>A), R95S (618C>T 619G>C 620A>T), E97N (624G>A 626A>T), E98D (629A>C), P99T (630C>A 632A>C), L100M (633C>A 635T>G), S109C (661C>G 662G>T), R113K (672C>A 673G>A 674T>G), K117A (684A>G 685A>C 686A>C), A122H (699G>C 700C>A 701A>C), I123V (702A>G), R125A (708A>G 709G>C 710G>A), D127S (714G>T 715A>C), E128D (719A>T), A144S (765G>T 767T>A), V146* (771G>T 772T>A 773T>G), L147I (774T>A 776G>T), K148A (777A>G 778A>C 779A>G), S154K (796G>A 797T>G), E159Q (810G>C), P166S (831C>T 833T>A), G169D (841G>A 842G>T), L172R (850T>G 851A>T), E176G (862A>G), I178M (869T>G), F182* (880T>G 881C>A), H184K (885C>A 887T>G), Q186R (892A>G), F192H (909T>C 910T>A), R194Q (916G>A), G195S (918G>T 919G>C), M198L (927A>C 929G>T), I200K (934T>A 935T>G), S203N (942T>A 943C>A), Q205K (948C>A), N214L (975A>C 976A>T 977T>C), S220R (993T>C 994C>G 995A>G), Y221F (997A>T 998C>T), G222Q (999G>C 1000G>A 1001T>A), L223Y (1003T>A 1004A>T), A224S (1005G>T 1007C>G), K228V (1017A>G 1018A>T 1019A>T), Q231G (1026C>G 1027A>G), E232A (1030A>C 1031G>A), S234A (1035A>G 1036G>C), F237V (1044T>G 1046C>T), F239E (1050T>G 1051T>A 1052T>A), P240H (1054C>A 1055A>C), L250I (1083C>A), W255L (1099G>T), T257V (1104A>G 1105C>T 1106A>G), I260M (1115T>G), F263I (1122T>A 1124C>T), M272I (1151G>T), K276Q (1161A>C), S277M (1164T>A 1165C>T 1166T>G), F279L (1172C>A), F280Y (1174T>A 1175T>C), L282F (1181A>T), R284S (1185A>T 1186G>C 1187G>T), A285N (1188G>A 1189C>A), A286N (1191G>A 1192C>A 1193A>T), E288del (1197_1199delGAA), L290F (1205A>T), L293E (1212T>G 1213T>A), E296D (1223A>T), H297L (1224C>T 1225A>T 1226C>A), S301Q (1236A>C 1237G>A 1238T>A), W305L (1248T>C 1249G>T 1250G>T), T308S (1258C>G), S311G (1266A>G 1268T>G), P312A (1269C>G 1271T>C), R314Q (1276G>A), D352I (1389G>A 1390A>T), K360S (1414A>G 1415A>T), I361L (1416A>C 1418A>T), D368N (1437G>A), K371R (1446A>C 1447A>G 1448A>T), L374Q (1455T>C 1456T>A), A375Y (1458G>T 1459C>A), K377R (1465A>G 1466G>A), Q378L (1468A>T 1469A>T), N380R (1474A>G 1475C>A), F381S (1477T>C 1478T>G), N383S (1482A>T 1483A>C), C384S (1486G>C), Y385F (1489A>T 1490T>C), L389F (1502A>C), M401I (1538G>A), L404F (1545C>T 1547T>C), M407C (1554A>T 1555T>G 1556G>C), I408M (1559A>G), P411S (1566C>A 1567C>G), S420del (1593_1597delAGCTC), I427F (1614A>T), V429I (1620G>A 1622T>C), N430V (1623A>G 1624A>T), K431P (1626A>C 1627A>C 1628A>T), L433M (1632T>A 1634A>G), T434V (1635A>G 1636C>T 1637A>C), Y438S (1647T>A 1648A>G), F450Y (1684T>A), H451Y (1686C>T), A455S (1698G>T 1700A>C), S458A (1707T>G), I464M (1727A>G), S470K (1744G>A 1745C>A), K471Y (1746A>T 1748A>T), G475T (1758G>A 1759G>C 1760A>G), C476S (1761T>A), N477T (1765A>C 1766C>T), S478T (1767T>A 1769T>C), G481A (1777G>C 1778G>T), S495T (1819G>C 1820T>A), K496E (1821A>G 1823A>G), N499R (1830A>C 1831A>G 1832T>G), E500Q (1833G>C), K502R (1840A>G), N509G (1860A>G 1861A>G), K511T (1867A>C), P515A (1878C>G 1880A>T), L518R (1887T>A 1888T>G 1889G>A), D521E (1898T>G), S535D (1938T>G 1939C>A 1940A>C), D539Y (1950G>T 1952T>C), L549T (1980T>A 1981T>C 1982A>T), A559Q (2010G>C 2011C>A 2012C>A), I562T (2020T>C), D565S (2028G>T 2029A>C 2030T>G), N567S (2034A>T 2035A>C 2036C>A), I568T (2038T>C), V570I (2043G>A), Q572P (2050A>C 2051A>C), R573I (2053G>T 2054A>T), L574F (2057G>T), G575W (2058G>T), Y579F (2071A>T), K580R (2074A>G 2075G>A), A583S (2082G>T 2084G>T), F586M (2091T>A 2093C>G), T587S (2094A>T), H600A (2133C>G 2134A>C 2135T>A), L601S (2136C>A 2137T>G 2138T>C), V604H (2145G>C 2146T>A 2147T>C), Y611L (2166T>C 2167A>T), W613Y (2173G>A 2174G>T), G618F (2187G>T 2188G>T), M619K (2191T>A 2192G>A), Y624F (2206A>T), V630I (2223G>A), S631D (2226A>G 2227G>A), L634R (2236T>G 2237T>G), K638T (2248A>C), V639P (2250G>C 2251T>C), L642I (2259T>A), Q643R (2263A>G 2264G>A), I644L (2265A>C), K647N (2276G>T), D648E (2279T>G), V655R (2298G>A 2299T>G 2300T>A), R664L (2326G>T), Q665R (2328C>A 2329A>G), E668K (2337G>A), L674V (2355C>G), N675S (2358A>T 2359A>C 2360C>T), I680V (2373A>G 2375C>T), C681A (2376T>G 2377G>C 2378T>A), M682A (2379A>G 2380T>C 2381G>A), N683S (2382A>T 2383A>C 2384C>A), P684A (2385C>G), D685E (2390T>A), S687T (2394T>A), K689D (2400A>G 2402G>T), S690Q (2403A>C 2404G>A 2405T>G), I692M (2411C>G), D693E (2414C>A), S694V (2415T>G 2416C>T 2417C>T), I696M (2423A>G), A698S (2427G>A 2428C>G 2429A>T), T699K (2431C>A), W701* (2438G>A), L704S (2445C>T 2446T>C 2447C>G), E705N (2448G>A 2450A>T), T706E (2451A>G 2452C>A 2453T>A), F708L (2457T>C 2459T>G), D709K (2460G>A 2462T>A), I711V (2466A>G), N727D (2514A>G), P728L (2518C>T 2519C>G), Q733R (2532C>A 2533A>G), W734Y (2536G>A 2537G>T), F736L (2541T>C 2543T>G), N739S (2551A>G 2552C>T), V741S (2556G>T 2557T>C), N745D (2568A>G), K746R (2571A>C 2572A>G) | | | | | | | | |

| | Begin | End | Coverage | Score | Concordance | Matches | Identities | I/D/M/F* | Stop Codons |
|---|---|---|---|---|---|---|---|---|---|
| **NT** | **428** | **2593** | **75.7%** | **1494** | **36.1%** | **2089 (99.6%)** | **1427 (68.0%)** | **0/9** | |

Codon mutations:

AAA34AAG (437A>G), ATC36CTT (441A>C 443C>T), CTT38TTT (447C>T), CTA42TTA (459C>T), GCC44GTC (466C>T), TCC45TCT (470C>T), GCT46ATG (471G>A 472C>T 473T>G), TTG48ATG (477T>A), ACA49CAT (480A>C 481C>A 482A>T), AAA50AAG (485A>G), ATC52GTC (489A>G), GCT53GCC (494T>C), CTT54CTA (497T>A), GCT55GCG (500T>G), GGA56GGC (503A>C), TGT57TGC (506T>C), GAC58GAT (509C>T), TTT59TTC (512T>C), GTA60ATA (513G>A), TCT61AAA (516T>A 517C>A 518T>A), AAA62ACA (520A>C), GTG63GTT (524G>T), ACT64GTG (525A>G 526C>T 527T>G), ACC65AAA (529C>A 530C>A), TTA66CTG (531T>C 533A>G), CGC67AAA (534C>A 535G>A 536C>A), CGG68CGA (539G>A), AAG69CGT (540A>C 541A>G 542G>T), TCG70TCA (545G>A), GGC71GGT (548C>T), TAT72TTT (550A>T), TTC74TAT (556T>A 557C>T), ACA75ACT (560A>T), TAC78TAT (569C>T), AAG80AAA (575G>A), TGC82TGT (581C>T), GCA83ACA (582G>A), GTA84GTT (587A>T), TGC85TCC (589G>C), TTA86CTT (591T>C 593A>T), CGA88CGT (599A>T), TAC89TAT (602C>T), TCC93AGT (612T>A 613C>G 614C>T), TTT94TAT (616T>A), CGA95TCT (618C>T 619G>C 620A>T), GAA97AAT (624G>A 626A>T), GAA98GAC (629A>C), CCA99ACC (630C>A 632A>C), CTT100ATG (633C>A 635T>G), TCG101TCT (638G>T), GTG102GTT (641G>T), CCT103CCA (644T>A), GTG104GTT (647G>T), CTC106TTG (651C>T 653C>G), ACA107ACC (656A>C), AGA108CGA (657A>C), TCG109TGT (661C>G 662G>T), GGT110GGA (665T>A), ATC111ATT (668C>T), CGT113AAG (672C>A 673G>A 674T>G), ATC115ATA (680C>A), CCC116CCT (683C>T), AAA117GCC (684A>G 685A>C 686A>C), TTG119TTA (692G>A), CGT120CGA (695T>A), GCA122CAC (699G>C 700C>A 701A>C), ATA123GTA (702A>G), CGT124AGG (705C>A 707T>G), AGG125GCA (708A>G 709G>C 710G>A), AAA126AAG (713A>G), GAT127TCT (714G>T 715A>C), GAA128GAT (719A>T), CAC129CAT (722C>T), GTT134GTC (737T>C), TAC137TAT (746C>T), TCA139TCT (752A>T), GGG142GGA (761G>A), GCT144TCA (765G>T 767T>A), AAG145AAA (770G>A), GTT146TAG (771G>T 772T>A 773T>G), TTG147ATT (774T>A 776G>T), AAA148GCG (777A>G 778A>C 779A>G), GTT149GTG (782T>G), AGT154AAG (796G>A 797T>G), GCA156GCC (803A>C), TTT158TTC (809T>C), GAA159CAA (810G>C), TCC160TCA (815C>A), ATA161ATT (818A>T), ACT162ACG (821T>G), ACC163ACT (824C>T), CCA164CCT (827A>T), CCT166TCA (831C>T 833T>A), GGG169GAT (841G>A 842G>T), CTA172CGT (850T>G 851A>T), GTG174GTT (857G>T), CTG175CTT (860G>T), GAA176GGA (862A>G), ATT178ATG (869T>G), ACG180ACT (875G>T), TTC182TGA (880T>G 881C>A), CAT184AAG (885C>A 887T>G), CAA186CGA (892A>G), CTA190CTT (905A>T), CCC191CCA (908C>A), TTT192CAT (909T>C 910T>A), CGA194CAA (916G>A), GGC195TCC (918G>T 919G>C), ATT196ATA (923T>A), CCA197CCT (926A>T), ATG198CTT (927A>C 929G>T), ATT200AAG (934T>A 935T>G), GGG201GGT (938G>T), TCT203AAT (942T>A 943C>A), CAA205AAA (948C>A), CCC212CCA (971C>A), CTA213TTA (972C>T), AAT214CTC (975A>C 976A>T 977T>C), AAT216AAC (983T>C), AGG219CGC (990A>C 992G>C), TCA220CGG (993T>C 994C>G 995A>G), TAC221TTT (997A>T 998C>T), GGT222CAA (999G>C 1000G>A 1001T>A), TTA223TAT (1003T>A 1004A>T), GCC224TCG (1005G>T 1007C>G), GTA225GTG (1010A>G), GAC227GAT (1016C>T), AAA228GTT (1017A>G 1018A>T 1019A>T), GTT229GTA (1022T>A), GAT230GAC (1025T>C), CAA231GGA (1026C>G 1027A>G), GAG232GCA (1030A>C 1031G>A), GCT233GCG (1034T>G), AGT234GCT (1035A>G 1036G>C), TTC237GTT (1044T>G 1046C>T), AAG238AAA (1049G>A), TTT239GAA (1050T>G 1051T>A 1052T>A), CCA240CAC (1054C>A 1055A>C), ATT242ATC (1061T>C), GTT244GTG (1067T>G), TTG246TTA (1073G>A), AAA247AAG (1076A>G), CTA250ATA (1083C>A), GCG251GCT (1088G>T), GCC252GCG (1091C>G), TGG255TTG (1099G>T), ACA257GTG (1104A>G 1105C>T 1106A>G), ATT260ATG (1115T>G), TCA262TCC (1121A>C), TTC263ATT (1122T>A 1124C>T), CAG264CAA (1127G>A), GAC265GAT (1130C>T), GGC266GGG (1133C>G), TCA269TCC (1142A>C), CCC270CCT (1145C>T), GGG271GGT (1148G>T), ATG272ATT (1151G>T), TTG273CTT (1152T>C 1154G>T), TCC275TCT (1160C>T), AAA276CAA (1161A>C), TCT277ATG (1164T>A 1165C>T 1166T>G), GTC278GTT (1169C>T), TTC279TTA (1172C>A), TTT280TAC (1174T>A 1175T>C), CCT281CCA (1178T>A), TTA282TTT (1181A>T), GAC283GAT (1184C>T), AGG284TCT (1185A>T 1186G>C 1187G>T), GCT285AAT (1188G>A 1189C>A), GCA286AAT (1191G>A 1192C>A 1193A>T), ACT287ACC (1196T>C), GAA288del (1197_1199delGAA), TTA290TTT (1205A>T), AAC292AAT (1211C>T), TTA293GAA (1212T>G 1213T>A), TCT294TCC (1217T>C), CTC295CTT (1220C>T), GAA296GAT (1223A>T), CAC297TTA (1224C>T 1225A>T 1226C>A), GAG299GAA (1232G>A), AGT301CAA (1236A>C 1237G>A 1238T>A), GTC302GTA (1241C>A), CCA304CCC (1247A>C), TGG305CTT (1248T>C 1249G>T 1250G>T), TTT306TTC (1253T>C), TCA307TCT (1256A>T), ACC308AGC (1258C>G), TTG310TTA (1265G>A), AGT311GGG (1266A>G 1268T>G), CCT312GCC (1269C>G 1271T>C), TAT313TAC (1274T>C), CGA314CAA (1276G>A), GGG315GGA (1280G>A), GTG316GTT (1283G>T), GTT319GTG (1292T>G), CTC347CTG (1376C>G), CAC348CAT (1379C>T), CCT349CCC (1382T>C), GTT350GTC (1385T>C), CAC351CAT (1388C>T), GAT352ATT (1389G>A 1390A>T), ACA356ACT (1403A>T), GTC357GTA (1406C>A), CTG358CTA (1409G>A), TCT359TCA (1412T>A), AAA360AGT (1414A>G 1415A>T), ATA361CTT (1416A>C 1418A>T), ACA363ACT (1424A>T), GAC364GAT (1427C>T), GGG365GGT (1430G>T), ACT366ACA (1433T>A), GAC368AAC (1437G>A), GAG370GAA (1445G>A), AAA371CGT (1446A>C 1447A>G 1448A>T), CCT372CCG (1451T>G), TTG374CAG (1455T>C 1456T>A), GCT375TAT (1458G>T 1459C>A), TTA376CTA (1461T>C), AAG377AGA (1465A>G 1466G>A), CAA378CTT (1468A>T 1469A>T), AAG379AAA (1472G>A), AAC380AGA (1474A>G 1475C>A), TTT381TCG (1477T>C 1478T>G), AAA382AAG (1481A>G), AAT383TCT (1482A>T 1483A>C), TGT384TCT (1486G>C), TAT385TTC (1489A>T 1490T>C), TTA389TTC (1502A>C), AAG390AAA (1505G>A), TCA391TCG (1508A>G), CCG397CCA (1526G>A), ATG401ATA (1538G>A), TAT402TAC (1541T>C), CTT404TTC (1545C>T 1547T>C), TCT406TCC (1553T>C), ATG407TGC (1554A>T 1555T>G 1556G>C), ATA408ATG (1559A>G), CCT411AGT (1566C>A 1567C>G), ACC412ACA (1571C>A), TTG413CTG (1572T>C), TCC415TCA (1580C>A), GTT418GTC (1589T>C), AGC420del (1593_1597delAGCTC), TCT421--A (1593_1597delAGCTC 1598T>A), CTT424CTC (1607T>C), ACG426ACT (1613G>T), ATC427TTC (1614A>T), CTT428TTG (1617C>T 1619T>G), GTT429ATC (1620G>A 1622T>C), AAC430GTC (1623A>G 1624A>T), AAA431CCT (1626A>C 1627A>C 1628A>T), TTA433ATG (1632T>A 1634A>G), ACA434GTC (1635A>G 1636C>T 1637A>C), AGA436AGG (1643A>G), TAT438AGT (1647T>A 1648A>G), ATA440ATT (1655A>T), TTT442TTC (1661T>C), CTG443TTA (1662C>T 1664G>A), ACT444ACA (1667T>A), GGG445GGA (1670G>A), CCT447CCG (1676T>G), TTG448TTA (1679G>A), GGC449GGT (1682C>T), TTC450TAC (1684T>A), CAC451TAC (1686C>T), GGT452GGA (1691T>A), TCT453TCA (1694T>A), GCA455TCC (1698G>T 1700A>C), CTT456CTA (1703T>A), TTC457TTT (1706C>T), TCG458GCG (1707T>G), TTG459CTT (1710T>C 1712G>T), TCT460TCC (1715T>C), CAC462CAT (1721C>T), ATA464ATG (1727A>G), TTA467CTA (1734T>C), GCA468GCT (1739A>T), AGC470AAA (1744G>A 1745C>A), AAA471TAT (1746A>T 1748A>T), TAT473TAC (1754T>C), GGA475ACG (1758G>A 1759G>C 1760A>G), TGT476AGT (1761T>A), AAC477ACT (1765A>C 1766C>T), TCT478ACC (1767T>A 1769T>C), CCA479CCT (1772A>T), GGG481GCT (1777G>C 1778G>T), GCT484GCC (1787T>C), TTA485CTG (1788T>C 1790A>G), CTA486TTA (1791C>T), AGT495ACA (1819G>C 1820T>A), AAA496GAG (1821A>G 1823A>G), GCC498GCG (1829C>G), AAT499CGG (1830A>C 1831A>G 1832T>G), GAG500CAG (1833G>C), AAG502AGG (1840A>G), TTA504TTG (1847A>G), TTG505CTG (1848T>C), AGA507AGG (1856A>G), CTC508CTT (1859C>T), AAT509GGT (1860A>G 1861A>G), GTC510GTT (1865C>T), AAA511ACA (1867A>C), ATT512ATC (1871T>C), TCG513TCT (1874G>T), ATA514-TA (1875delA), CCA515GCT (1878C>G 1880A>T), TCT517TCG (1886T>G), TTG518AGA (1887T>A 1888T>G 1889G>A), TCT520TCC (1895T>C), GAT521GAG (1898T>G), AAT522AAC (1901T>C), ATT525ATC (1910T>C), GCG528GCT (1919G>T), AGG530AGA (1925G>A), ACT533ACC (1934T>C), AAA534AAG (1937A>G), TCA535GAC (1938T>G 1939C>A 1940A>C), GAT539TAC (1950G>T 1952T>C), TTA540CTT (1953T>C 1955A>T), ATT543ATC (1964T>C), TCC544TCT (1967C>T), GCT547GCA (1976T>A), TTA549ACT (1980T>A 1981T>C 1982A>T), TCT550TCC (1985T>C), CGT552AGG (1989C>A 1991T>G), ACT553ACA (1994T>A), GTT555GTG (2000T>G), GGA556GGT (2003A>T), CTT557CTG (2006T>G), TGC558TGT (2009C>T), GCC559CAA (2010G>C 2011C>A 2012C>A), ATC562ACC (2020T>C), GAT565TCG (2028G>T 2029A>C 2030T>G), AAC567TCA (2034A>T 2035A>C 2036C>A), ATA568ACA (2038T>C), TCA569TCG (2042A>G), GTT570ATT (2043G>A), CAA572CCC (2050A>C 2051A>C), AGA573ATT (2053G>T 2054A>T), TTG574TTT (2057G>T), GGG575TGG (2058G>T), GGT576GGG (2063T>G), GGG578GGA (2069G>A), TAC579TTC (2071A>T), AAG580AGA (2074A>G 2075G>A), GTA581GTT (2078A>T), GCG583TCT (2082G>T 2084G>T), CGC584CGT (2087C>T), TTA585TTG (2090A>G), TTC586ATG (2091T>A 2093C>G), ACT587TCT (2094A>T), ACC588ACT (2099C>T), AGC590TCT (2103A>T 2104G>C 2105C>T), AAG591AAA (2108G>A), GAG594GAA (2117G>A), CGT595AGA (2118C>A 2120T>A), CTA596CTG (2123A>G), GCT598GCG (2129T>G), CAT600GCA (2133C>G 2134A>C 2135T>A), CTT601AGC (2136C>A 2137T>G 2138T>C), AAG602AAA (2141G>A), GTT604CAC (2145G>C 2146T>A 2147T>C), CGG605AGG (2148C>A), CAC607CAT (2156C>T), TTA608CTA (2157T>C), CTA609TTA (2160C>T), TAC611CTC (2166T>C 2167A>T), TGG613TAT (2173G>A 2174G>T), GGA616GGG (2183A>G), AGG617AGA (2186G>A), GGT618TTT (2187G>T 2188G>T), ATG619AAA (2191T>A 2192G>A), CCC620CCT (2195C>T), CCC623CCT (2204C>T), TAT624TTT (2206A>T), AAA626AAG (2213A>G), GGG627GGA (2216G>A), GTA630ATA (2223G>A), AGT631GAT (2226A>G 2227G>A), TAC632TAT (2231C>T), CTT634CGG (2236T>G 2237T>G), AAA635AAG (2240A>G), AAA638ACA (2248A>C), GTT639CCT (2250G>C 2251T>C), TTA642ATA (2259T>A), CAG643CGA (2263A>G 2264G>A), ATC644CTC (2265A>C), AAG647AAT (2276G>T), GAT648GAG (2279T>G), CTG649TTG (2280C>T), TTC651TTT (2288C>T), GAA654GAG (2297A>G), GTT655AGA (2298G>A 2299T>G 2300T>A), ATA657ATC (2306A>C), CTA658CTT (2309A>T), CGT660CGC (2315T>C), CTA663TTA (2322C>T), CGC664CTC (2326G>T), CAA665AGA (2328C>A 2329A>G), GTA667GTG (2336A>G), GAA668AAA (2337G>A), TTA671CTG (2346T>C 2348A>G), AAG672AAA (2351G>A), CTC674GTC (2355C>G), AAC675TCT (2358A>T 2359A>C 2360C>T), ACG679ACT (2372G>T), ATC680GTT (2373A>G 2375C>T), TGT681GCA (2376T>G 2377G>C 2378T>A), ATG682GCA (2379A>G 2380T>C 2381G>A), AAC683TCA (2382A>T 2383A>C 2384C>A), CCA684GCA (2385C>G), GAT685GAA (2390T>A), TCC687ACC (2394T>A), AAG689GAT (2400A>G 2402G>T), AGT690CAG (2403A>C 2404G>A 2405T>G), CTC691TTG (2406C>T 2408C>G), ATC692ATG (2411C>G), GAC693GAA (2414C>A), TCC694GTT (2415T>G 2416C>T 2417C>T), ATA696ATG (2423A>G), TGC697TGT (2426C>T), GCA698AGT (2427G>A 2428C>G 2429A>T), ACA699AAA (2431C>A), TGG701TGA (2438G>A), AAA702AAG (2441A>G), AGG703CGA (2442A>C 2444G>A), CTC704TCG (2445C>T 2446T>C 2447C>G), GAA705AAT (2448G>A 2450A>T), ACT706GAA (2451A>G 2452C>A 2453T>A), GAT707GAC (2456T>C), TTT708CTG (2457T>C 2459T>G), GAT709AAA (2460G>A 2462T>A), TTA710TTG (2465A>G), ATC711GTC (2466A>G), GGC714GGT (2477C>T), CTA715TTA (2478C>T), AAA718AAG (2489A>G), TGT719TGC (2492T>C), TAC720TAT (2495C>T), GGT723GGA (2504T>A), GGT725GGC (2510T>C), AAT727GAT (2514A>G), CCC728CTG (2518C>T 2519C>G), ACC730ACT (2525C>T), ACT731ACA (2528T>A), CAG733AGG (2532C>A 2533A>G), TGG734TAT (2536G>A 2537G>T), TTT736CTG (2541T>C 2543T>G), AAC739AGT (2551A>G 2552C>T), ACT740ACC (2555T>C), GTT741TCT (2556G>T 2557T>C), ATT742ATC (2561T>C), AAT745GAT (2568A>G), AAA746CGA (2571A>C 2572A>G), TTA749TTG (2582A>G), GGG750GGA (2585G>A)

**Proteins**

| | Begin | End | Coverage | Score | Concordance | Matches | Identities | I/D/M/F* | Stop Codons |
|---|---|---|---|---|---|---|---|---|---|
| RNA-dependent RNA polymerase (YP_010798874.1) | 32 | 753 | 90.0% | 3210 | 65.4% | 697 (99.7%) | 482 (69.0%) | 0/2/2/2 | 3 |

| | Begin | End | Coverage | Score | Concordance | Matches | Identities | I/D/M/F* | Stop Codons |
|---|---|---|---|---|---|---|---|---|---|
| **NT** | **428** | **2593** | **75.7%** | **1494** | **36.1%** | **2089 (99.6%)** | **1427 (68.0%)** | **0/9** | |

Protein mutations:

I36L (441A>C 443C>T), L38F (447C>T), A44V (466C>T), A46M (471G>A 472C>T 473T>G), L48M (477T>A), T49H (480A>C 481C>A 482A>T), I52V (489A>G), V60I (513G>A), S61K (516T>A 517C>A 518T>A), K62T (520A>C), T64V (525A>G 526C>T 527T>G), T65K (529C>A 530C>A), R67K (534C>A 535G>A 536C>A), K69R (540A>C 541A>G 542G>T), Y72F (550A>T), F74Y (556T>A 557C>T), A83T (582G>A), C85S (589G>C), F94Y (616T>A), R95S (618C>T 619G>C 620A>T), E97N (624G>A 626A>T), E98D (629A>C), P99T (630C>A 632A>C), L100M (633C>A 635T>G), S109C (661C>G 662G>T), R113K (672C>A 673G>A 674T>G), K117A (684A>G 685A>C 686A>C), A122H (699G>C 700C>A 701A>C), I123V (702A>G), R125A (708A>G 709G>C 710G>A), D127S (714G>T 715A>C), E128D (719A>T), A144S (765G>T 767T>A), V146* (771G>T 772T>A 773T>G), L147I (774T>A 776G>T), K148A (777A>G 778A>C 779A>G), S154K (796G>A 797T>G), E159Q (810G>C), P166S (831C>T 833T>A), G169D (841G>A 842G>T), L172R (850T>G 851A>T), E176G (862A>G), I178M (869T>G), F182* (880T>G 881C>A), H184K (885C>A 887T>G), Q186R (892A>G), F192H (909T>C 910T>A), R194Q (916G>A), G195S (918G>T 919G>C), M198L (927A>C 929G>T), I200K (934T>A 935T>G), S203N (942T>A 943C>A), Q205K (948C>A), N214L (975A>C 976A>T 977T>C), S220R (993T>C 994C>G 995A>G), Y221F (997A>T 998C>T), G222Q (999G>C 1000G>A 1001T>A), L223Y (1003T>A 1004A>T), A224S (1005G>T 1007C>G), K228V (1017A>G 1018A>T 1019A>T), Q231G (1026C>G 1027A>G), E232A (1030A>C 1031G>A), S234A (1035A>G 1036G>C), F237V (1044T>G 1046C>T), F239E (1050T>G 1051T>A 1052T>A), P240H (1054C>A 1055A>C), L250I (1083C>A), W255L (1099G>T), T257V (1104A>G 1105C>T 1106A>G), I260M (1115T>G), F263I (1122T>A 1124C>T), M272I (1151G>T), K276Q (1161A>C), S277M (1164T>A 1165C>T 1166T>G), F279L (1172C>A), F280Y (1174T>A 1175T>C), L282F (1181A>T), R284S (1185A>T 1186G>C 1187G>T), A285N (1188G>A 1189C>A), A286N (1191G>A 1192C>A 1193A>T), E288del (1197_1199delGAA), L290F (1205A>T), L293E (1212T>G 1213T>A), E296D (1223A>T), H297L (1224C>T 1225A>T 1226C>A), S301Q (1236A>C 1237G>A 1238T>A), W305L (1248T>C 1249G>T 1250G>T), T308S (1258C>G), S311G (1266A>G 1268T>G), P312A (1269C>G 1271T>C), R314Q (1276G>A), D352I (1389G>A 1390A>T), K360S (1414A>G 1415A>T), I361L (1416A>C 1418A>T), D368N (1437G>A), K371R (1446A>C 1447A>G 1448A>T), L374Q (1455T>C 1456T>A), A375Y (1458G>T 1459C>A), K377R (1465A>G 1466G>A), Q378L (1468A>T 1469A>T), N380R (1474A>G 1475C>A), F381S (1477T>C 1478T>G), N383S (1482A>T 1483A>C), C384S (1486G>C), Y385F (1489A>T 1490T>C), L389F (1502A>C), M401I (1538G>A), L404F (1545C>T 1547T>C), M407C (1554A>T 1555T>G 1556G>C), I408M (1559A>G), P411S (1566C>A 1567C>G), S420del (1593_1597delAGCTC), I427F (1614A>T), V429I (1620G>A 1622T>C), N430V (1623A>G 1624A>T), K431P (1626A>C 1627A>C 1628A>T), L433M (1632T>A 1634A>G), T434V (1635A>G 1636C>T 1637A>C), Y438S (1647T>A 1648A>G), F450Y (1684T>A), H451Y (1686C>T), A455S (1698G>T 1700A>C), S458A (1707T>G), I464M (1727A>G), S470K (1744G>A 1745C>A), K471Y (1746A>T 1748A>T), G475T (1758G>A 1759G>C 1760A>G), C476S (1761T>A), N477T (1765A>C 1766C>T), S478T (1767T>A 1769T>C), G481A (1777G>C 1778G>T), S495T (1819G>C 1820T>A), K496E (1821A>G 1823A>G), N499R (1830A>C 1831A>G 1832T>G), E500Q (1833G>C), K502R (1840A>G), N509G (1860A>G 1861A>G), K511T (1867A>C), P515A (1878C>G 1880A>T), L518R (1887T>A 1888T>G 1889G>A), D521E (1898T>G), S535D (1938T>G 1939C>A 1940A>C), D539Y (1950G>T 1952T>C), L549T (1980T>A 1981T>C 1982A>T), A559Q (2010G>C 2011C>A 2012C>A), I562T (2020T>C), D565S (2028G>T 2029A>C 2030T>G), N567S (2034A>T 2035A>C 2036C>A), I568T (2038T>C), V570I (2043G>A), Q572P (2050A>C 2051A>C), R573I (2053G>T 2054A>T), L574F (2057G>T), G575W (2058G>T), Y579F (2071A>T), K580R (2074A>G 2075G>A), A583S (2082G>T 2084G>T), F586M (2091T>A 2093C>G), T587S (2094A>T), H600A (2133C>G 2134A>C 2135T>A), L601S (2136C>A 2137T>G 2138T>C), V604H (2145G>C 2146T>A 2147T>C), Y611L (2166T>C 2167A>T), W613Y (2173G>A 2174G>T), G618F (2187G>T 2188G>T), M619K (2191T>A 2192G>A), Y624F (2206A>T), V630I (2223G>A), S631D (2226A>G 2227G>A), L634R (2236T>G 2237T>G), K638T (2248A>C), V639P (2250G>C 2251T>C), L642I (2259T>A), Q643R (2263A>G 2264G>A), I644L (2265A>C), K647N (2276G>T), D648E (2279T>G), V655R (2298G>A 2299T>G 2300T>A), R664L (2326G>T), Q665R (2328C>A 2329A>G), E668K (2337G>A), L674V (2355C>G), N675S (2358A>T 2359A>C 2360C>T), I680V (2373A>G 2375C>T), C681A (2376T>G 2377G>C 2378T>A), M682A (2379A>G 2380T>C 2381G>A), N683S (2382A>T 2383A>C 2384C>A), P684A (2385C>G), D685E (2390T>A), S687T (2394T>A), K689D (2400A>G 2402G>T), S690Q (2403A>C 2404G>A 2405T>G), I692M (2411C>G), D693E (2414C>A), S694V (2415T>G 2416C>T 2417C>T), I696M (2423A>G), A698S (2427G>A 2428C>G 2429A>T), T699K (2431C>A), W701* (2438G>A), L704S (2445C>T 2446T>C 2447C>G), E705N (2448G>A 2450A>T), T706E (2451A>G 2452C>A 2453T>A), F708L (2457T>C 2459T>G), D709K (2460G>A 2462T>A), I711V (2466A>G), N727D (2514A>G), P728L (2518C>T 2519C>G), Q733R (2532C>A 2533A>G), W734Y (2536G>A 2537G>T), F736L (2541T>C 2543T>G), N739S (2551A>G 2552C>T), V741S (2556G>T 2557T>C), N745D (2568A>G), K746R (2571A>C 2572A>G)

Codon mutations:

AAA34AAG (437A>G), ATC36CTT (441A>C 443C>T), CTT38TTT (447C>T), CTA42TTA (459C>T), GCC44GTC (466C>T), TCC45TCT (470C>T), GCT46ATG (471G>A 472C>T 473T>G), TTG48ATG (477T>A), ACA49CAT (480A>C 481C>A 482A>T), AAA50AAG (485A>G), ATC52GTC (489A>G), GCT53GCC (494T>C), CTT54CTA (497T>A), GCT55GCG (500T>G), GGA56GGC (503A>C), TGT57TGC (506T>C), GAC58GAT (509C>T), TTT59TTC (512T>C), GTA60ATA (513G>A), TCT61AAA (516T>A 517C>A 518T>A), AAA62ACA (520A>C), GTG63GTT (524G>T), ACT64GTG (525A>G 526C>T 527T>G), ACC65AAA (529C>A 530C>A), TTA66CTG (531T>C 533A>G), CGC67AAA (534C>A 535G>A 536C>A), CGG68CGA (539G>A), AAG69CGT (540A>C 541A>G 542G>T), TCG70TCA (545G>A), GGC71GGT (548C>T), TAT72TTT (550A>T), TTC74TAT (556T>A 557C>T), ACA75ACT (560A>T), TAC78TAT (569C>T), AAG80AAA (575G>A), TGC82TGT (581C>T), GCA83ACA (582G>A), GTA84GTT (587A>T), TGC85TCC (589G>C), TTA86CTT (591T>C 593A>T), CGA88CGT (599A>T), TAC89TAT (602C>T), TCC93AGT (612T>A 613C>G 614C>T), TTT94TAT (616T>A), CGA95TCT (618C>T 619G>C 620A>T), GAA97AAT (624G>A 626A>T), GAA98GAC (629A>C), CCA99ACC (630C>A 632A>C), CTT100ATG (633C>A 635T>G), TCG101TCT (638G>T), GTG102GTT (641G>T), CCT103CCA (644T>A), GTG104GTT (647G>T), CTC106TTG (651C>T 653C>G), ACA107ACC (656A>C), AGA108CGA (657A>C), TCG109TGT (661C>G 662G>T), GGT110GGA (665T>A), ATC111ATT (668C>T), CGT113AAG (672C>A 673G>A 674T>G), ATC115ATA (680C>A), CCC116CCT (683C>T), AAA117GCC (684A>G 685A>C 686A>C), TTG119TTA (692G>A), CGT120CGA (695T>A), GCA122CAC (699G>C 700C>A 701A>C), ATA123GTA (702A>G), CGT124AGG (705C>A 707T>G), AGG125GCA (708A>G 709G>C 710G>A), AAA126AAG (713A>G), GAT127TCT (714G>T 715A>C), GAA128GAT (719A>T), CAC129CAT (722C>T), GTT134GTC (737T>C), TAC137TAT (746C>T), TCA139TCT (752A>T), GGG142GGA (761G>A), GCT144TCA (765G>T 767T>A), AAG145AAA (770G>A), GTT146TAG (771G>T 772T>A 773T>G), TTG147ATT (774T>A 776G>T), AAA148GCG (777A>G 778A>C 779A>G), GTT149GTG (782T>G), AGT154AAG (796G>A 797T>G), GCA156GCC (803A>C), TTT158TTC (809T>C), GAA159CAA (810G>C), TCC160TCA (815C>A), ATA161ATT (818A>T), ACT162ACG (821T>G), ACC163ACT (824C>T), CCA164CCT (827A>T), CCT166TCA (831C>T 833T>A), GGG169GAT (841G>A 842G>T), CTA172CGT (850T>G 851A>T), GTG174GTT (857G>T), CTG175CTT (860G>T), GAA176GGA (862A>G), ATT178ATG (869T>G), ACG180ACT (875G>T), TTC182TGA (880T>G 881C>A), CAT184AAG (885C>A 887T>G), CAA186CGA (892A>G), CTA190CTT (905A>T), CCC191CCA (908C>A), TTT192CAT (909T>C 910T>A), CGA194CAA (916G>A), GGC195TCC (918G>T 919G>C), ATT196ATA (923T>A), CCA197CCT (926A>T), ATG198CTT (927A>C 929G>T), ATT200AAG (934T>A 935T>G), GGG201GGT (938G>T), TCT203AAT (942T>A 943C>A), CAA205AAA (948C>A), CCC212CCA (971C>A), CTA213TTA (972C>T), AAT214CTC (975A>C 976A>T 977T>C), AAT216AAC (983T>C), AGG219CGC (990A>C 992G>C), TCA220CGG (993T>C 994C>G 995A>G), TAC221TTT (997A>T 998C>T), GGT222CAA (999G>C 1000G>A 1001T>A), TTA223TAT (1003T>A 1004A>T), GCC224TCG (1005G>T 1007C>G), GTA225GTG (1010A>G), GAC227GAT (1016C>T), AAA228GTT (1017A>G 1018A>T 1019A>T), GTT229GTA (1022T>A), GAT230GAC (1025T>C), CAA231GGA (1026C>G 1027A>G), GAG232GCA (1030A>C 1031G>A), GCT233GCG (1034T>G), AGT234GCT (1035A>G 1036G>C), TTC237GTT (1044T>G 1046C>T), AAG238AAA (1049G>A), TTT239GAA (1050T>G 1051T>A 1052T>A), CCA240CAC (1054C>A 1055A>C), ATT242ATC (1061T>C), GTT244GTG (1067T>G), TTG246TTA (1073G>A), AAA247AAG (1076A>G), CTA250ATA (1083C>A), GCG251GCT (1088G>T), GCC252GCG (1091C>G), TGG255TTG (1099G>T), ACA257GTG (1104A>G 1105C>T 1106A>G), ATT260ATG (1115T>G), TCA262TCC (1121A>C), TTC263ATT (1122T>A 1124C>T), CAG264CAA (1127G>A), GAC265GAT (1130C>T), GGC266GGG (1133C>G), TCA269TCC (1142A>C), CCC270CCT (1145C>T), GGG271GGT (1148G>T), ATG272ATT (1151G>T), TTG273CTT (1152T>C 1154G>T), TCC275TCT (1160C>T), AAA276CAA (1161A>C), TCT277ATG (1164T>A 1165C>T 1166T>G), GTC278GTT (1169C>T), TTC279TTA (1172C>A), TTT280TAC (1174T>A 1175T>C), CCT281CCA (1178T>A), TTA282TTT (1181A>T), GAC283GAT (1184C>T), AGG284TCT (1185A>T 1186G>C 1187G>T), GCT285AAT (1188G>A 1189C>A), GCA286AAT (1191G>A 1192C>A 1193A>T), ACT287ACC (1196T>C), GAA288del (1197_1199delGAA), TTA290TTT (1205A>T), AAC292AAT (1211C>T), TTA293GAA (1212T>G 1213T>A), TCT294TCC (1217T>C), CTC295CTT (1220C>T), GAA296GAT (1223A>T), CAC297TTA (1224C>T 1225A>T 1226C>A), GAG299GAA (1232G>A), AGT301CAA (1236A>C 1237G>A 1238T>A), GTC302GTA (1241C>A), CCA304CCC (1247A>C), TGG305CTT (1248T>C 1249G>T 1250G>T), TTT306TTC (1253T>C), TCA307TCT (1256A>T), ACC308AGC (1258C>G), TTG310TTA (1265G>A), AGT311GGG (1266A>G 1268T>G), CCT312GCC (1269C>G 1271T>C), TAT313TAC (1274T>C), CGA314CAA (1276G>A), GGG315GGA (1280G>A), GTG316GTT (1283G>T), GTT319GTG (1292T>G), CTC347CTG (1376C>G), CAC348CAT (1379C>T), CCT349CCC (1382T>C), GTT350GTC (1385T>C), CAC351CAT (1388C>T), GAT352ATT (1389G>A 1390A>T), ACA356ACT (1403A>T), GTC357GTA (1406C>A), CTG358CTA (1409G>A), TCT359TCA (1412T>A), AAA360AGT (1414A>G 1415A>T), ATA361CTT (1416A>C 1418A>T), ACA363ACT (1424A>T), GAC364GAT (1427C>T), GGG365GGT (1430G>T), ACT366ACA (1433T>A), GAC368AAC (1437G>A), GAG370GAA (1445G>A), AAA371CGT (1446A>C 1447A>G 1448A>T), CCT372CCG (1451T>G), TTG374CAG (1455T>C 1456T>A), GCT375TAT (1458G>T 1459C>A), TTA376CTA (1461T>C), AAG377AGA (1465A>G 1466G>A), CAA378CTT (1468A>T 1469A>T), AAG379AAA (1472G>A), AAC380AGA (1474A>G 1475C>A), TTT381TCG (1477T>C 1478T>G), AAA382AAG (1481A>G), AAT383TCT (1482A>T 1483A>C), TGT384TCT (1486G>C), TAT385TTC (1489A>T 1490T>C), TTA389TTC (1502A>C), AAG390AAA (1505G>A), TCA391TCG (1508A>G), CCG397CCA (1526G>A), ATG401ATA (1538G>A), TAT402TAC (1541T>C), CTT404TTC (1545C>T 1547T>C), TCT406TCC (1553T>C), ATG407TGC (1554A>T 1555T>G 1556G>C), ATA408ATG (1559A>G), CCT411AGT (1566C>A 1567C>G), ACC412ACA (1571C>A), TTG413CTG (1572T>C), TCC415TCA (1580C>A), GTT418GTC (1589T>C), AGC420del (1593_1597delAGCTC), TCT421--A (1593_1597delAGCTC 1598T>A), CTT424CTC (1607T>C), ACG426ACT (1613G>T), ATC427TTC (1614A>T), CTT428TTG (1617C>T 1619T>G), GTT429ATC (1620G>A 1622T>C), AAC430GTC (1623A>G 1624A>T), AAA431CCT (1626A>C 1627A>C 1628A>T), TTA433ATG (1632T>A 1634A>G), ACA434GTC (1635A>G 1636C>T 1637A>C), AGA436AGG (1643A>G), TAT438AGT (1647T>A 1648A>G), ATA440ATT (1655A>T), TTT442TTC (1661T>C), CTG443TTA (1662C>T 1664G>A), ACT444ACA (1667T>A), GGG445GGA (1670G>A), CCT447CCG (1676T>G), TTG448TTA (1679G>A), GGC449GGT (1682C>T), TTC450TAC (1684T>A), CAC451TAC (1686C>T), GGT452GGA (1691T>A), TCT453TCA (1694T>A), GCA455TCC (1698G>T 1700A>C), CTT456CTA (1703T>A), TTC457TTT (1706C>T), TCG458GCG (1707T>G), TTG459CTT (1710T>C 1712G>T), TCT460TCC (1715T>C), CAC462CAT (1721C>T), ATA464ATG (1727A>G), TTA467CTA (1734T>C), GCA468GCT (1739A>T), AGC470AAA (1744G>A 1745C>A), AAA471TAT (1746A>T 1748A>T), TAT473TAC (1754T>C), GGA475ACG (1758G>A 1759G>C 1760A>G), TGT476AGT (1761T>A), AAC477ACT (1765A>C 1766C>T), TCT478ACC (1767T>A 1769T>C), CCA479CCT (1772A>T), GGG481GCT (1777G>C 1778G>T), GCT484GCC (1787T>C), TTA485CTG (1788T>C 1790A>G), CTA486TTA (1791C>T), AGT495ACA (1819G>C 1820T>A), AAA496GAG (1821A>G 1823A>G), GCC498GCG (1829C>G), AAT499CGG (1830A>C 1831A>G 1832T>G), GAG500CAG (1833G>C), AAG502AGG (1840A>G), TTA504TTG (1847A>G), TTG505CTG (1848T>C), AGA507AGG (1856A>G), CTC508CTT (1859C>T), AAT509GGT (1860A>G 1861A>G), GTC510GTT (1865C>T), AAA511ACA (1867A>C), ATT512ATC (1871T>C), TCG513TCT (1874G>T), ATA514-TA (1875delA), CCA515GCT (1878C>G 1880A>T), TCT517TCG (1886T>G), TTG518AGA (1887T>A 1888T>G 1889G>A), TCT520TCC (1895T>C), GAT521GAG (1898T>G), AAT522AAC (1901T>C), ATT525ATC (1910T>C), GCG528GCT (1919G>T), AGG530AGA (1925G>A), ACT533ACC (1934T>C), AAA534AAG (1937A>G), TCA535GAC (1938T>G 1939C>A 1940A>C), GAT539TAC (1950G>T 1952T>C), TTA540CTT (1953T>C 1955A>T), ATT543ATC (1964T>C), TCC544TCT (1967C>T), GCT547GCA (1976T>A), TTA549ACT (1980T>A 1981T>C 1982A>T), TCT550TCC (1985T>C), CGT552AGG (1989C>A 1991T>G), ACT553ACA (1994T>A), GTT555GTG (2000T>G), GGA556GGT (2003A>T), CTT557CTG (2006T>G), TGC558TGT (2009C>T), GCC559CAA (2010G>C 2011C>A 2012C>A), ATC562ACC (2020T>C), GAT565TCG (2028G>T 2029A>C 2030T>G), AAC567TCA (2034A>T 2035A>C 2036C>A), ATA568ACA (2038T>C), TCA569TCG (2042A>G), GTT570ATT (2043G>A), CAA572CCC (2050A>C 2051A>C), AGA573ATT (2053G>T 2054A>T), TTG574TTT (2057G>T), GGG575TGG (2058G>T), GGT576GGG (2063T>G), GGG578GGA (2069G>A), TAC579TTC (2071A>T), AAG580AGA (2074A>G 2075G>A), GTA581GTT (2078A>T), GCG583TCT (2082G>T 2084G>T), CGC584CGT (2087C>T), TTA585TTG (2090A>G), TTC586ATG (2091T>A 2093C>G), ACT587TCT (2094A>T), ACC588ACT (2099C>T), AGC590TCT (2103A>T 2104G>C 2105C>T), AAG591AAA (2108G>A), GAG594GAA (2117G>A), CGT595AGA (2118C>A 2120T>A), CTA596CTG (2123A>G), GCT598GCG (2129T>G), CAT600GCA (2133C>G 2134A>C 2135T>A), CTT601AGC (2136C>A 2137T>G 2138T>C), AAG602AAA (2141G>A), GTT604CAC (2145G>C 2146T>A 2147T>C), CGG605AGG (2148C>A), CAC607CAT (2156C>T), TTA608CTA (2157T>C), CTA609TTA (2160C>T), TAC611CTC (2166T>C 2167A>T), TGG613TAT (2173G>A 2174G>T), GGA616GGG (2183A>G), AGG617AGA (2186G>A), GGT618TTT (2187G>T 2188G>T), ATG619AAA (2191T>A 2192G>A), CCC620CCT (2195C>T), CCC623CCT (2204C>T), TAT624TTT (2206A>T), AAA626AAG (2213A>G), GGG627GGA (2216G>A), GTA630ATA (2223G>A), AGT631GAT (2226A>G 2227G>A), TAC632TAT (2231C>T), CTT634CGG (2236T>G 2237T>G), AAA635AAG (2240A>G), AAA638ACA (2248A>C), GTT639CCT (2250G>C 2251T>C), TTA642ATA (2259T>A), CAG643CGA (2263A>G 2264G>A), ATC644CTC (2265A>C), AAG647AAT (2276G>T), GAT648GAG (2279T>G), CTG649TTG (2280C>T), TTC651TTT (2288C>T), GAA654GAG (2297A>G), GTT655AGA (2298G>A 2299T>G 2300T>A), ATA657ATC (2306A>C), CTA658CTT (2309A>T), CGT660CGC (2315T>C), CTA663TTA (2322C>T), CGC664CTC (2326G>T), CAA665AGA (2328C>A 2329A>G), GTA667GTG (2336A>G), GAA668AAA (2337G>A), TTA671CTG (2346T>C 2348A>G), AAG672AAA (2351G>A), CTC674GTC (2355C>G), AAC675TCT (2358A>T 2359A>C 2360C>T), ACG679ACT (2372G>T), ATC680GTT (2373A>G 2375C>T), TGT681GCA (2376T>G 2377G>C 2378T>A), ATG682GCA (2379A>G 2380T>C 2381G>A), AAC683TCA (2382A>T 2383A>C 2384C>A), CCA684GCA (2385C>G), GAT685GAA (2390T>A), TCC687ACC (2394T>A), AAG689GAT (2400A>G 2402G>T), AGT690CAG (2403A>C 2404G>A 2405T>G), CTC691TTG (2406C>T 2408C>G), ATC692ATG (2411C>G), GAC693GAA (2414C>A), TCC694GTT (2415T>G 2416C>T 2417C>T), ATA696ATG (2423A>G), TGC697TGT (2426C>T), GCA698AGT (2427G>A 2428C>G 2429A>T), ACA699AAA (2431C>A), TGG701TGA (2438G>A), AAA702AAG (2441A>G), AGG703CGA (2442A>C 2444G>A), CTC704TCG (2445C>T 2446T>C 2447C>G), GAA705AAT (2448G>A 2450A>T), ACT706GAA (2451A>G 2452C>A 2453T>A), GAT707GAC (2456T>C), TTT708CTG (2457T>C 2459T>G), GAT709AAA (2460G>A 2462T>A), TTA710TTG (2465A>G), ATC711GTC (2466A>G), GGC714GGT (2477C>T), CTA715TTA (2478C>T), AAA718AAG (2489A>G), TGT719TGC (2492T>C), TAC720TAT (2495C>T), GGT723GGA (2504T>A), GGT725GGC (2510T>C), AAT727GAT (2514A>G), CCC728CTG (2518C>T 2519C>G), ACC730ACT (2525C>T), ACT731ACA (2528T>A), CAG733AGG (2532C>A 2533A>G), TGG734TAT (2536G>A 2537G>T), TTT736CTG (2541T>C 2543T>G), AAC739AGT (2551A>G 2552C>T), ACT740ACC (2555T>C), GTT741TCT (2556G>T 2557T>C), ATT742ATC (2561T>C), AAT745GAT (2568A>G), AAA746CGA (2571A>C 2572A>G), TTA749TTG (2582A>G), GGG750GGA (2585G>A)

*: Inserts / Deletes / Misaligned / Frameshifts

#### Analysis details

This analysis was performed with panviral2.64

#### NGS Details (UN62): Solendovirus venanicotianae

##### Assembly

| | |
|---|---|
| **Coverage Length** | 1561 (5 contig(s)) |
| **Depth Of Coverage** | 8.5 |
| **Number Of Reads** | 124 |
| **Reads Per Million** | 2.87 rpm (after QC) |
| **Ambiguities** | 10 |
| **Assembly Method** | de novo + reference guided assembly |
| **Consensus Caller** | Bcf Tools |

##### Coverage Map

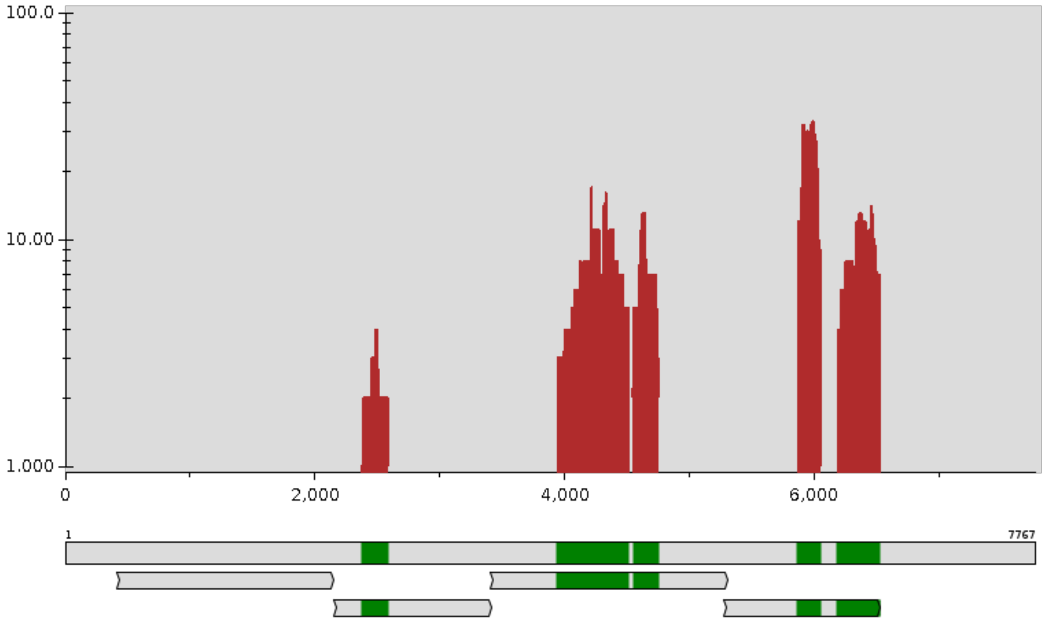

##### Assignment

| | |
|---|---|
| **Type** | Solendovirus venanicotianae (Taxonomy ID: 3048371) |
| **Reference Genome** | NC_003378.1 |
| **NT Identity (%)** | 76.9529 |
| **AA Identity (%)** | 73.5409 |
| **Number Of Stop Codons** | 2 |
| **Number Of CDS** | 4 |

##### Alignment

| | |
|---|---|
| **Alignment Score** | 1669.0 (NT) + 2601.0 (AA) = 4270.0 |
| **Concordance (%)** | 64.6284 |

| Alignment Method | Global, seeded, nucleotide + amino acids (AGA) |
|---|---|

#### Genome Region

Sequence starts at position 2371 and ends at position 6532 relative to NC_003378.1 reference sequence.

#### Alignment Detailed Statistics

| | Begin | End | Coverage | Score | Concordance | Matches | Identities | I/D/M/F* | Stop Codons |
|---|---|---|---|---|---|---|---|---|---|
| **NT** | **2371** | **6532** | **20.1%** | **1669** | **54.2%** | **1549 (99.2%)** | **1192 (76.4%)** | **0/12** | |
| Mutations: | 2373T>A, 2376T>C, 2379C>T, 2388A>T, 2389G>A, 2390G>A, 2394T>A, 2397G>A, 2400C>T, 2426T>C, 2438A>G, 2442A>T, 2454G>A, 2472G>A, 2475T>C, 2482T>C, 2493T>A, 2511A>T, 2514T>C, 2535C>A, 2551C>T, 2564G>C, 2565A>G, 2571C>A, 3935A>T, 3939G>C, 3944A>G, 3946T>C, 3947A>T, 3951A>G, 3952A>G, 3953T>A, 3956T>A, 3959C>T, 3962T>C, 3966C>G, 3969G>A, 3970G>A, 3977G>T, 3981C>A, 3982A>G, 3984A>C, 3986A>C, 3987A>G, 3988C>A, 3994T>G, 3997A>T, 3998A>T, 3999A>T, 4002G>A, 4003A>T, 4004A>C, 4007A>T, 4009T>A, 4010T>A, 4013A>T, 4014G>C, 4015A>T, 4016G>T, 4018A>C, 4019T>G, 4021G>T, 4030C>G, 4031A>G, 4032C>G, 4033A>T, 4036A>G, 4040T>A, 4041C>A, 4043A>G, 4044A>T, 4045A>T, 4052C>T, 4053T>G, 4054T>A, 4055T>A, 4058T>A, 4062T>A, 4063T>A, 4065A>T, 4067A>T, 4068G>A, 4073T>C, 4075A>G, 4079A>T, 4082T>G, 4085A>T, 4086A>C, 4091T>A, 4092T>C, 4094A>T, 4095T>C, 4097G>C, 4098G>A, 4100T>A, 4101A>T, 4102A>T, 4103T>A, 4109T>A, 4113C>A, 4114A>G, 4124T>G, 4127C>T, 4128A>C, 4129A>C, 4135C>G, 4136A>C, 4142A>T, 4151A>G, 4152G>A, 4156A>G, 4157T>A, 4160A>T, 4164A>G, 4165G>C, 4166T>G, 4184A>G, 4185A>G, 4186G>C, 4187C>T, 4188A>C, 4190G>A, 4196C>T, 4200G>A, 4207G>A, 4209A>G, 4211T>G, 4212T>C, 4214A>T, 4228A>T, 4235T>C, 4238T>A, 4241T>C, 4250C>T, 4253T>C, 4260T>C, 4262A>T, 4265G>A, 4274A>G, 4313G>A, 4325T>C, 4328T>C, 4329T>C, 4331G>A, 4335C>T, 4340G>A, 4343T>A, 4364T>A, 4376A>C, 4400T>A, 4401T>C, 4406A>T, 4430A>T, 4444T>G, 4451C>T, 4462A>G, 4463T>C, 4484A>T, 4500C>T, 4503C>T, 4513G>A, 4550C>T, 4552C>T, 4553A>G, 4554C>A, 4559T>A, 4560A>T, 4562A>T, 4566A>T, 4567A>C, 4568C>T, 4571T>A, 4573G>A, 4574T>A, 4580C>T, 4586C>T, 4592C>A, 4595A>G, 4596G>T, 4604T>A, 4612A>C, 4613T>A, 4616G>A, 4648A>G, 4649T>C, 4662C>T, 4665A>T, 4676A>G, 4688C>T, 4691T>C, 4692C>T, 4693T>C, 4694T>A, 4695G>A, 4697T>C, 4701A>C, 4703C>A, 4706T>A, 4715A>G, 4718G>A, 4738G>T, 4740A>T, 4748C>T, 5865C>T, 5867A>G, 5868T>C, 5871A>G, 5872T>C, 5874G>A, 5877A>T, 5880T>A, 5881T>C, 5882T>A, 5886A>T, 5889T>A, 5892A>C, 5894G>A, 5901A>C, 5902T>C, 5908G>A, 5909C>A, 5910C>A, 5913C>A, 5916C>T, 5921A>G, 5922C>Y, 5925C>T, 5928A>W, 5929C>Y, 5931A>R, 5932C>Y, 5935G>A, 5936A>R, 5937T>Y, 5940T>C, 5950C>T, 5952G>T, 5953A>G, 5954C>A, 5955A>Y, 5959C>G, 5964G>A, 5968G>T, 5970C>R, 5972A>C, 5973T>Y, 5979T>A, 5991G>A, 5997C>T, 5999T>C, 6000G>A, 6001C>A, 6002A>C, 6009G>A, 6012G>A, 6015A>T, 6017C>T, 6018T>A, 6023G>A, 6024G>C, 6026C>A, 6027A>C, 6033C>T, 6036C>T, 6037G>A, 6040C>A, 6041G>A, 6042A>G, 6045C>T, 6181G>A, 6183A>T, 6186C>T, 6195C>A, 6201G>A, 6204A>T, 6209G>A, 6211A>C, 6214T>G, 6215C>T, 6219T>G, 6223C>A, 6225A>C, 6232C>T, 6246T>C, 6247C>A, 6248T>C, 6253C>G, 6254A>G, 6258T>C, 6259G>C, 6264T>G, 6269G>A, 6270C>T, 6271T>C, 6277T>A, 6279A>C, 6282G>A, 6285A>C, 6290C>T, 6300T>C, 6304A>C, 6307T>A, 6308G>T, 6310_6321delAGAGATATAAGA, 6325G>A, 6326C>G, 6340C>G, 6341T>C, 6344G>T, 6345A>G, 6352G>A, 6354C>A, 6355T>A, 6357A>G, 6359G>C, 6360T>A, 6361C>T, 6363T>G, 6369A>T, 6375A>G, 6376C>A, 6380C>A, 6381T>A, 6382A>C, 6384A>T, 6386C>T, 6387C>T, 6401G>A, 6402G>A, 6404G>A, 6407A>G, 6408T>A, 6411C>T, 6414C>T, 6417C>G, 6418C>A, 6419C>A, 6423T>A, 6424C>T, 6426G>A, 6427T>C, 6444G>A, 6446T>A, 6451A>G, 6453T>G, 6456C>A, 6462T>C, 6471C>T, 6477C>T, 6489A>C, 6502A>G, 6504T>A, 6505G>A, 6507T>C, 6513C>T, 6522G>A, 6525A>C, 6526T>G | | | | | | | | |

##### CDS

| | Begin | End | Coverage | Score | Concordance | Matches | Identities | I/D/M/F* | Stop Codons |
|---|---|---|---|---|---|---|---|---|---|
| Tvvgp2 | 73 | 146 | 17.6% | 474 | 93.3% | 74 (100%) | 70 (94.6%) | 0/0/0/0 | 0 |
| Protein mutations: | G79K (2389G>A 2390G>A), I91T (2426T>C), E95G (2438A>G), R137T (2564G>C 2565A>G) | | | | | | | | |
| Codon mutations: | ATT73ATA (2373T>A), GAT74GAC (2376T>C), TGC75TGT (2379C>T), ACA78ACT (2388A>T), GGA79AAA (2389G>A 2390G>A), GGT80GGA (2394T>A), AAG81AAA (2397G>A), GCC82GCT (2400C>T), ATA91ACA (2426T>C), GAA95GGA (2438A>G), ATA96ATT (2442A>T), AAG100AAA (2454G>A), AAG106AAA (2472G>A), TAT107TAC (2475T>C), TTA110CTA (2482T>C), ACT113ACA (2493T>A), GCA119GCT (2511A>T), TGT120TGC (2514T>C), ACC127ACA (2535C>A), CTA133TTA (2551C>T), AGA137ACG (2564G>C 2565A>G), ATC139ATA (2571C>A) | | | | | | | | |
| Tvvgp3 | 176 | 449 | 41.7% | 1324 | 72.1% | 265 (100%) | 194 (73.2%) | 0/0/0/0 | 2 |
| Protein mutations: | E177Q (3939G>C), L179S (3946T>C 3947A>T), N181G (3951A>G 3952A>G 3953T>A), D182E (3956T>A), Q186E (3966C>G), G187K (3969G>A 3970G>A), E189D (3977G>T), H191S (3981C>A 3982A>G), K192H (3984A>C 3986A>C), T193E (3987A>G 3988C>A), V195G (3994T>G), K196I (3997A>T 3998A>T), I197L (3999A>T), E198I (4002G>A 4003A>T 4004A>C), I200K (4009T>A 4010T>A), E201D (4013A>T), E202L (4014G>C 4015A>T 4016G>T), N203T (4018A>C 4019T>G), S204I (4021G>T), T207R (4030C>G 4031A>G), Q208V (4032C>G 4033A>T), K209R (4036A>G), L211M (4041C>A 4043A>G), K212L (4044A>T 4045A>T), F215E (4053T>G 4054T>A 4055T>A), N216K (4058T>A), L218K (4062T>A 4063T>A), T219S (4065A>T 4067A>T), E220K (4068G>A), K222R (4075A>G), H224Q (4082T>G), K226Q (4086A>C), D227E (4091T>A), D230K (4098G>A 4100T>A), N231L (4101A>T 4102A>T 4103T>A), Y233* (4109T>A), Q235R (4113C>A 4114A>G), N238K (4124T>G), K240P (4128A>C 4129A>C), T242S (4135C>G 4136A>C), I247M (4151A>G), V248I (4152G>A), N249R (4156A>G 4157T>A), K250N (4160A>T), S252A (4164A>G 4165G>C 4166T>G), S259A (4185A>G 4186G>C 4187C>T), D264N (4200G>A), R266K (4207G>A), N267E (4209A>G 4211T>G), K273I (4228A>T), N276K (4238T>A), D311E (4343T>A), F345C (4444T>G), N351S (4462A>G 4463T>C), R368K (4513G>A), A381V (4552C>T 4553A>G), H382N (4554C>A), I384F (4560A>T 4562A>T), N386S (4566A>T 4567A>C 4568C>T), G388E (4573G>A 4574T>A), A396S (4596G>T), N401T (4612A>C 4613T>A), N413S (4648A>G 4649T>C), Q418* (4662C>T), T419S (4665A>T), L428S (4692C>T 4693T>C 4694T>A), D429N (4695G>A 4697T>C), N431Q (4701A>C 4703C>A), G443V (4738G>T), I444L (4740A>T) | | | | | | | | |
| Codon mutations: | AAA175.AT (3935A>T), GAA177CAA (3939G>C), AAA178AAG (3944A>G), TTA179TCT (3946T>C 3947A>T), AAT181GGA (3951A>G 3952A>G 3953T>A), GAT182GAA (3956T>A), AAC183AAT (3959C>T), CCT184CCC (3962T>C), CAA186GAA (3966C>G), GGA187AAA (3969G>A 3970G>A), GAG189GAT (3977G>T), CAT191AGT (3981C>A 3982A>G), AAA192CAC (3984A>C 3986A>C), ACA193GAA (3987A>G 3988C>A), GTA195GGA (3994T>G), AAA196ATT (3997A>T 3998A>T), ATA197TTA (3999A>T), GAA198ATC (4002G>A 4003A>T 4004A>C), CTA199CTT (4007A>T), ATT200AAA (4009T>A 4010T>A), GAA201GAT (4013A>T), GAG202CTT (4014G>C 4015A>T 4016G>T), AAT203ACG (4018A>C 4019T>G), AGT204ATT (4021G>T), ACA207AGG (4030C>G 4031A>G), CAA208GTA (4032C>G 4033A>T), AAA209AGA (4036A>G), CCT210CCA (4040T>A), CTA211ATG (4041C>A 4043A>G), AAA212TTA (4044A>T 4045A>T), AAC214AAT (4052C>T), TTT215GAA (4053T>G 4054T>A 4055T>A), AAT216AAA (4058T>A), TTA218AAA (4062T>A 4063T>A), ACA219TCT (4065A>T 4067A>T), GAA220AAA (4068G>A), TTT221TTC (4073T>C), AAA222AGA (4075A>G), ATA223ATT (4079A>T), CAT224CAG (4082T>G), ATA225ATT (4085A>T), AAA226CAA (4086A>C), GAT227GAA (4091T>A), TTA228CTT (4092T>C 4094A>T), TTG229CTC (4095T>C 4097G>C), GAT230AAA (4098G>A 4100T>A), AAT231TTA (4101A>T 4102A>T 4103T>A), TAT233TAA (4109T>A), CAA235AGA (4113C>A 4114A>G), AAT238AAG (4124T>G), AGC239AGT (4127C>T), AAA240CCA (4128A>C 4129A>C), ACA242AGC (4135C>G 4136A>C), CCA244CCT (4142A>T), ATA247ATG (4151A>G), GTA248ATA (4152G>A), AAT249AGA (4156A>G 4157T>A), AAA250AAT (4160A>T), AGT252GCG (4164A>G 4165G>C 4166T>G), AAA258AAG (4184A>G), AGC259GCT (4185A>G 4186G>C 4187C>T), AGG260CGA (4188A>C 4190G>A), GTC262GTT (4196C>T), GAC264AAC (4200G>A), AGA266AAA (4207G>A), AAT267GAG (4209A>G 4211T>G), TTA268CTT (4212T>C 4214A>T), AAA273ATA (4228A>T), TAT275TAC (4235T>C), AAT276AAA (4238T>A), TAT277TAC (4241T>C), CCC280CCT (4250C>T), AAT281AAC (4253T>C), TTA284CTT (4260T>C 4262A>T), AAG285AAA (4265G>A), CAA288CAG (4274A>G), AAG301AAA (4313G>A), TAT305TAC (4325T>C), CAT306CAC (4328T>C), TTG307CTA (4329T>C 4331G>A), CTA309TTA (4335C>T), GAG310GAA (4340G>A), GAT311GAA (4343T>A), GCT318GCA (4364T>A), CCA322CCC (4376A>C), GTT330GTA (4400T>A), TTA331CTA (4401T>C), CCA332CCT (4406A>T), GGA340GGT (4430A>T), TTT345TGT (4444T>G), GAC347GAT (4451C>T), AAT351AGC (4462A>G 4463T>C), GTA358GTT (4484A>T), CTA364TTA (4500C>T), CTA365TTA (4503C>T), AGA368AAA (4513G>A), TTC380TTT (4550C>T), GCA381GTG (4552C>T 4553A>G), CAT382AAT (4554C>A), ATT383ATA (4559T>A), ATA384TTT (4560A>T 4562A>T), AAC386TCT (4566A>T 4567A>C 4568C>T), TCT387TCA (4571T>A), GGT388GAA (4573G>A 4574T>A), AGC390AGT (4580C>T), AGC392AGT (4586C>T), ACC394ACA (4592C>A), AAA395AAG (4595A>G), GCA396TCA (4596G>T), ATT398ATA (4604T>A), AAT401ACA (4612A>C 4613T>A), CAG402CAA (4616G>A), AAT413AGC (4648A>G 4649T>C), CAA418TAA (4662C>T), ACT419TCT (4665A>T), GTA422GTG (4676A>G), ATC426ATT (4688C>T), AAT427AAC (4691T>C), CTT428TCA (4692C>T 4693T>C 4694T>A), GAT429AAC (4695G>A 4697T>C), AAC431CAA (4701A>C 4703C>A), ATT432ATA (4706T>A), AAA435AAG (4715A>G), AAG436AAA (4718G>A), GGA443GTA (4738G>T), ATA444TTA (4740A>T), AAC446AAT (4748C>T) | | | | | | | | |
| Tvvgp4 | 197 | 418 | 42.8% | 803 | 65.8% | 175 (97.8%) | 114 (63.7%) | 0/4/1/0 | 0 |
| Protein mutations: | N198S (5867A>G 5868T>C), L203Q (5881T>C 5882T>A), N205K (5889T>A), S207N (5894G>A), K209N (5901A>C), A212K (5908G>A 5909C>A 5910C>A), N216S (5921A>G 5922C>Y), Q226Y (5950C>T 5952G>T), T227D (5953A>G 5954C>A 5955A>Y), Q229E (5959C>G), A232S (5968G>T 5970C>R), N233T (5972A>C 5973T>Y), M242T (5999T>C 6000G>A), Q243T (6001C>A 6002A>C), T248I (6017C>T 6018T>A), G250D (6023G>A 6024G>C), T251N (6026C>A 6027A>C), V255I (6037G>A), R256K (6040C>A 6041G>A 6042A>G), E303N (6181G>A 6183A>T), N307K (6195C>A), G312E (6209G>A), I313L (6211A>C), S314V (6214T>G 6215C>T), L317I (6223C>A 6225A>C), L325T (6247C>A 6248T>C), Q327G (6253C>G 6254A>G), A329P (6259G>C), I330M (6264T>G), S332N (6269G>A 6270C>T), Y333H (6271T>C), S335T (6277T>A 6279A>C), E337D (6285A>C), T339I (6290C>T), N344H (6304A>C), C345I (6307T>A 6308G>T), R346_R349del (6310_6321delAGAGATATAAGA), A351R (6325G>A 6326C>G), L356A (6340C>G 6341T>C), R357M (6344G>T 6345A>G), V360I (6352G>A 6354C>A), L361M (6355T>A 6357A>G), S362T (6359G>C 6360T>A), K365N (6369A>T), Q368K (6376C>A), P369Q (6380C>A 6381T>A), T370P (6382A>C 6384A>T), T371I (6386C>T 6387C>T), R376K (6401G>A 6402G>A), R377K (6404G>A), N378R (6407A>G 6408T>A), P382K (6418C>A 6419C>A), D383E (6423T>A), L391Q (6446T>A), S393G (6451A>G 6453T>G), H394Q (6456C>A), K405N (6489A>C), I410V (6502A>G 6504T>A), V411I (6505G>A 6507T>C), E417D (6525A>C), *418E (6526T>G) | | | | | | | | |

| | Begin | End | Coverage | Score | Concordance | Matches | Identities | I/D/M/F* | Stop Codons |
|---|---|---|---|---|---|---|---|---|---|
| **NT** | **2371** | **6532** | **20.1%** | **1669** | **54.2%** | **1549 (99.2%)** | **1192 (76.4%)** | **0/12** | |
| Codon mutations: | TAC197TAT (5865C>T), AAT198AGC (5867A>G 5868T>C), AAA199AAG (5871A>G), TTG200CTA (5872T>C 5874G>A), ATA201ATT (5877A>T), GCT202GCA (5880T>A), TTA203CAA (5881T>C 5882T>A), CCA204CCT (5886A>T), AAT205AAA (5889T>A), ACA206ACC (5892A>C), AGT207AAT (5894G>A), AAA209AAC (5901A>C), TTA210CTA (5902T>C), GCC212AAA (5908G>A 5909C>A 5910C>A), ACC213ACA (5913C>A), TGC214TGT (5916C>T), AAC216AGY (5921A>G 5922C>Y), TAC217TAT (5925C>T), GGA218GGW (5928A>W), CTA219YTR (5929C>Y 5931A>R), CTA220YTA (5932C>Y), GAT221ARY (5935G>A 5936A>R 5937T>Y), ACT222ACC (5940T>C), CAG226TAT (5950C>T 5952G>T), ACA227GAY (5953A>G 5954C>A 5955A>Y), CAA229GAA (5959C>G), GAG230GAA (5964G>A), GCC232TCR (5968G>T 5970C>R), AAT233ACY (5972A>C 5973T>Y), CCT235CCA (5979T>A), AAG239AAA (5991G>A), TTC241TTT (5997C>T), ATG242ACA (5999T>C 6000G>A), CAA243ACA (6001C>A 6002A>C), AAG245AAA (6009G>A), AGG246AGA (6012G>A), ATA247ATT (6015A>T), ACT248ATA (6017C>T 6018T>A), GGG250GAC (6023G>A 6024G>C), ACA251AAC (6026C>A 6027A>C), TTC253TTT (6033C>T), TAC254TAT (6036C>T), GTA255ATA (6037G>A), CGA256AAG (6040C>A 6041G>A 6042A>G), TTC257TTT (6045C>T), GAA303AAT (6181G>A 6183A>T), TTC304TTT (6186C>T), AAC307AAA (6195C>A), AGG309AGA (6201G>A), ATA310ATT (6204A>T), GGA312GAA (6209G>A), ATA313CTA (6211A>C), TCT314GTT (6214T>G 6215C>T), ACT315ACG (6219T>G), CTA317ATC (6223C>A 6225A>C), CTA320TTA (6232C>T), TAT324TAC (6246T>C), CTA325ACA (6247C>A 6248T>C), CAG327GGG (6253C>G 6254A>G), AAT328AAC (6258T>C), GCA329CCA (6259G>C), ATT330ATG (6264T>G), AGC332AAT (6269G>A 6270C>T), TAT333CAT (6271T>C), TCA335ACC (6277T>A 6279A>C), AGG336AGA (6282G>A), GAA337GAC (6285A>C), ACA339ATA (6290C>T), TAT342TAC (6300T>C), AAT344CAT (6304A>C), TGC345ATC (6307T>A 6308G>T), AGA346_AGA349del (6310_6321delAGAGATATAAGA), GCA351AGA (6325G>A 6326C>G), CTA356GCA (6340C>G 6341T>C), AGA357ATG (6344G>T 6345A>G), GTC360ATA (6352G>A 6354C>A), TTA361ATG (6355T>A 6357A>G), AGT362ACA (6359G>C 6360T>A), CTT363TTG (6361C>T 6363T>G), AAA365AAT (6369A>T), GAA367GAG (6375A>G), CAA368AAA (6376C>A), CCT369CAA (6380C>A 6381T>A), ACA370CCT (6382A>C 6384A>T), ACC371ATT (6386C>T 6387C>T), AGG376AAA (6401G>A 6402G>A), AGG377AAG (6404G>A), AAT378AGA (6407A>G 6408T>A), TTC379TTT (6411C>T), ATC380ATT (6414C>T), TCC381TCG (6417C>G), CCA382AAA (6418C>A 6419C>A), GAT383GAA (6423T>A), CTG384TTA (6424C>T 6426G>A), TTA385CTA (6427T>C), AAG390AAA (6444G>A), CTA391CAA (6446T>A), AGT393GGG (6451A>G 6453T>G), CAC394CAA (6456C>A), TAT396TAC (6462T>C), CAC399CAT (6471C>T), TGC401TGT (6477C>T), AAA405AAC (6489A>C), ATT410GTA (6502A>G 6504T>A), GTT411ATC (6505G>A 6507T>C), GAC413GAT (6513C>T), CTG416CTA (6522G>A), GAA417GAC (6525A>C), TAA418GAA (6526T>G) | | | | | | | | |
| **Proteins** | | | | | | | | | |
| putative cell-to-cell movement protein (NP_569140.1) | 73 | 146 | 17.6% | 474 | 93.3% | 74 (100%) | 70 (94.6%) | 0/0/0/0 | 0 |
| Protein mutations: | G79K (2389G>A 2390G>A), I91T (2426T>C), E95G (2438A>G), R137T (2564G>C 2565A>G) | | | | | | | | |
| Codon mutations: | ATT73ATA (2373T>A), GAT74GAC (2376T>C), TGC75TGT (2379C>T), ACA78ACT (2388A>T), GGA79AAA (2389G>A 2390G>A), GGT80GGA (2394T>A), AAG81AAA (2397G>A), GCC82GCT (2400C>T), ATA91ACA (2426T>C), GAA95GGA (2438A>G), ATA96ATT (2442A>T), AAG100AAA (2454G>A), AAG106AAA (2472G>A), TAT107TAC (2475T>C), TTA110CTA (2482T>C), ACT113ACA (2493T>A), GCA119GCT (2511A>T), TGT120TGC (2514T>C), ACC127ACA (2535C>A), CTA133TTA (2551C>T), AGA137ACG (2564G>C 2565A>G), ATC139ATA (2571C>A) | | | | | | | | |
| polyprotein (NP_569141.1) | 176 | 449 | 41.7% | 1324 | 72.1% | 265 (100%) | 194 (73.2%) | 0/0/0/0 | 2 |
| Protein mutations: | E177Q (3939G>C), L179S (3946T>C 3947A>T), N181G (3951A>G 3952A>G 3953T>A), D182E (3956T>A), Q186E (3966C>G), G187K (3969G>A 3970G>A), E189D (3977G>T), H191S (3981C>A 3982A>G), K192H (3984A>C 3986A>C), T193E (3987A>G 3988C>A), V195G (3994T>G), K196I (3997A>T 3998A>T), I197L (3999A>T), E198I (4002G>A 4003A>T 4004A>C), I200K (4009T>A 4010T>A), E201D (4013A>T), E202L (4014G>C 4015A>T 4016G>T), N203T (4018A>C 4019T>G), S204I (4021G>T), T207R (4030C>G 4031A>G), Q208V (4032C>G 4033A>T), K209R (4036A>G), L211M (4041C>A 4043A>G), K212L (4044A>T 4045A>T), F215E (4053T>G 4054T>A 4055T>A), N216K (4058T>A), L218K (4062T>A 4063T>A), T219S (4065A>T 4067A>T), E220K (4068G>A), K222R (4075A>G), H224Q (4082T>G), K226Q (4086A>C), D227E (4091T>A), D230K (4098G>A 4100T>A), N231L (4101A>T 4102A>T 4103T>A), Y233* (4109T>A), Q235R (4113C>A 4114A>G), N238K (4124T>G), K240P (4128A>C 4129A>C), T242S (4135C>G 4136A>C), I247M (4151A>G), V248I (4152G>A), N249R (4156A>G 4157T>A), K250N (4160A>T), S252A (4164A>G 4165G>C 4166T>G), S259A (4185A>G 4186G>C 4187C>T), D264N (4200G>A), R266K (4207G>A), N267E (4209A>G 4211T>G), K273I (4228A>T), N276K (4238T>A), D311E (4343T>A), F345C (4444T>G), N351S (4462A>G 4463T>C), R368K (4513G>A), A381V (4552C>T 4553A>G), H382N (4554C>A), I384F (4560A>T 4562A>T), N386S (4566A>T 4567A>C 4568C>T), G388E (4573G>A 4574T>A), A396S (4596G>T), N401T (4612A>C 4613T>A), N413S (4648A>G 4649T>C), Q418* (4662C>T), T419S (4665A>T), L428S (4692C>T 4693T>C 4694T>A), D429N (4695G>A 4697T>C), N431Q (4701A>C 4703C>A), G443V (4738G>T), I444L (4740A>T) | | | | | | | | |
| Codon mutations: | AAA175.AT (3935A>T), GAA177CAA (3939G>C), AAA178AAG (3944A>G), TTA179TCT (3946T>C 3947A>T), AAT181GGA (3951A>G 3952A>G 3953T>A), GAT182GAA (3956T>A), AAC183AAT (3959C>T), CCT184CCC (3962T>C), CAA186GAA (3966C>G), GGA187AAA (3969G>A 3970G>A), GAG189GAT (3977G>T), CAT191AGT (3981C>A 3982A>G), AAA192CAC (3984A>C 3986A>C), ACA193GAA (3987A>G 3988C>A), GTA195GGA (3994T>G), AAA196ATT (3997A>T 3998A>T), ATA197TTA (3999A>T), GAA198ATC (4002G>A 4003A>T 4004A>C), CTA199CTT (4007A>T), ATT200AAA (4009T>A 4010T>A), GAA201GAT (4013A>T), GAG202CTT (4014G>C 4015A>T 4016G>T), AAT203ACG (4018A>C 4019T>G), AGT204ATT (4021G>T), ACA207AGG (4030C>G 4031A>G), CAA208GTA (4032C>G 4033A>T), AAA209AGA (4036A>G), CCT210CCA (4040T>A), CTA211ATG (4041C>A 4043A>G), AAA212TTA (4044A>T 4045A>T), AAC214AAT (4052C>T), TTT215GAA (4053T>G 4054T>A 4055T>A), AAT216AAA (4058T>A), TTA218AAA (4062T>A 4063T>A), ACA219TCT (4065A>T 4067A>T), GAA220AAA (4068G>A), TTT221TTC (4073T>C), AAA222AGA (4075A>G), ATA223ATT (4079A>T), CAT224CAG (4082T>G), ATA225ATT (4085A>T), AAA226CAA (4086A>C), GAT227GAA (4091T>A), TTA228CTT (4092T>C 4094A>T), TTG229CTC (4095T>C 4097G>C), GAT230AAA (4098G>A 4100T>A), AAT231TTA (4101A>T 4102A>T 4103T>A), TAT233TAA (4109T>A), CAA235AGA (4113C>A 4114A>G), AAT238AAG (4124T>G), AGC239AGT (4127C>T), AAA240CCA (4128A>C 4129A>C), ACA242AGC (4135C>G 4136A>C), CCA244CCT (4142A>T), ATA247ATG (4151A>G), GTA248ATA (4152G>A), AAT249AGA (4156A>G 4157T>A), AAA250AAT (4160A>T), AGT252GCG (4164A>G 4165G>C 4166T>G), AAA258AAG (4184A>G), AGC259GCT (4185A>G 4186G>C 4187C>T), AGG260CGA (4188A>C 4190G>A), GTC262GTT (4196C>T), GAC264AAC (4200G>A), AGA266AAA (4207G>A), AAT267GAG (4209A>G 4211T>G), TTA268CTT (4212T>C 4214A>T), AAA273ATA (4228A>T), TAT275TAC (4235T>C), AAT276AAA (4238T>A), TAT277TAC (4241T>C), CCC280CCT (4250C>T), AAT281AAC (4253T>C), TTA284CTT (4260T>C 4262A>T), AAG285AAA (4265G>A), CAA288CAG (4274A>G), AAG301AAA (4313G>A), TAT305TAC (4325T>C), CAT306CAC (4328T>C), TTG307CTA (4329T>C 4331G>A), CTA309TTA (4335C>T), GAG310GAA (4340G>A), GAT311GAA (4343T>A), GCT318GCA (4364T>A), CCA322CCC (4376A>C), GTT330GTA (4400T>A), TTA331CTA (4401T>C), CCA332CCT (4406A>T), GGA340GGT (4430A>T), TTT345TGT (4444T>G), GAC347GAT (4451C>T), AAT351AGC (4462A>G 4463T>C), GTA358GTT (4484A>T), CTA364TTA (4500C>T), CTA365TTA (4503C>T), AGA368AAA (4513G>A), TTC380TTT (4550C>T), GCA381GTG (4552C>T 4553A>G), CAT382AAT (4554C>A), ATT383ATA (4559T>A), ATA384TTT (4560A>T 4562A>T), AAC386TCT (4566A>T 4567A>C 4568C>T), TCT387TCA (4571T>A), GGT388GAA (4573G>A 4574T>A), AGC390AGT (4580C>T), AGC392AGT (4586C>T), ACC394ACA (4592C>A), AAA395AAG (4595A>G), GCA396TCA (4596G>T), ATT398ATA (4604T>A), AAT401ACA (4612A>C 4613T>A), CAG402CAA (4616G>A), AAT413AGC (4648A>G 4649T>C), CAA418TAA (4662C>T), ACT419TCT (4665A>T), GTA422GTG (4676A>G), ATC426ATT (4688C>T), AAT427AAC (4691T>C), CTT428TCA (4692C>T 4693T>C 4694T>A), GAT429AAC (4695G>A 4697T>C), AAC431CAA (4701A>C 4703C>A), ATT432ATA (4706T>A), AAA435AAG (4715A>G), AAG436AAA (4718G>A), GGA443GTA (4738G>T), ATA444TTA (4740A>T), AAC446AAT (4748C>T) | | | | | | | | |
| putative transactivator factor (NP_569142.1) | 197 | 418 | 42.8% | 803 | 65.8% | 175 (97.8%) | 114 (63.7%) | 0/4/1/0 | 0 |
| Protein mutations: | N198S (5867A>G 5868T>C), L203Q (5881T>C 5882T>A), N205K (5889T>A), S207N (5894G>A), K209N (5901A>C), A212K (5908G>A 5909C>A 5910C>A), N216S (5921A>G 5922C>Y), Q226Y (5950C>T 5952G>T), T227D (5953A>G 5954C>A 5955A>Y), Q229E (5959C>G), A232S (5968G>T 5970C>R), N233T (5972A>C 5973T>Y), M242T (5999T>C 6000G>A), Q243T (6001C>A 6002A>C), T248I (6017C>T 6018T>A), G250D (6023G>A 6024G>C), T251N (6026C>A 6027A>C), V255I (6037G>A), R256K (6040C>A 6041G>A 6042A>G), E303N (6181G>A 6183A>T), N307K (6195C>A), G312E (6209G>A), I313L (6211A>C), S314V (6214T>G 6215C>T), L317I (6223C>A 6225A>C), L325T (6247C>A 6248T>C), Q327G (6253C>G 6254A>G), A329P (6259G>C), I330M (6264T>G), S332N (6269G>A 6270C>T), Y333H (6271T>C), S335T (6277T>A 6279A>C), E337D (6285A>C), T339I (6290C>T), N344H (6304A>C), C345I (6307T>A 6308G>T), R346_R349del (6310_6321delAGAGATATAAGA), A351R (6325G>A 6326C>G), L356A (6340C>G 6341T>C), R357M (6344G>T 6345A>G), V360I (6352G>A 6354C>A), L361M (6355T>A 6357A>G), S362T (6359G>C 6360T>A), K365N (6369A>T), Q368K (6376C>A), P369Q (6380C>A 6381T>A), T370P (6382A>C 6384A>T), T371I (6386C>T 6387C>T), R376K (6401G>A 6402G>A), R377K (6404G>A), N378R (6407A>G 6408T>A), P382K (6418C>A 6419C>A), D383E (6423T>A), L391Q (6446T>A), S393G (6451A>G 6453T>G), H394Q (6456C>A), K405N (6489A>C), I410V (6502A>G 6504T>A), V411I (6505G>A 6507T>C), E417D (6525A>C), *418E (6526T>G) | | | | | | | | |
| Codon mutations: | TAC197TAT (5865C>T), AAT198AGC (5867A>G 5868T>C), AAA199AAG (5871A>G), TTG200CTA (5872T>C 5874G>A), ATA201ATT (5877A>T), GCT202GCA (5880T>A), TTA203CAA (5881T>C 5882T>A), CCA204CCT (5886A>T), AAT205AAA (5889T>A), ACA206ACC (5892A>C), AGT207AAT (5894G>A), AAA209AAC (5901A>C), TTA210CTA (5902T>C), GCC212AAA (5908G>A 5909C>A 5910C>A), ACC213ACA (5913C>A), TGC214TGT (5916C>T), AAC216AGY (5921A>G 5922C>Y), TAC217TAT (5925C>T), GGA218GGW (5928A>W), CTA219YTR (5929C>Y 5931A>R), CTA220YTA (5932C>Y), GAT221ARY (5935G>A 5936A>R 5937T>Y), ACT222ACC (5940T>C), CAG226TAT (5950C>T 5952G>T), ACA227GAY (5953A>G 5954C>A 5955A>Y), CAA229GAA (5959C>G), GAG230GAA (5964G>A), GCC232TCR (5968G>T 5970C>R), AAT233ACY (5972A>C 5973T>Y), CCT235CCA (5979T>A), AAG239AAA (5991G>A), TTC241TTT (5997C>T), ATG242ACA (5999T>C 6000G>A), CAA243ACA (6001C>A 6002A>C), AAG245AAA (6009G>A), AGG246AGA (6012G>A), ATA247ATT (6015A>T), ACT248ATA (6017C>T 6018T>A), GGG250GAC (6023G>A 6024G>C), ACA251AAC (6026C>A 6027A>C), TTC253TTT (6033C>T), TAC254TAT (6036C>T), GTA255ATA (6037G>A), CGA256AAG (6040C>A 6041G>A 6042A>G), TTC257TTT (6045C>T), GAA303AAT (6181G>A 6183A>T), TTC304TTT (6186C>T), AAC307AAA (6195C>A), AGG309AGA (6201G>A), ATA310ATT (6204A>T), GGA312GAA (6209G>A), ATA313CTA (6211A>C), TCT314GTT (6214T>G 6215C>T), ACT315ACG (6219T>G), CTA317ATC (6223C>A 6225A>C), CTA320TTA (6232C>T), TAT324TAC (6246T>C), CTA325ACA (6247C>A 6248T>C), CAG327GGG (6253C>G 6254A>G), AAT328AAC (6258T>C), GCA329CCA (6259G>C), ATT330ATG (6264T>G), AGC332AAT (6269G>A 6270C>T), TAT333CAT (6271T>C), TCA335ACC (6277T>A 6279A>C), AGG336AGA (6282G>A), GAA337GAC (6285A>C), ACA339ATA (6290C>T), TAT342TAC (6300T>C), AAT344CAT (6304A>C), TGC345ATC (6307T>A 6308G>T), AGA346_AGA349del (6310_6321delAGAGATATAAGA), GCA351AGA (6325G>A 6326C>G), CTA356GCA (6340C>G 6341T>C), AGA357ATG (6344G>T 6345A>G), GTC360ATA (6352G>A 6354C>A), TTA361ATG (6355T>A 6357A>G), AGT362ACA (6359G>C 6360T>A), CTT363TTG (6361C>T 6363T>G), AAA365AAT (6369A>T), GAA367GAG (6375A>G), CAA368AAA (6376C>A), CCT369CAA (6380C>A 6381T>A), ACA370CCT (6382A>C 6384A>T), ACC371ATT (6386C>T 6387C>T), AGG376AAA (6401G>A 6402G>A), AGG377AAG (6404G>A), AAT378AGA (6407A>G 6408T>A), TTC379TTT (6411C>T), ATC380ATT (6414C>T), TCC381TCG (6417C>G), CCA382AAA (6418C>A 6419C>A), GAT383GAA (6423T>A), CTG384TTA (6424C>T 6426G>A), TTA385CTA (6427T>C), AAG390AAA (6444G>A), CTA391CAA (6446T>A), AGT393GGG (6451A>G 6453T>G), CAC394CAA (6456C>A), TAT396TAC (6462T>C), CAC399CAT (6471C>T), TGC401TGT (6477C>T), AAA405AAC (6489A>C), ATT410GTA (6502A>G 6504T>A), GTT411ATC (6505G>A 6507T>C), GAC413GAT (6513C>T), CTG416CTA (6522G>A), GAA417GAC (6525A>C), TAA418GAA (6526T>G) | | | | | | | | |

*: Inserts / Deletes / Misaligned / Frameshifts

#### Analysis details

This analysis was performed with panviral2.64

### NGS Details (UN62): Tomato chocolate spot virus (segment RNA2)

#### Assembly

| | |
|---|---|
| **Coverage Length** | 409 (1 contig(s)) |
| **Depth Of Coverage** | 5209.1 |
| **Number Of Reads** | 18591 |
| **Reads Per Million** | 430.96 rpm (after QC) |
| **Ambiguities** | 0 |
| **Assembly Method** | de novo + reference guided assembly |
| **Consensus Caller** | Bcf Tools |

#### Coverage Map

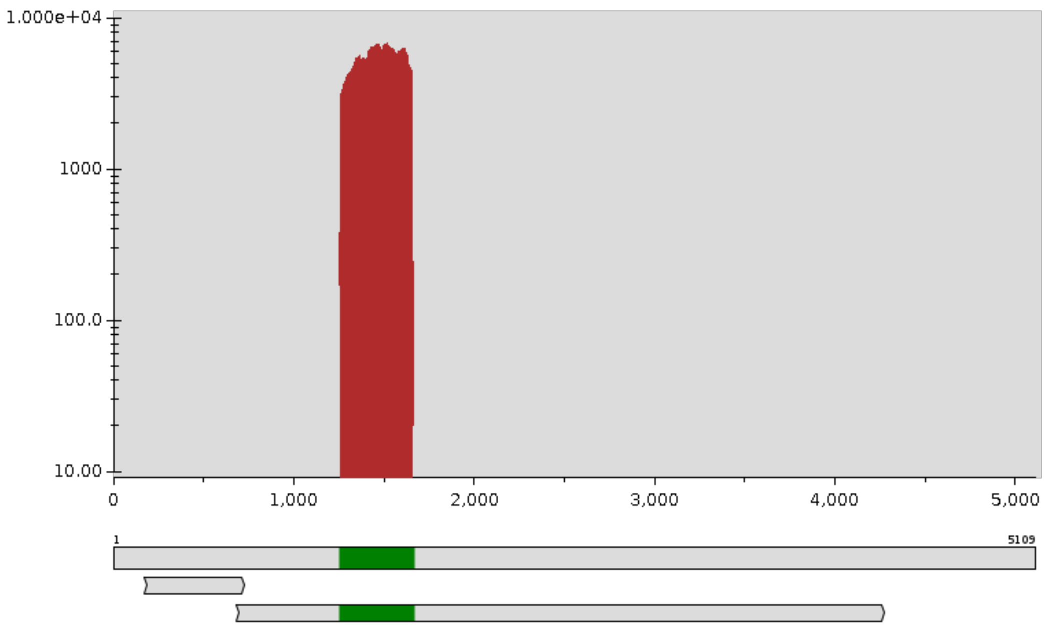

#### Assignment

| | |
|---|---|
| **Type** | Tomato chocolate spot virus (Taxonomy ID: 661101) |
| **Reference Genome** | NC_013076.1 |
| **NT Identity (%)** | 66.988 |
| **AA Identity (%)** | 70.5036 |
| **Number Of Stop Codons** | 0 |
| **Number Of CDS** | 2 |

#### Alignment

| | |
|---|---|
| **Alignment Score** | 279.0 (NT) + 653.0 (AA) = 932.0 |
| **Concordance (%)** | 53.8728 |

| **Alignment Method** | Global, seeded, nucleotide + amino acids (AGA) |
|---|---|

#### Genome Region

Sequence starts at position 1256 and ends at position 1664 relative to NC_013076.1 reference sequence.

#### Alignment Detailed Statistics

| | Begin | End | Coverage | Score | Concordance | Matches | Identities | I/D/M/F* | Stop Codons |
|---|---|---|---|---|---|---|---|---|---|
| **NT** | **1256** | **1664** | **8.0%** | **279** | **34.1%** | **409 (98.6%)** | **278 (67.0%)** | **6/0** | |
| Mutations: | 1267G>A, 1273T>A, 1277T>A, 1278C>G, 1285C>A, 1291A>G, 1294A>T, 1295T>C, 1297G>C, 1300C>T, 1306T>A, 1309A>G, 1312T>C, 1315C>T, 1316C>T, 1317A>C, 1318A>T, 1327G>A, 1333T>A, 1336C>A, 1351A>T, 1354A>C, 1360T>G, 1366G>A, 1367C>G, 1369A>G, 1372T>G, 1373T>G, 1375C>G, 1376C>T, 1377A>C, 1378G>T, 1379A>C, 1380T>A, 1381G>A, 1385T>C, 1387G>A, 1394C>A, 1396T>A, 1397G>C, 1405T>A, 1408A>T, 1410A>C, 1411G>A, 1412A>C, 1413G>T, 1414G>T, 1423T>C, 1426A>T, 1428G>A, 1429A>C, 1433G>A, 1438A>T, 1440T>C, 1441G>C, 1442A>C, 1443G>A, 1444T>G, 1448A>T, 1449G>C, 1450C>T, 1453C>A, 1454A>T, 1456G>A, 1458C>T, 1459T>G, 1463A>G, 1464G>C, 1465C>G, 1468T>A, 1471C>G, 1474G>T, 1477A>C, 1483G>A, 1489C>A, 1493G>A, 1495G>C, 1501T>C, 1503G>A, 1505C>A, 1513A>T, 1516C>G, 1522A>G, 1525A>G, 1531C>T, 1532G>T, 1534G>T, 1535C>T, 1537T>A, 1539A>G, 1540C>G, 1541A>G, 1547C>T, 1548G>C, 1551T>C, 1552A>G, 1555_1556insGTAGGT, 1556A>G, 1559C>A, 1572C>G, 1573T>G, 1574C>G, 1575A>T, 1576C>A, 1582A>T, 1585C>T, 1586A>T, 1588T>C, 1589G>A, 1590G>A, 1592G>T, 1593G>T, 1595T>C, 1603A>C, 1606T>C, 1610A>G, 1618A>G, 1620G>C, 1622T>A, 1624C>T, 1625A>G, 1627C>T, 1628A>T, 1630G>T, 1633T>C, 1634G>C, 1635T>A, 1636T>G, 1645T>C, 1649C>T, 1650T>A, 1651G>T | | | | | | | | |

##### CDS

| | Begin | End | Coverage | Score | Concordance | Matches | Identities | I/D/M/F* | Stop Codons |
|---|---|---|---|---|---|---|---|---|---|
| ToChV_s2_gp2 | 191 | 327 | 11.4% | 653 | 70.7% | 137 (98.6%) | 98 (70.5%) | 2/0/0/0 | 0 |
| Protein mutations: | Q211S (1316C>T 1317A>C 1318A>T), Q228E (1367C>G 1369A>G), S229R (1372T>G), S230A (1373T>G 1375C>G), Q231S (1376C>T 1377A>C 1378G>T), M232Q (1379A>C 1380T>A 1381G>A), A238P (1397G>C), E242A (1410A>C 1411G>A), R243L (1412A>C 1413G>T 1414G>T), R248H (1428G>A 1429A>C), D250N (1433G>A), L252P (1440T>C 1441G>C), S253Q (1442A>C 1443G>A 1444T>G), M257L (1454A>T 1456G>A), T258M (1458C>T 1459T>G), S260A (1463A>G 1464G>C 1465C>G), V270I (1493G>A 1495G>C), R273K (1503G>A), V283F (1532G>T 1534G>T), P284S (1535C>T 1537T>A), N285R (1539A>G 1540C>G), T286A (1541A>G), R288S (1547C>T 1548G>C), L289S (1551T>C 1552A>G), T290_M291insVG (1555_1556insGTAGGT), M291V (1556A>G), A296G (1572C>G 1573T>G), H297V (1574C>G 1575A>T 1576C>A), E299D (1582A>T), T301S (1586A>T 1588T>C), G302N (1589G>A 1590G>A), G303L (1592G>T 1593G>T), K309E (1610A>G), G312A (1620G>C), F313I (1622T>A 1624C>T), N314D (1625A>G 1627C>T), M315F (1628A>T 1630G>T), V317Q (1634G>C 1635T>A 1636T>G), L322Y (1649C>T 1650T>A 1651G>T) | | | | | | | | |
| Codon mutations: | CAG194CAA (1267G>A), ACT196ACA (1273T>A), TCT198AGT (1277T>A 1278C>G), GCC200GCA (1285C>A), GAA202GAG (1291A>G), GCA203GCT (1294A>T), TTG204CTC (1295T>C 1297G>C), AAC205AAT (1300C>T), CCT207CCA (1306T>A), CAA208CAG (1309A>G), TTT209TTC (1312T>C), TTC210TTT (1315C>T), CAA211TCT (1316C>T 1317A>C 1318A>T), CAG214CAA (1327G>A), GCT216GCA (1333T>A), GCC217GCA (1336C>A), TCA222TCT (1351A>T), GTA223GTC (1354A>C), TCT225TCG (1360T>G), GGG227GGA (1366G>A), CAA228GAG (1367C>G 1369A>G), AGT229AGG (1372T>G), TCC230GCG (1373T>G 1375C>G), CAG231TCT (1376C>T 1377A>C 1378G>T), ATG232CAA (1379A>C 1380T>A 1381G>A), TTG234CTA (1385T>C 1387G>A), CGT237AGA (1394C>A 1396T>A), GCT238CCT (1397G>C), GTT240GTA (1405T>A), GGA241GGT (1408A>T), GAG242GCA (1410A>C 1411G>A), AGG243CTT (1412A>C 1413G>T 1414G>T), AAT246AAC (1423T>C), ACA247ACT (1426A>T), CGA248CAC (1428G>A 1429A>C), GAT250AAT (1433G>A), GTA251GTT (1438A>T), CTG252CCC (1440T>C 1441G>C), AGT253CAG (1442A>C 1443G>A 1444T>G), AGC255TCT (1448A>T 1449G>C 1450C>T), TCC256TCA (1453C>A), ATG257TTA (1454A>T 1456G>A), ACT258ATG (1458C>T 1459T>G), AGC260GCG (1463A>G 1464G>C 1465C>G), TCT261TCA (1468T>A), TCC262TCG (1471C>G), CTG263CTT (1474G>T), CGA264CGC (1477A>C), CCG266CCA (1483G>A), CTC268CTA (1489C>A), GTG270ATC (1493G>A 1495G>C), AGT272AGC (1501T>C), AGG273AAG (1503G>A), CGG274AGG (1505C>A), CCA276CCT (1513A>T), TCC277TCG (1516C>G), GAA279GAG (1522A>G), CTA280CTG (1525A>G), GAC282GAT (1531C>T), GTG283TTT (1532G>T 1534G>T), CCT284TCA (1535C>T 1537T>A), AAC285AGG (1539A>G 1540C>G), ACC286GCC (1541A>G), CGA288TCA (1547C>T 1548G>C), TTA289TCG (1551T>C 1552A>G), ACA290_ATG291insGTAGGT (1555_1556insGTAGGT), ATG291GTG (1556A>G), CGA292AGA (1559C>A), GCT296GGG (1572C>G 1573T>G), CAC297GTA (1574C>G 1575A>T 1576C>A), GAA299GAT (1582A>T), GAC300GAT (1585C>T), ACT301TCC (1586A>T 1588T>C), GGT302AAT (1589G>A 1590G>A), GGA303TTA (1592G>T 1593G>T), TTG304CTG (1595T>C), CCA306CCC (1603A>C), GTT307GTC (1606T>C), AAG309GAG (1610A>G), CAA311CAG (1618A>G), GGC312GCC (1620G>C), TTC313ATT (1622T>A 1624C>T), AAC314GAT (1625A>G 1627C>T), ATG315TTT (1628A>T 1630G>T), GGT316GGC (1633T>C), GTT317CAG (1634G>C 1635T>A 1636T>G), GAT320GAC (1645T>C), CTG322TAT (1649C>T 1650T>A 1651G>T) | | | | | | | | |

##### Proteins

| | Begin | End | Coverage | Score | Concordance | Matches | Identities | I/D/M/F* | Stop Codons |
|---|---|---|---|---|---|---|---|---|---|
| polyprotein (YP_003097231.1) | 191 | 327 | 11.4% | 653 | 70.7% | 137 (98.6%) | 98 (70.5%) | 2/0/0/0 | 0 |
| Protein mutations: | Q211S (1316C>T 1317A>C 1318A>T), Q228E (1367C>G 1369A>G), S229R (1372T>G), S230A (1373T>G 1375C>G), Q231S (1376C>T 1377A>C 1378G>T), M232Q (1379A>C 1380T>A 1381G>A), A238P (1397G>C), E242A (1410A>C 1411G>A), R243L (1412A>C 1413G>T 1414G>T), R248H (1428G>A 1429A>C), D250N (1433G>A), L252P (1440T>C 1441G>C), S253Q (1442A>C 1443G>A 1444T>G), M257L (1454A>T 1456G>A), T258M (1458C>T 1459T>G), S260A (1463A>G 1464G>C 1465C>G), V270I (1493G>A 1495G>C), R273K (1503G>A), V283F (1532G>T 1534G>T), P284S (1535C>T 1537T>A), N285R (1539A>G 1540C>G), T286A (1541A>G), R288S (1547C>T 1548G>C), L289S (1551T>C 1552A>G), T290_M291insVG (1555_1556insGTAGGT), M291V (1556A>G), A296G (1572C>G 1573T>G), H297V (1574C>G 1575A>T 1576C>A), E299D (1582A>T), T301S (1586A>T 1588T>C), G302N (1589G>A 1590G>A), G303L (1592G>T 1593G>T), K309E (1610A>G), G312A (1620G>C), F313I (1622T>A 1624C>T), N314D (1625A>G 1627C>T), M315F (1628A>T 1630G>T), V317Q (1634G>C 1635T>A 1636T>G), L322Y (1649C>T 1650T>A 1651G>T) | | | | | | | | |
| Codon mutations: | CAG194CAA (1267G>A), ACT196ACA (1273T>A), TCT198AGT (1277T>A 1278C>G), GCC200GCA (1285C>A), GAA202GAG (1291A>G), GCA203GCT (1294A>T), TTG204CTC (1295T>C 1297G>C), AAC205AAT (1300C>T), CCT207CCA (1306T>A), CAA208CAG (1309A>G), TTT209TTC (1312T>C), TTC210TTT (1315C>T), CAA211TCT (1316C>T 1317A>C 1318A>T), CAG214CAA (1327G>A), GCT216GCA (1333T>A), GCC217GCA (1336C>A), TCA222TCT (1351A>T), GTA223GTC (1354A>C), TCT225TCG (1360T>G), GGG227GGA (1366G>A), CAA228GAG (1367C>G 1369A>G), AGT229AGG (1372T>G), TCC230GCG (1373T>G 1375C>G), CAG231TCT (1376C>T 1377A>C 1378G>T), ATG232CAA (1379A>C 1380T>A 1381G>A), TTG234CTA (1385T>C 1387G>A), CGT237AGA (1394C>A 1396T>A), GCT238CCT (1397G>C), GTT240GTA (1405T>A), GGA241GGT (1408A>T), GAG242GCA (1410A>C 1411G>A), AGG243CTT (1412A>C 1413G>T 1414G>T), AAT246AAC (1423T>C), ACA247ACT (1426A>T), CGA248CAC (1428G>A 1429A>C), GAT250AAT (1433G>A), GTA251GTT (1438A>T), CTG252CCC (1440T>C 1441G>C), AGT253CAG (1442A>C 1443G>A 1444T>G), AGC255TCT (1448A>T 1449G>C 1450C>T), TCC256TCA (1453C>A), ATG257TTA (1454A>T 1456G>A), ACT258ATG (1458C>T 1459T>G), AGC260GCG (1463A>G 1464G>C 1465C>G), TCT261TCA (1468T>A), TCC262TCG (1471C>G), CTG263CTT (1474G>T), CGA264CGC (1477A>C), CCG266CCA (1483G>A), CTC268CTA (1489C>A), GTG270ATC (1493G>A 1495G>C), AGT272AGC (1501T>C), AGG273AAG (1503G>A), CGG274AGG (1505C>A), CCA276CCT (1513A>T), TCC277TCG (1516C>G), GAA279GAG (1522A>G), CTA280CTG (1525A>G), GAC282GAT (1531C>T), GTG283TTT (1532G>T 1534G>T), CCT284TCA (1535C>T 1537T>A), AAC285AGG (1539A>G 1540C>G), ACC286GCC (1541A>G), CGA288TCA (1547C>T 1548G>C), TTA289TCG (1551T>C 1552A>G), ACA290_ATG291insGTAGGT (1555_1556insGTAGGT), ATG291GTG (1556A>G), CGA292AGA (1559C>A), GCT296GGG (1572C>G 1573T>G), CAC297GTA (1574C>G 1575A>T 1576C>A), GAA299GAT (1582A>T), GAC300GAT (1585C>T), ACT301TCC (1586A>T 1588T>C), GGT302AAT (1589G>A 1590G>A), GGA303TTA (1592G>T 1593G>T), TTG304CTG (1595T>C), CCA306CCC (1603A>C), GTT307GTC (1606T>C), AAG309GAG (1610A>G), CAA311CAG (1618A>G), GGC312GCC (1620G>C), TTC313ATT (1622T>A 1624C>T), AAC314GAT (1625A>G 1627C>T), ATG315TTT (1628A>T 1630G>T), GGT316GGC (1633T>C), GTT317CAG (1634G>C 1635T>A 1636T>G), GAT320GAC (1645T>C), CTG322TAT (1649C>T 1650T>A 1651G>T) | | | | | | | | |

*: Inserts / Deletes / Misaligned / Frameshifts

#### Analysis details

This analysis was performed with panviral2.64

### NGS Details (UN62): Tomato chocolate spot virus (segment RNA 1)

#### Assembly

| | |
|---|---|
| **Coverage Length** | 278 (1 contig(s)) |
| **Depth Of Coverage** | 24274.9 |
| **Number Of Reads** | 61095 |
| **Reads Per Million** | 1416.24 rpm (after QC) |
| **Ambiguities** | 0 |
| **Assembly Method** | de novo + reference guided assembly |
| **Consensus Caller** | Bcf Tools |

#### Coverage Map

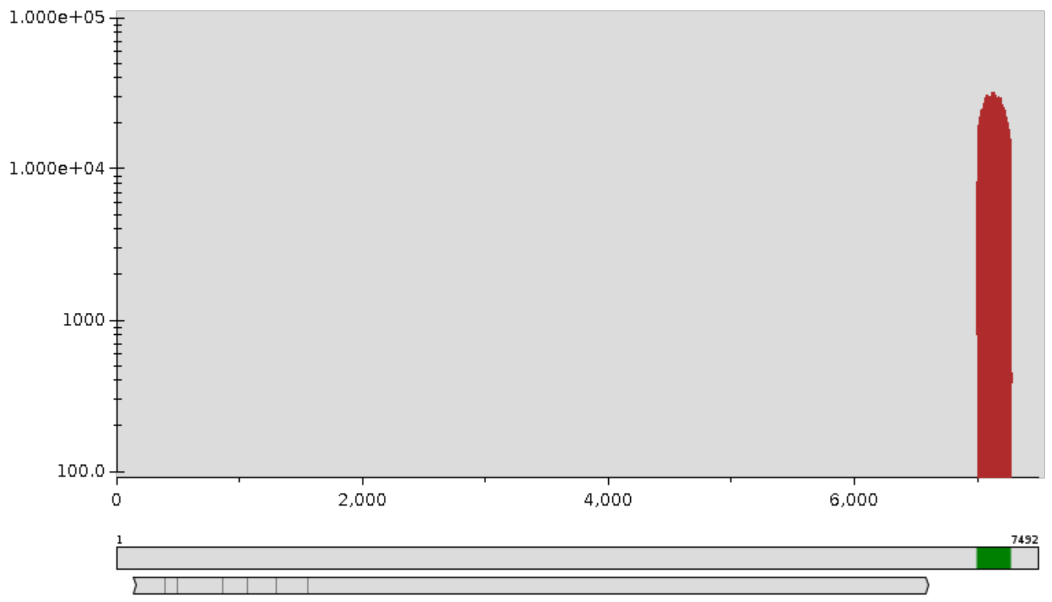

#### Assignment

| | |
|---|---|
| **Type** | Tomato chocolate spot virus (Taxonomy ID: 661101) |
| **Reference Genome** | NC_013075.1 |
| **NT Identity (%)** | 82.0789 |
| **AA Identity (%)** | 0.0 |
| **Number Of Stop Codons** | 0 |
| **Number Of CDS** | 1 |

#### Alignment

| | |
|---|---|
| **Alignment Score** | 350.0 (NT) + 0.0 (AA) = 350.0 |
| **Concordance (%)** | 62.9496 |

| Alignment Method | Global, seeded, nucleotide + amino acids (AGA) |
|---|---|

#### Genome Region

Sequence starts at position 6998 and ends at position 7275 relative to NC_013075.1 reference sequence.

#### Alignment Detailed Statistics

| | Begin | End | Coverage | Score | Concordance | Matches | Identities | I/D/M/F* | Stop Codons |
|---|---|---|---|---|---|---|---|---|---|
| **NT** | **6998** | **7275** | **3.7%** | **350** | **62.9%** | **278 (99.6%)** | **229 (82.1%)** | **1/0** | |
| Mutations: | 7007T>C, 7012T>G, 7018C>T, 7019A>G, 7020A>G, 7020_7021insT, 7023C>A, 7042G>T, 7044A>T, 7047G>A, 7048C>G, 7066G>A, 7082G>A, 7083T>C, 7098C>T, 7099T>C, 7104T>A, 7107G>A, 7116C>G, 7121G>A, 7127A>C, 7140C>T, 7151A>T, 7152T>C, 7183C>T, 7184T>C, 7185G>T, 7186T>G, 7187T>C, 7188G>A, 7196C>T, 7197A>G, 7198A>C, 7199C>A, 7200A>G, 7201G>A, 7207A>T, 7208A>T, 7209T>G, 7210C>T, 7212T>A, 7219T>A, 7224T>C, 7242A>T, 7255A>T, 7257G>A, 7258G>C, 7260T>C, 7262C>G, 7264T>C | | | | | | | | |

**CDS**

**Proteins**

*: Inserts / Deletes / Misaligned / Frameshifts

#### Analysis details

This analysis was performed with panviral2.64

### NGS Details (UN62): Noumeavirus

#### Assembly

| | |
|---|---|
| **Coverage Length** | 217 (1 contig(s)) |
| **Depth Of Coverage** | 10898.8 |
| **Number Of Reads** | 24245 |
| **Reads Per Million** | 562.02 rpm (after QC) |
| **Ambiguities** | 0 |
| **Assembly Method** | de novo + reference guided assembly |
| **Consensus Caller** | Bcf Tools |

#### Coverage Map

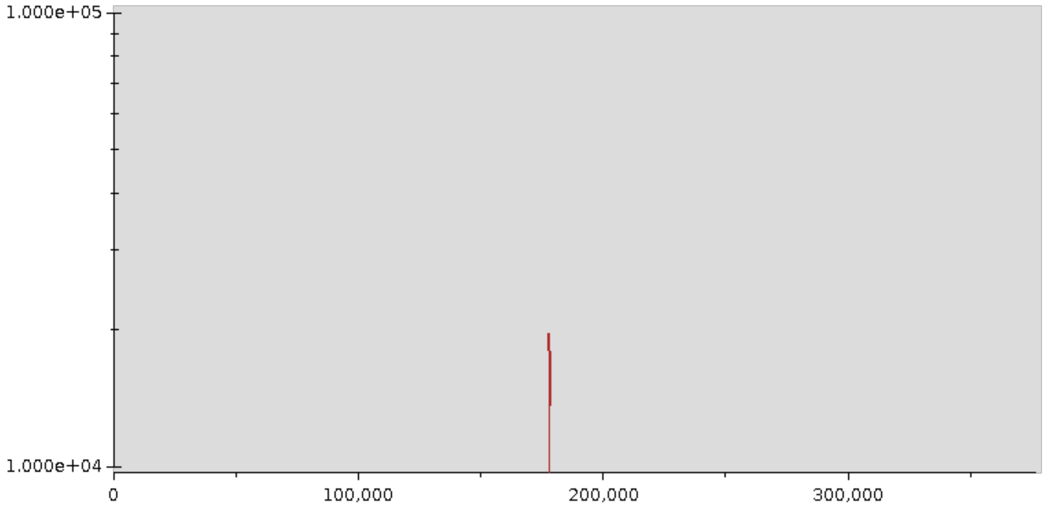

#### Assignment

| | |
|---|---|
| **Type** | Noumeavirus (Taxonomy ID: 1955558) |
| **Reference Genome** | NC_033775.1 |
| **NT Identity (%)** | 82.0276 |
| **AA Identity (%)** | 94.4444 |
| **Number Of Stop Codons** | 0 |
| **Number Of CDS** | 452 |

#### Alignment

| | |
|---|---|
| **Alignment Score** | 278.0 (NT) + 437.0 (AA) = 715.0 |
| **Concordance (%)** | 80.8824 |
| **Alignment Method** | Local, heuristic, nucleotide (BLASTN) |

#### Genome Region

Sequence starts at position 177922 and ends at position 178138 relative to NC_033775.1 reference sequence.

#### Alignment Detailed Statistics

| | Begin | End | Coverage | Score | Concordance | Matches | Identities | I/D/M/F* | Stop Codons |
|---|---|---|---|---|---|---|---|---|---|
| **NT** | **177922** | **178138** | **0.1%** | **278** | **64.1%** | **217 (100%)** | **178 (82.0%)** | **0/0** | |

Mutations: 177944C>G, 177956G>C, 177965C>T, 177974A>G, 177977C>T, 177980A>T, 177992G>C, 177995C>T, 177998C>G, 178002C>G, 178003A>C, 178004A>T, 178019A>G, 178025A>G, 178034G>A, 178043A>G, 178044C>A, 178046A>G, 178049T>G, 178058C>T, 178061A>T, 178064A>G, 178073A>G, 178079A>C, 178080A>C, 178082G>C, 178091C>G, 178094T>C, 178097T>C, 178100C>T, 178101G>A, 178106A>G, 178109A>G, 178112A>G, 178115A>G, 178118A>C, 178119C>T, 178130T>G, 178136A>G

*: Inserts / Deletes / Misaligned / Frameshifts

#### Analysis details

This analysis was performed with panviral2.64

### NGS Details (UN62): Tomato necrotic dwarf virus (segment RNA1)

#### Assembly

| | |
|---|---|
| **Coverage Length** | 749 (2 contig(s)) |
| **Depth Of Coverage** | 356.8 |
| **Number Of Reads** | 2020 |
| **Reads Per Million** | 46.83 rpm (after QC) |
| **Ambiguities** | 0 |
| **Assembly Method** | de novo + reference guided assembly |
| **Consensus Caller** | Bcf Tools |

#### Coverage Map

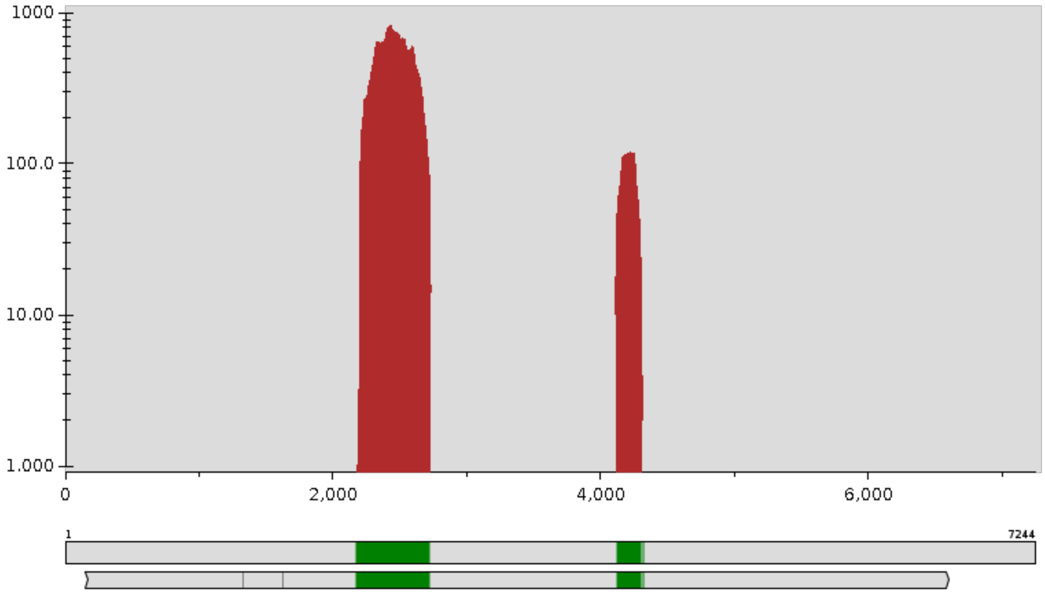

#### Assignment

| | |
|---|---|
| **Type** | Tomato necrotic dwarf virus (Taxonomy ID: 1481465) |
| **Reference Genome** | NC_027926.1 |
| **NT Identity (%)** | 64.7606 |
| **AA Identity (%)** | 63.2 |
| **Number Of Stop Codons** | 0 |
| **Number Of CDS** | 1 |

#### Alignment

| | |
|---|---|
| **Alignment Score** | 438.0 (NT) + 1167.0 (AA) = 1605.0 |
| **Concordance (%)** | 49.0076 |

| | |
|---|---|
| **Alignment Method** | Global, seeded, nucleotide + amino acids (AGA) |

#### Genome Region

Sequence starts at position 2174 and ends at position 4311 relative to NC_027926.1 reference sequence.

#### Alignment Detailed Statistics

| | Begin | End | Coverage | Score | Concordance | Matches | Identities | I/D/M/F* | Stop Codons |
|---|---|---|---|---|---|---|---|---|---|
| **NT** | **2174** | **4311** | **10.3%** | **438** | **29.2%** | **749 (99.6%)** | **487 (64.8%)** | **3/0** | |
| Mutations: | 2174C>A, 2178C>T, 2179A>T, 2182A>T, 2184C>G, 2185T>G, 2186A>T, 2193T>A, 2194G>T, 2196C>A, 2198A>G, 2199T>A, 2200G>C, 2201C>A, 2203G>C, 2205A>G, 2208A>G, 2211C>T, 2212A>G, 2213G>C, 2214G>T, 2225A>G, 2229T>C, 2245C>A, 2256T>C, 2259A>G, 2262C>T, 2268C>T, 2272G>A, 2273C>G, 2274T>C, 2275T>A, 2277C>A, 2283C>T, 2285G>C, 2290A>G, 2293A>C, 2294A>T, 2300C>A, 2304A>C, 2308A>C, 2309C>G, 2310A>C, 2311G>A, 2313G>T, 2314A>C, 2316G>T, 2317C>G, 2318G>A, 2319C>G, 2320A>C, 2321C>A, 2325A>G, 2326T>A, 2327T>A, 2328T>A, 2331A>G, 2332C>T, 2337T>G, 2338T>G, 2339G>C, 2340T>G, 2341A>C, 2344G>C, 2346T>G, 2354T>C, 2355T>A, 2362T>C, 2364G>T, 2366T>C, 2367T>G, 2370C>T, 2373C>T, 2374C>G, 2376G>A, 2389T>A, 2390T>G, 2391T>A, 2394G>A, 2400T>C, 2401T>C, 2409A>T, 2412C>T, 2421T>C, 2424T>C, 2432C>G, 2433C>G, 2440T>G, 2442C>A, 2442_2443insGGT, 2444A>C, 2445A>T, 2448G>C, 2449G>A, 2450C>A, 2451C>G, 2454A>G, 2455G>A, 2456A>G, 2457C>A, 2458T>A, 2459C>A, 2464G>A, 2466C>T, 2468A>T, 2469G>C, 2470A>G, 2471T>A, 2475T>C, 2476T>A, 2477A>T, 2478C>G, 2479C>A, 2480A>T, 2481C>T, 2482A>G, 2483A>G, 2484A>T, 2487T>C, 2488G>A, 2489T>A, 2493T>C, 2494T>A, 2496C>A, 2501A>T, 2502A>C, 2503G>C, 2505A>G, 2507C>T, 2508T>G, 2512T>A, 2513G>A, 2514G>T, 2515A>G, 2520A>G, 2523A>C, 2525A>T, 2526T>C, 2527G>A, 2528C>T, 2529A>C, 2530C>T, 2534G>A, 2553C>A, 2556A>G, 2560A>T, 2562C>G, 2565G>A, 2568G>A, 2577A>C, 2578T>A, 2579T>C, 2580G>C, 2581T>A, 2583T>G, 2584G>A, 2585G>A, 2586G>T, 2588G>C, 2589A>C, 2590A>G, 2592A>T, 2595A>T, 2601T>C, 2602C>A, 2605G>A, 2610T>C, 2611T>C, 2612G>T, 2613G>C, 2614C>A, 2616A>G, 2617A>G, 2618G>C, 2619C>A, 2620A>G, 2621A>C, 2622T>A, 2623T>A, 2627T>C, 2628T>C, 2631A>T, 2632A>T, 2633G>T, 2634T>A, 2638A>G, 2639G>A, 2643A>C, 2649T>G, 2652T>C, 2655C>A, 2658A>C, 2673C>T, 2679T>C, 2685A>G, 2688G>A, 2691C>T, 2694G>A, 2697C>G, 2698C>A, 2699C>A, 2700C>G, 2701T>A, 2705G>A, 2707C>T, 2709G>C, 2710A>G, 2711A>C, 2712G>A, 2715T>A, 2717T>C, 2718G>C, 2722G>C, 2724G>A, 4116T>C, 4119G>A, 4122A>C, 4137C>T, 4149C>G, 4150A>C, 4161T>C, 4173C>T, 4176C>T, 4177A>T, 4178C>G, 4179C>T, 4188A>T, 4189T>G, 4190T>C, 4197T>C, 4200C>A, 4201A>C, 4206A>T, 4209T>C, 4215T>A, 4218A>T, 4221A>C, 4222G>A, 4230C>G, 4233T>C, 4234T>C, 4235C>A, 4243T>A, 4245T>A, 4246C>G, 4248T>G, 4251T>G, 4254C>T, 4256A>G, 4257C>T, 4259A>G, 4263C>T, 4267G>T, 4269T>A, 4279A>G, 4280C>A, 4281T>G, 4284G>A, 4287T>C, 4293T>C, 4302A>G, 4305T>C, 4311C>T | | | | | | | | |

##### CDS

| | Begin | End | Coverage | Score | Concordance | Matches | Identities | I/D/M/F* | Stop Codons |
|---|---|---|---|---|---|---|---|---|---|
| APL33_sRA11gp1 | 676 | 1387 | 11.6% | 1167 | 64.8% | 249 (99.6%) | 158 (63.2%) | 1/0/0/0 | 0 |
| Protein mutations: | M677L (2179A>T), I678L (2182A>T 2184C>G), Y679V (2185T>G 2186A>T), A682S (2194G>T 2196C>A), N683R (2198A>G 2199T>A), A684H (2200G>C 2201C>A), E685Q (2203G>C 2205A>G), R688A (2212A>G 2213G>C 2214G>T), Q692R (2225A>G), L699I (2245C>A), A708S (2272G>A 2273C>G 2274T>C), S709T (2275T>A 2277C>A), C712S (2285G>C), K714E (2290A>G), K715L (2293A>C 2294A>T), A717D (2300C>A), T720R (2308A>C 2309C>G 2310A>C), E721N (2311G>A 2313G>T), M722L (2314A>C 2316G>T), R723E (2317C>G 2318G>A 2319C>G), T724Q (2320A>C 2321C>A), F726K (2326T>A 2327T>A 2328T>A), H728Y (2332C>T), N729K (2337T>G), C730A (2338T>G 2339G>C 2340T>G), I731L (2341A>C), D732Q (2344G>C 2346T>G), V735A (2354T>C 2355T>A), I739T (2366T>C 2367T>G), Q742E (2374C>G 2376G>A), F747R (2389T>A 2390T>G 2391T>A), T761R (2432C>G 2433C>G), S764A (2440T>G 2442C>A), S764_K765insG (2442_2443insGGT), K765T (2444A>C 2445A>T), E766D (2448G>C), A767K (2449G>A 2450C>A 2451C>G), D769R (2455G>A 2456A>G 2457C>A), S770N (2458T>A 2459C>A), V772I (2464G>A 2466C>T), K773I (2468A>T 2469G>C), I774E (2470A>G 2471T>A), Y776M (2476T>A 2477A>T 2478C>G), H777I (2479C>A 2480A>T 2481C>T), K778G (2482A>G 2483A>G 2484A>T), V780K (2488G>A 2489T>A), Y782K (2494T>A 2496C>A), K784I (2501A>T 2502A>C), E785Q (2503G>C 2505A>G), T786M (2507C>T 2508T>G), W788N (2512T>A 2513G>A 2514G>T), N789D (2515A>G), Y792F (2525A>T 2526T>C), A793I (2527G>A 2528C>T 2529A>C), R795K (2534G>A), I804L (2560A>T 2562C>G), M805I (2565G>A), L810T (2578T>A 2579T>C 2580G>C), F811M (2581T>A 2583T>G), G812N (2584G>A 2585G>A 2586G>T), G813A (2588G>C 2589A>C), I814V (2590A>G 2592A>T), L818I (2602C>A), A819T (2605G>A), W821L (2611T>C 2612G>T 2613G>C), Q822K (2614C>A 2616A>G), S823A (2617A>G 2618G>C 2619C>A), N824A (2620A>G 2621A>C 2622T>A), S825T (2623T>A), F826S (2627T>C 2628T>C), S828L (2632A>T 2633G>T 2634T>A), S830D (2638A>G 2639G>A), P850K (2698C>A 2699C>A 2700C>G), S851T (2701T>A), R852H (2705G>A), L853F (2707C>T 2709G>C), K854A (2710A>G 2711A>C 2712G>A), N855K (2715T>A), L856S (2717T>C 2718G>C), V858L (2722G>C 2724G>A), T1343C (4177A>T 4178C>G 4179C>T), L1347A (4189T>G 4190T>C), I1351L (4201A>C), D1358N (4222G>A), S1362H (4234T>C 4235C>A), S1365T (4243T>A 4245T>A), L1366V (4246C>G 4248T>G), N1369S (4256A>G 4257C>T), N1370S (4259A>G), A1373S (4267G>T 4269T>A), T1377E (4279A>G 4280C>A 4281T>G) | | | | | | | | |
| Codon mutations: | ACT675.AT (2174C>A), GAC676GAT (2178C>T), ATG677TTG (2179A>T), ATC678TTG (2182A>T 2184C>G), TAC679GTC (2185T>G 2186A>T), GGT681GGA (2193T>A), GCC682TCA (2194G>T 2196C>A), AAT683AGA (2198A>G 2199T>A), GCT684CAT (2200G>C 2201C>A), GAA685CAG (2203G>C 2205A>G), CAA686CAG (2208A>G), TAC687TAT (2211C>T), AGG688GCT (2212A>G 2213G>C 2214G>T), CAA692CGA (2225A>G), AAT693AAC (2229T>C), CTC699ATC (2245C>A), TTT702TTC (2256T>C), AAA703AAG (2259A>G), AGC704AGT (2262C>T), ATC706ATT (2268C>T), GCT708AGC (2272G>A 2273C>G 2274T>C), TCC709ACA (2275T>A 2277C>A), ATC711ATT (2283C>T), TGT712TCT (2285G>C), AAG714GAG (2290A>G), AAG715CTG (2293A>C 2294A>T), GCT717GAT (2300C>A), GTA718GTC (2304A>C), ACA720CGC (2308A>C 2309C>G 2310A>C), GAG721AAT (2311G>A 2313G>T), ATG722CTT (2314A>C 2316G>T), CGC723GAG (2317C>G 2318G>A 2319C>G), ACA724CAA (2320A>C 2321C>A), GGA725GGG (2325A>G), TTT726AAA (2326T>A 2327T>A 2328T>A), GCA727GCG (2331A>G), CAT728TAT (2332C>T), AAT729AAG (2337T>G), TGT730GCG (2338T>G 2339G>C 2340T>G), ATT731CTT (2341A>C), GAT732CAG (2344G>C 2346T>G), GTT735GCA (2354T>C 2355T>A), TTG738CTT (2362T>C 2364G>T), ATT739ACG (2366T>C 2367T>G), TGC740TGT (2370C>T), ATC741ATT (2373C>T), CAG742GAA (2374C>G 2376G>A), TTT747AGA (2389T>A 2390T>G 2391T>A), GTG748GTA (2394G>A), TTT750TTC (2400T>C), TTG751CTG (2401T>C), ACA753ACT (2409A>T), GCC754GCT (2412C>T), CGT757CGC (2421T>C), TAT758TAC (2424T>C), ACC761AGG (2432C>G 2433C>G), TCC764GCA (2440T>G 2442C>A), TCC764_AAA765insGGT (2442_2443insGGT), AAA765ACT (2444A>C 2445A>T), GAG766GAC (2448G>C), GCC767AAG (2449G>A 2450C>A 2451C>G), AAA768AAG (2454A>G), GAC769AGA (2455G>A 2456A>G 2457C>A), TCT770AAT (2458T>A 2459C>A), GTC772ATT (2464G>A 2466C>T), AAG773ATC (2468A>T 2469G>C), ATA774GAA (2470A>G 2471T>A), TGT775TGC (2475T>C), TAC776ATG (2476T>A 2477A>T 2478C>G), CAC777ATT (2479C>A 2480A>T 2481C>T), AAA778GGT (2482A>G 2483A>G 2484A>T), ATT779ATC (2487T>C), GTG780AAG (2488G>A 2489T>A), GAT781GAC (2493T>C), TAC782AAA (2494T>A 2496C>A), AAA784ATC (2501A>T 2502A>C), GAA785CAG (2503G>C 2505A>G), ACT786ATG (2507C>T 2508T>G), TGG788AAT (2512T>A 2513G>A 2514G>T), AAC789GAC (2515A>G), TTA790TTG (2520A>G), CCA791CCC (2523A>C), TAT792TTC (2525A>T 2526T>C), GCA793ATC (2527G>A 2528C>T 2529A>C), CTG794TTG (2530C>T), AGG795AAG (2534G>A), GGC801GGA (2553C>A), CTA802CTG (2556A>G), ATC804TTG (2560A>T 2562C>G), ATG805ATA (2565G>A), GCG806GCA (2568G>A), GGA809GGC (2577A>C), TTG810ACC (2578T>A 2579T>C 2580G>C), TTT811ATG (2581T>A 2583T>G), GGG812AAT (2584G>A 2585G>A 2586G>T), GGA813GCC (2588G>C 2589A>C), ATA814GTT (2590A>G 2592A>T), ACA815ACT (2595A>T), TTT817TTC (2601T>C), CTT818ATT (2602C>A), GCT819ACT (2605G>A), TGT820TGC (2610T>C), TGG821CTC (2611T>C 2612G>T 2613G>C), CAA822AAG (2614C>A 2616A>G), AGC823GCA (2617A>G 2618G>C 2619C>A), AAT824GCA (2620A>G 2621A>C 2622T>A), TCT825ACT (2623T>A), TTT826TCC (2627T>C 2628T>C), CCA827CCT (2631A>T), AGT828TTA (2632A>T 2633G>T 2634T>A), AGT830GAT (2638A>G 2639G>A), GGA831GGC (2643A>C), GGT833GGG (2649T>G), GGT834GGC (2652T>C), GTC835GTA (2655C>A), ACA836ACC (2658A>C), AGC841AGT (2673C>T), TCT843TCC (2679T>C), AAA845AAG (2685A>G), AAG846AAA (2688G>A), AAC847AAT (2691C>T), AAG848AAA (2694G>A), GGC849GGG (2697C>G), CCC850AAG (2698C>A 2699C>A 2700C>G), TCT851ACT (2701T>A), CGT852CAT (2705G>A), CTG853TTC (2707C>T 2709G>C), AAG854GCA (2710A>G 2711A>C 2712G>A), AAT855AAA (2715T>A), TTG856TCC (2717T>C 2718G>C), GTG858CTA (2722G>C 2724G>A), TTT1322.TC (4116T>C), GAG1323GAA (4119G>A), ATA1324ATC (4122A>C), TAC1329TAT (4137C>T), GTC1333GTG (4149C>G), AGG1334CGG (4150A>C), TTT1337TTC (4161T>C), TCC1341TCT (4173C>T), GCC1342GCT (4176C>T), ACC1343TGT (4177A>T 4178C>G 4179C>T), GCA1346GCT (4188A>T), TTA1347GCA (4189T>G 4190T>C), AAT1349AAC (4197T>C), GTC1350GTA (4200C>A), ATC1351CTC (4201A>C), CCA1352CCT (4206A>T), TGT1353TGC (4209T>C), GTT1355GTA (4215T>A), GGA1356GGT (4218A>T), ATA1357ATC (4221A>C), GAT1358AAT (4222G>A), ACC1360ACG (4230C>G), AGT1361AGC (4233T>C), TCT1362CAT (4234T>C 4235C>A), TCT1365ACA (4243T>A 4245T>A), CTT1366GTG (4246C>G 4248T>G), CTT1367CTG (4251T>G), GCC1368GCT (4254C>T), AAC1369AGT (4256A>G 4257C>T), AAC1370AGC (4259A>G), TTC1371TTT (4263C>T), GCT1373TCA (4267G>T 4269T>A), ACT1377GAG (4279A>G 4280C>A 4281T>G), GGG1378GGA (4284G>A), TTT1379TTC (4287T>C), GCT1381GCC (4293T>C), TCA1384TCG (4302A>G), AGT1385AGC (4305T>C), GAC1387..T (4311C>T) | | | | | | | | |

##### Proteins

| | Begin | End | Coverage | Score | Concordance | Matches | Identities | I/D/M/F* | Stop Codons |
|---|---|---|---|---|---|---|---|---|---|
| polyprotein (YP_009165993.1) | 676 | 1387 | 11.6% | 1167 | 64.8% | 249 (99.6%) | 158 (63.2%) | 1/0/0/0 | 0 |
| Protein mutations: | M677L (2179A>T), I678L (2182A>T 2184C>G), Y679V (2185T>G 2186A>T), A682S (2194G>T 2196C>A), N683R (2198A>G 2199T>A), A684H (2200G>C 2201C>A), E685Q (2203G>C 2205A>G), R688A (2212A>G 2213G>C 2214G>T), Q692R (2225A>G), L699I (2245C>A), A708S (2272G>A 2273C>G 2274T>C), S709T (2275T>A 2277C>A), C712S (2285G>C), K714E (2290A>G), K715L (2293A>C 2294A>T), A717D (2300C>A), T720R (2308A>C 2309C>G 2310A>C), E721N (2311G>A 2313G>T), M722L (2314A>C 2316G>T), R723E (2317C>G 2318G>A 2319C>G), T724Q (2320A>C 2321C>A), F726K (2326T>A 2327T>A 2328T>A), H728Y (2332C>T), N729K (2337T>G), C730A (2338T>G 2339G>C 2340T>G), I731L (2341A>C), D732Q (2344G>C 2346T>G), V735A (2354T>C 2355T>A), I739T (2366T>C 2367T>G), Q742E (2374C>G 2376G>A), F747R (2389T>A 2390T>G 2391T>A), T761R (2432C>G 2433C>G), S764A (2440T>G 2442C>A), S764_K765insG (2442_2443insGGT), K765T (2444A>C 2445A>T), E766D (2448G>C), A767K (2449G>A 2450C>A 2451C>G), D769R (2455G>A 2456A>G 2457C>A), S770N (2458T>A 2459C>A), V772I (2464G>A 2466C>T), K773I (2468A>T 2469G>C), I774E (2470A>G 2471T>A), Y776M (2476T>A 2477A>T 2478C>G), H777I (2479C>A 2480A>T 2481C>T), K778G (2482A>G 2483A>G 2484A>T), V780K (2488G>A 2489T>A), Y782K (2494T>A 2496C>A), K784I (2501A>T 2502A>C), E785Q (2503G>C 2505A>G), T786M (2507C>T 2508T>G), W788N (2512T>A 2513G>A 2514G>T), N789D (2515A>G), Y792F (2525A>T 2526T>C), A793I (2527G>A 2528C>T 2529A>C), R795K (2534G>A), I804L (2560A>T 2562C>G), M805I (2565G>A), L810T (2578T>A 2579T>C 2580G>C), F811M (2581T>A 2583T>G), G812N (2584G>A 2585G>A 2586G>T), G813A (2588G>C 2589A>C), I814V (2590A>G 2592A>T), L818I (2602C>A), A819T (2605G>A), W821L (2611T>C 2612G>T 2613G>C), Q822K (2614C>A 2616A>G), S823A (2617A>G 2618G>C 2619C>A), N824A (2620A>G 2621A>C 2622T>A), S825T (2623T>A), F826S (2627T>C 2628T>C), S828L (2632A>T 2633G>T 2634T>A), S830D (2638A>G 2639G>A), P850K (2698C>A 2699C>A 2700C>G), S851T (2701T>A), R852H (2705G>A), L853F (2707C>T 2709G>C), K854A (2710A>G 2711A>C 2712G>A), N855K (2715T>A), L856S (2717T>C 2718G>C), V858L (2722G>C 2724G>A), T1343C (4177A>T 4178C>G 4179C>T), L1347A (4189T>G 4190T>C), I1351L (4201A>C), D1358N (4222G>A), S1362H (4234T>C 4235C>A), S1365T (4243T>A 4245T>A), L1366V (4246C>G 4248T>G), N1369S (4256A>G 4257C>T), N1370S (4259A>G), A1373S (4267G>T 4269T>A), T1377E (4279A>G 4280C>A 4281T>G) | | | | | | | | |

| | Begin | End | Coverage | Score | Concordance | Matches | Identities | I/D/M/F* | Stop Codons |
|---|---|---|---|---|---|---|---|---|---|
| **NT** | **2174** | **4311** | **10.3%** | **438** | **29.2%** | **749 (99.6%)** | **487 (64.8%)** | **3/0** | |
| Codon mutations: | ACT675.AT (2174C>A), GAC676GAT (2178C>T), ATG677TTG (2179A>T), ATC678TTG (2182A>T 2184C>G), TAC679GTC (2185T>G 2186A>T), GGT681GGA (2193T>A), GCC682TCA (2194G>T 2196C>A), AAT683AGA (2198A>G 2199T>A), GCT684CAT (2200G>C 2201C>A), GAA685CAG (2203G>C 2205A>G), CAA686CAG (2208A>G), TAC687TAT (2211C>T), AGG688GCT (2212A>G 2213G>C 2214G>T), CAA692CGA (2225A>G), AAT693AAC (2229T>C), CTC699ATC (2245C>A), TTT702TTC (2256T>C), AAA703AAG (2259A>G), AGC704AGT (2262C>T), ATC706ATT (2268C>T), GCT708AGC (2272G>A 2273C>G 2274T>C), TCC709ACA (2275T>A 2277C>A), ATC711ATT (2283C>T), TGT712TCT (2285G>C), AAG714GAG (2290A>G), AAG715CTG (2293A>C 2294A>T), GCT717GAT (2300C>A), GTA718GTC (2304A>C), ACA720CGC (2308A>C 2309C>G 2310A>C), GAG721AAT (2311G>A 2313G>T), ATG722CTT (2314A>C 2316G>T), CGC723GAG (2317C>G 2318G>A 2319C>G), ACA724CAA (2320A>C 2321C>A), GGA725GGG (2325A>G), TTT726AAA (2326T>A 2327T>A 2328T>A), GCA727GCG (2331A>G), CAT728TAT (2332C>T), AAT729AAG (2337T>G), TGT730GCG (2338T>G 2339G>C 2340T>G), ATT731CTT (2341A>C), GAT732CAG (2344G>C 2346T>G), GTT735GCA (2354T>C 2355T>A), TTG738CTT (2362T>C 2364G>T), ATT739ACG (2366T>C 2367T>G), TGC740TGT (2370C>T), ATC741ATT (2373C>T), CAG742GAA (2374C>G 2376G>A), TTT747AGA (2389T>A 2390T>G 2391T>A), GTG748GTA (2394G>A), TTT750TTC (2400T>C), TTG751CTG (2401T>C), ACA753ACT (2409A>T), GCC754GCT (2412C>T), CGT757CGC (2421T>C), TAT758TAC (2424T>C), ACC761AGG (2432C>G 2433C>G), TCC764GCA (2440T>G 2442C>A), TCC764_AAA765insGGT (2442_2443insGGT), AAA765ACT (2444A>C 2445A>T), GAG766GAC (2448G>C), GCC767AAG (2449G>A 2450C>A 2451C>G), AAA768AAG (2454A>G), GAC769AGA (2455G>A 2456A>G 2457C>A), TCT770AAT (2458T>A 2459C>A), GTC772ATT (2464G>A 2466C>T), AAG773ATC (2468A>T 2469G>C), ATA774GAA (2470A>G 2471T>A), TGT775TGC (2475T>C), TAC776ATG (2476T>A 2477A>T 2478C>G), CAC777ATT (2479C>A 2480A>T 2481C>T), AAA778GGT (2482A>G 2483A>G 2484A>T), ATT779ATC (2487T>C), GTG780AAG (2488G>A 2489T>A), GAT781GAC (2493T>C), TAC782AAA (2494T>A 2496C>A), AAA784ATC (2501A>T 2502A>C), GAA785CAG (2503G>C 2505A>G), ACT786ATG (2507C>T 2508T>G), TGG788AAT (2512T>A 2513G>A 2514G>T), AAC789GAC (2515A>G), TTA790TTG (2520A>G), CCA791CCC (2523A>C), TAT792TTC (2525A>T 2526T>C), GCA793ATC (2527G>A 2528C>T 2529A>C), CTG794TTG (2530C>T), AGG795AAG (2534G>A), GGC801GGA (2553C>A), CTA802CTG (2556A>G), ATC804TTG (2560A>T 2562C>G), ATG805ATA (2565G>A), GCG806GCA (2568G>A), GGA809GGC (2577A>C), TTG810ACC (2578T>A 2579T>C 2580G>C), TTT811ATG (2581T>A 2583T>G), GGG812AAT (2584G>A 2585G>A 2586G>T), GGA813GCC (2588G>C 2589A>C), ATA814GTT (2590A>G 2592A>T), ACA815ACT (2595A>T), TTT817TTC (2601T>C), CTT818ATT (2602C>A), GCT819ACT (2605G>A), TGT820TGC (2610T>C), TGG821CTC (2611T>C 2612G>T 2613G>C), CAA822AAG (2614C>A 2616A>G), AGC823GCA (2617A>G 2618G>C 2619C>A), AAT824GCA (2620A>G 2621A>C 2622T>A), TCT825ACT (2623T>A), TTT826TCC (2627T>C 2628T>C), CCA827CCT (2631A>T), AGT828TTA (2632A>T 2633G>T 2634T>A), AGT830GAT (2638A>G 2639G>A), GGA831GGC (2643A>C), GGT833GGG (2649T>G), GGT834GGC (2652T>C), GTC835GTA (2655C>A), ACA836ACC (2658A>C), AGC841AGT (2673C>T), TCT843TCC (2679T>C), AAA845AAG (2685A>G), AAG846AAA (2688G>A), AAC847AAT (2691C>T), AAG848AAA (2694G>A), GGC849GGG (2697C>G), CCC850AAG (2698C>A 2699C>A 2700C>G), TCT851ACT (2701T>A), CGT852CAT (2705G>A), CTG853TTC (2707C>T 2709G>C), AAG854GCA (2710A>G 2711A>C 2712G>A), AAT855AAA (2715T>A), TTG856TCC (2717T>C 2718G>C), GTG858CTA (2722G>C 2724G>A), TTT1322.TC (4116T>C), GAG1323GAA (4119G>A), ATA1324ATC (4122A>C), TAC1329TAT (4137C>T), GTC1333GTG (4149C>G), AGG1334CGG (4150A>C), TTT1337TTC (4161T>C), TCC1341TCT (4173C>T), GCC1342GCT (4176C>T), ACC1343TGT (4177A>T 4178C>G 4179C>T), GCA1346GCT (4188A>T), TTA1347GCA (4189T>G 4190T>C), AAT1349AAC (4197T>C), GTC1350GTA (4200C>A), ATC1351CTC (4201A>C), CCA1352CCT (4206A>T), TGT1353TGC (4209T>C), GTT1355GTA (4215T>A), GGA1356GGT (4218A>T), ATA1357ATC (4221A>C), GAT1358AAT (4222G>A), ACC1360ACG (4230C>G), AGT1361AGC (4233T>C), TCT1362CAT (4234T>C 4235C>A), TCT1365ACA (4243T>A 4245T>A), CTT1366GTG (4246C>G 4248T>G), CTT1367CTG (4251T>G), GCC1368GCT (4254C>T), AAC1369AGT (4256A>G 4257C>T), AAC1370AGC (4259A>G), TTC1371TTT (4263C>T), GCT1373TCA (4267G>T 4269T>A), ACT1377GAG (4279A>G 4280C>A 4281T>G), GGG1378GGA (4284G>A), TTT1379TTC (4287T>C), GCT1381GCC (4293T>C), TCA1384TCG (4302A>G), AGT1385AGC (4305T>C), GAC1387..T (4311C>T) | | | | | | | | |

*: Inserts / Deletes / Misaligned / Frameshifts

#### Analysis details

This analysis was performed with panviral2.64

### NGS Details (UN62): Cladosporium fulvum T-1 virus

#### Assembly

| | |
|---|---|
| **Coverage Length** | 1370 (3 contig(s)) |
| **Depth Of Coverage** | 39.2 |
| **Number Of Reads** | 511 |
| **Reads Per Million** | 11.85 rpm (after QC) |
| **Ambiguities** | 0 |
| **Assembly Method** | de novo + reference guided assembly |
| **Consensus Caller** | Bcf Tools |

#### Coverage Map

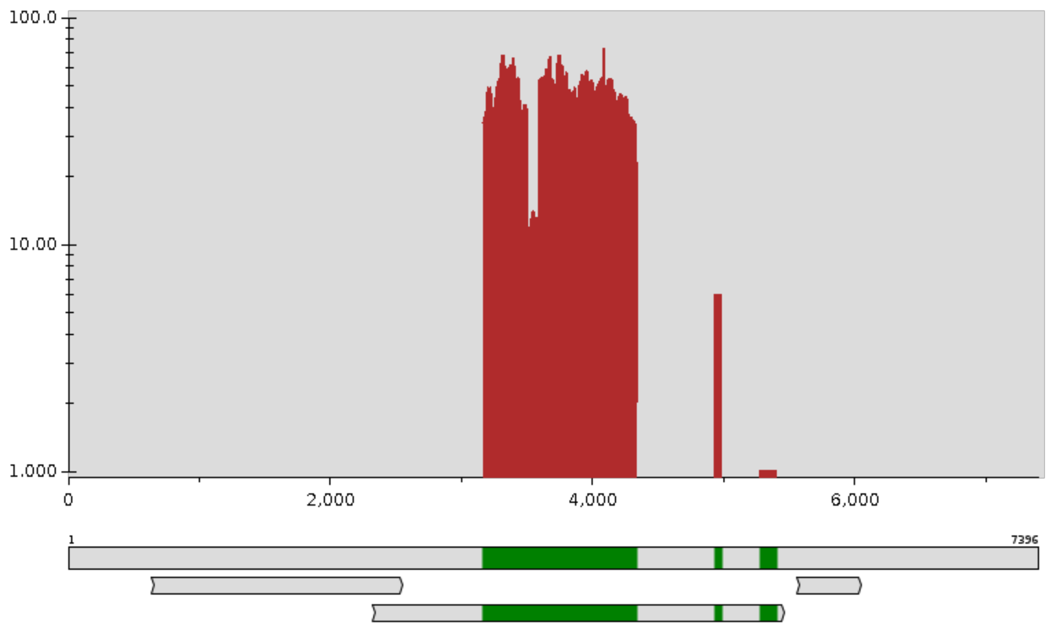

#### Assignment

| | |
|---|---|
| **Type** | Cladosporium fulvum T-1 virus (Taxonomy ID: 2052899) |
| **Reference Genome** | NC_043491.1 |
| **NT Identity (%)** | 52.9758 |
| **AA Identity (%)** | 47.2406 |
| **Number Of Stop Codons** | 0 |
| **Number Of CDS** | 3 |

#### Alignment

| | |
|---|---|
| **Alignment Score** | 117.0 (NT) + 1494.0 (AA) = 1611.0 |
| **Concordance (%)** | 27.8431 |

| Alignment Method | Global, seeded, nucleotide + amino acids (AGA) |
|---|---|

#### Genome Region

Sequence starts at position 3165 and ends at position 5404 relative to NC_043491.1 reference sequence.

#### Alignment Detailed Statistics

| | Begin | End | Coverage | Score | Concordance | Matches | Identities | I/D/M/F* | Stop Codons |
|---|---|---|---|---|---|---|---|---|---|
| **NT** | **3165** | **5404** | **18.5%** | **117** | **4.4%** | **1352 (98.0%)** | **721 (52.3%)** | **9/18** | |
| Mutations: | 3165A>G, 3167T>G, 3170A>G, 3171G>A, 3172T>A, 3175T>A, 3178C>A, 3179A>G, 3181G>C, 3182G>T, 3183A>G, 3184G>T, 3185T>G, 3186A>T, 3187T>G, 3188C>G, 3189G>C, 3190C>A, 3191A>G, 3194T>C, 3195G>T, 3197T>C, 3198C>T, 3200C>A, 3201G>A, 3204T>A, 3205C>G, 3207T>A, 3208T>C, 3210A>C, 3211A>T, 3214G>T, 3216A>T, 3217A>G, 3219G>T, 3220A>G, 3221G>C, 3222G>C, 3223C>A, 3224A>C, 3225A>C, 3226G>T, 3229C>G, 3230G>T, 3231C>T, 3232C>G, 3233T>C, 3234T>C, 3236C>A, 3237C>A, 3238T>G, 3243A>T, 3244C>G, 3246A>C, 3251T>A, 3253_3254insAGGGATATT, 3261A>G, 3262G>A, 3265A>T, 3266A>G, 3268C>G, 3269A>T, 3271T>G, 3272C>A, 3273A>T, 3280G>T, 3281A>T, 3282A>C, 3283A>T, 3284G>A, 3285A>C, 3289T>G, 3292A>T, 3293T>G, 3294G>C, 3295G>A, 3296G>C, 3297G>A, 3299C>G, 3301C>T, 3303T>C, 3304A>T, 3307T>C, 3308C>A, 3309A>G, 3314T>G, 3317G>C, 3318A>C, 3319G>T, 3326C>T, 3328A>G, 3329C>G, 3330A>C, 3331G>T, 3332A>G, 3333C>A, 3334C>A, 3337A>G, 3338C>A, 3340A>G, 3341G>A, 3343A>G, 3344T>C, 3345G>A, 3352G>C, 3355G>A, 3356A>T, 3357A>T, 3361A>T, 3363C>A, 3364C>T, 3365A>T, 3366A>C, 3371T>C, 3372G>T, 3376A>T, 3378G>A, 3379A>G, 3381G>C, 3382A>T, 3385C>T, 3387C>A, 3388C>G, 3389T>G, 3391A>T, 3392A>C, 3393G>C, 3394T>C, 3395G>T, 3396C>A, 3397A>T, 3400A>T, 3401A>G, 3403T>A, 3406A>T, 3407T>G, 3408G>T, 3409C>T, 3410A>T, 3412G>A, 3415C>T, 3416G>C, 3417T>A, 3418T>G, 3419C>A, 3420C>A, 3425G>C, 3426C>A, 3427A>G, 3428A>G, 3430C>T, 3434A>T, 3435A>C, 3437C>T, 3440C>A, 3442A>G, 3443C>A, 3445C>G, 3446G>T, 3447T>G, 3448A>C, 3449C>A, 3450A>T, 3451A>G, 3454C>T, 3457C>T, 3460A>G, 3461A>G, 3462A>C, 3464T>C, 3470G>A, 3473A>G, 3479A>G, 3481C>T, 3487C>T, 3488C>A, 3489G>A, 3490A>G, 3493T>C, 3497C>A, 3499A>C, 3502C>A, 3503A>C, 3504A>T, 3508C>T, 3510A>C, 3511A>T, 3514A>T, 3515G>C, 3516C>T, 3517A>C, 3518C>T, 3519A>T, 3520A>T, 3523C>T, 3524A>C, 3525G>A, 3527T>C, 3529A>T, 3530A>G, 3531C>G, 3533G>C, 3534G>A, 3536T>G, 3539G>C, 3540A>G, 3541C>T, 3546A>T, 3550G>C, 3554A>T, 3556T>A, 3559C>T, 3562A>T, 3565A>G, 3566G>T, 3567A>C, 3568C>T, 3570C>G, 3571C>A, 3573T>A, 3578G>C, 3579C>A, 3580T>A, 3581A>G, 3583C>G, 3584C>A, 3589G>C, 3598A>G, 3601A>T, 3604A>G, 3605T>C, 3606G>C, 3607G>T, 3610A>G, 3613C>T, 3614G>A, 3618T>G, 3619C>T, 3620A>G, 3621G>T, 3622G>A, 3625A>T, 3628A>G, 3631C>T, 3634A>G, 3635C>T, 3636T>C, 3637C>A, 3640C>T, 3647T>C, 3649G>T, 3652C>A, 3658A>G, 3659A>T, 3661G>T, 3664A>G, 3670C>T, 3673C>T, 3676A>C, 3679C>G, 3682A>T, 3683T>A, 3687G>T, 3688C>T, 3689C>T, 3690A>G, 3691G>C, 3692G>A, 3694C>T, 3697T>A, 3698G>A, 3700C>G, 3703C>T, 3705A>G, 3706A>C, 3707A>G, 3708C>T, 3712T>G, 3713A>T, 3714G>A, 3715A>T, 3716G>C, 3717A>C, 3719C>T, 3720T>A, 3721A>C, 3724C>T, 3728G>A, 3729T>A, 3730G>T, 3732G>T, 3733C>T, 3736C>A, 3739T>A, 3741C>T, 3742T>C, 3745C>T, 3746A>C, 3748G>A, 3751C>T, 3757A>T, 3758C>G, 3760G>C, 3761G>A, 3763C>A, 3766C>T, 3768C>G, 3769A>T, 3770A>C, 3774G>A, 3775A>T, 3776T>A, 3778C>G, 3779C>T, 3781C>G, 3782C>G, 3784G>T, 3791A>G, 3792C>T, 3793C>T, 3795A>G, 3796G>C, 3799A>T, 3800G>T, 3802T>G, 3803C>A, 3804A>G, 3805A>T, 3806G>T, 3807A>T, 3808T>G, 3811G>A, 3814C>A, 3815G>T, 3816A>C, 3817A>T, 3820A>T, 3821C>T, 3823C>A, 3825C>G, 3827A>C, 3829G>A, 3831C>A, 3833G>A, 3834G>C, 3835A>T, 3836T>C, 3839A>T, 3841G>T, 3842A>G, 3843C>T, 3844A>T, 3845G>A, 3846C>A, 3848C>A, 3849C>T, 3850C>G, 3856A>G, 3859C>T, 3862A>G, 3865C>T, 3866C>G, 3867A>C, 3869A>C, 3871G>A, 3877A>G, 3878G>A, 3880C>A, 3886T>C, 3887T>C, 3892C>G, 3893T>C, 3894T>A, 3896A>C, 3898C>A, 3899A>G, 3901C>T, 3905A>G, 3906C>A, 3909C>A, 3910A>C, 3911G>C, 3912G>A, 3913G>A, 3914A>G, 3918C>G, 3922C>G, 3925C>T, 3929G>A, 3930C>A, 3931A>G, 3934G>A, 3935A>G, 3936C>T, 3937A>G, 3940G>A, 3941T>G, 3943A>C, 3946C>T, 3947A>G, 3948G>T, 3949A>T, 3952A>T, 3957C>A, 3960A>C, 3964G>C, 3967G>A, 3970A>T, 3973C>G, 3980G>C, 3982A>G, 3983C>A, 3984A>G, 3988A>T, 3992C>T, 3994T>G, 3997A>G, 4000C>A, 4003C>T, 4010A>T, 4012C>T, 4025A>G, 4026A>C, 4027G>T, 4029A>G, 4030C>A, 4033T>C, 4041C>A, 4048A>T, 4049C>G, 4052A>C, 4057G>T, 4058A>G, 4059T>A, 4060G>T, 4061C>T, 4063T>G, 4064A>C, 4065C>T, 4066A>G, 4068G>A, 4069A>G, 4072A>G, 4075C>A, 4081C>G, 4085A>G, 4086A>T, 4091G>A, 4092G>C, 4093A>C, 4094A>G, 4099A>G, 4100C>T, 4101A>G, 4102G>T, 4103A>G, 4104C>A, 4105C>G, 4107A>C, 4108A>G, 4109G>A, 4110C>G, 4111G>C, 4114C>T, 4115A>C, 4119G>A, 4120A>G, 4129A>C, 4130C>T, 4131A>T, 4132G>A, 4134G>T, 4135C>G, 4136G>A, 4147A>T, 4149C>T, 4150G>T, 4153T>C, 4154C>G, 4155G>C, 4157C>T, 4160T>C, 4161T>C, 4165T>G, 4167G>A, 4168T>G, 4169A>C, 4170G>C, 4171C>T, 4172A>T, 4173A>T, 4174G>T, 4177A>G, 4180C>A, 4184A>G, 4186C>G, 4187G>C, 4192C>T, 4195C>T, 4201T>G, 4204T>C, 4206T>A, 4207G>A, 4210A>T, 4211A>G, 4213A>G, 4216C>A, 4218C>G, 4219A>C, 4220T>G, 4221G>T, 4222T>A, 4223C>T, 4225A>G, 4226A>G, 4227C>T, 4228A>G, 4231G>A, 4232_4237delACACAC, 4240T>A, 4243G>T, 4244_4249delAAAAGA, 4252C>T, 4255A>G, 4263A>T, 4265T>G, 4267T>A, 4268T>A, 4269C>G, 4271C>A, 4277A>T, 4279G>A, 4281C>A, 4282C>T, 4283A>G, 4284C>A, 4285A>T, 4288G>T, 4291A>G, 4296A>G, 4297C>G, 4301G>T, 4302A>C, 4303C>T, 4305T>C, 4306C>T, 4312C>A, 4319C>A, 4321T>G, 4322C>G, 4324A>T, 4327C>A, 4328A>G, 4330T>G, 4333T>G, 4334G>C, 4923A>G, 4924A>T, 4933C>G, 4934G>A, 4935A>G, 4936G>T, 4942G>T, 4954T>A, 4957G>T, 4958A>T, 4959A>G, 4960A>C, 4963C>G, 4966G>A, 4968G>A, 4972A>C, 4975G>C, 4976G>A, 4978T>G, 5276C>T, 5277A>G, 5278A>C, 5284A>T, 5285C>A, 5292A>T, 5294A>G, 5296C>G, 5298A>G, 5301A>C, 5302C>T, 5303G>C, 5304C>A, 5305A>G, 5309G>A, 5311C>G, 5312A>G, 5314C>T, 5319T>C, 5321T>A, 5322C>A, 5323A>G, 5326A>G, 5329G>A, 5330C>G, 5331C>A, 5332A>T, 5333A>T, 5338G>A, 5339C>G, 5341G>A, 5342A>T, 5345G>C, 5348C>T, 5349T>A, 5350G>T, 5354A>T, 5355A>T, 5356C>A, 5359C>G, 5360A>T, 5361A>G, 5362A>G, 5363T>A, 5364C>G, 5365A>T, 5368G>A, 5369A>G, 5370C>T, 5371A>C, 5374T>C, 5375T>G, 5376C>G, 5377G>A, 5379C>G, 5383G>A, 5390A>G, 5391T>A, 5393_5398delCGAACT, 5404A>C | | | | | | | | |
| **CDS** | | | | | | | | | |
| homologue_of_retroviral_POL_genes | 281 | 1026 | 43.6% | 1494 | 47.5% | 450 (98.0%) | 214 (46.6%) | 3/6/0/0 | 0 |

| | Begin | End | Coverage | Score | Concordance | Matches | Identities | I/D/M/F* | Stop Codons |
|---|---|---|---|---|---|---|---|---|---|
| **NT** | **3165** | **5404** | **18.5%** | **117** | **4.4%** | **1352 (98.0%)** | **721 (52.3%)** | **9/18** | |
| Protein mutations: | L281V (3167T>G), S282E (3170A>G 3171G>A 3172T>A), K285D (3179A>G 3181G>C), E286C (3182G>T 3183A>G 3184G>T), Y287V (3185T>G 3186A>T 3187T>G), R288A (3188C>G 3189G>C 3190C>A), K289E (3191A>G), W290L (3194T>C 3195G>T), S291L (3197T>C 3198C>T), R292K (3200C>A 3201G>A), L293Q (3204T>A 3205C>G), F294Y (3207T>A 3208T>C), E295A (3210A>C 3211A>T), E296D (3214G>T), E297V (3216A>T 3217A>G), R298M (3219G>T 3220A>G), G299P (3221G>C 3222G>C 3223C>A), K300P (3224A>C 3225A>C 3226G>T), D301E (3229C>G), A302L (3230G>T 3231C>T 3232C>G), L303P (3233T>C 3234T>C), P304K (3236C>A 3237C>A 3238T>G), H306L (3243A>T 3244C>G), Q307P (3246A>C), W309R (3251T>A), W309_D310insRDI (3253_3254insAGGGATATT), K312R (3261A>G 3262G>A), N314E (3266A>G 3268C>G), I315L (3269A>T 3271T>G), Q316M (3272C>A 3273A>T), K319S (3281A>T 3282A>C 3283A>T), E320T (3284G>A 3285A>C), W323A (3293T>G 3294G>C 3295G>A), G324Q (3296G>C 3297G>A), P325A (3299C>G 3301C>T), L326P (3303T>C 3304A>T), Q328R (3308C>A 3309A>G), S330A (3314T>G), E331P (3317G>C 3318A>C 3319G>T), Q335A (3329C>G 3330A>C 3331G>T), T336E (3332A>G 3333C>A 3334C>A), E339K (3341G>A 3343A>G), W340Q (3344T>C 3345G>A), K342N (3352G>C), K344L (3356A>T 3357A>T), A346D (3363C>A 3364C>T), K347S (3365A>T 3366A>C), W349L (3371T>C 3372G>T), R351Q (3378G>A 3379A>G), R352P (3381G>C 3382A>T), T354K (3387C>A 3388C>G), S355A (3389T>G 3391A>T), S356P (3392A>C 3393G>C 3394T>C), A357Y (3395G>T 3396C>A 3397A>T), T359A (3401A>G 3403T>A), C361V (3407T>G 3408G>T 3409C>T), M362L (3410A>T 3412G>A), V364Q (3416G>C 3417T>A 3418T>G), P365K (3419C>A 3420C>A), A367Q (3425G>C 3426C>A 3427A>G), N368D (3428A>G 3430C>T), K370S (3434A>T 3435A>C), L373M (3443C>A 3445C>G), V374C (3446G>T 3447T>G 3448A>C), Q375M (3449C>A 3450A>T 3451A>G), K379A (3461A>G 3462A>C), E382K (3470G>A), I383V (3473A>G), I385V (3479A>G 3481C>T), R388K (3488C>A 3489G>A 3490A>G), L391I (3497C>A 3499A>C), N393L (3503A>C 3504A>T), E395A (3510A>C 3511A>T), E396D (3514A>T), A397L (3515G>C 3516C>T 3517A>C), Q398F (3518C>T 3519A>T 3520A>T), R400Q (3524A>C 3525G>A), T402G (3530A>G 3531C>G), G403Q (3533G>C 3534G>A), S404A (3536T>G), D405R (3539G>C 3540A>G 3541C>T), Y407F (3546A>T), I410L (3554A>T 3556T>A), D414S (3566G>T 3567A>C 3568C>T), A415G (3570C>G 3571C>A), F416Y (3573T>A), A418Q (3578G>C 3579C>A 3580T>A), I419V (3581A>G 3583C>G), M421I (3589G>C), E425D (3601A>T), W427P (3605T>C 3606G>C 3607G>T), A430T (3614G>A), F431C (3618T>G 3619C>T), R432V (3620A>G 3621G>T 3622G>A), L437S (3635C>T 3636T>C 3637C>A), M445F (3659A>T 3661G>T), S453T (3683T>A), C454F (3687G>T 3688C>T), Q455C (3689C>T 3690A>G 3691G>C), D456N (3692G>A 3694C>T), V458M (3698G>A 3700C>G), E460G (3705A>G 3706A>C), T461V (3707A>G 3708C>T), R463Y (3713A>T 3714G>A 3715A>T), D464P (3716G>C 3717A>C), L465Y (3719C>T 3720T>A 3721A>C), V468N (3728G>A 3729T>A 3730G>T), C469F (3732G>T 3733C>T), A472V (3741C>T 3742T>C), M474L (3746A>C 3748G>A), L478V (3758C>G 3760G>C), V479I (3761G>A 3763C>A), T481S (3768C>G 3769A>T), K482Q (3770A>C), G483D (3774G>A 3775A>T), S484T (3776T>A 3778C>G), Q486D (3782C>G 3784G>T), T489V (3791A>G 3792C>T 3793C>T), K490S (3795A>G 3796G>C), Q491H (3799A>T), V492L (3800G>T 3802T>G), Q493S (3803C>A 3804A>G 3805A>T), D494L (3806G>T 3807A>T 3808T>G), F496L (3814C>A), E497S (3815G>T 3816A>C 3817A>T), T500R (3825C>G), K501Q (3827A>C 3829G>A), S502Y (3831C>A), G503T (3833G>A 3834G>C 3835A>T), F504L (3836T>C), K505Y (3839A>T 3841G>T), T506V (3842A>G 3843C>T 3844A>T), A507K (3845G>A 3846C>A), P508M (3848C>A 3849C>T 3850C>G), H514A (3866C>G 3867A>C), K515Q (3869A>C 3871G>A), V518I (3878G>A 3880C>A), F523H (3893T>C 3894T>A), I524L (3896A>C 3898C>A), I525V (3899A>G 3901C>T), T527E (3905A>G 3906C>A), T528N (3909C>A 3910A>C), G529Q (3911G>C 3912G>A 3913G>A), I530V (3914A>G), T531R (3918C>G), I532M (3922C>G), A535K (3929G>A 3930C>A 3931A>G), T537V (3935A>G 3936C>T 3937A>G), S539A (3941T>G 3943A>C), R541V (3947A>G 3948G>T 3949A>T), E542D (3952A>T), P544Q (3957C>A), E545A (3960A>C), V552L (3980G>C 3982A>G), Q553R (3983C>A 3984A>G), N562Y (4010A>T 4012C>T), K567A (4025A>G 4026A>C 4027G>T), D568G (4029A>G 4030C>A), T572K (4041C>A), P575A (4049C>G), M576L (4052A>C), M578D (4058A>G 4059T>A 4060G>T), T580L (4064A>C 4065C>T 4066A>G), R581K (4068G>A 4069A>G), D583E (4075C>A), N585K (4081C>G), K587V (4085A>G 4086A>T), G589T (4091G>A 4092G>C 4093A>C), K590E (4094A>G), Q592C (4100C>T 4101A>G 4102G>T), T593E (4103A>G 4104C>A 4105C>G), E594A (4107A>C 4108A>G), A595S (4109G>A 4110C>G 4111G>C), K597Q (4115A>C), R598K (4119G>A 4120A>G), E601D (4129A>C), Q602L (4130C>T 4131A>T 4132G>A), C603L (4134G>T 4135C>G), A604T (4136G>A), T608I (4149C>T 4150G>T), R610A (4154C>G 4155G>C), F612P (4160T>C 4161T>C), D613E (4165T>G), G614E (4167G>A 4168T>G), S615P (4169A>C 4170G>C 4171C>T), K616F (4172A>T 4173A>T 4174G>T), I620V (4184A>G 4186C>G), E621Q (4187G>C), M627K (4206T>A 4207G>A), I629V (4211A>G 4213A>G), A631G (4218C>G 4219A>C), C632V (4220T>G 4221G>T 4222T>A), T634V (4226A>G 4227C>T 4228A>G), T636_H637del (4232_4237delACACAC), D638E (4240T>A), K640_R641del (4244_4249delAAAAGA), Y646F (4263A>T), Y647E (4265T>G 4267T>A), M651L (4277A>T 4279G>A), T652N (4281C>A 4282C>T), T653D (4283A>G 4284C>A 4285A>T), N657R (4296A>G 4297C>G), D659S (4301G>T 4302A>C 4303C>T), I660T (4305T>C 4306C>T), D662E (4312C>A), L665M (4319C>A 4321T>G), L666V (4322C>G 4324A>T), I668V (4328A>G 4330T>G), A670P (4334G>C), D869E (4933C>G), E870S (4934G>A 4935A>G 4936G>T), K878C (4958A>T 4959A>G 4960A>C), R881K (4968G>A), K883N (4975G>C), D884K (4976G>A 4978T>G), Q984C (5276C>T 5277A>G 5278A>C), Y989F (5292A>T), I990V (5294A>G 5296C>G), N991S (5298A>G), Y992S (5301A>C 5302C>T), A993Q (5303G>C 5304C>A 5305A>G), D995K (5309G>A 5311C>G), N996D (5312A>G 5314C>T), V998A (5319T>C), S999K (5321T>A 5322C>A 5323A>G), P1002D (5330C>G 5331C>A 5332A>T), M1003L (5333A>T), Q1005E (5339C>G 5341G>A), I1006F (5342A>T), A1007P (5345G>C), L1008Y (5348C>T 5349T>A 5350G>T), N1010L (5354A>T 5355A>T 5356C>A), H1011Q (5359C>G), K1012W (5360A>T 5361A>G 5362A>G), T1015V (5369A>G 5370C>T 5371A>C), S1017G (5375T>G 5376C>G 5377G>A), T1018R (5379C>G), M1022E (5390A>G 5391T>A), R1023_T1024del (5393_5398delCGAACT) | | | | | | | | |

| | Begin | End | Coverage | Score | Concordance | Matches | Identities | I/D/M/F* | Stop Codons |
|---|---|---|---|---|---|---|---|---|---|
| **NT** | **3165** | **5404** | **18.5%** | **117** | **4.4%** | **1352 (98.0%)** | **721 (52.3%)** | **9/18** | |
| Codon mutations: | GAG280.GG (3165A>G), TTG281GTG (3167T>G), AGT282GAA (3170A>G 3171G>A 3172T>A), ATT283ATA (3175T>A), CCC284CCA (3178C>A), AAG285GAC (3179A>G 3181G>C), GAG286TGT (3182G>T 3183A>G 3184G>T), TAT287GTG (3185T>G 3186A>T 3187T>G), CGC288GCA (3188C>G 3189G>C 3190C>A), AAG289GAG (3191A>G), TGG290CTG (3194T>C 3195G>T), TCG291CTG (3197T>C 3198C>T), CGA292AAA (3200C>A 3201G>A), CTC293CAG (3204T>A 3205C>G), TTT294TAC (3207T>A 3208T>C), GAA295GCT (3210A>C 3211A>T), GAG296GAT (3214G>T), GAA297GTG (3216A>T 3217A>G), AGA298ATG (3219G>T 3220A>G), GGC299CCA (3221G>C 3222G>C 3223C>A), AAG300CCT (3224A>C 3225A>C 3226G>T), GAC301GAG (3229C>G), GCC302TTG (3230G>T 3231C>T 3232C>G), TTA303CCA (3233T>C 3234T>C), CCT304AAG (3236C>A 3237C>A 3238T>G), CAC306CTG (3243A>T 3244C>G), CAA307CCA (3246A>C), TGG309AGG (3251T>A), TGG309_GAT310insAGGGATATT (3253_3254insAGGGATATT), AAG312AGA (3261A>G 3262G>A), ATA313ATT (3265A>T), AAC314GAG (3266A>G 3268C>G), ATT315TTG (3269A>T 3271T>G), CAG316ATG (3272C>A 3273A>T), GGG318GGT (3280G>T), AAA319TCT (3281A>T 3282A>C 3283A>T), GAG320ACG (3284G>A 3285A>C), CCT321CCG (3289T>G), CCA322CCT (3292A>T), TGG323GCA (3293T>G 3294G>C 3295G>A), GGA324CAA (3296G>C 3297G>A), CCC325GCT (3299C>G 3301C>T), CTA326CCT (3303T>C 3304A>T), TAT327TAC (3307T>C), CAA328AGA (3308C>A 3309A>G), TCT330GCT (3314T>G), GAG331CCT (3317G>C 3318A>C 3319G>T), CTA334TTG (3326C>T 3328A>G), CAG335GCT (3329C>G 3330A>C 3331G>T), ACC336GAA (3332A>G 3333C>A 3334C>A), CTA337CTG (3337A>G), CGA338AGG (3338C>A 3340A>G), GAA339AAG (3341G>A 3343A>G), TGG340CAG (3344T>C 3345G>A), AAG342AAC (3352G>C), GAG343GAA (3355G>A), AAG344TTG (3356A>T 3357A>T), CTA345CTT (3361A>T), GCC346GAT (3363C>A 3364C>T), AAA347TCA (3365A>T 3366A>C), TGG349CTG (3371T>C 3372G>T), ATA350ATT (3376A>T), CGA351CAG (3378G>A 3379A>G), CGA352CCT (3381G>C 3382A>T), TCC353TCT (3385C>T), ACC354AAG (3387C>A 3388C>G), TCA355GCT (3389T>G 3391A>T), AGT356CCC (3392A>C 3393G>C 3394T>C), GCA357TAT (3395G>T 3396C>A 3397A>T), GGA358GGT (3400A>T), ACT359GCA (3401A>G 3403T>A), CCA360CCT (3406A>T), TGC361GTT (3407T>G 3408G>T 3409C>T), ATG362TTA (3410A>T 3412G>A), TTC363TTT (3415C>T), GTT364CAG (3416G>C 3417T>A 3418T>G), CCA365AAA (3419C>A 3420C>A), GCA367CAG (3425G>C 3426C>A 3427A>G), AAC368GAT (3428A>G 3430C>T), AAA370TCA (3434A>T 3435A>C), CTA371TTA (3437C>T), CGA372AGG (3440C>A 3442A>G), CTC373ATG (3443C>A 3445C>G), GTA374TGC (3446G>T 3447T>G 3448A>C), CAA375ATG (3449C>A 3450A>T 3451A>G), GAC376GAT (3454C>T), TAC377TAT (3457C>T), CGA378CGG (3460A>G), AAG379GCG (3461A>G 3462A>C), TTG380CTG (3464T>C), GAG382AAG (3470G>A), ATC383GTC (3473A>G), ATC385GTT (3479A>G 3481C>T), AAC387AAT (3487C>T), CGA388AAG (3488C>A 3489G>A 3490A>G), TAT389TAC (3493T>C), CTA391ATC (3497C>A 3499A>C), CCC392CCA (3502C>A), AAC393CTC (3503A>C 3504A>T), ATC394ATT (3508C>T), GAA395GCT (3510A>C 3511A>T), GAA396GAT (3514A>T), GCA397CTC (3515G>C 3516C>T 3517A>C), CAA398TTT (3518C>T 3519A>T 3520A>T), GAC399GAT (3523C>T), AGA400CAA (3524A>C 3525G>A), TTA401CTT (3527T>C 3529A>T), ACC402GGC (3530A>G 3531C>G), GGA403CAA (3533G>C 3534G>A), TCA404GCA (3536T>G), GAC405CGT (3539G>C 3540A>G 3541C>T), TAC407TTC (3546A>T), ACG408ACC (3550G>C), ATT410TTA (3554A>T 3556T>A), GAC411GAT (3559C>T), CTA412CTT (3562A>T), CGA413CGG (3565A>G), GAC414TCT (3566G>T 3567A>C 3568C>T), GCC415GGA (3570C>G 3571C>A), TTC416TAC (3573T>A), GCT418CAA (3578G>C 3579C>A 3580T>A), ATC419GTG (3581A>G 3583C>G), CGA420AGA (3584C>A), ATG421ATC (3589G>C), GGA424GGG (3598A>G), GAA425GAT (3601A>T), GAA426GAG (3604A>G), TGG427CCT (3605T>C 3606G>C 3607G>T), AAA428AAG (3610A>G), ACC429ACT (3613C>T), GCT430ACT (3614G>A), TTC431TGT (3618T>G 3619C>T), AGG432GTA (3620A>G 3621G>T 3622G>A), ACA433ACT (3625A>T), AGA434AGG (3628A>G), TAC435TAT (3631C>T), GGA436GGG (3634A>G), CTC437TCA (3635C>T 3636T>C 3637C>A), TAC438TAT (3640C>T), TTG441CTT (3647T>C 3649G>T), GTC442GTA (3652C>A), CCA444CCG (3658A>G), ATG445TTT (3659A>T 3661G>T), GGA446GGG (3664A>G), ACC448ACT (3670C>T), AAC449AAT (3673C>T), GCA450GCC (3676A>C), CCC451CCG (3679C>G), GCA452GCT (3682A>T), TCC453ACC (3683T>A), TGC454TTT (3687G>T 3688C>T), CAG455TGC (3689C>T 3690A>G 3691G>C), GAC456AAT (3692G>A 3694C>T), CTT457CTA (3697T>A), GTC458ATG (3698G>A 3700C>G), AAC459AAT (3703C>T), GAA460GGC (3705A>G 3706A>C), ACA461GTA (3707A>G 3708C>T), CTT462CTG (3712T>G), AGA463TAT (3713A>T 3714G>A 3715A>T), GAC464CCC (3716G>C 3717A>C), CTA465TAC (3719C>T 3720T>A 3721A>C), CTC466CTT (3724C>T), GTG468AAT (3728G>A 3729T>A 3730G>T), TGC469TTT (3732G>T 3733C>T), GTC470GTA (3736C>A), GTT471GTA (3739T>A), GCT472GTC (3741C>T 3742T>C), TAC473TAT (3745C>T), ATG474CTA (3746A>C 3748G>A), GAC475GAT (3751C>T), ATA477ATT (3757A>T), CTG478GTC (3758C>G 3760G>C), GTC479ATA (3761G>A 3763C>A), TAC480TAT (3766C>T), ACA481AGT (3768C>G 3769A>T), AAA482CAA (3770A>C), GGA483GAT (3774G>A 3775A>T), TCC484ACG (3776T>A 3778C>G), CTC485TTG (3779C>T 3781C>G), CAG486GAT (3782C>G 3784G>T), ACC489GTT (3791A>G 3792C>T 3793C>T), AAG490AGC (3795A>G 3796G>C), CAA491CAT (3799A>T), GTT492TTG (3800G>T 3802T>G), CAA493AGT (3803C>A 3804A>G 3805A>T), GAT494TTG (3806G>T 3807A>T 3808T>G), GTG495GTA (3811G>A), TTC496TTA (3814C>A), GAA497TCT (3815G>T 3816A>C 3817A>T), CGA498CGT (3820A>T), CTC499TTA (3821C>T 3823C>A), ACG500AGG (3825C>G), AAG501CAA (3827A>C 3829G>A), TCC502TAC (3831C>A), GGA503ACT (3833G>A 3834G>C 3835A>T), TTC504CTC (3836T>C), AAG505TAT (3839A>T 3841G>T), ACA506GTT (3842A>G 3843C>T 3844A>T), GCA507AAA (3845G>A 3846C>A), CCC508ATG (3848C>A 3849C>T 3850C>G), AAA510AAG (3856A>G), TGC511TGT (3859C>T), GAA512GAG (3862A>G), TTC513TTT (3865C>T), CAC514GCC (3866C>G 3867A>C), AAG515CAA (3869A>C 3871G>A), GAA517GAG (3877A>G), GTC518ATA (3878G>A 3880C>A), TTT520TTC (3886T>C), TTA521CTA (3887T>C), GGC522GGG (3892C>G), TTT523CAT (3893T>C 3894T>A), ATC524CTA (3896A>C 3898C>A), ATC525GTT (3899A>G 3901C>T), ACA527GAA (3905A>G 3906C>A), ACA528AAC (3909C>A 3910A>C), GGG529CAA (3911G>C 3912G>A 3913G>A), ATA530GTA (3914A>G), ACG531AGG (3918C>G), ATC532ATG (3922C>G), GAC533GAT (3925C>T), GCA535AAG (3929G>A 3930C>A 3931A>G), AAG536AAA (3934G>A), ACA537GTG (3935A>G 3936C>T 3937A>G), CAG538CAA (3940G>A), TCA539GCC (3941T>G 3943A>C), ATC540ATT (3946C>T), AGA541GTT (3947A>G 3948G>T 3949A>T), GAA542GAT (3952A>T), CCA544CAA (3957C>A), GAA545GCA (3960A>C), CCG546CCC (3964G>C), AAG547AAA (3967G>A), ACA548ACT (3970A>T), GTC549GTG (3973C>G), GTA552CTG (3980G>C 3982A>G), CAG553AGG (3983C>A 3984A>G), TCA554TCT (3988A>T), CTT556TTG (3992C>T 3994T>G), GGA557GGG (3997A>G), CTC558CTA (4000C>A), GCC559GCT (4003C>T), AAC562TAT (4010A>T 4012C>T), AAG567GCT (4025A>G 4026A>C 4027G>T), GAC568GGA (4029A>G 4030C>A), TAT569TAC (4033T>C), ACA572AAA (4041C>A), GCA574GCT (4048A>T), CCA575GCA (4049C>G), ATG576CTG (4052A>C), ACG577ACT (4057G>T), ATG578GAT (4058A>G 4059T>A 4060G>T), CTT579TTG (4061C>T 4063T>G), ACA580CTG (4064A>C 4065C>T 4066A>G), AGA581AAG (4068G>A 4069A>G), AAA582AAG (4072A>G), GAC583GAA (4075C>A), AAC585AAG (4081C>G), AAA587GTA (4085A>G 4086A>T), GGA589ACC (4091G>A 4092G>C 4093A>C), AAA590GAA (4094A>G), GAA591GAG (4099A>G), CAG592TGT (4100C>T 4101A>G 4102G>T), ACC593GAG (4103A>G 4104C>A 4105C>G), GAA594GCG (4107A>C 4108A>G), GCG595AGC (4109G>A 4110C>G 4111G>C), TTC596TTT (4114C>T), AAA597CAA (4115A>C), AGA598AAG (4119G>A 4120A>G), GAA601GAC (4129A>C), CAG602TTA (4130C>T 4131A>T 4132G>A), TGC603TTG (4134G>T 4135C>G), GCT604ACT (4136G>A), CCA607CCT (4147A>T), ACG608ATT (4149C>T 4150G>T), CTT609CTC (4153T>C), CGA610GCA (4154C>G 4155G>C), CTA611TTA (4157C>T), TTC612CCC (4160T>C 4161T>C), GAT613GAG (4165T>G), GGT614GAG (4167G>A 4168T>G), AGC615CCT (4169A>C 4170G>C 4171C>T), AAG616TTT (4172A>T 4173A>T 4174G>T), GAA617GAG (4177A>G), GTC618GTA (4180C>A), ATC620GTG (4184A>G 4186C>G), GAG621CAG (4187G>C), ACC622ACT (4192C>T), GAC623GAT (4195C>T), TCT625TCG (4201T>G), GAT626GAC (4204T>C), ATG627AAA (4206T>A 4207G>A), GCA628GCT (4210A>T), ATA629GTG (4211A>G 4213A>G), GGC630GGA (4216C>A), GCA631GGC (4218C>G 4219A>C), TGT632GTA (4220T>G 4221G>T 4222T>A), CTA633TTG (4223C>T 4225A>G), ACA634GTG (4226A>G 4227C>T 4228A>G), CAG635CAA (4231G>A), ACA636_CAC637del (4232_4237delACACAC), GAT638GAA (4240T>A), GGG639GGT (4243G>T), AAA640_AGA641del (4244_4249delAAAAGA), CAC642CAT (4252C>T), CCA643CCG (4255A>G), TAT646TTT (4263A>T), TAT647GAA (4265T>G 4267T>A), TCC648AGC (4268T>A 4269C>G), CGG649AGG (4271C>A), ATG651TTA (4277A>T 4279G>A), ACC652AAT (4281C>A 4282C>T), ACA653GAT (4283A>G 4284C>A 4285A>T), GCG654GCT (4288G>T), GAA655GAG (4291A>G), AAC657AGG (4296A>G 4297C>G), GAC659TCT (4301G>T 4302A>C 4303C>T), ATC660ACT (4305T>C 4306C>T), GAC662GAA (4312C>A), CTT665ATG (4319C>A 4321T>G), CTA666GTT (4322C>G 4324A>T), GCC667GCA (4327C>A), ATT668GTG (4328A>G 4330T>G), GTT669GTG (4333T>G), GCC670CCC (4334G>C), AAA866.GT (4923A>G 4924A>T), GAC869GAG (4933C>G), GAG870AGT (4934G>A 4935A>G 4936G>T), ACG872ACT (4942G>T), ATT876ATA (4954T>A), ACG877ACT (4957G>T), AAA878TGC (4958A>T 4959A>G 4960A>C), CTC879CTG (4963C>G), CCG880CCA (4966G>A), AGG881AAG (4968G>A), TCA882TCC (4972A>C), AAG883AAC (4975G>C), GAT884AAG (4976G>A 4978T>G), CAA984TGC (5276C>T 5277A>G 5278A>C), CTA986CTT (5284A>T), CGG987AGG (5285C>A), TAC989TTC (5292A>T), ATC990GTG (5294A>G 5296C>G), AAC991AGC (5298A>G), TAC992TCT (5301A>C 5302C>T), GCA993CAG (5303G>C 5304C>A 5305A>G), GAC995AAG (5309G>A 5311C>G), AAC996GAT (5312A>G 5314C>T), GTT998GCT (5319T>C), TCA999AAG (5321T>A 5322C>A 5323A>G), TTA1000TTG (5326A>G), CTG1001CTA (5329G>A), CCA1002GAT (5330C>G 5331C>A 5332A>T), ATG1003TTG (5333A>T), GCG1004GCA (5338G>A), CAG1005GAA (5339C>G 5341G>A), ATC1006TTC (5342A>T), GCA1007CCA (5345G>C), CTG1008TAT (5348C>T 5349T>A 5350G>T), AAC1010TTA (5354A>T 5355A>T 5356C>A), CAC1011CAG (5359C>G), AAA1012TGG (5360A>T 5361A>G 5362A>G), TCA1013AGT (5363T>A 5364C>G 5365A>T), GAG1014GAA (5368G>A), ACA1015GTC (5369A>G 5370C>T 5371A>C), ACT1016ACC (5374T>C), TCG1017GGA (5375T>G 5376C>G 5377G>A), ACG1018AGG (5379C>G), ACG1019ACA (5383G>A), ATG1022GAG (5390A>G 5391T>A), CGA1023_ACT1024del (5393_5398delCGAACT), GCA1026GCC (5404A>C) | | | | | | | | |

**Proteins**

| | Begin | End | Coverage | Score | Concordance | Matches | Identities | I/D/M/F* | Stop Codons |
|---|---|---|---|---|---|---|---|---|---|
| Reverse Transcriptase (YP_009666308.1) | 281 | 1026 | 43.6% | 1494 | 47.5% | 450 (98.0%) | 214 (46.6%) | 3/6/0/0 | 0 |

| | Begin | End | Coverage | Score | Concordance | Matches | Identities | I/D/M/F* | Stop Codons |
|---|---|---|---|---|---|---|---|---|---|
| **NT** | **3165** | **5404** | **18.5%** | **117** | **4.4%** | **1352 (98.0%)** | **721 (52.3%)** | **9/18** | |
| Protein mutations: | L281V (3167T>G), S282E (3170A>G 3171G>A 3172T>A), K285D (3179A>G 3181G>C), E286C (3182G>T 3183A>G 3184G>T), Y287V (3185T>G 3186A>T 3187T>G), R288A (3188C>G 3189G>C 3190C>A), K289E (3191A>G), W290L (3194T>C 3195G>T), S291L (3197T>C 3198C>T), R292K (3200C>A 3201G>A), L293Q (3204T>A 3205C>G), F294Y (3207T>A 3208T>C), E295A (3210A>C 3211A>T), E296D (3214G>T), E297V (3216A>T 3217A>G), R298M (3219G>T 3220A>G), G299P (3221G>C 3222G>C 3223C>A), K300P (3224A>C 3225A>C 3226G>T), D301E (3229C>G), A302L (3230G>T 3231C>T 3232C>G), L303P (3233T>C 3234T>C), P304K (3236C>A 3237C>A 3238T>G), H306L (3243A>T 3244C>G), Q307P (3246A>C), W309R (3251T>A), W309_D310insRDI (3253_3254insAGGGATATT), K312R (3261A>G 3262G>A), N314E (3266A>G 3268C>G), I315L (3269A>T 3271T>G), Q316M (3272C>A 3273A>T), K319S (3281A>T 3282A>C 3283A>T), E320T (3284G>A 3285A>C), W323A (3293T>G 3294G>C 3295G>A), G324Q (3296G>C 3297G>A), P325A (3299C>G 3301C>T), L326P (3303T>C 3304A>T), Q328R (3308C>A 3309A>G), S330A (3314T>G), E331P (3317G>C 3318A>C 3319G>T), Q335A (3329C>G 3330A>C 3331G>T), T336E (3332A>G 3333C>A 3334C>A), E339K (3341G>A 3343A>G), W340Q (3344T>C 3345G>A), K342N (3352G>C), K344L (3356A>T 3357A>T), A346D (3363C>A 3364C>T), K347S (3365A>T 3366A>C), W349L (3371T>C 3372G>T), R351Q (3378G>A 3379A>G), R352P (3381G>C 3382A>T), T354K (3387C>A 3388C>G), S355A (3389T>G 3391A>T), S356P (3392A>C 3393G>C 3394T>C), A357Y (3395G>T 3396C>A 3397A>T), T359A (3401A>G 3403T>A), C361V (3407T>G 3408G>T 3409C>T), M362L (3410A>T 3412G>A), V364Q (3416G>C 3417T>A 3418T>G), P365K (3419C>A 3420C>A), A367Q (3425G>C 3426C>A 3427A>G), N368D (3428A>G 3430C>T), K370S (3434A>T 3435A>C), L373M (3443C>A 3445C>G), V374C (3446G>T 3447T>G 3448A>C), Q375M (3449C>A 3450A>T 3451A>G), K379A (3461A>G 3462A>C), E382K (3470G>A), I383V (3473A>G), I385V (3479A>G 3481C>T), R388K (3488C>A 3489G>A 3490A>G), L391I (3497C>A 3499A>C), N393L (3503A>C 3504A>T), E395A (3510A>C 3511A>T), E396D (3514A>T), A397L (3515G>C 3516C>T 3517A>C), Q398F (3518C>T 3519A>T 3520A>T), R400Q (3524A>C 3525G>A), T402G (3530A>G 3531C>G), G403Q (3533G>C 3534G>A), S404A (3536T>G), D405R (3539G>C 3540A>G 3541C>T), Y407F (3546A>T), I410L (3554A>T 3556T>A), D414S (3566G>T 3567A>C 3568C>T), A415G (3570C>G 3571C>A), F416Y (3573T>A), A418Q (3578G>C 3579C>A 3580T>A), I419V (3581A>G 3583C>G), M421I (3589G>C), E425D (3601A>T), W427P (3605T>C 3606G>C 3607G>T), A430T (3614G>A), F431C (3618T>G 3619C>T), R432V (3620A>G 3621G>T 3622G>A), L437S (3635C>T 3636T>C 3637C>A), M445F (3659A>T 3661G>T), S453T (3683T>A), C454F (3687G>T 3688C>T), Q455C (3689C>T 3690A>G 3691G>C), D456N (3692G>A 3694C>T), V458M (3698G>A 3700C>G), E460G (3705A>G 3706A>C), T461V (3707A>G 3708C>T), R463Y (3713A>T 3714G>A 3715A>T), D464P (3716G>C 3717A>C), L465Y (3719C>T 3720T>A 3721A>C), V468N (3728G>A 3729T>A 3730G>T), C469F (3732G>T 3733C>T), A472V (3741C>T 3742T>C), M474L (3746A>C 3748G>A), L478V (3758C>G 3760G>C), V479I (3761G>A 3763C>A), T481S (3768C>G 3769A>T), K482Q (3770A>C), G483D (3774G>A 3775A>T), S484T (3776T>A 3778C>G), Q486D (3782C>G 3784G>T), T489V (3791A>G 3792C>T 3793C>T), K490S (3795A>G 3796G>C), Q491H (3799A>T), V492L (3800G>T 3802T>G), Q493S (3803C>A 3804A>G 3805A>T), D494L (3806G>T 3807A>T 3808T>G), F496L (3814C>A), E497S (3815G>T 3816A>C 3817A>T), T500R (3825C>G), K501Q (3827A>C 3829G>A), S502Y (3831C>A), G503T (3833G>A 3834G>C 3835A>T), F504L (3836T>C), K505Y (3839A>T 3841G>T), T506V (3842A>G 3843C>T 3844A>T), A507K (3845G>A 3846C>A), P508M (3848C>A 3849C>T 3850C>G), H514A (3866C>G 3867A>C), K515Q (3869A>C 3871G>A), V518I (3878G>A 3880C>A), F523H (3893T>C 3894T>A), I524L (3896A>C 3898C>A), I525V (3899A>G 3901C>T), T527E (3905A>G 3906C>A), T528N (3909C>A 3910A>C), G529Q (3911G>C 3912G>A 3913G>A), I530V (3914A>G), T531R (3918C>G), I532M (3922C>G), A535K (3929G>A 3930C>A 3931A>G), T537V (3935A>G 3936C>T 3937A>G), S539A (3941T>G 3943A>C), R541V (3947A>G 3948G>T 3949A>T), E542D (3952A>T), P544Q (3957C>A), E545A (3960A>C), V552L (3980G>C 3982A>G), Q553R (3983C>A 3984A>G), N562Y (4010A>T 4012C>T), K567A (4025A>G 4026A>C 4027G>T), D568G (4029A>G 4030C>A), T572K (4041C>A), P575A (4049C>G), M576L (4052A>C), M578D (4058A>G 4059T>A 4060G>T), T580L (4064A>C 4065C>T 4066A>G), R581K (4068G>A 4069A>G), D583E (4075C>A), N585K (4081C>G), K587V (4085A>G 4086A>T), G589T (4091G>A 4092G>C 4093A>C), K590E (4094A>G), Q592C (4100C>T 4101A>G 4102G>T), T593E (4103A>G 4104C>A 4105C>G), E594A (4107A>C 4108A>G), A595S (4109G>A 4110C>G 4111G>C), K597Q (4115A>C), R598K (4119G>A 4120A>G), E601D (4129A>C), Q602L (4130C>T 4131A>T 4132G>A), C603L (4134G>T 4135C>G), A604T (4136G>A), T608I (4149C>T 4150G>T), R610A (4154C>G 4155G>C), F612P (4160T>C 4161T>C), D613E (4165T>G), G614E (4167G>A 4168T>G), S615P (4169A>C 4170G>C 4171C>T), K616F (4172A>T 4173A>T 4174G>T), I620V (4184A>G 4186C>G), E621Q (4187G>C), M627K (4206T>A 4207G>A), I629V (4211A>G 4213A>G), A631G (4218C>G 4219A>C), C632V (4220T>G 4221G>T 4222T>A), T634V (4226A>G 4227C>T 4228A>G), T636_H637del (4232_4237delACACAC), D638E (4240T>A), K640_R641del (4244_4249delAAAAGA), Y646F (4263A>T), Y647E (4265T>G 4267T>A), M651L (4277A>T 4279G>A), T652N (4281C>A 4282C>T), T653D (4283A>G 4284C>A 4285A>T), N657R (4296A>G 4297C>G), D659S (4301G>T 4302A>C 4303C>T), I660T (4305T>C 4306C>T), D662E (4312C>A), L665M (4319C>A 4321T>G), L666V (4322C>G 4324A>T), I668V (4328A>G 4330T>G), A670P (4334G>C), D869E (4933C>G), E870S (4934G>A 4935A>G 4936G>T), K878C (4958A>T 4959A>G 4960A>C), R881K (4968G>A), K883N (4975G>C), D884K (4976G>A 4978T>G), Q984C (5276C>T 5277A>G 5278A>C), Y989F (5292A>T), I990V (5294A>G 5296C>G), N991S (5298A>G), Y992S (5301A>C 5302C>T), A993Q (5303G>C 5304C>A 5305A>G), D995K (5309G>A 5311C>G), N996D (5312A>G 5314C>T), V998A (5319T>C), S999K (5321T>A 5322C>A 5323A>G), P1002D (5330C>G 5331C>A 5332A>T), M1003L (5333A>T), Q1005E (5339C>G 5341G>A), I1006F (5342A>T), A1007P (5345G>C), L1008Y (5348C>T 5349T>A 5350G>T), N1010L (5354A>T 5355A>T 5356C>A), H1011Q (5359C>G), K1012W (5360A>T 5361A>G 5362A>G), T1015V (5369A>G 5370C>T 5371A>C), S1017G (5375T>G 5376C>G 5377G>A), T1018R (5379C>G), M1022E (5390A>G 5391T>A), R1023_T1024del (5393_5398delCGAACT) | | | | | | | | |

| | Begin | End | Coverage | Score | Concordance | Matches | Identities | I/D/M/F* | Stop Codons |
|---|---|---|---|---|---|---|---|---|---|
| **NT** | **3165** | **5404** | **18.5%** | **117** | **4.4%** | **1352 (98.0%)** | **721 (52.3%)** | **9/18** | |
| Codon mutations: | GAG280.GG (3165A>G), TTG281GTG (3167T>G), AGT282GAA (3170A>G 3171G>A 3172T>A), ATT283ATA (3175T>A), CCC284CCA (3178C>A), AAG285GAC (3179A>G 3181G>C), GAG286TGT (3182G>T 3183A>G 3184G>T), TAT287GTG (3185T>G 3186A>T 3187T>G), CGC288GCA (3188C>G 3189G>C 3190C>A), AAG289GAG (3191A>G), TGG290CTG (3194T>C 3195G>T), TCG291CTG (3197T>C 3198C>T), CGA292AAA (3200C>A 3201G>A), CTC293CAG (3204T>A 3205C>G), TTT294TAC (3207T>A 3208T>C), GAA295GCT (3210A>C 3211A>T), GAG296GAT (3214G>T), GAA297GTG (3216A>T 3217A>G), AGA298ATG (3219G>T 3220A>G), GGC299CCA (3221G>C 3222G>C 3223C>A), AAG300CCT (3224A>C 3225A>C 3226G>T), GAC301GAG (3229C>G), GCC302TTG (3230G>T 3231C>T 3232C>G), TTA303CCA (3233T>C 3234T>C), CCT304AAG (3236C>A 3237C>A 3238T>G), CAC306CTG (3243A>T 3244C>G), CAA307CCA (3246A>C), TGG309AGG (3251T>A), TGG309_GAT310insAGGGATATT (3253_3254insAGGGATATT), AAG312AGA (3261A>G 3262G>A), ATA313ATT (3265A>T), AAC314GAG (3266A>G 3268C>G), ATT315TTG (3269A>T 3271T>G), CAG316ATG (3272C>A 3273A>T), GGG318GGT (3280G>T), AAA319TCT (3281A>T 3282A>C 3283A>T), GAG320ACG (3284G>A 3285A>C), CCT321CCG (3289T>G), CCA322CCT (3292A>T), TGG323GCA (3293T>G 3294G>C 3295G>A), GGA324CAA (3296G>C 3297G>A), CCC325GCT (3299C>G 3301C>T), CTA326CCT (3303T>C 3304A>T), TAT327TAC (3307T>C), CAA328AGA (3308C>A 3309A>G), TCT330GCT (3314T>G), GAG331CCT (3317G>C 3318A>C 3319G>T), CTA334TTG (3326C>T 3328A>G), CAG335GCT (3329C>G 3330A>C 3331G>T), ACC336GAA (3332A>G 3333C>A 3334C>A), CTA337CTG (3337A>G), CGA338AGG (3338C>A 3340A>G), GAA339AAG (3341G>A 3343A>G), TGG340CAG (3344T>C 3345G>A), AAG342AAC (3352G>C), GAG343GAA (3355G>A), AAG344TTG (3356A>T 3357A>T), CTA345CTT (3361A>T), GCC346GAT (3363C>A 3364C>T), AAA347TCA (3365A>T 3366A>C), TGG349CTG (3371T>C 3372G>T), ATA350ATT (3376A>T), CGA351CAG (3378G>A 3379A>G), CGA352CCT (3381G>C 3382A>T), TCC353TCT (3385C>T), ACC354AAG (3387C>A 3388C>G), TCA355GCT (3389T>G 3391A>T), AGT356CCC (3392A>C 3393G>C 3394T>C), GCA357TAT (3395G>T 3396C>A 3397A>T), GGA358GGT (3400A>T), ACT359GCA (3401A>G 3403T>A), CCA360CCT (3406A>T), TGC361GTT (3407T>G 3408G>T 3409C>T), ATG362TTA (3410A>T 3412G>A), TTC363TTT (3415C>T), GTT364CAG (3416G>C 3417T>A 3418T>G), CCA365AAA (3419C>A 3420C>A), GCA367CAG (3425G>C 3426C>A 3427A>G), AAC368GAT (3428A>G 3430C>T), AAA370TCA (3434A>T 3435A>C), CTA371TTA (3437C>T), CGA372AGG (3440C>A 3442A>G), CTC373ATG (3443C>A 3445C>G), GTA374TGC (3446G>T 3447T>G 3448A>C), CAA375ATG (3449C>A 3450A>T 3451A>G), GAC376GAT (3454C>T), TAC377TAT (3457C>T), CGA378CGG (3460A>G), AAG379GCG (3461A>G 3462A>C), TTG380CTG (3464T>C), GAG382AAG (3470G>A), ATC383GTC (3473A>G), ATC385GTT (3479A>G 3481C>T), AAC387AAT (3487C>T), CGA388AAG (3488C>A 3489G>A 3490A>G), TAT389TAC (3493T>C), CTA391ATC (3497C>A 3499A>C), CCC392CCA (3502C>A), AAC393CTC (3503A>C 3504A>T), ATC394ATT (3508C>T), GAA395GCT (3510A>C 3511A>T), GAA396GAT (3514A>T), GCA397CTC (3515G>C 3516C>T 3517A>C), CAA398TTT (3518C>T 3519A>T 3520A>T), GAC399GAT (3523C>T), AGA400CAA (3524A>C 3525G>A), TTA401CTT (3527T>C 3529A>T), ACC402GGC (3530A>G 3531C>G), GGA403CAA (3533G>C 3534G>A), TCA404GCA (3536T>G), GAC405CGT (3539G>C 3540A>G 3541C>T), TAC407TTC (3546A>T), ACG408ACC (3550G>C), ATT410TTA (3554A>T 3556T>A), GAC411GAT (3559C>T), CTA412CTT (3562A>T), CGA413CGG (3565A>G), GAC414TCT (3566G>T 3567A>C 3568C>T), GCC415GGA (3570C>G 3571C>A), TTC416TAC (3573T>A), GCT418CAA (3578G>C 3579C>A 3580T>A), ATC419GTG (3581A>G 3583C>G), CGA420AGA (3584C>A), ATG421ATC (3589G>C), GGA424GGG (3598A>G), GAA425GAT (3601A>T), GAA426GAG (3604A>G), TGG427CCT (3605T>C 3606G>C 3607G>T), AAA428AAG (3610A>G), ACC429ACT (3613C>T), GCT430ACT (3614G>A), TTC431TGT (3618T>G 3619C>T), AGG432GTA (3620A>G 3621G>T 3622G>A), ACA433ACT (3625A>T), AGA434AGG (3628A>G), TAC435TAT (3631C>T), GGA436GGG (3634A>G), CTC437TCA (3635C>T 3636T>C 3637C>A), TAC438TAT (3640C>T), TTG441CTT (3647T>C 3649G>T), GTC442GTA (3652C>A), CCA444CCG (3658A>G), ATG445TTT (3659A>T 3661G>T), GGA446GGG (3664A>G), ACC448ACT (3670C>T), AAC449AAT (3673C>T), GCA450GCC (3676A>C), CCC451CCG (3679C>G), GCA452GCT (3682A>T), TCC453ACC (3683T>A), TGC454TTT (3687G>T 3688C>T), CAG455TGC (3689C>T 3690A>G 3691G>C), GAC456AAT (3692G>A 3694C>T), CTT457CTA (3697T>A), GTC458ATG (3698G>A 3700C>G), AAC459AAT (3703C>T), GAA460GGC (3705A>G 3706A>C), ACA461GTA (3707A>G 3708C>T), CTT462CTG (3712T>G), AGA463TAT (3713A>T 3714G>A 3715A>T), GAC464CCC (3716G>C 3717A>C), CTA465TAC (3719C>T 3720T>A 3721A>C), CTC466CTT (3724C>T), GTG468AAT (3728G>A 3729T>A 3730G>T), TGC469TTT (3732G>T 3733C>T), GTC470GTA (3736C>A), GTT471GTA (3739T>A), GCT472GTC (3741C>T 3742T>C), TAC473TAT (3745C>T), ATG474CTA (3746A>C 3748G>A), GAC475GAT (3751C>T), ATA477ATT (3757A>T), CTG478GTC (3758C>G 3760G>C), GTC479ATA (3761G>A 3763C>A), TAC480TAT (3766C>T), ACA481AGT (3768C>G 3769A>T), AAA482CAA (3770A>C), GGA483GAT (3774G>A 3775A>T), TCC484ACG (3776T>A 3778C>G), CTC485TTG (3779C>T 3781C>G), CAG486GAT (3782C>G 3784G>T), ACC489GTT (3791A>G 3792C>T 3793C>T), AAG490AGC (3795A>G 3796G>C), CAA491CAT (3799A>T), GTT492TTG (3800G>T 3802T>G), CAA493AGT (3803C>A 3804A>G 3805A>T), GAT494TTG (3806G>T 3807A>T 3808T>G), GTG495GTA (3811G>A), TTC496TTA (3814C>A), GAA497TCT (3815G>T 3816A>C 3817A>T), CGA498CGT (3820A>T), CTC499TTA (3821C>T 3823C>A), ACG500AGG (3825C>G), AAG501CAA (3827A>C 3829G>A), TCC502TAC (3831C>A), GGA503ACT (3833G>A 3834G>C 3835A>T), TTC504CTC (3836T>C), AAG505TAT (3839A>T 3841G>T), ACA506GTT (3842A>G 3843C>T 3844A>T), GCA507AAA (3845G>A 3846C>A), CCC508ATG (3848C>A 3849C>T 3850C>G), AAA510AAG (3856A>G), TGC511TGT (3859C>T), GAA512GAG (3862A>G), TTC513TTT (3865C>T), CAC514GCC (3866C>G 3867A>C), AAG515CAA (3869A>C 3871G>A), GAA517GAG (3877A>G), GTC518ATA (3878G>A 3880C>A), TTT520TTC (3886T>C), TTA521CTA (3887T>C), GGC522GGG (3892C>G), TTT523CAT (3893T>C 3894T>A), ATC524CTA (3896A>C 3898C>A), ATC525GTT (3899A>G 3901C>T), ACA527GAA (3905A>G 3906C>A), ACA528AAC (3909C>A 3910A>C), GGG529CAA (3911G>C 3912G>A 3913G>A), ATA530GTA (3914A>G), ACG531AGG (3918C>G), ATC532ATG (3922C>G), GAC533GAT (3925C>T), GCA535AAG (3929G>A 3930C>A 3931A>G), AAG536AAA (3934G>A), ACA537GTG (3935A>G 3936C>T 3937A>G), CAG538CAA (3940G>A), TCA539GCC (3941T>G 3943A>C), ATC540ATT (3946C>T), AGA541GTT (3947A>G 3948G>T 3949A>T), GAA542GAT (3952A>T), CCA544CAA (3957C>A), GAA545GCA (3960A>C), CCG546CCC (3964G>C), AAG547AAA (3967G>A), ACA548ACT (3970A>T), GTC549GTG (3973C>G), GTA552CTG (3980G>C 3982A>G), CAG553AGG (3983C>A 3984A>G), TCA554TCT (3988A>T), CTT556TTG (3992C>T 3994T>G), GGA557GGG (3997A>G), CTC558CTA (4000C>A), GCC559GCT (4003C>T), AAC562TAT (4010A>T 4012C>T), AAG567GCT (4025A>G 4026A>C 4027G>T), GAC568GGA (4029A>G 4030C>A), TAT569TAC (4033T>C), ACA572AAA (4041C>A), GCA574GCT (4048A>T), CCA575GCA (4049C>G), ATG576CTG (4052A>C), ACG577ACT (4057G>T), ATG578GAT (4058A>G 4059T>A 4060G>T), CTT579TTG (4061C>T 4063T>G), ACA580CTG (4064A>C 4065C>T 4066A>G), AGA581AAG (4068G>A 4069A>G), AAA582AAG (4072A>G), GAC583GAA (4075C>A), AAC585AAG (4081C>G), AAA587GTA (4085A>G 4086A>T), GGA589ACC (4091G>A 4092G>C 4093A>C), AAA590GAA (4094A>G), GAA591GAG (4099A>G), CAG592TGT (4100C>T 4101A>G 4102G>T), ACC593GAG (4103A>G 4104C>A 4105C>G), GAA594GCG (4107A>C 4108A>G), GCG595AGC (4109G>A 4110C>G 4111G>C), TTC596TTT (4114C>T), AAA597CAA (4115A>C), AGA598AAG (4119G>A 4120A>G), GAA601GAC (4129A>C), CAG602TTA (4130C>T 4131A>T 4132G>A), TGC603TTG (4134G>T 4135C>G), GCT604ACT (4136G>A), CCA607CCT (4147A>T), ACG608ATT (4149C>T 4150G>T), CTT609CTC (4153T>C), CGA610GCA (4154C>G 4155G>C), CTA611TTA (4157C>T), TTC612CCC (4160T>C 4161T>C), GAT613GAG (4165T>G), GGT614GAG (4167G>A 4168T>G), AGC615CCT (4169A>C 4170G>C 4171C>T), AAG616TTT (4172A>T 4173A>T 4174G>T), GAA617GAG (4177A>G), GTC618GTA (4180C>A), ATC620GTG (4184A>G 4186C>G), GAG621CAG (4187G>C), ACC622ACT (4192C>T), GAC623GAT (4195C>T), TCT625TCG (4201T>G), GAT626GAC (4204T>C), ATG627AAA (4206T>A 4207G>A), GCA628GCT (4210A>T), ATA629GTG (4211A>G 4213A>G), GGC630GGA (4216C>A), GCA631GGC (4218C>G 4219A>C), TGT632GTA (4220T>G 4221G>T 4222T>A), CTA633TTG (4223C>T 4225A>G), ACA634GTG (4226A>G 4227C>T 4228A>G), CAG635CAA (4231G>A), ACA636_CAC637del (4232_4237delACACAC), GAT638GAA (4240T>A), GGG639GGT (4243G>T), AAA640_AGA641del (4244_4249delAAAAGA), CAC642CAT (4252C>T), CCA643CCG (4255A>G), TAT646TTT (4263A>T), TAT647GAA (4265T>G 4267T>A), TCC648AGC (4268T>A 4269C>G), CGG649AGG (4271C>A), ATG651TTA (4277A>T 4279G>A), ACC652AAT (4281C>A 4282C>T), ACA653GAT (4283A>G 4284C>A 4285A>T), GCG654GCT (4288G>T), GAA655GAG (4291A>G), AAC657AGG (4296A>G 4297C>G), GAC659TCT (4301G>T 4302A>C 4303C>T), ATC660ACT (4305T>C 4306C>T), GAC662GAA (4312C>A), CTT665ATG (4319C>A 4321T>G), CTA666GTT (4322C>G 4324A>T), GCC667GCA (4327C>A), ATT668GTG (4328A>G 4330T>G), GTT669GTG (4333T>G), GCC670CCC (4334G>C), AAA866.GT (4923A>G 4924A>T), GAC869GAG (4933C>G), GAG870AGT (4934G>A 4935A>G 4936G>T), ACG872ACT (4942G>T), ATT876ATA (4954T>A), ACG877ACT (4957G>T), AAA878TGC (4958A>T 4959A>G 4960A>C), CTC879CTG (4963C>G), CCG880CCA (4966G>A), AGG881AAG (4968G>A), TCA882TCC (4972A>C), AAG883AAC (4975G>C), GAT884AAG (4976G>A 4978T>G), CAA984TGC (5276C>T 5277A>G 5278A>C), CTA986CTT (5284A>T), CGG987AGG (5285C>A), TAC989TTC (5292A>T), ATC990GTG (5294A>G 5296C>G), AAC991AGC (5298A>G), TAC992TCT (5301A>C 5302C>T), GCA993CAG (5303G>C 5304C>A 5305A>G), GAC995AAG (5309G>A 5311C>G), AAC996GAT (5312A>G 5314C>T), GTT998GCT (5319T>C), TCA999AAG (5321T>A 5322C>A 5323A>G), TTA1000TTG (5326A>G), CTG1001CTA (5329G>A), CCA1002GAT (5330C>G 5331C>A 5332A>T), ATG1003TTG (5333A>T), GCG1004GCA (5338G>A), CAG1005GAA (5339C>G 5341G>A), ATC1006TTC (5342A>T), GCA1007CCA (5345G>C), CTG1008TAT (5348C>T 5349T>A 5350G>T), AAC1010TTA (5354A>T 5355A>T 5356C>A), CAC1011CAG (5359C>G), AAA1012TGG (5360A>T 5361A>G 5362A>G), TCA1013AGT (5363T>A 5364C>G 5365A>T), GAG1014GAA (5368G>A), ACA1015GTC (5369A>G 5370C>T 5371A>C), ACT1016ACC (5374T>C), TCG1017GGA (5375T>G 5376C>G 5377G>A), ACG1018AGG (5379C>G), ACG1019ACA (5383G>A), ATG1022GAG (5390A>G 5391T>A), CGA1023_ACT1024del (5393_5398delCGAACT), GCA1026GCC (5404A>C) | | | | | | | | |

*: Inserts / Deletes / Misaligned / Frameshifts

#### Analysis details

This analysis was performed with panviral2.64

### NGS Details (UN62): Bracoviriform congregatae (segment Circle 7)

#### Assembly

| | |
|---|---|
| **Coverage Length** | 143 (1 contig(s)) |
| **Depth Of Coverage** | 3.1 |
| **Number Of Reads** | 5 |
| **Reads Per Million** | 0.12 rpm (after QC) |
| **Ambiguities** | 0 |
| **Assembly Method** | de novo + reference guided assembly |
| **Consensus Caller** | Bcf Tools |

#### Coverage Map

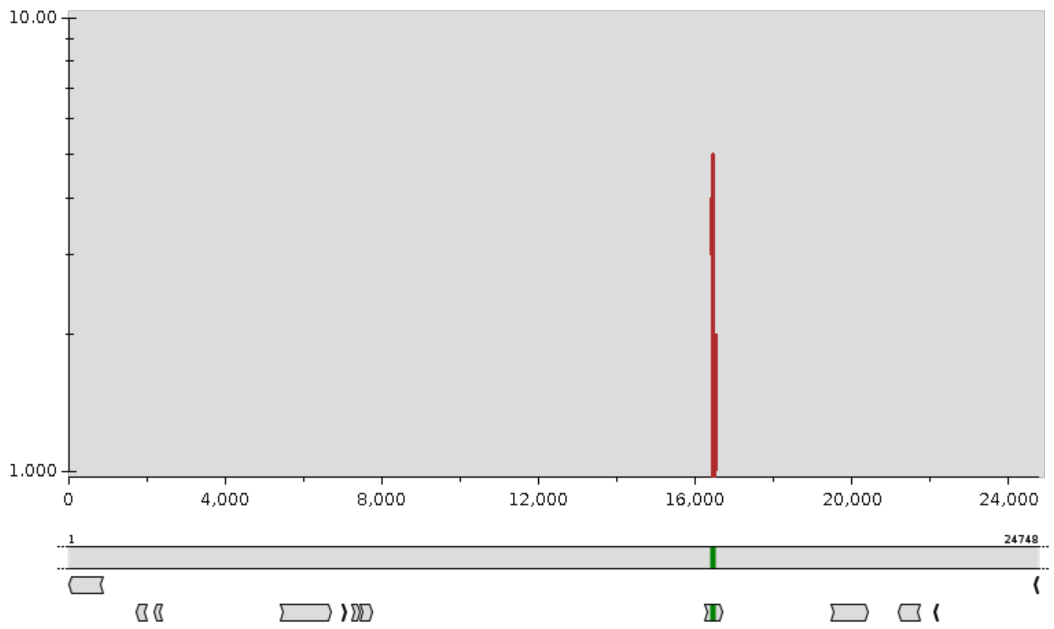

#### Assignment

| | |
|---|---|
| **Type** | Bracoviriform congregatae (Taxonomy ID: 39640) |
| **Reference Genome** | NC_006639.1 |
| **NT Identity (%)** | 75.5245 |
| **AA Identity (%)** | 85.4167 |
| **Number Of Stop Codons** | 0 |
| **Number Of CDS** | 5 |

#### Alignment

| | |
|---|---|
| **Alignment Score** | 146.0 (NT) + 284.0 (AA) = 430.0 |
| **Concordance (%)** | 72.3906 |

| Alignment Method | Global, seeded, nucleotide + amino acids (AGA) |
|---|---|

#### Genome Region

Sequence starts at position 16402 and ends at position 16544 relative to NC_006639.1 reference sequence.

#### Alignment Detailed Statistics

| | Begin | End | Coverage | Score | Concordance | Matches | Identities | I/D/M/F* | Stop Codons |
|---|---|---|---|---|---|---|---|---|---|
| **NT** | **16402** | **16544** | **0.6%** | **146** | **51.0%** | **143 (100%)** | **108 (75.5%)** | **0/0** | |
| Mutations: | 16402A>C, 16408A>T, 16411G>C, 16414A>G, 16417A>C, 16418T>C, 16420G>C, 16423A>C, 16426A>G, 16429G>C, 16432C>A, 16435T>C, 16444T>C, 16449G>A, 16450A>G, 16451T>A, 16452C>T, 16459A>C, 16462T>C, 16471A>G, 16474A>T, 16486A>C, 16489T>C, 16495T>C, 16501G>C, 16504T>C, 16516A>T, 16526G>A, 16531T>A, 16533G>C, 16534A>C, 16535T>A, 16538G>A, 16540C>T, 16543T>C | | | | | | | | |
| **CDS** | | | | | | | | | |
| Histone | 58 | 105 | 30.8% | 284 | 88.8% | 48 (100%) | 41 (85.4%) | 0/0/0/0 | 0 |
| Protein mutations: | R73K (16449G>A 16450A>G), S74I (16451T>A 16452C>T), V99I (16526G>A), G101A (16533G>C 16534A>C), L102M (16535T>A), V103I (16538G>A 16540C>T) | | | | | | | | |
| Codon mutations: | GTA57..C (16402A>C), GGA59GGT (16408A>T), GGG60GGC (16411G>C), AAA61AAG (16414A>G), GGA62GGC (16417A>C), TTG63CTC (16418T>C 16420G>C), GGA64GGC (16423A>C), AAA65AAG (16426A>G), GGG66GGC (16429G>C), GGC67GGA (16432C>A), GCT68GCC (16435T>C), CAT71CAC (16444T>C), AGA73AAG (16449G>A 16450A>G), TCT74ATT (16451T>A 16452C>T), CGA76CGC (16459A>C), GAT77GAC (16462T>C), CAA80CAG (16471A>G), GGA81GGT (16474A>T), CCA85CCC (16486A>C), GCT86GCC (16489T>C), CGT88CGC (16495T>C), CTG90CTC (16501G>C), GCT91GCC (16504T>C), GGA95GGT (16516A>T), GTC99ATC (16526G>A), TCT100TCA (16531T>A), GGA101GCC (16533G>C 16534A>C), TTG102ATG (16535T>A), GTC103ATT (16538G>A 16540C>T), TAT104TAC (16543T>C) | | | | | | | | |
| **Proteins** | | | | | | | | | |
| hypothetical protein (YP_184795.1) | 58 | 105 | 30.8% | 284 | 88.8% | 48 (100%) | 41 (85.4%) | 0/0/0/0 | 0 |
| Protein mutations: | R73K (16449G>A 16450A>G), S74I (16451T>A 16452C>T), V99I (16526G>A), G101A (16533G>C 16534A>C), L102M (16535T>A), V103I (16538G>A 16540C>T) | | | | | | | | |
| Codon mutations: | GTA57..C (16402A>C), GGA59GGT (16408A>T), GGG60GGC (16411G>C), AAA61AAG (16414A>G), GGA62GGC (16417A>C), TTG63CTC (16418T>C 16420G>C), GGA64GGC (16423A>C), AAA65AAG (16426A>G), GGG66GGC (16429G>C), GGC67GGA (16432C>A), GCT68GCC (16435T>C), CAT71CAC (16444T>C), AGA73AAG (16449G>A 16450A>G), TCT74ATT (16451T>A 16452C>T), CGA76CGC (16459A>C), GAT77GAC (16462T>C), CAA80CAG (16471A>G), GGA81GGT (16474A>T), CCA85CCC (16486A>C), GCT86GCC (16489T>C), CGT88CGC (16495T>C), CTG90CTC (16501G>C), GCT91GCC (16504T>C), GGA95GGT (16516A>T), GTC99ATC (16526G>A), TCT100TCA (16531T>A), GGA101GCC (16533G>C 16534A>C), TTG102ATG (16535T>A), GTC103ATT (16538G>A 16540C>T), TAT104TAC (16543T>C) | | | | | | | | |

*: Inserts / Deletes / Misaligned / Frameshifts

#### Analysis details

This analysis was performed with panviral2.64

### NGS Details (UN62): Bracoviriform congregatae (segment Circle 7)

#### Assembly

| | |
|---|---|
| **Coverage Length** | 112 (1 contig(s)) |
| **Depth Of Coverage** | 198.1 |
| **Number Of Reads** | 277 |
| **Reads Per Million** | 6.42 rpm (after QC) |
| **Ambiguities** | 0 |
| **Assembly Method** | de novo + reference guided assembly |
| **Consensus Caller** | Bcf Tools |

#### Coverage Map

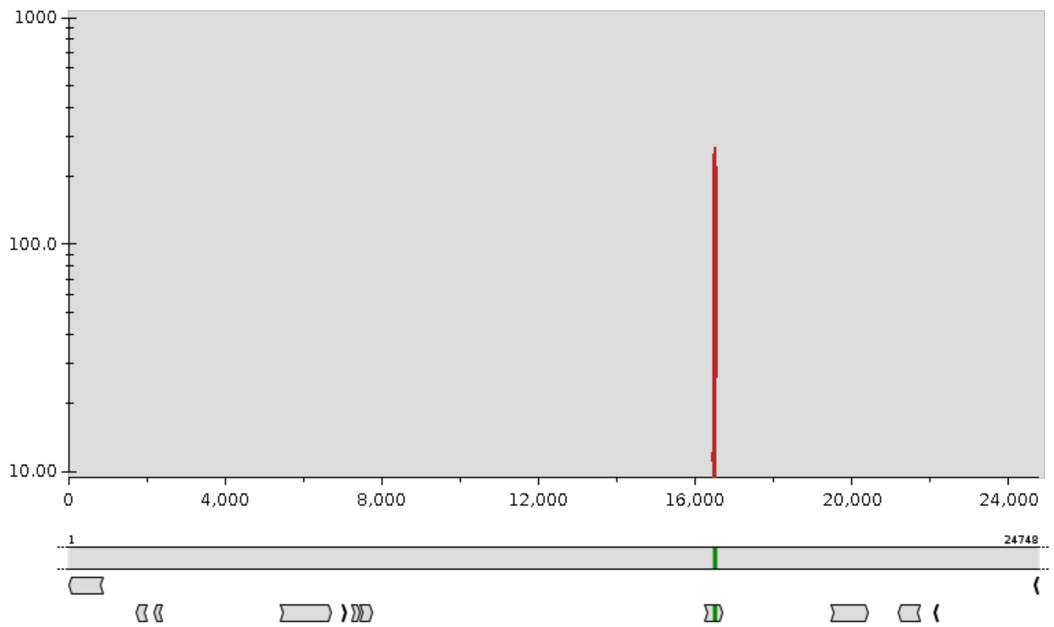

#### Assignment

| | |
|---|---|
| **Type** | Bracoviriform congregatae (Taxonomy ID: 39640) |
| **Reference Genome** | NC_006639.1 |
| **NT Identity (%)** | 78.5714 |
| **AA Identity (%)** | 86.8421 |
| **Number Of Stop Codons** | 0 |
| **Number Of CDS** | 5 |

#### Alignment

| | |
|---|---|
| **Alignment Score** | 128.0 (NT) + 221.0 (AA) = 349.0 |
| **Concordance (%)** | 75.378 |

| Alignment Method | Global, seeded, nucleotide + amino acids (AGA) |
|---|---|

#### Genome Region

Sequence starts at position 16454 and ends at position 16565 relative to NC_006639.1 reference sequence.

#### Alignment Detailed Statistics

| | Begin | End | Coverage | Score | Concordance | Matches | Identities | I/D/M/F* | Stop Codons |
|---|---|---|---|---|---|---|---|---|---|
| **NT** | **16454** | **16565** | **0.5%** | **128** | **57.1%** | **112 (100%)** | **88 (78.6%)** | **0/0** | |
| Mutations: | 16468C>T, 16474A>T, 16480C>A, 16486A>G, 16492C>T, 16493C>A, 16495T>G, 16499C>T, 16508C>A, 16510T>A, 16513T>A, 16519C>G, 16526G>A, 16528C>T, 16529T>A, 16530C>G, 16534A>T, 16535T>C, 16537G>T, 16538G>A, 16546A>G, 16551T>C, 16557A>G, 16558T>G | | | | | | | | |
| **CDS** | | | | | | | | | |
| Histone | 75 | 112 | 24.4% | 221 | 90.9% | 38 (100%) | 33 (86.8%) | 0/0/0/0 | 0 |
| Protein mutations: | V99I (16526G>A 16528C>T), V103I (16538G>A), I107T (16551T>C), D109G (16557A>G 16558T>G) | | | | | | | | |
| Codon mutations: | ATC79ATT (16468C>T), GGA81GGT (16474A>T), ACC83ACA (16480C>A), CCA85CCG (16486A>G), ATC87ATT (16492C>T), CGT88AGG (16493C>A 16495T>G), CTG90TTG (16499C>T), CGT93AGA (16508C>A 16510T>A), GGT94GGA (16513T>A), GTC96GTG (16519C>G), GTC99ATT (16526G>A 16528C>T), TCT100AGT (16529T>A 16530C>G), GGA101GGT (16534A>T), TTG102CTT (16535T>C 16537G>T), GTC103ATC (16538G>A), GAA105GAG (16546A>G), ATT107ACT (16551T>C), GAT109GGG (16557A>G 16558T>G) | | | | | | | | |
| **Proteins** | | | | | | | | | |
| hypothetical protein (YP_184795.1) | 75 | 112 | 24.4% | 221 | 90.9% | 38 (100%) | 33 (86.8%) | 0/0/0/0 | 0 |
| Protein mutations: | V99I (16526G>A 16528C>T), V103I (16538G>A), I107T (16551T>C), D109G (16557A>G 16558T>G) | | | | | | | | |
| Codon mutations: | ATC79ATT (16468C>T), GGA81GGT (16474A>T), ACC83ACA (16480C>A), CCA85CCG (16486A>G), ATC87ATT (16492C>T), CGT88AGG (16493C>A 16495T>G), CTG90TTG (16499C>T), CGT93AGA (16508C>A 16510T>A), GGT94GGA (16513T>A), GTC96GTG (16519C>G), GTC99ATT (16526G>A 16528C>T), TCT100AGT (16529T>A 16530C>G), GGA101GGT (16534A>T), TTG102CTT (16535T>C 16537G>T), GTC103ATC (16538G>A), GAA105GAG (16546A>G), ATT107ACT (16551T>C), GAT109GGG (16557A>G 16558T>G) | | | | | | | | |

*: Inserts / Deletes / Misaligned / Frameshifts

#### Analysis details

This analysis was performed with panviral2.64

### NGS Details (UN62): Bracoviriform congregatae (segment Circle 7)

#### Assembly

| | |
|---|---|
| **Coverage Length** | 180 (1 contig(s)) |
| **Depth Of Coverage** | 206.7 |
| **Number Of Reads** | 408 |
| **Reads Per Million** | 9.46 rpm (after QC) |
| **Ambiguities** | 0 |
| **Assembly Method** | de novo + reference guided assembly |
| **Consensus Caller** | Bcf Tools |

#### Coverage Map

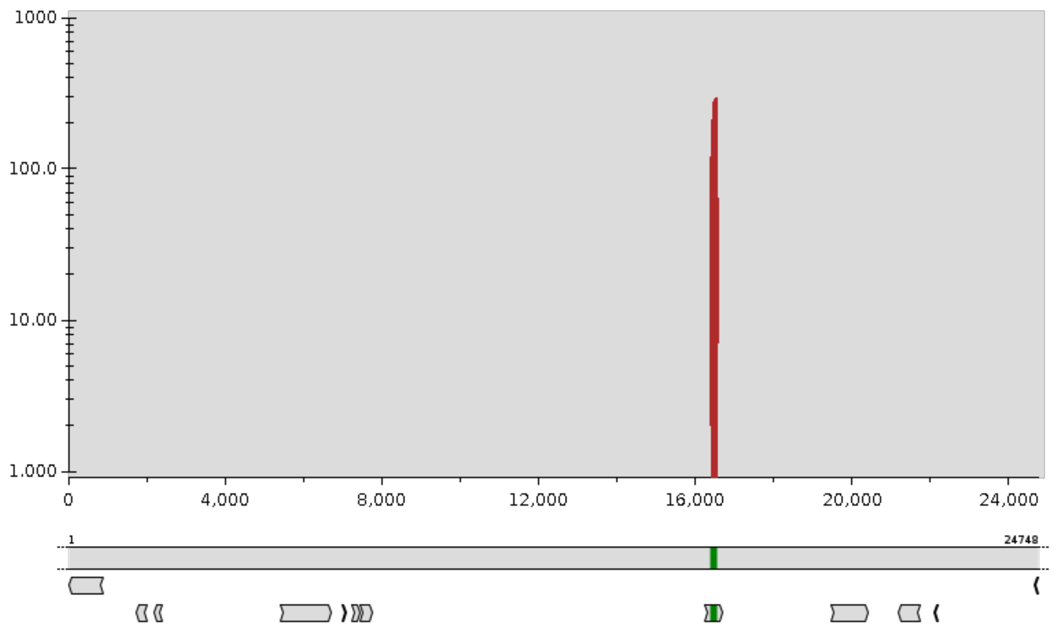

#### Assignment

| | |
|---|---|
| **Type** | Bracoviriform congregatae (Taxonomy ID: 39640) |
| **Reference Genome** | NC_006639.1 |
| **NT Identity (%)** | 78.8889 |
| **AA Identity (%)** | 90.0 |
| **Number Of Stop Codons** | 0 |
| **Number Of CDS** | 5 |

#### Alignment

| | |
|---|---|
| **Alignment Score** | 208.0 (NT) + 360.0 (AA) = 568.0 |
| **Concordance (%)** | 75.7333 |

| Alignment Method | Global, seeded, nucleotide + amino acids (AGA) |
| --- | --- |

#### Genome Region

Sequence starts at position 16394 and ends at position 16573 relative to NC_006639.1 reference sequence.

#### Alignment Detailed Statistics

| | Begin | End | Coverage | Score | Concordance | Matches | Identities | I/D/M/F* | Stop Codons |
| --- | --- | --- | --- | --- | --- | --- | --- | --- | --- |
| **NT** | **16394** | **16573** | **0.7%** | **208** | **57.8%** | **180 (100%)** | **142 (78.9%)** | **0/0** | |
| Mutations: | 16405A>G, 16411G>A, 16414A>G, 16417A>G, 16426A>G, 16429G>A, 16432C>A, 16444T>C, 16447C>T, 16449G>A, 16450A>G, 16451T>G, 16452C>T, 16456T>C, 16459A>C, 16471A>G, 16486A>T, 16492C>T, 16493C>A, 16495T>G, 16504T>G, 16508C>A, 16510T>A, 16513T>A, 16519C>G, 16526G>A, 16528C>T, 16529T>A, 16530C>G, 16534A>T, 16535T>C, 16537G>T, 16538G>A, 16546A>G, 16551T>C, 16557A>G, 16558T>G, 16562T>C | | | | | | | | |

##### CDS

| | Begin | End | Coverage | Score | Concordance | Matches | Identities | I/D/M/F* | Stop Codons |
| --- | --- | --- | --- | --- | --- | --- | --- | --- | --- |
| Histone | 55 | 114 | 38.5% | 360 | 92.3% | 60 (100%) | 54 (90.0%) | 0/0/0/0 | 0 |
| Protein mutations: | R73K (16449G>A 16450A>G), S74V (16451T>G 16452C>T), V99I (16526G>A 16528C>T), V103I (16538G>A), I107T (16551T>C), D109G (16557A>G 16558T>G) | | | | | | | | |
| Codon mutations: | AAA58AAG (16405A>G), GGG60GGA (16411G>A), AAA61AAG (16414A>G), GGA62GGG (16417A>G), AAA65AAG (16426A>G), GGG66GGA (16429G>A), GGC67GGA (16432C>A), CAT71CAC (16444T>C), CGC72CGT (16447C>T), AGA73AAG (16449G>A 16450A>G), TCT74GTT (16451T>G 16452C>T), CTT75CTC (16456T>C), CGA76CGC (16459A>C), CAA80CAG (16471A>G), CCA85CCT (16486A>T), ATC87ATT (16492C>T), CGT88AGG (16493C>A 16495T>G), GCT91GCG (16504T>G), CGT93AGA (16508C>A 16510T>A), GGT94GGA (16513T>A), GTC96GTG (16519C>G), GTC99ATT (16526G>A 16528C>T), TCT100AGT (16529T>A 16530C>G), GGA101GGT (16534A>T), TTG102CTT (16535T>C 16537G>T), GTC103ATC (16538G>A), GAA105GAG (16546A>G), ATT107ACT (16551T>C), GAT109GGG (16557A>G 16558T>G), TTG111CTG (16562T>C) | | | | | | | | |

##### Proteins

| | Begin | End | Coverage | Score | Concordance | Matches | Identities | I/D/M/F* | Stop Codons |
| --- | --- | --- | --- | --- | --- | --- | --- | --- | --- |
| hypothetical protein (YP_184795.1) | 55 | 114 | 38.5% | 360 | 92.3% | 60 (100%) | 54 (90.0%) | 0/0/0/0 | 0 |
| Protein mutations: | R73K (16449G>A 16450A>G), S74V (16451T>G 16452C>T), V99I (16526G>A 16528C>T), V103I (16538G>A), I107T (16551T>C), D109G (16557A>G 16558T>G) | | | | | | | | |
| Codon mutations: | AAA58AAG (16405A>G), GGG60GGA (16411G>A), AAA61AAG (16414A>G), GGA62GGG (16417A>G), AAA65AAG (16426A>G), GGG66GGA (16429G>A), GGC67GGA (16432C>A), CAT71CAC (16444T>C), CGC72CGT (16447C>T), AGA73AAG (16449G>A 16450A>G), TCT74GTT (16451T>G 16452C>T), CTT75CTC (16456T>C), CGA76CGC (16459A>C), CAA80CAG (16471A>G), CCA85CCT (16486A>T), ATC87ATT (16492C>T), CGT88AGG (16493C>A 16495T>G), GCT91GCG (16504T>G), CGT93AGA (16508C>A 16510T>A), GGT94GGA (16513T>A), GTC96GTG (16519C>G), GTC99ATT (16526G>A 16528C>T), TCT100AGT (16529T>A 16530C>G), GGA101GGT (16534A>T), TTG102CTT (16535T>C 16537G>T), GTC103ATC (16538G>A), GAA105GAG (16546A>G), ATT107ACT (16551T>C), GAT109GGG (16557A>G 16558T>G), TTG111CTG (16562T>C) | | | | | | | | |

*: Inserts / Deletes / Misaligned / Frameshifts

#### Analysis details

This analysis was performed with panviral2.64

### NGS Details (UN62): Cassava brown streak virus

#### Assembly

| | |
|---|---|
| **Coverage Length** | 526 (1 contig(s)) |
| **Depth Of Coverage** | 85.7 |
| **Number Of Reads** | 391 |
| **Reads Per Million** | 9.06 rpm (after QC) |
| **Ambiguities** | 0 |
| **Assembly Method** | de novo + reference guided assembly |
| **Consensus Caller** | Bcf Tools |

#### Coverage Map

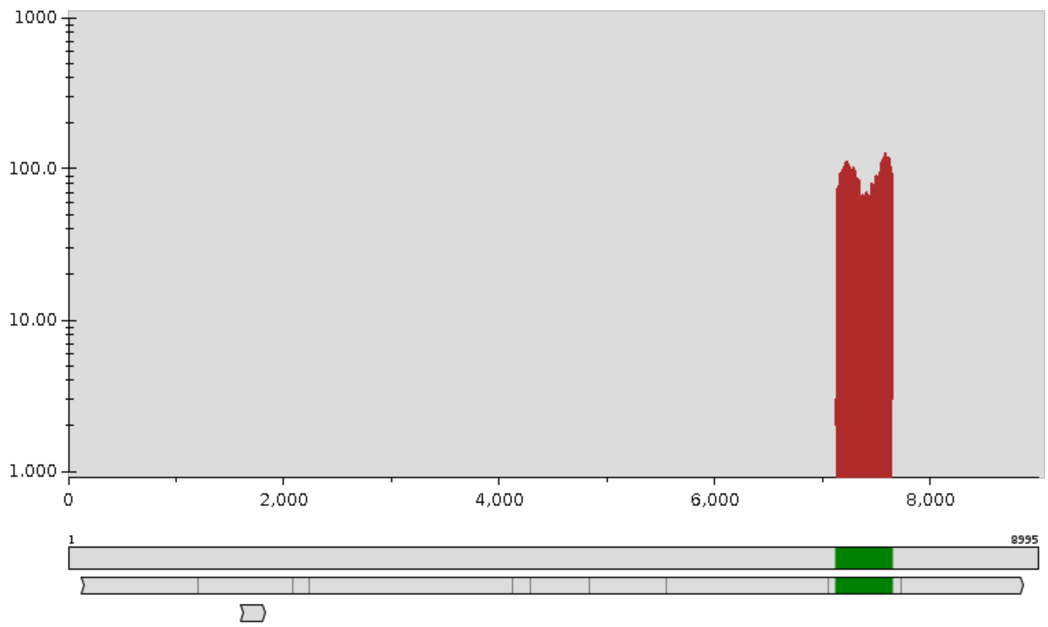

#### Assignment

| | |
|---|---|
| **Type** | Cassava brown streak virus (Taxonomy ID: 137758) |
| **Reference Genome** | NC_012698.2 |
| **NT Identity (%)** | 60.8365 |
| **AA Identity (%)** | 57.7143 |
| **Number Of Stop Codons** | 0 |
| **Number Of CDS** | 2 |

#### Alignment

| | |
|---|---|
| **Alignment Score** | 228.0 (NT) + 738.0 (AA) = 966.0 |
| **Concordance (%)** | 42.5738 |

| Alignment Method | Global, seeded, nucleotide + amino acids (AGA) |
|---|---|

#### Genome Region

Sequence starts at position 7123 and ends at position 7648 relative to NC_012698.2 reference sequence.

#### Alignment Detailed Statistics

| | Begin | End | Coverage | Score | Concordance | Matches | Identities | I/D/M/F* | Stop Codons |
|---|---|---|---|---|---|---|---|---|---|
| **NT** | **7123** | **7648** | **5.8%** | **228** | **21.7%** | **526 (100%)** | **320 (60.8%)** | **0/0** | |
| Mutations: | 7130A>G, 7133A>G, 7134A>G, 7136T>G, 7142C>T, 7145A>T, 7148A>C, 7151G>A, 7157A>T, 7158C>A, 7163A>G, 7167A>G, 7175A>T, 7177A>G, 7178A>G, 7179C>G, 7180A>C, 7184A>T, 7185T>C, 7190T>C, 7192C>A, 7193C>G, 7194A>T, 7202A>C, 7203A>T, 7205C>T, 7206A>C, 7207T>A, 7208C>G, 7211T>C, 7213G>T, 7214G>T, 7217A>G, 7218G>C, 7229A>T, 7234C>T, 7235A>T, 7241A>G, 7242A>G, 7243C>A, 7244A>G, 7245G>C, 7246T>C, 7250G>A, 7253A>T, 7256T>A, 7257A>T, 7258T>C, 7270T>C, 7271A>T, 7272C>A, 7273G>A, 7274T>A, 7275G>A, 7277A>T, 7280G>T, 7281T>G, 7283T>C, 7284G>A, 7286G>A, 7287T>G, 7288T>A, 7290A>G, 7292T>G, 7293G>A, 7294G>A, 7295A>T, 7298G>A, 7304T>G, 7307C>A, 7310A>T, 7313A>G, 7319A>T, 7320A>T, 7323T>C, 7325G>C, 7331T>C, 7332G>A, 7334T>C, 7337T>C, 7340C>T, 7343T>G, 7355A>G, 7358A>T, 7361C>T, 7364T>C, 7374A>T, 7377G>C, 7380G>A, 7381G>A, 7385T>C, 7388A>T, 7390T>A, 7391A>T, 7397A>T, 7398T>C, 7400A>T, 7401T>A, 7403T>C, 7406G>C, 7409G>A, 7410G>T, 7413G>A, 7414A>T, 7416C>G, 7418G>T, 7422C>G, 7424A>G, 7425A>G, 7432T>C, 7433G>A, 7436T>A, 7437A>T, 7438G>A, 7442T>C, 7443C>A, 7445C>G, 7448C>T, 7449G>A, 7451G>C, 7455G>T, 7457T>A, 7458T>C, 7462T>C, 7463A>T, 7464A>C, 7465A>T, 7467A>G, 7468A>G, 7469A>G, 7470G>C, 7471T>C, 7472T>A, 7473G>A, 7474G>A, 7476G>A, 7477A>C, 7481T>G, 7484T>G, 7487A>G, 7489T>C, 7490C>T, 7494A>G, 7495A>T, 7496G>T, 7499T>A, 7500G>A, 7501T>A, 7502G>A, 7503C>A, 7504T>C, 7505A>G, 7506A>C, 7507G>T, 7508A>G, 7511T>A, 7512G>A, 7517T>A, 7520A>G, 7521A>C, 7522T>C, 7523G>A, 7524C>G, 7526A>T, 7527C>A, 7529C>G, 7535A>C, 7539T>A, 7540C>A, 7541A>T, 7544T>C, 7547G>A, 7553C>T, 7556A>G, 7559T>A, 7562C>T, 7568A>T, 7570T>A, 7572A>G, 7573A>G, 7576G>A, 7577G>T, 7578A>G, 7579G>A, 7580A>C, 7581A>C, 7586A>T, 7588T>A, 7589T>C, 7599A>C, 7600T>C, 7601G>C, 7602G>A, 7603C>A, 7604C>G, 7607G>A, 7610G>A, 7616T>C, 7618T>A, 7622A>T, 7628T>C, 7631A>C, 7632T>G, 7633T>G, 7635C>A, 7636G>A | | | | | | | | |

##### CDS

| | Begin | End | Coverage | Score | Concordance | Matches | Identities | I/D/M/F* | Stop Codons |
|---|---|---|---|---|---|---|---|---|---|
| CBSV_gp1 | 2334 | 2508 | 6.0% | 738 | 60.1% | 175 (100%) | 101 (57.7%) | 0/0/0/0 | 0 |
| Protein mutations: | I2337V (7134A>G 7136T>G), Q2345K (7158C>A), K2348E (7167A>G), K2351R (7177A>G 7178A>G), Q2352A (7179C>G 7180A>C), F2354L (7185T>C), P2356Q (7192C>A 7193C>G), T2357S (7194A>T), I2360F (7203A>T 7205C>T), I2361Q (7206A>C 7207T>A 7208C>G), R2363L (7213G>T 7214G>T), V2365L (7218G>C), P2370L (7234C>T 7235A>T), T2373E (7242A>G 7243C>A 7244A>G), V2374P (7245G>C 7246T>C), E2376D (7253A>T), I2378S (7257A>T 7258T>C), V2382A (7270T>C 7271A>T), R2383K (7272C>A 7273G>A 7274T>A), V2384I (7275G>A 7277A>T), S2386A (7281T>G 7283T>C), E2387K (7284G>A 7286G>A), L2388E (7287T>G 7288T>A), I2389V (7290A>G 7292T>G), G2390N (7293G>A 7294G>A 7295A>T), S2399C (7320A>T), D2403N (7332G>A 7334T>C), N2406K (7343T>G), M2417L (7374A>T), E2418Q (7377G>C), G2419K (7380G>A 7381G>A), L2422H (7390T>A 7391A>T), Y2426N (7401T>A 7403T>C), K2427N (7406G>C), V2429L (7410G>T), E2430M (7413G>A 7414A>T), P2431A (7416C>G 7418G>T), Q2433E (7422C>G 7424A>G), N2434D (7425A>G), M2436T (7432T>C 7433G>A), S2438Y (7437A>T 7438G>A), L2440M (7443C>A 7445C>G), V2442I (7449G>A 7451G>C), A2444S (7455G>T 7457T>A), F2445L (7458T>C), V2446A (7462T>C 7463A>T), N2447L (7464A>C 7465A>T), K2448G (7467A>G 7468A>G 7469A>G), V2449P (7470G>C 7471T>C 7472T>A), G2450N (7473G>A 7474G>A), D2451T (7476G>A 7477A>C), D2452E (7481T>G), I2454M (7487A>G), I2455T (7489T>C 7490C>T), K2457V (7494A>G 7495A>T 7496G>T), V2459K (7500G>A 7501T>A 7502G>A), L2460T (7503C>A 7504T>C 7505A>G), R2461L (7506A>C 7507G>T 7508A>G), E2463K (7512G>A), M2466P (7521A>C 7522T>C 7523G>A), P2467A (7524C>G 7526A>T), S2472N (7539T>A 7540C>A 7541A>T), L2482H (7570T>A), N2483G (7572A>G 7573A>G), W2484Y (7576G>A 7577G>T), R2485D (7578A>G 7579G>A 7580A>C), K2486Q (7581A>C), F2488Y (7588T>A 7589T>C), M2492P (7599A>C 7600T>C 7601G>C), A2493K (7602G>A 7603C>A 7604C>G), M2498K (7618T>A), F2503G (7632T>G 7633T>G), R2504K (7635C>A 7636G>A) | | | | | | | | |
| Codon mutations: | GCA2335GCG (7130A>G), CCA2336CCG (7133A>G), ATT2337GTG (7134A>G 7136T>G), TTC2339TTT (7142C>T), GTA2340GTT (7145A>T), ACA2341ACC (7148A>C), GGG2342GGA (7151G>A), GCA2344GCT (7157A>T), CAA2345AAA (7158C>A), AAA2346AAG (7163A>G), AAG2348GAG (7167A>G), GTA2350GTT (7175A>T), AAA2351AGG (7177A>G 7178A>G), CAA2352GCA (7179C>G 7180A>C), ATA2353ATT (7184A>T), TTT2354CTT (7185T>C), GGT2355GGC (7190T>C), CCC2356CAG (7192C>A 7193C>G), ACT2357TCT (7194A>T), CCA2359CCC (7202A>C), ATC2360TTT (7203A>T 7205C>T), ATC2361CAG (7206A>C 7207T>A 7208C>G), TCT2362TCC (7211T>C), CGG2363CTT (7213G>T 7214G>T), AAA2364AAG (7217A>G), GTT2365CTT (7218G>C), CCA2368CCT (7229A>T), CCA2370CTT (7234C>T 7235A>T), GGA2372GGG (7241A>G), ACA2373GAG (7242A>G 7243C>A 7244A>G), GTT2374CCT (7245G>C 7246T>C), GAG2375GAA (7250G>A), GAA2376GAT (7253A>T), ATT2377ATA (7256T>A), ATC2378TCC (7257A>T 7258T>C), GTA2382GCT (7270T>C 7271A>T), CGT2383AAA (7272C>A 7273G>A 7274T>A), GTA2384ATT (7275G>A 7277A>T), GCG2385GCT (7280G>T), TCT2386GCC (7281T>G 7283T>C), GAG2387AAA (7284G>A 7286G>A), TTG2388GAG (7287T>G 7288T>A), ATT2389GTG (7290A>G 7292T>G), GGA2390AAT (7293G>A 7294G>A 7295A>T), GGG2391GGA (7298G>A), GTT2393GTG (7304T>G), CTC2394CTA (7307C>A), GTA2395GTT (7310A>T), GAA2396GAG (7313A>G), ACA2398ACT (7319A>T), AGT2399TGT (7320A>T), TTG2400CTC (7323T>C 7325G>C), TTT2402TTC (7331T>C), GAT2403AAC (7332G>A 7334T>C), GCT2404GCC (7337T>C), CTC2405CTT (7340C>T), AAT2406AAG (7343T>G), GGA2410GGG (7355A>G), CCA2411CCT (7358A>T), TAC2412TAT (7361C>T), ATT2413ATC (7364T>C), ATG2417TTG (7374A>T), GAA2418CAA (7377G>C), GGG2419AAG (7380G>A 7381G>A), ATT2420ATC (7385T>C), GGA2421GGT (7388A>T), CTA2422CAT (7390T>A 7391A>T), GGA2424GGT (7397A>T), TTA2425CTT (7398T>C 7400A>T), TAT2426AAC (7401T>A 7403T>C), AAG2427AAC (7406G>C), TTG2428TTA (7409G>A), GTG2429TTG (7410G>T), GAG2430ATG (7413G>A 7414A>T), CCG2431GCT (7416C>G 7418G>T), CAA2433GAG (7422C>G 7424A>G), AAT2434GAT (7425A>G), ATG2436ACA (7432T>C 7433G>A), GCT2437GCA (7436T>A), AGT2438TAT (7437A>T 7438G>A), GCT2439GCC (7442T>C), CTC2440ATG (7443C>A 7445C>G), TGC2441TGT (7448C>T), GTG2442ATC (7449G>A 7451G>C), GCT2444TCA (7455G>T 7457T>A), TTT2445CTT (7458T>C), GTA2446GCT (7462T>C 7463A>T), AAT2447CTT (7464A>C 7465A>T), AAA2448GGG (7467A>G 7468A>G 7469A>G), GTT2449CCA (7470G>C 7471T>C 7472T>A), GGT2450AAT (7473G>A 7474G>A), GAT2451ACT (7476G>A 7477A>C), GAT2452GAG (7481T>G), CCT2453CCG (7484T>G), ATA2454ATG (7487A>G), ATC2455ACT (7489T>C 7490C>T), AAG2457GTT (7494A>G 7495A>T 7496G>T), GGT2458GGA (7499T>A), GTG2459AAA (7500G>A 7501T>A 7502G>A), CTA2460ACG (7503C>A 7504T>C 7505A>G), AGA2461CTG (7506A>C 7507G>T 7508A>G), GGT2462GGA (7511T>A), GAG2463AAG (7512G>A), ATT2464ATA (7517T>A), GTA2465GTG (7520A>G), ATG2466CCA (7521A>C 7522T>C 7523G>A), CCA2467GCT (7524C>G 7526A>T), CGC2468AGG (7527C>A 7529C>G), CCA2470CCC (7535A>C), TCA2472AAT (7539T>A 7540C>A 7541A>T), TTT2473TTC (7544T>C), GGG2474GGA (7547G>A), GAC2476GAT (7553C>T), CCA2477CCG (7556A>G), ATT2478ATA (7559T>A), TTC2479TTT (7562C>T), CCA2481CCT (7568A>T), CTT2482CAT (7570T>A), AAC2483GGC (7572A>G 7573A>G), TGG2484TAT (7576G>A 7577G>T), AGA2485GAC (7578A>G 7579G>A 7580A>C), AAG2486CAG (7581A>C), ACA2487ACT (7586A>T), TTT2488TAC (7588T>A 7589T>C), ATG2492CCC (7599A>C 7600T>C 7601G>C), GCC2493AAG (7602G>A 7603C>A 7604C>G), GAG2494GAA (7607G>A), GAG2495GAA (7610G>A), AAT2497AAC (7616T>C), ATG2498AAG (7618T>A), ATA2499ATT (7622A>T), CAT2501CAC (7628T>C), CGA2502CGC (7631A>C), TTT2503GGT (7632T>G 7633T>G), CGA2504AAA (7635C>A 7636G>A) | | | | | | | | |

##### Proteins

| | Begin | End | Coverage | Score | Concordance | Matches | Identities | I/D/M/F* | Stop Codons |
|---|---|---|---|---|---|---|---|---|---|
| polyprotein (YP_007027011.1) | 2334 | 2508 | 6.0% | 738 | 60.1% | 175 (100%) | 101 (57.7%) | 0/0/0/0 | 0 |
| Protein mutations: | I2337V (7134A>G 7136T>G), Q2345K (7158C>A), K2348E (7167A>G), K2351R (7177A>G 7178A>G), Q2352A (7179C>G 7180A>C), F2354L (7185T>C), P2356Q (7192C>A 7193C>G), T2357S (7194A>T), I2360F (7203A>T 7205C>T), I2361Q (7206A>C 7207T>A 7208C>G), R2363L (7213G>T 7214G>T), V2365L (7218G>C), P2370L (7234C>T 7235A>T), T2373E (7242A>G 7243C>A 7244A>G), V2374P (7245G>C 7246T>C), E2376D (7253A>T), I2378S (7257A>T 7258T>C), V2382A (7270T>C 7271A>T), R2383K (7272C>A 7273G>A 7274T>A), V2384I (7275G>A 7277A>T), S2386A (7281T>G 7283T>C), E2387K (7284G>A 7286G>A), L2388E (7287T>G 7288T>A), I2389V (7290A>G 7292T>G), G2390N (7293G>A 7294G>A 7295A>T), S2399C (7320A>T), D2403N (7332G>A 7334T>C), N2406K (7343T>G), M2417L (7374A>T), E2418Q (7377G>C), G2419K (7380G>A 7381G>A), L2422H (7390T>A 7391A>T), Y2426N (7401T>A 7403T>C), K2427N (7406G>C), V2429L (7410G>T), E2430M (7413G>A 7414A>T), P2431A (7416C>G 7418G>T), Q2433E (7422C>G 7424A>G), N2434D (7425A>G), M2436T (7432T>C 7433G>A), S2438Y (7437A>T 7438G>A), L2440M (7443C>A 7445C>G), V2442I (7449G>A 7451G>C), A2444S (7455G>T 7457T>A), F2445L (7458T>C), V2446A (7462T>C 7463A>T), N2447L (7464A>C 7465A>T), K2448G (7467A>G 7468A>G 7469A>G), V2449P (7470G>C 7471T>C 7472T>A), G2450N (7473G>A 7474G>A), D2451T (7476G>A 7477A>C), D2452E (7481T>G), I2454M (7487A>G), I2455T (7489T>C 7490C>T), K2457V (7494A>G 7495A>T 7496G>T), V2459K (7500G>A 7501T>A 7502G>A), L2460T (7503C>A 7504T>C 7505A>G), R2461L (7506A>C 7507G>T 7508A>G), E2463K (7512G>A), M2466P (7521A>C 7522T>C 7523G>A), P2467A (7524C>G 7526A>T), S2472N (7539T>A 7540C>A 7541A>T), L2482H (7570T>A), N2483G (7572A>G 7573A>G), W2484Y (7576G>A 7577G>T), R2485D (7578A>G 7579G>A 7580A>C), K2486Q (7581A>C), F2488Y (7588T>A 7589T>C), M2492P (7599A>C 7600T>C 7601G>C), A2493K (7602G>A 7603C>A 7604C>G), M2498K (7618T>A), F2503G (7632T>G 7633T>G), R2504K (7635C>A 7636G>A) | | | | | | | | |

| | Begin | End | Coverage | Score | Concordance | Matches | Identities | I/D/M/F* | Stop Codons |
|---|---|---|---|---|---|---|---|---|---|
| **NT** | **7123** | **7648** | **5.8%** | **228** | **21.7%** | **526 (100%)** | **320 (60.8%)** | **0/0** | |
| Codon mutations: | GCA2335GCG (7130A>G), CCA2336CCG (7133A>G), ATT2337GTG (7134A>G 7136T>G), TTC2339TTT (7142C>T), GTA2340GTT (7145A>T), ACA2341ACC (7148A>C), GGG2342GGA (7151G>A), GCA2344GCT (7157A>T), CAA2345AAA (7158C>A), AAA2346AAG (7163A>G), AAG2348GAG (7167A>G), GTA2350GTT (7175A>T), AAA2351AGG (7177A>G 7178A>G), CAA2352GCA (7179C>G 7180A>C), ATA2353ATT (7184A>T), TTT2354CTT (7185T>C), GGT2355GGC (7190T>C), CCC2356CAG (7192C>A 7193C>G), ACT2357TCT (7194A>T), CCA2359CCC (7202A>C), ATC2360TTT (7203A>T 7205C>T), ATC2361CAG (7206A>C 7207T>A 7208C>G), TCT2362TCC (7211T>C), CGG2363CTT (7213G>T 7214G>T), AAA2364AAG (7217A>G), GTT2365CTT (7218G>C), CCA2368CCT (7229A>T), CCA2370CTT (7234C>T 7235A>T), GGA2372GGG (7241A>G), ACA2373GAG (7242A>G 7243C>A 7244A>G), GTT2374CCT (7245G>C 7246T>C), GAG2375GAA (7250G>A), GAA2376GAT (7253A>T), ATT2377ATA (7256T>A), ATC2378TCC (7257A>T 7258T>C), GTA2382GCT (7270T>C 7271A>T), CGT2383AAA (7272C>A 7273G>A 7274T>A), GTA2384ATT (7275G>A 7277A>T), GCG2385GCT (7280G>T), TCT2386GCC (7281T>G 7283T>C), GAG2387AAA (7284G>A 7286G>A), TTG2388GAG (7287T>G 7288T>A), ATT2389GTG (7290A>G 7292T>G), GGA2390AAT (7293G>A 7294G>A 7295A>T), GGG2391GGA (7298G>A), GTT2393GTG (7304T>G), CTC2394CTA (7307C>A), GTA2395GTT (7310A>T), GAA2396GAG (7313A>G), ACA2398ACT (7319A>T), AGT2399TGT (7320A>T), TTG2400CTC (7323T>C 7325G>C), TTT2402TTC (7331T>C), GAT2403AAC (7332G>A 7334T>C), GCT2404GCC (7337T>C), CTC2405CTT (7340C>T), AAT2406AAG (7343T>G), GGA2410GGG (7355A>G), CCA2411CCT (7358A>T), TAC2412TAT (7361C>T), ATT2413ATC (7364T>C), ATG2417TTG (7374A>T), GAA2418CAA (7377G>C), GGG2419AAG (7380G>A 7381G>A), ATT2420ATC (7385T>C), GGA2421GGT (7388A>T), CTA2422CAT (7390T>A 7391A>T), GGA2424GGT (7397A>T), TTA2425CTT (7398T>C 7400A>T), TAT2426AAC (7401T>A 7403T>C), AAG2427AAC (7406G>C), TTG2428TTA (7409G>A), GTG2429TTG (7410G>T), GAG2430ATG (7413G>A 7414A>T), CCG2431GCT (7416C>G 7418G>T), CAA2433GAG (7422C>G 7424A>G), AAT2434GAT (7425A>G), ATG2436ACA (7432T>C 7433G>A), GCT2437GCA (7436T>A), AGT2438TAT (7437A>T 7438G>A), GCT2439GCC (7442T>C), CTC2440ATG (7443C>A 7445C>G), TGC2441TGT (7448C>T), GTG2442ATC (7449G>A 7451G>C), GCT2444TCA (7455G>T 7457T>A), TTT2445CTT (7458T>C), GTA2446GCT (7462T>C 7463A>T), AAT2447CTT (7464A>C 7465A>T), AAA2448GGG (7467A>G 7468A>G 7469A>G), GTT2449CCA (7470G>C 7471T>C 7472T>A), GGT2450AAT (7473G>A 7474G>A), GAT2451ACT (7476G>A 7477A>C), GAT2452GAG (7481T>G), CCT2453CCG (7484T>G), ATA2454ATG (7487A>G), ATC2455ACT (7489T>C 7490C>T), AAG2457GTT (7494A>G 7495A>T 7496G>T), GGT2458GGA (7499T>A), GTG2459AAA (7500G>A 7501T>A 7502G>A), CTA2460ACG (7503C>A 7504T>C 7505A>G), AGA2461CTG (7506A>C 7507G>T 7508A>G), GGT2462GGA (7511T>A), GAG2463AAG (7512G>A), ATT2464ATA (7517T>A), GTA2465GTG (7520A>G), ATG2466CCA (7521A>C 7522T>C 7523G>A), CCA2467GCT (7524C>G 7526A>T), CGC2468AGG (7527C>A 7529C>G), CCA2470CCC (7535A>C), TCA2472AAT (7539T>A 7540C>A 7541A>T), TTT2473TTC (7544T>C), GGG2474GGA (7547G>A), GAC2476GAT (7553C>T), CCA2477CCG (7556A>G), ATT2478ATA (7559T>A), TTC2479TTT (7562C>T), CCA2481CCT (7568A>T), CTT2482CAT (7570T>A), AAC2483GGC (7572A>G 7573A>G), TGG2484TAT (7576G>A 7577G>T), AGA2485GAC (7578A>G 7579G>A 7580A>C), AAG2486CAG (7581A>C), ACA2487ACT (7586A>T), TTT2488TAC (7588T>A 7589T>C), ATG2492CCC (7599A>C 7600T>C 7601G>C), GCC2493AAG (7602G>A 7603C>A 7604C>G), GAG2494GAA (7607G>A), GAG2495GAA (7610G>A), AAT2497AAC (7616T>C), ATG2498AAG (7618T>A), ATA2499ATT (7622A>T), CAT2501CAC (7628T>C), CGA2502CGC (7631A>C), TTT2503GGT (7632T>G 7633T>G), CGA2504AAA (7635C>A 7636G>A) | | | | | | | | |
| HAM1-like protein (YP_007032446.1) | 26 | 200 | 77.4% | 738 | 60.1% | 175 (100%) | 101 (57.7%) | 0/0/0/0 | 0 |
| Protein mutations: | I29V (7134A>G 7136T>G), Q37K (7158C>A), K40E (7167A>G), K43R (7177A>G 7178A>G), Q44A (7179C>G 7180A>C), F46L (7185T>C), P48Q (7192C>A 7193C>G), T49S (7194A>T), I52F (7203A>T 7205C>T), I53Q (7206A>C 7207T>A 7208C>G), R55L (7213G>T 7214G>T), V57L (7218G>C), P62L (7234C>T 7235A>T), T65E (7242A>G 7243C>A 7244A>G), V66P (7245G>C 7246T>C), E68D (7253A>T), I70S (7257A>T 7258T>C), V74A (7270T>C 7271A>T), R75K (7272C>A 7273G>A 7274T>A), V76I (7275G>A 7277A>T), S78A (7281T>G 7283T>C), E79K (7284G>A 7286G>A), L80E (7287T>G 7288T>A), I81V (7290A>G 7292T>G), G82N (7293G>A 7294G>A 7295A>T), S91C (7320A>T), D95N (7332G>A 7334T>C), N98K (7343T>G), M109L (7374A>T), E110Q (7377G>C), G111K (7380G>A 7381G>A), L114H (7390T>A 7391A>T), Y118N (7401T>A 7403T>C), K119N (7406G>C), V121L (7410G>T), E122M (7413G>A 7414A>T), P123A (7416C>G 7418G>T), Q125E (7422C>G 7424A>G), N126D (7425A>G), M128T (7432T>C 7433G>A), S130Y (7437A>T 7438G>A), L132M (7443C>A 7445C>G), V134I (7449G>A 7451G>C), A136S (7455G>T 7457T>A), F137L (7458T>C), V138A (7462T>C 7463A>T), N139L (7464A>C 7465A>T), K140G (7467A>G 7468A>G 7469A>G), V141P (7470G>C 7471T>C 7472T>A), G142N (7473G>A 7474G>A), D143T (7476G>A 7477A>C), D144E (7481T>G), I146M (7487A>G), I147T (7489T>C 7490C>T), K149V (7494A>G 7495A>T 7496G>T), V151K (7500G>A 7501T>A 7502G>A), L152T (7503C>A 7504T>C 7505A>G), R153L (7506A>C 7507G>T 7508A>G), E155K (7512G>A), M158P (7521A>C 7522T>C 7523G>A), P159A (7524C>G 7526A>T), S164N (7539T>A 7540C>A 7541A>T), L174H (7570T>A), N175G (7572A>G 7573A>G), W176Y (7576G>A 7577G>T), R177D (7578A>G 7579G>A 7580A>C), K178Q (7581A>C), F180Y (7588T>A 7589T>C), M184P (7599A>C 7600T>C 7601G>C), A185K (7602G>A 7603C>A 7604C>G), M190K (7618T>A), F195G (7632T>G 7633T>G), R196K (7635C>A 7636G>A) | | | | | | | | |
| Codon mutations: | GCA27GCG (7130A>G), CCA28CCG (7133A>G), ATT29GTG (7134A>G 7136T>G), TTC31TTT (7142C>T), GTA32GTT (7145A>T), ACA33ACC (7148A>C), GGG34GGA (7151G>A), GCA36GCT (7157A>T), CAA37AAA (7158C>A), AAA38AAG (7163A>G), AAG40GAG (7167A>G), GTA42GTT (7175A>T), AAA43AGG (7177A>G 7178A>G), CAA44GCA (7179C>G 7180A>C), ATA45ATT (7184A>T), TTT46CTT (7185T>C), GGT47GGC (7190T>C), CCC48CAG (7192C>A 7193C>G), ACT49TCT (7194A>T), CCA51CCC (7202A>C), ATC52TTT (7203A>T 7205C>T), ATC53CAG (7206A>C 7207T>A 7208C>G), TCT54TCC (7211T>C), CGG55CTT (7213G>T 7214G>T), AAA56AAG (7217A>G), GTT57CTT (7218G>C), CCA60CCT (7229A>T), CCA62CTT (7234C>T 7235A>T), GGA64GGG (7241A>G), ACA65GAG (7242A>G 7243C>A 7244A>G), GTT66CCT (7245G>C 7246T>C), GAG67GAA (7250G>A), GAA68GAT (7253A>T), ATT69ATA (7256T>A), ATC70TCC (7257A>T 7258T>C), GTA74GCT (7270T>C 7271A>T), CGT75AAA (7272C>A 7273G>A 7274T>A), GTA76ATT (7275G>A 7277A>T), GCG77GCT (7280G>T), TCT78GCC (7281T>G 7283T>C), GAG79AAA (7284G>A 7286G>A), TTG80GAG (7287T>G 7288T>A), ATT81GTG (7290A>G 7292T>G), GGA82AAT (7293G>A 7294G>A 7295A>T), GGG83GGA (7298G>A), GTT85GTG (7304T>G), CTC86CTA (7307C>A), GTA87GTT (7310A>T), GAA88GAG (7313A>G), ACA90ACT (7319A>T), AGT91TGT (7320A>T), TTG92CTC (7323T>C 7325G>C), TTT94TTC (7331T>C), GAT95AAC (7332G>A 7334T>C), GCT96GCC (7337T>C), CTC97CTT (7340C>T), AAT98AAG (7343T>G), GGA102GGG (7355A>G), CCA103CCT (7358A>T), TAC104TAT (7361C>T), ATT105ATC (7364T>C), ATG109TTG (7374A>T), GAA110CAA (7377G>C), GGG111AAG (7380G>A 7381G>A), ATT112ATC (7385T>C), GGA113GGT (7388A>T), CTA114CAT (7390T>A 7391A>T), GGA116GGT (7397A>T), TTA117CTT (7398T>C 7400A>T), TAT118AAC (7401T>A 7403T>C), AAG119AAC (7406G>C), TTG120TTA (7409G>A), GTG121TTG (7410G>T), GAG122ATG (7413G>A 7414A>T), CCG123GCT (7416C>G 7418G>T), CAA125GAG (7422C>G 7424A>G), AAT126GAT (7425A>G), ATG128ACA (7432T>C 7433G>A), GCT129GCA (7436T>A), AGT130TAT (7437A>T 7438G>A), GCT131GCC (7442T>C), CTC132ATG (7443C>A 7445C>G), TGC133TGT (7448C>T), GTG134ATC (7449G>A 7451G>C), GCT136TCA (7455G>T 7457T>A), TTT137CTT (7458T>C), GTA138GCT (7462T>C 7463A>T), AAT139CTT (7464A>C 7465A>T), AAA140GGG (7467A>G 7468A>G 7469A>G), GTT141CCA (7470G>C 7471T>C 7472T>A), GGT142AAT (7473G>A 7474G>A), GAT143ACT (7476G>A 7477A>C), GAT144GAG (7481T>G), CCT145CCG (7484T>G), ATA146ATG (7487A>G), ATC147ACT (7489T>C 7490C>T), AAG149GTT (7494A>G 7495A>T 7496G>T), GGT150GGA (7499T>A), GTG151AAA (7500G>A 7501T>A 7502G>A), CTA152ACG (7503C>A 7504T>C 7505A>G), AGA153CTG (7506A>C 7507G>T 7508A>G), GGT154GGA (7511T>A), GAG155AAG (7512G>A), ATT156ATA (7517T>A), GTA157GTG (7520A>G), ATG158CCA (7521A>C 7522T>C 7523G>A), CCA159GCT (7524C>G 7526A>T), CGC160AGG (7527C>A 7529C>G), CCA162CCC (7535A>C), TCA164AAT (7539T>A 7540C>A 7541A>T), TTT165TTC (7544T>C), GGG166GGA (7547G>A), GAC168GAT (7553C>T), CCA169CCG (7556A>G), ATT170ATA (7559T>A), TTC171TTT (7562C>T), CCA173CCT (7568A>T), CTT174CAT (7570T>A), AAC175GGC (7572A>G 7573A>G), TGG176TAT (7576G>A 7577G>T), AGA177GAC (7578A>G 7579G>A 7580A>C), AAG178CAG (7581A>C), ACA179ACT (7586A>T), TTT180TAC (7588T>A 7589T>C), ATG184CCC (7599A>C 7600T>C 7601G>C), GCC185AAG (7602G>A 7603C>A 7604C>G), GAG186GAA (7607G>A), GAG187GAA (7610G>A), AAT189AAC (7616T>C), ATG190AAG (7618T>A), ATA191ATT (7622A>T), CAT193CAC (7628T>C), CGA194CGC (7631A>C), TTT195GGT (7632T>G 7633T>G), CGA196AAA (7635C>A 7636G>A) | | | | | | | | |

*: Inserts / Deletes / Misaligned / Frameshifts

#### Analysis details

This analysis was performed with panviral2.64

### NGS Details (UN62): Human gammaherpesvirus 8

#### Assembly

| | |
|---|---|
| **Coverage Length** | 135 (1 contig(s)) |
| **Depth Of Coverage** | 200.8 |
| **Number Of Reads** | 322 |
| **Reads Per Million** | 7.46 rpm (after QC) |
| **Ambiguities** | 0 |
| **Assembly Method** | de novo + reference guided assembly |
| **Consensus Caller** | Bcf Tools |

#### Coverage Map

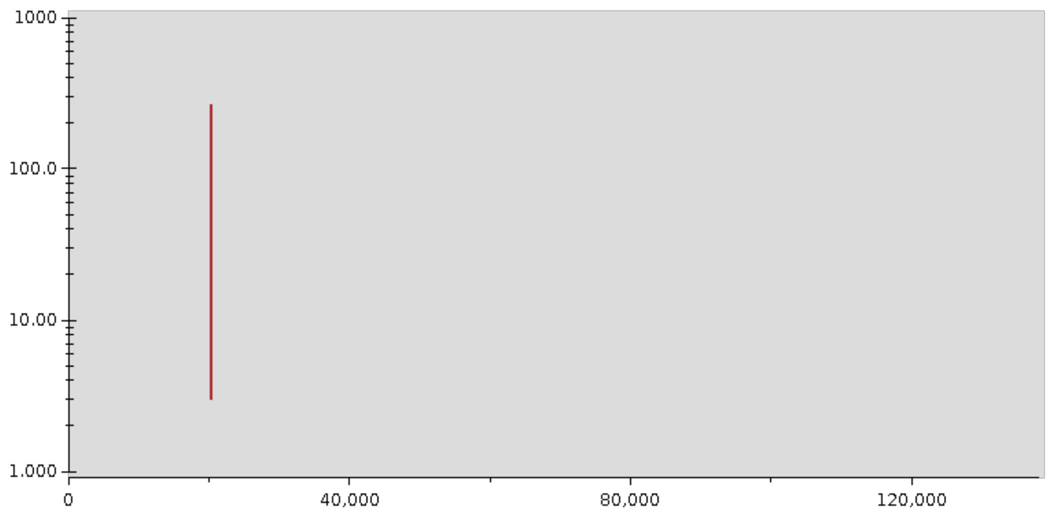

#### Assignment

| | |
|---|---|
| **Type** | Human gammaherpesvirus 8 (Taxonomy ID: 37296) |
| **Subtype** | Could not assign |
| **Reference Genome** | NC_009333.1 |
| **NT Identity (%)** | 83.7037 |
| **AA Identity (%)** | 84.4444 |
| **Number Of Stop Codons** | 0 |
| **Number Of CDS** | 86 |

#### Alignment

| | |
|---|---|
| **Alignment Score** | 182.0 (NT) + 287.0 (AA) = 469.0 |
| **Concordance (%)** | 78.2972 |
| **Alignment Method** | Local, heuristic, nucleotide (BLASTN) |

#### Genome Region

Sequence starts at position 20273 and ends at position 20407 relative to NC_009333.1 reference sequence.

#### Alignment Detailed Statistics

| | Begin | End | Coverage | Score | Concordance | Matches | Identities | I/D/M/F* | Stop Codons |
|---|---|---|---|---|---|---|---|---|---|
| **NT** | **20273** | **20407** | **0.1%** | **182** | **67.4%** | **135 (100%)** | **113 (83.7%)** | **0/0** | |

Mutations: 20293A>T, 20296A>C, 20311G>A, 20314T>G, 20315C>G, 20317C>A, 20320C>T, 20322T>G, 20331G>T, 20341A>C, 20347A>C, 20350G>A, 20352C>T, 20353A>G, 20369C>G, 20370A>C, 20371T>A, 20376A>T, 20383G>A, 20386C>T, 20389A>G, 20392C>T

*: Inserts / Deletes / Misaligned / Frameshifts

#### Analysis details

This analysis was performed with panviral2.64

### NGS Details (UN62): Hibiscus bacilliform virus GD1

#### Assembly

| | |
|---|---|
| **Coverage Length** | 332 (1 contig(s)) |
| **Depth Of Coverage** | 11.3 |
| **Number Of Reads** | 53 |
| **Reads Per Million** | 1.23 rpm (after QC) |
| **Ambiguities** | 0 |
| **Assembly Method** | de novo + reference guided assembly |
| **Consensus Caller** | Bcf Tools |

#### Coverage Map

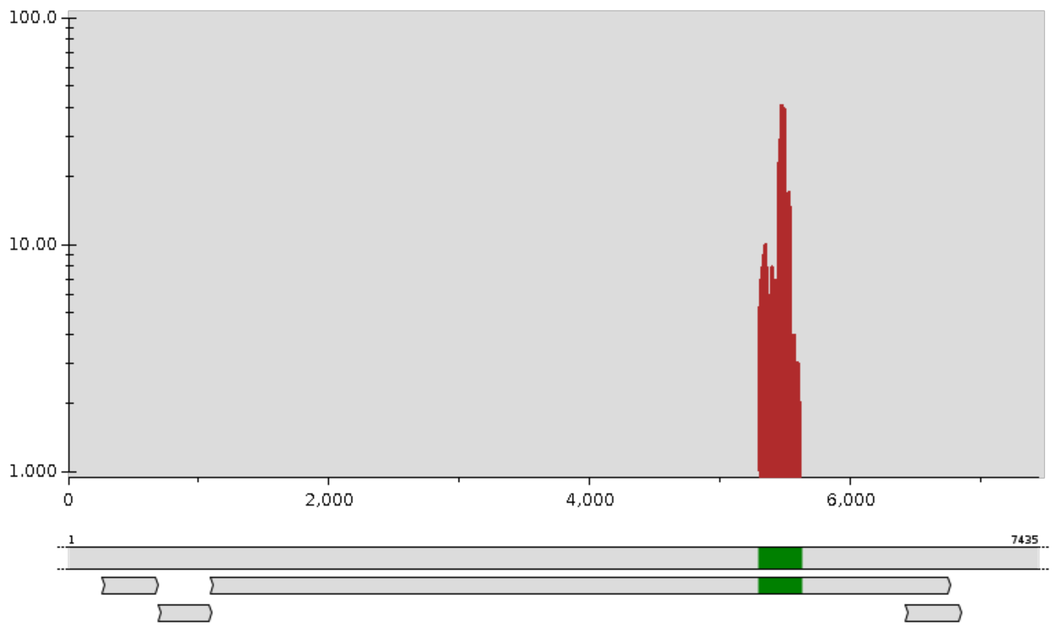

#### Assignment

| | |
|---|---|
| **Type** | Hibiscus bacilliform virus GD1 (Taxonomy ID: 1459800) |
| **Reference Genome** | NC_023485.1 |
| **NT Identity (%)** | 51.6418 |
| **AA Identity (%)** | 42.8571 |
| **Number Of Stop Codons** | 0 |
| **Number Of CDS** | 4 |

#### Alignment

| | |
|---|---|
| **Alignment Score** | 16.0 (NT) + 322.0 (AA) = 338.0 |
| **Concordance (%)** | 23.2782 |

| | |
|---|---|
| **Alignment Method** | Global, seeded, nucleotide + amino acids (AGA) |

#### Genome Region

Sequence starts at position 5295 and ends at position 5626 relative to NC_023485.1 reference sequence.

#### Alignment Detailed Statistics

| | Begin | End | Coverage | Score | Concordance | Matches | Identities | I/D/M/F* | Stop Codons |
|---|---|---|---|---|---|---|---|---|---|
| **NT** | **5295** | **5626** | **4.5%** | **16** | **2.4%** | **332 (99.1%)** | **173 (51.6%)** | **3/0** | |
| Mutations: | 5306A>G, 5307G>A, 5310C>A, 5315C>T, 5316A>C, 5317G>C, 5318T>A, 5319C>T, 5324A>T, 5325G>A, 5327C>A, 5330C>T, 5331A>G, 5333C>T, 5334A>G, 5335C>A, 5337A>C, 5339T>A, 5340A>T, 5342G>T, 5343A>G, 5345G>T, 5346A>C, 5347G>A, 5348G>A, 5349G>C, 5352G>C, 5353G>A, 5354C>A, 5355C>G, 5356A>G, 5357T>G, 5360T>A, 5362A>G, 5363G>T, 5364A>T, 5365T>A, 5366C>T, 5368A>T, 5369C>T, 5376T>G, 5384C>T, 5385A>T, 5386A>G, 5387G>T, 5388A>T, 5389G>C, 5393A>G, 5395T>A, 5399T>C, 5403G>T, 5405G>A, 5406G>T, 5407C>G, 5408T>G, 5409A>T, 5411G>A, 5412C>A, 5414C>G, 5415C>G, 5416C>A, 5417G>A, 5419A>G, 5420G>T, 5421T>G, 5422C>A, 5423T>C, 5426C>T, 5427A>C, 5428A>T, 5430T>A, 5431G>A, 5434C>T, 5435T>G, 5440T>A, 5442T>C, 5445G>A, 5447T>G, 5448C>A, 5449C>G, 5450A>G, 5451G>T, 5456A>G, 5457C>T, 5458T>A, 5459G>T, 5461A>T, 5467G>T, 5468G>T, 5471A>T, 5474T>C, 5478C>T, 5480A>C, 5486A>T, 5487C>T, 5489T>A, 5491A>C, 5492A>T, 5495C>T, 5502G>A, 5504A>T, 5507A>G, 5511C>A, 5512A>T, 5514C>A, 5515G>A, 5516A>T, 5517A>T, 5518A>T, 5519G>T, 5523G>A, 5525C>T, 5527A>G, 5528C>G, 5531C>T, 5537A>G, 5538G>C, 5539G>C, 5540_5541insTTC, 5541A>C, 5542C>T, 5546A>T, 5547G>A, 5548C>T, 5549C>G, 5552T>C, 5553A>G, 5554T>G, 5555C>G, 5557C>T, 5558T>G, 5563A>T, 5564C>T, 5568G>A, 5573C>T, 5574A>G, 5576A>G, 5577C>T, 5579C>G, 5580G>A, 5584T>A, 5585T>C, 5588C>T, 5590A>G, 5592T>A, 5593C>G, 5594T>C, 5595G>A, 5600G>A, 5603A>G, 5607A>G, 5608T>A, 5609C>A, 5613C>A, 5616A>T, 5618A>G, 5619C>A, 5622G>A, 5624C>A, 5625A>G | | | | | | | | |

##### CDS

| | Begin | End | Coverage | Score | Concordance | Matches | Identities | I/D/M/F* | Stop Codons |
|---|---|---|---|---|---|---|---|---|---|
| BU35_gp3 | 1402 | 1512 | 5.9% | 322 | 40.6% | 111 (99.1%) | 48 (42.9%) | 1/0/0/0 | 0 |
| Protein mutations: | D1406N (5307G>A), Q1407K (5310C>A), S1409P (5316A>C 5317G>C 5318T>A), G1412R (5325G>A 5327C>A), N1414D (5331A>G 5333C>T), T1415D (5334A>G 5335C>A), I1416L (5337A>C 5339T>A), M1417F (5340A>T 5342G>T), K1418D (5343A>G 5345G>T), R1419Q (5346A>C 5347G>A 5348G>A), V1420L (5349G>C), G1421Q (5352G>C 5353G>A 5354C>A), H1422G (5355C>G 5356A>G 5357T>G), K1424S (5362A>G 5363G>T), I1425Y (5364A>T 5365T>A 5366C>T), Y1426F (5368A>T 5369C>T), F1429V (5376T>G), K1432C (5385A>T 5386A>G 5387G>T), F1435Y (5395T>A), V1438L (5403G>T 5405G>A), A1439W (5406G>T 5407C>G 5408T>G), M1440L (5409A>T 5411G>A), H1441K (5412C>A 5414C>G), P1442E (5415C>G 5416C>A 5417G>A), E1443G (5419A>G 5420G>T), S1444D (5421T>G 5422C>A 5423T>C), K1446L (5427A>C 5428A>T), W1447K (5430T>A 5431G>A), T1448M (5434C>T 5435T>G), F1450Y (5440T>A), W1451R (5442T>C), V1452M (5445G>A 5447T>G), P1453R (5448C>A 5449C>G 5450A>G), D1454Y (5451G>T), L1456Y (5457C>T 5458T>A 5459G>T), Y1457F (5461A>T), W1459F (5467G>T 5468G>T), P1463S (5478C>T 5480A>C), K1467T (5491A>C 5492A>T), A1471T (5502G>A 5504A>T), Q1474M (5511C>A 5512A>T), R1475N (5514C>A 5515G>A 5516A>T), K1476F (5517A>T 5518A>T 5519G>T), D1478N (5523G>A 5525C>T), N1479R (5527A>G 5528C>G), G1483P (5538G>C 5539G>C), G1483_T1484insF (5540_5541insTTC), T1484L (5541A>C 5542C>T), E1485D (5546A>T), A1486M (5547G>A 5548C>T 5549C>G), I1488G (5553A>G 5554T>G 5555C>G), A1489V (5557C>T 5558T>G), Y1491F (5563A>T 5564C>T), D1493N (5568G>A), I1495V (5574A>G 5576A>G), V1497I (5580G>A), F1498Y (5584T>A 5585T>C), Q1500R (5590A>G), E1502K (5595G>A), I1506E (5607A>G 5608T>A 5609C>A), H1508N (5613C>A), I1509L (5616A>T 5618A>G), V1511I (5622G>A 5624C>A) | | | | | | | | |
| Codon mutations: | AAA1405AAG (5306A>G), GAT1406AAT (5307G>A), CAG1407AAG (5310C>A), TAC1408TAT (5315C>T), AGT1409CCA (5316A>C 5317G>C 5318T>A), CTG1410TTG (5319C>T), CCA1411CCT (5324A>T), GGC1412AGA (5325G>A 5327C>A), ATC1413ATT (5330C>T), AAC1414GAT (5331A>G 5333C>T), ACC1415GAC (5334A>G 5335C>A), ATT1416CTA (5337A>C 5339T>A), ATG1417TTT (5340A>T 5342G>T), AAG1418GAT (5343A>G 5345G>T), AGG1419CAA (5346A>C 5347G>A 5348G>A), GTT1420CTT (5349G>C), GGC1421CAA (5352G>C 5353G>A 5354C>A), CAT1422GGG (5355C>G 5356A>G 5357T>G), GCT1423GCA (5360T>A), AAG1424AGT (5362A>G 5363G>T), ATC1425TAT (5364A>T 5365T>A 5366C>T), TAC1426TTT (5368A>T 5369C>T), TTT1429GTT (5376T>G), CTC1431CTT (5384C>T), AAG1432TGT (5385A>T 5386A>G 5387G>T), AGT1433TCT (5388A>T 5389G>C), GGA1434GGG (5393A>G), TTT1435TAT (5395T>A), CAT1436CAC (5399T>C), GTG1438TTA (5403G>T 5405G>A), GCT1439TGG (5406G>T 5407C>G 5408T>G), ATG1440TTA (5409A>T 5411G>A), CAC1441AAG (5412C>A 5414C>G), CCG1442GAA (5415C>G 5416C>A 5417G>A), GAG1443GGT (5419A>G 5420G>T), TCT1444GAC (5421T>G 5422C>A 5423T>C), ATC1445ATT (5426C>T), AAG1446CTG (5427A>C 5428A>T), TGG1447AAG (5430T>A 5431G>A), ACT1448ATG (5434C>T 5435T>G), TTC1450TAC (5440T>A), TGG1451CGG (5442T>C), GTT1452ATG (5445G>A 5447T>G), CCA1453AGG (5448C>A 5449C>G 5450A>G), GAT1454TAT (5451G>T), GGA1455GGG (5456A>G), CTG1456TAT (5457C>T 5458T>A 5459G>T), TAT1457TTT (5461A>T), TGG1459TTT (5467G>T 5468G>T), CTA1460CTT (5471A>T), GTT1461GTC (5474T>C), CCA1463TCC (5478C>T 5480A>C), GGA1465GGT (5486A>T), CTT1466TTA (5487C>T 5489T>A), AAA1467ACT (5491A>C 5492A>T), AAC1468AAT (5495C>T), GCA1471ACT (5502G>A 5504A>T), GTA1472GTG (5507A>G), CAG1474ATG (5511C>A 5512A>T), CGA1475AAT (5514C>A 5515G>A 5516A>T), AAG1476TTT (5517A>T 5518A>T 5519G>T), GAC1478AAT (5523G>A 5525C>T), AAC1479AGG (5527A>G 5528C>G), GTC1480GTT (5531C>T), AAA1482AAG (5537A>G), GGT1483CCT (5538G>C 5539G>C), GGT1483_ACT1484insTTC (5540_5541insTTC), ACT1484CTT (5541A>C 5542C>T), GAA1485GAT (5546A>T), GCC1486ATG (5547G>A 5548C>T 5549C>G), TTT1487TTC (5552T>C), ATC1488GGG (5553A>G 5554T>G 5555C>G), GCT1489GTG (5557C>T 5558T>G), TAC1491TTT (5563A>T 5564C>T), GAT1493AAT (5568G>A), GAC1494GAT (5573C>T), ATA1495GTG (5574A>G 5576A>G), CTC1496TTG (5577C>T 5579C>G), GTA1497ATA (5580G>A), TTT1498TAC (5584T>A 5585T>C), TCC1499TCT (5588C>T), CAG1500CGG (5590A>G), TCT1501AGC (5592T>A 5593C>G 5594T>C), GAG1502AAG (5595G>A), GAG1503GAA (5600G>A), GAA1504GAG (5603A>G), ATC1506GAA (5607A>G 5608T>A 5609C>A), CAC1508AAC (5613C>A), ATA1509TTG (5616A>T 5618A>G), CGG1510AGG (5619C>A), GTC1511ATA (5622G>A 5624C>A), ATG1512GT. (5625A>G) | | | | | | | | |

##### Proteins

| | Begin | End | Coverage | Score | Concordance | Matches | Identities | I/D/M/F* | Stop Codons |
|---|---|---|---|---|---|---|---|---|---|
| polyprotein (YP_009002585.1) | 1402 | 1512 | 5.9% | 322 | 40.6% | 111 (99.1%) | 48 (42.9%) | 1/0/0/0 | 0 |
| Protein mutations: | D1406N (5307G>A), Q1407K (5310C>A), S1409P (5316A>C 5317G>C 5318T>A), G1412R (5325G>A 5327C>A), N1414D (5331A>G 5333C>T), T1415D (5334A>G 5335C>A), I1416L (5337A>C 5339T>A), M1417F (5340A>T 5342G>T), K1418D (5343A>G 5345G>T), R1419Q (5346A>C 5347G>A 5348G>A), V1420L (5349G>C), G1421Q (5352G>C 5353G>A 5354C>A), H1422G (5355C>G 5356A>G 5357T>G), K1424S (5362A>G 5363G>T), I1425Y (5364A>T 5365T>A 5366C>T), Y1426F (5368A>T 5369C>T), F1429V (5376T>G), K1432C (5385A>T 5386A>G 5387G>T), F1435Y (5395T>A), V1438L (5403G>T 5405G>A), A1439W (5406G>T 5407C>G 5408T>G), M1440L (5409A>T 5411G>A), H1441K (5412C>A 5414C>G), P1442E (5415C>G 5416C>A 5417G>A), E1443G (5419A>G 5420G>T), S1444D (5421T>G 5422C>A 5423T>C), K1446L (5427A>C 5428A>T), W1447K (5430T>A 5431G>A), T1448M (5434C>T 5435T>G), F1450Y (5440T>A), W1451R (5442T>C), V1452M (5445G>A 5447T>G), P1453R (5448C>A 5449C>G 5450A>G), D1454Y (5451G>T), L1456Y (5457C>T 5458T>A 5459G>T), Y1457F (5461A>T), W1459F (5467G>T 5468G>T), P1463S (5478C>T 5480A>C), K1467T (5491A>C 5492A>T), A1471T (5502G>A 5504A>T), Q1474M (5511C>A 5512A>T), R1475N (5514C>A 5515G>A 5516A>T), K1476F (5517A>T 5518A>T 5519G>T), D1478N (5523G>A 5525C>T), N1479R (5527A>G 5528C>G), G1483P (5538G>C 5539G>C), G1483_T1484insF (5540_5541insTTC), T1484L (5541A>C 5542C>T), E1485D (5546A>T), A1486M (5547G>A 5548C>T 5549C>G), I1488G (5553A>G 5554T>G 5555C>G), A1489V (5557C>T 5558T>G), Y1491F (5563A>T 5564C>T), D1493N (5568G>A), I1495V (5574A>G 5576A>G), V1497I (5580G>A), F1498Y (5584T>A 5585T>C), Q1500R (5590A>G), E1502K (5595G>A), I1506E (5607A>G 5608T>A 5609C>A), H1508N (5613C>A), I1509L (5616A>T 5618A>G), V1511I (5622G>A 5624C>A) | | | | | | | | |
| Codon mutations: | AAA1405AAG (5306A>G), GAT1406AAT (5307G>A), CAG1407AAG (5310C>A), TAC1408TAT (5315C>T), AGT1409CCA (5316A>C 5317G>C 5318T>A), CTG1410TTG (5319C>T), CCA1411CCT (5324A>T), GGC1412AGA (5325G>A 5327C>A), ATC1413ATT (5330C>T), AAC1414GAT (5331A>G 5333C>T), ACC1415GAC (5334A>G 5335C>A), ATT1416CTA (5337A>C 5339T>A), ATG1417TTT (5340A>T 5342G>T), AAG1418GAT (5343A>G 5345G>T), AGG1419CAA (5346A>C 5347G>A 5348G>A), GTT1420CTT (5349G>C), GGC1421CAA (5352G>C 5353G>A 5354C>A), CAT1422GGG (5355C>G 5356A>G 5357T>G), GCT1423GCA (5360T>A), AAG1424AGT (5362A>G 5363G>T), ATC1425TAT (5364A>T 5365T>A 5366C>T), TAC1426TTT (5368A>T 5369C>T), TTT1429GTT (5376T>G), CTC1431CTT (5384C>T), AAG1432TGT (5385A>T 5386A>G 5387G>T), AGT1433TCT (5388A>T 5389G>C), GGA1434GGG (5393A>G), TTT1435TAT (5395T>A), CAT1436CAC (5399T>C), GTG1438TTA (5403G>T 5405G>A), GCT1439TGG (5406G>T 5407C>G 5408T>G), ATG1440TTA (5409A>T 5411G>A), CAC1441AAG (5412C>A 5414C>G), CCG1442GAA (5415C>G 5416C>A 5417G>A), GAG1443GGT (5419A>G 5420G>T), TCT1444GAC (5421T>G 5422C>A 5423T>C), ATC1445ATT (5426C>T), AAG1446CTG (5427A>C 5428A>T), TGG1447AAG (5430T>A 5431G>A), ACT1448ATG (5434C>T 5435T>G), TTC1450TAC (5440T>A), TGG1451CGG (5442T>C), GTT1452ATG (5445G>A 5447T>G), CCA1453AGG (5448C>A 5449C>G 5450A>G), GAT1454TAT (5451G>T), GGA1455GGG (5456A>G), CTG1456TAT (5457C>T 5458T>A 5459G>T), TAT1457TTT (5461A>T), TGG1459TTT (5467G>T 5468G>T), CTA1460CTT (5471A>T), GTT1461GTC (5474T>C), CCA1463TCC (5478C>T 5480A>C), GGA1465GGT (5486A>T), CTT1466TTA (5487C>T 5489T>A), AAA1467ACT (5491A>C 5492A>T), AAC1468AAT (5495C>T), GCA1471ACT (5502G>A 5504A>T), GTA1472GTG (5507A>G), CAG1474ATG (5511C>A 5512A>T), CGA1475AAT (5514C>A 5515G>A 5516A>T), AAG1476TTT (5517A>T 5518A>T 5519G>T), GAC1478AAT (5523G>A 5525C>T), AAC1479AGG (5527A>G 5528C>G), GTC1480GTT (5531C>T), AAA1482AAG (5537A>G), GGT1483CCT (5538G>C 5539G>C), GGT1483_ACT1484insTTC (5540_5541insTTC), ACT1484CTT (5541A>C 5542C>T), GAA1485GAT (5546A>T), GCC1486ATG (5547G>A 5548C>T 5549C>G), TTT1487TTC (5552T>C), ATC1488GGG (5553A>G 5554T>G 5555C>G), GCT1489GTG (5557C>T 5558T>G), TAC1491TTT (5563A>T 5564C>T), GAT1493AAT (5568G>A), GAC1494GAT (5573C>T), ATA1495GTG (5574A>G 5576A>G), CTC1496TTG (5577C>T 5579C>G), GTA1497ATA (5580G>A), TTT1498TAC (5584T>A 5585T>C), TCC1499TCT (5588C>T), CAG1500CGG (5590A>G), TCT1501AGC (5592T>A 5593C>G 5594T>C), GAG1502AAG (5595G>A), GAG1503GAA (5600G>A), GAA1504GAG (5603A>G), ATC1506GAA (5607A>G 5608T>A 5609C>A), CAC1508AAC (5613C>A), ATA1509TTG (5616A>T 5618A>G), CGG1510AGG (5619C>A), GTC1511ATA (5622G>A 5624C>A), ATG1512GT. (5625A>G) | | | | | | | | |

*: Inserts / Deletes / Misaligned / Frameshifts

#### Analysis details

This analysis was performed with panviral2.64

### NGS Details (UN62): Caulimovirus venafragariae

#### Assembly

| | |
|---|---|
| **Coverage Length** | 876 (3 contig(s)) |
| **Depth Of Coverage** | 6.8 |
| **Number Of Reads** | 51 |
| **Reads Per Million** | 1.18 rpm (after QC) |
| **Ambiguities** | 0 |
| **Assembly Method** | de novo + reference guided assembly |
| **Consensus Caller** | Bcf Tools |

#### Coverage Map

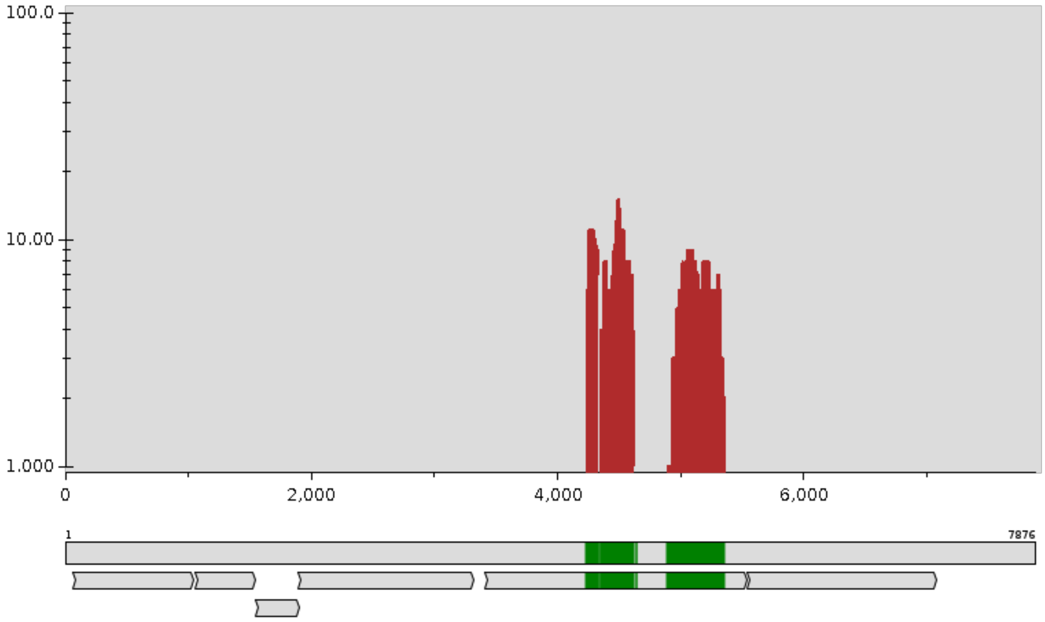

#### Assignment

| | |
|---|---|
| **Type** | Caulimovirus venafragariae (Taxonomy ID: 3048344) |
| **Reference Genome** | NC_001725.1 |
| **NT Identity (%)** | 57.6389 |
| **AA Identity (%)** | 51.7361 |
| **Number Of Stop Codons** | 0 |
| **Number Of CDS** | 6 |

#### Alignment

| | |
|---|---|
| **Alignment Score** | 243.0 (NT) + 1130.0 (AA) = 1373.0 |
| **Concordance (%)** | 37.2389 |

| Alignment Method | Global, seeded, nucleotide + amino acids (AGA) |
|---|---|

#### Genome Region

Sequence starts at position 4224 and ends at position 5362 relative to NC_001725.1 reference sequence.

#### Alignment Detailed Statistics

| | Begin | End | Coverage | Score | Concordance | Matches | Identities | I/D/M/F* | Stop Codons |
|---|---|---|---|---|---|---|---|---|---|
| **NT** | **4224** | **5362** | **11.1%** | **243** | **14.2%** | **864 (98.6%)** | **498 (56.8%)** | **0/12** | |

Mutations:

4224A>G, 4227C>T, 4228A>C, 4230T>C, 4232A>G, 4235A>T, 4236A>C, 4242T>A, 4244A>G, 4245G>A, 4246T>G, 4247G>T, 4254A>G, 4259A>T, 4260G>A, 4263T>C, 4264T>A, 4265C>A, 4267C>G, 4268C>A, 4269T>A, 4275C>T, 4276C>A, 4277G>A, 4279G>A, 4280A>C, 4281A>T, 4284G>A, 4292C>T, 4299C>T, 4300G>C, 4306C>T, 4309C>T, 4311C>A, 4314G>T, 4315C>A, 4317T>G, 4318G>A, 4319G>A, 4320T>A, 4321A>T, 4323C>A, 4326C>A, 4330C>A, 4331C>G, 4332T>A, 4335C>A, 4337A>C, 4338G>A, 4343C>G, 4347T>C, 4348T>A, 4349C>G, 4350C>A, 4354C>G, 4359C>A, 4371G>A, 4374T>C, 4378G>A, 4379C>G, 4383G>A, 4386C>A, 4387A>G, 4388A>T, 4390C>A, 4395C>A, 4413T>A, 4416C>T, 4424A>G, 4425G>A, 4426T>C, 4428A>T, 4434C>T, 4435C>A, 4440A>T, 4442A>G, 4443G>A, 4444G>A, 4445G>T, 4446A>G, 4451G>C, 4452C>T, 4455C>T, 4456C>A, 4457T>A, 4459C>T, 4461T>A, 4464T>A, 4465A>G, 4470G>A, 4471G>A, 4473G>T, 4474C>G, 4479G>A, 4480C>A, 4482T>A, 4483C>A, 4485A>T, 4491C>A, 4492G>C, 4493G>A, 4497T>A, 4500G>A, 4503C>A, 4508A>T, 4509C>T, 4510T>A, 4511C>G, 4513T>A, 4514C>A, 4515C>A, 4521C>T, 4530T>A, 4535T>A, 4542G>A, 4546C>A, 4547G>A, 4548C>A, 4549C>A, 4551T>G, 4552G>C, 4553C>A, 4554T>C, 4557A>T, 4560A>T, 4562C>G, 4563A>T, 4566T>A, 4567C>G, 4569G>A, 4570C>T, 4571T>G, 4572A>G, 4575C>A, 4576G>A, 4578T>A, 4581C>T, 4583G>C, 4584C>T, 4590C>A, 4591C>G, 4599C>T, 4601A>T, 4602C>T, 4605A>G, 4613T>C, 4614C>A, 4618C>T, 4620T>A, 4633C>A, 4635A>T, 4884G>A, 4886G>C, 4893C>A, 4894G>T, 4895G>C, 4897C>A, 4898T>A, 4900C>A, 4902C>G, 4903T>A, 4905A>C, 4906G>T, 4907T>G, 4910C>A, 4911T>A, 4921C>A, 4926T>A, 4928G>A, 4930G>A, 4931A>C, 4932T>A, 4934G>A, 4935G>A, 4936A>C, 4938T>A, 4940C>A, 4941C>A, 4942T>A, 4944A>C, 4950A>G, 4954T>C, 4956A>T, 4959C>A, 4960C>T, 4961T>G, 4962C>T, 4963C>T, 4971T>C, 4972A>G, 4973T>C, 4974C>A, 4975T>A, 4976C>G, 4977C>T, 4979C>A, 4980T>A, 4982A>T, 4984T>A, 4986T>A, 4990A>G, 4992A>C, 4993A>T, 5001C>A, 5002C>A, 5003T>A, 5004T>G, 5007G>A, 5008T>A, 5009C>A, 5010A>T, 5012C>T, 5019G>A, 5020G>A, 5021T>G, 5022T>G, 5023A>C, 5024A>T, 5025A>T, 5026C>A, 5029A>C, 5030A>G, 5031G>A, 5035G>T, 5036A>C, 5037G>C, 5039T>A, 5040A>C, 5041A>C, 5042C>G, 5043T>A, 5044T>A, 5045G>T, 5047T>G, 5048C>G, 5049T>A, 5055T>A, 5058G>T, 5059A>G, 5062G>C, 5064C>T, 5067G>T, 5068G>A, 5071A>T, 5072C>G, 5076C>T, 5077A>C, 5078G>A, 5081A>G, 5082A>C, 5083A>C, 5088G>A, 5090G>A, 5091T>A, 5092C>G, 5093T>A, 5095G>T, 5096T>G, 5097A>T, 5098A>T, 5099A>C, 5100A>C, 5102C>A, 5103T>A, 5110G>A, 5112C>G, 5115T>A, 5116T>A, 5117A>G, 5118C>A, 5119A>C, 5120A>T, 5124T>G, 5125T>G, 5126C>A, 5133G>A, 5136T>C, 5139G>C, 5140C>T, 5141C>T, 5142T>A, 5143A>G, 5145C>T, 5146A>T, 5148C>A, 5149G>C, 5152T>A, 5153G>C, 5154C>A, 5160A>T, 5167G>T, 5182A>T, 5184C>A, 5185C>T, 5187C>A, 5188A>C, 5191G>A, 5193A>T, 5194A>G, 5196A>C, 5200C>A, 5201C>A, 5202A>T, 5206_5217delGGTAAAGAGGTA, 5220A>T, 5223C>T, 5230G>A, 5232T>A, 5233T>A, 5234C>G, 5235A>T, 5238A>T, 5241C>A, 5247A>T, 5248C>G, 5249C>A, 5251G>A, 5253T>C, 5256G>A, 5258A>C, 5259G>T, 5262T>A, 5266C>T, 5267A>C, 5268T>A, 5270G>C, 5271T>A, 5280G>A, 5284A>T, 5286C>A, 5287C>T, 5289T>G, 5290T>G, 5295T>A, 5296A>G, 5298T>A, 5300A>G, 5301G>A, 5303C>G, 5304A>C, 5307C>A, 5308A>C, 5309A>G, 5311G>A, 5312C>A, 5313T>A, 5317A>T, 5318G>C, 5319A>T, 5324A>T, 5326A>C, 5328C>A, 5329C>T, 5331T>A, 5334T>A, 5335T>A, 5337T>G, 5338A>C, 5343T>C, 5344C>A, 5346T>C, 5347G>A, 5349A>C, 5352G>A, 5353A>T, 5355T>G, 5358T>C

##### CDS

| | Begin | End | Coverage | Score | Concordance | Matches | Identities | I/D/M/F* | Stop Codons |
|---|---|---|---|---|---|---|---|---|---|
| ORF_V | 274 | 653 | 41.2% | 1130 | 55.6% | 288 (98.6%) | 149 (51.0%) | 0/4/1/0 | 0 |

Protein mutations:

T275P (4228A>C 4230T>C), N276S (4232A>G), K277I (4235A>T 4236A>C), K280R (4244A>G 4245G>A), C281V (4246T>G 4247G>T), Q285L (4259A>T 4260G>A), S287K (4264T>A 4265C>A), P288E (4267C>G 4268C>A 4269T>A), R291K (4276C>A 4277G>A), E292T (4279G>A 4280A>C 4281A>T), T296I (4292C>T), E299Q (4300G>C), K303N (4314G>T), L304M (4315C>A 4317T>G), G305K (4318G>A 4319G>A 4320T>A), I306L (4321A>T 4323C>A), P309R (4330C>A 4331C>G 4332T>A), P313R (4343C>G), S315R (4348T>A 4349C>G 4350C>A), P317A (4354C>G), A325S (4378G>A 4379C>G), K328V (4387A>G 4388A>T), K340R (4424A>G 4425G>A), H344N (4435C>A), K346R (4442A>G 4443G>A), G347M (4444G>A 4445G>T 4446A>G), G349A (4451G>C 4452C>T), L351K (4456C>A 4457T>A), N354D (4465A>G), E356N (4471G>A 4473G>T), Q357E (4474C>G), L359I (4480C>A 4482T>A), Q360N (4483C>A 4485A>T), G363Q (4492G>C 4493G>A), Y368F (4508A>T 4509C>T), S370K (4513T>A 4514C>A 4515C>A), F377Y (4535T>A), R381K (4546C>A 4547G>A 4548C>A), L382M (4549C>A 4551T>G), A383H (4552G>C 4553C>A 4554T>C), E385D (4560A>T), T386S (4562C>G 4563A>T), Q388E (4567C>G 4569G>A), L389W (4570C>T 4571T>G 4572A>G), A391T (4576G>A 4578T>A), S393T (4583G>C 4584C>T), Q396E (4591C>G), Y399F (4601A>T 4602C>T), V403A (4613T>C 4614C>A), P405S (4618C>T 4620T>A), S494T (4886G>C), G497S (4894G>T 4895G>C), L498K (4897C>A 4898T>A), H499K (4900C>A 4902C>G), L500I (4903T>A 4905A>C), V501C (4906G>T 4907T>G), A502E (4910C>A 4911T>A), Q506K (4921C>A), S508N (4928G>A), D509T (4930G>A 4931A>C 4932T>A), R510K (4934G>A 4935G>A), N511Q (4936A>C 4938T>A), A512E (4940C>A 4941C>A), L513I (4942T>A 4944A>C), L519C (4960C>T 4961T>G 4962C>T), I523A (4972A>G 4973T>C 4974C>A), A525E (4979C>A 4980T>A), Y526F (4982A>T), F527I (4984T>A 4986T>A), K529D (4990A>G 4992A>C), I530L (4993A>T), N532K (5001C>A), L533K (5002C>A 5003T>A 5004T>G), S535N (5008T>A 5009C>A 5010A>T), P536L (5012C>T), V539R (5020G>A 5021T>G 5022T>G), K540L (5023A>C 5024A>T 5025A>T), L541I (5026C>A), K542R (5029A>C 5030A>G 5031G>A), E544S (5035G>T 5036A>C 5037G>C), I545N (5039T>A 5040A>C), T546R (5041A>C 5042C>G 5043T>A), W547M (5044T>A 5045G>T), S548G (5047T>G 5048C>G 5049T>A), E551D (5058G>T), K552E (5059A>G), D553H (5062G>C 5064C>T), E555K (5068G>A), T556C (5071A>T 5072C>G), R558Q (5077A>C 5078G>A), K559S (5081A>G 5082A>C), I560L (5083A>C), S562K (5090G>A 5091T>A), L563E (5092C>G 5093T>A), V564C (5095G>T 5096T>G 5097A>T), K565S (5098A>T 5099A>C 5100A>C), T566K (5102C>A 5103T>A), D569K (5110G>A 5112C>G), Y571R (5116T>A 5117A>G 5118C>A), N572L (5119A>C 5120A>T), S574E (5125T>G 5126C>A), K578N (5139G>C), P579L (5140C>T 5141C>T 5142T>A), I580V (5143A>G 5145C>T), I581L (5146A>T 5148C>A), E582Q (5149G>C), C583T (5152T>A 5153G>C 5154C>A), D588Y (5167G>T), I593L (5182A>T 5184C>A), K595Q (5188A>C), A596T (5191G>A 5193A>T), K597D (5194A>G 5196A>C), P599N (5200C>A 5201C>A 5202A>T), G601_V604del (5206_5217delGGTAAAGAGGTA), A609T (5230G>A 5232T>A), K614N (5247A>T), P615E (5248C>G 5249C>A), A616T (5251G>A 5253T>C), K618T (5258A>C 5259G>T), N619K (5262T>A), H621S (5266C>T 5267A>C 5268T>A), S622T (5270G>C 5271T>A), I627L (5284A>T 5286C>A), S629A (5290T>G), I631V (5296A>G 5298T>A), K632R (5300A>G 5301G>A), A633G (5303C>G 5304A>C), K635R (5308A>C 5309A>G), A636K (5311G>A 5312C>A 5313T>A), R638S (5317A>T 5318G>C 5319A>T), Y640F (5324A>T), I641L (5326A>C 5328C>A), Y644K (5335T>A 5337T>G), K645Q (5338A>C), L647I (5344C>A 5346T>C), V648I (5347G>A 5349A>C), T650S (5353A>T 5355T>G)

Codon mutations:

TTA273..G (4224A>G), GAC274GAT (4227C>T), ACT275CCC (4228A>C 4230T>C), AAC276AGC (4232A>G), AAA277ATC (4235A>T 4236A>C), ATT279ATA (4242T>A), AAG280AGA (4244A>G 4245G>A), TGT281GTT (4246T>G 4247G>T), CCA283CCG (4254A>G), CAG285CTA (4259A>T 4260G>A), TAT286TAC (4263T>C), TCA287AAA (4264T>A 4265C>A), CCT288GAA (4267C>G 4268C>A 4269T>A), GAC290GAT (4275C>T), CGA291AAA (4276C>A 4277G>A), GAA292ACT (4279G>A 4280A>C 4281A>T), GAG293GAA (4284G>A), ACT296ATT (4292C>T), ATC298ATT (4299C>T), GAA299CAA (4300G>C), CTA301TTA (4306C>T), CTC302TTA (4309C>T 4311C>A), AAG303AAT (4314G>T), CTT304ATG (4315C>A 4317T>G), GGT305AAA (4318G>A 4319G>A 4320T>A), ATC306TTA (4321A>T 4323C>A), ATC307ATA (4326C>A), CCT309AGA (4330C>A 4331C>G 4332T>A), AGC310..A (4335C>A), AAG311.CA (4337A>C 4338G>A), CCA313CGA (4343C>G), CAT314CAC (4347T>C), TCC315AGA (4348T>A 4349C>G 4350C>A), CCA317GCA (4354C>G), GCC318GCA (4359C>A), AGG322AGA (4371G>A), AAT323AAC (4374T>C), GCC325AGC (4378G>A 4379C>G), GAG326GAA (4383G>A), ATC327ATA (4386C>A), AAA328GTA (4387A>G 4388A>T), CGA329AGA (4390C>A), GGC330GGA (4395C>A), ATT336ATA (4413T>A), AAC337AAT (4416C>T), AAG340AGA (4424A>G 4425G>A), TTA341CTT (4426T>C 4428A>T), GAC343GAT (4434C>T), CAT344AAT (4435C>A), ACA345ACT (4440A>T), AAG346AGA (4442A>G 4443G>A), GGA347ATG (4444G>A 4445G>T 4446A>G), GGC349GCT (4451G>C 4452C>T), TAC350TAT (4455C>T), CTA351AAA (4456C>A 4457T>A), CTT352TTA (4459C>T 4461T>A), CCT353CCA (4464T>A), AAC354GAC (4465A>G), AAG355AAA (4470G>A), GAG356AAT (4471G>A 4473G>T), CAA357GAA (4474C>G), CTG358CTA (4479G>A), CTT359ATA (4480C>A 4482T>A), CAA360AAT (4483C>A 4485A>T), ATC362ATA (4491C>A), GGA363CAA (4492G>C 4493G>A), GGT364GGA (4497T>A), AAG365AAA (4500G>A), ACC366ACA (4503C>A), TAC368TTT (4508A>T 4509C>T), TCT369AGT (4510T>A 4511C>G), TCC370AAA (4513T>A 4514C>A 4515C>A), GAC372GAT (4521C>T), TCT375TCA (4530T>A), TTT377TAT (4535T>A), CAG379CAA (4542G>A), CGC381AAA (4546C>A 4547G>A 4548C>A), CTT382ATG (4549C>A 4551T>G), GCT383CAC (4552G>C 4553C>A 4554T>C), CCA384CCT (4557A>T), GAA385GAT (4560A>T), ACA386AGT (4562C>G 4563A>T), ATT387ATA (4566T>A), CAG388GAA (4567C>G 4569G>A), CTA389TGG (4570C>T 4571T>G 4572A>G), ACC390ACA (4575C>A), GCT391ACA (4576G>A 4578T>A), TTC392TTT (4581C>T), AGC393ACT (4583G>C 4584C>T), CCC395CCA (4590C>A), CAA396GAA (4591C>G), CAC398CAT (4599C>T), TAC399TTT (4601A>T 4602C>T), GAA400GAG (4605A>G), GTC403GCA (4613T>C 4614C>A), CCT405TCA (4618C>T 4620T>A), CAA410A.T (4633C>A 4635A>T), CAG493.AA (4884G>A), AGT494ACT (4886G>C), ATC496ATA (4893C>A), GGA497TCA (4894G>T 4895G>C), CTA498AAA (4897C>A 4898T>A), CAC499AAG (4900C>A 4902C>G), TTA500ATC (4903T>A 4905A>C), GTT501TGT (4906G>T 4907T>G), GCT502GAA (4910C>A 4911T>A), CAA506AAA (4921C>A), CTT507CTA (4926T>A), AGT508AAT (4928G>A), GAT509ACA (4930G>A 4931A>C 4932T>A), AGG510AAA (4934G>A 4935G>A), AAT511CAA (4936A>C 4938T>A), GCC512GAA (4940C>A 4941C>A), TTA513ATC (4942T>A 4944A>C), AGA515AGG (4950A>G), TTA517CTT (4954T>C 4956A>T), GGC518GGA (4959C>A), CTC519TGT (4960C>T 4961T>G 4962C>T), CTA520TTA (4963C>T), TAT522TAC (4971T>C), ATC523GCA (4972A>G 4973T>C 4974C>A), TCC524AGT (4975T>A 4976C>G 4977C>T), GCT525GAA (4979C>A 4980T>A), TAT526TTT (4982A>T), TTT527ATA (4984T>A 4986T>A), AAA529GAC (4990A>G 4992A>C), ATA530TTA (4993A>T), AAC532AAA (5001C>A), CTT533AAG (5002C>A 5003T>A 5004T>G), AGG534AGA (5007G>A), TCA535AAT (5008T>A 5009C>A 5010A>T), CCT536CTT (5012C>T), CAG538CAA (5019G>A), GTT539AGG (5020G>A 5021T>G 5022T>G), AAA540CTT (5023A>C 5024A>T 5025A>T), CTT541ATT (5026C>A), AAG542CGA (5029A>C 5030A>G 5031G>A), GAG544TCC (5035G>T 5036A>C 5037G>C), ATA545AAC (5039T>A 5040A>C), ACT546CGA (5041A>C 5042C>G 5043T>A), TGG547ATG (5044T>A 5045G>T), TCT548GGA (5047T>G 5048C>G 5049T>A), ACT550ACA (5055T>A), GAG551GAT (5058G>T), AAA552GAA (5059A>G), GAC553CAT (5062G>C 5064C>T), ACG554ACT (5067G>T), GAA555AAA (5068G>A), ACT556TGT (5071A>T 5072C>G), GTC557GTT (5076C>T), AGA558CAA (5077A>C 5078G>A), AAA559AGC (5081A>G 5082A>C), ATA560CTA (5083A>C), AAG561AAA (5088G>A), AGT562AAA (5090G>A 5091T>A), CTA563GAA (5092C>G 5093T>A), GTA564TGT (5095G>T 5096T>G 5097A>T), AAA565TCC (5098A>T 5099A>C 5100A>C), ACT566AAA (5102C>A 5103T>A), GAC569AAG (5110G>A 5112C>G), CTT570CTA (5115T>A), TAC571AGA (5116T>A 5117A>G 5118C>A), AAT572CTT (5119A>C 5120A>T), CCT573CCG (5124T>G), TCA574GAA (5125T>G 5126C>A), GAG576GAA (5133G>A), GAT577GAC (5136T>C), AAG578AAC (5139G>C), CCT579TTA (5140C>T 5141C>T 5142T>A), ATC580GTT (5143A>G 5145C>T), ATC581TTA (5146A>T 5148C>A), GAA582CAA (5149G>C), TGC583ACA (5152T>A 5153G>C 5154C>A), GCA585GCT (5160A>T), GAC588TAC (5167G>T), ATC593TTA (5182A>T 5184C>A), CTC594TTA (5185C>T 5187C>A), AAA595CAA (5188A>C), GCA596ACT (5191G>A 5193A>T), AAA597GAC (5194A>G 5196A>C), CCA599AAT (5200C>A 5201C>A 5202A>T), GGT601_GTA604del (5206_5217delGGTAAAGAGGTA), ATA605ATT (5220A>T), TGC606TGT (5223C>T), GCT609ACA (5230G>A 5232T>A), TCA610AGT (5233T>A 5234C>G 5235A>T), GGA611GGT (5238A>T), ACC612ACA (5241C>A), AAA614AAT (5247A>T), CCA615GAA (5248C>G 5249C>A), GCT616ACC (5251G>A 5253T>C), GAG617GAA (5256G>A), AAG618ACT (5258A>C 5259G>T), AAT619AAA (5262T>A), CAT621TCA (5266C>T 5267A>C 5268T>A), AGT622ACA (5270G>C 5271T>A), AAG625AAA (5280G>A), ATC627TTA (5284A>T 5286C>A), CTT628TTG (5287C>T 5289T>G), TCC629GCC (5290T>G), ATT630ATA (5295T>A), ATT631GTA (5296A>G 5298T>A), AAG632AGA (5300A>G 5301G>A), GCA633GGC (5303C>G 5304A>C), ATC634ATA (5307C>A), AAA635CGA (5308A>C 5309A>G), GCT636AAA (5311G>A 5312C>A 5313T>A), AGA638TCT (5317A>T 5318G>C 5319A>T), TAT640TTT (5324A>T), ATC641CTA (5326A>C 5328C>A), CTT642TTA (5329C>T 5331T>A), CCT643CCA (5334T>A), TAT644AAG (5335T>A 5337T>G), AAA645CAA (5338A>C), TTT646TTC (5343T>C), CTT647ATC (5344C>A 5346T>C), GTA648ATC (5347G>A 5349A>C), AGG649AGA (5352G>A), ACT650TCG (5353A>T 5355T>G), GAT651GAC (5358T>C)

##### Proteins

| | Begin | End | Coverage | Score | Concordance | Matches | Identities | I/D/M/F* | Stop Codons |
|---|---|---|---|---|---|---|---|---|---|
| hypothetical protein (NP_043933.1) | 274 | 653 | 41.2% | 1130 | 55.6% | 288 (98.6%) | 149 (51.0%) | 0/4/1/0 | 0 |

| | Begin | End | Coverage | Score | Concordance | Matches | Identities | I/D/M/F* | Stop Codons |
|---|---|---|---|---|---|---|---|---|---|
| **NT** | **4224** | **5362** | **11.1%** | **243** | **14.2%** | **864 (98.6%)** | **498 (56.8%)** | **0/12** | |

| | |
|---|---|
| Protein mutations: | T275P (4228A>C 4230T>C), N276S (4232A>G), K277I (4235A>T 4236A>C), K280R (4244A>G 4245G>A), C281V (4246T>G 4247G>T), Q285L (4259A>T 4260G>A), S287K (4264T>A 4265C>A), P288E (4267C>G 4268C>A 4269T>A), R291K (4276C>A 4277G>A), E292T (4279G>A 4280A>C 4281A>T), T296I (4292C>T), E299Q (4300G>C), K303N (4314G>T), L304M (4315C>A 4317T>G), G305K (4318G>A 4319G>A 4320T>A), I306L (4321A>T 4323C>A), P309R (4330C>A 4331C>G 4332T>A), P313R (4343C>G), S315R (4348T>A 4349C>G 4350C>A), P317A (4354C>G), A325S (4378G>A 4379C>G), K328V (4387A>G 4388A>T), K340R (4424A>G 4425G>A), H344N (4435C>A), K346R (4442A>G 4443G>A), G347M (4444G>A 4445G>T 4446A>G), G349A (4451G>C 4452C>T), L351K (4456C>A 4457T>A), N354D (4465A>G), E356N (4471G>A 4473G>T), Q357E (4474C>G), L359I (4480C>A 4482T>A), Q360N (4483C>A 4485A>T), G363Q (4492G>C 4493G>A), Y368F (4508A>T 4509C>T), S370K (4513T>A 4514C>A 4515C>A), F377Y (4535T>A), R381K (4546C>A 4547G>A 4548C>A), L382M (4549C>A 4551T>G), A383H (4552G>C 4553C>A 4554T>C), E385D (4560A>T), T386S (4562C>G 4563A>T), Q388E (4567C>G 4569G>A), L389W (4570C>T 4571T>G 4572A>G), A391T (4576G>A 4578T>A), S393T (4583G>C 4584C>T), Q396E (4591C>G), Y399F (4601A>T 4602C>T), V403A (4613T>C 4614C>A), P405S (4618C>T 4620T>A), S494T (4886G>C), G497S (4894G>T 4895G>C), L498K (4897C>A 4898T>A), H499K (4900C>A 4902C>G), L500I (4903T>A 4905A>C), V501C (4906G>T 4907T>G), A502E (4910C>A 4911T>A), Q506K (4921C>A), S508N (4928G>A), D509T (4930G>A 4931A>C 4932T>A), R510K (4934G>A 4935G>A), N511Q (4936A>C 4938T>A), A512E (4940C>A 4941C>A), L513I (4942T>A 4944A>C), L519C (4960C>T 4961T>G 4962C>T), I523A (4972A>G 4973T>C 4974C>A), A525E (4979C>A 4980T>A), Y526F (4982A>T), F527I (4984T>A 4986T>A), K529D (4990A>G 4992A>C), I530L (4993A>T), N532K (5001C>A), L533K (5002C>A 5003T>A 5004T>G), S535N (5008T>A 5009C>A 5010A>T), P536L (5012C>T), V539R (5020G>A 5021T>G 5022T>G), K540L (5023A>C 5024A>T 5025A>T), L541I (5026C>A), K542R (5029A>C 5030A>G 5031G>A), E544S (5035G>T 5036A>C 5037G>C), I545N (5039T>A 5040A>C), T546R (5041A>C 5042C>G 5043T>A), W547M (5044T>A 5045G>T), S548G (5047T>G 5048C>G 5049T>A), E551D (5058G>T), K552E (5059A>G), D553H (5062G>C 5064C>T), E555K (5068G>A), T556C (5071A>T 5072C>G), R558Q (5077A>C 5078G>A), K559S (5081A>G 5082A>C), I560L (5083A>C), S562K (5090G>A 5091T>A), L563E (5092C>G 5093T>A), V564C (5095G>T 5096T>G 5097A>T), K565S (5098A>T 5099A>C 5100A>C), T566K (5102C>A 5103T>A), D569K (5110G>A 5112C>G), Y571R (5116T>A 5117A>G 5118C>A), N572L (5119A>C 5120A>T), S574E (5125T>G 5126C>A), K578N (5139G>C), P579L (5140C>T 5141C>T 5142T>A), I580V (5143A>G 5145C>T), I581L (5146A>T 5148C>A), E582Q (5149G>C), C583T (5152T>A 5153G>C 5154C>A), D588Y (5167G>T), I593L (5182A>T 5184C>A), K595Q (5188A>C), A596T (5191G>A 5193A>T), K597D (5194A>G 5196A>C), P599N (5200C>A 5201C>A 5202A>T), G601_V604del (5206_5217delGGTAAAGAGGTA), A609T (5230G>A 5232T>A), K614N (5247A>T), P615E (5248C>G 5249C>A), A616T (5251G>A 5253T>C), K618T (5258A>C 5259G>T), N619K (5262T>A), H621S (5266C>T 5267A>C 5268T>A), S622T (5270G>C 5271T>A), I627L (5284A>T 5286C>A), S629A (5290T>G), I631V (5296A>G 5298T>A), K632R (5300A>G 5301G>A), A633G (5303C>G 5304A>C), K635R (5308A>C 5309A>G), A636K (5311G>A 5312C>A 5313T>A), R638S (5317A>T 5318G>C 5319A>T), Y640F (5324A>T), I641L (5326A>C 5328C>A), Y644K (5335T>A 5337T>G), K645Q (5338A>C), L647I (5344C>A 5346T>C), V648I (5347G>A 5349A>C), T650S (5353A>T 5355T>G) |
| Codon mutations: | TTA273..G (4224A>G), GAC274GAT (4227C>T), ACT275CCC (4228A>C 4230T>C), AAC276AGC (4232A>G), AAA277ATC (4235A>T 4236A>C), ATT279ATA (4242T>A), AAG280AGA (4244A>G 4245G>A), TGT281GTT (4246T>G 4247G>T), CCA283CCG (4254A>G), CAG285CTA (4259A>T 4260G>A), TAT286TAC (4263T>C), TCA287AAA (4264T>A 4265C>A), CCT288GAA (4267C>G 4268C>A 4269T>A), GAC290GAT (4275C>T), CGA291AAA (4276C>A 4277G>A), GAA292ACT (4279G>A 4280A>C 4281A>T), GAG293GAA (4284G>A), ACT296ATT (4292C>T), ATC298ATT (4299C>T), GAA299CAA (4300G>C), CTA301TTA (4306C>T), CTC302TTA (4309C>T 4311C>A), AAG303AAT (4314G>T), CTT304ATG (4315C>A 4317T>G), GGT305AAA (4318G>A 4319G>A 4320T>A), ATC306TTA (4321A>T 4323C>A), ATC307ATA (4326C>A), CCT309AGA (4330C>A 4331C>G 4332T>A), AGC310..A (4335C>A), AAG311.CA (4337A>C 4338G>A), CCA313CGA (4343C>G), CAT314CAC (4347T>C), TCC315AGA (4348T>A 4349C>G 4350C>A), CCA317GCA (4354C>G), GCC318GCA (4359C>A), AGG322AGA (4371G>A), AAT323AAC (4374T>C), GCC325AGC (4378G>A 4379C>G), GAG326GAA (4383G>A), ATC327ATA (4386C>A), AAA328GTA (4387A>G 4388A>T), CGA329AGA (4390C>A), GGC330GGA (4395C>A), ATT336ATA (4413T>A), AAC337AAT (4416C>T), AAG340AGA (4424A>G 4425G>A), TTA341CTT (4426T>C 4428A>T), GAC343GAT (4434C>T), CAT344AAT (4435C>A), ACA345ACT (4440A>T), AAG346AGA (4442A>G 4443G>A), GGA347ATG (4444G>A 4445G>T 4446A>G), GGC349GCT (4451G>C 4452C>T), TAC350TAT (4455C>T), CTA351AAA (4456C>A 4457T>A), CTT352TTA (4459C>T 4461T>A), CCT353CCA (4464T>A), AAC354GAC (4465A>G), AAG355AAA (4470G>A), GAG356AAT (4471G>A 4473G>T), CAA357GAA (4474C>G), CTG358CTA (4479G>A), CTT359ATA (4480C>A 4482T>A), CAA360AAT (4483C>A 4485A>T), ATC362ATA (4491C>A), GGA363CAA (4492G>C 4493G>A), GGT364GGA (4497T>A), AAG365AAA (4500G>A), ACC366ACA (4503C>A), TAC368TTT (4508A>T 4509C>T), TCT369AGT (4510T>A 4511C>G), TCC370AAA (4513T>A 4514C>A 4515C>A), GAC372GAT (4521C>T), TCT375TCA (4530T>A), TTT377TAT (4535T>A), CAG379CAA (4542G>A), CGC381AAA (4546C>A 4547G>A 4548C>A), CTT382ATG (4549C>A 4551T>G), GCT383CAC (4552G>C 4553C>A 4554T>C), CCA384CCT (4557A>T), GAA385GAT (4560A>T), ACA386AGT (4562C>G 4563A>T), ATT387ATA (4566T>A), CAG388GAA (4567C>G 4569G>A), CTA389TGG (4570C>T 4571T>G 4572A>G), ACC390ACA (4575C>A), GCT391ACA (4576G>A 4578T>A), TTC392TTT (4581C>T), AGC393ACT (4583G>C 4584C>T), CCC395CCA (4590C>A), CAA396GAA (4591C>G), CAC398CAT (4599C>T), TAC399TTT (4601A>T 4602C>T), GAA400GAG (4605A>G), GTC403GCA (4613T>C 4614C>A), CCT405TCA (4618C>T 4620T>A), CAA410A.T (4633C>A 4635A>T), CAG493.AA (4884G>A), AGT494ACT (4886G>C), ATC496ATA (4893C>A), GGA497TCA (4894G>T 4895G>C), CTA498AAA (4897C>A 4898T>A), CAC499AAG (4900C>A 4902C>G), TTA500ATC (4903T>A 4905A>C), GTT501TGT (4906G>T 4907T>G), GCT502GAA (4910C>A 4911T>A), CAA506AAA (4921C>A), CTT507CTA (4926T>A), AGT508AAT (4928G>A), GAT509ACA (4930G>A 4931A>C 4932T>A), AGG510AAA (4934G>A 4935G>A), AAT511CAA (4936A>C 4938T>A), GCC512GAA (4940C>A 4941C>A), TTA513ATC (4942T>A 4944A>C), AGA515AGG (4950A>G), TTA517CTT (4954T>C 4956A>T), GGC518GGA (4959C>A), CTC519TGT (4960C>T 4961T>G 4962C>T), CTA520TTA (4963C>T), TAT522TAC (4971T>C), ATC523GCA (4972A>G 4973T>C 4974C>A), TCC524AGT (4975T>A 4976C>G 4977C>T), GCT525GAA (4979C>A 4980T>A), TAT526TTT (4982A>T), TTT527ATA (4984T>A 4986T>A), AAA529GAC (4990A>G 4992A>C), ATA530TTA (4993A>T), AAC532AAA (5001C>A), CTT533AAG (5002C>A 5003T>A 5004T>G), AGG534AGA (5007G>A), TCA535AAT (5008T>A 5009C>A 5010A>T), CCT536CTT (5012C>T), CAG538CAA (5019G>A), GTT539AGG (5020G>A 5021T>G 5022T>G), AAA540CTT (5023A>C 5024A>T 5025A>T), CTT541ATT (5026C>A), AAG542CGA (5029A>C 5030A>G 5031G>A), GAG544TCC (5035G>T 5036A>C 5037G>C), ATA545AAC (5039T>A 5040A>C), ACT546CGA (5041A>C 5042C>G 5043T>A), TGG547ATG (5044T>A 5045G>T), TCT548GGA (5047T>G 5048C>G 5049T>A), ACT550ACA (5055T>A), GAG551GAT (5058G>T), AAA552GAA (5059A>G), GAC553CAT (5062G>C 5064C>T), ACG554ACT (5067G>T), GAA555AAA (5068G>A), ACT556TGT (5071A>T 5072C>G), GTC557GTT (5076C>T), AGA558CAA (5077A>C 5078G>A), AAA559AGC (5081A>G 5082A>C), ATA560CTA (5083A>C), AAG561AAA (5088G>A), AGT562AAA (5090G>A 5091T>A), CTA563GAA (5092C>G 5093T>A), GTA564TGT (5095G>T 5096T>G 5097A>T), AAA565TCC (5098A>T 5099A>C 5100A>C), ACT566AAA (5102C>A 5103T>A), GAC569AAG (5110G>A 5112C>G), CTT570CTA (5115T>A), TAC571AGA (5116T>A 5117A>G 5118C>A), AAT572CTT (5119A>C 5120A>T), CCT573CCG (5124T>G), TCA574GAA (5125T>G 5126C>A), GAG576GAA (5133G>A), GAT577GAC (5136T>C), AAG578AAC (5139G>C), CCT579TTA (5140C>T 5141C>T 5142T>A), ATC580GTT (5143A>G 5145C>T), ATC581TTA (5146A>T 5148C>A), GAA582CAA (5149G>C), TGC583ACA (5152T>A 5153G>C 5154C>A), GCA585GCT (5160A>T), GAC588TAC (5167G>T), ATC593TTA (5182A>T 5184C>A), CTC594TTA (5185C>T 5187C>A), AAA595CAA (5188A>C), GCA596ACT (5191G>A 5193A>T), AAA597GAC (5194A>G 5196A>C), CCA599AAT (5200C>A 5201C>A 5202A>T), GGT601_GTA604del (5206_5217delGGTAAAGAGGTA), ATA605ATT (5220A>T), TGC606TGT (5223C>T), GCT609ACA (5230G>A 5232T>A), TCA610AGT (5233T>A 5234C>G 5235A>T), GGA611GGT (5238A>T), ACC612ACA (5241C>A), AAA614AAT (5247A>T), CCA615GAA (5248C>G 5249C>A), GCT616ACC (5251G>A 5253T>C), GAG617GAA (5256G>A), AAG618ACT (5258A>C 5259G>T), AAT619AAA (5262T>A), CAT621TCA (5266C>T 5267A>C 5268T>A), AGT622ACA (5270G>C 5271T>A), AAG625AAA (5280G>A), ATC627TTA (5284A>T 5286C>A), CTT628TTG (5287C>T 5289T>G), TCC629GCC (5290T>G), ATT630ATA (5295T>A), ATT631GTA (5296A>G 5298T>A), AAG632AGA (5300A>G 5301G>A), GCA633GGC (5303C>G 5304A>C), ATC634ATA (5307C>A), AAA635CGA (5308A>C 5309A>G), GCT636AAA (5311G>A 5312C>A 5313T>A), AGA638TCT (5317A>T 5318G>C 5319A>T), TAT640TTT (5324A>T), ATC641CTA (5326A>C 5328C>A), CTT642TTA (5329C>T 5331T>A), CCT643CCA (5334T>A), TAT644AAG (5335T>A 5337T>G), AAA645CAA (5338A>C), TTT646TTC (5343T>C), CTT647ATC (5344C>A 5346T>C), GTA648ATC (5347G>A 5349A>C), AGG649AGA (5352G>A), ACT650TCG (5353A>T 5355T>G), GAT651GAC (5358T>C) |

*: Inserts / Deletes / Misaligned / Frameshifts

#### Analysis details

This analysis was performed with panviral2.64

### NGS Details (UN62): Badnavirus occultipomeae

#### Assembly

| | |
|---|---|
| **Coverage Length** | 429 (2 contig(s)) |
| **Depth Of Coverage** | 8.1 |
| **Number Of Reads** | 46 |
| **Reads Per Million** | 1.07 rpm (after QC) |
| **Ambiguities** | 0 |
| **Assembly Method** | de novo + reference guided assembly |
| **Consensus Caller** | Bcf Tools |

#### Coverage Map

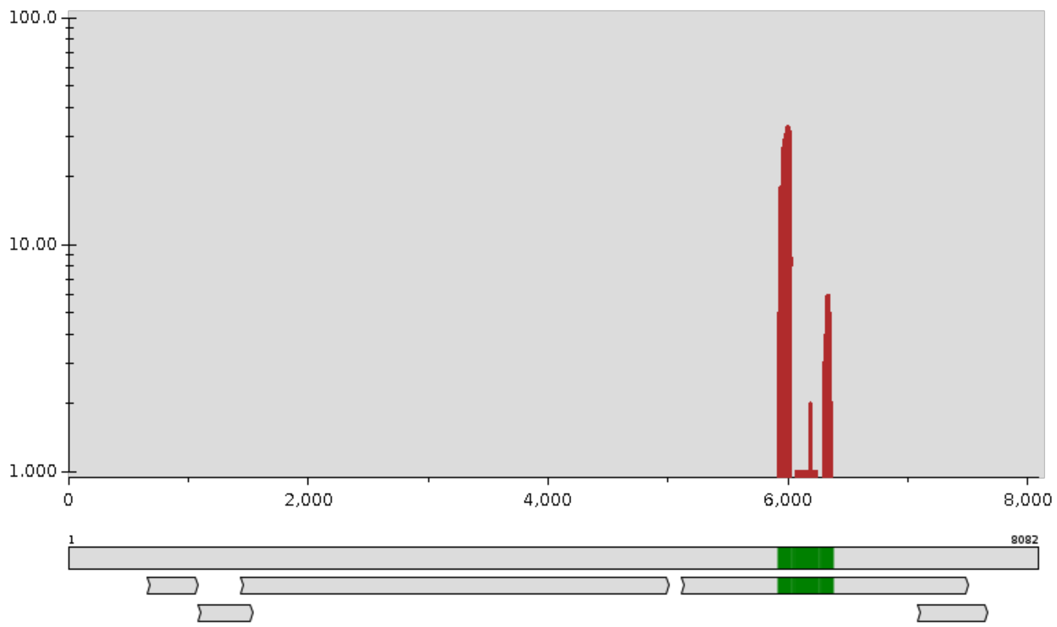

#### Assignment

| | |
|---|---|
| **Type** | Badnavirus occultipomeae (Taxonomy ID: 3048353) |
| **Reference Genome** | NC_015655.1 |
| **NT Identity (%)** | 49.537 |
| **AA Identity (%)** | 38.3562 |
| **Number Of Stop Codons** | 0 |
| **Number Of CDS** | 5 |

#### Alignment

| | |
|---|---|
| **Alignment Score** | -14.0 (NT) + 253.0 (AA) = 239.0 |
| **Concordance (%)** | 14.3371 |

| **Alignment Method** | Global, seeded, nucleotide + amino acids (AGA) |
|---|---|

#### Genome Region

Sequence starts at position 5911 and ends at position 6377 relative to NC_015655.1 reference sequence.

#### Alignment Detailed Statistics

| | Begin | End | Coverage | Score | Concordance | Matches | Identities | I/D/M/F* | Stop Codons |
|---|---|---|---|---|---|---|---|---|---|
| **NT** | **5911** | **6377** | **5.3%** | **-14** | **-1.6%** | **429 (99.3%)** | **214 (49.5%)** | **3/0** | |
| Mutations: | 5911G>A, 5915T>A, 5916T>G, 5917A>G, 5918G>A, 5919A>T, 5921G>A, 5922C>G, 5924A>G, 5925A>C, 5926C>A, 5927A>T, 5929A>T, 5935G>T, 5936T>G, 5938T>A, 5939A>T, 5940C>T, 5941A>G, 5948A>G, 5949G>T, 5951G>A, 5952C>A, 5959G>A, 5961C>G, 5962A>G, 5963A>T, 5964T>C, 5967A>T, 5968G>A, 5969A>T, 5973A>G, 5974G>A, 5977C>A, 5982T>C, 5983T>C, 5985C>G, 5986C>T, 5988A>G, 5991G>T, 5992G>A, 5997C>T, 5998A>G, 6000C>T, 6001A>G, 6002C>A, 6004A>T, 6006T>G, 6007G>T, 6009C>T, 6010A>G, 6011G>A, 6013A>C, 6014G>A, 6016A>C, 6018A>T, 6019T>C, 6020C>A, 6021T>A, 6025A>G, 6026A>C, 6027A>G, 6029A>C, 6030G>T, 6031A>G, 6033A>G, 6036C>T, 6039A>T, 6042A>G, 6043T>A, 6051C>G, 6053A>G, 6055G>T, 6057A>G, 6060C>T, 6062T>A, 6069G>A, 6070A>T, 6072A>G, 6075A>G, 6076A>G, 6078G>T, 6079G>A, 6080A>G, 6084A>G, 6085A>T, 6086A>C, 6087G>T, 6088T>G, 6089C>A, 6090C>T, 6092A>T, 6093A>C, 6096A>G, 6097T>A, 6098G>A, 6109T>C, 6116C>G, 6117A>G, 6118G>T, 6120A>T, 6123G>T, 6125T>A, 6135T>C, 6136G>C, 6137A>T, 6138A>G, 6141C>T, 6145C>T, 6153A>G, 6154C>T, 6156G>A, 6158T>C, 6159G>T, 6165A>C, 6168A>T, 6173A>C, 6178C>A, 6179A>T, 6181A>G, 6182G>A, 6183A>C, 6184A>T, 6185A>T, 6190G>A, 6192T>C, 6193A>C, 6194A>G, 6195T>G, 6197C>T, 6198A>G, 6201T>C, 6203G>A, 6205G>T, 6206G>C, 6207A>T, 6207_6208insTAT, 6208A>C, 6209C>T, 6210A>T, 6213T>C, 6214G>C, 6215C>A, 6216A>G, 6220A>G, 6222C>G, 6223G>A, 6224C>T, 6225A>T, 6228A>G, 6230A>T, 6231T>C, 6240C>T, 6243A>C, 6251T>A, 6255T>C, 6256G>C, 6260A>G, 6262G>A, 6264A>G, 6270G>A, 6276G>A, 6279T>A, 6282T>C, 6283C>T, 6285C>A, 6286T>A, 6287T>G, 6288A>G, 6289A>G, 6290A>T, 6292T>G, 6295G>C, 6296C>T, 6300A>G, 6302T>C, 6304G>T, 6310A>G, 6311G>A, 6312A>T, 6316G>C, 6317G>A, 6318G>A, 6322A>T, 6323T>G, 6325C>G, 6326T>C, 6327A>T, 6329G>A, 6330C>G, 6331C>T, 6332C>T, 6333A>C, 6334A>T, 6339A>G, 6340A>T, 6341T>A, 6342G>T, 6343A>G, 6345A>G, 6346A>T, 6349G>T, 6351A>G, 6352G>C, 6354C>A, 6355A>G, 6358G>T, 6359A>C, 6363G>A, 6364G>T, 6365A>C, 6366T>C, 6369C>T, 6375A>C, 6376A>C, 6377T>A | | | | | | | | |

##### CDS

| | Begin | End | Coverage | Score | Concordance | Matches | Identities | I/D/M/F* | Stop Codons |
|---|---|---|---|---|---|---|---|---|---|
| SPBVa_gp4 | 266 | 421 | 18.2% | 253 | 24.1% | 145 (99.3%) | 56 (38.4%) | 1/0/6/0 | 0 |
| Protein mutations: | E266K (5911G>A), I267K (5915T>A 5916T>G), R268D (5917A>G 5918G>A 5919A>T), G269E (5921G>A 5922C>G), K270S (5924A>G 5925A>C), Q271I (5926C>A 5927A>T), R272W (5929A>T), V274C (5935G>T 5936T>G), Y275I (5938T>A 5939A>T 5940C>T), N276D (5941A>G), K278S (5948A>G 5949G>T), R279Q (5951G>A 5952C>A), D282K (5959G>A 5961C>G), N283V (5962A>G 5963A>T 5964T>C), E285I (5968G>A 5969A>T), D287N (5974G>A), Q288K (5977C>A), S290P (5983T>C 5985C>G), G293R (5992G>A), N295D (5998A>G 6000C>T), T296D (6001A>G 6002C>A), I297L (6004A>T 6006T>G), V298F (6007G>T 6009C>T), S299D (6010A>G 6011G>A), R300Q (6013A>C 6014G>A), I301L (6016A>C 6018A>T), S302Q (6019T>C 6020C>A 6021T>A), K304A (6025A>G 6026A>C 6027A>G), K305T (6029A>C 6030G>T), F316Y (6062T>A), I319L (6070A>T 6072A>G), M321V (6076A>G 6078G>T), E322R (6079G>A 6080A>G), K324S (6085A>T 6086A>C 6087G>T), S325D (6088T>G 6089C>A 6090C>T), K326I (6092A>T 6093A>C), W328K (6097T>A 6098G>A), W332R (6109T>C), P334R (6116C>G 6117A>G), E335Y (6118G>T 6120A>T), L337H (6125T>A), E341L (6136G>C 6137A>T 6138A>G), P344S (6145C>T), M348T (6158T>C 6159G>T), D353A (6173A>C), Q355M (6178C>A 6179A>T), R356D (6181A>G 6182G>A 6183A>C), K357L (6184A>T 6185A>T), D359N (6190G>A 6192T>C), N360R (6193A>C 6194A>G 6195T>G), A361V (6197C>T 6198A>G), R363K (6203G>A), G364S (6205G>T 6206G>C 6207A>T), G364_T365insY (6207_6208insTAT), T365L (6208A>C 6209C>T 6210A>T), A367Q (6214G>C 6215C>A 6216A>G), I369V (6220A>G 6222C>G), A370I (6223G>A 6224C>T 6225A>T), Y372F (6230A>T 6231T>C), F379Y (6251T>A), L391R (6286T>A 6287T>G 6288A>G), N392V (6289A>G 6290A>T), L393V (6292T>G), A394L (6295G>C 6296C>T), I396T (6302T>C), V397L (6304G>T), R399D (6310A>G 6311G>A 6312A>T), G401Q (6316G>C 6317G>A 6318G>A), I403C (6322A>T 6323T>G), L404A (6325C>G 6326T>C 6327A>T), S405K (6329G>A 6330C>G), P406F (6331C>T 6332C>T 6333A>C), T407S (6334A>T), M409Y (6340A>T 6341T>A 6342G>T), K410E (6343A>G 6345A>G), I411F (6346A>T), G412W (6349G>T 6351A>G), V413L (6352G>C 6354C>A), K414E (6355A>G), E415S (6358G>T 6359A>C), D417S (6364G>T 6365A>C 6366T>C) | | | | | | | | |
| Codon mutations: | GAG266AAG (5911G>A), ATT267AAG (5915T>A 5916T>G), AGA268GAT (5917A>G 5918G>A 5919A>T), GGC269GAG (5921G>A 5922C>G), AAA270AGC (5924A>G 5925A>C), CAA271ATA (5926C>A 5927A>T), AGG272TGG (5929A>T), GTT274TGT (5935G>T 5936T>G), TAC275ATT (5938T>A 5939A>T 5940C>T), AAC276GAC (5941A>G), AAG278AGT (5948A>G 5949G>T), CGC279CAA (5951G>A 5952C>A), GAC282AAG (5959G>A 5961C>G), AAT283GTC (5962A>G 5963A>T 5964T>C), ACA284ACT (5967A>T), GAA285ATA (5968G>A 5969A>T), AAA286AAG (5973A>G), GAT287AAT (5974G>A), CAG288AAG (5977C>A), TAT289TAC (5982T>C), TCC290CCG (5983T>C 5985C>G), CTA291TTG (5986C>T 5988A>G), CCG292CCT (5991G>T), GGA293AGA (5992G>A), ATC294ATT (5997C>T), AAC295GAT (5998A>G 6000C>T), ACT296GAT (6001A>G 6002C>A), ATT297TTG (6004A>T 6006T>G), GTC298TTT (6007G>T 6009C>T), AGT299GAT (6010A>G 6011G>A), AGA300CAA (6013A>C 6014G>A), ATA301CTT (6016A>C 6018A>T), TCT302CAA (6019T>C 6020C>A 6021T>A), AAA304GCG (6025A>G 6026A>C 6027A>G), AAG305ACT (6029A>C 6030G>T), ATA306G.G (6031A>G 6033A>G), TTC307..T (6036C>T), TCA308..T (6039A>T), AAA309..G (6042A>G), TTT310A.. (6043T>A), CTC312..G (6051C>G), AAA313.G. (6053A>G), GCA314T.G (6055G>T 6057A>G), GGC315..T (6060C>T), TTT316TAT (6062T>A), CAG318CAA (6069G>A), ATA319TTG (6070A>T 6072A>G), AGA320AGG (6075A>G), ATG321GTT (6076A>G 6078G>T), GAG322AGG (6079G>A 6080A>G), GAA323GAG (6084A>G), AAG324TCT (6085A>T 6086A>C 6087G>T), TCC325GAT (6088T>G 6089C>A 6090C>T), AAA326ATC (6092A>T 6093A>C), CCA327CCG (6096A>G), TGG328AAG (6097T>A 6098G>A), TGG332CGG (6109T>C), CCA334CGG (6116C>G 6117A>G), GAA335TAT (6118G>T 6120A>T), GGG336GGT (6123G>T), CTC337CAC (6125T>A), TTT340TTC (6135T>C), GAA341CTG (6136G>C 6137A>T 6138A>G), GTC342GTT (6141C>T), CCC344TCC (6145C>T), GGA346GGG (6153A>G), CTG347TTA (6154C>T 6156G>A), ATG348ACT (6158T>C 6159G>T), GCA350GCC (6165A>C), CCA351CCT (6168A>T), GAT353GCT (6173A>C), CAG355ATG (6178C>A 6179A>T), AGA356GAC (6181A>G 6182G>A 6183A>C), AAG357TTG (6184A>T 6185A>T), GAT359AAC (6190G>A 6192T>C), AAT360CGG (6193A>C 6194A>G 6195T>G), GCA361GTG (6197C>T 6198A>G), TTT362TTC (6201T>C), AGG363AAG (6203G>A), GGA364TCT (6205G>T 6206G>C 6207A>T), GGA364_ACA365insTAT (6207_6208insTAT), ACA365CTT (6208A>C 6209C>T 6210A>T), GAT366GAC (6213T>C), GCA367CAG (6214G>C 6215C>A 6216A>G), ATC369GTG (6220A>G 6222C>G), GCA370ATT (6223G>A 6224C>T 6225A>T), GTA371GTG (6228A>G), TAT372TTC (6230A>T 6231T>C), GAC375GAT (6240C>T), ATA376ATC (6243A>C), TTC379TAC (6251T>A), TCT380TCC (6255T>C), GAA381C.. (6256G>C), AAT382.G. (6260A>G), GAA383A.G (6262G>A 6264A>G), GAG385..A (6270G>A), GAG387..A (6276G>A), GAT388..A (6279T>A), CAT389..C (6282T>C), CTC390TTA (6283C>T 6285C>A), TTA391AGG (6286T>A 6287T>G 6288A>G), AAT392GTT (6289A>G 6290A>T), TTG393GTG (6292T>G), GCC394CTC (6295G>C 6296C>T), CAA395CAG (6300A>G), ATT396ACT (6302T>C), GTG397TTG (6304G>T), AGA399GAT (6310A>G 6311G>A 6312A>T), GGG401CAA (6316G>C 6317G>A 6318G>A), ATT403TGT (6322A>T 6323T>G), CTA404GCT (6325C>G 6326T>C 6327A>T), AGC405AAG (6329G>A 6330C>G), CCA406TTC (6331C>T 6332C>T 6333A>C), ACA407TCA (6334A>T), AAA408AAG (6339A>G), ATG409TAT (6340A>T 6341T>A 6342G>T), AAA410GAG (6343A>G 6345A>G), ATT411TTT (6346A>T), GGA412TGG (6349G>T 6351A>G), GTC413CTA (6352G>C 6354C>A), AAA414GAA (6355A>G), GAA415TCA (6358G>T 6359A>C), GTG416GTA (6363G>A), GAT417TCC (6364G>T 6365A>C 6366T>C), TTC418TTT (6369C>T), GGA420GGC (6375A>C), ATA421CA. (6376A>C 6377T>A) | | | | | | | | |

##### Proteins

| | Begin | End | Coverage | Score | Concordance | Matches | Identities | I/D/M/F* | Stop Codons |
|---|---|---|---|---|---|---|---|---|---|
| RNaseH/reverse transcriptase (YP_004581513.1) | 266 | 421 | 18.2% | 253 | 24.1% | 145 (99.3%) | 56 (38.4%) | 1/0/6/0 | 0 |
| Protein mutations: | E266K (5911G>A), I267K (5915T>A 5916T>G), R268D (5917A>G 5918G>A 5919A>T), G269E (5921G>A 5922C>G), K270S (5924A>G 5925A>C), Q271I (5926C>A 5927A>T), R272W (5929A>T), V274C (5935G>T 5936T>G), Y275I (5938T>A 5939A>T 5940C>T), N276D (5941A>G), K278S (5948A>G 5949G>T), R279Q (5951G>A 5952C>A), D282K (5959G>A 5961C>G), N283V (5962A>G 5963A>T 5964T>C), E285I (5968G>A 5969A>T), D287N (5974G>A), Q288K (5977C>A), S290P (5983T>C 5985C>G), G293R (5992G>A), N295D (5998A>G 6000C>T), T296D (6001A>G 6002C>A), I297L (6004A>T 6006T>G), V298F (6007G>T 6009C>T), S299D (6010A>G 6011G>A), R300Q (6013A>C 6014G>A), I301L (6016A>C 6018A>T), S302Q (6019T>C 6020C>A 6021T>A), K304A (6025A>G 6026A>C 6027A>G), K305T (6029A>C 6030G>T), F316Y (6062T>A), I319L (6070A>T 6072A>G), M321V (6076A>G 6078G>T), E322R (6079G>A 6080A>G), K324S (6085A>T 6086A>C 6087G>T), S325D (6088T>G 6089C>A 6090C>T), K326I (6092A>T 6093A>C), W328K (6097T>A 6098G>A), W332R (6109T>C), P334R (6116C>G 6117A>G), E335Y (6118G>T 6120A>T), L337H (6125T>A), E341L (6136G>C 6137A>T 6138A>G), P344S (6145C>T), M348T (6158T>C 6159G>T), D353A (6173A>C), Q355M (6178C>A 6179A>T), R356D (6181A>G 6182G>A 6183A>C), K357L (6184A>T 6185A>T), D359N (6190G>A 6192T>C), N360R (6193A>C 6194A>G 6195T>G), A361V (6197C>T 6198A>G), R363K (6203G>A), G364S (6205G>T 6206G>C 6207A>T), G364_T365insY (6207_6208insTAT), T365L (6208A>C 6209C>T 6210A>T), A367Q (6214G>C 6215C>A 6216A>G), I369V (6220A>G 6222C>G), A370I (6223G>A 6224C>T 6225A>T), Y372F (6230A>T 6231T>C), F379Y (6251T>A), L391R (6286T>A 6287T>G 6288A>G), N392V (6289A>G 6290A>T), L393V (6292T>G), A394L (6295G>C 6296C>T), I396T (6302T>C), V397L (6304G>T), R399D (6310A>G 6311G>A 6312A>T), G401Q (6316G>C 6317G>A 6318G>A), I403C (6322A>T 6323T>G), L404A (6325C>G 6326T>C 6327A>T), S405K (6329G>A 6330C>G), P406F (6331C>T 6332C>T 6333A>C), T407S (6334A>T), M409Y (6340A>T 6341T>A 6342G>T), K410E (6343A>G 6345A>G), I411F (6346A>T), G412W (6349G>T 6351A>G), V413L (6352G>C 6354C>A), K414E (6355A>G), E415S (6358G>T 6359A>C), D417S (6364G>T 6365A>C 6366T>C) | | | | | | | | |

| | Begin | End | Coverage | Score | Concordance | Matches | Identities | I/D/M/F* | Stop Codons |
|---|---|---|---|---|---|---|---|---|---|
| **NT** | **5911** | **6377** | **5.3%** | **-14** | **-1.6%** | **429 (99.3%)** | **214 (49.5%)** | **3/0** | |
| Codon mutations: | GAG266AAG (5911G>A), ATT267AAG (5915T>A 5916T>G), AGA268GAT (5917A>G 5918G>A 5919A>T), GGC269GAG (5921G>A 5922C>G), AAA270AGC (5924A>G 5925A>C), CAA271ATA (5926C>A 5927A>T), AGG272TGG (5929A>T), GTT274TGT (5935G>T 5936T>G), TAC275ATT (5938T>A 5939A>T 5940C>T), AAC276GAC (5941A>G), AAG278AGT (5948A>G 5949G>T), CGC279CAA (5951G>A 5952C>A), GAC282AAG (5959G>A 5961C>G), AAT283GTC (5962A>G 5963A>T 5964T>C), ACA284ACT (5967A>T), GAA285ATA (5968G>A 5969A>T), AAA286AAG (5973A>G), GAT287AAT (5974G>A), CAG288AAG (5977C>A), TAT289TAC (5982T>C), TCC290CCG (5983T>C 5985C>G), CTA291TTG (5986C>T 5988A>G), CCG292CCT (5991G>T), GGA293AGA (5992G>A), ATC294ATT (5997C>T), AAC295GAT (5998A>G 6000C>T), ACT296GAT (6001A>G 6002C>A), ATT297TTG (6004A>T 6006T>G), GTC298TTT (6007G>T 6009C>T), AGT299GAT (6010A>G 6011G>A), AGA300CAA (6013A>C 6014G>A), ATA301CTT (6016A>C 6018A>T), TCT302CAA (6019T>C 6020C>A 6021T>A), AAA304GCG (6025A>G 6026A>C 6027A>G), AAG305ACT (6029A>C 6030G>T), ATA306G.G (6031A>G 6033A>G), TTC307..T (6036C>T), TCA308..T (6039A>T), AAA309..G (6042A>G), TTT310A.. (6043T>A), CTC312..G (6051C>G), AAA313.G. (6053A>G), GCA314T.G (6055G>T 6057A>G), GGC315..T (6060C>T), TTT316TAT (6062T>A), CAG318CAA (6069G>A), ATA319TTG (6070A>T 6072A>G), AGA320AGG (6075A>G), ATG321GTT (6076A>G 6078G>T), GAG322AGG (6079G>A 6080A>G), GAA323GAG (6084A>G), AAG324TCT (6085A>T 6086A>C 6087G>T), TCC325GAT (6088T>G 6089C>A 6090C>T), AAA326ATC (6092A>T 6093A>C), CCA327CCG (6096A>G), TGG328AAG (6097T>A 6098G>A), TGG332CGG (6109T>C), CCA334CGG (6116C>G 6117A>G), GAA335TAT (6118G>T 6120A>T), GGG336GGT (6123G>T), CTC337CAC (6125T>A), TTT340TTC (6135T>C), GAA341CTG (6136G>C 6137A>T 6138A>G), GTC342GTT (6141C>T), CCC344TCC (6145C>T), GGA346GGG (6153A>G), CTG347TTA (6154C>T 6156G>A), ATG348ACT (6158T>C 6159G>T), GCA350GCC (6165A>C), CCA351CCT (6168A>T), GAT353GCT (6173A>C), CAG355ATG (6178C>A 6179A>T), AGA356GAC (6181A>G 6182G>A 6183A>C), AAG357TTG (6184A>T 6185A>T), GAT359AAC (6190G>A 6192T>C), AAT360CGG (6193A>C 6194A>G 6195T>G), GCA361GTG (6197C>T 6198A>G), TTT362TTC (6201T>C), AGG363AAG (6203G>A), GGA364TCT (6205G>T 6206G>C 6207A>T), GGA364_ACA365insTAT (6207_6208insTAT), ACA365CTT (6208A>C 6209C>T 6210A>T), GAT366GAC (6213T>C), GCA367CAG (6214G>C 6215C>A 6216A>G), ATC369GTG (6220A>G 6222C>G), GCA370ATT (6223G>A 6224C>T 6225A>T), GTA371GTG (6228A>G), TAT372TTC (6230A>T 6231T>C), GAC375GAT (6240C>T), ATA376ATC (6243A>C), TTC379TAC (6251T>A), TCT380TCC (6255T>C), GAA381C.. (6256G>C), AAT382.G. (6260A>G), GAA383A.G (6262G>A 6264A>G), GAG385..A (6270G>A), GAG387..A (6276G>A), GAT388..A (6279T>A), CAT389..C (6282T>C), CTC390TTA (6283C>T 6285C>A), TTA391AGG (6286T>A 6287T>G 6288A>G), AAT392GTT (6289A>G 6290A>T), TTG393GTG (6292T>G), GCC394CTC (6295G>C 6296C>T), CAA395CAG (6300A>G), ATT396ACT (6302T>C), GTG397TTG (6304G>T), AGA399GAT (6310A>G 6311G>A 6312A>T), GGG401CAA (6316G>C 6317G>A 6318G>A), ATT403TGT (6322A>T 6323T>G), CTA404GCT (6325C>G 6326T>C 6327A>T), AGC405AAG (6329G>A 6330C>G), CCA406TTC (6331C>T 6332C>T 6333A>C), ACA407TCA (6334A>T), AAA408AAG (6339A>G), ATG409TAT (6340A>T 6341T>A 6342G>T), AAA410GAG (6343A>G 6345A>G), ATT411TTT (6346A>T), GGA412TGG (6349G>T 6351A>G), GTC413CTA (6352G>C 6354C>A), AAA414GAA (6355A>G), GAA415TCA (6358G>T 6359A>C), GTG416GTA (6363G>A), GAT417TCC (6364G>T 6365A>C 6366T>C), TTC418TTT (6369C>T), GGA420GGC (6375A>C), ATA421CA. (6376A>C 6377T>A) | | | | | | | | |

*: Inserts / Deletes / Misaligned / Frameshifts

#### Analysis details

This analysis was performed with panviral2.64

### NGS Details (UN62): Errantivirus

#### Assembly

| | |
|---|---|
| **Coverage Length** | 575 (3 contig(s)) |
| **Depth Of Coverage** | 8.9 |
| **Number Of Reads** | 38 |
| **Reads Per Million** | 0.88 rpm (after QC) |
| **Ambiguities** | 0 |
| **Assembly Method** | de novo + reference guided assembly |
| **Consensus Caller** | Bcf Tools |

#### Coverage Map

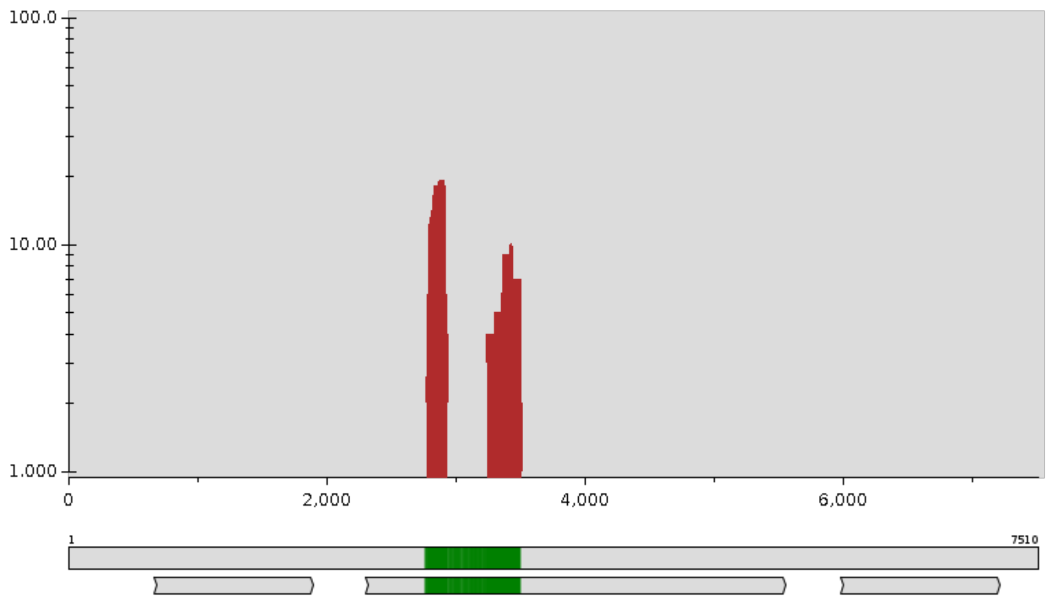

#### Assignment

| | |
|---|---|
| **Type** | Errantivirus (Taxonomy ID: 186666) |
| **Reference Genome** | NC_038512.1 |
| **NT Identity (%)** | 37.9433 |
| **AA Identity (%)** | 33.3333 |
| **Number Of Stop Codons** | 4 |
| **Number Of CDS** | 3 |

#### Alignment

| | |
|---|---|
| **Alignment Score** | -310.0 (NT) + -175.0 (AA) = -485.0 |
| **Concordance (%)** | -27.8351 |

| Alignment Method | Global, seeded, nucleotide + amino acids (AGA) |
|---|---|

#### Genome Region

Sequence starts at position 2767 and ends at position 3509 relative to NC_038512.1 reference sequence.

#### Alignment Detailed Statistics

| | Begin | End | Coverage | Score | Concordance | Matches | Identities | I/D/M/F* | Stop Codons |
|---|---|---|---|---|---|---|---|---|---|
| **NT** | **2767** | **3509** | **7.7%** | **-310** | **-28.3%** | **563 (97.7%)** | **214 (37.2%)** | **1/12** | |

Mutations: 2767T>A, 2769G>A, 2773C>A, 2777A>T, 2779C>A, 2781C>A, 2783A>G, 2785A>T, 2786A>T, 2788G>A, 2791G>A, 2794C>G, 2795C>A, 2806A>T, 2807A>C, 2809A>G, 2812A>T, 2813T>A, 2814C>G, 2816G>A, 2818C>A, 2821T>C, 2822G>C, 2824A>G, 2828A>G, 2830C>G, 2830_2831insT, 2831T>G, 2836C>G, 2837A>G, 2839A>G, 2841G>T, 2842G>A, 2843G>T, 2849C>A, 2850C>G, 2851C>A, 2857A>G, 2858_2860delATC, 2863C>T, 2865C>G, 2869T>C, 2870G>C, 2871G>T, 2872G>A, 2873_2881delAAACAAAAG, 2883G>T, 2885C>A, 2887T>A, 2888C>A, 2890C>G, 2891G>T, 2892T>G, 2893A>C, 2894G>A, 2896T>A, 2899C>T, 2901T>A, 2905T>G, 2906A>C, 2908G>A, 2909T>C, 2911G>C, 2915G>A, 2919A>T, 2920G>C, 2924A>G, 2926C>T, 2927G>A, 2929T>G, 2935A>G, 2937A>T, 2938C>T, 2941G>A, 2944A>T, 2949A>T, 2950C>T, 2953A>T, 2954A>G, 2955G>A, 2959C>G, 2960G>T, 2963C>T, 2968C>T, 2969A>G, 2974A>G, 2975G>C, 2976G>A, 2977T>A, 2978A>G, 2979A>G, 2981T>G, 2982G>C, 2983C>T, 2984C>A, 2986A>T, 2988A>T, 2989C>T, 2993A>T, 2995C>A, 2997C>A, 2998C>G, 2999T>C, 3001A>T, 3005T>C, 3007G>T, 3008G>C, 3009C>A, 3010A>T, 3011A>T, 3012G>C, 3016G>A, 3018T>A, 3020T>C, 3025G>A, 3026G>A, 3028G>T, 3029G>A, 3030A>G, 3032A>G, 3034G>T, 3035G>A, 3036A>G, 3037C>G, 3038C>G, 3039C>T, 3040T>A, 3041C>G, 3042A>C, 3043A>G, 3049A>T, 3050T>C, 3052G>C, 3058C>T, 3061G>T, 3066A>G, 3067C>G, 3068G>A, 3069T>C, 3071G>C, 3073A>C, 3074C>T, 3076C>T, 3079G>T, 3084T>A, 3091C>T, 3092C>T, 3094T>A, 3095C>G, 3096G>T, 3097A>C, 3103T>C, 3104A>T, 3106G>T, 3109A>T, 3112A>G, 3114A>C, 3115A>T, 3118C>T, 3119T>G, 3121A>G, 3124A>G, 3127T>C, 3130T>A, 3137A>G, 3140G>C, 3142T>G, 3146G>A, 3148C>T, 3149A>C, 3151T>G, 3154C>A, 3155C>T, 3157A>T, 3159G>A, 3160A>G, 3162G>C, 3163T>A, 3164C>T, 3165T>A, 3166C>T, 3168A>T, 3169A>T, 3171A>G, 3172T>G, 3175C>G, 3176A>T, 3178C>T, 3179T>G, 3180G>T, 3182C>T, 3184C>A, 3187C>T, 3189A>T, 3190C>T, 3191C>T, 3196C>T, 3202T>C, 3203A>T, 3205T>A, 3206G>A, 3208C>A, 3214T>C, 3216C>A, 3217T>G, 3218T>A, 3219C>G, 3220C>T, 3223A>G, 3224C>G, 3226G>A, 3229A>T, 3233C>T, 3236G>C, 3238G>A, 3239A>C, 3242C>T, 3245G>A, 3247A>G, 3248C>A, 3249G>T, 3250A>G, 3253T>G, 3256C>T, 3257C>G, 3259A>T, 3261G>T, 3262A>T, 3265T>G, 3272A>C, 3273G>A, 3275A>T, 3276A>C, 3278T>C, 3280C>A, 3281A>T, 3282A>T, 3283A>T, 3284A>G, 3285T>C, 3287C>A, 3289A>G, 3291T>G, 3292G>A, 3293G>T, 3294A>C, 3298G>C, 3300C>G, 3301C>A, 3302G>T, 3304A>T, 3307C>T, 3308T>G, 3309T>G, 3310G>A, 3311A>C, 3313G>A, 3315T>C, 3316C>A, 3317G>T, 3319A>G, 3321C>T, 3324C>A, 3325T>G, 3331T>A, 3337C>T, 3338A>G, 3340C>A, 3343A>T, 3344A>T, 3345G>C, 3346C>T, 3347A>G, 3348G>C, 3349G>T, 3352C>A, 3356A>G, 3358C>T, 3359A>T, 3361G>C, 3362C>T, 3363C>G, 3364T>A, 3365A>C, 3370T>A, 3371G>A, 3373T>G, 3378T>C, 3379T>G, 3380T>G, 3381C>A, 3382C>A, 3385T>A, 3386A>T, 3389C>A, 3390A>G, 3394A>T, 3396A>G, 3397T>G, 3398C>T, 3399T>C, 3400G>A, 3402T>A, 3403T>G, 3406A>C, 3407A>T, 3408A>T, 3409G>A, 3410A>T, 3415T>G, 3416A>G, 3417A>C, 3421A>T, 3424A>T, 3426A>G, 3428C>A, 3429A>G, 3430A>T, 3433T>C, 3436A>G, 3439C>G, 3442T>A, 3443C>G, 3444T>C, 3445C>G, 3448T>G, 3451T>C, 3454C>T, 3455C>A, 3459A>G, 3460A>G, 3463C>T, 3464A>G, 3466T>G, 3467C>G, 3468C>A, 3471A>G, 3472T>C, 3476G>T, 3479C>T, 3480G>C, 3482C>A, 3485A>T, 3487A>C, 3488A>T, 3489A>C, 3490A>T, 3493C>T, 3494C>T, 3496T>G, 3499A>C, 3500C>T

##### CDS

| | Begin | End | Coverage | Score | Concordance | Matches | Identities | I/D/M/F* | Stop Codons |
|---|---|---|---|---|---|---|---|---|---|
| D1R33_gp2 | 160 | 402 | 16.3% | -175 | -11.3% | 173 (97.2%) | 58 (32.6%) | 1/4/23/1 | 4 |

Protein mutations: K160D (2783A>G 2785A>T), M161L (2786A>T 2788G>A), D163E (2794C>G), Q164K (2795C>A), S170R (2813T>A 2814C>G), D171K (2816G>A 2818C>A), A173P (2822G>C 2824A>G), S175G (2828A>G 2830C>G), S175_S176insX (2830_2831insT), S176A (2831T>G), I178V (2837A>G 2839A>G), W179L (2841G>T 2842G>A), V180F (2843G>T), P182R (2849C>A 2850C>G 2851C>A), I185del (2858_2860delATC), A187G (2865C>G), G189L (2870G>C 2871G>T 2872G>A), K190_K192del (2873_2881delAAACAAAAG), W193L (2883G>T), L195M (2888C>A 2890C>G), V196C (2891G>T 2892T>G 2893A>C), V197I (2894G>A 2896T>A), F199Y (2901T>A), K201Q (2906A>C 2908G>A), E204K (2915G>A), K205I (2919A>T 2920G>C), I207V (2924A>G 2926C>T), D208K (2927G>A 2929T>G), Y211F (2937A>T 2938C>T), G224Q (2975G>C 2976G>A 2977T>A), C226A (2981T>G 2982G>C 2983C>T), A235H (3008G>C 3009C>A 3010A>T), D244R (3035G>A 3036A>G 3037C>G), P245V (3038C>G 3039C>T 3040T>A), Q246A (3041C>G 3042A>C 3043A>G), R264V (3095C>G 3096G>T 3097A>C), L287Y (3164C>T 3165T>A 3166C>T), E314K (3245G>A 3247A>G), R315M (3248C>A 3249G>T 3250A>G), Q318D (3257C>G 3259A>T), R319I (3261G>T 3262A>T), S323H (3272A>C 3273G>A), N324S (3275A>T 3276A>C), F325L (3278T>C 3280C>A), K326F (3281A>T 3282A>T 3283A>T), I327A (3284A>G 3285T>C), Q328K (3287C>A 3289A>G), M329R (3291T>G 3292G>A), D330S (3293G>T 3294A>C), K331N (3298G>C), S332* (3300C>G 3301C>A), E333Y (3302G>T 3304A>T), L335G (3308T>G 3309T>G 3310G>A), K336Q (3311A>C 3313G>A), L337P (3315T>C 3316C>A), E338* (3317G>T 3319A>G), T339I (3321C>T), A340E (3324C>A 3325T>G), I345V (3338A>G 3340C>A), R348A (3347A>G 3348G>C 3349G>T), D349E (3352C>A), I351V (3356A>G 3358C>T), K352Y (3359A>T 3361G>C), P353* (3362C>T 3363C>G 3364T>A), N354H (3365A>C), D356K (3371G>A 3373T>G), I358T (3378T>C 3379T>G), S359E (3380T>G 3381C>A 3382C>A), I361F (3386A>T), Q362R (3389C>A 3390A>G), K363N (3394A>T), Y364W (3396A>G 3397T>G), L365S (3398C>T 3399T>C 3400G>A), I366K (3402T>A 3403T>G), K368L (3407A>T 3408A>T 3409G>A), T369S (3410A>T), K371A (3416A>G 3417A>C), E372D (3421A>T), K374R (3426A>G), Q375S (3428C>A 3429A>G 3430A>T), L380A (3443C>G 3444T>C 3445C>G), K385R (3459A>G 3460A>G), I387V (3464A>G 3466T>G), P388E (3467C>G 3468C>A), D389G (3471A>G 3472T>C), A391S (3476G>T), R392S (3479C>T 3480G>C), L393I (3482C>A), T394S (3485A>T 3487A>C), K395S (3488A>T 3489A>C 3490A>T), Q399* (3500C>T)

Codon mutations: GTT154..A (2767T>A), AGG155.A. (2769G>A), GAC156..A (2773C>A), ATC158T.A (2777A>T 2779C>A), ACG159.AG (2781C>A), AAA160GAT (2783A>G 2785A>T), ATG161TTA (2786A>T 2788G>A), TTG162TTA (2791G>A), GAC163GAG (2794C>G), CAA164AAA (2795C>A), ATA167ATT (2806A>T), AGA168CGG (2807A>C 2809A>G), CCA169CCT (2812A>T), TCA170AGA (2813T>A 2814C>G), GAC171AAA (2816G>A 2818C>A), TCT172TCC (2821T>C), GCA173CCG (2822G>C 2824A>G), AGC175GGG (2828A>G 2830C>G), AGC175_TCA176insT-- (2830_2831insT), TCA176GCA (2831T>G), CCC177CCG (2836C>G), ATA178GTG (2837A>G 2839A>G), TGG179TTA (2841G>T 2842G>A), GTT180TTT (2843G>T), CCC182AGA (2849C>A 2850C>G 2851C>A), AAA184AAG (2857A>G), ATC185del (2858_2860delATC), GAC186GAT (2863C>T), GCT187GGT (2865C>G), TCT188TCC (2869T>C), GGG189CTA (2870G>C 2871G>T 2872G>A), AAA190_AAG192del (2873_2881delAAACAAAAG), TGG193TTG (2883G>T), CGT194AGA (2885C>A 2887T>A), CTC195ATG (2888C>A 2890C>G), GTA196TGC (2891G>T 2892T>G 2893A>C), GTT197ATA (2894G>A 2896T>A), GAC198GAT (2899C>T), TTC199TAC (2901T>A), CGT200CGG (2905T>G), AAG201CAA (2906A>C 2908G>A), TTG202CTC (2909T>C 2911G>C), GAG204AAG (2915G>A), AAG205ATC (2919A>T 2920G>C), ATC207GTT (2924A>G 2926C>T), GAT208AAG (2927G>A 2929T>G), AAA210AAG (2935A>G), TAC211TTT (2937A>T 2938C>T), CCG212..A (2941G>A), ATA213..T (2944A>T), AAC215.TT (2949A>T 2950C>T), ATA216..T (2953A>T), AGT217GA. (2954A>G 2955G>A), GAC218..G (2959C>G), GTA219T.. (2960G>T), CTT220T.. (2963C>T), GAC221..T (2968C>T), AAG222G.. (2969A>G), TTA223..G (2974A>G), GGT224CAA (2975G>C 2976G>A 2977T>A), AAG225GG. (2978A>G 2979A>G), TGC226GCT (2981T>G 2982G>C 2983C>T), CAA227A.T (2984C>A 2986A>T), TAC228.TT (2988A>T 2989C>T), ACC230T.A (2993A>T 2995C>A), ACC231.AG (2997C>A 2998C>G), TTA232C.T (2999T>C 3001A>T), TTG234C.T (3005T>C 3007G>T), GCA235CAT (3008G>C 3009C>A 3010A>T), AGT236TC. (3011A>T 3012G>C), GGG237..A (3016G>A), TTT238.A. (3018T>A), TAT239C.. (3020T>C), CAG240..A (3025G>A), GTG241A.T (3026G>A 3028G>T), GAG242AG. (3029G>A 3030A>G), ATG243G.T (3032A>G 3034G>T), GAC244AGG (3035G>A 3036A>G 3037C>G), CCT245GTA (3038C>G 3039C>T 3040T>A), CAA246GCG (3041C>G 3042A>C 3043A>G), ATA248..T (3049A>T), TCG249C.C (3050T>C 3052G>C), ACC251..T (3058C>T), GCG252..T (3061G>T), AAC254.GG (3066A>G 3067C>G), GTA255AC. (3068G>A 3069T>C), GAA256C.C (3071G>C 3073A>C), CAC257T.T (3074C>T 3076C>T), GGG258..T (3079G>T), TTT260.A. (3084T>A), TTC262..T (3091C>T), CTT263T.A (3092C>T 3094T>A), CGA264GTC (3095C>G 3096G>T 3097A>C), CCT266..C (3103T>C), ATG267T.T (3104A>T 3106G>T), GGA268..T (3109A>T), TTA269..G (3112A>G), AAA270.CT (3114A>C 3115A>T), AAC271..T (3118C>T), TCA272G.G (3119T>G 3121A>G), CCA273..G (3124A>G), TCT274..C (3127T>C), ACT275..A (3130T>A), AGA278G.. (3137A>G), GTT279C.G (3140G>C 3142T>G), GAC281A.T (3146G>A 3148C>T), AAT282C.G (3149A>C 3151T>G), GTC283..A (3154C>A), CTA284T.T (3155C>T 3157A>T), AGA285.AG (3159G>A 3160A>G), GGT286.CA (3162G>C 3163T>A), CTC287TAT (3164C>T 3165T>A 3166C>T), CAA288.TT (3168A>T 3169A>T), AAT289.GG (3171A>G 3172T>G), AAC290..G (3175C>G), ATC291T.T (3176A>T 3178C>T), TGT292GT. (3179T>G 3180G>T), CTC293T.A (3182C>T 3184C>A), GTC294..T (3187C>T), TAC295.TT (3189A>T 3190C>T), CTT296T.. (3191C>T), GAC297..T (3196C>T), ATT299..C (3202T>C), ATT300T.A (3203A>T 3205T>A), GTC301A.A (3206G>A 3208C>A), AGT303..C (3214T>C), ACT304.AG (3216C>A 3217T>G), TCC305AGT (3218T>A 3219C>G 3220C>T), CTA306..G (3223A>G), CAG307G.A (3224C>G 3226G>A), GAA308..T (3229A>T), CTG310T.. (3233C>T), GAG311C.A (3236G>C 3238G>A), AAC312C.C (3239A>C), CTG313TTG (3242C>T), GAA314AAG (3245G>A 3247A>G), CGA315ATG (3248C>A 3249G>T 3250A>G), GTT316GTG (3253T>G), TTC317TTT (3256C>T), CAA318GAT (3257C>G 3259A>T), AGA319ATT (3261G>T 3262A>T), CTT320CTG (3265T>G), AGT323CAT (3272A>C 3273G>A), AAC324TCC (3275A>T 3276A>C), TTC325CTA (3278T>C 3280C>A), AAA326TTT (3281A>T 3282A>T 3283A>T), ATT327GCT (3284A>G 3285T>C), CAA328AAG (3287C>A 3289A>G), ATG329AGA (3291T>G 3292G>A), GAC330TCC (3293G>T 3294A>C), AAG331AAC (3298G>C), TCC332TGA (3300C>G 3301C>A), GAA333TAT (3302G>T 3304A>T), TTC334TTT (3307C>T), TTG335GGA (3308T>G 3309T>G 3310G>A), AAG336CAA (3311A>C 3313G>A), CTC337CCA (3315T>C 3316C>A), GAA338TAG (3317G>T 3319A>G), ACT339ATT (3321C>T), GCT340GAG (3324C>A 3325T>G), CTT342CTA (3331T>A), CAC344CAT (3337C>T), ATC345GTA (3338A>G 3340C>A), ATA346ATT (3343A>T), AGC347TCT (3344A>T 3345G>C 3346C>T), AGG348GCT (3347A>G 3348G>C 3349G>T), GAC349GAA (3352C>A), ATC351GTT (3356A>G 3358C>T), AAG352TAC (3359A>T 3361G>C), CCT353TGA (3362C>T 3363C>G 3364T>A), AAC354CAC (3365A>C), CCT355CCA (3370T>A), GAT356AAG (3371G>A 3373T>G), ATT358ACG (3378T>C 3379T>G), TCC359GAA (3380T>G 3381C>A 3382C>A), GCT360GCA (3385T>A), ATT361TTT (3386A>T), CAA362AGA (3389C>A 3390A>G), AAA363AAT (3394A>T), TAT364TGG (3396A>G 3397T>G), CTG365TCA (3398C>T 3399T>C 3400G>A), ATT366AAG (3402T>A 3403T>G), CCA367CCC (3406A>C), AAG368TTA (3407A>T 3408A>T 3409G>A), ACC369TCC (3410A>T), CCT370CCG (3415T>G), AAG371GCG (3416A>G 3417A>C), GAA372GAT (3421A>T), ATA373ATT (3424A>T), AAA374AGA (3426A>G), CAA375AGT (3428C>A 3429A>G 3430A>T), TTT376TTC (3433T>C), TTA377TTG (3436A>G), GGC378GGG (3439C>G), CTT379CTA (3442T>A), CTC380GCG (3443C>G 3444T>C 3445C>G), GGT381GGG (3448T>G), TAT382TAC (3451T>C), TAC383TAT (3454C>T), CGA384AGA (3455C>A), AAA385AGG (3459A>G 3460A>G), TTC386TTT (3463C>T), ATT387GTG (3464A>G 3466T>G), CCA388GAA (3467C>G 3468C>A), GAT389GGC (3471A>G 3472T>C), GCA391TCA (3476G>T), CGA392TCA (3479C>T 3480G>C), CTC393ATC (3482C>A), ACA394TCC (3485A>T 3487A>C), AAA395TCT (3488A>T 3489A>C 3490A>T), CCC396CCT (3493C>T), CTT397TTG (3494C>T 3496T>G), ACA398ACC (3499A>C), CAG399TAG (3500C>T)

##### Proteins

| | Begin | End | Coverage | Score | Concordance | Matches | Identities | I/D/M/F* | Stop Codons |
|---|---|---|---|---|---|---|---|---|---|
| ORF B (YP_009507248.1) | 160 | 402 | 16.3% | -175 | -11.3% | 173 (97.2%) | 58 (32.6%) | 1/4/23/1 | 4 |

| | Begin | End | Coverage | Score | Concordance | Matches | Identities | I/D/M/F* | Stop Codons |
|---|---|---|---|---|---|---|---|---|---|
| **NT** | **2767** | **3509** | **7.7%** | **-310** | **-28.3%** | **563 (97.7%)** | **214 (37.2%)** | **1/12** | |
| Protein mutations: | K160D (2783A>G 2785A>T), M161L (2786A>T 2788G>A), D163E (2794C>G), Q164K (2795C>A), S170R (2813T>A 2814C>G), D171K (2816G>A 2818C>A), A173P (2822G>C 2824A>G), S175G (2828A>G 2830C>G), S175_S176insX (2830_2831insT), S176A (2831T>G), I178V (2837A>G 2839A>G), W179L (2841G>T 2842G>A), V180F (2843G>T), P182R (2849C>A 2850C>G 2851C>A), I185del (2858_2860delATC), A187G (2865C>G), G189L (2870G>C 2871G>T 2872G>A), K190_K192del (2873_2881delAAACAAAAG), W193L (2883G>T), L195M (2888C>A 2890C>G), V196C (2891G>T 2892T>G 2893A>C), V197I (2894G>A 2896T>A), F199Y (2901T>A), K201Q (2906A>C 2908G>A), E204K (2915G>A), K205I (2919A>T 2920G>C), I207V (2924A>G 2926C>T), D208K (2927G>A 2929T>G), Y211F (2937A>T 2938C>T), G224Q (2975G>C 2976G>A 2977T>A), C226A (2981T>G 2982G>C 2983C>T), A235H (3008G>C 3009C>A 3010A>T), D244R (3035G>A 3036A>G 3037C>G), P245V (3038C>G 3039C>T 3040T>A), Q246A (3041C>G 3042A>C 3043A>G), R264V (3095C>G 3096G>T 3097A>C), L287Y (3164C>T 3165T>A 3166C>T), E314K (3245G>A 3247A>G), R315M (3248C>A 3249G>T 3250A>G), Q318D (3257C>G 3259A>T), R319I (3261G>T 3262A>T), S323H (3272A>C 3273G>A), N324S (3275A>T 3276A>C), F325L (3278T>C 3280C>A), K326F (3281A>T 3282A>T 3283A>T), I327A (3284A>G 3285T>C), Q328K (3287C>A 3289A>G), M329R (3291T>G 3292G>A), D330S (3293G>T 3294A>C), K331N (3298G>C), S332* (3300C>G 3301C>A), E333Y (3302G>T 3304A>T), L335G (3308T>G 3309T>G 3310G>A), K336Q (3311A>C 3313G>A), L337P (3315T>C 3316C>A), E338* (3317G>T 3319A>G), T339I (3321C>T), A340E (3324C>A 3325T>G), I345V (3338A>G 3340C>A), R348A (3347A>G 3348G>C 3349G>T), D349E (3352C>A), I351V (3356A>G 3358C>T), K352Y (3359A>T 3361G>C), P353* (3362C>T 3363C>G 3364T>A), N354H (3365A>C), D356K (3371G>A 3373T>G), I358T (3378T>C 3379T>G), S359E (3380T>G 3381C>A 3382C>A), I361F (3386A>T), Q362R (3389C>A 3390A>G), K363N (3394A>T), Y364W (3396A>G 3397T>G), L365S (3398C>T 3399T>C 3400G>A), I366K (3402T>A 3403T>G), K368L (3407A>T 3408A>T 3409G>A), T369S (3410A>T), K371A (3416A>G 3417A>C), E372D (3421A>T), K374R (3426A>G), Q375S (3428C>A 3429A>G 3430A>T), L380A (3443C>G 3444T>C 3445C>G), K385R (3459A>G 3460A>G), I387V (3464A>G 3466T>G), P388E (3467C>G 3468C>A), D389G (3471A>G 3472T>C), A391S (3476G>T), R392S (3479C>T 3480G>C), L393I (3482C>A), T394S (3485A>T 3487A>C), K395S (3488A>T 3489A>C 3490A>T), Q399* (3500C>T) | | | | | | | | |
| Codon mutations: | GTT154..A (2767T>A), AGG155.A. (2769G>A), GAC156..A (2773C>A), ATC158T.A (2777A>T 2779C>A), ACG159.AG (2781C>A), AAA160GAT (2783A>G 2785A>T), ATG161TTA (2786A>T 2788G>A), TTG162TTA (2791G>A), GAC163GAG (2794C>G), CAA164AAA (2795C>A), ATA167ATT (2806A>T), AGA168CGG (2807A>C 2809A>G), CCA169CCT (2812A>T), TCA170AGA (2813T>A 2814C>G), GAC171AAA (2816G>A 2818C>A), TCT172TCC (2821T>C), GCA173CCG (2822G>C 2824A>G), AGC175GGG (2828A>G 2830C>G), AGC175_TCA176insT-- (2830_2831insT), TCA176GCA (2831T>G), CCC177CCG (2836C>G), ATA178GTG (2837A>G 2839A>G), TGG179TTA (2841G>T 2842G>A), GTT180TTT (2843G>T), CCC182AGA (2849C>A 2850C>G 2851C>A), AAA184AAG (2857A>G), ATC185del (2858_2860delATC), GAC186GAT (2863C>T), GCT187GGT (2865C>G), TCT188TCC (2869T>C), GGG189CTA (2870G>C 2871G>T 2872G>A), AAA190_AAG192del (2873_2881delAAACAAAAG), TGG193TTG (2883G>T), CGT194AGA (2885C>A 2887T>A), CTC195ATG (2888C>A 2890C>G), GTA196TGC (2891G>T 2892T>G 2893A>C), GTT197ATA (2894G>A 2896T>A), GAC198GAT (2899C>T), TTC199TAC (2901T>A), CGT200CGG (2905T>G), AAG201CAA (2906A>C 2908G>A), TTG202CTC (2909T>C 2911G>C), GAG204AAG (2915G>A), AAG205ATC (2919A>T 2920G>C), ATC207GTT (2924A>G 2926C>T), GAT208AAG (2927G>A 2929T>G), AAA210AAG (2935A>G), TAC211TTT (2937A>T 2938C>T), CCG212..A (2941G>A), ATA213..T (2944A>T), AAC215.TT (2949A>T 2950C>T), ATA216..T (2953A>T), AGT217GA. (2954A>G 2955G>A), GAC218..G (2959C>G), GTA219T.. (2960G>T), CTT220T.. (2963C>T), GAC221..T (2968C>T), AAG222G.. (2969A>G), TTA223..G (2974A>G), GGT224CAA (2975G>C 2976G>A 2977T>A), AAG225GG. (2978A>G 2979A>G), TGC226GCT (2981T>G 2982G>C 2983C>T), CAA227A.T (2984C>A 2986A>T), TAC228.TT (2988A>T 2989C>T), ACC230T.A (2993A>T 2995C>A), ACC231.AG (2997C>A 2998C>G), TTA232C.T (2999T>C 3001A>T), TTG234C.T (3005T>C 3007G>T), GCA235CAT (3008G>C 3009C>A 3010A>T), AGT236TC. (3011A>T 3012G>C), GGG237..A (3016G>A), TTT238.A. (3018T>A), TAT239C.. (3020T>C), CAG240..A (3025G>A), GTG241A.T (3026G>A 3028G>T), GAG242AG. (3029G>A 3030A>G), ATG243G.T (3032A>G 3034G>T), GAC244AGG (3035G>A 3036A>G 3037C>G), CCT245GTA (3038C>G 3039C>T 3040T>A), CAA246GCG (3041C>G 3042A>C 3043A>G), ATA248..T (3049A>T), TCG249C.C (3050T>C 3052G>C), ACC251..T (3058C>T), GCG252..T (3061G>T), AAC254.GG (3066A>G 3067C>G), GTA255AC. (3068G>A 3069T>C), GAA256C.C (3071G>C 3073A>C), CAC257T.T (3074C>T 3076C>T), GGG258..T (3079G>T), TTT260.A. (3084T>A), TTC262..T (3091C>T), CTT263T.A (3092C>T 3094T>A), CGA264GTC (3095C>G 3096G>T 3097A>C), CCT266..C (3103T>C), ATG267T.T (3104A>T 3106G>T), GGA268..T (3109A>T), TTA269..G (3112A>G), AAA270.CT (3114A>C 3115A>T), AAC271..T (3118C>T), TCA272G.G (3119T>G 3121A>G), CCA273..G (3124A>G), TCT274..C (3127T>C), ACT275..A (3130T>A), AGA278G.. (3137A>G), GTT279C.G (3140G>C 3142T>G), GAC281A.T (3146G>A 3148C>T), AAT282C.G (3149A>C 3151T>G), GTC283..A (3154C>A), CTA284T.T (3155C>T 3157A>T), AGA285.AG (3159G>A 3160A>G), GGT286.CA (3162G>C 3163T>A), CTC287TAT (3164C>T 3165T>A 3166C>T), CAA288.TT (3168A>T 3169A>T), AAT289.GG (3171A>G 3172T>G), AAC290..G (3175C>G), ATC291T.T (3176A>T 3178C>T), TGT292GT. (3179T>G 3180G>T), CTC293T.A (3182C>T 3184C>A), GTC294..T (3187C>T), TAC295.TT (3189A>T 3190C>T), CTT296T.. (3191C>T), GAC297..T (3196C>T), ATT299..C (3202T>C), ATT300T.A (3203A>T 3205T>A), GTC301A.A (3206G>A 3208C>A), AGT303..C (3214T>C), ACT304.AG (3216C>A 3217T>G), TCC305AGT (3218T>A 3219C>G 3220C>T), CTA306..G (3223A>G), CAG307G.A (3224C>G 3226G>A), GAA308..T (3229A>T), CTG310T.. (3233C>T), GAG311C.A (3236G>C 3238G>A), AAC312C.C (3239A>C), CTG313TTG (3242C>T), GAA314AAG (3245G>A 3247A>G), CGA315ATG (3248C>A 3249G>T 3250A>G), GTT316GTG (3253T>G), TTC317TTT (3256C>T), CAA318GAT (3257C>G 3259A>T), AGA319ATT (3261G>T 3262A>T), CTT320CTG (3265T>G), AGT323CAT (3272A>C 3273G>A), AAC324TCC (3275A>T 3276A>C), TTC325CTA (3278T>C 3280C>A), AAA326TTT (3281A>T 3282A>T 3283A>T), ATT327GCT (3284A>G 3285T>C), CAA328AAG (3287C>A 3289A>G), ATG329AGA (3291T>G 3292G>A), GAC330TCC (3293G>T 3294A>C), AAG331AAC (3298G>C), TCC332TGA (3300C>G 3301C>A), GAA333TAT (3302G>T 3304A>T), TTC334TTT (3307C>T), TTG335GGA (3308T>G 3309T>G 3310G>A), AAG336CAA (3311A>C 3313G>A), CTC337CCA (3315T>C 3316C>A), GAA338TAG (3317G>T 3319A>G), ACT339ATT (3321C>T), GCT340GAG (3324C>A 3325T>G), CTT342CTA (3331T>A), CAC344CAT (3337C>T), ATC345GTA (3338A>G 3340C>A), ATA346ATT (3343A>T), AGC347TCT (3344A>T 3345G>C 3346C>T), AGG348GCT (3347A>G 3348G>C 3349G>T), GAC349GAA (3352C>A), ATC351GTT (3356A>G 3358C>T), AAG352TAC (3359A>T 3361G>C), CCT353TGA (3362C>T 3363C>G 3364T>A), AAC354CAC (3365A>C), CCT355CCA (3370T>A), GAT356AAG (3371G>A 3373T>G), ATT358ACG (3378T>C 3379T>G), TCC359GAA (3380T>G 3381C>A 3382C>A), GCT360GCA (3385T>A), ATT361TTT (3386A>T), CAA362AGA (3389C>A 3390A>G), AAA363AAT (3394A>T), TAT364TGG (3396A>G 3397T>G), CTG365TCA (3398C>T 3399T>C 3400G>A), ATT366AAG (3402T>A 3403T>G), CCA367CCC (3406A>C), AAG368TTA (3407A>T 3408A>T 3409G>A), ACC369TCC (3410A>T), CCT370CCG (3415T>G), AAG371GCG (3416A>G 3417A>C), GAA372GAT (3421A>T), ATA373ATT (3424A>T), AAA374AGA (3426A>G), CAA375AGT (3428C>A 3429A>G 3430A>T), TTT376TTC (3433T>C), TTA377TTG (3436A>G), GGC378GGG (3439C>G), CTT379CTA (3442T>A), CTC380GCG (3443C>G 3444T>C 3445C>G), GGT381GGG (3448T>G), TAT382TAC (3451T>C), TAC383TAT (3454C>T), CGA384AGA (3455C>A), AAA385AGG (3459A>G 3460A>G), TTC386TTT (3463C>T), ATT387GTG (3464A>G 3466T>G), CCA388GAA (3467C>G 3468C>A), GAT389GGC (3471A>G 3472T>C), GCA391TCA (3476G>T), CGA392TCA (3479C>T 3480G>C), CTC393ATC (3482C>A), ACA394TCC (3485A>T 3487A>C), AAA395TCT (3488A>T 3489A>C 3490A>T), CCC396CCT (3493C>T), CTT397TTG (3494C>T 3496T>G), ACA398ACC (3499A>C), CAG399TAG (3500C>T) | | | | | | | | |

*: Inserts / Deletes / Misaligned / Frameshifts

#### Analysis details

This analysis was performed with panviral2.64

### NGS Details (UN62): Badnavirus volubetulae

#### Assembly

| | |
|---|---|
| **Coverage Length** | 339 (1 contig(s)) |
| **Depth Of Coverage** | 10.1 |
| **Number Of Reads** | 30 |
| **Reads Per Million** | 0.70 rpm (after QC) |
| **Ambiguities** | 0 |
| **Assembly Method** | de novo + reference guided assembly |
| **Consensus Caller** | Bcf Tools |

#### Coverage Map

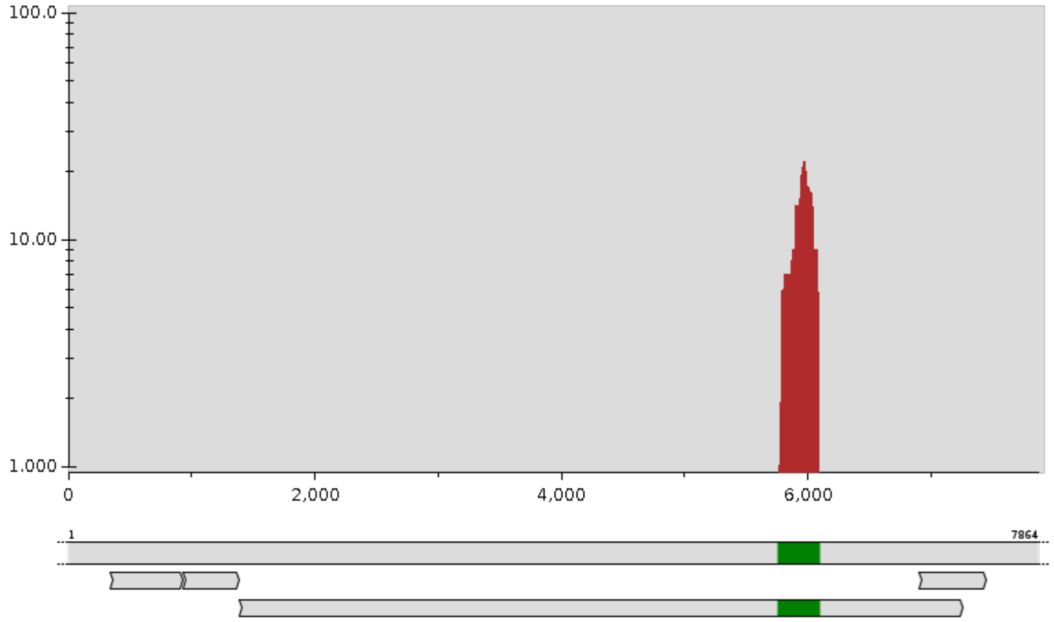

#### Assignment

| | |
|---|---|
| **Type** | Badnavirus volubetulae (Taxonomy ID: 3047683) |
| **Reference Genome** | NC_040635.1 |
| **NT Identity (%)** | 52.6316 |
| **AA Identity (%)** | 42.1053 |
| **Number Of Stop Codons** | 0 |
| **Number Of CDS** | 4 |

#### Alignment

| | |
|---|---|
| **Alignment Score** | 30.0 (NT) + 350.0 (AA) = 380.0 |
| **Concordance (%)** | 25.3502 |

| **Alignment Method** | Global, seeded, nucleotide + amino acids (AGA) |
|---|---|

#### Genome Region

Sequence starts at position 5758 and ends at position 6096 relative to NC_040635.1 reference sequence.

#### Alignment Detailed Statistics

| | Begin | End | Coverage | Score | Concordance | Matches | Identities | I/D/M/F* | Stop Codons |
|---|---|---|---|---|---|---|---|---|---|
| **NT** | **5758** | **6096** | **4.3%** | **30** | **4.4%** | **339 (99.1%)** | **180 (52.6%)** | **3/0** | |
| Mutations: | 5758C>A, 5760C>T, 5763T>C, 5770A>C, 5771G>A, 5772T>A, 5773C>A, 5775G>T, 5778C>T, 5779C>A, 5780A>G, 5782A>G, 5783A>T, 5788C>G, 5789A>T, 5790C>G, 5794G>A, 5797C>A, 5799A>G, 5803A>C, 5804G>C, 5808A>T, 5811A>C, 5812G>A, 5814A>G, 5817C>T, 5818A>G, 5820C>T, 5821A>G, 5822C>A, 5824A>T, 5826T>G, 5827G>T, 5829C>T, 5830C>G, 5832G>T, 5834G>A, 5835C>A, 5836A>C, 5839G>C, 5841T>A, 5843A>G, 5844T>G, 5849A>G, 5850G>C, 5851G>T, 5852T>A, 5853A>T, 5856C>T, 5859A>T, 5862A>G, 5863T>A, 5865C>T, 5872A>C, 5873A>G, 5874G>T, 5875A>T, 5880C>A, 5882T>A, 5886T>C, 5890G>C, 5892C>T, 5894C>G, 5896A>G, 5899G>A, 5901T>G, 5904A>G, 5905G>A, 5906A>G, 5907A>T, 5908T>G, 5909C>A, 5910A>T, 5911A>G, 5913A>T, 5916A>G, 5917T>A, 5918G>A, 5919G>A, 5921C>T, 5922A>G, 5925A>T, 5930T>G, 5931C>A, 5935C>A, 5936C>G, 5937A>G, 5938G>T, 5944T>C, 5945T>A, 5946G>T, 5948A>T, 5949C>T, 5954G>T, 5955G>C, 5958A>T, 5961T>C, 5965C>T, 5967A>C, 5974C>T, 5976T>G, 5978A>C, 5979A>T, 5986C>T, 5988A>G, 5989G>A, 5990C>A, 5991A>T, 5997C>T, 5998C>A, 5999A>T, 6000G>A, 6001A>G, 6002G>A, 6003A>T, 6004A>T, 6005A>T, 6006A>G, 6010G>A, 6014A>G, 6015T>G, 6016T>G, 6017G>T, 6018C>T, 6021T>C, 6024A>G, 6025G>C, 6026G>C, 6028A>T, 6029C>T, 6030G>C, 6031G>C, 6032A>T, 6036A>C, 6036_6037insATA, 6039C>T, 6040A>G, 6044C>T, 6045T>G, 6048A>G, 6050A>T, 6051C>T, 6054C>T, 6057T>C, 6061A>G, 6063T>A, 6067G>A, 6069C>A, 6071T>A, 6076G>C, 6077A>G, 6082C>A, 6083A>G, 6084T>G, 6085A>G, 6088C>G, 6093T>C, 6094G>T, 6095A>T, 6096A>G | | | | | | | | |

##### CDS

| | Begin | End | Coverage | Score | Concordance | Matches | Identities | I/D/M/F* | Stop Codons |
|---|---|---|---|---|---|---|---|---|---|
| BLRaV_gp3 | 1457 | 1569 | 5.8% | 350 | 42.6% | 113 (99.1%) | 48 (42.1%) | 1/0/0/0 | 0 |
| Protein mutations: | L1457I (5758C>A 5760C>T), S1461Q (5770A>C 5771G>A 5772T>A), L1462I (5773C>A 5775G>T), Q1464R (5779C>A 5780A>G), N1465V (5782A>G 5783A>T), H1467V (5788C>G 5789A>T 5790C>G), D1469N (5794G>A), Q1470K (5797C>A 5799A>G), S1472P (5803A>C 5804G>C), G1475R (5812G>A 5814A>G), N1477D (5818A>G 5820C>T), T1478D (5821A>G 5822C>A), I1479L (5824A>T 5826T>G), V1480F (5827G>T 5829C>T), Q1481D (5830C>G 5832G>T), R1482Q (5834G>A 5835C>A), I1483L (5836A>C), G1484R (5839G>C 5841T>A), N1485R (5843A>G 5844T>G), K1487S (5849A>G 5850G>C), V1488Y (5851G>T 5852T>A 5853A>T), F1492I (5863T>A 5865C>T), K1495R (5872A>C 5873A>G 5874G>T), S1496C (5875A>T), F1498Y (5882T>A), V1501L (5890G>C 5892C>T), T1502S (5894C>G), M1503V (5896A>G), D1504K (5899G>A 5901T>G), E1506S (5905G>A 5906A>G 5907A>T), S1507D (5908T>G 5909C>A 5910A>T), I1508V (5911A>G 5913A>T), W1510K (5917T>A 5918G>A 5919G>A), T1511M (5921C>T 5922A>G), L1514R (5930T>G 5931C>A), P1516R (5935C>A 5936C>G 5937A>G), D1517Y (5938G>T), L1519H (5944T>C 5945T>A 5946G>T), Y1520F (5948A>T 5949C>T), W1522F (5954G>T 5955G>C), P1526S (5965C>T 5967A>C), K1530T (5978A>C 5979A>T), P1533S (5986C>T 5988A>G), A1534N (5989G>A 5990C>A 5991A>T), Q1537I (5998C>A 5999A>T 6000G>A), R1538D (6001A>G 6002G>A 6003A>T), K1539L (6004A>T 6005A>T 6006A>G), D1541N (6010G>A), N1542R (6014A>G 6015T>G), C1543V (6016T>G 6017G>T 6018C>T), G1546P (6025G>C 6026G>C), T1547F (6028A>T 6029C>T 6030G>C), E1548L (6031G>C 6032A>T), E1549D (6036A>C), E1549_F1550insI (6036_6037insATA), I1551V (6040A>G), A1552V (6044C>T 6045T>G), Y1554F (6050A>T 6051C>T), I1558V (6061A>G 6063T>A), V1560I (6067G>A 6069C>A), F1561Y (6071T>A), E1563R (6076G>C 6077A>G), H1565R (6082C>A 6083A>G 6084T>G), K1566E (6085A>G), Q1567E (6088C>G), E1569L (6094G>T 6095A>T 6096A>G) | | | | | | | | |
| Codon mutations: | CTC1457ATT (5758C>A 5760C>T), GAT1458GAC (5763T>C), AGT1461CAA (5770A>C 5771G>A 5772T>A), CTG1462ATT (5773C>A 5775G>T), AAC1463AAT (5778C>T), CAG1464AGG (5779C>A 5780A>G), AAC1465GTC (5782A>G 5783A>T), CAC1467GTG (5788C>G 5789A>T 5790C>G), GAT1469AAT (5794G>A), CAA1470AAG (5797C>A 5799A>G), AGC1472CCC (5803A>C 5804G>C), CTA1473CTT (5808A>T), CCA1474CCC (5811A>C), GGA1475AGG (5812G>A 5814A>G), ATC1476ATT (5817C>T), AAC1477GAT (5818A>G 5820C>T), ACT1478GAT (5821A>G 5822C>A), ATT1479TTG (5824A>T 5826T>G), GTC1480TTT (5827G>T 5829C>T), CAG1481GAT (5830C>G 5832G>T), CGC1482CAA (5834G>A 5835C>A), ATA1483CTA (5836A>C), GGT1484CGA (5839G>C 5841T>A), AAT1485AGG (5843A>G 5844T>G), AAG1487AGC (5849A>G 5850G>C), GTA1488TAT (5851G>T 5852T>A 5853A>T), TTC1489TTT (5856C>T), TCA1490TCT (5859A>T), AAA1491AAG (5862A>G), TTC1492ATT (5863T>A 5865C>T), AAG1495CGT (5872A>C 5873A>G 5874G>T), AGT1496TGT (5875A>T), GGC1497GGA (5880C>A), TTT1498TAT (5882T>A), CAT1499CAC (5886T>C), GTC1501CTT (5890G>C 5892C>T), ACC1502AGC (5894C>G), ATG1503GTG (5896A>G), GAT1504AAG (5899G>A 5901T>G), GAA1505GAG (5904A>G), GAA1506AGT (5905G>A 5906A>G 5907A>T), TCA1507GAT (5908T>G 5909C>A 5910A>T), ATA1508GTT (5911A>G 5913A>T), CCA1509CCG (5916A>G), TGG1510AAA (5917T>A 5918G>A 5919G>A), ACA1511ATG (5921C>T 5922A>G), GCA1512GCT (5925A>T), CTC1514CGA (5930T>G 5931C>A), CCA1516AGG (5935C>A 5936C>G 5937A>G), GAT1517TAT (5938G>T), TTG1519CAT (5944T>C 5945T>A 5946G>T), TAC1520TTT (5948A>T 5949C>T), TGG1522TTC (5954G>T 5955G>C), CTA1523CTT (5958A>T), GTT1524GTC (5961T>C), CCA1526TCC (5965C>T 5967A>C), CTT1529TTG (5974C>T 5976T>G), AAA1530ACT (5978A>C 5979A>T), CCA1533TCG (5986C>T 5988A>G), GCA1534AAT (5989G>A 5990C>A 5991A>T), TTC1536TTT (5997C>T), CAG1537ATA (5998C>A 5999A>T 6000G>A), AGA1538GAT (6001A>G 6002G>A 6003A>T), AAA1539TTG (6004A>T 6005A>T 6006A>G), GAT1541AAT (6010G>A), AAT1542AGG (6014A>G 6015T>G), TGC1543GTT (6016T>G 6017G>T 6018C>T), TTT1544TTC (6021T>C), AAA1545AAG (6024A>G), GGT1546CCT (6025G>C 6026G>C), ACG1547TTC (6028A>T 6029C>T 6030G>C), GAA1548CTA (6031G>C 6032A>T), GAA1549GAC (6036A>C), GAA1549_TTC1550insATA (6036_6037insATA), TTC1550TTT (6039C>T), ATA1551GTA (6040A>G), GCT1552GTG (6044C>T 6045T>G), GTA1553GTG (6048A>G), TAC1554TTT (6050A>T 6051C>T), ATC1555ATT (6054C>T), GAT1556GAC (6057T>C), ATT1558GTA (6061A>G 6063T>A), GTC1560ATA (6067G>A 6069C>A), TTC1561TAC (6071T>A), GAA1563CGA (6076G>C 6077A>G), CAT1565AGG (6082C>A 6083A>G 6084T>G), AAA1566GAA (6085A>G), CAG1567GAG (6088C>G), CAT1568CAC (6093T>C), GAA1569TTG (6094G>T 6095A>T 6096A>G) | | | | | | | | |

##### Proteins

| | Begin | End | Coverage | Score | Concordance | Matches | Identities | I/D/M/F* | Stop Codons |
|---|---|---|---|---|---|---|---|---|---|
| polyprotein (YP_009552737.1) | 1457 | 1569 | 5.8% | 350 | 42.6% | 113 (99.1%) | 48 (42.1%) | 1/0/0/0 | 0 |
| Protein mutations: | L1457I (5758C>A 5760C>T), S1461Q (5770A>C 5771G>A 5772T>A), L1462I (5773C>A 5775G>T), Q1464R (5779C>A 5780A>G), N1465V (5782A>G 5783A>T), H1467V (5788C>G 5789A>T 5790C>G), D1469N (5794G>A), Q1470K (5797C>A 5799A>G), S1472P (5803A>C 5804G>C), G1475R (5812G>A 5814A>G), N1477D (5818A>G 5820C>T), T1478D (5821A>G 5822C>A), I1479L (5824A>T 5826T>G), V1480F (5827G>T 5829C>T), Q1481D (5830C>G 5832G>T), R1482Q (5834G>A 5835C>A), I1483L (5836A>C), G1484R (5839G>C 5841T>A), N1485R (5843A>G 5844T>G), K1487S (5849A>G 5850G>C), V1488Y (5851G>T 5852T>A 5853A>T), F1492I (5863T>A 5865C>T), K1495R (5872A>C 5873A>G 5874G>T), S1496C (5875A>T), F1498Y (5882T>A), V1501L (5890G>C 5892C>T), T1502S (5894C>G), M1503V (5896A>G), D1504K (5899G>A 5901T>G), E1506S (5905G>A 5906A>G 5907A>T), S1507D (5908T>G 5909C>A 5910A>T), I1508V (5911A>G 5913A>T), W1510K (5917T>A 5918G>A 5919G>A), T1511M (5921C>T 5922A>G), L1514R (5930T>G 5931C>A), P1516R (5935C>A 5936C>G 5937A>G), D1517Y (5938G>T), L1519H (5944T>C 5945T>A 5946G>T), Y1520F (5948A>T 5949C>T), W1522F (5954G>T 5955G>C), P1526S (5965C>T 5967A>C), K1530T (5978A>C 5979A>T), P1533S (5986C>T 5988A>G), A1534N (5989G>A 5990C>A 5991A>T), Q1537I (5998C>A 5999A>T 6000G>A), R1538D (6001A>G 6002G>A 6003A>T), K1539L (6004A>T 6005A>T 6006A>G), D1541N (6010G>A), N1542R (6014A>G 6015T>G), C1543V (6016T>G 6017G>T 6018C>T), G1546P (6025G>C 6026G>C), T1547F (6028A>T 6029C>T 6030G>C), E1548L (6031G>C 6032A>T), E1549D (6036A>C), E1549_F1550insI (6036_6037insATA), I1551V (6040A>G), A1552V (6044C>T 6045T>G), Y1554F (6050A>T 6051C>T), I1558V (6061A>G 6063T>A), V1560I (6067G>A 6069C>A), F1561Y (6071T>A), E1563R (6076G>C 6077A>G), H1565R (6082C>A 6083A>G 6084T>G), K1566E (6085A>G), Q1567E (6088C>G), E1569L (6094G>T 6095A>T 6096A>G) | | | | | | | | |
| Codon mutations: | CTC1457ATT (5758C>A 5760C>T), GAT1458GAC (5763T>C), AGT1461CAA (5770A>C 5771G>A 5772T>A), CTG1462ATT (5773C>A 5775G>T), AAC1463AAT (5778C>T), CAG1464AGG (5779C>A 5780A>G), AAC1465GTC (5782A>G 5783A>T), CAC1467GTG (5788C>G 5789A>T 5790C>G), GAT1469AAT (5794G>A), CAA1470AAG (5797C>A 5799A>G), AGC1472CCC (5803A>C 5804G>C), CTA1473CTT (5808A>T), CCA1474CCC (5811A>C), GGA1475AGG (5812G>A 5814A>G), ATC1476ATT (5817C>T), AAC1477GAT (5818A>G 5820C>T), ACT1478GAT (5821A>G 5822C>A), ATT1479TTG (5824A>T 5826T>G), GTC1480TTT (5827G>T 5829C>T), CAG1481GAT (5830C>G 5832G>T), CGC1482CAA (5834G>A 5835C>A), ATA1483CTA (5836A>C), GGT1484CGA (5839G>C 5841T>A), AAT1485AGG (5843A>G 5844T>G), AAG1487AGC (5849A>G 5850G>C), GTA1488TAT (5851G>T 5852T>A 5853A>T), TTC1489TTT (5856C>T), TCA1490TCT (5859A>T), AAA1491AAG (5862A>G), TTC1492ATT (5863T>A 5865C>T), AAG1495CGT (5872A>C 5873A>G 5874G>T), AGT1496TGT (5875A>T), GGC1497GGA (5880C>A), TTT1498TAT (5882T>A), CAT1499CAC (5886T>C), GTC1501CTT (5890G>C 5892C>T), ACC1502AGC (5894C>G), ATG1503GTG (5896A>G), GAT1504AAG (5899G>A 5901T>G), GAA1505GAG (5904A>G), GAA1506AGT (5905G>A 5906A>G 5907A>T), TCA1507GAT (5908T>G 5909C>A 5910A>T), ATA1508GTT (5911A>G 5913A>T), CCA1509CCG (5916A>G), TGG1510AAA (5917T>A 5918G>A 5919G>A), ACA1511ATG (5921C>T 5922A>G), GCA1512GCT (5925A>T), CTC1514CGA (5930T>G 5931C>A), CCA1516AGG (5935C>A 5936C>G 5937A>G), GAT1517TAT (5938G>T), TTG1519CAT (5944T>C 5945T>A 5946G>T), TAC1520TTT (5948A>T 5949C>T), TGG1522TTC (5954G>T 5955G>C), CTA1523CTT (5958A>T), GTT1524GTC (5961T>C), CCA1526TCC (5965C>T 5967A>C), CTT1529TTG (5974C>T 5976T>G), AAA1530ACT (5978A>C 5979A>T), CCA1533TCG (5986C>T 5988A>G), GCA1534AAT (5989G>A 5990C>A 5991A>T), TTC1536TTT (5997C>T), CAG1537ATA (5998C>A 5999A>T 6000G>A), AGA1538GAT (6001A>G 6002G>A 6003A>T), AAA1539TTG (6004A>T 6005A>T 6006A>G), GAT1541AAT (6010G>A), AAT1542AGG (6014A>G 6015T>G), TGC1543GTT (6016T>G 6017G>T 6018C>T), TTT1544TTC (6021T>C), AAA1545AAG (6024A>G), GGT1546CCT (6025G>C 6026G>C), ACG1547TTC (6028A>T 6029C>T 6030G>C), GAA1548CTA (6031G>C 6032A>T), GAA1549GAC (6036A>C), GAA1549_TTC1550insATA (6036_6037insATA), TTC1550TTT (6039C>T), ATA1551GTA (6040A>G), GCT1552GTG (6044C>T 6045T>G), GTA1553GTG (6048A>G), TAC1554TTT (6050A>T 6051C>T), ATC1555ATT (6054C>T), GAT1556GAC (6057T>C), ATT1558GTA (6061A>G 6063T>A), GTC1560ATA (6067G>A 6069C>A), TTC1561TAC (6071T>A), GAA1563CGA (6076G>C 6077A>G), CAT1565AGG (6082C>A 6083A>G 6084T>G), AAA1566GAA (6085A>G), CAG1567GAG (6088C>G), CAT1568CAC (6093T>C), GAA1569TTG (6094G>T 6095A>T 6096A>G) | | | | | | | | |

*: Inserts / Deletes / Misaligned / Frameshifts

#### Analysis details

This analysis was performed with panviral2.64

#### NGS Details (UN62): Badnavirus zetavirgamusae

##### Assembly

| | |
|---|---|
| **Coverage Length** | 471 (1 contig(s)) |
| **Depth Of Coverage** | 8.4 |
| **Number Of Reads** | 29 |
| **Reads Per Million** | 0.67 rpm (after QC) |
| **Ambiguities** | 0 |
| **Assembly Method** | de novo + reference guided assembly |
| **Consensus Caller** | Bcf Tools |

##### Coverage Map

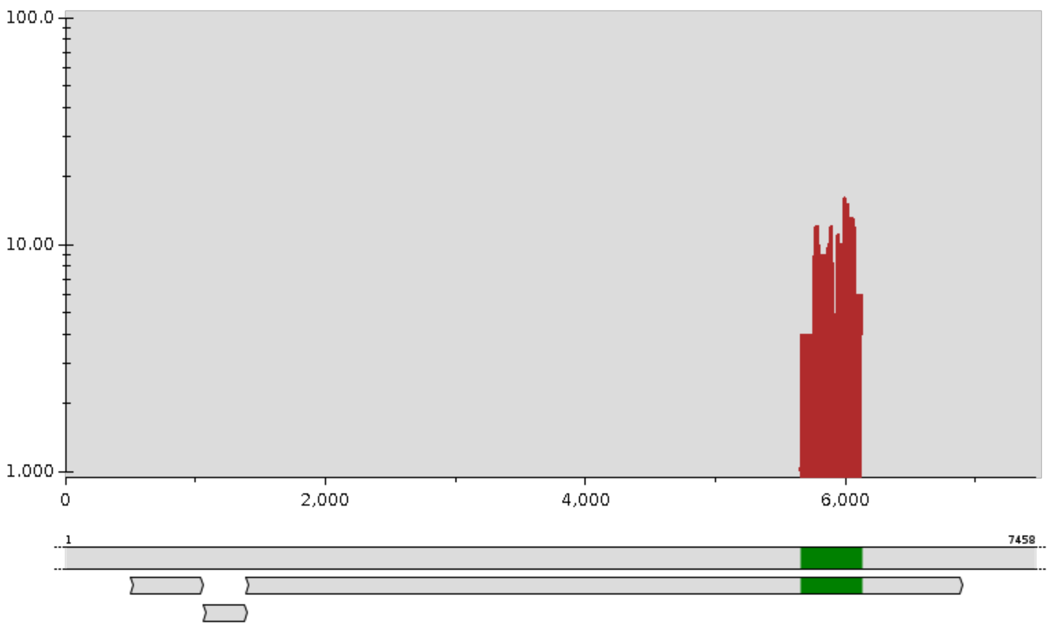

##### Assignment

| | |
|---|---|
| **Type** | Badnavirus zetavirgamusae (Taxonomy ID: 3047391) |
| **Reference Genome** | NC_015503.1 |
| **NT Identity (%)** | 53.8627 |
| **AA Identity (%)** | 40.3846 |
| **Number Of Stop Codons** | 3 |
| **Number Of CDS** | 3 |

##### Alignment

| | |
|---|---|
| **Alignment Score** | 40.0 (NT) + 133.0 (AA) = 173.0 |
| **Concordance (%)** | 11.9048 |

| Alignment Method | Global, seeded, nucleotide + amino acids (AGA) |
|---|---|

#### Genome Region

Sequence starts at position 5657 and ends at position 6127 relative to NC_015503.1 reference sequence.

#### Alignment Detailed Statistics

| | Begin | End | Coverage | Score | Concordance | Matches | Identities | I/D/M/F* | Stop Codons |
|---|---|---|---|---|---|---|---|---|---|
| **NT** | **5657** | **6127** | **6.3%** | **40** | **4.4%** | **466 (98.9%)** | **251 (53.3%)** | **0/5** | |
| Mutations: | 5657G>A, 5658T>C, 5660T>G, 5662A>C, 5664A>G, 5666G>A, 5667A>T, 5669G>T, 5670G>A, 5671A>G, 5673A>C, 5675C>A, 5677G>T, 5679A>G, 5680A>T, 5681G>A, 5683A>G, 5684G>T, 5690A>T, 5691T>A, 5696T>A, 5697C>G, 5699C>A, 5703C>A, 5704C>A, 5707C>A, 5708A>G, 5711G>A, 5717G>A, 5718A>T, 5720C>A, 5721G>T, 5723A>T, 5724G>A, 5725T>A, 5726T>A, 5728A>C, 5729G>C, 5730G>C, 5731T>A, 5732A>T, 5733G>A, 5736G>A, 5741T>C, 5742C>T, 5746G>A, 5747A>G, 5749G>T, 5750C>T, 5752C>T, 5753T>A, 5754G>T, 5756T>A, 5759T>A, 5763A>T, 5764A>G, 5765T>A, 5766C>A, 5768C>G, 5769C>A, 5771G>A, 5772G>A, 5774T>A, 5781delC, 5783A>C, 5789T>A, 5791T>C, 5792G>T, 5793C>A, 5799A>G, 5801A>C, 5802G>T, 5804G>A, 5807T>G, 5808T>A, 5810T>G, 5811G>C, 5812A>C, 5813T>A, 5814G>T, 5817G>A, 5818A>T, 5819A>T, 5821A>G, 5823_5825delCTG, 5828G>A, 5829A>C, 5830C>A, 5831A>C, 5834G>A, 5836A>G, 5838G>A, 5839G>T, 5841C>A, 5842T>C, 5843C>A, 5847T>A, 5848C>T, 5849C>T, 5850T>C, 5851G>T, 5852G>T, 5853C>A, 5855G>A, 5857C>G, 5858A>T, 5859A>T, 5861T>A, 5862C>A, 5864C>A, 5873C>T, 5876G>A, 5877G>A, 5878G>A, 5879A>T, 5880C>T, 5881A>T, 5882C>T, 5885C>A, 5889A>G, 5892A>T, 5894G>A, 5903C>A, 5904C>G, 5909T>A, 5915A>C, 5919C>T, 5921C>T, 5924A>G, 5927T>A, 5928T>G, 5929C>G, 5930G>A, 5931G>A, 5932A>G, 5933A>T, 5939G>A, 5940G>T, 5942G>A, 5945A>G, 5946A>T, 5947G>A, 5948A>T, 5951G>T, 5955T>A, 5956C>A, 5963C>T, 5964T>A, 5965G>T, 5966G>C, 5970A>T, 5973A>G, 5976A>C, 5977G>A, 5984G>C, 5987G>A, 5990A>T, 5991G>A, 5996C>T, 5998G>A, 6001G>A, 6002G>A, 6005G>C, 6006delC, 6007C>T, 6008C>A, 6009A>G, 6011A>T, 6012A>T, 6017A>T, 6023C>T, 6025C>T, 6029G>A, 6030G>T, 6031A>C, 6032T>C, 6034C>G, 6035A>T, 6038C>T, 6039A>T, 6042G>A, 6044C>A, 6053G>A, 6059C>G, 6060T>A, 6061C>G, 6062T>C, 6063A>T, 6064T>C, 6065G>C, 6067A>C, 6068T>A, 6070G>A, 6079G>C, 6083G>T, 6085G>T, 6086T>G, 6087T>A, 6089T>A, 6090T>G, 6091G>T, 6092G>C, 6095G>A, 6096A>C, 6097A>C, 6101C>T, 6105G>T, 6106C>A, 6108G>A, 6109A>G, 6111C>A, 6112C>A, 6113A>T, 6115G>A, 6117A>G, 6118G>A, 6125G>A, 6126C>A | | | | | | | | |

##### CDS

| | Begin | End | Coverage | Score | Concordance | Matches | Identities | I/D/M/F* | Stop Codons |
|---|---|---|---|---|---|---|---|---|---|
| BaSUIV_gp3 | 1422 | 1578 | 8.5% | 133 | 12.5% | 156 (99.4%) | 63 (40.1%) | 0/1/2/2 | 3 |
| Protein mutations: | Y1422Q (5658T>C 5660T>G), K1423T (5662A>C), M1424V (5664A>G 5666G>A), M1425F (5667A>T 5669G>T), E1426R (5670G>A 5671A>G), I1427L (5673A>C 5675C>A), C1428F (5677G>T), K1429V (5679A>G 5680A>T 5681G>A), K1430S (5683A>G 5684G>T), L1433I (5691T>A), L1435V (5697C>G 5699C>A), P1437K (5703C>A 5704C>A), T1438K (5707C>A 5708A>G), I1442L (5718A>T 5720C>A), G1443C (5721G>T 5723A>T), V1444K (5724G>A 5725T>A 5726T>A), K1445T (5728A>C 5729G>C), V1446H (5730G>C 5731T>A 5732A>T), V1447I (5733G>A), D1448N (5736G>A), G1451E (5746G>A 5747A>G), S1452I (5749G>T 5750C>T), T1453I (5752C>T 5753T>A), V1454L (5754G>T 5756T>A), N1457* (5763A>T 5764A>G 5765T>A), H1458K (5766C>A 5768C>G), L1459I (5769C>A 5771G>A), D1460K (5772G>A 5774T>A), V1466A (5791T>C 5792G>T), Q1467K (5793C>A), I1469V (5799A>G 5801A>C), V1470L (5802G>T 5804G>A), D1471E (5807T>G), F1472M (5808T>A 5810T>G), D1473P (5811G>C 5812A>C 5813T>A), E1474* (5814G>T), E1475I (5817G>A 5818A>T 5819A>T), K1476R (5821A>G), L1477del (5823_5825delCTG), T1479H (5829A>C 5830C>A 5831A>C), K1481R (5836A>G), G1482I (5838G>A 5839G>T), L1483T (5841C>A 5842T>C 5843C>A), S1485I (5847T>A 5848C>T 5849C>T), W1486L (5850T>C 5851G>T 5852G>T), L1487I (5853C>A 5855G>A), A1488G (5857C>G 5858A>T), I1489L (5859A>T 5861T>A), L1490I (5862C>A 5864C>A), G1495N (5877G>A 5878G>A 5879A>T), H1496F (5880C>T 5881A>T 5882C>T), N1499D (5889A>G), M1500L (5892A>T 5894G>A), L1504V (5904C>G), P1509S (5919C>T 5921C>T), S1512G (5928T>G 5929C>G 5930G>A), E1513S (5931G>A 5932A>G 5933A>T), E1516* (5940G>T 5942G>A), R1518Y (5946A>T 5947G>A 5948A>T), L1519F (5951G>T), S1521N (5955T>A 5956C>A), W1524I (5964T>A 5965G>T 5966G>C), I1526L (5970A>T), I1527V (5973A>G), R1528Q (5976A>C 5977G>A), M1530I (5984G>C), E1532D (5990A>T), E1533K (5991G>A), R1535K (5998G>A), R1536K (6001G>A 6002G>A), L1537F (6005G>C), K1539D (6009A>G 6011A>T), M1540L (6012A>T), P1544L (6025C>T), D1546S (6030G>T 6031A>C 6032T>C), A1547G (6034C>G 6035A>T), I1549L (6039A>T), V1550I (6042G>A 6044C>A), M1557S (6063A>T 6064T>C 6065G>C), N1558T (6067A>C 6068T>A), G1559E (6070G>A), G1562A (6079G>C), C1564L (6085G>T 6086T>G), Y1565K (6087T>A 6089T>A), W1566V (6090T>G 6091G>T 6092G>C), K1568P (6096A>C 6097A>C), A1571Y (6105G>T 6106C>A), D1572S (6108G>A 6109A>G), P1573N (6111C>A 6112C>A 6113A>T), R1574K (6115G>A), S1575D (6117A>G 6118G>A) | | | | | | | | |
| Codon mutations: | TTG1421..A (5657G>A), TAT1422CAG (5658T>C 5660T>G), AAA1423ACA (5662A>C), ATG1424GTA (5664A>G 5666G>A), ATG1425TTT (5667A>T 5669G>T), GAA1426AGA (5670G>A 5671A>G), ATC1427CTA (5673A>C 5675C>A), TGT1428TTT (5677G>T), AAG1429GTA (5679A>G 5680A>T 5681G>A), AAG1430AGT (5683A>G 5684G>T), GGA1432GGT (5690A>T), TTA1433ATA (5691T>A), ATT1434ATA (5696T>A), CTC1435GTA (5697C>G 5699C>A), CCA1437AAA (5703C>A 5704C>A), ACA1438AAG (5707C>A 5708A>G), AAG1439AAA (5711G>A), AAG1441AAA (5717G>A), ATC1442TTA (5718A>T 5720C>A), GGA1443TGT (5721G>T 5723A>T), GTT1444AAA (5724G>A 5725T>A 5726T>A), AAG1445ACC (5728A>C 5729G>C), GTA1446CAT (5730G>C 5731T>A 5732A>T), GTA1447ATA (5733G>A), GAT1448AAT (5736G>A), TTT1449TTC (5741T>C), CTA1450TTA (5742C>T), GGA1451GAG (5746G>A 5747A>G), AGC1452ATT (5749G>T 5750C>T), ACT1453ATA (5752C>T 5753T>A), GTT1454TTA (5754G>T 5756T>A), GGT1455GGA (5759T>A), AAT1457TGA (5763A>T 5764A>G 5765T>A), CAC1458AAG (5766C>A 5768C>G), CTG1459ATA (5769C>A 5771G>A), GAT1460AAA (5772G>A 5774T>A), CCA1463-CC (5781delC 5783A>C), ATT1465ATA (5789T>A), GTG1466GCT (5791T>C 5792G>T), CAA1467AAA (5793C>A), ATA1469GTC (5799A>G 5801A>C), GTG1470TTA (5802G>T 5804G>A), GAT1471GAG (5807T>G), TTT1472ATG (5808T>A 5810T>G), GAT1473CCA (5811G>C 5812A>C 5813T>A), GAA1474TAA (5814G>T), GAA1475ATT (5817G>A 5818A>T 5819A>T), AAA1476AGA (5821A>G), CTG1477del (5823_5825delCTG), AAG1478AAA (5828G>A), ACA1479CAC (5829A>C 5830C>A 5831A>C), AAG1480AAA (5834G>A), AAA1481AGA (5836A>G), GGT1482ATT (5838G>A 5839G>T), CTC1483ACA (5841C>A 5842T>C 5843C>A), TCC1485ATT (5847T>A 5848C>T 5849C>T), TGG1486CTT (5850T>C 5851G>T 5852G>T), CTG1487ATA (5853C>A 5855G>A), GCA1488GGT (5857C>G 5858A>T), ATT1489TTA (5859A>T 5861T>A), CTC1490ATA (5862C>A 5864C>A), GCC1493GCT (5873C>T), AGG1494AGA (5876G>A), GGA1495AAT (5877G>A 5878G>A 5879A>T), CAC1496TTT (5880C>T 5881A>T 5882C>T), ATC1497ATA (5885C>A), AAC1499GAC (5889A>G), ATG1500TTA (5892A>T 5894G>A), ATC1503ATA (5903C>A), CTA1504GTA (5904C>G), GGT1505GGA (5909T>A), CTA1507CTC (5915A>C), CCC1509TCT (5919C>T 5921C>T), AAA1510AAG (5924A>G), ACT1511ACA (5927T>A), TCG1512GGA (5928T>G 5929C>G 5930G>A), GAA1513AGT (5931G>A 5932A>G 5933A>T), GGG1515GGA (5939G>A), GAG1516TAA (5940G>T 5942G>A), AAA1517AAG (5945A>G), AGA1518TAT (5946A>T 5947G>A 5948A>T), TTG1519TTT (5951G>T), TCT1521AAT (5955T>A 5956C>A), GAC1523GAT (5963C>T), TGG1524ATC (5964T>A 5965G>T 5966G>C), ATA1526TTA (5970A>T), ATC1527GTC (5973A>G), AGA1528CAA (5976A>C 5977G>A), ATG1530ATC (5984G>C), AAG1531AAA (5987G>A), GAA1532GAT (5990A>T), GAA1533AAA (5991G>A), GTC1534GTT (5996C>T), AGA1535AAA (5998G>A), AGG1536AAA (6001G>A 6002G>A), TTG1537TTC (6005G>C), CCC1538-TA (6006delC 6007C>T 6008C>A), AAA1539GAT (6009A>G 6011A>T), ATG1540TTG (6012A>T), GCA1541GCT (6017A>T), CCC1543CCT (6023C>T), CCA1544CTA (6025C>T), GAG1545GAA (6029G>A), GAT1546TCC (6030G>T 6031A>C 6032T>C), GCA1547GGT (6034C>G 6035A>T), TAC1548TAT (6038C>T), ATA1549TTA (6039A>T), GTC1550ATA (6042G>A 6044C>A), ACG1553ACA (6053G>A), GGC1555GGG (6059C>G), TCT1556AGC (6060T>A 6061C>G 6062T>C), ATG1557TCC (6063A>T 6064T>C 6065G>C), AAT1558ACA (6067A>C 6068T>A), GGA1559GAA (6070G>A), GGA1562GCA (6079G>C), GTG1563GTT (6083G>T), TGT1564TTG (6085G>T 6086T>G), TAT1565AAA (6087T>A 6089T>A), TGG1566GTC (6090T>G 6091G>T 6092G>C), AAG1567AAA (6095G>A), AAA1568CCA (6096A>C 6097A>C), AGC1569AGT (6101C>T), GCT1571TAT (6105G>T 6106C>A), GAT1572AGT (6108G>A 6109A>G), CCA1573AAT (6111C>A 6112C>A 6113A>T), AGA1574AAA (6115G>A), AGT1575GAT (6117A>G 6118G>A), GAG1577GAA (6125G>A), CAA1578AA. (6126C>A) | | | | | | | | |

##### Proteins

| | Begin | End | Coverage | Score | Concordance | Matches | Identities | I/D/M/F* | Stop Codons |
|---|---|---|---|---|---|---|---|---|---|
| polyprotein (YP_004442827.1) | 1422 | 1578 | 8.5% | 133 | 12.5% | 156 (99.4%) | 63 (40.1%) | 0/1/2/2 | 3 |
| Protein mutations: | Y1422Q (5658T>C 5660T>G), K1423T (5662A>C), M1424V (5664A>G 5666G>A), M1425F (5667A>T 5669G>T), E1426R (5670G>A 5671A>G), I1427L (5673A>C 5675C>A), C1428F (5677G>T), K1429V (5679A>G 5680A>T 5681G>A), K1430S (5683A>G 5684G>T), L1433I (5691T>A), L1435V (5697C>G 5699C>A), P1437K (5703C>A 5704C>A), T1438K (5707C>A 5708A>G), I1442L (5718A>T 5720C>A), G1443C (5721G>T 5723A>T), V1444K (5724G>A 5725T>A 5726T>A), K1445T (5728A>C 5729G>C), V1446H (5730G>C 5731T>A 5732A>T), V1447I (5733G>A), D1448N (5736G>A), G1451E (5746G>A 5747A>G), S1452I (5749G>T 5750C>T), T1453I (5752C>T 5753T>A), V1454L (5754G>T 5756T>A), N1457* (5763A>T 5764A>G 5765T>A), H1458K (5766C>A 5768C>G), L1459I (5769C>A 5771G>A), D1460K (5772G>A 5774T>A), V1466A (5791T>C 5792G>T), Q1467K (5793C>A), I1469V (5799A>G 5801A>C), V1470L (5802G>T 5804G>A), D1471E (5807T>G), F1472M (5808T>A 5810T>G), D1473P (5811G>C 5812A>C 5813T>A), E1474* (5814G>T), E1475I (5817G>A 5818A>T 5819A>T), K1476R (5821A>G), L1477del (5823_5825delCTG), T1479H (5829A>C 5830C>A 5831A>C), K1481R (5836A>G), G1482I (5838G>A 5839G>T), L1483T (5841C>A 5842T>C 5843C>A), S1485I (5847T>A 5848C>T 5849C>T), W1486L (5850T>C 5851G>T 5852G>T), L1487I (5853C>A 5855G>A), A1488G (5857C>G 5858A>T), I1489L (5859A>T 5861T>A), L1490I (5862C>A 5864C>A), G1495N (5877G>A 5878G>A 5879A>T), H1496F (5880C>T 5881A>T 5882C>T), N1499D (5889A>G), M1500L (5892A>T 5894G>A), L1504V (5904C>G), P1509S (5919C>T 5921C>T), S1512G (5928T>G 5929C>G 5930G>A), E1513S (5931G>A 5932A>G 5933A>T), E1516* (5940G>T 5942G>A), R1518Y (5946A>T 5947G>A 5948A>T), L1519F (5951G>T), S1521N (5955T>A 5956C>A), W1524I (5964T>A 5965G>T 5966G>C), I1526L (5970A>T), I1527V (5973A>G), R1528Q (5976A>C 5977G>A), M1530I (5984G>C), E1532D (5990A>T), E1533K (5991G>A), R1535K (5998G>A), R1536K (6001G>A 6002G>A), L1537F (6005G>C), K1539D (6009A>G 6011A>T), M1540L (6012A>T), P1544L (6025C>T), D1546S (6030G>T 6031A>C 6032T>C), A1547G (6034C>G 6035A>T), I1549L (6039A>T), V1550I (6042G>A 6044C>A), M1557S (6063A>T 6064T>C 6065G>C), N1558T (6067A>C 6068T>A), G1559E (6070G>A), G1562A (6079G>C), C1564L (6085G>T 6086T>G), Y1565K (6087T>A 6089T>A), W1566V (6090T>G 6091G>T 6092G>C), K1568P (6096A>C 6097A>C), A1571Y (6105G>T 6106C>A), D1572S (6108G>A 6109A>G), P1573N (6111C>A 6112C>A 6113A>T), R1574K (6115G>A), S1575D (6117A>G 6118G>A) | | | | | | | | |

| | Begin | End | Coverage | Score | Concordance | Matches | Identities | I/D/M/F* | Stop Codons |
|---|---|---|---|---|---|---|---|---|---|
| **NT** | **5657** | **6127** | **6.3%** | **40** | **4.4%** | **466 (98.9%)** | **251 (53.3%)** | **0/5** | |
| Codon mutations: | TTG1421..A (5657G>A), TAT1422CAG (5658T>C 5660T>G), AAA1423ACA (5662A>C), ATG1424GTA (5664A>G 5666G>A), ATG1425TTT (5667A>T 5669G>T), GAA1426AGA (5670G>A 5671A>G), ATC1427CTA (5673A>C 5675C>A), TGT1428TTT (5677G>T), AAG1429GTA (5679A>G 5680A>T 5681G>A), AAG1430AGT (5683A>G 5684G>T), GGA1432GGT (5690A>T), TTA1433ATA (5691T>A), ATT1434ATA (5696T>A), CTC1435GTA (5697C>G 5699C>A), CCA1437AAA (5703C>A 5704C>A), ACA1438AAG (5707C>A 5708A>G), AAG1439AAA (5711G>A), AAG1441AAA (5717G>A), ATC1442TTA (5718A>T 5720C>A), GGA1443TGT (5721G>T 5723A>T), GTT1444AAA (5724G>A 5725T>A 5726T>A), AAG1445ACC (5728A>C 5729G>C), GTA1446CAT (5730G>C 5731T>A 5732A>T), GTA1447ATA (5733G>A), GAT1448AAT (5736G>A), TTT1449TTC (5741T>C), CTA1450TTA (5742C>T), GGA1451GAG (5746G>A 5747A>G), AGC1452ATT (5749G>T 5750C>T), ACT1453ATA (5752C>T 5753T>A), GTT1454TTA (5754G>T 5756T>A), GGT1455GGA (5759T>A), AAT1457TGA (5763A>T 5764A>G 5765T>A), CAC1458AAG (5766C>A 5768C>G), CTG1459ATA (5769C>A 5771G>A), GAT1460AAA (5772G>A 5774T>A), CCA1463-CC (5781delC 5783A>C), ATT1465ATA (5789T>A), GTG1466GCT (5791T>C 5792G>T), CAA1467AAA (5793C>A), ATA1469GTC (5799A>G 5801A>C), GTG1470TTA (5802G>T 5804G>A), GAT1471GAG (5807T>G), TTT1472ATG (5808T>A 5810T>G), GAT1473CCA (5811G>C 5812A>C 5813T>A), GAA1474TAA (5814G>T), GAA1475ATT (5817G>A 5818A>T 5819A>T), AAA1476AGA (5821A>G), CTG1477del (5823_5825delCTG), AAG1478AAA (5828G>A), ACA1479CAC (5829A>C 5830C>A 5831A>C), AAG1480AAA (5834G>A), AAA1481AGA (5836A>G), GGT1482ATT (5838G>A 5839G>T), CTC1483ACA (5841C>A 5842T>C 5843C>A), TCC1485ATT (5847T>A 5848C>T 5849C>T), TGG1486CTT (5850T>C 5851G>T 5852G>T), CTG1487ATA (5853C>A 5855G>A), GCA1488GGT (5857C>G 5858A>T), ATT1489TTA (5859A>T 5861T>A), CTC1490ATA (5862C>A 5864C>A), GCC1493GCT (5873C>T), AGG1494AGA (5876G>A), GGA1495AAT (5877G>A 5878G>A 5879A>T), CAC1496TTT (5880C>T 5881A>T 5882C>T), ATC1497ATA (5885C>A), AAC1499GAC (5889A>G), ATG1500TTA (5892A>T 5894G>A), ATC1503ATA (5903C>A), CTA1504GTA (5904C>G), GGT1505GGA (5909T>A), CTA1507CTC (5915A>C), CCC1509TCT (5919C>T 5921C>T), AAA1510AAG (5924A>G), ACT1511ACA (5927T>A), TCG1512GGA (5928T>G 5929C>G 5930G>A), GAA1513AGT (5931G>A 5932A>G 5933A>T), GGG1515GGA (5939G>A), GAG1516TAA (5940G>T 5942G>A), AAA1517AAG (5945A>G), AGA1518TAT (5946A>T 5947G>A 5948A>T), TTG1519TTT (5951G>T), TCT1521AAT (5955T>A 5956C>A), GAC1523GAT (5963C>T), TGG1524ATC (5964T>A 5965G>T 5966G>C), ATA1526TTA (5970A>T), ATC1527GTC (5973A>G), AGA1528CAA (5976A>C 5977G>A), ATG1530ATC (5984G>C), AAG1531AAA (5987G>A), GAA1532GAT (5990A>T), GAA1533AAA (5991G>A), GTC1534GTT (5996C>T), AGA1535AAA (5998G>A), AGG1536AAA (6001G>A 6002G>A), TTG1537TTC (6005G>C), CCC1538-TA (6006delC 6007C>T 6008C>A), AAA1539GAT (6009A>G 6011A>T), ATG1540TTG (6012A>T), GCA1541GCT (6017A>T), CCC1543CCT (6023C>T), CCA1544CTA (6025C>T), GAG1545GAA (6029G>A), GAT1546TCC (6030G>T 6031A>C 6032T>C), GCA1547GGT (6034C>G 6035A>T), TAC1548TAT (6038C>T), ATA1549TTA (6039A>T), GTC1550ATA (6042G>A 6044C>A), ACG1553ACA (6053G>A), GGC1555GGG (6059C>G), TCT1556AGC (6060T>A 6061C>G 6062T>C), ATG1557TCC (6063A>T 6064T>C 6065G>C), AAT1558ACA (6067A>C 6068T>A), GGA1559GAA (6070G>A), GGA1562GCA (6079G>C), GTG1563GTT (6083G>T), TGT1564TTG (6085G>T 6086T>G), TAT1565AAA (6087T>A 6089T>A), TGG1566GTC (6090T>G 6091G>T 6092G>C), AAG1567AAA (6095G>A), AAA1568CCA (6096A>C 6097A>C), AGC1569AGT (6101C>T), GCT1571TAT (6105G>T 6106C>A), GAT1572AGT (6108G>A 6109A>G), CCA1573AAT (6111C>A 6112C>A 6113A>T), AGA1574AAA (6115G>A), AGT1575GAT (6117A>G 6118G>A), GAG1577GAA (6125G>A), CAA1578AA. (6126C>A) | | | | | | | | |

*: Inserts / Deletes / Misaligned / Frameshifts

#### Analysis details

This analysis was performed with panviral2.64

### NGS Details (UN62): Badnavirus maculasmallanthi

#### Assembly

| | |
|---|---|
| **Coverage Length** | 245 (1 contig(s)) |
| **Depth Of Coverage** | 6.2 |
| **Number Of Reads** | 15 |
| **Reads Per Million** | 0.35 rpm (after QC) |
| **Ambiguities** | 0 |
| **Assembly Method** | de novo + reference guided assembly |
| **Consensus Caller** | Bcf Tools |

#### Coverage Map

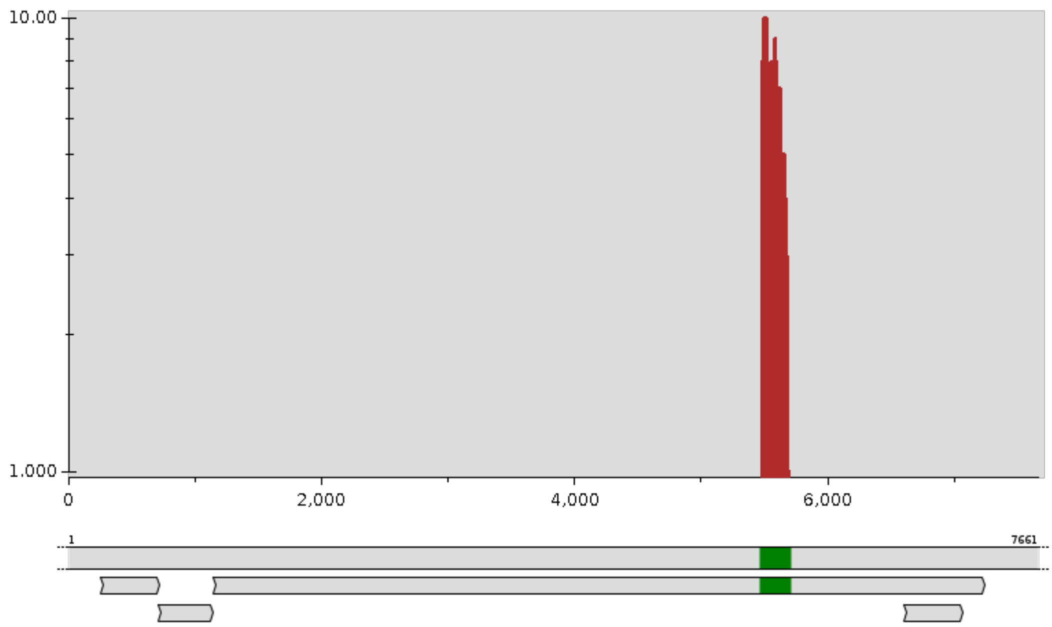

#### Assignment

| | |
|---|---|
| **Type** | Badnavirus maculasmallanthi (Taxonomy ID: 3048453) |
| **Reference Genome** | NC_026472.1 |
| **NT Identity (%)** | 57.1429 |
| **AA Identity (%)** | 48.7805 |
| **Number Of Stop Codons** | 2 |
| **Number Of CDS** | 4 |

#### Alignment

| | |
|---|---|
| **Alignment Score** | 70.0 (NT) + 301.0 (AA) = 371.0 |
| **Concordance (%)** | 34.9012 |

| Alignment Method | Global, seeded, nucleotide + amino acids (AGA) |
|---|---|

#### Genome Region

Sequence starts at position 5469 and ends at position 5713 relative to NC_026472.1 reference sequence.

#### Alignment Detailed Statistics

| | Begin | End | Coverage | Score | Concordance | Matches | Identities | I/D/M/F* | Stop Codons |
|---|---|---|---|---|---|---|---|---|---|
| **NT** | **5469** | **5713** | **3.2%** | **70** | **14.3%** | **245 (100%)** | **140 (57.1%)** | **0/0** | |
| Mutations: | 5476A>G, 5478T>C, 5481C>T, 5485A>G, 5487A>T, 5488C>T, 5490C>G, 5493T>C, 5494G>A, 5496C>A, 5497A>G, 5499T>C, 5503T>G, 5504A>T, 5505C>T, 5512C>A, 5516A>T, 5518T>C, 5520C>A, 5521C>A, 5527G>A, 5528G>T, 5529G>T, 5532A>C, 5533A>G, 5535C>A, 5536A>G, 5537C>A, 5538C>G, 5545A>G, 5547A>T, 5548A>G, 5549G>A, 5551A>T, 5553A>G, 5556G>T, 5557A>G, 5558A>G, 5559T>C, 5560G>T, 5562A>T, 5565G>A, 5566A>G, 5571C>T, 5574A>T, 5578T>A, 5583T>C, 5584C>T, 5587A>C, 5588A>G, 5589G>A, 5590T>G, 5591C>T, 5592T>A, 5595A>C, 5597T>A, 5598T>C, 5605G>A, 5608G>T, 5609C>G, 5610C>A, 5615A>C, 5616A>C, 5617G>C, 5618A>C, 5619A>G, 5620G>A, 5621A>C, 5623T>G, 5624C>A, 5626A>G, 5632T>A, 5633G>A, 5637G>A, 5638G>T, 5640C>A, 5645T>G, 5648T>C, 5649C>T, 5651C>A, 5652A>T, 5653G>T, 5654G>C, 5655A>C, 5658A>G, 5659T>A, 5660T>A, 5661A>C, 5664C>T, 5667A>G, 5669G>A, 5670G>T, 5671C>T, 5673T>A, 5676C>G, 5682A>C, 5688A>G, 5691C>A, 5693A>C, 5694A>T, 5698G>A, 5703C>T, 5704G>T, 5708T>G, 5709T>C | | | | | | | | |

##### CDS

| | Begin | End | Coverage | Score | Concordance | Matches | Identities | I/D/M/F* | Stop Codons |
|---|---|---|---|---|---|---|---|---|---|
| UF61_gp3 | 1443 | 1524 | 4.0% | 301 | 51.3% | 82 (100%) | 40 (48.8%) | 0/0/0/0 | 2 |
| Protein mutations: | N1445D (5476A>G 5478T>C), T1448A (5485A>G 5487A>T), D1451K (5494G>A 5496C>A), N1452D (5497A>G 5499T>C), Y1454V (5503T>G 5504A>T 5505C>T), Q1457K (5512C>A), Y1458F (5516A>T), S1459P (5518T>C 5520C>A), L1460I (5521C>A), G1462I (5527G>A 5528G>T 5529G>T), N1464E (5533A>G 5535C>A), T1465E (5536A>G 5537C>A 5538C>G), K1468D (5545A>G 5547A>T), R1469E (5548A>G 5549G>A), I1470L (5551A>T 5553A>G), N1472G (5557A>G 5558A>G 5559T>C), A1473S (5560G>T 5562A>T), I1475V (5566A>G), F1479I (5578T>A), K1482R (5587A>C 5588A>G 5589G>A), S1483V (5590T>G 5591C>T 5592T>A), F1485Y (5597T>A 5598T>C), V1488I (5605G>A), A1489* (5608G>T 5609C>G 5610C>A), E1491A (5615A>C 5616A>C), E1492P (5617G>C 5618A>C 5619A>G), E1493T (5620G>A 5621A>C), S1494D (5623T>G 5624C>A), I1495V (5626A>G), W1497K (5632T>A 5633G>A), A1499S (5638G>T 5640C>A), L1501* (5645T>G), I1502T (5648T>C 5649C>T), P1503H (5651C>A 5652A>T), G1504S (5653G>T 5654G>C 5655A>C), L1506N (5659T>A 5660T>A 5661A>C), W1509Y (5669G>A 5670G>T), K1517T (5693A>C 5694A>T), A1519T (5698G>A), A1521S (5704G>T), I1522S (5708T>G 5709T>C) | | | | | | | | |
| Codon mutations: | AAT1445GAC (5476A>G 5478T>C), TAC1446TAT (5481C>T), ACA1448GCT (5485A>G 5487A>T), CTC1449TTG (5488C>T 5490C>G), AAT1450AAC (5493T>C), GAC1451AAA (5494G>A 5496C>A), AAT1452GAC (5497A>G 5499T>C), TAC1454GTT (5503T>G 5504A>T 5505C>T), CAG1457AAG (5512C>A), TAT1458TTT (5516A>T), TCC1459CCA (5518T>C 5520C>A), CTT1460ATT (5521C>A), GGG1462ATT (5527G>A 5528G>T 5529G>T), ATA1463ATC (5532A>C), AAC1464GAA (5533A>G 5535C>A), ACC1465GAG (5536A>G 5537C>A 5538C>G), AAA1468GAT (5545A>G 5547A>T), AGA1469GAA (5548A>G 5549G>A), ATA1470TTG (5551A>T 5553A>G), GGG1471GGT (5556G>T), AAT1472GGC (5557A>G 5558A>G 5559T>C), GCA1473TCT (5560G>T 5562A>T), AAG1474AAA (5565G>A), ATA1475GTA (5566A>G), TTC1476TTT (5571C>T), TCA1477TCT (5574A>T), TTT1479ATT (5578T>A), GAT1480GAC (5583T>C), CTG1481TTG (5584C>T), AAG1482CGA (5587A>C 5588A>G 5589G>A), TCT1483GTA (5590T>G 5591C>T 5592T>A), GGA1484GGC (5595A>C), TTT1485TAC (5597T>A 5598T>C), GTT1488ATT (5605G>A), GCC1489TGA (5608G>T 5609C>G 5610C>A), GAA1491GCC (5615A>C 5616A>C), GAA1492CCG (5617G>C 5618A>C 5619A>G), GAA1493ACA (5620G>A 5621A>C), TCC1494GAC (5623T>G 5624C>A), ATA1495GTA (5626A>G), TGG1497AAG (5632T>A 5633G>A), ACG1498ACA (5637G>A), GCC1499TCA (5638G>T 5640C>A), TTA1501TGA (5645T>G), ATC1502ACT (5648T>C 5649C>T), CCA1503CAT (5651C>A 5652A>T), GGA1504TCC (5653G>T 5654G>C 5655A>C), GGA1505GGG (5658A>G), TTA1506AAC (5659T>A 5660T>A 5661A>C), TAC1507TAT (5664C>T), GAA1508GAG (5667A>G), TGG1509TAT (5669G>A 5670G>T), CTT1510TTA (5671C>T 5673T>A), GTC1511GTG (5676C>G), CCA1513CCC (5682A>C), GGA1515GGG (5688A>G), CTC1516CTA (5691C>A), AAA1517ACT (5693A>C 5694A>T), GCA1519ACA (5698G>A), CCC1520CCT (5703C>T), GCT1521TCT (5704G>T), ATT1522AGC (5708T>G 5709T>C) | | | | | | | | |

##### Proteins

| | Begin | End | Coverage | Score | Concordance | Matches | Identities | I/D/M/F* | Stop Codons |
|---|---|---|---|---|---|---|---|---|---|
| ORF3 (YP_009121747.1) | 1443 | 1524 | 4.0% | 301 | 51.3% | 82 (100%) | 40 (48.8%) | 0/0/0/0 | 2 |
| Protein mutations: | N1445D (5476A>G 5478T>C), T1448A (5485A>G 5487A>T), D1451K (5494G>A 5496C>A), N1452D (5497A>G 5499T>C), Y1454V (5503T>G 5504A>T 5505C>T), Q1457K (5512C>A), Y1458F (5516A>T), S1459P (5518T>C 5520C>A), L1460I (5521C>A), G1462I (5527G>A 5528G>T 5529G>T), N1464E (5533A>G 5535C>A), T1465E (5536A>G 5537C>A 5538C>G), K1468D (5545A>G 5547A>T), R1469E (5548A>G 5549G>A), I1470L (5551A>T 5553A>G), N1472G (5557A>G 5558A>G 5559T>C), A1473S (5560G>T 5562A>T), I1475V (5566A>G), F1479I (5578T>A), K1482R (5587A>C 5588A>G 5589G>A), S1483V (5590T>G 5591C>T 5592T>A), F1485Y (5597T>A 5598T>C), V1488I (5605G>A), A1489* (5608G>T 5609C>G 5610C>A), E1491A (5615A>C 5616A>C), E1492P (5617G>C 5618A>C 5619A>G), E1493T (5620G>A 5621A>C), S1494D (5623T>G 5624C>A), I1495V (5626A>G), W1497K (5632T>A 5633G>A), A1499S (5638G>T 5640C>A), L1501* (5645T>G), I1502T (5648T>C 5649C>T), P1503H (5651C>A 5652A>T), G1504S (5653G>T 5654G>C 5655A>C), L1506N (5659T>A 5660T>A 5661A>C), W1509Y (5669G>A 5670G>T), K1517T (5693A>C 5694A>T), A1519T (5698G>A), A1521S (5704G>T), I1522S (5708T>G 5709T>C) | | | | | | | | |
| Codon mutations: | AAT1445GAC (5476A>G 5478T>C), TAC1446TAT (5481C>T), ACA1448GCT (5485A>G 5487A>T), CTC1449TTG (5488C>T 5490C>G), AAT1450AAC (5493T>C), GAC1451AAA (5494G>A 5496C>A), AAT1452GAC (5497A>G 5499T>C), TAC1454GTT (5503T>G 5504A>T 5505C>T), CAG1457AAG (5512C>A), TAT1458TTT (5516A>T), TCC1459CCA (5518T>C 5520C>A), CTT1460ATT (5521C>A), GGG1462ATT (5527G>A 5528G>T 5529G>T), ATA1463ATC (5532A>C), AAC1464GAA (5533A>G 5535C>A), ACC1465GAG (5536A>G 5537C>A 5538C>G), AAA1468GAT (5545A>G 5547A>T), AGA1469GAA (5548A>G 5549G>A), ATA1470TTG (5551A>T 5553A>G), GGG1471GGT (5556G>T), AAT1472GGC (5557A>G 5558A>G 5559T>C), GCA1473TCT (5560G>T 5562A>T), AAG1474AAA (5565G>A), ATA1475GTA (5566A>G), TTC1476TTT (5571C>T), TCA1477TCT (5574A>T), TTT1479ATT (5578T>A), GAT1480GAC (5583T>C), CTG1481TTG (5584C>T), AAG1482CGA (5587A>C 5588A>G 5589G>A), TCT1483GTA (5590T>G 5591C>T 5592T>A), GGA1484GGC (5595A>C), TTT1485TAC (5597T>A 5598T>C), GTT1488ATT (5605G>A), GCC1489TGA (5608G>T 5609C>G 5610C>A), GAA1491GCC (5615A>C 5616A>C), GAA1492CCG (5617G>C 5618A>C 5619A>G), GAA1493ACA (5620G>A 5621A>C), TCC1494GAC (5623T>G 5624C>A), ATA1495GTA (5626A>G), TGG1497AAG (5632T>A 5633G>A), ACG1498ACA (5637G>A), GCC1499TCA (5638G>T 5640C>A), TTA1501TGA (5645T>G), ATC1502ACT (5648T>C 5649C>T), CCA1503CAT (5651C>A 5652A>T), GGA1504TCC (5653G>T 5654G>C 5655A>C), GGA1505GGG (5658A>G), TTA1506AAC (5659T>A 5660T>A 5661A>C), TAC1507TAT (5664C>T), GAA1508GAG (5667A>G), TGG1509TAT (5669G>A 5670G>T), CTT1510TTA (5671C>T 5673T>A), GTC1511GTG (5676C>G), CCA1513CCC (5682A>C), GGA1515GGG (5688A>G), CTC1516CTA (5691C>A), AAA1517ACT (5693A>C 5694A>T), GCA1519ACA (5698G>A), CCC1520CCT (5703C>T), GCT1521TCT (5704G>T), ATT1522AGC (5708T>G 5709T>C) | | | | | | | | |

*: Inserts / Deletes / Misaligned / Frameshifts

#### Analysis details

This analysis was performed with panviral2.64

### NGS Details (UN62): Chinaberry tree badnavirus 1

#### Assembly

| | |
|---|---|
| **Coverage Length** | 249 (1 contig(s)) |
| **Depth Of Coverage** | 6.5 |
| **Number Of Reads** | 13 |
| **Reads Per Million** | 0.30 rpm (after QC) |
| **Ambiguities** | 0 |
| **Assembly Method** | de novo + reference guided assembly |
| **Consensus Caller** | Bcf Tools |

#### Coverage Map

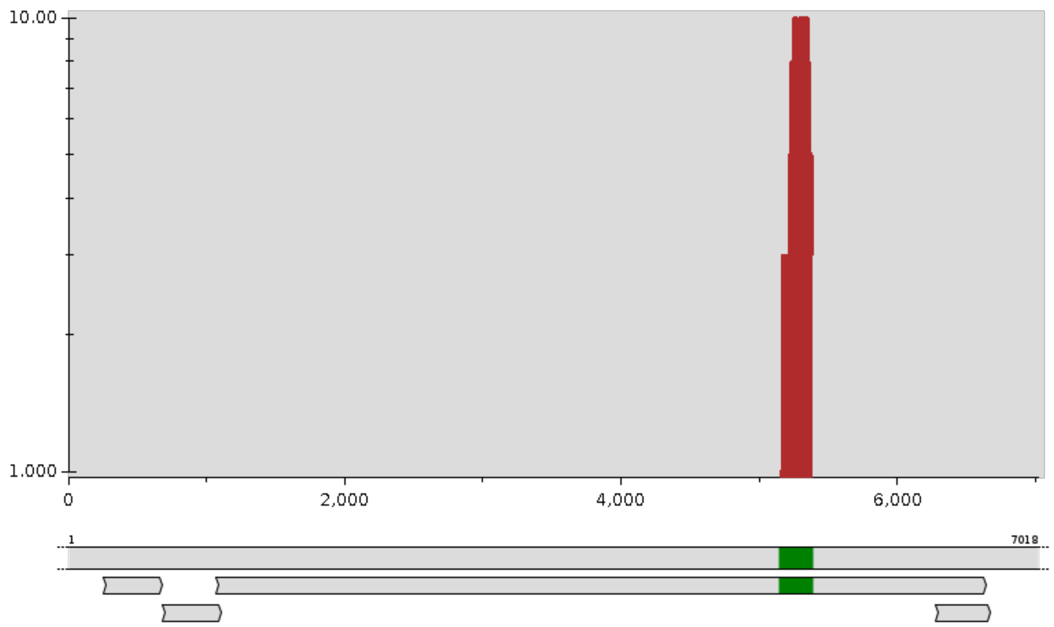

#### Assignment

| | |
|---|---|
| **Type** | Chinaberry tree badnavirus 1 (Taxonomy ID: 2908099) |
| **Reference Genome** | NC_077165.1 |
| **NT Identity (%)** | 55.4217 |
| **AA Identity (%)** | 45.7831 |
| **Number Of Stop Codons** | 1 |
| **Number Of CDS** | 4 |

#### Alignment

| | |
|---|---|
| **Alignment Score** | 54.0 (NT) + 275.0 (AA) = 329.0 |
| **Concordance (%)** | 29.5597 |

| Alignment Method | Global, seeded, nucleotide + amino acids (AGA) |
|---|---|

#### Genome Region

Sequence starts at position 5144 and ends at position 5392 relative to NC_077165.1 reference sequence.

#### Alignment Detailed Statistics

| | Begin | End | Coverage | Score | Concordance | Matches | Identities | I/D/M/F* | Stop Codons |
|---|---|---|---|---|---|---|---|---|---|
| **NT** | **5144** | **5392** | **3.5%** | **54** | **10.8%** | **249 (100%)** | **138 (55.4%)** | **0/0** | |
| Mutations: | 5144A>G, 5145A>C, 5147G>T, 5148G>T, 5149T>G, 5150A>T, 5151T>G, 5153C>T, 5154A>G, 5156C>T, 5163A>G, 5164G>A, 5168G>C, 5172G>C, 5175C>T, 5180C>A, 5181C>G, 5182A>T, 5183C>G, 5190C>T, 5192G>A, 5194A>T, 5195T>C, 5196A>C, 5197G>C, 5198T>A, 5199C>A, 5201A>T, 5204A>T, 5205G>A, 5206G>T, 5207A>G, 5210C>A, 5211C>G, 5213C>G, 5214A>G, 5215C>G, 5217A>C, 5219C>A, 5220A>T, 5223A>G, 5224G>A, 5225G>T, 5226A>G, 5227G>A, 5228A>G, 5229A>T, 5231T>G, 5234C>A, 5237A>G, 5240C>T, 5242A>G, 5243G>A, 5244A>G, 5249T>C, 5252C>T, 5256T>A, 5264G>T, 5266A>G, 5268T>G, 5270G>T, 5273C>T, 5275T>A, 5282A>G, 5283G>T, 5285A>G, 5286G>A, 5287C>G, 5288C>A, 5293A>C, 5294C>A, 5295C>A, 5297G>A, 5298G>C, 5299A>C, 5300G>A, 5301T>G, 5302C>A, 5306A>G, 5307G>C, 5309A>T, 5310T>A, 5311G>A, 5312G>A, 5316G>A, 5322T>A, 5323G>A, 5324G>A, 5325G>A, 5326T>C, 5329C>A, 5330A>T, 5333T>A, 5336G>C, 5338T>A, 5345A>G, 5347G>T, 5348G>T, 5349C>T, 5357T>G, 5360A>T, 5363C>T, 5366G>C, 5367T>C, 5371A>C, 5372A>T, 5378T>A, 5381A>T, 5382G>T, 5385A>T, 5386T>C | | | | | | | | |

##### CDS

| | Begin | End | Coverage | Score | Concordance | Matches | Identities | I/D/M/F* | Stop Codons |
|---|---|---|---|---|---|---|---|---|---|
| QKV46_gp3 | 1359 | 1441 | 4.5% | 275 | 43.6% | 83 (100%) | 38 (45.8%) | 0/0/0/0 | 1 |
| Protein mutations: | M1359L (5145A>C 5147G>T), V1360C (5148G>T 5149T>G 5150A>T), F1361V (5151T>G 5153C>T), N1362D (5154A>G 5156C>T), R1365E (5163A>G 5164G>A), D1368H (5172G>C), H1371V (5181C>G 5182A>T 5183C>G), Q1374* (5190C>T 5192G>A), Y1375F (5194A>T 5195T>C), S1376P (5196A>C 5197G>C 5198T>A), L1377I (5199C>A 5201A>T), G1379M (5205G>A 5206G>T 5207A>G), H1381E (5211C>G 5213C>G), T1382G (5214A>G 5215C>G), I1383L (5217A>C 5219C>A), M1384L (5220A>T), R1385D (5223A>G 5224G>A 5225G>T), R1386E (5226A>G 5227G>A 5228A>G), I1387L (5229A>T 5231T>G), Q1391R (5242A>G 5243G>A), I1392V (5244A>G), F1396I (5256T>A), K1399R (5266A>G), S1400A (5268T>G 5270G>T), F1402Y (5275T>A), V1405L (5283G>T 5285A>G), A1406R (5286G>A 5287C>G 5288C>A), D1408A (5293A>C 5294C>A), P1409T (5295C>A 5297G>A), E1410P (5298G>C 5299A>C 5300G>A), S1411D (5301T>G 5302C>A), E1413H (5307G>C 5309A>T), W1414K (5310T>A 5311G>A 5312G>A), A1416T (5316G>A), W1418K (5322T>A 5323G>A 5324G>A), V1419T (5325G>A 5326T>C), P1420H (5329C>A 5330A>T), D1421E (5333T>A), L1423H (5338T>A), W1426F (5347G>T 5348G>T), I1429M (5357T>G), K1434T (5371A>C 5372A>T), A1438S (5382G>T), I1439S (5385A>T 5386T>C) | | | | | | | | |
| Codon mutations: | AGA1358..G (5144A>G), ATG1359CTT (5145A>C 5147G>T), GTA1360TGT (5148G>T 5149T>G 5150A>T), TTC1361GTT (5151T>G 5153C>T), AAC1362GAT (5154A>G 5156C>T), AGG1365GAG (5163A>G 5164G>A), CTG1366CTC (5168G>C), GAT1368CAT (5172G>C), CTA1369TTA (5175C>T), ACC1370ACA (5180C>A), CAC1371GTG (5181C>G 5182A>T 5183C>G), CAG1374TAA (5190C>T 5192G>A), TAT1375TTC (5194A>T 5195T>C), AGT1376CCA (5196A>C 5197G>C 5198T>A), CTA1377ATT (5199C>A 5201A>T), CCA1378CCT (5204A>T), GGA1379ATG (5205G>A 5206G>T 5207A>G), ATC1380ATA (5210C>A), CAC1381GAG (5211C>G 5213C>G), ACC1382GGC (5214A>G 5215C>G), ATC1383CTA (5217A>C 5219C>A), ATG1384TTG (5220A>T), AGG1385GAT (5223A>G 5224G>A 5225G>T), AGA1386GAG (5226A>G 5227G>A 5228A>G), ATT1387TTG (5229A>T 5231T>G), GGC1388GGA (5234C>A), CAA1389CAG (5237A>G), GCC1390GCT (5240C>T), CAG1391CGA (5242A>G 5243G>A), ATA1392GTA (5244A>G), TTT1393TTC (5249T>C), TCC1394TCT (5252C>T), TTT1396ATT (5256T>A), CTG1398CTT (5264G>T), AAA1399AGA (5266A>G), TCG1400GCT (5268T>G 5270G>T), GGC1401GGT (5273C>T), TTT1402TAT (5275T>A), CAA1404CAG (5282A>G), GTA1405TTG (5283G>T 5285A>G), GCC1406AGA (5286G>A 5287C>G 5288C>A), GAC1408GCA (5293A>C 5294C>A), CCG1409ACA (5295C>A 5297G>A), GAG1410CCA (5298G>C 5299A>C 5300G>A), TCT1411GAT (5301T>G 5302C>A), GTA1412GTG (5306A>G), GAA1413CAT (5307G>C 5309A>T), TGG1414AAA (5310T>A 5311G>A 5312G>A), GCA1416ACA (5316G>A), TGG1418AAA (5322T>A 5323G>A 5324G>A), GTT1419ACT (5325G>A 5326T>C), CCA1420CAT (5329C>A 5330A>T), GAT1421GAA (5333T>A), GGG1422GGC (5336G>C), CTT1423CAT (5338T>A), GAA1425GAG (5345A>G), TGG1426TTT (5347G>T 5348G>T), CTG1427TTG (5349C>T), ATT1429ATG (5357T>G), CCA1430CCT (5360A>T), TTC1431TTT (5363C>T), GGG1432GGC (5366G>C), TTA1433CTA (5367T>C), AAA1434ACT (5371A>C 5372A>T), GCT1436GCA (5378T>A), CCA1437CCT (5381A>T), GCT1438TCT (5382G>T), ATA1439TCA (5385A>T 5386T>C) | | | | | | | | |

##### Proteins

| | Begin | End | Coverage | Score | Concordance | Matches | Identities | I/D/M/F* | Stop Codons |
|---|---|---|---|---|---|---|---|---|---|
| ORF3 (YP_010805662.1) | 1359 | 1441 | 4.5% | 275 | 43.6% | 83 (100%) | 38 (45.8%) | 0/0/0/0 | 1 |
| Protein mutations: | M1359L (5145A>C 5147G>T), V1360C (5148G>T 5149T>G 5150A>T), F1361V (5151T>G 5153C>T), N1362D (5154A>G 5156C>T), R1365E (5163A>G 5164G>A), D1368H (5172G>C), H1371V (5181C>G 5182A>T 5183C>G), Q1374* (5190C>T 5192G>A), Y1375F (5194A>T 5195T>C), S1376P (5196A>C 5197G>C 5198T>A), L1377I (5199C>A 5201A>T), G1379M (5205G>A 5206G>T 5207A>G), H1381E (5211C>G 5213C>G), T1382G (5214A>G 5215C>G), I1383L (5217A>C 5219C>A), M1384L (5220A>T), R1385D (5223A>G 5224G>A 5225G>T), R1386E (5226A>G 5227G>A 5228A>G), I1387L (5229A>T 5231T>G), Q1391R (5242A>G 5243G>A), I1392V (5244A>G), F1396I (5256T>A), K1399R (5266A>G), S1400A (5268T>G 5270G>T), F1402Y (5275T>A), V1405L (5283G>T 5285A>G), A1406R (5286G>A 5287C>G 5288C>A), D1408A (5293A>C 5294C>A), P1409T (5295C>A 5297G>A), E1410P (5298G>C 5299A>C 5300G>A), S1411D (5301T>G 5302C>A), E1413H (5307G>C 5309A>T), W1414K (5310T>A 5311G>A 5312G>A), A1416T (5316G>A), W1418K (5322T>A 5323G>A 5324G>A), V1419T (5325G>A 5326T>C), P1420H (5329C>A 5330A>T), D1421E (5333T>A), L1423H (5338T>A), W1426F (5347G>T 5348G>T), I1429M (5357T>G), K1434T (5371A>C 5372A>T), A1438S (5382G>T), I1439S (5385A>T 5386T>C) | | | | | | | | |
| Codon mutations: | AGA1358..G (5144A>G), ATG1359CTT (5145A>C 5147G>T), GTA1360TGT (5148G>T 5149T>G 5150A>T), TTC1361GTT (5151T>G 5153C>T), AAC1362GAT (5154A>G 5156C>T), AGG1365GAG (5163A>G 5164G>A), CTG1366CTC (5168G>C), GAT1368CAT (5172G>C), CTA1369TTA (5175C>T), ACC1370ACA (5180C>A), CAC1371GTG (5181C>G 5182A>T 5183C>G), CAG1374TAA (5190C>T 5192G>A), TAT1375TTC (5194A>T 5195T>C), AGT1376CCA (5196A>C 5197G>C 5198T>A), CTA1377ATT (5199C>A 5201A>T), CCA1378CCT (5204A>T), GGA1379ATG (5205G>A 5206G>T 5207A>G), ATC1380ATA (5210C>A), CAC1381GAG (5211C>G 5213C>G), ACC1382GGC (5214A>G 5215C>G), ATC1383CTA (5217A>C 5219C>A), ATG1384TTG (5220A>T), AGG1385GAT (5223A>G 5224G>A 5225G>T), AGA1386GAG (5226A>G 5227G>A 5228A>G), ATT1387TTG (5229A>T 5231T>G), GGC1388GGA (5234C>A), CAA1389CAG (5237A>G), GCC1390GCT (5240C>T), CAG1391CGA (5242A>G 5243G>A), ATA1392GTA (5244A>G), TTT1393TTC (5249T>C), TCC1394TCT (5252C>T), TTT1396ATT (5256T>A), CTG1398CTT (5264G>T), AAA1399AGA (5266A>G), TCG1400GCT (5268T>G 5270G>T), GGC1401GGT (5273C>T), TTT1402TAT (5275T>A), CAA1404CAG (5282A>G), GTA1405TTG (5283G>T 5285A>G), GCC1406AGA (5286G>A 5287C>G 5288C>A), GAC1408GCA (5293A>C 5294C>A), CCG1409ACA (5295C>A 5297G>A), GAG1410CCA (5298G>C 5299A>C 5300G>A), TCT1411GAT (5301T>G 5302C>A), GTA1412GTG (5306A>G), GAA1413CAT (5307G>C 5309A>T), TGG1414AAA (5310T>A 5311G>A 5312G>A), GCA1416ACA (5316G>A), TGG1418AAA (5322T>A 5323G>A 5324G>A), GTT1419ACT (5325G>A 5326T>C), CCA1420CAT (5329C>A 5330A>T), GAT1421GAA (5333T>A), GGG1422GGC (5336G>C), CTT1423CAT (5338T>A), GAA1425GAG (5345A>G), TGG1426TTT (5347G>T 5348G>T), CTG1427TTG (5349C>T), ATT1429ATG (5357T>G), CCA1430CCT (5360A>T), TTC1431TTT (5363C>T), GGG1432GGC (5366G>C), TTA1433CTA (5367T>C), AAA1434ACT (5371A>C 5372A>T), GCT1436GCA (5378T>A), CCA1437CCT (5381A>T), GCT1438TCT (5382G>T), ATA1439TCA (5385A>T 5386T>C) | | | | | | | | |

*: Inserts / Deletes / Misaligned / Frameshifts

#### Analysis details

This analysis was performed with panviral2.64

### NGS Details (UN62): Pinus nigra virus 1

#### Assembly

| | |
|---|---|
| **Coverage Length** | 291 (1 contig(s)) |
| **Depth Of Coverage** | 4.5 |
| **Number Of Reads** | 12 |
| **Reads Per Million** | 0.28 rpm (after QC) |
| **Ambiguities** | 0 |
| **Assembly Method** | de novo + reference guided assembly |
| **Consensus Caller** | Bcf Tools |

#### Coverage Map

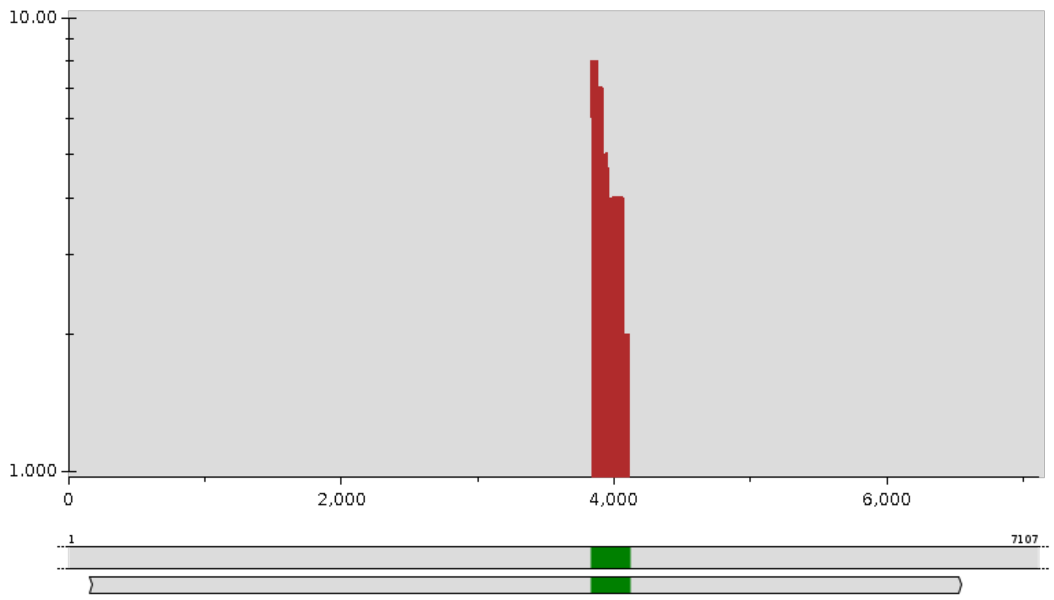

#### Assignment

| | |
|---|---|
| **Type** | Pinus nigra virus 1 (Taxonomy ID: 2267679) |
| **Reference Genome** | NC_040841.1 |
| **NT Identity (%)** | 55.3265 |
| **AA Identity (%)** | 47.4227 |
| **Number Of Stop Codons** | 0 |
| **Number Of CDS** | 1 |

#### Alignment

| | |
|---|---|
| **Alignment Score** | 62.0 (NT) + 360.0 (AA) = 422.0 |
| **Concordance (%)** | 32.3372 |

| **Alignment Method** | Global, seeded, nucleotide + amino acids (AGA) |
|---|---|

#### Genome Region

Sequence starts at position 3830 and ends at position 4120 relative to NC_040841.1 reference sequence.

#### Alignment Detailed Statistics

| | Begin | End | Coverage | Score | Concordance | Matches | Identities | I/D/M/F* | Stop Codons |
|---|---|---|---|---|---|---|---|---|---|
| **NT** | **3830** | **4120** | **4.1%** | **62** | **10.7%** | **291 (100%)** | **161 (55.3%)** | **0/0** | |
| Mutations: | 3830C>A, 3835A>T, 3836G>A, 3837A>G, 3842A>T, 3843G>C, 3845G>A, 3850C>T, 3852A>C, 3853C>T, 3855A>G, 3856T>G, 3857G>A, 3858C>G, 3859C>T, 3865T>G, 3868A>C, 3873T>A, 3874A>T, 3877A>T, 3879G>A, 3880G>T, 3883T>C, 3884C>G, 3885A>T, 3887T>G, 3888C>T, 3889T>G, 3894T>A, 3897T>A, 3904T>C, 3905A>T, 3906A>C, 3907A>T, 3913A>G, 3914A>T, 3916G>A, 3919A>T, 3922A>T, 3925T>C, 3928T>C, 3931A>G, 3932C>T, 3934T>G, 3935C>T, 3937T>G, 3941G>T, 3944A>G, 3946C>T, 3947A>T, 3948C>T, 3949T>A, 3950C>A, 3952G>A, 3953A>T, 3954A>G, 3955T>G, 3956T>A, 3958T>A, 3960A>G, 3961T>A, 3964C>T, 3968T>A, 3970G>T, 3973A>T, 3975G>A, 3976A>T, 3977C>A, 3978C>A, 3980G>A, 3982A>G, 3984T>A, 3985G>T, 3986A>C, 3989A>T, 3991G>A, 3992C>A, 3995A>C, 3996A>G, 3999T>C, 4001C>T, 4002A>T, 4003G>T, 4004C>A, 4006T>A, 4007A>G, 4008G>C, 4009T>C, 4012G>T, 4013G>A, 4015G>T, 4017T>A, 4021C>T, 4024G>A, 4036G>A, 4042G>T, 4044A>T, 4045T>C, 4047A>G, 4048T>G, 4051G>A, 4054A>T, 4061C>T, 4062A>C, 4063G>T, 4064C>G, 4065C>A, 4067G>A, 4069A>G, 4072T>C, 4074G>A, 4075A>G, 4076C>T, 4084A>T, 4092T>A, 4093A>T, 4094T>G, 4095G>T, 4096T>C, 4099A>C, 4100G>T, 4101C>T, 4102A>T, 4106T>C, 4107T>A, 4108C>A, 4111T>C, 4112C>G, 4114G>A, 4120A>T | | | | | | | | |

##### CDS

| | Begin | End | Coverage | Score | Concordance | Matches | Identities | I/D/M/F* | Stop Codons |
|---|---|---|---|---|---|---|---|---|---|
| EXL67_gp1 | 1226 | 1322 | 4.6% | 360 | 49.8% | 97 (100%) | 46 (47.4%) | 0/0/0/0 | 0 |
| Protein mutations: | L1226I (3830C>A), E1228R (3836G>A 3837A>G), E1231K (3845G>A), H1233P (3852A>C 3853C>T), Y1234W (3855A>G 3856T>G), A1235S (3857G>A 3858C>G 3859C>T), L1240Y (3873T>A 3874A>T), R1242N (3879G>A 3880G>T), H1244V (3884C>G 3885A>T), S1245V (3887T>G 3888C>T 3889T>G), I1247K (3894T>A), V1248E (3897T>A), K1251S (3905A>T 3906A>C 3907A>T), M1254L (3914A>T 3916G>A), P1260S (3932C>T 3934T>G), A1263S (3941G>T), I1264V (3944A>G 3946C>T), T1265L (3947A>T 3948C>T 3949T>A), Q1266K (3950C>A 3952G>A), N1267W (3953A>T 3954A>G 3955T>G), F1268I (3956T>A 3958T>A), N1269R (3960A>G 3961T>A), L1272I (3968T>A 3970G>T), R1274N (3975G>A 3976A>T), P1275K (3977C>A 3978C>A), E1276K (3980G>A 3982A>G), V1277D (3984T>A 3985G>T), I1278L (3986A>C), M1279L (3989A>T 3991G>A), Q1280K (3992C>A), K1281R (3995A>C 3996A>G), I1282T (3999T>C), Q1283F (4001C>T 4002A>T 4003G>T), H1284K (4004C>A 4006T>A), S1285A (4007A>G 4008G>C 4009T>C), K1286N (4012G>T), V1287I (4013G>A 4015G>T), F1288Y (4017T>A), Y1297F (4044A>T 4045T>C), Y1298W (4047A>G 4048T>G), Q1303S (4061C>T 4062A>C 4063G>T), P1304E (4064C>G 4065C>A), E1305K (4067G>A 4069A>G), R1307K (4074G>A 4075A>G), H1308Y (4076C>T), I1313N (4092T>A 4093A>T), C1314V (4094T>G 4095G>T 4096T>C), A1316F (4100G>T 4101C>T 4102A>T), F1318Q (4106T>C 4107T>A 4108C>A), Q1320E (4112C>G 4114G>A), K1322N (4120A>T) | | | | | | | | |
| Codon mutations: | CTT1226ATT (3830C>A), ATA1227ATT (3835A>T), GAG1228AGG (3836G>A 3837A>G), AGT1230TCT (3842A>T 3843G>C), GAA1231AAA (3845G>A), TCC1232TCT (3850C>T), CAC1233CCT (3852A>C 3853C>T), TAT1234TGG (3855A>G 3856T>G), GCC1235AGT (3857G>A 3858C>G 3859C>T), TCT1237TCG (3865T>G), GCA1238GCC (3868A>C), TTA1240TAT (3873T>A 3874A>T), GTA1241GTT (3877A>T), AGG1242AAT (3879G>A 3880G>T), AAT1243AAC (3883T>C), CAT1244GTT (3884C>G 3885A>T), TCT1245GTG (3887T>G 3888C>T 3889T>G), ATA1247AAA (3894T>A), GTA1248GAA (3897T>A), GGT1250GGC (3904T>C), AAA1251TCT (3905A>T 3906A>C 3907A>T), AGA1253AGG (3913A>G), ATG1254TTA (3914A>T 3916G>A), GTA1255GTT (3919A>T), ATA1256ATT (3922A>T), AAT1257AAC (3925T>C), TAT1258TAC (3928T>C), AAA1259AAG (3931A>G), CCT1260TCG (3932C>T 3934T>G), CTT1261TTG (3935C>T 3937T>G), GCC1263TCC (3941G>T), ATC1264GTT (3944A>G 3946C>T), ACT1265TTA (3947A>T 3948C>T 3949T>A), CAG1266AAA (3950C>A 3952G>A), AAT1267TGG (3953A>T 3954A>G 3955T>G), TTT1268ATA (3956T>A 3958T>A), AAT1269AGA (3960A>G 3961T>A), TAC1270TAT (3964C>T), TTG1272ATT (3968T>A 3970G>T), CCA1273CCT (3973A>T), AGA1274AAT (3975G>A 3976A>T), CCA1275AAA (3977C>A 3978C>A), GAA1276AAG (3980G>A 3982A>G), GTG1277GAT (3984T>A 3985G>T), ATT1278CTT (3986A>C), ATG1279TTA (3989A>T 3991G>A), CAA1280AAA (3992C>A), AAA1281CGA (3995A>C 3996A>G), ATT1282ACT (3999T>C), CAG1283TTT (4001C>T 4002A>T 4003G>T), CAT1284AAA (4004C>A 4006T>A), AGT1285GCC (4007A>G 4008G>C 4009T>C), AAG1286AAT (4012G>T), GTG1287ATT (4013G>A 4015G>T), TTC1288TAC (4017T>A), AGC1289AGT (4021C>T), AAG1290AAA (4024G>A), AAG1294AAA (4036G>A), GGG1296GGT (4042G>T), TAT1297TTC (4044A>T 4045T>C), TAT1298TGG (4047A>G 4048T>G), CAG1299CAA (4051G>A), ATA1300ATT (4054A>T), CAG1303TCT (4061C>T 4062A>C 4063G>T), CCA1304GAA (4064C>G 4065C>A), GAA1305AAG (4067G>A 4069A>G), GAT1306GAC (4072T>C), AGA1307AAG (4074G>A 4075A>G), CAC1308TAC (4076C>T), ACA1310ACT (4084A>T), ATA1313AAT (4092T>A 4093A>T), TGT1314GTC (4094T>G 4095G>T 4096T>C), CCA1315CCC (4099A>C), GCA1316TTT (4100G>T 4101C>T 4102A>T), TTC1318CAA (4106T>C 4107T>A 4108C>A), TAT1319TAC (4111T>C), CAG1320GAA (4112C>G 4114G>A), AAA1322AAT (4120A>T) | | | | | | | | |

##### Proteins

| | Begin | End | Coverage | Score | Concordance | Matches | Identities | I/D/M/F* | Stop Codons |
|---|---|---|---|---|---|---|---|---|---|
| polyprotein (YP_009553669.1) | 1226 | 1322 | 4.6% | 360 | 49.8% | 97 (100%) | 46 (47.4%) | 0/0/0/0 | 0 |
| Protein mutations: | L1226I (3830C>A), E1228R (3836G>A 3837A>G), E1231K (3845G>A), H1233P (3852A>C 3853C>T), Y1234W (3855A>G 3856T>G), A1235S (3857G>A 3858C>G 3859C>T), L1240Y (3873T>A 3874A>T), R1242N (3879G>A 3880G>T), H1244V (3884C>G 3885A>T), S1245V (3887T>G 3888C>T 3889T>G), I1247K (3894T>A), V1248E (3897T>A), K1251S (3905A>T 3906A>C 3907A>T), M1254L (3914A>T 3916G>A), P1260S (3932C>T 3934T>G), A1263S (3941G>T), I1264V (3944A>G 3946C>T), T1265L (3947A>T 3948C>T 3949T>A), Q1266K (3950C>A 3952G>A), N1267W (3953A>T 3954A>G 3955T>G), F1268I (3956T>A 3958T>A), N1269R (3960A>G 3961T>A), L1272I (3968T>A 3970G>T), R1274N (3975G>A 3976A>T), P1275K (3977C>A 3978C>A), E1276K (3980G>A 3982A>G), V1277D (3984T>A 3985G>T), I1278L (3986A>C), M1279L (3989A>T 3991G>A), Q1280K (3992C>A), K1281R (3995A>C 3996A>G), I1282T (3999T>C), Q1283F (4001C>T 4002A>T 4003G>T), H1284K (4004C>A 4006T>A), S1285A (4007A>G 4008G>C 4009T>C), K1286N (4012G>T), V1287I (4013G>A 4015G>T), F1288Y (4017T>A), Y1297F (4044A>T 4045T>C), Y1298W (4047A>G 4048T>G), Q1303S (4061C>T 4062A>C 4063G>T), P1304E (4064C>G 4065C>A), E1305K (4067G>A 4069A>G), R1307K (4074G>A 4075A>G), H1308Y (4076C>T), I1313N (4092T>A 4093A>T), C1314V (4094T>G 4095G>T 4096T>C), A1316F (4100G>T 4101C>T 4102A>T), F1318Q (4106T>C 4107T>A 4108C>A), Q1320E (4112C>G 4114G>A), K1322N (4120A>T) | | | | | | | | |
| Codon mutations: | CTT1226ATT (3830C>A), ATA1227ATT (3835A>T), GAG1228AGG (3836G>A 3837A>G), AGT1230TCT (3842A>T 3843G>C), GAA1231AAA (3845G>A), TCC1232TCT (3850C>T), CAC1233CCT (3852A>C 3853C>T), TAT1234TGG (3855A>G 3856T>G), GCC1235AGT (3857G>A 3858C>G 3859C>T), TCT1237TCG (3865T>G), GCA1238GCC (3868A>C), TTA1240TAT (3873T>A 3874A>T), GTA1241GTT (3877A>T), AGG1242AAT (3879G>A 3880G>T), AAT1243AAC (3883T>C), CAT1244GTT (3884C>G 3885A>T), TCT1245GTG (3887T>G 3888C>T 3889T>G), ATA1247AAA (3894T>A), GTA1248GAA (3897T>A), GGT1250GGC (3904T>C), AAA1251TCT (3905A>T 3906A>C 3907A>T), AGA1253AGG (3913A>G), ATG1254TTA (3914A>T 3916G>A), GTA1255GTT (3919A>T), ATA1256ATT (3922A>T), AAT1257AAC (3925T>C), TAT1258TAC (3928T>C), AAA1259AAG (3931A>G), CCT1260TCG (3932C>T 3934T>G), CTT1261TTG (3935C>T 3937T>G), GCC1263TCC (3941G>T), ATC1264GTT (3944A>G 3946C>T), ACT1265TTA (3947A>T 3948C>T 3949T>A), CAG1266AAA (3950C>A 3952G>A), AAT1267TGG (3953A>T 3954A>G 3955T>G), TTT1268ATA (3956T>A 3958T>A), AAT1269AGA (3960A>G 3961T>A), TAC1270TAT (3964C>T), TTG1272ATT (3968T>A 3970G>T), CCA1273CCT (3973A>T), AGA1274AAT (3975G>A 3976A>T), CCA1275AAA (3977C>A 3978C>A), GAA1276AAG (3980G>A 3982A>G), GTG1277GAT (3984T>A 3985G>T), ATT1278CTT (3986A>C), ATG1279TTA (3989A>T 3991G>A), CAA1280AAA (3992C>A), AAA1281CGA (3995A>C 3996A>G), ATT1282ACT (3999T>C), CAG1283TTT (4001C>T 4002A>T 4003G>T), CAT1284AAA (4004C>A 4006T>A), AGT1285GCC (4007A>G 4008G>C 4009T>C), AAG1286AAT (4012G>T), GTG1287ATT (4013G>A 4015G>T), TTC1288TAC (4017T>A), AGC1289AGT (4021C>T), AAG1290AAA (4024G>A), AAG1294AAA (4036G>A), GGG1296GGT (4042G>T), TAT1297TTC (4044A>T 4045T>C), TAT1298TGG (4047A>G 4048T>G), CAG1299CAA (4051G>A), ATA1300ATT (4054A>T), CAG1303TCT (4061C>T 4062A>C 4063G>T), CCA1304GAA (4064C>G 4065C>A), GAA1305AAG (4067G>A 4069A>G), GAT1306GAC (4072T>C), AGA1307AAG (4074G>A 4075A>G), CAC1308TAC (4076C>T), ACA1310ACT (4084A>T), ATA1313AAT (4092T>A 4093A>T), TGT1314GTC (4094T>G 4095G>T 4096T>C), CCA1315CCC (4099A>C), GCA1316TTT (4100G>T 4101C>T 4102A>T), TTC1318CAA (4106T>C 4107T>A 4108C>A), TAT1319TAC (4111T>C), CAG1320GAA (4112C>G 4114G>A), AAA1322AAT (4120A>T) | | | | | | | | |

*: Inserts / Deletes / Misaligned / Frameshifts

#### Analysis details

This analysis was performed with panviral2.64

### NGS Details (UN62): Dioscovirus dioscoreae

#### Assembly

| | |
|---|---|
| **Coverage Length** | 434 (1 contig(s)) |
| **Depth Of Coverage** | 2.7 |
| **Number Of Reads** | 11 |
| **Reads Per Million** | 0.25 rpm (after QC) |
| **Ambiguities** | 0 |
| **Assembly Method** | de novo + reference guided assembly |
| **Consensus Caller** | Bcf Tools |

#### Coverage Map

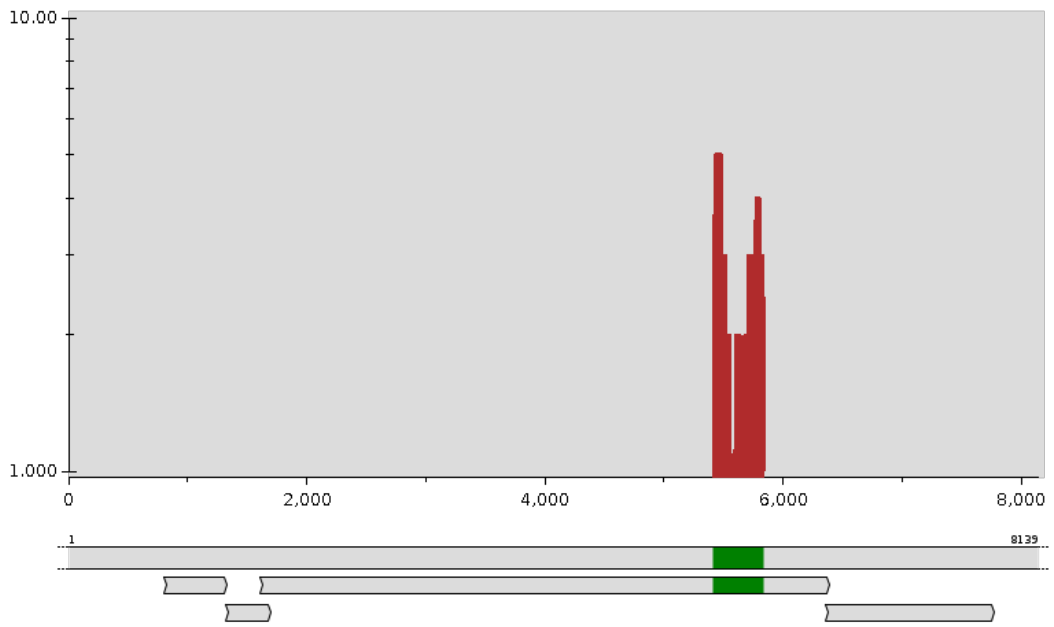

#### Assignment

| | |
|---|---|
| **Type** | Dioscovirus dioscoreae (Taxonomy ID: 3052184) |
| **Reference Genome** | NC_040712.1 |
| **NT Identity (%)** | 61.1241 |
| **AA Identity (%)** | 54.9296 |
| **Number Of Stop Codons** | 0 |
| **Number Of CDS** | 4 |

#### Alignment

| | |
|---|---|
| **Alignment Score** | 156.0 (NT) + 442.0 (AA) = 598.0 |
| **Concordance (%)** | 35.1765 |

| Alignment Method | Global, seeded, nucleotide + amino acids (AGA) |
|---|---|

#### Genome Region

Sequence starts at position 5410 and ends at position 5843 relative to NC_040712.1 reference sequence.

#### Alignment Detailed Statistics

| | Begin | End | Coverage | Score | Concordance | Matches | Identities | I/D/M/F* | Stop Codons |
|---|---|---|---|---|---|---|---|---|---|
| **NT** | **5410** | **5843** | **5.3%** | **156** | **19.0%** | **427 (98.4%)** | **261 (60.1%)** | **0/7** | |
| Mutations: | 5410G>C, 5416A>C, 5417A>T, 5419G>T, 5429T>C, 5430G>A, 5431T>G, 5432C>T, 5433T>A, 5434G>C, 5435C>A, 5439C>A, 5441G>T, 5444T>A, 5449T>A, 5457A>T, 5468A>T, 5471C>T, 5475C>T, 5477C>A, 5486A>C, 5490T>C, 5491C>A, 5493G>A, 5495A>C, 5498C>T, 5502C>A, 5514A>G, 5516T>A, 5522C>T, 5523A>T, 5524A>C, 5525A>T, 5526A>G, 5529T>A, 5531T>G, 5532_5534delTCA, 5535G>A, 5537A>T, 5540T>C, 5541G>C, 5543C>A, 5544T>A, 5545G>T, 5549T>C, 5567G>A, 5572T>G, 5576T>A, 5577G>T, 5578A>C, 5581G>C, 5582T>A, 5583A>T, 5584T>A, 5585A>C, 5589C>G, 5595G>C, 5598C>A, 5599A>C, 5604T>C, 5606A>T, 5607T>A, 5608T>A, 5610C>G, 5612G>A, 5615C>T, 5617T>C, 5618C>A, 5619C>A, 5622G>A, 5624A>T, 5628A>G, 5629A>T, 5630G>T, 5633A>C, 5634G>A, 5636A>T, 5639G>A, 5640C>A, 5643A>G, 5649T>A, 5650C>G, 5651A>C, 5652A>C, 5655delA, 5656C>A, 5660A>G, 5661T>G, 5662T>C, 5663G>T, 5670G>A, 5671G>A, 5672A>T, 5673G>A, 5674T>A, 5676G>A, 5677C>T, 5678A>T, 5679A>G, 5681T>A, 5690C>T, 5696C>A, 5697C>A, 5699A>G, 5700G>A, 5701A>T, 5702A>T, 5705A>T, 5706G>T, 5707G>C, 5709G>A, 5712G>A, 5713G>A, 5714A>T, 5715A>G, 5716A>G, 5717A>G, 5718G>A, 5720T>A, 5721C>G, 5724C>T, 5726C>A, 5729G>A, 5730C>A, 5732C>T, 5738A>T, 5740T>C, 5744A>G, 5747T>A, 5751T>A, 5752T>C, 5753A>G, 5759T>C, 5760C>T, 5762T>G, 5763_5765delGAA, 5768C>T, 5769C>A, 5775G>A, 5776A>C, 5781T>A, 5782T>A, 5783A>G, 5784A>C, 5786G>A, 5788G>A, 5789T>A, 5790C>A, 5795G>A, 5798G>A, 5801C>T, 5802T>C, 5804A>T, 5807G>T, 5808A>T, 5809T>G, 5810A>T, 5811T>C, 5813A>G, 5816C>T, 5822C>A, 5823A>G, 5825A>T, 5826A>G, 5828T>G, 5830A>T, 5834C>T, 5835C>T, 5837A>G | | | | | | | | |

##### CDS

| | Begin | End | Coverage | Score | Concordance | Matches | Identities | I/D/M/F* | Stop Codons |
|---|---|---|---|---|---|---|---|---|---|
| EXK67_gp3 | 1268 | 1411 | 9.0% | 442 | 43.8% | 142 (98.6%) | 78 (54.2%) | 0/2/1/1 | 0 |
| Protein mutations: | E1269A (5416A>C 5417A>T), W1270L (5419G>T), V1274S (5430G>A 5431T>G 5432C>T), C1275T (5433T>A 5434G>C 5435C>A), Q1277N (5439C>A 5441G>T), F1280Y (5449T>A), I1283L (5457A>T), S1294Q (5490T>C 5491C>A), V1295I (5493G>A 5495A>C), N1302E (5514A>G 5516T>A), K1305S (5523A>T 5524A>C 5525A>T), K1306E (5526A>G), Y1307K (5529T>A 5531T>G), S1308del (5532_5534delTCA), E1309N (5535G>A 5537A>T), V1311L (5541G>C 5543C>A), C1312I (5544T>A 5545G>T), F1321C (5572T>G), E1323S (5577G>T 5578A>C), S1324T (5581G>C 5582T>A), I1325Y (5583A>T 5584T>A 5585A>C), Q1327E (5589C>G), V1329L (5595G>C), Q1330T (5598C>A 5599A>C), L1333K (5607T>A 5608T>A), Q1334E (5610C>G 5612G>A), F1336S (5617T>C 5618C>A), Q1337K (5619C>A), V1338I (5622G>A 5624A>T), K1340V (5628A>G 5629A>T 5630G>T), E1341D (5633A>C), E1342N (5634G>A 5636A>T), L1344I (5640C>A), I1345V (5643A>G), K1348Q (5652A>C), L1351A (5661T>G 5662T>C 5663G>T), G1354N (5670G>A 5671G>A 5672A>T), V1355K (5673G>A 5674T>A), A1356I (5676G>A 5677C>T 5678A>T), N1357E (5679A>G 5681T>A), L1363M (5697C>A 5699A>G), E1364I (5700G>A 5701A>T 5702A>T), G1366S (5706G>T 5707G>C), E1367K (5709G>A), G1368N (5712G>A 5713G>A 5714A>T), K1369G (5715A>G 5716A>G 5717A>G), V1370I (5718G>A 5720T>A), Q1371E (5721C>G), P1374T (5730C>A 5732C>T), L1377S (5740T>C), N1379K (5747T>A), L1381T (5751T>A 5752T>C 5753A>G), P1384S (5760C>T 5762T>G), E1385del (5763_5765delGAA), Q1387K (5769C>A), E1389T (5775G>A 5776A>C), L1391K (5781T>A 5782T>A 5783A>G), K1392Q (5784A>C 5786G>A), G1393E (5788G>A 5789T>A), L1394I (5790C>A), I1400C (5808A>T 5809T>G 5810A>T), R1405G (5823A>G 5825A>T), N1406E (5826A>G 5828T>G), Y1407F (5830A>T), P1409S (5835C>T 5837A>G) | | | | | | | | |
| Codon mutations: | AGT1267.CT (5410G>C), GAA1269GCT (5416A>C 5417A>T), TGG1270TTG (5419G>T), TTT1273TTC (5429T>C), GTC1274AGT (5430G>A 5431T>G 5432C>T), TGC1275ACA (5433T>A 5434G>C 5435C>A), CAG1277AAT (5439C>A 5441G>T), GGT1278GGA (5444T>A), TTT1280TAT (5449T>A), ATA1283TTA (5457A>T), CCA1286CCT (5468A>T), TTC1287TTT (5471C>T), CTC1289TTA (5475C>T 5477C>A), GCA1292GCC (5486A>C), TCA1294CAA (5490T>C 5491C>A), GTA1295ATC (5493G>A 5495A>C), TTC1296TTT (5498C>T), CGA1298AGA (5502C>A), AAT1302GAA (5514A>G 5516T>A), TTC1304TTT (5522C>T), AAA1305TCT (5523A>T 5524A>C 5525A>T), AAA1306GAA (5526A>G), TAT1307AAG (5529T>A 5531T>G), TCA1308del (5532_5534delTCA), GAA1309AAT (5535G>A 5537A>T), TTT1310TTC (5540T>C), GTC1311CTA (5541G>C 5543C>A), TGT1312ATT (5544T>A 5545G>T), GTT1313GTC (5549T>C), TTG1319TTA (5567G>A), TTT1321TGT (5572T>G), TCT1322TCA (5576T>A), GAA1323TCA (5577G>T 5578A>C), AGT1324ACA (5581G>C 5582T>A), ATA1325TAC (5583A>T 5584T>A 5585A>C), CAA1327GAA (5589C>G), GTA1329CTA (5595G>C), CAA1330ACA (5598C>A 5599A>C), TTA1332CTT (5604T>C 5606A>T), TTA1333AAA (5607T>A 5608T>A), CAG1334GAA (5610C>G 5612G>A), TTC1335TTT (5615C>T), TTC1336TCA (5617T>C 5618C>A), CAA1337AAA (5619C>A), GTA1338ATT (5622G>A 5624A>T), AAG1340GTT (5628A>G 5629A>T 5630G>T), GAA1341GAC (5633A>C), GAA1342AAT (5634G>A 5636A>T), GGG1343GGA (5639G>A), CTT1344ATT (5640C>A), ATT1345GTT (5643A>G), TCA1347AGC (5649T>A 5650C>G 5651A>C), AAA1348CAA (5652A>C), ACA1349-AA (5655delA 5656C>A), AAA1350AAG (5660A>G), TTG1351GCT (5661T>G 5662T>C 5663G>T), GGA1354AAT (5670G>A 5671G>A 5672A>T), GTA1355AAA (5673G>A 5674T>A), GCA1356ATT (5676G>A 5677C>T 5678A>T), AAT1357GAA (5679A>G 5681T>A), TTC1360TTT (5690C>T), GGC1362GGA (5696C>A), CTA1363ATG (5697C>A 5699A>G), GAA1364ATT (5700G>A 5701A>T 5702A>T), ATA1365ATT (5705A>T), GGA1366TCA (5706G>T 5707G>C), GAA1367AAA (5709G>A), GGA1368AAT (5712G>A 5713G>A 5714A>T), AAA1369GGG (5715A>G 5716A>G 5717A>G), GTT1370ATA (5718G>A 5720T>A), CAA1371GAA (5721C>G), CTC1372TTA (5724C>T 5726C>A), CAG1373CAA (5729G>A), CCC1374ACT (5730C>A 5732C>T), ATA1376ATT (5738A>T), TTA1377TCA (5740T>C), AAA1378AAG (5744A>G), AAT1379AAA (5747T>A), TTA1381ACG (5751T>A 5752T>C 5753A>G), TTT1383TTC (5759T>C), CCT1384TCG (5760C>T 5762T>G), GAA1385del (5763_5765delGAA), GAC1386GAT (5768C>T), CAA1387AAA (5769C>A), GAA1389ACA (5775G>A 5776A>C), TTA1391AAG (5781T>A 5782T>A 5783A>G), AAG1392CAA (5784A>C 5786G>A), GGT1393GAA (5788G>A 5789T>A), CTA1394ATA (5790C>A), CAG1395CAA (5795G>A), AAG1396AAA (5798G>A), TTC1397TTT (5801C>T), TTA1398CTT (5802T>C 5804A>T), GGG1399GGT (5807G>T), ATA1400TGT (5808A>T 5809T>G 5810A>T), TTA1401CTG (5811T>C 5813A>G), AAC1402AAT (5816C>T), GCC1404GCA (5822C>A), AGA1405GGT (5823A>G 5825A>T), AAT1406GAG (5826A>G 5828T>G), TAT1407TTT (5830A>T), ATC1408ATT (5834C>T), CCA1409TCG (5835C>T 5837A>G) | | | | | | | | |

##### Proteins

| | Begin | End | Coverage | Score | Concordance | Matches | Identities | I/D/M/F* | Stop Codons |
|---|---|---|---|---|---|---|---|---|---|
| ORF3 (YP_009553219.1) | 1268 | 1411 | 9.0% | 442 | 43.8% | 142 (98.6%) | 78 (54.2%) | 0/2/1/1 | 0 |
| Protein mutations: | E1269A (5416A>C 5417A>T), W1270L (5419G>T), V1274S (5430G>A 5431T>G 5432C>T), C1275T (5433T>A 5434G>C 5435C>A), Q1277N (5439C>A 5441G>T), F1280Y (5449T>A), I1283L (5457A>T), S1294Q (5490T>C 5491C>A), V1295I (5493G>A 5495A>C), N1302E (5514A>G 5516T>A), K1305S (5523A>T 5524A>C 5525A>T), K1306E (5526A>G), Y1307K (5529T>A 5531T>G), S1308del (5532_5534delTCA), E1309N (5535G>A 5537A>T), V1311L (5541G>C 5543C>A), C1312I (5544T>A 5545G>T), F1321C (5572T>G), E1323S (5577G>T 5578A>C), S1324T (5581G>C 5582T>A), I1325Y (5583A>T 5584T>A 5585A>C), Q1327E (5589C>G), V1329L (5595G>C), Q1330T (5598C>A 5599A>C), L1333K (5607T>A 5608T>A), Q1334E (5610C>G 5612G>A), F1336S (5617T>C 5618C>A), Q1337K (5619C>A), V1338I (5622G>A 5624A>T), K1340V (5628A>G 5629A>T 5630G>T), E1341D (5633A>C), E1342N (5634G>A 5636A>T), L1344I (5640C>A), I1345V (5643A>G), K1348Q (5652A>C), L1351A (5661T>G 5662T>C 5663G>T), G1354N (5670G>A 5671G>A 5672A>T), V1355K (5673G>A 5674T>A), A1356I (5676G>A 5677C>T 5678A>T), N1357E (5679A>G 5681T>A), L1363M (5697C>A 5699A>G), E1364I (5700G>A 5701A>T 5702A>T), G1366S (5706G>T 5707G>C), E1367K (5709G>A), G1368N (5712G>A 5713G>A 5714A>T), K1369G (5715A>G 5716A>G 5717A>G), V1370I (5718G>A 5720T>A), Q1371E (5721C>G), P1374T (5730C>A 5732C>T), L1377S (5740T>C), N1379K (5747T>A), L1381T (5751T>A 5752T>C 5753A>G), P1384S (5760C>T 5762T>G), E1385del (5763_5765delGAA), Q1387K (5769C>A), E1389T (5775G>A 5776A>C), L1391K (5781T>A 5782T>A 5783A>G), K1392Q (5784A>C 5786G>A), G1393E (5788G>A 5789T>A), L1394I (5790C>A), I1400C (5808A>T 5809T>G 5810A>T), R1405G (5823A>G 5825A>T), N1406E (5826A>G 5828T>G), Y1407F (5830A>T), P1409S (5835C>T 5837A>G) | | | | | | | | |
| Codon mutations: | AGT1267.CT (5410G>C), GAA1269GCT (5416A>C 5417A>T), TGG1270TTG (5419G>T), TTT1273TTC (5429T>C), GTC1274AGT (5430G>A 5431T>G 5432C>T), TGC1275ACA (5433T>A 5434G>C 5435C>A), CAG1277AAT (5439C>A 5441G>T), GGT1278GGA (5444T>A), TTT1280TAT (5449T>A), ATA1283TTA (5457A>T), CCA1286CCT (5468A>T), TTC1287TTT (5471C>T), CTC1289TTA (5475C>T 5477C>A), GCA1292GCC (5486A>C), TCA1294CAA (5490T>C 5491C>A), GTA1295ATC (5493G>A 5495A>C), TTC1296TTT (5498C>T), CGA1298AGA (5502C>A), AAT1302GAA (5514A>G 5516T>A), TTC1304TTT (5522C>T), AAA1305TCT (5523A>T 5524A>C 5525A>T), AAA1306GAA (5526A>G), TAT1307AAG (5529T>A 5531T>G), TCA1308del (5532_5534delTCA), GAA1309AAT (5535G>A 5537A>T), TTT1310TTC (5540T>C), GTC1311CTA (5541G>C 5543C>A), TGT1312ATT (5544T>A 5545G>T), GTT1313GTC (5549T>C), TTG1319TTA (5567G>A), TTT1321TGT (5572T>G), TCT1322TCA (5576T>A), GAA1323TCA (5577G>T 5578A>C), AGT1324ACA (5581G>C 5582T>A), ATA1325TAC (5583A>T 5584T>A 5585A>C), CAA1327GAA (5589C>G), GTA1329CTA (5595G>C), CAA1330ACA (5598C>A 5599A>C), TTA1332CTT (5604T>C 5606A>T), TTA1333AAA (5607T>A 5608T>A), CAG1334GAA (5610C>G 5612G>A), TTC1335TTT (5615C>T), TTC1336TCA (5617T>C 5618C>A), CAA1337AAA (5619C>A), GTA1338ATT (5622G>A 5624A>T), AAG1340GTT (5628A>G 5629A>T 5630G>T), GAA1341GAC (5633A>C), GAA1342AAT (5634G>A 5636A>T), GGG1343GGA (5639G>A), CTT1344ATT (5640C>A), ATT1345GTT (5643A>G), TCA1347AGC (5649T>A 5650C>G 5651A>C), AAA1348CAA (5652A>C), ACA1349-AA (5655delA 5656C>A), AAA1350AAG (5660A>G), TTG1351GCT (5661T>G 5662T>C 5663G>T), GGA1354AAT (5670G>A 5671G>A 5672A>T), GTA1355AAA (5673G>A 5674T>A), GCA1356ATT (5676G>A 5677C>T 5678A>T), AAT1357GAA (5679A>G 5681T>A), TTC1360TTT (5690C>T), GGC1362GGA (5696C>A), CTA1363ATG (5697C>A 5699A>G), GAA1364ATT (5700G>A 5701A>T 5702A>T), ATA1365ATT (5705A>T), GGA1366TCA (5706G>T 5707G>C), GAA1367AAA (5709G>A), GGA1368AAT (5712G>A 5713G>A 5714A>T), AAA1369GGG (5715A>G 5716A>G 5717A>G), GTT1370ATA (5718G>A 5720T>A), CAA1371GAA (5721C>G), CTC1372TTA (5724C>T 5726C>A), CAG1373CAA (5729G>A), CCC1374ACT (5730C>A 5732C>T), ATA1376ATT (5738A>T), TTA1377TCA (5740T>C), AAA1378AAG (5744A>G), AAT1379AAA (5747T>A), TTA1381ACG (5751T>A 5752T>C 5753A>G), TTT1383TTC (5759T>C), CCT1384TCG (5760C>T 5762T>G), GAA1385del (5763_5765delGAA), GAC1386GAT (5768C>T), CAA1387AAA (5769C>A), GAA1389ACA (5775G>A 5776A>C), TTA1391AAG (5781T>A 5782T>A 5783A>G), AAG1392CAA (5784A>C 5786G>A), GGT1393GAA (5788G>A 5789T>A), CTA1394ATA (5790C>A), CAG1395CAA (5795G>A), AAG1396AAA (5798G>A), TTC1397TTT (5801C>T), TTA1398CTT (5802T>C 5804A>T), GGG1399GGT (5807G>T), ATA1400TGT (5808A>T 5809T>G 5810A>T), TTA1401CTG (5811T>C 5813A>G), AAC1402AAT (5816C>T), GCC1404GCA (5822C>A), AGA1405GGT (5823A>G 5825A>T), AAT1406GAG (5826A>G 5828T>G), TAT1407TTT (5830A>T), ATC1408ATT (5834C>T), CCA1409TCG (5835C>T 5837A>G) | | | | | | | | |

*: Inserts / Deletes / Misaligned / Frameshifts

#### Analysis details

This analysis was performed with panviral2.64

### NGS Details (UN62): Duamitovirus peex1

#### Assembly

| | |
|---|---|
| **Coverage Length** | 263 (1 contig(s)) |
| **Depth Of Coverage** | 5.1 |
| **Number Of Reads** | 10 |
| **Reads Per Million** | 0.23 rpm (after QC) |
| **Ambiguities** | 0 |
| **Assembly Method** | de novo + reference guided assembly |
| **Consensus Caller** | Bcf Tools |

#### Coverage Map

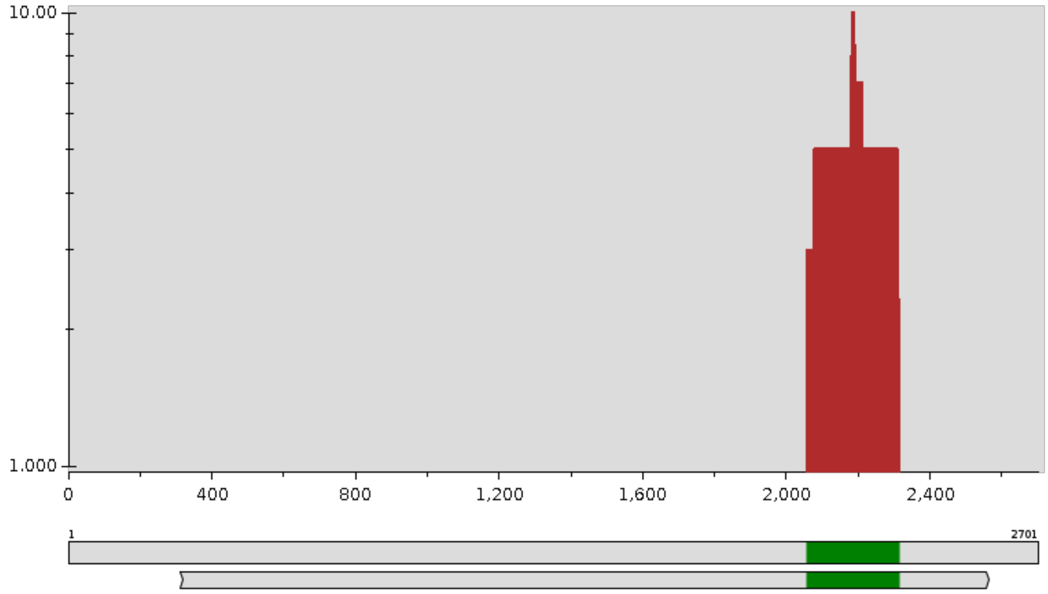

#### Assignment

| | |
|---|---|
| **Type** | Duamitovirus peex1 (Taxonomy ID: 2955799) |
| **Reference Genome** | NC_076525.1 |
| **NT Identity (%)** | 73.384 |
| **AA Identity (%)** | 64.3678 |
| **Number Of Stop Codons** | 0 |
| **Number Of CDS** | 1 |

#### Alignment

| | |
|---|---|
| **Alignment Score** | 246.0 (NT) + 438.0 (AA) = 684.0 |
| **Concordance (%)** | 57.4307 |

| Alignment Method | Global, seeded, nucleotide + amino acids (AGA) |
|---|---|

#### Genome Region

Sequence starts at position 2056 and ends at position 2318 relative to NC_076525.1 reference sequence.

#### Alignment Detailed Statistics

| | Begin | End | Coverage | Score | Concordance | Matches | Identities | I/D/M/F* | Stop Codons |
|---|---|---|---|---|---|---|---|---|---|
| **NT** | **2056** | **2318** | **9.7%** | **246** | **46.8%** | **263 (100%)** | **193 (73.4%)** | **0/0** | |
| Mutations: | 2057C>A, 2066G>A, 2067A>C, 2069T>A, 2075T>G, 2077T>C, 2088T>C, 2090T>A, 2091T>A, 2095T>G, 2096A>G, 2098C>T, 2102G>T, 2104G>T, 2105G>A, 2109T>C, 2119G>T, 2120G>T, 2133G>T, 2134G>T, 2135A>T, 2136A>G, 2137A>G, 2138G>A, 2139C>A, 2145G>A, 2156A>G, 2158G>A, 2159A>G, 2161G>C, 2162T>G, 2165G>A, 2171C>A, 2173C>A, 2180T>G, 2184A>G, 2188A>C, 2189G>T, 2190T>G, 2195A>G, 2202G>C, 2210G>A, 2211T>A, 2213G>C, 2214A>T, 2216A>T, 2223C>G, 2225A>T, 2226C>T, 2231G>A, 2234A>T, 2240T>A, 2245A>G, 2246T>G, 2249G>A, 2255A>T, 2258A>G, 2261C>G, 2268G>A, 2270A>C, 2272G>T, 2274C>A, 2283G>A, 2292C>T, 2294T>G, 2295C>A, 2296G>C, 2297A>T, 2306A>T, 2315C>T | | | | | | | | |
| **CDS** | | | | | | | | | |
| RdRp | 583 | 669 | 11.6% | 438 | 65.1% | 87 (100%) | 56 (64.4%) | 0/0/0/0 | 0 |
| Protein mutations: | I586L (2067A>C 2069T>A), V589A (2077T>C), S593P (2088T>C 2090T>A), Y594N (2091T>A), I595R (2095T>G 2096A>G), S596L (2098C>T), Q597H (2102G>T), R598L (2104G>T 2105G>A), S600P (2109T>C), W603F (2119G>T 2120G>T), G608F (2133G>T 2134G>T 2135A>T), K609G (2136A>G 2137A>G 2138G>A), P610T (2139C>A), D612N (2145G>A), R616K (2158G>A 2159A>G), G617A (2161G>C 2162T>G), A621D (2173C>A), K625E (2184A>G), E626A (2188A>C 2189G>T), L627V (2190T>G), E631Q (2202G>C), L634I (2211T>A 2213G>C), I635F (2214A>T 2216A>T), Q638D (2223C>G 2225A>T), L641F (2234A>T), H645R (2245A>G 2246T>G), V653I (2268G>A 2270A>C), R654L (2272G>T), H655N (2274C>A), E658K (2283G>A), R662T (2295C>A 2296G>C 2297A>T) | | | | | | | | |
| Codon mutations: | CGC582.GA (2057C>A), AGG585AGA (2066G>A), ATT586CTA (2067A>C 2069T>A), GCT588GCG (2075T>G), GTT589GCT (2077T>C), TCT593CCA (2088T>C 2090T>A), TAC594AAC (2091T>A), ATA595AGG (2095T>G 2096A>G), TCA596TTA (2098C>T), CAG597CAT (2102G>T), CGG598CTA (2104G>T 2105G>A), TCT600CCT (2109T>C), TGG603TTT (2119G>T 2120G>T), GGA608TTT (2133G>T 2134G>T 2135A>T), AAG609GGA (2136A>G 2137A>G 2138G>A), CCC610ACC (2139C>A), GAT612AAT (2145G>A), TTA615TTG (2156A>G), AGA616AAG (2158G>A 2159A>G), GGT617GCG (2161G>C 2162T>G), AAG618AAA (2165G>A), ATC620ATA (2171C>A), GCT621GAT (2173C>A), CTT623CTG (2180T>G), AAG625GAG (2184A>G), GAG626GCT (2188A>C 2189G>T), TTA627GTA (2190T>G), AAA628AAG (2195A>G), GAG631CAG (2202G>C), CAG633CAA (2210G>A), TTG634ATC (2211T>A 2213G>C), ATA635TTT (2214A>T 2216A>T), CAA638GAT (2223C>G 2225A>T), CTG639TTG (2226C>T), GTG640GTA (2231G>A), TTA641TTT (2234A>T), GGT643GGA (2240T>A), CAT645CGG (2245A>G 2246T>G), GAG646GAA (2249G>A), CTA648CTT (2255A>T), GAA649GAG (2258A>G), CGC650CGG (2261C>G), GTA653ATC (2268G>A 2270A>C), CGT654CTT (2272G>T), CAT655AAT (2274C>A), GAA658AAA (2283G>A), CTT661TTG (2292C>T 2294T>G), CGA662ACT (2295C>A 2296G>C 2297A>T), TCA665TCT (2306A>T), CAC668CAT (2315C>T) | | | | | | | | |
| **Proteins** | | | | | | | | | |
| RNA-dependent RNA polymerase (YP_010798875.1) | 583 | 669 | 11.6% | 438 | 65.1% | 87 (100%) | 56 (64.4%) | 0/0/0/0 | 0 |
| Protein mutations: | I586L (2067A>C 2069T>A), V589A (2077T>C), S593P (2088T>C 2090T>A), Y594N (2091T>A), I595R (2095T>G 2096A>G), S596L (2098C>T), Q597H (2102G>T), R598L (2104G>T 2105G>A), S600P (2109T>C), W603F (2119G>T 2120G>T), G608F (2133G>T 2134G>T 2135A>T), K609G (2136A>G 2137A>G 2138G>A), P610T (2139C>A), D612N (2145G>A), R616K (2158G>A 2159A>G), G617A (2161G>C 2162T>G), A621D (2173C>A), K625E (2184A>G), E626A (2188A>C 2189G>T), L627V (2190T>G), E631Q (2202G>C), L634I (2211T>A 2213G>C), I635F (2214A>T 2216A>T), Q638D (2223C>G 2225A>T), L641F (2234A>T), H645R (2245A>G 2246T>G), V653I (2268G>A 2270A>C), R654L (2272G>T), H655N (2274C>A), E658K (2283G>A), R662T (2295C>A 2296G>C 2297A>T) | | | | | | | | |
| Codon mutations: | CGC582.GA (2057C>A), AGG585AGA (2066G>A), ATT586CTA (2067A>C 2069T>A), GCT588GCG (2075T>G), GTT589GCT (2077T>C), TCT593CCA (2088T>C 2090T>A), TAC594AAC (2091T>A), ATA595AGG (2095T>G 2096A>G), TCA596TTA (2098C>T), CAG597CAT (2102G>T), CGG598CTA (2104G>T 2105G>A), TCT600CCT (2109T>C), TGG603TTT (2119G>T 2120G>T), GGA608TTT (2133G>T 2134G>T 2135A>T), AAG609GGA (2136A>G 2137A>G 2138G>A), CCC610ACC (2139C>A), GAT612AAT (2145G>A), TTA615TTG (2156A>G), AGA616AAG (2158G>A 2159A>G), GGT617GCG (2161G>C 2162T>G), AAG618AAA (2165G>A), ATC620ATA (2171C>A), GCT621GAT (2173C>A), CTT623CTG (2180T>G), AAG625GAG (2184A>G), GAG626GCT (2188A>C 2189G>T), TTA627GTA (2190T>G), AAA628AAG (2195A>G), GAG631CAG (2202G>C), CAG633CAA (2210G>A), TTG634ATC (2211T>A 2213G>C), ATA635TTT (2214A>T 2216A>T), CAA638GAT (2223C>G 2225A>T), CTG639TTG (2226C>T), GTG640GTA (2231G>A), TTA641TTT (2234A>T), GGT643GGA (2240T>A), CAT645CGG (2245A>G 2246T>G), GAG646GAA (2249G>A), CTA648CTT (2255A>T), GAA649GAG (2258A>G), CGC650CGG (2261C>G), GTA653ATC (2268G>A 2270A>C), CGT654CTT (2272G>T), CAT655AAT (2274C>A), GAA658AAA (2283G>A), CTT661TTG (2292C>T 2294T>G), CGA662ACT (2295C>A 2296G>C 2297A>T), TCA665TCT (2306A>T), CAC668CAT (2315C>T) | | | | | | | | |

*: Inserts / Deletes / Misaligned / Frameshifts

#### Analysis details

This analysis was performed with panviral2.64

### NGS Details (UN62): Cacao swollen shoot Ghana J virus

#### Assembly

| | |
|---|---|
| **Coverage Length** | 305 (1 contig(s)) |
| **Depth Of Coverage** | 4.1 |
| **Number Of Reads** | 10 |
| **Reads Per Million** | 0.23 rpm (after QC) |
| **Ambiguities** | 0 |
| **Assembly Method** | de novo + reference guided assembly |
| **Consensus Caller** | Bcf Tools |

#### Coverage Map

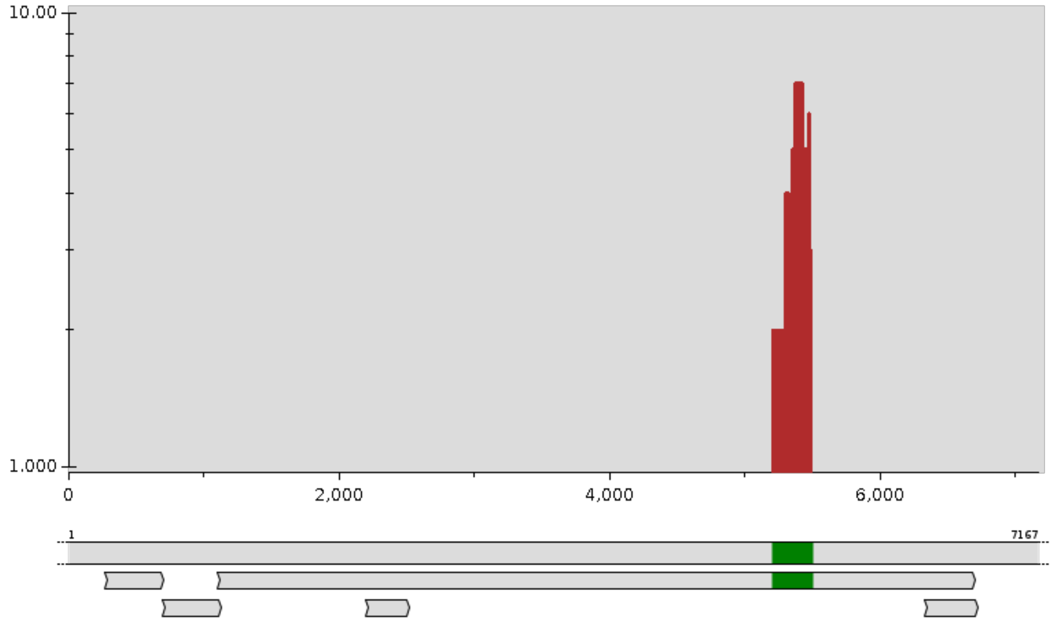

#### Assignment

| | |
|---|---|
| **Type** | Cacao swollen shoot Ghana J virus (Taxonomy ID: 2056880) |
| **Reference Genome** | NC_040807.1 |
| **NT Identity (%)** | 58.0328 |
| **AA Identity (%)** | 49.505 |
| **Number Of Stop Codons** | 0 |
| **Number Of CDS** | 5 |

#### Alignment

| | |
|---|---|
| **Alignment Score** | 98.0 (NT) + 346.0 (AA) = 444.0 |
| **Concordance (%)** | 32.9866 |

| Alignment Method | Global, seeded, nucleotide + amino acids (AGA) |
|---|---|

#### Genome Region

Sequence starts at position 5200 and ends at position 5504 relative to NC_040807.1 reference sequence.

#### Alignment Detailed Statistics

| | Begin | End | Coverage | Score | Concordance | Matches | Identities | I/D/M/F* | Stop Codons |
|---|---|---|---|---|---|---|---|---|---|
| **NT** | **5200** | **5504** | **4.3%** | **98** | **16.1%** | **305 (100%)** | **177 (58.0%)** | **0/0** | |
| Mutations: | 5201G>A, 5204G>A, 5205G>T, 5210T>A, 5220A>C, 5221G>A, 5222A>T, 5223C>T, 5225C>A, 5229G>A, 5231C>A, 5232A>T, 5236C>T, 5237T>G, 5238G>A, 5241A>T, 5242A>G, 5243A>G, 5244G>A, 5245A>T, 5246C>T, 5247C>A, 5248A>G, 5253T>A, 5256C>A, 5258A>T, 5261A>T, 5262G>A, 5263G>A, 5264A>T, 5268C>A, 5271A>G, 5272C>A, 5274A>T, 5276C>G, 5277C>T, 5279G>A, 5280A>T, 5281A>C, 5282G>T, 5284A>G, 5285G>A, 5286G>C, 5288C>G, 5290G>A, 5292A>G, 5294C>T, 5295A>G, 5296T>C, 5297G>T, 5300G>T, 5303T>A, 5306C>T, 5307A>T, 5308G>C, 5309C>A, 5312G>A, 5319C>T, 5327T>A, 5330T>A, 5332T>A, 5333C>T, 5334C>T, 5335A>G, 5336C>G, 5339G>A, 5340G>A, 5342T>A, 5343G>C, 5344C>A, 5345T>G, 5348G>T, 5349G>T, 5350C>T, 5352G>A, 5357A>G, 5358T>C, 5359C>A, 5362T>C, 5364C>T, 5365C>A, 5366A>T, 5367T>A, 5369G>A, 5378C>T, 5379T>A, 5380G>A, 5381G>T, 5384A>C, 5388C>T, 5389A>T, 5390G>T, 5394T>C, 5395T>A, 5398T>A, 5402G>A, 5406C>A, 5407T>A, 5408C>T, 5424T>C, 5426G>A, 5439G>T, 5441A>T, 5443T>A, 5444C>A, 5447C>T, 5450A>G, 5451C>A, 5452G>A, 5453G>A, 5455A>T, 5456G>T, 5460G>A, 5463C>G, 5465A>T, 5466T>A, 5467G>A, 5474A>T, 5475G>C, 5476G>C, 5478A>T, 5479C>A, 5481G>A, 5482A>T, 5483A>G, 5489T>C, 5493G>A, 5494C>T | | | | | | | | |

##### CDS

| | Begin | End | Coverage | Score | Concordance | Matches | Identities | I/D/M/F* | Stop Codons |
|---|---|---|---|---|---|---|---|---|---|
| EXL33_gp3 | 1369 | 1469 | 5.4% | 346 | 46.5% | 101 (100%) | 50 (49.5%) | 0/0/0/0 | 0 |
| Protein mutations: | V1370F (5205G>T), F1371L (5210T>A), R1375H (5220A>C 5221G>A 5222A>T), D1378K (5229G>A 5231C>A), N1379Y (5232A>T), T1380M (5236C>T 5237T>G), E1381K (5238G>A), K1382W (5241A>T 5242A>G 5243A>G), D1383I (5244G>A 5245A>T 5246C>T), Q1384R (5247C>A 5248A>G), S1386T (5253T>A), L1387I (5256C>A 5258A>T), G1389N (5262G>A 5263G>A 5264A>T), Q1391K (5268C>A), T1392D (5271A>G 5272C>A), I1393L (5274A>T 5276C>G), K1395S (5280A>T 5281A>C 5282G>T), K1396R (5284A>G 5285G>A), V1397L (5286G>C 5288C>G), C1398Y (5290G>A), N1399D (5292A>G 5294C>T), M1400A (5295A>G 5296T>C 5297G>T), K1401N (5300G>T), F1412Y (5332T>A 5333C>T), H1413W (5334C>T 5335A>G 5336C>G), V1415I (5340G>A 5342T>A), A1416Q (5343G>C 5344C>A 5345T>G), M1417I (5348G>T), A1418F (5349G>T 5350C>T), E1419K (5352G>A), S1421H (5358T>C 5359C>A), I1422T (5362T>C), P1423Y (5364C>T 5365C>A 5366A>T), W1424R (5367T>A 5369G>A), W1428N (5379T>A 5380G>A 5381G>T), Q1431F (5388C>T 5389A>T 5390G>T), L1433Q (5394T>C 5395T>A), F1434Y (5398T>A), L1437N (5406C>A 5407T>A 5408C>T), A1448S (5439G>T 5441A>T), V1449E (5443T>A 5444C>A), R1452K (5451C>A 5452G>A 5453G>A), K1453I (5455A>T 5456G>T), D1455N (5460G>A), Q1456D (5463C>G 5465A>T), C1457N (5466T>A 5467G>A), K1459N (5474A>T), G1460P (5475G>C 5476G>C), T1461Y (5478A>T 5479C>A), E1462M (5481G>A 5482A>T 5483A>G), A1466I (5493G>A 5494C>T) | | | | | | | | |
| Codon mutations: | AGG1368.GA (5201G>A), CTG1369CTA (5204G>A), GTT1370TTT (5205G>T), TTT1371TTA (5210T>A), AGA1375CAT (5220A>C 5221G>A 5222A>T), CTC1376TTA (5223C>T 5225C>A), GAC1378AAA (5229G>A 5231C>A), AAT1379TAT (5232A>T), ACT1380ATG (5236C>T 5237T>G), GAA1381AAA (5238G>A), AAA1382TGG (5241A>T 5242A>G 5243A>G), GAC1383ATT (5244G>A 5245A>T 5246C>T), CAG1384AGG (5247C>A 5248A>G), TCT1386ACT (5253T>A), CTA1387ATT (5256C>A 5258A>T), CCA1388CCT (5261A>T), GGA1389AAT (5262G>A 5263G>A 5264A>T), CAG1391AAG (5268C>A), ACT1392GAT (5271A>G 5272C>A), ATC1393TTG (5274A>T 5276C>G), CTG1394TTA (5277C>T 5279G>A), AAG1395TCT (5280A>T 5281A>C 5282G>T), AAG1396AGA (5284A>G 5285G>A), GTC1397CTG (5286G>C 5288C>G), TGT1398TAT (5290G>A), AAC1399GAT (5292A>G 5294C>T), ATG1400GCT (5295A>G 5296T>C 5297G>T), AAG1401AAT (5300G>T), ATT1402ATA (5303T>A), TTC1403TTT (5306C>T), AGC1404TCA (5307A>T 5308G>C 5309C>A), AAG1405AAA (5312G>A), CTA1408TTA (5319C>T), TCT1410TCA (5327T>A), GGT1411GGA (5330T>A), TTC1412TAT (5332T>A 5333C>T), CAC1413TGG (5334C>T 5335A>G 5336C>G), CAG1414CAA (5339G>A), GTT1415ATA (5340G>A 5342T>A), GCT1416CAG (5343G>C 5344C>A 5345T>G), ATG1417ATT (5348G>T), GCT1418TTT (5349G>T 5350C>T), GAA1419AAA (5352G>A), GAA1420GAG (5357A>G), TCT1421CAT (5358T>C 5359C>A), ATT1422ACT (5362T>C), CCA1423TAT (5364C>T 5365C>A 5366A>T), TGG1424AGA (5367T>A 5369G>A), TTC1427TTT (5378C>T), TGG1428AAT (5379T>A 5380G>A 5381G>T), GTA1429GTC (5384A>C), CAG1431TTT (5388C>T 5389A>T 5390G>T), TTA1433CAA (5394T>C 5395T>A), TTT1434TAT (5398T>A), GAG1435GAA (5402G>A), CTC1437AAT (5406C>A 5407T>A 5408C>T), TTG1443CTA (5424T>C 5426G>A), GCA1448TCT (5439G>T 5441A>T), GTC1449GAA (5443T>A 5444C>A), TTC1450TTT (5447C>T), CAA1451CAG (5450A>G), CGG1452AAA (5451C>A 5452G>A 5453G>A), AAG1453ATT (5455A>T 5456G>T), GAT1455AAT (5460G>A), CAA1456GAT (5463C>G 5465A>T), TGT1457AAT (5466T>A 5467G>A), AAA1459AAT (5474A>T), GGA1460CCA (5475G>C 5476G>C), ACT1461TAT (5478A>T 5479C>A), GAA1462ATG (5481G>A 5482A>T 5483A>G), TTT1464TTC (5489T>C), GCT1466ATT (5493G>A 5494C>T) | | | | | | | | |

##### Proteins

| | Begin | End | Coverage | Score | Concordance | Matches | Identities | I/D/M/F* | Stop Codons |
|---|---|---|---|---|---|---|---|---|---|
| ORF3 polyprotein (YP_009553538.1) | 1369 | 1469 | 5.4% | 346 | 46.5% | 101 (100%) | 50 (49.5%) | 0/0/0/0 | 0 |
| Protein mutations: | V1370F (5205G>T), F1371L (5210T>A), R1375H (5220A>C 5221G>A 5222A>T), D1378K (5229G>A 5231C>A), N1379Y (5232A>T), T1380M (5236C>T 5237T>G), E1381K (5238G>A), K1382W (5241A>T 5242A>G 5243A>G), D1383I (5244G>A 5245A>T 5246C>T), Q1384R (5247C>A 5248A>G), S1386T (5253T>A), L1387I (5256C>A 5258A>T), G1389N (5262G>A 5263G>A 5264A>T), Q1391K (5268C>A), T1392D (5271A>G 5272C>A), I1393L (5274A>T 5276C>G), K1395S (5280A>T 5281A>C 5282G>T), K1396R (5284A>G 5285G>A), V1397L (5286G>C 5288C>G), C1398Y (5290G>A), N1399D (5292A>G 5294C>T), M1400A (5295A>G 5296T>C 5297G>T), K1401N (5300G>T), F1412Y (5332T>A 5333C>T), H1413W (5334C>T 5335A>G 5336C>G), V1415I (5340G>A 5342T>A), A1416Q (5343G>C 5344C>A 5345T>G), M1417I (5348G>T), A1418F (5349G>T 5350C>T), E1419K (5352G>A), S1421H (5358T>C 5359C>A), I1422T (5362T>C), P1423Y (5364C>T 5365C>A 5366A>T), W1424R (5367T>A 5369G>A), W1428N (5379T>A 5380G>A 5381G>T), Q1431F (5388C>T 5389A>T 5390G>T), L1433Q (5394T>C 5395T>A), F1434Y (5398T>A), L1437N (5406C>A 5407T>A 5408C>T), A1448S (5439G>T 5441A>T), V1449E (5443T>A 5444C>A), R1452K (5451C>A 5452G>A 5453G>A), K1453I (5455A>T 5456G>T), D1455N (5460G>A), Q1456D (5463C>G 5465A>T), C1457N (5466T>A 5467G>A), K1459N (5474A>T), G1460P (5475G>C 5476G>C), T1461Y (5478A>T 5479C>A), E1462M (5481G>A 5482A>T 5483A>G), A1466I (5493G>A 5494C>T) | | | | | | | | |
| Codon mutations: | AGG1368.GA (5201G>A), CTG1369CTA (5204G>A), GTT1370TTT (5205G>T), TTT1371TTA (5210T>A), AGA1375CAT (5220A>C 5221G>A 5222A>T), CTC1376TTA (5223C>T 5225C>A), GAC1378AAA (5229G>A 5231C>A), AAT1379TAT (5232A>T), ACT1380ATG (5236C>T 5237T>G), GAA1381AAA (5238G>A), AAA1382TGG (5241A>T 5242A>G 5243A>G), GAC1383ATT (5244G>A 5245A>T 5246C>T), CAG1384AGG (5247C>A 5248A>G), TCT1386ACT (5253T>A), CTA1387ATT (5256C>A 5258A>T), CCA1388CCT (5261A>T), GGA1389AAT (5262G>A 5263G>A 5264A>T), CAG1391AAG (5268C>A), ACT1392GAT (5271A>G 5272C>A), ATC1393TTG (5274A>T 5276C>G), CTG1394TTA (5277C>T 5279G>A), AAG1395TCT (5280A>T 5281A>C 5282G>T), AAG1396AGA (5284A>G 5285G>A), GTC1397CTG (5286G>C 5288C>G), TGT1398TAT (5290G>A), AAC1399GAT (5292A>G 5294C>T), ATG1400GCT (5295A>G 5296T>C 5297G>T), AAG1401AAT (5300G>T), ATT1402ATA (5303T>A), TTC1403TTT (5306C>T), AGC1404TCA (5307A>T 5308G>C 5309C>A), AAG1405AAA (5312G>A), CTA1408TTA (5319C>T), TCT1410TCA (5327T>A), GGT1411GGA (5330T>A), TTC1412TAT (5332T>A 5333C>T), CAC1413TGG (5334C>T 5335A>G 5336C>G), CAG1414CAA (5339G>A), GTT1415ATA (5340G>A 5342T>A), GCT1416CAG (5343G>C 5344C>A 5345T>G), ATG1417ATT (5348G>T), GCT1418TTT (5349G>T 5350C>T), GAA1419AAA (5352G>A), GAA1420GAG (5357A>G), TCT1421CAT (5358T>C 5359C>A), ATT1422ACT (5362T>C), CCA1423TAT (5364C>T 5365C>A 5366A>T), TGG1424AGA (5367T>A 5369G>A), TTC1427TTT (5378C>T), TGG1428AAT (5379T>A 5380G>A 5381G>T), GTA1429GTC (5384A>C), CAG1431TTT (5388C>T 5389A>T 5390G>T), TTA1433CAA (5394T>C 5395T>A), TTT1434TAT (5398T>A), GAG1435GAA (5402G>A), CTC1437AAT (5406C>A 5407T>A 5408C>T), TTG1443CTA (5424T>C 5426G>A), GCA1448TCT (5439G>T 5441A>T), GTC1449GAA (5443T>A 5444C>A), TTC1450TTT (5447C>T), CAA1451CAG (5450A>G), CGG1452AAA (5451C>A 5452G>A 5453G>A), AAG1453ATT (5455A>T 5456G>T), GAT1455AAT (5460G>A), CAA1456GAT (5463C>G 5465A>T), TGT1457AAT (5466T>A 5467G>A), AAA1459AAT (5474A>T), GGA1460CCA (5475G>C 5476G>C), ACT1461TAT (5478A>T 5479C>A), GAA1462ATG (5481G>A 5482A>T 5483A>G), TTT1464TTC (5489T>C), GCT1466ATT (5493G>A 5494C>T) | | | | | | | | |

*: Inserts / Deletes / Misaligned / Frameshifts

#### Analysis details

This analysis was performed with panviral2.64

### NGS Details (UN62): Hibiscus bacilliform virus GD1

#### Assembly

| | |
|---|---|
| **Coverage Length** | 238 (1 contig(s)) |
| **Depth Of Coverage** | 5.5 |
| **Number Of Reads** | 10 |
| **Reads Per Million** | 0.23 rpm (after QC) |
| **Ambiguities** | 0 |
| **Assembly Method** | de novo + reference guided assembly |
| **Consensus Caller** | Bcf Tools |

#### Coverage Map

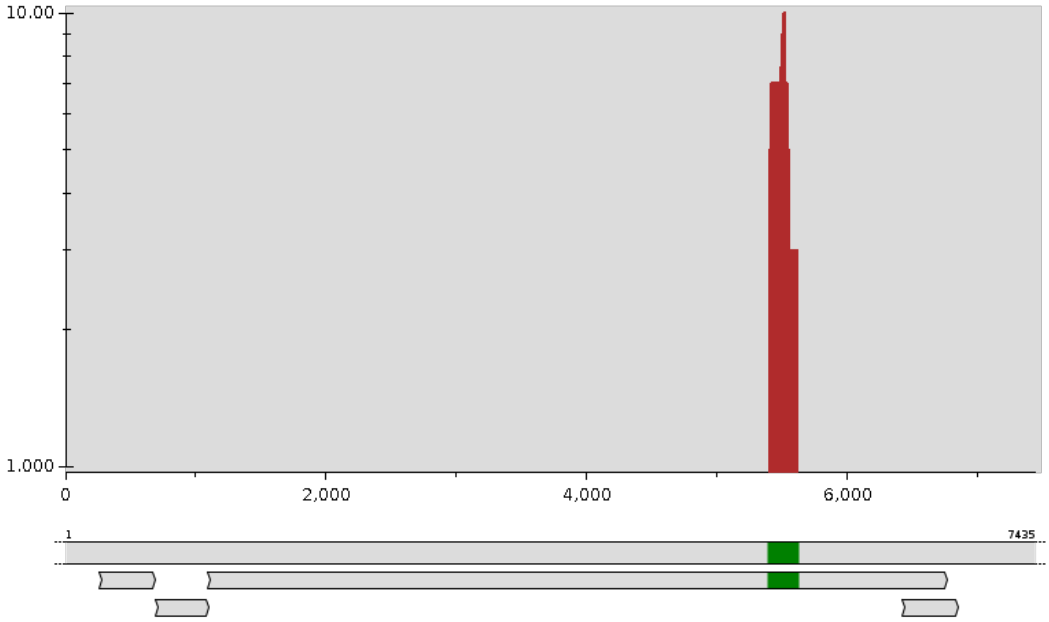

#### Assignment

| | |
|---|---|
| **Type** | Hibiscus bacilliform virus GD1 (Taxonomy ID: 1459800) |
| **Reference Genome** | NC_023485.1 |
| **NT Identity (%)** | 56.4315 |
| **AA Identity (%)** | 46.25 |
| **Number Of Stop Codons** | 1 |
| **Number Of CDS** | 4 |

#### Alignment

| | |
|---|---|
| **Alignment Score** | 56.0 (NT) + 268.0 (AA) = 324.0 |
| **Concordance (%)** | 30.5949 |

| **Alignment Method** | Global, seeded, nucleotide + amino acids (AGA) |
|---|---|

#### Genome Region

Sequence starts at position 5386 and ends at position 5623 relative to NC_023485.1 reference sequence.

#### Alignment Detailed Statistics

| | Begin | End | Coverage | Score | Concordance | Matches | Identities | I/D/M/F* | Stop Codons |
|---|---|---|---|---|---|---|---|---|---|
| **NT** | **5386** | **5623** | **3.2%** | **56** | **11.8%** | **238 (98.8%)** | **136 (56.4%)** | **3/0** | |
| Mutations: | 5390T>G, 5393A>G, 5395T>A, 5403G>T, 5406G>A, 5407C>G, 5408T>A, 5409A>G, 5411G>T, 5414C>G, 5415C>G, 5416C>A, 5418G>T, 5419A>C, 5420G>A, 5421T>G, 5422C>A, 5427A>C, 5428A>T, 5430T>A, 5431G>A, 5438T>A, 5441C>T, 5442T>C, 5445G>A, 5446T>C, 5447T>A, 5449C>G, 5450A>C, 5451G>T, 5456A>T, 5458T>A, 5459G>C, 5465A>G, 5467G>T, 5468G>C, 5469C>G, 5471A>G, 5478C>T, 5479C>A, 5480A>T, 5486A>G, 5491A>C, 5492A>T, 5495C>T, 5497C>T, 5501A>T, 5504A>T, 5506T>C, 5507A>T, 5511C>A, 5512A>T, 5514C>G, 5515G>A, 5517A>C, 5518A>T, 5523G>A, 5525C>T, 5526A>T, 5527A>G, 5528C>A, 5531C>G, 5532T>A, 5534C>T, 5537A>G, 5538G>T, 5539G>C, 5540_5541insTAC, 5541A>T, 5542C>T, 5543T>G, 5546A>T, 5547G>C, 5548C>A, 5549C>A, 5555C>G, 5556G>A, 5557C>T, 5558T>C, 5561A>G, 5563A>T, 5564C>T, 5567T>A, 5573C>T, 5576A>T, 5577C>T, 5579C>G, 5582A>T, 5585T>C, 5589C>A, 5592T>A, 5593C>G, 5598G>A, 5599A>G, 5600G>T, 5603A>T, 5607A>G, 5609C>G, 5612G>T, 5616A>T, 5618A>G, 5619C>A, 5621G>A | | | | | | | | |

##### CDS

| | Begin | End | Coverage | Score | Concordance | Matches | Identities | I/D/M/F* | Stop Codons |
|---|---|---|---|---|---|---|---|---|---|
| BU35_gp3 | 1433 | 1511 | 4.2% | 268 | 45.3% | 79 (98.8%) | 37 (46.3%) | 1/0/0/0 | 1 |
| Protein mutations: | S1433R (5390T>G), F1435Y (5395T>A), V1438L (5403G>T), A1439R (5406G>A 5407C>G 5408T>A), M1440V (5409A>G 5411G>T), H1441Q (5414C>G), P1442E (5415C>G 5416C>A), E1443S (5418G>T 5419A>C 5420G>A), S1444D (5421T>G 5422C>A), K1446L (5427A>C 5428A>T), W1447K (5430T>A 5431G>A), W1451R (5442T>C), V1452T (5445G>A 5446T>C 5447T>A), P1453R (5449C>G 5450A>C), D1454Y (5451G>T), L1456H (5458T>A 5459G>C), W1459F (5467G>T 5468G>C), L1460V (5469C>G 5471A>G), P1463Y (5478C>T 5479C>A 5480A>T), K1467T (5491A>C 5492A>T), A1469V (5497C>T), V1472A (5506T>C 5507A>T), Q1474M (5511C>A 5512A>T), R1475E (5514C>G 5515G>A), K1476L (5517A>C 5518A>T), D1478N (5523G>A 5525C>T), N1479* (5526A>T 5527A>G 5528C>A), F1481I (5532T>A 5534C>T), G1483S (5538G>T 5539G>C), G1483_T1484insY (5540_5541insTAC), T1484L (5541A>T 5542C>T 5543T>G), E1485D (5546A>T), A1486Q (5547G>C 5548C>A 5549C>A), I1488M (5555C>G), A1489I (5556G>A 5557C>T 5558T>C), Y1491F (5563A>T 5564C>T), Q1500K (5589C>A), E1503S (5598G>A 5599A>G 5600G>T), E1504D (5603A>T), I1506V (5607A>G 5609C>G), K1507N (5612G>T), I1509L (5616A>T 5618A>G) | | | | | | | | |
| Codon mutations: | AGT1433AGG (5390T>G), GGA1434GGG (5393A>G), TTT1435TAT (5395T>A), GTG1438TTG (5403G>T), GCT1439AGA (5406G>A 5407C>G 5408T>A), ATG1440GTT (5409A>G 5411G>T), CAC1441CAG (5414C>G), CCG1442GAG (5415C>G 5416C>A), GAG1443TCA (5418G>T 5419A>C 5420G>A), TCT1444GAT (5421T>G 5422C>A), AAG1446CTG (5427A>C 5428A>T), TGG1447AAG (5430T>A 5431G>A), GCT1449GCA (5438T>A), TTC1450TTT (5441C>T), TGG1451CGG (5442T>C), GTT1452ACA (5445G>A 5446T>C 5447T>A), CCA1453CGC (5449C>G 5450A>C), GAT1454TAT (5451G>T), GGA1455GGT (5456A>T), CTG1456CAC (5458T>A 5459G>C), GAA1458GAG (5465A>G), TGG1459TTC (5467G>T 5468G>C), CTA1460GTG (5469C>G 5471A>G), CCA1463TAT (5478C>T 5479C>A 5480A>T), GGA1465GGG (5486A>G), AAA1467ACT (5491A>C 5492A>T), AAC1468AAT (5495C>T), GCC1469GTC (5497C>T), CCA1470CCT (5501A>T), GCA1471GCT (5504A>T), GTA1472GCT (5506T>C 5507A>T), CAG1474ATG (5511C>A 5512A>T), CGA1475GAA (5514C>G 5515G>A), AAG1476CTG (5517A>C 5518A>T), GAC1478AAT (5523G>A 5525C>T), AAC1479TGA (5526A>T 5527A>G 5528C>A), GTC1480GTG (5531C>G), TTC1481ATT (5532T>A 5534C>T), AAA1482AAG (5537A>G), GGT1483TCT (5538G>T 5539G>C), GGT1483_ACT1484insTAC (5540_5541insTAC), ACT1484TTG (5541A>T 5542C>T 5543T>G), GAA1485GAT (5546A>T), GCC1486CAA (5547G>C 5548C>A 5549C>A), ATC1488ATG (5555C>G), GCT1489ATC (5556G>A 5557C>T 5558T>C), GTA1490GTG (5561A>G), TAC1491TTT (5563A>T 5564C>T), ATT1492ATA (5567T>A), GAC1494GAT (5573C>T), ATA1495ATT (5576A>T), CTC1496TTG (5577C>T 5579C>G), GTA1497GTT (5582A>T), TTT1498TTC (5585T>C), CAG1500AAG (5589C>A), TCT1501AGT (5592T>A 5593C>G), GAG1503AGT (5598G>A 5599A>G 5600G>T), GAA1504GAT (5603A>T), ATC1506GTG (5607A>G 5609C>G), AAG1507AAT (5612G>T), ATA1509TTG (5616A>T 5618A>G), CGG1510AGA (5619C>A 5621G>A) | | | | | | | | |

##### Proteins

| | Begin | End | Coverage | Score | Concordance | Matches | Identities | I/D/M/F* | Stop Codons |
|---|---|---|---|---|---|---|---|---|---|
| polyprotein (YP_009002585.1) | 1433 | 1511 | 4.2% | 268 | 45.3% | 79 (98.8%) | 37 (46.3%) | 1/0/0/0 | 1 |
| Protein mutations: | S1433R (5390T>G), F1435Y (5395T>A), V1438L (5403G>T), A1439R (5406G>A 5407C>G 5408T>A), M1440V (5409A>G 5411G>T), H1441Q (5414C>G), P1442E (5415C>G 5416C>A), E1443S (5418G>T 5419A>C 5420G>A), S1444D (5421T>G 5422C>A), K1446L (5427A>C 5428A>T), W1447K (5430T>A 5431G>A), W1451R (5442T>C), V1452T (5445G>A 5446T>C 5447T>A), P1453R (5449C>G 5450A>C), D1454Y (5451G>T), L1456H (5458T>A 5459G>C), W1459F (5467G>T 5468G>C), L1460V (5469C>G 5471A>G), P1463Y (5478C>T 5479C>A 5480A>T), K1467T (5491A>C 5492A>T), A1469V (5497C>T), V1472A (5506T>C 5507A>T), Q1474M (5511C>A 5512A>T), R1475E (5514C>G 5515G>A), K1476L (5517A>C 5518A>T), D1478N (5523G>A 5525C>T), N1479* (5526A>T 5527A>G 5528C>A), F1481I (5532T>A 5534C>T), G1483S (5538G>T 5539G>C), G1483_T1484insY (5540_5541insTAC), T1484L (5541A>T 5542C>T 5543T>G), E1485D (5546A>T), A1486Q (5547G>C 5548C>A 5549C>A), I1488M (5555C>G), A1489I (5556G>A 5557C>T 5558T>C), Y1491F (5563A>T 5564C>T), Q1500K (5589C>A), E1503S (5598G>A 5599A>G 5600G>T), E1504D (5603A>T), I1506V (5607A>G 5609C>G), K1507N (5612G>T), I1509L (5616A>T 5618A>G) | | | | | | | | |
| Codon mutations: | AGT1433AGG (5390T>G), GGA1434GGG (5393A>G), TTT1435TAT (5395T>A), GTG1438TTG (5403G>T), GCT1439AGA (5406G>A 5407C>G 5408T>A), ATG1440GTT (5409A>G 5411G>T), CAC1441CAG (5414C>G), CCG1442GAG (5415C>G 5416C>A), GAG1443TCA (5418G>T 5419A>C 5420G>A), TCT1444GAT (5421T>G 5422C>A), AAG1446CTG (5427A>C 5428A>T), TGG1447AAG (5430T>A 5431G>A), GCT1449GCA (5438T>A), TTC1450TTT (5441C>T), TGG1451CGG (5442T>C), GTT1452ACA (5445G>A 5446T>C 5447T>A), CCA1453CGC (5449C>G 5450A>C), GAT1454TAT (5451G>T), GGA1455GGT (5456A>T), CTG1456CAC (5458T>A 5459G>C), GAA1458GAG (5465A>G), TGG1459TTC (5467G>T 5468G>C), CTA1460GTG (5469C>G 5471A>G), CCA1463TAT (5478C>T 5479C>A 5480A>T), GGA1465GGG (5486A>G), AAA1467ACT (5491A>C 5492A>T), AAC1468AAT (5495C>T), GCC1469GTC (5497C>T), CCA1470CCT (5501A>T), GCA1471GCT (5504A>T), GTA1472GCT (5506T>C 5507A>T), CAG1474ATG (5511C>A 5512A>T), CGA1475GAA (5514C>G 5515G>A), AAG1476CTG (5517A>C 5518A>T), GAC1478AAT (5523G>A 5525C>T), AAC1479TGA (5526A>T 5527A>G 5528C>A), GTC1480GTG (5531C>G), TTC1481ATT (5532T>A 5534C>T), AAA1482AAG (5537A>G), GGT1483TCT (5538G>T 5539G>C), GGT1483_ACT1484insTAC (5540_5541insTAC), ACT1484TTG (5541A>T 5542C>T 5543T>G), GAA1485GAT (5546A>T), GCC1486CAA (5547G>C 5548C>A 5549C>A), ATC1488ATG (5555C>G), GCT1489ATC (5556G>A 5557C>T 5558T>C), GTA1490GTG (5561A>G), TAC1491TTT (5563A>T 5564C>T), ATT1492ATA (5567T>A), GAC1494GAT (5573C>T), ATA1495ATT (5576A>T), CTC1496TTG (5577C>T 5579C>G), GTA1497GTT (5582A>T), TTT1498TTC (5585T>C), CAG1500AAG (5589C>A), TCT1501AGT (5592T>A 5593C>G), GAG1503AGT (5598G>A 5599A>G 5600G>T), GAA1504GAT (5603A>T), ATC1506GTG (5607A>G 5609C>G), AAG1507AAT (5612G>T), ATA1509TTG (5616A>T 5618A>G), CGG1510AGA (5619C>A 5621G>A) | | | | | | | | |

*: Inserts / Deletes / Misaligned / Frameshifts

#### Analysis details

This analysis was performed with panviral2.64

### NGS Details (UN62): Badnavirus maculasmallanthi

#### Assembly

| | |
|---|---|
| **Coverage Length** | 422 (1 contig(s)) |
| **Depth Of Coverage** | 2.8 |
| **Number Of Reads** | 9 |
| **Reads Per Million** | 0.21 rpm (after QC) |
| **Ambiguities** | 0 |
| **Assembly Method** | de novo + reference guided assembly |
| **Consensus Caller** | Bcf Tools |

#### Coverage Map

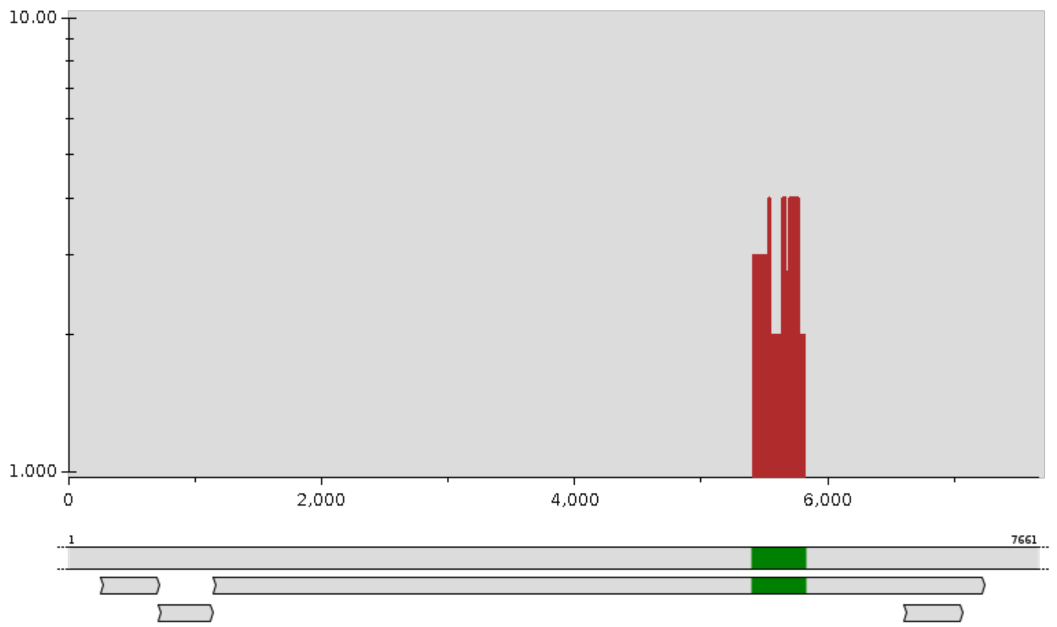

#### Assignment

| | |
|---|---|
| **Type** | Badnavirus maculasmallanthi (Taxonomy ID: 3048453) |
| **Reference Genome** | NC_026472.1 |
| **NT Identity (%)** | 51.79 |
| **AA Identity (%)** | 45.0 |
| **Number Of Stop Codons** | 0 |
| **Number Of CDS** | 4 |

#### Alignment

| | |
|---|---|
| **Alignment Score** | 9.0 (NT) + 415.0 (AA) = 424.0 |
| **Concordance (%)** | 24.0091 |

| Alignment Method | Global, seeded, nucleotide + amino acids (AGA) |
|---|---|

#### Genome Region

Sequence starts at position 5407 and ends at position 5828 relative to NC_026472.1 reference sequence.

#### Alignment Detailed Statistics

| | Begin | End | Coverage | Score | Concordance | Matches | Identities | I/D/M/F* | Stop Codons |
|---|---|---|---|---|---|---|---|---|---|
| **NT** | **5407** | **5828** | **5.5%** | **9** | **1.1%** | **416 (97.9%)** | **217 (51.1%)** | **3/6** | |
| Mutations: | 5407G>T, 5408T>C, 5409C>A, 5410A>C, 5411A>C, 5412C>T, 5414C>G, 5415T>G, 5419A>G, 5422A>C, 5427T>C, 5428_5433delGACCCA, 5434G>C, 5435C>T, 5437A>T, 5438C>T, 5439A>T, 5441G>T, 5442C>G, 5446G>A, 5448A>G, 5450C>A, 5451A>G, 5452A>G, 5454G>T, 5457A>G, 5458A>T, 5459A>C, 5460T>A, 5461G>A, 5462A>T, 5463T>G, 5470G>T, 5471T>G, 5472G>T, 5473T>A, 5475C>T, 5476A>G, 5481C>T, 5485A>C, 5486C>A, 5487A>G, 5488C>T, 5490C>G, 5493T>C, 5494G>A, 5496C>G, 5497A>G, 5498A>T, 5499T>G, 5502A>T, 5503T>G, 5504A>T, 5505C>A, 5508A>G, 5509G>A, 5511T>C, 5512C>A, 5514G>A, 5518T>C, 5523T>C, 5527G>C, 5529G>T, 5532A>C, 5533A>G, 5535C>T, 5537C>A, 5538C>T, 5541G>A, 5542C>T, 5544A>T, 5545A>G, 5547A>T, 5548A>C, 5549G>A, 5550A>G, 5551A>T, 5554G>C, 5555G>A, 5557A>G, 5558A>G, 5559T>A, 5563A>G, 5564A>C, 5565G>C, 5574A>T, 5578T>A, 5588A>G, 5589G>A, 5591C>T, 5592T>C, 5597T>A, 5604A>G, 5605G>T, 5607T>G, 5608G>A, 5609C>A, 5610C>G, 5613G>T, 5614G>C, 5616A>T, 5617G>C, 5618A>C, 5619A>T, 5620G>T, 5621A>C, 5622A>T, 5623T>G, 5624C>A, 5625C>T, 5628A>T, 5631C>T, 5632T>A, 5633G>A, 5637G>A, 5640C>T, 5643T>C, 5644T>C, 5645T>G, 5646A>G, 5648T>C, 5650C>A, 5651C>G, 5652A>G, 5653G>T, 5654G>A, 5655A>T, 5658A>C, 5659T>C, 5660T>A, 5661A>T, 5664C>T, 5667A>G, 5669G>T, 5670G>T, 5676C>T, 5680C>T, 5682A>G, 5685C>T, 5688A>G, 5689C>T, 5691C>G, 5693A>C, 5694A>T, 5700A>T, 5703C>G, 5704G>A, 5706T>A, 5707A>G, 5708T>C, 5709T>A, 5713C>A, 5714A>T, 5716A>G, 5717G>A, 5719A>T, 5720A>T, 5725G>A, 5728A>G, 5729A>G, 5730A>G, 5731T>G, 5732G>T, 5733C>G, 5736C>T, 5737A>C, 5739A>T, 5739_5740insCCG, 5740G>T, 5743A>T, 5744C>T, 5745A>G, 5748A>T, 5749G>T, 5750A>C, 5751G>A, 5755A>G, 5757C>G, 5758G>A, 5759C>T, 5760C>T, 5765A>T, 5766C>T, 5769C>T, 5772C>T, 5775C>T, 5778C>T, 5779C>T, 5784A>T, 5793C>A, 5795G>C, 5799A>G, 5803G>A, 5810A>T, 5813G>A, 5814G>A, 5817T>C, 5820A>G, 5822A>G, 5823G>A, 5827A>G | | | | | | | | |

##### CDS

| | | | | | | | | | |
|---|---|---|---|---|---|---|---|---|---|
| UF61_gp3 | 1422 | 1562 | 6.9% | 415 | 43.5% | 139 (97.9%) | 63 (44.4%) | 1/2/0/0 | 0 |
| Protein mutations: | V1422S (5407G>T 5408T>C 5409C>A), N1423P (5410A>C 5411A>C 5412C>T), S1424W (5414C>G 5415T>G), T1426A (5419A>G), T1427P (5422A>C), D1429_P1430del (5428_5433delGACCCA), A1431L (5434G>C 5435C>T), T1432F (5437A>T 5438C>T 5439A>T), G1433V (5441G>T 5442C>G), E1435K (5446G>A 5448A>G), T1436K (5450C>A 5451A>G), K1437D (5452A>G 5454G>T), N1439S (5458A>T 5459A>C 5460T>A), D1440M (5461G>A 5462A>T 5463T>G), V1443C (5470G>T 5471T>G 5472G>T), F1444I (5473T>A 5475C>T), N1445D (5476A>G), T1448Q (5485A>C 5486C>A 5487A>G), D1451K (5494G>A 5496C>G), N1452V (5497A>G 5498A>T 5499T>G), Y1454V (5503T>G 5504A>T 5505C>A), D1456N (5509G>A 5511T>C), Q1457K (5512C>A 5514G>A), S1459P (5518T>C), G1462R (5527G>C 5529G>T), N1464D (5533A>G 5535C>T), T1465N (5537C>A 5538C>T), L1467F (5542C>T 5544A>T), K1468D (5545A>G 5547A>T), R1469Q (5548A>C 5549G>A 5550A>G), I1470L (5551A>T), G1471Q (5554G>C 5555G>A), N1472G (5557A>G 5558A>G 5559T>A), K1474A (5563A>G 5564A>C 5565G>C), F1479I (5578T>A), K1482R (5588A>G 5589G>A), S1483F (5591C>T 5592T>C), F1485Y (5597T>A), V1488L (5605G>T 5607T>G), A1489K (5608G>A 5609C>A 5610C>G), M1490I (5613G>T), E1491H (5614G>C 5616A>T), E1492P (5617G>C 5618A>C 5619A>T), E1493S (5620G>T 5621A>C 5622A>T), S1494D (5623T>G 5624C>A 5625C>T), W1497K (5632T>A 5633G>A), L1501R (5644T>C 5645T>G 5646A>G), I1502T (5648T>C), P1503R (5650C>A 5651C>G 5652A>G), G1504Y (5653G>T 5654G>A 5655A>T), L1506H (5659T>C 5660T>A 5661A>T), W1509F (5669G>T 5670G>T), P1513S (5680C>T 5682A>G), K1517T (5693A>C 5694A>T), A1521T (5704G>A 5706T>A), I1522A (5707A>G 5708T>C 5709T>A), Q1524M (5713C>A 5714A>T), R1525E (5716A>G 5717G>A), K1526L (5719A>T 5720A>T), D1528N (5725G>A), K1529G (5728A>G 5729A>G 5730A>G), C1530V (5731T>G 5732G>T 5733C>G), K1532H (5737A>C 5739A>T), K1532_D1533insP (5739_5740insCCG), D1533Y (5740G>T), T1534L (5743A>T 5744C>T 5745A>G), E1535D (5748A>T), E1536S (5749G>T 5750A>C 5751G>A), I1538V (5755A>G 5757C>G), A1539I (5758G>A 5759C>T 5760C>T), Y1541F (5765A>T 5766C>T), N1550K (5793C>A), S1551T (5795G>C), D1554N (5803G>A), E1556V (5810A>T), R1557K (5813G>A 5814G>A), K1560R (5822A>G 5823G>A) | | | | | | | | |
| Codon mutations: | GTC1422TCA (5407G>T 5408T>C 5409C>A), AAC1423CCT (5410A>C 5411A>C 5412C>T), TCT1424TGG (5414C>G 5415T>G), ACC1426GCC (5419A>G), ACG1427CCG (5422A>C), GTT1428GTC (5427T>C), GAC1429_CCA1430del (5428_5433delGACCCA), GCA1431CTA (5434G>C 5435C>T), ACA1432TTT (5437A>T 5438C>T 5439A>T), GGC1433GTG (5441G>T 5442C>G), GAA1435AAG (5446G>A 5448A>G), ACA1436AAG (5450C>A 5451A>G), AAG1437GAT (5452A>G 5454G>T), GGA1438GGG (5457A>G), AAT1439TCA (5458A>T 5459A>C 5460T>A), GAT1440ATG (5461G>A 5462A>T 5463T>G), GTG1443TGT (5470G>T 5471T>G 5472G>T), TTC1444ATT (5473T>A 5475C>T), AAT1445GAT (5476A>G), TAC1446TAT (5481C>T), ACA1448CAG (5485A>C 5486C>A 5487A>G), CTC1449TTG (5488C>T 5490C>G), AAT1450AAC (5493T>C), GAC1451AAG (5494G>A 5496C>G), AAT1452GTG (5497A>G 5498A>T 5499T>G), ACA1453ACT (5502A>T), TAC1454GTA (5503T>G 5504A>T 5505C>A), AAA1455AAG (5508A>G), GAT1456AAC (5509G>A 5511T>C), CAG1457AAA (5512C>A 5514G>A), TCC1459CCC (5518T>C), CTT1460CTC (5523T>C), GGG1462CGT (5527G>C 5529G>T), ATA1463ATC (5532A>C), AAC1464GAT (5533A>G 5535C>T), ACC1465AAT (5537C>A 5538C>T), TTG1466TTA (5541G>A), CTA1467TTT (5542C>T 5544A>T), AAA1468GAT (5545A>G 5547A>T), AGA1469CAG (5548A>C 5549G>A 5550A>G), ATA1470TTA (5551A>T), GGG1471CAG (5554G>C 5555G>A), AAT1472GGA (5557A>G 5558A>G 5559T>A), AAG1474GCC (5563A>G 5564A>C 5565G>C), TCA1477TCT (5574A>T), TTT1479ATT (5578T>A), AAG1482AGA (5588A>G 5589G>A), TCT1483TTC (5591C>T 5592T>C), TTT1485TAT (5597T>A), CAA1487CAG (5604A>G), GTT1488TTG (5605G>T 5607T>G), GCC1489AAG (5608G>A 5609C>A 5610C>G), ATG1490ATT (5613G>T), GAA1491CAT (5614G>C 5616A>T), GAA1492CCT (5617G>C 5618A>C 5619A>T), GAA1493TCT (5620G>T 5621A>C 5622A>T), TCC1494GAT (5623T>G 5624C>A 5625C>T), ATA1495ATT (5628A>T), CCC1496CCT (5631C>T), TGG1497AAG (5632T>A 5633G>A), ACG1498ACA (5637G>A), GCC1499GCT (5640C>T), TTT1500TTC (5643T>C), TTA1501CGG (5644T>C 5645T>G 5646A>G), ATC1502ACC (5648T>C), CCA1503AGG (5650C>A 5651C>G 5652A>G), GGA1504TAT (5653G>T 5654G>A 5655A>T), GGA1505GGC (5658A>C), TTA1506CAT (5659T>C 5660T>A 5661A>T), TAC1507TAT (5664C>T), GAA1508GAG (5667A>G), TGG1509TTT (5669G>T 5670G>T), GTC1511GTT (5676C>T), CCA1513TCG (5680C>T 5682A>G), TTC1514TTT (5685C>T), GGA1515GGG (5688A>G), CTC1516TTG (5689C>T 5691C>G), AAA1517ACT (5693A>C 5694A>T), GCA1519GCT (5700A>T), CCC1520CCG (5703C>G), GCT1521ACA (5704G>A 5706T>A), ATT1522GCA (5707A>G 5708T>C 5709T>A), CAG1524ATG (5713C>A 5714A>T), AGA1525GAA (5716A>G 5717G>A), AAA1526TTA (5719A>T 5720A>T), GAT1528AAT (5725G>A), AAA1529GGG (5728A>G 5729A>G 5730A>G), TGC1530GTG (5731T>G 5732G>T 5733C>G), TTC1531TTT (5736C>T), AAA1532CAT (5737A>C 5739A>T), AAA1532_GAC1533insCCG (5739_5740insCCG), GAC1533TAC (5740G>T), ACA1534TTG (5743A>T 5744C>T 5745A>G), GAA1535GAT (5748A>T), GAG1536TCA (5749G>T 5750A>C 5751G>A), ATC1538GTG (5755A>G 5757C>G), GCC1539ATT (5758G>A 5759C>T 5760C>T), TAC1541TTT (5765A>T 5766C>T), ATC1542ATT (5769C>T), GAC1543GAT (5772C>T), GAC1544GAT (5775C>T), ATC1545ATT (5778C>T), CTG1546TTG (5779C>T), GTA1547GTT (5784A>T), AAC1550AAA (5793C>A), AGT1551ACT (5795G>C), GAA1552GAG (5799A>G), GAT1554AAT (5803G>A), GAA1556GTA (5810A>T), AGG1557AAA (5813G>A 5814G>A), CAT1558CAC (5817T>C), TTA1559TTG (5820A>G), AAG1560AGA (5822A>G 5823G>A), ATG1562GT. (5827A>G) | | | | | | | | |

##### Proteins

| | | | | | | | | | |
|---|---|---|---|---|---|---|---|---|---|
| ORF3 (YP_009121747.1) | 1422 | 1562 | 6.9% | 415 | 43.5% | 139 (97.9%) | 63 (44.4%) | 1/2/0/0 | 0 |
| Protein mutations: | V1422S (5407G>T 5408T>C 5409C>A), N1423P (5410A>C 5411A>C 5412C>T), S1424W (5414C>G 5415T>G), T1426A (5419A>G), T1427P (5422A>C), D1429_P1430del (5428_5433delGACCCA), A1431L (5434G>C 5435C>T), T1432F (5437A>T 5438C>T 5439A>T), G1433V (5441G>T 5442C>G), E1435K (5446G>A 5448A>G), T1436K (5450C>A 5451A>G), K1437D (5452A>G 5454G>T), N1439S (5458A>T 5459A>C 5460T>A), D1440M (5461G>A 5462A>T 5463T>G), V1443C (5470G>T 5471T>G 5472G>T), F1444I (5473T>A 5475C>T), N1445D (5476A>G), T1448Q (5485A>C 5486C>A 5487A>G), D1451K (5494G>A 5496C>G), N1452V (5497A>G 5498A>T 5499T>G), Y1454V (5503T>G 5504A>T 5505C>A), D1456N (5509G>A 5511T>C), Q1457K (5512C>A 5514G>A), S1459P (5518T>C), G1462R (5527G>C 5529G>T), N1464D (5533A>G 5535C>T), T1465N (5537C>A 5538C>T), L1467F (5542C>T 5544A>T), K1468D (5545A>G 5547A>T), R1469Q (5548A>C 5549G>A 5550A>G), I1470L (5551A>T), G1471Q (5554G>C 5555G>A), N1472G (5557A>G 5558A>G 5559T>A), K1474A (5563A>G 5564A>C 5565G>C), F1479I (5578T>A), K1482R (5588A>G 5589G>A), S1483F (5591C>T 5592T>C), F1485Y (5597T>A), V1488L (5605G>T 5607T>G), A1489K (5608G>A 5609C>A 5610C>G), M1490I (5613G>T), E1491H (5614G>C 5616A>T), E1492P (5617G>C 5618A>C 5619A>T), E1493S (5620G>T 5621A>C 5622A>T), S1494D (5623T>G 5624C>A 5625C>T), W1497K (5632T>A 5633G>A), L1501R (5644T>C 5645T>G 5646A>G), I1502T (5648T>C), P1503R (5650C>A 5651C>G 5652A>G), G1504Y (5653G>T 5654G>A 5655A>T), L1506H (5659T>C 5660T>A 5661A>T), W1509F (5669G>T 5670G>T), P1513S (5680C>T 5682A>G), K1517T (5693A>C 5694A>T), A1521T (5704G>A 5706T>A), I1522A (5707A>G 5708T>C 5709T>A), Q1524M (5713C>A 5714A>T), R1525E (5716A>G 5717G>A), K1526L (5719A>T 5720A>T), D1528N (5725G>A), K1529G (5728A>G 5729A>G 5730A>G), C1530V (5731T>G 5732G>T 5733C>G), K1532H (5737A>C 5739A>T), K1532_D1533insP (5739_5740insCCG), D1533Y (5740G>T), T1534L (5743A>T 5744C>T 5745A>G), E1535D (5748A>T), E1536S (5749G>T 5750A>C 5751G>A), I1538V (5755A>G 5757C>G), A1539I (5758G>A 5759C>T 5760C>T), Y1541F (5765A>T 5766C>T), N1550K (5793C>A), S1551T (5795G>C), D1554N (5803G>A), E1556V (5810A>T), R1557K (5813G>A 5814G>A), K1560R (5822A>G 5823G>A) | | | | | | | | |

| | Begin | End | Coverage | Score | Concordance | Matches | Identities | I/D/M/F* | Stop Codons |
|---|---|---|---|---|---|---|---|---|---|
| **NT** | **5407** | **5828** | **5.5%** | **9** | **1.1%** | **416 (97.9%)** | **217 (51.1%)** | **3/6** | |
| Codon mutations: | GTC1422TCA (5407G>T 5408T>C 5409C>A), AAC1423CCT (5410A>C 5411A>C 5412C>T), TCT1424TGG (5414C>G 5415T>G), ACC1426GCC (5419A>G), ACG1427CCG (5422A>C), GTT1428GTC (5427T>C), GAC1429_CCA1430del (5428_5433delGACCCA), GCA1431CTA (5434G>C 5435C>T), ACA1432TTT (5437A>T 5438C>T 5439A>T), GGC1433GTG (5441G>T 5442C>G), GAA1435AAG (5446G>A 5448A>G), ACA1436AAG (5450C>A 5451A>G), AAG1437GAT (5452A>G 5454G>T), GGA1438GGG (5457A>G), AAT1439TCA (5458A>T 5459A>C 5460T>A), GAT1440ATG (5461G>A 5462A>T 5463T>G), GTG1443TGT (5470G>T 5471T>G 5472G>T), TTC1444ATT (5473T>A 5475C>T), AAT1445GAT (5476A>G), TAC1446TAT (5481C>T), ACA1448CAG (5485A>C 5486C>A 5487A>G), CTC1449TTG (5488C>T 5490C>G), AAT1450AAC (5493T>C), GAC1451AAG (5494G>A 5496C>G), AAT1452GTG (5497A>G 5498A>T 5499T>G), ACA1453ACT (5502A>T), TAC1454GTA (5503T>G 5504A>T 5505C>A), AAA1455AAG (5508A>G), GAT1456AAC (5509G>A 5511T>C), CAG1457AAA (5512C>A 5514G>A), TCC1459CCC (5518T>C), CTT1460CTC (5523T>C), GGG1462CGT (5527G>C 5529G>T), ATA1463ATC (5532A>C), AAC1464GAT (5533A>G 5535C>T), ACC1465AAT (5537C>A 5538C>T), TTG1466TTA (5541G>A), CTA1467TTT (5542C>T 5544A>T), AAA1468GAT (5545A>G 5547A>T), AGA1469CAG (5548A>C 5549G>A 5550A>G), ATA1470TTA (5551A>T), GGG1471CAG (5554G>C 5555G>A), AAT1472GGA (5557A>G 5558A>G 5559T>A), AAG1474GCC (5563A>G 5564A>C 5565G>C), TCA1477TCT (5574A>T), TTT1479ATT (5578T>A), AAG1482AGA (5588A>G 5589G>A), TCT1483TTC (5591C>T 5592T>C), TTT1485TAT (5597T>A), CAA1487CAG (5604A>G), GTT1488TTG (5605G>T 5607T>G), GCC1489AAG (5608G>A 5609C>A 5610C>G), ATG1490ATT (5613G>T), GAA1491CAT (5614G>C 5616A>T), GAA1492CCT (5617G>C 5618A>C 5619A>T), GAA1493TCT (5620G>T 5621A>C 5622A>T), TCC1494GAT (5623T>G 5624C>A 5625C>T), ATA1495ATT (5628A>T), CCC1496CCT (5631C>T), TGG1497AAG (5632T>A 5633G>A), ACG1498ACA (5637G>A), GCC1499GCT (5640C>T), TTT1500TTC (5643T>C), TTA1501CGG (5644T>C 5645T>G 5646A>G), ATC1502ACC (5648T>C), CCA1503AGG (5650C>A 5651C>G 5652A>G), GGA1504TAT (5653G>T 5654G>A 5655A>T), GGA1505GGC (5658A>C), TTA1506CAT (5659T>C 5660T>A 5661A>T), TAC1507TAT (5664C>T), GAA1508GAG (5667A>G), TGG1509TTT (5669G>T 5670G>T), GTC1511GTT (5676C>T), CCA1513TCG (5680C>T 5682A>G), TTC1514TTT (5685C>T), GGA1515GGG (5688A>G), CTC1516TTG (5689C>T 5691C>G), AAA1517ACT (5693A>C 5694A>T), GCA1519GCT (5700A>T), CCC1520CCG (5703C>G), GCT1521ACA (5704G>A 5706T>A), ATT1522GCA (5707A>G 5708T>C 5709T>A), CAG1524ATG (5713C>A 5714A>T), AGA1525GAA (5716A>G 5717G>A), AAA1526TTA (5719A>T 5720A>T), GAT1528AAT (5725G>A), AAA1529GGG (5728A>G 5729A>G 5730A>G), TGC1530GTG (5731T>G 5732G>T 5733C>G), TTC1531TTT (5736C>T), AAA1532CAT (5737A>C 5739A>T), AAA1532_GAC1533insCCG (5739_5740insCCG), GAC1533TAC (5740G>T), ACA1534TTG (5743A>T 5744C>T 5745A>G), GAA1535GAT (5748A>T), GAG1536TCA (5749G>T 5750A>C 5751G>A), ATC1538GTG (5755A>G 5757C>G), GCC1539ATT (5758G>A 5759C>T 5760C>T), TAC1541TTT (5765A>T 5766C>T), ATC1542ATT (5769C>T), GAC1543GAT (5772C>T), GAC1544GAT (5775C>T), ATC1545ATT (5778C>T), CTG1546TTG (5779C>T), GTA1547GTT (5784A>T), AAC1550AAA (5793C>A), AGT1551ACT (5795G>C), GAA1552GAG (5799A>G), GAT1554AAT (5803G>A), GAA1556GTA (5810A>T), AGG1557AAA (5813G>A 5814G>A), CAT1558CAC (5817T>C), TTA1559TTG (5820A>G), AAG1560AGA (5822A>G 5823G>A), ATG1562GT. (5827A>G) | | | | | | | | |

*: Inserts / Deletes / Misaligned / Frameshifts

#### Analysis details

This analysis was performed with panviral2.64

### NGS Details (UN62): Badnavirus venazanthoxyli

#### Assembly

| | |
|---|---|
| **Coverage Length** | 278 (1 contig(s)) |
| **Depth Of Coverage** | 3.8 |
| **Number Of Reads** | 8 |
| **Reads Per Million** | 0.19 rpm (after QC) |
| **Ambiguities** | 0 |
| **Assembly Method** | de novo + reference guided assembly |
| **Consensus Caller** | Bcf Tools |

#### Coverage Map

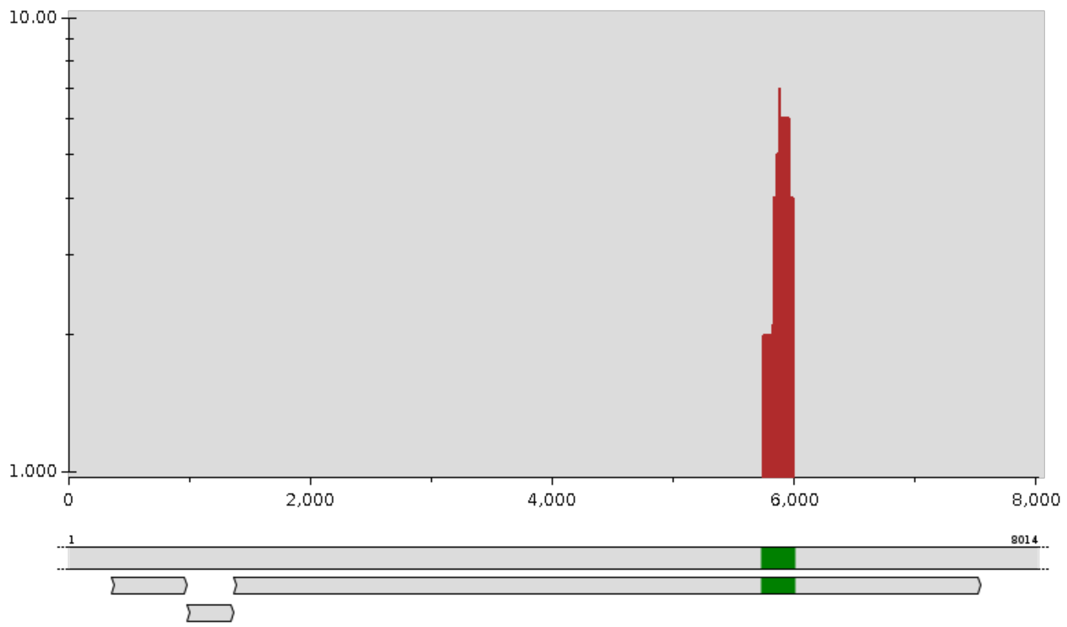

#### Assignment

| | |
|---|---|
| **Type** | Badnavirus venazanthoxyli (Taxonomy ID: 3047858) |
| **Reference Genome** | NC_076349.1 |
| **NT Identity (%)** | 58.363 |
| **AA Identity (%)** | 53.1915 |
| **Number Of Stop Codons** | 0 |
| **Number Of CDS** | 3 |

#### Alignment

| | |
|---|---|
| **Alignment Score** | 88.0 (NT) + 365.0 (AA) = 453.0 |
| **Concordance (%)** | 37.5622 |

| Alignment Method | Global, seeded, nucleotide + amino acids (AGA) |
|---|---|

#### Genome Region

Sequence starts at position 5732 and ends at position 6009 relative to NC_076349.1 reference sequence.

#### Alignment Detailed Statistics

| | Begin | End | Coverage | Score | Concordance | Matches | Identities | I/D/M/F* | Stop Codons |
|---|---|---|---|---|---|---|---|---|---|
| **NT** | **5732** | **6009** | **3.5%** | **88** | **15.8%** | **278 (98.9%)** | **164 (58.4%)** | **3/0** | |
| Mutations: | 5741C>G, 5744T>A, 5746G>C, 5750G>A, 5756A>G, 5757A>T, 5759A>G, 5763A>T, 5764G>C, 5768G>A, 5769T>G, 5771T>A, 5777A>G, 5779A>G, 5780G>A, 5783A>T, 5788T>A, 5795A>G, 5796G>A, 5798G>C, 5799G>A, 5800C>G, 5801C>A, 5806A>T, 5807T>G, 5809A>T, 5810A>G, 5814A>G, 5815G>A, 5819C>T, 5823T>A, 5824G>A, 5825G>A, 5836T>A, 5837C>A, 5839G>C, 5842C>A, 5843A>T, 5844G>A, 5845C>T, 5846A>G, 5849G>A, 5850T>C, 5851T>A, 5852G>T, 5858A>G, 5860G>T, 5861G>T, 5862C>T, 5873A>C, 5876C>T, 5879C>A, 5880C>T, 5882A>G, 5884A>C, 5885G>T, 5891T>A, 5894A>T, 5895G>T, 5897A>C, 5898G>A, 5899T>C, 5900C>T, 5906G>A, 5907C>T, 5909G>T, 5910A>C, 5911A>C, 5916G>A, 5919G>C, 5921A>T, 5922T>G, 5923G>T, 5924C>T, 5927C>T, 5928A>C, 5931G>C, 5932G>A, 5933_5934insTAC, 5935C>T, 5937G>A, 5938A>G, 5940G>A, 5945C>T, 5946A>G, 5948T>C, 5949G>T, 5950C>T, 5951C>A, 5956A>T, 5958A>T, 5960C>T, 5963C>T, 5966T>C, 5969A>T, 5972T>G, 5973G>A, 5977T>A, 5978C>T, 5979T>A, 5980C>G, 5981C>T, 5984C>G, 5985A>T, 5988A>T, 5990C>A, 5991A>C, 5992A>T, 5999T>C, 6000G>C, 6001A>T, 6002G>T, 6004A>T, 6005G>T, 6008C>T | | | | | | | | |

##### CDS

| | Begin | End | Coverage | Score | Concordance | Matches | Identities | I/D/M/F* | Stop Codons |
|---|---|---|---|---|---|---|---|---|---|
| QKN22_gp3 | 1457 | 1549 | 4.5% | 365 | 55.3% | 93 (98.9%) | 50 (53.2%) | 1/0/0/0 | 0 |
| Protein mutations: | G1461A (5746G>C), I1465L (5757A>T 5759A>G), F1469V (5769T>G 5771T>A), K1472R (5779A>G 5780G>A), F1475Y (5788T>A), V1478I (5796G>A 5798G>C), A1479R (5799G>A 5800C>G 5801C>A), D1481V (5806A>T 5807T>G), E1482V (5809A>T 5810A>G), S1484D (5814A>G 5815G>A), W1487K (5823T>A 5824G>A 5825G>A), I1491K (5836T>A 5837C>A), S1492T (5839G>C), P1493H (5842C>A 5843A>T), A1494M (5844G>A 5845C>T 5846A>G), L1496H (5850T>C 5851T>A 5852G>T), W1499F (5860G>T 5861G>T), K1507T (5884A>C 5885G>T), A1511S (5895G>T 5897A>C), V1512T (5898G>A 5899T>C 5900C>T), R1515C (5907C>T 5909G>T), K1516P (5910A>C 5911A>C), D1518N (5916G>A), E1519H (5919G>C 5921A>T), C1520V (5922T>G 5923G>T 5924C>T), K1522Q (5928A>C), G1523Q (5931G>C 5932G>A), G1523_T1524insY (5933_5934insTAC), T1524M (5935C>T), E1525R (5937G>A 5938A>G), E1526K (5940G>A), I1528V (5946A>G 5948T>C), A1529L (5949G>T 5950C>T 5951C>A), Y1531F (5956A>T), I1532F (5958A>T 5960C>T), V1537I (5973G>A), F1538Y (5977T>A 5978C>T), N1540K (5984C>G), T1541S (5985A>T), I1542L (5988A>T 5990C>A), K1543L (5991A>C 5992A>T), E1546L (6000G>C 6001A>T 6002G>T), K1547I (6004A>T 6005G>T) | | | | | | | | |
| Codon mutations: | TCC1459TCG (5741C>G), CTT1460CTA (5744T>A), GGA1461GCA (5746G>C), GGG1462GGA (5750G>A), AAA1464AAG (5756A>G), ATA1465TTG (5757A>T 5759A>G), AGC1467TCC (5763A>T 5764G>C), AAG1468AAA (5768G>A), TTT1469GTA (5769T>G 5771T>A), TTA1471TTG (5777A>G), AAG1472AGA (5779A>G 5780G>A), TCA1473TCT (5783A>T), TTC1475TAC (5788T>A), CAA1477CAG (5795A>G), GTG1478ATC (5796G>A 5798G>C), GCC1479AGA (5799G>A 5800C>G 5801C>A), GAT1481GTG (5806A>T 5807T>G), GAA1482GTG (5809A>T 5810A>G), AGC1484GAC (5814A>G 5815G>A), ATC1485ATT (5819C>T), TGG1487AAA (5823T>A 5824G>A 5825G>A), ATC1491AAA (5836T>A 5837C>A), AGT1492ACT (5839G>C), CCA1493CAT (5842C>A 5843A>T), GCA1494ATG (5844G>A 5845C>T 5846A>G), GGG1495GGA (5849G>A), TTG1496CAT (5850T>C 5851T>A 5852G>T), GAA1498GAG (5858A>G), TGG1499TTT (5860G>T 5861G>T), CTG1500TTG (5862C>T), CCA1503CCC (5873A>C), TTC1504TTT (5876C>T), GGC1505GGA (5879C>A), CTA1506TTG (5880C>T 5882A>G), AAG1507ACT (5884A>C 5885G>T), GCT1509GCA (5891T>A), CCA1510CCT (5894A>T), GCA1511TCC (5895G>T 5897A>C), GTC1512ACT (5898G>A 5899T>C 5900C>T), CAG1514CAA (5906G>A), CGG1515TGT (5907C>T 5909G>T), AAG1516CCG (5910A>C 5911A>C), GAT1518AAT (5916G>A), GAA1519CAT (5919G>C 5921A>T), TGC1520GTT (5922T>G 5923G>T 5924C>T), TTC1521TTT (5927C>T), AAA1522CAA (5928A>C), GGA1523CAA (5931G>C 5932G>A), GGA1523_ACG1524insTAC (5933_5934insTAC), ACG1524ATG (5935C>T), GAG1525AGG (5937G>A 5938A>G), GAG1526AAG (5940G>A), TTC1527TTT (5945C>T), ATT1528GTC (5946A>G 5948T>C), GCC1529TTA (5949G>T 5950C>T 5951C>A), TAT1531TTT (5956A>T), ATC1532TTT (5958A>T 5960C>T), GAC1533GAT (5963C>T), GAT1534GAC (5966T>C), ATA1535ATT (5969A>T), CTT1536CTG (5972T>G), GTC1537ATC (5973G>A), TTC1538TAT (5977T>A 5978C>T), TCC1539AGT (5979T>A 5980C>G 5981C>T), AAC1540AAG (5984C>G), ACC1541TCC (5985A>T), ATC1542TTA (5988A>T 5990C>A), AAG1543CTG (5991A>C 5992A>T), CAT1545CAC (5999T>C), GAG1546CTT (6000G>C 6001A>T 6002G>T), AAG1547ATT (6004A>T 6005G>T), CAC1548CAT (6008C>T) | | | | | | | | |

##### Proteins

| | Begin | End | Coverage | Score | Concordance | Matches | Identities | I/D/M/F* | Stop Codons |
|---|---|---|---|---|---|---|---|---|---|
| polyprotein (YP_010798182.1) | 1457 | 1549 | 4.5% | 365 | 55.3% | 93 (98.9%) | 50 (53.2%) | 1/0/0/0 | 0 |
| Protein mutations: | G1461A (5746G>C), I1465L (5757A>T 5759A>G), F1469V (5769T>G 5771T>A), K1472R (5779A>G 5780G>A), F1475Y (5788T>A), V1478I (5796G>A 5798G>C), A1479R (5799G>A 5800C>G 5801C>A), D1481V (5806A>T 5807T>G), E1482V (5809A>T 5810A>G), S1484D (5814A>G 5815G>A), W1487K (5823T>A 5824G>A 5825G>A), I1491K (5836T>A 5837C>A), S1492T (5839G>C), P1493H (5842C>A 5843A>T), A1494M (5844G>A 5845C>T 5846A>G), L1496H (5850T>C 5851T>A 5852G>T), W1499F (5860G>T 5861G>T), K1507T (5884A>C 5885G>T), A1511S (5895G>T 5897A>C), V1512T (5898G>A 5899T>C 5900C>T), R1515C (5907C>T 5909G>T), K1516P (5910A>C 5911A>C), D1518N (5916G>A), E1519H (5919G>C 5921A>T), C1520V (5922T>G 5923G>T 5924C>T), K1522Q (5928A>C), G1523Q (5931G>C 5932G>A), G1523_T1524insY (5933_5934insTAC), T1524M (5935C>T), E1525R (5937G>A 5938A>G), E1526K (5940G>A), I1528V (5946A>G 5948T>C), A1529L (5949G>T 5950C>T 5951C>A), Y1531F (5956A>T), I1532F (5958A>T 5960C>T), V1537I (5973G>A), F1538Y (5977T>A 5978C>T), N1540K (5984C>G), T1541S (5985A>T), I1542L (5988A>T 5990C>A), K1543L (5991A>C 5992A>T), E1546L (6000G>C 6001A>T 6002G>T), K1547I (6004A>T 6005G>T) | | | | | | | | |
| Codon mutations: | TCC1459TCG (5741C>G), CTT1460CTA (5744T>A), GGA1461GCA (5746G>C), GGG1462GGA (5750G>A), AAA1464AAG (5756A>G), ATA1465TTG (5757A>T 5759A>G), AGC1467TCC (5763A>T 5764G>C), AAG1468AAA (5768G>A), TTT1469GTA (5769T>G 5771T>A), TTA1471TTG (5777A>G), AAG1472AGA (5779A>G 5780G>A), TCA1473TCT (5783A>T), TTC1475TAC (5788T>A), CAA1477CAG (5795A>G), GTG1478ATC (5796G>A 5798G>C), GCC1479AGA (5799G>A 5800C>G 5801C>A), GAT1481GTG (5806A>T 5807T>G), GAA1482GTG (5809A>T 5810A>G), AGC1484GAC (5814A>G 5815G>A), ATC1485ATT (5819C>T), TGG1487AAA (5823T>A 5824G>A 5825G>A), ATC1491AAA (5836T>A 5837C>A), AGT1492ACT (5839G>C), CCA1493CAT (5842C>A 5843A>T), GCA1494ATG (5844G>A 5845C>T 5846A>G), GGG1495GGA (5849G>A), TTG1496CAT (5850T>C 5851T>A 5852G>T), GAA1498GAG (5858A>G), TGG1499TTT (5860G>T 5861G>T), CTG1500TTG (5862C>T), CCA1503CCC (5873A>C), TTC1504TTT (5876C>T), GGC1505GGA (5879C>A), CTA1506TTG (5880C>T 5882A>G), AAG1507ACT (5884A>C 5885G>T), GCT1509GCA (5891T>A), CCA1510CCT (5894A>T), GCA1511TCC (5895G>T 5897A>C), GTC1512ACT (5898G>A 5899T>C 5900C>T), CAG1514CAA (5906G>A), CGG1515TGT (5907C>T 5909G>T), AAG1516CCG (5910A>C 5911A>C), GAT1518AAT (5916G>A), GAA1519CAT (5919G>C 5921A>T), TGC1520GTT (5922T>G 5923G>T 5924C>T), TTC1521TTT (5927C>T), AAA1522CAA (5928A>C), GGA1523CAA (5931G>C 5932G>A), GGA1523_ACG1524insTAC (5933_5934insTAC), ACG1524ATG (5935C>T), GAG1525AGG (5937G>A 5938A>G), GAG1526AAG (5940G>A), TTC1527TTT (5945C>T), ATT1528GTC (5946A>G 5948T>C), GCC1529TTA (5949G>T 5950C>T 5951C>A), TAT1531TTT (5956A>T), ATC1532TTT (5958A>T 5960C>T), GAC1533GAT (5963C>T), GAT1534GAC (5966T>C), ATA1535ATT (5969A>T), CTT1536CTG (5972T>G), GTC1537ATC (5973G>A), TTC1538TAT (5977T>A 5978C>T), TCC1539AGT (5979T>A 5980C>G 5981C>T), AAC1540AAG (5984C>G), ACC1541TCC (5985A>T), ATC1542TTA (5988A>T 5990C>A), AAG1543CTG (5991A>C 5992A>T), CAT1545CAC (5999T>C), GAG1546CTT (6000G>C 6001A>T 6002G>T), AAG1547ATT (6004A>T 6005G>T), CAC1548CAT (6008C>T) | | | | | | | | |

*: Inserts / Deletes / Misaligned / Frameshifts

#### Analysis details

This analysis was performed with panviral2.64

### NGS Details (UN62): Dioscovirus dioscoreae

#### Assembly

| | |
|---|---|
| **Coverage Length** | 344 (1 contig(s)) |
| **Depth Of Coverage** | 3.1 |
| **Number Of Reads** | 8 |
| **Reads Per Million** | 0.19 rpm (after QC) |
| **Ambiguities** | 0 |
| **Assembly Method** | de novo + reference guided assembly |
| **Consensus Caller** | Bcf Tools |

#### Coverage Map

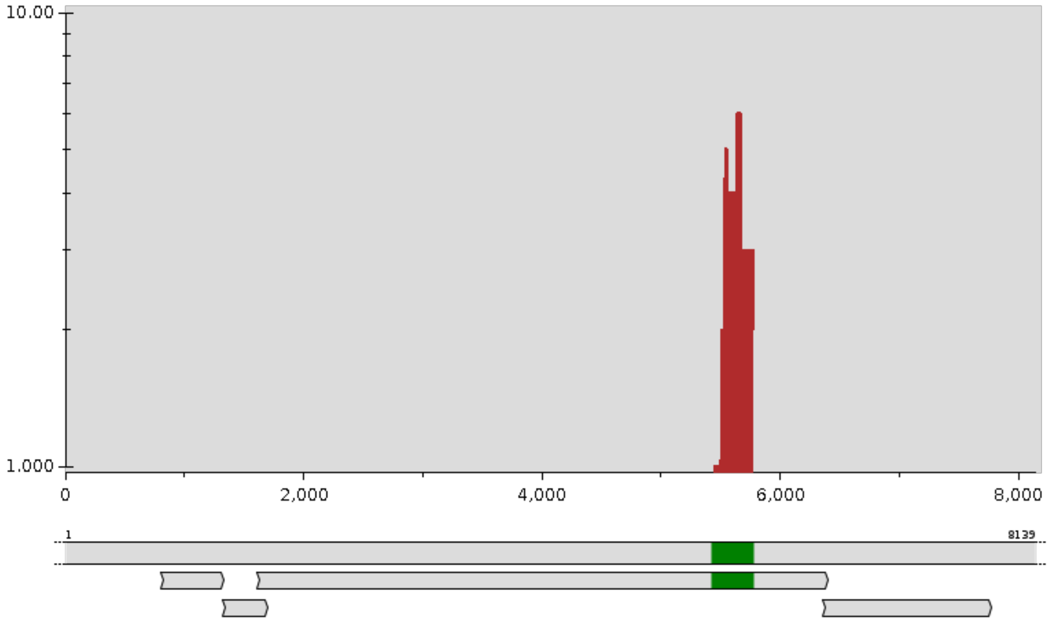

#### Assignment

| | |
|---|---|
| **Type** | Dioscovirus dioscoreae (Taxonomy ID: 3052184) |
| **Reference Genome** | NC_040712.1 |
| **NT Identity (%)** | 63.4286 |
| **AA Identity (%)** | 48.7179 |
| **Number Of Stop Codons** | 0 |
| **Number Of CDS** | 4 |

#### Alignment

| | |
|---|---|
| **Alignment Score** | 176.0 (NT) + 460.0 (AA) = 636.0 |
| **Concordance (%)** | 43.3243 |

| **Alignment Method** | Global, seeded, nucleotide + amino acids (AGA) |
|---|---|

#### Genome Region

Sequence starts at position 5436 and ends at position 5779 relative to NC_040712.1 reference sequence.

#### Alignment Detailed Statistics

| | Begin | End | Coverage | Score | Concordance | Matches | Identities | I/D/M/F* | Stop Codons |
|---|---|---|---|---|---|---|---|---|---|
| **NT** | **5436** | **5779** | **4.2%** | **176** | **26.3%** | **341 (96.6%)** | **222 (62.9%)** | **9/3** | |
| Mutations: | 5439C>G, 5441G>A, 5444T>A, 5452A>T, 5453A>G, 5457A>T, 5461T>C, 5466C>T, 5480A>G, 5482A>C, 5486A>C, 5489T>C, 5490T>C, 5492A>C, 5492_5493insCCCCCCCAA, 5493G>A, 5498C>T, 5502C>A, 5507A>G, 5514A>G, 5516T>A, 5523A>G, 5524A>G, 5526A>G, 5532T>A, 5533C>A, 5535G>A, 5538T>A, 5543C>A, 5544T>C, 5545G>T, 5546T>A, 5549T>C, 5553A>G, 5558C>T, 5561T>C, 5564T>C, 5565T>C, 5567G>A, 5568A>G, 5573T>C, 5574T>A, 5575C>G, 5577G>A, 5578A>G, 5580A>G, 5581G>A, 5585A>G, 5589C>G, 5595G>C, 5598C>G, 5607T>C, 5608T>A, 5610C>A, 5611A>C, 5612G>A, 5613T>A, 5615C>T, 5618C>T, 5619C>A, 5620A>G, 5621A>G, 5622G>T, 5626G>T, 5627T>C, 5628A>G, 5629A>T, 5630G>A, 5631G>A, 5632A>G, 5633A>T, 5634G>A, 5639G>A, 5640C>A, 5642T>A, 5645T>A, 5646T>A, 5648G>A, 5649T>A, 5651A>C, 5656C>A, 5657A>G, 5661T>A, 5664A>G, 5667A>T, 5670G>T, 5672A>T, 5673G>A, 5674T>A, 5676G>A, 5679A>C, 5685G>A, 5687A>C, 5691C>T, 5693T>A, 5696C>A, 5697C>G, 5700G>A, 5701A>C, 5702A>T, 5703A>T, 5711A>T, 5714A>G, 5717A>G, 5718G>A, 5720T>A, 5721C>A, 5726C>A, 5739T>G, 5740T>C, 5741A>T, 5748A>G, 5756A>T, 5757T>A, 5759T>G, 5762T>A, 5763_5765delGAA, 5768C>T, 5769C>A, 5777A>T, 5779C>A | | | | | | | | |
| **CDS** | | | | | | | | | |
| EXK67_gp3 | 1276 | 1390 | 7.2% | 460 | 57.3% | 114 (96.6%) | 57 (48.3%) | 3/1/0/0 | 0 |
| Protein mutations: | Q1277E (5439C>G 5441G>A), E1281V (5452A>T 5453A>G), I1283L (5457A>T), V1284A (5461T>C), P1286S (5466C>T), N1291T (5482A>C), S1294P (5490T>C 5492A>C), S1294_V1295insPPQ (5492_5493insCCCCCCCAA), V1295I (5493G>A), N1302E (5514A>G 5516T>A), K1305G (5523A>G 5524A>G), K1306E (5526A>G), S1308K (5532T>A 5533C>A), E1309K (5535G>A), F1310I (5538T>A), C1312L (5544T>C 5545G>T 5546T>A), I1315V (5553A>G), I1320V (5568A>G), E1323R (5577G>A 5578A>G), S1324D (5580A>G 5581G>A), I1325M (5585A>G), Q1327E (5589C>G), V1329L (5595G>C), Q1330E (5598C>G), L1333Q (5607T>C 5608T>A), Q1334T (5610C>A 5611A>C 5612G>A), F1335I (5613T>A 5615C>T), Q1337R (5619C>A 5620A>G 5621A>G), V1338L (5622G>T), C1339F (5626G>T 5627T>C), K1340V (5628A>G 5629A>T 5630G>A), E1341S (5631G>A 5632A>G 5633A>T), E1342K (5634G>A), L1344I (5640C>A 5642T>A), L1346I (5646T>A 5648G>A), S1347T (5649T>A 5651A>C), T1349K (5656C>A 5657A>G), L1351M (5661T>A), K1352E (5664A>G), I1353L (5667A>T), G1354C (5670G>T 5672A>T), V1355K (5673G>A 5674T>A), A1356T (5676G>A), N1357H (5679A>C), E1359N (5685G>A 5687A>C), L1363V (5697C>G), E1364T (5700G>A 5701A>C 5702A>T), I1365L (5703A>T), E1367D (5711A>T), V1370I (5718G>A 5720T>A), Q1371K (5721C>A), L1377A (5739T>G 5740T>C 5741A>T), I1380V (5748A>G), E1382D (5756A>T), F1383M (5757T>A 5759T>G), E1385del (5763_5765delGAA), Q1387K (5769C>A), E1389D (5777A>T) | | | | | | | | |
| Codon mutations: | CAG1277GAA (5439C>G 5441G>A), GGT1278GGA (5444T>A), GAA1281GTG (5452A>T 5453A>G), ATA1283TTA (5457A>T), GTA1284GCA (5461T>C), CCA1286TCA (5466C>T), AAA1290AAG (5480A>G), AAT1291ACT (5482A>C), GCA1292GCC (5486A>C), CCT1293CCC (5489T>C), TCA1294CCC (5490T>C 5492A>C), TCA1294_GTA1295insCCCCCCCAA (5492_5493insCCCCCCCAA), GTA1295ATA (5493G>A), TTC1296TTT (5498C>T), CGA1298AGA (5502C>A), AAA1299AAG (5507A>G), AAT1302GAA (5514A>G 5516T>A), AAA1305GGA (5523A>G 5524A>G), AAA1306GAA (5526A>G), TCA1308AAA (5532T>A 5533C>A), GAA1309AAA (5535G>A), TTT1310ATT (5538T>A), GTC1311GTA (5543C>A), TGT1312CTA (5544T>C 5545G>T 5546T>A), GTT1313GTC (5549T>C), ATA1315GTA (5553A>G), GAC1316GAT (5558C>T), GAT1317GAC (5561T>C), ATT1318ATC (5564T>C), TTG1319CTA (5565T>C 5567G>A), ATA1320GTA (5568A>G), TTT1321TTC (5573T>C), TCT1322AGT (5574T>A 5575C>G), GAA1323AGA (5577G>A 5578A>G), AGT1324GAT (5580A>G 5581G>A), ATA1325ATG (5585A>G), CAA1327GAA (5589C>G), GTA1329CTA (5595G>C), CAA1330GAA (5598C>G), TTA1333CAA (5607T>C 5608T>A), CAG1334ACA (5610C>A 5611A>C 5612G>A), TTC1335ATT (5613T>A 5615C>T), TTC1336TTT (5618C>T), CAA1337AGG (5619C>A 5620A>G 5621A>G), GTA1338TTA (5622G>T), TGT1339TTC (5626G>T 5627T>C), AAG1340GTA (5628A>G 5629A>T 5630G>A), GAA1341AGT (5631G>A 5632A>G 5633A>T), GAA1342AAA (5634G>A), GGG1343GGA (5639G>A), CTT1344ATA (5640C>A 5642T>A), ATT1345ATA (5645T>A), TTG1346ATA (5646T>A 5648G>A), TCA1347ACC (5649T>A 5651A>C), ACA1349AAG (5656C>A 5657A>G), TTG1351ATG (5661T>A), AAA1352GAA (5664A>G), ATA1353TTA (5667A>T), GGA1354TGT (5670G>T 5672A>T), GTA1355AAA (5673G>A 5674T>A), GCA1356ACA (5676G>A), AAT1357CAT (5679A>C), GAA1359AAC (5685G>A 5687A>C), CTT1361TTA (5691C>T 5693T>A), GGC1362GGA (5696C>A), CTA1363GTA (5697C>G), GAA1364ACT (5700G>A 5701A>C 5702A>T), ATA1365TTA (5703A>T), GAA1367GAT (5711A>T), GGA1368GGG (5714A>G), AAA1369AAG (5717A>G), GTT1370ATA (5718G>A 5720T>A), CAA1371AAA (5721C>A), CTC1372CTA (5726C>A), TTA1377GCT (5739T>G 5740T>C 5741A>T), ATA1380GTA (5748A>G), GAA1382GAT (5756A>T), TTT1383ATG (5757T>A 5759T>G), CCT1384CCA (5762T>A), GAA1385del (5763_5765delGAA), GAC1386GAT (5768C>T), CAA1387AAA (5769C>A), GAA1389GAT (5777A>T), ACA1390AA. (5779C>A) | | | | | | | | |
| **Proteins** | | | | | | | | | |
| ORF3 (YP_009553219.1) | 1276 | 1390 | 7.2% | 460 | 57.3% | 114 (96.6%) | 57 (48.3%) | 3/1/0/0 | 0 |
| Protein mutations: | Q1277E (5439C>G 5441G>A), E1281V (5452A>T 5453A>G), I1283L (5457A>T), V1284A (5461T>C), P1286S (5466C>T), N1291T (5482A>C), S1294P (5490T>C 5492A>C), S1294_V1295insPPQ (5492_5493insCCCCCCCAA), V1295I (5493G>A), N1302E (5514A>G 5516T>A), K1305G (5523A>G 5524A>G), K1306E (5526A>G), S1308K (5532T>A 5533C>A), E1309K (5535G>A), F1310I (5538T>A), C1312L (5544T>C 5545G>T 5546T>A), I1315V (5553A>G), I1320V (5568A>G), E1323R (5577G>A 5578A>G), S1324D (5580A>G 5581G>A), I1325M (5585A>G), Q1327E (5589C>G), V1329L (5595G>C), Q1330E (5598C>G), L1333Q (5607T>C 5608T>A), Q1334T (5610C>A 5611A>C 5612G>A), F1335I (5613T>A 5615C>T), Q1337R (5619C>A 5620A>G 5621A>G), V1338L (5622G>T), C1339F (5626G>T 5627T>C), K1340V (5628A>G 5629A>T 5630G>A), E1341S (5631G>A 5632A>G 5633A>T), E1342K (5634G>A), L1344I (5640C>A 5642T>A), L1346I (5646T>A 5648G>A), S1347T (5649T>A 5651A>C), T1349K (5656C>A 5657A>G), L1351M (5661T>A), K1352E (5664A>G), I1353L (5667A>T), G1354C (5670G>T 5672A>T), V1355K (5673G>A 5674T>A), A1356T (5676G>A), N1357H (5679A>C), E1359N (5685G>A 5687A>C), L1363V (5697C>G), E1364T (5700G>A 5701A>C 5702A>T), I1365L (5703A>T), E1367D (5711A>T), V1370I (5718G>A 5720T>A), Q1371K (5721C>A), L1377A (5739T>G 5740T>C 5741A>T), I1380V (5748A>G), E1382D (5756A>T), F1383M (5757T>A 5759T>G), E1385del (5763_5765delGAA), Q1387K (5769C>A), E1389D (5777A>T) | | | | | | | | |
| Codon mutations: | CAG1277GAA (5439C>G 5441G>A), GGT1278GGA (5444T>A), GAA1281GTG (5452A>T 5453A>G), ATA1283TTA (5457A>T), GTA1284GCA (5461T>C), CCA1286TCA (5466C>T), AAA1290AAG (5480A>G), AAT1291ACT (5482A>C), GCA1292GCC (5486A>C), CCT1293CCC (5489T>C), TCA1294CCC (5490T>C 5492A>C), TCA1294_GTA1295insCCCCCCCAA (5492_5493insCCCCCCCAA), GTA1295ATA (5493G>A), TTC1296TTT (5498C>T), CGA1298AGA (5502C>A), AAA1299AAG (5507A>G), AAT1302GAA (5514A>G 5516T>A), AAA1305GGA (5523A>G 5524A>G), AAA1306GAA (5526A>G), TCA1308AAA (5532T>A 5533C>A), GAA1309AAA (5535G>A), TTT1310ATT (5538T>A), GTC1311GTA (5543C>A), TGT1312CTA (5544T>C 5545G>T 5546T>A), GTT1313GTC (5549T>C), ATA1315GTA (5553A>G), GAC1316GAT (5558C>T), GAT1317GAC (5561T>C), ATT1318ATC (5564T>C), TTG1319CTA (5565T>C 5567G>A), ATA1320GTA (5568A>G), TTT1321TTC (5573T>C), TCT1322AGT (5574T>A 5575C>G), GAA1323AGA (5577G>A 5578A>G), AGT1324GAT (5580A>G 5581G>A), ATA1325ATG (5585A>G), CAA1327GAA (5589C>G), GTA1329CTA (5595G>C), CAA1330GAA (5598C>G), TTA1333CAA (5607T>C 5608T>A), CAG1334ACA (5610C>A 5611A>C 5612G>A), TTC1335ATT (5613T>A 5615C>T), TTC1336TTT (5618C>T), CAA1337AGG (5619C>A 5620A>G 5621A>G), GTA1338TTA (5622G>T), TGT1339TTC (5626G>T 5627T>C), AAG1340GTA (5628A>G 5629A>T 5630G>A), GAA1341AGT (5631G>A 5632A>G 5633A>T), GAA1342AAA (5634G>A), GGG1343GGA (5639G>A), CTT1344ATA (5640C>A 5642T>A), ATT1345ATA (5645T>A), TTG1346ATA (5646T>A 5648G>A), TCA1347ACC (5649T>A 5651A>C), ACA1349AAG (5656C>A 5657A>G), TTG1351ATG (5661T>A), AAA1352GAA (5664A>G), ATA1353TTA (5667A>T), GGA1354TGT (5670G>T 5672A>T), GTA1355AAA (5673G>A 5674T>A), GCA1356ACA (5676G>A), AAT1357CAT (5679A>C), GAA1359AAC (5685G>A 5687A>C), CTT1361TTA (5691C>T 5693T>A), GGC1362GGA (5696C>A), CTA1363GTA (5697C>G), GAA1364ACT (5700G>A 5701A>C 5702A>T), ATA1365TTA (5703A>T), GAA1367GAT (5711A>T), GGA1368GGG (5714A>G), AAA1369AAG (5717A>G), GTT1370ATA (5718G>A 5720T>A), CAA1371AAA (5721C>A), CTC1372CTA (5726C>A), TTA1377GCT (5739T>G 5740T>C 5741A>T), ATA1380GTA (5748A>G), GAA1382GAT (5756A>T), TTT1383ATG (5757T>A 5759T>G), CCT1384CCA (5762T>A), GAA1385del (5763_5765delGAA), GAC1386GAT (5768C>T), CAA1387AAA (5769C>A), GAA1389GAT (5777A>T), ACA1390AA. (5779C>A) | | | | | | | | |

*: Inserts / Deletes / Misaligned / Frameshifts

#### Analysis details

This analysis was performed with panviral2.64

### NGS Details (UN62): Duamitovirus soch1

#### Assembly

| | |
|---|---|
| **Coverage Length** | 357 (1 contig(s)) |
| **Depth Of Coverage** | 2.5 |
| **Number Of Reads** | 7 |
| **Reads Per Million** | 0.16 rpm (after QC) |
| **Ambiguities** | 0 |
| **Assembly Method** | de novo + reference guided assembly |
| **Consensus Caller** | Bcf Tools |

#### Coverage Map

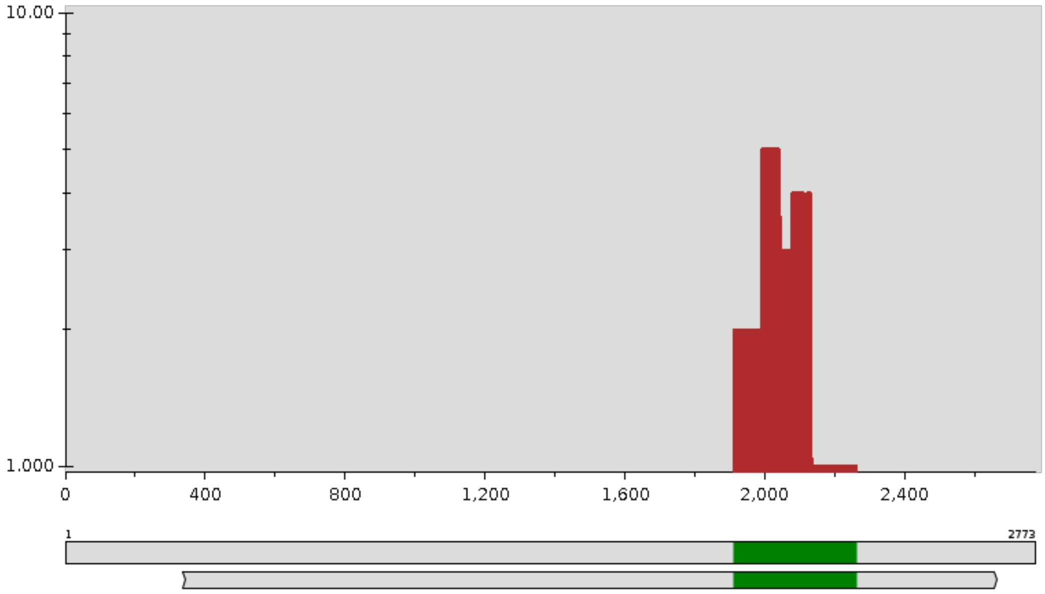

#### Assignment

| | |
|---|---|
| **Type** | Duamitovirus soch1 (Taxonomy ID: 2955838) |
| **Reference Genome** | NC_076524.1 |
| **NT Identity (%)** | 69.1643 |
| **AA Identity (%)** | 73.913 |
| **Number Of Stop Codons** | 0 |
| **Number Of CDS** | 1 |

#### Alignment

| | |
|---|---|
| **Alignment Score** | 247.0 (NT) + 469.0 (AA) = 716.0 |
| **Concordance (%)** | 54.2424 |

| Alignment Method | Global, seeded, nucleotide + amino acids (AGA) |
|---|---|

#### Genome Region

Sequence starts at position 1910 and ends at position 2266 relative to NC_076524.1 reference sequence.

#### Alignment Detailed Statistics

| | Begin | End | Coverage | Score | Concordance | Matches | Identities | I/D/M/F* | Stop Codons |
|---|---|---|---|---|---|---|---|---|---|
| **NT** | **1910** | **2266** | **12.9%** | **247** | **36.6%** | **347 (97.2%)** | **240 (67.2%)** | **0/10** | |
| Mutations: | 1919G>T, 1933C>T, 1934T>A, 1935A>C, 1936A>G, 1938T>G, 1939C>A, 1940A>T, 1948T>A, 1949T>A, 1953T>A, 1955A>C, 1961A>C, 1964T>C, 1967C>T, 1970T>C, 1972G>A, 1980T>A, 1981T>C, 1982A>C, 1985T>C, 1989C>A, 1991T>A, 1994T>G, 2000T>A, 2003A>C, 2006T>G, 2010G>C, 2011C>A, 2012C>A, 2013C>T, 2015C>G, 2016G>T, 2018A>C, 2019A>T, 2020T>G, 2022A>C, 2023A>G, 2024G>A, 2027T>C, 2028G>T, 2029A>C, 2030T>G, 2033T>A, 2034A>G, 2036C>A, 2042A>G, 2043G>A, 2044T>C, 2054A>G, 2055T>C, 2057G>A, 2059G>C, 2066C>A, 2069G>T, 2072C>T, 2078A>C, 2082G>T, 2084G>T, 2088T>C, 2090A>G, 2091T>A, 2093C>G, 2094A>T, 2099C>T, 2102A>G, 2103A>T, 2104G>C, 2105C>A, 2106A>C, 2107A>C, 2108G>A, 2111T>A, 2117G>A, 2118C>A, 2120T>A, 2124_2133delAAAGCTGCTC, 2135T>G, 2138T>C, 2141G>A, 2146T>G, 2148C>A, 2152C>T, 2158T>C, 2167A>T, 2168C>A, 2173G>T, 2174G>T, 2186G>A, 2191T>A, 2192G>A, 2195C>T, 2204C>T, 2208T>C, 2210G>A, 2213A>G, 2225A>G, 2230A>T, 2231C>T, 2234C>T, 2237T>G, 2239A>G, 2240A>G, 2243A>G, 2250G>C, 2251T>C, 2252T>C, 2264G>T | | | | | | | | |

##### CDS

| | Begin | End | Coverage | Score | Concordance | Matches | Identities | I/D/M/F* | Stop Codons |
|---|---|---|---|---|---|---|---|---|---|
| RdRp | 526 | 644 | 15.2% | 469 | 59.3% | 115 (97.5%) | 85 (72.0%) | 0/3/1/1 | 0 |
| Protein mutations: | T533I (1933C>T 1934T>A), K534R (1935A>C 1936A>G), S535D (1938T>G 1939C>A 1940A>T), I538K (1948T>A 1949T>A), L540I (1953T>A 1955A>C), R546K (1972G>A), L549T (1980T>A 1981T>C 1982A>C), A559Q (2010G>C 2011C>A 2012C>A), A561S (2016G>T 2018A>C), I562C (2019A>T 2020T>G), K563R (2022A>C 2023A>G 2024G>A), D565S (2028G>T 2029A>C 2030T>G), N567E (2034A>G 2036C>A), V570T (2043G>A 2044T>C), G575A (2059G>C), A583S (2082G>T 2084G>T), F586M (2091T>A 2093C>G), T587S (2094A>T), K591P (2106A>C 2107A>C 2108G>A), K597_A599del (2124_2133delAAAGCTGCTC), V604G (2146T>G), S606L (2152C>T), L608S (2158T>C), Y611L (2167A>T 2168C>A), W613F (2173G>T 2174G>T), M619K (2191T>A 2192G>A), Y632F (2230A>T 2231C>T), K635R (2239A>G 2240A>G), V639P (2250G>C 2251T>C 2252T>C), Q643H (2264G>T) | | | | | | | | |
| Codon mutations: | GCG528GCT (1919G>T), ACT533ATA (1933C>T 1934T>A), AAA534CGA (1935A>C 1936A>G), TCA535GAT (1938T>G 1939C>A 1940A>T), ATT538AAA (1948T>A 1949T>A), TTA540ATC (1953T>A 1955A>C), CCA542CCC (1961A>C), ATT543ATC (1964T>C), TCC544TCT (1967C>T), CTT545CTC (1970T>C), AGA546AAA (1972G>A), TTA549ACC (1980T>A 1981T>C 1982A>C), TCT550TCC (1985T>C), CGT552AGA (1989C>A 1991T>A), ACT553ACG (1994T>G), GTT555GTA (2000T>A), GGA556GGC (2003A>C), CTT557CTG (2006T>G), GCC559CAA (2010G>C 2011C>A 2012C>A), CTC560TTG (2013C>T 2015C>G), GCA561TCC (2016G>T 2018A>C), ATC562TGC (2019A>T 2020T>G), AAG563CGA (2022A>C 2023A>G 2024G>A), TAT564TAC (2027T>C), GAT565TCG (2028G>T 2029A>C 2030T>G), ATT566ATA (2033T>A), AAC567GAA (2034A>G 2036C>A), TCA569TCG (2042A>G), GTT570ACT (2043G>A 2044T>C), AGA573AGG (2054A>G), TTG574CTA (2055T>C 2057G>A), GGG575GCG (2059G>C), GCC577GCA (2066C>A), GGG578GGT (2069G>T), TAC579TAT (2072C>T), GTA581GTC (2078A>C), GCG583TCT (2082G>T 2084G>T), TTA585CTG (2088T>C 2090A>G), TTC586ATG (2091T>A 2093C>G), ACT587TCT (2094A>T), ACC588ACT (2099C>T), CAA589CAG (2102A>G), AGC590TCA (2103A>T 2104G>C 2105C>A), AAG591CCA (2106A>C 2107A>C 2108G>A), CGT592CGA (2111T>A), GAG594GAA (2117G>A), CGT595AGA (2118C>A 2120T>A), AAA597_GCT599del (2124_2133delAAAGCTGCTC), CAT600-AG (2124_2133delAAAGCTGCTC 2135T>G), CTT601CTC (2138T>C), AAG602AAA (2141G>A), GTT604GGT (2146T>G), CGG605AGG (2148C>A), TCA606TTA (2152C>T), TTA608TCA (2158T>C), TAC611TTA (2167A>T 2168C>A), TGG613TTT (2173G>T 2174G>T), AGG617AGA (2186G>A), ATG619AAA (2191T>A 2192G>A), CCC620CCT (2195C>T), CCC623CCT (2204C>T), TTG625CTA (2208T>C 2210G>A), AAA626AAG (2213A>G), GTA630GTG (2225A>G), TAC632TTT (2230A>T 2231C>T), CTC633CTT (2234C>T), CTT634CTG (2237T>G), AAA635AGG (2239A>G 2240A>G), GAA636GAG (2243A>G), GTT639CCC (2250G>C 2251T>C 2252T>C), CAG643CAT (2264G>T) | | | | | | | | |

##### Proteins

| | Begin | End | Coverage | Score | Concordance | Matches | Identities | I/D/M/F* | Stop Codons |
|---|---|---|---|---|---|---|---|---|---|
| RNA-dependent RNA polymerase (YP_010798874.1) | 526 | 644 | 15.2% | 469 | 59.3% | 115 (97.5%) | 85 (72.0%) | 0/3/1/1 | 0 |
| Protein mutations: | T533I (1933C>T 1934T>A), K534R (1935A>C 1936A>G), S535D (1938T>G 1939C>A 1940A>T), I538K (1948T>A 1949T>A), L540I (1953T>A 1955A>C), R546K (1972G>A), L549T (1980T>A 1981T>C 1982A>C), A559Q (2010G>C 2011C>A 2012C>A), A561S (2016G>T 2018A>C), I562C (2019A>T 2020T>G), K563R (2022A>C 2023A>G 2024G>A), D565S (2028G>T 2029A>C 2030T>G), N567E (2034A>G 2036C>A), V570T (2043G>A 2044T>C), G575A (2059G>C), A583S (2082G>T 2084G>T), F586M (2091T>A 2093C>G), T587S (2094A>T), K591P (2106A>C 2107A>C 2108G>A), K597_A599del (2124_2133delAAAGCTGCTC), V604G (2146T>G), S606L (2152C>T), L608S (2158T>C), Y611L (2167A>T 2168C>A), W613F (2173G>T 2174G>T), M619K (2191T>A 2192G>A), Y632F (2230A>T 2231C>T), K635R (2239A>G 2240A>G), V639P (2250G>C 2251T>C 2252T>C), Q643H (2264G>T) | | | | | | | | |
| Codon mutations: | GCG528GCT (1919G>T), ACT533ATA (1933C>T 1934T>A), AAA534CGA (1935A>C 1936A>G), TCA535GAT (1938T>G 1939C>A 1940A>T), ATT538AAA (1948T>A 1949T>A), TTA540ATC (1953T>A 1955A>C), CCA542CCC (1961A>C), ATT543ATC (1964T>C), TCC544TCT (1967C>T), CTT545CTC (1970T>C), AGA546AAA (1972G>A), TTA549ACC (1980T>A 1981T>C 1982A>C), TCT550TCC (1985T>C), CGT552AGA (1989C>A 1991T>A), ACT553ACG (1994T>G), GTT555GTA (2000T>A), GGA556GGC (2003A>C), CTT557CTG (2006T>G), GCC559CAA (2010G>C 2011C>A 2012C>A), CTC560TTG (2013C>T 2015C>G), GCA561TCC (2016G>T 2018A>C), ATC562TGC (2019A>T 2020T>G), AAG563CGA (2022A>C 2023A>G 2024G>A), TAT564TAC (2027T>C), GAT565TCG (2028G>T 2029A>C 2030T>G), ATT566ATA (2033T>A), AAC567GAA (2034A>G 2036C>A), TCA569TCG (2042A>G), GTT570ACT (2043G>A 2044T>C), AGA573AGG (2054A>G), TTG574CTA (2055T>C 2057G>A), GGG575GCG (2059G>C), GCC577GCA (2066C>A), GGG578GGT (2069G>T), TAC579TAT (2072C>T), GTA581GTC (2078A>C), GCG583TCT (2082G>T 2084G>T), TTA585CTG (2088T>C 2090A>G), TTC586ATG (2091T>A 2093C>G), ACT587TCT (2094A>T), ACC588ACT (2099C>T), CAA589CAG (2102A>G), AGC590TCA (2103A>T 2104G>C 2105C>A), AAG591CCA (2106A>C 2107A>C 2108G>A), CGT592CGA (2111T>A), GAG594GAA (2117G>A), CGT595AGA (2118C>A 2120T>A), AAA597_GCT599del (2124_2133delAAAGCTGCTC), CAT600-AG (2124_2133delAAAGCTGCTC 2135T>G), CTT601CTC (2138T>C), AAG602AAA (2141G>A), GTT604GGT (2146T>G), CGG605AGG (2148C>A), TCA606TTA (2152C>T), TTA608TCA (2158T>C), TAC611TTA (2167A>T 2168C>A), TGG613TTT (2173G>T 2174G>T), AGG617AGA (2186G>A), ATG619AAA (2191T>A 2192G>A), CCC620CCT (2195C>T), CCC623CCT (2204C>T), TTG625CTA (2208T>C 2210G>A), AAA626AAG (2213A>G), GTA630GTG (2225A>G), TAC632TTT (2230A>T 2231C>T), CTC633CTT (2234C>T), CTT634CTG (2237T>G), AAA635AGG (2239A>G 2240A>G), GAA636GAG (2243A>G), GTT639CCC (2250G>C 2251T>C 2252T>C), CAG643CAT (2264G>T) | | | | | | | | |

*: Inserts / Deletes / Misaligned / Frameshifts

#### Analysis details

This analysis was performed with panviral2.64

#### NGS Details (UN62): Escherichia virus DE3

##### Assembly

| | |
|---|---|
| **Coverage Length** | 226 (1 contig(s)) |
| **Depth Of Coverage** | 3.6 |
| **Number Of Reads** | 6 |
| **Reads Per Million** | 0.14 rpm (after QC) |
| **Ambiguities** | 0 |
| **Assembly Method** | de novo + reference guided assembly |
| **Consensus Caller** | Bcf Tools |

##### Coverage Map

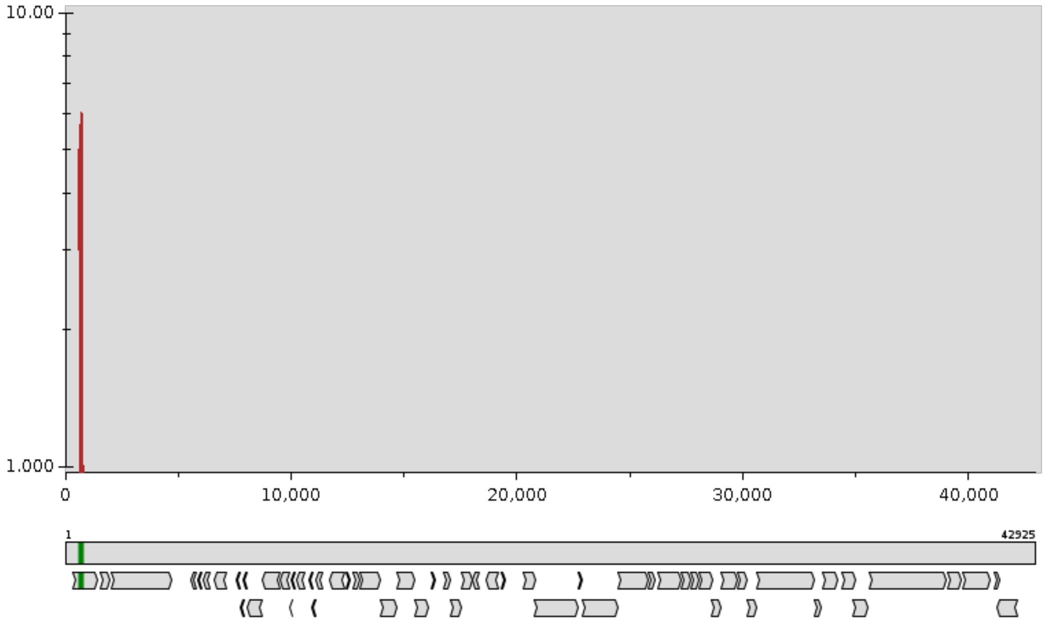

##### Assignment

| | |
|---|---|
| **Type** | Escherichia virus DE3 (Taxonomy ID: 2169967) |
| **Reference Genome** | NC_042057.1 |
| **NT Identity (%)** | 100.0 |
| **AA Identity (%)** | 100.0 |
| **Number Of Stop Codons** | 0 |
| **Number Of CDS** | 57 |

##### Alignment

| | |
|---|---|
| **Alignment Score** | 452.0 (NT) + 491.0 (AA) = 943.0 |
| **Concordance (%)** | 100.0 |

| Alignment Method | Local, heuristic, nucleotide (BLASTN) |
|---|---|

#### Genome Region

Sequence starts at position 607 and ends at position 832 relative to NC_042057.1 reference sequence.

#### Alignment Detailed Statistics

| | Begin | End | Coverage | Score | Concordance | Matches | Identities | I/D/M/F * | Stop Codons |
|---|---|---|---|---|---|---|---|---|---|
| **NT** | **607** | **832** | **0.5%** | **452** | **100%** | **226 (100%)** | **226 (100%)** | **0/0** | |
| Mutations: | *none* | | | | | | | | |
| **CDS** | | | | | | | | | |
| lacI | 90 | 164 | 20.8% | 491 | 98.4% | 75 (100%) | 75 (100%) | 0/0/0/0 | 0 |
| **Proteins** | | | | | | | | | |
| DNA-binding transcriptional repressor LacI (YP_009617199.1) | 90 | 164 | 20.8% | 491 | 98.4% | 75 (100%) | 75 (100%) | 0/0/0/0 | 0 |
| Protein mutations: | *none* | | | | | | | | |
| Codon mutations: | *none* | | | | | | | | |

*: Inserts / Deletes / Misaligned / Frameshifts

#### Analysis details

This analysis was performed with panviral2.64

### NGS Details (UN62): Potato virus X

#### Assembly

| | |
|---|---|
| **Coverage Length** | 225 (1 contig(s)) |
| **Depth Of Coverage** | 2.4 |
| **Number Of Reads** | 4 |
| **Reads Per Million** | 0.09 rpm (after QC) |
| **Ambiguities** | 0 |
| **Assembly Method** | de novo + reference guided assembly |
| **Consensus Caller** | Bcf Tools |

#### Coverage Map

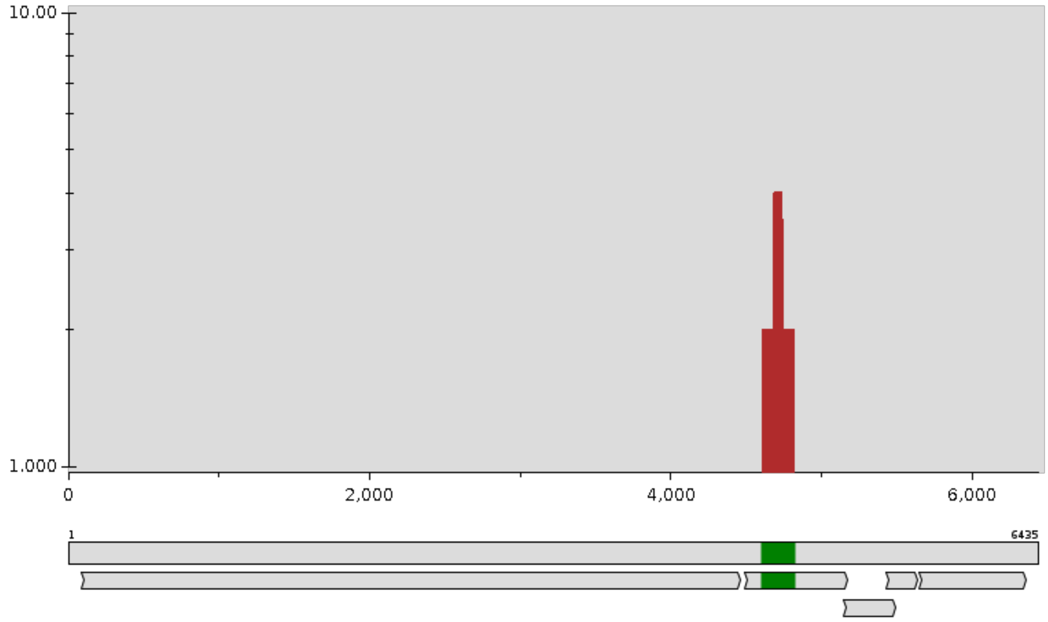

#### Assignment

| | |
|---|---|
| **Type** | Potato virus X (Taxonomy ID: 12183) |
| **Reference Genome** | NC_011620.1 |
| **NT Identity (%)** | 73.6607 |
| **AA Identity (%)** | 88.0 |
| **Number Of Stop Codons** | 0 |
| **Number Of CDS** | 5 |

#### Alignment

| | |
|---|---|
| **Alignment Score** | 202.0 (NT) + 365.0 (AA) = 567.0 |
| **Concordance (%)** | 67.3888 |

| Alignment Method | Global, seeded, nucleotide + amino acids (AGA) |
|---|---|

#### Genome Region

Sequence starts at position 4600 and ends at position 4824 relative to NC_011620.1 reference sequence.

#### Alignment Detailed Statistics

| | Begin | End | Coverage | Score | Concordance | Matches | Identities | I/D/M/F* | Stop Codons |
|---|---|---|---|---|---|---|---|---|---|
| **NT** | **4600** | **4824** | **3.5%** | **202** | **46.1%** | **224 (99.6%)** | **165 (73.3%)** | **0/1** | |
| Mutations: | 4602A>G, 4603A>C, 4605G>T, 4608G>A, 4611G>A, 4612A>C, 4614C>T, 4615C>G, 4616T>C, 4618A>C, 4624C>T, 4629A>T, 4635C>G, 4641T>C, 4644A>C, 4650T>A, 4656T>G, 4662G>A, 4663G>A, 4665G>C, 4668T>C, 4671C>T, 4674A>G, 4677T>A, 4683C>A, 4689G>A, 4690delA, 4695A>C, 4698A>G, 4701T>C, 4710G>A, 4716T>C, 4719C>T, 4722A>C, 4725C>T, 4728C>T, 4731T>C, 4737T>C, 4740T>G, 4741T>C, 4746C>T, 4747A>G, 4748A>C, 4749C>G, 4759A>G, 4761C>A, 4762T>G, 4767C>T, 4770G>A, 4773A>T, 4776T>A, 4782T>A, 4788T>G, 4797A>G, 4800T>G, 4803G>A, 4804T>C, 4809C>T, 4812A>T, 4815G>A | | | | | | | | |

##### CDS

| | Begin | End | Coverage | Score | Concordance | Matches | Identities | I/D/M/F* | Stop Codons |
|---|---|---|---|---|---|---|---|---|---|
| PVX_gp2 | 39 | 113 | 33.0% | 365 | 67.7% | 75 (100%) | 66 (88.0%) | 0/0/1/1 | 0 |
| Protein mutations: | I43L (4612A>C 4614C>T), L44A (4615C>G 4616T>C), P47S (4624C>T), V60I (4663G>A 4665G>C), N88A (4747A>G 4748A>C 4749C>G), N92E (4759A>G 4761C>A), S93A (4762T>G), F107L (4804T>C) | | | | | | | | |
| Codon mutations: | CTA39CTG (4602A>G), AGG40CGT (4603A>C 4605G>T), AAG41AAA (4608G>A), TTG42TTA (4611G>A), ATC43CTT (4612A>C 4614C>T), CTC44GCC (4615C>G 4616T>C), AGA45CGA (4618A>C), CCA47TCA (4624C>T), ACA48ACT (4629A>T), ACC50ACG (4635C>G), CAT52CAC (4641T>C), ACA53ACC (4644A>C), GGT55GGA (4650T>A), CCT57CCG (4656T>G), AAG59AAA (4662G>A), GTG60ATC (4663G>A 4665G>C), AGT61AGC (4668T>C), ATC62ATT (4671C>T), AGA63AGG (4674A>G), ACT64ACA (4677T>A), GGC66GGA (4683C>A), CAG68CAA (4689G>A), AAG69-AG (4690delA), CCA70CCC (4695A>C), GGA71GGG (4698A>G), CCT72CCC (4701T>C), GAG75GAA (4710G>A), AAT77AAC (4716T>C), TTC78TTT (4719C>T), GCA79GCC (4722A>C), ATC80ATT (4725C>T), CTC81CTT (4728C>T), GAT82GAC (4731T>C), TAT84TAC (4737T>C), ACT85ACG (4740T>G), TTG86CTG (4741T>C), GAC87GAT (4746C>T), AAC88GCG (4747A>G 4748A>C 4749C>G), AAC92GAA (4759A>G 4761C>A), TCA93GCA (4762T>G), TAC94TAT (4767C>T), CAG95CAA (4770G>A), GCA96GCT (4773A>T), CTT97CTA (4776T>A), GCT99GCA (4782T>A), CCT101CCG (4788T>G), GCA104GCG (4797A>G), CCT105CCG (4800T>G), GAG106GAA (4803G>A), TTT107CTT (4804T>C), AGC108AGT (4809C>T), CTA109CTT (4812A>T), GAG110GAA (4815G>A) | | | | | | | | |

##### Proteins

| | Begin | End | Coverage | Score | Concordance | Matches | Identities | I/D/M/F* | Stop Codons |
|---|---|---|---|---|---|---|---|---|---|
| 25K protein (YP_002332930.1) | 39 | 113 | 33.0% | 365 | 67.7% | 75 (100%) | 66 (88.0%) | 0/0/1/1 | 0 |
| Protein mutations: | I43L (4612A>C 4614C>T), L44A (4615C>G 4616T>C), P47S (4624C>T), V60I (4663G>A 4665G>C), N88A (4747A>G 4748A>C 4749C>G), N92E (4759A>G 4761C>A), S93A (4762T>G), F107L (4804T>C) | | | | | | | | |
| Codon mutations: | CTA39CTG (4602A>G), AGG40CGT (4603A>C 4605G>T), AAG41AAA (4608G>A), TTG42TTA (4611G>A), ATC43CTT (4612A>C 4614C>T), CTC44GCC (4615C>G 4616T>C), AGA45CGA (4618A>C), CCA47TCA (4624C>T), ACA48ACT (4629A>T), ACC50ACG (4635C>G), CAT52CAC (4641T>C), ACA53ACC (4644A>C), GGT55GGA (4650T>A), CCT57CCG (4656T>G), AAG59AAA (4662G>A), GTG60ATC (4663G>A 4665G>C), AGT61AGC (4668T>C), ATC62ATT (4671C>T), AGA63AGG (4674A>G), ACT64ACA (4677T>A), GGC66GGA (4683C>A), CAG68CAA (4689G>A), AAG69-AG (4690delA), CCA70CCC (4695A>C), GGA71GGG (4698A>G), CCT72CCC (4701T>C), GAG75GAA (4710G>A), AAT77AAC (4716T>C), TTC78TTT (4719C>T), GCA79GCC (4722A>C), ATC80ATT (4725C>T), CTC81CTT (4728C>T), GAT82GAC (4731T>C), TAT84TAC (4737T>C), ACT85ACG (4740T>G), TTG86CTG (4741T>C), GAC87GAT (4746C>T), AAC88GCG (4747A>G 4748A>C 4749C>G), AAC92GAA (4759A>G 4761C>A), TCA93GCA (4762T>G), TAC94TAT (4767C>T), CAG95CAA (4770G>A), GCA96GCT (4773A>T), CTT97CTA (4776T>A), GCT99GCA (4782T>A), CCT101CCG (4788T>G), GCA104GCG (4797A>G), CCT105CCG (4800T>G), GAG106GAA (4803G>A), TTT107CTT (4804T>C), AGC108AGT (4809C>T), CTA109CTT (4812A>T), GAG110GAA (4815G>A) | | | | | | | | |

*: Inserts / Deletes / Misaligned / Frameshifts

#### Analysis details

This analysis was performed with panviral2.64

### NGS Details (UN62): Pandoravirus salinus

#### Assembly

| | |
|---|---|
| **Coverage Length** | 122 (1 contig(s)) |
| **Depth Of Coverage** | 2.7 |
| **Number Of Reads** | 4 |
| **Reads Per Million** | 0.09 rpm (after QC) |
| **Ambiguities** | 0 |
| **Assembly Method** | de novo + reference guided assembly |
| **Consensus Caller** | Bcf Tools |

#### Coverage Map

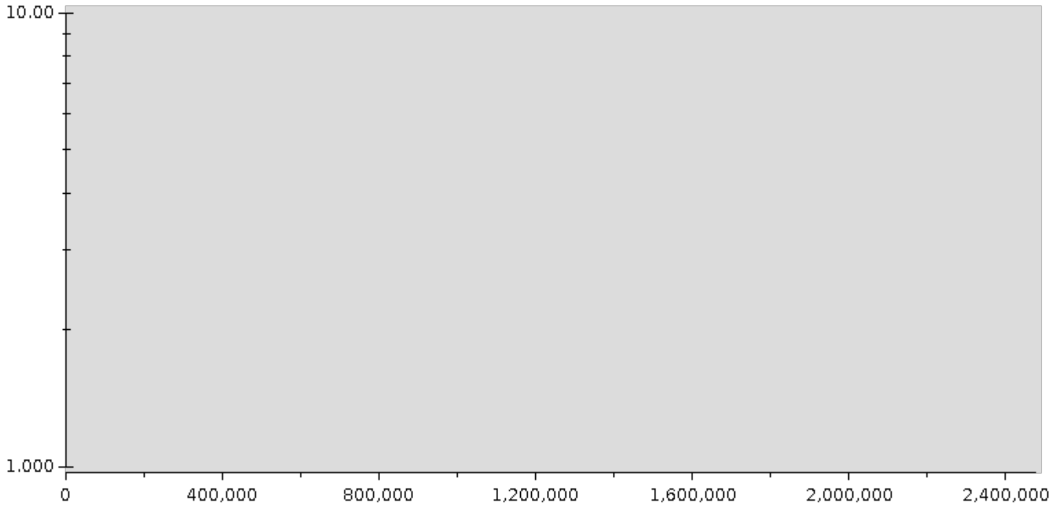

#### Assignment

| | |
|---|---|
| **Type** | Pandoravirus salinus (Taxonomy ID: 1349410) |
| **Reference Genome** | NC_022098.1 |
| **NT Identity (%)** | 75.4098 |
| **AA Identity (%)** | 80.4878 |
| **Number Of Stop Codons** | 0 |
| **Number Of CDS** | 1430 |

#### Alignment

| | |
|---|---|
| **Alignment Score** | 124.0 (NT) + 272.0 (AA) = 396.0 |
| **Concordance (%)** | 71.7391 |
| **Alignment Method** | Local, heuristic, nucleotide (BLASTN) |

#### Genome Region

Sequence starts at position 796297 and ends at position 796418 relative to NC_022098.1 reference sequence.

#### Alignment Detailed Statistics

| | Begin | End | Coverage | Score | Concordance | Matches | Identities | I/D/M/F* | Stop Codons |
|---|---|---|---|---|---|---|---|---|---|
| **NT** | **796297** | **796418** | **0.1%** | **124** | **50.8%** | **122 (100%)** | **92 (75.4%)** | **0/0** | |

Mutations: 796299G>A, 796302G>A, 796304C>T, 796305C>T, 796308C>T, 796314G>A, 796320A>G, 796323G>A, 796326G>A, 796329C>T, 796338T>C, 796347C>T, 796349C>G, 796353T>G, 796355T>C, 796356C>A, 796357C>T, 796358G>C, 796359T>A, 796365G>C, 796366C>T, 796367G>T, 796377C>G, 796380A>C, 796383A>G, 796389A>G, 796401G>T, 796402C>T, 796404T>G, 796410C>A

*: Inserts / Deletes / Misaligned / Frameshifts

#### Analysis details

This analysis was performed with panviral2.64

### NGS Details (UN62): Dioscorea bacilliform virus

#### Assembly

| | |
|---|---|
| **Coverage Length** | 315 (1 contig(s)) |
| **Depth Of Coverage** | 1.7 |
| **Number Of Reads** | 4 |
| **Reads Per Million** | 0.09 rpm (after QC) |
| **Ambiguities** | 0 |
| **Assembly Method** | de novo + reference guided assembly |
| **Consensus Caller** | Bcf Tools |

#### Coverage Map

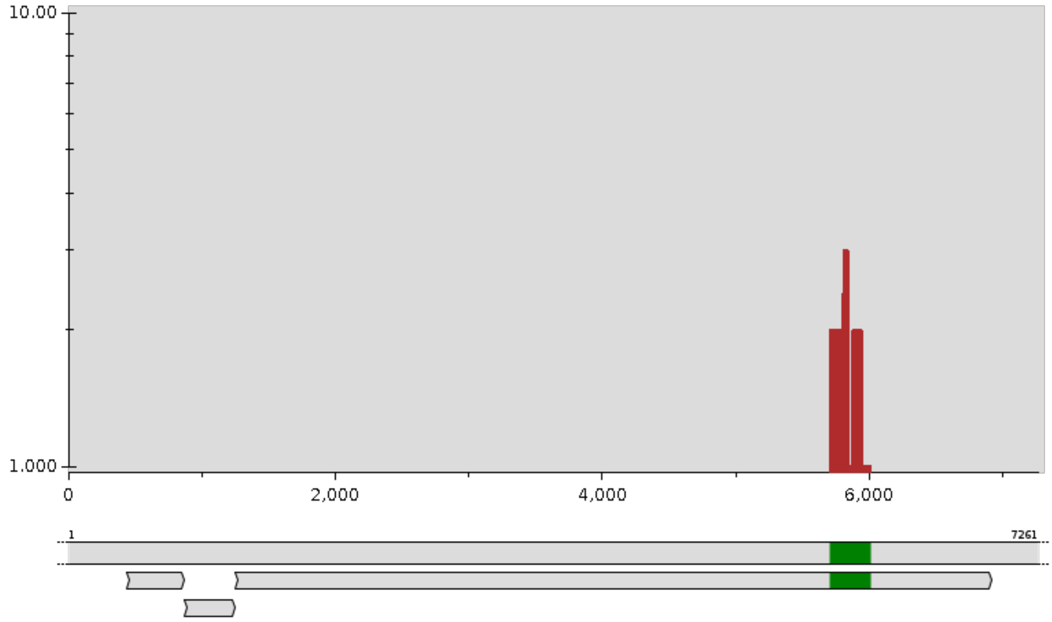

#### Assignment

| | |
|---|---|
| **Type** | Dioscorea bacilliform virus (Taxonomy ID: 52996) |
| **Reference Genome** | NC_009010.1 |
| **NT Identity (%)** | 52.9412 |
| **AA Identity (%)** | 47.0588 |
| **Number Of Stop Codons** | 2 |
| **Number Of CDS** | 3 |

#### Alignment

| | |
|---|---|
| **Alignment Score** | 0.0 (NT) + 313.0 (AA) = 313.0 |
| **Concordance (%)** | 24.6457 |

| **Alignment Method** | Global, seeded, nucleotide + amino acids (AGA) |
|---|---|

#### Genome Region

Sequence starts at position 5702 and ends at position 6016 relative to NC_009010.1 reference sequence.

#### Alignment Detailed Statistics

| | Begin | End | Coverage | Score | Concordance | Matches | Identities | I/D/M/F* | Stop Codons |
|---|---|---|---|---|---|---|---|---|---|
| **NT** | **5702** | **6016** | **4.3%** | **0** | **0%** | **306 (97.1%)** | **162 (51.4%)** | **0/9** | |
| Mutations: | 5702A>C, 5705G>C, 5706A>T, 5707G>T, 5709C>A, 5710G>T, 5712T>C, 5713C>T, 5715A>C, 5716A>T, 5717G>A, 5719A>G, 5721A>T, 5722_5724delATA, 5726A>T, 5729C>T, 5730A>C, 5734T>A, 5740T>C, 5742A>C, 5745G>A, 5746A>T, 5747G>C, 5749G>A, 5750G>A, 5751C>A, 5755C>T, 5756A>G, 5757T>G, 5761G>A, 5763G>A, 5764G>C, 5765C>A, 5769G>T, 5770G>A, 5771C>A, 5772G>C, 5775A>C, 5776C>A, 5777A>G, 5779T>G, 5780C>A, 5781A>T, 5782G>T, 5783T>G, 5784T>C, 5785G>T, 5787A>T, 5788T>A, 5789G>A, 5790G>A, 5793G>T, 5794G>T, 5795C>T, 5796G>T, 5800C>A, 5801T>C, 5802A>G, 5804C>T, 5805G>A, 5809G>T, 5810G>T, 5811A>T, 5816T>A, 5817T>G, 5820C>T, 5826G>A, 5827T>A, 5828T>A, 5829G>T, 5830G>T, 5832G>T, 5838A>C, 5841C>T, 5844A>G, 5847G>A, 5850G>A, 5853T>C, 5854G>A, 5856A>T, 5859A>T, 5860G>T, 5861C>T, 5862A>T, 5864T>A, 5865G>A, 5868C>T, 5871G>A, 5873G>A, 5874G>A, 5876A>T, 5877G>A, 5881G>A, 5883C>T, 5884A>G, 5885A>T, 5886C>T, 5887G>A, 5889C>T, 5892C>T, 5893C>A, 5894G>A, 5895A>T, 5896G>C, 5897G>C, 5899A>T, 5900C>T, 5902G>T, 5903A>C, 5905G>A, 5910T>C, 5914G>A, 5915C>T, 5916T>A, 5919A>C, 5922C>T, 5932A>G, 5934A>T, 5937G>A, 5940G>A, 5947G>A, 5948A>C, 5949A>T, 5952A>C, 5953G>A, 5954A>T, 5955A>T, 5958G>C, 5959G>C, 5967G>C, 5970G>A, 5976G>T, 5978G>A, 5981T>G, 5982A>T, 5985G>T, 5992_5994delATC, 5998C>T, 6000A>T, 6004_6006delCAC, 6009G>A, 6010C>A, 6011T>A, 6012T>A, 6013G>T, 6014T>G, 6015T>A | | | | | | | | |

##### CDS

| | Begin | End | Coverage | Score | Concordance | Matches | Identities | I/D/M/F* | Stop Codons |
|---|---|---|---|---|---|---|---|---|---|
| DBV_gp3 | 1487 | 1591 | 5.6% | 313 | 44.6% | 102 (97.1%) | 48 (45.7%) | 0/3/0/0 | 2 |
| Protein mutations: | R1487T (5705G>C 5706A>T), V1488L (5707G>T 5709C>A), G1489C (5710G>T 5712T>C), Q1490Y (5713C>T 5715A>C), S1491Y (5716A>T 5717G>A), K1492D (5719A>G 5721A>T), I1493del (5722_5724delATA), Y1494F (5726A>T), S1495F (5729C>T 5730A>C), F1497I (5734T>A), G1502K (5749G>A 5750G>A 5751C>A), H1504W (5755C>T 5756A>G 5757T>G), V1506I (5761G>A 5763G>A), A1507Q (5764G>C 5765C>A), M1508I (5769G>T), A1509N (5770G>A 5771C>A 5772G>C), Q1511R (5776C>A 5777A>G), S1512D (5779T>G 5780C>A 5781A>T), V1513C (5782G>T 5783T>G 5784T>C), E1514Y (5785G>T 5787A>T), W1515K (5788T>A 5789G>A 5790G>A), A1517F (5794G>T 5795C>T 5796G>T), L1519T (5800C>A 5801T>C 5802A>G), A1520V (5804C>T 5805G>A), G1522F (5809G>T 5810G>T 5811A>T), L1524Q (5816T>A 5817T>G), W1527* (5826G>A), L1528N (5827T>A 5828T>A 5829G>T), V1529F (5830G>T 5832G>T), A1537T (5854G>A 5856A>T), A1539F (5860G>T 5861C>T 5862A>T), V1540E (5864T>A 5865G>A), R1543K (5873G>A 5874G>A), K1544I (5876A>T 5877G>A), D1546N (5881G>A 5883C>T), N1547V (5884A>G 5885A>T 5886C>T), V1548I (5887G>A 5889C>T), R1550N (5893C>A 5894G>A 5895A>T), G1551P (5896G>C 5897G>C), T1552F (5899A>T 5900C>T), E1553S (5902G>T 5903A>C), E1554K (5905G>A), A1557I (5914G>A 5915C>T 5916T>A), I1563V (5932A>G 5934A>T), E1568T (5947G>A 5948A>C 5949A>T), E1570I (5953G>A 5954A>T 5955A>T), E1571D (5958G>C), E1572Q (5959G>C), L1574F (5967G>C), R1578K (5978G>A), I1579S (5981T>G 5982A>T), I1583del (5992_5994delATC), Q1585Y (5998C>T 6000A>T), H1587del (6004_6006delCAC), L1589K (6010C>A 6011T>A 6012T>A), V1590* (6013G>T 6014T>G 6015T>A) | | | | | | | | |
| Codon mutations: | AAG1486.CG (5702A>C), AGA1487ACT (5705G>C 5706A>T), GTC1488TTA (5707G>T 5709C>A), GGT1489TGC (5710G>T 5712T>C), CAA1490TAC (5713C>T 5715A>C), AGC1491TAC (5716A>T 5717G>A), AAA1492GAT (5719A>G 5721A>T), ATA1493del (5722_5724delATA), TAT1494TTT (5726A>T), TCA1495TTC (5729C>T 5730A>C), TTT1497ATT (5734T>A), TTA1499CTC (5740T>C 5742A>C), AAG1500AAA (5745G>A), AGC1501TCC (5746A>T 5747G>C), GGC1502AAA (5749G>A 5750G>A 5751C>A), CAT1504TGG (5755C>T 5756A>G 5757T>G), GTG1506ATA (5761G>A 5763G>A), GCA1507CAA (5764G>C 5765C>A), ATG1508ATT (5769G>T), GCG1509AAC (5770G>A 5771C>A 5772G>C), CCA1510CCC (5775A>C), CAA1511AGA (5776C>A 5777A>G), TCA1512GAT (5779T>G 5780C>A 5781A>T), GTT1513TGC (5782G>T 5783T>G 5784T>C), GAA1514TAT (5785G>T 5787A>T), TGG1515AAA (5788T>A 5789G>A 5790G>A), ACG1516ACT (5793G>T), GCG1517TTT (5794G>T 5795C>T 5796G>T), CTA1519ACG (5800C>A 5801T>C 5802A>G), GCG1520GTA (5804C>T 5805G>A), GGA1522TTT (5809G>T 5810G>T 5811A>T), CTT1524CAG (5816T>A 5817T>G), TAC1525TAT (5820C>T), TGG1527TGA (5826G>A), TTG1528AAT (5827T>A 5828T>A 5829G>T), GTG1529TTT (5830G>T 5832G>T), CCA1531CCC (5838A>C), TTC1532TTT (5841C>T), GGA1533GGG (5844A>G), CTG1534CTA (5847G>A), AAG1535AAA (5850G>A), AAT1536AAC (5853T>C), GCA1537ACT (5854G>A 5856A>T), CCA1538CCT (5859A>T), GCA1539TTT (5860G>T 5861C>T 5862A>T), GTG1540GAA (5864T>A 5865G>A), TTC1541TTT (5868C>T), CAG1542CAA (5871G>A), AGG1543AAA (5873G>A 5874G>A), AAG1544ATA (5876A>T 5877G>A), GAC1546AAT (5881G>A 5883C>T), AAC1547GTT (5884A>G 5885A>T 5886C>T), GTC1548ATT (5887G>A 5889C>T), TTC1549TTT (5892C>T), CGA1550AAT (5893C>A 5894G>A 5895A>T), GGT1551CCT (5896G>C 5897G>C), ACT1552TTT (5899A>T 5900C>T), GAA1553TCA (5902G>T 5903A>C), GAA1554AAA (5905G>A), TTT1555TTC (5910T>C), GCT1557ATA (5914G>A 5915C>T 5916T>A), GTA1558GTC (5919A>C), TAC1559TAT (5922C>T), ATA1563GTT (5932A>G 5934A>T), CTG1564CTA (5937G>A), GTG1565GTA (5940G>A), GAA1568ACT (5947G>A 5948A>C 5949A>T), ACA1569ACC (5952A>C), GAA1570ATT (5953G>A 5954A>T 5955A>T), GAG1571GAC (5958G>C), GAG1572CAG (5959G>C), TTG1574TTC (5967G>C), AAG1575AAA (5970G>A), CTG1577CTT (5976G>T), AGG1578AAG (5978G>A), ATA1579AGT (5981T>G 5982A>T), CTG1580CTT (5985G>T), ATC1583del (5992_5994delATC), CAA1585TAT (5998C>T 6000A>T), CAC1587del (6004_6006delCAC), GGG1588GGA (6009G>A), CTT1589AAA (6010C>A 6011T>A 6012T>A), GTT1590TGA (6013G>T 6014T>G 6015T>A) | | | | | | | | |

##### Proteins

| | Begin | End | Coverage | Score | Concordance | Matches | Identities | I/D/M/F* | Stop Codons |
|---|---|---|---|---|---|---|---|---|---|
| ORF3 protein (YP_001036293.1) | 1487 | 1591 | 5.6% | 313 | 44.6% | 102 (97.1%) | 48 (45.7%) | 0/3/0/0 | 2 |
| Protein mutations: | R1487T (5705G>C 5706A>T), V1488L (5707G>T 5709C>A), G1489C (5710G>T 5712T>C), Q1490Y (5713C>T 5715A>C), S1491Y (5716A>T 5717G>A), K1492D (5719A>G 5721A>T), I1493del (5722_5724delATA), Y1494F (5726A>T), S1495F (5729C>T 5730A>C), F1497I (5734T>A), G1502K (5749G>A 5750G>A 5751C>A), H1504W (5755C>T 5756A>G 5757T>G), V1506I (5761G>A 5763G>A), A1507Q (5764G>C 5765C>A), M1508I (5769G>T), A1509N (5770G>A 5771C>A 5772G>C), Q1511R (5776C>A 5777A>G), S1512D (5779T>G 5780C>A 5781A>T), V1513C (5782G>T 5783T>G 5784T>C), E1514Y (5785G>T 5787A>T), W1515K (5788T>A 5789G>A 5790G>A), A1517F (5794G>T 5795C>T 5796G>T), L1519T (5800C>A 5801T>C 5802A>G), A1520V (5804C>T 5805G>A), G1522F (5809G>T 5810G>T 5811A>T), L1524Q (5816T>A 5817T>G), W1527* (5826G>A), L1528N (5827T>A 5828T>A 5829G>T), V1529F (5830G>T 5832G>T), A1537T (5854G>A 5856A>T), A1539F (5860G>T 5861C>T 5862A>T), V1540E (5864T>A 5865G>A), R1543K (5873G>A 5874G>A), K1544I (5876A>T 5877G>A), D1546N (5881G>A 5883C>T), N1547V (5884A>G 5885A>T 5886C>T), V1548I (5887G>A 5889C>T), R1550N (5893C>A 5894G>A 5895A>T), G1551P (5896G>C 5897G>C), T1552F (5899A>T 5900C>T), E1553S (5902G>T 5903A>C), E1554K (5905G>A), A1557I (5914G>A 5915C>T 5916T>A), I1563V (5932A>G 5934A>T), E1568T (5947G>A 5948A>C 5949A>T), E1570I (5953G>A 5954A>T 5955A>T), E1571D (5958G>C), E1572Q (5959G>C), L1574F (5967G>C), R1578K (5978G>A), I1579S (5981T>G 5982A>T), I1583del (5992_5994delATC), Q1585Y (5998C>T 6000A>T), H1587del (6004_6006delCAC), L1589K (6010C>A 6011T>A 6012T>A), V1590* (6013G>T 6014T>G 6015T>A) | | | | | | | | |
| Codon mutations: | AAG1486.CG (5702A>C), AGA1487ACT (5705G>C 5706A>T), GTC1488TTA (5707G>T 5709C>A), GGT1489TGC (5710G>T 5712T>C), CAA1490TAC (5713C>T 5715A>C), AGC1491TAC (5716A>T 5717G>A), AAA1492GAT (5719A>G 5721A>T), ATA1493del (5722_5724delATA), TAT1494TTT (5726A>T), TCA1495TTC (5729C>T 5730A>C), TTT1497ATT (5734T>A), TTA1499CTC (5740T>C 5742A>C), AAG1500AAA (5745G>A), AGC1501TCC (5746A>T 5747G>C), GGC1502AAA (5749G>A 5750G>A 5751C>A), CAT1504TGG (5755C>T 5756A>G 5757T>G), GTG1506ATA (5761G>A 5763G>A), GCA1507CAA (5764G>C 5765C>A), ATG1508ATT (5769G>T), GCG1509AAC (5770G>A 5771C>A 5772G>C), CCA1510CCC (5775A>C), CAA1511AGA (5776C>A 5777A>G), TCA1512GAT (5779T>G 5780C>A 5781A>T), GTT1513TGC (5782G>T 5783T>G 5784T>C), GAA1514TAT (5785G>T 5787A>T), TGG1515AAA (5788T>A 5789G>A 5790G>A), ACG1516ACT (5793G>T), GCG1517TTT (5794G>T 5795C>T 5796G>T), CTA1519ACG (5800C>A 5801T>C 5802A>G), GCG1520GTA (5804C>T 5805G>A), GGA1522TTT (5809G>T 5810G>T 5811A>T), CTT1524CAG (5816T>A 5817T>G), TAC1525TAT (5820C>T), TGG1527TGA (5826G>A), TTG1528AAT (5827T>A 5828T>A 5829G>T), GTG1529TTT (5830G>T 5832G>T), CCA1531CCC (5838A>C), TTC1532TTT (5841C>T), GGA1533GGG (5844A>G), CTG1534CTA (5847G>A), AAG1535AAA (5850G>A), AAT1536AAC (5853T>C), GCA1537ACT (5854G>A 5856A>T), CCA1538CCT (5859A>T), GCA1539TTT (5860G>T 5861C>T 5862A>T), GTG1540GAA (5864T>A 5865G>A), TTC1541TTT (5868C>T), CAG1542CAA (5871G>A), AGG1543AAA (5873G>A 5874G>A), AAG1544ATA (5876A>T 5877G>A), GAC1546AAT (5881G>A 5883C>T), AAC1547GTT (5884A>G 5885A>T 5886C>T), GTC1548ATT (5887G>A 5889C>T), TTC1549TTT (5892C>T), CGA1550AAT (5893C>A 5894G>A 5895A>T), GGT1551CCT (5896G>C 5897G>C), ACT1552TTT (5899A>T 5900C>T), GAA1553TCA (5902G>T 5903A>C), GAA1554AAA (5905G>A), TTT1555TTC (5910T>C), GCT1557ATA (5914G>A 5915C>T 5916T>A), GTA1558GTC (5919A>C), TAC1559TAT (5922C>T), ATA1563GTT (5932A>G 5934A>T), CTG1564CTA (5937G>A), GTG1565GTA (5940G>A), GAA1568ACT (5947G>A 5948A>C 5949A>T), ACA1569ACC (5952A>C), GAA1570ATT (5953G>A 5954A>T 5955A>T), GAG1571GAC (5958G>C), GAG1572CAG (5959G>C), TTG1574TTC (5967G>C), AAG1575AAA (5970G>A), CTG1577CTT (5976G>T), AGG1578AAG (5978G>A), ATA1579AGT (5981T>G 5982A>T), CTG1580CTT (5985G>T), ATC1583del (5992_5994delATC), CAA1585TAT (5998C>T 6000A>T), CAC1587del (6004_6006delCAC), GGG1588GGA (6009G>A), CTT1589AAA (6010C>A 6011T>A 6012T>A), GTT1590TGA (6013G>T 6014T>G 6015T>A) | | | | | | | | |

*: Inserts / Deletes / Misaligned / Frameshifts

#### Analysis details

This analysis was performed with panviral2.64

### NGS Details (UN62): Rhizobium phage RHEph10

#### Assembly

| | |
|---|---|
| **Coverage Length** | 135 (1 contig(s)) |
| **Depth Of Coverage** | 2.9 |
| **Number Of Reads** | 3 |
| **Reads Per Million** | 0.07 rpm (after QC) |
| **Ambiguities** | 0 |
| **Assembly Method** | de novo + reference guided assembly |
| **Consensus Caller** | Bcf Tools |

#### Coverage Map

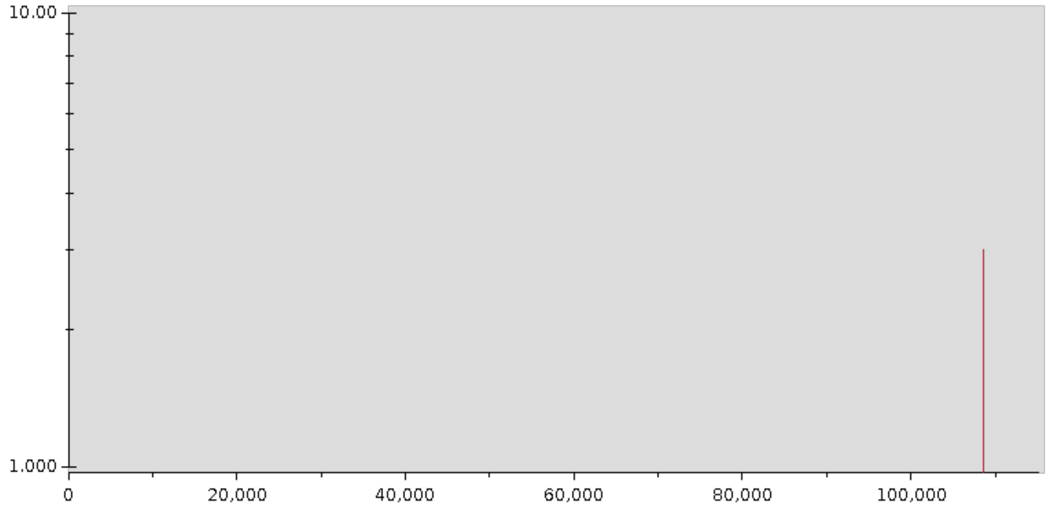

#### Assignment

| | |
|---|---|
| **Type** | Rhizobium phage RHEph10 (Taxonomy ID: 1220717) |
| **Reference Genome** | NC_034248.1 |
| **NT Identity (%)** | 80.0 |
| **AA Identity (%)** | 82.2222 |
| **Number Of Stop Codons** | 0 |
| **Number Of CDS** | 171 |

#### Alignment

| | |
|---|---|
| **Alignment Score** | 162.0 (NT) + 249.0 (AA) = 411.0 |
| **Concordance (%)** | 74.0541 |
| **Alignment Method** | Local, heuristic, nucleotide (BLASTN) |

#### Genome Region

Sequence starts at position 108586 and ends at position 108720 relative to NC_034248.1 reference sequence.

#### Alignment Detailed Statistics

| | Begin | End | Coverage | Score | Concordance | Matches | Identities | I/D/M/F* | Stop Codons |
|---|---|---|---|---|---|---|---|---|---|
| **NT** | **108586** | **108720** | **0.1%** | **162** | **60.0%** | **135 (100%)** | **108 (80.0%)** | **0/0** | |

Mutations: 108589T>C, 108602T>G, 108603C>G, 108604G>T, 108610C>G, 108613C>G, 108625T>C, 108628T>C, 108631T>C, 108632C>T, 108634G>C, 108635A>C, 108636A>G, 108637G>C, 108643A>G, 108647G>C, 108649C>G, 108650A>G, 108652G>C, 108656A>G, 108657C>G, 108660A>G, 108664C>T, 108670C>G, 108676C>G, 108703C>T, 108712A>C

*: Inserts / Deletes / Misaligned / Frameshifts

#### Analysis details

This analysis was performed with panviral2.64

### NGS Details (UN62): Melanoplus sanguinipes entomopoxvirus

#### Assembly

| | |
|---|---|
| **Coverage Length** | 135 (1 contig(s)) |
| **Depth Of Coverage** | 2.0 |
| **Number Of Reads** | 2 |
| **Reads Per Million** | 0.05 rpm (after QC) |
| **Ambiguities** | 0 |
| **Assembly Method** | de novo + reference guided assembly |
| **Consensus Caller** | Bcf Tools |

#### Coverage Map

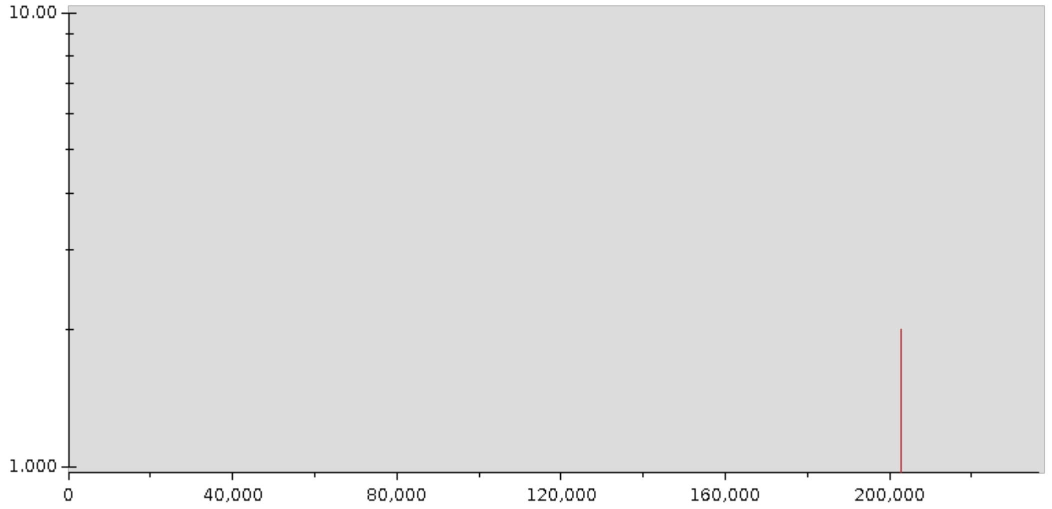

#### Assignment

| | |
|---|---|
| **Type** | Melanoplus sanguinipes entomopoxvirus (Taxonomy ID: 83191) |
| **Reference Genome** | NC_001993.1 |
| **NT Identity (%)** | 80.0 |
| **AA Identity (%)** | 80.0 |
| **Number Of Stop Codons** | 0 |
| **Number Of CDS** | 267 |

#### Alignment

| | |
|---|---|
| **Alignment Score** | 162.0 (NT) + 334.0 (AA) = 496.0 |
| **Concordance (%)** | 76.1905 |
| **Alignment Method** | Local, heuristic, nucleotide (BLASTN) |

#### Genome Region

Sequence starts at position 202763 and ends at position 202897 relative to NC_001993.1 reference sequence.

#### Alignment Detailed Statistics

| | Begin | End | Coverage | Score | Concordance | Matches | Identities | I/D/M/F* | Stop Codons |
|---|---|---|---|---|---|---|---|---|---|
| **NT** | **202763** | **202897** | **0.1%** | **162** | **60.0%** | **135 (100%)** | **108 (80.0%)** | **0/0** | |

Mutations: 202768T>C, 202771A>T, 202773G>T, 202774T>C, 202775C>T, 202777A>G, 202789A>G, 202790C>T, 202798T>C, 202807A>T, 202813T>A, 202817G>T, 202818G>C, 202826T>C, 202831T>C, 202832G>C, 202834T>C, 202835C>T, 202836T>G, 202852A>T, 202874T>A, 202876T>A, 202880T>C, 202885A>G, 202888A>G, 202891T>A, 202894A>G

*: Inserts / Deletes / Misaligned / Frameshifts

#### Analysis details

This analysis was performed with panviral2.64

#### NGS Details (UN62): Wadgaonvirus wv5004651

##### Assembly

| | |
|---|---|
| **Coverage Length** | 189 (1 contig(s)) |
| **Depth Of Coverage** | 1.4 |
| **Number Of Reads** | 2 |
| **Reads Per Million** | 0.05 rpm (after QC) |
| **Ambiguities** | 0 |
| **Assembly Method** | de novo + reference guided assembly |
| **Consensus Caller** | Bcf Tools |

##### Coverage Map

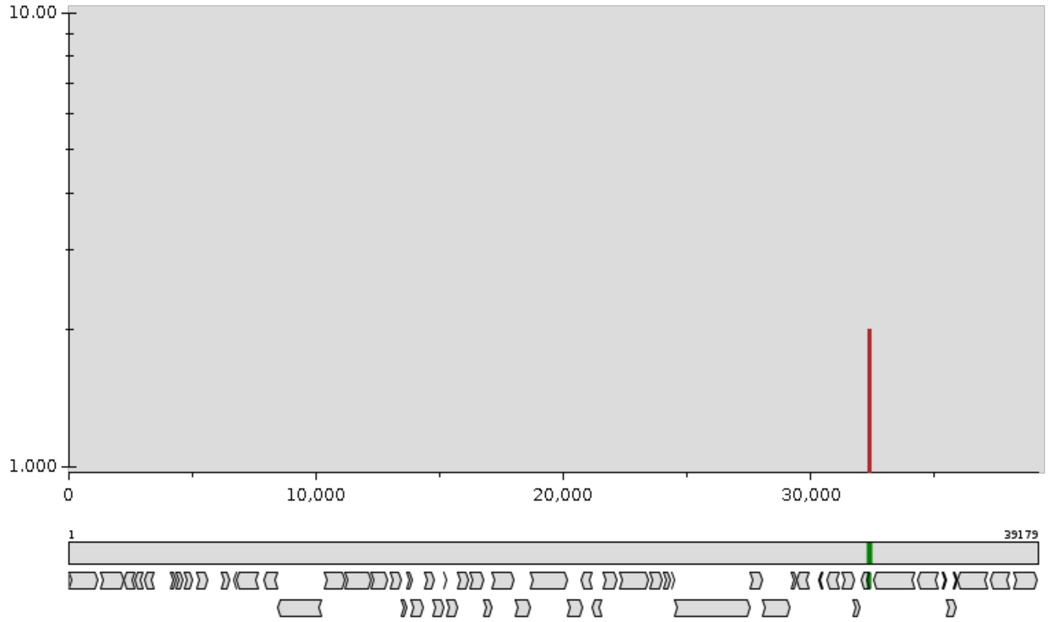

##### Assignment

| | |
|---|---|
| **Type** | Wadgaonvirus wv5004651 (Taxonomy ID: 2956672) |
| **Reference Genome** | NC_049342.1 |
| **NT Identity (%)** | 100.0 |
| **AA Identity (%)** | 100.0 |
| **Number Of Stop Codons** | 0 |
| **Number Of CDS** | 57 |

##### Alignment

| | |
|---|---|
| **Alignment Score** | 378.0 (NT) + 256.0 (AA) = 634.0 |
| **Concordance (%)** | 100.0 |

| Alignment Method | Local, heuristic, nucleotide (BLASTN) |
|---|---|

#### Genome Region

Sequence starts at position 32293 and ends at position 32481 relative to NC_049342.1 reference sequence.

#### Alignment Detailed Statistics

| | Begin | End | Coverage | Score | Concordance | Matches | Identities | I/D/M/F * | Stop Codons |
|---|---|---|---|---|---|---|---|---|---|
| **NT** | **32293** | **32481** | **0.5%** | **378** | **100%** | **189 (100%)** | **189 (100%)** | **0/0** | |
| Mutations: | *none* | | | | | | | | |
| **CDS** | | | | | | | | | |
| HYO67_gp49 | 1 | 40 | 35.1% | 256 | 100% | 40 (100%) | 40 (100%) | 0/0/0/0 | 0 |
| **Proteins** | | | | | | | | | |
| outer membrane protein assembly factor BamE (YP_009949248.1) | 1 | 40 | 35.1% | 256 | 100% | 40 (100%) | 40 (100%) | 0/0/0/0 | 0 |
| Protein mutations: | *none* | | | | | | | | |
| Codon mutations: | *none* | | | | | | | | |

*: Inserts / Deletes / Misaligned / Frameshifts

#### Analysis details

This analysis was performed with panviral2.64

### NGS Details (UN62): Enterobacteria phage P7

#### Assembly

| | |
|---|---|
| **Coverage Length** | 172 (1 contig(s)) |
| **Depth Of Coverage** | 1.6 |
| **Number Of Reads** | 2 |
| **Reads Per Million** | 0.05 rpm (after QC) |
| **Ambiguities** | 0 |
| **Assembly Method** | de novo + reference guided assembly |
| **Consensus Caller** | Bcf Tools |

#### Coverage Map

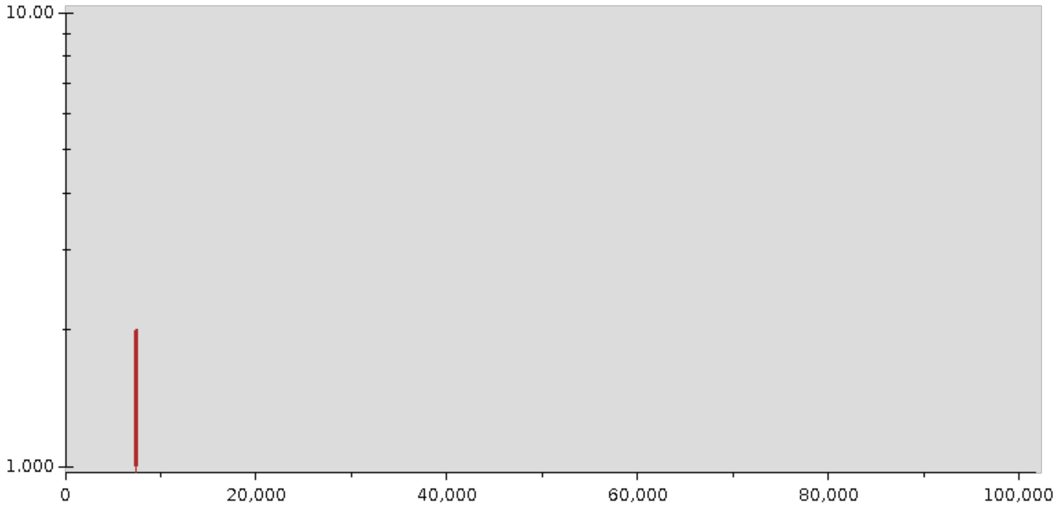

#### Assignment

| | |
|---|---|
| **Type** | Enterobacteria phage P7 (Taxonomy ID: 10682) |
| **Reference Genome** | NC_050152.1 |
| **NT Identity (%)** | 100.0 |
| **AA Identity (%)** | 100.0 |
| **Number Of Stop Codons** | 1 |
| **Number Of CDS** | 113 |

#### Alignment

| | |
|---|---|
| **Alignment Score** | 344.0 (NT) + 119.0 (AA) = 463.0 |
| **Concordance (%)** | 100.0 |
| **Alignment Method** | Local, heuristic, nucleotide (BLASTN) |

#### Genome Region

Sequence starts at position 7371 and ends at position 7542 relative to NC_050152.1 reference sequence.

#### Alignment Detailed Statistics

| | Begin | End | Coverage | Score | Concordance | Matches | Identities | I/D/M/F* | Stop Codons |
|---|---|---|---|---|---|---|---|---|---|
| **NT** | **7371** | **7542** | **0.2%** | **344** | **100%** | **172 (100%)** | **172 (100%)** | **0/0** | |

Mutations: *none*

*: Inserts / Deletes / Misaligned / Frameshifts

#### Analysis details

This analysis was performed with panviral2.64

### NGS Details (UN62): Orthomarburgvirus marburgense

#### Assembly

| | |
|---|---|
| **Coverage Length** | 270 (2 contig(s)) |
| **Depth Of Coverage** | 1.0 |
| **Number Of Reads** | 2 |
| **Reads Per Million** | 0.05 rpm (after QC) |
| **Ambiguities** | 0 |
| **Assembly Method** | de novo + reference guided assembly |
| **Consensus Caller** | Bcf Tools |

#### Coverage Map

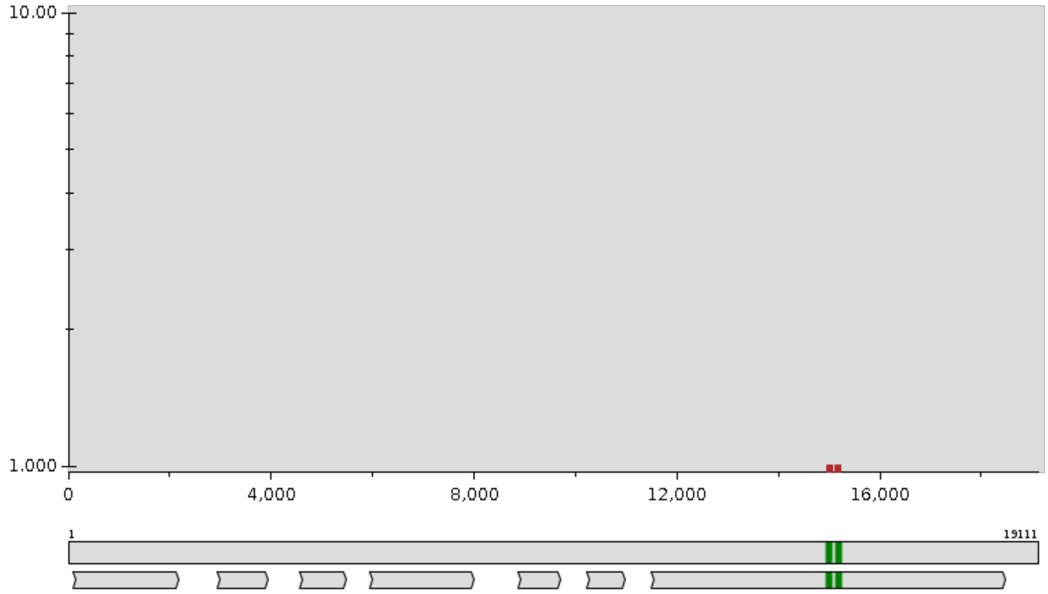

#### Assignment

| | |
|---|---|
| **Type** | Orthomarburgvirus marburgense (Taxonomy ID: 11269) |
| **Reference Genome** | NC_001608.3 |
| **NT Identity (%)** | 70.3704 |
| **AA Identity (%)** | 98.8889 |
| **Number Of Stop Codons** | 0 |
| **Number Of CDS** | 7 |

#### Alignment

| | |
|---|---|
| **Alignment Score** | 220.0 (NT) + 601.0 (AA) = 821.0 |
| **Concordance (%)** | 71.9544 |

| Alignment Method | Global, seeded, nucleotide + amino acids (AGA) |
|---|---|

#### Genome Region

Sequence starts at position 14942 and ends at position 15245 relative to NC_001608.3 reference sequence.

#### Alignment Detailed Statistics

| | Begin | End | Coverage | Score | Concordance | Matches | Identities | I/D/M/F* | Stop Codons |
|---|---|---|---|---|---|---|---|---|---|
| **NT** | **14942** | **15245** | **1.4%** | **220** | **40.7%** | **270 (100%)** | **190 (70.4%)** | **0/0** | |
| Mutations: | 14945A>G, 14948A>G, 14957A>G, 14958T>A, 14959C>G, 14960T>C, 14963A>C, 14966C>T, 14969A>C, 14972A>G, 14973T>A, 14974C>G, 14975A>C, 14978T>C, 14981A>C, 14984C>T, 14985T>C, 14988T>C, 14993G>C, 14996T>C, 14999C>T, 15008A>G, 15012C>A, 15014A>G, 15017T>G, 15020A>G, 15023A>C, 15029T>C, 15035A>C, 15039T>C, 15041A>G, 15047T>C, 15050C>G, 15053T>C, 15056T>C, 15059T>C, 15062T>G, 15066C>A, 15071A>G, 15074T>G, 15113G>C, 15116A>C, 15119A>C, 15122T>C, 15123C>A, 15125A>G, 15128A>C, 15131G>C, 15134T>C, 15137T>C, 15143A>C, 15144C>A, 15146G>A, 15149A>C, 15152A>G, 15155T>C, 15164T>C, 15167T>C, 15173C>T, 15174T>C, 15176A>G, 15182A>G, 15188C>T, 15191A>T, 15192T>A, 15193C>G, 15194A>C, 15197A>C, 15200A>C, 15203T>G, 15206A>G, 15209A>G, 15212T>C, 15215T>C, 15224T>G, 15227T>C, 15230A>G, 15231T>C, 15234T>C, 15236A>G | | | | | | | | |
| **CDS** | | | | | | | | | |
| L | 1155 | 1255 | 3.9% | 601 | 97.9% | 90 (100%) | 89 (98.9%) | 0/0/0/0 | 0 |
| **Proteins** | | | | | | | | | |
| RNA-dependent RNA polymerase (YP_001531159.1) | 1155 | 1255 | 3.9% | 601 | 97.9% | 90 (100%) | 89 (98.9%) | 0/0/0/0 | 0 |
| Protein mutations: | *none* | | | | | | | | |
| Codon mutations: | GAA1155GAG (14945A>G), CAA1156CAG (14948A>G), CTA1159CTG (14957A>G), TCT1160AGC (14958T>A 14959C>G 14960T>C), TCA1161TCC (14963A>C), GAC1162GAT (14966C>T), TCA1163TCC (14969A>C), AAA1164AAG (14972A>G), TCA1165AGC (14973T>A 14974C>G 14975A>C), ACT1166ACC (14978T>C), ATA1167ATC (14981A>C), AAC1168AAT (14984C>T), TTG1169CTG (14985T>C), TTG1170CTG (14988T>C), CCG1171CCC (14993G>C), TAT1172TAC (14996T>C), GAC1173GAT (14999C>T), GAA1176GAG (15008A>G), CGA1178AGG (15012C>A 15014A>G), CTT1179CTG (15017T>G), GAA1180GAG (15020A>G), GGA1181GGC (15023A>C), AAT1183AAC (15029T>C), ACA1185ACC (15035A>C), TTA1187CTG (15039T>C 15041A>G), TAT1189TAC (15047T>C), GTC1190GTG (15050C>G), AGT1191AGC (15053T>C), TGT1192TGC (15056T>C), GCT1193GCC (15059T>C), CTT1194CTG (15062T>G), CGG1196AGG (15066C>A), AAA1197AAG (15071A>G), GTT1198GTG (15074T>G), ACG1211ACC (15113G>C), ATA1212ATC (15116A>C), GGA1213GGC (15119A>C), AAT1214AAC (15122T>C), CGA1215AGG (15123C>A 15125A>G), GCA1216GCC (15128A>C), CCG1217CCC (15131G>C), TAT1218TAC (15134T>C), ATT1219ATC (15137T>C), TCA1221TCC (15143A>C), CGG1222AGA (15144C>A 15146G>A), ACA1223ACC (15149A>C), GAA1224GAG (15152A>G), GAT1225GAC (15155T>C), GGT1228GGC (15164T>C), TAT1229TAC (15167T>C), CCC1231CCT (15173C>T), TTA1232CTG (15174T>C 15176A>G), GTA1234GTG (15182A>G), TGC1236TGT (15188C>T), CCA1237CCT (15191A>T), TCA1238AGC (15192T>A 15193C>G 15194A>C), GCA1239GCC (15197A>C), GCA1240GCC (15200A>C), CTT1241CTG (15203T>G), AAA1242AAG (15206A>G), GAA1243GAG (15209A>G), GCT1244GCC (15212T>C), ATT1245ATC (15215T>C), GTT1248GTG (15224T>G), TCT1249TCC (15227T>C), AGA1250AGG (15230A>G), TTG1251CTG (15231T>C), TTA1252CTG (15234T>C 15236A>G) | | | | | | | | |

*: Inserts / Deletes / Misaligned / Frameshifts

#### Analysis details

This analysis was performed with panviral2.64

### NGS Details (UN62): Sclerotinia sclerotiorum deltaflexivirus 2

#### Assembly

| | |
|---|---|
| **Coverage Length** | 216 (1 contig(s)) |
| **Depth Of Coverage** | 1.2 |
| **Number Of Reads** | 2 |
| **Reads Per Million** | 0.05 rpm (after QC) |
| **Ambiguities** | 0 |
| **Assembly Method** | de novo + reference guided assembly |
| **Consensus Caller** | Bcf Tools |

#### Coverage Map

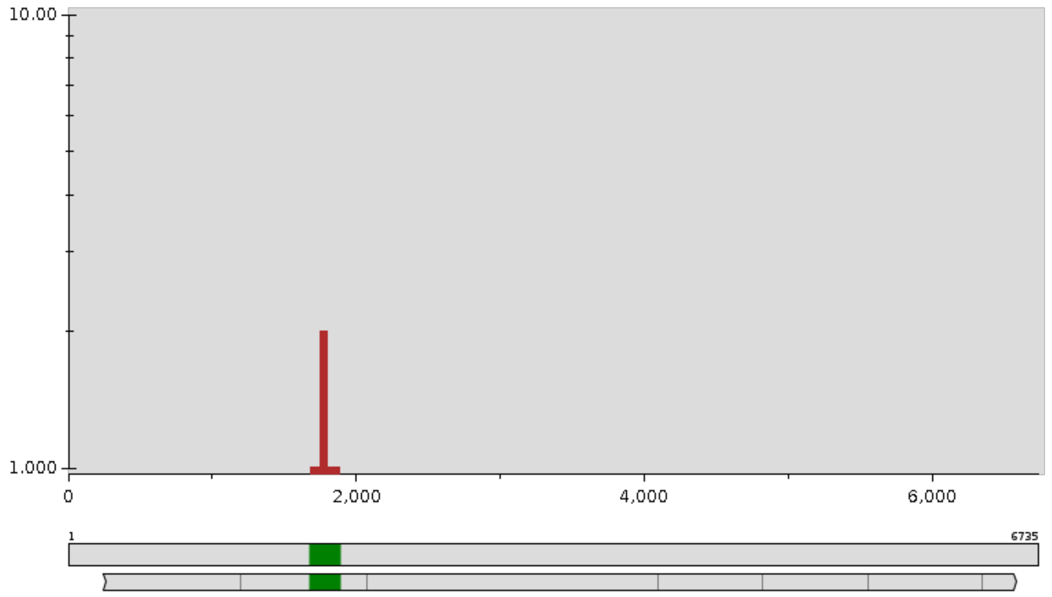

#### Assignment

| | |
|---|---|
| **Type** | Sclerotinia sclerotiorum deltaflexivirus 2 (Taxonomy ID: 2219092) |
| **Reference Genome** | NC_040649.1 |
| **NT Identity (%)** | 60.7306 |
| **AA Identity (%)** | 65.7534 |
| **Number Of Stop Codons** | 0 |
| **Number Of CDS** | 1 |

#### Alignment

| | |
|---|---|
| **Alignment Score** | 88.0 (NT) + 342.0 (AA) = 430.0 |
| **Concordance (%)** | 44.3756 |

| Alignment Method | Global, seeded, nucleotide + amino acids (AGA) |
|---|---|

#### Genome Region

Sequence starts at position 1675 and ends at position 1890 relative to NC_040649.1 reference sequence.

#### Alignment Detailed Statistics

| | Begin | End | Coverage | Score | Concordance | Matches | Identities | I/D/M/F* | Stop Codons |
|---|---|---|---|---|---|---|---|---|---|
| **NT** | **1675** | **1890** | **3.2%** | **88** | **20.4%** | **216 (98.6%)** | **133 (60.7%)** | **3/0** | |
| Mutations: | 1675C>T, 1678A>G, 1683T>C, 1686T>G, 1687T>G, 1688G>T, 1689C>T, 1692G>C, 1693A>G, 1695A>C, 1698A>C, 1704G>A, 1710C>T, 1712G>T, 1713G>C, 1716C>T, 1717C>T, 1719A>G, 1722T>G, 1725C>T, 1731C>T, 1735T>G, 1736C>A, 1737C>G, 1738C>T, 1743T>C, 1744G>A, 1746C>T, 1749C>T, 1750C>G, 1751G>A, 1752A>G, 1753T>A, 1754A>T, 1755C>T, 1756T>C, 1758C>T, 1759A>G, 1761A>T, 1762A>G, 1763A>G, 1770G>A, 1772T>C, 1773_1774insCTG, 1774A>G, 1776A>G, 1780T>A, 1782T>A, 1785T>A, 1788G>A, 1792G>A, 1797G>C, 1800C>G, 1804A>G, 1805G>C, 1806G>T, 1809C>T, 1810G>T, 1811A>T, 1812A>T, 1818T>G, 1819C>T, 1821C>G, 1827T>C, 1828C>A, 1831T>G, 1832C>A, 1833T>G, 1837C>G, 1839C>T, 1840A>C, 1841C>G, 1842T>G, 1845C>T, 1848C>A, 1849A>C, 1850A>G, 1854G>C, 1866C>A, 1867A>C, 1868A>G, 1881T>C, 1884C>T, 1887C>G | | | | | | | | |
| **CDS** | | | | | | | | | |
| EXK03_gp1 | 477 | 548 | 3.4% | 342 | 63.7% | 72 (98.6%) | 48 (65.8%) | 1/0/0/0 | 0 |
| Protein mutations: | I478V (1678A>G), C481V (1687T>G 1688G>T 1689C>T), I483V (1693A>G 1695A>C), W489F (1712G>T 1713G>C), S497E (1735T>G 1736C>A 1737C>G), D500N (1744G>A 1746C>T), R502E (1750C>G 1751G>A 1752A>G), Y503I (1753T>A 1754A>T 1755C>T), C504R (1756T>C 1758C>T), K505D (1759A>G 1761A>T), N506G (1762A>G 1763A>G), I509T (1772T>C), I509_I510insL (1773_1774insCTG), I510V (1774A>G 1776A>G), F512I (1780T>A 1782T>A), D516N (1792G>A), K517N (1797G>C), R520A (1804A>G 1805G>C 1806G>T), E522F (1810G>T 1811A>T 1812A>T), S529E (1831T>G 1832C>A 1833T>G), L531V (1837C>G 1839C>T), T532R (1840A>C 1841C>G 1842T>G), N534K (1848C>A), K535R (1849A>C 1850A>G), N541R (1867A>C 1868A>G) | | | | | | | | |
| Codon mutations: | CTG477TTG (1675C>T), ATT478GTT (1678A>G), GCT479GCC (1683T>C), ACT480ACG (1686T>G), TGC481GTT (1687T>G 1688G>T 1689C>T), GTG482GTC (1692G>C), ATA483GTC (1693A>G 1695A>C), CCA484CCC (1698A>C), GAG486GAA (1704G>A), CTC488CTT (1710C>T), TGG489TTC (1712G>T 1713G>C), GAC490GAT (1716C>T), CTA491TTG (1717C>T 1719A>G), CCT492CCG (1722T>G), GCC493GCT (1725C>T), ACC495ACT (1731C>T), TCC497GAG (1735T>G 1736C>A 1737C>G), CTG498TTG (1738C>T), TAT499TAC (1743T>C), GAC500AAT (1744G>A 1746C>T), TTC501TTT (1749C>T), CGA502GAG (1750C>G 1751G>A 1752A>G), TAC503ATT (1753T>A 1754A>T 1755C>T), TGC504CGT (1756T>C 1758C>T), AAA505GAT (1759A>G 1761A>T), AAC506GGC (1762A>G 1763A>G), AAG508AAA (1770G>A), ATC509ACC (1772T>C), ATC509_ATA510insCTG (1773_1774insCTG), ATA510GTG (1774A>G 1776A>G), TTT512ATA (1780T>A 1782T>A), CCT513CCA (1785T>A), GAG514GAA (1788G>A), GAC516AAC (1792G>A), AAG517AAC (1797G>C), GGC518GGG (1800C>G), AGG520GCT (1804A>G 1805G>C 1806G>T), TAC521TAT (1809C>T), GAA522TTT (1810G>T 1811A>T 1812A>T), CCT524CCG (1818T>G), CTC525TTG (1819C>T 1821C>G), GCT527GCC (1827T>C), CGG528AGG (1828C>A), TCT529GAG (1831T>G 1832C>A 1833T>G), CTC531GTT (1837C>G 1839C>T), ACT532CGG (1840A>C 1841C>G 1842T>G), GCC533GCT (1845C>T), AAC534AAA (1848C>A), AAG535CGG (1849A>C 1850A>G), CTG536CTC (1854G>C), GGC540GGA (1866C>A), AAT541CGT (1867A>C 1868A>G), CAT545CAC (1881T>C), GTC546GTT (1884C>T), TCC547TCG (1887C>G) | | | | | | | | |
| **Proteins** | | | | | | | | | |
| polyprotein (YP_009552771.1) | 477 | 548 | 3.4% | 342 | 63.7% | 72 (98.6%) | 48 (65.8%) | 1/0/0/0 | 0 |
| Protein mutations: | I478V (1678A>G), C481V (1687T>G 1688G>T 1689C>T), I483V (1693A>G 1695A>C), W489F (1712G>T 1713G>C), S497E (1735T>G 1736C>A 1737C>G), D500N (1744G>A 1746C>T), R502E (1750C>G 1751G>A 1752A>G), Y503I (1753T>A 1754A>T 1755C>T), C504R (1756T>C 1758C>T), K505D (1759A>G 1761A>T), N506G (1762A>G 1763A>G), I509T (1772T>C), I509_I510insL (1773_1774insCTG), I510V (1774A>G 1776A>G), F512I (1780T>A 1782T>A), D516N (1792G>A), K517N (1797G>C), R520A (1804A>G 1805G>C 1806G>T), E522F (1810G>T 1811A>T 1812A>T), S529E (1831T>G 1832C>A 1833T>G), L531V (1837C>G 1839C>T), T532R (1840A>C 1841C>G 1842T>G), N534K (1848C>A), K535R (1849A>C 1850A>G), N541R (1867A>C 1868A>G) | | | | | | | | |
| Codon mutations: | CTG477TTG (1675C>T), ATT478GTT (1678A>G), GCT479GCC (1683T>C), ACT480ACG (1686T>G), TGC481GTT (1687T>G 1688G>T 1689C>T), GTG482GTC (1692G>C), ATA483GTC (1693A>G 1695A>C), CCA484CCC (1698A>C), GAG486GAA (1704G>A), CTC488CTT (1710C>T), TGG489TTC (1712G>T 1713G>C), GAC490GAT (1716C>T), CTA491TTG (1717C>T 1719A>G), CCT492CCG (1722T>G), GCC493GCT (1725C>T), ACC495ACT (1731C>T), TCC497GAG (1735T>G 1736C>A 1737C>G), CTG498TTG (1738C>T), TAT499TAC (1743T>C), GAC500AAT (1744G>A 1746C>T), TTC501TTT (1749C>T), CGA502GAG (1750C>G 1751G>A 1752A>G), TAC503ATT (1753T>A 1754A>T 1755C>T), TGC504CGT (1756T>C 1758C>T), AAA505GAT (1759A>G 1761A>T), AAC506GGC (1762A>G 1763A>G), AAG508AAA (1770G>A), ATC509ACC (1772T>C), ATC509_ATA510insCTG (1773_1774insCTG), ATA510GTG (1774A>G 1776A>G), TTT512ATA (1780T>A 1782T>A), CCT513CCA (1785T>A), GAG514GAA (1788G>A), GAC516AAC (1792G>A), AAG517AAC (1797G>C), GGC518GGG (1800C>G), AGG520GCT (1804A>G 1805G>C 1806G>T), TAC521TAT (1809C>T), GAA522TTT (1810G>T 1811A>T 1812A>T), CCT524CCG (1818T>G), CTC525TTG (1819C>T 1821C>G), GCT527GCC (1827T>C), CGG528AGG (1828C>A), TCT529GAG (1831T>G 1832C>A 1833T>G), CTC531GTT (1837C>G 1839C>T), ACT532CGG (1840A>C 1841C>G 1842T>G), GCC533GCT (1845C>T), AAC534AAA (1848C>A), AAG535CGG (1849A>C 1850A>G), CTG536CTC (1854G>C), GGC540GGA (1866C>A), AAT541CGT (1867A>C 1868A>G), CAT545CAC (1881T>C), GTC546GTT (1884C>T), TCC547TCG (1887C>G) | | | | | | | | |
| viral methyltransferase (YP_009553706.1) | 159 | 230 | 24.7% | 342 | 63.7% | 72 (98.6%) | 48 (65.8%) | 1/0/0/0 | 0 |
| Protein mutations: | I160V (1678A>G), C163V (1687T>G 1688G>T 1689C>T), I165V (1693A>G 1695A>C), W171F (1712G>T 1713G>C), S179E (1735T>G 1736C>A 1737C>G), D182N (1744G>A 1746C>T), R184E (1750C>G 1751G>A 1752A>G), Y185I (1753T>A 1754A>T 1755C>T), C186R (1756T>C 1758C>T), K187D (1759A>G 1761A>T), N188G (1762A>G 1763A>G), I191T (1772T>C), I191_I192insL (1773_1774insCTG), I192V (1774A>G 1776A>G), F194I (1780T>A 1782T>A), D198N (1792G>A), K199N (1797G>C), R202A (1804A>G 1805G>C 1806G>T), E204F (1810G>T 1811A>T 1812A>T), S211E (1831T>G 1832C>A 1833T>G), L213V (1837C>G 1839C>T), T214R (1840A>C 1841C>G 1842T>G), N216K (1848C>A), K217R (1849A>C 1850A>G), N223R (1867A>C 1868A>G) | | | | | | | | |
| Codon mutations: | CTG159TTG (1675C>T), ATT160GTT (1678A>G), GCT161GCC (1683T>C), ACT162ACG (1686T>G), TGC163GTT (1687T>G 1688G>T 1689C>T), GTG164GTC (1692G>C), ATA165GTC (1693A>G 1695A>C), CCA166CCC (1698A>C), GAG168GAA (1704G>A), CTC170CTT (1710C>T), TGG171TTC (1712G>T 1713G>C), GAC172GAT (1716C>T), CTA173TTG (1717C>T 1719A>G), CCT174CCG (1722T>G), GCC175GCT (1725C>T), ACC177ACT (1731C>T), TCC179GAG (1735T>G 1736C>A 1737C>G), CTG180TTG (1738C>T), TAT181TAC (1743T>C), GAC182AAT (1744G>A 1746C>T), TTC183TTT (1749C>T), CGA184GAG (1750C>G 1751G>A 1752A>G), TAC185ATT (1753T>A 1754A>T 1755C>T), TGC186CGT (1756T>C 1758C>T), AAA187GAT (1759A>G 1761A>T), AAC188GGC (1762A>G 1763A>G), AAG190AAA (1770G>A), ATC191ACC (1772T>C), ATC191_ATA192insCTG (1773_1774insCTG), ATA192GTG (1774A>G 1776A>G), TTT194ATA (1780T>A 1782T>A), CCT195CCA (1785T>A), GAG196GAA (1788G>A), GAC198AAC (1792G>A), AAG199AAC (1797G>C), GGC200GGG (1800C>G), AGG202GCT (1804A>G 1805G>C 1806G>T), TAC203TAT (1809C>T), GAA204TTT (1810G>T 1811A>T 1812A>T), CCT206CCG (1818T>G), CTC207TTG (1819C>T 1821C>G), GCT209GCC (1827T>C), CGG210AGG (1828C>A), TCT211GAG (1831T>G 1832C>A 1833T>G), CTC213GTT (1837C>G 1839C>T), ACT214CGG (1840A>C 1841C>G 1842T>G), GCC215GCT (1845C>T), AAC216AAA (1848C>A), AAG217CGG (1849A>C 1850A>G), CTG218CTC (1854G>C), GGC222GGA (1866C>A), AAT223CGT (1867A>C 1868A>G), CAT227CAC (1881T>C), GTC228GTT (1884C>T), TCC229TCG (1887C>G) | | | | | | | | |

*: Inserts / Deletes / Misaligned / Frameshifts

#### Analysis details

This analysis was performed with panviral2.64

### NGS Details (UN62): Friend murine leukemia virus

#### Assembly

| | |
|---|---|
| **Coverage Length** | 135 (1 contig(s)) |
| **Depth Of Coverage** | 1.0 |
| **Number Of Reads** | 1 |
| **Reads Per Million** | 0.02 rpm (after QC) |
| **Ambiguities** | 0 |
| **Assembly Method** | de novo + reference guided assembly |
| **Consensus Caller** | Bcf Tools |

#### Coverage Map

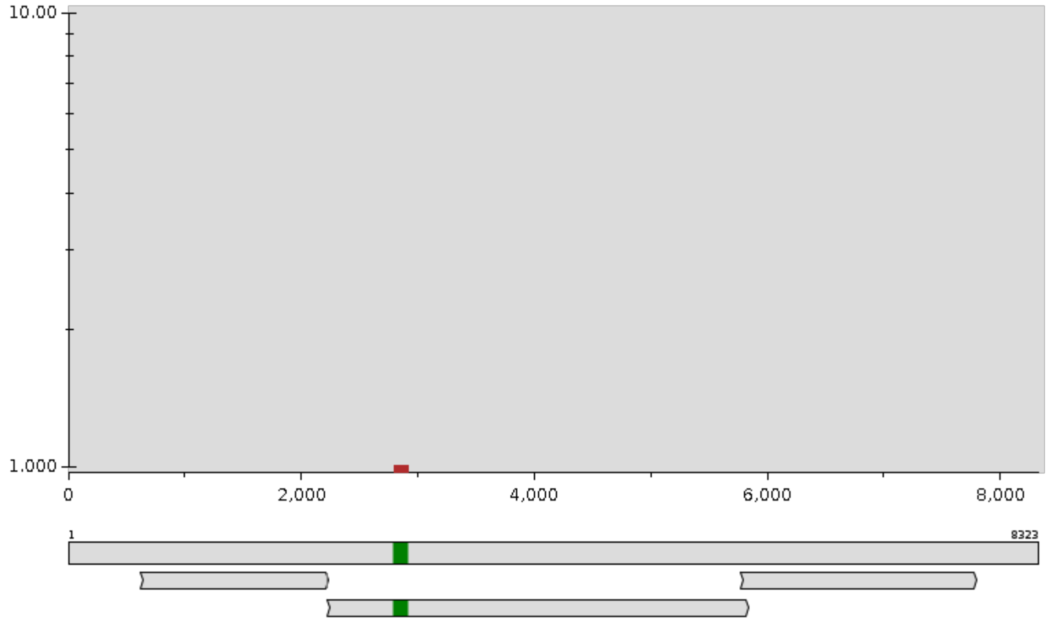

#### Assignment

| | |
|---|---|
| **Type** | Friend murine leukemia virus (Taxonomy ID: 11795) |
| **Reference Genome** | NC_001362.1 |
| **NT Identity (%)** | 77.037 |
| **AA Identity (%)** | 97.7778 |
| **Number Of Stop Codons** | 0 |
| **Number Of CDS** | 3 |

#### Alignment

| | |
|---|---|
| **Alignment Score** | 146.0 (NT) + 329.0 (AA) = 475.0 |
| **Concordance (%)** | 79.2988 |

| Alignment Method | Global, seeded, nucleotide + amino acids (AGA) |
|---|---|

#### Genome Region

Sequence starts at position 2789 and ends at position 2923 relative to NC_001362.1 reference sequence.

#### Alignment Detailed Statistics

| | Begin | End | Coverage | Score | Concordance | Matches | Identities | I/D/M/F* | Stop Codons |
|---|---|---|---|---|---|---|---|---|---|
| **NT** | **2789** | **2923** | **1.6%** | **146** | **54.1%** | **135 (100%)** | **104 (77.0%)** | **0/0** | |
| Mutations: | 2790C>G, 2794T>A, 2795C>G, 2796A>C, 2799A>G, 2805C>G, 2806A>C, 2808A>C, 2814G>C, 2817C>T, 2820G>A, 2823C>G, 2826C>T, 2829A>T, 2833A>C, 2835A>C, 2850A>C, 2859A>G, 2862C>G, 2869T>A, 2870C>G, 2874C>G, 2883G>C, 2886C>G, 2892A>G, 2895C>G, 2898T>G, 2901G>A, 2910G>C, 2913T>C, 2916T>C | | | | | | | | |
| **CDS** | | | | | | | | | |
| pol | 191 | 235 | 3.7% | 329 | 94.3% | 45 (100%) | 44 (97.8%) | 0/0/0/0 | 0 |
| **Proteins** | | | | | | | | | |
| RNA-dependent DNA polymerase (NP_040333.1) | 191 | 235 | 3.7% | 329 | 94.3% | 45 (100%) | 44 (97.8%) | 0/0/0/0 | 0 |
| Protein mutations: | *none* | | | | | | | | |
| Codon mutations: | CCC190.CG (2790C>G), TCA192AGC (2794T>A 2795C>G 2796A>C), CAA193CAG (2799A>G), GCC195GCG (2805C>G), AGA196CGC (2806A>C 2808A>C), GGG198GGC (2814G>C), ATC199ATT (2817C>T), AAG200AAA (2820G>A), CCC201CCG (2823C>G), CAC202CAT (2826C>T), ATA203ATT (2829A>T), AGA205CGC (2833A>C 2835A>C), GGA210GGC (2850A>C), GTA213GTG (2859A>G), CCC214CCG (2862C>G), TCC217AGC (2869T>A 2870C>G), CCC218CCG (2874C>G), ACG221ACC (2883G>C), CCC222CCG (2886C>G), CTA224CTG (2892A>G), CCC225CCG (2895C>G), GTT226GTG (2898T>G), AAG227AAA (2901G>A), GGG230GGC (2910G>C), ACT231ACC (2913T>C), AAT232AAC (2916T>C) | | | | | | | | |

*: Inserts / Deletes / Misaligned / Frameshifts

#### Analysis details

This analysis was performed with panviral2.64

### NGS Details (UN62): Moloney murine leukemia virus

#### Assembly

| | |
|---|---|
| **Coverage Length** | 135 (1 contig(s)) |
| **Depth Of Coverage** | 1.0 |
| **Number Of Reads** | 1 |
| **Reads Per Million** | 0.02 rpm (after QC) |
| **Ambiguities** | 0 |
| **Assembly Method** | de novo + reference guided assembly |
| **Consensus Caller** | Bcf Tools |

#### Coverage Map

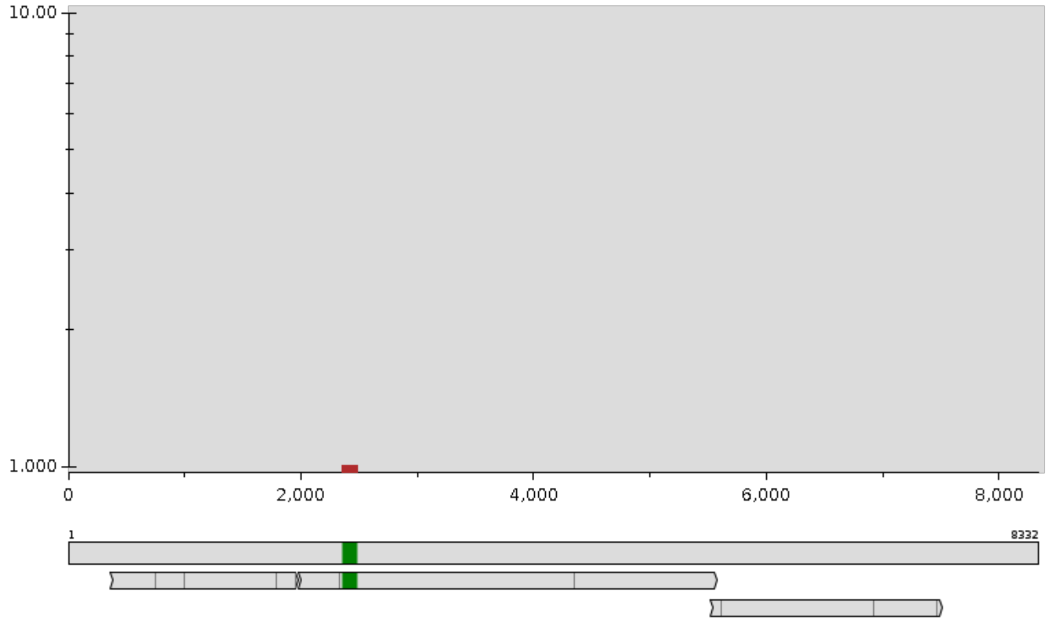

#### Assignment

| | |
|---|---|
| **Type** | Moloney murine leukemia virus (Taxonomy ID: 11801) |
| **Reference Genome** | NC_001501.1 |
| **NT Identity (%)** | 76.2963 |
| **AA Identity (%)** | 100.0 |
| **Number Of Stop Codons** | 0 |
| **Number Of CDS** | 3 |

#### Alignment

| | |
|---|---|
| **Alignment Score** | 142.0 (NT) + 327.0 (AA) = 469.0 |
| **Concordance (%)** | 78.5595 |

| Alignment Method | Global, seeded, nucleotide + amino acids (AGA) |
|---|---|

#### Genome Region

Sequence starts at position 2352 and ends at position 2486 relative to NC_001501.1 reference sequence.

#### Alignment Detailed Statistics

| | Begin | End | Coverage | Score | Concordance | Matches | Identities | I/D/M/F* | Stop Codons |
|---|---|---|---|---|---|---|---|---|---|
| **NT** | **2352** | **2486** | **1.6%** | **142** | **52.6%** | **135 (100%)** | **103 (76.3%)** | **0/0** | |
| Mutations: | 2354G>A, 2360G>C, 2363A>G, 2369G>A, 2373T>A, 2374C>G, 2375A>C, 2381G>A, 2384A>G, 2390T>G, 2391T>A, 2392C>G, 2393T>C, 2396A>G, 2399G>C, 2400T>A, 2401C>G, 2405A>C, 2412T>A, 2413C>G, 2414T>C, 2423T>G, 2429C>G, 2444G>C, 2453A>C, 2459A>G, 2462T>G, 2468A>G, 2471T>G, 2474T>G, 2480C>T, 2483A>T | | | | | | | | |
| **CDS** | | | | | | | | | |
| gag-pol | 665 | 709 | 2.6% | 327 | 100% | 45 (100%) | 45 (100%) | 0/0/0/0 | 0 |
| **Proteins** | | | | | | | | | |
| Pr180 (NP_057933.2) | 665 | 709 | 2.6% | 327 | 100% | 45 (100%) | 45 (100%) | 0/0/0/0 | 0 |
| Protein mutations: | *none* | | | | | | | | |
| Codon mutations: | GAG665GAA (2354G>A), CGG667CGC (2360G>C), CTA668CTG (2363A>G), GAG670GAA (2369G>A), TCA672AGC (2373T>A 2374C>G 2375A>C), GAG674GAA (2381G>A), CCA675CCG (2384A>G), GTT677GTG (2390T>G), TCT678AGC (2391T>A 2392C>G 2393T>C), CTA679CTG (2396A>G), GGG680GGC (2399G>C), TCC681AGC (2400T>A 2401C>G), ACA682ACC (2405A>C), TCT685AGC (2412T>A 2413C>G 2414T>C), CCT688CCG (2423T>G), GCC690GCG (2429C>G), GGG695GGC (2444G>C), GGA698GGC (2453A>C), GCA700GCG (2459A>G), GTT701GTG (2462T>G), CAA703CAG (2468A>G), GCT704GCG (2471T>G), CCT705CCG (2474T>G), ATC707ATT (2480C>T), ATA708ATT (2483A>T) | | | | | | | | |
| p80 RT (NP_955591.1) | 6 | 50 | 6.7% | 327 | 100% | 45 (100%) | 45 (100%) | 0/0/0/0 | 0 |
| Protein mutations: | *none* | | | | | | | | |
| Codon mutations: | GAG6GAA (2354G>A), CGG8CGC (2360G>C), CTA9CTG (2363A>G), GAG11GAA (2369G>A), TCA13AGC (2373T>A 2374C>G 2375A>C), GAG15GAA (2381G>A), CCA16CCG (2384A>G), GTT18GTG (2390T>G), TCT19AGC (2391T>A 2392C>G 2393T>C), CTA20CTG (2396A>G), GGG21GGC (2399G>C), TCC22AGC (2400T>A 2401C>G), ACA23ACC (2405A>C), TCT26AGC (2412T>A 2413C>G 2414T>C), CCT29CCG (2423T>G), GCC31GCG (2429C>G), GGG36GGC (2444G>C), GGA39GGC (2453A>C), GCA41GCG (2459A>G), GTT42GTG (2462T>G), CAA44CAG (2468A>G), GCT45GCG (2471T>G), CCT46CCG (2474T>G), ATC48ATT (2480C>T), ATA49ATT (2483A>T) | | | | | | | | |

*: Inserts / Deletes / Misaligned / Frameshifts

#### Analysis details

This analysis was performed with panviral2.64

### NGS Details (UN62): Ochrobactrum phage POA1180

#### Assembly

| | |
|---|---|
| **Coverage Length** | 135 (1 contig(s)) |
| **Depth Of Coverage** | 1.0 |
| **Number Of Reads** | 1 |
| **Reads Per Million** | 0.02 rpm (after QC) |
| **Ambiguities** | 0 |
| **Assembly Method** | de novo + reference guided assembly |
| **Consensus Caller** | Bcf Tools |

#### Coverage Map

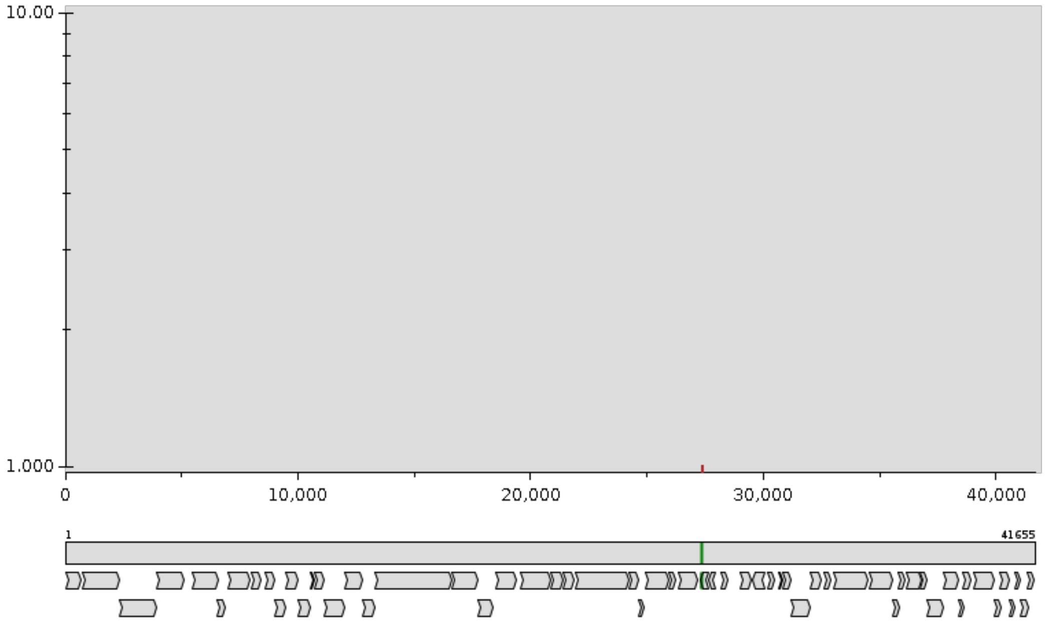

#### Assignment

| | |
|---|---|
| **Type** | Ochrobactrum phage POA1180 (Taxonomy ID: 1897640) |
| **Reference Genome** | NC_070931.1 |
| **NT Identity (%)** | 79.2593 |
| **AA Identity (%)** | 71.1111 |
| **Number Of Stop Codons** | 0 |
| **Number Of CDS** | 58 |

#### Alignment

| | |
|---|---|
| **Alignment Score** | 158.0 (NT) + 269.0 (AA) = 427.0 |
| **Concordance (%)** | 71.1667 |

| Alignment Method | Local, heuristic, nucleotide (BLASTN) |
|---|---|

#### Genome Region

Sequence starts at position 27298 and ends at position 27432 relative to NC_070931.1 reference sequence.

#### Alignment Detailed Statistics

| | Begin | End | Coverage | Score | Concordance | Matches | Identities | I/D/M/F* | Stop Codons |
|---|---|---|---|---|---|---|---|---|---|
| **NT** | **27298** | **27432** | **0.3%** | **158** | **58.5%** | **135 (100%)** | **107 (79.3%)** | **0/0** | |
| Mutations: | 27301A>C, 27309A>C, 27312A>G, 27314G>T, 27322T>C, 27323T>A, 27324C>G, 27325G>C, 27327C>G, 27328T>C, 27329T>G, 27335G>T, 27336A>C, 27352A>G, 27354G>C, 27357G>C, 27366T>G, 27372C>G, 27375T>C, 27378C>G, 27384G>C, 27386G>C, 27388G>A, 27393A>G, 27418G>C, 27419T>G, 27420G>C, 27425C>T | | | | | | | | |
| **CDS** | | | | | | | | | |
| PQB33_gp31 | 59 | 103 | 41.3% | 269 | 78.7% | 45 (100%) | 32 (71.1%) | 0/0/0/0 | 0 |
| Protein mutations: | G61S (27425C>T), D62E (27420G>C), T63R (27418G>C 27419T>G), T73M (27388G>A), L74V (27384G>C 27386G>C), I85T (27352A>G), L91M (27335G>T), K93R (27327C>G 27328T>C 27329T>G), T94S (27324C>G 27325G>C), K95W (27322T>C 27323T>A), L98I (27312A>G 27314G>T), I102S (27301A>C) | | | | | | | | |
| Codon mutations: | GGC61AGC (27425C>T), GAC62GAG (27420G>C), ACG63CGG (27418G>C 27419T>G), TAT71TAC (27393A>G), ACG73ATG (27388G>A), CTC74GTG (27384G>C 27386G>C), CCG76CCC (27378C>G), GGA77GGG (27375T>C), CCG78CCC (27372C>G), GCA80GCC (27366T>G), CTC83CTG (27357G>C), GCC84GCG (27354G>C), ATC85ACC (27352A>G), CTT90CTG (27336A>C), CTG91ATG (27335G>T), AAG93CGC (27327C>G 27328T>C 27329T>G), ACG94AGC (27324C>G 27325G>C), AAG95TGG (27322T>C 27323T>A), CTT98ATC (27312A>G 27314G>T), GTT99GTG (27309A>C), ATC102AGC (27301A>C) | | | | | | | | |
| **Proteins** | | | | | | | | | |
| chromate transport protein (YP_010665114.1) | 59 | 103 | 41.3% | 269 | 78.7% | 45 (100%) | 32 (71.1%) | 0/0/0/0 | 0 |
| Protein mutations: | G61S (27425C>T), D62E (27420G>C), T63R (27418G>C 27419T>G), T73M (27388G>A), L74V (27384G>C 27386G>C), I85T (27352A>G), L91M (27335G>T), K93R (27327C>G 27328T>C 27329T>G), T94S (27324C>G 27325G>C), K95W (27322T>C 27323T>A), L98I (27312A>G 27314G>T), I102S (27301A>C) | | | | | | | | |
| Codon mutations: | GGC61AGC (27425C>T), GAC62GAG (27420G>C), ACG63CGG (27418G>C 27419T>G), TAT71TAC (27393A>G), ACG73ATG (27388G>A), CTC74GTG (27384G>C 27386G>C), CCG76CCC (27378C>G), GGA77GGG (27375T>C), CCG78CCC (27372C>G), GCA80GCC (27366T>G), CTC83CTG (27357G>C), GCC84GCG (27354G>C), ATC85ACC (27352A>G), CTT90CTG (27336A>C), CTG91ATG (27335G>T), AAG93CGC (27327C>G 27328T>C 27329T>G), ACG94AGC (27324C>G 27325G>C), AAG95TGG (27322T>C 27323T>A), CTT98ATC (27312A>G 27314G>T), GTT99GTG (27309A>C), ATC102AGC (27301A>C) | | | | | | | | |

*: Inserts / Deletes / Misaligned / Frameshifts

#### Analysis details

This analysis was performed with panviral2.64

### Analysis of UN70

| | |
|---|---|
| **Technology** | Paired-end short reads |
| **Input Files** | UN70_R1.fq.gz (1.69 GB), UN70_R2.fq.gz (1.74 GB) |
| **Submitted On** | 2023-09-26 13:21:22 UTC |
| **Duration** | 3h 43m 14s |
| **Tool Version** | panviral2.64 |
| **Location** | https://www.genomedetective.com/db/ui/analysis/01dff0d7-3177-4836-8378-54a03e04e81d |

#### Statistics

| | |
|---|---|
| **Original Read Length** | 20 - 150 |
| **Trimmed Read Length** | 50 - 135 |

| | # Reads | % of Reads |
|---|---|---|
| **Input file** | 53127568 | 100.0% |
| **After QC** | 52752992 | 99.3% |
| **After filtering** | 6204606 | 11.7% |
| **Mapped back to contigs** | 3034086 | 5.7% |

#### Assignments

| Assignment | No. of Reads | Depth of Coverage | Identity | | Genome Coverage | |
|---|---|---|---|---|---|---|
| | | | NT | AA | | |
| Torradovirus lycopersici (2 segments out of 2) | 2444918 | 24879.5 | 89.1% | 95.2% | 99.7% | |
| Torradovirus lycopersici (segment RNA 2) | 1691517 | 40729.6 | 87.4% | 91.4% | 99.6% | 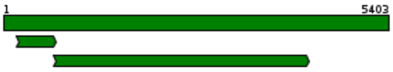 |
| Torradovirus lycopersici (segment RNA 1) | 753401 | 13912.1 | 90.3% | 97.6% | 99.7% | 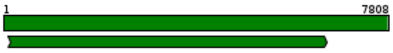 |
| Torradovirus marchitezum (3 segments out of 2) | 227889 | 4171.1 | 64.0% | 64.3% | 59.2% | |
| Torradovirus marchitezum (segment RNA 2) | 19430 | 8120.3 | 75.8% | 87.9% | 5.4% | 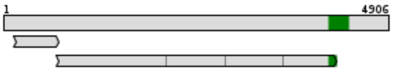 |
| Torradovirus marchitezum (segment RNA 2) | 13 | 7.0 | 77.4% | 84.6% | 3.9% | 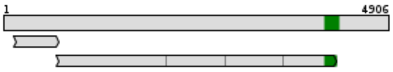 |
| Torradovirus marchitezum (segment RNA 1) | 208446 | 4133.7 | 63.2% | 63.5% | 93.0% | 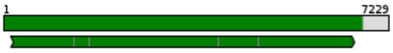 |
| Potato virus Y | 167938 | 2313.2 | 82.7% | 90.1% | 99.8% | 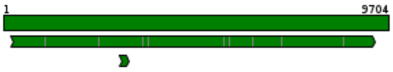 |

| Assignment | No. of Reads | Depth of Coverage | Identity | | Genome Coverage | |
|---|---|---|---|---|---|---|
| | | | NT | AA | | |
| Bracoviriform glomeratae (segment NC_043292.1) | 47201 | 17187.9 | 74.5% | 84.5% | 72.8% | 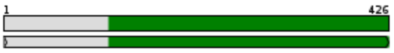 |
| Diachasmimorpha longicaudata entomopoxvirus (segment NC_043455.1) | 11204 | 1241.9 | 59.4% | 55.6% | 84.1% | 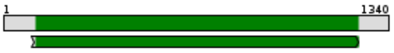 |
| Potato leafroll virus | 796 | 18.2 | 98.8% | 98.4% | 97.1% | 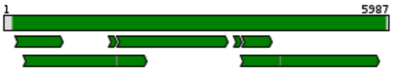 |
| Solendovirus venanicotianae | 234 | 12.1 | 77.6% | 74.2% | 27.1% | 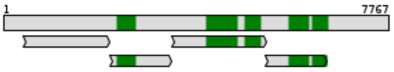 |
| Duamitovirus soch1 | 210 | 11.7 | 69.4% | 69.9% | 72.7% | 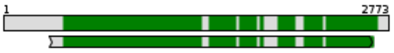 |

#### Discoveries

| Similar to | No. of Reads | Depth of Coverage | Identity | | Genome Coverage | |
|---|---|---|---|---|---|---|
| | | | NT | AA | | |
| Tomato chocolate spot virus (2 segments out of 2) | 93634 | 11276.0 | 69.8% | 80.2% | 10.1% | |
| Tomato chocolate spot virus (segment RNA 1) | 74446 | 13296.7 | 69.9% | 93.5% | 12.5% | 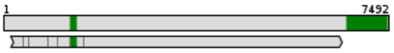 |
| Tomato chocolate spot virus (segment RNA2) | 19188 | 5827.1 | 69.7% | 75.0% | 6.8% | 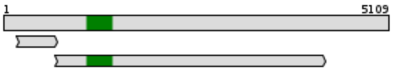 |
| Brazilian marseillevirus | 13004 | 5346.7 | 79.8% | 94.7% | 0.1% | 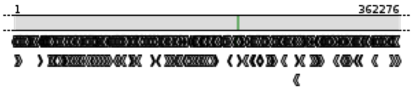 |
| Tokyovirus A1 | 12194 | 6107.3 | 80.6% | 93.1% | 0.0% | 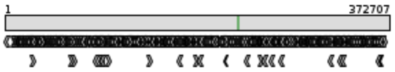 |
| Lausannevirus | 10606 | 4682.9 | 80.5% | 95.2% | 0.1% | 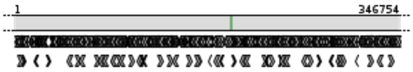 |
| Marseillevirus marseillevirus | 1457 | 1004.1 | 89.4% | 94.3% | 0.0% | 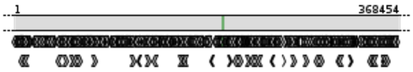 |
| Badnavirus occultipomeae | 541 | 139.6 | 53.1% | 45.5% | 5.3% | 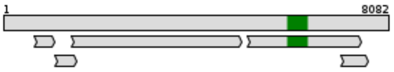 |
| Cassava brown streak virus | 521 | 121.8 | 60.8% | 57.7% | 5.8% | 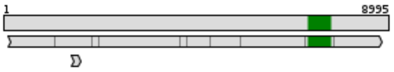 |
| Human gammaherpesvirus 8 (subtype: Could not assign) | 386 | 240.5 | 83.3% | 84.1% | 0.1% | 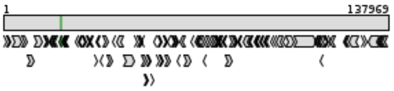 |

| Similar to | No. of Reads | Depth of Coverage | Identity | | Genome Coverage | |
|---|---|---|---|---|---|---|
| | | | NT | AA | | |
| Makelovirus prm1 | 316 | 40.8 | 76.4% | 87.3% | 0.7% | 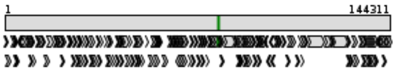 |
| Cladosporium fulvum T-1 virus | 132 | 14.0 | 54.4% | 50.6% | 13.3% | 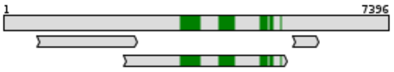 |
| Lowelvirus tuscon4d | 115 | 27.6 | 79.6% | 89.9% | 0.2% | 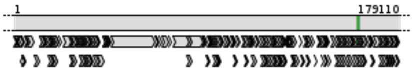 |
| Errantivirus | 112 | 13.9 | 51.7% | 43.4% | 11.5% | 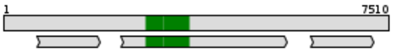 |
| Caulimovirus tesselloscrophulariae | 80 | 15.8 | 59.0% | 46.3% | 7.5% | 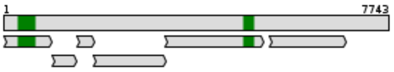 |
| Tomato necrotic dwarf virus (segment RNA1) | 79 | 21.2 | 69.4% | 74.5% | 6.0% | 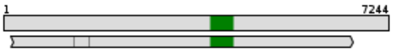 |
| Badnavirus maculaucubae | 77 | 24.5 | 54.7% | 45.0% | 3.5% | 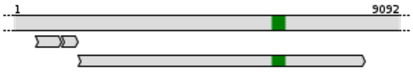 |
| Neptunevirus srim50 | 69 | 10.5 | 72.0% | 87.8% | 0.3% | 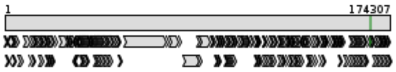 |
| Badnavirus alphacolocalasiae | 42 | 21.2 | 53.6% | 44.9% | 3.1% | 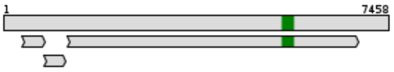 |
| Caulimovirus venafragariae | 36 | 7.0 | 59.9% | 59.1% | 8.3% | 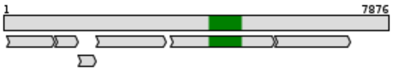 |
| Alisovirus socal22 | 28 | 19.1 | 79.9% | 85.4% | 0.1% | 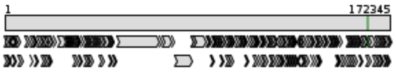 |
| Badnavirus maculakalanchoes | 27 | 10.4 | 55.2% | 48.9% | 3.7% | 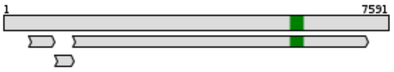 |
| Dioscovirus dioscoreae | 26 | 4.1 | 62.5% | 48.2% | 7.4% | 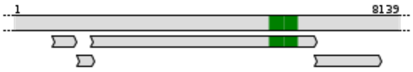 |
| Chinaberry tree badnavirus 1 | 26 | 7.5 | 53.9% | 43.8% | 6.6% | 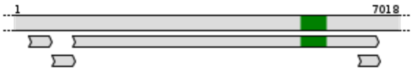 |
| Duamitovirus peex1 | 22 | 5.8 | 71.1% | 66.9% | 14.1% | 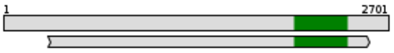 |
| Badnavirus deltainflatheobromae | 20 | 6.1 | 52.2% | 49.1% | 4.4% | 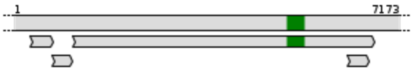 |
| Cavemovirus collusipomeae | 15 | 7.1 | 61.6% | 50.0% | 3.3% | 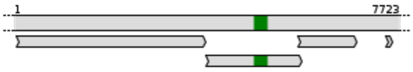 |

| Similar to | No. of Reads | Depth of Coverage | Identity | | Genome Coverage | |
|---|---|---|---|---|---|---|
| | | | NT | AA | | |
| Petuvirus venapetuniae | 14 | 2.8 | 64.0% | 57.9% | 8.5% | 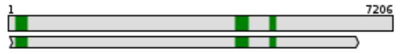 |
| Epiphyllum badnavirus 1 | 14 | 5.1 | 52.3% | 45.7% | 4.3% | 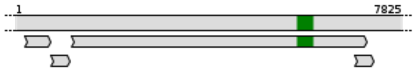 |
| Caulimovirus venafragariae | 11 | 3.0 | 64.3% | 59.3% | 6.4% | 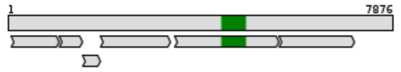 |
| Gihfavirus pelohabitans | 10 | 4.8 | 97.3% | 0.0% | 4.2% | 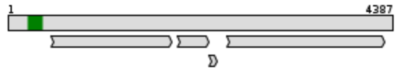 |
| Caulimovirus tessellomirabilis | 10 | 4.0 | 57.0% | 48.6% | 4.4% | 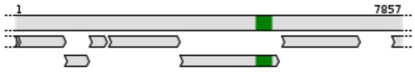 |
| Badnavirus venabougainvilleae | 10 | 3.5 | 51.7% | 39.7% | 4.3% | 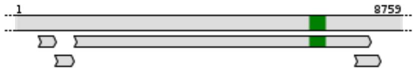 |
| Caulimovirus tessellomirabilis | 8 | 3.3 | 59.3% | 47.6% | 4.0% | 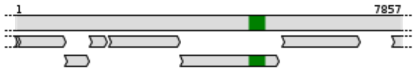 |
| Cavemovirus collusipomeae | 8 | 3.2 | 63.1% | 51.0% | 4.0% | 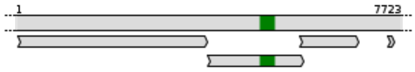 |
| Cavemovirus collusipomeae | 8 | 3.7 | 62.8% | 49.4% | 3.1% | 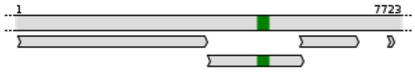 |
| Duamitovirus chqu1 | 8 | 6.4 | 68.1% | 75.6% | 4.9% | 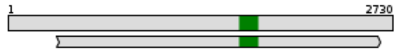 |
| Badnavirus tessellocastaneae | 7 | 3.4 | 55.6% | 55.4% | 3.5% | 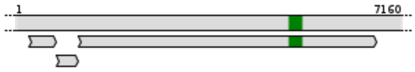 |
| Caulimovirus tessellomirabilis | 7 | 3.1 | 58.7% | 49.5% | 4.0% | 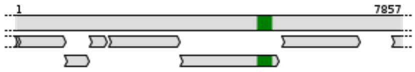 |
| Badnavirus alphacolocalasiae | 5 | 2.3 | 58.9% | 52.1% | 3.9% | 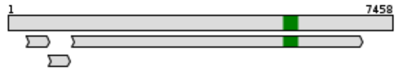 |
| Caulimovirus venafragariae | 4 | 1.6 | 62.6% | 60.9% | 3.3% | 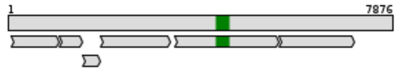 |
| Fragaria chiloensis cryptic virus (segment RNA 3) | 4 | 2.1 | 54.9% | 42.9% | 15.9% | 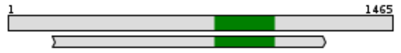 |
| Colombian datura virus | 2 | 1.3 | 99.0% | 100.0% | 2.2% | 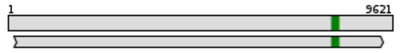 |
| Limeum africanum associated virus | 1 | 1.0 | 72.6% | 75.6% | 4.6% | 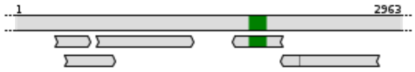 |

#### NGS Details (UN70): Torradovirus marchitezum (segment RNA 2)

##### Assembly

| | |
|---|---|
| **Coverage Length** | 190 (1 contig(s)) |
| **Depth Of Coverage** | 7.0 |
| **Number Of Reads** | 13 |
| **Reads Per Million** | 0.25 rpm (after QC) |
| **Ambiguities** | 0 |
| **Assembly Method** | de novo + reference guided assembly |
| **Consensus Caller** | Bcf Tools |

##### Coverage Map

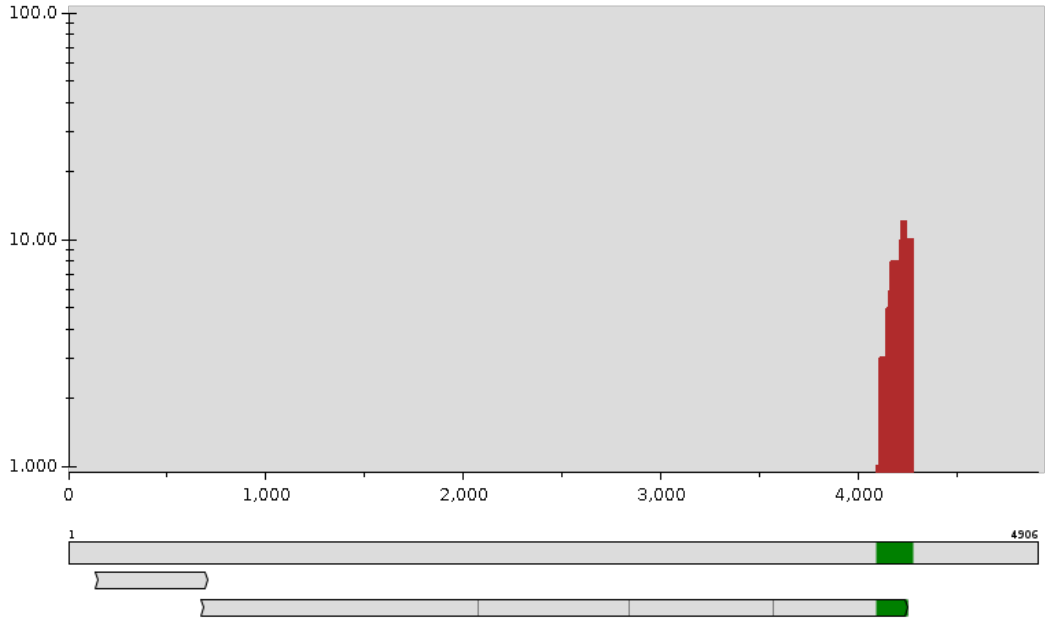

##### Assignment

| | |
|---|---|
| **Type** | Torradovirus marchitezum (Taxonomy ID: 3048376) |
| **Reference Genome** | NC_010988.1 |
| **NT Identity (%)** | 77.3684 |
| **AA Identity (%)** | 84.6154 |
| **Number Of Stop Codons** | 1 |
| **Number Of CDS** | 2 |

##### Alignment

| | |
|---|---|
| **Alignment Score** | 208.0 (NT) + 298.0 (AA) = 506.0 |
| **Concordance (%)** | 69.6011 |

| Alignment Method | Global, seeded, nucleotide + amino acids (AGA) |
|---|---|

#### Genome Region

Sequence starts at position 4089 and ends at position 4278 relative to NC_010988.1 reference sequence.

#### Alignment Detailed Statistics

| | Begin | End | Coverage | Score | Concordance | Matches | Identities | I/D/M/F* | Stop Codons |
|---|---|---|---|---|---|---|---|---|---|
| **NT** | **4089** | **4278** | **3.9%** | **208** | **54.7%** | **190 (100%)** | **147 (77.4%)** | **0/0** | |
| Mutations: | 4090G>T, 4096C>A, 4102G>A, 4105A>C, 4106C>A, 4108T>A, 4109A>G, 4111A>C, 4112T>A, 4113C>A, 4114T>G, 4117T>A, 4120A>T, 4122T>C, 4129C>T, 4132G>A, 4144C>T, 4150T>C, 4153T>A, 4154T>C, 4155C>T, 4159C>A, 4163C>A, 4168A>G, 4171A>G, 4177T>C, 4180C>A, 4181A>G, 4189A>G, 4192T>G, 4195C>T, 4198C>T, 4210C>A, 4213A>T, 4228A>G, 4242T>A, 4247A>G, 4248A>C, 4259C>T, 4261A>G, 4267T>A, 4274T>C, 4276G>A | | | | | | | | |
| **CDS** | | | | | | | | | |
| ToMarV_RNA2gp2 | 1141 | 1192 | 4.4% | 298 | 84.7% | 52 (100%) | 44 (84.6%) | 0/0/0/0 | 1 |
| Protein mutations: | L1146I (4106C>A 4108T>A), I1147V (4109A>G 4111A>C), S1148K (4112T>A 4113C>A 4114T>G), L1151P (4122T>C), S1162L (4154T>C 4155C>T), H1165N (4163C>A), N1171D (4181A>G), I1191N (4242T>A) | | | | | | | | |
| Codon mutations: | GTG1140.TT (4090G>T), TCC1142TCA (4096C>A), GAG1144GAA (4102G>A), ATA1145ATC (4105A>C), CTT1146ATA (4106C>A 4108T>A), ATA1147GTC (4109A>G 4111A>C), TCT1148AAG (4112T>A 4113C>A 4114T>G), CCT1149CCA (4117T>A), GGA1150GGT (4120A>T), CTT1151CCT (4122T>C), TTC1153TTT (4129C>T), AAG1154AAA (4132G>A), GCC1158GCT (4144C>T), CCT1160CCC (4150T>C), CCT1161CCA (4153T>A), TCT1162CTT (4154T>C 4155C>T), GCC1163GCA (4159C>A), CAT1165AAT (4163C>A), GAA1166GAG (4168A>G), AAA1167AAG (4171A>G), CTT1169CTC (4177T>C), GGC1170GGA (4180C>A), AAT1171GAT (4181A>G), CAA1173CAG (4189A>G), ACT1174ACG (4192T>G), CAC1175CAT (4195C>T), ACC1176ACT (4198C>T), ACC1180ACA (4210C>A), TCA1181TCT (4213A>T), CTA1186CTG (4228A>G), ATT1191AAT (4242T>A) | | | | | | | | |
| **Proteins** | | | | | | | | | |
| polyprotein (YP_001976149.1) | 1141 | 1192 | 4.4% | 298 | 84.7% | 52 (100%) | 44 (84.6%) | 0/0/0/0 | 1 |
| Protein mutations: | L1146I (4106C>A 4108T>A), I1147V (4109A>G 4111A>C), S1148K (4112T>A 4113C>A 4114T>G), L1151P (4122T>C), S1162L (4154T>C 4155C>T), H1165N (4163C>A), N1171D (4181A>G), I1191N (4242T>A) | | | | | | | | |
| Codon mutations: | GTG1140.TT (4090G>T), TCC1142TCA (4096C>A), GAG1144GAA (4102G>A), ATA1145ATC (4105A>C), CTT1146ATA (4106C>A 4108T>A), ATA1147GTC (4109A>G 4111A>C), TCT1148AAG (4112T>A 4113C>A 4114T>G), CCT1149CCA (4117T>A), GGA1150GGT (4120A>T), CTT1151CCT (4122T>C), TTC1153TTT (4129C>T), AAG1154AAA (4132G>A), GCC1158GCT (4144C>T), CCT1160CCC (4150T>C), CCT1161CCA (4153T>A), TCT1162CTT (4154T>C 4155C>T), GCC1163GCA (4159C>A), CAT1165AAT (4163C>A), GAA1166GAG (4168A>G), AAA1167AAG (4171A>G), CTT1169CTC (4177T>C), GGC1170GGA (4180C>A), AAT1171GAT (4181A>G), CAA1173CAG (4189A>G), ACT1174ACG (4192T>G), CAC1175CAT (4195C>T), ACC1176ACT (4198C>T), ACC1180ACA (4210C>A), TCA1181TCT (4213A>T), CTA1186CTG (4228A>G), ATT1191AAT (4242T>A) | | | | | | | | |
| Coat protein C (YP_001976157.1) | 175 | 225 | 22.7% | 297 | 84.6% | 51 (100%) | 43 (84.3%) | 0/0/0/0 | 0 |
| Protein mutations: | L180I (4106C>A 4108T>A), I181V (4109A>G 4111A>C), S182K (4112T>A 4113C>A 4114T>G), L185P (4122T>C), S196L (4154T>C 4155C>T), H199N (4163C>A), N205D (4181A>G), I225N (4242T>A) | | | | | | | | |
| Codon mutations: | GTG174.TT (4090G>T), TCC176TCA (4096C>A), GAG178GAA (4102G>A), ATA179ATC (4105A>C), CTT180ATA (4106C>A 4108T>A), ATA181GTC (4109A>G 4111A>C), TCT182AAG (4112T>A 4113C>A 4114T>G), CCT183CCA (4117T>A), GGA184GGT (4120A>T), CTT185CCT (4122T>C), TTC187TTT (4129C>T), AAG188AAA (4132G>A), GCC192GCT (4144C>T), CCT194CCC (4150T>C), CCT195CCA (4153T>A), TCT196CTT (4154T>C 4155C>T), GCC197GCA (4159C>A), CAT199AAT (4163C>A), GAA200GAG (4168A>G), AAA201AAG (4171A>G), CTT203CTC (4177T>C), GGC204GGA (4180C>A), AAT205GAT (4181A>G), CAA207CAG (4189A>G), ACT208ACG (4192T>G), CAC209CAT (4195C>T), ACC210ACT (4198C>T), ACC214ACA (4210C>A), TCA215TCT (4213A>T), CTA220CTG (4228A>G), ATT225AAT (4242T>A) | | | | | | | | |

*: Inserts / Deletes / Misaligned / Frameshifts

#### Analysis details

This analysis was performed with panviral2.64

### NGS Details (UN70): Torradovirus marchitezum (segment RNA 2)

#### Assembly

| | |
|---|---|
| **Coverage Length** | 264 (1 contig(s)) |
| **Depth Of Coverage** | 8120.3 |
| **Number Of Reads** | 19430 |
| **Reads Per Million** | 368.32 rpm (after QC) |
| **Ambiguities** | 0 |
| **Assembly Method** | de novo + reference guided assembly |
| **Consensus Caller** | Bcf Tools |

#### Coverage Map

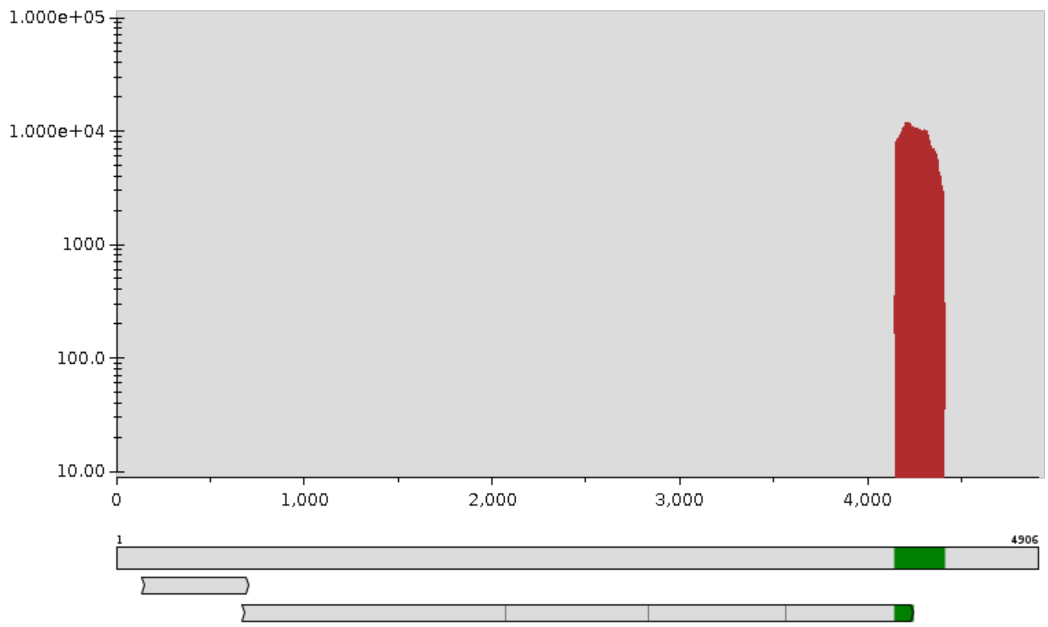

#### Assignment

| | |
|---|---|
| **Type** | Torradovirus marchitezum (Taxonomy ID: 3048376) |
| **Reference Genome** | NC_010988.1 |
| **NT Identity (%)** | 75.7692 |
| **AA Identity (%)** | 87.8788 |
| **Number Of Stop Codons** | 1 |
| **Number Of CDS** | 2 |

#### Alignment

| | |
|---|---|
| **Alignment Score** | 219.0 (NT) + 91.0 (AA) = 310.0 |
| **Concordance (%)** | 52.8846 |

| Alignment Method | Global, seeded, nucleotide + amino acids (AGA) |
|---|---|

#### Genome Region

Sequence starts at position 4147 and ends at position 4410 relative to NC_010988.1 reference sequence.

#### Alignment Detailed Statistics

| | Begin | End | Coverage | Score | Concordance | Matches | Identities | I/D/M/F* | Stop Codons |
|---|---|---|---|---|---|---|---|---|---|
| **NT** | **4147** | **4410** | **5.4%** | **219** | **44.8%** | **257 (96.3%)** | **197 (73.8%)** | **3/7** | |
| Mutations: | 4156T>G, 4159C>T, 4162C>T, 4180C>T, 4181A>G, 4187_4188delCA, 4189A>C, 4192T>G, 4195C>T, 4198C>T, 4201T>C, 4204G>A, 4213A>T, 4219T>C, 4222T>A, 4225G>A, 4228A>G, 4229C>A, 4231T>A, 4234T>C, 4237T>C, 4238G>T, 4242T>A, 4243T>C, 4248A>C, 4252T>A, 4279A>T, 4279_4280insT, 4280A>T, 4280_4281insC, 4283A>G, 4286C>T, 4287T>A, 4288T>C, 4291C>A, 4294C>T, 4306G>A, 4309G>A, 4312G>T, 4313C>A, 4313_4314insG, 4315A>G, 4316A>T, 4318T>C, 4319G>T, 4320G>A, 4323T>G, 4324T>A, 4328T>C, 4330G>A, 4333A>G, 4335G>T, 4336A>C, 4338A>G, 4339_4343delAGGGA, 4345A>T, 4346G>C, 4349A>C, 4364C>T, 4365T>C, 4369A>G, 4372A>G, 4373G>A, 4392T>C, 4393C>T | | | | | | | | |
| **CDS** | | | | | | | | | |
| ToMarV_RNA2gp2 | 1160 | 1192 | 2.8% | 91 | 37.4% | 33 (100%) | 29 (87.9%) | 0/0/1/1 | 1 |
| Protein mutations: | N1171D (4181A>G), A1190S (4238G>T), I1191N (4242T>A 4243T>C) | | | | | | | | |
| Codon mutations: | TCT1162TCG (4156T>G), GCC1163GCT (4159C>T), AAC1164AAT (4162C>T), GGC1170GGT (4180C>T), AAT1171GAT (4181A>G), CAA1173--C (4187_4188delCA 4189A>C), ACT1174ACG (4192T>G), CAC1175CAT (4195C>T), ACC1176ACT (4198C>T), TAT1177TAC (4201T>C), CAG1178CAA (4204G>A), TCA1181TCT (4213A>T), TTT1183TTC (4219T>C), TCT1184TCA (4222T>A), GAG1185GAA (4225G>A), CTA1186CTG (4228A>G), CGT1187AGA (4229C>A 4231T>A), GAT1188GAC (4234T>C), TTT1189TTC (4237T>C), GCG1190TCG (4238G>T), ATT1191AAC (4242T>A 4243T>C) | | | | | | | | |
| **Proteins** | | | | | | | | | |
| polyprotein (YP_001976149.1) | 1160 | 1192 | 2.8% | 91 | 37.4% | 33 (100%) | 29 (87.9%) | 0/0/1/1 | 1 |
| Protein mutations: | N1171D (4181A>G), A1190S (4238G>T), I1191N (4242T>A 4243T>C) | | | | | | | | |
| Codon mutations: | TCT1162TCG (4156T>G), GCC1163GCT (4159C>T), AAC1164AAT (4162C>T), GGC1170GGT (4180C>T), AAT1171GAT (4181A>G), CAA1173--C (4187_4188delCA 4189A>C), ACT1174ACG (4192T>G), CAC1175CAT (4195C>T), ACC1176ACT (4198C>T), TAT1177TAC (4201T>C), CAG1178CAA (4204G>A), TCA1181TCT (4213A>T), TTT1183TTC (4219T>C), TCT1184TCA (4222T>A), GAG1185GAA (4225G>A), CTA1186CTG (4228A>G), CGT1187AGA (4229C>A 4231T>A), GAT1188GAC (4234T>C), TTT1189TTC (4237T>C), GCG1190TCG (4238G>T), ATT1191AAC (4242T>A 4243T>C) | | | | | | | | |
| Coat protein C (YP_001976157.1) | 194 | 225 | 14.2% | 90 | 37.2% | 32 (100%) | 28 (87.5%) | 0/0/1/1 | 0 |
| Protein mutations: | N205D (4181A>G), A224S (4238G>T), I225N (4242T>A 4243T>C) | | | | | | | | |
| Codon mutations: | TCT196TCG (4156T>G), GCC197GCT (4159C>T), AAC198AAT (4162C>T), GGC204GGT (4180C>T), AAT205GAT (4181A>G), CAA207--C (4187_4188delCA 4189A>C), ACT208ACG (4192T>G), CAC209CAT (4195C>T), ACC210ACT (4198C>T), TAT211TAC (4201T>C), CAG212CAA (4204G>A), TCA215TCT (4213A>T), TTT217TTC (4219T>C), TCT218TCA (4222T>A), GAG219GAA (4225G>A), CTA220CTG (4228A>G), CGT221AGA (4229C>A 4231T>A), GAT222GAC (4234T>C), TTT223TTC (4237T>C), GCG224TCG (4238G>T), ATT225AAC (4242T>A 4243T>C) | | | | | | | | |

*: Inserts / Deletes / Misaligned / Frameshifts

#### Analysis details

This analysis was performed with panviral2.64

### NGS Details (UN70): Torradovirus marchitezum (segment RNA 1)

#### Assembly

| | |
|---|---|
| **Coverage Length** | 6726 (1 contig(s)) |
| **Depth Of Coverage** | 4133.7 |
| **Number Of Reads** | 208446 |
| **Reads Per Million** | 3951.36 rpm (after QC) |
| **Ambiguities** | 0 |
| **Assembly Method** | de novo + reference guided assembly |
| **Consensus Caller** | Bcf Tools |

#### Coverage Map

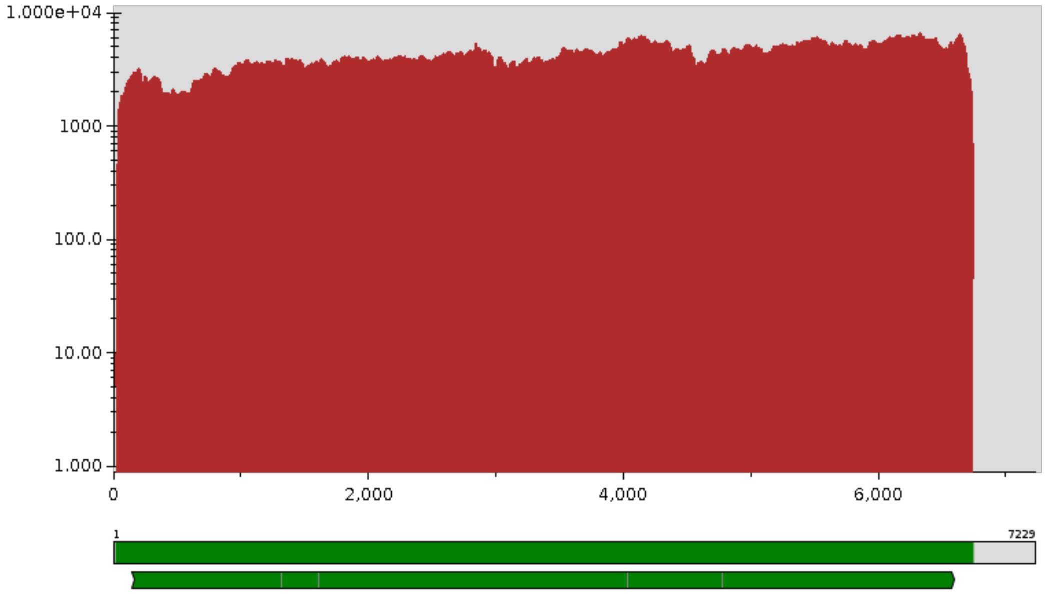

#### Assignment

| | |
|---|---|
| **Type** | Torradovirus marchitezum (Taxonomy ID: 3048376) |
| **Reference Genome** | NC_010987.1 |
| **NT Identity (%)** | 63.1901 |
| **AA Identity (%)** | 63.4669 |
| **Number Of Stop Codons** | 0 |
| **Number Of CDS** | 1 |

#### Alignment

| | |
|---|---|
| **Alignment Score** | 3341.0 (NT) + 9871.0 (AA) = 13212.0 |
| **Concordance (%)** | 47.1941 |

| Alignment Method | Global, seeded, nucleotide + amino acids (AGA) |
|---|---|

#### Genome Region

Sequence starts at position 22 and ends at position 6747 relative to NC_010987.1 reference sequence.

#### Alignment Detailed Statistics

| | Begin | End | Coverage | Score | Concordance | Matches | Identities | I/D/M/F* | Stop Codons |
|---|---|---|---|---|---|---|---|---|---|
| **NT** | **22** | **6747** | **93.0%** | **3341** | **25.4%** | **6659 (98.4%)** | **4235 (62.6%)** | **43/67** | |

| | Begin | End | Coverage | Score | Concordance | Matches | Identities | I/D/M/F* | Stop Codons |
|---|---|---|---|---|---|---|---|---|---|
| NT | 22 | 6747 | 93.0% | 3341 | 25.4% | 6659 (98.4%) | 4235 (62.6%) | 43/67 | |

Mutations:

32T>C, 37G>A, 40C>T, 59A>C, 61T>C, 62T>A, 63G>T, 66A>C, 67_78delTTTGATTCTTGT, 81A>T, 82C>A, 83T>C, 87C>G, 89A>T, 90A>T, 91C>T, 94C>T, 99T>A, 100_101insC, 108A>T, 112G>C, 114C>T, 115C>T, 116A>C, 117T>C, 118T>C, 119A>T, 122T>G, 125A>T, 128C>A, 129C>A, 130T>A, 133_134delAT, 136T>A, 146T>A, 160T>G, 161C>G, 162T>A, 163C>G, 164C>T, 165_188delAAGGTTGCTGGTGTTGATGACGCC, 189A>C, 190G>C, 192T>C, 195T>G, 200C>T, 201A>T, 203C>A, 204A>C, 217G>C, 218T>A, 222C>G, 227A>T, 228A>T, 229A>C, 234A>T, 239C>T, 240A>T, 241A>C, 242A>T, 243T>C, 251A>G, 252G>A, 254G>T, 255G>C, 257T>C, 258A>T, 259G>C, 260T>C, 264A>G, 268C>A, 269C>T, 271C>A, 272T>A, 273A>G, 275G>T, 283A>C, 284G>C, 288T>G, 289C>T, 296T>C, 297C>A, 299T>A, 306C>G, 308G>C, 311A>G, 314A>G, 315T>C, 317T>A, 320G>T, 322C>T, 323T>G, 325A>C, 326T>A, 328T>C, 329G>A, 330T>G, 332C>A, 338T>A, 343T>C, 344G>C, 345G>T, 347C>T, 350T>A, 353T>C, 355G>A, 357T>G, 359A>T, 360C>G, 362T>A, 363A>G, 365T>A, 367A>G, 374G>A, 377C>T, 380A>G, 385G>C, 386T>A, 387G>A, 388C>A, 390C>A, 392T>A, 395A>G, 398T>A, 401A>C, 404T>C, 405C>A, 407T>C, 410A>G, 412C>G, 413G>T, 416C>T, 417T>G, 418C>A, 422C>T, 423T>G, 425G>T, 426T>A, 427C>A, 428C>A, 431A>T, 437C>T, 441T>A, 446C>A, 447A>G, 448A>T, 449G>A, 453A>C, 455G>T, 458T>A, 459T>C, 461A>T, 464A>G, 465T>A, 468T>G, 471G>T, 472A>T, 473G>A, 477T>G, 479G>C, 480T>A, 481C>A, 482C>G, 483A>G, 489G>T, 491T>C, 493A>T, 494A>C, 495T>C, 502T>A, 503T>C, 505C>A, 506T>A, 510G>T, 512G>A, 516A>G, 518T>A, 522A>G, 523G>T, 527A>G, 528G>T, 529A>C, 530T>A, 531G>T, 542T>G, 543G>T, 545A>T, 546T>A, 547A>T, 548T>A, 550G>C, 555C>T, 557T>A, 560A>T, 567A>T, 569A>G, 570A>G, 572A>C, 573T>A, 575G>T, 578A>G, 579G>C, 581G>A, 584T>A, 587A>G, 589T>A, 593T>A, 596T>C, 600A>G, 601G>T, 602T>A, 611T>A, 612G>A, 613A>C, 615G>C, 616G>C, 618G>T, 621T>A, 622C>A, 626C>T, 627A>G, 628C>T, 629T>A, 632C>T, 633A>G, 634A>T, 635A>T, 637A>C, 639A>T, 641C>G, 646T>C, 647G>A, 648A>G, 650A>T, 653C>T, 659C>T, 660A>T, 662G>T, 665T>C, 668A>T, 669A>T, 670G>C, 671T>A, 672C>T, 674A>G, 677C>T, 697A>C, 702G>T, 704C>T, 706A>G, 707A>G, 708C>A, 709G>A, 710A>G, 713C>T, 714A>T, 717A>C, 718T>A, 720G>C, 725C>G, 731T>A, 732A>G, 733G>C, 734T>A, 735A>G, 737G>A, 746G>T, 750T>A, 751C>G, 752A>T, 753G>T, 755T>A, 756T>A, 758A>G, 770G>T, 771T>A, 772C>G, 773T>C, 776G>A, 777T>C, 779G>T, 780A>T, 782G>A, 786A>T, 789A>T, 791A>G, 803G>T, 806T>C, 809G>A, 812G>T, 815G>A, 818G>A, 821C>T, 822A>G, 827A>T, 830G>T, 836T>G, 848C>T, 849T>C, 851G>T, 852A>T, 853G>C, 854C>A, 861T>C, 866C>A, 872T>C, 875C>G, 878T>C, 881G>T, 882A>G, 884A>T, 896T>G, 899G>A, 911A>G, 914A>G, 918A>C, 919C>A, 920T>G, 921C>T, 923T>G, 927C>A, 929T>G, 932T>C, 938A>G, 944C>T, 947C>A, 950C>T, 953G>A, 956T>C, 974A>G, 978A>G, 980A>G, 983C>T, 984G>A, 988C>T, 989C>T, 995A>C, 998C>T, 1010G>A, 1019T>G, 1024G>A, 1028A>T, 1031G>C, 1032C>T, 1034G>A, 1046G>T, 1050G>C, 1059C>A, 1060A>C, 1064G>A, 1068G>T, 1069A>C, 1070T>G, 1072C>A, 1073A>G, 1076G>T, 1077T>G, 1078G>C, 1082T>A, 1085T>A, 1086G>T, 1088A>T, 1090C>T, 1100G>A, 1103C>T, 1109C>A, 1113A>C, 1114A>C, 1115G>T, 1118T>A, 1121C>T, 1124T>A, 1126C>T, 1127C>T, 1130A>G, 1134A>C, 1136C>A, 1140C>T, 1142G>A, 1144G>A, 1145G>A, 1148A>G, 1149G>C, 1151A>G, 1154T>C, 1155C>G, 1160G>A, 1163C>A, 1178A>C, 1190T>C, 1193G>A, 1199C>T, 1200A>T, 1215A>T, 1216G>C, 1226T>A, 1227G>A, 1233T>C, 1238A>T, 1242G>C, 1244T>G, 1245T>G, 1246A>T, 1253A>G, 1254A>G, 1255G>C, 1256T>C, 1257T>A, 1259T>C, 1260A>C, 1261G>A, 1262G>A, 1265C>T, 1266A>T, 1267T>C, 1268G>T, 1271C>T, 1280G>A, 1284C>A, 1286T>G, 1292A>G, 1298T>C, 1304A>G, 1313C>T, 1316G>A, 1317T>G, 1319C>T, 1325G>A, 1337T>A, 1340A>G, 1342C>G, 1343C>T, 1350T>C, 1352A>T, 1355T>C, 1358T>C, 1367G>T, 1368T>C, 1370G>C, 1373C>T, 1374A>G, 1375G>C, 1385G>A, 1391A>G, 1394C>G, 1397T>C, 1400G>C, 1401C>T, 1403A>G, 1409A>G, 1418G>A, 1421G>A, 1445T>A, 1448C>T, 1457T>A, 1458A>G, 1460C>G, 1463G>T, 1472C>T, 1478A>T, 1481A>T, 1482T>C, 1484G>T, 1487A>G, 1493G>A, 1496T>A, 1505C>T, 1508A>G, 1511T>C, 1514T>A, 1517A>G, 1529T>C, 1538A>T, 1541A>C, 1544A>G, 1547A>T, 1553C>T, 1554T>C, 1556A>T, 1565C>T, 1571T>C, 1577C>T, 1583C>A, 1589T>C, 1592C>T, 1595C>A, 1598C>A, 1604T>C, 1607A>C, 1616A>C, 1623C>T, 1628C>A, 1631A>T, 1637T>A, 1640C>T, 1643A>G, 1649T>G, 1652T>A, 1655A>G, 1661A>T, 1667T>A, 1673G>A, 1674C>A, 1679A>G, 1682G>A, 1685C>T, 1688C>T, 1689C>T, 1691G>A, 1694G>T, 1700T>C, 1702T>G, 1703G>A, 1708A>T, 1715G>T, 1719T>G, 1720C>A, 1721C>T, 1722A>C, 1724T>A, 1725G>A, 1727A>T, 1728G>A, 1729A>C, 1733T>A, 1736G>A, 1738G>C, 1739A>C, 1745C>A, 1746A>G, 1747G>T, 1748A>T, 1752A>G, 1754C>T, 1755A>G, 1757A>G, 1763G>A, 1775T>C, 1781T>C, 1787T>A, 1790C>T, 1793G>A, 1794A>G, 1795A>C, 1799G>A, 1800A>T, 1806A>G, 1807G>A, 1811T>C, 1814T>C, 1818G>C, 1820G>A, 1829A>G, 1832C>A, 1835A>G, 1838T>C, 1840A>G, 1844A>G, 1847T>C, 1848G>T, 1849C>A, 1850A>T, 1853A>G, 1856A>G, 1861A>G, 1862G>C, 1866T>C, 1868G>T, 1870T>A, 1871T>C, 1872G>A, 1874A>T, 1875G>A, 1878G>A, 1879A>C, 1883T>G, 1886C>A, 1887A>T, 1889A>G, 1890A>G, 1891G>C, 1893C>G, 1894C>A, 1895T>C, 1899A>G, 1901A>T, 1910T>C, 1911C>A, 1913A>T, 1917T>G, 1921C>T, 1922A>T, 1928T>A, 1932A>C, 1934A>G, 1936C>A, 1937G>T, 1941G>T, 1943G>C, 1944G>C, 1946T>A, 1948G>A, 1949G>T, 1952T>C, 1953_1955delAGA, 1958G>A, 1960G>A, 1961C>A, 1962A>G, 1963C>T, 1964C>T, 1965A>T, 1966A>T, 1969A>C, 1970A>T, 1971G>C, 1972A>C, 1976A>G, 1980A>G, 1981T>C, 1983T>G, 1984T>C, 1985G>T, 1986G>T, 1987T>A, 1988G>T, 1989C>G, 1990C>A, 1991C>G, 1992A>C, 1993C>A, 1994T>G, 1995G>A, 1996T>C, 1997T>A, 1998G>A, 2000G>T, 2002G>C, 2003T>A, 2007A>G, 2008A>C, 2012T>A, 2014G>A, 2016G>A, 2018G>T, 2019A>G, 2021C>T, 2024T>C, 2025G>T, 2027A>C, 2028A>C, 2029G>A, 2030A>G, 2033G>A, 2039G>A, 2042A>T, 2048A>C, 2049T>C, 2051G>C, 2060A>T, 2066G>A, 2069C>G, 2073T>A, 2076G>A, 2079C>A, 2080C>A, 2081T>A, 2082G>T, 2083A>T, 2084G>T, 2087A>G, 2088C>T, 2093T>C, 2096T>A, 2104A>T, 2105T>C, 2106C>T, 2107A>C, 2108T>C, 2117C>T, 2120G>C, 2127T>G, 2128C>T, 2129T>G, 2132T>A, 2133G>T, 2134A>C, 2136G>T, 2137A>C, 2141A>T, 2142C>G, 2143A>C, 2144T>C, 2148G>C, 2152T>A, 2156C>T, 2157A>G, 2159C>A, 2161A>T, 2162A>G, 2166A>G, 2167G>T, 2169G>A, 2170T>A, 2171C>T, 2172A>G, 2174C>T, 2175A>T, 2181T>G, 2182A>T, 2183T>C, 2185G>A, 2193G>A, 2194A>G, 2196G>C, 2199G>C, 2201A>G, 2208C>G, 2209G>C, 2210A>T, 2219A>G, 2221A>G, 2237C>T, 2243A>C, 2249C>T, 2252T>C, 2268G>A, 2269C>G, 2270A>C, 2271T>A, 2279C>T, 2285G>A, 2286A>G, 2289A>C, 2290A>T, 2291A>G, 2294A>G, 2296C>A, 2297A>T, 2300G>C, 2303G>A, 2304A>C, 2305C>G, 2306A>C, 2307G>A, 2309G>T, 2310A>C, 2312G>T, 2313A>G, 2314G>A, 2316A>C, 2317C>A, 2318T>A, 2321A>G, 2322T>A, 2323T>A, 2324T>A, 2327A>G, 2328C>T, 2331T>A, 2332C>A, 2333T>G, 2334T>G, 2335G>C, 2336C>G, 2337A>C, 2340G>C, 2342T>G, 2345A>T, 2350T>C, 2351T>A, 2354C>T, 2357C>A, 2360C>T, 2362T>C, 2363A>G, 2369C>T, 2372G>A, 2385T>A, 2386T>G, 2387T>A, 2390T>A, 2396T>C, 2397T>C, 2399A>G, 2405A>T, 2408A>T, 2415A>C, 2417A>C, 2426C>T, 2428A>G, 2429C>G, 2435A>T, 2436T>G, 2438_2439insGGT, 2440A>C, 2441A>T, 2444G>C, 2445C>A, 2446C>A, 2447A>G, 2451G>A, 2452A>G, 2454T>A, 2455C>A, 2456A>T, 2462C>T, 2464A>T, 2465A>C, 2466A>G, 2467T>A, 2468T>A, 2472T>A, 2473A>T, 2474C>G, 2475C>A, 2476A>T, 2478A>G, 2479A>G, 2480A>T, 2484G>A, 2485T>A, 2486T>G, 2489A>C, 2490T>A, 2492T>A, 2495C>G, 2497A>T, 2498G>C, 2499G>C, 2501T>G, 2503C>T, 2504A>G, 2508T>A, 2509G>A, 2510G>T, 2511A>G, 2512G>A, 2514C>T, 2516T>G, 2521A>T, 2522T>C, 2523G>A, 2524C>T, 2525A>C, 2526C>T, 2530G>A, 2531A>G, 2537C>T, 2546A>C, 2549T>A, 2556A>T, 2558A>G, 2561G>A, 2564T>A, 2573A>C, 2574G>A, 2575T>C, 2576T>C, 2577T>A, 2579T>G, 2580G>A, 2581G>A, 2582A>T, 2584G>C, 2586A>G, 2591A>T, 2594A>T, 2597T>C, 2598T>A, 2600G>T, 2601G>A, 2603G>T, 2607T>C, 2608G>T, 2609G>C, 2610C>A, 2613A>G, 2614G>C, 2615T>A, 2616A>G, 2617A>C, 2618C>A, 2619T>A, 2623T>C, 2627C>T, 2628A>T, 2629A>T, 2630T>A, 2633A>T, 2634T>G, 2635C>A, 2636G>T, 2645A>G, 2648T>C, 2651G>A, 2660A>G, 2672A>C, 2678T>C, 2684G>A, 2687C>T, 2690G>A, 2693T>G, 2696T>G, 2698A>C, 2699G>T, 2699_2700insCAT, 2700C>T, 2703A>G, 2704G>C, 2708T>A, 2709C>T, 2710T>C, 2711T>C, 2712C>T, 2714C>G, 2715G>C, 2717T>A, 2720T>G, 2724A>G, 2725G>C, 2726T>G, 2727T>G, 2729T>A, 2730C>G, 2732A>T, 2733T>A, 2734C>A, 2735A>C, 2736T>A, 2737T>A, 2738G>C, 2739G>C, 2740C>A, 2741A>T, 2742C>G, 2743A>T, 2744A>G, 2747T>C, 2747_2748insCGG, 2753T>A, 2755C>A, 2756T>A, 2757G>C, 2759A>G, 2771A>T, 2775C>G, 2777G>A, 2780C>T, 2783C>G, 2784A>C, 2786G>C, 2792T>C, 2798G>T, 2801G>T, 2804A>G, 2808C>A, 2812G>C, 2822G>C, 2825C>T, 2828T>C, 2834C>T, 2837C>G, 2840C>T, 2843G>T, 2858A>T, 2859A>C, 2861G>C, 2864C>T, 2867A>G, 2870T>A, 2876A>C, 2879C>T, 2880C>A, 2881A>G, 2882C>A, 2885T>C, 2886G>A, 2887C>T, 2888T>G, 2889C>A, 2891A>G, 2897T>C, 2903A>T, 2907A>T, 2909A>C, 2915C>T, 2920C>A, 2921C>G, 2922A>C, 2927T>C, 2928C>A, 2929G>T, 2930A>T, 2932C>G, 2936A>T, 2939T>G, 2942A>G, 2944A>T, 2948C>T, 2953G>C, 2954A>C, 2955G>C, 2962G>A, 2964A>T, 2966T>G, 2969T>A, 2975C>T, 2981C>T, 2982A>G, 2984C>G, 2988C>T, 2994C>A, 2996C>G, 2999T>A, 3002T>G, 3017A>C, 3020G>A, 3021C>T, 3023T>A, 3026C>T, 3029C>A, 3030C>A, 3032T>A, 3038C>T, 3044A>T, 3045A>C, 3047A>C, 3050G>A, 3056G>T, 3062C>T, 3064T>A, 3066A>T, 3068A>T, 3078C>A, 3079A>T, 3080A>G, 3083A>C, 3090T>G, 3091G>T, 3092T>G, 3095A>G, 3101T>A, 3104G>T, 3107T>A, 3109C>A, 3110A>G, 3116A>C, 3117C>A, 3118G>A, 3119T>G, 3120C>A, 3121T>A, 3122T>A, 3123G>T, 3124G>C, 3131A>G, 3135T>G, 3140C>T, 3143G>A, 3144A>T, 3145T>A, 3146G>T, 3167A>C, 3173A>C, 3175T>A, 3176T>A, 3182G>C, 3183G>A, 3184A>C, 3190A>G, 3192C>T, 3194G>A, 3195A>G, 3197C>T, 3200C>G, 3201T>A, 3202C>A, 3203C>T, 3204A>G, 3205G>C, 3206T>A, 3210A>G, 3211A>C, 3212G>T, 3213A>C, 3216T>A, 3221T>A, 3225C>G, 3226A>T, 3227A>T, 3228T>G, 3230T>A, 3236A>G, 3245A>T, 3249G>A, 3251G>T, 3254A>G, 3260T>A, 3263C>T, 3269C>T, 3272G>T, 3273T>C, 3275G>T, 3278A>C, 3279C>T, 3281G>A, 3284A>G, 3289G>C, 3293T>G, 3299T>C, 3305A>G, 3308T>G, 3309G>T, 3310T>G, 3311G>C, 3320C>A, 3323C>A, 3326C>T, 3335A>T, 3343G>A, 3344C>T, 3345G>A, 3346T>C, 3347A>C, 3350A>T, 3353T>A, 3356A>G, 3357C>A, 3358G>A, 3359T>G, 3362A>T, 3364T>A, 3365T>C, 3366A>G, 3368C>G, 3369T>G, 3377T>C, 3380T>A, 3386T>A, 3387A>G, 3389C>G, 3391A>T, 3392C>T, 3395G>A, 3398A>T, 3402C>T, 3404A>G, 3407G>A, 3416A>C, 3419T>G, 3422G>A, 3425G>C, 3426G>A, 3428C>T, 3433C>T, 3434A>G, 3437A>T, 3443A>G, 3445G>A, 3452T>A, 3453A>C, 3455T>G, 3456C>T, 3462T>C, 3463C>A, 3464A>C, 3465A>C, 3467T>G, 3471A>T, 3472G>C, 3475G>A, 3476A>G, 3479A>G, 3480A>G, 3481C>A, 3483C>A, 3484A>G, 3485G>A, 3486C>T, 3488T>C, 3497A>G, 3500C>A, 3503A>G, 3507T>G, 3508A>C, 3509C>T, 3511A>T, 3513C>T, 3515T>G, 3518A>C, 3521A>T, 3524T>A, 3527A>G, 3528C>T, 3529A>T, 3533G>A, 3534T>A, 3539C>T, 3542C>T, 3548T>G, 3551C>G, 3557G>A, 3560C>A, 3563T>C, 3566G>T, 3567C>A, 3568G>A, 3569G>A, 3570A>C, 3571A>G, 3572A>C, 3575C>T, 3577A>C, 3578A>C, 3579T>C, 3587A>G, 3590A>T, 3593C>T, 3596A>G, 3597C>G, 3598C>A, 3599T>C, 3602A>T, 3603G>C, 3608A>T, 3611A>C, 3624A>T, 3626A>T, 3628T>C, 3629T>C, 3630C>A, 3632T>A, 3641A>G, 3644T>A, 3647T>A, 3650T>G, 3656C>T, 3671G>T, 3674A>C, 3675A>G, 3676T>A, 3680A>G, 3683C>T, 3697G>C, 3698T>C, 3699G>A, 3700G>A, 3701A>C, 3707T>G, 3710T>G, 3711G>C, 3712A>C, 3713C>A, 3719C>A, 3722A>G, 3725A>G, 3728A>C, 3731C>T, 3734A>G, 3735A>G, 3736A>G, 3740C>T, 3743G>A, 3744T>G, 3745G>C, 3746C>A, 3749G>C, 3750C>A, 3752G>T, 3755A>T, 3759A>G, 3761C>G, 3762A>C, 3764G>T, 3765C>A, 3766G>A, 3767T>G, 3769G>T, 3770C>G, 3774A>C, 3775A>G, 3776G>A, 3777A>G, 3779C>G, 3782T>C, 3789C>A, 3790G>A, 3791T>A, 3794A>T, 3795T>G, 3796G>T, 3797T>A, 3798G>A, 3799A>T, 3800A>C, 3803G>A, 3804G>A, 3827A>G, 3838G>T, 3845C>T, 3847A>T, 3853G>C, 3854C>A, 3860G>A, 3861C>T, 3869A>G, 3872T>A, 3875C>T, 3878A>T, 3884C>T, 3887C>T, 3894A>T, 3899G>A, 3900A>C, 3902G>T, 3905G>T, 3906A>T, 3907T>C, 3908G>T, 3914G>A, 3917A>G, 3927C>G, 3930T>G, 3933A>C, 3934C>A, 3935T>A, 3936G>C, 3937G>C, 3938C>A, 3939G>A, 3941A>C, 3942G>T, 3943G>C, 3944A>G, 3945G>C, 3950A>G, 3953G>T, 3954G>C, 3955A>T, 3956A>T, 3965A>G, 3966G>A, 3971T>A, 3977A>C, 3983A>T, 3996C>T, 3998A>G, 4001T>C, 4002G>T, 4003T>C, 4008A>G, 4010T>A, 4014A>C, 4021T>C, 4022C>A, 4023C>T, 4024A>T, 4028G>T, 4031T>C, 4034G>C, 4040C>A, 4046A>G, 4049C>T, 4052C>G, 4064C>G, 4065C>T, 4067C>G, 4079A>G, 4082C>T, 4088C>G, 4094A>G, 4101A>C, 4103G>C, 4106C>G, 4109T>C, 4112G>A, 4115A>C, 4116C>T, 4118T>G, 4130C>T, 4137T>C, 4142C>G, 4143A>C, 4148G>A, 4151T>C, 4154T>C, 4160T>C, 4163C>T, 4166A>T, 4169C>T, 4170A>T, 4171C>G, 4172A>T, 4182T>G, 4183T>C, 4184G>A, 4190T>C, 4192C>T, 4193T>A, 4194A>C, 4196A>C, 4199A>T, 4205A>G, 4208T>A, 4214T>C, 4215G>A, 4220T>A, 4223A>G, 4226T>C, 4227T>C, 4228C>A, 4239T>G, 4242T>C, 4247A>T, 4249A>G, 4251G>A, 4253G>C, 4259A>G, 4260G>T, 4262T>A, 4265G>A, 4268A>T, 4271C>T, 4273T>A, 4280T>C, 4286T>C, 4292T>C, 4295C>G, 4316T>C, 4317G>A, 4319T>C, 4325T>A, 4328T>C, 4331G>A, 4344T>C, 4354A>C, 4355A>T, 4357A>T, 4358C>T, 4361C>T, 4364A>T, 4365T>G, 4366C>A, 4367A>T, 4368A>G, 4376C>G, 4385T>G, 4388C>A, 4395C>A, 4397A>G, 4400T>C, 4401G>A, 4404C>T, 4406T>A, 4418A>C, 4424T>C, 4430C>T, 4433A>C, 4436G>C, 4439C>T, 4443G>A, 4445T>C, 4448A>G, 4449G>T, 4455A>C, 4457G>C, 4458T>C, 4460G>A, 4463T>G, 4466G>T, 4469C>A, 4475A>C, 4478A>T, 4481A>T, 4487A>T, 4488C>T, 4490A>G, 4493G>T, 4499C>A, 4502C>T, 4508T>A, 4509C>T, 4511C>G, 4514C>A, 4530C>A, 4532A>G, 4535T>C, 4538C>G, 4548T>C, 4553A>G, 4556A>G, 4562C>A, 4566A>G, 4568T>C, 4569G>A, 4570C>A, 4571T>A, 4576G>A, 4577G>A, 4578G>T, 4581A>G, 4583A>G, 4586A>C, 4587G>A, 4592A>G, 4597G>A, 4601T>C, 4611C>T, 4613A>G, 4622T>A, 4625T>A, 4631A>T, 4637C>T, 4640T>C, 4641C>T, 4643T>G, 4649T>G, 4652A>T, 4656T>A, 4658A>G, 4662C>T, 4664C>G, 4665C>A, 4666A>G, 4673T>C, 4674T>G, 4675C>A, 4679G>A, 4683A>T, 4684A>C, 4685T>C, 4688A>T, 4692C>A, 4693A>G, 4694T>A, 4697G>A, 4698T>C, 4700A>G, 4704C>G, 4705T>C, 4706G>T, 4709C>T, 4712T>C, 4715A>C, 4716A>G, 4731C>T, 4733T>G, 4739A>G, 4742G>A, 4751T>A, 4752A>G, 4754T>G, 4755G>A, 4758G>A, 4759T>C, 4761C>A, 4768A>T, 4769A>G, 4784A>C, 4787T>C, 4793G>C, 4799A>G, 4805T>C, 4808A>G, 4817T>A, 4818C>T, 4820T>G, 4826C>T, 4829C>A, 4835T>A, 4838T>A, 4844G>A, 4854A>G, 4859T>C, 4868G>T, 4877A>T, 4880A>G, 4883T>A, 4886C>A, 4887C>G, 4888C>T, 4889A>G, 4892T>A, 4901C>T, 4902A>G, 4903C>T, 4904T>A, 4907G>A, 4910C>T, 4914A>T, 4915T>A, 4916G>T, 4923G>A, 4925G>T, 4934A>C, 4937A>C, 4938C>A, 4940T>A, 4946T>C, 4955C>T, 4958T>G, 4960G>A, 4961T>G, 4963T>C, 4964G>C, 4965G>A, 4973C>A, 4975A>G, 4976G>T, 4977C>T, 4982C>A, 4983T>A, 4984T>A, 4985A>T, 4992C>T, 4993A>T, 4994G>T, 4995A>G, 4996C>G, 4997G>A, 5000G>T, 5001T>C, 5003G>C, 5004G>A, 5006T>G, 5007G>C, 5009A>G, 5012G>A, 5013G>A, 5014G>A, 5016T>C, 5019A>C, 5020G>C, 5021A>C, 5022G>A, 5023C>T, 5024T>C, 5025G>A, 5031T>A, 5033T>G, 5036T>C, 5037T>A, 5039C>T, 5042T>C, 5044T>A, 5045G>T, 5049G>C, 5051G>T, 5055C>A, 5056A>G, 5057G>T, 5058G>C, 5064A>T, 5066T>G, 5067G>A, 5070A>G, 5075A>C, 5081G>A, 5085T>G, 5086C>A, 5087A>G, 5090T>C, 5091G>A, 5092C>T, 5093C>T, 5094C>G, 5095T>G, 5097A>G, 5098C>T, 5099A>T, 5100A>T, 5102T>G, 5103T>A, 5104G>T, 5105T>G, 5108A>G, 5109A>C, 5111G>A, 5112G>T, 5115A>G, 5117A>T, 5120A>T, 5121G>A, 5122C>A, 5123T>A, 5125T>A, 5127T>A, 5129C>A, 5132T>C, 5135G>C, 5138A>T, 5141G>A, 5142G>C, 5143A>C, 5144C>T, 5147A>T, 5150T>G, 5153G>A, 5159C>T, 5160_5162delACA, 5172C>G, 5173G>A, 5174T>G, 5176A>G, 5178C>G, 5180G>C, 5181A>C, 5184T>G, 5185G>A, 5186G>A, 5191C>T, 5192A>G, 5195A>T, 5196A>G, 5198C>T, 5199A>G, 5201T>A, 5202G>T, 5203G>T, 5207T>C, 5211T>A, 5212C>G, 5213A>T, 5216A>T, 5219T>G, 5226G>T, 5228C>A, 5229C>A, 5232T>C, 5234C>A, 5235A>C, 5236C>A, 5237G>A, 5241G>T, 5242A>C, 5243T>A, 5244G>T, 5245A>C, 5249G>T, 5250T>A, 5255T>C, 5256A>C, 5258T>G, 5264C>T, 5267T>C, 5270A>T, 5273T>A, 5276A>C, 5277A>C, 5278A>G, 5279G>C, 5281A>G, 5282G>A, 5283T>A, 5285C>A, 5287A>C, 5288T>A, 5294T>G, 5300T>G, 5303A>T, 5309C>T, 5333G>A, 5339G>C, 5346A>T, 5349T>G, 5354T>A, 5357A>C, 5358A>C, 5359T>A, 5364G>C, 5365C>A, 5366C>T, 5370A>C, 5371A>G, 5372G>C, 5375T>G, 5376G>A, 5377T>A, 5378G>T, 5379T>C, 5383C>A, 5386G>T, 5387A>T, 5390A>T, 5391C>G, 5395T>G, 5396A>G, 5399A>T, 5403A>T, 5404A>C, 5409G>A, 5410T>C, 5411T>C, 5417G>T, 5423A>T, 5429A>T, 5435T>A, 5436G>A, 5438C>A, 5440G>C, 5441C>T, 5447T>C, 5456G>A, 5457C>T, 5460A>T, 5462G>T, 5465C>A, 5468A>G, 5471A>C, 5474A>T, 5475A>C, 5476G>A, 5477G>T, 5483T>C, 5484T>A, 5486T>G, 5487G>A, 5488T>C, 5489G>C, 5490G>A, 5491A>C, 5492G>A, 5496A>C, 5498A>T, 5501A>T, 5504T>G, 5505A>C, 5507A>C, 5508T>C, 5510G>A, 5513C>A, 5516A>C, 5519T>G, 5521G>C, 5522A>T, 5523A>G, 5524A>G, 5525T>G, 5531T>G, 5534C>T, 5537C>G, 5540T>C, 5546T>C, 5555C>T, 5560T>C, 5564G>A, 5567T>C, 5570T>G, 5573T>C, 5574C>A, 5582G>T, 5583C>G, 5584C>A, 5585T>A, 5586C>G, 5588T>C, 5588_5589insACC, 5589G>T, 5590G>C, 5592G>C, 5597A>G, 5598G>T, 5602C>G, 5603T>A, 5606A>G, 5612T>C, 5613G>A, 5617C>A, 5618G>T, 5619C>G, 5620T>G, 5622_5624delCAC, 5626G>C, 5627T>C, 5629G>C, 5635T>A, 5636G>C, 5642A>T, 5642_5643insCACAGT, 5644G>C, 5645G>A, 5649G>T, 5650A>C, 5652T>G, 5654G>T, 5657C>A, 5658A>C, 5659A>T, 5660C>T, 5663C>A, 5664C>A, 5666C>G, 5669T>A, 5670A>G, 5672A>T, 5674A>C, 5676G>T, 5677G>C, 5678T>C, 5681T>C, 5682T>C, 5684A>C, 5685G>T, 5687A>T, 5688A>G, 5691C>A, 5695C>T, 5696A>G, 5700A>G,

| | |
|---|---|
| | 5702G>T, 5705G>T, 5706A>T, 5714T>C, 5715G>A, 5716T>C, 5717G>A, 5718G>A, 5720A>T, 5723A>T, 5726A>T, 5727C>G, 5729T>A, 5735A>T, 5740G>C, 5741A>C, 5742A>C, 5744T>G, 5746T>G, 5747T>C, 5750C>A, 5753A>G, 5754C>A, 5757G>A, 5759T>C, 5762T>G, 5763G>A, 5766A>C, 5767G>A, 5769T>G, 5770T>C, 5771G>T, 5772C>T, 5774A>G, 5775C>G, 5777C>G, 5778T>A, 5779A>C, 5780T>A, 5784G>T, 5785A>C, 5786G>T, 5787C>G, 5788A>T, 5789G>C, 5790G>A, 5792A>T, 5798A>T, 5799G>A, 5800A>C, 5801G>T, 5802T>C, 5804A>T, 5805A>G, 5806T>C, 5807T>C, 5808C>G, 5809A>C, 5812A>G, 5813G>A, 5814A>G, 5815A>G, 5816T>A, 5817A>G, 5818C>T, 5819A>C, 5823C>G, 5825A>G, 5828T>A, 5829C>T, 5837C>T, 5843G>A, 5844C>T, 5846G>C, 5847A>T, 5848G>C, 5849T>C, 5850A>C, 5851T>A, 5852G>A, 5853G>C, 5855A>C, 5856A>G, 5859G>T, 5860T>G, 5861G>T, 5862C>G, 5865C>A, 5866A>T, 5867A>G, 5868C>A, 5870G>A, 5871A>C, 5873A>T, 5876T>G, 5880G>C, 5882T>G, 5883A>G, 5885G>A, 5886C>A, 5887T>G, 5891G>A, 5895T>G, 5897T>A, 5898T>G, 5900C>A, 5901C>T, 5902C>T, 5903A>C, 5905A>T, 5906A>G, 5908A>G, 5909A>G, 5911C>G, 5912T>C, 5913A>G, 5914C>T, 5915C>T, 5917T>A, 5918T>G, 5919C>A, 5921G>A, 5922A>C, 5924G>A, 5925C>A, 5927C>G, 5928A>T, 5929A>T, 5930A>G, 5931A>T, 5932A>C, 5933T>C, 5936T>C, 5937G>A, 5939T>G, 5940G>A, 5943G>C, 5944C>A, 5945T>A, 5946G>C, 5948A>G, 5949T>A, 5950G>T, 5951T>G, 5952C>T, 5953T>C, 5956C>A, 5958T>A, 5959T>C, 5961A>C, 5965C>T, 5966A>T, 5967A>G, 5968G>C, 5969G>C, 5972_5973insCCATGTTGTGTCACA, 5978G>T, 5979C>A, 5981C>A, 5983T>C, 5985G>T, 5986T>G, 5987T>G, 5988T>C, 5990C>A, 5992A>C, 5993A>T, 5997A>G, 5999A>C, 6000C>A, 6003T>G, 6005G>C, 6007G>T, 6008T>G, 6009T>A, 6011C>A, 6013C>A, 6015T>G, 6017G>T, 6020A>G, 6022T>A, 6024T>C, 6025G>C, 6026T>A, 6029T>A, 6030_6032delGAA, 6033A>T, 6034G>C, 6039_6050delGTTGCTTCAAGG, 6051T>C, 6056A>T, 6059G>A, 6061G>C, 6062T>A, 6063C>G, 6065A>G, 6068C>T, 6070C>T, 6074C>G, 6078T>C, 6079T>C, 6080G>T, 6083T>A, 6086T>A, 6087T>C, 6088C>T, 6089C>A, 6092A>T, 6093T>G, 6097A>G, 6102T>A, 6104A>G, 6108G>C, 6109A>G, 6110A>C, 6112G>A, 6113A>T, 6114C>A, 6115A>G, 6116C>T, 6117A>G, 6118C>G, 6122A>G, 6126A>T, 6127T>A, 6128G>C, 6129A>T, 6130G>C, 6131T>A, 6135C>A, 6136A>C, 6137G>A, 6139G>C, 6140T>A, 6141A>C, 6146G>A, 6153C>T, 6155G>T, 6157A>G, 6161A>G, 6164T>C, 6165C>A, 6167A>G, 6169T>G, 6170A>T, 6171A>C, 6173C>G, 6174T>G, 6175T>A, 6176C>A, 6177C>G, 6178A>G, 6179A>G, 6180G>T, 6181T>A, 6182A>T, 6183A>T, 6186A>G, 6187C>A, 6188A>T, 6190G>A, 6191T>A, 6193A>C, 6194G>C, 6198T>G, 6200A>G, 6201A>G, 6203G>C, 6204A>C, 6205A>G, 6206G>C, 6209T>C, 6210G>A, 6212A>T, 6215T>G, 6219T>A, 6221T>G, 6222C>G, 6223A>G, 6224G>C, 6226A>T, 6227G>T, 6228A>T, 6229A>G, 6230G>T, 6231A>G, 6233A>G, 6236C>G, 6237A>G, 6238C>A, 6239C>G, 6240A>C, 6241C>G, 6243C>A, 6245A>C, 6246C>T, 6248A>G, 6249A>G, 6251C>G, 6252A>C, 6253A>G, 6254A>C, 6255G>A, 6256T>C, 6258T>A, 6260C>T, 6261T>G, 6262T>C, 6263G>T, 6270G>T, 6271A>C, 6272G>C, 6273G>A, 6274A>C, 6276A>T, 6277G>C, 6278T>A, 6280A>T, 6281T>G, 6282C>A, 6284G>C, 6285A>G, 6286C>A, 6287T>G, 6290G>T, 6294T>A, 6295G>C, 6297A>G, 6299A>G, 6300G>T, 6302T>C, 6303T>A, 6305T>A, 6306C>G, 6308G>T, 6310T>G, 6314C>T, 6315C>T, 6316A>T, 6317G>T, 6318A>G, 6319G>A, 6320G>T, 6321_6323delATA, 6324A>C, 6326C>T, 6328C>T, 6330C>A, 6333A>G, 6334A>C, 6336T>G, 6338C>G, 6339A>G, 6340G>T, 6341A>T, 6342G>T, 6344G>C, 6345C>A, 6348G>C, 6349A>C, 6350A>T, 6353T>G, 6359T>C, 6361A>C, 6364T>G, 6366C>G, 6368T>A, 6369C>A, 6370C>T, 6371T>C, 6372G>A, 6374A>T, 6376G>A, 6377A>T, 6378A>G, 6379T>C, 6383C>T, 6392C>T, 6394T>C, 6402A>G, 6404A>G, 6405A>T, 6407A>G, 6411T>G, 6412C>A, 6413T>A, 6416G>C, 6420G>A, 6421A>G, 6423T>C, 6424C>A, 6427A>G, 6428T>G, 6429A>T, 6430G>C, 6431C>A, 6432C>A, 6433G>A, 6434A>G, 6435G>T, 6436A>C, 6437T>A, 6438A>C, 6440T>A, 6442A>G, 6444T>A, 6445C>A, 6446C>G, 6448A>T, 6449T>C, 6450T>A, 6453G>T, 6455G>T, 6456T>G, 6458A>C, 6460C>A, 6462C>G, 6463C>A, 6464C>A, 6465A>G, 6467C>T, 6468A>G, 6471C>A, 6473G>A, 6474G>T, 6475G>C, 6476G>A, 6477T>A, 6478G>A, 6479T>C, 6480T>C, 6482C>G, 6483A>C, 6488C>A, 6489C>G, 6490T>C, 6494C>T, 6495G>T, 6496T>A, 6497G>T, 6498C>T, 6503T>A, 6505G>A, 6507C>A, 6509A>G, 6510G>C, 6511G>T, 6514A>T, 6515G>A, 6517A>G, 6518A>T, 6519G>C, 6520A>T, 6521T>A, 6522C>G, 6524T>G, 6525C>T, 6526C>A, 6528G>T, 6529T>A, 6531C>G, 6532G>A, 6533G>T, 6534A>T, 6535G>C, 6536G>T, 6537T>A, 6539T>A, 6540T>G, 6541C>A, 6543A>T, 6545G>T, 6546G>T, 6548T>G, 6549T>G, 6550T>A, 6552C>T, 6554T>G, 6555A>T, 6556A>T, 6558A>C, 6560C>A, 6561T>A, 6562G>T, 6563T>A, 6566A>T, 6572T>A, 6576A>G, 6582T>G, 6583A>T, 6588G>T, 6590G>C, 6594T>G, 6596A>G, 6597G>T, 6598T>C, 6599_6600delCA, 6603C>G, 6606T>A, 6609G>A, 6610A>G, 6611C>A, 6612T>C, 6614A>C, 6620T>C, 6621G>A, 6624A>T, 6624_6625insG, 6626A>T, 6629C>T, 6643G>A, 6647_6648insACAAACAT, 6649T>A, 6652A>G, 6654C>T, 6655G>T, 6657T>G, 6658A>T, 6659G>A, 6660G>T, 6661A>C, 6663G>A, 6664G>C, 6667T>G, 6669G>A, 6672A>C, 6683C>A, 6687C>T, 6688T>C, 6692A>G, 6695A>G, 6696G>A, 6715T>C |

**CDS**

| | | | | | | | | | |
|---|---|---|---|---|---|---|---|---|---|
| ToMarV_RNA1gp1 | 1 | 2152 | 100% | 9871 | 66.5% | 2135 (98.7%) | 1362 (63.0%) | 11/17/0/0 | 0 |

| | Begin | End | Coverage | Score | Concordance | Matches | Identities | I/D/M/F* | Stop Codons |
|---|---|---|---|---|---|---|---|---|---|
| **NT** | **22** | **6747** | **93.0%** | **3341** | **25.4%** | **6659 (98.4%)** | **4235 (62.6%)** | **43/67** | |

Protein mutations:

F7W (160T>G 161C>G), K9_A16del (165_188delAAGGTTGCTGGTGTTGATGACGCC), S17P (189A>C 190G>C), S18P (192T>C), S19A (195T>G), T21S (201A>T 203C>A), T22P (204A>C), G26A (217G>C 218T>A), L28V (222C>G), N30S (228A>T 229A>C), I32F (234A>T), K34S (240A>T 241A>C 242A>T), F35L (243T>C), A38T (252G>A 254G>T), V39L (255G>C 257T>C), I42V (264A>G), T43N (268C>A 269C>T), T44K (271C>A 272T>A), M45V (273A>G 275G>T), K48T (283A>C 284G>C), S50V (288T>G 289C>T), L53I (297C>A 299T>A), L56V (306C>G 308G>C), S59P (315T>C 317T>A), T61M (322C>T 323T>G), D62A (325A>C 326T>A), V63A (328T>C 329G>A), S64A (330T>G 332C>A), V68A (343T>C 344G>C), A69S (345G>T 347C>T), S72N (355G>A), S73A (357T>G 359A>T), L74V (360C>G 362T>A), N75E (363A>G 365T>A), K76R (367A>G), G82A (385G>C 386T>A), A83K (387G>A 388C>A), L84I (390C>A 392T>A), I85M (395A>G), D86E (398T>A), L89I (405C>A 407T>C), A91G (412C>G 413G>T), S93D (417T>G 418C>A), L95V (423T>G 425G>T), S96K (426T>A 427C>A 428C>A), S101T (441T>A), K103V (447A>G 448A>T 449G>A), M105L (453A>C 455G>T), L109M (465T>A), F110V (468T>G), E111L (471G>T 472A>T 473G>A), L113V (477T>G 479G>C), S114K (480T>A 481C>A 482C>G), N115D (483A>G), V117F (489G>T 491T>C), K118I (493A>T 494A>C), F121Y (502T>A 503T>C), A122E (505C>A 506T>A), V124L (510G>T 512G>A), N126E (516A>G 518T>A), S128V (522A>G 523G>T), D130S (528G>T 529A>C 530T>A), V131L (531G>T), V135F (543G>T 545A>T), Y136I (546T>A 547A>T 548T>A), G137A (550G>C), I143L (567A>T 569A>G), T144A (570A>G 572A>C), L145I (573T>A 575G>T), V147L (579G>C 581G>A), F150Y (589T>A), S154V (600A>G 601G>T 602T>A), E158T (612G>A 613A>C), G159P (615G>C 616G>C), V160F (618G>T), S161N (621T>A 622C>A), T163V (627A>G 628C>T 629T>A), K165V (633A>G 634A>T 635A>T), N166T (637A>C), I167L (639A>T 641C>G), L169S (646T>C 647G>A), I170V (648A>G 650A>T), M174F (660A>T 662G>T), K186T (697A>C), A188S (702G>T 704C>T), E189G (706A>G 707A>G), R190K (708C>A 709G>A 710A>G), T192S (714A>T), M193Q (717A>C 718T>A), V194L (720G>C), S198A (732A>G 733G>C 734T>A), M199V (735A>G 737G>A), V205L (753G>T 755T>A), S206T (756T>A 758A>G), M214L (780A>T 782G>A), T216S (786A>T), I217L (789A>T 791A>G), I228V (822A>G), T248A (882A>G 884A>T), T260Q (918A>C 919C>A 920T>G), I280V (978A>G 980A>G), G282S (984G>A), T283I (988C>T 989C>T), S295N (1024G>A), E304Q (1050G>C), Q307T (1059C>A 1060A>C), D310S (1068G>T 1069A>C 1070T>G), A311E (1072C>A 1073A>G), C313A (1077T>G 1078G>C), D314E (1082T>A), A316S (1086G>T 1088A>T), T317M (1090C>T), K325P (1113A>C 1114A>C 1115G>T), T329I (1126C>T 1127C>T), N332Q (1134A>C 1136C>A), R335K (1144G>A 1145G>A), E337Q (1149G>C 1151A>G), L339V (1155C>G), M354L (1200A>T), A363T (1227G>A), E366D (1238A>T), V368L (1242G>C 1244T>G), Y369V (1245T>G 1246A>T), S372A (1254A>G 1255G>C 1256T>C), C373S (1257T>A 1259T>C), R374Q (1260A>C 1261G>A 1262G>A), M376S (1266A>T 1267T>C 1268G>T), S393A (1317T>G 1319C>T), S401C (1342C>G 1343C>T), R412A (1374A>G 1375G>C), I440V (1458A>G 1460C>G), M521R (1702T>G 1703G>A), E523V (1708A>T), E525D (1715G>T), S527D (1719T>G 1720C>A 1721C>T), T528P (1722A>C 1724T>A), E529N (1725G>A 1727A>T), D530T (1728G>A 1729A>C), G533A (1738G>C 1739A>C), R536V (1746A>G 1747G>T 1748A>T), N538D (1752A>G 1754C>T), I539V (1755A>G 1757A>G), N552A (1794A>G 1795A>C), T554S (1800A>T), S556D (1806A>G 1807G>A), E560Q (1818G>C 1820G>A), Q567R (1840A>G), A570Y (1848G>T 1849C>A 1850A>T), K574S (1861A>G 1862G>C), F577Y (1870T>A 1871T>C), E578N (1872G>A 1874A>T), A579T (1875G>A), E580T (1878G>A 1879A>C), I583L (1887A>T 1889A>G), S584A (1890A>G 1891G>C), P585D (1893C>G 1894C>A 1895T>C), I587V (1899A>G 1901A>T), Q591N (1911C>A 1913A>T), S593A (1917T>G), A594V (1921C>T 1922A>T), D596E (1928T>A), I598L (1932A>C 1934A>G), A599D (1936C>A 1937G>T), V601F (1941G>T 1943G>C), D602Q (1944G>C 1946T>A), G603D (1948G>A 1949G>T), R605del (1953_1955delAGA), R607Q (1960G>A 1961C>A), T608V (1962A>G 1963C>T 1964C>T), K609L (1965A>T 1966A>T), Q610P (1969A>C 1970A>T), D611P (1971G>C 1972A>C), I614A (1980A>G 1981T>C), L615A (1983T>G 1984T>C 1985G>T), V616Y (1986G>T 1987T>A 1988G>T), P617E (1989C>G 1990C>A 1991C>G), T618Q (1992A>C 1993C>A 1994T>G), V619T (1995G>A 1996T>C 1997T>A), V620I (1998G>A 2000G>T), S621T (2002G>C 2003T>A), N623A (2007A>G 2008A>C), D624E (2012T>A), R625K (2014G>A), V626I (2016G>A 2018G>T), I627V (2019A>G 2021C>T), A629S (2025G>T 2027A>C), R630Q (2028A>C 2029G>A 2030A>G), L645M (2073T>A), D646N (2076G>A), P647K (2079C>A 2080C>A 2081T>A), E648F (2082G>T 2083A>T 2084G>T), L650F (2088C>T), H655L (2104A>T 2105T>C), H656S (2106C>T 2107A>C 2108T>C), M660I (2120G>C), S663V (2127T>G 2128C>T 2129T>G), D665S (2133G>T 2134A>C), D666S (2136G>T 2137A>C), E667D (2141A>T), H668A (2142C>G 2143A>C 2144T>C), E670Q (2148G>C), I671N (2152T>A), N673E (2157A>G 2159C>A), K674M (2161A>T 2162A>G), R676V (2166A>G 2167G>T), V677N (2169G>A 2170T>A 2171C>T), N678D (2172A>G 2174C>T), M679L (2175A>T), Y681V (2181T>G 2182A>T 2183T>C), R682K (2185G>A), D685S (2193G>A 2194A>G), A686P (2196G>C), E687Q (2199G>C 2201A>G), R690A (2208C>G 2209G>C 2210A>T), Q694R (2221A>G), A710S (2268G>A 2269C>G 2270A>C), S711T (2271T>A), K716E (2286A>G), K717L (2289A>C 2290A>T 2291A>G), A719D (2296C>A 2297A>T), T722R (2304A>C 2305C>G 2306A>C), E723N (2307G>A 2309G>T), M724L (2310A>C 2312G>T), R725E (2313A>G 2314G>A), T726Q (2316A>C 2317C>A 2318T>A), F728K (2322T>A 2323T>A 2324T>A), H730Y (2328C>T), S731K (2331T>A 2332C>A 2333T>G), C732A (2334T>G 2335G>C 2336C>G), I733L (2337A>C), D734Q (2340G>C 2342T>G), V737A (2350T>C 2351T>A), I741T (2362T>C 2363A>G), F749R (2385T>A 2386T>G 2387T>A), N763R (2428A>G 2429C>G), S766A (2436T>G), S766_K767insG (2438_2439insGGT), K767T (2440A>C 2441A>T), E768D (2444G>C), P769K (2445C>A 2446C>A 2447A>G), E771R (2451G>A 2452A>G), S772N (2454T>A 2455C>A 2456A>T), K775I (2464A>T 2465A>C), I776E (2466A>G 2467T>A 2468T>A), Y778M (2472T>A 2473A>T 2474C>G), H779I (2475C>A 2476A>T), K780G (2478A>G 2479A>G 2480A>T), V782K (2484G>A 2485T>A 2486T>G), E783D (2489A>C), Y784K (2490T>A 2492T>A), I785M (2495C>G), K786I (2497A>T 2498G>C), D787Q (2499G>C 2501T>G), T788M (2503C>T 2504A>G), W790N (2508T>A 2509G>A 2510G>T), S791D (2511A>G 2512G>A), Y794F (2521A>T 2522T>C), A795I (2523G>A 2524C>T 2525A>C), R797K (2530G>A 2531A>G), I806L (2556A>T 2558A>G), M807I (2561G>A), V812T (2574G>A 2575T>C 2576T>C), F813M (2577T>A 2579T>G), G814N (2580G>A 2581G>A 2582A>T), G815A (2584G>C), I816V (2586A>G), L820I (2598T>A 2600G>T), A821T (2601G>A 2603G>T), W823L (2607T>C 2608G>T 2609G>C), Q824K (2610C>A), S825A (2613A>G 2614G>C 2615T>A), N826A (2616A>G 2617A>C 2618C>A), S827T (2619T>A), F828S (2623T>C), N830L (2628A>T 2629A>T 2630T>A), S832D (2634T>G 2635C>A 2636G>T), N852K (2696T>G), K853T (2698A>C 2699G>T), K853_L854insH (2699_2700insCAT), L854F (2700C>T), R855A (2703A>G 2704G>C), N856K (2708T>A), L857S (2709C>T 2710T>C 2711T>C), V859L (2715G>C 2717T>A), S862A (2724A>G 2725G>C 2726T>G), S863A (2727T>G 2729T>A), Q864D (2730C>G 2732A>T), S865N (2733T>A 2734C>A 2735A>C), L866N (2736T>A 2737T>A 2738G>C), A867H (2739G>C 2740C>A 2741A>T), Q868V (2742C>G 2743A>T 2744A>G), D869_W870insR (2747_2748insCGG), A872E (2755C>A 2756T>A), E873Q (2757G>C 2759A>G), Q879E (2775C>G 2777G>A), K882H (2784A>C 2786G>C), L890I (2808C>A), G891A (2812G>C), M907L (2859A>C 2861G>C), H914R (2880C>A 2881A>G 2882C>A), A916M (2886G>A 2887C>T 2888T>G), T923S (2907A>T 2909A>C), A927E (2920C>A 2921C>G), N928H (2922A>C), R930I (2928C>A 2929G>T 2930A>T), S931C (2932C>G), K932N (2936A>T), Y935F (2944A>T), R938T (2953G>C 2954A>C), E939Q (2955G>C), R941K (2962G>A), I942L (2964A>T 2966T>G), I948V (2982A>G 2984C>G), P961S (3021C>T 3023T>A), E972D (3056G>T), F975Y (3064T>A), T976S (3066A>T 3068A>T), Q980M (3078C>A 3079A>T 3080A>G), C984V (3090T>G 3091G>T 3092T>G), T990K (3109C>A 3110A>G), R993K (3117C>A 3118G>A 3119T>G), L994K (3120C>A 3121T>A 3122T>A), G995S (3123G>T 3124G>C), F999V (3135T>G), M1002Y (3144A>T 3145T>A 3146G>T), V1012E (3175T>A 3176T>A), E1015T (3183G>A 3184A>C), E1017G (3190A>G), N1019D (3195A>G 3197C>T), S1021N (3201T>A 3202C>A 3203C>T), S1022A (3204A>G 3205G>C 3206T>A), K1024A (3210A>G 3211A>C 3212G>T), I1025L (3213A>C), C1026S (3216T>A), Q1029V (3225C>G 3226A>T 3227A>T), S1030A (3228T>G 3230T>A), G1037S (3249G>A 3251G>T), S1050T (3289G>C), V1057C (3309G>T 3310T>G 3311G>C), G1068D (3343G>A 3344C>T), V1069T (3345G>A 3346T>C 3347A>C), R1073K (3357C>A 3358G>A 3359T>G), F1075Y (3364T>A 3365T>C), I1076V (3366A>G 3368C>G), S1077A (3369T>G), I1083V (3387A>G 3389C>G), Y1084F (3391A>T 3392C>T), E1086D (3398A>T), V1096I (3426G>A 3428C>T), A1098V (3433C>T 3434A>G), R1102H (3445G>A), D1104E (3452T>A), I1105L (3453A>C 3455T>G), S1108H (3462T>C 3463C>A 3464A>C), I1109L (3465A>C 3467T>G), R1112K (3475G>A 3476A>G), T1114E (3480A>G 3481C>A), Q1115R (3483C>A 3484A>G 3485G>A), L1116F (3486C>T 3488T>C), N1120K (3500C>A), Y1123A (3507T>G 3508A>C 3509C>T), Y1124F (3511A>T), Q1130L (3528C>T 3529A>T), L1132M (3534T>A), R1143K (3567C>A 3568G>A 3569G>A), K1144R (3570A>C 3571A>G 3572A>C), Q1146P (3577A>C 3578A>C), P1153D (3597C>G 3598C>A 3599T>C), E1155Q (3603G>C), T1162S (3624A>T 3626A>T), V1163A (3628T>C 3629T>C), H1164K (3630C>A 3632T>A), D1169E (3647T>A), M1179E (3675A>G 3676T>A), C1186S (3697G>C 3698T>C), G1187N (3699G>A 3700G>A 3701A>C), D1189E (3707T>G), D1190E (3710T>G), D1191P (3711G>C 3712A>C 3713C>A), N1199G (3735A>G 3736A>G), C1202A (3744T>G 3745G>C 3746C>A), L1204I (3750C>A 3752G>T), I1207V (3759A>G 3761C>G), M1208L (3762A>C 3764G>T), R1209K (3765C>A 3766G>A 3767T>G), S1210M (3769G>T 3770C>G), K1212R (3774A>C 3775A>G 3776G>A), I1213V (3777A>G 3779C>G), R1217K (3789C>A 3790G>A 3791T>A), C1219V (3795T>G 3796G>T 3797T>A), E1220I (3798G>A 3799A>T 3800A>C), D1222N (3804G>A), C1233F (3838G>T), Y1236F (3847A>T), G1238A (3853G>C 3854C>A), M1252L (3894A>T), M1254L (3900A>C 3902G>T), M1256S (3906A>T 3907T>C 3908G>T), Q1263E (3927C>G), S1264A (3930T>G), T1265Q (3933A>C 3934C>A 3935T>A), G1266P (3936G>C 3937G>C 3938C>A), E1267N (3939G>A 3941A>C), G1268S (3942G>T 3943G>C 3944A>G), E1269Q (3945G>C), E1272L (3954G>C 3955A>T 3956A>T), D1276N (3966G>A), E1281D (3983A>T), V1288S (4002G>T 4003T>C), I1290V (4008A>G 4010T>A), K1292Q (4014A>C), I1294T (4021T>C 4022C>A), H1295F (4023C>T 4024A>T), T1344C (4170A>T 4171C>G 4172A>T), L1348A (4182T>G 4183T>C 4184G>A), A1351V (4192C>T 4193T>A), I1352L (4194A>C 4196A>C), D1359N (4215G>A), S1363H (4227T>C 4228C>A), L1367V (4239T>G), N1370S (4249A>G), G1371S (4251G>A 4253G>C), A1374S (4260G>T 4262T>A), V1378E (4273T>A), V1393I (4317G>A 4319T>C), E1405A (4354A>C 4355A>T), Y1406F (4357A>T 4358C>T), S1409D (4365T>G 4366C>A 4367A>T), K1410E (4368A>G), D1415E (4385T>G), A1421T (4401G>A), D1435N (4443G>A 4445T>C), V1437L (4449G>T), I1476V (4566A>G 4568T>C), A1477K (4569G>A 4570C>A 4571T>A), R1479K (4576G>A 4577G>A), A1480S (4578G>T), I1481V (4581A>G 4583A>G), V1483I (4587G>A), S1486N (4597G>A), L1506M (4656T>A 4658A>G), Q1509R (4665C>A 4666A>G), S1512D (4674T>G 4675C>A), N1515S (4683A>T 4684A>C 4685T>C), H1518R (4692C>A 4693A>G 4694T>A), L1522A (4704C>G 4705T>C 4706G>T), I1526V (4716A>G), N1538E (4752A>G 4754T>G), E1539K (4755G>A), V1540T (4758G>A 4759T>C), Q1541K (4761C>A), Q1543L (4768A>T 4769A>G), H1563Q (4829C>A), I1572V (4854A>G), M1576I (4868G>T), P1583V (4887C>G 4888C>T 4889A>G), T1588V (4902A>G 4903C>T 4904T>A), M1592Y (4914A>T 4915T>A 4916G>T), V1595I (4923G>A 4925G>T), R1607Q (4960G>A 4961T>G), V1608A (4963T>C 4964G>C), A1609T (4965G>A), D1611E (4973C>A), K1612S (4975A>G 4976G>T), L1613F (4977C>T), L1615N (4983T>A 4984T>A 4985A>T), Q1618F (4992C>T 4993A>T 4994G>T), T1619G (4995A>G 4996C>G 4997G>A), D1622K (5004G>A 5006T>G), E1623Q (5007G>C 5009A>G), G1625N (5013G>A 5014G>A), F1626L (5016T>C), R1627P (5019A>C 5020G>C 5021A>C), A1628I (5022G>A 5023C>T 5024T>C), V1629M (5025G>A), F1631M (5031T>A 5033T>G), S1633T (5037T>A 5039C>T), L1635Y (5044T>A 5045G>T), V1637L (5049G>C 5051G>T), Q1639S (5055C>A 5056A>G 5057G>T), E1640Q (5058G>C), I1642L (5064A>T 5066T>G), A1643T (5067G>A), K1644E (5070A>G), S1649E (5085T>G 5086C>A 5087A>G), A1651I (5091G>A 5092C>T 5093C>T), L1652G (5094C>G 5095T>G), T1653V (5097A>G 5098C>T 5099A>T), I1654L (5100A>T 5102T>G), C1655M (5103T>A 5104G>T 5105T>G), M1657L (5109A>C 5111G>A), A1658S (5112G>T), K1659D (5115A>G 5117A>T), A1661K (5121G>A 5122C>A 5123T>A), I1662N (5125T>A), S1663T (5127T>A 5129C>A), D1668P (5142G>C 5143A>C 5144C>T), T1674del (5160_5162delACA), R1678E (5172C>G 5173G>A 5174T>G), D1679G (5176A>G), L1680V (5178C>G 5180G>C), N1681H (5181A>C), W1682E (5184T>G 5185G>A 5186G>A), A1684V (5191C>T 5192A>G), N1686D (5196A>G 5198C>T), I1687V (5199A>G 5201T>A), G1688F (5202G>T 5203G>T), A1696S (5226G>T 5228C>A), H1697N (5229C>A), Y1698Q (5232T>C 5234C>A), T1699Q (5235A>C 5236C>A 5237G>A), D1701S (5241G>T 5242A>C 5243T>A), E1702S (5244G>T 5245A>C), Y1704N (5250T>A), I1706L (5256A>C 5258T>G), K1713R (5277A>C 5278A>G 5279G>C), K1714R (5281A>G 5282G>A), S1715T (5283T>A 5285C>A), N1716T (5287A>C 5288T>A), M1731I (5333G>A), M1736L (5346A>T), S1737A (5349T>G), M1740Q (5358A>C 5359T>A), A1742H (5364G>C 5365C>A 5366C>T), K1744R (5370A>C 5371A>G 5372G>C), N1745K (5375T>G), V1746N (5376G>A 5377T>A 5378G>T), A1748E (5383C>A), G1749V (5386G>T 5387A>T), H1751D (5391C>G), V1752G (5395T>G 5396A>G), N1755S (5403A>T 5404A>C), V1757T (5409G>A 5410T>C 5411T>C), V1766I (5436G>A 5438C>A), G1767A (5440G>C 5441C>T), M1774F (5460A>T 5462G>T), R1779H (5475A>C 5476G>A 5477G>T), C1782R (5484T>A 5486T>G), V1783T (5487G>A 5488T>C 5489G>C), E1784T (5490G>A 5491A>C 5492G>A), I1786L (5496A>C 5498A>T), I1793M (5519T>G), G1794A (5521G>C 5522A>T), N1795G (5523A>G 5524A>G 5525T>G), V1807A (5560T>C), F1810L (5570T>G), Q1812K (5574C>A), P1815E (5583C>G 5584C>A 5585T>A), H1816D (5586C>G 5588T>C), H1816_G1817insT (5588_5589insACC), G1817S (5589G>T 5590G>C), D1818H (5592G>C), V1820L (5598G>T), A1821G (5602C>G 5603T>A), A1825T (5613G>A), S1826Y (5617C>A 5618G>T), L1827G (5619C>G 5620T>G), H1828del (5622_5624delCAC), S1829T (5626G>C 5627T>C), S1830T (5629G>C), M1832N (5635T>A 5636G>C), Q1834H (5642A>T), Q1834_G1835insHS (5642_5643insCACAGT), G1835A (5644G>C 5645G>A), E1837S (5649G>T 5650A>C), L1838V (5652T>G 5654G>T), N1840L (5658A>C 5659A>T 5660C>T), I1844V (5670A>G 5672A>T), H1845P (5674A>C), G1846S (5676G>T 5677G>C 5678T>C), A1849S (5685G>T 5687A>T), K1850E (5688A>G), Q1851K (5691C>A), T1852M (5695C>T 5696A>G), M1854V (5700A>G 5702G>T), M1856L (5706A>T), V1859T (5715G>A 5716T>C 5717G>A), V1860I (5718G>A 5720A>T), H1863E (5727C>G 5729T>A), G1867A (5740G>C 5741A>C), N1868Q (5742A>C 5744T>G), F1869C (5746T>G 5747T>C), Q1872K (5754C>A), D1873N (5757G>A 5759T>C), E1875K (5763G>A), S1876H (5766A>C 5767G>A), L1877A (5769T>G 5770T>C 5771G>T), H1879E (5775C>G 5777C>G), Y1880T (5778T>A 5779A>C 5780T>A), E1882S (5784G>T 5785A>C 5786G>T), Q1883V (5787C>G 5788A>T 5789G>C), G1884S (5790G>A 5792A>T), E1887T (5799G>A 5800A>C 5801G>T), I1889A (5805A>G 5806T>C 5807T>C), Q1890A (5808C>G 5809A>C), K1891R (5812A>G 5813G>A), N1892G (5814A>G 5815A>G 5816T>A), T1893V (5817A>G 5818C>T 5819A>C), L1895V (5823C>G 5825A>G), L1902F (5844C>T 5846G>C), M1904Q (5850A>C 5851T>A 5852G>A), E1905H (5853G>C 5855A>C), N1906D (5856A>G), V1907C (5859G>T 5860T>G 5861G>T), Q1908E (5862C>G), Q1909M (5865C>A 5866A>T 5867A>G), L1910I (5868C>A 5870G>A), I1911L (5871A>C 5873A>T), D1912E (5876T>G), V1914L (5880G>C 5882T>G), K1915E (5883A>G 5885G>A), L1916S (5886C>A 5887T>G), F1919V (5895T>G 5897T>A), F1920V (5898T>G 5900C>A), P1921F (5901C>T 5902C>T 5903A>C), K1922M (5905A>T 5906A>G), E1923G (5908A>G 5909A>G), T1924S (5911C>G 5912T>C), T1925V (5913A>G 5914C>T 5915C>T), I1926K (5917T>A 5918T>G), Q1927K (5919C>A 5921G>A), K1928Q (5922A>C 5924G>A), L1929M (5925C>A 5927C>G), K1930L (5928A>T 5929A>T 5930A>G), N1931S (5931A>T 5932A>C 5933T>C), V1933M (5937G>A 5939T>G), D1934N (5940G>A), A1935Q (5943G>C 5944C>A 5945T>A), E1936Q (5946G>C 5948A>G), C1937M (5949T>A 5950G>T 5951T>G), L1938S (5952C>T 5953T>C), T1939K (5956C>A), F1940T (5958T>A 5959T>C), K1941Q (5961A>C), A1942V (5965C>T 5966A>T), R1943A (5967A>G 5968G>C 5969G>C), S1944_F1945insPCCVT (5972_5973insCCATGTTGTGTCACA), R1946S (5978G>T), H1947K (5979C>A 5981C>A), V1948A (5983T>C), V1949W (5985G>T 5986T>G 5987T>G), F1950L (5988T>C 5990C>A), K1951T (5992A>C 5993A>T), K1953D (5997A>G 5999A>C), L1954I (6000C>A), L1955V (6003T>G 6005G>C), S1956M (6007G>T 6008T>G), S1957T (6009T>A 6011C>A), T1958K (6013C>A), W1959G (6015T>G 6017G>T), M1961K (6022T>A), C1962P (6024T>C 6025G>C 6026T>A), E1964del (6030_6032delGAA), V1967_R1970del (6039_6050delGTTGCTTCAAGG), S1971P (6051T>C), S1974T (6061G>C 6062T>A), L1975V (6063C>G 6065A>G), P1977L (6070C>T), L1980P (6078T>C 6079T>C 6080G>T), S1983L (6087T>C 6088C>T 6089C>A), L1985V (6093T>G), K1986R (6097A>G), L1988M (6102T>A 6104A>G), E1990R (6108G>C 6109A>G 6110A>C), R1991N (6112G>A 6113A>T), H1992S (6114C>A 6115A>G 6116C>T), T1993G (6117A>G 6118C>G), M1996Y

(6126A>T 6127T>A 6128G>C), Q1999T (6135C>A 6136A>C 6137G>A), S2000T (6139G>C 6140T>A), M2001L (6141A>C), L2005F (6153C>T 6155G>T), K2006R (6157A>G), I2007M (6161A>G), L2009M (6165C>A 6167A>G), L2010C (6169T>G 6170A>T), N2011Q (6171A>C 6173C>G), F2012E (6174T>G 6175T>A 6176C>A), Q2013G (6177C>G 6178A>G 6179A>G), V2014Y (6180G>T 6181T>A 6182A>T), I2015L (6183A>T), T2016D (6186A>G 6187C>A 6188A>T), S2017K (6190G>A 6191T>A), E2018A (6193A>C 6194G>C), L2020V (6198T>G 6200A>G), K2021D (6201A>G 6203G>C), K2022R (6204A>C 6205A>G 6206G>C), E2024N (6210G>A 6212A>T), D2025E (6215T>G), F2027M (6219T>A 6221T>G), Q2028G (6222C>G 6223A>G 6224G>C), E2029V (6226A>T 6227G>T), K2030C (6228A>T 6229A>G 6230G>T), I2031V (6231A>G 6233A>G), T2033E (6237A>G 6238C>A 6239C>G), T2034R (6240A>C 6241C>G), L2035I (6243C>A 6245A>C), I2037V (6249A>G 6251C>G), K2038R (6252A>C 6253A>G 6254A>C), V2039T (6255G>A 6256T>C), F2040I (6258T>A 6260C>T), L2041A (6261T>G 6262T>C 6263G>T), E2044S (6270G>T 6271A>C 6272G>C), E2045T (6273G>A 6274A>C), Y2047L (6280A>T 6281T>G), Q2048N (6282C>A 6284G>C), T2049E (6285A>G 6286C>A 6287T>G), E2050D (6290G>T), C2052T (6294T>A 6295G>C), I2053V (6297A>G 6299A>G), D2054Y (6300G>T 6302T>C), S2055T (6303T>A 6305T>A), Q2056D (6306C>G 6308G>T), V2057G (6310T>G), Q2059F (6315C>T 6316A>T 6317G>T), R2060D (6318A>G 6319G>A 6320G>T), I2061del (6321_6323delATA), N2062H (6324A>C 6326C>T), S2063F (6328C>T), Q2064K (6330C>A), K2065A (6333A>G 6334A>C), F2066V (6336T>G 6338C>G), R2067V (6339A>G 6340G>T 6341A>T), V2068F (6342G>T 6344G>C), Q2069K (6345C>A), E2070P (6348G>C 6349A>C 6350A>T), Y2074S (6361A>C), L2075W (6364T>G), H2076E (6366C>G 6368T>A), P2077I (6369C>A 6370C>T 6371T>C), E2078N (6372G>A 6374A>T), G2079D (6376G>A 6377A>T), I2080A (6378A>G 6379T>C), V2085A (6394T>C), I2088V (6402A>G 6404A>G), I2089L (6405A>T 6407A>G), S2091E (6411T>G 6412C>A 6413T>A), E2094R (6420G>A 6421A>G), S2095H (6423T>C 6424C>A), Y2096W (6427A>G 6428T>G), R2098K (6432C>A 6433G>A 6434A>G), D2099S (6435G>T 6436A>C 6437T>A), S2100R (6438A>C 6440T>A), N2101S (6442A>G), S2102K (6444T>A 6445C>A 6446C>G), Y2103F (6448A>T 6449T>C), S2104T (6450T>A), V2105F (6453G>T 6455G>T), S2106A (6456T>G 6458A>C), T2107N (6460C>A), P2108E (6462C>G 6463C>A 6464C>A), I2109V (6465A>G 6467C>T), K2110E (6468A>G), L2111I (6471C>A 6473G>A), G2112S (6474G>T 6475G>C 6476G>A), C2113N (6477T>A 6478G>A 6479T>C), F2114L (6480T>C 6482C>G), I2115L (6483A>C), F2116L (6488C>A), L2117A (6489C>G 6490T>C), V2119Y (6495G>T 6496T>A 6497G>T), D2121E (6503T>A), S2122N (6505G>A), Q2123K (6507C>A 6509A>G), G2124L (6510G>C 6511G>T), K2125I (6514A>T 6515G>A), E2126G (6517A>G 6518A>T), D2127L (6519G>C 6520A>T 6521T>A), H2128E (6522C>G 6524T>G), P2129Y (6525C>T 6526C>A), V2130Y (6528G>T 6529T>A), R2131D (6531C>G 6532G>A 6533G>T), R2132S (6534A>T 6535G>C 6536G>T), F2133I (6537T>A 6539T>A), S2134E (6540T>G 6541C>A), T2135S (6543A>T 6545G>T), V2136L (6546G>T 6548T>G), F2137D (6549T>G 6550T>A), K2139L (6555A>T 6556A>T), N2140Q (6558A>C 6560C>A), C2141I (6561T>A 6562G>T 6563T>A), E2142D (6566A>T), T2146A (6576A>G), Y2148V (6582T>G 6583A>T), E2150Y (6588G>T 6590G>C), *2152G (6594T>G 6596A>G)

| | Begin | End | Coverage | Score | Concordance | Matches | Identities | I/D/M/F* | Stop Codons |
|---|---|---|---|---|---|---|---|---|---|
| NT | 22 | 6747 | 93.0% | 3341 | 25.4% | 6659 (98.4%) | 4235 (62.6%) | 43/67 | |

GCT2GCA (146T>A), TTC7TGG (160T>G 161C>G), TCC8AGT (162T>A 163C>G 164C>T), AAG9_GCC16del (165_188delAAGGTTGCTGGTGTTGATGACGCC), AGT17CCT (189A>C 190G>C), TCT18CCT (192T>C), TCT19GCT (195T>G), TCC20TCT (200C>T), ACC21TCA (201A>T 203C>A), ACT22CCT (204A>C), GGT26GCA (217G>C 218T>A), CTT28GTT (222C>G), TCA29TCT (227A>T), AAT30TCT (228A>T 229A>C), ATT32TTT (234A>T), TCC33TCT (239C>T), AAA34TCT (240A>T 241A>C 242A>T), TTT35CTT (243T>C), AAA37AAG (251A>G), GCG38ACT (252G>A 254G>T), GTT39CTC (255G>C 257T>C), AGT40TCC (258A>T 259G>C 260T>C), ATT42GTT (264A>G), ACC43AAT (268C>A 269C>T), ACT44AAA (271C>A 272T>A), ATG45GTT (273A>G 275G>T), AAG48ACC (283A>C 284G>C), TCA50GTA (288T>G 289C>T), CAT52CAC (296T>C), CTT53ATA (297C>A 299T>A), CTG56GTC (306C>G 308G>C), AAA57AAG (311A>G), CCA58CCG (314A>G), TCT59CCA (315T>C 317T>A), GTG60GTT (320G>T), ACT61ATG (322C>T 323T>G), GAT62GCA (325A>C 326T>A), GTG63GCA (328T>C 329G>A), TCC64GCA (330T>G 332C>A), TCT66TCA (338T>A), GTG68GCC (343T>C 344G>C), GCC69TCT (345G>T 347C>T), ACT70ACA (350T>A), TGT71TGC (353T>C), AGT72AAT (355G>A), TCA73GCT (357T>G 359A>T), CTT74GTA (360C>G 362T>A), AAT75GAA (363A>G 365T>A), AAA76AGA (367A>G), TTG78TTA (374G>A), GAC79GAT (377C>T), AAA80AAG (380A>G), GGT82GCA (385G>C 386T>A), GCA83AAA (387G>A 388C>A), CTT84ATA (390C>A 392T>A), ATA85ATG (395A>G), GAT86GAA (398T>A), CCA87CCC (401A>C), TTT88TTC (404T>C), CTT89ATC (405C>A 407T>C), AAA90AAG (410A>G), GCG91GGT (412C>G 413G>T), TAC92TAT (416C>T), TCT93GAT (417T>G 418C>A), TTC94TTT (422C>T), TTG95GTT (423T>G 425G>T), TCC96AAA (426T>A 427C>A 428C>A), ACA97ACT (431A>T), TAC99TAT (437C>T), TCT101ACT (441T>A), ATC102ATA (446C>A), AAG103GTA (447A>G 448A>T 449G>A), ATG105CTT (453A>C 455G>T), GTT106GTA (458T>A), TTA107CTT (459T>C 461A>T), AAA108AAG (464A>G), TTG109ATG (465T>A), TTT110GTT (468T>G), GAG111TTA (471G>T 472A>T 473G>A), TTG113GTC (477T>G 479G>C), TCC114AAG (480T>A 481C>A 482C>G), AAT115GAT (483A>G), GTT117TTC (489G>T 491T>C), AAA118ATC (493A>T 494A>C), TTA119CTA (495T>C), TTT121TAC (502T>A 503T>C), GCT122GAA (505C>A 506T>A), GTG124TTA (510G>T 512G>A), AAT126GAA (516A>G 518T>A), AGT128GTT (522A>G 523G>T), GAA129GAG (527A>G), GAT130TCA (528G>T 529A>C 530T>A), GTA131TTA (531G>T), GTT134GTG (542T>G), GTA135TTT (543G>T 545A>T), TAT136ATA (546T>A 547A>T 548T>A), GGT137GCT (550G>C), CTT139TTA (555C>T 557T>A), GTA140GTT (560A>T), ATA143TTG (567A>T 569A>G), ACA144GCC (570A>G 572A>C), TTG145ATT (573T>A 575G>T), TTA146TTG (578A>G), GTG147CTA (579G>C 581G>A), CTT148CTA (584T>A), CTA149CTG (587A>G), TTT150TAT (589T>A), GTT151GTA (593T>A), TGT152TGC (596T>C), AGT154GTA (600A>G 601G>T 602T>A), GTT157GTA (611T>A), GAG158ACG (612G>A 613A>C), GGT159CCT (615G>C 616G>C), GTT160TTT (618G>T), TCT161AAT (621T>A 622C>A), AAC162AAT (626C>T), ACT163GTA (627A>G 628C>T 629T>A), GTC164GTT (632C>T), AAA165GTT (633A>G 634A>T 635A>T), AAT166ACT (637A>C), ATC167TTG (639A>T 641C>G), TTG169TCA (646T>C 647G>A), ATA170GTT (648A>G 650A>T), GTC171GTT (653C>T), AAC173AAT (659C>T), ATG174TTT (660A>T 662G>T), TTT175TTC (665T>C), TCA176TCT (668A>T), AGT177TCA (669A>T 670G>C 671T>A), CTA178TTG (672C>T 674A>G), TAC179TAT (677C>T), AAA186ACA (697A>C), GCC188TCT (702G>T 704C>T), GAA189GGG (706A>G 707A>G), CGA190AAG (708C>A 709G>A 710A>G), TTC191TTT (713C>T), ACC192TCC (714A>T), ATG193CAG (717A>C 718T>A), GTG194CTG (720G>C), GCC195GCG (725C>G), GCT197GCA (731T>A), AGT198GCA (732A>G 733G>C 734T>A), ATG199GTA (735A>G 737G>A), GGG202GGT (746G>T), TCA204AGT (750T>A 751C>G 752A>T), GTT205TTA (753G>T 755T>A), TCA206ACG (756T>A 758A>G), ACG210ACT (770G>T), TCT211AGC (771T>A 772C>G 773T>C), CAG212CAA (776G>A), TTG213CTT (777T>C 779G>T), ATG214TTA (780A>T 782G>A), ACA216TCA (786A>T), ATA217TTG (789A>T 791A>G), GGG221GGT (803G>T), ATT222ATC (806T>C), TCG223TCA (809G>A), ACG224ACT (812G>T), TTG225TTA (815G>A), GTG226GTA (818G>A), TTC227TTT (821C>T), ATT228GTT (822A>G), GCA229GCT (827A>T), GTG230GTT (830G>T), GGT232GGG (836T>G), GGC236GGT (848C>T), TTG237CTT (849T>C 851G>T), AGC238TCA (852A>T 853G>C 854C>A), TTG241CTG (861T>C), TCC242TCA (866C>A), ATT244ATC (872T>C), CTC245CTG (875C>G), TAT246TAC (878T>C), TCG247TCT (881G>T), ACA248GCT (882A>G 884A>T), GCT252GCG (896T>G), CAG253CAA (899G>A), CAA257CAG (911A>G), CTA258CTG (914A>G), ACT260CAG (918A>C 919C>A 920T>G), CTT261TTG (921C>T 923T>G), CGT263AGG (927C>A 929T>G), AAT264AAC (932T>C), AAA266AAG (938A>G), TGC268TGT (944C>T), ACC269ACA (947C>A), TCC270TCT (950C>T), CAG271CAA (953G>A), GCT272GCC (956T>C), GAA278GAG (974A>G), ATA280GTG (978A>G 980A>G), GTC281GTT (983C>T), GGC282AGC (984G>A), ACC283ATT (988C>T 989C>T), GGA285GGC (995A>C), TTC286TTT (998C>T), GTG290GTA (1010G>A), GCT293GCG (1019T>G), AGT295AAT (1024G>A), GCA296GCT (1028A>T), ACG297ACC (1031G>C), CTG298TTA (1032C>T 1034G>A), CTG302CTT (1046G>T), GAG304CAG (1050G>C), CAG307ACG (1059C>A 1060A>C), GAG308GAA (1064G>A), GAT310TCG (1068G>T 1069A>C 1070T>G), GCA311GAG (1072C>A 1073A>G), GTG312GTT (1076G>T), TGT313GCT (1077T>G 1078G>C), GAT314GAA (1082T>A), CCT315CCA (1085T>A), GCA316TCT (1086G>T 1088A>T), ACG317ATG (1090C>T), GAG320GAA (1100G>A), AAC321AAT (1103C>T), GCC323GCA (1109C>A), AAG325CCT (1113A>C 1114A>C 1115G>T), GCT326GCA (1118T>A), TTC327TTT (1121C>T), CCT328CCA (1124T>A), ACC329ATT (1126C>T 1127C>T), AAA330AAG (1130A>G), AAC332CAA (1134A>C 1136C>A), CTG334TTA (1140C>T 1142G>A), AGG335AAA (1144G>A 1145G>A), GAA336GAG (1148A>G), GAA337CAG (1149G>C 1151A>G), GCT338GCC (1154T>C), CTC339GTC (1155C>G), AAG340AAA (1160G>A), ATC341ATA (1163C>A), GCA346GCC (1178A>C), GTT350GTC (1190T>C), GCG351GCA (1193G>A), TTC353TTT (1199C>T), ATG354TTG (1200A>T), AGT359TCT (1215A>T 1216G>C), ATT362ATA (1226T>A), GCG363ACG (1227G>A), TTA365CTA (1233T>C), GAA366GAT (1238A>T), GTT368CTG (1242G>C 1244T>G), TAT369GTT (1245T>G 1246A>T), GAA371GAG (1253A>G), AGT372GCC (1254A>G 1255G>C 1256T>C), TGT373AGC (1257T>A 1259T>C), AGG374CAA (1260A>C 1261G>A 1262G>A), CAC375CAT (1265C>T), ATG376TCT (1266A>T 1267T>C 1268G>T), GGC377GGT (1271C>T), CAG380CAA (1280G>A), CGT382AGG (1284C>A 1286T>G), GAA384GAG (1292A>G), TTT386TTC (1298T>C), GTA388GTG (1304A>G), TTC391TTT (1313C>T), GGG392GGA (1316G>A), TCC393GCT (1317T>G 1319C>T), GGG395GGA (1325G>A), TCT399TCA (1337T>A), ACA400ACG (1340A>G), TCC401TGT (1342C>G 1343C>T), TTA404CTT (1350T>C 1352A>T), TTT405TTC (1355T>C), ATT406ATC (1358T>C), GTG409GTT (1367G>T), TTG410CTC (1368T>C 1370G>C), GAC411GAT (1373C>T), AGA412GCA (1374A>G 1375G>C), GAG415GAA (1385G>A), AAA417AAG (1391A>G), CTC418CTG (1394C>G), AAT419AAC (1397T>C), CGG420CGC (1400G>C), CTA421TTG (1401C>T 1403A>G), GCA423GCG (1409A>G), AAG426AAA (1418G>A), AGG427AGA (1421G>A), GCT435GCA (1445T>A), CAC436CAT (1448C>T), GCT439GCA (1457T>A), ATC440GTG (1458A>G 1460C>G), CTG441CTT (1463G>T), GAC444GAT (1472C>T), GGA446GGT (1478A>T), GCA447GCT (1481A>T), TTG448CTT (1482T>C 1484G>T), CGA449CGG (1487A>G), GGG451GGA (1493G>A), GCT452GCA (1496T>A), TGC455TGT (1505C>T), CAA456CAG (1508A>G), GAT457GAC (1511T>C), ATT458ATA (1514T>A), AAA459AAG (1517A>G), GAT463GAC (1529T>C), TCA466TCT (1538A>T), ACA467ACC (1541A>C), CAA468CAG (1544A>G), CCA469CCT (1547A>T), CCC471CCT (1553C>T), TTA472CTT (1554T>C 1556A>T), GCC475GCT (1565C>T), GTT477GTC (1571T>C), GAC479GAT (1577C>T), GGC481GGA (1583C>A), CAT483CAC (1589T>C), TTC484TTT (1592C>T), ACC485ACA (1595C>A), TCC486TCA (1598C>A), TAT488TAC (1604T>C), ATA489ATC (1607A>C), ACA492ACC (1616A>C), CTG495TTG (1623C>T), ATC496ATA (1628C>A), TCA497TCT (1631A>T), CCT499CCA (1637T>A), GCC500GCT (1640C>T), CAA501CAG (1643A>G), GGT503GGG (1649T>G), CTT504CTA (1652T>A), ACA505ACG (1655A>G), CCA507CCT (1661A>T), GCT509GCA (1667T>A), GAG511GAA (1673G>A), CGG512AGG (1674C>A), AGA513AGG (1679A>G), AGG514AGA (1682G>A), GAC515GAT (1685C>T), ATC516ATT (1688C>T), CTG517TTA (1689C>T 1691G>A), GTG518GTT (1694G>T), TGT520TGC (1700T>C), ATG521AGA (1702T>G 1703G>A), GAG523GTG (1708A>T), GAG525GAT (1715G>T), TCC527GAT (1719T>G 1720C>A 1721C>T), ACT528CCA (1722A>C 1724T>A), GAA529AAT (1725G>A 1727A>T), GAT530ACT (1728G>A 1729A>C), CCT531CCA (1733T>A), ACG532ACA (1736G>A), GGA533GCC (1738G>C 1739A>C), CTC535CTA (1745C>A), AGA536GTT (1746A>G 1747G>T 1748A>T), AAC538GAT (1752A>G 1754C>T), ATA539GTG (1755A>G 1757A>G), GAG541GAA (1763G>A), CCT545CCC (1775T>C), GCT547GCC (1781T>C), ACT549ACA (1787T>A), CAC550CAT (1790C>T), AGG551AGA (1793G>A), AAT552GCT (1794A>G 1795A>C), TTG553TTA (1799G>A), ACC554TCC (1800A>T), AGT556GAT (1806A>G 1807G>A), GAT557GAC (1811T>C), CTT558CTC (1814T>C), GAG560CAA (1818G>C 1820G>A), GTA563GTG (1829A>G), GCC564GCA (1832C>A), AAA565AAG (1835A>G), TGT566TGC (1838T>C), CAG567CGG (1840A>G), GTA568GTG (1844A>G), CAT569CAC (1847T>C), GCA570TAT (1848G>T 1849C>A 1850A>T), GAA571GAG (1853A>G), GTA572GTG (1856A>G), AAG574AGC (1861A>G 1862G>C), TTG576CTT (1866T>C 1868G>T), TTT577TAC (1870T>A 1871T>C), GAA578AAT (1872G>A 1874A>T), GCT579ACT (1875G>A), GAA580ACA (1878G>A 1879A>C), TCT581TCG (1883T>G), GGC582GGA (1886C>A), ATA583TTG (1887A>T 1889A>G), AGT584GCT (1890A>G 1891G>C), CCT585GAC (1893C>G 1894C>A 1895T>C), ATA587GTT (1899A>G 1901A>T), GTT590GTC (1910T>C), CAA591AAT (1911C>A 1913A>T), TCT593GCT (1917T>G), GCA594GTT (1921C>T 1922A>T), GAT596GAA (1928T>A), ATA598CTG (1932A>C 1934A>G), GCG599GAT (1936C>A 1937G>T), GTG601TTC (1941G>T 1943G>C), GAT602CAA (1944G>C 1946T>A), GGG603GAT (1948G>A 1949G>T), GCT604GCC (1952T>C), AGA605del (1953_1955delAGA), TTG606TTA (1958G>A), CGC607CAA (1960G>A 1961C>A), ACC608GTT (1962A>G 1963C>T 1964C>T), AAG609TTG (1965A>T 1966A>T), CAA610CCT (1969A>C 1970A>T), GAT611CCT (1971G>C 1972A>C), GAA612GAG (1976A>G), ATA614GCA (1980A>G 1981T>C), TTG615GCT (1983T>G 1984T>C 1985G>T), GTG616TAT (1986G>T 1987T>A 1988G>T), CCC617GAG (1989C>G 1990C>A 1991C>G), ACT618CAG (1992A>C 1993C>A 1994T>G), GTT619ACA (1995G>A 1996T>C 1997T>A), GTG620ATT (1998G>A 2000G>T), AGT621ACA (2002G>C 2003T>A), AAT623GCT (2007A>G 2008A>C), GAT624GAA (2012T>A), AGA625AAA (2014G>A), GTG626ATT (2016G>A 2018G>T), ATC627GTT (2019A>G 2021C>T), TAT628TAC (2024T>C), GCA629TCC (2025G>T 2027A>C), AGA630CAG (2028A>C 2029G>A 2030A>G), GAG631GAA (2033G>A), ACG633ACA (2039G>A), GTA634GTT (2042A>T), GCA636GCC (2048A>C), TTG637CTC (2049T>C 2051G>C), GCA640GCT (2060A>T), CAG642CAA (2066G>A), GGC643GGG (2069C>G), TTG645ATG (2073T>A), GAC646AAC (2076G>A), CCT647AAA (2079C>A 2080C>A 2081T>A), GAG648TTT (2082G>T 2083A>T 2084G>T), GAA649GAG (2087A>G), CTT650TTT (2088C>T), TTT651TTC (2093T>C), CCT652CCA (2096T>A), CAT655CTC (2104A>T 2105T>C), CAT656TCC (2106C>T 2107A>C 2108T>C), GCC659GCT (2117C>T), ATG660ATC (2120G>C), TCT663GTG (2127T>G 2128C>T 2129T>G), CTT664CTA (2132T>A), GAT665TCT (2133G>T 2134A>C), GAT666TCT (2136G>T 2137A>C), GAA667GAT (2141A>T), CAT668GCC (2142C>G 2143A>C 2144T>C), GAG670CAG (2148G>C), ATT671AAT (2152T>A), TTC672TTT (2156C>T), AAC673GAA (2157A>G 2159C>A), AAA674ATG (2161A>T 2162A>G), AGA676GTA (2166A>G 2167G>T), GTC677AAT (2169G>A 2170T>A 2171C>T), AAC678GAT (2172A>G 2174C>T), ATG679TTG (2175A>T), TAT681GTC (2181T>G 2182A>T 2183T>C), AGA682AAA (2185G>A), GAT685AGT (2193G>A 2194A>G), GCT686CCT (2196G>C), GAA687CAG (2199G>C 2201A>G), CGA690GCT (2208C>G 2209G>C 2210A>T), GTA693GTG (2219A>G), CAA694CGA (2221A>G), GAC699GAT (2237C>T), ATA701ATC (2243A>C), CAC703CAT (2249C>T), TTT704TTC (2252T>C), GCA710AGC (2268G>A 2269C>G 2270A>C), TCA711ACA (2271T>A), ATC713ATT (2279C>T), GAG715GAA (2285G>A), AAG716GAG (2286A>G), AAA717CTG (2289A>C 2290A>T 2291A>G), CTA718CTG (2294A>G), GCA719GAT (2296C>A 2297A>T), GTG720GTC (2300G>C), CAG721CAA (2303G>A), ACA722CGC (2304A>C 2305C>G 2306A>C), GAG723AAT (2307G>A 2309G>T), ATG724CTT (2310A>C 2312G>T), AGG725GAG (2313A>G 2314G>A), ACT726CAA (2316A>C 2317C>A 2318T>A), GGA727GGG (2321A>G), TTT728AAA (2322T>A 2323T>A 2324T>A), GCA729GCG (2327A>G), CAT730TAT (2328C>T), TCT731AAG (2331T>A 2332C>A 2333T>G), TGC732GCG (2334T>G 2335G>C 2336C>G), ATT733CTT (2337A>C), GAT734CAG (2340G>C 2342T>G), GCA735GCT (2345A>T), GTT737GCA (2350T>C 2351T>A), GAC738GAT (2354C>T), ACC739ACA (2357C>A), CTC740CTT (2360C>T), ATA741ACG (2362T>C 2363A>G), ATC743ATT (2369C>T), GAG744GAA (2372G>A), TTT749AGA (2385T>A 2386T>G 2387T>A), GTT750GTA (2390T>A), TTT752TTC (2396T>C), TTA753CTG (2397T>C 2399A>G), ACA755ACT (2405A>T), GCA756GCT (2408A>T), AGA759CGC (2415A>C 2417A>C), TGC762TGT (2426C>T), AAC763AGG (2428A>G 2429C>G), GTA765GTT (2435A>T), TCA766GCA (2436T>G), TCA766_AAA767insGGT (2438_2439insGGT), AAA767ACT (2440A>C 2441A>T), GAG768GAC (2444G>C), CCA769AAG (2445C>A 2446C>A 2447A>G), GAA771AGA (2451G>A 2452A>G), TCA772AAT (2454T>A 2455C>A 2456A>T), ATC774ATT (2462C>T), AAA775ATC (2464A>T 2465A>C), ATT776GAA (2466A>G 2467T>A 2468T>A), TAC778ATG (2472T>A 2473A>T 2474C>G), CAT779ATT (2475C>A 2476A>T), AAA780GGT (2478A>G 2479A>G 2480A>T), GTT782AAG (2484G>A 2485T>A 2486T>G), GAA783GAC (2489A>C), TAT784AAA (2490T>A 2492T>A), ATC785ATG (2495C>G), AAG786ATC (2497A>T 2498G>C), GAT787CAG (2499G>C 2501T>G), ACA788ATG (2503C>T 2504A>G), TGG790AAT (2508T>A 2509G>A 2510G>T), AGC791GAC (2511A>G 2512G>A), CTT792TTG (2514C>T 2516T>G), TAT794TTC (2521A>T 2522T>C), GCA795ATC (2523G>A 2524C>T 2525A>C), CTG796TTG (2526C>T), AGA797AAG (2530G>A 2531A>G), CTC799CTT (2537C>T), GCA802GCC (2546A>C), GGT803GGA (2549T>A), ATA806TTG (2556A>T 2558A>G), ATG807ATA (2561G>A), GCT808GCA (2564T>A), GGA811GGC (2573A>C), GTT812ACC (2574G>A 2575T>C 2576T>C), TTT813ATG (2577T>A 2579T>G), GGA814AAT (2580G>A 2581G>A 2582A>T), GGC815GCC (2584G>C), ATT816GTT (2586A>G), ACA817ACT (2591A>T), GCA818GCT (2594A>T), TTT819TTC (2597T>C), TTG820ATT (2598T>A 2600G>T), GCG821ACT (2601G>A 2603G>T), TGG823CTC (2607T>C 2608G>T 2609G>C), CAG824AAG (2610C>A), AGT825GCA (2613A>G 2614G>C 2615T>A), AAC826GCA (2616A>G 2617A>C 2618C>A), TCT827ACT (2619T>A), TTC828TCC (2623T>C), CCC829CCT (2627C>T), AAT830TTA (2628A>T 2629A>T 2630T>A), GCA831GCT (2633A>T), TCG832GAT (2634T>G 2635C>A 2636G>T), GGA835GGG (2645A>G), GGT836GGC (2648T>C), GTG837GTA (2651G>A), GAA840GAG (2660A>G), ATA844ATC (2672A>C), AGT846AGC (2678T>C), AAG848AAA (2684G>A), AAC849AAT (2687C>T), AAG850AAA (2690G>A), GGT851GGG (2693T>G), AAT852AAG (2696T>G), AAG853ACT (2698A>C 2699G>T), AAG853_CTC854insCAT (2699_2700insCAT), CTC854TTC (2700C>T), AGA855GCA (2703A>G 2704G>C), AAT856AAA (2708T>A), CTT857TCC (2709C>T 2710T>C 2711T>C), CTC858TTG (2712C>T 2714C>G), GTT859CTA (2715G>C 2717T>A), GGT860GGG (2720T>G), AGT862GCG (2724A>G 2725G>C 2726T>G), TCT863GCA (2727T>G 2729T>A), CAA864GAT (2730C>G 2732A>T), TCA865AAC (2733T>A 2734C>A 2735A>C), TTG866AAC (2736T>A 2737T>A 2738G>C), GCA867CAT (2739G>C 2740C>A 2741A>T), CAA868GTG (2742C>G 2743A>T 2744A>G), GAT869GAC (2747T>C), GAT869_TGG870insCGG (2747_2748insCGG), GCT871GCA (2753T>A), GCT872GAA (2755C>A 2756T>A), GAA873CAG (2757G>C 2759A>G), GTA877GTT (2771A>T), CAG879GAA (2775C>G 2777G>A), AGC880AGT (2780C>T), CTC881CTG (2783C>G), AAG882CAC (2784A>C 2786G>C), AAT884AAC (2792T>C), GTG886GTT (2798G>T), GTG887GTT (2801G>T), TTA888TTG (2804A>G), CTT890ATT (2808C>A), GGT891GCT (2812G>C), GTG894GTC (2822G>C), TAC895TAT (2825C>T), TTT896TTC (2828T>C), GGC898GGT (2834C>T), ACC899ACG (2837C>G), TAC900TAT (2840C>T), GTG901GTT (2843G>T), ATA906ATT (2858A>T), ATG907CTC (2859A>C 2861G>C), ACC908ACT (2864C>T), GTA909GTG (2867A>G), GCT910GCA (2870T>A), GCA912GCC (2876A>C), TTC913TTT (2879C>T), CAC914AGA (2880C>A 2881A>G 2882C>A), AAT915AAC (2885T>C), GCT916ATG (2886G>A 2887C>T 2888T>G), CGA917AGG (2889C>A 2891A>G), GGT919GGC (2897T>C), CCA921CCT (2903A>T), ACA923TCC (2907A>T 2909A>C), ATC925ATT (2915C>T), GCC927GAG (2920C>A 2921C>G), AAT928CAT (2922A>C), TCT929TCC (2927T>C), CGA930ATT (2928C>A 2929G>T 2930A>T), TCT931TGT (2932C>G), AAA932AAT (2936A>T), GTT933GTG (2939T>G), CAA934CAG (2942A>G), TAC935TTC (2944A>T), AAC936AAT (2948C>T), AGA938ACC (2953G>C 2954A>C), GAA939CAA (2955G>C), AGG941AAG (2962G>A), ATT942TTG (2964A>T 2966T>G), ATT943ATA (2969T>A), GGC945GGT (2975C>T), GAC947GAT (2981C>T), ATC948GTG (2982A>G 2984C>G), CTG950TTG (2988C>T), CGC952AGG (2994C>A 2996C>G), GTT953GTA (2999T>A), GGT954GGG (3002T>G), CCA959CCC (3017A>C), AAG960AAA (3020G>A), CCT961TCA (3021C>T 3023T>A), GAC962GAT (3026C>T), ATC963ATA (3029C>A), CGT964AGA (3030C>A 3032T>A), CAC966CAT (3038C>T), CCA968CCT (3044A>T), AGA969CGC (3045A>C 3047A>C),

Codon mutations:

AGG970AGA (3050G>A), GAG972GAT (3056G>T), TGC974TGT (3062C>T), TTC975TAC (3064T>A), ACA976TCT (3066A>T 3068A>T), CAA980ATG (3078C>A 3079A>T 3080A>G), GGA981GGC (3083A>C), TGT984GTG (3090T>G 3091G>T 3092T>G), AGA985AGG (3095A>G), GTT987GTA (3101T>A), GCG988GCT (3104G>T), TCT989TCA (3107T>A), ACA990AAG (3109C>A 3110A>G), CCA992CCC (3116A>C), CGT993AAG (3117C>A 3118G>A 3119T>G), CTT994AAA (3120C>A 3121T>A 3122T>A), GGC995TCC (3123G>T 3124G>C), TTA997TTG (3131A>G), TTT999GTT (3135T>G), CTC1000CTT (3140C>T), AAG1001AAA (3143G>A), ATG1002TAT (3144A>T 3145T>A 3146G>T), GGA1009GGC (3167A>C), ACA1011ACC (3173A>C), GTT1012GAA (3175T>A 3176T>A), GTG1014GTC (3182G>C), GAA1015ACA (3183G>A 3184A>C), GAA1017GGA (3190A>G), CTG1018TTA (3192C>T 3194G>A), AAC1019GAT (3195A>G 3197C>T), TCC1020TCG (3200C>G), TCC1021AAT (3201T>A 3202C>A 3203C>T), AGT1022GCA (3204A>G 3205G>C 3206T>A), AAG1024GCT (3210A>G 3211A>C 3212G>T), ATT1025CTT (3213A>C), TGC1026AGC (3216T>A), TCT1027TCA (3221T>A), CAA1029GTT (3225C>G 3226A>T 3227A>T), TCT1030GCA (3228T>G 3230T>A), GAA1032GAG (3236A>G), ATA1035ATT (3245A>T), GGG1037AGT (3249G>A 3251G>T), GAA1038GAG (3254A>G), GGT1040GGA (3260T>A), GAC1041GAT (3263C>T), GGC1043GGT (3269C>T), ACG1044ACT (3272G>T), TTG1045CTT (3273T>C 3275G>T), CTA1046CTC (3278A>C), CTG1047TTA (3279C>T 3281G>A), TTA1048TTG (3284A>G), AGT1050ACT (3289G>C), GTT1051GTG (3293T>G), AAT1053AAC (3299T>C), CAA1055CAG (3305A>G), CCT1056CCG (3308T>G), GTG1057TGC (3309G>T 3310T>G 3311G>C), GGC1060GGA (3320C>A), ATC1061ATA (3323C>A), CAC1062CAT (3326C>T), TCA1065TCT (3335A>T), GGC1068GAT (3343G>A 3344C>T), GTA1069ACC (3345G>A 3346T>C 3347A>C), GCA1070GCT (3350A>T), GCT1071GCA (3353T>A), GAA1072GAG (3356A>G), CGT1073AAG (3357C>A 3358G>A 3359T>G), GGA1074GGT (3362A>T), TTT1075TAC (3364T>A 3365T>C), ATC1076GTG (3366A>G 3368C>G), TCT1077GCT (3369T>G), AAT1079AAC (3377T>C), GCT1080GCA (3380T>A), GCT1082GCA (3386T>A), ATC1083GTG (3387A>G 3389C>G), TAC1084TTT (3391A>T 3392C>T), AGG1085AGA (3395G>A), GAA1086GAT (3398A>T), CTA1088TTG (3402C>T 3404A>G), GAG1089GAA (3407G>A), CCA1092CCC (3416A>C), ACT1093ACG (3419T>G), GGG1094GGA (3422G>A), CCG1095CCC (3425G>C), GTC1096ATT (3426G>A 3428C>T), GCA1098GTG (3433C>T 3434A>G), GCA1099GCT (3437A>T), GTA1101GTG (3443A>G), CGC1102CAC (3445G>A), GAT1104GAA (3452T>A), ATT1105CTG (3453A>C 3455T>G), CTG1106TTG (3456C>T), TCA1108CAC (3462T>C 3463C>A 3464A>C), ATT1109CTG (3465A>C 3467T>G), AGC1111TCC (3471A>T 3472G>C), AGA1112AAG (3475G>A 3476A>G), GAA1113GAG (3479A>G), ACA1114GAA (3480A>G 3481C>A), CAG1115AGA (3483C>A 3484A>G 3485G>A), CTT1116TTC (3486C>T 3488T>C), GAA1119GAG (3497A>G), AAC1120AAA (3500C>A), CAA1121CAG (3503A>G), TAC1123GCT (3507T>G 3508A>C 3509C>T), TAC1124TTC (3511A>T), CTT1125TTG (3513C>T 3515T>G), GGA1126GGC (3518A>C), ACA1127ACT (3521A>T), GTT1128GTA (3524T>A), CCA1129CCG (3527A>G), CAG1130TTG (3528C>T 3529A>T), GAG1131GAA (3533G>A), TTG1132ATG (3534T>A), GCC1133GCT (3539C>T), GCC1134GCT (3542C>T), GTT1136GTG (3548T>G), CCC1137CCG (3551C>G), AAG1139AAA (3557G>A), ACC1140ACA (3560C>A), ACT1141ACC (3563T>C), CTG1142CTT (3566G>T), CGG1143AAA (3567C>A 3568G>A 3569G>A), AAA1144CGC (3570A>C 3571A>G 3572A>C), AGC1145AGT (3575C>T), CAA1146CCC (3577A>C 3578A>C), TTG1147CTG (3579T>C), GAA1149GAG (3587A>G), GCA1150GCT (3590A>T), TTC1151TTT (3593C>T), GGA1152GGG (3596A>G), CCT1153GAC (3597C>G 3598C>A 3599T>C), GCA1154GCT (3602A>T), GAG1155CAG (3603G>C), ACA1156ACT (3608A>T), GCA1157GCC (3611A>C), ACA1162TCT (3624A>T 3626A>T), GTT1163GCC (3628T>C 3629T>C), CAT1164AAA (3630C>A 3632T>A), AGA1167AGG (3641A>G), GGT1168GGA (3644T>A), GAT1169GAA (3647T>A), GGT1170GGG (3650T>G), GAC1172GAT (3656C>T), GGG1177GGT (3671G>T), GTA1178GTC (3674A>C), ATG1179GAG (3675A>G 3676T>A), AAA1180AAG (3680A>G), TAC1181TAT (3683C>T), TGT1186TCC (3697G>C 3698T>C), GGA1187AAC (3699G>A 3700G>A 3701A>C), GAT1189GAG (3707T>G), GAT1190GAG (3710T>G), GAC1191CCA (3711G>C 3712A>C 3713C>A), GCC1193GCA (3719C>A), AAA1194AAG (3722A>G), CTA1195CTG (3725A>G), GCA1196GCC (3728A>C), TTC1197TTT (3731C>T), GAA1198GAG (3734A>G), AAT1199GGT (3735A>G 3736A>G), CTC1200CTT (3740C>T), AAG1201AAA (3743G>A), TGC1202GCA (3744T>G 3745G>C 3746C>A), TCG1203TCC (3749G>C), CTG1204ATT (3750C>A 3752G>T), CTA1205CTT (3755A>T), ATC1207GTG (3759A>G 3761C>G), ATG1208CTT (3762A>C 3764G>T), CGT1209AAG (3765C>A 3766G>A 3767T>G), AGC1210ATG (3769G>T 3770C>G), AAG1212CGA (3774A>C 3775A>G 3776G>A), ATC1213GTG (3777A>G 3779C>G), CCT1214CCC (3782T>C), CGT1217AAA (3789C>A 3790G>A 3791T>A), CCA1218CCT (3794A>T), TGT1219GTA (3795T>G 3796G>T 3797T>A), GAA1220ATC (3798G>A 3799A>T 3800A>C), AGG1221AGA (3803G>A), GAT1222AAT (3804G>A), GGA1229GGG (3827A>G), TGT1233TTT (3838G>T), TAC1235TAT (3845C>T), TAT1236TTT (3847A>T), GGC1238GCA (3853G>C 3854C>A), GAG1240GAA (3860G>A), CTG1241TTG (3861C>T), ACA1243ACG (3869A>G), TCT1244TCA (3872T>A), TGC1245TGT (3875C>T), GGA1246GGT (3878A>T), CCC1248CCT (3884C>T), TTC1249TTT (3887C>T), ATG1252TTG (3894A>T), GGG1253GGA (3899G>A), ATG1254CTT (3900A>C 3902G>T), GGG1255GGT (3905G>T), ATG1256TCT (3906A>T 3907T>C 3908G>T), AAG1258AAA (3914G>A), AGA1259AGG (3917A>G), CAA1263GAA (3927C>G), TCC1264GCC (3930T>G), ACT1265CAA (3933A>C 3934C>A 3935T>A), GGC1266CCA (3936G>C 3937G>C 3938C>A), GAA1267AAC (3939G>A 3941A>C), GGA1268TCG (3942G>T 3943G>C 3944A>G), GAG1269CAG (3945G>C), AGA1270AGG (3950A>G), GTG1271GTT (3953G>T), GAA1272CTT (3954G>C 3955A>T 3956A>T), AGA1275AGG (3965A>G), GAC1276AAC (3966G>A), ACT1277ACA (3971T>A), GTA1279GTC (3977A>C), GAA1281GAT (3983A>T), CTA1286TTG (3996C>T 3998A>G), GAT1287GAC (4001T>C), GTG1288TCG (4002G>T 4003T>C), ATT1290GTA (4008A>G 4010T>A), AAA1292CAA (4014A>C), ATC1294ACA (4021T>C 4022C>A), CAT1295TTT (4023C>T 4024A>T), GTG1296GTT (4028G>T), GAT1297GAC (4031T>C), CTG1298CTC (4034G>C), ACC1300ACA (4040C>A), CAA1302CAG (4046A>G), TGC1303TGT (4049C>T), GCC1304GCG (4052C>G), CGC1308CGG (4064C>G), CTC1309TTG (4065C>T 4067C>G), AAA1313AAG (4079A>G), ATC1314ATT (4082C>T), GGC1316GGG (4088C>G), AGA1318AGG (4094A>G), AGG1321CGC (4101A>C 4103G>C), CTC1322CTG (4106C>G), TTT1323TTC (4109T>C), GAG1324GAA (4112G>A), ATA1325ATC (4115A>C), CTT1326TTG (4116C>T 4118T>G), TAC1330TAT (4130C>T), TTG1333CTG (4137T>C), GTC1334GTG (4142C>G), AGG1335CGG (4143A>C), AAG1336AAA (4148G>A), TAT1337TAC (4151T>C), TTT1338TTC (4154T>C), GAT1340GAC (4160T>C), TTC1341TTT (4163C>T), TCA1342TCT (4166A>T), GCC1343GCT (4169C>T), ACA1344TGT (4170A>T 4171C>G 4172A>T), TTG1348GCA (4182T>G 4183T>C 4184G>A), AAT1350AAC (4190T>C), GCT1351GTA (4192C>T 4193T>A), ATA1352CTC (4194A>C 4196A>C), CCA1353CCT (4199A>T), AAA1355AAG (4205A>G), GTT1356GTA (4208T>A), ATT1358ATC (4214T>C), GAT1359AAT (4215G>A), CCT1360CCA (4220T>A), ACA1361ACG (4223A>G), AGT1362AGC (4226T>C), TCT1363CAT (4227T>C 4228C>A), TTG1367GTG (4239T>G), TTG1368CTG (4242T>C), GCA1369GCT (4247A>T), AAT1370AGT (4249A>G), GGG1371AGC (4251G>A 4253G>C), AGA1373AGG (4259A>G), GCT1374TCA (4260G>T 4262T>A), GTG1375GTA (4265G>A), TCA1376TCT (4268A>T), GAC1377GAT (4271C>T), GTG1378GAG (4273T>A), TTT1380TTC (4280T>C), GCT1382GCC (4286T>C), TAT1384TAC (4292T>C), TCC1385TCG (4295C>G), CCT1392CCC (4316T>C), GTT1393ATC (4317G>A 4319T>C), GCT1395GCA (4325T>A), TTT1396TTC (4328T>C), CAG1397CAA (4331G>A), TTG1402CTG (4344T>C), GAA1405GCT (4354A>C 4355A>T), TAC1406TTT (4357A>T 4358C>T), TAC1407TAT (4361C>T), GGA1408GGT (4364A>T), TCA1409GAT (4365T>G 4366C>A 4367A>T), AAG1410GAG (4368A>G), GGC1412GGG (4376C>G), GAT1415GAG (4385T>G), TCC1416TCA (4388C>A), CGA1419AGG (4395C>A 4397A>G), CAT1420CAC (4400T>C), GCA1421ACA (4401G>A), CTT1422TTA (4404C>T 4406T>A), GCA1426GCC (4418A>C), TGT1428TGC (4424T>C), TAC1430TAT (4430C>T), ACA1431ACC (4433A>C), CTG1432CTC (4436G>C), TGC1433TGT (4439C>T), GAT1435AAC (4443G>A 4445T>C), AAA1436AAG (4448A>G), GTG1437TTG (4449G>T), AGG1439CGC (4455A>C 4457G>C), TTG1440CTA (4458T>C 4460G>A), GTT1441GTG (4463T>G), GGG1442GGT (4466G>T), GGC1443GGA (4469C>A), CCA1445CCC (4475A>C), TCA1446TCT (4478A>T), GGA1447GGT (4481A>T), GCA1449GCT (4487A>T), CTA1450TTG (4488C>T 4490A>G), ACG1451ACT (4493G>T), ATC1453ATA (4499C>A), TTC1454TTT (4502C>T), TCT1456TCA (4508T>A), CTC1457TTG (4509C>T 4511C>G), CTC1458CTA (4514C>A), CGA1464AGG (4530C>A 4532A>G), TAT1465TAC (4535T>C), GCC1466GCG (4538C>G), TTG1470CTG (4548T>C), TTA1471TTG (4553A>G), AGA1472AGG (4556A>G), CCC1474CCA (4562C>A), ATT1476GTC (4566A>G 4568T>C), GCT1477AAA (4569G>A 4570C>A 4571T>A), AGG1479AAA (4576G>A 4577G>A), GCT1480TCT (4578G>T), ATA1481GTG (4581A>G 4583A>G), GGA1482GGC (4586A>C), GTT1483ATT (4587G>A), AAA1484AAG (4592A>G), AGT1486AAT (4597G>A), GAT1487GAC (4601T>C), CTA1491TTG (4611C>T 4613A>G), GCT1494GCA (4622T>A), GTT1495GTA (4625T>A), GGA1497GGT (4631A>T), GAC1499GAT (4637C>T), AAT1500AAC (4640T>C), CTT1501TTG (4641C>T 4643T>G), GCT1503GCG (4649T>G), GTA1504GTT (4652A>T), TTA1506ATG (4656T>A 4658A>G), CTC1508TTG (4662C>T 4664C>G), CAG1509AGG (4665C>A 4666A>G), TAT1511TAC (4673T>C), TCT1512GAT (4674T>G 4675C>A), CTG1513CTA (4679G>A), AAT1515TCC (4683A>T 4684A>C 4685T>C), ATA1516ATT (4688A>T), CAT1518AGA (4692C>A 4693A>G 4694T>A), GAG1519GAA (4697G>A), TTA1520CTG (4698T>C 4700A>G), CTG1522GCT (4704C>G 4705T>C 4706G>T), GTC1523GTT (4709C>T), AAT1524AAC (4712T>C), GTA1525GTC (4715A>C), ATC1526GTC (4716A>G), CTT1531TTG (4731C>T 4733T>G), AAA1533AAG (4739A>G), TCG1534TCA (4742G>A), GTT1537GTA (4751T>A), AAT1538GAG (4752A>G 4754T>G), GAG1539AAG (4755G>A), GTA1540ACA (4758G>A 4759T>C), CAA1541AAA (4761C>A), CAA1543CTG (4768A>T 4769A>G), CTA1548CTC (4784A>C), ACT1549ACC (4787T>C), CTG1551CTC (4793G>C), AGA1553AGG (4799A>G), TTT1555TTC (4805T>C), AAA1556AAG (4808A>G), GCT1559GCA (4817T>A), CTT1560TTG (4818C>T 4820T>G), TAC1562TAT (4826C>T), CAC1563CAA (4829C>A), GCT1565GCA (4835T>A), CCT1566CCA (4838T>A), AAG1568AAA (4844G>A), ATC1572GTC (4854A>G), ATT1573ATC (4859T>C), ATG1576ATT (4868G>T), ATA1579ATT (4877A>T), AGA1580AGG (4880A>G), CCT1581CCA (4883T>A), GCC1582GCA (4886C>A), CCA1583GTG (4887C>G 4888C>T 4889A>G), GGT1584GGA (4892T>A), GAC1587GAT (4901C>T), ACT1588GTA (4902A>G 4903C>T 4904T>A), CAG1589CAA (4907G>A), GCC1590GCT (4910C>T), ATG1592TAT (4914A>T 4915T>A 4916G>T), GTG1595ATT (4923G>A 4925G>T), GGA1598GGC (4934A>C), ATA1599ATC (4937A>C), CGT1600AGA (4938C>A 4940T>A), GCT1602GCC (4946T>C), CAC1605CAT (4955C>T), GGT1606GGG (4958T>G), CGT1607CAG (4960G>A 4961T>G), GTG1608GCC (4963T>C 4964G>C), GCT1609ACT (4965G>A), GAC1611GAA (4973C>A), AAG1612AGT (4975A>G 4976G>T), CTT1613TTT (4977C>T), GTC1614GTA (4982C>A), TTA1615AAT (4983T>A 4984T>A 4985A>T), CAG1618TTT (4992C>T 4993A>T 4994G>T), ACG1619GGA (4995A>G 4996C>G 4997G>A), GCG1620GCT (5000G>T), TTG1621CTC (5001T>C 5003G>C), GAT1622AAG (5004G>A 5006T>G), GAA1623CAG (5007G>C 5009A>G), AGG1624AGA (5012G>A), GGT1625AAT (5013G>A 5014G>A), TTC1626CTC (5016T>C), AGA1627CCC (5019A>C 5020G>C 5021A>C), GCT1628ATC (5022G>A 5023C>T 5024T>C), GTG1629ATG (5025G>A), TTT1631ATG (5031T>A 5033T>G), CCT1632CCC (5036T>C), TCC1633ACT (5037T>A 5039C>T), TAT1634TAC (5042T>C), TTG1635TAT (5044T>A 5045G>T), GTG1637CTT (5049G>C 5051G>T), CAG1639AGT (5055C>A 5056A>G 5057G>T), GAA1640CAA (5058G>C), ATT1642TTG (5064A>T 5066T>G), GCA1643ACA (5067G>A), AAG1644GAG (5070A>G), GTA1645GTC (5075A>C), GGG1647GGA (5081G>A), TCA1649GAG (5085T>G 5086C>A 5087A>G), AGT1650AGC (5090T>C), GCC1651ATT (5091G>A 5092C>T 5093C>T), CTG1652GGG (5094C>G 5095T>G), ACA1653GTT (5097A>G 5098C>T 5099A>T), ATT1654TTG (5100A>T 5102T>G), TGT1655ATG (5103T>A 5104G>T 5105T>G), GAA1656GAG (5108A>G), ATG1657CTA (5109A>C 5111G>A), GCA1658TCA (5112G>T), AAA1659GAT (5115A>G 5117A>T), GCA1660GCT (5120A>T), GCT1661AAA (5121G>A 5122C>A 5123T>A), ATT1662AAT (5125T>A), TCC1663ACA (5127T>A 5129C>A), TAT1664TAC (5132T>C), ACG1665ACC (5135G>C), CCA1666CCT (5138A>T), TTG1667TTA (5141G>A), GAC1668CCT (5142G>C 5143A>C 5144C>T), GCA1669GCT (5147A>T), GGT1670GGG (5150T>G), GAG1671GAA (5153G>A), ATC1673ATT (5159C>T), ACA1674del (5160_5162delACA), CGT1678GAG (5172C>G 5173G>A 5174T>G), GAT1679GGT (5176A>G), CTG1680GTC (5178C>G 5180G>C), AAT1681CAT (5181A>C), TGG1682GAA (5184T>G 5185G>A 5186G>A), GCA1684GTG (5191C>T 5192A>G), CCA1685CCT (5195A>T), AAC1686GAT (5196A>G 5198C>T), ATT1687GTA (5199A>G 5201T>A), GGT1688TTT (5202G>T 5203G>T), TTT1689TTC (5207T>C), TCA1691AGT (5211T>A 5212C>G 5213A>T), GCA1692GCT (5216A>T), CGT1693CGG (5219T>G), GCC1696TCA (5226G>T 5228C>A), CAC1697AAC (5229C>A), TAC1698CAA (5232T>C 5234C>A), ACG1699CAA (5235A>C 5236C>A 5237G>A), GAT1701TCA (5241G>T 5242A>C 5243T>A), GAA1702TCA (5244G>T 5245A>C), GGG1703GGT (5249G>T), TAC1704AAC (5250T>A), ATT1705ATC (5255T>C), ATT1706CTG (5256A>C 5258T>G), AAC1708AAT (5264C>T), TGT1709TGC (5267T>C), ACA1710ACT (5270A>T), GGT1711GGA (5273T>A), GCA1712GCC (5276A>C), AAG1713CGC (5277A>C 5278A>G 5279G>C), AAG1714AGA (5281A>G 5282G>A), TCC1715ACA (5283T>A 5285C>A), AAT1716ACA (5287A>C 5288T>A), GTT1718GTG (5294T>G), GGT1720GGG (5300T>G), CCA1721CCT (5303A>T), AAC1723AAT (5309C>T), ATG1731ATA (5333G>A), CCG1733CCC (5339G>C), ATG1736TTG (5346A>T), TCA1737GCA (5349T>G), GCT1738GCA (5354T>A), ATA1739ATC (5357A>C), ATG1740CAG (5358A>C 5359T>A), GCC1742CAT (5364G>C 5365C>A 5366C>T), AAG1744CGC (5370A>C 5371A>G 5372G>C), AAT1745AAG (5375T>G), GTG1746AAT (5376G>A 5377T>A 5378G>T), TTG1747CTG (5379T>C), GCA1748GAA (5383C>A), GGA1749GTT (5386G>T 5387A>T), GGA1750GGT (5390A>T), CAT1751GAT (5391C>G), GTA1752GGG (5395T>G 5396A>G), GCA1753GCT (5399A>T), AAT1755TCT (5403A>T 5404A>C), GTT1757ACC (5409G>A 5410T>C 5411T>C), GTG1759GTT (5417G>T), GGA1761GGT (5423A>T), GGA1763GGT (5429A>T), GCT1765GCA (5435T>A), GTC1766ATA (5436G>A 5438C>A), GGC1767GCT (5440G>C 5441C>T), ATT1769ATC (5447T>C), GCG1772GCA (5456G>A), CTG1773TTG (5457C>T), ATG1774TTT (5460A>T 5462G>T), GCC1775GCA (5465C>A), TTA1776TTG (5468A>G), GCA1777GCC (5471A>C), ACA1778ACT (5474A>T), AGG1779CAT (5475A>C 5476G>A 5477G>T), TAT1781TAC (5483T>C), TGT1782AGG (5484T>A 5486T>G), GTG1783ACC (5487G>A 5488T>C 5489G>C), GAG1784ACA (5490G>A 5491A>C 5492G>A), ATA1786CTT (5496A>C 5498A>T), ATA1787ATT (5501A>T), GTT1788GTG (5504T>G), AGA1789CGC (5505A>C 5507A>C), TTG1790CTA (5508T>C 5510G>A), CGC1791CGA (5513C>A), ACA1792ACC (5516A>C), ATT1793ATG (5519T>G), GGA1794GCT (5521G>C 5522A>T), AAT1795GGG (5523A>G 5524A>G 5525T>G), CTT1797CTG (5531T>G), GAC1798GAT (5534C>T), CTC1799CTG (5537C>G), AAT1800AAC (5540T>C), TAT1802TAC (5546T>C), GGC1805GGT (5555C>T), GTG1807GCG (5560T>C), CAG1808CAA (5564G>A), TAT1809TAC (5567T>C), TTT1810TTG (5570T>G), CTT1811CTC (5573T>C), CAA1812AAA (5574C>A), GTG1814GTT (5582G>T), CCT1815GAA (5583C>G 5584C>A 5585T>A), CAT1816GAC (5586C>G 5588T>C), CAT1816_GGA1817insACC (5588_5589insACC), GGA1817TCA (5589G>T 5590G>C), GAC1818CAC (5592G>C), AAA1819AAG (5597A>G), GTG1820TTG (5598G>T), GCT1821GGA (5602C>G 5603T>A), CAA1822CAG (5606A>G), GGT1824GGC (5612T>C), GCA1825ACA (5613G>A), TCG1826TAT (5617C>A 5618G>T), CTC1827GGC (5619C>G 5620T>G), CAC1828del (5622_5624delCAC), AGT1829ACC (5626G>C 5627T>C), AGT1830ACT (5629G>C), ATG1832AAC (5635T>A 5636G>C), CAA1834CAT (5642A>T), CAA1834_GGG1835insCACAGT (5642_5643insCACAGT), GGG1835GCA (5644G>C 5645G>A), GAA1837TCA (5649G>T 5650A>C), TTG1838GTT (5652T>G 5654G>T), GGC1839GGA (5657C>A), AAC1840CTT (5658A>C 5659A>T 5660C>T), CTC1841CTA (5663C>A), CGC1842AGG (5664C>A 5666C>G), ATT1843ATA (5669T>A), ATA1844GTT (5670A>G 5672A>T), CAT1845CCT (5674A>C), GGT1846TCC (5676G>T 5677G>C 5678T>C), GAT1847GAC (5681T>C), TTA1848CTC (5682T>C 5684A>C), GCA1849TCT (5685G>T 5687A>T), AAA1850GAA (5688A>G), CAG1851AAG (5691C>A), ACA1852ATG (5695C>T 5696A>G), ATG1854GTT (5700A>G 5702G>T), CGG1855CGT (5705G>T), ATG1856TTG (5706A>T), TAT1858TAC (5714T>C), GTG1859ACA (5715G>A 5716T>C 5717G>A), GTA1860ATT (5718G>A 5720A>T), GGA1861GGT (5723A>T), CCA1862CCT (5726A>T), CAT1863GAA (5727C>G 5729T>A), GGA1865GGT (5735A>T), GGA1867GCC (5740G>C 5741A>C), AAT1868CAG (5742A>C 5744T>G), TTT1869TGC (5746T>G 5747T>C), TCC1870TCA (5750C>A), ACA1871ACG (5753A>G), CAG1872AAG (5754C>A), GAT1873AAC (5757G>A 5759T>C), CTT1874CTG (5762T>G), GAG1875AAG (5763G>A), AGT1876CAT (5766A>C 5767G>A), TTG1877GCT (5769T>G 5770T>C 5771G>T), CTA1878TTG (5772C>T 5774A>G), CAC1879GAG (5775C>G 5777C>G), TAT1880ACA (5778T>A 5779A>C 5780T>A), GAG1882TCT (5784G>T 5785A>C 5786G>T), CAG1883GTC (5787C>G 5788A>T 5789G>C), GGA1884AGT (5790G>A 5792A>T), GCA1886GCT (5798A>T), GAG1887ACT (5799G>A 5800A>C 5801G>T), TTA1888CTT (5802T>C 5804A>T), ATT1889GCC (5805A>G 5806T>C 5807T>C), CAA1890GCA (5808C>G 5809A>C), AAG1891AGA (5812A>G 5813G>A), AAT1892GGA (5814A>G 5815A>G 5816T>A), ACA1893GTC (5817A>G 5818C>T 5819A>C), CTA1895GTG (5823C>G 5825A>G), ACT1896ACA (5828T>A), CTG1897TTG (5829C>T), TTC1899TTT (5837C>T), GAG1901GAA (5843G>A), CTG1902TTC (5844C>T 5846G>C), AGT1903TCC (5847A>T 5848G>C 5849T>C), ATG1904CAA (5850A>C 5851T>A 5852G>A), GAA1905CAC (5853G>C 5855A>C), AAT1906GAT (5856A>G), GTG1907TGT (5859G>T 5860T>G 5861G>T), CAA1908GAA (5862C>G), CAA1909ATG (5865C>A 5866A>T 5867A>G), CTG1910ATA (5868C>A 5870G>A), ATA1911CTT (5871A>C 5873A>T), GAT1912GAG (5876T>G), GTT1914CTG (5880G>C 5882T>G), AAG1915GAA (5883A>G 5885G>A), CTT1916AGT (5886C>A 5887T>G), CAG1917CAA (5891G>A), TTT1919GTA (5895T>G 5897T>A), TTC1920GTA (5898T>G 5900C>A), CCA1921TTC (5901C>T 5902C>T 5903A>C), AAA1922ATG (5905A>T 5906A>G), GAA1923GGG (5908A>G 5909A>G), ACT1924AGC

| | |
|---|---|
| | (5911C>G 5912T>C), ACC1925GTT (5913A>G 5914C>T 5915C>T), ATT1926AAG (5917T>A 5918T>G), CAG1927AAA (5919C>A 5921G>A), AAG1928CAA (5922A>C 5924G>A), CTC1929ATG (5925C>A 5927C>G), AAA1930TTG (5928A>T 5929A>T 5930A>G), AAT1931TCC (5931A>T 5932A>C 5933T>C), TTT1932TTC (5936T>C), GTT1933ATG (5937G>A 5939T>G), GAT1934AAT (5940G>A), GCT1935CAA (5943G>C 5944C>A 5945T>A), GAA1936CAG (5946G>C 5948A>G), TGT1937ATG (5949T>A 5950G>T 5951T>G), CTA1938TCA (5952C>T 5953T>C), ACA1939AAA (5956C>A), TTC1940ACC (5958T>A 5959T>C), AAA1941CAA (5961A>C), GCA1942GTT (5965C>T 5966A>T), AGG1943GCC (5967A>G 5968G>C 5969G>C), AGC1944_TTT1945insCCATGTTGTGTCACA (5972_5973insCCATGTTGTGTCACA), AGG1946AGT (5978G>T), CAC1947AAA (5979C>A 5981C>A), GTA1948GCA (5983T>C), GTT1949TGG (5985G>T 5986T>G 5987T>G), TTC1950CTA (5988T>C 5990C>A), AAA1951ACT (5992A>C 5993A>T), AAA1953GAC (5997A>G 5999A>C), CTT1954ATT (6000C>A), TTG1955GTC (6003T>G 6005G>C), AGT1956ATG (6007G>T 6008T>G), TCC1957ACA (6009T>A 6011C>A), ACA1958AAA (6013C>A), TGG1959GGT (6015T>G 6017G>T), AAA1960AAG (6020A>G), ATG1961AAG (6022T>A), TGT1962CCA (6024T>C 6025G>C 6026T>A), GGT1963GGA (6029T>A), GAA1964del (6030_6032delGAA), AGT1965TCT (6033A>T 6034G>C), GTT1967_AGG1970del (6039_6050delGTTGCTTCAAGG), TCT1971CCT (6051T>C), GCA1972GCT (6056A>T), GAG1973GAA (6059G>A), AGT1974ACA (6061G>C 6062T>A), CTA1975GTG (6063C>G 6065A>G), TTC1976TTT (6068C>T), CCT1977CTT (6070C>T), GGC1978GGG (6074C>G), TTG1980CCT (6078T>C 6079T>C 6080G>T), TCT1981TCA (6083T>A), GCT1982GCA (6086T>A), TCC1983CTA (6087T>C 6088C>T 6089C>A), GTA1984GTT (6092A>T), TTG1985GTG (6093T>G), AAA1986AGA (6097A>G), TTA1988ATG (6102T>A 6104A>G), GAA1990CGC (6108G>C 6109A>G 6110A>C), AGA1991AAT (6112G>A 6113A>T), CAC1992AGT (6114C>A 6115A>G 6116C>T), ACA1993GGA (6117A>G 6118C>G), AGA1994AGG (6122A>G), ATG1996TAC (6126A>T 6127T>A 6128G>C), AGT1997TCA (6129A>T 6130G>C 6131T>A), CAG1999ACA (6135C>A 6136A>C 6137G>A), AGT2000ACA (6139G>C 6140T>A), ATG2001CTG (6141A>C), GAG2002GAA (6146G>A), CTG2005TTT (6153C>T 6155G>T), AAA2006AGA (6157A>G), ATA2007ATG (6161A>G), TAT2008TAC (6164T>C), CTA2009ATG (6165C>A 6167A>G), TTA2010TGT (6169T>G 6170A>T), AAC2011CAG (6171A>C 6173C>G), TTC2012GAA (6174T>G 6175T>A 6176C>A), CAA2013GGG (6177C>G 6178A>G 6179A>G), GTA2014TAT (6180G>T 6181T>A 6182A>T), ATA2015TTA (6183A>T), ACA2016GAT (6186A>G 6187C>A 6188A>T), AGT2017AAA (6190G>A 6191T>A), GAG2018GCC (6193A>C 6194G>C), TTA2020GTG (6198T>G 6200A>G), AAG2021GAC (6201A>G 6203G>C), AAG2022CGC (6204A>C 6205A>G 6206G>C), TTT2023TTC (6209T>C), GAA2024AAT (6210G>A 6212A>T), GAT2025GAG (6215T>G), TTT2027ATG (6219T>A 6221T>G), CAG2028GGC (6222C>G 6223A>G 6224G>C), GAG2029GTT (6226A>T 6227G>T), AAG2030TGT (6228A>T 6229A>G 6230G>T), ATA2031GTG (6231A>G 6233A>G), TCC2032TCG (6236C>G), ACC2033GAG (6237A>G 6238C>A 6239C>G), ACC2034CGC (6240A>C 6241C>G), CTA2035ATC (6243C>A 6245A>C), CTA2036TTG (6246C>T 6248A>G), ATC2037GTG (6249A>G 6251C>G), AAA2038CGC (6252A>C 6253A>G 6254A>C), GTA2039ACA (6255G>A 6256T>C), TTC2040ATT (6258T>A 6260C>T), TTG2041GCT (6261T>G 6262T>C 6263G>T), GAG2044TCC (6270G>T 6271A>C 6272G>C), GAG2045ACG (6273G>A 6274A>C), AGT2046TCA (6276A>T 6277G>C 6278T>A), TAT2047TTG (6280A>T 6281T>G), CAG2048AAC (6282C>A 6284G>C), ACT2049GAG (6285A>G 6286C>A 6287T>G), GAG2050GAT (6290G>T), TGT2052ACT (6294T>A 6295G>C), ATA2053GTG (6297A>G 6299A>G), GAT2054TAC (6300G>T 6302T>C), TCT2055ACA (6303T>A 6305T>A), CAG2056GAT (6306C>G 6308G>T), GTG2057GGG (6310T>G), CTC2058CTT (6314C>T), CAG2059TTT (6315C>T 6316A>T 6317G>T), AGG2060GAT (6318A>G 6319G>A 6320G>T), ATA2061del (6321_6323delATA), AAC2062CAT (6324A>C 6326C>T), TCC2063TTC (6328C>T), CAG2064AAG (6330C>A), AAA2065GCA (6333A>G 6334A>C), TTC2066GTG (6336T>G 6338C>G), AGA2067GTT (6339A>G 6340G>T 6341A>T), GTG2068TTC (6342G>T 6344G>C), CAG2069AAG (6345C>A), GAA2070CCT (6348G>C 6349A>C 6350A>T), GGT2071GGG (6353T>G), TTT2073TTC (6359T>C), TAT2074TCT (6361A>C), TTG2075TGG (6364T>G), CAT2076GAA (6366C>G 6368T>A), CCT2077ATC (6369C>A 6370C>T 6371T>C), GAA2078AAT (6372G>A 6374A>T), GGA2079GAT (6376G>A 6377A>T), ATC2080GCC (6378A>G 6379T>C), AAC2081AAT (6383C>T), GCC2084GCT (6392C>T), GTA2085GCA (6394T>C), ATA2088GTG (6402A>G 6404A>G), ATA2089TTG (6405A>T 6407A>G), TCT2091GAA (6411T>G 6412C>A 6413T>A), CTG2092CTC (6416G>C), GAG2094AGG (6420G>A 6421A>G), TCC2095CAC (6423T>C 6424C>A), TAT2096TGG (6427A>G 6428T>G), AGC2097TCA (6429A>T 6430G>C 6431C>A), CGA2098AAG (6432C>A 6433G>A 6434A>G), GAT2099TCA (6435G>T 6436A>C 6437T>A), AGT2100CGA (6438A>C 6440T>A), AAC2101AGC (6442A>G), TCC2102AAG (6444T>A 6445C>A 6446C>G), TAT2103TTC (6448A>T 6449T>C), TCT2104ACT (6450T>A), GTG2105TTT (6453G>T 6455G>T), TCA2106GCC (6456T>G 6458A>C), ACT2107AAT (6460C>A), CCC2108GAA (6462C>G 6463C>A 6464C>A), ATC2109GTT (6465A>G 6467C>T), AAA2110GAA (6468A>G), CTG2111ATA (6471C>A 6473G>A), GGG2112TCA (6474G>T 6475G>C 6476G>A), TGT2113AAC (6477T>A 6478G>A 6479T>C), TTC2114CTG (6480T>C 6482C>G), ATT2115CTT (6483A>C), TTC2116TTA (6488C>A), CTT2117GCT (6489C>G 6490T>C), GTC2118GTT (6494C>T), GTG2119TAT (6495G>T 6496T>A 6497G>T), CTA2120TTA (6498C>T), GAT2121GAA (6503T>A), AGT2122AAT (6505G>A), CAA2123AAG (6507C>A 6509A>G), GGC2124CTC (6510G>C 6511G>T), AAG2125ATA (6514A>T 6515G>A), GAA2126GGT (6517A>G 6518A>T), GAT2127CTA (6519G>C 6520A>T 6521T>A), CAT2128GAG (6522C>G 6524T>G), CCT2129TAT (6525C>T 6526C>A), GTT2130TAT (6528G>T 6529T>A), CGG2131GAT (6531C>G 6532G>A 6533G>T), AGG2132TCT (6534A>T 6535G>C 6536G>T), TTT2133ATA (6537T>A 6539T>A), TCA2134GAA (6540T>G 6541C>A), ACG2135TCT (6543A>T 6545G>T), GTT2136TTG (6546G>T 6548T>G), TTC2137GAC (6549T>G 6550T>A), CTT2138TTG (6552C>T 6554T>G), AAA2139TTA (6555A>T 6556A>T), AAC2140CAA (6558A>C 6560C>A), TGT2141ATA (6561T>A 6562G>T 6563T>A), GAA2142GAT (6566A>T), ATT2144ATA (6572T>A), ACA2146GCA (6576A>G), TAC2148GTC (6582T>G 6583A>T), GAG2150TAC (6588G>T 6590G>C), TGA2152GGG (6594T>G 6596A>G) |

**Proteins**

| | | | | | | | | | |
|---|---|---|---|---|---|---|---|---|---|
| polyprotein (YP_001976147.1) | 1 | 2152 | 100% | 9871 | 66.5% | 2135 (98.7%) | 1362 (63.0%) | 11/17/0/0 | 0 |

| | Begin | End | Coverage | Score | Concordance | Matches | Identities | I/D/M/F* | Stop Codons |
|---|---|---|---|---|---|---|---|---|---|
| **NT** | **22** | **6747** | **93.0%** | **3341** | **25.4%** | **6659 (98.4%)** | **4235 (62.6%)** | **43/67** | |

Protein mutations:

F7W (160T>G 161C>G), K9_A16del (165_188delAAGGTTGCTGGTGTTGATGACGCC), S17P (189A>C 190G>C), S18P (192T>C), S19A (195T>G), T21S (201A>T 203C>A), T22P (204A>C), G26A (217G>C 218T>A), L28V (222C>G), N30S (228A>T 229A>C), I32F (234A>T), K34S (240A>T 241A>C 242A>T), F35L (243T>C), A38T (252G>A 254G>T), V39L (255G>C 257T>C), I42V (264A>G), T43N (268C>A 269C>T), T44K (271C>A 272T>A), M45V (273A>G 275G>T), K48T (283A>C 284G>C), S50V (288T>G 289C>T), L53I (297C>A 299T>A), L56V (306C>G 308G>C), S59P (315T>C 317T>A), T61M (322C>T 323T>G), D62A (325A>C 326T>A), V63A (328T>C 329G>A), S64A (330T>G 332C>A), V68A (343T>C 344G>C), A69S (345G>T 347C>T), S72N (355G>A), S73A (357T>G 359A>T), L74V (360C>G 362T>A), N75E (363A>G 365A>T), K76R (367A>G), G82A (385G>C 386T>A), A83K (387G>A 388C>A), L84I (390C>A 392T>A), I85M (395A>G), D86E (398T>A), L89I (405C>A 407T>C), A91G (412C>G 413G>T), S93D (417T>G 418C>A), L95V (423T>G 425G>T), S96K (426T>A 427C>A 428C>A), S101T (441T>A), K103V (447A>G 448A>T 449G>A), M105L (453A>C 455G>T), L109M (465T>A), F110V (468T>G), E111L (471G>T 472A>T 473G>A), L113V (477T>G 479G>C), S114K (480T>A 481C>A 482C>G), N115D (483A>G), V117F (489G>T 491T>C), K118I (493A>T 494A>C), F121Y (502T>A 503T>C), A122E (505C>A 506T>A), V124L (510G>T 512G>A), N126E (516A>G 518T>A), S128V (522A>G 523G>T), D130S (528G>T 529A>C 530T>A), V131L (531G>T), V135F (543G>T 545A>T), Y136I (546T>A 547A>T 548T>A), G137A (550G>C), I143L (567A>T 569A>G), T144A (570A>G 572A>C), L145I (573T>A 575G>T), V147L (579G>C 581G>A), F150Y (589T>A), S154V (600A>G 601G>T 602T>A), E158T (612G>A 613A>C), G159P (615G>C 616G>C), V160F (618G>T), S161N (621T>A 622C>A), T163V (627A>G 628C>T 629T>A), K165V (633A>G 634A>T 635A>T), N166T (637A>C), I167L (639A>T 641C>G), L169S (646T>C 647G>A), I170V (648A>G 650A>T), M174F (660A>T 662G>T), K186T (697A>C), A188S (702G>T 704C>T), E189G (706A>G 707A>G), R190K (708C>A 709G>A 710A>G), T192S (714A>T), M193Q (717A>C 718T>A), V194L (720G>C), S198A (732A>G 733G>C 734T>A), M199V (735A>G 737G>A), V205L (753G>T 755T>A), S206T (756T>A 758A>G), M214L (780A>T 782G>A), T216S (786A>T), I217L (789A>T 791A>G), I228V (822A>G), T248A (882A>G 884A>T), T260Q (918A>C 919C>A 920T>G), I280V (978A>G 980A>G), G282S (984G>A), T283I (988C>T 989C>T), S295N (1024G>A), E304Q (1050G>C), Q307T (1059C>A 1060A>C), D310S (1068G>T 1069A>C 1070T>G), A311E (1072C>A 1073A>G), C313A (1077T>G 1078G>C), D314E (1082T>A), A316S (1086G>T 1088A>T), T317M (1090C>T), K325P (1113A>C 1114A>C 1115G>T), T329I (1126C>T 1127C>T), N332Q (1134A>C 1136C>A), R335K (1144G>A 1145G>A), E337Q (1149G>C 1151A>G), L339V (1155C>G), M354L (1200A>T), A363T (1227G>A), E366D (1238A>T), V368L (1242G>C 1244T>G), Y369V (1245T>G 1246A>T), S372A (1254A>G 1255G>C 1256T>C), C373S (1257T>A 1259T>C), R374Q (1260A>C 1261G>A 1262G>A), M376S (1266A>T 1267T>C 1268G>T), S393A (1317T>G 1319C>T), S401C (1342C>G 1343C>T), R412A (1374A>G 1375G>C), I440V (1458A>G 1460C>G), M521R (1702T>G 1703G>A), E523V (1708A>T), E525D (1715G>T), S527D (1719T>G 1720C>A 1721C>T), T528P (1722A>C 1724T>A), E529N (1725G>A 1727A>T), D530T (1728G>A 1729A>C), G533A (1738G>C 1739A>C), R536V (1746A>G 1747G>T 1748A>T), N538D (1752A>G 1754C>T), I539V (1755A>G 1757A>G), N552A (1794A>G 1795A>C), T554S (1800A>T), S556D (1806A>G 1807G>A), E560Q (1818G>C 1820G>A), Q567R (1840A>G), A570Y (1848G>T 1849C>A 1850A>T), K574S (1861A>G 1862G>C), F577Y (1870T>A 1871T>C), E578N (1872G>A 1874A>T), A579T (1875G>A), E580T (1878G>A 1879A>C), I583L (1887A>T 1889A>G), S584A (1890A>G 1891G>C), P585D (1893C>G 1894C>A 1895T>C), I587V (1899A>G 1901A>T), Q591N (1911C>A 1913A>T), S593A (1917T>G), A594V (1921C>T 1922A>T), D596E (1928T>A), I598L (1932A>C 1934A>G), A599D (1936C>A 1937G>T), V601F (1941G>T 1943G>C), D602Q (1944G>C 1946T>A), G603D (1948G>A 1949G>T), R605del (1953_1955delAGA), R607Q (1960G>A 1961C>A), T608V (1962A>G 1963C>T 1964C>T), K609L (1965A>T 1966A>T), Q610P (1969A>C 1970A>T), D611P (1971G>C 1972A>C), I614A (1980A>G 1981T>C), L615A (1983T>G 1984T>C 1985G>T), V616Y (1986G>T 1987T>A 1988G>T), P617E (1989C>G 1990C>A 1991C>G), T618Q (1992A>C 1993C>A 1994T>G), V619T (1995G>A 1996T>C 1997T>A), V620I (1998G>A 2000G>T), S621T (2002G>C 2003T>A), N623A (2007A>G 2008A>C), D624E (2012T>A), R625K (2014G>A), V626I (2016G>A 2018G>T), I627V (2019A>G 2021C>T), A629S (2025G>T 2027A>C), R630Q (2028A>C 2029G>A 2030A>G), L645M (2073T>A), D646N (2076G>A), P647K (2079C>A 2080C>A 2081T>A), E648F (2082G>T 2083A>T 2084G>T), L650F (2088C>T), H655L (2104A>T 2105T>C), H656S (2106C>T 2107A>C 2108T>C), M660I (2120G>C), S663V (2127T>G 2128C>T 2129T>G), D665S (2133G>T 2134A>C), D666S (2136G>T 2137A>C), E667D (2141A>T), H668A (2142C>G 2143A>C 2144T>C), E670Q (2148G>C), I671N (2152T>A), N673E (2157A>G 2159C>A), K674M (2161A>T 2162A>G), R676V (2166A>G 2167G>T), V677N (2169G>A 2170T>A 2171C>T), N678D (2172A>G 2174C>T), M679L (2175A>T), Y681V (2181T>G 2182A>T 2183T>C), R682K (2185G>A), D685S (2193G>A 2194A>G), A686P (2196G>C), E687Q (2199G>C 2201A>G), R690A (2208C>G 2209G>C 2210A>T), Q694R (2221A>G), A710S (2268G>A 2269C>G 2270A>C), S711T (2271T>A), K716E (2286A>G), K717L (2289A>C 2290A>T 2291A>G), A719D (2296C>A 2297A>T), T722R (2304A>C 2305C>G 2306A>C), E723N (2307G>A 2309G>T), M724L (2310A>C 2312G>T), R725E (2313A>G 2314G>A), T726Q (2316A>C 2317C>A 2318T>A), F728K (2322T>A 2323T>A 2324T>A), H730Y (2328C>T), S731K (2331T>A 2332C>A 2333T>G), C732A (2334T>G 2335G>C 2336C>G), I733L (2337A>C), D734Q (2340G>C 2342T>G), V737A (2350T>C 2351T>A), I741T (2362T>C 2363A>G), F749R (2385T>A 2386T>G 2387T>A), N763R (2428A>G 2429C>G), S766A (2436T>G), S766_K767insG (2438_2439insGGT), K767T (2440A>C 2441A>T), E768D (2444G>C), P769K (2445C>A 2446C>A 2447A>G), E771R (2451G>A 2452A>G), S772N (2454T>A 2455C>A 2456A>T), K775I (2464A>T 2465A>C), I776E (2466A>G 2467T>A 2468T>A), Y778M (2472T>A 2473A>T 2474C>G), H779I (2475C>A 2476A>T), K780G (2478A>G 2479A>G 2480A>T), V782K (2484G>A 2485T>A 2486T>G), E783D (2489A>C), Y784K (2490T>A 2492T>A), I785M (2495C>G), K786I (2497A>T 2498G>C), D787Q (2499G>C 2501T>G), T788M (2503C>T 2504A>G), W790N (2508T>A 2509G>A 2510G>T), S791D (2511A>G 2512G>A), Y794F (2521A>T 2522T>C), A795I (2523G>A 2524C>T 2525A>C), R797K (2530G>A 2531A>G), I806L (2556A>T 2558A>G), M807I (2561G>A), V812T (2574G>A 2575T>C 2576T>C), F813M (2577T>A 2579T>G), G814N (2580G>A 2581G>A 2582A>T), G815A (2584G>C), I816V (2586A>G), L820I (2598T>A 2600G>T), A821T (2601G>A 2603G>T), W823L (2607T>C 2608G>T 2609G>C), Q824K (2610C>A), S825A (2613A>G 2614G>C 2615T>A), N826A (2616A>G 2617A>C 2618C>A), S827T (2619T>A), F828S (2623T>C), N830L (2628A>T 2629A>T 2630T>A), S832D (2634T>G 2635C>A 2636G>T), N852K (2696T>G), K853T (2698A>C 2699G>T), K853_L854insH (2699_2700insCAT), L854F (2700C>T), R855A (2703A>G 2704G>C), N856K (2708T>A), L857S (2709C>T 2710T>C 2711T>C), V859L (2715G>C 2717T>A), S862A (2724A>G 2725G>C 2726T>G), S863A (2727T>G 2729T>A), Q864D (2730C>G 2732A>T), S865N (2733T>A 2734C>A 2735A>C), L866N (2736T>A 2737T>A 2738G>C), A867H (2739G>C 2740C>A 2741A>T), Q868V (2742C>G 2743A>T 2744A>G), D869_W870insR (2747_2748insCGG), A872E (2755C>A 2756T>A), E873Q (2757G>C 2759A>G), Q879E (2775C>G 2777G>A), K882H (2784A>C 2786G>C), L890I (2808C>A), G891A (2812G>C), M907L (2859A>C 2861G>C), H914R (2880C>A 2881A>G 2882C>A), A916M (2886G>A 2887C>T 2888T>G), T923S (2907A>T 2909A>C), A927E (2920C>A 2921C>G), N928H (2922A>C), R930I (2928C>A 2929G>T 2930A>T), S931C (2932C>G), K932N (2936A>T), Y935F (2944A>T), R938T (2953G>C 2954A>C), E939Q (2955G>C), R941K (2962G>A), I942L (2964A>T 2966T>G), I948V (2982A>G 2984C>G), P961S (3021C>T 3023T>A), E972D (3056G>T), F975Y (3064T>A), T976S (3066A>T 3068A>T), Q980M (3078C>A 3079A>T 3080A>G), C984V (3090T>G 3091G>T 3092T>G), T990K (3109C>A 3110A>G), R993K (3117C>A 3118G>A 3119T>G), L994K (3120C>A 3121T>A 3122T>A), G995S (3123G>T 3124G>C), F999V (3135T>G), M1002Y (3144A>T 3145T>A 3146G>T), V1012E (3175T>A 3176T>A), E1015T (3183G>A 3184A>C), E1017G (3190A>G), N1019D (3195A>G 3197C>T), S1021N (3201T>A 3202C>A 3203C>T), S1022A (3204A>G 3205G>C 3206T>A), K1024A (3210A>G 3211A>C 3212G>T), I1025L (3213A>C), C1026S (3216T>A), Q1029V (3225C>G 3226A>T 3227A>T), S1030A (3228T>G 3230T>A), G1037S (3249G>A 3251G>T), S1050T (3289G>C), V1057C (3309G>T 3310T>G 3311G>C), G1068D (3343G>A 3344C>T), V1069T (3345G>A 3346T>C 3347A>C), R1073K (3357C>A 3358G>A 3359T>G), F1075Y (3364T>A 3365T>C), I1076V (3366A>G 3368C>G), S1077A (3369T>G), I1083V (3387A>G 3389C>G), Y1084F (3391A>T 3392C>T), E1086D (3398A>T), V1096I (3426G>A 3428C>T), A1098V (3433C>T 3434A>G), R1102H (3445G>A), D1104E (3452T>A), I1105L (3453A>C 3455T>G), S1108H (3462T>C 3463C>A 3464A>C), I1109L (3465A>C 3467T>G), R1112K (3475G>A 3476A>G), T1114E (3480A>G 3481C>A), Q1115R (3483C>A 3484A>G 3485G>A), L1116F (3486C>T 3488T>C), N1120K (3500C>A), Y1123A (3507T>G 3508A>C 3509C>T), Y1124F (3511A>T), Q1130L (3528C>T 3529A>T), L1132M (3534T>A), R1143K (3567C>A 3568G>A 3569G>A), K1144R (3570A>C 3571A>G 3572A>C), Q1146P (3577A>C 3578A>C), P1153D (3597C>G 3598C>A 3599T>C), E1155Q (3603G>C), T1162S (3624A>T 3626A>T), V1163A (3628T>C 3629T>C), H1164K (3630C>A 3632T>A), D1169E (3647T>A), M1179E (3675A>G 3676T>A), C1186S (3697G>C 3698T>C), G1187N (3699G>A 3700G>A 3701A>C), D1189E (3707T>G), D1190E (3710T>G), D1191P (3711G>C 3712A>C 3713C>A), N1199G (3735A>G 3736A>G), C1202A (3744T>G 3745G>C 3746C>A), L1204I (3750C>A 3752G>T), I1207V (3759A>G 3761C>G), M1208L (3762A>C 3764G>T), R1209K (3765C>A 3766G>A 3767T>G), S1210M (3769G>T 3770C>G), K1212R (3774A>C 3775A>G 3776G>A), I1213V (3777A>G 3779C>G), R1217K (3789C>A 3790G>A 3791T>A), C1219V (3795T>G 3796G>T 3797T>A), E1220I (3798G>A 3799A>T 3800A>C), D1222N (3804G>A), C1233F (3838G>T), Y1236F (3847A>T), G1238A (3853G>C 3854C>A), M1252L (3894A>T), M1254L (3900A>C 3902G>T), M1256S (3906A>T 3907T>C 3908G>T), Q1263E (3927C>G), S1264A (3930T>G), T1265Q (3933A>C 3934C>A 3935T>A), G1266P (3936G>C 3937G>C 3938C>A), E1267N (3939G>A 3941A>C), G1268S (3942G>T 3943G>C 3944A>G), E1269Q (3945G>C), E1272L (3954G>C 3955A>T 3956A>T), D1276N (3966G>A), E1281D (3983A>T), V1288S (4002G>T 4003T>C), I1290V (4008A>G 4010T>A), K1292Q (4014A>C), I1294T (4021T>C 4022C>A), H1295F (4023C>T 4024A>T), T1344C (4170A>T 4171C>G 4172A>T), L1348A (4182T>G 4183T>C 4184G>A), A1351V (4192C>T 4193T>A), I1352L (4194A>C 4196A>C), D1359N (4215G>A), S1363H (4227T>C 4228C>A), L1367V (4239T>G), N1370S (4249A>G), G1371S (4251G>A 4253G>C), A1374S (4260G>T 4262T>A), V1378E (4273T>A), V1393I (4317G>A 4319T>C), E1405A (4354A>C 4355A>T), Y1406F (4357A>T 4358C>T), S1409D (4365T>G 4366C>A 4367A>T), K1410E (4368A>G), D1415E (4385T>G), A1421T (4401G>A), D1435N (4443G>A 4445T>C), V1437L (4449G>T), I1476V (4566A>G 4568T>C), A1477K (4569G>A 4570C>A 4571T>A), R1479K (4576G>A 4577G>A), A1480S (4578G>T), I1481V (4581A>G 4583A>G), V1483I (4587G>A), S1486N (4597G>A), L1506M (4656T>A 4658A>G), Q1509R (4665C>A 4666A>G), S1512D (4674T>G 4675C>A), N1515S (4683A>T 4684A>C 4685T>C), H1518R (4692C>A 4693A>G 4694T>A), L1522A (4704C>G 4705T>C 4706G>T), I1526V (4716A>G), N1538E (4752A>G 4754T>G), E1539K (4755G>A), V1540T (4758G>A 4759T>C), Q1541K (4761C>A), Q1543L (4768A>T 4769A>G), H1563Q (4829C>A), I1572V (4854A>G), M1576I (4868G>T), P1583V (4887C>G 4888C>T 4889A>G), T1588V (4902A>G 4903C>T 4904T>A), M1592Y (4914A>T 4915T>A 4916G>T), V1595I (4923G>A 4925G>T), R1607Q (4960G>A 4961T>G), V1608A (4963T>C 4964G>C), A1609T (4965G>A), D1611E (4973C>A), K1612S (4975A>G 4976G>T), L1613F (4977C>T), L1615N (4983T>A 4984T>A 4985A>T), Q1618F (4992C>T 4993A>T 4994G>T), T1619G (4995A>G 4996C>G 4997G>A), D1622K (5004G>A 5006T>G), E1623Q (5007G>C 5009A>G), G1625N (5013G>A 5014G>A), F1626L (5016T>C), R1627P (5019A>C 5020G>C 5021A>C), A1628I (5022G>A 5023C>T 5024T>C), V1629M (5025G>A), F1631M (5031T>A 5033T>G), S1633T (5037T>A 5039C>T), L1635Y (5044T>A 5045G>T), V1637L (5049G>C 5051G>T), Q1639S (5055C>A 5056A>G 5057G>T), E1640Q (5058G>C), I1642L (5064A>T 5066T>G), A1643T (5067G>A), K1644E (5070A>G), S1649E (5085T>G 5086C>A 5087A>G), A1651I (5091G>A 5092C>T 5093C>T), L1652G (5094C>G 5095T>G), T1653V (5097A>G 5098C>T 5099A>T), I1654L (5100A>T 5102T>G), C1655M (5103T>A 5104G>T 5105T>G), M1657L (5109A>C 5111G>A), A1658S (5112G>T), K1659D (5115A>G 5117A>T), A1661K (5121G>A 5122C>A 5123T>A), I1662N (5125T>A), S1663T (5127T>A 5129C>A), D1668P (5142G>C 5143A>C 5144C>T), T1674del (5160_5162delACA), R1678E (5172C>G 5173G>A 5174T>G), D1679G (5176A>G), L1680V (5178C>G 5180G>C), N1681H (5181A>C), W1682E (5184T>G 5185G>A 5186G>A), A1684V (5191C>T 5192A>G), N1686D (5196A>G 5198C>T), I1687V (5199A>G 5201T>A), G1688F (5202G>T 5203G>T), A1696S (5226G>T 5228C>A), H1697N (5229C>A), Y1698Q (5232T>C 5234C>A), T1699Q (5235A>C 5236C>A 5237G>A), D1701S (5241G>T 5242A>C 5243T>A), E1702S (5244G>T 5245A>C), Y1704N (5250T>A), I1706L (5256A>C 5258T>G), K1713R (5277A>C 5278A>G 5279G>C), K1714R (5281A>G 5282G>A), S1715T (5283T>A 5285C>A), N1716T (5287A>C 5288T>A), M1731I (5333G>A), M1736L (5346A>T), S1737A (5349T>G), M1740Q (5358A>C 5359T>A), A1742H (5364G>C 5365C>A 5366C>T), K1744R (5370A>C 5371A>G 5372G>C), N1745K (5375T>G), V1746N (5376G>A 5377T>A 5378G>T), A1748E (5383C>A), G1749V (5386G>T 5387A>T), H1751D (5391C>G), V1752G (5395T>G 5396A>G), N1755S (5403A>T 5404A>C), V1757T (5409G>A 5410T>C 5411T>C), V1766I (5436G>A 5438C>A), G1767A (5440G>C 5441C>T), M1774F (5460A>T 5462G>T), R1779H (5475A>C 5476G>A 5477G>T), C1782R (5484T>A 5486T>G), V1783T (5487G>A 5488T>C 5489G>C), E1784T (5490G>A 5491A>C 5492G>A), I1786L (5496A>C 5498A>T), I1793M (5519T>G), G1794A (5521G>C 5522A>T), N1795G (5523A>G 5524A>G 5525T>G), V1807A (5560T>C), F1810L (5570T>G), Q1812K (5574C>A), P1815E (5583C>G 5584C>A 5585T>A), H1816D (5586C>G 5588T>C), H1816_G1817insT (5588_5589insACC), G1817S (5589G>T 5590G>C), D1818H (5592G>C), V1820L (5598G>T), A1821G (5602C>G 5603T>A), A1825T (5613G>A), S1826Y (5617C>A 5618G>T), L1827G (5619C>G 5620T>G), H1828del (5622_5624delCAC), S1829T (5626G>C 5627T>C), S1830T (5629G>C), M1832N (5635T>A 5636G>C), Q1834H (5642A>T), Q1834_G1835insHS (5642_5643insCACAGT), G1835A (5644G>C 5645G>A), E1837S (5649G>T 5650A>C), L1838V (5652T>G 5654G>T), N1840L (5658A>C 5659A>T 5660C>T), I1844V (5670A>G 5672A>T), H1845P (5674A>C), G1846S (5676G>T 5677G>C 5678T>C), A1849S (5685G>T 5687A>T), K1850E (5688A>G), Q1851K (5691C>A), T1852M (5695C>T 5696A>G), M1854V (5700A>G 5702G>T), M1856L (5706A>T), V1859T (5715G>A 5716T>C 5717G>A), V1860I (5718G>A 5720A>T), H1863E (5727C>G 5729T>A), G1867A (5740G>C 5741A>C), N1868Q (5742A>C 5744T>G), F1869C (5746T>G 5747T>C), Q1872K (5754C>A), D1873N (5757G>A 5759T>C), E1875K (5763G>A), S1876H (5766A>C 5767G>A), L1877A (5769T>G 5770T>C 5771G>T), H1879E (5775C>G 5777C>G), Y1880T (5778T>A 5779A>C 5780T>A), E1882S (5784G>T 5785A>C 5786G>T), Q1883V (5787C>G 5788A>T 5789G>C), G1884S (5790G>A 5792A>T), E1887T (5799G>A 5800A>C 5801G>T), I1889A (5805A>G 5806T>C 5807T>C), Q1890A (5808C>G 5809A>C), K1891R (5812A>G 5813G>A), N1892G (5814A>G 5815A>G 5816T>A), T1893V (5817A>G 5818C>T 5819A>C), L1895V (5823C>G 5825A>G), L1902F (5844C>T 5846G>C), M1904Q (5850A>C 5851T>A 5852G>A), E1905H (5853G>C 5855A>C), N1906D (5856A>G), V1907C (5859G>T 5860T>G 5861G>T), Q1908E (5862C>G), Q1909M (5865C>A 5866A>T 5867A>G), L1910I (5868C>A 5870G>A), I1911L (5871A>C 5873A>T), D1912E (5876T>G), V1914L (5880G>C 5882T>G), K1915E (5883A>G 5885G>A), L1916S (5886C>A 5887T>G), F1919V (5895T>G 5897T>A), F1920V (5898T>G 5900C>A), P1921F (5901C>T 5902C>T 5903A>C), K1922M (5905A>T 5906A>G), E1923G (5908A>G 5909A>G), T1924S (5911C>G 5912T>C), T1925V (5913A>G 5914C>T 5915C>T), I1926K (5917T>A 5918T>G), Q1927K (5919C>A 5921G>A), K1928Q (5922A>C 5924G>A), L1929M (5925C>A 5927C>G), K1930L (5928A>T 5929A>T 5930A>G), N1931S (5931A>T 5932A>C 5933T>C), V1933M (5937G>A 5939T>G), D1934N (5940G>A), A1935Q (5943G>C 5944C>A 5945T>A), E1936Q (5946G>C 5948A>G), C1937M (5949T>A 5950G>T 5951T>G), L1938S (5952C>T 5953T>C), T1939K (5956C>A), F1940T (5958T>A 5959T>C), K1941Q (5961A>C), A1942V (5965C>T 5966A>T), R1943A (5967A>G 5968G>C 5969G>C), S1944_F1945insPCCVT (5972_5973insCCATGTTGTGTCACA), R1946S (5978G>T), H1947K (5979C>A 5981C>A), V1948A (5983T>C), V1949W (5985G>T 5986T>G 5987T>G), F1950L (5988T>C 5990C>A), K1951T (5992A>C 5993A>T), K1953D (5997A>G 5999A>C), L1954I (6000C>A), L1955V (6003T>G 6005G>C), S1956M (6007G>T 6008T>G), S1957T (6009T>A 6011C>A), T1958K (6013C>A), W1959G (6015T>G 6017G>T), M1961K (6022T>A), C1962P (6024T>C 6025G>C 6026T>A), E1964del (6030_6032delGAA), V1967_R1970del (6039_6050delGTTGCTTCAAGG), S1971P (6051T>C), S1974T (6061G>C 6062T>A), L1975V (6063C>G 6065A>G), P1977L (6070C>T), L1980P (6078T>C 6079T>C 6080G>T), S1983L (6087T>C 6088C>T 6089C>A), L1985V (6093T>G), K1986R (6097A>G), L1988M (6102T>A 6104A>G), E1990R (6108G>C 6109A>G 6110A>C), R1991N (6112G>A 6113A>T), H1992S (6114C>A 6115A>G 6116C>T), T1993G (6117A>G 6118C>G), M1996Y

(6126A>T 6127T>A 6128G>C), Q1999T (6135C>A 6136A>C 6137G>A), S2000T (6139G>C 6140T>A), M2001L (6141A>C), L2005F (6153C>T 6155G>T), K2006R (6157A>G), I2007M (6161A>G), L2009M (6165C>A 6167A>G), L2010C (6169T>G 6170A>T), N2011Q (6171A>C 6173C>G), F2012E (6174T>G 6175T>A 6176C>A), Q2013G (6177C>G 6178A>G 6179A>G), V2014Y (6180G>T 6181T>A 6182A>T), I2015L (6183A>T), T2016D (6186A>G 6187C>A 6188A>T), S2017K (6190G>A 6191T>A), E2018A (6193A>C 6194G>C), L2020V (6198T>G 6200A>G), K2021D (6201A>G 6203G>C), K2022R (6204A>C 6205A>G 6206G>C), E2024N (6210G>A 6212A>T), D2025E (6215T>G), F2027M (6219T>A 6221T>G), Q2028G (6222C>G 6223A>G 6224G>C), E2029V (6226A>T 6227G>T), K2030C (6228A>T 6229A>G 6230G>T), I2031V (6231A>G 6233A>G), T2033E (6237A>G 6238C>A 6239C>G), T2034R (6240A>C 6241C>G), L2035I (6243C>A 6245A>C), I2037V (6249A>G 6251C>G), K2038R (6252A>C 6253A>G 6254A>C), V2039T (6255G>A 6256T>C), F2040I (6258T>A 6260C>T), L2041A (6261T>G 6262T>C 6263G>T), E2044S (6270G>T 6271A>C 6272G>C), E2045T (6273G>A 6274A>C), Y2047L (6280A>T 6281T>G), Q2048N (6282C>A 6284G>C), T2049E (6285A>G 6286C>A 6287T>G), E2050D (6290G>T), C2052T (6294T>A 6295G>C), I2053V (6297A>G 6299A>G), D2054Y (6300G>T 6302T>C), S2055T (6303T>A 6305T>A), Q2056D (6306C>G 6308G>T), V2057G (6310T>G), Q2059F (6315C>T 6316A>T 6317G>T), R2060D (6318A>G 6319G>A 6320G>T), I2061del (6321_6323delATA), N2062H (6324A>C 6326C>T), S2063F (6328C>T), Q2064K (6330C>A), K2065A (6333A>G 6334A>C), F2066V (6336T>G 6338C>G), R2067V (6339A>G 6340G>T 6341A>T), V2068F (6342G>T 6344G>C), Q2069K (6345C>A), E2070P (6348G>C 6349A>C 6350A>T), Y2074S (6361A>C), L2075W (6364T>G), H2076E (6366C>G 6368T>A), P2077I (6369C>A 6370C>T 6371T>C), E2078N (6372G>A 6374A>T), G2079D (6376G>A 6377A>T), I2080A (6378A>G 6379T>C), V2085A (6394T>C), I2088V (6402A>G 6404A>G), I2089L (6405A>T 6407A>G), S2091E (6411T>G 6412C>A 6413T>A), E2094R (6420G>A 6421A>G), S2095H (6423T>C 6424C>A), Y2096W (6427A>G 6428T>G), R2098K (6432C>A 6433G>A 6434A>G), D2099S (6435G>T 6436A>C 6437T>A), S2100R (6438A>C 6440T>A), N2101S (6442A>G), S2102K (6444T>A 6445C>A 6446C>G), Y2103F (6448A>T 6449T>C), S2104T (6450T>A), V2105F (6453G>T 6455G>T), S2106A (6456T>G 6458A>C), T2107N (6460C>A), P2108E (6462C>G 6463C>A 6464C>A), I2109V (6465A>G 6467C>T), K2110E (6468A>G), L2111I (6471C>A 6473G>A), G2112S (6474G>T 6475G>C 6476G>A), C2113N (6477T>A 6478G>A 6479T>C), F2114L (6480T>C 6482C>G), I2115L (6483A>C), F2116L (6488C>A), L2117A (6489C>G 6490T>C), V2119Y (6495G>T 6496T>A 6497G>T), D2121E (6503T>A), S2122N (6505G>A), Q2123K (6507C>A 6509A>G), G2124L (6510G>C 6511G>T), K2125I (6514A>T 6515G>A), E2126G (6517A>G 6518A>T), D2127L (6519G>C 6520A>T 6521T>A), H2128E (6522C>G 6524T>G), P2129Y (6525C>T 6526C>A), V2130Y (6528G>T 6529T>A), R2131D (6531C>G 6532G>A 6533G>T), R2132S (6534A>T 6535G>C 6536G>T), F2133I (6537T>A 6539T>A), S2134E (6540T>G 6541C>A), T2135S (6543A>T 6545G>T), V2136L (6546G>T 6548T>G), F2137D (6549T>G 6550T>A), K2139L (6555A>T 6556A>T), N2140Q (6558A>C 6560C>A), C2141I (6561T>A 6562G>T 6563T>A), E2142D (6566A>T), T2146A (6576A>G), Y2148V (6582T>G 6583A>T), E2150Y (6588G>T 6590G>C), *2152G (6594T>G 6596A>G)

| | Begin | End | Coverage | Score | Concordance | Matches | Identities | I/D/M/F* | Stop Codons |
|---|---|---|---|---|---|---|---|---|---|
| **NT** | **22** | **6747** | **93.0%** | **3341** | **25.4%** | **6659 (98.4%)** | **4235 (62.6%)** | **43/67** | |

GCT2GCA (146T>A), TTC7TGG (160T>G 161C>G), TCC8AGT (162T>A 163C>G 164C>T), AAG9_GCC16del (165_188delAAGGTTGCTGGTGTTGATGACGCC), AGT17CCT (189A>C 190G>C), TCT18CCT (192T>C), TCT19GCT (195T>G), TCC20TCT (200C>T), ACC21TCA (201A>T 203C>A), ACT22CCT (204A>C), GGT26GCA (217G>C 218T>A), CTT28GTT (222C>G), TCA29TCT (227A>T), AAT30TCT (228A>T 229A>C), ATT32TTT (234A>T), TCC33TCT (239C>T), AAA34TCT (240A>T 241A>C 242A>T), TTT35CTT (243T>C), AAA37AAG (251A>G), GCG38ACT (252G>A 254G>T), GTT39CTC (255G>C 257T>C), AGT40TCC (258A>T 259G>C 260T>C), ATT42GTT (264A>G), ACC43AAT (268C>A 269C>T), ACT44AAA (271C>A 272T>A), ATG45GTT (273A>G 275G>T), AAG48ACC (283A>C 284G>C), TCA50GTA (288T>G 289C>T), CAT52CAC (296T>C), CTT53ATA (297C>A 299T>A), CTG56GTC (306C>G 308G>C), AAA57AAG (311A>G), CCA58CCG (314A>G), TCT59CCA (315T>C 317T>A), GTG60GTT (320G>T), ACT61ATG (322C>T 323T>G), GAT62GCA (325A>C 326T>A), GTG63GCA (328T>C 329G>A), TCC64GCA (330T>G 332C>A), TCT66TCA (338T>A), GTG68GCC (343T>C 344G>C), GCC69TCT (345G>T 347C>T), ACT70ACA (350T>A), TGT71TGC (353T>C), AGT72AAT (355G>A), TCA73GCT (357T>G 359A>T), CTT74GTA (360C>G 362T>A), AAT75GAA (363A>G 365T>A), AAA76AGA (367A>G), TTG78TTA (374G>A), GAC79GAT (377C>T), AAA80AAG (380A>G), GGT82GCA (385G>C 386T>A), GCA83AAA (387G>A 388C>A), CTT84ATA (390C>A 392T>A), ATA85ATG (395A>G), GAT86GAA (398T>A), CCA87CCC (401A>C), TTT88TTC (404T>C), CTT89ATC (405C>A 407T>C), AAA90AAG (410A>G), GCG91GGT (412C>G 413G>T), TAC92TAT (416C>T), TCT93GAT (417T>G 418C>A), TTC94TTT (422C>T), TTG95GTT (423T>G 425G>T), TCC96AAA (426T>A 427C>A 428C>A), ACA97ACT (431A>T), TAC99TAT (437C>T), TCT101ACT (441T>A), ATC102ATA (446C>A), AAG103GTA (447A>G 448A>T 449G>A), ATG105CTT (453A>C 455G>T), GTT106GTA (458T>A), TTA107CTT (459T>C 461A>T), AAA108AAG (464A>G), TTG109ATG (465T>A), TTT110GTT (468T>G), GAG111TTA (471G>T 472A>T 473G>A), TTG113GTC (477T>G 479G>C), TCC114AAG (480T>A 481C>A 482C>G), AAT115GAT (483A>G), GTT117TTC (489G>T 491T>C), AAA118ATC (493A>T 494A>C), TTA119CTA (495T>C), TTT121TAC (502T>A 503T>C), GCT122GAA (505C>A 506T>A), GTG124TTA (510G>T 512G>A), AAT126GAA (516A>G 518T>A), AGT128GTT (522A>G 523G>T), GAA129GAG (527A>G), GAT130TCA (528G>T 529A>C 530T>A), GTA131TTA (531G>T), GTT134GTG (542T>G), GTA135TTT (543G>T 545A>T), TAT136ATA (546T>A 547A>T 548T>A), GGT137GCT (550G>C), CTT139TTA (555C>T 557T>A), GTA140GTT (560A>T), ATA143TTG (567A>T 569A>G), ACA144GCC (570A>G 572A>C), TTG145ATT (573T>A 575G>T), TTA146TTG (578A>G), GTG147CTA (579G>C 581G>A), CTT148CTA (584T>A), CTA149CTG (587A>G), TTT150TAT (589T>A), GTT151GTA (593T>A), TGT152TGC (596T>C), AGT154GTA (600A>G 601G>T 602T>A), GTT157GTA (611T>A), GAG158ACG (612G>A 613A>C), GGT159CCT (615G>C 616G>C), GTT160TTT (618G>T), TCT161AAT (621T>A 622C>A), AAC162AAT (626C>T), ACT163GTA (627A>G 628C>T 629T>A), GTC164GTT (632C>T), AAA165GTT (633A>G 634A>T 635A>T), AAT166ACT (637A>C), ATC167TTG (639A>T 641C>G), TTG169TCA (646T>C 647G>A), ATA170GTT (648A>G 650A>T), GTC171GTT (653C>T), AAC173AAT (659C>T), ATG174TTT (660A>T 662G>T), TTT175TTC (665T>C), TCA176TCT (668A>T), AGT177TCA (669A>T 670G>C 671T>A), CTA178TTG (672C>T 674A>G), TAC179TAT (677C>T), AAA186ACA (697A>C), GCC188TCT (702G>T 704C>T), GAA189GGG (706A>G 707A>G), CGA190AAG (708C>A 709G>A 710A>G), TTC191TTT (713C>T), ACC192TCC (714A>T), ATG193CAG (717A>C 718T>A), GTG194CTG (720G>C), GCC195GCG (725C>G), GCT197GCA (731T>A), AGT198GCA (732A>G 733G>C 734T>A), ATG199GTA (735A>G 737G>A), GGG202GGT (746G>T), TCA204AGT (750T>A 751C>G 752A>T), GTT205TTA (753G>T 755T>A), TCA206ACG (756T>A 758A>G), ACG210ACT (770G>T), TCT211AGC (771T>A 772C>G 773T>C), CAG212CAA (776G>A), TTG213CTT (777T>C 779G>T), ATG214TTA (780A>T 782G>A), ACA216TCA (786A>T), ATA217TTG (789A>T 791A>G), GGG221GGT (803G>T), ATT222ATC (806T>C), TCG223TCA (809G>A), ACG224ACT (812G>T), TTG225TTA (815G>A), GTG226GTA (818G>A), TTC227TTT (821C>T), ATT228GTT (822A>G), GCA229GCT (827A>T), GTG230GTT (830G>T), GGT232GGG (836T>G), GGC236GGT (848C>T), TTG237CTT (849T>C 851G>T), AGC238TCA (852A>T 853G>C 854C>A), TTG241CTG (861T>C), TCC242TCA (866C>A), ATT244ATC (872T>C), CTC245CTG (875C>G), TAT246TAC (878T>C), TCG247TCT (881G>T), ACA248GCT (882A>G 884A>T), GCT252GCG (896T>G), CAG253CAA (899G>A), CAA257CAG (911A>G), CTA258CTG (914A>G), ACT260CAG (918A>C 919C>A 920T>G), CTT261TTG (921C>T 923T>G), CGT263AGG (927C>A 929T>G), AAT264AAC (932T>C), AAA266AAG (938A>G), TGC268TGT (944C>T), ACC269ACA (947C>A), TCC270TCT (950C>T), CAG271CAA (953G>A), GCT272GCC (956T>C), GAA278GAG (974A>G), ATA280GTG (978A>G 980A>G), GTC281GTT (983C>T), GGC282AGC (984G>A), ACC283ATT (988C>T 989C>T), GGA285GGC (995A>C), TTC286TTT (998C>T), GTG290GTA (1010G>A), GCT293GCG (1019T>G), AGT295AAT (1024G>A), GCA296GCT (1028A>T), ACG297ACC (1031G>C), CTG298TTA (1032C>T 1034G>A), CTG302CTT (1046G>T), GAG304CAG (1050G>C), CAG307ACG (1059C>A 1060A>C), GAG308GAA (1064G>A), GAT310TCG (1068G>T 1069A>C 1070T>G), GCA311GAG (1072C>A 1073A>G), GTG312GTT (1076G>T), TGT313GCT (1077T>G 1078G>C), GAT314GAA (1082T>A), CCT315CCA (1085T>A), GCA316TCT (1086G>T 1088A>T), ACG317ATG (1090C>T), GAG320GAA (1100G>A), AAC321AAT (1103C>T), GCC323GCA (1109C>A), AAG325CCT (1113A>C 1114A>C 1115G>T), GCT326GCA (1118T>A), TTC327TTT (1121C>T), CCT328CCA (1124T>A), ACC329ATT (1126C>T 1127C>T), AAA330AAG (1130A>G), AAC332CAA (1134A>C 1136C>A), CTG334TTA (1140C>T 1142G>A), AGG335AAA (1144G>A 1145G>A), GAA336GAG (1148A>G), GAA337CAG (1149G>C 1151A>G), GCT338GCC (1154T>C), CTC339GTC (1155C>G), AAG340AAA (1160G>A), ATC341ATA (1163C>A), GCA346GCC (1178A>C), GTT350GTC (1190T>C), GCG351GCA (1193G>A), TTC353TTT (1199C>T), ATG354TTG (1200A>T), AGT359TCT (1215A>T 1216G>C), ATT362ATA (1226T>A), GCG363ACG (1227G>A), TTA365CTA (1233T>C), GAA366GAT (1238A>T), GTT368CTG (1242G>C 1244T>G), TAT369GTT (1245T>G 1246A>T), GAA371GAG (1253A>G), AGT372GCC (1254A>G 1255G>C 1256T>C), TGT373AGC (1257T>A 1259T>C), AGG374CAA (1260A>C 1261G>A 1262G>A), CAC375CAT (1265C>T), ATG376TCT (1266A>T 1267T>C 1268G>T), GGC377GGT (1271C>T), CAG380CAA (1280G>A), CGT382AGG (1284C>A 1286T>G), GAA384GAG (1292A>G), TTT386TTC (1298T>C), GTA388GTG (1304A>G), TTC391TTT (1313C>T), GGG392GGA (1316G>A), TCC393GCT (1317T>G 1319C>T), GGG395GGA (1325G>A), TCT399TCA (1337T>A), ACA400ACG (1340A>G), TCC401TGT (1342C>G 1343C>T), TTA404CTT (1350T>C 1352A>T), TTT405TTC (1355T>C), ATT406ATC (1358T>C), GTG409GTT (1367G>T), TTG410CTC (1368T>C 1370G>C), GAC411GAT (1373C>T), AGA412GCA (1374A>G 1375G>C), GAG415GAA (1385G>A), AAA417AAG (1391A>G), CTC418CTG (1394C>G), AAT419AAC (1397T>C), CGG420CGC (1400G>C), CTA421TTG (1401C>T 1403A>G), GCA423GCG (1409A>G), AAG426AAA (1418G>A), AGG427AGA (1421G>A), GCT435GCA (1445T>A), CAC436CAT (1448C>T), GCT439GCA (1457T>A), ATC440GTG (1458A>G 1460C>G), CTG441CTT (1463G>T), GAC444GAT (1472C>T), GGA446GGT (1478A>T), GCA447GCT (1481A>T), TTG448CTT (1482T>C 1484G>T), CGA449CGG (1487A>G), GGG451GGA (1493G>A), GCT452GCA (1496T>A), TGC455TGT (1505C>T), CAA456CAG (1508A>G), GAT457GAC (1511T>C), ATT458ATA (1514T>A), AAA459AAG (1517A>G), GAT463GAC (1529T>C), TCA466TCT (1538A>T), ACA467ACC (1541A>C), CAA468CAG (1544A>G), CCA469CCT (1547A>T), CCC471CCT (1553C>T), TTA472CTT (1554T>C 1556A>T), GCC475GCT (1565C>T), GTT477GTC (1571T>C), GAC479GAT (1577C>T), GGC481GGA (1583C>A), CAT483CAC (1589T>C), TTC484TTT (1592C>T), ACC485ACA (1595C>A), TCC486TCA (1598C>A), TAT488TAC (1604T>C), ATA489ATC (1607A>C), ACA492ACC (1616A>C), CTG495TTG (1623C>T), ATC496ATA (1628C>A), TCA497TCT (1631A>T), CCT499CCA (1637T>A), GCC500GCT (1640C>T), CAA501CAG (1643A>G), GGT503GGG (1649T>G), CTT504CTA (1652T>A), ACA505ACG (1655A>G), CCA507CCT (1661A>T), GCT509GCA (1667T>A), GAG511GAA (1673G>A), CGG512AGG (1674C>A), AGA513AGG (1679A>G), AGG514AGA (1682G>A), GAC515GAT (1685C>T), ATC516ATT (1688C>T), CTG517TTA (1689C>T 1691G>A), GTG518GTT (1694G>T), TGT520TGC (1700T>C), ATG521AGA (1702T>G 1703G>A), GAG523GTG (1708A>T), GAG525GAT (1715G>T), TCC527GAT (1719T>G 1720C>A 1721C>T), ACT528CCA (1722A>C 1724T>A), GAA529AAT (1725G>A 1727A>T), GAT530ACT (1728G>A 1729A>C), CCT531CCA (1733T>A), ACG532ACA (1736G>A), GGA533GCC (1738G>C 1739A>C), CTC535CTA (1745C>A), AGA536GTT (1746A>G 1747G>T 1748A>T), AAC538GAT (1752A>G 1754C>T), ATA539GTG (1755A>G 1757A>G), GAG541GAA (1763G>A), CCT545CCC (1775T>C), GCT547GCC (1781T>C), ACT549ACA (1787T>A), CAC550CAT (1790C>T), AGG551AGA (1793G>A), AAT552GCT (1794A>G 1795A>C), TTG553TTA (1799G>A), ACC554TCC (1800A>T), AGT556GAT (1806A>G 1807G>A), GAT557GAC (1811T>C), CTT558CTC (1814T>C), GAG560CAA (1818G>C 1820G>A), GTA563GTG (1829A>G), GCC564GCA (1832C>A), AAA565AAG (1835A>G), TGT566TGC (1838T>C), CAG567CGG (1840A>G), GTA568GTG (1844A>G), CAT569CAC (1847T>C), GCA570TAT (1848G>T 1849C>A 1850A>T), GAA571GAG (1853A>G), GTA572GTG (1856A>G), AAG574AGC (1861A>G 1862G>C), TTG576CTT (1866T>C 1868G>T), TTT577TAC (1870T>A 1871T>C), GAA578AAT (1872G>A 1874A>T), GCT579ACT (1875G>A), GAA580ACA (1878G>A 1879A>C), TCT581TCG (1883T>G), GGC582GGA (1886C>A), ATA583TTG (1887A>T 1889A>G), AGT584GCT (1890A>G 1891G>C), CCT585GAC (1893C>G 1894C>A 1895T>C), ATA587GTT (1899A>G 1901A>T), GTT590GTC (1910T>C), CAA591AAT (1911C>A 1913A>T), TCT593GCT (1917T>G), GCA594GTT (1921C>T 1922A>T), GAT596GAA (1928T>A), ATA598CTG (1932A>C 1934A>G), GCG599GAT (1936C>A 1937G>T), GTG601TTC (1941G>T 1943G>C), GAT602CAA (1944G>C 1946T>A), GGG603GAT (1948G>A 1949G>T), GCT604GCC (1952T>C), AGA605del (1953_1955delAGA), TTG606TTA (1958G>A), CGC607CAA (1960G>A 1961C>A), ACC608GTT (1962A>G 1963C>T 1964C>T), AAG609TTG (1965A>T 1966A>T), CAA610CCT (1969A>C 1970A>T), GAT611CCT (1971G>C 1972A>C), GAA612GAG (1976A>G), ATA614GCA (1980A>G 1981T>C), TTG615GCT (1983T>G 1984T>C 1985G>T), GTG616TAT (1986G>T 1987T>A 1988G>T), CCC617GAG (1989C>G 1990C>A 1991C>G), ACT618CAG (1992A>C 1993C>A 1994T>G), GTT619ACA (1995G>A 1996T>C 1997T>A), GTG620ATT (1998G>A 2000G>T), AGT621ACA (2002G>C 2003T>A), AAT623GCT (2007A>G 2008A>C), GAT624GAA (2012T>A), AGA625AAA (2014G>A), GTG626ATT (2016G>A 2018G>T), ATC627GTT (2019A>G 2021C>T), TAT628TAC (2024T>C), GCA629TCC (2025G>T 2027A>C), AGA630CAG (2028A>C 2029G>A 2030A>G), GAG631GAA (2033G>A), ACG633ACA (2039G>A), GTA634GTT (2042A>T), GCA636GCC (2048A>C), TTG637CTC (2049T>C 2051G>C), GCA640GCT (2060A>T), CAG642CAA (2066G>A), GGC643GGG (2069C>G), TTG645ATG (2073T>A), GAC646AAC (2076G>A), CCT647AAA (2079C>A 2080C>A 2081T>A), GAG648TTT (2082G>T 2083A>T 2084G>T), GAA649GAG (2087A>G), CTT650TTT (2088C>T), TTT651TTC (2093T>C), CCT652CCA (2096T>A), CAT655CTC (2104A>T 2105T>C), CAT656TCC (2106C>T 2107A>C 2108T>C), GCC659GCT (2117C>T), ATG660ATC (2120G>C), TCT663GTG (2127T>G 2128C>T 2129T>G), CTT664CTA (2132T>A), GAT665TCT (2133G>T 2134A>C), GAT666TCT (2136G>T 2137A>C), GAA667GAT (2141A>T), CAT668GCC (2142C>G 2143A>C 2144T>C), GAG670CAG (2148G>C), ATT671AAT (2152T>A), TTC672TTT (2156C>T), AAC673GAA (2157A>G 2159C>A), AAA674ATG (2161A>T 2162A>G), AGA676GTA (2166A>G 2167G>T), GTC677AAT (2169G>A 2170T>A 2171C>T), AAC678GAT (2172A>G 2174C>T), ATG679TTG (2175A>T), TAT681GTC (2181T>G 2182A>T 2183T>C), AGA682AAA (2185G>A), GAT685AGT (2193G>A 2194A>G), GCT686CCT (2196G>C), GAA687CAG (2199G>C 2201A>G), CGA690GCT (2208C>G 2209G>C 2210A>T), GTA693GTG (2219A>G), CAA694CGA (2221A>G), GAC699GAT (2237C>T), ATA701ATC (2243A>C), CAC703CAT (2249C>T), TTT704TTC (2252T>C), GCA710AGC (2268G>A 2269C>G 2270A>C), TCA711ACA (2271T>A), ATC713ATT (2279C>T), GAG715GAA (2285G>A), AAG716GAG (2286A>G), AAA717CTG (2289A>C 2290A>T 2291A>G), CTA718CTG (2294A>G), GCA719GAT (2296C>A 2297A>T), GTG720GTC (2300G>C), CAG721CAA (2303G>A), ACA722CGC (2304A>C 2305C>G 2306A>C), GAG723AAT (2307G>A 2309G>T), ATG724CTT (2310A>C 2312G>T), AGG725GAG (2313A>G 2314G>A), ACT726CAA (2316A>C 2317C>A 2318T>A), GGA727GGG (2321A>G), TTT728AAA (2322T>A 2323T>A 2324T>A), GCA729GCG (2327A>G), CAT730TAT (2328C>T), TCT731AAG (2331T>A 2332C>A 2333T>G), TGC732GCG (2334T>G 2335G>C 2336C>G), ATT733CTT (2337A>C), GAT734CAG (2340G>C 2342T>G), GCA735GCT (2345A>T), GTT737GCA (2350T>C 2351T>A), GAC738GAT (2354C>T), ACC739ACA (2357C>A), CTC740CTT (2360C>T), ATA741ACG (2362T>C 2363A>G), ATC743ATT (2369C>T), GAG744GAA (2372G>A), TTT749AGA (2385T>A 2386T>G 2387T>A), GTT750GTA (2390T>A), TTT752TTC (2396T>C), TTA753CTG (2397T>C 2399A>G), ACA755ACT (2405A>T), GCA756GCT (2408A>T), AGA759CGC (2415A>C 2417A>C), TGC762TGT (2426C>T), AAC763AGG (2428A>G 2429C>G), GTA765GTT (2435A>T), TCA766GCA (2436T>G), TCA766_AAA767insGGT (2438_2439insGGT), AAA767ACT (2440A>C 2441A>T), GAG768GAC (2444G>C), CCA769AAG (2445C>A 2446C>A 2447A>G), GAA771AGA (2451G>A 2452A>G), TCA772AAT (2454T>A 2455C>A 2456A>T), ATC774ATT (2462C>T), AAA775ATC (2464A>T 2465A>C), ATT776GAA (2466A>G 2467T>A 2468T>A), TAC778ATG (2472T>A 2473A>T 2474C>G), CAT779ATT (2475C>A 2476A>T), AAA780GGT (2478A>G 2479A>G 2480A>T), GTT782AAG (2484G>A 2485T>A 2486T>G), GAA783GAC (2489A>C), TAT784AAA (2490T>A 2492T>A), ATC785ATG (2495C>G), AAG786ATC (2497A>T 2498G>C), GAT787CAG (2499G>C 2501T>G), ACA788ATG (2503C>T 2504A>G), TGG790AAT (2508T>A 2509G>A 2510G>T), AGC791GAC (2511A>G 2512G>A), CTT792TTG (2514C>T 2516T>G), TAT794TTC (2521A>T 2522T>C), GCA795ATC (2523G>A 2524C>T 2525A>C), CTG796TTG (2526C>T), AGA797AAG (2530G>A 2531A>G), CTC799CTT (2537C>T), GCA802GCC (2546A>C), GGT803GGA (2549T>A), ATA806TTG (2556A>T 2558A>G), ATG807ATA (2561G>A), GCT808GCA (2564T>A), GGA811GGC (2573A>C), GTT812ACC (2574G>A 2575T>C 2576T>C), TTT813ATG (2577T>A 2579T>G), GGA814AAT (2580G>A 2581G>A 2582A>T), GGC815GCC (2584G>C), ATT816GTT (2586A>G), ACA817ACT (2591A>T), GCA818GCT (2594A>T), TTT819TTC (2597T>C), TTG820ATT (2598T>A 2600G>T), GCG821ACT (2601G>A 2603G>T), TGG823CTC (2607T>C 2608G>T 2609G>C), CAG824AAG (2610C>A), AGT825GCA (2613A>G 2614G>C 2615T>A), AAC826GCA (2616A>G 2617A>C 2618C>A), TCT827ACT (2619T>A), TTC828TCC (2623T>C), CCC829CCT (2627C>T), AAT830TTA (2628A>T 2629A>T 2630T>A), GCA831GCT (2633A>T), TCG832GAT (2634T>G 2635C>A 2636G>T), GGA835GGG (2645A>G), GGT836GGC (2648T>C), GTG837GTA (2651G>A), GAA840GAG (2660A>G), ATA844ATC (2672A>C), AGT846AGC (2678T>C), AAG848AAA (2684G>A), AAC849AAT (2687C>T), AAG850AAA (2690G>A), GGT851GGG (2693T>G), AAT852AAG (2696T>G), AAG853ACT (2698A>C 2699G>T), AAG853_CTC854insCAT (2699_2700insCAT), CTC854TTC (2700C>T), AGA855GCA (2703A>G 2704G>C), AAT856AAA (2708T>A), CTT857TCC (2709C>T 2710T>C 2711T>C), CTC858TTG (2712C>T 2714C>G), GTT859CTA (2715G>C 2717T>A), GGT860GGG (2720T>G), AGT862GCG (2724A>G 2725G>C 2726T>G), TCT863GCA (2727T>G 2729T>A), CAA864GAT (2730C>G 2732A>T), TCA865AAC (2733T>A 2734C>A 2735A>C), TTG866AAC (2736T>A 2737T>A 2738G>C), GCA867CAT (2739G>C 2740C>A 2741A>T), CAA868GTG (2742C>G 2743A>T 2744A>G), GAT869GAC (2747T>C), GAT869_TGG870insCGG (2747_2748insCGG), GCT871GCA (2753T>A), GCT872GAA (2755C>A 2756T>A), GAA873CAG (2757G>C 2759A>G), GTA877GTT (2771A>T), CAG879GAA (2775C>G 2777G>A), AGC880AGT (2780C>T), CTC881CTG (2783C>G), AAG882CAC (2784A>C 2786G>C), AAT884AAC (2792T>C), GTG886GTT (2798G>T), GTG887GTT (2801G>T), TTA888TTG (2804A>G), CTT890ATT (2808C>A), GGT891GCT (2812G>C), GTG894GTC (2822G>C), TAC895TAT (2825C>T), TTT896TTC (2828T>C), GGC898GGT (2834C>T), ACC899ACG (2837C>G), TAC900TAT (2840C>T), GTG901GTT (2843G>T), ATA906ATT (2858A>T), ATG907CTC (2859A>C 2861G>C), ACC908ACT (2864C>T), GTA909GTG (2867A>G), GCT910GCA (2870T>A), GCA912GCC (2876A>C), TTC913TTT (2879C>T), CAC914AGA (2880C>A 2881A>G 2882C>A), AAT915AAC (2885T>C), GCT916ATG (2886G>A 2887C>T 2888T>G), CGA917AGG (2889C>A 2891A>G), GGT919GGC (2897T>C), CCA921CCT (2903A>T), ACA923TCC (2907A>T 2909A>C), ATC925ATT (2915C>T), GCC927GAG (2920C>A 2921C>G), AAT928CAT (2922A>C), TCT929TCC (2927T>C), CGA930ATT (2928C>A 2929G>T 2930A>T), TCT931TGT (2932C>G), AAA932AAT (2936A>T), GTT933GTG (2939T>G), CAA934CAG (2942A>G), TAC935TTC (2944A>T), AAC936AAT (2948C>T), AGA938ACC (2953G>C 2954A>C), GAA939CAA (2955G>C), AGG941AAG (2962G>A), ATT942TTG (2964A>T 2966T>G), ATT943ATA (2969T>A), GGC945GGT (2975C>T), GAC947GAT (2981C>T), ATC948GTG (2982A>G 2984C>G), CTG950TTG (2988C>T), CGC952AGG (2994C>A 2996C>G), GTT953GTA (2999T>A), GGT954GGG (3002T>G), CCA959CCC (3017A>C), AAG960AAA (3020G>A), CCT961TCA (3021C>T 3023T>A), GAC962GAT (3026C>T), ATC963ATA (3029C>A), CGT964AGA (3030C>A 3032T>A), CAC966CAT (3038C>T), CCA968CCT (3044A>T), AGA969CGC (3045A>C 3047A>C),

| | |
|---|---|
| Codon mutations: | AGG970AGA (3050G>A), GAG972GAT (3056G>T), TGC974TGT (3062C>T), TTC975TAC (3064T>A), ACA976TCT (3066A>T 3068A>T), CAA980ATG (3078C>A 3079A>T 3080A>G), GGA981GGC (3083A>C), TGT984GTG (3090T>G 3091G>T 3092T>G), AGA985AGG (3095A>G), GTT987GTA (3101T>A), GCG988GCT (3104G>T), TCT989TCA (3107T>A), ACA990AAG (3109C>A 3110A>G), CCA992CCC (3116A>C), CGT993AAG (3117C>A 3118G>A 3119T>G), CTT994AAA (3120C>A 3121T>A 3122T>A), GGC995TCC (3123G>T 3124G>C), TTA997TTG (3131A>G), TTT999GTT (3135T>G), CTC1000CTT (3140C>T), AAG1001AAA (3143G>A), ATG1002TAT (3144A>T 3145T>A 3146G>T), GGA1009GGC (3167A>C), ACA1011ACC (3173A>C), GTT1012GAA (3175T>A 3176T>A), GTG1014GTC (3182G>C), GAA1015ACA (3183G>A 3184A>C), GAA1017GGA (3190A>G), CTG1018TTA (3192C>T 3194G>A), AAC1019GAT (3195A>G 3197C>T), TCC1020TCG (3200C>G), TCC1021AAT (3201T>A 3202C>A 3203C>T), AGT1022GCA (3204A>G 3205G>C 3206T>A), AAG1024GCT (3210A>G 3211A>C 3212G>T), ATT1025CTT (3213A>C), TGC1026AGC (3216T>A), TCT1027TCA (3221T>A), CAA1029GTT (3225C>G 3226A>T 3227A>T), TCT1030GCA (3228T>G 3230T>A), GAA1032GAG (3236A>G), ATA1035ATT (3245A>T), GGG1037AGT (3249G>A 3251G>T), GAA1038GAG (3254A>G), GGT1040GGA (3260T>A), GAC1041GAT (3263C>T), GGC1043GGT (3269C>T), ACG1044ACT (3272G>T), TTG1045CTT (3273T>C 3275G>T), CTA1046CTC (3278A>C), CTG1047TTA (3279C>T 3281G>A), TTA1048TTG (3284A>G), AGT1050ACT (3289G>C), GTT1051GTG (3293T>G), AAT1053AAC (3299T>C), CAA1055CAG (3305A>G), CCT1056CCG (3308T>G), GTG1057TGC (3309G>T 3310T>G 3311G>C), GGC1060GGA (3320C>A), ATC1061ATA (3323C>A), CAC1062CAT (3326C>T), TCA1065TCT (3335A>T), GGC1068GAT (3343G>A 3344C>T), GTA1069ACC (3345G>A 3346T>C 3347A>C), GCA1070GCT (3350A>T), GCT1071GCA (3353T>A), GAA1072GAG (3356A>G), CGT1073AAG (3357C>A 3358G>A 3359T>G), GGA1074GGT (3362A>T), TTT1075TAC (3364T>A 3365T>C), ATC1076GTG (3366A>G 3368C>G), TCT1077GCT (3369T>G), AAT1079AAC (3377T>C), GCT1080GCA (3380T>A), GCT1082GCA (3386T>A), ATC1083GTG (3387A>G 3389C>G), TAC1084TTT (3391A>T 3392C>T), AGG1085AGA (3395G>A), GAA1086GAT (3398A>T), CTA1088TTG (3402C>T 3404A>G), GAG1089GAA (3407G>A), CCA1092CCC (3416A>C), ACT1093ACG (3419T>G), GGG1094GGA (3422G>A), CCG1095CCC (3425G>C), GTC1096ATT (3426G>A 3428C>T), GCA1098GTG (3433C>T 3434A>G), GCA1099GCT (3437A>T), GTA1101GTG (3443A>G), CGC1102CAC (3445G>A), GAT1104GAA (3452T>A), ATT1105CTG (3453A>C 3455T>G), CTG1106TTG (3456C>T), TCA1108CAC (3462T>C 3463C>A 3464A>C), ATT1109CTG (3465A>C 3467T>G), AGC1111TCC (3471A>T 3472G>C), AGA1112AAG (3475G>A 3476A>G), GAA1113GAG (3479A>G), ACA1114GAA (3480A>G 3481C>A), CAG1115AGA (3483C>A 3484A>G 3485G>A), CTT1116TTC (3486C>T 3488T>C), GAA1119GAG (3497A>G), AAC1120AAA (3500C>A), CAA1121CAG (3503A>G), TAC1123GCT (3507T>G 3508A>C 3509C>T), TAC1124TTC (3511A>T), CTT1125TTG (3513C>T 3515T>G), GGA1126GGC (3518A>C), ACA1127ACT (3521A>T), GTT1128GTA (3524T>A), CCA1129CCG (3527A>G), CAG1130TTG (3528C>T 3529A>T), GAG1131GAA (3533G>A), TTG1132ATG (3534T>A), GCC1133GCT (3539C>T), GCC1134GCT (3542C>T), GTT1136GTG (3548T>G), CCC1137CCG (3551C>G), AAG1139AAA (3557G>A), ACC1140ACA (3560C>A), ACT1141ACC (3563T>C), CTG1142CTT (3566G>T), CGG1143AAA (3567C>A 3568G>A 3569G>A), AAA1144CGC (3570A>C 3571A>G 3572A>C), AGC1145AGT (3575C>T), CAA1146CCC (3577A>C 3578A>C), TTG1147CTG (3579T>C), GAA1149GAG (3587A>G), GCA1150GCT (3590A>T), TTC1151TTT (3593C>T), GGA1152GGG (3596A>G), CCT1153GAC (3597C>G 3598C>A 3599T>C), GCA1154GCT (3602A>T), GAG1155CAG (3603G>C), ACA1156ACT (3608A>T), GCA1157GCC (3611A>C), ACA1162TCT (3624A>T 3626A>T), GTT1163GCC (3628T>C 3629T>C), CAT1164AAA (3630C>A 3632T>A), AGA1167AGG (3641A>G), GGT1168GGA (3644T>A), GAT1169GAA (3647T>A), GGT1170GGG (3650T>G), GAC1172GAT (3656C>T), GGG1177GGT (3671G>T), GTA1178GTC (3674A>C), ATG1179GAG (3675A>G 3676T>A), AAA1180AAG (3680A>G), TAC1181TAT (3683C>T), TGT1186TCC (3697G>C 3698T>C), GGA1187AAC (3699G>A 3700G>A 3701A>C), GAT1189GAG (3707T>G), GAT1190GAG (3710T>G), GAC1191CCA (3711G>C 3712A>C 3713C>A), GCC1193GCA (3719C>A), AAA1194AAG (3722A>G), CTA1195CTG (3725A>G), GCA1196GCC (3728A>C), TTC1197TTT (3731C>T), GAA1198GAG (3734A>G), AAT1199GGT (3735A>G 3736A>G), CTC1200CTT (3740C>T), AAG1201AAA (3743G>A), TGC1202GCA (3744T>G 3745G>C 3746C>A), TCG1203TCC (3749G>C), CTG1204ATT (3750C>A 3752G>T), CTA1205CTT (3755A>T), ATC1207GTG (3759A>G 3761C>G), ATG1208CTT (3762A>C 3764G>T), CGT1209AAG (3765C>A 3766G>A 3767T>G), AGC1210ATG (3769G>T 3770C>G), AAG1212CGA (3774A>C 3775A>G 3776G>A), ATC1213GTG (3777A>G 3779C>G), CCT1214CCC (3782T>C), CGT1217AAA (3789C>A 3790G>A 3791T>A), CCA1218CCT (3794A>T), TGT1219GTA (3795T>G 3796G>T 3797T>A), GAA1220ATC (3798G>A 3799A>T 3800A>C), AGG1221AGA (3803G>A), GAT1222AAT (3804G>A), GGA1229GGG (3827A>G), TGT1233TTT (3838G>T), TAC1235TAT (3845C>T), TAT1236TTT (3847A>T), GGC1238GCA (3853G>C 3854C>A), GAG1240GAA (3860G>A), CTG1241TTG (3861C>T), ACA1243ACG (3869A>G), TCT1244TCA (3872T>A), TGC1245TGT (3875C>T), GGA1246GGT (3878A>T), CCC1248CCT (3884C>T), TTC1249TTT (3887C>T), ATG1252TTG (3894A>T), GGG1253GGA (3899G>A), ATG1254CTT (3900A>C 3902G>T), GGG1255GGT (3905G>T), ATG1256TCT (3906A>T 3907T>C 3908G>T), AAG1258AAA (3914G>A), AGA1259AGG (3917A>G), CAA1263GAA (3927C>G), TCC1264GCC (3930T>G), ACT1265CAA (3933A>C 3934C>A 3935T>A), GGC1266CCA (3936G>C 3937G>C 3938C>A), GAA1267AAC (3939G>A 3941A>C), GGA1268TCG (3942G>T 3943G>C 3944A>G), GAG1269CAG (3945G>C), AGA1270AGG (3950A>G), GTG1271GTT (3953G>T), GAA1272CTT (3954G>C 3955A>T 3956A>T), AGA1275AGG (3965A>G), GAC1276AAC (3966G>A), ACT1277ACA (3971T>A), GTA1279GTC (3977A>C), GAA1281GAT (3983A>T), CTA1286TTG (3996C>T 3998A>G), GAT1287GAC (4001T>C), GTG1288TCG (4002G>T 4003T>C), ATT1290GTA (4008A>G 4010T>A), AAA1292CAA (4014A>C), ATC1294ACA (4021T>C 4022C>A), CAT1295TTT (4023C>T 4024A>T), GTG1296GTT (4028G>T), GAT1297GAC (4031T>C), CTG1298CTC (4034G>C), ACC1300ACA (4040C>A), CAA1302CAG (4046A>G), TGC1303TGT (4049C>T), GCC1304GCG (4052C>G), CGC1308CGG (4064C>G), CTC1309TTG (4065C>T 4067C>G), AAA1313AAG (4079A>G), ATC1314ATT (4082C>T), GGC1316GGG (4088C>G), AGA1318AGG (4094A>G), AGG1321CGC (4101A>C 4103G>C), CTC1322CTG (4106C>G), TTT1323TTC (4109T>C), GAG1324GAA (4112G>A), ATA1325ATC (4115A>C), CTT1326TTG (4116C>T 4118T>G), TAC1330TAT (4130C>T), TTG1333CTG (4137T>C), GTC1334GTG (4142C>G), AGG1335CGG (4143A>C), AAG1336AAA (4148G>A), TAT1337TAC (4151T>C), TTT1338TTC (4154T>C), GAT1340GAC (4160T>C), TTC1341TTT (4163C>T), TCA1342TCT (4166A>T), GCC1343GCT (4169C>T), ACA1344TGT (4170A>T 4171C>G 4172A>T), TTG1348GCA (4182T>G 4183T>C 4184G>A), AAT1350AAC (4190T>C), GCT1351GTA (4192C>T 4193T>A), ATA1352CTC (4194A>C 4196A>C), CCA1353CCT (4199A>T), AAA1355AAG (4205A>G), GTT1356GTA (4208T>A), ATT1358ATC (4214T>C), GAT1359AAT (4215G>A), CCT1360CCA (4220T>A), ACA1361ACG (4223A>G), AGT1362AGC (4226T>C), TCT1363CAT (4227T>C 4228C>A), TTG1367GTG (4239T>G), TTG1368CTG (4242T>C), GCA1369GCT (4247A>T), AAT1370AGT (4249A>G), GGG1371AGC (4251G>A 4253G>C), AGA1373AGG (4259A>G), GCT1374TCA (4260G>T 4262T>A), GTG1375GTA (4265G>A), TCA1376TCT (4268A>T), GAC1377GAT (4271C>T), GTG1378GAG (4273T>A), TTT1380TTC (4280T>C), GCT1382GCC (4286T>C), TAT1384TAC (4292T>C), TCC1385TCG (4295C>G), CCT1392CCC (4316T>C), GTT1393ATC (4317G>A 4319T>C), GCT1395GCA (4325T>A), TTT1396TTC (4328T>C), CAG1397CAA (4331G>A), TTG1402CTG (4344T>C), GAA1405GCT (4354A>C 4355A>T), TAC1406TTT (4357A>T 4358C>T), TAC1407TAT (4361C>T), GGA1408GGT (4364A>T), TCA1409GAT (4365T>G 4366C>A 4367A>T), AAG1410GAG (4368A>G), GGC1412GGG (4376C>G), GAT1415GAG (4385T>G), TCC1416TCA (4388C>A), CGA1419AGG (4395C>A 4397A>G), CAT1420CAC (4400T>C), GCA1421ACA (4401G>A), CTT1422TTA (4404C>T 4406T>A), GCA1426GCC (4418A>C), TGT1428TGC (4424T>C), TAC1430TAT (4430C>T), ACA1431ACC (4433A>C), CTG1432CTC (4436G>C), TGC1433TGT (4439C>T), GAT1435AAC (4443G>A 4445T>C), AAA1436AAG (4448A>G), GTG1437TTG (4449G>T), AGG1439CGC (4455A>C 4457G>C), TTG1440CTA (4458T>C 4460G>A), GTT1441GTG (4463T>G), GGG1442GGT (4466G>T), GGC1443GGA (4469C>A), CCA1445CCC (4475A>C), TCA1446TCT (4478A>T), GGA1447GGT (4481A>T), GCA1449GCT (4487A>T), CTA1450TTG (4488C>T 4490A>G), ACG1451ACT (4493G>T), ATC1453ATA (4499C>A), TTC1454TTT (4502C>T), TCT1456TCA (4508T>A), CTC1457TTG (4509C>T 4511C>G), CTC1458CTA (4514C>A), CGA1464AGG (4530C>A 4532A>G), TAT1465TAC (4535T>C), GCC1466GCG (4538C>G), TTG1470CTG (4548T>C), TTA1471TTG (4553A>G), AGA1472AGG (4556A>G), CCC1474CCA (4562C>A), ATT1476GTC (4566A>G 4568T>C), GCT1477AAA (4569G>A 4570C>A 4571T>A), AGG1479AAA (4576G>A 4577G>A), GCT1480TCT (4578G>T), ATA1481GTG (4581A>G 4583A>G), GGA1482GGC (4586A>C), GTT1483ATT (4587G>A), AAA1484AAG (4592A>G), AGT1486AAT (4597G>A), GAT1487GAC (4601T>C), CTA1491TTG (4611C>T 4613A>G), GCT1494GCA (4622T>A), GTT1495GTA (4625T>A), GGA1497GGT (4631A>T), GAC1499GAT (4637C>T), AAT1500AAC (4640T>C), CTT1501TTG (4641C>T 4643T>G), GCT1503GCG (4649T>G), GTA1504GTT (4652A>T), TTA1506ATG (4656T>A 4658A>G), CTC1508TTG (4662C>T 4664C>G), CAG1509AGG (4665C>A 4666A>G), TAT1511TAC (4673T>C), TCT1512GAT (4674T>G 4675C>A), CTG1513CTA (4679G>A), AAT1515TCC (4683A>T 4684A>C 4685T>C), ATA1516ATT (4688A>T), CAT1518AGA (4692C>A 4693A>G 4694T>A), GAG1519GAA (4697G>A), TTA1520CTG (4698T>C 4700A>G), CTG1522GCT (4704C>G 4705T>C 4706G>T), GTC1523GTT (4709C>T), AAT1524AAC (4712T>C), GTA1525GTC (4715A>C), ATC1526GTC (4716A>G), CTT1531TTG (4731C>T 4733T>G), AAA1533AAG (4739A>G), TCG1534TCA (4742G>A), GTT1537GTA (4751T>A), AAT1538GAG (4752A>G 4754T>G), GAG1539AAG (4755G>A), GTA1540ACA (4758G>A 4759T>C), CAA1541AAA (4761C>A), CAA1543CTG (4768A>T 4769A>G), CTA1548CTC (4784A>C), ACT1549ACC (4787T>C), CTG1551CTC (4793G>C), AGA1553AGG (4799A>G), TTT1555TTC (4805T>C), AAA1556AAG (4808A>G), GCT1559GCA (4817T>A), CTT1560TTG (4818C>T 4820T>G), TAC1562TAT (4826C>T), CAC1563CAA (4829C>A), GCT1565GCA (4835T>A), CCT1566CCA (4838T>A), AAG1568AAA (4844G>A), ATC1572GTC (4854A>G), ATT1573ATC (4859T>C), ATG1576ATT (4868G>T), ATA1579ATT (4877A>T), AGA1580AGG (4880A>G), CCT1581CCA (4883T>A), GCC1582GCA (4886C>A), CCA1583GTG (4887C>G 4888C>T 4889A>G), GGT1584GGA (4892T>A), GAC1587GAT (4901C>T), ACT1588GTA (4902A>G 4903C>T 4904T>A), CAG1589CAA (4907G>A), GCC1590GCT (4910C>T), ATG1592TAT (4914A>T 4915T>A 4916G>T), GTG1595ATT (4923G>A 4925G>T), GGA1598GGC (4934A>C), ATA1599ATC (4937A>C), CGT1600AGA (4938C>A 4940T>A), GCT1602GCC (4946T>C), CAC1605CAT (4955C>T), GGT1606GGG (4958T>G), CGT1607CAG (4960G>A 4961T>G), GTG1608GCC (4963T>C 4964G>C), GCT1609ACT (4965G>A), GAC1611GAA (4973C>A), AAG1612AGT (4975A>G 4976G>T), CTT1613TTT (4977C>T), GTC1614GTA (4982C>A), TTA1615AAT (4983T>A 4984T>A 4985A>T), CAG1618TTT (4992C>T 4993A>T 4994G>T), ACG1619GGA (4995A>G 4996C>G 4997G>A), GCG1620GCT (5000G>T), TTG1621CTC (5001T>C 5003G>C), GAT1622AAG (5004G>A 5006T>G), GAA1623CAG (5007G>C 5009A>G), AGG1624AGA (5012G>A), GGT1625AAT (5013G>A 5014G>A), TTC1626CTC (5016T>C), AGA1627CCC (5019A>C 5020G>C 5021A>C), GCT1628ATC (5022G>A 5023C>T 5024T>C), GTG1629ATG (5025G>A), TTT1631ATG (5031T>A 5033T>G), CCT1632CCC (5036T>C), TCC1633ACT (5037T>A 5039C>T), TAT1634TAC (5042T>C), TTG1635TAT (5044T>A 5045G>T), GTG1637CTT (5049G>C 5051G>T), CAG1639AGT (5055C>A 5056A>G 5057G>T), GAA1640CAA (5058G>C), ATT1642TTG (5064A>T 5066T>G), GCA1643ACA (5067G>A), AAG1644GAG (5070A>G), GTA1645GTC (5075A>C), GGG1647GGA (5081G>A), TCA1649GAG (5085T>G 5086C>A 5087A>G), AGT1650AGC (5090T>C), GCC1651ATT (5091G>A 5092C>T 5093C>T), CTG1652GGG (5094C>G 5095T>G), ACA1653GTT (5097A>G 5098C>T 5099A>T), ATT1654TTG (5100A>T 5102T>G), TGT1655ATG (5103T>A 5104G>T 5105T>G), GAA1656GAG (5108A>G), ATG1657CTA (5109A>C 5111G>A), GCA1658TCA (5112G>T), AAA1659GAT (5115A>G 5117A>T), GCA1660GCT (5120A>T), GCT1661AAA (5121G>A 5122C>A 5123T>A), ATT1662AAT (5125T>A), TCC1663ACA (5127T>A 5129C>A), TAT1664TAC (5132T>C), ACG1665ACC (5135G>C), CCA1666CCT (5138A>T), TTG1667TTA (5141G>A), GAC1668CCT (5142G>C 5143A>C 5144C>T), GCA1669GCT (5147A>T), GGT1670GGG (5150T>G), GAG1671GAA (5153G>A), ATC1673ATT (5159C>T), ACA1674del (5160_5162delACA), CGT1678GAG (5172C>G 5173G>A 5174T>G), GAT1679GGT (5176A>G), CTG1680GTC (5178C>G 5180G>C), AAT1681CAT (5181A>C), TGG1682GAA (5184T>G 5185G>A 5186G>A), GCA1684GTG (5191C>T 5192A>G), CCA1685CCT (5195A>T), AAC1686GAT (5196A>G 5198C>T), ATT1687GTA (5199A>G 5201T>A), GGT1688TTT (5202G>T 5203G>T), TTT1689TTC (5207T>C), TCA1691AGT (5211T>A 5212C>G 5213A>T), GCA1692GCT (5216A>T), CGT1693CGG (5219T>G), GCC1696TCA (5226G>T 5228C>A), CAC1697AAC (5229C>A), TAC1698CAA (5232T>C 5234C>A), ACG1699CAA (5235A>C 5236C>A 5237G>A), GAT1701TCA (5241G>T 5242A>C 5243T>A), GAA1702TCA (5244G>T 5245A>C), GGG1703GGT (5249G>T), TAC1704AAC (5250T>A), ATT1705ATC (5255T>C), ATT1706CTG (5256A>C 5258T>G), AAC1708AAT (5264C>T), TGT1709TGC (5267T>C), ACA1710ACT (5270A>T), GGT1711GGA (5273T>A), GCA1712GCC (5276A>C), AAG1713CGC (5277A>C 5278A>G 5279G>C), AAG1714AGA (5281A>G 5282G>A), TCC1715ACA (5283T>A 5285C>A), AAT1716ACA (5287A>C 5288T>A), GTT1718GTG (5294T>G), GGT1720GGG (5300T>G), CCA1721CCT (5303A>T), AAC1723AAT (5309C>T), ATG1731ATA (5333G>A), CCG1733CCC (5339G>C), ATG1736TTG (5346A>T), TCA1737GCA (5349T>G), GCT1738GCA (5354T>A), ATA1739ATC (5357A>C), ATG1740CAG (5358A>C 5359T>A), GCC1742CAT (5364G>C 5365C>A 5366C>T), AAG1744CGC (5370A>C 5371A>G 5372G>C), AAT1745AAG (5375T>G), GTG1746AAT (5376G>A 5377T>A 5378G>T), TTG1747CTG (5379T>C), GCA1748GAA (5383C>A), GGA1749GTT (5386G>T 5387A>T), GGA1750GGT (5390A>T), CAT1751GAT (5391C>G), GTA1752GGG (5395T>G 5396A>G), GCA1753GCT (5399A>T), AAT1755TCT (5403A>T 5404A>C), GTT1757ACC (5409G>A 5410T>C 5411T>C), GTG1759GTT (5417G>T), GGA1761GGT (5423A>T), GGA1763GGT (5429A>T), GCT1765GCA (5435T>A), GTC1766ATA (5436G>A 5438C>A), GGC1767GCT (5440G>C 5441C>T), ATT1769ATC (5447T>C), GCG1772GCA (5456G>A), CTG1773TTG (5457C>T), ATG1774TTT (5460A>T 5462G>T), GCC1775GCA (5465C>A), TTA1776TTG (5468A>G), GCA1777GCC (5471A>C), ACA1778ACT (5474A>T), AGG1779CAT (5475A>C 5476G>A 5477G>T), TAT1781TAC (5483T>C), TGT1782AGG (5484T>A 5486T>G), GTG1783ACC (5487G>A 5488T>C 5489G>C), GAG1784ACA (5490G>A 5491A>C 5492G>A), ATA1786CTT (5496A>C 5498A>T), ATA1787ATT (5501A>T), GTT1788GTG (5504T>G), AGA1789CGC (5505A>C 5507A>C), TTG1790CTA (5508T>C 5510G>A), CGC1791CGA (5513C>A), ACA1792ACC (5516A>C), ATT1793ATG (5519T>G), GGA1794GCT (5521G>C 5522A>T), AAT1795GGG (5523A>G 5524A>G 5525T>G), CTT1797CTG (5531T>G), GAC1798GAT (5534C>T), CTC1799CTG (5537C>G), AAT1800AAC (5540T>C), TAT1802TAC (5546T>C), GGC1805GGT (5555C>T), GTG1807GCG (5560T>C), CAG1808CAA (5564G>A), TAT1809TAC (5567T>C), TTT1810TTG (5570T>G), CTT1811CTC (5573T>C), CAA1812AAA (5574C>A), GTG1814GTT (5582G>T), CCT1815GAA (5583C>G 5584C>A 5585T>A), CAT1816GAC (5586C>G 5588T>C), CAT1816_GGA1817insACC (5588_5589insACC), GGA1817TCA (5589G>T 5590G>C), GAC1818CAC (5592G>C), AAA1819AAG (5597A>G), GTG1820TTG (5598G>T), GCT1821GGA (5602C>G 5603T>A), CAA1822CAG (5606A>G), GGT1824GGC (5612T>C), GCA1825ACA (5613G>A), TCG1826TAT (5617C>A 5618G>T), CTC1827GGC (5619C>G 5620T>G), CAC1828del (5622_5624delCAC), AGT1829ACC (5626G>C 5627T>C), AGT1830ACT (5629G>C), ATG1832AAC (5635T>A 5636G>C), CAA1834CAT (5642A>T), CAA1834_GGG1835insCACAGT (5642_5643insCACAGT), GGG1835GCA (5644G>C 5645G>A), GAA1837TCA (5649G>T 5650A>C), TTG1838GTT (5652T>G 5654G>T), GGC1839GGA (5657C>A), AAC1840CTT (5658A>C 5659A>T 5660C>T), CTC1841CTA (5663C>A), CGC1842AGG (5664C>A 5666C>G), ATT1843ATA (5669T>A), ATA1844GTT (5670A>G 5672A>T), CAT1845CCT (5674A>C), GGT1846TCC (5676G>T 5677G>C 5678T>C), GAT1847GAC (5681T>C), TTA1848CTC (5682T>C 5684A>C), GCA1849TCT (5685G>T 5687A>T), AAA1850GAA (5688A>G), CAG1851AAG (5691C>A), ACA1852ATG (5695C>T 5696A>G), ATG1854GTT (5700A>G 5702G>T), CGG1855CGT (5705G>T), ATG1856TTG (5706A>T), TAT1858TAC (5714T>C), GTG1859ACA (5715G>A 5716T>C 5717G>A), GTA1860ATT (5718G>A 5720A>T), GGA1861GGT (5723A>T), CCA1862CCT (5726A>T), CAT1863GAA (5727C>G 5729T>A), GGA1865GGT (5735A>T), GGA1867GCC (5740G>C 5741A>C), AAT1868CAG (5742A>C 5744T>G), TTT1869TGC (5746T>G 5747T>C), TCC1870TCA (5750C>A), ACA1871ACG (5753A>G), CAG1872AAG (5754C>A), GAT1873AAC (5757G>A 5759T>C), CTT1874CTG (5762T>G), GAG1875AAG (5763G>A), AGT1876CAT (5766A>C 5767G>A), TTG1877GCT (5769T>G 5770T>C 5771G>T), CTA1878TTG (5772C>T 5774A>G), CAC1879GAG (5775C>G 5777C>G), TAT1880ACA (5778T>A 5779A>C 5780T>A), GAG1882TCT (5784G>T 5785A>C 5786G>T), CAG1883GTC (5787C>G 5788A>T 5789G>C), GGA1884AGT (5790G>A 5792A>T), GCA1886GCT (5798A>T), GAG1887ACT (5799G>A 5800A>C 5801G>T), TTA1888CTT (5802T>C 5804A>T), ATT1889GCC (5805A>G 5806T>C 5807T>C), CAA1890GCA (5808C>G 5809A>C), AAG1891AGA (5812A>G 5813G>A), AAT1892GGA (5814A>G 5815A>G 5816T>A), ACA1893GTC (5817A>G 5818C>T 5819A>C), CTA1895GTG (5823C>G 5825A>G), ACT1896ACA (5828T>A), CTG1897TTG (5829C>T), TTC1899TTT (5837C>T), GAG1901GAA (5843G>A), CTG1902TTC (5844C>T 5846G>C), AGT1903TCC (5847A>T 5848G>C 5849T>C), ATG1904CAA (5850A>C 5851T>A 5852G>A), GAA1905CAC (5853G>C 5855A>C), AAT1906GAT (5856A>G), GTG1907TGT (5859G>T 5860T>G 5861G>T), CAA1908GAA (5862C>G), CAA1909ATG (5865C>A 5866A>T 5867A>G), CTG1910ATA (5868C>A 5870G>A), ATA1911CTT (5871A>C 5873A>T), GAT1912GAG (5876T>G), GTT1914CTG (5880G>C 5882T>G), AAG1915GAA (5883A>G 5885G>A), CTT1916AGT (5886C>A 5887T>G), CAG1917CAA (5891G>A), TTT1919GTA (5895T>G 5897T>A), TTC1920GTA (5898T>G 5900C>A), CCA1921TTC (5901C>T 5902C>T 5903A>C), AAA1922ATG (5905A>T 5906A>G), GAA1923GGG (5908A>G 5909A>G), ACT1924AGC |

| | | | | | | | | | |
|---|---|---|---|---|---|---|---|---|---|
| | (5911C>G 5912T>C), ACC1925GTT (5913A>G 5914C>T 5915C>T), ATT1926AAG (5917T>A 5918T>G), CAG1927AAA (5919C>A 5921G>A), AAG1928CAA (5922A>C 5924G>A), CTC1929ATG (5925C>A 5927C>G), AAA1930TTG (5928A>T 5929A>T 5930A>G), AAT1931TCC (5931A>T 5932A>C 5933T>C), TTT1932TTC (5936T>C), GTT1933ATG (5937G>A 5939T>G), GAT1934AAT (5940G>A), GCT1935CAA (5943G>C 5944C>A 5945T>A), GAA1936CAG (5946G>C 5948A>G), TGT1937ATG (5949T>A 5950G>T 5951T>G), CTA1938TCA (5952C>T 5953T>C), ACA1939AAA (5956C>A), TTC1940ACC (5958T>A 5959T>C), AAA1941CAA (5961A>C), GCA1942GTT (5965C>T 5966A>T), AGG1943GCC (5967A>G 5968G>C 5969G>C), AGC1944_TTT1945insCCATGTTGTGTCACA (5972_5973insCCATGTTGTGTCACA), AGG1946AGT (5978G>T), CAC1947AAA (5979C>A 5981C>A), GTA1948GCA (5983T>C), GTT1949TGG (5985G>T 5986T>G 5987T>G), TTC1950CTA (5988T>C 5990C>A), AAA1951ACT (5992A>C 5993A>T), AAA1953GAC (5997A>G 5999A>C), CTT1954ATT (6000C>A), TTG1955GTC (6003T>G 6005G>C), AGT1956ATG (6007G>T 6008T>G), TCC1957ACA (6009T>A 6011C>A), ACA1958AAA (6013C>A), TGG1959GGT (6015T>G 6017G>T), AAA1960AAG (6020A>G), ATG1961AAG (6022T>A), TGT1962CCA (6024T>C 6025G>C 6026T>A), GGT1963GGA (6029T>A), GAA1964del (6030_6032delGAA), AGT1965TCT (6033A>T 6034G>C), GTT1967_AGG1970del (6039_6050delGTTGCTTCAAGG), TCT1971CCT (6051T>C), GCA1972GCT (6056A>T), GAG1973GAA (6059G>A), AGT1974ACA (6061G>C 6062T>A), CTA1975GTG (6063C>G 6065A>G), TTC1976TTT (6068C>T), CCT1977CTT (6070C>T), GGC1978GGG (6074C>G), TTG1980CCT (6078T>C 6079T>C 6080G>T), TCT1981TCA (6083T>A), GCT1982GCA (6086T>A), TCC1983CTA (6087T>C 6088C>T 6089C>A), GTA1984GTT (6092A>T), TTG1985GTG (6093T>G), AAA1986AGA (6097A>G), TTA1988ATG (6102T>A 6104A>G), GAA1990CGC (6108G>C 6109A>G 6110A>C), AGA1991AAT (6112G>A 6113A>T), CAC1992AGT (6114C>A 6115A>G 6116C>T), ACA1993GGA (6117A>G 6118C>G), AGA1994AGG (6122A>G), ATG1996TAC (6126A>T 6127T>A 6128G>C), AGT1997TCA (6129A>T 6130G>C 6131T>A), CAG1999ACA (6135C>A 6136A>C 6137G>A), AGT2000ACA (6139G>C 6140T>A), ATG2001CTG (6141A>C), GAG2002GAA (6146G>A), CTG2005TTT (6153C>T 6155G>T), AAA2006AGA (6157A>G), ATA2007ATG (6161A>G), TAT2008TAC (6164T>C), CTA2009ATG (6165C>A 6167A>G), TTA2010TGT (6169T>G 6170A>T), AAC2011CAG (6171A>C 6173C>G), TTC2012GAA (6174T>G 6175T>A 6176C>A), CAA2013GGG (6177C>G 6178A>G 6179A>G), GTA2014TAT (6180G>T 6181T>A 6182A>T), ATA2015TTA (6183A>T), ACA2016GAT (6186A>G 6187C>A 6188A>T), AGT2017AAA (6190G>A 6191T>A), GAG2018GCC (6193A>C 6194G>C), TTA2020GTG (6198T>G 6200A>G), AAG2021GAC (6201A>G 6203G>C), AAG2022CGC (6204A>C 6205A>G 6206G>C), TTT2023TTC (6209T>C), GAA2024AAT (6210G>A 6212A>T), GAT2025GAG (6215T>G), TTT2027ATG (6219T>A 6221T>G), CAG2028GGC (6222C>G 6223A>G 6224G>C), GAG2029GTT (6226A>T 6227G>T), AAG2030TGT (6228A>T 6229A>G 6230G>T), ATA2031GTG (6231A>G 6233A>G), TCC2032TCG (6236C>G), ACC2033GAG (6237A>G 6238C>A 6239C>G), ACC2034CGC (6240A>C 6241C>G), CTA2035ATC (6243C>A 6245A>C), CTA2036TTG (6246C>T 6248A>G), ATC2037GTG (6249A>G 6251C>G), AAA2038CGC (6252A>C 6253A>G 6254A>C), GTA2039ACA (6255G>A 6256T>C), TTC2040ATT (6258T>A 6260C>T), TTG2041GCT (6261T>G 6262T>C 6263G>T), GAG2044TCC (6270G>T 6271A>C 6272G>C), GAG2045ACG (6273G>A 6274A>C), AGT2046TCA (6276A>T 6277G>C 6278T>A), TAT2047TTG (6280A>T 6281T>G), CAG2048AAC (6282C>A 6284G>C), ACT2049GAG (6285A>G 6286C>A 6287T>G), GAG2050GAT (6290G>T), TGT2052ACT (6294T>A 6295G>C), ATA2053GTG (6297A>G 6299A>G), GAT2054TAC (6300G>T 6302T>C), TCT2055ACA (6303T>A 6305T>A), CAG2056GAT (6306C>G 6308G>T), GTG2057GGG (6310T>G), CTC2058CTT (6314C>T), CAG2059TTT (6315C>T 6316A>T 6317G>T), AGG2060GAT (6318A>G 6319G>A 6320G>T), ATA2061del (6321_6323delATA), AAC2062CAT (6324A>C 6326C>T), TCC2063TTC (6328C>T), CAG2064AAG (6330C>A), AAA2065GCA (6333A>G 6334A>C), TTC2066GTG (6336T>G 6338C>G), AGA2067GTT (6339A>G 6340G>T 6341A>T), GTG2068TTC (6342G>T 6344G>C), CAG2069AAG (6345C>A), GAA2070CCT (6348G>C 6349A>C 6350A>T), GGT2071GGG (6353T>G), TTT2073TTC (6359T>C), TAT2074TCT (6361A>C), TTG2075TGG (6364T>G), CAT2076GAA (6366C>G 6368T>A), CCT2077ATC (6369C>A 6370C>T 6371T>C), GAA2078AAT (6372G>A 6374A>T), GGA2079GAT (6376G>A 6377A>T), ATC2080GCC (6378A>G 6379T>C), AAC2081AAT (6383C>T), GCC2084GCT (6392C>T), GTA2085GCA (6394T>C), ATA2088GTG (6402A>G 6404A>G), ATA2089TTG (6405A>T 6407A>G), TCT2091GAA (6411T>G 6412C>A 6413T>A), CTG2092CTC (6416G>C), GAG2094AGG (6420G>A 6421A>G), TCC2095CAC (6423T>C 6424C>A), TAT2096TGG (6427A>G 6428T>G), AGC2097TCA (6429A>T 6430G>C 6431C>A), CGA2098AAG (6432C>A 6433G>A 6434A>G), GAT2099TCA (6435G>T 6436A>C 6437T>A), AGT2100CGA (6438A>C 6440T>A), AAC2101AGC (6442A>G), TCC2102AAG (6444T>A 6445C>A 6446C>G), TAT2103TTC (6448A>T 6449T>C), TCT2104ACT (6450T>A), GTG2105TTT (6453G>T 6455G>T), TCA2106GCC (6456T>G 6458A>C), ACT2107AAT (6460C>A), CCC2108GAA (6462C>G 6463C>A 6464C>A), ATC2109GTT (6465A>G 6467C>T), AAA2110GAA (6468A>G), CTG2111ATA (6471C>A 6473G>A), GGG2112TCA (6474G>T 6475G>C 6476G>A), TGT2113AAC (6477T>A 6478G>A 6479T>C), TTC2114CTG (6480T>C 6482C>G), ATT2115CTT (6483A>C), TTC2116TTA (6488C>A), CTT2117GCT (6489C>G 6490T>C), GTC2118GTT (6494C>T), GTG2119TAT (6495G>T 6496T>A 6497G>T), CTA2120TTA (6498C>T), GAT2121GAA (6503T>A), AGT2122AAT (6505G>A), CAA2123AAG (6507C>A 6509A>G), GGC2124CTC (6510G>C 6511G>T), AAG2125ATA (6514A>T 6515G>A), GAA2126GGT (6517A>G 6518A>T), GAT2127CTA (6519G>C 6520A>T 6521T>A), CAT2128GAG (6522C>G 6524T>G), CCT2129TAT (6525C>T 6526C>A), GTT2130TAT (6528G>T 6529T>A), CGG2131GAT (6531C>G 6532G>A 6533G>T), AGG2132TCT (6534A>T 6535G>C 6536G>T), TTT2133ATA (6537T>A 6539T>A), TCA2134GAA (6540T>G 6541C>A), ACG2135TCT (6543A>T 6545G>T), GTT2136TTG (6546G>T 6548T>G), TTC2137GAC (6549T>G 6550T>A), CTT2138TTG (6552C>T 6554T>G), AAA2139TTA (6555A>T 6556A>T), AAC2140CAA (6558A>C 6560C>A), TGT2141ATA (6561T>A 6562G>T 6563T>A), GAA2142GAT (6566A>T), ATT2144ATA (6572T>A), ACA2146GCA (6576A>G), TAC2148GTC (6582T>G 6583A>T), GAG2150TAC (6588G>T 6590G>C), TGA2152GGG (6594T>G 6596A>G) | | | | | | | | |
| helicase (YP_001976154.1) | 1 | 98 | 100% | 654 | 97.5% | 98 (100%) | 95 (96.9%) | 0/0/0/0 | 0 |
| Protein mutations: | S5C (1342C>G 1343C>T), R16A (1374A>G 1375G>C), I44V (1458A>G 1460C>G) | | | | | | | | |
| Codon mutations: | TCT3TCA (1337T>A), ACA4ACG (1340A>G), TCC5TGT (1342C>G 1343C>T), TTA8CTT (1350T>C 1352A>T), TTT9TTC (1355T>C), ATT10ATC (1358T>C), GTG13GTT (1367G>T), TTG14CTC (1368T>C 1370G>C), GAC15GAT (1373C>T), AGA16GCA (1374A>G 1375G>C), GAG19GAA (1385G>A), AAA21AAG (1391A>G), CTC22CTG (1394C>G), AAT23AAC (1397T>C), CGG24CGC (1400G>C), CTA25TTG (1401C>T 1403A>G), GCA27GCG (1409A>G), AAG30AAA (1418G>A), AGG31AGA (1421G>A), GCT39GCA (1445T>A), CAC40CAT (1448C>T), GCT43GCA (1457T>A), ATC44GTG (1458A>G 1460C>G), CTG45CTT (1463G>T), GAC48GAT (1472C>T), GGA50GGT (1478A>T), GCA51GCT (1481A>T), TTG52CTT (1482T>C 1484G>T), CGA53CGG (1487A>G), GGG55GGA (1493G>A), GCT56GCA (1496T>A), TGC59TGT (1505C>T), CAA60CAG (1508A>G), GAT61GAC (1511T>C), ATT62ATA (1514T>A), AAA63AAG (1517A>G), GAT67GAC (1529T>C), TCA70TCT (1538A>T), ACA71ACC (1541A>C), CAA72CAG (1544A>G), CCA73CCT (1547A>T), CCC75CCT (1553C>T), TTA76CTT (1554T>C 1556A>T), GCC79GCT (1565C>T), GTT81GTC (1571T>C), GAC83GAT (1577C>T), GGC85GGA (1583C>A), CAT87CAC (1589T>C), TTC88TTT (1592C>T), ACC89ACA (1595C>A), TCC90TCA (1598C>A), TAT92TAC (1604T>C), ATA93ATC (1607A>C), ACA96ACC (1616A>C) | | | | | | | | |
| RNA-dependent RNA polymerase (YP_001976153.1) | 1 | 249 | 100% | 1552 | 88.4% | 249 (100%) | 210 (84.3%) | 0/0/0/0 | 0 |
| Protein mutations: | T40C (4170A>T 4171C>G 4172A>T), L44A (4182T>G 4183T>C 4184G>A), A47V (4192C>T 4193T>A), I48L (4194A>C 4196A>C), D55N (4215G>A), S59H (4227T>C 4228C>A), L63V (4239T>G), N66S (4249A>G), G67S (4251G>A 4253G>C), A70S (4260G>T 4262T>A), V74E (4273T>A), V89I (4317G>A 4319T>C), E101A (4354A>C 4355A>T), Y102F (4357A>T 4358C>T), S105D (4365T>G 4366C>A 4367A>T), K106E (4368A>G), D111E (4385T>G), A117T (4401G>A), D131N (4443G>A 4445T>C), V133L (4449G>T), I172V (4566A>G 4568T>C), A173K (4569G>A 4570C>A 4571T>A), R175K (4576G>A 4577G>A), A176S (4578G>T), I177V (4581A>G 4583A>G), V179I (4587G>A), S182N (4597G>A), L202M (4656T>A 4658A>G), Q205R (4665C>A 4666A>G), S208D (4674T>G 4675C>A), N211S (4683A>T 4684A>C 4685T>C), H214R (4692C>A 4693A>G 4694T>A), L218A (4704C>G 4705T>C 4706G>T), I222V (4716A>G), N234E (4752A>G 4754T>G), E235K (4755G>A), V236T (4758G>A 4759T>C), Q237K (4761C>A), Q239L (4768A>T 4769A>G) | | | | | | | | |
| Codon mutations: | CGC4CGG (4064C>G), CTC5TTG (4065C>T 4067C>G), AAA9AAG (4079A>G), ATC10ATT (4082C>T), GGC12GGG (4088C>G), AGA14AGG (4094A>G), AGG17CGC (4101A>C 4103G>C), CTC18CTG (4106C>G), TTT19TTC (4109T>C), GAG20GAA (4112G>A), ATA21ATC (4115A>C), CTT22TTG (4116C>T 4118T>G), TAC26TAT (4130C>T), TTG29CTG (4137T>C), GTC30GTG (4142C>G), AGG31CGG (4143A>C), AAG32AAA (4148G>A), TAT33TAC (4151T>C), TTT34TTC (4154T>C), GAT36GAC (4160T>C), TTC37TTT (4163C>T), TCA38TCT (4166A>T), GCC39GCT (4169C>T), ACA40TGT (4170A>T 4171C>G 4172A>T), TTG44GCA (4182T>G 4183T>C 4184G>A), AAT46AAC (4190T>C), GCT47GTA (4192C>T 4193T>A), ATA48CTC (4194A>C 4196A>C), CCA49CCT (4199A>T), AAA51AAG (4205A>G), GTT52GTA (4208T>A), ATT54ATC (4214T>C), GAT55AAT (4215G>A), CCT56CCA (4220T>A), ACA57ACG (4223A>G), AGT58AGC (4226T>C), TCT59CAT (4227T>C 4228C>A), TTG63GTG (4239T>G), TTG64CTG (4242T>C), GCA65GCT (4247A>T), AAT66AGT (4249A>G), GGG67AGC (4251G>A 4253G>C), AGA69AGG (4259A>G), GCT70TCA (4260G>T 4262T>A), GTG71GTA (4265G>A), TCA72TCT (4268A>T), GAC73GAT (4271C>T), GTG74GAG (4273T>A), TTT76TTC (4280T>C), GCT78GCC (4286T>C), TAT80TAC (4292T>C), TCC81TCG (4295C>G), CCT88CCC (4316T>C), GTT89ATC (4317G>A 4319T>C), GCT91GCA (4325T>A), TTT92TTC (4328T>C), CAG93CAA (4331G>A), TTG98CTG (4344T>C), GAA101GCT (4354A>C 4355A>T), TAC102TTT (4357A>T 4358C>T), TAC103TAT (4361C>T), GGA104GGT (4364A>T), TCA105GAT (4365T>G 4366C>A 4367A>T), AAG106GAG (4368A>G), GGC108GGG (4376C>G), GAT111GAG (4385T>G), TCC112TCA (4388C>A), CGA115AGG (4395C>A 4397A>G), CAT116CAC (4400T>C), GCA117ACA (4401G>A), CTT118TTA (4404C>T 4406T>A), GCA122GCC (4418A>C), TGT124TGC (4424T>C), TAC126TAT (4430C>T), ACA127ACC (4433A>C), CTG128CTC (4436G>C), TGC129TGT (4439C>T), GAT131AAC (4443G>A 4445T>C), AAA132AAG (4448A>G), GTG133TTG (4449G>T), AGG135CGC (4455A>C 4457G>C), TTG136CTA (4458T>C 4460G>A), GTT137GTG (4463T>G), GGG138GGT (4466G>T), GGC139GGA (4469C>A), CCA141CCC (4475A>C), TCA142TCT (4478A>T), GGA143GGT (4481A>T), GCA145GCT (4487A>T), CTA146TTG (4488C>T 4490A>G), ACG147ACT (4493G>T), ATC149ATA (4499C>A), TTC150TTT (4502C>T), TCT152TCA (4508T>A), CTC153TTG (4509C>T 4511C>G), CTC154CTA (4514C>A), CGA160AGG (4530C>A 4532A>G), TAT161TAC (4535T>C), GCC162GCG (4538C>G), TTG166CTG (4548T>C), TTA167TTG (4553A>G), AGA168AGG (4556A>G), CCC170CCA (4562C>A), ATT172GTC (4566A>G 4568T>C), GCT173AAA (4569G>A 4570C>A 4571T>A), AGG175AAA (4576G>A 4577G>A), GCT176TCT (4578G>T), ATA177GTG (4581A>G 4583A>G), GGA178GGC (4586A>C), GTT179ATT (4587G>A), AAA180AAG (4592A>G), AGT182AAT (4597G>A), GAT183GAC (4601T>C), CTA187TTG (4611C>T 4613A>G), GCT190GCA (4622T>A), GTT191GTA (4625T>A), GGA193GGT (4631A>T), GAC195GAT (4637C>T), AAT196AAC (4640T>C), CTT197TTG (4641C>T 4643T>G), GCT199GCG (4649T>G), GTA200GTT (4652A>T), TTA202ATG (4656T>A 4658A>G), CTC204TTG (4662C>T 4664C>G), CAG205AGG (4665C>A 4666A>G), TAT207TAC (4673T>C), TCT208GAT (4674T>G 4675C>A), CTG209CTA (4679G>A), AAT211TCC (4683A>T 4684A>C 4685T>C), ATA212ATT (4688A>T), CAT214AGA (4692C>A 4693A>G 4694T>A), GAG215GAA (4697G>A), TTA216CTG (4698T>C 4700A>G), CTG218GCT (4704C>G 4705T>C 4706G>T), GTC219GTT (4709C>T), AAT220AAC (4712T>C), GTA221GTC (4715A>C), ATC222GTC (4716A>G), CTT227TTG (4731C>T 4733T>G), AAA229AAG (4739A>G), TCG230TCA (4742G>A), GTT233GTA (4751T>A), AAT234GAG (4752A>G 4754T>G), GAG235AAG (4755G>A), GTA236ACA (4758G>A 4759T>C), CAA237AAA (4761C>A), CAA239CTG (4768A>T 4769A>G), CTA244CTC (4784A>C), ACT245ACC (4787T>C), CTG247CTC (4793G>C), AGA249AGG (4799A>G) | | | | | | | | |

*: Inserts / Deletes / Misaligned / Frameshifts

#### Analysis details

This analysis was performed with panviral2.64

### NGS Details (UN70): Torradovirus lycopersici (segment RNA 1)

#### Assembly

| | |
|---|---|
| **Coverage Length** | 7781 (1 contig(s)) |
| **Depth Of Coverage** | 13912.1 |
| **Number Of Reads** | 753401 |
| **Reads Per Million** | 14281.67 rpm (after QC) |
| **Ambiguities** | 0 |
| **Assembly Method** | de novo + reference guided assembly |
| **Consensus Caller** | Bcf Tools |

#### Coverage Map

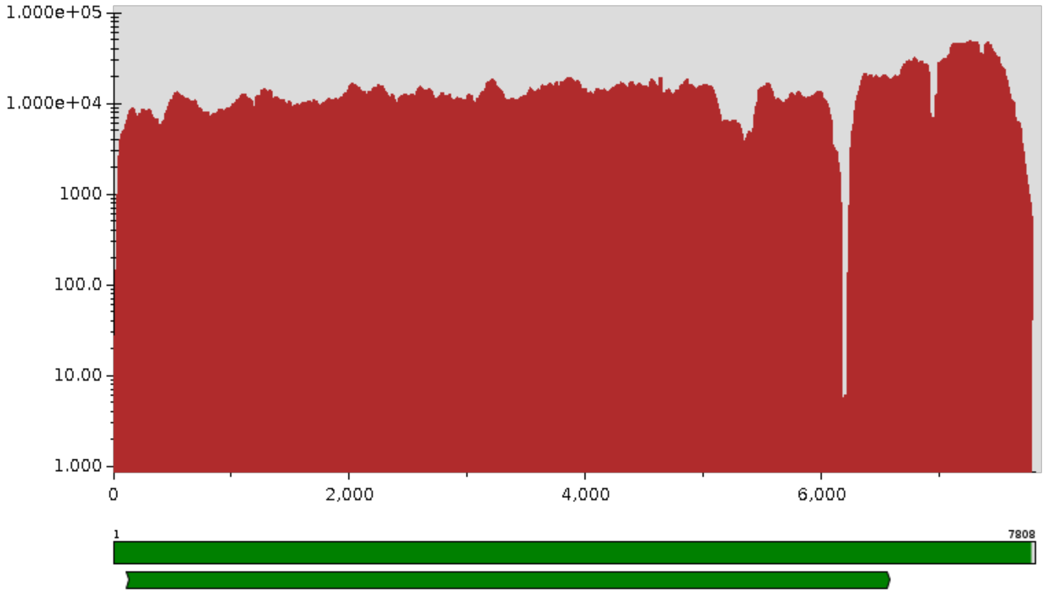

#### Assignment

| | |
|---|---|
| **Type** | Torradovirus lycopersici (Taxonomy ID: 3048378) |
| **Reference Genome** | NC_009013.1 |
| **NT Identity (%)** | 90.3062 |
| **AA Identity (%)** | 97.6378 |
| **Number Of Stop Codons** | 1 |
| **Number Of CDS** | 1 |

#### Alignment

| | |
|---|---|
| **Alignment Score** | 11522.0 (NT) + 14846.0 (AA) = 26368.0 |
| **Concordance (%)** | 89.2228 |

| Alignment Method | Global, seeded, nucleotide + amino acids (AGA) |
|---|---|

#### Genome Region

Sequence starts at position 1 and ends at position 7781 relative to NC_009013.1 reference sequence.

#### Alignment Detailed Statistics

| | Begin | End | Coverage | Score | Concordance | Matches | Identities | I/D/M/F* | Stop Codons |
|---|---|---|---|---|---|---|---|---|---|
| **NT** | **1** | **7781** | **99.7%** | **11522** | **79.6%** | **7469 (95.9%)** | **6754 (86.7%)** | **10/312** | |
| Mutations: | 2T>C, 5A>T, 9G>A, 12A>T, 13T>C, 14T>A, 15T>C, 16T>G, 27C>A, 69C>T, 76_77insT, 87C>T, 95A>G, 100T>C, 101G>A, 102G>A, 115T>C, 127C>T, 128C>T, 130C>T, 135T>C, 148T>C, 151A>G, 158G>T, 161A>G, 175T>C, 178T>C, 184T>C, 190G>A, 197G>A, 199T>A, 202T>C, 205T>A, 208T>C, 220C>T, 224C>T, 253A>G, 259T>C, 262T>A, 265C>T, 338T>C, 340G>A, 361A>G, 367A>G, 370C>T, 376C>T, 385C>T, 388T>C, 409A>G, 451A>G, 487C>T, 490A>G, 502G>A, 541T>C, 559C>T, 577A>G, 578C>T, 655C>T, 676A>G, 694A>G, 727A>G, 760C>T, 781C>T, 790C>T, 799A>G, 805T>C, 818C>T, 820A>G, 832A>T, 835A>C, 841T>C, 842C>T, 847C>T, 883G>A, 886T>C, 892G>A, 901T>C, 913T>C, 925C>T, 938C>T, 952A>G, 955C>T, 964T>G, 970G>A, 973T>C, 977G>A, 980T>C, 997C>T, 1006T>C, 1012C>T, 1013T>C, 1018C>T, 1033G>A, 1060T>C, 1063C>T, 1078A>T, 1081C>T, 1090C>T, 1093T>C, 1105A>G, 1108T>C, 1109C>T, 1165C>T, 1180G>A, 1196A>G, 1207T>C, 1231G>A, 1240C>T, 1255G>A, 1267T>C, 1300G>A, 1303A>G, 1319C>T, 1324C>T, 1351A>G, 1354G>A, 1411T>C, 1417T>C, 1429C>T, 1441C>T, 1450T>C, 1451C>T, 1480C>T, 1481A>G, 1489T>C, 1522C>T, 1552C>T, 1555C>T, 1570A>G, 1573T>C, 1582C>T, 1585A>C, 1591C>T, 1594T>C, 1607G>T, 1612G>A, 1618T>G, 1619C>T, 1624C>T, 1630C>T, 1639T>C, 1642G>A, 1645A>G, 1669C>T, 1672G>A, 1681T>A, 1684G>T, 1699T>C, 1705T>C, 1737A>G, 1738A>G, 1750A>T, 1756A>G, 1765T>G, 1781T>C, 1783G>A, 1784C>T, 1789A>G, 1810G>A, 1825C>T, 1834A>G, 1867G>A, 1879A>G, 1888A>G, 1910T>C, 1912G>A, 1916A>G, 1960A>G, 1962G>A, 1966A>G, 1984C>T, 1990G>A, 1993G>A, 2009T>C, 2023T>C, 2056T>C, 2062C>T, 2077C>A, 2082T>A, 2092A>G, 2107A>G, 2128T>C, 2138C>T, 2143T>C, 2149T>C, 2155T>C, 2174C>T, 2179A>C, 2181A>G, 2182G>A, 2200T>C, 2224C>T, 2236A>G, 2241A>G, 2250G>A, 2251C>T, 2257C>T, 2263G>A, 2275T>C, 2287G>A, 2290C>T, 2314C>T, 2323A>T, 2347C>T, 2356C>T, 2359G>A, 2377G>A, 2392A>G, 2409G>A, 2443T>C, 2446G>A, 2479G>A, 2485C>T, 2497G>A, 2500T>C, 2503G>A, 2506T>C, 2509C>T, 2512T>C, 2542G>A, 2548C>T, 2557C>T, 2620G>A, 2662C>T, 2665T>C, 2734C>T, 2743A>G, 2755C>T, 2773A>G, 2809C>T, 2812G>A, 2827A>G, 2860C>T, 2881T>C, 2914C>T, 2926G>A, 2929C>T, 2950C>T, 2951C>T, 2980C>T, 2983G>A, 2998A>G, 3007T>C, 3028T>C, 3031T>C, 3034G>A, 3040T>C, 3058A>G, 3064A>G, 3073G>A, 3079T>C, 3109T>C, 3115G>A, 3145A>G, 3151C>T, 3163T>C, 3169C>T, 3172T>C, 3202T>C, 3241T>C, 3253C>T, 3256G>A, 3262C>T, 3283T>C, 3289T>C, 3298T>C, 3307A>G, 3358A>T, 3406A>C, 3418C>A, 3421T>C, 3463A>G, 3466G>A, 3481C>T, 3484A>T, 3487G>A, 3492C>T, 3508G>A, 3559G>A, 3574T>C, 3577A>G, 3586T>C, 3591C>T, 3625C>T, 3628G>A, 3646C>T, 3661A>G, 3664T>C, 3673G>A, 3676C>T, 3686C>T, 3706T>C, 3716T>C, 3721T>A, 3739A>T, 3754G>A, 3784C>T, 3814C>T, 3868C>T, 3871T>C, 3883G>A, 3892G>A, 3898T>C, 3907A>T, 3916C>T, 3925A>G, 3928G>A, 3949C>T, 3955G>A, 3961G>A, 3970A>G, 3973T>C, 4018A>G, 4034T>C, 4036G>A, 4057G>A, 4066T>C, 4084C>T, 4118C>T, 4129C>T, 4138G>A, 4147C>A, 4159G>A, 4162A>T, 4168A>G, 4186A>T, 4195A>G, 4207G>A, 4210T>C, 4219C>T, 4228C>T, 4273A>G, 4282C>A, 4288C>T, 4291T>C, 4300T>C, 4318C>T, 4321C>T, 4324T>C, 4339T>C, 4348C>T, 4354T>C, 4360A>G, 4393T>C, 4417T>C, 4421T>C, 4459T>C, 4462C>T, 4465C>T, 4468C>T, 4471T>C, 4474T>C, 4486T>C, 4489T>C, 4591T>C, 4625T>C, 4636T>C, 4640C>T, 4651T>A, 4654C>A, 4657A>G, 4661T>C, 4663G>A, 4666G>A, 4672G>A, 4678T>C, 4681T>A, 4684A>C, 4687G>A, 4696C>A, 4699C>T, 4720T>A, 4759C>T, 4774G>A, 4783A>G, 4786A>G, 4789C>T, 4813A>G, 4816T>C, 4898G>A, 4906G>A, 4924T>C, 4946A>G, 4960C>T, 4969G>A, 4975G>A, 4993A>C, 4999T>C, 5005T>C, 5012T>C, 5017G>A, 5023C>T, 5044A>C, 5059G>A, 5065C>T, 5074T>C, 5077T>C, 5079T>C, 5102T>C, 5113C>T, 5119T>C, 5120G>A, 5146T>C, 5148C>A, 5167T>C, 5170C>T, 5176C>T, 5200G>A, 5218T>G, 5244G>A, 5266C>T, 5281C>T, 5290G>A, 5329C>T, 5344A>T, 5347C>A, 5354G>A, 5356G>T, 5365A>C, 5369G>A, 5371G>T, 5386C>G, 5392G>A, 5407A>C, 5413C>T, 5419A>T, 5429C>T, 5437T>C, 5440C>G, 5446T>C, 5461T>C, 5467G>T, 5471T>C, 5491T>C, 5492A>C, 5497T>C, 5533T>G, 5537A>C, 5575A>G, 5578T>C, 5581A>G, 5591G>T, 5599T>C, 5603A>C, 5611A>T, 5617A>T, 5641T>C, 5647T>C, 5674A>G, 5677G>T, 5686G>A, 5710T>A, 5716G>T, 5719A>C, 5731C>T, 5734T>C, 5737T>C, 5739G>A, 5746T>C, 5750T>C, 5752G>A, 5758C>T, 5773G>A, 5782A>G, 5784T>C, 5786T>C, 5797A>G, 5806C>T, 5836A>T, 5849T>C, 5850T>A, 5851G>A, 5869G>A, 5875A>G, 5878G>C, 5884T>C, 5887C>A, 5890C>T, 5902G>T, 5908C>T, 5909T>C, 5914T>C, 5923T>C, 5927C>T, 5944C>T, 5955A>G, 5965G>A, 5998A>G, 6004C>T, 6016A>G, 6019A>G, 6028C>T, 6031A>T, 6033C>T, 6035G>T, 6055T>A, 6067C>A, 6070T>C, 6091A>G, 6092C>T, 6100T>G, 6103G>A, 6106G>A, 6112A>G, 6118T>C, 6121A>T, 6139T>C, 6154C>G, 6163C>G, 6166C>T, 6184A>G, 6187T>G, 6193C>T, 6196A>G, 6202G>C, 6211A>G, 6217G>T, 6223A>T, 6230A>G, 6232A>G, 6235G>A, 6253A>C, 6256C>A, 6262A>T, 6265G>A, 6268G>T, 6274C>G, 6296G>C, 6298A>G, 6308T>C, 6313T>C, 6322G>A, 6327A>G, 6358T>C, 6484A>C, 6490A>G, 6493T>C, 6500A>T, 6517C>T, 6532C>T, 6545C>T, 6571C>T, 6587G>A, 6661T>C, 6670C>T, 6675C>T, 6685A>G, 6742A>T, 6766delC, 6786T>C, 6798_6799insA, 6806T>G, 6807A>T, 6809C>T, 6811_6832delGAGAGTACTGACTTTAACTAGT, 6835G>T, 6838_6850delGTCCGGCTCCATT, 6854A>T, 6858C>T, 6859C>A, 6860_6894delAATGAATTTACATTGGTTAAAGAGATTTGCACGCC, 6897T>A, 6905T>G, 6907T>A, 6908G>T, 6911T>C, 6912C>A, 6921T>G, 6923C>G, 6924T>A, 6927_6928delGT, 6931G>A, 6933A>C, 6935G>A, 6937T>G, 6938C>A, 6940T>A, 6941T>C, 6943A>G, 6947_6960delAGGTTTACAAAGTA, 6966A>T, 6968C>T, 6975T>A, 6977delG, 6984_6985insG, 6988G>A, 6994_7013delGCTATCCCATGATAGTCCTT, 7018T>A, 7020_7026delAAGGGTT, 7031T>A, 7033_7062delGGTTTGCATCTTTTGCCGTGATGAAAGATG, 7067A>T, 7068G>A, 7071T>A, 7074C>A, 7077T>C, 7083G>T, 7085_7089delAGGCT, 7091A>T, 7094C>A, 7096A>G, 7097C>T, 7099C>G, 7102A>G, 7106A>G, 7107G>A, 7111G>T, 7112T>G, 7114T>A, 7115G>T, 7117_7177delCTGAGTCCTATAATCAGTCCGTTTGAAATTCGATAATTTTCCGTAGCTTGCGTCAAGCTGC, 7182G>T, 7186G>A, 7187G>A, 7188_7225delGGTGTGAGTGAAGATGCGCCGTACCACGTCTTCCCCGG, 7231C>A, 7234G>C, 7235T>C, 7236G>C, 7238T>G, 7239T>G, 7240C>A, 7241A>T, 7243A>T, 7246G>T, 7247G>T, 7248G>T, 7249C>T, 7253C>A, 7254_7262delAGAATAGAG, 7268A>C, 7269A>T, 7271T>A, 7274_7320delTGTGATGGTGTATATCACGATAAAAGTGACACCCGGGTTGTGCTGCG, 7322C>A, 7323T>C, 7326T>G, 7329A>G, 7331A>T, 7333G>C, 7334A>C, 7335G>C, 7336C>T, 7338C>T, 7342T>A, 7343C>G, 7345C>G, 7346A>C, 7347C>T, 7348C>G, 7350T>C, 7353A>T, 7354G>A, 7358A>C, 7360G>A, 7361A>C, 7362A>C, 7364C>T, 7372A>T, 7375T>G, 7376T>G, 7377T>A, 7379A>T, 7380A>T, 7397G>A, 7398C>A, 7399G>A, 7400A>T, 7404C>T, 7407A>T, 7408G>T, 7423A>T, 7425G>A, 7435T>C, 7436G>A, 7437T>C, 7444C>T, 7445A>G, 7457A>G, 7458T>C, 7460G>A, 7503C>T, 7505A>G, 7506T>C, 7511_7512insT, 7513A>C, 7514A>T, 7515A>C, 7525_7526insG, 7526T>A, 7526_7527insG, 7545T>A, 7554A>G, 7569G>A, 7589T>A, 7600T>G, 7601T>A, 7602T>G, 7603_7604insC, 7610C>T, 7612G>A, 7615A>G, 7616A>C, 7617A>T, 7618C>T, 7625_7631delATTGCTT, 7637G>T, 7641T>A, 7642T>A, 7644G>T, 7647T>A, 7650T>A, 7652T>G, 7653A>T, 7654C>T, 7665G>T, 7666T>C, 7667G>T, 7678C>T, 7686T>G, 7687T>A, 7689G>T, 7698G>A, 7713G>A, 7717A>C, 7719T>A, 7739T>A, 7742_7743insC, 7749T>C, 7751T>C, 7752G>A, 7754_7755insG, 7756_7757insT, 7762C>T, 7763A>C, 7764A>T, 7767C>T, 7773G>T, 7775G>A, 7778T>A, 7781G>T | | | | | | | | |

##### CDS

| | Begin | End | Coverage | Score | Concordance | Matches | Identities | I/D/M/F* | Stop Codons |
|---|---|---|---|---|---|---|---|---|---|
| ToTV_sRNA1gp1 | 1 | 2159 | 100% | 14846 | 98.5% | 2159 (100%) | 2108 (97.6%) | 0/0/0/0 | 1 |
| Protein mutations: | P8S (128C>T 130C>T), F10S (135T>C), A18S (158G>T), T19A (161A>G), A31T (197G>A 199T>A), V291I (977G>A), T364A (1196A>G), I459V (1481A>G), A501S (1607G>T), E526D (1684G>T), K544R (1737A>G 1738A>G), I604V (1916A>G), S619N (1962G>A), F659Y (2082T>A), K692R (2181A>G 2182G>A), N712S (2241A>G), S715N (2250G>A 2251C>T), R768K (2409G>A), A1129V (3492C>T), A1162V (3591C>T), S1538R (4720T>A), V1598I (4898G>A), T1614A (4946A>G), I1658T (5079T>C), V1672I (5120G>A), T1681N (5148C>A), R1713K (5244G>A), V1750I (5354G>A 5356G>T), V1755I (5369G>A 5371G>T), T1796P (5492A>C), K1811Q (5537A>C), A1829S (5591G>T), T1833P (5603A>C), Q1857H (5677G>T), R1878K (5739G>A), I1893T (5784T>C), L1915Q (5849T>C 5850T>A 5851G>A), N1950S (5955A>G), T1976I (6033C>T), A1977S (6035G>T), L1996F (6092C>T), I2002M (6112A>G), M2032I (6202G>C), E2039D (6223A>T), I2042V (6230A>G 6232A>G), L2049F (6253A>C), S2056R (6274C>G), V2064L (6296G>C 6298A>G), Q2074R (6327A>G), I2132F (6500A>T), L2147F (6545C>T) | | | | | | | | |

| | Begin | End | Coverage | Score | Concordance | Matches | Identities | I/D/M/F* | Stop Codons |
|---|---|---|---|---|---|---|---|---|---|
| **NT** | **1** | **7781** | **99.7%** | **11522** | **79.6%** | **7469 (95.9%)** | **6754 (86.7%)** | **10/312** | |

Codon mutations:

TTT3TTC (115T>C), TTC7TTT (127C>T), CCC8TCT (128C>T 130C>T), TTC10TCC (135T>C), ACT14ACC (148T>C), GAA15GAG (151A>G), GCT18TCT (158G>T), ACC19GCC (161A>G), GGT23GGC (175T>C), TCT24TCC (178T>C), TTT26TTC (184T>C), GAG28GAA (190G>A), GCT31ACA (197G>A 199T>A), AGT32AGC (202T>C), ATT33ATA (205T>A), AGT34AGC (208T>C), CGC38CGT (220C>T), CTG40TTG (224C>T), CAA49CAG (253A>G), TCT51TCC (259T>C), TCT52TCA (262T>A), CAC53CAT (265C>T), TTG78CTA (338T>C 340G>A), TTA85TTG (361A>G), GAA87GAG (367A>G), CCC88CCT (370C>T), ATC90ATT (376C>T), TAC93TAT (385C>T), TCT94TCC (388T>C), AAA101AAG (409A>G), AAA115AAG (451A>G), GAC127GAT (487C>T), AAA128AAG (490A>G), GTG132GTA (502G>A), TCT145TCC (541T>C), TTC151TTT (559C>T), GTA157GTG (577A>G), CTA158TTA (578C>T), GAC183GAT (655C>T), CAA190CAG (676A>G), GCA196GCG (694A>G), TCA207TCG (727A>G), CTC218CTT (760C>T), ACC225ACT (781C>T), TTC228TTT (790C>T), GTA231GTG (799A>G), GGT233GGC (805T>C), CTA238TTG (818C>T 820A>G), CTA242CTT (832A>T), TCA243TCC (835A>C), ATT245ATC (841T>C), CTG246TTG (842C>T), TAC247TAT (847C>T), TTG259TTA (883G>A), TTT260TTC (886T>C), CTG262CTA (892G>A), AAT265AAC (901T>C), TGT269TGC (913T>C), GCC273GCT (925C>T), CTG278TTG (938C>T), GTA282GTG (952A>G), GAC283GAT (955C>T), GGT286GGG (964T>G), AAG288AAA (970G>A), AAT289AAC (973T>C), GTT291ATT (977G>A), TTG292CTG (980T>C), GCC297GCT (997C>T), TCT300TCC (1006T>C), GAC302GAT (1012C>T), TTA303CTA (1013T>C), TTC304TTT (1018C>T), GAG309GAA (1033G>A), CAT318CAC (1060T>C), CGC319CGT (1063C>T), GCA324GCT (1078A>T), AAC325AAT (1081C>T), TTC328TTT (1090C>T), ACT329ACC (1093T>C), CAA333CAG (1105A>G), CAT334CAC (1108T>C), CTA335TTA (1109C>T), GCC353GCT (1165C>T), AGG358AGA (1180G>A), ACC364GCC (1196A>G), GAT367GAC (1207T>C), CAG375CAA (1231G>A), GGC378GGT (1240C>T), AGG383AGA (1255G>A), TTT387TTC (1267T>C), GGG398GGA (1300G>A), AAA399AAG (1303A>G), CTG405TTG (1319C>T), TTC406TTT (1324C>T), GAA415GAG (1351A>G), GAG416GAA (1354G>A), TAT435TAC (1411T>C), CAT437CAC (1417T>C), ATC441ATT (1429C>T), GAC445GAT (1441C>T), GCT448GCC (1450T>C), CTG449TTG (1451C>T), GAC458GAT (1480C>T), ATT459GTT (1481A>G), GAT461GAC (1489T>C), CCC472CCT (1522C>T), GGC482GGT (1552C>T), CGC483CGT (1555C>T), CGA488CGG (1570A>G), TAT489TAC (1573T>C), GCC492GCT (1582C>T), ACA493ACC (1585A>C), AAC495AAT (1591C>T), CTT496CTC (1594T>C), GCT501TCT (1607G>T), CAG502CAA (1612G>A), GGT504GGG (1618T>G), CTA505TTA (1619C>T), ACC506ACT (1624C>T), CCC508CCT (1630C>T), TTT511TTC (1639T>C), GAG512GAA (1642G>A), AGA513AGG (1645A>G), TGC521TGT (1669C>T), AGG522AGA (1672G>A), GGT525GGA (1681T>A), GAG526GAT (1684G>T), GCT531GCC (1699T>C), ACT533ACC (1705T>C), AAA544AGG (1737A>G 1738A>G), GCA548GCT (1750A>T), ACA550ACG (1756A>G), GGT553GGG (1765T>G), TTG559CTA (1781T>C 1783G>A), CTG560TTG (1784C>T), GAA561GAG (1789A>G), AAG568AAA (1810G>A), ATC573ATT (1825C>T), AAA576AAG (1834A>G), CAG587CAA (1867G>A), TCA591TCG (1879A>G), GAA594GAG (1888A>G), TTG602CTA (1910T>C 1912G>A), ATC604GTC (1916A>G), GTA618GTG (1960A>G), AGT619AAT (1962G>A), GAA620GAG (1966A>G), TAC626TAT (1984C>T), CAG628CAA (1990G>A), GAG629GAA (1993G>A), TTG635CTG (2009T>C), TAT639TAC (2023T>C), CCT650CCC (2056T>C), GAC652GAT (2062C>T), GCC657GCA (2077C>A), TTT659TAT (2082T>A), CTA662CTG (2092A>G), AAA667AAG (2107A>G), GTT674GTC (2128T>C), CTG678TTG (2138C>T), TAT679TAC (2143T>C), GGT681GGC (2149T>C), TGT683TGC (2155T>C), CTG690TTG (2174C>T), ATA691ATC (2179A>C), AAG692AGA (2181A>G 2182G>A), TAT698TAC (2200T>C), ATC706ATT (2224C>T), ACA710ACG (2236A>G), AAT712AGT (2241A>G), AGC715AAT (2250G>A 2251C>T), GAC717GAT (2257C>T), CAG719CAA (2263G>A), CCT723CCC (2275T>C), GCG727GCA (2287G>A), CAC728CAT (2290C>T), GAC736GAT (2314C>T), ATA739ATT (2323A>T), CAC747CAT (2347C>T), TTC750TTT (2356C>T), TTG751TTA (2359G>A), AGG757AGA (2377G>A), AGA762AGG (2392A>G), AGG768AAG (2409G>A), ATT779ATC (2443T>C), GTG780GTA (2446G>A), CCG791CCA (2479G>A), ATC793ATT (2485C>T), CTG797CTA (2497G>A), ATT798ATC (2500T>C), AAG799AAA (2503G>A), GCT800GCC (2506T>C), GGC801GGT (2509C>T), CTT802CTC (2512T>C), GGG812GGA (2542G>A), GTC814GTT (2548C>T), TTC817TTT (2557C>T), GAG838GAA (2620G>A), ATC852ATT (2662C>T), TTT853TTC (2665T>C), ATC876ATT (2734C>T), GGA879GGG (2743A>G), AAC883AAT (2755C>T), CTA889CTG (2773A>G), TGC901TGT (2809C>T), TCG902TCA (2812G>A), ACA907ACG (2827A>G), GGC918GGT (2860C>T), CAT925CAC (2881T>C), GCC936GCT (2914C>T), CGG940CGA (2926G>A), TTC941TTT (2929C>T), GTC948GTT (2950C>T), CTG949TTG (2951C>T), CCC958CCT (2980C>T), AAG959AAA (2983G>A), AAA964AAG (2998A>G), CCT967CCC (3007T>C), TTT974TTC (3028T>C), TCT975TCC (3031T>C), AAG976AAA (3034G>A), ACT978ACC (3040T>C), AGA984AGG (3058A>G), GTA986GTG (3064A>G), AAG989AAA (3073G>A), GCT991GCC (3079T>C), TTT1001TTC (3109T>C), GTG1003GTA (3115G>A), GTA1013GTG (3145A>G), TAC1015TAT (3151C>T), TCT1019TCC (3163T>C), TCC1021TCT (3169C>T), TTT1022TTC (3172T>C), TAT1032TAC (3202T>C), CTT1045CTC (3241T>C), AAC1049AAT (3253C>T), GTG1050GTA (3256G>A), GAC1052GAT (3262C>T), GGT1059GGC (3283T>C), CAT1061CAC (3289T>C), TCT1064TCC (3298T>C), GAA1067GAG (3307A>G), CGA1084CGT (3358A>T), GTA1100GTC (3406A>C), CTC1104CTA (3418C>A), CTT1105CTC (3421T>C), AAA1119AAG (3463A>G), CAG1120CAA (3466G>A), GGC1125GGT (3481C>T), ACA1126ACT (3484A>T), TTG1127TTA (3487G>A), GCT1129GTT (3492C>T), ACG1134ACA (3508G>A), GGG1151GGA (3559G>A), GCT1156GCC (3574T>C), CCA1157CCG (3577A>G), ATT1160ATC (3586T>C), GCT1162GTT (3591C>T), TAC1173TAT (3625C>T), GTG1174GTA (3628G>A), TAC1180TAT (3646C>T), CAA1185CAG (3661A>G), AAT1186AAC (3664T>C), GAG1189GAA (3673G>A), GAC1190GAT (3676C>T), CTA1194TTA (3686C>T), CGT1200CGC (3706T>C), TTG1204CTG (3716T>C), CCT1205CCA (3721T>A), CGA1211CGT (3739A>T), AAG1216AAA (3754G>A), CTC1226CTT (3784C>T), GAC1236GAT (3814C>T), GGC1254GGT (3868C>T), ACT1255ACC (3871T>C), GAG1259GAA (3883G>A), GAG1262GAA (3892G>A), AGT1264AGC (3898T>C), GGA1267GGT (3907A>T), GTC1270GTT (3916C>T), AAA1273AAG (3925A>G), AGG1274AGA (3928G>A), GAC1281GAT (3949C>T), GAG1283GAA (3955G>A), TTG1285TTA (3961G>A), GAA1288GAG (3970A>G), ATT1289ATC (3973T>C), AAA1304AAG (4018A>G), TTG1310CTA (4034T>C 4036G>A), CGG1317CGA (4057G>A), CGT1320CGC (4066T>C), CCC1326CCT (4084C>T), CTG1338TTG (4118C>T), TCC1341TCT (4129C>T), CTG1344CTA (4138G>A), TCC1347TCA (4147C>A), CTG1351CTA (4159G>A), CCA1352CCT (4162A>T), AAA1354AAG (4168A>G), GGA1360GGT (4186A>T), GAA1363GAG (4195A>G), TTG1367TTA (4207G>A), GCT1368GCC (4210T>C), TTC1371TTT (4219C>T), GTC1374GTT (4228C>T), AGA1389AGG (4273A>G), ATC1392ATA (4282C>A), GCC1394GCT (4288C>T), TTT1395TTC (4291T>C), TTT1398TTC (4300T>C), GAC1404GAT (4318C>T), TAC1405TAT (4321C>T), TAT1406TAC (4324T>C), GGT1411GGC (4339T>C), GAC1414GAT (4348C>T), AAT1416AAC (4354T>C), AGA1418AGG (4360A>G), TAT1429TAC (4393T>C), TTT1437TTC (4417T>C), TTG1439CTG (4421T>C), GTT1451GTC (4459T>C), ATC1452ATT (4462C>T), TTC1453TTT (4465C>T), AAC1454AAT (4468C>T), TCT1455TCC (4471T>C), CTT1456CTC (4474T>C), TTT1460TTC (4486T>C), TAT1461TAC (4489T>C), TAT1495TAC (4591T>C), TTG1507CTG (4625T>C), TAT1510TAC (4636T>C), CTG1512TTG (4640C>T), ATT1515ATA (4651T>A), GCC1516GCA (4654C>A), CAA1517CAG (4657A>G), TTG1519CTA (4661T>C 4663G>A), GAG1520GAA (4666G>A), GTG1522GTA (4672G>A), GTT1524GTC (4678T>C), ATT1525ATA (4681T>A), ATA1526ATC (4684A>C), AAG1527AAA (4687G>A), ATC1530ATA (4696C>A), GAC1531GAT (4699C>T), AGT1538AGA (4720T>A), AGC1551AGT (4759C>T), AGG1556AGA (4774G>A), CTA1559CTG (4783A>G), GGA1560GGG (4786A>G), TAC1561TAT (4789C>T), GTA1569GTG (4813A>G), TCT1570TCC (4816T>C), GTT1598ATT (4898G>A), GAG1600GAA (4906G>A), CCT1606CCC (4924T>C), ACA1614GCA (4946A>G), AAC1618AAT (4960C>T), AAG1621AAA (4969G>A), CGG1623CGA (4975G>A), ACA1629ACC (4993A>C), CCT1631CCC (4999T>C), TAT1633TAC (5005T>C), TTG1636CTG (5012T>C), GAG1637GAA (5017G>A), GAC1639GAT (5023C>T), GGA1646GGC (5044A>C), GGG1651GGA (5059G>A), ATC1653ATT (5065C>T), CTT1656CTC (5074T>C), CAT1657CAC (5077T>C), ATT1658ACT (5079T>C), TTG1666CTG (5102T>C), GGC1669GGT (5113C>T), ACT1671ACC (5119T>C), GTT1672ATT (5120G>A), CAT1680CAC (5146T>C), ACT1681AAT (5148C>A), AGT1687AGC (5167T>C), TTC1688TTT (5170C>T), TCC1690TCT (5176C>T), CAG1698CAA (5200G>A), GTT1704GTG (5218T>G), AGA1713AAA (5244G>A), CCC1720CCT (5266C>T), GAC1725GAT (5281C>T), GGG1728GGA (5290G>A), CAC1741CAT (5329C>T), GTA1746GTT (5344A>T), ACC1747ACA (5347C>A), GTG1750ATT (5354G>A 5356G>T), CCA1753CCC (5365A>C), GTG1755ATT (5369G>A 5371G>T), GGC1760GGG (5386C>G), GGG1762GGA (5392G>A), CCA1767CCC (5407A>C), TGC1769TGT (5413C>T), GCA1771GCT (5419A>T), CTG1775TTG (5429C>T), ACT1777ACC (5437T>C), GGC1778GGG (5440C>G), TAT1780TAC (5446T>C), GTT1785GTC (5461T>C), GTG1787GTT (5467G>T), TTG1789CTG (5471T>C), GTT1795GTC (5491T>C), ACA1796CCA (5492A>C), GAT1797GAC (5497T>C), CTT1809CTG (5533T>G), AAA1811CAA (5537A>C), ACA1823ACG (5575A>G), TGT1824TGC (5578T>C), AAA1825AAG (5581A>G), GCC1829TCC (5591G>T), GGT1831GGC (5599T>C), ACA1833CCA (5603A>C), GCA1835GCT (5611A>T), ATA1837ATT (5617A>T), GGT1845GGC (5641T>C), TTT1847TTC (5647T>C), CTA1856CTG (5674A>G), CAG1857CAT (5677G>T), CAG1860CAA (5686G>A), CCT1868CCA (5710T>A), GGG1870GGT (5716G>T), GGA1871GGC (5719A>C), CAC1875CAT (5731C>T), ACT1876ACC (5734T>C), ACT1877ACC (5737T>C), AGA1878AAA (5739G>A), CTT1880CTC (5746T>C), TTG1882CTA (5750T>C 5752G>A), CTC1884CTT (5758C>T), AAG1889AAA (5773G>A), CAA1892CAG (5782A>G), ATT1893ACT (5784T>C), TTA1894CTA (5786T>C), AGA1897AGG (5797A>G), TTC1900TTT (5806C>T), TCA1910TCT (5836A>T), TTG1915CAA (5849T>C 5850T>A 5851G>A), AGG1921AGA (5869G>A), CAA1923CAG (5875A>G), GGG1924GGC (5878G>C), TTT1926TTC (5884T>C), CCC1927CCA (5887C>A), CGC1928CGT (5890C>T), GTG1932GTT (5902G>T), TAC1934TAT (5908C>T), TTG1935CTG (5909T>C), CTT1936CTC (5914T>C), TTT1939TTC (5923T>C), CTG1941TTG (5927C>T), CAC1946CAT (5944C>T), AAC1950AGC (5955A>G), ACG1953ACA (5965G>A), AAA1964AAG (5998A>G), ACC1966ACT (6004C>T), AAA1970AAG (6016A>G), GAA1971GAG (6019A>G), TCC1974TCT (6028C>T), GCA1975GCT (6031A>T), ACA1976ATA (6033C>T), GCT1977TCT (6035G>T), GGT1983GGA (6055T>A), TCC1987TCA (6067C>A), GCT1988GCC (6070T>C), CTA1995CTG (6091A>G), CTT1996TTT (6092C>T), CTT1998CTG (6100T>G), CAG1999CAA (6103G>A), AAG2000AAA (6106G>A), ATA2002ATG (6112A>G), TGT2004TGC (6118T>C), TCA2005TCT (6121A>T), TTT2011TTC (6139T>C), GTC2016GTG (6154C>G), GGC2019GGG (6163C>G), CAC2020CAT (6166C>T), GTA2026GTG (6184A>G), ACT2027ACG (6187T>G), TTC2029TTT (6193C>T), CAA2030CAG (6196A>G), ATG2032ATC (6202G>C), CAA2035CAG (6211A>G), CTG2037CTT (6217G>T), GAA2039GAT (6223A>T), ATA2042GTG (6230A>G 6232A>G), GAG2043GAA (6235G>A), TTA2049TTC (6253A>C), ACC2050ACA (6256C>A), ACA2052ACT (6262A>T), CTG2053CTA (6265G>A), TCG2054TCT (6268G>T), AGC2056AGG (6274C>G), GTA2064CTG (6296G>C 6298A>G), TTG2068CTG (6308T>C), GAT2069GAC (6313T>C), GAG2072GAA (6322G>A), CAG2074CGG (6327A>G), AAT2084AAC (6358T>C), GCA2126GCC (6484A>C), AGA2128AGG (6490A>G), CAT2129CAC (6493T>C), ATT2132TTT (6500A>T), TCC2137TCT (6517C>T), CAC2142CAT (6532C>T), CTT2147TTT (6545C>T), TAC2155TAT (6571C>T)

#### Proteins

| | Begin | End | Coverage | Score | Concordance | Matches | Identities | I/D/M/F* | Stop Codons |
|---|---|---|---|---|---|---|---|---|---|
| polyprotein (YP_001039627.1) | 1 | 2159 | 100% | 14846 | 98.5% | 2159 (100%) | 2108 (97.6%) | 0/0/0/0 | 1 |

Protein mutations:

P8S (128C>T 130C>T), F10S (135T>C), A18S (158G>T), T19A (161A>G), A31T (197G>A 199T>A), V291I (977G>A), T364A (1196A>G), I459V (1481A>G), A501S (1607G>T), E526D (1684G>T), K544R (1737A>G 1738A>G), I604V (1916A>G), S619N (1962G>A), F659Y (2082T>A), K692R (2181A>G 2182G>A), N712S (2241A>G), S715N (2250G>A 2251C>T), R768K (2409G>A), A1129V (3492C>T), A1162V (3591C>T), S1538R (4720T>A), V1598I (4898G>A), T1614A (4946A>G), I1658T (5079T>C), V1672I (5120G>A), T1681N (5148C>A), R1713K (5244G>A), V1750I (5354G>A 5356G>T), V1755I (5369G>A 5371G>T), T1796P (5492A>C), K1811Q (5537A>C), A1829S (5591G>T), T1833P (5603A>C), Q1857H (5677G>T), R1878K (5739G>A), I1893T (5784T>C), L1915Q (5849T>C 5850T>A 5851G>A), N1950S (5955A>G), T1976I (6033C>T), A1977S (6035G>T), L1996F (6092C>T), I2002M (6112A>G), M2032I (6202G>C), E2039D (6223A>T), I2042V (6230A>G 6232A>G), L2049F (6253A>C), S2056R (6274C>G), V2064L (6296G>C 6298A>G), Q2074R (6327A>G), I2132F (6500A>T), L2147F (6545C>T)

| | Begin | End | Coverage | Score | Concordance | Matches | Identities | I/D/M/F* | Stop Codons |
|---|---|---|---|---|---|---|---|---|---|
| **NT** | **1** | **7781** | **99.7%** | **11522** | **79.6%** | **7469 (95.9%)** | **6754 (86.7%)** | **10/312** | |
| Codon mutations: | TTT3TTC (115T>C), TTC7TTT (127C>T), CCC8TCT (128C>T 130C>T), TTC10TCC (135T>C), ACT14ACC (148T>C), GAA15GAG (151A>G), GCT18TCT (158G>T), ACC19GCC (161A>G), GGT23GGC (175T>C), TCT24TCC (178T>C), TTT26TTC (184T>C), GAG28GAA (190G>A), GCT31ACA (197G>A 199T>A), AGT32AGC (202T>C), ATT33ATA (205T>A), AGT34AGC (208T>C), CGC38CGT (220C>T), CTG40TTG (224C>T), CAA49CAG (253A>G), TCT51TCC (259T>C), TCT52TCA (262T>A), CAC53CAT (265C>T), TTG78CTA (338T>C 340G>A), TTA85TTG (361A>G), GAA87GAG (367A>G), CCC88CCT (370C>T), ATC90ATT (376C>T), TAC93TAT (385C>T), TCT94TCC (388T>C), AAA101AAG (409A>G), AAA115AAG (451A>G), GAC127GAT (487C>T), AAA128AAG (490A>G), GTG132GTA (502G>A), TCT145TCC (541T>C), TTC151TTT (559C>T), GTA157GTG (577A>G), CTA158TTA (578C>T), GAC183GAT (655C>T), CAA190CAG (676A>G), GCA196GCG (694A>G), TCA207TCG (727A>G), CTC218CTT (760C>T), ACC225ACT (781C>T), TTC228TTT (790C>T), GTA231GTG (799A>G), GGT233GGC (805T>C), CTA238TTG (818C>T 820A>G), CTA242CTT (832A>T), TCA243TCC (835A>C), ATT245ATC (841T>C), CTG246TTG (842C>T), TAC247TAT (847C>T), TTG259TTA (883G>A), TTT260TTC (886T>C), CTG262CTA (892G>A), AAT265AAC (901T>C), TGT269TGC (913T>C), GCC273GCT (925C>T), CTG278TTG (938C>T), GTA282GTG (952A>G), GAC283GAT (955C>T), GGT286GGG (964T>G), AAG288AAA (970G>A), AAT289AAC (973T>C), GTT291ATT (977G>A), TTG292CTG (980T>C), GCC297GCT (997C>T), TCT300TCC (1006T>C), GAC302GAT (1012C>T), TTA303CTA (1013T>C), TTC304TTT (1018C>T), GAG309GAA (1033G>A), CAT318CAC (1060T>C), CGC319CGT (1063C>T), GCA324GCT (1078A>T), AAC325AAT (1081C>T), TTC328TTT (1090C>T), ACT329ACC (1093T>C), CAA333CAG (1105A>G), CAT334CAC (1108T>C), CTA335TTA (1109C>T), GCC353GCT (1165C>T), AGG358AGA (1180G>A), ACC364GCC (1196A>G), GAT367GAC (1207T>C), CAG375CAA (1231G>A), GGC378GGT (1240C>T), AGG383AGA (1255G>A), TTT387TTC (1267T>C), GGG398GGA (1300G>A), AAA399AAG (1303A>G), CTG405TTG (1319C>T), TTC406TTT (1324C>T), GAA415GAG (1351A>G), GAG416GAA (1354G>A), TAT435TAC (1411T>C), CAT437CAC (1417T>C), ATC441ATT (1429C>T), GAC445GAT (1441C>T), GCT448GCC (1450T>C), CTG449TTG (1451C>T), GAC458GAT (1480C>T), ATT459GTT (1481A>G), GAT461GAC (1489T>C), CCC472CCT (1522C>T), GGC482GGT (1552C>T), CGC483CGT (1555C>T), CGA488CGG (1570A>G), TAT489TAC (1573T>C), GCC492GCT (1582C>T), ACA493ACC (1585A>C), AAC495AAT (1591C>T), CTT496CTC (1594T>C), GCT501TCT (1607G>T), CAG502CAA (1612G>A), GGT504GGG (1618T>G), CTA505TTA (1619C>T), ACC506ACT (1624C>T), CCC508CCT (1630C>T), TTT511TTC (1639T>C), GAG512GAA (1642G>A), AGA513AGG (1645A>G), TGC521TGT (1669C>T), AGG522AGA (1672G>A), GGT525GGA (1681T>A), GAG526GAT (1684G>T), GCT531GCC (1699T>C), ACT533ACC (1705T>C), AAA544AGG (1737A>G 1738A>G), GCA548GCT (1750A>T), ACA550ACG (1756A>G), GGT553GGG (1765T>G), TTG559CTA (1781T>C 1783G>A), CTG560TTG (1784C>T), GAA561GAG (1789A>G), AAG568AAA (1810G>A), ATC573ATT (1825C>T), AAA576AAG (1834A>G), CAG587CAA (1867G>A), TCA591TCG (1879A>G), GAA594GAG (1888A>G), TTG602CTA (1910T>C 1912G>A), ATC604GTC (1916A>G), GTA618GTG (1960A>G), AGT619AAT (1962G>A), GAA620GAG (1966A>G), TAC626TAT (1984C>T), CAG628CAA (1990G>A), GAG629GAA (1993G>A), TTG635CTG (2009T>C), TAT639TAC (2023T>C), CCT650CCC (2056T>C), GAC652GAT (2062C>T), GCC657GCA (2077C>A), TTT659TAT (2082T>A), CTA662CTG (2092A>G), AAA667AAG (2107A>G), GTT674GTC (2128T>C), CTG678TTG (2138C>T), TAT679TAC (2143T>C), GGT681GGC (2149T>C), TGT683TGC (2155T>C), CTG690TTG (2174C>T), ATA691ATC (2179A>C), AAG692AGA (2181A>G 2182G>A), TAT698TAC (2200T>C), ATC706ATT (2224C>T), ACA710ACG (2236A>G), AAT712AGT (2241A>G), AGC715AAT (2250G>A 2251C>T), GAC717GAT (2257C>T), CAG719CAA (2263G>A), CCT723CCC (2275T>C), GCG727GCA (2287G>A), CAC728CAT (2290C>T), GAC736GAT (2314C>T), ATA739ATT (2323A>T), CAC747CAT (2347C>T), TTC750TTT (2356C>T), TTG751TTA (2359G>A), AGG757AGA (2377G>A), AGA762AGG (2392A>G), AGG768AAG (2409G>A), ATT779ATC (2443T>C), GTG780GTA (2446G>A), CCG791CCA (2479G>A), ATC793ATT (2485C>T), CTG797CTA (2497G>A), ATT798ATC (2500T>C), AAG799AAA (2503G>A), GCT800GCC (2506T>C), GGC801GGT (2509C>T), CTT802CTC (2512T>C), GGG812GGA (2542G>A), GTC814GTT (2548C>T), TTC817TTT (2557C>T), GAG838GAA (2620G>A), ATC852ATT (2662C>T), TTT853TTC (2665T>C), ATC876ATT (2734C>T), GGA879GGG (2743A>G), AAC883AAT (2755C>T), CTA889CTG (2773A>G), TGC901TGT (2809C>T), TCG902TCA (2812G>A), ACA907ACG (2827A>G), GGC918GGT (2860C>T), CAT925CAC (2881T>C), GCC936GCT (2914C>T), CGG940CGA (2926G>A), TTC941TTT (2929C>T), GTC948GTT (2950C>T), CTG949TTG (2951C>T), CCC958CCT (2980C>T), AAG959AAA (2983G>A), AAA964AAG (2998A>G), CCT967CCC (3007T>C), TTT974TTC (3028T>C), TCT975TCC (3031T>C), AAG976AAA (3034G>A), ACT978ACC (3040T>C), AGA984AGG (3058A>G), GTA986GTG (3064A>G), AAG989AAA (3073G>A), GCT991GCC (3079T>C), TTT1001TTC (3109T>C), GTG1003GTA (3115G>A), GTA1013GTG (3145A>G), TAC1015TAT (3151C>T), TCT1019TCC (3163T>C), TCC1021TCT (3169C>T), TTT1022TTC (3172T>C), TAT1032TAC (3202T>C), CTT1045CTC (3241T>C), AAC1049AAT (3253C>T), GTG1050GTA (3256G>A), GAC1052GAT (3262C>T), GGT1059GGC (3283T>C), CAT1061CAC (3289T>C), TCT1064TCC (3298T>C), GAA1067GAG (3307A>G), CGA1084CGT (3358A>T), GTA1100GTC (3406A>C), CTC1104CTA (3418C>A), CTT1105CTC (3421T>C), AAA1119AAG (3463A>G), CAG1120CAA (3466G>A), GGC1125GGT (3481C>T), ACA1126ACT (3484A>T), TTG1127TTA (3487G>A), GCT1129GTT (3492C>T), ACG1134ACA (3508G>A), GGG1151GGA (3559G>A), GCT1156GCC (3574T>C), CCA1157CCG (3577A>G), ATT1160ATC (3586T>C), GCT1162GTT (3591C>T), TAC1173TAT (3625C>T), GTG1174GTA (3628G>A), TAC1180TAT (3646C>T), CAA1185CAG (3661A>G), AAT1186AAC (3664T>C), GAG1189GAA (3673G>A), GAC1190GAT (3676C>T), CTA1194TTA (3686C>T), CGT1200CGC (3706T>C), TTG1204CTG (3716T>C), CCT1205CCA (3721T>A), CGA1211CGT (3739A>T), AAG1216AAA (3754G>A), CTC1226CTT (3784C>T), GAC1236GAT (3814C>T), GGC1254GGT (3868C>T), ACT1255ACC (3871T>C), GAG1259GAA (3883G>A), GAG1262GAA (3892G>A), AGT1264AGC (3898T>C), GGA1267GGT (3907A>T), GTC1270GTT (3916C>T), AAA1273AAG (3925A>G), AGG1274AGA (3928G>A), GAC1281GAT (3949C>T), GAG1283GAA (3955G>A), TTG1285TTA (3961G>A), GAA1288GAG (3970A>G), ATT1289ATC (3973T>C), AAA1304AAG (4018A>G), TTG1310CTA (4034T>C 4036G>A), CGG1317CGA (4057G>A), CGT1320CGC (4066T>C), CCC1326CCT (4084C>T), CTG1338TTG (4118C>T), TCC1341TCT (4129C>T), CTG1344CTA (4138G>A), TCC1347TCA (4147C>A), CTG1351CTA (4159G>A), CCA1352CCT (4162A>T), AAA1354AAG (4168A>G), GGA1360GGT (4186A>T), GAA1363GAG (4195A>G), TTG1367TTA (4207G>A), GCT1368GCC (4210T>C), TTC1371TTT (4219C>T), GTC1374GTT (4228C>T), AGA1389AGG (4273A>G), ATC1392ATA (4282C>A), GCC1394GCT (4288C>T), TTT1395TTC (4291T>C), TTT1398TTC (4300T>C), GAC1404GAT (4318C>T), TAC1405TAT (4321C>T), TAT1406TAC (4324T>C), GGT1411GGC (4339T>C), GAC1414GAT (4348C>T), AAT1416AAC (4354T>C), AGA1418AGG (4360A>G), TAT1429TAC (4393T>C), TTT1437TTC (4417T>C), TTG1439CTG (4421T>C), GTT1451GTC (4459T>C), ATC1452ATT (4462C>T), TTC1453TTT (4465C>T), AAC1454AAT (4468C>T), TCT1455TCC (4471T>C), CTT1456CTC (4474T>C), TTT1460TTC (4486T>C), TAT1461TAC (4489T>C), TAT1495TAC (4591T>C), TTG1507CTG (4625T>C), TAT1510TAC (4636T>C), CTG1512TTG (4640C>T), ATT1515ATA (4651T>A), GCC1516GCA (4654C>A), CAA1517CAG (4657A>G), TTG1519CTA (4661T>C 4663G>A), GAG1520GAA (4666G>A), GTG1522GTA (4672G>A), GTT1524GTC (4678T>C), ATT1525ATA (4681T>A), ATA1526ATC (4684A>C), AAG1527AAA (4687G>A), ATC1530ATA (4696C>A), GAC1531GAT (4699C>T), AGT1538AGA (4720T>A), AGC1551AGT (4759C>T), AGG1556AGA (4774G>A), CTA1559CTG (4783A>G), GGA1560GGG (4786A>G), TAC1561TAT (4789C>T), GTA1569GTG (4813A>G), TCT1570TCC (4816T>C), GTT1598ATT (4898G>A), GAG1600GAA (4906G>A), CCT1606CCC (4924T>C), ACA1614GCA (4946A>G), AAC1618AAT (4960C>T), AAG1621AAA (4969G>A), CGG1623CGA (4975G>A), ACA1629ACC (4993A>C), CCT1631CCC (4999T>C), TAT1633TAC (5005T>C), TTG1636CTG (5012T>C), GAG1637GAA (5017G>A), GAC1639GAT (5023C>T), GGA1646GGC (5044A>C), GGG1651GGA (5059G>A), ATC1653ATT (5065C>T), CTT1656CTC (5074T>C), CAT1657CAC (5077T>C), ATT1658ACT (5079T>C), TTG1666CTG (5102T>C), GGC1669GGT (5113C>T), ACT1671ACC (5119T>C), GTT1672ATT (5120G>A), CAT1680CAC (5146T>C), ACT1681AAT (5148C>A), AGT1687AGC (5167T>C), TTC1688TTT (5170C>T), TCC1690TCT (5176C>T), CAG1698CAA (5200G>A), GTT1704GTG (5218T>G), AGA1713AAA (5244G>A), CCC1720CCT (5266C>T), GAC1725GAT (5281C>T), GGG1728GGA (5290G>A), CAC1741CAT (5329C>T), GTA1746GTT (5344A>T), ACC1747ACA (5347C>A), GTG1750ATT (5354G>A 5356G>T), CCA1753CCC (5365A>C), GTG1755ATT (5369G>A 5371G>T), GGC1760GGG (5386C>G), GGG1762GGA (5392G>A), CCA1767CCC (5407A>C), TGC1769TGT (5413C>T), GCA1771GCT (5419A>T), CTG1775TTG (5429C>T), ACT1777ACC (5437T>C), GGC1778GGG (5440C>G), TAT1780TAC (5446T>C), GTT1785GTC (5461T>C), GTG1787GTT (5467G>T), TTG1789CTG (5471T>C), GTT1795GTC (5491T>C), ACA1796CCA (5492A>C), GAT1797GAC (5497T>C), CTT1809CTG (5533T>G), AAA1811CAA (5537A>C), ACA1823ACG (5575A>G), TGT1824TGC (5578T>C), AAA1825AAG (5581A>G), GCC1829TCC (5591G>T), GGT1831GGC (5599T>C), ACA1833CCA (5603A>C), GCA1835GCT (5611A>T), ATA1837ATT (5617A>T), GGT1845GGC (5641T>C), TTT1847TTC (5647T>C), CTA1856CTG (5674A>G), CAG1857CAT (5677G>T), CAG1860CAA (5686G>A), CCT1868CCA (5710T>A), GGG1870GGT (5716G>T), GGA1871GGC (5719A>C), CAC1875CAT (5731C>T), ACT1876ACC (5734T>C), ACT1877ACC (5737T>C), AGA1878AAA (5739G>A), CTT1880CTC (5746T>C), TTG1882CTA (5750T>C 5752G>A), CTC1884CTT (5758C>T), AAG1889AAA (5773G>A), CAA1892CAG (5782A>G), ATT1893ACT (5784T>C), TTA1894CTA (5786T>C), AGA1897AGG (5797A>G), TTC1900TTT (5806C>T), TCA1910TCT (5836A>T), TTG1915CAA (5849T>C 5850T>A 5851G>A), AGG1921AGA (5869G>A), CAA1923CAG (5875A>G), GGG1924GGC (5878G>C), TTT1926TTC (5884T>C), CCC1927CCA (5887C>A), CGC1928CGT (5890C>T), GTG1932GTT (5902G>T), TAC1934TAT (5908C>T), TTG1935CTG (5909T>C), CTT1936CTC (5914T>C), TTT1939TTC (5923T>C), CTG1941TTG (5927C>T), CAC1946CAT (5944C>T), AAC1950AGC (5955A>G), ACG1953ACA (5965G>A), AAA1964AAG (5998A>G), ACC1966ACT (6004C>T), AAA1970AAG (6016A>G), GAA1971GAG (6019A>G), TCC1974TCT (6028C>T), GCA1975GCT (6031A>T), ACA1976ATA (6033C>T), GCT1977TCT (6035G>T), GGT1983GGA (6055T>A), TCC1987TCA (6067C>A), GCT1988GCC (6070T>C), CTA1995CTG (6091A>G), CTT1996TTT (6092C>T), CTT1998CTG (6100T>G), CAG1999CAA (6103G>A), AAG2000AAA (6106G>A), ATA2002ATG (6112A>G), TGT2004TGC (6118T>C), TCA2005TCT (6121A>T), TTT2011TTC (6139T>C), GTC2016GTG (6154C>G), GGC2019GGG (6163C>G), CAC2020CAT (6166C>T), GTA2026GTG (6184A>G), ACT2027ACG (6187T>G), TTC2029TTT (6193C>T), CAA2030CAG (6196A>G), ATG2032ATC (6202G>C), CAA2035CAG (6211A>G), CTG2037CTT (6217G>T), GAA2039GAT (6223A>T), ATA2042GTG (6230A>G 6232A>G), GAG2043GAA (6235G>A), TTA2049TTC (6253A>C), ACC2050ACA (6256C>A), ACA2052ACT (6262A>T), CTG2053CTA (6265G>A), TCG2054TCT (6268G>T), AGC2056AGG (6274C>G), GTA2064CTG (6296G>C 6298A>G), TTG2068CTG (6308T>C), GAT2069GAC (6313T>C), GAG2072GAA (6322G>A), CAG2074CGG (6327A>G), AAT2084AAC (6358T>C), GCA2126GCC (6484A>C), AGA2128AGG (6490A>G), CAT2129CAC (6493T>C), ATT2132TTT (6500A>T), TCC2137TCT (6517C>T), CAC2142CAT (6532C>T), CTT2147TTT (6545C>T), TAC2155TAT (6571C>T) | | | | | | | | |

*: Inserts / Deletes / Misaligned / Frameshifts

#### Analysis details

This analysis was performed with panviral2.64

### NGS Details (UN70): Torradovirus lycopersici (segment RNA 2)

#### Assembly

| | |
|---|---|
| **Coverage Length** | 5384 (1 contig(s)) |
| **Depth Of Coverage** | 40729.6 |
| **Number Of Reads** | 1691517 |
| **Reads Per Million** | 32064.85 rpm (after QC) |
| **Ambiguities** | 0 |
| **Assembly Method** | de novo + reference guided assembly |
| **Consensus Caller** | Bcf Tools |

#### Coverage Map

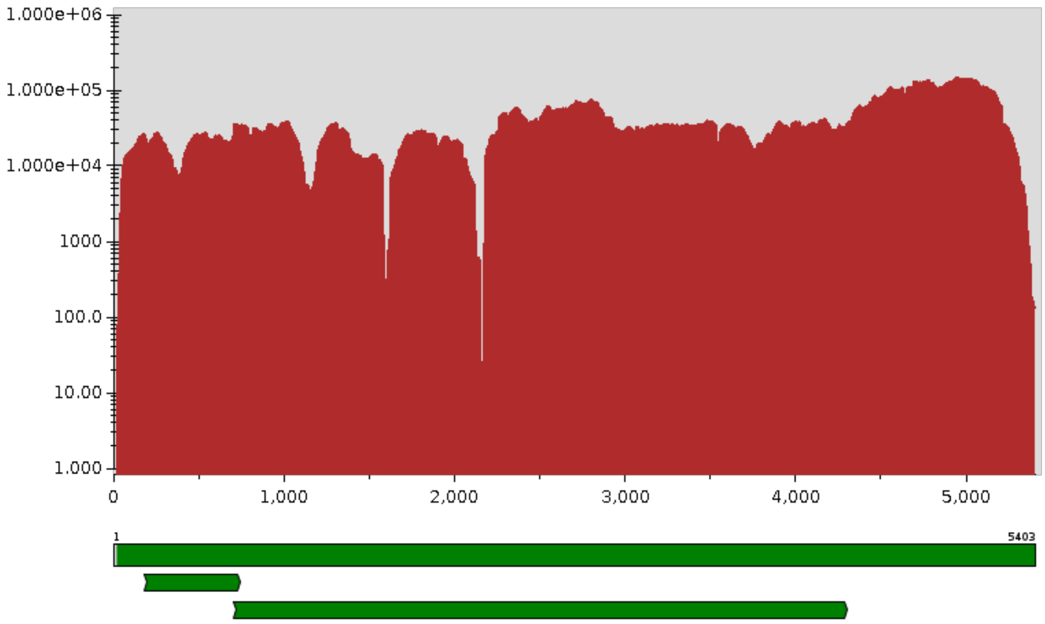

#### Assignment

| | |
|---|---|
| **Type** | Torradovirus lycopersici (Taxonomy ID: 3048378) |
| **Reference Genome** | NC_009032.1 |
| **NT Identity (%)** | 87.3561 |
| **AA Identity (%)** | 91.4203 |
| **Number Of Stop Codons** | 3 |
| **Number Of CDS** | 2 |

#### Alignment

| | |
|---|---|
| **Alignment Score** | 8032.0 (NT) + 8819.0 (AA) = 16851.0 |
| **Concordance (%)** | 83.1943 |

| Alignment Method | Global, seeded, nucleotide + amino acids (AGA) |
|---|---|

#### Genome Region

Sequence starts at position 20 and ends at position 5403 relative to NC_009032.1 reference sequence.

#### Alignment Detailed Statistics

| | Begin | End | Coverage | Score | Concordance | Matches | Identities | I/D/M/F* | Stop Codons |
|---|---|---|---|---|---|---|---|---|---|
| **NT** | **20** | **5403** | **99.6%** | **8032** | **74.6%** | **5384 (99.9%)** | **4705 (87.4%)** | **2/0** | |
| Mutations: | 20A>G, 78C>T, 89G>A, 90C>T, 97C>T, 107G>A, 111C>A, 137T>C, 144A>G, 277C>T, 280T>C, 283C>T, 295T>C, 301A>T, 313C>T, 343T>C, 370C>T, 372A>C, 376C>T, 388T>A, 400T>C, 401T>C, 418T>G, 433C>T, 448C>A, 460T>C, 481C>A, 487A>G, 507G>A, 511T>A, 565A>T, 568A>T, 576C>A, 583A>T, 586A>G, 587T>C, 589A>T, 603A>C, 607G>A, 613A>G, 622A>G, 644C>G, 654T>A, 679G>A, 696T>C, 721C>T, 733A>G, 748A>G, 750A>G, 797T>C, 806C>T, 833C>T, 844A>T, 911T>C, 929T>A, 932A>T, 968A>G, 986G>A, 995T>C, 1040A>C, 1043A>G, 1052A>G, 1061T>C, 1073C>A, 1083C>T, 1094G>A, 1095G>A, 1100G>A, 1109C>T, 1112G>T, 1115G>A, 1118A>C, 1128T>C, 1130A>G, 1139G>A, 1142C>T, 1151A>G, 1163A>G, 1169C>G, 1178C>T, 1187A>T, 1205T>A, 1220T>C, 1235C>A, 1247A>G, 1262T>C, 1268C>T, 1277A>G, 1313T>C, 1317C>A, 1319A>G, 1325A>G, 1328T>C, 1331C>T, 1343G>A, 1355G>A, 1361A>T, 1367A>T, 1370A>T, 1376T>A, 1383A>G, 1385C>T, 1388C>T, 1390C>G, 1394A>T, 1395A>T, 1397T>A, 1400T>C, 1403A>T, 1406G>A, 1409T>A, 1412G>T, 1418G>A, 1426A>C, 1427G>T, 1434G>A, 1439A>C, 1451G>T, 1454A>T, 1456G>C, 1457C>T, 1460A>G, 1461G>T, 1465G>A, 1472C>G, 1475G>C, 1478C>G, 1481C>T, 1487C>G, 1490G>A, 1491G>A, 1493G>A, 1496C>T, 1502C>T, 1514T>C, 1524T>C, 1526C>T, 1529A>T, 1538A>T, 1543C>T, 1545A>G, 1547C>T, 1553T>C, 1559C>A, 1560C>T, 1562C>A, 1568C>G, 1574G>A, 1575C>A, 1577T>G, 1589A>G, 1593C>G, 1595G>T, 1603G>A, 1605A>G, 1608T>C, 1610C>G, 1613A>G, 1616G>A, 1621G>C, 1622T>A, 1631G>A, 1634T>A, 1646T>C, 1651A>G, 1655T>A, 1661T>C, 1662C>T, 1663T>A, 1664G>T, 1667C>T, 1674C>T, 1676T>C, 1677G>A, 1679G>T, 1685T>C, 1688T>C, 1691T>G, 1695A>G, 1700G>A, 1721G>C, 1724C>T, 1727A>G, 1730C>T, 1740C>T, 1745G>A, 1748G>A, 1749A>C, 1754G>A, 1757A>G, 1760T>G, 1763T>A, 1766A>G, 1769A>T, 1778A>C, 1784A>C, 1790G>A, 1793T>A, 1794G>A, 1797T>G, 1799C>G, 1805C>T, 1811A>C, 1814T>C, 1817A>T, 1819T>A, 1820C>G, 1823C>T, 1832G>C, 1841C>T, 1843A>G, 1845T>C, 1847G>T, 1850G>A, 1856C>T, 1863C>T, 1865C>T, 1875G>A, 1881C>T, 1892C>T, 1898T>G, 1899G>A, 1900T>C, 1901G>A, 1906C>T, 1907C>G, 1913T>A, 1928A>C, 1931C>T, 1938A>G, 1941C>T, 1943T>G, 1946C>A, 1947T>C, 1952A>C, 1955G>A, 1958A>G, 1967G>A, 1970A>G, 1979A>G, 1986G>A, 1994T>C, 2002T>A, 2012C>T, 2024T>C, 2027T>A, 2049T>A, 2051A>T, 2054G>T, 2059A>G, 2060A>C, 2066C>G, 2074T>A, 2083T>C, 2084G>A, 2087A>C, 2094C>G, 2096G>C, 2098C>A, 2099G>A, 2100C>A, 2101G>T, 2102C>G, 2106A>G, 2107G>A, 2111T>G, 2114C>T, 2115A>C, 2116A>G, 2119A>C, 2120G>A, 2123G>T, 2125C>T, 2126C>T, 2127A>G, 2128G>T, 2130G>A, 2132T>C, 2135G>A, 2136G>C, 2138G>T, 2144C>G, 2159T>A, 2162T>C, 2164T>G, 2170A>T, 2171A>T, 2174A>T, 2177A>T, 2178T>G, 2183C>T, 2186A>C, 2189C>T, 2195A>C, 2198G>A, 2199G>A, 2200T>C, 2204C>T, 2205A>G, 2210G>A, 2216C>G, 2219T>C, 2225T>A, 2228C>T, 2229A>G, 2230T>A, 2231G>T, 2240T>A, 2243T>C, 2252A>C, 2256G>A, 2259A>C, 2261T>A, 2262G>C, 2270A>G, 2285T>C, 2303A>G, 2312T>C, 2321T>C, 2375C>T, 2393G>A, 2414T>C, 2420T>A, 2429G>A, 2432G>T, 2433C>T, 2436T>G, 2444G>T, 2447C>T, 2450T>C, 2453G>A, 2459C>G, 2465T>G, 2468A>G, 2469T>A, 2471C>A, 2474G>A, 2489A>C, 2492G>A, 2495A>G, 2501A>G, 2503G>A, 2507T>C, 2516C>T, 2523A>G, 2546T>C, 2561T>A, 2567T>A, 2573T>C, 2609C>G, 2627C>T, 2630C>A, 2645G>A, 2648T>C, 2651C>T, 2666G>A, 2669T>G, 2672A>C, 2675A>T, 2693G>A, 2702T>C, 2708A>G, 2717T>C, 2720C>A, 2729C>T, 2732T>G, 2747C>A, 2753G>T, 2774A>T, 2792C>T, 2795C>G, 2798G>A, 2799G>A, 2804T>C, 2814T>C, 2835A>G, 2837C>T, 2840T>C, 2841C>T, 2846G>A, 2852A>G, 2861G>A, 2864C>A, 2867C>T, 2870C>A, 2879T>A, 2882T>A, 2886T>C, 2888G>A, 2891A>G, 2894A>G, 2900A>T, 2909T>C, 2915A>G, 2921C>T, 2927A>G, 2936A>C, 2939A>G, 2948A>G, 2954G>A, 2957A>T, 2960G>A, 2966C>T, 2972T>C, 2981G>T, 2982G>A, 2996T>G, 2999T>A, 3003T>C, 3011C>T, 3012A>G, 3014C>T, 3020T>C, 3023G>C, 3029C>T, 3032A>T, 3044C>T, 3047T>A, 3050G>C, 3051C>T, 3059A>T, 3062C>T, 3063T>C, 3065G>A, 3071A>G, 3077A>T, 3083C>A, 3101T>G, 3104C>T, 3107T>G, 3113G>C, 3120A>G, 3122T>G, 3125T>C, 3143C>T, 3146T>C, 3152T>A, 3158G>A, 3161C>A, 3164T>C, 3167A>T, 3170G>T, 3173T>C, 3176A>C, 3179G>T, 3182C>T, 3187G>A, 3194C>T, 3195A>G, 3197C>T, 3200T>C, 3206C>T, 3227T>G, 3230T>A, 3236C>T, 3239T>G, 3242G>A, 3248A>C, 3251A>T, 3253A>G, 3265G>A, 3270A>T, 3272G>T, 3278C>T, 3290G>C, 3293A>G, 3296T>C, 3297C>T, 3299T>A, 3303A>T, 3305C>T, 3311T>A, 3324C>T, 3329G>T, 3350C>T, 3354A>C, 3359G>A, 3368A>G, 3371C>G, 3372T>C, 3383G>A, 3386A>T, 3395A>C, 3404A>G, 3410G>A, 3413C>T, 3428G>A, 3429C>T, 3431T>G, 3434A>G, 3437C>T, 3440A>T, 3447T>C, 3449G>A, 3452C>A, 3455T>A, 3458G>A, 3461C>T, 3470T>C, 3473T>C, 3479C>T, 3482T>C, 3485C>T, 3488A>G, 3491C>T, 3503A>G, 3508A>G, 3515G>T, 3522C>T, 3524A>G, 3527C>T, 3530T>A, 3533G>A, 3543T>A, 3551C>T, 3560T>C, 3564A>C, 3567T>A, 3569A>G, 3575G>A, 3578A>G, 3590A>C, 3605A>G, 3611G>A, 3617A>T, 3620G>A, 3623A>T, 3629G>C, 3631C>A, 3634T>C, 3638A>G, 3641C>T, 3650A>T, 3656G>T, 3659T>C, 3662A>G, 3674G>C, 3689C>T, 3698A>C, 3704G>A, 3707T>G, 3708T>C, 3713A>C, 3716C>T, 3722C>A, 3728C>G, 3737C>T, 3742A>C, 3743G>C, 3746G>A, 3749A>T, 3752G>A, 3755A>G, 3770A>C, 3773C>T, 3776A>T, 3779C>T, 3800C>T, 3815A>C, 3827C>T, 3833T>A, 3839A>G, 3842T>C, 3845A>G, 3848G>A, 3851G>A, 3852A>G, 3854T>G, 3857C>A, 3866A>C, 3869A>T, 3872A>T, 3878G>C, 3881C>T, 3882A>G, 3884C>A, 3885A>G, 3890A>G, 3893T>C, 3896A>T, 3905C>T, 3911T>C, 3914T>A, 3920A>T, 3923C>T, 3926A>C, 3927A>C, 3928A>G, 3929G>A, 3932C>A, 3933A>G, 3935C>T, 3941T>G, 3944C>A, 3947T>C, 3950T>A, 3953T>A, 3960G>A, 3971T>A, 3983T>A, 3984A>G, 3986G>T, 3988G>A, 3989C>T, 3990G>A, 3991A>G, 3995C>T, 3998C>T, 4001T>C, 4016T>G, 4025G>A, 4028T>C, 4031T>C, 4034T>G, 4035A>C, 4037A>T, 4040C>T, 4052C>T, 4059A>G, 4061A>T, 4067A>T, 4068A>G, 4069A>G, 4070T>C, 4072C>T, 4073T>G, 4077A>C, 4088T>A, 4091G>A, 4100A>C, 4103A>T, 4106A>C, 4112C>G, 4124A>G, 4133A>G, 4142A>C, 4145C>T, 4149C>A, 4151G>A, 4166G>A, 4181C>T, 4187G>A, 4188C>G, 4196T>A, 4208A>T, 4211A>T, 4223G>A, 4224C>T, 4227T>C, 4229G>T, 4244G>A, 4250C>G, 4253C>T, 4256T>G, 4262T>C, 4265T>A, 4274A>T, 4281A>C, 4284A>G, 4286T>A, 4289T>C, 4300T>C, 4302T>G, 4316T>C, 4354G>A, 4355T>G, 4457A>G, 4590G>A, 4749T>A, 4812C>T, 4814T>C, 4823A>G, 4974_4975insT, 5089T>C, 5206C>T, 5218C>T, 5225C>T, 5235_5236insT, 5241T>C, 5249A>G, 5262A>T | | | | | | | | |
| **CDS** | | | | | | | | | |
| ToTV_sRNA2gp1 | 1 | 188 | 100% | 1156 | 95.6% | 188 (100%) | 180 (95.7%) | 0/0/0/0 | 1 |
| Protein mutations: | N64T (372A>C), R109K (507G>A), Q128H (565A>T), T132K (576C>A), D141A (603A>C), L155V (644C>G), L158Q (654T>A), L172P (696T>C) | | | | | | | | |
| Codon mutations: | GGC32GGT (277C>T), TCT33TCC (280T>C), GGC34GGT (283C>T), AGT38AGC (295T>C), CCA40CCT (301A>T), TTC44TTT (313C>T), ATT54ATC (343T>C), AAC63AAT (370C>T), AAT64ACT (372A>C), AGC65AGT (376C>T), GTT69GTA (388T>A), TAT73TAC (400T>C), TTA74CTA (401T>C), ACT79ACG (418T>G), GGC84GGT (433C>T), GGC89GGA (448C>A), CAT93CAC (460T>C), ACC100ACA (481C>A), AGA102AGG (487A>G), AGG109AAG (507G>A), ATT110ATA (511T>A), CAA128CAT (565A>T), CCA129CCT (568A>T), ACG132AAG (576C>A), ACA134ACT (583A>T), CAA135CAG (586A>G), TTA136CTT (587T>C 589A>T), GAC141GCC (603A>C), GAG142GAA (607G>A), AAA144AAG (613A>G), TTA147TTG (622A>G), CTT155GTT (644C>G), CTA158CAA (654T>A), TTG166TTA (679G>A), CTA172CCA (696T>C), GGC180GGT (721C>T), GAA184GAG (733A>G) | | | | | | | | |
| ToTV_sRNA2gp2 | 1 | 1199 | 100% | 7663 | 92.6% | 1199 (100%) | 1088 (90.7%) | 0/0/0/0 | 2 |
| Protein mutations: | A7V (721C>T), K11R (733A>G), K16R (748A>G), S17G (750A>G), Y48F (844A>T), V132I (1095G>A), T228A (1383A>G 1385C>T), S230* (1390C>G), T232S (1395A>T 1397T>A), Q242P (1426A>C 1427G>T), D245N (1434G>A), S252T (1456G>C 1457C>T), A254S (1461G>T), R255K (1465G>A), I257M (1472C>G), V264I (1491G>A 1493G>A), S275P (1524T>C 1526C>T), S281F (1543C>T), T282A (1545A>G 1547C>T), H286Q (1559C>A), Q298D (1593C>G 1595G>T), S301N (1603G>A), R302G (1605A>G), Y303Q (1608T>C 1610C>G), G307A (1621G>C 1622T>A), K317R (1651A>G), L321Y (1662C>T 1663T>A 1664G>T), H325Y (1674C>T 1676T>C), V326I (1677G>A 1679G>T), I332V (1695A>G), V365I (1794G>A), C366G (1797T>G 1799C>G), K370N (1811A>C), I373K (1819T>A 1820C>G), K377N (1832G>C), H381R (1843A>G), H388Y (1863C>T 1865C>T), V392M (1875G>A), I399M (1898T>G), V400T (1899G>A 1900T>C 1901G>A), A402V (1906C>T 1907C>G), E409D (1928A>C), N413D (1938A>G), D429N (1986G>A), F434Y (2002T>A), L450I (2049T>A 2051A>T), K453S (2059A>G 2060A>C), L458Q (2074T>A), V461A (2083T>C 2084G>A), Q465D (2094C>G 2096G>C), A466E (2098C>A 2099G>A), R467M (2100C>A 2101G>T 2102C>G), R469E (2106A>G 2107G>A), F470L (2111T>G), K472R (2115A>C 2116A>G), E473A (2119A>C 2120G>A), A475V (2125C>T 2126C>T), R476V (2127A>G 2128G>T), A477T (2130G>A 2132T>C), V479L (2136G>C 2138G>T), N481K (2144C>G), V488G (2164T>G), E490V (2170A>T 2171A>T), S493A (2178T>G), V500T (2199G>A 2200T>C), M502V (2205A>G), D508E (2225T>A), M510D (2229A>G 2230T>A 2231G>T), E517D (2252A>C), A519T (2256G>A), N520Q (2259A>C 2261T>A), E521Q (2262G>C), S579A (2436T>G), S590T (2469T>A 2471C>A), R601K (2503G>A), I608V (2523A>G), A700T (2799G>A), I712V (2835A>G 2837C>T), A761T (2982G>A), I771V (3012A>G 3014C>T), I807V (3120A>G 3122T>G), S829N (3187G>A), T832A (3195A>G 3197C>T), K851R (3253A>G), R855K (3265G>A), T857S (3270A>T 3272G>T), T868S (3303A>T 3305C>T), N936S (3508A>G), S948T (3543T>A), K955Q (3564A>C), L956M (3567T>A 3569A>G), P977Q (3631C>A), L978P (3634T>C), Q1014P (3742A>C 3743G>C), T1051A (3852A>G 3854T>G), I1061V (3882A>G 3884C>A), T1062A (3885A>G), K1076R (3927A>C 3928A>G 3929G>A), T1078A (3933A>G 3935C>T), A1087T (3960G>A), M1095V (3984A>G 3986G>T), S1096N (3988G>A 3989C>T), D1097S (3990G>A 3991A>G), H1105Q (4016T>G), I1120V (4059A>G 4061A>T), N1123G (4068A>G 4069A>G 4070T>C), A1124V (4072C>T 4073T>G), L1150I (4149C>A 4151G>A), Q1163E (4188C>G), H1175Y (4224C>T), S1195G (4284A>G 4286T>A) | | | | | | | | |

| | Begin | End | Coverage | Score | Concordance | Matches | Identities | I/D/M/F* | Stop Codons |
|---|---|---|---|---|---|---|---|---|---|
| **NT** | **20** | **5403** | **99.6%** | **8032** | **74.6%** | **5384 (99.9%)** | **4705 (87.4%)** | **2/0** | |
| Codon mutations: | GCT7GTT (721C>T), AAG11AGG (733A>G), AAA16AGA (748A>G), AGT17GGT (750A>G), GTT32GTC (797T>C), GGC35GGT (806C>T), GGC44GGT (833C>T), TAT48TTT (844A>T), AAT70AAC (911T>C), TCT76TCA (929T>A), TCA77TCT (932A>T), GGA89GGG (968A>G), TTG95TTA (986G>A), GTT98GTC (995T>C), GTA113GTC (1040A>C), AAA114AAG (1043A>G), TTA117TTG (1052A>G), AGT120AGC (1061T>C), ACC124ACA (1073C>A), CTG128TTG (1083C>T), AAG131AAA (1094G>A), GTT132ATT (1095G>A), TTG133TTA (1100G>A), TCC136TCT (1109C>T), GGG137GGT (1112G>T), GGG138GGA (1115G>A), GCA139GCC (1118A>C), TTA143CTG (1128T>C 1130A>G), CCG146CCA (1139G>A), ACC147ACT (1142C>T), CTA150CTG (1151A>G), AAA154AAG (1163A>G), GGC156GGG (1169C>G), TTC159TTT (1178C>T), ACA162ACT (1187A>T), ACT168ACA (1205T>A), CCT173CCC (1220T>C), GCC178GCA (1235C>A), CAA182CAG (1247A>G), CGT187CGC (1262T>C), GCC189GCT (1268C>T), TCA192TCG (1277A>G), CTT204CTC (1313T>C), CGA206AGG (1317C>A 1319A>G), CAA208CAG (1325A>G), TTT209TTC (1328T>C), TTC210TTT (1331C>T), CAG214CAA (1343G>A), CAG218CAA (1355G>A), ATA220ATT (1361A>T), TCA222TCT (1367A>T), GTA223GTT (1370A>T), TCT225TCA (1376T>A), ACC228GCT (1383A>G 1385C>T), TCC229TCT (1388C>T), TCA230TGA (1390C>G), GCA231GCT (1394A>T), ACT232TCA (1395A>T 1397T>A), GAT233GAC (1400T>C), TCA234TCT (1403A>T), GTG235GTA (1406G>A), CTT236CTA (1409T>A), CCG237CCT (1412G>T), GTG239GTA (1418G>A), CAG242CCT (1426A>C 1427G>T), GAC245AAC (1434G>A), ACA246ACC (1439A>C), GTG250GTT (1451G>T), CCA251CCT (1454A>T), AGC252ACT (1456G>C 1457C>T), GAA253GAG (1460A>G), GCA254TCA (1461G>T), AGA255AAA (1465G>A), ATC257ATG (1472C>G), CGG258CGC (1475G>C), TCC259TCG (1478C>G), TCC260TCT (1481C>T), CTC262CTG (1487C>G), CGG263CGA (1490G>A), GTG264ATA (1491G>A 1493G>A), CCC265CCT (1496C>T), CTC267CTT (1502C>T), AGT271AGC (1514T>C), TCC275CCT (1524T>C 1526C>T), TCA276TCT (1529A>T), ATA279ATT (1538A>T), TCT281TTT (1543C>T), ACC282GCT (1545A>G 1547C>T), ACT284ACC (1553T>C), CAC286CAA (1559C>A), CTC287TTA (1560C>T 1562C>A), GGC289GGG (1568C>G), ACG291ACA (1574G>A), CGT292AGG (1575C>A 1577T>G), GTA296GTG (1589A>G), CAG298GAT (1593C>G 1595G>T), AGT301AAT (1603G>A), AGG302GGG (1605A>G), TAC303CAG (1608T>C 1610C>G), GAA304GAG (1613A>G), GAG305GAA (1616G>A), GGT307GCA (1621G>C 1622T>A), GGG310GGA (1631G>A), GTT311GTA (1634T>A), GTT315GTC (1646T>C), AAG317AGG (1651A>G), GCT318GCA (1655T>A), GGT320GGC (1661T>C), CTG321TAT (1662C>T 1663T>A 1664G>T), AAC322AAT (1667C>T), CAT325TAC (1674C>T 1676T>C), GTG326ATT (1677G>A 1679G>T), GAT328GAC (1685T>C), AAT329AAC (1688T>C), CTT330CTG (1691T>G), ATT332GTT (1695A>G), GAG333GAA (1700G>A), CTG340CTC (1721G>C), CCC341CCT (1724C>T), GAA342GAG (1727A>G), AAC343AAT (1730C>T), CTG347TTG (1740C>T), TCG348TCA (1745G>A), GTG349GTA (1748G>A), AGA350CGA (1749A>C), CAG351CAA (1754G>A), CAA352CAG (1757A>G), GTT353GTG (1760T>G), GCT354GCA (1763T>A), GCA355GCG (1766A>G), GCA356GCT (1769A>T), GCA359GCC (1778A>C), GCA361GCC (1784A>C), AAG363AAA (1790G>A), GGT364GGA (1793T>A), GTT365ATT (1794G>A), TGC366GGG (1797T>G 1799C>G), GTC368GTT (1805C>T), AAA370AAC (1811A>C), CAT371CAC (1814T>C), ATA372ATT (1817A>T), ATC373AAG (1819T>A 1820C>G), AAC374AAT (1823C>T), AAG377AAC (1832G>C), ATC380ATT (1841C>T), CAC381CGC (1843A>G), TTG382CTT (1845T>C 1847G>T), GAG383GAA (1850G>A), ATC385ATT (1856C>T), CAC388TAT (1863C>T 1865C>T), GTG392ATG (1875G>A), CTG394TTG (1881C>T), TAC397TAT (1892C>T), ATT399ATG (1898T>G), GTG400ACA (1899G>A 1900T>C 1901G>A), GCC402GTG (1906C>T 1907C>G), TCT404TCA (1913T>A), GAA409GAC (1928A>C), AAC410AAT (1931C>T), AAT413GAT (1938A>G), CTT414TTG (1941C>T 1943T>G), ACC415ACA (1946C>A), TTG416CTG (1947T>C), GCA417GCC (1952A>C), CAG418CAA (1955G>A), AGA419AGG (1958A>G), GAG422GAA (1967G>A), CTA423CTG (1970A>G), CAA426CAG (1979A>G), GAT429AAT (1986G>A), TAT431TAC (1994T>C), TTT434TAT (2002T>A), AAC437AAT (2012C>T), TAT441TAC (2024T>C), CCT442CCA (2027T>A), TTA450ATT (2049T>A 2051A>T), ACG451ACT (2054G>T), AAA453AGC (2059A>G 2060A>C), CGC455CGG (2066C>G), CTA458CAA (2074T>A), GTG461GCA (2083T>C 2084G>A), GCA462GCC (2087A>C), CAG465GAC (2094C>G 2096G>C), GCG466GAA (2098C>A 2099G>A), CGC467ATG (2100C>A 2101G>T 2102C>G), AGA469GAA (2106A>G 2107G>A), TTT470TTG (2111T>G), TAC471TAT (2114C>T), AAG472CGG (2115A>C 2116A>G), GAG473GCA (2119A>C 2120G>A), GCG474GCT (2123G>T), GCC475GTT (2125C>T 2126C>T), AGG476GTG (2127A>G 2128G>T), GCT477ACC (2130G>A 2132T>C), CAG478CAA (2135G>A), GTG479CTT (2136G>C 2138G>T), AAC481AAG (2144C>G), ACT486ACA (2159T>A), AGT487AGC (2162T>C), GTG488GGG (2164T>G), GAA490GTT (2170A>T 2171A>T), ATA491ATT (2174A>T), CCA492CCT (2177A>T), TCC493GCC (2178T>G), ACC494ACT (2183C>T), TCA495TCC (2186A>C), TTC496TTT (2189C>T), ACA498ACC (2195A>C), AAG499AAA (2198G>A), GTT500ACT (2199G>A 2200T>C), GCC501GCT (2204C>T), ATG502GTG (2205A>G), GAG503GAA (2210G>A), GGC505GGG (2216C>G), AGT506AGC (2219T>C), GAT508GAA (2225T>A), AGC509AGT (2228C>T), ATG510GAT (2229A>G 2230T>A 2231G>T), GCT513GCA (2240T>A), ATT514ATC (2243T>C), GAA517GAC (2252A>C), GCC519ACC (2256G>A), AAT520CAA (2259A>C 2261T>A), GAA521CAA (2262G>C), GTA523GTG (2270A>G), GTT528GTC (2285T>C), CAA534CAG (2303A>G), AAT537AAC (2312T>C), GGT540GGC (2321T>C), TTC558TTT (2375C>T), GAG564GAA (2393G>A), AGT571AGC (2414T>C), GTT573GTA (2420T>A), GAG576GAA (2429G>A), GTG577GTT (2432G>T), CTA578TTA (2433C>T), TCA579GCA (2436T>G), GCG581GCT (2444G>T), CTC582CTT (2447C>T), CCT583CCC (2450T>C), GCG584GCA (2453G>A), CTC586CTG (2459C>G), GGT588GGG (2465T>G), AAA589AAG (2468A>G), TCC590ACA (2469T>A 2471C>A), AAG591AAA (2474G>A), GGA596GGC (2489A>C), TCG597TCA (2492G>A), CAA598CAG (2495A>G), CTA600CTG (2501A>G), AGG601AAG (2503G>A), TAT602TAC (2507T>C), GCC605GCT (2516C>T), ATT608GTT (2523A>G), TAT615TAC (2546T>C), GGT620GGA (2561T>A), ATT622ATA (2567T>A), GGT624GGC (2573T>C), CTC636CTG (2609C>G), TTC642TTT (2627C>T), ATC643ATA (2630C>A), CTG648CTA (2645G>A), TAT649TAC (2648T>C), GCC650GCT (2651C>T), CTG655CTA (2666G>A), GTT656GTG (2669T>G), TCA657TCC (2672A>C), GCA658GCT (2675A>T), GGG664GGA (2693G>A), TGT667TGC (2702T>C), ACA669ACG (2708A>G), GGT672GGC (2717T>C), ATC673ATA (2720C>A), TTC676TTT (2729C>T), GTT677GTG (2732T>G), GCC682GCA (2747C>A), GGG684GGT (2753G>T), ATA691ATT (2774A>T), CAC697CAT (2792C>T), CCC698CCG (2795C>G), TTG699TTA (2798G>A), GCT700ACT (2799G>A), AGT701AGC (2804T>C), TTG705CTG (2814T>C), ATC712GTT (2835A>G 2837C>T), CAT713CAC (2840T>C), CTG714TTG (2841C>T), CAG715CAA (2846G>A), AAA717AAG (2852A>G), TCG720TCA (2861G>A), ACC721ACA (2864C>A), AAC722AAT (2867C>T), ATC723ATA (2870C>A), CCT726CCA (2879T>A), CCT727CCA (2882T>A), TTG729CTA (2886T>C 2888G>A), CGA730CGG (2891A>G), GCA731GCG (2894A>G), GCA733GCT (2900A>T), GGT736GGC (2909T>C), AAA738AAG (2915A>G), GAC740GAT (2921C>T), ACA742ACG (2927A>G), CCA745CCC (2936A>C), CGA746CGG (2939A>G), ACA749ACG (2948A>G), CAG751CAA (2954G>A), GTA752GTT (2957A>T), CTG753CTA (2960G>A), CAC755CAT (2966C>T), AAT757AAC (2972T>C), GTG760GTT (2981G>T), GCT761ACT (2982G>A), GGT765GGG (2996T>G), ACT766ACA (2999T>A), TTA768CTA (3003T>C), AGC770AGT (3011C>T), ATC771GTT (3012A>G 3014C>T), TCT773TCC (3020T>C), CCG774CCC (3023G>C), GGC776GGT (3029C>T), ATA777ATT (3032A>T), GAC781GAT (3044C>T), GGT782GGA (3047T>A), ACG783ACC (3050G>C), CTG784TTG (3051C>T), CCA786CCT (3059A>T), TCC787TCT (3062C>T), TTG788CTA (3063T>C 3065G>A), GGA790GGG (3071A>G), ATA792ATT (3077A>T), CGC794CGA (3083C>A), ACT800ACG (3101T>G), GGC801GGT (3104C>T), ACT802ACG (3107T>G), GTG804GTC (3113G>C), ATT807GTG (3120A>G 3122T>G), TGT808TGC (3125T>C), TTC814TTT (3143C>T), CAT815CAC (3146T>C), GGT817GGA (3152T>A), TTG819TTA (3158G>A), GCC820GCA (3161C>A), ATT821ATC (3164T>C), GGA822GGT (3167A>T), CTG823CTT (3170G>T), GGT824GGC (3173T>C), ACA825ACC (3176A>C), CTG826CTT (3179G>T), AAC827AAT (3182C>T), AGC829AAC (3187G>A), TCC831TCT (3194C>T), ACC832GCT (3195A>G 3197C>T), CCT833CCC (3200T>C), GAC835GAT (3206C>T), GTT842GTG (3227T>G), ATT843ATA (3230T>A), AAC845AAT (3236C>T), CTT846CTG (3239T>G), GAG847GAA (3242G>A), GGA849GGC (3248A>C), CGA850CGT (3251A>T), AAG851AGG (3253A>G), AGG855AAG (3265G>A), ACG857TCT (3270A>T 3272G>T), ACC859ACT (3278C>T), GGG863GGC (3290G>C), AAA864AAG (3293A>G), AAT865AAC (3296T>C), CTT866TTA (3297C>T 3299T>A), ACC868TCT (3303A>T 3305C>T), GGT870GGA (3311T>A), CTA875TTA (3324C>T), CCG876CCT (3329G>T), CAC883CAT (3350C>T), AGG885CGG (3354A>C), TTG886TTA (3359G>A), ACA889ACG (3368A>G), GTC890GTG (3371C>G), TTG891CTG (3372T>C), CTG894CTA (3383G>A), GTA895GTT (3386A>T), TCA898TCC (3395A>C), CTA901CTG (3404A>G), ACG903ACA (3410G>A), GTC904GTT (3413C>T), CAG909CAA (3428G>A), CTT910TTG (3429C>T 3431T>G), AAA911AAG (3434A>G), TGC912TGT (3437C>T), ATA913ATT (3440A>T), TTG916CTA (3447T>C 3449G>A), GTC917GTA (3452C>A), CTT918CTA (3455T>A), GGG919GGA (3458G>A), GGC920GGT (3461C>T), TCT923TCC (3470T>C), GTT924GTC (3473T>C), CCC926CCT (3479C>T), ATT927ATC (3482T>C), TAC928TAT (3485C>T), GGA929GGG (3488A>G), CAC930CAT (3491C>T), GGA934GGG (3503A>G), AAT936AGT (3508A>G), GTG938GTT (3515G>T), CTA941TTG (3522C>T 3524A>G), TTC942TTT (3527C>T), TCT943TCA (3530T>A), GAG944GAA (3533G>A), TCT948ACT (3543T>A), CGC950CGT (3551C>T), ATT953ATC (3560T>C), AAA955CAA (3564A>C), TTA956ATG (3567T>A 3569A>G), AAG958AAA (3575G>A), GAA959GAG (3578A>G), ACA963ACC (3590A>C), GGA968GGG (3605A>G), AAG970AAA (3611G>A), CCA972CCT (3617A>T), CAG973CAA (3620G>A), GTA974GTT (3623A>T), GTG976GTC (3629G>C), CCG977CAG (3631C>A), CTC978CCC (3634T>C), AGA979AGG (3638A>G), GAC980GAT (3641C>T), TCA983TCT (3650A>T), GGG985GGT (3656G>T), GCT986GCC (3659T>C), GTA987GTG (3662A>G), GTG991GTC (3674G>C), GAC996GAT (3689C>T), CGA999CGC (3698A>C), TTG1001TTA (3704G>A), GTT1002GTG (3707T>G), TTA1003CTA (3708T>C), CCA1004CCC (3713A>C), TGC1005TGT (3716C>T), CCC1007CCA (3722C>A), TCC1009TCG (3728C>G), TTC1012TTT (3737C>T), CAG1014CCC (3742A>C 3743G>C), GGG1015GGA (3746G>A), GCA1016GCT (3749A>T), CTG1017CTA (3752G>A), GTA1018GTG (3755A>G), ACA1023ACC (3770A>C), TGC1024TGT (3773C>T), CCA1025CCT (3776A>T), TTC1026TTT (3779C>T), TTC1033TTT (3800C>T), GGA1038GGC (3815A>C), TAC1042TAT (3827C>T), ATT1044ATA (3833T>A), GTA1046GTG (3839A>G), CAT1047CAC (3842T>C), AGA1048AGG (3845A>G), GTG1049GTA (3848G>A), CAG1050CAA (3851G>A), ACT1051GCG (3852A>G 3854T>G), TCC1052TCA (3857C>A), ATA1055ATC (3866A>C), GGA1056GGT (3869A>T), GGA1057GGT (3872A>T), CTG1059CTC (3878G>C), AAC1060AAT (3881C>T), ATC1061GTA (3882A>G 3884C>A), ACC1062GCC (3885A>G), TTA1063TTG (3890A>G), GAT1064GAC (3893T>C), TCA1065TCT (3896A>T), TAC1068TAT (3905C>T), TTT1070TTC (3911T>C), CCT1071CCA (3914T>A), GGA1073GGT (3920A>T), ATC1074ATT (3923C>T), TCA1075TCC (3926A>C), AAG1076CGA (3927A>C 3928A>G 3929G>A), GGC1077GGA (3932C>A), ACC1078GCT (3933A>G 3935C>T), GTT1080GTG (3941T>G), GTC1081GTA (3944C>A), TCT1082TCC (3947T>C), GCT1083GCA (3950T>A), GGT1084GGA (3953T>A), GCA1087ACA (3960G>A), GCT1090GCA (3971T>A), GGT1094GGA (3983T>A), ATG1095GTT (3984A>G 3986G>T), AGC1096AAT (3988G>A 3989C>T), GAC1097AGC (3990G>A 3991A>G), AAC1098AAT (3995C>T), ATC1099ATT (3998C>T), TTT1100TTC (4001T>C), CAT1105CAG (4016T>G), GAG1108GAA (4025G>A), TTT1109TTC (4028T>C), TTT1110TTC (4031T>C), CCT1111CCG (4034T>G), AGA1112CGT (4035A>C 4037A>T), CGC1113CGT (4040C>T), GCC1117GCT (4052C>T), ATA1120GTT (4059A>G 4061A>T), CCA1122CCT (4067A>T), AAT1123GGC (4068A>G 4069A>G 4070T>C), GCT1124GTG (4072C>T 4073T>G), AGA1126CGA (4077A>C), ACT1129ACA (4088T>A), CTG1130CTA (4091G>A), CGA1133CGC (4100A>C), CTA1134CTT (4103A>T), GGA1135GGC (4106A>C), CTC1137CTG (4112C>G), TTA1141TTG (4124A>G), AAA1144AAG (4133A>G), ATA1147ATC (4142A>C), AGC1148AGT (4145C>T), CTG1150ATA (4149C>A 4151G>A), AAG1155AAA (4166G>A), TTC1160TTT (4181C>T), TTG1162TTA (4187G>A), CAA1163GAA (4188C>G), GCT1165GCA (4196T>A), TCA1169TCT (4208A>T), GCA1170GCT (4211A>T), AAG1174AAA (4223G>A), CAT1175TAT (4224C>T), TTG1176CTT (4227T>C 4229G>T), ACG1181ACA (4244G>A), ACC1183ACG (4250C>G), TAC1184TAT (4253C>T), CTT1185CTG (4256T>G), ACT1187ACC (4262T>C), CCT1188CCA (4265T>A), TCA1191TCT (4274A>T), AGG1194CGG (4281A>C), AGT1195GGA (4284A>G 4286T>A), TTT1196TTC (4289T>C) | | | | | | | | |

#### Proteins

| | | | | | | | | | |
|---|---|---|---|---|---|---|---|---|---|
| Orf1 (YP_001040017.1) | 1 | 188 | 100% | 1156 | 95.6% | 188 (100%) | 180 (95.7%) | 0/0/0/0 | 1 |
| Protein mutations: | N64T (372A>C), R109K (507G>A), Q128H (565A>T), T132K (576C>A), D141A (603A>C), L155V (644C>G), L158Q (654T>A), L172P (696T>C) | | | | | | | | |
| Codon mutations: | GGC32GGT (277C>T), TCT33TCC (280T>C), GGC34GGT (283C>T), AGT38AGC (295T>C), CCA40CCT (301A>T), TTC44TTT (313C>T), ATT54ATC (343T>C), AAC63AAT (370C>T), AAT64ACT (372A>C), AGC65AGT (376C>T), GTT69GTA (388T>A), TAT73TAC (400T>C), TTA74CTA (401T>C), ACT79ACG (418T>G), GGC84GGT (433C>T), GGC89GGA (448C>A), CAT93CAC (460T>C), ACC100ACA (481C>A), AGA102AGG (487A>G), AGG109AAG (507G>A), ATT110ATA (511T>A), CAA128CAT (565A>T), CCA129CCT (568A>T), ACG132AAG (576C>A), ACA134ACT (583A>T), CAA135CAG (586A>G), TTA136CTT (587T>C 589A>T), GAC141GCC (603A>C), GAG142GAA (607G>A), AAA144AAG (613A>G), TTA147TTG (622A>G), CTT155GTT (644C>G), CTA158CAA (654T>A), TTG166TTA (679G>A), CTA172CCA (696T>C), GGC180GGT (721C>T), GAA184GAG (733A>G) | | | | | | | | |
| polyprotein (YP_001040018.1) | 1 | 1199 | 100% | 7663 | 92.6% | 1199 (100%) | 1088 (90.7%) | 0/0/0/0 | 2 |

| | Begin | End | Coverage | Score | Concordance | Matches | Identities | I/D/M/F* | Stop Codons |
|---|---|---|---|---|---|---|---|---|---|
| **NT** | **20** | **5403** | **99.6%** | **8032** | **74.6%** | **5384 (99.9%)** | **4705 (87.4%)** | **2/0** | |

| | |
|---|---|
| Protein mutations: | A7V (721C>T), K11R (733A>G), K16R (748A>G), S17G (750A>G), Y48F (844A>T), V132I (1095G>A), T228A (1383A>G 1385C>T), S230* (1390C>G), T232S (1395A>T 1397T>A), Q242P (1426A>C 1427G>T), D245N (1434G>A), S252T (1456G>C 1457C>T), A254S (1461G>T), R255K (1465G>A), I257M (1472C>G), V264I (1491G>A 1493G>A), S275P (1524T>C 1526C>T), S281F (1543C>T), T282A (1545A>G 1547C>T), H286Q (1559C>A), Q298D (1593C>G 1595G>T), S301N (1603G>A), R302G (1605A>G), Y303Q (1608T>C 1610C>G), G307A (1621G>C 1622T>A), K317R (1651A>G), L321Y (1662C>T 1663T>A 1664G>T), H325Y (1674C>T 1676T>C), V326I (1677G>A 1679G>T), I332V (1695A>G), V365I (1794G>A), C366G (1797T>G 1799C>G), K370N (1811A>C), I373K (1819T>A 1820C>G), K377N (1832G>C), H381R (1843A>G), H388Y (1863C>T 1865C>T), V392M (1875G>A), I399M (1898T>G), V400T (1899G>A 1900T>C 1901G>A), A402V (1906C>T 1907C>G), E409D (1928A>C), N413D (1938A>G), D429N (1986G>A), F434Y (2002T>A), L450I (2049T>A 2051A>T), K453S (2059A>G 2060A>C), L458Q (2074T>A), V461A (2083T>C 2084G>A), Q465D (2094C>G 2096G>C), A466E (2098C>A 2099G>A), R467M (2100C>A 2101G>T 2102C>G), R469E (2106A>G 2107G>A), F470L (2111T>G), K472R (2115A>C 2116A>G), E473A (2119A>C 2120G>A), A475V (2125C>T 2126C>T), R476V (2127A>G 2128G>T), A477T (2130G>A 2132T>C), V479L (2136G>C 2138G>T), N481K (2144C>G), V488G (2164T>G), E490V (2170A>T 2171A>T), S493A (2178T>G), V500T (2199G>A 2200T>C), M502V (2205A>G), D508E (2225T>A), M510D (2229A>G 2230T>A 2231G>T), E517D (2252A>C), A519T (2256G>A), N520Q (2259A>C 2261T>A), E521Q (2262G>C), S579A (2436T>G), S590T (2469T>A 2471C>A), R601K (2503G>A), I608V (2523A>G), A700T (2799G>A), I712V (2835A>G 2837C>T), A761T (2982G>A), I771V (3012A>G 3014C>T), I807V (3120A>G 3122T>G), S829N (3187G>A), T832A (3195A>G 3197C>T), K851R (3253A>G), R855K (3265G>A), T857S (3270A>T 3272G>T), T868S (3303A>T 3305C>T), N936S (3508A>G), S948T (3543T>A), K955Q (3564A>C), L956M (3567T>A 3569A>G), P977Q (3631C>A), L978P (3634T>C), Q1014P (3742A>C 3743G>C), T1051A (3852A>G 3854T>G), I1061V (3882A>G 3884C>A), T1062A (3885A>G), K1076R (3927A>C 3928A>G 3929G>A), T1078A (3933A>G 3935C>T), A1087T (3960G>A), M1095V (3984A>G 3986G>T), S1096N (3988G>A 3989C>T), D1097S (3990G>A 3991A>G), H1105Q (4016T>G), I1120V (4059A>G 4061A>T), N1123G (4068A>G 4069A>G 4070T>C), A1124V (4072C>T 4073T>G), L1150I (4149C>A 4151G>A), Q1163E (4188C>G), H1175Y (4224C>T), S1195G (4284A>G 4286T>A) |
| Codon mutations: | GCT7GTT (721C>T), AAG11AGG (733A>G), AAA16AGA (748A>G), AGT17GGT (750A>G), GTT32GTC (797T>C), GGC35GGT (806C>T), GGC44GGT (833C>T), TAT48TTT (844A>T), AAT70AAC (911T>C), TCT76TCA (929T>A), TCA77TCT (932A>T), GGA89GGG (968A>G), TTG95TTA (986G>A), GTT98GTC (995T>C), GTA113GTC (1040A>C), AAA114AAG (1043A>G), TTA117TTG (1052A>G), AGT120AGC (1061T>C), ACC124ACA (1073C>A), CTG128TTG (1083C>T), AAG131AAA (1094G>A), GTT132ATT (1095G>A), TTG133TTA (1100G>A), TCC136TCT (1109C>T), GGG137GGT (1112G>T), GGG138GGA (1115G>A), GCA139GCC (1118A>C), TTA143CTG (1128T>C 1130A>G), CCG146CCA (1139G>A), ACC147ACT (1142C>T), CTA150CTG (1151A>G), AAA154AAG (1163A>G), GGC156GGG (1169C>G), TTC159TTT (1178C>T), ACA162ACT (1187A>T), ACT168ACA (1205T>A), CCT173CCC (1220T>C), GCC178GCA (1235C>A), CAA182CAG (1247A>G), CGT187CGC (1262T>C), GCC189GCT (1268C>T), TCA192TCG (1277A>G), CTT204CTC (1313T>C), CGA206AGG (1317C>A 1319A>G), CAA208CAG (1325A>G), TTT209TTC (1328T>C), TTC210TTT (1331C>T), CAG214CAA (1343G>A), CAG218CAA (1355G>A), ATA220ATT (1361A>T), TCA222TCT (1367A>T), GTA223GTT (1370A>T), TCT225TCA (1376T>A), ACC228GCT (1383A>G 1385C>T), TCC229TCT (1388C>T), TCA230TGA (1390C>G), GCA231GCT (1394A>T), ACT232TCA (1395A>T 1397T>A), GAT233GAC (1400T>C), TCA234TCT (1403A>T), GTG235GTA (1406G>A), CTT236CTA (1409T>A), CCG237CCT (1412G>T), GTG239GTA (1418G>A), CAG242CCT (1426A>C 1427G>T), GAC245AAC (1434G>A), ACA246ACC (1439A>C), GTG250GTT (1451G>T), CCA251CCT (1454A>T), AGC252ACT (1456G>C 1457C>T), GAA253GAG (1460A>G), GCA254TCA (1461G>T), AGA255AAA (1465G>A), ATC257ATG (1472C>G), CGG258CGC (1475G>C), TCC259TCG (1478C>G), TCC260TCT (1481C>T), CTC262CTG (1487C>G), CGG263CGA (1490G>A), GTG264ATA (1491G>A 1493G>A), CCC265CCT (1496C>T), CTC267CTT (1502C>T), AGT271AGC (1514T>C), TCC275CCT (1524T>C 1526C>T), TCA276TCT (1529A>T), ATA279ATT (1538A>T), TCT281TTT (1543C>T), ACC282GCT (1545A>G 1547C>T), ACT284ACC (1553T>C), CAC286CAA (1559C>A), CTC287TTA (1560C>T 1562C>A), GGC289GGG (1568C>G), ACG291ACA (1574G>A), CGT292AGG (1575C>A 1577T>G), GTA296GTG (1589A>G), CAG298GAT (1593C>G 1595G>T), AGT301AAT (1603G>A), AGG302GGG (1605A>G), TAC303CAG (1608T>C 1610C>G), GAA304GAG (1613A>G), GAG305GAA (1616G>A), GGT307GCA (1621G>C 1622T>A), GGG310GGA (1631G>A), GTT311GTA (1634T>A), GTT315GTC (1646T>C), AAG317AGG (1651A>G), GCT318GCA (1655T>A), GGT320GGC (1661T>C), CTG321TAT (1662C>T 1663T>A 1664G>T), AAC322AAT (1667C>T), CAT325TAC (1674C>T 1676T>C), GTG326ATT (1677G>A 1679G>T), GAT328GAC (1685T>C), AAT329AAC (1688T>C), CTT330CTG (1691T>G), ATT332GTT (1695A>G), GAG333GAA (1700G>A), CTG340CTC (1721G>C), CCC341CCT (1724C>T), GAA342GAG (1727A>G), AAC343AAT (1730C>T), CTG347TTG (1740C>T), TCG348TCA (1745G>A), GTG349GTA (1748G>A), AGA350CGA (1749A>C), CAG351CAA (1754G>A), CAA352CAG (1757A>G), GTT353GTG (1760T>G), GCT354GCA (1763T>A), GCA355GCG (1766A>G), GCA356GCT (1769A>T), GCA359GCC (1778A>C), GCA361GCC (1784A>C), AAG363AAA (1790G>A), GGT364GGA (1793T>A), GTT365ATT (1794G>A), TGC366GGG (1797T>G 1799C>G), GTC368GTT (1805C>T), AAA370AAC (1811A>C), CAT371CAC (1814T>C), ATA372ATT (1817A>T), ATC373AAG (1819T>A 1820C>G), AAC374AAT (1823C>T), AAG377AAC (1832G>C), ATC380ATT (1841C>T), CAC381CGC (1843A>G), TTG382CTT (1845T>C 1847G>T), GAG383GAA (1850G>A), ATC385ATT (1856C>T), CAC388TAT (1863C>T 1865C>T), GTG392ATG (1875G>A), CTG394TTG (1881C>T), TAC397TAT (1892C>T), ATT399ATG (1898T>G), GTG400ACA (1899G>A 1900T>C 1901G>A), GCC402GTG (1906C>T 1907C>G), TCT404TCA (1913T>A), GAA409GAC (1928A>C), AAC410AAT (1931C>T), AAT413GAT (1938A>G), CTT414TTG (1941C>T 1943T>G), ACC415ACA (1946C>A), TTG416CTG (1947T>C), GCA417GCC (1952A>C), CAG418CAA (1955G>A), AGA419AGG (1958A>G), GAG422GAA (1967G>A), CTA423CTG (1970A>G), CAA426CAG (1979A>G), GAT429AAT (1986G>A), TAT431TAC (1994T>C), TTT434TAT (2002T>A), AAC437AAT (2012C>T), TAT441TAC (2024T>C), CCT442CCA (2027T>A), TTA450ATT (2049T>A 2051A>T), ACG451ACT (2054G>T), AAA453AGC (2059A>G 2060A>C), CGC455CGG (2066C>G), CTA458CAA (2074T>A), GTG461GCA (2083T>C 2084G>A), GCA462GCC (2087A>C), CAG465GAC (2094C>G 2096G>C), GCG466GAA (2098C>A 2099G>A), CGC467ATG (2100C>A 2101G>T 2102C>G), AGA469GAA (2106A>G 2107G>A), TTT470TTG (2111T>G), TAC471TAT (2114C>T), AAG472CGG (2115A>C 2116A>G), GAG473GCA (2119A>C 2120G>A), GCG474GCT (2123G>T), GCC475GTT (2125C>T 2126C>T), AGG476GTG (2127A>G 2128G>T), GCT477ACC (2130G>A 2132T>C), CAG478CAA (2135G>A), GTG479CTT (2136G>C 2138G>T), AAC481AAG (2144C>G), ACT486ACA (2159T>A), AGT487AGC (2162T>C), GTG488GGG (2164T>G), GAA490GTT (2170A>T 2171A>T), ATA491ATT (2174A>T), CCA492CCT (2177A>T), TCC493GCC (2178T>G), ACC494ACT (2183C>T), TCA495TCC (2186A>C), TTC496TTT (2189C>T), ACA498ACC (2195A>C), AAG499AAA (2198G>A), GTT500ACT (2199G>A 2200T>C), GCC501GCT (2204C>T), ATG502GTG (2205A>G), GAG503GAA (2210G>A), GGC505GGG (2216C>G), AGT506AGC (2219T>C), GAT508GAA (2225T>A), AGC509AGT (2228C>T), ATG510GAT (2229A>G 2230T>A 2231G>T), GCT513GCA (2240T>A), ATT514ATC (2243T>C), GAA517GAC (2252A>C), GCC519ACC (2256G>A), AAT520CAA (2259A>C 2261T>A), GAA521CAA (2262G>C), GTA523GTG (2270A>G), GTT528GTC (2285T>C), CAA534CAG (2303A>G), AAT537AAC (2312T>C), GGT540GGC (2321T>C), TTC558TTT (2375C>T), GAG564GAA (2393G>A), AGT571AGC (2414T>C), GTT573GTA (2420T>A), GAG576GAA (2429G>A), GTG577GTT (2432G>T), CTA578TTA (2433C>T), TCA579GCA (2436T>G), GCG581GCT (2444G>T), CTC582CTT (2447C>T), CCT583CCC (2450T>C), GCG584GCA (2453G>A), CTC586CTG (2459C>G), GGT588GGG (2465T>G), AAA589AAG (2468A>G), TCC590ACA (2469T>A 2471C>A), AAG591AAA (2474G>A), GGA596GGC (2489A>C), TCG597TCA (2492G>A), CAA598CAG (2495A>G), CTA600CTG (2501A>G), AGG601AAG (2503G>A), TAT602TAC (2507T>C), GCC605GCT (2516C>T), ATT608GTT (2523A>G), TAT615TAC (2546T>C), GGT620GGA (2561T>A), ATT622ATA (2567T>A), GGT624GGC (2573T>C), CTC636CTG (2609C>G), TTC642TTT (2627C>T), ATC643ATA (2630C>A), CTG648CTA (2645G>A), TAT649TAC (2648T>C), GCC650GCT (2651C>T), CTG655CTA (2666G>A), GTT656GTG (2669T>G), TCA657TCC (2672A>C), GCA658GCT (2675A>T), GGG664GGA (2693G>A), TGT667TGC (2702T>C), ACA669ACG (2708A>G), GGT672GGC (2717T>C), ATC673ATA (2720C>A), TTC676TTT (2729C>T), GTT677GTG (2732T>G), GCC682GCA (2747C>A), GGG684GGT (2753G>T), ATA691ATT (2774A>T), CAC697CAT (2792C>T), CCC698CCG (2795C>G), TTG699TTA (2798G>A), GCT700ACT (2799G>A), AGT701AGC (2804T>C), TTG705CTG (2814T>C), ATC712GTT (2835A>G 2837C>T), CAT713CAC (2840T>C), CTG714TTG (2841C>T), CAG715CAA (2846G>A), AAA717AAG (2852A>G), TCG720TCA (2861G>A), ACC721ACA (2864C>A), AAC722AAT (2867C>T), ATC723ATA (2870C>A), CCT726CCA (2879T>A), CCT727CCA (2882T>A), TTG729CTA (2886T>C 2888G>A), CGA730CGG (2891A>G), GCA731GCG (2894A>G), GCA733GCT (2900A>T), GGT736GGC (2909T>C), AAA738AAG (2915A>G), GAC740GAT (2921C>T), ACA742ACG (2927A>G), CCA745CCC (2936A>C), CGA746CGG (2939A>G), ACA749ACG (2948A>G), CAG751CAA (2954G>A), GTA752GTT (2957A>T), CTG753CTA (2960G>A), CAC755CAT (2966C>T), AAT757AAC (2972T>C), GTG760GTT (2981G>T), GCT761ACT (2982G>A), GGT765GGG (2996T>G), ACT766ACA (2999T>A), TTA768CTA (3003T>C), AGC770AGT (3011C>T), ATC771GTT (3012A>G 3014C>T), TCT773TCC (3020T>C), CCG774CCC (3023G>C), GGC776GGT (3029C>T), ATA777ATT (3032A>T), GAC781GAT (3044C>T), GGT782GGA (3047T>A), ACG783ACC (3050G>C), CTG784TTG (3051C>T), CCA786CCT (3059A>T), TCC787TCT (3062C>T), TTG788CTA (3063T>C 3065G>A), GGA790GGG (3071A>G), ATA792ATT (3077A>T), CGC794CGA (3083C>A), ACT800ACG (3101T>G), GGC801GGT (3104C>T), ACT802ACG (3107T>G), GTG804GTC (3113G>C), ATT807GTG (3120A>G 3122T>G), TGT808TGC (3125T>C), TTC814TTT (3143C>T), CAT815CAC (3146T>C), GGT817GGA (3152T>A), TTG819TTA (3158G>A), GCC820GCA (3161C>A), ATT821ATC (3164T>C), GGA822GGT (3167A>T), CTG823CTT (3170G>T), GGT824GGC (3173T>C), ACA825ACC (3176A>C), CTG826CTT (3179G>T), AAC827AAT (3182C>T), AGC829AAC (3187G>A), TCC831TCT (3194C>T), ACC832GCT (3195A>G 3197C>T), CCT833CCC (3200T>C), GAC835GAT (3206C>T), GTT842GTG (3227T>G), ATT843ATA (3230T>A), AAC845AAT (3236C>T), CTT846CTG (3239T>G), GAG847GAA (3242G>A), GGA849GGC (3248A>C), CGA850CGT (3251A>T), AAG851AGG (3253A>G), AGG855AAG (3265G>A), ACG857TCT (3270A>T 3272G>T), ACC859ACT (3278C>T), GGG863GGC (3290G>C), AAA864AAG (3293A>G), AAT865AAC (3296T>C), CTT866TTA (3297C>T 3299T>A), ACC868TCT (3303A>T 3305C>T), GGT870GGA (3311T>A), CTA875TTA (3324C>T), CCG876CCT (3329G>T), CAC883CAT (3350C>T), AGG885CGG (3354A>C), TTG886TTA (3359G>A), ACA889ACG (3368A>G), GTC890GTG (3371C>G), TTG891CTG (3372T>C), CTG894CTA (3383G>A), GTA895GTT (3386A>T), TCA898TCC (3395A>C), CTA901CTG (3404A>G), ACG903ACA (3410G>A), GTC904GTT (3413C>T), CAG909CAA (3428G>A), CTT910TTG (3429C>T 3431T>G), AAA911AAG (3434A>G), TGC912TGT (3437C>T), ATA913ATT (3440A>T), TTG916CTA (3447T>C 3449G>A), GTC917GTA (3452C>A), CTT918CTA (3455T>A), GGG919GGA (3458G>A), GGC920GGT (3461C>T), TCT923TCC (3470T>C), GTT924GTC (3473T>C), CCC926CCT (3479C>T), ATT927ATC (3482T>C), TAC928TAT (3485C>T), GGA929GGG (3488A>G), CAC930CAT (3491C>T), GGA934GGG (3503A>G), AAT936AGT (3508A>G), GTG938GTT (3515G>T), CTA941TTG (3522C>T 3524A>G), TTC942TTT (3527C>T), TCT943TCA (3530T>A), GAG944GAA (3533G>A), TCT948ACT (3543T>A), CGC950CGT (3551C>T), ATT953ATC (3560T>C), AAA955CAA (3564A>C), TTA956ATG (3567T>A 3569A>G), AAG958AAA (3575G>A), GAA959GAG (3578A>G), ACA963ACC (3590A>C), GGA968GGG (3605A>G), AAG970AAA (3611G>A), CCA972CCT (3617A>T), CAG973CAA (3620G>A), GTA974GTT (3623A>T), GTG976GTC (3629G>C), CCG977CAG (3631C>A), CTC978CCC (3634T>C), AGA979AGG (3638A>G), GAC980GAT (3641C>T), TCA983TCT (3650A>T), GGG985GGT (3656G>T), GCT986GCC (3659T>C), GTA987GTG (3662A>G), GTG991GTC (3674G>C), GAC996GAT (3689C>T), CGA999CGC (3698A>C), TTG1001TTA (3704G>A), GTT1002GTG (3707T>G), TTA1003CTA (3708T>C), CCA1004CCC (3713A>C), TGC1005TGT (3716C>T), CCC1007CCA (3722C>A), TCC1009TCG (3728C>G), TTC1012TTT (3737C>T), CAG1014CCC (3742A>C 3743G>C), GGG1015GGA (3746G>A), GCA1016GCT (3749A>T), CTG1017CTA (3752G>A), GTA1018GTG (3755A>G), ACA1023ACC (3770A>C), TGC1024TGT (3773C>T), CCA1025CCT (3776A>T), TTC1026TTT (3779C>T), TTC1033TTT (3800C>T), GGA1038GGC (3815A>C), TAC1042TAT (3827C>T), ATT1044ATA (3833T>A), GTA1046GTG (3839A>G), CAT1047CAC (3842T>C), AGA1048AGG (3845A>G), GTG1049GTA (3848G>A), CAG1050CAA (3851G>A), ACT1051GCG (3852A>G 3854T>G), TCC1052TCA (3857C>A), ATA1055ATC (3866A>C), GGA1056GGT (3869A>T), GGA1057GGT (3872A>T), CTG1059CTC (3878G>C), AAC1060AAT (3881C>T), ATC1061GTA (3882A>G 3884C>A), ACC1062GCC (3885A>G), TTA1063TTG (3890A>G), GAT1064GAC (3893T>C), TCA1065TCT (3896A>T), TAC1068TAT (3905C>T), TTT1070TTC (3911T>C), CCT1071CCA (3914T>A), GGA1073GGT (3920A>T), ATC1074ATT (3923C>T), TCA1075TCC (3926A>C), AAG1076CGA (3927A>C 3928A>G 3929G>A), GGC1077GGA (3932C>A), ACC1078GCT (3933A>G 3935C>T), GTT1080GTG (3941T>G), GTC1081GTA (3944C>A), TCT1082TCC (3947T>C), GCT1083GCA (3950T>A), GGT1084GGA (3953T>A), GCA1087ACA (3960G>A), GCT1090GCA (3971T>A), GGT1094GGA (3983T>A), ATG1095GTT (3984A>G 3986G>T), AGC1096AAT (3988G>A 3989C>T), GAC1097AGC (3990G>A 3991A>G), AAC1098AAT (3995C>T), ATC1099ATT (3998C>T), TTT1100TTC (4001T>C), CAT1105CAG (4016T>G), GAG1108GAA (4025G>A), TTT1109TTC (4028T>C), TTT1110TTC (4031T>C), CCT1111CCG (4034T>G), AGA1112CGT (4035A>C 4037A>T), CGC1113CGT (4040C>T), GCC1117GCT (4052C>T), ATA1120GTT (4059A>G 4061A>T), CCA1122CCT (4067A>T), AAT1123GGC (4068A>G 4069A>G 4070T>C), GCT1124GTG (4072C>T 4073T>G), AGA1126CGA (4077A>C), ACT1129ACA (4088T>A), CTG1130CTA (4091G>A), CGA1133CGC (4100A>C), CTA1134CTT (4103A>T), GGA1135GGC (4106A>C), CTC1137CTG (4112C>G), TTA1141TTG (4124A>G), AAA1144AAG (4133A>G), ATA1147ATC (4142A>C), AGC1148AGT (4145C>T), CTG1150ATA (4149C>A 4151G>A), AAG1155AAA (4166G>A), TTC1160TTT (4181C>T), TTG1162TTA (4187G>A), CAA1163GAA (4188C>G), GCT1165GCA (4196T>A), TCA1169TCT (4208A>T), GCA1170GCT (4211A>T), AAG1174AAA (4223G>A), CAT1175TAT (4224C>T), TTG1176CTT (4227T>C 4229G>T), ACG1181ACA (4244G>A), ACC1183ACG (4250C>G), TAC1184TAT (4253C>T), CTT1185CTG (4256T>G), ACT1187ACC (4262T>C), CCT1188CCA (4265T>A), TCA1191TCT (4274A>T), AGG1194CGG (4281A>C), AGT1195GGA (4284A>G 4286T>A), TTT1196TTC (4289T>C) |

*: Inserts / Deletes / Misaligned / Frameshifts

#### Analysis details

This analysis was performed with panviral2.64

### NGS Details (UN70): Potato virus Y

#### Assembly

| | |
|---|---|
| **Coverage Length** | 9686 (1 contig(s)) |
| **Depth Of Coverage** | 2313.2 |
| **Number Of Reads** | 167938 |
| **Reads Per Million** | 3183.48 rpm (after QC) |
| **Ambiguities** | 0 |
| **Assembly Method** | de novo + reference guided assembly |
| **Consensus Caller** | Bcf Tools |

#### Coverage Map

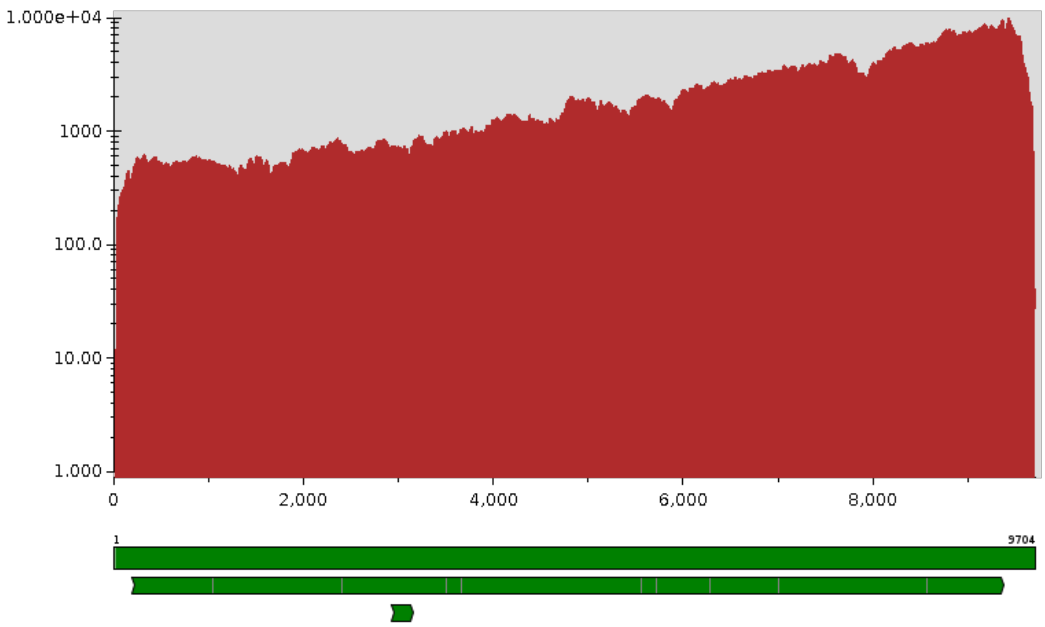

#### Assignment

| | |
|---|---|
| **Type** | Potato virus Y (Taxonomy ID: 12216) |
| **Reference Genome** | NC_001616.1 |
| **NT Identity (%)** | 82.6814 |
| **AA Identity (%)** | 90.1274 |
| **Number Of Stop Codons** | 3 |
| **Number Of CDS** | 2 |

#### Alignment

| | |
|---|---|
| **Alignment Score** | 12534.0 (NT) + 19051.0 (AA) = 31585.0 |
| **Concordance (%)** | 78.2307 |

| Alignment Method | Global, seeded, nucleotide + amino acids (AGA) |
|---|---|

#### Genome Region

Sequence starts at position 19 and ends at position 9704 relative to NC_001616.1 reference sequence.

#### Alignment Detailed Statistics

| | Begin | End | Coverage | Score | Concordance | Matches | Identities | I/D/M/F* | Stop Codons |
|---|---|---|---|---|---|---|---|---|---|
| NT | 19 | 9704 | 99.8% | 12534 | 65.0% | 9674 (99.7%) | 8011 (82.6%) | 15/12 | |

Mutations:

19C>T, 23A>T, 32A>T, 42_43insGAAATAT, 45A>T, 47T>A, 50T>G, 53A>G, 54C>T, 55G>T, 56C>T, 58C>T, 60T>A, 60_61insA, 75C>T, 79C>T, 83G>A, 85T>C, 87C>T, 89G>A, 90T>C, 93A>T, 94A>G, 95A>T, 100T>C, 101T>A, 102C>T, 104T>C, 107C>T, 110T>G, 112C>T, 114C>T, 115T>C, 116A>T, 117G>C, 119A>G, 120C>T, 121A>T, 125T>G, 126T>C, 127G>A, 128G>T, 131A>G, 132C>T, 139A>T, 139_140insG, 141C>T, 143C>T, 146C>T, 150C>T, 150_151insT, 154T>C, 156C>A, 158T>A, 160A>C, 164C>T, 168C>A, 169C>G, 171A>C, 172T>C, 173T>A, 174T>G, 177G>C, 178A>G, 179T>C, 180C>T, 181C>A, 182T>G, 183C>T, 193T>C, 194T>C, 196C>A, 202A>G, 208C>T, 209T>C, 210G>T, 211T>G, 220G>A, 221T>A, 223T>G, 226A>G, 235A>T, 241C>T, 247A>T, 251T>C, 255G>T, 256C>T, 258A>G, 262T>C, 263A>G, 265T>G, 267T>C, 273A>T, 277A>T, 280A>G, 283G>T, 286G>A, 287G>A, 289T>C, 293G>A, 294T>C, 301T>C, 304C>T, 305G>A, 306C>T, 309A>G, 311C>T, 316A>T, 317A>G, 318C>T, 319A>G, 322A>G, 331A>T, 339T>A, 340C>A, 342A>G, 347A>G, 348A>C, 349A>G, 350T>C, 355T>A, 356A>G, 357C>T, 364T>C, 365G>A, 366T>C, 367G>A, 368C>T, 369T>C, 373A>G, 376C>T, 383T>C, 384T>G, 388G>A, 391C>T, 394A>G, 397C>T, 400G>A, 406T>C, 408C>T, 409C>T, 412G>A, 415A>T, 416A>G, 417T>A, 424T>C, 427G>A, 433A>G, 436G>C, 437G>A, 440A>G, 441G>A, 442G>A, 443A>C, 444A>G, 445G>A, 448T>A, 454A>G, 456A>C, 460T>C, 461C>A, 469A>G, 475C>T, 481T>A, 502T>C, 505A>T, 511C>T, 517T>A, 520T>A, 523A>T, 531C>T, 535G>A, 536C>T, 539C>T, 540A>G, 541A>G, 549G>A, 550A>G, 556C>T, 559T>C, 562T>C, 568T>C, 571A>G, 580T>C, 585T>C, 586C>G, 593C>T, 594G>A, 595C>T, 597C>G, 599A>G, 601A>G, 602A>C, 603T>C, 604A>G, 613A>G, 616A>G, 619C>G, 622G>A, 625G>A, 628T>C, 631C>T, 635A>G, 637T>C, 644A>G, 655T>C, 661G>A, 663A>C, 664G>C, 667A>G, 673G>C, 679C>T, 682C>A, 683T>C, 691T>C, 699C>G, 703T>A, 706T>A, 719A>G, 721G>A, 722A>G, 724A>G, 725C>T, 727T>G, 730T>A, 731G>T, 734T>C, 735A>C, 736C>T, 737T>C, 738C>G, 739C>T, 744C>T, 745G>C, 749C>T, 751A>C, 754T>C, 765T>A, 770T>C, 773C>A, 774G>A, 775A>G, 776C>A, 790C>T, 801T>A, 813G>C, 814A>G, 816T>A, 818T>C, 823A>G, 825G>A, 826T>A, 833C>A, 835G>A, 841C>T, 853T>C, 859C>T, 860C>T, 862C>T, 863A>G, 865T>C, 867T>C, 869A>G, 871C>T, 872A>T, 874A>G, 879G>A, 886T>C, 898C>T, 903A>G, 906C>A, 907A>C, 909A>C, 910A>C, 912G>A, 913C>T, 922C>A, 931T>A, 938T>C, 939C>T, 940A>T, 943A>G, 952C>T, 967A>G, 973A>G, 976G>C, 980T>A, 982G>C, 991A>G, 994T>C, 999G>A, 1003T>C, 1007C>A, 1008A>G, 1009G>A, 1010A>G, 1012T>G, 1013A>G, 1018A>G, 1019A>C, 1021C>G, 1030C>T, 1039G>A, 1042T>C, 1045C>A, 1048C>A, 1051T>C, 1060G>A, 1063T>A, 1069C>T, 1083G>A, 1093T>C, 1097T>G, 1099G>A, 1105C>T, 1114A>G, 1117T>G, 1123A>G, 1126T>A, 1129C>T, 1143G>A, 1147A>T, 1150T>A, 1154T>A, 1156G>A, 1165C>T, 1168T>C, 1172C>T, 1174T>A, 1201C>T, 1221G>A, 1229G>A, 1230T>C, 1231T>C, 1233G>A, 1234C>T, 1240G>A, 1246G>A, 1250T>C, 1252G>A, 1255T>C, 1257A>C, 1258A>G, 1267A>G, 1273T>C, 1276G>A, 1278A>G, 1283T>C, 1290C>T, 1291G>A, 1298G>A, 1299A>C, 1300C>T, 1306T>C, 1307A>G, 1309A>G, 1312T>C, 1316A>G, 1318T>A, 1327G>A, 1329T>C, 1333G>T, 1336A>G, 1343C>T, 1345A>G, 1354G>A, 1357G>T, 1360C>T, 1367C>T, 1369C>G, 1373C>A, 1378C>T, 1384G>A, 1385A>G, 1393A>G, 1396C>T, 1405G>T, 1408A>G, 1411G>A, 1415G>T, 1417A>T, 1420G>C, 1429T>C, 1432A>G, 1436G>A, 1438C>T, 1439T>C, 1441A>G, 1444T>C, 1453C>T, 1456G>T, 1468A>G, 1499T>C, 1501G>A, 1504T>C, 1507G>A, 1510C>T, 1513A>G, 1516A>G, 1519A>T, 1522G>A, 1525C>T, 1528G>A, 1534C>T, 1540T>A, 1546C>T, 1555A>G, 1561T>C, 1564A>C, 1567T>A, 1573C>T, 1576A>G, 1583T>C, 1588T>A, 1594G>A, 1600C>T, 1609G>A, 1618G>A, 1621C>T, 1624C>T, 1627C>T, 1631C>T, 1639A>G, 1651C>T, 1654C>G, 1675T>C, 1693A>G, 1702T>C, 1707A>G, 1718G>A, 1726A>G, 1729C>T, 1732A>C, 1738T>C, 1742A>G, 1744C>T, 1747C>T, 1753T>C, 1758G>A, 1768G>A, 1786C>T, 1789A>G, 1790G>A, 1795C>T, 1805T>C, 1807A>G, 1813G>A, 1819G>A, 1825G>A, 1837C>T, 1840T>C, 1849A>G, 1858C>G, 1864A>G, 1870C>T, 1876T>C, 1882A>G, 1888T>A, 1891T>C, 1906T>C, 1915A>T, 1930T>C, 1936C>T, 1937A>G, 1939T>C, 1945A>G, 1948A>G, 1951C>T, 1960A>T, 1978A>G, 1981T>A, 1984C>T, 1987T>C, 1996C>T, 2004T>A, 2020A>G, 2023G>A, 2026T>C, 2029G>T, 2032G>A, 2059T>C, 2068T>G, 2071C>T, 2072G>A, 2074G>A, 2087C>T, 2089G>A, 2101C>T, 2104G>A, 2107G>A, 2116G>A, 2122C>T, 2137C>T, 2158T>C, 2164C>T, 2170A>G, 2182T>C, 2188G>A, 2191C>A, 2208G>A, 2218T>C, 2224C>T, 2227A>T, 2242G>A, 2245C>T, 2254G>A, 2257T>C, 2260C>T, 2266C>G, 2272A>G, 2275G>A, 2293A>G, 2299C>T, 2302G>A, 2305G>A, 2314A>T, 2317T>C, 2323T>C, 2326A>G, 2341G>A, 2347A>G, 2350T>C, 2356G>A, 2365T>C, 2371A>G, 2374A>G, 2383T>C, 2386A>T, 2389A>G, 2392T>C, 2395T>C, 2414A>G, 2415A>G, 2416T>G, 2419T>A, 2420A>T, 2440A>T, 2467T>C, 2476T>C, 2482G>A, 2491G>A, 2497G>T, 2503G>A, 2518A>G, 2521T>C, 2523A>G, 2533A>G, 2539G>A, 2545A>G, 2551G>A, 2554T>A, 2558C>T, 2561T>C, 2563G>C, 2569A>G, 2572A>G, 2584T>C, 2587C>T, 2593G>C, 2599T>C, 2608T>C, 2620T>C, 2629G>A, 2632G>A, 2634G>A, 2638G>A, 2644T>A, 2653A>G, 2665T>C, 2678C>T, 2683A>G, 2689T>C, 2698A>C, 2707A>G, 2710G>A, 2716A>G, 2719A>G, 2722C>T, 2723G>A, 2728A>G, 2734G>A, 2737T>A, 2746T>C, 2747G>A, 2749T>A, 2752A>T, 2755T>A, 2761C>T, 2764C>T, 2779T>C, 2785G>T, 2791C>T, 2795C>A, 2806C>T, 2816T>C, 2818G>C, 2822G>A, 2824G>A, 2827G>A, 2836T>C, 2839G>A, 2842T>C, 2863C>A, 2864C>T, 2882C>A, 2885A>C, 2886G>A, 2893C>T, 2895C>T, 2902C>T, 2905A>G, 2911T>C, 2931A>G, 2940A>G, 2947T>C, 2971_2972insA, 2989T>C, 2991C>T, 2994A>G, 2999G>A, 3000A>G, 3016delC, 3021G>A, 3024A>T, 3025C>T, 3036C>T, 3039A>G, 3040A>C, 3042A>C, 3043G>A, 3052G>A, 3061A>G, 3082G>A, 3091C>T, 3099C>T, 3100C>A, 3104G>A, 3105T>G, 3106G>A, 3109C>T, 3112A>G, 3118T>C, 3121C>T, 3127A>G, 3132G>A, 3137C>A, 3138G>A, 3139A>G, 3151T>C, 3152T>A, 3154C>T, 3165G>A, 3169A>G, 3199C>T, 3203A>C, 3208T>C, 3214A>G, 3217C>T, 3218T>C, 3223C>A, 3230G>A, 3235C>T, 3257T>C, 3262C>T, 3265C>T, 3266G>A, 3268A>G, 3271G>A, 3275G>A, 3284G>A, 3301G>A, 3315G>A, 3316A>G, 3318A>C, 3325G>A, 3328G>A, 3337A>T, 3349G>A, 3355C>T, 3358C>T, 3359A>G, 3367A>G, 3370T>C, 3377T>C, 3379A>G, 3394A>G, 3397C>T, 3400T>G, 3406A>G, 3418C>T, 3421T>A, 3427T>C, 3433G>A, 3436A>T, 3439A>G, 3442C>T, 3445T>C, 3457G>A, 3460T>C, 3463T>A, 3469G>T, 3472G>A, 3496C>T, 3499G>A, 3505C>T, 3513T>G, 3520A>G, 3529A>G, 3532A>G, 3545A>G, 3548G>A, 3551T>C, 3554G>A, 3556C>T, 3583T>C, 3586T>C, 3589C>T, 3595C>T, 3607T>C, 3613T>C, 3616G>A, 3619T>C, 3623C>G, 3628C>T, 3632C>A, 3637T>C, 3638T>C, 3643A>G, 3661A>G, 3670G>A, 3682T>C, 3688G>A, 3697G>A, 3698A>G, 3706T>C, 3709T>C, 3713T>C, 3715G>A, 3717G>A, 3721G>A, 3731C>A, 3732G>A, 3736T>A, 3746C>T, 3748A>G, 3754A>G, 3763T>C, 3778A>G, 3781A>G, 3784C>T, 3796T>C, 3799A>G, 3805A>T, 3811C>T, 3814A>G, 3829C>T, 3838T>C, 3844A>G, 3847A>C, 3856T>A, 3862G>T, 3871C>T, 3877C>A, 3883C>T, 3886A>G, 3892A>G, 3895C>T, 3898T>C, 3904A>G, 3905C>A, 3907G>A, 3913T>C, 3928G>A, 3931A>T, 3937G>A, 3946T>C, 3955T>C, 3958G>A, 3961C>T, 3964A>G, 3967T>C, 3976A>G, 3979T>C, 3985A>T, 3997A>T, 4000G>A, 4006C>T, 4009T>G, 4012C>T, 4015A>G, 4024T>C, 4030A>G, 4033A>T, 4042G>A, 4052C>T, 4054G>A, 4057T>C, 4063T>A, 4066A>G, 4073A>G, 4078T>C, 4081C>T, 4084T>C, 4096C>T, 4108C>T, 4111A>T, 4117G>A, 4118C>T, 4120A>G, 4126C>T, 4128T>A, 4129C>T, 4132C>T, 4135T>C, 4147A>G, 4148T>C, 4151G>T, 4162C>T, 4166G>A, 4171A>C, 4180G>A, 4190C>T, 4198T>C, 4201C>T, 4210G>T, 4213C>T, 4219T>C, 4220C>T, 4223C>A, 4225G>A, 4231T>G, 4237T>C, 4252A>G, 4253T>C, 4255A>G, 4258A>G, 4270T>C, 4282A>G, 4285G>A, 4291A>G, 4303G>A, 4312C>T, 4319A>G, 4321A>G, 4324G>T, 4327G>A, 4336G>T, 4339T>C, 4348A>T, 4360A>G, 4366T>C, 4369T>A, 4375T>C, 4387T>C, 4395T>A, 4406G>A, 4411T>C, 4438T>C, 4444C>A, 4450C>T, 4456C>T, 4465C>T, 4475A>G, 4477G>T, 4480C>T, 4481A>T, 4495C>T, 4507G>A, 4510C>T, 4513T>G, 4517C>T, 4528C>T, 4534A>G, 4540C>T, 4544G>T, 4545C>T, 4546G>A, 4548G>A, 4558T>C, 4564A>T, 4570C>T, 4576A>T, 4597G>A, 4600C>T, 4606C>T, 4609G>A, 4615G>A, 4618C>T, 4625T>C, 4627G>T, 4648C>T, 4651T>C, 4660G>A, 4663C>T, 4669T>A, 4679G>A, 4681G>C, 4684T>C, 4687T>G, 4699G>A, 4702A>G, 4708A>G, 4711G>A, 4712C>T, 4738A>G, 4744A>T, 4747A>G, 4748T>C, 4765T>C, 4796G>A, 4798C>T, 4803C>G, 4804T>C, 4807G>A, 4810G>T, 4813T>G, 4816T>C, 4831T>C, 4834C>T, 4840A>T, 4852A>G, 4855C>T, 4867T>C, 4870G>C, 4873T>A, 4882T>C, 4903A>T, 4909G>A, 4915T>C, 4918A>G, 4919T>C, 4921G>A, 4924T>C, 4927C>G, 4933T>C, 4936C>A, 4939G>A, 4942T>C, 4945C>T, 4963A>G, 4972T>C, 4990T>C, 5002A>G, 5023T>C, 5024T>C, 5029C>T, 5032T>C, 5038T>C, 5044A>T, 5053A>T, 5056G>A, 5057A>G, 5060A>G, 5071G>T, 5090C>A, 5095A>G, 5098G>T, 5100C>T, 5104A>G, 5107A>T, 5110T>C, 5113A>G, 5116G>A, 5119A>G, 5120G>T, 5122C>T, 5126A>G, 5137T>C, 5140C>T, 5143A>G, 5146G>A, 5161C>A, 5167A>G, 5173T>G, 5182G>A, 5183G>A, 5188C>T, 5191G>A, 5197A>G, 5200C>T, 5206C>T, 5209A>G, 5218C>T, 5230G>A, 5236C>T, 5239C>T, 5245C>T, 5263A>T, 5269C>T, 5272C>T, 5275C>A, 5278C>T, 5281A>C, 5284A>G, 5287T>C, 5288C>T, 5293A>T, 5294T>C, 5296G>A, 5302G>A, 5311T>A, 5314A>G, 5323A>T, 5347T>C, 5353C>T, 5365C>T, 5386T>C, 5395C>A, 5422A>G, 5425T>C, 5431C>T, 5437G>A, 5452T>C, 5458G>A, 5467T>C, 5470A>G, 5471C>T, 5473A>G, 5482C>T, 5485A>C, 5488T>C, 5489T>C, 5491G>A, 5497T>C, 5506T>C, 5509G>A, 5512G>A, 5530T>C, 5533G>A, 5536G>T, 5539G>A, 5548T>G, 5554C>T, 5560T>A, 5561G>A, 5563G>A, 5566G>C, 5569A>G, 5584C>G, 5593G>A, 5596G>A, 5597A>G, 5605C>A, 5617G>T, 5619C>T, 5620T>G, 5629G>A, 5630A>C, 5641C>T, 5650A>T, 5662A>T, 5665A>G, 5689A>G, 5693G>A, 5698G>A, 5713A>G, 5716G>C, 5719A>G, 5743C>A, 5755C>A, 5758T>C, 5764T>C, 5770A>G, 5779C>T, 5782T>C, 5791C>T, 5803C>T, 5836G>A, 5839A>G, 5845A>G, 5848A>G, 5853A>G, 5857T>C, 5860C>T, 5863A>T, 5869T>A, 5875T>C, 5884C>T, 5893C>T, 5894A>G, 5896C>G, 5899C>T, 5908G>A, 5911T>C, 5923G>A, 5938A>G, 5941C>T, 5944T>C, 5947T>C, 5959G>A, 5960C>G, 5961G>C, 5962G>C, 5974A>G, 5986T>A, 5989C>T, 5990A>G, 5998T>C, 6008A>C, 6010A>T, 6019A>T, 6022G>T, 6025A>T, 6030A>C, 6049C>A, 6058G>A, 6061A>G, 6064C>A, 6070T>C, 6072G>A, 6073T>C, 6074A>C, 6079G>A, 6081C>A, 6085A>T, 6091A>T, 6094C>T, 6111G>C, 6112T>C, 6118A>G, 6124G>A, 6127G>A, 6128A>G, 6130T>C, 6133T>C, 6136A>G, 6148C>T, 6157A>G, 6158G>A, 6163T>C, 6169A>G, 6170A>T, 6172A>G, 6178C>T, 6187A>G, 6190T>C, 6196G>A, 6203C>T, 6205C>T, 6208A>G, 6209C>T, 6211A>G, 6217G>A, 6229T>A, 6230G>A, 6238C>T, 6247G>A, 6253A>T, 6257G>A, 6258C>A, 6274T>C, 6282A>G, 6286G>A, 6298C>T, 6301G>A, 6304A>G, 6307C>T, 6313C>T, 6322C>T, 6328A>T, 6331T>G, 6334T>C, 6337G>A, 6338C>T, 6358T>C, 6361G>A, 6362G>A, 6367A>T, 6370G>A, 6376C>T, 6379T>G, 6385A>C, 6391A>T, 6394A>G, 6397C>T, 6401G>A, 6405C>T, 6412C>T, 6418A>G, 6421T>C, 6429A>T, 6436T>A, 6445G>A, 6450A>G, 6454C>A, 6460C>T, 6463T>A, 6469C>T, 6472G>A, 6475G>A, 6481T>C, 6485C>A, 6487C>T, 6490T>C, 6496C>T, 6502G>A, 6514T>C, 6517G>A, 6527C>A, 6532C>A, 6541G>A, 6544A>G, 6556C>T, 6559T>C, 6565G>A, 6569T>C, 6571G>A, 6586T>A, 6588C>T, 6598A>G, 6601A>G, 6604T>C, 6607T>C, 6616A>G, 6622C>T, 6631G>A, 6634G>A, 6635T>C, 6640T>A, 6646G>C, 6647A>G, 6652C>T, 6664C>T, 6670T>G, 6677A>G, 6679A>G, 6685C>T, 6691A>T, 6694C>T, 6715A>G, 6716G>A, 6719A>G, 6724A>G, 6734C>T, 6742G>A, 6745G>A, 6754C>T, 6760A>G, 6764A>T, 6766A>G, 6775T>A, 6782C>T, 6787A>T, 6790C>T, 6795C>T, 6796A>G, 6799C>A, 6802C>A, 6808C>T, 6811C>T, 6814C>T, 6823C>T, 6829A>G, 6843A>G, 6847C>T, 6853A>C, 6857A>G, 6862G>A, 6871A>G, 6875G>A, 6876T>C, 6877C>A, 6880G>A, 6883T>A, 6889T>G, 6901C>T, 6916C>T, 6919G>A, 6925A>G, 6928T>C, 6934C>A, 6949G>C, 6955C>T, 6958A>G, 6976A>G, 6980C>T, 6985C>T, 6997A>T, 7000G>T, 7001G>A, 7003G>A, 7012T>C, 7015G>A, 7018T>C, 7032T>A, 7039C>T, 7040T>C, 7042G>A, 7048A>G, 7060T>A, 7063C>A, 7078C>T, 7082T>C, 7084A>G, 7087A>G, 7090C>A, 7096T>C, 7099A>G, 7102T>C, 7111G>A, 7117A>G, 7125C>A, 7129A>G, 7132T>C, 7133C>T, 7135G>A, 7141G>A, 7144T>C, 7145G>T, 7147A>G, 7150G>A, 7156G>A, 7159A>T, 7160_7165delGAGGCA, 7171C>T, 7189G>T, 7198A>G, 7201C>T, 7205C>T, 7210T>C, 7216T>A, 7219G>A, 7225C>A, 7228G>A, 7240G>A, 7243T>C, 7249A>G, 7252T>G, 7258T>A, 7261T>C, 7264T>A, 7267C>G, 7270G>A, 7273_7274insT, 7274C>G, 7275G>T, 7281A>T, 7284T>C, 7296C>T, 7311C>T, 7314C>T, 7317G>A, 7319delA, 7321T>A, 7327C>T, 7330C>T, 7333C>T, 7334A>C, 7342T>C, 7343G>A, 7344C>A, 7345A>C, 7354T>C, 7363A>G, 7372C>T, 7378G>T, 7381C>T, 7396A>T, 7399C>T, 7405C>T, 7407G>T, 7408T>G, 7414G>T, 7415T>G, 7420A>G, 7429C>T, 7432T>C, 7441T>C, 7459G>A, 7462A>G, 7468A>T, 7471C>A, 7477A>G, 7480C>T, 7486G>T, 7495T>C, 7498A>G, 7501T>C, 7503T>C, 7504G>A, 7513T>A, 7519C>T, 7522A>G, 7531G>A, 7537G>A, 7540C>T, 7546T>C, 7570G>A, 7573G>A, 7579G>A, 7588T>C, 7595C>T, 7604T>C, 7606G>A, 7615T>A, 7624T>C, 7633C>T, 7636C>T, 7639T>C, 7648T>C, 7666A>G, 7672T>C, 7678A>T, 7684G>A, 7690T>C, 7696T>C, 7702T>G, 7709G>A, 7713A>G, 7714A>G, 7715C>T, 7726T>C, 7735G>A, 7744A>G, 7747C>T, 7762C>T, 7768G>A, 7771T>C, 7780A>G, 7783A>C, 7786T>C, 7793C>T, 7795A>G, 7798C>T, 7804T>C, 7812C>T, 7816C>T, 7834A>G, 7837C>T, 7844G>T, 7846G>A, 7855G>A, 7861G>A, 7868T>C, 7870A>G, 7873C>T, 7876T>A, 7879G>A, 7882T>A, 7883G>A, 7888C>T, 7894A>T, 7897T>C, 7921C>T, 7927G>A, 7936T>C, 7939T>C, 7942T>C, 7966G>A, 7969T>C, 7975T>C, 7987C>T, 7990T>C, 8002C>T, 8005T>C, 8032A>G, 8038C>T, 8041C>T, 8043G>A, 8080G>A, 8089G>A, 8095G>A, 8098G>A, 8107C>T, 8110T>C, 8116T>C, 8128A>G, 8146T>C, 8151A>G, 8152C>T, 8164G>A, 8178G>A, 8182G>A, 8191G>A, 8197C>T, 8203C>T, 8208G>A, 8212C>G, 8213C>T, 8218A>G, 8221C>T, 8224G>A, 8239A>G, 8260T>C, 8263A>G, 8281T>C, 8284A>G, 8294C>G, 8296A>G, 8299G>A, 8302C>T, 8317T>C, 8320C>T, 8326T>A, 8335G>A, 8338C>T, 8349C>T, 8353A>G, 8362C>T, 8365A>G, 8380C>T, 8383A>G, 8389A>G, 8404T>C, 8408G>T, 8413A>G, 8419G>T, 8434T>C, 8440T>C, 8452G>A, 8458A>G, 8460G>A, 8464A>G, 8467G>A, 8480G>A, 8482T>A, 8497G>A, 8498C>T, 8500A>G, 8502G>A, 8503A>G, 8506C>T, 8515A>T, 8524C>T, 8527A>C, 8528T>C, 8530A>G, 8539G>C, 8546C>T, 8547T>G, 8552T>A, 8563A>G, 8574C>G, 8585A>G, 8598G>A, 8602C>T, 8603A>T, 8604A>C, 8605C>T, 8616C>T, 8620A>G, 8622C>A, 8625A>G, 8626G>A, 8629G>A, 8632C>T, 8638C>T, 8641G>A, 8647C>T, 8648C>T, 8649C>T, 8650G>C, 8653C>T, 8656A>G, 8658G>A, 8664A>C, 8665T>G, 8674T>G, 8679C>T, 8680A>T, 8683C>A, 8692G>A, 8695A>T, 8707G>A, 8708A>C, 8716G>A, 8722C>T, 8725G>A, 8731A>G, 8737A>G, 8743C>T, 8745C>A, 8746A>G, 8749C>T, 8752G>A, 8755A>T, 8770C>T, 8771T>C, 8776A>G, 8779T>C, 8782G>A, 8791T>C, 8797A>G, 8800A>G, 8803A>G, 8809T>C, 8812T>C, 8818T>C, 8824G>A, 8827A>G, 8848G>A, 8854T>C, 8856A>G, 8857G>A, 8863G>A, 8865G>A, 8867A>C, 8869G>T, 8872A>G, 8883G>A, 8888A>T, 8929C>T, 8954G>A, 8959C>T, 8962A>T, 8983G>A, 8984A>G, 8986T>C, 8988A>T, 8998G>A, 9004G>A, 9007G>A, 9016C>T, 9037C>A, 9085A>T, 9097C>T, 9136A>T, 9145G>T, 9149A>G, 9150T>G, 9151G>A, 9152G>A, 9157A>G, 9160G>T, 9163T>C, 9184G>A, 9196A>G, 9214G>A, 9217A>G, 9220G>A, 9244A>T, 9247G>A, 9256C>T, 9260C>T, 9262T>C, 9274G>A, 9280C>T, 9289C>T, 9304G>A, 9334C>T, 9364C>G, 9377_9378insT, 9387C>T, 9404G>A, 9411C>T, 9412A>G, 9414A>T, 9437C>T, 9454A>G, 9458T>C, 9467G>A, 9470_9471insT, 9482G>A, 9483C>T, 9498T>C, 9516T>G, 9517delA, 9533G>A, 9534G>T, 9548A>T, 9551C>T, 9553_9554delTA, 9556A>T, 9557T>A, 9558A>T, 9574T>A, 9581G>T, 9582T>G, 9583G>C, 9584_9585insG, 9590C>T, 9591T>C, 9607C>T, 9610T>C, 9615delC, 9620A>G, 9648A>G, 9690C>T

| | Begin | End | Coverage | Score | Concordance | Matches | Identities | I/D/M/F* | Stop Codons |
|---|---|---|---|---|---|---|---|---|---|
| **NT** | **19** | **9704** | **99.8%** | **12534** | **65.0%** | **9674 (99.7%)** | **8011 (82.6%)** | **15/12** | |
| **CDS** | | | | | | | | | |
| PVYgp1 | 1 | 3064 | 100% | 18956 | 89.8% | 3062 (99.9%) | 2776 (90.5%) | 2/2/4/4 | 1 |
| Protein mutations: | Y4Q (194T>C 196C>A), C9L (209T>C 210G>T 211T>G), F13M (221T>A 223T>G), S23P (251T>C), C24F (255G>T 256C>T), E25G (258A>G), I27V (263A>G 265T>G), V28A (267T>C), E30V (273A>T), A35T (287G>A 289T>C), V37T (293G>A 294T>C), A41I (305G>A 306C>T), D42G (309A>G), E44D (316A>T), T45V (317A>G 318C>T 319A>G), L52Q (339T>A 340C>A), K53R (342A>G), K55A (347A>G 348A>C 349A>G), Y56H (350T>C), T58V (356A>G 357C>T), V61T (365G>A 366T>C 367G>A), L62S (368C>T 369T>C), F67R (383T>C 384T>G), A75V (408C>T 409C>T), M78E (416A>G 417T>A), E85K (437G>A), R86E (440A>G 441G>A 442G>A), K87R (443A>C 444A>G 445G>A), D88E (448T>A), E91A (456A>C), H93N (461C>A), D111E (517T>A), S116F (531C>T), P118S (536C>T), Q119W (539C>T 540A>G 541A>G), R122K (549G>A 550A>G), V134A (585T>C 586C>G), R137Y (593C>T 594G>A 595C>T), P138R (597C>G), I139V (599A>G 601A>G), I140P (602A>C 603T>C 604A>G), M147I (625G>A), I151V (635A>G 637T>C), I154V (644A>G), E160A (663A>C 664G>C), H166Q (682C>A), T172S (699C>G), H174Q (706T>A), K179E (719A>G 721G>A), I180V (722A>G 724A>G), A183S (731G>T), Y184P (734T>C 735A>C 736C>T), S185R (737T>C 738C>G 739C>T), A187V (744C>T 745G>C), R189C (749C>T 751A>C), M194K (765T>A), R197K (773C>A 774G>A 775A>G), M206K (801T>A), G210A (813G>C 814A>G), L211H (816T>A), R214Q (825G>A 826T>A), R226C (860C>T 862C>T), T227A (863A>G 865T>C), I228T (867T>C), N229D (869A>G 871C>T), I230L (872A>T 874A>G), R232K (879G>A), N240S (903A>G), T241N (906C>A 907A>C), K242T (909A>C 910A>C), S243N (912G>A 913C>T), S252L (938T>C 939C>T 940A>T), L266I (980T>A 982G>C), R272K (999G>A), Q275R (1007C>A 1008A>G 1009G>A), S276G (1010A>G 1012T>G), I277V (1013A>G), N279Q (1019A>C 1021C>G), D288E (1048C>A), R300Q (1083G>A), S305A (1097T>G 1099G>A), R320K (1143G>A), L324I (1154T>A 1156G>A), S346N (1221G>A), V349T (1229G>A 1230T>C 1231T>C), S350N (1233G>A 1234C>T), K358T (1257A>C 1258A>G), N365S (1278A>G), A369V (1290C>T 1291G>A), D372T (1298G>A 1299A>C 1300C>T), I375V (1307A>G 1309A>G), N378E (1316A>G 1318T>A), I382T (1329T>C), L397I (1373C>A), I401V (1385A>G), E407D (1405G>T), A411S (1415G>T 1417A>T), V418I (1436G>A 1438C>T), E508G (1707A>G), A512T (1718G>A), I520V (1742A>G 1744C>T), S525N (1758G>A), V536I (1790G>A), I585V (1937A>G 1939T>C), F607Y (2004T>A), I628M (2068T>G), V630I (2072G>A 2074G>A), R675K (2208G>A), D694E (2266C>G), N744G (2414A>G 2415A>G 2416T>G), S746C (2420A>T), K780R (2523A>G), R817K (2634G>A), V847I (2723G>A), A855T (2747G>A 2749T>A), H871N (2795C>A), V880I (2822G>A 2824G>A), P900T (2882C>A), S901H (2885A>C 2886G>A), T904M (2895C>T), N916S (2931A>G), N919S (2940A>G), E929_N930insX (2971_2972insA), T936I (2991C>T), H937R (2994A>G), E939R (2999G>A 3000A>G), R946Q (3021G>A), Y947F (3024A>T 3025C>T), T951I (3036C>T), E952G (3039A>G 3040A>C), K953T (3042A>C 3043G>A), A972V (3099C>T 3100C>A), V974R (3104G>A 3105T>G 3106G>A), S983N (3132G>A), R985K (3137C>A 3138G>A 3139A>G), F990I (3152T>A 3154C>T), C994Y (3165G>A), F1012L (3218T>C), V1016I (3230G>A), V1028M (3266G>A 3268A>G), V1031M (3275G>A), A1034T (3284G>A), R1044K (3315G>A 3316A>G), E1045A (3318A>C), M1059V (3359A>G), D1072E (3400T>G), V1110G (3513T>G), M1121V (3545A>G), A1122T (3548G>A), V1124I (3554G>A 3556C>T), L1147V (3623C>G), L1150M (3632C>A), Y1152H (3638T>C), I1172V (3698A>G), S1178N (3717G>A), R1183K (3731C>A 3732G>A), I1297V (4073A>G), F1315Y (4128T>A 4129C>T), A1323S (4151G>T), V1328I (4166G>A), L1347I (4223C>A 4225G>A), I1379V (4319A>G 4321A>G), F1404Y (4395T>A), V1408I (4406G>A), M1431V (4475A>G 4477G>T), T1433S (4481A>T), A1454L (4544G>T 4545C>T 4546G>A), R1455K (4548G>A), V1499I (4679G>A 4681G>C), V1538I (4796G>A 4798C>T), T1540S (4803C>G 4804T>C), S1625G (5057A>G), T1626A (5060A>G), L1636I (5090C>A), A1639V (5100C>T), E1641D (5107A>T), V1646F (5120G>T 5122C>T), I1648V (5126A>G), V1667I (5183G>A), A1793T (5561G>A 5563G>A), I1805V (5597A>G), N1807K (5605C>A), A1812V (5619C>T 5620T>G), I1816L (5630A>C), V1837I (5693G>A), K1890R (5853A>G), I1904V (5894A>G 5896C>G), R1926A (5960C>G 5961G>C 5962G>C), I1936V (5990A>G), E1945D (6019A>T), K1949T (6030A>C), D1955E (6049C>A), M1958I (6058G>A), S1963N (6072G>A 6073T>C), N1964H (6074A>C), T1966N (6081C>A), C1976S (6111G>C 6112T>C), I1982V (6128A>G 6130T>C), V1992I (6158G>A), T1996S (6170A>T 6172A>G), L2007F (6203C>T 6205C>T), V2016I (6230G>A), A2025K (6257G>A 6258C>A), K2033R (6282A>G), A2060T (6362G>A), V2073I (6401G>A), A2074V (6405C>T), Y2082F (6429A>T), Q2089R (6450A>G), H2101N (6485C>A 6487C>T), L2115I (6527C>A), T2135I (6588C>T), Y2151H (6635T>C), I2155V (6647A>G), I2165V (6677A>G 6679A>G), D2178N (6716G>A), N2179D (6719A>G), I2194L (6764A>T 6766A>G), A2204V (6795C>T 6796A>G), H2205Q (6799C>A), K2220R (6843A>G), N2225D (6857A>G), V2231T (6875G>A 6876T>C 6877C>A), D2250E (6934C>A), V2273I (7001G>A 7003G>A), F2283Y (7032T>A), T2314K (7125C>A), A2321S (7145G>T 7147A>G), E2326_A2327del (7160_7165delGAGGCA), D2344E (7216T>A), D2358E (7258T>A), D2363_R2364insX (7273_7274insT), R2364V (7274C>G 7275G>T), H2366L (7281A>T), L2367S (7284T>C), S2371L (7296C>T), S2376F (7311C>T), T2377I (7314C>T), C2378Y (7317G>A), K2384Q (7334A>C), A2387N (7343G>A 7344C>A 7345A>C), S2408M (7407G>T 7408T>G), C2411G (7415T>G), L2440S (7503T>C 7504G>A), D2509N (7709G>A), K2510R (7713A>G 7714A>G), T2543I (7812C>T), V2554L (7844G>T 7846G>A), V2567I (7883G>A), S2620N (8043G>A), N2656S (8151A>G 8152C>T), R2665K (8178G>A), R2675K (8208G>A), P2704A (8294C>G 8296A>G), S2722F (8349C>T), A2742S (8408G>T), M2756I (8452G>A), R2759K (8460G>A), A2766T (8480G>A 8482T>A), R2773K (8502G>A 8503A>G), E2777D (8515A>T), E2785D (8539G>C), L2788C (8546C>T 8547T>G), S2790T (8552T>A), A2797G (8574C>G), I2801V (8585A>G), G2805E (8598G>A), N2807S (8603A>T 8604A>C 8605C>T), A2811V (8616C>T), P2813Q (8622C>A), E2814G (8625A>G 8626G>A), P2822F (8648C>T 8649C>T 8650G>C), G2825E (8658G>A), D2827A (8664A>C 8665T>G), A2832V (8679C>T 8680A>T), T2854K (8745C>A 8746A>G), E2891G (8856A>G 8857G>A), R2894Q (8865G>A), M2895L (8867A>C 8869G>T), G2900E (8883G>A), T2902S (8888A>T), V2924I (8954G>A), N2934D (8984A>G 8986T>C), E2935V (8988A>T), M2989G (9149A>G 9150T>G 9151G>A), G2990S (9152G>A), P3026S (9260C>T 9262T>C) | | | | | | | | |

| | Begin | End | Coverage | Score | Concordance | Matches | Identities | I/D/M/F* | Stop Codons |
|---|---|---|---|---|---|---|---|---|---|
| NT | 19 | 9704 | 99.8% | 12534 | 65.0% | 9674 (99.7%) | 8011 (82.6%) | 15/12 | |

Codon mutations:

ACT3ACC (193T>C), TAC4CAA (194T>C 196C>A), TCA6TCG (202A>G), ATC8ATT (208C>T), TGT9CTG (209T>C 210G>T 211T>G), TCG12TCA (220G>A), TTT13ATG (221T>A 223T>G), GAA14GAG (226A>G), CTA17CTT (235A>T), TAC19TAT (241C>T), CCA21CCT (247A>T), TCT23CCT (251T>C), TGC24TTT (255G>T 256C>T), GAG25GGG (258A>G), CAT26CAC (262T>C), ATT27GTG (263A>G 265T>G), GTG28GCG (267T>C), GAA30GTA (273A>T), CGA31CGT (277A>T), GAA32GAG (280A>G), GTG33GTT (283G>T), CCG34CCA (286G>A), GCT35ACC (287G>A 289T>C), GTT37ACT (293G>A 294T>C), CCT39CCC (301T>C), TTC40TTT (304C>T), GCA41ATA (305G>A 306C>T), GAT42GGT (309A>G), CTG43TTG (311C>T), GAA44GAT (316A>T), ACA45GTG (317A>G 318C>T 319A>G), CAA46CAG (322A>G), GCA49GCT (331A>T), CTC52CAA (339T>A 340C>A), AAG53AGG (342A>G), AAA55GCG (347A>G 348A>C 349A>G), TAT56CAT (350T>C), GCT57GCA (355T>A), ACT58GTT (356A>G 357C>T), CGT60CGC (364T>C), GTG61ACA (365G>A 366T>C 367G>A), CTC62TCC (368C>T 369T>C), AAA63AAG (373A>G), AAC64AAT (376C>T), TTT67CGT (383T>C 384T>G), ACG68ACA (388G>A), TAC69TAT (391C>T), CGA70CGG (394A>G), TAC71TAT (397C>T), AAG72AAA (400G>A), GAT74GAC (406T>C), GCC75GTT (408C>T 409C>T), CAG76CAA (412G>A), ATA77ATT (415A>T), ATG78GAG (416A>G 417T>A), ATT80ATC (424T>C), CAG81CAA (427G>A), AAA83AAG (433A>G), CTG84CTC (436G>C), GAG85AAG (437G>A), AGG86GAA (440A>G 441G>A 442G>A), AAG87CGA (443A>C 444A>G 445G>A), GAT88GAA (448T>A), GAA90GAG (454A>G), GAA91GCA (456A>C), TAT92TAC (460T>C), CAC93AAC (461C>A), CAA95CAG (469A>G), GCC97GCT (475C>T), CCT99CCA (481T>A), ACT106ACC (502T>C), ATA107ATT (505A>T), GGC109GGT (511C>T), GAT111GAA (517T>A), CCT112CCA (520T>A), CCA113CCT (523A>T), TCT116TTT (531C>T), GAG117GAA (535G>A), CCA118TCA (536C>T), CAA119TGG (539C>T 540A>G 541A>G), AGA122AAG (549G>A 550A>G), ATC124ATT (556C>T), ATT125ATC (559T>C), CAT126CAC (562T>C), ACT128ACC (568T>C), CCA129CCG (571A>G), CGT132CGC (580T>C), GTC134GCG (585T>C 586C>G), CGC137TAT (593C>T 594G>A 595C>T), CCC138CGC (597C>G), ATA139GTG (599A>G 601A>G), ATA140CCG (602A>C 603T>C 604A>G), ACA143ACG (613A>G), GAA144GAG (616A>G), GGC145GGG (619C>G), CAG146CAA (622G>A), ATG147ATA (625G>A), AAT148AAC (628T>C), CAC149CAT (631C>T), ATT151GTC (635A>G 637T>C), ATA154GTA (644A>G), ATT157ATC (655T>C), TCG159TCA (661G>A), GAG160GCC (663A>C 664G>C), AAA161AAG (667A>G), GGG163GGC (673G>C), GTC165GTT (679C>T), CAC166CAA (682C>A), TTA167CTA (683T>C), AGT169AGC (691T>C), ACC172AGC (699C>G), ACT173ACA (703T>A), CAT174CAA (706T>A), AAG179GAA (719A>G 721G>A), ATA180GTG (722A>G 724A>G), CTT181TTG (725C>T 727T>G), GGT182GGA (730T>A), GCA183TCA (731G>T), TAC184CCT (734T>C 735A>C 736C>T), TCC185CGT (737T>C 738C>G 739C>T), GCG187GTC (744C>T 745G>C), CGA189TGC (749C>T 751A>C), ACT190ACC (754T>C), ATG194AAG (765T>A), TTG196CTG (770T>C), CGA197AAG (773C>A 774G>A 775A>G), CGG198AGG (776C>A), TTC202TTT (790C>T), ATG206AAG (801T>A), GGA210GCG (813G>C 814A>G), CTT211CAT (816T>A), TTG212CTG (818T>C), CAA213CAG (823A>G), CGT214CAA (825G>A 826T>A), CGG217AGA (833C>A 835G>A), GAC219GAT (841C>T), AAT223AAC (853T>C), GTC225GTT (859C>T), CGC226TGT (860C>T 862C>T), ACT227GCC (863A>G 865T>C), ATC228ACC (867T>C), AAC229GAT (869A>G 871C>T), ATA230TTG (872A>T 874A>G), AGG232AAG (879G>A), GAT234GAC (886T>C), ATC238ATT (898C>T), AAC240AGC (903A>G), ACA241AAC (906C>A 907A>C), AAA242ACC (909A>C 910A>C), AGC243AAT (912G>A 913C>T), GGC246GGA (922C>A), GGT249GGA (931T>A), TCA252CTT (938T>C 939C>T 940A>T), GGA253GGG (943A>G), TTC256TTT (952C>T), TCA261TCG (967A>G), GAA263GAG (973A>G), GGG264GGC (976G>C), TTG266ATC (980T>A 982G>C), GCA269GCG (991A>G), CGT270CGC (994T>C), AGA272AAA (999G>A), GTT273GTC (1003T>C), CAG275AGA (1007C>A 1008A>G 1009G>A), AGT276GGG (1010A>G 1012T>G), ATT277GTT (1013A>G), TTA278TTG (1018A>G), AAC279CAG (1019A>C 1021C>G), ATC282ATT (1030C>T), TCG285TCA (1039G>A), AAT286AAC (1042T>C), GCC287GCA (1045C>A), GAC288GAA (1048C>A), AAT289AAC (1051T>C), AAG292AAA (1060G>A), GGT293GGA (1063T>A), GAC295GAT (1069C>T), CGA300CAA (1083G>A), TAT303TAC (1093T>C), TCG305GCA (1097T>G 1099G>A), CAC307CAT (1105C>T), GTA310GTG (1114A>G), GCT311GCG (1117T>G), TTA313TTG (1123A>G), CCT314CCA (1126T>A), GTC315GTT (1129C>T), AGG320AAG (1143G>A), GTA321GTT (1147A>T), GCT322GCA (1150T>A), TTG324ATA (1154T>A 1156G>A), CAC327CAT (1165C>T), AGT328AGC (1168T>C), CTT330TTA (1172C>T 1174T>A), ACC339ACT (1201C>T), AGC346AAC (1221G>A), GTT349ACC (1229G>A 1230T>C 1231T>C), AGC350AAT (1233G>A 1234C>T), CTG352CTA (1240G>A), AAG354AAA (1246G>A), TTG356CTA (1250T>C 1252G>A), CAT357CAC (1255T>C), AAA358ACG (1257A>C 1258A>G), AGA361AGG (1267A>G), GGT363GGC (1273T>C), TTG364TTA (1276G>A), AAT365AGT (1278A>G), TTG367CTG (1283T>C), GCG369GTA (1290C>T 1291G>A), GAC372ACT (1298G>A 1299A>C 1300C>T), TTT374TTC (1306T>C), ATA375GTG (1307A>G 1309A>G), CAT376CAC (1312T>C), AAT378GAA (1316A>G 1318T>A), TTG381TTA (1327G>A), ATA382ACA (1329T>C), GCG383GCT (1333G>T), TTA384TTG (1336A>G), CTA387TTG (1343C>T 1345A>G), CCG390CCA (1354G>A), GTG391GTT (1357G>T), GAC392GAT (1360C>T), CTC395TTG (1367C>T 1369C>G), CTT397ATT (1373C>A), TTC398TTT (1378C>T), GAG400GAA (1384G>A), ATA401GTA (1385A>G), AAA403AAG (1393A>G), TCC404TCT (1396C>T), GAG407GAT (1405G>T), AAA408AAG (1408A>G), CAG409CAA (1411G>A), GCA411TCT (1415G>T 1417A>T), CCG412CCC (1420G>C), AAT415AAC (1429T>C), TTA416TTG (1432A>G), GTC418ATT (1436G>A 1438C>T), TTA419CTG (1439T>C 1441A>G), AAT420AAC (1444T>C), TTC423TTT (1453C>T), CTG424CTT (1456G>T), GAA428GAG (1468A>G), TTG439CTA (1499T>C 1501G>A), AGT440AGC (1504T>C), TTG441TTA (1507G>A), CTC442CTT (1510C>T), GAA443GAG (1513A>G), TTA444TTG (1516A>G), GCA445GCT (1519A>T), AGG446AGA (1522G>A), TTC447TTT (1525C>T), CAG448CAA (1528G>A), AAC450AAT (1534C>T), ACT452ACA (1540T>A), AAC454AAT (1546C>T), AAA457AAG (1555A>G), GAT459GAC (1561T>C), ATA460ATC (1564A>C), TCT461TCA (1567T>A), TTC463TTT (1573C>T), AGA464AGG (1576A>G), TTA467CTA (1583T>C), TCT468TCA (1588T>A), AAG470AAA (1594G>A), AAC472AAT (1600C>T), CTG475CTA (1609G>A), TCG478TCA (1618G>A), TGC479TGT (1621C>T), GAC480GAT (1624C>T), AAC481AAT (1627C>T), CTG483TTG (1631C>T), AAA485AAG (1639A>G), TTC489TTT (1651C>T), CTC490CTG (1654C>G), CAT497CAC (1675T>C), TCA503TCG (1693A>G), TTT506TTC (1702T>C), GAA508GGA (1707A>G), GCA512ACA (1718G>A), GGA514GGG (1726A>G), TAC515TAT (1729C>T), TCA516TCC (1732A>C), TAT518TAC (1738T>C), ATC520GTT (1742A>G 1744C>T), CGC521CGT (1747C>T), CAT523CAC (1753T>C), AGT525AAT (1758G>A), AGG528AGA (1768G>A), AAC534AAT (1786C>T), TTA535TTG (1789A>G), GTT536ATT (1790G>A), GTC537GTT (1795C>T), TTA541CTG (1805T>C 1807A>G), GAG543GAA (1813G>A), AGG545AGA (1819G>A), AAG547AAA (1825G>A), GAC551GAT (1837C>T), TAT552TAC (1840T>C), CAA555CAG (1849A>G), GTC558GTG (1858C>G), AAA560AAG (1864A>G), TGC562TGT (1870C>T), AGT564AGC (1876T>C), AAA566AAG (1882A>G), GGT568GGA (1888T>A), AAT569AAC (1891T>C), TGT574TGC (1906T>C), ACA577ACT (1915A>T), GGT582GGC (1930T>C), GCC584GCT (1936C>T), ATT585GTC (1937A>G 1939T>C), TCA587TCG (1945A>G), ACA588ACG (1948A>G), TTC589TTT (1951C>T), CCA592CCT (1960A>T), GTA598GTG (1978A>G), ATT599ATA (1981T>A), GGC600GGT (1984C>T), AAT601AAC (1987T>C), GAC604GAT (1996C>T), TTT607TAT (2004T>A), AAA612AAG (2020A>G), GGG613GGA (2023G>A), GAT614GAC (2026T>C), TCG615TCT (2029G>T), GAG616GAA (2032G>A), TAT625TAC (2059T>C), ATT628ATG (2068T>G), AAC629AAT (2071C>T), GTG630ATA (2072G>A 2074G>A), CTG635TTA (2087C>T 2089G>A), AGC639AGT (2101C>T), GAG640GAA (2104G>A), GAG641GAA (2107G>A), AAG644AAA (2116G>A), TTC646TTT (2122C>T), CGC651CGT (2137C>T), CTT658CTC (2158T>C), ACC660ACT (2164C>T), CCA662CCG (2170A>G), GAT666GAC (2182T>C), GCG668GCA (2188G>A), ACC669ACA (2191C>A), AGA675AAA (2208G>A), TAT678TAC (2218T>C), GAC680GAT (2224C>T), GTA681GTT (2227A>T), TTG686TTA (2242G>A), CCC687CCT (2245C>T), TTG690TTA (2254G>A), GTT691GTC (2257T>C), GAC692GAT (2260C>T), GAC694GAG (2266C>G), CAA696CAG (2272A>G), ACG697ACA (2275G>A), TCA703TCG (2293A>G), GGC705GGT (2299C>T), TCG706TCA (2302G>A), CAG707CAA (2305G>A), GGA710GGT (2314A>T), TAT711TAC (2317T>C), ATT713ATC (2323T>C), CTA714CTG (2326A>G), GTG719GTA (2341G>A), CAA721CAG (2347A>G), CTT722CTC (2350T>C), TTG724TTA (2356G>A), AAT727AAC (2365T>C), GAA729GAG (2371A>G), TTA730TTG (2374A>G), GAT733GAC (2383T>C), ATA734ATT (2386A>T), AAA735AAG (2389A>G), CAT736CAC (2392T>C), TAT737TAC (2395T>C), AAT744GGG (2414A>G 2415A>G 2416T>G), GCT745GCA (2419T>A), AGC746TGC (2420A>T), ACA752ACT (2440A>T), GTT761GTC (2467T>C), TCT764TCC (2476T>C), TCG766TCA (2482G>A), CTG769CTA (2491G>A), CTG771CTT (2497G>T), TTG773TTA (2503G>A), AGA778AGG (2518A>G), CCT779CCC (2521T>C), AAG780AGG (2523A>G), AGA783AGG (2533A>G), TTG785TTA (2539G>A), TTA787TTG (2545A>G), GAG789GAA (2551G>A), CCT790CCA (2554T>A), CTG792TTG (2558C>T), TTG793CTC (2561T>C 2563G>C), CTA795CTG (2569A>G), TCA796TCG (2572A>G), CCT800CCC (2584T>C), GGC801GGT (2587C>T), CTG803CTC (2593G>C), GCT805GCC (2599T>C), AAT808AAC (2608T>C), TTT812TTC (2620T>C), GCG815GCA (2629G>A), GTG816GTA (2632G>A), AGG817AAG (2634G>A), TTG818TTA (2638G>A), ATT820ATA (2644T>A), AAA823AAG (2653A>G), GCT827GCC (2665T>C), CTA832TTA (2678C>T), CTA833CTG (2683A>G), GCT835GCC (2689T>C), CTA838CTC (2698A>C), TCA841TCG (2707A>G), GCG842GCA (2710G>A), GAA844GAG (2716A>G), ACA845ACG (2719A>G), CTC846CTT (2722C>T), GTC847ATC (2723G>A), GCA848GCG (2728A>G), AGG850AGA (2734G>A), ATT851ATA (2737T>A), GAT854GAC (2746T>C), GCT855ACA (2747G>A 2749T>A), GCA856GCT (2752A>T), GCT857GCA (2755T>A), GAC859GAT (2761C>T), CTC860CTT (2764C>T), TGT865TGC (2779T>C), GGG867GGT (2785G>T), AAC869AAT (2791C>T), CAT871AAT (2795C>A), TAC874TAT (2806C>T), TTG878CTC (2816T>C 2818G>C), GTG880ATA (2822G>A 2824G>A), TTG881TTA (2827G>A), GTT884GTC (2836T>C), AAG885AAA (2839G>A), AAT886AAC (2842T>C), ACC893ACA (2863C>A), CTA894TTA (2864C>T), CCA900ACA (2882C>A), AGT901CAT (2885A>C 2886G>A), AAC903AAT (2893C>T), ACG904ATG (2895C>T), GTC906GTT (2902C>T), GTA907GTG (2905A>G), ATT909ATC (2911T>C), AAT916AGT (2931A>G), AAC919AGC (2940A>G), GCT921GCC (2947T>C), GAA929_AAT930insA-- (2971_2972insA), GGT935GGC (2989T>C), ACT936ATT (2991C>T), CAT937CGT (2994A>G), GAG939AGG (2999G>A 3000A>G), TCC944TC- (3016delC), CGG946CAG (3021G>A), TAC947TTT (3024A>T 3025C>T), ACA951ATA (3036C>T), GAA952GGC (3039A>G 3040A>C), AAG953ACA (3042A>C 3043G>A), TTG956TTA (3052G>A), TTA959TTG (3061A>G), GCG966GCA (3082G>A), GGC969GGT (3091C>T), GCC972GTA (3099C>T 3100C>A), GTG974AGA (3104G>A 3105T>G 3106G>A), GTC975GTT (3109C>T), AAA976AAG (3112A>G), ACT978ACC (3118T>C), GCC979GCT (3121C>T), GGA981GGG (3127A>G), AGC983AAC (3132G>A), CGA985AAG (3137C>A 3138G>A 3139A>G), TAT989TAC (3151T>C), TTC990ATT (3152T>A 3154C>T), TGT994TAT (3165G>A), GTA995GTG (3169A>G), ATC1005ATT (3199C>T), AGG1007CGG (3203A>C), CGT1008CGC (3208T>C), CCA1010CCG (3214A>G), ACC1011ACT (3217C>T), TTT1012CTT (3218T>C), GTC1013GTA (3223C>A), GTT1016ATT (3230G>A), AAC1017AAT (3235C>T), TTA1025CTA (3257T>C), ACC1026ACT (3262C>T), AGC1027AGT (3265C>T), GTA1028ATG (3266G>A 3268A>G), GTG1029GTA (3271G>A), GTG1031ATG (3275G>A), GCA1034ACA (3284G>A), CAG1039CAA (3301G>A), AGA1044AAG (3315G>A 3316A>G), GAA1045GCA (3318A>C), GAG1047GAA (3325G>A), TTG1048TTA (3328G>A), ATA1051ATT (3337A>T), GAG1055GAA (3349G>A), GTC1057GTT (3355C>T), TGC1058TGT (3358C>T), ATG1059GTG (3359A>G), CTA1061CTG (3367A>G), TAT1062TAC (3370T>C), TTA1065CTG (3377T>C 3379A>G), GAA1070GAG (3394A>G), CGC1071CGT (3397C>T), GAT1072GAG (3400T>G), ACA1074ACG (3406A>G), TAC1078TAT (3418C>T), ATT1079ATA (3421T>A), TAT1081TAC (3427T>C), AAG1083AAA (3433G>A), TCA1084TCT (3436A>T), GTA1085GTG (3439A>G), AAC1086AAT (3442C>T), CCT1087CCC (3445T>C), CAG1091CAA (3457G>A), TTT1092TTC (3460T>C), GCT1093GCA (3463T>A), GCG1095GCT (3469G>T), CAG1096CAA (3472G>A), CAC1104CAT (3496C>T), CAG1105CAA (3499G>A), TCC1107TCT (3505C>T), GTT1110GGT (3513T>G), AAA1112AAG (3520A>G), GAA1115GAG (3529A>G), CAA1116CAG (3532A>G), ATG1121GTG (3545A>G), GCT1122ACT (3548G>A), TTA1123CTA (3551T>C), GTC1124ATT (3554G>A 3556C>T), AGT1133AGC (3583T>C), GAT1134GAC (3586T>C), TGC1135TGT (3589C>T), TTC1137TTT (3595C>T), AAT1141AAC (3607T>C), TTT1143TTC (3613T>C), AAG1144AAA (3616G>A), GGT1145GGC (3619T>C), CTT1147GTT (3623C>G), TCC1148TCT (3628C>T), CTG1150ATG (3632C>A), GAT1151GAC (3637T>C), TAT1152CAT (3638T>C), GAA1153GAG (3643A>G), TTA1159TTG (3661A>G), GTG1162GTA (3670G>A), TTT1166TTC (3682T>C), GAG1168GAA (3688G>A), GAG1171GAA (3697G>A), ATT1172GTT (3698A>G), GAT1174GAC (3706T>C), TTT1175TTC (3709T>C), TTG1177CTA (3713T>C 3715G>A), AGT1178AAT (3717G>A), GAG1179GAA (3721G>A), CGA1183AAA (3731C>A 3732G>A), ACT1184ACA (3736T>A), CTA1188TTG (3746C>T 3748A>G), ACA1190ACG (3754A>G), AGT1193AGC (3763T>C), CGA1198CGG (3778A>G), CAA1199CAG (3781A>G), ATC1200ATT (3784C>T), CAT1204CAC (3796T>C), ACA1205ACG (3799A>G), CCA1207CCT (3805A>T), TAC1209TAT (3811C>T), AGA1210AGG (3814A>G), TTC1215TTT (3829C>T), TTT1218TTC (3838T>C), AGA1220AGG (3844A>G), GCA1221GCC (3847A>C), GTT1224GTA (3856T>A), GTG1226GTT (3862G>T), GAC1229GAT (3871C>T), GCC1231GCA (3877C>A), AGC1233AGT (3883C>T), GAA1234GAG (3886A>G), CTA1236CTG (3892A>G), GAC1237GAT (3895C>T), TTT1238TTC (3898T>C), GTA1240GTG (3904A>G), CGG1241AGA (3905C>A 3907G>A), GCT1243GCC (3913T>C), AAG1248AAA (3928G>A), TCA1249TCT (3931A>T), GGG1251GGA (3937G>A), GTT1254GTC (3946T>C), AGT1257AGC (3955T>C), GTG1258GTA (3958G>A), GCC1259GCT (3961C>T), GGA1260GGG (3964A>G), TCT1261TCC (3967T>C), TTA1264TTG (3976A>G), ATT1265ATC (3979T>C), CCA1267CCT (3985A>T), CTA1271CTT (3997A>T), GCG1272GCA (4000G>A), AAC1274AAT (4006C>T), GTT1275GTG (4009T>G), TTC1276TTT (4012C>T), AAA1277AAG (4015A>G), TCT1280TCC (4024T>C), GAA1282GAG (4030A>G), CCA1283CCT (4033A>T), AAG1286AAA (4042G>A), CTG1290TTA (4052C>T 4054G>A), CGT1291CGC (4057T>C), CGT1293CGA (4063T>A), GGA1294GGG (4066A>G), ATA1297GTA (4073A>G), TTT1298TTC (4078T>C), GGC1299GGT (4081C>T), TCT1300TCC (4084T>C), TCC1304TCT (4096C>T), AGC1308AGT (4108C>T), GGA1309GGT (4111A>T), GCG1311GCA (4117G>A), CTA1312TTG (4118C>T 4120A>G), TAC1314TAT (4126C>T), TTC1315TAT (4128T>A 4129C>T), GCC1316GCT (4132C>T), AAT1317AAC (4135T>C), CAA1321CAG (4147A>G), TTA1322CTA (4148T>C), GCT1323TCT (4151G>T), AAC1326AAT (4162C>T), GTA1328ATA (4166G>A), ATA1329ATC (4171A>C), GAG1332GAA (4180G>A), CTG1336TTG (4190C>T), CCT1338CCC (4198T>C), TCC1339TCT (4201C>T), GCG1342GCT (4210G>T), TTC1343TTT (4213C>T), AGT1345AGC (4219T>C), CTG1346TTG (4220C>T), CTG1347ATA (4223C>A 4225G>A), GTT1349GTG (4231T>G), CAT1351CAC (4237T>C), GTA1356GTG (4252A>G), TTA1357CTG (4253T>C 4255A>G), AAA1358AAG (4258A>G), ACT1362ACC (4270T>C), AGA1366AGG (4282A>G), GAG1367GAA (4285G>A), GAA1369GAG (4291A>G), CAG1373CAA (4303G>A), GTC1376GTT (4312C>T), ATA1379GTG (4319A>G 4321A>G), GTG1380GTT (4324G>T), GAG1381GAA (4327G>A), CTG1384CTT (4336G>T), TCT1385TCC (4339T>C), TCA1388TCT (4348A>T), GCA1392GCG (4360A>G), GGT1394GGC (4366T>C), TCT1395TCA (4369T>A), ACT1397ACC (4375T>C), GTT1401GTC (4387T>C), TTT1404TAT (4395T>A), GTA1408ATA (4406G>A), CTT1409CTC (4411T>C), GTT1418GTC (4438T>C), ACC1420ACA (4444C>A), GCC1422GCT (4450C>T), CTC1424CTT (4456C>T), GAC1427GAT (4465C>T), ATG1431GTT (4475A>G 4477G>T), GTC1432GTT (4480C>T), ACA1433TCA (4481A>T), GGC1437GGT (4495C>T), AAG1441AAA (4507G>A), CAC1442CAT (4510C>T), GGT1443GGG (4513T>G), CTA1445TTA (4517C>T), GTC1448GTT (4528C>T), AAA1450AAG (4534A>G), ACC1452ACT (4540C>T), GCG1454TTA (4544G>T 4545C>T 4546G>A), AGA1455AAA (4548G>A), TTT1458TTC (4558T>C), GTA1460GTT (4564A>T), ACC1462ACT (4570C>T), ATA1464ATT (4576A>T), TTG1471TTA (4597G>A), GAC1472GAT (4600C>T), GAC1474GAT (4606C>T), GTG1475GTA (4609G>A), GTG1477GTA (4615G>A), GAC1478GAT (4618C>T), TTG1481CTT (4625T>C 4627G>T), GAC1488GAT (4648C>T), ATT1489ATC (4651T>C), AGG1492AGA (4660G>A), AGC1493AGT (4663C>T), GCT1495GCA (4669T>A), GTG1499ATC (4679G>A 4681G>C), AGT1500AGC (4684T>C), GTT1501GTG (4687T>G), GAG1505GAA (4699G>A), AGA1506AGG (4702A>G), CAA1508CAG (4708A>G), AGG1509AGA (4711G>A), CTG1510TTG (4712C>T), AAA1518AAG (4738A>G), GTA1520GTT (4744A>T), GCA1521GCG (4747A>G), TTG1522CTG (4748T>C), ACT1527ACC (4765T>C), GTC1538ATT (4796G>A 4798C>T), ACT1540AGC (4803C>G 4804T>C), GAG1541GAA (4807G>A), GCG1542GCT (4810G>T), GCT1543GCG (4813T>G), CTT1544CTC (4816T>C), TAT1549TAC (4831T>C), AAC1550AAT (4834C>T), CCA1552CCT (4840A>T),

| | | | | | | | | | |
|---|---|---|---|---|---|---|---|---|---|
| | GGA1556GGG (4852A>G), GGC1557GGT (4855C>T), AGT1561AGC (4867T>C), CTG1562CTC (4870G>C), ATT1563ATA (4873T>A), TGT1566TGC (4882T>C), ACA1573ACT (4903A>T), CAG1575CAA (4909G>A), TTT1577TTC (4915T>C), GAA1578GAG (4918A>G), TTG1579CTA (4919T>C 4921G>A), AGT1580AGC (4924T>C), CCC1581CCG (4927C>G), TTT1583TTC (4933T>C), ATC1584ATA (4936C>A), CAG1585CAA (4939G>A), AAT1586AAC (4942T>C), TTC1587TTT (4945C>T), TCA1593TCG (4963A>G), CCT1596CCC (4972T>C), CTT1602CTC (4990T>C), AAA1606AAG (5002A>G), CCT1613CCC (5023T>C), TTG1614CTG (5024T>C), TGC1615TGT (5029C>T), GAT1616GAC (5032T>C), TCT1618TCC (5038T>C), CCA1620CCT (5044A>T), GCA1623GCT (5053A>T), TCG1624TCA (5056G>A), AGT1625GGT (5057A>G), ACT1626GCT (5060A>G), TCG1629TCT (5071G>T), CTT1636ATT (5090C>A), GGA1637GGG (5095A>G), GTG1638GTT (5098G>T), GCC1639GTC (5100C>T), TTA1640TTG (5104A>G), GAA1641GAT (5107A>T), ATT1642ATC (5110T>C), CCA1643CCG (5113A>G), AAG1644AAA (5116G>A), CAA1645CAG (5119A>G), GTC1646TTT (5120G>T 5122C>T), ATT1648GTT (5126A>G), CAT1651CAC (5137T>C), ATC1652ATT (5140C>T), AAA1653AAG (5143A>G), GAG1654GAA (5146G>A), CTC1659CTA (5161C>A), GAA1661GAG (5167A>G), CTT1663CTG (5173T>G), ACG1666ACA (5182G>A), GTT1667ATT (5183G>A), GTC1668GTT (5188C>T), AAG1669AAA (5191G>A), AAA1671AAG (5197A>G), GAC1672GAT (5200C>T), TGC1674TGT (5206C>T), TTA1675TTG (5209A>G), AGC1678AGT (5218C>T), TCG1682TCA (5230G>A), ATC1684ATT (5236C>T), AGC1685AGT (5239C>T), ATC1687ATT (5245C>T), ACA1693ACT (5263A>T), CTC1695CTT (5269C>T), TTC1696TTT (5272C>T), GCC1697GCA (5275C>A), ATC1698ATT (5278C>T), CCA1699CCC (5281A>C), AGA1700AGG (5284A>G), ACT1701ACC (5287T>C), CTA1702TTA (5288C>T), ATA1703ATT (5293A>T), TTG1704CTA (5294T>C 5296G>A), GAG1706GAA (5302G>A), CTT1709CTA (5311T>A), GAA1710GAG (5314A>G), CGA1713CGT (5323A>T), AGT1721AGC (5347T>C), ATC1723ATT (5353C>T), TGC1727TGT (5365C>T), GTT1734GTC (5386T>C), ACC1737ACA (5395C>A), AAA1746AAG (5422A>G), GAT1747GAC (5425T>C), ACC1749ACT (5431C>T), GAG1751GAA (5437G>A), CTT1756CTC (5452T>C), AAG1758AAA (5458G>A), AGT1761AGC (5467T>C), CAA1762CAG (5470A>G), CTA1763TTG (5471C>T 5473A>G), TTC1766TTT (5482C>T), TCA1767TCC (5485A>C), AAT1768AAC (5488T>C), TTG1769CTA (5489T>C 5491G>A), GGT1771GGC (5497T>C), TGT1774TGC (5506T>C), GAG1775GAA (5509G>A), GAG1776GAA (5512G>A), TAT1782TAC (5530T>C), GAG1783GAA (5533G>A), TCG1784TCT (5536G>T), TTG1785TTA (5539G>A), GTT1788GTG (5548T>G), CAC1790CAT (5554C>T), GCT1792GCA (5560T>A), GCG1793ACA (5561G>A 5563G>A), ACG1794ACC (5566G>C), TCA1795TCG (5569A>G), CTC1800CTG (5584C>G), AAG1803AAA (5593G>A), GGG1804GGA (5596G>A), ATT1805GTT (5597A>G), AAC1807AAA (5605C>A), GTG1811GTT (5617G>T), GCT1812GTG (5619C>T 5620T>G), TTG1815TTA (5629G>A), ATC1816CTC (5630A>C), GGC1819GGT (5641C>T), GCA1822GCT (5650A>T), ATA1826ATT (5662A>T), GGA1827GGG (5665A>G), CAA1835CAG (5689A>G), GTT1837ATT (5693G>A), GAG1838GAA (5698G>A), CAA1843CAG (5713A>G), GGG1844GGC (5716G>C), AAA1845AAG (5719A>G), GCC1853GCA (5743C>A), CGC1857CGA (5755C>A), CAT1858CAC (5758T>C), CGT1860CGC (5764T>C), AAA1862AAG (5770A>G), GGC1865GGT (5779C>T), TTT1866TTC (5782T>C), GAC1869GAT (5791C>T), GAC1873GAT (5803C>T), AGG1884AGA (5836G>A), AAA1885AAG (5839A>G), GGA1887GGG (5845A>G), AAA1888AAG (5848A>G), AAA1890AGA (5853A>G), GGT1891GGC (5857T>C), ACC1892ACT (5860C>T), ACA1893ACT (5863A>T), GGT1895GGA (5869T>A), GGT1897GGC (5875T>C), AGC1900AGT (5884C>T), TTC1903TTT (5893C>T), ATC1904GTG (5894A>G 5896C>G), AAC1905AAT (5899C>T), GGG1908GGA (5908G>A), TTT1909TTC (5911T>C), GAG1913GAA (5923G>A), CAA1918CAG (5938A>G), TTC1919TTT (5941C>T), GTT1920GTC (5944T>C), GAT1921GAC (5947T>C), GGG1925GGA (5959G>A), CGG1926GCC (5960C>G 5961G>C 5962G>C), GAA1930GAG (5974A>G), GCT1934GCA (5986T>A), GAC1935GAT (5989C>T), ATT1936GTT (5990A>G), GAT1938GAC (5998T>C), AGA1942CGT (6008A>C 6010A>T), GAA1945GAT (6019A>T), GTG1946GTT (6022G>T), CGA1947CGT (6025A>T), AAA1949ACA (6030A>C), GAC1955GAA (6049C>A), ATG1958ATA (6058G>A), CAA1959CAG (6061A>G), GCC1960GCA (6064C>A), GGT1962GGC (6070T>C), AGT1963AAC (6072G>A 6073T>C), AAC1964CAC (6074A>C), ACG1965ACA (6079G>A), ACC1966AAC (6081C>A), ATA1967ATT (6085A>T), GCA1969GCT (6091A>T), TAC1970TAT (6094C>T), TGT1976TCC (6111G>C 6112T>C), AAA1978AAG (6118A>G), TTG1980TTA (6124G>A), AAG1981AAA (6127G>A), ATT1982GTC (6128A>G 6130T>C), GAT1983GAC (6133T>C), TTA1984TTG (6136A>G), AAC1988AAT (6148C>T), AAA1991AAG (6157A>G), GTT1992ATT (6158G>A), TGT1993TGC (6163T>C), AAA1995AAG (6169A>G), ACA1996TCG (6170A>T 6172A>G), GGC1998GGT (6178C>T), AAA2001AAG (6187A>G), TTT2002TTC (6190T>C), GAG2004GAA (6196G>A), CTC2007TTT (6203C>T 6205C>T), GAA2008GAG (6208A>G), CTA2009TTG (6209C>T 6211A>G), CAG2011CAA (6217G>A), GCT2015GCA (6229T>A), GTA2016ATA (6230G>A), GTC2018GTT (6238C>T), AAG2021AAA (6247G>A), ATA2023ATT (6253A>T), GCA2025AAA (6257G>A 6258C>A), CAT2030CAC (6274T>C), AAA2033AGA (6282A>G), TCG2034TCA (6286G>A), GGC2038GGT (6298C>T), TTG2039TTA (6301G>A), AGA2040AGG (6304A>G), GAC2041GAT (6307C>T), AAC2043AAT (6313C>T), GCC2046GCT (6322C>T), ACA2048ACT (6328A>T), GTT2049GTG (6331T>G), TGT2050TGC (6334T>C), AGG2051AGA (6337G>A), CTG2052TTG (6338C>T), TAT2058TAC (6358T>C), GGG2059GGA (6361G>A), GCA2060ACA (6362G>A), TCA2061TCT (6367A>T), GAG2062GAA (6370G>A), TAC2064TAT (6376C>T), GGT2065GGG (6379T>G), GGA2067GGC (6385A>C), GGA2069GGT (6391A>T), GCA2070GCG (6394A>G), TAC2071TAT (6397C>T), GTA2073ATA (6401G>A), GCG2074GTG (6405C>T), CAC2076CAT (6412C>T), TTA2078TTG (6418A>G), TTT2079TTC (6421T>C), TAC2082TTC (6429A>T), GGT2084GGA (6436T>A), GAG2087GAA (6445G>A), CAA2089CGA (6450A>G), TCC2090TCA (6454C>A), CAC2092CAT (6460C>T), GGT2093GGA (6463T>A), TTC2095TTT (6469C>T), AGG2096AGA (6472G>A), GTG2097GTA (6475G>A), AAT2099AAC (6481T>C), CAC2101AAT (6485C>A 6487C>T), AGT2102AGC (6490T>C), AGC2104AGT (6496C>T), CTG2106CTA (6502G>A), GGT2110GGC (6514T>C), AGG2111AGA (6517G>A), CTC2115ATC (6527C>A), ATC2116ATA (6532C>A), CCG2119CCA (6541G>A), AAA2120AAG (6544A>G), GTC2124GTT (6556C>T), TTT2125TTC (6559T>C), CAG2127CAA (6565G>A), TTG2129CTA (6569T>C 6571G>A), CCT2134CCA (6586T>A), ACA2135ATA (6588C>T), GAA2138GAG (6598A>G), AGA2139AGG (6601A>G), ATT2140ATC (6604T>C), TGT2141TGC (6607T>C), GGA2144GGG (6616A>G), AAC2146AAT (6622C>T), GAG2149GAA (6631G>A), AAG2150AAA (6634G>A), TAT2151CAT (6635T>C), GCT2152GCA (6640T>A), TCG2154TCC (6646G>C), ATC2155GTC (6647A>G), ATC2156ATT (6652C>T), AGC2160AGT (6664C>T), ACT2162ACG (6670T>G), ATA2165GTG (6677A>G 6679A>G), GGC2167GGT (6685C>T), ACA2169ACT (6691A>T), TTC2170TTT (6694C>T), ACA2177ACG (6715A>G), GAT2178AAT (6716G>A), AAT2179GAT (6719A>G), GGA2180GGG (6724A>G), CTA2184TTA (6734C>T), GTG2186GTA (6742G>A), GTG2187GTA (6745G>A), GCC2190GCT (6754C>T), GGA2192GGG (6760A>G), ATA2194TTG (6764A>T 6766A>G), ATT2197ATA (6775T>A), CTG2200TTG (6782C>T), GCA2201GCT (6787A>T), AAC2202AAT (6790C>T), GCA2204GTG (6795C>T 6796A>G), CAC2205CAA (6799C>A), ACC2206ACA (6802C>A), AAC2208AAT (6808C>T), TAC2209TAT (6811C>T), TAC2210TAT (6814C>T), TTC2213TTT (6823C>T), GAA2215GAG (6829A>G), AAG2220AGG (6843A>G), TAC2221TAT (6847C>T), CGA2223CGC (6853A>C), AAT2225GAT (6857A>G), GAG2226GAA (6862G>A), GAA2229GAG (6871A>G), GTC2231ACA (6875G>A 6876T>C 6877C>A), AAG2232AAA (6880G>A), TCT2233TCA (6883T>A), GTT2235GTG (6889T>G), GAC2239GAT (6901C>T), GGC2244GGT (6916C>T), CCG2245CCA (6919G>A), AAA2247AAG (6925A>G), CTT2248CTC (6928T>C), GAC2250GAA (6934C>A), GGG2255GGC (6949G>C), TTC2257TTT (6955C>T), AAA2258AAG (6958A>G), CAA2264CAG (6976A>G), CTA2266TTA (6980C>T), ATC2267ATT (6985C>T), GTA2271GTT (6997A>T), GTG2272GTT (7000G>T), GTG2273ATA (7001G>A 7003G>A), GCT2276GCC (7012T>C), AAG2277AAA (7015G>A), CAT2278CAC (7018T>C), TTT2283TAT (7032T>A), GCC2285GCT (7039C>T), TTG2286CTA (7040T>C 7042G>A), GGA2288GGG (7048A>G), GCT2292GCA (7060T>A), GTC2293GTA (7063C>A), AGC2298AGT (7078C>T), TTA2300CTG (7082T>C 7084A>G), GTA2301GTG (7087A>G), ACC2302ACA (7090C>A), CAT2304CAC (7096T>C), GTA2305GTG (7099A>G), GTT2306GTC (7102T>C), GAG2309GAA (7111G>A), CGA2311CGG (7117A>G), ACA2314AAA (7125C>A), GAA2315GAG (7129A>G), TTT2316TTC (7132T>C), CTG2317TTA (7133C>T 7135G>A), GTG2319GTA (7141G>A), GAT2320GAC (7144T>C), GCA2321TCG (7145G>T 7147A>G), GAG2322GAA (7150G>A), GAG2324GAA (7156G>A), GCA2325GCT (7159A>T), GAG2326_GCA2327del (7160_7165delGAGGCA), TTC2329TTT (7171C>T), GCG2335GCT (7189G>T), AAA2338AAG (7198A>G), AGC2339AGT (7201C>T), CTA2341TTA (7205C>T), AAT2342AAC (7210T>C), GAT2344GAA (7216T>A), GCG2345GCA (7219G>A), ATC2347ATA (7225C>A), AAG2348AAA (7228G>A), AAG2352AAA (7240G>A), TAT2353TAC (7243T>C), AAA2355AAG (7249A>G), CCT2356CCG (7252T>G), GAT2358GAA (7258T>A), GTT2359GTC (7261T>C), GGT2360GGA (7264T>A), GTC2361GTG (7267C>G), GTG2362GTA (7270G>A), GAT2363_CGG2364insT-- (7273_7274insT), CGG2364GTG (7274C>G 7275G>T), CAT2366CTT (7281A>T), TTG2367TCG (7284T>C), TCA2371TTA (7296C>T), TCT2376TTT (7311C>T), ACC2377ATC (7314C>T), TGC2378TAC (7317G>A), AAT2379-AA (7319delA 7321T>A), CAC2381CAT (7327C>T), GGC2382GGT (7330C>T), TTC2383TTT (7333C>T), AAG2384CAG (7334A>C), TGT2386TGC (7342T>C), GCA2387AAC (7343G>A 7344C>A 7345A>C), ACT2390ACC (7354T>C), CAA2393CAG (7363A>G), TTC2396TTT (7372C>T), GCG2398GCT (7378G>T), CTC2399CTT (7381C>T), GCA2404GCT (7396A>T), GTC2405GTT (7399C>T), GCC2407GCT (7405C>T), AGT2408ATG (7407G>T 7408T>G), GGG2410GGT (7414G>T), TGC2411GGC (7415T>G), AAA2412AAG (7420A>G), GAC2415GAT (7429C>T), TAT2416TAC (7432T>C), CAT2419CAC (7441T>C), AAG2425AAA (7459G>A), GAA2426GAG (7462A>G), ATA2428ATT (7468A>T), GTC2429GTA (7471C>A), CAA2431CAG (7477A>G), AGC2432AGT (7480C>T), CTG2434CTT (7486G>T), TAT2437TAC (7495T>C), AAA2438AAG (7498A>G), GGT2439GGC (7501T>C), TTG2440TCA (7503T>C 7504G>A), ATT2443ATA (7513T>A), AAC2445AAT (7519C>T), GGA2446GGG (7522A>G), AAG2449AAA (7531G>A), GAG2451GAA (7537G>A), CTC2452CTT (7540C>T), TGT2454TGC (7546T>C), AAG2462AAA (7570G>A), ACG2463ACA (7573G>A), ACG2465ACA (7579G>A), GCT2468GCC (7588T>C), CTA2471TTA (7595C>T), TTG2474CTA (7604T>C 7606G>A), GGT2477GGA (7615T>A), TGT2480TGC (7624T>C), GAC2483GAT (7633C>T), TTC2484TTT (7636C>T), AAT2485AAC (7639T>C), TTT2488TTC (7648T>C), GAA2494GAG (7666A>G), TGT2496TGC (7672T>C), ACA2498ACT (7678A>T), GGG2500GGA (7684G>A), ACT2502ACC (7690T>C), TTT2504TTC (7696T>C), GGT2506GGG (7702T>G), GAT2509AAT (7709G>A), AAA2510AGG (7713A>G 7714A>G), CTG2511TTG (7715C>T), CGT2514CGC (7726T>C), GAG2517GAA (7735G>A), GTA2520GTG (7744A>G), TAC2521TAT (7747C>T), GGC2526GGT (7762C>T), CAG2528CAA (7768G>A), TTT2529TTC (7771T>C), TCA2532TCG (7780A>G), CTA2533CTC (7783A>C), ACT2534ACC (7786T>C), CTA2537TTG (7793C>T 7795A>G), ATC2538ATT (7798C>T), GCT2540GCC (7804T>C), ACC2543ATC (7812C>T), ATC2544ATT (7816C>T), GAA2550GAG (7834A>G), GAC2551GAT (7837C>T), GTG2554TTA (7844G>T 7846G>A), CAG2557CAA (7855G>A), CTG2559CTA (7861G>A), TTA2562CTG (7868T>C 7870A>G), TAC2563TAT (7873C>T), ACT2564ACA (7876T>A), GAG2565GAA (7879G>A), ATT2566ATA (7882T>A), GTT2567ATT (7883G>A), TAC2568TAT (7888C>T), CCA2570CCT (7894A>T), ATT2571ATC (7897T>C), GTC2579GTT (7921C>T), AAG2581AAA (7927G>A), GGT2584GGC (7936T>C), AAT2585AAC (7939T>C), AAT2586AAC (7942T>C), GTG2594GTA (7966G>A), GAT2595GAC (7969T>C), TCT2597TCC (7975T>C), GTC2601GTT (7987C>T), CTT2602CTC (7990T>C), TAC2606TAT (8002C>T), GCT2607GCC (8005T>C), GAA2616GAG (8032A>G), ATC2618ATT (8038C>T), GAC2619GAT (8041C>T), AGC2620AAC (8043G>A), TTG2632TTA (8080G>A), GTG2635GTA (8089G>A), CCG2637CCA (8095G>A), GAG2638GAA (8098G>A), AGC2641AGT (8107C>T), ATT2642ATC (8110T>C), GAT2644GAC (8116T>C), CAA2648CAG (8128A>G), GGT2654GGC (8146T>C), AAC2656AGT (8151A>G 8152C>T), TCG2660TCA (8164G>A), AGG2665AAG (8178G>A), AAG2666AAA (8182G>A), TTG2669TTA (8191G>A), TTC2671TTT (8197C>T), TCC2673TCT (8203C>T), AGA2675AAA (8208G>A), GGC2676GGG (8212C>G), CTG2677TTG (8213C>T), CTA2678CTG (8218A>G), ATC2679ATT (8221C>T), GAG2680GAA (8224G>A), CCA2685CCG (8239A>G), ATT2692ATC (8260T>C), GTA2693GTG (8263A>G), GAT2699GAC (8281T>C), AGA2700AGG (8284A>G), CCA2704GCG (8294C>G 8296A>G), GAG2705GAA (8299G>A), CAC2706CAT (8302C>T), ATT2711ATC (8317T>C), TGC2712TGT (8320C>T), GCT2714GCA (8326T>A), GAG2717GAA (8335G>A), TCC2718TCT (8338C>T), TCT2722TTT (8349C>T), GAA2723GAG (8353A>G), CAC2726CAT (8362C>T), CAA2727CAG (8365A>G), TAC2732TAT (8380C>T), TCA2733TCG (8383A>G), TTA2735TTG (8389A>G), CCT2740CCC (8404T>C), GCA2742TCA (8408G>T), ACA2743ACG (8413A>G), GCG2745GCT (8419G>T), GCT2750GCC (8434T>C), TAT2752TAC (8440T>C), ATG2756ATA (8452G>A), CTA2758CTG (8458A>G), AGG2759AAG (8460G>A), AAA2760AAG (8464A>G), CTG2761CTA (8467G>A), GCT2766ACA (8480G>A 8482T>A), GAG2771GAA (8497G>A), CTA2772TTG (8498C>T 8500A>G), AGA2773AAG (8502G>A 8503A>G), GCC2774GCT (8506C>T), GAA2777GAT (8515A>T), GTC2780GTT (8524C>T), GCA2781GCC (8527A>C), TTA2782CTG (8528T>C 8530A>G), GAG2785GAC (8539G>C), CTT2788TGT (8546C>T 8547T>G), TCT2790ACT (8552T>A), GTA2793GTG (8563A>G), GCA2797GGA (8574C>G), ATT2801GTT (8585A>G), GGA2805GAA (8598G>A), AGC2806AGT (8602C>T), AAC2807TCT (8603A>T 8604A>C 8605C>T), GCA2811GTA (8616C>T), AAA2812AAG (8620A>G), CCA2813CAA (8622C>A), GAG2814GGA (8625A>G 8626G>A), CAG2815CAA (8629G>A), GGC2816GGT (8632C>T), ATC2818ATT (8638C>T), CAG2819CAA (8641G>A), AAC2821AAT (8647C>T), CCG2822TTC (8648C>T 8649C>T 8650G>C), AAC2823AAT (8653C>T), AAA2824AAG (8656A>G), GGA2825GAA (8658G>A), GAT2827GCG (8664A>C 8665T>G), GTT2830GTG (8674T>G), GCA2832GTT (8679C>T 8680A>T), GGC2833GGA (8683C>A), GGG2836GGA (8692G>A), ACA2837ACT (8695A>T), CCG2841CCA (8707G>A), AGA2842CGA (8708A>C), AAG2844AAA (8716G>A), ATC2846ATT (8722C>T), ACG2847ACA (8725G>A), AAA2849AAG (8731A>G), AGA2851AGG (8737A>G), CCC2853CCT (8743C>T), ACA2854AAG (8745C>A 8746A>G), AGC2855AGT (8749C>T), AAG2856AAA (8752G>A), GGA2857GGT (8755A>T), AAC2862AAT (8770C>T), TTA2863CTA (8771T>C), GAA2864GAG (8776A>G), CAT2865CAC (8779T>C), TTG2866TTA (8782G>A), TAT2869TAC (8791T>C), CCA2871CCG (8797A>G), CAA2872CAG (8800A>G), CAA2873CAG (8803A>G), GAT2875GAC (8809T>C), ATT2876ATC (8812T>C), AAT2878AAC (8818T>C), CGG2880CGA (8824G>A), GCA2881GCG (8827A>G), ACG2888ACA (8848G>A), TAT2890TAC (8854T>C), GAG2891GGA (8856A>G 8857G>A), GTG2893GTA (8863G>A), CGG2894CAG (8865G>A), ATG2895CTT (8867A>C 8869G>T), GCA2896GCG (8872A>G), GGA2900GAA (8883G>A), ACT2902TCT (8888A>T), TGC2915TGT (8929C>T), GTC2924ATC (8954G>A), AAC2925AAT (8959C>T), GGA2926GGT (8962A>T), GGG2933GGA (8983G>A), AAT2934GAC (8984A>G 8986T>C), GAA2935GTA (8988A>T), GAG2938GAA (8998G>A), CCG2940CCA (9004G>A), TTG2941TTA (9007G>A), ATC2944ATT (9016C>T), ACC2951ACA (9037C>A), ATA2967ATT (9085A>T), AAC2971AAT (9097C>T), CGA2984CGT (9136A>T), CGG2987CGT (9145G>T), ATG2989GGA (9149A>G 9150T>G 9151G>A), GGT2990AGT (9152G>A), TTA2991TTG (9157A>G), GCG2992GCT (9160G>T), CGT2993CGC (9163T>C), GAG3000GAA (9184G>A), CGA3004CGG (9196A>G), AGG3010AGA (9214G>A), GAA3011GAG (9217A>G), GCG3012GCA (9220G>A), GCA3020GCT (9244A>T), TTG3021TTA (9247G>A), GCC3024GCT (9256C>T), CCT3026TCC (9260C>T 9262T>C), GGG3030GGA (9274G>A), GAC3032GAT (9280C>T), ATC3035ATT (9289C>T), GAG3040GAA (9304G>A), GTC3050GTT (9334C>T), GTC3060GTG (9364C>G) | | | | | | | | |
| PVYgp2 | 1 | 75 | 100% | 95 | 18.1% | 75 (98.7%) | 54 (71.1%) | 1/0/3/2 | 2 |
| Protein mutations: | R17_K17insX (2971_2972insA), Y23H (2989T>C 2991C>T), R26K (2999G>A 3000A>G), H35Y (3025C>T), K40H (3040A>C 3042A>C), G41S (3043G>A), E44K (3052G>A), I47V (3061A>G), V54I (3082G>A), P57S (3091C>T), P60T (3100C>A), G61E (3104G>A 3105T>G), G62S (3106G>A), Q63* (3109C>T), R64G (3112A>G), C66R (3118T>C), L67F (3121C>T), I69V (3127A>G), A72E (3137C>A 3138G>A), I73V (3139A>G) | | | | | | | | |
| Codon mutations: | AAA4AAG (2931A>G), GAA7GAG (2940A>G), TTG10CTG (2947T>C), AGA17_AAA17insAA- (2971_2972insA), TAC23CAT (2989T>C 2991C>T), TCA24TCG (2994A>G), AGA26AAG (2999G>A 3000A>G), CAC32-AC (3016delC), TCG33TCA (3021G>A), GTA34GTT (3024A>T), CAT35TAT (3025C>T), CAC38CAT (3036C>T), AGA39AGG (3039A>G), AAA40CAC (3040A>C 3042A>C), GGC41AGC (3043G>A), GAA44AAA (3052G>A), ATA47GTA (3061A>G), GTT54ATT (3082G>A), CCG57TCG (3091C>T), CGC59CGT (3099C>T), CCA60ACA (3100C>A), GGT61GAG (3104G>A 3105T>G), GGT62AGT (3106G>A), CAA63TAA (3109C>T), AGG64GGG (3112A>G), TGC66CGC (3118T>C), CTC67TTC (3121C>T), ATT69GTT (3127A>G), GAG70GAA (3132G>A), GCG72GAA (3137C>A 3138G>A), ATT73GTT (3139A>G) | | | | | | | | |

#### Proteins

| | Begin | End | Coverage | Score | Concordance | Matches | Identities | I/D/M/F* | Stop Codons |
|---|---|---|---|---|---|---|---|---|---|
| **NT** | **19** | **9704** | **99.8%** | **12534** | **65.0%** | **9674 (99.7%)** | **8011 (82.6%)** | **15/12** | |
| polyprotein (NP_056759.1) | 1 | 3064 | 100% | 18956 | 89.8% | 3062 (99.9%) | 2776 (90.5%) | 2/2/4/4 | 1 |
| Protein mutations: | Y4Q (194T>C 196C>A), C9L (209T>C 210G>T 211T>G), F13M (221T>A 223T>G), S23P (251T>C), C24F (255G>T 256C>T), E25G (258A>G), I27V (263A>G 265T>G), V28A (267T>C), E30V (273A>T), A35T (287G>A 289T>C), V37T (293G>A 294T>C), A41I (305G>A 306C>T), D42G (309A>G), E44D (316A>T), T45V (317A>G 318C>T 319A>G), L52Q (339T>A 340C>A), K53R (342A>G), K55A (347A>G 348A>C 349A>G), Y56H (350T>C), T58V (356A>G 357C>T), V61T (365G>A 366T>C 367G>A), L62S (368C>T 369T>C), F67R (383T>C 384T>G), A75V (408C>T 409C>T), M78E (416A>G 417T>A), E85K (437G>A), R86E (440A>G 441G>A 442G>A), K87R (443A>C 444A>G 445G>A), D88E (448T>A), E91A (456A>C), H93N (461C>A), D111E (517T>A), S116F (531C>T), P118S (536C>T), Q119W (539C>T 540A>G 541A>G), R122K (549G>A 550A>G), V134A (585T>C 586C>G), R137Y (593C>T 594G>A 595C>T), P138R (597C>G), I139V (599A>G 601A>G), I140P (602A>C 603T>C 604A>G), M147I (625G>A), I151V (635A>G 637T>C), I154V (644A>G), E160A (663A>C 664G>C), H166Q (682C>A), T172S (699C>G), H174Q (706T>A), K179E (719A>G 721G>A), I180V (722A>G 724A>G), A183S (731G>T), Y184P (734T>C 735A>C 736C>T), S185R (737T>C 738C>G 739C>T), A187V (744C>T 745G>C), R189C (749C>T 751A>C), M194K (765T>A), R197K (773C>A 774G>A 775A>G), M206K (801T>A), G210A (813G>C 814A>G), L211H (816T>A), R214Q (825G>A 826T>A), R226C (860C>T 862C>T), T227A (863A>G 865T>C), I228T (867T>C), N229D (869A>G 871C>T), I230L (872A>T 874A>G), R232K (879G>A), N240S (903A>G), T241N (906C>A 907A>C), K242T (909A>C 910A>C), S243N (912G>A 913C>T), S252L (938T>C 939C>T 940A>T), L266I (980T>A 982G>C), R272K (999G>A), Q275R (1007C>A 1008A>G 1009G>A), S276G (1010A>G 1012T>G), I277V (1013A>G), N279Q (1019A>C 1021C>G), D288E (1048C>A), R300Q (1083G>A), S305A (1097T>G 1099G>A), R320K (1143G>A), L324I (1154T>A 1156G>A), S346N (1221G>A), V349T (1229G>A 1230T>C 1231T>C), S350N (1233G>A 1234C>T), K358T (1257A>C 1258A>G), N365S (1278A>G), A369V (1290C>T 1291G>A), D372T (1298G>A 1299A>C 1300C>T), I375V (1307A>G 1309A>G), N378E (1316A>G 1318T>A), I382T (1329T>C), L397I (1373C>A), I401V (1385A>G), E407D (1405G>T), A411S (1415G>T 1417A>T), V418I (1436G>A 1438C>T), E508G (1707A>G), A512T (1718G>A), I520V (1742A>G 1744C>T), S525N (1758G>A), V536I (1790G>A), I585V (1937A>G 1939T>C), F607Y (2004T>A), I628M (2068T>G), V630I (2072G>A 2074G>A), R675K (2208G>A), D694E (2266C>G), N744G (2414A>G 2415A>G 2416T>G), S746C (2420A>T), K780R (2523A>G), R817K (2634G>A), V847I (2723G>A), A855T (2747G>A 2749T>A), H871N (2795C>A), V880I (2822G>A 2824G>A), P900T (2882C>A), S901H (2885A>C 2886G>A), T904M (2895C>T), N916S (2931A>G), N919S (2940A>G), E929_N930insX (2971_2972insA), T936I (2991C>T), H937R (2994A>G), E939R (2999G>A 3000A>G), R946Q (3021G>A), Y947F (3024A>T 3025C>T), T951I (3036C>T), E952G (3039A>G 3040A>C), K953T (3042A>C 3043G>A), A972V (3099C>T 3100C>A), V974R (3104G>A 3105T>G 3106G>A), S983N (3132G>A), R985K (3137C>A 3138G>A 3139A>G), F990I (3152T>A 3154C>T), C994Y (3165G>A), F1012L (3218T>C), V1016I (3230G>A), V1028M (3266G>A 3268A>G), V1031M (3275G>A), A1034T (3284G>A), R1044K (3315G>A 3316A>G), E1045A (3318A>C), M1059V (3359A>G), D1072E (3400T>G), V1110G (3513T>G), M1121V (3545A>G), A1122T (3548G>A), V1124I (3554G>A 3556C>T), L1147V (3623C>G), L1150M (3632C>A), Y1152H (3638T>C), I1172V (3698A>G), S1178N (3717G>A), R1183K (3731C>A 3732G>A), I1297V (4073A>G), F1315Y (4128T>A 4129C>T), A1323S (4151G>T), V1328I (4166G>A), L1347I (4223C>A 4225G>A), I1379V (4319A>G 4321A>G), F1404Y (4395T>A), V1408I (4406G>A), M1431V (4475A>G 4477G>T), T1433S (4481A>T), A1454L (4544G>T 4545C>T 4546G>A), R1455K (4548G>A), V1499I (4679G>A 4681G>C), V1538I (4796G>A 4798C>T), T1540S (4803C>G 4804T>C), S1625G (5057A>G), T1626A (5060A>G), L1636I (5090C>A), A1639V (5100C>T), E1641D (5107A>T), V1646F (5120G>T 5122C>T), I1648V (5126A>G), V1667I (5183G>A), A1793T (5561G>A 5563G>A), I1805V (5597A>G), N1807K (5605C>A), A1812V (5619C>T 5620T>G), I1816L (5630A>C), V1837I (5693G>A), K1890R (5853A>G), I1904V (5894A>G 5896C>G), R1926A (5960C>G 5961G>C 5962G>C), I1936V (5990A>G), E1945D (6019A>T), K1949T (6030A>C), D1955E (6049C>A), M1958I (6058G>A), S1963N (6072G>A 6073T>C), N1964H (6074A>C), T1966N (6081C>A), C1976S (6111G>C 6112T>C), I1982V (6128A>G 6130T>C), V1992I (6158G>A), T1996S (6170A>T 6172A>G), L2007F (6203C>T 6205C>T), V2016I (6230G>A), A2025K (6257G>A 6258C>A), K2033R (6282A>G), A2060T (6362G>A), V2073I (6401G>A), A2074V (6405C>T), Y2082F (6429A>T), Q2089R (6450A>G), H2101N (6485C>A 6487C>T), L2115I (6527C>A), T2135I (6588C>T), Y2151H (6635T>C), I2155V (6647A>G), I2165V (6677A>G 6679A>G), D2178N (6716G>A), N2179D (6719A>G), I2194L (6764A>T 6766A>G), A2204V (6795C>T 6796A>G), H2205Q (6799C>A), K2220R (6843A>G), N2225D (6857A>G), V2231T (6875G>A 6876T>C 6877C>A), D2250E (6934C>A), V2273I (7001G>A 7003G>A), F2283Y (7032T>A), T2314K (7125C>A), A2321S (7145G>T 7147A>G), E2326_A2327del (7160_7165delGAGGCA), D2344E (7216T>A), D2358E (7258T>A), D2363_R2364insX (7273_7274insT), R2364V (7274C>G 7275G>T), H2366L (7281A>T), L2367S (7284T>C), S2371L (7296C>T), S2376F (7311C>T), T2377I (7314C>T), C2378Y (7317G>A), K2384Q (7334A>C), A2387N (7343G>A 7344C>A 7345A>C), S2408M (7407G>T 7408T>G), C2411G (7415T>G), L2440S (7503T>C 7504G>A), D2509N (7709G>A), K2510R (7713A>G 7714A>G), T2543I (7812C>T), V2554L (7844G>T 7846G>A), V2567I (7883G>A), S2620N (8043G>A), N2656S (8151A>G 8152C>T), R2665K (8178G>A), R2675K (8208G>A), P2704A (8294C>G 8296A>G), S2722F (8349C>T), A2742S (8408G>T), M2756I (8452G>A), R2759K (8460G>A), A2766T (8480G>A 8482T>A), R2773K (8502G>A 8503A>G), E2777D (8515A>T), E2785D (8539G>C), L2788C (8546C>T 8547T>G), S2790T (8552T>A), A2797G (8574C>G), I2801V (8585A>G), G2805E (8598G>A), N2807S (8603A>T 8604A>C 8605C>T), A2811V (8616C>T), P2813Q (8622C>A), E2814G (8625A>G 8626G>A), P2822F (8648C>T 8649C>T 8650G>C), G2825E (8658G>A), D2827A (8664A>C 8665T>G), A2832V (8679C>T 8680A>T), T2854K (8745C>A 8746A>G), E2891G (8856A>G 8857G>A), R2894Q (8865G>A), M2895L (8867A>C 8869G>T), G2900E (8883G>A), T2902S (8888A>T), V2924I (8954G>A), N2934D (8984A>G 8986T>C), E2935V (8988A>T), M2989G (9149A>G 9150T>G 9151G>A), G2990S (9152G>A), P3026S (9260C>T 9262T>C) | | | | | | | | |

| | Begin | End | Coverage | Score | Concordance | Matches | Identities | I/D/M/F* | Stop Codons |
|---|---|---|---|---|---|---|---|---|---|
| **NT** | **19** | **9704** | **99.8%** | **12534** | **65.0%** | **9674 (99.7%)** | **8011 (82.6%)** | **15/12** | |

Codon mutations:

ACT3ACC (193T>C), TAC4CAA (194T>C 196C>A), TCA6TCG (202A>G), ATC8ATT (208C>T), TGT9CTG (209T>C 210G>T 211T>G), TCG12TCA (220G>A), TTT13ATG (221T>A 223T>G), GAA14GAG (226A>G), CTA17CTT (235A>T), TAC19TAT (241C>T), CCA21CCT (247A>T), TCT23CCT (251T>C), TGC24TTT (255G>T 256C>T), GAG25GGG (258A>G), CAT26CAC (262T>C), ATT27GTG (263A>G 265T>G), GTG28GCG (267T>C), GAA30GTA (273A>T), CGA31CGT (277A>T), GAA32GAG (280A>G), GTG33GTT (283G>T), CCG34CCA (286G>A), GCT35ACC (287G>A 289T>C), GTT37ACT (293G>A 294T>C), CCT39CCC (301T>C), TTC40TTT (304C>T), GCA41ATA (305G>A 306C>T), GAT42GGT (309A>G), CTG43TTG (311C>T), GAA44GAT (316A>T), ACA45GTG (317A>G 318C>T 319A>G), CAA46CAG (322A>G), GCA49GCT (331A>T), CTC52CAA (339T>A 340C>A), AAG53AGG (342A>G), AAA55GCG (347A>G 348A>C 349A>G), TAT56CAT (350T>C), GCT57GCA (355T>A), ACT58GTT (356A>G 357C>T), CGT60CGC (364T>C), GTG61ACA (365G>A 366T>C 367G>A), CTC62TCC (368C>T 369T>C), AAA63AAG (373A>G), AAC64AAT (376C>T), TTT67CGT (383T>C 384T>G), ACG68ACA (388G>A), TAC69TAT (391C>T), CGA70CGG (394A>G), TAC71TAT (397C>T), AAG72AAA (400G>A), GAT74GAC (406T>C), GCC75GTT (408C>T 409C>T), CAG76CAA (412G>A), ATA77ATT (415A>T), ATG78GAG (416A>G 417T>A), ATT80ATC (424T>C), CAG81CAA (427G>A), AAA83AAG (433A>G), CTG84CTC (436G>C), GAG85AAG (437G>A), AGG86GAA (440A>G 441G>A 442G>A), AAG87CGA (443A>C 444A>G 445G>A), GAT88GAA (448T>A), GAA90GAG (454A>G), GAA91GCA (456A>C), TAT92TAC (460T>C), CAC93AAC (461C>A), CAA95CAG (469A>G), GCC97GCT (475C>T), CCT99CCA (481T>A), ACT106ACC (502T>C), ATA107ATT (505A>T), GGC109GGT (511C>T), GAT111GAA (517T>A), CCT112CCA (520T>A), CCA113CCT (523A>T), TCT116TTT (531C>T), GAG117GAA (535G>A), CCA118TCA (536C>T), CAA119TGG (539C>T 540A>G 541A>G), AGA122AAG (549G>A 550A>G), ATC124ATT (556C>T), ATT125ATC (559T>C), CAT126CAC (562T>C), ACT128ACC (568T>C), CCA129CCG (571A>G), CGT132CGC (580T>C), GTC134GCG (585T>C 586C>G), CGC137TAT (593C>T 594G>A 595C>T), CCC138CGC (597C>G), ATA139GTG (599A>G 601A>G), ATA140CCG (602A>C 603T>C 604A>G), ACA143ACG (613A>G), GAA144GAG (616A>G), GGC145GGG (619C>G), CAG146CAA (622G>A), ATG147ATA (625G>A), AAT148AAC (628T>C), CAC149CAT (631C>T), ATT151GTC (635A>G 637T>C), ATA154GTA (644A>G), ATT157ATC (655T>C), TCG159TCA (661G>A), GAG160GCC (663A>C 664G>C), AAA161AAG (667A>G), GGG163GGC (673G>C), GTC165GTT (679C>T), CAC166CAA (682C>A), TTA167CTA (683T>C), AGT169AGC (691T>C), ACC172AGC (699C>G), ACT173ACA (703T>A), CAT174CAA (706T>A), AAG179GAA (719A>G 721G>A), ATA180GTG (722A>G 724A>G), CTT181TTG (725C>T 727T>G), GGT182GGA (730T>A), GCA183TCA (731G>T), TAC184CCT (734T>C 735A>C 736C>T), TCC185CGT (737T>C 738C>G 739C>T), GCG187GTC (744C>T 745G>C), CGA189TGC (749C>T 751A>C), ACT190ACC (754T>C), ATG194AAG (765T>A), TTG196CTG (770T>C), CGA197AAG (773C>A 774G>A 775A>G), CGG198AGG (776C>A), TTC202TTT (790C>T), ATG206AAG (801T>A), GGA210GCG (813G>C 814A>G), CTT211CAT (816T>A), TTG212CTG (818T>C), CAA213CAG (823A>G), CGT214CAA (825G>A 826T>A), CGG217AGA (833C>A 835G>A), GAC219GAT (841C>T), AAT223AAC (853T>C), GTC225GTT (859C>T), CGC226TGT (860C>T 862C>T), ACT227GCC (863A>G 865T>C), ATC228ACC (867T>C), AAC229GAT (869A>G 871C>T), ATA230TTG (872A>T 874A>G), AGG232AAG (879G>A), GAT234GAC (886T>C), ATC238ATT (898C>T), AAC240AGC (903A>G), ACA241AAC (906C>A 907A>C), AAA242ACC (909A>C 910A>C), AGC243AAT (912G>A 913C>T), GGC246GGA (922C>A), GGT249GGA (931T>A), TCA252CTT (938T>C 939C>T 940A>T), GGA253GGG (943A>G), TTC256TTT (952C>T), TCA261TCG (967A>G), GAA263GAG (973A>G), GGG264GGC (976G>C), TTG266ATC (980T>A 982G>C), GCA269GCG (991A>G), CGT270CGC (994T>C), AGA272AAA (999G>A), GTT273GTC (1003T>C), CAG275AGA (1007C>A 1008A>G 1009G>A), AGT276GGG (1010A>G 1012T>G), ATT277GTT (1013A>G), TTA278TTG (1018A>G), AAC279CAG (1019A>C 1021C>G), ATC282ATT (1030C>T), TCG285TCA (1039G>A), AAT286AAC (1042T>C), GCC287GCA (1045C>A), GAC288GAA (1048C>A), AAT289AAC (1051T>C), AAG292AAA (1060G>A), GGT293GGA (1063T>A), GAC295GAT (1069C>T), CGA300CAA (1083G>A), TAT303TAC (1093T>C), TCG305GCA (1097T>G 1099G>A), CAC307CAT (1105C>T), GTA310GTG (1114A>G), GCT311GCG (1117T>G), TTA313TTG (1123A>G), CCT314CCA (1126T>A), GTC315GTT (1129C>T), AGG320AAG (1143G>A), GTA321GTT (1147A>T), GCT322GCA (1150T>A), TTG324ATA (1154T>A 1156G>A), CAC327CAT (1165C>T), AGT328AGC (1168T>C), CTT330TTA (1172C>T 1174T>A), ACC339ACT (1201C>T), AGC346AAC (1221G>A), GTT349ACC (1229G>A 1230T>C 1231T>C), AGC350AAT (1233G>A 1234C>T), CTG352CTA (1240G>A), AAG354AAA (1246G>A), TTG356CTA (1250T>C 1252G>A), CAT357CAC (1255T>C), AAA358ACG (1257A>C 1258A>G), AGA361AGG (1267A>G), GGT363GGC (1273T>C), TTG364TTA (1276G>A), AAT365AGT (1278A>G), TTG367CTG (1283T>C), GCG369GTA (1290C>T 1291G>A), GAC372ACT (1298G>A 1299A>C 1300C>T), TTT374TTC (1306T>C), ATA375GTG (1307A>G 1309A>G), CAT376CAC (1312T>C), AAT378GAA (1316A>G 1318T>A), TTG381TTA (1327G>A), ATA382ACA (1329T>C), GCG383GCT (1333G>T), TTA384TTG (1336A>G), CTA387TTG (1343C>T 1345A>G), CCG390CCA (1354G>A), GTG391GTT (1357G>T), GAC392GAT (1360C>T), CTC395TTG (1367C>T 1369C>G), CTT397ATT (1373C>A), TTC398TTT (1378C>T), GAG400GAA (1384G>A), ATA401GTA (1385A>G), AAA403AAG (1393A>G), TCC404TCT (1396C>T), GAG407GAT (1405G>T), AAA408AAG (1408A>G), CAG409CAA (1411G>A), GCA411TCT (1415G>T 1417A>T), CCG412CCC (1420G>C), AAT415AAC (1429T>C), TTA416TTG (1432A>G), GTC418ATT (1436G>A 1438C>T), TTA419CTG (1439T>C 1441A>G), AAT420AAC (1444T>C), TTC423TTT (1453C>T), CTG424CTT (1456G>T), GAA428GAG (1468A>G), TTG439CTA (1499T>C 1501G>A), AGT440AGC (1504T>C), TTG441TTA (1507G>A), CTC442CTT (1510C>T), GAA443GAG (1513A>G), TTA444TTG (1516A>G), GCA445GCT (1519A>T), AGG446AGA (1522G>A), TTC447TTT (1525C>T), CAG448CAA (1528G>A), AAC450AAT (1534C>T), ACT452ACA (1540T>A), AAC454AAT (1546C>T), AAA457AAG (1555A>G), GAT459GAC (1561T>C), ATA460ATC (1564A>C), TCT461TCA (1567T>A), TTC463TTT (1573C>T), AGA464AGG (1576A>G), TTA467CTA (1583T>C), TCT468TCA (1588T>A), AAG470AAA (1594G>A), AAC472AAT (1600C>T), CTG475CTA (1609G>A), TCG478TCA (1618G>A), TGC479TGT (1621C>T), GAC480GAT (1624C>T), AAC481AAT (1627C>T), CTG483TTG (1631C>T), AAA485AAG (1639A>G), TTC489TTT (1651C>T), CTC490CTG (1654C>G), CAT497CAC (1675T>C), TCA503TCG (1693A>G), TTT506TTC (1702T>C), GAA508GGA (1707A>G), GCA512ACA (1718G>A), GGA514GGG (1726A>G), TAC515TAT (1729C>T), TCA516TCC (1732A>C), TAT518TAC (1738T>C), ATC520GTT (1742A>G 1744C>T), CGC521CGT (1747C>T), CAT523CAC (1753T>C), AGT525AAT (1758G>A), AGG528AGA (1768G>A), AAC534AAT (1786C>T), TTA535TTG (1789A>G), GTT536ATT (1790G>A), GTC537GTT (1795C>T), TTA541CTG (1805T>C 1807A>G), GAG543GAA (1813G>A), AGG545AGA (1819G>A), AAG547AAA (1825G>A), GAC551GAT (1837C>T), TAT552TAC (1840T>C), CAA555CAG (1849A>G), GTC558GTG (1858C>G), AAA560AAG (1864A>G), TGC562TGT (1870C>T), AGT564AGC (1876T>C), AAA566AAG (1882A>G), GGT568GGA (1888T>A), AAT569AAC (1891T>C), TGT574TGC (1906T>C), ACA577ACT (1915A>T), GGT582GGC (1930T>C), GCC584GCT (1936C>T), ATT585GTC (1937A>G 1939T>C), TCA587TCG (1945A>G), ACA588ACG (1948A>G), TTC589TTT (1951C>T), CCA592CCT (1960A>T), GTA598GTG (1978A>G), ATT599ATA (1981T>A), GGC600GGT (1984C>T), AAT601AAC (1987T>C), GAC604GAT (1996C>T), TTT607TAT (2004T>A), AAA612AAG (2020A>G), GGG613GGA (2023G>A), GAT614GAC (2026T>C), TCG615TCT (2029G>T), GAG616GAA (2032G>A), TAT625TAC (2059T>C), ATT628ATG (2068T>G), AAC629AAT (2071C>T), GTG630ATA (2072G>A 2074G>A), CTG635TTA (2087C>T 2089G>A), AGC639AGT (2101C>T), GAG640GAA (2104G>A), GAG641GAA (2107G>A), AAG644AAA (2116G>A), TTC646TTT (2122C>T), CGC651CGT (2137C>T), CTT658CTC (2158T>C), ACC660ACT (2164C>T), CCA662CCG (2170A>G), GAT666GAC (2182T>C), GCG668GCA (2188G>A), ACC669ACA (2191C>A), AGA675AAA (2208G>A), TAT678TAC (2218T>C), GAC680GAT (2224C>T), GTA681GTT (2227A>T), TTG686TTA (2242G>A), CCC687CCT (2245C>T), TTG690TTA (2254G>A), GTT691GTC (2257T>C), GAC692GAT (2260C>T), GAC694GAG (2266C>G), CAA696CAG (2272A>G), ACG697ACA (2275G>A), TCA703TCG (2293A>G), GGC705GGT (2299C>T), TCG706TCA (2302G>A), CAG707CAA (2305G>A), GGA710GGT (2314A>T), TAT711TAC (2317T>C), ATT713ATC (2323T>C), CTA714CTG (2326A>G), GTG719GTA (2341G>A), CAA721CAG (2347A>G), CTT722CTC (2350T>C), TTG724TTA (2356G>A), AAT727AAC (2365T>C), GAA729GAG (2371A>G), TTA730TTG (2374A>G), GAT733GAC (2383T>C), ATA734ATT (2386A>T), AAA735AAG (2389A>G), CAT736CAC (2392T>C), TAT737TAC (2395T>C), AAT744GGG (2414A>G 2415A>G 2416T>G), GCT745GCA (2419T>A), AGC746TGC (2420A>T), ACA752ACT (2440A>T), GTT761GTC (2467T>C), TCT764TCC (2476T>C), TCG766TCA (2482G>A), CTG769CTA (2491G>A), CTG771CTT (2497G>T), TTG773TTA (2503G>A), AGA778AGG (2518A>G), CCT779CCC (2521T>C), AAG780AGG (2523A>G), AGA783AGG (2533A>G), TTG785TTA (2539G>A), TTA787TTG (2545A>G), GAG789GAA (2551G>A), CCT790CCA (2554T>A), CTG792TTG (2558C>T), TTG793CTC (2561T>C 2563G>C), CTA795CTG (2569A>G), TCA796TCG (2572A>G), CCT800CCC (2584T>C), GGC801GGT (2587C>T), CTG803CTC (2593G>C), GCT805GCC (2599T>C), AAT808AAC (2608T>C), TTT812TTC (2620T>C), GCG815GCA (2629G>A), GTG816GTA (2632G>A), AGG817AAG (2634G>A), TTG818TTA (2638G>A), ATT820ATA (2644T>A), AAA823AAG (2653A>G), GCT827GCC (2665T>C), CTA832TTA (2678C>T), CTA833CTG (2683A>G), GCT835GCC (2689T>C), CTA838CTC (2698A>C), TCA841TCG (2707A>G), GCG842GCA (2710G>A), GAA844GAG (2716A>G), ACA845ACG (2719A>G), CTC846CTT (2722C>T), GTC847ATC (2723G>A), GCA848GCG (2728A>G), AGG850AGA (2734G>A), ATT851ATA (2737T>A), GAT854GAC (2746T>C), GCT855ACA (2747G>A 2749T>A), GCA856GCT (2752A>T), GCT857GCA (2755T>A), GAC859GAT (2761C>T), CTC860CTT (2764C>T), TGT865TGC (2779T>C), GGG867GGT (2785G>T), AAC869AAT (2791C>T), CAT871AAT (2795C>A), TAC874TAT (2806C>T), TTG878CTC (2816T>C 2818G>C), GTG880ATA (2822G>A 2824G>A), TTG881TTA (2827G>A), GTT884GTC (2836T>C), AAG885AAA (2839G>A), AAT886AAC (2842T>C), ACC893ACA (2863C>A), CTA894TTA (2864C>T), CCA900ACA (2882C>A), AGT901CAT (2885A>C 2886G>A), AAC903AAT (2893C>T), ACG904ATG (2895C>T), GTC906GTT (2902C>T), GTA907GTG (2905A>G), ATT909ATC (2911T>C), AAT916AGT (2931A>G), AAC919AGC (2940A>G), GCT921GCC (2947T>C), GAA929_AAT930insA-- (2971_2972insA), GGT935GGC (2989T>C), ACT936ATT (2991C>T), CAT937CGT (2994A>G), GAG939AGG (2999G>A 3000A>G), TCC944TC- (3016delC), CGG946CAG (3021G>A), TAC947TTT (3024A>T 3025C>T), ACA951ATA (3036C>T), GAA952GGC (3039A>G 3040A>C), AAG953ACA (3042A>C 3043G>A), TTG956TTA (3052G>A), TTA959TTG (3061A>G), GCG966GCA (3082G>A), GGC969GGT (3091C>T), GCC972GTA (3099C>T 3100C>A), GTG974AGA (3104G>A 3105T>G 3106G>A), GTC975GTT (3109C>T), AAA976AAG (3112A>G), ACT978ACC (3118T>C), GCC979GCT (3121C>T), GGA981GGG (3127A>G), AGC983AAC (3132G>A), CGA985AAG (3137C>A 3138G>A 3139A>G), TAT989TAC (3151T>C), TTC990ATT (3152T>A 3154C>T), TGT994TAT (3165G>A), GTA995GTG (3169A>G), ATC1005ATT (3199C>T), AGG1007CGG (3203A>C), CGT1008CGC (3208T>C), CCA1010CCG (3214A>G), ACC1011ACT (3217C>T), TTT1012CTT (3218T>C), GTC1013GTA (3223C>A), GTT1016ATT (3230G>A), AAC1017AAT (3235C>T), TTA1025CTA (3257T>C), ACC1026ACT (3262C>T), AGC1027AGT (3265C>T), GTA1028ATG (3266G>A 3268A>G), GTG1029GTA (3271G>A), GTG1031ATG (3275G>A), GCA1034ACA (3284G>A), CAG1039CAA (3301G>A), AGA1044AAG (3315G>A 3316A>G), GAA1045GCA (3318A>C), GAG1047GAA (3325G>A), TTG1048TTA (3328G>A), ATA1051ATT (3337A>T), GAG1055GAA (3349G>A), GTC1057GTT (3355C>T), TGC1058TGT (3358C>T), ATG1059GTG (3359A>G), CTA1061CTG (3367A>G), TAT1062TAC (3370T>C), TTA1065CTG (3377T>C 3379A>G), GAA1070GAG (3394A>G), CGC1071CGT (3397C>T), GAT1072GAG (3400T>G), ACA1074ACG (3406A>G), TAC1078TAT (3418C>T), ATT1079ATA (3421T>A), TAT1081TAC (3427T>C), AAG1083AAA (3433G>A), TCA1084TCT (3436A>T), GTA1085GTG (3439A>G), AAC1086AAT (3442C>T), CCT1087CCC (3445T>C), CAG1091CAA (3457G>A), TTT1092TTC (3460T>C), GCT1093GCA (3463T>A), GCG1095GCT (3469G>T), CAG1096CAA (3472G>A), CAC1104CAT (3496C>T), CAG1105CAA (3499G>A), TCC1107TCT (3505C>T), GTT1110GGT (3513T>G), AAA1112AAG (3520A>G), GAA1115GAG (3529A>G), CAA1116CAG (3532A>G), ATG1121GTG (3545A>G), GCT1122ACT (3548G>A), TTA1123CTA (3551T>C), GTC1124ATT (3554G>A 3556C>T), AGT1133AGC (3583T>C), GAT1134GAC (3586T>C), TGC1135TGT (3589C>T), TTC1137TTT (3595C>T), AAT1141AAC (3607T>C), TTT1143TTC (3613T>C), AAG1144AAA (3616G>A), GGT1145GGC (3619T>C), CTT1147GTT (3623C>G), TCC1148TCT (3628C>T), CTG1150ATG (3632C>A), GAT1151GAC (3637T>C), TAT1152CAT (3638T>C), GAA1153GAG (3643A>G), TTA1159TTG (3661A>G), GTG1162GTA (3670G>A), TTT1166TTC (3682T>C), GAG1168GAA (3688G>A), GAG1171GAA (3697G>A), ATT1172GTT (3698A>G), GAT1174GAC (3706T>C), TTT1175TTC (3709T>C), TTG1177CTA (3713T>C 3715G>A), AGT1178AAT (3717G>A), GAG1179GAA (3721G>A), CGA1183AAA (3731C>A 3732G>A), ACT1184ACA (3736T>A), CTA1188TTG (3746C>T 3748A>G), ACA1190ACG (3754A>G), AGT1193AGC (3763T>C), CGA1198CGG (3778A>G), CAA1199CAG (3781A>G), ATC1200ATT (3784C>T), CAT1204CAC (3796T>C), ACA1205ACG (3799A>G), CCA1207CCT (3805A>T), TAC1209TAT (3811C>T), AGA1210AGG (3814A>G), TTC1215TTT (3829C>T), TTT1218TTC (3838T>C), AGA1220AGG (3844A>G), GCA1221GCC (3847A>C), GTT1224GTA (3856T>A), GTG1226GTT (3862G>T), GAC1229GAT (3871C>T), GCC1231GCA (3877C>A), AGC1233AGT (3883C>T), GAA1234GAG (3886A>G), CTA1236CTG (3892A>G), GAC1237GAT (3895C>T), TTT1238TTC (3898T>C), GTA1240GTG (3904A>G), CGG1241AGA (3905C>A 3907G>A), GCT1243GCC (3913T>C), AAG1248AAA (3928G>A), TCA1249TCT (3931A>T), GGG1251GGA (3937G>A), GTT1254GTC (3946T>C), AGT1257AGC (3955T>C), GTG1258GTA (3958G>A), GCC1259GCT (3961C>T), GGA1260GGG (3964A>G), TCT1261TCC (3967T>C), TTA1264TTG (3976A>G), ATT1265ATC (3979T>C), CCA1267CCT (3985A>T), CTA1271CTT (3997A>T), GCG1272GCA (4000G>A), AAC1274AAT (4006C>T), GTT1275GTG (4009T>G), TTC1276TTT (4012C>T), AAA1277AAG (4015A>G), TCT1280TCC (4024T>C), GAA1282GAG (4030A>G), CCA1283CCT (4033A>T), AAG1286AAA (4042G>A), CTG1290TTA (4052C>T 4054G>A), CGT1291CGC (4057T>C), CGT1293CGA (4063T>A), GGA1294GGG (4066A>G), ATA1297GTA (4073A>G), TTT1298TTC (4078T>C), GGC1299GGT (4081C>T), TCT1300TCC (4084T>C), TCC1304TCT (4096C>T), AGC1308AGT (4108C>T), GGA1309GGT (4111A>T), GCG1311GCA (4117G>A), CTA1312TTG (4118C>T 4120A>G), TAC1314TAT (4126C>T), TTC1315TAT (4128T>A 4129C>T), GCC1316GCT (4132C>T), AAT1317AAC (4135T>C), CAA1321CAG (4147A>G), TTA1322CTA (4148T>C), GCT1323TCT (4151G>T), AAC1326AAT (4162C>T), GTA1328ATA (4166G>A), ATA1329ATC (4171A>C), GAG1332GAA (4180G>A), CTG1336TTG (4190C>T), CCT1338CCC (4198T>C), TCC1339TCT (4201C>T), GCG1342GCT (4210G>T), TTC1343TTT (4213C>T), AGT1345AGC (4219T>C), CTG1346TTG (4220C>T), CTG1347ATA (4223C>A 4225G>A), GTT1349GTG (4231T>G), CAT1351CAC (4237T>C), GTA1356GTG (4252A>G), TTA1357CTG (4253T>C 4255A>G), AAA1358AAG (4258A>G), ACT1362ACC (4270T>C), AGA1366AGG (4282A>G), GAG1367GAA (4285G>A), GAA1369GAG (4291A>G), CAG1373CAA (4303G>A), GTC1376GTT (4312C>T), ATA1379GTG (4319A>G 4321A>G), GTG1380GTT (4324G>T), GAG1381GAA (4327G>A), CTG1384CTT (4336G>T), TCT1385TCC (4339T>C), TCA1388TCT (4348A>T), GCA1392GCG (4360A>G), GGT1394GGC (4366T>C), TCT1395TCA (4369T>A), ACT1397ACC (4375T>C), GTT1401GTC (4387T>C), TTT1404TAT (4395T>A), GTA1408ATA (4406G>A), CTT1409CTC (4411T>C), GTT1418GTC (4438T>C), ACC1420ACA (4444C>A), GCC1422GCT (4450C>T), CTC1424CTT (4456C>T), GAC1427GAT (4465C>T), ATG1431GTT (4475A>G 4477G>T), GTC1432GTT (4480C>T), ACA1433TCA (4481A>T), GGC1437GGT (4495C>T), AAG1441AAA (4507G>A), CAC1442CAT (4510C>T), GGT1443GGG (4513T>G), CTA1445TTA (4517C>T), GTC1448GTT (4528C>T), AAA1450AAG (4534A>G), ACC1452ACT (4540C>T), GCG1454TTA (4544G>T 4545C>T 4546G>A), AGA1455AAA (4548G>A), TTT1458TTC (4558T>C), GTA1460GTT (4564A>T), ACC1462ACT (4570C>T), ATA1464ATT (4576A>T), TTG1471TTA (4597G>A), GAC1472GAT (4600C>T), GAC1474GAT (4606C>T), GTG1475GTA (4609G>A), GTG1477GTA (4615G>A), GAC1478GAT (4618C>T), TTG1481CTT (4625T>C 4627G>T), GAC1488GAT (4648C>T), ATT1489ATC (4651T>C), AGG1492AGA (4660G>A), AGC1493AGT (4663C>T), GCT1495GCA (4669T>A), GTG1499ATC (4679G>A 4681G>C), AGT1500AGC (4684T>C), GTT1501GTG (4687T>G), GAG1505GAA (4699G>A), AGA1506AGG (4702A>G), CAA1508CAG (4708A>G), AGG1509AGA (4711G>A), CTG1510TTG (4712C>T), AAA1518AAG (4738A>G), GTA1520GTT (4744A>T), GCA1521GCG (4747A>G), TTG1522CTG (4748T>C), ACT1527ACC (4765T>C), GTC1538ATT (4796G>A 4798C>T), ACT1540AGC (4803C>G 4804T>C), GAG1541GAA (4807G>A), GCG1542GCT (4810G>T), GCT1543GCG (4813T>G), CTT1544CTC (4816T>C), TAT1549TAC (4831T>C), AAC1550AAT (4834C>T), CCA1552CCT (4840A>T),

| | | | | | | | | | |
|---|---|---|---|---|---|---|---|---|---|
| | GGA1556GGG (4852A>G), GGC1557GGT (4855C>T), AGT1561AGC (4867T>C), CTG1562CTC (4870G>C), ATT1563ATA (4873T>A), TGT1566TGC (4882T>C), ACA1573ACT (4903A>T), CAG1575CAA (4909G>A), TTT1577TTC (4915T>C), GAA1578GAG (4918A>G), TTG1579CTA (4919T>C 4921G>A), AGT1580AGC (4924T>C), CCC1581CCG (4927C>G), TTT1583TTC (4933T>C), ATC1584ATA (4936C>A), CAG1585CAA (4939G>A), AAT1586AAC (4942T>C), TTC1587TTT (4945C>T), TCA1593TCG (4963A>G), CCT1596CCC (4972T>C), CTT1602CTC (4990T>C), AAA1606AAG (5002A>G), CCT1613CCC (5023T>C), TTG1614CTG (5024T>C), TGC1615TGT (5029C>T), GAT1616GAC (5032T>C), TCT1618TCC (5038T>C), CCA1620CCT (5044A>T), GCA1623GCT (5053A>T), TCG1624TCA (5056G>A), AGT1625GGT (5057A>G), ACT1626GCT (5060A>G), TCG1629TCT (5071G>T), CTT1636ATT (5090C>A), GGA1637GGG (5095A>G), GTG1638GTT (5098G>T), GCC1639GTC (5100C>T), TTA1640TTG (5104A>G), GAA1641GAT (5107A>T), ATT1642ATC (5110T>C), CCA1643CCG (5113A>G), AAG1644AAA (5116G>A), CAA1645CAG (5119A>G), GTC1646TTT (5120G>T 5122C>T), ATT1648GTT (5126A>G), CAT1651CAC (5137T>C), ATC1652ATT (5140C>T), AAA1653AAG (5143A>G), GAG1654GAA (5146G>A), CTC1659CTA (5161C>A), GAA1661GAG (5167A>G), CTT1663CTG (5173T>G), ACG1666ACA (5182G>A), GTT1667ATT (5183G>A), GTC1668GTT (5188C>T), AAG1669AAA (5191G>A), AAA1671AAG (5197A>G), GAC1672GAT (5200C>T), TGC1674TGT (5206C>T), TTA1675TTG (5209A>G), AGC1678AGT (5218C>T), TCG1682TCA (5230G>A), ATC1684ATT (5236C>T), AGC1685AGT (5239C>T), ATC1687ATT (5245C>T), ACA1693ACT (5263A>T), CTC1695CTT (5269C>T), TTC1696TTT (5272C>T), GCC1697GCA (5275C>A), ATC1698ATT (5278C>T), CCA1699CCC (5281A>C), AGA1700AGG (5284A>G), ACT1701ACC (5287T>C), CTA1702TTA (5288C>T), ATA1703ATT (5293A>T), TTG1704CTA (5294T>C 5296G>A), GAG1706GAA (5302G>A), CTT1709CTA (5311T>A), GAA1710GAG (5314A>G), CGA1713CGT (5323A>T), AGT1721AGC (5347T>C), ATC1723ATT (5353C>T), TGC1727TGT (5365C>T), GTT1734GTC (5386T>C), ACC1737ACA (5395C>A), AAA1746AAG (5422A>G), GAT1747GAC (5425T>C), ACC1749ACT (5431C>T), GAG1751GAA (5437G>A), CTT1756CTC (5452T>C), AAG1758AAA (5458G>A), AGT1761AGC (5467T>C), CAA1762CAG (5470A>G), CTA1763TTG (5471C>T 5473A>G), TTC1766TTT (5482C>T), TCA1767TCC (5485A>C), AAT1768AAC (5488T>C), TTG1769CTA (5489T>C 5491G>A), GGT1771GGC (5497T>C), TGT1774TGC (5506T>C), GAG1775GAA (5509G>A), GAG1776GAA (5512G>A), TAT1782TAC (5530T>C), GAG1783GAA (5533G>A), TCG1784TCT (5536G>T), TTG1785TTA (5539G>A), GTT1788GTG (5548T>G), CAC1790CAT (5554C>T), GCT1792GCA (5560T>A), GCG1793ACA (5561G>A 5563G>A), ACG1794ACC (5566G>C), TCA1795TCG (5569A>G), CTC1800CTG (5584C>G), AAG1803AAA (5593G>A), GGG1804GGA (5596G>A), ATT1805GTT (5597A>G), AAC1807AAA (5605C>A), GTG1811GTT (5617G>T), GCT1812GTG (5619C>T 5620T>G), TTG1815TTA (5629G>A), ATC1816CTC (5630A>C), GGC1819GGT (5641C>T), GCA1822GCT (5650A>T), ATA1826ATT (5662A>T), GGA1827GGG (5665A>G), CAA1835CAG (5689A>G), GTT1837ATT (5693G>A), GAG1838GAA (5698G>A), CAA1843CAG (5713A>G), GGG1844GGC (5716G>C), AAA1845AAG (5719A>G), GCC1853GCA (5743C>A), CGC1857CGA (5755C>A), CAT1858CAC (5758T>C), CGT1860CGC (5764T>C), AAA1862AAG (5770A>G), GGC1865GGT (5779C>T), TTT1866TTC (5782T>C), GAC1869GAT (5791C>T), GAC1873GAT (5803C>T), AGG1884AGA (5836G>A), AAA1885AAG (5839A>G), GGA1887GGG (5845A>G), AAA1888AAG (5848A>G), AAA1890AGA (5853A>G), GGT1891GGC (5857T>C), ACC1892ACT (5860C>T), ACA1893ACT (5863A>T), GGT1895GGA (5869T>A), GGT1897GGC (5875T>C), AGC1900AGT (5884C>T), TTC1903TTT (5893C>T), ATC1904GTG (5894A>G 5896C>G), AAC1905AAT (5899C>T), GGG1908GGA (5908G>A), TTT1909TTC (5911T>C), GAG1913GAA (5923G>A), CAA1918CAG (5938A>G), TTC1919TTT (5941C>T), GTT1920GTC (5944T>C), GAT1921GAC (5947T>C), GGG1925GGA (5959G>A), CGG1926GCC (5960C>G 5961G>C 5962G>C), GAA1930GAG (5974A>G), GCT1934GCA (5986T>A), GAC1935GAT (5989C>T), ATT1936GTT (5990A>G), GAT1938GAC (5998T>C), AGA1942CGT (6008A>C 6010A>T), GAA1945GAT (6019A>T), GTG1946GTT (6022G>T), CGA1947CGT (6025A>T), AAA1949ACA (6030A>C), GAC1955GAA (6049C>A), ATG1958ATA (6058G>A), CAA1959CAG (6061A>G), GCC1960GCA (6064C>A), GGT1962GGC (6070T>C), AGT1963AAC (6072G>A 6073T>C), AAC1964CAC (6074A>C), ACG1965ACA (6079G>A), ACC1966AAC (6081C>A), ATA1967ATT (6085A>T), GCA1969GCT (6091A>T), TAC1970TAT (6094C>T), TGT1976TCC (6111G>C 6112T>C), AAA1978AAG (6118A>G), TTG1980TTA (6124G>A), AAG1981AAA (6127G>A), ATT1982GTC (6128A>G 6130T>C), GAT1983GAC (6133T>C), TTA1984TTG (6136A>G), AAC1988AAT (6148C>T), AAA1991AAG (6157A>G), GTT1992ATT (6158G>A), TGT1993TGC (6163T>C), AAA1995AAG (6169A>G), ACA1996TCG (6170A>T 6172A>G), GGC1998GGT (6178C>T), AAA2001AAG (6187A>G), TTT2002TTC (6190T>C), GAG2004GAA (6196G>A), CTC2007TTT (6203C>T 6205C>T), GAA2008GAG (6208A>G), CTA2009TTG (6209C>T 6211A>G), CAG2011CAA (6217G>A), GCT2015GCA (6229T>A), GTA2016ATA (6230G>A), GTC2018GTT (6238C>T), AAG2021AAA (6247G>A), ATA2023ATT (6253A>T), GCA2025AAA (6257G>A 6258C>A), CAT2030CAC (6274T>C), AAA2033AGA (6282A>G), TCG2034TCA (6286G>A), GGC2038GGT (6298C>T), TTG2039TTA (6301G>A), AGA2040AGG (6304A>G), GAC2041GAT (6307C>T), AAC2043AAT (6313C>T), GCC2046GCT (6322C>T), ACA2048ACT (6328A>T), GTT2049GTG (6331T>G), TGT2050TGC (6334T>C), AGG2051AGA (6337G>A), CTG2052TTG (6338C>T), TAT2058TAC (6358T>C), GGG2059GGA (6361G>A), GCA2060ACA (6362G>A), TCA2061TCT (6367A>T), GAG2062GAA (6370G>A), TAC2064TAT (6376C>T), GGT2065GGG (6379T>G), GGA2067GGC (6385A>C), GGA2069GGT (6391A>T), GCA2070GCG (6394A>G), TAC2071TAT (6397C>T), GTA2073ATA (6401G>A), GCG2074GTG (6405C>T), CAC2076CAT (6412C>T), TTA2078TTG (6418A>G), TTT2079TTC (6421T>C), TAC2082TTC (6429A>T), GGT2084GGA (6436T>A), GAG2087GAA (6445G>A), CAA2089CGA (6450A>G), TCC2090TCA (6454C>A), CAC2092CAT (6460C>T), GGT2093GGA (6463T>A), TTC2095TTT (6469C>T), AGG2096AGA (6472G>A), GTG2097GTA (6475G>A), AAT2099AAC (6481T>C), CAC2101AAT (6485C>A 6487C>T), AGT2102AGC (6490T>C), AGC2104AGT (6496C>T), CTG2106CTA (6502G>A), GGT2110GGC (6514T>C), AGG2111AGA (6517G>A), CTC2115ATC (6527C>A), ATC2116ATA (6532C>A), CCG2119CCA (6541G>A), AAA2120AAG (6544A>G), GTC2124GTT (6556C>T), TTT2125TTC (6559T>C), CAG2127CAA (6565G>A), TTG2129CTA (6569T>C 6571G>A), CCT2134CCA (6586T>A), ACA2135ATA (6588C>T), GAA2138GAG (6598A>G), AGA2139AGG (6601A>G), ATT2140ATC (6604T>C), TGT2141TGC (6607T>C), GGA2144GGG (6616A>G), AAC2146AAT (6622C>T), GAG2149GAA (6631G>A), AAG2150AAA (6634G>A), TAT2151CAT (6635T>C), GCT2152GCA (6640T>A), TCG2154TCC (6646G>C), ATC2155GTC (6647A>G), ATC2156ATT (6652C>T), AGC2160AGT (6664C>T), ACT2162ACG (6670T>G), ATA2165GTG (6677A>G 6679A>G), GGC2167GGT (6685C>T), ACA2169ACT (6691A>T), TTC2170TTT (6694C>T), ACA2177ACG (6715A>G), GAT2178AAT (6716G>A), AAT2179GAT (6719A>G), GGA2180GGG (6724A>G), CTA2184TTA (6734C>T), GTG2186GTA (6742G>A), GTG2187GTA (6745G>A), GCC2190GCT (6754C>T), GGA2192GGG (6760A>G), ATA2194TTG (6764A>T 6766A>G), ATT2197ATA (6775T>A), CTG2200TTG (6782C>T), GCA2201GCT (6787A>T), AAC2202AAT (6790C>T), GCA2204GTG (6795C>T 6796A>G), CAC2205CAA (6799C>A), ACC2206ACA (6802C>A), AAC2208AAT (6808C>T), TAC2209TAT (6811C>T), TAC2210TAT (6814C>T), TTC2213TTT (6823C>T), GAA2215GAG (6829A>G), AAG2220AGG (6843A>G), TAC2221TAT (6847C>T), CGA2223CGC (6853A>C), AAT2225GAT (6857A>G), GAG2226GAA (6862G>A), GAA2229GAG (6871A>G), GTC2231ACA (6875G>A 6876T>C 6877C>A), AAG2232AAA (6880G>A), TCT2233TCA (6883T>A), GTT2235GTG (6889T>G), GAC2239GAT (6901C>T), GGC2244GGT (6916C>T), CCG2245CCA (6919G>A), AAA2247AAG (6925A>G), CTT2248CTC (6928T>C), GAC2250GAA (6934C>A), GGG2255GGC (6949G>C), TTC2257TTT (6955C>T), AAA2258AAG (6958A>G), CAA2264CAG (6976A>G), CTA2266TTA (6980C>T), ATC2267ATT (6985C>T), GTA2271GTT (6997A>T), GTG2272GTT (7000G>T), GTG2273ATA (7001G>A 7003G>A), GCT2276GCC (7012T>C), AAG2277AAA (7015G>A), CAT2278CAC (7018T>C), TTT2283TAT (7032T>A), GCC2285GCT (7039C>T), TTG2286CTA (7040T>C 7042G>A), GGA2288GGG (7048A>G), GCT2292GCA (7060T>A), GTC2293GTA (7063C>A), AGC2298AGT (7078C>T), TTA2300CTG (7082T>C 7084A>G), GTA2301GTG (7087A>G), ACC2302ACA (7090C>A), CAT2304CAC (7096T>C), GTA2305GTG (7099A>G), GTT2306GTC (7102T>C), GAG2309GAA (7111G>A), CGA2311CGG (7117A>G), ACA2314AAA (7125C>A), GAA2315GAG (7129A>G), TTT2316TTC (7132T>C), CTG2317TTA (7133C>T 7135G>A), GTG2319GTA (7141G>A), GAT2320GAC (7144T>C), GCA2321TCG (7145G>T 7147A>G), GAG2322GAA (7150G>A), GAG2324GAA (7156G>A), GCA2325GCT (7159A>T), GAG2326_GCA2327del (7160_7165delGAGGCA), TTC2329TTT (7171C>T), GCG2335GCT (7189G>T), AAA2338AAG (7198A>G), AGC2339AGT (7201C>T), CTA2341TTA (7205C>T), AAT2342AAC (7210T>C), GAT2344GAA (7216T>A), GCG2345GCA (7219G>A), ATC2347ATA (7225C>A), AAG2348AAA (7228G>A), AAG2352AAA (7240G>A), TAT2353TAC (7243T>C), AAA2355AAG (7249A>G), CCT2356CCG (7252T>G), GAT2358GAA (7258T>A), GTT2359GTC (7261T>C), GGT2360GGA (7264T>A), GTC2361GTG (7267C>G), GTG2362GTA (7270G>A), GAT2363_CGG2364insT-- (7273_7274insT), CGG2364GTG (7274C>G 7275G>T), CAT2366CTT (7281A>T), TTG2367TCG (7284T>C), TCA2371TTA (7296C>T), TCT2376TTT (7311C>T), ACC2377ATC (7314C>T), TGC2378TAC (7317G>A), AAT2379-AA (7319delA 7321T>A), CAC2381CAT (7327C>T), GGC2382GGT (7330C>T), TTC2383TTT (7333C>T), AAG2384CAG (7334A>C), TGT2386TGC (7342T>C), GCA2387AAC (7343G>A 7344C>A 7345A>C), ACT2390ACC (7354T>C), CAA2393CAG (7363A>G), TTC2396TTT (7372C>T), GCG2398GCT (7378G>T), CTC2399CTT (7381C>T), GCA2404GCT (7396A>T), GTC2405GTT (7399C>T), GCC2407GCT (7405C>T), AGT2408ATG (7407G>T 7408T>G), GGG2410GGT (7414G>T), TGC2411GGC (7415T>G), AAA2412AAG (7420A>G), GAC2415GAT (7429C>T), TAT2416TAC (7432T>C), CAT2419CAC (7441T>C), AAG2425AAA (7459G>A), GAA2426GAG (7462A>G), ATA2428ATT (7468A>T), GTC2429GTA (7471C>A), CAA2431CAG (7477A>G), AGC2432AGT (7480C>T), CTG2434CTT (7486G>T), TAT2437TAC (7495T>C), AAA2438AAG (7498A>G), GGT2439GGC (7501T>C), TTG2440TCA (7503T>C 7504G>A), ATT2443ATA (7513T>A), AAC2445AAT (7519C>T), GGA2446GGG (7522A>G), AAG2449AAA (7531G>A), GAG2451GAA (7537G>A), CTC2452CTT (7540C>T), TGT2454TGC (7546T>C), AAG2462AAA (7570G>A), ACG2463ACA (7573G>A), ACG2465ACA (7579G>A), GCT2468GCC (7588T>C), CTA2471TTA (7595C>T), TTG2474CTA (7604T>C 7606G>A), GGT2477GGA (7615T>A), TGT2480TGC (7624T>C), GAC2483GAT (7633C>T), TTC2484TTT (7636C>T), AAT2485AAC (7639T>C), TTT2488TTC (7648T>C), GAA2494GAG (7666A>G), TGT2496TGC (7672T>C), ACA2498ACT (7678A>T), GGG2500GGA (7684G>A), ACT2502ACC (7690T>C), TTT2504TTC (7696T>C), GGT2506GGG (7702T>G), GAT2509AAT (7709G>A), AAA2510AGG (7713A>G 7714A>G), CTG2511TTG (7715C>T), CGT2514CGC (7726T>C), GAG2517GAA (7735G>A), GTA2520GTG (7744A>G), TAC2521TAT (7747C>T), GGC2526GGT (7762C>T), CAG2528CAA (7768G>A), TTT2529TTC (7771T>C), TCA2532TCG (7780A>G), CTA2533CTC (7783A>C), ACT2534ACC (7786T>C), CTA2537TTG (7793C>T 7795A>G), ATC2538ATT (7798C>T), GCT2540GCC (7804T>C), ACC2543ATC (7812C>T), ATC2544ATT (7816C>T), GAA2550GAG (7834A>G), GAC2551GAT (7837C>T), GTG2554TTA (7844G>T 7846G>A), CAG2557CAA (7855G>A), CTG2559CTA (7861G>A), TTA2562CTG (7868T>C 7870A>G), TAC2563TAT (7873C>T), ACT2564ACA (7876T>A), GAG2565GAA (7879G>A), ATT2566ATA (7882T>A), GTT2567ATT (7883G>A), TAC2568TAT (7888C>T), CCA2570CCT (7894A>T), ATT2571ATC (7897T>C), GTC2579GTT (7921C>T), AAG2581AAA (7927G>A), GGT2584GGC (7936T>C), AAT2585AAC (7939T>C), AAT2586AAC (7942T>C), GTG2594GTA (7966G>A), GAT2595GAC (7969T>C), TCT2597TCC (7975T>C), GTC2601GTT (7987C>T), CTT2602CTC (7990T>C), TAC2606TAT (8002C>T), GCT2607GCC (8005T>C), GAA2616GAG (8032A>G), ATC2618ATT (8038C>T), GAC2619GAT (8041C>T), AGC2620AAC (8043G>A), TTG2632TTA (8080G>A), GTG2635GTA (8089G>A), CCG2637CCA (8095G>A), GAG2638GAA (8098G>A), AGC2641AGT (8107C>T), ATT2642ATC (8110T>C), GAT2644GAC (8116T>C), CAA2648CAG (8128A>G), GGT2654GGC (8146T>C), AAC2656AGT (8151A>G 8152C>T), TCG2660TCA (8164G>A), AGG2665AAG (8178G>A), AAG2666AAA (8182G>A), TTG2669TTA (8191G>A), TTC2671TTT (8197C>T), TCC2673TCT (8203C>T), AGA2675AAA (8208G>A), GGC2676GGG (8212C>G), CTG2677TTG (8213C>T), CTA2678CTG (8218A>G), ATC2679ATT (8221C>T), GAG2680GAA (8224G>A), CCA2685CCG (8239A>G), ATT2692ATC (8260T>C), GTA2693GTG (8263A>G), GAT2699GAC (8281T>C), AGA2700AGG (8284A>G), CCA2704GCG (8294C>G 8296A>G), GAG2705GAA (8299G>A), CAC2706CAT (8302C>T), ATT2711ATC (8317T>C), TGC2712TGT (8320C>T), GCT2714GCA (8326T>A), GAG2717GAA (8335G>A), TCC2718TCT (8338C>T), TCT2722TTT (8349C>T), GAA2723GAG (8353A>G), CAC2726CAT (8362C>T), CAA2727CAG (8365A>G), TAC2732TAT (8380C>T), TCA2733TCG (8383A>G), TTA2735TTG (8389A>G), CCT2740CCC (8404T>C), GCA2742TCA (8408G>T), ACA2743ACG (8413A>G), GCG2745GCT (8419G>T), GCT2750GCC (8434T>C), TAT2752TAC (8440T>C), ATG2756ATA (8452G>A), CTA2758CTG (8458A>G), AGG2759AAG (8460G>A), AAA2760AAG (8464A>G), CTG2761CTA (8467G>A), GCT2766ACA (8480G>A 8482T>A), GAG2771GAA (8497G>A), CTA2772TTG (8498C>T 8500A>G), AGA2773AAG (8502G>A 8503A>G), GCC2774GCT (8506C>T), GAA2777GAT (8515A>T), GTC2780GTT (8524C>T), GCA2781GCC (8527A>C), TTA2782CTG (8528T>C 8530A>G), GAG2785GAC (8539G>C), CTT2788TGT (8546C>T 8547T>G), TCT2790ACT (8552T>A), GTA2793GTG (8563A>G), GCA2797GGA (8574C>G), ATT2801GTT (8585A>G), GGA2805GAA (8598G>A), AGC2806AGT (8602C>T), AAC2807TCT (8603A>T 8604A>C 8605C>T), GCA2811GTA (8616C>T), AAA2812AAG (8620A>G), CCA2813CAA (8622C>A), GAG2814GGA (8625A>G 8626G>A), CAG2815CAA (8629G>A), GGC2816GGT (8632C>T), ATC2818ATT (8638C>T), CAG2819CAA (8641G>A), AAC2821AAT (8647C>T), CCG2822TTC (8648C>T 8649C>T 8650G>C), AAC2823AAT (8653C>T), AAA2824AAG (8656A>G), GGA2825GAA (8658G>A), GAT2827GCG (8664A>C 8665T>G), GTT2830GTG (8674T>G), GCA2832GTT (8679C>T 8680A>T), GGC2833GGA (8683C>A), GGG2836GGA (8692G>A), ACA2837ACT (8695A>T), CCG2841CCA (8707G>A), AGA2842CGA (8708A>C), AAG2844AAA (8716G>A), ATC2846ATT (8722C>T), ACG2847ACA (8725G>A), AAA2849AAG (8731A>G), AGA2851AGG (8737A>G), CCC2853CCT (8743C>T), ACA2854AAG (8745C>A 8746A>G), AGC2855AGT (8749C>T), AAG2856AAA (8752G>A), GGA2857GGT (8755A>T), AAC2862AAT (8770C>T), TTA2863CTA (8771T>C), GAA2864GAG (8776A>G), CAT2865CAC (8779T>C), TTG2866TTA (8782G>A), TAT2869TAC (8791T>C), CCA2871CCG (8797A>G), CAA2872CAG (8800A>G), CAA2873CAG (8803A>G), GAT2875GAC (8809T>C), ATT2876ATC (8812T>C), AAT2878AAC (8818T>C), CGG2880CGA (8824G>A), GCA2881GCG (8827A>G), ACG2888ACA (8848G>A), TAT2890TAC (8854T>C), GAG2891GGA (8856A>G 8857G>A), GTG2893GTA (8863G>A), CGG2894CAG (8865G>A), ATG2895CTT (8867A>C 8869G>T), GCA2896GCG (8872A>G), GGA2900GAA (8883G>A), ACT2902TCT (8888A>T), TGC2915TGT (8929C>T), GTC2924ATC (8954G>A), AAC2925AAT (8959C>T), GGA2926GGT (8962A>T), GGG2933GGA (8983G>A), AAT2934GAC (8984A>G 8986T>C), GAA2935GTA (8988A>T), GAG2938GAA (8998G>A), CCG2940CCA (9004G>A), TTG2941TTA (9007G>A), ATC2944ATT (9016C>T), ACC2951ACA (9037C>A), ATA2967ATT (9085A>T), AAC2971AAT (9097C>T), CGA2984CGT (9136A>T), CGG2987CGT (9145G>T), ATG2989GGA (9149A>G 9150T>G 9151G>A), GGT2990AGT (9152G>A), TTA2991TTG (9157A>G), GCG2992GCT (9160G>T), CGT2993CGC (9163T>C), GAG3000GAA (9184G>A), CGA3004CGG (9196A>G), AGG3010AGA (9214G>A), GAA3011GAG (9217A>G), GCG3012GCA (9220G>A), GCA3020GCT (9244A>T), TTG3021TTA (9247G>A), GCC3024GCT (9256C>T), CCT3026TCC (9260C>T 9262T>C), GGG3030GGA (9274G>A), GAC3032GAT (9280C>T), ATC3035ATT (9289C>T), GAG3040GAA (9304G>A), GTC3050GTT (9334C>T), GTC3060GTG (9364C>G) | | | | | | | | |
| P1 protein (NP_734243.1) | 1 | 284 | 100% | 1414 | 73.7% | 284 (100%) | 206 (72.5%) | 0/0/0/0 | 0 |

| | Begin | End | Coverage | Score | Concordance | Matches | Identities | I/D/M/F* | Stop Codons |
|---|---|---|---|---|---|---|---|---|---|
| **NT** | **19** | **9704** | **99.8%** | **12534** | **65.0%** | **9674 (99.7%)** | **8011 (82.6%)** | **15/12** | |
| Protein mutations: | Y4Q (194T>C 196C>A), C9L (209T>C 210G>T 211T>G), F13M (221T>A 223T>G), S23P (251T>C), C24F (255G>T 256C>T), E25G (258A>G), I27V (263A>G 265T>G), V28A (267T>C), E30V (273A>T), A35T (287G>A 289T>C), V37T (293G>A 294T>C), A41I (305G>A 306C>T), D42G (309A>G), E44D (316A>T), T45V (317A>G 318C>T 319A>G), L52Q (339T>A 340C>A), K53R (342A>G), K55A (347A>G 348A>C 349A>G), Y56H (350T>C), T58V (356A>G 357C>T), V61T (365G>A 366T>C 367G>A), L62S (368C>T 369T>C), F67R (383T>C 384T>G), A75V (408C>T 409C>T), M78E (416A>G 417T>A), E85K (437G>A), R86E (440A>G 441G>A 442G>A), K87R (443A>C 444A>G 445G>A), D88E (448T>A), E91A (456A>C), H93N (461C>A), D111E (517T>A), S116F (531C>T), P118S (536C>T), Q119W (539C>T 540A>G 541A>G), R122K (549G>A 550A>G), V134A (585T>C 586C>G), R137Y (593C>T 594G>A 595C>T), P138R (597C>G), I139V (599A>G 601A>G), I140P (602A>C 603T>C 604A>G), M147I (625G>A), I151V (635A>G 637T>C), I154V (644A>G), E160A (663A>C 664G>C), H166Q (682C>A), T172S (699C>G), H174Q (706T>A), K179E (719A>G 721G>A), I180V (722A>G 724A>G), A183S (731G>T), Y184P (734T>C 735A>C 736C>T), S185R (737T>C 738C>G 739C>T), A187V (744C>T 745G>C), R189C (749C>T 751A>C), M194K (765T>A), R197K (773C>A 774G>A 775A>G), M206K (801T>A), G210A (813G>C 814A>G), L211H (816T>A), R214Q (825G>A 826T>A), R226C (860C>T 862C>T), T227A (863A>G 865T>C), I228T (867T>C), N229D (869A>G 871C>T), I230L (872A>T 874A>G), R232K (879G>A), N240S (903A>G), T241N (906C>A 907A>C), K242T (909A>C 910A>C), S243N (912G>A 913C>T), S252L (938T>C 939C>T 940A>T), L266I (980T>A 982G>C), R272K (999G>A), Q275R (1007C>A 1008A>G 1009G>A), S276G (1010A>G 1012T>G), I277V (1013A>G), N279Q (1019A>C 1021C>G) | | | | | | | | |
| Codon mutations: | ACT3ACC (193T>C), TAC4CAA (194T>C 196C>A), TCA6TCG (202A>G), ATC8ATT (208C>T), TGT9CTG (209T>C 210G>T 211T>G), TCG12TCA (220G>A), TTT13ATG (221T>A 223T>G), GAA14GAG (226A>G), CTA17CTT (235A>T), TAC19TAT (241C>T), CCA21CCT (247A>T), TCT23CCT (251T>C), TGC24TTT (255G>T 256C>T), GAG25GGG (258A>G), CAT26CAC (262T>C), ATT27GTG (263A>G 265T>G), GTG28GCG (267T>C), GAA30GTA (273A>T), CGA31CGT (277A>T), GAA32GAG (280A>G), GTG33GTT (283G>T), CCG34CCA (286G>A), GCT35ACC (287G>A 289T>C), GTT37ACT (293G>A 294T>C), CCT39CCC (301T>C), TTC40TTT (304C>T), GCA41ATA (305G>A 306C>T), GAT42GGT (309A>G), CTG43TTG (311C>T), GAA44GAT (316A>T), ACA45GTG (317A>G 318C>T 319A>G), CAA46CAG (322A>G), GCA49GCT (331A>T), CTC52CAA (339T>A 340C>A), AAG53AGG (342A>G), AAA55GCG (347A>G 348A>C 349A>G), TAT56CAT (350T>C), GCT57GCA (355T>A), ACT58GTT (356A>G 357C>T), CGT60CGC (364T>C), GTG61ACA (365G>A 366T>C 367G>A), CTC62TCC (368C>T 369T>C), AAA63AAG (373A>G), AAC64AAT (376C>T), TTT67CGT (383T>C 384T>G), ACG68ACA (388G>A), TAC69TAT (391C>T), CGA70CGG (394A>G), TAC71TAT (397C>T), AAG72AAA (400G>A), GAT74GAC (406T>C), GCC75GTT (408C>T 409C>T), CAG76CAA (412G>A), ATA77ATT (415A>T), ATG78GAG (416A>G 417T>A), ATT80ATC (424T>C), CAG81CAA (427G>A), AAA83AAG (433A>G), CTG84CTC (436G>C), GAG85AAG (437G>A), AGG86GAA (440A>G 441G>A 442G>A), AAG87CGA (443A>C 444A>G 445G>A), GAT88GAA (448T>A), GAA90GAG (454A>G), GAA91GCA (456A>C), TAT92TAC (460T>C), CAC93AAC (461C>A), CAA95CAG (469A>G), GCC97GCT (475C>T), CCT99CCA (481T>A), ACT106ACC (502T>C), ATA107ATT (505A>T), GGC109GGT (511C>T), GAT111GAA (517T>A), CCT112CCA (520T>A), CCA113CCT (523A>T), TCT116TTT (531C>T), GAG117GAA (535G>A), CCA118TCA (536C>T), CAA119TGG (539C>T 540A>G 541A>G), AGA122AAG (549G>A 550A>G), ATC124ATT (556C>T), ATT125ATC (559T>C), CAT126CAC (562T>C), ACT128ACC (568T>C), CCA129CCG (571A>G), CGT132CGC (580T>C), GTC134GCG (585T>C 586C>G), CGC137TAT (593C>T 594G>A 595C>T), CCC138CGC (597C>G), ATA139GTG (599A>G 601A>G), ATA140CCG (602A>C 603T>C 604A>G), ACA143ACG (613A>G), GAA144GAG (616A>G), GGC145GGG (619C>G), CAG146CAA (622G>A), ATG147ATA (625G>A), AAT148AAC (628T>C), CAC149CAT (631C>T), ATT151GTC (635A>G 637T>C), ATA154GTA (644A>G), ATT157ATC (655T>C), TCG159TCA (661G>A), GAG160GCC (663A>C 664G>C), AAA161AAG (667A>G), GGG163GGC (673G>C), GTC165GTT (679C>T), CAC166CAA (682C>A), TTA167CTA (683T>C), AGT169AGC (691T>C), ACC172AGC (699C>G), ACT173ACA (703T>A), CAT174CAA (706T>A), AAG179GAA (719A>G 721G>A), ATA180GTG (722A>G 724A>G), CTT181TTG (725C>T 727T>G), GGT182GGA (730T>A), GCA183TCA (731G>T), TAC184CCT (734T>C 735A>C 736C>T), TCC185CGT (737T>C 738C>G 739C>T), GCG187GTC (744C>T 745G>C), CGA189TGC (749C>T 751A>C), ACT190ACC (754T>C), ATG194AAG (765T>A), TTG196CTG (770T>C), CGA197AAG (773C>A 774G>A 775A>G), CGG198AGG (776C>A), TTC202TTT (790C>T), ATG206AAG (801T>A), GGA210GCG (813G>C 814A>G), CTT211CAT (816T>A), TTG212CTG (818T>C), CAA213CAG (823A>G), CGT214CAA (825G>A 826T>A), CGG217AGA (833C>A 835G>A), GAC219GAT (841C>T), AAT223AAC (853T>C), GTC225GTT (859C>T), CGC226TGT (860C>T 862C>T), ACT227GCC (863A>G 865T>C), ATC228ACC (867T>C), AAC229GAT (869A>G 871C>T), ATA230TTG (872A>T 874A>G), AGG232AAG (879G>A), GAT234GAC (886T>C), ATC238ATT (898C>T), AAC240AGC (903A>G), ACA241AAC (906C>A 907A>C), AAA242ACC (909A>C 910A>C), AGC243AAT (912G>A 913C>T), GGC246GGA (922C>A), GGT249GGA (931T>A), TCA252CTT (938T>C 939C>T 940A>T), GGA253GGG (943A>G), TTC256TTT (952C>T), TCA261TCG (967A>G), GAA263GAG (973A>G), GGG264GGC (976G>C), TTG266ATC (980T>A 982G>C), GCA269GCG (991A>G), CGT270CGC (994T>C), AGA272AAA (999G>A), GTT273GTC (1003T>C), CAG275AGA (1007C>A 1008A>G 1009G>A), AGT276GGG (1010A>G 1012T>G), ATT277GTT (1013A>G), TTA278TTG (1018A>G), AAC279CAG (1019A>C 1021C>G), ATC282ATT (1030C>T) | | | | | | | | |
| HC-Pro protein (NP_734244.1) | 1 | 456 | 100% | 3144 | 95.7% | 456 (100%) | 425 (93.2%) | 0/0/0/0 | 0 |
| Protein mutations: | D4E (1048C>A), R16Q (1083G>A), S21A (1097T>G 1099G>A), R36K (1143G>A), L40I (1154T>A 1156G>A), S62N (1221G>A), V65T (1229G>A 1230T>C 1231T>C), S66N (1233G>A 1234C>T), K74T (1257A>C 1258A>G), N81S (1278A>G), A85V (1290C>T 1291G>A), D88T (1298G>A 1299A>C 1300C>T), I91V (1307A>G 1309A>G), N94E (1316A>G 1318T>A), I98T (1329T>C), L113I (1373C>A), I117V (1385A>G), E123D (1405G>T), A127S (1415G>T 1417A>T), V134I (1436G>A 1438C>T), E224G (1707A>G), A228T (1718G>A), I236V (1742A>G 1744C>T), S241N (1758G>A), V252I (1790G>A), I301V (1937A>G 1939T>C), F323Y (2004T>A), I344M (2068T>G), V346I (2072G>A 2074G>A), R391K (2208G>A), D410E (2266C>G) | | | | | | | | |
| Codon mutations: | TCG1TCA (1039G>A), AAT2AAC (1042T>C), GCC3GCA (1045C>A), GAC4GAA (1048C>A), AAT5AAC (1051T>C), AAG8AAA (1060G>A), GGT9GGA (1063T>A), GAC11GAT (1069C>T), CGA16CAA (1083G>A), TAT19TAC (1093T>C), TCG21GCA (1097T>G 1099G>A), CAC23CAT (1105C>T), GTA26GTG (1114A>G), GCT27GCG (1117T>G), TTA29TTG (1123A>G), CCT30CCA (1126T>A), GTC31GTT (1129C>T), AGG36AAG (1143G>A), GTA37GTT (1147A>T), GCT38GCA (1150T>A), TTG40ATA (1154T>A 1156G>A), CAC43CAT (1165C>T), AGT44AGC (1168T>C), CTT46TTA (1172C>T 1174T>A), ACC55ACT (1201C>T), AGC62AAC (1221G>A), GTT65ACC (1229G>A 1230T>C 1231T>C), AGC66AAT (1233G>A 1234C>T), CTG68CTA (1240G>A), AAG70AAA (1246G>A), TTG72CTA (1250T>C 1252G>A), CAT73CAC (1255T>C), AAA74ACG (1257A>C 1258A>G), AGA77AGG (1267A>G), GGT79GGC (1273T>C), TTG80TTA (1276G>A), AAT81AGT (1278A>G), TTG83CTG (1283T>C), GCG85GTA (1290C>T 1291G>A), GAC88ACT (1298G>A 1299A>C 1300C>T), TTT90TTC (1306T>C), ATA91GTG (1307A>G 1309A>G), CAT92CAC (1312T>C), AAT94GAA (1316A>G 1318T>A), TTG97TTA (1327G>A), ATA98ACA (1329T>C), GCG99GCT (1333G>T), TTA100TTG (1336A>G), CTA103TTG (1343C>T 1345A>G), CCG106CCA (1354G>A), GTG107GTT (1357G>T), GAC108GAT (1360C>T), CTC111TTG (1367C>T 1369C>G), CTT113ATT (1373C>A), TTC114TTT (1378C>T), GAG116GAA (1384G>A), ATA117GTA (1385A>G), AAA119AAG (1393A>G), TCC120TCT (1396C>T), GAG123GAT (1405G>T), AAA124AAG (1408A>G), CAG125CAA (1411G>A), GCA127TCT (1415G>T 1417A>T), CCG128CCC (1420G>C), AAT131AAC (1429T>C), TTA132TTG (1432A>G), GTC134ATT (1436G>A 1438C>T), TTA135CTG (1439T>C 1441A>G), AAT136AAC (1444T>C), TTC139TTT (1453C>T), CTG140CTT (1456G>T), GAA144GAG (1468A>G), TTG155CTA (1499T>C 1501G>A), AGT156AGC (1504T>C), TTG157TTA (1507G>A), CTC158CTT (1510C>T), GAA159GAG (1513A>G), TTA160TTG (1516A>G), GCA161GCT (1519A>T), AGG162AGA (1522G>A), TTC163TTT (1525C>T), CAG164CAA (1528G>A), AAC166AAT (1534C>T), ACT168ACA (1540T>A), AAC170AAT (1546C>T), AAA173AAG (1555A>G), GAT175GAC (1561T>C), ATA176ATC (1564A>C), TCT177TCA (1567T>A), TTC179TTT (1573C>T), AGA180AGG (1576A>G), TTA183CTA (1583T>C), TCT184TCA (1588T>A), AAG186AAA (1594G>A), AAC188AAT (1600C>T), CTG191CTA (1609G>A), TCG194TCA (1618G>A), TGC195TGT (1621C>T), GAC196GAT (1624C>T), AAC197AAT (1627C>T), CTG199TTG (1631C>T), AAA201AAG (1639A>G), TTC205TTT (1651C>T), CTC206CTG (1654C>G), CAT213CAC (1675T>C), TCA219TCG (1693A>G), TTT222TTC (1702T>C), GAA224GGA (1707A>G), GCA228ACA (1718G>A), GGA230GGG (1726A>G), TAC231TAT (1729C>T), TCA232TCC (1732A>C), TAT234TAC (1738T>C), ATC236GTT (1742A>G 1744C>T), CGC237CGT (1747C>T), CAT239CAC (1753T>C), AGT241AAT (1758G>A), AGG244AGA (1768G>A), AAC250AAT (1786C>T), TTA251TTG (1789A>G), GTT252ATT (1790G>A), GTC253GTT (1795C>T), TTA257CTG (1805T>C 1807A>G), GAG259GAA (1813G>A), AGG261AGA (1819G>A), AAG263AAA (1825G>A), GAC267GAT (1837C>T), TAT268TAC (1840T>C), CAA271CAG (1849A>G), GTC274GTG (1858C>G), AAA276AAG (1864A>G), TGC278TGT (1870C>T), AGT280AGC (1876T>C), AAA282AAG (1882A>G), GGT284GGA (1888T>A), AAT285AAC (1891T>C), TGT290TGC (1906T>C), ACA293ACT (1915A>T), GGT298GGC (1930T>C), GCC300GCT (1936C>T), ATT301GTC (1937A>G 1939T>C), TCA303TCG (1945A>G), ACA304ACG (1948A>G), TTC305TTT (1951C>T), CCA308CCT (1960A>T), GTA314GTG (1978A>G), ATT315ATA (1981T>A), GGC316GGT (1984C>T), AAT317AAC (1987T>C), GAC320GAT (1996C>T), TTT323TAT (2004T>A), AAA328AAG (2020A>G), GGG329GGA (2023G>A), GAT330GAC (2026T>C), TCG331TCT (2029G>T), GAG332GAA (2032G>A), TAT341TAC (2059T>C), ATT344ATG (2068T>G), AAC345AAT (2071C>T), GTG346ATA (2072G>A 2074G>A), CTG351TTA (2087C>T 2089G>A), AGC355AGT (2101C>T), GAG356GAA (2104G>A), GAG357GAA (2107G>A), AAG360AAA (2116G>A), TTC362TTT (2122C>T), CGC367CGT (2137C>T), CTT374CTC (2158T>C), ACC376ACT (2164C>T), CCA378CCG (2170A>G), GAT382GAC (2182T>C), GCG384GCA (2188G>A), ACC385ACA (2191C>A), AGA391AAA (2208G>A), TAT394TAC (2218T>C), GAC396GAT (2224C>T), GTA397GTT (2227A>T), TTG402TTA (2242G>A), CCC403CCT (2245C>T), TTG406TTA (2254G>A), GTT407GTC (2257T>C), GAC408GAT (2260C>T), GAC410GAG (2266C>G), CAA412CAG (2272A>G), ACG413ACA (2275G>A), TCA419TCG (2293A>G), GGC421GGT (2299C>T), TCG422TCA (2302G>A), CAG423CAA (2305G>A), GGA426GGT (2314A>T), TAT427TAC (2317T>C), ATT429ATC (2323T>C), CTA430CTG (2326A>G), GTG435GTA (2341G>A), CAA437CAG (2347A>G), CTT438CTC (2350T>C), TTG440TTA (2356G>A), AAT443AAC (2365T>C), GAA445GAG (2371A>G), TTA446TTG (2374A>G), GAT449GAC (2383T>C), ATA450ATT (2386A>T), AAA451AAG (2389A>G), CAT452CAC (2392T>C), TAT453TAC (2395T>C) | | | | | | | | |
| P3 protein (NP_734251.1) | 1 | 365 | 100% | 1928 | 79.8% | 365 (99.7%) | 328 (89.6%) | 1/0/2/2 | 0 |
| Protein mutations: | N4G (2414A>G 2415A>G 2416T>G), S6C (2420A>T), K40R (2523A>G), R77K (2634G>A), V107I (2723G>A), A115T (2747G>A 2749T>A), H131N (2795C>A), V140I (2822G>A 2824G>A), P160T (2882C>A), S161H (2885A>C 2886G>A), T164M (2895C>T), N176S (2931A>G), N179S (2940A>G), E189_N190insX (2971_2972insA), T196I (2991C>T), H197R (2994A>G), E199R (2999G>A 3000A>G), R206Q (3021G>A), Y207F (3024A>T 3025C>T), T211I (3036C>T), E212G (3039A>G 3040A>C), K213T (3042A>C 3043G>A), A232V (3099C>T 3100C>A), V234R (3104G>A 3105T>G 3106G>A), S243N (3132G>A), R245K (3137C>A 3138G>A 3139A>G), F250I (3152T>A 3154C>T), C254Y (3165G>A), F272L (3218T>C), V276I (3230G>A), V288M (3266G>A 3268A>G), V291M (3275G>A), A294T (3284G>A), R304K (3315G>A 3316A>G), E305A (3318A>C), M319V (3359A>G), D332E (3400T>G) | | | | | | | | |

| | Begin | End | Coverage | Score | Concordance | Matches | Identities | I/D/M/F* | Stop Codons |
|---|---|---|---|---|---|---|---|---|---|
| **NT** | **19** | **9704** | **99.8%** | **12534** | **65.0%** | **9674 (99.7%)** | **8011 (82.6%)** | **15/12** | |
| Codon mutations: | AAT4GGG (2414A>G 2415A>G 2416T>G), GCT5GCA (2419T>A), AGC6TGC (2420A>T), ACA12ACT (2440A>T), GTT21GTC (2467T>C), TCT24TCC (2476T>C), TCG26TCA (2482G>A), CTG29CTA (2491G>A), CTG31CTT (2497G>T), TTG33TTA (2503G>A), AGA38AGG (2518A>G), CCT39CCC (2521T>C), AAG40AGG (2523A>G), AGA43AGG (2533A>G), TTG45TTA (2539G>A), TTA47TTG (2545A>G), GAG49GAA (2551G>A), CCT50CCA (2554T>A), CTG52TTG (2558C>T), TTG53CTC (2561T>C 2563G>C), CTA55CTG (2569A>G), TCA56TCG (2572A>G), CCT60CCC (2584T>C), GGC61GGT (2587C>T), CTG63CTC (2593G>C), GCT65GCC (2599T>C), AAT68AAC (2608T>C), TTT72TTC (2620T>C), GCG75GCA (2629G>A), GTG76GTA (2632G>A), AGG77AAG (2634G>A), TTG78TTA (2638G>A), ATT80ATA (2644T>A), AAA83AAG (2653A>G), GCT87GCC (2665T>C), CTA92TTA (2678C>T), CTA93CTG (2683A>G), GCT95GCC (2689T>C), CTA98CTC (2698A>C), TCA101TCG (2707A>G), GCG102GCA (2710G>A), GAA104GAG (2716A>G), ACA105ACG (2719A>G), CTC106CTT (2722C>T), GTC107ATC (2723G>A), GCA108GCG (2728A>G), AGG110AGA (2734G>A), ATT111ATA (2737T>A), GAT114GAC (2746T>C), GCT115ACA (2747G>A 2749T>A), GCA116GCT (2752A>T), GCT117GCA (2755T>A), GAC119GAT (2761C>T), CTC120CTT (2764C>T), TGT125TGC (2779T>C), GGG127GGT (2785G>T), AAC129AAT (2791C>T), CAT131AAT (2795C>A), TAC134TAT (2806C>T), TTG138CTC (2816T>C 2818G>C), GTG140ATA (2822G>A 2824G>A), TTG141TTA (2827G>A), GTT144GTC (2836T>C), AAG145AAA (2839G>A), AAT146AAC (2842T>C), ACC153ACA (2863C>A), CTA154TTA (2864C>T), CCA160ACA (2882C>A), AGT161CAT (2885A>C 2886G>A), AAC163AAT (2893C>T), ACG164ATG (2895C>T), GTC166GTT (2902C>T), GTA167GTG (2905A>G), ATT169ATC (2911T>C), AAT176AGT (2931A>G), AAC179AGC (2940A>G), GCT181GCC (2947T>C), GAA189_AAT190insA-- (2971_2972insA), GGT195GGC (2989T>C), ACT196ATT (2991C>T), CAT197CGT (2994A>G), GAG199AGG (2999G>A 3000A>G), TCC204TC- (3016delC), CGG206CAG (3021G>A), TAC207TTT (3024A>T 3025C>T), ACA211ATA (3036C>T), GAA212GGC (3039A>G 3040A>C), AAG213ACA (3042A>C 3043G>A), TTG216TTA (3052G>A), TTA219TTG (3061A>G), GCG226GCA (3082G>A), GGC229GGT (3091C>T), GCC232GTA (3099C>T 3100C>A), GTG234AGA (3104G>A 3105T>G 3106G>A), GTC235GTT (3109C>T), AAA236AAG (3112A>G), ACT238ACC (3118T>C), GCC239GCT (3121C>T), GGA241GGG (3127A>G), AGC243AAC (3132G>A), CGA245AAG (3137C>A 3138G>A 3139A>G), TAT249TAC (3151T>C), TTC250ATT (3152T>A 3154C>T), TGT254TAT (3165G>A), GTA255GTG (3169A>G), ATC265ATT (3199C>T), AGG267CGG (3203A>C), CGT268CGC (3208T>C), CCA270CCG (3214A>G), ACC271ACT (3217C>T), TTT272CTT (3218T>C), GTC273GTA (3223C>A), GTT276ATT (3230G>A), AAC277AAT (3235C>T), TTA285CTA (3257T>C), ACC286ACT (3262C>T), AGC287AGT (3265C>T), GTA288ATG (3266G>A 3268A>G), GTG289GTA (3271G>A), GTG291ATG (3275G>A), GCA294ACA (3284G>A), CAG299CAA (3301G>A), AGA304AAG (3315G>A 3316A>G), GAA305GCA (3318A>C), GAG307GAA (3325G>A), TTG308TTA (3328G>A), ATA311ATT (3337A>T), GAG315GAA (3349G>A), GTC317GTT (3355C>T), TGC318TGT (3358C>T), ATG319GTG (3359A>G), CTA321CTG (3367A>G), TAT322TAC (3370T>C), TTA325CTG (3377T>C 3379A>G), GAA330GAG (3394A>G), CGC331CGT (3397C>T), GAT332GAG (3400T>G), ACA334ACG (3406A>G), TAC338TAT (3418C>T), ATT339ATA (3421T>A), TAT341TAC (3427T>C), AAG343AAA (3433G>A), TCA344TCT (3436A>T), GTA345GTG (3439A>G), AAC346AAT (3442C>T), CCT347CCC (3445T>C), CAG351CAA (3457G>A), TTT352TTC (3460T>C), GCT353GCA (3463T>A), GCG355GCT (3469G>T), CAG356CAA (3472G>A), CAC364CAT (3496C>T), CAG365CAA (3499G>A) | | | | | | | | |
| 6K1 protein (NP_734245.1) | 1 | 52 | 100% | 305 | 90.5% | 52 (100%) | 45 (86.5%) | 0/0/0/0 | 0 |
| Protein mutations: | V5G (3513T>G), M16V (3545A>G), A17T (3548G>A), V19I (3554G>A 3556C>T), L42V (3623C>G), L45M (3632C>A), Y47H (3638T>C) | | | | | | | | |
| Codon mutations: | TCC2TCT (3505C>T), GTT5GGT (3513T>G), AAA7AAG (3520A>G), GAA10GAG (3529A>G), CAA11CAG (3532A>G), ATG16GTG (3545A>G), GCT17ACT (3548G>A), TTA18CTA (3551T>C), GTC19ATT (3554G>A 3556C>T), AGT28AGC (3583T>C), GAT29GAC (3586T>C), TGC30TGT (3589C>T), TTC32TTT (3595C>T), AAT36AAC (3607T>C), TTT38TTC (3613T>C), AAG39AAA (3616G>A), GGT40GGC (3619T>C), CTT42GTT (3623C>G), TCC43TCT (3628C>T), CTG45ATG (3632C>A), GAT46GAC (3637T>C), TAT47CAT (3638T>C), GAA48GAG (3643A>G) | | | | | | | | |
| CI protein (NP_734246.1) | 1 | 634 | 100% | 4190 | 97.9% | 634 (100%) | 608 (95.9%) | 0/0/0/0 | 0 |
| Protein mutations: | I15V (3698A>G), S21N (3717G>A), R26K (3731C>A 3732G>A), I140V (4073A>G), F158Y (4128T>A 4129C>T), A166S (4151G>T), V171I (4166G>A), L190I (4223C>A 4225G>A), I222V (4319A>G 4321A>G), F247Y (4395T>A), V251I (4406G>A), M274V (4475A>G 4477G>T), T276S (4481A>T), A297L (4544G>T 4545C>T 4546G>A), R298K (4548G>A), V342I (4679G>A 4681G>C), V381I (4796G>A 4798C>T), T383S (4803C>G 4804T>C), S468G (5057A>G), T469A (5060A>G), L479I (5090C>A), A482V (5100C>T), E484D (5107A>T), V489F (5120G>T 5122C>T), I491V (5126A>G), V510I (5183G>A) | | | | | | | | |
| Codon mutations: | TTA2TTG (3661A>G), GTG5GTA (3670G>A), TTT9TTC (3682T>C), GAG11GAA (3688G>A), GAG14GAA (3697G>A), ATT15GTT (3698A>G), GAT17GAC (3706T>C), TTT18TTC (3709T>C), TTG20CTA (3713T>C 3715G>A), AGT21AAT (3717G>A), GAG22GAA (3721G>A), CGA26AAA (3731C>A 3732G>A), ACT27ACA (3736T>A), CTA31TTG (3746C>T 3748A>G), ACA33ACG (3754A>G), AGT36AGC (3763T>C), CGA41CGG (3778A>G), CAA42CAG (3781A>G), ATC43ATT (3784C>T), CAT47CAC (3796T>C), ACA48ACG (3799A>G), CCA50CCT (3805A>T), TAC52TAT (3811C>T), AGA53AGG (3814A>G), TTC58TTT (3829C>T), TTT61TTC (3838T>C), AGA63AGG (3844A>G), GCA64GCC (3847A>C), GTT67GTA (3856T>A), GTG69GTT (3862G>T), GAC72GAT (3871C>T), GCC74GCA (3877C>A), AGC76AGT (3883C>T), GAA77GAG (3886A>G), CTA79CTG (3892A>G), GAC80GAT (3895C>T), TTT81TTC (3898T>C), GTA83GTG (3904A>G), CGG84AGA (3905C>A 3907G>A), GCT86GCC (3913T>C), AAG91AAA (3928G>A), TCA92TCT (3931A>T), GGG94GGA (3937G>A), GTT97GTC (3946T>C), AGT100AGC (3955T>C), GTG101GTA (3958G>A), GCC102GCT (3961C>T), GGA103GGG (3964A>G), TCT104TCC (3967T>C), TTA107TTG (3976A>G), ATT108ATC (3979T>C), CCA110CCT (3985A>T), CTA114CTT (3997A>T), GCG115GCA (4000G>A), AAC117AAT (4006C>T), GTT118GTG (4009T>G), TTC119TTT (4012C>T), AAA120AAG (4015A>G), TCT123TCC (4024T>C), GAA125GAG (4030A>G), CCA126CCT (4033A>T), AAG129AAA (4042G>A), CTG133TTA (4052C>T 4054G>A), CGT134CGC (4057T>C), CGT136CGA (4063T>A), GGA137GGG (4066A>G), ATA140GTA (4073A>G), TTT141TTC (4078T>C), GGC142GGT (4081C>T), TCT143TCC (4084T>C), TCC147TCT (4096C>T), AGC151AGT (4108C>T), GGA152GGT (4111A>T), GCG154GCA (4117G>A), CTA155TTG (4118C>T 4120A>G), TAC157TAT (4126C>T), TTC158TAT (4128T>A 4129C>T), GCC159GCT (4132C>T), AAT160AAC (4135T>C), CAA164CAG (4147A>G), TTA165CTA (4148T>C), GCT166TCT (4151G>T), AAC169AAT (4162C>T), GTA171ATA (4166G>A), ATA172ATC (4171A>C), GAG175GAA (4180G>A), CTG179TTG (4190C>T), CCT181CCC (4198T>C), TCC182TCT (4201C>T), GCG185GCT (4210G>T), TTC186TTT (4213C>T), AGT188AGC (4219T>C), CTG189TTG (4220C>T), CTG190ATA (4223C>A 4225G>A), GTT192GTG (4231T>G), CAT194CAC (4237T>C), GTA199GTG (4252A>G), TTA200CTG (4253T>C 4255A>G), AAA201AAG (4258A>G), ACT205ACC (4270T>C), AGA209AGG (4282A>G), GAG210GAA (4285G>A), GAA212GAG (4291A>G), CAG216CAA (4303G>A), GTC219GTT (4312C>T), ATA222GTG (4319A>G 4321A>G), GTG223GTT (4324G>T), GAG224GAA (4327G>A), CTG227CTT (4336G>T), TCT228TCC (4339T>C), TCA231TCT (4348A>T), GCA235GCG (4360A>G), GGT237GGC (4366T>C), TCT238TCA (4369T>A), ACT240ACC (4375T>C), GTT244GTC (4387T>C), TTT247TAT (4395T>A), GTA251ATA (4406G>A), CTT252CTC (4411T>C), GTT261GTC (4438T>C), ACC263ACA (4444C>A), GCC265GCT (4450C>T), CTC267CTT (4456C>T), GAC270GAT (4465C>T), ATG274GTT (4475A>G 4477G>T), GTC275GTT (4480C>T), ACA276TCA (4481A>T), GGC280GGT (4495C>T), AAG284AAA (4507G>A), CAC285CAT (4510C>T), GGT286GGG (4513T>G), CTA288TTA (4517C>T), GTC291GTT (4528C>T), AAA293AAG (4534A>G), ACC295ACT (4540C>T), GCG297TTA (4544G>T 4545C>T 4546G>A), AGA298AAA (4548G>A), TTT301TTC (4558T>C), GTA303GTT (4564A>T), ACC305ACT (4570C>T), ATA307ATT (4576A>T), TTG314TTA (4597G>A), GAC315GAT (4600C>T), GAC317GAT (4606C>T), GTG318GTA (4609G>A), GTG320GTA (4615G>A), GAC321GAT (4618C>T), TTG324CTT (4625T>C 4627G>T), GAC331GAT (4648C>T), ATT332ATC (4651T>C), AGG335AGA (4660G>A), AGC336AGT (4663C>T), GCT338GCA (4669T>A), GTG342ATC (4679G>A 4681G>C), AGT343AGC (4684T>C), GTT344GTG (4687T>G), GAG348GAA (4699G>A), AGA349AGG (4702A>G), CAA351CAG (4708A>G), AGG352AGA (4711G>A), CTG353TTG (4712C>T), AAA361AAG (4738A>G), GTA363GTT (4744A>T), GCA364GCG (4747A>G), TTG365CTG (4748T>C), ACT370ACC (4765T>C), GTC381ATT (4796G>A 4798C>T), ACT383AGC (4803C>G 4804T>C), GAG384GAA (4807G>A), GCG385GCT (4810G>T), GCT386GCG (4813T>G), CTT387CTC (4816T>C), TAT392TAC (4831T>C), AAC393AAT (4834C>T), CCA395CCT (4840A>T), GGA399GGG (4852A>G), GGC400GGT (4855C>T), AGT404AGC (4867T>C), CTG405CTC (4870G>C), ATT406ATA (4873T>A), TGT409TGC (4882T>C), ACA416ACT (4903A>T), CAG418CAA (4909G>A), TTT420TTC (4915T>C), GAA421GAG (4918A>G), TTG422CTA (4919T>C 4921G>A), AGT423AGC (4924T>C), CCC424CCG (4927C>G), TTT426TTC (4933T>C), ATC427ATA (4936C>A), CAG428CAA (4939G>A), AAT429AAC (4942T>C), TTC430TTT (4945C>T), TCA436TCG (4963A>G), CCT439CCC (4972T>C), CTT445CTC (4990T>C), AAA449AAG (5002A>G), CCT456CCC (5023T>C), TTG457CTG (5024T>C), TGC458TGT (5029C>T), GAT459GAC (5032T>C), TCT461TCC (5038T>C), CCA463CCT (5044A>T), GCA466GCT (5053A>T), TCG467TCA (5056G>A), AGT468GGT (5057A>G), ACT469GCT (5060A>G), TCG472TCT (5071G>T), CTT479ATT (5090C>A), GGA480GGG (5095A>G), GTG481GTT (5098G>T), GCC482GTC (5100C>T), TTA483TTG (5104A>G), GAA484GAT (5107A>T), ATT485ATC (5110T>C), CCA486CCG (5113A>G), AAG487AAA (5116G>A), CAA488CAG (5119A>G), GTC489TTT (5120G>T 5122C>T), ATT491GTT (5126A>G), CAT494CAC (5137T>C), ATC495ATT (5140C>T), AAA496AAG (5143A>G), GAG497GAA (5146G>A), CTC502CTA (5161C>A), GAA504GAG (5167A>G), CTT506CTG (5173T>G), ACG509ACA (5182G>A), GTT510ATT (5183G>A), GTC511GTT (5188C>T), AAG512AAA (5191G>A), AAA514AAG (5197A>G), GAC515GAT (5200C>T), TGC517TGT (5206C>T), TTA518TTG (5209A>G), AGC521AGT (5218C>T), TCG525TCA (5230G>A), ATC527ATT (5236C>T), AGC528AGT (5239C>T), ATC530ATT (5245C>T), ACA536ACT (5263A>T), CTC538CTT (5269C>T), TTC539TTT (5272C>T), GCC540GCA (5275C>A), ATC541ATT (5278C>T), CCA542CCC (5281A>C), AGA543AGG (5284A>G), ACT544ACC (5287T>C), CTA545TTA (5288C>T), ATA546ATT (5293A>T), TTG547CTA (5294T>C 5296G>A), GAG549GAA (5302G>A), CTT552CTA (5311T>A), GAA553GAG (5314A>G), CGA556CGT (5323A>T), AGT564AGC (5347T>C), ATC566ATT (5353C>T), TGC570TGT (5365C>T), GTT577GTC (5386T>C), ACC580ACA (5395C>A), AAA589AAG (5422A>G), GAT590GAC (5425T>C), ACC592ACT (5431C>T), GAG594GAA (5437G>A), CTT599CTC (5452T>C), AAG601AAA (5458G>A), AGT604AGC (5467T>C), CAA605CAG (5470A>G), CTA606TTG (5471C>T 5473A>G), TTC609TTT (5482C>T), TCA610TCC (5485A>C), AAT611AAC (5488T>C), TTG612CTA (5489T>C 5491G>A), GGT614GGC (5497T>C), TGT617TGC (5506T>C), GAG618GAA (5509G>A), GAG619GAA (5512G>A), TAT625TAC (5530T>C), GAG626GAA (5533G>A), TCG627TCT (5536G>T), TTG628TTA (5539G>A), GTT631GTG (5548T>G), CAC633CAT (5554C>T) | | | | | | | | |
| 6K2 protein (NP_734247.1) | 1 | 52 | 100% | 303 | 93.5% | 52 (100%) | 46 (88.5%) | 0/0/0/0 | 0 |
| Protein mutations: | A2T (5561G>A 5563G>A), I14V (5597A>G), N16K (5605C>A), A21V (5619C>T 5620T>G), I25L (5630A>C), V46I (5693G>A) | | | | | | | | |
| Codon mutations: | GCT1GCA (5560T>A), GCG2ACA (5561G>A 5563G>A), ACG3ACC (5566G>C), TCA4TCG (5569A>G), CTC9CTG (5584C>G), AAG12AAA (5593G>A), GGG13GGA (5596G>A), ATT14GTT (5597A>G), AAC16AAA (5605C>A), GTG20GTT (5617G>T), GCT21GTG (5619C>T 5620T>G), TTG24TTA (5629G>A), ATC25CTC (5630A>C), GGC28GGT (5641C>T), GCA31GCT (5650A>T), ATA35ATT (5662A>T), GGA36GGG (5665A>G), CAA44CAG (5689A>G), GTT46ATT (5693G>A), GAG47GAA (5698G>A), CAA52CAG (5713A>G) | | | | | | | | |
| NIa-VPg protein (NP_734252.1) | 1 | 188 | 100% | 1227 | 93.3% | 188 (100%) | 170 (90.4%) | 0/0/0/0 | 0 |
| Protein mutations: | K47R (5853A>G), I61V (5894A>G 5896C>G), R83A (5960C>G 5961G>C 5962G>C), I93V (5990A>G), E102D (6019A>T), K106T (6030A>C), D112E (6049C>A), M115I (6058G>A), S120N (6072G>A 6073T>C), N121H (6074A>C), T123N (6081C>A), C133S (6111G>C 6112T>C), I139V (6128A>G 6130T>C), V149I (6158G>A), T153S (6170A>T 6172A>G), L164F (6203C>T 6205C>T), V173I (6230G>A), A182K (6257G>A 6258C>A) | | | | | | | | |

| | Begin | End | Coverage | Score | Concordance | Matches | Identities | I/D/M/F* | Stop Codons |
|---|---|---|---|---|---|---|---|---|---|
| **NT** | **19** | **9704** | **99.8%** | **12534** | **65.0%** | **9674 (99.7%)** | **8011 (82.6%)** | **15/12** | |
| Codon mutations: | GGG1GGC (5716G>C), AAA2AAG (5719A>G), GCC10GCA (5743C>A), CGC14CGA (5755C>A), CAT15CAC (5758T>C), CGT17CGC (5764T>C), AAA19AAG (5770A>G), GGC22GGT (5779C>T), TTT23TTC (5782T>C), GAC26GAT (5791C>T), GAC30GAT (5803C>T), AGG41AGA (5836G>A), AAA42AAG (5839A>G), GGA44GGG (5845A>G), AAA45AAG (5848A>G), AAA47AGA (5853A>G), GGT48GGC (5857T>C), ACC49ACT (5860C>T), ACA50ACT (5863A>T), GGT52GGA (5869T>A), GGT54GGC (5875T>C), AGC57AGT (5884C>T), TTC60TTT (5893C>T), ATC61GTG (5894A>G 5896C>G), AAC62AAT (5899C>T), GGG65GGA (5908G>A), TTT66TTC (5911T>C), GAG70GAA (5923G>A), CAA75CAG (5938A>G), TTC76TTT (5941C>T), GTT77GTC (5944T>C), GAT78GAC (5947T>C), GGG82GGA (5959G>A), CGG83GCC (5960C>G 5961G>C 5962G>C), GAA87GAG (5974A>G), GCT91GCA (5986T>A), GAC92GAT (5989C>T), ATT93GTT (5990A>G), GAT95GAC (5998T>C), AGA99CGT (6008A>C 6010A>T), GAA102GAT (6019A>T), GTG103GTT (6022G>T), CGA104CGT (6025A>T), AAA106ACA (6030A>C), GAC112GAA (6049C>A), ATG115ATA (6058G>A), CAA116CAG (6061A>G), GCC117GCA (6064C>A), GGT119GGC (6070T>C), AGT120AAC (6072G>A 6073T>C), AAC121CAC (6074A>C), ACG122ACA (6079G>A), ACC123AAC (6081C>A), ATA124ATT (6085A>T), GCA126GCT (6091A>T), TAC127TAT (6094C>T), TGT133TCC (6111G>C 6112T>C), AAA135AAG (6118A>G), TTG137TTA (6124G>A), AAG138AAA (6127G>A), ATT139GTC (6128A>G 6130T>C), GAT140GAC (6133T>C), TTA141TTG (6136A>G), AAC145AAT (6148C>T), AAA148AAG (6157A>G), GTT149ATT (6158G>A), TGT150TGC (6163T>C), AAA152AAG (6169A>G), ACA153TCG (6170A>T 6172A>G), GGC155GGT (6178C>T), AAA158AAG (6187A>G), TTT159TTC (6190T>C), GAG161GAA (6196G>A), CTC164TTT (6203C>T 6205C>T), GAA165GAG (6208A>G), CTA166TTG (6209C>T 6211A>G), CAG168CAA (6217G>A), GCT172GCA (6229T>A), GTA173ATA (6230G>A), GTC175GTT (6238C>T), AAG178AAA (6247G>A), ATA180ATT (6253A>T), GCA182AAA (6257G>A 6258C>A), CAT187CAC (6274T>C) | | | | | | | | |
| NIa-Pro protein (NP_734248.1) | 1 | 244 | 100% | 1651 | 93.5% | 244 (100%) | 222 (91.0%) | 0/0/0/0 | 0 |
| Protein mutations: | K2R (6282A>G), A29T (6362G>A), V42I (6401G>A), A43V (6405C>T), Y51F (6429A>T), Q58R (6450A>G), H70N (6485C>A 6487C>T), L84I (6527C>A), T104I (6588C>T), Y120H (6635T>C), I124V (6647A>G), I134V (6677A>G 6679A>G), D147N (6716G>A), N148D (6719A>G), I163L (6764A>T 6766A>G), A173V (6795C>T 6796A>G), H174Q (6799C>A), K189R (6843A>G), N194D (6857A>G), V200T (6875G>A 6876T>C 6877C>A), D219E (6934C>A), V242I (7001G>A 7003G>A) | | | | | | | | |
| Codon mutations: | AAA2AGA (6282A>G), TCG3TCA (6286G>A), GGC7GGT (6298C>T), TTG8TTA (6301G>A), AGA9AGG (6304A>G), GAC10GAT (6307C>T), AAC12AAT (6313C>T), GCC15GCT (6322C>T), ACA17ACT (6328A>T), GTT18GTG (6331T>G), TGT19TGC (6334T>C), AGG20AGA (6337G>A), CTG21TTG (6338C>T), TAT27TAC (6358T>C), GGG28GGA (6361G>A), GCA29ACA (6362G>A), TCA30TCT (6367A>T), GAG31GAA (6370G>A), TAC33TAT (6376C>T), GGT34GGG (6379T>G), GGA36GGC (6385A>C), GGA38GGT (6391A>T), GCA39GCG (6394A>G), TAC40TAT (6397C>T), GTA42ATA (6401G>A), GCG43GTG (6405C>T), CAC45CAT (6412C>T), TTA47TTG (6418A>G), TTT48TTC (6421T>C), TAC51TTC (6429A>T), GGT53GGA (6436T>A), GAG56GAA (6445G>A), CAA58CGA (6450A>G), TCC59TCA (6454C>A), CAC61CAT (6460C>T), GGT62GGA (6463T>A), TTC64TTT (6469C>T), AGG65AGA (6472G>A), GTG66GTA (6475G>A), AAT68AAC (6481T>C), CAC70AAT (6485C>A 6487C>T), AGT71AGC (6490T>C), AGC73AGT (6496C>T), CTG75CTA (6502G>A), GGT79GGC (6514T>C), AGG80AGA (6517G>A), CTC84ATC (6527C>A), ATC85ATA (6532C>A), CCG88CCA (6541G>A), AAA89AAG (6544A>G), GTC93GTT (6556C>T), TTT94TTC (6559T>C), CAG96CAA (6565G>A), TTG98CTA (6569T>C 6571G>A), CCT103CCA (6586T>A), ACA104ATA (6588C>T), GAA107GAG (6598A>G), AGA108AGG (6601A>G), ATT109ATC (6604T>C), TGT110TGC (6607T>C), GGA113GGG (6616A>G), AAC115AAT (6622C>T), GAG118GAA (6631G>A), AAG119AAA (6634G>A), TAT120CAT (6635T>C), GCT121GCA (6640T>A), TCG123TCC (6646G>C), ATC124GTC (6647A>G), ATC125ATT (6652C>T), AGC129AGT (6664C>T), ACT131ACG (6670T>G), ATA134GTG (6677A>G 6679A>G), GGC136GGT (6685C>T), ACA138ACT (6691A>T), TTC139TTT (6694C>T), ACA146ACG (6715A>G), GAT147AAT (6716G>A), AAT148GAT (6719A>G), GGA149GGG (6724A>G), CTA153TTA (6734C>T), GTG155GTA (6742G>A), GTG156GTA (6745G>A), GCC159GCT (6754C>T), GGA161GGG (6760A>G), ATA163TTG (6764A>T 6766A>G), ATT166ATA (6775T>A), CTG169TTG (6782C>T), GCA170GCT (6787A>T), AAC171AAT (6790C>T), GCA173GTG (6795C>T 6796A>G), CAC174CAA (6799C>A), ACC175ACA (6802C>A), AAC177AAT (6808C>T), TAC178TAT (6811C>T), TAC179TAT (6814C>T), TTC182TTT (6823C>T), GAA184GAG (6829A>G), AAG189AGG (6843A>G), TAC190TAT (6847C>T), CGA192CGC (6853A>C), AAT194GAT (6857A>G), GAG195GAA (6862G>A), GAA198GAG (6871A>G), GTC200ACA (6875G>A 6876T>C 6877C>A), AAG201AAA (6880G>A), TCT202TCA (6883T>A), GTT204GTG (6889T>G), GAC208GAT (6901C>T), GGC213GGT (6916C>T), CCG214CCA (6919G>A), AAA216AAG (6925A>G), CTT217CTC (6928T>C), GAC219GAA (6934C>A), GGG224GGC (6949G>C), TTC226TTT (6955C>T), AAA227AAG (6958A>G), CAA233CAG (6976A>G), CTA235TTA (6980C>T), ATC236ATT (6985C>T), GTA240GTT (6997A>T), GTG241GTT (7000G>T), GTG242ATA (7001G>A 7003G>A) | | | | | | | | |
| NIb protein (NP_734249.1) | 1 | 521 | 100% | 3134 | 86.2% | 519 (99.4%) | 481 (92.1%) | 1/2/2/2 | 0 |
| Protein mutations: | F8Y (7032T>A), T39K (7125C>A), A46S (7145G>T 7147A>G), E51_A52del (7160_7165delGAGGCA), D69E (7216T>A), D83E (7258T>A), D88_R89insX (7273_7274insT), R89V (7274C>G 7275G>T), H91L (7281A>T), L92S (7284T>C), S96L (7296C>T), S101F (7311C>T), T102I (7314C>T), C103Y (7317G>A), K109Q (7334A>C), A112N (7343G>A 7344C>A 7345A>C), S133M (7407G>T 7408T>G), C136G (7415T>G), L165S (7503T>C 7504G>A), D234N (7709G>A), K235R (7713A>G 7714A>G), T268I (7812C>T), V279L (7844G>T 7846G>A), V292I (7883G>A), S345N (8043G>A), N381S (8151A>G 8152C>T), R390K (8178G>A), R400K (8208G>A), P429A (8294C>G 8296A>G), S447F (8349C>T), A467S (8408G>T), M481I (8452G>A), R484K (8460G>A), A491T (8480G>A 8482T>A), R498K (8502G>A 8503A>G), E502D (8515A>T), E510D (8539G>C), L513C (8546C>T 8547T>G), S515T (8552T>A) | | | | | | | | |
| Codon mutations: | GCT1GCC (7012T>C), AAG2AAA (7015G>A), CAT3CAC (7018T>C), TTT8TAT (7032T>A), GCC10GCT (7039C>T), TTG11CTA (7040T>C 7042G>A), GGA13GGG (7048A>G), GCT17GCA (7060T>A), GTC18GTA (7063C>A), AGC23AGT (7078C>T), TTA25CTG (7082T>C 7084A>G), GTA26GTG (7087A>G), ACC27ACA (7090C>A), CAT29CAC (7096T>C), GTA30GTG (7099A>G), GTT31GTC (7102T>C), GAG34GAA (7111G>A), CGA36CGG (7117A>G), ACA39AAA (7125C>A), GAA40GAG (7129A>G), TTT41TTC (7132T>C), CTG42TTA (7133C>T 7135G>A), GTG44GTA (7141G>A), GAT45GAC (7144T>C), GCA46TCG (7145G>T 7147A>G), GAG47GAA (7150G>A), GAG49GAA (7156G>A), GCA50GCT (7159A>T), GAG51_GCA52del (7160_7165delGAGGCA), TTC54TTT (7171C>T), GCG60GCT (7189G>T), AAA63AAG (7198A>G), AGC64AGT (7201C>T), CTA66TTA (7205C>T), AAT67AAC (7210T>C), GAT69GAA (7216T>A), GCG70GCA (7219G>A), ATC72ATA (7225C>A), AAG73AAA (7228G>A), AAG77AAA (7240G>A), TAT78TAC (7243T>C), AAA80AAG (7249A>G), CCT81CCG (7252T>G), GAT83GAA (7258T>A), GTT84GTC (7261T>C), GGT85GGA (7264T>A), GTC86GTG (7267C>G), GTG87GTA (7270G>A), GAT88_CGG89insT-- (7273_7274insT), CGG89GTG (7274C>G 7275G>T), CAT91CTT (7281A>T), TTG92TCG (7284T>C), TCA96TTA (7296C>T), TCT101TTT (7311C>T), ACC102ATC (7314C>T), TGC103TAC (7317G>A), AAT104-AA (7319delA 7321T>A), CAC106CAT (7327C>T), GGC107GGT (7330C>T), TTC108TTT (7333C>T), AAG109CAG (7334A>C), TGT111TGC (7342T>C), GCA112AAC (7343G>A 7344C>A 7345A>C), ACT115ACC (7354T>C), CAA118CAG (7363A>G), TTC121TTT (7372C>T), GCG123GCT (7378G>T), CTC124CTT (7381C>T), GCA129GCT (7396A>T), GTC130GTT (7399C>T), GCC132GCT (7405C>T), AGT133ATG (7407G>T 7408T>G), GGG135GGT (7414G>T), TGC136GGC (7415T>G), AAA137AAG (7420A>G), GAC140GAT (7429C>T), TAT141TAC (7432T>C), CAT144CAC (7441T>C), AAG150AAA (7459G>A), GAA151GAG (7462A>G), ATA153ATT (7468A>T), GTC154GTA (7471C>A), CAA156CAG (7477A>G), AGC157AGT (7480C>T), CTG159CTT (7486G>T), TAT162TAC (7495T>C), AAA163AAG (7498A>G), GGT164GGC (7501T>C), TTG165TCA (7503T>C 7504G>A), ATT168ATA (7513T>A), AAC170AAT (7519C>T), GGA171GGG (7522A>G), AAG174AAA (7531G>A), GAG176GAA (7537G>A), CTC177CTT (7540C>T), TGT179TGC (7546T>C), AAG187AAA (7570G>A), ACG188ACA (7573G>A), ACG190ACA (7579G>A), GCT193GCC (7588T>C), CTA196TTA (7595C>T), TTG199CTA (7604T>C 7606G>A), GGT202GGA (7615T>A), TGT205TGC (7624T>C), GAC208GAT (7633C>T), TTC209TTT (7636C>T), AAT210AAC (7639T>C), TTT213TTC (7648T>C), GAA219GAG (7666A>G), TGT221TGC (7672T>C), ACA223ACT (7678A>T), GGG225GGA (7684G>A), ACT227ACC (7690T>C), TTT229TTC (7696T>C), GGT231GGG (7702T>G), GAT234AAT (7709G>A), AAA235AGG (7713A>G 7714A>G), CTG236TTG (7715C>T), CGT239CGC (7726T>C), GAG242GAA (7735G>A), GTA245GTG (7744A>G), TAC246TAT (7747C>T), GGC251GGT (7762C>T), CAG253CAA (7768G>A), TTT254TTC (7771T>C), TCA257TCG (7780A>G), CTA258CTC (7783A>C), ACT259ACC (7786T>C), CTA262TTG (7793C>T 7795A>G), ATC263ATT (7798C>T), GCT265GCC (7804T>C), ACC268ATC (7812C>T), ATC269ATT (7816C>T), GAA275GAG (7834A>G), GAC276GAT (7837C>T), GTG279TTA (7844G>T 7846G>A), CAG282CAA (7855G>A), CTG284CTA (7861G>A), TTA287CTG (7868T>C 7870A>G), TAC288TAT (7873C>T), ACT289ACA (7876T>A), GAG290GAA (7879G>A), ATT291ATA (7882T>A), GTT292ATT (7883G>A), TAC293TAT (7888C>T), CCA295CCT (7894A>T), ATT296ATC (7897T>C), GTC304GTT (7921C>T), AAG306AAA (7927G>A), GGT309GGC (7936T>C), AAT310AAC (7939T>C), AAT311AAC (7942T>C), GTG319GTA (7966G>A), GAT320GAC (7969T>C), TCT322TCC (7975T>C), GTC326GTT (7987C>T), CTT327CTC (7990T>C), TAC331TAT (8002C>T), GCT332GCC (8005T>C), GAA341GAG (8032A>G), ATC343ATT (8038C>T), GAC344GAT (8041C>T), AGC345AAC (8043G>A), TTG357TTA (8080G>A), GTG360GTA (8089G>A), CCG362CCA (8095G>A), GAG363GAA (8098G>A), AGC366AGT (8107C>T), ATT367ATC (8110T>C), GAT369GAC (8116T>C), CAA373CAG (8128A>G), GGT379GGC (8146T>C), AAC381AGT (8151A>G 8152C>T), TCG385TCA (8164G>A), AGG390AAG (8178G>A), AAG391AAA (8182G>A), TTG394TTA (8191G>A), TTC396TTT (8197C>T), TCC398TCT (8203C>T), AGA400AAA (8208G>A), GGC401GGG (8212C>G), CTG402TTG (8213C>T), CTA403CTG (8218A>G), ATC404ATT (8221C>T), GAG405GAA (8224G>A), CCA410CCG (8239A>G), ATT417ATC (8260T>C), GTA418GTG (8263A>G), GAT424GAC (8281T>C), AGA425AGG (8284A>G), CCA429GCG (8294C>G 8296A>G), GAG430GAA (8299G>A), CAC431CAT (8302C>T), ATT436ATC (8317T>C), TGC437TGT (8320C>T), GCT439GCA (8326T>A), GAG442GAA (8335G>A), TCC443TCT (8338C>T), TCT447TTT (8349C>T), GAA448GAG (8353A>G), CAC451CAT (8362C>T), CAA452CAG (8365A>G), TAC457TAT (8380C>T), TCA458TCG (8383A>G), TTA460TTG (8389A>G), CCT465CCC (8404T>C), GCA467TCA (8408G>T), ACA468ACG (8413A>G), GCG470GCT (8419G>T), GCT475GCC (8434T>C), TAT477TAC (8440T>C), ATG481ATA (8452G>A), CTA483CTG (8458A>G), AGG484AAG (8460G>A), AAA485AAG (8464A>G), CTG486CTA (8467G>A), GCT491ACA (8480G>A 8482T>A), GAG496GAA (8497G>A), CTA497TTG (8498C>T 8500A>G), AGA498AAG (8502G>A 8503A>G), GCC499GCT (8506C>T), GAA502GAT (8515A>T), GTC505GTT (8524C>T), GCA506GCC (8527A>C), TTA507CTG (8528T>C 8530A>G), GAG510GAC (8539G>C), CTT513TGT (8546C>T 8547T>G), TCT515ACT (8552T>A), GTA518GTG (8563A>G) | | | | | | | | |
| coat protein (NP_734250.1) | 1 | 267 | 100% | 1659 | 91.0% | 267 (100%) | 244 (91.4%) | 0/0/0/0 | 0 |
| Protein mutations: | A1G (8574C>G), I5V (8585A>G), G9E (8598G>A), N11S (8603A>T 8604A>C 8605C>T), A15V (8616C>T), P17Q (8622C>A), E18G (8625A>G 8626G>A), P26F (8648C>T 8649C>T 8650G>C), G29E (8658G>A), D31A (8664A>C 8665T>G), A36V (8679C>T 8680A>T), T58K (8745C>A 8746A>G), E95G (8856A>G 8857G>A), R98Q (8865G>A), M99L (8867A>C 8869G>T), G104E (8883G>A), T106S (8888A>T), V128I (8954G>A), N138D (8984A>G 8986T>C), E139V (8988A>T), M193G (9149A>G 9150T>G 9151G>A), G194S (9152G>A), P230S (9260C>T 9262T>C) | | | | | | | | |
| Codon mutations: | GCA1GGA (8574C>G), ATT5GTT (8585A>G), GGA9GAA (8598G>A), AGC10AGT (8602C>T), AAC11TCT (8603A>T 8604A>C 8605C>T), GCA15GTA (8616C>T), AAA16AAG (8620A>G), CCA17CAA (8622C>A), GAG18GGA (8625A>G 8626G>A), CAG19CAA (8629G>A), GGC20GGT (8632C>T), ATC22ATT (8638C>T), CAG23CAA (8641G>A), AAC25AAT (8647C>T), CCG26TTC (8648C>T 8649C>T 8650G>C), AAC27AAT (8653C>T), AAA28AAG (8656A>G), GGA29GAA (8658G>A), GAT31GCG (8664A>C 8665T>G), GTT34GTG (8674T>G), GCA36GTT (8679C>T 8680A>T), GGC37GGA (8683C>A), GGG40GGA (8692G>A), ACA41ACT (8695A>T), CCG45CCA (8707G>A), AGA46CGA (8708A>C), AAG48AAA (8716G>A), ATC50ATT (8722C>T), ACG51ACA (8725G>A), AAA53AAG (8731A>G), AGA55AGG (8737A>G), CCC57CCT (8743C>T), ACA58AAG (8745C>A 8746A>G), AGC59AGT (8749C>T), AAG60AAA (8752G>A), GGA61GGT (8755A>T), AAC66AAT (8770C>T), TTA67CTA (8771T>C), GAA68GAG (8776A>G), CAT69CAC (8779T>C), TTG70TTA (8782G>A), TAT73TAC (8791T>C), CCA75CCG (8797A>G), CAA76CAG (8800A>G), CAA77CAG (8803A>G), GAT79GAC (8809T>C), ATT80ATC (8812T>C), AAT82AAC (8818T>C), CGG84CGA (8824G>A), GCA85GCG (8827A>G), ACG92ACA (8848G>A), TAT94TAC (8854T>C), GAG95GGA (8856A>G 8857G>A), GTG97GTA (8863G>A), CGG98CAG (8865G>A), ATG99CTT (8867A>C 8869G>T), GCA100GCG (8872A>G), GGA104GAA (8883G>A), ACT106TCT (8888A>T), TGC119TGT (8929C>T), GTC128ATC (8954G>A), AAC129AAT (8959C>T), GGA130GGT (8962A>T), GGG137GGA (8983G>A), AAT138GAC (8984A>G 8986T>C), GAA139GTA (8988A>T), GAG142GAA (8998G>A), CCG144CCA (9004G>A), TTG145TTA (9007G>A), ATC148ATT (9016C>T), ACC155ACA (9037C>A), ATA171ATT (9085A>T), AAC175AAT (9097C>T), CGA188CGT (9136A>T), CGG191CGT (9145G>T), ATG193GGA (9149A>G 9150T>G 9151G>A), GGT194AGT (9152G>A), TTA195TTG (9157A>G), GCG196GCT (9160G>T), CGT197CGC (9163T>C), GAG204GAA (9184G>A), CGA208CGG (9196A>G), AGG214AGA (9214G>A), GAA215GAG (9217A>G), GCG216GCA (9220G>A), GCA224GCT (9244A>T), TTG225TTA (9247G>A), GCC228GCT (9256C>T), CCT230TCC (9260C>T 9262T>C), GGG234GGA (9274G>A), GAC236GAT (9280C>T), ATC239ATT (9289C>T), GAG244GAA (9304G>A), GTC254GTT (9334C>T), GTC264GTG (9364C>G) | | | | | | | | |

| | Begin | End | Coverage | Score | Concordance | Matches | Identities | I/D/M/F* | Stop Codons |
|---|---|---|---|---|---|---|---|---|---|
| **NT** | **19** | **9704** | **99.8%** | **12534** | **65.0%** | **9674 (99.7%)** | **8011 (82.6%)** | **15/12** | |
| PIPO (YP_006393460.1) | 1 | 75 | 100% | 95 | 18.1% | 75 (98.7%) | 54 (71.1%) | 1/0/3/2 | 2 |
| Protein mutations: | R17_K17insX (2971_2972insA), Y23H (2989T>C 2991C>T), R26K (2999G>A 3000A>G), H35Y (3025C>T), K40H (3040A>C 3042A>C), G41S (3043G>A), E44K (3052G>A), I47V (3061A>G), V54I (3082G>A), P57S (3091C>T), P60T (3100C>A), G61E (3104G>A 3105T>G), G62S (3106G>A), Q63* (3109C>T), R64G (3112A>G), C66R (3118T>C), L67F (3121C>T), I69V (3127A>G), A72E (3137C>A 3138G>A), I73V (3139A>G) | | | | | | | | |
| Codon mutations: | AAA4AAG (2931A>G), GAA7GAG (2940A>G), TTG10CTG (2947T>C), AGA17_AAA17insAA- (2971_2972insA), TAC23CAT (2989T>C 2991C>T), TCA24TCG (2994A>G), AGA26AAG (2999G>A 3000A>G), CAC32-AC (3016delC), TCG33TCA (3021G>A), GTA34GTT (3024A>T), CAT35TAT (3025C>T), CAC38CAT (3036C>T), AGA39AGG (3039A>G), AAA40CAC (3040A>C 3042A>C), GGC41AGC (3043G>A), GAA44AAA (3052G>A), ATA47GTA (3061A>G), GTT54ATT (3082G>A), CCG57TCG (3091C>T), CGC59CGT (3099C>T), CCA60ACA (3100C>A), GGT61GAG (3104G>A 3105T>G), GGT62AGT (3106G>A), CAA63TAA (3109C>T), AGG64GGG (3112A>G), TGC66CGC (3118T>C), CTC67TTC (3121C>T), ATT69GTT (3127A>G), GAG70GAA (3132G>A), GCG72GAA (3137C>A 3138G>A), ATT73GTT (3139A>G) | | | | | | | | |

*: Inserts / Deletes / Misaligned / Frameshifts

#### Analysis details

This analysis was performed with panviral2.64

### NGS Details (UN70): Bracoviriform glomeratae (segment NC_043292.1)

#### Assembly

| | |
|---|---|
| **Coverage Length** | 310 (1 contig(s)) |
| **Depth Of Coverage** | 17187.9 |
| **Number Of Reads** | 47201 |
| **Reads Per Million** | 894.75 rpm (after QC) |
| **Ambiguities** | 0 |
| **Assembly Method** | de novo + reference guided assembly |
| **Consensus Caller** | Bcf Tools |

#### Coverage Map

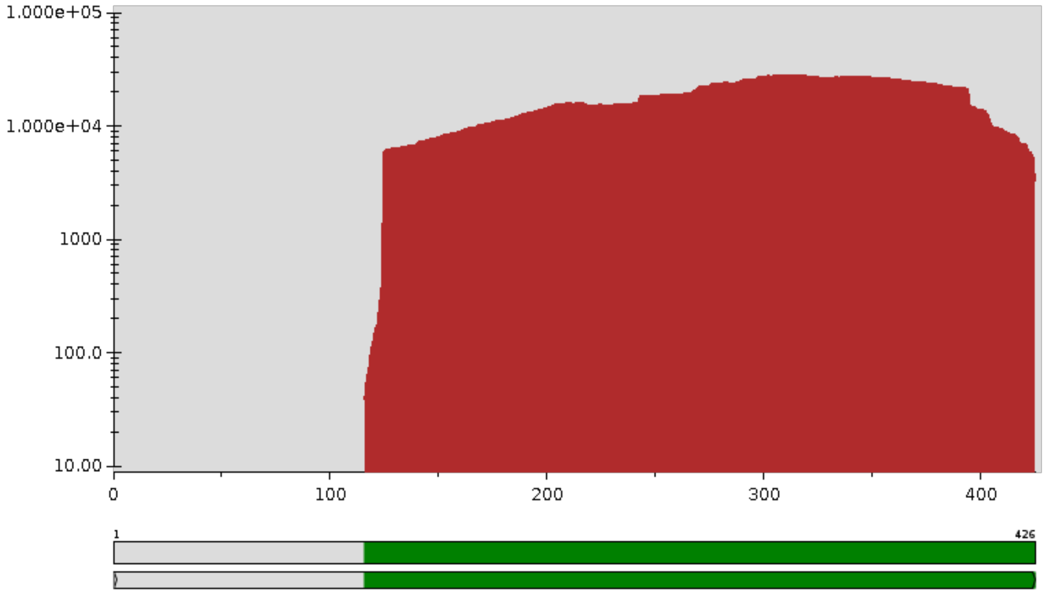

#### Assignment

| | |
|---|---|
| **Type** | Bracoviriform glomeratae (Taxonomy ID: 257816) |
| **Reference Genome** | NC_043292.1 |
| **NT Identity (%)** | 74.5161 |
| **AA Identity (%)** | 84.466 |
| **Number Of Stop Codons** | 1 |
| **Number Of CDS** | 1 |

#### Alignment

| | |
|---|---|
| **Alignment Score** | 304.0 (NT) + 567.0 (AA) = 871.0 |
| **Concordance (%)** | 68.2067 |

| | |
|---|---|
| **Alignment Method** | Global, seeded, nucleotide + amino acids (AGA) |

#### Genome Region

Sequence starts at position 117 and ends at position 426 relative to NC_043292.1 reference sequence.

#### Alignment Detailed Statistics

| | Begin | End | Coverage | Score | Concordance | Matches | Identities | I/D/M/F* | Stop Codons |
|---|---|---|---|---|---|---|---|---|---|
| **NT** | **117** | **426** | **72.8%** | **304** | **49.0%** | **310 (100%)** | **231 (74.5%)** | **0/0** | |
| Mutations: | 126G>T, 129C>A, 132A>G, 136A>G, 137C>G, 138G>C, 153A>G, 155C>G, 156G>A, 159C>A, 166C>A, 168T>G, 169T>C, 171T>C, 172C>A, 174C>G, 177A>G, 183T>A, 184C>A, 189T>C, 195C>T, 198A>G, 201A>C, 207C>T, 214A>G, 219C>T, 222T>C, 223C>A, 225T>G, 228G>T, 235C>A, 237T>G, 243A>T, 246C>G, 261A>G, 264A>G, 265G>A, 273A>G, 278T>C, 279T>C, 284A>G, 285T>A, 288T>G, 289T>C, 300C>T, 301C>T, 303C>G, 309C>T, 312T>G, 315C>T, 322G>T, 329T>C, 333T>C, 336C>T, 339A>G, 345A>C, 347A>G, 348A>G, 349C>A, 351T>A, 360C>T, 363C>T, 366C>T, 375C>T, 384A>C, 385T>C, 387G>C, 390A>G, 391C>A, 392A>G, 393T>A, 394A>C, 400C>A, 402T>G, 404T>C, 405G>T, 406A>T, 408C>G, 416A>T | | | | | | | | |
| **CDS** | | | | | | | | | |
| FK954_p501 | 40 | 142 | 72.5% | 567 | 85.3% | 103 (100%) | 87 (84.5%) | 0/0/0/0 | 1 |
| Protein mutations: | L42F (126G>T), T46G (136A>G 137C>G 138G>C), A52G (155C>G 156G>A), Y57H (169T>C 171T>C), T72A (214A>G), V89I (265G>A), I93T (278T>C 279T>C), D95G (284A>G 285T>A), A108S (322G>T), I110T (329T>C), K116R (347A>G 348A>G), H131R (391C>A 392A>G 393T>A), K132Q (394A>C), M135T (404T>C 405G>T), I136L (406A>T 408C>G), Y139F (416A>T) | | | | | | | | |
| Codon mutations: | TTG42TTT (126G>T), GGC43GGA (129C>A), AAA44AAG (132A>G), ACG46GGC (136A>G 137C>G 138G>C), AAA51AAG (153A>G), GCG52GGA (155C>G 156G>A), GGC53GGA (159C>A), CGT56AGG (166C>A 168T>G), TAT57CAC (169T>C 171T>C), CGC58AGG (172C>A 174C>G), AAA59AAG (177A>G), CTT61CTA (183T>A), CGA62AGA (184C>A), GAT63GAC (189T>C), ATC65ATT (195C>T), CAA66CAG (198A>G), GGA67GGC (201A>C), ACC69ACT (207C>T), ACT72GCT (214A>G), ATC73ATT (219C>T), CGT74CGC (222T>C), CGT75AGG (223C>A 225T>G), CTG76CTT (228G>T), CGT79AGG (235C>A 237T>G), GGA81GGT (243A>T), GTC82GTG (246C>G), GGA87GGG (261A>G), TTA88TTG (264A>G), GTC89ATC (265G>A), GAA91GAG (273A>G), ATT93ACC (278T>C 279T>C), GAT95GGA (284A>G 285T>A), GTT96GTG (288T>G), TTG97CTG (289T>C), TTC100TTT (300C>T), CTC101TTG (301C>T 303C>G), AAC103AAT (309C>T), GTT104GTG (312T>G), ATC105ATT (315C>T), GCT108TCT (322G>T), ATC110ACC (329T>C), TAT111TAC (333T>C), ACC112ACT (336C>T), GAA113GAG (339A>G), GCA115GCC (345A>C), AAA116AGG (347A>G 348A>G), CGT117AGA (349C>A 351T>A), GTC120GTT (360C>T), ACC121ACT (363C>T), GCC122GCT (366C>T), GTC125GTT (375C>T), GCA128GCC (384A>C), TTG129CTC (385T>C 387G>C), AAA130AAG (390A>G), CAT131AGA (391C>A 392A>G 393T>A), AAA132CAA (394A>C), CGT134AGG (400C>A 402T>G), ATG135ACT (404T>C 405G>T), ATC136TTG (406A>T 408C>G), TAT139TTT (416A>T) | | | | | | | | |
| **Proteins** | | | | | | | | | |
| putative histone 4 (YP_009665791.1) | 40 | 142 | 72.5% | 567 | 85.3% | 103 (100%) | 87 (84.5%) | 0/0/0/0 | 1 |
| Protein mutations: | L42F (126G>T), T46G (136A>G 137C>G 138G>C), A52G (155C>G 156G>A), Y57H (169T>C 171T>C), T72A (214A>G), V89I (265G>A), I93T (278T>C 279T>C), D95G (284A>G 285T>A), A108S (322G>T), I110T (329T>C), K116R (347A>G 348A>G), H131R (391C>A 392A>G 393T>A), K132Q (394A>C), M135T (404T>C 405G>T), I136L (406A>T 408C>G), Y139F (416A>T) | | | | | | | | |
| Codon mutations: | TTG42TTT (126G>T), GGC43GGA (129C>A), AAA44AAG (132A>G), ACG46GGC (136A>G 137C>G 138G>C), AAA51AAG (153A>G), GCG52GGA (155C>G 156G>A), GGC53GGA (159C>A), CGT56AGG (166C>A 168T>G), TAT57CAC (169T>C 171T>C), CGC58AGG (172C>A 174C>G), AAA59AAG (177A>G), CTT61CTA (183T>A), CGA62AGA (184C>A), GAT63GAC (189T>C), ATC65ATT (195C>T), CAA66CAG (198A>G), GGA67GGC (201A>C), ACC69ACT (207C>T), ACT72GCT (214A>G), ATC73ATT (219C>T), CGT74CGC (222T>C), CGT75AGG (223C>A 225T>G), CTG76CTT (228G>T), CGT79AGG (235C>A 237T>G), GGA81GGT (243A>T), GTC82GTG (246C>G), GGA87GGG (261A>G), TTA88TTG (264A>G), GTC89ATC (265G>A), GAA91GAG (273A>G), ATT93ACC (278T>C 279T>C), GAT95GGA (284A>G 285T>A), GTT96GTG (288T>G), TTG97CTG (289T>C), TTC100TTT (300C>T), CTC101TTG (301C>T 303C>G), AAC103AAT (309C>T), GTT104GTG (312T>G), ATC105ATT (315C>T), GCT108TCT (322G>T), ATC110ACC (329T>C), TAT111TAC (333T>C), ACC112ACT (336C>T), GAA113GAG (339A>G), GCA115GCC (345A>C), AAA116AGG (347A>G 348A>G), CGT117AGA (349C>A 351T>A), GTC120GTT (360C>T), ACC121ACT (363C>T), GCC122GCT (366C>T), GTC125GTT (375C>T), GCA128GCC (384A>C), TTG129CTC (385T>C 387G>C), AAA130AAG (390A>G), CAT131AGA (391C>A 392A>G 393T>A), AAA132CAA (394A>C), CGT134AGG (400C>A 402T>G), ATG135ACT (404T>C 405G>T), ATC136TTG (406A>T 408C>G), TAT139TTT (416A>T) | | | | | | | | |

*: Inserts / Deletes / Misaligned / Frameshifts

#### Analysis details

This analysis was performed with panviral2.64

### NGS Details (UN70): Diachasmimorpha longicaudata entomopoxvirus (segment NC_043455.1)

#### Assembly

| | |
|---|---|
| **Coverage Length** | 1127 (1 contig(s)) |
| **Depth Of Coverage** | 1241.9 |
| **Number Of Reads** | 11204 |
| **Reads Per Million** | 212.39 rpm (after QC) |
| **Ambiguities** | 0 |
| **Assembly Method** | de novo + reference guided assembly |
| **Consensus Caller** | Bcf Tools |

#### Coverage Map

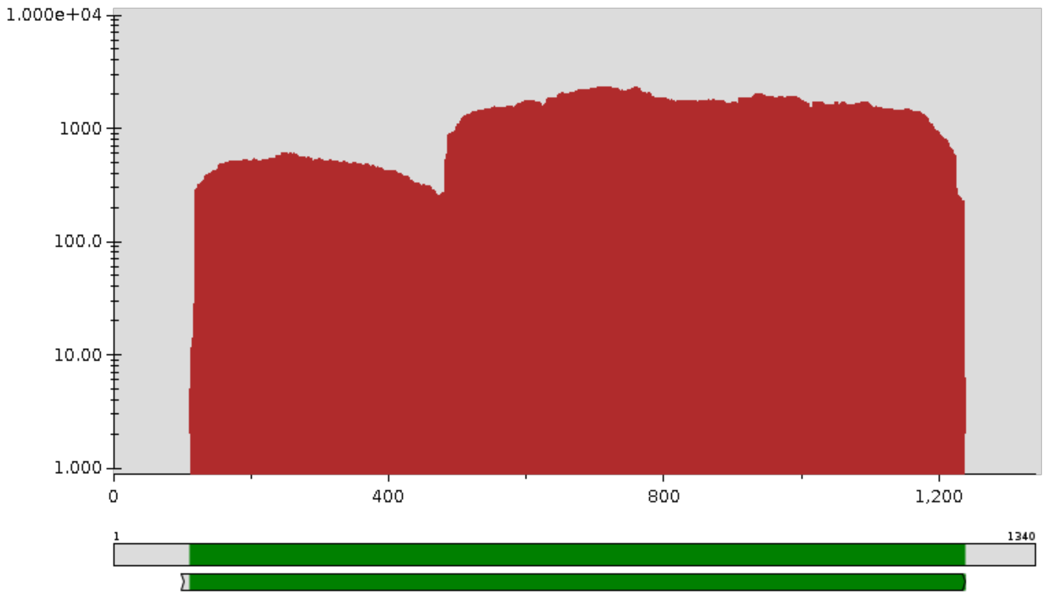

#### Assignment

| | |
|---|---|
| **Type** | Diachasmimorpha longicaudata entomopoxvirus (Taxonomy ID: 109981) |
| **Reference Genome** | NC_043455.1 |
| **NT Identity (%)** | 59.3611 |
| **AA Identity (%)** | 55.5851 |
| **Number Of Stop Codons** | 0 |
| **Number Of CDS** | 1 |

#### Alignment

| | |
|---|---|
| **Alignment Score** | 404.0 (NT) + 1470.0 (AA) = 1874.0 |
| **Concordance (%)** | 39.7961 |

| Alignment Method | Global, seeded, nucleotide + amino acids (AGA) |
|---|---|

#### Genome Region

Sequence starts at position 112 and ends at position 1238 relative to NC_043455.1 reference sequence.

#### Alignment Detailed Statistics

| | Begin | End | Coverage | Score | Concordance | Matches | Identities | I/D/M/F* | Stop Codons |
|---|---|---|---|---|---|---|---|---|---|
| **NT** | **112** | **1238** | **84.1%** | **404** | **18.1%** | **1124 (99.5%)** | **669 (59.2%)** | **3/3** | |

Mutations: 121A>T, 126A>C, 127C>T, 128C>G, 130T>G, 136C>G, 137C>A, 139C>T, 142G>A, 143A>G, 145A>T, 146A>G, 148T>C, 149A>C, 151C>T, 154A>T, 156A>G, 157G>A, 160T>G, 163A>T, 166T>C, 167T>C, 168C>A, 169T>G, 171G>A, 172T>C, 175A>G, 178T>C, 187T>A, 190A>T, 191A>G, 193T>C, 203A>C, 204A>G, 208C>T, 209A>G, 211T>A, 214T>G, 217A>T, 218T>A, 219G>T, 223T>A, 226T>G, 230A>C, 231A>G, 232A>T, 238G>C, 241A>T, 243T>C, 244C>T, 247A>G, 256G>C, 259A>T, 262A>C, 271A>C, 272G>T, 274A>C, 276C>T, 277A>G, 278T>A, 279A>T, 283A>T, 284A>C, 286C>T, 288G>C, 289T>C, 294T>G, 295A>C, 298G>A, 299C>G, 300A>T, 301A>T, 302A>G, 304A>G, 310T>C, 311T>A, 312C>A, 313T>A, 320A>G, 322T>G, 323A>G, 325C>A, 331T>A, 337C>A, 341A>T, 343C>T, 346A>T, 352T>G, 355A>G, 356T>C, 358G>T, 361T>A, 362C>G, 363T>C, 364C>T, 368G>A, 370T>G, 371C>G, 374C>A, 375G>A, 376T>G, 379A>G, 380T>A, 382G>T, 383C>T, 384A>T, 385A>G, 386A>G, 394G>A, 395A>G, 400T>C, 401C>A, 403A>T, 404_406delTAT, 409T>C, 410T>A, 412T>A, 413A>C, 415A>G, 416T>G, 417G>C, 418T>A, 421A>T, 423T>C, 424T>G, 438C>A, 442T>C, 443A>G, 445C>G, 446A>G, 447A>G, 448G>T, 452A>G, 453G>A, 454C>T, 455C>A, 456A>T, 457G>C, 458G>A, 459A>G, 462C>A, 463T>G, 467A>G, 469_470insCAT, 470A>G, 471A>G, 474C>T, 475C>A, 478G>A, 482T>G, 484G>C, 485A>G, 487T>C, 490A>T, 499A>C, 502A>T, 503A>G, 505G>A, 506A>T, 511T>C, 512C>A, 514A>G, 515C>T, 517C>G, 518A>C, 519C>G, 521C>A, 523A>G, 524A>C, 526A>G, 527A>T, 528G>C, 530A>C, 532C>T, 533G>C, 534A>G, 535T>C, 536A>C, 538C>T, 539A>G, 541A>C, 542G>C, 543C>A, 544A>C, 547T>C, 550A>G, 553A>G, 554G>T, 559A>T, 560A>C, 562A>G, 571T>C, 577G>A, 581T>C, 583G>C, 584A>T, 585T>C, 587G>A, 588A>G, 589T>A, 590A>G, 591A>G, 595T>C, 596T>A, 597T>A, 602A>C, 604A>G, 607A>T, 608C>T, 610A>T, 620G>C, 622A>G, 623T>C, 626T>C, 628T>G, 629G>C, 630A>C, 632A>C, 633G>C, 634T>A, 635C>A, 637T>G, 638G>A, 640C>T, 647A>G, 648T>G, 650T>G, 652A>T, 655A>C, 658A>T, 661A>T, 664C>T, 665G>A, 667T>G, 671T>C, 674A>G, 675G>A, 678T>C, 679T>C, 680A>C, 682C>T, 683A>G, 685T>A, 687C>T, 688A>T, 689T>A, 692C>A, 693A>G, 695G>A, 696T>A, 697C>G, 700T>C, 705G>A, 706A>C, 707G>A, 709T>A, 712C>T, 715A>G, 717A>G, 718A>G, 724G>T, 727A>G, 730A>G, 731A>C, 732A>G, 735C>A, 739A>G, 740T>C, 742G>C, 745A>T, 748C>T, 757T>C, 759G>A, 765A>T, 770A>G, 772A>C, 778A>T, 779A>G, 781G>C, 784A>G, 785A>G, 787T>A, 790T>G, 792T>G, 793T>G, 796A>G, 797G>C, 798C>T, 799A>T, 805A>T, 810T>G, 811T>C, 820T>C, 823C>G, 824C>A, 825A>C, 830A>G, 831G>C, 833C>A, 838T>C, 841A>G, 843C>G, 844A>T, 845C>G, 847A>C, 850A>C, 854T>G, 855G>T, 856T>G, 859T>C, 862A>C, 863C>A, 865T>G, 866A>C, 868A>T, 869C>A, 878G>T, 879T>G, 880A>G, 885T>C, 886G>T, 889A>T, 890T>A, 891G>A, 892T>G, 893T>A, 895A>G, 896A>C, 897C>G, 898A>C, 900A>G, 901T>C, 902A>C, 904A>T, 905A>G, 908T>C, 909T>A, 915C>T, 919A>T, 920A>G, 921G>C, 922C>A, 924T>C, 925C>T, 934T>C, 938T>G, 939C>A, 943A>G, 944C>A, 946A>C, 947G>A, 948A>C, 949A>T, 950C>A, 952T>A, 956T>A, 958T>C, 965A>C, 966A>G, 970A>G, 973T>C, 974A>C, 977G>T, 978A>C, 982T>G, 983A>T, 984A>C, 986A>T, 989A>C, 991A>T, 992A>G, 994A>G, 997T>C, 998C>A, 1000T>C, 1001T>A, 1003A>T, 1007A>G, 1012A>T, 1015A>G, 1019A>C, 1021A>T, 1024C>T, 1033T>A, 1036A>G, 1040A>G, 1042A>C, 1045T>C, 1046T>C, 1048A>T, 1051T>C, 1054C>T, 1057T>C, 1063T>C, 1064T>C, 1070C>A, 1071A>C, 1073A>C, 1075T>A, 1076A>C, 1077G>C, 1084T>C, 1087T>C, 1088A>C, 1090A>G, 1103A>C, 1105A>T, 1111T>G, 1112A>C, 1120C>A, 1121C>A, 1123A>G, 1126A>G, 1129G>T, 1130A>G, 1132T>A, 1141T>C, 1145A>G, 1147A>C, 1150T>A, 1151G>A, 1153A>G, 1156C>T, 1159A>T, 1160T>G, 1161T>A, 1163C>A, 1164C>G, 1165A>G, 1167A>T, 1172G>T, 1173C>T, 1177A>C, 1178T>A, 1180A>T, 1181G>C, 1183A>G, 1185C>A, 1192T>C, 1196A>G, 1197C>T, 1198C>T, 1199A>G, 1200A>T, 1204A>C, 1207T>G, 1210A>G, 1211A>C, 1218A>C, 1219G>C, 1223A>G, 1225T>G, 1227T>C, 1232T>C, 1234G>C, 1235A>C, 1237A>G

##### CDS

| | Begin | End | Coverage | Score | Concordance | Matches | Identities | I/D/M/F* | Stop Codons |
|---|---|---|---|---|---|---|---|---|---|
| FLA14_p101 | 5 | 380 | 98.9% | 1470 | 59.1% | 375 (99.5%) | 209 (55.4%) | 1/1/0/0 | 0 |

Protein mutations: E7D (121A>T), D9A (126A>C 127C>T), H10E (128C>G 130T>G), L13I (137C>A 139C>T), K15D (143A>G 145A>T), N16D (146A>G 148T>C), I17L (149A>C 151C>T), K19R (156A>G 157G>A), S23Q (167T>C 168C>A 169T>G), C24Y (171G>A 172T>C), T31A (191A>G 193T>C), K35R (203A>C 204A>G), I37V (209A>G 211T>A), F38L (214T>G), C40I (218T>A 219G>T), K44R (230A>C 231A>G 232A>T), V48A (243T>C 244C>T), A58S (272G>T 274A>C), T59M (276C>T 277A>G), Y60I (278T>A 279A>T), I62L (284A>C 286C>T), S63T (288G>C 289T>C), L65C (294T>G 295A>C), Q67V (299C>G 300A>T 301A>T), I68V (302A>G 304A>G), S71K (311T>A 312C>A 313T>A), N74E (320A>G 322T>G), I75V (323A>G 325C>A), T81S (341A>T 343C>T), L88A (362C>G 363T>C 364C>T), A90T (368G>A 370T>G), Q91E (371C>G), R92K (374C>A 375G>A 376T>G), L94I (380T>A 382G>T), Q95L (383C>T 384A>T 385A>G), T96A (386A>G), N99D (395A>G), L101I (401C>A 403A>T), Y102del (404_406delTAT), F104I (410T>A 412T>A), K105Q (413A>C 415A>G), C106A (416T>G 417G>C 418T>A), Q107H (421A>T), V108A (423T>C 424T>G), T113K (438C>A), I115V (443A>G 445C>G), K116G (446A>G 447A>G 448G>T), S118D (452A>G 453G>A 454C>T), Q119I (455C>A 456A>T 457G>C), E120R (458G>A 459A>G), T121K (462C>A 463T>G), K123E (467A>G), K123_K124insH (469_470insCAT), K124G (470A>G 471A>G), A125V (474C>T 475C>A), L128V (482T>G 484G>C), I129V (485A>G 487T>C), M135V (503A>G 505G>A), I136F (506A>T), L138M (512C>A 514A>G), T140R (518A>C 519C>G), K142Q (524A>C 526A>G), I144L (530A>C 532C>T), D145R (533G>C 534A>G 535T>C), T146P (536A>C 538C>T), K147D (539A>G 541A>C), A148H (542G>C 543C>A 544A>C), I151M (553A>G), V152F (554G>T), I154L (560A>C 562A>G), I162S (584A>T 585T>C), D163R (587G>A 588A>G 589T>A), N164G (590A>G 591A>G), L166K (596T>A 597T>A), K168Q (602A>C 604A>G), Q170Y (608C>T 610A>T), E174Q (620G>C 622A>G), F175L (623T>C), F176L (626T>C 628T>G), E177P (629G>C 630A>C), S178P (632A>C 633G>C 634T>A), H179K (635C>A 637T>G), V180I (638G>A 640C>T), I183G (647A>G 648T>G), L184V (650T>G 652A>T), L185F (655A>C), V189M (665G>A 667T>G), S191P (671T>C), R192E (674A>G 675G>A), V193A (678T>C 679T>C), I194L (680A>C 682C>T), N195E (683A>G 685T>A), T196I (687C>T 688A>T), S197T (689T>A), Q198R (692C>A 693A>G), V199K (695G>A 696T>A 697C>G), R202N (705G>A 706A>C), D203K (707G>A 709T>A), K206R (717A>G 718A>G), N211R (731A>C 732A>G), A212D (735C>A), R220K (759G>A), Y222F (765A>T), I224V (770A>G 772A>C), K227D (779A>G 781G>C), N229E (785A>G 787T>A), D230E (790T>G), F231W (792T>G 793T>G), A233L (797G>C 798C>T 799A>T), F237C (810T>G 811T>C), D241E (823C>G), H242T (824C>A 825A>C), S244A (830A>G 831G>C), L245I (833C>A), T248S (843C>G 844A>T), L249V (845C>G 847A>C), C252V (854T>G 855G>T 856T>G), Q257K (869C>A), V260W (878G>T 879T>G 880A>G), M262T (885T>C 886G>T), E263D (889A>T), C264K (890T>A 891G>A 892T>G), L265M (893T>A 895A>G), T266R (896A>C 897C>G 898A>C), N267S (900A>G 901T>C), N269D (905A>G), F270H (908T>C 909T>A), A272V (915C>T), S274A (920A>G 921G>C 922C>A), I275T (924T>C 925C>T), S280D (938T>G 939C>A), Q282N (944C>A 946A>C), E283T (947G>A 948A>C 949A>T), F286I (956T>A 958T>C), K289R (965A>C 966A>G), D293S (977G>T 978A>C), K295S (983A>T 984A>C), T296S (986A>T), I298V (992A>G 994A>G), L300I (998C>A 1000T>C), S301T (1001T>A 1003A>T), N303D (1007A>G), I314V (1040A>G 1042A>C), H324T (1070C>A 1071A>C), N325Q (1073A>C 1075T>A), R326P (1076A>C 1077G>C), I330L (1088A>C 1090A>G), I344V (1130A>G 1132T>A), I349V (1145A>G 1147A>C), E351K (1151G>A 1153A>G), E353D (1159A>T), L354E (1160T>G 1161T>A), P355R (1163C>A 1164C>G 1165A>G), K356M (1167A>T), A358F (1172G>T 1173C>T), E359D (1177A>C), L360I (1178T>A 1180A>T), E361Q (1181G>C 1183A>G), T362K (1185C>A), T366V (1196A>G 1197C>T 1198C>T), K367V (1199A>G 1200A>T), D369E (1207T>G), M371L (1211A>C), E373A (1218A>C 1219G>C), I375V (1223A>G 1225T>G), V376A (1227T>C), I379L (1235A>C 1237A>G)

| | Begin | End | Coverage | Score | Concordance | Matches | Identities | I/D/M/F* | Stop Codons |
|---|---|---|---|---|---|---|---|---|---|
| **NT** | **112** | **1238** | **84.1%** | **404** | **18.1%** | **1124 (99.5%)** | **669 (59.2%)** | **3/3** | |
| Codon mutations: | GAA7GAT (121A>T), GAC9GCT (126A>C 127C>T), CAT10GAG (128C>G 130T>G), GGC12GGG (136C>G), CTC13ATT (137C>A 139C>T), AAG14AAA (142G>A), AAA15GAT (143A>G 145A>T), AAT16GAC (146A>G 148T>C), ATC17CTT (149A>C 151C>T), CTA18CTT (154A>T), AAG19AGA (156A>G 157G>A), GGT20GGG (160T>G), ATA21ATT (163A>T), TAT22TAC (166T>C), TCT23CAG (167T>C 168C>A 169T>G), TGT24TAC (171G>A 172T>C), GGA25GGG (175A>G), TTT26TTC (178T>C), CCT29CCA (187T>A), TCA30TCT (190A>T), ACT31GCC (191A>G 193T>C), AAA35CGA (203A>C 204A>G), GCC36GCT (208C>T), ATT37GTA (209A>G 211T>A), TTT38TTG (214T>G), CCA39CCT (217A>T), TGT40ATT (218T>A 219G>T), ATT41ATA (223T>A), TCT42TCG (226T>G), AAA44CGT (230A>C 231A>G 232A>T), GTG46GTC (238G>C), ATA47ATT (241A>T), GTC48GCT (243T>C 244C>T), CAA49CAG (247A>G), TCG52TCC (256G>C), GGA53GGT (259A>T), ACA54ACC (262A>C), ACA57ACC (271A>C), GCA58TCC (272G>T 274A>C), ACA59ATG (276C>T 277A>G), TAT60ATT (278T>A 279A>T), GCA61GCT (283A>T), ATC62CTT (284A>C 286C>T), AGT63ACC (288G>C 289T>C), TTA65TGC (294T>G 295A>C), CAG66CAA (298G>A), CAA67GTT (299C>G 300A>T 301A>T), ATA68GTG (302A>G 304A>G), ACT70ACC (310T>C), TCT71AAA (311T>A 312C>A 313T>A), AAT74GAG (320A>G 322T>G), ATC75GTA (323A>G 325C>A), GCT77GCA (331T>A), ATC79ATA (337C>A), ACC81TCT (341A>T 343C>T), CCA82CCT (346A>T), CGT84CGG (352T>G), GAA85GAG (355A>G), TTG86CTT (356T>C 358G>T), GCT87GCA (361T>A), CTC88GCT (362C>G 363T>C 364C>T), GCT90ACG (368G>A 370T>G), CAA91GAA (371C>G), CGT92AAG (374C>A 375G>A 376T>G), GTA93GTG (379A>G), TTG94ATT (380T>A 382G>T), CAA95TTG (383C>T 384A>T 385A>G), ACA96GCA (386A>G), GGG98GGA (394G>A), AAT99GAT (395A>G), TAT100TAC (400T>C), CTA101ATT (401C>A 403A>T), TAT102del (404_406delTAT), AAT103AAC (409T>C), TTT104ATA (410T>A 412T>A), AAA105CAG (413A>C 415A>G), TGT106GCA (416T>G 417G>C 418T>A), CAA107CAT (421A>T), GTT108GCG (423T>C 424T>G), ACA113AAA (438C>A), AGT114AGC (442T>C), ATC115GTG (443A>G 445C>G), AAG116GGT (446A>G 447A>G 448G>T), AGC118GAT (452A>G 453G>A 454C>T), CAG119ATC (455C>A 456A>T 457G>C), GAA120AGA (458G>A 459A>G), ACT121AAG (462C>A 463T>G), AAG123GAG (467A>G), AAG123_AAA124insCAT (469_470insCAT), AAA124GGA (470A>G 471A>G), GCC125GTA (474C>T 475C>A), CAG126CAA (478G>A), TTG128GTC (482T>G 484G>C), ATT129GTC (485A>G 487T>C), GGA130GGT (490A>T), GGA133GGC (499A>C), CGA134CGT (502A>T), ATG135GTA (503A>G 505G>A), ATT136TTT (506A>T), GAT137GAC (511T>C), CTA138ATG (512C>A 514A>G), CTC139TTG (515C>T 517C>G), ACC140CGC (518A>C 519C>G), CGA141AGG (521C>A 523A>G), AAA142CAG (524A>C 526A>G), AGT143TCT (527A>T 528G>C), ATC144CTT (530A>C 532C>T), GAT145CGC (533G>C 534A>G 535T>C), ACC146CCT (536A>C 538C>T), AAA147GAC (539A>G 541A>C), GCA148CAC (542G>C 543C>A 544A>C), ATT149ATC (547T>C), AAA150AAG (550A>G), ATA151ATG (553A>G), GTT152TTT (554G>T), GTA153GTT (559A>T), ATA154CTG (560A>C 562A>G), GCT157GCC (571T>C), GAG159GAA (577G>A), TTG161CTC (581T>C 583G>C), ATA162TCA (584A>T 585T>C), GAT163AGA (587G>A 588A>G 589T>A), AAT164GGT (590A>G 591A>G), TTT165TTC (595T>C), TTG166AAG (596T>A 597T>A), AAA168CAG (602A>C 604A>G), ATA169ATT (607A>T), CAA170TAT (608C>T 610A>T), GAA174CAG (620G>C 622A>G), TTT175CTT (623T>C), TTT176CTG (626T>C 628T>G), GAA177CCA (629G>C 630A>C), AGT178CCA (632A>C 633G>C 634T>A), CAT179AAG (635C>A 637T>G), GTC180ATT (638G>A 640C>T), ATT183GGT (647A>G 648T>G), TTA184GTT (650T>G 652A>T), TTA185TTC (655A>C), TCA186TCT (658A>T), GCA187GCT (661A>T), ACC188ACT (664C>T), GTT189ATG (665G>A 667T>G), TCA191CCA (671T>C), AGG192GAG (674A>G 675G>A), GTT193GCC (678T>C 679T>C), ATC194CTT (680A>C 682C>T), AAT195GAA (683A>G 685T>A), ACA196ATT (687C>T 688A>T), TCT197ACT (689T>A), CAG198AGG (692C>A 693A>G), GTC199AAG (695G>A 696T>A 697C>G), TTT200TTC (700T>C), AGA202AAC (705G>A 706A>C), GAT203AAA (707G>A 709T>A), CCC204CCT (712C>T), GTA205GTG (715A>G), AAA206AGG (717A>G 718A>G), CTG208CTT (724G>T), GTA209GTG (727A>G), AAA210AAG (730A>G), AAT211CGT (731A>C 732A>G), GCT212GAT (735C>A), GAA213GAG (739A>G), TTG214CTC (740T>C 742G>C), ACA215ACT (745A>T), CTC216CTT (748C>T), ATT219ATC (757T>C), AGG220AAG (759G>A), TAT222TTT (765A>T), ATA224GTC (770A>G 772A>C), GTA226GTT (778A>T), AAG227GAC (779A>G 781G>C), AAA228AAG (784A>G), AAT229GAA (785A>G 787T>A), GAT230GAG (790T>G), TTT231TGG (792T>G 793T>G), AAA232AAG (796A>G), GCA233CTT (797G>C 798C>T 799A>T), ACA235ACT (805A>T), TTT237TGC (810T>G 811T>C), TAT240TAC (820T>C), GAC241GAG (823C>G), CAC242ACC (824C>A 825A>C), AGC244GCC (830A>G 831G>C), CTC245ATC (833C>A), ACT246ACC (838T>C), CAA247CAG (841A>G), ACA248AGT (843C>G 844A>T), CTA249GTC (845C>G 847A>C), ATA250ATC (850A>C), TGT252GTG (854T>G 855G>T 856T>G), AAT253AAC (859T>C), ACA254ACC (862A>C), CGT255AGG (863C>A 865T>G), AGA256CGT (866A>C 868A>T), CAA257AAA (869C>A), GTA260TGG (878G>T 879T>G 880A>G), ATG262ACT (885T>C 886G>T), GAA263GAT (889A>T), TGT264AAG (890T>A 891G>A 892T>G), TTA265ATG (893T>A 895A>G), ACA266CGC (896A>C 897C>G 898A>C), AAT267AGC (900A>G 901T>C), AGA268CGT (902A>C 904A>T), AAT269GAT (905A>G), TTC270CAC (908T>C 909T>A), GCA272GTA (915C>T), TCA273TCT (919A>T), AGC274GCA (920A>G 921G>C 922C>A), ATC275ACT (924T>C 925C>T), GAT278GAC (934T>C), TCC280GAC (938T>G 939C>A), CAA281CAG (943A>G), CAA282AAC (944C>A 946A>C), GAA283ACT (947G>A 948A>C 949A>T), CGT284AGA (950C>A 952T>A), TTT286ATC (956T>A 958T>C), AAA289CGA (965A>C 966A>G), GAA290GAG (970A>G), TTT291TTC (973T>C), AGA292CGA (974A>C), GAT293TCT (977G>T 978A>C), GGT294GGG (982T>G), AAA295TCA (983A>T 984A>C), ACT296TCT (986A>T), AGA297CGT (989A>C 991A>T), ATA298GTG (992A>G 994A>G), CTT299CTC (997T>C), CTT300ATC (998C>A 1000T>C), TCA301ACT (1001T>A 1003A>T), AAT303GAT (1007A>G), CTA304CTT (1012A>T), TTA305TTG (1015A>G), AGA307CGT (1019A>C 1021A>T), GGC308GGT (1024C>T), GTT311GTA (1033T>A), CAA312CAG (1036A>G), ATA314GTC (1040A>G 1042A>C), TCT315TCC (1045T>C), TTA316CTT (1046T>C 1048A>T), GTT317GTC (1051T>C), ATC318ATT (1054C>T), AAT319AAC (1057T>C), GAT321GAC (1063T>C), TTG322CTG (1064T>C), CAT324ACT (1070C>A 1071A>C), AAT325CAA (1073A>C 1075T>A), AGA326CCA (1076A>C 1077G>C), AAT328AAC (1084T>C), TAT329TAC (1087T>C), ATA330CTG (1088A>C 1090A>G), AGA335CGT (1103A>C 1105A>T), GGT337GGG (1111T>G), AGA338CGA (1112A>C), GGC340GGA (1120C>A), CGA341AGG (1121C>A 1123A>G), AAA342AAG (1126A>G), GGG343GGT (1129G>T), ATT344GTA (1130A>G 1132T>A), AAT347AAC (1141T>C), ATA349GTC (1145A>G 1147A>C), ACT350ACA (1150T>A), GAA351AAG (1151G>A 1153A>G), GAC352GAT (1156C>T), GAA353GAT (1159A>T), TTA354GAA (1160T>G 1161T>A), CCA355AGG (1163C>A 1164C>G 1165A>G), AAG356ATG (1167A>T), GCT358TTT (1172G>T 1173C>T), GAA359GAC (1177A>C), TTA360ATT (1178T>A 1180A>T), GAA361CAG (1181G>C 1183A>G), ACG362AAG (1185C>A), TAT364TAC (1192T>C), ACC366GTT (1196A>G 1197C>T 1198C>T), AAA367GTA (1199A>G 1200A>T), ATA368ATC (1204A>C), GAT369GAG (1207T>G), GAA370GAG (1210A>G), ATG371CTG (1211A>C), GAG373GCC (1218A>C 1219G>C), ATT375GTG (1223A>G 1225T>G), GTT376GCT (1227T>C), TTG378CTC (1232T>C 1234G>C), ATA379CTG (1235A>C 1237A>G) | | | | | | | | |

#### Proteins

| | Begin | End | Coverage | Score | Concordance | Matches | Identities | I/D/M/F* | Stop Codons |
|---|---|---|---|---|---|---|---|---|---|
| hypothetical protein (YP_009666148.1) | 5 | 380 | 98.9% | 1470 | 59.1% | 375 (99.5%) | 209 (55.4%) | 1/1/0/0 | 0 |
| Protein mutations: | E7D (121A>T), D9A (126A>C 127C>T), H10E (128C>G 130T>G), L13I (137C>A 139C>T), K15D (143A>G 145A>T), N16D (146A>G 148T>C), I17L (149A>C 151C>T), K19R (156A>G 157G>A), S23Q (167T>C 168C>A 169T>G), C24Y (171G>A 172T>C), T31A (191A>G 193T>C), K35R (203A>C 204A>G), I37V (209A>G 211T>A), F38L (214T>G), C40I (218T>A 219G>T), K44R (230A>C 231A>G 232A>T), V48A (243T>C 244C>T), A58S (272G>T 274A>C), T59M (276C>T 277A>G), Y60I (278T>A 279A>T), I62L (284A>C 286C>T), S63T (288G>C 289T>C), L65C (294T>G 295A>C), Q67V (299C>G 300A>T 301A>T), I68V (302A>G 304A>G), S71K (311T>A 312C>A 313T>A), N74E (320A>G 322T>G), I75V (323A>G 325C>A), T81S (341A>T 343C>T), L88A (362C>G 363T>C 364C>T), A90T (368G>A 370T>G), Q91E (371C>G), R92K (374C>A 375G>A 376T>G), L94I (380T>A 382G>T), Q95L (383C>T 384A>T 385A>G), T96A (386A>G), N99D (395A>G), L101I (401C>A 403A>T), Y102del (404_406delTAT), F104I (410T>A 412T>A), K105Q (413A>C 415A>G), C106A (416T>G 417G>C 418T>A), Q107H (421A>T), V108A (423T>C 424T>G), T113K (438C>A), I115V (443A>G 445C>G), K116G (446A>G 447A>G 448G>T), S118D (452A>G 453G>A 454C>T), Q119I (455C>A 456A>T 457G>C), E120R (458G>A 459A>G), T121K (462C>A 463T>G), K123E (467A>G), K123_K124insH (469_470insCAT), K124G (470A>G 471A>G), A125V (474C>T 475C>A), L128V (482T>G 484G>C), I129V (485A>G 487T>C), M135V (503A>G 505G>A), I136F (506A>T), L138M (512C>A 514A>G), T140R (518A>C 519C>G), K142Q (524A>C 526A>G), I144L (530A>C 532C>T), D145R (533G>C 534A>G 535T>C), T146P (536A>C 538C>T), K147D (539A>G 541A>C), A148H (542G>C 543C>A 544A>C), I151M (553A>G), V152F (554G>T), I154L (560A>C 562A>G), I162S (584A>T 585T>C), D163R (587G>A 588A>G 589T>A), N164G (590A>G 591A>G), L166K (596T>A 597T>A), K168Q (602A>C 604A>G), Q170Y (608C>T 610A>T), E174Q (620G>C 622A>G), F175L (623T>C), F176L (626T>C 628T>G), E177P (629G>C 630A>C), S178P (632A>C 633G>C 634T>A), H179K (635C>A 637T>G), V180I (638G>A 640C>T), I183G (647A>G 648T>G), L184V (650T>G 652A>T), L185F (655A>C), V189M (665G>A 667T>G), S191P (671T>C), R192E (674A>G 675G>A), V193A (678T>C 679T>C), I194L (680A>C 682C>T), N195E (683A>G 685T>A), T196I (687C>T 688A>T), S197T (689T>A), Q198R (692C>A 693A>G), V199K (695G>A 696T>A 697C>G), R202N (705G>A 706A>C), D203K (707G>A 709T>A), K206R (717A>G 718A>G), N211R (731A>C 732A>G), A212D (735C>A), R220K (759G>A), Y222F (765A>T), I224V (770A>G 772A>C), K227D (779A>G 781G>C), N229E (785A>G 787T>A), D230E (790T>G), F231W (792T>G 793T>G), A233L (797G>C 798C>T 799A>T), F237C (810T>G 811T>C), D241E (823C>G), H242T (824C>A 825A>C), S244A (830A>G 831G>C), L245I (833C>A), T248S (843C>G 844A>T), L249V (845C>G 847A>C), C252V (854T>G 855G>T 856T>G), Q257K (869C>A), V260W (878G>T 879T>G 880A>G), M262T (885T>C 886G>T), E263D (889A>T), C264K (890T>A 891G>A 892T>G), L265M (893T>A 895A>G), T266R (896A>C 897C>G 898A>C), N267S (900A>G 901T>C), N269D (905A>G), F270H (908T>C 909T>A), A272V (915C>T), S274A (920A>G 921G>C 922C>A), I275T (924T>C 925C>T), S280D (938T>G 939C>A), Q282N (944C>A 946A>C), E283T (947G>A 948A>C 949A>T), F286I (956T>A 958T>C), K289R (965A>C 966A>G), D293S (977G>T 978A>C), K295S (983A>T 984A>C), T296S (986A>T), I298V (992A>G 994A>G), L300I (998C>A 1000T>C), S301T (1001T>A 1003A>T), N303D (1007A>G), I314V (1040A>G 1042A>C), H324T (1070C>A 1071A>C), N325Q (1073A>C 1075T>A), R326P (1076A>C 1077G>C), I330L (1088A>C 1090A>G), I344V (1130A>G 1132T>A), I349V (1145A>G 1147A>C), E351K (1151G>A 1153A>G), E353D (1159A>T), L354E (1160T>G 1161T>A), P355R (1163C>A 1164C>G 1165A>G), K356M (1167A>T), A358F (1172G>T 1173C>T), E359D (1177A>C), L360I (1178T>A 1180A>T), E361Q (1181G>C 1183A>G), T362K (1185C>A), T366V (1196A>G 1197C>T 1198C>T), K367V (1199A>G 1200A>T), D369E (1207T>G), M371L (1211A>C), E373A (1218A>C 1219G>C), I375V (1223A>G 1225T>G), V376A (1227T>C), I379L (1235A>C 1237A>G) | | | | | | | | |

| | Begin | End | Coverage | Score | Concordance | Matches | Identities | I/D/M/F* | Stop Codons |
|---|---|---|---|---|---|---|---|---|---|
| **NT** | **112** | **1238** | **84.1%** | **404** | **18.1%** | **1124 (99.5%)** | **669 (59.2%)** | **3/3** | |
| Codon mutations: | GAA7GAT (121A>T), GAC9GCT (126A>C 127C>T), CAT10GAG (128C>G 130T>G), GGC12GGG (136C>G), CTC13ATT (137C>A 139C>T), AAG14AAA (142G>A), AAA15GAT (143A>G 145A>T), AAT16GAC (146A>G 148T>C), ATC17CTT (149A>C 151C>T), CTA18CTT (154A>T), AAG19AGA (156A>G 157G>A), GGT20GGG (160T>G), ATA21ATT (163A>T), TAT22TAC (166T>C), TCT23CAG (167T>C 168C>A 169T>G), TGT24TAC (171G>A 172T>C), GGA25GGG (175A>G), TTT26TTC (178T>C), CCT29CCA (187T>A), TCA30TCT (190A>T), ACT31GCC (191A>G 193T>C), AAA35CGA (203A>C 204A>G), GCC36GCT (208C>T), ATT37GTA (209A>G 211T>A), TTT38TTG (214T>G), CCA39CCT (217A>T), TGT40ATT (218T>A 219G>T), ATT41ATA (223T>A), TCT42TCG (226T>G), AAA44CGT (230A>C 231A>G 232A>T), GTG46GTC (238G>C), ATA47ATT (241A>T), GTC48GCT (243T>C 244C>T), CAA49CAG (247A>G), TCG52TCC (256G>C), GGA53GGT (259A>T), ACA54ACC (262A>C), ACA57ACC (271A>C), GCA58TCC (272G>T 274A>C), ACA59ATG (276C>T 277A>G), TAT60ATT (278T>A 279A>T), GCA61GCT (283A>T), ATC62CTT (284A>C 286C>T), AGT63ACC (288G>C 289T>C), TTA65TGC (294T>G 295A>C), CAG66CAA (298G>A), CAA67GTT (299C>G 300A>T 301A>T), ATA68GTG (302A>G 304A>G), ACT70ACC (310T>C), TCT71AAA (311T>A 312C>A 313T>A), AAT74GAG (320A>G 322T>G), ATC75GTA (323A>G 325C>A), GCT77GCA (331T>A), ATC79ATA (337C>A), ACC81TCT (341A>T 343C>T), CCA82CCT (346A>T), CGT84CGG (352T>G), GAA85GAG (355A>G), TTG86CTT (356T>C 358G>T), GCT87GCA (361T>A), CTC88GCT (362C>G 363T>C 364C>T), GCT90ACG (368G>A 370T>G), CAA91GAA (371C>G), CGT92AAG (374C>A 375G>A 376T>G), GTA93GTG (379A>G), TTG94ATT (380T>A 382G>T), CAA95TTG (383C>T 384A>T 385A>G), ACA96GCA (386A>G), GGG98GGA (394G>A), AAT99GAT (395A>G), TAT100TAC (400T>C), CTA101ATT (401C>A 403A>T), TAT102del (404_406delTAT), AAT103AAC (409T>C), TTT104ATA (410T>A 412T>A), AAA105CAG (413A>C 415A>G), TGT106GCA (416T>G 417G>C 418T>A), CAA107CAT (421A>T), GTT108GCG (423T>C 424T>G), ACA113AAA (438C>A), AGT114AGC (442T>C), ATC115GTG (443A>G 445C>G), AAG116GGT (446A>G 447A>G 448G>T), AGC118GAT (452A>G 453G>A 454C>T), CAG119ATC (455C>A 456A>T 457G>C), GAA120AGA (458G>A 459A>G), ACT121AAG (462C>A 463T>G), AAG123GAG (467A>G), AAG123_AAA124insCAT (469_470insCAT), AAA124GGA (470A>G 471A>G), GCC125GTA (474C>T 475C>A), CAG126CAA (478G>A), TTG128GTC (482T>G 484G>C), ATT129GTC (485A>G 487T>C), GGA130GGT (490A>T), GGA133GGC (499A>C), CGA134CGT (502A>T), ATG135GTA (503A>G 505G>A), ATT136TTT (506A>T), GAT137GAC (511T>C), CTA138ATG (512C>A 514A>G), CTC139TTG (515C>T 517C>G), ACC140CGC (518A>C 519C>G), CGA141AGG (521C>A 523A>G), AAA142CAG (524A>C 526A>G), AGT143TCT (527A>T 528G>C), ATC144CTT (530A>C 532C>T), GAT145CGC (533G>C 534A>G 535T>C), ACC146CCT (536A>C 538C>T), AAA147GAC (539A>G 541A>C), GCA148CAC (542G>C 543C>A 544A>C), ATT149ATC (547T>C), AAA150AAG (550A>G), ATA151ATG (553A>G), GTT152TTT (554G>T), GTA153GTT (559A>T), ATA154CTG (560A>C 562A>G), GCT157GCC (571T>C), GAG159GAA (577G>A), TTG161CTC (581T>C 583G>C), ATA162TCA (584A>T 585T>C), GAT163AGA (587G>A 588A>G 589T>A), AAT164GGT (590A>G 591A>G), TTT165TTC (595T>C), TTG166AAG (596T>A 597T>A), AAA168CAG (602A>C 604A>G), ATA169ATT (607A>T), CAA170TAT (608C>T 610A>T), GAA174CAG (620G>C 622A>G), TTT175CTT (623T>C), TTT176CTG (626T>C 628T>G), GAA177CCA (629G>C 630A>C), AGT178CCA (632A>C 633G>C 634T>A), CAT179AAG (635C>A 637T>G), GTC180ATT (638G>A 640C>T), ATT183GGT (647A>G 648T>G), TTA184GTT (650T>G 652A>T), TTA185TTC (655A>C), TCA186TCT (658A>T), GCA187GCT (661A>T), ACC188ACT (664C>T), GTT189ATG (665G>A 667T>G), TCA191CCA (671T>C), AGG192GAG (674A>G 675G>A), GTT193GCC (678T>C 679T>C), ATC194CTT (680A>C 682C>T), AAT195GAA (683A>G 685T>A), ACA196ATT (687C>T 688A>T), TCT197ACT (689T>A), CAG198AGG (692C>A 693A>G), GTC199AAG (695G>A 696T>A 697C>G), TTT200TTC (700T>C), AGA202AAC (705G>A 706A>C), GAT203AAA (707G>A 709T>A), CCC204CCT (712C>T), GTA205GTG (715A>G), AAA206AGG (717A>G 718A>G), CTG208CTT (724G>T), GTA209GTG (727A>G), AAA210AAG (730A>G), AAT211CGT (731A>C 732A>G), GCT212GAT (735C>A), GAA213GAG (739A>G), TTG214CTC (740T>C 742G>C), ACA215ACT (745A>T), CTC216CTT (748C>T), ATT219ATC (757T>C), AGG220AAG (759G>A), TAT222TTT (765A>T), ATA224GTC (770A>G 772A>C), GTA226GTT (778A>T), AAG227GAC (779A>G 781G>C), AAA228AAG (784A>G), AAT229GAA (785A>G 787T>A), GAT230GAG (790T>G), TTT231TGG (792T>G 793T>G), AAA232AAG (796A>G), GCA233CTT (797G>C 798C>T 799A>T), ACA235ACT (805A>T), TTT237TGC (810T>G 811T>C), TAT240TAC (820T>C), GAC241GAG (823C>G), CAC242ACC (824C>A 825A>C), AGC244GCC (830A>G 831G>C), CTC245ATC (833C>A), ACT246ACC (838T>C), CAA247CAG (841A>G), ACA248AGT (843C>G 844A>T), CTA249GTC (845C>G 847A>C), ATA250ATC (850A>C), TGT252GTG (854T>G 855G>T 856T>G), AAT253AAC (859T>C), ACA254ACC (862A>C), CGT255AGG (863C>A 865T>G), AGA256CGT (866A>C 868A>T), CAA257AAA (869C>A), GTA260TGG (878G>T 879T>G 880A>G), ATG262ACT (885T>C 886G>T), GAA263GAT (889A>T), TGT264AAG (890T>A 891G>A 892T>G), TTA265ATG (893T>A 895A>G), ACA266CGC (896A>C 897C>G 898A>C), AAT267AGC (900A>G 901T>C), AGA268CGT (902A>C 904A>T), AAT269GAT (905A>G), TTC270CAC (908T>C 909T>A), GCA272GTA (915C>T), TCA273TCT (919A>T), AGC274GCA (920A>G 921G>C 922C>A), ATC275ACT (924T>C 925C>T), GAT278GAC (934T>C), TCC280GAC (938T>G 939C>A), CAA281CAG (943A>G), CAA282AAC (944C>A 946A>C), GAA283ACT (947G>A 948A>C 949A>T), CGT284AGA (950C>A 952T>A), TTT286ATC (956T>A 958T>C), AAA289CGA (965A>C 966A>G), GAA290GAG (970A>G), TTT291TTC (973T>C), AGA292CGA (974A>C), GAT293TCT (977G>T 978A>C), GGT294GGG (982T>G), AAA295TCA (983A>T 984A>C), ACT296TCT (986A>T), AGA297CGT (989A>C 991A>T), ATA298GTG (992A>G 994A>G), CTT299CTC (997T>C), CTT300ATC (998C>A 1000T>C), TCA301ACT (1001T>A 1003A>T), AAT303GAT (1007A>G), CTA304CTT (1012A>T), TTA305TTG (1015A>G), AGA307CGT (1019A>C 1021A>T), GGC308GGT (1024C>T), GTT311GTA (1033T>A), CAA312CAG (1036A>G), ATA314GTC (1040A>G 1042A>C), TCT315TCC (1045T>C), TTA316CTT (1046T>C 1048A>T), GTT317GTC (1051T>C), ATC318ATT (1054C>T), AAT319AAC (1057T>C), GAT321GAC (1063T>C), TTG322CTG (1064T>C), CAT324ACT (1070C>A 1071A>C), AAT325CAA (1073A>C 1075T>A), AGA326CCA (1076A>C 1077G>C), AAT328AAC (1084T>C), TAT329TAC (1087T>C), ATA330CTG (1088A>C 1090A>G), AGA335CGT (1103A>C 1105A>T), GGT337GGG (1111T>G), AGA338CGA (1112A>C), GGC340GGA (1120C>A), CGA341AGG (1121C>A 1123A>G), AAA342AAG (1126A>G), GGG343GGT (1129G>T), ATT344GTA (1130A>G 1132T>A), AAT347AAC (1141T>C), ATA349GTC (1145A>G 1147A>C), ACT350ACA (1150T>A), GAA351AAG (1151G>A 1153A>G), GAC352GAT (1156C>T), GAA353GAT (1159A>T), TTA354GAA (1160T>G 1161T>A), CCA355AGG (1163C>A 1164C>G 1165A>G), AAG356ATG (1167A>T), GCT358TTT (1172G>T 1173C>T), GAA359GAC (1177A>C), TTA360ATT (1178T>A 1180A>T), GAA361CAG (1181G>C 1183A>G), ACG362AAG (1185C>A), TAT364TAC (1192T>C), ACC366GTT (1196A>G 1197C>T 1198C>T), AAA367GTA (1199A>G 1200A>T), ATA368ATC (1204A>C), GAT369GAG (1207T>G), GAA370GAG (1210A>G), ATG371CTG (1211A>C), GAG373GCC (1218A>C 1219G>C), ATT375GTG (1223A>G 1225T>G), GTT376GCT (1227T>C), TTG378CTC (1232T>C 1234G>C), ATA379CTG (1235A>C 1237A>G) | | | | | | | | |

*: Inserts / Deletes / Misaligned / Frameshifts

#### Analysis details

This analysis was performed with panviral2.64

### NGS Details (UN70): Potato leafroll virus

#### Assembly

| | |
|---|---|
| **Coverage Length** | 5812 (1 contig(s)) |
| **Depth Of Coverage** | 18.2 |
| **Number Of Reads** | 796 |
| **Reads Per Million** | 15.09 rpm (after QC) |
| **Ambiguities** | 0 |
| **Assembly Method** | read mapping against reference + variant calling |
| **Consensus Caller** | Bcf Tools |

#### Coverage Map

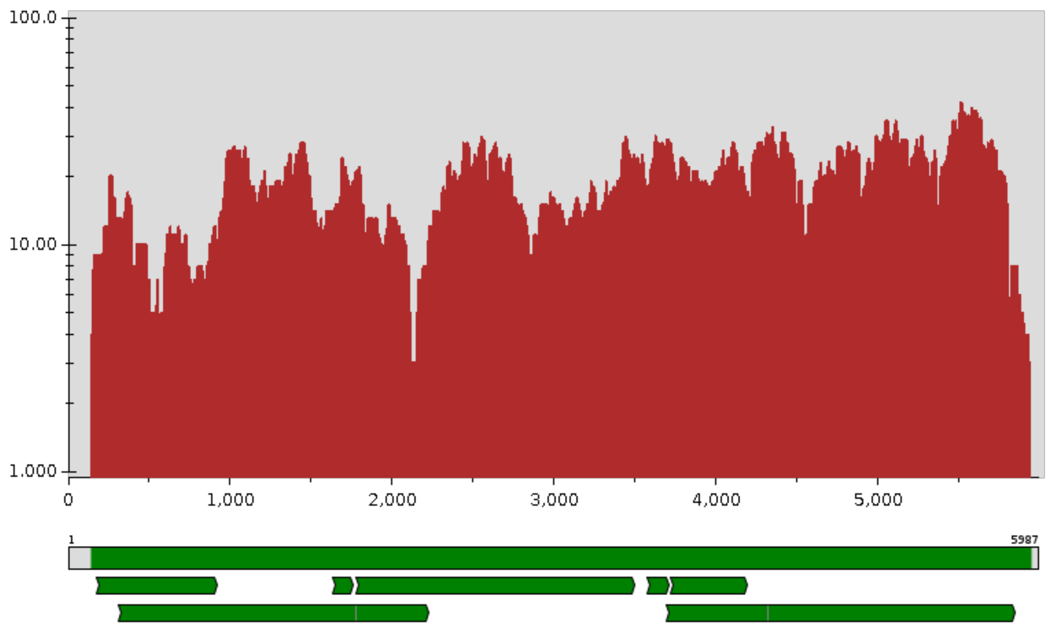

#### Assignment

| | |
|---|---|
| **Type** | Potato leafroll virus (Taxonomy ID: 12045) |
| **Reference Genome** | NC_001747.1 |
| **NT Identity (%)** | 98.7612 |
| **AA Identity (%)** | 98.399 |
| **Number Of Stop Codons** | 9 |
| **Number Of CDS** | 8 |

#### Alignment

| | |
|---|---|
| **Alignment Score** | 11336.0 (NT) + 21364.0 (AA) = 32700.0 |
| **Concordance (%)** | 98.2897 |

| Alignment Method | Global, seeded, nucleotide + amino acids (AGA) |
|---|---|

#### Genome Region

Sequence starts at position 137 and ends at position 5948 relative to NC_001747.1 reference sequence.

#### Alignment Detailed Statistics

| | Begin | End | Coverage | Score | Concordance | Matches | Identities | I/D/M/F* | Stop Codons |
|---|---|---|---|---|---|---|---|---|---|
| **NT** | **137** | **5948** | **97.1%** | **11336** | **97.5%** | **5812 (100%)** | **5740 (98.8%)** | **0/0** | |
| Mutations: | 561C>T, 666A>C, 715C>T, 825T>G, 844A>G, 864C>T, 921G>A, 934A>G, 958G>A, 986C>T, 1094A>G, 1242A>G, 1294G>A, 1356G>A, 1394A>T, 1399G>A, 1700C>A, 1717G>A, 1754C>T, 1780A>C, 1830C>G, 1894C>A, 1951G>A, 1989A>G, 2007T>C, 2064A>G, 2079C>T, 2316T>G, 2340T>C, 2361T>G, 2373G>A, 2470G>A, 2550G>A, 2568T>C, 2586A>G, 2610T>C, 2649C>T, 2729A>G, 2796T>C, 2806A>G, 2820G>A, 2865G>A, 2898T>C, 2916C>T, 3000T>C, 3036C>T, 3042G>A, 3066T>C, 3186T>C, 3333C>T, 3378G>A, 3379A>G, 3788T>G, 3830C>T, 3939A>T, 4031A>G, 4034C>T, 4094G>A, 4113G>A, 4172T>C, 4214T>G, 4658G>A, 4661T>C, 4733C>G, 4817T>C, 4883C>T, 5111C>T, 5129G>A, 5189C>T, 5199C>T, 5216A>C, 5268A>G | | | | | | | | |

##### CDS

| | Begin | End | Coverage | Score | Concordance | Matches | Identities | I/D/M/F* | Stop Codons |
|---|---|---|---|---|---|---|---|---|---|
| PLRVgp1 | 1 | 248 | 100% | 1720 | 99.3% | 248 (100%) | 246 (99.2%) | 0/0/0/0 | 1 |
| Protein mutations: | Q164H (666A>C), T224A (844A>G) | | | | | | | | |
| Codon mutations: | TAC129TAT (561C>T), CAA164CAC (666A>C), CTA181TTA (715C>T), GCT217GCG (825T>G), ACC224GCC (844A>G), GTC230GTT (864C>T) | | | | | | | | |
| PLRVgp2 | 1 | 1063 | 100% | 7356 | 98.6% | 1063 (100%) | 1045 (98.3%) | 0/0/0/0 | 1 |
| Protein mutations: | T85I (561C>T), K120T (666A>C), L173R (825T>G), S186F (864C>T), R205K (921G>A), T263A (1094A>G), K312R (1242A>G), M329I (1294G>A), G350E (1356G>A), N363Y (1394A>T), Q465K (1700C>A), S492R (1780A>C), Q530K (1894C>A), E549K (1951G>A), A722T (2470G>A), N808S (2729A>G), T834A (2806A>G), I1025V (3379A>G) | | | | | | | | |
| Codon mutations: | ACA85ATA (561C>T), AAA120ACA (666A>C), CTC136CTT (715C>T), CTA173CGA (825T>G), TTA179TTG (844A>G), TCC186TTC (864C>T), AGG205AAG (921G>A), GGA209GGG (934A>G), CAG217CAA (958G>A), CTA227TTA (986C>T), ACG263GCG (1094A>G), AAA312AGA (1242A>G), ATG329ATA (1294G>A), GGA350GAA (1356G>A), AAT363TAT (1394A>T), CTG364CTA (1399G>A), CAA465AAA (1700C>A), GAG470GAA (1717G>A), CTA483TTA (1754C>T), AGC492CGC (1780A>C), CGC508CGG (1830C>G), CAA530AAA (1894C>A), GAA549AAA (1951G>A), GAA561GAG (1989A>G), TTT567TTC (2007T>C), CCA586CCG (2064A>G), TGC591TGT (2079C>T), GTT670GTG (2316T>G), ACT678ACC (2340T>C), CTT685CTG (2361T>G), CTG689CTA (2373G>A), GCA722ACA (2470G>A), TCG748TCA (2550G>A), GAT754GAC (2568T>C), CTA760CTG (2586A>G), CCT768CCC (2610T>C), AGC781AGT (2649C>T), AAT808AGT (2729A>G), ACT830ACC (2796T>C), ACT834GCT (2806A>G), TTG838TTA (2820G>A), TTG853TTA (2865G>A), ACT864ACC (2898T>C), GAC870GAT (2916C>T), CGT898CGC (3000T>C), TCC910TCT (3036C>T), CTG912CTA (3042G>A), CTT920CTC (3066T>C), GAT960GAC (3186T>C), TAC1009TAT (3333C>T), GTG1024GTA (3378G>A), ATT1025GTT (3379A>G) | | | | | | | | |
| PLRVgp3 | 1 | 640 | 100% | 4193 | 97.7% | 640 (100%) | 623 (97.3%) | 0/0/0/0 | 1 |
| Protein mutations: | T85I (561C>T), K120T (666A>C), L173R (825T>G), S186F (864C>T), R205K (921G>A), T263A (1094A>G), K312R (1242A>G), M329I (1294G>A), G350E (1356G>A), N363Y (1394A>T), Q465K (1700C>A), Q491H (1780A>C), A508G (1830C>G), K561R (1989A>G), L567S (2007T>C), Q586R (2064A>G), A591V (2079C>T) | | | | | | | | |
| Codon mutations: | ACA85ATA (561C>T), AAA120ACA (666A>C), CTC136CTT (715C>T), CTA173CGA (825T>G), TTA179TTG (844A>G), TCC186TTC (864C>T), AGG205AAG (921G>A), GGA209GGG (934A>G), CAG217CAA (958G>A), CTA227TTA (986C>T), ACG263GCG (1094A>G), AAA312AGA (1242A>G), ATG329ATA (1294G>A), GGA350GAA (1356G>A), AAT363TAT (1394A>T), CTG364CTA (1399G>A), CAA465AAA (1700C>A), GAG470GAA (1717G>A), CTA483TTA (1754C>T), CAA491CAC (1780A>C), GCT508GGT (1830C>G), ATC529ATA (1894C>A), CAG548CAA (1951G>A), AAG561AGG (1989A>G), TTA567TCA (2007T>C), CAG586CGG (2064A>G), GCG591GTG (2079C>T) | | | | | | | | |
| Rap1 | 1 | 42 | 100% | 280 | 97.6% | 42 (100%) | 41 (97.6%) | 0/0/0/0 | 1 |
| Protein mutations: | R29K (1717G>A) | | | | | | | | |
| Codon mutations: | CTC23CTA (1700C>A), AGA29AAA (1717G>A), CTC41CTT (1754C>T) | | | | | | | | |
| ORF3a | 1 | 46 | 100% | 299 | 100% | 46 (100%) | 46 (100%) | 0/0/0/0 | 1 |
| PLRVgp4 | 1 | 718 | 100% | 5018 | 99.6% | 718 (100%) | 714 (99.4%) | 0/0/0/0 | 2 |
| Protein mutations: | T83S (3939A>T), V141I (4113G>A), P503S (5199C>T), T526A (5268A>G) | | | | | | | | |
| Codon mutations: | GTT32GTG (3788T>G), CGC46CGT (3830C>T), ACC83TCC (3939A>T), ACA113ACG (4031A>G), AGC114AGT (4034C>T), GAG134GAA (4094G>A), GTA141ATA (4113G>A), ACT160ACC (4172T>C), TCT174TCG (4214T>G), CGG322CGA (4658G>A), GGT323GGC (4661T>C), GTC347GTG (4733C>G), GGT375GGC (4817T>C), TTC397TTT (4883C>T), GAC473GAT (5111C>T), ACG479ACA (5129G>A), AAC499AAT (5189C>T), CCA503TCA (5199C>T), CGA508CGC (5216A>C), ACA526GCA (5268A>G) | | | | | | | | |
| PLRVgp5 | 1 | 209 | 100% | 1431 | 99.7% | 209 (100%) | 207 (99.0%) | 0/0/0/0 | 1 |
| Protein mutations: | T83S (3939A>T), V141I (4113G>A) | | | | | | | | |
| Codon mutations: | GTT32GTG (3788T>G), CGC46CGT (3830C>T), ACC83TCC (3939A>T), ACA113ACG (4031A>G), AGC114AGT (4034C>T), GAG134GAA (4094G>A), GTA141ATA (4113G>A), ACT160ACC (4172T>C), TCT174TCG (4214T>G) | | | | | | | | |
| PLRVgp6 | 1 | 157 | 100% | 1067 | 96.8% | 157 (100%) | 151 (96.2%) | 0/0/0/0 | 1 |
| Protein mutations: | F24C (3788T>G), A38V (3830C>T), Q105R (4031A>G), A106V (4034C>T), S126N (4094G>A), L152P (4172T>C) | | | | | | | | |
| Codon mutations: | TTC24TGC (3788T>G), GCC38GTC (3830C>T), ACA74ACT (3939A>T), CAA105CGA (4031A>G), GCA106GTA (4034C>T), AGT126AAT (4094G>A), AAG132AAA (4113G>A), CTT152CCT (4172T>C) | | | | | | | | |

##### Proteins

| | Begin | End | Coverage | Score | Concordance | Matches | Identities | I/D/M/F* | Stop Codons |
|---|---|---|---|---|---|---|---|---|---|
| P0 protein (NP_056746.1) | 1 | 248 | 100% | 1720 | 99.3% | 248 (100%) | 246 (99.2%) | 0/0/0/0 | 1 |
| Protein mutations: | Q164H (666A>C), T224A (844A>G) | | | | | | | | |
| Codon mutations: | TAC129TAT (561C>T), CAA164CAC (666A>C), CTA181TTA (715C>T), GCT217GCG (825T>G), ACC224GCC (844A>G), GTC230GTT (864C>T) | | | | | | | | |
| RNA-dependent RNA polymerase (NP_056748.3) | 1 | 1063 | 100% | 7356 | 98.6% | 1063 (100%) | 1045 (98.3%) | 0/0/0/0 | 1 |
| Protein mutations: | T85I (561C>T), K120T (666A>C), L173R (825T>G), S186F (864C>T), R205K (921G>A), T263A (1094A>G), K312R (1242A>G), M329I (1294G>A), G350E (1356G>A), N363Y (1394A>T), Q465K (1700C>A), S492R (1780A>C), Q530K (1894C>A), E549K (1951G>A), A722T (2470G>A), N808S (2729A>G), T834A (2806A>G), I1025V (3379A>G) | | | | | | | | |
| Codon mutations: | ACA85ATA (561C>T), AAA120ACA (666A>C), CTC136CTT (715C>T), CTA173CGA (825T>G), TTA179TTG (844A>G), TCC186TTC (864C>T), AGG205AAG (921G>A), GGA209GGG (934A>G), CAG217CAA (958G>A), CTA227TTA (986C>T), ACG263GCG (1094A>G), AAA312AGA (1242A>G), ATG329ATA (1294G>A), GGA350GAA (1356G>A), AAT363TAT (1394A>T), CTG364CTA (1399G>A), CAA465AAA (1700C>A), GAG470GAA (1717G>A), CTA483TTA (1754C>T), AGC492CGC (1780A>C), CGC508CGG (1830C>G), CAA530AAA (1894C>A), GAA549AAA (1951G>A), GAA561GAG (1989A>G), TTT567TTC (2007T>C), CCA586CCG (2064A>G), TGC591TGT (2079C>T), GTT670GTG (2316T>G), ACT678ACC (2340T>C), CTT685CTG (2361T>G), CTG689CTA (2373G>A), GCA722ACA (2470G>A), TCG748TCA (2550G>A), GAT754GAC (2568T>C), CTA760CTG (2586A>G), CCT768CCC (2610T>C), AGC781AGT (2649C>T), AAT808AGT (2729A>G), ACT830ACC (2796T>C), ACT834GCT (2806A>G), TTG838TTA (2820G>A), TTG853TTA (2865G>A), ACT864ACC (2898T>C), GAC870GAT (2916C>T), CGT898CGC (3000T>C), TCC910TCT (3036C>T), CTG912CTA (3042G>A), CTT920CTC (3066T>C), GAT960GAC (3186T>C), TAC1009TAT (3333C>T), GTG1024GTA (3378G>A), ATT1025GTT (3379A>G) | | | | | | | | |

| | Begin | End | Coverage | Score | Concordance | Matches | Identities | I/D/M/F* | Stop Codons |
|---|---|---|---|---|---|---|---|---|---|
| **NT** | **137** | **5948** | **97.1%** | **11336** | **97.5%** | **5812 (100%)** | **5740 (98.8%)** | **0/0** | |
| P1 protein (NP_056747.1) | 1 | 640 | 100% | 4193 | 97.7% | 640 (100%) | 623 (97.3%) | 0/0/0/0 | 1 |
| Protein mutations: | T85I (561C>T), K120T (666A>C), L173R (825T>G), S186F (864C>T), R205K (921G>A), T263A (1094A>G), K312R (1242A>G), M329I (1294G>A), G350E (1356G>A), N363Y (1394A>T), Q465K (1700C>A), Q491H (1780A>C), A508G (1830C>G), K561R (1989A>G), L567S (2007T>C), Q586R (2064A>G), A591V (2079C>T) | | | | | | | | |
| Codon mutations: | ACA85ATA (561C>T), AAA120ACA (666A>C), CTC136CTT (715C>T), CTA173CGA (825T>G), TTA179TTG (844A>G), TCC186TTC (864C>T), AGG205AAG (921G>A), GGA209GGG (934A>G), CAG217CAA (958G>A), CTA227TTA (986C>T), ACG263GCG (1094A>G), AAA312AGA (1242A>G), ATG329ATA (1294G>A), GGA350GAA (1356G>A), AAT363TAT (1394A>T), CTG364CTA (1399G>A), CAA465AAA (1700C>A), GAG470GAA (1717G>A), CTA483TTA (1754C>T), CAA491CAC (1780A>C), GCT508GGT (1830C>G), ATC529ATA (1894C>A), CAG548CAA (1951G>A), AAG561AGG (1989A>G), TTA567TCA (2007T>C), CAG586CGG (2064A>G), GCG591GTG (2079C>T) | | | | | | | | |
| Replication-associated protein (YP_006355442.1) | 1 | 42 | 100% | 280 | 97.6% | 42 (100%) | 41 (97.6%) | 0/0/0/0 | 1 |
| Protein mutations: | R29K (1717G>A) | | | | | | | | |
| Codon mutations: | CTC23CTA (1700C>A), AGA29AAA (1717G>A), CTC41CTT (1754C>T) | | | | | | | | |
| protein 3a (YP_009179365.2) | 1 | 46 | 100% | 299 | 100% | 46 (100%) | 46 (100%) | 0/0/0/0 | 1 |
| Protein mutations: | *none* | | | | | | | | |
| Codon mutations: | *none* | | | | | | | | |
| CP read-through protein (NP_056751.2) | 1 | 718 | 100% | 5018 | 99.6% | 718 (100%) | 714 (99.4%) | 0/0/0/0 | 2 |
| Protein mutations: | T83S (3939A>T), V141I (4113G>A), P503S (5199C>T), T526A (5268A>G) | | | | | | | | |
| Codon mutations: | GTT32GTG (3788T>G), CGC46CGT (3830C>T), ACC83TCC (3939A>T), ACA113ACG (4031A>G), AGC114AGT (4034C>T), GAG134GAA (4094G>A), GTA141ATA (4113G>A), ACT160ACC (4172T>C), TCT174TCG (4214T>G), CGG322CGA (4658G>A), GGT323GGC (4661T>C), GTC347GTG (4733C>G), GGT375GGC (4817T>C), TTC397TTT (4883C>T), GAC473GAT (5111C>T), ACG479ACA (5129G>A), AAC499AAT (5189C>T), CCA503TCA (5199C>T), CGA508CGC (5216A>C), ACA526GCA (5268A>G) | | | | | | | | |
| coat protein (NP_056749.1) | 1 | 209 | 100% | 1431 | 99.7% | 209 (100%) | 207 (99.0%) | 0/0/0/0 | 1 |
| Protein mutations: | T83S (3939A>T), V141I (4113G>A) | | | | | | | | |
| Codon mutations: | GTT32GTG (3788T>G), CGC46CGT (3830C>T), ACC83TCC (3939A>T), ACA113ACG (4031A>G), AGC114AGT (4034C>T), GAG134GAA (4094G>A), GTA141ATA (4113G>A), ACT160ACC (4172T>C), TCT174TCG (4214T>G) | | | | | | | | |
| movement protein (NP_056750.1) | 1 | 157 | 100% | 1067 | 96.8% | 157 (100%) | 151 (96.2%) | 0/0/0/0 | 1 |
| Protein mutations: | F24C (3788T>G), A38V (3830C>T), Q105R (4031A>G), A106V (4034C>T), S126N (4094G>A), L152P (4172T>C) | | | | | | | | |
| Codon mutations: | TTC24TGC (3788T>G), GCC38GTC (3830C>T), ACA74ACT (3939A>T), CAA105CGA (4031A>G), GCA106GTA (4034C>T), AGT126AAT (4094G>A), AAG132AAA (4113G>A), CTT152CCT (4172T>C) | | | | | | | | |

*: Inserts / Deletes / Misaligned / Frameshifts

#### Analysis details

This analysis was performed with panviral2.64

### NGS Details (UN70): Solendovirus venanicotianae

#### Assembly

| | |
|---|---|
| **Coverage Length** | 2106 (5 contig(s)) |
| **Depth Of Coverage** | 12.1 |
| **Number Of Reads** | 234 |
| **Reads Per Million** | 4.44 rpm (after QC) |
| **Ambiguities** | 12 |
| **Assembly Method** | de novo + reference guided assembly |
| **Consensus Caller** | Bcf Tools |

#### Coverage Map

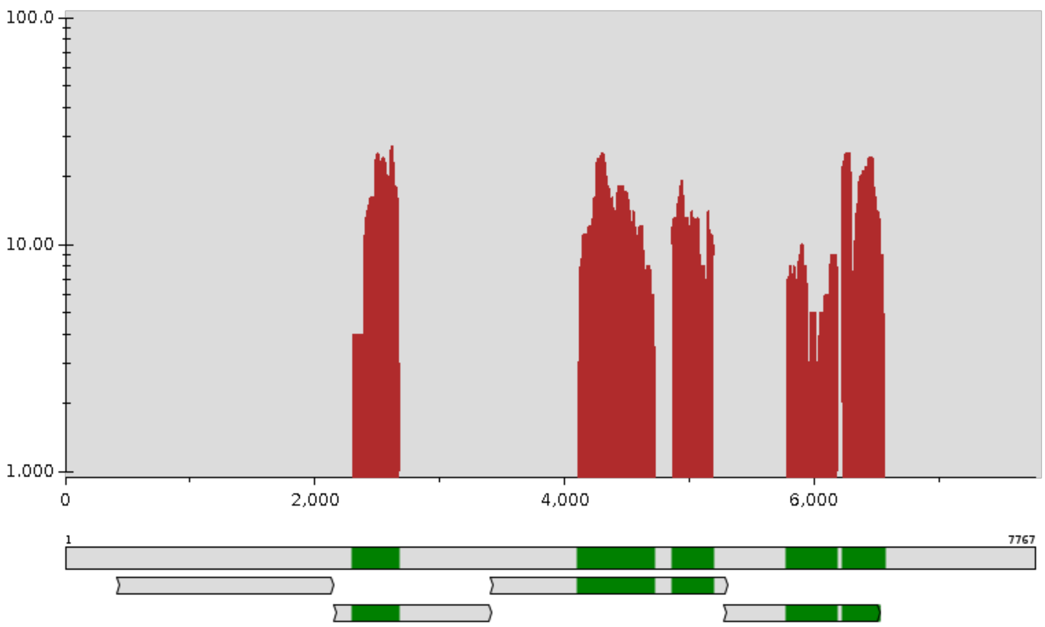

#### Assignment

| | |
|---|---|
| **Type** | Solendovirus venanicotianae (Taxonomy ID: 3048371) |
| **Reference Genome** | NC_003378.1 |
| **NT Identity (%)** | 77.55 |
| **AA Identity (%)** | 74.2313 |
| **Number Of Stop Codons** | 3 |
| **Number Of CDS** | 4 |

#### Alignment

| | |
|---|---|
| **Alignment Score** | 2301.0 (NT) + 3423.0 (AA) = 5724.0 |
| **Concordance (%)** | 65.2828 |

| Alignment Method | Global, seeded, nucleotide + amino acids (AGA) |
|---|---|

#### Genome Region

Sequence starts at position 2300 and ends at position 6573 relative to NC_003378.1 reference sequence.

#### Alignment Detailed Statistics

| | Begin | End | Coverage | Score | Concordance | Matches | Identities | I/D/M/F* | Stop Codons |
|---|---|---|---|---|---|---|---|---|---|
| **NT** | **2300** | **6573** | **27.1%** | **2301** | **55.2%** | **2097 (99.5%)** | **1627 (77.2%)** | **1/9** | |

Mutations: 2308A>T, 2313A>C, 2314A>T, 2316T>C, 2322C>T, 2328A>G, 2330A>C, 2332A>G, 2333A>T, 2343C>T, 2347G>A, 2350G>A, 2352T>A, 2355C>T, 2356C>T, 2368A>G, 2370A>G, 2373T>A, 2376T>Y, 2379C>Y, 2383T>Y, 2384A>W, 2385T>Y, 2388A>Y, 2389G>A, 2390G>A, 2391A>R, 2394T>A, 2396A>G, 2397G>A, 2399C>T, 2400C>T, 2402A>G, 2410A>C, 2417A>G, 2418C>T, 2420A>G, 2422A>C, 2425A>T, 2426T>C, 2430A>C, 2434A>C, 2443C>A, 2445A>G, 2446G>A, 2448T>G, 2454G>A, 2455G>C, 2457A>T, 2461A>G, 2463C>A, 2469T>A, 2472G>A, 2481C>T, 2489G>C, 2493T>C, 2500T>C, 2503A>G, 2535C>A, 2538T>A, 2545A>C, 2551C>T, 2555C>T, 2559T>C, 2562T>C, 2565A>G, 2568A>T, 2569A>T, 2571C>A, 2572C>G, 2589C>T, 2590A>G, 2595A>C, 2597G>C, 2598T>A, 2602A>G, 2606A>G, 2609G>A, 2613T>C, 2622T>C, 2628G>A, 2637T>C, 2638A>G, 2643A>C, 2646C>T, 2647G>C, 2649C>A, 2665C>T, 4105A>G, 4115A>G, 4116G>A, 4121T>C, 4122A>G, 4124T>A, 4125A>G, 4127C>T, 4139T>C, 4181A>T, 4190G>A, 4196C>T, 4199A>T, 4202C>T, 4235T>C, 4241T>C, 4250C>A, 4256A>G, 4260T>C, 4262A>T, 4265G>A, 4284T>A, 4286T>C, 4292C>T, 4298T>C, 4307C>T, 4313G>A, 4325T>C, 4328T>C, 4329T>C, 4331G>A, 4340G>A, 4343T>A, 4362G>A, 4364T>A, 4374C>G, 4400T>A, 4421T>C, 4423C>A, 4430A>T, 4436C>T, 4451C>T, 4453A>G, 4454C>T, 4478T>C, 4484A>T, 4500C>T, 4503C>T, 4508T>C, 4513G>A, 4517A>T, 4518C>A, 4523T>A, 4530A>T, 4545A>C, 4550C>T, 4551G>A, 4552C>T, 4554C>G, 4568C>T, 4571T>A, 4574T>A, 4580C>T, 4586C>T, 4589A>G, 4591C>A, 4592C>A, 4596G>T, 4604T>A, 4631A>G, 4648A>G, 4661G>R, 4662C>Y, 4665A>T, 4670T>Y, 4674G>R, 4676A>R, 4688C>T, 4692C>T, 4693T>C, 4694T>A, 4695G>A, 4701A>C, 4703C>A, 4706T>A, 4717A>G, 4718G>A, 4725C>T, 4859G>A, 4865T>A, 4868G>A, 4869A>T, 4872T>C, 4874G>T, 4897A>G, 4898C>T, 4907T>C, 4913C>T, 4919A>T, 4924C>T, 4925C>A, 4931T>A, 4938A>G, 4944T>G, 4950A>G, 4952T>G, 4953T>C, 4954A>G, 4955T>A, 4961T>C, 4963G>A, 4970T>A, 4971C>T, 4973T>A, 4976G>A, 4978A>C, 4979T>C, 4982A>G, 4986A>G, 5012C>T, 5018C>T, 5027A>T, 5028T>A, 5030T>A, 5032A>T, 5033T>C, 5034A>T, 5039A>C, 5040C>T, 5042A>G, 5043C>G, 5046G>A, 5047A>T, 5054G>A, 5057A>G, 5065G>A, 5076T>C, 5077T>A, 5079G>T, 5081A>G, 5094C>T, 5097T>C, 5102A>C, 5117A>T, 5119T>C, 5120T>G, 5125C>A, 5126C>T, 5128G>A, 5132T>C, 5135T>A, 5136G>A, 5140G>A, 5145G>A, 5156A>G, 5174T>A, 5180A>G, 5189T>C, 5195C>T, 5772A>T, 5775T>C, 5781A>C, 5784G>A, 5786A>C, 5787T>A, 5789T>A, 5796T>C, 5797A>T, 5798A>T, 5802T>A, 5805A>C, 5807G>A, 5811C>A, 5812C>A, 5814A>T, 5815T>A, 5816C>T, 5817T>C, 5820C>T, 5821C>A, 5824A>G, 5826T>A, 5831A>C, 5832C>A, 5833G>C, 5835A>G, 5838C>T, 5850C>A, 5851C>A, 5853T>A, 5857A>C, 5858C>A, 5872T>C, 5874G>A, 5877A>T, 5880T>A, 5881T>C, 5882T>A, 5889T>G, 5892A>T, 5894G>A, 5898A>T, 5901A>C, 5907A>C, 5908G>A, 5909C>A, 5910C>A, 5913C>A, 5916C>T, 5921A>G, 5922C>T, 5925C>T, 5935G>A, 5936A>G, 5940T>C, 5950C>T, 5952G>C, 5953A>G, 5954C>A, 5955A>T, 5959C>G, 5964G>A, 5965A>T, 5968G>A, 5970C>A, 5972A>C, 5979T>A, 5982G>A, 5988C>T, 5991G>A, 5997C>T, 5999T>C, 6001C>A, 6002A>C, 6003A>C, 6006T>C, 6009G>A, 6010A>C, 6012G>A, 6013A>G, 6015A>T, 6018T>A, 6024G>A, 6026C>A, 6027A>T, 6033C>T, 6036C>T, 6037G>A, 6040C>A, 6041G>A, 6045C>T, 6049T>A, 6050C>G, 6051G>A, 6054T>A, 6055A>C, 6067C>T, 6072T>C, 6075A>T, 6078G>A, 6084G>A, 6087T>A, 6088A>G, 6090C>G, 6093A>C, 6099T>C, 6102C>T, 6105A>G, 6108T>A, 6114C>G, 6117A>T, 6120G>A, 6123A>T, 6124A>G, 6129C>A, 6139A>G, 6146A>T, 6147A>G, 6154G>C, 6155A>C, 6163G>C, 6164A>C, 6167A>G, 6169G>A, 6170T>C, 6171A>T, 6172A>G, 6174T>G, 6177T>A, 6232C>T, 6246T>C, 6247C>A, 6248T>C, 6253C>G, 6254A>G, 6258T>C, 6259G>C, 6264T>G, 6269G>A, 6270C>T, 6271T>C, 6277_6283delTCAAGGG, 6285A>C, 6290C>T, 6301T>G, 6308G>C, 6309C>A, 6311G>A, 6315T>A, 6318A>C, 6319A>C, 6321A>C, 6325G>A, 6326C>G, 6340C>G, 6341T>C, 6352G>A, 6354C>A, 6355T>A, 6357A>G, 6359G>C, 6360T>A, 6361C>T, 6363T>G, 6369A>T, 6375A>G, 6376C>A, 6380C>A, 6381T>A, 6382A>C, 6384A>T, 6386C>T, 6387C>T, 6389C>T, 6401G>A, 6402G>A, 6404G>A, 6405G>T, 6407A>G, 6408T>A, 6411C>T, 6414C>T, 6417C>G, 6418C>G, 6419C>A, 6420A>C, 6423T>A, 6424C>T, 6426G>A, 6427T>C, 6446T>A, 6451A>G, 6453T>G, 6456C>A, 6462T>C, 6471C>T, 6477C>T, 6480A>G, 6489A>C, 6502A>G, 6504T>A, 6505G>A, 6507T>C, 6513C>T, 6522G>A, 6525A>C, 6526T>G, 6530T>A, 6531C>A, 6532_6533insA, 6537G>T, 6542A>T, 6545_6546delCG, 6551A>C, 6553C>G, 6555A>T, 6556C>A, 6557A>C, 6558A>T, 6561T>C, 6563T>A, 6564G>C, 6566C>A, 6570A>G, 6572G>T

##### CDS

| | Begin | End | Coverage | Score | Concordance | Matches | Identities | I/D/M/F* | Stop Codons |
|---|---|---|---|---|---|---|---|---|---|
| Tvvgp2 | 50 | 174 | 29.8% | 632 | 73.7% | 125 (100%) | 88 (70.4%) | 0/0/1/0 | 1 |

Protein mutations: I52L (2308A>T), K53N (2313A>C), N54Y (2314A>T 2316T>C), Q59P (2330A>C), K60V (2332A>G 2333A>T), V65I (2347G>A), V66I (2350G>A 2352T>A), Q68* (2356C>T), I72V (2368A>G 2370A>G), G79K (2389G>A 2390G>A 2391A>R), K81R (2396A>G 2397G>A), A82V (2399C>T 2400C>T), K83R (2402A>G), I86L (2410A>C), N88S (2417A>G 2418C>T), K89R (2420A>G), I91S (2425A>T 2426T>C), K94Q (2434A>C), Q97K (2443C>A 2445A>G), D98K (2446G>A 2448T>G), A101P (2455G>C 2457A>T), N103E (2461A>G 2463C>A), G112A (2489G>C), I117V (2503A>G), I131L (2545A>C), A134V (2555C>T), I139L (2569A>T 2571C>A), Q140E (2572C>G), I146V (2590A>G), S148T (2597G>C 2598T>A), I150V (2602A>G), K151R (2606A>G), G152E (2609G>A), I162V (2638A>G), A165P (2647G>C 2649C>A)

Codon mutations: ATA52TTA (2308A>T), AAA53AAC (2313A>C), AAT54TAC (2314A>T 2316T>C), TTC56TTT (2322C>T), AGA58AGG (2328A>G), CAA59CCA (2330A>C), AAA60GTA (2332A>G 2333A>T), TAC63TAT (2343C>T), GTA65ATA (2347G>A), GTT66ATA (2350G>A 2352T>A), AGC67AGT (2355C>T), CAA68TAA (2356C>T), ATA72GTG (2368A>G 2370A>G), ATT73ATA (2373T>A), GAT74GAY (2376T>Y), TGC75TGY (2379C>Y), TAT77YWY (2383T>Y 2384A>W 2385T>Y), ACA78ACY (2388A>Y), GGA79AAR (2389G>A 2390G>A 2391A>R), GGT80GGA (2394T>A), AAG81AGA (2396A>G 2397G>A), GCC82GTT (2399C>T 2400C>T), AAA83AGA (2402A>G), ATA86CTA (2410A>C), AAC88AGT (2417A>G 2418C>T), AAA89AGA (2420A>G), AGA90CGA (2422A>C), ATA91TCA (2425A>T 2426T>C), ATA92ATC (2430A>C), AAA94CAA (2434A>C), CAA97AAG (2443C>A 2445A>G), GAT98AAG (2446G>A 2448T>G), AAG100AAA (2454G>A), GCA101CCT (2455G>C 2457A>T), AAC103GAA (2461A>G 2463C>A), ATT105ATA (2469T>A), AAG106AAA (2472G>A), CAC109CAT (2481C>T), GGA112GCA (2489G>C), ACT113ACC (2493T>C), TTA116CTA (2500T>C), ATA117GTA (2503A>G), ACC127ACA (2535C>A), CCT128CCA (2538T>A), ATA131CTA (2545A>C), CTA133TTA (2551C>T), GCA134GTA (2555C>T), GAT135GAC (2559T>C), GAT136GAC (2562T>C), AGA137AGG (2565A>G), ATA138ATT (2568A>T), ATC139TTA (2569A>T 2571C>A), CAA140GAA (2572C>G), AGC145AGT (2589C>T), ATA146GTA (2590A>G), ATA147ATC (2595A>C), AGT148ACA (2597G>C 2598T>A), ATA150GTA (2602A>G), AAA151AGA (2606A>G), GGA152GAA (2609G>A), AAT153AAC (2613T>C), TAT156TAC (2622T>C), AAG158AAA (2628G>A), TTT161TTC (2637T>C), ATA162GTA (2638A>G), ATA163ATC (2643A>C), AGC164AGT (2646C>T), GCC165CCA (2647G>C 2649C>A), CTA171TTA (2665C>T)

| | Begin | End | Coverage | Score | Concordance | Matches | Identities | I/D/M/F* | Stop Codons |
|---|---|---|---|---|---|---|---|---|---|
| Tvvgp3 | 233 | 596 | 50.5% | 1789 | 79.1% | 321 (100%) | 264 (82.2%) | 0/0/3/0 | 2 |

Protein mutations: E236K (4116G>A), N238E (4122A>G 4124T>A), S239G (4125A>G 4127C>T), Y292N (4284T>A 4286T>C), D311E (4343T>A), A318T (4362G>A 4364T>A), P322A (4374C>G), A338E (4423C>A), N348S (4453A>G 4454C>T), R368K (4513G>A), Q370K (4518C>A), D371E (4523T>A), I374L (4530A>T), K379Q (4545A>C), A381I (4551G>A 4552C>T), H382D (4554C>G), T394K (4591C>A 4592C>A), A396S (4596G>T), N413S (4648A>G), T419S (4665A>T), L428S (4692C>T 4693T>C 4694T>A), D429N (4695G>A), N431Q (4701A>C 4703C>A), K436R (4717A>G 4718G>A), Q439* (4725C>T), I487L (4869A>T), Y496C (4897A>G 4898C>T), K503N (4919A>T), T505I (4924C>T 4925C>A), T510A (4938A>G), S512A (4944T>G), N514E (4950A>G 4952T>G), Y515R (4953T>C 4954A>G 4955T>A), G518E (4963G>A), Y523S (4978A>C 4979T>C), N526D (4986A>G), S540T (5028T>A 5030T>A), Y541F (5032A>T 5033T>C), T542S (5034A>T), E543D (5039A>C), P544S (5040C>T 5042A>G), Q545E (5043C>G), E546I (5046G>A 5047A>T), W548* (5054G>A), R552K (5065G>A), F556H (5076T>C 5077T>A), A557S (5079G>T 5081A>G), V570A (5119T>C 5120T>G), T572N (5125C>A 5126C>T), R573K (5128G>A), V576I (5136G>A), R577K (5140G>A), D579N (5145G>A)

Codon mutations: AAA232.GA (4105A>G), CAA235CAG (4115A>G), GAA236AAA (4116G>A), AGT237AGC (4121T>C), AAT238GAA (4122A>G 4124T>A), AGC239GGT (4125A>G 4127C>T), AGT243AGC (4139T>C), GGA257GGT (4181A>T), AGG260AGA (4190G>A), GTC262GTT (4196C>T), ATA263ATT (4199A>T), GAC264GAT (4202C>T), TAT275TAC (4235T>C), TAT277TAC (4241T>C), CCC280CCA (4250C>A), AAA282AAG (4256A>G), TTA284CTT (4260T>C 4262A>T), AAG285AAA (4265G>A), TAT292AAC (4284T>A 4286T>C), TAC294TAT (4292C>T), AGT296AGC (4298T>C), GAC299GAT (4307C>T), AAG301AAA (4313G>A), TAT305TAC (4325T>C), CAT306CAC (4328T>C), TTG307CTA (4329T>C 4331G>A), GAG310GAA (4340G>A), GAT311GAA (4343T>A), GCT318ACA (4362G>A 4364T>A), CCA322GCA (4374C>G), GTT330GTA (4400T>A), AAT337AAC (4421T>C), GCA338GAA (4423C>A), GGA340GGT (4430A>T), TAC342TAT (4436C>T), GAC347GAT (4451C>T), AAC348AGT (4453A>G 4454C>T), TGT356TGC (4478T>C), GTA358GTT (4484A>T), CTA364TTA (4500C>T), CTA365TTA (4503C>T), TAT366TAC (4508T>C), AGA368AAA (4513G>A), ACA369ACT (4517A>T), CAA370AAA (4518C>A), GAT371GAA (4523T>A), ATA374TTA (4530A>T), AAA379CAA (4545A>C), TTC380TTT (4550C>T), GCA381ATA (4551G>A 4552C>T), CAT382GAT (4554C>G), AAC386AAT (4568C>T), TCT387TCA (4571T>A), GGT388GGA (4574T>A), AGC390AGT (4580C>T), AGC392AGT (4586C>T), AAA393AAG (4589A>G), ACC394AAA (4591C>A 4592C>A), GCA396TCA (4596G>T), ATT398ATA (4604T>A), GGA407GGG (4631A>G), AAT413AGT (4648A>G), ATG417ATR (4661G>R), CAA418YAA (4662C>Y), ACT419TCT (4665A>T), CAT420CAY (4670T>Y), GTA422RTR (4674G>R 4676A>R), ATC426ATT (4688C>T), CTT428TCA (4692C>T 4693T>C 4694T>A), GAT429AAT (4695G>A), AAC431CAA (4701A>C 4703C>A), ATT432ATA (4706T>A), AAG436AGA (4717A>G 4718G>A), CAA439TAA (4725C>T), AAG483..A (4859G>A), ATT485ATA (4865T>A), AAG486AAA (4868G>A), ATA487TTA (4869A>T), TTG488CTT (4872T>C 4874G>T), TAC496TGT (4897A>G 4898C>T), GAT499GAC (4907T>C), AAC501AAT (4913C>T), AAA503AAT (4919A>T), ACC505ATA (4924C>T 4925C>A), ATT507ATA (4931T>A), ACA510GCA (4938A>G), TCA512GCA (4944T>G), AAT514GAG (4950A>G 4952T>G), TAT515CGA (4953T>C 4954A>G 4955T>A), TAT517TAC (4961T>C), GGA518GAA (4963G>A), GTT520GTA (4970T>A), CTT521TTA (4971C>T 4973T>A), AAG522AAA (4976G>A), TAT523TCC (4978A>C 4979T>C), AGA524AGG (4982A>G), AAT526GAT (4986A>G), TGC534TGT (5012C>T), TAC536TAT (5018C>T), GGA539GGT (5027A>T), TCT540ACA (5028T>A 5030T>A), TAT541TTC (5032A>T 5033T>C), ACA542TCA (5034A>T), GAA543GAC (5039A>C), CCA544TCG (5040C>T 5042A>G), CAA545GAA (5043C>G), GAA546ATA (5046G>A 5047A>T), TGG548TGA (5054G>A), GAA549GAG (5057A>G), AGA552AAA (5065G>A), TTT556CAT (5076T>C 5077T>A), GCA557TCG (5079G>T 5081A>G), CTA562TTA (5094C>T), TTA563CTA (5097T>C), GCA564GCC (5102A>C), ATA569ATT (5117A>T), GTT570GCG (5119T>C 5120T>G), ACC572AAT (5125C>A 5126C>T), AGA573AAA (5128G>A), TTT574TTC (5132T>C), ATT575ATA (5135T>A), GTA576ATA (5136G>A), AGA577AAA (5140G>A), GAC579AAC (5145G>A), CAA582CAG (5156A>G), ACT588ACA (5174T>A), AAA590AAG (5180A>G), GAT593GAC (5189T>C), GTC595GTT (5195C>T)

| | Begin | End | Coverage | Score | Concordance | Matches | Identities | I/D/M/F* | Stop Codons |
|---|---|---|---|---|---|---|---|---|---|
| Tvvgp4 | 167 | 418 | 57.2% | 1002 | 60.7% | 237 (99.2%) | 155 (64.9%) | 0/2/1/1 | 0 |

Protein mutations: N171T (5786A>C 5787T>A), F172Y (5789T>A), K175L (5797A>T 5798A>T), N176K (5802T>A), R178K (5807G>A), Q180N (5812C>A 5814A>T), S181I (5815T>A 5816C>T 5817T>C), Q183K (5821C>A), N184E (5824A>G 5826T>A), N186T (5831A>C 5832C>A), E187Q (5833G>C 5835A>G), H192Q (5850C>A), H193K (5851C>A 5853T>A), T195Q (5857A>C 5858C>A), L203Q (5881T>C 5882T>A), N205K (5889T>G), S207N (5894G>A), K209N (5901A>C), A212K (5908G>A 5909C>A 5910C>A), N216S (5921A>G 5922C>T), D221S (5935G>A 5936A>G), Q226Y (5950C>T 5952G>C), T227D (5953A>G 5954C>A 5955A>T), Q229E (5959C>G), I231L (5965A>T), A232T (5968G>A 5970C>A), N233T (5972A>C), M242T (5999T>C), Q243T (6001C>A 6002A>C 6003A>C), I247V (6013A>G 6015A>T), T251N (6026C>A 6027A>T), V255I (6037G>A), R256K (6040C>A 6041G>A), S259R (6049T>A 6050C>G 6051G>A), T261P (6055A>C), E267D (6075A>T), I272V (6088A>G 6090C>G), E283D (6123A>T), M284V (6124A>G), K289E (6139A>G), E291V (6146A>T 6147A>G), E294P (6154G>C 6155A>C), E297P (6163G>C 6164A>C), K298R (6167A>G), V299T (6169G>A 6170T>C 6171A>T), N300E (6172A>G 6174T>G), L325T (6247C>A 6248T>C), Q327G (6253C>G 6254A>G), A329P (6259G>C), I330M (6264T>G), S332N (6269G>A 6270C>T), Y333H (6271T>C), S335_R336del (6277_6283delTCAAGGG), T339I (6290C>T), S343A (6301T>G), C345S (6308G>C 6309C>A), R346K (6311G>A), D347E (6315T>A), A351R (6325G>A 6326C>G), L356A (6340C>G 6341T>C), V360I (6352G>A 6354C>A), L361M (6355T>A 6357A>G), S362T (6359G>C 6360T>A), K365N (6369A>T), Q368K (6376C>A), P369Q (6380C>A 6381T>A), T370P (6382A>C 6384A>T), T371I (6386C>T 6387C>T), T372I (6389C>T), R376K (6401G>A 6402G>A), R377N (6404G>A 6405G>T), N378R (6407A>G 6408T>A), P382D (6418C>G 6419C>A 6420A>C), D383E (6423T>A), L391Q (6446T>A), S393G (6451A>G 6453T>G), H394Q (6456C>A), K405N (6489A>C), I410V (6502A>G 6504T>A), V411I (6505G>A 6507T>C), E417D (6525A>C), *418E (6526T>G)

| | Begin | End | Coverage | Score | Concordance | Matches | Identities | I/D/M/F* | Stop Codons |
|---|---|---|---|---|---|---|---|---|---|
| **NT** | **2300** | **6573** | **27.1%** | **2301** | **55.2%** | **2097 (99.5%)** | **1627 (77.2%)** | **1/9** | |
| Codon mutations: | ATA166.TT (5772A>T), CAT167CAC (5775T>C), ATA169ATC (5781A>C), CAG170CAA (5784G>A), AAT171ACA (5786A>C 5787T>A), TTT172TAT (5789T>A), AAT174AAC (5796T>C), AAA175TTA (5797A>T 5798A>T), AAT176AAA (5802T>A), CCA177CCC (5805A>C), AGA178AAA (5807G>A), TCC179TCA (5811C>A), CAA180AAT (5812C>A 5814A>T), TCT181ATC (5815T>A 5816C>T 5817T>C), ACC182ACT (5820C>T), CAA183AAA (5821C>A), AAT184GAA (5824A>G 5826T>A), AAC186ACA (5831A>C 5832C>A), GAA187CAG (5833G>C 5835A>G), GAC188GAT (5838C>T), CAC192CAA (5850C>A), CAT193AAA (5851C>A 5853T>A), ACA195CAA (5857A>C 5858C>A), TTG200CTA (5872T>C 5874G>A), ATA201ATT (5877A>T), GCT202GCA (5880T>A), TTA203CAA (5881T>C 5882T>A), AAT205AAG (5889T>G), ACA206ACT (5892A>T), AGT207AAT (5894G>A), GCA208GCT (5898A>T), AAA209AAC (5901A>C), GTA211GTC (5907A>C), GCC212AAA (5908G>A 5909C>A 5910C>A), ACC213ACA (5913C>A), TGC214TGT (5916C>T), AAC216AGT (5921A>G 5922C>T), TAC217TAT (5925C>T), GAT221AGT (5935G>A 5936A>G), ACT222ACC (5940T>C), CAG226TAC (5950C>T 5952G>C), ACA227GAT (5953A>G 5954C>A 5955A>T), CAA229GAA (5959C>G), GAG230GAA (5964G>A), ATA231TTA (5965A>T), GCC232ACA (5968G>A 5970C>A), AAT233ACT (5972A>C), CCT235CCA (5979T>A), GAG236GAA (5982G>A), TAC238TAT (5988C>T), AAG239AAA (5991G>A), TTC241TTT (5997C>T), ATG242ACG (5999T>C), CAA243ACC (6001C>A 6002A>C 6003A>C), TAT244TAC (6006T>C), AAG245AAA (6009G>A), AGG246CGA (6010A>C 6012G>A), ATA247GTT (6013A>G 6015A>T), ACT248ACA (6018T>A), GGG250GGA (6024G>A), ACA251AAT (6026C>A 6027A>T), TTC253TTT (6033C>T), TAC254TAT (6036C>T), GTA255ATA (6037G>A), CGA256AAA (6040C>A 6041G>A), TTC257TTT (6045C>T), TCG259AGA (6049T>A 6050C>G 6051G>A), GCT260GCA (6054T>A), ACA261CCA (6055A>C), CTA265TTA (6067C>T), TAT266TAC (6072T>C), GAA267GAT (6075A>T), GAG268GAA (6078G>A), AAG270AAA (6084G>A), CCT271CCA (6087T>A), ATC272GTG (6088A>G 6090C>G), ATA273ATC (6093A>C), GTT275GTC (6099T>C), ATC276ATT (6102C>T), AAA277AAG (6105A>G), ATT278ATA (6108T>A), CTC280CTG (6114C>G), ACA281ACT (6117A>T), AGG282AGA (6120G>A), GAA283GAT (6123A>T), ATG284GTG (6124A>G), ATC285ATA (6129C>A), AAG289GAG (6139A>G), GAA291GTG (6146A>T 6147A>G), GAA294CCA (6154G>C 6155A>C), GAA297CCA (6163G>C 6164A>C), AAA298AGA (6167A>G), GTA299ACT (6169G>A 6170T>C 6171A>T), AAT300GAG (6172A>G 6174T>G), ATT301ATA (6177T>A), CTA320TTA (6232C>T), TAT324TAC (6246T>C), CTA325ACA (6247C>A 6248T>C), CAG327GGG (6253C>G 6254A>G), AAT328AAC (6258T>C), GCA329CCA (6259G>C), ATT330ATG (6264T>G), AGC332AAT (6269G>A 6270C>T), TAT333CAT (6271T>C), TCA335_AGG336del (6277_6283delTCAAGGG), GAA337-AC (6277_6283delTCAAGGG 6285A>C), ACA339ATA (6290C>T), TCA343GCA (6301T>G), TGC345TCA (6308G>C 6309C>A), AGA346AAA (6311G>A), GAT347GAA (6315T>A), ATA348ATC (6318A>C), AGA349CGC (6319A>C 6321A>C), GCA351AGA (6325G>A 6326C>G), CTA356GCA (6340C>G 6341T>C), GTC360ATA (6352G>A 6354C>A), TTA361ATG (6355T>A 6357A>G), AGT362ACA (6359G>C 6360T>A), CTT363TTG (6361C>T 6363T>G), AAA365AAT (6369A>T), GAA367GAG (6375A>G), CAA368AAA (6376C>A), CCT369CAA (6380C>A 6381T>A), ACA370CCT (6382A>C 6384A>T), ACC371ATT (6386C>T 6387C>T), ACA372ATA (6389C>T), AGG376AAA (6401G>A 6402G>A), AGG377AAT (6404G>A 6405G>T), AAT378AGA (6407A>G 6408T>A), TTC379TTT (6411C>T), ATC380ATT (6414C>T), TCC381TCG (6417C>G), CCA382GAC (6418C>G 6419C>A 6420A>C), GAT383GAA (6423T>A), CTG384TTA (6424C>T 6426G>A), TTA385CTA (6427T>C), CTA391CAA (6446T>A), AGT393GGG (6451A>G 6453T>G), CAC394CAA (6456C>A), TAT396TAC (6462T>C), CAC399CAT (6471C>T), TGC401TGT (6477C>T), TCA402TCG (6480A>G), AAA405AAC (6489A>C), ATT410GTA (6502A>G 6504T>A), GTT411ATC (6505G>A 6507T>C), GAC413GAT (6513C>T), CTG416CTA (6522G>A), GAA417GAC (6525A>C), TAA418GAA (6526T>G) | | | | | | | | |

**Proteins**

| | Begin | End | Coverage | Score | Concordance | Matches | Identities | I/D/M/F* | Stop Codons |
|---|---|---|---|---|---|---|---|---|---|
| putative cell-to-cell movement protein (NP_569140.1) | 50 | 174 | 29.8% | 632 | 73.7% | 125 (100%) | 88 (70.4%) | 0/0/1/0 | 1 |
| Protein mutations: | I52L (2308A>T), K53N (2313A>C), N54Y (2314A>T 2316T>C), Q59P (2330A>C), K60V (2332A>G 2333A>T), V65I (2347G>A), V66I (2350G>A 2352T>A), Q68* (2356C>T), I72V (2368A>G 2370A>G), G79K (2389G>A 2390G>A 2391A>R), K81R (2396A>G 2397G>A), A82V (2399C>T 2400C>T), K83R (2402A>G), I86L (2410A>C), N88S (2417A>G 2418C>T), K89R (2420A>G), I91S (2425A>T 2426T>C), K94Q (2434A>C), Q97K (2443C>A 2445A>G), D98K (2446G>A 2448T>G), A101P (2455G>C 2457A>T), N103E (2461A>G 2463C>A), G112A (2489G>C), I117V (2503A>G), I131L (2545A>C), A134V (2555C>T), I139L (2569A>T 2571C>A), Q140E (2572C>G), I146V (2590A>G), S148T (2597G>C 2598T>A), I150V (2602A>G), K151R (2606A>G), G152E (2609G>A), I162V (2638A>G), A165P (2647G>C 2649C>A) | | | | | | | | |
| Codon mutations: | ATA52TTA (2308A>T), AAA53AAC (2313A>C), AAT54TAC (2314A>T 2316T>C), TTC56TTT (2322C>T), AGA58AGG (2328A>G), CAA59CCA (2330A>C), AAA60GTA (2332A>G 2333A>T), TAC63TAT (2343C>T), GTA65ATA (2347G>A), GTT66ATA (2350G>A 2352T>A), AGC67AGT (2355C>T), CAA68TAA (2356C>T), ATA72GTG (2368A>G 2370A>G), ATT73ATA (2373T>A), GAT74GAY (2376T>Y), TGC75TGY (2379C>Y), TAT77YWY (2383T>Y 2384A>W 2385T>Y), ACA78ACY (2388A>Y), GGA79AAR (2389G>A 2390G>A 2391A>R), GGT80GGA (2394T>A), AAG81AGA (2396A>G 2397G>A), GCC82GTT (2399C>T 2400C>T), AAA83AGA (2402A>G), ATA86CTA (2410A>C), AAC88AGT (2417A>G 2418C>T), AAA89AGA (2420A>G), AGA90CGA (2422A>C), ATA91TCA (2425A>T 2426T>C), ATA92ATC (2430A>C), AAA94CAA (2434A>C), CAA97AAG (2443C>A 2445A>G), GAT98AAG (2446G>A 2448T>G), AAG100AAA (2454G>A), GCA101CCT (2455G>C 2457A>T), AAC103GAA (2461A>G 2463C>A), ATT105ATA (2469T>A), AAG106AAA (2472G>A), CAC109CAT (2481C>T), GGA112GCA (2489G>C), ACT113ACC (2493T>C), TTA116CTA (2500T>C), ATA117GTA (2503A>G), ACC127ACA (2535C>A), CCT128CCA (2538T>A), ATA131CTA (2545A>C), CTA133TTA (2551C>T), GCA134GTA (2555C>T), GAT135GAC (2559T>C), GAT136GAC (2562T>C), AGA137AGG (2565A>G), ATA138ATT (2568A>T), ATC139TTA (2569A>T 2571C>A), CAA140GAA (2572C>G), AGC145AGT (2589C>T), ATA146GTA (2590A>G), ATA147ATC (2595A>C), AGT148ACA (2597G>C 2598T>A), ATA150GTA (2602A>G), AAA151AGA (2606A>G), GGA152GAA (2609G>A), AAT153AAC (2613T>C), TAT156TAC (2622T>C), AAG158AAA (2628G>A), TTT161TTC (2637T>C), ATA162GTA (2638A>G), ATA163ATC (2643A>C), AGC164AGT (2646C>T), GCC165CCA (2647G>C 2649C>A), CTA171TTA (2665C>T) | | | | | | | | |
| polyprotein (NP_569141.1) | 233 | 596 | 50.5% | 1789 | 79.1% | 321 (100%) | 264 (82.2%) | 0/0/3/0 | 2 |
| Protein mutations: | E236K (4116G>A), N238E (4122A>G 4124T>A), S239G (4125A>G 4127C>T), Y292N (4284T>A 4286T>C), D311E (4343T>A), A318T (4362G>A 4364T>A), P322A (4374C>G), A338E (4423C>A), N348S (4453A>G 4454C>T), R368K (4513G>A), Q370K (4518C>A), D371E (4523T>A), I374L (4530A>T), K379Q (4545A>C), A381I (4551G>A 4552C>T), H382D (4554C>G), T394K (4591C>A 4592C>A), A396S (4596G>T), N413S (4648A>G), T419S (4665A>T), L428S (4692C>T 4693T>C 4694T>A), D429N (4695G>A), N431Q (4701A>C 4703C>A), K436R (4717A>G 4718G>A), Q439* (4725C>T), I487L (4869A>T), Y496C (4897A>G 4898C>T), K503N (4919A>T), T505I (4924C>T 4925C>A), T510A (4938A>G), S512A (4944T>G), N514E (4950A>G 4952T>G), Y515R (4953T>C 4954A>G 4955T>A), G518E (4963G>A), Y523S (4978A>C 4979T>C), N526D (4986A>G), S540T (5028T>A 5030T>A), Y541F (5032A>T 5033T>C), T542S (5034A>T), E543D (5039A>C), P544S (5040C>T 5042A>G), Q545E (5043C>G), E546I (5046G>A 5047A>T), W548* (5054G>A), R552K (5065G>A), F556H (5076T>C 5077T>A), A557S (5079G>T 5081A>G), V570A (5119T>C 5120T>G), T572N (5125C>A 5126C>T), R573K (5128G>A), V576I (5136G>A), R577K (5140G>A), D579N (5145G>A) | | | | | | | | |
| Codon mutations: | AAA232.GA (4105A>G), CAA235CAG (4115A>G), GAA236AAA (4116G>A), AGT237AGC (4121T>C), AAT238GAA (4122A>G 4124T>A), AGC239GGT (4125A>G 4127C>T), AGT243AGC (4139T>C), GGA257GGT (4181A>T), AGG260AGA (4190G>A), GTC262GTT (4196C>T), ATA263ATT (4199A>T), GAC264GAT (4202C>T), TAT275TAC (4235T>C), TAT277TAC (4241T>C), CCC280CCA (4250C>A), AAA282AAG (4256A>G), TTA284CTT (4260T>C 4262A>T), AAG285AAA (4265G>A), TAT292AAC (4284T>A 4286T>C), TAC294TAT (4292C>T), AGT296AGC (4298T>C), GAC299GAT (4307C>T), AAG301AAA (4313G>A), TAT305TAC (4325T>C), CAT306CAC (4328T>C), TTG307CTA (4329T>C 4331G>A), GAG310GAA (4340G>A), GAT311GAA (4343T>A), GCT318ACA (4362G>A 4364T>A), CCA322GCA (4374C>G), GTT330GTA (4400T>A), AAT337AAC (4421T>C), GCA338GAA (4423C>A), GGA340GGT (4430A>T), TAC342TAT (4436C>T), GAC347GAT (4451C>T), AAC348AGT (4453A>G 4454C>T), TGT356TGC (4478T>C), GTA358GTT (4484A>T), CTA364TTA (4500C>T), CTA365TTA (4503C>T), TAT366TAC (4508T>C), AGA368AAA (4513G>A), ACA369ACT (4517A>T), CAA370AAA (4518C>A), GAT371GAA (4523T>A), ATA374TTA (4530A>T), AAA379CAA (4545A>C), TTC380TTT (4550C>T), GCA381ATA (4551G>A 4552C>T), CAT382GAT (4554C>G), AAC386AAT (4568C>T), TCT387TCA (4571T>A), GGT388GGA (4574T>A), AGC390AGT (4580C>T), AGC392AGT (4586C>T), AAA393AAG (4589A>G), ACC394AAA (4591C>A 4592C>A), GCA396TCA (4596G>T), ATT398ATA (4604T>A), GGA407GGG (4631A>G), AAT413AGT (4648A>G), ATG417ATR (4661G>R), CAA418YAA (4662C>Y), ACT419TCT (4665A>T), CAT420CAY (4670T>Y), GTA422RTR (4674G>R 4676A>R), ATC426ATT (4688C>T), CTT428TCA (4692C>T 4693T>C 4694T>A), GAT429AAT (4695G>A), AAC431CAA (4701A>C 4703C>A), ATT432ATA (4706T>A), AAG436AGA (4717A>G 4718G>A), CAA439TAA (4725C>T), AAG483..A (4859G>A), ATT485ATA (4865T>A), AAG486AAA (4868G>A), ATA487TTA (4869A>T), TTG488CTT (4872T>C 4874G>T), TAC496TGT (4897A>G 4898C>T), GAT499GAC (4907T>C), AAC501AAT (4913C>T), AAA503AAT (4919A>T), ACC505ATA (4924C>T 4925C>A), ATT507ATA (4931T>A), ACA510GCA (4938A>G), TCA512GCA (4944T>G), AAT514GAG (4950A>G 4952T>G), TAT515CGA (4953T>C 4954A>G 4955T>A), TAT517TAC (4961T>C), GGA518GAA (4963G>A), GTT520GTA (4970T>A), CTT521TTA (4971C>T 4973T>A), AAG522AAA (4976G>A), TAT523TCC (4978A>C 4979T>C), AGA524AGG (4982A>G), AAT526GAT (4986A>G), TGC534TGT (5012C>T), TAC536TAT (5018C>T), GGA539GGT (5027A>T), TCT540ACA (5028T>A 5030T>A), TAT541TTC (5032A>T 5033T>C), ACA542TCA (5034A>T), GAA543GAC (5039A>C), CCA544TCG (5040C>T 5042A>G), CAA545GAA (5043C>G), GAA546ATA (5046G>A 5047A>T), TGG548TGA (5054G>A), GAA549GAG (5057A>G), AGA552AAA (5065G>A), TTT556CAT (5076T>C 5077T>A), GCA557TCG (5079G>T 5081A>G), CTA562TTA (5094C>T), TTA563CTA (5097T>C), GCA564GCC (5102A>C), ATA569ATT (5117A>T), GTT570GCG (5119T>C 5120T>G), ACC572AAT (5125C>A 5126C>T), AGA573AAA (5128G>A), TTT574TTC (5132T>C), ATT575ATA (5135T>A), GTA576ATA (5136G>A), AGA577AAA (5140G>A), GAC579AAC (5145G>A), CAA582CAG (5156A>G), ACT588ACA (5174T>A), AAA590AAG (5180A>G), GAT593GAC (5189T>C), GTC595GTT (5195C>T) | | | | | | | | |
| putative transactivator factor (NP_569142.1) | 167 | 418 | 57.2% | 1002 | 60.7% | 237 (99.2%) | 155 (64.9%) | 0/2/1/1 | 0 |
| Protein mutations: | N171T (5786A>C 5787T>A), F172Y (5789T>A), K175L (5797A>T 5798A>T), N176K (5802T>A), R178K (5807G>A), Q180N (5812C>A 5814A>T), S181I (5815T>A 5816C>T 5817T>C), Q183K (5821C>A), N184E (5824A>G 5826T>A), N186T (5831A>C 5832C>A), E187Q (5833G>C 5835A>G), H192Q (5850C>A), H193K (5851C>A 5853T>A), T195Q (5857A>C 5858C>A), L203Q (5881T>C 5882T>A), N205K (5889T>G), S207N (5894G>A), K209N (5901A>C), A212K (5908G>A 5909C>A 5910C>A), N216S (5921A>G 5922C>T), D221S (5935G>A 5936A>G), Q226Y (5950C>T 5952G>C), T227D (5953A>G 5954C>A 5955A>T), Q229E (5959C>G), I231L (5965A>T), A232T (5968G>A 5970C>A), N233T (5972A>C), M242T (5999T>C), Q243T (6001C>A 6002A>C 6003A>C), I247V (6013A>G 6015A>T), T251N (6026C>A 6027A>T), V255I (6037G>A), R256K (6040C>A 6041G>A), S259R (6049T>A 6050C>G 6051G>A), T261P (6055A>C), E267D (6075A>T), I272V (6088A>G 6090C>G), E283D (6123A>T), M284V (6124A>G), K289E (6139A>G), E291V (6146A>T 6147A>G), E294P (6154G>C 6155A>C), E297P (6163G>C 6164A>C), K298R (6167A>G), V299T (6169G>A 6170T>C 6171A>T), N300E (6172A>G 6174T>G), L325T (6247C>A 6248T>C), Q327G (6253C>G 6254A>G), A329P (6259G>C), I330M (6264T>G), S332N (6269G>A 6270C>T), Y333H (6271T>C), S335_R336del (6277_6283delTCAAGGG), T339I (6290C>T), S343A (6301T>G), C345S (6308G>C 6309C>A), R346K (6311G>A), D347E (6315T>A), A351R (6325G>A 6326C>G), L356A (6340C>G 6341T>C), V360I (6352G>A 6354C>A), L361M (6355T>A 6357A>G), S362T (6359G>C 6360T>A), K365N (6369A>T), Q368K (6376C>A), P369Q (6380C>A 6381T>A), T370P (6382A>C 6384A>T), T371I (6386C>T 6387C>T), T372I (6389C>T), R376K (6401G>A 6402G>A), R377N (6404G>A 6405G>T), N378R (6407A>G 6408T>A), P382D (6418C>G 6419C>A 6420A>C), D383E (6423T>A), L391Q (6446T>A), S393G (6451A>G 6453T>G), H394Q (6456C>A), K405N (6489A>C), I410V (6502A>G 6504T>A), V411I (6505G>A 6507T>C), E417D (6525A>C), *418E (6526T>G) | | | | | | | | |

| | Begin | End | Coverage | Score | Concordance | Matches | Identities | I/D/M/F* | Stop Codons |
|---|---|---|---|---|---|---|---|---|---|
| **NT** | **2300** | **6573** | **27.1%** | **2301** | **55.2%** | **2097 (99.5%)** | **1627 (77.2%)** | **1/9** | |

| | |
|---|---|
| Codon mutations: | ATA166.TT (5772A>T), CAT167CAC (5775T>C), ATA169ATC (5781A>C), CAG170CAA (5784G>A), AAT171ACA (5786A>C 5787T>A), TTT172TAT (5789T>A), AAT174AAC (5796T>C), AAA175TTA (5797A>T 5798A>T), AAT176AAA (5802T>A), CCA177CCC (5805A>C), AGA178AAA (5807G>A), TCC179TCA (5811C>A), CAA180AAT (5812C>A 5814A>T), TCT181ATC (5815T>A 5816C>T 5817T>C), ACC182ACT (5820C>T), CAA183AAA (5821C>A), AAT184GAA (5824A>G 5826T>A), AAC186ACA (5831A>C 5832C>A), GAA187CAG (5833G>C 5835A>G), GAC188GAT (5838C>T), CAC192CAA (5850C>A), CAT193AAA (5851C>A 5853T>A), ACA195CAA (5857A>C 5858C>A), TTG200CTA (5872T>C 5874G>A), ATA201ATT (5877A>T), GCT202GCA (5880T>A), TTA203CAA (5881T>C 5882T>A), AAT205AAG (5889T>G), ACA206ACT (5892A>T), AGT207AAT (5894G>A), GCA208GCT (5898A>T), AAA209AAC (5901A>C), GTA211GTC (5907A>C), GCC212AAA (5908G>A 5909C>A 5910C>A), ACC213ACA (5913C>A), TGC214TGT (5916C>T), AAC216AGT (5921A>G 5922C>T), TAC217TAT (5925C>T), GAT221AGT (5935G>A 5936A>G), ACT222ACC (5940T>C), CAG226TAC (5950C>T 5952G>C), ACA227GAT (5953A>G 5954C>A 5955A>T), CAA229GAA (5959C>G), GAG230GAA (5964G>A), ATA231TTA (5965A>T), GCC232ACA (5968G>A 5970C>A), AAT233ACT (5972A>C), CCT235CCA (5979T>A), GAG236GAA (5982G>A), TAC238TAT (5988C>T), AAG239AAA (5991G>A), TTC241TTT (5997C>T), ATG242ACG (5999T>C), CAA243ACC (6001C>A 6002A>C 6003A>C), TAT244TAC (6006T>C), AAG245AAA (6009G>A), AGG246CGA (6010A>C 6012G>A), ATA247GTT (6013A>G 6015A>T), ACT248ACA (6018T>A), GGG250GGA (6024G>A), ACA251AAT (6026C>A 6027A>T), TTC253TTT (6033C>T), TAC254TAT (6036C>T), GTA255ATA (6037G>A), CGA256AAA (6040C>A 6041G>A), TTC257TTT (6045C>T), TCG259AGA (6049T>A 6050C>G 6051G>A), GCT260GCA (6054T>A), ACA261CCA (6055A>C), CTA265TTA (6067C>T), TAT266TAC (6072T>C), GAA267GAT (6075A>T), GAG268GAA (6078G>A), AAG270AAA (6084G>A), CCT271CCA (6087T>A), ATC272GTG (6088A>G 6090C>G), ATA273ATC (6093A>C), GTT275GTC (6099T>C), ATC276ATT (6102C>T), AAA277AAG (6105A>G), ATT278ATA (6108T>A), CTC280CTG (6114C>G), ACA281ACT (6117A>T), AGG282AGA (6120G>A), GAA283GAT (6123A>T), ATG284GTG (6124A>G), ATC285ATA (6129C>A), AAG289GAG (6139A>G), GAA291GTG (6146A>T 6147A>G), GAA294CCA (6154G>C 6155A>C), GAA297CCA (6163G>C 6164A>C), AAA298AGA (6167A>G), GTA299ACT (6169G>A 6170T>C 6171A>T), AAT300GAG (6172A>G 6174T>G), ATT301ATA (6177T>A), CTA320TTA (6232C>T), TAT324TAC (6246T>C), CTA325ACA (6247C>A 6248T>C), CAG327GGG (6253C>G 6254A>G), AAT328AAC (6258T>C), GCA329CCA (6259G>C), ATT330ATG (6264T>G), AGC332AAT (6269G>A 6270C>T), TAT333CAT (6271T>C), TCA335_AGG336del (6277_6283delTCAAGGG), GAA337-AC (6277_6283delTCAAGGG 6285A>C), ACA339ATA (6290C>T), TCA343GCA (6301T>G), TGC345TCA (6308G>C 6309C>A), AGA346AAA (6311G>A), GAT347GAA (6315T>A), ATA348ATC (6318A>C), AGA349CGC (6319A>C 6321A>C), GCA351AGA (6325G>A 6326C>G), CTA356GCA (6340C>G 6341T>C), GTC360ATA (6352G>A 6354C>A), TTA361ATG (6355T>A 6357A>G), AGT362ACA (6359G>C 6360T>A), CTT363TTG (6361C>T 6363T>G), AAA365AAT (6369A>T), GAA367GAG (6375A>G), CAA368AAA (6376C>A), CCT369CAA (6380C>A 6381T>A), ACA370CCT (6382A>C 6384A>T), ACC371ATT (6386C>T 6387C>T), ACA372ATA (6389C>T), AGG376AAA (6401G>A 6402G>A), AGG377AAT (6404G>A 6405G>T), AAT378AGA (6407A>G 6408T>A), TTC379TTT (6411C>T), ATC380ATT (6414C>T), TCC381TCG (6417C>G), CCA382GAC (6418C>G 6419C>A 6420A>C), GAT383GAA (6423T>A), CTG384TTA (6424C>T 6426G>A), TTA385CTA (6427T>C), CTA391CAA (6446T>A), AGT393GGG (6451A>G 6453T>G), CAC394CAA (6456C>A), TAT396TAC (6462T>C), CAC399CAT (6471C>T), TGC401TGT (6477C>T), TCA402TCG (6480A>G), AAA405AAC (6489A>C), ATT410GTA (6502A>G 6504T>A), GTT411ATC (6505G>A 6507T>C), GAC413GAT (6513C>T), CTG416CTA (6522G>A), GAA417GAC (6525A>C), TAA418GAA (6526T>G) |

*: Inserts / Deletes / Misaligned / Frameshifts

#### Analysis details

This analysis was performed with panviral2.64

### NGS Details (UN70): Duamitovirus soch1

#### Assembly

| | |
|---|---|
| **Coverage Length** | 2016 (7 contig(s)) |
| **Depth Of Coverage** | 11.7 |
| **Number Of Reads** | 210 |
| **Reads Per Million** | 3.98 rpm (after QC) |
| **Ambiguities** | 0 |
| **Assembly Method** | de novo + reference guided assembly |
| **Consensus Caller** | Bcf Tools |

#### Coverage Map

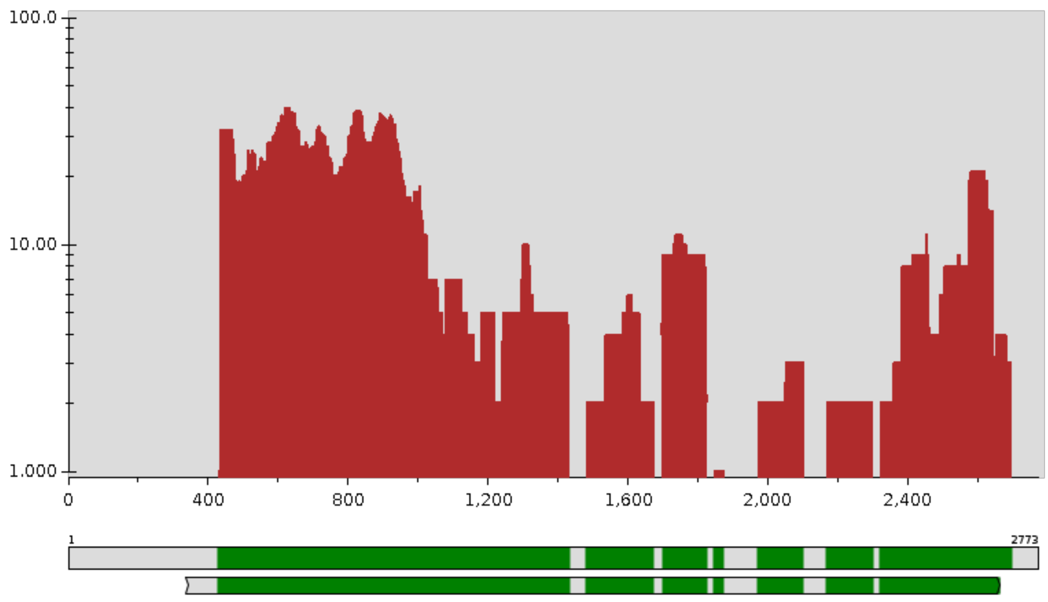

#### Assignment

| | |
|---|---|
| **Type** | Duamitovirus soch1 (Taxonomy ID: 2955838) |
| **Reference Genome** | NC_076524.1 |
| **NT Identity (%)** | 69.4071 |
| **AA Identity (%)** | 69.863 |
| **Number Of Stop Codons** | 3 |
| **Number Of CDS** | 1 |

#### Alignment

| | |
|---|---|
| **Alignment Score** | 1522.0 (NT) + 3215.0 (AA) = 4737.0 |
| **Concordance (%)** | 56.1056 |

| Alignment Method | Global, seeded, nucleotide + amino acids (AGA) |
|---|---|

#### Genome Region

Sequence starts at position 428 and ends at position 2700 relative to NC_076524.1 reference sequence.

#### Alignment Detailed Statistics

| | Begin | End | Coverage | Score | Concordance | Matches | Identities | I/D/M/F* | Stop Codons |
|---|---|---|---|---|---|---|---|---|---|
| **NT** | **428** | **2700** | **72.7%** | **1522** | **38.3%** | **2007 (99.6%)** | **1393 (69.1%)** | **0/9** | |
| Mutations: | 437A>G, 441A>C, 443C>T, 447C>T, 459C>T, 466C>T, 470C>T, 471G>A, 472C>T, 473T>G, 477T>A, 480A>C, 481C>A, 482A>T, 485A>G, 489A>G, 494T>C, 497T>A, 500T>G, 503A>C, 506T>C, 509C>T, 512T>C, 513G>A, 516T>A, 517C>A, 518T>A, 520A>C, 524G>T, 525A>G, 526C>T, 527T>G, 529C>A, 530C>A, 531T>C, 533A>G, 534C>A, 535G>A, 536C>A, 539G>A, 540A>C, 541A>G, 542G>T, 545G>A, 548C>T, 550A>T, 556T>A, 557C>T, 560A>T, 569C>T, 575G>A, 581C>T, 582G>A, 587A>T, 589G>C, 591T>C, 593A>T, 599A>T, 602C>T, 612T>A, 613C>G, 614C>T, 616T>A, 618C>T, 619G>C, 620A>T, 624G>A, 626A>T, 629A>C, 630C>A, 632A>C, 633C>A, 635T>G, 638G>T, 641G>T, 644T>A, 647G>T, 651C>T, 653C>G, 656A>C, 657A>C, 661C>G, 662G>T, 665T>A, 668C>T, 672C>A, 673G>A, 674T>G, 680C>A, 683C>T, 684A>G, 685A>C, 686A>C, 692G>A, 695T>A, 699G>C, 700C>A, 701A>C, 702A>G, 705C>A, 707T>G, 708A>G, 709G>C, 710G>A, 713A>G, 714G>T, 715A>C, 719A>T, 722C>T, 737T>C, 746C>T, 752A>T, 761G>A, 765G>T, 767T>A, 770G>A, 771G>T, 772T>A, 773T>G, 774T>A, 776G>T, 777A>G, 778A>C, 779A>G, 782T>G, 796G>A, 797T>G, 803A>C, 809T>C, 810G>C, 815C>A, 818A>T, 821T>G, 824C>T, 827A>T, 831C>T, 833T>A, 841G>A, 842G>T, 850T>G, 851A>T, 857G>T, 860G>T, 862A>G, 869T>G, 875G>T, 880T>G, 881C>A, 885C>A, 887T>G, 892A>G, 905A>T, 908C>A, 909T>C, 910T>A, 916G>A, 918G>T, 919G>C, 923T>A, 926A>T, 927A>C, 929G>T, 934T>A, 935T>G, 938G>T, 942T>A, 943C>A, 948C>A, 971C>A, 972C>T, 975A>C, 976A>T, 977T>C, 983T>C, 990A>C, 992G>C, 993T>C, 994C>G, 995A>G, 997A>T, 998C>T, 999G>C, 1000G>A, 1001T>A, 1003T>A, 1004A>T, 1005G>T, 1007C>G, 1010A>G, 1016C>T, 1017A>G, 1018A>T, 1019A>T, 1022T>A, 1025T>C, 1026C>G, 1027A>G, 1030A>C, 1031G>A, 1034T>G, 1035A>G, 1036G>C, 1044T>G, 1046C>T, 1049G>A, 1050T>G, 1051T>A, 1052T>A, 1054C>A, 1055A>C, 1061T>C, 1067T>G, 1073G>A, 1076A>G, 1083C>A, 1088G>T, 1091C>G, 1099G>T, 1104A>G, 1105C>T, 1106A>G, 1115T>G, 1121A>C, 1122T>A, 1124C>T, 1127G>A, 1130C>T, 1133C>G, 1142A>C, 1145C>T, 1148G>T, 1151G>T, 1152T>C, 1154G>T, 1160C>T, 1161A>C, 1164T>A, 1165C>T, 1166T>G, 1169C>T, 1172C>A, 1174T>A, 1175T>C, 1178T>A, 1181A>T, 1184C>T, 1185A>T, 1186G>C, 1187G>T, 1188G>A, 1189C>A, 1191G>A, 1192C>A, 1193A>T, 1196T>C, 1197_1199delGAA, 1205A>T, 1211C>T, 1212T>G, 1213T>A, 1217T>C, 1220C>T, 1223A>T, 1224C>T, 1225A>T, 1226C>A, 1232G>A, 1236A>C, 1237G>A, 1238T>A, 1241C>A, 1247A>C, 1248T>C, 1249G>T, 1250G>T, 1253T>C, 1256A>T, 1258C>G, 1265G>A, 1266A>G, 1268T>G, 1269C>G, 1271T>C, 1274T>C, 1276G>A, 1280G>A, 1283G>T, 1292T>G, 1301G>A, 1302C>T, 1304T>G, 1306G>C, 1308G>C, 1309C>A, 1310A>G, 1312T>C, 1313G>C, 1316A>T, 1319G>A, 1322C>T, 1328A>G, 1332A>C, 1334A>G, 1338G>A, 1340A>C, 1345T>C, 1346A>T, 1349G>A, 1355C>T, 1357A>C, 1358C>T, 1376C>G, 1379C>T, 1382T>C, 1385T>C, 1388C>T, 1389G>A, 1390A>T, 1403A>T, 1406C>A, 1409G>A, 1412T>A, 1414A>G, 1415A>T, 1416A>C, 1418A>T, 1424A>T, 1427C>T, 1430G>T, 1433T>A, 1481A>T, 1486G>C, 1489A>T, 1490T>C, 1502A>C, 1505G>A, 1508A>G, 1526G>A, 1533G>T, 1538G>A, 1541T>C, 1545C>T, 1547T>C, 1553T>C, 1554A>T, 1555T>G, 1556G>C, 1559A>G, 1566C>A, 1567C>G, 1571C>A, 1572T>C, 1580C>A, 1589T>C, 1593_1597delAGCTC, 1598T>A, 1607T>C, 1613G>T, 1614A>T, 1617C>T, 1619T>G, 1620G>A, 1622T>C, 1623A>G, 1624A>T, 1626A>C, 1627A>C, 1628A>T, 1632T>A, 1634A>C, 1644G>A, 1664G>A, 1670G>A, 1676T>A, 1700A>C, 1703T>A, 1706C>T, 1707T>G, 1710T>C, 1712G>T, 1715T>C, 1721C>T, 1727A>G, 1734T>C, 1739A>T, 1744G>A, 1745C>A, 1746A>T, 1748A>T, 1754T>C, 1758G>A, 1759G>C, 1760A>G, 1761T>A, 1765A>C, 1766C>T, 1767T>A, 1769T>C, 1772A>T, 1777G>C, 1778G>T, 1787T>C, 1788T>C, 1790A>G, 1791C>T, 1819G>C, 1820T>A, 1821A>G, 1823A>G, 1856A>G, 1859C>T, 1860A>G, 1861A>G, 1865C>T, 1867A>C, 1868A>C, 1874G>T, 1875A>T, 1972G>A, 1973A>G, 1976T>A, 1980T>A, 1981T>C, 1984C>T, 1985T>C, 1991T>C, 2000T>G, 2003A>T, 2006T>G, 2010G>A, 2011C>A, 2013C>A, 2015C>T, 2020T>C, 2030T>C, 2036C>T, 2039A>G, 2052A>C, 2054A>G, 2060G>T, 2063T>A, 2066C>T, 2071A>T, 2074A>G, 2083C>A, 2084G>C, 2087C>T, 2088T>C, 2090A>T, 2093C>T, 2102A>G, 2103A>T, 2104G>C, 2105C>A, 2168C>A, 2173G>T, 2174G>T, 2186G>A, 2191T>A, 2192G>A, 2195C>T, 2204C>T, 2208T>C, 2210G>A, 2213A>G, 2225A>G, 2230A>T, 2231C>T, 2234C>T, 2237T>G, 2239A>G, 2240A>G, 2243A>G, 2250G>C, 2251T>C, 2252T>C, 2264G>T, 2273A>T, 2274A>G, 2276G>A, 2280C>A, 2285G>A, 2288C>T, 2322C>T, 2326G>T, 2328C>A, 2330A>T, 2336A>G, 2337G>A, 2346T>C, 2348A>T, 2349A>C, 2350A>G, 2355C>G, 2357C>T, 2358A>T, 2359A>C, 2360C>T, 2369T>C, 2372G>T, 2377G>A, 2378T>G, 2382A>T, 2383A>C, 2384C>T, 2385C>T, 2386C>T, 2387A>T, 2390T>A, 2396C>T, 2399T>C, 2400A>G, 2402G>A, 2403A>C, 2404G>A, 2405T>A, 2406C>T, 2408C>G, 2411C>G, 2414C>T, 2415T>G, 2417C>T, 2420T>C, 2423A>T, 2431C>A, 2432A>G, 2435C>A, 2441A>G, 2442A>C, 2444G>A, 2445C>T, 2446T>C, 2447C>G, 2448G>A, 2450A>T, 2451A>G, 2452C>A, 2453T>A, 2456T>C, 2457T>C, 2459T>G, 2462T>A, 2465A>G, 2466A>G, 2477C>T, 2478C>T, 2489A>G, 2492T>C, 2495C>T, 2504T>A, 2510T>C, 2514A>G, 2518C>T, 2519C>G, 2524C>T, 2525C>T, 2528T>A, 2532C>A, 2533A>G, 2536G>A, 2537G>T, 2543T>G, 2551A>G, 2552C>T, 2555T>C, 2556G>T, 2557T>C, 2561T>C, 2568A>G, 2571A>C, 2572A>G, 2582A>T, 2585G>A, 2597G>A, 2598A>G, 2599A>C, 2600C>T, 2609T>A, 2613T>G, 2615T>A, 2618T>A, 2622G>T, 2624A>T, 2626A>T, 2627C>A, 2629A>C, 2631A>T, 2634A>C, 2635C>T, 2638G>T, 2639A>G, 2644A>G, 2646G>T, 2647A>G, 2648A>T, 2649G>A, 2650T>G, 2652G>T, 2653G>C, 2654C>A, 2656T>A, 2657A>T, 2658A>G, 2659G>A, 2660A>C, 2661C>T, 2664T>A, 2665A>G, 2667delG, 2674T>A, 2678C>T, 2681G>A, 2686T>A, 2689T>C, 2690G>C, 2694A>G, 2695T>A | | | | | | | | |

##### CDS

| | Begin | End | Coverage | Score | Concordance | Matches | Identities | I/D/M/F* | Stop Codons |
|---|---|---|---|---|---|---|---|---|---|
| RdRp | 32 | 777 | 84.8% | 3215 | 69.2% | 657 (99.7%) | 459 (69.7%) | 0/2/1/1 | 3 |
| Protein mutations: | I36L (441A>C 443C>T), L38F (447C>T), A44V (466C>T), A46M (471G>A 472C>T 473T>G), L48M (477T>A), T49H (480A>C 481C>A 482A>T), I52V (489A>G), V60I (513G>A), S61K (516T>A 517C>A 518T>A), K62T (520A>C), T64V (525A>G 526C>T 527T>G), T65K (529C>A 530C>A), R67K (534C>A 535G>A 536C>A), K69R (540A>C 541A>G 542G>T), Y72F (550A>T), F74Y (556T>A 557C>T), A83T (582G>A), C85S (589G>C), F94Y (616T>A), R95S (618C>T 619G>C 620A>T), E97N (624G>A 626A>T), E98D (629A>C), P99T (630C>A 632A>C), L100M (633C>A 635T>G), S109C (661C>G 662G>T), R113K (672C>A 673G>A 674T>G), K117A (684A>G 685A>C 686A>C), A122H (699G>C 700C>A 701A>C), I123V (702A>G), R125A (708A>G 709G>C 710G>A), D127S (714G>T 715A>C), E128D (719A>T), A144S (765G>T 767T>A), V146* (771G>T 772T>A 773T>G), L147I (774T>A 776G>T), K148A (777A>G 778A>C 779A>G), S154K (796G>A 797T>G), E159Q (810G>C), P166S (831C>T 833T>A), G169D (841G>A 842G>T), L172R (850T>G 851A>T), E176G (862A>G), I178M (869T>G), F182* (880T>G 881C>A), H184K (885C>A 887T>G), Q186R (892A>G), F192H (909T>C 910T>A), R194Q (916G>A), G195S (918G>T 919G>C), M198L (927A>C 929G>T), I200K (934T>A 935T>G), S203N (942T>A 943C>A), Q205K (948C>A), N214L (975A>C 976A>T 977T>C), S220R (993T>C 994C>G 995A>G), Y221F (997A>T 998C>T), G222Q (999G>C 1000G>A 1001T>A), L223Y (1003T>A 1004A>T), A224S (1005G>T 1007C>G), K228V (1017A>G 1018A>T 1019A>T), Q231G (1026C>G 1027A>G), E232A (1030A>C 1031G>A), S234A (1035A>G 1036G>C), F237V (1044T>G 1046C>T), F239E (1050T>G 1051T>A 1052T>A), P240H (1054C>A 1055A>C), L250I (1083C>A), W255L (1099G>T), T257V (1104A>G 1105C>T 1106A>G), I260M (1115T>G), F263I (1122T>A 1124C>T), M272I (1151G>T), K276Q (1161A>C), S277M (1164T>A 1165C>T 1166T>G), F279L (1172C>A), F280Y (1174T>A 1175T>C), L282F (1181A>T), R284S (1185A>T 1186G>C 1187G>T), A285N (1188G>A 1189C>A), A286N (1191G>A 1192C>A 1193A>T), E288del (1197_1199delGAA), L290F (1205A>T), L293E (1212T>G 1213T>A), E296D (1223A>T), H297L (1224C>T 1225A>T 1226C>A), S301Q (1236A>C 1237G>A 1238T>A), W305L (1248T>C 1249G>T 1250G>T), T308S (1258C>G), S311G (1266A>G 1268T>G), P312A (1269C>G 1271T>C), R314Q (1276G>A), G324A (1306G>C), A325Q (1308G>C 1309C>A 1310A>G), V326A (1312T>C 1313G>C), V335I (1338G>A 1340A>C), V337A (1345T>C 1346A>T), M338I (1349G>A), Y341S (1357A>C 1358C>T), D352I (1389G>A 1390A>T), K360S (1414A>G 1415A>T), I361L (1416A>C 1418A>T), C384S (1486G>C), Y385F (1489A>T 1490T>C), L389F (1502A>C), V400F (1533G>T), M401I (1538G>A), L404F (1545C>T 1547T>C), M407C (1554A>T 1555T>G 1556G>C), I408M (1559A>G), P411S (1566C>A 1567C>G), S420del (1593_1597delAGCTC), I427F (1614A>T), V429I (1620G>A 1622T>C), N430V (1623A>G 1624A>T), K431P (1626A>C 1627A>C 1628A>T), L433I (1632T>A 1634A>C), V437M (1644G>A), S458A (1707T>G), I464M (1727A>G), S470K (1744G>A 1745C>A), K471Y (1746A>T 1748A>T), G475T (1758G>A 1759G>C 1760A>G), C476S (1761T>A), N477T (1765A>C 1766C>T), S478T (1767T>A 1769T>C), G481A (1777G>C 1778G>T), S495T (1819G>C 1820T>A), K496E (1821A>G 1823A>G), N509G (1860A>G 1861A>G), K511T (1867A>C 1868A>C), I514L (1875A>T), L549T (1980T>A 1981T>C), S550F (1984C>T 1985T>C), A559N (2010G>A 2011C>A), L560I (2013C>A 2015C>T), I562T (2020T>C), I568M (2039A>G), Y579F (2071A>T), K580R (2074A>G), A583D (2083C>A 2084G>C), W613F (2173G>T 2174G>T), M619K (2191T>A 2192G>A), Y632F (2230A>T 2231C>T), K635R (2239A>G 2240A>G), V639P (2250G>C 2251T>C 2252T>C), Q643H (2264G>T), K647E (2274A>G 2276G>A), L649M (2280C>A), R664L (2326G>T), Q665N (2328C>A 2330A>T), E668K (2337G>A), K672R (2349A>C 2350A>G), L674V (2355C>G 2357C>T), N675S (2358A>T 2359A>C 2360C>T), C681* (2377G>A 2378T>G), N683S (2382A>T 2383A>C 2384C>T), P684F (2385C>T 2386C>T 2387A>T), D685E (2390T>A), K689E (2400A>G 2402G>A), S690Q (2403A>C 2404G>A 2405T>A), I692M (2411C>G), S694A (2415T>G 2417C>T), T699K (2431C>A 2432A>G), L704S (2445C>T 2446T>C 2447C>G), E705N (2448G>A 2450A>T), T706E (2451A>G 2452C>A 2453T>A), F708L (2457T>C 2459T>G), D709E (2462T>A), I711V (2466A>G), N727D (2514A>G), P728L (2518C>T 2519C>G), T730I (2524C>T 2525C>T), Q733R (2532C>A 2533A>G), W734Y (2536G>A 2537G>T), F736L (2543T>G), N739S (2551A>G 2552C>T), V741S (2556G>T 2557T>C), N745D (2568A>G), K746R (2571A>C 2572A>G), L749F (2582A>T), N755A (2598A>G 2599A>C 2600C>T), S760A (2613T>G 2615T>A), E763Y (2622G>T 2624A>T), Y764L (2626A>T 2627C>A), Q765P (2629A>C), T766S (2631A>T), T767L (2634A>C 2635C>T), G768V (2638G>T 2639A>G), E770G (2644A>G), E771C (2646G>T 2647A>G 2648A>T), V772R (2649G>A 2650T>G), G773S (2652G>T 2653G>C 2654C>A), L774Y (2656T>A 2657A>T), R775D (2658A>G 2659G>A 2660A>C), L776F (2661C>T), *777R (2664T>A 2665A>G) | | | | | | | | |

| | Begin | End | Coverage | Score | Concordance | Matches | Identities | I/D/M/F* | Stop Codons |
|---|---|---|---|---|---|---|---|---|---|
| **NT** | **428** | **2700** | **72.7%** | **1522** | **38.3%** | **2007 (99.6%)** | **1393 (69.1%)** | **0/9** | |

Codon mutations: AAA34AAG (437A>G), ATC36CTT (441A>C 443C>T), CTT38TTT (447C>T), CTA42TTA (459C>T), GCC44GTC (466C>T), TCC45TCT (470C>T), GCT46ATG (471G>A 472C>T 473T>G), TTG48ATG (477T>A), ACA49CAT (480A>C 481C>A 482A>T), AAA50AAG (485A>G), ATC52GTC (489A>G), GCT53GCC (494T>C), CTT54CTA (497T>A), GCT55GCG (500T>G), GGA56GGC (503A>C), TGT57TGC (506T>C), GAC58GAT (509C>T), TTT59TTC (512T>C), GTA60ATA (513G>A), TCT61AAA (516T>A 517C>A 518T>A), AAA62ACA (520A>C), GTG63GTT (524G>T), ACT64GTG (525A>G 526C>T 527T>G), ACC65AAA (529C>A 530C>A), TTA66CTG (531T>C 533A>G), CGC67AAA (534C>A 535G>A 536C>A), CGG68CGA (539G>A), AAG69CGT (540A>C 541A>G 542G>T), TCG70TCA (545G>A), GGC71GGT (548C>T), TAT72TTT (550A>T), TTC74TAT (556T>A 557C>T), ACA75ACT (560A>T), TAC78TAT (569C>T), AAG80AAA (575G>A), TGC82TGT (581C>T), GCA83ACA (582G>A), GTA84GTT (587A>T), TGC85TCC (589G>C), TTA86CTT (591T>C 593A>T), CGA88CGT (599A>T), TAC89TAT (602C>T), TCC93AGT (612T>A 613C>G 614C>T), TTT94TAT (616T>A), CGA95TCT (618C>T 619G>C 620A>T), GAA97AAT (624G>A 626A>T), GAA98GAC (629A>C), CCA99ACC (630C>A 632A>C), CTT100ATG (633C>A 635T>G), TCG101TCT (638G>T), GTG102GTT (641G>T), CCT103CCA (644T>A), GTG104GTT (647G>T), CTC106TTG (651C>T 653C>G), ACA107ACC (656A>C), AGA108CGA (657A>C), TCG109TGT (661C>G 662G>T), GGT110GGA (665T>A), ATC111ATT (668C>T), CGT113AAG (672C>A 673G>A 674T>G), ATC115ATA (680C>A), CCC116CCT (683C>T), AAA117GCC (684A>G 685A>C 686A>C), TTG119TTA (692G>A), CGT120CGA (695T>A), GCA122CAC (699G>C 700C>A 701A>C), ATA123GTA (702A>G), CGT124AGG (705C>A 707T>G), AGG125GCA (708A>G 709G>C 710G>A), AAA126AAG (713A>G), GAT127TCT (714G>T 715A>C), GAA128GAT (719A>T), CAC129CAT (722C>T), GTT134GTC (737T>C), TAC137TAT (746C>T), TCA139TCT (752A>T), GGG142GGA (761G>A), GCT144TCA (765G>T 767T>A), AAG145AAA (770G>A), GTT146TAG (771G>T 772T>A 773T>G), TTG147ATT (774T>A 776G>T), AAA148GCG (777A>G 778A>C 779A>G), GTT149GTG (782T>G), AGT154AAG (796G>A 797T>G), GCA156GCC (803A>C), TTT158TTC (809T>C), GAA159CAA (810G>C), TCC160TCA (815C>A), ATA161ATT (818A>T), ACT162ACG (821T>G), ACC163ACT (824C>T), CCA164CCT (827A>T), CCT166TCA (831C>T 833T>A), GGG169GAT (841G>A 842G>T), CTA172CGT (850T>G 851A>T), GTG174GTT (857G>T), CTG175CTT (860G>T), GAA176GGA (862A>G), ATT178ATG (869T>G), ACG180ACT (875G>T), TTC182TGA (880T>G 881C>A), CAT184AAG (885C>A 887T>G), CAA186CGA (892A>G), CTA190CTT (905A>T), CCC191CCA (908C>A), TTT192CAT (909T>C 910T>A), CGA194CAA (916G>A), GGC195TCC (918G>T 919G>C), ATT196ATA (923T>A), CCA197CCT (926A>T), ATG198CTT (927A>C 929G>T), ATT200AAG (934T>A 935T>G), GGG201GGT (938G>T), TCT203AAT (942T>A 943C>A), CAA205AAA (948C>A), CCC212CCA (971C>A), CTA213TTA (972C>T), AAT214CTC (975A>C 976A>T 977T>C), AAT216AAC (983T>C), AGG219CGC (990A>C 992G>C), TCA220CGG (993T>C 994C>G 995A>G), TAC221TTT (997A>T 998C>T), GGT222CAA (999G>C 1000G>A 1001T>A), TTA223TAT (1003T>A 1004A>T), GCC224TCG (1005G>T 1007C>G), GTA225GTG (1010A>G), GAC227GAT (1016C>T), AAA228GTT (1017A>G 1018A>T 1019A>T), GTT229GTA (1022T>A), GAT230GAC (1025T>C), CAA231GGA (1026C>G 1027A>G), GAG232GCA (1030A>C 1031G>A), GCT233GCG (1034T>G), AGT234GCT (1035A>G 1036G>C), TTC237GTT (1044T>G 1046C>T), AAG238AAA (1049G>A), TTT239GAA (1050T>G 1051T>A 1052T>A), CCA240CAC (1054C>A 1055A>C), ATT242ATC (1061T>C), GTT244GTG (1067T>G), TTG246TTA (1073G>A), AAA247AAG (1076A>G), CTA250ATA (1083C>A), GCG251GCT (1088G>T), GCC252GCG (1091C>G), TGG255TTG (1099G>T), ACA257GTG (1104A>G 1105C>T 1106A>G), ATT260ATG (1115T>G), TCA262TCC (1121A>C), TTC263ATT (1122T>A 1124C>T), CAG264CAA (1127G>A), GAC265GAT (1130C>T), GGC266GGG (1133C>G), TCA269TCC (1142A>C), CCC270CCT (1145C>T), GGG271GGT (1148G>T), ATG272ATT (1151G>T), TTG273CTT (1152T>C 1154G>T), TCC275TCT (1160C>T), AAA276CAA (1161A>C), TCT277ATG (1164T>A 1165C>T 1166T>G), GTC278GTT (1169C>T), TTC279TTA (1172C>A), TTT280TAC (1174T>A 1175T>C), CCT281CCA (1178T>A), TTA282TTT (1181A>T), GAC283GAT (1184C>T), AGG284TCT (1185A>T 1186G>C 1187G>T), GCT285AAT (1188G>A 1189C>A), GCA286AAT (1191G>A 1192C>A 1193A>T), ACT287ACC (1196T>C), GAA288del (1197_1199delGAA), TTA290TTT (1205A>T), AAC292AAT (1211C>T), TTA293GAA (1212T>G 1213T>A), TCT294TCC (1217T>C), CTC295CTT (1220C>T), GAA296GAT (1223A>T), CAC297TTA (1224C>T 1225A>T 1226C>A), GAG299GAA (1232G>A), AGT301CAA (1236A>C 1237G>A 1238T>A), GTC302GTA (1241C>A), CCA304CCC (1247A>C), TGG305CTT (1248T>C 1249G>T 1250G>T), TTT306TTC (1253T>C), TCA307TCT (1256A>T), ACC308AGC (1258C>G), TTG310TTA (1265G>A), AGT311GGG (1266A>G 1268T>G), CCT312GCC (1269C>G 1271T>C), TAT313TAC (1274T>C), CGA314CAA (1276G>A), GGG315GGA (1280G>A), GTG316GTT (1283G>T), GTT319GTG (1292T>G), AGG322AGA (1301G>A), CTT323TTG (1302C>T 1304T>G), GGT324GCT (1306G>C), GCA325CAG (1308G>C 1309C>A 1310A>G), GTG326GCC (1312T>C 1313G>C), ATA327ATT (1316A>T), GAG328GAA (1319G>A), GGC329GGT (1322C>T), GGA331GGG (1328A>G), AGA333CGG (1332A>C 1334A>G), GTA335ATC (1338G>A 1340A>C), GTA337GCT (1345T>C 1346A>T), ATG338ATA (1349G>A), AAC340AAT (1355C>T), TAC341TCT (1357A>C 1358C>T), CTC347CTG (1376C>G), CAC348CAT (1379C>T), CCT349CCC (1382T>C), GTT350GTC (1385T>C), CAC351CAT (1388C>T), GAT352ATT (1389G>A 1390A>T), ACA356ACT (1403A>T), GTC357GTA (1406C>A), CTG358CTA (1409G>A), TCT359TCA (1412T>A), AAA360AGT (1414A>G 1415A>T), ATA361CTT (1416A>C 1418A>T), ACA363ACT (1424A>T), GAC364GAT (1427C>T), GGG365GGT (1430G>T), ACT366ACA (1433T>A), AAA382..T (1481A>T), TGT384TCT (1486G>C), TAT385TTC (1489A>T 1490T>C), TTA389TTC (1502A>C), AAG390AAA (1505G>A), TCA391TCG (1508A>G), CCG397CCA (1526G>A), GTC400TTC (1533G>T), ATG401ATA (1538G>A), TAT402TAC (1541T>C), CTT404TTC (1545C>T 1547T>C), TCT406TCC (1553T>C), ATG407TGC (1554A>T 1555T>G 1556G>C), ATA408ATG (1559A>G), CCT411AGT (1566C>A 1567C>G), ACC412ACA (1571C>A), TTG413CTG (1572T>C), TCC415TCA (1580C>A), GTT418GTC (1589T>C), AGC420del (1593_1597delAGCTC), TCT421--A (1593_1597delAGCTC 1598T>A), CTT424CTC (1607T>C), ACG426ACT (1613G>T), ATC427TTC (1614A>T), CTT428TTG (1617C>T 1619T>G), GTT429ATC (1620G>A 1622T>C), AAC430GTC (1623A>G 1624A>T), AAA431CCT (1626A>C 1627A>C 1628A>T), TTA433ATC (1632T>A 1634A>C), GTG437ATG (1644G>A), CTG443CTA (1664G>A), GGG445GGA (1670G>A), CCT447CCA (1676T>A), GCA455.CC (1700A>C), CTT456CTA (1703T>A), TTC457TTT (1706C>T), TCG458GCG (1707T>G), TTG459CTT (1710T>C 1712G>T), TCT460TCC (1715T>C), CAC462CAT (1721C>T), ATA464ATG (1727A>G), TTA467CTA (1734T>C), GCA468GCT (1739A>T), AGC470AAA (1744G>A 1745C>A), AAA471TAT (1746A>T 1748A>T), TAT473TAC (1754T>C), GGA475ACG (1758G>A 1759G>C 1760A>G), TGT476AGT (1761T>A), AAC477ACT (1765A>C 1766C>T), TCT478ACC (1767T>A 1769T>C), CCA479CCT (1772A>T), GGG481GCT (1777G>C 1778G>T), GCT484GCC (1787T>C), TTA485CTG (1788T>C 1790A>G), CTA486TTA (1791C>T), AGT495ACA (1819G>C 1820T>A), AAA496GAG (1821A>G 1823A>G), AGA507AGG (1856A>G), CTC508CTT (1859C>T), AAT509GGT (1860A>G 1861A>G), GTC510GTT (1865C>T), AAA511ACC (1867A>C 1868A>C), TCG513TCT (1874G>T), ATA514TTA (1875A>T), AGA546.AG (1972G>A 1973A>G), GCT547GCA (1976T>A), TTA549ACA (1980T>A 1981T>C), TCT550TTC (1984C>T 1985T>C), CGT552CGC (1991T>C), GTT555GTG (2000T>G), GGA556GGT (2003A>T), CTT557CTG (2006T>G), GCC559AAC (2010G>A 2011C>A), CTC560ATT (2013C>A 2015C>T), ATC562ACC (2020T>C), GAT565GAC (2030T>C), AAC567AAT (2036C>T), ATA568ATG (2039A>G), AGA573CGG (2052A>C 2054A>G), GGG575GGT (2060G>T), GGT576GGA (2063T>A), GCC577GCT (2066C>T), TAC579TTC (2071A>T), AAG580AGG (2074A>G), GCG583GAC (2083C>A 2084G>C), CGC584CGT (2087C>T), TTA585CTT (2088T>C 2090A>T), TTC586TTT (2093C>T), CAA589CAG (2102A>G), AGC590TCA (2103A>T 2104G>C 2105C>A), TAC611..A (2168C>A), TGG613TTT (2173G>T 2174G>T), AGG617AGA (2186G>A), ATG619AAA (2191T>A 2192G>A), CCC620CCT (2195C>T), CCC623CCT (2204C>T), TTG625CTA (2208T>C 2210G>A), AAA626AAG (2213A>G), GTA630GTG (2225A>G), TAC632TTT (2230A>T 2231C>T), CTC633CTT (2234C>T), CTT634CTG (2237T>G), AAA635AGG (2239A>G 2240A>G), GAA636GAG (2243A>G), GTT639CCC (2250G>C 2251T>C 2252T>C), CAG643CAT (2264G>T), CCA646CCT (2273A>T), AAG647GAA (2274A>G 2276G>A), CTG649ATG (2280C>A), GTG650GTA (2285G>A), TTC651TTT (2288C>T), CTA663TTA (2322C>T), CGC664CTC (2326G>T), CAA665AAT (2328C>A 2330A>T), GTA667GTG (2336A>G), GAA668AAA (2337G>A), TTA671CTT (2346T>C 2348A>T), AAG672CGG (2349A>C 2350A>G), CTC674GTT (2355C>G 2357C>T), AAC675TCT (2358A>T 2359A>C 2360C>T), CAT678CAC (2369T>C), ACG679ACT (2372G>T), TGT681TAG (2377G>A 2378T>G), AAC683TCT (2382A>T 2383A>C 2384C>T), CCA684TTT (2385C>T 2386C>T 2387A>T), GAT685GAA (2390T>A), TCC687TCT (2396C>T), ATT688ATC (2399T>C), AAG689GAA (2400A>G 2402G>A), AGT690CAA (2403A>C 2404G>A 2405T>A), CTC691TTG (2406C>T 2408C>G), ATC692ATG (2411C>G), GAC693GAT (2414C>T), TCC694GCT (2415T>G 2417C>T), CCT695CCC (2420T>C), ATA696ATT (2423A>T), ACA699AAG (2431C>A 2432A>G), TCC700TCA (2435C>A), AAA702AAG (2441A>G), AGG703CGA (2442A>C 2444G>A), CTC704TCG (2445C>T 2446T>C 2447C>G), GAA705AAT (2448G>A 2450A>T), ACT706GAA (2451A>G 2452C>A 2453T>A), GAT707GAC (2456T>C), TTT708CTG (2457T>C 2459T>G), GAT709GAA (2462T>A), TTA710TTG (2465A>G), ATC711GTC (2466A>G), GGC714GGT (2477C>T), CTA715TTA (2478C>T), AAA718AAG (2489A>G), TGT719TGC (2492T>C), TAC720TAT (2495C>T), GGT723GGA (2504T>A), GGT725GGC (2510T>C), AAT727GAT (2514A>G), CCC728CTG (2518C>T 2519C>G), ACC730ATT (2524C>T 2525C>T), ACT731ACA (2528T>A), CAG733AGG (2532C>A 2533A>G), TGG734TAT (2536G>A 2537G>T), TTT736TTG (2543T>G), AAC739AGT (2551A>G 2552C>T), ACT740ACC (2555T>C), GTT741TCT (2556G>T 2557T>C), ATT742ATC (2561T>C), AAT745GAT (2568A>G), AAA746CGA (2571A>C 2572A>G), TTA749TTT (2582A>T), GGG750GGA (2585G>A), GGG754GGA (2597G>A), AAC755GCT (2598A>G 2599A>C 2600C>T), ATT758ATA (2609T>A), TCT760GCA (2613T>G 2615T>A), CCT761CCA (2618T>A), GAA763TAT (2622G>T 2624A>T), TAC764TTA (2626A>T 2627C>A), CAA765CCA (2629A>C), ACT766TCT (2631A>T), ACA767CTA (2634A>C 2635C>T), GGA768GTG (2638G>T 2639A>G), GAG770GGG (2644A>G), GAA771TGT (2646G>T 2647A>G 2648A>T), GTG772AGG (2649G>A 2650T>G), GGC773TCA (2652G>T 2653G>C 2654C>A), TTA774TAT (2656T>A 2657A>T), AGA775GAC (2658A>G 2659G>A 2660A>C), CTT776TTT (2661C>T), TAG777AGG (2664T>A 2665A>G)

#### Proteins

| | Begin | End | Coverage | Score | Concordance | Matches | Identities | I/D/M/F* | Stop Codons |
|---|---|---|---|---|---|---|---|---|---|
| RNA-dependent RNA polymerase (YP_010798874.1) | 32 | 777 | 84.8% | 3215 | 69.2% | 657 (99.7%) | 459 (69.7%) | 0/2/1/1 | 3 |

Protein mutations: I36L (441A>C 443C>T), L38F (447C>T), A44V (466C>T), A46M (471G>A 472C>T 473T>G), L48M (477T>A), T49H (480A>C 481C>A 482A>T), I52V (489A>G), V60I (513G>A), S61K (516T>A 517C>A 518T>A), K62T (520A>C), T64V (525A>G 526C>T 527T>G), T65K (529C>A 530C>A), R67K (534C>A 535G>A 536C>A), K69R (540A>C 541A>G 542G>T), Y72F (550A>T), F74Y (556T>A 557C>T), A83T (582G>A), C85S (589G>C), F94Y (616T>A), R95S (618C>T 619G>C 620A>T), E97N (624G>A 626A>T), E98D (629A>C), P99T (630C>A 632A>C), L100M (633C>A 635T>G), S109C (661C>G 662G>T), R113K (672C>A 673G>A 674T>G), K117A (684A>G 685A>C 686A>C), A122H (699G>C 700C>A 701A>C), I123V (702A>G), R125A (708A>G 709G>C 710G>A), D127S (714G>T 715A>C), E128D (719A>T), A144S (765G>T 767T>A), V146* (771G>T 772T>A 773T>G), L147I (774T>A 776G>T), K148A (777A>G 778A>C 779A>G), S154K (796G>A 797T>G), E159Q (810G>C), P166S (831C>T 833T>A), G169D (841G>A 842G>T), L172R (850T>G 851A>T), E176G (862A>G), I178M (869T>G), F182* (880T>G 881C>A), H184K (885C>A 887T>G), Q186R (892A>G), F192H (909T>C 910T>A), R194Q (916G>A), G195S (918G>T 919G>C), M198L (927A>C 929G>T), I200K (934T>A 935T>G), S203N (942T>A 943C>A), Q205K (948C>A), N214L (975A>C 976A>T 977T>C), S220R (993T>C 994C>G 995A>G), Y221F (997A>T 998C>T), G222Q (999G>C 1000G>A 1001T>A), L223Y (1003T>A 1004A>T), A224S (1005G>T 1007C>G), K228V (1017A>G 1018A>T 1019A>T), Q231G (1026C>G 1027A>G), E232A (1030A>C 1031G>A), S234A (1035A>G 1036G>C), F237V (1044T>G 1046C>T), F239E (1050T>G 1051T>A 1052T>A), P240H (1054C>A 1055A>C), L250I (1083C>A), W255L (1099G>T), T257V (1104A>G 1105C>T 1106A>G), I260M (1115T>G), F263I (1122T>A 1124C>T), M272I (1151G>T), K276Q (1161A>C), S277M (1164T>A 1165C>T 1166T>G), F279L (1172C>A), F280Y (1174T>A 1175T>C), L282F (1181A>T), R284S (1185A>T 1186G>C 1187G>T), A285N (1188G>A 1189C>A), A286N (1191G>A 1192C>A 1193A>T), E288del (1197_1199delGAA), L290F (1205A>T), L293E (1212T>G 1213T>A), E296D (1223A>T), H297L (1224C>T 1225A>T 1226C>A), S301Q (1236A>C 1237G>A 1238T>A), W305L (1248T>C 1249G>T 1250G>T), T308S (1258C>G), S311G (1266A>G 1268T>G), P312A (1269C>G 1271T>C), R314Q (1276G>A), G324A (1306G>C), A325Q (1308G>C 1309C>A 1310A>G), V326A (1312T>C 1313G>C), V335I (1338G>A 1340A>C), V337A (1345T>C 1346A>T), M338I (1349G>A), Y341S (1357A>C 1358C>T), D352I (1389G>A 1390A>T), K360S (1414A>G 1415A>T), I361L (1416A>C 1418A>T), C384S (1486G>C), Y385F (1489A>T 1490T>C), L389F (1502A>C), V400F (1533G>T), M401I (1538G>A), L404F (1545C>T 1547T>C), M407C (1554A>T 1555T>G 1556G>C), I408M (1559A>G), P411S (1566C>A 1567C>G), S420del (1593_1597delAGCTC), I427F (1614A>T), V429I (1620G>A 1622T>C), N430V (1623A>G 1624A>T), K431P (1626A>C 1627A>C 1628A>T), L433I (1632T>A 1634A>C), V437M (1644G>A), S458A (1707T>G), I464M (1727A>G), S470K (1744G>A 1745C>A), K471Y (1746A>T 1748A>T), G475T (1758G>A 1759G>C 1760A>G), C476S (1761T>A), N477T (1765A>C 1766C>T), S478T (1767T>A 1769T>C), G481A (1777G>C 1778G>T), S495T (1819G>C 1820T>A), K496E (1821A>G 1823A>G), N509G (1860A>G 1861A>G), K511T (1867A>C 1868A>C), I514L (1875A>T), L549T (1980T>A 1981T>C), S550F (1984C>T 1985T>C), A559N (2010G>A 2011C>A), L560I (2013C>A 2015C>T), I562T (2020T>C), I568M (2039A>G), Y579F (2071A>T), K580R (2074A>G), A583D (2083C>A 2084G>C), W613F (2173G>T 2174G>T), M619K (2191T>A 2192G>A), Y632F (2230A>T 2231C>T), K635R (2239A>G 2240A>G), V639P (2250G>C 2251T>C 2252T>C), Q643H (2264G>T), K647E (2274A>G 2276G>A), L649M (2280C>A), R664L (2326G>T), Q665N (2328C>A 2330A>T), E668K (2337G>A), K672R (2349A>C 2350A>G), L674V (2355C>G 2357C>T), N675S (2358A>T 2359A>C 2360C>T), C681* (2377G>A 2378T>G), N683S (2382A>T 2383A>C 2384C>T), P684F (2385C>T 2386C>T 2387A>T), D685E (2390T>A), K689E (2400A>G 2402G>A), S690Q (2403A>C 2404G>A 2405T>A), I692M (2411C>G), S694A (2415T>G 2417C>T), T699K (2431C>A 2432A>G), L704S (2445C>T 2446T>C 2447C>G), E705N (2448G>A 2450A>T), T706E (2451A>G 2452C>A 2453T>A), F708L (2457T>C 2459T>G), D709E (2462T>A), I711V (2466A>G), N727D (2514A>G), P728L (2518C>T 2519C>G), T730I (2524C>T 2525C>T), Q733R (2532C>A 2533A>G), W734Y (2536G>A 2537G>T), F736L (2543T>G), N739S (2551A>G 2552C>T), V741S (2556G>T 2557T>C), N745D (2568A>G), K746R (2571A>C 2572A>G), L749F (2582A>T), N755A (2598A>G 2599A>C 2600C>T), S760A (2613T>G 2615T>A), E763Y (2622G>T 2624A>T), Y764L (2626A>T 2627C>A), Q765P (2629A>C), T766S (2631A>T), T767L (2634A>C 2635C>T), G768V (2638G>T 2639A>G), E770G (2644A>G), E771C (2646G>T 2647A>G 2648A>T), V772R (2649G>A 2650T>G), G773S (2652G>T 2653G>C 2654C>A), L774Y (2656T>A 2657A>T), R775D (2658A>G 2659G>A 2660A>C), L776F (2661C>T), *777R (2664T>A 2665A>G)

| | Begin | End | Coverage | Score | Concordance | Matches | Identities | I/D/M/F* | Stop Codons |
|---|---|---|---|---|---|---|---|---|---|
| **NT** | **428** | **2700** | **72.7%** | **1522** | **38.3%** | **2007 (99.6%)** | **1393 (69.1%)** | **0/9** | |

Codon mutations:

AAA34AAG (437A>G), ATC36CTT (441A>C 443C>T), CTT38TTT (447C>T), CTA42TTA (459C>T), GCC44GTC (466C>T), TCC45TCT (470C>T), GCT46ATG (471G>A 472C>T 473T>G), TTG48ATG (477T>A), ACA49CAT (480A>C 481C>A 482A>T), AAA50AAG (485A>G), ATC52GTC (489A>G), GCT53GCC (494T>C), CTT54CTA (497T>A), GCT55GCG (500T>G), GGA56GGC (503A>C), TGT57TGC (506T>C), GAC58GAT (509C>T), TTT59TTC (512T>C), GTA60ATA (513G>A), TCT61AAA (516T>A 517C>A 518T>A), AAA62ACA (520A>C), GTG63GTT (524G>T), ACT64GTG (525A>G 526C>T 527T>G), ACC65AAA (529C>A 530C>A), TTA66CTG (531T>C 533A>G), CGC67AAA (534C>A 535G>A 536C>A), CGG68CGA (539G>A), AAG69CGT (540A>C 541A>G 542G>T), TCG70TCA (545G>A), GGC71GGT (548C>T), TAT72TTT (550A>T), TTC74TAT (556T>A 557C>T), ACA75ACT (560A>T), TAC78TAT (569C>T), AAG80AAA (575G>A), TGC82TGT (581C>T), GCA83ACA (582G>A), GTA84GTT (587A>T), TGC85TCC (589G>C), TTA86CTT (591T>C 593A>T), CGA88CGT (599A>T), TAC89TAT (602C>T), TCC93AGT (612T>A 613C>G 614C>T), TTT94TAT (616T>A), CGA95TCT (618C>T 619G>C 620A>T), GAA97AAT (624G>A 626A>T), GAA98GAC (629A>C), CCA99ACC (630C>A 632A>C), CTT100ATG (633C>A 635T>G), TCG101TCT (638G>T), GTG102GTT (641G>T), CCT103CCA (644T>A), GTG104GTT (647G>T), CTC106TTG (651C>T 653C>G), ACA107ACC (656A>C), AGA108CGA (657A>C), TCG109TGT (661C>G 662G>T), GGT110GGA (665T>A), ATC111ATT (668C>T), CGT113AAG (672C>A 673G>A 674T>G), ATC115ATA (680C>A), CCC116CCT (683C>T), AAA117GCC (684A>G 685A>C 686A>C), TTG119TTA (692G>A), CGT120CGA (695T>A), GCA122CAC (699G>C 700C>A 701A>C), ATA123GTA (702A>G), CGT124AGG (705C>A 707T>G), AGG125GCA (708A>G 709G>C 710G>A), AAA126AAG (713A>G), GAT127TCT (714G>T 715A>C), GAA128GAT (719A>T), CAC129CAT (722C>T), GTT134GTC (737T>C), TAC137TAT (746C>T), TCA139TCT (752A>T), GGG142GGA (761G>A), GCT144TCA (765G>T 767T>A), AAG145AAA (770G>A), GTT146TAG (771G>T 772T>A 773T>G), TTG147ATT (774T>A 776G>T), AAA148GCG (777A>G 778A>C 779A>G), GTT149GTG (782T>G), AGT154AAG (796G>A 797T>G), GCA156GCC (803A>C), TTT158TTC (809T>C), GAA159CAA (810G>C), TCC160TCA (815C>A), ATA161ATT (818A>T), ACT162ACG (821T>G), ACC163ACT (824C>T), CCA164CCT (827A>T), CCT166TCA (831C>T 833T>A), GGG169GAT (841G>A 842G>T), CTA172CGT (850T>G 851A>T), GTG174GTT (857G>T), CTG175CTT (860G>T), GAA176GGA (862A>G), ATT178ATG (869T>G), ACG180ACT (875G>T), TTC182TGA (880T>G 881C>A), CAT184AAG (885C>A 887T>G), CAA186CGA (892A>G), CTA190CTT (905A>T), CCC191CCA (908C>A), TTT192CAT (909T>C 910T>A), CGA194CAA (916G>A), GGC195TCC (918G>T 919G>C), ATT196ATA (923T>A), CCA197CCT (926A>T), ATG198CTT (927A>C 929G>T), ATT200AAG (934T>A 935T>G), GGG201GGT (938G>T), TCT203AAT (942T>A 943C>A), CAA205AAA (948C>A), CCC212CCA (971C>A), CTA213TTA (972C>T), AAT214CTC (975A>C 976A>T 977T>C), AAT216AAC (983T>C), AGG219CGC (990A>C 992G>C), TCA220CGG (993T>C 994C>G 995A>G), TAC221TTT (997A>T 998C>T), GGT222CAA (999G>C 1000G>A 1001T>A), TTA223TAT (1003T>A 1004A>T), GCC224TCG (1005G>T 1007C>G), GTA225GTG (1010A>G), GAC227GAT (1016C>T), AAA228GTT (1017A>G 1018A>T 1019A>T), GTT229GTA (1022T>A), GAT230GAC (1025T>C), CAA231GGA (1026C>G 1027A>G), GAG232GCA (1030A>C 1031G>A), GCT233GCG (1034T>G), AGT234GCT (1035A>G 1036G>C), TTC237GTT (1044T>G 1046C>T), AAG238AAA (1049G>A), TTT239GAA (1050T>G 1051T>A 1052T>A), CCA240CAC (1054C>A 1055A>C), ATT242ATC (1061T>C), GTT244GTG (1067T>G), TTG246TTA (1073G>A), AAA247AAG (1076A>G), CTA250ATA (1083C>A), GCG251GCT (1088G>T), GCC252GCG (1091C>G), TGG255TTG (1099G>T), ACA257GTG (1104A>G 1105C>T 1106A>G), ATT260ATG (1115T>G), TCA262TCC (1121A>C), TTC263ATT (1122T>A 1124C>T), CAG264CAA (1127G>A), GAC265GAT (1130C>T), GGC266GGG (1133C>G), TCA269TCC (1142A>C), CCC270CCT (1145C>T), GGG271GGT (1148G>T), ATG272ATT (1151G>T), TTG273CTT (1152T>C 1154G>T), TCC275TCT (1160C>T), AAA276CAA (1161A>C), TCT277ATG (1164T>A 1165C>T 1166T>G), GTC278GTT (1169C>T), TTC279TTA (1172C>A), TTT280TAC (1174T>A 1175T>C), CCT281CCA (1178T>A), TTA282TTT (1181A>T), GAC283GAT (1184C>T), AGG284TCT (1185A>T 1186G>C 1187G>T), GCT285AAT (1188G>A 1189C>A), GCA286AAT (1191G>A 1192C>A 1193A>T), ACT287ACC (1196T>C), GAA288del (1197_1199delGAA), TTA290TTT (1205A>T), AAC292AAT (1211C>T), TTA293GAA (1212T>G 1213T>A), TCT294TCC (1217T>C), CTC295CTT (1220C>T), GAA296GAT (1223A>T), CAC297TTA (1224C>T 1225A>T 1226C>A), GAG299GAA (1232G>A), AGT301CAA (1236A>C 1237G>A 1238T>A), GTC302GTA (1241C>A), CCA304CCC (1247A>C), TGG305CTT (1248T>C 1249G>T 1250G>T), TTT306TTC (1253T>C), TCA307TCT (1256A>T), ACC308AGC (1258C>G), TTG310TTA (1265G>A), AGT311GGG (1266A>G 1268T>G), CCT312GCC (1269C>G 1271T>C), TAT313TAC (1274T>C), CGA314CAA (1276G>A), GGG315GGA (1280G>A), GTG316GTT (1283G>T), GTT319GTG (1292T>G), AGG322AGA (1301G>A), CTT323TTG (1302C>T 1304T>G), GGT324GCT (1306G>C), GCA325CAG (1308G>C 1309C>A 1310A>G), GTG326GCC (1312T>C 1313G>C), ATA327ATT (1316A>T), GAG328GAA (1319G>A), GGC329GGT (1322C>T), GGA331GGG (1328A>G), AGA333CGG (1332A>C 1334A>G), GTA335ATC (1338G>A 1340A>C), GTA337GCT (1345T>C 1346A>T), ATG338ATA (1349G>A), AAC340AAT (1355C>T), TAC341TCT (1357A>C 1358C>T), CTC347CTG (1376C>G), CAC348CAT (1379C>T), CCT349CCC (1382T>C), GTT350GTC (1385T>C), CAC351CAT (1388C>T), GAT352ATT (1389G>A 1390A>T), ACA356ACT (1403A>T), GTC357GTA (1406C>A), CTG358CTA (1409G>A), TCT359TCA (1412T>A), AAA360AGT (1414A>G 1415A>T), ATA361CTT (1416A>C 1418A>T), ACA363ACT (1424A>T), GAC364GAT (1427C>T), GGG365GGT (1430G>T), ACT366ACA (1433T>A), AAA382..T (1481A>T), TGT384TCT (1486G>C), TAT385TTC (1489A>T 1490T>C), TTA389TTC (1502A>C), AAG390AAA (1505G>A), TCA391TCG (1508A>G), CCG397CCA (1526G>A), GTC400TTC (1533G>T), ATG401ATA (1538G>A), TAT402TAC (1541T>C), CTT404TTC (1545C>T 1547T>C), TCT406TCC (1553T>C), ATG407TGC (1554A>T 1555T>G 1556G>C), ATA408ATG (1559A>G), CCT411AGT (1566C>A 1567C>G), ACC412ACA (1571C>A), TTG413CTG (1572T>C), TCC415TCA (1580C>A), GTT418GTC (1589T>C), AGC420del (1593_1597delAGCTC), TCT421--A (1593_1597delAGCTC 1598T>A), CTT424CTC (1607T>C), ACG426ACT (1613G>T), ATC427TTC (1614A>T), CTT428TTG (1617C>T 1619T>G), GTT429ATC (1620G>A 1622T>C), AAC430GTC (1623A>G 1624A>T), AAA431CCT (1626A>C 1627A>C 1628A>T), TTA433ATC (1632T>A 1634A>C), GTG437ATG (1644G>A), CTG443CTA (1664G>A), GGG445GGA (1670G>A), CCT447CCA (1676T>A), GCA455.CC (1700A>C), CTT456CTA (1703T>A), TTC457TTT (1706C>T), TCG458GCG (1707T>G), TTG459CTT (1710T>C 1712G>T), TCT460TCC (1715T>C), CAC462CAT (1721C>T), ATA464ATG (1727A>G), TTA467CTA (1734T>C), GCA468GCT (1739A>T), AGC470AAA (1744G>A 1745C>A), AAA471TAT (1746A>T 1748A>T), TAT473TAC (1754T>C), GGA475ACG (1758G>A 1759G>C 1760A>G), TGT476AGT (1761T>A), AAC477ACT (1765A>C 1766C>T), TCT478ACC (1767T>A 1769T>C), CCA479CCT (1772A>T), GGG481GCT (1777G>C 1778G>T), GCT484GCC (1787T>C), TTA485CTG (1788T>C 1790A>G), CTA486TTA (1791C>T), AGT495ACA (1819G>C 1820T>A), AAA496GAG (1821A>G 1823A>G), AGA507AGG (1856A>G), CTC508CTT (1859C>T), AAT509GGT (1860A>G 1861A>G), GTC510GTT (1865C>T), AAA511ACC (1867A>C 1868A>C), TCG513TCT (1874G>T), ATA514TTA (1875A>T), AGA546.AG (1972G>A 1973A>G), GCT547GCA (1976T>A), TTA549ACA (1980T>A 1981T>C), TCT550TTC (1984C>T 1985T>C), CGT552CGC (1991T>C), GTT555GTG (2000T>G), GGA556GGT (2003A>T), CTT557CTG (2006T>G), GCC559AAC (2010G>A 2011C>A), CTC560ATT (2013C>A 2015C>T), ATC562ACC (2020T>C), GAT565GAC (2030T>C), AAC567AAT (2036C>T), ATA568ATG (2039A>G), AGA573CGG (2052A>C 2054A>G), GGG575GGT (2060G>T), GGT576GGA (2063T>A), GCC577GCT (2066C>T), TAC579TTC (2071A>T), AAG580AGG (2074A>G), GCG583GAC (2083C>A 2084G>C), CGC584CGT (2087C>T), TTA585CTT (2088T>C 2090A>T), TTC586TTT (2093C>T), CAA589CAG (2102A>G), AGC590TCA (2103A>T 2104G>C 2105C>A), TAC611..A (2168C>A), TGG613TTT (2173G>T 2174G>T), AGG617AGA (2186G>A), ATG619AAA (2191T>A 2192G>A), CCC620CCT (2195C>T), CCC623CCT (2204C>T), TTG625CTA (2208T>C 2210G>A), AAA626AAG (2213A>G), GTA630GTG (2225A>G), TAC632TTT (2230A>T 2231C>T), CTC633CTT (2234C>T), CTT634CTG (2237T>G), AAA635AGG (2239A>G 2240A>G), GAA636GAG (2243A>G), GTT639CCC (2250G>C 2251T>C 2252T>C), CAG643CAT (2264G>T), CCA646CCT (2273A>T), AAG647GAA (2274A>G 2276G>A), CTG649ATG (2280C>A), GTG650GTA (2285G>A), TTC651TTT (2288C>T), CTA663TTA (2322C>T), CGC664CTC (2326G>T), CAA665AAT (2328C>A 2330A>T), GTA667GTG (2336A>G), GAA668AAA (2337G>A), TTA671CTT (2346T>C 2348A>T), AAG672CGG (2349A>C 2350A>G), CTC674GTT (2355C>G 2357C>T), AAC675TCT (2358A>T 2359A>C 2360C>T), CAT678CAC (2369T>C), ACG679ACT (2372G>T), TGT681TAG (2377G>A 2378T>G), AAC683TCT (2382A>T 2383A>C 2384C>T), CCA684TTT (2385C>T 2386C>T 2387A>T), GAT685GAA (2390T>A), TCC687TCT (2396C>T), ATT688ATC (2399T>C), AAG689GAA (2400A>G 2402G>A), AGT690CAA (2403A>C 2404G>A 2405T>A), CTC691TTG (2406C>T 2408C>G), ATC692ATG (2411C>G), GAC693GAT (2414C>T), TCC694GCT (2415T>G 2417C>T), CCT695CCC (2420T>C), ATA696ATT (2423A>T), ACA699AAG (2431C>A 2432A>G), TCC700TCA (2435C>A), AAA702AAG (2441A>G), AGG703CGA (2442A>C 2444G>A), CTC704TCG (2445C>T 2446T>C 2447C>G), GAA705AAT (2448G>A 2450A>T), ACT706GAA (2451A>G 2452C>A 2453T>A), GAT707GAC (2456T>C), TTT708CTG (2457T>C 2459T>G), GAT709GAA (2462T>A), TTA710TTG (2465A>G), ATC711GTC (2466A>G), GGC714GGT (2477C>T), CTA715TTA (2478C>T), AAA718AAG (2489A>G), TGT719TGC (2492T>C), TAC720TAT (2495C>T), GGT723GGA (2504T>A), GGT725GGC (2510T>C), AAT727GAT (2514A>G), CCC728CTG (2518C>T 2519C>G), ACC730ATT (2524C>T 2525C>T), ACT731ACA (2528T>A), CAG733AGG (2532C>A 2533A>G), TGG734TAT (2536G>A 2537G>T), TTT736TTG (2543T>G), AAC739AGT (2551A>G 2552C>T), ACT740ACC (2555T>C), GTT741TCT (2556G>T 2557T>C), ATT742ATC (2561T>C), AAT745GAT (2568A>G), AAA746CGA (2571A>C 2572A>G), TTA749TTT (2582A>T), GGG750GGA (2585G>A), GGG754GGA (2597G>A), AAC755GCT (2598A>G 2599A>C 2600C>T), ATT758ATA (2609T>A), TCT760GCA (2613T>G 2615T>A), CCT761CCA (2618T>A), GAA763TAT (2622G>T 2624A>T), TAC764TTA (2626A>T 2627C>A), CAA765CCA (2629A>C), ACT766TCT (2631A>T), ACA767CTA (2634A>C 2635C>T), GGA768GTG (2638G>T 2639A>G), GAG770GGG (2644A>G), GAA771TGT (2646G>T 2647A>G 2648A>T), GTG772AGG (2649G>A 2650T>G), GGC773TCA (2652G>T 2653G>C 2654C>A), TTA774TAT (2656T>A 2657A>T), AGA775GAC (2658A>G 2659G>A 2660A>C), CTT776TTT (2661C>T), TAG777AGG (2664T>A 2665A>G)

*: Inserts / Deletes / Misaligned / Frameshifts

#### Analysis details

This analysis was performed with panviral2.64

#### NGS Details (UN70): Tomato chocolate spot virus (segment RNA2)

##### Assembly

| | |
|---|---|
| **Coverage Length** | 346 (1 contig(s)) |
| **Depth Of Coverage** | 5827.1 |
| **Number Of Reads** | 19188 |
| **Reads Per Million** | 363.73 rpm (after QC) |
| **Ambiguities** | 0 |
| **Assembly Method** | de novo + reference guided assembly |
| **Consensus Caller** | Bcf Tools |

##### Coverage Map

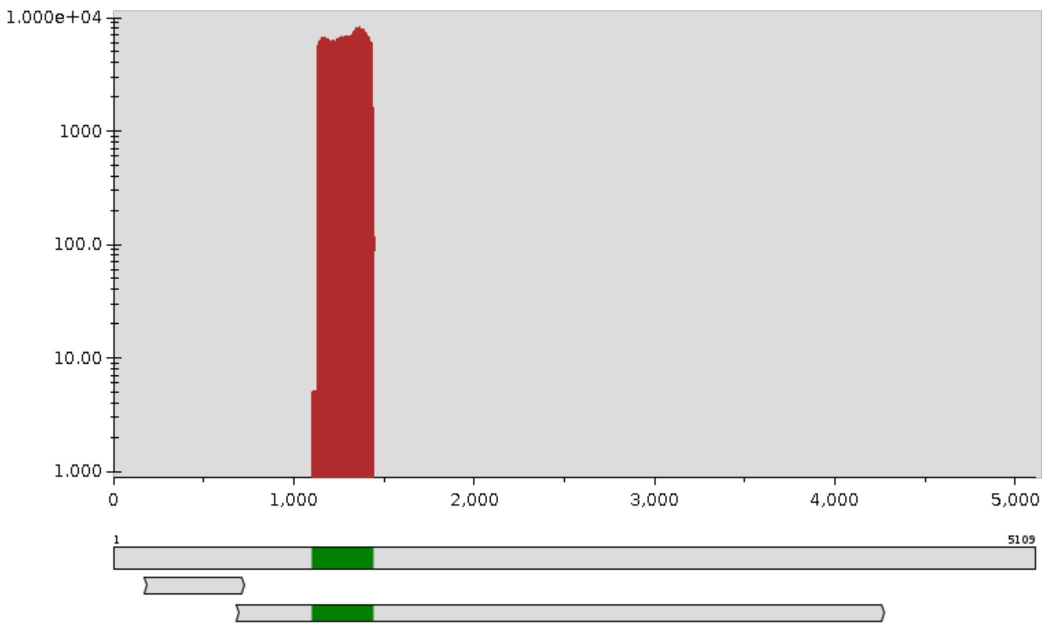

##### Assignment

| | |
|---|---|
| **Type** | Tomato chocolate spot virus (Taxonomy ID: 661101) |
| **Reference Genome** | NC_013076.1 |
| **NT Identity (%)** | 69.6532 |
| **AA Identity (%)** | 75.0 |
| **Number Of Stop Codons** | 1 |
| **Number Of CDS** | 2 |

##### Alignment

| | |
|---|---|
| **Alignment Score** | 272.0 (NT) + 591.0 (AA) = 863.0 |
| **Concordance (%)** | 58.6676 |

| **Alignment Method** | Global, seeded, nucleotide + amino acids (AGA) |
|---|---|

#### Genome Region

Sequence starts at position 1100 and ends at position 1445 relative to NC_013076.1 reference sequence.

#### Alignment Detailed Statistics

| | Begin | End | Coverage | Score | Concordance | Matches | Identities | I/D/M/F* | Stop Codons |
|---|---|---|---|---|---|---|---|---|---|
| **NT** | **1100** | **1445** | **6.8%** | **272** | **39.3%** | **346 (100%)** | **241 (69.7%)** | **0/0** | |
| Mutations: | 1103C>A, 1107C>G, 1108C>T, 1109G>C, 1111G>A, 1112C>T, 1115A>T, 1116T>C, 1117G>T, 1120T>G, 1121C>G, 1124A>T, 1126T>A, 1127A>G, 1128T>A, 1130A>G, 1132A>C, 1133T>C, 1138C>T, 1142C>G, 1144C>T, 1153A>G, 1156A>C, 1162C>T, 1181G>C, 1182A>T, 1183G>T, 1186T>C, 1192T>G, 1195A>T, 1197A>T, 1198G>A, 1201A>T, 1204T>A, 1210C>T, 1213G>T, 1216A>G, 1219G>A, 1222A>T, 1228A>G, 1231A>G, 1238G>A, 1239A>G, 1240T>C, 1241A>C, 1246G>A, 1252C>T, 1255A>T, 1258C>T, 1259T>A, 1261T>G, 1262C>G, 1263A>C, 1267G>A, 1273T>A, 1277T>A, 1278C>G, 1285C>A, 1291A>G, 1294A>T, 1295T>C, 1297G>C, 1300C>T, 1306T>A, 1309A>G, 1312T>C, 1315C>T, 1316C>T, 1317A>C, 1318A>T, 1327G>A, 1333T>A, 1336C>A, 1351A>T, 1354A>C, 1360T>G, 1366G>A, 1367C>G, 1369A>G, 1372T>G, 1373T>G, 1375C>G, 1376C>T, 1377A>C, 1378G>T, 1379A>C, 1380T>A, 1381G>A, 1385T>C, 1387G>A, 1394C>A, 1396T>A, 1397G>C, 1405T>A, 1408A>T, 1410A>C, 1411G>A, 1412A>C, 1413G>T, 1414G>T, 1423T>C, 1426A>T, 1428G>A, 1429A>C, 1433G>A | | | | | | | | |

##### CDS

| | Begin | End | Coverage | Score | Concordance | Matches | Identities | I/D/M/F* | Stop Codons |
|---|---|---|---|---|---|---|---|---|---|
| ToChV_s2_gp2 | 139 | 254 | 9.7% | 591 | 75.2% | 116 (100%) | 87 (75.0%) | 0/0/0/0 | 1 |
| Protein mutations: | Q140K (1103C>A), A141G (1107C>G 1108C>T), V142L (1109G>C 1111G>A), L143F (1112C>T), M144S (1115A>T 1116T>C 1117G>T), S145R (1120T>G), P146A (1121C>G), S147* (1124A>T 1126T>A), I148D (1127A>G 1128T>A), T149A (1130A>G 1132A>C), L153V (1142C>G 1144C>T), E166L (1181G>C 1182A>T 1183G>T), Q171L (1197A>T 1198G>A), D185S (1238G>A 1239A>G 1240T>C), T186P (1241A>C), S192T (1259T>A 1261T>G), Q193A (1262C>G 1263A>C), Q211S (1316C>T 1317A>C 1318A>T), Q228E (1367C>G 1369A>G), S229R (1372T>G), S230A (1373T>G 1375C>G), Q231S (1376C>T 1377A>C 1378G>T), M232Q (1379A>C 1380T>A 1381G>A), A238P (1397G>C), E242A (1410A>C 1411G>A), R243L (1412A>C 1413G>T 1414G>T), R248H (1428G>A 1429A>C), D250N (1433G>A) | | | | | | | | |
| Codon mutations: | CAA140AAA (1103C>A), GCC141GGT (1107C>G 1108C>T), GTG142CTA (1109G>C 1111G>A), CTC143TTC (1112C>T), ATG144TCT (1115A>T 1116T>C 1117G>T), AGT145AGG (1120T>G), CCT146GCT (1121C>G), AGT147TGA (1124A>T 1126T>A), ATC148GAC (1127A>G 1128T>A), ACA149GCC (1130A>G 1132A>C), TTG150CTG (1133T>C), CCC151CCT (1138C>T), CTC153GTT (1142C>G 1144C>T), GGA156GGG (1153A>G), CCA157CCC (1156A>C), TTC159TTT (1162C>T), GAG166CTT (1181G>C 1182A>T 1183G>T), AAT167AAC (1186T>C), CGT169CGG (1192T>G), GCA170GCT (1195A>T), CAG171CTA (1197A>T 1198G>A), ATA172ATT (1201A>T), CCT173CCA (1204T>A), TCC175TCT (1210C>T), GTG176GTT (1213G>T), GTA177GTG (1216A>G), GCG178GCA (1219G>A), ATA179ATT (1222A>T), AAA181AAG (1228A>G), CAA182CAG (1231A>G), GAT185AGC (1238G>A 1239A>G 1240T>C), ACA186CCA (1241A>C), AGG187AGA (1246G>A), GCC189GCT (1252C>T), GTA190GTT (1255A>T), TAC191TAT (1258C>T), TCT192ACG (1259T>A 1261T>G), CAA193GCA (1262C>G 1263A>C), CAG194CAA (1267G>A), ACT196ACA (1273T>A), TCT198AGT (1277T>A 1278C>G), GCC200GCA (1285C>A), GAA202GAG (1291A>G), GCA203GCT (1294A>T), TTG204CTC (1295T>C 1297G>C), AAC205AAT (1300C>T), CCT207CCA (1306T>A), CAA208CAG (1309A>G), TTT209TTC (1312T>C), TTC210TTT (1315C>T), CAA211TCT (1316C>T 1317A>C 1318A>T), CAG214CAA (1327G>A), GCT216GCA (1333T>A), GCC217GCA (1336C>A), TCA222TCT (1351A>T), GTA223GTC (1354A>C), TCT225TCG (1360T>G), GGG227GGA (1366G>A), CAA228GAG (1367C>G 1369A>G), AGT229AGG (1372T>G), TCC230GCG (1373T>G 1375C>G), CAG231TCT (1376C>T 1377A>C 1378G>T), ATG232CAA (1379A>C 1380T>A 1381G>A), TTG234CTA (1385T>C 1387G>A), CGT237AGA (1394C>A 1396T>A), GCT238CCT (1397G>C), GTT240GTA (1405T>A), GGA241GGT (1408A>T), GAG242GCA (1410A>C 1411G>A), AGG243CTT (1412A>C 1413G>T 1414G>T), AAT246AAC (1423T>C), ACA247ACT (1426A>T), CGA248CAC (1428G>A 1429A>C), GAT250AAT (1433G>A) | | | | | | | | |

##### Proteins

| | Begin | End | Coverage | Score | Concordance | Matches | Identities | I/D/M/F* | Stop Codons |
|---|---|---|---|---|---|---|---|---|---|
| polyprotein (YP_003097231.1) | 139 | 254 | 9.7% | 591 | 75.2% | 116 (100%) | 87 (75.0%) | 0/0/0/0 | 1 |
| Protein mutations: | Q140K (1103C>A), A141G (1107C>G 1108C>T), V142L (1109G>C 1111G>A), L143F (1112C>T), M144S (1115A>T 1116T>C 1117G>T), S145R (1120T>G), P146A (1121C>G), S147* (1124A>T 1126T>A), I148D (1127A>G 1128T>A), T149A (1130A>G 1132A>C), L153V (1142C>G 1144C>T), E166L (1181G>C 1182A>T 1183G>T), Q171L (1197A>T 1198G>A), D185S (1238G>A 1239A>G 1240T>C), T186P (1241A>C), S192T (1259T>A 1261T>G), Q193A (1262C>G 1263A>C), Q211S (1316C>T 1317A>C 1318A>T), Q228E (1367C>G 1369A>G), S229R (1372T>G), S230A (1373T>G 1375C>G), Q231S (1376C>T 1377A>C 1378G>T), M232Q (1379A>C 1380T>A 1381G>A), A238P (1397G>C), E242A (1410A>C 1411G>A), R243L (1412A>C 1413G>T 1414G>T), R248H (1428G>A 1429A>C), D250N (1433G>A) | | | | | | | | |
| Codon mutations: | CAA140AAA (1103C>A), GCC141GGT (1107C>G 1108C>T), GTG142CTA (1109G>C 1111G>A), CTC143TTC (1112C>T), ATG144TCT (1115A>T 1116T>C 1117G>T), AGT145AGG (1120T>G), CCT146GCT (1121C>G), AGT147TGA (1124A>T 1126T>A), ATC148GAC (1127A>G 1128T>A), ACA149GCC (1130A>G 1132A>C), TTG150CTG (1133T>C), CCC151CCT (1138C>T), CTC153GTT (1142C>G 1144C>T), GGA156GGG (1153A>G), CCA157CCC (1156A>C), TTC159TTT (1162C>T), GAG166CTT (1181G>C 1182A>T 1183G>T), AAT167AAC (1186T>C), CGT169CGG (1192T>G), GCA170GCT (1195A>T), CAG171CTA (1197A>T 1198G>A), ATA172ATT (1201A>T), CCT173CCA (1204T>A), TCC175TCT (1210C>T), GTG176GTT (1213G>T), GTA177GTG (1216A>G), GCG178GCA (1219G>A), ATA179ATT (1222A>T), AAA181AAG (1228A>G), CAA182CAG (1231A>G), GAT185AGC (1238G>A 1239A>G 1240T>C), ACA186CCA (1241A>C), AGG187AGA (1246G>A), GCC189GCT (1252C>T), GTA190GTT (1255A>T), TAC191TAT (1258C>T), TCT192ACG (1259T>A 1261T>G), CAA193GCA (1262C>G 1263A>C), CAG194CAA (1267G>A), ACT196ACA (1273T>A), TCT198AGT (1277T>A 1278C>G), GCC200GCA (1285C>A), GAA202GAG (1291A>G), GCA203GCT (1294A>T), TTG204CTC (1295T>C 1297G>C), AAC205AAT (1300C>T), CCT207CCA (1306T>A), CAA208CAG (1309A>G), TTT209TTC (1312T>C), TTC210TTT (1315C>T), CAA211TCT (1316C>T 1317A>C 1318A>T), CAG214CAA (1327G>A), GCT216GCA (1333T>A), GCC217GCA (1336C>A), TCA222TCT (1351A>T), GTA223GTC (1354A>C), TCT225TCG (1360T>G), GGG227GGA (1366G>A), CAA228GAG (1367C>G 1369A>G), AGT229AGG (1372T>G), TCC230GCG (1373T>G 1375C>G), CAG231TCT (1376C>T 1377A>C 1378G>T), ATG232CAA (1379A>C 1380T>A 1381G>A), TTG234CTA (1385T>C 1387G>A), CGT237AGA (1394C>A 1396T>A), GCT238CCT (1397G>C), GTT240GTA (1405T>A), GGA241GGT (1408A>T), GAG242GCA (1410A>C 1411G>A), AGG243CTT (1412A>C 1413G>T 1414G>T), AAT246AAC (1423T>C), ACA247ACT (1426A>T), CGA248CAC (1428G>A 1429A>C), GAT250AAT (1433G>A) | | | | | | | | |

*: Inserts / Deletes / Misaligned / Frameshifts

#### Analysis details

This analysis was performed with panviral2.64

### NGS Details (UN70): Tomato chocolate spot virus (segment RNA 1)

#### Assembly

| | |
|---|---|
| **Coverage Length** | 933 (2 contig(s)) |
| **Depth Of Coverage** | 13296.7 |
| **Number Of Reads** | 74446 |
| **Reads Per Million** | 1411.22 rpm (after QC) |
| **Ambiguities** | 0 |
| **Assembly Method** | de novo + reference guided assembly |
| **Consensus Caller** | Bcf Tools |

#### Coverage Map

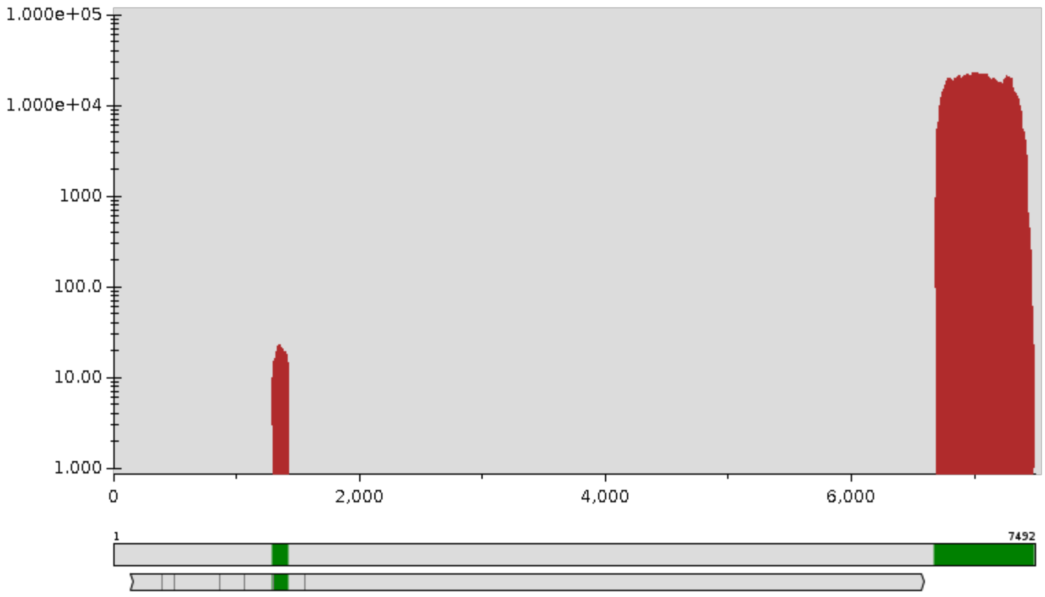

#### Assignment

| | |
|---|---|
| **Type** | Tomato chocolate spot virus (Taxonomy ID: 661101) |
| **Reference Genome** | NC_013075.1 |
| **NT Identity (%)** | 69.9115 |
| **AA Identity (%)** | 93.4783 |
| **Number Of Stop Codons** | 0 |
| **Number Of CDS** | 1 |

#### Alignment

| | |
|---|---|
| **Alignment Score** | 191.0 (NT) + 306.0 (AA) = 497.0 |
| **Concordance (%)** | 34.0878 |

| Alignment Method | Global, seeded, nucleotide + amino acids (AGA) |
|---|---|

#### Genome Region

Sequence starts at position 1291 and ends at position 7484 relative to NC_013075.1 reference sequence.

#### Alignment Detailed Statistics

| | Begin | End | Coverage | Score | Concordance | Matches | Identities | I/D/M/F* | Stop Codons |
|---|---|---|---|---|---|---|---|---|---|
| **NT** | **1291** | **7484** | **12.5%** | **191** | **16.8%** | **771 (80.9%)** | **553 (58.0%)** | **20/162** | |
| Mutations: | 1296C>T, 1299T>C, 1326C>A, 1332C>T, 1335G>A, 1341A>G, 1343C>G, 1344C>T, 1348A>C, 1350A>C, 1351T>C, 1353G>T, 1356T>C, 1359A>C, 1368G>T, 1375A>G, 1376G>C, 1383A>T, 1386G>A, 1389T>A, 1392A>G, 1395C>G, 1401T>C, 1402C>T, 1404C>G, 1410A>G, 1413G>A, 1419G>A, 6706A>G, 6707T>A, 6721G>C, 6722G>C, 6729T>C, 6731G>A, 6735G>A, 6736T>G, 6740C>A, 6741G>A, 6753G>T, 6757T>G, 6758_6759insT, 6759_6760insG, 6766G>A, 6772C>T, 6777T>C, 6795A>G, 6806C>T, 6814C>T, 6815_6834delAATTGTTTGGTGTTGTCAAC, 6837G>T, 6840G>A, 6842C>A, 6843T>C, 6845T>G, 6847_6850delGCCA, 6853A>G, 6855A>T, 6857C>A, 6859C>T, 6860C>G, 6861C>G, 6862C>T, 6863T>C, 6867_6868delGC, 6870T>G, 6872A>G, 6873T>A, 6878A>G, 6879A>T, 6881G>A, 6885A>C, 6886G>T, 6887A>T, 6889_6894delGTGGAG, 6897G>A, 6903C>A, 6906G>C, 6908A>C, 6909A>G, 6911C>G, 6912A>G, 6914A>G, 6915A>T, 6918A>T, 6919_6921delGAG, 6923C>T, 6924T>C, 6926T>C, 6930_6937delGGTCGGGC, 6942G>C, 6944_6946delTGG, 6949G>A, 6950C>G, 6951C>T, 6953G>C, 6955A>G, 6956T>G, 6958A>T, 6960A>T, 6961G>T, 6962C>T, 6964T>C, 6965T>C, 6967G>A, 6969G>A, 6970T>G, 6972T>A, 6973G>A, 6976T>A, 6978T>A, 6982_6985delCATC, 6987G>T, 6989T>A, 6991T>C, 6996A>C, 6998C>T, 7001_7012delCTCGTTTGGAGT, 7014T>G, 7016A>G, 7018C>G, 7020A>T, 7022delT, 7026G>T, 7030_7035delTGGTGT, 7040A>C, 7041C>G, 7042G>A, 7047_7067delGCAATGACACCCGGGGTGTGC, 7073C>G, 7077T>C, 7079A>T, 7080A>G, 7084_7088delAAGTA, 7091G>A, 7093T>C, 7097delT, 7100C>G, 7102T>A, 7103A>T, 7108_7116delAACTAGAGC, 7118T>G, 7121_7129delGTCTCAACT, 7132A>G, 7134G>T, 7135A>G, 7136G>A, 7140_7145delCCCGTT, 7148A>T, 7151A>C, 7152T>A, 7154G>A, 7155A>C, 7156T>C, 7158A>C, 7159A>T, 7161T>C, 7161_7162insC, 7165G>C, 7169C>G, 7172delG, 7177A>C, 7180_7183delCTGC, 7186T>G, 7188G>T, 7190T>C, 7193G>A, 7195_7218delGCAACAGTGAAGAATCATCACACC, 7220C>G, 7223C>A, 7224T>C, 7225T>C, 7227C>T, 7228C>G, 7230A>C, 7231G>T, 7234T>G, 7235C>G, 7238C>T, 7241G>C, 7244G>A, 7245T>A, 7247_7248delTC, 7251A>T, 7259_7260delCT, 7262C>G, 7264T>A, 7266G>A, 7270A>C, 7273_7274delAG, 7288G>C, 7289T>C, 7291T>G, 7292T>G, 7293G>A, 7294C>T, 7296_7297insGCTTCTCCTAGTTAA, 7298G>A, 7299A>C, 7300C>G, 7302A>G, 7303T>C, 7304G>A, 7308C>T, 7310T>A, 7311A>T, 7314G>C, 7315T>C, 7316T>A, 7317T>G, 7318G>T, 7319G>A, 7321T>A, 7325C>A, 7327T>A, 7328A>T, 7330A>T, 7335_7336insT, 7342T>G, 7346A>T, 7348A>T, 7350G>C, 7351G>T, 7352_7353insC, 7356A>G, 7358C>G, 7367C>T, 7369G>A, 7372G>T, 7373delA, 7375T>C, 7376G>C, 7378A>C, 7381C>G, 7385C>T, 7387T>G, 7388T>C, 7389C>A, 7390A>T, 7393_7396delTATT, 7407G>T, 7410delG, 7425A>C, 7428delC, 7430T>C, 7431C>A, 7447A>C, 7453A>G, 7454G>A, 7455A>T, 7459T>C, 7460C>G | | | | | | | | |

##### CDS

| | Begin | End | Coverage | Score | Concordance | Matches | Identities | I/D/M/F* | Stop Codons |
|---|---|---|---|---|---|---|---|---|---|
| ToChSV_s1_gp1 | 385 | 430 | 2.1% | 306 | 94.2% | 46 (100%) | 43 (93.5%) | 0/0/0/0 | 0 |
| Protein mutations: | S402C (1343C>G 1344C>T), R413A (1375A>G 1376G>C) | | | | | | | | |
| Codon mutations: | CCC386CCT (1296C>T), TTT387TTC (1299T>C), GGC396GGA (1326C>A), GGC398GGT (1332C>T), AAG399AAA (1335G>A), ACA401ACG (1341A>G), TCC402TGT (1343C>G 1344C>T), AGA404CGC (1348A>C 1350A>C), TTG405CTT (1351T>C 1353G>T), TTT406TTC (1356T>C), ATA407ATC (1359A>C), GTG410GTT (1368G>T), AGA413GCA (1375A>G 1376G>C), GGA415GGT (1383A>T), GAG416GAA (1386G>A), CCT417CCA (1389T>A), AAA418AAG (1392A>G), CTC419CTG (1395C>G), CGT421CGC (1401T>C), CTC422TTG (1402C>T 1404C>G), GCA424GCG (1410A>G), GTG425GTA (1413G>A), AAG427AAA (1419G>A) | | | | | | | | |

##### Proteins

| | Begin | End | Coverage | Score | Concordance | Matches | Identities | I/D/M/F* | Stop Codons |
|---|---|---|---|---|---|---|---|---|---|
| polyprotein (YP_003097229.1) | 385 | 430 | 2.1% | 306 | 94.2% | 46 (100%) | 43 (93.5%) | 0/0/0/0 | 0 |
| Protein mutations: | S402C (1343C>G 1344C>T), R413A (1375A>G 1376G>C) | | | | | | | | |
| Codon mutations: | CCC386CCT (1296C>T), TTT387TTC (1299T>C), GGC396GGA (1326C>A), GGC398GGT (1332C>T), AAG399AAA (1335G>A), ACA401ACG (1341A>G), TCC402TGT (1343C>G 1344C>T), AGA404CGC (1348A>C 1350A>C), TTG405CTT (1351T>C 1353G>T), TTT406TTC (1356T>C), ATA407ATC (1359A>C), GTG410GTT (1368G>T), AGA413GCA (1375A>G 1376G>C), GGA415GGT (1383A>T), GAG416GAA (1386G>A), CCT417CCA (1389T>A), AAA418AAG (1392A>G), CTC419CTG (1395C>G), CGT421CGC (1401T>C), CTC422TTG (1402C>T 1404C>G), GCA424GCG (1410A>G), GTG425GTA (1413G>A), AAG427AAA (1419G>A) | | | | | | | | |
| RNA-dependent RNA polymerase (YP_003097234.1) | 1 | 41 | 48.8% | 268 | 93.4% | 41 (100%) | 38 (92.7%) | 0/0/0/0 | 0 |
| Protein mutations: | S13C (1343C>G 1344C>T), R24A (1375A>G 1376G>C) | | | | | | | | |
| Codon mutations: | GGC7GGA (1326C>A), GGC9GGT (1332C>T), AAG10AAA (1335G>A), ACA12ACG (1341A>G), TCC13TGT (1343C>G 1344C>T), AGA15CGC (1348A>C 1350A>C), TTG16CTT (1351T>C 1353G>T), TTT17TTC (1356T>C), ATA18ATC (1359A>C), GTG21GTT (1368G>T), AGA24GCA (1375A>G 1376G>C), GGA26GGT (1383A>T), GAG27GAA (1386G>A), CCT28CCA (1389T>A), AAA29AAG (1392A>G), CTC30CTG (1395C>G), CGT32CGC (1401T>C), CTC33TTG (1402C>T 1404C>G), GCA35GCG (1410A>G), GTG36GTA (1413G>A), AAG38AAA (1419G>A) | | | | | | | | |

*: Inserts / Deletes / Misaligned / Frameshifts

#### Analysis details

This analysis was performed with panviral2.64

### NGS Details (UN70): Brazilian marseillevirus

#### Assembly

| | |
|---|---|
| **Coverage Length** | 233 (1 contig(s)) |
| **Depth Of Coverage** | 5346.7 |
| **Number Of Reads** | 13004 |
| **Reads Per Million** | 246.51 rpm (after QC) |
| **Ambiguities** | 0 |
| **Assembly Method** | de novo + reference guided assembly |
| **Consensus Caller** | Bcf Tools |

#### Coverage Map

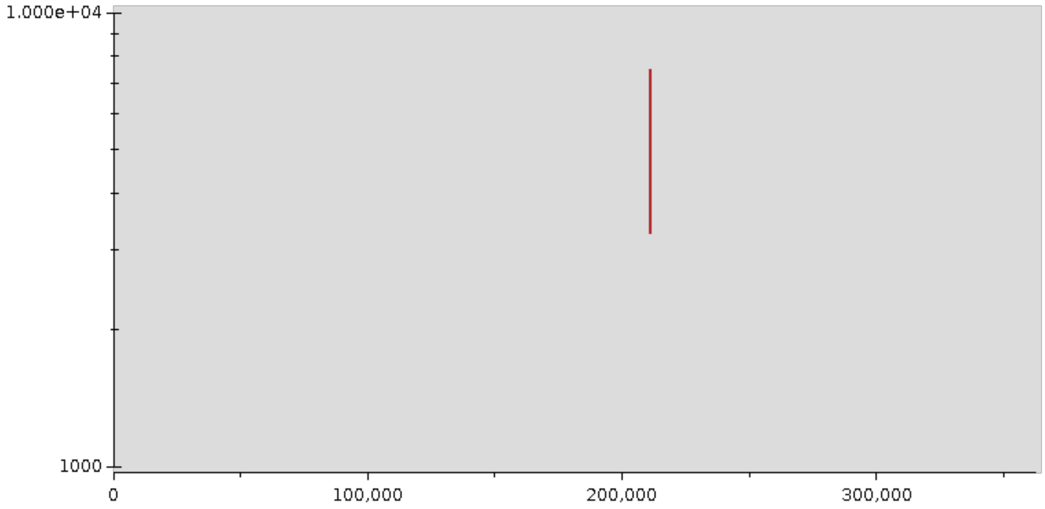

#### Assignment

| | |
|---|---|
| **Type** | Brazilian marseillevirus (Taxonomy ID: 1813599) |
| **Reference Genome** | NC_029692.1 |
| **NT Identity (%)** | 79.8283 |
| **AA Identity (%)** | 94.6667 |
| **Number Of Stop Codons** | 1 |
| **Number Of CDS** | 491 |

#### Alignment

| | |
|---|---|
| **Alignment Score** | 278.0 (NT) + 454.0 (AA) = 732.0 |
| **Concordance (%)** | 78.3726 |
| **Alignment Method** | Local, heuristic, nucleotide (BLASTN) |

#### Genome Region

Sequence starts at position 210884 and ends at position 211116 relative to NC_029692.1 reference sequence.

#### Alignment Detailed Statistics

| | Begin | End | Coverage | Score | Concordance | Matches | Identities | I/D/M/F* | Stop Codons |
|---|---|---|---|---|---|---|---|---|---|
| **NT** | **210884** | **211116** | **0.1%** | **278** | **59.7%** | **233 (100%)** | **186 (79.8%)** | **0/0** | |

Mutations: 210899T>C, 210901T>G, 210902T>A, 210908A>C, 210910G>A, 210917A>G, 210920G>T, 210926T>C, 210929T>C, 210938A>G, 210944C>A, 210955T>G, 210956T>G, 210962T>C, 210965C>A, 210974G>C, 210977T>G, 210980G>A, 210986A>C, 210988G>A, 210989G>T, 210991G>T, 210995T>C, 211001A>T, 211004A>G, 211009C>T, 211010T>G, 211022C>T, 211031T>G, 211032T>G, 211033G>C, 211037C>A, 211043T>G, 211046G>A, 211055A>G, 211067C>T, 211070G>A, 211073A>G, 211079C>T, 211088C>A, 211091G>C, 211093G>A, 211100T>C, 211103A>G, 211106C>A, 211108C>T, 211109C>T

*: Inserts / Deletes / Misaligned / Frameshifts

#### Analysis details

This analysis was performed with panviral2.64

### NGS Details (UN70): Tokyovirus A1

#### Assembly

| | |
|---|---|
| **Coverage Length** | 180 (1 contig(s)) |
| **Depth Of Coverage** | 6107.3 |
| **Number Of Reads** | 12194 |
| **Reads Per Million** | 231.15 rpm (after QC) |
| **Ambiguities** | 0 |
| **Assembly Method** | de novo + reference guided assembly |
| **Consensus Caller** | Bcf Tools |

#### Coverage Map

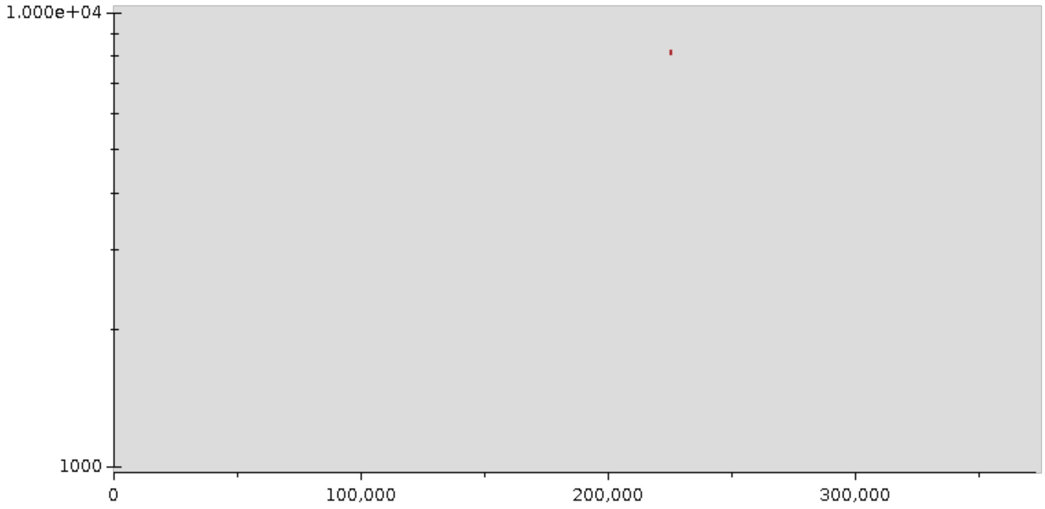

#### Assignment

| | |
|---|---|
| **Type** | Tokyovirus A1 (Taxonomy ID: 1826170) |
| **Reference Genome** | NC_030230.1 |
| **NT Identity (%)** | 80.5556 |
| **AA Identity (%)** | 93.1034 |
| **Number Of Stop Codons** | 1 |
| **Number Of CDS** | 470 |

#### Alignment

| | |
|---|---|
| **Alignment Score** | 220.0 (NT) + 353.0 (AA) = 573.0 |
| **Concordance (%)** | 78.4932 |
| **Alignment Method** | Local, heuristic, nucleotide (BLASTN) |

#### Genome Region

Sequence starts at position 225483 and ends at position 225662 relative to NC_030230.1 reference sequence.

#### Alignment Detailed Statistics

| | Begin | End | Coverage | Score | Concordance | Matches | Identities | I/D/M/F* | Stop Codons |
|---|---|---|---|---|---|---|---|---|---|
| **NT** | **225483** | **225662** | **0.1%** | **220** | **61.1%** | **180 (100%)** | **145 (80.6%)** | **0/0** | |
| Mutations: | 225498T>C, 225504G>C, 225507A>C, 225510G>A, 225513G>T, 225519G>T, 225528T>C, 225533C>T, 225534A>G, 225543G>C, 225549A>G, 225552T>G, 225554T>G, 225555C>G, 225564T>G, 225573C>A, 225579G>A, 225582G>A, 225585A>C, 225590G>T, 225594T>C, 225600G>C, 225603C>T, 225609A>T, 225621T>C, 225624G>T, 225630T>G, 225631T>G, 225632G>C, 225636C>G, 225640G>T, 225641C>T, 225642T>G, 225648G>T, 225651G>A | | | | | | | | |

*: Inserts / Deletes / Misaligned / Frameshifts

#### Analysis details

This analysis was performed with panviral2.64

### NGS Details (UN70): Lausannevirus

#### Assembly

| | |
|---|---|
| **Coverage Length** | 195 (1 contig(s)) |
| **Depth Of Coverage** | 4682.9 |
| **Number Of Reads** | 10606 |
| **Reads Per Million** | 201.05 rpm (after QC) |
| **Ambiguities** | 1 |
| **Assembly Method** | de novo + reference guided assembly |
| **Consensus Caller** | Bcf Tools |

#### Coverage Map

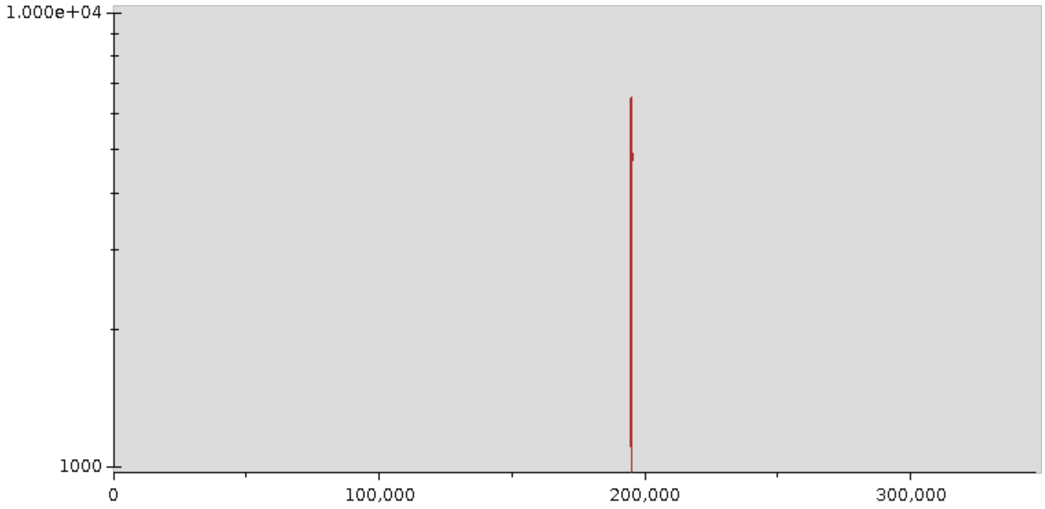

#### Assignment

| | |
|---|---|
| **Type** | Lausannevirus (Taxonomy ID: 999883) |
| **Reference Genome** | NC_015326.1 |
| **NT Identity (%)** | 80.5128 |
| **AA Identity (%)** | 95.2381 |
| **Number Of Stop Codons** | 1 |
| **Number Of CDS** | 444 |

#### Alignment

| | |
|---|---|
| **Alignment Score** | 240.0 (NT) + 358.0 (AA) = 598.0 |
| **Concordance (%)** | 75.7921 |
| **Alignment Method** | Local, heuristic, nucleotide (BLASTN) |

#### Genome Region

Sequence starts at position 194911 and ends at position 195105 relative to NC_015326.1 reference sequence.

#### Alignment Detailed Statistics

| | Begin | End | Coverage | Score | Concordance | Matches | Identities | I/D/M/F* | Stop Codons |
|---|---|---|---|---|---|---|---|---|---|
| **NT** | **194911** | **195105** | **0.1%** | **240** | **61.5%** | **195 (100%)** | **157 (80.5%)** | **0/0** | |

Mutations: 194925T>M, 194931A>C, 194937G>A, 194943A>G, 194946G>C, 194952T>C, 194955T>C, 194958A>G, 194967A>G, 194970G>C, 194979T>G, 194981T>G, 194982C>G, 194988T>C, 195003T>A, 195012A>C, 195015T>C, 195017G>T, 195018T>C, 195021T>C, 195033A>G, 195036G>C, 195042T>C, 195048T>C, 195054T>C, 195057T>A, 195058T>G, 195059G>C, 195060C>T, 195063G>C, 195066G>A, 195069C>G, 195072A>G, 195081C>A, 195087T>C, 195093T>C, 195096A>C, 195098G>A

*: Inserts / Deletes / Misaligned / Frameshifts

#### Analysis details

This analysis was performed with panviral2.64

### NGS Details (UN70): Marseillevirus marseillevirus

#### Assembly

| | |
|---|---|
| **Coverage Length** | 104 (1 contig(s)) |
| **Depth Of Coverage** | 1004.1 |
| **Number Of Reads** | 1457 |
| **Reads Per Million** | 27.62 rpm (after QC) |
| **Ambiguities** | 0 |
| **Assembly Method** | de novo + reference guided assembly |
| **Consensus Caller** | Bcf Tools |

#### Coverage Map

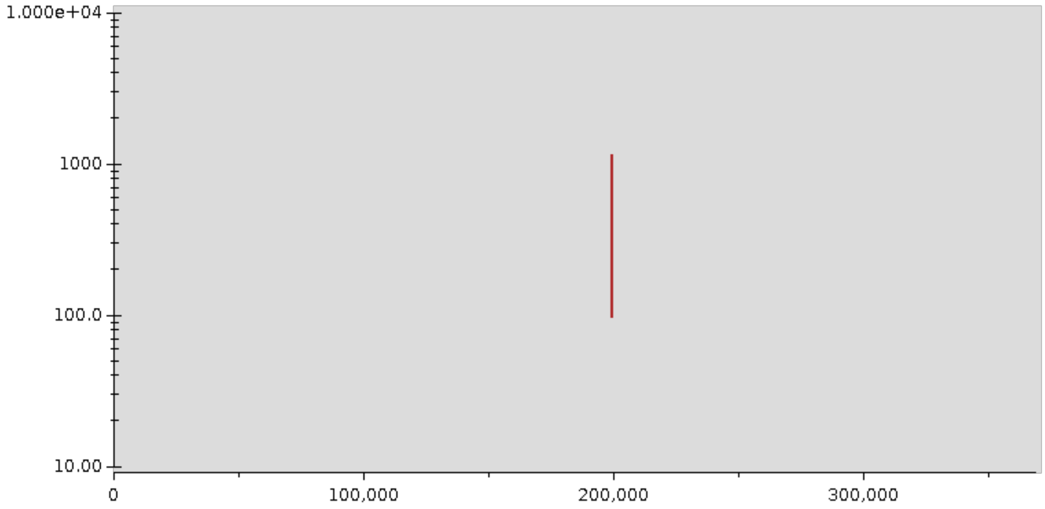

#### Assignment

| | |
|---|---|
| **Type** | Marseillevirus marseillevirus (Taxonomy ID: 694581) |
| **Reference Genome** | NC_013756.1 |
| **NT Identity (%)** | 89.4231 |
| **AA Identity (%)** | 94.2857 |
| **Number Of Stop Codons** | 0 |
| **Number Of CDS** | 428 |

#### Alignment

| | |
|---|---|
| **Alignment Score** | 164.0 (NT) + 235.0 (AA) = 399.0 |
| **Concordance (%)** | 89.8649 |
| **Alignment Method** | Local, heuristic, nucleotide (BLASTN) |

#### Genome Region

Sequence starts at position 199424 and ends at position 199527 relative to NC_013756.1 reference sequence.

#### Alignment Detailed Statistics

| | Begin | End | Coverage | Score | Concordance | Matches | Identities | I/D/M/F* | Stop Codons |
|---|---|---|---|---|---|---|---|---|---|
| NT | 199424 | 199527 | 0.1% | 164 | 78.8% | 104 (100%) | 93 (89.4%) | 0/0 | |

Mutations: 199435C>T, 199449C>T, 199459A>T, 199464A>G, 199489T>C, 199492G>T, 199495G>A, 199501G>C, 199504A>C, 199506G>T, 199516C>T

*: Inserts / Deletes / Misaligned / Frameshifts

#### Analysis details

This analysis was performed with panviral2.64

### NGS Details (UN70): Badnavirus occultipomeae

#### Assembly

| | |
|---|---|
| **Coverage Length** | 432 (1 contig(s)) |
| **Depth Of Coverage** | 139.6 |
| **Number Of Reads** | 541 |
| **Reads Per Million** | 10.26 rpm (after QC) |
| **Ambiguities** | 0 |
| **Assembly Method** | de novo + reference guided assembly |
| **Consensus Caller** | Bcf Tools |

#### Coverage Map

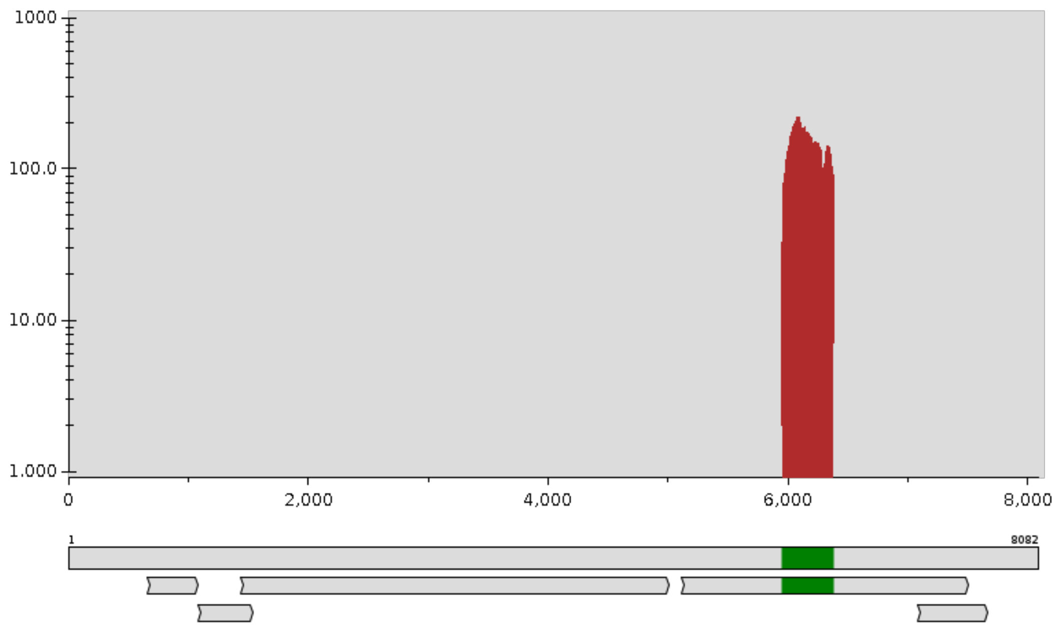

#### Assignment

| | |
|---|---|
| **Type** | Badnavirus occultipomeae (Taxonomy ID: 3048353) |
| **Reference Genome** | NC_015655.1 |
| **NT Identity (%)** | 53.1034 |
| **AA Identity (%)** | 45.5172 |
| **Number Of Stop Codons** | 0 |
| **Number Of CDS** | 5 |

#### Alignment

| | |
|---|---|
| **Alignment Score** | 48.0 (NT) + 460.0 (AA) = 508.0 |
| **Concordance (%)** | 27.3707 |

| Alignment Method | Global, seeded, nucleotide + amino acids (AGA) |
|---|---|

#### Genome Region

Sequence starts at position 5950 and ends at position 6381 relative to NC_015655.1 reference sequence.

#### Alignment Detailed Statistics

| | Begin | End | Coverage | Score | Concordance | Matches | Identities | I/D/M/F* | Stop Codons |
|---|---|---|---|---|---|---|---|---|---|
| **NT** | **5950** | **6381** | **5.3%** | **48** | **5.6%** | **432 (99.3%)** | **231 (53.1%)** | **3/0** | |
| Mutations: | 5958T>C, 5959G>A, 5961C>A, 5962A>G, 5963A>T, 5964T>G, 5967A>T, 5968G>A, 5969A>T, 5973A>G, 5974G>A, 5977C>A, 5983T>C, 5986C>T, 5988A>G, 5992G>C, 5994A>T, 5997C>T, 5998A>G, 6000C>T, 6001A>G, 6002C>A, 6004A>T, 6006T>G, 6007G>T, 6009C>T, 6010A>G, 6011G>A, 6013A>C, 6014G>A, 6015A>G, 6016A>T, 6019T>C, 6020C>A, 6021T>G, 6025A>G, 6026A>C, 6027A>T, 6028A>T, 6029A>C, 6030G>T, 6031A>G, 6033A>G, 6039A>T, 6042A>G, 6043T>A, 6048T>C, 6049C>T, 6051C>G, 6053A>G, 6055G>T, 6057A>G, 6060C>T, 6062T>A, 6070A>T, 6072A>G, 6076A>G, 6078G>T, 6079G>C, 6080A>G, 6084A>G, 6085A>T, 6086A>C, 6087G>A, 6088T>G, 6089C>A, 6090C>T, 6092A>T, 6093A>C, 6096A>G, 6097T>A, 6098G>A, 6105T>G, 6108T>C, 6109T>A, 6114T>C, 6116C>G, 6117A>C, 6118G>T, 6120A>T, 6123G>C, 6125T>A, 6126C>T, 6132A>G, 6137A>T, 6138A>G, 6141C>T, 6145C>T, 6147C>T, 6153A>G, 6154C>T, 6158T>C, 6159G>T, 6165A>C, 6168A>T, 6171A>T, 6173A>C, 6178C>A, 6179A>T, 6181A>G, 6182G>A, 6183A>G, 6184A>T, 6185A>T, 6190G>A, 6192T>C, 6193A>C, 6194A>G, 6195T>A, 6197C>T, 6198A>G, 6201T>C, 6203G>A, 6204G>A, 6205G>T, 6206G>C, 6207A>T, 6207_6208insTAT, 6208A>C, 6209C>T, 6210A>T, 6214G>C, 6215C>A, 6216A>G, 6220A>G, 6222C>G, 6223G>A, 6224C>T, 6225A>T, 6228A>G, 6230A>T, 6231T>C, 6234T>A, 6240C>T, 6243A>C, 6249A>T, 6251T>A, 6252C>T, 6255T>C, 6256G>A, 6258A>G, 6260A>G, 6264A>G, 6265G>A, 6266A>G, 6267A>T, 6270G>T, 6275A>T, 6277G>A, 6279T>A, 6282T>C, 6285C>G, 6286T>A, 6287T>G, 6288A>G, 6290A>T, 6292T>G, 6295G>T, 6296C>T, 6297C>G, 6300A>G, 6302T>G, 6303T>A, 6304G>T, 6310A>G, 6311G>A, 6315A>G, 6316G>A, 6317G>A, 6319C>T, 6321T>A, 6322A>T, 6323T>A, 6325C>G, 6326T>C, 6327A>C, 6329G>A, 6330C>G, 6331C>T, 6332C>T, 6333A>C, 6334A>T, 6336A>C, 6339A>G, 6340A>T, 6341T>G, 6342G>T, 6343A>G, 6345A>G, 6346A>T, 6349G>T, 6351A>G, 6352G>C, 6354C>A, 6355A>C, 6357A>G, 6358G>T, 6359A>C, 6363G>A, 6364G>T, 6365A>C, 6369C>T | | | | | | | | |

##### CDS

| | Begin | End | Coverage | Score | Concordance | Matches | Identities | I/D/M/F* | Stop Codons |
|---|---|---|---|---|---|---|---|---|---|
| SPBVa_gp4 | 279 | 422 | 18.1% | 460 | 46.4% | 144 (99.3%) | 66 (45.5%) | 1/0/0/0 | 0 |
| Protein mutations: | D282K (5959G>A 5961C>A), N283V (5962A>G 5963A>T 5964T>G), E285I (5968G>A 5969A>T), D287N (5974G>A), Q288K (5977C>A), S290P (5983T>C), G293R (5992G>C 5994A>T), N295D (5998A>G 6000C>T), T296D (6001A>G 6002C>A), I297L (6004A>T 6006T>G), V298F (6007G>T 6009C>T), S299D (6010A>G 6011G>A), R300Q (6013A>C 6014G>A 6015A>G), I301L (6016A>T), S302Q (6019T>C 6020C>A 6021T>G), K304A (6025A>G 6026A>C 6027A>T), K305S (6028A>T 6029A>C 6030G>T), I306V (6031A>G 6033A>G), F310I (6043T>A), K313R (6053A>G), A314S (6055G>T 6057A>G), F316Y (6062T>A), I319L (6070A>T 6072A>G), M321V (6076A>G 6078G>T), E322R (6079G>C 6080A>G), K324S (6085A>T 6086A>C 6087G>A), S325D (6088T>G 6089C>A 6090C>T), K326I (6092A>T 6093A>C), W328K (6097T>A 6098G>A), W332R (6109T>A), P334R (6116C>G 6117A>C), E335Y (6118G>T 6120A>T), L337H (6125T>A 6126C>T), E341V (6137A>T 6138A>G), P344S (6145C>T 6147C>T), M348T (6158T>C 6159G>T), D353A (6173A>C), Q355M (6178C>A 6179A>T), R356E (6181A>G 6182G>A 6183A>G), K357L (6184A>T 6185A>T), D359N (6190G>A 6192T>C), N360R (6193A>C 6194A>G 6195T>A), A361V (6197C>T 6198A>G), R363K (6203G>A 6204G>A), G364S (6205G>T 6206G>C 6207A>T), G364_T365insY (6207_6208insTAT), T365L (6208A>C 6209C>T 6210A>T), A367Q (6214G>C 6215C>A 6216A>G), I369V (6220A>G 6222C>G), A370I (6223G>A 6224C>T 6225A>T), Y372F (6230A>T 6231T>C), F379Y (6251T>A 6252C>T), E381K (6256G>A 6258A>G), N382S (6260A>G), E384S (6265G>A 6266A>G 6267A>T), E385D (6270G>T), E387V (6275A>T), D388K (6277G>A 6279T>A), L391R (6286T>A 6287T>G 6288A>G), N392I (6290A>T), L393V (6292T>G), A394L (6295G>T 6296C>T 6297C>G), I396R (6302T>G 6303T>A), V397L (6304G>T), R399E (6310A>G 6311G>A), G401K (6316G>A 6317G>A), I403Y (6322A>T 6323T>A), L404A (6325C>G 6326T>C 6327A>C), S405K (6329G>A 6330C>G), P406F (6331C>T 6332C>T 6333A>C), T407S (6334A>T 6336A>C), M409C (6340A>T 6341T>G 6342G>T), K410E (6343A>G 6345A>G), I411F (6346A>T), G412W (6349G>T 6351A>G), V413L (6352G>C 6354C>A), K414Q (6355A>C 6357A>G), E415S (6358G>T 6359A>C), D417S (6364G>T 6365A>C) | | | | | | | | |
| Codon mutations: | AAT281AAC (5958T>C), GAC282AAA (5959G>A 5961C>A), AAT283GTG (5962A>G 5963A>T 5964T>G), ACA284ACT (5967A>T), GAA285ATA (5968G>A 5969A>T), AAA286AAG (5973A>G), GAT287AAT (5974G>A), CAG288AAG (5977C>A), TCC290CCC (5983T>C), CTA291TTG (5986C>T 5988A>G), GGA293CGT (5992G>C 5994A>T), ATC294ATT (5997C>T), AAC295GAT (5998A>G 6000C>T), ACT296GAT (6001A>G 6002C>A), ATT297TTG (6004A>T 6006T>G), GTC298TTT (6007G>T 6009C>T), AGT299GAT (6010A>G 6011G>A), AGA300CAG (6013A>C 6014G>A 6015A>G), ATA301TTA (6016A>T), TCT302CAG (6019T>C 6020C>A 6021T>G), AAA304GCT (6025A>G 6026A>C 6027A>T), AAG305TCT (6028A>T 6029A>C 6030G>T), ATA306GTG (6031A>G 6033A>G), TCA308TCT (6039A>T), AAA309AAG (6042A>G), TTT310ATT (6043T>A), GAT311GAC (6048T>C), CTC312TTG (6049C>T 6051C>G), AAA313AGA (6053A>G), GCA314TCG (6055G>T 6057A>G), GGC315GGT (6060C>T), TTT316TAT (6062T>A), ATA319TTG (6070A>T 6072A>G), ATG321GTT (6076A>G 6078G>T), GAG322CGG (6079G>C 6080A>G), GAA323GAG (6084A>G), AAG324TCA (6085A>T 6086A>C 6087G>A), TCC325GAT (6088T>G 6089C>A 6090C>T), AAA326ATC (6092A>T 6093A>C), CCA327CCG (6096A>G), TGG328AAG (6097T>A 6098G>A), GCT330GCG (6105T>G), TTT331TTC (6108T>C), TGG332AGG (6109T>A), ACT333ACC (6114T>C), CCA334CGC (6116C>G 6117A>C), GAA335TAT (6118G>T 6120A>T), GGG336GGC (6123G>C), CTC337CAT (6125T>A 6126C>T), GAA339GAG (6132A>G), GAA341GTG (6137A>T 6138A>G), GTC342GTT (6141C>T), CCC344TCT (6145C>T 6147C>T), GGA346GGG (6153A>G), CTG347TTG (6154C>T), ATG348ACT (6158T>C 6159G>T), GCA350GCC (6165A>C), CCA351CCT (6168A>T), GCA352GCT (6171A>T), GAT353GCT (6173A>C), CAG355ATG (6178C>A 6179A>T), AGA356GAG (6181A>G 6182G>A 6183A>G), AAG357TTG (6184A>T 6185A>T), GAT359AAC (6190G>A 6192T>C), AAT360CGA (6193A>C 6194A>G 6195T>A), GCA361GTG (6197C>T 6198A>G), TTT362TTC (6201T>C), AGG363AAA (6203G>A 6204G>A), GGA364TCT (6205G>T 6206G>C 6207A>T), GGA364_ACA365insTAT (6207_6208insTAT), ACA365CTT (6208A>C 6209C>T 6210A>T), GCA367CAG (6214G>C 6215C>A 6216A>G), ATC369GTG (6220A>G 6222C>G), GCA370ATT (6223G>A 6224C>T 6225A>T), GTA371GTG (6228A>G), TAT372TTC (6230A>T 6231T>C), ATT373ATA (6234T>A), GAC375GAT (6240C>T), ATA376ATC (6243A>C), GTA378GTT (6249A>T), TTC379TAT (6251T>A 6252C>T), TCT380TCC (6255T>C), GAA381AAG (6256G>A 6258A>G), AAT382AGT (6260A>G), GAA383GAG (6264A>G), GAA384AGT (6265G>A 6266A>G 6267A>T), GAG385GAT (6270G>T), GAG387GTG (6275A>T), GAT388AAA (6277G>A 6279T>A), CAT389CAC (6282T>C), CTC390CTG (6285C>G), TTA391AGG (6286T>A 6287T>G 6288A>G), AAT392ATT (6290A>T), TTG393GTG (6292T>G), GCC394TTG (6295G>T 6296C>T 6297C>G), CAA395CAG (6300A>G), ATT396AGA (6302T>G 6303T>A), GTG397TTG (6304G>T), AGA399GAA (6310A>G 6311G>A), GAA400GAG (6315A>G), GGG401AAG (6316G>A 6317G>A), CTT402TTA (6319C>T 6321T>A), ATT403TAT (6322A>T 6323T>A), CTA404GCC (6325C>G 6326T>C 6327A>C), AGC405AAG (6329G>A 6330C>G), CCA406TTC (6331C>T 6332C>T 6333A>C), ACA407TCC (6334A>T 6336A>C), AAA408AAG (6339A>G), ATG409TGT (6340A>T 6341T>G 6342G>T), AAA410GAG (6343A>G 6345A>G), ATT411TTT (6346A>T), GGA412TGG (6349G>T 6351A>G), GTC413CTA (6352G>C 6354C>A), AAA414CAG (6355A>C 6357A>G), GAA415TCA (6358G>T 6359A>C), GTG416GTA (6363G>A), GAT417TCT (6364G>T 6365A>C), TTC418TTT (6369C>T) | | | | | | | | |

##### Proteins

| | Begin | End | Coverage | Score | Concordance | Matches | Identities | I/D/M/F* | Stop Codons |
|---|---|---|---|---|---|---|---|---|---|
| RNaseH/reverse transcriptase (YP_004581513.1) | 279 | 422 | 18.1% | 460 | 46.4% | 144 (99.3%) | 66 (45.5%) | 1/0/0/0 | 0 |
| Protein mutations: | D282K (5959G>A 5961C>A), N283V (5962A>G 5963A>T 5964T>G), E285I (5968G>A 5969A>T), D287N (5974G>A), Q288K (5977C>A), S290P (5983T>C), G293R (5992G>C 5994A>T), N295D (5998A>G 6000C>T), T296D (6001A>G 6002C>A), I297L (6004A>T 6006T>G), V298F (6007G>T 6009C>T), S299D (6010A>G 6011G>A), R300Q (6013A>C 6014G>A 6015A>G), I301L (6016A>T), S302Q (6019T>C 6020C>A 6021T>G), K304A (6025A>G 6026A>C 6027A>T), K305S (6028A>T 6029A>C 6030G>T), I306V (6031A>G 6033A>G), F310I (6043T>A), K313R (6053A>G), A314S (6055G>T 6057A>G), F316Y (6062T>A), I319L (6070A>T 6072A>G), M321V (6076A>G 6078G>T), E322R (6079G>C 6080A>G), K324S (6085A>T 6086A>C 6087G>A), S325D (6088T>G 6089C>A 6090C>T), K326I (6092A>T 6093A>C), W328K (6097T>A 6098G>A), W332R (6109T>A), P334R (6116C>G 6117A>C), E335Y (6118G>T 6120A>T), L337H (6125T>A 6126C>T), E341V (6137A>T 6138A>G), P344S (6145C>T 6147C>T), M348T (6158T>C 6159G>T), D353A (6173A>C), Q355M (6178C>A 6179A>T), R356E (6181A>G 6182G>A 6183A>G), K357L (6184A>T 6185A>T), D359N (6190G>A 6192T>C), N360R (6193A>C 6194A>G 6195T>A), A361V (6197C>T 6198A>G), R363K (6203G>A 6204G>A), G364S (6205G>T 6206G>C 6207A>T), G364_T365insY (6207_6208insTAT), T365L (6208A>C 6209C>T 6210A>T), A367Q (6214G>C 6215C>A 6216A>G), I369V (6220A>G 6222C>G), A370I (6223G>A 6224C>T 6225A>T), Y372F (6230A>T 6231T>C), F379Y (6251T>A 6252C>T), E381K (6256G>A 6258A>G), N382S (6260A>G), E384S (6265G>A 6266A>G 6267A>T), E385D (6270G>T), E387V (6275A>T), D388K (6277G>A 6279T>A), L391R (6286T>A 6287T>G 6288A>G), N392I (6290A>T), L393V (6292T>G), A394L (6295G>T 6296C>T 6297C>G), I396R (6302T>G 6303T>A), V397L (6304G>T), R399E (6310A>G 6311G>A), G401K (6316G>A 6317G>A), I403Y (6322A>T 6323T>A), L404A (6325C>G 6326T>C 6327A>C), S405K (6329G>A 6330C>G), P406F (6331C>T 6332C>T 6333A>C), T407S (6334A>T 6336A>C), M409C (6340A>T 6341T>G 6342G>T), K410E (6343A>G 6345A>G), I411F (6346A>T), G412W (6349G>T 6351A>G), V413L (6352G>C 6354C>A), K414Q (6355A>C 6357A>G), E415S (6358G>T 6359A>C), D417S (6364G>T 6365A>C) | | | | | | | | |

| | Begin | End | Coverage | Score | Concordance | Matches | Identities | I/D/M/F* | Stop Codons |
|---|---|---|---|---|---|---|---|---|---|
| **NT** | **5950** | **6381** | **5.3%** | **48** | **5.6%** | **432 (99.3%)** | **231 (53.1%)** | **3/0** | |
| Codon mutations: | AAT281AAC (5958T>C), GAC282AAA (5959G>A 5961C>A), AAT283GTG (5962A>G 5963A>T 5964T>G), ACA284ACT (5967A>T), GAA285ATA (5968G>A 5969A>T), AAA286AAG (5973A>G), GAT287AAT (5974G>A), CAG288AAG (5977C>A), TCC290CCC (5983T>C), CTA291TTG (5986C>T 5988A>G), GGA293CGT (5992G>C 5994A>T), ATC294ATT (5997C>T), AAC295GAT (5998A>G 6000C>T), ACT296GAT (6001A>G 6002C>A), ATT297TTG (6004A>T 6006T>G), GTC298TTT (6007G>T 6009C>T), AGT299GAT (6010A>G 6011G>A), AGA300CAG (6013A>C 6014G>A 6015A>G), ATA301TTA (6016A>T), TCT302CAG (6019T>C 6020C>A 6021T>G), AAA304GCT (6025A>G 6026A>C 6027A>T), AAG305TCT (6028A>T 6029A>C 6030G>T), ATA306GTG (6031A>G 6033A>G), TCA308TCT (6039A>T), AAA309AAG (6042A>G), TTT310ATT (6043T>A), GAT311GAC (6048T>C), CTC312TTG (6049C>T 6051C>G), AAA313AGA (6053A>G), GCA314TCG (6055G>T 6057A>G), GGC315GGT (6060C>T), TTT316TAT (6062T>A), ATA319TTG (6070A>T 6072A>G), ATG321GTT (6076A>G 6078G>T), GAG322CGG (6079G>C 6080A>G), GAA323GAG (6084A>G), AAG324TCA (6085A>T 6086A>C 6087G>A), TCC325GAT (6088T>G 6089C>A 6090C>T), AAA326ATC (6092A>T 6093A>C), CCA327CCG (6096A>G), TGG328AAG (6097T>A 6098G>A), GCT330GCG (6105T>G), TTT331TTC (6108T>C), TGG332AGG (6109T>A), ACT333ACC (6114T>C), CCA334CGC (6116C>G 6117A>C), GAA335TAT (6118G>T 6120A>T), GGG336GGC (6123G>C), CTC337CAT (6125T>A 6126C>T), GAA339GAG (6132A>G), GAA341GTG (6137A>T 6138A>G), GTC342GTT (6141C>T), CCC344TCT (6145C>T 6147C>T), GGA346GGG (6153A>G), CTG347TTG (6154C>T), ATG348ACT (6158T>C 6159G>T), GCA350GCC (6165A>C), CCA351CCT (6168A>T), GCA352GCT (6171A>T), GAT353GCT (6173A>C), CAG355ATG (6178C>A 6179A>T), AGA356GAG (6181A>G 6182G>A 6183A>G), AAG357TTG (6184A>T 6185A>T), GAT359AAC (6190G>A 6192T>C), AAT360CGA (6193A>C 6194A>G 6195T>A), GCA361GTG (6197C>T 6198A>G), TTT362TTC (6201T>C), AGG363AAA (6203G>A 6204G>A), GGA364TCT (6205G>T 6206G>C 6207A>T), GGA364_ACA365insTAT (6207_6208insTAT), ACA365CTT (6208A>C 6209C>T 6210A>T), GCA367CAG (6214G>C 6215C>A 6216A>G), ATC369GTG (6220A>G 6222C>G), GCA370ATT (6223G>A 6224C>T 6225A>T), GTA371GTG (6228A>G), TAT372TTC (6230A>T 6231T>C), ATT373ATA (6234T>A), GAC375GAT (6240C>T), ATA376ATC (6243A>C), GTA378GTT (6249A>T), TTC379TAT (6251T>A 6252C>T), TCT380TCC (6255T>C), GAA381AAG (6256G>A 6258A>G), AAT382AGT (6260A>G), GAA383GAG (6264A>G), GAA384AGT (6265G>A 6266A>G 6267A>T), GAG385GAT (6270G>T), GAG387GTG (6275A>T), GAT388AAA (6277G>A 6279T>A), CAT389CAC (6282T>C), CTC390CTG (6285C>G), TTA391AGG (6286T>A 6287T>G 6288A>G), AAT392ATT (6290A>T), TTG393GTG (6292T>G), GCC394TTG (6295G>T 6296C>T 6297C>G), CAA395CAG (6300A>G), ATT396AGA (6302T>G 6303T>A), GTG397TTG (6304G>T), AGA399GAA (6310A>G 6311G>A), GAA400GAG (6315A>G), GGG401AAG (6316G>A 6317G>A), CTT402TTA (6319C>T 6321T>A), ATT403TAT (6322A>T 6323T>A), CTA404GCC (6325C>G 6326T>C 6327A>C), AGC405AAG (6329G>A 6330C>G), CCA406TTC (6331C>T 6332C>T 6333A>C), ACA407TCC (6334A>T 6336A>C), AAA408AAG (6339A>G), ATG409TGT (6340A>T 6341T>G 6342G>T), AAA410GAG (6343A>G 6345A>G), ATT411TTT (6346A>T), GGA412TGG (6349G>T 6351A>G), GTC413CTA (6352G>C 6354C>A), AAA414CAG (6355A>C 6357A>G), GAA415TCA (6358G>T 6359A>C), GTG416GTA (6363G>A), GAT417TCT (6364G>T 6365A>C), TTC418TTT (6369C>T) | | | | | | | | |

*: Inserts / Deletes / Misaligned / Frameshifts

#### Analysis details

This analysis was performed with panviral2.64

### NGS Details (UN70): Cassava brown streak virus

#### Assembly

| | |
|---|---|
| **Coverage Length** | 525 (1 contig(s)) |
| **Depth Of Coverage** | 121.8 |
| **Number Of Reads** | 521 |
| **Reads Per Million** | 9.88 rpm (after QC) |
| **Ambiguities** | 0 |
| **Assembly Method** | de novo + reference guided assembly |
| **Consensus Caller** | Bcf Tools |

#### Coverage Map

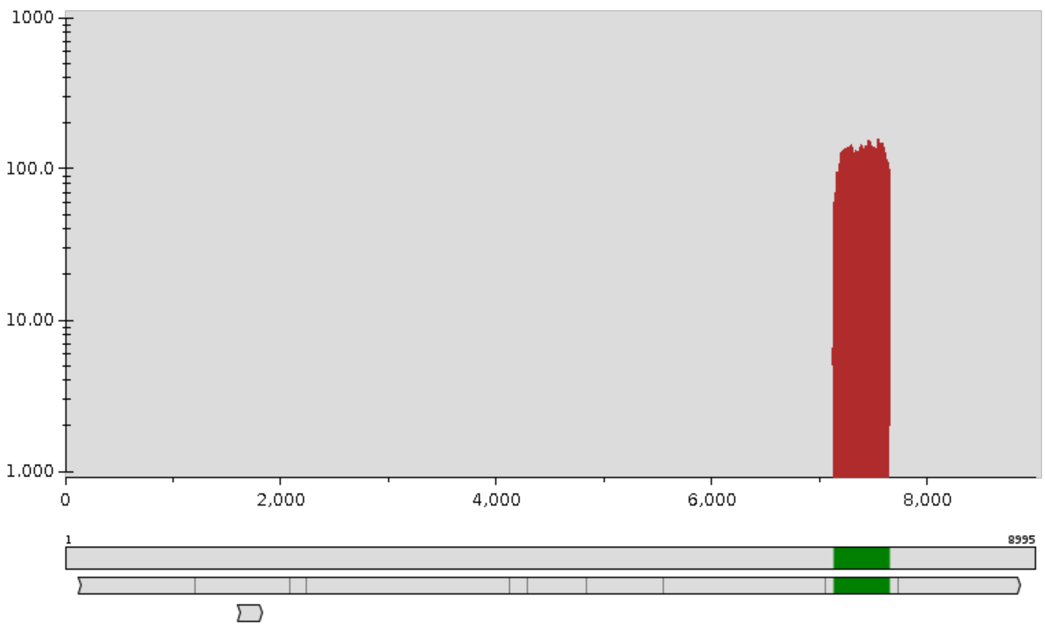

#### Assignment

| | |
|---|---|
| **Type** | Cassava brown streak virus (Taxonomy ID: 137758) |
| **Reference Genome** | NC_012698.2 |
| **NT Identity (%)** | 60.7619 |
| **AA Identity (%)** | 57.7143 |
| **Number Of Stop Codons** | 0 |
| **Number Of CDS** | 2 |

#### Alignment

| | |
|---|---|
| **Alignment Score** | 226.0 (NT) + 736.0 (AA) = 962.0 |
| **Concordance (%)** | 42.4724 |

| Alignment Method | Global, seeded, nucleotide + amino acids (AGA) |
|---|---|

#### Genome Region

Sequence starts at position 7126 and ends at position 7650 relative to NC_012698.2 reference sequence.

#### Alignment Detailed Statistics

| | Begin | End | Coverage | Score | Concordance | Matches | Identities | I/D/M/F* | Stop Codons |
|---|---|---|---|---|---|---|---|---|---|
| **NT** | **7126** | **7650** | **5.8%** | **226** | **21.5%** | **525 (100%)** | **319 (60.8%)** | **0/0** | |
| Mutations: | 7130A>G, 7133A>G, 7134A>G, 7136T>G, 7142C>T, 7145A>T, 7148A>C, 7151G>A, 7157A>T, 7158C>A, 7163A>G, 7167A>G, 7175A>T, 7177A>G, 7178A>G, 7179C>G, 7180A>C, 7184A>T, 7185T>C, 7190T>C, 7192C>A, 7193C>G, 7194A>T, 7202A>C, 7203A>T, 7205C>T, 7206A>C, 7207T>A, 7208C>G, 7211T>C, 7213G>T, 7214G>T, 7217A>G, 7218G>C, 7229A>T, 7234C>T, 7235A>T, 7241A>G, 7242A>G, 7243C>A, 7244A>G, 7245G>C, 7246T>C, 7250G>A, 7253A>T, 7256T>A, 7257A>T, 7258T>C, 7270T>C, 7271A>T, 7272C>A, 7273G>A, 7274T>A, 7275G>A, 7277A>T, 7280G>T, 7281T>G, 7283T>C, 7284G>A, 7286G>A, 7287T>G, 7288T>A, 7290A>G, 7292T>G, 7293G>A, 7294G>A, 7295A>T, 7298G>A, 7304T>G, 7307C>A, 7310A>T, 7313A>G, 7319A>T, 7320A>T, 7323T>C, 7325G>C, 7331T>C, 7332G>A, 7334T>C, 7337T>C, 7340C>T, 7343T>G, 7355A>G, 7358A>T, 7361C>T, 7364T>C, 7374A>T, 7377G>C, 7380G>A, 7381G>A, 7385T>C, 7388A>T, 7390T>A, 7391A>T, 7397A>T, 7398T>C, 7400A>T, 7401T>A, 7403T>C, 7406G>C, 7409G>A, 7410G>T, 7413G>A, 7414A>T, 7416C>G, 7418G>T, 7422C>G, 7424A>G, 7425A>G, 7432T>C, 7433G>A, 7436T>A, 7437A>T, 7438G>A, 7442T>C, 7443C>A, 7445C>G, 7448C>T, 7449G>A, 7451G>C, 7455G>T, 7457T>A, 7458T>C, 7462T>C, 7463A>T, 7464A>C, 7465A>T, 7467A>G, 7468A>G, 7469A>G, 7470G>C, 7471T>C, 7472T>A, 7473G>A, 7474G>A, 7476G>A, 7477A>C, 7481T>G, 7484T>G, 7487A>G, 7489T>C, 7490C>T, 7494A>G, 7495A>T, 7496G>T, 7499T>A, 7500G>A, 7501T>A, 7502G>A, 7503C>A, 7504T>C, 7505A>G, 7506A>C, 7507G>T, 7508A>G, 7511T>A, 7512G>A, 7517T>A, 7520A>G, 7521A>C, 7522T>C, 7523G>A, 7524C>G, 7526A>T, 7527C>A, 7529C>G, 7535A>C, 7539T>A, 7540C>A, 7541A>T, 7544T>C, 7547G>A, 7553C>T, 7556A>G, 7559T>A, 7562C>T, 7568A>T, 7570T>A, 7572A>G, 7573A>G, 7576G>A, 7577G>T, 7578A>G, 7579G>A, 7580A>C, 7581A>C, 7586A>T, 7588T>A, 7589T>C, 7599A>C, 7600T>C, 7601G>C, 7602G>A, 7603C>A, 7604C>G, 7607G>A, 7610G>A, 7616T>C, 7618T>A, 7622A>T, 7628T>C, 7631A>C, 7632T>G, 7633T>G, 7635C>A, 7636G>A | | | | | | | | |

##### CDS

| | Begin | End | Coverage | Score | Concordance | Matches | Identities | I/D/M/F* | Stop Codons |
|---|---|---|---|---|---|---|---|---|---|
| CBSV_gp1 | 2335 | 2509 | 6.0% | 736 | 60.0% | 175 (100%) | 101 (57.7%) | 0/0/0/0 | 0 |
| Protein mutations: | I2337V (7134A>G 7136T>G), Q2345K (7158C>A), K2348E (7167A>G), K2351R (7177A>G 7178A>G), Q2352A (7179C>G 7180A>C), F2354L (7185T>C), P2356Q (7192C>A 7193C>G), T2357S (7194A>T), I2360F (7203A>T 7205C>T), I2361Q (7206A>C 7207T>A 7208C>G), R2363L (7213G>T 7214G>T), V2365L (7218G>C), P2370L (7234C>T 7235A>T), T2373E (7242A>G 7243C>A 7244A>G), V2374P (7245G>C 7246T>C), E2376D (7253A>T), I2378S (7257A>T 7258T>C), V2382A (7270T>C 7271A>T), R2383K (7272C>A 7273G>A 7274T>A), V2384I (7275G>A 7277A>T), S2386A (7281T>G 7283T>C), E2387K (7284G>A 7286G>A), L2388E (7287T>G 7288T>A), I2389V (7290A>G 7292T>G), G2390N (7293G>A 7294G>A 7295A>T), S2399C (7320A>T), D2403N (7332G>A 7334T>C), N2406K (7343T>G), M2417L (7374A>T), E2418Q (7377G>C), G2419K (7380G>A 7381G>A), L2422H (7390T>A 7391A>T), Y2426N (7401T>A 7403T>C), K2427N (7406G>C), V2429L (7410G>T), E2430M (7413G>A 7414A>T), P2431A (7416C>G 7418G>T), Q2433E (7422C>G 7424A>G), N2434D (7425A>G), M2436T (7432T>C 7433G>A), S2438Y (7437A>T 7438G>A), L2440M (7443C>A 7445C>G), V2442I (7449G>A 7451G>C), A2444S (7455G>T 7457T>A), F2445L (7458T>C), V2446A (7462T>C 7463A>T), N2447L (7464A>C 7465A>T), K2448G (7467A>G 7468A>G 7469A>G), V2449P (7470G>C 7471T>C 7472T>A), G2450N (7473G>A 7474G>A), D2451T (7476G>A 7477A>C), D2452E (7481T>G), I2454M (7487A>G), I2455T (7489T>C 7490C>T), K2457V (7494A>G 7495A>T 7496G>T), V2459K (7500G>A 7501T>A 7502G>A), L2460T (7503C>A 7504T>C 7505A>G), R2461L (7506A>C 7507G>T 7508A>G), E2463K (7512G>A), M2466P (7521A>C 7522T>C 7523G>A), P2467A (7524C>G 7526A>T), S2472N (7539T>A 7540C>A 7541A>T), L2482H (7570T>A), N2483G (7572A>G 7573A>G), W2484Y (7576G>A 7577G>T), R2485D (7578A>G 7579G>A 7580A>C), K2486Q (7581A>C), F2488Y (7588T>A 7589T>C), M2492P (7599A>C 7600T>C 7601G>C), A2493K (7602G>A 7603C>A 7604C>G), M2498K (7618T>A), F2503G (7632T>G 7633T>G), R2504K (7635C>A 7636G>A) | | | | | | | | |
| Codon mutations: | GCA2335GCG (7130A>G), CCA2336CCG (7133A>G), ATT2337GTG (7134A>G 7136T>G), TTC2339TTT (7142C>T), GTA2340GTT (7145A>T), ACA2341ACC (7148A>C), GGG2342GGA (7151G>A), GCA2344GCT (7157A>T), CAA2345AAA (7158C>A), AAA2346AAG (7163A>G), AAG2348GAG (7167A>G), GTA2350GTT (7175A>T), AAA2351AGG (7177A>G 7178A>G), CAA2352GCA (7179C>G 7180A>C), ATA2353ATT (7184A>T), TTT2354CTT (7185T>C), GGT2355GGC (7190T>C), CCC2356CAG (7192C>A 7193C>G), ACT2357TCT (7194A>T), CCA2359CCC (7202A>C), ATC2360TTT (7203A>T 7205C>T), ATC2361CAG (7206A>C 7207T>A 7208C>G), TCT2362TCC (7211T>C), CGG2363CTT (7213G>T 7214G>T), AAA2364AAG (7217A>G), GTT2365CTT (7218G>C), CCA2368CCT (7229A>T), CCA2370CTT (7234C>T 7235A>T), GGA2372GGG (7241A>G), ACA2373GAG (7242A>G 7243C>A 7244A>G), GTT2374CCT (7245G>C 7246T>C), GAG2375GAA (7250G>A), GAA2376GAT (7253A>T), ATT2377ATA (7256T>A), ATC2378TCC (7257A>T 7258T>C), GTA2382GCT (7270T>C 7271A>T), CGT2383AAA (7272C>A 7273G>A 7274T>A), GTA2384ATT (7275G>A 7277A>T), GCG2385GCT (7280G>T), TCT2386GCC (7281T>G 7283T>C), GAG2387AAA (7284G>A 7286G>A), TTG2388GAG (7287T>G 7288T>A), ATT2389GTG (7290A>G 7292T>G), GGA2390AAT (7293G>A 7294G>A 7295A>T), GGG2391GGA (7298G>A), GTT2393GTG (7304T>G), CTC2394CTA (7307C>A), GTA2395GTT (7310A>T), GAA2396GAG (7313A>G), ACA2398ACT (7319A>T), AGT2399TGT (7320A>T), TTG2400CTC (7323T>C 7325G>C), TTT2402TTC (7331T>C), GAT2403AAC (7332G>A 7334T>C), GCT2404GCC (7337T>C), CTC2405CTT (7340C>T), AAT2406AAG (7343T>G), GGA2410GGG (7355A>G), CCA2411CCT (7358A>T), TAC2412TAT (7361C>T), ATT2413ATC (7364T>C), ATG2417TTG (7374A>T), GAA2418CAA (7377G>C), GGG2419AAG (7380G>A 7381G>A), ATT2420ATC (7385T>C), GGA2421GGT (7388A>T), CTA2422CAT (7390T>A 7391A>T), GGA2424GGT (7397A>T), TTA2425CTT (7398T>C 7400A>T), TAT2426AAC (7401T>A 7403T>C), AAG2427AAC (7406G>C), TTG2428TTA (7409G>A), GTG2429TTG (7410G>T), GAG2430ATG (7413G>A 7414A>T), CCG2431GCT (7416C>G 7418G>T), CAA2433GAG (7422C>G 7424A>G), AAT2434GAT (7425A>G), ATG2436ACA (7432T>C 7433G>A), GCT2437GCA (7436T>A), AGT2438TAT (7437A>T 7438G>A), GCT2439GCC (7442T>C), CTC2440ATG (7443C>A 7445C>G), TGC2441TGT (7448C>T), GTG2442ATC (7449G>A 7451G>C), GCT2444TCA (7455G>T 7457T>A), TTT2445CTT (7458T>C), GTA2446GCT (7462T>C 7463A>T), AAT2447CTT (7464A>C 7465A>T), AAA2448GGG (7467A>G 7468A>G 7469A>G), GTT2449CCA (7470G>C 7471T>C 7472T>A), GGT2450AAT (7473G>A 7474G>A), GAT2451ACT (7476G>A 7477A>C), GAT2452GAG (7481T>G), CCT2453CCG (7484T>G), ATA2454ATG (7487A>G), ATC2455ACT (7489T>C 7490C>T), AAG2457GTT (7494A>G 7495A>T 7496G>T), GGT2458GGA (7499T>A), GTG2459AAA (7500G>A 7501T>A 7502G>A), CTA2460ACG (7503C>A 7504T>C 7505A>G), AGA2461CTG (7506A>C 7507G>T 7508A>G), GGT2462GGA (7511T>A), GAG2463AAG (7512G>A), ATT2464ATA (7517T>A), GTA2465GTG (7520A>G), ATG2466CCA (7521A>C 7522T>C 7523G>A), CCA2467GCT (7524C>G 7526A>T), CGC2468AGG (7527C>A 7529C>G), CCA2470CCC (7535A>C), TCA2472AAT (7539T>A 7540C>A 7541A>T), TTT2473TTC (7544T>C), GGG2474GGA (7547G>A), GAC2476GAT (7553C>T), CCA2477CCG (7556A>G), ATT2478ATA (7559T>A), TTC2479TTT (7562C>T), CCA2481CCT (7568A>T), CTT2482CAT (7570T>A), AAC2483GGC (7572A>G 7573A>G), TGG2484TAT (7576G>A 7577G>T), AGA2485GAC (7578A>G 7579G>A 7580A>C), AAG2486CAG (7581A>C), ACA2487ACT (7586A>T), TTT2488TAC (7588T>A 7589T>C), ATG2492CCC (7599A>C 7600T>C 7601G>C), GCC2493AAG (7602G>A 7603C>A 7604C>G), GAG2494GAA (7607G>A), GAG2495GAA (7610G>A), AAT2497AAC (7616T>C), ATG2498AAG (7618T>A), ATA2499ATT (7622A>T), CAT2501CAC (7628T>C), CGA2502CGC (7631A>C), TTT2503GGT (7632T>G 7633T>G), CGA2504AAA (7635C>A 7636G>A) | | | | | | | | |

##### Proteins

| | Begin | End | Coverage | Score | Concordance | Matches | Identities | I/D/M/F* | Stop Codons |
|---|---|---|---|---|---|---|---|---|---|
| polyprotein (YP_007027011.1) | 2335 | 2509 | 6.0% | 736 | 60.0% | 175 (100%) | 101 (57.7%) | 0/0/0/0 | 0 |
| Protein mutations: | I2337V (7134A>G 7136T>G), Q2345K (7158C>A), K2348E (7167A>G), K2351R (7177A>G 7178A>G), Q2352A (7179C>G 7180A>C), F2354L (7185T>C), P2356Q (7192C>A 7193C>G), T2357S (7194A>T), I2360F (7203A>T 7205C>T), I2361Q (7206A>C 7207T>A 7208C>G), R2363L (7213G>T 7214G>T), V2365L (7218G>C), P2370L (7234C>T 7235A>T), T2373E (7242A>G 7243C>A 7244A>G), V2374P (7245G>C 7246T>C), E2376D (7253A>T), I2378S (7257A>T 7258T>C), V2382A (7270T>C 7271A>T), R2383K (7272C>A 7273G>A 7274T>A), V2384I (7275G>A 7277A>T), S2386A (7281T>G 7283T>C), E2387K (7284G>A 7286G>A), L2388E (7287T>G 7288T>A), I2389V (7290A>G 7292T>G), G2390N (7293G>A 7294G>A 7295A>T), S2399C (7320A>T), D2403N (7332G>A 7334T>C), N2406K (7343T>G), M2417L (7374A>T), E2418Q (7377G>C), G2419K (7380G>A 7381G>A), L2422H (7390T>A 7391A>T), Y2426N (7401T>A 7403T>C), K2427N (7406G>C), V2429L (7410G>T), E2430M (7413G>A 7414A>T), P2431A (7416C>G 7418G>T), Q2433E (7422C>G 7424A>G), N2434D (7425A>G), M2436T (7432T>C 7433G>A), S2438Y (7437A>T 7438G>A), L2440M (7443C>A 7445C>G), V2442I (7449G>A 7451G>C), A2444S (7455G>T 7457T>A), F2445L (7458T>C), V2446A (7462T>C 7463A>T), N2447L (7464A>C 7465A>T), K2448G (7467A>G 7468A>G 7469A>G), V2449P (7470G>C 7471T>C 7472T>A), G2450N (7473G>A 7474G>A), D2451T (7476G>A 7477A>C), D2452E (7481T>G), I2454M (7487A>G), I2455T (7489T>C 7490C>T), K2457V (7494A>G 7495A>T 7496G>T), V2459K (7500G>A 7501T>A 7502G>A), L2460T (7503C>A 7504T>C 7505A>G), R2461L (7506A>C 7507G>T 7508A>G), E2463K (7512G>A), M2466P (7521A>C 7522T>C 7523G>A), P2467A (7524C>G 7526A>T), S2472N (7539T>A 7540C>A 7541A>T), L2482H (7570T>A), N2483G (7572A>G 7573A>G), W2484Y (7576G>A 7577G>T), R2485D (7578A>G 7579G>A 7580A>C), K2486Q (7581A>C), F2488Y (7588T>A 7589T>C), M2492P (7599A>C 7600T>C 7601G>C), A2493K (7602G>A 7603C>A 7604C>G), M2498K (7618T>A), F2503G (7632T>G 7633T>G), R2504K (7635C>A 7636G>A) | | | | | | | | |

| | Begin | End | Coverage | Score | Concordance | Matches | Identities | I/D/M/F* | Stop Codons |
|---|---|---|---|---|---|---|---|---|---|
| **NT** | **7126** | **7650** | **5.8%** | **226** | **21.5%** | **525 (100%)** | **319 (60.8%)** | **0/0** | |
| Codon mutations: | GCA2335GCG (7130A>G), CCA2336CCG (7133A>G), ATT2337GTG (7134A>G 7136T>G), TTC2339TTT (7142C>T), GTA2340GTT (7145A>T), ACA2341ACC (7148A>C), GGG2342GGA (7151G>A), GCA2344GCT (7157A>T), CAA2345AAA (7158C>A), AAA2346AAG (7163A>G), AAG2348GAG (7167A>G), GTA2350GTT (7175A>T), AAA2351AGG (7177A>G 7178A>G), CAA2352GCA (7179C>G 7180A>C), ATA2353ATT (7184A>T), TTT2354CTT (7185T>C), GGT2355GGC (7190T>C), CCC2356CAG (7192C>A 7193C>G), ACT2357TCT (7194A>T), CCA2359CCC (7202A>C), ATC2360TTT (7203A>T 7205C>T), ATC2361CAG (7206A>C 7207T>A 7208C>G), TCT2362TCC (7211T>C), CGG2363CTT (7213G>T 7214G>T), AAA2364AAG (7217A>G), GTT2365CTT (7218G>C), CCA2368CCT (7229A>T), CCA2370CTT (7234C>T 7235A>T), GGA2372GGG (7241A>G), ACA2373GAG (7242A>G 7243C>A 7244A>G), GTT2374CCT (7245G>C 7246T>C), GAG2375GAA (7250G>A), GAA2376GAT (7253A>T), ATT2377ATA (7256T>A), ATC2378TCC (7257A>T 7258T>C), GTA2382GCT (7270T>C 7271A>T), CGT2383AAA (7272C>A 7273G>A 7274T>A), GTA2384ATT (7275G>A 7277A>T), GCG2385GCT (7280G>T), TCT2386GCC (7281T>G 7283T>C), GAG2387AAA (7284G>A 7286G>A), TTG2388GAG (7287T>G 7288T>A), ATT2389GTG (7290A>G 7292T>G), GGA2390AAT (7293G>A 7294G>A 7295A>T), GGG2391GGA (7298G>A), GTT2393GTG (7304T>G), CTC2394CTA (7307C>A), GTA2395GTT (7310A>T), GAA2396GAG (7313A>G), ACA2398ACT (7319A>T), AGT2399TGT (7320A>T), TTG2400CTC (7323T>C 7325G>C), TTT2402TTC (7331T>C), GAT2403AAC (7332G>A 7334T>C), GCT2404GCC (7337T>C), CTC2405CTT (7340C>T), AAT2406AAG (7343T>G), GGA2410GGG (7355A>G), CCA2411CCT (7358A>T), TAC2412TAT (7361C>T), ATT2413ATC (7364T>C), ATG2417TTG (7374A>T), GAA2418CAA (7377G>C), GGG2419AAG (7380G>A 7381G>A), ATT2420ATC (7385T>C), GGA2421GGT (7388A>T), CTA2422CAT (7390T>A 7391A>T), GGA2424GGT (7397A>T), TTA2425CTT (7398T>C 7400A>T), TAT2426AAC (7401T>A 7403T>C), AAG2427AAC (7406G>C), TTG2428TTA (7409G>A), GTG2429TTG (7410G>T), GAG2430ATG (7413G>A 7414A>T), CCG2431GCT (7416C>G 7418G>T), CAA2433GAG (7422C>G 7424A>G), AAT2434GAT (7425A>G), ATG2436ACA (7432T>C 7433G>A), GCT2437GCA (7436T>A), AGT2438TAT (7437A>T 7438G>A), GCT2439GCC (7442T>C), CTC2440ATG (7443C>A 7445C>G), TGC2441TGT (7448C>T), GTG2442ATC (7449G>A 7451G>C), GCT2444TCA (7455G>T 7457T>A), TTT2445CTT (7458T>C), GTA2446GCT (7462T>C 7463A>T), AAT2447CTT (7464A>C 7465A>T), AAA2448GGG (7467A>G 7468A>G 7469A>G), GTT2449CCA (7470G>C 7471T>C 7472T>A), GGT2450AAT (7473G>A 7474G>A), GAT2451ACT (7476G>A 7477A>C), GAT2452GAG (7481T>G), CCT2453CCG (7484T>G), ATA2454ATG (7487A>G), ATC2455ACT (7489T>C 7490C>T), AAG2457GTT (7494A>G 7495A>T 7496G>T), GGT2458GGA (7499T>A), GTG2459AAA (7500G>A 7501T>A 7502G>A), CTA2460ACG (7503C>A 7504T>C 7505A>G), AGA2461CTG (7506A>C 7507G>T 7508A>G), GGT2462GGA (7511T>A), GAG2463AAG (7512G>A), ATT2464ATA (7517T>A), GTA2465GTG (7520A>G), ATG2466CCA (7521A>C 7522T>C 7523G>A), CCA2467GCT (7524C>G 7526A>T), CGC2468AGG (7527C>A 7529C>G), CCA2470CCC (7535A>C), TCA2472AAT (7539T>A 7540C>A 7541A>T), TTT2473TTC (7544T>C), GGG2474GGA (7547G>A), GAC2476GAT (7553C>T), CCA2477CCG (7556A>G), ATT2478ATA (7559T>A), TTC2479TTT (7562C>T), CCA2481CCT (7568A>T), CTT2482CAT (7570T>A), AAC2483GGC (7572A>G 7573A>G), TGG2484TAT (7576G>A 7577G>T), AGA2485GAC (7578A>G 7579G>A 7580A>C), AAG2486CAG (7581A>C), ACA2487ACT (7586A>T), TTT2488TAC (7588T>A 7589T>C), ATG2492CCC (7599A>C 7600T>C 7601G>C), GCC2493AAG (7602G>A 7603C>A 7604C>G), GAG2494GAA (7607G>A), GAG2495GAA (7610G>A), AAT2497AAC (7616T>C), ATG2498AAG (7618T>A), ATA2499ATT (7622A>T), CAT2501CAC (7628T>C), CGA2502CGC (7631A>C), TTT2503GGT (7632T>G 7633T>G), CGA2504AAA (7635C>A 7636G>A) | | | | | | | | |
| HAM1-like protein (YP_007032446.1) | 27 | 201 | 77.4% | 736 | 60.0% | 175 (100%) | 101 (57.7%) | 0/0/0/0 | 0 |
| Protein mutations: | I29V (7134A>G 7136T>G), Q37K (7158C>A), K40E (7167A>G), K43R (7177A>G 7178A>G), Q44A (7179C>G 7180A>C), F46L (7185T>C), P48Q (7192C>A 7193C>G), T49S (7194A>T), I52F (7203A>T 7205C>T), I53Q (7206A>C 7207T>A 7208C>G), R55L (7213G>T 7214G>T), V57L (7218G>C), P62L (7234C>T 7235A>T), T65E (7242A>G 7243C>A 7244A>G), V66P (7245G>C 7246T>C), E68D (7253A>T), I70S (7257A>T 7258T>C), V74A (7270T>C 7271A>T), R75K (7272C>A 7273G>A 7274T>A), V76I (7275G>A 7277A>T), S78A (7281T>G 7283T>C), E79K (7284G>A 7286G>A), L80E (7287T>G 7288T>A), I81V (7290A>G 7292T>G), G82N (7293G>A 7294G>A 7295A>T), S91C (7320A>T), D95N (7332G>A 7334T>C), N98K (7343T>G), M109L (7374A>T), E110Q (7377G>C), G111K (7380G>A 7381G>A), L114H (7390T>A 7391A>T), Y118N (7401T>A 7403T>C), K119N (7406G>C), V121L (7410G>T), E122M (7413G>A 7414A>T), P123A (7416C>G 7418G>T), Q125E (7422C>G 7424A>G), N126D (7425A>G), M128T (7432T>C 7433G>A), S130Y (7437A>T 7438G>A), L132M (7443C>A 7445C>G), V134I (7449G>A 7451G>C), A136S (7455G>T 7457T>A), F137L (7458T>C), V138A (7462T>C 7463A>T), N139L (7464A>C 7465A>T), K140G (7467A>G 7468A>G 7469A>G), V141P (7470G>C 7471T>C 7472T>A), G142N (7473G>A 7474G>A), D143T (7476G>A 7477A>C), D144E (7481T>G), I146M (7487A>G), I147T (7489T>C 7490C>T), K149V (7494A>G 7495A>T 7496G>T), V151K (7500G>A 7501T>A 7502G>A), L152T (7503C>A 7504T>C 7505A>G), R153L (7506A>C 7507G>T 7508A>G), E155K (7512G>A), M158P (7521A>C 7522T>C 7523G>A), P159A (7524C>G 7526A>T), S164N (7539T>A 7540C>A 7541A>T), L174H (7570T>A), N175G (7572A>G 7573A>G), W176Y (7576G>A 7577G>T), R177D (7578A>G 7579G>A 7580A>C), K178Q (7581A>C), F180Y (7588T>A 7589T>C), M184P (7599A>C 7600T>C 7601G>C), A185K (7602G>A 7603C>A 7604C>G), M190K (7618T>A), F195G (7632T>G 7633T>G), R196K (7635C>A 7636G>A) | | | | | | | | |
| Codon mutations: | GCA27GCG (7130A>G), CCA28CCG (7133A>G), ATT29GTG (7134A>G 7136T>G), TTC31TTT (7142C>T), GTA32GTT (7145A>T), ACA33ACC (7148A>C), GGG34GGA (7151G>A), GCA36GCT (7157A>T), CAA37AAA (7158C>A), AAA38AAG (7163A>G), AAG40GAG (7167A>G), GTA42GTT (7175A>T), AAA43AGG (7177A>G 7178A>G), CAA44GCA (7179C>G 7180A>C), ATA45ATT (7184A>T), TTT46CTT (7185T>C), GGT47GGC (7190T>C), CCC48CAG (7192C>A 7193C>G), ACT49TCT (7194A>T), CCA51CCC (7202A>C), ATC52TTT (7203A>T 7205C>T), ATC53CAG (7206A>C 7207T>A 7208C>G), TCT54TCC (7211T>C), CGG55CTT (7213G>T 7214G>T), AAA56AAG (7217A>G), GTT57CTT (7218G>C), CCA60CCT (7229A>T), CCA62CTT (7234C>T 7235A>T), GGA64GGG (7241A>G), ACA65GAG (7242A>G 7243C>A 7244A>G), GTT66CCT (7245G>C 7246T>C), GAG67GAA (7250G>A), GAA68GAT (7253A>T), ATT69ATA (7256T>A), ATC70TCC (7257A>T 7258T>C), GTA74GCT (7270T>C 7271A>T), CGT75AAA (7272C>A 7273G>A 7274T>A), GTA76ATT (7275G>A 7277A>T), GCG77GCT (7280G>T), TCT78GCC (7281T>G 7283T>C), GAG79AAA (7284G>A 7286G>A), TTG80GAG (7287T>G 7288T>A), ATT81GTG (7290A>G 7292T>G), GGA82AAT (7293G>A 7294G>A 7295A>T), GGG83GGA (7298G>A), GTT85GTG (7304T>G), CTC86CTA (7307C>A), GTA87GTT (7310A>T), GAA88GAG (7313A>G), ACA90ACT (7319A>T), AGT91TGT (7320A>T), TTG92CTC (7323T>C 7325G>C), TTT94TTC (7331T>C), GAT95AAC (7332G>A 7334T>C), GCT96GCC (7337T>C), CTC97CTT (7340C>T), AAT98AAG (7343T>G), GGA102GGG (7355A>G), CCA103CCT (7358A>T), TAC104TAT (7361C>T), ATT105ATC (7364T>C), ATG109TTG (7374A>T), GAA110CAA (7377G>C), GGG111AAG (7380G>A 7381G>A), ATT112ATC (7385T>C), GGA113GGT (7388A>T), CTA114CAT (7390T>A 7391A>T), GGA116GGT (7397A>T), TTA117CTT (7398T>C 7400A>T), TAT118AAC (7401T>A 7403T>C), AAG119AAC (7406G>C), TTG120TTA (7409G>A), GTG121TTG (7410G>T), GAG122ATG (7413G>A 7414A>T), CCG123GCT (7416C>G 7418G>T), CAA125GAG (7422C>G 7424A>G), AAT126GAT (7425A>G), ATG128ACA (7432T>C 7433G>A), GCT129GCA (7436T>A), AGT130TAT (7437A>T 7438G>A), GCT131GCC (7442T>C), CTC132ATG (7443C>A 7445C>G), TGC133TGT (7448C>T), GTG134ATC (7449G>A 7451G>C), GCT136TCA (7455G>T 7457T>A), TTT137CTT (7458T>C), GTA138GCT (7462T>C 7463A>T), AAT139CTT (7464A>C 7465A>T), AAA140GGG (7467A>G 7468A>G 7469A>G), GTT141CCA (7470G>C 7471T>C 7472T>A), GGT142AAT (7473G>A 7474G>A), GAT143ACT (7476G>A 7477A>C), GAT144GAG (7481T>G), CCT145CCG (7484T>G), ATA146ATG (7487A>G), ATC147ACT (7489T>C 7490C>T), AAG149GTT (7494A>G 7495A>T 7496G>T), GGT150GGA (7499T>A), GTG151AAA (7500G>A 7501T>A 7502G>A), CTA152ACG (7503C>A 7504T>C 7505A>G), AGA153CTG (7506A>C 7507G>T 7508A>G), GGT154GGA (7511T>A), GAG155AAG (7512G>A), ATT156ATA (7517T>A), GTA157GTG (7520A>G), ATG158CCA (7521A>C 7522T>C 7523G>A), CCA159GCT (7524C>G 7526A>T), CGC160AGG (7527C>A 7529C>G), CCA162CCC (7535A>C), TCA164AAT (7539T>A 7540C>A 7541A>T), TTT165TTC (7544T>C), GGG166GGA (7547G>A), GAC168GAT (7553C>T), CCA169CCG (7556A>G), ATT170ATA (7559T>A), TTC171TTT (7562C>T), CCA173CCT (7568A>T), CTT174CAT (7570T>A), AAC175GGC (7572A>G 7573A>G), TGG176TAT (7576G>A 7577G>T), AGA177GAC (7578A>G 7579G>A 7580A>C), AAG178CAG (7581A>C), ACA179ACT (7586A>T), TTT180TAC (7588T>A 7589T>C), ATG184CCC (7599A>C 7600T>C 7601G>C), GCC185AAG (7602G>A 7603C>A 7604C>G), GAG186GAA (7607G>A), GAG187GAA (7610G>A), AAT189AAC (7616T>C), ATG190AAG (7618T>A), ATA191ATT (7622A>T), CAT193CAC (7628T>C), CGA194CGC (7631A>C), TTT195GGT (7632T>G 7633T>G), CGA196AAA (7635C>A 7636G>A) | | | | | | | | |

*: Inserts / Deletes / Misaligned / Frameshifts

#### Analysis details

This analysis was performed with panviral2.64

### NGS Details (UN70): Human gammaherpesvirus 8

#### Assembly

| | |
|---|---|
| **Coverage Length** | 132 (1 contig(s)) |
| **Depth Of Coverage** | 240.5 |
| **Number Of Reads** | 386 |
| **Reads Per Million** | 7.32 rpm (after QC) |
| **Ambiguities** | 0 |
| **Assembly Method** | de novo + reference guided assembly |
| **Consensus Caller** | Bcf Tools |

#### Coverage Map

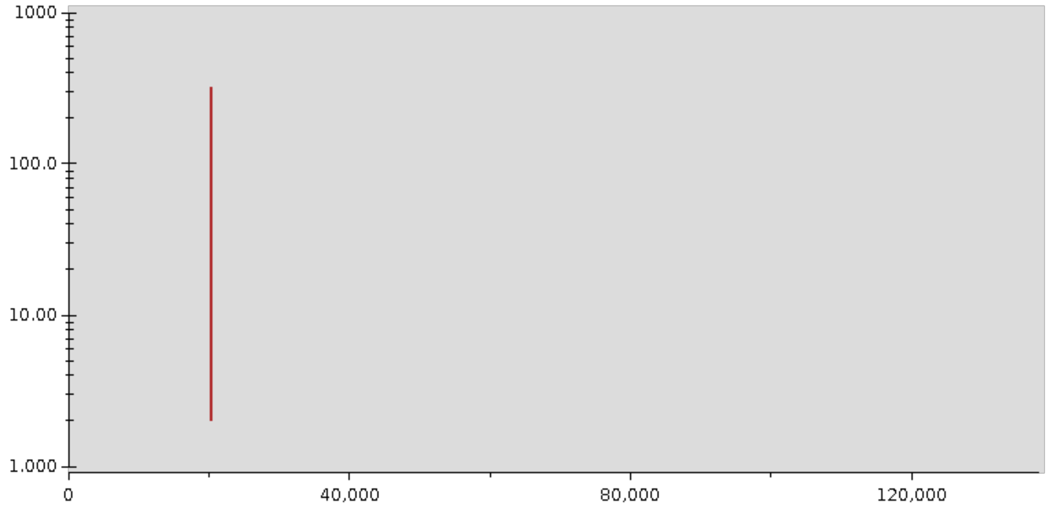

#### Assignment

| | |
|---|---|
| **Type** | Human gammaherpesvirus 8 (Taxonomy ID: 37296) |
| **Subtype** | Could not assign |
| **Reference Genome** | NC_009333.1 |
| **NT Identity (%)** | 83.3333 |
| **AA Identity (%)** | 84.0909 |
| **Number Of Stop Codons** | 0 |
| **Number Of CDS** | 86 |

#### Alignment

| | |
|---|---|
| **Alignment Score** | 176.0 (NT) + 283.0 (AA) = 459.0 |
| **Concordance (%)** | 77.9287 |
| **Alignment Method** | Local, heuristic, nucleotide (BLASTN) |

#### Genome Region

Sequence starts at position 20273 and ends at position 20404 relative to NC_009333.1 reference sequence.

#### Alignment Detailed Statistics

| | Begin | End | Coverage | Score | Concordance | Matches | Identities | I/D/M/F* | Stop Codons |
|---|---|---|---|---|---|---|---|---|---|
| **NT** | **20273** | **20404** | **0.1%** | **176** | **66.7%** | **132 (100%)** | **110 (83.3%)** | **0/0** | |

Mutations: 20293A>T, 20296A>C, 20311G>A, 20314T>G, 20315C>G, 20317C>A, 20320C>T, 20322T>G, 20331G>T, 20341A>C, 20347A>C, 20350G>A, 20352C>T, 20353A>G, 20369C>G, 20370A>C, 20371T>A, 20376A>T, 20383G>A, 20386C>T, 20389A>G, 20392C>T

*: Inserts / Deletes / Misaligned / Frameshifts

#### Analysis details

This analysis was performed with panviral2.64

### NGS Details (UN70): Makelovirus prm1

#### Assembly

| | |
|---|---|
| **Coverage Length** | 996 (1 contig(s)) |
| **Depth Of Coverage** | 40.8 |
| **Number Of Reads** | 316 |
| **Reads Per Million** | 5.99 rpm (after QC) |
| **Ambiguities** | 0 |
| **Assembly Method** | de novo + reference guided assembly |
| **Consensus Caller** | Bcf Tools |

#### Coverage Map

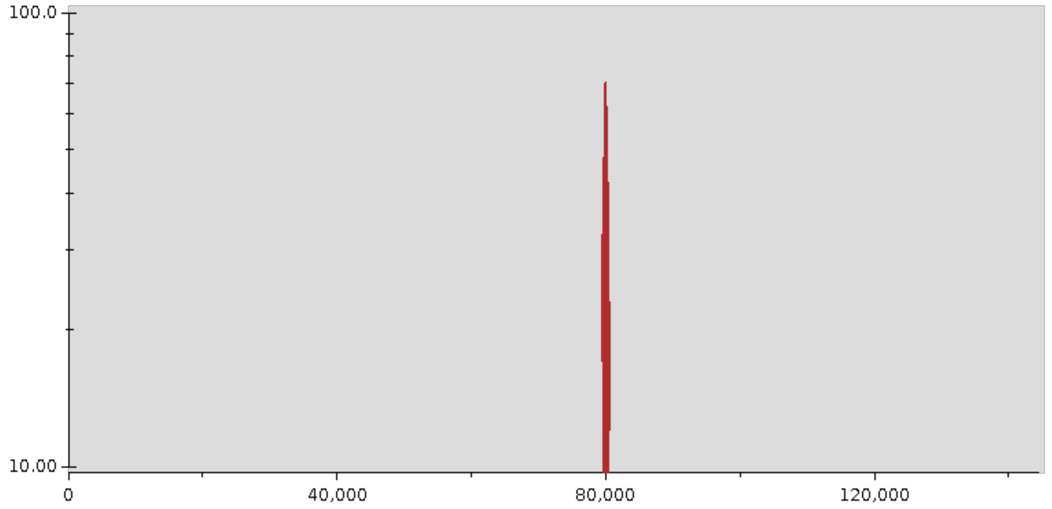

#### Assignment

| | |
|---|---|
| **Type** | Makelovirus prm1 (Taxonomy ID: 2956181) |
| **Reference Genome** | NC_055761.1 |
| **NT Identity (%)** | 76.4056 |
| **AA Identity (%)** | 87.3494 |
| **Number Of Stop Codons** | 0 |
| **Number Of CDS** | 190 |

#### Alignment

| | |
|---|---|
| **Alignment Score** | 1052.0 (NT) + 2227.0 (AA) = 3279.0 |
| **Concordance (%)** | 74.0682 |
| **Alignment Method** | Local, heuristic, nucleotide (BLASTN) |

#### Genome Region

Sequence starts at position 79515 and ends at position 80510 relative to NC_055761.1 reference sequence.

#### Alignment Detailed Statistics

| | Begin | End | Coverage | Score | Concordance | Matches | Identities | I/D/M/F* | Stop Codons |
|---|---|---|---|---|---|---|---|---|---|
| **NT** | **79515** | **80510** | **0.7%** | **1052** | **52.8%** | **996 (100%)** | **761 (76.4%)** | **0/0** | |

Mutations: 79519A>G, 79520A>T, 79522A>G, 79523A>C, 79529C>T, 79530G>A, 79532G>C, 79536G>A, 79541G>T, 79542T>A, 79543C>G, 79544T>C, 79547A>T, 79548A>G, 79550C>A, 79563G>A, 79565C>T, 79568T>A, 79574C>T, 79580G>T, 79581C>T, 79592A>T, 79595T>C, 79596C>T, 79598G>A, 79602G>A, 79604A>G, 79610C>T, 79611G>T, 79612T>C, 79613C>T, 79614T>G, 79615G>T, 79616C>A, 79619C>T, 79624C>T, 79628G>C, 79632G>A, 79643C>T, 79646T>A, 79649G>A, 79655C>T, 79658C>T, 79664C>T, 79673A>T, 79676C>T, 79677G>T, 79679T>A, 79682T>G, 79688G>A, 79689A>C, 79691G>T, 79694T>C, 79703C>T, 79712T>C, 79718T>C, 79719G>A, 79722G>A, 79724A>T, 79728T>A, 79733C>T, 79734A>G, 79735A>C, 79736C>A, 79739A>T, 79742T>A, 79746C>T, 79748T>A, 79751C>T, 79754C>T, 79760C>A, 79763T>C, 79772T>G, 79779C>G, 79791C>T, 79793C>A, 79799T>C, 79811C>T, 79812C>G, 79817G>A, 79818G>A, 79820C>T, 79823A>T, 79824T>C, 79826T>A, 79833C>T, 79835G>A, 79836A>C, 79838C>T, 79842A>G, 79844C>A, 79845T>G, 79846T>C, 79847C>T, 79850C>T, 79871A>G, 79874C>T, 79875T>A, 79876C>G, 79879A>T, 79880T>C, 79895T>A, 79898C>T, 79904C>T, 79905T>G, 79906G>C, 79907C>T, 79916C>T, 79919T>A, 79922A>T, 79931T>A, 79937A>T, 79938T>A, 79940T>C, 79946C>T, 79950C>T, 79953G>A, 79958T>C, 79961T>A, 79962T>A, 79973T>A, 79974T>A, 79975C>G, 79976G>T, 79979C>T, 79985C>T, 79994C>T, 79997C>A, 80000C>A, 80018C>T, 80025C>A, 80030C>A, 80039A>T, 80042A>G, 80054G>T, 80063C>A, 80066C>T, 80069T>C, 80073C>T, 80075T>A, 80078T>C, 80090C>A, 80096T>C, 80099T>C, 80105G>A, 80114A>T, 80120C>T, 80126T>C, 80127C>T, 80132T>A, 80139T>A, 80140C>G, 80141G>T, 80142C>T, 80145G>A, 80147T>C, 80148C>A, 80150T>G, 80159C>A, 80162G>A, 80165C>T, 80168G>A, 80171C>T, 80172C>G, 80173A>C, 80174G>T, 80177C>T, 80178T>G, 80180T>A, 80188A>G, 80189G>A, 80201A>G, 80207G>A, 80210C>T, 80213C>T, 80216C>T, 80222G>A, 80225T>C, 80237C>T, 80240C>T, 80243T>C, 80246C>A, 80247C>T, 80261C>T, 80265T>A, 80266C>G, 80267C>T, 80279C>T, 80286C>T, 80288G>A, 80291T>C, 80300T>A, 80306A>T, 80315T>A, 80318C>A, 80333T>C, 80336A>T, 80337C>T, 80339T>A, 80343G>A, 80346T>A, 80347C>G, 80351C>T, 80357A>T, 80366G>A, 80369C>T, 80378C>T, 80387G>A, 80390C>T, 80391A>G, 80393C>A, 80394A>G, 80396G>T, 80399T>C, 80400G>A, 80409A>C, 80410A>G, 80411A>T, 80414C>A, 80415C>A, 80417C>T, 80433G>A, 80435T>C, 80436C>A, 80438A>C, 80447C>T, 80453G>T, 80456G>T, 80462G>A, 80465A>T, 80471C>T, 80474G>A, 80486C>T, 80498G>A, 80504G>A

*: Inserts / Deletes / Misaligned / Frameshifts

#### Analysis details

This analysis was performed with panviral2.64

### NGS Details (UN70): Cladosporium fulvum T-1 virus

#### Assembly

| | |
|---|---|
| **Coverage Length** | 983 (5 contig(s)) |
| **Depth Of Coverage** | 14.0 |
| **Number Of Reads** | 132 |
| **Reads Per Million** | 2.50 rpm (after QC) |
| **Ambiguities** | 0 |
| **Assembly Method** | de novo + reference guided assembly |
| **Consensus Caller** | Bcf Tools |

#### Coverage Map

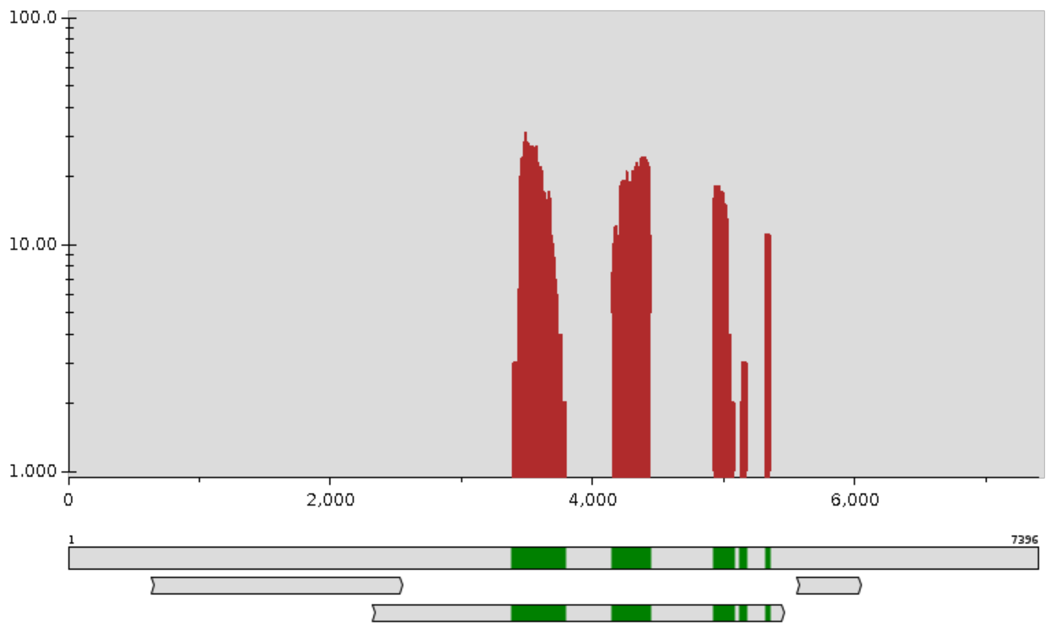

#### Assignment

| | |
|---|---|
| **Type** | Cladosporium fulvum T-1 virus (Taxonomy ID: 2052899) |
| **Reference Genome** | NC_043491.1 |
| **NT Identity (%)** | 54.4317 |
| **AA Identity (%)** | 50.625 |
| **Number Of Stop Codons** | 0 |
| **Number Of CDS** | 3 |

#### Alignment

| | |
|---|---|
| **Alignment Score** | 101.0 (NT) + 1177.0 (AA) = 1278.0 |
| **Concordance (%)** | 32.1752 |

| Alignment Method | Global, seeded, nucleotide + amino acids (AGA) |
|---|---|

#### Genome Region

Sequence starts at position 3387 and ends at position 5357 relative to NC_043491.1 reference sequence.

#### Alignment Detailed Statistics

| | Begin | End | Coverage | Score | Concordance | Matches | Identities | I/D/M/F* | Stop Codons |
|---|---|---|---|---|---|---|---|---|---|
| **NT** | **3387** | **5357** | **13.3%** | **101** | **5.5%** | **950 (95.8%)** | **522 (52.6%)** | **9/33** | |
| Mutations: | 3387C>A, 3388C>A, 3389T>G, 3391A>T, 3392A>C, 3393G>C, 3394T>A, 3395G>T, 3396C>A, 3397A>T, 3401A>G, 3403T>A, 3406A>T, 3407T>G, 3408G>T, 3410A>T, 3415C>T, 3416G>C, 3417T>A, 3418T>G, 3419C>A, 3420C>A, 3421A>G, 3424A>G, 3425G>A, 3426C>A, 3428A>G, 3430C>T, 3434A>T, 3435A>C, 3442A>C, 3443C>A, 3445C>G, 3446G>T, 3447T>G, 3448A>C, 3449C>A, 3450A>T, 3451A>G, 3454C>T, 3457C>T, 3460A>G, 3461A>G, 3462A>C, 3464T>C, 3470G>A, 3473A>G, 3475C>A, 3478G>T, 3479A>G, 3481C>G, 3488C>A, 3489G>A, 3490A>G, 3493T>C, 3496G>A, 3497C>G, 3499A>T, 3502C>T, 3503A>T, 3504A>T, 3505C>G, 3506A>G, 3508C>G, 3509G>C, 3511A>G, 3514A>T, 3515G>C, 3516C>T, 3517A>G, 3518C>A, 3519A>T, 3520A>G, 3526A>G, 3529A>G, 3531C>G, 3533G>A, 3534G>A, 3536T>G, 3538A>G, 3539G>T, 3540A>G, 3546A>T, 3550G>T, 3554A>C, 3556T>A, 3559C>T, 3562A>T, 3563C>A, 3565A>G, 3566G>T, 3567A>C, 3568C>A, 3570C>G, 3573T>A, 3574C>T, 3576A>G, 3577T>G, 3578G>C, 3579C>A, 3580T>G, 3581A>G, 3583C>T, 3584C>A, 3586A>G, 3589G>A, 3598A>G, 3601A>T, 3604A>G, 3605T>C, 3606G>C, 3607G>T, 3610A>G, 3613C>T, 3614G>A, 3618T>G, 3619C>T, 3620A>G, 3621G>T, 3622G>A, 3625A>T, 3628A>G, 3631C>T, 3634A>G, 3635C>T, 3636T>C, 3637C>A, 3640C>T, 3647T>C, 3649G>T, 3652C>A, 3658A>G, 3659A>T, 3661G>T, 3664A>G, 3670C>T, 3673C>T, 3676A>C, 3679C>G, 3682A>T, 3683T>A, 3687G>T, 3688C>T, 3689C>T, 3690A>G, 3691G>C, 3692G>A, 3695C>T, 3697T>A, 3698G>A, 3700C>G, 3703C>T, 3706A>T, 3707A>G, 3708C>T, 3709A>C, 3712T>G, 3713A>T, 3714G>A, 3715A>T, 3719C>T, 3720T>A, 3721A>T, 3724C>T, 3727C>T, 3729T>A, 3732G>T, 3733C>T, 3736C>A, 3739T>A, 3741C>T, 3742T>C, 3746A>T, 3748G>A, 3751C>T, 3757A>T, 3758C>G, 3760G>C, 3761G>A, 3763C>A, 3766C>T, 3768C>G, 3769A>T, 3773G>A, 3775A>T, 3776T>A, 3778C>G, 3779C>T, 3781C>G, 3782C>G, 3784G>T, 3791A>G, 3792C>T, 3793C>G, 3796G>C, 3799A>T, 4146C>G, 4150G>C, 4150_4151insAATATTGCG, 4151C>G, 4154C>G, 4155G>C, 4156A>T, 4157C>G, 4158T>A, 4159A>C, 4162C>T, 4166G>T, 4167G>T, 4168T>A, 4169A>C, 4170G>C, 4171C>A, 4172A>T, 4173A>T, 4174G>T, 4180C>G, 4186C>T, 4187G>A, 4189G>T, 4195C>T, 4198G>C, 4204T>C, 4205A>T, 4207G>C, 4210A>T, 4213A>T, 4216C>G, 4218C>G, 4219A>T, 4220T>G, 4221G>T, 4222T>G, 4225A>T, 4227C>T, 4228A>T, 4231G>A, 4233C>A, 4234A>T, 4235C>G, 4238_4249delGATGGGAAAAGA, 4252C>T, 4255A>T, 4258G>T, 4261C>A, 4263A>T, 4265T>G, 4267T>G, 4268T>A, 4269C>G, 4271C>A, 4277A>T, 4281C>A, 4282C>T, 4283A>C, 4285A>G, 4286G>A, 4294G>A, 4296A>G, 4297C>A, 4300C>T, 4301G>T, 4302A>C, 4303C>T, 4304A>G, 4305T>C, 4306C>A, 4312C>A, 4319C>A, 4321T>G, 4322C>G, 4324A>G, 4327C>A, 4328A>T, 4330T>G, 4331G>C, 4333T>C, 4334G>C, 4335C>A, 4336C>T, 4337G>T, 4338C>G, 4339C>T, 4340A>C, 4342G>T, 4346C>A, 4347A>C, 4348T>A, 4354A>G, 4360C>T, 4361G>C, 4364G>T, 4365A>T, 4366G>A, 4369C>A, 4370C>A, 4373_4375delCCG, 4381A>T, 4382A>G, 4383C>T, 4384G>A, 4385A>G, 4387T>A, 4388C>A, 4389T>A, 4390T>G, 4391T>A, 4393A>G, 4397C>A, 4399C>T, 4400A>G, 4401A>T, 4403A>G, 4404A>C, 4406C>A, 4407T>A, 4408C>T, 4411G>T, 4419C>A, 4420G>A, 4423G>T, 4424A>C, 4425C>A, 4426G>A, 4427A>C, 4430G>A, 4433C>T, 4435T>G, 4437C>G, 4438C>T, 4439C>T, 4440G>T, 4441A>G, 4443G>A, 4444A>G, 4924A>T, 4933C>G, 4934G>A, 4935A>G, 4936G>T, 4937G>A, 4939T>C, 4954T>A, 4957G>C, 4958A>T, 4959A>G, 4960A>T, 4961C>T, 4963C>G, 4966G>A, 4968G>A, 4972A>C, 4973_4975delAAG, 4979_4993delCGAGTCACAGGACAA, 4995C>G, 4996C>G, 5002C>A, 5004T>C, 5005G>C, 5009C>T, 5011A>G, 5014C>T, 5015A>G, 5020C>A, 5023C>T, 5027C>T, 5029C>G, 5038T>C, 5041A>C, 5044C>T, 5047C>T, 5050T>G, 5053T>C, 5055C>T, 5056A>C, 5057T>A, 5058C>A, 5061A>C, 5062A>T, 5063A>G, 5064T>C, 5065A>T, 5067A>G, 5074A>C, 5075G>A, 5078C>G, 5079A>C, 5081C>G, 5082T>C, 5083C>T, 5085G>C, 5122A>G, 5123T>C, 5125C>G, 5129G>A, 5130A>G, 5131A>G, 5132G>C, 5133T>C, 5134A>C, 5135T>A, 5137C>T, 5143A>T, 5146C>T, 5147A>C, 5153A>C, 5154A>C, 5157T>G, 5161C>T, 5164A>T, 5165T>G, 5166C>G, 5172A>T, 5319T>C, 5320T>G, 5321T>A, 5322C>A, 5323A>G, 5324T>C, 5329G>A, 5330C>G, 5331C>A, 5332A>C, 5333A>G, 5338G>C, 5341G>A, 5342A>T, 5344C>T, 5345G>T, 5348C>T, 5349T>A, 5350G>T, 5353C>T | | | | | | | | |

**CDS**

| | Begin | End | Coverage | Score | Concordance | Matches | Identities | I/D/M/F* | Stop Codons |
|---|---|---|---|---|---|---|---|---|---|
| homologue_of_retroviral_POL_genes | 355 | 1011 | 31.4% | 1177 | 53.8% | 317 (95.8%) | 162 (48.9%) | 3/11/0/0 | 0 |
| Protein mutations: | S355A (3389T>G 3391A>T), S356P (3392A>C 3393G>C 3394T>A), A357Y (3395G>T 3396C>A 3397A>T), T359A (3401A>G 3403T>A), C361V (3407T>G 3408G>T), M362L (3410A>T), V364Q (3416G>C 3417T>A 3418T>G), P365K (3419C>A 3420C>A 3421A>G), A367K (3425G>A 3426C>A), N368D (3428A>G 3430C>T), K370S (3434A>T 3435A>C), L373M (3443C>A 3445C>G), V374C (3446G>T 3447T>G 3448A>C), Q375M (3449C>A 3450A>T 3451A>G), K379A (3461A>G 3462A>C), E382K (3470G>A), I383V (3473A>G 3475C>A), I385V (3479A>G 3481C>G), R388K (3488C>A 3489G>A 3490A>G), L391V (3497C>G 3499A>T), N393L (3503A>T 3504A>T 3505C>G), I394V (3506A>G 3508C>G), E395Q (3509G>C 3511A>G), E396D (3514A>T), A397L (3515G>C 3516C>T 3517A>G), Q398M (3518C>A 3519A>T 3520A>G), T402S (3531C>G), G403K (3533G>A 3534G>A), S404A (3536T>G 3538A>G), D405C (3539G>T 3540A>G), Y407F (3546A>T), I410L (3554A>C 3556T>A), D414S (3566G>T 3567A>C 3568C>A), A415G (3570C>G), F416Y (3573T>A 3574C>T), Y417W (3576A>G 3577T>G), A418Q (3578G>C 3579C>A 3580T>G), I419V (3581A>G 3583C>T), M421I (3589G>A), E425D (3601A>T), W427P (3605T>C 3606G>C 3607G>T), A430T (3614G>A), F431C (3618T>G 3619C>T), R432V (3620A>G 3621G>T 3622G>A), L437S (3635C>T 3636T>C 3637C>A), M445F (3659A>T 3661G>T), S453T (3683T>A), C454F (3687G>T 3688C>T), Q455C (3689C>T 3690A>G 3691G>C), D456N (3692G>A), V458M (3698G>A 3700C>G), E460D (3706A>T), T461V (3707A>G 3708C>T 3709A>C), R463Y (3713A>T 3714G>A 3715A>T), L465Y (3719C>T 3720T>A 3721A>T), V468E (3729T>A), C469F (3732G>T 3733C>T), A472V (3741C>T 3742T>C), M474L (3746A>T 3748G>A), L478V (3758C>G 3760G>C), V479I (3761G>A 3763C>A), T481S (3768C>G 3769A>T), G483S (3773G>A 3775A>T), S484T (3776T>A 3778C>G), Q486D (3782C>G 3784G>T), T489V (3791A>G 3792C>T 3793C>G), K490N (3796G>C), Q491H (3799A>T), T608_L609insNIA (4150_4151insAATATTGCG), L609V (4151C>G), R610A (4154C>G 4155G>C 4156A>T), L611D (4157C>G 4158T>A 4159A>C), G614L (4166G>T 4167G>T 4168T>A), S615P (4169A>C 4170G>C 4171C>A), K616F (4172A>T 4173A>T 4174G>T), E621N (4187G>A 4189G>T), M627F (4205A>T 4207G>C), A631G (4218C>G 4219A>T), C632V (4220T>G 4221G>T 4222T>G), T634I (4227C>T 4228A>T), T636N (4233C>A 4234A>T), H637D (4235C>G), D638_R641del (4238_4249delGATGGGAAAAGA), Y646F (4263A>T), Y647E (4265T>G 4267T>G), M651L (4277A>T), T652N (4281C>A 4282C>T), T653P (4283A>C 4285A>G), A654T (4286G>A), N657R (4296A>G 4297C>A), D659S (4301G>T 4302A>C 4303C>T), I660A (4304A>G 4305T>C 4306C>A), D662E (4312C>A), L665M (4319C>A 4321T>G), L666V (4322C>G 4324A>G), I668L (4328A>T 4330T>G), V669L (4331G>C 4333T>C), A670H (4334G>C 4335C>A 4336C>T), A671C (4337G>T 4338C>G 4339C>T), M672L (4340A>C 4342G>T), H674T (4346C>A 4347A>C 4348T>A), V679L (4361G>C), E680L (4364G>T 4365A>T 4366G>A), P682T (4370C>A), P683del (4373_4375delCCG), L685F (4381A>T), T686V (4382A>G 4383C>T 4384G>A), I687V (4385A>G 4387T>A), L688K (4388C>A 4389T>A 4390T>G), S689T (4391T>A 4393A>G), H691N (4397C>A 4399C>T), K692V (4400A>G 4401A>T), N693A (4403A>G 4404A>C), L694N (4406C>A 4407T>A 4408C>T), T698K (4419C>A 4420G>A), T700Q (4424A>C 4425C>A 4426G>A), K701Q (4427A>C), E702K (4430G>A), T704S (4437C>G 4438C>T), R705L (4439C>T 4440G>T 4441A>G), R706K (4443G>A 4444A>G), K866N (4924A>T), D869E (4933C>G), E870S (4934G>A 4935A>G 4936G>T), V871I (4937G>A 4939T>C), K878C (4958A>T 4959A>G 4960A>T), R881K (4968G>A), K883del (4973_4975delAAG), R885_Q889del (4979_4993delCGAGTCACAGGACAA), A890G (4995C>G 4996C>G), D892E (5002C>A), M893T (5004T>C 5005G>C), M897V (5015A>G), I908M (5050T>G), A910V (5055C>T 5056A>C), S911K (5057T>A 5058C>A), E912A (5061A>C 5062A>T), I913A (5063A>G 5064T>C 5065A>T), Y914C (5067A>G), E917K (5075G>A), Q918A (5078C>G 5079A>C), L919A (5081C>G 5082T>C 5083C>T), F933L (5123T>C 5125C>G), E935R (5129G>A 5130A>G 5131A>G), V936P (5132G>C 5133T>C 5134A>C), F937I (5135T>A 5137C>T), K943P (5153A>C 5154A>C), L944R (5157T>G), S947G (5165T>G 5166C>G), Y949F (5172A>T), V998A (5319T>C 5320T>G), S999K (5321T>A 5322C>A 5323A>G), P1002D (5330C>G 5331C>A 5332A>C), M1003V (5333A>G), I1006F (5342A>T 5344C>T), A1007S (5345G>T), L1008Y (5348C>T 5349T>A 5350G>T) | | | | | | | | |

| | Begin | End | Coverage | Score | Concordance | Matches | Identities | I/D/M/F* | Stop Codons |
|---|---|---|---|---|---|---|---|---|---|
| **NT** | **3387** | **5357** | **13.3%** | **101** | **5.5%** | **950 (95.8%)** | **522 (52.6%)** | **9/33** | |
| Codon mutations: | ACC354.AA (3387C>A 3388C>A), TCA355GCT (3389T>G 3391A>T), AGT356CCA (3392A>C 3393G>C 3394T>A), GCA357TAT (3395G>T 3396C>A 3397A>T), ACT359GCA (3401A>G 3403T>A), CCA360CCT (3406A>T), TGC361GTC (3407T>G 3408G>T), ATG362TTG (3410A>T), TTC363TTT (3415C>T), GTT364CAG (3416G>C 3417T>A 3418T>G), CCA365AAG (3419C>A 3420C>A 3421A>G), AAA366AAG (3424A>G), GCA367AAA (3425G>A 3426C>A), AAC368GAT (3428A>G 3430C>T), AAA370TCA (3434A>T 3435A>C), CGA372CGC (3442A>C), CTC373ATG (3443C>A 3445C>G), GTA374TGC (3446G>T 3447T>G 3448A>C), CAA375ATG (3449C>A 3450A>T 3451A>G), GAC376GAT (3454C>T), TAC377TAT (3457C>T), CGA378CGG (3460A>G), AAG379GCG (3461A>G 3462A>C), TTG380CTG (3464T>C), GAG382AAG (3470G>A), ATC383GTA (3473A>G 3475C>A), ACG384ACT (3478G>T), ATC385GTG (3479A>G 3481C>G), CGA388AAG (3488C>A 3489G>A 3490A>G), TAT389TAC (3493T>C), CCG390CCA (3496G>A), CTA391GTT (3497C>G 3499A>T), CCC392CCT (3502C>T), AAC393TTG (3503A>T 3504A>T 3505C>G), ATC394GTG (3506A>G 3508C>G), GAA395CAG (3509G>C 3511A>G), GAA396GAT (3514A>T), GCA397CTG (3515G>C 3516C>T 3517A>G), CAA398ATG (3518C>A 3519A>T 3520A>G), AGA400AGG (3526A>G), TTA401TTG (3529A>G), ACC402AGC (3531C>G), GGA403AAA (3533G>A 3534G>A), TCA404GCG (3536T>G 3538A>G), GAC405TGC (3539G>T 3540A>G), TAC407TTC (3546A>T), ACG408ACT (3550G>T), ATT410CTA (3554A>C 3556T>A), GAC411GAT (3559C>T), CTA412CTT (3562A>T), CGA413AGG (3563C>A 3565A>G), GAC414TCA (3566G>T 3567A>C 3568C>A), GCC415GGC (3570C>G), TTC416TAT (3573T>A 3574C>T), TAT417TGG (3576A>G 3577T>G), GCT418CAG (3578G>C 3579C>A 3580T>G), ATC419GTT (3581A>G 3583C>T), CGA420AGG (3584C>A 3586A>G), ATG421ATA (3589G>A), GGA424GGG (3598A>G), GAA425GAT (3601A>T), GAA426GAG (3604A>G), TGG427CCT (3605T>C 3606G>C 3607G>T), AAA428AAG (3610A>G), ACC429ACT (3613C>T), GCT430ACT (3614G>A), TTC431TGT (3618T>G 3619C>T), AGG432GTA (3620A>G 3621G>T 3622G>A), ACA433ACT (3625A>T), AGA434AGG (3628A>G), TAC435TAT (3631C>T), GGA436GGG (3634A>G), CTC437TCA (3635C>T 3636T>C 3637C>A), TAC438TAT (3640C>T), TTG441CTT (3647T>C 3649G>T), GTC442GTA (3652C>A), CCA444CCG (3658A>G), ATG445TTT (3659A>T 3661G>T), GGA446GGG (3664A>G), ACC448ACT (3670C>T), AAC449AAT (3673C>T), GCA450GCC (3676A>C), CCC451CCG (3679C>G), GCA452GCT (3682A>T), TCC453ACC (3683T>A), TGC454TTT (3687G>T 3688C>T), CAG455TGC (3689C>T 3690A>G 3691G>C), GAC456AAC (3692G>A), CTT457TTA (3695C>T 3697T>A), GTC458ATG (3698G>A 3700C>G), AAC459AAT (3703C>T), GAA460GAT (3706A>T), ACA461GTC (3707A>G 3708C>T 3709A>C), CTT462CTG (3712T>G), AGA463TAT (3713A>T 3714G>A 3715A>T), CTA465TAT (3719C>T 3720T>A 3721A>T), CTC466CTT (3724C>T), GAC467GAT (3727C>T), GTG468GAG (3729T>A), TGC469TTT (3732G>T 3733C>T), GTC470GTA (3736C>A), GTT471GTA (3739T>A), GCT472GTC (3741C>T 3742T>C), ATG474TTA (3746A>T 3748G>A), GAC475GAT (3751C>T), ATA477ATT (3757A>T), CTG478GTC (3758C>G 3760G>C), GTC479ATA (3761G>A 3763C>A), TAC480TAT (3766C>T), ACA481AGT (3768C>G 3769A>T), GGA483AGT (3773G>A 3775A>T), TCC484ACG (3776T>A 3778C>G), CTC485TTG (3779C>T 3781C>G), CAG486GAT (3782C>G 3784G>T), ACC489GTG (3791A>G 3792C>T 3793C>G), AAG490AAC (3796G>C), CAA491CAT (3799A>T), CCA607.GA (4146C>G), ACG608ACC (4150G>C), ACG608_CTT609insAATATTGCG (4150_4151insAATATTGCG), CTT609GTT (4151C>G), CGA610GCT (4154C>G 4155G>C 4156A>T), CTA611GAC (4157C>G 4158T>A 4159A>C), TTC612TTT (4162C>T), GGT614TTA (4166G>T 4167G>T 4168T>A), AGC615CCA (4169A>C 4170G>C 4171C>A), AAG616TTT (4172A>T 4173A>T 4174G>T), GTC618GTG (4180C>G), ATC620ATT (4186C>T), GAG621AAT (4187G>A 4189G>T), GAC623GAT (4195C>T), GCG624GCC (4198G>C), GAT626GAC (4204T>C), ATG627TTC (4205A>T 4207G>C), GCA628GCT (4210A>T), ATA629ATT (4213A>T), GGC630GGG (4216C>G), GCA631GGT (4218C>G 4219A>T), TGT632GTG (4220T>G 4221G>T 4222T>G), CTA633CTT (4225A>T), ACA634ATT (4227C>T 4228A>T), CAG635CAA (4231G>A), ACA636AAT (4233C>A 4234A>T), CAC637GAC (4235C>G), GAT638_AGA641del (4238_4249delGATGGGAAAAGA), CAC642CAT (4252C>T), CCA643CCT (4255A>T), GTG644GTT (4258G>T), GCC645GCA (4261C>A), TAT646TTT (4263A>T), TAT647GAG (4265T>G 4267T>G), TCC648AGC (4268T>A 4269C>G), CGG649AGG (4271C>A), ATG651TTG (4277A>T), ACC652AAT (4281C>A 4282C>T), ACA653CCG (4283A>C 4285A>G), GCG654ACG (4286G>A), CAG656CAA (4294G>A), AAC657AGA (4296A>G 4297C>A), TAC658TAT (4300C>T), GAC659TCT (4301G>T 4302A>C 4303C>T), ATC660GCA (4304A>G 4305T>C 4306C>A), GAC662GAA (4312C>A), CTT665ATG (4319C>A 4321T>G), CTA666GTG (4322C>G 4324A>G), GCC667GCA (4327C>A), ATT668TTG (4328A>T 4330T>G), GTT669CTC (4331G>C 4333T>C), GCC670CAT (4334G>C 4335C>A 4336C>T), GCC671TGT (4337G>T 4338C>G 4339C>T), ATG672CTT (4340A>C 4342G>T), CAT674ACA (4346C>A 4347A>C 4348T>A), AGA676AGG (4354A>G), TAC678TAT (4360C>T), GTC679CTC (4361G>C), GAG680TTA (4364G>T 4365A>T 4366G>A), GGC681GGA (4369C>A), CCA682ACA (4370C>A), CCG683del (4373_4375delCCG), TTA685TTT (4381A>T), ACG686GTA (4382A>G 4383C>T 4384G>A), ATT687GTA (4385A>G 4387T>A), CTT688AAG (4388C>A 4389T>A 4390T>G), TCA689ACG (4391T>A 4393A>G), CAC691AAT (4397C>A 4399C>T), AAG692GTG (4400A>G 4401A>T), AAT693GCT (4403A>G 4404A>C), CTC694AAT (4406C>A 4407T>A 4408C>T), ACG695ACT (4411G>T), ACG698AAA (4419C>A 4420G>A), ACG699ACT (4423G>T), ACG700CAA (4424A>C 4425C>A 4426G>A), AAG701CAG (4427A>C), GAA702AAA (4430G>A), CTT703TTG (4433C>T 4435T>G), ACC704AGT (4437C>G 4438C>T), CGA705TTG (4439C>T 4440G>T 4441A>G), AGA706AAG (4443G>A 4444A>G), AAA866AAT (4924A>T), GAC869GAG (4933C>G), GAG870AGT (4934G>A 4935A>G 4936G>T), GTT871ATC (4937G>A 4939T>C), ATT876ATA (4954T>A), ACG877ACC (4957G>C), AAA878TGT (4958A>T 4959A>G 4960A>T), CTC879TTG (4961C>T 4963C>G), CCG880CCA (4966G>A), AGG881AAG (4968G>A), TCA882TCC (4972A>C), AAG883del (4973_4975delAAG), CGA885_CAA889del (4979_4993delCGAGTCACAGGACAA), GCC890GGG (4995C>G 4996C>G), GAC892GAA (5002C>A), ATG893ACC (5004T>C 5005G>C), CTA895TTG (5009C>T 5011A>G), GTC896GTT (5014C>T), ATG897GTG (5015A>G), GTC898GTA (5020C>A), GAC899GAT (5023C>T), CTC901TTG (5027C>T 5029C>G), TAT904TAC (5038T>C), GCA905GCC (5041A>C), CAC906CAT (5044C>T), TTC907TTT (5047C>T), ATT908ATG (5050T>G), CCT909CCC (5053T>C), GCA910GTC (5055C>T 5056A>C), TCA911AAA (5057T>A 5058C>A), GAA912GCT (5061A>C 5062A>T), ATA913GCT (5063A>G 5064T>C 5065A>T), TAC914TGC (5067A>G), GCA916GCC (5074A>C), GAG917AAG (5075G>A), CAG918GCG (5078C>G 5079A>C), CTC919GCT (5081C>G 5082T>C 5083C>T), GGA920GC. (5085G>C), GGA932GGG (5122A>G), TTC933CTG (5123T>C 5125C>G), GAA935AGG (5129G>A 5130A>G 5131A>G), GTA936CCC (5132G>C 5133T>C 5134A>C), TTC937ATT (5135T>A 5137C>T), ACA939ACT (5143A>T), GAC940GAT (5146C>T), AGA941CGA (5147A>C), AAG943CCG (5153A>C 5154A>C), CTC944CGC (5157T>G), TTC945TTT (5161C>T), ACA946ACT (5164A>T), TCA947GGA (5165T>G 5166C>G), TAC949TTC (5172A>T), GTT998GCG (5319T>C 5320T>G), TCA999AAG (5321T>A 5322C>A 5323A>G), TTA1000CTA (5324T>C), CTG1001CTA (5329G>A), CCA1002GAC (5330C>G 5331C>A 5332A>C), ATG1003GTG (5333A>G), GCG1004GCC (5338G>C), CAG1005CAA (5341G>A), ATC1006TTT (5342A>T 5344C>T), GCA1007TCA (5345G>T), CTG1008TAT (5348C>T 5349T>A 5350G>T), AAC1009AAT (5353C>T) | | | | | | | | |

#### Proteins

| | Begin | End | Coverage | Score | Concordance | Matches | Identities | I/D/M/F* | Stop Codons |
|---|---|---|---|---|---|---|---|---|---|
| Reverse Transcriptase (YP_009666308.1) | 355 | 1011 | 31.4% | 1177 | 53.8% | 317 (95.8%) | 162 (48.9%) | 3/11/0/0 | 0 |
| Protein mutations: | S355A (3389T>G 3391A>T), S356P (3392A>C 3393G>C 3394T>A), A357Y (3395G>T 3396C>A 3397A>T), T359A (3401A>G 3403T>A), C361V (3407T>G 3408G>T), M362L (3410A>T), V364Q (3416G>C 3417T>A 3418T>G), P365K (3419C>A 3420C>A 3421A>G), A367K (3425G>A 3426C>A), N368D (3428A>G 3430C>T), K370S (3434A>T 3435A>C), L373M (3443C>A 3445C>G), V374C (3446G>T 3447T>G 3448A>C), Q375M (3449C>A 3450A>T 3451A>G), K379A (3461A>G 3462A>C), E382K (3470G>A), I383V (3473A>G 3475C>A), I385V (3479A>G 3481C>G), R388K (3488C>A 3489G>A 3490A>G), L391V (3497C>G 3499A>T), N393L (3503A>T 3504A>T 3505C>G), I394V (3506A>G 3508C>G), E395Q (3509G>C 3511A>G), E396D (3514A>T), A397L (3515G>C 3516C>T 3517A>G), Q398M (3518C>A 3519A>T 3520A>G), T402S (3531C>G), G403K (3533G>A 3534G>A), S404A (3536T>G 3538A>G), D405C (3539G>T 3540A>G), Y407F (3546A>T), I410L (3554A>C 3556T>A), D414S (3566G>T 3567A>C 3568C>A), A415G (3570C>G), F416Y (3573T>A 3574C>T), Y417W (3576A>G 3577T>G), A418Q (3578G>C 3579C>A 3580T>G), I419V (3581A>G 3583C>T), M421I (3589G>A), E425D (3601A>T), W427P (3605T>C 3606G>C 3607G>T), A430T (3614G>A), F431C (3618T>G 3619C>T), R432V (3620A>G 3621G>T 3622G>A), L437S (3635C>T 3636T>C 3637C>A), M445F (3659A>T 3661G>T), S453T (3683T>A), C454F (3687G>T 3688C>T), Q455C (3689C>T 3690A>G 3691G>C), D456N (3692G>A), V458M (3698G>A 3700C>G), E460D (3706A>T), T461V (3707A>G 3708C>T 3709A>C), R463Y (3713A>T 3714G>A 3715A>T), L465Y (3719C>T 3720T>A 3721A>T), V468E (3729T>A), C469F (3732G>T 3733C>T), A472V (3741C>T 3742T>C), M474L (3746A>T 3748G>A), L478V (3758C>G 3760G>C), V479I (3761G>A 3763C>A), T481S (3768C>G 3769A>T), G483S (3773G>A 3775A>T), S484T (3776T>A 3778C>G), Q486D (3782C>G 3784G>T), T489V (3791A>G 3792C>T 3793C>G), K490N (3796G>C), Q491H (3799A>T), T608_L609insNIA (4150_4151insAATATTGCG), L609V (4151C>G), R610A (4154C>G 4155G>C 4156A>T), L611D (4157C>G 4158T>A 4159A>C), G614L (4166G>T 4167G>T 4168T>A), S615P (4169A>C 4170G>C 4171C>A), K616F (4172A>T 4173A>T 4174G>T), E621N (4187G>A 4189G>T), M627F (4205A>T 4207G>C), A631G (4218C>G 4219A>T), C632V (4220T>G 4221G>T 4222T>G), T634I (4227C>T 4228A>T), T636N (4233C>A 4234A>T), H637D (4235C>G), D638_R641del (4238_4249delGATGGGAAAAGA), Y646F (4263A>T), Y647E (4265T>G 4267T>G), M651L (4277A>T), T652N (4281C>A 4282C>T), T653P (4283A>C 4285A>G), A654T (4286G>A), N657R (4296A>G 4297C>A), D659S (4301G>T 4302A>C 4303C>T), I660A (4304A>G 4305T>C 4306C>A), D662E (4312C>A), L665M (4319C>A 4321T>G), L666V (4322C>G 4324A>G), I668L (4328A>T 4330T>G), V669L (4331G>C 4333T>C), A670H (4334G>C 4335C>A 4336C>T), A671C (4337G>T 4338C>G 4339C>T), M672L (4340A>C 4342G>T), H674T (4346C>A 4347A>C 4348T>A), V679L (4361G>C), E680L (4364G>T 4365A>T 4366G>A), P682T (4370C>A), P683del (4373_4375delCCG), L685F (4381A>T), T686V (4382A>G 4383C>T 4384G>A), I687V (4385A>G 4387T>A), L688K (4388C>A 4389T>A 4390T>G), S689T (4391T>A 4393A>G), H691N (4397C>A 4399C>T), K692V (4400A>G 4401A>T), N693A (4403A>G 4404A>C), L694N (4406C>A 4407T>A 4408C>T), T698K (4419C>A 4420G>A), T700Q (4424A>C 4425C>A 4426G>A), K701Q (4427A>C), E702K (4430G>A), T704S (4437C>G 4438C>T), R705L (4439C>T 4440G>T 4441A>G), R706K (4443G>A 4444A>G), K866N (4924A>T), D869E (4933C>G), E870S (4934G>A 4935A>G 4936G>T), V871I (4937G>A 4939T>C), K878C (4958A>T 4959A>G 4960A>T), R881K (4968G>A), K883del (4973_4975delAAG), R885_Q889del (4979_4993delCGAGTCACAGGACAA), A890G (4995C>G 4996C>G), D892E (5002C>A), M893T (5004T>C 5005G>C), M897V (5015A>G), I908M (5050T>G), A910V (5055C>T 5056A>C), S911K (5057T>A 5058C>A), E912A (5061A>C 5062A>T), I913A (5063A>G 5064T>C 5065A>T), Y914C (5067A>G), E917K (5075G>A), Q918A (5078C>G 5079A>C), L919A (5081C>G 5082T>C 5083C>T), F933L (5123T>C 5125C>G), E935R (5129G>A 5130A>G 5131A>G), V936P (5132G>C 5133T>C 5134A>C), F937I (5135T>A 5137C>T), K943P (5153A>C 5154A>C), L944R (5157T>G), S947G (5165T>G 5166C>G), Y949F (5172A>T), V998A (5319T>C 5320T>G), S999K (5321T>A 5322C>A 5323A>G), P1002D (5330C>G 5331C>A 5332A>C), M1003V (5333A>G), I1006F (5342A>T 5344C>T), A1007S (5345G>T), L1008Y (5348C>T 5349T>A 5350G>T) | | | | | | | | |

| | Begin | End | Coverage | Score | Concordance | Matches | Identities | I/D/M/F* | Stop Codons |
|---|---|---|---|---|---|---|---|---|---|
| **NT** | **3387** | **5357** | **13.3%** | **101** | **5.5%** | **950 (95.8%)** | **522 (52.6%)** | **9/33** | |
| Codon mutations: | ACC354.AA (3387C>A 3388C>A), TCA355GCT (3389T>G 3391A>T), AGT356CCA (3392A>C 3393G>C 3394T>A), GCA357TAT (3395G>T 3396C>A 3397A>T), ACT359GCA (3401A>G 3403T>A), CCA360CCT (3406A>T), TGC361GTC (3407T>G 3408G>T), ATG362TTG (3410A>T), TTC363TTT (3415C>T), GTT364CAG (3416G>C 3417T>A 3418T>G), CCA365AAG (3419C>A 3420C>A 3421A>G), AAA366AAG (3424A>G), GCA367AAA (3425G>A 3426C>A), AAC368GAT (3428A>G 3430C>T), AAA370TCA (3434A>T 3435A>C), CGA372CGC (3442A>C), CTC373ATG (3443C>A 3445C>G), GTA374TGC (3446G>T 3447T>G 3448A>C), CAA375ATG (3449C>A 3450A>T 3451A>G), GAC376GAT (3454C>T), TAC377TAT (3457C>T), CGA378CGG (3460A>G), AAG379GCG (3461A>G 3462A>C), TTG380CTG (3464T>C), GAG382AAG (3470G>A), ATC383GTA (3473A>G 3475C>A), ACG384ACT (3478G>T), ATC385GTG (3479A>G 3481C>G), CGA388AAG (3488C>A 3489G>A 3490A>G), TAT389TAC (3493T>C), CCG390CCA (3496G>A), CTA391GTT (3497C>G 3499A>T), CCC392CCT (3502C>T), AAC393TTG (3503A>T 3504A>T 3505C>G), ATC394GTG (3506A>G 3508C>G), GAA395CAG (3509G>C 3511A>G), GAA396GAT (3514A>T), GCA397CTG (3515G>C 3516C>T 3517A>G), CAA398ATG (3518C>A 3519A>T 3520A>G), AGA400AGG (3526A>G), TTA401TTG (3529A>G), ACC402AGC (3531C>G), GGA403AAA (3533G>A 3534G>A), TCA404GCG (3536T>G 3538A>G), GAC405TGC (3539G>T 3540A>G), TAC407TTC (3546A>T), ACG408ACT (3550G>T), ATT410CTA (3554A>C 3556T>A), GAC411GAT (3559C>T), CTA412CTT (3562A>T), CGA413AGG (3563C>A 3565A>G), GAC414TCA (3566G>T 3567A>C 3568C>A), GCC415GGC (3570C>G), TTC416TAT (3573T>A 3574C>T), TAT417TGG (3576A>G 3577T>G), GCT418CAG (3578G>C 3579C>A 3580T>G), ATC419GTT (3581A>G 3583C>T), CGA420AGG (3584C>A 3586A>G), ATG421ATA (3589G>A), GGA424GGG (3598A>G), GAA425GAT (3601A>T), GAA426GAG (3604A>G), TGG427CCT (3605T>C 3606G>C 3607G>T), AAA428AAG (3610A>G), ACC429ACT (3613C>T), GCT430ACT (3614G>A), TTC431TGT (3618T>G 3619C>T), AGG432GTA (3620A>G 3621G>T 3622G>A), ACA433ACT (3625A>T), AGA434AGG (3628A>G), TAC435TAT (3631C>T), GGA436GGG (3634A>G), CTC437TCA (3635C>T 3636T>C 3637C>A), TAC438TAT (3640C>T), TTG441CTT (3647T>C 3649G>T), GTC442GTA (3652C>A), CCA444CCG (3658A>G), ATG445TTT (3659A>T 3661G>T), GGA446GGG (3664A>G), ACC448ACT (3670C>T), AAC449AAT (3673C>T), GCA450GCC (3676A>C), CCC451CCG (3679C>G), GCA452GCT (3682A>T), TCC453ACC (3683T>A), TGC454TTT (3687G>T 3688C>T), CAG455TGC (3689C>T 3690A>G 3691G>C), GAC456AAC (3692G>A), CTT457TTA (3695C>T 3697T>A), GTC458ATG (3698G>A 3700C>G), AAC459AAT (3703C>T), GAA460GAT (3706A>T), ACA461GTC (3707A>G 3708C>T 3709A>C), CTT462CTG (3712T>G), AGA463TAT (3713A>T 3714G>A 3715A>T), CTA465TAT (3719C>T 3720T>A 3721A>T), CTC466CTT (3724C>T), GAC467GAT (3727C>T), GTG468GAG (3729T>A), TGC469TTT (3732G>T 3733C>T), GTC470GTA (3736C>A), GTT471GTA (3739T>A), GCT472GTC (3741C>T 3742T>C), ATG474TTA (3746A>T 3748G>A), GAC475GAT (3751C>T), ATA477ATT (3757A>T), CTG478GTC (3758C>G 3760G>C), GTC479ATA (3761G>A 3763C>A), TAC480TAT (3766C>T), ACA481AGT (3768C>G 3769A>T), GGA483AGT (3773G>A 3775A>T), TCC484ACG (3776T>A 3778C>G), CTC485TTG (3779C>T 3781C>G), CAG486GAT (3782C>G 3784G>T), ACC489GTG (3791A>G 3792C>T 3793C>G), AAG490AAC (3796G>C), CAA491CAT (3799A>T), CCA607.GA (4146C>G), ACG608ACC (4150G>C), ACG608_CTT609insAATATTGCG (4150_4151insAATATTGCG), CTT609GTT (4151C>G), CGA610GCT (4154C>G 4155G>C 4156A>T), CTA611GAC (4157C>G 4158T>A 4159A>C), TTC612TTT (4162C>T), GGT614TTA (4166G>T 4167G>T 4168T>A), AGC615CCA (4169A>C 4170G>C 4171C>A), AAG616TTT (4172A>T 4173A>T 4174G>T), GTC618GTG (4180C>G), ATC620ATT (4186C>T), GAG621AAT (4187G>A 4189G>T), GAC623GAT (4195C>T), GCG624GCC (4198G>C), GAT626GAC (4204T>C), ATG627TTC (4205A>T 4207G>C), GCA628GCT (4210A>T), ATA629ATT (4213A>T), GGC630GGG (4216C>G), GCA631GGT (4218C>G 4219A>T), TGT632GTG (4220T>G 4221G>T 4222T>G), CTA633CTT (4225A>T), ACA634ATT (4227C>T 4228A>T), CAG635CAA (4231G>A), ACA636AAT (4233C>A 4234A>T), CAC637GAC (4235C>G), GAT638_AGA641del (4238_4249delGATGGGAAAAGA), CAC642CAT (4252C>T), CCA643CCT (4255A>T), GTG644GTT (4258G>T), GCC645GCA (4261C>A), TAT646TTT (4263A>T), TAT647GAG (4265T>G 4267T>G), TCC648AGC (4268T>A 4269C>G), CGG649AGG (4271C>A), ATG651TTG (4277A>T), ACC652AAT (4281C>A 4282C>T), ACA653CCG (4283A>C 4285A>G), GCG654ACG (4286G>A), CAG656CAA (4294G>A), AAC657AGA (4296A>G 4297C>A), TAC658TAT (4300C>T), GAC659TCT (4301G>T 4302A>C 4303C>T), ATC660GCA (4304A>G 4305T>C 4306C>A), GAC662GAA (4312C>A), CTT665ATG (4319C>A 4321T>G), CTA666GTG (4322C>G 4324A>G), GCC667GCA (4327C>A), ATT668TTG (4328A>T 4330T>G), GTT669CTC (4331G>C 4333T>C), GCC670CAT (4334G>C 4335C>A 4336C>T), GCC671TGT (4337G>T 4338C>G 4339C>T), ATG672CTT (4340A>C 4342G>T), CAT674ACA (4346C>A 4347A>C 4348T>A), AGA676AGG (4354A>G), TAC678TAT (4360C>T), GTC679CTC (4361G>C), GAG680TTA (4364G>T 4365A>T 4366G>A), GGC681GGA (4369C>A), CCA682ACA (4370C>A), CCG683del (4373_4375delCCG), TTA685TTT (4381A>T), ACG686GTA (4382A>G 4383C>T 4384G>A), ATT687GTA (4385A>G 4387T>A), CTT688AAG (4388C>A 4389T>A 4390T>G), TCA689ACG (4391T>A 4393A>G), CAC691AAT (4397C>A 4399C>T), AAG692GTG (4400A>G 4401A>T), AAT693GCT (4403A>G 4404A>C), CTC694AAT (4406C>A 4407T>A 4408C>T), ACG695ACT (4411G>T), ACG698AAA (4419C>A 4420G>A), ACG699ACT (4423G>T), ACG700CAA (4424A>C 4425C>A 4426G>A), AAG701CAG (4427A>C), GAA702AAA (4430G>A), CTT703TTG (4433C>T 4435T>G), ACC704AGT (4437C>G 4438C>T), CGA705TTG (4439C>T 4440G>T 4441A>G), AGA706AAG (4443G>A 4444A>G), AAA866AAT (4924A>T), GAC869GAG (4933C>G), GAG870AGT (4934G>A 4935A>G 4936G>T), GTT871ATC (4937G>A 4939T>C), ATT876ATA (4954T>A), ACG877ACC (4957G>C), AAA878TGT (4958A>T 4959A>G 4960A>T), CTC879TTG (4961C>T 4963C>G), CCG880CCA (4966G>A), AGG881AAG (4968G>A), TCA882TCC (4972A>C), AAG883del (4973_4975delAAG), CGA885_CAA889del (4979_4993delCGAGTCACAGGACAA), GCC890GGG (4995C>G 4996C>G), GAC892GAA (5002C>A), ATG893ACC (5004T>C 5005G>C), CTA895TTG (5009C>T 5011A>G), GTC896GTT (5014C>T), ATG897GTG (5015A>G), GTC898GTA (5020C>A), GAC899GAT (5023C>T), CTC901TTG (5027C>T 5029C>G), TAT904TAC (5038T>C), GCA905GCC (5041A>C), CAC906CAT (5044C>T), TTC907TTT (5047C>T), ATT908ATG (5050T>G), CCT909CCC (5053T>C), GCA910GTC (5055C>T 5056A>C), TCA911AAA (5057T>A 5058C>A), GAA912GCT (5061A>C 5062A>T), ATA913GCT (5063A>G 5064T>C 5065A>T), TAC914TGC (5067A>G), GCA916GCC (5074A>C), GAG917AAG (5075G>A), CAG918GCG (5078C>G 5079A>C), CTC919GCT (5081C>G 5082T>C 5083C>T), GGA920GC. (5085G>C), GGA932GGG (5122A>G), TTC933CTG (5123T>C 5125C>G), GAA935AGG (5129G>A 5130A>G 5131A>G), GTA936CCC (5132G>C 5133T>C 5134A>C), TTC937ATT (5135T>A 5137C>T), ACA939ACT (5143A>T), GAC940GAT (5146C>T), AGA941CGA (5147A>C), AAG943CCG (5153A>C 5154A>C), CTC944CGC (5157T>G), TTC945TTT (5161C>T), ACA946ACT (5164A>T), TCA947GGA (5165T>G 5166C>G), TAC949TTC (5172A>T), GTT998GCG (5319T>C 5320T>G), TCA999AAG (5321T>A 5322C>A 5323A>G), TTA1000CTA (5324T>C), CTG1001CTA (5329G>A), CCA1002GAC (5330C>G 5331C>A 5332A>C), ATG1003GTG (5333A>G), GCG1004GCC (5338G>C), CAG1005CAA (5341G>A), ATC1006TTT (5342A>T 5344C>T), GCA1007TCA (5345G>T), CTG1008TAT (5348C>T 5349T>A 5350G>T), AAC1009AAT (5353C>T) | | | | | | | | |

*: Inserts / Deletes / Misaligned / Frameshifts

#### Analysis details

This analysis was performed with panviral2.64

### NGS Details (UN70): Lowelvirus tuscon4d

#### Assembly

| | |
|---|---|
| **Coverage Length** | 397 (2 contig(s)) |
| **Depth Of Coverage** | 27.6 |
| **Number Of Reads** | 115 |
| **Reads Per Million** | 2.18 rpm (after QC) |
| **Ambiguities** | 1 |
| **Assembly Method** | de novo + reference guided assembly |
| **Consensus Caller** | Bcf Tools |

#### Coverage Map

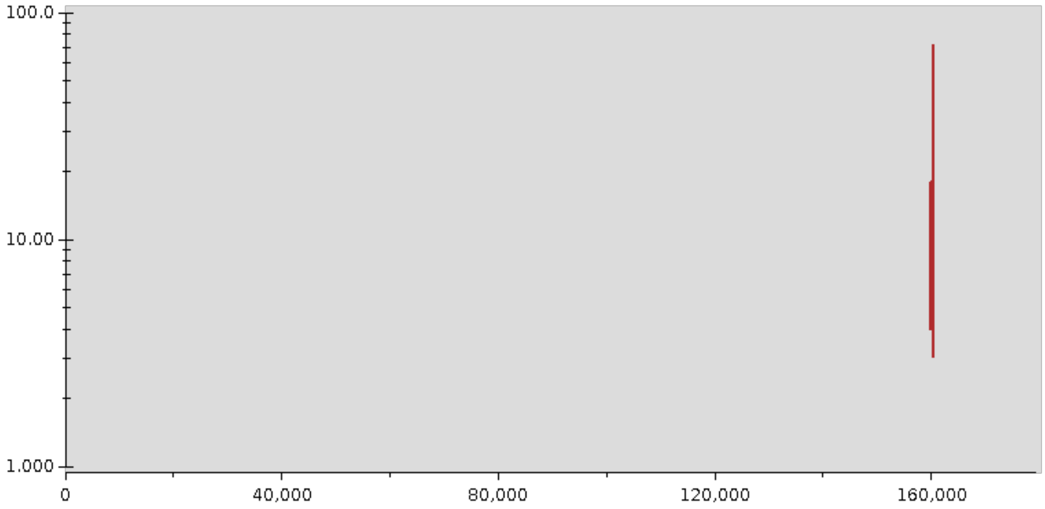

#### Assignment

| | |
|---|---|
| **Type** | Lowelvirus tuscon4d (Taxonomy ID: 2956131) |
| **Reference Genome** | NC_026923.1 |
| **NT Identity (%)** | 79.597 |
| **AA Identity (%)** | 89.9225 |
| **Number Of Stop Codons** | 1 |
| **Number Of CDS** | 218 |

#### Alignment

| | |
|---|---|
| **Alignment Score** | 472.0 (NT) + 855.0 (AA) = 1327.0 |
| **Concordance (%)** | 76.794 |
| **Alignment Method** | Local, heuristic, nucleotide (BLASTN) |

#### Genome Region

Sequence starts at position 159734 and ends at position 160404 relative to NC_026923.1 reference sequence.

#### Alignment Detailed Statistics

| | Begin | End | Coverage | Score | Concordance | Matches | Identities | I/D/M/F* | Stop Codons |
|---|---|---|---|---|---|---|---|---|---|
| **NT** | **159734** | **160404** | **0.2%** | **472** | **59.4%** | **397 (100%)** | **316 (79.6%)** | **0/0** | |

Mutations: 159738T>A, 159739C>T, 159740T>C, 159741A>T, 159745G>C, 159746G>A, 159747T>A, 159748A>T, 159750C>G, 159751C>A, 159753T>A, 159759C>T, 159765T>A, 159768T>C, 159771G>A, 159783T>A, 159798C>T, 159799A>T, 159800G>C, 159801C>T, 159807C>T, 159810T>G, 159813C>T, 159822G>A, 159832T>G, 159833C>G, 159834A>T, 159855C>T, 159858G>T, 159867A>T, 159870C>A, 159874A>C, 159880C>A, 159882G>C, 159883T>C, 159885G>A, 159888C>T, 159889C>T, 159894G>A, 159897T>G, 159900C>T, 159903C>T, 159906C>T, 159913C>T, 159915C>G, 159924C>T, 159942T>C, 159946A>G, 159949C>T, 159951A>G, 160246A>C, 160248A>C, 160254A>G, 160270G>A, 160275C>T, 160287G>A, 160290C>W, 160293C>A, 160296A>C, 160299G>A, 160302T>C, 160308A>T, 160309C>A, 160326A>G, 160327C>G, 160330G>C, 160331C>A, 160332T>G, 160338A>G, 160341A>T, 160350C>T, 160356C>A, 160359C>T, 160368A>G, 160371A>T, 160377T>A, 160378A>C, 160380A>T, 160383T>A, 160389A>T, 160392G>T

*: Inserts / Deletes / Misaligned / Frameshifts

#### Analysis details

This analysis was performed with panviral2.64

### NGS Details (UN70): Errantivirus

#### Assembly

| | |
|---|---|
| **Coverage Length** | 860 (1 contig(s)) |
| **Depth Of Coverage** | 13.9 |
| **Number Of Reads** | 112 |
| **Reads Per Million** | 2.12 rpm (after QC) |
| **Ambiguities** | 0 |
| **Assembly Method** | de novo + reference guided assembly |
| **Consensus Caller** | Bcf Tools |

#### Coverage Map

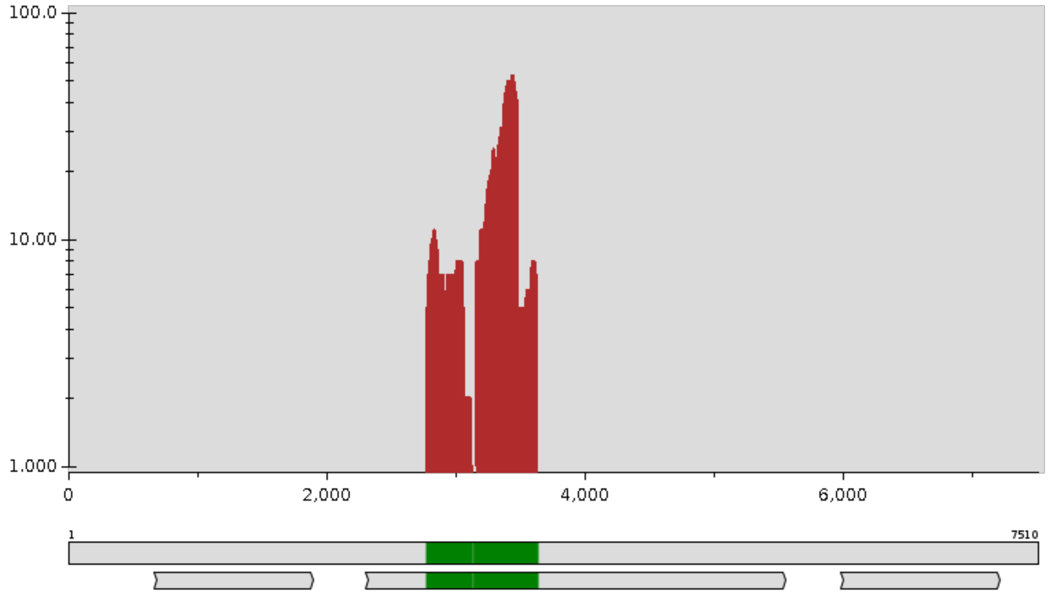

#### Assignment

| | |
|---|---|
| **Type** | Errantivirus (Taxonomy ID: 186666) |
| **Reference Genome** | NC_038512.1 |
| **NT Identity (%)** | 51.6627 |
| **AA Identity (%)** | 43.3692 |
| **Number Of Stop Codons** | 0 |
| **Number Of CDS** | 3 |

#### Alignment

| | |
|---|---|
| **Alignment Score** | 11.0 (NT) + 856.0 (AA) = 867.0 |
| **Concordance (%)** | 24.6097 |

| Alignment Method | Global, seeded, nucleotide + amino acids (AGA) |
|---|---|

#### Genome Region

Sequence starts at position 2769 and ends at position 3638 relative to NC_038512.1 reference sequence.

#### Alignment Detailed Statistics

| | Begin | End | Coverage | Score | Concordance | Matches | Identities | I/D/M/F* | Stop Codons |
|---|---|---|---|---|---|---|---|---|---|
| **NT** | **2769** | **3638** | **11.5%** | **11** | **0.7%** | **842 (97.9%)** | **435 (50.6%)** | **0/18** | |
| Mutations: | 2769G>A, 2771G>A, 2773C>G, 2778T>G, 2779C>G, 2781C>A, 2782G>A, 2783A>G, 2789T>C, 2791G>A, 2792G>C, 2794C>G, 2801A>G, 2803T>C, 2807A>C, 2808G>A, 2813T>A, 2814C>G, 2815A>T, 2816G>C, 2822G>C, 2826G>A, 2827G>C, 2828A>T, 2829G>C, 2833A>T, 2836C>A, 2837A>G, 2839A>G, 2841G>T, 2843G>T, 2845T>A, 2849_2851delCCC, 2859T>A, 2860C>A, 2863C>T, 2865C>G, 2866T>G, 2867T>A, 2870_2881delGGGAAACAAAAG, 2885C>A, 2887T>A, 2888C>T, 2891G>T, 2892T>A, 2893A>T, 2894G>A, 2901T>A, 2903C>A, 2905T>A, 2906A>G, 2908G>T, 2914C>T, 2915G>A, 2918A>T, 2919A>T, 2927G>A, 2929T>A, 2932C>T, 2937A>T, 2938C>T, 2939C>A, 2941G>A, 2944A>T, 2947A>C, 2949A>T, 2953A>T, 2954A>G, 2955G>A, 2956T>G, 2959C>G, 2960G>T, 2965T>G, 2968C>T, 2969A>G, 2971G>A, 2974A>G, 2977T>A, 2978A>G, 2979A>G, 2980G>T, 2982G>C, 2986A>G, 2987T>A, 2988A>T, 2997C>A, 2998C>G, 2999T>C, 3001A>T, 3005T>C, 3008G>A, 3009C>G, 3011A>T, 3012G>C, 3013T>A, 3016G>T, 3018T>A, 3020T>C, 3025G>A, 3026G>A, 3028G>T, 3029G>A, 3030A>G, 3035G>A, 3036A>C, 3037C>A, 3038C>A, 3039C>A, 3040T>G, 3041C>G, 3043A>G, 3047A>G, 3049A>C, 3050T>C, 3051C>A, 3058C>T, 3059G>A, 3061G>A, 3066A>G, 3067C>G, 3068G>A, 3069T>C, 3070A>T, 3071G>C, 3073A>T, 3074C>T, 3075A>C, 3076C>T, 3079G>C, 3082T>C, 3084T>A, 3088A>G, 3090T>A, 3094T>A, 3095C>G, 3096G>T, 3103T>C, 3104A>T, 3106G>T, 3109A>C, 3110T>C, 3112A>C, 3114A>C, 3115A>T, 3118C>T, 3119T>G, 3124A>T, 3127T>A, 3129C>G, 3137A>T, 3139A>T, 3140G>C, 3142T>A, 3146G>A, 3148C>T, 3149A>C, 3151T>G, 3154C>T, 3155C>T, 3157A>T, 3158A>C, 3159G>A, 3160A>G, 3161G>C, 3162G>C, 3164C>T, 3168A>T, 3169A>C, 3170A>G, 3172T>C, 3174A>T, 3175C>A, 3176A>T, 3178C>T, 3179T>G, 3180G>T, 3181T>G, 3182C>G, 3184C>A, 3189A>T, 3191C>A, 3200A>G, 3202T>A, 3203A>C, 3205T>G, 3206G>A, 3208C>A, 3211T>C, 3212A>T, 3213G>C, 3215A>C, 3216C>G, 3217T>A, 3218T>A, 3219C>G, 3220C>T, 3221C>A, 3222T>A, 3223A>G, 3224C>G, 3226G>A, 3232C>T, 3233C>G, 3234T>A, 3235G>A, 3238G>A, 3239A>T, 3241C>T, 3242C>T, 3245G>A, 3246A>G, 3247A>G, 3248C>A, 3249G>T, 3253T>A, 3254T>C, 3256C>G, 3259A>G, 3261G>C, 3263C>T, 3265T>G, 3268A>G, 3271A>T, 3272A>G, 3273G>A, 3274T>G, 3277C>G, 3280C>G, 3281A>T, 3283A>T, 3284A>G, 3285T>C, 3286T>A, 3287C>A, 3289A>G, 3290A>T, 3292G>T, 3300C>G, 3307C>T, 3309T>G, 3311A>T, 3312A>T, 3314C>A, 3315T>A, 3316C>T, 3320A>G, 3321C>T, 3325T>C, 3327A>T, 3334T>G, 3337C>T, 3338A>G, 3340C>A, 3341A>G, 3343A>T, 3344A>T, 3345G>C, 3346C>G, 3347A>G, 3349G>T, 3352C>T, 3355T>G, 3362C>G, 3363C>T, 3364T>G, 3365A>G, 3370T>G, 3371G>A, 3373T>G, 3376G>A, 3378T>C, 3380T>G, 3381C>A, 3382C>G, 3385T>A, 3386A>G, 3388T>C, 3389C>A, 3390A>G, 3394A>T, 3396A>G, 3397T>G, 3399T>C, 3400G>A, 3402T>G, 3403T>G, 3406A>T, 3407A>T, 3408A>T, 3409G>A, 3410A>T, 3412C>T, 3415T>G, 3417A>C, 3418G>T, 3421A>T, 3424A>T, 3426A>G, 3428C>A, 3429A>G, 3430A>C, 3433T>C, 3436A>G, 3439C>A, 3440C>T, 3442T>G, 3443C>G, 3444T>C, 3445C>A, 3451T>C, 3454C>T, 3455C>A, 3457A>G, 3459A>G, 3464A>G, 3466T>A, 3467C>G, 3468C>A, 3471A>G, 3476G>T, 3478A>T, 3479C>A, 3480G>T, 3481A>T, 3482C>A, 3484C>T, 3486C>G, 3487A>T, 3489A>G, 3490A>G, 3493C>A, 3498C>T, 3499A>G, 3500C>G, 3502G>T, 3504G>T, 3505C>G, 3506T>C, 3508A>G, 3514A>G, 3516G>A, 3517T>C, 3518_3520delAGT, 3523A>C, 3524G>T, 3526A>C, 3529T>A, 3530C>T, 3531T>G, 3532T>G, 3533A>T, 3534G>C, 3536C>G, 3537C>A, 3538C>A, 3539G>A, 3540A>T, 3541A>T, 3542T>G, 3543A>C, 3545G>A, 3546T>C, 3547A>T, 3548A>C, 3550T>A, 3551G>C, 3557G>C, 3559A>T, 3560C>A, 3564G>T, 3565T>G, 3568A>G, 3569A>G, 3570C>T, 3571T>A, 3572C>G, 3573T>C, 3574G>A, 3580C>A, 3581A>T, 3582A>C, 3583C>T, 3585A>C, 3586C>T, 3589A>T, 3590A>G, 3592A>C, 3593T>C, 3596C>G, 3597A>C, 3600A>T, 3601C>A, 3604A>T, 3605G>A, 3607C>T, 3609T>A, 3611A>T, 3615G>T, 3616A>T, 3617G>C, 3618A>C, 3619A>T, 3623A>G, 3624A>T, 3625C>T, 3626C>G, 3628C>T, 3629A>G, 3630C>A, 3637C>T | | | | | | | | |

**CDS**

| | Begin | End | Coverage | Score | Concordance | Matches | Identities | I/D/M/F* | Stop Codons |
|---|---|---|---|---|---|---|---|---|---|
| D1R33_gp2 | 156 | 445 | 26.3% | 856 | 43.8% | 279 (97.9%) | 121 (42.5%) | 0/6/1/0 | 0 |
| Protein mutations: | D156K (2771G>A 2773C>G), I158R (2778T>G 2779C>G), T159K (2781C>A 2782G>A), K160E (2783A>G), D163Q (2792G>C 2794C>G), I166V (2801A>G 2803T>C), R168Q (2807A>C 2808G>A), D171H (2816G>C), A173P (2822G>C), W174Y (2826G>A 2827G>C), I178V (2837A>G 2839A>G), W179L (2841G>T), V180L (2843G>T 2845T>A), P182del (2849_2851delCCC), I185K (2859T>A 2860C>A), A187G (2865C>G 2866T>G), S188T (2867T>A), G189_K192del (2870_2881delGGGAAACAAAAG), L195F (2888C>T), V196Y (2891G>T 2892T>A 2893A>T), V197I (2894G>A), F199Y (2901T>A), K201D (2906A>G 2908G>T), E204K (2915G>A), K205L (2918A>T 2919A>T), D208K (2927G>A 2929T>A), Y211F (2937A>T 2938C>T), P212T (2939C>A 2941G>A), N215I (2949A>T), S217E (2954A>G 2955G>A 2956T>G), D218E (2959C>G), V219L (2960G>T), K222E (2969A>G 2971G>A), K225G (2978A>G 2979A>G 2980G>T), C226S (2982G>C), Y228I (2987T>A 2988A>T), T231K (2997C>A 2998C>G), A235R (3008G>A 3009C>G), F238Y (3018T>A), Y239H (3020T>C), V241I (3026G>A 3028G>T), E242R (3029G>A 3030A>G), D244T (3035G>A 3036A>C 3037C>A), P245K (3038C>A 3039C>A 3040T>G), Q246E (3041C>G 3043A>G), I248V (3047A>G 3049A>C), S249Q (3050T>C 3051C>A), A252T (3059G>A 3061G>A), N254R (3066A>G 3067C>G), V255T (3068G>A 3069T>C 3070A>T), E256H (3071G>C 3073A>T), H257S (3074C>T 3075A>C 3076C>T), F260Y (3084T>A), F262Y (3090T>A), R264V (3095C>G 3096G>T), M267F (3104A>T 3106G>T), K270T (3114A>C 3115A>T), S272A (3119T>G), T275S (3129C>G), R285Q (3158A>C 3159G>A 3160A>G), G286P (3161G>C 3162G>C), L287F (3164C>T), Q288L (3168A>T 3169A>C), N289D (3170A>G 3172T>C), N290I (3174A>T 3175C>A), I291F (3176A>T 3178C>T), C292V (3179T>G 3180G>T 3181T>G), L293V (3182C>G 3184C>A), Y295F (3189A>T), L296I (3191C>A), I299V (3200A>G 3202T>A), I300L (3203A>C 3205T>G), V301I (3206G>A 3208C>A), T304R (3215A>C 3216C>G 3217T>A), L306K (3221C>A 3222T>A 3223A>G), Q307E (3224C>G 3226G>A), L310E (3233C>G 3234T>A 3235G>A), N312Y (3239A>T 3241C>T), E314R (3245G>A 3246A>G 3247A>G), R315I (3248C>A 3249G>T), F317L (3254T>C 3256C>G), R319T (3261G>C), E322D (3271A>T), S323E (3272A>G 3273G>A 3274T>G), N324K (3277C>G), F325L (3280C>G), K326Y (3281A>T 3283A>T), I327A (3284A>G 3285T>C 3286T>A), Q328K (3287C>A 3289A>G), M329F (3290A>T 3292G>T), S332C (3300C>G), L335W (3309T>G), K336L (3311A>T 3312A>T), L337N (3314C>A 3315T>A 3316C>T), T339V (3320A>G 3321C>T), Y341F (3327A>T), I345V (3338A>G 3340C>A), I346V (3341A>G 3343A>T), R348G (3347A>G 3349G>T), P353V (3362C>G 3363C>T 3364T>G), N354D (3365A>G), D356K (3371G>A 3373T>G), I358T (3378T>C), S359E (3380T>G 3381C>A 3382C>G), I361V (3386A>G 3388T>C), Q362R (3389C>A 3390A>G), K363N (3394A>T), Y364W (3396A>G 3397T>G), L365P (3399T>C 3400G>A), I366R (3402T>G 3403T>G), K368L (3407A>T 3408A>T 3409G>A), T369S (3410A>T 3412C>T), K371T (3417A>C 3418G>T), E372D (3421A>T), K374R (3426A>G), Q375S (3428C>A 3429A>G 3430A>C), L380A (3443C>G 3444T>C 3445C>A), K385R (3459A>G), I387V (3464A>G 3466T>A), P388E (3467C>G 3468C>A), D389G (3471A>G), A391S (3476G>T 3478A>T), R392I (3479C>A 3480G>T 3481A>T), L393I (3482C>A 3484C>T), T394S (3486C>G 3487A>T), K395R (3489A>G 3490A>G), T398M (3498C>T 3499A>G), Q399D (3500C>G 3502G>T), C400L (3504G>T 3505C>G), G404D (3516G>A 3517T>C), S405del (3518_3520delAGT), K406N (3523A>C), V407F (3524G>T 3526A>C), L409W (3530C>T 3531T>G 3532T>G), P411E (3536C>G 3537C>A 3538C>A), E412I (3539G>A 3540A>T 3541A>T), Y413A (3542T>G 3543A>C), V414T (3545G>A 3546T>C 3547A>T), N415Q (3548A>C 3550T>A), A416P (3551G>C), E418H (3557G>C 3559A>T), H419N (3560C>A), C420L (3564G>T 3565T>G), T422V (3569A>G 3570C>T 3571T>A), L423A (3572C>G 3573T>C 3574G>A), N426S (3581A>T 3582A>C 3583C>T), D427A (3585A>C 3586C>T), I429V (3590A>G 3592A>C), Q431A (3596C>G 3597A>C), Y432L (3600A>T 3601C>A), D434N (3605G>A 3607C>T), F435Y (3609T>A), T436S (3611A>T), R437I (3615G>T 3616A>T), E438P (3617G>C 3618A>C 3619A>T), N440V (3623A>G 3624A>T 3625C>T), L441V (3626C>G 3628C>T), T442E (3629A>G 3630C>A) | | | | | | | | |

| | Begin | End | Coverage | Score | Concordance | Matches | Identities | I/D/M/F* | Stop Codons |
|---|---|---|---|---|---|---|---|---|---|
| **NT** | **2769** | **3638** | **11.5%** | **11** | **0.7%** | **842 (97.9%)** | **435 (50.6%)** | **0/18** | |
| Codon mutations: | AGG155.AG (2769G>A), GAC156AAG (2771G>A 2773C>G), ATC158AGG (2778T>G 2779C>G), ACG159AAA (2781C>A 2782G>A), AAA160GAA (2783A>G), TTG162CTA (2789T>C 2791G>A), GAC163CAG (2792G>C 2794C>G), ATT166GTC (2801A>G 2803T>C), AGA168CAA (2807A>C 2808G>A), TCA170AGT (2813T>A 2814C>G 2815A>T), GAC171CAC (2816G>C), GCA173CCA (2822G>C), TGG174TAC (2826G>A 2827G>C), AGC175TCC (2828A>T 2829G>C), TCA176TCT (2833A>T), CCC177CCA (2836C>A), ATA178GTG (2837A>G 2839A>G), TGG179TTG (2841G>T), GTT180TTA (2843G>T 2845T>A), CCC182del (2849_2851delCCC), ATC185AAA (2859T>A 2860C>A), GAC186GAT (2863C>T), GCT187GGG (2865C>G 2866T>G), TCT188ACT (2867T>A), GGG189_AAG192del (2870_2881delGGGAAACAAAAG), CGT194AGA (2885C>A 2887T>A), CTC195TTC (2888C>T), GTA196TAT (2891G>T 2892T>A 2893A>T), GTT197ATT (2894G>A), TTC199TAC (2901T>A), CGT200AGA (2903C>A 2905T>A), AAG201GAT (2906A>G 2908G>T), AAC203AAT (2914C>T), GAG204AAG (2915G>A), AAG205TTG (2918A>T 2919A>T), GAT208AAA (2927G>A 2929T>A), GAC209GAT (2932C>T), TAC211TTT (2937A>T 2938C>T), CCG212ACA (2939C>A 2941G>A), ATA213ATT (2944A>T), CCA214CCC (2947A>C), AAC215ATC (2949A>T), ATA216ATT (2953A>T), AGT217GAG (2954A>G 2955G>A 2956T>G), GAC218GAG (2959C>G), GTA219TTA (2960G>T), CTT220CTG (2965T>G), GAC221GAT (2968C>T), AAG222GAA (2969A>G 2971G>A), TTA223TTG (2974A>G), GGT224GGA (2977T>A), AAG225GGT (2978A>G 2979A>G 2980G>T), TGC226TCC (2982G>C), CAA227CAG (2986A>G), TAC228ATC (2987T>A 2988A>T), ACC231AAG (2997C>A 2998C>G), TTA232CTT (2999T>C 3001A>T), TTG234CTG (3005T>C), GCA235AGA (3008G>A 3009C>G), AGT236TCA (3011A>T 3012G>C 3013T>A), GGG237GGT (3016G>T), TTT238TAT (3018T>A), TAT239CAT (3020T>C), CAG240CAA (3025G>A), GTG241ATT (3026G>A 3028G>T), GAG242AGG (3029G>A 3030A>G), GAC244ACA (3035G>A 3036A>C 3037C>A), CCT245AAG (3038C>A 3039C>A 3040T>G), CAA246GAG (3041C>G 3043A>G), ATA248GTC (3047A>G 3049A>C), TCG249CAG (3050T>C 3051C>A), ACC251ACT (3058C>T), GCG252ACA (3059G>A 3061G>A), AAC254AGG (3066A>G 3067C>G), GTA255ACT (3068G>A 3069T>C 3070A>T), GAA256CAT (3071G>C 3073A>T), CAC257TCT (3074C>T 3075A>C 3076C>T), GGG258GGC (3079G>C), CAT259CAC (3082T>C), TTT260TAT (3084T>A), GAA261GAG (3088A>G), TTC262TAC (3090T>A), CTT263CTA (3094T>A), CGA264GTA (3095C>G 3096G>T), CCT266CCC (3103T>C), ATG267TTT (3104A>T 3106G>T), GGA268GGC (3109A>C), TTA269CTC (3110T>C 3112A>C), AAA270ACT (3114A>C 3115A>T), AAC271AAT (3118C>T), TCA272GCA (3119T>G), CCA273CCT (3124A>T), TCT274TCA (3127T>A), ACT275AGT (3129C>G), AGA278T.T (3137A>T 3139A>T), GTT279C.A (3140G>C 3142T>A), GAC281A.T (3146G>A 3148C>T), AAT282C.G (3149A>C 3151T>G), GTC283..T (3154C>T), CTA284T.T (3155C>T 3157A>T), AGA285CAG (3158A>C 3159G>A 3160A>G), GGT286CCT (3161G>C 3162G>C), CTC287TTC (3164C>T), CAA288CTC (3168A>T 3169A>C), AAT289GAC (3170A>G 3172T>C), AAC290ATA (3174A>T 3175C>A), ATC291TTT (3176A>T 3178C>T), TGT292GTG (3179T>G 3180G>T 3181T>G), CTC293GTA (3182C>G 3184C>A), TAC295TTC (3189A>T), CTT296ATT (3191C>A), ATT299GTA (3200A>G 3202T>A), ATT300CTG (3203A>C 3205T>G), GTC301ATA (3206G>A 3208C>A), TAT302TAC (3211T>C), AGT303TCT (3212A>T 3213G>C), ACT304CGA (3215A>C 3216C>G 3217T>A), TCC305AGT (3218T>A 3219C>G 3220C>T), CTA306AAG (3221C>A 3222T>A 3223A>G), CAG307GAA (3224C>G 3226G>A), CAC309CAT (3232C>T), CTG310GAA (3233C>G 3234T>A 3235G>A), GAG311GAA (3238G>A), AAC312TAT (3239A>T 3241C>T), CTG313TTG (3242C>T), GAA314AGG (3245G>A 3246A>G 3247A>G), CGA315ATA (3248C>A 3249G>T), GTT316GTA (3253T>A), TTC317CTG (3254T>C 3256C>G), CAA318CAG (3259A>G), AGA319ACA (3261G>C), CTT320TTG (3263C>T 3265T>G), AGA321AGG (3268A>G), GAA322GAT (3271A>T), AGT323GAG (3272A>G 3273G>A 3274T>G), AAC324AAG (3277C>G), TTC325TTG (3280C>G), AAA326TAT (3281A>T 3283A>T), ATT327GCA (3284A>G 3285T>C 3286T>A), CAA328AAG (3287C>A 3289A>G), ATG329TTT (3290A>T 3292G>T), TCC332TGC (3300C>G), TTC334TTT (3307C>T), TTG335TGG (3309T>G), AAG336TTG (3311A>T 3312A>T), CTC337AAT (3314C>A 3315T>A 3316C>T), ACT339GTT (3320A>G 3321C>T), GCT340GCC (3325T>C), TAT341TTT (3327A>T), GGT343GGG (3334T>G), CAC344CAT (3337C>T), ATC345GTA (3338A>G 3340C>A), ATA346GTT (3341A>G 3343A>T), AGC347TCG (3344A>T 3345G>C 3346C>G), AGG348GGT (3347A>G 3349G>T), GAC349GAT (3352C>T), GGT350GGG (3355T>G), CCT353GTG (3362C>G 3363C>T 3364T>G), AAC354GAC (3365A>G), CCT355CCG (3370T>G), GAT356AAG (3371G>A 3373T>G), AAG357AAA (3376G>A), ATT358ACT (3378T>C), TCC359GAG (3380T>G 3381C>A 3382C>G), GCT360GCA (3385T>A), ATT361GTC (3386A>G 3388T>C), CAA362AGA (3389C>A 3390A>G), AAA363AAT (3394A>T), TAT364TGG (3396A>G 3397T>G), CTG365CCA (3399T>C 3400G>A), ATT366AGG (3402T>G 3403T>G), CCA367CCT (3406A>T), AAG368TTA (3407A>T 3408A>T 3409G>A), ACC369TCT (3410A>T 3412C>T), CCT370CCG (3415T>G), AAG371ACT (3417A>C 3418G>T), GAA372GAT (3421A>T), ATA373ATT (3424A>T), AAA374AGA (3426A>G), CAA375AGC (3428C>A 3429A>G 3430A>C), TTT376TTC (3433T>C), TTA377TTG (3436A>G), GGC378GGA (3439C>A), CTT379TTG (3440C>T 3442T>G), CTC380GCA (3443C>G 3444T>C 3445C>A), TAT382TAC (3451T>C), TAC383TAT (3454C>T), CGA384AGG (3455C>A 3457A>G), AAA385AGA (3459A>G), ATT387GTA (3464A>G 3466T>A), CCA388GAA (3467C>G 3468C>A), GAT389GGT (3471A>G), GCA391TCT (3476G>T 3478A>T), CGA392ATT (3479C>A 3480G>T 3481A>T), CTC393ATT (3482C>A 3484C>T), ACA394AGT (3486C>G 3487A>T), AAA395AGG (3489A>G 3490A>G), CCC396CCA (3493C>A), ACA398ATG (3498C>T 3499A>G), CAG399GAT (3500C>G 3502G>T), TGC400TTG (3504G>T 3505C>G), TTA401CTG (3506T>C 3508A>G), AAA403AAG (3514A>G), GGT404GAC (3516G>A 3517T>C), AGT405del (3518_3520delAGT), AAA406AAC (3523A>C), GTA407TTC (3524G>T 3526A>C), ACT408ACA (3529T>A), CTT409TGG (3530C>T 3531T>G 3532T>G), AGT410TCT (3533A>T 3534G>C), CCC411GAA (3536C>G 3537C>A 3538C>A), GAA412ATT (3539G>A 3540A>T 3541A>T), TAT413GCT (3542T>G 3543A>C), GTA414ACT (3545G>A 3546T>C 3547A>T), AAT415CAA (3548A>C 3550T>A), GCT416CCT (3551G>C), GAA418CAT (3557G>C 3559A>T), CAC419AAC (3560C>A), TGT420TTG (3564G>T 3565T>G), AAA421AAG (3568A>G), ACT422GTA (3569A>G 3570C>T 3571T>A), CTG423GCA (3572C>G 3573T>C 3574G>A), ACC425ACA (3580C>A), AAC426TCT (3581A>T 3582A>C 3583C>T), GAC427GCT (3585A>C 3586C>T), CCA428CCT (3589A>T), ATA429GTC (3590A>G 3592A>C), TTA430CTA (3593T>C), CAA431GCA (3596C>G 3597A>C), TAC432TTA (3600A>T 3601C>A), CCA433CCT (3604A>T), GAC434AAT (3605G>A 3607C>T), TTT435TAT (3609T>A), ACC436TCC (3611A>T), AGA437ATT (3615G>T 3616A>T), GAA438CCT (3617G>C 3618A>C 3619A>T), AAC440GTT (3623A>G 3624A>T 3625C>T), CTC441GTT (3626C>G 3628C>T), ACG442GAG (3629A>G 3630C>A), GAC444GAT (3637C>T) | | | | | | | | |

#### Proteins

| | Begin | End | Coverage | Score | Concordance | Matches | Identities | I/D/M/F* | Stop Codons |
|---|---|---|---|---|---|---|---|---|---|
| ORF B (YP_009507248.1) | 156 | 445 | 26.3% | 856 | 43.8% | 279 (97.9%) | 121 (42.5%) | 0/6/1/0 | 0 |
| Protein mutations: | D156K (2771G>A 2773C>G), I158R (2778T>G 2779C>G), T159K (2781C>A 2782G>A), K160E (2783A>G), D163Q (2792G>C 2794C>G), I166V (2801A>G 2803T>C), R168Q (2807A>C 2808G>A), D171H (2816G>C), A173P (2822G>C), W174Y (2826G>A 2827G>C), I178V (2837A>G 2839A>G), W179L (2841G>T), V180L (2843G>T 2845T>A), P182del (2849_2851delCCC), I185K (2859T>A 2860C>A), A187G (2865C>G 2866T>G), S188T (2867T>A), G189_K192del (2870_2881delGGGAAACAAAAG), L195F (2888C>T), V196Y (2891G>T 2892T>A 2893A>T), V197I (2894G>A), F199Y (2901T>A), K201D (2906A>G 2908G>T), E204K (2915G>A), K205L (2918A>T 2919A>T), D208K (2927G>A 2929T>A), Y211F (2937A>T 2938C>T), P212T (2939C>A 2941G>A), N215I (2949A>T), S217E (2954A>G 2955G>A 2956T>G), D218E (2959C>G), V219L (2960G>T), K222E (2969A>G 2971G>A), K225G (2978A>G 2979A>G 2980G>T), C226S (2982G>C), Y228I (2987T>A 2988A>T), T231K (2997C>A 2998C>G), A235R (3008G>A 3009C>G), F238Y (3018T>A), Y239H (3020T>C), V241I (3026G>A 3028G>T), E242R (3029G>A 3030A>G), D244T (3035G>A 3036A>C 3037C>A), P245K (3038C>A 3039C>A 3040T>G), Q246E (3041C>G 3043A>G), I248V (3047A>G 3049A>C), S249Q (3050T>C 3051C>A), A252T (3059G>A 3061G>A), N254R (3066A>G 3067C>G), V255T (3068G>A 3069T>C 3070A>T), E256H (3071G>C 3073A>T), H257S (3074C>T 3075A>C 3076C>T), F260Y (3084T>A), F262Y (3090T>A), R264V (3095C>G 3096G>T), M267F (3104A>T 3106G>T), K270T (3114A>C 3115A>T), S272A (3119T>G), T275S (3129C>G), R285Q (3158A>C 3159G>A 3160A>G), G286P (3161G>C 3162G>C), L287F (3164C>T), Q288L (3168A>T 3169A>C), N289D (3170A>G 3172T>C), N290I (3174A>T 3175C>A), I291F (3176A>T 3178C>T), C292V (3179T>G 3180G>T 3181T>G), L293V (3182C>G 3184C>A), Y295F (3189A>T), L296I (3191C>A), I299V (3200A>G 3202T>A), I300L (3203A>C 3205T>G), V301I (3206G>A 3208C>A), T304R (3215A>C 3216C>G 3217T>A), L306K (3221C>A 3222T>A 3223A>G), Q307E (3224C>G 3226G>A), L310E (3233C>G 3234T>A 3235G>A), N312Y (3239A>T 3241C>T), E314R (3245G>A 3246A>G 3247A>G), R315I (3248C>A 3249G>T), F317L (3254T>C 3256C>G), R319T (3261G>C), E322D (3271A>T), S323E (3272A>G 3273G>A 3274T>G), N324K (3277C>G), F325L (3280C>G), K326Y (3281A>T 3283A>T), I327A (3284A>G 3285T>C 3286T>A), Q328K (3287C>A 3289A>G), M329F (3290A>T 3292G>T), S332C (3300C>G), L335W (3309T>G), K336L (3311A>T 3312A>T), L337N (3314C>A 3315T>A 3316C>T), T339V (3320A>G 3321C>T), Y341F (3327A>T), I345V (3338A>G 3340C>A), I346V (3341A>G 3343A>T), R348G (3347A>G 3349G>T), P353V (3362C>G 3363C>T 3364T>G), N354D (3365A>G), D356K (3371G>A 3373T>G), I358T (3378T>C), S359E (3380T>G 3381C>A 3382C>G), I361V (3386A>G 3388T>C), Q362R (3389C>A 3390A>G), K363N (3394A>T), Y364W (3396A>G 3397T>G), L365P (3399T>C 3400G>A), I366R (3402T>G 3403T>G), K368L (3407A>T 3408A>T 3409G>A), T369S (3410A>T 3412C>T), K371T (3417A>C 3418G>T), E372D (3421A>T), K374R (3426A>G), Q375S (3428C>A 3429A>G 3430A>C), L380A (3443C>G 3444T>C 3445C>A), K385R (3459A>G), I387V (3464A>G 3466T>A), P388E (3467C>G 3468C>A), D389G (3471A>G), A391S (3476G>T 3478A>T), R392I (3479C>A 3480G>T 3481A>T), L393I (3482C>A 3484C>T), T394S (3486C>G 3487A>T), K395R (3489A>G 3490A>G), T398M (3498C>T 3499A>G), Q399D (3500C>G 3502G>T), C400L (3504G>T 3505C>G), G404D (3516G>A 3517T>C), S405del (3518_3520delAGT), K406N (3523A>C), V407F (3524G>T 3526A>C), L409W (3530C>T 3531T>G 3532T>G), P411E (3536C>G 3537C>A 3538C>A), E412I (3539G>A 3540A>T 3541A>T), Y413A (3542T>G 3543A>C), V414T (3545G>A 3546T>C 3547A>T), N415Q (3548A>C 3550T>A), A416P (3551G>C), E418H (3557G>C 3559A>T), H419N (3560C>A), C420L (3564G>T 3565T>G), T422V (3569A>G 3570C>T 3571T>A), L423A (3572C>G 3573T>C 3574G>A), N426S (3581A>T 3582A>C 3583C>T), D427A (3585A>C 3586C>T), I429V (3590A>G 3592A>C), Q431A (3596C>G 3597A>C), Y432L (3600A>T 3601C>A), D434N (3605G>A 3607C>T), F435Y (3609T>A), T436S (3611A>T), R437I (3615G>T 3616A>T), E438P (3617G>C 3618A>C 3619A>T), N440V (3623A>G 3624A>T 3625C>T), L441V (3626C>G 3628C>T), T442E (3629A>G 3630C>A) | | | | | | | | |

| | Begin | End | Coverage | Score | Concordance | Matches | Identities | I/D/M/F* | Stop Codons |
|---|---|---|---|---|---|---|---|---|---|
| NT | 2769 | 3638 | 11.5% | 11 | 0.7% | 842 (97.9%) | 435 (50.6%) | 0/18 | |

Codon mutations: AGG155.AG (2769G>A), GAC156AAG (2771G>A 2773C>G), ATC158AGG (2778T>G 2779C>G), ACG159AAA (2781C>A 2782G>A), AAA160GAA (2783A>G), TTG162CTA (2789T>C 2791G>A), GAC163CAG (2792G>C 2794C>G), ATT166GTC (2801A>G 2803T>C), AGA168CAA (2807A>C 2808G>A), TCA170AGT (2813T>A 2814C>G 2815A>T), GAC171CAC (2816G>C), GCA173CCA (2822G>C), TGG174TAC (2826G>A 2827G>C), AGC175TCC (2828A>T 2829G>C), TCA176TCT (2833A>T), CCC177CCA (2836C>A), ATA178GTG (2837A>G 2839A>G), TGG179TTG (2841G>T), GTT180TTA (2843G>T 2845T>A), CCC182del (2849_2851delCCC), ATC185AAA (2859T>A 2860C>A), GAC186GAT (2863C>T), GCT187GGG (2865C>G 2866T>G), TCT188ACT (2867T>A), GGG189_AAG192del (2870_2881delGGGAAACAAAAG), CGT194AGA (2885C>A 2887T>A), CTC195TTC (2888C>T), GTA196TAT (2891G>T 2892T>A 2893A>T), GTT197ATT (2894G>A), TTC199TAC (2901T>A), CGT200AGA (2903C>A 2905T>A), AAG201GAT (2906A>G 2908G>T), AAC203AAT (2914C>T), GAG204AAG (2915G>A), AAG205TTG (2918A>T 2919A>T), GAT208AAA (2927G>A 2929T>A), GAC209GAT (2932C>T), TAC211TTT (2937A>T 2938C>T), CCG212ACA (2939C>A 2941G>A), ATA213ATT (2944A>T), CCA214CCC (2947A>C), AAC215ATC (2949A>T), ATA216ATT (2953A>T), AGT217GAG (2954A>G 2955G>A 2956T>G), GAC218GAG (2959C>G), GTA219TTA (2960G>T), CTT220CTG (2965T>G), GAC221GAT (2968C>T), AAG222GAA (2969A>G 2971G>A), TTA223TTG (2974A>G), GGT224GGA (2977T>A), AAG225GGT (2978A>G 2979A>G 2980G>T), TGC226TCC (2982G>C), CAA227CAG (2986A>G), TAC228ATC (2987T>A 2988A>T), ACC231AAG (2997C>A 2998C>G), TTA232CTT (2999T>C 3001A>T), TTG234CTG (3005T>C), GCA235AGA (3008G>A 3009C>G), AGT236TCA (3011A>T 3012G>C 3013T>A), GGG237GGT (3016G>T), TTT238TAT (3018T>A), TAT239CAT (3020T>C), CAG240CAA (3025G>A), GTG241ATT (3026G>A 3028G>T), GAG242AGG (3029G>A 3030A>G), GAC244ACA (3035G>A 3036A>C 3037C>A), CCT245AAG (3038C>A 3039C>A 3040T>G), CAA246GAG (3041C>G 3043A>G), ATA248GTC (3047A>G 3049A>C), TCG249CAG (3050T>C 3051C>A), ACC251ACT (3058C>T), GCG252ACA (3059G>A 3061G>A), AAC254AGG (3066A>G 3067C>G), GTA255ACT (3068G>A 3069T>C 3070A>T), GAA256CAT (3071G>C 3073A>T), CAC257TCT (3074C>T 3075A>C 3076C>T), GGG258GGC (3079G>C), CAT259CAC (3082T>C), TTT260TAT (3084T>A), GAA261GAG (3088A>G), TTC262TAC (3090T>A), CTT263CTA (3094T>A), CGA264GTA (3095C>G 3096G>T), CCT266CCC (3103T>C), ATG267TTT (3104A>T 3106G>T), GGA268GGC (3109A>C), TTA269CTC (3110T>C 3112A>C), AAA270ACT (3114A>C 3115A>T), AAC271AAT (3118C>T), TCA272GCA (3119T>G), CCA273CCT (3124A>T), TCT274TCA (3127T>A), ACT275AGT (3129C>G), AGA278T.T (3137A>T 3139A>T), GTT279C.A (3140G>C 3142T>A), GAC281A.T (3146G>A 3148C>T), AAT282C.G (3149A>C 3151T>G), GTC283..T (3154C>T), CTA284T.T (3155C>T 3157A>T), AGA285CAG (3158A>C 3159G>A 3160A>G), GGT286CCT (3161G>C 3162G>C), CTC287TTC (3164C>T), CAA288CTC (3168A>T 3169A>C), AAT289GAC (3170A>G 3172T>C), AAC290ATA (3174A>T 3175C>A), ATC291TTT (3176A>T 3178C>T), TGT292GTG (3179T>G 3180G>T 3181T>G), CTC293GTA (3182C>G 3184C>A), TAC295TTC (3189A>T), CTT296ATT (3191C>A), ATT299GTA (3200A>G 3202T>A), ATT300CTG (3203A>C 3205T>G), GTC301ATA (3206G>A 3208C>A), TAT302TAC (3211T>C), AGT303TCT (3212A>T 3213G>C), ACT304CGA (3215A>C 3216C>G 3217T>A), TCC305AGT (3218T>A 3219C>G 3220C>T), CTA306AAG (3221C>A 3222T>A 3223A>G), CAG307GAA (3224C>G 3226G>A), CAC309CAT (3232C>T), CTG310GAA (3233C>G 3234T>A 3235G>A), GAG311GAA (3238G>A), AAC312TAT (3239A>T 3241C>T), CTG313TTG (3242C>T), GAA314AGG (3245G>A 3246A>G 3247A>G), CGA315ATA (3248C>A 3249G>T), GTT316GTA (3253T>A), TTC317CTG (3254T>C 3256C>G), CAA318CAG (3259A>G), AGA319ACA (3261G>C), CTT320TTG (3263C>T 3265T>G), AGA321AGG (3268A>G), GAA322GAT (3271A>T), AGT323GAG (3272A>G 3273G>A 3274T>G), AAC324AAG (3277C>G), TTC325TTG (3280C>G), AAA326TAT (3281A>T 3283A>T), ATT327GCA (3284A>G 3285T>C 3286T>A), CAA328AAG (3287C>A 3289A>G), ATG329TTT (3290A>T 3292G>T), TCC332TGC (3300C>G), TTC334TTT (3307C>T), TTG335TGG (3309T>G), AAG336TTG (3311A>T 3312A>T), CTC337AAT (3314C>A 3315T>A 3316C>T), ACT339GTT (3320A>G 3321C>T), GCT340GCC (3325T>C), TAT341TTT (3327A>T), GGT343GGG (3334T>G), CAC344CAT (3337C>T), ATC345GTA (3338A>G 3340C>A), ATA346GTT (3341A>G 3343A>T), AGC347TCG (3344A>T 3345G>C 3346C>G), AGG348GGT (3347A>G 3349G>T), GAC349GAT (3352C>T), GGT350GGG (3355T>G), CCT353GTG (3362C>G 3363C>T 3364T>G), AAC354GAC (3365A>G), CCT355CCG (3370T>G), GAT356AAG (3371G>A 3373T>G), AAG357AAA (3376G>A), ATT358ACT (3378T>C), TCC359GAG (3380T>G 3381C>A 3382C>G), GCT360GCA (3385T>A), ATT361GTC (3386A>G 3388T>C), CAA362AGA (3389C>A 3390A>G), AAA363AAT (3394A>T), TAT364TGG (3396A>G 3397T>G), CTG365CCA (3399T>C 3400G>A), ATT366AGG (3402T>G 3403T>G), CCA367CCT (3406A>T), AAG368TTA (3407A>T 3408A>T 3409G>A), ACC369TCT (3410A>T 3412C>T), CCT370CCG (3415T>G), AAG371ACT (3417A>C 3418G>T), GAA372GAT (3421A>T), ATA373ATT (3424A>T), AAA374AGA (3426A>G), CAA375AGC (3428C>A 3429A>G 3430A>C), TTT376TTC (3433T>C), TTA377TTG (3436A>G), GGC378GGA (3439C>A), CTT379TTG (3440C>T 3442T>G), CTC380GCA (3443C>G 3444T>C 3445C>A), TAT382TAC (3451T>C), TAC383TAT (3454C>T), CGA384AGG (3455C>A 3457A>G), AAA385AGA (3459A>G), ATT387GTA (3464A>G 3466T>A), CCA388GAA (3467C>G 3468C>A), GAT389GGT (3471A>G), GCA391TCT (3476G>T 3478A>T), CGA392ATT (3479C>A 3480G>T 3481A>T), CTC393ATT (3482C>A 3484C>T), ACA394AGT (3486C>G 3487A>T), AAA395AGG (3489A>G 3490A>G), CCC396CCA (3493C>A), ACA398ATG (3498C>T 3499A>G), CAG399GAT (3500C>G 3502G>T), TGC400TTG (3504G>T 3505C>G), TTA401CTG (3506T>C 3508A>G), AAA403AAG (3514A>G), GGT404GAC (3516G>A 3517T>C), AGT405del (3518_3520delAGT), AAA406AAC (3523A>C), GTA407TTC (3524G>T 3526A>C), ACT408ACA (3529T>A), CTT409TGG (3530C>T 3531T>G 3532T>G), AGT410TCT (3533A>T 3534G>C), CCC411GAA (3536C>G 3537C>A 3538C>A), GAA412ATT (3539G>A 3540A>T 3541A>T), TAT413GCT (3542T>G 3543A>C), GTA414ACT (3545G>A 3546T>C 3547A>T), AAT415CAA (3548A>C 3550T>A), GCT416CCT (3551G>C), GAA418CAT (3557G>C 3559A>T), CAC419AAC (3560C>A), TGT420TTG (3564G>T 3565T>G), AAA421AAG (3568A>G), ACT422GTA (3569A>G 3570C>T 3571T>A), CTG423GCA (3572C>G 3573T>C 3574G>A), ACC425ACA (3580C>A), AAC426TCT (3581A>T 3582A>C 3583C>T), GAC427GCT (3585A>C 3586C>T), CCA428CCT (3589A>T), ATA429GTC (3590A>G 3592A>C), TTA430CTA (3593T>C), CAA431GCA (3596C>G 3597A>C), TAC432TTA (3600A>T 3601C>A), CCA433CCT (3604A>T), GAC434AAT (3605G>A 3607C>T), TTT435TAT (3609T>A), ACC436TCC (3611A>T), AGA437ATT (3615G>T 3616A>T), GAA438CCT (3617G>C 3618A>C 3619A>T), AAC440GTT (3623A>G 3624A>T 3625C>T), CTC441GTT (3626C>G 3628C>T), ACG442GAG (3629A>G 3630C>A), GAC444GAT (3637C>T)

*: Inserts / Deletes / Misaligned / Frameshifts

#### Analysis details

This analysis was performed with panviral2.64

#### NGS Details (UN70): Caulimovirus tesselloscrophulariae

##### Assembly

| | |
|---|---|
| **Coverage Length** | 583 (2 contig(s)) |
| **Depth Of Coverage** | 15.8 |
| **Number Of Reads** | 80 |
| **Reads Per Million** | 1.52 rpm (after QC) |
| **Ambiguities** | 0 |
| **Assembly Method** | de novo + reference guided assembly |
| **Consensus Caller** | Bcf Tools |

##### Coverage Map

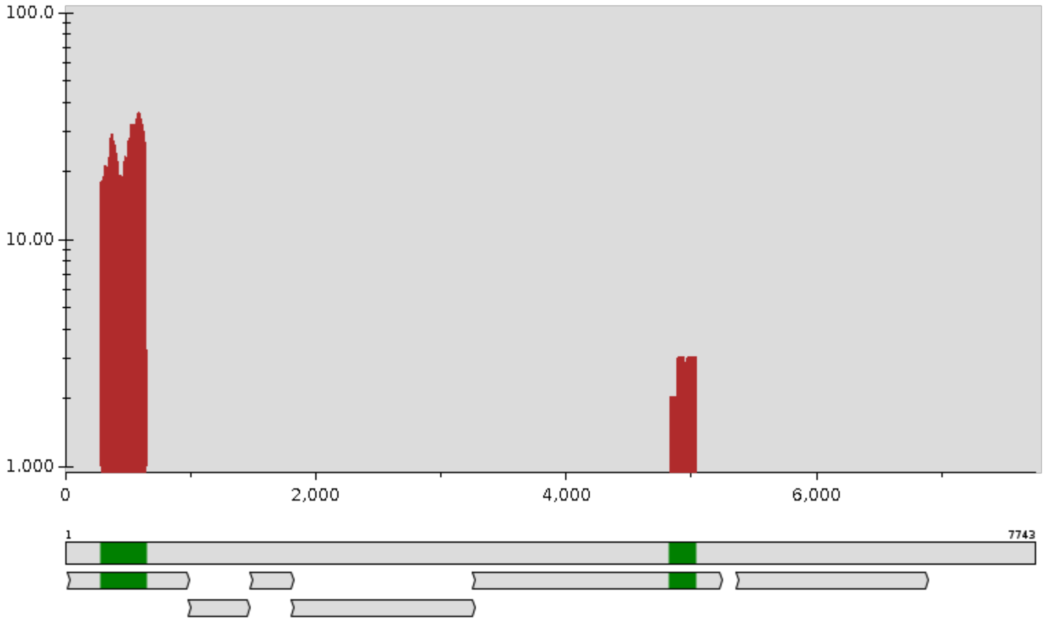

##### Assignment

| | |
|---|---|
| **Type** | Caulimovirus tesselloscrophulariae (Taxonomy ID: 3047834) |
| **Reference Genome** | NC_003554.1 |
| **NT Identity (%)** | 59.0193 |
| **AA Identity (%)** | 46.3158 |
| **Number Of Stop Codons** | 0 |
| **Number Of CDS** | 6 |

##### Alignment

| | |
|---|---|
| **Alignment Score** | 176.0 (NT) + 588.0 (AA) = 764.0 |
| **Concordance (%)** | 32.9168 |

| Alignment Method | Global, seeded, nucleotide + amino acids (AGA) |
|---|---|

#### Genome Region

Sequence starts at position 286 and ends at position 5043 relative to NC_003554.1 reference sequence.

#### Alignment Detailed Statistics

| | Begin | End | Coverage | Score | Concordance | Matches | Identities | I/D/M/F* | Stop Codons |
|---|---|---|---|---|---|---|---|---|---|
| **NT** | **286** | **5043** | **7.5%** | **176** | **15.8%** | **571 (97.9%)** | **337 (57.8%)** | **0/12** | |
| Mutations: | 296C>A, 299G>C, 300T>A, 302T>A, 304T>A, 305C>T, 307T>A, 308C>A, 309C>A, 311C>T, 313T>A, 314A>T, 322G>A, 324A>C, 325G>A, 326G>A, 327A>T, 328A>G, 331A>T, 332C>A, 334A>T, 339A>T, 340G>C, 343G>A, 344A>C, 345T>A, 346G>A, 348A>G, 353G>C, 354A>C, 355T>A, 356C>G, 358A>C, 359T>A, 360C>G, 361A>C, 362G>A, 363T>A, 368A>C, 369G>A, 373A>G, 376C>G, 377T>A, 378C>A, 379A>T, 380A>T, 381T>A, 382G>T, 383A>G, 385C>T, 388C>T, 389C>T, 391T>G, 396C>G, 398G>A, 401A>C, 403G>A, 406T>C, 407C>A, 409A>G, 410C>G, 412T>G, 414C>A, 416G>T, 417C>T, 419C>A, 420A>C, 421A>T, 425C>A, 426G>A, 427G>A, 428C>G, 433A>T, 436T>A, 437G>A, 440A>T, 441C>G, 442G>T, 443T>C, 445A>T, 446G>A, 448T>A, 450A>T, 454G>A, 455G>A, 456C>A, 457A>C, 461A>T, 462T>C, 463C>T, 469T>A, 473A>T, 475C>T, 476G>A, 478C>T, 482C>A, 483G>C, 484A>T, 486A>G, 490T>A, 491A>G, 493T>A, 494C>A, 495T>A, 496G>T, 499C>T, 503G>A, 504C>T, 507C>T, 508A>T, 509A>G, 510G>A, 518C>T, 520C>A, 526T>C, 527G>A, 528G>C, 535T>A, 536A>C, 538G>T, 544C>A, 545G>T, 546T>A, 550C>T, 553T>C, 555A>G, 558T>A, 560G>T, 561C>G, 562A>T, 563T>A, 565G>A, 572C>A, 575A>G, 576G>A, 577C>T, 578A>G, 579A>C, 580G>T, 583T>C, 586A>T, 587G>A, 590A>G, 593A>G, 598A>G, 601C>T, 604T>G, 605A>C, 606T>A, 608C>T, 609A>T, 613A>G, 614T>G, 616T>C, 617G>A, 628C>T, 632A>T, 634G>C, 637A>G, 638A>C, 4825A>G, 4828T>A, 4834G>A, 4838T>A, 4839A>G, 4840C>A, 4844C>T, 4846T>A, 4847A>G, 4859C>A, 4861T>C, 4864C>A, 4865A>G, 4867T>A, 4868A>T, 4870T>A, 4871G>C, 4876A>T, 4882A>T, 4889A>T, 4890G>A, 4892T>C, 4893T>A, 4900C>A, 4902G>C, 4903T>A, 4904G>T, 4906C>A, 4907C>T, 4910A>C, 4913G>A, 4916_4924delAGAGCGCTT, 4927C>T, 4928G>C, 4929G>T, 4930T>A, 4931G>A, 4932T>A, 4937_4939delTTA, 4945C>T, 4948G>A, 4953G>C, 4955T>A, 4956C>G, 4957A>T, 4962G>C, 4966C>T, 4969A>T, 4970C>G, 4973G>A, 4980A>C, 4981G>T, 4983A>G, 4984T>G, 4987T>C, 4988C>T, 4989A>T, 4990C>A, 4992G>C, 4993C>A, 4999T>A, 5014A>T, 5015G>A, 5017C>A, 5018A>G, 5019A>T, 5020G>A, 5021C>A, 5022A>G, 5025T>G, 5026A>C, 5029C>A, 5030A>C, 5031C>G, 5032C>A, 5041A>T | | | | | | | | |
| **CDS** | | | | | | | | | |
| FMVgp1 | 92 | 212 | 37.3% | 334 | 43.5% | 121 (100%) | 51 (42.1%) | 0/0/0/0 | 0 |
| Protein mutations: | Q95K (296C>A), V96H (299G>C 300T>A), Y97K (302T>A 304T>A), P99N (308C>A 309C>A), I101L (314A>T), Q104P (324A>C 325G>A), E105M (326G>A 327A>T 328A>G), Q107N (332C>A 334A>T), K109I (339A>T 340G>C), M111Q (344A>C 345T>A 346G>A), K112R (348A>G), D114P (353G>C 354A>C 355T>A), P115A (356C>G 358A>C), V117K (362G>A 363T>A), S119H (368A>C 369G>A), I121M (376C>G), S122N (377T>A 378C>A 379A>T), M123Y (380A>T 381T>A 382G>T), I124V (383A>G 385C>T), A128G (396C>G), V129I (398G>A), K130Q (401A>C 403G>A), L132M (407C>A 409A>G), L133V (410C>G 412T>G), T134K (414C>A), A135F (416G>T 417C>T), Q136T (419C>A 420A>C 421A>T), R138K (425C>A 426G>A 427G>A), Q139E (428C>G), D142N (437G>A), T143C (440A>T 441C>G 442G>T), S144P (443T>C 445A>T), V145I (446G>A 448T>A), K146I (450A>T), M147I (454G>A), A148N (455G>A 456C>A 457A>C), I150S (461A>T 462T>C 463C>T), D152E (469T>A), I154F (473A>T 475C>T), V155I (476G>A 478C>T), R157T (482C>A 483G>C 484A>T), K158R (486A>G), D159E (490T>A), S160G (491A>G 493T>A), L161N (494C>A 495T>A 496G>T), A164I (503G>A 504C>T), A165V (507C>T 508A>T), R166E (509A>G 510G>A), G172T (527G>A 528G>C), F174L (535T>A), M175L (536A>C 538G>T), V178Y (545G>T 546T>A), K181R (555A>G), F182Y (558T>A), A183C (560G>T 561C>G 562A>T), L184I (563T>A 565G>A), Q187K (572C>A), S188D (575A>G 576G>A 577C>T), K189A (578A>G 579A>C 580G>T), L191F (586A>T), D192N (587G>A), K193E (590A>G), T194A (593A>G), F197L (604T>G), I198H (605A>C 606T>A), H199F (608C>T 609A>T), F201V (614T>G 616T>C), E202K (617G>A), M207F (632A>T 634G>C), T209P (638A>C) | | | | | | | | |
| Codon mutations: | CAA95AAA (296C>A), GTT96CAT (299G>C 300T>A), TAT97AAA (302T>A 304T>A), CTT98TTA (305C>T 307T>A), CCT99AAT (308C>A 309C>A), CTT100TTA (311C>T 313T>A), ATA101TTA (314A>T), AAG103AAA (322G>A), CAG104CCA (324A>C 325G>A), GAA105ATG (326G>A 327A>T 328A>G), ATA106ATT (331A>T), CAA107AAT (332C>A 334A>T), AAG109ATC (339A>T 340G>C), CTG110CTA (343G>A), ATG111CAA (344A>C 345T>A 346G>A), AAA112AGA (348A>G), GAT114CCA (353G>C 354A>C 355T>A), CCA115GCC (356C>G 358A>C), TCA116AGC (359T>A 360C>G 361A>C), GTA117AAA (362G>A 363T>A), AGT119CAT (368A>C 369G>A), AAA120AAG (373A>G), ATC121ATG (376C>G), TCA122AAT (377T>A 378C>A 379A>T), ATG123TAT (380A>T 381T>A 382G>T), ATC124GTT (383A>G 385C>T), CAC125CAT (388C>T), CTT126TTG (389C>T 391T>G), GCA128GGA (396C>G), GTA129ATA (398G>A), AAG130CAA (401A>C 403G>A), ATT131ATC (406T>C), CTA132ATG (407C>A 409A>G), CTT133GTG (410C>G 412T>G), ACA134AAA (414C>A), GCC135TTC (416G>T 417C>T), CAA136ACT (419C>A 420A>C 421A>T), CGG138AAA (425C>A 426G>A 427G>A), CAA139GAA (428C>G), GGA140GGT (433A>T), ATT141ATA (436T>A), GAT142AAT (437G>A), ACG143TGT (440A>T 441C>G 442G>T), TCA144CCT (443T>C 445A>T), GTT145ATA (446G>A 448T>A), AAA146ATA (450A>T), ATG147ATA (454G>A), GCA148AAC (455G>A 456C>A 457A>C), ATC150TCT (461A>T 462T>C 463C>T), GAT152GAA (469T>A), ATC154TTT (473A>T 475C>T), GTC155ATT (476G>A 478C>T), CGA157ACT (482C>A 483G>C 484A>T), AAA158AGA (486A>G), GAT159GAA (490T>A), AGT160GGA (491A>G 493T>A), CTG161AAT (494C>A 495T>A 496G>T), CTC162CTT (499C>T), GCA164ATA (503G>A 504C>T), GCA165GTT (507C>T 508A>T), AGA166GAA (509A>G 510G>A), CTC169TTA (518C>T 520C>A), TAT171TAC (526T>C), GGT172ACT (527G>A 528G>C), TTT174TTA (535T>A), ATG175CTT (536A>C 538G>T), ACC177ACA (544C>A), GTT178TAT (545G>T 546T>A), TAC179TAT (550C>T), CCT180CCC (553T>C), AAA181AGA (555A>G), TTT182TAT (558T>A), GCA183TGT (560G>T 561C>G 562A>T), TTG184ATA (563T>A 565G>A), CAA187AAA (572C>A), AGC188GAT (575A>G 576G>A 577C>T), AAG189GCT (578A>G 579A>C 580G>T), AAT190AAC (583T>C), TTA191TTT (586A>T), GAT192AAT (587G>A), AAA193GAA (590A>G), ACA194GCA (593A>G), TTA195TTG (598A>G), AGC196AGT (601C>T), TTT197TTG (604T>G), ATT198CAT (605A>C 606T>A), CAT199TTT (608C>T 609A>T), CAA200CAG (613A>G), TTT201GTC (614T>G 616T>C), GAA202AAA (617G>A), GAC205GAT (628C>T), ATG207TTC (632A>T 634G>C), AAA208AAG (637A>G), ACA209CCA (638A>C) | | | | | | | | |
| FMVgp5 | 527 | 599 | 10.9% | 254 | 55.5% | 69 (94.5%) | 37 (50.7%) | 0/4/0/0 | 0 |
| Protein mutations: | F527L (4828T>A), Y531R (4838T>A 4839A>G 4840C>A), P533S (4844C>T 4846T>A), K534E (4847A>G), H538N (4859C>A 4861T>C), I540V (4865A>G 4867T>A), I541L (4868A>T 4870T>A), E542Q (4871G>C), S548Y (4889A>T 4890G>A), F549H (4892T>C 4893T>A), G552A (4902G>C 4903T>A), V553L (4904G>T 4906C>A), K555Q (4910A>C), A556T (4913G>A), R557_L559del (4916_4924delAGAGCGCTT), G561L (4928G>C 4929G>T 4930T>A), V562N (4931G>A 4932T>A), L564del (4937_4939delTTA), S569T (4953G>C), S572T (4962G>C), K574N (4969A>T), Q575E (4970C>G), A576T (4973G>A), K578T (4980A>C 4981G>T), N579R (4983A>G 4984T>G), H581L (4988C>T 4989A>T 4990C>A), S582T (4992G>C 4993C>A), D584E (4999T>A), V590I (5015G>A 5017C>A), K591V (5018A>G 5019A>T 5020G>A), Q592R (5021C>A 5022A>G), V593G (5025T>G 5026A>C), T595R (5030A>C 5031C>G 5032C>A) | | | | | | | | |
| Codon mutations: | TCA526..G (4825A>G), TTT527TTA (4828T>A), AAG529AAA (4834G>A), TAC531AGA (4838T>A 4839A>G 4840C>A), CCT533TCA (4844C>T 4846T>A), AAA534GAA (4847A>G), CAT538AAC (4859C>A 4861T>C), CTC539CTA (4864C>A), ATT540GTA (4865A>G 4867T>A), ATT541TTA (4868A>T 4870T>A), GAA542CAA (4871G>C), ACA543ACT (4876A>T), GCA545GCT (4882A>T), AGT548TAT (4889A>T 4890G>A), TTT549CAT (4892T>C 4893T>A), GGC551GGA (4900C>A), GGT552GCA (4902G>C 4903T>A), GTC553TTA (4904G>T 4906C>A), CTA554TTA (4907C>T), AAA555CAA (4910A>C), GCA556ACA (4913G>A), AGA557_CTT559del (4916_4924delAGAGCGCTT), GAC560GAT (4927C>T), GGT561CTA (4928G>C 4929G>T 4930T>A), GTT562AAT (4931G>A 4932T>A), TTA564del (4937_4939delTTA), TGC566TGT (4945C>T), AGG567AGA (4948G>A), AGT569ACT (4953G>C), TCA570AGT (4955T>A 4956C>G 4957A>T), AGC572ACC (4962G>C), TTC573TTT (4966C>T), AAA574AAT (4969A>T), CAA575GAA (4970C>G), GCA576ACA (4973G>A), AAG578ACT (4980A>C 4981G>T), AAT579AGG (4983A>G 4984T>G), TAT580TAC (4987T>C), CAC581TTA (4988C>T 4989A>T 4990C>A), AGC582ACA (4992G>C 4993C>A), GAT584GAA (4999T>A), GCA589GCT (5014A>T), GTC590ATA (5015G>A 5017C>A), AAG591GTA (5018A>G 5019A>T 5020G>A), CAA592AGA (5021C>A 5022A>G), GTA593GGC (5025T>G 5026A>C), ATC594ATA (5029C>A), ACC595CGA (5030A>C 5031C>G 5032C>A), TCA598TCT (5041A>T) | | | | | | | | |
| **Proteins** | | | | | | | | | |
| hypothetical protein (NP_619544.1) | 92 | 212 | 37.3% | 334 | 43.5% | 121 (100%) | 51 (42.1%) | 0/0/0/0 | 0 |
| Protein mutations: | Q95K (296C>A), V96H (299G>C 300T>A), Y97K (302T>A 304T>A), P99N (308C>A 309C>A), I101L (314A>T), Q104P (324A>C 325G>A), E105M (326G>A 327A>T 328A>G), Q107N (332C>A 334A>T), K109I (339A>T 340G>C), M111Q (344A>C 345T>A 346G>A), K112R (348A>G), D114P (353G>C 354A>C 355T>A), P115A (356C>G 358A>C), V117K (362G>A 363T>A), S119H (368A>C 369G>A), I121M (376C>G), S122N (377T>A 378C>A 379A>T), M123Y (380A>T 381T>A 382G>T), I124V (383A>G 385C>T), A128G (396C>G), V129I (398G>A), K130Q (401A>C 403G>A), L132M (407C>A 409A>G), L133V (410C>G 412T>G), T134K (414C>A), A135F (416G>T 417C>T), Q136T (419C>A 420A>C 421A>T), R138K (425C>A 426G>A 427G>A), Q139E (428C>G), D142N (437G>A), T143C (440A>T 441C>G 442G>T), S144P (443T>C 445A>T), V145I (446G>A 448T>A), K146I (450A>T), M147I (454G>A), A148N (455G>A 456C>A 457A>C), I150S (461A>T 462T>C 463C>T), D152E (469T>A), I154F (473A>T 475C>T), V155I (476G>A 478C>T), R157T (482C>A 483G>C 484A>T), K158R (486A>G), D159E (490T>A), S160G (491A>G 493T>A), L161N (494C>A 495T>A 496G>T), A164I (503G>A 504C>T), A165V (507C>T 508A>T), R166E (509A>G 510G>A), G172T (527G>A 528G>C), F174L (535T>A), M175L (536A>C 538G>T), V178Y (545G>T 546T>A), K181R (555A>G), F182Y (558T>A), A183C (560G>T 561C>G 562A>T), L184I (563T>A 565G>A), Q187K (572C>A), S188D (575A>G 576G>A 577C>T), K189A (578A>G 579A>C 580G>T), L191F (586A>T), D192N (587G>A), K193E (590A>G), T194A (593A>G), F197L (604T>G), I198H (605A>C 606T>A), H199F (608C>T 609A>T), F201V (614T>G 616T>C), E202K (617G>A), M207F (632A>T 634G>C), T209P (638A>C) | | | | | | | | |

| | Begin | End | Coverage | Score | Concordance | Matches | Identities | I/D/M/F* | Stop Codons |
|---|---|---|---|---|---|---|---|---|---|
| **NT** | **286** | **5043** | **7.5%** | **176** | **15.8%** | **571 (97.9%)** | **337 (57.8%)** | **0/12** | |
| Codon mutations: | CAA95AAA (296C>A), GTT96CAT (299G>C 300T>A), TAT97AAA (302T>A 304T>A), CTT98TTA (305C>T 307T>A), CCT99AAT (308C>A 309C>A), CTT100TTA (311C>T 313T>A), ATA101TTA (314A>T), AAG103AAA (322G>A), CAG104CCA (324A>C 325G>A), GAA105ATG (326G>A 327A>T 328A>G), ATA106ATT (331A>T), CAA107AAT (332C>A 334A>T), AAG109ATC (339A>T 340G>C), CTG110CTA (343G>A), ATG111CAA (344A>C 345T>A 346G>A), AAA112AGA (348A>G), GAT114CCA (353G>C 354A>C 355T>A), CCA115GCC (356C>G 358A>C), TCA116AGC (359T>A 360C>G 361A>C), GTA117AAA (362G>A 363T>A), AGT119CAT (368A>C 369G>A), AAA120AAG (373A>G), ATC121ATG (376C>G), TCA122AAT (377T>A 378C>A 379A>T), ATG123TAT (380A>T 381T>A 382G>T), ATC124GTT (383A>G 385C>T), CAC125CAT (388C>T), CTT126TTG (389C>T 391T>G), GCA128GGA (396C>G), GTA129ATA (398G>A), AAG130CAA (401A>C 403G>A), ATT131ATC (406T>C), CTA132ATG (407C>A 409A>G), CTT133GTG (410C>G 412T>G), ACA134AAA (414C>A), GCC135TTC (416G>T 417C>T), CAA136ACT (419C>A 420A>C 421A>T), CGG138AAA (425C>A 426G>A 427G>A), CAA139GAA (428C>G), GGA140GGT (433A>T), ATT141ATA (436T>A), GAT142AAT (437G>A), ACG143TGT (440A>T 441C>G 442G>T), TCA144CCT (443T>C 445A>T), GTT145ATA (446G>A 448T>A), AAA146ATA (450A>T), ATG147ATA (454G>A), GCA148AAC (455G>A 456C>A 457A>C), ATC150TCT (461A>T 462T>C 463C>T), GAT152GAA (469T>A), ATC154TTT (473A>T 475C>T), GTC155ATT (476G>A 478C>T), CGA157ACT (482C>A 483G>C 484A>T), AAA158AGA (486A>G), GAT159GAA (490T>A), AGT160GGA (491A>G 493T>A), CTG161AAT (494C>A 495T>A 496G>T), CTC162CTT (499C>T), GCA164ATA (503G>A 504C>T), GCA165GTT (507C>T 508A>T), AGA166GAA (509A>G 510G>A), CTC169TTA (518C>T 520C>A), TAT171TAC (526T>C), GGT172ACT (527G>A 528G>C), TTT174TTA (535T>A), ATG175CTT (536A>C 538G>T), ACC177ACA (544C>A), GTT178TAT (545G>T 546T>A), TAC179TAT (550C>T), CCT180CCC (553T>C), AAA181AGA (555A>G), TTT182TAT (558T>A), GCA183TGT (560G>T 561C>G 562A>T), TTG184ATA (563T>A 565G>A), CAA187AAA (572C>A), AGC188GAT (575A>G 576G>A 577C>T), AAG189GCT (578A>G 579A>C 580G>T), AAT190AAC (583T>C), TTA191TTT (586A>T), GAT192AAT (587G>A), AAA193GAA (590A>G), ACA194GCA (593A>G), TTA195TTG (598A>G), AGC196AGT (601C>T), TTT197TTG (604T>G), ATT198CAT (605A>C 606T>A), CAT199TTT (608C>T 609A>T), CAA200CAG (613A>G), TTT201GTC (614T>G 616T>C), GAA202AAA (617G>A), GAC205GAT (628C>T), ATG207TTC (632A>T 634G>C), AAA208AAG (637A>G), ACA209CCA (638A>C) | | | | | | | | |
| hypothetical protein (NP_619548.1) | 527 | 599 | 10.9% | 254 | 55.5% | 69 (94.5%) | 37 (50.7%) | 0/4/0/0 | 0 |
| Protein mutations: | F527L (4828T>A), Y531R (4838T>A 4839A>G 4840C>A), P533S (4844C>T 4846T>A), K534E (4847A>G), H538N (4859C>A 4861T>C), I540V (4865A>G 4867T>A), I541L (4868A>T 4870T>A), E542Q (4871G>C), S548Y (4889A>T 4890G>A), F549H (4892T>C 4893T>A), G552A (4902G>C 4903T>A), V553L (4904G>T 4906C>A), K555Q (4910A>C), A556T (4913G>A), R557_L559del (4916_4924delAGAGCGCTT), G561L (4928G>C 4929G>T 4930T>A), V562N (4931G>A 4932T>A), L564del (4937_4939delTTA), S569T (4953G>C), S572T (4962G>C), K574N (4969A>T), Q575E (4970C>G), A576T (4973G>A), K578T (4980A>C 4981G>T), N579R (4983A>G 4984T>G), H581L (4988C>T 4989A>T 4990C>A), S582T (4992G>C 4993C>A), D584E (4999T>A), V590I (5015G>A 5017C>A), K591V (5018A>G 5019A>T 5020G>A), Q592R (5021C>A 5022A>G), V593G (5025T>G 5026A>C), T595R (5030A>C 5031C>G 5032C>A) | | | | | | | | |
| Codon mutations: | TCA526..G (4825A>G), TTT527TTA (4828T>A), AAG529AAA (4834G>A), TAC531AGA (4838T>A 4839A>G 4840C>A), CCT533TCA (4844C>T 4846T>A), AAA534GAA (4847A>G), CAT538AAC (4859C>A 4861T>C), CTC539CTA (4864C>A), ATT540GTA (4865A>G 4867T>A), ATT541TTA (4868A>T 4870T>A), GAA542CAA (4871G>C), ACA543ACT (4876A>T), GCA545GCT (4882A>T), AGT548TAT (4889A>T 4890G>A), TTT549CAT (4892T>C 4893T>A), GGC551GGA (4900C>A), GGT552GCA (4902G>C 4903T>A), GTC553TTA (4904G>T 4906C>A), CTA554TTA (4907C>T), AAA555CAA (4910A>C), GCA556ACA (4913G>A), AGA557_CTT559del (4916_4924delAGAGCGCTT), GAC560GAT (4927C>T), GGT561CTA (4928G>C 4929G>T 4930T>A), GTT562AAT (4931G>A 4932T>A), TTA564del (4937_4939delTTA), TGC566TGT (4945C>T), AGG567AGA (4948G>A), AGT569ACT (4953G>C), TCA570AGT (4955T>A 4956C>G 4957A>T), AGC572ACC (4962G>C), TTC573TTT (4966C>T), AAA574AAT (4969A>T), CAA575GAA (4970C>G), GCA576ACA (4973G>A), AAG578ACT (4980A>C 4981G>T), AAT579AGG (4983A>G 4984T>G), TAT580TAC (4987T>C), CAC581TTA (4988C>T 4989A>T 4990C>A), AGC582ACA (4992G>C 4993C>A), GAT584GAA (4999T>A), GCA589GCT (5014A>T), GTC590ATA (5015G>A 5017C>A), AAG591GTA (5018A>G 5019A>T 5020G>A), CAA592AGA (5021C>A 5022A>G), GTA593GGC (5025T>G 5026A>C), ATC594ATA (5029C>A), ACC595CGA (5030A>C 5031C>G 5032C>A), TCA598TCT (5041A>T) | | | | | | | | |

*: Inserts / Deletes / Misaligned / Frameshifts

#### Analysis details

This analysis was performed with panviral2.64

### NGS Details (UN70): Tomato necrotic dwarf virus (segment RNA1)

#### Assembly

| | |
|---|---|
| **Coverage Length** | 435 (1 contig(s)) |
| **Depth Of Coverage** | 21.2 |
| **Number Of Reads** | 79 |
| **Reads Per Million** | 1.50 rpm (after QC) |
| **Ambiguities** | 0 |
| **Assembly Method** | de novo + reference guided assembly |
| **Consensus Caller** | Bcf Tools |

#### Coverage Map

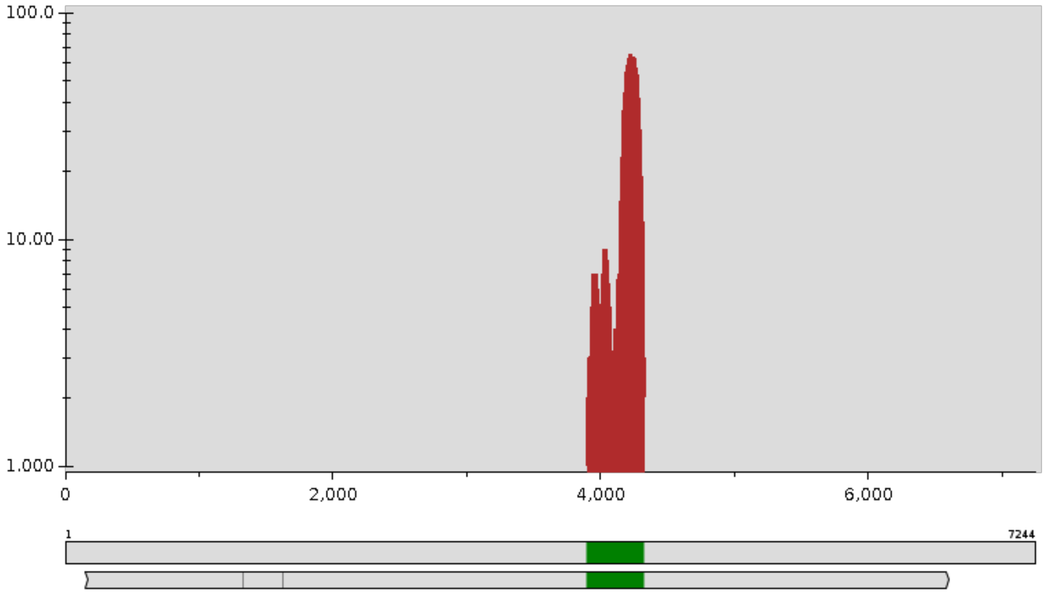

#### Assignment

| | |
|---|---|
| **Type** | Tomato necrotic dwarf virus (Taxonomy ID: 1481465) |
| **Reference Genome** | NC_027926.1 |
| **NT Identity (%)** | 69.4253 |
| **AA Identity (%)** | 74.4828 |
| **Number Of Stop Codons** | 0 |
| **Number Of CDS** | 1 |

#### Alignment

| | |
|---|---|
| **Alignment Score** | 338.0 (NT) + 788.0 (AA) = 1126.0 |
| **Concordance (%)** | 59.4823 |

| **Alignment Method** | Global, seeded, nucleotide + amino acids (AGA) |
|---|---|

#### Genome Region

Sequence starts at position 3895 and ends at position 4329 relative to NC_027926.1 reference sequence.

#### Alignment Detailed Statistics

| | Begin | End | Coverage | Score | Concordance | Matches | Identities | I/D/M/F* | Stop Codons |
|---|---|---|---|---|---|---|---|---|---|
| **NT** | **3895** | **4329** | **6.0%** | **338** | **38.9%** | **435 (100%)** | **302 (69.4%)** | **0/0** | |
| Mutations: | 3901A>T, 3906C>A, 3907A>C, 3909G>T, 3912A>T, 3913A>T, 3914T>C, 3915G>T, 3921G>A, 3922C>A, 3927G>A, 3933C>G, 3935T>A, 3937C>G, 3939A>C, 3940A>C, 3941C>A, 3943G>C, 3944G>C, 3946G>A, 3948A>C, 3949G>T, 3950G>C, 3951T>G, 3952G>C, 3954T>G, 3956A>G, 3957A>G, 3960A>T, 3961G>C, 3962A>T, 3963G>T, 3966T>C, 3968G>A, 3973G>A, 3981A>T, 3984G>C, 3990A>T, 3991G>C, 3993T>A, 3999G>A, 4002A>G, 4003C>T, 4009A>T, 4010T>C, 4011T>G, 4014A>G, 4015A>G, 4017T>A, 4023G>A, 4028T>C, 4030C>T, 4031A>T, 4035G>T, 4038T>C, 4041A>C, 4042A>G, 4047C>A, 4050A>C, 4053A>G, 4059C>G, 4062G>A, 4068G>A, 4071C>G, 4072C>T, 4074T>G, 4076C>A, 4077T>A, 4083A>T, 4093G>C, 4104A>G, 4107A>C, 4108A>C, 4110A>C, 4111T>C, 4113A>G, 4116T>C, 4119G>A, 4122A>C, 4137C>T, 4141A>C, 4149C>G, 4150A>C, 4161T>C, 4173C>T, 4176C>T, 4177A>T, 4178C>G, 4179C>T, 4188A>T, 4189T>G, 4190T>C, 4197T>C, 4200C>A, 4201A>C, 4206A>T, 4209T>C, 4215T>A, 4218A>T, 4221A>C, 4222G>A, 4230C>G, 4233T>C, 4234T>C, 4235C>A, 4243T>A, 4245T>A, 4246C>G, 4248T>G, 4251T>G, 4254C>T, 4256A>G, 4257C>T, 4259A>G, 4263C>T, 4267G>T, 4269T>A, 4279A>G, 4280C>A, 4281T>G, 4284G>A, 4287T>C, 4293T>C, 4302A>G, 4305T>C, 4311C>T, 4314G>A, 4315C>A, 4317C>A, 4323A>C, 4324G>A, 4326G>C, 4329C>T | | | | | | | | |

##### CDS

| | Begin | End | Coverage | Score | Concordance | Matches | Identities | I/D/M/F* | Stop Codons |
|---|---|---|---|---|---|---|---|---|---|
| APL33_sRA11gp1 | 1249 | 1393 | 6.7% | 788 | 77.0% | 145 (100%) | 108 (74.5%) | 0/0/0/0 | 0 |
| Protein mutations: | M1251L (3901A>T), M1253L (3907A>C 3909G>T), M1255S (3913A>T 3914T>C 3915G>T), V1262E (3935T>A), P1263A (3937C>G 3939A>C), T1264Q (3940A>C 3941C>A), G1265P (3943G>C 3944G>C), E1266N (3946G>A 3948A>C), G1267S (3949G>T 3950G>C 3951T>G), D1268Q (3952G>C 3954T>G), K1269R (3956A>G 3957A>G), E1271L (3961G>C 3962A>T 3963G>T), R1273K (3968G>A), D1275N (3973G>A), E1280D (3990A>T), A1281P (3991G>C 3993T>A), I1287S (4009A>T 4010T>C 4011T>G), I1289V (4015A>G 4017T>A), I1293T (4028T>C), H1294F (4030C>T 4031A>T), I1298V (4042A>G), P1309Q (4076C>A 4077T>A), E1311D (4083A>T), G1315R (4093G>C), M1331L (4141A>C), T1343C (4177A>T 4178C>G 4179C>T), L1347A (4189T>G 4190T>C), I1351L (4201A>C), D1358N (4222G>A), S1362H (4234T>C 4235C>A), S1365T (4243T>A 4245T>A), L1366V (4246C>G 4248T>G), N1369S (4256A>G 4257C>T), N1370S (4259A>G), A1373S (4267G>T 4269T>A), T1377E (4279A>G 4280C>A 4281T>G), V1392I (4324G>A 4326G>C) | | | | | | | | |
| Codon mutations: | ATG1251TTG (3901A>T), GGC1252GGA (3906C>A), ATG1253CTT (3907A>C 3909G>T), GGA1254GGT (3912A>T), ATG1255TCT (3913A>T 3914T>C 3915G>T), AAG1257AAA (3921G>A), CGG1258AGG (3922C>A), GAG1259GAA (3927G>A), GTC1261GTG (3933C>G), GTA1262GAA (3935T>A), CCA1263GCC (3937C>G 3939A>C), ACA1264CAA (3940A>C 3941C>A), GGA1265CCA (3943G>C 3944G>C), GAA1266AAC (3946G>A 3948A>C), GGT1267TCG (3949G>T 3950G>C 3951T>G), GAT1268CAG (3952G>C 3954T>G), AAA1269AGG (3956A>G 3957A>G), GTA1270GTT (3960A>T), GAG1271CTT (3961G>C 3962A>T 3963G>T), CTT1272CTC (3966T>C), AGA1273AAA (3968G>A), GAC1275AAC (3973G>A), CCA1277CCT (3981A>T), GTG1278GTC (3984G>C), GAA1280GAT (3990A>T), GCT1281CCA (3991G>C 3993T>A), GAG1283GAA (3999G>A), GAA1284GAG (4002A>G), CTG1285TTG (4003C>T), ATT1287TCG (4009A>T 4010T>C 4011T>G), CAA1288CAG (4014A>G), ATT1289GTA (4015A>G 4017T>A), CAG1291CAA (4023G>A), ATA1293ACA (4028T>C), CAT1294TTT (4030C>T 4031A>T), GTG1295GTT (4035G>T), GAT1296GAC (4038T>C), CTA1297CTC (4041A>C), ATC1298GTC (4042A>G), ACC1299ACA (4047C>A), ACA1300ACC (4050A>C), CAA1301CAG (4053A>G), GCC1303GCG (4059C>G), AAG1304AAA (4062G>A), GAG1306GAA (4068G>A), CGC1307CGG (4071C>G), CTT1308TTG (4072C>T 4074T>G), CCT1309CAA (4076C>A 4077T>A), GAA1311GAT (4083A>T), GGG1315CGG (4093G>C), AAA1318AAG (4104A>G), ACA1319ACC (4107A>C), AGA1320CGC (4108A>C 4110A>C), TTA1321CTG (4111T>C 4113A>G), TTT1322TTC (4116T>C), GAG1323GAA (4119G>A), ATA1324ATC (4122A>C), TAC1329TAT (4137C>T), ATG1331CTG (4141A>C), GTC1333GTG (4149C>G), AGG1334CGG (4150A>C), TTT1337TTC (4161T>C), TCC1341TCT (4173C>T), GCC1342GCT (4176C>T), ACC1343TGT (4177A>T 4178C>G 4179C>T), GCA1346GCT (4188A>T), TTA1347GCA (4189T>G 4190T>C), AAT1349AAC (4197T>C), GTC1350GTA (4200C>A), ATC1351CTC (4201A>C), CCA1352CCT (4206A>T), TGT1353TGC (4209T>C), GTT1355GTA (4215T>A), GGA1356GGT (4218A>T), ATA1357ATC (4221A>C), GAT1358AAT (4222G>A), ACC1360ACG (4230C>G), AGT1361AGC (4233T>C), TCT1362CAT (4234T>C 4235C>A), TCT1365ACA (4243T>A 4245T>A), CTT1366GTG (4246C>G 4248T>G), CTT1367CTG (4251T>G), GCC1368GCT (4254C>T), AAC1369AGT (4256A>G 4257C>T), AAC1370AGC (4259A>G), TTC1371TTT (4263C>T), GCT1373TCA (4267G>T 4269T>A), ACT1377GAG (4279A>G 4280C>A 4281T>G), GGG1378GGA (4284G>A), TTT1379TTC (4287T>C), GCT1381GCC (4293T>C), TCA1384TCG (4302A>G), AGT1385AGC (4305T>C), GAC1387GAT (4311C>T), GGG1388GGA (4314G>A), CGC1389AGA (4315C>A 4317C>A), CCA1391CCC (4323A>C), GTG1392ATC (4324G>A 4326G>C), TTC1393TTT (4329C>T) | | | | | | | | |

##### Proteins

| | Begin | End | Coverage | Score | Concordance | Matches | Identities | I/D/M/F* | Stop Codons |
|---|---|---|---|---|---|---|---|---|---|
| polyprotein (YP_009165993.1) | 1249 | 1393 | 6.7% | 788 | 77.0% | 145 (100%) | 108 (74.5%) | 0/0/0/0 | 0 |
| Protein mutations: | M1251L (3901A>T), M1253L (3907A>C 3909G>T), M1255S (3913A>T 3914T>C 3915G>T), V1262E (3935T>A), P1263A (3937C>G 3939A>C), T1264Q (3940A>C 3941C>A), G1265P (3943G>C 3944G>C), E1266N (3946G>A 3948A>C), G1267S (3949G>T 3950G>C 3951T>G), D1268Q (3952G>C 3954T>G), K1269R (3956A>G 3957A>G), E1271L (3961G>C 3962A>T 3963G>T), R1273K (3968G>A), D1275N (3973G>A), E1280D (3990A>T), A1281P (3991G>C 3993T>A), I1287S (4009A>T 4010T>C 4011T>G), I1289V (4015A>G 4017T>A), I1293T (4028T>C), H1294F (4030C>T 4031A>T), I1298V (4042A>G), P1309Q (4076C>A 4077T>A), E1311D (4083A>T), G1315R (4093G>C), M1331L (4141A>C), T1343C (4177A>T 4178C>G 4179C>T), L1347A (4189T>G 4190T>C), I1351L (4201A>C), D1358N (4222G>A), S1362H (4234T>C 4235C>A), S1365T (4243T>A 4245T>A), L1366V (4246C>G 4248T>G), N1369S (4256A>G 4257C>T), N1370S (4259A>G), A1373S (4267G>T 4269T>A), T1377E (4279A>G 4280C>A 4281T>G), V1392I (4324G>A 4326G>C) | | | | | | | | |
| Codon mutations: | ATG1251TTG (3901A>T), GGC1252GGA (3906C>A), ATG1253CTT (3907A>C 3909G>T), GGA1254GGT (3912A>T), ATG1255TCT (3913A>T 3914T>C 3915G>T), AAG1257AAA (3921G>A), CGG1258AGG (3922C>A), GAG1259GAA (3927G>A), GTC1261GTG (3933C>G), GTA1262GAA (3935T>A), CCA1263GCC (3937C>G 3939A>C), ACA1264CAA (3940A>C 3941C>A), GGA1265CCA (3943G>C 3944G>C), GAA1266AAC (3946G>A 3948A>C), GGT1267TCG (3949G>T 3950G>C 3951T>G), GAT1268CAG (3952G>C 3954T>G), AAA1269AGG (3956A>G 3957A>G), GTA1270GTT (3960A>T), GAG1271CTT (3961G>C 3962A>T 3963G>T), CTT1272CTC (3966T>C), AGA1273AAA (3968G>A), GAC1275AAC (3973G>A), CCA1277CCT (3981A>T), GTG1278GTC (3984G>C), GAA1280GAT (3990A>T), GCT1281CCA (3991G>C 3993T>A), GAG1283GAA (3999G>A), GAA1284GAG (4002A>G), CTG1285TTG (4003C>T), ATT1287TCG (4009A>T 4010T>C 4011T>G), CAA1288CAG (4014A>G), ATT1289GTA (4015A>G 4017T>A), CAG1291CAA (4023G>A), ATA1293ACA (4028T>C), CAT1294TTT (4030C>T 4031A>T), GTG1295GTT (4035G>T), GAT1296GAC (4038T>C), CTA1297CTC (4041A>C), ATC1298GTC (4042A>G), ACC1299ACA (4047C>A), ACA1300ACC (4050A>C), CAA1301CAG (4053A>G), GCC1303GCG (4059C>G), AAG1304AAA (4062G>A), GAG1306GAA (4068G>A), CGC1307CGG (4071C>G), CTT1308TTG (4072C>T 4074T>G), CCT1309CAA (4076C>A 4077T>A), GAA1311GAT (4083A>T), GGG1315CGG (4093G>C), AAA1318AAG (4104A>G), ACA1319ACC (4107A>C), AGA1320CGC (4108A>C 4110A>C), TTA1321CTG (4111T>C 4113A>G), TTT1322TTC (4116T>C), GAG1323GAA (4119G>A), ATA1324ATC (4122A>C), TAC1329TAT (4137C>T), ATG1331CTG (4141A>C), GTC1333GTG (4149C>G), AGG1334CGG (4150A>C), TTT1337TTC (4161T>C), TCC1341TCT (4173C>T), GCC1342GCT (4176C>T), ACC1343TGT (4177A>T 4178C>G 4179C>T), GCA1346GCT (4188A>T), TTA1347GCA (4189T>G 4190T>C), AAT1349AAC (4197T>C), GTC1350GTA (4200C>A), ATC1351CTC (4201A>C), CCA1352CCT (4206A>T), TGT1353TGC (4209T>C), GTT1355GTA (4215T>A), GGA1356GGT (4218A>T), ATA1357ATC (4221A>C), GAT1358AAT (4222G>A), ACC1360ACG (4230C>G), AGT1361AGC (4233T>C), TCT1362CAT (4234T>C 4235C>A), TCT1365ACA (4243T>A 4245T>A), CTT1366GTG (4246C>G 4248T>G), CTT1367CTG (4251T>G), GCC1368GCT (4254C>T), AAC1369AGT (4256A>G 4257C>T), AAC1370AGC (4259A>G), TTC1371TTT (4263C>T), GCT1373TCA (4267G>T 4269T>A), ACT1377GAG (4279A>G 4280C>A 4281T>G), GGG1378GGA (4284G>A), TTT1379TTC (4287T>C), GCT1381GCC (4293T>C), TCA1384TCG (4302A>G), AGT1385AGC (4305T>C), GAC1387GAT (4311C>T), GGG1388GGA (4314G>A), CGC1389AGA (4315C>A 4317C>A), CCA1391CCC (4323A>C), GTG1392ATC (4324G>A 4326G>C), TTC1393TTT (4329C>T) | | | | | | | | |

*: Inserts / Deletes / Misaligned / Frameshifts

#### Analysis details

This analysis was performed with panviral2.64

### NGS Details (UN70): Badnavirus maculaucubae

#### Assembly

| | |
|---|---|
| **Coverage Length** | 321 (1 contig(s)) |
| **Depth Of Coverage** | 24.5 |
| **Number Of Reads** | 77 |
| **Reads Per Million** | 1.46 rpm (after QC) |
| **Ambiguities** | 0 |
| **Assembly Method** | de novo + reference guided assembly |
| **Consensus Caller** | Bcf Tools |

#### Coverage Map

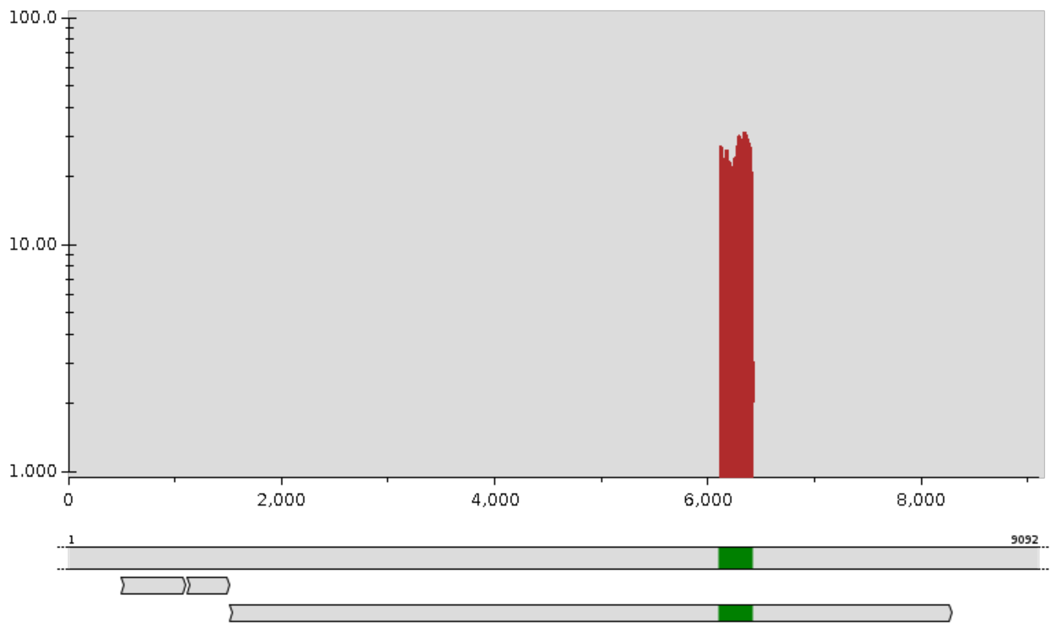

#### Assignment

| | |
|---|---|
| **Type** | Badnavirus maculaucubae (Taxonomy ID: 3051986) |
| **Reference Genome** | NC_076606.1 |
| **NT Identity (%)** | 54.6828 |
| **AA Identity (%)** | 45.045 |
| **Number Of Stop Codons** | 1 |
| **Number Of CDS** | 3 |

#### Alignment

| | |
|---|---|
| **Alignment Score** | 45.0 (NT) + 220.0 (AA) = 265.0 |
| **Concordance (%)** | 22.0497 |

| Alignment Method | Global, seeded, nucleotide + amino acids (AGA) |
|---|---|

#### Genome Region

Sequence starts at position 6102 and ends at position 6422 relative to NC_076606.1 reference sequence.

#### Alignment Detailed Statistics

| | Begin | End | Coverage | Score | Concordance | Matches | Identities | I/D/M/F* | Stop Codons |
|---|---|---|---|---|---|---|---|---|---|
| **NT** | **6102** | **6422** | **3.5%** | **45** | **7.2%** | **319 (95.8%)** | **181 (54.4%)** | **12/2** | |

| | |
|---|---|
| Mutations: | 6102C>T, 6105A>G, 6106T>A, 6112T>C, 6114G>T, 6115A>C, 6116A>G, 6120T>C, 6122G>A, 6125T>A, 6132G>A, 6133G>T, 6136G>A, 6137C>G, 6138A>G, 6139A>G, 6141G>A, 6142G>A, 6147A>G, 6149C>G, 6150A>T, 6151_6152delAG, 6153C>G, 6156T>C, 6159A>G, 6160T>A, 6161G>A, 6165G>T, 6171C>T, 6173T>G, 6179C>A, 6181G>T, 6184G>A, 6186A>T, 6188T>A, 6189A>C, 6191A>T, 6192C>T, 6197G>T, 6198G>T, 6199C>T, 6201A>G, 6204A>T, 6207G>A, 6208C>T, 6210A>T, 6216T>A, 6221A>C, 6222G>A, 6228A>T, 6231T>C, 6235G>A, 6236T>C, 6237G>T, 6240C>T, 6241C>A, 6242A>T, 6243A>G, 6244A>G, 6245G>A, 6246G>C, 6247A>T, 6248A>T, 6253G>A, 6257A>G, 6258C>G, 6259T>G, 6260G>T, 6261T>A, 6266G>A, 6268G>C, 6269G>C, 6270T>G, 6271A>T, 6272C>A, 6273A>T, 6274G>T, 6275A>T, 6276A>G, 6279T>C, 6279_6280insATG, 6282C>T, 6283A>G, 6285C>A, 6287C>T, 6288T>G, 6291T>C, 6292T>C, 6293A>T, 6297T>A, 6303T>C, 6306T>A, 6307C>T, 6312A>T, 6314T>A, 6319G>C, 6320A>G, 6321A>G, 6325C>G, 6326T>C, 6327T>C, 6328C>A, 6330A>G, 6333G>A, 6336C>T, 6342G>T, 6345T>C, 6346C>T, 6348C>G, 6350A>G, 6351A>G, 6354A>T, 6355A>G, 6358C>T, 6361G>C, 6365T>C, 6366A>T, 6371G>A, 6372G>A, 6373A>C, 6374A>T, 6375G>T, 6376A>T, 6377A>G, 6378T>A, 6382C>A, 6383T>C, 6384A>C, 6386T>A, 6387_6388insAATTGTTCG, 6390C>A, 6393C>T, 6394C>T, 6397A>G, 6398C>T, 6402A>T, 6403A>G, 6408A>T, 6409A>T, 6411A>T, 6413C>G |

##### CDS

| | Begin | End | Coverage | Score | Concordance | Matches | Identities | I/D/M/F* | Stop Codons |
|---|---|---|---|---|---|---|---|---|---|
| QKP79_gp3 | 1532 | 1638 | 4.7% | 220 | 28.5% | 107 (96.4%) | 50 (45.0%) | 4/0/1/1 | 1 |

| | |
|---|---|
| Protein mutations: | F1533I (6106T>A), K1536R (6115A>C 6116A>G), G1538D (6122G>A), F1539Y (6125T>A), V1542L (6133G>T), A1543R (6136G>A 6137C>G 6138A>G), M1544V (6139A>G 6141G>A), E1545K (6142G>A), S1547C (6149C>G 6150A>T), W1551K (6160T>A 6161G>A), I1555R (6173T>G), P1557Q (6179C>A), D1558Y (6181G>T), G1559S (6184G>A 6186A>T), L1560H (6188T>A 6189A>C), Y1561F (6191A>T 6192C>T), W1563F (6197G>T 6198G>T), M1566I (6207G>A), P1567S (6208C>T 6210A>T), K1571T (6221A>C 6222G>A), V1576T (6235G>A 6236T>C 6237G>T), Q1578M (6241C>A 6242A>T 6243A>G), R1579D (6244A>G 6245G>A 6246G>C), K1580L (6247A>T 6248A>T), D1582N (6253G>A), N1583R (6257A>G 6258C>G), C1584V (6259T>G 6260G>T 6261T>A), R1586K (6266G>A), G1587P (6268G>C 6269G>C 6270T>G), T1588Y (6271A>T 6272C>A 6273A>T), E1589L (6274G>T 6275A>T 6276A>G), D1590_F1591insM (6279_6280insATG), I1592V (6283A>G 6285C>A), A1593V (6287C>T 6288T>G), Y1595L (6292T>C 6293A>T), F1602Y (6314T>A), E1604R (6319G>C 6320A>G 6321A>G), L1606A (6325C>G 6326T>C 6327T>C), Q1607K (6328C>A 6330A>G), E1611D (6342G>T), K1614R (6350A>G 6351A>G), M1616V (6355A>G), E1618Q (6361G>C), I1619T (6365T>C 6366A>T), R1621K (6371G>A 6372G>A), K1622L (6373A>C 6374A>T 6375G>T), N1623* (6376A>T 6377A>G 6378T>A), L1625T (6382C>A 6383T>C 6384A>C), V1626D (6386T>A), V1626_L1627insNCS (6387_6388insAATTGTTCG), P1629S (6394C>T), T1630V (6397A>G 6398C>T), K1631N (6402A>T), M1632V (6403A>G), K1633N (6408A>T), I1634F (6409A>T 6411A>T), A1635G (6413C>G) |
| Codon mutations: | AGC1531..T (6102C>T), AAA1532AAG (6105A>G), TTT1533ATT (6106T>A), TTG1535CTT (6112T>C 6114G>T), AAA1536CGA (6115A>C 6116A>G), TCT1537TCC (6120T>C), GGT1538GAT (6122G>A), TTT1539TAT (6125T>A), CAG1541CAA (6132G>A), GTG1542TTG (6133G>T), GCA1543AGG (6136G>A 6137C>G 6138A>G), ATG1544GTA (6139A>G 6141G>A), GAA1545AAA (6142G>A), GAA1546GAG (6147A>G), TCA1547TGT (6149C>G 6150A>T), AGC1548--G (6151_6152delAG 6153C>G), ATT1549ATC (6156T>C), CCA1550CCG (6159A>G), TGG1551AAG (6160T>A 6161G>A), ACG1552ACT (6165G>T), TTC1554TTT (6171C>T), ATA1555AGA (6173T>G), CCA1557CAA (6179C>A), GAT1558TAT (6181G>T), GGA1559AGT (6184G>A 6186A>T), CTA1560CAC (6188T>A 6189A>C), TAC1561TTT (6191A>T 6192C>T), TGG1563TTT (6197G>T 6198G>T), CTA1564TTG (6199C>T 6201A>G), GTA1565GTT (6204A>T), ATG1566ATA (6207G>A), CCA1567TCT (6208C>T 6210A>T), GGT1569GGA (6216T>A), AAG1571ACA (6221A>C 6222G>A), GCA1573GCT (6228A>T), CCT1574CCC (6231T>C), GTG1576ACT (6235G>A 6236T>C 6237G>T), TTC1577TTT (6240C>T), CAA1578ATG (6241C>A 6242A>T 6243A>G), AGG1579GAC (6244A>G 6245G>A 6246G>C), AAA1580TTA (6247A>T 6248A>T), GAC1582AAC (6253G>A), AAC1583AGG (6257A>G 6258C>G), TGT1584GTA (6259T>G 6260G>T 6261T>A), AGG1586AAG (6266G>A), GGT1587CCG (6268G>C 6269G>C 6270T>G), ACA1588TAT (6271A>T 6272C>A 6273A>T), GAA1589TTG (6274G>T 6275A>T 6276A>G), GAT1590GAC (6279T>C), GAT1590_TTC1591insATG (6279_6280insATG), TTC1591TTT (6282C>T), ATC1592GTA (6283A>G 6285C>A), GCT1593GTG (6287C>T 6288T>G), GTT1594GTC (6291T>C), TAC1595CTC (6292T>C 6293A>T), ATT1596ATA (6297T>A), GAT1598GAC (6303T>C), ATT1599ATA (6306T>A), CTG1600TTG (6307C>T), GTA1601GTT (6312A>T), TTC1602TAC (6314T>A), GAA1604CGG (6319G>C 6320A>G 6321A>G), CTT1606GCC (6325C>G 6326T>C 6327T>C), CAA1607AAG (6328C>A 6330A>G), GAG1608GAA (6333G>A), CAC1609CAT (6336C>T), GAG1611GAT (6342G>T), CAT1612CAC (6345T>C), CTC1613TTG (6346C>T 6348C>G), AAA1614AGG (6350A>G 6351A>G), ATA1615ATT (6354A>T), ATG1616GTG (6355A>G), CTG1617TTG (6358C>T), GAA1618CAA (6361G>C), ATA1619ACT (6365T>C 6366A>T), AGG1621AAA (6371G>A 6372G>A), AAG1622CTT (6373A>C 6374A>T 6375G>T), AAT1623TGA (6376A>T 6377A>G 6378T>A), CTA1625ACC (6382C>A 6383T>C 6384A>C), GTC1626GAC (6386T>A), GTC1626_CTC1627insAATTGTTCG (6387_6388insAATTGTTCG), CTC1627CTA (6390C>A), AGC1628AGT (6393C>T), CCA1629TCA (6394C>T), ACA1630GTA (6397A>G 6398C>T), AAA1631AAT (6402A>T), ATG1632GTG (6403A>G), AAA1633AAT (6408A>T), ATA1634TTT (6409A>T 6411A>T), GCT1635GGT (6413C>G) |

##### Proteins

| | Begin | End | Coverage | Score | Concordance | Matches | Identities | I/D/M/F* | Stop Codons |
|---|---|---|---|---|---|---|---|---|---|
| hypothetical protein (YP_010799265.1) | 1532 | 1638 | 4.7% | 220 | 28.5% | 107 (96.4%) | 50 (45.0%) | 4/0/1/1 | 1 |

| | |
|---|---|
| Protein mutations: | F1533I (6106T>A), K1536R (6115A>C 6116A>G), G1538D (6122G>A), F1539Y (6125T>A), V1542L (6133G>T), A1543R (6136G>A 6137C>G 6138A>G), M1544V (6139A>G 6141G>A), E1545K (6142G>A), S1547C (6149C>G 6150A>T), W1551K (6160T>A 6161G>A), I1555R (6173T>G), P1557Q (6179C>A), D1558Y (6181G>T), G1559S (6184G>A 6186A>T), L1560H (6188T>A 6189A>C), Y1561F (6191A>T 6192C>T), W1563F (6197G>T 6198G>T), M1566I (6207G>A), P1567S (6208C>T 6210A>T), K1571T (6221A>C 6222G>A), V1576T (6235G>A 6236T>C 6237G>T), Q1578M (6241C>A 6242A>T 6243A>G), R1579D (6244A>G 6245G>A 6246G>C), K1580L (6247A>T 6248A>T), D1582N (6253G>A), N1583R (6257A>G 6258C>G), C1584V (6259T>G 6260G>T 6261T>A), R1586K (6266G>A), G1587P (6268G>C 6269G>C 6270T>G), T1588Y (6271A>T 6272C>A 6273A>T), E1589L (6274G>T 6275A>T 6276A>G), D1590_F1591insM (6279_6280insATG), I1592V (6283A>G 6285C>A), A1593V (6287C>T 6288T>G), Y1595L (6292T>C 6293A>T), F1602Y (6314T>A), E1604R (6319G>C 6320A>G 6321A>G), L1606A (6325C>G 6326T>C 6327T>C), Q1607K (6328C>A 6330A>G), E1611D (6342G>T), K1614R (6350A>G 6351A>G), M1616V (6355A>G), E1618Q (6361G>C), I1619T (6365T>C 6366A>T), R1621K (6371G>A 6372G>A), K1622L (6373A>C 6374A>T 6375G>T), N1623* (6376A>T 6377A>G 6378T>A), L1625T (6382C>A 6383T>C 6384A>C), V1626D (6386T>A), V1626_L1627insNCS (6387_6388insAATTGTTCG), P1629S (6394C>T), T1630V (6397A>G 6398C>T), K1631N (6402A>T), M1632V (6403A>G), K1633N (6408A>T), I1634F (6409A>T 6411A>T), A1635G (6413C>G) |
| Codon mutations: | AGC1531..T (6102C>T), AAA1532AAG (6105A>G), TTT1533ATT (6106T>A), TTG1535CTT (6112T>C 6114G>T), AAA1536CGA (6115A>C 6116A>G), TCT1537TCC (6120T>C), GGT1538GAT (6122G>A), TTT1539TAT (6125T>A), CAG1541CAA (6132G>A), GTG1542TTG (6133G>T), GCA1543AGG (6136G>A 6137C>G 6138A>G), ATG1544GTA (6139A>G 6141G>A), GAA1545AAA (6142G>A), GAA1546GAG (6147A>G), TCA1547TGT (6149C>G 6150A>T), AGC1548--G (6151_6152delAG 6153C>G), ATT1549ATC (6156T>C), CCA1550CCG (6159A>G), TGG1551AAG (6160T>A 6161G>A), ACG1552ACT (6165G>T), TTC1554TTT (6171C>T), ATA1555AGA (6173T>G), CCA1557CAA (6179C>A), GAT1558TAT (6181G>T), GGA1559AGT (6184G>A 6186A>T), CTA1560CAC (6188T>A 6189A>C), TAC1561TTT (6191A>T 6192C>T), TGG1563TTT (6197G>T 6198G>T), CTA1564TTG (6199C>T 6201A>G), GTA1565GTT (6204A>T), ATG1566ATA (6207G>A), CCA1567TCT (6208C>T 6210A>T), GGT1569GGA (6216T>A), AAG1571ACA (6221A>C 6222G>A), GCA1573GCT (6228A>T), CCT1574CCC (6231T>C), GTG1576ACT (6235G>A 6236T>C 6237G>T), TTC1577TTT (6240C>T), CAA1578ATG (6241C>A 6242A>T 6243A>G), AGG1579GAC (6244A>G 6245G>A 6246G>C), AAA1580TTA (6247A>T 6248A>T), GAC1582AAC (6253G>A), AAC1583AGG (6257A>G 6258C>G), TGT1584GTA (6259T>G 6260G>T 6261T>A), AGG1586AAG (6266G>A), GGT1587CCG (6268G>C 6269G>C 6270T>G), ACA1588TAT (6271A>T 6272C>A 6273A>T), GAA1589TTG (6274G>T 6275A>T 6276A>G), GAT1590GAC (6279T>C), GAT1590_TTC1591insATG (6279_6280insATG), TTC1591TTT (6282C>T), ATC1592GTA (6283A>G 6285C>A), GCT1593GTG (6287C>T 6288T>G), GTT1594GTC (6291T>C), TAC1595CTC (6292T>C 6293A>T), ATT1596ATA (6297T>A), GAT1598GAC (6303T>C), ATT1599ATA (6306T>A), CTG1600TTG (6307C>T), GTA1601GTT (6312A>T), TTC1602TAC (6314T>A), GAA1604CGG (6319G>C 6320A>G 6321A>G), CTT1606GCC (6325C>G 6326T>C 6327T>C), CAA1607AAG (6328C>A 6330A>G), GAG1608GAA (6333G>A), CAC1609CAT (6336C>T), GAG1611GAT (6342G>T), CAT1612CAC (6345T>C), CTC1613TTG (6346C>T 6348C>G), AAA1614AGG (6350A>G 6351A>G), ATA1615ATT (6354A>T), ATG1616GTG (6355A>G), CTG1617TTG (6358C>T), GAA1618CAA (6361G>C), ATA1619ACT (6365T>C 6366A>T), AGG1621AAA (6371G>A 6372G>A), AAG1622CTT (6373A>C 6374A>T 6375G>T), AAT1623TGA (6376A>T 6377A>G 6378T>A), CTA1625ACC (6382C>A 6383T>C 6384A>C), GTC1626GAC (6386T>A), GTC1626_CTC1627insAATTGTTCG (6387_6388insAATTGTTCG), CTC1627CTA (6390C>A), AGC1628AGT (6393C>T), CCA1629TCA (6394C>T), ACA1630GTA (6397A>G 6398C>T), AAA1631AAT (6402A>T), ATG1632GTG (6403A>G), AAA1633AAT (6408A>T), ATA1634TTT (6409A>T 6411A>T), GCT1635GGT (6413C>G) |

*: Inserts / Deletes / Misaligned / Frameshifts

#### Analysis details

This analysis was performed with panviral2.64

### NGS Details (UN70): Neptunevirus srim50

#### Assembly

| | |
|---|---|
| **Coverage Length** | 564 (1 contig(s)) |
| **Depth Of Coverage** | 10.5 |
| **Number Of Reads** | 69 |
| **Reads Per Million** | 1.31 rpm (after QC) |
| **Ambiguities** | 0 |
| **Assembly Method** | de novo + reference guided assembly |
| **Consensus Caller** | Bcf Tools |

#### Coverage Map

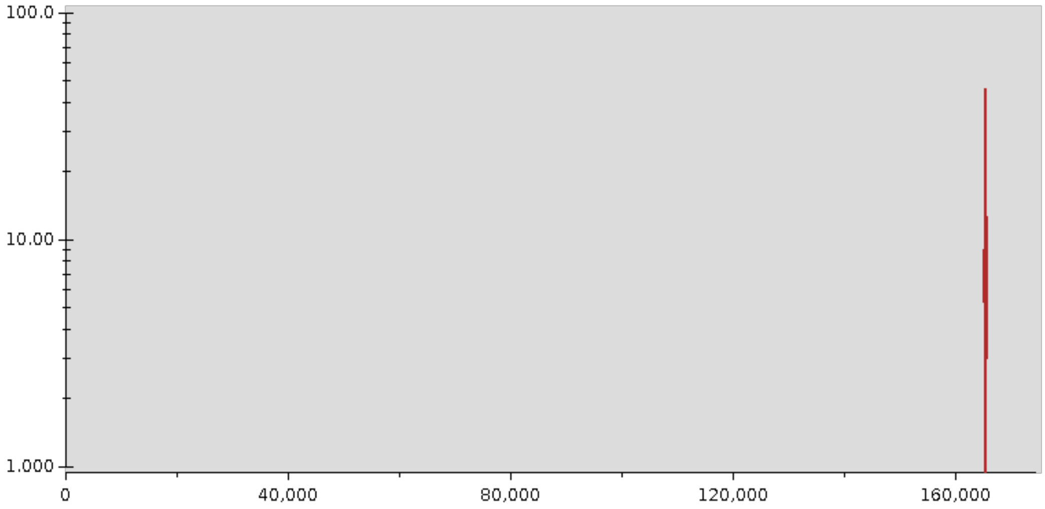

#### Assignment

| | |
|---|---|
| **Type** | Neptunevirus srim50 (Taxonomy ID: 2734122) |
| **Reference Genome** | NC_031242.1 |
| **NT Identity (%)** | 71.9858 |
| **AA Identity (%)** | 87.766 |
| **Number Of Stop Codons** | 0 |
| **Number Of CDS** | 227 |

#### Alignment

| | |
|---|---|
| **Alignment Score** | 496.0 (NT) + 1298.0 (AA) = 1794.0 |
| **Concordance (%)** | 71.1622 |
| **Alignment Method** | Local, heuristic, nucleotide (BLASTN) |

#### Genome Region

Sequence starts at position 165152 and ends at position 165715 relative to NC_031242.1 reference sequence.

#### Alignment Detailed Statistics

| | Begin | End | Coverage | Score | Concordance | Matches | Identities | I/D/M/F* | Stop Codons |
|---|---|---|---|---|---|---|---|---|---|
| **NT** | **165152** | **165715** | **0.3%** | **496** | **44.0%** | **564 (100%)** | **406 (72.0%)** | **0/0** | |
| Mutations: | 165155C>T, 165159G>A, 165161C>T, 165162C>A, 165164T>G, 165167C>T, 165174C>T, 165176T>A, 165177A>C, 165178A>G, 165179A>G, 165182A>G, 165200T>A, 165209T>C, 165212A>T, 165215T>A, 165216C>T, 165218T>G, 165221T>C, 165227C>T, 165228A>T, 165229C>G, 165233T>C, 165237C>T, 165239T>C, 165245T>A, 165255C>T, 165257T>C, 165260T>A, 165263G>T, 165266T>A, 165269T>C, 165272C>T, 165275C>A, 165278G>T, 165279A>T, 165280G>C, 165281T>A, 165290T>C, 165293C>T, 165297C>T, 165299T>G, 165302G>C, 165308C>T, 165312C>T, 165314T>G, 165317G>A, 165320T>C, 165321G>T, 165322C>G, 165323A>C, 165326C>T, 165329T>C, 165330C>T, 165332T>A, 165338G>C, 165341A>G, 165353A>T, 165356A>T, 165357G>A, 165359C>T, 165360G>A, 165361C>G, 165363A>T, 165365G>A, 165367G>C, 165368T>A, 165374T>G, 165375C>T, 165377T>G, 165378C>T, 165381C>T, 165383T>A, 165398G>A, 165399T>G, 165401T>A, 165407G>A, 165410C>T, 165413C>T, 165415T>C, 165416C>T, 165419C>T, 165425C>T, 165429C>T, 165431T>G, 165437A>T, 165440C>G, 165452G>T, 165455G>T, 165461C>T, 165467C>T, 165472C>G, 165473A>C, 165476T>A, 165489C>T, 165491T>A, 165494C>T, 165497G>A, 165500T>C, 165503A>G, 165506G>T, 165509A>T, 165513C>T, 165514T>C, 165516A>G, 165518C>T, 165519G>C, 165520G>A, 165521T>A, 165522A>T, 165524T>G, 165525C>A, 165527T>A, 165533C>T, 165539T>A, 165542T>C, 165545T>A, 165557T>A, 165566A>T, 165569C>T, 165573A>T, 165574G>C, 165578T>A, 165584T>G, 165587G>T, 165593T>A, 165596G>A, 165599A>T, 165602A>G, 165605G>T, 165606T>G, 165607C>G, 165608C>T, 165617C>T, 165623G>T, 165624T>A, 165625C>G, 165626G>T, 165629C>T, 165632G>T, 165635C>A, 165641A>T, 165644C>A, 165647C>T, 165654C>A, 165656G>C, 165659C>A, 165662C>T, 165663C>T, 165665C>T, 165674C>T, 165680C>T, 165686G>C, 165687C>T, 165692T>C, 165701C>T, 165710C>A, 165713A>T | | | | | | | | |

*: Inserts / Deletes / Misaligned / Frameshifts

#### Analysis details

This analysis was performed with panviral2.64

### NGS Details (UN70): Badnavirus alphacolocalasiae

#### Assembly

| | |
|---|---|
| **Coverage Length** | 232 (1 contig(s)) |
| **Depth Of Coverage** | 21.2 |
| **Number Of Reads** | 42 |
| **Reads Per Million** | 0.80 rpm (after QC) |
| **Ambiguities** | 0 |
| **Assembly Method** | de novo + reference guided assembly |
| **Consensus Caller** | Bcf Tools |

#### Coverage Map

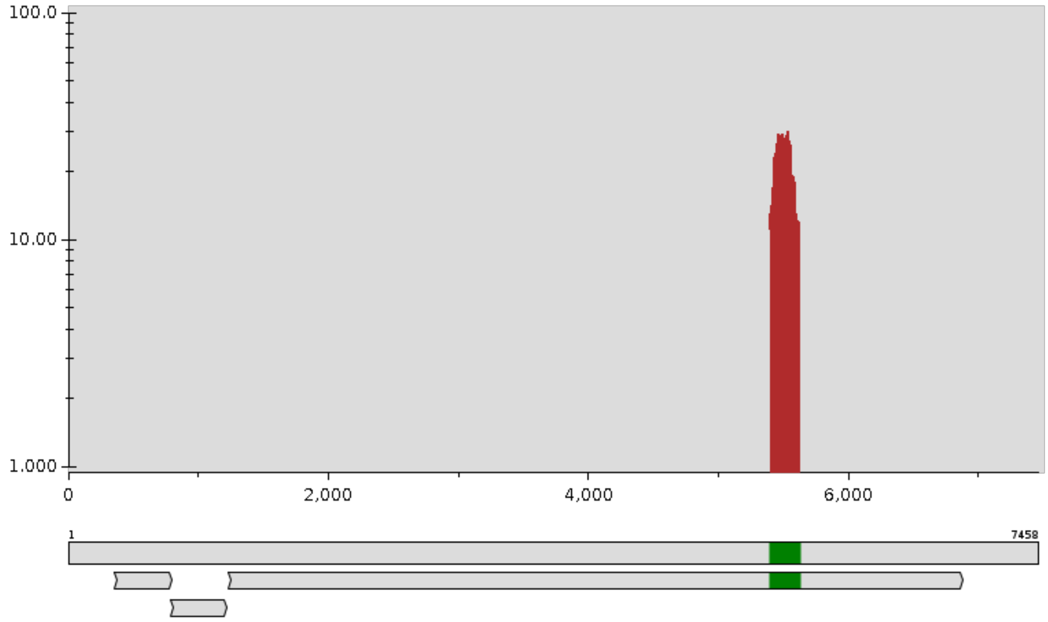

#### Assignment

| | |
|---|---|
| **Type** | Badnavirus alphacolocalasiae (Taxonomy ID: 3048356) |
| **Reference Genome** | NC_004450.1 |
| **NT Identity (%)** | 53.617 |
| **AA Identity (%)** | 44.8718 |
| **Number Of Stop Codons** | 0 |
| **Number Of CDS** | 3 |

#### Alignment

| | |
|---|---|
| **Alignment Score** | 28.0 (NT) + 255.0 (AA) = 283.0 |
| **Concordance (%)** | 27.5024 |

| Alignment Method | Global, seeded, nucleotide + amino acids (AGA) |
|---|---|

#### Genome Region

Sequence starts at position 5399 and ends at position 5630 relative to NC_004450.1 reference sequence.

#### Alignment Detailed Statistics

| | Begin | End | Coverage | Score | Concordance | Matches | Identities | I/D/M/F* | Stop Codons |
|---|---|---|---|---|---|---|---|---|---|
| **NT** | **5399** | **5630** | **3.1%** | **28** | **6.0%** | **232 (98.7%)** | **126 (53.6%)** | **3/0** | |
| Mutations: | 5399A>T, 5405T>C, 5407T>A, 5409C>A, 5414A>G, 5415G>A, 5417T>C, 5418G>T, 5419C>G, 5420T>C, 5424G>T, 5425A>C, 5426T>A, 5429C>A, 5430G>A, 5432T>G, 5433T>G, 5434C>A, 5437T>A, 5438G>A, 5442T>C, 5443G>T, 5444G>C, 5447A>T, 5450T>A, 5453C>T, 5455T>G, 5456A>C, 5457G>A, 5458T>C, 5459T>A, 5462A>T, 5463C>A, 5469T>A, 5471A>T, 5475G>T, 5476A>G, 5477A>C, 5479G>A, 5480G>C, 5481C>A, 5482T>A, 5485C>T, 5486A>G, 5498C>T, 5499C>T, 5501C>A, 5504A>G, 5511C>G, 5512C>G, 5513A>T, 5517G>A, 5518T>C, 5521T>A, 5528A>G, 5529A>G, 5530A>C, 5531A>T, 5535G>C, 5537T>A, 5538G>A, 5539C>A, 5540A>T, 5541G>A, 5546C>T, 5547A>G, 5549A>T, 5551G>A, 5552G>C, 5554G>T, 5555T>A, 5555_5556insCTC, 5556G>C, 5558A>C, 5562T>A, 5563T>A, 5564C>T, 5565C>G, 5567C>T, 5569C>A, 5571G>T, 5572T>G, 5573C>T, 5577A>G, 5585T>C, 5586A>T, 5588T>G, 5589C>G, 5591G>A, 5594A>G, 5595T>A, 5596T>A, 5597T>G, 5603C>G, 5606T>G, 5607G>A, 5608A>G, 5610G>A, 5611A>G, 5612A>T, 5615T>C, 5619G>T, 5620C>T, 5621A>G, 5625C>G, 5628C>T, 5630G>A | | | | | | | | |
| **CDS** | | | | | | | | | |
| TaBVgp3 | 1392 | 1468 | 4.1% | 255 | 44.8% | 77 (98.7%) | 35 (44.9%) | 1/0/0/0 | 0 |
| Protein mutations: | F1394Y (5407T>A), H1395N (5409C>A), V1397I (5415G>A 5417T>C), A1398C (5418G>T 5419C>G 5420T>C), D1400S (5424G>T 5425A>C 5426T>A), D1402K (5430G>A 5432T>G), S1403D (5433T>G 5434C>A), V1404E (5437T>A 5438G>A), W1406L (5442T>C 5443G>T 5444G>C), L1410R (5455T>G 5456A>C), V1411T (5457G>A 5458T>C 5459T>A), Q1413K (5463C>A), L1415I (5469T>A 5471A>T), E1417C (5475G>T 5476A>G 5477A>C), W1418Y (5479G>A 5480G>C), L1419K (5481C>A 5482T>A), A1420V (5485C>T 5486A>G), P1429G (5511C>G 5512C>G 5513A>T), V1431T (5517G>A 5518T>C), F1432Y (5521T>A), K1435A (5529A>G 5530A>C 5531A>T), D1437Q (5535G>C 5537T>A), A1438N (5538G>A 5539C>A 5540A>T), V1439I (5541G>A), K1441D (5547A>G 5549A>T), G1442D (5551G>A 5552G>C), C1443L (5554G>T 5555T>A), C1443_E1444insL (5555_5556insCTC), E1444H (5556G>C 5558A>C), F1446N (5562T>A 5563T>A 5564C>T), L1447V (5565C>G 5567C>T), A1448E (5569C>A), V1449C (5571G>T 5572T>G 5573C>T), I1451V (5577A>G), I1454L (5586A>T 5588T>G), L1455V (5589C>G 5591G>A), F1457K (5595T>A 5596T>A 5597T>G), N1459K (5603C>G), N1460K (5606T>G), E1461R (5607G>A 5608A>G), E1462S (5610G>A 5611A>G 5612A>T), A1465L (5619G>T 5620C>T 5621A>G), H1467D (5625C>G) | | | | | | | | |
| Codon mutations: | AAA1391..T (5399A>T), GGT1393GGC (5405T>C), TTT1394TAT (5407T>A), CAC1395AAC (5409C>A), CAA1396CAG (5414A>G), GTT1397ATC (5415G>A 5417T>C), GCT1398TGC (5418G>T 5419C>G 5420T>C), GAT1400TCA (5424G>T 5425A>C 5426T>A), CCC1401CCA (5429C>A), GAT1402AAG (5430G>A 5432T>G), TCT1403GAT (5433T>G 5434C>A), GTG1404GAA (5437T>A 5438G>A), TGG1406CTC (5442T>C 5443G>T 5444G>C), ACA1407ACT (5447A>T), GCT1408GCA (5450T>A), TTC1409TTT (5453C>T), CTA1410CGC (5455T>G 5456A>C), GTT1411ACA (5457G>A 5458T>C 5459T>A), CCA1412CCT (5462A>T), CAA1413AAA (5463C>A), TTA1415ATT (5469T>A 5471A>T), GAA1417TGC (5475G>T 5476A>G 5477A>C), TGG1418TAC (5479G>A 5480G>C), CTG1419AAG (5481C>A 5482T>A), GCA1420GTG (5485C>T 5486A>G), GGC1424GGT (5498C>T), CTC1425TTA (5499C>T 5501C>A), AAA1426AAG (5504A>G), CCA1429GGT (5511C>G 5512C>G 5513A>T), GTA1431ACA (5517G>A 5518T>C), TTT1432TAT (5521T>A), AGA1434AGG (5528A>G), AAA1435GCT (5529A>G 5530A>C 5531A>T), GAT1437CAA (5535G>C 5537T>A), GCA1438AAT (5538G>A 5539C>A 5540A>T), GTA1439ATA (5541G>A), TTC1440TTT (5546C>T), AAA1441GAT (5547A>G 5549A>T), GGG1442GAC (5551G>A 5552G>C), TGT1443TTA (5554G>T 5555T>A), TGT1443_GAA1444insCTC (5555_5556insCTC), GAA1444CAC (5556G>C 5558A>C), TTC1446AAT (5562T>A 5563T>A 5564C>T), CTC1447GTT (5565C>G 5567C>T), GCA1448GAA (5569C>A), GTC1449TGT (5571G>T 5572T>G 5573C>T), ATT1451GTT (5577A>G), GAT1453GAC (5585T>C), ATT1454TTG (5586A>T 5588T>G), CTG1455GTA (5589C>G 5591G>A), GTA1456GTG (5594A>G), TTT1457AAG (5595T>A 5596T>A 5597T>G), AAC1459AAG (5603C>G), AAT1460AAG (5606T>G), GAG1461AGG (5607G>A 5608A>G), GAA1462AGT (5610G>A 5611A>G 5612A>T), GAT1463GAC (5615T>C), GCA1465TTG (5619G>T 5620C>T 5621A>G), CAC1467GAC (5625C>G), CTG1468TTA (5628C>T 5630G>A) | | | | | | | | |
| **Proteins** | | | | | | | | | |
| polyprotein (NP_758808.1) | 1392 | 1468 | 4.1% | 255 | 44.8% | 77 (98.7%) | 35 (44.9%) | 1/0/0/0 | 0 |
| Protein mutations: | F1394Y (5407T>A), H1395N (5409C>A), V1397I (5415G>A 5417T>C), A1398C (5418G>T 5419C>G 5420T>C), D1400S (5424G>T 5425A>C 5426T>A), D1402K (5430G>A 5432T>G), S1403D (5433T>G 5434C>A), V1404E (5437T>A 5438G>A), W1406L (5442T>C 5443G>T 5444G>C), L1410R (5455T>G 5456A>C), V1411T (5457G>A 5458T>C 5459T>A), Q1413K (5463C>A), L1415I (5469T>A 5471A>T), E1417C (5475G>T 5476A>G 5477A>C), W1418Y (5479G>A 5480G>C), L1419K (5481C>A 5482T>A), A1420V (5485C>T 5486A>G), P1429G (5511C>G 5512C>G 5513A>T), V1431T (5517G>A 5518T>C), F1432Y (5521T>A), K1435A (5529A>G 5530A>C 5531A>T), D1437Q (5535G>C 5537T>A), A1438N (5538G>A 5539C>A 5540A>T), V1439I (5541G>A), K1441D (5547A>G 5549A>T), G1442D (5551G>A 5552G>C), C1443L (5554G>T 5555T>A), C1443_E1444insL (5555_5556insCTC), E1444H (5556G>C 5558A>C), F1446N (5562T>A 5563T>A 5564C>T), L1447V (5565C>G 5567C>T), A1448E (5569C>A), V1449C (5571G>T 5572T>G 5573C>T), I1451V (5577A>G), I1454L (5586A>T 5588T>G), L1455V (5589C>G 5591G>A), F1457K (5595T>A 5596T>A 5597T>G), N1459K (5603C>G), N1460K (5606T>G), E1461R (5607G>A 5608A>G), E1462S (5610G>A 5611A>G 5612A>T), A1465L (5619G>T 5620C>T 5621A>G), H1467D (5625C>G) | | | | | | | | |
| Codon mutations: | AAA1391..T (5399A>T), GGT1393GGC (5405T>C), TTT1394TAT (5407T>A), CAC1395AAC (5409C>A), CAA1396CAG (5414A>G), GTT1397ATC (5415G>A 5417T>C), GCT1398TGC (5418G>T 5419C>G 5420T>C), GAT1400TCA (5424G>T 5425A>C 5426T>A), CCC1401CCA (5429C>A), GAT1402AAG (5430G>A 5432T>G), TCT1403GAT (5433T>G 5434C>A), GTG1404GAA (5437T>A 5438G>A), TGG1406CTC (5442T>C 5443G>T 5444G>C), ACA1407ACT (5447A>T), GCT1408GCA (5450T>A), TTC1409TTT (5453C>T), CTA1410CGC (5455T>G 5456A>C), GTT1411ACA (5457G>A 5458T>C 5459T>A), CCA1412CCT (5462A>T), CAA1413AAA (5463C>A), TTA1415ATT (5469T>A 5471A>T), GAA1417TGC (5475G>T 5476A>G 5477A>C), TGG1418TAC (5479G>A 5480G>C), CTG1419AAG (5481C>A 5482T>A), GCA1420GTG (5485C>T 5486A>G), GGC1424GGT (5498C>T), CTC1425TTA (5499C>T 5501C>A), AAA1426AAG (5504A>G), CCA1429GGT (5511C>G 5512C>G 5513A>T), GTA1431ACA (5517G>A 5518T>C), TTT1432TAT (5521T>A), AGA1434AGG (5528A>G), AAA1435GCT (5529A>G 5530A>C 5531A>T), GAT1437CAA (5535G>C 5537T>A), GCA1438AAT (5538G>A 5539C>A 5540A>T), GTA1439ATA (5541G>A), TTC1440TTT (5546C>T), AAA1441GAT (5547A>G 5549A>T), GGG1442GAC (5551G>A 5552G>C), TGT1443TTA (5554G>T 5555T>A), TGT1443_GAA1444insCTC (5555_5556insCTC), GAA1444CAC (5556G>C 5558A>C), TTC1446AAT (5562T>A 5563T>A 5564C>T), CTC1447GTT (5565C>G 5567C>T), GCA1448GAA (5569C>A), GTC1449TGT (5571G>T 5572T>G 5573C>T), ATT1451GTT (5577A>G), GAT1453GAC (5585T>C), ATT1454TTG (5586A>T 5588T>G), CTG1455GTA (5589C>G 5591G>A), GTA1456GTG (5594A>G), TTT1457AAG (5595T>A 5596T>A 5597T>G), AAC1459AAG (5603C>G), AAT1460AAG (5606T>G), GAG1461AGG (5607G>A 5608A>G), GAA1462AGT (5610G>A 5611A>G 5612A>T), GAT1463GAC (5615T>C), GCA1465TTG (5619G>T 5620C>T 5621A>G), CAC1467GAC (5625C>G), CTG1468TTA (5628C>T 5630G>A) | | | | | | | | |

*: Inserts / Deletes / Misaligned / Frameshifts

#### Analysis details

This analysis was performed with panviral2.64

### NGS Details (UN70): Caulimovirus venafragariae

#### Assembly

| | |
|---|---|
| **Coverage Length** | 654 (1 contig(s)) |
| **Depth Of Coverage** | 7.0 |
| **Number Of Reads** | 36 |
| **Reads Per Million** | 0.68 rpm (after QC) |
| **Ambiguities** | 0 |
| **Assembly Method** | de novo + reference guided assembly |
| **Consensus Caller** | Bcf Tools |

#### Coverage Map

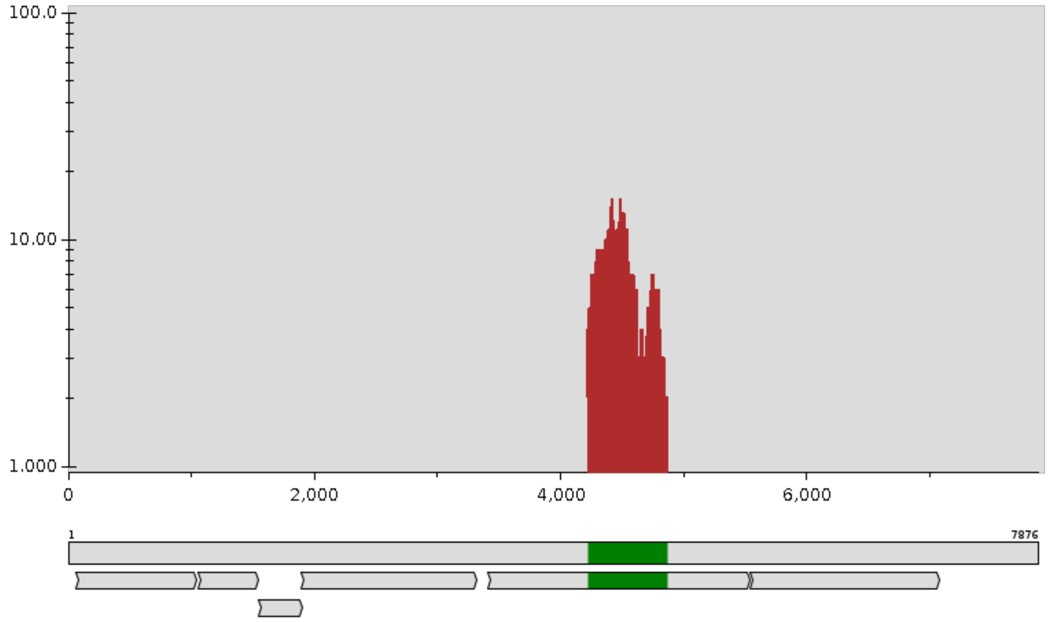

#### Assignment

| | |
|---|---|
| **Type** | Caulimovirus venafragariae (Taxonomy ID: 3048344) |
| **Reference Genome** | NC_001725.1 |
| **NT Identity (%)** | 59.9379 |
| **AA Identity (%)** | 59.0698 |
| **Number Of Stop Codons** | 0 |
| **Number Of CDS** | 6 |

#### Alignment

| | |
|---|---|
| **Alignment Score** | 219.0 (NT) + 779.0 (AA) = 998.0 |
| **Concordance (%)** | 38.4732 |

| **Alignment Method** | Global, seeded, nucleotide + amino acids (AGA) |
|---|---|

#### Genome Region

Sequence starts at position 4219 and ends at position 4872 relative to NC_001725.1 reference sequence.

#### Alignment Detailed Statistics

| | Begin | End | Coverage | Score | Concordance | Matches | Identities | I/D/M/F* | Stop Codons |
|---|---|---|---|---|---|---|---|---|---|
| **NT** | **4219** | **4872** | **8.3%** | **219** | **17.5%** | **644 (98.5%)** | **386 (59.0%)** | **0/10** | |
| Mutations: | 4219T>C, 4222T>A, 4223T>A, 4228A>C, 4230T>C, 4232A>G, 4235A>T, 4236A>C, 4242T>A, 4244A>G, 4245G>A, 4246T>G, 4247G>T, 4254A>T, 4259A>T, 4260G>A, 4263T>C, 4264T>A, 4265C>A, 4266A>T, 4267C>G, 4268C>A, 4269T>A, 4275C>T, 4276C>A, 4277G>A, 4279G>A, 4280A>C, 4281A>T, 4284G>A, 4292C>T, 4299C>T, 4300G>C, 4306C>T, 4309C>T, 4311C>A, 4314G>T, 4315C>A, 4317T>G, 4318G>A, 4319G>A, 4320T>A, 4321A>T, 4323C>A, 4326C>A, 4331C>A, 4335C>T, 4338G>A, 4348T>A, 4349C>G, 4350C>T, 4351T>A, 4352C>G, 4353A>C, 4359C>A, 4362C>T, 4371G>A, 4374T>C, 4379C>G, 4383G>A, 4384A>C, 4385T>A, 4386C>A, 4390delC, 4391G>C, 4392A>T, 4395C>A, 4402A>C, 4413T>A, 4423A>G, 4426T>C, 4428A>T, 4432G>A, 4434C>A, 4435C>A, 4438A>T, 4439C>G, 4440A>T, 4441A>G, 4442A>T, 4443G>C, 4444G>T, 4445G>T, 4446A>C, 4449T>C, 4452C>A, 4455C>T, 4456C>T, 4458A>C, 4459C>A, 4461T>C, 4464T>A, 4467C>T, 4470G>A, 4471G>A, 4473G>T, 4474C>A, 4476A>C, 4479G>T, 4480C>A, 4482T>A, 4483C>A, 4485A>T, 4486A>C, 4487G>T, 4488A>T, 4489A>G, 4490T>C, 4492G>A, 4493G>A, 4497T>A, 4500G>A, 4505T>A, 4506T>C, 4508A>T, 4509C>T, 4512T>A, 4513T>A, 4514C>A, 4515C>A, 4521C>T, 4524T>C, 4528T>A, 4529C>G, 4542G>A, 4543G>A, 4546C>A, 4547G>A, 4548C>A, 4551T>A, 4553C>A, 4554T>A, 4555C>G, 4556C>A, 4558G>A, 4560A>T, 4561A>T, 4563A>T, 4566T>A, 4567C>G, 4568A>C, 4569G>T, 4570C>T, 4575C>A, 4578T>A, 4584C>T, 4585T>A, 4586G>C, 4587T>A, 4590C>A, 4591C>A, 4593A>T, 4614C>G, 4633C>A, 4635A>T, 4642G>C, 4643C>A, 4644T>A, 4647T>C, 4650C>T, 4656G>A, 4657C>A, 4659C>A, 4665C>T, 4670G>T, 4671T>A, 4672C>T, 4674C>T, 4675A>T, 4676G>C, 4677C>T, 4678A>G, 4680C>A, 4682T>A, 4684T>A, 4687_4692delCCACAG, 4696T>C, 4697G>T, 4698T>A, 4699G>A, 4700C>T, 4701T>C, 4704A>C, 4708G>A, 4710T>A, 4716C>T, 4719C>A, 4720A>C, 4722C>A, 4723G>A, 4725C>A, 4727T>G, 4728C>T, 4729A>T, 4730G>C, 4731C>T, 4734A>T, 4735A>T, 4737T>A, 4738G>T, 4740A>T, 4741G>C, 4746A>G, 4753G>A, 4754G>C, 4758T>C, 4759G>C, 4761C>T, 4765A>C, 4766T>A, 4767T>A, 4768G>T, 4771C>T, 4772T>C, 4773A>T, 4774A>C, 4776C>A, 4777A>T, 4778G>T, 4782C>T, 4783A>G, 4784A>T, 4785A>T, 4787C>A, 4788C>A, 4789C>A, 4790T>A, 4791A>T, 4797A>T, 4801C>T, 4803T>A, 4807A>C, 4809A>G, 4812G>A, 4815A>G, 4817C>A, 4818C>G, 4819C>G, 4821G>T, 4822T>A, 4824G>A, 4825T>A, 4827C>T, 4832C>A, 4833A>T, 4834A>G, 4835C>A, 4836T>A, 4839C>T, 4840A>G, 4842T>A, 4851T>A, 4852C>A, 4854A>G, 4855G>A, 4860A>T, 4861G>T, 4862A>C, 4863A>C, 4865G>A, 4867_4869delGGA | | | | | | | | |

##### CDS

| | | | | | | | | | |
|---|---|---|---|---|---|---|---|---|---|
| ORF_V | 272 | 489 | 30.7% | 779 | 52.1% | 215 (98.6%) | 127 (58.3%) | 0/3/1/1 | 0 |
| Protein mutations: | L273K (4222T>A 4223T>A), T275P (4228A>C 4230T>C), N276S (4232A>G), K277I (4235A>T 4236A>C), K280R (4244A>G 4245G>A), C281V (4246T>G 4247G>T), Q285L (4259A>T 4260G>A), S287N (4264T>A 4265C>A 4266A>T), P288E (4267C>G 4268C>A 4269T>A), R291K (4276C>A 4277G>A), E292T (4279G>A 4280A>C 4281A>T), T296I (4292C>T), E299Q (4300G>C), K303N (4314G>T), L304M (4315C>A 4317T>G), G305K (4318G>A 4319G>A 4320T>A), I306L (4321A>T 4323C>A), P309H (4331C>A), A325G (4379C>G), I327Q (4384A>C 4385T>A 4386C>A), K340E (4423A>G), D343K (4432G>A 4434C>A), H344N (4435C>A), T345C (4438A>T 4439C>G 4440A>T), K346V (4441A>G 4442A>T 4443G>C), G347F (4444G>T 4445G>T 4446A>C), L351F (4456C>T 4458A>C), L352I (4459C>A 4461T>C), E356N (4471G>A 4473G>T), Q357N (4474C>A 4476A>C), L359I (4480C>A 4482T>A), Q360N (4483C>A 4485A>T), R361L (4486A>C 4487G>T 4488A>T), I362A (4489A>G 4490T>C), G363K (4492G>A 4493G>A), F367Y (4505T>A 4506T>C), Y368F (4508A>T 4509C>T), S370K (4513T>A 4514C>A 4515C>A), V380I (4543G>A), R381K (4546C>A 4547G>A 4548C>A), A383E (4553C>A 4554T>A), P384E (4555C>G 4556C>A), E385N (4558G>A 4560A>T), T386S (4561A>T 4563A>T), Q388A (4567C>G 4568A>C 4569G>T), C394T (4585T>A 4586G>C 4587T>A), Q396N (4591C>A 4593A>T), Q410N (4633C>A 4635A>T), A413Q (4642G>C 4643C>A 4644T>A), H418K (4657C>A 4659C>A), S422I (4670G>T 4671T>A), L423F (4672C>T 4674C>T), N425E (4678A>G 4680C>A), M426K (4682T>A), Y427N (4684T>A), P428_Q429del (4687_4692delCCACAG), C431L (4696T>C 4697G>T 4698T>A), A432I (4699G>A 4700C>T 4701T>C), V435I (4708G>A 4710T>A), I439L (4720A>C 4722C>A), V440I (4723G>A 4725C>A), F441C (4727T>G 4728C>T), K443N (4734A>T), T444S (4735A>T 4737T>A), E445Y (4738G>T 4740A>T), E446Q (4741G>C), G450T (4753G>A 4754G>C), V452L (4759G>C 4761C>T), I454Q (4765A>C 4766T>A 4767T>A), V455F (4768G>T), L456S (4771C>T 4772T>C 4773A>T), N457Q (4774A>C 4776C>A), R458L (4777A>T 4778G>T), K460V (4783A>G 4784A>T 4785A>T), A461E (4787C>A 4788C>A), L462N (4789C>A 4790T>A 4791A>T), K468Q (4807A>C 4809A>G), A471E (4817C>A 4818C>G), Q472D (4819C>G 4821G>T), L473I (4822T>A 4824G>A), C474S (4825T>A 4827C>T), T476N (4832C>A 4833A>T), T477E (4834A>G 4835C>A 4836T>A), N479E (4840A>G 4842T>A), L483M (4852C>A 4854A>G), V484I (4855G>A), E486S (4861G>T 4862A>C 4863A>C), R487K (4865G>A), G488del (4867_4869delGGA) | | | | | | | | |
| Codon mutations: | TTA272CTA (4219T>C), TTA273AAA (4222T>A 4223T>A), ACT275CCC (4228A>C 4230T>C), AAC276AGC (4232A>G), AAA277ATC (4235A>T 4236A>C), ATT279ATA (4242T>A), AAG280AGA (4244A>G 4245G>A), TGT281GTT (4246T>G 4247G>T), CCA283CCT (4254A>T), CAG285CTA (4259A>T 4260G>A), TAT286TAC (4263T>C), TCA287AAT (4264T>A 4265C>A 4266A>T), CCT288GAA (4267C>G 4268C>A 4269T>A), GAC290GAT (4275C>T), CGA291AAA (4276C>A 4277G>A), GAA292ACT (4279G>A 4280A>C 4281A>T), GAG293GAA (4284G>A), ACT296ATT (4292C>T), ATC298ATT (4299C>T), GAA299CAA (4300G>C), CTA301TTA (4306C>T), CTC302TTA (4309C>T 4311C>A), AAG303AAT (4314G>T), CTT304ATG (4315C>A 4317T>G), GGT305AAA (4318G>A 4319G>A 4320T>A), ATC306TTA (4321A>T 4323C>A), ATC307ATA (4326C>A), CCT309CAT (4331C>A), AGC310AGT (4335C>T), AAG311AAA (4338G>A), TCC315AGT (4348T>A 4349C>G 4350C>T), TCA316AGC (4351T>A 4352C>G 4353A>C), GCC318GCA (4359C>A), TTC319TTT (4362C>T), AGG322AGA (4371G>A), AAT323AAC (4374T>C), GCC325GGC (4379C>G), GAG326GAA (4383G>A), ATC327CAA (4384A>C 4385T>A 4386C>A), CGA329-CT (4390delC 4391G>C 4392A>T), GGC330GGA (4395C>A), AGA333CGA (4402A>C), ATT336ATA (4413T>A), AAG340GAG (4423A>G), TTA341CTT (4426T>C 4428A>T), GAC343AAA (4432G>A 4434C>A), CAT344AAT (4435C>A), ACA345TGT (4438A>T 4439C>G 4440A>T), AAG346GTC (4441A>G 4442A>T 4443G>C), GGA347TTC (4444G>T 4445G>T 4446A>C), GAT348GAC (4449T>C), GGC349GGA (4452C>A), TAC350TAT (4455C>T), CTA351TTC (4456C>T 4458A>C), CTT352ATC (4459C>A 4461T>C), CCT353CCA (4464T>A), AAC354AAT (4467C>T), AAG355AAA (4470G>A), GAG356AAT (4471G>A 4473G>T), CAA357AAC (4474C>A 4476A>C), CTG358CTT (4479G>T), CTT359ATA (4480C>A 4482T>A), CAA360AAT (4483C>A 4485A>T), AGA361CTT (4486A>C 4487G>T 4488A>T), ATC362GCC (4489A>G 4490T>C), GGA363AAA (4492G>A 4493G>A), GGT364GGA (4497T>A), AAG365AAA (4500G>A), TTT367TAC (4505T>A 4506T>C), TAC368TTT (4508A>T 4509C>T), TCT369TCA (4512T>A), TCC370AAA (4513T>A 4514C>A 4515C>A), GAC372GAT (4521C>T), TGT373TGC (4524T>C), TCT375AGT (4528T>A 4529C>G), CAG379CAA (4542G>A), GTA380ATA (4543G>A), CGC381AAA (4546C>A 4547G>A 4548C>A), CTT382CTA (4551T>A), GCT383GAA (4553C>A 4554T>A), CCA384GAA (4555C>G 4556C>A), GAA385AAT (4558G>A 4560A>T), ACA386TCT (4561A>T 4563A>T), ATT387ATA (4566T>A), CAG388GCT (4567C>G 4568A>C 4569G>T), CTA389TTA (4570C>T), ACC390ACA (4575C>A), GCT391GCA (4578T>A), AGC393AGT (4584C>T), TGT394ACA (4585T>A 4586G>C 4587T>A), CCC395CCA (4590C>A), CAA396AAT (4591C>A 4593A>T), GTC403GTG (4614C>G), CAA410AAT (4633C>A 4635A>T), GCT413CAA (4642G>C 4643C>A 4644T>A), ATT414ATC (4647T>C), TTC415TTT (4650C>T), AGG417AGA (4656G>A), CAC418AAA (4657C>A 4659C>A), GAC420GAT (4665C>T), AGT422ATA (4670G>T 4671T>A), CTC423TTT (4672C>T 4674C>T), AGC424TCT (4675A>T 4676G>C 4677C>T), AAC425GAA (4678A>G 4680C>A), ATG426AAG (4682T>A), TAT427AAT (4684T>A), CCA428_CAG429del (4687_4692delCCACAG), TGT431CTA (4696T>C 4697G>T 4698T>A), GCT432ATC (4699G>A 4700C>T 4701T>C), GTA433GTC (4704A>C), GTT435ATA (4708G>A 4710T>A), GAC437GAT (4716C>T), ATC438ATA (4719C>A), ATC439CTA (4720A>C 4722C>A), GTC440ATA (4723G>A 4725C>A), TTC441TGT (4727T>G 4728C>T), AGC442TCT (4729A>T 4730G>C 4731C>T), AAA443AAT (4734A>T), ACT444TCA (4735A>T 4737T>A), GAA445TAT (4738G>T 4740A>T), GAA446CAA (4741G>C), GAA447GAG (4746A>G), GGA450ACA (4753G>A 4754G>C), CAT451CAC (4758T>C), GTC452CTT (4759G>C 4761C>T), ATT454CAA (4765A>C 4766T>A 4767T>A), GTT455TTT (4768G>T), CTA456TCT (4771C>T 4772T>C 4773A>T), AAC457CAA (4774A>C 4776C>A), AGA458TTA (4777A>T 4778G>T), TGC459TGT (4782C>T), AAA460GTT (4783A>G 4784A>T 4785A>T), GCC461GAA (4787C>A 4788C>A), CTA462AAT (4789C>A 4790T>A 4791A>T), ATA464ATT (4797A>T), CTT466TTA (4801C>T 4803T>A), AAA468CAG (4807A>C 4809A>G), AAG469AAA (4812G>A), AAA470AAG (4815A>G), GCC471GAG (4817C>A 4818C>G), CAG472GAT (4819C>G 4821G>T), TTG473ATA (4822T>A 4824G>A), TGC474AGT (4825T>A 4827C>T), ACA476AAT (4832C>A 4833A>T), ACT477GAA (4834A>G 4835C>A 4836T>A), ATC478ATT (4839C>T), AAT479GAA (4840A>G 4842T>A), GGT482GGA (4851T>A), CTA483ATG (4852C>A 4854A>G), GTA484ATA (4855G>A), ATA485ATT (4860A>T), GAA486TCC (4861G>T 4862A>C 4863A>C), AGA487AAA (4865G>A), GGA488del (4867_4869delGGA) | | | | | | | | |

##### Proteins

| | | | | | | | | | |
|---|---|---|---|---|---|---|---|---|---|
| hypothetical protein (NP_043933.1) | 272 | 489 | 30.7% | 779 | 52.1% | 215 (98.6%) | 127 (58.3%) | 0/3/1/1 | 0 |
| Protein mutations: | L273K (4222T>A 4223T>A), T275P (4228A>C 4230T>C), N276S (4232A>G), K277I (4235A>T 4236A>C), K280R (4244A>G 4245G>A), C281V (4246T>G 4247G>T), Q285L (4259A>T 4260G>A), S287N (4264T>A 4265C>A 4266A>T), P288E (4267C>G 4268C>A 4269T>A), R291K (4276C>A 4277G>A), E292T (4279G>A 4280A>C 4281A>T), T296I (4292C>T), E299Q (4300G>C), K303N (4314G>T), L304M (4315C>A 4317T>G), G305K (4318G>A 4319G>A 4320T>A), I306L (4321A>T 4323C>A), P309H (4331C>A), A325G (4379C>G), I327Q (4384A>C 4385T>A 4386C>A), K340E (4423A>G), D343K (4432G>A 4434C>A), H344N (4435C>A), T345C (4438A>T 4439C>G 4440A>T), K346V (4441A>G 4442A>T 4443G>C), G347F (4444G>T 4445G>T 4446A>C), L351F (4456C>T 4458A>C), L352I (4459C>A 4461T>C), E356N (4471G>A 4473G>T), Q357N (4474C>A 4476A>C), L359I (4480C>A 4482T>A), Q360N (4483C>A 4485A>T), R361L (4486A>C 4487G>T 4488A>T), I362A (4489A>G 4490T>C), G363K (4492G>A 4493G>A), F367Y (4505T>A 4506T>C), Y368F (4508A>T 4509C>T), S370K (4513T>A 4514C>A 4515C>A), V380I (4543G>A), R381K (4546C>A 4547G>A 4548C>A), A383E (4553C>A 4554T>A), P384E (4555C>G 4556C>A), E385N (4558G>A 4560A>T), T386S (4561A>T 4563A>T), Q388A (4567C>G 4568A>C 4569G>T), C394T (4585T>A 4586G>C 4587T>A), Q396N (4591C>A 4593A>T), Q410N (4633C>A 4635A>T), A413Q (4642G>C 4643C>A 4644T>A), H418K (4657C>A 4659C>A), S422I (4670G>T 4671T>A), L423F (4672C>T 4674C>T), N425E (4678A>G 4680C>A), M426K (4682T>A), Y427N (4684T>A), P428_Q429del (4687_4692delCCACAG), C431L (4696T>C 4697G>T 4698T>A), A432I (4699G>A 4700C>T 4701T>C), V435I (4708G>A 4710T>A), I439L (4720A>C 4722C>A), V440I (4723G>A 4725C>A), F441C (4727T>G 4728C>T), K443N (4734A>T), T444S (4735A>T 4737T>A), E445Y (4738G>T 4740A>T), E446Q (4741G>C), G450T (4753G>A 4754G>C), V452L (4759G>C 4761C>T), I454Q (4765A>C 4766T>A 4767T>A), V455F (4768G>T), L456S (4771C>T 4772T>C 4773A>T), N457Q (4774A>C 4776C>A), R458L (4777A>T 4778G>T), K460V (4783A>G 4784A>T 4785A>T), A461E (4787C>A 4788C>A), L462N (4789C>A 4790T>A 4791A>T), K468Q (4807A>C 4809A>G), A471E (4817C>A 4818C>G), Q472D (4819C>G 4821G>T), L473I (4822T>A 4824G>A), C474S (4825T>A 4827C>T), T476N (4832C>A 4833A>T), T477E (4834A>G 4835C>A 4836T>A), N479E (4840A>G 4842T>A), L483M (4852C>A 4854A>G), V484I (4855G>A), E486S (4861G>T 4862A>C 4863A>C), R487K (4865G>A), G488del (4867_4869delGGA) | | | | | | | | |

| | Begin | End | Coverage | Score | Concordance | Matches | Identities | I/D/M/F* | Stop Codons |
|---|---|---|---|---|---|---|---|---|---|
| **NT** | **4219** | **4872** | **8.3%** | **219** | **17.5%** | **644 (98.5%)** | **386 (59.0%)** | **0/10** | |
| Codon mutations: | TTA272CTA (4219T>C), TTA273AAA (4222T>A 4223T>A), ACT275CCC (4228A>C 4230T>C), AAC276AGC (4232A>G), AAA277ATC (4235A>T 4236A>C), ATT279ATA (4242T>A), AAG280AGA (4244A>G 4245G>A), TGT281GTT (4246T>G 4247G>T), CCA283CCT (4254A>T), CAG285CTA (4259A>T 4260G>A), TAT286TAC (4263T>C), TCA287AAT (4264T>A 4265C>A 4266A>T), CCT288GAA (4267C>G 4268C>A 4269T>A), GAC290GAT (4275C>T), CGA291AAA (4276C>A 4277G>A), GAA292ACT (4279G>A 4280A>C 4281A>T), GAG293GAA (4284G>A), ACT296ATT (4292C>T), ATC298ATT (4299C>T), GAA299CAA (4300G>C), CTA301TTA (4306C>T), CTC302TTA (4309C>T 4311C>A), AAG303AAT (4314G>T), CTT304ATG (4315C>A 4317T>G), GGT305AAA (4318G>A 4319G>A 4320T>A), ATC306TTA (4321A>T 4323C>A), ATC307ATA (4326C>A), CCT309CAT (4331C>A), AGC310AGT (4335C>T), AAG311AAA (4338G>A), TCC315AGT (4348T>A 4349C>G 4350C>T), TCA316AGC (4351T>A 4352C>G 4353A>C), GCC318GCA (4359C>A), TTC319TTT (4362C>T), AGG322AGA (4371G>A), AAT323AAC (4374T>C), GCC325GGC (4379C>G), GAG326GAA (4383G>A), ATC327CAA (4384A>C 4385T>A 4386C>A), CGA329-CT (4390delC 4391G>C 4392A>T), GGC330GGA (4395C>A), AGA333CGA (4402A>C), ATT336ATA (4413T>A), AAG340GAG (4423A>G), TTA341CTT (4426T>C 4428A>T), GAC343AAA (4432G>A 4434C>A), CAT344AAT (4435C>A), ACA345TGT (4438A>T 4439C>G 4440A>T), AAG346GTC (4441A>G 4442A>T 4443G>C), GGA347TTC (4444G>T 4445G>T 4446A>C), GAT348GAC (4449T>C), GGC349GGA (4452C>A), TAC350TAT (4455C>T), CTA351TTC (4456C>T 4458A>C), CTT352ATC (4459C>A 4461T>C), CCT353CCA (4464T>A), AAC354AAT (4467C>T), AAG355AAA (4470G>A), GAG356AAT (4471G>A 4473G>T), CAA357AAC (4474C>A 4476A>C), CTG358CTT (4479G>T), CTT359ATA (4480C>A 4482T>A), CAA360AAT (4483C>A 4485A>T), AGA361CTT (4486A>C 4487G>T 4488A>T), ATC362GCC (4489A>G 4490T>C), GGA363AAA (4492G>A 4493G>A), GGT364GGA (4497T>A), AAG365AAA (4500G>A), TTT367TAC (4505T>A 4506T>C), TAC368TTT (4508A>T 4509C>T), TCT369TCA (4512T>A), TCC370AAA (4513T>A 4514C>A 4515C>A), GAC372GAT (4521C>T), TGT373TGC (4524T>C), TCT375AGT (4528T>A 4529C>G), CAG379CAA (4542G>A), GTA380ATA (4543G>A), CGC381AAA (4546C>A 4547G>A 4548C>A), CTT382CTA (4551T>A), GCT383GAA (4553C>A 4554T>A), CCA384GAA (4555C>G 4556C>A), GAA385AAT (4558G>A 4560A>T), ACA386TCT (4561A>T 4563A>T), ATT387ATA (4566T>A), CAG388GCT (4567C>G 4568A>C 4569G>T), CTA389TTA (4570C>T), ACC390ACA (4575C>A), GCT391GCA (4578T>A), AGC393AGT (4584C>T), TGT394ACA (4585T>A 4586G>C 4587T>A), CCC395CCA (4590C>A), CAA396AAT (4591C>A 4593A>T), GTC403GTG (4614C>G), CAA410AAT (4633C>A 4635A>T), GCT413CAA (4642G>C 4643C>A 4644T>A), ATT414ATC (4647T>C), TTC415TTT (4650C>T), AGG417AGA (4656G>A), CAC418AAA (4657C>A 4659C>A), GAC420GAT (4665C>T), AGT422ATA (4670G>T 4671T>A), CTC423TTT (4672C>T 4674C>T), AGC424TCT (4675A>T 4676G>C 4677C>T), AAC425GAA (4678A>G 4680C>A), ATG426AAG (4682T>A), TAT427AAT (4684T>A), CCA428_CAG429del (4687_4692delCCACAG), TGT431CTA (4696T>C 4697G>T 4698T>A), GCT432ATC (4699G>A 4700C>T 4701T>C), GTA433GTC (4704A>C), GTT435ATA (4708G>A 4710T>A), GAC437GAT (4716C>T), ATC438ATA (4719C>A), ATC439CTA (4720A>C 4722C>A), GTC440ATA (4723G>A 4725C>A), TTC441TGT (4727T>G 4728C>T), AGC442TCT (4729A>T 4730G>C 4731C>T), AAA443AAT (4734A>T), ACT444TCA (4735A>T 4737T>A), GAA445TAT (4738G>T 4740A>T), GAA446CAA (4741G>C), GAA447GAG (4746A>G), GGA450ACA (4753G>A 4754G>C), CAT451CAC (4758T>C), GTC452CTT (4759G>C 4761C>T), ATT454CAA (4765A>C 4766T>A 4767T>A), GTT455TTT (4768G>T), CTA456TCT (4771C>T 4772T>C 4773A>T), AAC457CAA (4774A>C 4776C>A), AGA458TTA (4777A>T 4778G>T), TGC459TGT (4782C>T), AAA460GTT (4783A>G 4784A>T 4785A>T), GCC461GAA (4787C>A 4788C>A), CTA462AAT (4789C>A 4790T>A 4791A>T), ATA464ATT (4797A>T), CTT466TTA (4801C>T 4803T>A), AAA468CAG (4807A>C 4809A>G), AAG469AAA (4812G>A), AAA470AAG (4815A>G), GCC471GAG (4817C>A 4818C>G), CAG472GAT (4819C>G 4821G>T), TTG473ATA (4822T>A 4824G>A), TGC474AGT (4825T>A 4827C>T), ACA476AAT (4832C>A 4833A>T), ACT477GAA (4834A>G 4835C>A 4836T>A), ATC478ATT (4839C>T), AAT479GAA (4840A>G 4842T>A), GGT482GGA (4851T>A), CTA483ATG (4852C>A 4854A>G), GTA484ATA (4855G>A), ATA485ATT (4860A>T), GAA486TCC (4861G>T 4862A>C 4863A>C), AGA487AAA (4865G>A), GGA488del (4867_4869delGGA) | | | | | | | | |

*: Inserts / Deletes / Misaligned / Frameshifts

#### Analysis details

This analysis was performed with panviral2.64

### NGS Details (UN70): Alisovirus socal22

#### Assembly

| | |
|---|---|
| **Coverage Length** | 144 (1 contig(s)) |
| **Depth Of Coverage** | 19.1 |
| **Number Of Reads** | 28 |
| **Reads Per Million** | 0.53 rpm (after QC) |
| **Ambiguities** | 0 |
| **Assembly Method** | de novo + reference guided assembly |
| **Consensus Caller** | Bcf Tools |

#### Coverage Map

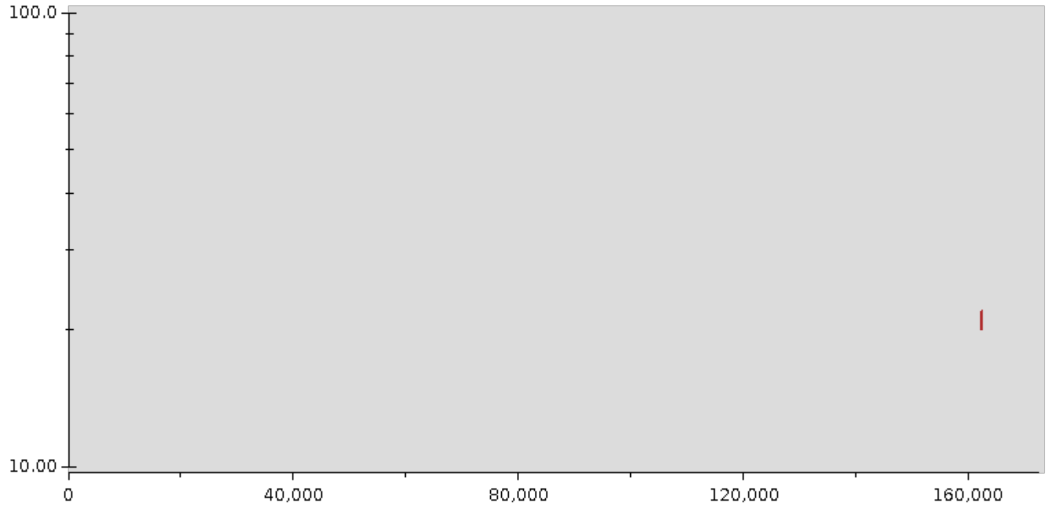

#### Assignment

| | |
|---|---|
| **Type** | Alisovirus socal22 (Taxonomy ID: 2955237) |
| **Reference Genome** | NC_031903.1 |
| **NT Identity (%)** | 79.8611 |
| **AA Identity (%)** | 85.4167 |
| **Number Of Stop Codons** | 1 |
| **Number Of CDS** | 214 |

#### Alignment

| | |
|---|---|
| **Alignment Score** | 172.0 (NT) + 277.0 (AA) = 449.0 |
| **Concordance (%)** | 75.3356 |
| **Alignment Method** | Local, heuristic, nucleotide (BLASTN) |

#### Genome Region

Sequence starts at position 162305 and ends at position 162448 relative to NC_031903.1 reference sequence.

#### Alignment Detailed Statistics

| | Begin | End | Coverage | Score | Concordance | Matches | Identities | I/D/M/F* | Stop Codons |
|---|---|---|---|---|---|---|---|---|---|
| **NT** | **162305** | **162448** | **0.1%** | **172** | **59.7%** | **144 (100%)** | **115 (79.9%)** | **0/0** | |

Mutations: 162314C>T, 162315A>G, 162319T>G, 162322A>C, 162324C>T, 162327T>A, 162328A>G, 162330T>A, 162331A>G, 162333T>C, 162336A>T, 162339C>A, 162345T>G, 162348T>C, 162351T>A, 162357T>C, 162360C>T, 162366T>C, 162369A>T, 162373C>T, 162378G>A, 162384T>C, 162393G>A, 162408A>G, 162411C>T, 162423G>A, 162429C>T, 162435G>A, 162438T>C

*: Inserts / Deletes / Misaligned / Frameshifts

#### Analysis details

This analysis was performed with panviral2.64

#### NGS Details (UN70): Badnavirus maculakalanchoes

##### Assembly

| | |
|---|---|
| **Coverage Length** | 278 (1 contig(s)) |
| **Depth Of Coverage** | 10.4 |
| **Number Of Reads** | 27 |
| **Reads Per Million** | 0.51 rpm (after QC) |
| **Ambiguities** | 0 |
| **Assembly Method** | de novo + reference guided assembly |
| **Consensus Caller** | Bcf Tools |

##### Coverage Map

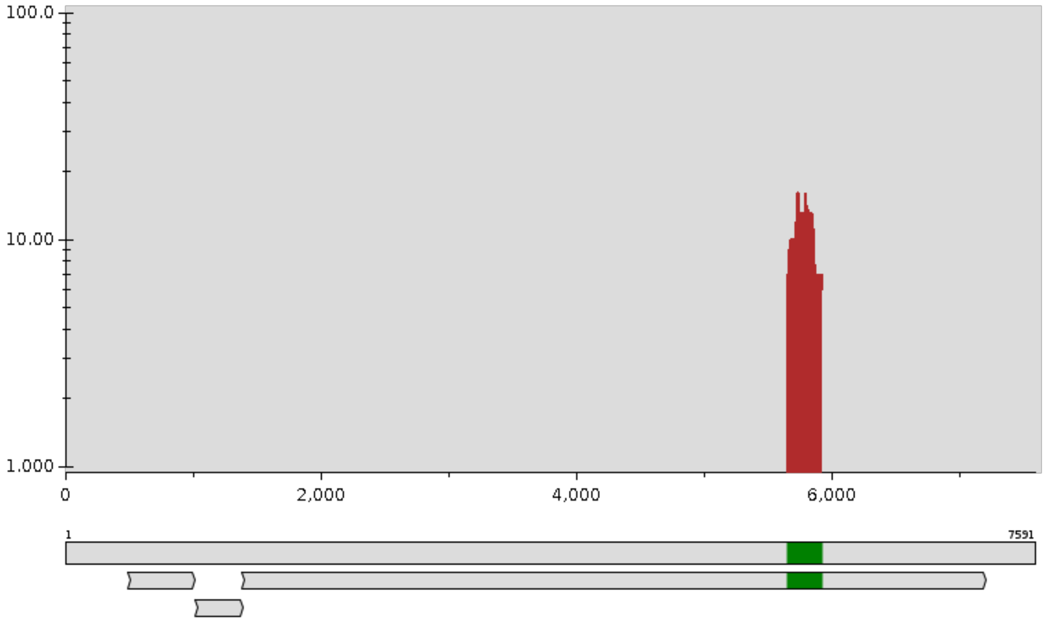

##### Assignment

| | |
|---|---|
| **Type** | Badnavirus maculakalanchoes (Taxonomy ID: 218387) |
| **Reference Genome** | NC_004540.1 |
| **NT Identity (%)** | 55.1601 |
| **AA Identity (%)** | 48.9362 |
| **Number Of Stop Codons** | 0 |
| **Number Of CDS** | 3 |

##### Alignment

| | |
|---|---|
| **Alignment Score** | 52.0 (NT) + 341.0 (AA) = 393.0 |
| **Concordance (%)** | 31.7447 |

| Alignment Method | Global, seeded, nucleotide + amino acids (AGA) |
|---|---|

#### Genome Region

Sequence starts at position 5649 and ends at position 5926 relative to NC_004540.1 reference sequence.

#### Alignment Detailed Statistics

| | Begin | End | Coverage | Score | Concordance | Matches | Identities | I/D/M/F* | Stop Codons |
|---|---|---|---|---|---|---|---|---|---|
| **NT** | **5649** | **5926** | **3.7%** | **52** | **9.4%** | **278 (98.9%)** | **155 (55.2%)** | **3/0** | |
| Mutations: | 5649A>T, 5656G>A, 5660A>T, 5661T>C, 5662A>T, 5663G>C, 5664C>T, 5668T>A, 5670T>A, 5673C>T, 5674C>T, 5676C>A, 5678A>G, 5680A>T, 5681G>C, 5682T>G, 5687T>A, 5688C>T, 5691C>T, 5694G>A, 5695G>C, 5698G>A, 5699C>G, 5700C>G, 5706G>A, 5707G>A, 5708A>T, 5712A>G, 5713T>A, 5714C>A, 5715A>T, 5716A>G, 5722T>A, 5723G>A, 5726C>A, 5727G>T, 5730C>T, 5734T>A, 5737G>A, 5739G>T, 5740A>C, 5741T>A, 5742A>T, 5743G>T, 5744A>C, 5745C>T, 5750T>A, 5751C>T, 5753A>T, 5754C>T, 5757A>G, 5759G>A, 5760G>T, 5763A>G, 5766A>G, 5775C>T, 5778T>G, 5781T>C, 5782A>T, 5783A>C, 5793T>A, 5796T>A, 5798T>C, 5799A>T, 5802C>T, 5806C>G, 5809A>T, 5810A>T, 5815G>A, 5818G>T, 5819A>C, 5820G>T, 5821T>G, 5822G>T, 5826C>T, 5828G>A, 5829A>C, 5830G>T, 5832A>T, 5832_5833insTAT, 5833A>C, 5836G>A, 5837A>G, 5839G>A, 5841G>A, 5842T>C, 5843T>A, 5844C>T, 5845A>G, 5847A>G, 5848G>T, 5849C>T, 5855A>T, 5857G>T, 5859G>T, 5868A>C, 5869C>T, 5871G>A, 5874C>A, 5876T>A, 5877C>T, 5878T>A, 5879C>G, 5881A>C, 5885G>C, 5886C>T, 5887G>T, 5889C>G, 5890C>G, 5895C>T, 5898T>C, 5899A>G, 5900A>T, 5903G>A, 5904G>A, 5908C>T, 5911C>A, 5913G>A, 5914A>T, 5915G>C, 5916G>T, 5917T>G, 5922C>T, 5925A>G | | | | | | | | |
| **CDS** | | | | | | | | | |
| KTSVgp3 | 1423 | 1515 | 4.8% | 341 | 49.3% | 93 (98.9%) | 46 (48.9%) | 1/0/0/0 | 0 |
| Protein mutations: | V1425I (5656G>A), Y1426F (5660A>T 5661T>C), F1429I (5668T>A 5670T>A), K1432R (5678A>G), F1435Y (5687T>A 5688C>T), V1438L (5695G>C), A1439R (5698G>A 5699C>G 5700C>G), E1442I (5707G>A 5708A>T), S1444N (5713T>A 5714C>A 5715A>T), I1445V (5716A>G), W1447K (5722T>A 5723G>A), T1448N (5726C>A 5727G>T), W1451R (5734T>A), A1452T (5737G>A 5739G>T), I1453H (5740A>C 5741T>A 5742A>T), D1454S (5743G>T 5744A>C 5745C>T), L1456H (5750T>A 5751C>T), Y1457F (5753A>T 5754C>T), W1459Y (5759G>A 5760G>T), K1467S (5782A>T 5783A>C), I1472T (5798T>C 5799A>T), R1475G (5806C>G), K1476L (5809A>T 5810A>T), D1478N (5815G>A), E1479S (5818G>T 5819A>C 5820G>T), C1480V (5821T>G 5822G>T), R1482H (5828G>A 5829A>C), G1483C (5830G>T 5832A>T), G1483_M1484insY (5832_5833insTAT), M1484L (5833A>C), E1485R (5836G>A 5837A>G), E1486K (5839G>A 5841G>A), F1487H (5842T>C 5843T>A 5844C>T), I1488V (5845A>G 5847A>G), A1489L (5848G>T 5849C>T), Y1491F (5855A>T), V1492F (5857G>T 5859G>T), F1498Y (5876T>A 5877C>T), N1500H (5881A>C), S1501T (5885G>C 5886C>T), V1502L (5887G>T 5889C>G), Q1503E (5890C>G), K1506V (5899A>G 5900A>T), R1507K (5903G>A 5904G>A), Q1510K (5911C>A 5913G>A), R1511S (5914A>T 5915G>C 5916G>T), F1512V (5917T>G) | | | | | | | | |
| Codon mutations: | ACA1422..T (5649A>T), GTC1425ATC (5656G>A), TAT1426TTC (5660A>T 5661T>C), AGC1427TCT (5662A>T 5663G>C 5664C>T), TTT1429ATA (5668T>A 5670T>A), GAC1430GAT (5673C>T), CTC1431TTA (5674C>T 5676C>A), AAG1432AGG (5678A>G), AGT1433TCG (5680A>T 5681G>C 5682T>G), TTC1435TAT (5687T>A 5688C>T), CAC1436CAT (5691C>T), CAG1437CAA (5694G>A), GTG1438CTG (5695G>C), GCC1439AGG (5698G>A 5699C>G 5700C>G), GCG1441GCA (5706G>A), GAA1442ATA (5707G>A 5708A>T), GAA1443GAG (5712A>G), TCA1444AAT (5713T>A 5714C>A 5715A>T), ATC1445GTC (5716A>G), TGG1447AAG (5722T>A 5723G>A), ACG1448AAT (5726C>A 5727G>T), GCC1449GCT (5730C>T), TGG1451AGG (5734T>A), GCG1452ACT (5737G>A 5739G>T), ATA1453CAT (5740A>C 5741T>A 5742A>T), GAC1454TCT (5743G>T 5744A>C 5745C>T), CTC1456CAT (5750T>A 5751C>T), TAC1457TTT (5753A>T 5754C>T), GAA1458GAG (5757A>G), TGG1459TAT (5759G>A 5760G>T), TTA1460TTG (5763A>G), GTA1461GTG (5766A>G), TTC1464TTT (5775C>T), GGT1465GGG (5778T>G), CTT1466CTC (5781T>C), AAA1467TCA (5782A>T 5783A>C), CCT1470CCA (5793T>A), GCT1471GCA (5796T>A), ATA1472ACT (5798T>C 5799A>T), TTC1473TTT (5802C>T), CGA1475GGA (5806C>G), AAA1476TTA (5809A>T 5810A>T), GAT1478AAT (5815G>A), GAG1479TCT (5818G>T 5819A>C 5820G>T), TGT1480GTT (5821T>G 5822G>T), TTC1481TTT (5826C>T), CGA1482CAC (5828G>A 5829A>C), GGA1483TGT (5830G>T 5832A>T), GGA1483_ATG1484insTAT (5832_5833insTAT), ATG1484CTG (5833A>C), GAG1485AGG (5836G>A 5837A>G), GAG1486AAA (5839G>A 5841G>A), TTC1487CAT (5842T>C 5843T>A 5844C>T), ATA1488GTG (5845A>G 5847A>G), GCA1489TTA (5848G>T 5849C>T), TAT1491TTT (5855A>T), GTG1492TTT (5857G>T 5859G>T), ATA1495ATC (5868A>C), CTG1496TTA (5869C>T 5871G>A), ATC1497ATA (5874C>A), TTC1498TAT (5876T>A 5877C>T), TCC1499AGC (5878T>A 5879C>G), AAC1500CAC (5881A>C), AGC1501ACT (5885G>C 5886C>T), GTC1502TTG (5887G>T 5889C>G), CAA1503GAA (5890C>G), GAC1504GAT (5895C>T), CAT1505CAC (5898T>C), AAG1506GTG (5899A>G 5900A>T), AGG1507AAA (5903G>A 5904G>A), CTG1509TTG (5908C>T), CAG1510AAA (5911C>A 5913G>A), AGG1511TCT (5914A>T 5915G>C 5916G>T), TTT1512GTT (5917T>G), TTC1513TTT (5922C>T), GAA1514GAG (5925A>G) | | | | | | | | |
| **Proteins** | | | | | | | | | |
| putative polyprotein (NP_777317.1) | 1423 | 1515 | 4.8% | 341 | 49.3% | 93 (98.9%) | 46 (48.9%) | 1/0/0/0 | 0 |
| Protein mutations: | V1425I (5656G>A), Y1426F (5660A>T 5661T>C), F1429I (5668T>A 5670T>A), K1432R (5678A>G), F1435Y (5687T>A 5688C>T), V1438L (5695G>C), A1439R (5698G>A 5699C>G 5700C>G), E1442I (5707G>A 5708A>T), S1444N (5713T>A 5714C>A 5715A>T), I1445V (5716A>G), W1447K (5722T>A 5723G>A), T1448N (5726C>A 5727G>T), W1451R (5734T>A), A1452T (5737G>A 5739G>T), I1453H (5740A>C 5741T>A 5742A>T), D1454S (5743G>T 5744A>C 5745C>T), L1456H (5750T>A 5751C>T), Y1457F (5753A>T 5754C>T), W1459Y (5759G>A 5760G>T), K1467S (5782A>T 5783A>C), I1472T (5798T>C 5799A>T), R1475G (5806C>G), K1476L (5809A>T 5810A>T), D1478N (5815G>A), E1479S (5818G>T 5819A>C 5820G>T), C1480V (5821T>G 5822G>T), R1482H (5828G>A 5829A>C), G1483C (5830G>T 5832A>T), G1483_M1484insY (5832_5833insTAT), M1484L (5833A>C), E1485R (5836G>A 5837A>G), E1486K (5839G>A 5841G>A), F1487H (5842T>C 5843T>A 5844C>T), I1488V (5845A>G 5847A>G), A1489L (5848G>T 5849C>T), Y1491F (5855A>T), V1492F (5857G>T 5859G>T), F1498Y (5876T>A 5877C>T), N1500H (5881A>C), S1501T (5885G>C 5886C>T), V1502L (5887G>T 5889C>G), Q1503E (5890C>G), K1506V (5899A>G 5900A>T), R1507K (5903G>A 5904G>A), Q1510K (5911C>A 5913G>A), R1511S (5914A>T 5915G>C 5916G>T), F1512V (5917T>G) | | | | | | | | |
| Codon mutations: | ACA1422..T (5649A>T), GTC1425ATC (5656G>A), TAT1426TTC (5660A>T 5661T>C), AGC1427TCT (5662A>T 5663G>C 5664C>T), TTT1429ATA (5668T>A 5670T>A), GAC1430GAT (5673C>T), CTC1431TTA (5674C>T 5676C>A), AAG1432AGG (5678A>G), AGT1433TCG (5680A>T 5681G>C 5682T>G), TTC1435TAT (5687T>A 5688C>T), CAC1436CAT (5691C>T), CAG1437CAA (5694G>A), GTG1438CTG (5695G>C), GCC1439AGG (5698G>A 5699C>G 5700C>G), GCG1441GCA (5706G>A), GAA1442ATA (5707G>A 5708A>T), GAA1443GAG (5712A>G), TCA1444AAT (5713T>A 5714C>A 5715A>T), ATC1445GTC (5716A>G), TGG1447AAG (5722T>A 5723G>A), ACG1448AAT (5726C>A 5727G>T), GCC1449GCT (5730C>T), TGG1451AGG (5734T>A), GCG1452ACT (5737G>A 5739G>T), ATA1453CAT (5740A>C 5741T>A 5742A>T), GAC1454TCT (5743G>T 5744A>C 5745C>T), CTC1456CAT (5750T>A 5751C>T), TAC1457TTT (5753A>T 5754C>T), GAA1458GAG (5757A>G), TGG1459TAT (5759G>A 5760G>T), TTA1460TTG (5763A>G), GTA1461GTG (5766A>G), TTC1464TTT (5775C>T), GGT1465GGG (5778T>G), CTT1466CTC (5781T>C), AAA1467TCA (5782A>T 5783A>C), CCT1470CCA (5793T>A), GCT1471GCA (5796T>A), ATA1472ACT (5798T>C 5799A>T), TTC1473TTT (5802C>T), CGA1475GGA (5806C>G), AAA1476TTA (5809A>T 5810A>T), GAT1478AAT (5815G>A), GAG1479TCT (5818G>T 5819A>C 5820G>T), TGT1480GTT (5821T>G 5822G>T), TTC1481TTT (5826C>T), CGA1482CAC (5828G>A 5829A>C), GGA1483TGT (5830G>T 5832A>T), GGA1483_ATG1484insTAT (5832_5833insTAT), ATG1484CTG (5833A>C), GAG1485AGG (5836G>A 5837A>G), GAG1486AAA (5839G>A 5841G>A), TTC1487CAT (5842T>C 5843T>A 5844C>T), ATA1488GTG (5845A>G 5847A>G), GCA1489TTA (5848G>T 5849C>T), TAT1491TTT (5855A>T), GTG1492TTT (5857G>T 5859G>T), ATA1495ATC (5868A>C), CTG1496TTA (5869C>T 5871G>A), ATC1497ATA (5874C>A), TTC1498TAT (5876T>A 5877C>T), TCC1499AGC (5878T>A 5879C>G), AAC1500CAC (5881A>C), AGC1501ACT (5885G>C 5886C>T), GTC1502TTG (5887G>T 5889C>G), CAA1503GAA (5890C>G), GAC1504GAT (5895C>T), CAT1505CAC (5898T>C), AAG1506GTG (5899A>G 5900A>T), AGG1507AAA (5903G>A 5904G>A), CTG1509TTG (5908C>T), CAG1510AAA (5911C>A 5913G>A), AGG1511TCT (5914A>T 5915G>C 5916G>T), TTT1512GTT (5917T>G), TTC1513TTT (5922C>T), GAA1514GAG (5925A>G) | | | | | | | | |

*: Inserts / Deletes / Misaligned / Frameshifts

#### Analysis details

This analysis was performed with panviral2.64

### NGS Details (UN70): Dioscovirus dioscoreae

#### Assembly

| | |
|---|---|
| **Coverage Length** | 603 (2 contig(s)) |
| **Depth Of Coverage** | 4.1 |
| **Number Of Reads** | 26 |
| **Reads Per Million** | 0.49 rpm (after QC) |
| **Ambiguities** | 0 |
| **Assembly Method** | de novo + reference guided assembly |
| **Consensus Caller** | Bcf Tools |

#### Coverage Map

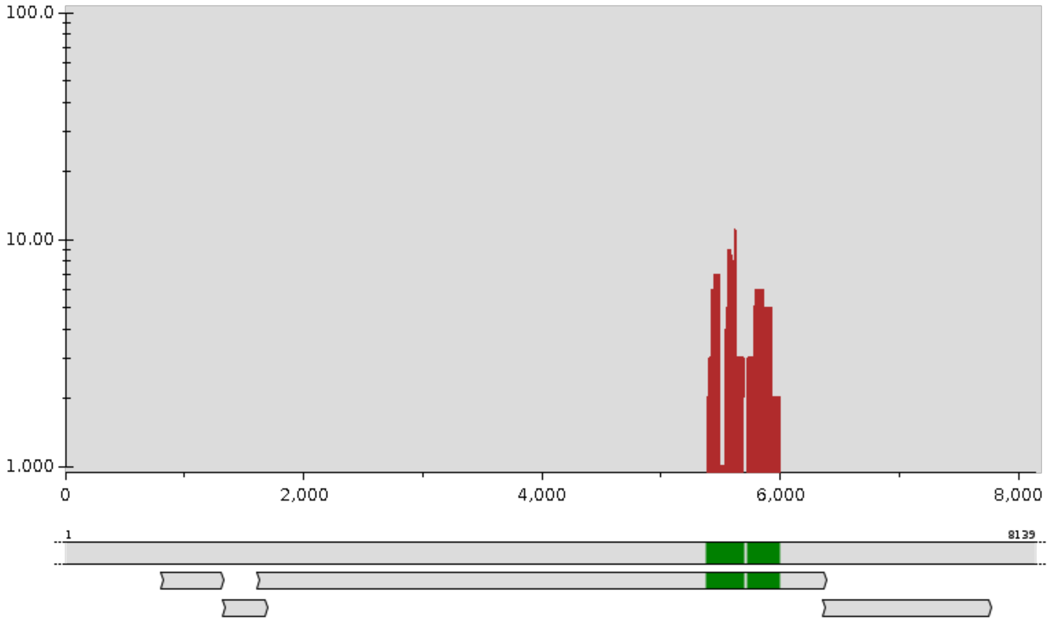

#### Assignment

| | |
|---|---|
| **Type** | Dioscovirus dioscoreae (Taxonomy ID: 3052184) |
| **Reference Genome** | NC_040712.1 |
| **NT Identity (%)** | 62.5 |
| **AA Identity (%)** | 48.2412 |
| **Number Of Stop Codons** | 1 |
| **Number Of CDS** | 4 |

#### Alignment

| | |
|---|---|
| **Alignment Score** | 288.0 (NT) + 708.0 (AA) = 996.0 |
| **Concordance (%)** | 38.3372 |

| Alignment Method | Global, seeded, nucleotide + amino acids (AGA) |
|---|---|

#### Genome Region

Sequence starts at position 5383 and ends at position 6002 relative to NC_040712.1 reference sequence.

#### Alignment Detailed Statistics

| | Begin | End | Coverage | Score | Concordance | Matches | Identities | I/D/M/F* | Stop Codons |
|---|---|---|---|---|---|---|---|---|---|
| **NT** | **5383** | **6002** | **7.4%** | **288** | **24.2%** | **600 (99.5%)** | **375 (62.2%)** | **0/3** | |
| Mutations: | 5383T>A, 5387C>T, 5394A>C, 5395T>G, 5396G>A, 5400G>A, 5403C>T, 5405A>T, 5406C>G, 5407A>G, 5408G>A, 5409A>G, 5410G>A, 5412A>G, 5413T>A, 5415G>T, 5416A>G, 5418T>A, 5419G>A, 5420G>A, 5423A>T, 5424G>A, 5426A>T, 5430G>A, 5431T>A, 5432C>G, 5433T>A, 5434G>T, 5436C>A, 5437C>A, 5439C>T, 5441G>T, 5443G>A, 5444T>G, 5446A>T, 5449T>A, 5453A>G, 5457A>T, 5462A>T, 5468A>C, 5471C>T, 5474A>C, 5475C>T, 5477C>G, 5479A>C, 5480A>T, 5486A>G, 5489T>C, 5490T>A, 5491C>A, 5493G>A, 5502C>A, 5505A>T, 5506A>T, 5511G>A, 5514A>G, 5519A>T, 5522C>T, 5524A>G, 5526A>G, 5530A>T, 5532T>G, 5533C>G, 5535G>A, 5541G>T, 5543C>T, 5544T>C, 5545G>T, 5553A>T, 5555A>T, 5558C>T, 5561T>C, 5564T>C, 5568A>G, 5577G>A, 5579A>G, 5581G>A, 5583A>C, 5585A>G, 5586C>G, 5588A>T, 5589C>G, 5591A>G, 5595G>A, 5598C>G, 5603T>C, 5604T>C, 5607T>A, 5608T>A, 5612G>A, 5613T>G, 5615C>T, 5618C>T, 5619C>G, 5626G>T, 5627T>C, 5628A>G, 5629A>T, 5630G>A, 5631G>A, 5632A>G, 5633A>T, 5634G>A, 5639G>A, 5640C>A, 5642T>A, 5645T>A, 5646T>A, 5648G>A, 5649T>A, 5651A>C, 5656C>A, 5657A>G, 5661T>A, 5664A>G, 5667A>T, 5670G>T, 5672A>T, 5673G>A, 5674T>A, 5676G>A, 5679A>C, 5685G>A, 5687A>C, 5691C>T, 5693T>A, 5721C>A, 5726C>A, 5739T>G, 5740T>C, 5741A>T, 5748A>G, 5756A>T, 5757T>A, 5759T>G, 5762T>A, 5763_5765delGAA, 5768C>T, 5769C>A, 5777A>T, 5779C>A, 5781T>A, 5782T>C, 5786G>A, 5788G>A, 5790C>T, 5795G>A, 5798G>A, 5807G>A, 5808A>T, 5811T>G, 5813A>G, 5816C>T, 5822C>T, 5828T>C, 5830A>T, 5834C>A, 5836C>A, 5843G>A, 5844A>G, 5846T>A, 5850T>A, 5851A>T, 5853A>G, 5855T>A, 5856A>G, 5859A>C, 5860T>C, 5864T>A, 5868A>G, 5869A>C, 5870T>A, 5873G>A, 5874T>A, 5875G>T, 5876T>A, 5877T>G, 5878C>G, 5879C>A, 5882C>T, 5889G>C, 5893G>A, 5895A>C, 5897A>C, 5903T>C, 5904T>A, 5906T>A, 5907C>G, 5910G>A, 5912C>T, 5913T>A, 5914G>T, 5915G>T, 5918G>A, 5919A>C, 5924T>A, 5925A>C, 5926G>A, 5928A>C, 5933C>T, 5940G>A, 5941T>A, 5945T>A, 5947C>A, 5948C>A, 5951A>C, 5952T>A, 5957C>A, 5958C>G, 5959C>A, 5960T>C, 5963G>A, 5965T>G, 5966T>C, 5967T>A, 5973A>C, 5974A>T, 5976C>G, 5977C>A, 5979C>T, 5980A>C, 5982A>G, 5983G>A, 5988A>C, 5990T>A, 5999G>A | | | | | | | | |

##### CDS

| | Begin | End | Coverage | Score | Concordance | Matches | Identities | I/D/M/F* | Stop Codons |
|---|---|---|---|---|---|---|---|---|---|
| EXK67_gp3 | 1259 | 1464 | 12.5% | 708 | 49.7% | 199 (99.5%) | 96 (48.0%) | 0/1/0/0 | 1 |
| Protein mutations: | M1262R (5394A>C 5395T>G 5396G>A), D1264N (5400G>A), P1265S (5403C>T 5405A>T), Q1266G (5406C>G 5407A>G 5408G>A), S1267D (5409A>G 5410G>A), I1268E (5412A>G 5413T>A), E1269* (5415G>T 5416A>G), W1270K (5418T>A 5419G>A 5420G>A), A1272T (5424G>A 5426A>T), V1274K (5430G>A 5431T>A 5432C>G), C1275I (5433T>A 5434G>T), P1276K (5436C>A 5437C>A), Q1277Y (5439C>T 5441G>T), G1278E (5443G>A 5444T>G), H1279L (5446A>T), F1280Y (5449T>A), I1283L (5457A>T), K1290T (5479A>C 5480A>T), S1294K (5490T>A 5491C>A), V1295I (5493G>A), K1299L (5505A>T 5506A>T), D1301N (5511G>A), N1302D (5514A>G), K1305R (5524A>G), K1306E (5526A>G), Y1307F (5530A>T), S1308G (5532T>G 5533C>G), E1309K (5535G>A), V1311F (5541G>T 5543C>T), C1312L (5544T>C 5545G>T), I1315F (5553A>T 5555A>T), I1320V (5568A>G), E1323K (5577G>A 5579A>G), S1324N (5581G>A), I1325L (5583A>C 5585A>G), Q1326D (5586C>G 5588A>T), Q1327E (5589C>G 5591A>G), V1329I (5595G>A), Q1330E (5598C>G), L1333K (5607T>A 5608T>A), F1335V (5613T>G 5615C>T), Q1337E (5619C>G), C1339F (5626G>T 5627T>C), K1340V (5628A>G 5629A>T 5630G>A), E1341S (5631G>A 5632A>G 5633A>T), E1342K (5634G>A), L1344I (5640C>A 5642T>A), L1346I (5646T>A 5648G>A), S1347T (5649T>A 5651A>C), T1349K (5656C>A 5657A>G), L1351M (5661T>A), K1352E (5664A>G), I1353L (5667A>T), G1354C (5670G>T 5672A>T), V1355K (5673G>A 5674T>A), A1356T (5676G>A), N1357H (5679A>C), E1359N (5685G>A 5687A>C), Q1371K (5721C>A), L1377A (5739T>G 5740T>C 5741A>T), I1380V (5748A>G), E1382D (5756A>T), F1383M (5757T>A 5759T>G), E1385del (5763_5765delGAA), Q1387K (5769C>A), E1389D (5777A>T), T1390K (5779C>A), L1391T (5781T>A 5782T>C), G1393D (5788G>A), I1400L (5808A>T), L1401V (5811T>G 5813A>G), Y1407F (5830A>T), P1409Q (5836C>A), S1412G (5844A>G 5846T>A), Y1414I (5850T>A 5851A>T), T1415A (5853A>G 5855T>A), R1416G (5856A>G), I1417P (5859A>C 5860T>C), F1418L (5864T>A), N1420A (5868A>G 5869A>C 5870T>A), C1422I (5874T>A 5875G>T 5876T>A), S1423G (5877T>G 5878C>G 5879C>A), E1427Q (5889G>C), R1428K (5893G>A), K1429H (5895A>C 5897A>C), S1432T (5904T>A 5906T>A), Q1433E (5907C>G), D1434N (5910G>A 5912C>T), W1435I (5913T>A 5914G>T 5915G>T), M1437L (5919A>C), R1439Q (5925A>C 5926G>A), K1440Q (5928A>C), V1444K (5940G>A 5941T>A), T1446K (5947C>A 5948C>A), K1447N (5951A>C), L1448I (5952T>A), P1450D (5958C>G 5959C>A 5960T>C), I1452S (5965T>G 5966T>C), L1453I (5967T>A), K1455L (5973A>C 5974A>T), P1456E (5976C>G 5977C>A), Q1457S (5979C>T 5980A>C), S1458D (5982A>G 5983G>A), I1460L (5988A>C 5990T>A) | | | | | | | | |
| Codon mutations: | TTC1258.AC (5383T>A), CAC1259CAT (5387C>T), ATG1262CGA (5394A>C 5395T>G 5396G>A), GAT1264AAT (5400G>A), CCA1265TCT (5403C>T 5405A>T), CAG1266GGA (5406C>G 5407A>G 5408G>A), AGT1267GAT (5409A>G 5410G>A), ATA1268GAA (5412A>G 5413T>A), GAA1269TGA (5415G>T 5416A>G), TGG1270AAA (5418T>A 5419G>A 5420G>A), ACA1271ACT (5423A>T), GCA1272ACT (5424G>A 5426A>T), GTC1274AAG (5430G>A 5431T>A 5432C>G), TGC1275ATC (5433T>A 5434G>T), CCA1276AAA (5436C>A 5437C>A), CAG1277TAT (5439C>T 5441G>T), GGT1278GAG (5443G>A 5444T>G), CAT1279CTT (5446A>T), TTT1280TAT (5449T>A), GAA1281GAG (5453A>G), ATA1283TTA (5457A>T), GTA1284GTT (5462A>T), CCA1286CCC (5468A>C), TTC1287TTT (5471C>T), GGA1288GGC (5474A>C), CTC1289TTG (5475C>T 5477C>G), AAA1290ACT (5479A>C 5480A>T), GCA1292GCG (5486A>G), CCT1293CCC (5489T>C), TCA1294AAA (5490T>A 5491C>A), GTA1295ATA (5493G>A), CGA1298AGA (5502C>A), AAA1299TTA (5505A>T 5506A>T), GAT1301AAT (5511G>A), AAT1302GAT (5514A>G), ATA1303ATT (5519A>T), TTC1304TTT (5522C>T), AAA1305AGA (5524A>G), AAA1306GAA (5526A>G), TAT1307TTT (5530A>T), TCA1308GGA (5532T>G 5533C>G), GAA1309AAA (5535G>A), GTC1311TTT (5541G>T 5543C>T), TGT1312CTT (5544T>C 5545G>T), ATA1315TTT (5553A>T 5555A>T), GAC1316GAT (5558C>T), GAT1317GAC (5561T>C), ATT1318ATC (5564T>C), ATA1320GTA (5568A>G), GAA1323AAG (5577G>A 5579A>G), AGT1324AAT (5581G>A), ATA1325CTG (5583A>C 5585A>G), CAA1326GAT (5586C>G 5588A>T), CAA1327GAG (5589C>G 5591A>G), GTA1329ATA (5595G>A), CAA1330GAA (5598C>G), CAT1331CAC (5603T>C), TTA1332CTA (5604T>C), TTA1333AAA (5607T>A 5608T>A), CAG1334CAA (5612G>A), TTC1335GTT (5613T>G 5615C>T), TTC1336TTT (5618C>T), CAA1337GAA (5619C>G), TGT1339TTC (5626G>T 5627T>C), AAG1340GTA (5628A>G 5629A>T 5630G>A), GAA1341AGT (5631G>A 5632A>G 5633A>T), GAA1342AAA (5634G>A), GGG1343GGA (5639G>A), CTT1344ATA (5640C>A 5642T>A), ATT1345ATA (5645T>A), TTG1346ATA (5646T>A 5648G>A), TCA1347ACC (5649T>A 5651A>C), ACA1349AAG (5656C>A 5657A>G), TTG1351ATG (5661T>A), AAA1352GAA (5664A>G), ATA1353TTA (5667A>T), GGA1354TGT (5670G>T 5672A>T), GTA1355AAA (5673G>A 5674T>A), GCA1356ACA (5676G>A), AAT1357CAT (5679A>C), GAA1359AAC (5685G>A 5687A>C), CTT1361TTA (5691C>T 5693T>A), CAA1371AAA (5721C>A), CTC1372CTA (5726C>A), TTA1377GCT (5739T>G 5740T>C 5741A>T), ATA1380GTA (5748A>G), GAA1382GAT (5756A>T), TTT1383ATG (5757T>A 5759T>G), CCT1384CCA (5762T>A), GAA1385del (5763_5765delGAA), GAC1386GAT (5768C>T), CAA1387AAA (5769C>A), GAA1389GAT (5777A>T), ACA1390AAA (5779C>A), TTA1391ACA (5781T>A 5782T>C), AAG1392AAA (5786G>A), GGT1393GAT (5788G>A), CTA1394TTA (5790C>T), CAG1395CAA (5795G>A), AAG1396AAA (5798G>A), GGG1399GGA (5807G>A), ATA1400TTA (5808A>T), TTA1401GTG (5811T>G 5813A>G), AAC1402AAT (5816C>T), GCC1404GCT (5822C>T), AAT1406AAC (5828T>C), TAT1407TTT (5830A>T), ATC1408ATA (5834C>A), CCA1409CAA (5836C>A), TTG1411TTA (5843G>A), AGT1412GGA (5844A>G 5846T>A), TAT1414ATT (5850T>A 5851A>T), ACT1415GCA (5853A>G 5855T>A), AGA1416GGA (5856A>G), ATA1417CCA (5859A>C 5860T>C), TTT1418TTA (5864T>A), AAT1420GCA (5868A>G 5869A>C 5870T>A), AAG1421AAA (5873G>A), TGT1422ATA (5874T>A 5875G>T 5876T>A), TCC1423GGA (5877T>G 5878C>G 5879C>A), AGC1424AGT (5882C>T), GAA1427CAA (5889G>C), AGA1428AAA (5893G>A), AAA1429CAC (5895A>C 5897A>C), AAT1431AAC (5903T>C), TCT1432ACA (5904T>A 5906T>A), CAA1433GAA (5907C>G), GAC1434AAT (5910G>A 5912C>T), TGG1435ATT (5913T>A 5914G>T 5915G>T), AAG1436AAA (5918G>A), ATG1437CTG (5919A>C), GTT1438GTA (5924T>A), AGA1439CAA (5925A>C 5926G>A), AAA1440CAA (5928A>C), ATC1441ATT (5933C>T), GTA1444AAA (5940G>A 5941T>A), ATT1445ATA (5945T>A), ACC1446AAA (5947C>A 5948C>A), AAA1447AAC (5951A>C), TTA1448ATA (5952T>A), CCC1449CCA (5957C>A), CCT1450GAC (5958C>G 5959C>A 5960T>C), TTG1451TTA (5963G>A), ATT1452AGC (5965T>G 5966T>C), TTA1453ATA (5967T>A), AAA1455CTA (5973A>C 5974A>T), CCA1456GAA (5976C>G 5977C>A), CAA1457TCA (5979C>T 5980A>C), AGT1458GAT (5982A>G 5983G>A), ATT1460CTA (5988A>C 5990T>A), GAG1463GAA (5999G>A) | | | | | | | | |

##### Proteins

| | Begin | End | Coverage | Score | Concordance | Matches | Identities | I/D/M/F* | Stop Codons |
|---|---|---|---|---|---|---|---|---|---|
| ORF3 (YP_009553219.1) | 1259 | 1464 | 12.5% | 708 | 49.7% | 199 (99.5%) | 96 (48.0%) | 0/1/0/0 | 1 |
| Protein mutations: | M1262R (5394A>C 5395T>G 5396G>A), D1264N (5400G>A), P1265S (5403C>T 5405A>T), Q1266G (5406C>G 5407A>G 5408G>A), S1267D (5409A>G 5410G>A), I1268E (5412A>G 5413T>A), E1269* (5415G>T 5416A>G), W1270K (5418T>A 5419G>A 5420G>A), A1272T (5424G>A 5426A>T), V1274K (5430G>A 5431T>A 5432C>G), C1275I (5433T>A 5434G>T), P1276K (5436C>A 5437C>A), Q1277Y (5439C>T 5441G>T), G1278E (5443G>A 5444T>G), H1279L (5446A>T), F1280Y (5449T>A), I1283L (5457A>T), K1290T (5479A>C 5480A>T), S1294K (5490T>A 5491C>A), V1295I (5493G>A), K1299L (5505A>T 5506A>T), D1301N (5511G>A), N1302D (5514A>G), K1305R (5524A>G), K1306E (5526A>G), Y1307F (5530A>T), S1308G (5532T>G 5533C>G), E1309K (5535G>A), V1311F (5541G>T 5543C>T), C1312L (5544T>C 5545G>T), I1315F (5553A>T 5555A>T), I1320V (5568A>G), E1323K (5577G>A 5579A>G), S1324N (5581G>A), I1325L (5583A>C 5585A>G), Q1326D (5586C>G 5588A>T), Q1327E (5589C>G 5591A>G), V1329I (5595G>A), Q1330E (5598C>G), L1333K (5607T>A 5608T>A), F1335V (5613T>G 5615C>T), Q1337E (5619C>G), C1339F (5626G>T 5627T>C), K1340V (5628A>G 5629A>T 5630G>A), E1341S (5631G>A 5632A>G 5633A>T), E1342K (5634G>A), L1344I (5640C>A 5642T>A), L1346I (5646T>A 5648G>A), S1347T (5649T>A 5651A>C), T1349K (5656C>A 5657A>G), L1351M (5661T>A), K1352E (5664A>G), I1353L (5667A>T), G1354C (5670G>T 5672A>T), V1355K (5673G>A 5674T>A), A1356T (5676G>A), N1357H (5679A>C), E1359N (5685G>A 5687A>C), Q1371K (5721C>A), L1377A (5739T>G 5740T>C 5741A>T), I1380V (5748A>G), E1382D (5756A>T), F1383M (5757T>A 5759T>G), E1385del (5763_5765delGAA), Q1387K (5769C>A), E1389D (5777A>T), T1390K (5779C>A), L1391T (5781T>A 5782T>C), G1393D (5788G>A), I1400L (5808A>T), L1401V (5811T>G 5813A>G), Y1407F (5830A>T), P1409Q (5836C>A), S1412G (5844A>G 5846T>A), Y1414I (5850T>A 5851A>T), T1415A (5853A>G 5855T>A), R1416G (5856A>G), I1417P (5859A>C 5860T>C), F1418L (5864T>A), N1420A (5868A>G 5869A>C 5870T>A), C1422I (5874T>A 5875G>T 5876T>A), S1423G (5877T>G 5878C>G 5879C>A), E1427Q (5889G>C), R1428K (5893G>A), K1429H (5895A>C 5897A>C), S1432T (5904T>A 5906T>A), Q1433E (5907C>G), D1434N (5910G>A 5912C>T), W1435I (5913T>A 5914G>T 5915G>T), M1437L (5919A>C), R1439Q (5925A>C 5926G>A), K1440Q (5928A>C), V1444K (5940G>A 5941T>A), T1446K (5947C>A 5948C>A), K1447N (5951A>C), L1448I (5952T>A), P1450D (5958C>G 5959C>A 5960T>C), I1452S (5965T>G 5966T>C), L1453I (5967T>A), K1455L (5973A>C 5974A>T), P1456E (5976C>G 5977C>A), Q1457S (5979C>T 5980A>C), S1458D (5982A>G 5983G>A), I1460L (5988A>C 5990T>A) | | | | | | | | |

| | Begin | End | Coverage | Score | Concordance | Matches | Identities | I/D/M/F* | Stop Codons |
|---|---|---|---|---|---|---|---|---|---|
| **NT** | **5383** | **6002** | **7.4%** | **288** | **24.2%** | **600 (99.5%)** | **375 (62.2%)** | **0/3** | |
| Codon mutations: | TTC1258.AC (5383T>A), CAC1259CAT (5387C>T), ATG1262CGA (5394A>C 5395T>G 5396G>A), GAT1264AAT (5400G>A), CCA1265TCT (5403C>T 5405A>T), CAG1266GGA (5406C>G 5407A>G 5408G>A), AGT1267GAT (5409A>G 5410G>A), ATA1268GAA (5412A>G 5413T>A), GAA1269TGA (5415G>T 5416A>G), TGG1270AAA (5418T>A 5419G>A 5420G>A), ACA1271ACT (5423A>T), GCA1272ACT (5424G>A 5426A>T), GTC1274AAG (5430G>A 5431T>A 5432C>G), TGC1275ATC (5433T>A 5434G>T), CCA1276AAA (5436C>A 5437C>A), CAG1277TAT (5439C>T 5441G>T), GGT1278GAG (5443G>A 5444T>G), CAT1279CTT (5446A>T), TTT1280TAT (5449T>A), GAA1281GAG (5453A>G), ATA1283TTA (5457A>T), GTA1284GTT (5462A>T), CCA1286CCC (5468A>C), TTC1287TTT (5471C>T), GGA1288GGC (5474A>C), CTC1289TTG (5475C>T 5477C>G), AAA1290ACT (5479A>C 5480A>T), GCA1292GCG (5486A>G), CCT1293CCC (5489T>C), TCA1294AAA (5490T>A 5491C>A), GTA1295ATA (5493G>A), CGA1298AGA (5502C>A), AAA1299TTA (5505A>T 5506A>T), GAT1301AAT (5511G>A), AAT1302GAT (5514A>G), ATA1303ATT (5519A>T), TTC1304TTT (5522C>T), AAA1305AGA (5524A>G), AAA1306GAA (5526A>G), TAT1307TTT (5530A>T), TCA1308GGA (5532T>G 5533C>G), GAA1309AAA (5535G>A), GTC1311TTT (5541G>T 5543C>T), TGT1312CTT (5544T>C 5545G>T), ATA1315TTT (5553A>T 5555A>T), GAC1316GAT (5558C>T), GAT1317GAC (5561T>C), ATT1318ATC (5564T>C), ATA1320GTA (5568A>G), GAA1323AAG (5577G>A 5579A>G), AGT1324AAT (5581G>A), ATA1325CTG (5583A>C 5585A>G), CAA1326GAT (5586C>G 5588A>T), CAA1327GAG (5589C>G 5591A>G), GTA1329ATA (5595G>A), CAA1330GAA (5598C>G), CAT1331CAC (5603T>C), TTA1332CTA (5604T>C), TTA1333AAA (5607T>A 5608T>A), CAG1334CAA (5612G>A), TTC1335GTT (5613T>G 5615C>T), TTC1336TTT (5618C>T), CAA1337GAA (5619C>G), TGT1339TTC (5626G>T 5627T>C), AAG1340GTA (5628A>G 5629A>T 5630G>A), GAA1341AGT (5631G>A 5632A>G 5633A>T), GAA1342AAA (5634G>A), GGG1343GGA (5639G>A), CTT1344ATA (5640C>A 5642T>A), ATT1345ATA (5645T>A), TTG1346ATA (5646T>A 5648G>A), TCA1347ACC (5649T>A 5651A>C), ACA1349AAG (5656C>A 5657A>G), TTG1351ATG (5661T>A), AAA1352GAA (5664A>G), ATA1353TTA (5667A>T), GGA1354TGT (5670G>T 5672A>T), GTA1355AAA (5673G>A 5674T>A), GCA1356ACA (5676G>A), AAT1357CAT (5679A>C), GAA1359AAC (5685G>A 5687A>C), CTT1361TTA (5691C>T 5693T>A), CAA1371AAA (5721C>A), CTC1372CTA (5726C>A), TTA1377GCT (5739T>G 5740T>C 5741A>T), ATA1380GTA (5748A>G), GAA1382GAT (5756A>T), TTT1383ATG (5757T>A 5759T>G), CCT1384CCA (5762T>A), GAA1385del (5763_5765delGAA), GAC1386GAT (5768C>T), CAA1387AAA (5769C>A), GAA1389GAT (5777A>T), ACA1390AAA (5779C>A), TTA1391ACA (5781T>A 5782T>C), AAG1392AAA (5786G>A), GGT1393GAT (5788G>A), CTA1394TTA (5790C>T), CAG1395CAA (5795G>A), AAG1396AAA (5798G>A), GGG1399GGA (5807G>A), ATA1400TTA (5808A>T), TTA1401GTG (5811T>G 5813A>G), AAC1402AAT (5816C>T), GCC1404GCT (5822C>T), AAT1406AAC (5828T>C), TAT1407TTT (5830A>T), ATC1408ATA (5834C>A), CCA1409CAA (5836C>A), TTG1411TTA (5843G>A), AGT1412GGA (5844A>G 5846T>A), TAT1414ATT (5850T>A 5851A>T), ACT1415GCA (5853A>G 5855T>A), AGA1416GGA (5856A>G), ATA1417CCA (5859A>C 5860T>C), TTT1418TTA (5864T>A), AAT1420GCA (5868A>G 5869A>C 5870T>A), AAG1421AAA (5873G>A), TGT1422ATA (5874T>A 5875G>T 5876T>A), TCC1423GGA (5877T>G 5878C>G 5879C>A), AGC1424AGT (5882C>T), GAA1427CAA (5889G>C), AGA1428AAA (5893G>A), AAA1429CAC (5895A>C 5897A>C), AAT1431AAC (5903T>C), TCT1432ACA (5904T>A 5906T>A), CAA1433GAA (5907C>G), GAC1434AAT (5910G>A 5912C>T), TGG1435ATT (5913T>A 5914G>T 5915G>T), AAG1436AAA (5918G>A), ATG1437CTG (5919A>C), GTT1438GTA (5924T>A), AGA1439CAA (5925A>C 5926G>A), AAA1440CAA (5928A>C), ATC1441ATT (5933C>T), GTA1444AAA (5940G>A 5941T>A), ATT1445ATA (5945T>A), ACC1446AAA (5947C>A 5948C>A), AAA1447AAC (5951A>C), TTA1448ATA (5952T>A), CCC1449CCA (5957C>A), CCT1450GAC (5958C>G 5959C>A 5960T>C), TTG1451TTA (5963G>A), ATT1452AGC (5965T>G 5966T>C), TTA1453ATA (5967T>A), AAA1455CTA (5973A>C 5974A>T), CCA1456GAA (5976C>G 5977C>A), CAA1457TCA (5979C>T 5980A>C), AGT1458GAT (5982A>G 5983G>A), ATT1460CTA (5988A>C 5990T>A), GAG1463GAA (5999G>A) | | | | | | | | |

*: Inserts / Deletes / Misaligned / Frameshifts

#### Analysis details

This analysis was performed with panviral2.64

### NGS Details (UN70): Chinaberry tree badnavirus 1

#### Assembly

| | |
|---|---|
| **Coverage Length** | 463 (1 contig(s)) |
| **Depth Of Coverage** | 7.5 |
| **Number Of Reads** | 26 |
| **Reads Per Million** | 0.49 rpm (after QC) |
| **Ambiguities** | 0 |
| **Assembly Method** | de novo + reference guided assembly |
| **Consensus Caller** | Bcf Tools |

#### Coverage Map

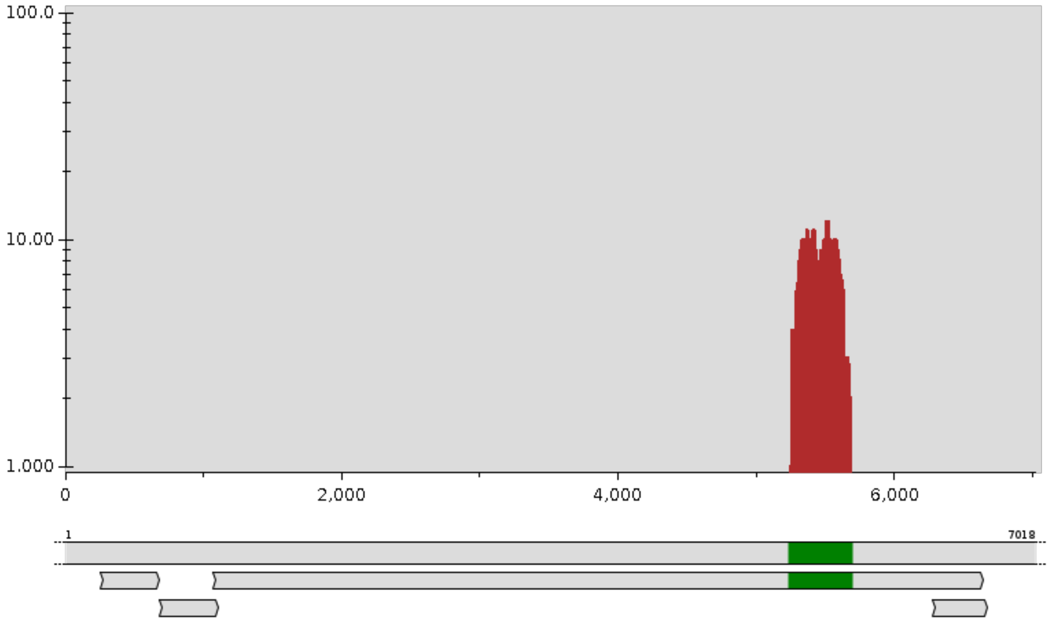

#### Assignment

| | |
|---|---|
| **Type** | Chinaberry tree badnavirus 1 (Taxonomy ID: 2908099) |
| **Reference Genome** | NC_077165.1 |
| **NT Identity (%)** | 53.9301 |
| **AA Identity (%)** | 43.7908 |
| **Number Of Stop Codons** | 0 |
| **Number Of CDS** | 4 |

#### Alignment

| | |
|---|---|
| **Alignment Score** | 22.0 (NT) + 211.0 (AA) = 233.0 |
| **Concordance (%)** | 16.1121 |

| Alignment Method | Global, seeded, nucleotide + amino acids (AGA) |
|---|---|

#### Genome Region

Sequence starts at position 5240 and ends at position 5702 relative to NC_077165.1 reference sequence.

#### Alignment Detailed Statistics

| | Begin | End | Coverage | Score | Concordance | Matches | Identities | I/D/M/F* | Stop Codons |
|---|---|---|---|---|---|---|---|---|---|
| **NT** | **5240** | **5702** | **6.6%** | **22** | **2.5%** | **455 (97.6%)** | **247 (53.0%)** | **3/8** | |
| Mutations: | 5246A>C, 5249T>C, 5250T>A, 5256T>C, 5266A>G, 5270G>A, 5273C>T, 5275T>A, 5279C>T, 5283G>A, 5285A>T, 5286G>A, 5287C>G, 5288C>G, 5292G>A, 5293A>C, 5294C>A, 5295C>A, 5296C>A, 5301T>G, 5302C>A, 5306A>C, 5307G>C, 5309A>G, 5310T>A, 5311G>A, 5312G>A, 5315A>T, 5316G>A, 5322T>A, 5325G>A, 5326T>C, 5329C>A, 5330A>T, 5331G>T, 5332A>C, 5336G>C, 5338T>A, 5339T>C, 5345A>G, 5347G>A, 5348G>C, 5351G>A, 5354T>A, 5357T>G, 5360A>C, 5363C>T, 5366G>C, 5367T>C, 5369A>C, 5371A>C, 5372A>T, 5378T>A, 5381A>T, 5382G>T, 5384T>A, 5386T>G, 5387A>T, 5390C>T, 5394A>G, 5396A>G, 5397A>C, 5398A>T, 5403G>A, 5405C>T, 5408A>T, 5409T>G, 5410G>T, 5411C>T, 5414C>T, 5419G>A, 5420_5421insCAC, 5421A>C, 5422C>T, 5423A>T, 5424G>A, 5425A>G, 5436G>T, 5437C>T, 5441C>T, 5443A>T, 5444C>T, 5445delA, 5447A>T, 5456C>T, 5457T>C, 5460G>A, 5462C>A, 5464T>A, 5465T>C, 5466T>A, 5467C>G, 5473A>G, 5474C>T, 5475G>C, 5476A>T, 5480G>A, 5483G>T, 5484C>G, 5486T>C, 5487G>T, 5488C>T, 5489A>G, 5490G>C, 5491G>A, 5492C>A, 5495T>C, 5496G>C, 5498A>G, 5501A>G, 5502A>G, 5503A>T, 5504A>T, 5506T>C, 5507G>T, 5510G>T, 5511G>A, 5513A>C, 5514A>T, 5517G>T, 5519A>G, 5521A>T, 5522G>C, 5523T>A, 5524C>A, 5525T>A, 5526A>C, 5529G>A, 5534A>G, 5535A>T, 5537C>G, 5538C>A, 5540A>C, 5542G>A, 5543C>G, 5544C>A, 5545C>A, 5546A>G, 5547A>T, 5553A>T, 5554T>G, 5555G>T, 5557G>T, 5558C>T, 5559C>T, 5563C>A, 5564C>G, 5567A>T, 5569G>A, 5571delT, 5572C>G, 5581T>A, 5582T>C, 5585A>T, 5588G>T, 5589G>C, 5590C>A, 5591A>C, 5592A>T, 5594T>C, 5597C>A, 5598A>T, 5600A>T, 5602G>C, 5605G>A, 5606A>G, 5607A>G, 5608A>G, 5609G>A, 5610C>G, 5612C>G, 5613C>G, 5614A>C, 5615A>T, 5616T>A, 5617T>C, 5619C>G, 5621G>T, 5624C>T, 5625C>A, 5627C>G, 5629T>A, 5630C>G, 5633C>T, 5634A>G, 5635A>C, 5636G>A, 5637_5639delAAG, 5641T>C, 5642G>A, 5645C>A, 5648G>A, 5649_5651delTTC, 5654T>A, 5655A>T, 5656A>G, 5660C>A, 5662G>T, 5663C>T, 5664T>C, 5665T>C, 5666A>T, 5667G>C, 5672C>T, 5674C>T, 5675T>A, 5678A>G, 5680G>A, 5681A>G, 5682C>T, 5687A>G, 5700C>T, 5702C>G | | | | | | | | |

##### CDS

| | | | | | | | | | |
|---|---|---|---|---|---|---|---|---|---|
| QKV46_gp3 | 1391 | 1544 | 8.3% | 211 | 20.1% | 152 (98.1%) | 67 (43.2%) | 1/2/2/2 | 0 |
| Protein mutations: | S1394T (5250T>A), F1396L (5256T>C), K1399R (5266A>G), F1402Y (5275T>A), V1405I (5283G>A 5285A>T), A1406R (5286G>A 5287C>G 5288C>G), D1408T (5292G>A 5293A>C 5294C>A), P1409K (5295C>A 5296C>A), S1411D (5301T>G 5302C>A), E1413Q (5307G>C 5309A>G), W1414K (5310T>A 5311G>A 5312G>A), A1416T (5316G>A), W1418R (5322T>A), V1419T (5325G>A 5326T>C), P1420H (5329C>A 5330A>T), D1421S (5331G>T 5332A>C), L1423H (5338T>A 5339T>C), W1426Y (5347G>A 5348G>C), I1429M (5357T>G), K1434T (5371A>C 5372A>T), A1438S (5382G>T 5384T>A), I1439S (5386T>G 5387A>T), R1442G (5394A>G 5396A>G), K1443L (5397A>C 5398A>T), D1445N (5403G>A 5405C>T), Q1446H (5408A>T), C1447V (5409T>G 5410G>T 5411C>T), G1450E (5419G>A), G1450_T1451insH (5420_5421insCAC), T1451L (5421A>C 5422C>T 5423A>T), E1452R (5424G>A 5425A>G), A1456L (5436G>T 5437C>T), Y1458F (5443A>T 5444C>T), V1464I (5460G>A 5462C>A), F1465Y (5464T>A 5465T>C), N1468S (5473A>G 5474C>T), E1469L (5475G>C 5476A>T), E1471D (5483G>T), H1472D (5484C>G 5486T>C), A1473L (5487G>T 5488C>T 5489A>G), G1474Q (5490G>C 5491G>A 5492C>A), V1476L (5496G>C 5498A>G), K1478V (5502A>G 5503A>T 5504A>T), M1479T (5506T>C 5507G>T), L1480F (5510G>T), E1481N (5511G>A 5513A>C), I1482L (5514A>T), V1483L (5517G>T 5519A>G), K1484I (5521A>T 5522G>C), S1485K (5523T>A 5524C>A 5525T>A), N1486H (5526A>C), G1487R (5529G>A), I1489L (5535A>T 5537C>G), L1490I (5538C>A 5540A>C), S1491K (5542G>A 5543C>G), P1492K (5544C>A 5545C>A 5546A>G), T1493S (5547A>T), M1495C (5553A>T 5554T>G 5555G>T), C1496F (5557G>T 5558C>T), A1498E (5563C>A 5564C>G), Q1499H (5567A>T), R1500Q (5569G>A), F1504Y (5581T>A 5582T>C), A1507H (5589G>C 5590C>A 5591A>C), I1508F (5592A>T 5594T>C), T1510S (5598A>T 5600A>T), G1511A (5602G>C), G1512E (5605G>A 5606A>G), K1513G (5607A>G 5608A>G 5609G>A), L1514V (5610C>G 5612C>G), Q1515A (5613C>G 5614A>C 5615A>T), L1516T (5616T>A 5617T>C), Q1517D (5619C>G 5621G>T), H1519K (5625C>A 5627C>G), I1520K (5629T>A 5630C>G), K1522A (5634A>G 5635A>C 5636G>A), K1523del (5637_5639delAAG), V1524A (5641T>C 5642G>A), F1527del (5649_5651delTTC), D1528E (5654T>A), K1529W (5655A>T 5656A>G), S1531I (5662G>T 5663C>T), L1532P (5664T>C 5665T>C 5666A>T), E1533Q (5667G>C), T1535I (5674C>T 5675T>A), G1537E (5680G>A 5681A>G) | | | | | | | | |
| Codon mutations: | ATA1392ATC (5246A>C), TTT1393TTC (5249T>C), TCC1394ACC (5250T>A), TTT1396CTT (5256T>C), AAA1399AGA (5266A>G), TCG1400TCA (5270G>A), GGC1401GGT (5273C>T), TTT1402TAT (5275T>A), CAC1403CAT (5279C>T), GTA1405ATT (5283G>A 5285A>T), GCC1406AGG (5286G>A 5287C>G 5288C>G), GAC1408ACA (5292G>A 5293A>C 5294C>A), CCG1409AAG (5295C>A 5296C>A), TCT1411GAT (5301T>G 5302C>A), GTA1412GTC (5306A>C), GAA1413CAG (5307G>C 5309A>G), TGG1414AAA (5310T>A 5311G>A 5312G>A), ACA1415ACT (5315A>T), GCA1416ACA (5316G>A), TGG1418AGG (5322T>A), GTT1419ACT (5325G>A 5326T>C), CCA1420CAT (5329C>A 5330A>T), GAT1421TCT (5331G>T 5332A>C), GGG1422GGC (5336G>C), CTT1423CAC (5338T>A 5339T>C), GAA1425GAG (5345A>G), TGG1426TAC (5347G>A 5348G>C), CTG1427CTA (5351G>A), GTT1428GTA (5354T>A), ATT1429ATG (5357T>G), CCA1430CCC (5360A>C), TTC1431TTT (5363C>T), GGG1432GGC (5366G>C), TTA1433CTC (5367T>C 5369A>C), AAA1434ACT (5371A>C 5372A>T), GCT1436GCA (5378T>A), CCA1437CCT (5381A>T), GCT1438TCA (5382G>T 5384T>A), ATA1439AGT (5386T>G 5387A>T), TTC1440TTT (5390C>T), AGA1442GGG (5394A>G 5396A>G), AAG1443CTG (5397A>C 5398A>T), GAC1445AAT (5403G>A 5405C>T), CAA1446CAT (5408A>T), TGC1447GTT (5409T>G 5410G>T 5411C>T), TTC1448TTT (5414C>T), GGA1450GAA (5419G>A), GGA1450_ACA1451insCAC (5420_5421insCAC), ACA1451CTT (5421A>C 5422C>T 5423A>T), GAG1452AGG (5424G>A 5425A>G), GCG1456TTG (5436G>T 5437C>T), GTC1457GTT (5441C>T), TAC1458TTT (5443A>T 5444C>T), ATA1459-TT (5445delA 5447A>T), ATC1462ATT (5456C>T), TTG1463CTG (5457T>C), GTC1464ATA (5460G>A 5462C>A), TTT1465TAC (5464T>A 5465T>C), TCC1466AGC (5466T>A 5467C>G), AAC1468AGT (5473A>G 5474C>T), GAA1469CTA (5475G>C 5476A>T), GAG1470GAA (5480G>A), GAG1471GAT (5483G>T), CAT1472GAC (5484C>G 5486T>C), GCA1473TTG (5487G>T 5488C>T 5489A>G), GGC1474CAA (5490G>C 5491G>A 5492C>A), CAT1475CAC (5495T>C), GTA1476CTG (5496G>C 5498A>G), AGA1477AGG (5501A>G), AAA1478GTT (5502A>G 5503A>T 5504A>T), ATG1479ACT (5506T>C 5507G>T), TTG1480TTT (5510G>T), GAA1481AAC (5511G>A 5513A>C), ATA1482TTA (5514A>T), GTA1483TTG (5517G>T 5519A>G), AAG1484ATC (5521A>T 5522G>C), TCT1485AAA (5523T>A 5524C>A 5525T>A), AAT1486CAT (5526A>C), GGA1487AGA (5529G>A), CTA1488CTG (5534A>G), ATC1489TTG (5535A>T 5537C>G), CTA1490ATC (5538C>A 5540A>C), AGC1491AAG (5542G>A 5543C>G), CCA1492AAG (5544C>A 5545C>A 5546A>G), ACA1493TCA (5547A>T), ATG1495TGT (5553A>T 5554T>G 5555G>T), TGC1496TTT (5557G>T 5558C>T), CTG1497TTG (5559C>T), GCC1498GAG (5563C>A 5564C>G), CAA1499CAT (5567A>T), CGA1500CAA (5569G>A), TCG1501-GG (5571delT 5572C>G), TTT1504TAC (5581T>A 5582T>C), CTA1505CTT (5585A>T), GGG1506GGT (5588G>T), GCA1507CAC (5589G>C 5590C>A 5591A>C), ATT1508TTC (5592A>T 5594T>C), ATC1509ATA (5597C>A), ACA1510TCT (5598A>T 5600A>T), GGA1511GCA (5602G>C), GGA1512GAG (5605G>A 5606A>G), AAG1513GGA (5607A>G 5608A>G 5609G>A), CTC1514GTG (5610C>G 5612C>G), CAA1515GCT (5613C>G 5614A>C 5615A>T), TTA1516ACA (5616T>A 5617T>C), CAG1517GAT (5619C>G 5621G>T), CCC1518CCT (5624C>T), CAC1519AAG (5625C>A 5627C>G), ATC1520AAG (5629T>A 5630C>G), ATC1521ATT (5633C>T), AAG1522GCA (5634A>G 5635A>C 5636G>A), AAG1523del (5637_5639delAAG), GTG1524GCA (5641T>C 5642G>A), GTC1525GTA (5645C>A), CAG1526CAA (5648G>A), TTC1527del (5649_5651delTTC), GAT1528GAA (5654T>A), AAG1529TGG (5655A>T 5656A>G), CCC1530CCA (5660C>A), AGC1531ATT (5662G>T 5663C>T), TTA1532CCT (5664T>C 5665T>C 5666A>T), GAG1533CAG (5667G>C), ACC1534ACT (5672C>T), ACT1535ATA (5674C>T 5675T>A), AAA1536AAG (5678A>G), GGA1537GAG (5680G>A 5681A>G), CTA1538TTA (5682C>T), AGA1539AGG (5687A>G), CTC1544TTG (5700C>T 5702C>G) | | | | | | | | |

##### Proteins

| | | | | | | | | | |
|---|---|---|---|---|---|---|---|---|---|
| ORF3 (YP_010805662.1) | 1391 | 1544 | 8.3% | 211 | 20.1% | 152 (98.1%) | 67 (43.2%) | 1/2/2/2 | 0 |
| Protein mutations: | S1394T (5250T>A), F1396L (5256T>C), K1399R (5266A>G), F1402Y (5275T>A), V1405I (5283G>A 5285A>T), A1406R (5286G>A 5287C>G 5288C>G), D1408T (5292G>A 5293A>C 5294C>A), P1409K (5295C>A 5296C>A), S1411D (5301T>G 5302C>A), E1413Q (5307G>C 5309A>G), W1414K (5310T>A 5311G>A 5312G>A), A1416T (5316G>A), W1418R (5322T>A), V1419T (5325G>A 5326T>C), P1420H (5329C>A 5330A>T), D1421S (5331G>T 5332A>C), L1423H (5338T>A 5339T>C), W1426Y (5347G>A 5348G>C), I1429M (5357T>G), K1434T (5371A>C 5372A>T), A1438S (5382G>T 5384T>A), I1439S (5386T>G 5387A>T), R1442G (5394A>G 5396A>G), K1443L (5397A>C 5398A>T), D1445N (5403G>A 5405C>T), Q1446H (5408A>T), C1447V (5409T>G 5410G>T 5411C>T), G1450E (5419G>A), G1450_T1451insH (5420_5421insCAC), T1451L (5421A>C 5422C>T 5423A>T), E1452R (5424G>A 5425A>G), A1456L (5436G>T 5437C>T), Y1458F (5443A>T 5444C>T), V1464I (5460G>A 5462C>A), F1465Y (5464T>A 5465T>C), N1468S (5473A>G 5474C>T), E1469L (5475G>C 5476A>T), E1471D (5483G>T), H1472D (5484C>G 5486T>C), A1473L (5487G>T 5488C>T 5489A>G), G1474Q (5490G>C 5491G>A 5492C>A), V1476L (5496G>C 5498A>G), K1478V (5502A>G 5503A>T 5504A>T), M1479T (5506T>C 5507G>T), L1480F (5510G>T), E1481N (5511G>A 5513A>C), I1482L (5514A>T), V1483L (5517G>T 5519A>G), K1484I (5521A>T 5522G>C), S1485K (5523T>A 5524C>A 5525T>A), N1486H (5526A>C), G1487R (5529G>A), I1489L (5535A>T 5537C>G), L1490I (5538C>A 5540A>C), S1491K (5542G>A 5543C>G), P1492K (5544C>A 5545C>A 5546A>G), T1493S (5547A>T), M1495C (5553A>T 5554T>G 5555G>T), C1496F (5557G>T 5558C>T), A1498E (5563C>A 5564C>G), Q1499H (5567A>T), R1500Q (5569G>A), F1504Y (5581T>A 5582T>C), A1507H (5589G>C 5590C>A 5591A>C), I1508F (5592A>T 5594T>C), T1510S (5598A>T 5600A>T), G1511A (5602G>C), G1512E (5605G>A 5606A>G), K1513G (5607A>G 5608A>G 5609G>A), L1514V (5610C>G 5612C>G), Q1515A (5613C>G 5614A>C 5615A>T), L1516T (5616T>A 5617T>C), Q1517D (5619C>G 5621G>T), H1519K (5625C>A 5627C>G), I1520K (5629T>A 5630C>G), K1522A (5634A>G 5635A>C 5636G>A), K1523del (5637_5639delAAG), V1524A (5641T>C 5642G>A), F1527del (5649_5651delTTC), D1528E (5654T>A), K1529W (5655A>T 5656A>G), S1531I (5662G>T 5663C>T), L1532P (5664T>C 5665T>C 5666A>T), E1533Q (5667G>C), T1535I (5674C>T 5675T>A), G1537E (5680G>A 5681A>G) | | | | | | | | |

| | Begin | End | Coverage | Score | Concordance | Matches | Identities | I/D/M/F* | Stop Codons |
|---|---|---|---|---|---|---|---|---|---|
| **NT** | **5240** | **5702** | **6.6%** | **22** | **2.5%** | **455 (97.6%)** | **247 (53.0%)** | **3/8** | |
| Codon mutations: | ATA1392ATC (5246A>C), TTT1393TTC (5249T>C), TCC1394ACC (5250T>A), TTT1396CTT (5256T>C), AAA1399AGA (5266A>G), TCG1400TCA (5270G>A), GGC1401GGT (5273C>T), TTT1402TAT (5275T>A), CAC1403CAT (5279C>T), GTA1405ATT (5283G>A 5285A>T), GCC1406AGG (5286G>A 5287C>G 5288C>G), GAC1408ACA (5292G>A 5293A>C 5294C>A), CCG1409AAG (5295C>A 5296C>A), TCT1411GAT (5301T>G 5302C>A), GTA1412GTC (5306A>C), GAA1413CAG (5307G>C 5309A>G), TGG1414AAA (5310T>A 5311G>A 5312G>A), ACA1415ACT (5315A>T), GCA1416ACA (5316G>A), TGG1418AGG (5322T>A), GTT1419ACT (5325G>A 5326T>C), CCA1420CAT (5329C>A 5330A>T), GAT1421TCT (5331G>T 5332A>C), GGG1422GGC (5336G>C), CTT1423CAC (5338T>A 5339T>C), GAA1425GAG (5345A>G), TGG1426TAC (5347G>A 5348G>C), CTG1427CTA (5351G>A), GTT1428GTA (5354T>A), ATT1429ATG (5357T>G), CCA1430CCC (5360A>C), TTC1431TTT (5363C>T), GGG1432GGC (5366G>C), TTA1433CTC (5367T>C 5369A>C), AAA1434ACT (5371A>C 5372A>T), GCT1436GCA (5378T>A), CCA1437CCT (5381A>T), GCT1438TCA (5382G>T 5384T>A), ATA1439AGT (5386T>G 5387A>T), TTC1440TTT (5390C>T), AGA1442GGG (5394A>G 5396A>G), AAG1443CTG (5397A>C 5398A>T), GAC1445AAT (5403G>A 5405C>T), CAA1446CAT (5408A>T), TGC1447GTT (5409T>G 5410G>T 5411C>T), TTC1448TTT (5414C>T), GGA1450GAA (5419G>A), GGA1450_ACA1451insCAC (5420_5421insCAC), ACA1451CTT (5421A>C 5422C>T 5423A>T), GAG1452AGG (5424G>A 5425A>G), GCG1456TTG (5436G>T 5437C>T), GTC1457GTT (5441C>T), TAC1458TTT (5443A>T 5444C>T), ATA1459-TT (5445delA 5447A>T), ATC1462ATT (5456C>T), TTG1463CTG (5457T>C), GTC1464ATA (5460G>A 5462C>A), TTT1465TAC (5464T>A 5465T>C), TCC1466AGC (5466T>A 5467C>G), AAC1468AGT (5473A>G 5474C>T), GAA1469CTA (5475G>C 5476A>T), GAG1470GAA (5480G>A), GAG1471GAT (5483G>T), CAT1472GAC (5484C>G 5486T>C), GCA1473TTG (5487G>T 5488C>T 5489A>G), GGC1474CAA (5490G>C 5491G>A 5492C>A), CAT1475CAC (5495T>C), GTA1476CTG (5496G>C 5498A>G), AGA1477AGG (5501A>G), AAA1478GTT (5502A>G 5503A>T 5504A>T), ATG1479ACT (5506T>C 5507G>T), TTG1480TTT (5510G>T), GAA1481AAC (5511G>A 5513A>C), ATA1482TTA (5514A>T), GTA1483TTG (5517G>T 5519A>G), AAG1484ATC (5521A>T 5522G>C), TCT1485AAA (5523T>A 5524C>A 5525T>A), AAT1486CAT (5526A>C), GGA1487AGA (5529G>A), CTA1488CTG (5534A>G), ATC1489TTG (5535A>T 5537C>G), CTA1490ATC (5538C>A 5540A>C), AGC1491AAG (5542G>A 5543C>G), CCA1492AAG (5544C>A 5545C>A 5546A>G), ACA1493TCA (5547A>T), ATG1495TGT (5553A>T 5554T>G 5555G>T), TGC1496TTT (5557G>T 5558C>T), CTG1497TTG (5559C>T), GCC1498GAG (5563C>A 5564C>G), CAA1499CAT (5567A>T), CGA1500CAA (5569G>A), TCG1501-GG (5571delT 5572C>G), TTT1504TAC (5581T>A 5582T>C), CTA1505CTT (5585A>T), GGG1506GGT (5588G>T), GCA1507CAC (5589G>C 5590C>A 5591A>C), ATT1508TTC (5592A>T 5594T>C), ATC1509ATA (5597C>A), ACA1510TCT (5598A>T 5600A>T), GGA1511GCA (5602G>C), GGA1512GAG (5605G>A 5606A>G), AAG1513GGA (5607A>G 5608A>G 5609G>A), CTC1514GTG (5610C>G 5612C>G), CAA1515GCT (5613C>G 5614A>C 5615A>T), TTA1516ACA (5616T>A 5617T>C), CAG1517GAT (5619C>G 5621G>T), CCC1518CCT (5624C>T), CAC1519AAG (5625C>A 5627C>G), ATC1520AAG (5629T>A 5630C>G), ATC1521ATT (5633C>T), AAG1522GCA (5634A>G 5635A>C 5636G>A), AAG1523del (5637_5639delAAG), GTG1524GCA (5641T>C 5642G>A), GTC1525GTA (5645C>A), CAG1526CAA (5648G>A), TTC1527del (5649_5651delTTC), GAT1528GAA (5654T>A), AAG1529TGG (5655A>T 5656A>G), CCC1530CCA (5660C>A), AGC1531ATT (5662G>T 5663C>T), TTA1532CCT (5664T>C 5665T>C 5666A>T), GAG1533CAG (5667G>C), ACC1534ACT (5672C>T), ACT1535ATA (5674C>T 5675T>A), AAA1536AAG (5678A>G), GGA1537GAG (5680G>A 5681A>G), CTA1538TTA (5682C>T), AGA1539AGG (5687A>G), CTC1544TTG (5700C>T 5702C>G) | | | | | | | | |

*: Inserts / Deletes / Misaligned / Frameshifts

#### Analysis details

This analysis was performed with panviral2.64

### NGS Details (UN70): Duamitovirus peex1

#### Assembly

| | |
|---|---|
| **Coverage Length** | 380 (1 contig(s)) |
| **Depth Of Coverage** | 5.8 |
| **Number Of Reads** | 22 |
| **Reads Per Million** | 0.42 rpm (after QC) |
| **Ambiguities** | 0 |
| **Assembly Method** | de novo + reference guided assembly |
| **Consensus Caller** | Bcf Tools |

#### Coverage Map

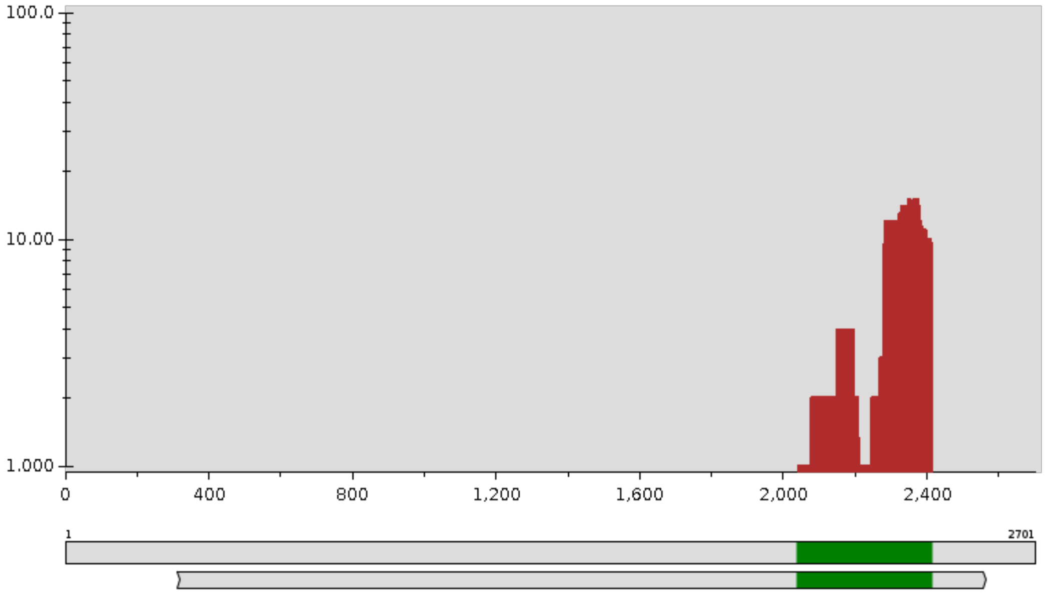

#### Assignment

| | |
|---|---|
| **Type** | Duamitovirus peex1 (Taxonomy ID: 2955799) |
| **Reference Genome** | NC_076525.1 |
| **NT Identity (%)** | 71.1286 |
| **AA Identity (%)** | 66.9291 |
| **Number Of Stop Codons** | 0 |
| **Number Of CDS** | 1 |

#### Alignment

| | |
|---|---|
| **Alignment Score** | 314.0 (NT) + 503.0 (AA) = 817.0 |
| **Concordance (%)** | 49.9408 |

| Alignment Method | Global, seeded, nucleotide + amino acids (AGA) |
|---|---|

#### Genome Region

Sequence starts at position 2038 and ends at position 2417 relative to NC_076525.1 reference sequence.

#### Alignment Detailed Statistics

| | Begin | End | Coverage | Score | Concordance | Matches | Identities | I/D/M/F* | Stop Codons |
|---|---|---|---|---|---|---|---|---|---|
| **NT** | **2038** | **2417** | **14.1%** | **314** | **41.3%** | **380 (99.7%)** | **271 (71.1%)** | **1/0** | |
| Mutations: | 2040A>T, 2042G>T, 2048G>A, 2057C>T, 2066G>A, 2067A>C, 2069T>G, 2072G>A, 2075T>G, 2077T>C, 2081T>A, 2082T>A, 2084T>C, 2088T>C, 2090T>A, 2091T>C, 2095T>G, 2096A>G, 2102G>T, 2104G>T, 2105G>A, 2109T>C, 2114T>C, 2119G>A, 2120G>T, 2129A>G, 2132G>A, 2133G>T, 2134G>T, 2135A>T, 2138G>A, 2141C>T, 2145G>A, 2152A>T, 2153_2154insT, 2156A>G, 2158G>A, 2159A>G, 2162T>A, 2171C>A, 2173C>A, 2180T>C, 2183T>G, 2189G>A, 2194A>C, 2209A>G, 2210G>A, 2211T>C, 2213G>C, 2214A>T, 2216A>T, 2222A>T, 2223C>G, 2225A>G, 2226C>T, 2234A>T, 2243A>G, 2244C>A, 2245A>G, 2246T>A, 2249G>A, 2255A>T, 2268G>T, 2272G>T, 2273T>C, 2274C>A, 2275A>G, 2276T>A, 2280A>G, 2283G>A, 2294T>G, 2295C>A, 2296G>A, 2301C>G, 2303T>C, 2306A>T, 2311A>G, 2312T>C, 2315C>T, 2321G>T, 2325A>G, 2326A>C, 2327T>A, 2330C>A, 2331C>G, 2333G>T, 2339G>A, 2345A>T, 2348C>T, 2351A>G, 2352C>T, 2354T>C, 2363G>T, 2373G>A, 2374C>G, 2375G>T, 2377C>A, 2378T>A, 2379T>A, 2388C>A, 2390T>A, 2393T>A, 2394C>A, 2396G>C, 2398C>A, 2399T>A, 2406C>G, 2411A>G, 2412G>A, 2414G>C | | | | | | | | |
| **CDS** | | | | | | | | | |
| RdRp | 577 | 702 | 16.8% | 503 | 53.9% | 126 (99.2%) | 85 (66.9%) | 1/0/1/1 | 0 |
| Protein mutations: | T577S (2040A>T 2042G>T), I586L (2067A>C 2069T>G), V589A (2077T>C), C591S (2082T>A 2084T>C), S593P (2088T>C 2090T>A), Y594H (2091T>C), I595R (2095T>G 2096A>G), Q597H (2102G>T), R598L (2104G>T 2105G>A), S600P (2109T>C), W603Y (2119G>A 2120G>T), G608F (2133G>T 2134G>T 2135A>T), D612N (2145G>A), Y614F (2152A>T), Y614_L615insX (2153_2154insT), R616K (2158G>A 2159A>G), A621D (2173C>A), K628T (2194A>C), Q633R (2209A>G 2210G>A), I635F (2214A>T 2216A>T), K637N (2222A>T), Q638E (2223C>G 2225A>G), L641F (2234A>T), H645R (2244C>A 2245A>G 2246T>A), V653L (2268G>T), R654L (2272G>T 2273T>C), H655R (2274C>A 2275A>G 2276T>A), M657V (2280A>G), E658K (2283G>A), R662K (2295C>A 2296G>A), L664V (2301C>G 2303T>C), Y667C (2311A>G 2312T>C), N672A (2325A>G 2326A>C 2327T>A), P674A (2331C>G 2333G>T), L681F (2352C>T 2354T>C), A688S (2373G>A 2374C>G 2375G>T), T689K (2377C>A 2378T>A), S690T (2379T>A), Q695N (2394C>A 2396G>C), T696K (2398C>A 2399T>A), H699D (2406C>G), V701I (2412G>A 2414G>C) | | | | | | | | |
| Codon mutations: | ACG577TCT (2040A>T 2042G>T), CAG579CAA (2048G>A), CGC582CGT (2057C>T), AGG585AGA (2066G>A), ATT586CTG (2067A>C 2069T>G), AAG587AAA (2072G>A), GCT588GCG (2075T>G), GTT589GCT (2077T>C), GCT590GCA (2081T>A), TGT591AGC (2082T>A 2084T>C), TCT593CCA (2088T>C 2090T>A), TAC594CAC (2091T>C), ATA595AGG (2095T>G 2096A>G), CAG597CAT (2102G>T), CGG598CTA (2104G>T 2105G>A), TCT600CCT (2109T>C), CTT601CTC (2114T>C), TGG603TAT (2119G>A 2120G>T), GGA606GGG (2129A>G), AGG607AGA (2132G>A), GGA608TTT (2133G>T 2134G>T 2135A>T), AAG609AAA (2138G>A), CCC610CCT (2141C>T), GAT612AAT (2145G>A), TAT614TTT (2152A>T), TAT614_TTA615insT-- (2153_2154insT), TTA615TTG (2156A>G), AGA616AAG (2158G>A 2159A>G), GGT617GGA (2162T>A), ATC620ATA (2171C>A), GCT621GAT (2173C>A), CTT623CTC (2180T>C), CGT624CGG (2183T>G), GAG626GAA (2189G>A), AAA628ACA (2194A>C), CAG633CGA (2209A>G 2210G>A), TTG634CTC (2211T>C 2213G>C), ATA635TTT (2214A>T 2216A>T), AAA637AAT (2222A>T), CAA638GAG (2223C>G 2225A>G), CTG639TTG (2226C>T), TTA641TTT (2234A>T), GAA644GAG (2243A>G), CAT645AGA (2244C>A 2245A>G 2246T>A), GAG646GAA (2249G>A), CTA648CTT (2255A>T), GTA653TTA (2268G>T), CGT654CTC (2272G>T 2273T>C), CAT655AGA (2274C>A 2275A>G 2276T>A), ATG657GTG (2280A>G), GAA658AAA (2283G>A), CTT661CTG (2294T>G), CGA662AAA (2295C>A 2296G>A), CTT664GTC (2301C>G 2303T>C), TCA665TCT (2306A>T), TAT667TGC (2311A>G 2312T>C), CAC668CAT (2315C>T), GTG670GTT (2321G>T), AAT672GCA (2325A>G 2326A>C 2327T>A), TCC673TCA (2330C>A), CCG674GCT (2331C>G 2333G>T), GTG676GTA (2339G>A), ATA678ATT (2345A>T), GAC679GAT (2348C>T), CAA680CAG (2351A>G), CTT681TTC (2352C>T 2354T>C), GTG684GTT (2363G>T), GCG688AGT (2373G>A 2374C>G 2375G>T), ACT689AAA (2377C>A 2378T>A), TCT690ACT (2379T>A), CGT693AGA (2388C>A 2390T>A), ACT694ACA (2393T>A), CAG695AAC (2394C>A 2396G>C), ACT696AAA (2398C>A 2399T>A), CAT699GAT (2406C>G), TTA700TTG (2411A>G), GTG701ATC (2412G>A 2414G>C) | | | | | | | | |
| **Proteins** | | | | | | | | | |
| RNA-dependent RNA polymerase (YP_010798875.1) | 577 | 702 | 16.8% | 503 | 53.9% | 126 (99.2%) | 85 (66.9%) | 1/0/1/1 | 0 |
| Protein mutations: | T577S (2040A>T 2042G>T), I586L (2067A>C 2069T>G), V589A (2077T>C), C591S (2082T>A 2084T>C), S593P (2088T>C 2090T>A), Y594H (2091T>C), I595R (2095T>G 2096A>G), Q597H (2102G>T), R598L (2104G>T 2105G>A), S600P (2109T>C), W603Y (2119G>A 2120G>T), G608F (2133G>T 2134G>T 2135A>T), D612N (2145G>A), Y614F (2152A>T), Y614_L615insX (2153_2154insT), R616K (2158G>A 2159A>G), A621D (2173C>A), K628T (2194A>C), Q633R (2209A>G 2210G>A), I635F (2214A>T 2216A>T), K637N (2222A>T), Q638E (2223C>G 2225A>G), L641F (2234A>T), H645R (2244C>A 2245A>G 2246T>A), V653L (2268G>T), R654L (2272G>T 2273T>C), H655R (2274C>A 2275A>G 2276T>A), M657V (2280A>G), E658K (2283G>A), R662K (2295C>A 2296G>A), L664V (2301C>G 2303T>C), Y667C (2311A>G 2312T>C), N672A (2325A>G 2326A>C 2327T>A), P674A (2331C>G 2333G>T), L681F (2352C>T 2354T>C), A688S (2373G>A 2374C>G 2375G>T), T689K (2377C>A 2378T>A), S690T (2379T>A), Q695N (2394C>A 2396G>C), T696K (2398C>A 2399T>A), H699D (2406C>G), V701I (2412G>A 2414G>C) | | | | | | | | |
| Codon mutations: | ACG577TCT (2040A>T 2042G>T), CAG579CAA (2048G>A), CGC582CGT (2057C>T), AGG585AGA (2066G>A), ATT586CTG (2067A>C 2069T>G), AAG587AAA (2072G>A), GCT588GCG (2075T>G), GTT589GCT (2077T>C), GCT590GCA (2081T>A), TGT591AGC (2082T>A 2084T>C), TCT593CCA (2088T>C 2090T>A), TAC594CAC (2091T>C), ATA595AGG (2095T>G 2096A>G), CAG597CAT (2102G>T), CGG598CTA (2104G>T 2105G>A), TCT600CCT (2109T>C), CTT601CTC (2114T>C), TGG603TAT (2119G>A 2120G>T), GGA606GGG (2129A>G), AGG607AGA (2132G>A), GGA608TTT (2133G>T 2134G>T 2135A>T), AAG609AAA (2138G>A), CCC610CCT (2141C>T), GAT612AAT (2145G>A), TAT614TTT (2152A>T), TAT614_TTA615insT-- (2153_2154insT), TTA615TTG (2156A>G), AGA616AAG (2158G>A 2159A>G), GGT617GGA (2162T>A), ATC620ATA (2171C>A), GCT621GAT (2173C>A), CTT623CTC (2180T>C), CGT624CGG (2183T>G), GAG626GAA (2189G>A), AAA628ACA (2194A>C), CAG633CGA (2209A>G 2210G>A), TTG634CTC (2211T>C 2213G>C), ATA635TTT (2214A>T 2216A>T), AAA637AAT (2222A>T), CAA638GAG (2223C>G 2225A>G), CTG639TTG (2226C>T), TTA641TTT (2234A>T), GAA644GAG (2243A>G), CAT645AGA (2244C>A 2245A>G 2246T>A), GAG646GAA (2249G>A), CTA648CTT (2255A>T), GTA653TTA (2268G>T), CGT654CTC (2272G>T 2273T>C), CAT655AGA (2274C>A 2275A>G 2276T>A), ATG657GTG (2280A>G), GAA658AAA (2283G>A), CTT661CTG (2294T>G), CGA662AAA (2295C>A 2296G>A), CTT664GTC (2301C>G 2303T>C), TCA665TCT (2306A>T), TAT667TGC (2311A>G 2312T>C), CAC668CAT (2315C>T), GTG670GTT (2321G>T), AAT672GCA (2325A>G 2326A>C 2327T>A), TCC673TCA (2330C>A), CCG674GCT (2331C>G 2333G>T), GTG676GTA (2339G>A), ATA678ATT (2345A>T), GAC679GAT (2348C>T), CAA680CAG (2351A>G), CTT681TTC (2352C>T 2354T>C), GTG684GTT (2363G>T), GCG688AGT (2373G>A 2374C>G 2375G>T), ACT689AAA (2377C>A 2378T>A), TCT690ACT (2379T>A), CGT693AGA (2388C>A 2390T>A), ACT694ACA (2393T>A), CAG695AAC (2394C>A 2396G>C), ACT696AAA (2398C>A 2399T>A), CAT699GAT (2406C>G), TTA700TTG (2411A>G), GTG701ATC (2412G>A 2414G>C) | | | | | | | | |

*: Inserts / Deletes / Misaligned / Frameshifts

#### Analysis details

This analysis was performed with panviral2.64

#### NGS Details (UN70): Badnavirus deltainflatheobromae

##### Assembly

| | |
|---|---|
| **Coverage Length** | 315 (1 contig(s)) |
| **Depth Of Coverage** | 6.1 |
| **Number Of Reads** | 20 |
| **Reads Per Million** | 0.38 rpm (after QC) |
| **Ambiguities** | 0 |
| **Assembly Method** | de novo + reference guided assembly |
| **Consensus Caller** | Bcf Tools |

##### Coverage Map

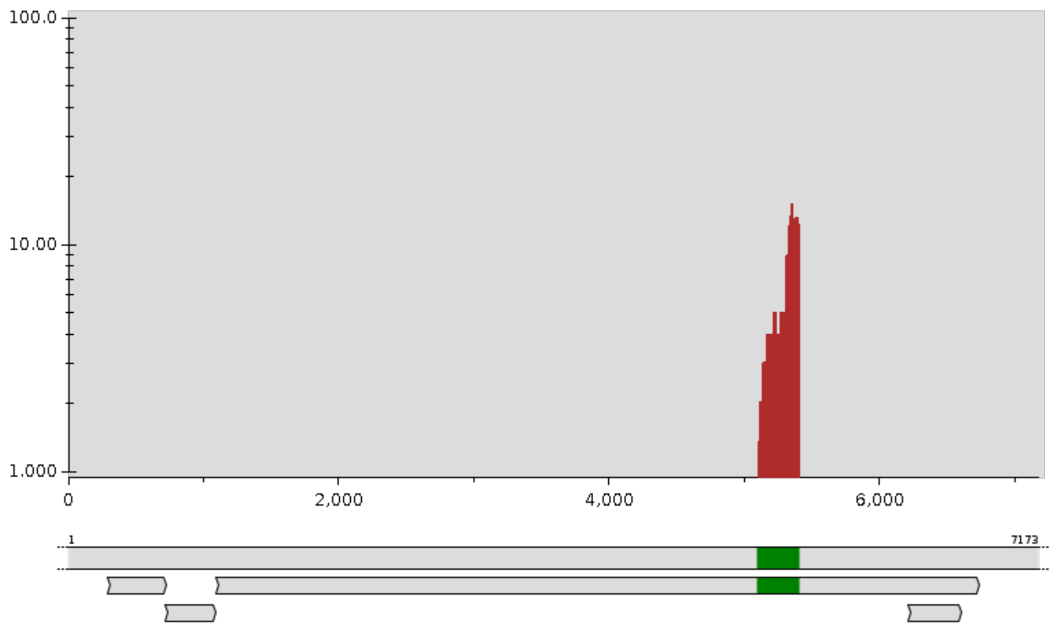

##### Assignment

| | |
|---|---|
| **Type** | Badnavirus deltainflatheobromae (Taxonomy ID: 3047710) |
| **Reference Genome** | NC_040622.1 |
| **NT Identity (%)** | 52.2013 |
| **AA Identity (%)** | 49.0566 |
| **Number Of Stop Codons** | 0 |
| **Number Of CDS** | 4 |

##### Alignment

| | |
|---|---|
| **Alignment Score** | 22.0 (NT) + 381.0 (AA) = 403.0 |
| **Concordance (%)** | 29.1606 |

| Alignment Method | Global, seeded, nucleotide + amino acids (AGA) |
|---|---|

#### Genome Region

Sequence starts at position 5098 and ends at position 5412 relative to NC_040622.1 reference sequence.

#### Alignment Detailed Statistics

| | Begin | End | Coverage | Score | Concordance | Matches | Identities | I/D/M/F* | Stop Codons |
|---|---|---|---|---|---|---|---|---|---|
| **NT** | **5098** | **5412** | **4.4%** | **22** | **3.5%** | **315 (99.1%)** | **166 (52.2%)** | **3/0** | |
| Mutations: | 5098A>G, 5099A>G, 5100A>G, 5101G>A, 5105G>A, 5109G>A, 5110A>C, 5114T>C, 5117G>T, 5119A>T, 5120A>C, 5123A>G, 5126C>T, 5127C>A, 5131A>T, 5133T>C, 5135C>A, 5136C>A, 5143G>T, 5144A>T, 5147C>T, 5148C>A, 5150A>G, 5151A>G, 5152C>A, 5153T>A, 5154A>T, 5157C>T, 5159C>A, 5160A>G, 5162G>T, 5163A>G, 5165A>G, 5166A>T, 5168T>G, 5169G>C, 5170G>A, 5171C>T, 5172A>G, 5173A>G, 5174C>A, 5175A>G, 5176A>C, 5177A>C, 5178C>A, 5181G>T, 5182T>A, 5186C>T, 5187A>T, 5188G>C, 5193T>C, 5195T>A, 5198T>C, 5199C>T, 5203A>G, 5207C>T, 5210G>T, 5212T>A, 5213T>C, 5216T>C, 5219A>G, 5220G>A, 5222T>A, 5223G>A, 5224C>G, 5225T>A, 5229G>T, 5230C>T, 5231A>T, 5235G>T, 5236A>C, 5237G>T, 5238T>A, 5239C>A, 5240C>T, 5243C>A, 5246C>T, 5247T>A, 5248G>A, 5249G>A, 5252T>G, 5253G>A, 5255C>A, 5258T>C, 5259T>A, 5261G>A, 5262G>A, 5263T>C, 5264T>C, 5266C>A, 5267A>T, 5273C>G, 5275T>A, 5276A>C, 5282A>G, 5284G>T, 5285G>C, 5286C>T, 5288T>A, 5291T>G, 5297A>C, 5303G>C, 5308A>C, 5309G>C, 5315A>C, 5318A>C, 5319G>T, 5321A>C, 5323T>C, 5324C>A, 5327C>T, 5330A>G, 5333A>C, 5334A>C, 5335A>T, 5336A>C, 5340G>A, 5345G>A, 5346T>A, 5347G>T, 5348C>A, 5351T>C, 5354G>T, 5356G>C, 5357A>T, 5357_5358insCAT, 5358A>C, 5359C>T, 5360A>T, 5361G>A, 5362A>G, 5364G>A, 5366T>A, 5369C>T, 5373G>C, 5374C>T, 5375A>G, 5378C>T, 5380A>T, 5382A>T, 5384C>T, 5393C>A, 5394T>C, 5396G>T, 5397G>A, 5399C>A, 5401T>A, 5404G>A, 5411C>T, 5412A>T | | | | | | | | |

##### CDS

| | Begin | End | Coverage | Score | Concordance | Matches | Identities | I/D/M/F* | Stop Codons |
|---|---|---|---|---|---|---|---|---|---|
| EXJ76_gp3 | 1337 | 1441 | 5.6% | 381 | 50.0% | 105 (99.1%) | 52 (49.1%) | 1/0/0/0 | 0 |
| Protein mutations: | R1337E (5100A>G 5101G>A), D1340T (5109G>A 5110A>C), E1343V (5119A>T 5120A>C), Q1346K (5127C>A), Y1347F (5131A>T), S1348P (5133T>C 5135C>A), L1349I (5136C>A), G1351V (5143G>T 5144A>T), Q1353K (5148C>A 5150A>G), T1354E (5151A>G 5152C>A 5153T>A), I1355L (5154A>T), K1357D (5160A>G 5162G>T), K1358E (5163A>G 5165A>G), I1359L (5166A>T 5168T>G), G1360H (5169G>C 5170G>A 5171C>T), N1361G (5172A>G 5173A>G 5174C>A), K1362A (5175A>G 5176A>C 5177A>C), Q1363K (5178C>A), V1364Y (5181G>T 5182T>A), F1368L (5193T>C 5195T>A), K1371R (5203A>G), F1374Y (5212T>A 5213T>C), V1377I (5220G>A 5222T>A), A1378R (5223G>A 5224C>G 5225T>A), A1380F (5229G>T 5230C>T 5231A>T), E1382S (5235G>T 5236A>C 5237G>T), S1383N (5238T>A 5239C>A 5240C>T), W1386K (5247T>A 5248G>A 5249G>A), A1388T (5253G>A 5255C>A), W1390R (5259T>A 5261G>A), V1391T (5262G>A 5263T>C 5264T>C), P1392H (5266C>A 5267A>T), L1395H (5275T>A 5276A>C), W1398F (5284G>T 5285G>C), K1406T (5308A>C 5309G>C), A1410S (5319G>T 5321A>C), I1411T (5323T>C 5324C>A), R1414S (5333A>C), K1415L (5334A>C 5335A>T 5336A>C), D1417N (5340G>A), C1419I (5346T>A 5347G>T 5348C>A), K1421N (5354G>T), G1422A (5356G>C 5357A>T), G1422_T1423insH (5357_5358insCAT), T1423L (5358A>C 5359C>T 5360A>T), E1424R (5361G>A 5362A>G), D1425K (5364G>A 5366T>A), A1428L (5373G>C 5374C>T 5375A>G), Y1430F (5380A>T), I1431F (5382A>T 5384C>T), V1436I (5397G>A 5399C>A), F1437Y (5401T>A), S1438N (5404G>A) | | | | | | | | |
| Codon mutations: | AAA1336.GG (5098A>G 5099A>G), AGA1337GAA (5100A>G 5101G>A), TTG1338TTA (5105G>A), GAC1340ACC (5109G>A 5110A>C), AAT1341AAC (5114T>C), ACG1342ACT (5117G>T), GAA1343GTC (5119A>T 5120A>C), AAA1344AAG (5123A>G), GAC1345GAT (5126C>T), CAG1346AAG (5127C>A), TAT1347TTT (5131A>T), TCC1348CCA (5133T>C 5135C>A), CTT1349ATT (5136C>A), GGA1351GTT (5143G>T 5144A>T), ATC1352ATT (5147C>T), CAA1353AAG (5148C>A 5150A>G), ACT1354GAA (5151A>G 5152C>A 5153T>A), ATA1355TTA (5154A>T), CTC1356TTA (5157C>T 5159C>A), AAG1357GAT (5160A>G 5162G>T), AAA1358GAG (5163A>G 5165A>G), ATT1359TTG (5166A>T 5168T>G), GGC1360CAT (5169G>C 5170G>A 5171C>T), AAC1361GGA (5172A>G 5173A>G 5174C>A), AAA1362GCC (5175A>G 5176A>C 5177A>C), CAG1363AAG (5178C>A), GTT1364TAT (5181G>T 5182T>A), TTC1365TTT (5186C>T), AGT1366TCT (5187A>T 5188G>C), TTT1368CTA (5193T>C 5195T>A), GAT1369GAC (5198T>C), CTG1370TTG (5199C>T), AAA1371AGA (5203A>G), TCC1372TCT (5207C>T), GGG1373GGT (5210G>T), TTT1374TAC (5212T>A 5213T>C), CAT1375CAC (5216T>C), CAA1376CAG (5219A>G), GTT1377ATA (5220G>A 5222T>A), GCT1378AGA (5223G>A 5224C>G 5225T>A), GCA1380TTT (5229G>T 5230C>T 5231A>T), GAG1382TCT (5235G>T 5236A>C 5237G>T), TCC1383AAT (5238T>A 5239C>A 5240C>T), ATC1384ATA (5243C>A), CCC1385CCT (5246C>T), TGG1386AAA (5247T>A 5248G>A 5249G>A), ACT1387ACG (5252T>G), GCC1388ACA (5253G>A 5255C>A), TTT1389TTC (5258T>C), TGG1390AGA (5259T>A 5261G>A), GTT1391ACC (5262G>A 5263T>C 5264T>C), CCA1392CAT (5266C>A 5267A>T), GGC1394GGG (5273C>G), CTA1395CAC (5275T>A 5276A>C), GAA1397GAG (5282A>G), TGG1398TTC (5284G>T 5285G>C), CTT1399TTA (5286C>T 5288T>A), GTT1400GTG (5291T>G), CCA1402CCC (5297A>C), GGG1404GGC (5303G>C), AAG1406ACC (5308A>C 5309G>C), GCA1408GCC (5315A>C), CCA1409CCC (5318A>C), GCA1410TCC (5319G>T 5321A>C), ATC1411ACA (5323T>C 5324C>A), TTC1412TTT (5327C>T), CAA1413CAG (5330A>G), AGA1414AGC (5333A>C), AAA1415CTC (5334A>C 5335A>T 5336A>C), GAT1417AAT (5340G>A), CAG1418CAA (5345G>A), TGC1419ATA (5346T>A 5347G>T 5348C>A), TTT1420TTC (5351T>C), AAG1421AAT (5354G>T), GGA1422GCT (5356G>C 5357A>T), GGA1422_ACA1423insCAT (5357_5358insCAT), ACA1423CTT (5358A>C 5359C>T 5360A>T), GAA1424AGA (5361G>A 5362A>G), GAT1425AAA (5364G>A 5366T>A), TTC1426TTT (5369C>T), GCA1428CTG (5373G>C 5374C>T 5375A>G), GTC1429GTT (5378C>T), TAC1430TTC (5380A>T), ATC1431TTT (5382A>T 5384C>T), ATC1434ATA (5393C>A), TTG1435CTT (5394T>C 5396G>T), GTC1436ATA (5397G>A 5399C>A), TTC1437TAC (5401T>A), AGC1438AAC (5404G>A), AAC1440AAT (5411C>T), ATG1441T.. (5412A>T) | | | | | | | | |

##### Proteins

| | Begin | End | Coverage | Score | Concordance | Matches | Identities | I/D/M/F* | Stop Codons |
|---|---|---|---|---|---|---|---|---|---|
| ORF3 polyprotein (YP_009552693.1) | 1337 | 1441 | 5.6% | 381 | 50.0% | 105 (99.1%) | 52 (49.1%) | 1/0/0/0 | 0 |
| Protein mutations: | R1337E (5100A>G 5101G>A), D1340T (5109G>A 5110A>C), E1343V (5119A>T 5120A>C), Q1346K (5127C>A), Y1347F (5131A>T), S1348P (5133T>C 5135C>A), L1349I (5136C>A), G1351V (5143G>T 5144A>T), Q1353K (5148C>A 5150A>G), T1354E (5151A>G 5152C>A 5153T>A), I1355L (5154A>T), K1357D (5160A>G 5162G>T), K1358E (5163A>G 5165A>G), I1359L (5166A>T 5168T>G), G1360H (5169G>C 5170G>A 5171C>T), N1361G (5172A>G 5173A>G 5174C>A), K1362A (5175A>G 5176A>C 5177A>C), Q1363K (5178C>A), V1364Y (5181G>T 5182T>A), F1368L (5193T>C 5195T>A), K1371R (5203A>G), F1374Y (5212T>A 5213T>C), V1377I (5220G>A 5222T>A), A1378R (5223G>A 5224C>G 5225T>A), A1380F (5229G>T 5230C>T 5231A>T), E1382S (5235G>T 5236A>C 5237G>T), S1383N (5238T>A 5239C>A 5240C>T), W1386K (5247T>A 5248G>A 5249G>A), A1388T (5253G>A 5255C>A), W1390R (5259T>A 5261G>A), V1391T (5262G>A 5263T>C 5264T>C), P1392H (5266C>A 5267A>T), L1395H (5275T>A 5276A>C), W1398F (5284G>T 5285G>C), K1406T (5308A>C 5309G>C), A1410S (5319G>T 5321A>C), I1411T (5323T>C 5324C>A), R1414S (5333A>C), K1415L (5334A>C 5335A>T 5336A>C), D1417N (5340G>A), C1419I (5346T>A 5347G>T 5348C>A), K1421N (5354G>T), G1422A (5356G>C 5357A>T), G1422_T1423insH (5357_5358insCAT), T1423L (5358A>C 5359C>T 5360A>T), E1424R (5361G>A 5362A>G), D1425K (5364G>A 5366T>A), A1428L (5373G>C 5374C>T 5375A>G), Y1430F (5380A>T), I1431F (5382A>T 5384C>T), V1436I (5397G>A 5399C>A), F1437Y (5401T>A), S1438N (5404G>A) | | | | | | | | |
| Codon mutations: | AAA1336.GG (5098A>G 5099A>G), AGA1337GAA (5100A>G 5101G>A), TTG1338TTA (5105G>A), GAC1340ACC (5109G>A 5110A>C), AAT1341AAC (5114T>C), ACG1342ACT (5117G>T), GAA1343GTC (5119A>T 5120A>C), AAA1344AAG (5123A>G), GAC1345GAT (5126C>T), CAG1346AAG (5127C>A), TAT1347TTT (5131A>T), TCC1348CCA (5133T>C 5135C>A), CTT1349ATT (5136C>A), GGA1351GTT (5143G>T 5144A>T), ATC1352ATT (5147C>T), CAA1353AAG (5148C>A 5150A>G), ACT1354GAA (5151A>G 5152C>A 5153T>A), ATA1355TTA (5154A>T), CTC1356TTA (5157C>T 5159C>A), AAG1357GAT (5160A>G 5162G>T), AAA1358GAG (5163A>G 5165A>G), ATT1359TTG (5166A>T 5168T>G), GGC1360CAT (5169G>C 5170G>A 5171C>T), AAC1361GGA (5172A>G 5173A>G 5174C>A), AAA1362GCC (5175A>G 5176A>C 5177A>C), CAG1363AAG (5178C>A), GTT1364TAT (5181G>T 5182T>A), TTC1365TTT (5186C>T), AGT1366TCT (5187A>T 5188G>C), TTT1368CTA (5193T>C 5195T>A), GAT1369GAC (5198T>C), CTG1370TTG (5199C>T), AAA1371AGA (5203A>G), TCC1372TCT (5207C>T), GGG1373GGT (5210G>T), TTT1374TAC (5212T>A 5213T>C), CAT1375CAC (5216T>C), CAA1376CAG (5219A>G), GTT1377ATA (5220G>A 5222T>A), GCT1378AGA (5223G>A 5224C>G 5225T>A), GCA1380TTT (5229G>T 5230C>T 5231A>T), GAG1382TCT (5235G>T 5236A>C 5237G>T), TCC1383AAT (5238T>A 5239C>A 5240C>T), ATC1384ATA (5243C>A), CCC1385CCT (5246C>T), TGG1386AAA (5247T>A 5248G>A 5249G>A), ACT1387ACG (5252T>G), GCC1388ACA (5253G>A 5255C>A), TTT1389TTC (5258T>C), TGG1390AGA (5259T>A 5261G>A), GTT1391ACC (5262G>A 5263T>C 5264T>C), CCA1392CAT (5266C>A 5267A>T), GGC1394GGG (5273C>G), CTA1395CAC (5275T>A 5276A>C), GAA1397GAG (5282A>G), TGG1398TTC (5284G>T 5285G>C), CTT1399TTA (5286C>T 5288T>A), GTT1400GTG (5291T>G), CCA1402CCC (5297A>C), GGG1404GGC (5303G>C), AAG1406ACC (5308A>C 5309G>C), GCA1408GCC (5315A>C), CCA1409CCC (5318A>C), GCA1410TCC (5319G>T 5321A>C), ATC1411ACA (5323T>C 5324C>A), TTC1412TTT (5327C>T), CAA1413CAG (5330A>G), AGA1414AGC (5333A>C), AAA1415CTC (5334A>C 5335A>T 5336A>C), GAT1417AAT (5340G>A), CAG1418CAA (5345G>A), TGC1419ATA (5346T>A 5347G>T 5348C>A), TTT1420TTC (5351T>C), AAG1421AAT (5354G>T), GGA1422GCT (5356G>C 5357A>T), GGA1422_ACA1423insCAT (5357_5358insCAT), ACA1423CTT (5358A>C 5359C>T 5360A>T), GAA1424AGA (5361G>A 5362A>G), GAT1425AAA (5364G>A 5366T>A), TTC1426TTT (5369C>T), GCA1428CTG (5373G>C 5374C>T 5375A>G), GTC1429GTT (5378C>T), TAC1430TTC (5380A>T), ATC1431TTT (5382A>T 5384C>T), ATC1434ATA (5393C>A), TTG1435CTT (5394T>C 5396G>T), GTC1436ATA (5397G>A 5399C>A), TTC1437TAC (5401T>A), AGC1438AAC (5404G>A), AAC1440AAT (5411C>T), ATG1441T.. (5412A>T) | | | | | | | | |

*: Inserts / Deletes / Misaligned / Frameshifts

#### Analysis details

This analysis was performed with panviral2.64

### NGS Details (UN70): Cavemovirus collusipomeae

#### Assembly

| | |
|---|---|
| **Coverage Length** | 258 (1 contig(s)) |
| **Depth Of Coverage** | 7.1 |
| **Number Of Reads** | 15 |
| **Reads Per Million** | 0.28 rpm (after QC) |
| **Ambiguities** | 0 |
| **Assembly Method** | de novo + reference guided assembly |
| **Consensus Caller** | Bcf Tools |

#### Coverage Map

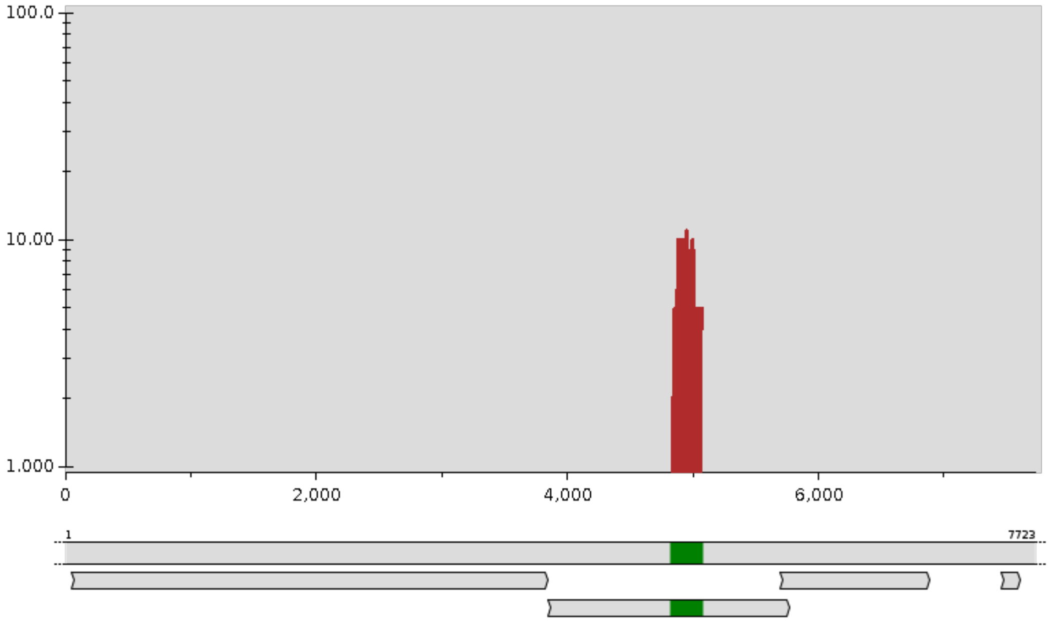

#### Assignment

| | |
|---|---|
| **Type** | Cavemovirus collusipomeae (Taxonomy ID: 3048352) |
| **Reference Genome** | NC_015328.1 |
| **NT Identity (%)** | 61.6279 |
| **AA Identity (%)** | 50.0 |
| **Number Of Stop Codons** | 0 |
| **Number Of CDS** | 4 |

#### Alignment

| | |
|---|---|
| **Alignment Score** | 120.0 (NT) + 326.0 (AA) = 446.0 |
| **Concordance (%)** | 39.9284 |

| Alignment Method | Global, seeded, nucleotide + amino acids (AGA) |
|---|---|

#### Genome Region

Sequence starts at position 4825 and ends at position 5082 relative to NC_015328.1 reference sequence.

#### Alignment Detailed Statistics

| | Begin | End | Coverage | Score | Concordance | Matches | Identities | I/D/M/F* | Stop Codons |
|---|---|---|---|---|---|---|---|---|---|
| **NT** | **4825** | **5082** | **3.3%** | **120** | **23.3%** | **258 (100%)** | **159 (61.6%)** | **0/0** | |
| Mutations: | 4826T>A, 4828A>G, 4831T>A, 4838T>C, 4839A>T, 4840T>A, 4841C>A, 4843T>A, 4847T>G, 4849A>T, 4852A>T, 4853A>T, 4854G>C, 4856A>G, 4857T>A, 4860A>T, 4864A>G, 4865C>A, 4868T>A, 4870T>C, 4874G>A, 4878G>A, 4879A>T, 4882A>T, 4887G>A, 4888A>T, 4896A>T, 4897T>A, 4906A>C, 4907C>A, 4912A>T, 4918A>T, 4924T>C, 4927A>T, 4930A>G, 4933A>T, 4935T>A, 4939G>T, 4945A>G, 4947A>T, 4948C>T, 4952G>A, 4953A>T, 4954A>G, 4959A>T, 4960A>C, 4962T>A, 4967T>C, 4969G>T, 4972A>T, 4975A>C, 4978T>C, 4980G>A, 4984T>A, 4987A>T, 4988A>G, 4990A>T, 4994A>C, 4997C>A, 5001G>A, 5002A>T, 5003A>T, 5005A>G, 5009T>A, 5011A>T, 5012A>T, 5013G>C, 5017G>A, 5018A>C, 5019A>C, 5024G>A, 5025C>T, 5026A>G, 5027G>A, 5028A>G, 5029G>T, 5030A>T, 5033G>T, 5034G>T, 5035A>T, 5036A>C, 5039C>A, 5040A>C, 5042A>C, 5044A>T, 5045A>G, 5047A>C, 5048G>C, 5049A>G, 5060G>C, 5061T>A, 5065G>T, 5068A>T, 5069G>T, 5070A>G, 5071A>C, 5076G>A, 5078G>A, 5079G>A | | | | | | | | |
| **CDS** | | | | | | | | | |
| SPCV_gp2 | 329 | 414 | 13.4% | 326 | 53.3% | 86 (100%) | 43 (50.0%) | 0/0/0/0 | 0 |
| Protein mutations: | L329M (4826T>A 4828A>G), Y333L (4838T>C 4839A>T 4840T>A), H334K (4841C>A 4843T>A), S336A (4847T>G 4849A>T), I339E (4856A>G 4857T>A), Y340F (4860A>T), Q342K (4865C>A), F343I (4868T>A 4870T>C), D345N (4874G>A), G346D (4878G>A 4879A>T), R349N (4887G>A 4888A>T), Y352L (4896A>T 4897T>A), L356I (4907C>A), F365Y (4935T>A), Y369F (4947A>T 4948C>T), E371M (4952G>A 4953A>T 4954A>G), K373I (4959A>T 4960A>C), I374K (4962T>A), E377D (4972A>T), R380K (4980G>A), N381K (4984T>A), I383V (4988A>G 4990A>T), K385Q (4994A>C), H386N (4997C>A), G387D (5001G>A 5002A>T), I388L (5003A>T 5005A>G), L390I (5009T>A 5011A>T), K393P (5018A>C 5019A>C), A395M (5024G>A 5025C>T 5026A>G), E396S (5027G>A 5028A>G 5029G>T), I397L (5030A>T), G398F (5033G>T 5034G>T 5035A>T), K399Q (5036A>C), Q400T (5039C>A 5040A>C), K401H (5042A>C 5044A>T), I402V (5045A>G 5047A>C), E403R (5048G>C 5049A>G), V407H (5060G>C 5061T>A), K408N (5065G>T), E410C (5069G>T 5070A>G 5071A>C), G412E (5076G>A), G413K (5078G>A 5079G>A) | | | | | | | | |
| Codon mutations: | TTA329ATG (4826T>A 4828A>G), CCT330CCA (4831T>A), TAT333CTA (4838T>C 4839A>T 4840T>A), CAT334AAA (4841C>A 4843T>A), TCA336GCT (4847T>G 4849A>T), CCA337CCT (4852A>T), AGT338TCT (4853A>T 4854G>C), ATA339GAA (4856A>G 4857T>A), TAT340TTT (4860A>T), CAA341CAG (4864A>G), CAA342AAA (4865C>A), TTT343ATC (4868T>A 4870T>C), GAT345AAT (4874G>A), GGA346GAT (4878G>A 4879A>T), ATA347ATT (4882A>T), AGA349AAT (4887G>A 4888A>T), TAT352TTA (4896A>T 4897T>A), ATA355ATC (4906A>C), CTT356ATT (4907C>A), GTA357GTT (4912A>T), ATA359ATT (4918A>T), GAT361GAC (4924T>C), ATA362ATT (4927A>T), TTA363TTG (4930A>G), ATA364ATT (4933A>T), TTT365TAT (4935T>A), TCG366TCT (4939G>T), ACA368ACG (4945A>G), TAC369TTT (4947A>T 4948C>T), GAA371ATG (4952G>A 4953A>T 4954A>G), AAA373ATC (4959A>T 4960A>C), ATA374AAA (4962T>A), TTG376CTT (4967T>C 4969G>T), GAA377GAT (4972A>T), ATA378ATC (4975A>C), TTT379TTC (4978T>C), AGA380AAA (4980G>A), AAT381AAA (4984T>A), ATA382ATT (4987A>T), ATA383GTT (4988A>G 4990A>T), AAA385CAA (4994A>C), CAT386AAT (4997C>A), GGA387GAT (5001G>A 5002A>T), ATA388TTG (5003A>T 5005A>G), TTA390ATT (5009T>A 5011A>T), AGT391TCT (5012A>T 5013G>C), AAG392AAA (5017G>A), AAA393CCA (5018A>C 5019A>C), GCA395ATG (5024G>A 5025C>T 5026A>G), GAG396AGT (5027G>A 5028A>G 5029G>T), ATA397TTA (5030A>T), GGA398TTT (5033G>T 5034G>T 5035A>T), AAA399CAA (5036A>C), CAA400ACA (5039C>A 5040A>C), AAA401CAT (5042A>C 5044A>T), ATA402GTC (5045A>G 5047A>C), GAA403CGA (5048G>C 5049A>G), GTT407CAT (5060G>C 5061T>A), AAG408AAT (5065G>T), ATA409ATT (5068A>T), GAA410TGC (5069G>T 5070A>G 5071A>C), GGA412GAA (5076G>A), GGA413AAA (5078G>A 5079G>A) | | | | | | | | |
| **Proteins** | | | | | | | | | |
| replicase (YP_004347415.1) | 329 | 414 | 13.4% | 326 | 53.3% | 86 (100%) | 43 (50.0%) | 0/0/0/0 | 0 |
| Protein mutations: | L329M (4826T>A 4828A>G), Y333L (4838T>C 4839A>T 4840T>A), H334K (4841C>A 4843T>A), S336A (4847T>G 4849A>T), I339E (4856A>G 4857T>A), Y340F (4860A>T), Q342K (4865C>A), F343I (4868T>A 4870T>C), D345N (4874G>A), G346D (4878G>A 4879A>T), R349N (4887G>A 4888A>T), Y352L (4896A>T 4897T>A), L356I (4907C>A), F365Y (4935T>A), Y369F (4947A>T 4948C>T), E371M (4952G>A 4953A>T 4954A>G), K373I (4959A>T 4960A>C), I374K (4962T>A), E377D (4972A>T), R380K (4980G>A), N381K (4984T>A), I383V (4988A>G 4990A>T), K385Q (4994A>C), H386N (4997C>A), G387D (5001G>A 5002A>T), I388L (5003A>T 5005A>G), L390I (5009T>A 5011A>T), K393P (5018A>C 5019A>C), A395M (5024G>A 5025C>T 5026A>G), E396S (5027G>A 5028A>G 5029G>T), I397L (5030A>T), G398F (5033G>T 5034G>T 5035A>T), K399Q (5036A>C), Q400T (5039C>A 5040A>C), K401H (5042A>C 5044A>T), I402V (5045A>G 5047A>C), E403R (5048G>C 5049A>G), V407H (5060G>C 5061T>A), K408N (5065G>T), E410C (5069G>T 5070A>G 5071A>C), G412E (5076G>A), G413K (5078G>A 5079G>A) | | | | | | | | |
| Codon mutations: | TTA329ATG (4826T>A 4828A>G), CCT330CCA (4831T>A), TAT333CTA (4838T>C 4839A>T 4840T>A), CAT334AAA (4841C>A 4843T>A), TCA336GCT (4847T>G 4849A>T), CCA337CCT (4852A>T), AGT338TCT (4853A>T 4854G>C), ATA339GAA (4856A>G 4857T>A), TAT340TTT (4860A>T), CAA341CAG (4864A>G), CAA342AAA (4865C>A), TTT343ATC (4868T>A 4870T>C), GAT345AAT (4874G>A), GGA346GAT (4878G>A 4879A>T), ATA347ATT (4882A>T), AGA349AAT (4887G>A 4888A>T), TAT352TTA (4896A>T 4897T>A), ATA355ATC (4906A>C), CTT356ATT (4907C>A), GTA357GTT (4912A>T), ATA359ATT (4918A>T), GAT361GAC (4924T>C), ATA362ATT (4927A>T), TTA363TTG (4930A>G), ATA364ATT (4933A>T), TTT365TAT (4935T>A), TCG366TCT (4939G>T), ACA368ACG (4945A>G), TAC369TTT (4947A>T 4948C>T), GAA371ATG (4952G>A 4953A>T 4954A>G), AAA373ATC (4959A>T 4960A>C), ATA374AAA (4962T>A), TTG376CTT (4967T>C 4969G>T), GAA377GAT (4972A>T), ATA378ATC (4975A>C), TTT379TTC (4978T>C), AGA380AAA (4980G>A), AAT381AAA (4984T>A), ATA382ATT (4987A>T), ATA383GTT (4988A>G 4990A>T), AAA385CAA (4994A>C), CAT386AAT (4997C>A), GGA387GAT (5001G>A 5002A>T), ATA388TTG (5003A>T 5005A>G), TTA390ATT (5009T>A 5011A>T), AGT391TCT (5012A>T 5013G>C), AAG392AAA (5017G>A), AAA393CCA (5018A>C 5019A>C), GCA395ATG (5024G>A 5025C>T 5026A>G), GAG396AGT (5027G>A 5028A>G 5029G>T), ATA397TTA (5030A>T), GGA398TTT (5033G>T 5034G>T 5035A>T), AAA399CAA (5036A>C), CAA400ACA (5039C>A 5040A>C), AAA401CAT (5042A>C 5044A>T), ATA402GTC (5045A>G 5047A>C), GAA403CGA (5048G>C 5049A>G), GTT407CAT (5060G>C 5061T>A), AAG408AAT (5065G>T), ATA409ATT (5068A>T), GAA410TGC (5069G>T 5070A>G 5071A>C), GGA412GAA (5076G>A), GGA413AAA (5078G>A 5079G>A) | | | | | | | | |

*: Inserts / Deletes / Misaligned / Frameshifts

#### Analysis details

This analysis was performed with panviral2.64

### NGS Details (UN70): Petuvirus venapetuniae

#### Assembly

| | |
|---|---|
| **Coverage Length** | 609 (3 contig(s)) |
| **Depth Of Coverage** | 2.8 |
| **Number Of Reads** | 14 |
| **Reads Per Million** | 0.27 rpm (after QC) |
| **Ambiguities** | 0 |
| **Assembly Method** | de novo + reference guided assembly |
| **Consensus Caller** | Bcf Tools |

#### Coverage Map

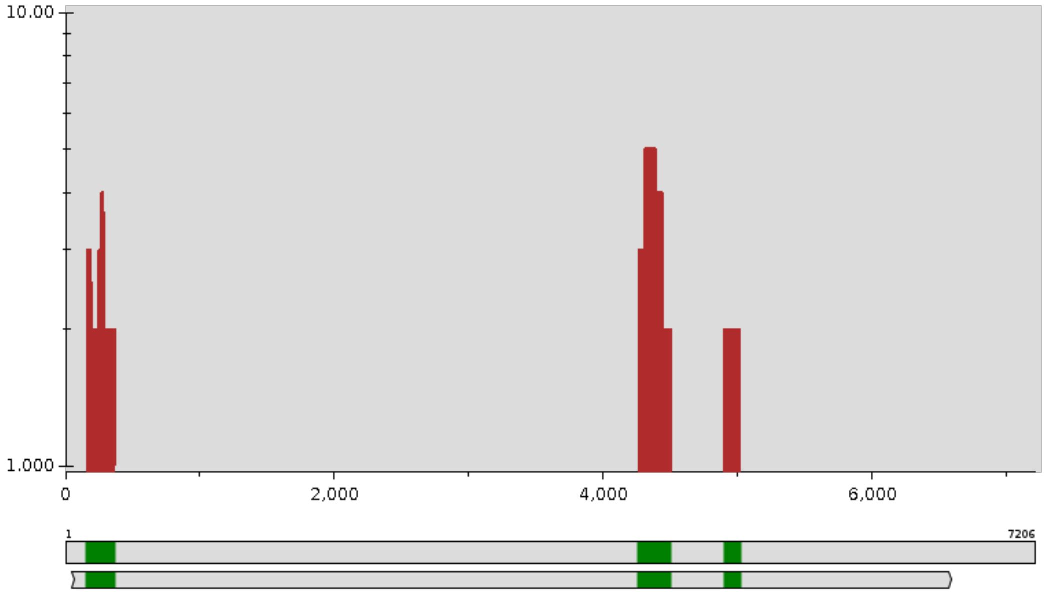

#### Assignment

| | |
|---|---|
| **Type** | Petuvirus venapetuniae (Taxonomy ID: 3048265) |
| **Reference Genome** | NC_001839.2 |
| **NT Identity (%)** | 64.0394 |
| **AA Identity (%)** | 57.9208 |
| **Number Of Stop Codons** | 2 |
| **Number Of CDS** | 1 |

#### Alignment

| | |
|---|---|
| **Alignment Score** | 342.0 (NT) + 831.0 (AA) = 1173.0 |
| **Concordance (%)** | 44.3814 |

| **Alignment Method** | Global, seeded, nucleotide + amino acids (AGA) |
|---|---|

#### Genome Region

Sequence starts at position 151 and ends at position 5024 relative to NC_001839.2 reference sequence.

#### Alignment Detailed Statistics

| | Begin | End | Coverage | Score | Concordance | Matches | Identities | I/D/M/F* | Stop Codons |
|---|---|---|---|---|---|---|---|---|---|
| **NT** | **151** | **5024** | **8.5%** | **342** | **28.1%** | **609 (100%)** | **390 (64.0%)** | **0/0** | |
| Mutations: | 152T>G, 159G>A, 161A>T, 165A>G, 170T>A, 171A>C, 172A>C, 173A>C, 177A>G, 179C>G, 186A>G, 187T>C, 192G>T, 193A>C, 194T>C, 197C>G, 199A>C, 200T>C, 201C>T, 203T>A, 207T>C, 209G>T, 210T>A, 221C>T, 233A>C, 238G>A, 239A>T, 241C>A, 242C>A, 243G>T, 245T>A, 246A>T, 248C>G, 251T>A, 255T>A, 256C>T, 257T>C, 260G>T, 263C>T, 265G>A, 269A>C, 278G>A, 279T>A, 280C>A, 281T>C, 286C>T, 287T>C, 288C>A, 289G>T, 290C>A, 291C>G, 292A>G, 293T>A, 296G>A, 298A>G, 310C>T, 317C>A, 321T>C, 323A>C, 324G>A, 329G>A, 333C>T, 336G>A, 338C>A, 339T>C, 344T>A, 347A>T, 348G>C, 349G>C, 359G>A, 362T>C, 363G>T, 366A>T, 4253A>C, 4255T>A, 4259G>A, 4262A>G, 4263T>A, 4264G>T, 4269G>A, 4271G>T, 4274T>C, 4275C>T, 4276A>T, 4281T>A, 4282T>A, 4283T>G, 4285A>G, 4287C>T, 4289A>G, 4292C>A, 4293G>A, 4298C>G, 4301T>C, 4302G>A, 4304T>A, 4309A>C, 4310A>T, 4314G>T, 4316A>T, 4320G>A, 4321A>C, 4325T>C, 4334C>T, 4337C>T, 4340G>C, 4341A>G, 4342G>C, 4343G>A, 4344T>G, 4345C>G, 4346A>T, 4349G>A, 4350C>A, 4352A>G, 4354T>A, 4355G>A, 4361A>T, 4362A>G, 4363A>C, 4364A>T, 4365T>C, 4366T>C, 4367A>T, 4368A>C, 4370G>A, 4371T>C, 4376C>T, 4386C>A, 4391A>T, 4397C>T, 4398C>A, 4400T>A, 4402T>A, 4403C>T, 4404C>T, 4406C>G, 4407C>A, 4410G>T, 4411A>G, 4412T>G, 4413G>A, 4414A>T, 4415C>T, 4417A>G, 4420T>A, 4421T>C, 4424C>T, 4430A>T, 4436G>A, 4437C>A, 4438T>G, 4439C>G, 4440A>G, 4441C>A, 4442A>T, 4443C>T, 4445C>A, 4448C>A, 4452C>A, 4453A>G, 4454C>A, 4459C>A, 4460C>T, 4461A>G, 4463A>T, 4466A>C, 4469G>T, 4470C>A, 4472C>A, 4481G>T, 4488C>T, 4490T>A, 4493G>A, 4501T>A, 4894C>T, 4896G>A, 4898A>C, 4904G>A, 4905A>C, 4907A>C, 4908C>T, 4910A>G, 4916T>C, 4922G>T, 4923A>G, 4925T>G, 4926G>A, 4935A>G, 4938A>C, 4939G>A, 4952A>T, 4953G>T, 4955G>T, 4956G>A, 4958T>C, 4959A>T, 4961T>G, 4962G>A, 4965C>T, 4973C>T, 4976T>A, 4981G>T, 4982T>G, 4983G>A, 4985T>G, 4988G>T, 4992A>C, 4994G>A, 5001C>T, 5002C>A, 5003A>T, 5006G>T, 5008C>T, 5009A>T, 5015A>G, 5018A>G, 5019T>G, 5020G>A | | | | | | | | |
| **CDS** | | | | | | | | | |
| PVCVgp1 | 35 | 1658 | 9.3% | 831 | 57.7% | 202 (100%) | 117 (57.9%) | 0/0/0/0 | 2 |
| Protein mutations: | V37I (159G>A 161A>T), I39V (165A>G), K41P (171A>C 172A>C 173A>C), N43E (177A>G 179C>G), I46A (186A>G 187T>C), D48S (192G>T 193A>C 194T>C), Y50S (199A>C 200T>C), L54I (210T>A), R63N (238G>A 239A>T), S64* (241C>A 242C>A), V65L (243G>T 245T>A), T66S (246A>T 248C>G), S69I (255T>A 256C>T 257T>C), Q70H (260G>T), R72H (265G>A), S77N (279T>A 280C>A 281T>C), P79L (286C>T 287T>C), R80I (288C>A 289G>T 290C>A), H81G (291C>G 292A>G 293T>A), K83R (298A>G), A87V (310C>T), D92N (324G>A), P95S (333C>T), V96I (336G>A 338C>A), F97L (339T>C), G100P (348G>C 349G>C), V105F (363G>T), T106S (366A>T), I1402K (4255T>A), C1405I (4263T>A 4264G>T), E1407N (4269G>A 4271G>T), Q1409L (4275C>T 4276A>T), F1411K (4281T>A 4282T>A 4283T>G), D1412G (4285A>G), E1415K (4293G>A), D1418K (4302G>A 4304T>A), Q1420P (4309A>C 4310A>T), A1422S (4314G>T 4316A>T), E1424T (4320G>A 4321A>C), K1430N (4340G>C), R1431A (4341A>G 4342G>C 4343G>A), S1432G (4344T>G 4345C>G 4346A>T), Q1434K (4350C>A 4352A>G), V1435E (4354T>A 4355G>A), K1438A (4362A>G 4363A>C 4364A>T), L1439P (4365T>C 4366T>C 4367A>T), Q1446K (4386C>A), H1450K (4398C>A 4400T>A), F1451Y (4402T>A 4403C>T), Q1453K (4407C>A), D1454W (4410G>T 4411A>G 4412T>G), D1455I (4413G>A 4414A>T 4415C>T), K1456R (4417A>G), F1457Y (4420T>A 4421T>C), L1463R (4437C>A 4438T>G 4439C>G), T1464D (4440A>G 4441C>A 4442A>T), F1466L (4448C>A), H1468R (4452C>A 4453A>G 4454C>A), S1470Y (4459C>A 4460C>T), K1471D (4461A>G 4463A>T), K1473N (4469G>T), L1474I (4470C>A 4472C>A), K1477N (4481G>T), F1484Y (4501T>A), S1615L (4894C>T), V1616I (4896G>A 4898A>C), I1619L (4905A>C 4907A>C), Q1620* (4908C>T 4910A>G), I1625V (4923A>G 4925T>G), V1626I (4926G>A), I1629V (4935A>G), R1630Q (4938A>C 4939G>A), E1635Y (4953G>T 4955G>T), V1636I (4956G>A 4958T>C), T1637S (4959A>T 4961T>G), E1638K (4962G>A), H1639Y (4965C>T), S1644M (4981G>T 4982T>G), D1645K (4983G>A 4985T>G), M1646I (4988G>T), K1648Q (4992A>C 4994G>A), P1651Y (5001C>T 5002C>A 5003A>T), A1653V (5008C>T 5009A>T), C1657D (5019T>G 5020G>A) | | | | | | | | |
| Codon mutations: | CTT34.TG (152T>G), GTA37ATT (159G>A 161A>T), ATC39GTC (165A>G), CTT40CTA (170T>A), AAA41CCC (171A>C 172A>C 173A>C), AAC43GAG (177A>G 179C>G), ATA46GCA (186A>G 187T>C), GAT48TCC (192G>T 193A>C 194T>C), ACC49ACG (197C>G), TAT50TCC (199A>C 200T>C), CTT51TTA (201C>T 203T>A), TTG53CTT (207T>C 209G>T), TTA54ATA (210T>A), TAC57TAT (221C>T), GCA61GCC (233A>C), AGA63AAT (238G>A 239A>T), TCC64TAA (241C>A 242C>A), GTT65TTA (243G>T 245T>A), ACC66TCG (246A>T 248C>G), CCT67CCA (251T>A), TCT69ATC (255T>A 256C>T 257T>C), CAG70CAT (260G>T), ATC71ATT (263C>T), CGT72CAT (265G>A), TCA73TCC (269A>C), CAG76CAA (278G>A), TCT77AAC (279T>A 280C>A 281T>C), CCT79CTC (286C>T 287T>C), CGC80ATA (288C>A 289G>T 290C>A), CAT81GGA (291C>G 292A>G 293T>A), GTG82GTA (296G>A), AAA83AGA (298A>G), GCT87GTT (310C>T), TCC89TCA (317C>A), TTA91CTC (321T>C 323A>C), GAT92AAT (324G>A), CAG93CAA (329G>A), CCT95TCT (333C>T), GTC96ATA (336G>A 338C>A), TTT97CTT (339T>C), GCT98GCA (344T>A), ACA99ACT (347A>T), GGA100CCA (348G>C 349G>C), CAG103CAA (359G>A), TTT104TTC (362T>C), GTT105TTT (363G>T), ACT106TCT (366A>T), GCA1401.CC (4253A>C), ATA1402AAA (4255T>A), AAG1403AAA (4259G>A), GAA1404GAG (4262A>G), TGT1405ATT (4263T>A 4264G>T), GAG1407AAT (4269G>A 4271G>T), CTT1408CTC (4274T>C), CAA1409TTA (4275C>T 4276A>T), TTT1411AAG (4281T>A 4282T>A 4283T>G), GAT1412GGT (4285A>G), CTA1413TTG (4287C>T 4289A>G), ATC1414ATA (4292C>A), GAA1415AAA (4293G>A), CCC1416CCG (4298C>G), TCT1417TCC (4301T>C), GAT1418AAA (4302G>A 4304T>A), CAA1420CCT (4309A>C 4310A>T), GCA1422TCT (4314G>T 4316A>T), GAA1424ACA (4320G>A 4321A>C), GCT1425GCC (4325T>C), GTC1428GTT (4334C>T), AAC1429AAT (4337C>T), AAG1430AAC (4340G>C), AGG1431GCA (4341A>G 4342G>C 4343G>A), TCA1432GGT (4344T>G 4345C>G 4346A>T), GAG1433GAA (4349G>A), CAA1434AAG (4350C>A 4352A>G), GTG1435GAA (4354T>A 4355G>A), GGA1437GGT (4361A>T), AAA1438GCT (4362A>G 4363A>C 4364A>T), TTA1439CCT (4365T>C 4366T>C 4367A>T), AGG1440CGA (4368A>C 4370G>A), TTA1441CTA (4371T>C), GTC1442GTT (4376C>T), CAA1446AAA (4386C>A), CCA1447CCT (4391A>T), AAC1449AAT (4397C>T), CAT1450AAA (4398C>A 4400T>A), TTC1451TAT (4402T>A 4403C>T), CTC1452TTG (4404C>T 4406C>G), CAA1453AAA (4407C>A), GAT1454TGG (4410G>T 4411A>G 4412T>G), GAC1455ATT (4413G>A 4414A>T 4415C>T), AAG1456AGG (4417A>G), TTT1457TAC (4420T>A 4421T>C), CCC1458CCT (4424C>T), CCA1460CCT (4430A>T), AAG1462AAA (4436G>A), CTC1463AGG (4437C>A 4438T>G 4439C>G), ACA1464GAT (4440A>G 4441C>A 4442A>T), CTC1465TTA (4443C>T 4445C>A), TTC1466TTA (4448C>A), CAC1468AGA (4452C>A 4453A>G 4454C>A), TCC1470TAT (4459C>A 4460C>T), AAA1471GAT (4461A>G 4463A>T), GCA1472GCC (4466A>C), AAG1473AAT (4469G>T), CTC1474ATA (4470C>A 4472C>A), AAG1477AAT (4481G>T), CTT1480TTA (4488C>T 4490T>A), AAG1481AAA (4493G>A), TTT1484TAT (4501T>A), TCG1615TTG (4894C>T), GTA1616ATC (4896G>A 4898A>C), CAG1618CAA (4904G>A), ATA1619CTC (4905A>C 4907A>C), CAA1620TAG (4908C>T 4910A>G), TTT1622TTC (4916T>C), GGG1624GGT (4922G>T), ATT1625GTG (4923A>G 4925T>G), GTT1626ATT (4926G>A), ATC1629GTC (4935A>G), AGA1630CAA (4938A>C 4939G>A), CCA1634CCT (4952A>T), GAG1635TAT (4953G>T 4955G>T), GTT1636ATC (4956G>A 4958T>C), ACT1637TCG (4959A>T 4961T>G), GAG1638AAG (4962G>A), CAT1639TAT (4965C>T), TCC1641TCT (4973C>T), CCT1642CCA (4976T>A), AGT1644ATG (4981G>T 4982T>G), GAT1645AAG (4983G>A 4985T>G), ATG1646ATT (4988G>T), AAG1648CAA (4992A>C 4994G>A), CCA1651TAT (5001C>T 5002C>A 5003A>T), CCG1652CCT (5006G>T), GCA1653GTT (5008C>T 5009A>T), GGA1655GGG (5015A>G), AAA1656AAG (5018A>G), TGT1657GAT (5019T>G 5020G>A) | | | | | | | | |
| **Proteins** | | | | | | | | | |
| ORF I polyprotein (NP_127504.1) | 35 | 1658 | 9.3% | 831 | 57.7% | 202 (100%) | 117 (57.9%) | 0/0/0/0 | 2 |
| Protein mutations: | V37I (159G>A 161A>T), I39V (165A>G), K41P (171A>C 172A>C 173A>C), N43E (177A>G 179C>G), I46A (186A>G 187T>C), D48S (192G>T 193A>C 194T>C), Y50S (199A>C 200T>C), L54I (210T>A), R63N (238G>A 239A>T), S64* (241C>A 242C>A), V65L (243G>T 245T>A), T66S (246A>T 248C>G), S69I (255T>A 256C>T 257T>C), Q70H (260G>T), R72H (265G>A), S77N (279T>A 280C>A 281T>C), P79L (286C>T 287T>C), R80I (288C>A 289G>T 290C>A), H81G (291C>G 292A>G 293T>A), K83R (298A>G), A87V (310C>T), D92N (324G>A), P95S (333C>T), V96I (336G>A 338C>A), F97L (339T>C), G100P (348G>C 349G>C), V105F (363G>T), T106S (366A>T), I1402K (4255T>A), C1405I (4263T>A 4264G>T), E1407N (4269G>A 4271G>T), Q1409L (4275C>T 4276A>T), F1411K (4281T>A 4282T>A 4283T>G), D1412G (4285A>G), E1415K (4293G>A), D1418K (4302G>A 4304T>A), Q1420P (4309A>C 4310A>T), A1422S (4314G>T 4316A>T), E1424T (4320G>A 4321A>C), K1430N (4340G>C), R1431A (4341A>G 4342G>C 4343G>A), S1432G (4344T>G 4345C>G 4346A>T), Q1434K (4350C>A 4352A>G), V1435E (4354T>A 4355G>A), K1438A (4362A>G 4363A>C 4364A>T), L1439P (4365T>C 4366T>C 4367A>T), Q1446K (4386C>A), H1450K (4398C>A 4400T>A), F1451Y (4402T>A 4403C>T), Q1453K (4407C>A), D1454W (4410G>T 4411A>G 4412T>G), D1455I (4413G>A 4414A>T 4415C>T), K1456R (4417A>G), F1457Y (4420T>A 4421T>C), L1463R (4437C>A 4438T>G 4439C>G), T1464D (4440A>G 4441C>A 4442A>T), F1466L (4448C>A), H1468R (4452C>A 4453A>G 4454C>A), S1470Y (4459C>A 4460C>T), K1471D (4461A>G 4463A>T), K1473N (4469G>T), L1474I (4470C>A 4472C>A), K1477N (4481G>T), F1484Y (4501T>A), S1615L (4894C>T), V1616I (4896G>A 4898A>C), I1619L (4905A>C 4907A>C), Q1620* (4908C>T 4910A>G), I1625V (4923A>G 4925T>G), V1626I (4926G>A), I1629V (4935A>G), R1630Q (4938A>C 4939G>A), E1635Y (4953G>T 4955G>T), V1636I (4956G>A 4958T>C), T1637S (4959A>T 4961T>G), E1638K (4962G>A), H1639Y (4965C>T), S1644M (4981G>T 4982T>G), D1645K (4983G>A 4985T>G), M1646I (4988G>T), K1648Q (4992A>C 4994G>A), P1651Y (5001C>T 5002C>A 5003A>T), A1653V (5008C>T 5009A>T), C1657D (5019T>G 5020G>A) | | | | | | | | |

| | Begin | End | Coverage | Score | Concordance | Matches | Identities | I/D/M/F* | Stop Codons |
|---|---|---|---|---|---|---|---|---|---|
| **NT** | **151** | **5024** | **8.5%** | **342** | **28.1%** | **609 (100%)** | **390 (64.0%)** | **0/0** | |

Codon mutations: CTT34.TG (152T>G), GTA37ATT (159G>A 161A>T), ATC39GTC (165A>G), CTT40CTA (170T>A), AAA41CCC (171A>C 172A>C 173A>C), AAC43GAG (177A>G 179C>G), ATA46GCA (186A>G 187T>C), GAT48TCC (192G>T 193A>C 194T>C), ACC49ACG (197C>G), TAT50TCC (199A>C 200T>C), CTT51TTA (201C>T 203T>A), TTG53CTT (207T>C 209G>T), TTA54ATA (210T>A), TAC57TAT (221C>T), GCA61GCC (233A>C), AGA63AAT (238G>A 239A>T), TCC64TAA (241C>A 242C>A), GTT65TTA (243G>T 245T>A), ACC66TCG (246A>T 248C>G), CCT67CCA (251T>A), TCT69ATC (255T>A 256C>T 257T>C), CAG70CAT (260G>T), ATC71ATT (263C>T), CGT72CAT (265G>A), TCA73TCC (269A>C), CAG76CAA (278G>A), TCT77AAC (279T>A 280C>A 281T>C), CCT79CTC (286C>T 287T>C), CGC80ATA (288C>A 289G>T 290C>A), CAT81GGA (291C>G 292A>G 293T>A), GTG82GTA (296G>A), AAA83AGA (298A>G), GCT87GTT (310C>T), TCC89TCA (317C>A), TTA91CTC (321T>C 323A>C), GAT92AAT (324G>A), CAG93CAA (329G>A), CCT95TCT (333C>T), GTC96ATA (336G>A 338C>A), TTT97CTT (339T>C), GCT98GCA (344T>A), ACA99ACT (347A>T), GGA100CCA (348G>C 349G>C), CAG103CAA (359G>A), TTT104TTC (362T>C), GTT105TTT (363G>T), ACT106TCT (366A>T), GCA1401.CC (4253A>C), ATA1402AAA (4255T>A), AAG1403AAA (4259G>A), GAA1404GAG (4262A>G), TGT1405ATT (4263T>A 4264G>T), GAG1407AAT (4269G>A 4271G>T), CTT1408CTC (4274T>C), CAA1409TTA (4275C>T 4276A>T), TTT1411AAG (4281T>A 4282T>A 4283T>G), GAT1412GGT (4285A>G), CTA1413TTG (4287C>T 4289A>G), ATC1414ATA (4292C>A), GAA1415AAA (4293G>A), CCC1416CCG (4298C>G), TCT1417TCC (4301T>C), GAT1418AAA (4302G>A 4304T>A), CAA1420CCT (4309A>C 4310A>T), GCA1422TCT (4314G>T 4316A>T), GAA1424ACA (4320G>A 4321A>C), GCT1425GCC (4325T>C), GTC1428GTT (4334C>T), AAC1429AAT (4337C>T), AAG1430AAC (4340G>C), AGG1431GCA (4341A>G 4342G>C 4343G>A), TCA1432GGT (4344T>G 4345C>G 4346A>T), GAG1433GAA (4349G>A), CAA1434AAG (4350C>A 4352A>G), GTG1435GAA (4354T>A 4355G>A), GGA1437GGT (4361A>T), AAA1438GCT (4362A>G 4363A>C 4364A>T), TTA1439CCT (4365T>C 4366T>C 4367A>T), AGG1440CGA (4368A>C 4370G>A), TTA1441CTA (4371T>C), GTC1442GTT (4376C>T), CAA1446AAA (4386C>A), CCA1447CCT (4391A>T), AAC1449AAT (4397C>T), CAT1450AAA (4398C>A 4400T>A), TTC1451TAT (4402T>A 4403C>T), CTC1452TTG (4404C>T 4406C>G), CAA1453AAA (4407C>A), GAT1454TGG (4410G>T 4411A>G 4412T>G), GAC1455ATT (4413G>A 4414A>T 4415C>T), AAG1456AGG (4417A>G), TTT1457TAC (4420T>A 4421T>C), CCC1458CCT (4424C>T), CCA1460CCT (4430A>T), AAG1462AAA (4436G>A), CTC1463AGG (4437C>A 4438T>G 4439C>G), ACA1464GAT (4440A>G 4441C>A 4442A>T), CTC1465TTA (4443C>T 4445C>A), TTC1466TTA (4448C>A), CAC1468AGA (4452C>A 4453A>G 4454C>A), TCC1470TAT (4459C>A 4460C>T), AAA1471GAT (4461A>G 4463A>T), GCA1472GCC (4466A>C), AAG1473AAT (4469G>T), CTC1474ATA (4470C>A 4472C>A), AAG1477AAT (4481G>T), CTT1480TTA (4488C>T 4490T>A), AAG1481AAA (4493G>A), TTT1484TAT (4501T>A), TCG1615TTG (4894C>T), GTA1616ATC (4896G>A 4898A>C), CAG1618CAA (4904G>A), ATA1619CTC (4905A>C 4907A>C), CAA1620TAG (4908C>T 4910A>G), TTT1622TTC (4916T>C), GGG1624GGT (4922G>T), ATT1625GTG (4923A>G 4925T>G), GTT1626ATT (4926G>A), ATC1629GTC (4935A>G), AGA1630CAA (4938A>C 4939G>A), CCA1634CCT (4952A>T), GAG1635TAT (4953G>T 4955G>T), GTT1636ATC (4956G>A 4958T>C), ACT1637TCG (4959A>T 4961T>G), GAG1638AAG (4962G>A), CAT1639TAT (4965C>T), TCC1641TCT (4973C>T), CCT1642CCA (4976T>A), AGT1644ATG (4981G>T 4982T>G), GAT1645AAG (4983G>A 4985T>G), ATG1646ATT (4988G>T), AAG1648CAA (4992A>C 4994G>A), CCA1651TAT (5001C>T 5002C>A 5003A>T), CCG1652CCT (5006G>T), GCA1653GTT (5008C>T 5009A>T), GGA1655GGG (5015A>G), AAA1656AAG (5018A>G), TGT1657GAT (5019T>G 5020G>A)

*: Inserts / Deletes / Misaligned / Frameshifts

#### Analysis details

This analysis was performed with panviral2.64

### NGS Details (UN70): Epiphyllum badnavirus 1

#### Assembly

| | |
|---|---|
| **Coverage Length** | 338 (1 contig(s)) |
| **Depth Of Coverage** | 5.1 |
| **Number Of Reads** | 14 |
| **Reads Per Million** | 0.27 rpm (after QC) |
| **Ambiguities** | 0 |
| **Assembly Method** | de novo + reference guided assembly |
| **Consensus Caller** | Bcf Tools |

#### Coverage Map

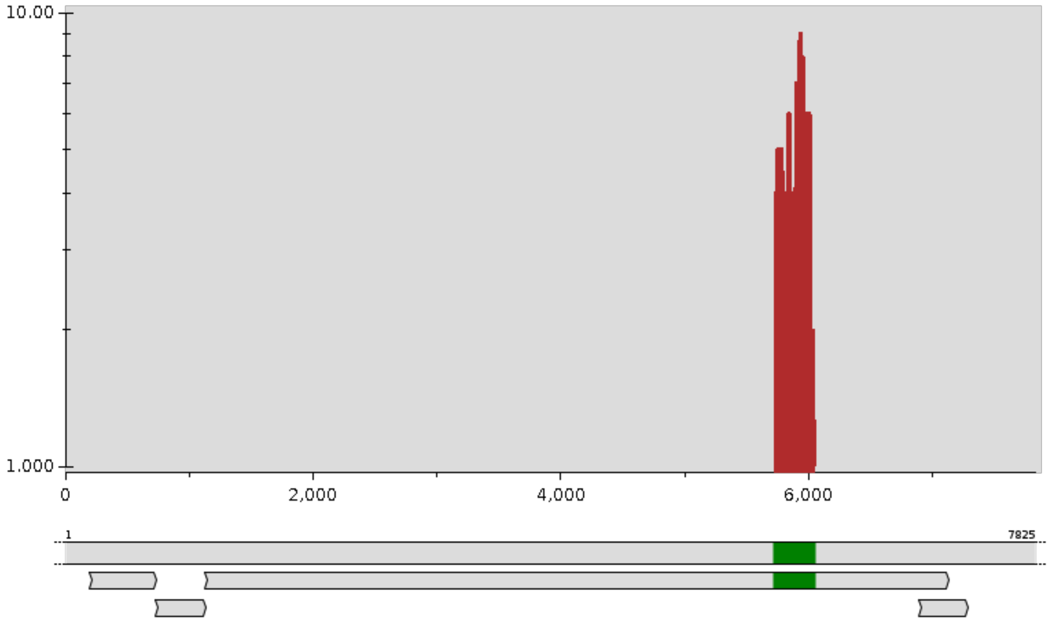

#### Assignment

| | |
|---|---|
| **Type** | Epiphyllum badnavirus 1 (Taxonomy ID: 2518008) |
| **Reference Genome** | NC_076247.1 |
| **NT Identity (%)** | 52.3121 |
| **AA Identity (%)** | 45.6897 |
| **Number Of Stop Codons** | 0 |
| **Number Of CDS** | 4 |

#### Alignment

| | |
|---|---|
| **Alignment Score** | 22.0 (NT) + 230.0 (AA) = 252.0 |
| **Concordance (%)** | 17.027 |

| Alignment Method | Global, seeded, nucleotide + amino acids (AGA) |
|---|---|

#### Genome Region

Sequence starts at position 5715 and ends at position 6052 relative to NC_076247.1 reference sequence.

#### Alignment Detailed Statistics

| | Begin | End | Coverage | Score | Concordance | Matches | Identities | I/D/M/F* | Stop Codons |
|---|---|---|---|---|---|---|---|---|---|
| **NT** | **5715** | **6052** | **4.3%** | **22** | **3.3%** | **338 (97.7%)** | **181 (52.3%)** | **8/0** | |
| Mutations: | 5715A>G, 5716G>A, 5718A>G, 5719G>A, 5724G>A, 5725A>G, 5727A>T, 5729G>A, 5730A>C, 5731A>T, 5732A>C, 5734G>T, 5735A>G, 5740A>C, 5742G>T, 5743G>T, 5744T>G, 5745C>T, 5746T>G, 5748C>A, 5749A>G, 5751C>T, 5754C>T, 5756A>G, 5757A>G, 5758A>G, 5759C>A, 5760C>G, 5761T>C, 5766C>T, 5767G>A, 5769C>T, 5770A>T, 5771A>G, 5775G>A, 5776T>G, 5784T>C, 5785C>A, 5787G>A, 5789A>T, 5790C>T, 5791T>C, 5793T>A, 5794C>A, 5799T>A, 5801G>T, 5806A>G, 5808C>A, 5809A>G, 5810C>A, 5811A>G, 5812A>C, 5814C>A, 5815A>T, 5817A>G, 5818C>G, 5820G>T, 5821A>G, 5823G>A, 5824G>C, 5826C>A, 5827T>C, 5828G>A, 5830A>G, 5831A>G, 5835C>A, 5838T>G, 5839G>T, 5840T>A, 5844C>T, 5847C>T, 5853C>G, 5856T>C, 5859G>T, 5861A>G, 5862G>A, 5863A>T, 5864G>C, 5865C>T, 5868A>T, 5870T>G, 5871C>T, 5877G>A, 5880T>C, 5882T>G, 5887G>T, 5888A>T, 5889C>T, 5892A>G, 5893G>C, 5894A>C, 5895A>T, 5896T>G, 5897C>A, 5901C>T, 5905T>A, 5906G>A, 5910C>T, 5913C>T, 5916C>T, 5917C>A, 5918T>G, 5922C>A, 5924C>A, 5925_5926insTGCAT, 5926G>C, 5931A>G, 5933T>A, 5937C>T, 5942G>T, 5943G>C, 5944T>C, 5949T>C, 5955A>T, 5958C>T, 5961A>T, 5966A>C, 5967G>T, 5976T>A, 5977G>T, 5978C>A, 5979T>C, 5980G>A, 5981T>C, 5982G>A, 5985C>T, 5988A>G, 5989C>A, 5991A>C, 5992A>C, 5993A>T, 5994G>A, 5998G>A, 6003C>G, 6004T>A, 6005G>T, 6009C>T, 6010A>C, 6011G>A, 6012A>T, 6015C>T, 6015_6016insCAT, 6018G>C, 6019T>A, 6020C>G, 6021A>G, 6022G>A, 6023G>A, 6024C>G, 6027T>C, 6028G>A, 6030A>T, 6031G>T, 6032C>T, 6033A>G, 6036T>A, 6038A>T, 6040A>T, 6051C>A | | | | | | | | |

##### CDS

| | Begin | End | Coverage | Score | Concordance | Matches | Identities | I/D/M/F* | Stop Codons |
|---|---|---|---|---|---|---|---|---|---|
| QKM20_gp3 | 1530 | 1642 | 5.6% | 230 | 28.2% | 113 (97.4%) | 53 (45.7%) | 3/0/1/1 | 0 |
| Protein mutations: | E1530K (5716G>A 5718A>G), E1531K (5719G>A), K1533D (5725A>G 5727A>T), G1534D (5729G>A 5730A>C), K1535S (5731A>T 5732A>C), E1536W (5734G>T 5735A>G), M1538L (5740A>C 5742G>T), V1539C (5743G>T 5744T>G 5745C>T), F1540V (5746T>G 5748C>A), N1541D (5749A>G 5751C>T), K1543R (5756A>G 5757A>G), T1544E (5758A>G 5759C>A 5760C>G), D1547N (5767G>A 5769C>T), N1548C (5770A>T 5771A>G), F1550V (5776T>G), Q1553K (5785C>A 5787G>A), Y1554F (5789A>T 5790C>T), S1555P (5791T>C 5793T>A), L1556I (5794C>A), G1558V (5801G>T), N1560E (5806A>G 5808C>A), T1561E (5809A>G 5810C>A 5811A>G), I1562L (5812A>C 5814C>A), I1563L (5815A>T 5817A>G), Q1564D (5818C>G 5820G>T), K1565E (5821A>G 5823G>A), V1566L (5824G>C 5826C>A), C1567H (5827T>C 5828G>A), N1568G (5830A>G 5831A>G), S1570R (5838T>G), V1571Y (5839G>T 5840T>A), F1575L (5853C>G), K1578R (5861A>G 5862G>A), F1581C (5870T>G 5871C>T), M1585R (5882T>G), D1587F (5887G>T 5888A>T 5889C>T), E1589P (5893G>C 5894A>C 5895A>T), S1590D (5896T>G 5897C>A), W1593K (5905T>A 5906G>A), L1597R (5917C>A 5918T>G), P1599Q (5924C>A), P1599_D1600insCX (5925_5926insTGCAT), D1600H (5926G>C), L1602H (5933T>A), W1605F (5942G>T 5943G>C), K1613T (5966A>C 5967G>T), A1617Y (5977G>T 5978C>A 5979T>C), V1618T (5980G>A 5981T>C 5982G>A), R1621S (5989C>A 5991A>C), K1622L (5992A>C 5993A>T 5994G>A), D1624N (5998G>A), N1625K (6003C>G), C1626I (6004T>A 6005G>T), R1628H (6010A>C 6011G>A 6012A>T), D1629_L1630insH (6015_6016insCAT), S1631R (6019T>A 6020C>G 6021A>G), G1632K (6022G>A 6023G>A 6024C>G), V1634I (6028G>A 6030A>T), A1635L (6031G>T 6032C>T 6033A>G), Y1637F (6038A>T), I1638F (6040A>T) | | | | | | | | |
| Codon mutations: | GGA1529..G (5715A>G), GAA1530AAG (5716G>A 5718A>G), GAG1531AAG (5719G>A), AAG1532AAA (5724G>A), AAA1533GAT (5725A>G 5727A>T), GGA1534GAC (5729G>A 5730A>C), AAG1535TCG (5731A>T 5732A>C), GAG1536TGG (5734G>T 5735A>G), ATG1538CTT (5740A>C 5742G>T), GTC1539TGT (5743G>T 5744T>G 5745C>T), TTC1540GTA (5746T>G 5748C>A), AAC1541GAT (5749A>G 5751C>T), TAC1542TAT (5754C>T), AAA1543AGG (5756A>G 5757A>G), ACC1544GAG (5758A>G 5759C>A 5760C>G), TTG1545CTG (5761T>C), AAC1546AAT (5766C>T), GAC1547AAT (5767G>A 5769C>T), AAC1548TGC (5770A>T 5771A>G), ACG1549ACA (5775G>A), TTC1550GTC (5776T>G), GAT1552GAC (5784T>C), CAG1553AAA (5785C>A 5787G>A), TAC1554TTT (5789A>T 5790C>T), TCT1555CCA (5791T>C 5793T>A), CTA1556ATA (5794C>A), CCT1557CCA (5799T>A), GGC1558GTC (5801G>T), AAC1560GAA (5806A>G 5808C>A), ACA1561GAG (5809A>G 5810C>A 5811A>G), ATC1562CTA (5812A>C 5814C>A), ATA1563TTG (5815A>T 5817A>G), CAG1564GAT (5818C>G 5820G>T), AAG1565GAA (5821A>G 5823G>A), GTC1566CTA (5824G>C 5826C>A), TGT1567CAT (5827T>C 5828G>A), AAT1568GGT (5830A>G 5831A>G), GCC1569GCA (5835C>A), AGT1570AGG (5838T>G), GTC1571TAC (5839G>T 5840T>A), TTC1572TTT (5844C>T), TCC1573TCT (5847C>T), TTC1575TTG (5853C>G), GAT1576GAC (5856T>C), CTG1577CTT (5859G>T), AAG1578AGA (5861A>G 5862G>A), AGC1579TCT (5863A>T 5864G>C 5865C>T), GGA1580GGT (5868A>T), TTC1581TGT (5870T>G 5871C>T), CAG1583CAA (5877G>A), ATT1584ATC (5880T>C), ATG1585AGG (5882T>G), GAC1587TTT (5887G>T 5888A>T 5889C>T), GAA1588GAG (5892A>G), GAA1589CCT (5893G>C 5894A>C 5895A>T), TCC1590GAC (5896T>G 5897C>A), ATC1591ATT (5901C>T), TGG1593AAG (5905T>A 5906G>A), ACC1594ACT (5910C>T), GCC1595GCT (5913C>T), TTC1596TTT (5916C>T), CTG1597AGG (5917C>A 5918T>G), ACC1598ACA (5922C>A), CCA1599CAA (5924C>A), CCA1599_GAT1600insTGCAT- (5925_5926insTGCAT), GAT1600CAT (5926G>C), GGA1601GGG (5931A>G), CTT1602CAT (5933T>A), TAC1603TAT (5937C>T), TGG1605TTC (5942G>T 5943G>C), TTA1606CTA (5944T>C), GTT1607GTC (5949T>C), CCA1609CCT (5955A>T), TTC1610TTT (5958C>T), GGA1611GGT (5961A>T), AAG1613ACT (5966A>C 5967G>T), CCT1616CCA (5976T>A), GCT1617TAC (5977G>T 5978C>A 5979T>C), GTG1618ACA (5980G>A 5981T>C 5982G>A), TTC1619TTT (5985C>T), CAA1620CAG (5988A>G), CGA1621AGC (5989C>A 5991A>C), AAG1622CTA (5992A>C 5993A>T 5994G>A), GAC1624AAC (5998G>A), AAC1625AAG (6003C>G), TGT1626ATT (6004T>A 6005G>T), TTC1627TTT (6009C>T), AGA1628CAT (6010A>C 6011G>A 6012A>T), GAC1629GAT (6015C>T), GAC1629_CTG1630insCAT (6015_6016insCAT), CTG1630CTC (6018G>C), TCA1631AGG (6019T>A 6020C>G 6021A>G), GGC1632AAG (6022G>A 6023G>A 6024C>G), TTT1633TTC (6027T>C), GTA1634ATT (6028G>A 6030A>T), GCA1635TTG (6031G>T 6032C>T 6033A>G), GTT1636GTA (6036T>A), TAC1637TTC (6038A>T), ATT1638TTT (6040A>T), ATC1641ATA (6051C>A) | | | | | | | | |

##### Proteins

| | Begin | End | Coverage | Score | Concordance | Matches | Identities | I/D/M/F* | Stop Codons |
|---|---|---|---|---|---|---|---|---|---|
| polyprotein (YP_010797894.1) | 1530 | 1642 | 5.6% | 230 | 28.2% | 113 (97.4%) | 53 (45.7%) | 3/0/1/1 | 0 |
| Protein mutations: | E1530K (5716G>A 5718A>G), E1531K (5719G>A), K1533D (5725A>G 5727A>T), G1534D (5729G>A 5730A>C), K1535S (5731A>T 5732A>C), E1536W (5734G>T 5735A>G), M1538L (5740A>C 5742G>T), V1539C (5743G>T 5744T>G 5745C>T), F1540V (5746T>G 5748C>A), N1541D (5749A>G 5751C>T), K1543R (5756A>G 5757A>G), T1544E (5758A>G 5759C>A 5760C>G), D1547N (5767G>A 5769C>T), N1548C (5770A>T 5771A>G), F1550V (5776T>G), Q1553K (5785C>A 5787G>A), Y1554F (5789A>T 5790C>T), S1555P (5791T>C 5793T>A), L1556I (5794C>A), G1558V (5801G>T), N1560E (5806A>G 5808C>A), T1561E (5809A>G 5810C>A 5811A>G), I1562L (5812A>C 5814C>A), I1563L (5815A>T 5817A>G), Q1564D (5818C>G 5820G>T), K1565E (5821A>G 5823G>A), V1566L (5824G>C 5826C>A), C1567H (5827T>C 5828G>A), N1568G (5830A>G 5831A>G), S1570R (5838T>G), V1571Y (5839G>T 5840T>A), F1575L (5853C>G), K1578R (5861A>G 5862G>A), F1581C (5870T>G 5871C>T), M1585R (5882T>G), D1587F (5887G>T 5888A>T 5889C>T), E1589P (5893G>C 5894A>C 5895A>T), S1590D (5896T>G 5897C>A), W1593K (5905T>A 5906G>A), L1597R (5917C>A 5918T>G), P1599Q (5924C>A), P1599_D1600insCX (5925_5926insTGCAT), D1600H (5926G>C), L1602H (5933T>A), W1605F (5942G>T 5943G>C), K1613T (5966A>C 5967G>T), A1617Y (5977G>T 5978C>A 5979T>C), V1618T (5980G>A 5981T>C 5982G>A), R1621S (5989C>A 5991A>C), K1622L (5992A>C 5993A>T 5994G>A), D1624N (5998G>A), N1625K (6003C>G), C1626I (6004T>A 6005G>T), R1628H (6010A>C 6011G>A 6012A>T), D1629_L1630insH (6015_6016insCAT), S1631R (6019T>A 6020C>G 6021A>G), G1632K (6022G>A 6023G>A 6024C>G), V1634I (6028G>A 6030A>T), A1635L (6031G>T 6032C>T 6033A>G), Y1637F (6038A>T), I1638F (6040A>T) | | | | | | | | |
| Codon mutations: | GGA1529..G (5715A>G), GAA1530AAG (5716G>A 5718A>G), GAG1531AAG (5719G>A), AAG1532AAA (5724G>A), AAA1533GAT (5725A>G 5727A>T), GGA1534GAC (5729G>A 5730A>C), AAG1535TCG (5731A>T 5732A>C), GAG1536TGG (5734G>T 5735A>G), ATG1538CTT (5740A>C 5742G>T), GTC1539TGT (5743G>T 5744T>G 5745C>T), TTC1540GTA (5746T>G 5748C>A), AAC1541GAT (5749A>G 5751C>T), TAC1542TAT (5754C>T), AAA1543AGG (5756A>G 5757A>G), ACC1544GAG (5758A>G 5759C>A 5760C>G), TTG1545CTG (5761T>C), AAC1546AAT (5766C>T), GAC1547AAT (5767G>A 5769C>T), AAC1548TGC (5770A>T 5771A>G), ACG1549ACA (5775G>A), TTC1550GTC (5776T>G), GAT1552GAC (5784T>C), CAG1553AAA (5785C>A 5787G>A), TAC1554TTT (5789A>T 5790C>T), TCT1555CCA (5791T>C 5793T>A), CTA1556ATA (5794C>A), CCT1557CCA (5799T>A), GGC1558GTC (5801G>T), AAC1560GAA (5806A>G 5808C>A), ACA1561GAG (5809A>G 5810C>A 5811A>G), ATC1562CTA (5812A>C 5814C>A), ATA1563TTG (5815A>T 5817A>G), CAG1564GAT (5818C>G 5820G>T), AAG1565GAA (5821A>G 5823G>A), GTC1566CTA (5824G>C 5826C>A), TGT1567CAT (5827T>C 5828G>A), AAT1568GGT (5830A>G 5831A>G), GCC1569GCA (5835C>A), AGT1570AGG (5838T>G), GTC1571TAC (5839G>T 5840T>A), TTC1572TTT (5844C>T), TCC1573TCT (5847C>T), TTC1575TTG (5853C>G), GAT1576GAC (5856T>C), CTG1577CTT (5859G>T), AAG1578AGA (5861A>G 5862G>A), AGC1579TCT (5863A>T 5864G>C 5865C>T), GGA1580GGT (5868A>T), TTC1581TGT (5870T>G 5871C>T), CAG1583CAA (5877G>A), ATT1584ATC (5880T>C), ATG1585AGG (5882T>G), GAC1587TTT (5887G>T 5888A>T 5889C>T), GAA1588GAG (5892A>G), GAA1589CCT (5893G>C 5894A>C 5895A>T), TCC1590GAC (5896T>G 5897C>A), ATC1591ATT (5901C>T), TGG1593AAG (5905T>A 5906G>A), ACC1594ACT (5910C>T), GCC1595GCT (5913C>T), TTC1596TTT (5916C>T), CTG1597AGG (5917C>A 5918T>G), ACC1598ACA (5922C>A), CCA1599CAA (5924C>A), CCA1599_GAT1600insTGCAT- (5925_5926insTGCAT), GAT1600CAT (5926G>C), GGA1601GGG (5931A>G), CTT1602CAT (5933T>A), TAC1603TAT (5937C>T), TGG1605TTC (5942G>T 5943G>C), TTA1606CTA (5944T>C), GTT1607GTC (5949T>C), CCA1609CCT (5955A>T), TTC1610TTT (5958C>T), GGA1611GGT (5961A>T), AAG1613ACT (5966A>C 5967G>T), CCT1616CCA (5976T>A), GCT1617TAC (5977G>T 5978C>A 5979T>C), GTG1618ACA (5980G>A 5981T>C 5982G>A), TTC1619TTT (5985C>T), CAA1620CAG (5988A>G), CGA1621AGC (5989C>A 5991A>C), AAG1622CTA (5992A>C 5993A>T 5994G>A), GAC1624AAC (5998G>A), AAC1625AAG (6003C>G), TGT1626ATT (6004T>A 6005G>T), TTC1627TTT (6009C>T), AGA1628CAT (6010A>C 6011G>A 6012A>T), GAC1629GAT (6015C>T), GAC1629_CTG1630insCAT (6015_6016insCAT), CTG1630CTC (6018G>C), TCA1631AGG (6019T>A 6020C>G 6021A>G), GGC1632AAG (6022G>A 6023G>A 6024C>G), TTT1633TTC (6027T>C), GTA1634ATT (6028G>A 6030A>T), GCA1635TTG (6031G>T 6032C>T 6033A>G), GTT1636GTA (6036T>A), TAC1637TTC (6038A>T), ATT1638TTT (6040A>T), ATC1641ATA (6051C>A) | | | | | | | | |

*: Inserts / Deletes / Misaligned / Frameshifts

#### Analysis details

This analysis was performed with panviral2.64

### NGS Details (UN70): Caulimovirus venafragariae

#### Assembly

| | |
|---|---|
| **Coverage Length** | 504 (1 contig(s)) |
| **Depth Of Coverage** | 3.0 |
| **Number Of Reads** | 11 |
| **Reads Per Million** | 0.21 rpm (after QC) |
| **Ambiguities** | 0 |
| **Assembly Method** | de novo + reference guided assembly |
| **Consensus Caller** | Bcf Tools |

#### Coverage Map

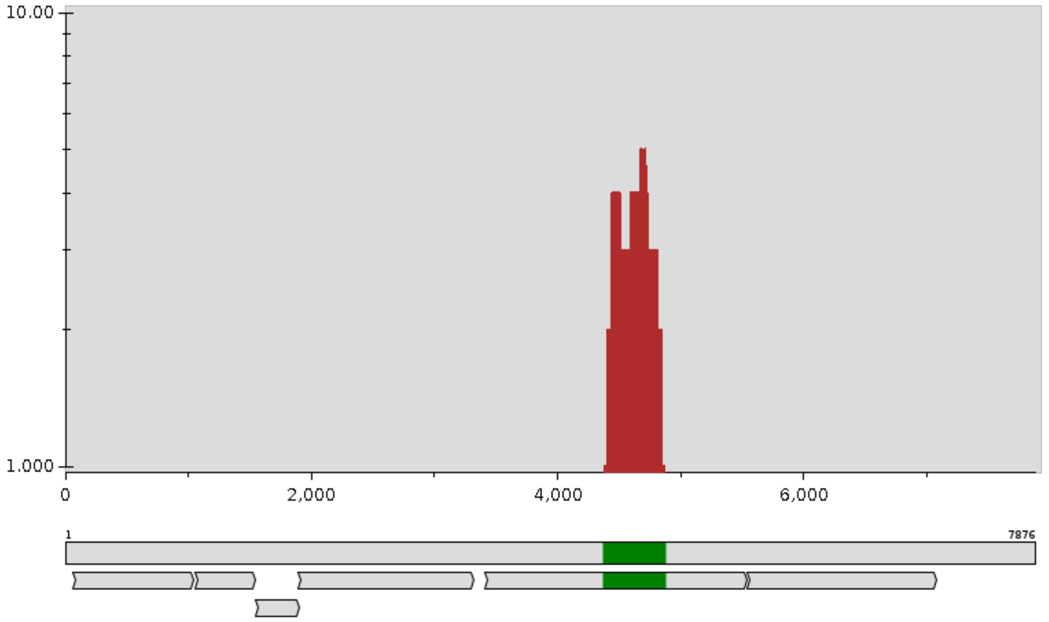

#### Assignment

| | |
|---|---|
| **Type** | Caulimovirus venafragariae (Taxonomy ID: 3048344) |
| **Reference Genome** | NC_001725.1 |
| **NT Identity (%)** | 64.2715 |
| **AA Identity (%)** | 59.2814 |
| **Number Of Stop Codons** | 2 |
| **Number Of CDS** | 6 |

#### Alignment

| | |
|---|---|
| **Alignment Score** | 274.0 (NT) + 730.0 (AA) = 1004.0 |
| **Concordance (%)** | 46.4385 |

| **Alignment Method** | Global, seeded, nucleotide + amino acids (AGA) |
|---|---|

#### Genome Region

Sequence starts at position 4373 and ends at position 4876 relative to NC_001725.1 reference sequence.

#### Alignment Detailed Statistics

| | Begin | End | Coverage | Score | Concordance | Matches | Identities | I/D/M/F* | Stop Codons |
|---|---|---|---|---|---|---|---|---|---|
| **NT** | **4373** | **4876** | **6.4%** | **274** | **27.7%** | **501 (99.4%)** | **322 (63.9%)** | **0/3** | |
| Mutations: | 4378G>A, 4379C>A, 4380C>T, 4383G>A, 4384A>C, 4386C>A, 4388A>T, 4389A>T, 4390C>A, 4395C>A, 4401A>C, 4410A>C, 4416C>T, 4424A>G, 4425G>A, 4434C>T, 4435C>A, 4441A>G, 4443G>A, 4444G>A, 4445G>T, 4446A>G, 4449T>C, 4452C>A, 4455C>T, 4456C>A, 4457T>A, 4461T>A, 4464T>A, 4465A>G, 4470G>A, 4471G>A, 4472A>C, 4473G>A, 4474C>G, 4477C>T, 4479G>A, 4480C>A, 4482T>A, 4483C>A, 4485A>T, 4491C>A, 4492G>C, 4493G>A, 4494A>G, 4497T>A, 4500G>A, 4502C>G, 4503C>A, 4504T>A, 4508A>T, 4509C>T, 4510T>A, 4511C>G, 4512T>C, 4513T>A, 4514C>A, 4515C>G, 4521C>T, 4527A>G, 4530T>A, 4533A>C, 4535T>A, 4542G>A, 4545A>T, 4546C>A, 4547G>A, 4548C>A, 4549C>A, 4551T>G, 4552G>C, 4553C>A, 4560A>C, 4562C>G, 4563A>T, 4567C>G, 4570C>T, 4571T>G, 4575C>A, 4578T>C, 4581C>T, 4582A>T, 4583G>A, 4584C>T, 4587T>C, 4590C>A, 4591C>G, 4599C>T, 4601A>T, 4607G>A, 4613T>C, 4614C>A, 4620T>A, 4633C>A, 4634A>C, 4635A>T, 4638C>T, 4641T>A, 4642G>C, 4644T>A, 4650C>T, 4653A>G, 4656G>A, 4657C>A, 4659C>A, 4665C>T, 4666G>A, 4668A>T, 4670G>T, 4672C>T, 4674C>T, 4675A>G, 4677C>A, 4678_4680delAAC, 4682T>G, 4683G>A, 4687C>A, 4688C>A, 4692G>T, 4695T>C, 4696T>A, 4697G>T, 4699G>T, 4700C>T, 4701T>A, 4704A>T, 4708G>A, 4710T>A, 4719C>T, 4720A>C, 4722C>A, 4725C>T, 4728C>T, 4731C>T, 4734A>T, 4737T>C, 4738G>A, 4739A>T, 4740A>G, 4741G>C, 4759G>C, 4762A>C, 4766T>A, 4767T>G, 4771C>T, 4773A>T, 4776C>T, 4778G>T, 4781G>T, 4782C>T, 4784A>T, 4787C>A, 4789C>A, 4790T>A, 4791A>T, 4794A>T, 4795A>T, 4798G>A, 4800G>T, 4801C>A, 4803T>A, 4806C>T, 4816G>A, 4817C>T, 4818C>A, 4819C>G, 4824G>A, 4834A>T, 4835C>A, 4839C>T, 4846C>T, 4848T>A, 4851T>A, 4852C>A, 4855G>A, 4856T>C, 4857A>C, 4862A>G, 4864A>G, 4865G>A | | | | | | | | |
| **CDS** | | | | | | | | | |
| ORF_V | 324 | 491 | 23.7% | 730 | 61.7% | 167 (99.4%) | 99 (58.9%) | 0/1/0/0 | 2 |
| Protein mutations: | A325N (4378G>A 4379C>A 4380C>T), I327L (4384A>C 4386C>A), K328I (4388A>T 4389A>T), K340R (4424A>G 4425G>A), H344N (4435C>A), K346E (4441A>G 4443G>A), G347M (4444G>A 4445G>T 4446A>G), L351K (4456C>A 4457T>A), N354D (4465A>G), E356T (4471G>A 4472A>C 4473G>A), Q357E (4474C>G), L359I (4480C>A 4482T>A), Q360N (4483C>A 4485A>T), G363Q (4492G>C 4493G>A 4494A>G), T366R (4502C>G 4503C>A), F367I (4504T>A), Y368F (4508A>T 4509C>T), S370K (4513T>A 4514C>A 4515C>G), F377Y (4535T>A), R381K (4546C>A 4547G>A 4548C>A), L382M (4549C>A 4551T>G), A383H (4552G>C 4553C>A), E385D (4560A>C), T386S (4562C>G 4563A>T), Q388E (4567C>G), L389* (4570C>T 4571T>G), S393Y (4582A>T 4583G>A 4584C>T), Q396E (4591C>G), Y399F (4601A>T), W401* (4607G>A), V403A (4613T>C 4614C>A), Q410T (4633C>A 4634A>C 4635A>T), A413P (4642G>C 4644T>A), H418K (4657C>A 4659C>A), E421N (4666G>A 4668A>T), S422I (4670G>T), L423F (4672C>T 4674C>T), S424G (4675A>G 4677C>A), N425del (4678_4680delAAC), M426R (4682T>G 4683G>A), P428K (4687C>A 4688C>A), Q429H (4692G>T), C431I (4696T>A 4697G>T), A432L (4699G>T 4700C>T 4701T>A), V435I (4708G>A 4710T>A), I439L (4720A>C 4722C>A), K443N (4734A>T), E445M (4738G>A 4739A>T 4740A>G), E446Q (4741G>C), V452L (4759G>C), K453Q (4762A>C), I454K (4766T>A 4767T>G), L456F (4771C>T 4773A>T), R458I (4778G>T), C459F (4781G>T 4782C>T), K460I (4784A>T), A461D (4787C>A), L462N (4789C>A 4790T>A 4791A>T), I464L (4795A>T), V465I (4798G>A 4800G>T), L466I (4801C>A 4803T>A), A471I (4816G>A 4817C>T 4818C>A), Q472E (4819C>G), T477Y (4834A>T 4835C>A), L483I (4852C>A), V484T (4855G>A 4856T>C 4857A>C), E486G (4862A>G), R487E (4864A>G 4865G>A) | | | | | | | | |
| Codon mutations: | GCC325AAT (4378G>A 4379C>A 4380C>T), GAG326GAA (4383G>A), ATC327CTA (4384A>C 4386C>A), AAA328ATT (4388A>T 4389A>T), CGA329AGA (4390C>A), GGC330GGA (4395C>A), GCA332GCC (4401A>C), GTA335GTC (4410A>C), AAC337AAT (4416C>T), AAG340AGA (4424A>G 4425G>A), GAC343GAT (4434C>T), CAT344AAT (4435C>A), AAG346GAA (4441A>G 4443G>A), GGA347ATG (4444G>A 4445G>T 4446A>G), GAT348GAC (4449T>C), GGC349GGA (4452C>A), TAC350TAT (4455C>T), CTA351AAA (4456C>A 4457T>A), CTT352CTA (4461T>A), CCT353CCA (4464T>A), AAC354GAC (4465A>G), AAG355AAA (4470G>A), GAG356ACA (4471G>A 4472A>C 4473G>A), CAA357GAA (4474C>G), CTG358TTA (4477C>T 4479G>A), CTT359ATA (4480C>A 4482T>A), CAA360AAT (4483C>A 4485A>T), ATC362ATA (4491C>A), GGA363CAG (4492G>C 4493G>A 4494A>G), GGT364GGA (4497T>A), AAG365AAA (4500G>A), ACC366AGA (4502C>G 4503C>A), TTT367ATT (4504T>A), TAC368TTT (4508A>T 4509C>T), TCT369AGC (4510T>A 4511C>G 4512T>C), TCC370AAG (4513T>A 4514C>A 4515C>G), GAC372GAT (4521C>T), AAA374AAG (4527A>G), TCT375TCA (4530T>A), GGA376GGC (4533A>C), TTT377TAT (4535T>A), CAG379CAA (4542G>A), GTA380GTT (4545A>T), CGC381AAA (4546C>A 4547G>A 4548C>A), CTT382ATG (4549C>A 4551T>G), GCT383CAT (4552G>C 4553C>A), GAA385GAC (4560A>C), ACA386AGT (4562C>G 4563A>T), CAG388GAG (4567C>G), CTA389TGA (4570C>T 4571T>G), ACC390ACA (4575C>A), GCT391GCC (4578T>C), TTC392TTT (4581C>T), AGC393TAT (4582A>T 4583G>A 4584C>T), TGT394TGC (4587T>C), CCC395CCA (4590C>A), CAA396GAA (4591C>G), CAC398CAT (4599C>T), TAC399TTC (4601A>T), TGG401TAG (4607G>A), GTC403GCA (4613T>C 4614C>A), CCT405CCA (4620T>A), CAA410ACT (4633C>A 4634A>C 4635A>T), GCC411GCT (4638C>T), CCT412CCA (4641T>A), GCT413CCA (4642G>C 4644T>A), TTC415TTT (4650C>T), CAA416CAG (4653A>G), AGG417AGA (4656G>A), CAC418AAA (4657C>A 4659C>A), GAC420GAT (4665C>T), GAA421AAT (4666G>A 4668A>T), AGT422ATT (4670G>T), CTC423TTT (4672C>T 4674C>T), AGC424GGA (4675A>G 4677C>A), AAC425del (4678_4680delAAC), ATG426AGA (4682T>G 4683G>A), CCA428AAA (4687C>A 4688C>A), CAG429CAT (4692G>T), TTT430TTC (4695T>C), TGT431ATT (4696T>A 4697G>T), GCT432TTA (4699G>T 4700C>T 4701T>A), GTA433GTT (4704A>T), GTT435ATA (4708G>A 4710T>A), ATC438ATT (4719C>T), ATC439CTA (4720A>C 4722C>A), GTC440GTT (4725C>T), TTC441TTT (4728C>T), AGC442AGT (4731C>T), AAA443AAT (4734A>T), ACT444ACC (4737T>C), GAA445ATG (4738G>A 4739A>T 4740A>G), GAA446CAA (4741G>C), GTC452CTC (4759G>C), AAA453CAA (4762A>C), ATT454AAG (4766T>A 4767T>G), CTA456TTT (4771C>T 4773A>T), AAC457AAT (4776C>T), AGA458ATA (4778G>T), TGC459TTT (4781G>T 4782C>T), AAA460ATA (4784A>T), GCC461GAC (4787C>A), CTA462AAT (4789C>A 4790T>A 4791A>T), GGA463GGT (4794A>T), ATA464TTA (4795A>T), GTG465ATT (4798G>A 4800G>T), CTT466ATA (4801C>A 4803T>A), AGC467AGT (4806C>T), GCC471ATA (4816G>A 4817C>T 4818C>A), CAG472GAG (4819C>G), TTG473TTA (4824G>A), ACT477TAT (4834A>T 4835C>A), ATC478ATT (4839C>T), CTT481TTA (4846C>T 4848T>A), GGT482GGA (4851T>A), CTA483ATA (4852C>A), GTA484ACC (4855G>A 4856T>C 4857A>C), GAA486GGA (4862A>G), AGA487GAA (4864A>G 4865G>A) | | | | | | | | |
| **Proteins** | | | | | | | | | |
| hypothetical protein (NP_043933.1) | 324 | 491 | 23.7% | 730 | 61.7% | 167 (99.4%) | 99 (58.9%) | 0/1/0/0 | 2 |
| Protein mutations: | A325N (4378G>A 4379C>A 4380C>T), I327L (4384A>C 4386C>A), K328I (4388A>T 4389A>T), K340R (4424A>G 4425G>A), H344N (4435C>A), K346E (4441A>G 4443G>A), G347M (4444G>A 4445G>T 4446A>G), L351K (4456C>A 4457T>A), N354D (4465A>G), E356T (4471G>A 4472A>C 4473G>A), Q357E (4474C>G), L359I (4480C>A 4482T>A), Q360N (4483C>A 4485A>T), G363Q (4492G>C 4493G>A 4494A>G), T366R (4502C>G 4503C>A), F367I (4504T>A), Y368F (4508A>T 4509C>T), S370K (4513T>A 4514C>A 4515C>G), F377Y (4535T>A), R381K (4546C>A 4547G>A 4548C>A), L382M (4549C>A 4551T>G), A383H (4552G>C 4553C>A), E385D (4560A>C), T386S (4562C>G 4563A>T), Q388E (4567C>G), L389* (4570C>T 4571T>G), S393Y (4582A>T 4583G>A 4584C>T), Q396E (4591C>G), Y399F (4601A>T), W401* (4607G>A), V403A (4613T>C 4614C>A), Q410T (4633C>A 4634A>C 4635A>T), A413P (4642G>C 4644T>A), H418K (4657C>A 4659C>A), E421N (4666G>A 4668A>T), S422I (4670G>T), L423F (4672C>T 4674C>T), S424G (4675A>G 4677C>A), N425del (4678_4680delAAC), M426R (4682T>G 4683G>A), P428K (4687C>A 4688C>A), Q429H (4692G>T), C431I (4696T>A 4697G>T), A432L (4699G>T 4700C>T 4701T>A), V435I (4708G>A 4710T>A), I439L (4720A>C 4722C>A), K443N (4734A>T), E445M (4738G>A 4739A>T 4740A>G), E446Q (4741G>C), V452L (4759G>C), K453Q (4762A>C), I454K (4766T>A 4767T>G), L456F (4771C>T 4773A>T), R458I (4778G>T), C459F (4781G>T 4782C>T), K460I (4784A>T), A461D (4787C>A), L462N (4789C>A 4790T>A 4791A>T), I464L (4795A>T), V465I (4798G>A 4800G>T), L466I (4801C>A 4803T>A), A471I (4816G>A 4817C>T 4818C>A), Q472E (4819C>G), T477Y (4834A>T 4835C>A), L483I (4852C>A), V484T (4855G>A 4856T>C 4857A>C), E486G (4862A>G), R487E (4864A>G 4865G>A) | | | | | | | | |
| Codon mutations: | GCC325AAT (4378G>A 4379C>A 4380C>T), GAG326GAA (4383G>A), ATC327CTA (4384A>C 4386C>A), AAA328ATT (4388A>T 4389A>T), CGA329AGA (4390C>A), GGC330GGA (4395C>A), GCA332GCC (4401A>C), GTA335GTC (4410A>C), AAC337AAT (4416C>T), AAG340AGA (4424A>G 4425G>A), GAC343GAT (4434C>T), CAT344AAT (4435C>A), AAG346GAA (4441A>G 4443G>A), GGA347ATG (4444G>A 4445G>T 4446A>G), GAT348GAC (4449T>C), GGC349GGA (4452C>A), TAC350TAT (4455C>T), CTA351AAA (4456C>A 4457T>A), CTT352CTA (4461T>A), CCT353CCA (4464T>A), AAC354GAC (4465A>G), AAG355AAA (4470G>A), GAG356ACA (4471G>A 4472A>C 4473G>A), CAA357GAA (4474C>G), CTG358TTA (4477C>T 4479G>A), CTT359ATA (4480C>A 4482T>A), CAA360AAT (4483C>A 4485A>T), ATC362ATA (4491C>A), GGA363CAG (4492G>C 4493G>A 4494A>G), GGT364GGA (4497T>A), AAG365AAA (4500G>A), ACC366AGA (4502C>G 4503C>A), TTT367ATT (4504T>A), TAC368TTT (4508A>T 4509C>T), TCT369AGC (4510T>A 4511C>G 4512T>C), TCC370AAG (4513T>A 4514C>A 4515C>G), GAC372GAT (4521C>T), AAA374AAG (4527A>G), TCT375TCA (4530T>A), GGA376GGC (4533A>C), TTT377TAT (4535T>A), CAG379CAA (4542G>A), GTA380GTT (4545A>T), CGC381AAA (4546C>A 4547G>A 4548C>A), CTT382ATG (4549C>A 4551T>G), GCT383CAT (4552G>C 4553C>A), GAA385GAC (4560A>C), ACA386AGT (4562C>G 4563A>T), CAG388GAG (4567C>G), CTA389TGA (4570C>T 4571T>G), ACC390ACA (4575C>A), GCT391GCC (4578T>C), TTC392TTT (4581C>T), AGC393TAT (4582A>T 4583G>A 4584C>T), TGT394TGC (4587T>C), CCC395CCA (4590C>A), CAA396GAA (4591C>G), CAC398CAT (4599C>T), TAC399TTC (4601A>T), TGG401TAG (4607G>A), GTC403GCA (4613T>C 4614C>A), CCT405CCA (4620T>A), CAA410ACT (4633C>A 4634A>C 4635A>T), GCC411GCT (4638C>T), CCT412CCA (4641T>A), GCT413CCA (4642G>C 4644T>A), TTC415TTT (4650C>T), CAA416CAG (4653A>G), AGG417AGA (4656G>A), CAC418AAA (4657C>A 4659C>A), GAC420GAT (4665C>T), GAA421AAT (4666G>A 4668A>T), AGT422ATT (4670G>T), CTC423TTT (4672C>T 4674C>T), AGC424GGA (4675A>G 4677C>A), AAC425del (4678_4680delAAC), ATG426AGA (4682T>G 4683G>A), CCA428AAA (4687C>A 4688C>A), CAG429CAT (4692G>T), TTT430TTC (4695T>C), TGT431ATT (4696T>A 4697G>T), GCT432TTA (4699G>T 4700C>T 4701T>A), GTA433GTT (4704A>T), GTT435ATA (4708G>A 4710T>A), ATC438ATT (4719C>T), ATC439CTA (4720A>C 4722C>A), GTC440GTT (4725C>T), TTC441TTT (4728C>T), AGC442AGT (4731C>T), AAA443AAT (4734A>T), ACT444ACC (4737T>C), GAA445ATG (4738G>A 4739A>T 4740A>G), GAA446CAA (4741G>C), GTC452CTC (4759G>C), AAA453CAA (4762A>C), ATT454AAG (4766T>A 4767T>G), CTA456TTT (4771C>T 4773A>T), AAC457AAT (4776C>T), AGA458ATA (4778G>T), TGC459TTT (4781G>T 4782C>T), AAA460ATA (4784A>T), GCC461GAC (4787C>A), CTA462AAT (4789C>A 4790T>A 4791A>T), GGA463GGT (4794A>T), ATA464TTA (4795A>T), GTG465ATT (4798G>A 4800G>T), CTT466ATA (4801C>A 4803T>A), AGC467AGT (4806C>T), GCC471ATA (4816G>A 4817C>T 4818C>A), CAG472GAG (4819C>G), TTG473TTA (4824G>A), ACT477TAT (4834A>T 4835C>A), ATC478ATT (4839C>T), CTT481TTA (4846C>T 4848T>A), GGT482GGA (4851T>A), CTA483ATA (4852C>A), GTA484ACC (4855G>A 4856T>C 4857A>C), GAA486GGA (4862A>G), AGA487GAA (4864A>G 4865G>A) | | | | | | | | |

*: Inserts / Deletes / Misaligned / Frameshifts

#### Analysis details

This analysis was performed with panviral2.64

### NGS Details (UN70): Gihfavirus pelohabitans

#### Assembly

| | |
|---|---|
| **Coverage Length** | 183 (1 contig(s)) |
| **Depth Of Coverage** | 4.8 |
| **Number Of Reads** | 10 |
| **Reads Per Million** | 0.19 rpm (after QC) |
| **Ambiguities** | 0 |
| **Assembly Method** | de novo + reference guided assembly |
| **Consensus Caller** | Bcf Tools |

#### Coverage Map

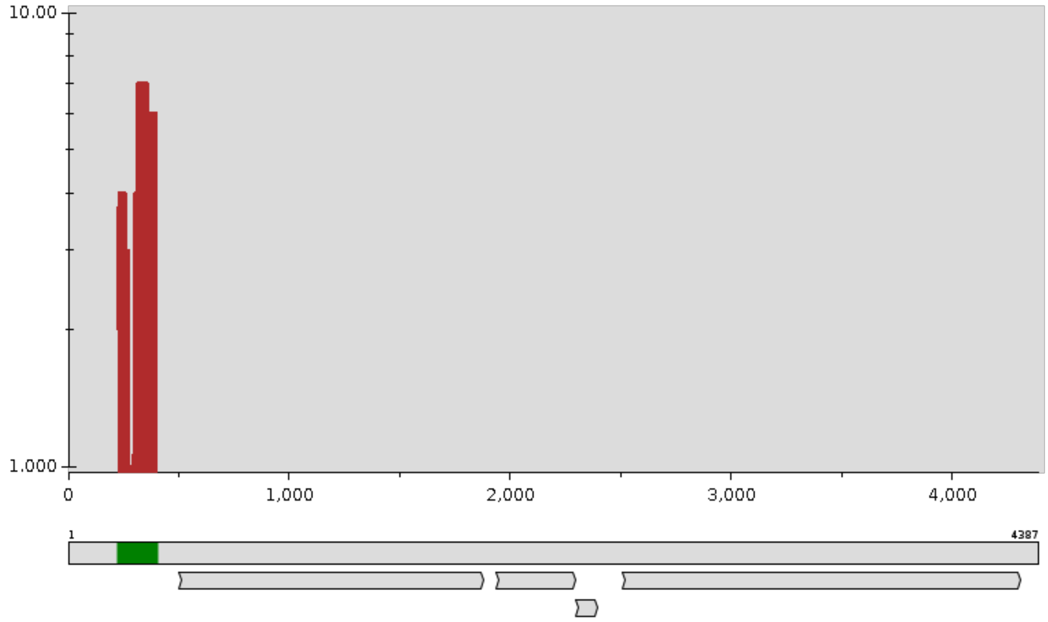

#### Assignment

| | |
|---|---|
| **Type** | Gihfavirus pelohabitans (Taxonomy ID: 2844652) |
| **Reference Genome** | NC_074583.1 |
| **NT Identity (%)** | 97.2678 |
| **AA Identity (%)** | 0.0 |
| **Number Of Stop Codons** | 0 |
| **Number Of CDS** | 4 |

#### Alignment

| | |
|---|---|
| **Alignment Score** | 346.0 (NT) + 0.0 (AA) = 346.0 |
| **Concordance (%)** | 94.5355 |

| Alignment Method | Global, seeded, nucleotide + amino acids (AGA) |
|---|---|

#### Genome Region

Sequence starts at position 222 and ends at position 404 relative to NC_074583.1 reference sequence.

#### Alignment Detailed Statistics

| | Begin | End | Coverage | Score | Concordance | Matches | Identities | I/D/M/F* | Stop Codons |
|---|---|---|---|---|---|---|---|---|---|
| **NT** | **222** | **404** | **4.2%** | **346** | **94.5%** | **183 (100%)** | **178 (97.3%)** | **0/0** | |
| Mutations: | 276G>T, 277C>T, 278A>C, 279A>G, 288A>T | | | | | | | | |

**CDS**

**Proteins**

*: Inserts / Deletes / Misaligned / Frameshifts

#### Analysis details

This analysis was performed with panviral2.64

### NGS Details (UN70): Caulimovirus tessellomirabilis

#### Assembly

| | |
|---|---|
| **Coverage Length** | 343 (1 contig(s)) |
| **Depth Of Coverage** | 4.0 |
| **Number Of Reads** | 10 |
| **Reads Per Million** | 0.19 rpm (after QC) |
| **Ambiguities** | 0 |
| **Assembly Method** | de novo + reference guided assembly |
| **Consensus Caller** | Bcf Tools |

#### Coverage Map

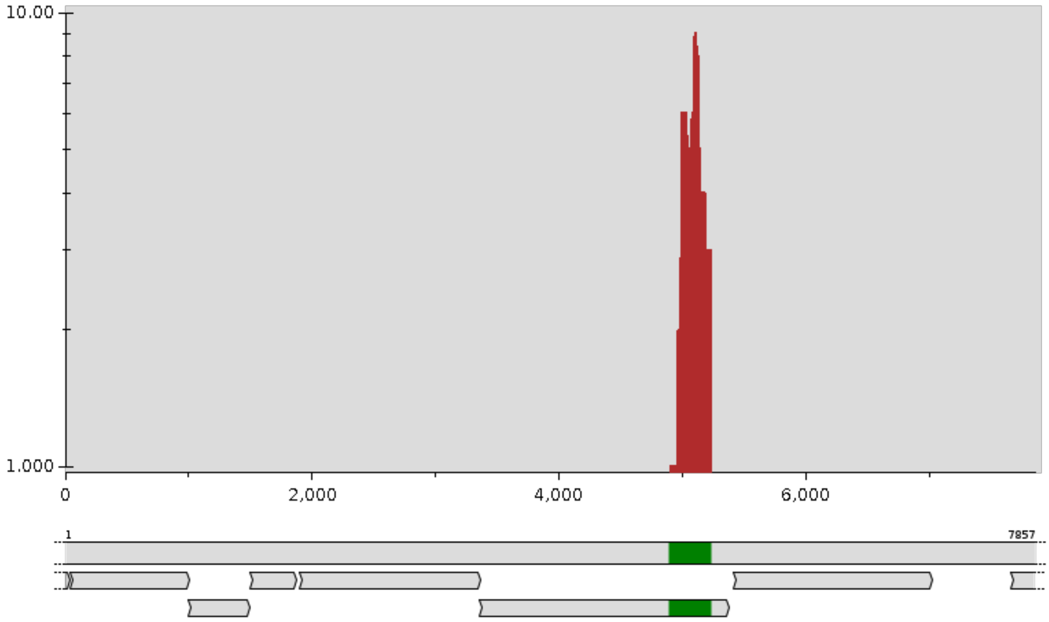

#### Assignment

| | |
|---|---|
| **Type** | Caulimovirus tessellomirabilis (Taxonomy ID: 3048203) |
| **Reference Genome** | NC_004036.1 |
| **NT Identity (%)** | 57.0122 |
| **AA Identity (%)** | 48.6239 |
| **Number Of Stop Codons** | 0 |
| **Number Of CDS** | 7 |

#### Alignment

| | |
|---|---|
| **Alignment Score** | 50.0 (NT) + 365.0 (AA) = 415.0 |
| **Concordance (%)** | 31.4394 |

| Alignment Method | Global, seeded, nucleotide + amino acids (AGA) |
|---|---|

#### Genome Region

Sequence starts at position 4896 and ends at position 5238 relative to NC_004036.1 reference sequence.

#### Alignment Detailed Statistics

| | Begin | End | Coverage | Score | Concordance | Matches | Identities | I/D/M/F* | Stop Codons |
|---|---|---|---|---|---|---|---|---|---|
| **NT** | **4896** | **5238** | **4.4%** | **50** | **8.1%** | **328 (95.6%)** | **187 (54.5%)** | **0/15** | |
| Mutations: | 4903C>T, 4907A>C, 4911A>G, 4912G>C, 4913A>C, 4918G>A, 4921A>G, 4923G>A, 4926T>G, 4927A>T, 4928A>G, 4929T>C, 4930T>C, 4933T>G, 4934T>C, 4936T>C, 4939A>G, 4942A>G, 4946T>A, 4947A>G, 4948T>G, 4955A>G, 4957G>A, 4958A>C, 4959A>C, 4966C>T, 4968G>A, 4969T>C, 4970T>C, 4975T>A, 4976A>T, 4978T>A, 4979G>C, 4984T>G, 4990T>C, 4993T>C, 4996T>C, 4997C>T, 5000T>C, 5001T>A, 5002C>T, 5008T>A, 5010G>C, 5012G>T, 5014C>G, 5015C>T, 5017T>A, 5018_5023delAAGGCC, 5029C>A, 5030A>G, 5031C>A, 5033G>T, 5034A>T, 5035A>G, 5036G>A, 5037G>A, 5042_5047delGAATTA, 5050C>T, 5053C>T, 5056G>A, 5060T>A, 5063T>A, 5064C>G, 5065A>T, 5071A>T, 5074C>T, 5077A>T, 5078C>G, 5079C>A, 5087T>A, 5088T>C, 5089G>T, 5091A>G, 5092T>A, 5095C>T, 5096C>T, 5097A>C, 5100G>C, 5101T>C, 5107G>A, 5110A>T, 5117C>T, 5122G>A, 5123G>A, 5125A>T, 5126A>G, 5127A>T, 5128A>T, 5130A>G, 5131G>A, 5133T>G, 5134G>A, 5137T>C, 5139C>G, 5140T>A, 5147A>T, 5148G>C, 5149T>A, 5150A>G, 5151T>C, 5154A>T, 5155T>C, 5159A>C, 5160C>T, 5161C>T, 5164T>A, 5165G>A, 5166T>A, 5167T>A, 5168T>C, 5169G>A, 5170T>A, 5173T>C, 5175C>T, 5176A>C, 5177G>A, 5180A>C, 5181G>A, 5182G>A, 5183A>T, 5188T>C, 5192_5194delGTA, 5196A>C, 5197T>C, 5199T>A, 5200T>G, 5201T>G, 5204A>T, 5205A>C, 5209A>T, 5212T>G, 5215G>A, 5216A>T, 5217A>T, 5218T>C, 5221A>T, 5224G>A, 5225A>C, 5228A>C, 5231G>T, 5232G>C, 5236C>A, 5237A>C, 5238G>C | | | | | | | | |

##### CDS

| | Begin | End | Coverage | Score | Concordance | Matches | Identities | I/D/M/F* | Stop Codons |
|---|---|---|---|---|---|---|---|---|---|
| MiMVgp6 | 515 | 628 | 16.9% | 365 | 51.0% | 109 (95.6%) | 53 (46.5%) | 0/5/0/0 | 0 |
| Protein mutations: | K518Q (4907A>C), K519S (4911A>G 4912G>C), I520L (4913A>C), G523E (4923G>A), L524C (4926T>G 4927A>T), I525A (4928A>G 4929T>C 4930T>C), N526K (4933T>G), F527L (4934T>C 4936T>C), Y531R (4946T>A 4947A>G 4948T>G), K534E (4955A>G 4957G>A), K535P (4958A>C 4959A>C), S538N (4968G>A 4969T>C), I541L (4976A>T 4978T>A), E542Q (4979G>C), H548Y (4997C>T), F549H (5000T>C 5001T>A 5002C>T), G552A (5010G>C), V553L (5012G>T 5014C>G), K555_A556del (5018_5023delAAGGCC), T559D (5030A>G 5031C>A), E560L (5033G>T 5034A>T 5035A>G), G561N (5036G>A 5037G>A), E563_L564del (5042_5047delGAATTA), S569T (5060T>A), K574N (5077A>T), P575E (5078C>G 5079C>A), L578T (5087T>A 5088T>C 5089G>T), N579R (5091A>G 5092T>A), H581S (5096C>T 5097A>C), S582T (5100G>C 5101T>C), K585N (5110A>T), V590I (5123G>A 5125A>T), K591V (5126A>G 5127A>T 5128A>T), Q592R (5130A>G 5131G>A), V593G (5133T>G 5134G>A), T595R (5139C>G 5140T>A), I599A (5150A>G 5151T>C), Y600F (5154A>T 5155T>C), T602L (5159A>C 5160C>T 5161C>T), V604K (5165G>A 5166T>A 5167T>A), C605Q (5168T>C 5169G>A 5170T>A), T607I (5175C>T 5176A>C), V608I (5177G>A), R609Q (5180A>C 5181G>A 5182G>A), T610S (5183A>T), V613del (5192_5194delGTA), N614T (5196A>C 5197T>C), L615Q (5199T>A 5200T>G), L616V (5201T>G), K617S (5204A>T 5205A>C), F619L (5212T>G), M620I (5215G>A), N621F (5216A>T 5217A>T 5218T>C), K622N (5221A>T), I624L (5225A>C), T625P (5228A>C), G626S (5231G>T 5232G>C), D627E (5236C>A) | | | | | | | | |
| Codon mutations: | TAC516TAT (4903C>T), AAA518CAA (4907A>C), AAG519AGC (4911A>G 4912G>C), ATA520CTA (4913A>C), AAG521AAA (4918G>A), AAA522AAG (4921A>G), GGA523GAA (4923G>A), TTA524TGT (4926T>G 4927A>T), ATT525GCC (4928A>G 4929T>C 4930T>C), AAT526AAG (4933T>G), TTT527CTC (4934T>C 4936T>C), CCA528CCG (4939A>G), AAA529AAG (4942A>G), TAT531AGG (4946T>A 4947A>G 4948T>G), AAG534GAA (4955A>G 4957G>A), AAA535CCA (4958A>C 4959A>C), GAC537GAT (4966C>T), AGT538AAC (4968G>A 4969T>C), TTA539CTA (4970T>C), ATT540ATA (4975T>A), ATT541TTA (4976A>T 4978T>A), GAA542CAA (4979G>C), ACT543ACG (4984T>G), GCT545GCC (4990T>C), TCT546TCC (4993T>C), GAT547GAC (4996T>C), CAC548TAC (4997C>T), TTC549CAT (5000T>C 5001T>A 5002C>T), GGT551GGA (5008T>A), GGA552GCA (5010G>C), GTC553TTG (5012G>T 5014C>G), CTT554TTA (5015C>T 5017T>A), AAG555_GCC556del (5018_5023delAAGGCC), ACC558ACA (5029C>A), ACT559GAT (5030A>G 5031C>A), GAA560TTG (5033G>T 5034A>T 5035A>G), GGT561AAT (5036G>A 5037G>A), GAA563_TTA564del (5042_5047delGAATTA), ATC565ATT (5050C>T), TGC566TGT (5053C>T), AGG567AGA (5056G>A), TCT569ACT (5060T>A), TCA570AGT (5063T>A 5064C>G 5065A>T), ACA572ACT (5071A>T), TTC573TTT (5074C>T), AAA574AAT (5077A>T), CCA575GAA (5078C>G 5079C>A), TTG578ACT (5087T>A 5088T>C 5089G>T), AAT579AGA (5091A>G 5092T>A), TAC580TAT (5095C>T), CAT581TCT (5096C>T 5097A>C), AGT582ACC (5100G>C 5101T>C), GAG584GAA (5107G>A), AAA585AAT (5110A>T), CTA588TTA (5117C>T), GCG589GCA (5122G>A), GTA590ATT (5123G>A 5125A>T), AAA591GTT (5126A>G 5127A>T 5128A>T), CAG592CGA (5130A>G 5131G>A), GTG593GGA (5133T>G 5134G>A), ATT594ATC (5137T>C), ACT595AGA (5139C>G 5140T>A), AGT598TCA (5147A>T 5148G>C 5149T>A), ATT599GCT (5150A>G 5151T>C), TAT600TTC (5154A>T 5155T>C), ACC602CTT (5159A>C 5160C>T 5161C>T), CCT603CCA (5164T>A), GTT604AAA (5165G>A 5166T>A 5167T>A), TGT605CAA (5168T>C 5169G>A 5170T>A), TTT606TTC (5173T>C), ACA607ATC (5175C>T 5176A>C), GTC608ATC (5177G>A), AGG609CAA (5180A>C 5181G>A 5182G>A), ACA610TCA (5183A>T), GAT611GAC (5188T>C), GTA613del (5192_5194delGTA), AAT614ACC (5196A>C 5197T>C), CTT615CAG (5199T>A 5200T>G), TTA616GTA (5201T>G), AAA617TCA (5204A>T 5205A>C), GGA618GGT (5209A>T), TTT619TTG (5212T>G), ATG620ATA (5215G>A), AAT621TTC (5216A>T 5217A>T 5218T>C), AAA622AAT (5221A>T), AAG623AAA (5224G>A), ATT624CTT (5225A>C), ACT625CCT (5228A>C), GGT626TCT (5231G>T 5232G>C), GAC627GAA (5236C>A), AGT628CC. (5237A>C 5238G>C) | | | | | | | | |

##### Proteins

| | Begin | End | Coverage | Score | Concordance | Matches | Identities | I/D/M/F* | Stop Codons |
|---|---|---|---|---|---|---|---|---|---|
| hypothetical protein (NP_659397.1) | 515 | 628 | 16.9% | 365 | 51.0% | 109 (95.6%) | 53 (46.5%) | 0/5/0/0 | 0 |
| Protein mutations: | K518Q (4907A>C), K519S (4911A>G 4912G>C), I520L (4913A>C), G523E (4923G>A), L524C (4926T>G 4927A>T), I525A (4928A>G 4929T>C 4930T>C), N526K (4933T>G), F527L (4934T>C 4936T>C), Y531R (4946T>A 4947A>G 4948T>G), K534E (4955A>G 4957G>A), K535P (4958A>C 4959A>C), S538N (4968G>A 4969T>C), I541L (4976A>T 4978T>A), E542Q (4979G>C), H548Y (4997C>T), F549H (5000T>C 5001T>A 5002C>T), G552A (5010G>C), V553L (5012G>T 5014C>G), K555_A556del (5018_5023delAAGGCC), T559D (5030A>G 5031C>A), E560L (5033G>T 5034A>T 5035A>G), G561N (5036G>A 5037G>A), E563_L564del (5042_5047delGAATTA), S569T (5060T>A), K574N (5077A>T), P575E (5078C>G 5079C>A), L578T (5087T>A 5088T>C 5089G>T), N579R (5091A>G 5092T>A), H581S (5096C>T 5097A>C), S582T (5100G>C 5101T>C), K585N (5110A>T), V590I (5123G>A 5125A>T), K591V (5126A>G 5127A>T 5128A>T), Q592R (5130A>G 5131G>A), V593G (5133T>G 5134G>A), T595R (5139C>G 5140T>A), I599A (5150A>G 5151T>C), Y600F (5154A>T 5155T>C), T602L (5159A>C 5160C>T 5161C>T), V604K (5165G>A 5166T>A 5167T>A), C605Q (5168T>C 5169G>A 5170T>A), T607I (5175C>T 5176A>C), V608I (5177G>A), R609Q (5180A>C 5181G>A 5182G>A), T610S (5183A>T), V613del (5192_5194delGTA), N614T (5196A>C 5197T>C), L615Q (5199T>A 5200T>G), L616V (5201T>G), K617S (5204A>T 5205A>C), F619L (5212T>G), M620I (5215G>A), N621F (5216A>T 5217A>T 5218T>C), K622N (5221A>T), I624L (5225A>C), T625P (5228A>C), G626S (5231G>T 5232G>C), D627E (5236C>A) | | | | | | | | |
| Codon mutations: | TAC516TAT (4903C>T), AAA518CAA (4907A>C), AAG519AGC (4911A>G 4912G>C), ATA520CTA (4913A>C), AAG521AAA (4918G>A), AAA522AAG (4921A>G), GGA523GAA (4923G>A), TTA524TGT (4926T>G 4927A>T), ATT525GCC (4928A>G 4929T>C 4930T>C), AAT526AAG (4933T>G), TTT527CTC (4934T>C 4936T>C), CCA528CCG (4939A>G), AAA529AAG (4942A>G), TAT531AGG (4946T>A 4947A>G 4948T>G), AAG534GAA (4955A>G 4957G>A), AAA535CCA (4958A>C 4959A>C), GAC537GAT (4966C>T), AGT538AAC (4968G>A 4969T>C), TTA539CTA (4970T>C), ATT540ATA (4975T>A), ATT541TTA (4976A>T 4978T>A), GAA542CAA (4979G>C), ACT543ACG (4984T>G), GCT545GCC (4990T>C), TCT546TCC (4993T>C), GAT547GAC (4996T>C), CAC548TAC (4997C>T), TTC549CAT (5000T>C 5001T>A 5002C>T), GGT551GGA (5008T>A), GGA552GCA (5010G>C), GTC553TTG (5012G>T 5014C>G), CTT554TTA (5015C>T 5017T>A), AAG555_GCC556del (5018_5023delAAGGCC), ACC558ACA (5029C>A), ACT559GAT (5030A>G 5031C>A), GAA560TTG (5033G>T 5034A>T 5035A>G), GGT561AAT (5036G>A 5037G>A), GAA563_TTA564del (5042_5047delGAATTA), ATC565ATT (5050C>T), TGC566TGT (5053C>T), AGG567AGA (5056G>A), TCT569ACT (5060T>A), TCA570AGT (5063T>A 5064C>G 5065A>T), ACA572ACT (5071A>T), TTC573TTT (5074C>T), AAA574AAT (5077A>T), CCA575GAA (5078C>G 5079C>A), TTG578ACT (5087T>A 5088T>C 5089G>T), AAT579AGA (5091A>G 5092T>A), TAC580TAT (5095C>T), CAT581TCT (5096C>T 5097A>C), AGT582ACC (5100G>C 5101T>C), GAG584GAA (5107G>A), AAA585AAT (5110A>T), CTA588TTA (5117C>T), GCG589GCA (5122G>A), GTA590ATT (5123G>A 5125A>T), AAA591GTT (5126A>G 5127A>T 5128A>T), CAG592CGA (5130A>G 5131G>A), GTG593GGA (5133T>G 5134G>A), ATT594ATC (5137T>C), ACT595AGA (5139C>G 5140T>A), AGT598TCA (5147A>T 5148G>C 5149T>A), ATT599GCT (5150A>G 5151T>C), TAT600TTC (5154A>T 5155T>C), ACC602CTT (5159A>C 5160C>T 5161C>T), CCT603CCA (5164T>A), GTT604AAA (5165G>A 5166T>A 5167T>A), TGT605CAA (5168T>C 5169G>A 5170T>A), TTT606TTC (5173T>C), ACA607ATC (5175C>T 5176A>C), GTC608ATC (5177G>A), AGG609CAA (5180A>C 5181G>A 5182G>A), ACA610TCA (5183A>T), GAT611GAC (5188T>C), GTA613del (5192_5194delGTA), AAT614ACC (5196A>C 5197T>C), CTT615CAG (5199T>A 5200T>G), TTA616GTA (5201T>G), AAA617TCA (5204A>T 5205A>C), GGA618GGT (5209A>T), TTT619TTG (5212T>G), ATG620ATA (5215G>A), AAT621TTC (5216A>T 5217A>T 5218T>C), AAA622AAT (5221A>T), AAG623AAA (5224G>A), ATT624CTT (5225A>C), ACT625CCT (5228A>C), GGT626TCT (5231G>T 5232G>C), GAC627GAA (5236C>A), AGT628CC. (5237A>C 5238G>C) | | | | | | | | |

*: Inserts / Deletes / Misaligned / Frameshifts

#### Analysis details

This analysis was performed with panviral2.64

### NGS Details (UN70): Badnavirus venabougainvilleae

#### Assembly

| | |
|---|---|
| **Coverage Length** | 374 (1 contig(s)) |
| **Depth Of Coverage** | 3.5 |
| **Number Of Reads** | 10 |
| **Reads Per Million** | 0.19 rpm (after QC) |
| **Ambiguities** | 0 |
| **Assembly Method** | de novo + reference guided assembly |
| **Consensus Caller** | Bcf Tools |

#### Coverage Map

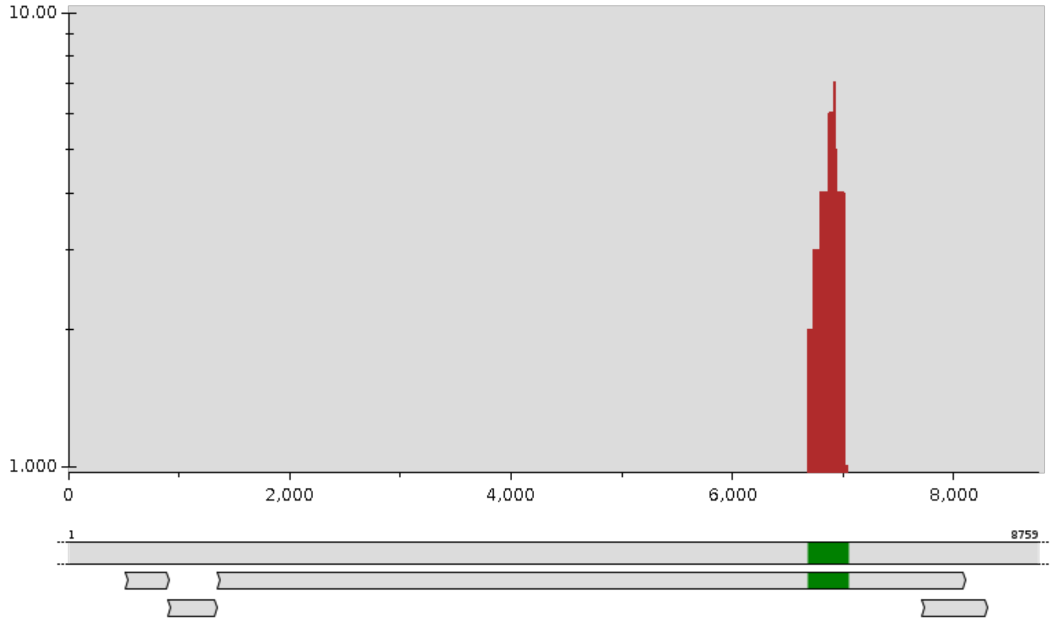

#### Assignment

| | |
|---|---|
| **Type** | Badnavirus venabougainvilleae (Taxonomy ID: 3047697) |
| **Reference Genome** | NC_011592.1 |
| **NT Identity (%)** | 51.7241 |
| **AA Identity (%)** | 39.6825 |
| **Number Of Stop Codons** | 1 |
| **Number Of CDS** | 4 |

#### Alignment

| | |
|---|---|
| **Alignment Score** | 20.0 (NT) + 330.0 (AA) = 350.0 |
| **Concordance (%)** | 21.4067 |

| Alignment Method | Global, seeded, nucleotide + amino acids (AGA) |
|---|---|

#### Genome Region

Sequence starts at position 6681 and ends at position 7054 relative to NC_011592.1 reference sequence.

#### Alignment Detailed Statistics

| | Begin | End | Coverage | Score | Concordance | Matches | Identities | I/D/M/F* | Stop Codons |
|---|---|---|---|---|---|---|---|---|---|
| **NT** | **6681** | **7054** | **4.3%** | **20** | **2.7%** | **374 (99.2%)** | **195 (51.7%)** | **3/0** | |
| Mutations: | 6681A>C, 6684C>G, 6686A>T, 6690A>C, 6693A>G, 6694T>G, 6697G>T, 6700C>T, 6704A>G, 6705G>A, 6706A>T, 6707G>C, 6713T>A, 6720G>T, 6721G>C, 6723T>G, 6724G>A, 6725C>G, 6726T>A, 6729G>T, 6730G>A, 6731A>G, 6732T>G, 6733C>G, 6735A>T, 6736G>T, 6737A>T, 6738A>T, 6739T>G, 6740C>A, 6744A>T, 6747A>T, 6748T>A, 6749G>A, 6753G>A, 6754G>A, 6756C>T, 6759C>T, 6761T>G, 6762T>G, 6766C>A, 6767C>G, 6768A>G, 6769C>T, 6771A>T, 6775T>C, 6776T>A, 6777G>C, 6779T>A, 6783A>G, 6785G>C, 6786G>C, 6789A>T, 6792C>G, 6796C>T, 6798A>C, 6804T>A, 6807A>G, 6809A>C, 6810G>T, 6816A>T, 6818C>T, 6822T>C, 6823A>G, 6824T>C, 6825C>A, 6828C>T, 6829C>A, 6830A>T, 6831A>G, 6832A>G, 6833G>A, 6835A>T, 6836A>T, 6841G>A, 6843C>T, 6844A>G, 6845A>T, 6846C>T, 6847T>G, 6848G>T, 6849C>A, 6853T>C, 6854C>T, 6857G>C, 6861C>T, 6862A>T, 6863G>T, 6864C>G, 6867_6868insTTA, 6870C>T, 6871A>G, 6874G>A, 6875C>T, 6879A>C, 6881A>T, 6885T>C, 6891C>T, 6900G>A, 6902T>A, 6903T>C, 6906T>C, 6907G>A, 6911G>C, 6912T>C, 6913G>A, 6916A>G, 6917G>A, 6918A>G, 6921C>A, 6925G>A, 6928A>C, 6930G>A, 6933T>C, 6934C>T, 6936C>G, 6939A>G, 6940G>A, 6943A>G, 6945G>A, 6948T>C, 6951A>G, 6952G>T, 6953T>G, 6954T>A, 6955T>C, 6956G>T, 6958C>A, 6959A>G, 6963A>G, 6966T>G, 6967G>A, 6968G>A, 6969C>G, 6973G>T, 6974T>A, 6976C>G, 6977T>C, 6979T>A, 6980C>A, 6982C>T, 6983C>T, 6984C>T, 6985A>T, 6990A>G, 6991A>T, 6992T>G, 6993G>C, 6997G>T, 7000G>T, 7001C>G, 7003G>C, 7005C>G, 7007A>G, 7008A>T, 7009A>T, 7011T>G, 7012A>G, 7014T>G, 7016A>C, 7017G>A, 7020C>T, 7023T>A, 7026A>C, 7027G>T, 7028C>A, 7029A>T, 7033C>G, 7036G>T, 7037G>C, 7038G>T, 7039A>G, 7040A>T, 7041T>A, 7042C>A, 7044A>G, 7045T>G, 7048G>A, 7052A>G, 7054T>G | | | | | | | | |

##### CDS

| | Begin | End | Coverage | Score | Concordance | Matches | Identities | I/D/M/F* | Stop Codons |
|---|---|---|---|---|---|---|---|---|---|
| BSCVV_gp3 | 1780 | 1904 | 5.5% | 330 | 36.9% | 125 (99.2%) | 50 (39.7%) | 1/0/0/0 | 1 |
| Protein mutations: | I1780M (6684C>G), Y1781F (6686A>T), F1784V (6694T>G), D1785Y (6697G>T), K1787R (6704A>G 6705G>A), F1790Y (6713T>A), Q1792H (6720G>T), V1793L (6721G>C 6723T>G), A1794R (6724G>A 6725C>G 6726T>A), M1795I (6729G>T), D1796R (6730G>A 6731A>G 6732T>G), P1797A (6733C>G 6735A>T), E1798F (6736G>T 6737A>T 6738A>T), S1799D (6739T>G 6740C>A), W1802K (6748T>A 6749G>A), A1804T (6754G>A 6756C>T), L1806R (6761T>G 6762T>G), P1808R (6766C>A 6767C>G 6768A>G), Q1809Y (6769C>T 6771A>T), L1811H (6775T>C 6776T>A 6777G>C), F1812Y (6779T>A), W1814S (6785G>C 6786G>C), P1818S (6796C>T 6798A>C), K1822T (6809A>C 6810G>T), P1825L (6818C>T), I1827A (6823A>G 6824T>C 6825C>A), Q1829M (6829C>A 6830A>T 6831A>G), R1830E (6832A>G 6833G>A), K1831L (6835A>T 6836A>T), D1833N (6841G>A 6843C>T), N1834V (6844A>G 6845A>T 6846C>T), C1835V (6847T>G 6848G>T 6849C>A), S1837L (6853T>C 6854C>T), R1838P (6857G>C), S1840L (6862A>T 6863G>T 6864C>G), D1841_F1842insL (6867_6868insTTA), I1843V (6871A>G), A1844I (6874G>A 6875C>T), Y1846F (6881A>T), F1853Y (6902T>A 6903T>C), E1855K (6907G>A), S1856T (6911G>C 6912T>C), E1857K (6913G>A), R1858E (6916A>G 6917G>A 6918A>G), D1859E (6921C>A), E1861K (6925G>A), K1862Q (6928A>C 6930G>A), V1866I (6940G>A), M1867V (6943A>G 6945G>A), V1870* (6952G>T 6953T>G 6954T>A), C1871L (6955T>C 6956G>T), Q1872R (6958C>A 6959A>G), N1874K (6966T>G), G1875K (6967G>A 6968G>A 6969C>G), V1877Y (6973G>T 6974T>A), L1878A (6976C>G 6977T>C), S1879N (6979T>A 6980C>A), P1880F (6982C>T 6983C>T 6984C>T), T1881S (6985A>T), M1883C (6991A>T 6992T>G 6993G>C), V1885F (6997G>T), A1886W (7000G>T 7001C>G), V1887L (7003G>C 7005C>G), K1888S (7007A>G 7008A>T), T1889S (7009A>T 7011T>G), I1890V (7012A>G 7014T>G), E1891A (7016A>C 7017G>A), A1895Y (7027G>T 7028C>A 7029A>T), L1897V (7033C>G), G1898S (7036G>T 7037G>C 7038G>T), N1899V (7039A>G 7040A>T 7041T>A), Q1900K (7042C>A 7044A>G), C1901G (7045T>G), V1902I (7048G>A), Q1903R (7052A>G) | | | | | | | | |
| Codon mutations: | AAA1779..C (6681A>C), ATC1780ATG (6684C>G), TAC1781TTC (6686A>T), TCA1782TCC (6690A>C), AAA1783AAG (6693A>G), TTT1784GTT (6694T>G), GAT1785TAT (6697G>T), CTG1786TTG (6700C>T), AAG1787AGA (6704A>G 6705G>A), AGT1788TCT (6706A>T 6707G>C), TTT1790TAT (6713T>A), CAG1792CAT (6720G>T), GTT1793CTG (6721G>C 6723T>G), GCT1794AGA (6724G>A 6725C>G 6726T>A), ATG1795ATT (6729G>T), GAT1796AGG (6730G>A 6731A>G 6732T>G), CCA1797GCT (6733C>G 6735A>T), GAA1798TTT (6736G>T 6737A>T 6738A>T), TCC1799GAC (6739T>G 6740C>A), ATA1800ATT (6744A>T), CCA1801CCT (6747A>T), TGG1802AAG (6748T>A 6749G>A), ACG1803ACA (6753G>A), GCC1804ACT (6754G>A 6756C>T), TTC1805TTT (6759C>T), CTT1806CGG (6761T>G 6762T>G), CCA1808AGG (6766C>A 6767C>G 6768A>G), CAA1809TAT (6769C>T 6771A>T), TTG1811CAC (6775T>C 6776T>A 6777G>C), TTT1812TAT (6779T>A), GAA1813GAG (6783A>G), TGG1814TCC (6785G>C 6786G>C), CTA1815CTT (6789A>T), GTC1816GTG (6792C>G), CCA1818TCC (6796C>T 6798A>C), GGT1820GGA (6804T>A), CTA1821CTG (6807A>G), AAG1822ACT (6809A>C 6810G>T), GCA1824GCT (6816A>T), CCA1825CTA (6818C>T), GCT1826GCC (6822T>C), ATC1827GCA (6823A>G 6824T>C 6825C>A), TTC1828TTT (6828C>T), CAA1829ATG (6829C>A 6830A>T 6831A>G), AGG1830GAG (6832A>G 6833G>A), AAG1831TTG (6835A>T 6836A>T), GAC1833AAT (6841G>A 6843C>T), AAC1834GTT (6844A>G 6845A>T 6846C>T), TGC1835GTA (6847T>G 6848G>T 6849C>A), TCA1837CTA (6853T>C 6854C>T), CGA1838CCA (6857G>C), TAC1839TAT (6861C>T), AGC1840TTG (6862A>T 6863G>T 6864C>G), GAT1841_TTC1842insTTA (6867_6868insTTA), TTC1842TTT (6870C>T), ATT1843GTT (6871A>G), GCT1844ATT (6874G>A 6875C>T), GTA1845GTC (6879A>C), TAC1846TTC (6881A>T), ATT1847ATC (6885T>C), GAC1849GAT (6891C>T), GTG1852GTA (6900G>A), TTT1853TAC (6902T>A 6903T>C), TCT1854TCC (6906T>C), GAG1855AAG (6907G>A), AGT1856ACC (6911G>C 6912T>C), GAA1857AAA (6913G>A), AGA1858GAG (6916A>G 6917G>A 6918A>G), GAC1859GAA (6921C>A), GAG1861AAG (6925G>A), AAG1862CAA (6928A>C 6930G>A), CAT1863CAC (6933T>C), CTC1864TTG (6934C>T 6936C>G), AGA1865AGG (6939A>G), GTT1866ATT (6940G>A), ATG1867GTA (6943A>G 6945G>A), CTT1868CTC (6948T>C), CAA1869CAG (6951A>G), GTT1870TGA (6952G>T 6953T>G 6954T>A), TGT1871CTT (6955T>C 6956G>T), CAA1872AGA (6958C>A 6959A>G), GAA1873GAG (6963A>G), AAT1874AAG (6966T>G), GGC1875AAG (6967G>A 6968G>A 6969C>G), GTT1877TAT (6973G>T 6974T>A), CTA1878GCA (6976C>G 6977T>C), TCC1879AAC (6979T>A 6980C>A), CCC1880TTT (6982C>T 6983C>T 6984C>T), ACT1881TCT (6985A>T), AAA1882AAG (6990A>G), ATG1883TGC (6991A>T 6992T>G 6993G>C), GTT1885TTT (6997G>T), GCG1886TGG (7000G>T 7001C>G), GTC1887CTG (7003G>C 7005C>G), AAA1888AGT (7007A>G 7008A>T), ACT1889TCG (7009A>T 7011T>G), ATT1890GTG (7012A>G 7014T>G), GAG1891GCA (7016A>C 7017G>A), TTC1892TTT (7020C>T), CTT1893CTA (7023T>A), GGA1894GGC (7026A>C), GCA1895TAT (7027G>T 7028C>A 7029A>T), CTA1897GTA (7033C>G), GGG1898TCT (7036G>T 7037G>C 7038G>T), AAT1899GTA (7039A>G 7040A>T 7041T>A), CAA1900AAG (7042C>A 7044A>G), TGC1901GGC (7045T>G), GTT1902ATT (7048G>A), CAG1903CGG (7052A>G), TTA1904G.. (7054T>G) | | | | | | | | |

##### Proteins

| | Begin | End | Coverage | Score | Concordance | Matches | Identities | I/D/M/F* | Stop Codons |
|---|---|---|---|---|---|---|---|---|---|
| polyprotein (YP_002321513.1) | 1780 | 1904 | 5.5% | 330 | 36.9% | 125 (99.2%) | 50 (39.7%) | 1/0/0/0 | 1 |
| Protein mutations: | I1780M (6684C>G), Y1781F (6686A>T), F1784V (6694T>G), D1785Y (6697G>T), K1787R (6704A>G 6705G>A), F1790Y (6713T>A), Q1792H (6720G>T), V1793L (6721G>C 6723T>G), A1794R (6724G>A 6725C>G 6726T>A), M1795I (6729G>T), D1796R (6730G>A 6731A>G 6732T>G), P1797A (6733C>G 6735A>T), E1798F (6736G>T 6737A>T 6738A>T), S1799D (6739T>G 6740C>A), W1802K (6748T>A 6749G>A), A1804T (6754G>A 6756C>T), L1806R (6761T>G 6762T>G), P1808R (6766C>A 6767C>G 6768A>G), Q1809Y (6769C>T 6771A>T), L1811H (6775T>C 6776T>A 6777G>C), F1812Y (6779T>A), W1814S (6785G>C 6786G>C), P1818S (6796C>T 6798A>C), K1822T (6809A>C 6810G>T), P1825L (6818C>T), I1827A (6823A>G 6824T>C 6825C>A), Q1829M (6829C>A 6830A>T 6831A>G), R1830E (6832A>G 6833G>A), K1831L (6835A>T 6836A>T), D1833N (6841G>A 6843C>T), N1834V (6844A>G 6845A>T 6846C>T), C1835V (6847T>G 6848G>T 6849C>A), S1837L (6853T>C 6854C>T), R1838P (6857G>C), S1840L (6862A>T 6863G>T 6864C>G), D1841_F1842insL (6867_6868insTTA), I1843V (6871A>G), A1844I (6874G>A 6875C>T), Y1846F (6881A>T), F1853Y (6902T>A 6903T>C), E1855K (6907G>A), S1856T (6911G>C 6912T>C), E1857K (6913G>A), R1858E (6916A>G 6917G>A 6918A>G), D1859E (6921C>A), E1861K (6925G>A), K1862Q (6928A>C 6930G>A), V1866I (6940G>A), M1867V (6943A>G 6945G>A), V1870* (6952G>T 6953T>G 6954T>A), C1871L (6955T>C 6956G>T), Q1872R (6958C>A 6959A>G), N1874K (6966T>G), G1875K (6967G>A 6968G>A 6969C>G), V1877Y (6973G>T 6974T>A), L1878A (6976C>G 6977T>C), S1879N (6979T>A 6980C>A), P1880F (6982C>T 6983C>T 6984C>T), T1881S (6985A>T), M1883C (6991A>T 6992T>G 6993G>C), V1885F (6997G>T), A1886W (7000G>T 7001C>G), V1887L (7003G>C 7005C>G), K1888S (7007A>G 7008A>T), T1889S (7009A>T 7011T>G), I1890V (7012A>G 7014T>G), E1891A (7016A>C 7017G>A), A1895Y (7027G>T 7028C>A 7029A>T), L1897V (7033C>G), G1898S (7036G>T 7037G>C 7038G>T), N1899V (7039A>G 7040A>T 7041T>A), Q1900K (7042C>A 7044A>G), C1901G (7045T>G), V1902I (7048G>A), Q1903R (7052A>G) | | | | | | | | |
| Codon mutations: | AAA1779..C (6681A>C), ATC1780ATG (6684C>G), TAC1781TTC (6686A>T), TCA1782TCC (6690A>C), AAA1783AAG (6693A>G), TTT1784GTT (6694T>G), GAT1785TAT (6697G>T), CTG1786TTG (6700C>T), AAG1787AGA (6704A>G 6705G>A), AGT1788TCT (6706A>T 6707G>C), TTT1790TAT (6713T>A), CAG1792CAT (6720G>T), GTT1793CTG (6721G>C 6723T>G), GCT1794AGA (6724G>A 6725C>G 6726T>A), ATG1795ATT (6729G>T), GAT1796AGG (6730G>A 6731A>G 6732T>G), CCA1797GCT (6733C>G 6735A>T), GAA1798TTT (6736G>T 6737A>T 6738A>T), TCC1799GAC (6739T>G 6740C>A), ATA1800ATT (6744A>T), CCA1801CCT (6747A>T), TGG1802AAG (6748T>A 6749G>A), ACG1803ACA (6753G>A), GCC1804ACT (6754G>A 6756C>T), TTC1805TTT (6759C>T), CTT1806CGG (6761T>G 6762T>G), CCA1808AGG (6766C>A 6767C>G 6768A>G), CAA1809TAT (6769C>T 6771A>T), TTG1811CAC (6775T>C 6776T>A 6777G>C), TTT1812TAT (6779T>A), GAA1813GAG (6783A>G), TGG1814TCC (6785G>C 6786G>C), CTA1815CTT (6789A>T), GTC1816GTG (6792C>G), CCA1818TCC (6796C>T 6798A>C), GGT1820GGA (6804T>A), CTA1821CTG (6807A>G), AAG1822ACT (6809A>C 6810G>T), GCA1824GCT (6816A>T), CCA1825CTA (6818C>T), GCT1826GCC (6822T>C), ATC1827GCA (6823A>G 6824T>C 6825C>A), TTC1828TTT (6828C>T), CAA1829ATG (6829C>A 6830A>T 6831A>G), AGG1830GAG (6832A>G 6833G>A), AAG1831TTG (6835A>T 6836A>T), GAC1833AAT (6841G>A 6843C>T), AAC1834GTT (6844A>G 6845A>T 6846C>T), TGC1835GTA (6847T>G 6848G>T 6849C>A), TCA1837CTA (6853T>C 6854C>T), CGA1838CCA (6857G>C), TAC1839TAT (6861C>T), AGC1840TTG (6862A>T 6863G>T 6864C>G), GAT1841_TTC1842insTTA (6867_6868insTTA), TTC1842TTT (6870C>T), ATT1843GTT (6871A>G), GCT1844ATT (6874G>A 6875C>T), GTA1845GTC (6879A>C), TAC1846TTC (6881A>T), ATT1847ATC (6885T>C), GAC1849GAT (6891C>T), GTG1852GTA (6900G>A), TTT1853TAC (6902T>A 6903T>C), TCT1854TCC (6906T>C), GAG1855AAG (6907G>A), AGT1856ACC (6911G>C 6912T>C), GAA1857AAA (6913G>A), AGA1858GAG (6916A>G 6917G>A 6918A>G), GAC1859GAA (6921C>A), GAG1861AAG (6925G>A), AAG1862CAA (6928A>C 6930G>A), CAT1863CAC (6933T>C), CTC1864TTG (6934C>T 6936C>G), AGA1865AGG (6939A>G), GTT1866ATT (6940G>A), ATG1867GTA (6943A>G 6945G>A), CTT1868CTC (6948T>C), CAA1869CAG (6951A>G), GTT1870TGA (6952G>T 6953T>G 6954T>A), TGT1871CTT (6955T>C 6956G>T), CAA1872AGA (6958C>A 6959A>G), GAA1873GAG (6963A>G), AAT1874AAG (6966T>G), GGC1875AAG (6967G>A 6968G>A 6969C>G), GTT1877TAT (6973G>T 6974T>A), CTA1878GCA (6976C>G 6977T>C), TCC1879AAC (6979T>A 6980C>A), CCC1880TTT (6982C>T 6983C>T 6984C>T), ACT1881TCT (6985A>T), AAA1882AAG (6990A>G), ATG1883TGC (6991A>T 6992T>G 6993G>C), GTT1885TTT (6997G>T), GCG1886TGG (7000G>T 7001C>G), GTC1887CTG (7003G>C 7005C>G), AAA1888AGT (7007A>G 7008A>T), ACT1889TCG (7009A>T 7011T>G), ATT1890GTG (7012A>G 7014T>G), GAG1891GCA (7016A>C 7017G>A), TTC1892TTT (7020C>T), CTT1893CTA (7023T>A), GGA1894GGC (7026A>C), GCA1895TAT (7027G>T 7028C>A 7029A>T), CTA1897GTA (7033C>G), GGG1898TCT (7036G>T 7037G>C 7038G>T), AAT1899GTA (7039A>G 7040A>T 7041T>A), CAA1900AAG (7042C>A 7044A>G), TGC1901GGC (7045T>G), GTT1902ATT (7048G>A), CAG1903CGG (7052A>G), TTA1904G.. (7054T>G) | | | | | | | | |

*: Inserts / Deletes / Misaligned / Frameshifts

#### Analysis details

This analysis was performed with panviral2.64

### NGS Details (UN70): Caulimovirus tessellomirabilis

#### Assembly

| | |
|---|---|
| **Coverage Length** | 316 (1 contig(s)) |
| **Depth Of Coverage** | 3.3 |
| **Number Of Reads** | 8 |
| **Reads Per Million** | 0.15 rpm (after QC) |
| **Ambiguities** | 0 |
| **Assembly Method** | de novo + reference guided assembly |
| **Consensus Caller** | Bcf Tools |

#### Coverage Map

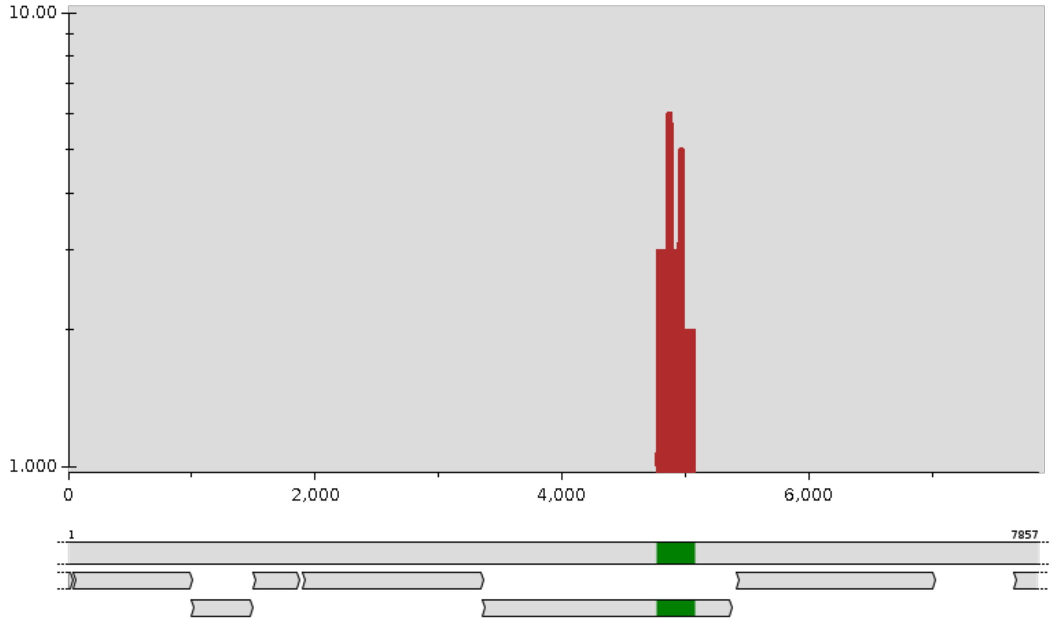

#### Assignment

| | |
|---|---|
| **Type** | Caulimovirus tessellomirabilis (Taxonomy ID: 3048203) |
| **Reference Genome** | NC_004036.1 |
| **NT Identity (%)** | 59.3443 |
| **AA Identity (%)** | 47.5728 |
| **Number Of Stop Codons** | 0 |
| **Number Of CDS** | 7 |

#### Alignment

| | |
|---|---|
| **Alignment Score** | 76.0 (NT) + 208.0 (AA) = 284.0 |
| **Concordance (%)** | 24.7012 |

| **Alignment Method** | Global, seeded, nucleotide + amino acids (AGA) |
|---|---|

#### Genome Region

Sequence starts at position 4769 and ends at position 5084 relative to NC_004036.1 reference sequence.

#### Alignment Detailed Statistics

| | Begin | End | Coverage | Score | Concordance | Matches | Identities | I/D/M/F* | Stop Codons |
|---|---|---|---|---|---|---|---|---|---|
| **NT** | **4769** | **5084** | **4.0%** | **76** | **13.1%** | **304 (95.9%)** | **181 (57.1%)** | **1/12** | |
| Mutations: | 4769C>G, 4772C>A, 4774T>A, 4789A>T, 4790G>T, 4791T>G, 4793C>T, 4797C>A, 4798A>T, 4806A>G, 4807T>A, 4808A>G, 4809G>A, 4810C>T, 4812A>T, 4813T>C, 4820A>G, 4822G>T, 4829G>A, 4833A>G, 4834A>G, 4835C>A, 4837G>A, 4840A>T, 4842C>T, 4843G>T, 4849G>A, 4850G>A, 4851T>A, 4853A>C, 4854A>T, 4855G>C, 4856T>A, 4858G>T, 4860A>G, 4861G>A, 4864A>G, 4865G>T, 4866A>C, 4867T>A, 4868C>A, 4870G>T, 4871G>C, 4872T>G, 4873A>T, 4874T>C, 4875G>T, 4876G>A, 4877A>G, 4878T>G, 4879C>A, 4885G>T, 4886C>G, 4889T>G, 4890C>A, 4891A>G, 4892G>C, 4898G>C, 4900T>A, 4902A>T, 4903C>G, 4903_4904insT, 4906T>A, 4907A>C, 4913A>C, 4915A>C, 4918G>A, 4919A>G, 4921A>G, 4923G>A, 4926T>G, 4927A>T, 4928A>G, 4929T>C, 4934T>C, 4942A>G, 4946T>A, 4947A>G, 4948T>A, 4954A>T, 4955A>G, 4957G>A, 4958A>C, 4959A>C, 4960A>C, 4966C>T, 4968G>A, 4973A>G, 4975T>C, 4976A>T, 4978T>A, 4979G>C, 4982A>T, 4984T>A, 4997C>T, 4999C>T, 5000T>C, 5001T>A, 5002C>T, 5004G>T, 5008T>A, 5010G>C, 5011A>T, 5012G>C, 5014C>A, 5015C>T, 5017T>A, 5018_5023delAAGGCC, 5029C>G, 5030A>G, 5031C>A, 5033G>T, 5034A>T, 5036G>A, 5037G>A, 5042_5047delGAATTA, 5053C>T, 5056G>A, 5060T>A, 5063T>A, 5064C>G, 5065A>T, 5071A>T, 5077A>C, 5078C>G, 5079C>A | | | | | | | | |
| **CDS** | | | | | | | | | |
| MiMVgp6 | 472 | 577 | 15.7% | 208 | 30.4% | 102 (95.3%) | 49 (45.8%) | 1/4/1/1 | 0 |
| Protein mutations: | Q472E (4769C>G), L473I (4772C>A 4774T>A), V479C (4790G>T 4791T>G), T481N (4797C>A 4798A>T), D484G (4806A>G 4807T>A), S485D (4808A>G 4809G>A 4810C>T), Y486F (4812A>T 4813T>C), K489D (4820A>G 4822G>T), E492K (4829G>A), K493R (4833A>G 4834A>G), K495N (4840A>T), P496L (4842C>T 4843G>T), V499K (4850G>A 4851T>A), K500L (4853A>C 4854A>T 4855G>C), L501I (4856T>A 4858G>T), K502R (4860A>G 4861G>A), D504S (4865G>T 4866A>C 4867T>A), Q505N (4868C>A 4870G>T), V506R (4871G>C 4872T>G 4873A>T), W507L (4874T>C 4875G>T 4876G>A), I508G (4877A>G 4878T>G 4879C>A), Q511E (4886C>G), S512E (4889T>G 4890C>A 4891A>G), D513H (4892G>C), D515Q (4898G>C 4900T>A), Y516L (4902A>T 4903C>G), Y516_V517insX (4903_4904insT), K518Q (4907A>C), I520L (4913A>C 4915A>C), K522E (4919A>G 4921A>G), G523E (4923G>A), L524C (4926T>G 4927A>T), I525A (4928A>G 4929T>C), F527L (4934T>C), Y531R (4946T>A 4947A>G 4948T>A), K534E (4955A>G 4957G>A), K535P (4958A>C 4959A>C 4960A>C), S538N (4968G>A), I540V (4973A>G 4975T>C), I541L (4976A>T 4978T>A), E542Q (4979G>C), T543S (4982A>T 4984T>A), H548Y (4997C>T 4999C>T), F549H (5000T>C 5001T>A 5002C>T), W550L (5004G>T), G552A (5010G>C 5011A>T), V553L (5012G>C 5014C>A), K555_A556del (5018_5023delAAGGCC), T559D (5030A>G 5031C>A), E560L (5033G>T 5034A>T), G561N (5036G>A 5037G>A), E563_L564del (5042_5047delGAATTA), S569T (5060T>A), K574N (5077A>C), P575E (5078C>G 5079C>A) | | | | | | | | |
| Codon mutations: | CAA472GAA (4769C>G), CTT473ATA (4772C>A 4774T>A), GGA478GGT (4789A>T), GTT479TGT (4790G>T 4791T>G), CTA480TTA (4793C>T), ACA481AAT (4797C>A 4798A>T), GAT484GGA (4806A>G 4807T>A), AGC485GAT (4808A>G 4809G>A 4810C>T), TAT486TTC (4812A>T 4813T>C), AAG489GAT (4820A>G 4822G>T), GAA492AAA (4829G>A), AAA493AGG (4833A>G 4834A>G), CGG494AGA (4835C>A 4837G>A), AAA495AAT (4840A>T), CCG496CTT (4842C>T 4843G>T), CAG498CAA (4849G>A), GTA499AAA (4850G>A 4851T>A), AAG500CTC (4853A>C 4854A>T 4855G>C), TTG501ATT (4856T>A 4858G>T), AAG502AGA (4860A>G 4861G>A), AAA503AAG (4864A>G), GAT504TCA (4865G>T 4866A>C 4867T>A), CAG505AAT (4868C>A 4870G>T), GTA506CGT (4871G>C 4872T>G 4873A>T), TGG507CTA (4874T>C 4875G>T 4876G>A), ATC508GGA (4877A>G 4878T>G 4879C>A), ACG510ACT (4885G>T), CAA511GAA (4886C>G), TCA512GAG (4889T>G 4890C>A 4891A>G), GAT513CAT (4892G>C), GAT515CAA (4898G>C 4900T>A), TAC516TTG (4902A>T 4903C>G), TAC516_GTT517insT-- (4903_4904insT), GTT517GTA (4906T>A), AAA518CAA (4907A>C), ATA520CTC (4913A>C 4915A>C), AAG521AAA (4918G>A), AAA522GAG (4919A>G 4921A>G), GGA523GAA (4923G>A), TTA524TGT (4926T>G 4927A>T), ATT525GCT (4928A>G 4929T>C), TTT527CTT (4934T>C), AAA529AAG (4942A>G), TAT531AGA (4946T>A 4947A>G 4948T>A), CCA533CCT (4954A>T), AAG534GAA (4955A>G 4957G>A), AAA535CCC (4958A>C 4959A>C 4960A>C), GAC537GAT (4966C>T), AGT538AAT (4968G>A), ATT540GTC (4973A>G 4975T>C), ATT541TTA (4976A>T 4978T>A), GAA542CAA (4979G>C), ACT543TCA (4982A>T 4984T>A), CAC548TAT (4997C>T 4999C>T), TTC549CAT (5000T>C 5001T>A 5002C>T), TGG550TTG (5004G>T), GGT551GGA (5008T>A), GGA552GCT (5010G>C 5011A>T), GTC553CTA (5012G>C 5014C>A), CTT554TTA (5015C>T 5017T>A), AAG555_GCC556del (5018_5023delAAGGCC), ACC558ACG (5029C>G), ACT559GAT (5030A>G 5031C>A), GAA560TTA (5033G>T 5034A>T), GGT561AAT (5036G>A 5037G>A), GAA563_TTA564del (5042_5047delGAATTA), TGC566TGT (5053C>T), AGG567AGA (5056G>A), TCT569ACT (5060T>A), TCA570AGT (5063T>A 5064C>G 5065A>T), ACA572ACT (5071A>T), AAA574AAC (5077A>C), CCA575GAA (5078C>G 5079C>A) | | | | | | | | |
| **Proteins** | | | | | | | | | |
| hypothetical protein (NP_659397.1) | 472 | 577 | 15.7% | 208 | 30.4% | 102 (95.3%) | 49 (45.8%) | 1/4/1/1 | 0 |
| Protein mutations: | Q472E (4769C>G), L473I (4772C>A 4774T>A), V479C (4790G>T 4791T>G), T481N (4797C>A 4798A>T), D484G (4806A>G 4807T>A), S485D (4808A>G 4809G>A 4810C>T), Y486F (4812A>T 4813T>C), K489D (4820A>G 4822G>T), E492K (4829G>A), K493R (4833A>G 4834A>G), K495N (4840A>T), P496L (4842C>T 4843G>T), V499K (4850G>A 4851T>A), K500L (4853A>C 4854A>T 4855G>C), L501I (4856T>A 4858G>T), K502R (4860A>G 4861G>A), D504S (4865G>T 4866A>C 4867T>A), Q505N (4868C>A 4870G>T), V506R (4871G>C 4872T>G 4873A>T), W507L (4874T>C 4875G>T 4876G>A), I508G (4877A>G 4878T>G 4879C>A), Q511E (4886C>G), S512E (4889T>G 4890C>A 4891A>G), D513H (4892G>C), D515Q (4898G>C 4900T>A), Y516L (4902A>T 4903C>G), Y516_V517insX (4903_4904insT), K518Q (4907A>C), I520L (4913A>C 4915A>C), K522E (4919A>G 4921A>G), G523E (4923G>A), L524C (4926T>G 4927A>T), I525A (4928A>G 4929T>C), F527L (4934T>C), Y531R (4946T>A 4947A>G 4948T>A), K534E (4955A>G 4957G>A), K535P (4958A>C 4959A>C 4960A>C), S538N (4968G>A), I540V (4973A>G 4975T>C), I541L (4976A>T 4978T>A), E542Q (4979G>C), T543S (4982A>T 4984T>A), H548Y (4997C>T 4999C>T), F549H (5000T>C 5001T>A 5002C>T), W550L (5004G>T), G552A (5010G>C 5011A>T), V553L (5012G>C 5014C>A), K555_A556del (5018_5023delAAGGCC), T559D (5030A>G 5031C>A), E560L (5033G>T 5034A>T), G561N (5036G>A 5037G>A), E563_L564del (5042_5047delGAATTA), S569T (5060T>A), K574N (5077A>C), P575E (5078C>G 5079C>A) | | | | | | | | |
| Codon mutations: | CAA472GAA (4769C>G), CTT473ATA (4772C>A 4774T>A), GGA478GGT (4789A>T), GTT479TGT (4790G>T 4791T>G), CTA480TTA (4793C>T), ACA481AAT (4797C>A 4798A>T), GAT484GGA (4806A>G 4807T>A), AGC485GAT (4808A>G 4809G>A 4810C>T), TAT486TTC (4812A>T 4813T>C), AAG489GAT (4820A>G 4822G>T), GAA492AAA (4829G>A), AAA493AGG (4833A>G 4834A>G), CGG494AGA (4835C>A 4837G>A), AAA495AAT (4840A>T), CCG496CTT (4842C>T 4843G>T), CAG498CAA (4849G>A), GTA499AAA (4850G>A 4851T>A), AAG500CTC (4853A>C 4854A>T 4855G>C), TTG501ATT (4856T>A 4858G>T), AAG502AGA (4860A>G 4861G>A), AAA503AAG (4864A>G), GAT504TCA (4865G>T 4866A>C 4867T>A), CAG505AAT (4868C>A 4870G>T), GTA506CGT (4871G>C 4872T>G 4873A>T), TGG507CTA (4874T>C 4875G>T 4876G>A), ATC508GGA (4877A>G 4878T>G 4879C>A), ACG510ACT (4885G>T), CAA511GAA (4886C>G), TCA512GAG (4889T>G 4890C>A 4891A>G), GAT513CAT (4892G>C), GAT515CAA (4898G>C 4900T>A), TAC516TTG (4902A>T 4903C>G), TAC516_GTT517insT-- (4903_4904insT), GTT517GTA (4906T>A), AAA518CAA (4907A>C), ATA520CTC (4913A>C 4915A>C), AAG521AAA (4918G>A), AAA522GAG (4919A>G 4921A>G), GGA523GAA (4923G>A), TTA524TGT (4926T>G 4927A>T), ATT525GCT (4928A>G 4929T>C), TTT527CTT (4934T>C), AAA529AAG (4942A>G), TAT531AGA (4946T>A 4947A>G 4948T>A), CCA533CCT (4954A>T), AAG534GAA (4955A>G 4957G>A), AAA535CCC (4958A>C 4959A>C 4960A>C), GAC537GAT (4966C>T), AGT538AAT (4968G>A), ATT540GTC (4973A>G 4975T>C), ATT541TTA (4976A>T 4978T>A), GAA542CAA (4979G>C), ACT543TCA (4982A>T 4984T>A), CAC548TAT (4997C>T 4999C>T), TTC549CAT (5000T>C 5001T>A 5002C>T), TGG550TTG (5004G>T), GGT551GGA (5008T>A), GGA552GCT (5010G>C 5011A>T), GTC553CTA (5012G>C 5014C>A), CTT554TTA (5015C>T 5017T>A), AAG555_GCC556del (5018_5023delAAGGCC), ACC558ACG (5029C>G), ACT559GAT (5030A>G 5031C>A), GAA560TTA (5033G>T 5034A>T), GGT561AAT (5036G>A 5037G>A), GAA563_TTA564del (5042_5047delGAATTA), TGC566TGT (5053C>T), AGG567AGA (5056G>A), TCT569ACT (5060T>A), TCA570AGT (5063T>A 5064C>G 5065A>T), ACA572ACT (5071A>T), AAA574AAC (5077A>C), CCA575GAA (5078C>G 5079C>A) | | | | | | | | |

*: Inserts / Deletes / Misaligned / Frameshifts

#### Analysis details

This analysis was performed with panviral2.64

### NGS Details (UN70): Cavemovirus collusipomeae

#### Assembly

| | |
|---|---|
| **Coverage Length** | 312 (1 contig(s)) |
| **Depth Of Coverage** | 3.2 |
| **Number Of Reads** | 8 |
| **Reads Per Million** | 0.15 rpm (after QC) |
| **Ambiguities** | 0 |
| **Assembly Method** | de novo + reference guided assembly |
| **Consensus Caller** | Bcf Tools |

#### Coverage Map

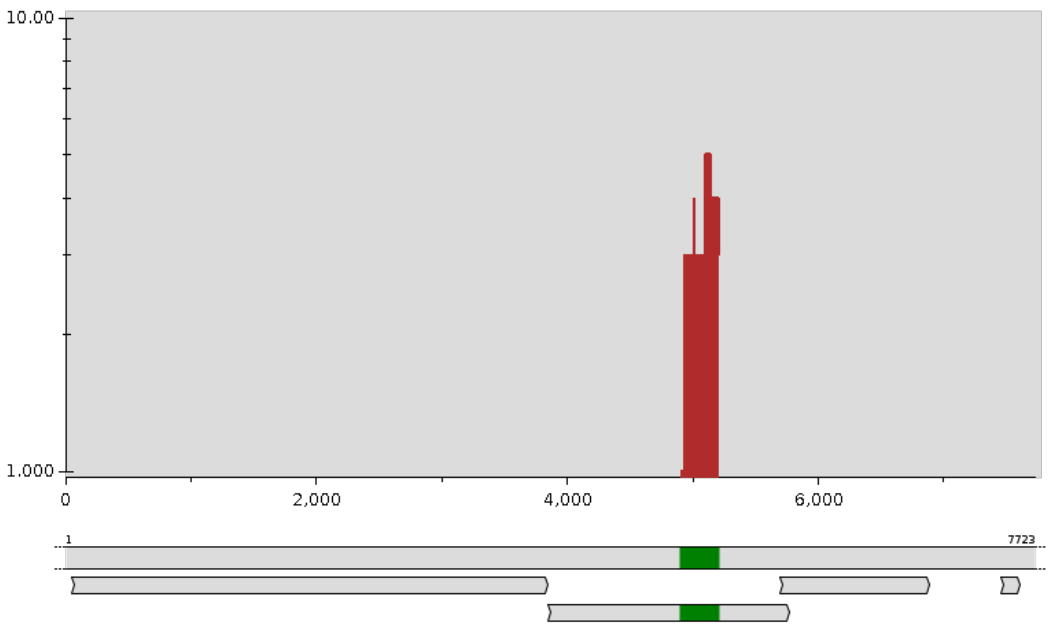

#### Assignment

| | |
|---|---|
| **Type** | Cavemovirus collusipomeae (Taxonomy ID: 3048352) |
| **Reference Genome** | NC_015328.1 |
| **NT Identity (%)** | 63.141 |
| **AA Identity (%)** | 50.9615 |
| **Number Of Stop Codons** | 0 |
| **Number Of CDS** | 4 |

#### Alignment

| | |
|---|---|
| **Alignment Score** | 164.0 (NT) + 356.0 (AA) = 520.0 |
| **Concordance (%)** | 40.0925 |

| Alignment Method | Global, seeded, nucleotide + amino acids (AGA) |
|---|---|

#### Genome Region

Sequence starts at position 4898 and ends at position 5209 relative to NC_015328.1 reference sequence.

#### Alignment Detailed Statistics

| | Begin | End | Coverage | Score | Concordance | Matches | Identities | I/D/M/F* | Stop Codons |
|---|---|---|---|---|---|---|---|---|---|
| **NT** | **4898** | **5209** | **4.0%** | **164** | **26.3%** | **312 (100%)** | **197 (63.1%)** | **0/0** | |
| Mutations: | 4898G>A, 4904A>C, 4907C>A, 4909T>C, 4912A>C, 4915T>C, 4935T>G, 4939G>T, 4942A>T, 4943A>T, 4949G>C, 4958A>T, 4959A>T, 4962T>C, 4963A>T, 4967T>C, 4969G>T, 4970G>A, 4973A>C, 4974T>A, 4978T>C, 4979A>T, 4980G>C, 4981A>T, 4982A>C, 4984T>A, 4985A>T, 4988A>T, 4989T>G, 4990A>T, 4991A>T, 4992T>G, 4993A>C, 4994A>G, 4997C>A, 5005A>T, 5008A>G, 5014T>C, 5015A>C, 5017G>A, 5023A>G, 5029G>C, 5033G>A, 5034G>A, 5035A>T, 5039C>A, 5041A>T, 5042A>G, 5047A>T, 5056A>C, 5060G>A, 5062T>G, 5064A>T, 5065G>A, 5068A>T, 5069G>T, 5070A>C, 5072C>A, 5074A>G, 5075G>A, 5076G>A, 5077A>T, 5080A>C, 5083A>T, 5092G>A, 5093C>A, 5102A>T, 5103T>C, 5105G>A, 5107T>A, 5110A>G, 5113A>C, 5115T>G, 5116A>T, 5120A>T, 5121A>T, 5122A>T, 5124A>C, 5125T>A, 5126A>G, 5127T>A, 5128A>T, 5132A>C, 5137A>C, 5139G>C, 5140T>A, 5144A>C, 5150T>A, 5152A>C, 5156T>A, 5157C>G, 5158A>G, 5159A>T, 5161A>T, 5162T>C, 5164A>T, 5169T>G, 5170A>T, 5171G>T, 5176C>T, 5177C>T, 5179A>C, 5180A>G, 5181T>C, 5182T>A, 5185A>T, 5186A>G, 5188T>A, 5191C>T, 5192T>A, 5198A>G, 5200T>C, 5204A>G, 5205G>C, 5206T>A | | | | | | | | |

##### CDS

| | Begin | End | Coverage | Score | Concordance | Matches | Identities | I/D/M/F* | Stop Codons |
|---|---|---|---|---|---|---|---|---|---|
| SPCV_gp2 | 353 | 456 | 16.2% | 356 | 52.9% | 104 (100%) | 53 (51.0%) | 0/0/0/0 | 0 |
| Protein mutations: | D353N (4898G>A), I355L (4904A>C), L356I (4907C>A 4909T>C), F365C (4935T>G), K367N (4942A>T), T368S (4943A>T), E370Q (4949G>C), K373L (4958A>T 4959A>T), I374T (4962T>C 4963A>T), E377K (4970G>A), I378Q (4973A>C 4974T>A), R380S (4979A>T 4980G>C 4981A>T), N381Q (4982A>C 4984T>A), I382L (4985A>T), I383C (4988A>T 4989T>G 4990A>T), I384C (4991A>T 4992T>G 4993A>C), K385E (4994A>G), H386N (4997C>A), K392Q (5015A>C 5017G>A), E396D (5029G>C), G398N (5033G>A 5034G>A 5035A>T), Q400N (5039C>A 5041A>T), K401E (5042A>G), V407M (5060G>A 5062T>G), K408I (5064A>T 5065G>A), E410S (5069G>T 5070A>C), Q411K (5072C>A 5074A>G), G412N (5075G>A 5076G>A 5077A>T), P418T (5093C>A), I421S (5102A>T 5103T>C), D422K (5105G>A 5107T>A), L425C (5115T>G 5116A>T), K427F (5120A>T 5121A>T 5122A>T), H428P (5124A>C 5125T>A), I429D (5126A>G 5127T>A 5128A>T), I431L (5132A>C), K432N (5137A>C), S433T (5139G>C 5140T>A), K435Q (5144A>C), L437I (5150T>A 5152A>C), S439R (5156T>A 5157C>G 5158A>G), I440F (5159A>T 5161A>T), L443C (5169T>G 5170A>T), V444L (5171G>T), Q446Y (5177C>T 5179A>C), I447A (5180A>G 5181T>C 5182T>A), R448S (5185A>T), N449E (5186A>G 5188T>A), L451I (5192T>A), N453D (5198A>G 5200T>C), S455A (5204A>G 5205G>C 5206T>A) | | | | | | | | |
| Codon mutations: | GAT353AAT (4898G>A), ATA355CTA (4904A>C), CTT356ATC (4907C>A 4909T>C), GTA357GTC (4912A>C), TAT358TAC (4915T>C), TTT365TGT (4935T>G), TCG366TCT (4939G>T), AAA367AAT (4942A>T), ACA368TCA (4943A>T), GAA370CAA (4949G>C), AAA373TTA (4958A>T 4959A>T), ATA374ACT (4962T>C 4963A>T), TTG376CTT (4967T>C 4969G>T), GAA377AAA (4970G>A), ATA378CAA (4973A>C 4974T>A), TTT379TTC (4978T>C), AGA380TCT (4979A>T 4980G>C 4981A>T), AAT381CAA (4982A>C 4984T>A), ATA382TTA (4985A>T), ATA383TGT (4988A>T 4989T>G 4990A>T), ATA384TGC (4991A>T 4992T>G 4993A>C), AAA385GAA (4994A>G), CAT386AAT (4997C>A), ATA388ATT (5005A>T), GTA389GTG (5008A>G), AGT391AGC (5014T>C), AAG392CAA (5015A>C 5017G>A), AAA394AAG (5023A>G), GAG396GAC (5029G>C), GGA398AAT (5033G>A 5034G>A 5035A>T), CAA400AAT (5039C>A 5041A>T), AAA401GAA (5042A>G), ATA402ATT (5047A>T), CTA405CTC (5056A>C), GTT407ATG (5060G>A 5062T>G), AAG408ATA (5064A>T 5065G>A), ATA409ATT (5068A>T), GAA410TCA (5069G>T 5070A>C), CAA411AAG (5072C>A 5074A>G), GGA412AAT (5075G>A 5076G>A 5077A>T), GGA413GGC (5080A>C), ATA414ATT (5083A>T), CAG417CAA (5092G>A), CCT418ACT (5093C>A), ATA421TCA (5102A>T 5103T>C), GAT422AAA (5105G>A 5107T>A), AAA423AAG (5110A>G), ATA424ATC (5113A>C), TTA425TGT (5115T>G 5116A>T), AAA427TTT (5120A>T 5121A>T 5122A>T), CAT428CCA (5124A>C 5125T>A), ATA429GAT (5126A>G 5127T>A 5128A>T), ATA431CTA (5132A>C), AAA432AAC (5137A>C), AGT433ACA (5139G>C 5140T>A), AAA435CAA (5144A>C), TTA437ATC (5150T>A 5152A>C), TCA439AGG (5156T>A 5157C>G 5158A>G), ATA440TTT (5159A>T 5161A>T), TTA441CTT (5162T>C 5164A>T), TTA443TGT (5169T>G 5170A>T), GTA444TTA (5171G>T), AAC445AAT (5176C>T), CAA446TAC (5177C>T 5179A>C), ATT447GCA (5180A>G 5181T>C 5182T>A), AGA448AGT (5185A>T), AAT449GAA (5186A>G 5188T>A), TTC450TTT (5191C>T), TTA451ATA (5192T>A), AAT453GAC (5198A>G 5200T>C), AGT455GCA (5204A>G 5205G>C 5206T>A) | | | | | | | | |

##### Proteins

| | Begin | End | Coverage | Score | Concordance | Matches | Identities | I/D/M/F* | Stop Codons |
|---|---|---|---|---|---|---|---|---|---|
| replicase (YP_004347415.1) | 353 | 456 | 16.2% | 356 | 52.9% | 104 (100%) | 53 (51.0%) | 0/0/0/0 | 0 |
| Protein mutations: | D353N (4898G>A), I355L (4904A>C), L356I (4907C>A 4909T>C), F365C (4935T>G), K367N (4942A>T), T368S (4943A>T), E370Q (4949G>C), K373L (4958A>T 4959A>T), I374T (4962T>C 4963A>T), E377K (4970G>A), I378Q (4973A>C 4974T>A), R380S (4979A>T 4980G>C 4981A>T), N381Q (4982A>C 4984T>A), I382L (4985A>T), I383C (4988A>T 4989T>G 4990A>T), I384C (4991A>T 4992T>G 4993A>C), K385E (4994A>G), H386N (4997C>A), K392Q (5015A>C 5017G>A), E396D (5029G>C), G398N (5033G>A 5034G>A 5035A>T), Q400N (5039C>A 5041A>T), K401E (5042A>G), V407M (5060G>A 5062T>G), K408I (5064A>T 5065G>A), E410S (5069G>T 5070A>C), Q411K (5072C>A 5074A>G), G412N (5075G>A 5076G>A 5077A>T), P418T (5093C>A), I421S (5102A>T 5103T>C), D422K (5105G>A 5107T>A), L425C (5115T>G 5116A>T), K427F (5120A>T 5121A>T 5122A>T), H428P (5124A>C 5125T>A), I429D (5126A>G 5127T>A 5128A>T), I431L (5132A>C), K432N (5137A>C), S433T (5139G>C 5140T>A), K435Q (5144A>C), L437I (5150T>A 5152A>C), S439R (5156T>A 5157C>G 5158A>G), I440F (5159A>T 5161A>T), L443C (5169T>G 5170A>T), V444L (5171G>T), Q446Y (5177C>T 5179A>C), I447A (5180A>G 5181T>C 5182T>A), R448S (5185A>T), N449E (5186A>G 5188T>A), L451I (5192T>A), N453D (5198A>G 5200T>C), S455A (5204A>G 5205G>C 5206T>A) | | | | | | | | |
| Codon mutations: | GAT353AAT (4898G>A), ATA355CTA (4904A>C), CTT356ATC (4907C>A 4909T>C), GTA357GTC (4912A>C), TAT358TAC (4915T>C), TTT365TGT (4935T>G), TCG366TCT (4939G>T), AAA367AAT (4942A>T), ACA368TCA (4943A>T), GAA370CAA (4949G>C), AAA373TTA (4958A>T 4959A>T), ATA374ACT (4962T>C 4963A>T), TTG376CTT (4967T>C 4969G>T), GAA377AAA (4970G>A), ATA378CAA (4973A>C 4974T>A), TTT379TTC (4978T>C), AGA380TCT (4979A>T 4980G>C 4981A>T), AAT381CAA (4982A>C 4984T>A), ATA382TTA (4985A>T), ATA383TGT (4988A>T 4989T>G 4990A>T), ATA384TGC (4991A>T 4992T>G 4993A>C), AAA385GAA (4994A>G), CAT386AAT (4997C>A), ATA388ATT (5005A>T), GTA389GTG (5008A>G), AGT391AGC (5014T>C), AAG392CAA (5015A>C 5017G>A), AAA394AAG (5023A>G), GAG396GAC (5029G>C), GGA398AAT (5033G>A 5034G>A 5035A>T), CAA400AAT (5039C>A 5041A>T), AAA401GAA (5042A>G), ATA402ATT (5047A>T), CTA405CTC (5056A>C), GTT407ATG (5060G>A 5062T>G), AAG408ATA (5064A>T 5065G>A), ATA409ATT (5068A>T), GAA410TCA (5069G>T 5070A>C), CAA411AAG (5072C>A 5074A>G), GGA412AAT (5075G>A 5076G>A 5077A>T), GGA413GGC (5080A>C), ATA414ATT (5083A>T), CAG417CAA (5092G>A), CCT418ACT (5093C>A), ATA421TCA (5102A>T 5103T>C), GAT422AAA (5105G>A 5107T>A), AAA423AAG (5110A>G), ATA424ATC (5113A>C), TTA425TGT (5115T>G 5116A>T), AAA427TTT (5120A>T 5121A>T 5122A>T), CAT428CCA (5124A>C 5125T>A), ATA429GAT (5126A>G 5127T>A 5128A>T), ATA431CTA (5132A>C), AAA432AAC (5137A>C), AGT433ACA (5139G>C 5140T>A), AAA435CAA (5144A>C), TTA437ATC (5150T>A 5152A>C), TCA439AGG (5156T>A 5157C>G 5158A>G), ATA440TTT (5159A>T 5161A>T), TTA441CTT (5162T>C 5164A>T), TTA443TGT (5169T>G 5170A>T), GTA444TTA (5171G>T), AAC445AAT (5176C>T), CAA446TAC (5177C>T 5179A>C), ATT447GCA (5180A>G 5181T>C 5182T>A), AGA448AGT (5185A>T), AAT449GAA (5186A>G 5188T>A), TTC450TTT (5191C>T), TTA451ATA (5192T>A), AAT453GAC (5198A>G 5200T>C), AGT455GCA (5204A>G 5205G>C 5206T>A) | | | | | | | | |

*: Inserts / Deletes / Misaligned / Frameshifts

#### Analysis details

This analysis was performed with panviral2.64

### NGS Details (UN70): Cavemovirus collusipomeae

#### Assembly

| | |
|---|---|
| **Coverage Length** | 239 (1 contig(s)) |
| **Depth Of Coverage** | 3.7 |
| **Number Of Reads** | 8 |
| **Reads Per Million** | 0.15 rpm (after QC) |
| **Ambiguities** | 0 |
| **Assembly Method** | de novo + reference guided assembly |
| **Consensus Caller** | Bcf Tools |

#### Coverage Map

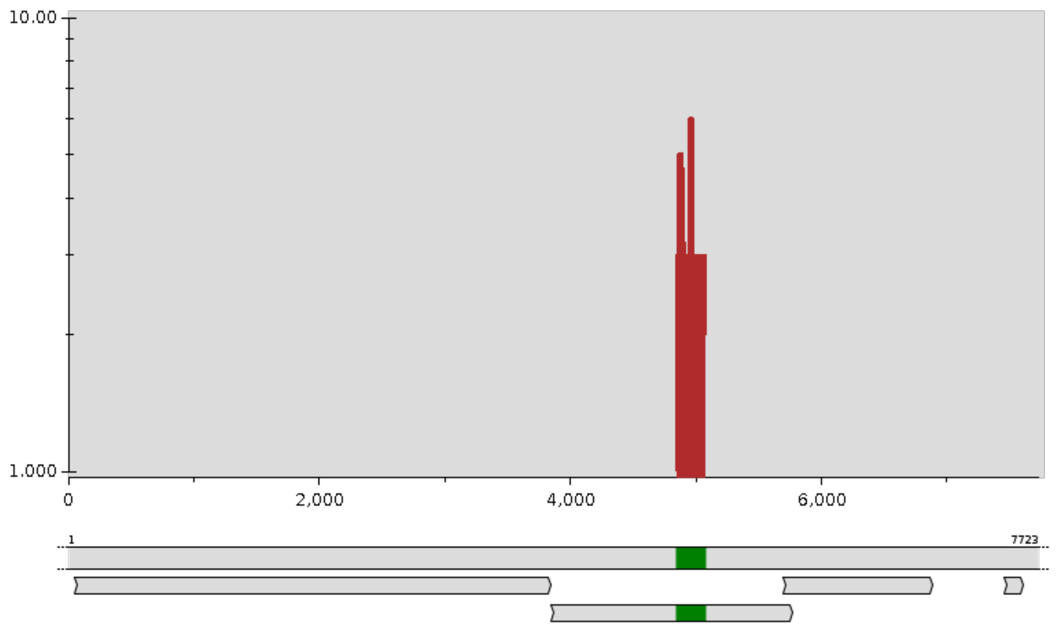

#### Assignment

| | |
|---|---|
| **Type** | Cavemovirus collusipomeae (Taxonomy ID: 3048352) |
| **Reference Genome** | NC_015328.1 |
| **NT Identity (%)** | 62.7615 |
| **AA Identity (%)** | 49.3671 |
| **Number Of Stop Codons** | 0 |
| **Number Of CDS** | 4 |

#### Alignment

| | |
|---|---|
| **Alignment Score** | 122.0 (NT) + 298.0 (AA) = 420.0 |
| **Concordance (%)** | 41.0557 |

| Alignment Method | Global, seeded, nucleotide + amino acids (AGA) |
|---|---|

#### Genome Region

Sequence starts at position 4842 and ends at position 5080 relative to NC_015328.1 reference sequence.

#### Alignment Detailed Statistics

| | Begin | End | Coverage | Score | Concordance | Matches | Identities | I/D/M/F* | Stop Codons |
|---|---|---|---|---|---|---|---|---|---|
| **NT** | **4842** | **5080** | **3.1%** | **122** | **25.5%** | **239 (100%)** | **150 (62.8%)** | **0/0** | |
| Mutations: | 4843T>A, 4847T>G, 4849A>C, 4852A>T, 4853A>T, 4854G>C, 4856A>G, 4857T>A, 4860A>T, 4865C>A, 4868T>A, 4874G>A, 4878G>A, 4879A>T, 4882A>T, 4887G>A, 4888A>T, 4896A>T, 4897T>G, 4906A>T, 4907C>A, 4912A>C, 4914A>G, 4915T>C, 4918A>T, 4924T>C, 4927A>C, 4930A>G, 4931A>G, 4935T>A, 4938C>T, 4939G>C, 4944C>T, 4947A>T, 4948C>T, 4952G>A, 4953A>T, 4954A>G, 4959A>T, 4960A>T, 4962T>A, 4963A>G, 4967T>C, 4969G>T, 4972A>C, 4975A>T, 4980G>A, 4981A>G, 4984T>A, 4987A>T, 4988A>G, 4990A>G, 4994A>C, 4997C>A, 5002A>T, 5003A>T, 5005A>G, 5008A>G, 5009T>A, 5012A>T, 5013G>C, 5017G>A, 5018A>C, 5019A>C, 5024G>A, 5025C>T, 5027G>A, 5028A>G, 5029G>T, 5030A>T, 5032A>G, 5033G>T, 5034G>T, 5035A>T, 5036A>C, 5039C>A, 5040A>C, 5042A>G, 5045A>G, 5047A>C, 5048G>C, 5049A>G, 5054C>T, 5060G>C, 5061T>A, 5065G>T, 5069G>T, 5070A>G, 5071A>T | | | | | | | | |

##### CDS

| | Begin | End | Coverage | Score | Concordance | Matches | Identities | I/D/M/F* | Stop Codons |
|---|---|---|---|---|---|---|---|---|---|
| SPCV_gp2 | 335 | 413 | 12.3% | 298 | 53.3% | 79 (100%) | 39 (49.4%) | 0/0/0/0 | 0 |
| Protein mutations: | S336A (4847T>G 4849A>C), I339E (4856A>G 4857T>A), Y340F (4860A>T), Q342K (4865C>A), F343I (4868T>A), D345N (4874G>A), G346D (4878G>A 4879A>T), R349N (4887G>A 4888A>T), Y352L (4896A>T 4897T>G), L356I (4907C>A), Y358C (4914A>G 4915T>C), I364V (4931A>G), F365Y (4935T>A), S366F (4938C>T 4939G>C), T368I (4944C>T), Y369F (4947A>T 4948C>T), E371M (4952G>A 4953A>T 4954A>G), K373I (4959A>T 4960A>T), I374K (4962T>A 4963A>G), E377D (4972A>C), R380K (4980G>A 4981A>G), N381K (4984T>A), I383V (4988A>G 4990A>G), K385Q (4994A>C), H386N (4997C>A), I388L (5003A>T 5005A>G), L390I (5009T>A), K393P (5018A>C 5019A>C), A395I (5024G>A 5025C>T), E396S (5027G>A 5028A>G 5029G>T), I397L (5030A>T 5032A>G), G398F (5033G>T 5034G>T 5035A>T), K399Q (5036A>C), Q400T (5039C>A 5040A>C), K401E (5042A>G), I402V (5045A>G 5047A>C), E403R (5048G>C 5049A>G), V407H (5060G>C 5061T>A), K408N (5065G>T), E410C (5069G>T 5070A>G 5071A>T) | | | | | | | | |
| Codon mutations: | CAT334.AA (4843T>A), TCA336GCC (4847T>G 4849A>C), CCA337CCT (4852A>T), AGT338TCT (4853A>T 4854G>C), ATA339GAA (4856A>G 4857T>A), TAT340TTT (4860A>T), CAA342AAA (4865C>A), TTT343ATT (4868T>A), GAT345AAT (4874G>A), GGA346GAT (4878G>A 4879A>T), ATA347ATT (4882A>T), AGA349AAT (4887G>A 4888A>T), TAT352TTG (4896A>T 4897T>G), ATA355ATT (4906A>T), CTT356ATT (4907C>A), GTA357GTC (4912A>C), TAT358TGC (4914A>G 4915T>C), ATA359ATT (4918A>T), GAT361GAC (4924T>C), ATA362ATC (4927A>C), TTA363TTG (4930A>G), ATA364GTA (4931A>G), TTT365TAT (4935T>A), TCG366TTC (4938C>T 4939G>C), ACA368ATA (4944C>T), TAC369TTT (4947A>T 4948C>T), GAA371ATG (4952G>A 4953A>T 4954A>G), AAA373ATT (4959A>T 4960A>T), ATA374AAG (4962T>A 4963A>G), TTG376CTT (4967T>C 4969G>T), GAA377GAC (4972A>C), ATA378ATT (4975A>T), AGA380AAG (4980G>A 4981A>G), AAT381AAA (4984T>A), ATA382ATT (4987A>T), ATA383GTG (4988A>G 4990A>G), AAA385CAA (4994A>C), CAT386AAT (4997C>A), GGA387GGT (5002A>T), ATA388TTG (5003A>T 5005A>G), GTA389GTG (5008A>G), TTA390ATA (5009T>A), AGT391TCT (5012A>T 5013G>C), AAG392AAA (5017G>A), AAA393CCA (5018A>C 5019A>C), GCA395ATA (5024G>A 5025C>T), GAG396AGT (5027G>A 5028A>G 5029G>T), ATA397TTG (5030A>T 5032A>G), GGA398TTT (5033G>T 5034G>T 5035A>T), AAA399CAA (5036A>C), CAA400ACA (5039C>A 5040A>C), AAA401GAA (5042A>G), ATA402GTC (5045A>G 5047A>C), GAA403CGA (5048G>C 5049A>G), CTA405TTA (5054C>T), GTT407CAT (5060G>C 5061T>A), AAG408AAT (5065G>T), GAA410TGT (5069G>T 5070A>G 5071A>T) | | | | | | | | |

##### Proteins

| | Begin | End | Coverage | Score | Concordance | Matches | Identities | I/D/M/F* | Stop Codons |
|---|---|---|---|---|---|---|---|---|---|
| replicase (YP_004347415.1) | 335 | 413 | 12.3% | 298 | 53.3% | 79 (100%) | 39 (49.4%) | 0/0/0/0 | 0 |
| Protein mutations: | S336A (4847T>G 4849A>C), I339E (4856A>G 4857T>A), Y340F (4860A>T), Q342K (4865C>A), F343I (4868T>A), D345N (4874G>A), G346D (4878G>A 4879A>T), R349N (4887G>A 4888A>T), Y352L (4896A>T 4897T>G), L356I (4907C>A), Y358C (4914A>G 4915T>C), I364V (4931A>G), F365Y (4935T>A), S366F (4938C>T 4939G>C), T368I (4944C>T), Y369F (4947A>T 4948C>T), E371M (4952G>A 4953A>T 4954A>G), K373I (4959A>T 4960A>T), I374K (4962T>A 4963A>G), E377D (4972A>C), R380K (4980G>A 4981A>G), N381K (4984T>A), I383V (4988A>G 4990A>G), K385Q (4994A>C), H386N (4997C>A), I388L (5003A>T 5005A>G), L390I (5009T>A), K393P (5018A>C 5019A>C), A395I (5024G>A 5025C>T), E396S (5027G>A 5028A>G 5029G>T), I397L (5030A>T 5032A>G), G398F (5033G>T 5034G>T 5035A>T), K399Q (5036A>C), Q400T (5039C>A 5040A>C), K401E (5042A>G), I402V (5045A>G 5047A>C), E403R (5048G>C 5049A>G), V407H (5060G>C 5061T>A), K408N (5065G>T), E410C (5069G>T 5070A>G 5071A>T) | | | | | | | | |
| Codon mutations: | CAT334.AA (4843T>A), TCA336GCC (4847T>G 4849A>C), CCA337CCT (4852A>T), AGT338TCT (4853A>T 4854G>C), ATA339GAA (4856A>G 4857T>A), TAT340TTT (4860A>T), CAA342AAA (4865C>A), TTT343ATT (4868T>A), GAT345AAT (4874G>A), GGA346GAT (4878G>A 4879A>T), ATA347ATT (4882A>T), AGA349AAT (4887G>A 4888A>T), TAT352TTG (4896A>T 4897T>G), ATA355ATT (4906A>T), CTT356ATT (4907C>A), GTA357GTC (4912A>C), TAT358TGC (4914A>G 4915T>C), ATA359ATT (4918A>T), GAT361GAC (4924T>C), ATA362ATC (4927A>C), TTA363TTG (4930A>G), ATA364GTA (4931A>G), TTT365TAT (4935T>A), TCG366TTC (4938C>T 4939G>C), ACA368ATA (4944C>T), TAC369TTT (4947A>T 4948C>T), GAA371ATG (4952G>A 4953A>T 4954A>G), AAA373ATT (4959A>T 4960A>T), ATA374AAG (4962T>A 4963A>G), TTG376CTT (4967T>C 4969G>T), GAA377GAC (4972A>C), ATA378ATT (4975A>T), AGA380AAG (4980G>A 4981A>G), AAT381AAA (4984T>A), ATA382ATT (4987A>T), ATA383GTG (4988A>G 4990A>G), AAA385CAA (4994A>C), CAT386AAT (4997C>A), GGA387GGT (5002A>T), ATA388TTG (5003A>T 5005A>G), GTA389GTG (5008A>G), TTA390ATA (5009T>A), AGT391TCT (5012A>T 5013G>C), AAG392AAA (5017G>A), AAA393CCA (5018A>C 5019A>C), GCA395ATA (5024G>A 5025C>T), GAG396AGT (5027G>A 5028A>G 5029G>T), ATA397TTG (5030A>T 5032A>G), GGA398TTT (5033G>T 5034G>T 5035A>T), AAA399CAA (5036A>C), CAA400ACA (5039C>A 5040A>C), AAA401GAA (5042A>G), ATA402GTC (5045A>G 5047A>C), GAA403CGA (5048G>C 5049A>G), CTA405TTA (5054C>T), GTT407CAT (5060G>C 5061T>A), AAG408AAT (5065G>T), GAA410TGT (5069G>T 5070A>G 5071A>T) | | | | | | | | |

*: Inserts / Deletes / Misaligned / Frameshifts

#### Analysis details

This analysis was performed with panviral2.64

#### NGS Details (UN70): Duamitovirus chqu1

##### Assembly

| | |
|---|---|
| **Coverage Length** | 135 (1 contig(s)) |
| **Depth Of Coverage** | 6.4 |
| **Number Of Reads** | 8 |
| **Reads Per Million** | 0.15 rpm (after QC) |
| **Ambiguities** | 0 |
| **Assembly Method** | de novo + reference guided assembly |
| **Consensus Caller** | Bcf Tools |

##### Coverage Map

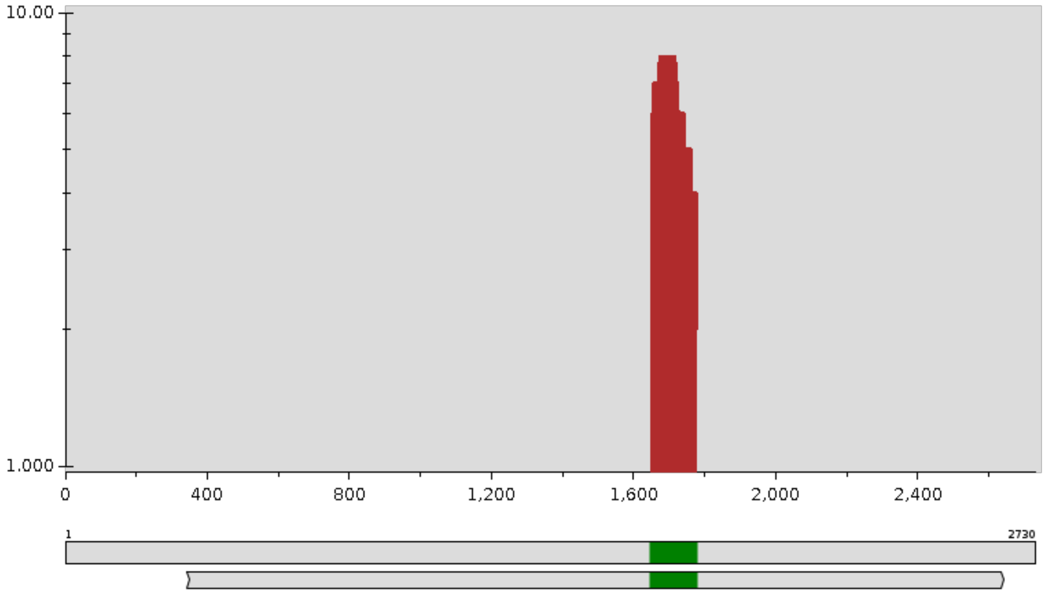

##### Assignment

| | |
|---|---|
| **Type** | Duamitovirus chqu1 (Taxonomy ID: 2955775) |
| **Reference Genome** | NC_040543.1 |
| **NT Identity (%)** | 68.1481 |
| **AA Identity (%)** | 75.5556 |
| **Number Of Stop Codons** | 1 |
| **Number Of CDS** | 1 |

##### Alignment

| | |
|---|---|
| **Alignment Score** | 98.0 (NT) + 265.0 (AA) = 363.0 |
| **Concordance (%)** | 59.8023 |

| Alignment Method | Global, seeded, nucleotide + amino acids (AGA) |
|---|---|

#### Genome Region

Sequence starts at position 1647 and ends at position 1781 relative to NC_040543.1 reference sequence.

#### Alignment Detailed Statistics

| | Begin | End | Coverage | Score | Concordance | Matches | Identities | I/D/M/F* | Stop Codons |
|---|---|---|---|---|---|---|---|---|---|
| **NT** | **1647** | **1781** | **4.9%** | **98** | **36.3%** | **135 (100%)** | **92 (68.1%)** | **0/0** | |
| Mutations: | 1651T>A, 1654G>A, 1657A>G, 1660A>G, 1661C>T, 1663C>A, 1669T>C, 1672T>C, 1678C>A, 1684A>C, 1690C>T, 1696A>T, 1699A>C, 1706C>T, 1707T>A, 1708G>T, 1709G>A, 1711T>G, 1718T>C, 1723G>T, 1726T>G, 1727G>A, 1729T>A, 1731T>A, 1732A>T, 1735C>A, 1741T>A, 1743G>C, 1744C>G, 1747A>T, 1748G>A, 1749A>C, 1750A>T, 1753T>C, 1756G>T, 1760A>G, 1761G>C, 1762G>T, 1765C>T, 1768T>C, 1771T>C, 1772G>C, 1777G>A | | | | | | | | |
| **CDS** | | | | | | | | | |
| EXI96_gp1 | 436 | 480 | 5.9% | 265 | 74.6% | 45 (100%) | 34 (75.6%) | 0/0/0/0 | 1 |
| Protein mutations: | C436* (1651T>A), L455Y (1706C>T 1707T>A 1708G>T), V456M (1709G>A 1711T>G), D462K (1727G>A 1729T>A), L463Y (1731T>A 1732A>T), S467T (1743G>C 1744C>G), R468S (1747A>T), E469T (1748G>A 1749A>C 1750A>T), R473A (1760A>G 1761G>C 1762G>T), V477L (1772G>C) | | | | | | | | |
| Codon mutations: | TGT436TGA (1651T>A), GGG437GGA (1654G>A), CAA438CAG (1657A>G), CCA439CCG (1660A>G), CTC440TTA (1661C>T 1663C>A), TAT442TAC (1669T>C), TAT443TAC (1672T>C), TCC445TCA (1678C>A), TCA447TCC (1684A>C), TTC449TTT (1690C>T), CTA451CTT (1696A>T), TCA452TCC (1699A>C), CTG455TAT (1706C>T 1707T>A 1708G>T), GTT456ATG (1709G>A 1711T>G), TTA459CTA (1718T>C), GCG460GCT (1723G>T), GCT461GCG (1726T>G), GAT462AAA (1727G>A 1729T>A), TTA463TAT (1731T>A 1732A>T), GCC464GCA (1735C>A), CCT466CCA (1741T>A), AGC467ACG (1743G>C 1744C>G), AGA468AGT (1747A>T), GAA469ACT (1748G>A 1749A>C 1750A>T), ACT470ACC (1753T>C), CCG471CCT (1756G>T), AGG473GCT (1760A>G 1761G>C 1762G>T), GAC474GAT (1765C>T), TAT475TAC (1768T>C), GCT476GCC (1771T>C), GTG477CTG (1772G>C), TTG478TTA (1777G>A) | | | | | | | | |
| **Proteins** | | | | | | | | | |
| RNA dependent RNA polymerase (YP_009551903.1) | 436 | 480 | 5.9% | 265 | 74.6% | 45 (100%) | 34 (75.6%) | 0/0/0/0 | 1 |
| Protein mutations: | C436* (1651T>A), L455Y (1706C>T 1707T>A 1708G>T), V456M (1709G>A 1711T>G), D462K (1727G>A 1729T>A), L463Y (1731T>A 1732A>T), S467T (1743G>C 1744C>G), R468S (1747A>T), E469T (1748G>A 1749A>C 1750A>T), R473A (1760A>G 1761G>C 1762G>T), V477L (1772G>C) | | | | | | | | |
| Codon mutations: | TGT436TGA (1651T>A), GGG437GGA (1654G>A), CAA438CAG (1657A>G), CCA439CCG (1660A>G), CTC440TTA (1661C>T 1663C>A), TAT442TAC (1669T>C), TAT443TAC (1672T>C), TCC445TCA (1678C>A), TCA447TCC (1684A>C), TTC449TTT (1690C>T), CTA451CTT (1696A>T), TCA452TCC (1699A>C), CTG455TAT (1706C>T 1707T>A 1708G>T), GTT456ATG (1709G>A 1711T>G), TTA459CTA (1718T>C), GCG460GCT (1723G>T), GCT461GCG (1726T>G), GAT462AAA (1727G>A 1729T>A), TTA463TAT (1731T>A 1732A>T), GCC464GCA (1735C>A), CCT466CCA (1741T>A), AGC467ACG (1743G>C 1744C>G), AGA468AGT (1747A>T), GAA469ACT (1748G>A 1749A>C 1750A>T), ACT470ACC (1753T>C), CCG471CCT (1756G>T), AGG473GCT (1760A>G 1761G>C 1762G>T), GAC474GAT (1765C>T), TAT475TAC (1768T>C), GCT476GCC (1771T>C), GTG477CTG (1772G>C), TTG478TTA (1777G>A) | | | | | | | | |

*: Inserts / Deletes / Misaligned / Frameshifts

#### Analysis details

This analysis was performed with panviral2.64

### NGS Details (UN70): Badnavirus tessellocastaneae

#### Assembly

| | |
|---|---|
| **Coverage Length** | 248 (1 contig(s)) |
| **Depth Of Coverage** | 3.4 |
| **Number Of Reads** | 7 |
| **Reads Per Million** | 0.13 rpm (after QC) |
| **Ambiguities** | 0 |
| **Assembly Method** | de novo + reference guided assembly |
| **Consensus Caller** | Bcf Tools |

#### Coverage Map

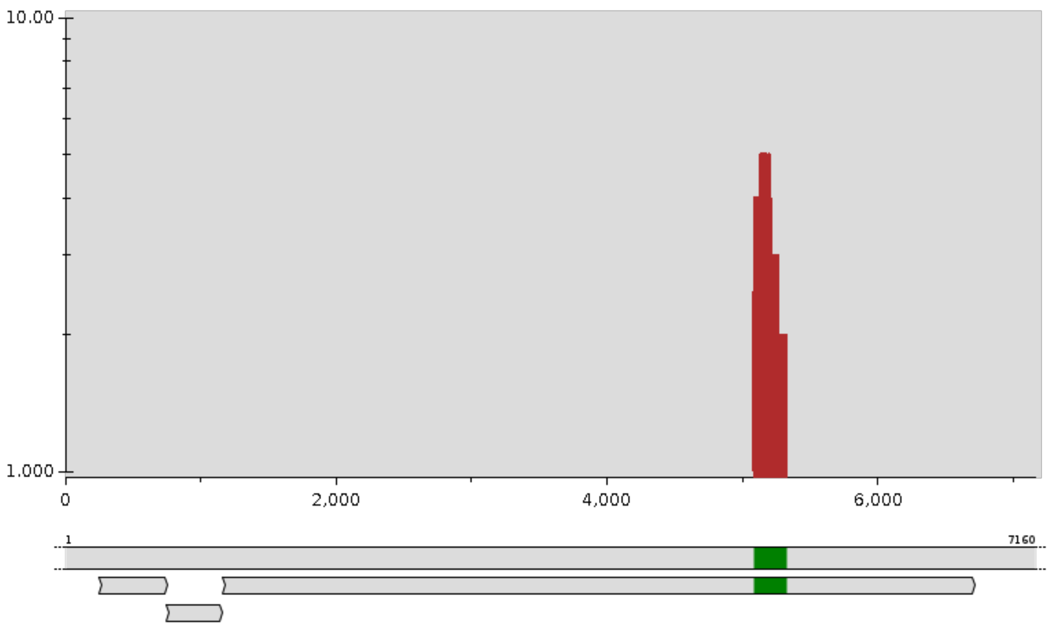

#### Assignment

| | |
|---|---|
| **Type** | Badnavirus tessellocastaneae (Taxonomy ID: 3051987) |
| **Reference Genome** | NC_076885.1 |
| **NT Identity (%)** | 55.6452 |
| **AA Identity (%)** | 55.4217 |
| **Number Of Stop Codons** | 0 |
| **Number Of CDS** | 3 |

#### Alignment

| | |
|---|---|
| **Alignment Score** | 56.0 (NT) + 329.0 (AA) = 385.0 |
| **Concordance (%)** | 35.9813 |

| Alignment Method | Global, seeded, nucleotide + amino acids (AGA) |
|---|---|

#### Genome Region

Sequence starts at position 5084 and ends at position 5331 relative to NC_076885.1 reference sequence.

#### Alignment Detailed Statistics

| | Begin | End | Coverage | Score | Concordance | Matches | Identities | I/D/M/F* | Stop Codons |
|---|---|---|---|---|---|---|---|---|---|
| **NT** | **5084** | **5331** | **3.5%** | **56** | **11.3%** | **248 (100%)** | **138 (55.6%)** | **0/0** | |
| Mutations: | 5087C>G, 5092T>C, 5098A>G, 5100C>A, 5102C>T, 5104C>A, 5109T>C, 5110G>C, 5113T>C, 5116C>T, 5117C>G, 5118A>T, 5122A>G, 5125C>T, 5126C>A, 5130A>T, 5132T>C, 5134C>A, 5135C>A, 5137C>T, 5142G>T, 5143A>T, 5146A>T, 5148C>A, 5149A>G, 5150T>G, 5151C>A, 5152C>A, 5153A>T, 5155C>A, 5156A>T, 5158C>A, 5159A>G, 5161G>T, 5162A>G, 5163G>A, 5164A>G, 5165G>T, 5167T>G, 5168G>C, 5169C>A, 5170A>T, 5174T>G, 5176A>C, 5180G>T, 5181T>A, 5182C>T, 5188C>T, 5192T>C, 5194T>A, 5197T>C, 5200A>G, 5202A>G, 5203G>A, 5204A>T, 5205G>C, 5209G>T, 5211T>A, 5215T>C, 5219G>A, 5221G>A, 5222G>A, 5223T>G, 5228G>T, 5229A>T, 5230A>T, 5234G>T, 5235A>C, 5236G>T, 5237T>A, 5238C>A, 5239A>T, 5245A>T, 5246T>A, 5247G>A, 5248G>A, 5251A>G, 5252G>A, 5254G>A, 5258G>A, 5259T>G, 5260G>A, 5261G>A, 5262T>C, 5263A>C, 5265C>A, 5266A>T, 5267G>C, 5268G>A, 5272A>G, 5273T>C, 5274T>A, 5275A>C, 5278C>T, 5283G>T, 5284G>C, 5285C>T, 5287G>A, 5290T>G, 5296A>C, 5302G>C, 5305A>T, 5307G>C, 5308A>C, 5314A>C, 5317A>C, 5320A>C, 5323C>A, 5326C>T, 5329A>G | | | | | | | | |

##### CDS

| | Begin | End | Coverage | Score | Concordance | Matches | Identities | I/D/M/F* | Stop Codons |
|---|---|---|---|---|---|---|---|---|---|
| ORF3 | 1308 | 1390 | 4.5% | 329 | 56.4% | 83 (100%) | 46 (55.4%) | 0/0/0/0 | 0 |
| Protein mutations: | L1309V (5087C>G), A1313E (5100C>A), M1316T (5109T>C 5110G>C), H1319V (5117C>G 5118A>T), Q1322K (5126C>A), Y1323F (5130A>T), S1324P (5132T>C 5134C>A), L1325I (5135C>A 5137C>T), G1327V (5142G>T 5143A>T), T1329K (5148C>A 5149A>G), S1330E (5150T>G 5151C>A 5152C>A), I1331L (5153A>T 5155C>A), I1332L (5156A>T 5158C>A), K1333D (5159A>G 5161G>T), R1334E (5162A>G 5163G>A 5164A>G), V1335L (5165G>T 5167T>G), A1336H (5168G>C 5169C>A 5170A>T), S1338A (5174T>G 5176A>C), V1340Y (5180G>T 5181T>A 5182C>T), F1344L (5192T>C 5194T>A), K1347R (5202A>G 5203G>A), F1350Y (5211T>A), V1353I (5219G>A 5221G>A), V1354R (5222G>A 5223T>G), E1356F (5228G>T 5229A>T 5230A>T), E1358S (5234G>T 5235A>C 5236G>T), S1359N (5237T>A 5238C>A 5239A>T), W1362K (5246T>A 5247G>A 5248G>A), A1364T (5252G>A 5254G>A), V1366R (5258G>A 5259T>G 5260G>A), V1367T (5261G>A 5262T>C 5263A>C), P1368H (5265C>A 5266A>T), G1369Q (5267G>C 5268G>A), L1371H (5273T>C 5274T>A 5275A>C), W1374F (5283G>T 5284G>C), R1382T (5307G>C 5308A>C) | | | | | | | | |
| Codon mutations: | CTA1309GTA (5087C>G), GAT1310GAC (5092T>C), AGA1312AGG (5098A>G), GCA1313GAA (5100C>A), CTC1314TTA (5102C>T 5104C>A), ATG1316ACC (5109T>C 5110G>C), AAT1317AAC (5113T>C), ACC1318ACT (5116C>T), CAC1319GTC (5117C>G 5118A>T), AAA1320AAG (5122A>G), GAC1321GAT (5125C>T), CAG1322AAG (5126C>A), TAT1323TTT (5130A>T), TCC1324CCA (5132T>C 5134C>A), CTC1325ATT (5135C>A 5137C>T), GGA1327GTT (5142G>T 5143A>T), ATA1328ATT (5146A>T), ACA1329AAG (5148C>A 5149A>G), TCC1330GAA (5150T>G 5151C>A 5152C>A), ATC1331TTA (5153A>T 5155C>A), ATC1332TTA (5156A>T 5158C>A), AAG1333GAT (5159A>G 5161G>T), AGA1334GAG (5162A>G 5163G>A 5164A>G), GTT1335TTG (5165G>T 5167T>G), GCA1336CAT (5168G>C 5169C>A 5170A>T), TCA1338GCC (5174T>G 5176A>C), GTC1340TAT (5180G>T 5181T>A 5182C>T), TCC1342TCT (5188C>T), TTT1344CTA (5192T>C 5194T>A), GAT1345GAC (5197T>C), TTA1346TTG (5200A>G), AAG1347AGA (5202A>G 5203G>A), AGT1348TCT (5204A>T 5205G>C), GGG1349GGT (5209G>T), TTC1350TAC (5211T>A), CAT1351CAC (5215T>C), GTG1353ATA (5219G>A 5221G>A), GTA1354AGA (5222G>A 5223T>G), GAA1356TTT (5228G>T 5229A>T 5230A>T), GAG1358TCT (5234G>T 5235A>C 5236G>T), TCA1359AAT (5237T>A 5238C>A 5239A>T), CCA1361CCT (5245A>T), TGG1362AAA (5246T>A 5247G>A 5248G>A), ACA1363ACG (5251A>G), GCG1364ACA (5252G>A 5254G>A), GTG1366AGA (5258G>A 5259T>G 5260G>A), GTA1367ACC (5261G>A 5262T>C 5263A>C), CCA1368CAT (5265C>A 5266A>T), GGA1369CAA (5267G>C 5268G>A), GGA1370GGG (5272A>G), TTA1371CAC (5273T>C 5274T>A 5275A>C), TAC1372TAT (5278C>T), TGG1374TTC (5283G>T 5284G>C), CTG1375TTA (5285C>T 5287G>A), GTT1376GTG (5290T>G), CCA1378CCC (5296A>C), GGG1380GGC (5302G>C), CTA1381CTT (5305A>T), AGA1382ACC (5307G>C 5308A>C), GCA1384GCC (5314A>C), CCA1385CCC (5317A>C), TCA1386TCC (5320A>C), ACC1387ACA (5323C>A), TTC1388TTT (5326C>T), CAA1389CAG (5329A>G) | | | | | | | | |

##### Proteins

| | Begin | End | Coverage | Score | Concordance | Matches | Identities | I/D/M/F* | Stop Codons |
|---|---|---|---|---|---|---|---|---|---|
| polyprotein (YP_010800602.1) | 1308 | 1390 | 4.5% | 329 | 56.4% | 83 (100%) | 46 (55.4%) | 0/0/0/0 | 0 |
| Protein mutations: | L1309V (5087C>G), A1313E (5100C>A), M1316T (5109T>C 5110G>C), H1319V (5117C>G 5118A>T), Q1322K (5126C>A), Y1323F (5130A>T), S1324P (5132T>C 5134C>A), L1325I (5135C>A 5137C>T), G1327V (5142G>T 5143A>T), T1329K (5148C>A 5149A>G), S1330E (5150T>G 5151C>A 5152C>A), I1331L (5153A>T 5155C>A), I1332L (5156A>T 5158C>A), K1333D (5159A>G 5161G>T), R1334E (5162A>G 5163G>A 5164A>G), V1335L (5165G>T 5167T>G), A1336H (5168G>C 5169C>A 5170A>T), S1338A (5174T>G 5176A>C), V1340Y (5180G>T 5181T>A 5182C>T), F1344L (5192T>C 5194T>A), K1347R (5202A>G 5203G>A), F1350Y (5211T>A), V1353I (5219G>A 5221G>A), V1354R (5222G>A 5223T>G), E1356F (5228G>T 5229A>T 5230A>T), E1358S (5234G>T 5235A>C 5236G>T), S1359N (5237T>A 5238C>A 5239A>T), W1362K (5246T>A 5247G>A 5248G>A), A1364T (5252G>A 5254G>A), V1366R (5258G>A 5259T>G 5260G>A), V1367T (5261G>A 5262T>C 5263A>C), P1368H (5265C>A 5266A>T), G1369Q (5267G>C 5268G>A), L1371H (5273T>C 5274T>A 5275A>C), W1374F (5283G>T 5284G>C), R1382T (5307G>C 5308A>C) | | | | | | | | |
| Codon mutations: | CTA1309GTA (5087C>G), GAT1310GAC (5092T>C), AGA1312AGG (5098A>G), GCA1313GAA (5100C>A), CTC1314TTA (5102C>T 5104C>A), ATG1316ACC (5109T>C 5110G>C), AAT1317AAC (5113T>C), ACC1318ACT (5116C>T), CAC1319GTC (5117C>G 5118A>T), AAA1320AAG (5122A>G), GAC1321GAT (5125C>T), CAG1322AAG (5126C>A), TAT1323TTT (5130A>T), TCC1324CCA (5132T>C 5134C>A), CTC1325ATT (5135C>A 5137C>T), GGA1327GTT (5142G>T 5143A>T), ATA1328ATT (5146A>T), ACA1329AAG (5148C>A 5149A>G), TCC1330GAA (5150T>G 5151C>A 5152C>A), ATC1331TTA (5153A>T 5155C>A), ATC1332TTA (5156A>T 5158C>A), AAG1333GAT (5159A>G 5161G>T), AGA1334GAG (5162A>G 5163G>A 5164A>G), GTT1335TTG (5165G>T 5167T>G), GCA1336CAT (5168G>C 5169C>A 5170A>T), TCA1338GCC (5174T>G 5176A>C), GTC1340TAT (5180G>T 5181T>A 5182C>T), TCC1342TCT (5188C>T), TTT1344CTA (5192T>C 5194T>A), GAT1345GAC (5197T>C), TTA1346TTG (5200A>G), AAG1347AGA (5202A>G 5203G>A), AGT1348TCT (5204A>T 5205G>C), GGG1349GGT (5209G>T), TTC1350TAC (5211T>A), CAT1351CAC (5215T>C), GTG1353ATA (5219G>A 5221G>A), GTA1354AGA (5222G>A 5223T>G), GAA1356TTT (5228G>T 5229A>T 5230A>T), GAG1358TCT (5234G>T 5235A>C 5236G>T), TCA1359AAT (5237T>A 5238C>A 5239A>T), CCA1361CCT (5245A>T), TGG1362AAA (5246T>A 5247G>A 5248G>A), ACA1363ACG (5251A>G), GCG1364ACA (5252G>A 5254G>A), GTG1366AGA (5258G>A 5259T>G 5260G>A), GTA1367ACC (5261G>A 5262T>C 5263A>C), CCA1368CAT (5265C>A 5266A>T), GGA1369CAA (5267G>C 5268G>A), GGA1370GGG (5272A>G), TTA1371CAC (5273T>C 5274T>A 5275A>C), TAC1372TAT (5278C>T), TGG1374TTC (5283G>T 5284G>C), CTG1375TTA (5285C>T 5287G>A), GTT1376GTG (5290T>G), CCA1378CCC (5296A>C), GGG1380GGC (5302G>C), CTA1381CTT (5305A>T), AGA1382ACC (5307G>C 5308A>C), GCA1384GCC (5314A>C), CCA1385CCC (5317A>C), TCA1386TCC (5320A>C), ACC1387ACA (5323C>A), TTC1388TTT (5326C>T), CAA1389CAG (5329A>G) | | | | | | | | |

*: Inserts / Deletes / Misaligned / Frameshifts

#### Analysis details

This analysis was performed with panviral2.64

### NGS Details (UN70): Caulimovirus tessellomirabilis

#### Assembly

| | |
|---|---|
| **Coverage Length** | 318 (1 contig(s)) |
| **Depth Of Coverage** | 3.1 |
| **Number Of Reads** | 7 |
| **Reads Per Million** | 0.13 rpm (after QC) |
| **Ambiguities** | 0 |
| **Assembly Method** | de novo + reference guided assembly |
| **Consensus Caller** | Bcf Tools |

#### Coverage Map

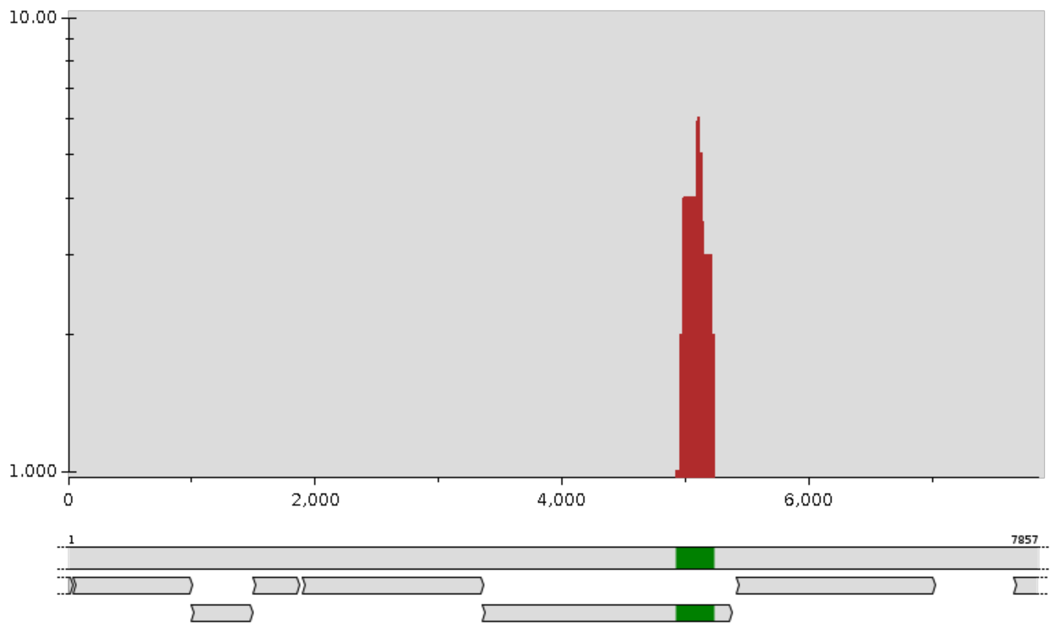

#### Assignment

| | |
|---|---|
| **Type** | Caulimovirus tessellomirabilis (Taxonomy ID: 3048203) |
| **Reference Genome** | NC_004036.1 |
| **NT Identity (%)** | 58.7459 |
| **AA Identity (%)** | 49.505 |
| **Number Of Stop Codons** | 0 |
| **Number Of CDS** | 7 |

#### Alignment

| | |
|---|---|
| **Alignment Score** | 64.0 (NT) + 343.0 (AA) = 407.0 |
| **Concordance (%)** | 33.1164 |

| **Alignment Method** | Global, seeded, nucleotide + amino acids (AGA) |
|---|---|

#### Genome Region

Sequence starts at position 4922 and ends at position 5239 relative to NC_004036.1 reference sequence.

#### Alignment Detailed Statistics

| | Begin | End | Coverage | Score | Concordance | Matches | Identities | I/D/M/F* | Stop Codons |
|---|---|---|---|---|---|---|---|---|---|
| **NT** | **4922** | **5239** | **4.0%** | **64** | **11.3%** | **303 (95.3%)** | **178 (56.0%)** | **0/15** | |
| Mutations: | 4928A>G, 4933T>G, 4934T>C, 4942A>G, 4946T>A, 4947A>G, 4948T>G, 4955A>G, 4957G>T, 4958A>C, 4959A>C, 4968G>A, 4970T>C, 4972A>G, 4973A>G, 4975T>A, 4976A>T, 4978T>A, 4979G>C, 4984T>A, 4985G>A, 4989C>T, 4990T>C, 4993T>C, 4997C>T, 4999C>T, 5000T>C, 5001T>A, 5002C>T, 5008T>A, 5010G>C, 5012G>T, 5014C>G, 5017T>A, 5018_5023delAAGGCC, 5029C>A, 5030A>G, 5031C>A, 5032T>C, 5033G>C, 5034A>T, 5036G>A, 5037G>A, 5038T>C, 5042_5047delGAATTA, 5050C>T, 5053C>T, 5060T>A, 5062T>A, 5063T>A, 5064C>G, 5065A>T, 5071A>T, 5074C>T, 5077A>C, 5078C>G, 5079C>A, 5084G>A, 5087T>G, 5089G>T, 5095C>T, 5096C>T, 5097A>C, 5098T>C, 5100G>C, 5101T>A, 5107G>A, 5117C>T, 5122G>A, 5126A>G, 5127A>T, 5129C>A, 5130A>G, 5131G>A, 5133T>G, 5134G>A, 5137T>A, 5138A>C, 5139C>A, 5140T>A, 5147A>T, 5148G>C, 5149T>G, 5150A>G, 5151T>C, 5154A>G, 5155T>C, 5156C>T, 5159A>T, 5160C>T, 5161C>A, 5164T>A, 5165G>A, 5166T>A, 5167T>A, 5169G>C, 5170T>A, 5174A>G, 5175C>T, 5176A>T, 5177G>A, 5179C>T, 5181G>A, 5182G>A, 5192_5194delGTA, 5196A>C, 5199T>A, 5200T>A, 5201T>A, 5204A>T, 5205A>C, 5209A>G, 5212T>G, 5215G>A, 5216A>T, 5217A>T, 5221A>C, 5224G>A, 5225A>C, 5227T>C, 5228A>C, 5231G>T, 5232G>C, 5233T>C, 5236C>A, 5237A>C, 5238G>C, 5239T>C | | | | | | | | |

##### CDS

| | Begin | End | Coverage | Score | Concordance | Matches | Identities | I/D/M/F* | Stop Codons |
|---|---|---|---|---|---|---|---|---|---|
| MiMVgp6 | 523 | 628 | 15.7% | 343 | 51.6% | 101 (95.3%) | 50 (47.2%) | 0/5/0/0 | 0 |
| Protein mutations: | I525V (4928A>G), N526K (4933T>G), F527L (4934T>C), Y531R (4946T>A 4947A>G 4948T>G), K534D (4955A>G 4957G>T), K535P (4958A>C 4959A>C), S538N (4968G>A), I540V (4973A>G 4975T>A), I541L (4976A>T 4978T>A), E542Q (4979G>C), D544N (4985G>A), A545V (4989C>T 4990T>C), H548Y (4997C>T 4999C>T), F549H (5000T>C 5001T>A 5002C>T), G552A (5010G>C), V553L (5012G>T 5014C>G), K555_A556del (5018_5023delAAGGCC), T559D (5030A>G 5031C>A 5032T>C), E560L (5033G>C 5034A>T), G561N (5036G>A 5037G>A 5038T>C), E563_L564del (5042_5047delGAATTA), S569T (5060T>A 5062T>A), K574N (5077A>C), P575E (5078C>G 5079C>A), E577K (5084G>A), L578V (5087T>G 5089G>T), H581S (5096C>T 5097A>C 5098T>C), S582T (5100G>C 5101T>A), K591V (5126A>G 5127A>T), Q592R (5129C>A 5130A>G 5131G>A), V593G (5133T>G 5134G>A), T595Q (5138A>C 5139C>A 5140T>A), I599A (5150A>G 5151T>C), Y600C (5154A>G 5155T>C), T602L (5159A>T 5160C>T 5161C>A), V604K (5165G>A 5166T>A 5167T>A), C605S (5169G>C 5170T>A), T607V (5174A>G 5175C>T 5176A>T), V608I (5177G>A 5179C>T), R609K (5181G>A 5182G>A), V613del (5192_5194delGTA), N614T (5196A>C), L615Q (5199T>A 5200T>A), L616I (5201T>A), K617S (5204A>T 5205A>C), F619L (5212T>G), M620I (5215G>A), N621F (5216A>T 5217A>T), K622N (5221A>C), I624L (5225A>C 5227T>C), T625P (5228A>C), G626S (5231G>T 5232G>C 5233T>C), D627E (5236C>A), S628P (5237A>C 5238G>C 5239T>C) | | | | | | | | |
| Codon mutations: | ATT525GTT (4928A>G), AAT526AAG (4933T>G), TTT527CTT (4934T>C), AAA529AAG (4942A>G), TAT531AGG (4946T>A 4947A>G 4948T>G), AAG534GAT (4955A>G 4957G>T), AAA535CCA (4958A>C 4959A>C), AGT538AAT (4968G>A), TTA539CTG (4970T>C 4972A>G), ATT540GTA (4973A>G 4975T>A), ATT541TTA (4976A>T 4978T>A), GAA542CAA (4979G>C), ACT543ACA (4984T>A), GAT544AAT (4985G>A), GCT545GTC (4989C>T 4990T>C), TCT546TCC (4993T>C), CAC548TAT (4997C>T 4999C>T), TTC549CAT (5000T>C 5001T>A 5002C>T), GGT551GGA (5008T>A), GGA552GCA (5010G>C), GTC553TTG (5012G>T 5014C>G), CTT554CTA (5017T>A), AAG555_GCC556del (5018_5023delAAGGCC), ACC558ACA (5029C>A), ACT559GAC (5030A>G 5031C>A 5032T>C), GAA560CTA (5033G>C 5034A>T), GGT561AAC (5036G>A 5037G>A 5038T>C), GAA563_TTA564del (5042_5047delGAATTA), ATC565ATT (5050C>T), TGC566TGT (5053C>T), TCT569ACA (5060T>A 5062T>A), TCA570AGT (5063T>A 5064C>G 5065A>T), ACA572ACT (5071A>T), TTC573TTT (5074C>T), AAA574AAC (5077A>C), CCA575GAA (5078C>G 5079C>A), GAA577AAA (5084G>A), TTG578GTT (5087T>G 5089G>T), TAC580TAT (5095C>T), CAT581TCC (5096C>T 5097A>C 5098T>C), AGT582ACA (5100G>C 5101T>A), GAG584GAA (5107G>A), CTA588TTA (5117C>T), GCG589GCA (5122G>A), AAA591GTA (5126A>G 5127A>T), CAG592AGA (5129C>A 5130A>G 5131G>A), GTG593GGA (5133T>G 5134G>A), ATT594ATA (5137T>A), ACT595CAA (5138A>C 5139C>A 5140T>A), AGT598TCG (5147A>T 5148G>C 5149T>G), ATT599GCT (5150A>G 5151T>C), TAT600TGC (5154A>G 5155T>C), CTA601TTA (5156C>T), ACC602TTA (5159A>T 5160C>T 5161C>A), CCT603CCA (5164T>A), GTT604AAA (5165G>A 5166T>A 5167T>A), TGT605TCA (5169G>C 5170T>A), ACA607GTT (5174A>G 5175C>T 5176A>T), GTC608ATT (5177G>A 5179C>T), AGG609AAA (5181G>A 5182G>A), GTA613del (5192_5194delGTA), AAT614ACT (5196A>C), CTT615CAA (5199T>A 5200T>A), TTA616ATA (5201T>A), AAA617TCA (5204A>T 5205A>C), GGA618GGG (5209A>G), TTT619TTG (5212T>G), ATG620ATA (5215G>A), AAT621TTT (5216A>T 5217A>T), AAA622AAC (5221A>C), AAG623AAA (5224G>A), ATT624CTC (5225A>C 5227T>C), ACT625CCT (5228A>C), GGT626TCC (5231G>T 5232G>C 5233T>C), GAC627GAA (5236C>A), AGT628CCC (5237A>C 5238G>C 5239T>C) | | | | | | | | |

##### Proteins

| | Begin | End | Coverage | Score | Concordance | Matches | Identities | I/D/M/F* | Stop Codons |
|---|---|---|---|---|---|---|---|---|---|
| hypothetical protein (NP_659397.1) | 523 | 628 | 15.7% | 343 | 51.6% | 101 (95.3%) | 50 (47.2%) | 0/5/0/0 | 0 |
| Protein mutations: | I525V (4928A>G), N526K (4933T>G), F527L (4934T>C), Y531R (4946T>A 4947A>G 4948T>G), K534D (4955A>G 4957G>T), K535P (4958A>C 4959A>C), S538N (4968G>A), I540V (4973A>G 4975T>A), I541L (4976A>T 4978T>A), E542Q (4979G>C), D544N (4985G>A), A545V (4989C>T 4990T>C), H548Y (4997C>T 4999C>T), F549H (5000T>C 5001T>A 5002C>T), G552A (5010G>C), V553L (5012G>T 5014C>G), K555_A556del (5018_5023delAAGGCC), T559D (5030A>G 5031C>A 5032T>C), E560L (5033G>C 5034A>T), G561N (5036G>A 5037G>A 5038T>C), E563_L564del (5042_5047delGAATTA), S569T (5060T>A 5062T>A), K574N (5077A>C), P575E (5078C>G 5079C>A), E577K (5084G>A), L578V (5087T>G 5089G>T), H581S (5096C>T 5097A>C 5098T>C), S582T (5100G>C 5101T>A), K591V (5126A>G 5127A>T), Q592R (5129C>A 5130A>G 5131G>A), V593G (5133T>G 5134G>A), T595Q (5138A>C 5139C>A 5140T>A), I599A (5150A>G 5151T>C), Y600C (5154A>G 5155T>C), T602L (5159A>T 5160C>T 5161C>A), V604K (5165G>A 5166T>A 5167T>A), C605S (5169G>C 5170T>A), T607V (5174A>G 5175C>T 5176A>T), V608I (5177G>A 5179C>T), R609K (5181G>A 5182G>A), V613del (5192_5194delGTA), N614T (5196A>C), L615Q (5199T>A 5200T>A), L616I (5201T>A), K617S (5204A>T 5205A>C), F619L (5212T>G), M620I (5215G>A), N621F (5216A>T 5217A>T), K622N (5221A>C), I624L (5225A>C 5227T>C), T625P (5228A>C), G626S (5231G>T 5232G>C 5233T>C), D627E (5236C>A), S628P (5237A>C 5238G>C 5239T>C) | | | | | | | | |
| Codon mutations: | ATT525GTT (4928A>G), AAT526AAG (4933T>G), TTT527CTT (4934T>C), AAA529AAG (4942A>G), TAT531AGG (4946T>A 4947A>G 4948T>G), AAG534GAT (4955A>G 4957G>T), AAA535CCA (4958A>C 4959A>C), AGT538AAT (4968G>A), TTA539CTG (4970T>C 4972A>G), ATT540GTA (4973A>G 4975T>A), ATT541TTA (4976A>T 4978T>A), GAA542CAA (4979G>C), ACT543ACA (4984T>A), GAT544AAT (4985G>A), GCT545GTC (4989C>T 4990T>C), TCT546TCC (4993T>C), CAC548TAT (4997C>T 4999C>T), TTC549CAT (5000T>C 5001T>A 5002C>T), GGT551GGA (5008T>A), GGA552GCA (5010G>C), GTC553TTG (5012G>T 5014C>G), CTT554CTA (5017T>A), AAG555_GCC556del (5018_5023delAAGGCC), ACC558ACA (5029C>A), ACT559GAC (5030A>G 5031C>A 5032T>C), GAA560CTA (5033G>C 5034A>T), GGT561AAC (5036G>A 5037G>A 5038T>C), GAA563_TTA564del (5042_5047delGAATTA), ATC565ATT (5050C>T), TGC566TGT (5053C>T), TCT569ACA (5060T>A 5062T>A), TCA570AGT (5063T>A 5064C>G 5065A>T), ACA572ACT (5071A>T), TTC573TTT (5074C>T), AAA574AAC (5077A>C), CCA575GAA (5078C>G 5079C>A), GAA577AAA (5084G>A), TTG578GTT (5087T>G 5089G>T), TAC580TAT (5095C>T), CAT581TCC (5096C>T 5097A>C 5098T>C), AGT582ACA (5100G>C 5101T>A), GAG584GAA (5107G>A), CTA588TTA (5117C>T), GCG589GCA (5122G>A), AAA591GTA (5126A>G 5127A>T), CAG592AGA (5129C>A 5130A>G 5131G>A), GTG593GGA (5133T>G 5134G>A), ATT594ATA (5137T>A), ACT595CAA (5138A>C 5139C>A 5140T>A), AGT598TCG (5147A>T 5148G>C 5149T>G), ATT599GCT (5150A>G 5151T>C), TAT600TGC (5154A>G 5155T>C), CTA601TTA (5156C>T), ACC602TTA (5159A>T 5160C>T 5161C>A), CCT603CCA (5164T>A), GTT604AAA (5165G>A 5166T>A 5167T>A), TGT605TCA (5169G>C 5170T>A), ACA607GTT (5174A>G 5175C>T 5176A>T), GTC608ATT (5177G>A 5179C>T), AGG609AAA (5181G>A 5182G>A), GTA613del (5192_5194delGTA), AAT614ACT (5196A>C), CTT615CAA (5199T>A 5200T>A), TTA616ATA (5201T>A), AAA617TCA (5204A>T 5205A>C), GGA618GGG (5209A>G), TTT619TTG (5212T>G), ATG620ATA (5215G>A), AAT621TTT (5216A>T 5217A>T), AAA622AAC (5221A>C), AAG623AAA (5224G>A), ATT624CTC (5225A>C 5227T>C), ACT625CCT (5228A>C), GGT626TCC (5231G>T 5232G>C 5233T>C), GAC627GAA (5236C>A), AGT628CCC (5237A>C 5238G>C 5239T>C) | | | | | | | | |

*: Inserts / Deletes / Misaligned / Frameshifts

#### Analysis details

This analysis was performed with panviral2.64

### NGS Details (UN70): Badnavirus alphacolocalasiae

#### Assembly

| | |
|---|---|
| **Coverage Length** | 288 (1 contig(s)) |
| **Depth Of Coverage** | 2.3 |
| **Number Of Reads** | 5 |
| **Reads Per Million** | 0.09 rpm (after QC) |
| **Ambiguities** | 0 |
| **Assembly Method** | de novo + reference guided assembly |
| **Consensus Caller** | Bcf Tools |

#### Coverage Map

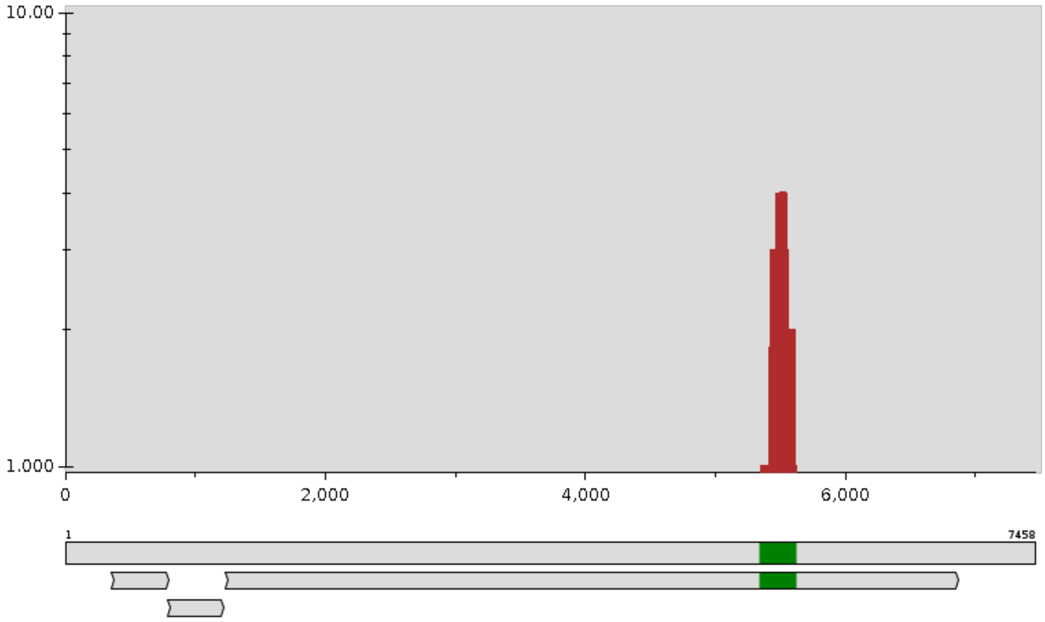

#### Assignment

| | |
|---|---|
| **Type** | Badnavirus alphacolocalasiae (Taxonomy ID: 3048356) |
| **Reference Genome** | NC_004450.1 |
| **NT Identity (%)** | 58.885 |
| **AA Identity (%)** | 52.0833 |
| **Number Of Stop Codons** | 0 |
| **Number Of CDS** | 3 |

#### Alignment

| | |
|---|---|
| **Alignment Score** | 92.0 (NT) + 244.0 (AA) = 336.0 |
| **Concordance (%)** | 31.3984 |

| **Alignment Method** | Global, seeded, nucleotide + amino acids (AGA) |
|---|---|

#### Genome Region

Sequence starts at position 5343 and ends at position 5630 relative to NC_004450.1 reference sequence.

#### Alignment Detailed Statistics

| | Begin | End | Coverage | Score | Concordance | Matches | Identities | I/D/M/F* | Stop Codons |
|---|---|---|---|---|---|---|---|---|---|
| **NT** | **5343** | **5630** | **3.9%** | **92** | **16.3%** | **287 (99.7%)** | **169 (58.7%)** | **0/1** | |
| Mutations: | 5347T>A, 5349A>C, 5359A>G, 5360G>A, 5361G>A, 5362T>C, 5363G>T, 5364A>T, 5366C>T, 5369T>A, 5371G>C, 5374C>A, 5378C>T, 5381T>C, 5382T>A, 5383C>G, 5394C>A, 5402T>A, 5408T>C, 5409C>T, 5410A>G, 5411C>G, 5415G>A, 5418G>C, 5419C>A, 5420T>A, 5423G>T, 5424G>T, 5425A>C, 5426T>G, 5427C>A, 5428C>A, 5429C>A, 5430G>A, 5432T>A, 5433T>G, 5434C>A, 5436G>A, 5437T>A, 5438G>A, 5439G>T, 5441A>T, 5442T>A, 5443G>A, 5444G>A, 5447A>T, 5454C>A, 5455T>A, 5456A>T, 5459T>C, 5462A>G, 5463C>T, 5464A>T, 5465A>T, 5468T>A, 5469T>C, 5470T>A, 5481C>A, 5482T>A, 5483G>T, 5486A>T, 5492T>A, 5498C>A, 5501C>A, 5507T>C, 5510T>C, 5513A>T, 5514G>T, 5515C>T, 5518T>A, 5519A>G, 5522T>C, 5527G>A, 5528A>C, 5530A>T, 5531A>C, 5534G>A, 5535G>A, 5538G>A, 5539C>A, 5540A>T, 5541G>A, 5543A>T, 5549A>C, 5550G>C, 5551G>A, 5552G>T, 5553T>A, 5554G>T, 5555T>C, 5557A>C, 5560A>T, 5564C>T, 5567C>A, 5568delG, 5569C>T, 5570A>C, 5573C>A, 5586A>G, 5588T>A, 5591G>A, 5592G>A, 5597T>C, 5600A>T, 5603C>G, 5604A>G, 5606T>C, 5607G>A, 5608A>T, 5609G>A, 5612A>T, 5613G>T, 5614A>C, 5618T>C, 5619G>T, 5620C>T, 5621A>T, 5627C>T, 5628C>T | | | | | | | | |
| **CDS** | | | | | | | | | |
| TaBVgp3 | 1373 | 1468 | 5.1% | 244 | 36.3% | 96 (100%) | 50 (52.1%) | 0/0/1/1 | 0 |
| Protein mutations: | L1374H (5347T>A), I1375L (5349A>C), K1378R (5359A>G 5360G>A), V1379T (5361G>A 5362T>C 5363G>T), I1380F (5364A>T 5366C>T), N1381K (5369T>A), S1382T (5371G>C), T1383N (5374C>A), L1390M (5394C>A), H1395W (5409C>T 5410A>G 5411C>G), V1397I (5415G>A), A1398Q (5418G>C 5419C>A 5420T>A), M1399I (5423G>T), D1400S (5424G>T 5425A>C 5426T>G), P1401K (5427C>A 5428C>A 5429C>A), D1402K (5430G>A 5432T>A), S1403D (5433T>G 5434C>A), V1404K (5436G>A 5437T>A 5438G>A), E1405Y (5439G>T 5441A>T), W1406K (5442T>A 5443G>A 5444G>A), L1410N (5454C>A 5455T>A 5456A>T), Q1413F (5463C>T 5464A>T 5465A>T), L1415Q (5469T>C 5470T>A), L1419N (5481C>A 5482T>A 5483G>T), A1430F (5514G>T 5515C>T), V1431E (5518T>A 5519A>G), R1434N (5527G>A 5528A>C), K1435I (5530A>T 5531A>C), M1436I (5534G>A), D1437N (5535G>A), A1438N (5538G>A 5539C>A 5540A>T), V1439I (5541G>A 5543A>T), K1441N (5549A>C), G1442H (5550G>C 5551G>A 5552G>T), C1443I (5553T>A 5554G>T 5555T>C), E1444A (5557A>C), K1445I (5560A>T), I1454V (5586A>G 5588T>A), V1456I (5592G>A), N1459K (5603C>G), N1460D (5604A>G 5606T>C), E1461I (5607G>A 5608A>T 5609G>A), E1462D (5612A>T), D1463S (5613G>T 5614A>C), A1465F (5619G>T 5620C>T 5621A>T) | | | | | | | | |
| Codon mutations: | CTT1374CAT (5347T>A), ATT1375CTT (5349A>C), AAG1378AGA (5359A>G 5360G>A), GTG1379ACT (5361G>A 5362T>C 5363G>T), ATC1380TTT (5364A>T 5366C>T), AAT1381AAA (5369T>A), AGC1382ACC (5371G>C), ACT1383AAT (5374C>A), ATC1384ATT (5378C>T), TAT1385TAC (5381T>C), TCT1386AGT (5382T>A 5383C>G), CTG1390ATG (5394C>A), TCT1392TCA (5402T>A), TTT1394TTC (5408T>C), CAC1395TGG (5409C>T 5410A>G 5411C>G), GTT1397ATT (5415G>A), GCT1398CAA (5418G>C 5419C>A 5420T>A), ATG1399ATT (5423G>T), GAT1400TCG (5424G>T 5425A>C 5426T>G), CCC1401AAA (5427C>A 5428C>A 5429C>A), GAT1402AAA (5430G>A 5432T>A), TCT1403GAT (5433T>G 5434C>A), GTG1404AAA (5436G>A 5437T>A 5438G>A), GAA1405TAT (5439G>T 5441A>T), TGG1406AAA (5442T>A 5443G>A 5444G>A), ACA1407ACT (5447A>T), CTA1410AAT (5454C>A 5455T>A 5456A>T), GTT1411GTC (5459T>C), CCA1412CCG (5462A>G), CAA1413TTT (5463C>T 5464A>T 5465A>T), GGT1414GGA (5468T>A), TTA1415CAA (5469T>C 5470T>A), CTG1419AAT (5481C>A 5482T>A 5483G>T), GCA1420GCT (5486A>T), CCT1422CCA (5492T>A), GGC1424GGA (5498C>A), CTC1425CTA (5501C>A), AAT1427AAC (5507T>C), GCT1428GCC (5510T>C), CCA1429CCT (5513A>T), GCC1430TTC (5514G>T 5515C>T), GTA1431GAG (5518T>A 5519A>G), TTT1432TTC (5522T>C), AGA1434AAC (5527G>A 5528A>C), AAA1435ATC (5530A>T 5531A>C), ATG1436ATA (5534G>A), GAT1437AAT (5535G>A), GCA1438AAT (5538G>A 5539C>A 5540A>T), GTA1439ATT (5541G>A 5543A>T), AAA1441AAC (5549A>C), GGG1442CAT (5550G>C 5551G>A 5552G>T), TGT1443ATC (5553T>A 5554G>T 5555T>C), GAA1444GCA (5557A>C), AAA1445ATA (5560A>T), TTC1446TTT (5564C>T), CTC1447CTA (5567C>A), GCA1448-TC (5568delG 5569C>T 5570A>C), GTC1449GTA (5573C>A), ATT1454GTA (5586A>G 5588T>A), CTG1455CTA (5591G>A), GTA1456ATA (5592G>A), TTT1457TTC (5597T>C), TCA1458TCT (5600A>T), AAC1459AAG (5603C>G), AAT1460GAC (5604A>G 5606T>C), GAG1461ATA (5607G>A 5608A>T 5609G>A), GAA1462GAT (5612A>T), GAT1463TCT (5613G>T 5614A>C), CAT1464CAC (5618T>C), GCA1465TTT (5619G>T 5620C>T 5621A>T), CAC1467CAT (5627C>T), CTG1468TTG (5628C>T) | | | | | | | | |
| **Proteins** | | | | | | | | | |
| polyprotein (NP_758808.1) | 1373 | 1468 | 5.1% | 244 | 36.3% | 96 (100%) | 50 (52.1%) | 0/0/1/1 | 0 |
| Protein mutations: | L1374H (5347T>A), I1375L (5349A>C), K1378R (5359A>G 5360G>A), V1379T (5361G>A 5362T>C 5363G>T), I1380F (5364A>T 5366C>T), N1381K (5369T>A), S1382T (5371G>C), T1383N (5374C>A), L1390M (5394C>A), H1395W (5409C>T 5410A>G 5411C>G), V1397I (5415G>A), A1398Q (5418G>C 5419C>A 5420T>A), M1399I (5423G>T), D1400S (5424G>T 5425A>C 5426T>G), P1401K (5427C>A 5428C>A 5429C>A), D1402K (5430G>A 5432T>A), S1403D (5433T>G 5434C>A), V1404K (5436G>A 5437T>A 5438G>A), E1405Y (5439G>T 5441A>T), W1406K (5442T>A 5443G>A 5444G>A), L1410N (5454C>A 5455T>A 5456A>T), Q1413F (5463C>T 5464A>T 5465A>T), L1415Q (5469T>C 5470T>A), L1419N (5481C>A 5482T>A 5483G>T), A1430F (5514G>T 5515C>T), V1431E (5518T>A 5519A>G), R1434N (5527G>A 5528A>C), K1435I (5530A>T 5531A>C), M1436I (5534G>A), D1437N (5535G>A), A1438N (5538G>A 5539C>A 5540A>T), V1439I (5541G>A 5543A>T), K1441N (5549A>C), G1442H (5550G>C 5551G>A 5552G>T), C1443I (5553T>A 5554G>T 5555T>C), E1444A (5557A>C), K1445I (5560A>T), I1454V (5586A>G 5588T>A), V1456I (5592G>A), N1459K (5603C>G), N1460D (5604A>G 5606T>C), E1461I (5607G>A 5608A>T 5609G>A), E1462D (5612A>T), D1463S (5613G>T 5614A>C), A1465F (5619G>T 5620C>T 5621A>T) | | | | | | | | |
| Codon mutations: | CTT1374CAT (5347T>A), ATT1375CTT (5349A>C), AAG1378AGA (5359A>G 5360G>A), GTG1379ACT (5361G>A 5362T>C 5363G>T), ATC1380TTT (5364A>T 5366C>T), AAT1381AAA (5369T>A), AGC1382ACC (5371G>C), ACT1383AAT (5374C>A), ATC1384ATT (5378C>T), TAT1385TAC (5381T>C), TCT1386AGT (5382T>A 5383C>G), CTG1390ATG (5394C>A), TCT1392TCA (5402T>A), TTT1394TTC (5408T>C), CAC1395TGG (5409C>T 5410A>G 5411C>G), GTT1397ATT (5415G>A), GCT1398CAA (5418G>C 5419C>A 5420T>A), ATG1399ATT (5423G>T), GAT1400TCG (5424G>T 5425A>C 5426T>G), CCC1401AAA (5427C>A 5428C>A 5429C>A), GAT1402AAA (5430G>A 5432T>A), TCT1403GAT (5433T>G 5434C>A), GTG1404AAA (5436G>A 5437T>A 5438G>A), GAA1405TAT (5439G>T 5441A>T), TGG1406AAA (5442T>A 5443G>A 5444G>A), ACA1407ACT (5447A>T), CTA1410AAT (5454C>A 5455T>A 5456A>T), GTT1411GTC (5459T>C), CCA1412CCG (5462A>G), CAA1413TTT (5463C>T 5464A>T 5465A>T), GGT1414GGA (5468T>A), TTA1415CAA (5469T>C 5470T>A), CTG1419AAT (5481C>A 5482T>A 5483G>T), GCA1420GCT (5486A>T), CCT1422CCA (5492T>A), GGC1424GGA (5498C>A), CTC1425CTA (5501C>A), AAT1427AAC (5507T>C), GCT1428GCC (5510T>C), CCA1429CCT (5513A>T), GCC1430TTC (5514G>T 5515C>T), GTA1431GAG (5518T>A 5519A>G), TTT1432TTC (5522T>C), AGA1434AAC (5527G>A 5528A>C), AAA1435ATC (5530A>T 5531A>C), ATG1436ATA (5534G>A), GAT1437AAT (5535G>A), GCA1438AAT (5538G>A 5539C>A 5540A>T), GTA1439ATT (5541G>A 5543A>T), AAA1441AAC (5549A>C), GGG1442CAT (5550G>C 5551G>A 5552G>T), TGT1443ATC (5553T>A 5554G>T 5555T>C), GAA1444GCA (5557A>C), AAA1445ATA (5560A>T), TTC1446TTT (5564C>T), CTC1447CTA (5567C>A), GCA1448-TC (5568delG 5569C>T 5570A>C), GTC1449GTA (5573C>A), ATT1454GTA (5586A>G 5588T>A), CTG1455CTA (5591G>A), GTA1456ATA (5592G>A), TTT1457TTC (5597T>C), TCA1458TCT (5600A>T), AAC1459AAG (5603C>G), AAT1460GAC (5604A>G 5606T>C), GAG1461ATA (5607G>A 5608A>T 5609G>A), GAA1462GAT (5612A>T), GAT1463TCT (5613G>T 5614A>C), CAT1464CAC (5618T>C), GCA1465TTT (5619G>T 5620C>T 5621A>T), CAC1467CAT (5627C>T), CTG1468TTG (5628C>T) | | | | | | | | |

*: Inserts / Deletes / Misaligned / Frameshifts

#### Analysis details

This analysis was performed with panviral2.64

### NGS Details (UN70): Caulimovirus venafragariae

#### Assembly

| | |
|---|---|
| **Coverage Length** | 262 (1 contig(s)) |
| **Depth Of Coverage** | 1.6 |
| **Number Of Reads** | 4 |
| **Reads Per Million** | 0.08 rpm (after QC) |
| **Ambiguities** | 0 |
| **Assembly Method** | de novo + reference guided assembly |
| **Consensus Caller** | Bcf Tools |

#### Coverage Map

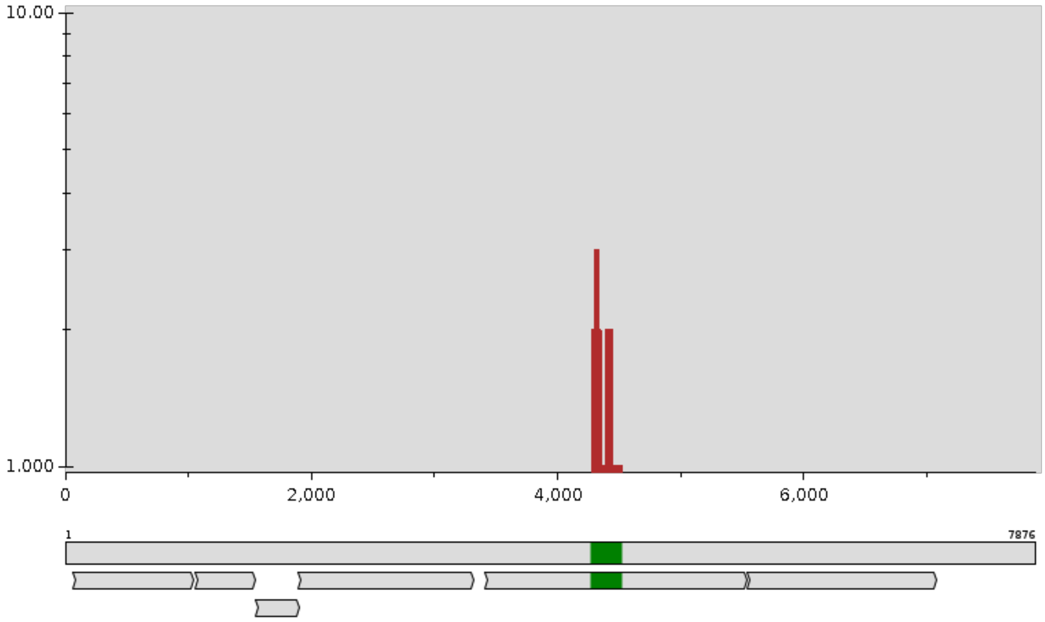

#### Assignment

| | |
|---|---|
| **Type** | Caulimovirus venafragariae (Taxonomy ID: 3048344) |
| **Reference Genome** | NC_001725.1 |
| **NT Identity (%)** | 62.5954 |
| **AA Identity (%)** | 60.9195 |
| **Number Of Stop Codons** | 0 |
| **Number Of CDS** | 6 |

#### Alignment

| | |
|---|---|
| **Alignment Score** | 132.0 (NT) + 378.0 (AA) = 510.0 |
| **Concordance (%)** | 45.9046 |

| Alignment Method | Global, seeded, nucleotide + amino acids (AGA) |
|---|---|

#### Genome Region

Sequence starts at position 4268 and ends at position 4529 relative to NC_001725.1 reference sequence.

#### Alignment Detailed Statistics

| | Begin | End | Coverage | Score | Concordance | Matches | Identities | I/D/M/F* | Stop Codons |
|---|---|---|---|---|---|---|---|---|---|
| **NT** | **4268** | **4529** | **3.3%** | **132** | **25.2%** | **262 (100%)** | **164 (62.6%)** | **0/0** | |
| Mutations: | 4268C>A, 4276C>G, 4277G>T, 4284G>T, 4289A>G, 4290A>C, 4292C>A, 4296G>A, 4299C>T, 4300G>A, 4311C>A, 4314G>C, 4315C>T, 4317T>A, 4320T>A, 4321A>G, 4323C>T, 4326C>A, 4330C>A, 4331C>G, 4332T>A, 4333A>T, 4334G>C, 4335C>T, 4337A>C, 4338G>A, 4339A>T, 4340G>C, 4341T>A, 4342C>A, 4343C>A, 4348T>A, 4349C>G, 4350C>A, 4354C>G, 4362C>T, 4371G>A, 4378G>A, 4379C>G, 4383G>A, 4386C>A, 4387A>G, 4388A>T, 4390C>A, 4395C>T, 4402A>C, 4403G>A, 4408G>A, 4413T>A, 4419T>C, 4424A>G, 4425G>A, 4426T>C, 4428A>T, 4434C>T, 4435C>A, 4437T>C, 4440A>T, 4442A>G, 4443G>A, 4444G>A, 4445G>C, 4446A>G, 4447G>T, 4451G>C, 4452C>T, 4455C>T, 4456C>A, 4457T>A, 4458A>G, 4461T>A, 4464T>A, 4465A>G, 4467C>T, 4470G>A, 4471G>A, 4472A>G, 4473G>T, 4474C>G, 4477C>T, 4479G>A, 4480C>A, 4482T>A, 4483C>A, 4485A>T, 4491C>A, 4492G>C, 4493G>A, 4503C>A, 4504T>G, 4506T>A, 4508A>T, 4510T>A, 4511C>A, 4513T>A, 4514C>A, 4515C>A, 4521C>T | | | | | | | | |

##### CDS

| | Begin | End | Coverage | Score | Concordance | Matches | Identities | I/D/M/F* | Stop Codons |
|---|---|---|---|---|---|---|---|---|---|
| ORF_V | 289 | 375 | 12.3% | 378 | 62.8% | 87 (100%) | 53 (60.9%) | 0/0/0/0 | 0 |
| Protein mutations: | R291V (4276C>G 4277G>T), E293D (4284G>T), K295S (4289A>G 4290A>C), T296N (4292C>A), E299K (4300G>A), K303N (4314G>C), I306V (4321A>G 4323C>T), P309R (4330C>A 4331C>G 4332T>A), K311T (4337A>C 4338G>A), P313K (4342C>A 4343C>A), S315R (4348T>A 4349C>G 4350C>A), P317A (4354C>G), A325S (4378G>A 4379C>G), K328V (4387A>G 4388A>T), R333Q (4402A>C 4403G>A), V335I (4408G>A), K340R (4424A>G 4425G>A), H344N (4435C>A 4437T>C), K346R (4442A>G 4443G>A), G347T (4444G>A 4445G>C 4446A>G), D348Y (4447G>T), G349A (4451G>C 4452C>T), L351K (4456C>A 4457T>A 4458A>G), N354D (4465A>G 4467C>T), E356S (4471G>A 4472A>G 4473G>T), Q357E (4474C>G), L359I (4480C>A 4482T>A), Q360N (4483C>A 4485A>T), G363Q (4492G>C 4493G>A), F367V (4504T>G 4506T>A), Y368F (4508A>T), S369N (4510T>A 4511C>A), S370K (4513T>A 4514C>A 4515C>A) | | | | | | | | |
| Codon mutations: | CCT288.AT (4268C>A), CGA291GTA (4276C>G 4277G>T), GAG293GAT (4284G>T), AAA295AGC (4289A>G 4290A>C), ACT296AAT (4292C>A), CAG297CAA (4296G>A), ATC298ATT (4299C>T), GAA299AAA (4300G>A), CTC302CTA (4311C>A), AAG303AAC (4314G>C), CTT304TTA (4315C>T 4317T>A), GGT305GGA (4320T>A), ATC306GTT (4321A>G 4323C>T), ATC307ATA (4326C>A), CCT309AGA (4330C>A 4331C>G 4332T>A), AGC310TCT (4333A>T 4334G>C 4335C>T), AAG311ACA (4337A>C 4338G>A), AGT312TCA (4339A>T 4340G>C 4341T>A), CCA313AAA (4342C>A 4343C>A), TCC315AGA (4348T>A 4349C>G 4350C>A), CCA317GCA (4354C>G), TTC319TTT (4362C>T), AGG322AGA (4371G>A), GCC325AGC (4378G>A 4379C>G), GAG326GAA (4383G>A), ATC327ATA (4386C>A), AAA328GTA (4387A>G 4388A>T), CGA329AGA (4390C>A), GGC330GGT (4395C>T), AGA333CAA (4402A>C 4403G>A), GTA335ATA (4408G>A), ATT336ATA (4413T>A), TAT338TAC (4419T>C), AAG340AGA (4424A>G 4425G>A), TTA341CTT (4426T>C 4428A>T), GAC343GAT (4434C>T), CAT344AAC (4435C>A 4437T>C), ACA345ACT (4440A>T), AAG346AGA (4442A>G 4443G>A), GGA347ACG (4444G>A 4445G>C 4446A>G), GAT348TAT (4447G>T), GGC349GCT (4451G>C 4452C>T), TAC350TAT (4455C>T), CTA351AAG (4456C>A 4457T>A 4458A>G), CTT352CTA (4461T>A), CCT353CCA (4464T>A), AAC354GAT (4465A>G 4467C>T), AAG355AAA (4470G>A), GAG356AGT (4471G>A 4472A>G 4473G>T), CAA357GAA (4474C>G), CTG358TTA (4477C>T 4479G>A), CTT359ATA (4480C>A 4482T>A), CAA360AAT (4483C>A 4485A>T), ATC362ATA (4491C>A), GGA363CAA (4492G>C 4493G>A), ACC366ACA (4503C>A), TTT367GTA (4504T>G 4506T>A), TAC368TTC (4508A>T), TCT369AAT (4510T>A 4511C>A), TCC370AAA (4513T>A 4514C>A 4515C>A), GAC372GAT (4521C>T) | | | | | | | | |

##### Proteins

| | Begin | End | Coverage | Score | Concordance | Matches | Identities | I/D/M/F* | Stop Codons |
|---|---|---|---|---|---|---|---|---|---|
| hypothetical protein (NP_043933.1) | 289 | 375 | 12.3% | 378 | 62.8% | 87 (100%) | 53 (60.9%) | 0/0/0/0 | 0 |
| Protein mutations: | R291V (4276C>G 4277G>T), E293D (4284G>T), K295S (4289A>G 4290A>C), T296N (4292C>A), E299K (4300G>A), K303N (4314G>C), I306V (4321A>G 4323C>T), P309R (4330C>A 4331C>G 4332T>A), K311T (4337A>C 4338G>A), P313K (4342C>A 4343C>A), S315R (4348T>A 4349C>G 4350C>A), P317A (4354C>G), A325S (4378G>A 4379C>G), K328V (4387A>G 4388A>T), R333Q (4402A>C 4403G>A), V335I (4408G>A), K340R (4424A>G 4425G>A), H344N (4435C>A 4437T>C), K346R (4442A>G 4443G>A), G347T (4444G>A 4445G>C 4446A>G), D348Y (4447G>T), G349A (4451G>C 4452C>T), L351K (4456C>A 4457T>A 4458A>G), N354D (4465A>G 4467C>T), E356S (4471G>A 4472A>G 4473G>T), Q357E (4474C>G), L359I (4480C>A 4482T>A), Q360N (4483C>A 4485A>T), G363Q (4492G>C 4493G>A), F367V (4504T>G 4506T>A), Y368F (4508A>T), S369N (4510T>A 4511C>A), S370K (4513T>A 4514C>A 4515C>A) | | | | | | | | |
| Codon mutations: | CCT288.AT (4268C>A), CGA291GTA (4276C>G 4277G>T), GAG293GAT (4284G>T), AAA295AGC (4289A>G 4290A>C), ACT296AAT (4292C>A), CAG297CAA (4296G>A), ATC298ATT (4299C>T), GAA299AAA (4300G>A), CTC302CTA (4311C>A), AAG303AAC (4314G>C), CTT304TTA (4315C>T 4317T>A), GGT305GGA (4320T>A), ATC306GTT (4321A>G 4323C>T), ATC307ATA (4326C>A), CCT309AGA (4330C>A 4331C>G 4332T>A), AGC310TCT (4333A>T 4334G>C 4335C>T), AAG311ACA (4337A>C 4338G>A), AGT312TCA (4339A>T 4340G>C 4341T>A), CCA313AAA (4342C>A 4343C>A), TCC315AGA (4348T>A 4349C>G 4350C>A), CCA317GCA (4354C>G), TTC319TTT (4362C>T), AGG322AGA (4371G>A), GCC325AGC (4378G>A 4379C>G), GAG326GAA (4383G>A), ATC327ATA (4386C>A), AAA328GTA (4387A>G 4388A>T), CGA329AGA (4390C>A), GGC330GGT (4395C>T), AGA333CAA (4402A>C 4403G>A), GTA335ATA (4408G>A), ATT336ATA (4413T>A), TAT338TAC (4419T>C), AAG340AGA (4424A>G 4425G>A), TTA341CTT (4426T>C 4428A>T), GAC343GAT (4434C>T), CAT344AAC (4435C>A 4437T>C), ACA345ACT (4440A>T), AAG346AGA (4442A>G 4443G>A), GGA347ACG (4444G>A 4445G>C 4446A>G), GAT348TAT (4447G>T), GGC349GCT (4451G>C 4452C>T), TAC350TAT (4455C>T), CTA351AAG (4456C>A 4457T>A 4458A>G), CTT352CTA (4461T>A), CCT353CCA (4464T>A), AAC354GAT (4465A>G 4467C>T), AAG355AAA (4470G>A), GAG356AGT (4471G>A 4472A>G 4473G>T), CAA357GAA (4474C>G), CTG358TTA (4477C>T 4479G>A), CTT359ATA (4480C>A 4482T>A), CAA360AAT (4483C>A 4485A>T), ATC362ATA (4491C>A), GGA363CAA (4492G>C 4493G>A), ACC366ACA (4503C>A), TTT367GTA (4504T>G 4506T>A), TAC368TTC (4508A>T), TCT369AAT (4510T>A 4511C>A), TCC370AAA (4513T>A 4514C>A 4515C>A), GAC372GAT (4521C>T) | | | | | | | | |

*: Inserts / Deletes / Misaligned / Frameshifts

#### Analysis details

This analysis was performed with panviral2.64

#### NGS Details (UN70): Fragaria chiloensis cryptic virus (segment RNA 3)

##### Assembly

| | |
|---|---|
| **Coverage Length** | 233 (1 contig(s)) |
| **Depth Of Coverage** | 2.1 |
| **Number Of Reads** | 4 |
| **Reads Per Million** | 0.08 rpm (after QC) |
| **Ambiguities** | 0 |
| **Assembly Method** | de novo + reference guided assembly |
| **Consensus Caller** | Bcf Tools |

##### Coverage Map

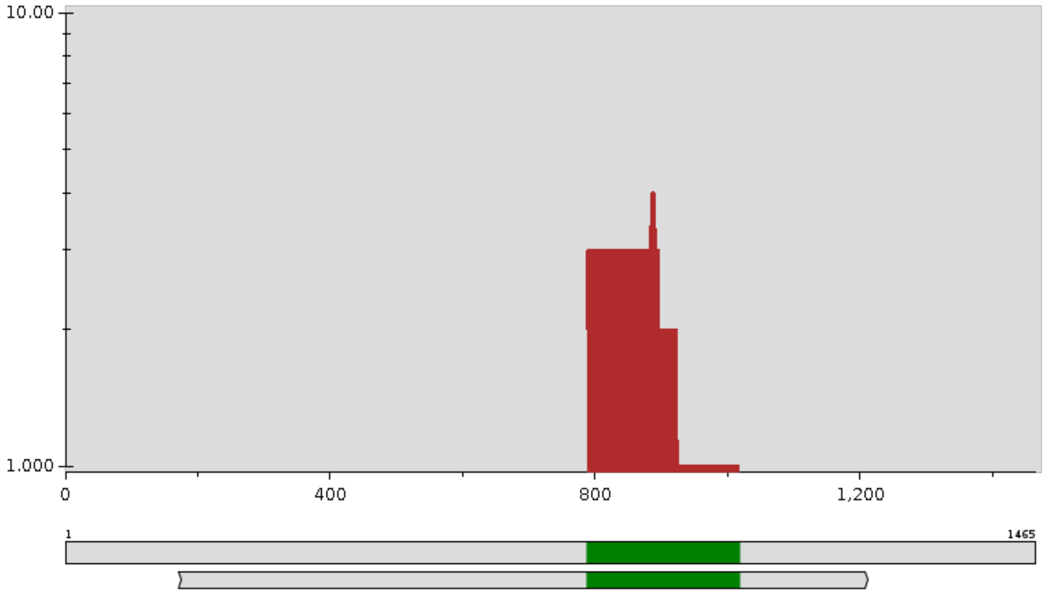

##### Assignment

| | |
|---|---|
| **Type** | Fragaria chiloensis cryptic virus (Taxonomy ID: 335204) |
| **Reference Genome** | NC_009520.1 |
| **NT Identity (%)** | 54.9356 |
| **AA Identity (%)** | 42.8571 |
| **Number Of Stop Codons** | 0 |
| **Number Of CDS** | 1 |

##### Alignment

| | |
|---|---|
| **Alignment Score** | 46.0 (NT) + 271.0 (AA) = 317.0 |
| **Concordance (%)** | 30.2193 |

| Alignment Method | Global, seeded, nucleotide + amino acids (AGA) |
|---|---|

#### Genome Region

Sequence starts at position 788 and ends at position 1020 relative to NC_009520.1 reference sequence.

#### Alignment Detailed Statistics

| | Begin | End | Coverage | Score | Concordance | Matches | Identities | I/D/M/F* | Stop Codons |
|---|---|---|---|---|---|---|---|---|---|
| **NT** | **788** | **1020** | **15.9%** | **46** | **9.9%** | **233 (100%)** | **128 (54.9%)** | **0/0** | |
| Mutations: | 788T>C, 792C>T, 794C>T, 795T>G, 798G>A, 801C>A, 804C>T, 807T>C, 812C>G, 813C>A, 816C>T, 817C>G, 819G>A, 820T>A, 828A>T, 829C>T, 831C>T, 832G>T, 834T>C, 835C>A, 836A>G, 837A>C, 838G>C, 840C>G, 842C>A, 843T>A, 847A>C, 848A>C, 849C>T, 852G>T, 853A>G, 854C>A, 855T>C, 856G>C, 857A>G, 858G>C, 861C>T, 864G>C, 868T>A, 869A>T, 876A>T, 877A>C, 878C>A, 879T>C, 880C>A, 882T>G, 885A>G, 886G>A, 889C>G, 890T>C, 894T>A, 897C>A, 900C>A, 901T>C, 902G>T, 903T>C, 904A>G, 906A>G, 909A>C, 910G>A, 912G>C, 915A>T, 924C>T, 926A>C, 927G>A, 929A>C, 933A>T, 936C>T, 947G>C, 950A>T, 955A>G, 957C>T, 958C>A, 960C>G, 961A>C, 963C>T, 965A>T, 967C>A, 969C>T, 970G>A, 971G>A, 973A>C, 974C>G, 977C>A, 978A>T, 980A>C, 981T>C, 982C>T, 984C>T, 986T>C, 987C>T, 988T>G, 990C>A, 991C>A, 993G>A, 994C>T, 1003A>G, 1004C>T, 1005A>T, 1007A>T, 1008C>G, 1014T>C, 1015C>A, 1017G>T, 1020T>G | | | | | | | | |

##### CDS

| | Begin | End | Coverage | Score | Concordance | Matches | Identities | I/D/M/F* | Stop Codons |
|---|---|---|---|---|---|---|---|---|---|
| FCCV_s3gp1 | 207 | 283 | 22.2% | 271 | 46.1% | 77 (100%) | 33 (42.9%) | 0/0/0/0 | 0 |
| Protein mutations: | A208V (794C>T 795T>G), A214G (812C>G 813C>A), Q216E (817C>G 819G>A), Y217N (820T>A), R220C (829C>T 831C>T), A221S (832G>T 834T>C), Q222S (835C>A 836A>G 837A>C), D223Q (838G>C 840C>G), A224E (842C>A 843T>A), N226P (847A>C 848A>C 849C>T), M227I (852G>T), T228D (853A>G 854C>A 855T>C), E229R (856G>C 857A>G 858G>C), Q231H (864G>C), Y233I (868T>A 869A>T), T236H (877A>C 878C>A 879T>C), L237M (880C>A 882T>G), D239N (886G>A), L240A (889C>G 890T>C), C244L (901T>C 902G>T 903T>C), K245E (904A>G 906A>G), V247I (910G>A 912G>C), E248D (915A>T), K252T (926A>C 927G>A), K253T (929A>C), S259T (947G>C), Y260F (950A>T), I262V (955A>G 957C>T), H263K (958C>A 960C>G), N264H (961A>C 963C>T), D265V (965A>T), H266N (967C>A 969C>T), G267N (970G>A 971G>A), T268R (973A>C 974C>G), S269Y (977C>A 978A>T), D270A (980A>C 981T>C), L271F (982C>T 984C>T), V272A (986T>C 987C>T), C273G (988T>G 990C>A), L274I (991C>A 993G>A), L275F (994C>T), T278V (1003A>G 1004C>T 1005A>T), N279M (1007A>T 1008C>G), P282T (1015C>A 1017G>T) | | | | | | | | |
| Codon mutations: | GTT206.CT (788T>C), ATC207ATT (792C>T), GCT208GTG (794C>T 795T>G), CCG209CCA (798G>A), ACC210ACA (801C>A), TAC211TAT (804C>T), CCT212CCC (807T>C), GCC214GGA (812C>G 813C>A), ACC215ACT (816C>T), CAG216GAA (817C>G 819G>A), TAT217AAT (820T>A), GGA219GGT (828A>T), CGC220TGT (829C>T 831C>T), GCT221TCC (832G>T 834T>C), CAA222AGC (835C>A 836A>G 837A>C), GAC223CAG (838G>C 840C>G), GCT224GAA (842C>A 843T>A), AAC226CCT (847A>C 848A>C 849C>T), ATG227ATT (852G>T), ACT228GAC (853A>G 854C>A 855T>C), GAG229CGC (856G>C 857A>G 858G>C), TAC230TAT (861C>T), CAG231CAC (864G>C), TAC233ATC (868T>A 869A>T), CCA235CCT (876A>T), ACT236CAC (877A>C 878C>A 879T>C), CTT237ATG (880C>A 882T>G), AAA238AAG (885A>G), GAT239AAT (886G>A), CTT240GCT (889C>G 890T>C), GGT241GGA (894T>A), ATC242ATA (897C>A), CCC243CCA (900C>A), TGT244CTC (901T>C 902G>T 903T>C), AAA245GAG (904A>G 906A>G), TCA246TCC (909A>C), GTG247ATC (910G>A 912G>C), GAA248GAT (915A>T), GAC251GAT (924C>T), AAG252ACA (926A>C 927G>A), AAG253ACG (929A>C), GGA254GGT (933A>T), ACC255ACT (936C>T), AGT259ACT (947G>C), TAC260TTC (950A>T), ATC262GTT (955A>G 957C>T), CAC263AAG (958C>A 960C>G), AAC264CAT (961A>C 963C>T), GAC265GTC (965A>T), CAC266AAT (967C>A 969C>T), GGT267AAT (970G>A 971G>A), ACA268CGA (973A>C 974C>G), TCA269TAT (977C>A 978A>T), GAT270GCC (980A>C 981T>C), CTC271TTT (982C>T 984C>T), GTC272GCT (986T>C 987C>T), TGC273GGA (988T>G 990C>A), CTG274ATA (991C>A 993G>A), CTT275TTT (994C>T), ACA278GTT (1003A>G 1004C>T 1005A>T), AAC279ATG (1007A>T 1008C>G), TCT281TCC (1014T>C), CCG282ACT (1015C>A 1017G>T), CTT283CTG (1020T>G) | | | | | | | | |

##### Proteins

| | Begin | End | Coverage | Score | Concordance | Matches | Identities | I/D/M/F* | Stop Codons |
|---|---|---|---|---|---|---|---|---|---|
| putative coat protein (YP_001274392.1) | 207 | 283 | 22.2% | 271 | 46.1% | 77 (100%) | 33 (42.9%) | 0/0/0/0 | 0 |
| Protein mutations: | A208V (794C>T 795T>G), A214G (812C>G 813C>A), Q216E (817C>G 819G>A), Y217N (820T>A), R220C (829C>T 831C>T), A221S (832G>T 834T>C), Q222S (835C>A 836A>G 837A>C), D223Q (838G>C 840C>G), A224E (842C>A 843T>A), N226P (847A>C 848A>C 849C>T), M227I (852G>T), T228D (853A>G 854C>A 855T>C), E229R (856G>C 857A>G 858G>C), Q231H (864G>C), Y233I (868T>A 869A>T), T236H (877A>C 878C>A 879T>C), L237M (880C>A 882T>G), D239N (886G>A), L240A (889C>G 890T>C), C244L (901T>C 902G>T 903T>C), K245E (904A>G 906A>G), V247I (910G>A 912G>C), E248D (915A>T), K252T (926A>C 927G>A), K253T (929A>C), S259T (947G>C), Y260F (950A>T), I262V (955A>G 957C>T), H263K (958C>A 960C>G), N264H (961A>C 963C>T), D265V (965A>T), H266N (967C>A 969C>T), G267N (970G>A 971G>A), T268R (973A>C 974C>G), S269Y (977C>A 978A>T), D270A (980A>C 981T>C), L271F (982C>T 984C>T), V272A (986T>C 987C>T), C273G (988T>G 990C>A), L274I (991C>A 993G>A), L275F (994C>T), T278V (1003A>G 1004C>T 1005A>T), N279M (1007A>T 1008C>G), P282T (1015C>A 1017G>T) | | | | | | | | |
| Codon mutations: | GTT206.CT (788T>C), ATC207ATT (792C>T), GCT208GTG (794C>T 795T>G), CCG209CCA (798G>A), ACC210ACA (801C>A), TAC211TAT (804C>T), CCT212CCC (807T>C), GCC214GGA (812C>G 813C>A), ACC215ACT (816C>T), CAG216GAA (817C>G 819G>A), TAT217AAT (820T>A), GGA219GGT (828A>T), CGC220TGT (829C>T 831C>T), GCT221TCC (832G>T 834T>C), CAA222AGC (835C>A 836A>G 837A>C), GAC223CAG (838G>C 840C>G), GCT224GAA (842C>A 843T>A), AAC226CCT (847A>C 848A>C 849C>T), ATG227ATT (852G>T), ACT228GAC (853A>G 854C>A 855T>C), GAG229CGC (856G>C 857A>G 858G>C), TAC230TAT (861C>T), CAG231CAC (864G>C), TAC233ATC (868T>A 869A>T), CCA235CCT (876A>T), ACT236CAC (877A>C 878C>A 879T>C), CTT237ATG (880C>A 882T>G), AAA238AAG (885A>G), GAT239AAT (886G>A), CTT240GCT (889C>G 890T>C), GGT241GGA (894T>A), ATC242ATA (897C>A), CCC243CCA (900C>A), TGT244CTC (901T>C 902G>T 903T>C), AAA245GAG (904A>G 906A>G), TCA246TCC (909A>C), GTG247ATC (910G>A 912G>C), GAA248GAT (915A>T), GAC251GAT (924C>T), AAG252ACA (926A>C 927G>A), AAG253ACG (929A>C), GGA254GGT (933A>T), ACC255ACT (936C>T), AGT259ACT (947G>C), TAC260TTC (950A>T), ATC262GTT (955A>G 957C>T), CAC263AAG (958C>A 960C>G), AAC264CAT (961A>C 963C>T), GAC265GTC (965A>T), CAC266AAT (967C>A 969C>T), GGT267AAT (970G>A 971G>A), ACA268CGA (973A>C 974C>G), TCA269TAT (977C>A 978A>T), GAT270GCC (980A>C 981T>C), CTC271TTT (982C>T 984C>T), GTC272GCT (986T>C 987C>T), TGC273GGA (988T>G 990C>A), CTG274ATA (991C>A 993G>A), CTT275TTT (994C>T), ACA278GTT (1003A>G 1004C>T 1005A>T), AAC279ATG (1007A>T 1008C>G), TCT281TCC (1014T>C), CCG282ACT (1015C>A 1017G>T), CTT283CTG (1020T>G) | | | | | | | | |

*: Inserts / Deletes / Misaligned / Frameshifts

#### Analysis details

This analysis was performed with panviral2.64

### NGS Details (UN70): Colombian datura virus

#### Assembly

| | |
|---|---|
| **Coverage Length** | 208 (1 contig(s)) |
| **Depth Of Coverage** | 1.3 |
| **Number Of Reads** | 2 |
| **Reads Per Million** | 0.04 rpm (after QC) |
| **Ambiguities** | 0 |
| **Assembly Method** | de novo + reference guided assembly |
| **Consensus Caller** | Bcf Tools |

#### Coverage Map

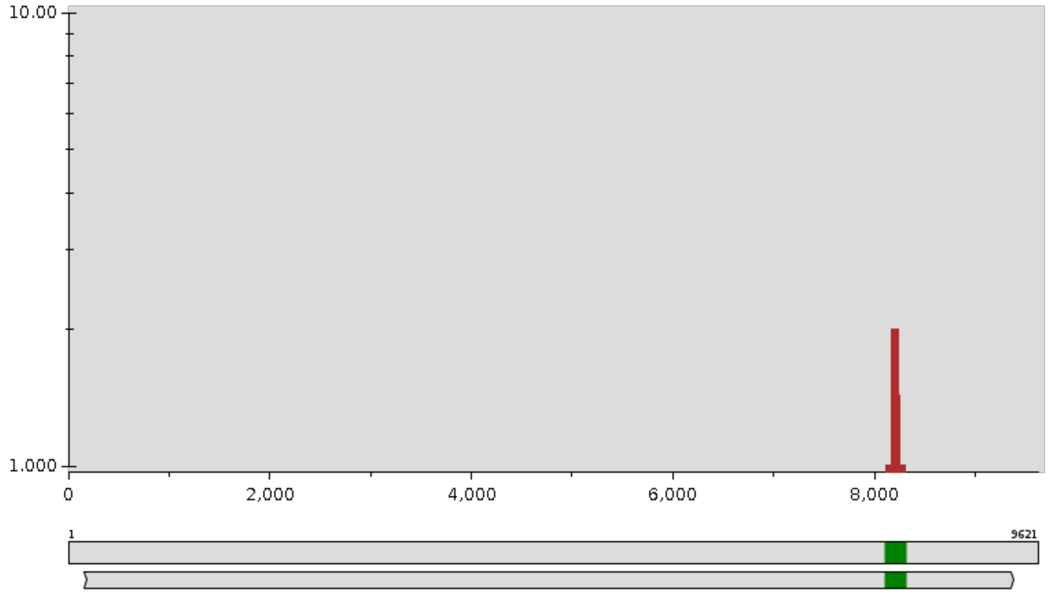

#### Assignment

| | |
|---|---|
| **Type** | Colombian datura virus (Taxonomy ID: 91613) |
| **Reference Genome** | NC_020072.1 |
| **NT Identity (%)** | 99.0385 |
| **AA Identity (%)** | 100.0 |
| **Number Of Stop Codons** | 0 |
| **Number Of CDS** | 1 |

#### Alignment

| | |
|---|---|
| **Alignment Score** | 408.0 (NT) + 478.0 (AA) = 886.0 |
| **Concordance (%)** | 99.1051 |

| Alignment Method | Global, seeded, nucleotide + amino acids (AGA) |
|---|---|

#### Genome Region

Sequence starts at position 8107 and ends at position 8314 relative to NC_020072.1 reference sequence.

#### Alignment Detailed Statistics

| | Begin | End | Coverage | Score | Concordance | Matches | Identities | I/D/M/F* | Stop Codons |
|---|---|---|---|---|---|---|---|---|---|
| **NT** | **8107** | **8314** | **2.2%** | **408** | **98.1%** | **208 (100%)** | **206 (99.0%)** | **0/0** | |
| Mutations: | 8107T>C, 8233T>A | | | | | | | | |
| **CDS** | | | | | | | | | |
| G357_gp1 | 2652 | 2720 | 2.2% | 478 | 97.2% | 69 (100%) | 69 (100%) | 0/0/0/0 | 0 |
| **Proteins** | | | | | | | | | |
| polyprotein (YP_007346986.1) | 2652 | 2720 | 2.2% | 478 | 97.2% | 69 (100%) | 69 (100%) | 0/0/0/0 | 0 |
| Protein mutations: | *none* | | | | | | | | |
| Codon mutations: | CAT2651..C (8107T>C), CTT2693CTA (8233T>A) | | | | | | | | |

*: Inserts / Deletes / Misaligned / Frameshifts

#### Analysis details

This analysis was performed with panviral2.64

### NGS Details (UN70): Limeum africanum associated virus

#### Assembly

| | |
|---|---|
| **Coverage Length** | 135 (1 contig(s)) |
| **Depth Of Coverage** | 1.0 |
| **Number Of Reads** | 1 |
| **Reads Per Million** | 0.02 rpm (after QC) |
| **Ambiguities** | 0 |
| **Assembly Method** | de novo + reference guided assembly |
| **Consensus Caller** | Bcf Tools |

#### Coverage Map

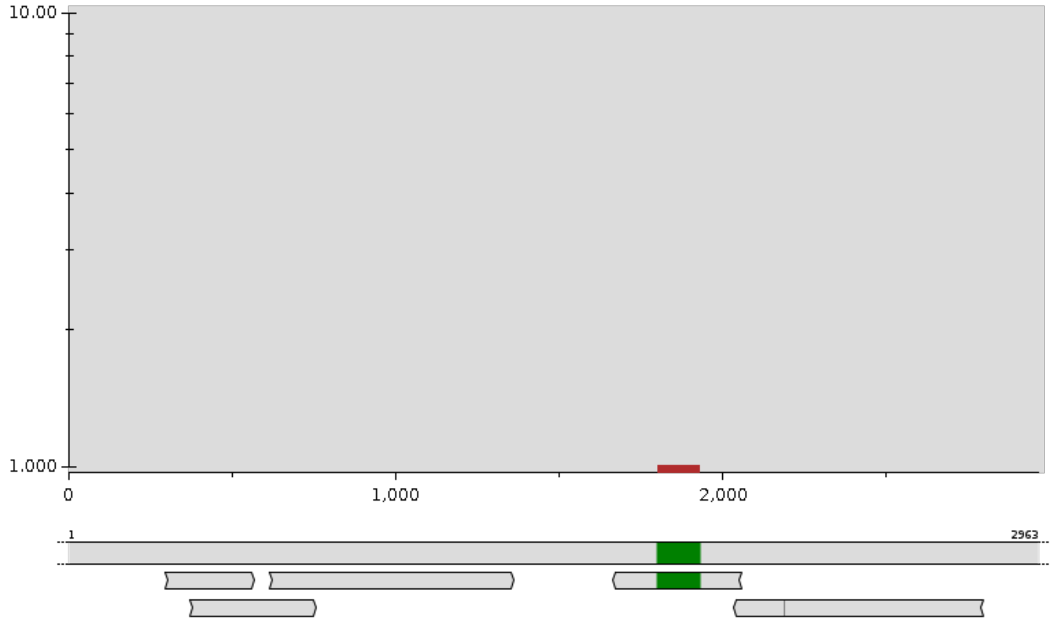

#### Assignment

| | |
|---|---|
| **Type** | Limeum africanum associated virus (Taxonomy ID: 2093276) |
| **Reference Genome** | NC_037063.1 |
| **NT Identity (%)** | 72.5926 |
| **AA Identity (%)** | 75.5556 |
| **Number Of Stop Codons** | 0 |
| **Number Of CDS** | 5 |

#### Alignment

| | |
|---|---|
| **Alignment Score** | 122.0 (NT) + 227.0 (AA) = 349.0 |
| **Concordance (%)** | 58.2638 |

| Alignment Method | Global, seeded, nucleotide + amino acids (AGA) |
|---|---|

#### Genome Region

Sequence starts at position 1799 and ends at position 1933 relative to NC_037063.1 reference sequence.

#### Alignment Detailed Statistics

| | Begin | End | Coverage | Score | Concordance | Matches | Identities | I/D/M/F* | Stop Codons |
|---|---|---|---|---|---|---|---|---|---|
| **NT** | **1799** | **1933** | **4.6%** | **122** | **45.2%** | **135 (100%)** | **98 (72.6%)** | **0/0** | |
| Mutations: | 1801T>C, 1810T>C, 1813C>T, 1816A>G, 1819C>T, 1828G>T, 1835A>T, 1838T>C, 1840A>G, 1841T>A, 1842G>A, 1846A>G, 1847G>C, 1854G>A, 1855T>C, 1857A>G, 1861T>C, 1864T>C, 1867A>G, 1868C>T, 1869A>T, 1877T>A, 1880T>A, 1881G>T, 1885A>C, 1888T>G, 1894A>G, 1909T>C, 1912A>T, 1914C>T, 1915A>G, 1916T>A, 1917A>T, 1921A>G, 1924C>A, 1925C>T, 1927A>G | | | | | | | | |
| **CDS** | | | | | | | | | |
| C5Y53_gp4 | 246 | 290 | 13.5% | 227 | 65.8% | 45 (100%) | 34 (75.6%) | 0/0/0/0 | 0 |
| Protein mutations: | W248Y (1924C>A 1925C>T), Y251I (1915A>G 1916T>A 1917A>T), A252T (1912A>T 1914C>T), Q263I (1880T>A 1881G>T), Y264F (1877T>A), C267N (1867A>G 1868C>T 1869A>T), S274C (1846A>G 1847G>C), H276F (1840A>G 1841T>A 1842G>A), D277G (1838T>C), F278Y (1835A>T) | | | | | | | | |
| Codon mutations: | AAT247AAC (1927A>G), TGG248TAT (1924C>A 1925C>T), GAT249GAC (1921A>G), TAT251ATC (1915A>G 1916T>A 1917A>T), GCT252ACA (1912A>T 1914C>T), TTA253TTG (1909T>C), GAT258GAC (1894A>G), ATA260ATC (1888T>G), CCT261CCG (1885A>C), CAA263ATA (1880T>A 1881G>T), TAT264TTT (1877T>A), TGT267AAC (1867A>G 1868C>T 1869A>T), AAA268AAG (1864T>C), AAA269AAG (1861T>C), TTA271CTG (1855T>C 1857A>G), CTA272TTA (1854G>A), TCT274TGC (1846A>G 1847G>C), CAT276TTC (1840A>G 1841T>A 1842G>A), GAC277GGC (1838T>C), TTT278TAT (1835A>T), GTC280GTA (1828G>T), AAG283AAA (1819C>T), TAT284TAC (1816A>G), AGG285AGA (1813C>T), AAA286AAG (1810T>C), AGA289AGG (1801T>C) | | | | | | | | |
| **Proteins** | | | | | | | | | |
| replication associated protein (YP_009465970.1) | 246 | 290 | 13.5% | 227 | 65.8% | 45 (100%) | 34 (75.6%) | 0/0/0/0 | 0 |
| Protein mutations: | W248Y (1924C>A 1925C>T), Y251I (1915A>G 1916T>A 1917A>T), A252T (1912A>T 1914C>T), Q263I (1880T>A 1881G>T), Y264F (1877T>A), C267N (1867A>G 1868C>T 1869A>T), S274C (1846A>G 1847G>C), H276F (1840A>G 1841T>A 1842G>A), D277G (1838T>C), F278Y (1835A>T) | | | | | | | | |
| Codon mutations: | AAT247AAC (1927A>G), TGG248TAT (1924C>A 1925C>T), GAT249GAC (1921A>G), TAT251ATC (1915A>G 1916T>A 1917A>T), GCT252ACA (1912A>T 1914C>T), TTA253TTG (1909T>C), GAT258GAC (1894A>G), ATA260ATC (1888T>G), CCT261CCG (1885A>C), CAA263ATA (1880T>A 1881G>T), TAT264TTT (1877T>A), TGT267AAC (1867A>G 1868C>T 1869A>T), AAA268AAG (1864T>C), AAA269AAG (1861T>C), TTA271CTG (1855T>C 1857A>G), CTA272TTA (1854G>A), TCT274TGC (1846A>G 1847G>C), CAT276TTC (1840A>G 1841T>A 1842G>A), GAC277GGC (1838T>C), TTT278TAT (1835A>T), GTC280GTA (1828G>T), AAG283AAA (1819C>T), TAT284TAC (1816A>G), AGG285AGA (1813C>T), AAA286AAG (1810T>C), AGA289AGG (1801T>C) | | | | | | | | |

*: Inserts / Deletes / Misaligned / Frameshifts

#### Analysis details

This analysis was performed with panviral2.64
